## Supplementary material for "A handle on mass coincidence errors in *de novo* sequencing of antibodies by bottom-up proteomics": Combined_077.html

Details Combined\_077 | Stitch OverviewUndefined

### Read Combined\_077

#### Sequence (length=15)

JSEVSDRPSGVSSRF

#### Spectrum 4862? Spectrum 4862 The raw spectrum of this peptide as annotated by Hecklib. The fragments are coloured according to ion type (see legend). Any peaks with a star '\*' as text can be hovered over to see the full details, first the ion type second the mass shift type. By hovering over the amino acids in the peptide or ions in the legend the corresponding peaks are highlighted. By toggling the 'Unassigned' label you can turn the background (unassigned) peaks on or off in the plot. By updating the slider in the Ion legend you can update the spectrum to only show the top X% of the peaks with labels. The top X% means any peak that is within X% of the highest intensity. By dragging in the spectrum you can zoom in to a specific part of the spectrum and use 'Zoom Out' to get back to the original zoom level. The annotation of the spectrum is based on the given sequence in the peptides file and is done with different software so inconsistencies are likely. The peaks are annotated based on the given sequence, with 20 ppm tolerance.

Copy Data

##### Spectrum 4862 (TSV)

###### Preview

```
Loading example...
```

*Click on the button to copy the data to your clipboard.*

Mz MinMz MaxIntensity Max

WidthHeightPeptide font sizePeptide stroke widthSpectrum font sizeSpectrum stroke widthCompact peptide

Ion legend

wxyz

abcd

OtherUnassignedIon^Charge^_Position_Show for top:%

JSEVSDRPSGVSSRF

04.55e+49.10e+41.37e+51.82e+5

Zoom Out

y^+1^_1_y^+1^_2_z^+1^_2_z^+3^_9_y^+1^_2_c^+1^_3_c^+3^_11_y^+1^_3_y^+3^_11_c^+2^_7_y^+1^_3_y^+2^_8_c^+1^_4_z^+1^_4_w^+1^_4_y^+3^_13_y^+1^_4_z^+1^_4_z^+2^_9_y^+1^_4_y^+2^_9_y^+3^_14_c^+1^_5_z^+2^_10_y^+2^_10_z^+1^_5_c^+2^_11_z^+1^_5_y^+2^_11_y^+2^_11_z^+2^_11_y^+1^_5_y^+2^_11_z^+1^_6_w^+2^_12_z^+1^_6_y^+2^_12_z^+2^_12_y^+2^_12_c^+1^_6_c^+2^_13_y^+1^_6_w^+2^_13_z^+2^_13_y^+2^_13_y^+2^_13_z^+2^_13_z^+1^_7_y^+2^_13_y^+1^_7_z^+1^_7_c^+2^_14_c^+2^_14_y^+1^_7_y^+2^_14_y^+2^_14_z^+2^_14_y^+2^_14_w^+1^_8_y^+1^_8_w^+1^_9_c^+1^_8_c^+1^_9_y^+1^_9_z^+1^_9_c^+1^_9_c^+1^_10_w^+1^_10_z^+1^_10_z^+1^_10_y^+1^_10_c^+1^_11_y^+1^_11_z^+1^_11_y^+1^_11_c^+1^_12_z^+1^_12_z^+1^_12_y^+1^_12_y^+1^_12_z^+1^_12_y^+1^_12_c^+1^_13_w^+1^_13_z^+1^_13_y^+1^_13_c^+1^_14_z^+1^_14_y^+1^_14_

0816163324493266

Fragment Matches Table

Show background peaks

| Position | Ion type | Intensity | mz Theoretical | mz Error (Th) | mz Error (ppm) | Charge | Series Number |
| --- | --- | --- | --- | --- | --- | --- | --- |
| - | - | 2410 | 120.1 | - | - | 0 | - |
| - | - | 477.2 | 134.7 | - | - | 0 | - |
| - | - | 417.8 | 136.4 | - | - | 0 | - |
| - | - | 474.6 | 148 | - | - | 0 | - |
| - | - | 587.2 | 148.9 | - | - | 0 | - |
| - | - | 1070 | 166.1 | - | - | 0 | - |
| 15 | y | 5126 | 166.1 | 0.0001555 | 0.9361 | +1 | 1 |
| - | - | 1.339E+04 | 173.1 | - | - | 0 | - |
| - | - | 1398 | 174.1 | - | - | 0 | - |
| - | - | 4716 | 183.1 | - | - | 0 | - |
| - | - | 545.8 | 184.1 | - | - | 0 | - |
| - | - | 647 | 191.1 | - | - | 0 | - |
| - | - | 1127 | 199.1 | - | - | 0 | - |
| - | - | 561.2 | 199.1 | - | - | 0 | - |
| - | - | 1316 | 200.1 | - | - | 0 | - |
| - | - | 584.7 | 200.6 | - | - | 0 | - |
| - | - | 2.187E+04 | 201.1 | - | - | 0 | - |
| - | - | 1.276E+04 | 201.1 | - | - | 0 | - |
| - | - | 3500 | 202.1 | - | - | 0 | - |
| - | - | 1360 | 202.1 | - | - | 0 | - |
| - | - | 2003 | 217.1 | - | - | 0 | - |
| - | - | 526.3 | 221.8 | - | - | 0 | - |
| - | - | 4025 | 223.1 | - | - | 0 | - |
| - | - | 780.8 | 229.1 | - | - | 0 | - |
| - | - | 2349 | 229.1 | - | - | 0 | - |
| - | - | 1732 | 244.1 | - | - | 0 | - |
| - | - | 1986 | 253.1 | - | - | 0 | - |
| - | - | 625.5 | 254.2 | - | - | 0 | - |
| - | - | 1529 | 261.1 | - | - | 0 | - |
| - | - | 2.664E+04 | 261.2 | - | - | 0 | - |
| - | - | 3050 | 262.2 | - | - | 0 | - |
| - | - | 1668 | 270.2 | - | - | 0 | - |
| - | - | 886.7 | 284.2 | - | - | 0 | - |
| - | - | 1128 | 287.2 | - | - | 0 | - |
| - | - | 2253 | 288.2 | - | - | 0 | - |
| - | - | 660 | 288.2 | - | - | 0 | - |
| - | - | 790.8 | 289.2 | - | - | 0 | - |
| - | - | 619.8 | 291.6 | - | - | 0 | - |
| - | - | 924.5 | 303.1 | - | - | 0 | - |
| 14 | y | 621.8 | 305.2 | 0.0009872 | 3.235 | +1 | 2 |
| 14 | z | 5304 | 306.2 | 0.0002117 | 0.6916 | +1 | 2 |
| - | - | 791.2 | 307.2 | - | - | 0 | - |
| - | - | 731.1 | 309.2 | - | - | 0 | - |
| - | - | 1813 | 310.1 | - | - | 0 | - |
| - | - | 1.106E+04 | 312.2 | - | - | 0 | - |
| - | - | 1003 | 313.2 | - | - | 0 | - |
| - | - | 583.4 | 313.5 | - | - | 0 | - |
| - | - | 1911 | 316.2 | - | - | 0 | - |
| 7 | z | 1347 | 320.2 | 0.001235 | 3.859 | +3 | 9 |
| - | - | 1035 | 321.2 | - | - | 0 | - |
| 14 | y | 6683 | 322.2 | 0.000317 | 0.984 | +1 | 2 |
| - | - | 1728 | 323.2 | - | - | 0 | - |
| - | - | 8212 | 330.2 | - | - | 0 | - |
| - | - | 1156 | 331.2 | - | - | 0 | - |
| - | - | 604.9 | 341.1 | - | - | 0 | - |
| 3 | c | 1169 | 347.2 | 0.0006652 | 1.916 | +1 | 3 |
| - | - | 3631 | 348.2 | - | - | 0 | - |
| - | - | 957.9 | 349.2 | - | - | 0 | - |
| - | - | 655.2 | 350.2 | - | - | 0 | - |
| - | - | 2135 | 350.6 | - | - | 0 | - |
| - | - | 1061 | 373.2 | - | - | 0 | - |
| 11 | c | 1.732E+04 | 376.2 | 0.001555 | 4.133 | +3 | 11 |
| - | - | 3328 | 377.2 | - | - | 0 | - |
| - | - | 1036 | 386.1 | - | - | 0 | - |
| - | - | 601.5 | 386.6 | - | - | 0 | - |
| - | - | 988.5 | 387.2 | - | - | 0 | - |
| - | - | 983.1 | 388.2 | - | - | 0 | - |
| - | - | 6538 | 391.2 | - | - | 0 | - |
| 13 | y | 2.857E+04 | 392.2 | 0.0002088 | 0.5324 | +1 | 3 |
| 5 | y | 1.188E+04 | 393.2 | 0.007093 | 18.04 | +3 | 11 |
| 7 | c | 3155 | 394.2 | 0.002929 | 7.43 | +2 | 7 |
| - | - | 908.3 | 394.7 | - | - | 0 | - |
| - | - | 2135 | 400.3 | - | - | 0 | - |
| - | - | 3720 | 401.2 | - | - | 0 | - |
| - | - | 729.5 | 403.3 | - | - | 0 | - |
| 13 | y | 6196 | 409.2 | 0.0008203 | 2.005 | +1 | 3 |
| - | - | 2050 | 410.2 | - | - | 0 | - |
| - | - | 9320 | 417.2 | - | - | 0 | - |
| - | - | 2297 | 418.2 | - | - | 0 | - |
| 8 | y | 1030 | 418.7 | 0.0008447 | 2.017 | +2 | 8 |
| - | - | 1575 | 428.2 | - | - | 0 | - |
| - | - | 691.5 | 429.2 | - | - | 0 | - |
| - | - | 5375 | 429.2 | - | - | 0 | - |
| - | - | 829 | 430.2 | - | - | 0 | - |
| - | - | 1619 | 430.2 | - | - | 0 | - |
| - | - | 1424 | 431.1 | - | - | 0 | - |
| - | - | 1036 | 431.6 | - | - | 0 | - |
| - | - | 2477 | 435.2 | - | - | 0 | - |
| - | - | 1225 | 436.2 | - | - | 0 | - |
| - | - | 814.1 | 436.2 | - | - | 0 | - |
| - | - | 4795 | 444.3 | - | - | 0 | - |
| - | - | 1490 | 445.2 | - | - | 0 | - |
| - | - | 4493 | 445.3 | - | - | 0 | - |
| 4 | c | 5339 | 446.3 | 0.0001217 | 0.2728 | +1 | 4 |
| - | - | 1686 | 447.3 | - | - | 0 | - |
| - | - | 629.4 | 450.6 | - | - | 0 | - |
| - | - | 731.3 | 451.1 | - | - | 0 | - |
| - | - | 1099 | 457.7 | - | - | 0 | - |
| - | - | 735.6 | 458.2 | - | - | 0 | - |
| - | - | 1232 | 459.7 | - | - | 0 | - |
| 12 | z | 1999 | 462.2 | 0.001193 | 2.582 | +1 | 4 |
| - | - | 780.1 | 462.9 | - | - | 0 | - |
| 12 | w | 1.976E+04 | 463.2 | 0.0007232 | 1.561 | +1 | 4 |
| - | - | 6221 | 464.2 | - | - | 0 | - |
| - | - | 1137 | 465.2 | - | - | 0 | - |
| - | - | 971.1 | 469.2 | - | - | 0 | - |
| - | - | 1578 | 472.7 | - | - | 0 | - |
| - | - | 1120 | 473.2 | - | - | 0 | - |
| - | - | 987.4 | 473.3 | - | - | 0 | - |
| - | - | 781.1 | 473.7 | - | - | 0 | - |
| - | - | 1280 | 474.2 | - | - | 0 | - |
| 3 | y | 1330 | 474.9 | 0.0006293 | 1.325 | +3 | 13 |
| - | - | 835.7 | 475.2 | - | - | 0 | - |
| - | - | 7173 | 478.2 | - | - | 0 | - |
| 12 | y | 7965 | 479.2 | 0.0005391 | 1.125 | +1 | 4 |
| - | - | 1490 | 480.2 | - | - | 0 | - |
| 12 | z | 1.42E+04 | 480.2 | 0.0003029 | 0.6307 | +1 | 4 |
| - | - | 3470 | 481.2 | - | - | 0 | - |
| - | - | 1011 | 481.7 | - | - | 0 | - |
| - | - | 894.9 | 486.2 | - | - | 0 | - |
| - | - | 941.6 | 486.7 | - | - | 0 | - |
| - | - | 841.8 | 487.7 | - | - | 0 | - |
| 7 | z | 1.423E+04 | 488.8 | 0.0008499 | 1.739 | +2 | 9 |
| - | - | 1596 | 489.2 | - | - | 0 | - |
| - | - | 7446 | 489.3 | - | - | 0 | - |
| - | - | 2120 | 489.7 | - | - | 0 | - |
| - | - | 2461 | 489.8 | - | - | 0 | - |
| - | - | 1008 | 490.2 | - | - | 0 | - |
| - | - | 2394 | 490.3 | - | - | 0 | - |
| - | - | 768.7 | 494.2 | - | - | 0 | - |
| - | - | 2898 | 495.2 | - | - | 0 | - |
| - | - | 1568 | 495.7 | - | - | 0 | - |
| - | - | 780.6 | 496.2 | - | - | 0 | - |
| 12 | y | 6177 | 496.3 | 0.0008659 | 1.745 | +1 | 4 |
| - | - | 4039 | 496.7 | - | - | 0 | - |
| 7 | y | 3676 | 496.8 | 0.0002464 | 0.4961 | +2 | 9 |
| - | - | 3196 | 497.3 | - | - | 0 | - |
| - | - | 1181 | 498.3 | - | - | 0 | - |
| - | - | 1806 | 499.3 | - | - | 0 | - |
| - | - | 985.6 | 503.7 | - | - | 0 | - |
| 2 | y | 1007 | 503.9 | 0.0005733 | 1.138 | +3 | 14 |
| - | - | 7420 | 504.2 | - | - | 0 | - |
| - | - | 843.9 | 504.3 | - | - | 0 | - |
| - | - | 1639 | 505.2 | - | - | 0 | - |
| - | - | 2139 | 509.7 | - | - | 0 | - |
| - | - | 1362 | 510.2 | - | - | 0 | - |
| - | - | 633.4 | 511.2 | - | - | 0 | - |
| - | - | 772.9 | 514.3 | - | - | 0 | - |
| - | - | 2925 | 516.3 | - | - | 0 | - |
| - | - | 1016 | 516.7 | - | - | 0 | - |
| - | - | 1021 | 518.2 | - | - | 0 | - |
| - | - | 1270 | 518.7 | - | - | 0 | - |
| - | - | 824.9 | 519.2 | - | - | 0 | - |
| - | - | 5605 | 524.3 | - | - | 0 | - |
| - | - | 3727 | 524.8 | - | - | 0 | - |
| - | - | 1303 | 525.3 | - | - | 0 | - |
| - | - | 906.2 | 531.3 | - | - | 0 | - |
| - | - | 4654 | 531.7 | - | - | 0 | - |
| - | - | 2074 | 532.2 | - | - | 0 | - |
| - | - | 9868 | 532.3 | - | - | 0 | - |
| - | - | 1406 | 532.7 | - | - | 0 | - |
| - | - | 768.9 | 532.8 | - | - | 0 | - |
| - | - | 1872 | 533.2 | - | - | 0 | - |
| 5 | c | 4.06E+04 | 533.3 | 0.0001985 | 0.3722 | +1 | 5 |
| - | - | 3333 | 533.7 | - | - | 0 | - |
| - | - | 1.091E+04 | 534.3 | - | - | 0 | - |
| - | - | 1944 | 535.3 | - | - | 0 | - |
| - | - | 4939 | 535.6 | - | - | 0 | - |
| - | - | 3810 | 535.9 | - | - | 0 | - |
| - | - | 1235 | 536.3 | - | - | 0 | - |
| - | - | 2384 | 540.3 | - | - | 0 | - |
| - | - | 3392 | 540.7 | - | - | 0 | - |
| - | - | 827.3 | 540.8 | - | - | 0 | - |
| - | - | 8085 | 541.2 | - | - | 0 | - |
| - | - | 4271 | 541.3 | - | - | 0 | - |
| - | - | 1.902E+04 | 541.6 | - | - | 0 | - |
| - | - | 2675 | 541.7 | - | - | 0 | - |
| - | - | 1.924E+04 | 541.9 | - | - | 0 | - |
| - | - | 1.614E+04 | 542.2 | - | - | 0 | - |
| - | - | 9874 | 542.3 | - | - | 0 | - |
| - | - | 1157 | 542.4 | - | - | 0 | - |
| - | - | 2843 | 542.6 | - | - | 0 | - |
| - | - | 1.949E+04 | 542.7 | - | - | 0 | - |
| - | - | 868 | 542.9 | - | - | 0 | - |
| - | - | 1467 | 543.2 | - | - | 0 | - |
| 6 | z | 2.325E+04 | 546.3 | 0.0004399 | 0.8053 | +2 | 10 |
| - | - | 2.466E+04 | 546.8 | - | - | 0 | - |
| - | - | 1.094E+04 | 547.3 | - | - | 0 | - |
| - | - | 3789 | 547.8 | - | - | 0 | - |
| - | - | 1335 | 549.3 | - | - | 0 | - |
| - | - | 1644 | 553.8 | - | - | 0 | - |
| 6 | y | 1.184E+04 | 554.3 | 0.0003552 | 0.6409 | +2 | 10 |
| - | - | 8144 | 554.8 | - | - | 0 | - |
| - | - | 2695 | 555.3 | - | - | 0 | - |
| - | - | 1269 | 556.3 | - | - | 0 | - |
| - | - | 1015 | 560.3 | - | - | 0 | - |
| 11 | z | 3491 | 561.3 | 0.0005896 | 1.05 | +1 | 5 |
| - | - | 2447 | 562.3 | - | - | 0 | - |
| - | - | 865.1 | 563.3 | - | - | 0 | - |
| 11 | c | 1.627E+04 | 564.3 | 0.009577 | 16.97 | +2 | 11 |
| - | - | 4460 | 565.3 | - | - | 0 | - |
| - | - | 1492 | 566.3 | - | - | 0 | - |
| - | - | 861.8 | 566.8 | - | - | 0 | - |
| - | - | 2096 | 567.8 | - | - | 0 | - |
| - | - | 989 | 568.3 | - | - | 0 | - |
| 11 | z | 8398 | 579.3 | 0.0007671 | 1.324 | +1 | 5 |
| - | - | 2868 | 580.3 | - | - | 0 | - |
| - | - | 710.5 | 581.8 | - | - | 0 | - |
| - | - | 1443 | 582.3 | - | - | 0 | - |
| - | - | 846.9 | 588.3 | - | - | 0 | - |
| 5 | y | 4919 | 588.8 | 0.009705 | 16.48 | +2 | 11 |
| 5 | y | 7299 | 589.3 | 4.049E-06 | 0.006871 | +2 | 11 |
| 5 | z | 2.363E+04 | 589.8 | 0.0001728 | 0.293 | +2 | 11 |
| - | - | 3.915E+04 | 590.3 | - | - | 0 | - |
| - | - | 2.096E+04 | 590.8 | - | - | 0 | - |
| - | - | 5952 | 591.3 | - | - | 0 | - |
| - | - | 2162 | 591.8 | - | - | 0 | - |
| - | - | 1.169E+04 | 592.3 | - | - | 0 | - |
| - | - | 4510 | 593.3 | - | - | 0 | - |
| 11 | y | 9535 | 595.3 | 0.000964 | 1.619 | +1 | 5 |
| - | - | 3271 | 596.3 | - | - | 0 | - |
| - | - | 6296 | 597.3 | - | - | 0 | - |
| 5 | y | 6.048E+04 | 597.8 | 0.0003933 | 0.6579 | +2 | 11 |
| - | - | 3.678E+04 | 598.3 | - | - | 0 | - |
| - | - | 1.533E+04 | 598.8 | - | - | 0 | - |
| - | - | 3252 | 599.3 | - | - | 0 | - |
| - | - | 1093 | 599.8 | - | - | 0 | - |
| - | - | 3628 | 601.8 | - | - | 0 | - |
| - | - | 2184 | 602.3 | - | - | 0 | - |
| - | - | 1181 | 611.3 | - | - | 0 | - |
| 10 | z | 674 | 618.3 | 0.003723 | 6.022 | +1 | 6 |
| - | - | 1652 | 619.3 | - | - | 0 | - |
| - | - | 990.5 | 620.3 | - | - | 0 | - |
| - | - | 1344 | 621.3 | - | - | 0 | - |
| - | - | 943.5 | 626.3 | - | - | 0 | - |
| - | - | 768 | 631.3 | - | - | 0 | - |
| 4 | w | 2022 | 631.8 | 0.001141 | 1.806 | +2 | 12 |
| - | - | 1512 | 632.3 | - | - | 0 | - |
| - | - | 930.6 | 632.8 | - | - | 0 | - |
| 10 | z | 3.791E+04 | 636.3 | 0.0007267 | 1.142 | +1 | 6 |
| - | - | 2.274E+04 | 637.3 | - | - | 0 | - |
| 4 | y | 6512 | 638.3 | 0.004428 | 6.936 | +2 | 12 |
| 4 | z | 3.919E+04 | 639.3 | 0.001061 | 1.66 | +2 | 12 |
| - | - | 2.743E+04 | 639.8 | - | - | 0 | - |
| - | - | 1.05E+04 | 640.3 | - | - | 0 | - |
| - | - | 3708 | 640.8 | - | - | 0 | - |
| - | - | 924.2 | 641.3 | - | - | 0 | - |
| - | - | 1684 | 646.8 | - | - | 0 | - |
| 4 | y | 5.838E+04 | 647.3 | 0.0007322 | 1.131 | +2 | 12 |
| - | - | 4.223E+04 | 647.8 | - | - | 0 | - |
| - | - | 1326 | 647.9 | - | - | 0 | - |
| 6 | c | 4.302E+04 | 648.3 | 0.004505 | 6.949 | +1 | 6 |
| - | - | 3153 | 648.8 | - | - | 0 | - |
| - | - | 1.098E+04 | 649.3 | - | - | 0 | - |
| - | - | 2576 | 650.3 | - | - | 0 | - |
| 13 | c | 1103 | 651.3 | 0.01101 | 16.9 | +2 | 13 |
| 10 | y | 3.928E+04 | 652.3 | 0.0007405 | 1.135 | +1 | 6 |
| - | - | 1.554E+04 | 653.3 | - | - | 0 | - |
| - | - | 2930 | 654.3 | - | - | 0 | - |
| 3 | w | 3.021E+04 | 674.3 | 0.000882 | 1.308 | +2 | 13 |
| - | - | 2.261E+04 | 674.8 | - | - | 0 | - |
| - | - | 1.073E+04 | 675.3 | - | - | 0 | - |
| - | - | 3206 | 675.8 | - | - | 0 | - |
| - | - | 1070 | 676.3 | - | - | 0 | - |
| - | - | 4665 | 679.4 | - | - | 0 | - |
| - | - | 1910 | 680.4 | - | - | 0 | - |
| - | - | 2097 | 681.8 | - | - | 0 | - |
| - | - | 888.5 | 682.8 | - | - | 0 | - |
| - | - | 765.8 | 688.4 | - | - | 0 | - |
| 3 | z | 923.5 | 694.8 | 0.001465 | 2.109 | +2 | 13 |
| - | - | 1399 | 696.3 | - | - | 0 | - |
| - | - | 1099 | 700.2 | - | - | 0 | - |
| 3 | y | 2742 | 702.8 | 0.0007556 | 1.075 | +2 | 13 |
| 3 | y | 2958 | 703.3 | 0.009922 | 14.11 | +2 | 13 |
| 3 | z | 2.405E+04 | 703.8 | 0.0009437 | 1.341 | +2 | 13 |
| - | - | 2.135E+04 | 704.3 | - | - | 0 | - |
| - | - | 1206 | 704.4 | - | - | 0 | - |
| - | - | 9625 | 704.8 | - | - | 0 | - |
| 9 | z | 1886 | 705.3 | 0.001907 | 2.704 | +1 | 7 |
| - | - | 2504 | 707.9 | - | - | 0 | - |
| - | - | 1234 | 708.4 | - | - | 0 | - |
| - | - | 803.6 | 708.9 | - | - | 0 | - |
| - | - | 3851 | 711.3 | - | - | 0 | - |
| 3 | y | 7.404E+04 | 711.9 | 0.000859 | 1.207 | +2 | 13 |
| - | - | 5.827E+04 | 712.4 | - | - | 0 | - |
| - | - | 2.497E+04 | 712.9 | - | - | 0 | - |
| - | - | 5469 | 713.4 | - | - | 0 | - |
| - | - | 907 | 713.9 | - | - | 0 | - |
| - | - | 6960 | 715.9 | - | - | 0 | - |
| - | - | 5373 | 716.4 | - | - | 0 | - |
| - | - | 1716 | 716.9 | - | - | 0 | - |
| - | - | 5006 | 717.9 | - | - | 0 | - |
| - | - | 3405 | 718.4 | - | - | 0 | - |
| - | - | 1079 | 718.9 | - | - | 0 | - |
| - | - | 1352 | 721.3 | - | - | 0 | - |
| 9 | y | 3849 | 722.3 | 0.001365 | 1.889 | +1 | 7 |
| 9 | z | 2.305E+04 | 723.4 | 0.0005587 | 0.7724 | +1 | 7 |
| - | - | 1.076E+04 | 724.4 | - | - | 0 | - |
| - | - | 6377 | 725.4 | - | - | 0 | - |
| - | - | 1611 | 725.9 | - | - | 0 | - |
| - | - | 789.1 | 726.4 | - | - | 0 | - |
| 14 | c | 1463 | 729.4 | 0.006773 | 9.286 | +2 | 14 |
| - | - | 1893 | 730.4 | - | - | 0 | - |
| - | - | 953.6 | 730.9 | - | - | 0 | - |
| - | - | 704.5 | 731.4 | - | - | 0 | - |
| - | - | 764.8 | 732.3 | - | - | 0 | - |
| 14 | c | 1.711E+05 | 737.9 | 0.0009489 | 1.286 | +2 | 14 |
| - | - | 1.384E+05 | 738.4 | - | - | 0 | - |
| - | - | 5.784E+04 | 738.9 | - | - | 0 | - |
| 9 | y | 2.232E+04 | 739.4 | 0.009728 | 13.16 | +1 | 7 |
| - | - | 4569 | 739.9 | - | - | 0 | - |
| - | - | 3543 | 740.4 | - | - | 0 | - |
| - | - | 1283 | 745.4 | - | - | 0 | - |
| 2 | y | 8510 | 746.4 | 0.001206 | 1.616 | +2 | 14 |
| 2 | y | 5877 | 746.9 | 0.006359 | 8.515 | +2 | 14 |
| 2 | z | 3.157E+04 | 747.4 | 0.001216 | 1.626 | +2 | 14 |
| - | - | 2.465E+04 | 747.9 | - | - | 0 | - |
| - | - | 1.165E+04 | 748.4 | - | - | 0 | - |
| - | - | 4520 | 748.9 | - | - | 0 | - |
| - | - | 1054 | 754.9 | - | - | 0 | - |
| 2 | y | 9.965E+04 | 755.4 | 0.000836 | 1.107 | +2 | 14 |
| - | - | 7.826E+04 | 755.9 | - | - | 0 | - |
| - | - | 3.718E+04 | 756.4 | - | - | 0 | - |
| - | - | 1.136E+04 | 756.9 | - | - | 0 | - |
| - | - | 2554 | 757.4 | - | - | 0 | - |
| - | - | 1180 | 759.4 | - | - | 0 | - |
| - | - | 1960 | 760.4 | - | - | 0 | - |
| - | - | 3352 | 760.4 | - | - | 0 | - |
| - | - | 3029 | 760.9 | - | - | 0 | - |
| - | - | 1447 | 761.4 | - | - | 0 | - |
| - | - | 1135 | 761.4 | - | - | 0 | - |
| - | - | 1792 | 761.9 | - | - | 0 | - |
| - | - | 864.6 | 767.9 | - | - | 0 | - |
| - | - | 1929 | 771.2 | - | - | 0 | - |
| - | - | 1442 | 772.2 | - | - | 0 | - |
| - | - | 1833 | 773.4 | - | - | 0 | - |
| - | - | 1251 | 773.9 | - | - | 0 | - |
| - | - | 4169 | 774.4 | - | - | 0 | - |
| - | - | 1634 | 774.9 | - | - | 0 | - |
| - | - | 2140 | 775.4 | - | - | 0 | - |
| - | - | 2514 | 775.9 | - | - | 0 | - |
| - | - | 883.2 | 776.4 | - | - | 0 | - |
| - | - | 1685 | 781.4 | - | - | 0 | - |
| - | - | 1.63E+04 | 781.9 | - | - | 0 | - |
| - | - | 3.235E+04 | 782.4 | - | - | 0 | - |
| - | - | 2.207E+04 | 782.9 | - | - | 0 | - |
| - | - | 1.305E+04 | 783.4 | - | - | 0 | - |
| - | - | 3808 | 783.9 | - | - | 0 | - |
| - | - | 3664 | 787.9 | - | - | 0 | - |
| - | - | 2842 | 788.4 | - | - | 0 | - |
| - | - | 3086 | 788.9 | - | - | 0 | - |
| - | - | 4.987E+04 | 789.4 | - | - | 0 | - |
| - | - | 3.778E+04 | 789.9 | - | - | 0 | - |
| - | - | 2.329E+04 | 790.4 | - | - | 0 | - |
| - | - | 4.608E+04 | 790.9 | - | - | 0 | - |
| - | - | 3.701E+04 | 791.4 | - | - | 0 | - |
| - | - | 1.625E+04 | 791.9 | - | - | 0 | - |
| - | - | 4936 | 792.4 | - | - | 0 | - |
| - | - | 1642 | 792.9 | - | - | 0 | - |
| 8 | w | 995.1 | 793.4 | 0.005633 | 7.1 | +1 | 8 |
| - | - | 5242 | 794.9 | - | - | 0 | - |
| - | - | 3759 | 795.4 | - | - | 0 | - |
| - | - | 5540 | 795.9 | - | - | 0 | - |
| - | - | 4566 | 796.4 | - | - | 0 | - |
| - | - | 5110 | 796.9 | - | - | 0 | - |
| - | - | 2396 | 797.4 | - | - | 0 | - |
| - | - | 899 | 797.9 | - | - | 0 | - |
| - | - | 2.088E+04 | 803.4 | - | - | 0 | - |
| - | - | 1.802E+05 | 803.9 | - | - | 0 | - |
| - | - | 1.441E+05 | 804.4 | - | - | 0 | - |
| - | - | 7.03E+04 | 804.9 | - | - | 0 | - |
| - | - | 2.303E+04 | 805.4 | - | - | 0 | - |
| - | - | 5035 | 805.9 | - | - | 0 | - |
| - | - | 1753 | 810.4 | - | - | 0 | - |
| - | - | 8.866E+04 | 811.9 | - | - | 0 | - |
| - | - | 1.24E+05 | 812.4 | - | - | 0 | - |
| - | - | 7.606E+04 | 812.9 | - | - | 0 | - |
| - | - | 3.364E+04 | 813.4 | - | - | 0 | - |
| - | - | 9674 | 813.9 | - | - | 0 | - |
| - | - | 1505 | 814.4 | - | - | 0 | - |
| - | - | 1634 | 829.4 | - | - | 0 | - |
| - | - | 1073 | 834.4 | - | - | 0 | - |
| - | - | 1124 | 835.4 | - | - | 0 | - |
| 8 | y | 5160 | 836.4 | 0.002069 | 2.473 | +1 | 8 |
| - | - | 2180 | 837.4 | - | - | 0 | - |
| - | - | 934.4 | 839.5 | - | - | 0 | - |
| - | - | 1205 | 840.5 | - | - | 0 | - |
| - | - | 2107 | 842.5 | - | - | 0 | - |
| - | - | 1377 | 843.5 | - | - | 0 | - |
| - | - | 1119 | 846.5 | - | - | 0 | - |
| - | - | 862.3 | 847.5 | - | - | 0 | - |
| - | - | 1.069E+04 | 857.5 | - | - | 0 | - |
| - | - | 1410 | 858.3 | - | - | 0 | - |
| - | - | 4026 | 858.5 | - | - | 0 | - |
| - | - | 2241 | 859.5 | - | - | 0 | - |
| - | - | 1143 | 865.4 | - | - | 0 | - |
| - | - | 1415 | 885.5 | - | - | 0 | - |
| - | - | 1646 | 886.5 | - | - | 0 | - |
| - | - | 892.7 | 887.5 | - | - | 0 | - |
| 7 | w | 2082 | 890.4 | 0.003735 | 4.195 | +1 | 9 |
| - | - | 1430 | 891.4 | - | - | 0 | - |
| - | - | 3551 | 900.5 | - | - | 0 | - |
| 8 | c | 4.57E+04 | 901.5 | 0.001448 | 1.607 | +1 | 8 |
| - | - | 2.184E+04 | 902.5 | - | - | 0 | - |
| - | - | 8499 | 903.5 | - | - | 0 | - |
| - | - | 1807 | 904.5 | - | - | 0 | - |
| - | - | 1315 | 916.5 | - | - | 0 | - |
| - | - | 1096 | 926.5 | - | - | 0 | - |
| - | - | 1925 | 927.5 | - | - | 0 | - |
| - | - | 975 | 928.5 | - | - | 0 | - |
| - | - | 2191 | 933.5 | - | - | 0 | - |
| - | - | 817.4 | 934.5 | - | - | 0 | - |
| - | - | 2.632E+04 | 944.5 | - | - | 0 | - |
| - | - | 1.342E+04 | 945.5 | - | - | 0 | - |
| - | - | 4696 | 946.5 | - | - | 0 | - |
| - | - | 918.1 | 947.5 | - | - | 0 | - |
| 9 | c | 794.3 | 971.5 | 0.001467 | 1.51 | +1 | 9 |
| - | - | 1601 | 972.5 | - | - | 0 | - |
| 7 | y | 815.4 | 975.5 | 0.002336 | 2.395 | +1 | 9 |
| 7 | z | 2.534E+04 | 976.5 | 0.0006755 | 0.6918 | +1 | 9 |
| - | - | 1.646E+04 | 977.5 | - | - | 0 | - |
| - | - | 6521 | 978.5 | - | - | 0 | - |
| - | - | 1688 | 979.5 | - | - | 0 | - |
| - | - | 6449 | 987.5 | - | - | 0 | - |
| 9 | c | 1.135E+05 | 988.5 | 0.001097 | 1.11 | +1 | 9 |
| - | - | 6.082E+04 | 989.5 | - | - | 0 | - |
| - | - | 706.7 | 990.4 | - | - | 0 | - |
| - | - | 1.848E+04 | 990.5 | - | - | 0 | - |
| - | - | 3503 | 991.5 | - | - | 0 | - |
| - | - | 3041 | 992.4 | - | - | 0 | - |
| - | - | 1800 | 993.4 | - | - | 0 | - |
| - | - | 760.6 | 995.5 | - | - | 0 | - |
| - | - | 6150 | 1002 | - | - | 0 | - |
| - | - | 4062 | 1003 | - | - | 0 | - |
| - | - | 1335 | 1004 | - | - | 0 | - |
| - | - | 990.5 | 1021 | - | - | 0 | - |
| - | - | 3400 | 1022 | - | - | 0 | - |
| - | - | 8509 | 1023 | - | - | 0 | - |
| - | - | 3432 | 1024 | - | - | 0 | - |
| - | - | 9192 | 1025 | - | - | 0 | - |
| - | - | 3.117E+04 | 1026 | - | - | 0 | - |
| - | - | 2670 | 1027 | - | - | 0 | - |
| - | - | 1501 | 1032 | - | - | 0 | - |
| 10 | c | 6.357E+04 | 1046 | 0.001545 | 1.478 | +1 | 10 |
| 6 | w | 3.485E+04 | 1047 | 0.00603 | 5.762 | +1 | 10 |
| - | - | 1.722E+04 | 1048 | - | - | 0 | - |
| - | - | 6921 | 1049 | - | - | 0 | - |
| - | - | 1955 | 1050 | - | - | 0 | - |
| - | - | 1057 | 1053 | - | - | 0 | - |
| - | - | 929.3 | 1060 | - | - | 0 | - |
| - | - | 3353 | 1064 | - | - | 0 | - |
| - | - | 1977 | 1065 | - | - | 0 | - |
| - | - | 3809 | 1066 | - | - | 0 | - |
| - | - | 2316 | 1067 | - | - | 0 | - |
| 6 | z | 932.1 | 1074 | 0.002519 | 2.347 | +1 | 10 |
| - | - | 5305 | 1080 | - | - | 0 | - |
| - | - | 936.2 | 1081 | - | - | 0 | - |
| - | - | 8.554E+04 | 1081 | - | - | 0 | - |
| - | - | 5.517E+04 | 1082 | - | - | 0 | - |
| - | - | 2.185E+04 | 1083 | - | - | 0 | - |
| - | - | 970.3 | 1084 | - | - | 0 | - |
| - | - | 1641 | 1084 | - | - | 0 | - |
| - | - | 3.012E+04 | 1084 | - | - | 0 | - |
| - | - | 1.136E+04 | 1085 | - | - | 0 | - |
| - | - | 1335 | 1086 | - | - | 0 | - |
| - | - | 2159 | 1086 | - | - | 0 | - |
| 6 | z | 2.235E+04 | 1092 | 0.0009542 | 0.8741 | +1 | 10 |
| - | - | 3.652E+04 | 1093 | - | - | 0 | - |
| - | - | 1.714E+04 | 1094 | - | - | 0 | - |
| - | - | 5460 | 1095 | - | - | 0 | - |
| - | - | 1087 | 1096 | - | - | 0 | - |
| - | - | 3519 | 1101 | - | - | 0 | - |
| - | - | 3163 | 1102 | - | - | 0 | - |
| - | - | 1498 | 1103 | - | - | 0 | - |
| - | - | 1915 | 1107 | - | - | 0 | - |
| 6 | y | 6298 | 1108 | 0.001412 | 1.275 | +1 | 10 |
| - | - | 3536 | 1109 | - | - | 0 | - |
| - | - | 1355 | 1110 | - | - | 0 | - |
| - | - | 837.8 | 1111 | - | - | 0 | - |
| - | - | 5124 | 1130 | - | - | 0 | - |
| - | - | 3264 | 1131 | - | - | 0 | - |
| - | - | 997.2 | 1132 | - | - | 0 | - |
| - | - | 1982 | 1144 | - | - | 0 | - |
| 11 | c | 5.317E+04 | 1145 | 0.0007582 | 0.6624 | +1 | 11 |
| - | - | 2.978E+04 | 1146 | - | - | 0 | - |
| - | - | 1.131E+04 | 1147 | - | - | 0 | - |
| - | - | 3205 | 1148 | - | - | 0 | - |
| - | - | 837.6 | 1149 | - | - | 0 | - |
| - | - | 1272 | 1173 | - | - | 0 | - |
| 5 | y | 983.1 | 1178 | 0.007136 | 6.06 | +1 | 11 |
| 5 | z | 4660 | 1179 | 6.84E-05 | 0.05803 | +1 | 11 |
| - | - | 1.787E+04 | 1180 | - | - | 0 | - |
| - | - | 1.053E+04 | 1181 | - | - | 0 | - |
| - | - | 4483 | 1182 | - | - | 0 | - |
| - | - | 1361 | 1183 | - | - | 0 | - |
| - | - | 923.9 | 1184 | - | - | 0 | - |
| - | - | 5304 | 1188 | - | - | 0 | - |
| - | - | 5052 | 1189 | - | - | 0 | - |
| - | - | 1865 | 1190 | - | - | 0 | - |
| - | - | 1160 | 1191 | - | - | 0 | - |
| - | - | 1125 | 1194 | - | - | 0 | - |
| 5 | y | 7421 | 1195 | 0.0003598 | 0.3012 | +1 | 11 |
| - | - | 4999 | 1196 | - | - | 0 | - |
| - | - | 2655 | 1197 | - | - | 0 | - |
| - | - | 827.2 | 1198 | - | - | 0 | - |
| - | - | 3463 | 1231 | - | - | 0 | - |
| 12 | c | 9.463E+04 | 1232 | 2.02E-05 | 0.0164 | +1 | 12 |
| - | - | 5.457E+04 | 1233 | - | - | 0 | - |
| - | - | 2.475E+04 | 1234 | - | - | 0 | - |
| - | - | 5651 | 1235 | - | - | 0 | - |
| - | - | 884.7 | 1236 | - | - | 0 | - |
| - | - | 3219 | 1259 | - | - | 0 | - |
| 4 | z | 2590 | 1260 | 0.005559 | 4.413 | +1 | 12 |
| 4 | z | 924.5 | 1261 | 0.01483 | 11.76 | +1 | 12 |
| - | - | 1254 | 1262 | - | - | 0 | - |
| - | - | 3993 | 1275 | - | - | 0 | - |
| 4 | y | 3618 | 1276 | 0.006244 | 4.895 | +1 | 12 |
| 4 | y | 1618 | 1277 | 0.02406 | 18.85 | +1 | 12 |
| 4 | z | 2969 | 1278 | 0.002196 | 1.719 | +1 | 12 |
| - | - | 2.204E+04 | 1279 | - | - | 0 | - |
| - | - | 1.356E+04 | 1280 | - | - | 0 | - |
| - | - | 6660 | 1281 | - | - | 0 | - |
| - | - | 1173 | 1282 | - | - | 0 | - |
| 4 | y | 6016 | 1294 | 0.0006842 | 0.5289 | +1 | 12 |
| - | - | 3875 | 1295 | - | - | 0 | - |
| - | - | 2807 | 1296 | - | - | 0 | - |
| 13 | c | 6.638E+04 | 1319 | 5.589E-05 | 0.04238 | +1 | 13 |
| - | - | 4.319E+04 | 1320 | - | - | 0 | - |
| - | - | 1.641E+04 | 1321 | - | - | 0 | - |
| - | - | 6070 | 1322 | - | - | 0 | - |
| - | - | 926.1 | 1323 | - | - | 0 | - |
| - | - | 1717 | 1345 | - | - | 0 | - |
| - | - | 6567 | 1346 | - | - | 0 | - |
| - | - | 4544 | 1347 | - | - | 0 | - |
| 3 | w | 1606 | 1348 | 0.002434 | 1.806 | +1 | 13 |
| - | - | 1562 | 1365 | - | - | 0 | - |
| - | - | 1053 | 1366 | - | - | 0 | - |
| - | - | 1518 | 1395 | - | - | 0 | - |
| - | - | 874.7 | 1396 | - | - | 0 | - |
| 3 | z | 4468 | 1407 | 0.000968 | 0.6881 | +1 | 13 |
| - | - | 3.436E+04 | 1408 | - | - | 0 | - |
| - | - | 2.793E+04 | 1409 | - | - | 0 | - |
| - | - | 1.182E+04 | 1410 | - | - | 0 | - |
| - | - | 3340 | 1411 | - | - | 0 | - |
| - | - | 4125 | 1417 | - | - | 0 | - |
| - | - | 2274 | 1418 | - | - | 0 | - |
| 3 | y | 1428 | 1423 | 0.01327 | 9.325 | +1 | 13 |
| - | - | 2033 | 1424 | - | - | 0 | - |
| - | - | 1200 | 1425 | - | - | 0 | - |
| - | - | 1384 | 1426 | - | - | 0 | - |
| - | - | 3517 | 1431 | - | - | 0 | - |
| - | - | 7797 | 1432 | - | - | 0 | - |
| - | - | 7049 | 1433 | - | - | 0 | - |
| - | - | 4144 | 1434 | - | - | 0 | - |
| - | - | 1063 | 1435 | - | - | 0 | - |
| - | - | 1747 | 1436 | - | - | 0 | - |
| - | - | 1169 | 1448 | - | - | 0 | - |
| - | - | 1347 | 1452 | - | - | 0 | - |
| - | - | 895.3 | 1453 | - | - | 0 | - |
| - | - | 7329 | 1459 | - | - | 0 | - |
| - | - | 5287 | 1460 | - | - | 0 | - |
| - | - | 2703 | 1461 | - | - | 0 | - |
| - | - | 1292 | 1462 | - | - | 0 | - |
| - | - | 933.5 | 1467 | - | - | 0 | - |
| - | - | 1000 | 1469 | - | - | 0 | - |
| 14 | c | 1.475E+04 | 1475 | 0.0008354 | 0.5665 | +1 | 14 |
| - | - | 4.53E+04 | 1476 | - | - | 0 | - |
| - | - | 3.162E+04 | 1477 | - | - | 0 | - |
| - | - | 1.292E+04 | 1478 | - | - | 0 | - |
| - | - | 3530 | 1479 | - | - | 0 | - |
| - | - | 1362 | 1480 | - | - | 0 | - |
| 2 | z | 3558 | 1494 | 0.002282 | 1.528 | +1 | 14 |
| - | - | 2.55E+04 | 1495 | - | - | 0 | - |
| - | - | 1.889E+04 | 1496 | - | - | 0 | - |
| - | - | 7708 | 1497 | - | - | 0 | - |
| - | - | 2842 | 1498 | - | - | 0 | - |
| - | - | 1819 | 1508 | - | - | 0 | - |
| - | - | 1661 | 1509 | - | - | 0 | - |
| 2 | y | 2371 | 1510 | 0.01847 | 12.23 | +1 | 14 |
| - | - | 1372 | 1511 | - | - | 0 | - |
| - | - | 1063 | 1521 | - | - | 0 | - |
| - | - | 1378 | 1522 | - | - | 0 | - |
| - | - | 7360 | 1523 | - | - | 0 | - |
| - | - | 7205 | 1524 | - | - | 0 | - |
| - | - | 3303 | 1525 | - | - | 0 | - |
| - | - | 1647 | 1526 | - | - | 0 | - |
| - | - | 1113 | 1536 | - | - | 0 | - |
| - | - | 1064 | 1537 | - | - | 0 | - |
| - | - | 1261 | 1538 | - | - | 0 | - |
| - | - | 2960 | 1548 | - | - | 0 | - |
| - | - | 4048 | 1549 | - | - | 0 | - |
| - | - | 1582 | 1550 | - | - | 0 | - |
| - | - | 1120 | 1551 | - | - | 0 | - |
| - | - | 2129 | 1564 | - | - | 0 | - |
| - | - | 3.671E+04 | 1565 | - | - | 0 | - |
| - | - | 5.063E+04 | 1566 | - | - | 0 | - |
| - | - | 2.902E+04 | 1567 | - | - | 0 | - |
| - | - | 1.206E+04 | 1568 | - | - | 0 | - |
| - | - | 3628 | 1569 | - | - | 0 | - |
| - | - | 4598 | 1579 | - | - | 0 | - |
| - | - | 8734 | 1580 | - | - | 0 | - |
| - | - | 1.015E+04 | 1581 | - | - | 0 | - |
| - | - | 1.253E+04 | 1582 | - | - | 0 | - |
| - | - | 7857 | 1583 | - | - | 0 | - |
| - | - | 3745 | 1584 | - | - | 0 | - |
| - | - | 1395 | 1585 | - | - | 0 | - |
| - | - | 2493 | 1590 | - | - | 0 | - |
| - | - | 1.188E+04 | 1591 | - | - | 0 | - |
| - | - | 1.024E+04 | 1592 | - | - | 0 | - |
| - | - | 4915 | 1593 | - | - | 0 | - |
| - | - | 1339 | 1594 | - | - | 0 | - |
| - | - | 3196 | 1596 | - | - | 0 | - |
| - | - | 1.254E+04 | 1597 | - | - | 0 | - |
| - | - | 9776 | 1598 | - | - | 0 | - |
| - | - | 4823 | 1599 | - | - | 0 | - |
| - | - | 1891 | 1600 | - | - | 0 | - |
| - | - | 1084 | 1606 | - | - | 0 | - |
| - | - | 2.785E+04 | 1607 | - | - | 0 | - |
| - | - | 1.255E+05 | 1608 | - | - | 0 | - |
| - | - | 9.651E+04 | 1609 | - | - | 0 | - |
| - | - | 4.868E+04 | 1610 | - | - | 0 | - |
| - | - | 1.691E+04 | 1611 | - | - | 0 | - |
| - | - | 3495 | 1612 | - | - | 0 | - |
| - | - | 6248 | 1623 | - | - | 0 | - |
| - | - | 3.136E+04 | 1624 | - | - | 0 | - |
| - | - | 1.252E+05 | 1625 | - | - | 0 | - |
| - | - | 9.546E+04 | 1626 | - | - | 0 | - |
| - | - | 4.829E+04 | 1627 | - | - | 0 | - |
| - | - | 1.436E+04 | 1628 | - | - | 0 | - |
| - | - | 3872 | 1629 | - | - | 0 | - |
| - | - | 927.2 | 2329 | - | - | 0 | - |
| - | - | 772.6 | 2341 | - | - | 0 | - |
| - | - | 694.4 | 2817 | - | - | 0 | - |
| - | - | 695.4 | 3233 | - | - | 0 | - |

m/z Charge Intensity FragmentType MassShift Position
120.08098602294922 0 2409.7646
134.6780548095703 0 477.16672
136.37803649902344 0 417.76437
148.0227813720703 0 474.5696
148.9468536376953 0 587.2074
166.0758056640625 0 1070.053
166.08641052246094 0 5125.613 y 14
173.12867736816406 0 13387.549
174.13243103027344 0 1397.5652
183.11300659179688 0 4715.552
184.1158447265625 0 545.77295
191.06893920898438 0 646.9546
199.07113647460938 0 1126.964
199.11903381347656 0 561.22736
200.11575317382812 0 1315.765
200.63851928710938 0 584.68396
201.12351989746094 0 21871.736
201.1349334716797 0 12759.266
202.12367248535156 0 3499.6086
202.1417694091797 0 1359.6514
217.081787109375 0 2002.7717
221.83541870117188 0 526.3405
223.1079559326172 0 4024.5503
229.11851501464844 0 780.80145
229.13006591796875 0 2348.5125
244.12933349609375 0 1731.7628
253.11865234375 0 1985.6665
254.1532440185547 0 625.5244
261.1233825683594 0 1529.4856
261.171142578125 0 26639.209
262.1742248535156 0 3049.788
270.15655517578125 0 1668.1823
284.1609802246094 0 886.6733
287.2196350097656 0 1127.5463
288.1672058105469 0 2253.2495
288.2023620605469 0 659.9696
289.1745910644531 0 790.75854
291.614990234375 0 619.8223
303.1458740234375 0 924.50195
305.16180419921875 0 621.7848 y Ammonia loss 13
306.1688537597656 0 5304.0093 z 13
307.1728210449219 0 791.168
309.20281982421875 0 731.0881
310.1402587890625 0 1812.831
312.1558837890625 0 11059.307
313.1587829589844 0 1003.3735
313.4654541015625 0 583.3973
316.1505432128906 0 1911.2222
320.17205810546875 0 1347.3113 z Water loss 6
321.1797790527344 0 1034.5223
322.18768310546875 0 6683.331 y 13
323.1903381347656 0 1727.6661
330.1666259765625 0 8212.09
331.17010498046875 0 1155.9683
341.1116943359375 0 604.89185
347.19317626953125 0 1169.4146 c 2
348.2033386230469 0 3631.1606
349.2091064453125 0 957.87866
350.1773986816406 0 655.1698
350.6006774902344 0 2135.1626
373.19757080078125 0 1060.7551
376.1983337402344 0 17322.895 c Water loss 10
377.2009582519531 0 3328.1223
386.11962890625 0 1036.2561
386.6213684082031 0 601.51416
387.23602294921875 0 988.54517
388.24041748046875 0 983.14667
391.1852111816406 0 6537.849
392.19305419921875 0 28567.441 y Ammonia loss 12
393.198486328125 0 11881.336 y Ammonia loss 4
394.20379638671875 0 3154.9802 c Ammonia loss 6
394.7047119140625 0 908.2902
400.2668151855469 0 2135.4766
401.2401428222656 0 3719.6777
403.25177001953125 0 729.4993
409.22021484375 0 6195.5654 y 12
410.2236633300781 0 2049.8909
417.1888732910156 0 9320.283
418.1912536621094 0 2296.893
418.717529296875 0 1029.8585 y 7
428.22674560546875 0 1575.3776
429.1884460449219 0 691.511
429.2349853515625 0 5374.8604
430.1515808105469 0 828.9689
430.2370910644531 0 1619.1088
431.1408386230469 0 1423.5406
431.6374206542969 0 1035.9484
435.23651123046875 0 2477.2197
436.1500244140625 0 1224.5199
436.2438659667969 0 814.10046
444.2569580078125 0 4794.9434
445.15704345703125 0 1490.2856
445.2554626464844 0 4492.6406
446.26080322265625 0 5339.202 c 3
447.26348876953125 0 1685.973
450.6422424316406 0 629.41455
451.14617919921875 0 731.27203
457.6884765625 0 1099.1761
458.18988037109375 0 735.5934
459.6512756347656 0 1231.9846
462.22332763671875 0 1998.97 z Water loss 11
462.8976745605469 0 780.11566
463.2306823730469 0 19755.568 w 11
464.2316589355469 0 6221.0884
465.23175048828125 0 1137.0339
469.1551208496094 0 971.12335
472.693359375 0 1577.5859
473.19842529296875 0 1120.1405
473.2732238769531 0 987.35455
473.7018737792969 0 781.0889
474.17742919921875 0 1279.6519
474.904541015625 0 1329.8533 y 2
475.2406005859375 0 835.682
478.2175598144531 0 7172.989
479.2243347167969 0 7964.898 y Ammonia loss 11
480.1826171875 0 1489.8966
480.2330017089844 0 14195.863 z 11
481.2364501953125 0 3469.75
481.69097900390625 0 1011.45825
486.1874084472656 0 894.8794
486.68853759765625 0 941.6384
487.6869812011719 0 841.7884
488.75872802734375 0 14226.878 z 6
489.1849060058594 0 1595.8098
489.2605285644531 0 7446.173
489.6859436035156 0 2120.052
489.7613220214844 0 2461.0554
490.1853332519531 0 1007.9525
490.28680419921875 0 2393.8486
494.236328125 0 768.6769
495.19189453125 0 2897.7664
495.6959533691406 0 1568.2012
496.1939392089844 0 780.5912
496.2522888183594 0 6177.474 y 11
496.6932373046875 0 4038.9102
496.7674865722656 0 3675.8801 y 6
497.26788330078125 0 3196.0063
498.25579833984375 0 1180.5162
499.2986145019531 0 1806.3606
503.68646240234375 0 985.5772
503.9151611328125 0 1007.24396 y 1
504.2208557128906 0 7419.755
504.2568664550781 0 843.8637
505.2254943847656 0 1638.5546
509.6855163574219 0 2138.6304
510.18701171875 0 1362.3035
511.1796875 0 633.38464
514.2728881835938 0 772.89075
516.2666625976562 0 2924.6855
516.6757202148438 0 1016.075
518.2396240234375 0 1020.73315
518.6887817382812 0 1270.3105
519.1939086914062 0 824.8846
524.27587890625 0 5605.464
524.7789916992188 0 3726.851
525.2798461914062 0 1302.5177
531.2885131835938 0 906.1911
531.6815185546875 0 4654.3096
532.1832275390625 0 2073.6501
532.2860107421875 0 9868.5
532.6825561523438 0 1405.7056
532.7763061523438 0 768.8537
533.1838989257812 0 1872.3481
533.2931518554688 0 40602.605 c 4
533.6834106445312 0 3333.221
534.2962646484375 0 10908.537
535.3065795898438 0 1943.7815
535.6067504882812 0 4938.7295
535.94091796875 0 3810.3057
536.2730102539062 0 1235.3882
540.3106079101562 0 2384.3357
540.6897583007812 0 3391.856
540.8170166015625 0 827.3015
541.18896484375 0 8084.6255
541.3468017578125 0 4271.1836
541.6101684570312 0 19017.674
541.6907958984375 0 2675.133
541.9442749023438 0 19244.264
542.1900634765625 0 16139.11
542.2782592773438 0 9874.275
542.3501586914062 0 1157.1158
542.6123657226562 0 2843.0947
542.6898193359375 0 19489.65
542.944091796875 0 867.9606
543.1902465820312 0 1466.9789
546.2717895507812 0 23251.71 z 5
546.7748413085938 0 24659.697
547.2763061523438 0 10941.12
547.7786865234375 0 3789.2205
549.2922973632812 0 1335.3668
553.77783203125 0 1643.6432
554.2810668945312 0 11844.056 y 5
554.78271484375 0 8144.1074
555.2852783203125 0 2695.3381
556.2847900390625 0 1268.8085
560.2809448242188 0 1015.36694
561.2911376953125 0 3490.8494 z Water loss 10
562.29736328125 0 2447.084
563.2999267578125 0 865.06396
564.2786254882812 0 16272.413 c Ammonia loss 10
565.2809448242188 0 4459.628
566.283447265625 0 1492.0746
566.7904052734375 0 861.7981
567.7872314453125 0 2095.893
568.2891845703125 0 989.0096
579.3018798828125 0 8397.529 z 10
580.3048095703125 0 2867.93
581.7799072265625 0 710.47015
582.2855834960938 0 1442.83
588.2880249023438 0 846.8902
588.78173828125 0 4919.274 y Water loss 4
589.283447265625 0 7298.787 y Ammonia loss 4
589.7875366210938 0 23630.885 z 4
590.2911987304688 0 39145.723
590.7926635742188 0 20959.145
591.2942504882812 0 5951.773
591.7935791015625 0 2162.2534
592.3333740234375 0 11685.457
593.3358154296875 0 4510.4365
595.32080078125 0 9535.0625 y 10
596.3232421875 0 3270.8618
597.2933349609375 0 6296.3276
597.797119140625 0 60475.57 y 4
598.2987670898438 0 36782.754
598.8003540039062 0 15334.367
599.302490234375 0 3251.9287
599.8069458007812 0 1093.0457
601.7809448242188 0 3627.8704
602.283447265625 0 2184.2073
611.2937622070312 0 1180.508
618.3082885742188 0 673.97217 z Water loss 9
619.3179321289062 0 1652.196
620.321044921875 0 990.4917
621.2999877929688 0 1344.0764
626.3063354492188 0 943.4954
631.294189453125 0 768.0399
631.8109741210938 0 2022.3789 w 3
632.3104248046875 0 1512.339
632.8197021484375 0 930.6254
636.3233032226562 0 37905.96 z 9
637.3283081054688 0 22739.043
638.330078125 0 6511.9033 y Water loss 3
639.3226318359375 0 39187.188 z 3
639.8241577148438 0 27430.371
640.3256225585938 0 10500.124
640.8267822265625 0 3708.1418
641.3257446289062 0 924.22675
646.8257446289062 0 1684.3834
647.3316650390625 0 58375.76 y 3
647.8330688476562 0 42226.043
647.8952026367188 0 1326.364
648.3244018554688 0 43017.95 c 5
648.8348388671875 0 3152.556
649.3250122070312 0 10983.4375
650.3265380859375 0 2576.4988
651.3312377929688 0 1102.8773 c Ammonia loss 12
652.342041015625 0 39275.574 y 9
653.3449096679688 0 15536.1
654.3483276367188 0 2930.222
674.3370971679688 0 30209.25 w 2
674.8382568359375 0 22610.057
675.3397827148438 0 10725.223
675.8414916992188 0 3206.0527
676.3424682617188 0 1069.6659
679.3653564453125 0 4665.4604
680.3674926757812 0 1909.7764
681.8406372070312 0 2097.4983
682.8201293945312 0 888.4952
688.3692626953125 0 765.7699
694.8390502929688 0 923.5364 z Water loss 2
696.3352661132812 0 1399.4111
700.1946411132812 0 1098.7737
702.84619140625 0 2741.6736 y Water loss 2
703.348876953125 0 2958.1736 y Ammonia loss 2
703.8438110351562 0 24047.807 z 2
704.34521484375 0 21347.475
704.4091796875 0 1206.2601
704.8463745117188 0 9624.685
705.345947265625 0 1885.8888 z Water loss 8
707.8707275390625 0 2503.8545
708.3657836914062 0 1233.544
708.8701171875 0 803.5938
711.3487548828125 0 3851.4136
711.8530883789062 0 74038.34 y 2
712.3544921875 0 58274.35
712.85595703125 0 24968.057
713.356201171875 0 5469.4937
713.8573608398438 0 907.0337
715.8782348632812 0 6960.0444
716.3782958984375 0 5373.178
716.880126953125 0 1715.6422
717.8548583984375 0 5005.743
718.3551635742188 0 3405.222
718.8588256835938 0 1078.6888
721.3411254882812 0 1352.3956
722.34814453125 0 3848.701 y Ammonia loss 8
723.3551635742188 0 23053.14 z 8
724.3583374023438 0 10755.542
725.3617553710938 0 6377.3447
725.8607177734375 0 1610.963
726.3619995117188 0 789.1136
729.364013671875 0 1463.1825 c Ammonia loss 13
730.374267578125 0 1893.0538
730.880615234375 0 953.6404
731.3712768554688 0 704.5306
732.3411254882812 0 764.7781
737.885009765625 0 171144.86 c 13
738.3861083984375 0 138440.39
738.8873901367188 0 57837.113
739.383056640625 0 22324.633 y 8
739.887939453125 0 4569.0864
740.37646484375 0 3543.3728
745.3972778320312 0 1282.6847
746.3617553710938 0 8510.003 y Water loss 1
746.861328125 0 5877.2085 y Ammonia loss 1
747.357666015625 0 31574.271 z 1
747.861328125 0 24646.266
748.3594360351562 0 11649.171
748.8628540039062 0 4520.304
754.8971557617188 0 1054.0673
755.3690795898438 0 99648.64 y 1
755.8706665039062 0 78260.52
756.371826171875 0 37176.773
756.8728637695312 0 11360.089
757.3743286132812 0 2554.2969
759.4014282226562 0 1180.1052
760.354248046875 0 1959.6682
760.4104614257812 0 3351.7227
760.8641357421875 0 3028.8596
761.35302734375 0 1446.5797
761.4200439453125 0 1134.7589
761.8670654296875 0 1792.4543
767.9083251953125 0 864.6356
771.2284545898438 0 1928.5468
772.2337036132812 0 1442.2972
773.3934326171875 0 1833.4181
773.900390625 0 1250.5269
774.3972778320312 0 4168.81
774.8939819335938 0 1633.5422
775.3870849609375 0 2140.0112
775.8818359375 0 2514.1768
776.3787231445312 0 883.18787
781.40869140625 0 1685.4261
781.90673828125 0 16295.097
782.4057006835938 0 32345.24
782.9042358398438 0 22068.406
783.40625 0 13053.719
783.9033203125 0 3807.5405
787.9007568359375 0 3663.8088
788.399169921875 0 2841.773
788.9002685546875 0 3086.3718
789.3845825195312 0 49869.96
789.8864135742188 0 37775.527
790.3887329101562 0 23285.47
790.8994750976562 0 46076.58
791.4016723632812 0 37009.652
791.9029541015625 0 16254.646
792.4050903320312 0 4935.525
792.9063720703125 0 1641.9719
793.3895263671875 0 995.14526 w 7
794.8983154296875 0 5242.106
795.4014892578125 0 3758.719
795.8985595703125 0 5540.3496
796.3917236328125 0 4565.6646
796.8934326171875 0 5110.4434
797.39599609375 0 2396.4192
797.897216796875 0 899.01373
803.4086303710938 0 20876
803.9029541015625 0 180206.39
804.4038696289062 0 144064.53
804.9049682617188 0 70299.6
805.4057006835938 0 23032.326
805.9080810546875 0 5034.694
810.379638671875 0 1753.037
811.9110717773438 0 88657.016
812.4137573242188 0 123986.016
812.9154052734375 0 76058.01
813.4164428710938 0 33637.414
813.91796875 0 9674.176
814.4191284179688 0 1505.4401
829.4446411132812 0 1634.3741
834.4141845703125 0 1072.9799
835.4077758789062 0 1123.683
836.4281616210938 0 5159.5747 y 7
837.4307861328125 0 2180.2024
839.5043334960938 0 934.39087
840.5040283203125 0 1205.3573
842.4509887695312 0 2106.822
843.4522705078125 0 1377.2578
846.4698486328125 0 1119.1901
847.4714965820312 0 862.2964
857.4607543945312 0 10690.259
858.2735595703125 0 1410.4862
858.4653930664062 0 4025.7327
859.4675903320312 0 2240.9253
865.4302978515625 0 1142.6925
885.4593505859375 0 1415.139
886.4580688476562 0 1646.0596
887.4628295898438 0 892.6518
890.4329223632812 0 2082.4631 w 6
891.4472045898438 0 1429.6571
900.4979858398438 0 3550.6462
901.4752197265625 0 45695.48 c 7
902.4776000976562 0 21841.611
903.4819946289062 0 8499.315
904.4789428710938 0 1807.0323
916.4739990234375 0 1315.0187
926.4776000976562 0 1095.8774
927.4876708984375 0 1925.0326
928.4874877929688 0 975.01636
933.5037841796875 0 2191.2932
934.49755859375 0 817.41095
944.4932861328125 0 26324.94
945.49609375 0 13417.878
946.4989013671875 0 4696.0303
947.5001831054688 0 918.073
971.477783203125 0 794.28345 c Ammonia loss 8
972.4823608398438 0 1600.5413
975.5029907226562 0 815.42523 y Ammonia loss 6
976.5091552734375 0 25337.996 z 6
977.5133666992188 0 16461.07
978.5162353515625 0 6521.0176
979.5181884765625 0 1687.9578
987.5000610351562 0 6448.8677
988.5068969726562 0 113542.625 c 8
989.5097045898438 0 60817.49
990.395263671875 0 706.69183
990.5126342773438 0 18483.953
991.5148315429688 0 3502.8748
992.386474609375 0 3040.8975
993.3894653320312 0 1800.3944
995.5191040039062 0 760.63556
1001.514892578125 0 6149.5166
1002.5184326171875 0 4061.8787
1003.5233764648438 0 1334.751
1021.3624877929688 0 990.5182
1022.3585815429688 0 3399.553
1023.3654174804688 0 8509.146
1024.369384765625 0 3431.5203
1025.3665771484375 0 9191.508
1026.3648681640625 0 31169.006
1027.367431640625 0 2669.777
1031.5185546875 0 1501.1654
1045.52880859375 0 63572.37 c 9
1046.53173828125 0 34847.543 w 5
1047.537841796875 0 17222.12
1048.544921875 0 6920.8706
1049.5447998046875 0 1954.8782
1052.6326904296875 0 1057.4847
1059.519287109375 0 929.3014
1064.349365234375 0 3352.8774
1065.34765625 0 1976.7499
1065.5352783203125 0 3809.4568
1066.5362548828125 0 2315.5752
1073.5223388671875 0 932.0608 z Water loss 5
1080.3726806640625 0 5304.7935
1080.623046875 0 936.22546
1081.3719482421875 0 85540.125
1082.374755859375 0 55166.28
1083.3746337890625 0 21848.305
1083.505615234375 0 970.32056
1083.62060546875 0 1641.1696
1084.37841796875 0 30116.572
1085.3818359375 0 11357.181
1085.5704345703125 0 1334.9303
1086.376708984375 0 2159.0828
1091.536376953125 0 22353.213 z 5
1092.542724609375 0 36523.254
1093.5458984375 0 17136.89
1094.54833984375 0 5459.953
1095.5487060546875 0 1086.7467
1100.5845947265625 0 3518.7598
1101.585693359375 0 3163.4846
1102.5926513671875 0 1497.9413
1106.5430908203125 0 1915.4266
1107.552734375 0 6297.7153 y 5
1108.5556640625 0 3535.7957
1109.559814453125 0 1355.3003
1110.5614013671875 0 837.77954
1129.5850830078125 0 5124.2764
1130.587646484375 0 3263.667
1131.593505859375 0 997.22485
1143.593994140625 0 1982.4948
1144.596435546875 0 53168.406 c 10
1145.599853515625 0 29784.525
1146.6031494140625 0 11312.13
1147.6053466796875 0 3204.9873
1148.61181640625 0 837.60376
1172.600341796875 0 1271.8016
1177.552490234375 0 983.0638 y Ammonia loss 4
1178.5673828125 0 4660.328 z 4
1179.5755615234375 0 17868.281
1180.5792236328125 0 10526.748
1181.58251953125 0 4482.954
1182.58056640625 0 1360.6445
1183.581787109375 0 923.9252
1187.6146240234375 0 5304.133
1188.6182861328125 0 5051.8564
1189.62158203125 0 1864.8724
1190.6312255859375 0 1159.6998
1193.584228515625 0 1124.9198
1194.5858154296875 0 7421.2344 y 4
1195.5908203125 0 4999.3584
1196.5936279296875 0 2654.8848
1197.6058349609375 0 827.18475
1230.619140625 0 3462.7664
1231.627685546875 0 94633.64 c 11
1232.6309814453125 0 54572.805
1233.6337890625 0 24745.115
1234.6370849609375 0 5650.7896
1235.6171875 0 884.6549
1258.628173828125 0 3218.7778
1259.630859375 0 2590.3733 z Water loss 3
1260.6241455078125 0 924.54456 z Ammonia loss 3
1261.623046875 0 1253.9661
1274.649658203125 0 3992.5166
1275.6502685546875 0 3617.703 y Water loss 3
1276.652099609375 0 1618.3389 y Ammonia loss 3
1277.6380615234375 0 2968.6396 z 3
1278.642822265625 0 22044.213
1279.64794921875 0 13563.671
1280.65185546875 0 6659.697
1281.6475830078125 0 1172.6207
1293.6552734375 0 6015.673 y 3
1294.65576171875 0 3874.8042
1295.660888671875 0 2807.189
1318.6597900390625 0 66381.55 c 12
1319.66259765625 0 43194.64
1320.6649169921875 0 16410.402
1321.6639404296875 0 6069.7964
1322.6632080078125 0 926.0597
1344.678955078125 0 1717.3793
1345.660888671875 0 6567.178
1346.666259765625 0 4543.5107
1347.6627197265625 0 1605.6786 w 2
1364.6651611328125 0 1561.588
1365.6536865234375 0 1052.7432
1394.632568359375 0 1518.4744
1395.64697265625 0 874.7486
1406.677490234375 0 4467.8027 z 2
1407.6849365234375 0 34358.28
1408.6883544921875 0 27928.994
1409.69091796875 0 11817.0205
1410.6927490234375 0 3339.776
1416.7244873046875 0 4124.6685
1417.7196044921875 0 2274.3792
1422.71044921875 0 1427.5037 y 2
1423.704345703125 0 2032.8385
1424.69482421875 0 1200.2372
1425.7022705078125 0 1383.5115
1430.7452392578125 0 3516.6116
1431.7567138671875 0 7796.5605
1432.7532958984375 0 7049.066
1433.7608642578125 0 4143.984
1434.733642578125 0 1063.2754
1435.6898193359375 0 1746.9269
1447.77490234375 0 1169.3842
1451.69970703125 0 1347.3445
1452.6820068359375 0 895.26996
1458.7401123046875 0 7329.3916
1459.7469482421875 0 5286.516
1460.7518310546875 0 2703.0881
1461.7601318359375 0 1291.5494
1466.7276611328125 0 933.5295
1468.730224609375 0 1000.2958
1474.760009765625 0 14748.88 c 13
1475.7672119140625 0 45295.277
1476.7684326171875 0 31616.63
1477.7659912109375 0 12923.005
1478.760498046875 0 3530.2336
1479.73291015625 0 1362.1868
1493.7127685546875 0 3557.662 z 1
1494.7169189453125 0 25503.74
1495.720947265625 0 18888.25
1496.7225341796875 0 7707.9404
1497.7288818359375 0 2841.7666
1507.766357421875 0 1818.539
1508.7242431640625 0 1660.6765
1509.7476806640625 0 2371.0837 y 1
1510.756591796875 0 1372.4906
1520.767822265625 0 1063.0831
1521.7763671875 0 1378.1907
1522.7276611328125 0 7360.4873
1523.730712890625 0 7205.1597
1524.73583984375 0 3302.9167
1525.7352294921875 0 1646.9152
1535.75341796875 0 1112.6969
1536.776123046875 0 1063.5596
1537.7730712890625 0 1260.9237
1547.77587890625 0 2960.17
1548.764404296875 0 4048.364
1549.760009765625 0 1582.1022
1550.75146484375 0 1119.7076
1563.80615234375 0 2128.5388
1564.7828369140625 0 36705.625
1565.783447265625 0 50629.88
1566.785888671875 0 29021.656
1567.7879638671875 0 12057.902
1568.7930908203125 0 3627.94
1578.8236083984375 0 4598.23
1579.8179931640625 0 8733.796
1580.810546875 0 10149.444
1581.8038330078125 0 12530.826
1582.804931640625 0 7857.4478
1583.8050537109375 0 3745.0146
1584.80029296875 0 1394.5516
1589.794189453125 0 2492.9463
1590.780029296875 0 11884.042
1591.7813720703125 0 10236.095
1592.781494140625 0 4915.1943
1593.786865234375 0 1338.8746
1595.828857421875 0 3196.2317
1596.8311767578125 0 12541.792
1597.835205078125 0 9775.871
1598.8404541015625 0 4822.913
1599.837646484375 0 1890.9503
1605.7908935546875 0 1083.5123
1606.80419921875 0 27853.58
1607.8026123046875 0 125540.83
1608.8040771484375 0 96509.74
1609.807373046875 0 48678.8
1610.8101806640625 0 16914.5
1611.815185546875 0 3495.1492
1622.8099365234375 0 6247.7793
1623.819091796875 0 31361.713
1624.8265380859375 0 125235.81
1625.8299560546875 0 95463.42
1626.83203125 0 48294.953
1627.833984375 0 14358.118
1628.8372802734375 0 3871.556
2328.557373046875 0 927.24
2341.298828125 0 772.56323
2817.16064453125 0 694.4061
3233.452880859375 0 695.36053

Spectrum Details

|  |  |
| --- | --- |
| Matched peaks? Matched peaks The total absolute number of peaks matched. Additionally in brackets the total fraction of peaks matched and the total number of peaks is shown. | 89 (14.04% of 634) |
| FDR? FDR The false discovery rate estimated for this peptide. It is calculated by matching all theoretical fragments with a non-integer shift with the raw peaks for this spectrum. This is done with 40 different shifts. The resulting percentage is the average number of annotated peaks over the number of annotated peaks with the correct spectrum. | 1.42% |
| Satellite FDR? Satellite FDR See the FDR for details on its calculation. This satellite ion specific FDR only contains the satellite ions (d/w) for I/L/J positions. | - |
| PSM Score? PSM Score The PSM Score as given by Hecklib to this annotated spectrum. It is shown with three significant figures. | 718 |

#### Spectrum 4731? Spectrum 4731 The raw spectrum of this peptide as annotated by Hecklib. The fragments are coloured according to ion type (see legend). Any peaks with a star '\*' as text can be hovered over to see the full details, first the ion type second the mass shift type. By hovering over the amino acids in the peptide or ions in the legend the corresponding peaks are highlighted. By toggling the 'Unassigned' label you can turn the background (unassigned) peaks on or off in the plot. By updating the slider in the Ion legend you can update the spectrum to only show the top X% of the peaks with labels. The top X% means any peak that is within X% of the highest intensity. By dragging in the spectrum you can zoom in to a specific part of the spectrum and use 'Zoom Out' to get back to the original zoom level. The annotation of the spectrum is based on the given sequence in the peptides file and is done with different software so inconsistencies are likely. The peaks are annotated based on the given sequence, with 20 ppm tolerance.

Copy Data

##### Spectrum 4731 (TSV)

###### Preview

```
Loading example...
```

*Click on the button to copy the data to your clipboard.*

Mz MinMz MaxIntensity Max

WidthHeightPeptide font sizePeptide stroke widthSpectrum font sizeSpectrum stroke widthCompact peptide

Ion legend

wxyz

abcd

OtherUnassignedIon^Charge^_Position_Show for top:%

JSEVSDRPSGVSSRF

08.47e+41.69e+52.54e+53.39e+5

Zoom Out

y^+2^_5_z^+1^_2_z^+3^_9_y^+1^_2_c^+3^_11_y^+1^_3_y^+3^_11_c^+2^_7_y^+1^_3_y^+2^_8_c^+1^_4_z^+1^_4_w^+1^_4_y^+3^_13_y^+3^_13_y^+3^_13_y^+1^_4_z^+1^_4_z^+2^_9_y^+1^_4_y^+2^_9_y^+3^_14_c^+1^_5_z^+2^_10_y^+2^_10_z^+1^_5_c^+2^_11_z^+1^_5_y^+2^_11_y^+2^_11_z^+2^_11_y^+1^_5_y^+2^_11_w^+2^_12_z^+1^_6_y^+2^_12_z^+2^_12_y^+2^_12_c^+1^_6_c^+2^_13_y^+1^_6_w^+2^_13_z^+2^_13_z^+2^_13_y^+2^_13_y^+2^_13_z^+2^_13_z^+1^_7_y^+2^_13_y^+1^_7_z^+1^_7_c^+2^_14_c^+2^_14_y^+1^_7_y^+2^_14_y^+2^_14_z^+2^_14_y^+2^_14_c^+1^_7_w^+1^_8_y^+1^_8_w^+1^_9_c^+1^_8_z^+1^_9_c^+1^_9_y^+1^_9_z^+1^_9_c^+1^_9_y^+1^_9_c^+1^_10_w^+1^_10_z^+1^_10_z^+1^_10_z^+1^_10_y^+1^_10_c^+1^_11_z^+1^_11_y^+1^_11_z^+1^_11_y^+1^_11_c^+1^_12_z^+1^_12_z^+1^_12_w^+1^_12_y^+1^_12_y^+1^_12_z^+1^_12_y^+1^_12_c^+1^_13_w^+1^_13_z^+1^_13_y^+1^_13_c^+1^_14_z^+1^_14_y^+1^_14_

0883176726503534

Fragment Matches Table

Show background peaks

| Position | Ion type | Intensity | mz Theoretical | mz Error (Th) | mz Error (ppm) | Charge | Series Number |
| --- | --- | --- | --- | --- | --- | --- | --- |
| - | - | 1242 | 120.1 | - | - | 0 | - |
| - | - | 670.5 | 121.1 | - | - | 0 | - |
| - | - | 940.6 | 129.1 | - | - | 0 | - |
| - | - | 964.6 | 149 | - | - | 0 | - |
| - | - | 868.5 | 172.4 | - | - | 0 | - |
| - | - | 2.66E+04 | 173.1 | - | - | 0 | - |
| - | - | 1206 | 173.4 | - | - | 0 | - |
| - | - | 2508 | 174.1 | - | - | 0 | - |
| - | - | 9968 | 183.1 | - | - | 0 | - |
| - | - | 1373 | 184.1 | - | - | 0 | - |
| - | - | 1092 | 184.8 | - | - | 0 | - |
| - | - | 896.9 | 191.8 | - | - | 0 | - |
| - | - | 2365 | 199.1 | - | - | 0 | - |
| - | - | 1014 | 199.1 | - | - | 0 | - |
| - | - | 5.038E+04 | 201.1 | - | - | 0 | - |
| - | - | 2.299E+04 | 201.1 | - | - | 0 | - |
| - | - | 6915 | 202.1 | - | - | 0 | - |
| - | - | 3511 | 202.1 | - | - | 0 | - |
| - | - | 1281 | 211.1 | - | - | 0 | - |
| - | - | 4954 | 217.1 | - | - | 0 | - |
| - | - | 1380 | 226.1 | - | - | 0 | - |
| - | - | 1736 | 229.1 | - | - | 0 | - |
| - | - | 3656 | 229.1 | - | - | 0 | - |
| - | - | 1536 | 242.1 | - | - | 0 | - |
| - | - | 3620 | 244.1 | - | - | 0 | - |
| - | - | 5406 | 261.1 | - | - | 0 | - |
| - | - | 5.47E+04 | 261.2 | - | - | 0 | - |
| - | - | 8797 | 262.2 | - | - | 0 | - |
| - | - | 2813 | 270.2 | - | - | 0 | - |
| - | - | 1046 | 284.2 | - | - | 0 | - |
| - | - | 3663 | 288.2 | - | - | 0 | - |
| - | - | 1153 | 289.2 | - | - | 0 | - |
| 11 | y | 1442 | 298.2 | 0.000959 | 3.216 | +2 | 5 |
| - | - | 1203 | 299.2 | - | - | 0 | - |
| - | - | 1058 | 303.1 | - | - | 0 | - |
| 14 | z | 1.105E+04 | 306.2 | 9.343E-05 | 0.3052 | +1 | 2 |
| - | - | 2546 | 307.2 | - | - | 0 | - |
| - | - | 2.653E+04 | 312.2 | - | - | 0 | - |
| - | - | 4077 | 313.2 | - | - | 0 | - |
| - | - | 4937 | 316.2 | - | - | 0 | - |
| 7 | z | 2035 | 320.2 | 0.001419 | 4.431 | +3 | 9 |
| - | - | 2585 | 321.2 | - | - | 0 | - |
| 14 | y | 1.371E+04 | 322.2 | 0.0004696 | 1.458 | +1 | 2 |
| - | - | 1288 | 323.2 | - | - | 0 | - |
| - | - | 1.566E+04 | 330.2 | - | - | 0 | - |
| - | - | 3691 | 331.2 | - | - | 0 | - |
| - | - | 984.1 | 347.9 | - | - | 0 | - |
| - | - | 8096 | 348.2 | - | - | 0 | - |
| - | - | 1838 | 349.2 | - | - | 0 | - |
| - | - | 4061 | 373.2 | - | - | 0 | - |
| 11 | c | 3.374E+04 | 376.2 | 0.001921 | 5.106 | +3 | 11 |
| - | - | 6255 | 377.2 | - | - | 0 | - |
| - | - | 1626 | 387.2 | - | - | 0 | - |
| - | - | 1.576E+04 | 391.2 | - | - | 0 | - |
| 13 | y | 5.973E+04 | 392.2 | 8.674E-05 | 0.2212 | +1 | 3 |
| 5 | y | 2.476E+04 | 393.2 | 0.007368 | 18.74 | +3 | 11 |
| 7 | c | 7195 | 394.2 | 0.00418 | 10.6 | +2 | 7 |
| - | - | 1936 | 395.2 | - | - | 0 | - |
| - | - | 7236 | 401.2 | - | - | 0 | - |
| - | - | 1645 | 403.2 | - | - | 0 | - |
| - | - | 1787 | 407.2 | - | - | 0 | - |
| 13 | y | 1.276E+04 | 409.2 | 0.0004236 | 1.035 | +1 | 3 |
| - | - | 1374 | 410.2 | - | - | 0 | - |
| - | - | 1907 | 411.2 | - | - | 0 | - |
| - | - | 1.918E+04 | 417.2 | - | - | 0 | - |
| - | - | 4102 | 418.2 | - | - | 0 | - |
| 8 | y | 2651 | 418.7 | 0.001363 | 3.256 | +2 | 8 |
| - | - | 1258 | 419.2 | - | - | 0 | - |
| - | - | 4726 | 428.2 | - | - | 0 | - |
| - | - | 1442 | 429.2 | - | - | 0 | - |
| - | - | 1.201E+04 | 429.2 | - | - | 0 | - |
| - | - | 3548 | 430.2 | - | - | 0 | - |
| - | - | 4407 | 435.2 | - | - | 0 | - |
| - | - | 3438 | 436.2 | - | - | 0 | - |
| - | - | 9475 | 444.3 | - | - | 0 | - |
| - | - | 8705 | 445.3 | - | - | 0 | - |
| 4 | c | 1.092E+04 | 446.3 | 0.0001828 | 0.4096 | +1 | 4 |
| - | - | 2286 | 447.3 | - | - | 0 | - |
| 12 | z | 3147 | 462.2 | 0.001163 | 2.516 | +1 | 4 |
| 12 | w | 4.057E+04 | 463.2 | 0.0003875 | 0.8365 | +1 | 4 |
| - | - | 1.306E+04 | 464.2 | - | - | 0 | - |
| - | - | 1945 | 465.2 | - | - | 0 | - |
| 3 | y | 1758 | 468.9 | 0.0009761 | 2.082 | +3 | 13 |
| 3 | y | 1559 | 469.2 | 0.006987 | 14.89 | +3 | 13 |
| - | - | 1326 | 473.3 | - | - | 0 | - |
| 3 | y | 1621 | 474.9 | 0.0005072 | 1.068 | +3 | 13 |
| - | - | 1795 | 475.2 | - | - | 0 | - |
| - | - | 1.517E+04 | 478.2 | - | - | 0 | - |
| 12 | y | 1.786E+04 | 479.2 | 0.0006001 | 1.252 | +1 | 4 |
| 12 | z | 2.918E+04 | 480.2 | 2.302E-06 | 0.004792 | +1 | 4 |
| - | - | 9212 | 481.2 | - | - | 0 | - |
| - | - | 1750 | 482.2 | - | - | 0 | - |
| 7 | z | 3.252E+04 | 488.8 | 0.0004227 | 0.8648 | +2 | 9 |
| - | - | 1.874E+04 | 489.3 | - | - | 0 | - |
| - | - | 7907 | 489.8 | - | - | 0 | - |
| - | - | 4380 | 490.3 | - | - | 0 | - |
| - | - | 2390 | 494.2 | - | - | 0 | - |
| - | - | 1320 | 495.2 | - | - | 0 | - |
| 12 | y | 1.202E+04 | 496.3 | 0.0009269 | 1.868 | +1 | 4 |
| 7 | y | 1.019E+04 | 496.8 | 0.0002464 | 0.4961 | +2 | 9 |
| - | - | 3992 | 497.3 | - | - | 0 | - |
| - | - | 3565 | 498.3 | - | - | 0 | - |
| - | - | 2461 | 499.3 | - | - | 0 | - |
| 2 | y | 4310 | 503.9 | 0.00097 | 1.925 | +3 | 14 |
| - | - | 1.727E+04 | 504.2 | - | - | 0 | - |
| - | - | 5104 | 505.2 | - | - | 0 | - |
| - | - | 1970 | 506.2 | - | - | 0 | - |
| - | - | 5543 | 516.3 | - | - | 0 | - |
| - | - | 1.066E+04 | 524.3 | - | - | 0 | - |
| - | - | 6893 | 524.8 | - | - | 0 | - |
| - | - | 3329 | 525.3 | - | - | 0 | - |
| - | - | 1330 | 529.9 | - | - | 0 | - |
| - | - | 1.811E+04 | 532.3 | - | - | 0 | - |
| 5 | c | 8.063E+04 | 533.3 | 1.538E-05 | 0.02884 | +1 | 5 |
| - | - | 2.101E+04 | 534.3 | - | - | 0 | - |
| - | - | 5336 | 535.3 | - | - | 0 | - |
| - | - | 1.125E+04 | 535.6 | - | - | 0 | - |
| - | - | 1.014E+04 | 535.9 | - | - | 0 | - |
| - | - | 4151 | 536.3 | - | - | 0 | - |
| - | - | 1419 | 536.6 | - | - | 0 | - |
| - | - | 1083 | 540.8 | - | - | 0 | - |
| - | - | 1769 | 541.3 | - | - | 0 | - |
| - | - | 9174 | 541.3 | - | - | 0 | - |
| - | - | 4.464E+04 | 541.6 | - | - | 0 | - |
| - | - | 1903 | 541.8 | - | - | 0 | - |
| - | - | 3.634E+04 | 541.9 | - | - | 0 | - |
| - | - | 2.562E+04 | 542.3 | - | - | 0 | - |
| - | - | 2606 | 542.4 | - | - | 0 | - |
| - | - | 7780 | 542.6 | - | - | 0 | - |
| 6 | z | 4.402E+04 | 546.3 | 0.0004399 | 0.8053 | +2 | 10 |
| - | - | 4.907E+04 | 546.8 | - | - | 0 | - |
| - | - | 2.079E+04 | 547.3 | - | - | 0 | - |
| - | - | 6389 | 547.8 | - | - | 0 | - |
| - | - | 2009 | 548.3 | - | - | 0 | - |
| - | - | 2108 | 549.3 | - | - | 0 | - |
| - | - | 4908 | 553.8 | - | - | 0 | - |
| 6 | y | 2.687E+04 | 554.3 | 0.0002551 | 0.4603 | +2 | 10 |
| - | - | 1.363E+04 | 554.8 | - | - | 0 | - |
| - | - | 3987 | 555.3 | - | - | 0 | - |
| - | - | 1987 | 556.3 | - | - | 0 | - |
| 11 | z | 6116 | 561.3 | 0.0007727 | 1.377 | +1 | 5 |
| - | - | 5043 | 562.3 | - | - | 0 | - |
| - | - | 1307 | 563.3 | - | - | 0 | - |
| 11 | c | 3.029E+04 | 564.3 | 0.01037 | 18.38 | +2 | 11 |
| - | - | 8274 | 565.3 | - | - | 0 | - |
| - | - | 2061 | 566.3 | - | - | 0 | - |
| - | - | 3108 | 567.8 | - | - | 0 | - |
| - | - | 3184 | 568.3 | - | - | 0 | - |
| - | - | 1356 | 568.8 | - | - | 0 | - |
| 11 | z | 1.802E+04 | 579.3 | 0.0002094 | 0.3615 | +1 | 5 |
| - | - | 5596 | 580.3 | - | - | 0 | - |
| - | - | 4091 | 581.8 | - | - | 0 | - |
| - | - | 1535 | 588.3 | - | - | 0 | - |
| 5 | y | 8247 | 588.8 | 0.008362 | 14.2 | +2 | 11 |
| 5 | y | 1.38E+04 | 589.3 | 0.0007975 | 1.353 | +2 | 11 |
| 5 | z | 4.535E+04 | 589.8 | 0.0001324 | 0.2245 | +2 | 11 |
| - | - | 7.754E+04 | 590.3 | - | - | 0 | - |
| - | - | 3.817E+04 | 590.8 | - | - | 0 | - |
| - | - | 1.128E+04 | 591.3 | - | - | 0 | - |
| - | - | 2859 | 591.8 | - | - | 0 | - |
| - | - | 2.47E+04 | 592.3 | - | - | 0 | - |
| - | - | 9133 | 593.3 | - | - | 0 | - |
| - | - | 1296 | 594.3 | - | - | 0 | - |
| 11 | y | 2.09E+04 | 595.3 | 0.001025 | 1.722 | +1 | 5 |
| - | - | 6261 | 596.3 | - | - | 0 | - |
| - | - | 1.106E+04 | 597.3 | - | - | 0 | - |
| 5 | y | 1.259E+05 | 597.8 | 0.0002102 | 0.3516 | +2 | 11 |
| - | - | 8.52E+04 | 598.3 | - | - | 0 | - |
| - | - | 3.314E+04 | 598.8 | - | - | 0 | - |
| - | - | 6992 | 599.3 | - | - | 0 | - |
| - | - | 1770 | 599.8 | - | - | 0 | - |
| - | - | 7390 | 601.8 | - | - | 0 | - |
| - | - | 3656 | 602.3 | - | - | 0 | - |
| - | - | 2356 | 611.3 | - | - | 0 | - |
| - | - | 1516 | 611.8 | - | - | 0 | - |
| - | - | 1547 | 612.3 | - | - | 0 | - |
| - | - | 1362 | 614.3 | - | - | 0 | - |
| - | - | 3016 | 619.3 | - | - | 0 | - |
| - | - | 1745 | 620.3 | - | - | 0 | - |
| - | - | 1887 | 621.3 | - | - | 0 | - |
| - | - | 1444 | 624.3 | - | - | 0 | - |
| - | - | 2528 | 631.3 | - | - | 0 | - |
| 4 | w | 3185 | 631.8 | 0.00108 | 1.709 | +2 | 12 |
| - | - | 3673 | 632.3 | - | - | 0 | - |
| - | - | 1315 | 632.4 | - | - | 0 | - |
| 10 | z | 7.223E+04 | 636.3 | 0.0003605 | 0.5666 | +1 | 6 |
| - | - | 4.28E+04 | 637.3 | - | - | 0 | - |
| 4 | y | 1.246E+04 | 638.3 | 0.004 | 6.267 | +2 | 12 |
| 4 | z | 7.082E+04 | 639.3 | 0.0007559 | 1.182 | +2 | 12 |
| - | - | 5.527E+04 | 639.8 | - | - | 0 | - |
| - | - | 2.171E+04 | 640.3 | - | - | 0 | - |
| - | - | 6704 | 640.8 | - | - | 0 | - |
| - | - | 2291 | 641.3 | - | - | 0 | - |
| - | - | 5565 | 646.8 | - | - | 0 | - |
| 4 | y | 1.221E+05 | 647.3 | 0.000305 | 0.4711 | +2 | 12 |
| - | - | 9.068E+04 | 647.8 | - | - | 0 | - |
| 6 | c | 8.88E+04 | 648.3 | 0.00475 | 7.326 | +1 | 6 |
| - | - | 1.004E+04 | 648.8 | - | - | 0 | - |
| - | - | 2.325E+04 | 649.3 | - | - | 0 | - |
| - | - | 4505 | 650.3 | - | - | 0 | - |
| 13 | c | 3262 | 651.3 | 0.01027 | 15.78 | +2 | 13 |
| - | - | 1836 | 651.8 | - | - | 0 | - |
| 10 | y | 8.223E+04 | 652.3 | 0.0002522 | 0.3866 | +1 | 6 |
| - | - | 2.772E+04 | 653.3 | - | - | 0 | - |
| - | - | 6115 | 654.3 | - | - | 0 | - |
| 3 | w | 6.55E+04 | 674.3 | 0.0002106 | 0.3123 | +2 | 13 |
| - | - | 4.961E+04 | 674.8 | - | - | 0 | - |
| - | - | 2.067E+04 | 675.3 | - | - | 0 | - |
| - | - | 8955 | 675.8 | - | - | 0 | - |
| - | - | 1.033E+04 | 679.4 | - | - | 0 | - |
| - | - | 3069 | 680.4 | - | - | 0 | - |
| - | - | 1210 | 680.8 | - | - | 0 | - |
| - | - | 2253 | 681.8 | - | - | 0 | - |
| - | - | 2736 | 682.4 | - | - | 0 | - |
| 3 | z | 1406 | 694.8 | 0.000732 | 1.053 | +2 | 13 |
| 3 | z | 1999 | 695.3 | 0.01025 | 14.74 | +2 | 13 |
| - | - | 2292 | 696.3 | - | - | 0 | - |
| 3 | y | 6580 | 702.8 | 0.003273 | 4.656 | +2 | 13 |
| 3 | y | 6679 | 703.3 | 0.009861 | 14.02 | +2 | 13 |
| 3 | z | 5.006E+04 | 703.8 | 0.0006996 | 0.9939 | +2 | 13 |
| - | - | 3.602E+04 | 704.3 | - | - | 0 | - |
| - | - | 1.863E+04 | 704.8 | - | - | 0 | - |
| 9 | z | 6027 | 705.3 | 0.003921 | 5.559 | +1 | 7 |
| - | - | 3620 | 707.9 | - | - | 0 | - |
| - | - | 3603 | 708.4 | - | - | 0 | - |
| - | - | 2844 | 708.9 | - | - | 0 | - |
| - | - | 1617 | 709.4 | - | - | 0 | - |
| - | - | 4352 | 711.3 | - | - | 0 | - |
| 3 | y | 1.514E+05 | 711.9 | 0.0003707 | 0.5208 | +2 | 13 |
| - | - | 1.166E+05 | 712.4 | - | - | 0 | - |
| - | - | 5.348E+04 | 712.9 | - | - | 0 | - |
| - | - | 1.524E+04 | 713.4 | - | - | 0 | - |
| - | - | 2990 | 713.9 | - | - | 0 | - |
| - | - | 1.261E+04 | 715.9 | - | - | 0 | - |
| - | - | 8873 | 716.4 | - | - | 0 | - |
| - | - | 6553 | 716.9 | - | - | 0 | - |
| - | - | 8898 | 717.9 | - | - | 0 | - |
| - | - | 7265 | 718.4 | - | - | 0 | - |
| - | - | 4783 | 718.9 | - | - | 0 | - |
| - | - | 4212 | 721.3 | - | - | 0 | - |
| 9 | y | 9504 | 722.3 | 0.0001002 | 0.1387 | +1 | 7 |
| 9 | z | 5.266E+04 | 723.4 | 0.0001925 | 0.2661 | +1 | 7 |
| - | - | 1.865E+04 | 724.4 | - | - | 0 | - |
| - | - | 1595 | 724.9 | - | - | 0 | - |
| - | - | 8823 | 725.4 | - | - | 0 | - |
| - | - | 3590 | 725.9 | - | - | 0 | - |
| - | - | 3114 | 726.4 | - | - | 0 | - |
| 14 | c | 3837 | 729.4 | 0.003049 | 4.181 | +2 | 14 |
| - | - | 3454 | 730.4 | - | - | 0 | - |
| - | - | 1913 | 731.4 | - | - | 0 | - |
| - | - | 1417 | 733.3 | - | - | 0 | - |
| - | - | 1872 | 734.4 | - | - | 0 | - |
| - | - | 1315 | 735.4 | - | - | 0 | - |
| - | - | 2311 | 737.4 | - | - | 0 | - |
| 14 | c | 3.286E+05 | 737.9 | 0.0005827 | 0.7897 | +2 | 14 |
| - | - | 2.574E+05 | 738.4 | - | - | 0 | - |
| - | - | 1.211E+05 | 738.9 | - | - | 0 | - |
| 9 | y | 4.719E+04 | 739.4 | 0.009789 | 13.24 | +1 | 7 |
| - | - | 7037 | 739.9 | - | - | 0 | - |
| - | - | 8309 | 740.4 | - | - | 0 | - |
| - | - | 1693 | 741.4 | - | - | 0 | - |
| - | - | 1896 | 744.4 | - | - | 0 | - |
| - | - | 2240 | 745.4 | - | - | 0 | - |
| 2 | y | 1.319E+04 | 746.4 | 0.002671 | 3.578 | +2 | 14 |
| 2 | y | 1.501E+04 | 746.9 | 0.005871 | 7.861 | +2 | 14 |
| 2 | z | 6.661E+04 | 747.4 | 0.001704 | 2.28 | +2 | 14 |
| - | - | 4.352E+04 | 747.9 | - | - | 0 | - |
| - | - | 2.348E+04 | 748.4 | - | - | 0 | - |
| - | - | 5106 | 748.9 | - | - | 0 | - |
| - | - | 2731 | 754.9 | - | - | 0 | - |
| 2 | y | 1.887E+05 | 755.4 | 0.0004088 | 0.5411 | +2 | 14 |
| - | - | 1.649E+05 | 755.9 | - | - | 0 | - |
| - | - | 7.135E+04 | 756.4 | - | - | 0 | - |
| - | - | 2.262E+04 | 756.9 | - | - | 0 | - |
| - | - | 5873 | 757.4 | - | - | 0 | - |
| - | - | 1722 | 759.4 | - | - | 0 | - |
| - | - | 3378 | 760.3 | - | - | 0 | - |
| - | - | 6390 | 760.4 | - | - | 0 | - |
| - | - | 5724 | 760.9 | - | - | 0 | - |
| - | - | 2826 | 761.4 | - | - | 0 | - |
| - | - | 1402 | 761.9 | - | - | 0 | - |
| - | - | 1656 | 762.4 | - | - | 0 | - |
| - | - | 1530 | 767.9 | - | - | 0 | - |
| - | - | 2979 | 768.4 | - | - | 0 | - |
| - | - | 3693 | 773.4 | - | - | 0 | - |
| - | - | 1499 | 773.9 | - | - | 0 | - |
| - | - | 7591 | 774.4 | - | - | 0 | - |
| - | - | 4574 | 774.9 | - | - | 0 | - |
| - | - | 4609 | 775.4 | - | - | 0 | - |
| - | - | 5774 | 775.9 | - | - | 0 | - |
| - | - | 3255 | 776.4 | - | - | 0 | - |
| - | - | 1931 | 776.9 | - | - | 0 | - |
| - | - | 1622 | 777.4 | - | - | 0 | - |
| - | - | 5308 | 781.4 | - | - | 0 | - |
| - | - | 3.272E+04 | 781.9 | - | - | 0 | - |
| - | - | 6.776E+04 | 782.4 | - | - | 0 | - |
| - | - | 5.363E+04 | 782.9 | - | - | 0 | - |
| - | - | 2.756E+04 | 783.4 | - | - | 0 | - |
| - | - | 8203 | 783.9 | - | - | 0 | - |
| 7 | c | 1760 | 787.4 | 0.002368 | 3.008 | +1 | 7 |
| - | - | 4571 | 787.9 | - | - | 0 | - |
| - | - | 7177 | 788.4 | - | - | 0 | - |
| - | - | 6570 | 788.9 | - | - | 0 | - |
| - | - | 9.854E+04 | 789.4 | - | - | 0 | - |
| - | - | 8.384E+04 | 789.9 | - | - | 0 | - |
| - | - | 4.352E+04 | 790.4 | - | - | 0 | - |
| - | - | 1.021E+05 | 790.9 | - | - | 0 | - |
| - | - | 8.033E+04 | 791.4 | - | - | 0 | - |
| - | - | 4.284E+04 | 791.9 | - | - | 0 | - |
| - | - | 1.082E+04 | 792.4 | - | - | 0 | - |
| - | - | 4387 | 792.9 | - | - | 0 | - |
| 8 | w | 1737 | 793.4 | 0.002581 | 3.253 | +1 | 8 |
| - | - | 1606 | 794.4 | - | - | 0 | - |
| - | - | 1.262E+04 | 794.9 | - | - | 0 | - |
| - | - | 1.069E+04 | 795.4 | - | - | 0 | - |
| - | - | 5468 | 795.9 | - | - | 0 | - |
| - | - | 1.125E+04 | 796.4 | - | - | 0 | - |
| - | - | 7207 | 796.9 | - | - | 0 | - |
| - | - | 3803 | 797.4 | - | - | 0 | - |
| - | - | 2128 | 797.9 | - | - | 0 | - |
| - | - | 2037 | 798.4 | - | - | 0 | - |
| - | - | 3.459E+04 | 803.4 | - | - | 0 | - |
| - | - | 3.355E+05 | 803.9 | - | - | 0 | - |
| - | - | 2.955E+05 | 804.4 | - | - | 0 | - |
| - | - | 1.414E+05 | 804.9 | - | - | 0 | - |
| - | - | 4.571E+04 | 805.4 | - | - | 0 | - |
| - | - | 1.23E+04 | 805.9 | - | - | 0 | - |
| - | - | 1457 | 811.4 | - | - | 0 | - |
| - | - | 1.644E+05 | 811.9 | - | - | 0 | - |
| - | - | 2.496E+05 | 812.4 | - | - | 0 | - |
| - | - | 1.539E+05 | 812.9 | - | - | 0 | - |
| - | - | 7.105E+04 | 813.4 | - | - | 0 | - |
| - | - | 2.369E+04 | 813.9 | - | - | 0 | - |
| - | - | 2659 | 814.4 | - | - | 0 | - |
| - | - | 3162 | 834.4 | - | - | 0 | - |
| - | - | 2094 | 835.4 | - | - | 0 | - |
| 8 | y | 1.292E+04 | 836.4 | 0.001397 | 1.671 | +1 | 8 |
| - | - | 4876 | 837.4 | - | - | 0 | - |
| - | - | 3607 | 842.5 | - | - | 0 | - |
| - | - | 1383 | 856.4 | - | - | 0 | - |
| - | - | 1.69E+04 | 857.5 | - | - | 0 | - |
| - | - | 7724 | 858.5 | - | - | 0 | - |
| - | - | 2426 | 859.5 | - | - | 0 | - |
| - | - | 1661 | 864.4 | - | - | 0 | - |
| - | - | 4229 | 885.5 | - | - | 0 | - |
| - | - | 4250 | 886.5 | - | - | 0 | - |
| 7 | w | 5776 | 890.4 | 0.002857 | 3.208 | +1 | 9 |
| - | - | 1788 | 891.4 | - | - | 0 | - |
| - | - | 1706 | 892.4 | - | - | 0 | - |
| - | - | 2114 | 893.5 | - | - | 0 | - |
| - | - | 4846 | 900.5 | - | - | 0 | - |
| 8 | c | 9.223E+04 | 901.5 | 0.000716 | 0.7943 | +1 | 8 |
| - | - | 4.232E+04 | 902.5 | - | - | 0 | - |
| - | - | 1.245E+04 | 903.5 | - | - | 0 | - |
| - | - | 2967 | 904.5 | - | - | 0 | - |
| - | - | 2048 | 916.5 | - | - | 0 | - |
| - | - | 2900 | 926.5 | - | - | 0 | - |
| - | - | 2430 | 927.5 | - | - | 0 | - |
| - | - | 2044 | 929.5 | - | - | 0 | - |
| - | - | 3995 | 933.5 | - | - | 0 | - |
| - | - | 5.345E+04 | 944.5 | - | - | 0 | - |
| - | - | 2.765E+04 | 945.5 | - | - | 0 | - |
| - | - | 8786 | 946.5 | - | - | 0 | - |
| - | - | 2920 | 947.5 | - | - | 0 | - |
| 7 | z | 2685 | 958.5 | 0.00355 | 3.703 | +1 | 9 |
| 9 | c | 1749 | 971.5 | 0.01037 | 10.68 | +1 | 9 |
| - | - | 3545 | 972.5 | - | - | 0 | - |
| 7 | y | 2112 | 975.5 | 0.007402 | 7.588 | +1 | 9 |
| 7 | z | 5.027E+04 | 976.5 | 0.0001262 | 0.1292 | +1 | 9 |
| - | - | 3.355E+04 | 977.5 | - | - | 0 | - |
| - | - | 1583 | 977.6 | - | - | 0 | - |
| - | - | 1.3E+04 | 978.5 | - | - | 0 | - |
| - | - | 1528 | 979.5 | - | - | 0 | - |
| - | - | 1.385E+04 | 987.5 | - | - | 0 | - |
| 9 | c | 2.247E+05 | 988.5 | 0.0007311 | 0.7396 | +1 | 9 |
| - | - | 1.187E+05 | 989.5 | - | - | 0 | - |
| - | - | 3.736E+04 | 990.5 | - | - | 0 | - |
| - | - | 7931 | 991.5 | - | - | 0 | - |
| 7 | y | 2071 | 992.5 | 0.01009 | 10.16 | +1 | 9 |
| - | - | 1.388E+04 | 1002 | - | - | 0 | - |
| - | - | 8170 | 1003 | - | - | 0 | - |
| - | - | 2899 | 1004 | - | - | 0 | - |
| - | - | 1491 | 1031 | - | - | 0 | - |
| - | - | 2133 | 1032 | - | - | 0 | - |
| - | - | 1808 | 1033 | - | - | 0 | - |
| 10 | c | 1.341E+05 | 1046 | 0.0008128 | 0.7774 | +1 | 10 |
| 6 | w | 7.383E+04 | 1047 | 0.006641 | 6.345 | +1 | 10 |
| - | - | 3.571E+04 | 1048 | - | - | 0 | - |
| - | - | 1.352E+04 | 1049 | - | - | 0 | - |
| - | - | 4149 | 1050 | - | - | 0 | - |
| - | - | 1713 | 1060 | - | - | 0 | - |
| 6 | z | 3457 | 1074 | 0.005293 | 4.931 | +1 | 10 |
| 6 | z | 2627 | 1075 | 0.01395 | 12.99 | +1 | 10 |
| - | - | 1746 | 1082 | - | - | 0 | - |
| - | - | 2780 | 1083 | - | - | 0 | - |
| - | - | 2043 | 1085 | - | - | 0 | - |
| 6 | z | 4.748E+04 | 1092 | 0.0005879 | 0.5386 | +1 | 10 |
| - | - | 7.805E+04 | 1093 | - | - | 0 | - |
| - | - | 3.511E+04 | 1094 | - | - | 0 | - |
| - | - | 9157 | 1095 | - | - | 0 | - |
| - | - | 1805 | 1096 | - | - | 0 | - |
| - | - | 1527 | 1100 | - | - | 0 | - |
| - | - | 7671 | 1101 | - | - | 0 | - |
| - | - | 4613 | 1102 | - | - | 0 | - |
| - | - | 2921 | 1103 | - | - | 0 | - |
| - | - | 1789 | 1104 | - | - | 0 | - |
| - | - | 3271 | 1107 | - | - | 0 | - |
| 6 | y | 1.091E+04 | 1108 | 0.0006801 | 0.614 | +1 | 10 |
| - | - | 7125 | 1109 | - | - | 0 | - |
| - | - | 3009 | 1110 | - | - | 0 | - |
| - | - | 9647 | 1130 | - | - | 0 | - |
| - | - | 5430 | 1131 | - | - | 0 | - |
| - | - | 2292 | 1132 | - | - | 0 | - |
| - | - | 2966 | 1144 | - | - | 0 | - |
| 11 | c | 9.735E+04 | 1145 | 0.0002699 | 0.2358 | +1 | 11 |
| - | - | 6.006E+04 | 1146 | - | - | 0 | - |
| - | - | 2.585E+04 | 1147 | - | - | 0 | - |
| - | - | 6873 | 1148 | - | - | 0 | - |
| - | - | 1670 | 1149 | - | - | 0 | - |
| 5 | z | 1714 | 1162 | 0.0008459 | 0.7283 | +1 | 11 |
| - | - | 2003 | 1173 | - | - | 0 | - |
| 5 | y | 2589 | 1178 | 0.002019 | 1.715 | +1 | 11 |
| 5 | z | 8848 | 1179 | 0.002388 | 2.026 | +1 | 11 |
| - | - | 3.643E+04 | 1180 | - | - | 0 | - |
| - | - | 2.117E+04 | 1181 | - | - | 0 | - |
| - | - | 9089 | 1182 | - | - | 0 | - |
| - | - | 1720 | 1183 | - | - | 0 | - |
| - | - | 1.024E+04 | 1188 | - | - | 0 | - |
| - | - | 6321 | 1189 | - | - | 0 | - |
| - | - | 4943 | 1190 | - | - | 0 | - |
| - | - | 2759 | 1191 | - | - | 0 | - |
| - | - | 1987 | 1194 | - | - | 0 | - |
| 5 | y | 1.463E+04 | 1195 | 0.000604 | 0.5056 | +1 | 11 |
| - | - | 1.047E+04 | 1196 | - | - | 0 | - |
| - | - | 5094 | 1197 | - | - | 0 | - |
| - | - | 1653 | 1198 | - | - | 0 | - |
| - | - | 8668 | 1231 | - | - | 0 | - |
| 12 | c | 1.883E+05 | 1232 | 0.0007526 | 0.6111 | +1 | 12 |
| - | - | 1.133E+05 | 1233 | - | - | 0 | - |
| - | - | 4.548E+04 | 1234 | - | - | 0 | - |
| - | - | 1.3E+04 | 1235 | - | - | 0 | - |
| - | - | 2631 | 1236 | - | - | 0 | - |
| - | - | 1457 | 1237 | - | - | 0 | - |
| - | - | 8913 | 1259 | - | - | 0 | - |
| 4 | z | 6774 | 1260 | 0.004338 | 3.444 | +1 | 12 |
| 4 | z | 2345 | 1261 | 0.01532 | 12.15 | +1 | 12 |
| 4 | w | 1771 | 1263 | 0.02006 | 15.88 | +1 | 12 |
| - | - | 7553 | 1275 | - | - | 0 | - |
| 4 | y | 4810 | 1276 | 0.007953 | 6.235 | +1 | 12 |
| 4 | y | 1929 | 1277 | 0.009289 | 7.276 | +1 | 12 |
| 4 | z | 8331 | 1278 | 0.0003671 | 0.2873 | +1 | 12 |
| - | - | 4.431E+04 | 1279 | - | - | 0 | - |
| - | - | 2.73E+04 | 1280 | - | - | 0 | - |
| - | - | 1.136E+04 | 1281 | - | - | 0 | - |
| - | - | 2211 | 1282 | - | - | 0 | - |
| 4 | y | 1.121E+04 | 1294 | 0.0001702 | 0.1316 | +1 | 12 |
| - | - | 9031 | 1295 | - | - | 0 | - |
| - | - | 4109 | 1296 | - | - | 0 | - |
| - | - | 1614 | 1297 | - | - | 0 | - |
| 13 | c | 1.333E+05 | 1319 | 0.0005545 | 0.4205 | +1 | 13 |
| - | - | 9.666E+04 | 1320 | - | - | 0 | - |
| - | - | 4.021E+04 | 1321 | - | - | 0 | - |
| - | - | 1.185E+04 | 1322 | - | - | 0 | - |
| - | - | 1625 | 1323 | - | - | 0 | - |
| - | - | 2940 | 1345 | - | - | 0 | - |
| - | - | 1.315E+04 | 1346 | - | - | 0 | - |
| - | - | 1.048E+04 | 1347 | - | - | 0 | - |
| 3 | w | 6284 | 1348 | 0.001458 | 1.082 | +1 | 13 |
| - | - | 1809 | 1349 | - | - | 0 | - |
| - | - | 2186 | 1365 | - | - | 0 | - |
| - | - | 2533 | 1395 | - | - | 0 | - |
| - | - | 2346 | 1396 | - | - | 0 | - |
| 3 | z | 9490 | 1407 | 0.00109 | 0.7749 | +1 | 13 |
| - | - | 7.371E+04 | 1408 | - | - | 0 | - |
| - | - | 5.384E+04 | 1409 | - | - | 0 | - |
| - | - | 2.323E+04 | 1410 | - | - | 0 | - |
| - | - | 7510 | 1411 | - | - | 0 | - |
| - | - | 2052 | 1412 | - | - | 0 | - |
| - | - | 2827 | 1415 | - | - | 0 | - |
| - | - | 1676 | 1416 | - | - | 0 | - |
| - | - | 5244 | 1417 | - | - | 0 | - |
| - | - | 4280 | 1418 | - | - | 0 | - |
| - | - | 2351 | 1419 | - | - | 0 | - |
| 3 | y | 2219 | 1423 | 0.007851 | 5.519 | +1 | 13 |
| - | - | 4349 | 1424 | - | - | 0 | - |
| - | - | 2810 | 1425 | - | - | 0 | - |
| - | - | 1456 | 1426 | - | - | 0 | - |
| - | - | 5134 | 1431 | - | - | 0 | - |
| - | - | 1.815E+04 | 1432 | - | - | 0 | - |
| - | - | 1.237E+04 | 1433 | - | - | 0 | - |
| - | - | 6047 | 1434 | - | - | 0 | - |
| - | - | 2174 | 1435 | - | - | 0 | - |
| - | - | 1568 | 1436 | - | - | 0 | - |
| - | - | 2130 | 1448 | - | - | 0 | - |
| - | - | 3067 | 1452 | - | - | 0 | - |
| - | - | 1952 | 1453 | - | - | 0 | - |
| - | - | 1.37E+04 | 1459 | - | - | 0 | - |
| - | - | 1.069E+04 | 1460 | - | - | 0 | - |
| - | - | 4485 | 1461 | - | - | 0 | - |
| - | - | 1818 | 1462 | - | - | 0 | - |
| - | - | 2343 | 1465 | - | - | 0 | - |
| 14 | c | 2.87E+04 | 1475 | 0.00108 | 0.732 | +1 | 14 |
| - | - | 8.568E+04 | 1476 | - | - | 0 | - |
| - | - | 6.054E+04 | 1477 | - | - | 0 | - |
| - | - | 2.836E+04 | 1478 | - | - | 0 | - |
| - | - | 1.025E+04 | 1479 | - | - | 0 | - |
| - | - | 1706 | 1480 | - | - | 0 | - |
| 2 | z | 7085 | 1494 | 0.003625 | 2.427 | +1 | 14 |
| - | - | 5.446E+04 | 1495 | - | - | 0 | - |
| - | - | 4.05E+04 | 1496 | - | - | 0 | - |
| - | - | 2.047E+04 | 1497 | - | - | 0 | - |
| - | - | 6769 | 1498 | - | - | 0 | - |
| - | - | 3978 | 1508 | - | - | 0 | - |
| 2 | y | 3463 | 1510 | 0.0192 | 12.72 | +1 | 14 |
| - | - | 2356 | 1511 | - | - | 0 | - |
| - | - | 2542 | 1522 | - | - | 0 | - |
| - | - | 1.704E+04 | 1523 | - | - | 0 | - |
| - | - | 1.482E+04 | 1524 | - | - | 0 | - |
| - | - | 8645 | 1525 | - | - | 0 | - |
| - | - | 3273 | 1526 | - | - | 0 | - |
| - | - | 2257 | 1536 | - | - | 0 | - |
| - | - | 2040 | 1537 | - | - | 0 | - |
| - | - | 2341 | 1538 | - | - | 0 | - |
| - | - | 2056 | 1539 | - | - | 0 | - |
| - | - | 2989 | 1547 | - | - | 0 | - |
| - | - | 5012 | 1548 | - | - | 0 | - |
| - | - | 5167 | 1549 | - | - | 0 | - |
| - | - | 4705 | 1550 | - | - | 0 | - |
| - | - | 1608 | 1551 | - | - | 0 | - |
| - | - | 1327 | 1554 | - | - | 0 | - |
| - | - | 2741 | 1563 | - | - | 0 | - |
| - | - | 6004 | 1564 | - | - | 0 | - |
| - | - | 7.664E+04 | 1565 | - | - | 0 | - |
| - | - | 9.555E+04 | 1566 | - | - | 0 | - |
| - | - | 5.857E+04 | 1567 | - | - | 0 | - |
| - | - | 2.224E+04 | 1568 | - | - | 0 | - |
| - | - | 7761 | 1569 | - | - | 0 | - |
| - | - | 2067 | 1570 | - | - | 0 | - |
| - | - | 2324 | 1578 | - | - | 0 | - |
| - | - | 1.101E+04 | 1579 | - | - | 0 | - |
| - | - | 1.844E+04 | 1580 | - | - | 0 | - |
| - | - | 2.048E+04 | 1581 | - | - | 0 | - |
| - | - | 2.524E+04 | 1582 | - | - | 0 | - |
| - | - | 1.506E+04 | 1583 | - | - | 0 | - |
| - | - | 7361 | 1584 | - | - | 0 | - |
| - | - | 1725 | 1585 | - | - | 0 | - |
| - | - | 6237 | 1590 | - | - | 0 | - |
| - | - | 2.385E+04 | 1591 | - | - | 0 | - |
| - | - | 1.878E+04 | 1592 | - | - | 0 | - |
| - | - | 8586 | 1593 | - | - | 0 | - |
| - | - | 3009 | 1594 | - | - | 0 | - |
| - | - | 4306 | 1596 | - | - | 0 | - |
| - | - | 1060 | 1597 | - | - | 0 | - |
| - | - | 2.607E+04 | 1597 | - | - | 0 | - |
| - | - | 1.94E+04 | 1598 | - | - | 0 | - |
| - | - | 9681 | 1599 | - | - | 0 | - |
| - | - | 2788 | 1600 | - | - | 0 | - |
| - | - | 1533 | 1601 | - | - | 0 | - |
| - | - | 4.824E+04 | 1607 | - | - | 0 | - |
| - | - | 2.428E+05 | 1608 | - | - | 0 | - |
| - | - | 2.005E+05 | 1609 | - | - | 0 | - |
| - | - | 1.012E+05 | 1610 | - | - | 0 | - |
| - | - | 3.133E+04 | 1611 | - | - | 0 | - |
| - | - | 9005 | 1612 | - | - | 0 | - |
| - | - | 1.073E+04 | 1623 | - | - | 0 | - |
| - | - | 2534 | 1624 | - | - | 0 | - |
| - | - | 6.22E+04 | 1624 | - | - | 0 | - |
| - | - | 2.418E+05 | 1625 | - | - | 0 | - |
| - | - | 1.824E+05 | 1626 | - | - | 0 | - |
| - | - | 9.279E+04 | 1627 | - | - | 0 | - |
| - | - | 2.985E+04 | 1628 | - | - | 0 | - |
| - | - | 7476 | 1629 | - | - | 0 | - |
| - | - | 1397 | 2228 | - | - | 0 | - |
| - | - | 1419 | 3243 | - | - | 0 | - |
| - | - | 1245 | 3499 | - | - | 0 | - |

m/z Charge Intensity FragmentType MassShift Position
120.0810775756836 0 1241.878
121.11397552490234 0 670.54156
129.1131591796875 0 940.5996
148.95449829101562 0 964.6495
172.3621063232422 0 868.50464
173.12852478027344 0 26596.58
173.4381866455078 0 1206.358
174.13197326660156 0 2508.421
183.1129913330078 0 9967.884
184.11654663085938 0 1372.9012
184.76817321777344 0 1091.6545
191.81692504882812 0 896.9326
199.07110595703125 0 2364.5417
199.1187744140625 0 1014.3264
201.12342834472656 0 50384.375
201.13482666015625 0 22988.736
202.12368774414062 0 6914.7705
202.14157104492188 0 3510.5361
211.10829162597656 0 1280.683
217.0821075439453 0 4953.6567
226.13026428222656 0 1379.9934
229.1182098388672 0 1735.9398
229.1297149658203 0 3655.5935
242.11351013183594 0 1536.0009
244.1446075439453 0 3619.5923
261.1231689453125 0 5406.0015
261.1709289550781 0 54701.668
262.1744689941406 0 8797.001
270.156494140625 0 2812.8145
284.16070556640625 0 1045.5378
288.16705322265625 0 3663.173
289.1727600097656 0 1153.3005
298.16259765625 0 1442.4598 y 10
299.1936950683594 0 1202.6056
303.0821228027344 0 1058.2239
306.1685485839844 0 11052.761 z 13
307.171630859375 0 2546.4155
312.1557312011719 0 26529.174
313.158935546875 0 4076.6047
316.1504211425781 0 4936.7334
320.1722412109375 0 2034.9874 z Water loss 6
321.1784362792969 0 2585.495
322.1878356933594 0 13710.989 y 13
323.1905822753906 0 1288.1307
330.1662292480469 0 15662.27
331.1694030761719 0 3690.5085
347.9123840332031 0 984.1126
348.203369140625 0 8096.489
349.2077941894531 0 1837.6614
373.1976013183594 0 4061.066
376.1979675292969 0 33737.836 c Water loss 10
377.199462890625 0 6255.281
387.2338562011719 0 1625.5309
391.1851806640625 0 15764.305
392.19293212890625 0 59731.223 y Ammonia loss 12
393.1987609863281 0 24757.422 y Ammonia loss 4
394.2050476074219 0 7195.4976 c Ammonia loss 6
395.2066650390625 0 1936.4436
401.23968505859375 0 7236.368
403.2481384277344 0 1644.7288
407.2034912109375 0 1787.3922
409.2198181152344 0 12756.199 y 12
410.2237854003906 0 1374.1735
411.22393798828125 0 1907.413
417.1884460449219 0 19184.191
418.1912536621094 0 4101.699
418.7180480957031 0 2651.283 y 7
419.2237854003906 0 1258.2627
428.226806640625 0 4726.1494
429.1836853027344 0 1442.0173
429.23486328125 0 12013.49
430.237548828125 0 3548.264
435.2359924316406 0 4407.207
436.2411193847656 0 3437.7312
444.25689697265625 0 9474.789
445.25616455078125 0 8705.449
446.2607421875 0 10922.758 c 3
447.26348876953125 0 2285.5833
462.2232971191406 0 3147.1602 z Water loss 11
463.2303466796875 0 40568.086 w 11
464.231201171875 0 13058.514
465.23150634765625 0 1944.6919
468.8994140625 0 1758.38 y Water loss 2
469.2353820800781 0 1559.3992 y Ammonia loss 2
473.27001953125 0 1326.0117
474.9044189453125 0 1620.9176 y 2
475.23822021484375 0 1795.0493
478.21759033203125 0 15168.891
479.2242736816406 0 17855.023 y Ammonia loss 11
480.2326965332031 0 29175.076 z 11
481.23626708984375 0 9211.759
482.2381591796875 0 1749.8184
488.75830078125 0 32517.557 z 6
489.260009765625 0 18740.74
489.76080322265625 0 7906.888
490.28668212890625 0 4380.338
494.237060546875 0 2390.2878
495.2337646484375 0 1320.2719
496.2523498535156 0 12018.73 y 11
496.7674865722656 0 10192.49 y 6
497.26678466796875 0 3992.3145
498.2558898925781 0 3564.5164
499.2995910644531 0 2460.5708
503.9155578613281 0 4310.4346 y 1
504.2207336425781 0 17274.998
505.2235107421875 0 5103.962
506.225830078125 0 1969.658
516.266357421875 0 5542.8105
524.2752685546875 0 10660.084
524.7786865234375 0 6893.1187
525.2789306640625 0 3329.4734
529.9357299804688 0 1329.9515
532.2852172851562 0 18111.572
533.29296875 0 80628.75 c 4
534.2966918945312 0 21014.807
535.3030395507812 0 5335.8555
535.6058959960938 0 11250.081
535.9398193359375 0 10137.911
536.2754516601562 0 4150.6045
536.6066284179688 0 1419.278
540.8155517578125 0 1082.9277
541.2805786132812 0 1768.916
541.3451538085938 0 9173.6455
541.6095581054688 0 44643.516
541.772705078125 0 1902.6074
541.9441528320312 0 36337.26
542.277587890625 0 25623.9
542.3500366210938 0 2606.3252
542.6129150390625 0 7779.795
546.2717895507812 0 44022.31 z 5
546.7743530273438 0 49072.25
547.2763061523438 0 20794.914
547.7774658203125 0 6389.4507
548.27880859375 0 2009.093
549.2911376953125 0 2108.4324
553.7772216796875 0 4908.358
554.2804565429688 0 26866.287 y 5
554.78173828125 0 13632.62
555.2857666015625 0 3986.653
556.289794921875 0 1986.5894
561.2913208007812 0 6115.891 z Water loss 10
562.2969970703125 0 5042.8994
563.3016967773438 0 1307.1537
564.27783203125 0 30292.65 c Ammonia loss 10
565.2808837890625 0 8274.178
566.2864990234375 0 2060.9937
567.7906494140625 0 3107.5015
568.2892456054688 0 3184.3591
568.7890625 0 1355.5104
579.3009033203125 0 18020.328 z 10
580.3045043945312 0 5595.8535
581.7823486328125 0 4090.693
588.2852172851562 0 1534.6766
588.7830810546875 0 8247.469 y Water loss 4
589.2826538085938 0 13797.899 y Ammonia loss 4
589.7872314453125 0 45349.008 z 4
590.2907104492188 0 77535.28
590.7926025390625 0 38171.094
591.2936401367188 0 11277.044
591.79345703125 0 2859.0195
592.3323974609375 0 24698.867
593.3343505859375 0 9132.7705
594.3327026367188 0 1295.6488
595.3208618164062 0 20904.805 y 10
596.3228759765625 0 6261.063
597.2928466796875 0 11063.24
597.7969360351562 0 125905.336 y 4
598.2984619140625 0 85201.47
598.7998657226562 0 33142.887
599.301025390625 0 6991.9673
599.797607421875 0 1770.3934
601.7809448242188 0 7390.2876
602.283203125 0 3656.0737
611.291748046875 0 2355.6963
611.7915649414062 0 1516.124
612.2954711914062 0 1547.0093
614.3218994140625 0 1362.3298
619.3212890625 0 3015.9482
620.3232421875 0 1745.0051
621.3006591796875 0 1886.7268
624.3192749023438 0 1444.2434
631.2987060546875 0 2528.2876
631.8109130859375 0 3185.21 w 3
632.3103637695312 0 3672.7878
632.3723754882812 0 1315.0615
636.3229370117188 0 72233.69 z 9
637.3277587890625 0 42803.43
638.3296508789062 0 12457.218 y Water loss 3
639.3223266601562 0 70820.04 z 3
639.8236083984375 0 55269.23
640.3248901367188 0 21710.256
640.8257446289062 0 6703.7065
641.3255004882812 0 2291.228
646.8277587890625 0 5565.247
647.3312377929688 0 122134.53 y 3
647.8327026367188 0 90683.35
648.3246459960938 0 88798.53 c 5
648.8355712890625 0 10042.193
649.3239135742188 0 23251.14
650.3267822265625 0 4505.3755
651.3305053710938 0 3262.4114 c Ammonia loss 12
651.8304443359375 0 1835.9602
652.341552734375 0 82228.9 y 9
653.343994140625 0 27723.166
654.3461303710938 0 6115.367
674.33642578125 0 65502.273 w 2
674.8380126953125 0 49609.715
675.3388671875 0 20666.494
675.8406372070312 0 8954.981
679.3650512695312 0 10327.7705
680.36767578125 0 3068.8086
680.795166015625 0 1209.9176
681.8424072265625 0 2252.791
682.3500366210938 0 2736.0964
694.8368530273438 0 1406.1244 z Water loss 2
695.33984375 0 1998.6581 z Ammonia loss 2
696.3387451171875 0 2292.224
702.8502197265625 0 6580.086 y Water loss 2
703.3488159179688 0 6678.5786 y Ammonia loss 2
703.8435668945312 0 50061.785 z 2
704.3445434570312 0 36016.65
704.8464965820312 0 18634.129
705.3479614257812 0 6027.004 z Water loss 8
707.8685913085938 0 3619.5747
708.36962890625 0 3603.356
708.866943359375 0 2843.5208
709.3551025390625 0 1616.5736
711.346435546875 0 4352.1587
711.8526000976562 0 151381.16 y 2
712.3541870117188 0 116610.664
712.855224609375 0 53483.945
713.3566284179688 0 15244.824
713.857421875 0 2990.303
715.8786010742188 0 12613.032
716.3766479492188 0 8873.401
716.8779296875 0 6553.09
717.852294921875 0 8897.708
718.3564453125 0 7265.4023
718.8555908203125 0 4783.337
721.3427124023438 0 4211.7036
722.3466796875 0 9504.191 y Ammonia loss 8
723.3547973632812 0 52659.652 z 8
724.3585205078125 0 18650.592
724.8651733398438 0 1594.7047
725.3616333007812 0 8823.295
725.8613891601562 0 3589.7625
726.3638916015625 0 3113.828
729.3677368164062 0 3836.9426 c Ammonia loss 13
730.3749389648438 0 3454.4749
731.3815307617188 0 1913.4935
733.3491821289062 0 1417.3523
734.3922729492188 0 1871.7574
735.3988037109375 0 1314.9918
737.3640747070312 0 2311.4924
737.8846435546875 0 328572.5 c 13
738.3856201171875 0 257351.75
738.8867797851562 0 121053.68
739.3831176757812 0 47193.61 y 8
739.8853149414062 0 7037.4595
740.3744506835938 0 8309.231
741.3780517578125 0 1692.8224
744.3915405273438 0 1896.2838
745.4005126953125 0 2239.5981
746.3602905273438 0 13189.079 y Water loss 1
746.86083984375 0 15013.956 y Ammonia loss 1
747.357177734375 0 66607.14 z 1
747.8604736328125 0 43524.258
748.359619140625 0 23484.855
748.8631591796875 0 5106.033
754.8945922851562 0 2730.6833
755.36865234375 0 188683.77 y 1
755.8702392578125 0 164851.47
756.3715209960938 0 71353.38
756.8721313476562 0 22618.791
757.374267578125 0 5873.4756
759.39990234375 0 1721.8113
760.349853515625 0 3378.468
760.40771484375 0 6389.958
760.8624267578125 0 5723.5903
761.4119873046875 0 2825.7039
761.8666381835938 0 1402.0789
762.36181640625 0 1655.6638
767.8992309570312 0 1529.5565
768.3949584960938 0 2979.0098
773.3931884765625 0 3693.1655
773.899658203125 0 1499.31
774.3939208984375 0 7590.7295
774.8976440429688 0 4573.8003
775.3858032226562 0 4608.9653
775.8785400390625 0 5774.37
776.3776245117188 0 3255.1467
776.8740234375 0 1931.1783
777.3787841796875 0 1621.7053
781.4050903320312 0 5308.2515
781.90673828125 0 32724.207
782.4052734375 0 67759.414
782.9049072265625 0 53627.832
783.405517578125 0 27560.15
783.9077758789062 0 8203.061
787.39208984375 0 1759.565 c Ammonia loss 6
787.9003295898438 0 4571.393
788.4008178710938 0 7176.611
788.900146484375 0 6570.408
789.3841552734375 0 98544.92
789.8861083984375 0 83840.75
790.388916015625 0 43517.32
790.8991088867188 0 102143.23
791.4009399414062 0 80334.51
791.9031982421875 0 42840.38
792.4033813476562 0 10817.698
792.9071044921875 0 4387.454
793.386474609375 0 1737.0095 w 7
794.3956298828125 0 1605.8391
794.89599609375 0 12622.141
795.3970947265625 0 10689.275
795.9010009765625 0 5468.1143
796.391357421875 0 11254.115
796.8919677734375 0 7207.2017
797.3937377929688 0 3802.8184
797.8973999023438 0 2127.7205
798.4080810546875 0 2037.3103
803.40869140625 0 34587.527
803.9024658203125 0 335483.25
804.4033203125 0 295493.5
804.9041137695312 0 141386.84
805.4052124023438 0 45706.26
805.9060668945312 0 12304.431
811.403076171875 0 1456.6101
811.9105834960938 0 164439.64
812.4132080078125 0 249602.67
812.9147338867188 0 153876.28
813.4163208007812 0 71053.414
813.9179077148438 0 23688.889
814.4182739257812 0 2659.0923
834.407958984375 0 3162.0938
835.412353515625 0 2093.8765
836.427490234375 0 12916.955 y 7
837.4293212890625 0 4875.9526
842.453125 0 3607.4324
856.4453125 0 1383.3729
857.4598388671875 0 16895.035
858.4644165039062 0 7724.059
859.4674072265625 0 2426.0068
864.4122924804688 0 1660.642
885.4583129882812 0 4229.358
886.4615478515625 0 4250.11
890.4395141601562 0 5776.36 w 6
891.44091796875 0 1787.7955
892.4481201171875 0 1706.3586
893.4727783203125 0 2113.714
900.4945678710938 0 4846.478
901.4744873046875 0 92227.7 c 7
902.47705078125 0 42323.73
903.4802856445312 0 12451.634
904.4859008789062 0 2966.9255
916.4803466796875 0 2047.924
926.4861450195312 0 2900.138
927.4808349609375 0 2430.4167
929.4863891601562 0 2044.0925
933.5025634765625 0 3995.1929
944.4931030273438 0 53448.35
945.4957885742188 0 27646.17
946.4996337890625 0 8785.883
947.5015869140625 0 2919.5325
958.50146484375 0 2685.1697 z Water loss 6
971.4896240234375 0 1749.1519 c Ammonia loss 8
972.4874877929688 0 3544.5483
975.508056640625 0 2111.5642 y Ammonia loss 6
976.5086059570312 0 50268.49 z 6
977.512939453125 0 33553.99
977.6270141601562 0 1583.1693
978.5148315429688 0 13000.766
979.5139770507812 0 1528.2083
987.4993286132812 0 13851.903
988.5065307617188 0 224717.61 c 8
989.5093383789062 0 118732.27
990.511474609375 0 37358.914
991.5152587890625 0 7931.006
992.5372924804688 0 2070.5315 y 6
1001.5142822265625 0 13884.038
1002.51806640625 0 8169.6675
1003.521484375 0 2899.0486
1030.5242919921875 0 1490.5101
1031.51806640625 0 2132.7563
1032.5206298828125 0 1807.8054
1045.528076171875 0 134088.34 c 9
1046.5311279296875 0 73826.03 w 5
1047.5377197265625 0 35706.66
1048.5452880859375 0 13517.8
1049.5479736328125 0 4149.116
1059.5186767578125 0 1712.9797
1073.5301513671875 0 3456.9285 z Water loss 5
1074.5228271484375 0 2627.3267 z Ammonia loss 5
1081.5533447265625 0 1745.5273
1082.5567626953125 0 2780.2375
1084.5726318359375 0 2042.5443
1091.5360107421875 0 47480.957 z 5
1092.541748046875 0 78050.05
1093.5455322265625 0 35113.086
1094.5498046875 0 9156.531
1095.542724609375 0 1804.7572
1099.57470703125 0 1527.3
1100.5841064453125 0 7671.0884
1101.58349609375 0 4612.7993
1102.5972900390625 0 2920.6428
1103.5975341796875 0 1789.0819
1106.544921875 0 3270.6204
1107.553466796875 0 10909.315 y 5
1108.5555419921875 0 7124.834
1109.55908203125 0 3009.2974
1129.5859375 0 9647.404
1130.593017578125 0 5430.094
1131.5963134765625 0 2291.9539
1143.592529296875 0 2966.3123
1144.595947265625 0 97352.38 c 10
1145.599609375 0 60056.453
1146.601806640625 0 25851.145
1147.6072998046875 0 6872.9443
1148.6109619140625 0 1669.8726
1161.541748046875 0 1713.5781 z Ammonia loss 4
1172.6033935546875 0 2003.404
1177.5616455078125 0 2589.2534 y Ammonia loss 4
1178.5650634765625 0 8848.464 z 4
1179.5745849609375 0 36433.19
1180.5772705078125 0 21171.697
1181.5816650390625 0 9089.013
1182.5899658203125 0 1720.0461
1187.61474609375 0 10243.369
1188.614501953125 0 6320.574
1189.6243896484375 0 4942.566
1190.62451171875 0 2759.0679
1193.5831298828125 0 1986.619
1194.5855712890625 0 14629.961 y 4
1195.58740234375 0 10472.612
1196.5955810546875 0 5094.3203
1197.6058349609375 0 1652.5232
1230.623291015625 0 8668.443
1231.626953125 0 188305.7 c 11
1232.6304931640625 0 113305.42
1233.6334228515625 0 45481.96
1234.6339111328125 0 12998.443
1235.6329345703125 0 2631.4165
1236.6282958984375 0 1457.0576
1258.6259765625 0 8913.1455
1259.629638671875 0 6774.0234 z Water loss 3
1260.6246337890625 0 2344.8115 z Ammonia loss 3
1262.6324462890625 0 1770.9358 w 3
1274.64453125 0 7552.8525
1275.6519775390625 0 4810.4507 y Water loss 3
1276.6373291015625 0 1928.7607 y Ammonia loss 3
1277.635498046875 0 8331.451 z 3
1278.642822265625 0 44305.293
1279.6458740234375 0 27297.137
1280.6475830078125 0 11360.802
1281.650146484375 0 2211.2808
1293.6544189453125 0 11213.69 y 3
1294.6558837890625 0 9031.419
1295.6614990234375 0 4109.421
1296.6746826171875 0 1614.4847
1318.6591796875 0 133328.42 c 12
1319.661865234375 0 96661.68
1320.6646728515625 0 40211.516
1321.6663818359375 0 11847.666
1322.6802978515625 0 1624.7611
1344.6724853515625 0 2940.084
1345.661376953125 0 13149.032
1346.6649169921875 0 10483.8545
1347.6636962890625 0 6284.23 w 2
1348.6556396484375 0 1809.3359
1364.657958984375 0 2186.353
1394.6304931640625 0 2533.4355
1395.6424560546875 0 2346.1145
1406.6773681640625 0 9489.942 z 2
1407.6842041015625 0 73710.38
1408.6878662109375 0 53836.156
1409.6903076171875 0 23228.447
1410.6900634765625 0 7510.0195
1411.6876220703125 0 2051.5583
1414.7366943359375 0 2826.9043
1415.732666015625 0 1676.1158
1416.716552734375 0 5243.787
1417.729248046875 0 4280.2373
1418.7325439453125 0 2350.5337
1422.6893310546875 0 2218.8599 y 2
1423.70361328125 0 4349.203
1424.7020263671875 0 2810.0483
1425.7037353515625 0 1456.2258
1430.748046875 0 5133.513
1431.7569580078125 0 18148.71
1432.753173828125 0 12369.229
1433.75341796875 0 6047.3022
1434.7442626953125 0 2173.849
1435.6644287109375 0 1567.9398
1447.7725830078125 0 2129.7278
1451.688232421875 0 3066.5417
1452.693115234375 0 1952.3373
1458.7398681640625 0 13704.999
1459.74462890625 0 10692.531
1460.7440185546875 0 4485.2266
1461.749755859375 0 1818.4917
1464.7318115234375 0 2342.9453
1474.759765625 0 28701.857 c 13
1475.76611328125 0 85678.45
1476.7674560546875 0 60535.895
1477.765380859375 0 28364.684
1478.762451171875 0 10246.328
1479.7691650390625 0 1706.349
1493.714111328125 0 7084.711 z 1
1494.715576171875 0 54459.47
1495.7200927734375 0 40503.582
1496.7237548828125 0 20468.395
1497.7222900390625 0 6768.53
1507.7523193359375 0 3978.1182
1509.7484130859375 0 3463.3992 y 1
1510.718505859375 0 2356.1892
1521.7852783203125 0 2541.6323
1522.72607421875 0 17041.865
1523.72900390625 0 14822.42
1524.7314453125 0 8645.176
1525.7392578125 0 3272.9795
1535.7578125 0 2257.1572
1536.7628173828125 0 2039.5233
1537.7716064453125 0 2340.5085
1538.7843017578125 0 2056.001
1546.765869140625 0 2988.5413
1547.76953125 0 5011.8643
1548.7657470703125 0 5167.4287
1549.7659912109375 0 4704.9116
1550.7691650390625 0 1608.3585
1553.7933349609375 0 1327.437
1562.820556640625 0 2740.5032
1563.79345703125 0 6003.7354
1564.78173828125 0 76644.27
1565.7822265625 0 95545.27
1566.7852783203125 0 58569.453
1567.7872314453125 0 22242.176
1568.7899169921875 0 7761.0977
1569.8016357421875 0 2067.453
1577.8009033203125 0 2324.0386
1578.8197021484375 0 11013.834
1579.81494140625 0 18436.066
1580.8084716796875 0 20482.04
1581.8018798828125 0 25238.629
1582.8038330078125 0 15058.432
1583.80712890625 0 7360.615
1584.7987060546875 0 1724.6045
1589.79345703125 0 6236.958
1590.77783203125 0 23847.43
1591.7822265625 0 18776.977
1592.7789306640625 0 8585.645
1593.7779541015625 0 3009.1777
1595.8221435546875 0 4306.018
1596.6185302734375 0 1060.3467
1596.831787109375 0 26067.14
1597.8345947265625 0 19398.414
1598.837890625 0 9680.62
1599.855224609375 0 2787.7239
1600.829833984375 0 1532.745
1606.8055419921875 0 48244.73
1607.8016357421875 0 242833.34
1608.803955078125 0 200475.06
1609.806396484375 0 101203.1
1610.81005859375 0 31327.635
1611.8111572265625 0 9004.998
1622.8140869140625 0 10729.342
1623.5738525390625 0 2534.1035
1623.8184814453125 0 62197.83
1624.82568359375 0 241784.62
1625.8287353515625 0 182444.33
1626.8310546875 0 92787.2
1627.833984375 0 29850.64
1628.837158203125 0 7476.441
2228.072509765625 0 1397.1338
3242.95458984375 0 1418.6028
3498.86474609375 0 1245.078

Spectrum Details

|  |  |
| --- | --- |
| Matched peaks? Matched peaks The total absolute number of peaks matched. Additionally in brackets the total fraction of peaks matched and the total number of peaks is shown. | 95 (16.52% of 575) |
| FDR? FDR The false discovery rate estimated for this peptide. It is calculated by matching all theoretical fragments with a non-integer shift with the raw peaks for this spectrum. This is done with 40 different shifts. The resulting percentage is the average number of annotated peaks over the number of annotated peaks with the correct spectrum. | 0.15% |
| Satellite FDR? Satellite FDR See the FDR for details on its calculation. This satellite ion specific FDR only contains the satellite ions (d/w) for I/L/J positions. | - |
| PSM Score? PSM Score The PSM Score as given by Hecklib to this annotated spectrum. It is shown with three significant figures. | 737 |

#### Spectrum 5111? Spectrum 5111 The raw spectrum of this peptide as annotated by Hecklib. The fragments are coloured according to ion type (see legend). Any peaks with a star '\*' as text can be hovered over to see the full details, first the ion type second the mass shift type. By hovering over the amino acids in the peptide or ions in the legend the corresponding peaks are highlighted. By toggling the 'Unassigned' label you can turn the background (unassigned) peaks on or off in the plot. By updating the slider in the Ion legend you can update the spectrum to only show the top X% of the peaks with labels. The top X% means any peak that is within X% of the highest intensity. By dragging in the spectrum you can zoom in to a specific part of the spectrum and use 'Zoom Out' to get back to the original zoom level. The annotation of the spectrum is based on the given sequence in the peptides file and is done with different software so inconsistencies are likely. The peaks are annotated based on the given sequence, with 20 ppm tolerance.

Copy Data

##### Spectrum 5111 (TSV)

###### Preview

```
Loading example...
```

*Click on the button to copy the data to your clipboard.*

Mz MinMz MaxIntensity Max

WidthHeightPeptide font sizePeptide stroke widthSpectrum font sizeSpectrum stroke widthCompact peptide

Ion legend

wxyz

abcd

OtherUnassignedIon^Charge^_Position_Show for top:%

JSEVSDRPSGVSSRF

02.24e+44.49e+46.73e+48.98e+4

Zoom Out

z^+1^_2_y^+1^_2_c^+3^_11_y^+1^_3_y^+3^_11_c^+2^_7_y^+1^_3_c^+1^_4_w^+1^_4_y^+1^_4_z^+1^_4_z^+2^_9_z^+3^_14_y^+1^_4_y^+2^_9_y^+3^_14_c^+1^_5_z^+2^_10_y^+2^_10_z^+1^_5_c^+2^_11_z^+1^_5_y^+2^_11_y^+2^_11_z^+2^_11_y^+1^_5_y^+2^_11_z^+1^_6_w^+2^_12_z^+1^_6_y^+2^_12_z^+2^_12_y^+2^_12_c^+1^_6_y^+1^_6_w^+2^_13_z^+2^_13_y^+2^_13_y^+2^_13_z^+2^_13_z^+1^_7_y^+2^_13_y^+1^_7_z^+1^_7_c^+2^_14_c^+2^_14_y^+1^_7_y^+2^_14_y^+2^_14_z^+2^_14_y^+2^_14_y^+1^_8_w^+1^_9_c^+1^_8_z^+1^_9_c^+1^_9_c^+1^_10_w^+1^_10_z^+1^_10_y^+1^_10_c^+1^_11_z^+1^_11_y^+1^_11_c^+1^_12_z^+1^_12_y^+1^_12_z^+1^_12_y^+1^_12_c^+1^_13_w^+1^_13_z^+1^_13_c^+1^_14_z^+1^_14_y^+1^_14_

0555111016652221

Fragment Matches Table

Show background peaks

| Position | Ion type | Intensity | mz Theoretical | mz Error (Th) | mz Error (ppm) | Charge | Series Number |
| --- | --- | --- | --- | --- | --- | --- | --- |
| - | - | 433.4 | 120.1 | - | - | 0 | - |
| - | - | 363.4 | 120.4 | - | - | 0 | - |
| - | - | 1069 | 142.1 | - | - | 0 | - |
| - | - | 928 | 148.9 | - | - | 0 | - |
| - | - | 469.2 | 155.1 | - | - | 0 | - |
| - | - | 455.3 | 159.4 | - | - | 0 | - |
| - | - | 440.4 | 160.1 | - | - | 0 | - |
| - | - | 435.9 | 163.3 | - | - | 0 | - |
| - | - | 6448 | 173.1 | - | - | 0 | - |
| - | - | 3092 | 173.5 | - | - | 0 | - |
| - | - | 2390 | 183.1 | - | - | 0 | - |
| - | - | 892.9 | 199.1 | - | - | 0 | - |
| - | - | 9292 | 201.1 | - | - | 0 | - |
| - | - | 6675 | 201.1 | - | - | 0 | - |
| - | - | 1321 | 202.1 | - | - | 0 | - |
| - | - | 519.7 | 208 | - | - | 0 | - |
| - | - | 671.6 | 217.1 | - | - | 0 | - |
| - | - | 767.8 | 229.1 | - | - | 0 | - |
| - | - | 1099 | 261.1 | - | - | 0 | - |
| - | - | 1.297E+04 | 261.2 | - | - | 0 | - |
| - | - | 2677 | 262.2 | - | - | 0 | - |
| - | - | 1015 | 270.2 | - | - | 0 | - |
| - | - | 502.5 | 285 | - | - | 0 | - |
| - | - | 1364 | 288.2 | - | - | 0 | - |
| - | - | 651.6 | 291.4 | - | - | 0 | - |
| 14 | z | 2761 | 306.2 | 1.878E-06 | 0.006133 | +1 | 2 |
| - | - | 604.8 | 307.2 | - | - | 0 | - |
| - | - | 4668 | 312.2 | - | - | 0 | - |
| - | - | 1208 | 313.2 | - | - | 0 | - |
| - | - | 957.4 | 316.1 | - | - | 0 | - |
| 14 | y | 3067 | 322.2 | 0.0002933 | 0.9104 | +1 | 2 |
| - | - | 740 | 323.2 | - | - | 0 | - |
| - | - | 3287 | 330.2 | - | - | 0 | - |
| - | - | 1837 | 348.2 | - | - | 0 | - |
| - | - | 552.5 | 362.8 | - | - | 0 | - |
| - | - | 607.4 | 373.2 | - | - | 0 | - |
| 11 | c | 8259 | 376.2 | 0.002562 | 6.81 | +3 | 11 |
| - | - | 582.1 | 377.2 | - | - | 0 | - |
| - | - | 1815 | 377.2 | - | - | 0 | - |
| - | - | 3714 | 391.2 | - | - | 0 | - |
| 13 | y | 1.493E+04 | 392.2 | 0.0003405 | 0.8682 | +1 | 3 |
| 5 | y | 5655 | 393.2 | 0.00746 | 18.97 | +3 | 11 |
| 7 | c | 2002 | 394.2 | 0.002197 | 5.572 | +2 | 7 |
| - | - | 631 | 394.9 | - | - | 0 | - |
| - | - | 1943 | 401.2 | - | - | 0 | - |
| 13 | y | 3315 | 409.2 | 0.0004919 | 1.202 | +1 | 3 |
| - | - | 1035 | 410.2 | - | - | 0 | - |
| - | - | 634.3 | 411.2 | - | - | 0 | - |
| - | - | 565.1 | 415.5 | - | - | 0 | - |
| - | - | 4883 | 417.2 | - | - | 0 | - |
| - | - | 1078 | 418.2 | - | - | 0 | - |
| - | - | 552.4 | 418.6 | - | - | 0 | - |
| - | - | 685.6 | 428.2 | - | - | 0 | - |
| - | - | 3762 | 429.2 | - | - | 0 | - |
| - | - | 1328 | 435.2 | - | - | 0 | - |
| - | - | 1156 | 436.2 | - | - | 0 | - |
| - | - | 3057 | 444.3 | - | - | 0 | - |
| - | - | 2573 | 445.3 | - | - | 0 | - |
| - | - | 702.1 | 445.6 | - | - | 0 | - |
| 4 | c | 2286 | 446.3 | 0.0008236 | 1.846 | +1 | 4 |
| 12 | w | 1.069E+04 | 463.2 | 0.0002654 | 0.573 | +1 | 4 |
| - | - | 2669 | 464.2 | - | - | 0 | - |
| - | - | 2046 | 478.1 | - | - | 0 | - |
| - | - | 4088 | 478.2 | - | - | 0 | - |
| 12 | y | 4441 | 479.2 | 0.001119 | 2.335 | +1 | 4 |
| 12 | z | 9081 | 480.2 | 0.000399 | 0.8309 | +1 | 4 |
| - | - | 2509 | 481.2 | - | - | 0 | - |
| 7 | z | 7176 | 488.8 | 0.0001572 | 0.3216 | +2 | 9 |
| - | - | 4459 | 489.3 | - | - | 0 | - |
| - | - | 2543 | 489.8 | - | - | 0 | - |
| 2 | z | 725.9 | 492.6 | 0.001903 | 3.864 | +3 | 14 |
| 12 | y | 2675 | 496.3 | 0.0001717 | 0.346 | +1 | 4 |
| 7 | y | 3310 | 496.8 | 0.0003944 | 0.794 | +2 | 9 |
| - | - | 1532 | 497.3 | - | - | 0 | - |
| 2 | y | 1361 | 497.9 | 0.0005744 | 1.154 | +3 | 14 |
| - | - | 687 | 498.3 | - | - | 0 | - |
| - | - | 3883 | 504.2 | - | - | 0 | - |
| - | - | 801.7 | 507.1 | - | - | 0 | - |
| - | - | 1516 | 516.3 | - | - | 0 | - |
| - | - | 3831 | 524.3 | - | - | 0 | - |
| - | - | 2098 | 524.8 | - | - | 0 | - |
| - | - | 1364 | 525.3 | - | - | 0 | - |
| - | - | 4196 | 532.3 | - | - | 0 | - |
| 5 | c | 1.988E+04 | 533.3 | 0.0007781 | 1.459 | +1 | 5 |
| - | - | 6084 | 534.3 | - | - | 0 | - |
| - | - | 1098 | 535.3 | - | - | 0 | - |
| - | - | 2180 | 535.6 | - | - | 0 | - |
| - | - | 2505 | 535.9 | - | - | 0 | - |
| - | - | 1231 | 536.3 | - | - | 0 | - |
| - | - | 717.3 | 539.8 | - | - | 0 | - |
| - | - | 753.4 | 540.3 | - | - | 0 | - |
| - | - | 1826 | 541.3 | - | - | 0 | - |
| - | - | 8391 | 541.6 | - | - | 0 | - |
| - | - | 5821 | 541.9 | - | - | 0 | - |
| - | - | 3198 | 542.3 | - | - | 0 | - |
| 6 | z | 1.165E+04 | 546.3 | 0.0004146 | 0.7589 | +2 | 10 |
| - | - | 1.184E+04 | 546.8 | - | - | 0 | - |
| - | - | 5246 | 547.3 | - | - | 0 | - |
| - | - | 1440 | 547.8 | - | - | 0 | - |
| 6 | y | 5203 | 554.3 | 0.0002332 | 0.4207 | +2 | 10 |
| - | - | 4269 | 554.8 | - | - | 0 | - |
| 11 | z | 1620 | 561.3 | 0.0009973 | 1.777 | +1 | 5 |
| - | - | 1955 | 562.3 | - | - | 0 | - |
| 11 | c | 9169 | 564.3 | 0.01098 | 19.46 | +2 | 11 |
| - | - | 2273 | 565.3 | - | - | 0 | - |
| - | - | 1080 | 567.8 | - | - | 0 | - |
| - | - | 783.5 | 568.3 | - | - | 0 | - |
| - | - | 786.4 | 568.8 | - | - | 0 | - |
| 11 | z | 4159 | 579.3 | 0.0003399 | 0.5867 | +1 | 5 |
| - | - | 1138 | 580.3 | - | - | 0 | - |
| - | - | 679.4 | 582.3 | - | - | 0 | - |
| 5 | y | 2374 | 588.8 | 0.01044 | 17.73 | +2 | 11 |
| 5 | y | 3303 | 589.3 | 0.0003011 | 0.511 | +2 | 11 |
| 5 | z | 1.35E+04 | 589.8 | 0.0004986 | 0.8454 | +2 | 11 |
| - | - | 2.173E+04 | 590.3 | - | - | 0 | - |
| - | - | 1.161E+04 | 590.8 | - | - | 0 | - |
| - | - | 2954 | 591.3 | - | - | 0 | - |
| - | - | 5374 | 592.3 | - | - | 0 | - |
| - | - | 1577 | 593.3 | - | - | 0 | - |
| 11 | y | 4952 | 595.3 | 0.0003536 | 0.594 | +1 | 5 |
| - | - | 970.6 | 596.3 | - | - | 0 | - |
| - | - | 2683 | 597.3 | - | - | 0 | - |
| 5 | y | 3.177E+04 | 597.8 | 0.0004002 | 0.6694 | +2 | 11 |
| - | - | 2.001E+04 | 598.3 | - | - | 0 | - |
| - | - | 4942 | 598.8 | - | - | 0 | - |
| - | - | 2083 | 599.3 | - | - | 0 | - |
| - | - | 1370 | 601.8 | - | - | 0 | - |
| - | - | 915.1 | 602.3 | - | - | 0 | - |
| 10 | z | 755.4 | 618.3 | 0.00415 | 6.712 | +1 | 6 |
| - | - | 964.1 | 619.3 | - | - | 0 | - |
| 4 | w | 815.6 | 631.8 | 0.0009343 | 1.479 | +2 | 12 |
| - | - | 1196 | 632.3 | - | - | 0 | - |
| 10 | z | 1.918E+04 | 636.3 | 0.0002498 | 0.3926 | +1 | 6 |
| - | - | 1.069E+04 | 637.3 | - | - | 0 | - |
| 4 | y | 3771 | 638.3 | 0.002169 | 3.398 | +2 | 12 |
| 4 | z | 2.153E+04 | 639.3 | 3.759E-05 | 0.05879 | +2 | 12 |
| - | - | 1.601E+04 | 639.8 | - | - | 0 | - |
| - | - | 5437 | 640.3 | - | - | 0 | - |
| - | - | 679.9 | 640.8 | - | - | 0 | - |
| - | - | 821.1 | 646.8 | - | - | 0 | - |
| 4 | y | 3.021E+04 | 647.3 | 0.0005495 | 0.8489 | +2 | 12 |
| - | - | 1.747E+04 | 647.8 | - | - | 0 | - |
| 6 | c | 2.081E+04 | 648.3 | 0.003773 | 5.82 | +1 | 6 |
| - | - | 2935 | 648.8 | - | - | 0 | - |
| - | - | 5384 | 649.3 | - | - | 0 | - |
| - | - | 1516 | 650.3 | - | - | 0 | - |
| 10 | y | 2.097E+04 | 652.3 | 0.0002971 | 0.4555 | +1 | 6 |
| - | - | 8003 | 653.3 | - | - | 0 | - |
| - | - | 1494 | 654.3 | - | - | 0 | - |
| 3 | w | 1.647E+04 | 674.3 | 0.0001556 | 0.2307 | +2 | 13 |
| - | - | 1.249E+04 | 674.8 | - | - | 0 | - |
| - | - | 5534 | 675.3 | - | - | 0 | - |
| - | - | 1956 | 675.8 | - | - | 0 | - |
| - | - | 2309 | 679.4 | - | - | 0 | - |
| - | - | 1561 | 680.4 | - | - | 0 | - |
| - | - | 896.5 | 681.8 | - | - | 0 | - |
| 3 | z | 857.2 | 694.8 | 0.005127 | 7.379 | +2 | 13 |
| 3 | y | 1093 | 702.8 | 0.001732 | 2.465 | +2 | 13 |
| 3 | y | 1896 | 703.3 | 0.01267 | 18.01 | +2 | 13 |
| 3 | z | 1.113E+04 | 703.8 | 0.0001549 | 0.2201 | +2 | 13 |
| - | - | 7806 | 704.3 | - | - | 0 | - |
| - | - | 4261 | 704.8 | - | - | 0 | - |
| 9 | z | 1100 | 705.3 | 0.003921 | 5.559 | +1 | 7 |
| - | - | 2529 | 708.4 | - | - | 0 | - |
| - | - | 1005 | 711.3 | - | - | 0 | - |
| 3 | y | 3.266E+04 | 711.9 | 0.0007889 | 1.108 | +2 | 13 |
| - | - | 2.531E+04 | 712.4 | - | - | 0 | - |
| - | - | 1.24E+04 | 712.9 | - | - | 0 | - |
| - | - | 3131 | 713.4 | - | - | 0 | - |
| - | - | 3193 | 715.9 | - | - | 0 | - |
| - | - | 2221 | 716.4 | - | - | 0 | - |
| - | - | 768.9 | 716.9 | - | - | 0 | - |
| - | - | 2003 | 717.9 | - | - | 0 | - |
| - | - | 1866 | 718.4 | - | - | 0 | - |
| - | - | 688.7 | 718.9 | - | - | 0 | - |
| - | - | 916.1 | 721.3 | - | - | 0 | - |
| 9 | y | 1989 | 722.3 | 0.002341 | 3.241 | +1 | 7 |
| 9 | z | 1.352E+04 | 723.4 | 0.000662 | 0.9152 | +1 | 7 |
| - | - | 6084 | 724.4 | - | - | 0 | - |
| - | - | 1906 | 725.4 | - | - | 0 | - |
| - | - | 1201 | 725.9 | - | - | 0 | - |
| - | - | 724.9 | 726.3 | - | - | 0 | - |
| 14 | c | 1323 | 729.4 | 0.01172 | 16.06 | +2 | 14 |
| - | - | 805.5 | 729.8 | - | - | 0 | - |
| - | - | 841.9 | 730.4 | - | - | 0 | - |
| - | - | 800.8 | 737.4 | - | - | 0 | - |
| 14 | c | 8.889E+04 | 737.9 | 0.0002107 | 0.2856 | +2 | 14 |
| - | - | 6.386E+04 | 738.4 | - | - | 0 | - |
| - | - | 3.269E+04 | 738.9 | - | - | 0 | - |
| 9 | y | 1.264E+04 | 739.4 | 0.008812 | 11.92 | +1 | 7 |
| - | - | 1603 | 739.9 | - | - | 0 | - |
| - | - | 1802 | 740.4 | - | - | 0 | - |
| - | - | 1095 | 744.4 | - | - | 0 | - |
| 2 | y | 2758 | 746.4 | 0.004624 | 6.195 | +2 | 14 |
| 2 | y | 3654 | 746.9 | 0.004833 | 6.471 | +2 | 14 |
| 2 | z | 1.514E+04 | 747.4 | 0.001887 | 2.525 | +2 | 14 |
| - | - | 1.249E+04 | 747.9 | - | - | 0 | - |
| - | - | 4805 | 748.4 | - | - | 0 | - |
| - | - | 2064 | 748.9 | - | - | 0 | - |
| - | - | 806.5 | 749.3 | - | - | 0 | - |
| 2 | y | 4.241E+04 | 755.4 | 0.0003847 | 0.5093 | +2 | 14 |
| - | - | 3.525E+04 | 755.9 | - | - | 0 | - |
| - | - | 1.568E+04 | 756.4 | - | - | 0 | - |
| - | - | 4395 | 756.9 | - | - | 0 | - |
| - | - | 2087 | 760.4 | - | - | 0 | - |
| - | - | 1216 | 760.9 | - | - | 0 | - |
| - | - | 1019 | 767.9 | - | - | 0 | - |
| - | - | 2464 | 774.4 | - | - | 0 | - |
| - | - | 1128 | 774.9 | - | - | 0 | - |
| - | - | 1232 | 775.9 | - | - | 0 | - |
| - | - | 838.4 | 781.4 | - | - | 0 | - |
| - | - | 7330 | 781.9 | - | - | 0 | - |
| - | - | 1.646E+04 | 782.4 | - | - | 0 | - |
| - | - | 1.206E+04 | 782.9 | - | - | 0 | - |
| - | - | 4801 | 783.4 | - | - | 0 | - |
| - | - | 1380 | 783.9 | - | - | 0 | - |
| - | - | 886.8 | 787.9 | - | - | 0 | - |
| - | - | 1882 | 788.4 | - | - | 0 | - |
| - | - | 1013 | 788.9 | - | - | 0 | - |
| - | - | 2.388E+04 | 789.4 | - | - | 0 | - |
| - | - | 2.551E+04 | 789.9 | - | - | 0 | - |
| - | - | 9823 | 790.4 | - | - | 0 | - |
| - | - | 2.467E+04 | 790.9 | - | - | 0 | - |
| - | - | 2.023E+04 | 791.4 | - | - | 0 | - |
| - | - | 8536 | 791.9 | - | - | 0 | - |
| - | - | 2977 | 792.4 | - | - | 0 | - |
| - | - | 760.1 | 792.9 | - | - | 0 | - |
| - | - | 3030 | 794.9 | - | - | 0 | - |
| - | - | 3055 | 795.4 | - | - | 0 | - |
| - | - | 2306 | 795.9 | - | - | 0 | - |
| - | - | 2620 | 796.4 | - | - | 0 | - |
| - | - | 1513 | 796.9 | - | - | 0 | - |
| - | - | 852.3 | 797.4 | - | - | 0 | - |
| - | - | 1.165E+04 | 803.4 | - | - | 0 | - |
| - | - | 8.172E+04 | 803.9 | - | - | 0 | - |
| - | - | 7.136E+04 | 804.4 | - | - | 0 | - |
| - | - | 3.771E+04 | 804.9 | - | - | 0 | - |
| - | - | 1.187E+04 | 805.4 | - | - | 0 | - |
| - | - | 3041 | 805.9 | - | - | 0 | - |
| - | - | 1086 | 810.4 | - | - | 0 | - |
| - | - | 4.555E+04 | 811.9 | - | - | 0 | - |
| - | - | 5.965E+04 | 812.4 | - | - | 0 | - |
| - | - | 4.3E+04 | 812.9 | - | - | 0 | - |
| - | - | 1.499E+04 | 813.4 | - | - | 0 | - |
| - | - | 4035 | 813.9 | - | - | 0 | - |
| 8 | y | 2711 | 836.4 | 0.0003116 | 0.3725 | +1 | 8 |
| - | - | 825.8 | 837.4 | - | - | 0 | - |
| - | - | 1019 | 842.5 | - | - | 0 | - |
| - | - | 4200 | 857.5 | - | - | 0 | - |
| - | - | 2249 | 858.5 | - | - | 0 | - |
| - | - | 772.8 | 859.5 | - | - | 0 | - |
| - | - | 1111 | 885.5 | - | - | 0 | - |
| - | - | 1137 | 886.5 | - | - | 0 | - |
| 7 | w | 1535 | 890.4 | 0.00127 | 1.426 | +1 | 9 |
| - | - | 1921 | 900.5 | - | - | 0 | - |
| 8 | c | 2.224E+04 | 901.5 | 0.0001667 | 0.1849 | +1 | 8 |
| - | - | 1.213E+04 | 902.5 | - | - | 0 | - |
| - | - | 4363 | 903.5 | - | - | 0 | - |
| - | - | 761.5 | 904.5 | - | - | 0 | - |
| - | - | 1140 | 926.5 | - | - | 0 | - |
| - | - | 899.3 | 927.5 | - | - | 0 | - |
| - | - | 1.279E+04 | 944.5 | - | - | 0 | - |
| - | - | 7842 | 945.5 | - | - | 0 | - |
| - | - | 2374 | 946.5 | - | - | 0 | - |
| 7 | z | 1.413E+04 | 976.5 | 0.0006062 | 0.6208 | +1 | 9 |
| - | - | 9517 | 977.5 | - | - | 0 | - |
| - | - | 3375 | 978.5 | - | - | 0 | - |
| - | - | 3677 | 987.5 | - | - | 0 | - |
| 9 | c | 6.179E+04 | 988.5 | 0.0006727 | 0.6806 | +1 | 9 |
| - | - | 2.759E+04 | 989.5 | - | - | 0 | - |
| - | - | 9016 | 990.5 | - | - | 0 | - |
| - | - | 2238 | 991.5 | - | - | 0 | - |
| - | - | 3280 | 1002 | - | - | 0 | - |
| - | - | 3165 | 1003 | - | - | 0 | - |
| - | - | 1112 | 1026 | - | - | 0 | - |
| 10 | c | 3.366E+04 | 1046 | 0.00053 | 0.5069 | +1 | 10 |
| 6 | w | 1.861E+04 | 1047 | 0.007495 | 7.162 | +1 | 10 |
| - | - | 9425 | 1048 | - | - | 0 | - |
| - | - | 3374 | 1049 | - | - | 0 | - |
| - | - | 701.1 | 1053 | - | - | 0 | - |
| - | - | 880.6 | 1075 | - | - | 0 | - |
| - | - | 2045 | 1081 | - | - | 0 | - |
| - | - | 1651 | 1082 | - | - | 0 | - |
| - | - | 651.4 | 1083 | - | - | 0 | - |
| - | - | 829.9 | 1083 | - | - | 0 | - |
| - | - | 929.3 | 1084 | - | - | 0 | - |
| - | - | 825.4 | 1086 | - | - | 0 | - |
| - | - | 944.7 | 1091 | - | - | 0 | - |
| 6 | z | 1.216E+04 | 1092 | 0.0005107 | 0.4679 | +1 | 10 |
| - | - | 2.075E+04 | 1093 | - | - | 0 | - |
| - | - | 8976 | 1094 | - | - | 0 | - |
| - | - | 2884 | 1095 | - | - | 0 | - |
| - | - | 1507 | 1101 | - | - | 0 | - |
| - | - | 1655 | 1102 | - | - | 0 | - |
| - | - | 1182 | 1107 | - | - | 0 | - |
| 6 | y | 2652 | 1108 | 0.002999 | 2.708 | +1 | 10 |
| - | - | 1709 | 1109 | - | - | 0 | - |
| - | - | 2377 | 1130 | - | - | 0 | - |
| - | - | 2208 | 1131 | - | - | 0 | - |
| 11 | c | 2.574E+04 | 1145 | 0.0008287 | 0.724 | +1 | 11 |
| - | - | 1.743E+04 | 1146 | - | - | 0 | - |
| - | - | 7350 | 1147 | - | - | 0 | - |
| - | - | 1599 | 1148 | - | - | 0 | - |
| 5 | z | 1884 | 1179 | 0.002388 | 2.026 | +1 | 11 |
| - | - | 9495 | 1180 | - | - | 0 | - |
| - | - | 5938 | 1181 | - | - | 0 | - |
| - | - | 2709 | 1182 | - | - | 0 | - |
| - | - | 1326 | 1183 | - | - | 0 | - |
| - | - | 2478 | 1188 | - | - | 0 | - |
| - | - | 1988 | 1189 | - | - | 0 | - |
| - | - | 1354 | 1190 | - | - | 0 | - |
| 5 | y | 5539 | 1195 | 0.0006167 | 0.5163 | +1 | 11 |
| - | - | 2755 | 1196 | - | - | 0 | - |
| - | - | 1233 | 1197 | - | - | 0 | - |
| - | - | 1882 | 1231 | - | - | 0 | - |
| 12 | c | 4.686E+04 | 1232 | 0.001973 | 1.602 | +1 | 12 |
| - | - | 3.041E+04 | 1233 | - | - | 0 | - |
| - | - | 1.229E+04 | 1234 | - | - | 0 | - |
| - | - | 2924 | 1235 | - | - | 0 | - |
| - | - | 844 | 1237 | - | - | 0 | - |
| - | - | 2880 | 1259 | - | - | 0 | - |
| 4 | z | 1518 | 1260 | 0.0005446 | 0.4323 | +1 | 12 |
| - | - | 2008 | 1275 | - | - | 0 | - |
| 4 | y | 1641 | 1276 | 0.002704 | 2.12 | +1 | 12 |
| 4 | z | 1922 | 1278 | 0.001222 | 0.9561 | +1 | 12 |
| - | - | 1.245E+04 | 1279 | - | - | 0 | - |
| - | - | 8638 | 1280 | - | - | 0 | - |
| - | - | 3865 | 1281 | - | - | 0 | - |
| - | - | 1459 | 1282 | - | - | 0 | - |
| 4 | y | 2965 | 1294 | 7.389E-05 | 0.05712 | +1 | 12 |
| - | - | 3912 | 1295 | - | - | 0 | - |
| - | - | 1357 | 1296 | - | - | 0 | - |
| 13 | c | 3.463E+04 | 1319 | 0.001897 | 1.439 | +1 | 13 |
| - | - | 2.414E+04 | 1320 | - | - | 0 | - |
| - | - | 9640 | 1321 | - | - | 0 | - |
| - | - | 2673 | 1322 | - | - | 0 | - |
| - | - | 816.4 | 1345 | - | - | 0 | - |
| - | - | 3582 | 1346 | - | - | 0 | - |
| - | - | 2361 | 1347 | - | - | 0 | - |
| 3 | w | 1571 | 1348 | 0.0028 | 2.078 | +1 | 13 |
| - | - | 1140 | 1365 | - | - | 0 | - |
| - | - | 1166 | 1396 | - | - | 0 | - |
| 3 | z | 3226 | 1407 | 0.0001306 | 0.09288 | +1 | 13 |
| - | - | 2.154E+04 | 1408 | - | - | 0 | - |
| - | - | 1.69E+04 | 1409 | - | - | 0 | - |
| - | - | 7300 | 1410 | - | - | 0 | - |
| - | - | 1649 | 1411 | - | - | 0 | - |
| - | - | 982.3 | 1417 | - | - | 0 | - |
| - | - | 1513 | 1418 | - | - | 0 | - |
| - | - | 896.7 | 1424 | - | - | 0 | - |
| - | - | 942.2 | 1426 | - | - | 0 | - |
| - | - | 1346 | 1431 | - | - | 0 | - |
| - | - | 6420 | 1432 | - | - | 0 | - |
| - | - | 4244 | 1433 | - | - | 0 | - |
| - | - | 2321 | 1434 | - | - | 0 | - |
| - | - | 4808 | 1459 | - | - | 0 | - |
| - | - | 2354 | 1460 | - | - | 0 | - |
| - | - | 1957 | 1461 | - | - | 0 | - |
| - | - | 749.9 | 1467 | - | - | 0 | - |
| 14 | c | 8545 | 1475 | 0.00169 | 1.146 | +1 | 14 |
| - | - | 2.517E+04 | 1476 | - | - | 0 | - |
| - | - | 1.794E+04 | 1477 | - | - | 0 | - |
| - | - | 6739 | 1478 | - | - | 0 | - |
| - | - | 3176 | 1479 | - | - | 0 | - |
| - | - | 1049 | 1480 | - | - | 0 | - |
| - | - | 728.7 | 1481 | - | - | 0 | - |
| 2 | z | 2043 | 1494 | 0.002282 | 1.528 | +1 | 14 |
| - | - | 1.487E+04 | 1495 | - | - | 0 | - |
| - | - | 1.322E+04 | 1496 | - | - | 0 | - |
| - | - | 5465 | 1497 | - | - | 0 | - |
| - | - | 1223 | 1498 | - | - | 0 | - |
| - | - | 1022 | 1508 | - | - | 0 | - |
| - | - | 685.4 | 1509 | - | - | 0 | - |
| 2 | y | 749.1 | 1510 | 0.008704 | 5.765 | +1 | 14 |
| - | - | 4271 | 1523 | - | - | 0 | - |
| - | - | 3396 | 1524 | - | - | 0 | - |
| - | - | 2276 | 1525 | - | - | 0 | - |
| - | - | 879 | 1526 | - | - | 0 | - |
| - | - | 807.1 | 1537 | - | - | 0 | - |
| - | - | 1876 | 1548 | - | - | 0 | - |
| - | - | 2006 | 1549 | - | - | 0 | - |
| - | - | 890.8 | 1550 | - | - | 0 | - |
| - | - | 1539 | 1564 | - | - | 0 | - |
| - | - | 2.165E+04 | 1565 | - | - | 0 | - |
| - | - | 2.77E+04 | 1566 | - | - | 0 | - |
| - | - | 1.72E+04 | 1567 | - | - | 0 | - |
| - | - | 6589 | 1568 | - | - | 0 | - |
| - | - | 2238 | 1569 | - | - | 0 | - |
| - | - | 3481 | 1579 | - | - | 0 | - |
| - | - | 4770 | 1580 | - | - | 0 | - |
| - | - | 5308 | 1581 | - | - | 0 | - |
| - | - | 7681 | 1582 | - | - | 0 | - |
| - | - | 4707 | 1583 | - | - | 0 | - |
| - | - | 1974 | 1584 | - | - | 0 | - |
| - | - | 2351 | 1590 | - | - | 0 | - |
| - | - | 7442 | 1591 | - | - | 0 | - |
| - | - | 5842 | 1592 | - | - | 0 | - |
| - | - | 2723 | 1593 | - | - | 0 | - |
| - | - | 1932 | 1594 | - | - | 0 | - |
| - | - | 1645 | 1596 | - | - | 0 | - |
| - | - | 8347 | 1597 | - | - | 0 | - |
| - | - | 6424 | 1598 | - | - | 0 | - |
| - | - | 3060 | 1599 | - | - | 0 | - |
| - | - | 1455 | 1600 | - | - | 0 | - |
| - | - | 1.659E+04 | 1607 | - | - | 0 | - |
| - | - | 7.655E+04 | 1608 | - | - | 0 | - |
| - | - | 5.999E+04 | 1609 | - | - | 0 | - |
| - | - | 2.776E+04 | 1610 | - | - | 0 | - |
| - | - | 9954 | 1611 | - | - | 0 | - |
| - | - | 1788 | 1612 | - | - | 0 | - |
| - | - | 752.4 | 1618 | - | - | 0 | - |
| - | - | 922.8 | 1621 | - | - | 0 | - |
| - | - | 3446 | 1623 | - | - | 0 | - |
| - | - | 2.019E+04 | 1624 | - | - | 0 | - |
| - | - | 7.755E+04 | 1625 | - | - | 0 | - |
| - | - | 6.082E+04 | 1626 | - | - | 0 | - |
| - | - | 2.674E+04 | 1627 | - | - | 0 | - |
| - | - | 513.3 | 1628 | - | - | 0 | - |
| - | - | 9322 | 1628 | - | - | 0 | - |
| - | - | 2652 | 1629 | - | - | 0 | - |
| - | - | 795.3 | 2199 | - | - | 0 | - |

m/z Charge Intensity FragmentType MassShift Position
120.08089447021484 0 433.39478
120.44309997558594 0 363.42078
142.09762573242188 0 1069.0251
148.94715881347656 0 927.99445
155.1189727783203 0 469.18625
159.4344940185547 0 455.2894
160.1071014404297 0 440.36774
163.25425720214844 0 435.9294
173.12831115722656 0 6447.975
173.45074462890625 0 3091.9885
183.11277770996094 0 2389.7559
199.07130432128906 0 892.9131
201.1231689453125 0 9292.402
201.1344757080078 0 6675.1313
202.1224822998047 0 1320.6711
208.02447509765625 0 519.73535
217.08128356933594 0 671.6179
229.12936401367188 0 767.77423
261.1232604980469 0 1099.1879
261.1705322265625 0 12973.371
262.1744689941406 0 2677.338
270.1565856933594 0 1015.28204
284.9512939453125 0 502.53336
288.1663513183594 0 1363.779
291.392822265625 0 651.6243
306.16864013671875 0 2760.794 z 13
307.17034912109375 0 604.8289
312.1551513671875 0 4668.385
313.1595764160156 0 1207.9907
316.1499328613281 0 957.4124
322.18707275390625 0 3067.4277 y 13
323.1906433105469 0 740.0338
330.166015625 0 3287.3025
348.2024230957031 0 1837.2844
362.7662658691406 0 552.5354
373.1972351074219 0 607.36285
376.19732666015625 0 8258.745 c Water loss 10
377.1571044921875 0 582.0677
377.1990661621094 0 1814.9974
391.1846618652344 0 3714.469
392.1925048828125 0 14932.934 y Ammonia loss 12
393.1988525390625 0 5654.522 y Ammonia loss 4
394.20306396484375 0 2002.1055 c Ammonia loss 6
394.87030029296875 0 630.95746
401.2398681640625 0 1942.9574
409.2189025878906 0 3315.112 y 12
410.2222900390625 0 1035.4618
411.22412109375 0 634.34424
415.4580383300781 0 565.09686
417.18792724609375 0 4883.134
418.1904296875 0 1078.3732
418.6407470703125 0 552.443
428.22552490234375 0 685.5529
429.2341613769531 0 3761.5813
435.23583984375 0 1327.9753
436.2406005859375 0 1155.7706
444.2562561035156 0 3056.7178
445.2560119628906 0 2573.4775
445.61822509765625 0 702.08594
446.2601013183594 0 2285.5552 c 3
463.230224609375 0 10688.579 w 11
464.2302551269531 0 2668.956
478.1136779785156 0 2045.9652
478.21697998046875 0 4087.623
479.2237548828125 0 4441.157 y Ammonia loss 11
480.2322998046875 0 9081.115 z 11
481.2350158691406 0 2508.5234
488.7577209472656 0 7176.0864 z 6
489.2595520019531 0 4458.567
489.7608337402344 0 2543.4268
492.5733947753906 0 725.9009 z Water loss 1
496.2512512207031 0 2675.0664 y 11
496.766845703125 0 3310.0867 y 6
497.2641906738281 0 1531.9784
497.9104919433594 0 1361.2384 y Water loss 1
498.25714111328125 0 687.01227
504.2202453613281 0 3882.7737
507.10345458984375 0 801.6681
516.2637329101562 0 1516.2107
524.27587890625 0 3830.874
524.77685546875 0 2098.3477
525.2814331054688 0 1363.8237
532.284912109375 0 4196.256
533.2921752929688 0 19879.076 c 4
534.29638671875 0 6083.583
535.298583984375 0 1098.0687
535.6050415039062 0 2180.3105
535.9405517578125 0 2505.2688
536.2710571289062 0 1230.5968
539.8134765625 0 717.33746
540.3116455078125 0 753.3513
541.3455200195312 0 1825.9597
541.6089477539062 0 8391.498
541.9436645507812 0 5820.538
542.2760620117188 0 3198.1907
546.2709350585938 0 11654.683 z 5
546.7734375 0 11835.699
547.2757568359375 0 5246.371
547.776123046875 0 1439.9222
554.2809448242188 0 5203.0894 y 5
554.7814331054688 0 4269.032
561.28955078125 0 1620.4915 z Water loss 10
562.2969970703125 0 1955.4026
564.2772216796875 0 9168.54 c Ammonia loss 10
565.2802124023438 0 2272.7043
567.7887573242188 0 1079.9666
568.2957153320312 0 783.5355
568.7959594726562 0 786.3733
579.3014526367188 0 4159.093 z 10
580.304931640625 0 1138.0687
582.2833251953125 0 679.40515
588.781005859375 0 2373.7092 y Water loss 4
589.2837524414062 0 3302.9326 y Ammonia loss 4
589.786865234375 0 13501.5625 z 4
590.2902221679688 0 21731.06
590.7918090820312 0 11609.846
591.2930908203125 0 2954.2346
592.3324584960938 0 5374.2705
593.3346557617188 0 1577.404
595.3201904296875 0 4951.7915 y 10
596.3209228515625 0 970.57715
597.2932739257812 0 2683.145
597.7963256835938 0 31767.074 y 4
598.2976684570312 0 20007.559
598.799072265625 0 4941.6245
599.3018798828125 0 2083.0383
601.7804565429688 0 1370.0374
602.2786865234375 0 915.1006
618.316162109375 0 755.4022 z Water loss 9
619.318359375 0 964.1217
631.8088989257812 0 815.63916 w 3
632.31201171875 0 1195.6329
636.3223266601562 0 19180.957 z 9
637.3264770507812 0 10694.775
638.3278198242188 0 3771.2385 y Water loss 3
639.321533203125 0 21532.963 z 3
639.8225708007812 0 16012.033
640.3243408203125 0 5436.8633
640.822021484375 0 679.8607
646.8274536132812 0 821.0615
647.3303833007812 0 30211.508 y 3
647.83203125 0 17467.996
648.3236694335938 0 20810.54 c 5
648.8345947265625 0 2934.6743
649.3238525390625 0 5384.361
650.3234252929688 0 1515.9932
652.3410034179688 0 20971.479 y 9
653.3436889648438 0 8003.2104
654.347412109375 0 1493.7091
674.3360595703125 0 16472.605 w 2
674.8370971679688 0 12487.563
675.3384399414062 0 5533.569
675.8394165039062 0 1955.615
679.3651123046875 0 2309.1829
680.3675537109375 0 1560.7612
681.8422241210938 0 896.4637
694.8427124023438 0 857.19025 z Water loss 2
702.84521484375 0 1093.2523 y Water loss 2
703.3516235351562 0 1896.3833 y Ammonia loss 2
703.8427124023438 0 11126.042 z 2
704.3435668945312 0 7805.796
704.8453369140625 0 4260.827
705.3479614257812 0 1099.545 z Water loss 8
708.3649291992188 0 2528.6743
711.3446655273438 0 1004.9121
711.8514404296875 0 32662.508 y 2
712.3532104492188 0 25312.951
712.85498046875 0 12403.334
713.35546875 0 3131.1914
715.87646484375 0 3192.9634
716.3775634765625 0 2221.3352
716.8834228515625 0 768.85376
717.8535766601562 0 2002.6414
718.3533325195312 0 1865.9324
718.8606567382812 0 688.7116
721.3386840820312 0 916.05554
722.34912109375 0 1989.0085 y Ammonia loss 8
723.3539428710938 0 13524.947 z 8
724.3572998046875 0 6084.401
725.361328125 0 1906.2668
725.8638916015625 0 1201.2347
726.3499145507812 0 724.85516
729.3590698242188 0 1322.6667 c Ammonia loss 13
729.8453369140625 0 805.5085
730.3532104492188 0 841.89105
737.363525390625 0 800.76984
737.8838500976562 0 88892.51 c 13
738.3849487304688 0 63862.21
738.885986328125 0 32692.404
739.3821411132812 0 12635.001 y 8
739.8873291015625 0 1602.6565
740.3717041015625 0 1801.658
744.387451171875 0 1095.0984
746.3583374023438 0 2758.187 y Water loss 1
746.8598022460938 0 3654.4473 y Ammonia loss 1
747.3569946289062 0 15135.454 z 1
747.8607177734375 0 12488.277
748.3588256835938 0 4805.3237
748.8631591796875 0 2064.0637
749.3486938476562 0 806.549
755.3678588867188 0 42411.395 y 1
755.8694458007812 0 35253.66
756.370361328125 0 15677.788
756.8712158203125 0 4394.5854
760.4059448242188 0 2087.0774
760.8584594726562 0 1216.077
767.884033203125 0 1019.22314
774.397705078125 0 2463.7014
774.8973388671875 0 1127.7333
775.8756713867188 0 1231.899
781.4080200195312 0 838.377
781.9064331054688 0 7330.017
782.4043579101562 0 16462.738
782.9046020507812 0 12059.958
783.4039916992188 0 4800.6167
783.905517578125 0 1380.278
787.9028930664062 0 886.78394
788.4005126953125 0 1881.6248
788.896484375 0 1012.7331
789.3832397460938 0 23875.57
789.884521484375 0 25507.195
790.3896484375 0 9822.501
790.898681640625 0 24670.15
791.4004516601562 0 20227.803
791.9020385742188 0 8535.517
792.4036254882812 0 2977.3354
792.9119873046875 0 760.14417
794.8956909179688 0 3029.6558
795.39599609375 0 3054.5742
795.89404296875 0 2305.6409
796.391357421875 0 2619.9578
796.8902587890625 0 1513.0034
797.3894653320312 0 852.25494
803.4072265625 0 11651.58
803.9014892578125 0 81720.35
804.4024658203125 0 71362.55
804.9035034179688 0 37707.684
805.40478515625 0 11873.379
805.9046630859375 0 3041.0747
810.3829956054688 0 1085.8228
811.9097290039062 0 45549.445
812.4122924804688 0 59654.543
812.9135131835938 0 42997.504
813.4149780273438 0 14992.651
813.9169311523438 0 4035.2517
836.42578125 0 2711.4646 y 7
837.4303588867188 0 825.79645
842.4546508789062 0 1019.1689
857.4606323242188 0 4199.923
858.4624633789062 0 2248.8706
859.4668579101562 0 772.76154
885.4583740234375 0 1110.6627
886.4589233398438 0 1136.9973
890.4379272460938 0 1534.537 w 6
900.4974975585938 0 1921.1366
901.4739379882812 0 22238.166 c 7
902.4757080078125 0 12125.572
903.4788818359375 0 4362.978
904.4852294921875 0 761.4779
926.4778442382812 0 1139.852
927.4881591796875 0 899.3076
944.4909057617188 0 12791.506
945.4954833984375 0 7841.847
946.4970703125 0 2373.578
976.5078735351562 0 14131.525 z 6
977.5126342773438 0 9517.097
978.515625 0 3374.947
987.497314453125 0 3676.9163
988.505126953125 0 61785.22 c 8
989.5086669921875 0 27594.213
990.5105590820312 0 9016.261
991.5126342773438 0 2238.2695
1001.5127563476562 0 3280.1003
1002.51904296875 0 3165.0168
1026.36181640625 0 1112.1393
1045.5267333984375 0 33662.574 c 9
1046.5302734375 0 18612.373 w 5
1047.5367431640625 0 9424.851
1048.54345703125 0 3373.9038
1052.615234375 0 701.1243
1074.53662109375 0 880.5935
1081.3717041015625 0 2044.5999
1082.3729248046875 0 1650.9631
1082.6304931640625 0 651.3961
1083.3602294921875 0 829.8693
1084.3792724609375 0 929.2605
1085.5831298828125 0 825.37115
1091.407958984375 0 944.7288
1091.534912109375 0 12162.476 z 5
1092.541259765625 0 20749.219
1093.544921875 0 8976.136
1094.5438232421875 0 2884.1094
1100.5806884765625 0 1506.6057
1101.5863037109375 0 1655.4086
1106.5489501953125 0 1181.9364
1107.5511474609375 0 2651.9414 y 5
1108.55517578125 0 1709.4232
1129.585205078125 0 2376.8088
1130.586669921875 0 2207.8625
1144.5948486328125 0 25744.389 c 10
1145.5987548828125 0 17433.123
1146.60302734375 0 7349.873
1147.604248046875 0 1599.4133
1178.5650634765625 0 1883.8245 z 4
1179.57421875 0 9495.179
1180.5738525390625 0 5938.4644
1181.580810546875 0 2709.2573
1182.594970703125 0 1326.4525
1187.6143798828125 0 2478.354
1188.6170654296875 0 1987.6536
1189.6295166015625 0 1354.2747
1194.5867919921875 0 5538.9395 y 4
1195.5843505859375 0 2755.1475
1196.5872802734375 0 1233.1091
1230.617919921875 0 1881.7654
1231.625732421875 0 46861.41 c 11
1232.62939453125 0 30414.334
1233.63232421875 0 12286.702
1234.635009765625 0 2924.3516
1236.618896484375 0 843.98413
1258.6246337890625 0 2879.9426
1259.624755859375 0 1517.9191 z Water loss 3
1274.643310546875 0 2008.1865
1275.646728515625 0 1641.1859 y Water loss 3
1277.6346435546875 0 1921.5828 z 3
1278.6402587890625 0 12452.01
1279.644775390625 0 8638.378
1280.647705078125 0 3864.8235
1281.6517333984375 0 1458.9976
1293.6546630859375 0 2965.3352 y 3
1294.6533203125 0 3911.8113
1295.6568603515625 0 1357.2657
1318.6578369140625 0 34626.16 c 12
1319.6612548828125 0 24139.17
1320.66259765625 0 9639.586
1321.665771484375 0 2672.718
1344.6884765625 0 816.3541
1345.65869140625 0 3581.7852
1346.6656494140625 0 2360.9885
1347.662353515625 0 1571.2402 w 2
1364.6505126953125 0 1139.9346
1395.6307373046875 0 1165.9586
1406.6785888671875 0 3225.8633 z 2
1407.682861328125 0 21536.91
1408.6865234375 0 16904.096
1409.6883544921875 0 7300.148
1410.6912841796875 0 1648.7781
1416.724365234375 0 982.34045
1417.7205810546875 0 1513.0212
1423.7039794921875 0 896.6903
1425.703857421875 0 942.2276
1430.75146484375 0 1346.3364
1431.755615234375 0 6420.1006
1432.75390625 0 4244.3027
1433.75439453125 0 2321.3855
1458.742919921875 0 4808.1025
1459.7471923828125 0 2354.0024
1460.7431640625 0 1957.0874
1466.712646484375 0 749.871
1474.7591552734375 0 8544.689 c 13
1475.765625 0 25169.307
1476.765380859375 0 17935.504
1477.7640380859375 0 6739.3896
1478.760009765625 0 3175.916
1479.7269287109375 0 1049.3715
1480.698974609375 0 728.7102
1493.7127685546875 0 2043.1318 z 1
1494.7161865234375 0 14869.272
1495.7181396484375 0 13217.28
1496.724853515625 0 5465.06
1497.725341796875 0 1223.0366
1507.7650146484375 0 1022.0754
1508.76123046875 0 685.4193
1509.7379150390625 0 749.0825 y 1
1522.7275390625 0 4270.764
1523.7288818359375 0 3395.848
1524.7388916015625 0 2275.8953
1525.74072265625 0 878.95154
1536.7476806640625 0 807.05457
1547.7713623046875 0 1875.9858
1548.766357421875 0 2005.5078
1549.7581787109375 0 890.8013
1563.7904052734375 0 1538.5532
1564.78076171875 0 21651.223
1565.7825927734375 0 27703.762
1566.7840576171875 0 17201.422
1567.7838134765625 0 6589.0996
1568.7911376953125 0 2238.0464
1578.82080078125 0 3481.115
1579.8154296875 0 4770.0234
1580.8056640625 0 5307.759
1581.8038330078125 0 7680.9033
1582.8017578125 0 4706.917
1583.8052978515625 0 1974.011
1589.794189453125 0 2351.124
1590.777587890625 0 7442.1357
1591.7818603515625 0 5841.7104
1592.7794189453125 0 2723.03
1593.7803955078125 0 1931.5994
1595.82275390625 0 1645.3531
1596.8289794921875 0 8346.943
1597.8323974609375 0 6424.478
1598.837158203125 0 3060.0232
1599.836669921875 0 1455.43
1606.8031005859375 0 16589.256
1607.8004150390625 0 76547.89
1608.8028564453125 0 59985.305
1609.8052978515625 0 27756.713
1610.808837890625 0 9954.114
1611.8104248046875 0 1788.3774
1617.93017578125 0 752.42224
1620.7723388671875 0 922.84576
1622.802490234375 0 3446.3096
1623.8160400390625 0 20189.137
1624.8243408203125 0 77552.24
1625.8270263671875 0 60817.633
1626.8310546875 0 26742.787
1627.601318359375 0 513.3101
1627.833251953125 0 9322.485
1628.833740234375 0 2651.7087
2198.592529296875 0 795.3209

Spectrum Details

|  |  |
| --- | --- |
| Matched peaks? Matched peaks The total absolute number of peaks matched. Additionally in brackets the total fraction of peaks matched and the total number of peaks is shown. | 74 (17.58% of 421) |
| FDR? FDR The false discovery rate estimated for this peptide. It is calculated by matching all theoretical fragments with a non-integer shift with the raw peaks for this spectrum. This is done with 40 different shifts. The resulting percentage is the average number of annotated peaks over the number of annotated peaks with the correct spectrum. | 0.29% |
| Satellite FDR? Satellite FDR See the FDR for details on its calculation. This satellite ion specific FDR only contains the satellite ions (d/w) for I/L/J positions. | - |
| PSM Score? PSM Score The PSM Score as given by Hecklib to this annotated spectrum. It is shown with three significant figures. | 515 |

#### Spectrum 4990? Spectrum 4990 The raw spectrum of this peptide as annotated by Hecklib. The fragments are coloured according to ion type (see legend). Any peaks with a star '\*' as text can be hovered over to see the full details, first the ion type second the mass shift type. By hovering over the amino acids in the peptide or ions in the legend the corresponding peaks are highlighted. By toggling the 'Unassigned' label you can turn the background (unassigned) peaks on or off in the plot. By updating the slider in the Ion legend you can update the spectrum to only show the top X% of the peaks with labels. The top X% means any peak that is within X% of the highest intensity. By dragging in the spectrum you can zoom in to a specific part of the spectrum and use 'Zoom Out' to get back to the original zoom level. The annotation of the spectrum is based on the given sequence in the peptides file and is done with different software so inconsistencies are likely. The peaks are annotated based on the given sequence, with 20 ppm tolerance.

Copy Data

##### Spectrum 4990 (TSV)

###### Preview

```
Loading example...
```

*Click on the button to copy the data to your clipboard.*

Mz MinMz MaxIntensity Max

WidthHeightPeptide font sizePeptide stroke widthSpectrum font sizeSpectrum stroke widthCompact peptide

Ion legend

wxyz

abcd

OtherUnassignedIon^Charge^_Position_Show for top:%

JSEVSDRPSGVSSRF

02.96e+45.93e+48.89e+41.19e+5

Zoom Out

y^+2^_4_z^+1^_2_y^+1^_2_c^+3^_11_y^+1^_3_y^+3^_11_c^+2^_7_y^+1^_3_y^+2^_8_c^+1^_4_z^+1^_4_w^+1^_4_y^+3^_13_y^+1^_4_z^+1^_4_z^+2^_9_y^+1^_4_y^+2^_9_y^+3^_14_c^+1^_5_z^+2^_10_z^+2^_10_y^+2^_10_z^+1^_5_c^+2^_11_z^+1^_5_z^+2^_11_z^+2^_11_y^+2^_11_y^+2^_11_z^+2^_11_y^+1^_5_y^+2^_11_w^+2^_12_z^+1^_6_y^+2^_12_z^+2^_12_y^+2^_12_c^+1^_6_y^+1^_6_w^+2^_13_y^+2^_13_y^+2^_13_z^+2^_13_z^+1^_7_y^+2^_13_y^+1^_7_z^+1^_7_c^+2^_14_c^+2^_14_y^+1^_7_y^+2^_14_y^+2^_14_z^+2^_14_y^+2^_14_y^+1^_8_w^+1^_9_c^+1^_8_z^+1^_9_z^+1^_9_c^+1^_9_c^+1^_10_w^+1^_10_z^+1^_10_z^+1^_10_z^+1^_10_y^+1^_10_c^+1^_11_z^+1^_11_y^+1^_11_c^+1^_12_z^+1^_12_w^+1^_12_y^+1^_12_y^+1^_12_z^+1^_12_y^+1^_12_c^+1^_13_w^+1^_13_z^+1^_13_y^+1^_13_c^+1^_14_z^+1^_14_

0758151522733030

Fragment Matches Table

Show background peaks

| Position | Ion type | Intensity | mz Theoretical | mz Error (Th) | mz Error (ppm) | Charge | Series Number |
| --- | --- | --- | --- | --- | --- | --- | --- |
| - | - | 1209 | 120.1 | - | - | 0 | - |
| - | - | 414.9 | 123.8 | - | - | 0 | - |
| - | - | 358.7 | 124.3 | - | - | 0 | - |
| - | - | 365.5 | 128.3 | - | - | 0 | - |
| - | - | 409.9 | 132.3 | - | - | 0 | - |
| - | - | 1345 | 142.1 | - | - | 0 | - |
| - | - | 406.7 | 156.6 | - | - | 0 | - |
| - | - | 463.3 | 160.1 | - | - | 0 | - |
| - | - | 475.7 | 171 | - | - | 0 | - |
| - | - | 8286 | 173.1 | - | - | 0 | - |
| - | - | 1736 | 173.5 | - | - | 0 | - |
| - | - | 743.5 | 174.1 | - | - | 0 | - |
| - | - | 2592 | 183.1 | - | - | 0 | - |
| - | - | 436 | 198.9 | - | - | 0 | - |
| - | - | 740 | 199.1 | - | - | 0 | - |
| - | - | 548.8 | 200.1 | - | - | 0 | - |
| - | - | 1.383E+04 | 201.1 | - | - | 0 | - |
| - | - | 8053 | 201.1 | - | - | 0 | - |
| - | - | 2702 | 202.1 | - | - | 0 | - |
| - | - | 926.2 | 202.1 | - | - | 0 | - |
| - | - | 1184 | 217.1 | - | - | 0 | - |
| - | - | 533.5 | 221.3 | - | - | 0 | - |
| - | - | 1556 | 223.1 | - | - | 0 | - |
| - | - | 698.7 | 229.1 | - | - | 0 | - |
| - | - | 1210 | 229.1 | - | - | 0 | - |
| - | - | 514.4 | 235.2 | - | - | 0 | - |
| - | - | 1108 | 244.1 | - | - | 0 | - |
| 12 | y | 560 | 248.6 | 9.063E-05 | 0.3645 | +2 | 4 |
| - | - | 573.7 | 249.1 | - | - | 0 | - |
| - | - | 1687 | 261.1 | - | - | 0 | - |
| - | - | 1.845E+04 | 261.2 | - | - | 0 | - |
| - | - | 2270 | 262.2 | - | - | 0 | - |
| - | - | 670.6 | 270.2 | - | - | 0 | - |
| - | - | 465.2 | 277 | - | - | 0 | - |
| - | - | 1370 | 288.2 | - | - | 0 | - |
| 14 | z | 3729 | 306.2 | 0.0001812 | 0.5919 | +1 | 2 |
| - | - | 751.1 | 307.2 | - | - | 0 | - |
| - | - | 858.9 | 310.1 | - | - | 0 | - |
| - | - | 6390 | 312.2 | - | - | 0 | - |
| - | - | 1185 | 313.2 | - | - | 0 | - |
| - | - | 1375 | 316.2 | - | - | 0 | - |
| - | - | 740.7 | 321.2 | - | - | 0 | - |
| 14 | y | 3990 | 322.2 | 1.185E-05 | 0.03677 | +1 | 2 |
| - | - | 710 | 323.2 | - | - | 0 | - |
| - | - | 3660 | 330.2 | - | - | 0 | - |
| - | - | 697.1 | 331.2 | - | - | 0 | - |
| - | - | 2202 | 348.2 | - | - | 0 | - |
| - | - | 1199 | 349.2 | - | - | 0 | - |
| - | - | 583.8 | 373.2 | - | - | 0 | - |
| 11 | c | 1.178E+04 | 376.2 | 0.002043 | 5.431 | +3 | 11 |
| - | - | 2824 | 377.2 | - | - | 0 | - |
| - | - | 1004 | 387.2 | - | - | 0 | - |
| - | - | 4511 | 391.2 | - | - | 0 | - |
| 13 | y | 1.775E+04 | 392.2 | 3.533E-05 | 0.09009 | +1 | 3 |
| 5 | y | 7844 | 393.2 | 0.007551 | 19.2 | +3 | 11 |
| 7 | c | 2303 | 394.2 | 0.004516 | 11.46 | +2 | 7 |
| - | - | 594.3 | 397.4 | - | - | 0 | - |
| - | - | 2579 | 401.2 | - | - | 0 | - |
| 13 | y | 4010 | 409.2 | 0.0005152 | 1.259 | +1 | 3 |
| - | - | 708.1 | 410.2 | - | - | 0 | - |
| - | - | 632.9 | 411.2 | - | - | 0 | - |
| - | - | 6250 | 417.2 | - | - | 0 | - |
| - | - | 1338 | 418.2 | - | - | 0 | - |
| 8 | y | 944.6 | 418.7 | 0.0004066 | 0.971 | +2 | 8 |
| - | - | 553.4 | 419 | - | - | 0 | - |
| - | - | 1234 | 428.2 | - | - | 0 | - |
| - | - | 4917 | 429.2 | - | - | 0 | - |
| - | - | 1382 | 430.2 | - | - | 0 | - |
| - | - | 1445 | 435.2 | - | - | 0 | - |
| - | - | 835.1 | 436.2 | - | - | 0 | - |
| - | - | 3857 | 444.3 | - | - | 0 | - |
| - | - | 3848 | 445.3 | - | - | 0 | - |
| 4 | c | 3011 | 446.3 | 0.001434 | 3.213 | +1 | 4 |
| - | - | 669.1 | 447.3 | - | - | 0 | - |
| 12 | z | 740.7 | 462.2 | 0.001553 | 3.36 | +1 | 4 |
| 12 | w | 1.211E+04 | 463.2 | 0.0003875 | 0.8365 | +1 | 4 |
| - | - | 4881 | 464.2 | - | - | 0 | - |
| 3 | y | 669.5 | 469.2 | 0.007444 | 15.87 | +3 | 13 |
| - | - | 5185 | 478.2 | - | - | 0 | - |
| 12 | y | 5932 | 479.2 | 0.000356 | 0.7428 | +1 | 4 |
| 12 | z | 1.006E+04 | 480.2 | 0.0001854 | 0.3861 | +1 | 4 |
| - | - | 2650 | 481.2 | - | - | 0 | - |
| 7 | z | 1.033E+04 | 488.8 | 0.0003616 | 0.7399 | +2 | 9 |
| - | - | 688.4 | 489.2 | - | - | 0 | - |
| - | - | 4640 | 489.3 | - | - | 0 | - |
| - | - | 2970 | 489.8 | - | - | 0 | - |
| - | - | 882.7 | 490.3 | - | - | 0 | - |
| 12 | y | 3728 | 496.3 | 0.0004997 | 1.007 | +1 | 4 |
| 7 | y | 2754 | 496.8 | 0.0001198 | 0.2411 | +2 | 9 |
| - | - | 2421 | 497.3 | - | - | 0 | - |
| - | - | 1017 | 498.2 | - | - | 0 | - |
| 2 | y | 926 | 503.9 | 0.0003728 | 0.7398 | +3 | 14 |
| - | - | 6538 | 504.2 | - | - | 0 | - |
| - | - | 2064 | 505.2 | - | - | 0 | - |
| - | - | 657.6 | 515.6 | - | - | 0 | - |
| - | - | 1140 | 516.3 | - | - | 0 | - |
| - | - | 668 | 523.2 | - | - | 0 | - |
| - | - | 3815 | 524.3 | - | - | 0 | - |
| - | - | 2362 | 524.8 | - | - | 0 | - |
| - | - | 1370 | 525.3 | - | - | 0 | - |
| - | - | 694.9 | 531.3 | - | - | 0 | - |
| - | - | 5246 | 532.3 | - | - | 0 | - |
| 5 | c | 2.712E+04 | 533.3 | 0.0002288 | 0.429 | +1 | 5 |
| - | - | 723.7 | 533.7 | - | - | 0 | - |
| - | - | 6775 | 534.3 | - | - | 0 | - |
| - | - | 1993 | 535.3 | - | - | 0 | - |
| - | - | 2531 | 535.6 | - | - | 0 | - |
| - | - | 2672 | 535.9 | - | - | 0 | - |
| - | - | 714.1 | 536.3 | - | - | 0 | - |
| 6 | z | 757.6 | 537.3 | 0.003403 | 6.334 | +2 | 10 |
| - | - | 1601 | 541.3 | - | - | 0 | - |
| - | - | 1.172E+04 | 541.6 | - | - | 0 | - |
| - | - | 9011 | 541.9 | - | - | 0 | - |
| - | - | 779.5 | 542.2 | - | - | 0 | - |
| - | - | 4844 | 542.3 | - | - | 0 | - |
| - | - | 2313 | 542.6 | - | - | 0 | - |
| - | - | 1308 | 542.7 | - | - | 0 | - |
| 6 | z | 1.472E+04 | 546.3 | 0.0001958 | 0.3584 | +2 | 10 |
| - | - | 1.645E+04 | 546.8 | - | - | 0 | - |
| - | - | 6060 | 547.3 | - | - | 0 | - |
| - | - | 1677 | 547.8 | - | - | 0 | - |
| - | - | 825 | 549.3 | - | - | 0 | - |
| - | - | 1033 | 550.3 | - | - | 0 | - |
| - | - | 1738 | 553.8 | - | - | 0 | - |
| 6 | y | 5921 | 554.3 | 0.0003772 | 0.6805 | +2 | 10 |
| - | - | 3868 | 554.8 | - | - | 0 | - |
| - | - | 2255 | 555.3 | - | - | 0 | - |
| - | - | 731.5 | 556.3 | - | - | 0 | - |
| 11 | z | 1816 | 561.3 | 0.00057 | 1.016 | +1 | 5 |
| - | - | 2333 | 562.3 | - | - | 0 | - |
| 11 | c | 1.181E+04 | 564.3 | 0.009882 | 17.51 | +2 | 11 |
| - | - | 4076 | 565.3 | - | - | 0 | - |
| - | - | 1152 | 567.8 | - | - | 0 | - |
| - | - | 1561 | 568.3 | - | - | 0 | - |
| 11 | z | 6428 | 579.3 | 0.000462 | 0.7974 | +1 | 5 |
| - | - | 2464 | 580.3 | - | - | 0 | - |
| 5 | z | 895.9 | 580.8 | 0.0006484 | 1.116 | +2 | 11 |
| 5 | z | 622.9 | 581.3 | 0.0004687 | 0.8063 | +2 | 11 |
| - | - | 984.4 | 581.8 | - | - | 0 | - |
| - | - | 830.6 | 582.3 | - | - | 0 | - |
| - | - | 782.9 | 586.3 | - | - | 0 | - |
| 5 | y | 1965 | 588.8 | 0.008729 | 14.82 | +2 | 11 |
| 5 | y | 4164 | 589.3 | 0.0004923 | 0.8355 | +2 | 11 |
| 5 | z | 1.279E+04 | 589.8 | 0.0001324 | 0.2245 | +2 | 11 |
| - | - | 2.477E+04 | 590.3 | - | - | 0 | - |
| - | - | 1.276E+04 | 590.8 | - | - | 0 | - |
| - | - | 4841 | 591.3 | - | - | 0 | - |
| - | - | 1538 | 591.8 | - | - | 0 | - |
| - | - | 9154 | 592.3 | - | - | 0 | - |
| - | - | 3345 | 593.3 | - | - | 0 | - |
| - | - | 788.6 | 594.3 | - | - | 0 | - |
| 11 | y | 6096 | 595.3 | 0.0005367 | 0.9016 | +1 | 5 |
| - | - | 2211 | 596.3 | - | - | 0 | - |
| - | - | 6192 | 597.3 | - | - | 0 | - |
| 5 | y | 4.2E+04 | 597.8 | 0.0001491 | 0.2495 | +2 | 11 |
| - | - | 2.02E+04 | 598.3 | - | - | 0 | - |
| - | - | 1.081E+04 | 598.8 | - | - | 0 | - |
| - | - | 1814 | 599.3 | - | - | 0 | - |
| - | - | 728.5 | 599.8 | - | - | 0 | - |
| - | - | 1920 | 601.8 | - | - | 0 | - |
| - | - | 1629 | 602.3 | - | - | 0 | - |
| - | - | 988.1 | 611.3 | - | - | 0 | - |
| - | - | 1092 | 619.3 | - | - | 0 | - |
| - | - | 806.9 | 620.3 | - | - | 0 | - |
| 4 | w | 1163 | 631.8 | 0.0007136 | 1.13 | +2 | 12 |
| - | - | 1479 | 632.3 | - | - | 0 | - |
| 10 | z | 2.684E+04 | 636.3 | 0.0002385 | 0.3748 | +1 | 6 |
| - | - | 1.415E+04 | 637.3 | - | - | 0 | - |
| 4 | y | 3903 | 638.3 | 0.003329 | 5.215 | +2 | 12 |
| 4 | z | 2.365E+04 | 639.3 | 0.0005117 | 0.8004 | +2 | 12 |
| - | - | 1.942E+04 | 639.8 | - | - | 0 | - |
| - | - | 7858 | 640.3 | - | - | 0 | - |
| - | - | 2566 | 640.8 | - | - | 0 | - |
| - | - | 1104 | 642.3 | - | - | 0 | - |
| - | - | 967.8 | 642.8 | - | - | 0 | - |
| - | - | 808.6 | 646.8 | - | - | 0 | - |
| 4 | y | 4.012E+04 | 647.3 | 0.0001829 | 0.2825 | +2 | 12 |
| - | - | 2.441E+04 | 647.8 | - | - | 0 | - |
| 6 | c | 2.742E+04 | 648.3 | 0.004689 | 7.232 | +1 | 6 |
| - | - | 2021 | 648.8 | - | - | 0 | - |
| - | - | 7691 | 649.3 | - | - | 0 | - |
| - | - | 1692 | 650.3 | - | - | 0 | - |
| 10 | y | 2.55E+04 | 652.3 | 0.0003743 | 0.5737 | +1 | 6 |
| - | - | 9816 | 653.3 | - | - | 0 | - |
| - | - | 1925 | 654.3 | - | - | 0 | - |
| - | - | 585.6 | 658.9 | - | - | 0 | - |
| 3 | w | 2.255E+04 | 674.3 | 0.0002716 | 0.4028 | +2 | 13 |
| - | - | 1.762E+04 | 674.8 | - | - | 0 | - |
| - | - | 6291 | 675.3 | - | - | 0 | - |
| - | - | 1743 | 675.8 | - | - | 0 | - |
| - | - | 2841 | 679.4 | - | - | 0 | - |
| - | - | 1114 | 680.4 | - | - | 0 | - |
| - | - | 1655 | 681.8 | - | - | 0 | - |
| - | - | 885.8 | 694.3 | - | - | 0 | - |
| 3 | y | 3168 | 702.8 | 0.001427 | 2.03 | +2 | 13 |
| 3 | y | 2065 | 703.3 | 0.00864 | 12.28 | +2 | 13 |
| 3 | z | 1.627E+04 | 703.8 | 0.0003333 | 0.4736 | +2 | 13 |
| - | - | 1.236E+04 | 704.3 | - | - | 0 | - |
| - | - | 871.9 | 704.4 | - | - | 0 | - |
| - | - | 4532 | 704.8 | - | - | 0 | - |
| 9 | z | 1547 | 705.3 | 0.0001981 | 0.2808 | +1 | 7 |
| - | - | 1445 | 707.9 | - | - | 0 | - |
| - | - | 1891 | 708.4 | - | - | 0 | - |
| - | - | 1535 | 711.3 | - | - | 0 | - |
| 3 | y | 4.515E+04 | 711.9 | 5.652E-05 | 0.0794 | +2 | 13 |
| - | - | 3.638E+04 | 712.4 | - | - | 0 | - |
| - | - | 1.555E+04 | 712.9 | - | - | 0 | - |
| - | - | 3550 | 713.4 | - | - | 0 | - |
| - | - | 1107 | 713.9 | - | - | 0 | - |
| - | - | 5380 | 715.9 | - | - | 0 | - |
| - | - | 3504 | 716.4 | - | - | 0 | - |
| - | - | 1075 | 716.9 | - | - | 0 | - |
| - | - | 697.1 | 717.4 | - | - | 0 | - |
| - | - | 4161 | 717.9 | - | - | 0 | - |
| - | - | 2199 | 718.4 | - | - | 0 | - |
| - | - | 2388 | 721.3 | - | - | 0 | - |
| - | - | 1347 | 721.7 | - | - | 0 | - |
| 9 | y | 2636 | 722.3 | 0.00126 | 1.744 | +1 | 7 |
| 9 | z | 1.824E+04 | 723.4 | 0.0001127 | 0.1558 | +1 | 7 |
| - | - | 7115 | 724.4 | - | - | 0 | - |
| - | - | 619.2 | 724.9 | - | - | 0 | - |
| - | - | 3376 | 725.4 | - | - | 0 | - |
| - | - | 1151 | 725.9 | - | - | 0 | - |
| - | - | 1496 | 726.4 | - | - | 0 | - |
| - | - | 590.2 | 726.9 | - | - | 0 | - |
| 14 | c | 776.1 | 729.4 | 0.004697 | 6.44 | +2 | 14 |
| - | - | 1088 | 730.4 | - | - | 0 | - |
| - | - | 653.1 | 732.3 | - | - | 0 | - |
| 14 | c | 1.173E+05 | 737.9 | 0.0002776 | 0.3762 | +2 | 14 |
| - | - | 8.899E+04 | 738.4 | - | - | 0 | - |
| - | - | 4.165E+04 | 738.9 | - | - | 0 | - |
| 9 | y | 1.577E+04 | 739.4 | 0.008141 | 11.01 | +1 | 7 |
| - | - | 1543 | 739.9 | - | - | 0 | - |
| - | - | 2739 | 740.4 | - | - | 0 | - |
| 2 | y | 5068 | 746.4 | 0.001206 | 1.616 | +2 | 14 |
| 2 | y | 5984 | 746.9 | 0.004772 | 6.39 | +2 | 14 |
| 2 | z | 2.334E+04 | 747.4 | 0.001826 | 2.443 | +2 | 14 |
| - | - | 1.359E+04 | 747.9 | - | - | 0 | - |
| - | - | 6826 | 748.4 | - | - | 0 | - |
| - | - | 2080 | 748.9 | - | - | 0 | - |
| 2 | y | 5.181E+04 | 755.4 | 0.0002257 | 0.2987 | +2 | 14 |
| - | - | 4.401E+04 | 755.9 | - | - | 0 | - |
| - | - | 2.419E+04 | 756.4 | - | - | 0 | - |
| - | - | 6683 | 756.9 | - | - | 0 | - |
| - | - | 2384 | 757.4 | - | - | 0 | - |
| - | - | 728.2 | 759.4 | - | - | 0 | - |
| - | - | 3072 | 760.4 | - | - | 0 | - |
| - | - | 889.5 | 760.9 | - | - | 0 | - |
| - | - | 768.3 | 761.4 | - | - | 0 | - |
| - | - | 869.6 | 761.9 | - | - | 0 | - |
| - | - | 1090 | 768.4 | - | - | 0 | - |
| - | - | 1170 | 773.4 | - | - | 0 | - |
| - | - | 2373 | 774.4 | - | - | 0 | - |
| - | - | 1149 | 774.9 | - | - | 0 | - |
| - | - | 977.8 | 775.4 | - | - | 0 | - |
| - | - | 2614 | 775.9 | - | - | 0 | - |
| - | - | 1417 | 781.4 | - | - | 0 | - |
| - | - | 1.055E+04 | 781.9 | - | - | 0 | - |
| - | - | 2.234E+04 | 782.4 | - | - | 0 | - |
| - | - | 1.648E+04 | 782.9 | - | - | 0 | - |
| - | - | 9119 | 783.4 | - | - | 0 | - |
| - | - | 2293 | 783.9 | - | - | 0 | - |
| - | - | 3423 | 788.4 | - | - | 0 | - |
| - | - | 2311 | 788.9 | - | - | 0 | - |
| - | - | 3.364E+04 | 789.4 | - | - | 0 | - |
| - | - | 3.112E+04 | 789.9 | - | - | 0 | - |
| - | - | 1.582E+04 | 790.4 | - | - | 0 | - |
| - | - | 3.364E+04 | 790.9 | - | - | 0 | - |
| - | - | 2.791E+04 | 791.4 | - | - | 0 | - |
| - | - | 1.291E+04 | 791.9 | - | - | 0 | - |
| - | - | 4378 | 792.4 | - | - | 0 | - |
| - | - | 892.1 | 794.4 | - | - | 0 | - |
| - | - | 3207 | 794.9 | - | - | 0 | - |
| - | - | 3780 | 795.4 | - | - | 0 | - |
| - | - | 3235 | 795.9 | - | - | 0 | - |
| - | - | 4229 | 796.4 | - | - | 0 | - |
| - | - | 2018 | 796.9 | - | - | 0 | - |
| - | - | 1525 | 797.4 | - | - | 0 | - |
| - | - | 1.423E+04 | 803.4 | - | - | 0 | - |
| - | - | 1.078E+05 | 803.9 | - | - | 0 | - |
| - | - | 9.655E+04 | 804.4 | - | - | 0 | - |
| - | - | 4.89E+04 | 804.9 | - | - | 0 | - |
| - | - | 1129 | 805 | - | - | 0 | - |
| - | - | 1.333E+04 | 805.4 | - | - | 0 | - |
| - | - | 4675 | 805.9 | - | - | 0 | - |
| - | - | 645.4 | 810.4 | - | - | 0 | - |
| - | - | 808.2 | 810.8 | - | - | 0 | - |
| - | - | 5.872E+04 | 811.9 | - | - | 0 | - |
| - | - | 8.035E+04 | 812.4 | - | - | 0 | - |
| - | - | 5.591E+04 | 812.9 | - | - | 0 | - |
| - | - | 2.262E+04 | 813.4 | - | - | 0 | - |
| - | - | 5134 | 813.9 | - | - | 0 | - |
| - | - | 1238 | 814.4 | - | - | 0 | - |
| 8 | y | 3544 | 836.4 | 0.001642 | 1.963 | +1 | 8 |
| - | - | 2032 | 837.4 | - | - | 0 | - |
| - | - | 973 | 838.4 | - | - | 0 | - |
| - | - | 944.1 | 842.5 | - | - | 0 | - |
| - | - | 7161 | 857.5 | - | - | 0 | - |
| - | - | 2845 | 858.5 | - | - | 0 | - |
| - | - | 1014 | 859.5 | - | - | 0 | - |
| - | - | 1203 | 862.4 | - | - | 0 | - |
| - | - | 797 | 871.8 | - | - | 0 | - |
| - | - | 1564 | 885.5 | - | - | 0 | - |
| - | - | 1247 | 886.5 | - | - | 0 | - |
| 7 | w | 1784 | 890.4 | 0.001355 | 1.521 | +1 | 9 |
| - | - | 2397 | 900.5 | - | - | 0 | - |
| 8 | c | 3.282E+04 | 901.5 | 0.0002888 | 0.3203 | +1 | 8 |
| - | - | 1.452E+04 | 902.5 | - | - | 0 | - |
| - | - | 5069 | 903.5 | - | - | 0 | - |
| - | - | 1150 | 904.5 | - | - | 0 | - |
| - | - | 744.6 | 919.4 | - | - | 0 | - |
| - | - | 1142 | 926.5 | - | - | 0 | - |
| - | - | 1467 | 927.5 | - | - | 0 | - |
| - | - | 808.7 | 936.4 | - | - | 0 | - |
| - | - | 1.615E+04 | 944.5 | - | - | 0 | - |
| - | - | 9845 | 945.5 | - | - | 0 | - |
| - | - | 3430 | 946.5 | - | - | 0 | - |
| - | - | 776.6 | 947.5 | - | - | 0 | - |
| 7 | z | 849.4 | 958.5 | 0.002024 | 2.112 | +1 | 9 |
| - | - | 767 | 961.5 | - | - | 0 | - |
| - | - | 738.6 | 973.5 | - | - | 0 | - |
| 7 | z | 1.63E+04 | 976.5 | 0.0005452 | 0.5583 | +1 | 9 |
| - | - | 1.077E+04 | 977.5 | - | - | 0 | - |
| - | - | 3481 | 978.5 | - | - | 0 | - |
| - | - | 699.1 | 979.5 | - | - | 0 | - |
| - | - | 6741 | 987.5 | - | - | 0 | - |
| 9 | c | 7.59E+04 | 988.5 | 0.0002428 | 0.2456 | +1 | 9 |
| - | - | 3.637E+04 | 989.5 | - | - | 0 | - |
| - | - | 1.252E+04 | 990.5 | - | - | 0 | - |
| - | - | 2358 | 991.5 | - | - | 0 | - |
| - | - | 962 | 992.5 | - | - | 0 | - |
| - | - | 4535 | 1002 | - | - | 0 | - |
| - | - | 1511 | 1003 | - | - | 0 | - |
| - | - | 954.1 | 1023 | - | - | 0 | - |
| - | - | 2325 | 1026 | - | - | 0 | - |
| - | - | 1107 | 1026 | - | - | 0 | - |
| - | - | 726.8 | 1031 | - | - | 0 | - |
| 10 | c | 4.288E+04 | 1046 | 0.0002024 | 0.1936 | +1 | 10 |
| 6 | w | 2.475E+04 | 1047 | 0.007007 | 6.695 | +1 | 10 |
| - | - | 1.327E+04 | 1048 | - | - | 0 | - |
| - | - | 4681 | 1049 | - | - | 0 | - |
| - | - | 1271 | 1050 | - | - | 0 | - |
| - | - | 1055 | 1061 | - | - | 0 | - |
| - | - | 1663 | 1068 | - | - | 0 | - |
| 6 | z | 952 | 1074 | 0.01045 | 9.738 | +1 | 10 |
| 6 | z | 1094 | 1075 | 0.003211 | 2.989 | +1 | 10 |
| - | - | 900.6 | 1076 | - | - | 0 | - |
| - | - | 734.6 | 1080 | - | - | 0 | - |
| - | - | 7120 | 1081 | - | - | 0 | - |
| - | - | 4862 | 1082 | - | - | 0 | - |
| - | - | 1780 | 1083 | - | - | 0 | - |
| - | - | 913.9 | 1083 | - | - | 0 | - |
| - | - | 2463 | 1083 | - | - | 0 | - |
| - | - | 2732 | 1084 | - | - | 0 | - |
| - | - | 1204 | 1086 | - | - | 0 | - |
| 6 | z | 1.587E+04 | 1092 | 0.0002217 | 0.2031 | +1 | 10 |
| - | - | 2.364E+04 | 1093 | - | - | 0 | - |
| - | - | 1.184E+04 | 1094 | - | - | 0 | - |
| - | - | 4668 | 1095 | - | - | 0 | - |
| - | - | 1099 | 1096 | - | - | 0 | - |
| - | - | 1927 | 1101 | - | - | 0 | - |
| - | - | 1785 | 1102 | - | - | 0 | - |
| - | - | 1494 | 1103 | - | - | 0 | - |
| - | - | 708.4 | 1107 | - | - | 0 | - |
| 6 | y | 3700 | 1108 | 0.001901 | 1.716 | +1 | 10 |
| - | - | 1045 | 1109 | - | - | 0 | - |
| - | - | 936 | 1110 | - | - | 0 | - |
| - | - | 3821 | 1130 | - | - | 0 | - |
| - | - | 2101 | 1131 | - | - | 0 | - |
| - | - | 999.1 | 1132 | - | - | 0 | - |
| - | - | 1422 | 1144 | - | - | 0 | - |
| 11 | c | 3.301E+04 | 1145 | 0.0003404 | 0.2974 | +1 | 11 |
| - | - | 1.999E+04 | 1146 | - | - | 0 | - |
| - | - | 7836 | 1147 | - | - | 0 | - |
| - | - | 1943 | 1148 | - | - | 0 | - |
| - | - | 964.1 | 1173 | - | - | 0 | - |
| 5 | z | 2498 | 1179 | 0.002266 | 1.922 | +1 | 11 |
| - | - | 1.322E+04 | 1180 | - | - | 0 | - |
| - | - | 6943 | 1181 | - | - | 0 | - |
| - | - | 3577 | 1182 | - | - | 0 | - |
| - | - | 1706 | 1183 | - | - | 0 | - |
| - | - | 3117 | 1188 | - | - | 0 | - |
| - | - | 3213 | 1189 | - | - | 0 | - |
| - | - | 1166 | 1190 | - | - | 0 | - |
| 5 | y | 5876 | 1195 | 0.001336 | 1.119 | +1 | 11 |
| - | - | 3439 | 1196 | - | - | 0 | - |
| - | - | 1760 | 1197 | - | - | 0 | - |
| - | - | 1869 | 1231 | - | - | 0 | - |
| 12 | c | 6.454E+04 | 1232 | 0.001241 | 1.008 | +1 | 12 |
| - | - | 4.249E+04 | 1233 | - | - | 0 | - |
| - | - | 1.577E+04 | 1234 | - | - | 0 | - |
| - | - | 4598 | 1235 | - | - | 0 | - |
| - | - | 1339 | 1236 | - | - | 0 | - |
| - | - | 755.2 | 1237 | - | - | 0 | - |
| - | - | 2761 | 1259 | - | - | 0 | - |
| 4 | z | 1635 | 1260 | 0.003728 | 2.96 | +1 | 12 |
| 4 | w | 857.9 | 1263 | 0.01774 | 14.05 | +1 | 12 |
| - | - | 2327 | 1275 | - | - | 0 | - |
| 4 | y | 1564 | 1276 | 0.01345 | 10.54 | +1 | 12 |
| 4 | y | 998.6 | 1277 | 0.0149 | 11.67 | +1 | 12 |
| 4 | z | 3952 | 1278 | 0.0004891 | 0.3829 | +1 | 12 |
| - | - | 1.627E+04 | 1279 | - | - | 0 | - |
| - | - | 1.2E+04 | 1280 | - | - | 0 | - |
| - | - | 4151 | 1281 | - | - | 0 | - |
| 4 | y | 4887 | 1294 | 0.002368 | 1.83 | +1 | 12 |
| - | - | 3431 | 1295 | - | - | 0 | - |
| - | - | 1540 | 1296 | - | - | 0 | - |
| 13 | c | 4.611E+04 | 1319 | 0.001165 | 0.8833 | +1 | 13 |
| - | - | 3.269E+04 | 1320 | - | - | 0 | - |
| - | - | 1.292E+04 | 1321 | - | - | 0 | - |
| - | - | 3612 | 1322 | - | - | 0 | - |
| - | - | 4819 | 1346 | - | - | 0 | - |
| - | - | 2575 | 1347 | - | - | 0 | - |
| 3 | w | 1651 | 1348 | 0.003045 | 2.259 | +1 | 13 |
| - | - | 969.1 | 1349 | - | - | 0 | - |
| - | - | 1014 | 1392 | - | - | 0 | - |
| 3 | z | 3481 | 1407 | 0.0002527 | 0.1797 | +1 | 13 |
| - | - | 2.763E+04 | 1408 | - | - | 0 | - |
| - | - | 1.755E+04 | 1409 | - | - | 0 | - |
| - | - | 7088 | 1410 | - | - | 0 | - |
| - | - | 2119 | 1411 | - | - | 0 | - |
| - | - | 1474 | 1412 | - | - | 0 | - |
| - | - | 1140 | 1415 | - | - | 0 | - |
| - | - | 1085 | 1416 | - | - | 0 | - |
| - | - | 2200 | 1417 | - | - | 0 | - |
| - | - | 2231 | 1418 | - | - | 0 | - |
| 3 | y | 810.9 | 1423 | 0.01371 | 9.637 | +1 | 13 |
| - | - | 1811 | 1424 | - | - | 0 | - |
| - | - | 1437 | 1425 | - | - | 0 | - |
| - | - | 1667 | 1431 | - | - | 0 | - |
| - | - | 6063 | 1432 | - | - | 0 | - |
| - | - | 5334 | 1433 | - | - | 0 | - |
| - | - | 2608 | 1434 | - | - | 0 | - |
| - | - | 964.6 | 1435 | - | - | 0 | - |
| - | - | 772.6 | 1438 | - | - | 0 | - |
| - | - | 767.5 | 1448 | - | - | 0 | - |
| - | - | 764.8 | 1449 | - | - | 0 | - |
| - | - | 1500 | 1452 | - | - | 0 | - |
| - | - | 5518 | 1459 | - | - | 0 | - |
| - | - | 3139 | 1460 | - | - | 0 | - |
| - | - | 2182 | 1461 | - | - | 0 | - |
| 14 | c | 1.069E+04 | 1475 | 0.0007133 | 0.4837 | +1 | 14 |
| - | - | 3.248E+04 | 1476 | - | - | 0 | - |
| - | - | 2.342E+04 | 1477 | - | - | 0 | - |
| - | - | 9771 | 1478 | - | - | 0 | - |
| - | - | 3048 | 1479 | - | - | 0 | - |
| - | - | 1040 | 1480 | - | - | 0 | - |
| 2 | z | 3031 | 1494 | 0.008385 | 5.614 | +1 | 14 |
| - | - | 2.001E+04 | 1495 | - | - | 0 | - |
| - | - | 1.365E+04 | 1496 | - | - | 0 | - |
| - | - | 6356 | 1497 | - | - | 0 | - |
| - | - | 2294 | 1498 | - | - | 0 | - |
| - | - | 2339 | 1508 | - | - | 0 | - |
| - | - | 770 | 1509 | - | - | 0 | - |
| - | - | 882.3 | 1510 | - | - | 0 | - |
| - | - | 934.9 | 1511 | - | - | 0 | - |
| - | - | 5441 | 1523 | - | - | 0 | - |
| - | - | 5446 | 1524 | - | - | 0 | - |
| - | - | 3256 | 1525 | - | - | 0 | - |
| - | - | 1152 | 1526 | - | - | 0 | - |
| - | - | 894.1 | 1527 | - | - | 0 | - |
| - | - | 1176 | 1537 | - | - | 0 | - |
| - | - | 1969 | 1548 | - | - | 0 | - |
| - | - | 3343 | 1549 | - | - | 0 | - |
| - | - | 1470 | 1550 | - | - | 0 | - |
| - | - | 905.2 | 1563 | - | - | 0 | - |
| - | - | 1849 | 1564 | - | - | 0 | - |
| - | - | 2.799E+04 | 1565 | - | - | 0 | - |
| - | - | 3.412E+04 | 1566 | - | - | 0 | - |
| - | - | 2.218E+04 | 1567 | - | - | 0 | - |
| - | - | 9748 | 1568 | - | - | 0 | - |
| - | - | 2225 | 1569 | - | - | 0 | - |
| - | - | 3900 | 1579 | - | - | 0 | - |
| - | - | 6453 | 1580 | - | - | 0 | - |
| - | - | 8029 | 1581 | - | - | 0 | - |
| - | - | 9539 | 1582 | - | - | 0 | - |
| - | - | 6372 | 1583 | - | - | 0 | - |
| - | - | 2774 | 1584 | - | - | 0 | - |
| - | - | 2501 | 1590 | - | - | 0 | - |
| - | - | 9432 | 1591 | - | - | 0 | - |
| - | - | 6136 | 1592 | - | - | 0 | - |
| - | - | 3525 | 1593 | - | - | 0 | - |
| - | - | 828.1 | 1594 | - | - | 0 | - |
| - | - | 1994 | 1596 | - | - | 0 | - |
| - | - | 8906 | 1597 | - | - | 0 | - |
| - | - | 7639 | 1598 | - | - | 0 | - |
| - | - | 4537 | 1599 | - | - | 0 | - |
| - | - | 1097 | 1600 | - | - | 0 | - |
| - | - | 1174 | 1606 | - | - | 0 | - |
| - | - | 1.994E+04 | 1607 | - | - | 0 | - |
| - | - | 9.432E+04 | 1608 | - | - | 0 | - |
| - | - | 7.447E+04 | 1609 | - | - | 0 | - |
| - | - | 3.492E+04 | 1610 | - | - | 0 | - |
| - | - | 1.19E+04 | 1611 | - | - | 0 | - |
| - | - | 2540 | 1612 | - | - | 0 | - |
| - | - | 5227 | 1623 | - | - | 0 | - |
| - | - | 476.6 | 1624 | - | - | 0 | - |
| - | - | 2.484E+04 | 1624 | - | - | 0 | - |
| - | - | 9.253E+04 | 1625 | - | - | 0 | - |
| - | - | 7.142E+04 | 1626 | - | - | 0 | - |
| - | - | 3.736E+04 | 1627 | - | - | 0 | - |
| - | - | 1.214E+04 | 1628 | - | - | 0 | - |
| - | - | 2550 | 1629 | - | - | 0 | - |
| - | - | 959.6 | 1635 | - | - | 0 | - |
| - | - | 762.1 | 3000 | - | - | 0 | - |

m/z Charge Intensity FragmentType MassShift Position
120.08097839355469 0 1209.2993
123.81603240966797 0 414.8634
124.33694458007812 0 358.74432
128.26171875 0 365.45526
132.25692749023438 0 409.87943
142.09767150878906 0 1344.5481
156.56663513183594 0 406.68774
160.1081085205078 0 463.25378
170.95933532714844 0 475.7021
173.1285400390625 0 8286.249
173.4515380859375 0 1735.6752
174.1321563720703 0 743.468
183.11289978027344 0 2592.475
198.90003967285156 0 436.04227
199.11891174316406 0 740.0266
200.11546325683594 0 548.8309
201.12344360351562 0 13827.84
201.13473510742188 0 8052.9385
202.12332153320312 0 2701.7104
202.1414337158203 0 926.159
217.08245849609375 0 1183.7742
221.34112548828125 0 533.4845
223.1083221435547 0 1555.6143
229.11862182617188 0 698.66064
229.1300048828125 0 1210.0651
235.22415161132812 0 514.40265
244.14410400390625 0 1107.6273
248.6294403076172 0 559.9879 y 11
249.13192749023438 0 573.6898
261.1231689453125 0 1686.8734
261.1709899902344 0 18452.826
262.1746520996094 0 2269.7996
270.15533447265625 0 670.639
277.0154113769531 0 465.15353
288.16693115234375 0 1370.4188
306.1688232421875 0 3728.9329 z 13
307.1722412109375 0 751.1409
310.13995361328125 0 858.8526
312.1556396484375 0 6390.0005
313.1579895019531 0 1184.9982
316.1501159667969 0 1374.6362
321.1783752441406 0 740.7361
322.1873779296875 0 3990.1443 y 13
323.18914794921875 0 709.96704
330.1667175292969 0 3659.9436
331.1686706542969 0 697.09
348.20330810546875 0 2202.0898
349.2094421386719 0 1199.2169
373.1975402832031 0 583.76184
376.1978454589844 0 11779.64 c Water loss 10
377.2010498046875 0 2824.2024
387.23516845703125 0 1004.29
391.1848449707031 0 4510.971
392.19281005859375 0 17748.69 y Ammonia loss 12
393.1989440917969 0 7844.053 y Ammonia loss 4
394.20538330078125 0 2303.1562 c Ammonia loss 6
397.44073486328125 0 594.3307
401.2392272949219 0 2579.386
409.21990966796875 0 4009.8425 y 12
410.2230224609375 0 708.1157
411.2275695800781 0 632.9095
417.1884460449219 0 6249.759
418.1920471191406 0 1338.173
418.7162780761719 0 944.60315 y 7
419.043212890625 0 553.40985
428.2255859375 0 1234.1655
429.2342224121094 0 4916.7817
430.2366943359375 0 1381.6357
435.2356262207031 0 1444.5912
436.2426452636719 0 835.1183
444.2567443847656 0 3857.1387
445.2555847167969 0 3848.2993
446.2594909667969 0 3011.431 c 3
447.26483154296875 0 669.078
462.2205810546875 0 740.7103 z Water loss 11
463.2303466796875 0 12109.174 w 11
464.2313537597656 0 4881.2534
469.23583984375 0 669.52454 y Ammonia loss 2
478.2173767089844 0 5184.756
479.2245178222656 0 5931.8623 y Ammonia loss 11
480.2325134277344 0 10056.347 z 11
481.237548828125 0 2650.3147
488.75823974609375 0 10332.052 z 6
489.1830749511719 0 688.4453
489.2602844238281 0 4640.1074
489.76171875 0 2970.0747
490.2889404296875 0 882.7218
496.2519226074219 0 3727.506 y 11
496.7671203613281 0 2754.4705 y 6
497.26715087890625 0 2421.222
498.2491149902344 0 1017.3942
503.9142150878906 0 926.02014 y 1
504.2206115722656 0 6537.534
505.22442626953125 0 2064.3853
515.5519409179688 0 657.59674
516.2637329101562 0 1140.1528
523.2393188476562 0 667.9725
524.2764282226562 0 3815.4417
524.7792358398438 0 2362.2034
525.281494140625 0 1369.7108
531.2857666015625 0 694.9258
532.2855834960938 0 5245.9863
533.292724609375 0 27118.852 c 4
533.73876953125 0 723.65894
534.2958374023438 0 6774.724
535.3043823242188 0 1992.9036
535.6060180664062 0 2530.7236
535.9403686523438 0 2671.5527
536.2730102539062 0 714.0763
537.2694702148438 0 757.5995 z Water loss 5
541.3474731445312 0 1600.8348
541.6095581054688 0 11716.474
541.9439086914062 0 9010.709
542.190185546875 0 779.46204
542.2786254882812 0 4844.4834
542.6130981445312 0 2312.9011
542.687255859375 0 1308.0585
546.2715454101562 0 14715.258 z 5
546.7742919921875 0 16447.111
547.2760620117188 0 6059.8057
547.7770385742188 0 1676.7645
549.2921752929688 0 825.0292
550.2906494140625 0 1033.4567
553.7760620117188 0 1738.1547
554.2803344726562 0 5920.5225 y 5
554.7822265625 0 3867.8413
555.2838745117188 0 2255.4084
556.28076171875 0 731.4549
561.2899780273438 0 1816.2377 z Water loss 10
562.2975463867188 0 2333.1975
564.2783203125 0 11805.923 c Ammonia loss 10
565.2811279296875 0 4076.235
567.7875366210938 0 1151.6653
568.2818603515625 0 1560.7488
579.3015747070312 0 6427.8945 z 10
580.30419921875 0 2464.4473
580.7814331054688 0 895.8678 z Water loss 4
581.2736206054688 0 622.9275 z Ammonia loss 4
581.7808227539062 0 984.4398
582.2861328125 0 830.62164
586.3347778320312 0 782.9321
588.78271484375 0 1965.0424 y Water loss 4
589.282958984375 0 4163.895 y Ammonia loss 4
589.7872314453125 0 12786.801 z 4
590.2904052734375 0 24771.102
590.7921142578125 0 12759.952
591.2939453125 0 4840.964
591.7913208007812 0 1538.1301
592.3328857421875 0 9154.105
593.3345336914062 0 3345.1213
594.3351440429688 0 788.587
595.3203735351562 0 6095.5605 y 10
596.32373046875 0 2210.91
597.293701171875 0 6191.812
597.796875 0 42003.332 y 4
598.2984619140625 0 20199.533
598.7994995117188 0 10808.916
599.3013305664062 0 1813.686
599.8024291992188 0 728.4597
601.7807006835938 0 1919.801
602.2840576171875 0 1629.1082
611.2929077148438 0 988.1011
619.3204345703125 0 1092.4128
620.3216552734375 0 806.87946
631.810546875 0 1163.3008 w 3
632.31201171875 0 1478.9495
636.3228149414062 0 26836.947 z 9
637.3282470703125 0 14149.142
638.3289794921875 0 3902.5352 y Water loss 3
639.3220825195312 0 23650.67 z 3
639.8233642578125 0 19418.502
640.3258056640625 0 7858.2363
640.8269653320312 0 2566.062
642.3145141601562 0 1103.9303
642.8110961914062 0 967.77716
646.8225708007812 0 808.62366
647.3311157226562 0 40119.688 y 3
647.8327026367188 0 24414.572
648.3245849609375 0 27423.023 c 5
648.8350830078125 0 2021.1517
649.3231201171875 0 7691.2085
650.3285522460938 0 1692.0134
652.3416748046875 0 25504.68 y 9
653.3441162109375 0 9816.221
654.3478393554688 0 1924.8472
658.8795776367188 0 585.62744
674.3364868164062 0 22550.463 w 2
674.8374633789062 0 17624.504
675.3399047851562 0 6290.779
675.8405151367188 0 1742.7815
679.3643798828125 0 2841.0405
680.3676147460938 0 1113.5405
681.8443603515625 0 1654.567
694.3349609375 0 885.7626
702.8455200195312 0 3168.3032 y Water loss 2
703.3475952148438 0 2064.734 y Ammonia loss 2
703.8432006835938 0 16269.338 z 2
704.3446655273438 0 12361.65
704.409912109375 0 871.91034
704.8460693359375 0 4531.9917
705.34423828125 0 1547.3232 z Water loss 8
707.8697509765625 0 1444.7488
708.3667602539062 0 1890.5355
711.349609375 0 1534.9542
711.8521728515625 0 45153.516 y 2
712.353759765625 0 36380.215
712.8551025390625 0 15550.837
713.3566284179688 0 3550.4253
713.8599853515625 0 1107.2455
715.8772583007812 0 5379.925
716.377685546875 0 3503.6646
716.882568359375 0 1074.8807
717.3784790039062 0 697.1321
717.8505859375 0 4161.2383
718.356689453125 0 2199.0571
721.338134765625 0 2387.9146
721.6715698242188 0 1346.6809
722.3455200195312 0 2635.7183 y Ammonia loss 8
723.3544921875 0 18238.729 z 8
724.3576049804688 0 7114.5156
724.8594970703125 0 619.17596
725.3623657226562 0 3376.413
725.8639526367188 0 1150.9711
726.36279296875 0 1496.3972
726.8755493164062 0 590.20917
729.3660888671875 0 776.0901 c Ammonia loss 13
730.3729248046875 0 1087.7328
732.330810546875 0 653.12036
737.8843383789062 0 117346.305 c 13
738.385498046875 0 88985.16
738.8865356445312 0 41652.324
739.3814697265625 0 15774.793 y 8
739.886962890625 0 1542.8988
740.3763427734375 0 2739.21
746.3617553710938 0 5068.1636 y Water loss 1
746.8597412109375 0 5983.849 y Ammonia loss 1
747.3570556640625 0 23335.973 z 1
747.8602294921875 0 13592.944
748.3588256835938 0 6825.7197
748.8613891601562 0 2080.4988
755.3684692382812 0 51812.38 y 1
755.8700561523438 0 44010.48
756.3707885742188 0 24186.242
756.8720092773438 0 6683.3345
757.3709106445312 0 2384.4077
759.39404296875 0 728.2406
760.4058227539062 0 3071.802
760.8629760742188 0 889.5037
761.40869140625 0 768.2903
761.86865234375 0 869.64215
768.3934326171875 0 1090.037
773.3884887695312 0 1169.8628
774.3967895507812 0 2372.7622
774.9000854492188 0 1149.1085
775.394287109375 0 977.7794
775.8792724609375 0 2614.0618
781.4097900390625 0 1416.8483
781.9058227539062 0 10547.2
782.4048461914062 0 22339.78
782.9044189453125 0 16483.564
783.4037475585938 0 9119.069
783.8953857421875 0 2292.8538
788.4000244140625 0 3423.0752
788.900390625 0 2311.198
789.383056640625 0 33644.52
789.8845825195312 0 31121.035
790.3882446289062 0 15816.111
790.8991088867188 0 33637.793
791.4005737304688 0 27906.807
791.9024658203125 0 12905.997
792.4024047851562 0 4378.2734
794.4033203125 0 892.12994
794.8965454101562 0 3207.0366
795.3980102539062 0 3779.7856
795.8969116210938 0 3235.4714
796.393798828125 0 4228.666
796.8903198242188 0 2017.6631
797.3948974609375 0 1524.7422
803.407958984375 0 14230.556
803.9020385742188 0 107772.8
804.4033813476562 0 96554.72
804.9039916992188 0 48898.9
804.994140625 0 1128.8525
805.4051513671875 0 13334.527
805.9067993164062 0 4674.9814
810.3814086914062 0 645.36804
810.8231811523438 0 808.246
811.9105224609375 0 58718
812.4131469726562 0 80351.625
812.914794921875 0 55911.03
813.4166259765625 0 22624.807
813.9190673828125 0 5133.9946
814.409912109375 0 1238.3372
836.427734375 0 3543.6797 y 7
837.4263305664062 0 2032.2336
838.4295043945312 0 973.0207
842.4507446289062 0 944.06573
857.460693359375 0 7161.4067
858.4654541015625 0 2845.244
859.4699096679688 0 1013.7365
862.3788452148438 0 1203.4172
871.7864990234375 0 796.9753
885.458740234375 0 1563.5569
886.458984375 0 1246.8438
890.435302734375 0 1784.3005 w 6
900.497314453125 0 2396.9255
901.4740600585938 0 32820.168 c 7
902.4769287109375 0 14523.204
903.4802856445312 0 5069.2363
904.48486328125 0 1149.917
919.3901977539062 0 744.59625
926.4801635742188 0 1141.6704
927.4893798828125 0 1466.6532
936.42138671875 0 808.705
944.4930419921875 0 16149.23
945.4956665039062 0 9845.211
946.4996337890625 0 3430.1235
947.49658203125 0 776.5914
958.4999389648438 0 849.4455 z Water loss 6
961.4954223632812 0 766.9664
973.48828125 0 738.615
976.5079345703125 0 16299.419 z 6
977.51318359375 0 10766.248
978.5131225585938 0 3481.4285
979.50732421875 0 699.1325
987.4967651367188 0 6740.5903
988.5060424804688 0 75904.14 c 8
989.5089721679688 0 36368.24
990.51171875 0 12520.071
991.512451171875 0 2358.2417
992.4503173828125 0 961.9981
1001.5137329101562 0 4535.208
1002.5191040039062 0 1511.418
1023.3635864257812 0 954.056
1026.363037109375 0 2324.6282
1026.4725341796875 0 1106.9506
1030.509765625 0 726.8109
1045.5274658203125 0 42875.484 c 9
1046.53076171875 0 24752.773 w 5
1047.536865234375 0 13268.546
1048.5419921875 0 4680.903
1049.5460205078125 0 1270.7367
1060.5029296875 0 1055.1522
1068.4617919921875 0 1662.8998
1073.514404296875 0 952.002 z Water loss 5
1074.5120849609375 0 1093.8042 z Ammonia loss 5
1075.5223388671875 0 900.6231
1080.3729248046875 0 734.5871
1081.370849609375 0 7120.14
1082.37255859375 0 4862.1875
1082.510009765625 0 1780.0319
1083.0140380859375 0 913.89166
1083.3717041015625 0 2462.8289
1084.3756103515625 0 2731.5571
1085.593017578125 0 1203.6376
1091.53564453125 0 15869.779 z 5
1092.54150390625 0 23642.236
1093.5447998046875 0 11843.212
1094.5491943359375 0 4667.783
1095.5504150390625 0 1098.9249
1100.57861328125 0 1926.7728
1101.587646484375 0 1785.3335
1102.591796875 0 1493.816
1106.5399169921875 0 708.41205
1107.55224609375 0 3699.869 y 5
1108.5560302734375 0 1045.084
1109.5567626953125 0 935.9836
1129.583740234375 0 3821.1868
1130.5860595703125 0 2100.5261
1131.5870361328125 0 999.069
1143.5860595703125 0 1421.5887
1144.5953369140625 0 33008.484 c 10
1145.5987548828125 0 19994.217
1146.6021728515625 0 7836.082
1147.5994873046875 0 1943.3954
1172.6014404296875 0 964.14496
1178.565185546875 0 2498.4858 z 4
1179.573974609375 0 13217.225
1180.5775146484375 0 6943.336
1181.5826416015625 0 3576.792
1182.572998046875 0 1706.125
1187.6092529296875 0 3116.8599
1188.615234375 0 3212.867
1189.63232421875 0 1165.9878
1194.5848388671875 0 5876.3213 y 4
1195.588134765625 0 3439.2385
1196.59521484375 0 1759.6241
1230.6226806640625 0 1868.6171
1231.62646484375 0 64542.938 c 11
1232.62939453125 0 42493.5
1233.6336669921875 0 15768.461
1234.6318359375 0 4597.917
1235.6329345703125 0 1338.9089
1236.625 0 755.1797
1258.62255859375 0 2761.4355
1259.6290283203125 0 1635.1206 z Water loss 3
1262.630126953125 0 857.8839 w 3
1274.6451416015625 0 2327.241
1275.657470703125 0 1564.4974 y Water loss 3
1276.6429443359375 0 998.62006 y Ammonia loss 3
1277.6353759765625 0 3952.0972 z 3
1278.64208984375 0 16271.48
1279.6458740234375 0 12004.186
1280.6495361328125 0 4151.0117
1293.6522216796875 0 4886.66 y 3
1294.6585693359375 0 3431.4075
1295.6639404296875 0 1540.089
1318.6585693359375 0 46111.406 c 12
1319.6611328125 0 32687.346
1320.66357421875 0 12923.301
1321.6673583984375 0 3612.3328
1345.6622314453125 0 4818.869
1346.6651611328125 0 2574.9624
1347.662109375 0 1651.3118 w 2
1348.6549072265625 0 969.1312
1391.679443359375 0 1014.0284
1406.6787109375 0 3480.5247 z 2
1407.6834716796875 0 27632.53
1408.6875 0 17551.846
1409.6883544921875 0 7087.5728
1410.68994140625 0 2118.9348
1411.695556640625 0 1474.1666
1414.727294921875 0 1140.3477
1415.72412109375 0 1085.0638
1416.716552734375 0 2199.7175
1417.7186279296875 0 2231.3374
1422.6834716796875 0 810.87616 y 2
1423.705322265625 0 1811.4937
1424.6988525390625 0 1436.7665
1430.7442626953125 0 1667.068
1431.75 0 6062.679
1432.751708984375 0 5334.076
1433.749755859375 0 2608.3506
1434.748779296875 0 964.5721
1437.6864013671875 0 772.6255
1447.7589111328125 0 767.459
1448.767578125 0 764.83105
1451.6888427734375 0 1499.9105
1458.74169921875 0 5517.89
1459.7415771484375 0 3139.3335
1460.7540283203125 0 2181.5188
1474.7601318359375 0 10685.53 c 13
1475.765380859375 0 32483.137
1476.765869140625 0 23416.059
1477.7640380859375 0 9770.834
1478.7506103515625 0 3048.0293
1479.733154296875 0 1040.4774
1493.7188720703125 0 3030.834 z 1
1494.714599609375 0 20009.562
1495.7193603515625 0 13654.843
1496.7232666015625 0 6356.48
1497.7236328125 0 2294.3535
1507.74853515625 0 2338.5232
1508.7393798828125 0 770.0171
1509.76416015625 0 882.2504
1510.7421875 0 934.9138
1522.7279052734375 0 5440.566
1523.7310791015625 0 5446.112
1524.7366943359375 0 3255.9639
1525.72802734375 0 1151.9873
1526.751220703125 0 894.06866
1536.8045654296875 0 1175.6965
1547.7762451171875 0 1969.1008
1548.763671875 0 3342.6592
1549.76123046875 0 1470.1454
1562.8096923828125 0 905.20624
1563.791259765625 0 1849.1007
1564.7822265625 0 27989.166
1565.7821044921875 0 34122.7
1566.7855224609375 0 22177.93
1567.7864990234375 0 9748.158
1568.7867431640625 0 2224.5908
1578.818603515625 0 3900.3208
1579.8123779296875 0 6453.047
1580.8079833984375 0 8029.287
1581.802001953125 0 9539.179
1582.7982177734375 0 6372.2944
1583.8013916015625 0 2773.5085
1589.7919921875 0 2501.4387
1590.77685546875 0 9431.849
1591.7818603515625 0 6135.5166
1592.7786865234375 0 3524.7996
1593.7802734375 0 828.14545
1595.8167724609375 0 1994.0103
1596.8289794921875 0 8905.558
1597.8355712890625 0 7639.012
1598.8369140625 0 4536.9795
1599.844482421875 0 1097.3119
1605.8077392578125 0 1173.6528
1606.8035888671875 0 19942.102
1607.8006591796875 0 94323.94
1608.8028564453125 0 74467.055
1609.8056640625 0 34922.727
1610.80908203125 0 11895.735
1611.8140869140625 0 2539.7434
1622.8087158203125 0 5227.2544
1623.611572265625 0 476.60498
1623.8177490234375 0 24840.443
1624.8251953125 0 92525.1
1625.828857421875 0 71424.99
1626.8310546875 0 37357.97
1627.833251953125 0 12137.389
1628.825439453125 0 2549.8215
1635.2923583984375 0 959.6025
3000.15673828125 0 762.0561

Spectrum Details

|  |  |
| --- | --- |
| Matched peaks? Matched peaks The total absolute number of peaks matched. Additionally in brackets the total fraction of peaks matched and the total number of peaks is shown. | 83 (16.44% of 505) |
| FDR? FDR The false discovery rate estimated for this peptide. It is calculated by matching all theoretical fragments with a non-integer shift with the raw peaks for this spectrum. This is done with 40 different shifts. The resulting percentage is the average number of annotated peaks over the number of annotated peaks with the correct spectrum. | 0.23% |
| Satellite FDR? Satellite FDR See the FDR for details on its calculation. This satellite ion specific FDR only contains the satellite ions (d/w) for I/L/J positions. | - |
| PSM Score? PSM Score The PSM Score as given by Hecklib to this annotated spectrum. It is shown with three significant figures. | 586 |

#### Spectrum 4533? Spectrum 4533 The raw spectrum of this peptide as annotated by Hecklib. The fragments are coloured according to ion type (see legend). Any peaks with a star '\*' as text can be hovered over to see the full details, first the ion type second the mass shift type. By hovering over the amino acids in the peptide or ions in the legend the corresponding peaks are highlighted. By toggling the 'Unassigned' label you can turn the background (unassigned) peaks on or off in the plot. By updating the slider in the Ion legend you can update the spectrum to only show the top X% of the peaks with labels. The top X% means any peak that is within X% of the highest intensity. By dragging in the spectrum you can zoom in to a specific part of the spectrum and use 'Zoom Out' to get back to the original zoom level. The annotation of the spectrum is based on the given sequence in the peptides file and is done with different software so inconsistencies are likely. The peaks are annotated based on the given sequence, with 20 ppm tolerance.

Copy Data

##### Spectrum 4533 (TSV)

###### Preview

```
Loading example...
```

*Click on the button to copy the data to your clipboard.*

Mz MinMz MaxIntensity Max

WidthHeightPeptide font sizePeptide stroke widthSpectrum font sizeSpectrum stroke widthCompact peptide

Ion legend

wxyz

abcd

OtherUnassignedIon^Charge^_Position_Show for top:%

JSEVSDRPSGVSSRF

04.24e+68.48e+61.27e+71.70e+7

Zoom Out

y^+3^_8_z^+1^_2_z^+3^_9_y^+1^_2_c^+3^_11_y^+1^_3_y^+3^_11_c^+2^_7_y^+1^_3_y^+2^_8_c^+1^_4_z^+1^_4_w^+1^_4_y^+3^_13_y^+3^_13_y^+1^_4_z^+1^_4_z^+2^_9_y^+1^_4_y^+2^_9_y^+3^_14_c^+1^_5_z^+2^_10_y^+2^_10_z^+1^_5_c^+2^_11_z^+1^_5_y^+2^_11_y^+2^_11_z^+2^_11_y^+1^_5_y^+2^_11_z^+1^_6_w^+2^_12_z^+1^_6_y^+2^_12_z^+2^_12_y^+2^_12_c^+1^_6_c^+2^_13_y^+1^_6_w^+2^_13_z^+2^_13_z^+2^_13_y^+2^_13_y^+2^_13_z^+2^_13_z^+1^_7_y^+2^_13_y^+1^_7_z^+1^_7_c^+2^_14_c^+2^_14_y^+1^_7_y^+2^_14_y^+2^_14_z^+2^_14_y^+2^_14_c^+1^_7_w^+1^_8_y^+1^_8_w^+1^_9_c^+1^_8_z^+1^_9_y^+1^_9_z^+1^_9_c^+1^_9_c^+1^_10_c^+1^_10_w^+1^_10_z^+1^_10_z^+1^_10_z^+1^_10_y^+1^_10_c^+1^_11_y^+1^_11_z^+1^_11_y^+1^_11_c^+1^_12_z^+1^_12_z^+1^_12_y^+1^_12_y^+1^_12_z^+1^_12_y^+1^_12_c^+1^_13_w^+1^_13_z^+1^_13_y^+1^_13_c^+1^_14_z^+1^_14_y^+1^_14_

041182312341645

Fragment Matches Table

Show background peaks

| Position | Ion type | Intensity | mz Theoretical | mz Error (Th) | mz Error (ppm) | Charge | Series Number |
| --- | --- | --- | --- | --- | --- | --- | --- |
| - | - | 5.094E+04 | 120.1 | - | - | 0 | - |
| - | - | 2.987E+04 | 124.2 | - | - | 0 | - |
| - | - | 3.26E+04 | 125 | - | - | 0 | - |
| - | - | 1.815E+05 | 142.1 | - | - | 0 | - |
| - | - | 5.362E+04 | 148.9 | - | - | 0 | - |
| - | - | 6.988E+04 | 173.1 | - | - | 0 | - |
| - | - | 1.344E+06 | 173.1 | - | - | 0 | - |
| - | - | 1.171E+05 | 173.4 | - | - | 0 | - |
| - | - | 1.096E+05 | 174.1 | - | - | 0 | - |
| - | - | 4.715E+04 | 176.4 | - | - | 0 | - |
| - | - | 5.384E+05 | 183.1 | - | - | 0 | - |
| - | - | 4.856E+04 | 183.6 | - | - | 0 | - |
| - | - | 9.198E+04 | 199.1 | - | - | 0 | - |
| - | - | 5.936E+04 | 199.1 | - | - | 0 | - |
| - | - | 1.034E+05 | 200.1 | - | - | 0 | - |
| - | - | 2.311E+06 | 201.1 | - | - | 0 | - |
| - | - | 1.18E+06 | 201.1 | - | - | 0 | - |
| - | - | 3.43E+05 | 202.1 | - | - | 0 | - |
| - | - | 1.472E+05 | 202.1 | - | - | 0 | - |
| - | - | 7.23E+04 | 211.1 | - | - | 0 | - |
| - | - | 2.157E+05 | 217.1 | - | - | 0 | - |
| - | - | 4.467E+04 | 226.7 | - | - | 0 | - |
| - | - | 7.321E+04 | 229.1 | - | - | 0 | - |
| - | - | 1.352E+05 | 229.1 | - | - | 0 | - |
| - | - | 1.969E+05 | 261.1 | - | - | 0 | - |
| - | - | 2.471E+06 | 261.2 | - | - | 0 | - |
| - | - | 4.41E+04 | 262.1 | - | - | 0 | - |
| - | - | 5.029E+04 | 262.1 | - | - | 0 | - |
| - | - | 7.514E+04 | 262.2 | - | - | 0 | - |
| - | - | 4.062E+05 | 262.2 | - | - | 0 | - |
| - | - | 4.571E+04 | 263.2 | - | - | 0 | - |
| - | - | 1.286E+05 | 270.2 | - | - | 0 | - |
| 8 | y | 4.512E+04 | 273.8 | 0.001057 | 3.859 | +3 | 8 |
| - | - | 9.406E+04 | 276.2 | - | - | 0 | - |
| - | - | 5.372E+04 | 277 | - | - | 0 | - |
| - | - | 9.689E+04 | 284.2 | - | - | 0 | - |
| - | - | 1.961E+05 | 288.2 | - | - | 0 | - |
| - | - | 4.692E+04 | 299.2 | - | - | 0 | - |
| - | - | 5.532E+04 | 303.1 | - | - | 0 | - |
| 14 | z | 4.99E+05 | 306.2 | 8.968E-05 | 0.2929 | +1 | 2 |
| - | - | 9.679E+04 | 307.2 | - | - | 0 | - |
| - | - | 9.754E+05 | 312.2 | - | - | 0 | - |
| - | - | 2.063E+05 | 313.2 | - | - | 0 | - |
| - | - | 1.744E+05 | 316.2 | - | - | 0 | - |
| 7 | z | 1.288E+05 | 320.2 | 0.001174 | 3.668 | +3 | 9 |
| - | - | 7.769E+04 | 321.2 | - | - | 0 | - |
| 14 | y | 6.855E+05 | 322.2 | 4.236E-05 | 0.1315 | +1 | 2 |
| - | - | 4.958E+04 | 327.2 | - | - | 0 | - |
| - | - | 8.449E+05 | 330.2 | - | - | 0 | - |
| - | - | 1.647E+05 | 331.2 | - | - | 0 | - |
| - | - | 3.472E+05 | 348.2 | - | - | 0 | - |
| - | - | 1.288E+05 | 349.2 | - | - | 0 | - |
| - | - | 1.543E+05 | 373.2 | - | - | 0 | - |
| - | - | 5.505E+04 | 374.2 | - | - | 0 | - |
| 11 | c | 1.57E+06 | 376.2 | 0.00189 | 5.025 | +3 | 11 |
| - | - | 3.728E+05 | 377.2 | - | - | 0 | - |
| - | - | 1.286E+05 | 387.2 | - | - | 0 | - |
| - | - | 5.817E+05 | 391.2 | - | - | 0 | - |
| 13 | y | 2.68E+06 | 392.2 | 3.533E-05 | 0.09009 | +1 | 3 |
| 5 | y | 1.122E+06 | 393.2 | 0.007337 | 18.66 | +3 | 11 |
| 7 | c | 3.61E+05 | 394.2 | 0.002837 | 7.198 | +2 | 7 |
| - | - | 3.044E+05 | 401.2 | - | - | 0 | - |
| - | - | 6.607E+04 | 407.2 | - | - | 0 | - |
| 13 | y | 5.162E+05 | 409.2 | 0.0004847 | 1.184 | +1 | 3 |
| - | - | 1.807E+05 | 410.2 | - | - | 0 | - |
| - | - | 6.035E+04 | 411.2 | - | - | 0 | - |
| - | - | 9.122E+05 | 417.2 | - | - | 0 | - |
| - | - | 2.028E+05 | 418.2 | - | - | 0 | - |
| 8 | y | 1.265E+05 | 418.7 | 0.0003564 | 0.8511 | +2 | 8 |
| - | - | 1.405E+05 | 428.2 | - | - | 0 | - |
| - | - | 6.868E+05 | 429.2 | - | - | 0 | - |
| - | - | 1.096E+05 | 430.2 | - | - | 0 | - |
| - | - | 2.554E+05 | 435.2 | - | - | 0 | - |
| - | - | 6.975E+04 | 436.2 | - | - | 0 | - |
| - | - | 1.458E+05 | 436.2 | - | - | 0 | - |
| - | - | 5.343E+04 | 439.2 | - | - | 0 | - |
| - | - | 4.338E+05 | 444.3 | - | - | 0 | - |
| - | - | 3.923E+05 | 445.3 | - | - | 0 | - |
| 4 | c | 5.024E+05 | 446.3 | 0.0004574 | 1.025 | +1 | 4 |
| - | - | 9.683E+04 | 447.3 | - | - | 0 | - |
| 12 | z | 1.741E+05 | 462.2 | 0.000278 | 0.6014 | +1 | 4 |
| - | - | 7.058E+04 | 462.9 | - | - | 0 | - |
| 12 | w | 1.934E+06 | 463.2 | 0.000418 | 0.9024 | +1 | 4 |
| - | - | 6.38E+05 | 464.2 | - | - | 0 | - |
| - | - | 2.039E+05 | 465.2 | - | - | 0 | - |
| 3 | y | 6.077E+04 | 468.9 | 0.002044 | 4.36 | +3 | 13 |
| 3 | y | 1.48E+05 | 474.9 | 0.0009576 | 2.016 | +3 | 13 |
| - | - | 9.638E+04 | 475.2 | - | - | 0 | - |
| - | - | 1.023E+05 | 475.6 | - | - | 0 | - |
| - | - | 7.111E+05 | 478.2 | - | - | 0 | - |
| 12 | y | 8.212E+05 | 479.2 | 0.0007832 | 1.634 | +1 | 4 |
| 12 | z | 1.582E+06 | 480.2 | 9.385E-05 | 0.1954 | +1 | 4 |
| - | - | 3.885E+05 | 481.2 | - | - | 0 | - |
| - | - | 6.07E+04 | 482.2 | - | - | 0 | - |
| 7 | z | 1.557E+06 | 488.8 | 0.0003616 | 0.7399 | +2 | 9 |
| - | - | 8.381E+05 | 489.3 | - | - | 0 | - |
| - | - | 3.229E+05 | 489.8 | - | - | 0 | - |
| - | - | 1.764E+05 | 490.3 | - | - | 0 | - |
| - | - | 8.104E+04 | 494.2 | - | - | 0 | - |
| 12 | y | 5.279E+05 | 496.3 | 0.0008049 | 1.622 | +1 | 4 |
| 7 | y | 4.995E+05 | 496.8 | 0.0005821 | 1.172 | +2 | 9 |
| - | - | 2.394E+05 | 497.3 | - | - | 0 | - |
| - | - | 7.985E+04 | 497.8 | - | - | 0 | - |
| - | - | 2.043E+05 | 498.3 | - | - | 0 | - |
| - | - | 1.321E+05 | 499.3 | - | - | 0 | - |
| - | - | 6.091E+04 | 500.3 | - | - | 0 | - |
| 2 | y | 2.442E+05 | 503.9 | 0.0006343 | 1.259 | +3 | 14 |
| - | - | 8.844E+05 | 504.2 | - | - | 0 | - |
| - | - | 7.032E+04 | 504.6 | - | - | 0 | - |
| - | - | 2.476E+05 | 505.2 | - | - | 0 | - |
| - | - | 6.032E+04 | 514.3 | - | - | 0 | - |
| - | - | 6.09E+04 | 515.3 | - | - | 0 | - |
| - | - | 2.058E+05 | 516.3 | - | - | 0 | - |
| - | - | 5.73E+05 | 524.3 | - | - | 0 | - |
| - | - | 4.264E+05 | 524.8 | - | - | 0 | - |
| - | - | 1.853E+05 | 525.3 | - | - | 0 | - |
| - | - | 8.773E+04 | 529.9 | - | - | 0 | - |
| - | - | 9.92E+04 | 531.3 | - | - | 0 | - |
| - | - | 8.458E+05 | 532.3 | - | - | 0 | - |
| 5 | c | 3.77E+06 | 533.3 | 4.565E-05 | 0.08561 | +1 | 5 |
| - | - | 1.138E+06 | 534.3 | - | - | 0 | - |
| - | - | 2.274E+05 | 535.3 | - | - | 0 | - |
| - | - | 5.726E+05 | 535.6 | - | - | 0 | - |
| - | - | 5.801E+05 | 535.9 | - | - | 0 | - |
| - | - | 2.324E+05 | 536.3 | - | - | 0 | - |
| - | - | 2.418E+06 | 541.6 | - | - | 0 | - |
| - | - | 1.713E+06 | 541.9 | - | - | 0 | - |
| - | - | 1.109E+06 | 542.3 | - | - | 0 | - |
| - | - | 2.559E+05 | 542.6 | - | - | 0 | - |
| - | - | 7.931E+04 | 542.9 | - | - | 0 | - |
| 6 | z | 2.114E+06 | 546.3 | 0.0001958 | 0.3584 | +2 | 10 |
| - | - | 2.306E+06 | 546.8 | - | - | 0 | - |
| - | - | 9.639E+05 | 547.3 | - | - | 0 | - |
| - | - | 2.759E+05 | 547.8 | - | - | 0 | - |
| - | - | 8.289E+04 | 548.3 | - | - | 0 | - |
| - | - | 1.261E+05 | 549.3 | - | - | 0 | - |
| - | - | 1.274E+05 | 553.8 | - | - | 0 | - |
| 6 | y | 1.202E+06 | 554.3 | 0.0005383 | 0.9712 | +2 | 10 |
| - | - | 6.473E+05 | 554.8 | - | - | 0 | - |
| - | - | 1.983E+05 | 555.3 | - | - | 0 | - |
| - | - | 1.011E+05 | 556.3 | - | - | 0 | - |
| 11 | z | 3.805E+05 | 561.3 | 0.0008142 | 1.451 | +1 | 5 |
| - | - | 3.493E+05 | 562.3 | - | - | 0 | - |
| 11 | c | 1.492E+06 | 564.3 | 0.01013 | 17.95 | +2 | 11 |
| - | - | 4.595E+05 | 565.3 | - | - | 0 | - |
| - | - | 1.095E+05 | 566.3 | - | - | 0 | - |
| - | - | 9.28E+04 | 566.8 | - | - | 0 | - |
| - | - | 8.434E+04 | 567.3 | - | - | 0 | - |
| - | - | 1.504E+05 | 567.8 | - | - | 0 | - |
| - | - | 1.949E+05 | 568.3 | - | - | 0 | - |
| 11 | z | 8.155E+05 | 579.3 | 0.0002178 | 0.376 | +1 | 5 |
| - | - | 2.835E+05 | 580.3 | - | - | 0 | - |
| - | - | 1.289E+05 | 581.8 | - | - | 0 | - |
| - | - | 5.576E+04 | 588.3 | - | - | 0 | - |
| 5 | y | 4.271E+05 | 588.8 | 0.01117 | 18.97 | +2 | 11 |
| 5 | y | 7.241E+05 | 589.3 | 0.0001872 | 0.3176 | +2 | 11 |
| 5 | z | 2.166E+06 | 589.8 | 5.072E-05 | 0.086 | +2 | 11 |
| - | - | 3.738E+06 | 590.3 | - | - | 0 | - |
| - | - | 1.887E+06 | 590.8 | - | - | 0 | - |
| - | - | 6.507E+05 | 591.3 | - | - | 0 | - |
| - | - | 1.302E+05 | 591.8 | - | - | 0 | - |
| - | - | 1.293E+06 | 592.3 | - | - | 0 | - |
| - | - | 3.869E+05 | 593.3 | - | - | 0 | - |
| 11 | y | 8.11E+05 | 595.3 | 0.0007809 | 1.312 | +1 | 5 |
| - | - | 3.032E+05 | 596.3 | - | - | 0 | - |
| - | - | 5.777E+05 | 597.3 | - | - | 0 | - |
| 5 | y | 5.996E+06 | 597.8 | 2.706E-05 | 0.04527 | +2 | 11 |
| - | - | 3.595E+06 | 598.3 | - | - | 0 | - |
| - | - | 1.62E+06 | 598.8 | - | - | 0 | - |
| - | - | 3.256E+05 | 599.3 | - | - | 0 | - |
| - | - | 3.776E+05 | 601.8 | - | - | 0 | - |
| - | - | 2.185E+05 | 602.3 | - | - | 0 | - |
| - | - | 5.956E+04 | 603.3 | - | - | 0 | - |
| - | - | 8.648E+04 | 605.3 | - | - | 0 | - |
| - | - | 1.272E+05 | 611.3 | - | - | 0 | - |
| - | - | 9.196E+04 | 611.8 | - | - | 0 | - |
| - | - | 6.374E+04 | 612.3 | - | - | 0 | - |
| 10 | z | 7.584E+04 | 618.3 | 0.002136 | 3.455 | +1 | 6 |
| - | - | 1.645E+05 | 619.3 | - | - | 0 | - |
| - | - | 8.144E+04 | 620.3 | - | - | 0 | - |
| - | - | 7.726E+04 | 621.3 | - | - | 0 | - |
| - | - | 8.586E+04 | 631.3 | - | - | 0 | - |
| 4 | w | 1.043E+05 | 631.8 | 0.001446 | 2.289 | +2 | 12 |
| - | - | 7.309E+04 | 632.3 | - | - | 0 | - |
| 10 | z | 3.528E+06 | 636.3 | 0.0003605 | 0.5666 | +1 | 6 |
| - | - | 2.203E+06 | 637.3 | - | - | 0 | - |
| 4 | y | 6.204E+05 | 638.3 | 0.004977 | 7.797 | +2 | 12 |
| 4 | z | 3.618E+06 | 639.3 | 0.0006338 | 0.9914 | +2 | 12 |
| - | - | 2.51E+06 | 639.8 | - | - | 0 | - |
| - | - | 1.062E+06 | 640.3 | - | - | 0 | - |
| - | - | 3.153E+05 | 640.8 | - | - | 0 | - |
| - | - | 1.194E+05 | 641.3 | - | - | 0 | - |
| - | - | 6.595E+04 | 642.3 | - | - | 0 | - |
| - | - | 7.112E+04 | 646.8 | - | - | 0 | - |
| 4 | y | 5.902E+06 | 647.3 | 0.0002439 | 0.3768 | +2 | 12 |
| - | - | 4.309E+06 | 647.8 | - | - | 0 | - |
| 6 | c | 4.235E+06 | 648.3 | 0.004994 | 7.703 | +1 | 6 |
| - | - | 4.476E+05 | 648.8 | - | - | 0 | - |
| - | - | 1.017E+06 | 649.3 | - | - | 0 | - |
| - | - | 2.259E+05 | 650.3 | - | - | 0 | - |
| 13 | c | 1.41E+05 | 651.3 | 0.009298 | 14.28 | +2 | 13 |
| 10 | y | 3.935E+06 | 652.3 | 0.0003743 | 0.5737 | +1 | 6 |
| - | - | 1.353E+06 | 653.3 | - | - | 0 | - |
| - | - | 3.706E+05 | 654.3 | - | - | 0 | - |
| - | - | 5.44E+04 | 655.3 | - | - | 0 | - |
| - | - | 7.543E+04 | 657.3 | - | - | 0 | - |
| 3 | w | 3.104E+06 | 674.3 | 0.0003327 | 0.4934 | +2 | 13 |
| - | - | 2.27E+06 | 674.8 | - | - | 0 | - |
| - | - | 1.094E+06 | 675.3 | - | - | 0 | - |
| - | - | 2.398E+05 | 675.8 | - | - | 0 | - |
| - | - | 4.13E+05 | 679.4 | - | - | 0 | - |
| - | - | 2.144E+05 | 680.4 | - | - | 0 | - |
| - | - | 2.031E+05 | 681.8 | - | - | 0 | - |
| - | - | 5.965E+04 | 690.3 | - | - | 0 | - |
| 3 | z | 7.442E+04 | 694.8 | 0.0001836 | 0.2642 | +2 | 13 |
| 3 | z | 1.482E+05 | 695.3 | 0.01147 | 16.5 | +2 | 13 |
| - | - | 6.356E+04 | 695.8 | - | - | 0 | - |
| - | - | 8.81E+04 | 696.3 | - | - | 0 | - |
| - | - | 9.595E+04 | 696.8 | - | - | 0 | - |
| - | - | 6.143E+04 | 701.4 | - | - | 0 | - |
| - | - | 6.937E+04 | 702.3 | - | - | 0 | - |
| 3 | y | 4.058E+05 | 702.8 | 0.0005872 | 0.8354 | +2 | 13 |
| 3 | y | 3.069E+05 | 703.3 | 0.009373 | 13.33 | +2 | 13 |
| 3 | z | 2.041E+06 | 703.8 | 0.0005775 | 0.8205 | +2 | 13 |
| - | - | 1.694E+06 | 704.3 | - | - | 0 | - |
| - | - | 8.27E+05 | 704.8 | - | - | 0 | - |
| 9 | z | 2.47E+05 | 705.3 | 0.003738 | 5.3 | +1 | 7 |
| - | - | 8.03E+04 | 705.9 | - | - | 0 | - |
| - | - | 2.584E+05 | 707.9 | - | - | 0 | - |
| - | - | 1.82E+05 | 708.4 | - | - | 0 | - |
| - | - | 6.366E+04 | 708.9 | - | - | 0 | - |
| - | - | 2.781E+05 | 711.3 | - | - | 0 | - |
| 3 | y | 8.29E+06 | 711.9 | 0.0001876 | 0.2636 | +2 | 13 |
| - | - | 6.135E+06 | 712.4 | - | - | 0 | - |
| - | - | 2.333E+06 | 712.9 | - | - | 0 | - |
| - | - | 6.813E+05 | 713.4 | - | - | 0 | - |
| - | - | 2.185E+05 | 713.9 | - | - | 0 | - |
| - | - | 5.484E+04 | 713.9 | - | - | 0 | - |
| - | - | 6.664E+05 | 715.9 | - | - | 0 | - |
| - | - | 7.012E+05 | 716.4 | - | - | 0 | - |
| - | - | 1.551E+05 | 716.9 | - | - | 0 | - |
| - | - | 5.437E+05 | 717.9 | - | - | 0 | - |
| - | - | 3.575E+05 | 718.4 | - | - | 0 | - |
| - | - | 1.372E+05 | 718.9 | - | - | 0 | - |
| - | - | 2.425E+05 | 721.3 | - | - | 0 | - |
| 9 | y | 2.945E+05 | 722.3 | 0.001016 | 1.406 | +1 | 7 |
| 9 | z | 2.395E+06 | 723.4 | 0.0001925 | 0.2661 | +1 | 7 |
| - | - | 1.143E+06 | 724.4 | - | - | 0 | - |
| - | - | 4.479E+05 | 725.4 | - | - | 0 | - |
| - | - | 2.082E+05 | 725.9 | - | - | 0 | - |
| - | - | 1.738E+05 | 726.4 | - | - | 0 | - |
| 14 | c | 9.9E+04 | 729.4 | 0.004392 | 6.022 | +2 | 14 |
| - | - | 1.45E+05 | 730.4 | - | - | 0 | - |
| - | - | 7.397E+04 | 730.9 | - | - | 0 | - |
| - | - | 7.566E+04 | 732.3 | - | - | 0 | - |
| - | - | 5.549E+04 | 733.3 | - | - | 0 | - |
| 14 | c | 1.629E+07 | 737.9 | 0.0005217 | 0.707 | +2 | 14 |
| - | - | 1.249E+07 | 738.4 | - | - | 0 | - |
| - | - | 5.731E+06 | 738.9 | - | - | 0 | - |
| 9 | y | 2.156E+06 | 739.4 | 0.009667 | 13.07 | +1 | 7 |
| - | - | 3.419E+05 | 739.9 | - | - | 0 | - |
| - | - | 3.365E+05 | 740.4 | - | - | 0 | - |
| - | - | 1.23E+05 | 741.4 | - | - | 0 | - |
| - | - | 1.158E+05 | 744.4 | - | - | 0 | - |
| 2 | y | 8.108E+05 | 746.4 | 0.001328 | 1.779 | +2 | 14 |
| 2 | y | 8.841E+05 | 746.9 | 0.005749 | 7.697 | +2 | 14 |
| 2 | z | 3.18E+06 | 747.4 | 0.00207 | 2.77 | +2 | 14 |
| - | - | 2.048E+06 | 747.9 | - | - | 0 | - |
| - | - | 1.105E+06 | 748.4 | - | - | 0 | - |
| - | - | 3.51E+05 | 748.9 | - | - | 0 | - |
| 2 | y | 9.389E+06 | 755.4 | 0.0003477 | 0.4603 | +2 | 14 |
| - | - | 7.6E+06 | 755.9 | - | - | 0 | - |
| - | - | 3.803E+06 | 756.4 | - | - | 0 | - |
| - | - | 1.051E+06 | 756.9 | - | - | 0 | - |
| - | - | 2.743E+05 | 757.4 | - | - | 0 | - |
| - | - | 1.061E+05 | 759.4 | - | - | 0 | - |
| - | - | 2.025E+05 | 760.4 | - | - | 0 | - |
| - | - | 3.221E+05 | 760.4 | - | - | 0 | - |
| - | - | 2.441E+05 | 760.9 | - | - | 0 | - |
| - | - | 1.049E+05 | 761.4 | - | - | 0 | - |
| - | - | 1.346E+05 | 761.9 | - | - | 0 | - |
| - | - | 6.53E+04 | 762.4 | - | - | 0 | - |
| - | - | 8.271E+04 | 767.9 | - | - | 0 | - |
| - | - | 8.712E+04 | 768.4 | - | - | 0 | - |
| - | - | 1.373E+05 | 773.4 | - | - | 0 | - |
| - | - | 2.108E+05 | 773.9 | - | - | 0 | - |
| - | - | 2.79E+05 | 774.4 | - | - | 0 | - |
| - | - | 1.446E+05 | 774.9 | - | - | 0 | - |
| - | - | 1.798E+05 | 775.4 | - | - | 0 | - |
| - | - | 1.909E+05 | 775.9 | - | - | 0 | - |
| - | - | 1.678E+05 | 776.4 | - | - | 0 | - |
| - | - | 8.489E+04 | 776.9 | - | - | 0 | - |
| - | - | 2.488E+05 | 781.4 | - | - | 0 | - |
| - | - | 1.28E+06 | 781.9 | - | - | 0 | - |
| - | - | 3.051E+06 | 782.4 | - | - | 0 | - |
| - | - | 2.588E+06 | 782.9 | - | - | 0 | - |
| - | - | 1.122E+06 | 783.4 | - | - | 0 | - |
| - | - | 3.304E+05 | 783.9 | - | - | 0 | - |
| - | - | 9.481E+04 | 784.4 | - | - | 0 | - |
| 7 | c | 1.614E+05 | 787.4 | 0.001026 | 1.303 | +1 | 7 |
| - | - | 2.991E+05 | 787.9 | - | - | 0 | - |
| - | - | 3.575E+05 | 788.4 | - | - | 0 | - |
| - | - | 3.581E+05 | 788.9 | - | - | 0 | - |
| - | - | 4.661E+06 | 789.4 | - | - | 0 | - |
| - | - | 4.259E+06 | 789.9 | - | - | 0 | - |
| - | - | 2.181E+06 | 790.4 | - | - | 0 | - |
| - | - | 4.728E+06 | 790.9 | - | - | 0 | - |
| - | - | 4.129E+06 | 791.4 | - | - | 0 | - |
| - | - | 2.135E+06 | 791.9 | - | - | 0 | - |
| - | - | 6.807E+05 | 792.4 | - | - | 0 | - |
| - | - | 2.2E+05 | 792.9 | - | - | 0 | - |
| 8 | w | 6.906E+04 | 793.4 | 0.009295 | 11.72 | +1 | 8 |
| - | - | 1.787E+05 | 794.4 | - | - | 0 | - |
| - | - | 5.433E+05 | 794.9 | - | - | 0 | - |
| - | - | 6.427E+05 | 795.4 | - | - | 0 | - |
| - | - | 2.739E+05 | 795.9 | - | - | 0 | - |
| - | - | 4.736E+05 | 796.4 | - | - | 0 | - |
| - | - | 4.96E+05 | 796.9 | - | - | 0 | - |
| - | - | 1.766E+05 | 797.4 | - | - | 0 | - |
| - | - | 1.233E+05 | 797.9 | - | - | 0 | - |
| - | - | 9.52E+04 | 798.4 | - | - | 0 | - |
| - | - | 7.4E+04 | 799.4 | - | - | 0 | - |
| - | - | 6.566E+04 | 800.5 | - | - | 0 | - |
| - | - | 1.839E+06 | 803.4 | - | - | 0 | - |
| - | - | 1.68E+07 | 803.9 | - | - | 0 | - |
| - | - | 1.471E+07 | 804.4 | - | - | 0 | - |
| - | - | 7.159E+06 | 804.9 | - | - | 0 | - |
| - | - | 2.209E+06 | 805.4 | - | - | 0 | - |
| - | - | 6.126E+05 | 805.9 | - | - | 0 | - |
| - | - | 7.736E+06 | 811.9 | - | - | 0 | - |
| - | - | 1.173E+07 | 812.4 | - | - | 0 | - |
| - | - | 8.147E+06 | 812.9 | - | - | 0 | - |
| - | - | 3.449E+06 | 813.4 | - | - | 0 | - |
| - | - | 9.236E+05 | 813.9 | - | - | 0 | - |
| - | - | 9.988E+04 | 814.4 | - | - | 0 | - |
| - | - | 7.221E+04 | 831.4 | - | - | 0 | - |
| - | - | 2.161E+05 | 834.4 | - | - | 0 | - |
| - | - | 1.193E+05 | 835.4 | - | - | 0 | - |
| 8 | y | 5.436E+05 | 836.4 | 0.0002505 | 0.2995 | +1 | 8 |
| - | - | 2.282E+05 | 837.4 | - | - | 0 | - |
| - | - | 9.583E+04 | 838.4 | - | - | 0 | - |
| - | - | 1.653E+05 | 842.5 | - | - | 0 | - |
| - | - | 9.977E+04 | 844.5 | - | - | 0 | - |
| - | - | 9.616E+05 | 857.5 | - | - | 0 | - |
| - | - | 3.781E+05 | 858.5 | - | - | 0 | - |
| - | - | 1.335E+05 | 859.5 | - | - | 0 | - |
| - | - | 1.919E+05 | 885.5 | - | - | 0 | - |
| - | - | 1.698E+05 | 886.5 | - | - | 0 | - |
| - | - | 6.038E+04 | 887.5 | - | - | 0 | - |
| 7 | w | 1.001E+05 | 890.4 | 0.003613 | 4.058 | +1 | 9 |
| - | - | 1.043E+05 | 891.4 | - | - | 0 | - |
| - | - | 2.705E+05 | 900.5 | - | - | 0 | - |
| 8 | c | 4.662E+06 | 901.5 | 0.0008381 | 0.9297 | +1 | 8 |
| - | - | 2.041E+06 | 902.5 | - | - | 0 | - |
| - | - | 6.664E+05 | 903.5 | - | - | 0 | - |
| - | - | 1.481E+05 | 904.5 | - | - | 0 | - |
| - | - | 1.74E+05 | 926.5 | - | - | 0 | - |
| - | - | 1.986E+05 | 927.5 | - | - | 0 | - |
| - | - | 9.067E+04 | 928.5 | - | - | 0 | - |
| - | - | 6.624E+04 | 943.5 | - | - | 0 | - |
| - | - | 2.606E+06 | 944.5 | - | - | 0 | - |
| - | - | 1.193E+06 | 945.5 | - | - | 0 | - |
| - | - | 4.183E+05 | 946.5 | - | - | 0 | - |
| - | - | 7.798E+04 | 947.5 | - | - | 0 | - |
| 7 | z | 1.111E+05 | 958.5 | 0.00123 | 1.284 | +1 | 9 |
| - | - | 9.465E+04 | 959.5 | - | - | 0 | - |
| - | - | 1.175E+05 | 972.5 | - | - | 0 | - |
| 7 | y | 8.156E+04 | 975.5 | 0.003523 | 3.612 | +1 | 9 |
| 7 | z | 2.573E+06 | 976.5 | 0.000179 | 0.1833 | +1 | 9 |
| - | - | 1.777E+06 | 977.5 | - | - | 0 | - |
| - | - | 8.893E+04 | 977.6 | - | - | 0 | - |
| - | - | 6.524E+05 | 978.5 | - | - | 0 | - |
| - | - | 1.29E+05 | 979.5 | - | - | 0 | - |
| - | - | 7.561E+05 | 987.5 | - | - | 0 | - |
| 9 | c | 1.086E+07 | 988.5 | 0.000548 | 0.5543 | +1 | 9 |
| - | - | 5.694E+06 | 989.5 | - | - | 0 | - |
| - | - | 1.759E+06 | 990.5 | - | - | 0 | - |
| - | - | 4.998E+05 | 991.5 | - | - | 0 | - |
| - | - | 1.293E+05 | 992.5 | - | - | 0 | - |
| - | - | 6.89E+05 | 1002 | - | - | 0 | - |
| - | - | 3.388E+05 | 1003 | - | - | 0 | - |
| - | - | 1.618E+05 | 1004 | - | - | 0 | - |
| 10 | c | 7.506E+04 | 1029 | 0.002093 | 2.035 | +1 | 10 |
| - | - | 8.634E+04 | 1031 | - | - | 0 | - |
| - | - | 9.314E+04 | 1032 | - | - | 0 | - |
| 10 | c | 6.481E+06 | 1046 | 0.0006907 | 0.6606 | +1 | 10 |
| 6 | w | 3.695E+06 | 1047 | 0.006641 | 6.345 | +1 | 10 |
| - | - | 1.891E+06 | 1048 | - | - | 0 | - |
| - | - | 7.109E+05 | 1049 | - | - | 0 | - |
| - | - | 1.584E+05 | 1050 | - | - | 0 | - |
| 6 | z | 1.465E+05 | 1074 | 0.004195 | 3.907 | +1 | 10 |
| 6 | z | 9.803E+04 | 1075 | 0.02018 | 18.78 | +1 | 10 |
| - | - | 8.9E+04 | 1076 | - | - | 0 | - |
| - | - | 6.492E+04 | 1085 | - | - | 0 | - |
| - | - | 1.085E+05 | 1086 | - | - | 0 | - |
| - | - | 6.84E+04 | 1086 | - | - | 0 | - |
| - | - | 1.062E+05 | 1087 | - | - | 0 | - |
| 6 | z | 2.112E+06 | 1092 | 0.0002217 | 0.2031 | +1 | 10 |
| - | - | 3.759E+06 | 1093 | - | - | 0 | - |
| - | - | 1.691E+06 | 1094 | - | - | 0 | - |
| - | - | 5.035E+05 | 1095 | - | - | 0 | - |
| - | - | 7.204E+04 | 1096 | - | - | 0 | - |
| - | - | 9.976E+04 | 1100 | - | - | 0 | - |
| - | - | 3.184E+05 | 1101 | - | - | 0 | - |
| - | - | 2.205E+05 | 1102 | - | - | 0 | - |
| - | - | 2.54E+05 | 1103 | - | - | 0 | - |
| - | - | 7.892E+04 | 1104 | - | - | 0 | - |
| - | - | 1.636E+05 | 1107 | - | - | 0 | - |
| 6 | y | 6.496E+05 | 1108 | 0.0003139 | 0.2834 | +1 | 10 |
| - | - | 3.135E+05 | 1109 | - | - | 0 | - |
| - | - | 1.428E+05 | 1110 | - | - | 0 | - |
| - | - | 4.434E+05 | 1130 | - | - | 0 | - |
| - | - | 2.782E+05 | 1131 | - | - | 0 | - |
| - | - | 1.02E+05 | 1132 | - | - | 0 | - |
| - | - | 1.633E+05 | 1144 | - | - | 0 | - |
| 11 | c | 4.76E+06 | 1145 | 0.0001479 | 0.1292 | +1 | 11 |
| - | - | 3.022E+06 | 1146 | - | - | 0 | - |
| - | - | 1.209E+06 | 1147 | - | - | 0 | - |
| - | - | 3.078E+05 | 1148 | - | - | 0 | - |
| - | - | 7.958E+04 | 1163 | - | - | 0 | - |
| - | - | 1.091E+05 | 1173 | - | - | 0 | - |
| - | - | 7.585E+04 | 1174 | - | - | 0 | - |
| 5 | y | 1.113E+05 | 1178 | 0.00678 | 5.758 | +1 | 11 |
| 5 | z | 4.441E+05 | 1179 | 0.0005567 | 0.4723 | +1 | 11 |
| - | - | 1.757E+06 | 1180 | - | - | 0 | - |
| - | - | 1.159E+06 | 1181 | - | - | 0 | - |
| - | - | 4.758E+05 | 1182 | - | - | 0 | - |
| - | - | 1.307E+05 | 1183 | - | - | 0 | - |
| - | - | 5.319E+05 | 1188 | - | - | 0 | - |
| - | - | 3.498E+05 | 1189 | - | - | 0 | - |
| - | - | 2.366E+05 | 1190 | - | - | 0 | - |
| - | - | 1.557E+05 | 1191 | - | - | 0 | - |
| 5 | y | 7.129E+05 | 1195 | 0.0004819 | 0.4034 | +1 | 11 |
| - | - | 5.116E+05 | 1196 | - | - | 0 | - |
| - | - | 2.068E+05 | 1197 | - | - | 0 | - |
| - | - | 3.807E+05 | 1231 | - | - | 0 | - |
| 12 | c | 9.092E+06 | 1232 | 0.0008747 | 0.7102 | +1 | 12 |
| - | - | 6.054E+06 | 1233 | - | - | 0 | - |
| - | - | 2.327E+06 | 1234 | - | - | 0 | - |
| - | - | 4.903E+05 | 1235 | - | - | 0 | - |
| - | - | 1.999E+05 | 1236 | - | - | 0 | - |
| - | - | 3.566E+05 | 1259 | - | - | 0 | - |
| 4 | z | 2.134E+05 | 1260 | 0.005925 | 4.704 | +1 | 12 |
| 4 | z | 1.096E+05 | 1261 | 0.01666 | 13.22 | +1 | 12 |
| - | - | 3.614E+05 | 1275 | - | - | 0 | - |
| 4 | y | 2.919E+05 | 1276 | 0.006366 | 4.991 | +1 | 12 |
| 4 | y | 1.429E+05 | 1277 | 0.01966 | 15.4 | +1 | 12 |
| 4 | z | 4.159E+05 | 1278 | 0.001832 | 1.434 | +1 | 12 |
| - | - | 2.29E+06 | 1279 | - | - | 0 | - |
| - | - | 1.484E+06 | 1280 | - | - | 0 | - |
| - | - | 6.208E+05 | 1281 | - | - | 0 | - |
| - | - | 1.451E+05 | 1282 | - | - | 0 | - |
| 4 | y | 5.215E+05 | 1294 | 0.0005622 | 0.4346 | +1 | 12 |
| - | - | 4.279E+05 | 1295 | - | - | 0 | - |
| - | - | 3.245E+05 | 1296 | - | - | 0 | - |
| - | - | 1.181E+05 | 1297 | - | - | 0 | - |
| - | - | 9.999E+04 | 1308 | - | - | 0 | - |
| 13 | c | 6.48E+06 | 1319 | 0.0009207 | 0.6982 | +1 | 13 |
| - | - | 4.552E+06 | 1320 | - | - | 0 | - |
| - | - | 2.005E+06 | 1321 | - | - | 0 | - |
| - | - | 5.602E+05 | 1322 | - | - | 0 | - |
| - | - | 6.233E+04 | 1323 | - | - | 0 | - |
| - | - | 1.475E+05 | 1345 | - | - | 0 | - |
| - | - | 7.362E+05 | 1346 | - | - | 0 | - |
| - | - | 3.95E+05 | 1347 | - | - | 0 | - |
| 3 | w | 2.443E+05 | 1348 | 0.002312 | 1.716 | +1 | 13 |
| - | - | 1.079E+05 | 1349 | - | - | 0 | - |
| - | - | 7.053E+04 | 1367 | - | - | 0 | - |
| - | - | 5.961E+04 | 1391 | - | - | 0 | - |
| - | - | 1.511E+05 | 1395 | - | - | 0 | - |
| - | - | 1.033E+05 | 1396 | - | - | 0 | - |
| 3 | z | 5.069E+05 | 1407 | 0.002067 | 1.469 | +1 | 13 |
| - | - | 3.541E+06 | 1408 | - | - | 0 | - |
| - | - | 2.684E+06 | 1409 | - | - | 0 | - |
| - | - | 1.375E+06 | 1410 | - | - | 0 | - |
| - | - | 2.833E+05 | 1411 | - | - | 0 | - |
| - | - | 2.784E+05 | 1417 | - | - | 0 | - |
| - | - | 1.332E+05 | 1418 | - | - | 0 | - |
| - | - | 8.289E+04 | 1419 | - | - | 0 | - |
| 3 | y | 9.256E+04 | 1423 | 0.00309 | 2.172 | +1 | 13 |
| - | - | 2.206E+05 | 1424 | - | - | 0 | - |
| - | - | 1.372E+05 | 1425 | - | - | 0 | - |
| - | - | 2.014E+05 | 1431 | - | - | 0 | - |
| - | - | 7.612E+05 | 1432 | - | - | 0 | - |
| - | - | 7.75E+05 | 1433 | - | - | 0 | - |
| - | - | 2.984E+05 | 1434 | - | - | 0 | - |
| - | - | 1.452E+05 | 1435 | - | - | 0 | - |
| - | - | 1.033E+05 | 1448 | - | - | 0 | - |
| - | - | 7.794E+04 | 1449 | - | - | 0 | - |
| - | - | 1.725E+05 | 1452 | - | - | 0 | - |
| - | - | 1.026E+05 | 1453 | - | - | 0 | - |
| - | - | 6.509E+05 | 1459 | - | - | 0 | - |
| - | - | 5.424E+05 | 1460 | - | - | 0 | - |
| - | - | 3.38E+05 | 1461 | - | - | 0 | - |
| - | - | 1.228E+05 | 1462 | - | - | 0 | - |
| - | - | 1.089E+05 | 1465 | - | - | 0 | - |
| - | - | 7.172E+04 | 1467 | - | - | 0 | - |
| 14 | c | 1.265E+06 | 1475 | 0.0007133 | 0.4837 | +1 | 14 |
| - | - | 4.117E+06 | 1476 | - | - | 0 | - |
| - | - | 2.893E+06 | 1477 | - | - | 0 | - |
| - | - | 1.191E+06 | 1478 | - | - | 0 | - |
| - | - | 4.024E+05 | 1479 | - | - | 0 | - |
| - | - | 9.965E+04 | 1480 | - | - | 0 | - |
| 2 | z | 4.355E+05 | 1494 | 0.002892 | 1.936 | +1 | 14 |
| - | - | 2.387E+06 | 1495 | - | - | 0 | - |
| - | - | 1.967E+06 | 1496 | - | - | 0 | - |
| - | - | 7.358E+05 | 1497 | - | - | 0 | - |
| - | - | 1.708E+05 | 1498 | - | - | 0 | - |
| - | - | 2.048E+05 | 1508 | - | - | 0 | - |
| - | - | 1.401E+05 | 1509 | - | - | 0 | - |
| 2 | y | 1.651E+05 | 1510 | 0.01212 | 8.029 | +1 | 14 |
| - | - | 9.191E+04 | 1511 | - | - | 0 | - |
| - | - | 8.654E+04 | 1512 | - | - | 0 | - |
| - | - | 1.299E+05 | 1522 | - | - | 0 | - |
| - | - | 7.838E+05 | 1523 | - | - | 0 | - |
| - | - | 7.014E+05 | 1524 | - | - | 0 | - |
| - | - | 4.056E+05 | 1525 | - | - | 0 | - |
| - | - | 1.462E+05 | 1526 | - | - | 0 | - |
| - | - | 7.226E+04 | 1527 | - | - | 0 | - |
| - | - | 1.178E+05 | 1536 | - | - | 0 | - |
| - | - | 1.015E+05 | 1538 | - | - | 0 | - |
| - | - | 2.368E+05 | 1548 | - | - | 0 | - |
| - | - | 3.043E+05 | 1549 | - | - | 0 | - |
| - | - | 1.607E+05 | 1550 | - | - | 0 | - |
| - | - | 1.432E+05 | 1551 | - | - | 0 | - |
| - | - | 1.162E+05 | 1554 | - | - | 0 | - |
| - | - | 8.453E+04 | 1555 | - | - | 0 | - |
| - | - | 2.566E+05 | 1564 | - | - | 0 | - |
| - | - | 3.933E+06 | 1565 | - | - | 0 | - |
| - | - | 4.699E+06 | 1566 | - | - | 0 | - |
| - | - | 3.058E+06 | 1567 | - | - | 0 | - |
| - | - | 1.221E+06 | 1568 | - | - | 0 | - |
| - | - | 3.476E+05 | 1569 | - | - | 0 | - |
| - | - | 7.809E+04 | 1578 | - | - | 0 | - |
| - | - | 4.88E+05 | 1579 | - | - | 0 | - |
| - | - | 7.885E+05 | 1580 | - | - | 0 | - |
| - | - | 8.424E+05 | 1581 | - | - | 0 | - |
| - | - | 1.148E+06 | 1582 | - | - | 0 | - |
| - | - | 7.292E+05 | 1583 | - | - | 0 | - |
| - | - | 3.329E+05 | 1584 | - | - | 0 | - |
| - | - | 8.172E+04 | 1585 | - | - | 0 | - |
| - | - | 3E+05 | 1590 | - | - | 0 | - |
| - | - | 1.238E+06 | 1591 | - | - | 0 | - |
| - | - | 1.033E+06 | 1592 | - | - | 0 | - |
| - | - | 4.488E+05 | 1593 | - | - | 0 | - |
| - | - | 1.47E+05 | 1594 | - | - | 0 | - |
| - | - | 1.848E+05 | 1596 | - | - | 0 | - |
| - | - | 1.253E+06 | 1597 | - | - | 0 | - |
| - | - | 1.035E+06 | 1598 | - | - | 0 | - |
| - | - | 5.816E+05 | 1599 | - | - | 0 | - |
| - | - | 1.217E+05 | 1600 | - | - | 0 | - |
| - | - | 2.308E+06 | 1607 | - | - | 0 | - |
| - | - | 1.194E+07 | 1608 | - | - | 0 | - |
| - | - | 1.016E+07 | 1609 | - | - | 0 | - |
| - | - | 5.055E+06 | 1610 | - | - | 0 | - |
| - | - | 1.42E+06 | 1611 | - | - | 0 | - |
| - | - | 3.238E+05 | 1612 | - | - | 0 | - |
| - | - | 4.489E+05 | 1623 | - | - | 0 | - |
| - | - | 2.863E+06 | 1624 | - | - | 0 | - |
| - | - | 1.156E+07 | 1625 | - | - | 0 | - |
| - | - | 9.736E+06 | 1626 | - | - | 0 | - |
| - | - | 4.822E+06 | 1627 | - | - | 0 | - |
| - | - | 1.307E+06 | 1628 | - | - | 0 | - |
| - | - | 3.805E+05 | 1629 | - | - | 0 | - |

m/z Charge Intensity FragmentType MassShift Position
120.08104705810547 0 50944.21
124.2380599975586 0 29873.926
124.98079681396484 0 32603.008
142.0975341796875 0 181549.7
148.94776916503906 0 53621.953
173.12022399902344 0 69884.66
173.12852478027344 0 1344375.2
173.4398193359375 0 117107.06
174.132080078125 0 109618.484
176.39341735839844 0 47146.9
183.11289978027344 0 538359.6
183.6128692626953 0 48563.29
199.0712127685547 0 91983.55
199.11932373046875 0 59357.156
200.11575317382812 0 103415.66
201.12342834472656 0 2310823.5
201.13487243652344 0 1179603.2
202.12344360351562 0 342950.66
202.14173889160156 0 147197.77
211.1078643798828 0 72300.91
217.08216857910156 0 215702.56
226.65248107910156 0 44673.64
229.11862182617188 0 73211.375
229.13021850585938 0 135177.86
261.1231994628906 0 196860.58
261.1708984375 0 2471337.2
262.06707763671875 0 44101.508
262.12689208984375 0 50288.395
262.1513366699219 0 75138.8
262.17425537109375 0 406207.28
263.1757507324219 0 45709.953
270.1567077636719 0 128595.34
273.8057556152344 0 45123.88 y Ammonia loss 7
276.18170166015625 0 94062.23
277.0317687988281 0 53720.04
284.1600036621094 0 96894.625
288.1669921875 0 196084.11
299.19830322265625 0 46920.113
303.14581298828125 0 55320.4
306.1687316894531 0 498968.6 z 13
307.1714172363281 0 96788
312.1556091308594 0 975402.44
313.15863037109375 0 206270.66
316.1507263183594 0 174422.05
320.1719970703125 0 128849.44 z Water loss 6
321.1807861328125 0 77686.02
322.1874084472656 0 685527.75 y 13
327.1902160644531 0 49575.01
330.16632080078125 0 844850.4
331.1696472167969 0 164654.6
348.2032775878906 0 347207.97
349.2090759277344 0 128839.62
373.1976623535156 0 154335.27
374.1996154785156 0 55051.246
376.197998046875 0 1570302.8 c Water loss 10
377.1988830566406 0 372810.03
387.2355651855469 0 128614.72
391.1849365234375 0 581657.44
392.19281005859375 0 2680293 y Ammonia loss 12
393.19873046875 0 1122097.9 y Ammonia loss 4
394.2037048339844 0 361011.06 c Ammonia loss 6
401.23974609375 0 304435.9
407.20318603515625 0 66066.69
409.2198791503906 0 516189.4 y 12
410.2231140136719 0 180693.05
411.2242431640625 0 60345.465
417.18853759765625 0 912241.6
418.1922912597656 0 202839.48
418.717041015625 0 126472.59 y 7
428.22589111328125 0 140512.14
429.2347717285156 0 686824.56
430.23638916015625 0 109644.03
435.2354431152344 0 255409.25
436.20941162109375 0 69750.52
436.24139404296875 0 145790.81
439.2213134765625 0 53430.363
444.2566833496094 0 433785.25
445.2551574707031 0 392317.16
446.2604675292969 0 502377 c 3
447.2650146484375 0 96830.31
462.222412109375 0 174133.47 z Water loss 11
462.8960266113281 0 70581.484
463.2303771972656 0 1934282.2 w 11
464.2310791015625 0 638020.7
465.23193359375 0 203945.94
468.8983459472656 0 60772.867 y Water loss 2
474.9029541015625 0 148012.25 y 2
475.23773193359375 0 96382
475.57244873046875 0 102349.56
478.2174987792969 0 711082.2
479.2240905761719 0 821240.6 y Ammonia loss 11
480.23260498046875 0 1581953 z 11
481.23699951171875 0 388521.03
482.2351989746094 0 60697.797
488.75823974609375 0 1556652.6 z 6
489.2597961425781 0 838077
489.7611389160156 0 322919.28
490.2898254394531 0 176440.5
494.23468017578125 0 81043.305
496.2522277832031 0 527853.75 y 11
496.767822265625 0 499525.94 y 6
497.2659606933594 0 239444.31
497.77001953125 0 79852.89
498.2519226074219 0 204310.89
499.3001403808594 0 132105.42
500.3038635253906 0 60914.945
503.91522216796875 0 244226.39 y 1
504.2203674316406 0 884369.7
504.5845642089844 0 70318.17
505.22308349609375 0 247598.23
514.2731323242188 0 60323.918
515.265380859375 0 60901.535
516.2676391601562 0 205807.72
524.276123046875 0 572959
524.7780151367188 0 426393.94
525.279052734375 0 185288.73
529.9361572265625 0 87731.586
531.2850341796875 0 99197.586
532.2857055664062 0 845755.44
533.2929077148438 0 3770227.2 c 4
534.2962036132812 0 1138058.2
535.3020629882812 0 227377.78
535.60546875 0 572572.1
535.9402465820312 0 580070.25
536.2755126953125 0 232395.22
541.6096801757812 0 2418162.5
541.94384765625 0 1713457.5
542.2777709960938 0 1108745.2
542.6129150390625 0 255882.33
542.9439697265625 0 79311.18
546.2715454101562 0 2113950.2 z 5
546.7742919921875 0 2306284.5
547.2759399414062 0 963891.4
547.7764282226562 0 275947.1
548.2765502929688 0 82885.52
549.293212890625 0 126075.8
553.7767333984375 0 127375.305
554.28125 0 1202279.5 y 5
554.7821655273438 0 647336.75
555.2841186523438 0 198280.22
556.2821044921875 0 101137.414
561.2897338867188 0 380489.8 z Water loss 10
562.2978515625 0 349310.06
564.278076171875 0 1491635.5 c Ammonia loss 10
565.2810668945312 0 459498.66
566.2848510742188 0 109534.336
566.7825927734375 0 92800.78
567.286865234375 0 84336.555
567.7898559570312 0 150391.48
568.2893676757812 0 194854.2
579.3013305664062 0 815465.5 z 10
580.3041381835938 0 283507.7
581.7846069335938 0 128940.445
588.29150390625 0 55758.05
588.7802734375 0 427055.53 y Water loss 4
589.2832641601562 0 724057.75 y Ammonia loss 4
589.7874145507812 0 2166146 z 4
590.2906494140625 0 3737826.2
590.7926025390625 0 1887418.2
591.2942504882812 0 650702.94
591.7943725585938 0 130168.9
592.3329467773438 0 1293313.4
593.3367919921875 0 386927.1
595.3206176757812 0 810982.06 y 10
596.3232421875 0 303152.88
597.2931518554688 0 577663.1
597.7967529296875 0 5995968.5 y 4
598.2984008789062 0 3594501.8
598.7998657226562 0 1620426.9
599.3018798828125 0 325582.16
601.7809448242188 0 377631.62
602.28271484375 0 218457.33
603.2952270507812 0 59557.227
605.3106689453125 0 86482.26
611.2922973632812 0 127153.42
611.7897338867188 0 91964.875
612.2979125976562 0 63739.305
618.3098754882812 0 75837.99 z Water loss 9
619.3211669921875 0 164508.25
620.3240966796875 0 81439.27
621.2979125976562 0 77258.555
631.2982177734375 0 85864.516
631.811279296875 0 104314.125 w 3
632.3092651367188 0 73094.414
636.3229370117188 0 3527536.8 z 9
637.3280029296875 0 2202975.8
638.3306274414062 0 620445.7 y Water loss 3
639.3222045898438 0 3618263.8 z 3
639.8235473632812 0 2509715.5
640.324951171875 0 1062417.9
640.8260498046875 0 315325.7
641.3297729492188 0 119404.27
642.3314208984375 0 65946.93
646.82666015625 0 71122.836
647.3311767578125 0 5902284 y 3
647.8328857421875 0 4309109.5
648.3248901367188 0 4234801.5 c 5
648.8357543945312 0 447585
649.32373046875 0 1016972.25
650.3264770507812 0 225910.95
651.3295288085938 0 140974.28 c Ammonia loss 12
652.3416748046875 0 3934988.5 y 9
653.3444213867188 0 1353449.5
654.3469848632812 0 370639.5
655.3160400390625 0 54398.71
657.3439331054688 0 75433.875
674.3365478515625 0 3103628 w 2
674.8377685546875 0 2270091.2
675.3392333984375 0 1094157.1
675.8422241210938 0 239772.89
679.364501953125 0 413020.75
680.3690795898438 0 214372.12
681.8452758789062 0 203056.42
690.3328247070312 0 59650.242
694.8377685546875 0 74422.086 z Water loss 2
695.341064453125 0 148188.12 z Ammonia loss 2
695.8366088867188 0 63556.11
696.33447265625 0 88098.42
696.83837890625 0 95949.32
701.3606567382812 0 61430.633
702.3441162109375 0 69373.35
702.8475341796875 0 405804.16 y Water loss 2
703.3483276367188 0 306921.12 y Ammonia loss 2
703.8434448242188 0 2040627.4 z 2
704.3447265625 0 1693584.6
704.8460693359375 0 826956.3
705.3477783203125 0 246978.14 z Water loss 8
705.8504638671875 0 80295.3
707.8701782226562 0 258412.16
708.36865234375 0 182026.2
708.869384765625 0 63658.99
711.3480834960938 0 278122.3
711.8524169921875 0 8290079.5 y 2
712.3540649414062 0 6135434.5
712.8551025390625 0 2332872.2
713.3563232421875 0 681321.94
713.8588256835938 0 218514.66
713.9227294921875 0 54837.62
715.8768310546875 0 666408
716.3786010742188 0 701161.94
716.8776245117188 0 155149.45
717.853271484375 0 543727.6
718.3538818359375 0 357525.03
718.854248046875 0 137185.16
721.3414306640625 0 242489.97
722.3457641601562 0 294455.1 y Ammonia loss 8
723.3547973632812 0 2395377 z 8
724.3580322265625 0 1143144.6
725.3627319335938 0 447896.5
725.8611450195312 0 208164.98
726.3640747070312 0 173782.86
729.3663940429688 0 99004.266 c Ammonia loss 13
730.3734130859375 0 145034.81
730.87939453125 0 73972.11
732.3331909179688 0 75662.28
733.3397216796875 0 55490.992
737.8845825195312 0 16291817 c 13
738.3856811523438 0 12493216
738.8869018554688 0 5731322.5
739.3829956054688 0 2155511.2 y 8
739.8853149414062 0 341921.12
740.3770141601562 0 336509.56
741.3755493164062 0 123036.06
744.38720703125 0 115756.21
746.3616333007812 0 810789.3 y Water loss 1
746.8607177734375 0 884078.06 y Ammonia loss 1
747.3568115234375 0 3179751.8 z 1
747.8602905273438 0 2047962.1
748.3596801757812 0 1105195.4
748.8628540039062 0 351021.44
755.3685913085938 0 9389440 y 1
755.8699951171875 0 7599737.5
756.371337890625 0 3802739.8
756.87255859375 0 1051177.2
757.3750610351562 0 274272.7
759.4003295898438 0 106115.19
760.35302734375 0 202507.81
760.4083251953125 0 322079.5
760.8631591796875 0 244113.34
761.4154052734375 0 104880.29
761.8692016601562 0 134587
762.3704833984375 0 65302.2
767.888916015625 0 82709.47
768.3865966796875 0 87118.47
773.3984985351562 0 137304.02
773.8934936523438 0 210791.83
774.3963623046875 0 278987.3
774.8924560546875 0 144636.14
775.3855590820312 0 179804.86
775.8812866210938 0 190906.64
776.3754272460938 0 167793.19
776.875732421875 0 84892.41
781.4059448242188 0 248818.56
781.9063720703125 0 1279573
782.4053344726562 0 3050676.5
782.904541015625 0 2588486.8
783.404296875 0 1122020.4
783.8998413085938 0 330359.03
784.4069213867188 0 94810.74
787.3934326171875 0 161367.81 c Ammonia loss 6
787.900390625 0 299136.9
788.4010009765625 0 357537.88
788.8988037109375 0 358101.7
789.3837280273438 0 4661109.5
789.88525390625 0 4259434
790.3880004882812 0 2180539
790.8989868164062 0 4728001.5
791.4012451171875 0 4129472.2
791.9026489257812 0 2134639.2
792.4042358398438 0 680686.44
792.9047241210938 0 220007.45
793.3931884765625 0 69055.69 w 7
794.3977661132812 0 178679.98
794.897216796875 0 543254.8
795.39892578125 0 642722.4
795.8995361328125 0 273850.4
796.3929443359375 0 473630.66
796.893310546875 0 496009.47
797.3975219726562 0 176645.14
797.8947143554688 0 123343.32
798.4174194335938 0 95197.8
799.4493408203125 0 73998.734
800.458740234375 0 65656.234
803.4085693359375 0 1838878.4
803.9024047851562 0 16796246
804.4034423828125 0 14705096
804.9041137695312 0 7159419
805.4055786132812 0 2209382.2
805.90673828125 0 612570.44
811.9105834960938 0 7736423.5
812.4132080078125 0 11725250
812.9149169921875 0 8147171.5
813.416259765625 0 3448719.8
813.9171142578125 0 923642.6
814.41943359375 0 99884.34
831.4332275390625 0 72208.125
834.410400390625 0 216102.72
835.4131469726562 0 119303.59
836.4258422851562 0 543628.56 y 7
837.42822265625 0 228199.3
838.4277954101562 0 95828.97
842.4506225585938 0 165266.17
844.4500122070312 0 99773.35
857.460693359375 0 961614.5
858.4627075195312 0 378147.2
859.4692993164062 0 133470.81
885.457763671875 0 191902.72
886.4605712890625 0 169804.16
887.4594116210938 0 60381.64
890.4330444335938 0 100094.66 w 6
891.4403686523438 0 104304.18
900.4965209960938 0 270496.53
901.474609375 0 4662369.5 c 7
902.476806640625 0 2040822.5
903.4813232421875 0 666446.06
904.4871826171875 0 148129.88
926.48046875 0 174005.64
927.487060546875 0 198594.39
928.487548828125 0 90671.76
943.4898071289062 0 66243.69
944.4929809570312 0 2605730.5
945.4954833984375 0 1193174.4
946.4996337890625 0 418338.6
947.5104370117188 0 77977.38
958.4991455078125 0 111137.55 z Water loss 6
959.5032348632812 0 94646.55
972.4852905273438 0 117517.64
975.4971313476562 0 81561.27 y Ammonia loss 6
976.50830078125 0 2573189.2 z 6
977.5126953125 0 1777386.4
977.62841796875 0 88927.45
978.5152587890625 0 652449.7
979.5177612304688 0 128988.31
987.4981079101562 0 756148.8
988.50634765625 0 10864469 c 8
989.50927734375 0 5693565.5
990.5120849609375 0 1759116.4
991.5139770507812 0 499799.4
992.4514770507812 0 129273.47
1001.514404296875 0 688950.75
1002.5185546875 0 338764.8
1003.5250854492188 0 161805.16
1028.5028076171875 0 75059.07 c Ammonia loss 9
1030.518310546875 0 86342.91
1031.515625 0 93142.445
1045.5279541015625 0 6480596 c 9
1046.5311279296875 0 3694716.5 w 5
1047.537109375 0 1891358.1
1048.544189453125 0 710852.1
1049.54638671875 0 158420.75
1073.529052734375 0 146469.05 z Water loss 5
1074.529052734375 0 98028.484 z Ammonia loss 5
1075.537109375 0 89001.12
1084.5809326171875 0 64924.49
1085.578369140625 0 108453.57
1086.1712646484375 0 68395.25
1086.5682373046875 0 106238.03
1091.53564453125 0 2112164.8 z 5
1092.5419921875 0 3759344.8
1093.544921875 0 1690730.4
1094.54638671875 0 503453.22
1095.5517578125 0 72041.46
1099.569091796875 0 99758.78
1100.581787109375 0 318377.22
1101.5828857421875 0 220540.53
1102.593017578125 0 254016.3
1103.5968017578125 0 78915.91
1106.5516357421875 0 163593.17
1107.5538330078125 0 649648.5 y 5
1108.5592041015625 0 313526.44
1109.56494140625 0 142774
1129.5848388671875 0 443381.3
1130.5875244140625 0 278174.94
1131.5975341796875 0 101960.87
1143.5927734375 0 163305.33
1144.5958251953125 0 4760340.5 c 10
1145.59912109375 0 3021920.8
1146.6026611328125 0 1208902.2
1147.608154296875 0 307841.47
1162.5595703125 0 79576.586
1172.6033935546875 0 109086.87
1173.6082763671875 0 75853.266
1177.56640625 0 111286.07 y Ammonia loss 4
1178.56689453125 0 444053.84 z 4
1179.574462890625 0 1757278.8
1180.5777587890625 0 1159264.6
1181.582275390625 0 475780.06
1182.584228515625 0 130738.37
1187.6143798828125 0 531924.3
1188.6192626953125 0 349778.78
1189.6243896484375 0 236583.45
1190.6297607421875 0 155730.56
1194.585693359375 0 712901.7 y 4
1195.58837890625 0 511638.1
1196.5889892578125 0 206821.61
1230.6192626953125 0 380743.72
1231.6268310546875 0 9092113 c 11
1232.6304931640625 0 6053539
1233.633056640625 0 2327013.5
1234.6361083984375 0 490269.53
1235.6300048828125 0 199917.98
1258.6259765625 0 356639.7
1259.6312255859375 0 213372.45 z Water loss 3
1260.6259765625 0 109579.89 z Ammonia loss 3
1274.643798828125 0 361351.78
1275.650390625 0 291850.34 y Water loss 3
1276.647705078125 0 142911.25 y Ammonia loss 3
1277.634033203125 0 415878.66 z 3
1278.642822265625 0 2290071.8
1279.646484375 0 1484402
1280.6485595703125 0 620832.9
1281.650146484375 0 145124.86
1293.6551513671875 0 521548.8 y 3
1294.656005859375 0 427928.62
1295.661865234375 0 324525.7
1296.66357421875 0 118135.81
1307.608154296875 0 99992.8
1318.6588134765625 0 6480107 c 12
1319.6619873046875 0 4551667
1320.6646728515625 0 2005083.4
1321.6673583984375 0 560175.06
1322.6734619140625 0 62328.05
1344.6741943359375 0 147480.61
1345.6610107421875 0 736183.25
1346.6644287109375 0 395040.78
1347.662841796875 0 244297.98 w 2
1348.6605224609375 0 107940.68
1366.6544189453125 0 70527.6
1390.6727294921875 0 59607.49
1394.632568359375 0 151138.81
1395.6474609375 0 103298.82
1406.6763916015625 0 506923.75 z 2
1407.683837890625 0 3541477.2
1408.687744140625 0 2683866.5
1409.6904296875 0 1375184.8
1410.6925048828125 0 283305.6
1416.72412109375 0 278436.1
1417.720703125 0 133192.97
1418.72802734375 0 82886.64
1422.694091796875 0 92556.23 y 2
1423.7005615234375 0 220593.11
1424.70751953125 0 137153.75
1430.74560546875 0 201423.38
1431.7564697265625 0 761209.3
1432.7518310546875 0 775010
1433.75341796875 0 298389.88
1434.756103515625 0 145189.66
1447.775146484375 0 103301.39
1448.757568359375 0 77944.47
1451.6910400390625 0 172530.45
1452.689208984375 0 102591.016
1458.7410888671875 0 650920.75
1459.7435302734375 0 542423.4
1460.7451171875 0 338008.88
1461.7479248046875 0 122849.08
1464.72119140625 0 108884.92
1466.7415771484375 0 71718.18
1474.7601318359375 0 1264709.4 c 13
1475.766357421875 0 4116561
1476.7662353515625 0 2892843.8
1477.7655029296875 0 1191050.2
1478.758056640625 0 402401.3
1479.7313232421875 0 99654.26
1493.71337890625 0 435542.47 z 1
1494.7161865234375 0 2387403.2
1495.7200927734375 0 1967398.1
1496.72509765625 0 735808.6
1497.731201171875 0 170819.42
1507.728759765625 0 204804.88
1508.75830078125 0 140142.12
1509.7413330078125 0 165076.47 y 1
1510.73046875 0 91913.734
1511.7396240234375 0 86541.09
1521.775634765625 0 129915.51
1522.7266845703125 0 783795
1523.72998046875 0 701426.25
1524.7373046875 0 405587.53
1525.7435302734375 0 146224.67
1526.75537109375 0 72258.47
1535.72998046875 0 117791.836
1537.779541015625 0 101487.49
1547.770263671875 0 236790.77
1548.7747802734375 0 304323.94
1549.759521484375 0 160712.1
1550.7630615234375 0 143208.92
1553.8106689453125 0 116183.766
1554.7998046875 0 84528.414
1563.7921142578125 0 256632.84
1564.781494140625 0 3933327.5
1565.782958984375 0 4698881
1566.78564453125 0 3058347.5
1567.7861328125 0 1220639.4
1568.7962646484375 0 347636.38
1577.811279296875 0 78092.46
1578.8175048828125 0 488031.3
1579.815185546875 0 788472.5
1580.8094482421875 0 842436.7
1581.80517578125 0 1147648.5
1582.8033447265625 0 729171.75
1583.805419921875 0 332867.25
1584.8062744140625 0 81723.05
1589.7947998046875 0 299958.06
1590.778076171875 0 1237748.9
1591.7789306640625 0 1033396.06
1592.779296875 0 448844.34
1593.7830810546875 0 147035.27
1595.8187255859375 0 184782.81
1596.8302001953125 0 1253222.5
1597.8341064453125 0 1034655.3
1598.8372802734375 0 581585.5
1599.839111328125 0 121652.664
1606.8048095703125 0 2308325.8
1607.8016357421875 0 11935695
1608.8035888671875 0 10155174
1609.806884765625 0 5055113
1610.8094482421875 0 1420134
1611.8106689453125 0 323848.94
1622.815185546875 0 448868.34
1623.8182373046875 0 2863463.2
1624.82568359375 0 11560090
1625.8287353515625 0 9735951
1626.831787109375 0 4821626.5
1627.8338623046875 0 1307477.5
1628.8380126953125 0 380546.78

Spectrum Details

|  |  |
| --- | --- |
| Matched peaks? Matched peaks The total absolute number of peaks matched. Additionally in brackets the total fraction of peaks matched and the total number of peaks is shown. | 92 (16.31% of 564) |
| FDR? FDR The false discovery rate estimated for this peptide. It is calculated by matching all theoretical fragments with a non-integer shift with the raw peaks for this spectrum. This is done with 40 different shifts. The resulting percentage is the average number of annotated peaks over the number of annotated peaks with the correct spectrum. | 0.05% |
| Satellite FDR? Satellite FDR See the FDR for details on its calculation. This satellite ion specific FDR only contains the satellite ions (d/w) for I/L/J positions. | - |
| PSM Score? PSM Score The PSM Score as given by Hecklib to this annotated spectrum. It is shown with three significant figures. | 718 |

#### Spectrum 5048? Spectrum 5048 The raw spectrum of this peptide as annotated by Hecklib. The fragments are coloured according to ion type (see legend). Any peaks with a star '\*' as text can be hovered over to see the full details, first the ion type second the mass shift type. By hovering over the amino acids in the peptide or ions in the legend the corresponding peaks are highlighted. By toggling the 'Unassigned' label you can turn the background (unassigned) peaks on or off in the plot. By updating the slider in the Ion legend you can update the spectrum to only show the top X% of the peaks with labels. The top X% means any peak that is within X% of the highest intensity. By dragging in the spectrum you can zoom in to a specific part of the spectrum and use 'Zoom Out' to get back to the original zoom level. The annotation of the spectrum is based on the given sequence in the peptides file and is done with different software so inconsistencies are likely. The peaks are annotated based on the given sequence, with 20 ppm tolerance.

Copy Data

##### Spectrum 5048 (TSV)

###### Preview

```
Loading example...
```

*Click on the button to copy the data to your clipboard.*

Mz MinMz MaxIntensity Max

WidthHeightPeptide font sizePeptide stroke widthSpectrum font sizeSpectrum stroke widthCompact peptide

Ion legend

wxyz

abcd

OtherUnassignedIon^Charge^_Position_Show for top:%

JSEVSDRPSGVSSRF

02.43e+44.86e+47.29e+49.73e+4

Zoom Out

y^+2^_4_z^+1^_2_z^+3^_9_y^+1^_2_c^+3^_11_y^+1^_3_y^+3^_11_c^+2^_7_y^+1^_3_c^+1^_4_w^+1^_4_y^+1^_4_z^+1^_4_z^+2^_9_y^+1^_4_y^+2^_9_y^+3^_14_y^+3^_14_c^+1^_5_z^+2^_10_y^+2^_10_z^+1^_5_c^+2^_11_z^+1^_5_y^+2^_11_y^+2^_11_z^+2^_11_y^+1^_5_y^+2^_11_z^+1^_6_z^+1^_6_y^+2^_12_y^+2^_12_z^+2^_12_y^+2^_12_c^+1^_6_y^+1^_6_w^+2^_13_z^+2^_13_y^+2^_13_y^+2^_13_z^+2^_13_z^+1^_7_y^+2^_13_y^+1^_7_z^+1^_7_c^+2^_14_c^+2^_14_y^+1^_7_y^+2^_14_y^+2^_14_z^+2^_14_y^+2^_14_y^+1^_8_w^+1^_9_c^+1^_8_z^+1^_9_z^+1^_9_c^+1^_9_c^+1^_10_w^+1^_10_z^+1^_10_y^+1^_10_c^+1^_11_z^+1^_11_y^+1^_11_c^+1^_12_z^+1^_12_z^+1^_12_y^+1^_12_z^+1^_12_y^+1^_12_c^+1^_13_w^+1^_13_z^+1^_13_c^+1^_14_z^+1^_14_y^+1^_14_

0757151422713028

Fragment Matches Table

Show background peaks

| Position | Ion type | Intensity | mz Theoretical | mz Error (Th) | mz Error (ppm) | Charge | Series Number |
| --- | --- | --- | --- | --- | --- | --- | --- |
| - | - | 580.5 | 120.1 | - | - | 0 | - |
| - | - | 414.9 | 123.3 | - | - | 0 | - |
| - | - | 400.4 | 130.4 | - | - | 0 | - |
| - | - | 692.3 | 142.1 | - | - | 0 | - |
| - | - | 528.6 | 149 | - | - | 0 | - |
| - | - | 568.7 | 153.1 | - | - | 0 | - |
| - | - | 460.2 | 157.4 | - | - | 0 | - |
| - | - | 394.3 | 159.1 | - | - | 0 | - |
| - | - | 6207 | 173.1 | - | - | 0 | - |
| - | - | 702.2 | 174.1 | - | - | 0 | - |
| - | - | 2263 | 183.1 | - | - | 0 | - |
| - | - | 874.3 | 199.1 | - | - | 0 | - |
| - | - | 570.5 | 199.8 | - | - | 0 | - |
| - | - | 543.4 | 200.2 | - | - | 0 | - |
| - | - | 9938 | 201.1 | - | - | 0 | - |
| - | - | 7265 | 201.1 | - | - | 0 | - |
| - | - | 2483 | 202.1 | - | - | 0 | - |
| - | - | 663.7 | 202.1 | - | - | 0 | - |
| - | - | 706 | 217.1 | - | - | 0 | - |
| - | - | 595.8 | 246.1 | - | - | 0 | - |
| 12 | y | 585.4 | 248.6 | 0.000289 | 1.162 | +2 | 4 |
| - | - | 497.7 | 258.4 | - | - | 0 | - |
| - | - | 924.2 | 261.1 | - | - | 0 | - |
| - | - | 1.682E+04 | 261.2 | - | - | 0 | - |
| - | - | 518 | 262.2 | - | - | 0 | - |
| - | - | 2320 | 262.2 | - | - | 0 | - |
| - | - | 1102 | 288.2 | - | - | 0 | - |
| - | - | 590.5 | 298.1 | - | - | 0 | - |
| - | - | 562.3 | 303.1 | - | - | 0 | - |
| 14 | z | 2594 | 306.2 | 6.291E-05 | 0.2055 | +1 | 2 |
| - | - | 632 | 307.2 | - | - | 0 | - |
| - | - | 6347 | 312.2 | - | - | 0 | - |
| - | - | 854.4 | 313.2 | - | - | 0 | - |
| - | - | 1450 | 316.2 | - | - | 0 | - |
| 7 | z | 894.4 | 320.2 | 0.001724 | 5.384 | +3 | 9 |
| 14 | y | 4754 | 322.2 | 0.0004154 | 1.289 | +1 | 2 |
| - | - | 930.1 | 323.2 | - | - | 0 | - |
| - | - | 3536 | 330.2 | - | - | 0 | - |
| - | - | 844.2 | 331.2 | - | - | 0 | - |
| - | - | 2005 | 348.2 | - | - | 0 | - |
| - | - | 623.2 | 349.2 | - | - | 0 | - |
| - | - | 490.5 | 357.6 | - | - | 0 | - |
| - | - | 1230 | 373.2 | - | - | 0 | - |
| 11 | c | 8934 | 376.2 | 0.002013 | 5.35 | +3 | 11 |
| - | - | 1943 | 377.2 | - | - | 0 | - |
| - | - | 692.4 | 378.2 | - | - | 0 | - |
| - | - | 3975 | 391.2 | - | - | 0 | - |
| 13 | y | 1.599E+04 | 392.2 | 0.0001879 | 0.4792 | +1 | 3 |
| 5 | y | 7359 | 393.2 | 0.007246 | 18.43 | +3 | 11 |
| 7 | c | 2608 | 394.2 | 0.003509 | 8.901 | +2 | 7 |
| - | - | 1977 | 401.2 | - | - | 0 | - |
| - | - | 639.6 | 407.2 | - | - | 0 | - |
| 13 | y | 3764 | 409.2 | 0.0001257 | 0.3072 | +1 | 3 |
| - | - | 1034 | 410.2 | - | - | 0 | - |
| - | - | 621.7 | 411.2 | - | - | 0 | - |
| - | - | 4592 | 417.2 | - | - | 0 | - |
| - | - | 1120 | 418.2 | - | - | 0 | - |
| - | - | 974.2 | 428.2 | - | - | 0 | - |
| - | - | 3623 | 429.2 | - | - | 0 | - |
| - | - | 656.9 | 430.2 | - | - | 0 | - |
| - | - | 1557 | 435.2 | - | - | 0 | - |
| - | - | 1075 | 436.2 | - | - | 0 | - |
| - | - | 3472 | 444.3 | - | - | 0 | - |
| - | - | 2771 | 445.3 | - | - | 0 | - |
| 4 | c | 2659 | 446.3 | 0.0007633 | 1.71 | +1 | 4 |
| - | - | 684.4 | 450.2 | - | - | 0 | - |
| - | - | 668.7 | 457.8 | - | - | 0 | - |
| 12 | w | 1.055E+04 | 463.2 | 0.0002044 | 0.4412 | +1 | 4 |
| - | - | 4093 | 464.2 | - | - | 0 | - |
| - | - | 615.5 | 465.2 | - | - | 0 | - |
| - | - | 621.2 | 473.3 | - | - | 0 | - |
| - | - | 600.2 | 474.3 | - | - | 0 | - |
| - | - | 1499 | 478.1 | - | - | 0 | - |
| - | - | 4093 | 478.2 | - | - | 0 | - |
| 12 | y | 5171 | 479.2 | 0.0008443 | 1.762 | +1 | 4 |
| 12 | z | 9613 | 480.2 | 0.0002464 | 0.5132 | +1 | 4 |
| - | - | 744.8 | 480.6 | - | - | 0 | - |
| - | - | 3270 | 481.2 | - | - | 0 | - |
| 7 | z | 8295 | 488.8 | 4.574E-06 | 0.009359 | +2 | 9 |
| - | - | 5040 | 489.3 | - | - | 0 | - |
| - | - | 1688 | 489.8 | - | - | 0 | - |
| - | - | 1199 | 490.3 | - | - | 0 | - |
| 12 | y | 3300 | 496.3 | 1.91E-05 | 0.03848 | +1 | 4 |
| 7 | y | 3736 | 496.8 | 0.0004906 | 0.9875 | +2 | 9 |
| - | - | 1492 | 497.3 | - | - | 0 | - |
| - | - | 746.9 | 497.8 | - | - | 0 | - |
| 2 | y | 587.4 | 497.9 | 0.001318 | 2.647 | +3 | 14 |
| 2 | y | 1492 | 503.9 | 0.0004949 | 0.982 | +3 | 14 |
| - | - | 5060 | 504.2 | - | - | 0 | - |
| - | - | 1656 | 505.2 | - | - | 0 | - |
| - | - | 969.8 | 507.1 | - | - | 0 | - |
| - | - | 1854 | 516.3 | - | - | 0 | - |
| - | - | 784.5 | 517.3 | - | - | 0 | - |
| - | - | 1570 | 524.3 | - | - | 0 | - |
| - | - | 3094 | 524.8 | - | - | 0 | - |
| - | - | 5077 | 532.3 | - | - | 0 | - |
| 5 | c | 2.36E+04 | 533.3 | 0.0003508 | 0.6579 | +1 | 5 |
| - | - | 6614 | 534.3 | - | - | 0 | - |
| - | - | 1554 | 535.3 | - | - | 0 | - |
| - | - | 3931 | 535.6 | - | - | 0 | - |
| - | - | 3623 | 535.9 | - | - | 0 | - |
| - | - | 1627 | 536.3 | - | - | 0 | - |
| - | - | 974.6 | 536.6 | - | - | 0 | - |
| - | - | 625.4 | 540.8 | - | - | 0 | - |
| - | - | 936 | 541.2 | - | - | 0 | - |
| - | - | 1959 | 541.3 | - | - | 0 | - |
| - | - | 9195 | 541.6 | - | - | 0 | - |
| - | - | 7980 | 541.9 | - | - | 0 | - |
| - | - | 802.8 | 542.2 | - | - | 0 | - |
| - | - | 3996 | 542.3 | - | - | 0 | - |
| - | - | 687.4 | 542.3 | - | - | 0 | - |
| - | - | 856.1 | 542.6 | - | - | 0 | - |
| - | - | 1001 | 542.9 | - | - | 0 | - |
| 6 | z | 1.116E+04 | 546.3 | 7.371E-05 | 0.1349 | +2 | 10 |
| - | - | 1.395E+04 | 546.8 | - | - | 0 | - |
| - | - | 6051 | 547.3 | - | - | 0 | - |
| - | - | 2404 | 547.8 | - | - | 0 | - |
| - | - | 1221 | 553.8 | - | - | 0 | - |
| 6 | y | 5202 | 554.3 | 0.0002942 | 0.5308 | +2 | 10 |
| - | - | 5135 | 554.8 | - | - | 0 | - |
| - | - | 1410 | 555.3 | - | - | 0 | - |
| 11 | z | 2138 | 561.3 | 8.176E-05 | 0.1457 | +1 | 5 |
| - | - | 1990 | 562.3 | - | - | 0 | - |
| - | - | 698.7 | 563.3 | - | - | 0 | - |
| 11 | c | 9524 | 564.3 | 0.01092 | 19.35 | +2 | 11 |
| - | - | 3091 | 565.3 | - | - | 0 | - |
| - | - | 899.1 | 567.3 | - | - | 0 | - |
| - | - | 822.6 | 567.8 | - | - | 0 | - |
| - | - | 951.9 | 568.3 | - | - | 0 | - |
| 11 | z | 5127 | 579.3 | 8.736E-05 | 0.1508 | +1 | 5 |
| - | - | 1508 | 580.3 | - | - | 0 | - |
| - | - | 766.1 | 581.8 | - | - | 0 | - |
| 5 | y | 2544 | 588.8 | 0.01093 | 18.56 | +2 | 11 |
| 5 | y | 3530 | 589.3 | 0.0009196 | 1.56 | +2 | 11 |
| 5 | z | 1.335E+04 | 589.8 | 0.0005596 | 0.9489 | +2 | 11 |
| - | - | 2.121E+04 | 590.3 | - | - | 0 | - |
| - | - | 9976 | 590.8 | - | - | 0 | - |
| - | - | 2501 | 591.3 | - | - | 0 | - |
| - | - | 730.7 | 591.8 | - | - | 0 | - |
| - | - | 7578 | 592.3 | - | - | 0 | - |
| - | - | 2454 | 593.3 | - | - | 0 | - |
| - | - | 1125 | 594.3 | - | - | 0 | - |
| - | - | 666.7 | 594.3 | - | - | 0 | - |
| 11 | y | 5996 | 595.3 | 0.0001095 | 0.1839 | +1 | 5 |
| - | - | 1635 | 596.3 | - | - | 0 | - |
| - | - | 3026 | 597.3 | - | - | 0 | - |
| 5 | y | 3.708E+04 | 597.8 | 0.000156 | 0.261 | +2 | 11 |
| - | - | 2.281E+04 | 598.3 | - | - | 0 | - |
| - | - | 8251 | 598.8 | - | - | 0 | - |
| - | - | 1633 | 599.3 | - | - | 0 | - |
| - | - | 1504 | 601.8 | - | - | 0 | - |
| - | - | 1625 | 602.3 | - | - | 0 | - |
| - | - | 629.8 | 603.3 | - | - | 0 | - |
| - | - | 1107 | 611.3 | - | - | 0 | - |
| - | - | 783.2 | 613.3 | - | - | 0 | - |
| 10 | z | 709.9 | 618.3 | 0.002075 | 3.356 | +1 | 6 |
| - | - | 1547 | 632.3 | - | - | 0 | - |
| - | - | 776.6 | 632.4 | - | - | 0 | - |
| 10 | z | 2.046E+04 | 636.3 | 5.536E-05 | 0.087 | +1 | 6 |
| - | - | 1.121E+04 | 637.3 | - | - | 0 | - |
| 4 | y | 3827 | 638.3 | 0.003451 | 5.406 | +2 | 12 |
| 4 | y | 661.9 | 638.8 | 0.003326 | 5.206 | +2 | 12 |
| 4 | z | 2.07E+04 | 639.3 | 0.0003897 | 0.6095 | +2 | 12 |
| - | - | 1.53E+04 | 639.8 | - | - | 0 | - |
| - | - | 800 | 639.9 | - | - | 0 | - |
| - | - | 6188 | 640.3 | - | - | 0 | - |
| - | - | 717.9 | 646.8 | - | - | 0 | - |
| 4 | y | 3.18E+04 | 647.3 | 0.0004275 | 0.6603 | +2 | 12 |
| - | - | 2.021E+04 | 647.8 | - | - | 0 | - |
| 6 | c | 2.434E+04 | 648.3 | 0.003834 | 5.914 | +1 | 6 |
| - | - | 2275 | 648.8 | - | - | 0 | - |
| - | - | 5501 | 649.3 | - | - | 0 | - |
| 10 | y | 2.239E+04 | 652.3 | 0.000114 | 0.1748 | +1 | 6 |
| - | - | 7001 | 653.3 | - | - | 0 | - |
| - | - | 1624 | 654.3 | - | - | 0 | - |
| - | - | 889.6 | 657.3 | - | - | 0 | - |
| - | - | 737.3 | 663.9 | - | - | 0 | - |
| - | - | 712.7 | 670.4 | - | - | 0 | - |
| - | - | 1121 | 674.3 | - | - | 0 | - |
| 3 | w | 1.717E+04 | 674.3 | 0.0002166 | 0.3213 | +2 | 13 |
| - | - | 1.384E+04 | 674.8 | - | - | 0 | - |
| - | - | 7895 | 675.3 | - | - | 0 | - |
| - | - | 1179 | 675.8 | - | - | 0 | - |
| - | - | 2465 | 679.4 | - | - | 0 | - |
| - | - | 590 | 680.4 | - | - | 0 | - |
| - | - | 1198 | 681.8 | - | - | 0 | - |
| - | - | 956 | 682.3 | - | - | 0 | - |
| 3 | z | 819.9 | 695.3 | 0.009274 | 13.34 | +2 | 13 |
| - | - | 730.9 | 695.8 | - | - | 0 | - |
| - | - | 625.5 | 698.8 | - | - | 0 | - |
| 3 | y | 1896 | 702.8 | 0.003944 | 5.612 | +2 | 13 |
| 3 | y | 1822 | 703.3 | 0.006138 | 8.727 | +2 | 13 |
| 3 | z | 1.192E+04 | 703.8 | 0.0005211 | 0.7404 | +2 | 13 |
| - | - | 1.059E+04 | 704.3 | - | - | 0 | - |
| - | - | 5090 | 704.8 | - | - | 0 | - |
| 9 | z | 2031 | 705.3 | 0.003677 | 5.213 | +1 | 7 |
| - | - | 1567 | 707.9 | - | - | 0 | - |
| - | - | 1800 | 708.4 | - | - | 0 | - |
| - | - | 961.5 | 708.9 | - | - | 0 | - |
| - | - | 736.5 | 709.4 | - | - | 0 | - |
| - | - | 2313 | 711.3 | - | - | 0 | - |
| 3 | y | 3.965E+04 | 711.9 | 0.0006058 | 0.8511 | +2 | 13 |
| - | - | 2.675E+04 | 712.4 | - | - | 0 | - |
| - | - | 1.431E+04 | 712.9 | - | - | 0 | - |
| - | - | 3155 | 713.4 | - | - | 0 | - |
| - | - | 1294 | 713.9 | - | - | 0 | - |
| - | - | 3775 | 715.9 | - | - | 0 | - |
| - | - | 2421 | 716.4 | - | - | 0 | - |
| - | - | 2417 | 716.9 | - | - | 0 | - |
| - | - | 3124 | 717.9 | - | - | 0 | - |
| - | - | 1490 | 718.4 | - | - | 0 | - |
| - | - | 1675 | 718.9 | - | - | 0 | - |
| - | - | 622.6 | 720.9 | - | - | 0 | - |
| - | - | 1200 | 721.3 | - | - | 0 | - |
| 9 | y | 2397 | 722.3 | 0.001626 | 2.251 | +1 | 7 |
| 9 | z | 1.269E+04 | 723.4 | 0.0003568 | 0.4933 | +1 | 7 |
| - | - | 7049 | 724.4 | - | - | 0 | - |
| - | - | 2339 | 725.4 | - | - | 0 | - |
| - | - | 1002 | 725.9 | - | - | 0 | - |
| 14 | c | 732.7 | 729.4 | 0.005186 | 7.11 | +2 | 14 |
| - | - | 1279 | 730.4 | - | - | 0 | - |
| - | - | 887.2 | 730.9 | - | - | 0 | - |
| - | - | 869.6 | 734.4 | - | - | 0 | - |
| 14 | c | 9.007E+04 | 737.9 | 8.865E-05 | 0.1201 | +2 | 14 |
| - | - | 7.324E+04 | 738.4 | - | - | 0 | - |
| - | - | 3.386E+04 | 738.9 | - | - | 0 | - |
| 9 | y | 1.391E+04 | 739.4 | 0.008751 | 11.84 | +1 | 7 |
| - | - | 2704 | 739.9 | - | - | 0 | - |
| - | - | 2041 | 740.4 | - | - | 0 | - |
| 2 | y | 4905 | 746.4 | 0.002671 | 3.578 | +2 | 14 |
| 2 | y | 4836 | 746.9 | 0.005322 | 7.125 | +2 | 14 |
| 2 | z | 1.948E+04 | 747.4 | 0.001399 | 1.871 | +2 | 14 |
| - | - | 1.33E+04 | 747.9 | - | - | 0 | - |
| - | - | 6581 | 748.4 | - | - | 0 | - |
| 2 | y | 4.865E+04 | 755.4 | 0.0002626 | 0.3477 | +2 | 14 |
| - | - | 4.316E+04 | 755.9 | - | - | 0 | - |
| - | - | 1.61E+04 | 756.4 | - | - | 0 | - |
| - | - | 4858 | 756.9 | - | - | 0 | - |
| - | - | 1468 | 757.4 | - | - | 0 | - |
| - | - | 2276 | 760.4 | - | - | 0 | - |
| - | - | 1846 | 760.9 | - | - | 0 | - |
| - | - | 743.5 | 772.8 | - | - | 0 | - |
| - | - | 1486 | 774.4 | - | - | 0 | - |
| - | - | 1202 | 774.9 | - | - | 0 | - |
| - | - | 1159 | 775.9 | - | - | 0 | - |
| - | - | 2017 | 781.4 | - | - | 0 | - |
| - | - | 7942 | 781.9 | - | - | 0 | - |
| - | - | 1.778E+04 | 782.4 | - | - | 0 | - |
| - | - | 1.499E+04 | 782.9 | - | - | 0 | - |
| - | - | 7380 | 783.4 | - | - | 0 | - |
| - | - | 1322 | 783.9 | - | - | 0 | - |
| - | - | 2374 | 787.9 | - | - | 0 | - |
| - | - | 3017 | 788.4 | - | - | 0 | - |
| - | - | 2244 | 788.9 | - | - | 0 | - |
| - | - | 2.925E+04 | 789.4 | - | - | 0 | - |
| - | - | 2.139E+04 | 789.9 | - | - | 0 | - |
| - | - | 1.113E+04 | 790.4 | - | - | 0 | - |
| - | - | 2.516E+04 | 790.9 | - | - | 0 | - |
| - | - | 2.234E+04 | 791.4 | - | - | 0 | - |
| - | - | 1.246E+04 | 791.9 | - | - | 0 | - |
| - | - | 2571 | 792.4 | - | - | 0 | - |
| - | - | 1324 | 792.9 | - | - | 0 | - |
| - | - | 4467 | 794.9 | - | - | 0 | - |
| - | - | 3984 | 795.4 | - | - | 0 | - |
| - | - | 2853 | 795.9 | - | - | 0 | - |
| - | - | 2860 | 796.4 | - | - | 0 | - |
| - | - | 1741 | 796.9 | - | - | 0 | - |
| - | - | 868.4 | 797.4 | - | - | 0 | - |
| - | - | 1.006E+04 | 803.4 | - | - | 0 | - |
| - | - | 9.629E+04 | 803.9 | - | - | 0 | - |
| - | - | 7.44E+04 | 804.4 | - | - | 0 | - |
| - | - | 4.067E+04 | 804.9 | - | - | 0 | - |
| - | - | 1.369E+04 | 805.4 | - | - | 0 | - |
| - | - | 2636 | 805.9 | - | - | 0 | - |
| - | - | 957.2 | 810.4 | - | - | 0 | - |
| - | - | 4.96E+04 | 811.9 | - | - | 0 | - |
| - | - | 6.431E+04 | 812.4 | - | - | 0 | - |
| - | - | 4.326E+04 | 812.9 | - | - | 0 | - |
| - | - | 1.715E+04 | 813.4 | - | - | 0 | - |
| - | - | 4885 | 813.9 | - | - | 0 | - |
| - | - | 1187 | 814.4 | - | - | 0 | - |
| - | - | 1157 | 835.4 | - | - | 0 | - |
| 8 | y | 3347 | 836.4 | 0.0007998 | 0.9563 | +1 | 8 |
| - | - | 1090 | 837.4 | - | - | 0 | - |
| - | - | 1258 | 842.5 | - | - | 0 | - |
| - | - | 5244 | 857.5 | - | - | 0 | - |
| - | - | 1745 | 858.5 | - | - | 0 | - |
| - | - | 1686 | 859.5 | - | - | 0 | - |
| - | - | 854.5 | 885.5 | - | - | 0 | - |
| - | - | 717.2 | 886.5 | - | - | 0 | - |
| 7 | w | 1496 | 890.4 | 0.002514 | 2.824 | +1 | 9 |
| - | - | 788.4 | 891.4 | - | - | 0 | - |
| - | - | 2416 | 900.5 | - | - | 0 | - |
| 8 | c | 2.592E+04 | 901.5 | 4.464E-05 | 0.04952 | +1 | 8 |
| - | - | 1.277E+04 | 902.5 | - | - | 0 | - |
| - | - | 3789 | 903.5 | - | - | 0 | - |
| - | - | 1208 | 904.5 | - | - | 0 | - |
| - | - | 835.5 | 916.5 | - | - | 0 | - |
| - | - | 1010 | 927.5 | - | - | 0 | - |
| - | - | 810.5 | 938.6 | - | - | 0 | - |
| - | - | 1.363E+04 | 944.5 | - | - | 0 | - |
| - | - | 8966 | 945.5 | - | - | 0 | - |
| - | - | 2550 | 946.5 | - | - | 0 | - |
| 7 | z | 921.2 | 958.5 | 0.0007837 | 0.8177 | +1 | 9 |
| - | - | 666.6 | 972.5 | - | - | 0 | - |
| 7 | z | 1.369E+04 | 976.5 | 0.0009114 | 0.9333 | +1 | 9 |
| - | - | 9796 | 977.5 | - | - | 0 | - |
| - | - | 4060 | 978.5 | - | - | 0 | - |
| - | - | 1119 | 979.5 | - | - | 0 | - |
| - | - | 4635 | 987.5 | - | - | 0 | - |
| 9 | c | 6.65E+04 | 988.5 | 0.0004896 | 0.4953 | +1 | 9 |
| - | - | 3.174E+04 | 989.5 | - | - | 0 | - |
| - | - | 1.096E+04 | 990.5 | - | - | 0 | - |
| - | - | 2411 | 991.5 | - | - | 0 | - |
| - | - | 5515 | 1002 | - | - | 0 | - |
| - | - | 1668 | 1003 | - | - | 0 | - |
| - | - | 1537 | 1026 | - | - | 0 | - |
| - | - | 892.9 | 1031 | - | - | 0 | - |
| - | - | 757.2 | 1031 | - | - | 0 | - |
| 10 | c | 3.821E+04 | 1046 | 0.00053 | 0.5069 | +1 | 10 |
| 6 | w | 2.29E+04 | 1047 | 0.007129 | 6.812 | +1 | 10 |
| - | - | 1.014E+04 | 1048 | - | - | 0 | - |
| - | - | 4281 | 1049 | - | - | 0 | - |
| - | - | 1236 | 1050 | - | - | 0 | - |
| - | - | 982.7 | 1080 | - | - | 0 | - |
| - | - | 3997 | 1081 | - | - | 0 | - |
| - | - | 2423 | 1082 | - | - | 0 | - |
| - | - | 850.7 | 1084 | - | - | 0 | - |
| 6 | z | 1.214E+04 | 1092 | 0.0002665 | 0.2442 | +1 | 10 |
| - | - | 1.929E+04 | 1093 | - | - | 0 | - |
| - | - | 1.053E+04 | 1094 | - | - | 0 | - |
| - | - | 3287 | 1095 | - | - | 0 | - |
| - | - | 2060 | 1101 | - | - | 0 | - |
| - | - | 1310 | 1102 | - | - | 0 | - |
| - | - | 1120 | 1103 | - | - | 0 | - |
| - | - | 1519 | 1107 | - | - | 0 | - |
| 6 | y | 3616 | 1108 | 0.001657 | 1.496 | +1 | 10 |
| - | - | 1275 | 1109 | - | - | 0 | - |
| - | - | 812.9 | 1110 | - | - | 0 | - |
| - | - | 1784 | 1130 | - | - | 0 | - |
| - | - | 1485 | 1131 | - | - | 0 | - |
| - | - | 669.4 | 1144 | - | - | 0 | - |
| 11 | c | 2.846E+04 | 1145 | 0.001317 | 1.151 | +1 | 11 |
| - | - | 1.721E+04 | 1146 | - | - | 0 | - |
| - | - | 6264 | 1147 | - | - | 0 | - |
| - | - | 1310 | 1148 | - | - | 0 | - |
| - | - | 967.8 | 1174 | - | - | 0 | - |
| 5 | z | 2162 | 1179 | 0.001167 | 0.9902 | +1 | 11 |
| - | - | 9638 | 1180 | - | - | 0 | - |
| - | - | 6310 | 1181 | - | - | 0 | - |
| - | - | 2349 | 1182 | - | - | 0 | - |
| - | - | 2512 | 1188 | - | - | 0 | - |
| - | - | 2127 | 1189 | - | - | 0 | - |
| 5 | y | 5553 | 1195 | 6.362E-06 | 0.005326 | +1 | 11 |
| - | - | 3605 | 1196 | - | - | 0 | - |
| - | - | 1415 | 1197 | - | - | 0 | - |
| - | - | 2043 | 1231 | - | - | 0 | - |
| 12 | c | 5.438E+04 | 1232 | 0.001973 | 1.602 | +1 | 12 |
| - | - | 3.154E+04 | 1233 | - | - | 0 | - |
| - | - | 1.358E+04 | 1234 | - | - | 0 | - |
| - | - | 3543 | 1235 | - | - | 0 | - |
| - | - | 764.4 | 1236 | - | - | 0 | - |
| - | - | 2587 | 1259 | - | - | 0 | - |
| 4 | z | 1510 | 1260 | 0.0007982 | 0.6337 | +1 | 12 |
| 4 | z | 1253 | 1261 | 0.01971 | 15.64 | +1 | 12 |
| - | - | 1722 | 1275 | - | - | 0 | - |
| 4 | y | 1456 | 1276 | 0.001361 | 1.067 | +1 | 12 |
| 4 | z | 2486 | 1278 | 0.00464 | 3.631 | +1 | 12 |
| - | - | 1.467E+04 | 1279 | - | - | 0 | - |
| - | - | 8940 | 1280 | - | - | 0 | - |
| - | - | 2920 | 1281 | - | - | 0 | - |
| 4 | y | 3686 | 1294 | 0.0031 | 2.396 | +1 | 12 |
| - | - | 3496 | 1295 | - | - | 0 | - |
| - | - | 1545 | 1296 | - | - | 0 | - |
| 13 | c | 3.677E+04 | 1319 | 0.001775 | 1.346 | +1 | 13 |
| - | - | 2.662E+04 | 1320 | - | - | 0 | - |
| - | - | 1.105E+04 | 1321 | - | - | 0 | - |
| - | - | 1974 | 1322 | - | - | 0 | - |
| - | - | 4093 | 1346 | - | - | 0 | - |
| - | - | 3199 | 1347 | - | - | 0 | - |
| 3 | w | 1480 | 1348 | 0.003045 | 2.259 | +1 | 13 |
| - | - | 708.1 | 1350 | - | - | 0 | - |
| - | - | 906.1 | 1365 | - | - | 0 | - |
| - | - | 735.3 | 1391 | - | - | 0 | - |
| - | - | 918.9 | 1396 | - | - | 0 | - |
| 3 | z | 1911 | 1407 | 0.002067 | 1.469 | +1 | 13 |
| - | - | 2.388E+04 | 1408 | - | - | 0 | - |
| - | - | 1.551E+04 | 1409 | - | - | 0 | - |
| - | - | 7175 | 1410 | - | - | 0 | - |
| - | - | 2498 | 1411 | - | - | 0 | - |
| - | - | 687.7 | 1416 | - | - | 0 | - |
| - | - | 1573 | 1417 | - | - | 0 | - |
| - | - | 1446 | 1418 | - | - | 0 | - |
| - | - | 1778 | 1424 | - | - | 0 | - |
| - | - | 1516 | 1431 | - | - | 0 | - |
| - | - | 4910 | 1432 | - | - | 0 | - |
| - | - | 4188 | 1433 | - | - | 0 | - |
| - | - | 1933 | 1434 | - | - | 0 | - |
| - | - | 4241 | 1459 | - | - | 0 | - |
| - | - | 3119 | 1460 | - | - | 0 | - |
| - | - | 1106 | 1461 | - | - | 0 | - |
| - | - | 956.9 | 1462 | - | - | 0 | - |
| 14 | c | 9276 | 1475 | 0.001324 | 0.8976 | +1 | 14 |
| - | - | 2.604E+04 | 1476 | - | - | 0 | - |
| - | - | 2.121E+04 | 1477 | - | - | 0 | - |
| - | - | 9001 | 1478 | - | - | 0 | - |
| - | - | 1987 | 1479 | - | - | 0 | - |
| 2 | z | 2558 | 1494 | 0.003211 | 2.15 | +1 | 14 |
| - | - | 1.711E+04 | 1495 | - | - | 0 | - |
| - | - | 1.105E+04 | 1496 | - | - | 0 | - |
| - | - | 4879 | 1497 | - | - | 0 | - |
| - | - | 1183 | 1498 | - | - | 0 | - |
| - | - | 922.1 | 1508 | - | - | 0 | - |
| - | - | 837.7 | 1509 | - | - | 0 | - |
| 2 | y | 843.6 | 1510 | 0.004554 | 3.016 | +1 | 14 |
| - | - | 660.4 | 1511 | - | - | 0 | - |
| - | - | 4370 | 1523 | - | - | 0 | - |
| - | - | 4169 | 1524 | - | - | 0 | - |
| - | - | 2619 | 1525 | - | - | 0 | - |
| - | - | 761 | 1526 | - | - | 0 | - |
| - | - | 1702 | 1548 | - | - | 0 | - |
| - | - | 1603 | 1549 | - | - | 0 | - |
| - | - | 1493 | 1550 | - | - | 0 | - |
| - | - | 1161 | 1551 | - | - | 0 | - |
| - | - | 754.3 | 1554 | - | - | 0 | - |
| - | - | 966 | 1564 | - | - | 0 | - |
| - | - | 2.133E+04 | 1565 | - | - | 0 | - |
| - | - | 2.835E+04 | 1566 | - | - | 0 | - |
| - | - | 1.736E+04 | 1567 | - | - | 0 | - |
| - | - | 7507 | 1568 | - | - | 0 | - |
| - | - | 2225 | 1569 | - | - | 0 | - |
| - | - | 908.5 | 1578 | - | - | 0 | - |
| - | - | 2930 | 1579 | - | - | 0 | - |
| - | - | 5453 | 1580 | - | - | 0 | - |
| - | - | 7614 | 1581 | - | - | 0 | - |
| - | - | 7430 | 1582 | - | - | 0 | - |
| - | - | 5302 | 1583 | - | - | 0 | - |
| - | - | 1987 | 1584 | - | - | 0 | - |
| - | - | 2318 | 1590 | - | - | 0 | - |
| - | - | 8315 | 1591 | - | - | 0 | - |
| - | - | 8063 | 1592 | - | - | 0 | - |
| - | - | 3493 | 1593 | - | - | 0 | - |
| - | - | 1205 | 1594 | - | - | 0 | - |
| - | - | 1804 | 1596 | - | - | 0 | - |
| - | - | 8868 | 1597 | - | - | 0 | - |
| - | - | 7048 | 1598 | - | - | 0 | - |
| - | - | 3625 | 1599 | - | - | 0 | - |
| - | - | 1499 | 1600 | - | - | 0 | - |
| - | - | 1.339E+04 | 1607 | - | - | 0 | - |
| - | - | 7.739E+04 | 1608 | - | - | 0 | - |
| - | - | 6.63E+04 | 1609 | - | - | 0 | - |
| - | - | 3.151E+04 | 1610 | - | - | 0 | - |
| - | - | 1.118E+04 | 1611 | - | - | 0 | - |
| - | - | 2786 | 1612 | - | - | 0 | - |
| - | - | 3924 | 1623 | - | - | 0 | - |
| - | - | 2.002E+04 | 1624 | - | - | 0 | - |
| - | - | 7.371E+04 | 1625 | - | - | 0 | - |
| - | - | 6.145E+04 | 1626 | - | - | 0 | - |
| - | - | 3.042E+04 | 1627 | - | - | 0 | - |
| - | - | 1.049E+04 | 1628 | - | - | 0 | - |
| - | - | 2609 | 1629 | - | - | 0 | - |
| - | - | 856.3 | 2998 | - | - | 0 | - |

m/z Charge Intensity FragmentType MassShift Position
120.080810546875 0 580.5389
123.28770446777344 0 414.87704
130.38687133789062 0 400.3686
142.09735107421875 0 692.30896
148.95494079589844 0 528.62885
153.10263061523438 0 568.6725
157.38742065429688 0 460.1912
159.12498474121094 0 394.30853
173.12852478027344 0 6207.1836
174.1320037841797 0 702.2168
183.11293029785156 0 2263.387
199.0714569091797 0 874.3185
199.82870483398438 0 570.4752
200.15106201171875 0 543.3829
201.1233367919922 0 9937.637
201.13467407226562 0 7264.634
202.12330627441406 0 2483.3538
202.1407928466797 0 663.65137
217.0813446044922 0 706.04944
246.14353942871094 0 595.78674
248.629638671875 0 585.3882 y 11
258.4145202636719 0 497.7038
261.12396240234375 0 924.22174
261.1708679199219 0 16822.582
262.1500244140625 0 518.0327
262.1743469238281 0 2320.3943
288.16693115234375 0 1101.9526
298.14031982421875 0 590.5071
303.14483642578125 0 562.3353
306.1685791015625 0 2594.0134 z 13
307.17193603515625 0 632.02576
312.15545654296875 0 6347.41
313.1597900390625 0 854.42285
316.1506652832031 0 1450.0907
320.17254638671875 0 894.4119 z Water loss 6
322.18695068359375 0 4754.356 y 13
323.1922302246094 0 930.08826
330.1661682128906 0 3535.7595
331.1687316894531 0 844.2119
348.2036437988281 0 2004.633
349.2084045410156 0 623.2165
357.6324462890625 0 490.50885
373.1976013183594 0 1230.1077
376.1978759765625 0 8933.99 c Water loss 10
377.2001037597656 0 1942.7051
378.2033996582031 0 692.4427
391.18475341796875 0 3975.3774
392.1926574707031 0 15990.945 y Ammonia loss 12
393.1986389160156 0 7358.9946 y Ammonia loss 4
394.2043762207031 0 2607.6755 c Ammonia loss 6
401.2391357421875 0 1977.219
407.20587158203125 0 639.641
409.2192687988281 0 3764.3962 y 12
410.2215576171875 0 1033.6991
411.2232666015625 0 621.6926
417.1883850097656 0 4592.016
418.1917419433594 0 1119.8326
428.22613525390625 0 974.19745
429.2342529296875 0 3623.0261
430.2356872558594 0 656.8725
435.23516845703125 0 1556.575
436.2400817871094 0 1075.0968
444.25701904296875 0 3471.9321
445.2564697265625 0 2770.6995
446.2616882324219 0 2659.4824 c 3
450.22039794921875 0 684.3743
457.78399658203125 0 668.68054
463.23016357421875 0 10551.583 w 11
464.23150634765625 0 4092.8528
465.2344055175781 0 615.45667
473.2752685546875 0 621.1861
474.27294921875 0 600.2079
478.1142883300781 0 1498.8824
478.2177734375 0 4093.4753
479.2240295410156 0 5171.4414 y Ammonia loss 11
480.2324523925781 0 9612.684 z 11
480.5985412597656 0 744.83655
481.23712158203125 0 3269.9631
488.75787353515625 0 8295.105 z 6
489.2603759765625 0 5039.553
489.7603759765625 0 1687.8417
490.2846374511719 0 1199.1011
496.25140380859375 0 3299.9597 y 11
496.7677307128906 0 3736.1965 y 6
497.2655944824219 0 1492.2357
497.77056884765625 0 746.9017
497.9123840332031 0 587.4269 y Water loss 1
503.9140930175781 0 1491.9152 y 1
504.2205505371094 0 5059.5967
505.2234191894531 0 1656.4315
507.1050720214844 0 969.8444
516.2666625976562 0 1853.8418
517.2694091796875 0 784.54034
524.276123046875 0 1570.3553
524.77685546875 0 3093.7407
532.2855224609375 0 5076.581
533.2926025390625 0 23596.486 c 4
534.2962646484375 0 6613.7236
535.3036499023438 0 1553.5579
535.605224609375 0 3930.6187
535.9392700195312 0 3623.2612
536.2727661132812 0 1626.7338
536.6101684570312 0 974.5694
540.8138427734375 0 625.38983
541.1911010742188 0 936.03204
541.3451538085938 0 1958.6168
541.6094360351562 0 9195.139
541.9436645507812 0 7979.5244
542.2271118164062 0 802.80426
542.2769775390625 0 3996.4287
542.316162109375 0 687.3565
542.6112670898438 0 856.0515
542.945556640625 0 1001.2549
546.2714233398438 0 11158.727 z 5
546.7739868164062 0 13950.685
547.2755737304688 0 6051.3696
547.777587890625 0 2403.5786
553.7769775390625 0 1221.2173
554.281005859375 0 5201.61 y 5
554.7819213867188 0 5134.69
555.28173828125 0 1409.5063
561.2904663085938 0 2137.8164 z Water loss 10
562.296875 0 1990.4347
563.2990112304688 0 698.7149
564.2772827148438 0 9523.982 c Ammonia loss 10
565.2819213867188 0 3091.4595
567.2892456054688 0 899.0729
567.7935180664062 0 822.6106
568.2929077148438 0 951.88367
579.301025390625 0 5126.842 z 10
580.3040771484375 0 1507.9065
581.7796630859375 0 766.1021
588.780517578125 0 2543.8286 y Water loss 4
589.2825317382812 0 3530.378 y Ammonia loss 4
589.7868041992188 0 13353.682 z 4
590.2900390625 0 21208.75
590.7921142578125 0 9976.181
591.2942504882812 0 2501.443
591.7935180664062 0 730.6882
592.332763671875 0 7577.8076
593.3362426757812 0 2453.5881
594.2748413085938 0 1125.0321
594.3341064453125 0 666.662
595.3199462890625 0 5996.2666 y 10
596.3236694335938 0 1635.0712
597.2930297851562 0 3025.7603
597.7965698242188 0 37075.453 y 4
598.2982177734375 0 22807.791
598.799072265625 0 8251.342
599.2984008789062 0 1633.0165
601.781494140625 0 1503.91
602.2841186523438 0 1625.2903
603.294921875 0 629.7558
611.2938842773438 0 1106.818
613.3185424804688 0 783.2135
618.3099365234375 0 709.859 z Water loss 9
632.310546875 0 1547.1144
632.36865234375 0 776.58734
636.3226318359375 0 20456.646 z 9
637.3273315429688 0 11213.923
638.3291015625 0 3826.9124 y Water loss 3
638.8209838867188 0 661.9183 y Ammonia loss 3
639.3219604492188 0 20697.71 z 3
639.8231811523438 0 15295.538
639.8860473632812 0 800.0364
640.3251342773438 0 6187.6216
646.8242797851562 0 717.9069
647.3305053710938 0 31796.049 y 3
647.8324584960938 0 20210.658
648.32373046875 0 24343.188 c 5
648.833984375 0 2275.061
649.3236083984375 0 5500.543
652.3411865234375 0 22385.783 y 9
653.3452758789062 0 7000.8877
654.34765625 0 1624.0209
657.3426513671875 0 889.60114
663.8553466796875 0 737.25977
670.3865356445312 0 712.6821
674.272705078125 0 1121.043
674.3359985351562 0 17171.496 w 2
674.8372802734375 0 13837.32
675.3388671875 0 7894.6353
675.837158203125 0 1179.1215
679.3638305664062 0 2464.8435
680.3569946289062 0 589.99176
681.8433227539062 0 1198.024
682.345703125 0 956.00085
695.3388671875 0 819.8722 z Ammonia loss 2
695.8426513671875 0 730.9423
698.8474731445312 0 625.505
702.8508911132812 0 1895.5719 y Water loss 2
703.3450927734375 0 1821.6337 y Ammonia loss 2
703.8423461914062 0 11921.523 z 2
704.3444213867188 0 10591.86
704.845703125 0 5090.153
705.3477172851562 0 2030.5542 z Water loss 8
707.8690795898438 0 1567.3748
708.3660278320312 0 1799.597
708.8634033203125 0 961.47406
709.427001953125 0 736.5191
711.3489990234375 0 2313.4692
711.8516235351562 0 39648.113 y 2
712.353759765625 0 26747.836
712.8551025390625 0 14314.969
713.356201171875 0 3155.104
713.8574829101562 0 1293.8127
715.8780517578125 0 3775.3547
716.3762817382812 0 2420.8618
716.8778076171875 0 2416.721
717.8519897460938 0 3123.5518
718.3546752929688 0 1490.4791
718.8545532226562 0 1674.9773
720.885986328125 0 622.6168
721.34033203125 0 1199.6775
722.3451538085938 0 2396.783 y Ammonia loss 8
723.354248046875 0 12686.87 z 8
724.35693359375 0 7048.5654
725.3624877929688 0 2339.2014
725.858642578125 0 1001.7108
729.3656005859375 0 732.6692 c Ammonia loss 13
730.3779296875 0 1279.2267
730.884521484375 0 887.23486
734.3985595703125 0 869.555
737.8839721679688 0 90073.99 c 13
738.3849487304688 0 73244.87
738.8863525390625 0 33861.125
739.382080078125 0 13910.927 y 8
739.8892211914062 0 2703.688
740.3721313476562 0 2040.6895
746.3602905273438 0 4904.8022 y Water loss 1
746.8602905273438 0 4836.454 y Ammonia loss 1
747.3574829101562 0 19481.615 z 1
747.8609008789062 0 13298.416
748.3574829101562 0 6580.649
755.3679809570312 0 48654.367 y 1
755.8694458007812 0 43159.17
756.3700561523438 0 16100.187
756.8724365234375 0 4858.0996
757.372802734375 0 1468.0243
760.4052124023438 0 2276.0156
760.86083984375 0 1846.2549
772.8167724609375 0 743.48206
774.3956298828125 0 1486.3906
774.8953247070312 0 1202.1116
775.879150390625 0 1158.8998
781.4033813476562 0 2016.7313
781.9060668945312 0 7941.95
782.4041748046875 0 17782.113
782.904052734375 0 14989.382
783.4048461914062 0 7379.9487
783.9020385742188 0 1321.6687
787.8994750976562 0 2374.0256
788.3970947265625 0 3017.2852
788.8984985351562 0 2243.8147
789.383056640625 0 29251.781
789.8842163085938 0 21385.135
790.3865966796875 0 11129.194
790.89794921875 0 25164.633
791.4006958007812 0 22336.291
791.90185546875 0 12460.719
792.4022216796875 0 2570.9468
792.9125366210938 0 1324.3699
794.8948974609375 0 4467.1685
795.3976440429688 0 3983.8914
795.8951416015625 0 2853.4377
796.392822265625 0 2860.0881
796.8924560546875 0 1741.4062
797.3948974609375 0 868.3663
803.4076538085938 0 10064.675
803.9019775390625 0 96290.88
804.4025268554688 0 74397.44
804.9034423828125 0 40665.46
805.404052734375 0 13691.745
805.9077758789062 0 2635.9417
810.3802490234375 0 957.2164
811.9098510742188 0 49595.09
812.4124755859375 0 64314.77
812.9142456054688 0 43256.78
813.4156494140625 0 17145.074
813.9180908203125 0 4885.236
814.4172973632812 0 1187.3606
835.4189453125 0 1157.434
836.42529296875 0 3346.859 y 7
837.4310913085938 0 1090.3518
842.453125 0 1257.9619
857.4606323242188 0 5244.2783
858.4607543945312 0 1745.0746
859.4701538085938 0 1685.8057
885.4554443359375 0 854.5457
886.4658813476562 0 717.2301
890.4341430664062 0 1496.4078 w 6
891.4407958984375 0 788.4398
900.4984741210938 0 2415.957
901.4738159179688 0 25924.676 c 7
902.4766845703125 0 12774.166
903.4810180664062 0 3789.1602
904.4849853515625 0 1208.1528
916.4801635742188 0 835.54535
927.487548828125 0 1010.2404
938.5607299804688 0 810.5485
944.4927368164062 0 13633.181
945.4950561523438 0 8966.037
946.4981689453125 0 2550.4749
958.4971313476562 0 921.2168 z Water loss 6
972.4838256835938 0 666.5911
976.507568359375 0 13686.602 z 6
977.5122680664062 0 9796.412
978.5155639648438 0 4060.278
979.5162353515625 0 1118.9174
987.499755859375 0 4635.28
988.5053100585938 0 66504.48 c 8
989.5081787109375 0 31743.494
990.5111083984375 0 10964.31
991.5104370117188 0 2410.647
1001.5130004882812 0 5514.8477
1002.5164794921875 0 1667.9519
1026.36962890625 0 1536.9363
1030.5150146484375 0 892.85724
1031.25927734375 0 757.1764
1045.5267333984375 0 38207.883 c 9
1046.5306396484375 0 22896.389 w 5
1047.5367431640625 0 10143.089
1048.543212890625 0 4281.253
1049.542236328125 0 1235.7593
1080.49755859375 0 982.6739
1081.3721923828125 0 3996.6157
1082.369140625 0 2422.9526
1084.38427734375 0 850.70306
1091.53515625 0 12138.138 z 5
1092.5411376953125 0 19293.275
1093.543212890625 0 10531.673
1094.5458984375 0 3286.7283
1100.5799560546875 0 2059.6301
1101.584228515625 0 1309.6586
1102.58203125 0 1119.5342
1106.5506591796875 0 1519.32
1107.552490234375 0 3615.844 y 5
1108.55908203125 0 1274.6627
1109.548828125 0 812.85376
1129.583251953125 0 1784.2954
1130.5843505859375 0 1485.0701
1143.590576171875 0 669.43854
1144.5943603515625 0 28463.887 c 10
1145.597900390625 0 17210.852
1146.6004638671875 0 6263.6445
1147.6064453125 0 1310.3665
1173.6070556640625 0 967.82104
1178.5662841796875 0 2161.5938 z 4
1179.5728759765625 0 9638.386
1180.574951171875 0 6310.1426
1181.5814208984375 0 2348.6624
1187.6141357421875 0 2512.4775
1188.61572265625 0 2127.262
1194.586181640625 0 5553.366 y 4
1195.583984375 0 3605.1982
1196.5966796875 0 1415.0089
1230.6123046875 0 2043.3811
1231.625732421875 0 54384.633 c 11
1232.628662109375 0 31542.154
1233.631591796875 0 13578.707
1234.6339111328125 0 3542.6016
1235.626953125 0 764.3743
1258.6270751953125 0 2586.875
1259.6260986328125 0 1509.921 z Water loss 3
1260.6290283203125 0 1252.6066 z Ammonia loss 3
1274.6455078125 0 1722.3884
1275.6453857421875 0 1456.0999 y Water loss 3
1277.6312255859375 0 2485.879 z 3
1278.64111328125 0 14671.281
1279.642333984375 0 8940.449
1280.6468505859375 0 2919.7332
1293.6514892578125 0 3686.337 y 3
1294.6541748046875 0 3495.8582
1295.6636962890625 0 1545.0527
1318.657958984375 0 36767.85 c 12
1319.6605224609375 0 26620.334
1320.6612548828125 0 11052.984
1321.666748046875 0 1974.0693
1345.660888671875 0 4093.2466
1346.6646728515625 0 3199.3796
1347.662109375 0 1480.3752 w 2
1349.6529541015625 0 708.0749
1364.660400390625 0 906.05066
1390.6748046875 0 735.3321
1395.628173828125 0 918.9311
1406.6763916015625 0 1911.2736 z 2
1407.6822509765625 0 23880.375
1408.6868896484375 0 15511.726
1409.6876220703125 0 7174.8975
1410.692138671875 0 2498.3782
1415.731201171875 0 687.7288
1416.7117919921875 0 1572.9718
1417.7025146484375 0 1445.6118
1423.6998291015625 0 1777.609
1430.7491455078125 0 1515.9232
1431.7528076171875 0 4909.911
1432.75244140625 0 4188.4785
1433.7523193359375 0 1933.1652
1458.736572265625 0 4240.7046
1459.74169921875 0 3118.7224
1460.74658203125 0 1106.0317
1461.7275390625 0 956.89124
1474.759521484375 0 9276.086 c 13
1475.7650146484375 0 26036.012
1476.7647705078125 0 21213.709
1477.7642822265625 0 9000.73
1478.7486572265625 0 1987.1029
1493.707275390625 0 2558.1296 z 1
1494.714111328125 0 17112.432
1495.7174072265625 0 11047.421
1496.718017578125 0 4879.351
1497.7254638671875 0 1183.4282
1507.7420654296875 0 922.1394
1508.7625732421875 0 837.72363
1509.7337646484375 0 843.551 y 1
1510.7415771484375 0 660.3724
1522.72509765625 0 4369.6353
1523.727294921875 0 4168.9424
1524.7332763671875 0 2619.4622
1525.7113037109375 0 761.0498
1547.7679443359375 0 1701.7716
1548.77099609375 0 1602.5133
1549.7625732421875 0 1493.1401
1550.7640380859375 0 1160.7532
1553.7845458984375 0 754.3459
1563.7957763671875 0 966.0289
1564.7813720703125 0 21327.086
1565.781982421875 0 28350.115
1566.7838134765625 0 17358.543
1567.7845458984375 0 7506.9395
1568.7889404296875 0 2224.5808
1577.8050537109375 0 908.46204
1578.8121337890625 0 2929.7158
1579.812744140625 0 5452.6235
1580.807861328125 0 7614.1562
1581.79833984375 0 7430.481
1582.7994384765625 0 5302.1978
1583.802490234375 0 1986.9589
1589.7890625 0 2318.477
1590.77978515625 0 8315.33
1591.77685546875 0 8063.1816
1592.7757568359375 0 3492.6836
1593.77734375 0 1204.7305
1595.8248291015625 0 1803.5502
1596.8292236328125 0 8867.9
1597.8349609375 0 7048.138
1598.83251953125 0 3624.988
1599.8389892578125 0 1498.5657
1606.801513671875 0 13391.565
1607.7999267578125 0 77387.164
1608.8026123046875 0 66296.5
1609.805908203125 0 31514.1
1610.80810546875 0 11178.782
1611.8095703125 0 2785.9058
1622.810546875 0 3923.828
1623.815673828125 0 20018.072
1624.8240966796875 0 73712.49
1625.828125 0 61453.527
1626.8316650390625 0 30419.422
1627.8321533203125 0 10486.73
1628.8404541015625 0 2608.696
2997.613037109375 0 856.2784

Spectrum Details

|  |  |
| --- | --- |
| Matched peaks? Matched peaks The total absolute number of peaks matched. Additionally in brackets the total fraction of peaks matched and the total number of peaks is shown. | 78 (16.88% of 462) |
| FDR? FDR The false discovery rate estimated for this peptide. It is calculated by matching all theoretical fragments with a non-integer shift with the raw peaks for this spectrum. This is done with 40 different shifts. The resulting percentage is the average number of annotated peaks over the number of annotated peaks with the correct spectrum. | 0.15% |
| Satellite FDR? Satellite FDR See the FDR for details on its calculation. This satellite ion specific FDR only contains the satellite ions (d/w) for I/L/J positions. | - |
| PSM Score? PSM Score The PSM Score as given by Hecklib to this annotated spectrum. It is shown with three significant figures. | 550 |

#### Spectrum 4796? Spectrum 4796 The raw spectrum of this peptide as annotated by Hecklib. The fragments are coloured according to ion type (see legend). Any peaks with a star '\*' as text can be hovered over to see the full details, first the ion type second the mass shift type. By hovering over the amino acids in the peptide or ions in the legend the corresponding peaks are highlighted. By toggling the 'Unassigned' label you can turn the background (unassigned) peaks on or off in the plot. By updating the slider in the Ion legend you can update the spectrum to only show the top X% of the peaks with labels. The top X% means any peak that is within X% of the highest intensity. By dragging in the spectrum you can zoom in to a specific part of the spectrum and use 'Zoom Out' to get back to the original zoom level. The annotation of the spectrum is based on the given sequence in the peptides file and is done with different software so inconsistencies are likely. The peaks are annotated based on the given sequence, with 20 ppm tolerance.

Copy Data

##### Spectrum 4796 (TSV)

###### Preview

```
Loading example...
```

*Click on the button to copy the data to your clipboard.*

Mz MinMz MaxIntensity Max

WidthHeightPeptide font sizePeptide stroke widthSpectrum font sizeSpectrum stroke widthCompact peptide

Ion legend

wxyz

abcd

OtherUnassignedIon^Charge^_Position_Show for top:%

JSEVSDRPSGVSSRF

06.43e+41.29e+51.93e+52.57e+5

Zoom Out

y^+1^_2_z^+1^_2_y^+1^_2_c^+1^_3_c^+3^_11_y^+1^_3_y^+3^_11_c^+2^_7_y^+1^_3_y^+2^_8_c^+1^_4_z^+1^_4_w^+1^_4_y^+1^_4_z^+1^_4_z^+2^_9_y^+1^_4_y^+2^_9_y^+3^_14_y^+3^_14_c^+1^_5_z^+2^_10_y^+2^_10_z^+1^_5_c^+2^_11_z^+1^_5_y^+2^_11_y^+2^_11_z^+2^_11_y^+1^_5_y^+2^_11_w^+2^_12_z^+1^_6_y^+2^_12_y^+2^_12_z^+2^_12_y^+2^_12_c^+1^_6_c^+2^_13_y^+1^_6_w^+2^_13_z^+2^_13_y^+2^_13_y^+2^_13_z^+2^_13_z^+1^_7_y^+2^_13_y^+1^_7_z^+1^_7_c^+2^_14_c^+2^_14_y^+1^_7_y^+2^_14_y^+2^_14_z^+2^_14_y^+2^_14_y^+1^_8_w^+1^_9_c^+1^_8_z^+1^_9_y^+1^_9_z^+1^_9_c^+1^_9_c^+1^_10_w^+1^_10_z^+1^_10_z^+1^_10_y^+1^_10_z^+1^_10_y^+1^_10_c^+1^_11_z^+1^_11_y^+1^_11_c^+1^_12_z^+1^_12_y^+1^_12_y^+1^_12_z^+1^_12_y^+1^_12_c^+1^_13_w^+1^_13_z^+1^_13_y^+1^_13_c^+1^_14_z^+1^_14_y^+1^_14_

0778155723353113

Fragment Matches Table

Show background peaks

| Position | Ion type | Intensity | mz Theoretical | mz Error (Th) | mz Error (ppm) | Charge | Series Number |
| --- | --- | --- | --- | --- | --- | --- | --- |
| - | - | 829.5 | 120.1 | - | - | 0 | - |
| - | - | 851.1 | 128.5 | - | - | 0 | - |
| - | - | 1192 | 148.9 | - | - | 0 | - |
| - | - | 832.2 | 169 | - | - | 0 | - |
| - | - | 1.548E+04 | 173.1 | - | - | 0 | - |
| - | - | 2403 | 173.5 | - | - | 0 | - |
| - | - | 6789 | 183.1 | - | - | 0 | - |
| - | - | 1324 | 199.1 | - | - | 0 | - |
| - | - | 971.6 | 199.1 | - | - | 0 | - |
| - | - | 1295 | 200.1 | - | - | 0 | - |
| - | - | 3.455E+04 | 201.1 | - | - | 0 | - |
| - | - | 1.747E+04 | 201.1 | - | - | 0 | - |
| - | - | 4872 | 202.1 | - | - | 0 | - |
| - | - | 1341 | 202.1 | - | - | 0 | - |
| - | - | 2916 | 217.1 | - | - | 0 | - |
| - | - | 973.5 | 229.1 | - | - | 0 | - |
| - | - | 3823 | 229.1 | - | - | 0 | - |
| - | - | 1.638E+04 | 244.1 | - | - | 0 | - |
| - | - | 1387 | 245.1 | - | - | 0 | - |
| - | - | 2297 | 261.1 | - | - | 0 | - |
| - | - | 4.181E+04 | 261.2 | - | - | 0 | - |
| - | - | 7028 | 262.2 | - | - | 0 | - |
| - | - | 3354 | 270.2 | - | - | 0 | - |
| - | - | 1040 | 281.5 | - | - | 0 | - |
| - | - | 1402 | 284.2 | - | - | 0 | - |
| - | - | 1775 | 288.2 | - | - | 0 | - |
| 14 | y | 1052 | 305.2 | 8.09E-05 | 0.2651 | +1 | 2 |
| 14 | z | 8206 | 306.2 | 0.0003033 | 0.9906 | +1 | 2 |
| - | - | 1791 | 307.2 | - | - | 0 | - |
| - | - | 1.806E+04 | 312.2 | - | - | 0 | - |
| - | - | 3093 | 313.2 | - | - | 0 | - |
| - | - | 3484 | 316.2 | - | - | 0 | - |
| 14 | y | 9921 | 322.2 | 0.0005001 | 1.552 | +1 | 2 |
| - | - | 1450 | 323.2 | - | - | 0 | - |
| - | - | 1.088E+04 | 330.2 | - | - | 0 | - |
| - | - | 1882 | 331.2 | - | - | 0 | - |
| 3 | c | 1260 | 347.2 | 3.668E-05 | 0.1056 | +1 | 3 |
| - | - | 5779 | 348.2 | - | - | 0 | - |
| - | - | 937.1 | 348.7 | - | - | 0 | - |
| - | - | 2366 | 349.2 | - | - | 0 | - |
| - | - | 2023 | 373.2 | - | - | 0 | - |
| 11 | c | 2.481E+04 | 376.2 | 0.001799 | 4.782 | +3 | 11 |
| - | - | 5959 | 377.2 | - | - | 0 | - |
| - | - | 992 | 385.2 | - | - | 0 | - |
| - | - | 1518 | 387.2 | - | - | 0 | - |
| - | - | 1.06E+04 | 391.2 | - | - | 0 | - |
| 13 | y | 4.408E+04 | 392.2 | 5.622E-05 | 0.1433 | +1 | 3 |
| 5 | y | 1.756E+04 | 393.2 | 0.007429 | 18.89 | +3 | 11 |
| 7 | c | 5470 | 394.2 | 0.004455 | 11.3 | +2 | 7 |
| - | - | 6536 | 401.2 | - | - | 0 | - |
| - | - | 1231 | 403.3 | - | - | 0 | - |
| - | - | 972.6 | 407.2 | - | - | 0 | - |
| 13 | y | 8241 | 409.2 | 0.0003321 | 0.8115 | +1 | 3 |
| - | - | 1886 | 410.2 | - | - | 0 | - |
| - | - | 1.451E+04 | 417.2 | - | - | 0 | - |
| - | - | 2612 | 418.2 | - | - | 0 | - |
| 8 | y | 1220 | 418.7 | 0.00115 | 2.746 | +2 | 8 |
| - | - | 2259 | 428.2 | - | - | 0 | - |
| - | - | 9125 | 429.2 | - | - | 0 | - |
| - | - | 2141 | 430.2 | - | - | 0 | - |
| - | - | 3519 | 435.2 | - | - | 0 | - |
| - | - | 2927 | 436.2 | - | - | 0 | - |
| - | - | 7375 | 444.3 | - | - | 0 | - |
| - | - | 6959 | 445.3 | - | - | 0 | - |
| 4 | c | 7393 | 446.3 | 6.137E-05 | 0.1375 | +1 | 4 |
| - | - | 981.3 | 455.2 | - | - | 0 | - |
| - | - | 1167 | 460.2 | - | - | 0 | - |
| 12 | z | 1928 | 462.2 | 0.001132 | 2.45 | +1 | 4 |
| 12 | w | 3.136E+04 | 463.2 | 0.0005096 | 1.1 | +1 | 4 |
| - | - | 1.01E+04 | 464.2 | - | - | 0 | - |
| - | - | 1391 | 465.2 | - | - | 0 | - |
| - | - | 1.169E+04 | 478.2 | - | - | 0 | - |
| 12 | y | 1.12E+04 | 479.2 | 0.0005391 | 1.125 | +1 | 4 |
| 12 | z | 2.178E+04 | 480.2 | 0.0002113 | 0.44 | +1 | 4 |
| - | - | 5200 | 481.2 | - | - | 0 | - |
| - | - | 1272 | 482.2 | - | - | 0 | - |
| 7 | z | 2.483E+04 | 488.8 | 0.0003616 | 0.7399 | +2 | 9 |
| - | - | 9969 | 489.3 | - | - | 0 | - |
| - | - | 5252 | 489.8 | - | - | 0 | - |
| - | - | 1676 | 490.3 | - | - | 0 | - |
| - | - | 1055 | 494.2 | - | - | 0 | - |
| 12 | y | 6382 | 496.3 | 0.0008354 | 1.683 | +1 | 4 |
| 7 | y | 6611 | 496.8 | 0.0007652 | 1.54 | +2 | 9 |
| - | - | 4122 | 497.3 | - | - | 0 | - |
| - | - | 1944 | 497.8 | - | - | 0 | - |
| 2 | y | 1495 | 497.9 | 0.0004523 | 0.9084 | +3 | 14 |
| - | - | 2537 | 498.3 | - | - | 0 | - |
| - | - | 1164 | 500.3 | - | - | 0 | - |
| 2 | y | 3585 | 503.9 | 0.0006038 | 1.198 | +3 | 14 |
| - | - | 1.392E+04 | 504.2 | - | - | 0 | - |
| - | - | 1068 | 504.3 | - | - | 0 | - |
| - | - | 3220 | 505.2 | - | - | 0 | - |
| - | - | 989.3 | 506.2 | - | - | 0 | - |
| - | - | 2484 | 513.4 | - | - | 0 | - |
| - | - | 3529 | 516.3 | - | - | 0 | - |
| - | - | 8243 | 524.3 | - | - | 0 | - |
| - | - | 4793 | 524.8 | - | - | 0 | - |
| - | - | 1406 | 525.3 | - | - | 0 | - |
| - | - | 1.168E+04 | 532.3 | - | - | 0 | - |
| 5 | c | 5.444E+04 | 533.3 | 0.0001375 | 0.2577 | +1 | 5 |
| - | - | 1.452E+04 | 534.3 | - | - | 0 | - |
| - | - | 4238 | 535.3 | - | - | 0 | - |
| - | - | 1.08E+04 | 535.6 | - | - | 0 | - |
| - | - | 6308 | 535.9 | - | - | 0 | - |
| - | - | 3692 | 536.3 | - | - | 0 | - |
| - | - | 2.105E+04 | 541.3 | - | - | 0 | - |
| - | - | 2.473E+04 | 541.6 | - | - | 0 | - |
| - | - | 2.131E+04 | 541.9 | - | - | 0 | - |
| - | - | 1.282E+04 | 542.3 | - | - | 0 | - |
| - | - | 5485 | 542.3 | - | - | 0 | - |
| - | - | 2067 | 542.6 | - | - | 0 | - |
| 6 | z | 3.505E+04 | 546.3 | 0.0001347 | 0.2467 | +2 | 10 |
| - | - | 3.516E+04 | 546.8 | - | - | 0 | - |
| - | - | 1.492E+04 | 547.3 | - | - | 0 | - |
| - | - | 1743 | 547.3 | - | - | 0 | - |
| - | - | 3839 | 547.8 | - | - | 0 | - |
| - | - | 2065 | 549.3 | - | - | 0 | - |
| - | - | 5064 | 553.8 | - | - | 0 | - |
| 6 | y | 1.973E+04 | 554.3 | 0.0001721 | 0.3105 | +2 | 10 |
| - | - | 8326 | 554.8 | - | - | 0 | - |
| - | - | 4361 | 555.3 | - | - | 0 | - |
| 11 | z | 5980 | 561.3 | 0.0006507 | 1.159 | +1 | 5 |
| - | - | 4652 | 562.3 | - | - | 0 | - |
| 11 | c | 2.002E+04 | 564.3 | 0.009821 | 17.4 | +2 | 11 |
| - | - | 5360 | 565.3 | - | - | 0 | - |
| - | - | 1175 | 566.3 | - | - | 0 | - |
| - | - | 1314 | 567.3 | - | - | 0 | - |
| - | - | 1618 | 567.8 | - | - | 0 | - |
| - | - | 2711 | 568.3 | - | - | 0 | - |
| 11 | z | 1.256E+04 | 579.3 | 3.471E-05 | 0.05992 | +1 | 5 |
| - | - | 4924 | 580.3 | - | - | 0 | - |
| - | - | 1403 | 581.8 | - | - | 0 | - |
| - | - | 1284 | 582.3 | - | - | 0 | - |
| 5 | y | 8517 | 588.8 | 0.01044 | 17.73 | +2 | 11 |
| 5 | y | 1.142E+04 | 589.3 | 5.699E-05 | 0.0967 | +2 | 11 |
| 5 | z | 3.306E+04 | 589.8 | 0.0001324 | 0.2245 | +2 | 11 |
| - | - | 5.858E+04 | 590.3 | - | - | 0 | - |
| - | - | 2.822E+04 | 590.8 | - | - | 0 | - |
| - | - | 9716 | 591.3 | - | - | 0 | - |
| - | - | 1778 | 591.8 | - | - | 0 | - |
| - | - | 1.902E+04 | 592.3 | - | - | 0 | - |
| - | - | 6625 | 593.3 | - | - | 0 | - |
| - | - | 1273 | 594.3 | - | - | 0 | - |
| 11 | y | 1.45E+04 | 595.3 | 0.0007809 | 1.312 | +1 | 5 |
| - | - | 3074 | 596.3 | - | - | 0 | - |
| - | - | 8072 | 597.3 | - | - | 0 | - |
| 5 | y | 8.187E+04 | 597.8 | 0.0002712 | 0.4537 | +2 | 11 |
| - | - | 5.987E+04 | 598.3 | - | - | 0 | - |
| - | - | 1.974E+04 | 598.8 | - | - | 0 | - |
| - | - | 1240 | 598.9 | - | - | 0 | - |
| - | - | 5732 | 599.3 | - | - | 0 | - |
| - | - | 4721 | 601.8 | - | - | 0 | - |
| - | - | 5046 | 602.3 | - | - | 0 | - |
| - | - | 1208 | 602.8 | - | - | 0 | - |
| - | - | 1618 | 605.3 | - | - | 0 | - |
| - | - | 1340 | 611.3 | - | - | 0 | - |
| - | - | 1238 | 617.3 | - | - | 0 | - |
| - | - | 2946 | 619.3 | - | - | 0 | - |
| - | - | 1060 | 621.3 | - | - | 0 | - |
| - | - | 1083 | 631.3 | - | - | 0 | - |
| 4 | w | 3380 | 631.8 | 0.001507 | 2.385 | +2 | 12 |
| - | - | 2738 | 632.3 | - | - | 0 | - |
| - | - | 1518 | 632.4 | - | - | 0 | - |
| 10 | z | 5.382E+04 | 636.3 | 0.0004216 | 0.6625 | +1 | 6 |
| - | - | 3.168E+04 | 637.3 | - | - | 0 | - |
| 4 | y | 9702 | 638.3 | 0.004122 | 6.458 | +2 | 12 |
| 4 | y | 1540 | 638.8 | 0.004607 | 7.212 | +2 | 12 |
| 4 | z | 5.406E+04 | 639.3 | 0.0007559 | 1.182 | +2 | 12 |
| - | - | 4.642E+04 | 639.8 | - | - | 0 | - |
| - | - | 1.362E+04 | 640.3 | - | - | 0 | - |
| - | - | 5492 | 640.8 | - | - | 0 | - |
| - | - | 1322 | 643.4 | - | - | 0 | - |
| - | - | 2863 | 646.8 | - | - | 0 | - |
| 4 | y | 8.682E+04 | 647.3 | 0.000305 | 0.4711 | +2 | 12 |
| - | - | 722.4 | 647.8 | - | - | 0 | - |
| - | - | 5.558E+04 | 647.8 | - | - | 0 | - |
| 6 | c | 6.033E+04 | 648.3 | 0.004689 | 7.232 | +1 | 6 |
| - | - | 6612 | 648.8 | - | - | 0 | - |
| - | - | 1.432E+04 | 649.3 | - | - | 0 | - |
| - | - | 3232 | 650.3 | - | - | 0 | - |
| 13 | c | 1541 | 651.3 | 0.01247 | 19.15 | +2 | 13 |
| 10 | y | 6.068E+04 | 652.3 | 0.0004963 | 0.7608 | +1 | 6 |
| - | - | 2.097E+04 | 653.3 | - | - | 0 | - |
| - | - | 4030 | 654.3 | - | - | 0 | - |
| - | - | 1153 | 668.8 | - | - | 0 | - |
| 3 | w | 4.9E+04 | 674.3 | 0.000882 | 1.308 | +2 | 13 |
| - | - | 3.493E+04 | 674.8 | - | - | 0 | - |
| - | - | 1.458E+04 | 675.3 | - | - | 0 | - |
| - | - | 5320 | 675.8 | - | - | 0 | - |
| - | - | 7668 | 679.4 | - | - | 0 | - |
| - | - | 2668 | 680.4 | - | - | 0 | - |
| - | - | 3091 | 681.8 | - | - | 0 | - |
| - | - | 1122 | 682.3 | - | - | 0 | - |
| 3 | z | 2559 | 695.3 | 0.01208 | 17.38 | +2 | 13 |
| - | - | 2000 | 696.3 | - | - | 0 | - |
| - | - | 1306 | 696.8 | - | - | 0 | - |
| 3 | y | 4541 | 702.8 | 0.0004651 | 0.6617 | +2 | 13 |
| 3 | y | 2051 | 703.3 | 0.006321 | 8.987 | +2 | 13 |
| 3 | z | 3.654E+04 | 703.8 | 0.0008216 | 1.167 | +2 | 13 |
| - | - | 2.487E+04 | 704.3 | - | - | 0 | - |
| - | - | 1.334E+04 | 704.8 | - | - | 0 | - |
| 9 | z | 4850 | 705.3 | 0.00325 | 4.607 | +1 | 7 |
| - | - | 4366 | 707.9 | - | - | 0 | - |
| - | - | 3450 | 708.4 | - | - | 0 | - |
| - | - | 1287 | 708.9 | - | - | 0 | - |
| - | - | 2147 | 711.3 | - | - | 0 | - |
| 3 | y | 1.05E+05 | 711.9 | 0.0003097 | 0.435 | +2 | 13 |
| - | - | 7.994E+04 | 712.4 | - | - | 0 | - |
| - | - | 3.367E+04 | 712.9 | - | - | 0 | - |
| - | - | 9974 | 713.4 | - | - | 0 | - |
| - | - | 1192 | 713.9 | - | - | 0 | - |
| - | - | 7672 | 715.9 | - | - | 0 | - |
| - | - | 7116 | 716.4 | - | - | 0 | - |
| - | - | 3081 | 716.9 | - | - | 0 | - |
| - | - | 5105 | 717.9 | - | - | 0 | - |
| - | - | 5844 | 718.4 | - | - | 0 | - |
| - | - | 3484 | 718.9 | - | - | 0 | - |
| - | - | 2446 | 721.3 | - | - | 0 | - |
| 9 | y | 6594 | 722.3 | 0.00187 | 2.589 | +1 | 7 |
| 9 | z | 3.799E+04 | 723.4 | 0.0004366 | 0.6036 | +1 | 7 |
| - | - | 1.588E+04 | 724.4 | - | - | 0 | - |
| - | - | 8131 | 725.4 | - | - | 0 | - |
| - | - | 3298 | 725.9 | - | - | 0 | - |
| - | - | 1801 | 726.4 | - | - | 0 | - |
| 14 | c | 2429 | 729.4 | 0.01025 | 14.06 | +2 | 14 |
| - | - | 1440 | 730.4 | - | - | 0 | - |
| - | - | 1113 | 730.9 | - | - | 0 | - |
| - | - | 1293 | 732.3 | - | - | 0 | - |
| 14 | c | 2.449E+05 | 737.9 | 0.0005217 | 0.707 | +2 | 14 |
| - | - | 1.992E+05 | 738.4 | - | - | 0 | - |
| - | - | 9.128E+04 | 738.9 | - | - | 0 | - |
| 9 | y | 3.257E+04 | 739.4 | 0.009789 | 13.24 | +1 | 7 |
| - | - | 4993 | 739.9 | - | - | 0 | - |
| - | - | 5391 | 740.4 | - | - | 0 | - |
| - | - | 1537 | 744.4 | - | - | 0 | - |
| 2 | y | 1.131E+04 | 746.4 | 0.001694 | 2.27 | +2 | 14 |
| 2 | y | 1.03E+04 | 746.9 | 0.006176 | 8.269 | +2 | 14 |
| 2 | z | 4.541E+04 | 747.4 | 0.0007272 | 0.9731 | +2 | 14 |
| - | - | 3.129E+04 | 747.9 | - | - | 0 | - |
| - | - | 1.63E+04 | 748.4 | - | - | 0 | - |
| - | - | 3817 | 748.9 | - | - | 0 | - |
| - | - | 2208 | 749.4 | - | - | 0 | - |
| - | - | 2151 | 754.9 | - | - | 0 | - |
| 2 | y | 1.288E+05 | 755.4 | 0.0005308 | 0.7028 | +2 | 14 |
| - | - | 1.075E+05 | 755.9 | - | - | 0 | - |
| - | - | 5.019E+04 | 756.4 | - | - | 0 | - |
| - | - | 1.502E+04 | 756.9 | - | - | 0 | - |
| - | - | 4272 | 757.4 | - | - | 0 | - |
| - | - | 2413 | 757.5 | - | - | 0 | - |
| - | - | 1903 | 759.4 | - | - | 0 | - |
| - | - | 5961 | 760.4 | - | - | 0 | - |
| - | - | 3393 | 760.9 | - | - | 0 | - |
| - | - | 2806 | 761.4 | - | - | 0 | - |
| - | - | 1750 | 761.9 | - | - | 0 | - |
| - | - | 2571 | 764.5 | - | - | 0 | - |
| - | - | 2436 | 767.9 | - | - | 0 | - |
| - | - | 1643 | 768.4 | - | - | 0 | - |
| - | - | 2947 | 773.4 | - | - | 0 | - |
| - | - | 2152 | 773.9 | - | - | 0 | - |
| - | - | 4489 | 774.4 | - | - | 0 | - |
| - | - | 2491 | 774.9 | - | - | 0 | - |
| - | - | 3245 | 775.4 | - | - | 0 | - |
| - | - | 1334 | 775.9 | - | - | 0 | - |
| - | - | 2171 | 776.4 | - | - | 0 | - |
| - | - | 4259 | 781.4 | - | - | 0 | - |
| - | - | 2.197E+04 | 781.9 | - | - | 0 | - |
| - | - | 4.919E+04 | 782.4 | - | - | 0 | - |
| - | - | 3.714E+04 | 782.9 | - | - | 0 | - |
| - | - | 1.612E+04 | 783.4 | - | - | 0 | - |
| - | - | 4923 | 783.9 | - | - | 0 | - |
| - | - | 1521 | 784.4 | - | - | 0 | - |
| - | - | 3222 | 787.9 | - | - | 0 | - |
| - | - | 4145 | 788.4 | - | - | 0 | - |
| - | - | 6033 | 788.9 | - | - | 0 | - |
| - | - | 6.561E+04 | 789.4 | - | - | 0 | - |
| - | - | 6.464E+04 | 789.9 | - | - | 0 | - |
| - | - | 3.003E+04 | 790.4 | - | - | 0 | - |
| - | - | 6.962E+04 | 790.9 | - | - | 0 | - |
| - | - | 6.008E+04 | 791.4 | - | - | 0 | - |
| - | - | 2.612E+04 | 791.9 | - | - | 0 | - |
| - | - | 8060 | 792.4 | - | - | 0 | - |
| - | - | 2713 | 792.9 | - | - | 0 | - |
| - | - | 2265 | 794.4 | - | - | 0 | - |
| - | - | 6066 | 794.9 | - | - | 0 | - |
| - | - | 6495 | 795.4 | - | - | 0 | - |
| - | - | 6326 | 795.9 | - | - | 0 | - |
| - | - | 7390 | 796.4 | - | - | 0 | - |
| - | - | 5522 | 796.9 | - | - | 0 | - |
| - | - | 2664 | 797.4 | - | - | 0 | - |
| - | - | 1306 | 797.9 | - | - | 0 | - |
| - | - | 2682 | 799.5 | - | - | 0 | - |
| - | - | 2.704E+04 | 803.4 | - | - | 0 | - |
| - | - | 2.546E+05 | 803.9 | - | - | 0 | - |
| - | - | 2.013E+05 | 804.4 | - | - | 0 | - |
| - | - | 1.069E+05 | 804.9 | - | - | 0 | - |
| - | - | 1706 | 805 | - | - | 0 | - |
| - | - | 3.031E+04 | 805.4 | - | - | 0 | - |
| - | - | 8870 | 805.9 | - | - | 0 | - |
| - | - | 1518 | 810.4 | - | - | 0 | - |
| - | - | 1568 | 811.4 | - | - | 0 | - |
| - | - | 1.262E+05 | 811.9 | - | - | 0 | - |
| - | - | 1.755E+05 | 812.4 | - | - | 0 | - |
| - | - | 1.108E+05 | 812.9 | - | - | 0 | - |
| - | - | 5.609E+04 | 813.4 | - | - | 0 | - |
| - | - | 1.133E+04 | 813.9 | - | - | 0 | - |
| - | - | 3291 | 814.4 | - | - | 0 | - |
| - | - | 3247 | 834.4 | - | - | 0 | - |
| 8 | y | 8250 | 836.4 | 0.0003116 | 0.3725 | +1 | 8 |
| - | - | 2863 | 837.4 | - | - | 0 | - |
| - | - | 1.357E+04 | 839.5 | - | - | 0 | - |
| - | - | 5063 | 840.5 | - | - | 0 | - |
| - | - | 3538 | 842.4 | - | - | 0 | - |
| - | - | 1469 | 843.4 | - | - | 0 | - |
| - | - | 1.577E+04 | 857.5 | - | - | 0 | - |
| - | - | 5204 | 858.5 | - | - | 0 | - |
| - | - | 3208 | 859.5 | - | - | 0 | - |
| - | - | 1176 | 878.4 | - | - | 0 | - |
| - | - | 2935 | 885.5 | - | - | 0 | - |
| - | - | 3722 | 886.5 | - | - | 0 | - |
| 7 | w | 3149 | 890.4 | 0.0009646 | 1.083 | +1 | 9 |
| - | - | 1351 | 891.4 | - | - | 0 | - |
| - | - | 4546 | 900.5 | - | - | 0 | - |
| 8 | c | 6.972E+04 | 901.5 | 0.0009602 | 1.065 | +1 | 8 |
| - | - | 3.45E+04 | 902.5 | - | - | 0 | - |
| - | - | 1.058E+04 | 903.5 | - | - | 0 | - |
| - | - | 3667 | 904.5 | - | - | 0 | - |
| - | - | 3105 | 927.5 | - | - | 0 | - |
| - | - | 1415 | 928.5 | - | - | 0 | - |
| - | - | 1161 | 935.5 | - | - | 0 | - |
| - | - | 3.879E+04 | 944.5 | - | - | 0 | - |
| - | - | 2.088E+04 | 945.5 | - | - | 0 | - |
| - | - | 5695 | 946.5 | - | - | 0 | - |
| - | - | 2490 | 947.5 | - | - | 0 | - |
| - | - | 5009 | 953.6 | - | - | 0 | - |
| - | - | 3417 | 954.6 | - | - | 0 | - |
| 7 | z | 2381 | 958.5 | 0.01271 | 13.26 | +1 | 9 |
| - | - | 1935 | 972.5 | - | - | 0 | - |
| 7 | y | 1439 | 975.5 | 7.769E-05 | 0.07964 | +1 | 9 |
| 7 | z | 3.492E+04 | 976.5 | 5.691E-05 | 0.05828 | +1 | 9 |
| - | - | 2.656E+04 | 977.5 | - | - | 0 | - |
| - | - | 7824 | 978.5 | - | - | 0 | - |
| - | - | 2012 | 979.5 | - | - | 0 | - |
| - | - | 1.35E+04 | 987.5 | - | - | 0 | - |
| 9 | c | 1.674E+05 | 988.5 | 0.0007921 | 0.8013 | +1 | 9 |
| - | - | 8.739E+04 | 989.5 | - | - | 0 | - |
| - | - | 2.792E+04 | 990.5 | - | - | 0 | - |
| - | - | 5828 | 991.5 | - | - | 0 | - |
| - | - | 1.069E+04 | 1002 | - | - | 0 | - |
| - | - | 5823 | 1003 | - | - | 0 | - |
| - | - | 2183 | 1004 | - | - | 0 | - |
| - | - | 1719 | 1011 | - | - | 0 | - |
| - | - | 2193 | 1031 | - | - | 0 | - |
| - | - | 2188 | 1032 | - | - | 0 | - |
| 10 | c | 9.871E+04 | 1046 | 0.0006907 | 0.6606 | +1 | 10 |
| 6 | w | 5.472E+04 | 1047 | 0.006152 | 5.879 | +1 | 10 |
| - | - | 2.337E+04 | 1048 | - | - | 0 | - |
| - | - | 7748 | 1049 | - | - | 0 | - |
| - | - | 2654 | 1050 | - | - | 0 | - |
| - | - | 1459 | 1060 | - | - | 0 | - |
| - | - | 4224 | 1067 | - | - | 0 | - |
| - | - | 2809 | 1068 | - | - | 0 | - |
| 6 | z | 2748 | 1074 | 0.0008103 | 0.7548 | +1 | 10 |
| 6 | z | 1717 | 1075 | 0.01835 | 17.08 | +1 | 10 |
| - | - | 3285 | 1083 | - | - | 0 | - |
| - | - | 1.064E+04 | 1084 | - | - | 0 | - |
| - | - | 4473 | 1085 | - | - | 0 | - |
| - | - | 2059 | 1086 | - | - | 0 | - |
| 6 | y | 1801 | 1091 | 0.0008446 | 0.7745 | +1 | 10 |
| 6 | z | 3.474E+04 | 1092 | 0.0009542 | 0.8741 | +1 | 10 |
| - | - | 5.134E+04 | 1093 | - | - | 0 | - |
| - | - | 2.564E+04 | 1094 | - | - | 0 | - |
| - | - | 8796 | 1095 | - | - | 0 | - |
| - | - | 2137 | 1096 | - | - | 0 | - |
| - | - | 1704 | 1100 | - | - | 0 | - |
| - | - | 3908 | 1101 | - | - | 0 | - |
| - | - | 3237 | 1102 | - | - | 0 | - |
| - | - | 2486 | 1103 | - | - | 0 | - |
| - | - | 3310 | 1107 | - | - | 0 | - |
| 6 | y | 8888 | 1108 | 0.0006627 | 0.5983 | +1 | 10 |
| - | - | 4661 | 1109 | - | - | 0 | - |
| - | - | 1465 | 1110 | - | - | 0 | - |
| - | - | 5531 | 1130 | - | - | 0 | - |
| - | - | 3670 | 1131 | - | - | 0 | - |
| - | - | 2290 | 1132 | - | - | 0 | - |
| - | - | 2589 | 1144 | - | - | 0 | - |
| 11 | c | 7.265E+04 | 1145 | 0.0002699 | 0.2358 | +1 | 11 |
| - | - | 4.828E+04 | 1146 | - | - | 0 | - |
| - | - | 2.001E+04 | 1147 | - | - | 0 | - |
| - | - | 4217 | 1148 | - | - | 0 | - |
| - | - | 1686 | 1172 | - | - | 0 | - |
| 5 | z | 6841 | 1179 | 0.001777 | 1.508 | +1 | 11 |
| - | - | 2.842E+04 | 1180 | - | - | 0 | - |
| - | - | 1.71E+04 | 1181 | - | - | 0 | - |
| - | - | 8205 | 1182 | - | - | 0 | - |
| - | - | 1780 | 1183 | - | - | 0 | - |
| - | - | 6737 | 1188 | - | - | 0 | - |
| - | - | 6270 | 1189 | - | - | 0 | - |
| - | - | 4145 | 1190 | - | - | 0 | - |
| - | - | 1416 | 1191 | - | - | 0 | - |
| 5 | y | 1.178E+04 | 1195 | 0.000604 | 0.5056 | +1 | 11 |
| - | - | 8585 | 1196 | - | - | 0 | - |
| - | - | 3402 | 1197 | - | - | 0 | - |
| - | - | 1805 | 1198 | - | - | 0 | - |
| - | - | 4579 | 1231 | - | - | 0 | - |
| 12 | c | 1.302E+05 | 1232 | 0.0006305 | 0.512 | +1 | 12 |
| - | - | 8.823E+04 | 1233 | - | - | 0 | - |
| - | - | 3.609E+04 | 1234 | - | - | 0 | - |
| - | - | 8056 | 1235 | - | - | 0 | - |
| - | - | 2909 | 1236 | - | - | 0 | - |
| - | - | 1408 | 1237 | - | - | 0 | - |
| - | - | 6175 | 1259 | - | - | 0 | - |
| 4 | z | 3414 | 1260 | 0.003362 | 2.669 | +1 | 12 |
| - | - | 5983 | 1275 | - | - | 0 | - |
| 4 | y | 2768 | 1276 | 0.002216 | 1.737 | +1 | 12 |
| 4 | y | 3043 | 1277 | 0.01503 | 11.77 | +1 | 12 |
| 4 | z | 6454 | 1278 | 0.0008554 | 0.6695 | +1 | 12 |
| - | - | 3.246E+04 | 1279 | - | - | 0 | - |
| - | - | 2.289E+04 | 1280 | - | - | 0 | - |
| - | - | 8447 | 1281 | - | - | 0 | - |
| - | - | 2524 | 1282 | - | - | 0 | - |
| - | - | 1302 | 1283 | - | - | 0 | - |
| 4 | y | 1.092E+04 | 1294 | 0.0009284 | 0.7176 | +1 | 12 |
| - | - | 7563 | 1295 | - | - | 0 | - |
| - | - | 4910 | 1296 | - | - | 0 | - |
| - | - | 1672 | 1297 | - | - | 0 | - |
| 13 | c | 9.737E+04 | 1319 | 0.0005545 | 0.4205 | +1 | 13 |
| - | - | 7.167E+04 | 1320 | - | - | 0 | - |
| - | - | 3.15E+04 | 1321 | - | - | 0 | - |
| - | - | 7819 | 1322 | - | - | 0 | - |
| - | - | 1996 | 1323 | - | - | 0 | - |
| - | - | 1925 | 1345 | - | - | 0 | - |
| - | - | 1.079E+04 | 1346 | - | - | 0 | - |
| - | - | 6473 | 1347 | - | - | 0 | - |
| 3 | w | 3426 | 1348 | 0.001106 | 0.8206 | +1 | 13 |
| - | - | 1316 | 1349 | - | - | 0 | - |
| - | - | 1459 | 1365 | - | - | 0 | - |
| - | - | 1250 | 1391 | - | - | 0 | - |
| - | - | 1452 | 1392 | - | - | 0 | - |
| - | - | 1723 | 1395 | - | - | 0 | - |
| - | - | 2061 | 1396 | - | - | 0 | - |
| 3 | z | 7400 | 1407 | 0.003409 | 2.424 | +1 | 13 |
| - | - | 5.427E+04 | 1408 | - | - | 0 | - |
| - | - | 4.014E+04 | 1409 | - | - | 0 | - |
| - | - | 1.858E+04 | 1410 | - | - | 0 | - |
| - | - | 5790 | 1411 | - | - | 0 | - |
| - | - | 1537 | 1415 | - | - | 0 | - |
| - | - | 4188 | 1417 | - | - | 0 | - |
| - | - | 3397 | 1418 | - | - | 0 | - |
| 3 | y | 1758 | 1423 | 0.005088 | 3.576 | +1 | 13 |
| - | - | 3974 | 1424 | - | - | 0 | - |
| - | - | 2040 | 1425 | - | - | 0 | - |
| - | - | 2860 | 1431 | - | - | 0 | - |
| - | - | 1.359E+04 | 1432 | - | - | 0 | - |
| - | - | 1.311E+04 | 1433 | - | - | 0 | - |
| - | - | 6121 | 1434 | - | - | 0 | - |
| - | - | 1925 | 1435 | - | - | 0 | - |
| - | - | 1993 | 1452 | - | - | 0 | - |
| - | - | 1757 | 1453 | - | - | 0 | - |
| - | - | 1294 | 1454 | - | - | 0 | - |
| - | - | 1.075E+04 | 1459 | - | - | 0 | - |
| - | - | 9649 | 1460 | - | - | 0 | - |
| - | - | 4590 | 1461 | - | - | 0 | - |
| - | - | 1555 | 1462 | - | - | 0 | - |
| 14 | c | 1.946E+04 | 1475 | 0.0008354 | 0.5665 | +1 | 14 |
| - | - | 6.922E+04 | 1476 | - | - | 0 | - |
| - | - | 4.532E+04 | 1477 | - | - | 0 | - |
| - | - | 1.999E+04 | 1478 | - | - | 0 | - |
| - | - | 5580 | 1479 | - | - | 0 | - |
| - | - | 1818 | 1480 | - | - | 0 | - |
| 2 | z | 5128 | 1494 | 0.003381 | 2.263 | +1 | 14 |
| - | - | 4.061E+04 | 1495 | - | - | 0 | - |
| - | - | 2.955E+04 | 1496 | - | - | 0 | - |
| - | - | 1.487E+04 | 1497 | - | - | 0 | - |
| - | - | 4909 | 1498 | - | - | 0 | - |
| - | - | 1633 | 1508 | - | - | 0 | - |
| - | - | 2670 | 1509 | - | - | 0 | - |
| 2 | y | 2802 | 1510 | 0.02018 | 13.37 | +1 | 14 |
| - | - | 2313 | 1511 | - | - | 0 | - |
| - | - | 1.155E+04 | 1523 | - | - | 0 | - |
| - | - | 1.041E+04 | 1524 | - | - | 0 | - |
| - | - | 7825 | 1525 | - | - | 0 | - |
| - | - | 1962 | 1526 | - | - | 0 | - |
| - | - | 2765 | 1538 | - | - | 0 | - |
| - | - | 1255 | 1539 | - | - | 0 | - |
| - | - | 1396 | 1547 | - | - | 0 | - |
| - | - | 2004 | 1548 | - | - | 0 | - |
| - | - | 4680 | 1549 | - | - | 0 | - |
| - | - | 3075 | 1550 | - | - | 0 | - |
| - | - | 1406 | 1551 | - | - | 0 | - |
| - | - | 1606 | 1563 | - | - | 0 | - |
| - | - | 3774 | 1564 | - | - | 0 | - |
| - | - | 5.521E+04 | 1565 | - | - | 0 | - |
| - | - | 7.029E+04 | 1566 | - | - | 0 | - |
| - | - | 4.776E+04 | 1567 | - | - | 0 | - |
| - | - | 2.083E+04 | 1568 | - | - | 0 | - |
| - | - | 7771 | 1569 | - | - | 0 | - |
| - | - | 1309 | 1570 | - | - | 0 | - |
| - | - | 1999 | 1578 | - | - | 0 | - |
| - | - | 9054 | 1579 | - | - | 0 | - |
| - | - | 1.367E+04 | 1580 | - | - | 0 | - |
| - | - | 1.518E+04 | 1581 | - | - | 0 | - |
| - | - | 2.033E+04 | 1582 | - | - | 0 | - |
| - | - | 1.154E+04 | 1583 | - | - | 0 | - |
| - | - | 4731 | 1584 | - | - | 0 | - |
| - | - | 1951 | 1585 | - | - | 0 | - |
| - | - | 5606 | 1590 | - | - | 0 | - |
| - | - | 2.142E+04 | 1591 | - | - | 0 | - |
| - | - | 1.335E+04 | 1592 | - | - | 0 | - |
| - | - | 7228 | 1593 | - | - | 0 | - |
| - | - | 3815 | 1594 | - | - | 0 | - |
| - | - | 3679 | 1596 | - | - | 0 | - |
| - | - | 2.039E+04 | 1597 | - | - | 0 | - |
| - | - | 1.568E+04 | 1598 | - | - | 0 | - |
| - | - | 6715 | 1599 | - | - | 0 | - |
| - | - | 2039 | 1600 | - | - | 0 | - |
| - | - | 1486 | 1606 | - | - | 0 | - |
| - | - | 3.639E+04 | 1607 | - | - | 0 | - |
| - | - | 1.97E+05 | 1608 | - | - | 0 | - |
| - | - | 1.558E+05 | 1609 | - | - | 0 | - |
| - | - | 7.758E+04 | 1610 | - | - | 0 | - |
| - | - | 2.298E+04 | 1611 | - | - | 0 | - |
| - | - | 6385 | 1612 | - | - | 0 | - |
| - | - | 9638 | 1623 | - | - | 0 | - |
| - | - | 2139 | 1624 | - | - | 0 | - |
| - | - | 5.407E+04 | 1624 | - | - | 0 | - |
| - | - | 1.875E+05 | 1625 | - | - | 0 | - |
| - | - | 1.58E+05 | 1626 | - | - | 0 | - |
| - | - | 7.56E+04 | 1627 | - | - | 0 | - |
| - | - | 2.303E+04 | 1628 | - | - | 0 | - |
| - | - | 7158 | 1629 | - | - | 0 | - |
| - | - | 1290 | 3083 | - | - | 0 | - |

m/z Charge Intensity FragmentType MassShift Position
120.08124542236328 0 829.49475
128.52626037597656 0 851.06757
148.94735717773438 0 1192.0934
169.01412963867188 0 832.2026
173.1286163330078 0 15480.082
173.4523468017578 0 2402.5786
183.11300659179688 0 6788.797
199.07130432128906 0 1323.9896
199.11903381347656 0 971.5926
200.115234375 0 1295.0222
201.12347412109375 0 34546.492
201.13491821289062 0 17465.486
202.1236114501953 0 4872.285
202.1414031982422 0 1340.6511
217.0821533203125 0 2915.5754
229.1181640625 0 973.4988
229.1299285888672 0 3822.6707
244.12939453125 0 16378.965
245.13253784179688 0 1387.4911
261.1236267089844 0 2297.0466
261.17108154296875 0 41811.137
262.1748962402344 0 7027.926
270.15594482421875 0 3354.3914
281.51220703125 0 1040.1528
284.1601257324219 0 1402.1696
288.16693115234375 0 1775.33
305.1607360839844 0 1052.2942 y Ammonia loss 13
306.1689453125 0 8205.981 z 13
307.172607421875 0 1790.7372
312.1557312011719 0 18058.621
313.1587829589844 0 3092.5896
316.1507568359375 0 3484.0796
322.1878662109375 0 9920.995 y 13
323.1917724609375 0 1449.9525
330.1662902832031 0 10884.627
331.16986083984375 0 1882.4261
347.1924743652344 0 1260.2686 c 2
348.2034912109375 0 5779.1836
348.6514892578125 0 937.0604
349.2093505859375 0 2365.9573
373.1982116699219 0 2022.8815
376.1980895996094 0 24814.049 c Water loss 10
377.2009582519531 0 5959.3315
385.19512939453125 0 992.02716
387.2359924316406 0 1517.9349
391.1850891113281 0 10601.08
392.1929016113281 0 44082.836 y Ammonia loss 12
393.1988220214844 0 17555.41 y Ammonia loss 4
394.205322265625 0 5469.8315 c Ammonia loss 6
401.23895263671875 0 6535.6973
403.2557373046875 0 1231.1605
407.201171875 0 972.60345
409.2197265625 0 8240.502 y 12
410.2240295410156 0 1885.897
417.18853759765625 0 14514.699
418.19207763671875 0 2611.9497
418.71783447265625 0 1219.5884 y 7
428.2275390625 0 2259.2888
429.2344055175781 0 9125.313
430.2377624511719 0 2141.2827
435.2353515625 0 3518.56
436.24029541015625 0 2926.6064
444.2568054199219 0 7374.651
445.2564392089844 0 6959.3984
446.260986328125 0 7393.2896 c 3
455.2264404296875 0 981.28845
460.20831298828125 0 1166.5195
462.2232666015625 0 1927.9137 z Water loss 11
463.23046875 0 31362.33 w 11
464.2304992675781 0 10103.832
465.23529052734375 0 1390.8298
478.21759033203125 0 11686.896
479.2243347167969 0 11196.902 y Ammonia loss 11
480.23291015625 0 21781.44 z 11
481.2356262207031 0 5199.8403
482.2400207519531 0 1271.5516
488.75823974609375 0 24828.832 z 6
489.2598876953125 0 9968.99
489.76202392578125 0 5252.238
490.2904968261719 0 1675.8558
494.233642578125 0 1054.7938
496.25225830078125 0 6382.3013 y 11
496.76800537109375 0 6610.594 y 6
497.2655029296875 0 4122.36
497.77142333984375 0 1944.039
497.9106140136719 0 1494.7153 y Water loss 1
498.2547912597656 0 2536.9463
500.30169677734375 0 1163.5142
503.9151916503906 0 3585.4182 y 1
504.22088623046875 0 13922.752
504.2566833496094 0 1068.3534
505.2244567871094 0 3220.0557
506.227294921875 0 989.3066
513.3509521484375 0 2483.5862
516.2674560546875 0 3529.153
524.2759399414062 0 8243.316
524.7796630859375 0 4792.917
525.2802124023438 0 1405.8333
532.2857666015625 0 11675.512
533.2930908203125 0 54437.96 c 4
534.2965087890625 0 14522.448
535.3012084960938 0 4238.1294
535.60546875 0 10803.316
535.9407348632812 0 6308.0166
536.2740478515625 0 3692.011
541.3458862304688 0 21045.375
541.6098022460938 0 24729.812
541.9440307617188 0 21311.207
542.2779541015625 0 12820.147
542.3494262695312 0 5485.1714
542.6091918945312 0 2066.56
546.271484375 0 35054.03 z 5
546.7745361328125 0 35161.367
547.2760620117188 0 14916.837
547.3203125 0 1742.7406
547.7771606445312 0 3838.5212
549.2899780273438 0 2064.5168
553.7776489257812 0 5063.516
554.2808837890625 0 19726.004 y 5
554.7825317382812 0 8326.041
555.284423828125 0 4361.4683
561.2911987304688 0 5979.888 z Water loss 10
562.2974243164062 0 4652.059
564.2783813476562 0 20020.729 c Ammonia loss 10
565.2824096679688 0 5359.985
566.288818359375 0 1174.5797
567.288818359375 0 1314.0956
567.7848510742188 0 1618.3335
568.2821655273438 0 2710.6982
579.3011474609375 0 12558.081 z 10
580.3049926757812 0 4924.4985
581.78271484375 0 1402.7703
582.2827758789062 0 1284.4246
588.781005859375 0 8516.618 y Water loss 4
589.2835083007812 0 11419.117 y Ammonia loss 4
589.7872314453125 0 33062.12 z 4
590.2908325195312 0 58579.117
590.7925415039062 0 28223.361
591.2937622070312 0 9715.58
591.7935180664062 0 1778.0564
592.3329467773438 0 19019.344
593.336669921875 0 6624.7686
594.27197265625 0 1273.179
595.3206176757812 0 14497.878 y 10
596.323486328125 0 3073.821
597.2930908203125 0 8072.138
597.7969970703125 0 81868.97 y 4
598.2985229492188 0 59867.367
598.7997436523438 0 19736.068
598.8527221679688 0 1239.9374
599.3016357421875 0 5732.2197
601.7816162109375 0 4720.955
602.2822875976562 0 5046.1484
602.7836303710938 0 1208.0951
605.3070068359375 0 1617.826
611.2935791015625 0 1339.8508
617.3274536132812 0 1238.4515
619.3192138671875 0 2945.5928
621.298095703125 0 1059.684
631.294677734375 0 1083.0104
631.8113403320312 0 3379.967 w 3
632.3114013671875 0 2737.6545
632.3718872070312 0 1517.7104
636.322998046875 0 53821.234 z 9
637.328125 0 31676.838
638.3297729492188 0 9702.394 y Water loss 3
638.822265625 0 1539.7826 y Ammonia loss 3
639.3223266601562 0 54055.25 z 3
639.8235473632812 0 46415.457
640.3253173828125 0 13615.817
640.8253784179688 0 5491.933
643.3767700195312 0 1321.8892
646.826416015625 0 2863.3164
647.3312377929688 0 86823.05 y 3
647.7839965820312 0 722.4463
647.8328247070312 0 55580.977
648.3245849609375 0 60331.19 c 5
648.8353881835938 0 6612.4165
649.323974609375 0 14318.769
650.3272705078125 0 3231.7366
651.3327026367188 0 1541.1788 c Ammonia loss 12
652.341796875 0 60684.113 y 9
653.3446655273438 0 20972.96
654.346435546875 0 4029.7104
668.80908203125 0 1152.8895
674.3370971679688 0 49004.098 w 2
674.837890625 0 34929.043
675.339111328125 0 14578.7
675.8408813476562 0 5320.3853
679.3645629882812 0 7668.1553
680.3661499023438 0 2668.4756
681.8457641601562 0 3091.3655
682.3405151367188 0 1122.0264
695.3416748046875 0 2559.0525 z Ammonia loss 2
696.3395385742188 0 2000.0065
696.8380737304688 0 1305.6923
702.847412109375 0 4541.2285 y Water loss 2
703.3452758789062 0 2050.8179 y Ammonia loss 2
703.8436889648438 0 36542.32 z 2
704.3446044921875 0 24867.016
704.845458984375 0 13342.355
705.3472900390625 0 4850.0054 z Water loss 8
707.8703002929688 0 4366.229
708.3660888671875 0 3449.5864
708.86474609375 0 1286.645
711.3489379882812 0 2146.7922
711.8525390625 0 104997.12 y 2
712.3541870117188 0 79937.625
712.8555297851562 0 33669.605
713.3565063476562 0 9973.868
713.8526000976562 0 1192.1663
715.8779296875 0 7672.3438
716.378173828125 0 7116.488
716.8804931640625 0 3080.755
717.8539428710938 0 5104.617
718.353271484375 0 5844.0396
718.8549194335938 0 3484.2246
721.33935546875 0 2446.1284
722.3449096679688 0 6593.761 y Ammonia loss 8
723.3550415039062 0 37988.793 z 8
724.35791015625 0 15876.905
725.36328125 0 8130.5435
725.8636474609375 0 3297.7236
726.3616943359375 0 1801.307
729.3605346679688 0 2428.509 c Ammonia loss 13
730.3753662109375 0 1440.1344
730.8795166015625 0 1112.66
732.329833984375 0 1292.7688
737.8845825195312 0 244880.22 c 13
738.3857421875 0 199160.42
738.8869018554688 0 91278.81
739.3831176757812 0 32570.832 y 8
739.8856201171875 0 4993.4546
740.3721313476562 0 5390.93
744.3888549804688 0 1536.978
746.3612670898438 0 11309.784 y Water loss 1
746.8611450195312 0 10299.168 y Ammonia loss 1
747.358154296875 0 45406.14 z 1
747.8609619140625 0 31294.967
748.35986328125 0 16303.937
748.8626708984375 0 3817.4758
749.3612060546875 0 2208.3647
754.897705078125 0 2151.0632
755.3687744140625 0 128801.72 y 1
755.8702392578125 0 107491.88
756.3713989257812 0 50189.344
756.8723754882812 0 15023.596
757.373779296875 0 4271.5713
757.457275390625 0 2413.0076
759.3967895507812 0 1903.2827
760.4091186523438 0 5961.2524
760.8633422851562 0 3392.9148
761.414794921875 0 2806.2153
761.8645629882812 0 1749.7716
764.4778442382812 0 2571.2617
767.8917236328125 0 2435.9185
768.3912353515625 0 1643.3184
773.3925170898438 0 2946.591
773.8978271484375 0 2152.0166
774.3933715820312 0 4488.8604
774.8971557617188 0 2490.7917
775.3881225585938 0 3244.636
775.8876342773438 0 1333.956
776.370361328125 0 2171.0786
781.40966796875 0 4259.351
781.9060668945312 0 21973.568
782.405517578125 0 49191.95
782.9049682617188 0 37143.117
783.4042358398438 0 16122.856
783.9019775390625 0 4922.9653
784.3938598632812 0 1520.5525
787.8984375 0 3221.8022
788.3993530273438 0 4145.1104
788.9015502929688 0 6033.477
789.3840942382812 0 65613.78
789.88525390625 0 64644.523
790.3878784179688 0 30032.424
790.8992309570312 0 69624.41
791.4013671875 0 60084.07
791.902587890625 0 26119.45
792.4046630859375 0 8060.049
792.9090576171875 0 2712.8142
794.3956909179688 0 2264.8645
794.897705078125 0 6065.8047
795.3995971679688 0 6494.582
795.8987426757812 0 6326.3774
796.3922119140625 0 7390.265
796.8905029296875 0 5522.0503
797.3960571289062 0 2664.421
797.9021606445312 0 1305.7878
799.451416015625 0 2681.9282
803.4093627929688 0 27035.63
803.9026489257812 0 254638.42
804.4035034179688 0 201293.9
804.9044189453125 0 106880.28
804.9946899414062 0 1706.1887
805.40576171875 0 30305.922
805.908203125 0 8870.3955
810.3743286132812 0 1518.014
811.4061889648438 0 1567.7886
811.9107666015625 0 126221.35
812.413330078125 0 175500.86
812.9149169921875 0 110767.664
813.4163818359375 0 56089.293
813.9179077148438 0 11333.631
814.4181518554688 0 3290.8223
834.412841796875 0 3247.3193
836.42578125 0 8250.224 y 7
837.4282836914062 0 2863.3257
839.4988403320312 0 13574.042
840.50341796875 0 5063.094
842.4482421875 0 3538.2903
843.446533203125 0 1469.3461
857.4608764648438 0 15773.689
858.4646606445312 0 5204.206
859.467041015625 0 3207.9373
878.4295654296875 0 1175.6838
885.4609375 0 2935.2566
886.4616088867188 0 3721.768
890.4376220703125 0 3149.1873 w 6
891.436279296875 0 1351.1992
900.4953002929688 0 4545.657
901.4747314453125 0 69722.125 c 7
902.4774780273438 0 34504.438
903.4800415039062 0 10583.129
904.4855346679688 0 3667.2317
927.4877319335938 0 3105.4155
928.489501953125 0 1414.7188
935.4639892578125 0 1161.0605
944.4928588867188 0 38793.81
945.4963989257812 0 20884.793
946.4991455078125 0 5694.9478
947.5030517578125 0 2489.6216
953.5773315429688 0 5009.41
954.5795288085938 0 3416.6028
958.5106201171875 0 2380.9158 z Water loss 6
972.4885864257812 0 1935.2136
975.500732421875 0 1439.2334 y Ammonia loss 6
976.5084228515625 0 34921.89 z 6
977.5128173828125 0 26556.465
978.5165405273438 0 7824.069
979.5308837890625 0 2012.4304
987.49951171875 0 13502.803
988.506591796875 0 167366.62 c 8
989.50927734375 0 87386.18
990.5120849609375 0 27919.191
991.51513671875 0 5828.4165
1001.51416015625 0 10687.662
1002.517333984375 0 5823.472
1003.5182495117188 0 2183.3616
1010.5377807617188 0 1719.0299
1030.5157470703125 0 2193.085
1031.513916015625 0 2187.5845
1045.5279541015625 0 98712.695 c 9
1046.5316162109375 0 54718.277 w 5
1047.5382080078125 0 23371.516
1048.5439453125 0 7748.1807
1049.550048828125 0 2653.8474
1059.5262451171875 0 1458.8613
1066.6011962890625 0 4224.4883
1067.5972900390625 0 2809.2139
1073.5240478515625 0 2747.7537 z Water loss 5
1074.5272216796875 0 1716.8552 z Ammonia loss 5
1082.6256103515625 0 3284.8677
1083.6268310546875 0 10635.708
1084.6258544921875 0 4472.944
1085.556884765625 0 2059.434
1090.5284423828125 0 1800.5482 y Ammonia loss 5
1091.536376953125 0 34738.69 z 5
1092.5421142578125 0 51342.55
1093.5445556640625 0 25638.232
1094.546142578125 0 8796.226
1095.5501708984375 0 2136.902
1099.5771484375 0 1704.4143
1100.583251953125 0 3907.7666
1101.5887451171875 0 3237.3745
1102.59912109375 0 2486.4753
1106.55078125 0 3310.151
1107.5548095703125 0 8888.377 y 5
1108.5572509765625 0 4660.6797
1109.5555419921875 0 1464.5853
1129.583251953125 0 5531.0977
1130.582763671875 0 3670.4585
1131.5899658203125 0 2290.1165
1143.594970703125 0 2589.36
1144.595947265625 0 72652.42 c 10
1145.5992431640625 0 48281.207
1146.60302734375 0 20007.336
1147.6065673828125 0 4216.902
1171.6171875 0 1685.743
1178.565673828125 0 6840.6865 z 4
1179.5750732421875 0 28423.791
1180.5782470703125 0 17095.2
1181.5830078125 0 8204.892
1182.5814208984375 0 1780.4551
1187.61181640625 0 6736.944
1188.6170654296875 0 6269.989
1189.625244140625 0 4145.0522
1190.630615234375 0 1415.836
1194.5855712890625 0 11777.168 y 4
1195.5882568359375 0 8585.005
1196.59228515625 0 3401.607
1197.604736328125 0 1805.0085
1230.6204833984375 0 4579.3413
1231.6270751953125 0 130217.586 c 11
1232.630615234375 0 88231.36
1233.633544921875 0 36089.19
1234.6357421875 0 8056.1855
1235.6246337890625 0 2908.9277
1236.6278076171875 0 1408.2644
1258.6224365234375 0 6174.7437
1259.628662109375 0 3414.3643 z Water loss 3
1274.645751953125 0 5983.2847
1275.646240234375 0 2767.7788 y Water loss 3
1276.64306640625 0 3042.9478 y Ammonia loss 3
1277.635009765625 0 6453.5347 z 3
1278.6427001953125 0 32457.299
1279.6451416015625 0 22891.328
1280.650390625 0 8446.835
1281.6444091796875 0 2524.1318
1282.649169921875 0 1301.778
1293.655517578125 0 10918.5 y 3
1294.6588134765625 0 7562.9844
1295.6607666015625 0 4909.8438
1296.661376953125 0 1671.6934
1318.6591796875 0 97374.95 c 12
1319.66162109375 0 71666.984
1320.664794921875 0 31495.379
1321.6661376953125 0 7819.334
1322.65771484375 0 1995.8531
1344.6783447265625 0 1925.4854
1345.6611328125 0 10793.464
1346.6646728515625 0 6473.0386
1347.666259765625 0 3426.4478 w 2
1348.66015625 0 1316.34
1364.65771484375 0 1458.6029
1390.673828125 0 1250.4
1391.656982421875 0 1451.5265
1394.6304931640625 0 1722.8629
1395.63525390625 0 2060.5593
1406.675048828125 0 7399.5093 z 2
1407.6849365234375 0 54270.1
1408.6878662109375 0 40140.85
1409.6915283203125 0 18582.832
1410.6951904296875 0 5790.2085
1414.734619140625 0 1537.3743
1416.7230224609375 0 4187.872
1417.7275390625 0 3396.9485
1422.7022705078125 0 1757.6989 y 2
1423.700927734375 0 3974.3267
1424.703125 0 2039.8063
1430.7572021484375 0 2859.6262
1431.756103515625 0 13586.13
1432.7542724609375 0 13112.27
1433.751953125 0 6120.588
1434.7613525390625 0 1925.1234
1451.6873779296875 0 1993.3685
1452.694091796875 0 1756.5562
1453.692138671875 0 1293.8502
1458.739990234375 0 10749.342
1459.74658203125 0 9649.332
1460.7484130859375 0 4590.029
1461.746826171875 0 1554.8435
1474.760009765625 0 19461.01 c 13
1475.766845703125 0 69216.94
1476.766357421875 0 45319.184
1477.75927734375 0 19991.955
1478.7542724609375 0 5579.532
1479.7529296875 0 1818.2195
1493.7138671875 0 5127.9243 z 1
1494.716552734375 0 40606.652
1495.72119140625 0 29548.1
1496.7225341796875 0 14872.84
1497.72265625 0 4909.2314
1507.73388671875 0 1632.6941
1508.748291015625 0 2669.634
1509.7493896484375 0 2802.0037 y 1
1510.7464599609375 0 2312.855
1522.7286376953125 0 11547.26
1523.728759765625 0 10412.15
1524.7386474609375 0 7825.367
1525.75048828125 0 1962.3224
1537.7843017578125 0 2765.231
1538.775634765625 0 1255.352
1546.74658203125 0 1396.2743
1547.773193359375 0 2003.8998
1548.767578125 0 4679.946
1549.7607421875 0 3074.7476
1550.7689208984375 0 1406.412
1562.806884765625 0 1605.727
1563.79638671875 0 3773.8904
1564.7828369140625 0 55213.25
1565.784912109375 0 70292.87
1566.7860107421875 0 47760.973
1567.78662109375 0 20830.186
1568.790771484375 0 7771.2607
1569.787353515625 0 1308.8467
1577.791259765625 0 1999.4608
1578.821044921875 0 9053.765
1579.8165283203125 0 13673.745
1580.8094482421875 0 15184.005
1581.8011474609375 0 20328.012
1582.8026123046875 0 11544.853
1583.807861328125 0 4730.5967
1584.8076171875 0 1951.1404
1589.79443359375 0 5606.48
1590.7799072265625 0 21415.361
1591.779541015625 0 13346.335
1592.7777099609375 0 7228.1714
1593.788818359375 0 3814.7324
1595.8309326171875 0 3678.8716
1596.830078125 0 20394.43
1597.83447265625 0 15681.658
1598.8387451171875 0 6715.4526
1599.8309326171875 0 2038.9149
1605.788330078125 0 1485.6619
1606.8031005859375 0 36393.383
1607.8021240234375 0 196965.39
1608.804443359375 0 155789.98
1609.8062744140625 0 77582.54
1610.810546875 0 22984.07
1611.8115234375 0 6385.1997
1622.8133544921875 0 9638.444
1623.5699462890625 0 2139.4553
1623.8182373046875 0 54074.082
1624.82568359375 0 187534.69
1625.8289794921875 0 157989.8
1626.832275390625 0 75597.81
1627.8330078125 0 23025.225
1628.8397216796875 0 7157.8027
3082.639404296875 0 1289.8507

Spectrum Details

|  |  |
| --- | --- |
| Matched peaks? Matched peaks The total absolute number of peaks matched. Additionally in brackets the total fraction of peaks matched and the total number of peaks is shown. | 86 (16.20% of 531) |
| FDR? FDR The false discovery rate estimated for this peptide. It is calculated by matching all theoretical fragments with a non-integer shift with the raw peaks for this spectrum. This is done with 40 different shifts. The resulting percentage is the average number of annotated peaks over the number of annotated peaks with the correct spectrum. | 0.69% |
| Satellite FDR? Satellite FDR See the FDR for details on its calculation. This satellite ion specific FDR only contains the satellite ions (d/w) for I/L/J positions. | - |
| PSM Score? PSM Score The PSM Score as given by Hecklib to this annotated spectrum. It is shown with three significant figures. | 680 |

#### Spectrum 4665? Spectrum 4665 The raw spectrum of this peptide as annotated by Hecklib. The fragments are coloured according to ion type (see legend). Any peaks with a star '\*' as text can be hovered over to see the full details, first the ion type second the mass shift type. By hovering over the amino acids in the peptide or ions in the legend the corresponding peaks are highlighted. By toggling the 'Unassigned' label you can turn the background (unassigned) peaks on or off in the plot. By updating the slider in the Ion legend you can update the spectrum to only show the top X% of the peaks with labels. The top X% means any peak that is within X% of the highest intensity. By dragging in the spectrum you can zoom in to a specific part of the spectrum and use 'Zoom Out' to get back to the original zoom level. The annotation of the spectrum is based on the given sequence in the peptides file and is done with different software so inconsistencies are likely. The peaks are annotated based on the given sequence, with 20 ppm tolerance.

Copy Data

##### Spectrum 4665 (TSV)

###### Preview

```
Loading example...
```

*Click on the button to copy the data to your clipboard.*

Mz MinMz MaxIntensity Max

WidthHeightPeptide font sizePeptide stroke widthSpectrum font sizeSpectrum stroke widthCompact peptide

Ion legend

wxyz

abcd

OtherUnassignedIon^Charge^_Position_Show for top:%

JSEVSDRPSGVSSRF

06.83e+51.37e+62.05e+62.73e+6

Zoom Out

a^+1^_2_d^+1^_2_y^+1^_1_a^+1^_2_b^+1^_2_y^+3^_5_b^+1^_2_y^+2^_4_b^+3^_7_a^+1^_3_y^+2^_5_a^+1^_3_y^+1^_2_b^+1^_3_b^+2^_6_y^+1^_2_b^+1^_3_y^+1^_3_y^+1^_3_b^+2^_7_y^+3^_11_y^+1^_3_y^+2^_8_b^+1^_4_y^+2^_8_b^+1^_4_y^+3^_12_y^+3^_13_y^+3^_13_y^+3^_13_y^+1^_4_y^+1^_4_b^+3^_14_y^+2^_9_y^+2^_9_y^+1^_4_y^+2^_9_y^+3^_14_y^+3^_14_b^+1^_5_\*\*\*y^+2^_10_y^+2^_10_y^+2^_10_b^+2^_11_b^+2^_11_y^+2^_11_y^+2^_11_y^+1^_5_y^+2^_11_b^+2^_12_b^+2^_12_b^+1^_6_y^+1^_6_y^+2^_12_y^+2^_12_b^+2^_13_y^+2^_12_y^+1^_6_y^+2^_13_y^+2^_13_y^+2^_13_y^+1^_7_y^+1^_7_y^+1^_7_y^+2^_14_y^+2^_14_y^+2^_14_b^+1^_7_b^+1^_7_y^+1^_8_y^+1^_8_y^+1^_8_b^+1^_8_b^+1^_9_y^+1^_9_b^+1^_10_y^+1^_10_y^+1^_11_y^+1^_12_

037875611331511

Fragment Matches Table

Show background peaks

| Position | Ion type | Intensity | mz Theoretical | mz Error (Th) | mz Error (ppm) | Charge | Series Number |
| --- | --- | --- | --- | --- | --- | --- | --- |
| - | - | 6.351E+04 | 120.1 | - | - | 0 | - |
| - | - | 4835 | 121.1 | - | - | 0 | - |
| - | - | 1574 | 125.1 | - | - | 0 | - |
| - | - | 2.366E+04 | 125.1 | - | - | 0 | - |
| - | - | 2.774E+04 | 126.1 | - | - | 0 | - |
| - | - | 1941 | 126.4 | - | - | 0 | - |
| - | - | 2353 | 127.1 | - | - | 0 | - |
| - | - | 1.816E+04 | 128.1 | - | - | 0 | - |
| - | - | 5988 | 129.1 | - | - | 0 | - |
| - | - | 1.725E+04 | 129.1 | - | - | 0 | - |
| - | - | 2166 | 129.1 | - | - | 0 | - |
| - | - | 4763 | 130.1 | - | - | 0 | - |
| - | - | 5876 | 130.1 | - | - | 0 | - |
| - | - | 5056 | 130.1 | - | - | 0 | - |
| - | - | 3324 | 131.1 | - | - | 0 | - |
| - | - | 4508 | 132.1 | - | - | 0 | - |
| - | - | 1980 | 132.7 | - | - | 0 | - |
| - | - | 3145 | 133.1 | - | - | 0 | - |
| - | - | 2561 | 138.1 | - | - | 0 | - |
| - | - | 1.018E+04 | 138.1 | - | - | 0 | - |
| - | - | 2003 | 139 | - | - | 0 | - |
| - | - | 3383 | 139.1 | - | - | 0 | - |
| - | - | 9680 | 140.1 | - | - | 0 | - |
| - | - | 2588 | 141.1 | - | - | 0 | - |
| - | - | 9625 | 144.1 | - | - | 0 | - |
| - | - | 2746 | 145.1 | - | - | 0 | - |
| - | - | 1928 | 147.1 | - | - | 0 | - |
| - | - | 1.274E+04 | 147.1 | - | - | 0 | - |
| - | - | 2495 | 149 | - | - | 0 | - |
| - | - | 4042 | 153.1 | - | - | 0 | - |
| - | - | 3.727E+04 | 154.1 | - | - | 0 | - |
| - | - | 3852 | 155.1 | - | - | 0 | - |
| 2 | a | 3.249E+04 | 155.1 | 0.0004424 | 2.852 | +1 | 2 |
| - | - | 1.424E+04 | 156.1 | - | - | 0 | - |
| - | - | 2.684E+04 | 157.1 | - | - | 0 | - |
| - | - | 3.067E+04 | 157.1 | - | - | 0 | - |
| - | - | 1.891E+04 | 157.1 | - | - | 0 | - |
| 2 | d | 2806 | 157.1 | 0.0007072 | 4.501 | +1 | 2 |
| - | - | 2295 | 158.1 | - | - | 0 | - |
| - | - | 3.52E+04 | 158.1 | - | - | 0 | - |
| - | - | 4078 | 158.1 | - | - | 0 | - |
| - | - | 2017 | 158.1 | - | - | 0 | - |
| - | - | 1.972E+04 | 159.1 | - | - | 0 | - |
| - | - | 1992 | 162.4 | - | - | 0 | - |
| - | - | 2718 | 165.1 | - | - | 0 | - |
| 15 | y | 3.927E+04 | 166.1 | 0.0004149 | 2.498 | +1 | 1 |
| - | - | 2356 | 166.1 | - | - | 0 | - |
| - | - | 3153 | 166.1 | - | - | 0 | - |
| - | - | 8686 | 167.1 | - | - | 0 | - |
| - | - | 4596 | 167.1 | - | - | 0 | - |
| - | - | 1819 | 169 | - | - | 0 | - |
| - | - | 7429 | 169.1 | - | - | 0 | - |
| - | - | 2061 | 169.1 | - | - | 0 | - |
| - | - | 2268 | 170 | - | - | 0 | - |
| - | - | 4.386E+04 | 171.1 | - | - | 0 | - |
| - | - | 6049 | 172.1 | - | - | 0 | - |
| - | - | 4078 | 172.1 | - | - | 0 | - |
| 2 | a | 4.709E+05 | 173.1 | 0.000452 | 2.611 | +1 | 2 |
| - | - | 5743 | 173.4 | - | - | 0 | - |
| - | - | 4544 | 174.1 | - | - | 0 | - |
| - | - | 4.579E+04 | 174.1 | - | - | 0 | - |
| - | - | 2230 | 174.6 | - | - | 0 | - |
| - | - | 5.272E+04 | 175.1 | - | - | 0 | - |
| - | - | 5.125E+04 | 175.1 | - | - | 0 | - |
| - | - | 3004 | 176.1 | - | - | 0 | - |
| - | - | 3243 | 176.1 | - | - | 0 | - |
| - | - | 3647 | 181.1 | - | - | 0 | - |
| - | - | 2659 | 181.1 | - | - | 0 | - |
| - | - | 6974 | 182.1 | - | - | 0 | - |
| 2 | b | 3.053E+05 | 183.1 | 0.0004618 | 2.522 | +1 | 2 |
| - | - | 5306 | 184.1 | - | - | 0 | - |
| - | - | 2.943E+04 | 184.1 | - | - | 0 | - |
| - | - | 3247 | 185.1 | - | - | 0 | - |
| - | - | 3.255E+04 | 185.1 | - | - | 0 | - |
| - | - | 2175 | 185.1 | - | - | 0 | - |
| - | - | 3485 | 187.1 | - | - | 0 | - |
| - | - | 3.482E+04 | 187.1 | - | - | 0 | - |
| - | - | 2518 | 188.1 | - | - | 0 | - |
| - | - | 1.895E+05 | 189.1 | - | - | 0 | - |
| - | - | 1.505E+04 | 190.1 | - | - | 0 | - |
| - | - | 2620 | 191.1 | - | - | 0 | - |
| - | - | 4538 | 192.1 | - | - | 0 | - |
| - | - | 6281 | 194.1 | - | - | 0 | - |
| - | - | 2560 | 195.1 | - | - | 0 | - |
| - | - | 4000 | 195.1 | - | - | 0 | - |
| - | - | 2123 | 197.1 | - | - | 0 | - |
| - | - | 2235 | 197.1 | - | - | 0 | - |
| - | - | 7.725E+04 | 199.1 | - | - | 0 | - |
| 11 | y | 4833 | 199.1 | 0.003843 | 19.3 | +3 | 5 |
| - | - | 2102 | 199.1 | - | - | 0 | - |
| - | - | 5421 | 200.1 | - | - | 0 | - |
| 2 | b | 6.117E+05 | 201.1 | 0.0004257 | 2.117 | +1 | 2 |
| - | - | 1.379E+04 | 202.1 | - | - | 0 | - |
| - | - | 5.56E+04 | 202.1 | - | - | 0 | - |
| - | - | 5.459E+04 | 203.1 | - | - | 0 | - |
| - | - | 5017 | 203.1 | - | - | 0 | - |
| - | - | 4431 | 203.1 | - | - | 0 | - |
| - | - | 3138 | 204.1 | - | - | 0 | - |
| - | - | 3512 | 210.1 | - | - | 0 | - |
| - | - | 3.139E+04 | 211.1 | - | - | 0 | - |
| - | - | 4249 | 211.1 | - | - | 0 | - |
| - | - | 1.043E+04 | 211.2 | - | - | 0 | - |
| - | - | 1.883E+04 | 212.1 | - | - | 0 | - |
| - | - | 2597 | 213.1 | - | - | 0 | - |
| - | - | 2675 | 216.1 | - | - | 0 | - |
| - | - | 3.314E+05 | 217.1 | - | - | 0 | - |
| - | - | 3.014E+04 | 218.1 | - | - | 0 | - |
| - | - | 4293 | 219.1 | - | - | 0 | - |
| - | - | 3541 | 220.1 | - | - | 0 | - |
| - | - | 2429 | 220.1 | - | - | 0 | - |
| - | - | 3323 | 223.1 | - | - | 0 | - |
| - | - | 2.737E+04 | 224.1 | - | - | 0 | - |
| - | - | 7527 | 226.1 | - | - | 0 | - |
| - | - | 6215 | 227.1 | - | - | 0 | - |
| - | - | 4519 | 227.1 | - | - | 0 | - |
| - | - | 5.715E+04 | 227.1 | - | - | 0 | - |
| - | - | 4877 | 228.1 | - | - | 0 | - |
| - | - | 3.48E+04 | 229.1 | - | - | 0 | - |
| - | - | 1.298E+04 | 230.1 | - | - | 0 | - |
| - | - | 2571 | 236.1 | - | - | 0 | - |
| - | - | 8833 | 237.1 | - | - | 0 | - |
| - | - | 1.145E+04 | 237.1 | - | - | 0 | - |
| - | - | 5.104E+04 | 242.1 | - | - | 0 | - |
| - | - | 3981 | 242.1 | - | - | 0 | - |
| - | - | 5107 | 243.1 | - | - | 0 | - |
| - | - | 1.615E+04 | 244.1 | - | - | 0 | - |
| - | - | 3566 | 244.1 | - | - | 0 | - |
| - | - | 1.776E+04 | 245.1 | - | - | 0 | - |
| 12 | y | 9766 | 248.6 | 0.0004721 | 1.899 | +2 | 4 |
| - | - | 2788 | 250.1 | - | - | 0 | - |
| - | - | 6294 | 254.1 | - | - | 0 | - |
| - | - | 4276 | 254.2 | - | - | 0 | - |
| - | - | 4604 | 255.1 | - | - | 0 | - |
| - | - | 6.289E+04 | 255.1 | - | - | 0 | - |
| - | - | 5342 | 256.1 | - | - | 0 | - |
| - | - | 8241 | 259.1 | - | - | 0 | - |
| - | - | 4601 | 259.2 | - | - | 0 | - |
| - | - | 2.296E+04 | 262.2 | - | - | 0 | - |
| 7 | b | 1.127E+04 | 263.1 | 0.003067 | 11.66 | +3 | 7 |
| - | - | 3143 | 263.2 | - | - | 0 | - |
| - | - | 2587 | 266.1 | - | - | 0 | - |
| - | - | 3624 | 267.1 | - | - | 0 | - |
| - | - | 2355 | 268.1 | - | - | 0 | - |
| - | - | 1.108E+04 | 270.1 | - | - | 0 | - |
| - | - | 3451 | 271.1 | - | - | 0 | - |
| - | - | 2668 | 271.1 | - | - | 0 | - |
| - | - | 1.077E+04 | 272.1 | - | - | 0 | - |
| - | - | 2785 | 278.1 | - | - | 0 | - |
| - | - | 7215 | 279.1 | - | - | 0 | - |
| - | - | 8083 | 280.1 | - | - | 0 | - |
| - | - | 4736 | 281.1 | - | - | 0 | - |
| - | - | 1.151E+04 | 284.1 | - | - | 0 | - |
| 3 | a | 2.357E+04 | 284.2 | 0.0004671 | 1.644 | +1 | 3 |
| - | - | 4107 | 285.2 | - | - | 0 | - |
| - | - | 2887 | 286.2 | - | - | 0 | - |
| - | - | 2603 | 288.1 | - | - | 0 | - |
| - | - | 1.279E+04 | 288.2 | - | - | 0 | - |
| 11 | y | 3474 | 289.2 | 0.005259 | 18.19 | +2 | 5 |
| - | - | 2992 | 290.1 | - | - | 0 | - |
| - | - | 2783 | 294.1 | - | - | 0 | - |
| - | - | 5625 | 295.2 | - | - | 0 | - |
| - | - | 1.474E+04 | 296.1 | - | - | 0 | - |
| - | - | 3356 | 297.1 | - | - | 0 | - |
| - | - | 3.153E+04 | 298.1 | - | - | 0 | - |
| - | - | 1.333E+04 | 299.1 | - | - | 0 | - |
| - | - | 4604 | 302.1 | - | - | 0 | - |
| 3 | a | 5288 | 302.2 | 0.0002478 | 0.8202 | +1 | 3 |
| 14 | y | 1.001E+05 | 305.2 | 0.0006515 | 2.135 | +1 | 2 |
| - | - | 2820 | 306.1 | - | - | 0 | - |
| - | - | 1.889E+04 | 306.2 | - | - | 0 | - |
| - | - | 2834 | 307.1 | - | - | 0 | - |
| - | - | 2465 | 308 | - | - | 0 | - |
| 3 | b | 2.97E+05 | 312.2 | 0.0006696 | 2.145 | +1 | 3 |
| - | - | 2404 | 313.1 | - | - | 0 | - |
| - | - | 5.028E+04 | 313.2 | - | - | 0 | - |
| - | - | 6.517E+04 | 313.2 | - | - | 0 | - |
| - | - | 3.902E+04 | 314.1 | - | - | 0 | - |
| - | - | 1.118E+04 | 314.2 | - | - | 0 | - |
| - | - | 8410 | 315.1 | - | - | 0 | - |
| 6 | b | 6.821E+04 | 316.2 | 0.000567 | 1.794 | +2 | 6 |
| - | - | 1.015E+04 | 317.2 | - | - | 0 | - |
| 14 | y | 2.084E+04 | 322.2 | 0.0009884 | 3.068 | +1 | 2 |
| - | - | 1.329E+04 | 324.1 | - | - | 0 | - |
| - | - | 3311 | 324.2 | - | - | 0 | - |
| - | - | 2421 | 325.1 | - | - | 0 | - |
| - | - | 3310 | 326.1 | - | - | 0 | - |
| 3 | b | 1.475E+05 | 330.2 | 0.0007251 | 2.196 | +1 | 3 |
| - | - | 3.466E+04 | 331.2 | - | - | 0 | - |
| - | - | 1.43E+04 | 332.2 | - | - | 0 | - |
| - | - | 2402 | 333.2 | - | - | 0 | - |
| - | - | 8142 | 336.2 | - | - | 0 | - |
| - | - | 2805 | 341.2 | - | - | 0 | - |
| - | - | 3.652E+04 | 341.2 | - | - | 0 | - |
| - | - | 7.917E+04 | 342.1 | - | - | 0 | - |
| - | - | 6890 | 342.2 | - | - | 0 | - |
| - | - | 1.554E+04 | 343.1 | - | - | 0 | - |
| - | - | 3517 | 344.1 | - | - | 0 | - |
| - | - | 9704 | 345.2 | - | - | 0 | - |
| - | - | 5230 | 345.7 | - | - | 0 | - |
| - | - | 2.711E+04 | 349.2 | - | - | 0 | - |
| - | - | 2721 | 350.1 | - | - | 0 | - |
| - | - | 2983 | 350.2 | - | - | 0 | - |
| - | - | 5151 | 357.2 | - | - | 0 | - |
| - | - | 4.912E+04 | 359.2 | - | - | 0 | - |
| - | - | 8769 | 360.2 | - | - | 0 | - |
| - | - | 3610 | 365.2 | - | - | 0 | - |
| - | - | 6991 | 367.2 | - | - | 0 | - |
| - | - | 4961 | 375.2 | - | - | 0 | - |
| - | - | 5618 | 380.2 | - | - | 0 | - |
| - | - | 4570 | 381.2 | - | - | 0 | - |
| - | - | 2785 | 382.2 | - | - | 0 | - |
| - | - | 5529 | 383.2 | - | - | 0 | - |
| - | - | 1.169E+04 | 385.2 | - | - | 0 | - |
| - | - | 2943 | 386.2 | - | - | 0 | - |
| 13 | y | 3393 | 391.2 | 0.001406 | 3.593 | +1 | 3 |
| 13 | y | 1.779E+04 | 392.2 | 0.0007581 | 1.933 | +1 | 3 |
| - | - | 6178 | 393.7 | - | - | 0 | - |
| 7 | b | 6893 | 394.2 | 0.002075 | 5.263 | +2 | 7 |
| - | - | 3962 | 394.7 | - | - | 0 | - |
| - | - | 6622 | 395.2 | - | - | 0 | - |
| - | - | 2810 | 395.2 | - | - | 0 | - |
| - | - | 4862 | 396.2 | - | - | 0 | - |
| - | - | 7504 | 396.7 | - | - | 0 | - |
| - | - | 7892 | 398.2 | - | - | 0 | - |
| 5 | y | 3480 | 398.9 | 0.001278 | 3.205 | +3 | 11 |
| - | - | 2847 | 399.7 | - | - | 0 | - |
| - | - | 8681 | 400.2 | - | - | 0 | - |
| - | - | 5.065E+04 | 401.2 | - | - | 0 | - |
| - | - | 1.147E+04 | 402.2 | - | - | 0 | - |
| - | - | 8723 | 403.2 | - | - | 0 | - |
| - | - | 2724 | 404.2 | - | - | 0 | - |
| - | - | 3178 | 405.7 | - | - | 0 | - |
| - | - | 3567 | 408.2 | - | - | 0 | - |
| 13 | y | 3.65E+04 | 409.2 | 0.0009424 | 2.303 | +1 | 3 |
| 8 | y | 9191 | 410.2 | 0.001485 | 3.62 | +2 | 8 |
| 4 | b | 1.065E+04 | 411.2 | 0.0003404 | 0.8279 | +1 | 4 |
| - | - | 3768 | 412.2 | - | - | 0 | - |
| - | - | 2647 | 413.2 | - | - | 0 | - |
| - | - | 5007 | 413.2 | - | - | 0 | - |
| - | - | 2753 | 414.2 | - | - | 0 | - |
| - | - | 1.968E+04 | 414.2 | - | - | 0 | - |
| - | - | 3388 | 414.3 | - | - | 0 | - |
| - | - | 3.034E+04 | 414.7 | - | - | 0 | - |
| - | - | 1.133E+04 | 415.2 | - | - | 0 | - |
| 8 | y | 3.463E+04 | 418.7 | 0.001058 | 2.527 | +2 | 8 |
| - | - | 1.286E+04 | 419.2 | - | - | 0 | - |
| - | - | 6288 | 419.7 | - | - | 0 | - |
| - | - | 1.123E+05 | 423.2 | - | - | 0 | - |
| - | - | 4.88E+04 | 423.7 | - | - | 0 | - |
| - | - | 1.367E+04 | 424.2 | - | - | 0 | - |
| - | - | 1.742E+04 | 428.2 | - | - | 0 | - |
| 4 | b | 4.366E+04 | 429.2 | 0.0003654 | 0.8512 | +1 | 4 |
| - | - | 2.359E+04 | 430.2 | - | - | 0 | - |
| 4 | y | 3369 | 431.9 | 0.003056 | 7.077 | +3 | 12 |
| - | - | 3634 | 432.2 | - | - | 0 | - |
| - | - | 4365 | 439.2 | - | - | 0 | - |
| - | - | 2.042E+04 | 441.2 | - | - | 0 | - |
| - | - | 2672 | 441.7 | - | - | 0 | - |
| - | - | 4419 | 442.2 | - | - | 0 | - |
| - | - | 1.195E+04 | 442.3 | - | - | 0 | - |
| - | - | 3362 | 443.2 | - | - | 0 | - |
| - | - | 7220 | 448.3 | - | - | 0 | - |
| - | - | 3239 | 450.7 | - | - | 0 | - |
| - | - | 1.243E+04 | 452.3 | - | - | 0 | - |
| - | - | 2994 | 454.2 | - | - | 0 | - |
| - | - | 3288 | 455.2 | - | - | 0 | - |
| - | - | 3925 | 456.2 | - | - | 0 | - |
| - | - | 3.375E+04 | 458.2 | - | - | 0 | - |
| - | - | 7306 | 459.2 | - | - | 0 | - |
| - | - | 3794 | 461.2 | - | - | 0 | - |
| - | - | 3558 | 463.2 | - | - | 0 | - |
| - | - | 2739 | 463.6 | - | - | 0 | - |
| - | - | 8173 | 467.2 | - | - | 0 | - |
| - | - | 3907 | 467.7 | - | - | 0 | - |
| 3 | y | 1.008E+04 | 468.9 | 0.0001216 | 0.2594 | +3 | 13 |
| 3 | y | 8792 | 469.2 | 0.008146 | 17.36 | +3 | 13 |
| - | - | 1.466E+04 | 469.3 | - | - | 0 | - |
| - | - | 3435 | 469.6 | - | - | 0 | - |
| - | - | 3353 | 470.2 | - | - | 0 | - |
| 3 | y | 6175 | 474.9 | 0.0003778 | 0.7955 | +3 | 13 |
| - | - | 5539 | 475.2 | - | - | 0 | - |
| - | - | 3997 | 478.2 | - | - | 0 | - |
| 12 | y | 4314 | 478.2 | 0.0008375 | 1.751 | +1 | 4 |
| 12 | y | 2.203E+04 | 479.2 | 0.001078 | 2.25 | +1 | 4 |
| - | - | 1.282E+04 | 480.7 | - | - | 0 | - |
| - | - | 9780 | 481.3 | - | - | 0 | - |
| - | - | 4308 | 481.8 | - | - | 0 | - |
| - | - | 3333 | 482.6 | - | - | 0 | - |
| - | - | 3647 | 482.9 | - | - | 0 | - |
| 14 | b | 3697 | 486.6 | 0.0005464 | 1.123 | +3 | 14 |
| - | - | 5513 | 487.3 | - | - | 0 | - |
| 7 | y | 1.24E+04 | 487.8 | 0.001348 | 2.763 | +2 | 9 |
| 7 | y | 3.005E+04 | 488.3 | 0.001589 | 3.254 | +2 | 9 |
| - | - | 1.481E+04 | 488.8 | - | - | 0 | - |
| - | - | 5363 | 489.3 | - | - | 0 | - |
| - | - | 2901 | 489.8 | - | - | 0 | - |
| - | - | 4016 | 496.2 | - | - | 0 | - |
| 12 | y | 1.407E+05 | 496.3 | 0.0009575 | 1.929 | +1 | 4 |
| 7 | y | 3.612E+05 | 496.8 | 0.0006737 | 1.356 | +2 | 9 |
| - | - | 2.118E+05 | 497.3 | - | - | 0 | - |
| - | - | 5.538E+04 | 497.8 | - | - | 0 | - |
| 2 | y | 6023 | 497.9 | 0.0006049 | 1.215 | +3 | 14 |
| - | - | 1.162E+04 | 498.3 | - | - | 0 | - |
| - | - | 4277 | 498.7 | - | - | 0 | - |
| - | - | 5327 | 500.2 | - | - | 0 | - |
| - | - | 3393 | 501.6 | - | - | 0 | - |
| 2 | y | 1.482E+04 | 503.9 | 0.0003291 | 0.6531 | +3 | 14 |
| - | - | 2.443E+04 | 504.2 | - | - | 0 | - |
| - | - | 9408 | 504.6 | - | - | 0 | - |
| - | - | 2.03E+04 | 505.3 | - | - | 0 | - |
| - | - | 4703 | 506.3 | - | - | 0 | - |
| - | - | 3136 | 511.3 | - | - | 0 | - |
| - | - | 3355 | 513.2 | - | - | 0 | - |
| - | - | 5.162E+04 | 513.4 | - | - | 0 | - |
| - | - | 1.27E+04 | 514.4 | - | - | 0 | - |
| - | - | 2.262E+04 | 515.3 | - | - | 0 | - |
| - | - | 6747 | 515.8 | - | - | 0 | - |
| 5 | b | 1.204E+04 | 516.3 | 0.002244 | 4.347 | +1 | 5 |
| - | - | 4629 | 518.2 | - | - | 0 | - |
| - | - | 1.097E+05 | 524.3 | - | - | 0 | - |
| - | - | 5.712E+04 | 524.8 | - | - | 0 | - |
| - | - | 1.794E+04 | 525.3 | - | - | 0 | - |
| - | - | 5312 | 525.8 | - | - | 0 | - |
| - | - | 4741 | 526.3 | - | - | 0 | - |
| - | - | 8351 | 529.6 | - | - | 0 | - |
| - | - | 5485 | 529.9 | - | - | 0 | - |
| - | - | 3140 | 530.3 | - | - | 0 | - |
| - | - | 7533 | 532.3 | - | - | 0 | - |
| 0 | Precursor | 7.177E+04 | 535.6 | 0.0005689 | 1.062 | +3 | -1 |
| 0 | Precursor | 5.978E+04 | 535.9 | 0.006792 | 12.67 | +3 | -1 |
| - | - | 3.125E+04 | 536.3 | - | - | 0 | - |
| - | - | 1.309E+04 | 536.6 | - | - | 0 | - |
| - | - | 4673 | 536.9 | - | - | 0 | - |
| - | - | 2859 | 537.8 | - | - | 0 | - |
| - | - | 4978 | 539.3 | - | - | 0 | - |
| - | - | 7.294E+04 | 541.3 | - | - | 0 | - |
| 0 | Precursor | 1.45E+05 | 541.6 | 0.0007095 | 1.31 | +3 | -1 |
| - | - | 1.438E+05 | 541.9 | - | - | 0 | - |
| - | - | 8.305E+04 | 542.3 | - | - | 0 | - |
| - | - | 2.052E+04 | 542.3 | - | - | 0 | - |
| - | - | 2.166E+04 | 542.6 | - | - | 0 | - |
| - | - | 8219 | 542.9 | - | - | 0 | - |
| - | - | 6123 | 543.3 | - | - | 0 | - |
| 6 | y | 4707 | 545.3 | 0.001565 | 2.869 | +2 | 10 |
| 6 | y | 9310 | 545.8 | 0.004963 | 9.093 | +2 | 10 |
| - | - | 5313 | 546.3 | - | - | 0 | - |
| - | - | 5342 | 547.2 | - | - | 0 | - |
| - | - | 4928 | 550.8 | - | - | 0 | - |
| 6 | y | 1.463E+05 | 554.3 | 0.0003552 | 0.6409 | +2 | 10 |
| - | - | 9.299E+04 | 554.8 | - | - | 0 | - |
| 11 | b | 2.951E+04 | 555.3 | 0.001016 | 1.829 | +2 | 11 |
| - | - | 4768 | 555.8 | - | - | 0 | - |
| - | - | 5421 | 556.3 | - | - | 0 | - |
| - | - | 3950 | 556.3 | - | - | 0 | - |
| - | - | 4211 | 557.3 | - | - | 0 | - |
| - | - | 3423 | 557.3 | - | - | 0 | - |
| - | - | 3962 | 558.8 | - | - | 0 | - |
| - | - | 8683 | 559.3 | - | - | 0 | - |
| 11 | b | 4268 | 564.3 | 6.67E-05 | 0.1182 | +2 | 11 |
| - | - | 2.926E+04 | 564.8 | - | - | 0 | - |
| - | - | 4413 | 565.2 | - | - | 0 | - |
| - | - | 1.171E+04 | 565.3 | - | - | 0 | - |
| - | - | 6897 | 565.8 | - | - | 0 | - |
| - | - | 4727 | 566.3 | - | - | 0 | - |
| - | - | 4193 | 567.3 | - | - | 0 | - |
| - | - | 4294 | 568.8 | - | - | 0 | - |
| - | - | 2.411E+04 | 569.3 | - | - | 0 | - |
| - | - | 3580 | 570.3 | - | - | 0 | - |
| - | - | 6119 | 572.3 | - | - | 0 | - |
| - | - | 1.103E+05 | 573.8 | - | - | 0 | - |
| - | - | 8.214E+04 | 574.3 | - | - | 0 | - |
| - | - | 1.792E+04 | 574.8 | - | - | 0 | - |
| - | - | 8124 | 575.3 | - | - | 0 | - |
| - | - | 8613 | 582.8 | - | - | 0 | - |
| - | - | 4905 | 583.2 | - | - | 0 | - |
| - | - | 6350 | 583.3 | - | - | 0 | - |
| - | - | 1.283E+04 | 584.3 | - | - | 0 | - |
| - | - | 5154 | 585.3 | - | - | 0 | - |
| - | - | 2.939E+04 | 587.3 | - | - | 0 | - |
| - | - | 8934 | 588.3 | - | - | 0 | - |
| 5 | y | 1.672E+04 | 588.8 | 0.0007318 | 1.243 | +2 | 11 |
| 5 | y | 2.041E+04 | 589.3 | 0.005855 | 9.936 | +2 | 11 |
| - | - | 9942 | 589.8 | - | - | 0 | - |
| - | - | 3.229E+04 | 592.3 | - | - | 0 | - |
| - | - | 1.338E+04 | 593.3 | - | - | 0 | - |
| 11 | y | 2.391E+04 | 595.3 | 0.0007809 | 1.312 | +1 | 5 |
| - | - | 5951 | 596.3 | - | - | 0 | - |
| 5 | y | 1.162E+06 | 597.8 | 0.0005764 | 0.9642 | +2 | 11 |
| - | - | 7.282E+05 | 598.3 | - | - | 0 | - |
| 12 | b | 2.808E+05 | 598.8 | 0.0009926 | 1.658 | +2 | 12 |
| 12 | b | 6.87E+04 | 599.3 | 0.01014 | 16.93 | +2 | 12 |
| - | - | 1.447E+04 | 599.8 | - | - | 0 | - |
| - | - | 2.238E+04 | 600.3 | - | - | 0 | - |
| - | - | 1.036E+04 | 601.3 | - | - | 0 | - |
| - | - | 3642 | 611.3 | - | - | 0 | - |
| - | - | 3138 | 611.8 | - | - | 0 | - |
| - | - | 3086 | 612.3 | - | - | 0 | - |
| - | - | 3923 | 617.3 | - | - | 0 | - |
| - | - | 4264 | 618.3 | - | - | 0 | - |
| - | - | 3972 | 620.3 | - | - | 0 | - |
| - | - | 6288 | 620.8 | - | - | 0 | - |
| - | - | 3818 | 623.3 | - | - | 0 | - |
| - | - | 5517 | 624.3 | - | - | 0 | - |
| - | - | 3893 | 625.3 | - | - | 0 | - |
| - | - | 3233 | 625.8 | - | - | 0 | - |
| - | - | 9871 | 626.3 | - | - | 0 | - |
| - | - | 4562 | 627.3 | - | - | 0 | - |
| - | - | 5.139E+04 | 629.3 | - | - | 0 | - |
| - | - | 2.546E+04 | 629.8 | - | - | 0 | - |
| - | - | 1.979E+04 | 630.3 | - | - | 0 | - |
| - | - | 6175 | 630.8 | - | - | 0 | - |
| 6 | b | 3102 | 631.3 | 0.008472 | 13.42 | +1 | 6 |
| - | - | 3723 | 631.8 | - | - | 0 | - |
| - | - | 6815 | 632.3 | - | - | 0 | - |
| - | - | 5103 | 632.8 | - | - | 0 | - |
| - | - | 4365 | 633.3 | - | - | 0 | - |
| 10 | y | 3250 | 634.3 | 0.0007798 | 1.229 | +1 | 6 |
| - | - | 8397 | 636.3 | - | - | 0 | - |
| 4 | y | 1.673E+05 | 638.3 | 0.005826 | 9.127 | +2 | 12 |
| 4 | y | 1.221E+05 | 638.8 | 0.002776 | 4.346 | +2 | 12 |
| - | - | 3805 | 639.3 | - | - | 0 | - |
| - | - | 5.481E+04 | 639.3 | - | - | 0 | - |
| - | - | 1.751E+04 | 639.8 | - | - | 0 | - |
| - | - | 4594 | 640.3 | - | - | 0 | - |
| - | - | 4999 | 641.3 | - | - | 0 | - |
| 13 | b | 7239 | 642.3 | 0.006768 | 10.54 | +2 | 13 |
| - | - | 1.359E+04 | 643.4 | - | - | 0 | - |
| - | - | 4334 | 644.4 | - | - | 0 | - |
| - | - | 1.69E+04 | 646.3 | - | - | 0 | - |
| - | - | 5471 | 646.8 | - | - | 0 | - |
| 4 | y | 1.038E+06 | 647.3 | 0.0006712 | 1.037 | +2 | 12 |
| - | - | 7.21E+05 | 647.8 | - | - | 0 | - |
| - | - | 2.98E+05 | 648.3 | - | - | 0 | - |
| - | - | 9.029E+04 | 648.8 | - | - | 0 | - |
| - | - | 1.403E+04 | 649.3 | - | - | 0 | - |
| 10 | y | 6.359E+04 | 652.3 | 0.0004353 | 0.6673 | +1 | 6 |
| - | - | 3.257E+04 | 653.3 | - | - | 0 | - |
| - | - | 1.732E+04 | 654.3 | - | - | 0 | - |
| - | - | 8886 | 655.3 | - | - | 0 | - |
| - | - | 7843 | 656.3 | - | - | 0 | - |
| - | - | 4151 | 657.3 | - | - | 0 | - |
| - | - | 3774 | 658.8 | - | - | 0 | - |
| - | - | 9780 | 659.3 | - | - | 0 | - |
| - | - | 3367 | 659.8 | - | - | 0 | - |
| - | - | 3617 | 660.4 | - | - | 0 | - |
| - | - | 3492 | 662.3 | - | - | 0 | - |
| - | - | 9997 | 663.8 | - | - | 0 | - |
| - | - | 1.213E+04 | 664.3 | - | - | 0 | - |
| - | - | 4406 | 664.8 | - | - | 0 | - |
| - | - | 6835 | 666.8 | - | - | 0 | - |
| - | - | 7047 | 667.3 | - | - | 0 | - |
| - | - | 3162 | 670.4 | - | - | 0 | - |
| - | - | 7.98E+04 | 671.3 | - | - | 0 | - |
| - | - | 3.278E+04 | 672.3 | - | - | 0 | - |
| - | - | 6.055E+04 | 672.8 | - | - | 0 | - |
| - | - | 4.951E+04 | 673.3 | - | - | 0 | - |
| - | - | 2.564E+04 | 673.8 | - | - | 0 | - |
| - | - | 5.544E+04 | 674.3 | - | - | 0 | - |
| - | - | 1.893E+04 | 675.3 | - | - | 0 | - |
| - | - | 3238 | 678.9 | - | - | 0 | - |
| - | - | 6848 | 681.3 | - | - | 0 | - |
| - | - | 3.014E+05 | 681.8 | - | - | 0 | - |
| - | - | 1.964E+05 | 682.3 | - | - | 0 | - |
| - | - | 9.479E+04 | 682.8 | - | - | 0 | - |
| - | - | 2.822E+04 | 683.3 | - | - | 0 | - |
| - | - | 5377 | 683.8 | - | - | 0 | - |
| - | - | 9465 | 686.8 | - | - | 0 | - |
| - | - | 6124 | 687.3 | - | - | 0 | - |
| - | - | 3102 | 687.8 | - | - | 0 | - |
| - | - | 6908 | 688.4 | - | - | 0 | - |
| - | - | 1.602E+05 | 689.4 | - | - | 0 | - |
| - | - | 5.617E+04 | 690.4 | - | - | 0 | - |
| - | - | 4879 | 690.8 | - | - | 0 | - |
| - | - | 1.029E+04 | 691.4 | - | - | 0 | - |
| - | - | 8423 | 694.3 | - | - | 0 | - |
| - | - | 5759 | 694.8 | - | - | 0 | - |
| - | - | 3129 | 695.3 | - | - | 0 | - |
| - | - | 4244 | 696.8 | - | - | 0 | - |
| - | - | 4703 | 697.3 | - | - | 0 | - |
| - | - | 3299 | 697.8 | - | - | 0 | - |
| - | - | 4.092E+04 | 699.3 | - | - | 0 | - |
| - | - | 1.076E+04 | 700.3 | - | - | 0 | - |
| - | - | 3907 | 701.3 | - | - | 0 | - |
| - | - | 3592 | 702.3 | - | - | 0 | - |
| 3 | y | 1.522E+05 | 702.8 | 0.0005261 | 0.7485 | +2 | 13 |
| 3 | y | 1.384E+05 | 703.3 | 0.009251 | 13.15 | +2 | 13 |
| - | - | 4.591E+04 | 703.8 | - | - | 0 | - |
| - | - | 2.365E+04 | 704.3 | - | - | 0 | - |
| - | - | 7917 | 704.9 | - | - | 0 | - |
| - | - | 4481 | 711.3 | - | - | 0 | - |
| 3 | y | 1.533E+06 | 711.9 | 0.0006759 | 0.9495 | +2 | 13 |
| - | - | 1.219E+06 | 712.4 | - | - | 0 | - |
| - | - | 5.476E+05 | 712.9 | - | - | 0 | - |
| - | - | 1.621E+05 | 713.4 | - | - | 0 | - |
| - | - | 3.796E+04 | 713.9 | - | - | 0 | - |
| - | - | 4585 | 716.9 | - | - | 0 | - |
| - | - | 3301 | 717.9 | - | - | 0 | - |
| 9 | y | 8240 | 721.4 | 0.002657 | 3.683 | +1 | 7 |
| 9 | y | 1.613E+04 | 722.3 | 0.00759 | 10.51 | +1 | 7 |
| - | - | 4343 | 722.8 | - | - | 0 | - |
| - | - | 9495 | 723.4 | - | - | 0 | - |
| - | - | 3438 | 725.4 | - | - | 0 | - |
| - | - | 7916 | 725.8 | - | - | 0 | - |
| - | - | 6860 | 726.4 | - | - | 0 | - |
| - | - | 1.26E+04 | 728.4 | - | - | 0 | - |
| - | - | 1.032E+04 | 728.9 | - | - | 0 | - |
| - | - | 4770 | 729.4 | - | - | 0 | - |
| - | - | 9446 | 731.4 | - | - | 0 | - |
| - | - | 4967 | 731.9 | - | - | 0 | - |
| - | - | 4159 | 732.4 | - | - | 0 | - |
| - | - | 4288 | 733.8 | - | - | 0 | - |
| - | - | 3110 | 734.4 | - | - | 0 | - |
| - | - | 5221 | 734.9 | - | - | 0 | - |
| - | - | 2.627E+04 | 737.4 | - | - | 0 | - |
| - | - | 2.798E+04 | 737.9 | - | - | 0 | - |
| - | - | 1.402E+04 | 738.4 | - | - | 0 | - |
| 9 | y | 6.412E+04 | 739.4 | 3.794E-05 | 0.05131 | +1 | 7 |
| - | - | 4794 | 739.9 | - | - | 0 | - |
| - | - | 4.728E+04 | 740.4 | - | - | 0 | - |
| - | - | 1.945E+04 | 740.9 | - | - | 0 | - |
| - | - | 1.731E+04 | 741.4 | - | - | 0 | - |
| - | - | 3522 | 741.9 | - | - | 0 | - |
| - | - | 4170 | 743.4 | - | - | 0 | - |
| - | - | 4415 | 745.4 | - | - | 0 | - |
| - | - | 5046 | 745.9 | - | - | 0 | - |
| 2 | y | 2.47E+05 | 746.4 | 0.00032 | 0.4288 | +2 | 14 |
| 2 | y | 1.941E+05 | 746.9 | 0.009594 | 12.85 | +2 | 14 |
| - | - | 9.616E+04 | 747.4 | - | - | 0 | - |
| - | - | 3.348E+04 | 747.9 | - | - | 0 | - |
| - | - | 9221 | 748.4 | - | - | 0 | - |
| - | - | 6276 | 749.4 | - | - | 0 | - |
| - | - | 8989 | 751.4 | - | - | 0 | - |
| - | - | 7461 | 751.9 | - | - | 0 | - |
| - | - | 5051 | 752.4 | - | - | 0 | - |
| - | - | 4653 | 753.4 | - | - | 0 | - |
| - | - | 4508 | 754.4 | - | - | 0 | - |
| 2 | y | 2.705E+06 | 755.4 | 0.0005919 | 0.7836 | +2 | 14 |
| - | - | 2.176E+06 | 755.9 | - | - | 0 | - |
| - | - | 1.052E+06 | 756.4 | - | - | 0 | - |
| - | - | 3.148E+05 | 756.9 | - | - | 0 | - |
| - | - | 6.562E+04 | 757.4 | - | - | 0 | - |
| - | - | 1.354E+04 | 758.4 | - | - | 0 | - |
| - | - | 9.206E+04 | 760.4 | - | - | 0 | - |
| - | - | 8.17E+04 | 760.9 | - | - | 0 | - |
| - | - | 4.16E+04 | 761.4 | - | - | 0 | - |
| - | - | 1.732E+04 | 761.9 | - | - | 0 | - |
| - | - | 5074 | 762.4 | - | - | 0 | - |
| - | - | 8447 | 768.4 | - | - | 0 | - |
| 7 | b | 5568 | 769.4 | 0.01268 | 16.48 | +1 | 7 |
| - | - | 2.409E+04 | 770.4 | - | - | 0 | - |
| - | - | 4006 | 771.4 | - | - | 0 | - |
| - | - | 6402 | 771.4 | - | - | 0 | - |
| - | - | 3902 | 781.4 | - | - | 0 | - |
| - | - | 4897 | 782.4 | - | - | 0 | - |
| - | - | 4292 | 785.4 | - | - | 0 | - |
| - | - | 1.024E+04 | 786.4 | - | - | 0 | - |
| 7 | b | 1.694E+04 | 787.4 | 0.004566 | 5.799 | +1 | 7 |
| - | - | 1.058E+04 | 788.4 | - | - | 0 | - |
| - | - | 3768 | 789.4 | - | - | 0 | - |
| - | - | 3811 | 792.4 | - | - | 0 | - |
| - | - | 3427 | 794.4 | - | - | 0 | - |
| - | - | 1.302E+04 | 798.4 | - | - | 0 | - |
| - | - | 4663 | 799.4 | - | - | 0 | - |
| - | - | 7176 | 800.4 | - | - | 0 | - |
| - | - | 5548 | 801.4 | - | - | 0 | - |
| - | - | 6732 | 806.4 | - | - | 0 | - |
| - | - | 3204 | 807.4 | - | - | 0 | - |
| - | - | 3.732E+04 | 810.4 | - | - | 0 | - |
| - | - | 1.14E+04 | 811.4 | - | - | 0 | - |
| - | - | 4078 | 812.4 | - | - | 0 | - |
| 8 | y | 2.417E+04 | 818.4 | 0.0001213 | 0.1482 | +1 | 8 |
| 8 | y | 2.392E+04 | 819.4 | 0.005486 | 6.695 | +1 | 8 |
| - | - | 1.143E+04 | 820.4 | - | - | 0 | - |
| - | - | 3685 | 821.4 | - | - | 0 | - |
| - | - | 2734 | 828.4 | - | - | 0 | - |
| - | - | 1.341E+04 | 828.4 | - | - | 0 | - |
| - | - | 5456 | 829.4 | - | - | 0 | - |
| - | - | 6948 | 830.4 | - | - | 0 | - |
| 8 | y | 3.571E+05 | 836.4 | 0.0002378 | 0.2843 | +1 | 8 |
| - | - | 1.464E+05 | 837.4 | - | - | 0 | - |
| - | - | 3.091E+04 | 838.4 | - | - | 0 | - |
| - | - | 7300 | 839.4 | - | - | 0 | - |
| - | - | 7960 | 845.4 | - | - | 0 | - |
| - | - | 5509 | 857.5 | - | - | 0 | - |
| - | - | 6842 | 858.4 | - | - | 0 | - |
| - | - | 4807 | 859.4 | - | - | 0 | - |
| 8 | b | 6510 | 867.4 | 0.0106 | 12.22 | +1 | 8 |
| - | - | 3589 | 872.4 | - | - | 0 | - |
| - | - | 8522 | 873.4 | - | - | 0 | - |
| - | - | 6539 | 881.4 | - | - | 0 | - |
| - | - | 5531 | 882.4 | - | - | 0 | - |
| - | - | 7753 | 885.4 | - | - | 0 | - |
| - | - | 1.089E+04 | 887.4 | - | - | 0 | - |
| - | - | 5859 | 888.4 | - | - | 0 | - |
| - | - | 8114 | 890.4 | - | - | 0 | - |
| - | - | 8328 | 891.4 | - | - | 0 | - |
| - | - | 3593 | 892.4 | - | - | 0 | - |
| - | - | 4967 | 897.4 | - | - | 0 | - |
| - | - | 1.197E+04 | 899.5 | - | - | 0 | - |
| - | - | 1.062E+04 | 900.5 | - | - | 0 | - |
| - | - | 3168 | 901.5 | - | - | 0 | - |
| - | - | 1.232E+04 | 909.4 | - | - | 0 | - |
| - | - | 7125 | 910.4 | - | - | 0 | - |
| - | - | 2.263E+04 | 915.4 | - | - | 0 | - |
| - | - | 9775 | 916.4 | - | - | 0 | - |
| - | - | 3848 | 917.4 | - | - | 0 | - |
| - | - | 5337 | 925.5 | - | - | 0 | - |
| - | - | 3489 | 926.5 | - | - | 0 | - |
| - | - | 1.547E+04 | 927.5 | - | - | 0 | - |
| - | - | 6392 | 928.5 | - | - | 0 | - |
| - | - | 8036 | 933.5 | - | - | 0 | - |
| - | - | 8894 | 943.5 | - | - | 0 | - |
| - | - | 1.667E+04 | 944.5 | - | - | 0 | - |
| - | - | 5240 | 945.5 | - | - | 0 | - |
| - | - | 4840 | 950.5 | - | - | 0 | - |
| - | - | 6021 | 957.5 | - | - | 0 | - |
| - | - | 1.193E+04 | 958.5 | - | - | 0 | - |
| - | - | 6510 | 959.5 | - | - | 0 | - |
| - | - | 7628 | 968.5 | - | - | 0 | - |
| - | - | 1.037E+04 | 969.5 | - | - | 0 | - |
| 9 | b | 4806 | 971.5 | 0.002317 | 2.385 | +1 | 9 |
| - | - | 6736 | 972.5 | - | - | 0 | - |
| 7 | y | 3.688E+04 | 975.5 | 0.001082 | 1.109 | +1 | 9 |
| - | - | 1.792E+04 | 976.5 | - | - | 0 | - |
| - | - | 6566 | 977.5 | - | - | 0 | - |
| - | - | 8035 | 984.5 | - | - | 0 | - |
| - | - | 4217 | 985.5 | - | - | 0 | - |
| - | - | 3.599E+04 | 986.5 | - | - | 0 | - |
| - | - | 3.372E+04 | 987.5 | - | - | 0 | - |
| - | - | 1.587E+04 | 988.5 | - | - | 0 | - |
| - | - | 9571 | 989.5 | - | - | 0 | - |
| - | - | 6182 | 990.5 | - | - | 0 | - |
| - | - | 4273 | 994.5 | - | - | 0 | - |
| - | - | 4432 | 995.5 | - | - | 0 | - |
| - | - | 1.125E+04 | 996.5 | - | - | 0 | - |
| - | - | 4646 | 997.5 | - | - | 0 | - |
| - | - | 3663 | 998.5 | - | - | 0 | - |
| - | - | 2.271E+04 | 1012 | - | - | 0 | - |
| - | - | 1.573E+04 | 1013 | - | - | 0 | - |
| - | - | 3.304E+04 | 1014 | - | - | 0 | - |
| - | - | 2.491E+04 | 1015 | - | - | 0 | - |
| - | - | 6288 | 1016 | - | - | 0 | - |
| 10 | b | 6495 | 1029 | 0.001569 | 1.525 | +1 | 10 |
| - | - | 4076 | 1030 | - | - | 0 | - |
| - | - | 3.64E+04 | 1030 | - | - | 0 | - |
| - | - | 1.776E+04 | 1031 | - | - | 0 | - |
| - | - | 7770 | 1032 | - | - | 0 | - |
| - | - | 4421 | 1034 | - | - | 0 | - |
| - | - | 7923 | 1056 | - | - | 0 | - |
| - | - | 3872 | 1058 | - | - | 0 | - |
| - | - | 2.04E+04 | 1074 | - | - | 0 | - |
| - | - | 9336 | 1075 | - | - | 0 | - |
| - | - | 1.349E+04 | 1084 | - | - | 0 | - |
| - | - | 7254 | 1085 | - | - | 0 | - |
| 6 | y | 2.589E+04 | 1091 | 0.0001319 | 0.121 | +1 | 10 |
| - | - | 1.626E+04 | 1092 | - | - | 0 | - |
| - | - | 8001 | 1093 | - | - | 0 | - |
| - | - | 6810 | 1101 | - | - | 0 | - |
| - | - | 2.112E+04 | 1102 | - | - | 0 | - |
| - | - | 1.065E+04 | 1103 | - | - | 0 | - |
| - | - | 3903 | 1104 | - | - | 0 | - |
| - | - | 6109 | 1112 | - | - | 0 | - |
| - | - | 5440 | 1113 | - | - | 0 | - |
| - | - | 9095 | 1119 | - | - | 0 | - |
| - | - | 6679 | 1120 | - | - | 0 | - |
| - | - | 5063 | 1130 | - | - | 0 | - |
| - | - | 3849 | 1132 | - | - | 0 | - |
| - | - | 5927 | 1135 | - | - | 0 | - |
| - | - | 5141 | 1143 | - | - | 0 | - |
| - | - | 8108 | 1160 | - | - | 0 | - |
| - | - | 1.768E+04 | 1161 | - | - | 0 | - |
| - | - | 1.06E+04 | 1162 | - | - | 0 | - |
| - | - | 4519 | 1171 | - | - | 0 | - |
| 5 | y | 7.853E+04 | 1178 | 0.0003 | 0.2548 | +1 | 11 |
| - | - | 4.726E+04 | 1179 | - | - | 0 | - |
| - | - | 1.889E+04 | 1180 | - | - | 0 | - |
| - | - | 5244 | 1181 | - | - | 0 | - |
| - | - | 6190 | 1188 | - | - | 0 | - |
| - | - | 1.518E+04 | 1189 | - | - | 0 | - |
| - | - | 8659 | 1190 | - | - | 0 | - |
| - | - | 3525 | 1197 | - | - | 0 | - |
| - | - | 1.042E+04 | 1206 | - | - | 0 | - |
| - | - | 9510 | 1207 | - | - | 0 | - |
| - | - | 3536 | 1208 | - | - | 0 | - |
| - | - | 4453 | 1241 | - | - | 0 | - |
| 4 | y | 1.749E+04 | 1277 | 0.001453 | 1.138 | +1 | 12 |
| - | - | 7376 | 1278 | - | - | 0 | - |
| - | - | 3947 | 1279 | - | - | 0 | - |
| - | - | 3540 | 1496 | - | - | 0 | - |

m/z Charge Intensity FragmentType MassShift Position
120.0811996459961 0 63507.98
121.08452606201172 0 4835.391
125.07117462158203 0 1574.209
125.10773468017578 0 23660.238
126.05535125732422 0 27744.771
126.39140319824219 0 1941.395
127.0583724975586 0 2352.8726
128.10739135742188 0 18164.867
129.06631469726562 0 5988.357
129.10260009765625 0 17245.326
129.1107635498047 0 2165.7578
130.05029296875 0 4763.0923
130.06167602539062 0 5876.0747
130.097900390625 0 5055.6978
131.0820770263672 0 3324.4119
132.10227966308594 0 4508.246
132.67587280273438 0 1980.1697
133.06103515625 0 3145.0654
138.06640625 0 2561.0496
138.09170532226562 0 10184.072
139.0499267578125 0 2002.7366
139.08709716796875 0 3382.5095
140.082275390625 0 9680.361
141.10296630859375 0 2587.5378
144.0659942626953 0 9624.802
145.06126403808594 0 2745.774
147.07135009765625 0 1927.5776
147.07688903808594 0 12743.242
148.95538330078125 0 2495.1213
153.06646728515625 0 4042.182
154.05026245117188 0 37267.453
155.05386352539062 0 3852.3733
155.1183319091797 0 32491.076 a Water loss 1
156.10235595703125 0 14239.072
157.0612030029297 0 26840.469
157.09756469726562 0 30673.395
157.10877990722656 0 18912.963
157.13424682617188 0 2806.0796 d 1
158.0648651123047 0 2294.8376
158.0928497314453 0 35202.758
158.09967041015625 0 4077.969
158.1124267578125 0 2017.1384
159.1132049560547 0 19720.059
162.377685546875 0 1991.5602
165.11399841308594 0 2717.8403
166.086669921875 0 39268.164 y 14
166.09530639648438 0 2355.6345
166.09808349609375 0 3152.5273
167.08192443847656 0 8686.425
167.0894317626953 0 4595.746
169.04441833496094 0 1818.974
169.09765625 0 7429.1445
169.13392639160156 0 2060.7988
170.04452514648438 0 2267.774
171.0768585205078 0 43863.598
172.06089782714844 0 6049.418
172.0802764892578 0 4078.2563
173.12890625 0 470866.06 a 1
173.43882751464844 0 5742.8096
174.12498474121094 0 4544.1187
174.13233947753906 0 45787.797
174.59544372558594 0 2230.2627
175.07177734375 0 52723.65
175.11940002441406 0 51245.965
176.07476806640625 0 3004.1394
176.1227264404297 0 3243.174
181.061279296875 0 3647.3838
181.1088409423828 0 2658.9084
182.0929718017578 0 6973.8076
183.11326599121094 0 305301.72 b Water loss 1
184.10838317871094 0 5305.8647
184.11669921875 0 29433.084
185.05618286132812 0 3247.4592
185.0924530029297 0 32548.422
185.1194610595703 0 2174.8677
187.09963989257812 0 3485.4963
187.10813903808594 0 34822.56
188.11155700683594 0 2518.183
189.0874481201172 0 189457.11
190.0909423828125 0 15048.405
191.09237670898438 0 2619.8933
192.11370849609375 0 4538.331
194.0926513671875 0 6281.183
195.07723999023438 0 2559.7412
195.1134796142578 0 4000.3594
197.10386657714844 0 2123.432
197.12832641601562 0 2234.5178
199.07174682617188 0 77250.14
199.1076202392578 0 4832.6064 y 10
199.118896484375 0 2101.618
200.075439453125 0 5420.558
201.12379455566406 0 611717.44 b 1
202.11898803710938 0 13786.321
202.12722778320312 0 55603.164
203.06668090820312 0 54586.676
203.1033477783203 0 5017.398
203.1300048828125 0 4430.619
204.07044982910156 0 3138.0835
210.08787536621094 0 3511.9087
211.1082000732422 0 31392.85
211.11880493164062 0 4249.3213
211.15553283691406 0 10425.535
212.1033935546875 0 18829.312
213.08737182617188 0 2596.5796
216.13380432128906 0 2675.4565
217.0823516845703 0 331380.66
218.08570861816406 0 30143.707
219.08767700195312 0 4293.258
220.0925750732422 0 3540.8784
220.1084747314453 0 2428.9534
223.10752868652344 0 3322.7039
224.1034393310547 0 27372.22
226.11892700195312 0 7527.3296
227.06680297851562 0 6215.0874
227.1021270751953 0 4518.5825
227.11436462402344 0 57146.594
228.11802673339844 0 4877.485
229.1188201904297 0 34798.574
230.11383056640625 0 12983.777
236.1031951904297 0 2571.3557
237.09873962402344 0 8833.172
237.13519287109375 0 11449.776
242.11402893066406 0 51035.16
242.1277618408203 0 3981.1885
243.11766052246094 0 5106.6133
244.129638671875 0 16148.332
244.1400909423828 0 3565.8025
245.12489318847656 0 17757.643
248.62982177734375 0 9766.19 y 11
250.13058471679688 0 2787.9448
254.114013671875 0 6294.2373
254.1614990234375 0 4275.846
255.09510803222656 0 4603.775
255.10935974121094 0 62888.418
256.1127014160156 0 5342.0933
259.14031982421875 0 8241.065
259.1549072265625 0 4601.294
262.1529235839844 0 22957.537
263.139404296875 0 11270.645 b 6
263.155029296875 0 3142.5913
266.11297607421875 0 2586.6934
267.1334533691406 0 3623.6091
268.12908935546875 0 2354.939
270.145751953125 0 11077.54
271.12823486328125 0 3450.9224
271.1430969238281 0 2667.7136
272.13568115234375 0 10771.485
278.12481689453125 0 2784.6465
279.1093444824219 0 7215.3784
280.1295471191406 0 8083.4844
281.1255798339844 0 4736.392
284.12451171875 0 11511.922
284.16094970703125 0 23570.518 a Water loss 2
285.16461181640625 0 4106.951
286.1527404785156 0 2887.226
288.1379089355469 0 2603.4482
288.1560363769531 0 12786.81
289.15301513671875 0 3474.499 y Water loss 10
290.1352233886719 0 2991.9072
294.1443786621094 0 2783.4395
295.1758728027344 0 5625.1855
296.1363220214844 0 14735.074
297.12127685546875 0 3355.5725
298.1405029296875 0 31525.254
299.13629150390625 0 13334.859
302.1347961425781 0 4604.198
302.1712951660156 0 5288.238 a 2
305.1614685058594 0 100068.71 y Ammonia loss 13
306.1201477050781 0 2820.3936
306.1646423339844 0 18891.035
307.1034240722656 0 2833.763
308.0495300292969 0 2465.3608
312.15606689453125 0 296995.53 b Water loss 2
313.13330078125 0 2403.647
313.1585998535156 0 50280.195
313.1877136230469 0 65171.8
314.14678955078125 0 39024.39
314.1909484863281 0 11177.989
315.1496276855469 0 8410.285
316.15087890625 0 68208.96 b 5
317.1516418457031 0 10153.465
322.1883544921875 0 20839.834 y 13
324.1312561035156 0 13286.748
324.1679992675781 0 3311.419
325.1365661621094 0 2421.113
326.1341552734375 0 3309.6316
330.16668701171875 0 147517.47 b 2
331.1707763671875 0 34663.72
332.1578674316406 0 14296.686
333.16094970703125 0 2401.5725
336.1781005859375 0 8141.9375
341.15887451171875 0 2804.5964
341.1828308105469 0 36515.223
342.1415100097656 0 79173.78
342.1849060058594 0 6890.361
343.14434814453125 0 15540.148
344.1480407714844 0 3517.049
345.1832275390625 0 9703.648
345.6850280761719 0 5229.668
349.18359375 0 27105.143
350.13482666015625 0 2721.0469
350.18658447265625 0 2983.101
357.1570739746094 0 5151.4043
359.1678161621094 0 49118.113
360.1706848144531 0 8768.801
365.1831970214844 0 3609.982
367.1611633300781 0 6991.3
375.1662902832031 0 4960.528
380.1565856933594 0 5617.7056
381.1892395019531 0 4569.9375
382.1707458496094 0 2784.9695
383.22918701171875 0 5529.0396
385.17181396484375 0 11692.764
386.1750793457031 0 2942.9734
391.2102355957031 0 3392.8335 y Water loss 12
392.193603515625 0 17789.775 y Ammonia loss 12
393.7096862792969 0 6178.2285
394.20294189453125 0 6893.0522 b 6
394.70440673828125 0 3962.2869
395.1567077636719 0 6622.035
395.2035827636719 0 2810.1985
396.1878356933594 0 4861.974
396.6850891113281 0 7504.054
398.2149963378906 0 7891.5806
398.8656311035156 0 3480.0525 y 4
399.7190856933594 0 2846.8167
400.2203063964844 0 8681.267
401.24017333984375 0 50652.996
402.2437744140625 0 11466.953
403.1827087402344 0 8723.211
404.1875305175781 0 2724.4634
405.7171936035156 0 3177.9185
408.2287902832031 0 3567.0845
409.2203369140625 0 36497.004 y 12
410.20489501953125 0 9191.077 y Ammonia loss 7
411.2241516113281 0 10654.722 b Water loss 3
412.2275695800781 0 3767.9216
413.1673278808594 0 2647.4336
413.2136535644531 0 5006.6943
414.17144775390625 0 2753.2715
414.2281494140625 0 19682.914
414.2823181152344 0 3387.7249
414.72186279296875 0 30338.748
415.2232360839844 0 11330.763
418.7177429199219 0 34632.992 y 7
419.21844482421875 0 12855.203
419.7193298339844 0 6288.438
423.23388671875 0 112331.97
423.7352600097656 0 48795.387
424.2366027832031 0 13668.014
428.2159423828125 0 17419.707
429.2347412109375 0 43655.473 b 3
430.2400817871094 0 23591.541
431.88665771484375 0 3369.0518 y 3
432.22515869140625 0 3634.0735
439.1954650878906 0 4364.6553
441.2101745605469 0 20420.56
441.7275390625 0 2671.8792
442.2101745605469 0 4419.1597
442.27764892578125 0 11951.526
443.2055358886719 0 3362.457
448.2521667480469 0 7219.5522
450.7344665527344 0 3239.2559
452.262451171875 0 12430.845
454.2281799316406 0 2994.0383
455.2256164550781 0 3288.3376
456.2203369140625 0 3924.6252
458.2368469238281 0 33747.81
459.2381591796875 0 7305.898
461.2113037109375 0 3794.3188
463.2310485839844 0 3558.288
463.5677185058594 0 2739.353
467.2443542480469 0 8172.5024
467.74383544921875 0 3907.358
468.9002685546875 0 10077.682 y Water loss 2
469.2365417480469 0 8791.898 y Ammonia loss 2
469.288818359375 0 14660.275
469.5716857910156 0 3435.0657
470.2366638183594 0 3353.3545
474.9035339355469 0 6174.9937 y 2
475.2379455566406 0 5538.724
478.2045593261719 0 3996.9226
478.2400207519531 0 4314.133 y Water loss 11
479.2259521484375 0 22030.344 y Ammonia loss 11
480.7478332519531 0 12818.768
481.25006103515625 0 9780.397
481.76190185546875 0 4307.648
482.567626953125 0 3332.6685
482.8996887207031 0 3647.1326
486.58349609375 0 3697.0723 b 13
487.2611389160156 0 5513.122
487.7633056640625 0 12399.319 y Water loss 6
488.25555419921875 0 30045.264 y Ammonia loss 6
488.75726318359375 0 14812.6875
489.25335693359375 0 5363.217
489.75897216796875 0 2900.7212
496.218505859375 0 4015.9255
496.25238037109375 0 140700.89 y 11
496.7679138183594 0 361198.47 y 6
497.26873779296875 0 211829.61
497.77020263671875 0 55378.156
497.91046142578125 0 6023.322 y Water loss 1
498.2685241699219 0 11615.919
498.74053955078125 0 4276.623
500.19879150390625 0 5327.0703
501.5736083984375 0 3393.3386
503.9149169921875 0 14815.333 y 1
504.249267578125 0 24432.264
504.583984375 0 9408.044
505.27301025390625 0 20301.18
506.27777099609375 0 4702.745
511.3345031738281 0 3136.4639
513.238525390625 0 3355.2236
513.351318359375 0 51624.594
514.3543090820312 0 12702.355
515.2572631835938 0 22620.268
515.7572631835938 0 6746.741
516.26416015625 0 12039.065 b 4
518.2091674804688 0 4629.3345
524.2630615234375 0 109712.16
524.7648315429688 0 57122.934
525.2648315429688 0 17939.941
525.7658081054688 0 5312.0425
526.267578125 0 4740.8447
529.6033935546875 0 8351.235
529.93408203125 0 5484.746
530.2698974609375 0 3140.431
532.2725830078125 0 7533
535.6063232421875 0 71771.62 Precursor Water loss
535.9405517578125 0 59784.387 Precursor Ammonia loss
536.274169921875 0 31245.416
536.6087036132812 0 13090.525
536.9404296875 0 4673.158
537.7620239257812 0 2859.0764
539.2954711914062 0 4978.4883
541.34619140625 0 72935.836
541.6099853515625 0 144962.94 Precursor
541.944091796875 0 143763.1
542.2779541015625 0 83053.95
542.3494873046875 0 20524.451
542.6124267578125 0 21661.504
542.9448852539062 0 8218.78
543.2503662109375 0 6123.4453
545.2738647460938 0 4707.425 y Water loss 5
545.7723999023438 0 9309.981 y Ammonia loss 5
546.2705078125 0 5312.7
547.2279052734375 0 5341.5107
550.7922973632812 0 4927.5312
554.2810668945312 0 146273.94 y 5
554.7828979492188 0 92986.84
555.283935546875 0 29513.904 b Water loss 10
555.7882080078125 0 4767.606
556.2808227539062 0 5420.668
556.323974609375 0 3949.8577
557.2671508789062 0 4211.2227
557.3222045898438 0 3422.9607
558.7930908203125 0 3961.9922
559.2830200195312 0 8683.332
564.2882690429688 0 4267.792 b 10
564.79150390625 0 29262.514
565.2383422851562 0 4413.175
565.29052734375 0 11708.646
565.7918701171875 0 6896.757
566.3057250976562 0 4726.925
567.2911376953125 0 4192.5273
568.7748413085938 0 4294.332
569.268798828125 0 24107.92
570.266357421875 0 3579.891
572.2792358398438 0 6118.8335
573.797119140625 0 110337.46
574.2982788085938 0 82137.695
574.8001708984375 0 17917.436
575.2982177734375 0 8123.847
582.7930908203125 0 8612.969
583.2474975585938 0 4905.0415
583.2938842773438 0 6350.408
584.316650390625 0 12829.35
585.3155517578125 0 5153.8594
587.27880859375 0 29394.271
588.2806396484375 0 8933.694
588.7921752929688 0 16725 y Water loss 4
589.289306640625 0 20413.182 y Ammonia loss 4
589.7888793945312 0 9942.481
592.3059692382812 0 32288.477
593.3072509765625 0 13382.938
595.3206176757812 0 23907.557 y 10
596.3206787109375 0 5951.4897
597.7973022460938 0 1161671.9 y 4
598.2986450195312 0 728188.44
598.7999267578125 0 280843.9 b Water loss 11
599.3010864257812 0 68696.09 b Ammonia loss 11
599.8025512695312 0 14467.349
600.275146484375 0 22382.008
601.2771606445312 0 10363.72
611.3139038085938 0 3641.7097
611.798828125 0 3138.2239
612.3042602539062 0 3085.6995
617.3054809570312 0 3923.4468
618.305419921875 0 4263.962
620.3062133789062 0 3972.3052
620.806640625 0 6288.152
623.3173217773438 0 3817.6672
624.313720703125 0 5516.9155
625.33740234375 0 3893.0576
625.8043212890625 0 3232.6787
626.3270874023438 0 9871.026
627.3258666992188 0 4562.4463
629.3143310546875 0 51388.223
629.8143310546875 0 25458.225
630.3157958984375 0 19790.031
630.8139038085938 0 6175.41
631.3018188476562 0 3102.1707 b 5
631.8236694335938 0 3722.97
632.3274536132812 0 6814.9434
632.8280029296875 0 5102.5244
633.3274536132812 0 4364.552
634.3299560546875 0 3249.839 y Water loss 9
636.3135375976562 0 8397.122
638.31982421875 0 167293.78 y Water loss 3
638.8204345703125 0 122108.164 y Ammonia loss 3
639.265625 0 3805.044
639.322021484375 0 54809.652
639.8236083984375 0 17514.344
640.3224487304688 0 4593.8857
641.3369140625 0 4999.125
642.3217163085938 0 7238.8247 b Water loss 12
643.3515625 0 13592.394
644.355224609375 0 4333.783
646.314697265625 0 16899.926
646.826171875 0 5471.3965
647.3316040039062 0 1038192.75 y 3
647.8329467773438 0 721017.44
648.3339233398438 0 298024.12
648.8353881835938 0 90288.28
649.3375244140625 0 14026.24
652.3417358398438 0 63592.44 y 9
653.3408813476562 0 32571.588
654.3231811523438 0 17316.193
655.3218383789062 0 8885.673
656.302734375 0 7843.422
657.2879028320312 0 4150.9507
658.8312377929688 0 3773.8542
659.3448486328125 0 9780.11
659.83642578125 0 3367.477
660.3502807617188 0 3616.515
662.3270263671875 0 3491.664
663.8250732421875 0 9996.552
664.3173828125 0 12125.431
664.8224487304688 0 4406.122
666.8292236328125 0 6834.815
667.3287353515625 0 7047.056
670.3624877929688 0 3161.7056
671.3473510742188 0 79798.38
672.3430786132812 0 32779.953
672.8291015625 0 60545.31
673.3320922851562 0 49514.008
673.831787109375 0 25638.809
674.311767578125 0 55435.87
675.313720703125 0 18934.44
678.9251708984375 0 3237.9912
681.3322143554688 0 6848.45
681.8344116210938 0 301447.88
682.3348999023438 0 196419.05
682.8369750976562 0 94792.266
683.3377075195312 0 28220.822
683.8417358398438 0 5376.7524
686.8268432617188 0 9464.758
687.3277587890625 0 6124.465
687.8309936523438 0 3101.8662
688.3709106445312 0 6907.988
689.3583374023438 0 160233.69
690.3612670898438 0 56170.875
690.8351440429688 0 4878.9595
691.3631591796875 0 10290.2
694.335693359375 0 8423.374
694.84423828125 0 5759.109
695.3378295898438 0 3129.0996
696.8493041992188 0 4243.7754
697.347900390625 0 4703.389
697.8467407226562 0 3298.7534
699.3425903320312 0 40924.35
700.3431396484375 0 10758.643
701.3472290039062 0 3906.6467
702.3407592773438 0 3591.5378
702.8474731445312 0 152236.55 y Water loss 2
703.3482055664062 0 138358.48 y Ammonia loss 2
703.8485107421875 0 45913.195
704.3468627929688 0 23651.318
704.8511962890625 0 7917.4736
711.3389282226562 0 4481.0684
711.8529052734375 0 1533259.5 y 2
712.3543090820312 0 1218876.1
712.8554077148438 0 547644.25
713.3565673828125 0 162107.05
713.8580932617188 0 37963.69
716.850341796875 0 4585.437
717.8605346679688 0 3300.9766
721.360107421875 0 8240.018 y Water loss 8
722.3543701171875 0 16131.113 y Ammonia loss 8
722.8499145507812 0 4342.8667
723.3552856445312 0 9495.465
725.3507080078125 0 3438.2908
725.8469848632812 0 7915.663
726.3524169921875 0 6860.4785
728.35302734375 0 12603.055
728.8536376953125 0 10323.519
729.3558959960938 0 4770.03
731.356201171875 0 9446.175
731.8595581054688 0 4966.8955
732.3618774414062 0 4158.8003
733.838134765625 0 4288.106
734.3514404296875 0 3109.978
734.8540649414062 0 5220.6357
737.3576049804688 0 26267.748
737.8573608398438 0 27977.965
738.3602294921875 0 14021.301
739.373291015625 0 64121.453 y 8
739.8635864257812 0 4794.005
740.3699340820312 0 47283.004
740.8644409179688 0 19446.22
741.3702392578125 0 17311.67
741.8630981445312 0 3522.3906
743.36767578125 0 4170.3696
745.3562622070312 0 4414.9907
745.8598022460938 0 5046.1016
746.36328125 0 247040.3 y Water loss 1
746.8645629882812 0 194098.8 y Ammonia loss 1
747.3649291992188 0 96156.45
747.8662719726562 0 33479.36
748.36181640625 0 9221.23
749.3583984375 0 6275.835
751.353759765625 0 8988.916
751.85791015625 0 7460.63
752.351806640625 0 5051.2876
753.3502807617188 0 4653.438
754.3568725585938 0 4507.6543
755.3688354492188 0 2705405 y 1
755.8701782226562 0 2176059
756.3713989257812 0 1052109.8
756.8724975585938 0 314826.3
757.3743286132812 0 65623.26
758.381591796875 0 13539.923
760.3607788085938 0 92060.34
760.8622436523438 0 81700.05
761.3634033203125 0 41595.42
761.86572265625 0 17321.346
762.3659057617188 0 5074.0254
768.365478515625 0 8447.078
769.3712158203125 0 5567.7295 b Water loss 6
770.4140625 0 24094.256
771.3570556640625 0 4005.603
771.4241943359375 0 6402.227
781.388916015625 0 3901.797
782.3761596679688 0 4897.2993
785.42626953125 0 4292.443
786.3760375976562 0 10242.875
787.389892578125 0 16939.818 b 6
788.39990234375 0 10581.282
789.4011840820312 0 3768.0413
792.3621215820312 0 3810.9045
794.4081420898438 0 3426.7244
798.4097900390625 0 13020.182
799.4134521484375 0 4662.506
800.4000854492188 0 7175.641
801.392333984375 0 5547.589
806.41650390625 0 6731.606
807.4178466796875 0 3204.3184
810.375 0 37321.574
811.3800659179688 0 11395.105
812.384765625 0 4077.8027
818.4156494140625 0 24173.025 y Water loss 7
819.405029296875 0 23917.615 y Ammonia loss 7
820.4048461914062 0 11425.297
821.4071044921875 0 3685.4436
828.3670043945312 0 2734.0984
828.4314575195312 0 13414.512
829.4373779296875 0 5456.026
830.4046630859375 0 6947.7354
836.4263305664062 0 357061.72 y 7
837.4293823242188 0 146354.89
838.4307250976562 0 30906.055
839.432373046875 0 7300.3257
845.40966796875 0 7959.9595
857.4557495117188 0 5509.137
858.3948364257812 0 6842.114
859.3974609375 0 4806.6504
867.4312744140625 0 6509.7393 b Ammonia loss 7
872.4186401367188 0 3589.129
873.4078979492188 0 8521.704
881.4445190429688 0 6539.313
882.4486694335938 0 5531.1045
885.4398193359375 0 7753.251
887.421142578125 0 10885.596
888.423095703125 0 5858.6064
890.4297485351562 0 8114.113
891.4265747070312 0 8328.455
892.4274291992188 0 3592.6782
897.4024047851562 0 4967.428
899.4563598632812 0 11973.8545
900.460205078125 0 10617.815
901.4573364257812 0 3167.932
909.4400024414062 0 12319.302
910.4432983398438 0 7124.745
915.4169311523438 0 22633.77
916.4225463867188 0 9775.225
917.423583984375 0 3847.77
925.4537963867188 0 5337.287
926.4600830078125 0 3488.6064
927.4548950195312 0 15469.421
928.4560546875 0 6391.657
933.4797973632812 0 8036.0767
943.459228515625 0 8893.549
944.470947265625 0 16671.549
945.4844970703125 0 5240.047
950.5091552734375 0 4840.232
957.4854736328125 0 6021.294
958.47802734375 0 11929.5
959.4808959960938 0 6510.3623
968.4786376953125 0 7627.6177
969.4766845703125 0 10374.263
971.4815673828125 0 4805.8345 b 8
972.4754638671875 0 6735.615
975.4995727539062 0 36878.195 y Ammonia loss 6
976.5009155273438 0 17920.762
977.50146484375 0 6565.746
984.4826049804688 0 8034.5146
985.4894409179688 0 4217.233
986.4896240234375 0 35990.133
987.4903564453125 0 33718.9
988.4847412109375 0 15869.235
989.4970092773438 0 9571.255
990.4874877929688 0 6182.475
994.4713134765625 0 4272.6577
995.4713134765625 0 4432.3228
996.4747924804688 0 11249.888
997.4727783203125 0 4646.366
998.4755249023438 0 3662.804
1012.4800415039062 0 22713.502
1013.4838256835938 0 15732.705
1014.4845581054688 0 33035.902
1015.4874877929688 0 24907.504
1016.4892578125 0 6287.62
1028.4991455078125 0 6495.3687 b 9
1029.5087890625 0 4076.0825
1030.4915771484375 0 36399.355
1031.49267578125 0 17762.918
1032.497802734375 0 7770.3
1033.5068359375 0 4420.8237
1055.5115966796875 0 7922.712
1057.5218505859375 0 3871.9578
1073.5186767578125 0 20398.072
1074.521484375 0 9336.473
1083.513916015625 0 13493.835
1084.5079345703125 0 7253.705
1090.5274658203125 0 25886.912 y Ammonia loss 5
1091.5321044921875 0 16259.519
1092.5335693359375 0 8000.5093
1100.5269775390625 0 6810.0127
1101.5191650390625 0 21116.65
1102.5198974609375 0 10654.511
1103.515869140625 0 3903.397
1111.5458984375 0 6109.454
1112.552978515625 0 5439.596
1118.5419921875 0 9094.963
1119.5355224609375 0 6679.207
1129.5592041015625 0 5063.351
1131.556396484375 0 3848.6763
1134.5518798828125 0 5926.8345
1142.5478515625 0 5141.47
1159.54931640625 0 8107.562
1160.551513671875 0 17681.414
1161.55615234375 0 10595.557
1170.5499267578125 0 4519.333
1177.559326171875 0 78531.22 y Ammonia loss 4
1178.561279296875 0 47256.22
1179.5626220703125 0 18886.615
1180.5711669921875 0 5243.718
1187.560546875 0 6190.3716
1188.5474853515625 0 15183.774
1189.55029296875 0 8659.007
1197.4320068359375 0 3524.872
1205.5736083984375 0 10423.348
1206.572998046875 0 9510.172
1207.5704345703125 0 3536.0583
1240.5933837890625 0 4452.8677
1276.6265869140625 0 17492.863 y Ammonia loss 3
1277.6328125 0 7375.86
1278.635009765625 0 3946.723
1496.2984619140625 0 3539.9507

Spectrum Details

|  |  |
| --- | --- |
| Matched peaks? Matched peaks The total absolute number of peaks matched. Additionally in brackets the total fraction of peaks matched and the total number of peaks is shown. | 82 (11.90% of 689) |
| FDR? FDR The false discovery rate estimated for this peptide. It is calculated by matching all theoretical fragments with a non-integer shift with the raw peaks for this spectrum. This is done with 40 different shifts. The resulting percentage is the average number of annotated peaks over the number of annotated peaks with the correct spectrum. | 0.26% |
| Satellite FDR? Satellite FDR See the FDR for details on its calculation. This satellite ion specific FDR only contains the satellite ions (d/w) for I/L/J positions. | - |
| PSM Score? PSM Score The PSM Score as given by Hecklib to this annotated spectrum. It is shown with three significant figures. | 552 |

#### Spectrum 4596? Spectrum 4596 The raw spectrum of this peptide as annotated by Hecklib. The fragments are coloured according to ion type (see legend). Any peaks with a star '\*' as text can be hovered over to see the full details, first the ion type second the mass shift type. By hovering over the amino acids in the peptide or ions in the legend the corresponding peaks are highlighted. By toggling the 'Unassigned' label you can turn the background (unassigned) peaks on or off in the plot. By updating the slider in the Ion legend you can update the spectrum to only show the top X% of the peaks with labels. The top X% means any peak that is within X% of the highest intensity. By dragging in the spectrum you can zoom in to a specific part of the spectrum and use 'Zoom Out' to get back to the original zoom level. The annotation of the spectrum is based on the given sequence in the peptides file and is done with different software so inconsistencies are likely. The peaks are annotated based on the given sequence, with 20 ppm tolerance.

Copy Data

##### Spectrum 4596 (TSV)

###### Preview

```
Loading example...
```

*Click on the button to copy the data to your clipboard.*

Mz MinMz MaxIntensity Max

WidthHeightPeptide font sizePeptide stroke widthSpectrum font sizeSpectrum stroke widthCompact peptide

Ion legend

wxyz

abcd

OtherUnassignedIon^Charge^_Position_Show for top:%

JSEVSDRPSGVSSRF

03.10e+66.20e+69.30e+61.24e+7

Zoom Out

a^+1^_2_y^+1^_1_a^+1^_2_b^+1^_2_y^+2^_3_y^+3^_5_b^+1^_2_y^+2^_4_b^+3^_7_a^+1^_3_y^+2^_5_a^+1^_3_y^+1^_2_b^+1^_3_b^+2^_6_y^+1^_2_y^+2^_6_b^+1^_3_y^+1^_3_y^+1^_3_b^+2^_7_y^+3^_11_y^+1^_3_y^+2^_8_b^+1^_4_y^+2^_8_b^+1^_4_y^+3^_13_y^+3^_13_y^+3^_13_y^+1^_4_y^+1^_4_y^+2^_9_y^+2^_9_y^+1^_4_y^+2^_9_y^+3^_14_y^+3^_14_b^+1^_5_\*\*\*y^+2^_10_y^+2^_10_b^+2^_11_b^+2^_11_y^+2^_11_y^+2^_11_y^+1^_5_y^+2^_11_b^+2^_12_b^+2^_12_y^+1^_6_y^+2^_12_y^+2^_12_b^+2^_13_y^+2^_12_y^+1^_6_y^+2^_13_y^+2^_13_y^+2^_13_y^+1^_7_y^+1^_7_y^+1^_7_y^+2^_14_y^+2^_14_y^+2^_14_b^+1^_7_y^+1^_8_y^+1^_8_y^+1^_8_b^+1^_8_b^+1^_9_b^+1^_9_y^+1^_9_b^+1^_10_y^+1^_10_y^+1^_11_y^+1^_12_

0855171025653420

Fragment Matches Table

Show background peaks

| Position | Ion type | Intensity | mz Theoretical | mz Error (Th) | mz Error (ppm) | Charge | Series Number |
| --- | --- | --- | --- | --- | --- | --- | --- |
| - | - | 3.026E+05 | 120.1 | - | - | 0 | - |
| - | - | 3.445E+04 | 121.1 | - | - | 0 | - |
| - | - | 9.883E+04 | 125.1 | - | - | 0 | - |
| - | - | 1.639E+05 | 126.1 | - | - | 0 | - |
| - | - | 9.555E+04 | 128.1 | - | - | 0 | - |
| - | - | 2.78E+04 | 129.1 | - | - | 0 | - |
| - | - | 5.751E+04 | 129.1 | - | - | 0 | - |
| - | - | 3.452E+04 | 130.1 | - | - | 0 | - |
| - | - | 1.908E+04 | 130.1 | - | - | 0 | - |
| - | - | 1.969E+04 | 130.1 | - | - | 0 | - |
| - | - | 1.526E+04 | 131.1 | - | - | 0 | - |
| - | - | 9412 | 131.2 | - | - | 0 | - |
| - | - | 1.155E+04 | 133.1 | - | - | 0 | - |
| - | - | 4.438E+04 | 138.1 | - | - | 0 | - |
| - | - | 1.557E+04 | 139.1 | - | - | 0 | - |
| - | - | 5.087E+04 | 140.1 | - | - | 0 | - |
| - | - | 1.463E+04 | 141.1 | - | - | 0 | - |
| - | - | 2.816E+04 | 144.1 | - | - | 0 | - |
| - | - | 9367 | 145.1 | - | - | 0 | - |
| - | - | 7.226E+04 | 147.1 | - | - | 0 | - |
| - | - | 2.125E+04 | 153.1 | - | - | 0 | - |
| - | - | 1.807E+05 | 154.1 | - | - | 0 | - |
| - | - | 1.251E+04 | 155.1 | - | - | 0 | - |
| 2 | a | 1.152E+05 | 155.1 | 0.0002288 | 1.475 | +1 | 2 |
| - | - | 1.014E+05 | 156.1 | - | - | 0 | - |
| - | - | 1.333E+04 | 156.1 | - | - | 0 | - |
| - | - | 1.483E+05 | 157.1 | - | - | 0 | - |
| - | - | 1.404E+05 | 157.1 | - | - | 0 | - |
| - | - | 9.828E+04 | 157.1 | - | - | 0 | - |
| - | - | 1.423E+05 | 158.1 | - | - | 0 | - |
| - | - | 1.726E+04 | 158.1 | - | - | 0 | - |
| - | - | 1.357E+04 | 159.1 | - | - | 0 | - |
| - | - | 1.025E+05 | 159.1 | - | - | 0 | - |
| - | - | 1.409E+04 | 160.1 | - | - | 0 | - |
| - | - | 1.652E+04 | 165.1 | - | - | 0 | - |
| 15 | y | 1.981E+05 | 166.1 | 0.0002775 | 1.671 | +1 | 1 |
| - | - | 1.291E+04 | 166.1 | - | - | 0 | - |
| - | - | 1.469E+04 | 166.1 | - | - | 0 | - |
| - | - | 4.009E+04 | 167.1 | - | - | 0 | - |
| - | - | 1.62E+04 | 167.1 | - | - | 0 | - |
| - | - | 2.503E+04 | 169.1 | - | - | 0 | - |
| - | - | 2.302E+05 | 171.1 | - | - | 0 | - |
| - | - | 1.043E+04 | 171.1 | - | - | 0 | - |
| - | - | 2.881E+04 | 172.1 | - | - | 0 | - |
| 2 | a | 2.161E+06 | 173.1 | 0.00033 | 1.906 | +1 | 2 |
| - | - | 4.664E+04 | 173.4 | - | - | 0 | - |
| - | - | 1.772E+05 | 174.1 | - | - | 0 | - |
| - | - | 2.141E+05 | 175.1 | - | - | 0 | - |
| - | - | 2.645E+05 | 175.1 | - | - | 0 | - |
| - | - | 1.257E+04 | 176.1 | - | - | 0 | - |
| - | - | 2.236E+04 | 182.1 | - | - | 0 | - |
| 2 | b | 1.397E+06 | 183.1 | 0.0002787 | 1.522 | +1 | 2 |
| - | - | 1.196E+04 | 184.1 | - | - | 0 | - |
| - | - | 2.114E+04 | 184.1 | - | - | 0 | - |
| - | - | 1.456E+05 | 184.1 | - | - | 0 | - |
| - | - | 2.085E+04 | 185.1 | - | - | 0 | - |
| - | - | 1.37E+05 | 185.1 | - | - | 0 | - |
| - | - | 1.811E+04 | 185.1 | - | - | 0 | - |
| - | - | 1.08E+04 | 187.1 | - | - | 0 | - |
| - | - | 1.65E+05 | 187.1 | - | - | 0 | - |
| - | - | 1.012E+06 | 189.1 | - | - | 0 | - |
| - | - | 6.382E+04 | 190.1 | - | - | 0 | - |
| - | - | 1.663E+04 | 192.1 | - | - | 0 | - |
| - | - | 2.2E+04 | 192.1 | - | - | 0 | - |
| - | - | 3.338E+04 | 194.1 | - | - | 0 | - |
| - | - | 1.449E+04 | 194.1 | - | - | 0 | - |
| - | - | 1.489E+04 | 195.1 | - | - | 0 | - |
| - | - | 1.294E+04 | 195.1 | - | - | 0 | - |
| 13 | y | 1.556E+04 | 196.1 | 0.0004939 | 2.519 | +2 | 3 |
| - | - | 3.243E+05 | 199.1 | - | - | 0 | - |
| - | - | 1.61E+04 | 199.1 | - | - | 0 | - |
| 11 | y | 1.149E+04 | 199.1 | 0.003629 | 18.23 | +3 | 5 |
| - | - | 1.202E+04 | 199.1 | - | - | 0 | - |
| - | - | 2.101E+04 | 200.1 | - | - | 0 | - |
| 2 | b | 2.701E+06 | 201.1 | 0.0002579 | 1.282 | +1 | 2 |
| - | - | 1.696E+04 | 202.1 | - | - | 0 | - |
| - | - | 6.504E+04 | 202.1 | - | - | 0 | - |
| - | - | 2.51E+05 | 202.1 | - | - | 0 | - |
| - | - | 2.516E+05 | 203.1 | - | - | 0 | - |
| - | - | 2.239E+04 | 203.1 | - | - | 0 | - |
| - | - | 1.621E+04 | 203.1 | - | - | 0 | - |
| - | - | 1.403E+04 | 204.1 | - | - | 0 | - |
| - | - | 1.865E+04 | 210.1 | - | - | 0 | - |
| - | - | 1.172E+05 | 211.1 | - | - | 0 | - |
| - | - | 1.079E+05 | 212.1 | - | - | 0 | - |
| - | - | 1.643E+04 | 213.1 | - | - | 0 | - |
| - | - | 2.132E+04 | 216.1 | - | - | 0 | - |
| - | - | 1.461E+06 | 217.1 | - | - | 0 | - |
| - | - | 1.398E+05 | 218.1 | - | - | 0 | - |
| - | - | 2.096E+04 | 219.1 | - | - | 0 | - |
| - | - | 2.848E+04 | 220.1 | - | - | 0 | - |
| - | - | 1.115E+05 | 224.1 | - | - | 0 | - |
| - | - | 1.792E+04 | 226.1 | - | - | 0 | - |
| - | - | 3.671E+04 | 227.1 | - | - | 0 | - |
| - | - | 3.017E+05 | 227.1 | - | - | 0 | - |
| - | - | 1.994E+04 | 228.1 | - | - | 0 | - |
| - | - | 1.34E+05 | 229.1 | - | - | 0 | - |
| - | - | 5.36E+04 | 230.1 | - | - | 0 | - |
| - | - | 4.569E+04 | 237.1 | - | - | 0 | - |
| - | - | 5.467E+04 | 237.1 | - | - | 0 | - |
| - | - | 1.203E+04 | 238.1 | - | - | 0 | - |
| - | - | 2.677E+05 | 242.1 | - | - | 0 | - |
| - | - | 2.364E+04 | 242.1 | - | - | 0 | - |
| - | - | 2.533E+04 | 243.1 | - | - | 0 | - |
| - | - | 1.329E+04 | 243.1 | - | - | 0 | - |
| - | - | 9.649E+04 | 244.1 | - | - | 0 | - |
| - | - | 1.545E+04 | 244.1 | - | - | 0 | - |
| - | - | 6.973E+04 | 245.1 | - | - | 0 | - |
| 12 | y | 4.316E+04 | 248.6 | 0.0003043 | 1.224 | +2 | 4 |
| - | - | 1.713E+04 | 252.1 | - | - | 0 | - |
| - | - | 3.075E+04 | 254.1 | - | - | 0 | - |
| - | - | 2.372E+04 | 254.2 | - | - | 0 | - |
| - | - | 3.018E+05 | 255.1 | - | - | 0 | - |
| - | - | 3.497E+04 | 256.1 | - | - | 0 | - |
| - | - | 1.623E+04 | 256.1 | - | - | 0 | - |
| - | - | 1.162E+04 | 256.7 | - | - | 0 | - |
| - | - | 3.778E+04 | 259.1 | - | - | 0 | - |
| - | - | 2.217E+04 | 259.2 | - | - | 0 | - |
| - | - | 9.993E+04 | 262.2 | - | - | 0 | - |
| 7 | b | 5.719E+04 | 263.1 | 0.002487 | 9.453 | +3 | 7 |
| - | - | 1.154E+04 | 263.2 | - | - | 0 | - |
| - | - | 1.403E+04 | 266.1 | - | - | 0 | - |
| - | - | 1.752E+04 | 267.1 | - | - | 0 | - |
| - | - | 3.314E+04 | 270.1 | - | - | 0 | - |
| - | - | 4.805E+04 | 272.1 | - | - | 0 | - |
| - | - | 2.337E+04 | 278.1 | - | - | 0 | - |
| - | - | 2.697E+04 | 279.1 | - | - | 0 | - |
| - | - | 2.659E+04 | 280.1 | - | - | 0 | - |
| - | - | 2.113E+04 | 281.1 | - | - | 0 | - |
| - | - | 6.462E+04 | 284.1 | - | - | 0 | - |
| 3 | a | 9.605E+04 | 284.2 | 0.0006807 | 2.395 | +1 | 3 |
| - | - | 2.646E+04 | 285.2 | - | - | 0 | - |
| - | - | 2.548E+04 | 288.1 | - | - | 0 | - |
| - | - | 5.119E+04 | 288.2 | - | - | 0 | - |
| - | - | 1.929E+04 | 289.2 | - | - | 0 | - |
| - | - | 1.979E+04 | 294.1 | - | - | 0 | - |
| - | - | 1.624E+04 | 295.1 | - | - | 0 | - |
| - | - | 1.557E+04 | 295.2 | - | - | 0 | - |
| - | - | 8.097E+04 | 296.1 | - | - | 0 | - |
| - | - | 2.072E+04 | 296.2 | - | - | 0 | - |
| - | - | 1.808E+04 | 297.1 | - | - | 0 | - |
| - | - | 1.397E+04 | 297.4 | - | - | 0 | - |
| - | - | 1.615E+05 | 298.1 | - | - | 0 | - |
| 11 | y | 2.322E+04 | 298.2 | 0.001051 | 3.523 | +2 | 5 |
| - | - | 5.575E+04 | 299.1 | - | - | 0 | - |
| - | - | 1.197E+04 | 301.2 | - | - | 0 | - |
| - | - | 4.79E+04 | 302.1 | - | - | 0 | - |
| 3 | a | 2.766E+04 | 302.2 | 0.0003394 | 1.123 | +1 | 3 |
| 14 | y | 4.795E+05 | 305.2 | 0.0004989 | 1.635 | +1 | 2 |
| - | - | 1.337E+04 | 306.1 | - | - | 0 | - |
| - | - | 7.836E+04 | 306.2 | - | - | 0 | - |
| 3 | b | 1.399E+06 | 312.2 | 0.0004255 | 1.363 | +1 | 3 |
| - | - | 1.998E+05 | 313.2 | - | - | 0 | - |
| - | - | 3.278E+05 | 313.2 | - | - | 0 | - |
| - | - | 2.264E+05 | 314.1 | - | - | 0 | - |
| - | - | 1.428E+04 | 314.2 | - | - | 0 | - |
| - | - | 4.942E+04 | 314.2 | - | - | 0 | - |
| - | - | 3.506E+04 | 315.2 | - | - | 0 | - |
| 6 | b | 3.345E+05 | 316.2 | 0.000506 | 1.6 | +2 | 6 |
| - | - | 5.909E+04 | 317.1 | - | - | 0 | - |
| - | - | 1.416E+04 | 319.2 | - | - | 0 | - |
| 14 | y | 9.346E+04 | 322.2 | 0.000195 | 0.6051 | +1 | 2 |
| - | - | 2.327E+04 | 323.2 | - | - | 0 | - |
| - | - | 6.908E+04 | 324.1 | - | - | 0 | - |
| - | - | 2.243E+04 | 325.1 | - | - | 0 | - |
| 10 | y | 1.061E+04 | 326.7 | 0.002566 | 7.855 | +2 | 6 |
| 3 | b | 6.631E+05 | 330.2 | 0.0004809 | 1.457 | +1 | 3 |
| - | - | 1.88E+05 | 331.2 | - | - | 0 | - |
| - | - | 8.345E+04 | 332.2 | - | - | 0 | - |
| - | - | 1.589E+04 | 332.2 | - | - | 0 | - |
| - | - | 2.542E+04 | 336.2 | - | - | 0 | - |
| - | - | 2.128E+04 | 336.7 | - | - | 0 | - |
| - | - | 1.976E+05 | 341.2 | - | - | 0 | - |
| - | - | 3.784E+05 | 342.1 | - | - | 0 | - |
| - | - | 3.333E+04 | 342.2 | - | - | 0 | - |
| - | - | 6.643E+04 | 343.1 | - | - | 0 | - |
| - | - | 2.025E+04 | 344.1 | - | - | 0 | - |
| - | - | 5.736E+04 | 345.2 | - | - | 0 | - |
| - | - | 2.195E+04 | 349.2 | - | - | 0 | - |
| - | - | 1.336E+05 | 349.2 | - | - | 0 | - |
| - | - | 1.439E+04 | 350.2 | - | - | 0 | - |
| - | - | 1.506E+04 | 352.2 | - | - | 0 | - |
| - | - | 1.93E+04 | 357.2 | - | - | 0 | - |
| - | - | 1.491E+04 | 358.2 | - | - | 0 | - |
| - | - | 2.681E+05 | 359.2 | - | - | 0 | - |
| - | - | 1.366E+04 | 359.2 | - | - | 0 | - |
| - | - | 2.897E+04 | 360.2 | - | - | 0 | - |
| - | - | 2.129E+04 | 365.2 | - | - | 0 | - |
| - | - | 2.368E+04 | 367.2 | - | - | 0 | - |
| - | - | 1.184E+04 | 373.6 | - | - | 0 | - |
| - | - | 2.129E+04 | 374.2 | - | - | 0 | - |
| - | - | 2.333E+04 | 375.2 | - | - | 0 | - |
| - | - | 2.046E+04 | 380.2 | - | - | 0 | - |
| - | - | 1.775E+04 | 383.2 | - | - | 0 | - |
| - | - | 6.493E+04 | 385.2 | - | - | 0 | - |
| - | - | 1.318E+04 | 389.9 | - | - | 0 | - |
| 13 | y | 1.474E+04 | 391.2 | 2.854E-05 | 0.07295 | +1 | 3 |
| - | - | 1.494E+04 | 392.2 | - | - | 0 | - |
| 13 | y | 1.003E+05 | 392.2 | 0.0001478 | 0.3768 | +1 | 3 |
| - | - | 2.653E+04 | 393.7 | - | - | 0 | - |
| 7 | b | 2.227E+04 | 394.2 | 0.0001533 | 0.3888 | +2 | 7 |
| - | - | 2.382E+04 | 395.2 | - | - | 0 | - |
| - | - | 2.239E+04 | 396.2 | - | - | 0 | - |
| - | - | 3.95E+04 | 398.2 | - | - | 0 | - |
| 5 | y | 1.368E+04 | 398.9 | 0.001553 | 3.893 | +3 | 11 |
| - | - | 1.797E+04 | 399.7 | - | - | 0 | - |
| - | - | 1.395E+04 | 400.2 | - | - | 0 | - |
| - | - | 3.671E+04 | 400.2 | - | - | 0 | - |
| - | - | 2.288E+05 | 401.2 | - | - | 0 | - |
| - | - | 5.466E+04 | 402.2 | - | - | 0 | - |
| - | - | 4.095E+04 | 403.2 | - | - | 0 | - |
| - | - | 1.592E+04 | 405.7 | - | - | 0 | - |
| - | - | 3.67E+04 | 405.7 | - | - | 0 | - |
| - | - | 1.779E+04 | 408.2 | - | - | 0 | - |
| 13 | y | 1.475E+05 | 409.2 | 0.0004541 | 1.11 | +1 | 3 |
| 8 | y | 3.966E+04 | 410.2 | 0.0004168 | 1.016 | +2 | 8 |
| 4 | b | 2.687E+04 | 411.2 | 0.0009108 | 2.215 | +1 | 4 |
| - | - | 3.035E+04 | 413.2 | - | - | 0 | - |
| - | - | 2.245E+04 | 413.2 | - | - | 0 | - |
| - | - | 8.538E+04 | 414.2 | - | - | 0 | - |
| - | - | 1.544E+05 | 414.7 | - | - | 0 | - |
| - | - | 6.278E+04 | 415.2 | - | - | 0 | - |
| - | - | 2.355E+04 | 415.7 | - | - | 0 | - |
| 8 | y | 1.883E+05 | 418.7 | 0.0004785 | 1.143 | +2 | 8 |
| - | - | 7.426E+04 | 419.2 | - | - | 0 | - |
| - | - | 2.373E+04 | 419.7 | - | - | 0 | - |
| - | - | 5.819E+05 | 423.2 | - | - | 0 | - |
| - | - | 2.408E+05 | 423.7 | - | - | 0 | - |
| - | - | 4.442E+04 | 424.2 | - | - | 0 | - |
| - | - | 1.514E+04 | 424.7 | - | - | 0 | - |
| - | - | 7.202E+04 | 428.2 | - | - | 0 | - |
| 4 | b | 2.221E+05 | 429.2 | 0.0003348 | 0.7801 | +1 | 4 |
| - | - | 1.059E+05 | 430.2 | - | - | 0 | - |
| - | - | 1.989E+04 | 431.2 | - | - | 0 | - |
| - | - | 2.414E+04 | 439.2 | - | - | 0 | - |
| - | - | 8.712E+04 | 441.2 | - | - | 0 | - |
| - | - | 2.136E+04 | 442.2 | - | - | 0 | - |
| - | - | 1.508E+04 | 443.2 | - | - | 0 | - |
| - | - | 1.646E+04 | 445.2 | - | - | 0 | - |
| - | - | 2.811E+04 | 448.3 | - | - | 0 | - |
| - | - | 5.491E+04 | 452.3 | - | - | 0 | - |
| - | - | 1.621E+04 | 453.3 | - | - | 0 | - |
| - | - | 1.559E+04 | 455.2 | - | - | 0 | - |
| - | - | 1.413E+04 | 455.7 | - | - | 0 | - |
| - | - | 1.701E+05 | 458.2 | - | - | 0 | - |
| - | - | 4.192E+04 | 459.2 | - | - | 0 | - |
| - | - | 2.061E+04 | 461.2 | - | - | 0 | - |
| - | - | 1.798E+04 | 462.9 | - | - | 0 | - |
| - | - | 3.079E+04 | 463.2 | - | - | 0 | - |
| - | - | 4.387E+04 | 467.2 | - | - | 0 | - |
| - | - | 2.69E+04 | 467.7 | - | - | 0 | - |
| - | - | 1.701E+04 | 468.2 | - | - | 0 | - |
| 3 | y | 1.736E+04 | 468.9 | 0.001281 | 2.733 | +3 | 13 |
| 3 | y | 2.859E+04 | 469.2 | 0.00598 | 12.74 | +3 | 13 |
| - | - | 5.737E+04 | 469.3 | - | - | 0 | - |
| - | - | 2.728E+04 | 470.2 | - | - | 0 | - |
| - | - | 1.577E+04 | 470.3 | - | - | 0 | - |
| - | - | 1.872E+04 | 471.7 | - | - | 0 | - |
| - | - | 1.266E+04 | 472.2 | - | - | 0 | - |
| 3 | y | 3.523E+04 | 474.9 | 0.001179 | 2.482 | +3 | 13 |
| - | - | 2.447E+04 | 475.2 | - | - | 0 | - |
| - | - | 1.534E+04 | 475.6 | - | - | 0 | - |
| - | - | 1.535E+04 | 478.2 | - | - | 0 | - |
| 12 | y | 3.285E+04 | 478.2 | 0.0005663 | 1.184 | +1 | 4 |
| 12 | y | 9.281E+04 | 479.2 | 0.0007121 | 1.486 | +1 | 4 |
| - | - | 1.444E+04 | 480.3 | - | - | 0 | - |
| - | - | 6.976E+04 | 480.7 | - | - | 0 | - |
| - | - | 4.002E+04 | 481.2 | - | - | 0 | - |
| - | - | 1.554E+04 | 481.8 | - | - | 0 | - |
| - | - | 2.797E+04 | 487.3 | - | - | 0 | - |
| 7 | y | 4.056E+04 | 487.8 | 0.0002085 | 0.4275 | +2 | 9 |
| 7 | y | 1.331E+05 | 488.3 | 0.002962 | 6.066 | +2 | 9 |
| - | - | 6.146E+04 | 488.8 | - | - | 0 | - |
| - | - | 3.179E+04 | 489.3 | - | - | 0 | - |
| - | - | 1.913E+04 | 491.9 | - | - | 0 | - |
| - | - | 1.383E+04 | 494.7 | - | - | 0 | - |
| - | - | 1.291E+04 | 496.2 | - | - | 0 | - |
| 12 | y | 6.64E+05 | 496.3 | 0.0003166 | 0.638 | +1 | 4 |
| 7 | y | 1.79E+06 | 496.8 | 0.000399 | 0.8032 | +2 | 9 |
| - | - | 8.671E+05 | 497.3 | - | - | 0 | - |
| - | - | 2.972E+05 | 497.8 | - | - | 0 | - |
| 2 | y | 5.796E+04 | 497.9 | 0.0001471 | 0.2955 | +3 | 14 |
| - | - | 6.067E+04 | 498.3 | - | - | 0 | - |
| - | - | 1.609E+04 | 498.9 | - | - | 0 | - |
| - | - | 2.189E+04 | 500.2 | - | - | 0 | - |
| - | - | 1.675E+04 | 501.8 | - | - | 0 | - |
| 2 | y | 7.16E+04 | 503.9 | 6.76E-05 | 0.1342 | +3 | 14 |
| - | - | 9.324E+04 | 504.2 | - | - | 0 | - |
| - | - | 3.195E+04 | 504.6 | - | - | 0 | - |
| - | - | 1.598E+04 | 504.9 | - | - | 0 | - |
| - | - | 1.303E+05 | 505.3 | - | - | 0 | - |
| - | - | 1.446E+04 | 506.3 | - | - | 0 | - |
| - | - | 2.185E+04 | 506.7 | - | - | 0 | - |
| - | - | 1.235E+04 | 507.2 | - | - | 0 | - |
| - | - | 1.522E+04 | 507.7 | - | - | 0 | - |
| - | - | 1.831E+04 | 513.2 | - | - | 0 | - |
| - | - | 1.636E+04 | 514.3 | - | - | 0 | - |
| - | - | 1.356E+05 | 515.3 | - | - | 0 | - |
| - | - | 3.794E+04 | 515.8 | - | - | 0 | - |
| 5 | b | 2.536E+04 | 516.3 | 0.0008403 | 1.628 | +1 | 5 |
| - | - | 1.604E+04 | 517.3 | - | - | 0 | - |
| - | - | 2.572E+04 | 518.2 | - | - | 0 | - |
| - | - | 5.211E+05 | 524.3 | - | - | 0 | - |
| - | - | 2.33E+05 | 524.8 | - | - | 0 | - |
| - | - | 7.351E+04 | 525.3 | - | - | 0 | - |
| - | - | 1.601E+04 | 526.2 | - | - | 0 | - |
| - | - | 2.816E+04 | 529.6 | - | - | 0 | - |
| - | - | 3.916E+04 | 529.9 | - | - | 0 | - |
| - | - | 2.387E+04 | 530.3 | - | - | 0 | - |
| - | - | 2.134E+04 | 532.3 | - | - | 0 | - |
| - | - | 1.505E+04 | 533.3 | - | - | 0 | - |
| 0 | Precursor | 2.895E+05 | 535.6 | 0.0002027 | 0.3785 | +3 | -1 |
| 0 | Precursor | 2.538E+05 | 535.9 | 0.006548 | 12.22 | +3 | -1 |
| - | - | 1.55E+05 | 536.3 | - | - | 0 | - |
| - | - | 5.169E+04 | 536.6 | - | - | 0 | - |
| - | - | 1.702E+04 | 539.3 | - | - | 0 | - |
| - | - | 1.679E+04 | 541.3 | - | - | 0 | - |
| 0 | Precursor | 6.759E+05 | 541.6 | 9.914E-05 | 0.183 | +3 | -1 |
| - | - | 6.196E+05 | 541.9 | - | - | 0 | - |
| - | - | 3.223E+05 | 542.3 | - | - | 0 | - |
| - | - | 7.857E+04 | 542.6 | - | - | 0 | - |
| - | - | 3.686E+04 | 543.3 | - | - | 0 | - |
| 6 | y | 2.077E+04 | 545.3 | 0.0008768 | 1.608 | +2 | 10 |
| - | - | 1.51E+04 | 546.3 | - | - | 0 | - |
| - | - | 1.75E+04 | 547.3 | - | - | 0 | - |
| - | - | 1.7E+04 | 550.3 | - | - | 0 | - |
| - | - | 2.676E+04 | 550.8 | - | - | 0 | - |
| - | - | 1.757E+04 | 551.3 | - | - | 0 | - |
| 6 | y | 7.169E+05 | 554.3 | 1.098E-05 | 0.0198 | +2 | 10 |
| - | - | 4.021E+05 | 554.8 | - | - | 0 | - |
| 11 | b | 1.378E+05 | 555.3 | 0.0008324 | 1.499 | +2 | 11 |
| 11 | b | 3.397E+04 | 555.8 | 0.01035 | 18.62 | +2 | 11 |
| - | - | 2.611E+04 | 556.3 | - | - | 0 | - |
| - | - | 2.282E+04 | 556.3 | - | - | 0 | - |
| - | - | 1.958E+04 | 556.8 | - | - | 0 | - |
| - | - | 3.406E+04 | 558.8 | - | - | 0 | - |
| - | - | 3.731E+04 | 559.3 | - | - | 0 | - |
| - | - | 1.276E+05 | 564.8 | - | - | 0 | - |
| - | - | 4.286E+04 | 565.2 | - | - | 0 | - |
| - | - | 7.06E+04 | 565.3 | - | - | 0 | - |
| - | - | 4.727E+04 | 565.8 | - | - | 0 | - |
| - | - | 4.275E+04 | 566.3 | - | - | 0 | - |
| - | - | 2.76E+04 | 567.3 | - | - | 0 | - |
| - | - | 1.88E+04 | 568.3 | - | - | 0 | - |
| - | - | 1.062E+05 | 569.3 | - | - | 0 | - |
| - | - | 4.052E+04 | 570.3 | - | - | 0 | - |
| - | - | 4.1E+04 | 572.3 | - | - | 0 | - |
| - | - | 1.848E+04 | 573.3 | - | - | 0 | - |
| - | - | 7.111E+05 | 573.8 | - | - | 0 | - |
| - | - | 3.349E+05 | 574.3 | - | - | 0 | - |
| - | - | 1.254E+05 | 574.8 | - | - | 0 | - |
| - | - | 5.823E+04 | 575.3 | - | - | 0 | - |
| - | - | 1.275E+04 | 576.3 | - | - | 0 | - |
| - | - | 4.437E+04 | 582.8 | - | - | 0 | - |
| - | - | 1.78E+04 | 583.2 | - | - | 0 | - |
| - | - | 5.793E+04 | 584.3 | - | - | 0 | - |
| - | - | 2.585E+04 | 585.3 | - | - | 0 | - |
| - | - | 1.322E+05 | 587.3 | - | - | 0 | - |
| - | - | 5.568E+04 | 588.3 | - | - | 0 | - |
| 5 | y | 1.038E+05 | 588.8 | 0.0002435 | 0.4135 | +2 | 11 |
| 5 | y | 1E+05 | 589.3 | 0.004879 | 8.279 | +2 | 11 |
| - | - | 3.658E+04 | 589.8 | - | - | 0 | - |
| - | - | 2.301E+04 | 590.3 | - | - | 0 | - |
| - | - | 1.752E+05 | 592.3 | - | - | 0 | - |
| - | - | 4.63E+04 | 593.3 | - | - | 0 | - |
| - | - | 1.443E+04 | 594.3 | - | - | 0 | - |
| 11 | y | 9.091E+04 | 595.3 | 0.0005367 | 0.9016 | +1 | 5 |
| - | - | 3.89E+04 | 596.3 | - | - | 0 | - |
| 5 | y | 5.144E+06 | 597.8 | 8.81E-05 | 0.1474 | +2 | 11 |
| - | - | 3.331E+06 | 598.3 | - | - | 0 | - |
| 12 | b | 1.213E+06 | 598.8 | 0.0005653 | 0.9441 | +2 | 12 |
| 12 | b | 3.218E+05 | 599.3 | 0.009534 | 15.91 | +2 | 12 |
| - | - | 7.245E+04 | 599.8 | - | - | 0 | - |
| - | - | 9.796E+04 | 600.3 | - | - | 0 | - |
| - | - | 2.284E+04 | 601.3 | - | - | 0 | - |
| - | - | 2.089E+04 | 602.8 | - | - | 0 | - |
| - | - | 1.646E+04 | 609.3 | - | - | 0 | - |
| - | - | 2.303E+04 | 615.3 | - | - | 0 | - |
| - | - | 2.552E+04 | 617.3 | - | - | 0 | - |
| - | - | 3.616E+04 | 617.8 | - | - | 0 | - |
| - | - | 1.613E+04 | 618.3 | - | - | 0 | - |
| - | - | 2.682E+04 | 620.3 | - | - | 0 | - |
| - | - | 3.789E+04 | 620.8 | - | - | 0 | - |
| - | - | 1.716E+04 | 621.3 | - | - | 0 | - |
| - | - | 2.669E+04 | 623.3 | - | - | 0 | - |
| - | - | 2.765E+04 | 624.3 | - | - | 0 | - |
| - | - | 6.365E+04 | 626.3 | - | - | 0 | - |
| - | - | 2.764E+05 | 629.3 | - | - | 0 | - |
| - | - | 1.798E+05 | 629.8 | - | - | 0 | - |
| - | - | 8.906E+04 | 630.3 | - | - | 0 | - |
| - | - | 1.825E+04 | 630.8 | - | - | 0 | - |
| - | - | 2.121E+04 | 632.8 | - | - | 0 | - |
| 10 | y | 3.22E+04 | 635.3 | 0.005561 | 8.753 | +1 | 6 |
| - | - | 4.258E+04 | 636.3 | - | - | 0 | - |
| 4 | y | 7.929E+05 | 638.3 | 0.006376 | 9.988 | +2 | 12 |
| 4 | y | 6.56E+05 | 638.8 | 0.002532 | 3.964 | +2 | 12 |
| - | - | 1.951E+04 | 639.3 | - | - | 0 | - |
| - | - | 2.748E+05 | 639.3 | - | - | 0 | - |
| - | - | 8.26E+04 | 639.8 | - | - | 0 | - |
| - | - | 3.629E+04 | 641.3 | - | - | 0 | - |
| 13 | b | 3.786E+04 | 642.3 | 0.009026 | 14.05 | +2 | 13 |
| - | - | 4.1E+04 | 643.4 | - | - | 0 | - |
| - | - | 2.496E+04 | 644.4 | - | - | 0 | - |
| - | - | 1.883E+04 | 645.3 | - | - | 0 | - |
| - | - | 9.663E+04 | 646.3 | - | - | 0 | - |
| 4 | y | 4.827E+06 | 647.3 | 0.0001219 | 0.1883 | +2 | 12 |
| - | - | 3.543E+06 | 647.8 | - | - | 0 | - |
| - | - | 1.513E+06 | 648.3 | - | - | 0 | - |
| - | - | 2.52E+04 | 648.4 | - | - | 0 | - |
| - | - | 3.38E+05 | 648.8 | - | - | 0 | - |
| - | - | 8.801E+04 | 649.3 | - | - | 0 | - |
| - | - | 1.88E+04 | 651.8 | - | - | 0 | - |
| 10 | y | 2.729E+05 | 652.3 | 0.0007854 | 1.204 | +1 | 6 |
| - | - | 1.431E+05 | 653.3 | - | - | 0 | - |
| - | - | 8.17E+04 | 654.3 | - | - | 0 | - |
| - | - | 4.937E+04 | 655.3 | - | - | 0 | - |
| - | - | 2.385E+04 | 656.3 | - | - | 0 | - |
| - | - | 2.014E+04 | 657.3 | - | - | 0 | - |
| - | - | 2.527E+04 | 658.8 | - | - | 0 | - |
| - | - | 5.126E+04 | 659.3 | - | - | 0 | - |
| - | - | 3.604E+04 | 663.8 | - | - | 0 | - |
| - | - | 4.637E+04 | 664.3 | - | - | 0 | - |
| - | - | 2.364E+04 | 664.8 | - | - | 0 | - |
| - | - | 6.778E+04 | 666.8 | - | - | 0 | - |
| - | - | 4.236E+04 | 667.3 | - | - | 0 | - |
| - | - | 3.442E+05 | 671.3 | - | - | 0 | - |
| - | - | 1.59E+05 | 672.3 | - | - | 0 | - |
| - | - | 2.767E+05 | 672.8 | - | - | 0 | - |
| - | - | 2.59E+05 | 673.3 | - | - | 0 | - |
| - | - | 7.639E+04 | 673.8 | - | - | 0 | - |
| - | - | 2.564E+05 | 674.3 | - | - | 0 | - |
| - | - | 1.721E+04 | 674.8 | - | - | 0 | - |
| - | - | 7.881E+04 | 675.3 | - | - | 0 | - |
| - | - | 2.848E+04 | 681.3 | - | - | 0 | - |
| - | - | 1.456E+06 | 681.8 | - | - | 0 | - |
| - | - | 1.062E+06 | 682.3 | - | - | 0 | - |
| - | - | 3.89E+05 | 682.8 | - | - | 0 | - |
| - | - | 9.403E+04 | 683.3 | - | - | 0 | - |
| - | - | 4.002E+04 | 686.8 | - | - | 0 | - |
| - | - | 3.439E+04 | 687.3 | - | - | 0 | - |
| - | - | 2.947E+04 | 687.8 | - | - | 0 | - |
| - | - | 2.321E+04 | 688.4 | - | - | 0 | - |
| - | - | 8.269E+05 | 689.4 | - | - | 0 | - |
| - | - | 2.863E+05 | 690.4 | - | - | 0 | - |
| - | - | 1.786E+04 | 690.8 | - | - | 0 | - |
| - | - | 7.791E+04 | 691.4 | - | - | 0 | - |
| - | - | 1.651E+04 | 693.8 | - | - | 0 | - |
| - | - | 5.146E+04 | 694.3 | - | - | 0 | - |
| - | - | 2.042E+04 | 694.8 | - | - | 0 | - |
| - | - | 3.996E+04 | 696.8 | - | - | 0 | - |
| - | - | 1.948E+04 | 697.3 | - | - | 0 | - |
| - | - | 1.644E+05 | 699.3 | - | - | 0 | - |
| - | - | 5.29E+04 | 700.3 | - | - | 0 | - |
| 3 | y | 7.741E+05 | 702.8 | 0.0001599 | 0.2275 | +2 | 13 |
| 3 | y | 6.552E+05 | 703.3 | 0.00919 | 13.07 | +2 | 13 |
| - | - | 2.478E+05 | 703.8 | - | - | 0 | - |
| - | - | 1.123E+05 | 704.3 | - | - | 0 | - |
| - | - | 3.017E+04 | 704.9 | - | - | 0 | - |
| 3 | y | 7.181E+06 | 711.9 | 0.0001176 | 0.1651 | +2 | 13 |
| - | - | 5.524E+06 | 712.4 | - | - | 0 | - |
| - | - | 3.987E+04 | 712.5 | - | - | 0 | - |
| - | - | 2.424E+06 | 712.9 | - | - | 0 | - |
| - | - | 6.8E+05 | 713.4 | - | - | 0 | - |
| - | - | 1.21E+05 | 713.9 | - | - | 0 | - |
| - | - | 2.736E+04 | 716.8 | - | - | 0 | - |
| - | - | 1.582E+04 | 717.4 | - | - | 0 | - |
| 9 | y | 4.321E+04 | 721.4 | 0.002043 | 2.832 | +1 | 7 |
| 9 | y | 5.477E+04 | 722.3 | 0.005027 | 6.959 | +1 | 7 |
| - | - | 3.305E+04 | 722.9 | - | - | 0 | - |
| - | - | 2.59E+04 | 723.4 | - | - | 0 | - |
| - | - | 1.84E+04 | 723.9 | - | - | 0 | - |
| - | - | 2.061E+04 | 725.8 | - | - | 0 | - |
| - | - | 3.437E+04 | 726.3 | - | - | 0 | - |
| - | - | 6.448E+04 | 728.4 | - | - | 0 | - |
| - | - | 5.512E+04 | 728.9 | - | - | 0 | - |
| - | - | 4.177E+04 | 729.4 | - | - | 0 | - |
| - | - | 5.817E+04 | 731.4 | - | - | 0 | - |
| - | - | 3.162E+04 | 731.9 | - | - | 0 | - |
| - | - | 2.358E+04 | 732.4 | - | - | 0 | - |
| - | - | 2.463E+04 | 732.9 | - | - | 0 | - |
| - | - | 4.477E+04 | 734.4 | - | - | 0 | - |
| - | - | 3.201E+04 | 734.9 | - | - | 0 | - |
| - | - | 1.389E+05 | 737.4 | - | - | 0 | - |
| - | - | 1.177E+05 | 737.9 | - | - | 0 | - |
| - | - | 5.747E+04 | 738.4 | - | - | 0 | - |
| - | - | 1.578E+04 | 738.9 | - | - | 0 | - |
| 9 | y | 3.42E+05 | 739.4 | 0.0006483 | 0.8768 | +1 | 7 |
| - | - | 2.515E+05 | 740.4 | - | - | 0 | - |
| - | - | 6.378E+04 | 740.9 | - | - | 0 | - |
| - | - | 7.785E+04 | 741.4 | - | - | 0 | - |
| - | - | 2.586E+04 | 745.4 | - | - | 0 | - |
| 2 | y | 1.072E+06 | 746.4 | 0.0001683 | 0.2254 | +2 | 14 |
| 2 | y | 8.458E+05 | 746.9 | 0.008739 | 11.7 | +2 | 14 |
| - | - | 4.394E+05 | 747.4 | - | - | 0 | - |
| - | - | 1.691E+05 | 747.9 | - | - | 0 | - |
| - | - | 4.054E+04 | 748.4 | - | - | 0 | - |
| - | - | 4.303E+04 | 751.4 | - | - | 0 | - |
| - | - | 4.104E+04 | 751.9 | - | - | 0 | - |
| - | - | 2.514E+04 | 753.4 | - | - | 0 | - |
| 2 | y | 1.228E+07 | 755.4 | 4.256E-05 | 0.05634 | +2 | 14 |
| - | - | 1.022E+07 | 755.9 | - | - | 0 | - |
| - | - | 4.748E+06 | 756.4 | - | - | 0 | - |
| - | - | 1.389E+06 | 756.9 | - | - | 0 | - |
| - | - | 3.341E+05 | 757.4 | - | - | 0 | - |
| - | - | 5.548E+04 | 758.4 | - | - | 0 | - |
| - | - | 2.436E+04 | 759.4 | - | - | 0 | - |
| - | - | 4.533E+05 | 760.4 | - | - | 0 | - |
| - | - | 3.731E+05 | 760.9 | - | - | 0 | - |
| - | - | 1.664E+05 | 761.4 | - | - | 0 | - |
| - | - | 5.598E+04 | 761.9 | - | - | 0 | - |
| - | - | 3.49E+04 | 768.4 | - | - | 0 | - |
| - | - | 3.095E+04 | 768.4 | - | - | 0 | - |
| - | - | 1.859E+04 | 769.3 | - | - | 0 | - |
| - | - | 9.919E+04 | 770.4 | - | - | 0 | - |
| - | - | 3.556E+04 | 771.4 | - | - | 0 | - |
| - | - | 1.654E+04 | 783.4 | - | - | 0 | - |
| - | - | 2.174E+04 | 785.4 | - | - | 0 | - |
| - | - | 6.004E+04 | 786.4 | - | - | 0 | - |
| 7 | b | 6.986E+04 | 787.4 | 0.003284 | 4.171 | +1 | 7 |
| - | - | 5.968E+04 | 788.4 | - | - | 0 | - |
| - | - | 1.952E+04 | 789.4 | - | - | 0 | - |
| - | - | 6.525E+04 | 798.4 | - | - | 0 | - |
| - | - | 3.744E+04 | 799.4 | - | - | 0 | - |
| - | - | 4.958E+04 | 800.4 | - | - | 0 | - |
| - | - | 2.14E+04 | 801.4 | - | - | 0 | - |
| - | - | 4.823E+04 | 806.4 | - | - | 0 | - |
| - | - | 1.659E+05 | 810.4 | - | - | 0 | - |
| - | - | 7.516E+04 | 811.4 | - | - | 0 | - |
| - | - | 1.974E+04 | 812.4 | - | - | 0 | - |
| 8 | y | 1.56E+05 | 818.4 | 0.0006722 | 0.8213 | +1 | 8 |
| 8 | y | 1.364E+05 | 819.4 | 0.005913 | 7.216 | +1 | 8 |
| - | - | 7.545E+04 | 820.4 | - | - | 0 | - |
| - | - | 2.942E+04 | 828.4 | - | - | 0 | - |
| - | - | 7.991E+04 | 828.4 | - | - | 0 | - |
| - | - | 4.956E+04 | 829.4 | - | - | 0 | - |
| - | - | 2.246E+04 | 830.4 | - | - | 0 | - |
| 8 | y | 1.646E+06 | 836.4 | 0.0006167 | 0.7373 | +1 | 8 |
| - | - | 7.4E+05 | 837.4 | - | - | 0 | - |
| - | - | 1.897E+05 | 838.4 | - | - | 0 | - |
| - | - | 3.537E+04 | 839.4 | - | - | 0 | - |
| - | - | 4.086E+04 | 845.4 | - | - | 0 | - |
| - | - | 3.892E+04 | 855.4 | - | - | 0 | - |
| - | - | 4.492E+04 | 857.4 | - | - | 0 | - |
| - | - | 2.419E+04 | 858.4 | - | - | 0 | - |
| 8 | b | 3.842E+04 | 867.4 | 0.0106 | 12.22 | +1 | 8 |
| - | - | 4.142E+04 | 873.4 | - | - | 0 | - |
| - | - | 3.437E+04 | 881.4 | - | - | 0 | - |
| - | - | 4.521E+04 | 885.4 | - | - | 0 | - |
| - | - | 6.987E+04 | 887.4 | - | - | 0 | - |
| - | - | 3.388E+04 | 888.4 | - | - | 0 | - |
| - | - | 2.685E+04 | 890.4 | - | - | 0 | - |
| - | - | 3.127E+04 | 891.4 | - | - | 0 | - |
| - | - | 6.415E+04 | 899.5 | - | - | 0 | - |
| - | - | 4.566E+04 | 900.5 | - | - | 0 | - |
| - | - | 2.189E+04 | 901.5 | - | - | 0 | - |
| - | - | 5.46E+04 | 909.4 | - | - | 0 | - |
| - | - | 2.057E+04 | 910.4 | - | - | 0 | - |
| - | - | 9.939E+04 | 915.4 | - | - | 0 | - |
| - | - | 3.774E+04 | 916.4 | - | - | 0 | - |
| - | - | 2.052E+04 | 917.4 | - | - | 0 | - |
| - | - | 1.834E+04 | 925.4 | - | - | 0 | - |
| - | - | 3.327E+04 | 926.4 | - | - | 0 | - |
| - | - | 9.669E+04 | 927.5 | - | - | 0 | - |
| - | - | 5.076E+04 | 928.5 | - | - | 0 | - |
| - | - | 1.734E+04 | 929.5 | - | - | 0 | - |
| - | - | 2.329E+04 | 933.5 | - | - | 0 | - |
| - | - | 6.956E+04 | 943.5 | - | - | 0 | - |
| - | - | 5.05E+04 | 944.5 | - | - | 0 | - |
| - | - | 4.28E+04 | 945.5 | - | - | 0 | - |
| - | - | 2.213E+04 | 951.5 | - | - | 0 | - |
| 9 | b | 2.695E+04 | 954.5 | 0.01117 | 11.7 | +1 | 9 |
| - | - | 3.057E+04 | 957.5 | - | - | 0 | - |
| - | - | 8.227E+04 | 958.5 | - | - | 0 | - |
| - | - | 2.758E+04 | 959.5 | - | - | 0 | - |
| - | - | 3.957E+04 | 968.5 | - | - | 0 | - |
| - | - | 2.789E+04 | 969.5 | - | - | 0 | - |
| - | - | 2.17E+04 | 970.5 | - | - | 0 | - |
| 9 | b | 2.51E+04 | 971.5 | 0.01147 | 11.81 | +1 | 9 |
| - | - | 4.15E+04 | 972.5 | - | - | 0 | - |
| 7 | y | 1.709E+05 | 975.5 | 0.0004716 | 0.4835 | +1 | 9 |
| - | - | 7.732E+04 | 976.5 | - | - | 0 | - |
| - | - | 4.334E+04 | 977.5 | - | - | 0 | - |
| - | - | 2.763E+04 | 984.5 | - | - | 0 | - |
| - | - | 2.339E+04 | 985.5 | - | - | 0 | - |
| - | - | 2.012E+05 | 986.5 | - | - | 0 | - |
| - | - | 1.585E+05 | 987.5 | - | - | 0 | - |
| - | - | 7.488E+04 | 988.5 | - | - | 0 | - |
| - | - | 5.066E+04 | 989.5 | - | - | 0 | - |
| - | - | 3.972E+04 | 990.5 | - | - | 0 | - |
| - | - | 1.803E+04 | 991.5 | - | - | 0 | - |
| - | - | 3.393E+04 | 996.5 | - | - | 0 | - |
| - | - | 3.532E+04 | 997.5 | - | - | 0 | - |
| - | - | 1.722E+04 | 1000 | - | - | 0 | - |
| - | - | 1.226E+05 | 1012 | - | - | 0 | - |
| - | - | 7.476E+04 | 1013 | - | - | 0 | - |
| - | - | 2.059E+05 | 1014 | - | - | 0 | - |
| - | - | 9.518E+04 | 1015 | - | - | 0 | - |
| - | - | 2.285E+04 | 1016 | - | - | 0 | - |
| 10 | b | 3.29E+04 | 1029 | 0.003278 | 3.187 | +1 | 10 |
| - | - | 2.137E+04 | 1030 | - | - | 0 | - |
| - | - | 1.824E+05 | 1030 | - | - | 0 | - |
| - | - | 1.137E+05 | 1031 | - | - | 0 | - |
| - | - | 3.528E+04 | 1033 | - | - | 0 | - |
| - | - | 3.152E+04 | 1056 | - | - | 0 | - |
| - | - | 1.718E+04 | 1057 | - | - | 0 | - |
| - | - | 1.85E+04 | 1058 | - | - | 0 | - |
| - | - | 1.883E+04 | 1073 | - | - | 0 | - |
| - | - | 1.047E+05 | 1074 | - | - | 0 | - |
| - | - | 5.232E+04 | 1075 | - | - | 0 | - |
| - | - | 1.919E+04 | 1076 | - | - | 0 | - |
| - | - | 6.066E+04 | 1084 | - | - | 0 | - |
| - | - | 2.537E+04 | 1085 | - | - | 0 | - |
| - | - | 3.635E+04 | 1087 | - | - | 0 | - |
| 6 | y | 1.161E+05 | 1091 | 0.00294 | 2.696 | +1 | 10 |
| - | - | 7.897E+04 | 1092 | - | - | 0 | - |
| - | - | 3.378E+04 | 1093 | - | - | 0 | - |
| - | - | 2.423E+04 | 1094 | - | - | 0 | - |
| - | - | 1.889E+04 | 1101 | - | - | 0 | - |
| - | - | 1.21E+05 | 1102 | - | - | 0 | - |
| - | - | 5.073E+04 | 1103 | - | - | 0 | - |
| - | - | 3.44E+04 | 1104 | - | - | 0 | - |
| - | - | 4.396E+04 | 1112 | - | - | 0 | - |
| - | - | 3.111E+04 | 1113 | - | - | 0 | - |
| - | - | 1.786E+04 | 1114 | - | - | 0 | - |
| - | - | 1.772E+04 | 1119 | - | - | 0 | - |
| - | - | 3.707E+04 | 1120 | - | - | 0 | - |
| - | - | 2.645E+04 | 1121 | - | - | 0 | - |
| - | - | 3.222E+04 | 1130 | - | - | 0 | - |
| - | - | 2.81E+04 | 1131 | - | - | 0 | - |
| - | - | 2.479E+04 | 1135 | - | - | 0 | - |
| - | - | 2.512E+04 | 1136 | - | - | 0 | - |
| - | - | 3.097E+04 | 1143 | - | - | 0 | - |
| - | - | 3.217E+04 | 1144 | - | - | 0 | - |
| - | - | 2.389E+04 | 1153 | - | - | 0 | - |
| - | - | 5.976E+04 | 1160 | - | - | 0 | - |
| - | - | 9.22E+04 | 1161 | - | - | 0 | - |
| - | - | 5.006E+04 | 1162 | - | - | 0 | - |
| - | - | 2.045E+04 | 1171 | - | - | 0 | - |
| 5 | y | 4.023E+05 | 1178 | 0.002009 | 1.706 | +1 | 11 |
| - | - | 2.517E+05 | 1179 | - | - | 0 | - |
| - | - | 8.448E+04 | 1180 | - | - | 0 | - |
| - | - | 1.8E+04 | 1181 | - | - | 0 | - |
| - | - | 3.151E+04 | 1188 | - | - | 0 | - |
| - | - | 6.937E+04 | 1189 | - | - | 0 | - |
| - | - | 3.272E+04 | 1190 | - | - | 0 | - |
| - | - | 5.811E+04 | 1206 | - | - | 0 | - |
| - | - | 5.5E+04 | 1207 | - | - | 0 | - |
| - | - | 2.452E+04 | 1208 | - | - | 0 | - |
| 4 | y | 7.275E+04 | 1277 | 0.002796 | 2.19 | +1 | 12 |
| - | - | 2.66E+04 | 1278 | - | - | 0 | - |
| - | - | 2.617E+04 | 1300 | - | - | 0 | - |
| - | - | 1.549E+04 | 1301 | - | - | 0 | - |
| - | - | 1.282E+04 | 2146 | - | - | 0 | - |
| - | - | 1.471E+04 | 2444 | - | - | 0 | - |
| - | - | 1.791E+04 | 3386 | - | - | 0 | - |

m/z Charge Intensity FragmentType MassShift Position
120.08108520507812 0 302605.75
121.08446502685547 0 34454.39
125.10758209228516 0 98828.31
126.05526733398438 0 163892.64
128.10731506347656 0 95551.63
129.06625366210938 0 27796.062
129.1025390625 0 57514.426
130.05029296875 0 34518.39
130.06141662597656 0 19080.096
130.0979461669922 0 19690.113
131.0816192626953 0 15256.945
131.17135620117188 0 9412.251
133.09686279296875 0 11549.892
138.09164428710938 0 44384.832
139.08668518066406 0 15573.013
140.08209228515625 0 50866.434
141.1025390625 0 14630.64
144.06602478027344 0 28162.75
145.06060791015625 0 9366.787
147.07669067382812 0 72263.96
153.0663299560547 0 21246.56
154.0501708984375 0 180673.89
155.05328369140625 0 12509.767
155.1181182861328 0 115235.51 a Water loss 1
156.1022491455078 0 101376.18
156.1212921142578 0 13334.874
157.0609893798828 0 148258.95
157.0974884033203 0 140408.98
157.108642578125 0 98278.45
158.09280395507812 0 142338.23
158.10031127929688 0 17259.52
159.09652709960938 0 13566.138
159.11300659179688 0 102457.05
160.11656188964844 0 14085.678
165.1138153076172 0 16524.926
166.08653259277344 0 198061.86 y 14
166.09530639648438 0 12905.895
166.0977783203125 0 14691.3955
167.0816192626953 0 40085.695
167.08889770507812 0 16198.294
169.0973358154297 0 25031.229
171.07669067382812 0 230231.36
171.11456298828125 0 10430.673
172.0607147216797 0 28809.746
173.1287841796875 0 2161401.5 a 1
173.44024658203125 0 46635.223
174.1321563720703 0 177157.84
175.0716552734375 0 214120.44
175.11924743652344 0 264509.03
176.07530212402344 0 12569.029
182.09304809570312 0 22358.281
183.1130828857422 0 1396720.5 b Water loss 1
184.097412109375 0 11961.281
184.108642578125 0 21143.742
184.11654663085938 0 145582.38
185.0558319091797 0 20845.062
185.09231567382812 0 136999.78
185.1016845703125 0 18110.734
187.07196044921875 0 10800.969
187.10801696777344 0 165003.02
189.0872802734375 0 1011935.6
190.09071350097656 0 63824.977
192.097900390625 0 16625.61
192.11312866210938 0 21999.803
194.0924530029297 0 33384.008
194.12933349609375 0 14494.285
195.07705688476562 0 14886.44
195.1129150390625 0 12943.982
196.10755920410156 0 15563.94 y Water loss 12
199.0715789794922 0 324336.94
199.08209228515625 0 16096.324
199.1078338623047 0 11492.484 y 10
199.11891174316406 0 12016.152
200.07472229003906 0 21006.219
201.12362670898438 0 2700712.2 b 1
202.0822296142578 0 16964.404
202.1184539794922 0 65038.62
202.12713623046875 0 251012.22
203.06651306152344 0 251601.38
203.10289001464844 0 22388.473
203.12869262695312 0 16214.445
204.0696563720703 0 14025.982
210.087646484375 0 18653.027
211.10794067382812 0 117156.47
212.1031951904297 0 107906.98
213.08702087402344 0 16433.914
216.13461303710938 0 21321.676
217.08218383789062 0 1461092
218.08560180664062 0 139803.36
219.08712768554688 0 20959.545
220.0928497314453 0 28479.168
224.103271484375 0 111488.484
226.11851501464844 0 17922.824
227.06651306152344 0 36713.473
227.1142120361328 0 301715.47
228.1175079345703 0 19942.107
229.1185302734375 0 134013.56
230.11376953125 0 53602.72
237.09860229492188 0 45687.54
237.1348419189453 0 54672.23
238.0826416015625 0 12025.02
242.11380004882812 0 267687.03
242.127685546875 0 23636.906
243.11729431152344 0 25327.248
243.1321563720703 0 13287.308
244.1295166015625 0 96492.98
244.14039611816406 0 15445.961
245.1250762939453 0 69727.07
248.62965393066406 0 43163.008 y 11
252.1344451904297 0 17132.389
254.1135711669922 0 30745.459
254.16224670410156 0 23717.457
255.10903930664062 0 301777.78
256.1123962402344 0 34969.07
256.128662109375 0 16233.603
256.6573486328125 0 11618.749
259.140625 0 37784.78
259.1559753417969 0 22172.732
262.1524963378906 0 99929.83
263.1388244628906 0 57185.156 b 6
263.1529235839844 0 11544.964
266.1142272949219 0 14034.057
267.133544921875 0 17520.326
270.14501953125 0 33136.53
272.13568115234375 0 48047.902
278.12469482421875 0 23366.576
279.1097106933594 0 26970.518
280.1297912597656 0 26592.035
281.1256103515625 0 21130.762
284.1241455078125 0 64616.754
284.1611633300781 0 96052.22 a Water loss 2
285.1649475097656 0 26461.39
288.13555908203125 0 25483.238
288.1555480957031 0 51189.832
289.1515197753906 0 19287.254
294.1455383300781 0 19785.672
295.1416015625 0 16243.731
295.17718505859375 0 15570.877
296.1356506347656 0 80970.28
296.1719665527344 0 20723.957
297.12005615234375 0 18078.068
297.4077453613281 0 13969.939
298.1402893066406 0 161503.28
298.1625061035156 0 23221.816 y 10
299.13653564453125 0 55753.266
301.1510925292969 0 11971.147
302.13507080078125 0 47902.125
302.17138671875 0 27662.42 a 2
305.16131591796875 0 479455.88 y Ammonia loss 13
306.12091064453125 0 13367.5
306.1642761230469 0 78362.21
312.15582275390625 0 1399401.6 b Water loss 2
313.1583557128906 0 199804.92
313.1875305175781 0 327767.1
314.1462707519531 0 226358.83
314.16290283203125 0 14278.426
314.1911315917969 0 49415.383
315.1500244140625 0 35058.438
316.15081787109375 0 334457.34 b 5
317.148681640625 0 59085.766
319.16204833984375 0 14159.276
322.18756103515625 0 93458.64 y 13
323.1717224121094 0 23265.523
324.130859375 0 69084.56
325.11358642578125 0 22430.818
326.6717224121094 0 10610.452 y 9
330.16644287109375 0 663139.2 b 2
331.17071533203125 0 187956.95
332.15753173828125 0 83445.984
332.177001953125 0 15892.428
336.17681884765625 0 25417.537
336.67889404296875 0 21283.31
341.1824951171875 0 197607.42
342.14129638671875 0 378437.5
342.1853942871094 0 33331.94
343.14459228515625 0 66431.08
344.1451721191406 0 20246.652
345.1827697753906 0 57364.914
349.1518249511719 0 21947.117
349.18341064453125 0 133603.64
350.1724548339844 0 14388.485
352.1617736816406 0 15057.775
357.1558532714844 0 19300.361
358.1606140136719 0 14913.157
359.1673583984375 0 268054.72
359.1893615722656 0 13663.481
360.1707763671875 0 28967.438
365.1819763183594 0 21291.53
367.16033935546875 0 23678.482
373.5972595214844 0 11836.668
374.18310546875 0 21291.514
375.16754150390625 0 23326.541
380.1562194824219 0 20459.512
383.2297058105469 0 17750.012
385.17181396484375 0 64925.645
389.9110412597656 0 13178.561
391.20880126953125 0 14744.275 y Water loss 12
392.160888671875 0 14938.896
392.1929931640625 0 100294.305 y Ammonia loss 12
393.70941162109375 0 26528.955
394.2007141113281 0 22266.998 b 6
395.1575012207031 0 23823.773
396.18939208984375 0 22385.936
398.2138366699219 0 39501.1
398.8653564453125 0 13681.223 y 4
399.7134094238281 0 17974.932
400.1865539550781 0 13948.406
400.2193908691406 0 36711.047
401.23974609375 0 228845.45
402.24334716796875 0 54657.402
403.1824035644531 0 40951.375
405.6882019042969 0 15921.901
405.7156066894531 0 36702.547
408.23016357421875 0 17787.979
409.2198486328125 0 147460.92 y 12
410.2038269042969 0 39656.676 y Ammonia loss 7
411.222900390625 0 26870.139 b Water loss 3
413.1662902832031 0 30349.695
413.2154846191406 0 22449.934
414.22802734375 0 85380.39
414.72100830078125 0 154381.38
415.2220458984375 0 62777.258
415.7211608886719 0 23552.15
418.7171630859375 0 188304.77 y 7
419.2181701660156 0 74263.734
419.7198486328125 0 23726.646
423.2335205078125 0 581910.7
423.7347412109375 0 240783.3
424.236572265625 0 44419.742
424.7406005859375 0 15139.094
428.21514892578125 0 72019.32
429.2347106933594 0 222094.11 b 3
430.2393493652344 0 105893.32
431.244140625 0 19889.86
439.1946105957031 0 24140.166
441.21063232421875 0 87122.234
442.21002197265625 0 21356.338
443.2034606933594 0 15083.327
445.2418518066406 0 16455.04
448.251953125 0 28109.387
452.26165771484375 0 54911.12
453.2660827636719 0 16213.752
455.2255554199219 0 15592.913
455.7284240722656 0 14129.941
458.23638916015625 0 170123.34
459.2369689941406 0 41916.445
461.2135314941406 0 20608.812
462.8973693847656 0 17982.5
463.23297119140625 0 30794.28
467.2424011230469 0 43869.92
467.7459411621094 0 26904.695
468.2464294433594 0 17005.652
468.89910888671875 0 17355.691 y Water loss 2
469.234375 0 28588.162 y Ammonia loss 2
469.2882995605469 0 57370.45
470.23675537109375 0 27278.064
470.291259765625 0 15772.038
471.7439880371094 0 18724.521
472.24688720703125 0 12655.884
474.90509033203125 0 35228.258 y 2
475.23748779296875 0 24470.92
475.5740661621094 0 15339.543
478.2038879394531 0 15349.642
478.2414245605469 0 32854.13 y Water loss 11
479.2255859375 0 92806.336 y Ammonia loss 11
480.2577209472656 0 14444.927
480.74676513671875 0 69758.516
481.2491455078125 0 40023.96
481.7582092285156 0 15539.324
487.25689697265625 0 27965.412
487.7617492675781 0 40555.13 y Water loss 6
488.2569274902344 0 133073.98 y Ammonia loss 6
488.7577209472656 0 61456.51
489.2592468261719 0 31791.04
491.911376953125 0 19132.65
494.7375183105469 0 13827.539
496.2185363769531 0 12912.839
496.2517395019531 0 664020.3 y 11
496.76763916015625 0 1790347.4 y 6
497.26806640625 0 867148.1
497.770263671875 0 297184.7
497.9109191894531 0 57959.473 y Water loss 1
498.2684326171875 0 60673.363
498.912841796875 0 16092.171
500.1963195800781 0 21885.785
501.7606201171875 0 16754.3
503.9145202636719 0 71602.82 y 1
504.2491760253906 0 93238.46
504.584228515625 0 31948.174
504.9169006347656 0 15977.412
505.2731628417969 0 130344.25
506.2780456542969 0 14457.278
506.74359130859375 0 21847.713
507.2386779785156 0 12347.969
507.749267578125 0 15216.843
513.2423706054688 0 18309.836
514.2615966796875 0 16363.732
515.2564086914062 0 135632.92
515.755615234375 0 37938.496
516.2655639648438 0 25357.805 b 4
517.2682495117188 0 16039.993
518.2106323242188 0 25721.322
524.2625122070312 0 521076.28
524.763916015625 0 233044.1
525.2650146484375 0 73508.27
526.2272338867188 0 16011.373
529.6023559570312 0 28157.566
529.9349365234375 0 39157.83
530.2686767578125 0 23866.076
532.2750244140625 0 21335.922
533.2750244140625 0 15054.688
535.60595703125 0 289479.9 Precursor Water loss
535.9403076171875 0 253836.69 Precursor Ammonia loss
536.2744750976562 0 155040.1
536.607421875 0 51689.695
539.293701171875 0 17022.746
541.2734375 0 16788.947
541.609375 0 675903.44 Precursor
541.9437866210938 0 619645.3
542.2778930664062 0 322326.28
542.6123046875 0 78566.78
543.2537231445312 0 36859.97
545.2763061523438 0 20765.734 y Water loss 5
546.2688598632812 0 15095.743
547.2736206054688 0 17495.49
550.2862548828125 0 16995.137
550.7947998046875 0 26759.762
551.2564086914062 0 17569.746
554.2807006835938 0 716936.2 y 5
554.781982421875 0 402116.5
555.2837524414062 0 137828.6 b Water loss 10
555.7852783203125 0 33974.953 b Ammonia loss 10
556.2789306640625 0 26110.332
556.322509765625 0 22818.684
556.7830200195312 0 19576.076
558.7904052734375 0 34058.438
559.282470703125 0 37307.777
564.7911376953125 0 127599.44
565.2368774414062 0 42862.824
565.2919311523438 0 70602.086
565.7909545898438 0 47270.18
566.3031616210938 0 42748.434
567.2891235351562 0 27602.945
568.2743530273438 0 18796.969
569.267578125 0 106167.97
570.2649536132812 0 40516.93
572.2783203125 0 40999.27
573.280029296875 0 18475.941
573.7966918945312 0 711062.44
574.2980346679688 0 334917
574.7998657226562 0 125352.36
575.29736328125 0 58227.01
576.2737426757812 0 12745.395
582.7919921875 0 44369.133
583.24658203125 0 17795.074
584.3145141601562 0 57932.613
585.31640625 0 25852.127
587.2786254882812 0 132234.9
588.2809448242188 0 55679.09
588.7916870117188 0 103765.01 y Water loss 4
589.288330078125 0 100021 y Ammonia loss 4
589.7855224609375 0 36580.645
590.2930297851562 0 23010.012
592.30517578125 0 175180.84
593.3056640625 0 46295.023
594.309814453125 0 14428.366
595.3203735351562 0 90905.19 y 10
596.3212890625 0 38904.047
597.7968139648438 0 5144282 y 4
598.2982788085938 0 3331106
598.7994995117188 0 1213189.1 b Water loss 11
599.3004760742188 0 321795.06 b Ammonia loss 11
599.8024291992188 0 72449.96
600.2747192382812 0 97959.44
601.27587890625 0 22841.533
602.790283203125 0 20886.932
609.3015747070312 0 16456.775
615.3153686523438 0 23032.188
617.303466796875 0 25523.268
617.80859375 0 36157.312
618.3057250976562 0 16134.543
620.31005859375 0 26820.703
620.8068237304688 0 37892.42
621.3086547851562 0 17161.432
623.321533203125 0 26687.5
624.316162109375 0 27646.37
626.3251953125 0 63646.355
629.314453125 0 276395.2
629.81396484375 0 179774.81
630.315185546875 0 89058.19
630.8089599609375 0 18254.408
632.8271484375 0 21211.896
635.3203125 0 32201.2 y Ammonia loss 9
636.3099975585938 0 42577.617
638.3192749023438 0 792862.5 y Water loss 3
638.8201904296875 0 656022.8 y Ammonia loss 3
639.2661743164062 0 19510.84
639.3215942382812 0 274818.16
639.822021484375 0 82604.38
641.3365478515625 0 36290.38
642.323974609375 0 37863.86 b Water loss 12
643.3505859375 0 41001.043
644.35986328125 0 24958.725
645.305908203125 0 18830.285
646.315185546875 0 96629.55
647.3310546875 0 4826691 y 3
647.83251953125 0 3542568.2
648.3334350585938 0 1513009.9
648.3997192382812 0 25201.932
648.8348999023438 0 337959.38
649.3353271484375 0 88007.734
651.8155517578125 0 18801.674
652.3405151367188 0 272851.56 y 9
653.3409423828125 0 143051.12
654.3228759765625 0 81696.37
655.3220825195312 0 49374.53
656.2965087890625 0 23854.05
657.2914428710938 0 20141.76
658.8296508789062 0 25273.225
659.345947265625 0 51257.957
663.823486328125 0 36044.332
664.3143920898438 0 46370.242
664.8220825195312 0 23641.107
666.8284301757812 0 67775.6
667.3268432617188 0 42356.215
671.3469848632812 0 344151.06
672.3426513671875 0 159003.12
672.828857421875 0 276699.47
673.3302612304688 0 258967.33
673.8297729492188 0 76385.81
674.3116455078125 0 256408.55
674.8278198242188 0 17208.287
675.3135986328125 0 78812.52
681.3375244140625 0 28477.18
681.8338012695312 0 1455802.8
682.3343505859375 0 1061719.8
682.83642578125 0 389040.94
683.3351440429688 0 94032.14
686.8255004882812 0 40018.223
687.326416015625 0 34392.63
687.829833984375 0 29472.441
688.3740844726562 0 23206.547
689.3579711914062 0 826923.8
690.3604125976562 0 286345.5
690.832275390625 0 17861.832
691.3612670898438 0 77905.85
693.836181640625 0 16510.941
694.3409423828125 0 51462.93
694.8345947265625 0 20417.723
696.8464965820312 0 39956.707
697.3480834960938 0 19480.104
699.3419799804688 0 164447.38
700.3438720703125 0 52904.625
702.8471069335938 0 774144.25 y Water loss 2
703.34814453125 0 655150.5 y Ammonia loss 2
703.8486938476562 0 247801.11
704.3472900390625 0 112318.22
704.8512573242188 0 30166.01
711.8521118164062 0 7180893.5 y 2
712.3536987304688 0 5524490
712.498779296875 0 39873.715
712.8548583984375 0 2424133.8
713.3561401367188 0 680040.06
713.8574829101562 0 120957.69
716.8433837890625 0 27362.941
717.3518676757812 0 15824.983
721.3648071289062 0 43206.508 y Water loss 8
722.351806640625 0 54773.64 y Ammonia loss 8
722.85302734375 0 33054.65
723.3561401367188 0 25901.094
723.8529052734375 0 18395.934
725.8475952148438 0 20614.26
726.3477783203125 0 34368.453
728.3513793945312 0 64480.715
728.8550415039062 0 55116.074
729.3531494140625 0 41768.37
731.3558959960938 0 58173.258
731.8597412109375 0 31617.793
732.360595703125 0 23575.652
732.8700561523438 0 24634.502
734.3516845703125 0 44766.395
734.8510131835938 0 32007.53
737.3570556640625 0 138907.47
737.8572387695312 0 117723.54
738.3577880859375 0 57466.855
738.8580322265625 0 15784.88
739.3726806640625 0 341987.4 y 8
740.3707275390625 0 251505.58
740.86328125 0 63783.277
741.3688354492188 0 77852.44
745.3569946289062 0 25862.795
746.36279296875 0 1072161.6 y Water loss 1
746.8637084960938 0 845786.25 y Ammonia loss 1
747.3639526367188 0 439403.8
747.8651123046875 0 169121.36
748.3645629882812 0 40536.062
751.3568725585938 0 43032.68
751.8556518554688 0 41038.984
753.3526000976562 0 25141.582
755.3682861328125 0 12283336 y 1
755.8696899414062 0 10224948
756.3707275390625 0 4748330
756.871826171875 0 1389350.1
757.3734741210938 0 334067.84
758.3788452148438 0 55477.316
759.3814697265625 0 24360.48
760.3599243164062 0 453304.97
760.8612060546875 0 373066.66
761.3623046875 0 166352.89
761.864013671875 0 55981.207
768.3583984375 0 34902.953
768.4066772460938 0 30951.686
769.349853515625 0 18590.037
770.4129028320312 0 99189.67
771.415283203125 0 35563.984
783.3814086914062 0 16542.783
785.4271240234375 0 21736.361
786.3795166015625 0 60037.848
787.3911743164062 0 69855 b 6
788.397216796875 0 59681.555
789.4024047851562 0 19517.477
798.4107666015625 0 65251.008
799.4157104492188 0 37441.55
800.3975830078125 0 49579.957
801.3956298828125 0 21399.584
806.4149169921875 0 48229.07
810.3743286132812 0 165885.38
811.3857421875 0 75156.59
812.3948364257812 0 19741.145
818.4148559570312 0 156044.61 y Water loss 7
819.4054565429688 0 136413.97 y Ammonia loss 7
820.4049072265625 0 75450.445
828.3734741210938 0 29421.48
828.4323120117188 0 79907.09
829.4306030273438 0 49556.97
830.4028930664062 0 22457.695
836.4254760742188 0 1645983 y 7
837.4286499023438 0 740001.9
838.4310302734375 0 189655.16
839.4325561523438 0 35366.082
845.4105834960938 0 40858.312
855.401123046875 0 38919.137
857.4456176757812 0 44917.64
858.3892211914062 0 24188.645
867.4312744140625 0 38421.668 b Ammonia loss 7
873.40283203125 0 41418.414
881.4486694335938 0 34372.35
885.439208984375 0 45205.51
887.4254760742188 0 69869.3
888.4213256835938 0 33883.96
890.4254150390625 0 26853.957
891.4259643554688 0 31269.953
899.4550170898438 0 64151.36
900.4585571289062 0 45660.53
901.4616088867188 0 21894.965
909.4402465820312 0 54597.35
910.444091796875 0 20574.795
915.4164428710938 0 99392.08
916.422607421875 0 37742.363
917.4227905273438 0 20520.56
925.44580078125 0 18335.15
926.4481201171875 0 33268.027
927.45263671875 0 96691.11
928.4563598632812 0 50763.285
929.4623413085938 0 17340.396
933.4773559570312 0 23290.959
943.4618530273438 0 69560.86
944.4723510742188 0 50501.453
945.4796752929688 0 42798.484
951.4635009765625 0 22130.617
954.4638671875 0 26954.256 b Ammonia loss 8
957.4765625 0 30566.94
958.4761352539062 0 82266.54
959.4810180664062 0 27576.475
968.4798583984375 0 39571.64
969.4756469726562 0 27891.074
970.4729614257812 0 21700.072
971.49072265625 0 25103.283 b 8
972.479248046875 0 41504.81
975.5001831054688 0 170881.86 y Ammonia loss 6
976.5023193359375 0 77322.85
977.5004272460938 0 43340.684
984.474853515625 0 27631.309
985.4896240234375 0 23385.543
986.4884033203125 0 201200.58
987.4893798828125 0 158460.1
988.4900512695312 0 74884.76
989.4960327148438 0 50655.14
990.4970703125 0 39724.625
991.4901733398438 0 18029.266
996.4763793945312 0 33929.574
997.4682006835938 0 35321.035
1000.4789428710938 0 17218.994
1012.4791259765625 0 122620.66
1013.4862060546875 0 74756.65
1014.4830932617188 0 205855.23
1015.4871215820312 0 95184.65
1016.4856567382812 0 22852.6
1028.4974365234375 0 32904.89 b 9
1029.5067138671875 0 21372.215
1030.4908447265625 0 182397.38
1031.493896484375 0 113668.35
1032.5008544921875 0 35280.195
1055.5052490234375 0 31515.209
1056.5155029296875 0 17176.654
1057.5157470703125 0 18500.105
1072.515625 0 18834.361
1073.52001953125 0 104691.09
1074.5191650390625 0 52324.887
1075.52685546875 0 19191.717
1083.511474609375 0 60656.72
1084.5101318359375 0 25371.627
1086.5496826171875 0 36349.598
1090.524658203125 0 116066.414 y Ammonia loss 5
1091.529296875 0 78974.5
1092.5323486328125 0 33782.92
1093.53759765625 0 24230.355
1100.5361328125 0 18889.785
1101.5155029296875 0 121043.375
1102.5152587890625 0 50730.715
1103.5269775390625 0 34403.098
1111.544677734375 0 43960.75
1112.547119140625 0 31110.607
1113.553466796875 0 17862.564
1118.5306396484375 0 17723.555
1119.53466796875 0 37066.473
1120.5426025390625 0 26447.074
1129.5606689453125 0 32216.605
1130.53857421875 0 28099.596
1134.5552978515625 0 24790.89
1135.5411376953125 0 25121.139
1142.5291748046875 0 30965.846
1143.543212890625 0 32173.592
1152.542236328125 0 23885.395
1159.54736328125 0 59757.637
1160.5513916015625 0 92197.81
1161.552734375 0 50063.21
1170.54638671875 0 20450.174
1177.5576171875 0 402300.4 y Ammonia loss 4
1178.560302734375 0 251742.22
1179.561767578125 0 84481.13
1180.5657958984375 0 17996.668
1187.5567626953125 0 31510.498
1188.550537109375 0 69366.48
1189.5545654296875 0 32716.309
1205.57275390625 0 58110.875
1206.569580078125 0 54995.12
1207.5667724609375 0 24522.947
1276.625244140625 0 72745.914 y Ammonia loss 3
1277.6263427734375 0 26600.54
1299.631591796875 0 26173.66
1300.638427734375 0 15490.252
2145.9482421875 0 12823.079
2444.39794921875 0 14713.843
3385.8486328125 0 17908.816

Spectrum Details

|  |  |
| --- | --- |
| Matched peaks? Matched peaks The total absolute number of peaks matched. Additionally in brackets the total fraction of peaks matched and the total number of peaks is shown. | 79 (12.06% of 655) |
| FDR? FDR The false discovery rate estimated for this peptide. It is calculated by matching all theoretical fragments with a non-integer shift with the raw peaks for this spectrum. This is done with 40 different shifts. The resulting percentage is the average number of annotated peaks over the number of annotated peaks with the correct spectrum. | 0.06% |
| Satellite FDR? Satellite FDR See the FDR for details on its calculation. This satellite ion specific FDR only contains the satellite ions (d/w) for I/L/J positions. | - |
| PSM Score? PSM Score The PSM Score as given by Hecklib to this annotated spectrum. It is shown with three significant figures. | 532 |

#### Spectrum 4925? Spectrum 4925 The raw spectrum of this peptide as annotated by Hecklib. The fragments are coloured according to ion type (see legend). Any peaks with a star '\*' as text can be hovered over to see the full details, first the ion type second the mass shift type. By hovering over the amino acids in the peptide or ions in the legend the corresponding peaks are highlighted. By toggling the 'Unassigned' label you can turn the background (unassigned) peaks on or off in the plot. By updating the slider in the Ion legend you can update the spectrum to only show the top X% of the peaks with labels. The top X% means any peak that is within X% of the highest intensity. By dragging in the spectrum you can zoom in to a specific part of the spectrum and use 'Zoom Out' to get back to the original zoom level. The annotation of the spectrum is based on the given sequence in the peptides file and is done with different software so inconsistencies are likely. The peaks are annotated based on the given sequence, with 20 ppm tolerance.

Copy Data

##### Spectrum 4925 (TSV)

###### Preview

```
Loading example...
```

*Click on the button to copy the data to your clipboard.*

Mz MinMz MaxIntensity Max

WidthHeightPeptide font sizePeptide stroke widthSpectrum font sizeSpectrum stroke widthCompact peptide

Ion legend

wxyz

abcd

OtherUnassignedIon^Charge^_Position_Show for top:%

JSEVSDRPSGVSSRF

01.50e+52.99e+54.49e+55.99e+5

Zoom Out

a^+1^_2_y^+1^_1_a^+1^_2_b^+1^_2_y^+3^_5_b^+1^_2_y^+2^_4_b^+3^_7_a^+1^_3_y^+2^_5_a^+1^_3_y^+1^_2_b^+1^_3_b^+2^_6_y^+1^_2_b^+1^_3_y^+1^_3_y^+1^_3_y^+3^_11_b^+2^_7_y^+3^_11_y^+1^_3_y^+2^_8_b^+1^_4_y^+2^_8_b^+1^_4_y^+3^_12_y^+3^_13_y^+3^_13_y^+3^_13_y^+1^_4_y^+1^_4_y^+2^_9_y^+2^_9_y^+1^_4_y^+2^_9_y^+3^_14_y^+3^_14_y^+3^_14_b^+1^_5_\*\*\*y^+2^_10_y^+2^_10_b^+2^_11_y^+2^_11_y^+2^_11_y^+1^_5_y^+2^_11_b^+2^_12_b^+2^_12_y^+1^_6_y^+2^_12_y^+2^_12_y^+2^_12_y^+1^_6_y^+2^_13_y^+2^_13_y^+2^_13_y^+1^_7_y^+1^_7_y^+1^_7_y^+2^_14_y^+2^_14_y^+2^_14_b^+1^_7_b^+1^_7_y^+1^_8_y^+1^_8_y^+1^_8_b^+1^_9_y^+1^_9_b^+1^_10_y^+1^_10_b^+1^_11_y^+1^_11_y^+1^_12_

0686137120572743

Fragment Matches Table

Show background peaks

| Position | Ion type | Intensity | mz Theoretical | mz Error (Th) | mz Error (ppm) | Charge | Series Number |
| --- | --- | --- | --- | --- | --- | --- | --- |
| - | - | 2.757E+04 | 120.1 | - | - | 0 | - |
| - | - | 2683 | 121.1 | - | - | 0 | - |
| - | - | 517 | 125.1 | - | - | 0 | - |
| - | - | 4134 | 125.1 | - | - | 0 | - |
| - | - | 6987 | 126.1 | - | - | 0 | - |
| - | - | 950.3 | 127.1 | - | - | 0 | - |
| - | - | 335.8 | 127.1 | - | - | 0 | - |
| - | - | 448.1 | 128 | - | - | 0 | - |
| - | - | 3783 | 128.1 | - | - | 0 | - |
| - | - | 1479 | 129.1 | - | - | 0 | - |
| - | - | 8149 | 129.1 | - | - | 0 | - |
| - | - | 377 | 129.1 | - | - | 0 | - |
| - | - | 1723 | 130.1 | - | - | 0 | - |
| - | - | 1100 | 130.1 | - | - | 0 | - |
| - | - | 625.7 | 130.1 | - | - | 0 | - |
| - | - | 572 | 130.1 | - | - | 0 | - |
| - | - | 569.7 | 131.1 | - | - | 0 | - |
| - | - | 1066 | 132.1 | - | - | 0 | - |
| - | - | 678.3 | 133.1 | - | - | 0 | - |
| - | - | 3720 | 136.1 | - | - | 0 | - |
| - | - | 2316 | 138.1 | - | - | 0 | - |
| - | - | 902.1 | 139.1 | - | - | 0 | - |
| - | - | 2287 | 140.1 | - | - | 0 | - |
| - | - | 1074 | 141.1 | - | - | 0 | - |
| - | - | 852.9 | 143.1 | - | - | 0 | - |
| - | - | 1706 | 144.1 | - | - | 0 | - |
| - | - | 789.9 | 145.1 | - | - | 0 | - |
| - | - | 3138 | 147.1 | - | - | 0 | - |
| - | - | 514.3 | 148.9 | - | - | 0 | - |
| - | - | 731.6 | 153.1 | - | - | 0 | - |
| - | - | 6611 | 154.1 | - | - | 0 | - |
| - | - | 742.4 | 155.1 | - | - | 0 | - |
| - | - | 613.1 | 155.1 | - | - | 0 | - |
| 2 | a | 6599 | 155.1 | 0.0003966 | 2.557 | +1 | 2 |
| - | - | 1382 | 156.1 | - | - | 0 | - |
| - | - | 3505 | 156.1 | - | - | 0 | - |
| - | - | 641.3 | 156.1 | - | - | 0 | - |
| - | - | 6196 | 157.1 | - | - | 0 | - |
| - | - | 5665 | 157.1 | - | - | 0 | - |
| - | - | 5673 | 157.1 | - | - | 0 | - |
| - | - | 6970 | 158.1 | - | - | 0 | - |
| - | - | 834.1 | 158.1 | - | - | 0 | - |
| - | - | 719.6 | 159.1 | - | - | 0 | - |
| - | - | 5335 | 159.1 | - | - | 0 | - |
| - | - | 679 | 160.1 | - | - | 0 | - |
| - | - | 4042 | 161 | - | - | 0 | - |
| - | - | 504.8 | 163 | - | - | 0 | - |
| - | - | 671.1 | 164.1 | - | - | 0 | - |
| - | - | 1101 | 165.1 | - | - | 0 | - |
| - | - | 508.8 | 165.1 | - | - | 0 | - |
| 15 | y | 2.01E+04 | 166.1 | 0.0003843 | 2.314 | +1 | 1 |
| - | - | 771.1 | 166.1 | - | - | 0 | - |
| - | - | 1016 | 167.1 | - | - | 0 | - |
| - | - | 2003 | 167.1 | - | - | 0 | - |
| - | - | 459.8 | 169.1 | - | - | 0 | - |
| - | - | 1791 | 169.1 | - | - | 0 | - |
| - | - | 514.7 | 170 | - | - | 0 | - |
| - | - | 9848 | 171.1 | - | - | 0 | - |
| - | - | 1176 | 172.1 | - | - | 0 | - |
| - | - | 624.1 | 172.1 | - | - | 0 | - |
| - | - | 453 | 172.7 | - | - | 0 | - |
| 2 | a | 9.734E+04 | 173.1 | 0.0004673 | 2.699 | +1 | 2 |
| - | - | 817.8 | 173.4 | - | - | 0 | - |
| - | - | 929.5 | 174.1 | - | - | 0 | - |
| - | - | 9624 | 174.1 | - | - | 0 | - |
| - | - | 1.039E+04 | 175.1 | - | - | 0 | - |
| - | - | 1.128E+04 | 175.1 | - | - | 0 | - |
| - | - | 624.5 | 175.1 | - | - | 0 | - |
| - | - | 490.3 | 176.1 | - | - | 0 | - |
| - | - | 850.4 | 177.1 | - | - | 0 | - |
| - | - | 641 | 181.1 | - | - | 0 | - |
| - | - | 569.7 | 181.1 | - | - | 0 | - |
| - | - | 2070 | 182.1 | - | - | 0 | - |
| - | - | 1002 | 182.1 | - | - | 0 | - |
| 2 | b | 6.721E+04 | 183.1 | 0.0004618 | 2.522 | +1 | 2 |
| - | - | 1330 | 183.1 | - | - | 0 | - |
| - | - | 474.8 | 184.1 | - | - | 0 | - |
| - | - | 1579 | 184.1 | - | - | 0 | - |
| - | - | 6178 | 184.1 | - | - | 0 | - |
| - | - | 1082 | 185.1 | - | - | 0 | - |
| - | - | 6650 | 185.1 | - | - | 0 | - |
| - | - | 530.1 | 186.1 | - | - | 0 | - |
| - | - | 867.6 | 187.1 | - | - | 0 | - |
| - | - | 7785 | 187.1 | - | - | 0 | - |
| - | - | 497.2 | 187.6 | - | - | 0 | - |
| - | - | 531.1 | 188.1 | - | - | 0 | - |
| - | - | 4.138E+04 | 189.1 | - | - | 0 | - |
| - | - | 2941 | 190.1 | - | - | 0 | - |
| - | - | 514.5 | 191.1 | - | - | 0 | - |
| - | - | 480.8 | 192.1 | - | - | 0 | - |
| - | - | 932.4 | 192.1 | - | - | 0 | - |
| - | - | 614.5 | 194.1 | - | - | 0 | - |
| - | - | 547 | 194.1 | - | - | 0 | - |
| - | - | 721 | 195.1 | - | - | 0 | - |
| - | - | 639.4 | 195.1 | - | - | 0 | - |
| - | - | 1305 | 197.1 | - | - | 0 | - |
| - | - | 1.516E+04 | 199.1 | - | - | 0 | - |
| 11 | y | 1831 | 199.1 | 0.003095 | 15.55 | +3 | 5 |
| - | - | 1040 | 200.1 | - | - | 0 | - |
| - | - | 517.2 | 200.1 | - | - | 0 | - |
| 2 | b | 1.312E+05 | 201.1 | 0.0004105 | 2.041 | +1 | 2 |
| - | - | 567.2 | 202.1 | - | - | 0 | - |
| - | - | 2569 | 202.1 | - | - | 0 | - |
| - | - | 1.148E+04 | 202.1 | - | - | 0 | - |
| - | - | 1.129E+04 | 203.1 | - | - | 0 | - |
| - | - | 991.5 | 203.1 | - | - | 0 | - |
| - | - | 812.3 | 203.1 | - | - | 0 | - |
| - | - | 1116 | 207.1 | - | - | 0 | - |
| - | - | 590.6 | 208.1 | - | - | 0 | - |
| - | - | 1020 | 210.1 | - | - | 0 | - |
| - | - | 528.6 | 210.1 | - | - | 0 | - |
| - | - | 6495 | 211.1 | - | - | 0 | - |
| - | - | 3945 | 212.1 | - | - | 0 | - |
| - | - | 879.3 | 213.1 | - | - | 0 | - |
| - | - | 906.3 | 216.1 | - | - | 0 | - |
| - | - | 6.764E+04 | 217.1 | - | - | 0 | - |
| - | - | 5014 | 218.1 | - | - | 0 | - |
| - | - | 506.5 | 219.1 | - | - | 0 | - |
| - | - | 1069 | 220.1 | - | - | 0 | - |
| - | - | 2902 | 223.1 | - | - | 0 | - |
| - | - | 5392 | 224.1 | - | - | 0 | - |
| - | - | 522.8 | 225.1 | - | - | 0 | - |
| - | - | 1113 | 225.1 | - | - | 0 | - |
| - | - | 1143 | 225.2 | - | - | 0 | - |
| - | - | 2284 | 226.1 | - | - | 0 | - |
| - | - | 792 | 227.1 | - | - | 0 | - |
| - | - | 931.3 | 227.1 | - | - | 0 | - |
| - | - | 1.281E+04 | 227.1 | - | - | 0 | - |
| - | - | 669.6 | 228.1 | - | - | 0 | - |
| - | - | 1480 | 228.1 | - | - | 0 | - |
| - | - | 6947 | 229.1 | - | - | 0 | - |
| - | - | 2389 | 230.1 | - | - | 0 | - |
| - | - | 739.4 | 233.1 | - | - | 0 | - |
| - | - | 780.1 | 233.2 | - | - | 0 | - |
| - | - | 1100 | 235.1 | - | - | 0 | - |
| - | - | 1819 | 237.1 | - | - | 0 | - |
| - | - | 2553 | 237.1 | - | - | 0 | - |
| - | - | 532.7 | 238.1 | - | - | 0 | - |
| - | - | 494.1 | 240.1 | - | - | 0 | - |
| - | - | 637.8 | 240.1 | - | - | 0 | - |
| - | - | 1.165E+04 | 242.1 | - | - | 0 | - |
| - | - | 816.7 | 242.2 | - | - | 0 | - |
| - | - | 880.6 | 243.1 | - | - | 0 | - |
| - | - | 4436 | 244.1 | - | - | 0 | - |
| - | - | 1093 | 244.1 | - | - | 0 | - |
| - | - | 3713 | 245.1 | - | - | 0 | - |
| 12 | y | 2507 | 248.6 | 0.0008536 | 3.433 | +2 | 4 |
| - | - | 878.1 | 249.1 | - | - | 0 | - |
| - | - | 1642 | 252.1 | - | - | 0 | - |
| - | - | 1200 | 252.1 | - | - | 0 | - |
| - | - | 2894 | 253 | - | - | 0 | - |
| - | - | 4214 | 253.1 | - | - | 0 | - |
| - | - | 1058 | 254.1 | - | - | 0 | - |
| - | - | 574.7 | 254.2 | - | - | 0 | - |
| - | - | 1.34E+04 | 255.1 | - | - | 0 | - |
| - | - | 518.1 | 255.1 | - | - | 0 | - |
| - | - | 1011 | 256.1 | - | - | 0 | - |
| - | - | 1072 | 256.1 | - | - | 0 | - |
| - | - | 774 | 257.1 | - | - | 0 | - |
| - | - | 680.7 | 258.1 | - | - | 0 | - |
| - | - | 2007 | 259.1 | - | - | 0 | - |
| - | - | 1038 | 259.2 | - | - | 0 | - |
| - | - | 4424 | 262.2 | - | - | 0 | - |
| 7 | b | 2032 | 263.1 | 0.002793 | 10.61 | +3 | 7 |
| - | - | 600 | 263.6 | - | - | 0 | - |
| - | - | 686.5 | 266.1 | - | - | 0 | - |
| - | - | 784.1 | 267.1 | - | - | 0 | - |
| - | - | 1119 | 267.1 | - | - | 0 | - |
| - | - | 1253 | 269.1 | - | - | 0 | - |
| - | - | 2624 | 270.1 | - | - | 0 | - |
| - | - | 1484 | 270.2 | - | - | 0 | - |
| - | - | 1969 | 272.1 | - | - | 0 | - |
| - | - | 641.6 | 274.1 | - | - | 0 | - |
| - | - | 998.2 | 279.1 | - | - | 0 | - |
| - | - | 1337 | 280.1 | - | - | 0 | - |
| - | - | 2000 | 281 | - | - | 0 | - |
| - | - | 820.7 | 283.1 | - | - | 0 | - |
| - | - | 2574 | 284.1 | - | - | 0 | - |
| 3 | a | 5029 | 284.2 | 0.0006197 | 2.181 | +1 | 3 |
| - | - | 609.2 | 285.1 | - | - | 0 | - |
| - | - | 1014 | 285.2 | - | - | 0 | - |
| - | - | 669.3 | 287.2 | - | - | 0 | - |
| - | - | 1240 | 288.1 | - | - | 0 | - |
| - | - | 2423 | 288.2 | - | - | 0 | - |
| - | - | 977.5 | 289.2 | - | - | 0 | - |
| - | - | 651.2 | 294.1 | - | - | 0 | - |
| - | - | 952.1 | 295.2 | - | - | 0 | - |
| - | - | 3261 | 296.1 | - | - | 0 | - |
| - | - | 662.7 | 296.2 | - | - | 0 | - |
| - | - | 700.9 | 297.1 | - | - | 0 | - |
| - | - | 1749 | 297.1 | - | - | 0 | - |
| - | - | 6354 | 298.1 | - | - | 0 | - |
| 11 | y | 825.6 | 298.2 | 0.004652 | 15.6 | +2 | 5 |
| - | - | 567.5 | 299.1 | - | - | 0 | - |
| - | - | 2463 | 299.1 | - | - | 0 | - |
| - | - | 659.7 | 300.1 | - | - | 0 | - |
| - | - | 914.3 | 301.1 | - | - | 0 | - |
| - | - | 643.6 | 302.1 | - | - | 0 | - |
| - | - | 2131 | 302.1 | - | - | 0 | - |
| 3 | a | 1309 | 302.2 | 0.0003089 | 1.022 | +1 | 3 |
| - | - | 610.5 | 303.6 | - | - | 0 | - |
| 14 | y | 2.103E+04 | 305.2 | 0.0005905 | 1.935 | +1 | 2 |
| - | - | 3294 | 306.2 | - | - | 0 | - |
| - | - | 642.7 | 307.2 | - | - | 0 | - |
| - | - | 726.9 | 310.1 | - | - | 0 | - |
| 3 | b | 6.633E+04 | 312.2 | 0.0006391 | 2.047 | +1 | 3 |
| - | - | 9799 | 313.2 | - | - | 0 | - |
| - | - | 1.307E+04 | 313.2 | - | - | 0 | - |
| - | - | 9277 | 314.1 | - | - | 0 | - |
| - | - | 2092 | 314.2 | - | - | 0 | - |
| - | - | 1531 | 315.1 | - | - | 0 | - |
| 6 | b | 1.591E+04 | 316.2 | 0.0006586 | 2.083 | +2 | 6 |
| - | - | 2512 | 317.2 | - | - | 0 | - |
| - | - | 1115 | 319.1 | - | - | 0 | - |
| - | - | 793 | 319.2 | - | - | 0 | - |
| - | - | 545.2 | 320.1 | - | - | 0 | - |
| 14 | y | 4196 | 322.2 | 0.0004696 | 1.458 | +1 | 2 |
| - | - | 927.8 | 323.2 | - | - | 0 | - |
| - | - | 772.3 | 323.2 | - | - | 0 | - |
| - | - | 3859 | 324.1 | - | - | 0 | - |
| - | - | 904.2 | 324.2 | - | - | 0 | - |
| - | - | 810.7 | 325.1 | - | - | 0 | - |
| - | - | 1079 | 326.1 | - | - | 0 | - |
| - | - | 572.5 | 329.1 | - | - | 0 | - |
| 3 | b | 3.029E+04 | 330.2 | 0.0007251 | 2.196 | +1 | 3 |
| - | - | 7472 | 331.2 | - | - | 0 | - |
| - | - | 2979 | 332.2 | - | - | 0 | - |
| - | - | 1362 | 332.2 | - | - | 0 | - |
| - | - | 1036 | 336.2 | - | - | 0 | - |
| - | - | 8807 | 341.2 | - | - | 0 | - |
| - | - | 1.815E+04 | 342.1 | - | - | 0 | - |
| - | - | 1606 | 342.2 | - | - | 0 | - |
| - | - | 1879 | 343.1 | - | - | 0 | - |
| - | - | 959.4 | 344.1 | - | - | 0 | - |
| - | - | 2228 | 345.2 | - | - | 0 | - |
| - | - | 970.3 | 349.2 | - | - | 0 | - |
| - | - | 5023 | 349.2 | - | - | 0 | - |
| - | - | 851.4 | 350.6 | - | - | 0 | - |
| - | - | 627 | 352.1 | - | - | 0 | - |
| - | - | 1.217E+04 | 359.2 | - | - | 0 | - |
| - | - | 2175 | 360.2 | - | - | 0 | - |
| - | - | 981 | 367.2 | - | - | 0 | - |
| - | - | 727.6 | 370.2 | - | - | 0 | - |
| - | - | 565.5 | 374.1 | - | - | 0 | - |
| - | - | 904 | 377.2 | - | - | 0 | - |
| - | - | 1196 | 380.2 | - | - | 0 | - |
| - | - | 738.6 | 381.2 | - | - | 0 | - |
| - | - | 2223 | 385.2 | - | - | 0 | - |
| - | - | 835 | 385.7 | - | - | 0 | - |
| 13 | y | 820.1 | 391.2 | 0.0002766 | 0.7071 | +1 | 3 |
| 13 | y | 3464 | 392.2 | 0.0003919 | 0.9993 | +1 | 3 |
| 5 | y | 1326 | 393.2 | 0.004194 | 10.67 | +3 | 11 |
| 7 | b | 1833 | 394.2 | 0.001342 | 3.405 | +2 | 7 |
| - | - | 962.2 | 394.7 | - | - | 0 | - |
| - | - | 1197 | 395.2 | - | - | 0 | - |
| - | - | 1286 | 396.2 | - | - | 0 | - |
| - | - | 744.7 | 396.7 | - | - | 0 | - |
| - | - | 1877 | 398.2 | - | - | 0 | - |
| 5 | y | 736.8 | 398.9 | 0.0002407 | 0.6034 | +3 | 11 |
| - | - | 830.4 | 399.2 | - | - | 0 | - |
| - | - | 1348 | 400.2 | - | - | 0 | - |
| - | - | 1973 | 400.3 | - | - | 0 | - |
| - | - | 1.066E+04 | 401.2 | - | - | 0 | - |
| - | - | 1077 | 401.3 | - | - | 0 | - |
| - | - | 1896 | 402.2 | - | - | 0 | - |
| - | - | 2604 | 403.2 | - | - | 0 | - |
| - | - | 838.9 | 404.2 | - | - | 0 | - |
| - | - | 1283 | 408.2 | - | - | 0 | - |
| 13 | y | 8161 | 409.2 | 0.0008814 | 2.154 | +1 | 3 |
| 8 | y | 1437 | 410.2 | 0.0006818 | 1.662 | +2 | 8 |
| 4 | b | 1804 | 411.2 | 0.0009813 | 2.386 | +1 | 4 |
| - | - | 1196 | 413.2 | - | - | 0 | - |
| - | - | 1067 | 413.2 | - | - | 0 | - |
| - | - | 3168 | 414.2 | - | - | 0 | - |
| - | - | 6529 | 414.7 | - | - | 0 | - |
| - | - | 2404 | 415.2 | - | - | 0 | - |
| - | - | 963.4 | 415.7 | - | - | 0 | - |
| 8 | y | 7866 | 418.7 | 0.001058 | 2.527 | +2 | 8 |
| - | - | 2462 | 419.2 | - | - | 0 | - |
| - | - | 1280 | 419.7 | - | - | 0 | - |
| - | - | 720.8 | 423.2 | - | - | 0 | - |
| - | - | 2.141E+04 | 423.2 | - | - | 0 | - |
| - | - | 1.008E+04 | 423.7 | - | - | 0 | - |
| - | - | 2441 | 424.2 | - | - | 0 | - |
| - | - | 3561 | 428.2 | - | - | 0 | - |
| 4 | b | 1.234E+04 | 429.2 | 0.0006705 | 1.562 | +1 | 4 |
| - | - | 4916 | 430.2 | - | - | 0 | - |
| - | - | 763.5 | 431.2 | - | - | 0 | - |
| - | - | 806.4 | 431.2 | - | - | 0 | - |
| 4 | y | 1028 | 431.9 | 0.0002091 | 0.4841 | +3 | 12 |
| - | - | 885.5 | 432.2 | - | - | 0 | - |
| - | - | 719.6 | 433.2 | - | - | 0 | - |
| - | - | 1090 | 439.2 | - | - | 0 | - |
| - | - | 3831 | 441.2 | - | - | 0 | - |
| - | - | 1021 | 443.2 | - | - | 0 | - |
| - | - | 842 | 445.2 | - | - | 0 | - |
| - | - | 1572 | 448.3 | - | - | 0 | - |
| - | - | 919.6 | 451.2 | - | - | 0 | - |
| - | - | 635.9 | 452.2 | - | - | 0 | - |
| - | - | 2827 | 452.3 | - | - | 0 | - |
| - | - | 747.3 | 455.2 | - | - | 0 | - |
| - | - | 7578 | 458.2 | - | - | 0 | - |
| - | - | 1972 | 459.2 | - | - | 0 | - |
| - | - | 2247 | 463.2 | - | - | 0 | - |
| - | - | 897.3 | 467.2 | - | - | 0 | - |
| - | - | 1051 | 467.7 | - | - | 0 | - |
| 3 | y | 1630 | 468.9 | 0.001709 | 3.644 | +3 | 13 |
| 3 | y | 1215 | 469.2 | 0.007536 | 16.06 | +3 | 13 |
| - | - | 3143 | 469.3 | - | - | 0 | - |
| - | - | 1089 | 470.2 | - | - | 0 | - |
| - | - | 1296 | 470.3 | - | - | 0 | - |
| - | - | 849.2 | 471.7 | - | - | 0 | - |
| 3 | y | 2019 | 474.9 | 0.0009345 | 1.968 | +3 | 13 |
| - | - | 2211 | 475.2 | - | - | 0 | - |
| 12 | y | 683.1 | 478.2 | 0.0005969 | 1.248 | +1 | 4 |
| 12 | y | 4095 | 479.2 | 0.0007121 | 1.486 | +1 | 4 |
| - | - | 701.8 | 480.2 | - | - | 0 | - |
| - | - | 4158 | 480.7 | - | - | 0 | - |
| - | - | 1479 | 481.2 | - | - | 0 | - |
| - | - | 892.9 | 481.8 | - | - | 0 | - |
| - | - | 775.1 | 482.2 | - | - | 0 | - |
| - | - | 1106 | 487.3 | - | - | 0 | - |
| 7 | y | 2008 | 487.8 | 0.0005442 | 1.116 | +2 | 9 |
| 7 | y | 5926 | 488.3 | 0.001711 | 3.504 | +2 | 9 |
| - | - | 2871 | 488.8 | - | - | 0 | - |
| - | - | 927.8 | 489.2 | - | - | 0 | - |
| - | - | 701.3 | 492.2 | - | - | 0 | - |
| - | - | 694.4 | 494.2 | - | - | 0 | - |
| - | - | 658.9 | 494.7 | - | - | 0 | - |
| - | - | 786.7 | 496.2 | - | - | 0 | - |
| 12 | y | 3.04E+04 | 496.3 | 0.0008354 | 1.683 | +1 | 4 |
| 7 | y | 8.134E+04 | 496.8 | 0.0007347 | 1.479 | +2 | 9 |
| - | - | 4.08E+04 | 497.3 | - | - | 0 | - |
| - | - | 1.239E+04 | 497.8 | - | - | 0 | - |
| 2 | y | 2397 | 497.9 | 9.702E-05 | 0.1949 | +3 | 14 |
| 2 | y | 1122 | 498.2 | 0.002109 | 4.233 | +3 | 14 |
| - | - | 1904 | 498.3 | - | - | 0 | - |
| - | - | 756.9 | 498.6 | - | - | 0 | - |
| - | - | 889.9 | 499.2 | - | - | 0 | - |
| - | - | 1267 | 500.2 | - | - | 0 | - |
| - | - | 655 | 501.8 | - | - | 0 | - |
| 2 | y | 3960 | 503.9 | 0.0001765 | 0.3503 | +3 | 14 |
| - | - | 1664 | 504.2 | - | - | 0 | - |
| - | - | 1203 | 504.6 | - | - | 0 | - |
| - | - | 967.2 | 504.9 | - | - | 0 | - |
| - | - | 4704 | 505.3 | - | - | 0 | - |
| - | - | 2271 | 506.3 | - | - | 0 | - |
| - | - | 851.2 | 507.2 | - | - | 0 | - |
| - | - | 716.8 | 510.7 | - | - | 0 | - |
| - | - | 988.4 | 511.3 | - | - | 0 | - |
| - | - | 1101 | 513.4 | - | - | 0 | - |
| - | - | 4029 | 515.3 | - | - | 0 | - |
| - | - | 2087 | 515.8 | - | - | 0 | - |
| 5 | b | 2084 | 516.3 | 0.003526 | 6.83 | +1 | 5 |
| - | - | 3908 | 520.3 | - | - | 0 | - |
| - | - | 873.1 | 520.8 | - | - | 0 | - |
| - | - | 2.108E+04 | 524.3 | - | - | 0 | - |
| - | - | 1.199E+04 | 524.8 | - | - | 0 | - |
| - | - | 4926 | 525.3 | - | - | 0 | - |
| - | - | 1125 | 525.8 | - | - | 0 | - |
| - | - | 2889 | 529.6 | - | - | 0 | - |
| - | - | 2873 | 529.9 | - | - | 0 | - |
| - | - | 2213 | 530.3 | - | - | 0 | - |
| - | - | 1039 | 530.6 | - | - | 0 | - |
| - | - | 2200 | 532.2 | - | - | 0 | - |
| - | - | 2311 | 532.3 | - | - | 0 | - |
| - | - | 778.2 | 533.3 | - | - | 0 | - |
| - | - | 793 | 533.7 | - | - | 0 | - |
| 0 | Precursor | 1.562E+04 | 535.6 | 0.0005079 | 0.9483 | +3 | -1 |
| 0 | Precursor | 1.498E+04 | 535.9 | 0.006548 | 12.22 | +3 | -1 |
| - | - | 7567 | 536.3 | - | - | 0 | - |
| - | - | 1294 | 536.6 | - | - | 0 | - |
| - | - | 787.1 | 536.9 | - | - | 0 | - |
| - | - | 717 | 538 | - | - | 0 | - |
| - | - | 796 | 538.3 | - | - | 0 | - |
| - | - | 905.5 | 540.2 | - | - | 0 | - |
| - | - | 2465 | 540.3 | - | - | 0 | - |
| - | - | 1251 | 540.8 | - | - | 0 | - |
| - | - | 1150 | 540.9 | - | - | 0 | - |
| - | - | 1023 | 541.3 | - | - | 0 | - |
| - | - | 896.5 | 541.3 | - | - | 0 | - |
| 0 | Precursor | 3.843E+04 | 541.6 | 0.0007095 | 1.31 | +3 | -1 |
| - | - | 2.996E+04 | 541.9 | - | - | 0 | - |
| - | - | 2221 | 542.2 | - | - | 0 | - |
| - | - | 1.78E+04 | 542.3 | - | - | 0 | - |
| - | - | 5400 | 542.6 | - | - | 0 | - |
| - | - | 2952 | 542.7 | - | - | 0 | - |
| - | - | 1305 | 542.9 | - | - | 0 | - |
| - | - | 1472 | 543.3 | - | - | 0 | - |
| 6 | y | 1502 | 545.8 | 0.004475 | 8.199 | +2 | 10 |
| - | - | 1219 | 550.3 | - | - | 0 | - |
| 6 | y | 3.202E+04 | 554.3 | 0.0005994 | 1.081 | +2 | 10 |
| - | - | 1.739E+04 | 554.8 | - | - | 0 | - |
| 11 | b | 7424 | 555.3 | 0.001443 | 2.598 | +2 | 11 |
| - | - | 992.5 | 555.8 | - | - | 0 | - |
| - | - | 1100 | 556.3 | - | - | 0 | - |
| - | - | 742.3 | 556.3 | - | - | 0 | - |
| - | - | 720.7 | 557.3 | - | - | 0 | - |
| - | - | 1027 | 558.8 | - | - | 0 | - |
| - | - | 1090 | 559.3 | - | - | 0 | - |
| - | - | 761.9 | 560.3 | - | - | 0 | - |
| - | - | 5278 | 564.8 | - | - | 0 | - |
| - | - | 1288 | 565.2 | - | - | 0 | - |
| - | - | 2789 | 565.3 | - | - | 0 | - |
| - | - | 727.3 | 565.8 | - | - | 0 | - |
| - | - | 949.7 | 568.8 | - | - | 0 | - |
| - | - | 4306 | 569.3 | - | - | 0 | - |
| - | - | 880.8 | 570.3 | - | - | 0 | - |
| - | - | 928 | 572.3 | - | - | 0 | - |
| - | - | 2.269E+04 | 573.8 | - | - | 0 | - |
| - | - | 1.682E+04 | 574.3 | - | - | 0 | - |
| - | - | 4891 | 574.8 | - | - | 0 | - |
| - | - | 1716 | 575.3 | - | - | 0 | - |
| - | - | 1474 | 582.8 | - | - | 0 | - |
| - | - | 1143 | 583.2 | - | - | 0 | - |
| - | - | 1065 | 584.3 | - | - | 0 | - |
| - | - | 1915 | 584.3 | - | - | 0 | - |
| - | - | 6483 | 587.3 | - | - | 0 | - |
| - | - | 2556 | 588.3 | - | - | 0 | - |
| 5 | y | 5386 | 588.8 | 0.0007318 | 1.243 | +2 | 11 |
| 5 | y | 3974 | 589.3 | 0.007198 | 12.22 | +2 | 11 |
| - | - | 2491 | 589.8 | - | - | 0 | - |
| - | - | 858.2 | 590.3 | - | - | 0 | - |
| - | - | 7175 | 592.3 | - | - | 0 | - |
| - | - | 2267 | 593.3 | - | - | 0 | - |
| 11 | y | 4812 | 595.3 | 0.0005977 | 1.004 | +1 | 5 |
| - | - | 2095 | 596.3 | - | - | 0 | - |
| 5 | y | 2.47E+05 | 597.8 | 0.0005153 | 0.8621 | +2 | 11 |
| - | - | 1.579E+05 | 598.3 | - | - | 0 | - |
| 12 | b | 5.692E+04 | 598.8 | 0.001115 | 1.861 | +2 | 12 |
| 12 | b | 1.331E+04 | 599.3 | 0.009839 | 16.42 | +2 | 12 |
| - | - | 2257 | 599.8 | - | - | 0 | - |
| - | - | 4328 | 600.3 | - | - | 0 | - |
| - | - | 1266 | 601.3 | - | - | 0 | - |
| - | - | 894 | 602.3 | - | - | 0 | - |
| - | - | 1164 | 602.8 | - | - | 0 | - |
| - | - | 1499 | 612.3 | - | - | 0 | - |
| - | - | 1312 | 617.3 | - | - | 0 | - |
| - | - | 1191 | 617.8 | - | - | 0 | - |
| - | - | 1201 | 618.3 | - | - | 0 | - |
| - | - | 1242 | 620.8 | - | - | 0 | - |
| - | - | 1077 | 623.3 | - | - | 0 | - |
| - | - | 1127 | 624.3 | - | - | 0 | - |
| - | - | 1339 | 625.8 | - | - | 0 | - |
| - | - | 1036 | 626.3 | - | - | 0 | - |
| - | - | 9501 | 629.3 | - | - | 0 | - |
| - | - | 8854 | 629.8 | - | - | 0 | - |
| - | - | 3408 | 630.3 | - | - | 0 | - |
| - | - | 1086 | 630.8 | - | - | 0 | - |
| - | - | 2277 | 632.3 | - | - | 0 | - |
| - | - | 1053 | 632.8 | - | - | 0 | - |
| 10 | y | 1414 | 635.3 | 0.005073 | 7.985 | +1 | 6 |
| 4 | y | 3.656E+04 | 638.3 | 0.006009 | 9.414 | +2 | 12 |
| 4 | y | 2.732E+04 | 638.8 | 0.003204 | 5.015 | +2 | 12 |
| - | - | 1.092E+04 | 639.3 | - | - | 0 | - |
| - | - | 3470 | 639.8 | - | - | 0 | - |
| - | - | 1403 | 640.3 | - | - | 0 | - |
| - | - | 2034 | 641.3 | - | - | 0 | - |
| - | - | 2512 | 643.4 | - | - | 0 | - |
| - | - | 3479 | 646.3 | - | - | 0 | - |
| 4 | y | 2.226E+05 | 647.3 | 0.0005491 | 0.8483 | +2 | 12 |
| - | - | 1.647E+05 | 647.8 | - | - | 0 | - |
| - | - | 6.994E+04 | 648.3 | - | - | 0 | - |
| - | - | 1121 | 648.4 | - | - | 0 | - |
| - | - | 1.483E+04 | 648.8 | - | - | 0 | - |
| - | - | 2826 | 649.3 | - | - | 0 | - |
| 10 | y | 1.457E+04 | 652.3 | 6.908E-05 | 0.1059 | +1 | 6 |
| - | - | 7341 | 653.3 | - | - | 0 | - |
| - | - | 3479 | 654.3 | - | - | 0 | - |
| - | - | 1925 | 655.3 | - | - | 0 | - |
| - | - | 1205 | 656.3 | - | - | 0 | - |
| - | - | 1003 | 657.3 | - | - | 0 | - |
| - | - | 2072 | 658.8 | - | - | 0 | - |
| - | - | 2316 | 659.3 | - | - | 0 | - |
| - | - | 1965 | 663.8 | - | - | 0 | - |
| - | - | 1885 | 664.3 | - | - | 0 | - |
| - | - | 930.2 | 664.8 | - | - | 0 | - |
| - | - | 1813 | 666.8 | - | - | 0 | - |
| - | - | 1197 | 667.3 | - | - | 0 | - |
| - | - | 828.6 | 667.8 | - | - | 0 | - |
| - | - | 1813 | 669.3 | - | - | 0 | - |
| - | - | 3850 | 669.8 | - | - | 0 | - |
| - | - | 2177 | 670.3 | - | - | 0 | - |
| - | - | 1953 | 670.8 | - | - | 0 | - |
| - | - | 1.472E+04 | 671.3 | - | - | 0 | - |
| - | - | 7235 | 672.3 | - | - | 0 | - |
| - | - | 1.227E+04 | 672.8 | - | - | 0 | - |
| - | - | 1.192E+04 | 673.3 | - | - | 0 | - |
| - | - | 4168 | 673.8 | - | - | 0 | - |
| - | - | 1.147E+04 | 674.3 | - | - | 0 | - |
| - | - | 3766 | 675.3 | - | - | 0 | - |
| - | - | 1032 | 676.3 | - | - | 0 | - |
| - | - | 6.276E+04 | 681.8 | - | - | 0 | - |
| - | - | 4.793E+04 | 682.3 | - | - | 0 | - |
| - | - | 1.935E+04 | 682.8 | - | - | 0 | - |
| - | - | 4598 | 683.3 | - | - | 0 | - |
| - | - | 1085 | 683.8 | - | - | 0 | - |
| - | - | 982.3 | 684.3 | - | - | 0 | - |
| - | - | 1931 | 686.8 | - | - | 0 | - |
| - | - | 1144 | 687.3 | - | - | 0 | - |
| - | - | 1298 | 687.8 | - | - | 0 | - |
| - | - | 1995 | 688.4 | - | - | 0 | - |
| - | - | 3.259E+04 | 689.4 | - | - | 0 | - |
| - | - | 1.308E+04 | 690.4 | - | - | 0 | - |
| - | - | 1060 | 690.8 | - | - | 0 | - |
| - | - | 2735 | 691.4 | - | - | 0 | - |
| - | - | 1092 | 693.8 | - | - | 0 | - |
| - | - | 1249 | 694.3 | - | - | 0 | - |
| - | - | 1452 | 696.8 | - | - | 0 | - |
| - | - | 1891 | 697.4 | - | - | 0 | - |
| - | - | 1202 | 697.9 | - | - | 0 | - |
| - | - | 813.2 | 698.4 | - | - | 0 | - |
| - | - | 7103 | 699.3 | - | - | 0 | - |
| - | - | 3031 | 700.3 | - | - | 0 | - |
| 3 | y | 3.393E+04 | 702.8 | 0.0007703 | 1.096 | +2 | 13 |
| 3 | y | 2.555E+04 | 703.3 | 0.008274 | 11.76 | +2 | 13 |
| - | - | 1.188E+04 | 703.8 | - | - | 0 | - |
| - | - | 3387 | 704.3 | - | - | 0 | - |
| - | - | 1607 | 704.8 | - | - | 0 | - |
| - | - | 969.5 | 706.4 | - | - | 0 | - |
| - | - | 1119 | 711.3 | - | - | 0 | - |
| 3 | y | 3.405E+05 | 711.9 | 0.0005538 | 0.778 | +2 | 13 |
| - | - | 2.671E+05 | 712.4 | - | - | 0 | - |
| - | - | 1.22E+05 | 712.9 | - | - | 0 | - |
| - | - | 3.042E+04 | 713.4 | - | - | 0 | - |
| - | - | 8354 | 713.9 | - | - | 0 | - |
| - | - | 796.5 | 716.8 | - | - | 0 | - |
| - | - | 1139 | 717.3 | - | - | 0 | - |
| - | - | 871.2 | 719.3 | - | - | 0 | - |
| - | - | 998.9 | 720.3 | - | - | 0 | - |
| 9 | y | 1764 | 721.4 | 0.002165 | 3.001 | +1 | 7 |
| 9 | y | 1758 | 722.3 | 0.007285 | 10.09 | +1 | 7 |
| - | - | 1504 | 722.9 | - | - | 0 | - |
| - | - | 1365 | 723.4 | - | - | 0 | - |
| - | - | 2059 | 725.8 | - | - | 0 | - |
| - | - | 1761 | 726.3 | - | - | 0 | - |
| - | - | 3639 | 728.4 | - | - | 0 | - |
| - | - | 1556 | 728.9 | - | - | 0 | - |
| - | - | 777.7 | 729.4 | - | - | 0 | - |
| - | - | 859.6 | 729.9 | - | - | 0 | - |
| - | - | 1987 | 731.4 | - | - | 0 | - |
| - | - | 1445 | 733.8 | - | - | 0 | - |
| - | - | 1025 | 734.4 | - | - | 0 | - |
| - | - | 753.8 | 735.4 | - | - | 0 | - |
| - | - | 5446 | 737.4 | - | - | 0 | - |
| - | - | 6506 | 737.9 | - | - | 0 | - |
| - | - | 2527 | 738.4 | - | - | 0 | - |
| - | - | 761.3 | 738.9 | - | - | 0 | - |
| 9 | y | 1.422E+04 | 739.4 | 0.0003893 | 0.5265 | +1 | 7 |
| - | - | 9644 | 740.4 | - | - | 0 | - |
| - | - | 3743 | 740.9 | - | - | 0 | - |
| - | - | 2736 | 741.4 | - | - | 0 | - |
| - | - | 991.9 | 741.9 | - | - | 0 | - |
| - | - | 1297 | 745.4 | - | - | 0 | - |
| 2 | y | 4.605E+04 | 746.4 | 0.0001979 | 0.2652 | +2 | 14 |
| 2 | y | 4.192E+04 | 746.9 | 0.009045 | 12.11 | +2 | 14 |
| - | - | 1.875E+04 | 747.4 | - | - | 0 | - |
| - | - | 7201 | 747.9 | - | - | 0 | - |
| - | - | 1903 | 748.4 | - | - | 0 | - |
| - | - | 932 | 749.4 | - | - | 0 | - |
| - | - | 2722 | 751.4 | - | - | 0 | - |
| - | - | 1109 | 751.9 | - | - | 0 | - |
| - | - | 900.3 | 752.4 | - | - | 0 | - |
| - | - | 1812 | 753.4 | - | - | 0 | - |
| - | - | 1869 | 754.4 | - | - | 0 | - |
| 2 | y | 5.926E+05 | 755.4 | 0.0005308 | 0.7028 | +2 | 14 |
| - | - | 4.905E+05 | 755.9 | - | - | 0 | - |
| - | - | 2.269E+05 | 756.4 | - | - | 0 | - |
| - | - | 6.85E+04 | 756.9 | - | - | 0 | - |
| - | - | 1.591E+04 | 757.4 | - | - | 0 | - |
| - | - | 1967 | 758.4 | - | - | 0 | - |
| - | - | 2.034E+04 | 760.4 | - | - | 0 | - |
| - | - | 1.599E+04 | 760.9 | - | - | 0 | - |
| - | - | 9996 | 761.4 | - | - | 0 | - |
| - | - | 2809 | 761.9 | - | - | 0 | - |
| - | - | 2230 | 768.4 | - | - | 0 | - |
| 7 | b | 1609 | 769.4 | 0.004621 | 6.006 | +1 | 7 |
| - | - | 4016 | 770.4 | - | - | 0 | - |
| - | - | 1096 | 771.4 | - | - | 0 | - |
| - | - | 891.2 | 782.4 | - | - | 0 | - |
| - | - | 1064 | 783.4 | - | - | 0 | - |
| - | - | 2944 | 786.4 | - | - | 0 | - |
| 7 | b | 3746 | 787.4 | 0.003894 | 4.946 | +1 | 7 |
| - | - | 2198 | 788.4 | - | - | 0 | - |
| - | - | 2744 | 798.4 | - | - | 0 | - |
| - | - | 1624 | 799.4 | - | - | 0 | - |
| - | - | 1537 | 800.4 | - | - | 0 | - |
| - | - | 1289 | 801.4 | - | - | 0 | - |
| - | - | 2128 | 806.4 | - | - | 0 | - |
| - | - | 1028 | 807.4 | - | - | 0 | - |
| - | - | 8306 | 810.4 | - | - | 0 | - |
| - | - | 3249 | 811.4 | - | - | 0 | - |
| - | - | 2240 | 812.4 | - | - | 0 | - |
| - | - | 1566 | 813.4 | - | - | 0 | - |
| 8 | y | 5524 | 818.4 | 0.0006111 | 0.7467 | +1 | 8 |
| 8 | y | 6909 | 819.4 | 0.006767 | 8.259 | +1 | 8 |
| - | - | 2418 | 820.4 | - | - | 0 | - |
| - | - | 3393 | 828.4 | - | - | 0 | - |
| - | - | 3471 | 829.4 | - | - | 0 | - |
| - | - | 1596 | 830.4 | - | - | 0 | - |
| 8 | y | 7.082E+04 | 836.4 | 0.0004209 | 0.5032 | +1 | 8 |
| - | - | 3.12E+04 | 837.4 | - | - | 0 | - |
| - | - | 9406 | 838.4 | - | - | 0 | - |
| - | - | 1443 | 839.4 | - | - | 0 | - |
| - | - | 1749 | 845.4 | - | - | 0 | - |
| - | - | 1063 | 855.4 | - | - | 0 | - |
| - | - | 1767 | 857.4 | - | - | 0 | - |
| - | - | 1137 | 858.4 | - | - | 0 | - |
| - | - | 926.4 | 872.4 | - | - | 0 | - |
| - | - | 1558 | 873.4 | - | - | 0 | - |
| - | - | 2122 | 881.4 | - | - | 0 | - |
| - | - | 1051 | 882.5 | - | - | 0 | - |
| - | - | 740.8 | 883.4 | - | - | 0 | - |
| - | - | 791.2 | 884.4 | - | - | 0 | - |
| - | - | 1837 | 885.4 | - | - | 0 | - |
| - | - | 868.4 | 886.4 | - | - | 0 | - |
| - | - | 2058 | 887.4 | - | - | 0 | - |
| - | - | 1513 | 890.4 | - | - | 0 | - |
| - | - | 1456 | 891.4 | - | - | 0 | - |
| - | - | 1041 | 892.4 | - | - | 0 | - |
| - | - | 3096 | 899.5 | - | - | 0 | - |
| - | - | 1949 | 900.5 | - | - | 0 | - |
| - | - | 2531 | 909.4 | - | - | 0 | - |
| - | - | 914.7 | 910.4 | - | - | 0 | - |
| - | - | 6175 | 915.4 | - | - | 0 | - |
| - | - | 1133 | 916.5 | - | - | 0 | - |
| - | - | 1116 | 917.5 | - | - | 0 | - |
| - | - | 913.9 | 921.5 | - | - | 0 | - |
| - | - | 1891 | 925.4 | - | - | 0 | - |
| - | - | 2680 | 927.5 | - | - | 0 | - |
| - | - | 1747 | 928.5 | - | - | 0 | - |
| - | - | 2402 | 943.5 | - | - | 0 | - |
| - | - | 2455 | 944.5 | - | - | 0 | - |
| - | - | 1738 | 945.5 | - | - | 0 | - |
| - | - | 1221 | 950.5 | - | - | 0 | - |
| - | - | 699.3 | 953.4 | - | - | 0 | - |
| - | - | 960.7 | 957.5 | - | - | 0 | - |
| - | - | 2279 | 958.5 | - | - | 0 | - |
| - | - | 1248 | 959.5 | - | - | 0 | - |
| - | - | 1463 | 968.5 | - | - | 0 | - |
| - | - | 1446 | 969.5 | - | - | 0 | - |
| - | - | 976.4 | 970.5 | - | - | 0 | - |
| 9 | b | 1593 | 971.5 | 0.007688 | 7.914 | +1 | 9 |
| - | - | 1850 | 972.5 | - | - | 0 | - |
| 7 | y | 7607 | 975.5 | 0.0006547 | 0.6712 | +1 | 9 |
| - | - | 3896 | 976.5 | - | - | 0 | - |
| - | - | 975 | 977.5 | - | - | 0 | - |
| - | - | 967.4 | 984.5 | - | - | 0 | - |
| - | - | 1052 | 985.5 | - | - | 0 | - |
| - | - | 8829 | 986.5 | - | - | 0 | - |
| - | - | 6727 | 987.5 | - | - | 0 | - |
| - | - | 2452 | 988.5 | - | - | 0 | - |
| - | - | 2352 | 989.5 | - | - | 0 | - |
| - | - | 1012 | 990.5 | - | - | 0 | - |
| - | - | 1473 | 994.5 | - | - | 0 | - |
| - | - | 839.8 | 995.5 | - | - | 0 | - |
| - | - | 2395 | 996.5 | - | - | 0 | - |
| - | - | 1812 | 997.5 | - | - | 0 | - |
| - | - | 6091 | 1012 | - | - | 0 | - |
| - | - | 3821 | 1013 | - | - | 0 | - |
| - | - | 7887 | 1014 | - | - | 0 | - |
| - | - | 3220 | 1015 | - | - | 0 | - |
| - | - | 1877 | 1016 | - | - | 0 | - |
| - | - | 1123 | 1020 | - | - | 0 | - |
| 10 | b | 1531 | 1029 | 0.00307 | 2.985 | +1 | 10 |
| - | - | 1212 | 1030 | - | - | 0 | - |
| - | - | 8355 | 1030 | - | - | 0 | - |
| - | - | 4775 | 1031 | - | - | 0 | - |
| - | - | 839.4 | 1032 | - | - | 0 | - |
| - | - | 1036 | 1034 | - | - | 0 | - |
| - | - | 933.8 | 1046 | - | - | 0 | - |
| - | - | 1247 | 1056 | - | - | 0 | - |
| - | - | 4499 | 1074 | - | - | 0 | - |
| - | - | 1576 | 1075 | - | - | 0 | - |
| - | - | 2165 | 1084 | - | - | 0 | - |
| - | - | 2138 | 1085 | - | - | 0 | - |
| 6 | y | 6193 | 1091 | 0.0008446 | 0.7745 | +1 | 10 |
| - | - | 3049 | 1092 | - | - | 0 | - |
| - | - | 986.3 | 1093 | - | - | 0 | - |
| - | - | 904.9 | 1094 | - | - | 0 | - |
| - | - | 1631 | 1101 | - | - | 0 | - |
| - | - | 4716 | 1102 | - | - | 0 | - |
| - | - | 2322 | 1103 | - | - | 0 | - |
| - | - | 1604 | 1112 | - | - | 0 | - |
| - | - | 1175 | 1113 | - | - | 0 | - |
| - | - | 1445 | 1119 | - | - | 0 | - |
| - | - | 1841 | 1120 | - | - | 0 | - |
| 11 | b | 742.6 | 1128 | 0.005041 | 4.471 | +1 | 11 |
| - | - | 1244 | 1130 | - | - | 0 | - |
| - | - | 1307 | 1131 | - | - | 0 | - |
| - | - | 1032 | 1132 | - | - | 0 | - |
| - | - | 1313 | 1135 | - | - | 0 | - |
| - | - | 1511 | 1143 | - | - | 0 | - |
| - | - | 1065 | 1144 | - | - | 0 | - |
| - | - | 847.1 | 1150 | - | - | 0 | - |
| - | - | 766.9 | 1154 | - | - | 0 | - |
| - | - | 1870 | 1160 | - | - | 0 | - |
| - | - | 3276 | 1161 | - | - | 0 | - |
| - | - | 1811 | 1162 | - | - | 0 | - |
| - | - | 826.3 | 1163 | - | - | 0 | - |
| - | - | 1720 | 1171 | - | - | 0 | - |
| - | - | 1078 | 1177 | - | - | 0 | - |
| 5 | y | 1.631E+04 | 1178 | 0.0007883 | 0.6694 | +1 | 11 |
| - | - | 1.115E+04 | 1179 | - | - | 0 | - |
| - | - | 3563 | 1180 | - | - | 0 | - |
| - | - | 1046 | 1181 | - | - | 0 | - |
| - | - | 1725 | 1188 | - | - | 0 | - |
| - | - | 3130 | 1189 | - | - | 0 | - |
| - | - | 1636 | 1190 | - | - | 0 | - |
| - | - | 799.1 | 1191 | - | - | 0 | - |
| - | - | 2240 | 1206 | - | - | 0 | - |
| - | - | 1874 | 1207 | - | - | 0 | - |
| 4 | y | 2455 | 1277 | 0.0009882 | 0.7741 | +1 | 12 |
| - | - | 2190 | 1278 | - | - | 0 | - |
| - | - | 728.7 | 1279 | - | - | 0 | - |
| - | - | 879.6 | 1301 | - | - | 0 | - |
| - | - | 764.1 | 2716 | - | - | 0 | - |

m/z Charge Intensity FragmentType MassShift Position
120.0811996459961 0 27568.982
121.08453369140625 0 2683.1626
125.07099914550781 0 517.014
125.10768127441406 0 4133.619
126.05534362792969 0 6987.203
127.05057525634766 0 950.32605
127.08731842041016 0 335.78857
128.03904724121094 0 448.13553
128.10739135742188 0 3783.1318
129.06622314453125 0 1478.8223
129.1026153564453 0 8148.97
129.11416625976562 0 377.0223
130.05020141601562 0 1722.8103
130.06149291992188 0 1099.617
130.097900390625 0 625.70526
130.10617065429688 0 572.03503
131.08184814453125 0 569.7142
132.1023712158203 0 1066.3577
133.06094360351562 0 678.26904
136.0761260986328 0 3720.3972
138.0918426513672 0 2316.2212
139.08706665039062 0 902.0757
140.0822296142578 0 2286.805
141.10276794433594 0 1074.359
143.0819854736328 0 852.8689
144.06590270996094 0 1705.8042
145.0609893798828 0 789.92
147.07691955566406 0 3138.001
148.94691467285156 0 514.34753
153.06625366210938 0 731.63275
154.05030822753906 0 6610.564
155.0535430908203 0 742.38654
155.0814666748047 0 613.13666
155.1182861328125 0 6599.2744 a Water loss 1
156.0771026611328 0 1382.0413
156.1024169921875 0 3505.498
156.1217041015625 0 641.29474
157.06118774414062 0 6195.97
157.0975799560547 0 5664.9707
157.1088104248047 0 5672.7285
158.0928497314453 0 6969.5747
158.0997314453125 0 834.08496
159.07688903808594 0 719.62665
159.1132354736328 0 5334.866
160.11695861816406 0 679.00446
161.03839111328125 0 4041.8037
163.0341339111328 0 504.7574
164.0819854736328 0 671.05884
165.05531311035156 0 1100.9706
165.11386108398438 0 508.7933
166.08663940429688 0 20099.883 y 14
166.09840393066406 0 771.0659
167.08212280273438 0 1016.0492
167.09005737304688 0 2002.7126
169.0611572265625 0 459.7651
169.0975799560547 0 1791.3063
170.04486083984375 0 514.6586
171.07684326171875 0 9847.917
172.060546875 0 1176.2966
172.08001708984375 0 624.11304
172.68035888671875 0 452.95898
173.12892150878906 0 97341.64 a 1
173.4381561279297 0 817.7787
174.12474060058594 0 929.51715
174.13226318359375 0 9623.766
175.07179260253906 0 10390.653
175.11940002441406 0 11278.668
175.13330078125 0 624.5429
176.12294006347656 0 490.27475
177.1024932861328 0 850.35834
181.06179809570312 0 641.0485
181.09725952148438 0 569.687
182.08145141601562 0 2070.418
182.0927734375 0 1002.1881
183.11326599121094 0 67213.93 b Water loss 1
183.14952087402344 0 1330.1162
184.0972137451172 0 474.76523
184.1085205078125 0 1579.1013
184.11663818359375 0 6178.2837
185.05613708496094 0 1081.6079
185.09251403808594 0 6650.331
186.0958709716797 0 530.0952
187.07176208496094 0 867.58765
187.1082000732422 0 7784.9854
187.5689239501953 0 497.22495
188.11154174804688 0 531.0977
189.08743286132812 0 41384.84
190.09083557128906 0 2940.9895
191.0943603515625 0 514.47253
192.09808349609375 0 480.75525
192.11373901367188 0 932.41595
194.0935821533203 0 614.45526
194.1290740966797 0 547.01056
195.0770721435547 0 720.97
195.1139373779297 0 639.38165
197.12879943847656 0 1305.1168
199.07177734375 0 15164.475
199.10836791992188 0 1830.9103 y 10
200.07534790039062 0 1039.5916
200.140625 0 517.1926
201.123779296875 0 131191.58 b 1
202.083251953125 0 567.21893
202.11880493164062 0 2568.7754
202.1272735595703 0 11478.545
203.066650390625 0 11291.477
203.1028289794922 0 991.4635
203.12913513183594 0 812.29297
207.1132049560547 0 1115.6512
208.10836791992188 0 590.55225
210.0879364013672 0 1019.6801
210.1242218017578 0 528.58057
211.10821533203125 0 6494.7725
212.10340881347656 0 3945.1743
213.0876922607422 0 879.2775
216.13504028320312 0 906.26465
217.08233642578125 0 67640.05
218.0857391357422 0 5013.7563
219.08680725097656 0 506.48636
220.09292602539062 0 1069.1511
223.1081085205078 0 2901.7327
224.1033935546875 0 5392.013
225.10714721679688 0 522.809
225.12380981445312 0 1112.5978
225.1712646484375 0 1142.673
226.11915588378906 0 2283.8843
227.0665740966797 0 791.98956
227.1021728515625 0 931.3439
227.1143035888672 0 12809.071
228.09808349609375 0 669.623
228.1177978515625 0 1479.5256
229.11874389648438 0 6947.3745
230.114013671875 0 2389.329
233.12879943847656 0 739.3699
233.1648712158203 0 780.1162
235.10777282714844 0 1100.2041
237.09881591796875 0 1818.8911
237.135009765625 0 2552.753
238.11827087402344 0 532.73596
240.0966033935547 0 494.10718
240.1342010498047 0 637.82385
242.1140594482422 0 11652.664
242.19839477539062 0 816.73267
243.11691284179688 0 880.59644
244.12960815429688 0 4435.606
244.14126586914062 0 1093.3458
245.1250762939453 0 3712.6821
248.6302032470703 0 2507.021 y 11
249.13113403320312 0 878.1314
252.11012268066406 0 1641.5342
252.1348114013672 0 1200.4312
253.0320281982422 0 2893.7837
253.11866760253906 0 4213.9404
254.11328125 0 1058.3533
254.16165161132812 0 574.6566
255.10931396484375 0 13400.508
255.1454315185547 0 518.1413
256.11285400390625 0 1010.8164
256.1293029785156 0 1071.9008
257.12548828125 0 773.9823
258.144287109375 0 680.7203
259.1400451660156 0 2007.3511
259.15509033203125 0 1037.7594
262.1528625488281 0 4423.6855
263.1391296386719 0 2031.8335 b 6
263.5815124511719 0 600.0111
266.1131896972656 0 686.4742
267.1097717285156 0 784.1208
267.1340026855469 0 1118.8038
269.10711669921875 0 1252.7035
270.1454162597656 0 2624.3528
270.1929016113281 0 1483.5161
272.1357421875 0 1968.7091
274.084228515625 0 641.5525
279.1098937988281 0 998.1884
280.129638671875 0 1336.6672
281.0267028808594 0 2000.4799
283.1413269042969 0 820.66547
284.1241455078125 0 2573.5393
284.1611022949219 0 5029.081 a Water loss 2
285.12725830078125 0 609.2408
285.16412353515625 0 1013.9773
287.22003173828125 0 669.2955
288.13616943359375 0 1240.226
288.1558532714844 0 2422.7673
289.1516418457031 0 977.5189
294.1450500488281 0 651.2378
295.1769714355469 0 952.0608
296.1360778808594 0 3260.9526
296.1717834472656 0 662.72296
297.05731201171875 0 700.87103
297.1210021972656 0 1748.9806
298.140380859375 0 6353.9463
298.1589050292969 0 825.55365 y 10
299.0607604980469 0 567.4682
299.1365051269531 0 2462.98
300.1369323730469 0 659.71204
301.0592346191406 0 914.2816
302.0598449707031 0 643.6195
302.1351623535156 0 2130.6646
302.1713562011719 0 1308.924 a 2
303.6315002441406 0 610.535
305.1614074707031 0 21030.24 y Ammonia loss 13
306.16461181640625 0 3293.6655
307.16326904296875 0 642.7026
310.13922119140625 0 726.9173
312.1560363769531 0 66333.96 b Water loss 2
313.15850830078125 0 9799.051
313.1877746582031 0 13074.026
314.1466064453125 0 9276.942
314.1905517578125 0 2091.9292
315.1496887207031 0 1530.5372
316.1509704589844 0 15912.678 b 5
317.1514587402344 0 2512.0435
319.1408996582031 0 1114.5646
319.1602783203125 0 793.0321
320.1368408203125 0 545.19476
322.1878356933594 0 4196.393 y 13
323.17095947265625 0 927.75323
323.1906433105469 0 772.28296
324.131103515625 0 3859.0364
324.1673583984375 0 904.16284
325.1145935058594 0 810.73895
326.1357116699219 0 1078.5994
329.1433410644531 0 572.5471
330.16668701171875 0 30294.336 b 2
331.1708984375 0 7472.1997
332.1578369140625 0 2979.17
332.1766662597656 0 1362.2847
336.178955078125 0 1035.647
341.1827087402344 0 8806.639
342.1415100097656 0 18147.309
342.1848449707031 0 1606.2603
343.1445617675781 0 1878.5927
344.14691162109375 0 959.36316
345.1828308105469 0 2228.426
349.15203857421875 0 970.31525
349.1838684082031 0 5022.5703
350.6011657714844 0 851.42847
352.1252746582031 0 626.9567
359.16778564453125 0 12173.776
360.1706237792969 0 2175.2703
367.162353515625 0 981.026
370.2194519042969 0 727.56964
374.1400146484375 0 565.5289
377.1790771484375 0 904.0308
380.15631103515625 0 1196.3342
381.1882019042969 0 738.6139
385.1724853515625 0 2223.0125
385.71282958984375 0 835.02234
391.2091064453125 0 820.0634 y Water loss 12
392.1932373046875 0 3463.7332 y Ammonia loss 12
393.1955871582031 0 1325.7721 y Ammonia loss 4
394.20220947265625 0 1833.0033 b 6
394.7055358886719 0 962.2098
395.15740966796875 0 1197.2373
396.1883239746094 0 1285.8053
396.6841125488281 0 744.72876
398.2132568359375 0 1876.8278
398.8666687011719 0 736.7582 y 4
399.16192626953125 0 830.37427
400.2207946777344 0 1347.5355
400.26727294921875 0 1973.2688
401.23992919921875 0 10659.422
401.26910400390625 0 1077.1458
402.24371337890625 0 1895.6312
403.1829528808594 0 2604.215
404.1852111816406 0 838.9024
408.22967529296875 0 1283.0568
409.22027587890625 0 8160.5386 y 12
410.2027282714844 0 1436.6383 y Ammonia loss 7
411.22479248046875 0 1804.1814 b Water loss 3
413.168212890625 0 1196.1476
413.21453857421875 0 1067.0785
414.2277526855469 0 3168.177
414.7220458984375 0 6528.509
415.2229919433594 0 2403.736
415.7241516113281 0 963.39264
418.7177429199219 0 7865.9106 y 7
419.2186279296875 0 2462.1973
419.72003173828125 0 1280.2919
423.20086669921875 0 720.78796
423.2338562011719 0 21405.52
423.7350769042969 0 10079.164
424.236083984375 0 2441.0881
428.2158508300781 0 3561.4773
429.23504638671875 0 12340.548 b 3
430.2390441894531 0 4916.0884
431.1755676269531 0 763.5058
431.2475280761719 0 806.4401
431.8899230957031 0 1027.5231 y 3
432.2252502441406 0 885.5132
433.182373046875 0 719.64484
439.1934814453125 0 1089.6708
441.2105407714844 0 3831.4312
443.2034912109375 0 1021.1208
445.241455078125 0 841.9797
448.25189208984375 0 1571.7006
451.23028564453125 0 919.636
452.22821044921875 0 635.9338
452.2620849609375 0 2827.1519
455.226318359375 0 747.33795
458.2367248535156 0 7577.6675
459.2391662597656 0 1972.2667
463.2322082519531 0 2246.6624
467.2423400878906 0 897.30396
467.7463073730469 0 1050.6704
468.898681640625 0 1629.8792 y Water loss 2
469.2359313964844 0 1215.1536 y Ammonia loss 2
469.2887268066406 0 3143.2441
470.2337646484375 0 1088.541
470.29248046875 0 1295.8711
471.74310302734375 0 849.1695
474.90484619140625 0 2018.5654 y 2
475.23956298828125 0 2210.531
478.241455078125 0 683.13696 y Water loss 11
479.2255859375 0 4095.2678 y Ammonia loss 11
480.2243957519531 0 701.77924
480.7474060058594 0 4158.4673
481.2469482421875 0 1478.9032
481.75701904296875 0 892.9403
482.23309326171875 0 775.1298
487.25848388671875 0 1106.0769
487.76141357421875 0 2008.3208 y Water loss 6
488.25567626953125 0 5926.168 y Ammonia loss 6
488.75811767578125 0 2870.8872
489.1845703125 0 927.797
492.2424011230469 0 701.2915
494.2418518066406 0 694.42975
494.7416076660156 0 658.91113
496.217529296875 0 786.6991
496.25225830078125 0 30402.877 y 11
496.7679748535156 0 81340.42 y 6
497.2685852050781 0 40795.76
497.7705078125 0 12388.379
497.9111633300781 0 2396.833 y Water loss 1
498.2411804199219 0 1122.2659 y Ammonia loss 1
498.27203369140625 0 1903.6643
498.5794372558594 0 756.9214
499.2450256347656 0 889.86346
500.1999206542969 0 1266.7893
501.7579345703125 0 655.03937
503.9147644042969 0 3959.8643 y 1
504.2489013671875 0 1663.6929
504.5860290527344 0 1202.901
504.9185791015625 0 967.18494
505.2740173339844 0 4704.4395
506.2718200683594 0 2270.9204
507.2436828613281 0 851.15753
510.7351379394531 0 716.81537
511.26348876953125 0 988.4025
513.350341796875 0 1100.7921
515.2566528320312 0 4028.9119
515.759765625 0 2086.9924
516.2628784179688 0 2083.7466 b 4
520.2645263671875 0 3908.2986
520.7645874023438 0 873.10156
524.262939453125 0 21075.955
524.7643432617188 0 11990.882
525.2652587890625 0 4926.0483
525.7669067382812 0 1125.472
529.602783203125 0 2889.1094
529.9344482421875 0 2872.8516
530.2697143554688 0 2213.129
530.598876953125 0 1038.8501
532.1883544921875 0 2199.9216
532.2732543945312 0 2310.6538
533.2743530273438 0 778.1526
533.6882934570312 0 792.9865
535.6062622070312 0 15615.54 Precursor Water loss
535.9403076171875 0 14977.022 Precursor Ammonia loss
536.2744750976562 0 7566.816
536.6054077148438 0 1294.1486
536.9429931640625 0 787.1135
538.0419921875 0 717.03723
538.3118896484375 0 796.02185
540.2091064453125 0 905.54645
540.2964477539062 0 2464.9182
540.811279296875 0 1250.5848
540.8818969726562 0 1150.4963
541.2775268554688 0 1022.6457
541.346923828125 0 896.4751
541.6099853515625 0 38431.18 Precursor
541.9442138671875 0 29961.14
542.1889038085938 0 2220.7935
542.27734375 0 17804.994
542.6129150390625 0 5400.4824
542.6895141601562 0 2952.076
542.9465942382812 0 1304.6254
543.2539672851562 0 1472.1289
545.7719116210938 0 1501.8522 y Ammonia loss 5
550.29052734375 0 1219.2014
554.2813110351562 0 32021.947 y 5
554.7825317382812 0 17389.904
555.2843627929688 0 7424.391 b Water loss 10
555.7860717773438 0 992.49426
556.27734375 0 1100.4552
556.3248291015625 0 742.2975
557.2710571289062 0 720.74023
558.7907104492188 0 1027.2828
559.28369140625 0 1090.3702
560.2932739257812 0 761.903
564.7922973632812 0 5277.8813
565.2365112304688 0 1288.2229
565.2913208007812 0 2788.5464
565.7922973632812 0 727.26025
568.7716674804688 0 949.69696
569.26904296875 0 4305.6284
570.2750854492188 0 880.7748
572.2802734375 0 927.95825
573.7969970703125 0 22687.865
574.2981567382812 0 16816.408
574.7998657226562 0 4890.7964
575.2967529296875 0 1716.2487
582.7918701171875 0 1473.9819
583.2457275390625 0 1143.442
584.2529296875 0 1065.253
584.3177490234375 0 1915.2212
587.2785034179688 0 6483.3633
588.2797241210938 0 2555.8174
588.7921752929688 0 5386.4443 y Water loss 4
589.2906494140625 0 3974.435 y Ammonia loss 4
589.788330078125 0 2491.1382
590.2880249023438 0 858.185
592.3058471679688 0 7175.33
593.3076782226562 0 2266.741
595.3204345703125 0 4812.051 y 10
596.3228149414062 0 2095.3088
597.7972412109375 0 247036.42 y 4
598.2987060546875 0 157891.77
598.800048828125 0 56919.36 b Water loss 11
599.30078125 0 13312.321 b Ammonia loss 11
599.8013916015625 0 2257.388
600.274658203125 0 4328.325
601.2783203125 0 1266.0399
602.2850952148438 0 894.0185
602.7867431640625 0 1164.3207
612.3094482421875 0 1499.2964
617.310546875 0 1312.0042
617.8110961914062 0 1190.5493
618.305419921875 0 1201.2943
620.8056640625 0 1242.4464
623.320068359375 0 1077.3262
624.3148803710938 0 1127.4434
625.8010864257812 0 1339.125
626.3237915039062 0 1035.7229
629.31591796875 0 9501.184
629.8140869140625 0 8854.203
630.3160400390625 0 3408.2336
630.8154296875 0 1085.7948
632.3261108398438 0 2277.3486
632.82666015625 0 1052.7671
635.31982421875 0 1413.6078 y Ammonia loss 9
638.3196411132812 0 36558.1 y Water loss 3
638.8208618164062 0 27317.373 y Ammonia loss 3
639.3219604492188 0 10920.146
639.82275390625 0 3469.7449
640.3192138671875 0 1403.1781
641.3380126953125 0 2033.5615
643.3533325195312 0 2511.6423
646.3155517578125 0 3478.8206
647.3314819335938 0 222604.11 y 3
647.8328247070312 0 164710.36
648.3338012695312 0 69940.9
648.4002685546875 0 1120.9521
648.8353271484375 0 14834.049
649.334228515625 0 2825.9365
652.3413696289062 0 14569.982 y 9
653.342041015625 0 7341.039
654.3262939453125 0 3478.5203
655.3182983398438 0 1925.0354
656.30029296875 0 1204.5156
657.2976684570312 0 1002.57886
658.829833984375 0 2072.0522
659.3444213867188 0 2315.906
663.8232421875 0 1964.5481
664.3153076171875 0 1884.5123
664.8256225585938 0 930.1824
666.827392578125 0 1812.9286
667.33056640625 0 1197.3196
667.838623046875 0 828.6322
669.3068237304688 0 1813.2954
669.80810546875 0 3849.8984
670.3095092773438 0 2177.0674
670.8148193359375 0 1952.5425
671.3471069335938 0 14719.935
672.341552734375 0 7234.7793
672.8292846679688 0 12270.06
673.3309936523438 0 11916.566
673.8308715820312 0 4168.042
674.3121337890625 0 11469.146
675.3161010742188 0 3766.3281
676.3186645507812 0 1032.3195
681.83447265625 0 62764.758
682.3348388671875 0 47934.117
682.8367309570312 0 19351.564
683.3355102539062 0 4597.753
683.8390502929688 0 1084.6505
684.335693359375 0 982.3197
686.8272705078125 0 1931.2528
687.3224487304688 0 1144.2948
687.8321533203125 0 1297.523
688.371826171875 0 1995.042
689.3584594726562 0 32586.809
690.3610229492188 0 13083.273
690.8348999023438 0 1059.7905
691.357177734375 0 2735.032
693.84326171875 0 1091.9205
694.3375854492188 0 1249.3956
696.849365234375 0 1451.6141
697.3590087890625 0 1891.221
697.8516235351562 0 1201.8477
698.3619384765625 0 813.21423
699.3425903320312 0 7103.127
700.345703125 0 3031.2615
702.8477172851562 0 33933.383 y Water loss 2
703.3472290039062 0 25550.861 y Ammonia loss 2
703.8485717773438 0 11878.605
704.3463745117188 0 3387.1765
704.8478393554688 0 1606.771
706.3590087890625 0 969.49347
711.3433227539062 0 1118.8723
711.852783203125 0 340453.25 y 2
712.3543090820312 0 267078.66
712.8554077148438 0 122032.016
713.3565673828125 0 30415.994
713.8577880859375 0 8353.734
716.8360595703125 0 796.4925
717.3447265625 0 1139.4683
719.3448486328125 0 871.2353
720.346435546875 0 998.9355
721.3649291992188 0 1764.0234 y Water loss 8
722.3540649414062 0 1757.5028 y Ammonia loss 8
722.8583374023438 0 1504.3212
723.3533935546875 0 1364.827
725.8482055664062 0 2058.7007
726.347900390625 0 1760.5815
728.3545532226562 0 3639.4214
728.8524780273438 0 1556.3351
729.3524780273438 0 777.72833
729.8641357421875 0 859.5661
731.3587646484375 0 1987.2339
733.842041015625 0 1445.4395
734.3587646484375 0 1024.5289
735.3552856445312 0 753.80194
737.3574829101562 0 5446.214
737.8571166992188 0 6505.9673
738.36083984375 0 2527.0034
738.8590087890625 0 761.32837
739.3737182617188 0 14219.949 y 8
740.3709106445312 0 9644.155
740.8649291992188 0 3743.4658
741.3717041015625 0 2736.4028
741.8701782226562 0 991.8785
745.3547973632812 0 1296.6777
746.3631591796875 0 46046.168 y Water loss 1
746.864013671875 0 41917.9 y Ammonia loss 1
747.3651733398438 0 18751.793
747.8662109375 0 7200.6904
748.3656616210938 0 1903.0001
749.357177734375 0 931.97705
751.355712890625 0 2722.1558
751.8590698242188 0 1109.0311
752.3531494140625 0 900.32477
753.35498046875 0 1812.0038
754.3580932617188 0 1868.9955
755.3687744140625 0 592621.9 y 1
755.8701782226562 0 490505.88
756.3712768554688 0 226862.33
756.8722534179688 0 68500.9
757.3740844726562 0 15908.191
758.3860473632812 0 1966.9381
760.3604736328125 0 20339.55
760.8613891601562 0 15992.5205
761.3629150390625 0 9996.107
761.8651733398438 0 2808.525
768.369140625 0 2229.8464
769.3792724609375 0 1609.2511 b Water loss 6
770.4129638671875 0 4015.5134
771.359130859375 0 1095.6998
782.380859375 0 891.2174
783.3804931640625 0 1064.4001
786.378173828125 0 2944.388
787.3905639648438 0 3745.7046 b 6
788.3981323242188 0 2198.1018
798.4092407226562 0 2743.7817
799.4138793945312 0 1624.483
800.4022827148438 0 1536.6846
801.390380859375 0 1289.4677
806.4157104492188 0 2127.8313
807.4168090820312 0 1027.886
810.3744506835938 0 8305.542
811.3796997070312 0 3248.8562
812.395751953125 0 2239.597
813.404052734375 0 1566.309
818.4149169921875 0 5524.33 y Water loss 7
819.4063110351562 0 6909.313 y Ammonia loss 7
820.4047241210938 0 2417.8745
828.4317016601562 0 3392.8206
829.43896484375 0 3471.2766
830.398681640625 0 1596.1322
836.426513671875 0 70821.57 y 7
837.4290161132812 0 31202.752
838.431396484375 0 9405.521
839.4332275390625 0 1442.6617
845.4121704101562 0 1748.6785
855.3995971679688 0 1063.4156
857.4406127929688 0 1766.6245
858.3896484375 0 1137.2671
872.4188232421875 0 926.4379
873.4136352539062 0 1557.7023
881.4449462890625 0 2121.9844
882.450927734375 0 1050.7094
883.4116821289062 0 740.79297
884.4165649414062 0 791.2338
885.43896484375 0 1836.8826
886.4432373046875 0 868.38367
887.4247436523438 0 2058.1943
890.4334106445312 0 1512.5045
891.4180297851562 0 1455.6271
892.427734375 0 1040.6836
899.458251953125 0 3096.487
900.454833984375 0 1948.8381
909.4432373046875 0 2530.8672
910.4483032226562 0 914.65717
915.4171142578125 0 6175.36
916.478515625 0 1133.0471
917.4795532226562 0 1116.4077
921.4564819335938 0 913.873
925.446533203125 0 1891.309
927.453125 0 2680.367
928.4556274414062 0 1746.581
943.4612426757812 0 2402.0916
944.4699096679688 0 2455.2297
945.4765014648438 0 1737.7964
950.5021362304688 0 1221.3309
953.444091796875 0 699.33624
957.4812622070312 0 960.7032
958.4712524414062 0 2279.2673
959.4813842773438 0 1248.0555
968.4813232421875 0 1463.0052
969.4797973632812 0 1445.8792
970.4686279296875 0 976.41113
971.4869384765625 0 1593.2518 b 8
972.4782104492188 0 1850.492
975.5 0 7607.4126 y Ammonia loss 6
976.501708984375 0 3895.9536
977.5078125 0 975.03906
984.4888305664062 0 967.4459
985.4913330078125 0 1052.2029
986.4889526367188 0 8828.54
987.4887084960938 0 6727.028
988.4866943359375 0 2452.4612
989.4964599609375 0 2351.825
990.4935913085938 0 1012.0384
994.4702758789062 0 1472.6868
995.4736328125 0 839.7789
996.47607421875 0 2395.4812
997.4691772460938 0 1811.6539
1012.48095703125 0 6091.352
1013.4822998046875 0 3820.8008
1014.4844360351562 0 7887.278
1015.4893798828125 0 3219.9304
1016.4902954101562 0 1877.3282
1020.4737548828125 0 1122.6731
1028.5037841796875 0 1530.7272 b 9
1029.5126953125 0 1212.1665
1030.491455078125 0 8354.655
1031.4935302734375 0 4774.575
1032.4949951171875 0 839.4204
1033.50634765625 0 1035.5444
1045.5078125 0 933.82007
1055.5028076171875 0 1247.0176
1073.5186767578125 0 4498.5957
1074.51611328125 0 1575.9208
1083.512451171875 0 2165.0417
1084.5101318359375 0 2138.2058
1090.5284423828125 0 6193.391 y Ammonia loss 5
1091.52783203125 0 3048.6362
1092.5234375 0 986.27325
1093.538818359375 0 904.9375
1100.531982421875 0 1631.1118
1101.5194091796875 0 4716.2456
1102.5230712890625 0 2322.167
1111.5469970703125 0 1603.5144
1112.5521240234375 0 1175.2428
1118.538818359375 0 1444.5938
1119.534423828125 0 1840.6346
1127.5640869140625 0 742.5537 b 10
1129.5543212890625 0 1244.0972
1130.5577392578125 0 1306.7633
1131.5548095703125 0 1031.9359
1134.5516357421875 0 1312.7039
1142.53662109375 0 1511.1703
1143.5399169921875 0 1065.2837
1149.570556640625 0 847.12274
1153.54345703125 0 766.9482
1159.5526123046875 0 1870.0145
1160.54931640625 0 3275.5044
1161.556396484375 0 1811.2096
1162.5438232421875 0 826.3216
1170.5458984375 0 1719.6354
1177.4210205078125 0 1077.5757
1177.558837890625 0 16314.734 y Ammonia loss 4
1178.561767578125 0 11147.293
1179.5626220703125 0 3563.1394
1180.5596923828125 0 1045.6715
1187.557373046875 0 1724.8757
1188.5555419921875 0 3130.0054
1189.5518798828125 0 1635.5802
1190.5655517578125 0 799.1325
1205.5743408203125 0 2240.2004
1206.568359375 0 1873.9318
1276.6290283203125 0 2454.8752 y Ammonia loss 3
1277.629150390625 0 2190.49
1278.6370849609375 0 728.738
1300.630126953125 0 879.62213
2715.760498046875 0 764.0856

Spectrum Details

|  |  |
| --- | --- |
| Matched peaks? Matched peaks The total absolute number of peaks matched. Additionally in brackets the total fraction of peaks matched and the total number of peaks is shown. | 78 (10.88% of 717) |
| FDR? FDR The false discovery rate estimated for this peptide. It is calculated by matching all theoretical fragments with a non-integer shift with the raw peaks for this spectrum. This is done with 40 different shifts. The resulting percentage is the average number of annotated peaks over the number of annotated peaks with the correct spectrum. | 0.49% |
| Satellite FDR? Satellite FDR See the FDR for details on its calculation. This satellite ion specific FDR only contains the satellite ions (d/w) for I/L/J positions. | - |
| PSM Score? PSM Score The PSM Score as given by Hecklib to this annotated spectrum. It is shown with three significant figures. | 532 |

#### Spectrum 7338? Spectrum 7338 The raw spectrum of this peptide as annotated by Hecklib. The fragments are coloured according to ion type (see legend). Any peaks with a star '\*' as text can be hovered over to see the full details, first the ion type second the mass shift type. By hovering over the amino acids in the peptide or ions in the legend the corresponding peaks are highlighted. By toggling the 'Unassigned' label you can turn the background (unassigned) peaks on or off in the plot. By updating the slider in the Ion legend you can update the spectrum to only show the top X% of the peaks with labels. The top X% means any peak that is within X% of the highest intensity. By dragging in the spectrum you can zoom in to a specific part of the spectrum and use 'Zoom Out' to get back to the original zoom level. The annotation of the spectrum is based on the given sequence in the peptides file and is done with different software so inconsistencies are likely. The peaks are annotated based on the given sequence, with 20 ppm tolerance.

Copy Data

##### Spectrum 7338 (TSV)

###### Preview

```
Loading example...
```

*Click on the button to copy the data to your clipboard.*

Mz MinMz MaxIntensity Max

WidthHeightPeptide font sizePeptide stroke widthSpectrum font sizeSpectrum stroke widthCompact peptide

Ion legend

wxyz

abcd

OtherUnassignedIon^Charge^_Position_Show for top:%

JSEVSDRPSGVSSRF

03.39e+46.79e+41.02e+51.36e+5

Zoom Out

z^+1^_2_y^+1^_2_c^+3^_11_y^+1^_3_z^+1^_3_c^+1^_4_w^+1^_4_y^+1^_4_z^+1^_4_z^+2^_9_y^+1^_4_y^+2^_9_c^+1^_5_z^+2^_10_y^+2^_10_c^+2^_11_z^+1^_5_y^+2^_11_z^+2^_11_y^+1^_5_y^+2^_11_z^+1^_6_y^+2^_12_z^+2^_12_y^+2^_12_c^+1^_6_y^+1^_6_w^+2^_13_z^+2^_13_y^+2^_13_z^+2^_13_y^+2^_13_z^+1^_7_c^+2^_14_c^+2^_14_c^+2^_14_y^+1^_7_y^+2^_14_y^+2^_14_z^+2^_14_y^+2^_14_y^+1^_8_c^+1^_8_z^+1^_9_c^+1^_9_c^+1^_10_w^+1^_10_z^+1^_10_c^+1^_11_y^+1^_11_c^+1^_12_z^+1^_12_y^+1^_12_c^+1^_13_c^+1^_14_

0778155623333111

Fragment Matches Table

Show background peaks

| Position | Ion type | Intensity | mz Theoretical | mz Error (Th) | mz Error (ppm) | Charge | Series Number |
| --- | --- | --- | --- | --- | --- | --- | --- |
| - | - | 403.9 | 141.5 | - | - | 0 | - |
| - | - | 536.5 | 148.9 | - | - | 0 | - |
| - | - | 441 | 166.8 | - | - | 0 | - |
| - | - | 2145 | 173.1 | - | - | 0 | - |
| - | - | 482.7 | 175.2 | - | - | 0 | - |
| - | - | 834.5 | 183.1 | - | - | 0 | - |
| - | - | 492.8 | 189.8 | - | - | 0 | - |
| - | - | 3783 | 201.1 | - | - | 0 | - |
| - | - | 1356 | 201.1 | - | - | 0 | - |
| - | - | 551.4 | 203.7 | - | - | 0 | - |
| - | - | 577.3 | 205.1 | - | - | 0 | - |
| - | - | 477.6 | 214.5 | - | - | 0 | - |
| - | - | 1329 | 227.1 | - | - | 0 | - |
| - | - | 509.3 | 229.1 | - | - | 0 | - |
| - | - | 665.9 | 229.2 | - | - | 0 | - |
| - | - | 696 | 242.1 | - | - | 0 | - |
| - | - | 497.3 | 249.1 | - | - | 0 | - |
| - | - | 3572 | 261.2 | - | - | 0 | - |
| - | - | 675.8 | 280.1 | - | - | 0 | - |
| - | - | 495.9 | 289.5 | - | - | 0 | - |
| - | - | 1432 | 301.1 | - | - | 0 | - |
| - | - | 1084 | 302.1 | - | - | 0 | - |
| 14 | z | 887.3 | 306.2 | 0.0009479 | 3.096 | +1 | 2 |
| - | - | 1442 | 312.2 | - | - | 0 | - |
| - | - | 631.3 | 316.1 | - | - | 0 | - |
| 14 | y | 706 | 322.2 | 4.236E-05 | 0.1315 | +1 | 2 |
| - | - | 1045 | 330.2 | - | - | 0 | - |
| - | - | 1102 | 330.2 | - | - | 0 | - |
| - | - | 558.8 | 335.4 | - | - | 0 | - |
| - | - | 711.7 | 348.2 | - | - | 0 | - |
| 11 | c | 2332 | 376.2 | 0.002806 | 7.459 | +3 | 11 |
| - | - | 668.4 | 377.2 | - | - | 0 | - |
| - | - | 580 | 384.2 | - | - | 0 | - |
| - | - | 1026 | 391.2 | - | - | 0 | - |
| 13 | y | 4358 | 392.2 | 0.0003405 | 0.8682 | +1 | 3 |
| 13 | z | 1093 | 393.2 | 0.001208 | 3.071 | +1 | 3 |
| - | - | 689.7 | 394.2 | - | - | 0 | - |
| - | - | 1349 | 412.2 | - | - | 0 | - |
| - | - | 1030 | 417.2 | - | - | 0 | - |
| - | - | 678.9 | 429.2 | - | - | 0 | - |
| - | - | 598.4 | 442.3 | - | - | 0 | - |
| - | - | 1109 | 444.3 | - | - | 0 | - |
| - | - | 631.3 | 445.2 | - | - | 0 | - |
| 4 | c | 620.2 | 446.3 | 0.002044 | 4.581 | +1 | 4 |
| 12 | w | 2843 | 463.2 | 0.0001618 | 0.3493 | +1 | 4 |
| - | - | 1164 | 464.2 | - | - | 0 | - |
| - | - | 1230 | 468.3 | - | - | 0 | - |
| - | - | 1038 | 478.2 | - | - | 0 | - |
| 12 | y | 1533 | 479.2 | 0.00176 | 3.672 | +1 | 4 |
| 12 | z | 2550 | 480.2 | 0.00107 | 2.229 | +1 | 4 |
| - | - | 644.8 | 481.2 | - | - | 0 | - |
| 7 | z | 2008 | 488.8 | 0.000148 | 0.3028 | +2 | 9 |
| - | - | 1346 | 489.3 | - | - | 0 | - |
| 12 | y | 1669 | 496.3 | 0.001392 | 2.806 | +1 | 4 |
| 7 | y | 631.1 | 496.8 | 0.001585 | 3.19 | +2 | 9 |
| - | - | 743.6 | 497.3 | - | - | 0 | - |
| - | - | 2073 | 504.2 | - | - | 0 | - |
| - | - | 1367 | 507.1 | - | - | 0 | - |
| - | - | 1892 | 511.3 | - | - | 0 | - |
| - | - | 1.5E+04 | 513.4 | - | - | 0 | - |
| - | - | 4142 | 514.4 | - | - | 0 | - |
| - | - | 3963 | 520.2 | - | - | 0 | - |
| - | - | 1707 | 521.2 | - | - | 0 | - |
| - | - | 1412 | 523.3 | - | - | 0 | - |
| - | - | 745 | 524.3 | - | - | 0 | - |
| 5 | c | 4908 | 533.3 | 0.000717 | 1.345 | +1 | 5 |
| - | - | 1265 | 534.3 | - | - | 0 | - |
| - | - | 1167 | 535.6 | - | - | 0 | - |
| - | - | 1789 | 540.2 | - | - | 0 | - |
| - | - | 2849 | 540.3 | - | - | 0 | - |
| - | - | 1375 | 540.8 | - | - | 0 | - |
| - | - | 3064 | 541.2 | - | - | 0 | - |
| - | - | 1.344E+05 | 541.3 | - | - | 0 | - |
| - | - | 1443 | 541.6 | - | - | 0 | - |
| - | - | 1660 | 541.9 | - | - | 0 | - |
| - | - | 1.247E+04 | 542.2 | - | - | 0 | - |
| - | - | 1.557E+04 | 542.3 | - | - | 0 | - |
| - | - | 4.387E+04 | 542.3 | - | - | 0 | - |
| - | - | 5178 | 542.8 | - | - | 0 | - |
| 6 | z | 2208 | 546.3 | 1.268E-05 | 0.02321 | +2 | 10 |
| - | - | 3462 | 546.8 | - | - | 0 | - |
| - | - | 1912 | 547.3 | - | - | 0 | - |
| 6 | y | 1433 | 554.3 | 0.0005383 | 0.9712 | +2 | 10 |
| - | - | 758.1 | 554.8 | - | - | 0 | - |
| 11 | c | 2455 | 564.3 | 0.009882 | 17.51 | +2 | 11 |
| - | - | 668.9 | 565.3 | - | - | 0 | - |
| 11 | z | 870.6 | 579.3 | 0.001133 | 1.956 | +1 | 5 |
| 5 | y | 992.8 | 589.3 | 0.0007284 | 1.236 | +2 | 11 |
| 5 | z | 2591 | 589.8 | 0.001963 | 3.329 | +2 | 11 |
| - | - | 4597 | 590.3 | - | - | 0 | - |
| - | - | 2650 | 590.8 | - | - | 0 | - |
| - | - | 690.9 | 591.3 | - | - | 0 | - |
| - | - | 1412 | 592.3 | - | - | 0 | - |
| - | - | 697.1 | 593.3 | - | - | 0 | - |
| 11 | y | 1423 | 595.3 | 0.0004398 | 0.7388 | +1 | 5 |
| - | - | 1656 | 596.4 | - | - | 0 | - |
| - | - | 747.1 | 597.4 | - | - | 0 | - |
| 5 | y | 7741 | 597.8 | 0.0005833 | 0.9757 | +2 | 11 |
| - | - | 6917 | 598.3 | - | - | 0 | - |
| - | - | 1962 | 598.8 | - | - | 0 | - |
| - | - | 601.7 | 625.4 | - | - | 0 | - |
| 10 | z | 3753 | 636.3 | 0.0002498 | 0.3926 | +1 | 6 |
| - | - | 2646 | 637.3 | - | - | 0 | - |
| 4 | y | 717.1 | 638.3 | 0.005099 | 7.988 | +2 | 12 |
| 4 | z | 6215 | 639.3 | 0.000709 | 1.109 | +2 | 12 |
| - | - | 3015 | 639.8 | - | - | 0 | - |
| - | - | 807.5 | 640.3 | - | - | 0 | - |
| 4 | y | 6661 | 647.3 | 0.0002439 | 0.3768 | +2 | 12 |
| - | - | 5763 | 647.8 | - | - | 0 | - |
| 6 | c | 4968 | 648.3 | 0.005055 | 7.797 | +1 | 6 |
| - | - | 1523 | 649.3 | - | - | 0 | - |
| 10 | y | 5151 | 652.3 | 0.0002522 | 0.3866 | +1 | 6 |
| - | - | 1198 | 653.3 | - | - | 0 | - |
| - | - | 1872 | 663.8 | - | - | 0 | - |
| - | - | 886.9 | 664.3 | - | - | 0 | - |
| 3 | w | 3872 | 674.3 | 0.0002777 | 0.4118 | +2 | 13 |
| - | - | 3222 | 674.8 | - | - | 0 | - |
| - | - | 1804 | 675.3 | - | - | 0 | - |
| - | - | 865.7 | 679.4 | - | - | 0 | - |
| - | - | 1830 | 694.3 | - | - | 0 | - |
| 3 | z | 1077 | 695.3 | 0.009885 | 14.22 | +2 | 13 |
| 3 | y | 585.5 | 703.3 | 0.001377 | 1.958 | +2 | 13 |
| 3 | z | 2163 | 703.8 | 0.001131 | 1.608 | +2 | 13 |
| - | - | 2542 | 704.3 | - | - | 0 | - |
| - | - | 673.9 | 710.3 | - | - | 0 | - |
| - | - | 912.7 | 711.4 | - | - | 0 | - |
| 3 | y | 8380 | 711.9 | 0.000911 | 1.28 | +2 | 13 |
| - | - | 7182 | 712.4 | - | - | 0 | - |
| - | - | 3468 | 712.9 | - | - | 0 | - |
| - | - | 830.5 | 713.4 | - | - | 0 | - |
| 9 | z | 3697 | 723.4 | 0.0003145 | 0.4348 | +1 | 7 |
| - | - | 2529 | 724.4 | - | - | 0 | - |
| - | - | 751.8 | 725.4 | - | - | 0 | - |
| - | - | 1387 | 728.4 | - | - | 0 | - |
| 14 | c | 769.4 | 728.9 | 0.01434 | 19.67 | +2 | 14 |
| 14 | c | 1105 | 729.4 | 0.001585 | 2.173 | +2 | 14 |
| 14 | c | 2.33E+04 | 737.9 | 0.0005769 | 0.7819 | +2 | 14 |
| - | - | 1.878E+04 | 738.4 | - | - | 0 | - |
| - | - | 7548 | 738.9 | - | - | 0 | - |
| 9 | y | 3055 | 739.4 | 0.007591 | 10.27 | +1 | 7 |
| 2 | y | 945.5 | 746.4 | 0.0004124 | 0.5526 | +2 | 14 |
| 2 | y | 907.8 | 746.9 | 0.006725 | 9.005 | +2 | 14 |
| 2 | z | 4330 | 747.4 | 0.003962 | 5.301 | +2 | 14 |
| - | - | 1812 | 747.9 | - | - | 0 | - |
| - | - | 2564 | 748.4 | - | - | 0 | - |
| 2 | y | 1.078E+04 | 755.4 | 0.0003237 | 0.4285 | +2 | 14 |
| - | - | 8195 | 755.9 | - | - | 0 | - |
| - | - | 5247 | 756.4 | - | - | 0 | - |
| - | - | 1176 | 756.9 | - | - | 0 | - |
| - | - | 726.9 | 757.4 | - | - | 0 | - |
| - | - | 831.8 | 779.4 | - | - | 0 | - |
| - | - | 1974 | 781.9 | - | - | 0 | - |
| - | - | 4118 | 782.4 | - | - | 0 | - |
| - | - | 3065 | 782.9 | - | - | 0 | - |
| - | - | 1173 | 783.4 | - | - | 0 | - |
| - | - | 678.2 | 788.4 | - | - | 0 | - |
| - | - | 7420 | 789.4 | - | - | 0 | - |
| - | - | 5580 | 789.9 | - | - | 0 | - |
| - | - | 3564 | 790.4 | - | - | 0 | - |
| - | - | 7582 | 790.9 | - | - | 0 | - |
| - | - | 4837 | 791.4 | - | - | 0 | - |
| - | - | 2323 | 791.9 | - | - | 0 | - |
| - | - | 869.5 | 792.4 | - | - | 0 | - |
| - | - | 884.1 | 796.4 | - | - | 0 | - |
| - | - | 996.5 | 801.9 | - | - | 0 | - |
| - | - | 1019 | 802.9 | - | - | 0 | - |
| - | - | 3450 | 803.4 | - | - | 0 | - |
| - | - | 2.246E+04 | 803.9 | - | - | 0 | - |
| - | - | 1.673E+04 | 804.4 | - | - | 0 | - |
| - | - | 8257 | 804.9 | - | - | 0 | - |
| - | - | 2653 | 805.4 | - | - | 0 | - |
| - | - | 758.5 | 809.9 | - | - | 0 | - |
| - | - | 3385 | 810.4 | - | - | 0 | - |
| - | - | 4977 | 810.9 | - | - | 0 | - |
| - | - | 2770 | 811.4 | - | - | 0 | - |
| - | - | 1.165E+04 | 811.9 | - | - | 0 | - |
| - | - | 1.731E+04 | 812.4 | - | - | 0 | - |
| - | - | 1.1E+04 | 812.9 | - | - | 0 | - |
| - | - | 3961 | 813.4 | - | - | 0 | - |
| - | - | 1509 | 813.9 | - | - | 0 | - |
| 8 | y | 927.4 | 836.4 | 0.001031 | 1.233 | +1 | 8 |
| - | - | 1588 | 857.5 | - | - | 0 | - |
| - | - | 943 | 858.5 | - | - | 0 | - |
| - | - | 974.5 | 864.4 | - | - | 0 | - |
| - | - | 979.2 | 880.4 | - | - | 0 | - |
| - | - | 817 | 881.4 | - | - | 0 | - |
| - | - | 990.2 | 895.4 | - | - | 0 | - |
| - | - | 661.3 | 897.4 | - | - | 0 | - |
| 8 | c | 5284 | 901.5 | 0.0003826 | 0.4244 | +1 | 8 |
| - | - | 3035 | 902.5 | - | - | 0 | - |
| - | - | 792.3 | 903.5 | - | - | 0 | - |
| - | - | 768.8 | 904.5 | - | - | 0 | - |
| - | - | 3007 | 921.5 | - | - | 0 | - |
| - | - | 1298 | 922.5 | - | - | 0 | - |
| - | - | 3461 | 944.5 | - | - | 0 | - |
| - | - | 1335 | 945.5 | - | - | 0 | - |
| - | - | 746.9 | 946.5 | - | - | 0 | - |
| - | - | 652 | 968.7 | - | - | 0 | - |
| 7 | z | 2773 | 976.5 | 0.001888 | 1.933 | +1 | 9 |
| - | - | 1514 | 977.5 | - | - | 0 | - |
| - | - | 1207 | 987.5 | - | - | 0 | - |
| 9 | c | 1.499E+04 | 988.5 | 0.0007338 | 0.7423 | +1 | 9 |
| - | - | 8053 | 989.5 | - | - | 0 | - |
| - | - | 2796 | 990.5 | - | - | 0 | - |
| - | - | 754.3 | 1003 | - | - | 0 | - |
| - | - | 551.6 | 1045 | - | - | 0 | - |
| 10 | c | 8348 | 1046 | 0.0004079 | 0.3902 | +1 | 10 |
| 6 | w | 4661 | 1047 | 0.007739 | 7.395 | +1 | 10 |
| - | - | 1935 | 1048 | - | - | 0 | - |
| - | - | 761.2 | 1052 | - | - | 0 | - |
| - | - | 1928 | 1053 | - | - | 0 | - |
| - | - | 778.1 | 1054 | - | - | 0 | - |
| - | - | 690.6 | 1057 | - | - | 0 | - |
| - | - | 1110 | 1066 | - | - | 0 | - |
| - | - | 964 | 1080 | - | - | 0 | - |
| - | - | 1345 | 1081 | - | - | 0 | - |
| - | - | 640.6 | 1081 | - | - | 0 | - |
| - | - | 1020 | 1082 | - | - | 0 | - |
| - | - | 900.1 | 1082 | - | - | 0 | - |
| - | - | 1045 | 1084 | - | - | 0 | - |
| - | - | 1.063E+04 | 1085 | - | - | 0 | - |
| - | - | 4692 | 1086 | - | - | 0 | - |
| 6 | z | 2647 | 1092 | 0.001487 | 1.363 | +1 | 10 |
| - | - | 4918 | 1093 | - | - | 0 | - |
| - | - | 1925 | 1094 | - | - | 0 | - |
| 11 | c | 5843 | 1145 | 0.001927 | 1.684 | +1 | 11 |
| - | - | 3697 | 1146 | - | - | 0 | - |
| - | - | 2042 | 1147 | - | - | 0 | - |
| - | - | 657.3 | 1152 | - | - | 0 | - |
| - | - | 1051 | 1153 | - | - | 0 | - |
| - | - | 2537 | 1180 | - | - | 0 | - |
| - | - | 747.2 | 1181 | - | - | 0 | - |
| - | - | 899.5 | 1182 | - | - | 0 | - |
| - | - | 752.6 | 1188 | - | - | 0 | - |
| - | - | 641.1 | 1191 | - | - | 0 | - |
| 5 | y | 1124 | 1195 | 0.001703 | 1.425 | +1 | 11 |
| - | - | 670.6 | 1198 | - | - | 0 | - |
| - | - | 1413 | 1199 | - | - | 0 | - |
| - | - | 740.2 | 1231 | - | - | 0 | - |
| 12 | c | 1.178E+04 | 1232 | 0.002584 | 2.098 | +1 | 12 |
| - | - | 7954 | 1233 | - | - | 0 | - |
| - | - | 3001 | 1234 | - | - | 0 | - |
| 4 | z | 697.6 | 1278 | 0.0001212 | 0.09486 | +1 | 12 |
| - | - | 2412 | 1279 | - | - | 0 | - |
| - | - | 2406 | 1280 | - | - | 0 | - |
| 4 | y | 851.9 | 1294 | 0.009351 | 7.229 | +1 | 12 |
| - | - | 709.9 | 1295 | - | - | 0 | - |
| 13 | c | 9517 | 1319 | 0.001165 | 0.8833 | +1 | 13 |
| - | - | 5172 | 1320 | - | - | 0 | - |
| - | - | 2340 | 1321 | - | - | 0 | - |
| - | - | 640.3 | 1327 | - | - | 0 | - |
| - | - | 857.3 | 1346 | - | - | 0 | - |
| - | - | 5566 | 1408 | - | - | 0 | - |
| - | - | 4447 | 1409 | - | - | 0 | - |
| - | - | 2251 | 1410 | - | - | 0 | - |
| - | - | 1200 | 1432 | - | - | 0 | - |
| - | - | 1300 | 1433 | - | - | 0 | - |
| - | - | 1476 | 1459 | - | - | 0 | - |
| - | - | 1191 | 1460 | - | - | 0 | - |
| - | - | 992.9 | 1461 | - | - | 0 | - |
| 14 | c | 1925 | 1475 | 0.001484 | 1.006 | +1 | 14 |
| - | - | 7324 | 1476 | - | - | 0 | - |
| - | - | 5310 | 1477 | - | - | 0 | - |
| - | - | 2588 | 1478 | - | - | 0 | - |
| - | - | 4555 | 1495 | - | - | 0 | - |
| - | - | 2775 | 1496 | - | - | 0 | - |
| - | - | 1316 | 1497 | - | - | 0 | - |
| - | - | 1419 | 1523 | - | - | 0 | - |
| - | - | 790.6 | 1524 | - | - | 0 | - |
| - | - | 749.7 | 1549 | - | - | 0 | - |
| - | - | 5197 | 1565 | - | - | 0 | - |
| - | - | 6467 | 1566 | - | - | 0 | - |
| - | - | 4973 | 1567 | - | - | 0 | - |
| - | - | 1679 | 1568 | - | - | 0 | - |
| - | - | 1372 | 1569 | - | - | 0 | - |
| - | - | 698.6 | 1579 | - | - | 0 | - |
| - | - | 1877 | 1580 | - | - | 0 | - |
| - | - | 1353 | 1581 | - | - | 0 | - |
| - | - | 1710 | 1582 | - | - | 0 | - |
| - | - | 1041 | 1583 | - | - | 0 | - |
| - | - | 2872 | 1591 | - | - | 0 | - |
| - | - | 1368 | 1592 | - | - | 0 | - |
| - | - | 924 | 1593 | - | - | 0 | - |
| - | - | 761.8 | 1596 | - | - | 0 | - |
| - | - | 1402 | 1597 | - | - | 0 | - |
| - | - | 1393 | 1598 | - | - | 0 | - |
| - | - | 1133 | 1599 | - | - | 0 | - |
| - | - | 1669 | 1605 | - | - | 0 | - |
| - | - | 775.2 | 1606 | - | - | 0 | - |
| - | - | 4252 | 1607 | - | - | 0 | - |
| - | - | 2.008E+04 | 1608 | - | - | 0 | - |
| - | - | 1.604E+04 | 1609 | - | - | 0 | - |
| - | - | 8125 | 1610 | - | - | 0 | - |
| - | - | 2050 | 1611 | - | - | 0 | - |
| - | - | 1678 | 1621 | - | - | 0 | - |
| - | - | 2789 | 1622 | - | - | 0 | - |
| - | - | 2467 | 1623 | - | - | 0 | - |
| - | - | 6236 | 1624 | - | - | 0 | - |
| - | - | 2.032E+04 | 1625 | - | - | 0 | - |
| - | - | 1.41E+04 | 1626 | - | - | 0 | - |
| - | - | 9625 | 1627 | - | - | 0 | - |
| - | - | 2001 | 1628 | - | - | 0 | - |
| - | - | 756.3 | 2530 | - | - | 0 | - |
| - | - | 805.6 | 3080 | - | - | 0 | - |

m/z Charge Intensity FragmentType MassShift Position
141.45944213867188 0 403.8507
148.94627380371094 0 536.52673
166.76348876953125 0 440.98297
173.1282958984375 0 2145.0107
175.24884033203125 0 482.73578
183.1129150390625 0 834.4837
189.80914306640625 0 492.83676
201.12313842773438 0 3783.082
201.1343994140625 0 1356.0757
203.70164489746094 0 551.40027
205.09698486328125 0 577.28876
214.47178649902344 0 477.5835
227.10260009765625 0 1328.6355
229.1175994873047 0 509.3446
229.1545867919922 0 665.8581
242.1488494873047 0 695.957
249.11473083496094 0 497.25784
261.1707763671875 0 3572.2288
280.1297302246094 0 675.7681
289.51397705078125 0 495.90204
301.0592346191406 0 1432.0184
302.0603332519531 0 1083.6204
306.1676940917969 0 887.31384 z 13
312.1554260253906 0 1442.2456
316.1293029785156 0 631.278
322.1874084472656 0 705.9514 y 13
330.16571044921875 0 1045.1821
330.1856689453125 0 1102.3313
335.4117126464844 0 558.77094
348.2034606933594 0 711.71075
376.19708251953125 0 2331.9036 c Water loss 10
377.1980285644531 0 668.3838
384.2325744628906 0 579.98944
391.18585205078125 0 1026.1593
392.1925048828125 0 4358.014 y Ammonia loss 12
393.199462890625 0 1093.4891 z 12
394.2140197753906 0 689.6985
412.2261962890625 0 1348.6521
417.18743896484375 0 1029.7385
429.2326965332031 0 678.92896
442.2792663574219 0 598.35425
444.2553405761719 0 1109.3506
445.1722106933594 0 631.3083
446.2588806152344 0 620.20044 c 3
463.22979736328125 0 2843.3806 w 11
464.2327880859375 0 1163.663
468.2943115234375 0 1229.8635
478.2176513671875 0 1038.4802
479.2231140136719 0 1532.7429 y Ammonia loss 11
480.23162841796875 0 2550.361 z 11
481.2330322265625 0 644.84827
488.7580261230469 0 2007.6444 z 6
489.2591247558594 0 1346.3754
496.2500305175781 0 1669.1772 y 11
496.7656555175781 0 631.07153 y 6
497.2629699707031 0 743.64435
504.2210388183594 0 2073.467
507.10357666015625 0 1366.7034
511.3351135253906 0 1891.8069
513.3505859375 0 15000.259
514.3543701171875 0 4141.6646
520.1613159179688 0 3962.8843
521.1646118164062 0 1707.365
523.3327026367188 0 1411.9556
524.2737426757812 0 744.9765
533.292236328125 0 4908.414 c 4
534.29638671875 0 1264.8643
535.605712890625 0 1166.5365
540.1679077148438 0 1788.5375
540.30908203125 0 2849.0522
540.8148803710938 0 1374.7621
541.243408203125 0 3063.5122
541.3456420898438 0 134362.72
541.6094360351562 0 1442.8115
541.9432373046875 0 1660.4585
542.22509765625 0 12473.992
542.2808227539062 0 15570.916
542.3485107421875 0 43870.5
542.7825317382812 0 5178.2236
546.2713623046875 0 2207.9048 z 5
546.7737426757812 0 3462.3813
547.2744140625 0 1911.6051
554.28125 0 1432.7859 y 5
554.7822875976562 0 758.0975
564.2783203125 0 2454.5396 c Ammonia loss 10
565.2832641601562 0 668.8578
579.30224609375 0 870.56415 z 10
589.2841796875 0 992.82996 y Ammonia loss 4
589.785400390625 0 2590.6013 z 4
590.2899780273438 0 4596.659
590.7921142578125 0 2650.3196
591.294677734375 0 690.87915
592.3314819335938 0 1411.8728
593.335205078125 0 697.1326
595.3193969726562 0 1423.126 y 10
596.3509521484375 0 1655.8187
597.3568115234375 0 747.0915
597.796142578125 0 7741.303 y 4
598.2976684570312 0 6917.0083
598.8003540039062 0 1962.5
625.4359741210938 0 601.7269
636.3223266601562 0 3753.0366 z 9
637.3259887695312 0 2646.1667
638.3307495117188 0 717.06573 y Water loss 3
639.3208618164062 0 6214.806 z 3
639.822021484375 0 3014.8655
640.3245239257812 0 807.49603
647.3311767578125 0 6660.6855 y 3
647.8321533203125 0 5762.592
648.324951171875 0 4967.9556 c 5
649.3223266601562 0 1523.1101
652.341552734375 0 5150.934 y 9
653.3413696289062 0 1197.8002
663.84423828125 0 1871.737
664.3477172851562 0 886.86145
674.3359375 0 3871.9556 w 2
674.8374633789062 0 3221.9834
675.3370971679688 0 1804.4648
679.3674926757812 0 865.6968
694.3330688476562 0 1829.7625
695.3394775390625 0 1076.6185 z Ammonia loss 2
703.34033203125 0 585.5139 y Ammonia loss 2
703.8417358398438 0 2162.824 z 2
704.3429565429688 0 2542.1455
710.3317260742188 0 673.9453
711.3854370117188 0 912.744
711.851318359375 0 8379.67 y 2
712.3534545898438 0 7181.7246
712.856201171875 0 3467.6758
713.3551635742188 0 830.5147
723.3549194335938 0 3697.0532 z 8
724.3544311523438 0 2528.8955
725.3641967773438 0 751.7931
728.3653564453125 0 1387.1123
728.8644409179688 0 769.3851 c Water loss 13
729.3692016601562 0 1105.2467 c Ammonia loss 13
737.8834838867188 0 23304.137 c 13
738.3847045898438 0 18775.82
738.88623046875 0 7547.629
739.3809204101562 0 3054.5474 y 8
746.362548828125 0 945.45856 y Water loss 1
746.8616943359375 0 907.83136 y Ammonia loss 1
747.3549194335938 0 4330.3267 z 1
747.8592529296875 0 1811.9321
748.3602905273438 0 2564.4734
755.367919921875 0 10781.465 y 1
755.869384765625 0 8195.133
756.3703002929688 0 5247.1562
756.8703002929688 0 1175.8148
757.3756103515625 0 726.88007
779.3814086914062 0 831.8448
781.9071044921875 0 1973.8983
782.4042358398438 0 4117.502
782.9043579101562 0 3065.3223
783.4078369140625 0 1172.8572
788.40771484375 0 678.19415
789.3832397460938 0 7420.4775
789.88525390625 0 5580.3794
790.3876953125 0 3563.9788
790.8980102539062 0 7582.302
791.401123046875 0 4837.4165
791.902587890625 0 2322.5002
792.3983154296875 0 869.4662
796.39404296875 0 884.0598
801.8861083984375 0 996.4853
802.8858032226562 0 1018.6226
803.4068603515625 0 3450.2112
803.9015502929688 0 22461.459
804.401611328125 0 16727.123
804.9035034179688 0 8257.13
805.4035034179688 0 2652.8186
809.8846435546875 0 758.5257
810.3915405273438 0 3385.3447
810.8917846679688 0 4977.039
811.3942260742188 0 2769.9062
811.9075927734375 0 11648.966
812.4112548828125 0 17305.113
812.91357421875 0 10999.538
813.4124145507812 0 3960.976
813.9104614257812 0 1508.557
836.4271240234375 0 927.4368 y 7
857.4571533203125 0 1588.2637
858.465576171875 0 943.0264
864.4119873046875 0 974.4999
880.4237060546875 0 979.20483
881.4374389648438 0 816.9686
895.4223022460938 0 990.22626
897.4380493164062 0 661.319
901.473388671875 0 5284.491 c 7
902.4773559570312 0 3035.2483
903.4766845703125 0 792.2965
904.485595703125 0 768.7712
921.5003662109375 0 3007.3335
922.50244140625 0 1297.9141
944.4907836914062 0 3461.3127
945.4907836914062 0 1334.6173
946.4896850585938 0 746.9182
968.6566772460938 0 652.05
976.506591796875 0 2773.1965 z 6
977.5135498046875 0 1514.2021
987.49951171875 0 1207.0673
988.5050659179688 0 14986.635 c 8
989.5073852539062 0 8053.066
990.5108032226562 0 2796.2817
1002.52099609375 0 754.3066
1045.4052734375 0 551.563
1045.52685546875 0 8348.123 c 9
1046.530029296875 0 4660.887 w 5
1047.5311279296875 0 1934.9069
1051.6318359375 0 761.16284
1052.626953125 0 1927.795
1053.6314697265625 0 778.0809
1057.15673828125 0 690.5972
1066.4715576171875 0 1109.9553
1080.1094970703125 0 963.9776
1080.6268310546875 0 1345.1166
1080.9718017578125 0 640.61554
1081.5135498046875 0 1019.7774
1081.6373291015625 0 900.0564
1083.5089111328125 0 1044.5145
1084.5623779296875 0 10628.409
1085.564208984375 0 4692.3516
1091.533935546875 0 2647.183 z 5
1092.540771484375 0 4917.8955
1093.5418701171875 0 1925.0414
1144.59375 0 5842.768 c 10
1145.600341796875 0 3697.4617
1146.60302734375 0 2041.646
1152.484130859375 0 657.34326
1153.4989013671875 0 1051.1691
1179.5750732421875 0 2537.3936
1180.5706787109375 0 747.20966
1181.5888671875 0 899.51074
1187.614013671875 0 752.6029
1191.4886474609375 0 641.0972
1194.58447265625 0 1124.4716 y 4
1197.6065673828125 0 670.6172
1198.576416015625 0 1413.1737
1230.633056640625 0 740.17365
1231.6251220703125 0 11778.878 c 11
1232.6282958984375 0 7953.5957
1233.6312255859375 0 3000.684
1277.635986328125 0 697.64 z 3
1278.6405029296875 0 2412.2466
1279.64599609375 0 2405.58
1293.6639404296875 0 851.86957 y 3
1294.6607666015625 0 709.87024
1318.6585693359375 0 9516.647 c 12
1319.6612548828125 0 5171.837
1320.6658935546875 0 2339.5244
1326.6845703125 0 640.3125
1345.667724609375 0 857.28906
1407.6815185546875 0 5565.756
1408.685791015625 0 4446.517
1409.6923828125 0 2251.019
1431.7581787109375 0 1200.0563
1432.76611328125 0 1299.5348
1458.7318115234375 0 1476.0985
1459.7513427734375 0 1190.5817
1460.751953125 0 992.88885
1474.7623291015625 0 1924.6154 c 13
1475.7659912109375 0 7323.8374
1476.76806640625 0 5310.2915
1477.7637939453125 0 2588.0325
1494.7156982421875 0 4554.891
1495.718994140625 0 2775.2654
1496.7216796875 0 1315.5186
1522.7392578125 0 1418.8531
1523.732666015625 0 790.63324
1548.75244140625 0 749.7004
1564.78076171875 0 5197.272
1565.7818603515625 0 6466.5913
1566.7855224609375 0 4973.4067
1567.787353515625 0 1679.2428
1568.7960205078125 0 1371.9603
1578.8177490234375 0 698.55536
1579.804931640625 0 1876.8209
1580.7979736328125 0 1352.9982
1581.8057861328125 0 1709.9097
1582.7918701171875 0 1040.8687
1590.7830810546875 0 2872.3267
1591.77392578125 0 1368.302
1592.7635498046875 0 924.0242
1595.8232421875 0 761.7994
1596.8299560546875 0 1401.6919
1597.8287353515625 0 1392.5751
1598.8289794921875 0 1132.749
1604.7554931640625 0 1668.771
1605.7662353515625 0 775.24225
1606.80322265625 0 4251.942
1607.8004150390625 0 20077.482
1608.80224609375 0 16038.618
1609.8067626953125 0 8124.7026
1610.8038330078125 0 2050.452
1620.7823486328125 0 1678.475
1621.78466796875 0 2788.6416
1622.7999267578125 0 2466.7144
1623.8153076171875 0 6235.9897
1624.8233642578125 0 20318.47
1625.8267822265625 0 14103.447
1626.8314208984375 0 9624.52
1627.83837890625 0 2000.8405
2530.368408203125 0 756.27795
3080.402099609375 0 805.5598

Spectrum Details

|  |  |
| --- | --- |
| Matched peaks? Matched peaks The total absolute number of peaks matched. Additionally in brackets the total fraction of peaks matched and the total number of peaks is shown. | 55 (18.09% of 304) |
| FDR? FDR The false discovery rate estimated for this peptide. It is calculated by matching all theoretical fragments with a non-integer shift with the raw peaks for this spectrum. This is done with 40 different shifts. The resulting percentage is the average number of annotated peaks over the number of annotated peaks with the correct spectrum. | 1.60% |
| Satellite FDR? Satellite FDR See the FDR for details on its calculation. This satellite ion specific FDR only contains the satellite ions (d/w) for I/L/J positions. | - |
| PSM Score? PSM Score The PSM Score as given by Hecklib to this annotated spectrum. It is shown with three significant figures. | 316 |

#### Spectrum 6589? Spectrum 6589 The raw spectrum of this peptide as annotated by Hecklib. The fragments are coloured according to ion type (see legend). Any peaks with a star '\*' as text can be hovered over to see the full details, first the ion type second the mass shift type. By hovering over the amino acids in the peptide or ions in the legend the corresponding peaks are highlighted. By toggling the 'Unassigned' label you can turn the background (unassigned) peaks on or off in the plot. By updating the slider in the Ion legend you can update the spectrum to only show the top X% of the peaks with labels. The top X% means any peak that is within X% of the highest intensity. By dragging in the spectrum you can zoom in to a specific part of the spectrum and use 'Zoom Out' to get back to the original zoom level. The annotation of the spectrum is based on the given sequence in the peptides file and is done with different software so inconsistencies are likely. The peaks are annotated based on the given sequence, with 20 ppm tolerance.

Copy Data

##### Spectrum 6589 (TSV)

###### Preview

```
Loading example...
```

*Click on the button to copy the data to your clipboard.*

Mz MinMz MaxIntensity Max

WidthHeightPeptide font sizePeptide stroke widthSpectrum font sizeSpectrum stroke widthCompact peptide

Ion legend

wxyz

abcd

OtherUnassignedIon^Charge^_Position_Show for top:%

JSEVSDRPSGVSSRF

05.17e+41.03e+51.55e+52.07e+5

Zoom Out

y^+1^_1_c^+2^_5_z^+1^_2_y^+1^_2_c^+3^_11_y^+1^_3_y^+3^_11_y^+1^_3_w^+1^_4_y^+1^_4_z^+1^_4_z^+2^_9_y^+1^_4_c^+1^_5_z^+2^_10_y^+2^_10_c^+2^_11_z^+1^_5_y^+2^_11_y^+2^_11_z^+2^_11_y^+1^_5_y^+2^_11_z^+1^_6_z^+2^_12_y^+2^_12_c^+1^_6_y^+1^_6_w^+2^_13_y^+2^_13_z^+2^_13_y^+2^_13_y^+1^_7_y^+1^_7_z^+1^_7_c^+2^_14_c^+2^_14_c^+2^_14_y^+1^_7_y^+2^_14_y^+2^_14_z^+2^_14_y^+2^_14_w^+1^_8_z^+1^_8_z^+1^_8_y^+1^_8_c^+1^_8_z^+1^_9_c^+1^_9_c^+1^_10_w^+1^_10_z^+1^_10_y^+1^_10_c^+1^_11_z^+1^_11_y^+1^_11_c^+1^_12_y^+1^_12_c^+1^_13_y^+1^_13_z^+1^_13_c^+1^_14_c^+1^_14_c^+1^_14_

0778155523333111

Fragment Matches Table

Show background peaks

| Position | Ion type | Intensity | mz Theoretical | mz Error (Th) | mz Error (ppm) | Charge | Series Number |
| --- | --- | --- | --- | --- | --- | --- | --- |
| - | - | 907.1 | 120.1 | - | - | 0 | - |
| - | - | 412 | 129.1 | - | - | 0 | - |
| - | - | 681.6 | 129.1 | - | - | 0 | - |
| - | - | 376.3 | 133.1 | - | - | 0 | - |
| - | - | 377.8 | 134.3 | - | - | 0 | - |
| - | - | 516.3 | 134.5 | - | - | 0 | - |
| - | - | 427.3 | 138.3 | - | - | 0 | - |
| - | - | 829 | 149 | - | - | 0 | - |
| - | - | 2898 | 155.1 | - | - | 0 | - |
| 15 | y | 720.2 | 166.1 | 0.0001955 | 1.177 | +1 | 1 |
| - | - | 423.6 | 168.7 | - | - | 0 | - |
| - | - | 7502 | 173.1 | - | - | 0 | - |
| - | - | 430.4 | 173.3 | - | - | 0 | - |
| - | - | 1715 | 173.5 | - | - | 0 | - |
| - | - | 749.7 | 174.1 | - | - | 0 | - |
| - | - | 551.1 | 175.8 | - | - | 0 | - |
| - | - | 469.5 | 179.7 | - | - | 0 | - |
| - | - | 1145 | 182.1 | - | - | 0 | - |
| - | - | 468.5 | 186.1 | - | - | 0 | - |
| - | - | 538 | 190 | - | - | 0 | - |
| - | - | 447.7 | 198.2 | - | - | 0 | - |
| - | - | 6763 | 201.1 | - | - | 0 | - |
| - | - | 2115 | 201.1 | - | - | 0 | - |
| - | - | 450.2 | 202.1 | - | - | 0 | - |
| - | - | 1943 | 215.1 | - | - | 0 | - |
| - | - | 527.7 | 220.9 | - | - | 0 | - |
| - | - | 586.7 | 223.1 | - | - | 0 | - |
| - | - | 670.6 | 226.1 | - | - | 0 | - |
| - | - | 689.2 | 245.1 | - | - | 0 | - |
| - | - | 955.9 | 251.2 | - | - | 0 | - |
| 5 | c | 2439 | 258.1 | 0.0002403 | 0.9308 | +2 | 5 |
| - | - | 5121 | 261.2 | - | - | 0 | - |
| - | - | 1609 | 268.2 | - | - | 0 | - |
| - | - | 546.3 | 284.9 | - | - | 0 | - |
| - | - | 781.6 | 288.2 | - | - | 0 | - |
| - | - | 586.4 | 293.1 | - | - | 0 | - |
| - | - | 1242 | 301.1 | - | - | 0 | - |
| - | - | 598.9 | 303.1 | - | - | 0 | - |
| 14 | z | 969.8 | 306.2 | 0.0002765 | 0.9032 | +1 | 2 |
| - | - | 1408 | 312.2 | - | - | 0 | - |
| - | - | 738.3 | 315.2 | - | - | 0 | - |
| - | - | 2493 | 317.2 | - | - | 0 | - |
| 14 | y | 980.8 | 322.2 | 0.0004764 | 1.479 | +1 | 2 |
| - | - | 690.4 | 330.2 | - | - | 0 | - |
| - | - | 2179 | 331.2 | - | - | 0 | - |
| - | - | 548.5 | 341.1 | - | - | 0 | - |
| 11 | c | 3103 | 376.2 | 0.001738 | 4.62 | +3 | 11 |
| - | - | 637.7 | 377.2 | - | - | 0 | - |
| - | - | 601.2 | 378.2 | - | - | 0 | - |
| - | - | 705.7 | 389.2 | - | - | 0 | - |
| - | - | 1228 | 391.2 | - | - | 0 | - |
| 13 | y | 4164 | 392.2 | 0.0001574 | 0.4013 | +1 | 3 |
| 5 | y | 2196 | 393.2 | 0.007643 | 19.44 | +3 | 11 |
| 13 | y | 1112 | 409.2 | 0.002201 | 5.378 | +1 | 3 |
| - | - | 1535 | 417.2 | - | - | 0 | - |
| - | - | 639.6 | 423.3 | - | - | 0 | - |
| - | - | 1283 | 429.2 | - | - | 0 | - |
| - | - | 1037 | 434.7 | - | - | 0 | - |
| - | - | 1104 | 439.7 | - | - | 0 | - |
| - | - | 915.4 | 442.3 | - | - | 0 | - |
| - | - | 946.6 | 444.3 | - | - | 0 | - |
| - | - | 708 | 445.3 | - | - | 0 | - |
| - | - | 1263 | 448.7 | - | - | 0 | - |
| - | - | 777 | 449.2 | - | - | 0 | - |
| - | - | 3776 | 451.3 | - | - | 0 | - |
| 12 | w | 3438 | 463.2 | 0.0002044 | 0.4412 | +1 | 4 |
| - | - | 1490 | 464.2 | - | - | 0 | - |
| - | - | 1.049E+04 | 468.3 | - | - | 0 | - |
| - | - | 3248 | 469.3 | - | - | 0 | - |
| - | - | 808.1 | 478.2 | - | - | 0 | - |
| 12 | y | 2092 | 479.2 | 0.0008137 | 1.698 | +1 | 4 |
| 12 | z | 3024 | 480.2 | 0.001192 | 2.483 | +1 | 4 |
| - | - | 1209 | 481.2 | - | - | 0 | - |
| 7 | z | 1957 | 488.8 | 0.000676 | 1.383 | +2 | 9 |
| - | - | 961.3 | 489.3 | - | - | 0 | - |
| - | - | 636.8 | 492.3 | - | - | 0 | - |
| - | - | 835.3 | 494.2 | - | - | 0 | - |
| - | - | 1330 | 494.3 | - | - | 0 | - |
| 12 | y | 3168 | 496.3 | 0.008015 | 16.15 | +1 | 4 |
| - | - | 1172 | 504.2 | - | - | 0 | - |
| - | - | 850.6 | 507.1 | - | - | 0 | - |
| - | - | 1181 | 507.3 | - | - | 0 | - |
| - | - | 3755 | 511.3 | - | - | 0 | - |
| - | - | 1020 | 512.3 | - | - | 0 | - |
| - | - | 641.2 | 513.1 | - | - | 0 | - |
| - | - | 2.036E+04 | 513.4 | - | - | 0 | - |
| - | - | 6257 | 514.4 | - | - | 0 | - |
| - | - | 2233 | 523.3 | - | - | 0 | - |
| - | - | 827.1 | 524.3 | - | - | 0 | - |
| - | - | 1007 | 524.8 | - | - | 0 | - |
| - | - | 760 | 532.3 | - | - | 0 | - |
| 5 | c | 4537 | 533.3 | 0.0003508 | 0.6579 | +1 | 5 |
| - | - | 1704 | 534.3 | - | - | 0 | - |
| - | - | 748.6 | 535.9 | - | - | 0 | - |
| - | - | 1125 | 538.3 | - | - | 0 | - |
| - | - | 887.5 | 539.8 | - | - | 0 | - |
| - | - | 5543 | 540.3 | - | - | 0 | - |
| - | - | 1465 | 540.8 | - | - | 0 | - |
| - | - | 2.047E+05 | 541.3 | - | - | 0 | - |
| - | - | 1752 | 541.6 | - | - | 0 | - |
| - | - | 1223 | 541.9 | - | - | 0 | - |
| - | - | 6.507E+04 | 542.3 | - | - | 0 | - |
| - | - | 1170 | 542.6 | - | - | 0 | - |
| - | - | 977.4 | 542.6 | - | - | 0 | - |
| 6 | z | 4058 | 546.3 | 0.0001958 | 0.3584 | +2 | 10 |
| - | - | 3267 | 546.8 | - | - | 0 | - |
| - | - | 1008 | 547.3 | - | - | 0 | - |
| - | - | 788.2 | 547.8 | - | - | 0 | - |
| - | - | 1236 | 550.3 | - | - | 0 | - |
| - | - | 931.8 | 551.3 | - | - | 0 | - |
| 6 | y | 2878 | 554.3 | 0.0001721 | 0.3105 | +2 | 10 |
| - | - | 780.7 | 554.8 | - | - | 0 | - |
| 11 | c | 2739 | 564.3 | 0.009272 | 16.43 | +2 | 11 |
| - | - | 824.2 | 565.3 | - | - | 0 | - |
| 11 | z | 1745 | 579.3 | 0.001805 | 3.115 | +1 | 5 |
| 5 | y | 725.3 | 588.8 | 0.01117 | 18.97 | +2 | 11 |
| 5 | y | 926.7 | 589.3 | 0.001957 | 3.321 | +2 | 11 |
| 5 | z | 3908 | 589.8 | 0.0002949 | 0.4999 | +2 | 11 |
| - | - | 4620 | 590.3 | - | - | 0 | - |
| - | - | 3734 | 590.8 | - | - | 0 | - |
| - | - | 749 | 591.3 | - | - | 0 | - |
| - | - | 2240 | 592.3 | - | - | 0 | - |
| - | - | 915.1 | 593.3 | - | - | 0 | - |
| - | - | 997.3 | 593.3 | - | - | 0 | - |
| - | - | 1196 | 595.2 | - | - | 0 | - |
| 11 | y | 1859 | 595.3 | 0.0002567 | 0.4313 | +1 | 5 |
| - | - | 607.4 | 596.3 | - | - | 0 | - |
| - | - | 1.338E+04 | 596.4 | - | - | 0 | - |
| - | - | 845.5 | 597.3 | - | - | 0 | - |
| - | - | 4361 | 597.4 | - | - | 0 | - |
| 5 | y | 8628 | 597.8 | 0.0003391 | 0.5673 | +2 | 11 |
| - | - | 5116 | 598.3 | - | - | 0 | - |
| - | - | 1025 | 598.4 | - | - | 0 | - |
| - | - | 2187 | 598.8 | - | - | 0 | - |
| - | - | 672.3 | 602.3 | - | - | 0 | - |
| - | - | 1151 | 610.3 | - | - | 0 | - |
| - | - | 4694 | 611.3 | - | - | 0 | - |
| - | - | 1331 | 612.3 | - | - | 0 | - |
| - | - | 1270 | 627.3 | - | - | 0 | - |
| 10 | z | 6235 | 636.3 | 5.536E-05 | 0.087 | +1 | 6 |
| - | - | 3732 | 637.3 | - | - | 0 | - |
| 4 | z | 6400 | 639.3 | 0.0003897 | 0.6095 | +2 | 12 |
| - | - | 5851 | 639.8 | - | - | 0 | - |
| - | - | 2159 | 640.3 | - | - | 0 | - |
| - | - | 613.2 | 640.8 | - | - | 0 | - |
| - | - | 678.4 | 642.3 | - | - | 0 | - |
| - | - | 749.4 | 646.8 | - | - | 0 | - |
| 4 | y | 7988 | 647.3 | 0.0001833 | 0.2832 | +2 | 12 |
| - | - | 5958 | 647.8 | - | - | 0 | - |
| 6 | c | 6457 | 648.3 | 0.003346 | 5.161 | +1 | 6 |
| - | - | 1976 | 649.3 | - | - | 0 | - |
| 10 | y | 6301 | 652.3 | 0.0001912 | 0.293 | +1 | 6 |
| - | - | 1682 | 653.3 | - | - | 0 | - |
| - | - | 1469 | 655.4 | - | - | 0 | - |
| - | - | 2142 | 655.8 | - | - | 0 | - |
| - | - | 2166 | 656.3 | - | - | 0 | - |
| - | - | 3359 | 663.3 | - | - | 0 | - |
| - | - | 9623 | 663.8 | - | - | 0 | - |
| - | - | 7543 | 664.3 | - | - | 0 | - |
| - | - | 3113 | 664.8 | - | - | 0 | - |
| - | - | 1250 | 665.3 | - | - | 0 | - |
| - | - | 1037 | 665.3 | - | - | 0 | - |
| 3 | w | 3801 | 674.3 | 0.00101 | 1.498 | +2 | 13 |
| - | - | 2678 | 674.8 | - | - | 0 | - |
| - | - | 1402 | 675.3 | - | - | 0 | - |
| - | - | 1027 | 679.4 | - | - | 0 | - |
| - | - | 783.7 | 680.4 | - | - | 0 | - |
| - | - | 1529 | 680.8 | - | - | 0 | - |
| - | - | 829.5 | 681.3 | - | - | 0 | - |
| - | - | 1080 | 692.8 | - | - | 0 | - |
| - | - | 806.7 | 693.1 | - | - | 0 | - |
| - | - | 1013 | 693.4 | - | - | 0 | - |
| - | - | 4205 | 694.4 | - | - | 0 | - |
| - | - | 1547 | 695.3 | - | - | 0 | - |
| - | - | 1153 | 696.4 | - | - | 0 | - |
| - | - | 999.9 | 697.4 | - | - | 0 | - |
| - | - | 664.2 | 701.3 | - | - | 0 | - |
| - | - | 1095 | 701.8 | - | - | 0 | - |
| - | - | 1758 | 702.3 | - | - | 0 | - |
| 3 | y | 1919 | 702.8 | 0.01388 | 19.75 | +2 | 13 |
| - | - | 1507 | 703.4 | - | - | 0 | - |
| - | - | 1632 | 703.7 | - | - | 0 | - |
| 3 | z | 2632 | 703.8 | 3.286E-05 | 0.04669 | +2 | 13 |
| - | - | 866.2 | 704.1 | - | - | 0 | - |
| - | - | 2164 | 704.3 | - | - | 0 | - |
| - | - | 892.7 | 704.4 | - | - | 0 | - |
| - | - | 1550 | 704.8 | - | - | 0 | - |
| - | - | 827.5 | 709.3 | - | - | 0 | - |
| - | - | 1935 | 709.8 | - | - | 0 | - |
| - | - | 3536 | 710.3 | - | - | 0 | - |
| - | - | 2531 | 710.8 | - | - | 0 | - |
| - | - | 9618 | 711.4 | - | - | 0 | - |
| 3 | y | 1.095E+04 | 711.9 | 0.0007279 | 1.023 | +2 | 13 |
| - | - | 9299 | 712.4 | - | - | 0 | - |
| - | - | 2788 | 712.9 | - | - | 0 | - |
| - | - | 1035 | 713.4 | - | - | 0 | - |
| - | - | 1833 | 715.9 | - | - | 0 | - |
| - | - | 709 | 716.9 | - | - | 0 | - |
| - | - | 1273 | 717.9 | - | - | 0 | - |
| - | - | 1005 | 718.4 | - | - | 0 | - |
| - | - | 2836 | 719.9 | - | - | 0 | - |
| - | - | 4062 | 720.4 | - | - | 0 | - |
| - | - | 3561 | 720.9 | - | - | 0 | - |
| 9 | y | 1887 | 721.4 | 0.005034 | 6.978 | +1 | 7 |
| - | - | 1415 | 721.9 | - | - | 0 | - |
| 9 | y | 1089 | 722.3 | 0.001121 | 1.551 | +1 | 7 |
| - | - | 680 | 722.7 | - | - | 0 | - |
| - | - | 4683 | 723.1 | - | - | 0 | - |
| 9 | z | 4552 | 723.4 | 0.00231 | 3.193 | +1 | 7 |
| - | - | 2942 | 723.4 | - | - | 0 | - |
| - | - | 1506 | 723.8 | - | - | 0 | - |
| - | - | 842.7 | 724.1 | - | - | 0 | - |
| - | - | 3445 | 724.4 | - | - | 0 | - |
| - | - | 1610 | 725.4 | - | - | 0 | - |
| - | - | 697.5 | 726.4 | - | - | 0 | - |
| - | - | 1773 | 727.9 | - | - | 0 | - |
| - | - | 9659 | 728.4 | - | - | 0 | - |
| 14 | c | 8530 | 728.9 | 0.01135 | 15.57 | +2 | 14 |
| 14 | c | 5117 | 729.4 | 0.002683 | 3.679 | +2 | 14 |
| - | - | 2818 | 729.9 | - | - | 0 | - |
| - | - | 1450 | 730.4 | - | - | 0 | - |
| - | - | 1584 | 737.4 | - | - | 0 | - |
| 14 | c | 2.386E+04 | 737.9 | 0.0003938 | 0.5337 | +2 | 14 |
| - | - | 1.902E+04 | 738.4 | - | - | 0 | - |
| - | - | 8231 | 738.9 | - | - | 0 | - |
| 9 | y | 3766 | 739.4 | 0.005455 | 7.378 | +1 | 7 |
| - | - | 761.5 | 744.4 | - | - | 0 | - |
| 2 | y | 1798 | 746.4 | 0.00145 | 1.943 | +2 | 14 |
| 2 | y | 783.1 | 746.9 | 0.006298 | 8.433 | +2 | 14 |
| 2 | z | 5950 | 747.4 | 0.001093 | 1.463 | +2 | 14 |
| - | - | 2904 | 747.9 | - | - | 0 | - |
| - | - | 1534 | 748.4 | - | - | 0 | - |
| - | - | 1132 | 749.4 | - | - | 0 | - |
| - | - | 819 | 751.4 | - | - | 0 | - |
| - | - | 973.9 | 752.4 | - | - | 0 | - |
| 2 | y | 1.374E+04 | 755.4 | 0.0001036 | 0.1371 | +2 | 14 |
| - | - | 1.132E+04 | 755.9 | - | - | 0 | - |
| - | - | 5360 | 756.4 | - | - | 0 | - |
| - | - | 2157 | 756.9 | - | - | 0 | - |
| - | - | 1166 | 757.4 | - | - | 0 | - |
| - | - | 738.3 | 760.4 | - | - | 0 | - |
| - | - | 3218 | 777.4 | - | - | 0 | - |
| - | - | 6272 | 778.4 | - | - | 0 | - |
| - | - | 2795 | 779.4 | - | - | 0 | - |
| - | - | 1697 | 779.9 | - | - | 0 | - |
| - | - | 1751 | 780.4 | - | - | 0 | - |
| - | - | 3126 | 780.9 | - | - | 0 | - |
| - | - | 3376 | 781.4 | - | - | 0 | - |
| - | - | 2617 | 781.9 | - | - | 0 | - |
| - | - | 4720 | 782.4 | - | - | 0 | - |
| - | - | 3646 | 782.9 | - | - | 0 | - |
| - | - | 917.7 | 783.4 | - | - | 0 | - |
| - | - | 826.3 | 783.9 | - | - | 0 | - |
| - | - | 2191 | 787.9 | - | - | 0 | - |
| - | - | 2628 | 788.4 | - | - | 0 | - |
| - | - | 1494 | 788.9 | - | - | 0 | - |
| - | - | 9819 | 789.4 | - | - | 0 | - |
| - | - | 7548 | 789.9 | - | - | 0 | - |
| - | - | 4057 | 790.4 | - | - | 0 | - |
| - | - | 8843 | 790.9 | - | - | 0 | - |
| - | - | 5788 | 791.4 | - | - | 0 | - |
| - | - | 3470 | 791.9 | - | - | 0 | - |
| 8 | w | 976.9 | 793.4 | 0.0002877 | 0.3626 | +1 | 8 |
| - | - | 1192 | 793.9 | - | - | 0 | - |
| - | - | 1109 | 794.4 | - | - | 0 | - |
| - | - | 1845 | 794.9 | - | - | 0 | - |
| - | - | 1141 | 795.4 | - | - | 0 | - |
| - | - | 1441 | 796.4 | - | - | 0 | - |
| - | - | 1113 | 796.9 | - | - | 0 | - |
| - | - | 1974 | 801.4 | - | - | 0 | - |
| - | - | 5447 | 801.9 | - | - | 0 | - |
| 8 | z | 9150 | 802.4 | 0.01198 | 14.93 | +1 | 8 |
| - | - | 7062 | 802.9 | - | - | 0 | - |
| 8 | z | 4139 | 803.4 | 0.01542 | 19.19 | +1 | 8 |
| - | - | 2.655E+04 | 803.9 | - | - | 0 | - |
| - | - | 2.12E+04 | 804.4 | - | - | 0 | - |
| - | - | 1.215E+04 | 804.9 | - | - | 0 | - |
| - | - | 2200 | 805.4 | - | - | 0 | - |
| - | - | 5165 | 809.9 | - | - | 0 | - |
| - | - | 2.709E+04 | 810.4 | - | - | 0 | - |
| - | - | 3.961E+04 | 810.9 | - | - | 0 | - |
| - | - | 2.454E+04 | 811.4 | - | - | 0 | - |
| - | - | 2.307E+04 | 811.9 | - | - | 0 | - |
| - | - | 2.312E+04 | 812.4 | - | - | 0 | - |
| - | - | 1.416E+04 | 812.9 | - | - | 0 | - |
| - | - | 5301 | 813.4 | - | - | 0 | - |
| - | - | 1512 | 813.9 | - | - | 0 | - |
| 8 | y | 873.4 | 836.4 | 0.00611 | 7.305 | +1 | 8 |
| - | - | 1328 | 857.5 | - | - | 0 | - |
| - | - | 985.8 | 858.5 | - | - | 0 | - |
| - | - | 1202 | 880.4 | - | - | 0 | - |
| - | - | 2929 | 896.4 | - | - | 0 | - |
| - | - | 2865 | 897.4 | - | - | 0 | - |
| - | - | 1302 | 898.5 | - | - | 0 | - |
| 8 | c | 7807 | 901.5 | 0.0004108 | 0.4558 | +1 | 8 |
| - | - | 3172 | 902.5 | - | - | 0 | - |
| - | - | 1797 | 903.5 | - | - | 0 | - |
| - | - | 959.3 | 906.4 | - | - | 0 | - |
| - | - | 2508 | 907.4 | - | - | 0 | - |
| - | - | 2230 | 908.4 | - | - | 0 | - |
| - | - | 2072 | 909.4 | - | - | 0 | - |
| - | - | 1379 | 910.4 | - | - | 0 | - |
| - | - | 4422 | 913.5 | - | - | 0 | - |
| - | - | 2791 | 914.5 | - | - | 0 | - |
| - | - | 783.5 | 915.5 | - | - | 0 | - |
| - | - | 956.5 | 925.4 | - | - | 0 | - |
| - | - | 3747 | 944.5 | - | - | 0 | - |
| - | - | 2637 | 945.5 | - | - | 0 | - |
| - | - | 1407 | 946.5 | - | - | 0 | - |
| 7 | z | 3683 | 976.5 | 0.001644 | 1.683 | +1 | 9 |
| - | - | 2262 | 977.5 | - | - | 0 | - |
| - | - | 1101 | 978.5 | - | - | 0 | - |
| - | - | 1497 | 987.5 | - | - | 0 | - |
| 9 | c | 1.696E+04 | 988.5 | 0.0007921 | 0.8013 | +1 | 9 |
| - | - | 7252 | 989.5 | - | - | 0 | - |
| - | - | 2577 | 990.5 | - | - | 0 | - |
| - | - | 1211 | 991.5 | - | - | 0 | - |
| - | - | 1013 | 1002 | - | - | 0 | - |
| - | - | 2462 | 1011 | - | - | 0 | - |
| - | - | 1169 | 1012 | - | - | 0 | - |
| - | - | 733.9 | 1023 | - | - | 0 | - |
| - | - | 4132 | 1024 | - | - | 0 | - |
| - | - | 8271 | 1025 | - | - | 0 | - |
| - | - | 3080 | 1026 | - | - | 0 | - |
| - | - | 5488 | 1027 | - | - | 0 | - |
| - | - | 1325 | 1027 | - | - | 0 | - |
| - | - | 3248 | 1028 | - | - | 0 | - |
| - | - | 1491 | 1029 | - | - | 0 | - |
| - | - | 1131 | 1039 | - | - | 0 | - |
| - | - | 871.9 | 1040 | - | - | 0 | - |
| - | - | 1551 | 1041 | - | - | 0 | - |
| - | - | 1040 | 1042 | - | - | 0 | - |
| 10 | c | 1.167E+04 | 1046 | 0.001667 | 1.595 | +1 | 10 |
| 6 | w | 6064 | 1047 | 0.007251 | 6.929 | +1 | 10 |
| - | - | 2820 | 1048 | - | - | 0 | - |
| - | - | 711.8 | 1050 | - | - | 0 | - |
| - | - | 2503 | 1053 | - | - | 0 | - |
| - | - | 1347 | 1055 | - | - | 0 | - |
| - | - | 1374 | 1056 | - | - | 0 | - |
| - | - | 1201 | 1064 | - | - | 0 | - |
| - | - | 1625 | 1071 | - | - | 0 | - |
| - | - | 764.6 | 1072 | - | - | 0 | - |
| - | - | 1316 | 1076 | - | - | 0 | - |
| - | - | 1309 | 1076 | - | - | 0 | - |
| - | - | 1058 | 1076 | - | - | 0 | - |
| - | - | 1193 | 1081 | - | - | 0 | - |
| - | - | 1235 | 1081 | - | - | 0 | - |
| - | - | 919.9 | 1082 | - | - | 0 | - |
| - | - | 1589 | 1084 | - | - | 0 | - |
| - | - | 2876 | 1084 | - | - | 0 | - |
| - | - | 1.444E+04 | 1085 | - | - | 0 | - |
| - | - | 1.356E+04 | 1085 | - | - | 0 | - |
| - | - | 9176 | 1086 | - | - | 0 | - |
| - | - | 4322 | 1086 | - | - | 0 | - |
| 6 | z | 2925 | 1092 | 0.002053 | 1.881 | +1 | 10 |
| - | - | 6038 | 1093 | - | - | 0 | - |
| - | - | 3249 | 1094 | - | - | 0 | - |
| - | - | 910.4 | 1095 | - | - | 0 | - |
| - | - | 1051 | 1095 | - | - | 0 | - |
| 6 | y | 738.1 | 1108 | 0.002511 | 2.267 | +1 | 10 |
| - | - | 727.4 | 1125 | - | - | 0 | - |
| - | - | 2420 | 1142 | - | - | 0 | - |
| - | - | 1100 | 1143 | - | - | 0 | - |
| - | - | 1145 | 1144 | - | - | 0 | - |
| 11 | c | 7263 | 1145 | 0.0005846 | 0.5107 | +1 | 11 |
| - | - | 5873 | 1146 | - | - | 0 | - |
| - | - | 1447 | 1147 | - | - | 0 | - |
| - | - | 1419 | 1151 | - | - | 0 | - |
| - | - | 3175 | 1153 | - | - | 0 | - |
| - | - | 1.564E+04 | 1153 | - | - | 0 | - |
| - | - | 8242 | 1154 | - | - | 0 | - |
| - | - | 3031 | 1156 | - | - | 0 | - |
| - | - | 1174 | 1156 | - | - | 0 | - |
| - | - | 1710 | 1170 | - | - | 0 | - |
| - | - | 717.9 | 1171 | - | - | 0 | - |
| 5 | z | 1072 | 1179 | 0.001885 | 1.599 | +1 | 11 |
| - | - | 3311 | 1180 | - | - | 0 | - |
| - | - | 2708 | 1181 | - | - | 0 | - |
| - | - | 1708 | 1182 | - | - | 0 | - |
| - | - | 3384 | 1183 | - | - | 0 | - |
| - | - | 2415 | 1184 | - | - | 0 | - |
| - | - | 1226 | 1185 | - | - | 0 | - |
| 5 | y | 1073 | 1195 | 0.004401 | 3.684 | +1 | 11 |
| - | - | 812.6 | 1196 | - | - | 0 | - |
| - | - | 3366 | 1198 | - | - | 0 | - |
| - | - | 8438 | 1199 | - | - | 0 | - |
| - | - | 6366 | 1200 | - | - | 0 | - |
| - | - | 2920 | 1201 | - | - | 0 | - |
| - | - | 892.4 | 1202 | - | - | 0 | - |
| 12 | c | 1.396E+04 | 1232 | 0.0007526 | 0.6111 | +1 | 12 |
| - | - | 8334 | 1233 | - | - | 0 | - |
| - | - | 1055 | 1233 | - | - | 0 | - |
| - | - | 3514 | 1234 | - | - | 0 | - |
| - | - | 672.1 | 1259 | - | - | 0 | - |
| - | - | 4002 | 1279 | - | - | 0 | - |
| - | - | 2594 | 1280 | - | - | 0 | - |
| - | - | 1048 | 1281 | - | - | 0 | - |
| - | - | 753.5 | 1284 | - | - | 0 | - |
| - | - | 659.1 | 1289 | - | - | 0 | - |
| - | - | 1000 | 1290 | - | - | 0 | - |
| - | - | 3897 | 1291 | - | - | 0 | - |
| - | - | 3511 | 1292 | - | - | 0 | - |
| - | - | 1994 | 1293 | - | - | 0 | - |
| 4 | y | 773.8 | 1294 | 0.00752 | 5.813 | +1 | 12 |
| - | - | 1364 | 1295 | - | - | 0 | - |
| - | - | 1024 | 1310 | - | - | 0 | - |
| - | - | 1393 | 1311 | - | - | 0 | - |
| 13 | c | 1.008E+04 | 1319 | 6.618E-05 | 0.05019 | +1 | 13 |
| - | - | 7504 | 1320 | - | - | 0 | - |
| - | - | 3880 | 1321 | - | - | 0 | - |
| - | - | 1430 | 1326 | - | - | 0 | - |
| - | - | 4832 | 1327 | - | - | 0 | - |
| - | - | 4421 | 1328 | - | - | 0 | - |
| - | - | 1355 | 1329 | - | - | 0 | - |
| - | - | 1639 | 1346 | - | - | 0 | - |
| - | - | 921.7 | 1347 | - | - | 0 | - |
| - | - | 2602 | 1404 | - | - | 0 | - |
| - | - | 2893 | 1405 | - | - | 0 | - |
| 3 | y | 1206 | 1406 | 0.01756 | 12.49 | +1 | 13 |
| 3 | z | 1520 | 1407 | 0.01305 | 9.279 | +1 | 13 |
| - | - | 7213 | 1408 | - | - | 0 | - |
| - | - | 4276 | 1409 | - | - | 0 | - |
| - | - | 1769 | 1410 | - | - | 0 | - |
| - | - | 735.4 | 1412 | - | - | 0 | - |
| - | - | 1150 | 1414 | - | - | 0 | - |
| - | - | 825.9 | 1417 | - | - | 0 | - |
| - | - | 788.4 | 1425 | - | - | 0 | - |
| - | - | 1536 | 1432 | - | - | 0 | - |
| - | - | 1552 | 1433 | - | - | 0 | - |
| - | - | 808.3 | 1439 | - | - | 0 | - |
| - | - | 1574 | 1440 | - | - | 0 | - |
| - | - | 1667 | 1441 | - | - | 0 | - |
| - | - | 1605 | 1442 | - | - | 0 | - |
| - | - | 807.6 | 1455 | - | - | 0 | - |
| - | - | 2102 | 1456 | - | - | 0 | - |
| 14 | c | 2218 | 1457 | 0.01261 | 8.656 | +1 | 14 |
| 14 | c | 1593 | 1458 | 0.004351 | 2.985 | +1 | 14 |
| - | - | 2106 | 1459 | - | - | 0 | - |
| - | - | 1578 | 1460 | - | - | 0 | - |
| 14 | c | 3117 | 1475 | 0.0006294 | 0.4268 | +1 | 14 |
| - | - | 8674 | 1476 | - | - | 0 | - |
| - | - | 3721 | 1477 | - | - | 0 | - |
| - | - | 1943 | 1478 | - | - | 0 | - |
| - | - | 4980 | 1495 | - | - | 0 | - |
| - | - | 3835 | 1496 | - | - | 0 | - |
| - | - | 1316 | 1497 | - | - | 0 | - |
| - | - | 1180 | 1504 | - | - | 0 | - |
| - | - | 903.8 | 1505 | - | - | 0 | - |
| - | - | 1128 | 1523 | - | - | 0 | - |
| - | - | 1109 | 1524 | - | - | 0 | - |
| - | - | 1006 | 1525 | - | - | 0 | - |
| - | - | 1693 | 1561 | - | - | 0 | - |
| - | - | 2641 | 1562 | - | - | 0 | - |
| - | - | 2242 | 1563 | - | - | 0 | - |
| - | - | 2116 | 1564 | - | - | 0 | - |
| - | - | 5126 | 1565 | - | - | 0 | - |
| - | - | 8065 | 1566 | - | - | 0 | - |
| - | - | 5735 | 1567 | - | - | 0 | - |
| - | - | 2180 | 1568 | - | - | 0 | - |
| - | - | 707.8 | 1575 | - | - | 0 | - |
| - | - | 1697 | 1576 | - | - | 0 | - |
| - | - | 2811 | 1577 | - | - | 0 | - |
| - | - | 1810 | 1578 | - | - | 0 | - |
| - | - | 1305 | 1579 | - | - | 0 | - |
| - | - | 2142 | 1580 | - | - | 0 | - |
| - | - | 2076 | 1581 | - | - | 0 | - |
| - | - | 2157 | 1582 | - | - | 0 | - |
| - | - | 2223 | 1583 | - | - | 0 | - |
| - | - | 882.5 | 1584 | - | - | 0 | - |
| - | - | 760 | 1587 | - | - | 0 | - |
| - | - | 827 | 1590 | - | - | 0 | - |
| - | - | 1357 | 1591 | - | - | 0 | - |
| - | - | 2093 | 1592 | - | - | 0 | - |
| - | - | 1991 | 1593 | - | - | 0 | - |
| - | - | 2138 | 1594 | - | - | 0 | - |
| - | - | 838.6 | 1595 | - | - | 0 | - |
| - | - | 1052 | 1596 | - | - | 0 | - |
| - | - | 1935 | 1597 | - | - | 0 | - |
| - | - | 1613 | 1598 | - | - | 0 | - |
| - | - | 929.1 | 1599 | - | - | 0 | - |
| - | - | 2932 | 1603 | - | - | 0 | - |
| - | - | 8875 | 1604 | - | - | 0 | - |
| - | - | 1.082E+04 | 1605 | - | - | 0 | - |
| - | - | 9355 | 1606 | - | - | 0 | - |
| - | - | 8507 | 1607 | - | - | 0 | - |
| - | - | 2.501E+04 | 1608 | - | - | 0 | - |
| - | - | 1.728E+04 | 1609 | - | - | 0 | - |
| - | - | 9462 | 1610 | - | - | 0 | - |
| - | - | 2449 | 1611 | - | - | 0 | - |
| - | - | 3265 | 1620 | - | - | 0 | - |
| - | - | 1.099E+04 | 1621 | - | - | 0 | - |
| - | - | 1.737E+04 | 1622 | - | - | 0 | - |
| - | - | 1.287E+04 | 1623 | - | - | 0 | - |
| - | - | 1.018E+04 | 1624 | - | - | 0 | - |
| - | - | 2.496E+04 | 1625 | - | - | 0 | - |
| - | - | 1.859E+04 | 1626 | - | - | 0 | - |
| - | - | 8556 | 1627 | - | - | 0 | - |
| - | - | 2698 | 1628 | - | - | 0 | - |
| - | - | 838.6 | 1629 | - | - | 0 | - |
| - | - | 667.4 | 1676 | - | - | 0 | - |
| - | - | 811.4 | 2078 | - | - | 0 | - |
| - | - | 1664 | 2168 | - | - | 0 | - |
| - | - | 3985 | 2169 | - | - | 0 | - |
| - | - | 3327 | 2170 | - | - | 0 | - |
| - | - | 1408 | 2171 | - | - | 0 | - |
| - | - | 831.7 | 3080 | - | - | 0 | - |

m/z Charge Intensity FragmentType MassShift Position
120.08088684082031 0 907.07294
129.06570434570312 0 411.9714
129.10238647460938 0 681.6077
133.106201171875 0 376.3443
134.30123901367188 0 377.8495
134.53672790527344 0 516.28436
138.2952423095703 0 427.30862
148.9541015625 0 828.9607
155.09266662597656 0 2897.6978
166.0860595703125 0 720.1718 y 14
168.7227325439453 0 423.5715
173.12835693359375 0 7501.63
173.25082397460938 0 430.40146
173.4516143798828 0 1714.7161
174.13156127929688 0 749.6558
175.82078552246094 0 551.11633
179.74412536621094 0 469.4662
182.08145141601562 0 1144.7744
186.12339782714844 0 468.48434
189.9710693359375 0 537.95355
198.1524200439453 0 447.7161
201.12319946289062 0 6763.299
201.1347198486328 0 2114.7417
202.12368774414062 0 450.16476
215.13864135742188 0 1942.7009
220.8910675048828 0 527.6782
223.10763549804688 0 586.7223
226.10850524902344 0 670.60266
245.13673400878906 0 689.20557
251.15013122558594 0 955.9176
258.14459228515625 0 2439.07 c Water loss 4
261.1708068847656 0 5120.839
268.1764221191406 0 1609.3293
284.88250732421875 0 546.26135
288.1668395996094 0 781.58545
293.1118469238281 0 586.44336
301.05865478515625 0 1241.8407
303.05560302734375 0 598.9018
306.1683654785156 0 969.77966 z 13
312.1552429199219 0 1408.056
315.16485595703125 0 738.2869
317.1582336425781 0 2492.7844
322.1868896484375 0 980.7904 y 13
330.1666564941406 0 690.38416
331.2341003417969 0 2179.4258
341.14520263671875 0 548.535
376.1981506347656 0 3103.1965 c Water loss 10
377.1963195800781 0 637.7462
378.21356201171875 0 601.21484
389.19085693359375 0 705.66547
391.1844787597656 0 1227.5309
392.19268798828125 0 4163.7827 y Ammonia loss 12
393.19903564453125 0 2195.7073 y Ammonia loss 4
409.2171936035156 0 1111.7286 y 12
417.1880187988281 0 1535.3025
423.2691345214844 0 639.59924
429.2347717285156 0 1282.8458
434.72088623046875 0 1036.8972
439.71380615234375 0 1103.9641
442.27716064453125 0 915.4189
444.2568359375 0 946.58704
445.2552795410156 0 707.9873
448.7168884277344 0 1262.7106
449.2182922363281 0 777.0329
451.2664489746094 0 3775.6023
463.23016357421875 0 3438.309 w 11
464.2274169921875 0 1490.1763
468.2931823730469 0 10487.443
469.2955627441406 0 3247.9468
478.2175598144531 0 808.11237
479.22406005859375 0 2092.3962 y Ammonia loss 11
480.23150634765625 0 3023.952 z 11
481.23297119140625 0 1208.7944
488.7572021484375 0 1957.0369 z 6
489.2593994140625 0 961.3129
492.2820129394531 0 636.8365
494.2361755371094 0 835.3129
494.3069763183594 0 1330.3257
496.243408203125 0 3167.7043 y 11
504.219970703125 0 1171.6931
507.1060791015625 0 850.61945
507.29132080078125 0 1180.5269
511.3352355957031 0 3755.3726
512.338134765625 0 1020.16534
513.1412353515625 0 641.17255
513.3509521484375 0 20355.668
514.3538818359375 0 6256.9233
523.3355712890625 0 2232.7131
524.3374633789062 0 827.11847
524.7780151367188 0 1006.8757
532.2831420898438 0 760.0278
533.2926025390625 0 4536.8164 c 4
534.2982177734375 0 1703.5204
535.9405517578125 0 748.5975
538.2520141601562 0 1124.5522
539.8108520507812 0 887.5105
540.310791015625 0 5542.6587
540.811279296875 0 1465.2075
541.3460083007812 0 204722
541.6110229492188 0 1752.2722
541.943359375 0 1222.5482
542.3486328125 0 65072.938
542.5609741210938 0 1169.9653
542.6080932617188 0 977.4486
546.2715454101562 0 4058.1016 z 5
546.7730712890625 0 3266.5203
547.275634765625 0 1007.7062
547.7788696289062 0 788.18335
550.2501220703125 0 1236.1448
551.2596435546875 0 931.79333
554.2808837890625 0 2878.0315 y 5
554.7822265625 0 780.74603
564.2789306640625 0 2738.7852 c Ammonia loss 10
565.27978515625 0 824.2309
579.3029174804688 0 1745.121 z 10
588.7802734375 0 725.34863 y Water loss 4
589.281494140625 0 926.6729 y Ammonia loss 4
589.7876586914062 0 3907.6675 z 4
590.2896728515625 0 4620.365
590.7922973632812 0 3733.9277
591.2950439453125 0 748.9745
592.3311157226562 0 2240.466
593.25634765625 0 915.07764
593.3383178710938 0 997.2832
595.2483520507812 0 1196.145
595.319580078125 0 1859.0211 y 10
596.2525024414062 0 607.42285
596.3514404296875 0 13378.375
597.2986450195312 0 845.4595
597.3553466796875 0 4360.548
597.79638671875 0 8627.886 y 4
598.29833984375 0 5115.795
598.3538818359375 0 1024.7146
598.8001708984375 0 2187.0593
602.2866821289062 0 672.2727
610.283447265625 0 1150.655
611.2679443359375 0 4694.4727
612.2716674804688 0 1331.2101
627.260986328125 0 1270.03
636.3226318359375 0 6235.4814 z 9
637.328125 0 3731.9492
639.3219604492188 0 6400.303 z 3
639.8240966796875 0 5850.6064
640.3250122070312 0 2158.847
640.828857421875 0 613.1635
642.337158203125 0 678.3604
646.8323974609375 0 749.40594
647.3307495117188 0 7988.352 y 3
647.8319091796875 0 5958.0156
648.3232421875 0 6457.32 c 5
649.3247680664062 0 1976.0375
652.3414916992188 0 6301.0444 y 9
653.3440551757812 0 1681.9158
655.351806640625 0 1469.0952
655.8447875976562 0 2142.393
656.3499755859375 0 2165.7063
663.3460083007812 0 3359.2864
663.8449096679688 0 9622.943
664.3467407226562 0 7543.4395
664.8462524414062 0 3113.1023
665.2841186523438 0 1249.596
665.3480224609375 0 1037.425
674.335205078125 0 3801.4087 w 2
674.8377685546875 0 2678.4023
675.3384399414062 0 1401.7258
679.3538818359375 0 1027.1844
680.363525390625 0 783.74097
680.7962036132812 0 1528.5546
681.296630859375 0 829.51654
692.759033203125 0 1080.2661
693.0901489257812 0 806.736
693.4253540039062 0 1012.85486
694.3515625 0 4205.4014
695.34912109375 0 1546.9633
696.3656005859375 0 1152.8086
697.36328125 0 999.8809
701.32958984375 0 664.1622
701.8245849609375 0 1094.8794
702.322021484375 0 1757.7905
702.8330688476562 0 1919.1643 y Water loss 2
703.4180908203125 0 1507.1675
703.7376098632812 0 1631.5244
703.8428344726562 0 2632.4138 z 2
704.0742797851562 0 866.22546
704.3438720703125 0 2163.7864
704.4074096679688 0 892.74713
704.8447265625 0 1549.6294
709.3388671875 0 827.49475
709.8360595703125 0 1935.248
710.333251953125 0 3535.977
710.833740234375 0 2531.2292
711.3778076171875 0 9618.386
711.8515014648438 0 10946.7705 y 2
712.3595581054688 0 9299.126
712.8551025390625 0 2787.9583
713.3867797851562 0 1034.6294
715.8746948242188 0 1833.1123
716.8778686523438 0 709.022
717.8544921875 0 1273.3171
718.3540649414062 0 1004.9794
719.8701171875 0 2836.442
720.3726196289062 0 4061.8352
720.871826171875 0 3561.0996
721.3677978515625 0 1887.2866 y Water loss 8
721.8748168945312 0 1415.2498
722.347900390625 0 1089.0985 y Ammonia loss 8
722.7473754882812 0 680.04297
723.0845336914062 0 4683.0986
723.352294921875 0 4552.2217 z 8
723.4212036132812 0 2942.2156
723.7535400390625 0 1506.0599
724.0869140625 0 842.6545
724.352783203125 0 3445.0593
725.3574829101562 0 1609.9559
726.3590698242188 0 697.5493
727.8726196289062 0 1773.411
728.3685302734375 0 9658.804
728.867431640625 0 8530.039 c Water loss 13
729.3681030273438 0 5117.384 c Ammonia loss 13
729.8607788085938 0 2817.7876
730.3619384765625 0 1449.7242
737.3623657226562 0 1583.8937
737.8836669921875 0 23863.459 c 13
738.3851928710938 0 19016.959
738.8851928710938 0 8231.317
739.3787841796875 0 3766.3953 y 8
744.3795166015625 0 761.521
746.3615112304688 0 1798.2883 y Water loss 1
746.8612670898438 0 783.09784 y Ammonia loss 1
747.3577880859375 0 5949.7827 z 1
747.861572265625 0 2904.2297
748.3554077148438 0 1534.1774
749.3638305664062 0 1131.8666
751.3601684570312 0 819.0057
752.3599243164062 0 973.9194
755.3683471679688 0 13740.731 y 1
755.8703002929688 0 11321.074
756.3707275390625 0 5359.919
756.8716430664062 0 2157.007
757.3712768554688 0 1166.2974
760.4081420898438 0 738.30054
777.379150390625 0 3217.6172
778.36572265625 0 6271.801
779.36767578125 0 2795.3408
779.88330078125 0 1696.5405
780.3768310546875 0 1751.0236
780.8829956054688 0 3126.009
781.3829345703125 0 3375.615
781.8983764648438 0 2616.5046
782.4033203125 0 4720.2715
782.9024047851562 0 3646.0054
783.399169921875 0 917.65173
783.904296875 0 826.2661
787.8912353515625 0 2191.1155
788.392822265625 0 2627.748
788.8943481445312 0 1493.624
789.3839111328125 0 9818.844
789.8866577148438 0 7547.753
790.3870849609375 0 4057.0547
790.8994140625 0 8843.039
791.4010620117188 0 5788.3965
791.9020385742188 0 3469.7405
793.3836059570312 0 976.853 w 7
793.8799438476562 0 1191.5189
794.381103515625 0 1109.1954
794.8887329101562 0 1844.937
795.3936767578125 0 1141.4188
796.38818359375 0 1440.5221
796.8901977539062 0 1112.8545
801.3882446289062 0 1974.3424
801.885498046875 0 5446.8213
802.3848266601562 0 9149.885 z Water loss 7
802.885498046875 0 7061.6445
803.396240234375 0 4138.703 z Ammonia loss 7
803.90185546875 0 26554.852
804.4031982421875 0 21195.879
804.904052734375 0 12151.978
805.4090576171875 0 2199.929
809.892578125 0 5165.464
810.3927001953125 0 27091.52
810.8939208984375 0 39612.492
811.395751953125 0 24544.928
811.9041137695312 0 23066.121
812.4103393554688 0 23121.953
812.913818359375 0 14157.961
813.4161987304688 0 5300.5127
813.9178466796875 0 1512.4728
836.4199829101562 0 873.37274 y 7
857.4586181640625 0 1328.436
858.4620361328125 0 985.7716
880.4334716796875 0 1202.4315
896.4259033203125 0 2928.7083
897.4494018554688 0 2864.6426
898.4617919921875 0 1301.7844
901.4741821289062 0 7807.4756 c 7
902.4766845703125 0 3172.226
903.4835205078125 0 1797.2106
906.4223022460938 0 959.32623
907.4117431640625 0 2508.2004
908.4171142578125 0 2229.798
909.4232788085938 0 2071.7117
910.4139404296875 0 1379.0052
913.4529418945312 0 4421.6655
914.4571533203125 0 2791.0483
915.458984375 0 783.51764
925.435791015625 0 956.4648
944.490478515625 0 3747.2988
945.4933471679688 0 2636.6628
946.4935913085938 0 1407.1431
976.5068359375 0 3683.4004 z 6
977.5133666992188 0 2262.3135
978.5164794921875 0 1100.5676
987.4999389648438 0 1497.2749
988.506591796875 0 16956.482 c 8
989.509521484375 0 7251.783
990.509521484375 0 2576.6125
991.5128173828125 0 1210.5702
1001.5211181640625 0 1013.07184
1010.5416259765625 0 2461.6467
1011.5441284179688 0 1168.5585
1023.4224243164062 0 733.8747
1024.4481201171875 0 4131.6797
1025.43994140625 0 8271.231
1026.4375 0 3080.2058
1026.5396728515625 0 5488.2573
1027.43798828125 0 1325.4434
1027.542724609375 0 3248.3394
1028.5418701171875 0 1490.7303
1039.134521484375 0 1131.0961
1040.4715576171875 0 871.8877
1041.4571533203125 0 1550.9537
1042.4659423828125 0 1040.4219
1045.5289306640625 0 11665.368 c 9
1046.530517578125 0 6064.269 w 5
1047.53955078125 0 2820.0183
1049.5576171875 0 711.78125
1052.6278076171875 0 2503.1433
1055.1217041015625 0 1347.1682
1055.6207275390625 0 1373.6921
1063.599365234375 0 1201.1135
1070.6326904296875 0 1624.5997
1071.6181640625 0 764.5625
1075.6304931640625 0 1315.8523
1075.994873046875 0 1308.8387
1076.4986572265625 0 1058.1387
1080.623779296875 0 1193.1317
1081.452392578125 0 1234.8536
1081.5826416015625 0 919.90045
1083.54248046875 0 1588.5128
1084.122314453125 0 2875.6997
1084.627197265625 0 14439.7705
1085.12841796875 0 13559.997
1085.629150390625 0 9176.322
1086.1287841796875 0 4321.8027
1091.5374755859375 0 2924.6382 z 5
1092.5433349609375 0 6037.602
1093.5478515625 0 3248.8977
1094.5457763671875 0 910.3988
1095.4559326171875 0 1050.6343
1107.5516357421875 0 738.0779 y 5
1124.5867919921875 0 727.3977
1141.570556640625 0 2420.001
1142.5743408203125 0 1099.8524
1143.5731201171875 0 1144.6953
1144.5950927734375 0 7263.187 c 10
1145.5994873046875 0 5872.653
1146.6070556640625 0 1447.1802
1151.4805908203125 0 1418.5249
1152.5013427734375 0 3175.3916
1153.4949951171875 0 15636.202
1154.4962158203125 0 8242.099
1155.5013427734375 0 3031.184
1156.4991455078125 0 1173.9026
1169.508544921875 0 1710.0588
1170.5107421875 0 717.8559
1178.5693359375 0 1071.7493 z 4
1179.57568359375 0 3311.1611
1180.5784912109375 0 2708.395
1181.59375 0 1707.6125
1182.5943603515625 0 3384.1216
1183.5975341796875 0 2415.469
1184.601318359375 0 1226.3135
1194.590576171875 0 1073.4971 y 4
1195.5843505859375 0 812.5765
1197.598876953125 0 3366.108
1198.5885009765625 0 8438.145
1199.5931396484375 0 6366.2812
1200.5975341796875 0 2919.5513
1201.5872802734375 0 892.3921
1231.626953125 0 13964.347 c 11
1232.6300048828125 0 8333.796
1232.7808837890625 0 1054.9783
1233.6309814453125 0 3513.75
1258.6199951171875 0 672.1394
1278.6419677734375 0 4001.6182
1279.6502685546875 0 2593.5916
1280.6484375 0 1048.194
1283.6634521484375 0 753.48444
1288.5615234375 0 659.1229
1289.5517578125 0 999.99365
1290.555908203125 0 3896.5051
1291.5604248046875 0 3510.5977
1292.5562744140625 0 1994.2183
1293.662109375 0 773.76807 y 3
1294.657958984375 0 1364.3601
1309.70458984375 0 1023.64886
1310.693359375 0 1392.5106
1318.65966796875 0 10075.414 c 12
1319.66162109375 0 7503.7925
1320.6658935546875 0 3880.1216
1325.6982421875 0 1429.7177
1326.6854248046875 0 4832.356
1327.6875 0 4420.555
1328.6904296875 0 1354.8334
1345.66162109375 0 1639.4028
1346.662353515625 0 921.7276
1403.6435546875 0 2601.5244
1404.6527099609375 0 2892.7722
1405.653076171875 0 1206.2313 y Ammonia loss 2
1406.6654052734375 0 1520.4891 z 2
1407.68408203125 0 7212.841
1408.68896484375 0 4275.7993
1409.6929931640625 0 1768.6459
1411.7071533203125 0 735.3673
1413.7235107421875 0 1150.4083
1416.72802734375 0 825.93384
1424.705322265625 0 788.43427
1431.750244140625 0 1536.1758
1432.75830078125 0 1551.7915
1438.724609375 0 808.32043
1439.7373046875 0 1573.5476
1440.7310791015625 0 1666.6615
1441.7257080078125 0 1604.9292
1454.7288818359375 0 807.5858
1455.741455078125 0 2102.2095
1456.7376708984375 0 2217.9763 c Water loss 13
1457.7386474609375 0 1593.0756 c Ammonia loss 13
1458.7349853515625 0 2105.888
1459.74267578125 0 1578.4136
1474.761474609375 0 3116.8223 c 13
1475.7655029296875 0 8673.638
1476.767578125 0 3721.2239
1477.7637939453125 0 1943.4595
1494.718505859375 0 4979.925
1495.719482421875 0 3834.5212
1496.713623046875 0 1316.2294
1503.7303466796875 0 1180.0692
1504.7244873046875 0 903.7773
1522.7073974609375 0 1128.0956
1523.7353515625 0 1108.7452
1524.74755859375 0 1005.5547
1560.7777099609375 0 1693.2148
1561.7730712890625 0 2641.0833
1562.7696533203125 0 2241.6892
1563.77880859375 0 2115.675
1564.7816162109375 0 5125.941
1565.7818603515625 0 8064.8174
1566.7847900390625 0 5734.7144
1567.7879638671875 0 2180.207
1574.778076171875 0 707.7768
1575.7747802734375 0 1696.7938
1576.77587890625 0 2811.1973
1577.7822265625 0 1810.3356
1578.81884765625 0 1304.637
1579.805419921875 0 2142.3113
1580.8026123046875 0 2075.6794
1581.8043212890625 0 2157.3606
1582.8001708984375 0 2222.8267
1583.8006591796875 0 882.5453
1586.755615234375 0 760.0458
1589.788818359375 0 827.02496
1590.781494140625 0 1356.6997
1591.779541015625 0 2093.3481
1592.7889404296875 0 1990.9618
1593.7994384765625 0 2137.7598
1594.7889404296875 0 838.61554
1595.79833984375 0 1052.3389
1596.8330078125 0 1934.6771
1597.828857421875 0 1613.3813
1598.82958984375 0 929.13464
1602.7750244140625 0 2932.4875
1603.7696533203125 0 8874.605
1604.765625 0 10821.755
1605.767333984375 0 9354.872
1606.78857421875 0 8506.813
1607.800537109375 0 25010.4
1608.8043212890625 0 17281.111
1609.8065185546875 0 9462.372
1610.8072509765625 0 2449.144
1619.7801513671875 0 3264.5334
1620.78759765625 0 10989.641
1621.7879638671875 0 17370.219
1622.795166015625 0 12874.492
1623.8074951171875 0 10177.128
1624.824462890625 0 24961.09
1625.8284912109375 0 18593.256
1626.8331298828125 0 8556.276
1627.8330078125 0 2698.4429
1628.8277587890625 0 838.5996
1676.3380126953125 0 667.4098
2078.272216796875 0 811.37787
2168.225341796875 0 1664.3008
2169.249755859375 0 3984.9854
2170.25048828125 0 3327.412
2171.264404296875 0 1408.4551
3079.770263671875 0 831.67633

Spectrum Details

|  |  |
| --- | --- |
| Matched peaks? Matched peaks The total absolute number of peaks matched. Additionally in brackets the total fraction of peaks matched and the total number of peaks is shown. | 65 (12.85% of 506) |
| FDR? FDR The false discovery rate estimated for this peptide. It is calculated by matching all theoretical fragments with a non-integer shift with the raw peaks for this spectrum. This is done with 40 different shifts. The resulting percentage is the average number of annotated peaks over the number of annotated peaks with the correct spectrum. | 1.87% |
| Satellite FDR? Satellite FDR See the FDR for details on its calculation. This satellite ion specific FDR only contains the satellite ions (d/w) for I/L/J positions. | - |
| PSM Score? PSM Score The PSM Score as given by Hecklib to this annotated spectrum. It is shown with three significant figures. | 480 |

#### Reverse Lookup? Reverse Lookup All places where this read could be placed.

| Group | Segment | Template | Template Part | Read Part | Score | Unique |
| --- | --- | --- | --- | --- | --- | --- |
| Homo sapiens Light Chain | IGLV | IGLV2-14 | [49..64] | [0..15] | 97 | False |
| Homo sapiens Light Chain | IGLV | IGLV2-18 | [49..64] | [0..15] | 88 | False |

| Recombined | Template Part | Read Part | Score | Unique |
| --- | --- | --- | --- | --- |
| REC-0-1\_002 | [49..64] | [0..15] | 120 | True |

#### Meta Information from Multiple reads

##### Number of combined reads

12

##### Intensity

0.9904

##### TotalArea

9.625E+09

##### Changes to the peptide sequence

JSEVSDRPSGVSSRF

L→JNo support for either Leucine or Isoleucine based on side chain ions (Position: 1)

#### Positional Score

Copy Data

##### Positional Score (TSV)

###### Preview

```
Loading example...
```

*Click on the button to copy the data to your clipboard.*

1001234567891011121314

Label Value
"0" 0.8
"1" 0.788
"2" 0.818
"3" 0.81
"4" 0.811
"5" 0.819
"6" 0.782
"7" 0.785
"8" 0.821
"9" 0.787
"10" 0.79
"11" 0.824
"12" 0.828
"13" 0.83
"14" 0.833

#### Meta Information from PEAKS

##### Scan Identifier

F3:4862

##### Original sequence

L

S

E

V

S

D

R

P

S

G

V

S

S

R

F

##### Posttranslational Modifications

##### Source File

D:\separate\_stitch\_analyses\xle-disambiguation\raw\20210323\_F1\_UM1\_Peng0013\_SA\_F59\_ingel\_3ug\_chymo.raw

##### Fraction

3

##### Scan Feature

-

##### De Novo Score

98

##### ConfidenceScore

98

### m/z

541.6099

##### Mass

1621.8059

##### Charge

3

##### Retention Time

26.81

##### Predicted Retention Time

-

##### Area

0

##### Parts Per Million

1.3

##### Fragmentation mode

ETHCD

##### Originating file

01 D:\separate\_stitch\_analyses\xle-disambiguation\20210325\_F59\_3ug\_DENOVO\_12.csv

#### Meta Information from PEAKS

##### Scan Identifier

F3:4731

##### Original sequence

L

S

E

V

S

D

R

P

S

G

V

S

S

R

F

##### Posttranslational Modifications

##### Source File

D:\separate\_stitch\_analyses\xle-disambiguation\raw\20210323\_F1\_UM1\_Peng0013\_SA\_F59\_ingel\_3ug\_chymo.raw

##### Fraction

3

##### Scan Feature

F3:5951

##### De Novo Score

98

##### ConfidenceScore

98

### m/z

541.6105

##### Mass

1621.8059

##### Charge

3

##### Retention Time

25.03

##### Predicted Retention Time

-

##### Area

2.406E+09

##### Parts Per Million

2.4

##### Fragmentation mode

ETHCD

##### Originating file

01 D:\separate\_stitch\_analyses\xle-disambiguation\20210325\_F59\_3ug\_DENOVO\_12.csv

#### Meta Information from PEAKS

##### Scan Identifier

F3:5111

##### Original sequence

L

S

E

V

S

D

R

P

S

G

V

S

S

R

F

##### Posttranslational Modifications

##### Source File

D:\separate\_stitch\_analyses\xle-disambiguation\raw\20210323\_F1\_UM1\_Peng0013\_SA\_F59\_ingel\_3ug\_chymo.raw

##### Fraction

3

##### Scan Feature

-

##### De Novo Score

97

##### ConfidenceScore

97

### m/z

541.6102

##### Mass

1621.8059

##### Charge

3

##### Retention Time

28.26

##### Predicted Retention Time

-

##### Area

0

##### Parts Per Million

1.7

##### Fragmentation mode

ETHCD

##### Originating file

01 D:\separate\_stitch\_analyses\xle-disambiguation\20210325\_F59\_3ug\_DENOVO\_12.csv

#### Meta Information from PEAKS

##### Scan Identifier

F3:4990

##### Original sequence

L

S

E

V

S

D

R

P

S

G

V

S

S

R

F

##### Posttranslational Modifications

##### Source File

D:\separate\_stitch\_analyses\xle-disambiguation\raw\20210323\_F1\_UM1\_Peng0013\_SA\_F59\_ingel\_3ug\_chymo.raw

##### Fraction

3

##### Scan Feature

-

##### De Novo Score

97

##### ConfidenceScore

97

### m/z

541.6101

##### Mass

1621.8059

##### Charge

3

##### Retention Time

27.54

##### Predicted Retention Time

-

##### Area

0

##### Parts Per Million

1.6

##### Fragmentation mode

ETHCD

##### Originating file

01 D:\separate\_stitch\_analyses\xle-disambiguation\20210325\_F59\_3ug\_DENOVO\_12.csv

#### Meta Information from PEAKS

##### Scan Identifier

F3:4533

##### Original sequence

L

S

E

V

S

D

R

P

S

G

V

S

S

R

F

##### Posttranslational Modifications

##### Source File

D:\separate\_stitch\_analyses\xle-disambiguation\raw\20210323\_F1\_UM1\_Peng0013\_SA\_F59\_ingel\_3ug\_chymo.raw

##### Fraction

3

##### Scan Feature

F3:5951

##### De Novo Score

97

##### ConfidenceScore

97

### m/z

541.6105

##### Mass

1621.8059

##### Charge

3

##### Retention Time

25.03

##### Predicted Retention Time

-

##### Area

2.406E+09

##### Parts Per Million

2.4

##### Fragmentation mode

ETHCD

##### Originating file

01 D:\separate\_stitch\_analyses\xle-disambiguation\20210325\_F59\_3ug\_DENOVO\_12.csv

#### Meta Information from PEAKS

##### Scan Identifier

F3:5048

##### Original sequence

L

S

E

V

S

D

R

P

S

G

V

S

S

R

F

##### Posttranslational Modifications

##### Source File

D:\separate\_stitch\_analyses\xle-disambiguation\raw\20210323\_F1\_UM1\_Peng0013\_SA\_F59\_ingel\_3ug\_chymo.raw

##### Fraction

3

##### Scan Feature

-

##### De Novo Score

97

##### ConfidenceScore

97

### m/z

541.6101

##### Mass

1621.8059

##### Charge

3

##### Retention Time

27.89

##### Predicted Retention Time

-

##### Area

0

##### Parts Per Million

1.6

##### Fragmentation mode

ETHCD

##### Originating file

01 D:\separate\_stitch\_analyses\xle-disambiguation\20210325\_F59\_3ug\_DENOVO\_12.csv

#### Meta Information from PEAKS

##### Scan Identifier

F3:4796

##### Original sequence

L

S

E

V

S

D

R

P

S

G

V

S

S

R

F

##### Posttranslational Modifications

##### Source File

D:\separate\_stitch\_analyses\xle-disambiguation\raw\20210323\_F1\_UM1\_Peng0013\_SA\_F59\_ingel\_3ug\_chymo.raw

##### Fraction

3

##### Scan Feature

-

##### De Novo Score

97

##### ConfidenceScore

97

### m/z

541.6102

##### Mass

1621.8059

##### Charge

3

##### Retention Time

26.44

##### Predicted Retention Time

-

##### Area

0

##### Parts Per Million

1.8

##### Fragmentation mode

ETHCD

##### Originating file

01 D:\separate\_stitch\_analyses\xle-disambiguation\20210325\_F59\_3ug\_DENOVO\_12.csv

#### Meta Information from PEAKS

##### Scan Identifier

F3:4665

##### Original sequence

L

S

E

V

S

D

R

P

S

G

V

S

S

R

F

##### Posttranslational Modifications

##### Source File

D:\separate\_stitch\_analyses\xle-disambiguation\raw\20210323\_F1\_UM1\_Peng0013\_SA\_F59\_ingel\_3ug\_chymo.raw

##### Fraction

3

##### Scan Feature

F3:5951

##### De Novo Score

97

##### ConfidenceScore

97

### m/z

541.6105

##### Mass

1621.8059

##### Charge

3

##### Retention Time

25.03

##### Predicted Retention Time

-

##### Area

2.406E+09

##### Parts Per Million

2.4

##### Fragmentation mode

HCD

##### Originating file

01 D:\separate\_stitch\_analyses\xle-disambiguation\20210325\_F59\_3ug\_DENOVO\_12.csv

#### Meta Information from PEAKS

##### Scan Identifier

F3:4596

##### Original sequence

L

S

E

V

S

D

R

P

S

G

V

S

S

R

F

##### Posttranslational Modifications

##### Source File

D:\separate\_stitch\_analyses\xle-disambiguation\raw\20210323\_F1\_UM1\_Peng0013\_SA\_F59\_ingel\_3ug\_chymo.raw

##### Fraction

3

##### Scan Feature

F3:5951

##### De Novo Score

96

##### ConfidenceScore

96

### m/z

541.6105

##### Mass

1621.8059

##### Charge

3

##### Retention Time

25.03

##### Predicted Retention Time

-

##### Area

2.406E+09

##### Parts Per Million

2.4

##### Fragmentation mode

HCD

##### Originating file

01 D:\separate\_stitch\_analyses\xle-disambiguation\20210325\_F59\_3ug\_DENOVO\_12.csv

#### Meta Information from PEAKS

##### Scan Identifier

F3:4925

##### Original sequence

L

S

E

V

S

D

R

P

S

G

V

S

S

R

F

##### Posttranslational Modifications

##### Source File

D:\separate\_stitch\_analyses\xle-disambiguation\raw\20210323\_F1\_UM1\_Peng0013\_SA\_F59\_ingel\_3ug\_chymo.raw

##### Fraction

3

##### Scan Feature

-

##### De Novo Score

96

##### ConfidenceScore

96

### m/z

541.6101

##### Mass

1621.8059

##### Charge

3

##### Retention Time

27.17

##### Predicted Retention Time

-

##### Area

0

##### Parts Per Million

1.6

##### Fragmentation mode

HCD

##### Originating file

01 D:\separate\_stitch\_analyses\xle-disambiguation\20210325\_F59\_3ug\_DENOVO\_12.csv

#### Meta Information from PEAKS

##### Scan Identifier

F3:7338

##### Original sequence

L

S

E

V

S

D

R

P

S

G

V

S

S

R

F

##### Posttranslational Modifications

##### Source File

D:\separate\_stitch\_analyses\xle-disambiguation\raw\20210323\_F1\_UM1\_Peng0013\_SA\_F59\_ingel\_3ug\_chymo.raw

##### Fraction

3

##### Scan Feature

-

##### De Novo Score

95

##### ConfidenceScore

95

### m/z

541.6098

##### Mass

1621.8059

##### Charge

3

##### Retention Time

40.83

##### Predicted Retention Time

-

##### Area

0

##### Parts Per Million

1

##### Fragmentation mode

ETHCD

##### Originating file

01 D:\separate\_stitch\_analyses\xle-disambiguation\20210325\_F59\_3ug\_DENOVO\_12.csv

#### Meta Information from PEAKS

##### Scan Identifier

F3:6589

##### Original sequence

L

S

E

V

S

D

R

P

S

G

V

S

S

R

F

##### Posttranslational Modifications

##### Source File

D:\separate\_stitch\_analyses\xle-disambiguation\raw\20210323\_F1\_UM1\_Peng0013\_SA\_F59\_ingel\_3ug\_chymo.raw

##### Fraction

3

##### Scan Feature

-

##### De Novo Score

95

##### ConfidenceScore

95

### m/z

541.6094

##### Mass

1621.8059

##### Charge

3

##### Retention Time

36.58

##### Predicted Retention Time

-

##### Area

0

##### Parts Per Million

0.2

##### Fragmentation mode

ETHCD

##### Originating file

01 D:\separate\_stitch\_analyses\xle-disambiguation\20210325\_F59\_3ug\_DENOVO\_12.csv
