## Supplementary material for "A handle on mass coincidence errors in *de novo* sequencing of antibodies by bottom-up proteomics": Combined_078.html

Details Combined\_078 | Stitch OverviewUndefined

### Read Combined\_078

#### Sequence (length=7)

DTJMJSR

#### Spectrum 4447? Spectrum 4447 The raw spectrum of this peptide as annotated by Hecklib. The fragments are coloured according to ion type (see legend). Any peaks with a star '\*' as text can be hovered over to see the full details, first the ion type second the mass shift type. By hovering over the amino acids in the peptide or ions in the legend the corresponding peaks are highlighted. By toggling the 'Unassigned' label you can turn the background (unassigned) peaks on or off in the plot. By updating the slider in the Ion legend you can update the spectrum to only show the top X% of the peaks with labels. The top X% means any peak that is within X% of the highest intensity. By dragging in the spectrum you can zoom in to a specific part of the spectrum and use 'Zoom Out' to get back to the original zoom level. The annotation of the spectrum is based on the given sequence in the peptides file and is done with different software so inconsistencies are likely. The peaks are annotated based on the given sequence, with 20 ppm tolerance.

Copy Data

##### Spectrum 4447 (TSV)

###### Preview

```
Loading example...
```

*Click on the button to copy the data to your clipboard.*

Mz MinMz MaxIntensity Max

WidthHeightPeptide font sizePeptide stroke widthSpectrum font sizeSpectrum stroke widthCompact peptide

Ion legend

wxyz

abcd

OtherUnassignedIon^Charge^_Position_Show for top:%

DTJMJSR

02.36e+44.73e+47.09e+49.45e+4

Zoom Out

y^+1^_1_a^+1^_2_y^+1^_1_y^+2^_3_a^+1^_2_b^+1^_2_b^+1^_2_y^+1^_2_y^+1^_2_y^+2^_4_y^+2^_4_y^+1^_2_a^+1^_3_a^+1^_3_b^+1^_3_y^+2^_5_b^+1^_3_y^+1^_3_y^+1^_3_y^+2^_6_y^+2^_6_y^+1^_3_\*\*b^+1^_4_b^+1^_4_y^+1^_4_y^+1^_4_b^+1^_5_b^+1^_5_y^+1^_5_y^+1^_5_y^+1^_6_

0777155523323110

Fragment Matches Table

Show background peaks

| Position | Ion type | Intensity | mz Theoretical | mz Error (Th) | mz Error (ppm) | Charge | Series Number |
| --- | --- | --- | --- | --- | --- | --- | --- |
| - | - | 966.9 | 120.1 | - | - | 0 | - |
| - | - | 504.8 | 121.1 | - | - | 0 | - |
| - | - | 357.5 | 123.6 | - | - | 0 | - |
| - | - | 1620 | 124.1 | - | - | 0 | - |
| - | - | 958.2 | 125.1 | - | - | 0 | - |
| - | - | 580.1 | 125.1 | - | - | 0 | - |
| - | - | 1.685E+04 | 126.1 | - | - | 0 | - |
| - | - | 997.1 | 127.1 | - | - | 0 | - |
| - | - | 740.9 | 127.1 | - | - | 0 | - |
| - | - | 5713 | 129.1 | - | - | 0 | - |
| - | - | 2669 | 129.1 | - | - | 0 | - |
| - | - | 442.6 | 129.1 | - | - | 0 | - |
| - | - | 1032 | 130 | - | - | 0 | - |
| - | - | 5870 | 130.1 | - | - | 0 | - |
| - | - | 906.4 | 130.1 | - | - | 0 | - |
| - | - | 883.2 | 136 | - | - | 0 | - |
| - | - | 838.3 | 136.1 | - | - | 0 | - |
| - | - | 524.6 | 138.1 | - | - | 0 | - |
| - | - | 608.4 | 139.1 | - | - | 0 | - |
| - | - | 1098 | 139.1 | - | - | 0 | - |
| - | - | 1222 | 141.1 | - | - | 0 | - |
| - | - | 2233 | 141.1 | - | - | 0 | - |
| - | - | 729.3 | 142.1 | - | - | 0 | - |
| - | - | 1513 | 142.1 | - | - | 0 | - |
| - | - | 1385 | 143.1 | - | - | 0 | - |
| - | - | 473.3 | 143.5 | - | - | 0 | - |
| - | - | 6.326E+04 | 144.1 | - | - | 0 | - |
| - | - | 504.7 | 145.1 | - | - | 0 | - |
| - | - | 3501 | 145.1 | - | - | 0 | - |
| - | - | 1801 | 152.1 | - | - | 0 | - |
| - | - | 3844 | 153.1 | - | - | 0 | - |
| - | - | 2249 | 154.1 | - | - | 0 | - |
| - | - | 703.8 | 155.1 | - | - | 0 | - |
| - | - | 672.1 | 157.1 | - | - | 0 | - |
| - | - | 815.7 | 157.1 | - | - | 0 | - |
| - | - | 1701 | 157.1 | - | - | 0 | - |
| - | - | 1533 | 157.1 | - | - | 0 | - |
| 7 | y | 1.318E+04 | 158.1 | 0.000172 | 1.088 | +1 | 1 |
| - | - | 544.1 | 167.1 | - | - | 0 | - |
| - | - | 1.772E+04 | 169.1 | - | - | 0 | - |
| - | - | 4823 | 170.1 | - | - | 0 | - |
| - | - | 740.5 | 170.1 | - | - | 0 | - |
| - | - | 1420 | 170.1 | - | - | 0 | - |
| 2 | a | 1.035E+04 | 171.1 | 0.0001347 | 0.7873 | +1 | 2 |
| - | - | 1007 | 172.1 | - | - | 0 | - |
| - | - | 921.9 | 172.1 | - | - | 0 | - |
| - | - | 2849 | 173.1 | - | - | 0 | - |
| 7 | y | 3.887E+04 | 175.1 | 0.0002037 | 1.163 | +1 | 1 |
| - | - | 2271 | 176.1 | - | - | 0 | - |
| 5 | y | 715.8 | 179.1 | 0.002286 | 12.76 | +2 | 3 |
| - | - | 1.956E+04 | 180.1 | - | - | 0 | - |
| - | - | 1145 | 181.1 | - | - | 0 | - |
| - | - | 2142 | 181.1 | - | - | 0 | - |
| - | - | 742.2 | 182.1 | - | - | 0 | - |
| - | - | 958.1 | 183.1 | - | - | 0 | - |
| - | - | 742.4 | 183.1 | - | - | 0 | - |
| - | - | 616.9 | 184.1 | - | - | 0 | - |
| - | - | 1033 | 185.1 | - | - | 0 | - |
| - | - | 761.8 | 185.1 | - | - | 0 | - |
| - | - | 1960 | 185.1 | - | - | 0 | - |
| - | - | 1080 | 185.2 | - | - | 0 | - |
| - | - | 1.825E+04 | 187.1 | - | - | 0 | - |
| - | - | 1111 | 188.1 | - | - | 0 | - |
| 2 | a | 6.326E+04 | 189.1 | 0.0001749 | 0.9248 | +1 | 2 |
| - | - | 4673 | 190.1 | - | - | 0 | - |
| - | - | 576.1 | 195.1 | - | - | 0 | - |
| - | - | 1.94E+04 | 197.1 | - | - | 0 | - |
| - | - | 2374 | 198.1 | - | - | 0 | - |
| - | - | 1698 | 198.1 | - | - | 0 | - |
| 2 | b | 2.106E+04 | 199.1 | 0.0001542 | 0.7744 | +1 | 2 |
| - | - | 2041 | 200.1 | - | - | 0 | - |
| - | - | 2596 | 201.1 | - | - | 0 | - |
| - | - | 736.3 | 201.1 | - | - | 0 | - |
| - | - | 2808 | 202.1 | - | - | 0 | - |
| - | - | 1908 | 203.1 | - | - | 0 | - |
| - | - | 3084 | 203.1 | - | - | 0 | - |
| - | - | 1312 | 210.1 | - | - | 0 | - |
| - | - | 908.1 | 211.1 | - | - | 0 | - |
| - | - | 502.2 | 213 | - | - | 0 | - |
| - | - | 5656 | 215.1 | - | - | 0 | - |
| 2 | b | 3.701E+04 | 217.1 | 0.0001333 | 0.614 | +1 | 2 |
| - | - | 2731 | 218.1 | - | - | 0 | - |
| - | - | 1043 | 220.1 | - | - | 0 | - |
| - | - | 497.2 | 222.1 | - | - | 0 | - |
| - | - | 617.7 | 223.1 | - | - | 0 | - |
| - | - | 3497 | 225.1 | - | - | 0 | - |
| - | - | 2924 | 226.2 | - | - | 0 | - |
| - | - | 2677 | 227.1 | - | - | 0 | - |
| - | - | 1050 | 228.1 | - | - | 0 | - |
| - | - | 1872 | 229 | - | - | 0 | - |
| - | - | 622.4 | 229 | - | - | 0 | - |
| - | - | 5484 | 229.6 | - | - | 0 | - |
| - | - | 948.5 | 230.1 | - | - | 0 | - |
| - | - | 1611 | 230.1 | - | - | 0 | - |
| - | - | 727.2 | 232.1 | - | - | 0 | - |
| - | - | 4806 | 233.1 | - | - | 0 | - |
| - | - | 1474 | 238.2 | - | - | 0 | - |
| - | - | 1002 | 240.1 | - | - | 0 | - |
| - | - | 3325 | 243.2 | - | - | 0 | - |
| - | - | 718.5 | 243.6 | - | - | 0 | - |
| - | - | 2633 | 244.1 | - | - | 0 | - |
| 6 | y | 3056 | 244.1 | 0.0001939 | 0.794 | +1 | 2 |
| 6 | y | 2.234E+04 | 245.1 | 0.0002023 | 0.8253 | +1 | 2 |
| - | - | 2206 | 246.1 | - | - | 0 | - |
| - | - | 1295 | 249.1 | - | - | 0 | - |
| - | - | 1015 | 251.1 | - | - | 0 | - |
| 4 | y | 754.1 | 253.1 | 0.002452 | 9.685 | +2 | 4 |
| - | - | 554.6 | 258.1 | - | - | 0 | - |
| - | - | 8096 | 261.1 | - | - | 0 | - |
| 4 | y | 2750 | 261.6 | 0.002757 | 10.54 | +2 | 4 |
| 6 | y | 4.312E+04 | 262.2 | 5.092E-05 | 0.1942 | +1 | 2 |
| - | - | 4026 | 263.2 | - | - | 0 | - |
| - | - | 744.7 | 264.1 | - | - | 0 | - |
| - | - | 655.1 | 264.2 | - | - | 0 | - |
| - | - | 2767 | 266.2 | - | - | 0 | - |
| - | - | 2519 | 267.1 | - | - | 0 | - |
| - | - | 594.7 | 268.1 | - | - | 0 | - |
| - | - | 1364 | 271 | - | - | 0 | - |
| - | - | 915.9 | 272.1 | - | - | 0 | - |
| - | - | 844.5 | 277.1 | - | - | 0 | - |
| - | - | 640.7 | 279 | - | - | 0 | - |
| - | - | 674.9 | 279.1 | - | - | 0 | - |
| - | - | 651.7 | 282.2 | - | - | 0 | - |
| 3 | a | 1.831E+04 | 284.2 | 7.035E-05 | 0.2476 | +1 | 3 |
| - | - | 645.6 | 285.1 | - | - | 0 | - |
| - | - | 2124 | 285.2 | - | - | 0 | - |
| - | - | 743.3 | 286.1 | - | - | 0 | - |
| - | - | 1.399E+04 | 286.2 | - | - | 0 | - |
| - | - | 4700 | 286.7 | - | - | 0 | - |
| - | - | 2134 | 293.9 | - | - | 0 | - |
| - | - | 750.9 | 294.1 | - | - | 0 | - |
| - | - | 1381 | 294.9 | - | - | 0 | - |
| - | - | 889 | 295.1 | - | - | 0 | - |
| - | - | 3861 | 297.1 | - | - | 0 | - |
| - | - | 697 | 298.2 | - | - | 0 | - |
| - | - | 1123 | 299.1 | - | - | 0 | - |
| - | - | 1210 | 299.2 | - | - | 0 | - |
| - | - | 871 | 299.7 | - | - | 0 | - |
| 3 | a | 1067 | 302.2 | 0.000553 | 1.83 | +1 | 3 |
| - | - | 744.7 | 308.2 | - | - | 0 | - |
| - | - | 709.1 | 309.2 | - | - | 0 | - |
| - | - | 1605 | 310.2 | - | - | 0 | - |
| - | - | 600.4 | 311 | - | - | 0 | - |
| - | - | 3811 | 311.9 | - | - | 0 | - |
| 3 | b | 2.775E+04 | 312.2 | 0.0001203 | 0.3855 | +1 | 3 |
| - | - | 3784 | 313.2 | - | - | 0 | - |
| 3 | y | 3.565E+04 | 318.2 | 0.002595 | 8.157 | +2 | 5 |
| - | - | 7897 | 318.7 | - | - | 0 | - |
| - | - | 2884 | 319.2 | - | - | 0 | - |
| - | - | 1329 | 319.7 | - | - | 0 | - |
| 3 | b | 8758 | 330.2 | 0.0005114 | 1.549 | +1 | 3 |
| - | - | 1044 | 331.2 | - | - | 0 | - |
| - | - | 965 | 332.7 | - | - | 0 | - |
| - | - | 821.1 | 333.2 | - | - | 0 | - |
| - | - | 624.5 | 334.7 | - | - | 0 | - |
| - | - | 3019 | 336.7 | - | - | 0 | - |
| - | - | 1363 | 340.2 | - | - | 0 | - |
| - | - | 2262 | 341.7 | - | - | 0 | - |
| - | - | 1507 | 342.2 | - | - | 0 | - |
| - | - | 1586 | 346.1 | - | - | 0 | - |
| - | - | 709 | 348.2 | - | - | 0 | - |
| - | - | 724.8 | 349.1 | - | - | 0 | - |
| - | - | 621.9 | 350.2 | - | - | 0 | - |
| - | - | 645 | 353.9 | - | - | 0 | - |
| - | - | 1944 | 354.9 | - | - | 0 | - |
| - | - | 1405 | 355.7 | - | - | 0 | - |
| 5 | y | 4910 | 357.2 | 0.0002062 | 0.5772 | +1 | 3 |
| 5 | y | 4563 | 358.2 | 0.0005487 | 1.532 | +1 | 3 |
| - | - | 714.6 | 358.2 | - | - | 0 | - |
| - | - | 713.5 | 359.2 | - | - | 0 | - |
| 2 | y | 692.7 | 359.7 | 0.00357 | 9.925 | +2 | 6 |
| - | - | 855 | 362.2 | - | - | 0 | - |
| - | - | 1471 | 362.2 | - | - | 0 | - |
| - | - | 723.6 | 362.7 | - | - | 0 | - |
| - | - | 1094 | 363.2 | - | - | 0 | - |
| - | - | 1347 | 364.7 | - | - | 0 | - |
| - | - | 864.3 | 365.2 | - | - | 0 | - |
| - | - | 874.4 | 367.2 | - | - | 0 | - |
| - | - | 1582 | 367.7 | - | - | 0 | - |
| - | - | 832.1 | 368.2 | - | - | 0 | - |
| 2 | y | 4647 | 368.7 | 0.002072 | 5.619 | +2 | 6 |
| - | - | 830.8 | 369.2 | - | - | 0 | - |
| - | - | 2496 | 372.2 | - | - | 0 | - |
| - | - | 1693 | 373.7 | - | - | 0 | - |
| 5 | y | 7.494E+04 | 375.2 | 0.0001813 | 0.4831 | +1 | 3 |
| - | - | 1.187E+04 | 376.2 | - | - | 0 | - |
| - | - | 2021 | 377.2 | - | - | 0 | - |
| - | - | 1.237E+04 | 385.2 | - | - | 0 | - |
| - | - | 3708 | 385.7 | - | - | 0 | - |
| - | - | 1456 | 386.2 | - | - | 0 | - |
| - | - | 716.6 | 386.7 | - | - | 0 | - |
| - | - | 862.2 | 390.7 | - | - | 0 | - |
| - | - | 1.021E+04 | 394.2 | - | - | 0 | - |
| - | - | 3164 | 394.7 | - | - | 0 | - |
| - | - | 800.1 | 395.2 | - | - | 0 | - |
| - | - | 769.2 | 395.2 | - | - | 0 | - |
| - | - | 918 | 399.2 | - | - | 0 | - |
| - | - | 1177 | 399.7 | - | - | 0 | - |
| - | - | 695.8 | 400.2 | - | - | 0 | - |
| - | - | 1052 | 401.2 | - | - | 0 | - |
| - | - | 5733 | 402.2 | - | - | 0 | - |
| - | - | 858.8 | 408.7 | - | - | 0 | - |
| - | - | 916.5 | 413.2 | - | - | 0 | - |
| 0 | Precursor | 1.796E+04 | 417.2 | 0.002458 | 5.891 | +2 | -1 |
| - | - | 6795 | 417.7 | - | - | 0 | - |
| - | - | 2512 | 418.2 | - | - | 0 | - |
| - | - | 583.7 | 422.2 | - | - | 0 | - |
| - | - | 2384 | 423.2 | - | - | 0 | - |
| - | - | 611.8 | 424.2 | - | - | 0 | - |
| - | - | 2440 | 425.2 | - | - | 0 | - |
| 0 | Precursor | 3320 | 426.2 | 0.001936 | 4.543 | +2 | -1 |
| - | - | 2147 | 428.3 | - | - | 0 | - |
| - | - | 787.7 | 430.2 | - | - | 0 | - |
| - | - | 1.518E+04 | 431.2 | - | - | 0 | - |
| - | - | 3314 | 432.3 | - | - | 0 | - |
| - | - | 3096 | 440.3 | - | - | 0 | - |
| - | - | 8681 | 441.2 | - | - | 0 | - |
| - | - | 1689 | 442.2 | - | - | 0 | - |
| - | - | 664.5 | 443.3 | - | - | 0 | - |
| - | - | 612.5 | 444.3 | - | - | 0 | - |
| - | - | 9.353E+04 | 458.3 | - | - | 0 | - |
| 4 | b | 4134 | 459.2 | 0.005067 | 11.03 | +1 | 4 |
| - | - | 1.783E+04 | 459.3 | - | - | 0 | - |
| - | - | 3011 | 460.3 | - | - | 0 | - |
| - | - | 1583 | 468.3 | - | - | 0 | - |
| 4 | b | 8180 | 477.2 | 0.004939 | 10.35 | +1 | 4 |
| - | - | 1395 | 478.2 | - | - | 0 | - |
| - | - | 584.2 | 480.3 | - | - | 0 | - |
| - | - | 1041 | 484.3 | - | - | 0 | - |
| - | - | 719.5 | 485.3 | - | - | 0 | - |
| - | - | 2027 | 486.3 | - | - | 0 | - |
| - | - | 848 | 488.3 | - | - | 0 | - |
| - | - | 5497 | 501.3 | - | - | 0 | - |
| - | - | 773.6 | 502.3 | - | - | 0 | - |
| 4 | y | 1990 | 504.3 | 0.003794 | 7.524 | +1 | 4 |
| - | - | 2184 | 507.2 | - | - | 0 | - |
| - | - | 1903 | 515.3 | - | - | 0 | - |
| - | - | 678.3 | 516.3 | - | - | 0 | - |
| 4 | y | 9.357E+04 | 522.3 | 0.004551 | 8.715 | +1 | 4 |
| - | - | 2.258E+04 | 523.3 | - | - | 0 | - |
| - | - | 5750 | 524.3 | - | - | 0 | - |
| - | - | 1195 | 532.3 | - | - | 0 | - |
| - | - | 588.1 | 537.2 | - | - | 0 | - |
| - | - | 3267 | 541.3 | - | - | 0 | - |
| - | - | 1894 | 544.3 | - | - | 0 | - |
| - | - | 1865 | 553.3 | - | - | 0 | - |
| - | - | 1894 | 554.3 | - | - | 0 | - |
| - | - | 6.067E+04 | 571.4 | - | - | 0 | - |
| 5 | b | 919.1 | 572.3 | 0.008252 | 14.42 | +1 | 5 |
| - | - | 1.858E+04 | 572.4 | - | - | 0 | - |
| - | - | 3615 | 573.4 | - | - | 0 | - |
| 5 | b | 722.3 | 590.3 | 0.003486 | 5.906 | +1 | 5 |
| 3 | y | 793.5 | 617.3 | 0.004355 | 7.055 | +1 | 5 |
| - | - | 730.2 | 620.3 | - | - | 0 | - |
| 3 | y | 6.134E+04 | 635.3 | 0.004167 | 6.558 | +1 | 5 |
| - | - | 1.633E+04 | 636.4 | - | - | 0 | - |
| - | - | 6514 | 637.4 | - | - | 0 | - |
| - | - | 1191 | 638.4 | - | - | 0 | - |
| - | - | 568.4 | 640.4 | - | - | 0 | - |
| - | - | 3420 | 640.4 | - | - | 0 | - |
| - | - | 1445 | 672.4 | - | - | 0 | - |
| - | - | 4404 | 682.4 | - | - | 0 | - |
| - | - | 2117 | 683.4 | - | - | 0 | - |
| 2 | y | 1217 | 736.4 | 0.009345 | 12.69 | +1 | 6 |
| - | - | 1723 | 741.5 | - | - | 0 | - |
| - | - | 7635 | 746.4 | - | - | 0 | - |
| - | - | 2392 | 747.4 | - | - | 0 | - |
| - | - | 743 | 748.4 | - | - | 0 | - |
| - | - | 594.8 | 1043 | - | - | 0 | - |
| - | - | 583.3 | 1194 | - | - | 0 | - |
| - | - | 669.7 | 2846 | - | - | 0 | - |
| - | - | 676.9 | 3079 | - | - | 0 | - |

m/z Charge Intensity FragmentType MassShift Position
120.08086395263672 0 966.8918
121.10136413574219 0 504.8482
123.62602233886719 0 357.53146
124.11238098144531 0 1620.2594
125.07113647460938 0 958.248
125.10779571533203 0 580.053
126.05518341064453 0 16850.414
127.05033874511719 0 997.0597
127.0585708618164 0 740.90967
129.0660858154297 0 5713.287
129.10240173339844 0 2668.5493
129.1138153076172 0 442.55447
130.04986572265625 0 1032.2483
130.06134033203125 0 5870.3823
130.09788513183594 0 906.4383
136.039794921875 0 883.1559
136.07595825195312 0 838.3325
138.0552215576172 0 524.60834
139.05052185058594 0 608.4386
139.1119842529297 0 1098.2163
141.1023712158203 0 1221.7932
141.1387481689453 0 2232.6982
142.1103057861328 0 729.2805
142.12286376953125 0 1512.9491
143.11822509765625 0 1384.7246
143.5256805419922 0 473.2769
144.0657501220703 0 63257.043
145.06344604492188 0 504.7032
145.069091796875 0 3500.9595
152.10726928710938 0 1801.0154
153.06607055664062 0 3844.2766
154.0501251220703 0 2249.156
155.07061767578125 0 703.7518
157.06112670898438 0 672.0905
157.09730529785156 0 815.6955
157.10842895507812 0 1700.8046
157.1336669921875 0 1533.4098
158.0925750732422 0 13181.177 y Ammonia loss 6
167.11798095703125 0 544.08154
169.1337432861328 0 17717.81
170.09255981445312 0 4823.4307
170.1051025390625 0 740.4881
170.13693237304688 0 1419.6309
171.07655334472656 0 10348.434 a Water loss 1
172.0609588623047 0 1006.8311
172.0797882080078 0 921.9016
173.12869262695312 0 2848.5974
175.11915588378906 0 38868.62 y 6
176.1226348876953 0 2270.71
179.1181640625 0 715.8162 y Water loss 4
180.1020965576172 0 19563.102
181.06072998046875 0 1145.4716
181.1055450439453 0 2142.3237
182.0924072265625 0 742.2161
183.1128387451172 0 958.1273
183.14930725097656 0 742.3931
184.10865783691406 0 616.87427
185.09228515625 0 1033.4003
185.11770629882812 0 761.7642
185.13995361328125 0 1959.5529
185.16500854492188 0 1080.1714
187.144287109375 0 18246.346
188.14739990234375 0 1111.0612
189.087158203125 0 63259.277 a 1
190.090576171875 0 4672.811
195.1133270263672 0 576.14374
197.1285858154297 0 19398.559
198.08741760253906 0 2374.3342
198.1320037841797 0 1698.4402
199.0714874267578 0 21064.375 b Water loss 1
200.0748748779297 0 2041.2756
201.12344360351562 0 2595.9038
201.13360595703125 0 736.27203
202.11868286132812 0 2807.9585
203.10284423828125 0 1907.8356
203.12794494628906 0 3084.1982
210.08721923828125 0 1312.1254
211.14453125 0 908.09436
213.0373992919922 0 502.15607
215.13922119140625 0 5655.9536
217.08203125 0 37006.836 b 1
218.0853271484375 0 2731.1382
220.12953186035156 0 1042.5116
222.12367248535156 0 497.1887
223.10824584960938 0 617.67267
225.1236572265625 0 3497.4736
226.15516662597656 0 2923.6367
227.11398315429688 0 2677.1636
228.09828186035156 0 1049.8619
229.00140380859375 0 1872.4878
229.01283264160156 0 622.4278
229.6398468017578 0 5483.6855
230.1247100830078 0 948.4915
230.1409912109375 0 1610.7284
232.14064025878906 0 727.1935
233.13194274902344 0 4805.911
238.1549835205078 0 1473.5497
240.13400268554688 0 1001.9253
243.1819305419922 0 3325.0947
243.63702392578125 0 718.521
244.10008239746094 0 2633.2285
244.14060974121094 0 3055.5432 y Water loss 5
245.1246337890625 0 22338.176 y Ammonia loss 5
246.12905883789062 0 2205.9985
249.12290954589844 0 1294.8212
251.10214233398438 0 1014.70526
253.12557983398438 0 754.0968 y Ammonia loss 3
258.14501953125 0 554.56995
261.126708984375 0 8096.124
261.63916015625 0 2749.781 y 3
262.1510314941406 0 43118.473 y 5
263.1539306640625 0 4025.8318
264.1346740722656 0 744.70215
264.15643310546875 0 655.07513
266.1500549316406 0 2766.5684
267.1337890625 0 2519.365
268.12847900390625 0 594.6507
271.0273742675781 0 1364.3878
272.1356506347656 0 915.936
277.11883544921875 0 844.4681
279.0479736328125 0 640.7485
279.13446044921875 0 674.9192
282.2174987792969 0 651.67444
284.1605529785156 0 18310.158 a Water loss 2
285.1456298828125 0 645.61255
285.16357421875 0 2124.0566
286.142333984375 0 743.32654
286.1818542480469 0 13989.839
286.68341064453125 0 4700.186
293.93438720703125 0 2134.0417
294.1433410644531 0 750.8963
294.9186096191406 0 1381.3046
295.1293640136719 0 889.0034
297.0582275390625 0 3860.5608
298.1775207519531 0 696.9948
299.053955078125 0 1122.8206
299.17138671875 0 1209.5399
299.6729431152344 0 871.0385
302.1716003417969 0 1066.6986 a 2
308.1605224609375 0 744.65186
309.2032470703125 0 709.137
310.2127380371094 0 1604.9628
311.0110168457031 0 600.40784
311.9453125 0 3811.1045
312.155517578125 0 27754.57 b Water loss 2
313.1585388183594 0 3784.3145
318.1810302734375 0 35648.07 y 2
318.68231201171875 0 7896.7334
319.18145751953125 0 2884.1335
319.6842956542969 0 1329.4275
330.1664733886719 0 8758.213 b 2
331.1697998046875 0 1044.1926
332.6920471191406 0 964.9503
333.19378662109375 0 821.0559
334.68603515625 0 624.52545
336.70574951171875 0 3018.9863
340.1978454589844 0 1363.4448
341.6977844238281 0 2261.9631
342.1996154785156 0 1507.2866
346.14263916015625 0 1585.7235
348.1602478027344 0 708.9749
349.0736083984375 0 724.8089
350.2084655761719 0 621.8711
353.8996276855469 0 644.98096
354.8849792480469 0 1944.3068
355.6934814453125 0 1404.5273
357.2242736816406 0 4909.663 y Water loss 4
358.20794677734375 0 4563.394 y Ammonia loss 4
358.2289123535156 0 714.6447
359.2122497558594 0 713.4896
359.7005615234375 0 692.72626 y Water loss 1
362.1742858886719 0 855.0272
362.2114562988281 0 1471.2255
362.7109680175781 0 723.5997
363.2003173828125 0 1093.9829
364.6910400390625 0 1347.0288
365.19366455078125 0 864.3328
367.2060852050781 0 874.4362
367.6953125 0 1581.6083
368.1930847167969 0 832.0994
368.704345703125 0 4646.9453 y 1
369.2030944824219 0 830.76184
372.1583251953125 0 2496.066
373.6966247558594 0 1692.7946
375.23486328125 0 74939.55 y 4
376.23773193359375 0 11867.946
377.2393798828125 0 2021.092
385.2157897949219 0 12372.026
385.7144470214844 0 3707.5684
386.21966552734375 0 1456.2043
386.68670654296875 0 716.59955
390.68951416015625 0 862.1904
394.2169494628906 0 10211.942
394.7175598144531 0 3163.6755
395.19378662109375 0 800.13947
395.2229309082031 0 769.22473
399.20135498046875 0 917.9512
399.694091796875 0 1177.3461
400.1975402832031 0 695.82965
401.214111328125 0 1052.1506
402.2098083496094 0 5733.256
408.7032165527344 0 858.7657
413.20538330078125 0 916.48584
417.2129211425781 0 17959.271 Precursor Water loss
417.7144775390625 0 6795.1436
418.2141418457031 0 2511.8599
422.2452392578125 0 583.72736
423.23443603515625 0 2384.0173
424.15191650390625 0 611.772
425.214599609375 0 2440.3079
426.2176818847656 0 3320.1707 Precursor
428.2613830566406 0 2147.2893
430.2407531738281 0 787.74243
431.2486267089844 0 15178.538
432.2530822753906 0 3314.0222
440.26141357421875 0 3096.391
441.2455749511719 0 8680.908
442.24981689453125 0 1688.9518
443.25030517578125 0 664.522
444.25543212890625 0 612.501
458.272216796875 0 93525.07
459.1909484863281 0 4134.3555 b Water loss 3
459.2749938964844 0 17834.895
460.2776794433594 0 3010.725
468.25665283203125 0 1583.2731
477.2013854980469 0 8180.486 b 3
478.2045593261719 0 1394.9984
480.2507019042969 0 584.16406
484.2502746582031 0 1041.2471
485.25347900390625 0 719.53284
486.26641845703125 0 2026.949
488.2718811035156 0 847.9978
501.2779541015625 0 5497.343
502.2799377441406 0 773.5725
504.2587585449219 0 1990.4539 y Water loss 3
507.246337890625 0 2184.228
515.29443359375 0 1902.9432
516.2951049804688 0 678.3218
522.2700805664062 0 93571.57 y 3
523.2728881835938 0 22582.15
524.2706909179688 0 5750.0215
532.2579345703125 0 1195.1412
537.1521606445312 0 588.07916
541.3445434570312 0 3267.3972
544.3320922851562 0 1894.0297
553.3456420898438 0 1864.8862
554.3296508789062 0 1893.8519
571.3556518554688 0 60668.074
572.2781982421875 0 919.0944 b Water loss 4
572.3585815429688 0 18582.084
573.361328125 0 3615.4736
590.2839965820312 0 722.293 b 4
617.3433837890625 0 793.5449 y Water loss 2
620.3280029296875 0 730.1592
635.353759765625 0 61344.08 y 2
636.356201171875 0 16332.863
637.356201171875 0 6513.748
638.3629760742188 0 1190.9033
640.358642578125 0 568.40045
640.412353515625 0 3420.3694
672.4033203125 0 1445.1279
682.3878173828125 0 4403.508
683.3897705078125 0 2116.9026
736.4066162109375 0 1216.7415 y 1
741.4632568359375 0 1722.9398
746.385009765625 0 7635.43
747.3903198242188 0 2392.146
748.3861694335938 0 742.9522
1043.3638916015625 0 594.7686
1193.8070068359375 0 583.25574
2845.84521484375 0 669.66956
3078.7431640625 0 676.9442

Spectrum Details

|  |  |
| --- | --- |
| Matched peaks? Matched peaks The total absolute number of peaks matched. Additionally in brackets the total fraction of peaks matched and the total number of peaks is shown. | 33 (12.13% of 272) |
| FDR? FDR The false discovery rate estimated for this peptide. It is calculated by matching all theoretical fragments with a non-integer shift with the raw peaks for this spectrum. This is done with 40 different shifts. The resulting percentage is the average number of annotated peaks over the number of annotated peaks with the correct spectrum. | 0.29% |
| Satellite FDR? Satellite FDR See the FDR for details on its calculation. This satellite ion specific FDR only contains the satellite ions (d/w) for I/L/J positions. | - |
| PSM Score? PSM Score The PSM Score as given by Hecklib to this annotated spectrum. It is shown with three significant figures. | 423 |

#### Spectrum 4337? Spectrum 4337 The raw spectrum of this peptide as annotated by Hecklib. The fragments are coloured according to ion type (see legend). Any peaks with a star '\*' as text can be hovered over to see the full details, first the ion type second the mass shift type. By hovering over the amino acids in the peptide or ions in the legend the corresponding peaks are highlighted. By toggling the 'Unassigned' label you can turn the background (unassigned) peaks on or off in the plot. By updating the slider in the Ion legend you can update the spectrum to only show the top X% of the peaks with labels. The top X% means any peak that is within X% of the highest intensity. By dragging in the spectrum you can zoom in to a specific part of the spectrum and use 'Zoom Out' to get back to the original zoom level. The annotation of the spectrum is based on the given sequence in the peptides file and is done with different software so inconsistencies are likely. The peaks are annotated based on the given sequence, with 20 ppm tolerance.

Copy Data

##### Spectrum 4337 (TSV)

###### Preview

```
Loading example...
```

*Click on the button to copy the data to your clipboard.*

Mz MinMz MaxIntensity Max

WidthHeightPeptide font sizePeptide stroke widthSpectrum font sizeSpectrum stroke widthCompact peptide

Ion legend

wxyz

abcd

OtherUnassignedIon^Charge^_Position_Show for top:%

DTJMJSR

03.20e+46.40e+49.60e+41.28e+5

Zoom Out

z^+1^_1_y^+1^_1_y^+1^_2_z^+1^_2_y^+2^_4_y^+1^_2_y^+2^_5_w^+1^_3_y^+1^_3_z^+1^_3_y^+2^_6_y^+1^_3_z^+1^_4_y^+1^_4_w^+1^_5_c^+1^_5_z^+1^_5_y^+1^_5_c^+1^_6_w^+1^_6_y^+1^_6_z^+1^_6_

0777155523323109

Fragment Matches Table

Show background peaks

| Position | Ion type | Intensity | mz Theoretical | mz Error (Th) | mz Error (ppm) | Charge | Series Number |
| --- | --- | --- | --- | --- | --- | --- | --- |
| - | - | 501.4 | 126.1 | - | - | 0 | - |
| - | - | 371.8 | 126.7 | - | - | 0 | - |
| - | - | 488.3 | 138.3 | - | - | 0 | - |
| - | - | 2441 | 144.1 | - | - | 0 | - |
| - | - | 617.6 | 149 | - | - | 0 | - |
| - | - | 1095 | 155.1 | - | - | 0 | - |
| - | - | 489 | 156.5 | - | - | 0 | - |
| - | - | 560.3 | 157.1 | - | - | 0 | - |
| 7 | z | 9688 | 159.1 | 0.0001137 | 0.7146 | +1 | 1 |
| - | - | 1.721E+04 | 160.1 | - | - | 0 | - |
| - | - | 1019 | 161.1 | - | - | 0 | - |
| - | - | 449.5 | 170.1 | - | - | 0 | - |
| - | - | 663 | 171.1 | - | - | 0 | - |
| - | - | 502.9 | 171.1 | - | - | 0 | - |
| - | - | 599.4 | 173.4 | - | - | 0 | - |
| 7 | y | 3256 | 175.1 | 0.0001885 | 1.076 | +1 | 1 |
| - | - | 1762 | 180.1 | - | - | 0 | - |
| - | - | 1840 | 185.1 | - | - | 0 | - |
| - | - | 578.8 | 185.1 | - | - | 0 | - |
| - | - | 933.9 | 187.1 | - | - | 0 | - |
| - | - | 1.226E+04 | 189.1 | - | - | 0 | - |
| - | - | 752.2 | 190.1 | - | - | 0 | - |
| - | - | 703.6 | 197.1 | - | - | 0 | - |
| - | - | 2553 | 199.1 | - | - | 0 | - |
| - | - | 1245 | 200.1 | - | - | 0 | - |
| - | - | 2435 | 203.1 | - | - | 0 | - |
| - | - | 707.4 | 215.1 | - | - | 0 | - |
| - | - | 8150 | 217.1 | - | - | 0 | - |
| - | - | 611 | 218.1 | - | - | 0 | - |
| - | - | 733.5 | 225.1 | - | - | 0 | - |
| - | - | 709 | 229 | - | - | 0 | - |
| - | - | 533.1 | 244.1 | - | - | 0 | - |
| - | - | 928.8 | 244.1 | - | - | 0 | - |
| 6 | y | 8611 | 245.1 | 0.000126 | 0.5141 | +1 | 2 |
| 6 | z | 5327 | 246.1 | 0.0001612 | 0.6548 | +1 | 2 |
| - | - | 2.817E+04 | 247.1 | - | - | 0 | - |
| - | - | 2022 | 248.1 | - | - | 0 | - |
| - | - | 1113 | 261.1 | - | - | 0 | - |
| - | - | 1193 | 261.1 | - | - | 0 | - |
| 4 | y | 911.7 | 261.6 | 0.002422 | 9.256 | +2 | 4 |
| 6 | y | 1.217E+04 | 262.2 | 1.012E-05 | 0.0386 | +1 | 2 |
| - | - | 1112 | 263.2 | - | - | 0 | - |
| - | - | 519.1 | 264.9 | - | - | 0 | - |
| - | - | 875.2 | 270.1 | - | - | 0 | - |
| - | - | 953 | 271 | - | - | 0 | - |
| - | - | 565.3 | 272.2 | - | - | 0 | - |
| - | - | 703.4 | 279 | - | - | 0 | - |
| - | - | 2809 | 284.2 | - | - | 0 | - |
| - | - | 695.5 | 285.2 | - | - | 0 | - |
| - | - | 3269 | 286.2 | - | - | 0 | - |
| - | - | 2047 | 290.1 | - | - | 0 | - |
| - | - | 1227 | 293.9 | - | - | 0 | - |
| - | - | 650.6 | 294.1 | - | - | 0 | - |
| - | - | 4228 | 297.1 | - | - | 0 | - |
| - | - | 2159 | 299.1 | - | - | 0 | - |
| - | - | 1191 | 311.9 | - | - | 0 | - |
| - | - | 7627 | 312.2 | - | - | 0 | - |
| - | - | 644.1 | 313.2 | - | - | 0 | - |
| 3 | y | 8481 | 318.2 | 0.002412 | 7.582 | +2 | 5 |
| - | - | 4154 | 318.7 | - | - | 0 | - |
| - | - | 1285 | 319.2 | - | - | 0 | - |
| 5 | w | 6814 | 330.2 | 0.000407 | 1.233 | +1 | 3 |
| - | - | 820.6 | 331.2 | - | - | 0 | - |
| - | - | 804.6 | 337.2 | - | - | 0 | - |
| - | - | 715 | 349.1 | - | - | 0 | - |
| - | - | 1629 | 353.9 | - | - | 0 | - |
| - | - | 1298 | 354.9 | - | - | 0 | - |
| 5 | y | 618.2 | 358.2 | 0.00183 | 5.11 | +1 | 3 |
| 5 | z | 2.187E+04 | 359.2 | 7.293E-05 | 0.203 | +1 | 3 |
| - | - | 9554 | 360.2 | - | - | 0 | - |
| - | - | 1467 | 361.2 | - | - | 0 | - |
| 2 | y | 1856 | 368.7 | 0.001675 | 4.543 | +2 | 6 |
| - | - | 827.4 | 369.2 | - | - | 0 | - |
| - | - | 727.1 | 372.9 | - | - | 0 | - |
| - | - | 2259 | 374.2 | - | - | 0 | - |
| 5 | y | 1.722E+04 | 375.2 | 0.0003339 | 0.8897 | +1 | 3 |
| - | - | 2949 | 376.2 | - | - | 0 | - |
| - | - | 2528 | 385.2 | - | - | 0 | - |
| - | - | 1351 | 388.2 | - | - | 0 | - |
| - | - | 3304 | 394.2 | - | - | 0 | - |
| - | - | 1198 | 394.7 | - | - | 0 | - |
| - | - | 627.1 | 399.7 | - | - | 0 | - |
| - | - | 663.2 | 408.2 | - | - | 0 | - |
| - | - | 2323 | 415.2 | - | - | 0 | - |
| - | - | 1651 | 416.2 | - | - | 0 | - |
| - | - | 4687 | 417.2 | - | - | 0 | - |
| - | - | 2874 | 417.7 | - | - | 0 | - |
| - | - | 1101 | 418.2 | - | - | 0 | - |
| - | - | 2981 | 425.2 | - | - | 0 | - |
| - | - | 1333 | 426.2 | - | - | 0 | - |
| - | - | 1325 | 442.3 | - | - | 0 | - |
| - | - | 6728 | 443.3 | - | - | 0 | - |
| - | - | 6822 | 444.3 | - | - | 0 | - |
| - | - | 597.1 | 445.3 | - | - | 0 | - |
| - | - | 501.4 | 452.3 | - | - | 0 | - |
| - | - | 7470 | 458.3 | - | - | 0 | - |
| - | - | 519.3 | 458.9 | - | - | 0 | - |
| - | - | 1977 | 459.2 | - | - | 0 | - |
| - | - | 1074 | 459.3 | - | - | 0 | - |
| - | - | 2841 | 477.2 | - | - | 0 | - |
| 4 | z | 934.5 | 506.2 | 0.003347 | 6.612 | +1 | 4 |
| - | - | 655.8 | 507.3 | - | - | 0 | - |
| 4 | y | 2.884E+04 | 522.3 | 0.004551 | 8.715 | +1 | 4 |
| - | - | 6824 | 523.3 | - | - | 0 | - |
| - | - | 2206 | 524.3 | - | - | 0 | - |
| - | - | 5536 | 571.4 | - | - | 0 | - |
| - | - | 1583 | 572.4 | - | - | 0 | - |
| 3 | w | 7156 | 576.3 | 0.004485 | 7.782 | +1 | 5 |
| - | - | 1929 | 577.3 | - | - | 0 | - |
| - | - | 693.8 | 590.3 | - | - | 0 | - |
| - | - | 586.4 | 603.1 | - | - | 0 | - |
| 5 | c | 671.9 | 607.3 | 0.008187 | 13.48 | +1 | 5 |
| 3 | z | 1669 | 619.3 | 0.003848 | 6.213 | +1 | 5 |
| 3 | y | 1.864E+04 | 635.3 | 0.004167 | 6.558 | +1 | 5 |
| - | - | 4883 | 636.4 | - | - | 0 | - |
| - | - | 1180 | 637.4 | - | - | 0 | - |
| - | - | 1105 | 662.3 | - | - | 0 | - |
| - | - | 4218 | 664.3 | - | - | 0 | - |
| - | - | 1520 | 665.3 | - | - | 0 | - |
| - | - | 992.7 | 668.4 | - | - | 0 | - |
| - | - | 1409 | 677.3 | - | - | 0 | - |
| - | - | 824.5 | 677.4 | - | - | 0 | - |
| 6 | c | 5066 | 694.3 | 0.003991 | 5.747 | +1 | 6 |
| - | - | 1202 | 695.3 | - | - | 0 | - |
| - | - | 956.3 | 701.4 | - | - | 0 | - |
| - | - | 603.4 | 704.4 | - | - | 0 | - |
| 2 | w | 1164 | 705.4 | 0.002471 | 3.504 | +1 | 6 |
| - | - | 4303 | 718.4 | - | - | 0 | - |
| 2 | y | 1.317E+04 | 719.4 | 0.003301 | 4.589 | +1 | 6 |
| 2 | z | 1.779E+04 | 720.4 | 0.002434 | 3.379 | +1 | 6 |
| - | - | 8247 | 721.4 | - | - | 0 | - |
| - | - | 2732 | 722.4 | - | - | 0 | - |
| - | - | 2712 | 728.4 | - | - | 0 | - |
| - | - | 1610 | 729.4 | - | - | 0 | - |
| - | - | 703 | 730.4 | - | - | 0 | - |
| - | - | 9721 | 735.4 | - | - | 0 | - |
| - | - | 2436 | 736.4 | - | - | 0 | - |
| - | - | 854 | 737.4 | - | - | 0 | - |
| - | - | 2430 | 746.4 | - | - | 0 | - |
| - | - | 1848 | 747.4 | - | - | 0 | - |
| - | - | 2336 | 748.4 | - | - | 0 | - |
| - | - | 967.2 | 751.3 | - | - | 0 | - |
| - | - | 780.6 | 752.3 | - | - | 0 | - |
| - | - | 549.6 | 756.9 | - | - | 0 | - |
| - | - | 1534 | 760.4 | - | - | 0 | - |
| - | - | 1128 | 766.4 | - | - | 0 | - |
| - | - | 2837 | 772.4 | - | - | 0 | - |
| - | - | 5859 | 773.4 | - | - | 0 | - |
| - | - | 2761 | 774.4 | - | - | 0 | - |
| - | - | 857.3 | 775.4 | - | - | 0 | - |
| - | - | 1419 | 776.4 | - | - | 0 | - |
| - | - | 798.8 | 777.4 | - | - | 0 | - |
| - | - | 5829 | 778.4 | - | - | 0 | - |
| - | - | 6971 | 779.3 | - | - | 0 | - |
| - | - | 1253 | 780.3 | - | - | 0 | - |
| - | - | 1793 | 780.4 | - | - | 0 | - |
| - | - | 740.8 | 781.4 | - | - | 0 | - |
| - | - | 2818 | 788.4 | - | - | 0 | - |
| - | - | 1462 | 789.4 | - | - | 0 | - |
| - | - | 6512 | 790.4 | - | - | 0 | - |
| - | - | 9.038E+04 | 791.4 | - | - | 0 | - |
| - | - | 5.814E+04 | 792.4 | - | - | 0 | - |
| - | - | 2.247E+04 | 793.4 | - | - | 0 | - |
| - | - | 5502 | 794.4 | - | - | 0 | - |
| - | - | 6302 | 796.4 | - | - | 0 | - |
| - | - | 1892 | 797.4 | - | - | 0 | - |
| - | - | 829.4 | 798.4 | - | - | 0 | - |
| - | - | 1402 | 806.4 | - | - | 0 | - |
| - | - | 752.9 | 808.4 | - | - | 0 | - |
| - | - | 2.65E+04 | 809.4 | - | - | 0 | - |
| - | - | 1.109E+04 | 810.4 | - | - | 0 | - |
| - | - | 4010 | 811.4 | - | - | 0 | - |
| - | - | 4477 | 817.4 | - | - | 0 | - |
| - | - | 2064 | 818.4 | - | - | 0 | - |
| - | - | 1008 | 819.4 | - | - | 0 | - |
| - | - | 909.5 | 820.4 | - | - | 0 | - |
| - | - | 1610 | 823.4 | - | - | 0 | - |
| - | - | 1676 | 824.4 | - | - | 0 | - |
| - | - | 4043 | 834.4 | - | - | 0 | - |
| - | - | 9.25E+04 | 835.4 | - | - | 0 | - |
| - | - | 3.541E+04 | 836.4 | - | - | 0 | - |
| - | - | 1.759E+04 | 837.4 | - | - | 0 | - |
| - | - | 2502 | 838.4 | - | - | 0 | - |
| - | - | 3.699E+04 | 851.4 | - | - | 0 | - |
| - | - | 1.267E+05 | 852.4 | - | - | 0 | - |
| - | - | 4.853E+04 | 853.4 | - | - | 0 | - |
| - | - | 1.628E+04 | 854.4 | - | - | 0 | - |
| - | - | 1478 | 855.4 | - | - | 0 | - |
| - | - | 689.1 | 1207 | - | - | 0 | - |
| - | - | 706.8 | 2806 | - | - | 0 | - |
| - | - | 708.1 | 3060 | - | - | 0 | - |
| - | - | 814.1 | 3078 | - | - | 0 | - |

m/z Charge Intensity FragmentType MassShift Position
126.05513763427734 0 501.4261
126.6501693725586 0 371.78525
138.34707641601562 0 488.32245
144.06568908691406 0 2440.5413
148.9547882080078 0 617.6151
155.07046508789062 0 1094.7036
156.5141143798828 0 489.00406
157.13365173339844 0 560.3032
159.100341796875 0 9688.322 z 6
160.10816955566406 0 17209.95
161.11154174804688 0 1018.57007
170.09249877929688 0 449.50772
171.07656860351562 0 663.0407
171.1216583251953 0 502.9019
173.43820190429688 0 599.4497
175.119140625 0 3256.4067 y 6
180.10205078125 0 1762.3921
185.11744689941406 0 1839.8917
185.13980102539062 0 578.82544
187.14422607421875 0 933.8644
189.08709716796875 0 12259.403
190.09097290039062 0 752.203
197.1283721923828 0 703.6177
199.07150268554688 0 2552.8271
200.12701416015625 0 1244.5787
203.12796020507812 0 2434.7095
215.13897705078125 0 707.4188
217.08200073242188 0 8150.369
218.08633422851562 0 610.9713
225.1231689453125 0 733.50195
229.00125122070312 0 708.97925
244.10064697265625 0 533.05597
244.11666870117188 0 928.8283
245.1245574951172 0 8611.208 y Ammonia loss 5
246.13209533691406 0 5327.103 z 5
247.14015197753906 0 28168.873
248.14334106445312 0 2022.1199
261.1274108886719 0 1112.927
261.1429748535156 0 1193.373
261.6388244628906 0 911.6823 y 3
262.1509704589844 0 12168.056 y 5
263.1547546386719 0 1111.9553
264.8895568847656 0 519.118
270.1202087402344 0 875.221
271.0270690917969 0 953.0355
272.1597900390625 0 565.3031
279.0482177734375 0 703.43445
284.1603088378906 0 2809.1104
285.1640930175781 0 695.49036
286.1820373535156 0 3269.1147
290.145751953125 0 2047.1144
293.9345703125 0 1226.7646
294.1466064453125 0 650.56744
297.05780029296875 0 4227.559
299.0541076660156 0 2159.2566
311.9449462890625 0 1190.7585
312.155517578125 0 7626.7183
313.1586608886719 0 644.1173
318.18084716796875 0 8480.95 y 2
318.6824951171875 0 4153.713
319.18304443359375 0 1285.1102
330.1767883300781 0 6814.438 w 4
331.1821594238281 0 820.59576
337.2046203613281 0 804.61694
349.0745849609375 0 715.0266
353.9006652832031 0 1629.3423
354.88458251953125 0 1298.2494
358.2066650390625 0 618.21814 y Ammonia loss 4
359.21624755859375 0 21868.785 z 4
360.2218933105469 0 9553.701
361.22613525390625 0 1466.7852
368.7039489746094 0 1856.2817 y 1
369.2057189941406 0 827.41895
372.89605712890625 0 727.0522
374.2268371582031 0 2259.3523
375.2347106933594 0 17223.963 y 4
376.2370300292969 0 2949.2268
385.21368408203125 0 2528.1553
388.2061767578125 0 1350.757
394.2173767089844 0 3303.8042
394.71875 0 1197.695
399.6974792480469 0 627.0708
408.20928955078125 0 663.2341
415.22998046875 0 2323.4402
416.23638916015625 0 1651.2838
417.2129821777344 0 4686.8325
417.7138366699219 0 2874.3623
418.2161865234375 0 1101.4852
425.21441650390625 0 2980.9001
426.2198791503906 0 1333.2279
442.2535705566406 0 1324.7474
443.2605895996094 0 6727.6523
444.268310546875 0 6822.1016
445.2750244140625 0 597.05194
452.3002624511719 0 501.44067
458.2724914550781 0 7469.882
458.9259033203125 0 519.3232
459.1904602050781 0 1977.4164
459.27374267578125 0 1073.994
477.2031555175781 0 2840.9048
506.2501525878906 0 934.47955 z 3
507.25244140625 0 655.818
522.2700805664062 0 28836.035 y 3
523.2730102539062 0 6823.709
524.2716674804688 0 2205.8628
571.3553466796875 0 5535.8516
572.3584594726562 0 1582.871
576.2805786132812 0 7155.571 w 2
577.2830200195312 0 1928.6866
590.2841796875 0 693.8227
603.1265258789062 0 586.4403
607.3152465820312 0 671.8777 c 4
619.334716796875 0 1669.4794 z 2
635.353759765625 0 18644.715 y 2
636.3568725585938 0 4882.559
637.3549194335938 0 1179.8152
662.3026733398438 0 1105.4463
664.3199462890625 0 4218.0615
665.3230590820312 0 1519.6658
668.359619140625 0 992.73065
677.3222045898438 0 1409.2655
677.3798828125 0 824.48456
694.3430786132812 0 5065.7246 c 5
695.3469848632812 0 1202.3805
701.4057006835938 0 956.299
704.37841796875 0 603.4367
705.3575439453125 0 1164.4742 w 1
718.3670043945312 0 4302.5854
719.3740234375 0 13167.698 y Ammonia loss 1
720.3809814453125 0 17788.443 z 1
721.3858032226562 0 8247.226
722.388671875 0 2732.4934
728.427978515625 0 2712.2065
729.4331665039062 0 1609.5479
730.4260864257812 0 703.04956
735.3565673828125 0 9720.945
736.3577270507812 0 2435.7793
737.3583374023438 0 854.0107
746.38720703125 0 2430.442
747.389404296875 0 1847.6475
748.4124145507812 0 2335.862
751.3389282226562 0 967.1856
752.3414916992188 0 780.5512
756.8723754882812 0 549.5993
760.4080810546875 0 1534.0605
766.4078369140625 0 1127.8638
772.4197387695312 0 2836.9517
773.4121704101562 0 5858.503
774.41259765625 0 2761.084
775.4232788085938 0 857.33575
776.3994750976562 0 1418.9819
777.396728515625 0 798.7516
778.3642578125 0 5828.819
779.3486938476562 0 6970.708
780.3428955078125 0 1253.0295
780.409912109375 0 1792.7422
781.4120483398438 0 740.8317
788.4371948242188 0 2817.9875
789.4425048828125 0 1462.0796
790.4141235351562 0 6512.189
791.4188842773438 0 90380.64
792.4183349609375 0 58138.277
793.4165649414062 0 22467.098
794.4139404296875 0 5501.924
796.3729248046875 0 6301.8936
797.3723754882812 0 1892.083
798.3765869140625 0 829.4495
806.4310913085938 0 1402.4717
808.4467163085938 0 752.91583
809.4057006835938 0 26497.79
810.4083862304688 0 11088.779
811.4085083007812 0 4009.912
817.3974609375 0 4477.2397
818.4014892578125 0 2064.2896
819.4002685546875 0 1007.6509
820.3912963867188 0 909.4873
823.4069213867188 0 1610.3118
824.4362182617188 0 1676.3237
834.424560546875 0 4042.543
835.4140014648438 0 92500.77
836.4176025390625 0 35410.043
837.4163818359375 0 17592.234
838.417724609375 0 2502.136
851.4273071289062 0 36985.05
852.4346923828125 0 126724.53
853.4374389648438 0 48530.79
854.4379272460938 0 16284.818
855.4360961914062 0 1477.8884
1207.0101318359375 0 689.1014
2805.530517578125 0 706.79504
3059.7509765625 0 708.08655
3078.311767578125 0 814.1146

Spectrum Details

|  |  |
| --- | --- |
| Matched peaks? Matched peaks The total absolute number of peaks matched. Additionally in brackets the total fraction of peaks matched and the total number of peaks is shown. | 22 (11.46% of 192) |
| FDR? FDR The false discovery rate estimated for this peptide. It is calculated by matching all theoretical fragments with a non-integer shift with the raw peaks for this spectrum. This is done with 40 different shifts. The resulting percentage is the average number of annotated peaks over the number of annotated peaks with the correct spectrum. | 0.54% |
| Satellite FDR? Satellite FDR See the FDR for details on its calculation. This satellite ion specific FDR only contains the satellite ions (d/w) for I/L/J positions. | 0.00% |
| PSM Score? PSM Score The PSM Score as given by Hecklib to this annotated spectrum. It is shown with three significant figures. | 205 |

#### Spectrum 4286? Spectrum 4286 The raw spectrum of this peptide as annotated by Hecklib. The fragments are coloured according to ion type (see legend). Any peaks with a star '\*' as text can be hovered over to see the full details, first the ion type second the mass shift type. By hovering over the amino acids in the peptide or ions in the legend the corresponding peaks are highlighted. By toggling the 'Unassigned' label you can turn the background (unassigned) peaks on or off in the plot. By updating the slider in the Ion legend you can update the spectrum to only show the top X% of the peaks with labels. The top X% means any peak that is within X% of the highest intensity. By dragging in the spectrum you can zoom in to a specific part of the spectrum and use 'Zoom Out' to get back to the original zoom level. The annotation of the spectrum is based on the given sequence in the peptides file and is done with different software so inconsistencies are likely. The peaks are annotated based on the given sequence, with 20 ppm tolerance.

Copy Data

##### Spectrum 4286 (TSV)

###### Preview

```
Loading example...
```

*Click on the button to copy the data to your clipboard.*

Mz MinMz MaxIntensity Max

WidthHeightPeptide font sizePeptide stroke widthSpectrum font sizeSpectrum stroke widthCompact peptide

Ion legend

wxyz

abcd

OtherUnassignedIon^Charge^_Position_Show for top:%

DTJMJSR

03.39e+46.79e+41.02e+51.36e+5

Zoom Out

y^+1^_1_a^+1^_2_y^+1^_1_y^+2^_3_a^+1^_2_b^+1^_2_b^+1^_2_y^+1^_2_y^+1^_2_y^+2^_4_y^+1^_2_a^+1^_3_a^+1^_3_b^+1^_3_y^+2^_5_b^+1^_3_y^+1^_3_y^+1^_3_y^+2^_6_y^+2^_6_y^+1^_3_\*\*b^+1^_4_b^+1^_4_y^+1^_4_y^+1^_4_y^+1^_4_b^+1^_5_b^+1^_5_y^+1^_5_y^+1^_5_y^+1^_6_

0779155723363114

Fragment Matches Table

Show background peaks

| Position | Ion type | Intensity | mz Theoretical | mz Error (Th) | mz Error (ppm) | Charge | Series Number |
| --- | --- | --- | --- | --- | --- | --- | --- |
| - | - | 5314 | 120.1 | - | - | 0 | - |
| - | - | 2261 | 124.1 | - | - | 0 | - |
| - | - | 1159 | 125.1 | - | - | 0 | - |
| - | - | 434.1 | 125.4 | - | - | 0 | - |
| - | - | 2.202E+04 | 126.1 | - | - | 0 | - |
| - | - | 388.4 | 126.1 | - | - | 0 | - |
| - | - | 958.8 | 127.1 | - | - | 0 | - |
| - | - | 1478 | 127.1 | - | - | 0 | - |
| - | - | 8911 | 129.1 | - | - | 0 | - |
| - | - | 1277 | 129.1 | - | - | 0 | - |
| - | - | 550.3 | 129.1 | - | - | 0 | - |
| - | - | 1380 | 130 | - | - | 0 | - |
| - | - | 8096 | 130.1 | - | - | 0 | - |
| - | - | 405.4 | 130.1 | - | - | 0 | - |
| - | - | 2212 | 130.1 | - | - | 0 | - |
| - | - | 760.8 | 133.1 | - | - | 0 | - |
| - | - | 612.7 | 133.1 | - | - | 0 | - |
| - | - | 517.7 | 134.1 | - | - | 0 | - |
| - | - | 834.2 | 136 | - | - | 0 | - |
| - | - | 564.8 | 136.1 | - | - | 0 | - |
| - | - | 934.9 | 138.1 | - | - | 0 | - |
| - | - | 1098 | 139.1 | - | - | 0 | - |
| - | - | 639 | 139.1 | - | - | 0 | - |
| - | - | 504.4 | 140.3 | - | - | 0 | - |
| - | - | 1099 | 141.1 | - | - | 0 | - |
| - | - | 3125 | 141.1 | - | - | 0 | - |
| - | - | 1263 | 142.1 | - | - | 0 | - |
| - | - | 419.6 | 142.5 | - | - | 0 | - |
| - | - | 1634 | 143.1 | - | - | 0 | - |
| - | - | 428.1 | 143.9 | - | - | 0 | - |
| - | - | 8.638E+04 | 144.1 | - | - | 0 | - |
| - | - | 5440 | 145.1 | - | - | 0 | - |
| - | - | 437.4 | 146.2 | - | - | 0 | - |
| - | - | 528 | 148.8 | - | - | 0 | - |
| - | - | 478.1 | 148.8 | - | - | 0 | - |
| - | - | 546.6 | 148.9 | - | - | 0 | - |
| - | - | 575.1 | 148.9 | - | - | 0 | - |
| - | - | 637.4 | 148.9 | - | - | 0 | - |
| - | - | 779.1 | 148.9 | - | - | 0 | - |
| - | - | 1146 | 148.9 | - | - | 0 | - |
| - | - | 965.5 | 148.9 | - | - | 0 | - |
| - | - | 1541 | 148.9 | - | - | 0 | - |
| - | - | 3588 | 148.9 | - | - | 0 | - |
| - | - | 5754 | 149 | - | - | 0 | - |
| - | - | 3530 | 149 | - | - | 0 | - |
| - | - | 1332 | 149 | - | - | 0 | - |
| - | - | 1149 | 149 | - | - | 0 | - |
| - | - | 930 | 149 | - | - | 0 | - |
| - | - | 680 | 149 | - | - | 0 | - |
| - | - | 754.5 | 149 | - | - | 0 | - |
| - | - | 540.4 | 149 | - | - | 0 | - |
| - | - | 455.7 | 149 | - | - | 0 | - |
| - | - | 481.5 | 149.1 | - | - | 0 | - |
| - | - | 1904 | 152.1 | - | - | 0 | - |
| - | - | 5975 | 153.1 | - | - | 0 | - |
| - | - | 440.6 | 153.1 | - | - | 0 | - |
| - | - | 455.5 | 153.3 | - | - | 0 | - |
| - | - | 4235 | 154.1 | - | - | 0 | - |
| - | - | 455.3 | 154.1 | - | - | 0 | - |
| - | - | 547 | 155.1 | - | - | 0 | - |
| - | - | 694.7 | 157.1 | - | - | 0 | - |
| - | - | 1199 | 157.1 | - | - | 0 | - |
| - | - | 1045 | 157.1 | - | - | 0 | - |
| - | - | 439.3 | 157.9 | - | - | 0 | - |
| 7 | y | 1.751E+04 | 158.1 | 0.000111 | 0.7019 | +1 | 1 |
| - | - | 1195 | 166.1 | - | - | 0 | - |
| - | - | 602 | 169.1 | - | - | 0 | - |
| - | - | 2.243E+04 | 169.1 | - | - | 0 | - |
| - | - | 6547 | 170.1 | - | - | 0 | - |
| - | - | 791 | 170.1 | - | - | 0 | - |
| - | - | 1477 | 170.1 | - | - | 0 | - |
| 2 | a | 1.409E+04 | 171.1 | 8.892E-05 | 0.5198 | +1 | 2 |
| - | - | 774.1 | 171.1 | - | - | 0 | - |
| - | - | 1126 | 172.1 | - | - | 0 | - |
| - | - | 751 | 172.1 | - | - | 0 | - |
| - | - | 656.9 | 173.1 | - | - | 0 | - |
| - | - | 1687 | 173.1 | - | - | 0 | - |
| - | - | 785.9 | 175.1 | - | - | 0 | - |
| 7 | y | 5.265E+04 | 175.1 | 0.0001122 | 0.6405 | +1 | 1 |
| - | - | 3216 | 176.1 | - | - | 0 | - |
| 5 | y | 1260 | 179.1 | 0.002225 | 12.42 | +2 | 3 |
| - | - | 2.665E+04 | 180.1 | - | - | 0 | - |
| - | - | 1077 | 181.1 | - | - | 0 | - |
| - | - | 673.8 | 181.1 | - | - | 0 | - |
| - | - | 2113 | 181.1 | - | - | 0 | - |
| - | - | 1146 | 182.1 | - | - | 0 | - |
| - | - | 1089 | 183.1 | - | - | 0 | - |
| - | - | 1106 | 184.1 | - | - | 0 | - |
| - | - | 1410 | 185.1 | - | - | 0 | - |
| - | - | 509.8 | 185.1 | - | - | 0 | - |
| - | - | 2444 | 185.1 | - | - | 0 | - |
| - | - | 2.531E+04 | 187.1 | - | - | 0 | - |
| - | - | 1904 | 188.1 | - | - | 0 | - |
| 2 | a | 9.406E+04 | 189.1 | 6.805E-05 | 0.3599 | +1 | 2 |
| - | - | 5775 | 190.1 | - | - | 0 | - |
| - | - | 555.5 | 191.1 | - | - | 0 | - |
| - | - | 3377 | 191.1 | - | - | 0 | - |
| - | - | 858 | 193.1 | - | - | 0 | - |
| - | - | 994.2 | 195.1 | - | - | 0 | - |
| - | - | 518.9 | 195.1 | - | - | 0 | - |
| - | - | 2.636E+04 | 197.1 | - | - | 0 | - |
| - | - | 2821 | 198.1 | - | - | 0 | - |
| - | - | 2116 | 198.1 | - | - | 0 | - |
| 2 | b | 3.075E+04 | 199.1 | 3.208E-05 | 0.1612 | +1 | 2 |
| - | - | 542.4 | 199.1 | - | - | 0 | - |
| - | - | 745.2 | 200.1 | - | - | 0 | - |
| - | - | 2073 | 200.1 | - | - | 0 | - |
| - | - | 3189 | 201.1 | - | - | 0 | - |
| - | - | 695 | 201.1 | - | - | 0 | - |
| - | - | 4972 | 202.1 | - | - | 0 | - |
| - | - | 1666 | 203.1 | - | - | 0 | - |
| - | - | 1574 | 203.1 | - | - | 0 | - |
| - | - | 494.5 | 204.9 | - | - | 0 | - |
| - | - | 508.1 | 206 | - | - | 0 | - |
| - | - | 1299 | 207.1 | - | - | 0 | - |
| - | - | 1576 | 210.1 | - | - | 0 | - |
| - | - | 975.6 | 211.1 | - | - | 0 | - |
| - | - | 911.8 | 211.1 | - | - | 0 | - |
| - | - | 594 | 215.1 | - | - | 0 | - |
| - | - | 4737 | 215.1 | - | - | 0 | - |
| 2 | b | 5.281E+04 | 217.1 | 1.122E-05 | 0.05169 | +1 | 2 |
| - | - | 4497 | 218.1 | - | - | 0 | - |
| - | - | 671 | 219.1 | - | - | 0 | - |
| - | - | 1733 | 220.1 | - | - | 0 | - |
| - | - | 655.8 | 221.1 | - | - | 0 | - |
| - | - | 637.5 | 223.1 | - | - | 0 | - |
| - | - | 773.1 | 224.1 | - | - | 0 | - |
| - | - | 5923 | 225.1 | - | - | 0 | - |
| - | - | 526.5 | 225.1 | - | - | 0 | - |
| - | - | 993.2 | 226.1 | - | - | 0 | - |
| - | - | 3629 | 227.1 | - | - | 0 | - |
| - | - | 1027 | 228.1 | - | - | 0 | - |
| - | - | 1081 | 229 | - | - | 0 | - |
| - | - | 1018 | 229.1 | - | - | 0 | - |
| - | - | 1.045E+04 | 229.6 | - | - | 0 | - |
| - | - | 1211 | 230.1 | - | - | 0 | - |
| - | - | 1786 | 230.1 | - | - | 0 | - |
| - | - | 1220 | 232.1 | - | - | 0 | - |
| - | - | 7369 | 233.1 | - | - | 0 | - |
| - | - | 577.2 | 234.1 | - | - | 0 | - |
| - | - | 1686 | 238.2 | - | - | 0 | - |
| - | - | 1069 | 239.1 | - | - | 0 | - |
| - | - | 1451 | 240.1 | - | - | 0 | - |
| - | - | 525.1 | 242.1 | - | - | 0 | - |
| - | - | 626.7 | 243.6 | - | - | 0 | - |
| - | - | 3875 | 244.1 | - | - | 0 | - |
| 6 | y | 2922 | 244.1 | 0.0002181 | 0.8935 | +1 | 2 |
| 6 | y | 3.48E+04 | 245.1 | 4.973E-05 | 0.2029 | +1 | 2 |
| - | - | 512.3 | 246.1 | - | - | 0 | - |
| - | - | 4095 | 246.1 | - | - | 0 | - |
| - | - | 1950 | 249.1 | - | - | 0 | - |
| - | - | 1534 | 251.1 | - | - | 0 | - |
| - | - | 746.8 | 252.1 | - | - | 0 | - |
| - | - | 535.9 | 254.1 | - | - | 0 | - |
| - | - | 618.6 | 254.1 | - | - | 0 | - |
| - | - | 869.1 | 256.1 | - | - | 0 | - |
| - | - | 521.8 | 256.2 | - | - | 0 | - |
| - | - | 1.295E+04 | 261.1 | - | - | 0 | - |
| 4 | y | 3687 | 261.6 | 0.002422 | 9.256 | +2 | 4 |
| 6 | y | 5.949E+04 | 262.2 | 0.0001017 | 0.3878 | +1 | 2 |
| - | - | 475.7 | 263.1 | - | - | 0 | - |
| - | - | 5101 | 263.2 | - | - | 0 | - |
| - | - | 479.9 | 263.3 | - | - | 0 | - |
| - | - | 788 | 264.2 | - | - | 0 | - |
| - | - | 697.9 | 265.2 | - | - | 0 | - |
| - | - | 2422 | 266.1 | - | - | 0 | - |
| - | - | 3466 | 267.1 | - | - | 0 | - |
| - | - | 788.4 | 267.2 | - | - | 0 | - |
| - | - | 856.7 | 268.1 | - | - | 0 | - |
| - | - | 1053 | 270.1 | - | - | 0 | - |
| - | - | 1178 | 271 | - | - | 0 | - |
| - | - | 1601 | 272.1 | - | - | 0 | - |
| - | - | 722.1 | 277.1 | - | - | 0 | - |
| - | - | 730.3 | 277.2 | - | - | 0 | - |
| - | - | 730.3 | 277.7 | - | - | 0 | - |
| - | - | 841.9 | 280.2 | - | - | 0 | - |
| - | - | 1344 | 282.2 | - | - | 0 | - |
| 3 | a | 2.667E+04 | 284.2 | 0.0001128 | 0.3968 | +1 | 3 |
| - | - | 970.2 | 285.1 | - | - | 0 | - |
| - | - | 3271 | 285.2 | - | - | 0 | - |
| - | - | 663.9 | 286.1 | - | - | 0 | - |
| - | - | 2.17E+04 | 286.2 | - | - | 0 | - |
| - | - | 4532 | 286.7 | - | - | 0 | - |
| - | - | 1006 | 287.1 | - | - | 0 | - |
| - | - | 815.4 | 291.2 | - | - | 0 | - |
| - | - | 1926 | 293.9 | - | - | 0 | - |
| - | - | 1850 | 294.1 | - | - | 0 | - |
| - | - | 1486 | 294.9 | - | - | 0 | - |
| - | - | 1616 | 295.1 | - | - | 0 | - |
| - | - | 3864 | 297.1 | - | - | 0 | - |
| - | - | 1853 | 299.2 | - | - | 0 | - |
| - | - | 1373 | 299.2 | - | - | 0 | - |
| - | - | 1114 | 299.7 | - | - | 0 | - |
| 3 | a | 1431 | 302.2 | 0.0002404 | 0.7957 | +1 | 3 |
| - | - | 1155 | 308.2 | - | - | 0 | - |
| - | - | 713.9 | 309.2 | - | - | 0 | - |
| - | - | 1477 | 310.2 | - | - | 0 | - |
| - | - | 4200 | 311.9 | - | - | 0 | - |
| 3 | b | 3.939E+04 | 312.2 | 1.746E-06 | 0.005592 | +1 | 3 |
| - | - | 612.7 | 312.9 | - | - | 0 | - |
| - | - | 5453 | 313.2 | - | - | 0 | - |
| 3 | y | 4.432E+04 | 318.2 | 0.002412 | 7.582 | +2 | 5 |
| - | - | 1.553E+04 | 318.7 | - | - | 0 | - |
| - | - | 5884 | 319.2 | - | - | 0 | - |
| - | - | 1558 | 319.7 | - | - | 0 | - |
| - | - | 619.3 | 320.2 | - | - | 0 | - |
| - | - | 764.7 | 323.2 | - | - | 0 | - |
| - | - | 745 | 325.2 | - | - | 0 | - |
| - | - | 698 | 326.1 | - | - | 0 | - |
| - | - | 1242 | 327.7 | - | - | 0 | - |
| 3 | b | 1.214E+04 | 330.2 | 5.369E-05 | 0.1626 | +1 | 3 |
| - | - | 1984 | 331.2 | - | - | 0 | - |
| - | - | 1193 | 331.2 | - | - | 0 | - |
| - | - | 1702 | 332.7 | - | - | 0 | - |
| - | - | 3812 | 336.7 | - | - | 0 | - |
| - | - | 899.2 | 337.2 | - | - | 0 | - |
| - | - | 534.5 | 339.2 | - | - | 0 | - |
| - | - | 2325 | 340.2 | - | - | 0 | - |
| - | - | 3085 | 341.7 | - | - | 0 | - |
| - | - | 1028 | 342.2 | - | - | 0 | - |
| - | - | 665.2 | 345.2 | - | - | 0 | - |
| - | - | 2275 | 346.1 | - | - | 0 | - |
| - | - | 1068 | 348.2 | - | - | 0 | - |
| - | - | 768.6 | 353.9 | - | - | 0 | - |
| - | - | 1195 | 354.7 | - | - | 0 | - |
| - | - | 2306 | 354.9 | - | - | 0 | - |
| - | - | 1040 | 355.7 | - | - | 0 | - |
| - | - | 674.7 | 355.9 | - | - | 0 | - |
| 5 | y | 6842 | 357.2 | 0.0004503 | 1.261 | +1 | 3 |
| 5 | y | 6749 | 358.2 | 0.000274 | 0.765 | +1 | 3 |
| - | - | 998.1 | 358.2 | - | - | 0 | - |
| - | - | 655.2 | 359.2 | - | - | 0 | - |
| 2 | y | 2244 | 359.7 | 0.0008232 | 2.289 | +2 | 6 |
| - | - | 976.5 | 362.2 | - | - | 0 | - |
| - | - | 5073 | 362.2 | - | - | 0 | - |
| - | - | 1052 | 362.7 | - | - | 0 | - |
| - | - | 1139 | 363.2 | - | - | 0 | - |
| - | - | 2001 | 364.7 | - | - | 0 | - |
| - | - | 1126 | 365.2 | - | - | 0 | - |
| - | - | 723.2 | 367.2 | - | - | 0 | - |
| - | - | 1967 | 367.7 | - | - | 0 | - |
| - | - | 899.1 | 368.2 | - | - | 0 | - |
| 2 | y | 6700 | 368.7 | 0.001919 | 5.205 | +2 | 6 |
| - | - | 1419 | 369.2 | - | - | 0 | - |
| - | - | 1280 | 371.9 | - | - | 0 | - |
| - | - | 3491 | 372.2 | - | - | 0 | - |
| - | - | 1116 | 372.9 | - | - | 0 | - |
| - | - | 704.7 | 373.2 | - | - | 0 | - |
| - | - | 598.1 | 373.4 | - | - | 0 | - |
| - | - | 2282 | 373.7 | - | - | 0 | - |
| 5 | y | 1.092E+05 | 375.2 | 0.0003339 | 0.8897 | +1 | 3 |
| - | - | 707.2 | 376.2 | - | - | 0 | - |
| - | - | 1202 | 376.2 | - | - | 0 | - |
| - | - | 1.633E+04 | 376.2 | - | - | 0 | - |
| - | - | 800.2 | 376.7 | - | - | 0 | - |
| - | - | 2531 | 377.2 | - | - | 0 | - |
| - | - | 1.579E+04 | 385.2 | - | - | 0 | - |
| - | - | 5033 | 385.7 | - | - | 0 | - |
| - | - | 1956 | 386.2 | - | - | 0 | - |
| - | - | 1231 | 386.7 | - | - | 0 | - |
| - | - | 819.2 | 387.2 | - | - | 0 | - |
| - | - | 1552 | 390.7 | - | - | 0 | - |
| - | - | 1.7E+04 | 394.2 | - | - | 0 | - |
| - | - | 6739 | 394.7 | - | - | 0 | - |
| - | - | 617.4 | 395.2 | - | - | 0 | - |
| - | - | 1960 | 395.2 | - | - | 0 | - |
| - | - | 2446 | 399.7 | - | - | 0 | - |
| - | - | 1130 | 400.2 | - | - | 0 | - |
| - | - | 1103 | 401.2 | - | - | 0 | - |
| - | - | 8014 | 402.2 | - | - | 0 | - |
| - | - | 1646 | 403.2 | - | - | 0 | - |
| - | - | 1307 | 408.2 | - | - | 0 | - |
| - | - | 1564 | 408.7 | - | - | 0 | - |
| - | - | 802.6 | 413.2 | - | - | 0 | - |
| - | - | 1624 | 416.2 | - | - | 0 | - |
| 0 | Precursor | 2.576E+04 | 417.2 | 0.002305 | 5.526 | +2 | -1 |
| - | - | 9112 | 417.7 | - | - | 0 | - |
| - | - | 2712 | 418.2 | - | - | 0 | - |
| - | - | 3784 | 423.2 | - | - | 0 | - |
| - | - | 884.9 | 424.2 | - | - | 0 | - |
| - | - | 990 | 424.2 | - | - | 0 | - |
| - | - | 2695 | 425.2 | - | - | 0 | - |
| 0 | Precursor | 3025 | 426.2 | 0.002211 | 5.187 | +2 | -1 |
| - | - | 783.2 | 426.7 | - | - | 0 | - |
| - | - | 1011 | 427.2 | - | - | 0 | - |
| - | - | 712.9 | 428.3 | - | - | 0 | - |
| - | - | 1095 | 430.2 | - | - | 0 | - |
| - | - | 2.36E+04 | 431.2 | - | - | 0 | - |
| - | - | 5537 | 432.3 | - | - | 0 | - |
| - | - | 1193 | 433.3 | - | - | 0 | - |
| - | - | 4223 | 440.3 | - | - | 0 | - |
| - | - | 837.5 | 441.2 | - | - | 0 | - |
| - | - | 1.206E+04 | 441.2 | - | - | 0 | - |
| - | - | 3308 | 442.2 | - | - | 0 | - |
| - | - | 637.9 | 443.3 | - | - | 0 | - |
| - | - | 595.4 | 456.3 | - | - | 0 | - |
| - | - | 1.274E+05 | 458.3 | - | - | 0 | - |
| 4 | b | 6380 | 459.2 | 0.004792 | 10.44 | +1 | 4 |
| - | - | 2.866E+04 | 459.3 | - | - | 0 | - |
| - | - | 846.3 | 460.2 | - | - | 0 | - |
| - | - | 4835 | 460.3 | - | - | 0 | - |
| - | - | 1356 | 468.3 | - | - | 0 | - |
| 4 | b | 1.131E+04 | 477.2 | 0.004847 | 10.16 | +1 | 4 |
| - | - | 2850 | 478.2 | - | - | 0 | - |
| - | - | 739 | 479.2 | - | - | 0 | - |
| - | - | 900.9 | 484.3 | - | - | 0 | - |
| - | - | 882.2 | 485.2 | - | - | 0 | - |
| - | - | 2769 | 486.3 | - | - | 0 | - |
| - | - | 1861 | 488.3 | - | - | 0 | - |
| - | - | 8306 | 501.3 | - | - | 0 | - |
| - | - | 2108 | 502.3 | - | - | 0 | - |
| 4 | y | 3785 | 504.3 | 0.004191 | 8.311 | +1 | 4 |
| 4 | y | 679.5 | 505.2 | 0.007541 | 14.93 | +1 | 4 |
| - | - | 2437 | 507.2 | - | - | 0 | - |
| - | - | 592.1 | 508.2 | - | - | 0 | - |
| - | - | 2590 | 515.3 | - | - | 0 | - |
| - | - | 710.6 | 516.3 | - | - | 0 | - |
| - | - | 2238 | 521.2 | - | - | 0 | - |
| 4 | y | 1.344E+05 | 522.3 | 0.004307 | 8.247 | +1 | 4 |
| - | - | 3.132E+04 | 523.3 | - | - | 0 | - |
| - | - | 1.04E+04 | 524.3 | - | - | 0 | - |
| - | - | 1018 | 525.3 | - | - | 0 | - |
| - | - | 891.1 | 526.3 | - | - | 0 | - |
| - | - | 1446 | 528.3 | - | - | 0 | - |
| - | - | 1019 | 529.3 | - | - | 0 | - |
| - | - | 1339 | 532.3 | - | - | 0 | - |
| - | - | 1420 | 543.3 | - | - | 0 | - |
| - | - | 787.1 | 544.3 | - | - | 0 | - |
| - | - | 2344 | 544.3 | - | - | 0 | - |
| - | - | 622 | 545.3 | - | - | 0 | - |
| - | - | 3493 | 553.3 | - | - | 0 | - |
| - | - | 2056 | 554.3 | - | - | 0 | - |
| - | - | 623.5 | 555.3 | - | - | 0 | - |
| - | - | 8.796E+04 | 571.4 | - | - | 0 | - |
| 5 | b | 1184 | 572.3 | 0.006543 | 11.43 | +1 | 5 |
| - | - | 2.599E+04 | 572.4 | - | - | 0 | - |
| - | - | 5031 | 573.4 | - | - | 0 | - |
| - | - | 919.9 | 581.3 | - | - | 0 | - |
| - | - | 640.7 | 583.1 | - | - | 0 | - |
| 5 | b | 1203 | 590.3 | 0.00428 | 7.25 | +1 | 5 |
| - | - | 720.1 | 597.3 | - | - | 0 | - |
| 3 | y | 1581 | 617.3 | 0.004111 | 6.66 | +1 | 5 |
| - | - | 711.9 | 618.3 | - | - | 0 | - |
| - | - | 1375 | 620.3 | - | - | 0 | - |
| - | - | 955.5 | 621.3 | - | - | 0 | - |
| 3 | y | 8.817E+04 | 635.3 | 0.003739 | 5.886 | +1 | 5 |
| - | - | 5005 | 636.3 | - | - | 0 | - |
| - | - | 2.587E+04 | 636.4 | - | - | 0 | - |
| - | - | 9188 | 637.4 | - | - | 0 | - |
| - | - | 2872 | 638.4 | - | - | 0 | - |
| - | - | 949.8 | 639.4 | - | - | 0 | - |
| - | - | 845.6 | 654.4 | - | - | 0 | - |
| - | - | 719.9 | 664.4 | - | - | 0 | - |
| - | - | 1783 | 672.4 | - | - | 0 | - |
| - | - | 7981 | 682.4 | - | - | 0 | - |
| - | - | 2585 | 683.4 | - | - | 0 | - |
| - | - | 791.4 | 684.4 | - | - | 0 | - |
| - | - | 649.4 | 707.9 | - | - | 0 | - |
| 2 | y | 1118 | 736.4 | 0.005133 | 6.971 | +1 | 6 |
| - | - | 527.6 | 737.4 | - | - | 0 | - |
| - | - | 1.252E+04 | 746.4 | - | - | 0 | - |
| - | - | 4520 | 747.4 | - | - | 0 | - |
| - | - | 1374 | 748.4 | - | - | 0 | - |
| - | - | 650.3 | 769.3 | - | - | 0 | - |
| - | - | 774.6 | 783.3 | - | - | 0 | - |
| - | - | 706.3 | 3081 | - | - | 0 | - |
| - | - | 719.8 | 3083 | - | - | 0 | - |

m/z Charge Intensity FragmentType MassShift Position
120.08092498779297 0 5313.9824
124.11226654052734 0 2260.9197
125.07107543945312 0 1159.1686
125.42955017089844 0 434.08704
126.05512237548828 0 22018.695
126.12840270996094 0 388.35092
127.05046081542969 0 958.8166
127.0585708618164 0 1478.1494
129.0659942626953 0 8911.425
129.10256958007812 0 1277.2078
129.11375427246094 0 550.34
130.0498504638672 0 1379.8744
130.06121826171875 0 8096.4663
130.0858154296875 0 405.35263
130.09756469726562 0 2211.741
133.06082153320312 0 760.79297
133.09698486328125 0 612.7426
134.14166259765625 0 517.7275
136.03961181640625 0 834.1979
136.07568359375 0 564.7651
138.0551300048828 0 934.9225
139.05007934570312 0 1097.599
139.11187744140625 0 639.02783
140.2751922607422 0 504.44376
141.10235595703125 0 1098.5098
141.1387939453125 0 3125.4854
142.12277221679688 0 1263.1484
142.52206420898438 0 419.55325
143.11807250976562 0 1633.6589
143.88905334472656 0 428.13733
144.06568908691406 0 86383.56
145.06900024414062 0 5439.925
146.23619079589844 0 437.4075
148.8338165283203 0 528.0469
148.84048461914062 0 478.1197
148.88304138183594 0 546.5994
148.8977508544922 0 575.0952
148.9048309326172 0 637.43335
148.91184997558594 0 779.1293
148.9191131591797 0 1145.5148
148.92669677734375 0 965.5398
148.9329376220703 0 1540.7076
148.94049072265625 0 3588.1907
148.9568634033203 0 5754.093
148.96456909179688 0 3529.975
148.97198486328125 0 1331.6282
148.97901916503906 0 1149.3098
148.98599243164062 0 929.9739
148.9932403564453 0 679.999
149.00039672851562 0 754.5113
149.0146942138672 0 540.35504
149.0218963623047 0 455.70526
149.14256286621094 0 481.5399
152.10714721679688 0 1903.8416
153.0659942626953 0 5974.8936
153.13938903808594 0 440.56213
153.32797241210938 0 455.51773
154.050048828125 0 4234.5815
154.0694122314453 0 455.32233
155.0706787109375 0 546.9761
157.0976104736328 0 694.6791
157.10841369628906 0 1199.0044
157.13377380371094 0 1044.8097
157.86080932617188 0 439.27487
158.09251403808594 0 17506.826 y Ammonia loss 6
166.05308532714844 0 1195.2006
169.0973358154297 0 602.0343
169.1336669921875 0 22429.13
170.09243774414062 0 6546.789
170.10546875 0 790.95087
170.13702392578125 0 1476.5107
171.07650756835938 0 14088.4375 a Water loss 1
171.11338806152344 0 774.0883
172.06048583984375 0 1125.984
172.0795440673828 0 751.0234
173.0557861328125 0 656.8834
173.12872314453125 0 1687.4869
175.08644104003906 0 785.9066
175.1190643310547 0 52654.72 y 6
176.12232971191406 0 3216.2053
179.11810302734375 0 1260.4432 y Water loss 4
180.10195922851562 0 26650.094
181.06076049804688 0 1076.7604
181.09744262695312 0 673.7892
181.1055450439453 0 2113.2046
182.0923614501953 0 1145.9191
183.11329650878906 0 1089.2183
184.1080780029297 0 1106.2684
185.09176635742188 0 1409.8855
185.1179962158203 0 509.8338
185.13980102539062 0 2444.1558
187.1441650390625 0 25308.826
188.14756774902344 0 1904.06
189.08705139160156 0 94057.84 a 1
190.09039306640625 0 5775.4585
191.09129333496094 0 555.4937
191.11802673339844 0 3376.7288
193.09664916992188 0 858.0115
195.1126251220703 0 994.23596
195.14852905273438 0 518.85516
197.12850952148438 0 26363.363
198.08717346191406 0 2821.0208
198.13201904296875 0 2115.9993
199.0713653564453 0 30746.352 b Water loss 1
199.1187286376953 0 542.36
200.0555877685547 0 745.1596
200.0750274658203 0 2073.3198
201.12330627441406 0 3188.6926
201.1331024169922 0 695.0187
202.11871337890625 0 4971.8164
203.1026611328125 0 1666.3116
203.12754821777344 0 1573.5616
204.8888397216797 0 494.46408
205.96353149414062 0 508.12616
207.11192321777344 0 1299.3915
210.08775329589844 0 1575.5854
211.10800170898438 0 975.6496
211.1438751220703 0 911.7724
215.1215057373047 0 593.97485
215.1388702392578 0 4737.1934
217.0819091796875 0 52809.188 b 1
218.08535766601562 0 4497.113
219.11212158203125 0 671.04956
220.1290283203125 0 1733.2072
221.12916564941406 0 655.75336
223.107421875 0 637.48016
224.13929748535156 0 773.07324
225.1234130859375 0 5923.0034
225.1433563232422 0 526.4827
226.1292266845703 0 993.238
227.11376953125 0 3628.5693
228.09780883789062 0 1026.9769
229.0013427734375 0 1081.4713
229.1299591064453 0 1017.55383
229.63973999023438 0 10451.06
230.12490844726562 0 1210.6945
230.1412811279297 0 1785.6075
232.14019775390625 0 1219.9783
233.13182067871094 0 7369.281
234.13487243652344 0 577.1596
238.15499877929688 0 1686.167
239.13864135742188 0 1069.3632
240.13381958007812 0 1451.4338
242.12611389160156 0 525.1076
243.63645935058594 0 626.69135
244.10009765625 0 3874.7244
244.14019775390625 0 2921.9534 y Water loss 5
245.12448120117188 0 34797.82 y Ammonia loss 5
246.10911560058594 0 512.312
246.1294403076172 0 4094.5293
249.1233367919922 0 1950.3024
251.1024169921875 0 1534.2406
252.13519287109375 0 746.80756
254.12620544433594 0 535.8702
254.14920043945312 0 618.6305
256.1293640136719 0 869.12616
256.16534423828125 0 521.77246
261.1265869140625 0 12952.88
261.6388244628906 0 3686.9666 y 3
262.15087890625 0 59493.984 y 5
263.1231384277344 0 475.70847
263.1539611816406 0 5101.018
263.2652587890625 0 479.88242
264.1543884277344 0 787.9939
265.190673828125 0 697.87494
266.14947509765625 0 2421.7488
267.1339416503906 0 3466.0952
267.15093994140625 0 788.3711
268.1292419433594 0 856.7409
270.120361328125 0 1053.3718
271.02685546875 0 1177.7142
272.13494873046875 0 1600.9227
277.11962890625 0 722.11285
277.1768493652344 0 730.3499
277.6679382324219 0 730.31805
280.16485595703125 0 841.8701
282.21759033203125 0 1343.8608
284.1603698730469 0 26665.883 a Water loss 2
285.1463317871094 0 970.16943
285.16375732421875 0 3271.3545
286.1409912109375 0 663.851
286.1816711425781 0 21698.04
286.6831970214844 0 4532.2095
287.1052551269531 0 1006.0942
291.17425537109375 0 815.3624
293.93475341796875 0 1925.8959
294.1449890136719 0 1849.7734
294.9178771972656 0 1485.5923
295.12921142578125 0 1616.1632
297.05828857421875 0 3863.7083
299.171630859375 0 1852.5616
299.1904296875 0 1372.7095
299.67303466796875 0 1114.1344
302.1708068847656 0 1431.3079 a 2
308.1606750488281 0 1154.7119
309.2043151855469 0 713.9464
310.2113342285156 0 1477.3124
311.9451904296875 0 4200.1606
312.1553955078125 0 39391.91 b Water loss 2
312.9459533691406 0 612.6647
313.15850830078125 0 5452.8154
318.18084716796875 0 44321.07 y 2
318.6822204589844 0 15529.149
319.1818542480469 0 5884.0913
319.68438720703125 0 1558.2703
320.185546875 0 619.34314
323.1722106933594 0 764.73315
325.1877136230469 0 744.99567
326.14581298828125 0 698.0304
327.7003479003906 0 1241.956
330.166015625 0 12140.147 b 2
331.16864013671875 0 1983.8092
331.18865966796875 0 1193.0621
332.6920471191406 0 1701.7946
336.7052307128906 0 3812.3923
337.2065734863281 0 899.1623
339.2123107910156 0 534.526
340.1975402832031 0 2324.8745
341.6976013183594 0 3084.9377
342.198486328125 0 1027.5665
345.22332763671875 0 665.2448
346.1423034667969 0 2274.8062
348.15838623046875 0 1067.5616
353.9027404785156 0 768.5853
354.686279296875 0 1195.151
354.8851013183594 0 2305.9045
355.6937561035156 0 1040.4829
355.8843994140625 0 674.65485
357.2240295410156 0 6842.331 y Water loss 4
358.2082214355469 0 6749.2017 y Ammonia loss 4
358.2281494140625 0 998.11414
359.2127380371094 0 655.2299
359.69781494140625 0 2244.1692 y Water loss 1
362.1745300292969 0 976.4975
362.21063232421875 0 5072.8716
362.71209716796875 0 1051.6406
363.1984558105469 0 1138.7803
364.6911315917969 0 2001.0728
365.1920166015625 0 1126.3607
367.2054748535156 0 723.1787
367.6944274902344 0 1967.0103
368.192626953125 0 899.06757
368.7041931152344 0 6699.707 y 1
369.20623779296875 0 1418.5261
371.91015625 0 1280.0155
372.158203125 0 3491.3904
372.894775390625 0 1116.1055
373.1626892089844 0 704.6874
373.3951110839844 0 598.08356
373.6967468261719 0 2282.297
375.2347106933594 0 109166.41 y 4
376.153564453125 0 707.2188
376.208251953125 0 1201.7974
376.2374267578125 0 16332.129
376.7071228027344 0 800.1607
377.2394714355469 0 2531.266
385.2157897949219 0 15792.221
385.7147216796875 0 5032.895
386.2205810546875 0 1956.0979
386.6850891113281 0 1231.1503
387.1917419433594 0 819.2461
390.6875 0 1552.3018
394.2166442871094 0 16996.008
394.718505859375 0 6738.92
395.1915588378906 0 617.43317
395.21990966796875 0 1960.4326
399.6945495605469 0 2445.7815
400.1944274902344 0 1130.0432
401.2140808105469 0 1103.4763
402.2093811035156 0 8013.96
403.21246337890625 0 1645.6549
408.20684814453125 0 1306.8605
408.70379638671875 0 1563.9741
413.20294189453125 0 802.636
416.2259826660156 0 1624.1827
417.2127685546875 0 25759.492 Precursor Water loss
417.71435546875 0 9112.357
418.2140808105469 0 2712.4211
423.2344665527344 0 3783.654
424.15130615234375 0 884.8956
424.21502685546875 0 989.97363
425.2141418457031 0 2694.9485
426.21795654296875 0 3025.3513 Precursor
426.71856689453125 0 783.21875
427.2215576171875 0 1010.84155
428.2639465332031 0 712.9092
430.23992919921875 0 1095.1399
431.2483215332031 0 23598.232
432.2527160644531 0 5536.9067
433.2536315917969 0 1192.7628
440.2613220214844 0 4222.624
441.18109130859375 0 837.4578
441.2454833984375 0 12056.131
442.24847412109375 0 3308.3984
443.2506408691406 0 637.9191
456.2561950683594 0 595.3669
458.2720031738281 0 127365.45
459.190673828125 0 6379.9287 b Water loss 3
459.2748107910156 0 28656.47
460.1936950683594 0 846.2554
460.2770080566406 0 4835.345
468.2568359375 0 1356.0873
477.2012939453125 0 11312.011 b 3
478.2035217285156 0 2849.888
479.1997985839844 0 738.97296
484.25164794921875 0 900.91064
485.2463073730469 0 882.1659
486.2666320800781 0 2769.374
488.2691955566406 0 1861.2349
501.2771301269531 0 8305.507
502.2808532714844 0 2107.796
504.2591552734375 0 3785.1484 y Water loss 3
505.24652099609375 0 679.46185 y Ammonia loss 3
507.2464599609375 0 2437.0083
508.24560546875 0 592.11456
515.2926025390625 0 2590.4038
516.295654296875 0 710.5552
521.2465209960938 0 2238.0654
522.2698364257812 0 134370.75 y 3
523.2725830078125 0 31322.047
524.2701416015625 0 10400.7295
525.2720336914062 0 1017.6255
526.288818359375 0 891.1232
528.3126220703125 0 1446.4865
529.3123779296875 0 1019.3585
532.2531127929688 0 1338.7948
543.3241577148438 0 1419.6599
544.283935546875 0 787.09827
544.330322265625 0 2343.7012
545.3355712890625 0 621.95734
553.3443603515625 0 3492.7964
554.330322265625 0 2056.0996
555.3377685546875 0 623.49805
571.3552856445312 0 87955.34
572.2764892578125 0 1184.284 b Water loss 4
572.3585815429688 0 25993.775
573.3602905273438 0 5030.5283
581.3399658203125 0 919.9436
583.1469116210938 0 640.6718
590.2847900390625 0 1202.5955 b 4
597.3370971679688 0 720.05286
617.3431396484375 0 1580.8281 y Water loss 2
618.3369140625 0 711.8787
620.3311767578125 0 1374.9934
621.3329467773438 0 955.456
635.3533325195312 0 88167.98 y 2
636.27197265625 0 5005.047
636.3562622070312 0 25865.002
637.3553466796875 0 9187.542
638.359375 0 2872.1816
639.3614501953125 0 949.8049
654.3911743164062 0 845.55383
664.3738403320312 0 719.8546
672.4025268554688 0 1783.0693
682.3869018554688 0 7981.3003
683.3893432617188 0 2584.9717
684.392822265625 0 791.3661
707.8651123046875 0 649.37665
736.4024047851562 0 1117.587 y 1
737.4042358398438 0 527.58527
746.384765625 0 12520.391
747.3882446289062 0 4519.8604
748.3863525390625 0 1374.4004
769.3451538085938 0 650.32465
783.341796875 0 774.56085
3081.344970703125 0 706.3143
3083.234619140625 0 719.80914

Spectrum Details

|  |  |
| --- | --- |
| Matched peaks? Matched peaks The total absolute number of peaks matched. Additionally in brackets the total fraction of peaks matched and the total number of peaks is shown. | 33 (8.99% of 367) |
| FDR? FDR The false discovery rate estimated for this peptide. It is calculated by matching all theoretical fragments with a non-integer shift with the raw peaks for this spectrum. This is done with 40 different shifts. The resulting percentage is the average number of annotated peaks over the number of annotated peaks with the correct spectrum. | 0.43% |
| Satellite FDR? Satellite FDR See the FDR for details on its calculation. This satellite ion specific FDR only contains the satellite ions (d/w) for I/L/J positions. | - |
| PSM Score? PSM Score The PSM Score as given by Hecklib to this annotated spectrum. It is shown with three significant figures. | 448 |

#### Spectrum 4554? Spectrum 4554 The raw spectrum of this peptide as annotated by Hecklib. The fragments are coloured according to ion type (see legend). Any peaks with a star '\*' as text can be hovered over to see the full details, first the ion type second the mass shift type. By hovering over the amino acids in the peptide or ions in the legend the corresponding peaks are highlighted. By toggling the 'Unassigned' label you can turn the background (unassigned) peaks on or off in the plot. By updating the slider in the Ion legend you can update the spectrum to only show the top X% of the peaks with labels. The top X% means any peak that is within X% of the highest intensity. By dragging in the spectrum you can zoom in to a specific part of the spectrum and use 'Zoom Out' to get back to the original zoom level. The annotation of the spectrum is based on the given sequence in the peptides file and is done with different software so inconsistencies are likely. The peaks are annotated based on the given sequence, with 20 ppm tolerance.

Copy Data

##### Spectrum 4554 (TSV)

###### Preview

```
Loading example...
```

*Click on the button to copy the data to your clipboard.*

Mz MinMz MaxIntensity Max

WidthHeightPeptide font sizePeptide stroke widthSpectrum font sizeSpectrum stroke widthCompact peptide

Ion legend

wxyz

abcd

OtherUnassignedIon^Charge^_Position_Show for top:%

DTJMJSR

01.85e+43.71e+45.56e+47.42e+4

Zoom Out

y^+1^_1_a^+1^_2_y^+1^_1_y^+2^_3_a^+1^_2_b^+1^_2_b^+1^_2_y^+1^_2_y^+1^_2_y^+2^_4_y^+1^_2_a^+1^_3_a^+1^_3_b^+1^_3_y^+2^_5_b^+1^_3_y^+1^_3_y^+1^_3_y^+2^_6_y^+2^_6_y^+1^_3_\*\*b^+1^_4_b^+1^_4_y^+1^_4_y^+1^_4_y^+1^_4_y^+1^_5_y^+1^_6_

041282312351647

Fragment Matches Table

Show background peaks

| Position | Ion type | Intensity | mz Theoretical | mz Error (Th) | mz Error (ppm) | Charge | Series Number |
| --- | --- | --- | --- | --- | --- | --- | --- |
| - | - | 2372 | 120.1 | - | - | 0 | - |
| - | - | 349.3 | 121.1 | - | - | 0 | - |
| - | - | 1149 | 124.1 | - | - | 0 | - |
| - | - | 1241 | 125.1 | - | - | 0 | - |
| - | - | 343.3 | 125.2 | - | - | 0 | - |
| - | - | 1.437E+04 | 126.1 | - | - | 0 | - |
| - | - | 455.3 | 127.1 | - | - | 0 | - |
| - | - | 5131 | 129.1 | - | - | 0 | - |
| - | - | 1368 | 129.1 | - | - | 0 | - |
| - | - | 909.5 | 130.1 | - | - | 0 | - |
| - | - | 4990 | 130.1 | - | - | 0 | - |
| - | - | 1215 | 130.1 | - | - | 0 | - |
| - | - | 371.1 | 132.5 | - | - | 0 | - |
| - | - | 397.1 | 133.1 | - | - | 0 | - |
| - | - | 464 | 136 | - | - | 0 | - |
| - | - | 370.4 | 136.1 | - | - | 0 | - |
| - | - | 773 | 139.1 | - | - | 0 | - |
| - | - | 507.4 | 139.1 | - | - | 0 | - |
| - | - | 2423 | 141.1 | - | - | 0 | - |
| - | - | 1669 | 141.1 | - | - | 0 | - |
| - | - | 413.9 | 142.1 | - | - | 0 | - |
| - | - | 512.8 | 142.1 | - | - | 0 | - |
| - | - | 1843 | 143.1 | - | - | 0 | - |
| - | - | 4.942E+04 | 144.1 | - | - | 0 | - |
| - | - | 577.4 | 145.1 | - | - | 0 | - |
| - | - | 3455 | 145.1 | - | - | 0 | - |
| - | - | 450.9 | 145.3 | - | - | 0 | - |
| - | - | 944.8 | 148.9 | - | - | 0 | - |
| - | - | 595.6 | 152.1 | - | - | 0 | - |
| - | - | 3287 | 153.1 | - | - | 0 | - |
| - | - | 1550 | 154.1 | - | - | 0 | - |
| - | - | 1149 | 155.1 | - | - | 0 | - |
| - | - | 622.6 | 157.1 | - | - | 0 | - |
| - | - | 1125 | 157.1 | - | - | 0 | - |
| - | - | 1372 | 157.1 | - | - | 0 | - |
| 7 | y | 1.086E+04 | 158.1 | 0.0002636 | 1.667 | +1 | 1 |
| - | - | 434.4 | 164.3 | - | - | 0 | - |
| - | - | 475.3 | 164.9 | - | - | 0 | - |
| - | - | 488.8 | 166.1 | - | - | 0 | - |
| - | - | 1.402E+04 | 169.1 | - | - | 0 | - |
| - | - | 4389 | 170.1 | - | - | 0 | - |
| - | - | 1058 | 170.1 | - | - | 0 | - |
| 2 | a | 8530 | 171.1 | 0.000211 | 1.233 | +1 | 2 |
| - | - | 600.1 | 172.1 | - | - | 0 | - |
| - | - | 1718 | 173.1 | - | - | 0 | - |
| - | - | 953.5 | 173.5 | - | - | 0 | - |
| - | - | 727.9 | 175.1 | - | - | 0 | - |
| 7 | y | 3.047E+04 | 175.1 | 0.0002495 | 1.425 | +1 | 1 |
| - | - | 1411 | 176.1 | - | - | 0 | - |
| 5 | y | 661.9 | 179.1 | 0.002316 | 12.93 | +2 | 3 |
| - | - | 1.511E+04 | 180.1 | - | - | 0 | - |
| - | - | 1019 | 180.1 | - | - | 0 | - |
| - | - | 1142 | 181.1 | - | - | 0 | - |
| - | - | 745 | 181.1 | - | - | 0 | - |
| - | - | 1645 | 181.1 | - | - | 0 | - |
| - | - | 541.7 | 183.1 | - | - | 0 | - |
| - | - | 578.6 | 183.1 | - | - | 0 | - |
| - | - | 999.4 | 185.1 | - | - | 0 | - |
| - | - | 1358 | 185.1 | - | - | 0 | - |
| - | - | 3784 | 187.1 | - | - | 0 | - |
| - | - | 1.336E+04 | 187.1 | - | - | 0 | - |
| - | - | 1627 | 188.1 | - | - | 0 | - |
| 2 | a | 5.311E+04 | 189.1 | 0.0002206 | 1.167 | +1 | 2 |
| - | - | 3500 | 190.1 | - | - | 0 | - |
| - | - | 791.7 | 191.1 | - | - | 0 | - |
| - | - | 1494 | 191.1 | - | - | 0 | - |
| - | - | 516.7 | 193.1 | - | - | 0 | - |
| - | - | 1335 | 195.1 | - | - | 0 | - |
| - | - | 1.448E+04 | 197.1 | - | - | 0 | - |
| - | - | 820.7 | 197.1 | - | - | 0 | - |
| - | - | 2711 | 198.1 | - | - | 0 | - |
| - | - | 1382 | 198.1 | - | - | 0 | - |
| 2 | b | 1.79E+04 | 199.1 | 0.0001999 | 1.004 | +1 | 2 |
| - | - | 1237 | 200.1 | - | - | 0 | - |
| - | - | 2296 | 201.1 | - | - | 0 | - |
| - | - | 3009 | 202.1 | - | - | 0 | - |
| - | - | 1026 | 203.1 | - | - | 0 | - |
| - | - | 2779 | 203.1 | - | - | 0 | - |
| - | - | 939.5 | 210.1 | - | - | 0 | - |
| - | - | 557.3 | 211.1 | - | - | 0 | - |
| - | - | 579 | 211.1 | - | - | 0 | - |
| - | - | 493.8 | 212.1 | - | - | 0 | - |
| - | - | 2440 | 215.1 | - | - | 0 | - |
| - | - | 592.8 | 216.1 | - | - | 0 | - |
| 2 | b | 3.062E+04 | 217.1 | 0.0001638 | 0.7546 | +1 | 2 |
| - | - | 1814 | 218.1 | - | - | 0 | - |
| - | - | 638.1 | 220.1 | - | - | 0 | - |
| - | - | 1199 | 220.1 | - | - | 0 | - |
| - | - | 526.9 | 223.1 | - | - | 0 | - |
| - | - | 612.7 | 224.1 | - | - | 0 | - |
| - | - | 2799 | 225.1 | - | - | 0 | - |
| - | - | 657.4 | 227 | - | - | 0 | - |
| - | - | 1862 | 227.1 | - | - | 0 | - |
| - | - | 534 | 228.1 | - | - | 0 | - |
| - | - | 4985 | 229 | - | - | 0 | - |
| - | - | 1269 | 229 | - | - | 0 | - |
| - | - | 3675 | 229.6 | - | - | 0 | - |
| - | - | 848.3 | 230.1 | - | - | 0 | - |
| - | - | 725.8 | 230.1 | - | - | 0 | - |
| - | - | 4121 | 233.1 | - | - | 0 | - |
| - | - | 922.5 | 237.1 | - | - | 0 | - |
| - | - | 1253 | 238.2 | - | - | 0 | - |
| - | - | 1361 | 240.1 | - | - | 0 | - |
| - | - | 592 | 243.6 | - | - | 0 | - |
| - | - | 1693 | 244.1 | - | - | 0 | - |
| 6 | y | 1677 | 244.1 | 0.0003249 | 1.331 | +1 | 2 |
| 6 | y | 1.937E+04 | 245.1 | 0.0002481 | 1.012 | +1 | 2 |
| - | - | 1845 | 246.1 | - | - | 0 | - |
| - | - | 901.1 | 249.1 | - | - | 0 | - |
| - | - | 822.1 | 251.1 | - | - | 0 | - |
| - | - | 687.8 | 258.1 | - | - | 0 | - |
| - | - | 6570 | 261.1 | - | - | 0 | - |
| 4 | y | 3455 | 261.6 | 0.002757 | 10.54 | +2 | 4 |
| 6 | y | 3.735E+04 | 262.2 | 0.0001425 | 0.5435 | +1 | 2 |
| - | - | 3439 | 263.2 | - | - | 0 | - |
| - | - | 1541 | 266.2 | - | - | 0 | - |
| - | - | 2259 | 267.1 | - | - | 0 | - |
| - | - | 480.4 | 267.2 | - | - | 0 | - |
| - | - | 845.6 | 269.1 | - | - | 0 | - |
| - | - | 626.6 | 270.1 | - | - | 0 | - |
| - | - | 4354 | 271 | - | - | 0 | - |
| - | - | 876.5 | 272 | - | - | 0 | - |
| - | - | 681.8 | 272.1 | - | - | 0 | - |
| - | - | 542.6 | 274.3 | - | - | 0 | - |
| - | - | 1037 | 277.1 | - | - | 0 | - |
| - | - | 1042 | 279 | - | - | 0 | - |
| - | - | 568.1 | 280.2 | - | - | 0 | - |
| - | - | 599.3 | 282.2 | - | - | 0 | - |
| 3 | a | 1.533E+04 | 284.2 | 0.0001924 | 0.6771 | +1 | 3 |
| - | - | 1752 | 285.2 | - | - | 0 | - |
| - | - | 1.277E+04 | 286.2 | - | - | 0 | - |
| - | - | 3252 | 286.7 | - | - | 0 | - |
| - | - | 806.8 | 287.2 | - | - | 0 | - |
| - | - | 3014 | 293.9 | - | - | 0 | - |
| - | - | 1428 | 294.1 | - | - | 0 | - |
| - | - | 501.6 | 294.3 | - | - | 0 | - |
| - | - | 1397 | 294.9 | - | - | 0 | - |
| - | - | 703.2 | 295.1 | - | - | 0 | - |
| - | - | 626.9 | 296.1 | - | - | 0 | - |
| - | - | 8576 | 297.1 | - | - | 0 | - |
| - | - | 583.7 | 297.2 | - | - | 0 | - |
| - | - | 1471 | 299.1 | - | - | 0 | - |
| - | - | 1100 | 299.2 | - | - | 0 | - |
| - | - | 532.4 | 300.5 | - | - | 0 | - |
| 3 | a | 1069 | 302.2 | 2.681E-05 | 0.08872 | +1 | 3 |
| - | - | 981.2 | 308.2 | - | - | 0 | - |
| - | - | 703.7 | 309.2 | - | - | 0 | - |
| - | - | 1284 | 310.2 | - | - | 0 | - |
| - | - | 832.4 | 310.6 | - | - | 0 | - |
| - | - | 4078 | 311.9 | - | - | 0 | - |
| 3 | b | 2.38E+04 | 312.2 | 0.0002424 | 0.7765 | +1 | 3 |
| - | - | 3367 | 313.2 | - | - | 0 | - |
| 3 | y | 2.689E+04 | 318.2 | 0.002657 | 8.349 | +2 | 5 |
| - | - | 7749 | 318.7 | - | - | 0 | - |
| - | - | 2011 | 319.2 | - | - | 0 | - |
| - | - | 1220 | 319.7 | - | - | 0 | - |
| - | - | 1228 | 327.1 | - | - | 0 | - |
| 3 | b | 7028 | 330.2 | 0.0003589 | 1.087 | +1 | 3 |
| - | - | 1078 | 331.1 | - | - | 0 | - |
| - | - | 937.7 | 331.2 | - | - | 0 | - |
| - | - | 896.1 | 332.7 | - | - | 0 | - |
| - | - | 2992 | 336.7 | - | - | 0 | - |
| - | - | 829.1 | 337.2 | - | - | 0 | - |
| - | - | 1179 | 340.2 | - | - | 0 | - |
| - | - | 1599 | 341.7 | - | - | 0 | - |
| - | - | 849.5 | 342.2 | - | - | 0 | - |
| - | - | 803.5 | 348.2 | - | - | 0 | - |
| - | - | 1548 | 349.1 | - | - | 0 | - |
| - | - | 727.2 | 349.2 | - | - | 0 | - |
| - | - | 944.4 | 354.7 | - | - | 0 | - |
| - | - | 2843 | 354.9 | - | - | 0 | - |
| - | - | 571.6 | 355.7 | - | - | 0 | - |
| 5 | y | 3134 | 357.2 | 0.0005724 | 1.602 | +1 | 3 |
| 5 | y | 3265 | 358.2 | 0.000213 | 0.5946 | +1 | 3 |
| - | - | 953.2 | 359.2 | - | - | 0 | - |
| 2 | y | 1664 | 359.7 | 0.001983 | 5.513 | +2 | 6 |
| - | - | 610.1 | 360.2 | - | - | 0 | - |
| - | - | 2314 | 362.2 | - | - | 0 | - |
| - | - | 592.7 | 362.7 | - | - | 0 | - |
| - | - | 606.6 | 364.7 | - | - | 0 | - |
| - | - | 689.9 | 368.2 | - | - | 0 | - |
| 2 | y | 4901 | 368.7 | 0.001644 | 4.46 | +2 | 6 |
| - | - | 907 | 369.2 | - | - | 0 | - |
| - | - | 903.4 | 371.9 | - | - | 0 | - |
| - | - | 1946 | 372.2 | - | - | 0 | - |
| - | - | 1576 | 372.9 | - | - | 0 | - |
| - | - | 1791 | 373.7 | - | - | 0 | - |
| 5 | y | 6.314E+04 | 375.2 | 5.921E-05 | 0.1578 | +1 | 3 |
| - | - | 606 | 376.2 | - | - | 0 | - |
| - | - | 1.114E+04 | 376.2 | - | - | 0 | - |
| - | - | 1202 | 377.2 | - | - | 0 | - |
| - | - | 1.147E+04 | 385.2 | - | - | 0 | - |
| - | - | 2647 | 385.7 | - | - | 0 | - |
| - | - | 614.1 | 386.2 | - | - | 0 | - |
| - | - | 572.3 | 386.7 | - | - | 0 | - |
| - | - | 585.3 | 390.7 | - | - | 0 | - |
| - | - | 1.031E+04 | 394.2 | - | - | 0 | - |
| - | - | 2790 | 394.7 | - | - | 0 | - |
| - | - | 1015 | 395.2 | - | - | 0 | - |
| - | - | 1117 | 399.7 | - | - | 0 | - |
| - | - | 808.2 | 401.2 | - | - | 0 | - |
| - | - | 4527 | 402.2 | - | - | 0 | - |
| - | - | 707.7 | 408.7 | - | - | 0 | - |
| - | - | 719.1 | 413.2 | - | - | 0 | - |
| - | - | 800.7 | 416.2 | - | - | 0 | - |
| 0 | Precursor | 1.35E+04 | 417.2 | 0.002794 | 6.696 | +2 | -1 |
| - | - | 4712 | 417.7 | - | - | 0 | - |
| - | - | 2173 | 418.2 | - | - | 0 | - |
| - | - | 1761 | 423.2 | - | - | 0 | - |
| - | - | 992.5 | 424.2 | - | - | 0 | - |
| - | - | 3064 | 425.2 | - | - | 0 | - |
| 0 | Precursor | 3632 | 426.2 | 0.002394 | 5.617 | +2 | -1 |
| - | - | 1.225E+04 | 431.2 | - | - | 0 | - |
| - | - | 3146 | 432.3 | - | - | 0 | - |
| - | - | 2364 | 440.3 | - | - | 0 | - |
| - | - | 7608 | 441.2 | - | - | 0 | - |
| - | - | 1184 | 442.2 | - | - | 0 | - |
| - | - | 6.781E+04 | 458.3 | - | - | 0 | - |
| 4 | b | 4251 | 459.2 | 0.00525 | 11.43 | +1 | 4 |
| - | - | 1.712E+04 | 459.3 | - | - | 0 | - |
| - | - | 2616 | 460.3 | - | - | 0 | - |
| - | - | 693.6 | 468.3 | - | - | 0 | - |
| 4 | b | 6902 | 477.2 | 0.005183 | 10.86 | +1 | 4 |
| - | - | 1521 | 478.2 | - | - | 0 | - |
| - | - | 876.8 | 484.3 | - | - | 0 | - |
| - | - | 1188 | 486.3 | - | - | 0 | - |
| - | - | 1530 | 488.3 | - | - | 0 | - |
| - | - | 577.8 | 494.2 | - | - | 0 | - |
| - | - | 3785 | 501.3 | - | - | 0 | - |
| - | - | 1625 | 502.3 | - | - | 0 | - |
| 4 | y | 1688 | 504.3 | 0.005076 | 10.07 | +1 | 4 |
| 4 | y | 599.8 | 505.2 | 0.009433 | 18.67 | +1 | 4 |
| - | - | 1405 | 507.2 | - | - | 0 | - |
| - | - | 1687 | 515.3 | - | - | 0 | - |
| - | - | 574.6 | 521.2 | - | - | 0 | - |
| 4 | y | 7.345E+04 | 522.3 | 0.004735 | 9.065 | +1 | 4 |
| - | - | 1.674E+04 | 523.3 | - | - | 0 | - |
| - | - | 5319 | 524.3 | - | - | 0 | - |
| - | - | 869.2 | 528.3 | - | - | 0 | - |
| - | - | 761.3 | 532.3 | - | - | 0 | - |
| - | - | 620.4 | 543.3 | - | - | 0 | - |
| - | - | 1818 | 544.3 | - | - | 0 | - |
| - | - | 1905 | 553.3 | - | - | 0 | - |
| - | - | 1187 | 554.3 | - | - | 0 | - |
| - | - | 5.002E+04 | 571.4 | - | - | 0 | - |
| - | - | 1.413E+04 | 572.4 | - | - | 0 | - |
| - | - | 2721 | 573.4 | - | - | 0 | - |
| - | - | 631.6 | 575.2 | - | - | 0 | - |
| - | - | 596.9 | 575.7 | - | - | 0 | - |
| - | - | 670.1 | 581.3 | - | - | 0 | - |
| - | - | 570.4 | 608.1 | - | - | 0 | - |
| - | - | 841.4 | 618.3 | - | - | 0 | - |
| - | - | 683.2 | 620.3 | - | - | 0 | - |
| 3 | y | 4.771E+04 | 635.3 | 0.004167 | 6.558 | +1 | 5 |
| - | - | 1544 | 636.3 | - | - | 0 | - |
| - | - | 1.545E+04 | 636.4 | - | - | 0 | - |
| - | - | 4875 | 637.4 | - | - | 0 | - |
| - | - | 1516 | 638.4 | - | - | 0 | - |
| - | - | 599.7 | 639.4 | - | - | 0 | - |
| - | - | 886.1 | 672.4 | - | - | 0 | - |
| - | - | 563.4 | 673.4 | - | - | 0 | - |
| - | - | 4205 | 682.4 | - | - | 0 | - |
| - | - | 823.8 | 683.4 | - | - | 0 | - |
| 2 | y | 1324 | 736.4 | 0.002814 | 3.821 | +1 | 6 |
| - | - | 7177 | 746.4 | - | - | 0 | - |
| - | - | 2364 | 747.4 | - | - | 0 | - |
| - | - | 684.2 | 748.4 | - | - | 0 | - |
| - | - | 732.4 | 932 | - | - | 0 | - |
| - | - | 587.4 | 1625 | - | - | 0 | - |
| - | - | 561.8 | 1631 | - | - | 0 | - |

m/z Charge Intensity FragmentType MassShift Position
120.08103942871094 0 2372.0996
121.1012954711914 0 349.30148
124.11225891113281 0 1149.1473
125.0711898803711 0 1241.1005
125.21556091308594 0 343.32507
126.05521392822266 0 14365.626
127.05066680908203 0 455.344
129.0660400390625 0 5130.755
129.1025390625 0 1367.9795
130.0502471923828 0 909.4787
130.06137084960938 0 4989.5503
130.0977020263672 0 1214.5402
132.5079803466797 0 371.14648
133.09779357910156 0 397.13312
136.0395050048828 0 464.0025
136.07583618164062 0 370.3949
139.05032348632812 0 773.03
139.11209106445312 0 507.4287
141.1024627685547 0 2422.7197
141.13880920410156 0 1669.4471
142.11021423339844 0 413.8813
142.1427764892578 0 512.8118
143.1180419921875 0 1843.1294
144.06576538085938 0 49420.344
145.06341552734375 0 577.3607
145.06910705566406 0 3454.8667
145.3382568359375 0 450.89566
148.9469757080078 0 944.7664
152.10723876953125 0 595.59515
153.06605529785156 0 3286.8647
154.0501251220703 0 1549.5337
155.07032775878906 0 1149.1368
157.0975341796875 0 622.6042
157.10862731933594 0 1125.1974
157.1337432861328 0 1372.2142
158.09266662597656 0 10860.568 y Ammonia loss 6
164.3276824951172 0 434.36478
164.91368103027344 0 475.26645
166.05332946777344 0 488.8198
169.1337890625 0 14015.159
170.09262084960938 0 4389.4644
170.13720703125 0 1057.8679
171.07662963867188 0 8529.638 a Water loss 1
172.06080627441406 0 600.1432
173.12860107421875 0 1718.0701
173.4518585205078 0 953.51984
175.08694458007812 0 727.9239
175.11920166015625 0 30474.2 y 6
176.1222686767578 0 1411.0718
179.11819458007812 0 661.936 y Water loss 4
180.10215759277344 0 15114.6875
180.11163330078125 0 1018.83875
181.06097412109375 0 1141.5966
181.0975341796875 0 744.99915
181.10574340820312 0 1645.2075
183.1136932373047 0 541.6682
183.1490936279297 0 578.6103
185.09234619140625 0 999.38245
185.13966369628906 0 1358.1965
187.10781860351562 0 3784.3752
187.14439392089844 0 13357.86
188.14794921875 0 1626.7949
189.0872039794922 0 53111.863 a 1
190.090576171875 0 3499.61
191.09194946289062 0 791.68933
191.11819458007812 0 1493.7861
193.11390686035156 0 516.6809
195.11334228515625 0 1334.678
197.128662109375 0 14476.741
197.1387481689453 0 820.73566
198.08749389648438 0 2710.8928
198.1322479248047 0 1382.2363
199.071533203125 0 17900.58 b Water loss 1
200.0753936767578 0 1237.4707
201.12344360351562 0 2296.2266
202.11880493164062 0 3009.1824
203.10299682617188 0 1026.243
203.1279754638672 0 2778.5947
210.08738708496094 0 939.5442
211.1080780029297 0 557.3134
211.14462280273438 0 579.01917
212.103515625 0 493.79138
215.13937377929688 0 2439.6606
216.0983123779297 0 592.8303
217.08206176757812 0 30620.984 b 1
218.0857391357422 0 1813.7351
220.11868286132812 0 638.0875
220.1293182373047 0 1199.3119
223.10902404785156 0 526.94617
224.14019775390625 0 612.6983
225.1237335205078 0 2798.644
227.01663208007812 0 657.3856
227.11375427246094 0 1862.3143
228.09849548339844 0 534.02106
229.00167846679688 0 4985.16
229.0123748779297 0 1268.7216
229.6399688720703 0 3674.9678
230.12547302246094 0 848.29956
230.1414794921875 0 725.84155
233.1321258544922 0 4121.058
237.09130859375 0 922.45764
238.15499877929688 0 1253.1758
240.1345977783203 0 1361.2634
243.63804626464844 0 591.95807
244.09996032714844 0 1693.004
244.1400909423828 0 1677.2164 y Water loss 5
245.1246795654297 0 19365.389 y Ammonia loss 5
246.13002014160156 0 1844.7737
249.1240234375 0 901.1337
251.1033172607422 0 822.0501
258.1456604003906 0 687.758
261.1268005371094 0 6569.6323
261.63916015625 0 3454.898 y 3
262.151123046875 0 37347.555 y 5
263.1544494628906 0 3439.3696
266.1501159667969 0 1541.2991
267.1338195800781 0 2258.6377
267.1504211425781 0 480.35522
269.09161376953125 0 845.6176
270.1195983886719 0 626.59357
271.0271911621094 0 4353.548
272.0284118652344 0 876.5385
272.1355895996094 0 681.7959
274.2786560058594 0 542.63184
277.1194763183594 0 1037.1929
279.0467834472656 0 1042.2456
280.1656494140625 0 568.12134
282.2172546386719 0 599.3117
284.1606750488281 0 15326.061 a Water loss 2
285.1639099121094 0 1752.4839
286.18194580078125 0 12765.489
286.68341064453125 0 3251.7405
287.1846008300781 0 806.7732
293.93438720703125 0 3013.981
294.14508056640625 0 1428.4764
294.3099365234375 0 501.6484
294.9183349609375 0 1397.2456
295.1297607421875 0 703.15686
296.1097412109375 0 626.91504
297.0583190917969 0 8576.193
297.1731872558594 0 583.6773
299.0546569824219 0 1471.3094
299.17193603515625 0 1099.617
300.4967956542969 0 532.3644
302.1710205078125 0 1068.6858 a 2
308.159912109375 0 981.2343
309.2042236328125 0 703.688
310.2119445800781 0 1284.4594
310.60986328125 0 832.44635
311.9452819824219 0 4077.7188
312.1556396484375 0 23800.06 b Water loss 2
313.1589050292969 0 3367.1997
318.18109130859375 0 26887.521 y 2
318.6825256347656 0 7748.661
319.1817321777344 0 2011.2651
319.6855163574219 0 1220.3335
327.1378173828125 0 1228.4443
330.16632080078125 0 7028.319 b 2
331.1227111816406 0 1078.1416
331.1682434082031 0 937.72217
332.6932373046875 0 896.13873
336.7057189941406 0 2992.0818
337.2064514160156 0 829.05884
340.19830322265625 0 1179.0369
341.69732666015625 0 1599.4714
342.1997985839844 0 849.53314
348.1605529785156 0 803.47284
349.0738830566406 0 1548.3231
349.15252685546875 0 727.18115
354.68658447265625 0 944.4484
354.8857116699219 0 2842.6284
355.6946105957031 0 571.6327
357.2239074707031 0 3134.4624 y Water loss 4
358.2082824707031 0 3264.8896 y Ammonia loss 4
359.2109069824219 0 953.1514
359.698974609375 0 1664.233 y Water loss 1
360.2020568847656 0 610.1341
362.2110595703125 0 2314.2861
362.7110595703125 0 592.72906
364.6898193359375 0 606.617
368.1944580078125 0 689.85925
368.70391845703125 0 4900.9883 y 1
369.2048034667969 0 907.0008
371.912109375 0 903.4446
372.158447265625 0 1945.8002
372.8955078125 0 1576.3308
373.6974182128906 0 1791.0848
375.2349853515625 0 63143.47 y 4
376.1522521972656 0 605.9777
376.2379150390625 0 11140.783
377.2403869628906 0 1202.4993
385.2162170410156 0 11466.374
385.7146911621094 0 2646.8608
386.221435546875 0 614.14734
386.6888122558594 0 572.3024
390.68865966796875 0 585.3378
394.2177429199219 0 10313.476
394.7186279296875 0 2790.3828
395.2183532714844 0 1014.8303
399.6933898925781 0 1116.5669
401.21630859375 0 808.1953
402.209716796875 0 4526.9175
408.700439453125 0 707.6672
413.208740234375 0 719.10785
416.2252502441406 0 800.66876
417.2132568359375 0 13502.874 Precursor Water loss
417.714111328125 0 4711.52
418.2146911621094 0 2172.9048
423.23553466796875 0 1761.0352
424.1551208496094 0 992.5497
425.2154235839844 0 3064.3816
426.2181396484375 0 3632.1567 Precursor
431.24871826171875 0 12245.215
432.25311279296875 0 3145.9744
440.26116943359375 0 2363.763
441.2459716796875 0 7607.727
442.2491149902344 0 1183.9094
458.2723388671875 0 67808.67
459.1911315917969 0 4250.8447 b Water loss 3
459.2752380371094 0 17119.484
460.2771301269531 0 2616.0635
468.2580871582031 0 693.5615
477.2016296386719 0 6902.331 b 3
478.2041015625 0 1521.0637
484.2547302246094 0 876.8303
486.26708984375 0 1187.9989
488.2686462402344 0 1529.8522
494.18304443359375 0 577.7825
501.2784423828125 0 3784.6838
502.2785949707031 0 1625.2075
504.2600402832031 0 1688.3513 y Water loss 3
505.2484130859375 0 599.8301 y Ammonia loss 3
507.24627685546875 0 1405.1697
515.2928466796875 0 1687.0967
521.2463989257812 0 574.58954
522.270263671875 0 73453.53 y 3
523.2730102539062 0 16740.844
524.271484375 0 5319.056
528.3154907226562 0 869.16266
532.255126953125 0 761.28705
543.3198852539062 0 620.44025
544.3327026367188 0 1818.4636
553.3462524414062 0 1904.7224
554.32763671875 0 1187.1864
571.35595703125 0 50016.832
572.3590087890625 0 14133.605
573.3612060546875 0 2720.897
575.2200927734375 0 631.5944
575.6932373046875 0 596.9105
581.3404541015625 0 670.115
608.1271362304688 0 570.3927
618.343017578125 0 841.38727
620.3285522460938 0 683.20935
635.353759765625 0 47714.152 y 2
636.2708740234375 0 1543.7826
636.3567504882812 0 15448.887
637.3565673828125 0 4875.2734
638.361083984375 0 1515.883
639.3775634765625 0 599.68896
672.40576171875 0 886.14764
673.4053344726562 0 563.4112
682.3882446289062 0 4204.6577
683.3929443359375 0 823.7697
736.4000854492188 0 1324.0382 y 1
746.3853759765625 0 7176.82
747.3898315429688 0 2364.4502
748.3807373046875 0 684.15735
931.9945678710938 0 732.40906
1625 0 587.36017
1630.681884765625 0 561.75024

Spectrum Details

|  |  |
| --- | --- |
| Matched peaks? Matched peaks The total absolute number of peaks matched. Additionally in brackets the total fraction of peaks matched and the total number of peaks is shown. | 30 (11.11% of 270) |
| FDR? FDR The false discovery rate estimated for this peptide. It is calculated by matching all theoretical fragments with a non-integer shift with the raw peaks for this spectrum. This is done with 40 different shifts. The resulting percentage is the average number of annotated peaks over the number of annotated peaks with the correct spectrum. | 0.48% |
| Satellite FDR? Satellite FDR See the FDR for details on its calculation. This satellite ion specific FDR only contains the satellite ions (d/w) for I/L/J positions. | - |
| PSM Score? PSM Score The PSM Score as given by Hecklib to this annotated spectrum. It is shown with three significant figures. | 376 |

#### Spectrum 4616? Spectrum 4616 The raw spectrum of this peptide as annotated by Hecklib. The fragments are coloured according to ion type (see legend). Any peaks with a star '\*' as text can be hovered over to see the full details, first the ion type second the mass shift type. By hovering over the amino acids in the peptide or ions in the legend the corresponding peaks are highlighted. By toggling the 'Unassigned' label you can turn the background (unassigned) peaks on or off in the plot. By updating the slider in the Ion legend you can update the spectrum to only show the top X% of the peaks with labels. The top X% means any peak that is within X% of the highest intensity. By dragging in the spectrum you can zoom in to a specific part of the spectrum and use 'Zoom Out' to get back to the original zoom level. The annotation of the spectrum is based on the given sequence in the peptides file and is done with different software so inconsistencies are likely. The peaks are annotated based on the given sequence, with 20 ppm tolerance.

Copy Data

##### Spectrum 4616 (TSV)

###### Preview

```
Loading example...
```

*Click on the button to copy the data to your clipboard.*

Mz MinMz MaxIntensity Max

WidthHeightPeptide font sizePeptide stroke widthSpectrum font sizeSpectrum stroke widthCompact peptide

Ion legend

wxyz

abcd

OtherUnassignedIon^Charge^_Position_Show for top:%

DTJMJSR

01.73e+43.47e+45.20e+46.93e+4

Zoom Out

y^+1^_1_a^+1^_2_y^+1^_1_y^+2^_3_a^+1^_2_b^+1^_2_b^+1^_2_y^+1^_2_y^+1^_2_y^+2^_4_y^+2^_4_y^+1^_2_d^+1^_3_a^+1^_3_a^+1^_3_b^+1^_3_y^+2^_5_b^+1^_3_y^+1^_3_y^+1^_3_y^+2^_6_y^+2^_6_y^+1^_3_\*\*b^+1^_4_b^+1^_4_y^+1^_4_y^+1^_4_y^+1^_5_y^+1^_5_y^+1^_6_

0658131519732631

Fragment Matches Table

Show background peaks

| Position | Ion type | Intensity | mz Theoretical | mz Error (Th) | mz Error (ppm) | Charge | Series Number |
| --- | --- | --- | --- | --- | --- | --- | --- |
| - | - | 1709 | 120.1 | - | - | 0 | - |
| - | - | 335.1 | 122.4 | - | - | 0 | - |
| - | - | 375.6 | 123.1 | - | - | 0 | - |
| - | - | 1687 | 124.1 | - | - | 0 | - |
| - | - | 669.7 | 125.1 | - | - | 0 | - |
| - | - | 1.209E+04 | 126.1 | - | - | 0 | - |
| - | - | 1015 | 127.1 | - | - | 0 | - |
| - | - | 509.6 | 127.1 | - | - | 0 | - |
| - | - | 5028 | 129.1 | - | - | 0 | - |
| - | - | 1325 | 129.1 | - | - | 0 | - |
| - | - | 1027 | 130.1 | - | - | 0 | - |
| - | - | 4527 | 130.1 | - | - | 0 | - |
| - | - | 420.7 | 130.1 | - | - | 0 | - |
| - | - | 856 | 130.1 | - | - | 0 | - |
| - | - | 386.9 | 133.4 | - | - | 0 | - |
| - | - | 474.8 | 136 | - | - | 0 | - |
| - | - | 722.4 | 136.1 | - | - | 0 | - |
| - | - | 467.9 | 139.1 | - | - | 0 | - |
| - | - | 615.6 | 139.1 | - | - | 0 | - |
| - | - | 763.3 | 141.1 | - | - | 0 | - |
| - | - | 1402 | 141.1 | - | - | 0 | - |
| - | - | 1011 | 142.1 | - | - | 0 | - |
| - | - | 836.6 | 143.1 | - | - | 0 | - |
| - | - | 834.2 | 143.1 | - | - | 0 | - |
| - | - | 4.532E+04 | 144.1 | - | - | 0 | - |
| - | - | 569.9 | 145.1 | - | - | 0 | - |
| - | - | 3059 | 145.1 | - | - | 0 | - |
| - | - | 1340 | 152.1 | - | - | 0 | - |
| - | - | 4015 | 153.1 | - | - | 0 | - |
| - | - | 454.1 | 153.1 | - | - | 0 | - |
| - | - | 422.4 | 153.6 | - | - | 0 | - |
| - | - | 1392 | 154.1 | - | - | 0 | - |
| - | - | 1813 | 155.1 | - | - | 0 | - |
| - | - | 1066 | 157.1 | - | - | 0 | - |
| - | - | 880 | 157.1 | - | - | 0 | - |
| 7 | y | 9551 | 158.1 | 0.000233 | 1.474 | +1 | 1 |
| - | - | 542.8 | 167.1 | - | - | 0 | - |
| - | - | 1.18E+04 | 169.1 | - | - | 0 | - |
| - | - | 4281 | 170.1 | - | - | 0 | - |
| - | - | 1131 | 170.1 | - | - | 0 | - |
| 2 | a | 7509 | 171.1 | 0.0002262 | 1.322 | +1 | 2 |
| - | - | 574.3 | 172.1 | - | - | 0 | - |
| - | - | 1123 | 173.1 | - | - | 0 | - |
| - | - | 557.6 | 173.4 | - | - | 0 | - |
| 7 | y | 3.036E+04 | 175.1 | 0.00028 | 1.599 | +1 | 1 |
| - | - | 1454 | 176.1 | - | - | 0 | - |
| 5 | y | 870 | 179.1 | 0.002896 | 16.17 | +2 | 3 |
| - | - | 1.439E+04 | 180.1 | - | - | 0 | - |
| - | - | 618.8 | 181.1 | - | - | 0 | - |
| - | - | 1634 | 181.1 | - | - | 0 | - |
| - | - | 605.2 | 182 | - | - | 0 | - |
| - | - | 762.9 | 184.1 | - | - | 0 | - |
| - | - | 852.7 | 185.1 | - | - | 0 | - |
| - | - | 1345 | 185.1 | - | - | 0 | - |
| - | - | 1.24E+04 | 187.1 | - | - | 0 | - |
| - | - | 1173 | 188.1 | - | - | 0 | - |
| 2 | a | 4.977E+04 | 189.1 | 0.0002512 | 1.328 | +1 | 2 |
| - | - | 3091 | 190.1 | - | - | 0 | - |
| - | - | 460 | 190.4 | - | - | 0 | - |
| - | - | 536.6 | 190.7 | - | - | 0 | - |
| - | - | 960.7 | 191.1 | - | - | 0 | - |
| - | - | 762 | 195.1 | - | - | 0 | - |
| - | - | 1.53E+04 | 197.1 | - | - | 0 | - |
| - | - | 1367 | 198.1 | - | - | 0 | - |
| - | - | 1528 | 198.1 | - | - | 0 | - |
| 2 | b | 1.536E+04 | 199.1 | 0.0002152 | 1.081 | +1 | 2 |
| - | - | 586.8 | 199.1 | - | - | 0 | - |
| - | - | 1267 | 200.1 | - | - | 0 | - |
| - | - | 1658 | 201.1 | - | - | 0 | - |
| - | - | 2339 | 202.1 | - | - | 0 | - |
| - | - | 1210 | 203.1 | - | - | 0 | - |
| - | - | 2147 | 203.1 | - | - | 0 | - |
| - | - | 578.3 | 207.1 | - | - | 0 | - |
| - | - | 1118 | 210.1 | - | - | 0 | - |
| - | - | 525.3 | 211.1 | - | - | 0 | - |
| - | - | 2359 | 215.1 | - | - | 0 | - |
| - | - | 838.6 | 216.1 | - | - | 0 | - |
| 2 | b | 2.683E+04 | 217.1 | 0.0001943 | 0.8952 | +1 | 2 |
| - | - | 1596 | 218.1 | - | - | 0 | - |
| - | - | 799.5 | 220.1 | - | - | 0 | - |
| - | - | 1191 | 220.1 | - | - | 0 | - |
| - | - | 2212 | 225.1 | - | - | 0 | - |
| - | - | 626.3 | 227 | - | - | 0 | - |
| - | - | 1552 | 227.1 | - | - | 0 | - |
| - | - | 986.6 | 228.1 | - | - | 0 | - |
| - | - | 5569 | 229 | - | - | 0 | - |
| - | - | 868.7 | 229 | - | - | 0 | - |
| - | - | 4551 | 229.6 | - | - | 0 | - |
| - | - | 769.7 | 230 | - | - | 0 | - |
| - | - | 771.7 | 230.1 | - | - | 0 | - |
| - | - | 825.9 | 230.1 | - | - | 0 | - |
| - | - | 3827 | 233.1 | - | - | 0 | - |
| - | - | 656 | 237.1 | - | - | 0 | - |
| - | - | 1065 | 238.2 | - | - | 0 | - |
| - | - | 789.7 | 240.1 | - | - | 0 | - |
| - | - | 1611 | 244.1 | - | - | 0 | - |
| 6 | y | 1982 | 244.1 | 0.0002396 | 0.9815 | +1 | 2 |
| 6 | y | 1.853E+04 | 245.1 | 0.0003091 | 1.261 | +1 | 2 |
| - | - | 1784 | 246.1 | - | - | 0 | - |
| - | - | 1138 | 247 | - | - | 0 | - |
| 4 | y | 649.8 | 253.1 | 0.004369 | 17.26 | +2 | 4 |
| - | - | 545.8 | 255.2 | - | - | 0 | - |
| - | - | 6139 | 261.1 | - | - | 0 | - |
| 4 | y | 2475 | 261.6 | 0.002544 | 9.722 | +2 | 4 |
| 6 | y | 3.303E+04 | 262.2 | 0.000173 | 0.6599 | +1 | 2 |
| - | - | 964.4 | 263.1 | - | - | 0 | - |
| - | - | 3648 | 263.2 | - | - | 0 | - |
| - | - | 1494 | 266.1 | - | - | 0 | - |
| - | - | 1504 | 267.1 | - | - | 0 | - |
| - | - | 777.2 | 268.1 | - | - | 0 | - |
| - | - | 589.4 | 270.1 | - | - | 0 | - |
| - | - | 4043 | 271 | - | - | 0 | - |
| - | - | 589.5 | 272 | - | - | 0 | - |
| - | - | 731.2 | 272.1 | - | - | 0 | - |
| 3 | d | 511.2 | 274.1 | 0.004448 | 16.23 | +1 | 3 |
| - | - | 1422 | 279 | - | - | 0 | - |
| - | - | 1077 | 280.2 | - | - | 0 | - |
| - | - | 578.3 | 282.2 | - | - | 0 | - |
| 3 | a | 1.378E+04 | 284.2 | 0.0002535 | 0.8919 | +1 | 3 |
| - | - | 1800 | 285.2 | - | - | 0 | - |
| - | - | 1.014E+04 | 286.2 | - | - | 0 | - |
| - | - | 3460 | 286.7 | - | - | 0 | - |
| - | - | 757.5 | 289 | - | - | 0 | - |
| - | - | 2252 | 293.9 | - | - | 0 | - |
| - | - | 935.5 | 294.1 | - | - | 0 | - |
| - | - | 1113 | 294.9 | - | - | 0 | - |
| - | - | 568.7 | 295.1 | - | - | 0 | - |
| - | - | 641.8 | 295.8 | - | - | 0 | - |
| - | - | 6992 | 297.1 | - | - | 0 | - |
| - | - | 657 | 297.2 | - | - | 0 | - |
| - | - | 740.2 | 298.1 | - | - | 0 | - |
| - | - | 1671 | 299.1 | - | - | 0 | - |
| - | - | 1693 | 299.2 | - | - | 0 | - |
| - | - | 817.2 | 299.2 | - | - | 0 | - |
| 3 | a | 921.2 | 302.2 | 0.0003015 | 0.9977 | +1 | 3 |
| - | - | 983.8 | 309.2 | - | - | 0 | - |
| - | - | 3667 | 311.9 | - | - | 0 | - |
| 3 | b | 2.367E+04 | 312.2 | 0.0002729 | 0.8743 | +1 | 3 |
| - | - | 3902 | 313.2 | - | - | 0 | - |
| 3 | y | 2.276E+04 | 318.2 | 0.002657 | 8.349 | +2 | 5 |
| - | - | 7499 | 318.7 | - | - | 0 | - |
| - | - | 2423 | 319.2 | - | - | 0 | - |
| - | - | 1111 | 319.7 | - | - | 0 | - |
| - | - | 550.9 | 326.1 | - | - | 0 | - |
| - | - | 701.8 | 327.7 | - | - | 0 | - |
| - | - | 527.8 | 328.3 | - | - | 0 | - |
| - | - | 636.1 | 329.5 | - | - | 0 | - |
| - | - | 766.6 | 330 | - | - | 0 | - |
| 3 | b | 6233 | 330.2 | 0.000603 | 1.826 | +1 | 3 |
| - | - | 700.1 | 331.2 | - | - | 0 | - |
| - | - | 966.2 | 331.2 | - | - | 0 | - |
| - | - | 1172 | 332.7 | - | - | 0 | - |
| - | - | 2262 | 336.7 | - | - | 0 | - |
| - | - | 939.1 | 340.2 | - | - | 0 | - |
| - | - | 2068 | 341.7 | - | - | 0 | - |
| - | - | 1503 | 346.1 | - | - | 0 | - |
| - | - | 790.7 | 349.1 | - | - | 0 | - |
| - | - | 2671 | 354.9 | - | - | 0 | - |
| - | - | 886.7 | 355.7 | - | - | 0 | - |
| 5 | y | 3180 | 357.2 | 0.0004042 | 1.131 | +1 | 3 |
| 5 | y | 3579 | 358.2 | 0.0005487 | 1.532 | +1 | 3 |
| - | - | 760.6 | 359.2 | - | - | 0 | - |
| 2 | y | 1046 | 359.7 | 0.003295 | 9.161 | +2 | 6 |
| - | - | 1937 | 362.2 | - | - | 0 | - |
| - | - | 668.4 | 362.7 | - | - | 0 | - |
| - | - | 799.5 | 363.2 | - | - | 0 | - |
| 2 | y | 2708 | 368.7 | 0.002194 | 5.95 | +2 | 6 |
| - | - | 986.4 | 369.2 | - | - | 0 | - |
| - | - | 720 | 371.9 | - | - | 0 | - |
| - | - | 2306 | 372.2 | - | - | 0 | - |
| - | - | 1037 | 372.9 | - | - | 0 | - |
| - | - | 714.1 | 373.2 | - | - | 0 | - |
| - | - | 1252 | 373.7 | - | - | 0 | - |
| 5 | y | 5.753E+04 | 375.2 | 2.869E-05 | 0.07646 | +1 | 3 |
| - | - | 1.016E+04 | 376.2 | - | - | 0 | - |
| - | - | 666.1 | 376.8 | - | - | 0 | - |
| - | - | 875.3 | 377.2 | - | - | 0 | - |
| - | - | 1.011E+04 | 385.2 | - | - | 0 | - |
| - | - | 2108 | 385.7 | - | - | 0 | - |
| - | - | 1409 | 386.2 | - | - | 0 | - |
| - | - | 681.4 | 386.7 | - | - | 0 | - |
| - | - | 725.2 | 390.7 | - | - | 0 | - |
| - | - | 9442 | 394.2 | - | - | 0 | - |
| - | - | 2653 | 394.7 | - | - | 0 | - |
| - | - | 1143 | 395.2 | - | - | 0 | - |
| - | - | 1428 | 399.7 | - | - | 0 | - |
| - | - | 836.1 | 401.2 | - | - | 0 | - |
| - | - | 3356 | 402.2 | - | - | 0 | - |
| - | - | 875.2 | 408.2 | - | - | 0 | - |
| - | - | 1639 | 408.7 | - | - | 0 | - |
| 0 | Precursor | 1.357E+04 | 417.2 | 0.002702 | 6.477 | +2 | -1 |
| - | - | 5431 | 417.7 | - | - | 0 | - |
| - | - | 1562 | 418.2 | - | - | 0 | - |
| - | - | 2371 | 423.2 | - | - | 0 | - |
| - | - | 1985 | 425.2 | - | - | 0 | - |
| 0 | Precursor | 1867 | 426.2 | 0.003279 | 7.694 | +2 | -1 |
| - | - | 727.7 | 426.7 | - | - | 0 | - |
| - | - | 842.1 | 430.2 | - | - | 0 | - |
| - | - | 1.198E+04 | 431.2 | - | - | 0 | - |
| - | - | 2726 | 432.3 | - | - | 0 | - |
| - | - | 2809 | 440.3 | - | - | 0 | - |
| - | - | 6675 | 441.2 | - | - | 0 | - |
| - | - | 1063 | 442.3 | - | - | 0 | - |
| - | - | 556.3 | 444.3 | - | - | 0 | - |
| - | - | 559.1 | 455.9 | - | - | 0 | - |
| - | - | 6.557E+04 | 458.3 | - | - | 0 | - |
| 4 | b | 2906 | 459.2 | 0.005128 | 11.17 | +1 | 4 |
| - | - | 1.415E+04 | 459.3 | - | - | 0 | - |
| - | - | 1279 | 460.2 | - | - | 0 | - |
| - | - | 2231 | 460.3 | - | - | 0 | - |
| - | - | 822.5 | 468.3 | - | - | 0 | - |
| 4 | b | 5225 | 477.2 | 0.005519 | 11.57 | +1 | 4 |
| - | - | 1294 | 478.2 | - | - | 0 | - |
| - | - | 880.2 | 484.3 | - | - | 0 | - |
| - | - | 1573 | 486.3 | - | - | 0 | - |
| - | - | 670.3 | 488.3 | - | - | 0 | - |
| - | - | 3611 | 501.3 | - | - | 0 | - |
| - | - | 1099 | 502.3 | - | - | 0 | - |
| 4 | y | 1286 | 504.3 | 0.003153 | 6.253 | +1 | 4 |
| - | - | 543.6 | 505.2 | - | - | 0 | - |
| - | - | 876.3 | 507.2 | - | - | 0 | - |
| - | - | 543.3 | 513 | - | - | 0 | - |
| - | - | 1643 | 515.3 | - | - | 0 | - |
| - | - | 656.5 | 516.3 | - | - | 0 | - |
| - | - | 915.7 | 521.3 | - | - | 0 | - |
| 4 | y | 6.863E+04 | 522.3 | 0.004796 | 9.182 | +1 | 4 |
| - | - | 1.654E+04 | 523.3 | - | - | 0 | - |
| - | - | 5225 | 524.3 | - | - | 0 | - |
| - | - | 588 | 528.3 | - | - | 0 | - |
| - | - | 944.8 | 532.3 | - | - | 0 | - |
| - | - | 1620 | 544.3 | - | - | 0 | - |
| - | - | 1987 | 553.3 | - | - | 0 | - |
| - | - | 1332 | 554.3 | - | - | 0 | - |
| - | - | 4.774E+04 | 571.4 | - | - | 0 | - |
| - | - | 1.192E+04 | 572.4 | - | - | 0 | - |
| - | - | 1820 | 573.4 | - | - | 0 | - |
| 3 | y | 1514 | 617.3 | 0.004233 | 6.857 | +1 | 5 |
| - | - | 822.6 | 620.3 | - | - | 0 | - |
| 3 | y | 4.395E+04 | 635.3 | 0.004411 | 6.942 | +1 | 5 |
| - | - | 938.4 | 636.3 | - | - | 0 | - |
| - | - | 761.1 | 636.3 | - | - | 0 | - |
| - | - | 1.402E+04 | 636.4 | - | - | 0 | - |
| - | - | 4198 | 637.4 | - | - | 0 | - |
| - | - | 1397 | 638.4 | - | - | 0 | - |
| - | - | 4486 | 682.4 | - | - | 0 | - |
| - | - | 1627 | 683.4 | - | - | 0 | - |
| 2 | y | 910.8 | 736.4 | 0.003485 | 4.733 | +1 | 6 |
| - | - | 6063 | 746.4 | - | - | 0 | - |
| - | - | 1799 | 747.4 | - | - | 0 | - |
| - | - | 637.4 | 819.9 | - | - | 0 | - |
| - | - | 660.5 | 947.9 | - | - | 0 | - |
| - | - | 610.2 | 948.4 | - | - | 0 | - |
| - | - | 597.8 | 1048 | - | - | 0 | - |
| - | - | 715.6 | 1072 | - | - | 0 | - |
| - | - | 668.9 | 2240 | - | - | 0 | - |
| - | - | 730 | 2605 | - | - | 0 | - |

m/z Charge Intensity FragmentType MassShift Position
120.08110046386719 0 1708.8604
122.43797302246094 0 335.08096
123.08122253417969 0 375.56308
124.1125259399414 0 1686.52
125.07115173339844 0 669.67194
126.05522155761719 0 12089.54
127.05863952636719 0 1014.78955
127.0869369506836 0 509.63763
129.06613159179688 0 5027.659
129.1025848388672 0 1324.8413
130.05015563964844 0 1026.6029
130.06137084960938 0 4527.121
130.06924438476562 0 420.6744
130.09762573242188 0 855.97125
133.38034057617188 0 386.92688
136.0398712158203 0 474.76575
136.0761260986328 0 722.4247
139.05047607421875 0 467.91528
139.1119384765625 0 615.55695
141.1024932861328 0 763.27985
141.1388397216797 0 1401.9559
142.12277221679688 0 1010.72723
143.0820770263672 0 836.59784
143.1182098388672 0 834.2223
144.06581115722656 0 45324.883
145.06329345703125 0 569.9206
145.0692901611328 0 3058.8567
152.10707092285156 0 1340.0748
153.06610107421875 0 4015.0576
153.0832977294922 0 454.1136
153.61866760253906 0 422.35388
154.05018615722656 0 1392.2047
155.0706024169922 0 1813.0925
157.1090087890625 0 1065.6108
157.13377380371094 0 880.0311
158.09263610839844 0 9550.744 y Ammonia loss 6
167.1179656982422 0 542.81366
169.13381958007812 0 11796.436
170.09271240234375 0 4281.054
170.13729858398438 0 1131.1295
171.07664489746094 0 7509.4224 a Water loss 1
172.0608367919922 0 574.2792
173.12887573242188 0 1123.2522
173.4376220703125 0 557.567
175.11923217773438 0 30364.424 y 6
176.1227569580078 0 1454.294
179.1187744140625 0 869.9513 y Water loss 4
180.1021728515625 0 14390.971
181.06117248535156 0 618.83
181.105712890625 0 1634.0509
182.0447540283203 0 605.2224
184.10789489746094 0 762.8732
185.09197998046875 0 852.73566
185.14010620117188 0 1344.6256
187.1443634033203 0 12397.601
188.14776611328125 0 1173.465
189.0872344970703 0 49769.688 a 1
190.09060668945312 0 3091.0618
190.42982482910156 0 459.98026
190.74935913085938 0 536.6422
191.1182861328125 0 960.6669
195.11326599121094 0 761.9698
197.12867736816406 0 15295.289
198.0873565673828 0 1366.5919
198.13214111328125 0 1528.1598
199.07154846191406 0 15360.254 b Water loss 1
199.1195068359375 0 586.75964
200.07470703125 0 1267.3605
201.12374877929688 0 1657.8676
202.11892700195312 0 2339.1194
203.1027069091797 0 1209.7058
203.12826538085938 0 2147.179
207.11215209960938 0 578.28577
210.0870819091797 0 1118.1777
211.10739135742188 0 525.2539
215.13941955566406 0 2359.2205
216.09762573242188 0 838.5677
217.08209228515625 0 26826.973 b 1
218.08560180664062 0 1596.0961
220.1183319091797 0 799.5493
220.12940979003906 0 1190.712
225.12359619140625 0 2211.7747
227.01580810546875 0 626.2544
227.11412048339844 0 1551.9565
228.09837341308594 0 986.6298
229.00173950195312 0 5568.6777
229.01185607910156 0 868.6995
229.63978576660156 0 4551.4136
230.00172424316406 0 769.7489
230.1248321533203 0 771.7401
230.14230346679688 0 825.90735
233.1322479248047 0 3827.2766
237.0908660888672 0 655.9672
238.1555938720703 0 1064.5626
240.1341094970703 0 789.6729
244.10064697265625 0 1610.944
244.14065551757812 0 1981.9268 y Water loss 5
245.12474060058594 0 18531.955 y Ammonia loss 5
246.1293182373047 0 1784.268
247.01243591308594 0 1137.8385
253.11875915527344 0 649.8046 y Ammonia loss 3
255.2126922607422 0 545.8348
261.12677001953125 0 6138.6357
261.6389465332031 0 2474.997 y 3
262.1511535644531 0 33028.19 y 5
263.0696716308594 0 964.35236
263.1545104980469 0 3647.9478
266.1495666503906 0 1493.7354
267.1339416503906 0 1504.1049
268.129150390625 0 777.1558
270.1200866699219 0 589.41003
271.02734375 0 4042.77
272.0272521972656 0 589.52704
272.1362609863281 0 731.21265
274.1441955566406 0 511.1816 d 2
279.0482482910156 0 1422.4984
280.1655578613281 0 1076.911
282.2178955078125 0 578.34985
284.1607360839844 0 13781.005 a Water loss 2
285.1634521484375 0 1800.269
286.1820068359375 0 10142.942
286.6836853027344 0 3459.9902
289.0373840332031 0 757.4686
293.9348449707031 0 2251.6143
294.1449279785156 0 935.4521
294.9190368652344 0 1112.6742
295.1294250488281 0 568.66864
295.8275146484375 0 641.8217
297.0581359863281 0 6992.386
297.17437744140625 0 656.976
298.0599670410156 0 740.18005
299.0545654296875 0 1671.1184
299.17132568359375 0 1693.1052
299.189208984375 0 817.2109
302.1707458496094 0 921.2219 a 2
309.202880859375 0 983.7541
311.94549560546875 0 3666.8037
312.1556701660156 0 23668.58 b Water loss 2
313.1587829589844 0 3902.0237
318.18109130859375 0 22756.352 y 2
318.68280029296875 0 7499.4995
319.182861328125 0 2422.7595
319.6836242675781 0 1110.5209
326.1455078125 0 550.90015
327.70050048828125 0 701.8235
328.31341552734375 0 527.7575
329.4576416015625 0 636.1003
329.95452880859375 0 766.5595
330.16656494140625 0 6233.3247 b 2
331.1681823730469 0 700.0592
331.18768310546875 0 966.2158
332.69342041015625 0 1171.8563
336.7057800292969 0 2262.4424
340.1990966796875 0 939.13983
341.69793701171875 0 2067.9446
346.14361572265625 0 1502.6714
349.0760192871094 0 790.65656
354.8859558105469 0 2671.1685
355.693603515625 0 886.657
357.2248840332031 0 3180.2893 y Water loss 4
358.20794677734375 0 3578.9272 y Ammonia loss 4
359.2389831542969 0 760.56903
359.7002868652344 0 1045.7058 y Water loss 1
362.211181640625 0 1937.2946
362.7130432128906 0 668.36774
363.2023010253906 0 799.53485
368.7044677734375 0 2708.3862 y 1
369.2057800292969 0 986.3756
371.9121398925781 0 720.0016
372.1589050292969 0 2305.8557
372.8954162597656 0 1037.1986
373.2180480957031 0 714.14545
373.69677734375 0 1252.0759
375.2350158691406 0 57534.867 y 4
376.2379150390625 0 10155.697
376.8427429199219 0 666.0936
377.24053955078125 0 875.2791
385.21563720703125 0 10107.735
385.7147521972656 0 2107.6104
386.2227783203125 0 1408.6757
386.6888732910156 0 681.414
390.6883239746094 0 725.23016
394.21710205078125 0 9441.604
394.7192687988281 0 2652.9846
395.2200927734375 0 1142.6914
399.69512939453125 0 1428.3043
401.2156677246094 0 836.0697
402.2099304199219 0 3355.6187
408.20703125 0 875.16974
408.7005920410156 0 1638.7203
417.2131652832031 0 13573.59 Precursor Water loss
417.7143859863281 0 5431.2144
418.21417236328125 0 1562.2618
423.2354736328125 0 2370.9932
425.2141418457031 0 1984.606
426.2190246582031 0 1867.2404 Precursor
426.72210693359375 0 727.70605
430.24127197265625 0 842.1393
431.24871826171875 0 11979.621
432.2527770996094 0 2725.6257
440.2620544433594 0 2808.8652
441.24578857421875 0 6675.2505
442.2502746582031 0 1063.2999
444.2559814453125 0 556.328
455.88958740234375 0 559.0503
458.2724304199219 0 65571.29
459.1910095214844 0 2906.0972 b Water loss 3
459.2748718261719 0 14147.868
460.1957702636719 0 1278.5496
460.2784729003906 0 2230.6462
468.2561950683594 0 822.49457
477.20196533203125 0 5224.908 b 3
478.20587158203125 0 1294.4985
484.2510070800781 0 880.22833
486.2673034667969 0 1572.5403
488.2701416015625 0 670.2715
501.2789001464844 0 3610.597
502.27813720703125 0 1099.3213
504.25811767578125 0 1286.2484 y Water loss 3
505.2494812011719 0 543.64716
507.24664306640625 0 876.2516
513.021728515625 0 543.3486
515.2953491210938 0 1642.51
516.298095703125 0 656.47327
521.254638671875 0 915.6872
522.2703247070312 0 68634.72 y 3
523.2728881835938 0 16540.172
524.271728515625 0 5225.081
528.3137817382812 0 587.9985
532.2555541992188 0 944.7513
544.331787109375 0 1620.2073
553.345458984375 0 1987.2161
554.329345703125 0 1331.8805
571.3560791015625 0 47737.742
572.35888671875 0 11916.338
573.3600463867188 0 1820.3926
617.34326171875 0 1513.6019 y Water loss 2
620.3289184570312 0 822.559
635.35400390625 0 43951.46 y 2
636.2706298828125 0 938.41455
636.2943115234375 0 761.0998
636.3567504882812 0 14022.809
637.3572387695312 0 4198.3613
638.3582763671875 0 1396.916
682.3871459960938 0 4486.114
683.392333984375 0 1627.0106
736.4007568359375 0 910.83966 y 1
746.3853149414062 0 6063.101
747.3877563476562 0 1798.9237
819.8675537109375 0 637.42737
947.9387817382812 0 660.51855
948.3785400390625 0 610.17
1047.623779296875 0 597.7797
1071.91162109375 0 715.6163
2239.567626953125 0 668.9302
2604.949462890625 0 730.0372

Spectrum Details

|  |  |
| --- | --- |
| Matched peaks? Matched peaks The total absolute number of peaks matched. Additionally in brackets the total fraction of peaks matched and the total number of peaks is shown. | 32 (12.50% of 256) |
| FDR? FDR The false discovery rate estimated for this peptide. It is calculated by matching all theoretical fragments with a non-integer shift with the raw peaks for this spectrum. This is done with 40 different shifts. The resulting percentage is the average number of annotated peaks over the number of annotated peaks with the correct spectrum. | 0.37% |
| Satellite FDR? Satellite FDR See the FDR for details on its calculation. This satellite ion specific FDR only contains the satellite ions (d/w) for I/L/J positions. | 2.38% |
| PSM Score? PSM Score The PSM Score as given by Hecklib to this annotated spectrum. It is shown with three significant figures. | 376 |

#### Spectrum 4676? Spectrum 4676 The raw spectrum of this peptide as annotated by Hecklib. The fragments are coloured according to ion type (see legend). Any peaks with a star '\*' as text can be hovered over to see the full details, first the ion type second the mass shift type. By hovering over the amino acids in the peptide or ions in the legend the corresponding peaks are highlighted. By toggling the 'Unassigned' label you can turn the background (unassigned) peaks on or off in the plot. By updating the slider in the Ion legend you can update the spectrum to only show the top X% of the peaks with labels. The top X% means any peak that is within X% of the highest intensity. By dragging in the spectrum you can zoom in to a specific part of the spectrum and use 'Zoom Out' to get back to the original zoom level. The annotation of the spectrum is based on the given sequence in the peptides file and is done with different software so inconsistencies are likely. The peaks are annotated based on the given sequence, with 20 ppm tolerance.

Copy Data

##### Spectrum 4676 (TSV)

###### Preview

```
Loading example...
```

*Click on the button to copy the data to your clipboard.*

Mz MinMz MaxIntensity Max

WidthHeightPeptide font sizePeptide stroke widthSpectrum font sizeSpectrum stroke widthCompact peptide

Ion legend

wxyz

abcd

OtherUnassignedIon^Charge^_Position_Show for top:%

DTJMJSR

01.68e+43.36e+45.04e+46.72e+4

Zoom Out

y^+1^_1_a^+1^_2_y^+1^_1_a^+1^_2_b^+1^_2_b^+1^_2_y^+1^_2_y^+1^_2_y^+2^_4_y^+1^_2_a^+1^_3_a^+1^_3_b^+1^_3_y^+2^_5_b^+1^_3_y^+1^_3_y^+1^_3_y^+2^_6_y^+2^_6_y^+1^_3_\*\*b^+1^_4_b^+1^_4_y^+1^_4_y^+1^_4_y^+1^_5_y^+1^_6_y^+1^_6_

0865173025943459

Fragment Matches Table

Show background peaks

| Position | Ion type | Intensity | mz Theoretical | mz Error (Th) | mz Error (ppm) | Charge | Series Number |
| --- | --- | --- | --- | --- | --- | --- | --- |
| - | - | 674.5 | 120.1 | - | - | 0 | - |
| - | - | 443.4 | 121.1 | - | - | 0 | - |
| - | - | 363.2 | 123.1 | - | - | 0 | - |
| - | - | 1283 | 124.1 | - | - | 0 | - |
| - | - | 970.8 | 125.1 | - | - | 0 | - |
| - | - | 384.4 | 125.1 | - | - | 0 | - |
| - | - | 1.267E+04 | 126.1 | - | - | 0 | - |
| - | - | 406.7 | 126.6 | - | - | 0 | - |
| - | - | 589.2 | 127.1 | - | - | 0 | - |
| - | - | 470 | 127.1 | - | - | 0 | - |
| - | - | 5438 | 129.1 | - | - | 0 | - |
| - | - | 594.1 | 129.1 | - | - | 0 | - |
| - | - | 402.4 | 129.1 | - | - | 0 | - |
| - | - | 3350 | 129.1 | - | - | 0 | - |
| - | - | 601.2 | 130 | - | - | 0 | - |
| - | - | 1230 | 130.1 | - | - | 0 | - |
| - | - | 4111 | 130.1 | - | - | 0 | - |
| - | - | 1902 | 130.1 | - | - | 0 | - |
| - | - | 945.3 | 130.1 | - | - | 0 | - |
| - | - | 387.1 | 131.7 | - | - | 0 | - |
| - | - | 749.1 | 136 | - | - | 0 | - |
| - | - | 635.9 | 136.1 | - | - | 0 | - |
| - | - | 408.1 | 136.1 | - | - | 0 | - |
| - | - | 469 | 139.1 | - | - | 0 | - |
| - | - | 610.2 | 139.1 | - | - | 0 | - |
| - | - | 433.8 | 139.8 | - | - | 0 | - |
| - | - | 1098 | 141.1 | - | - | 0 | - |
| - | - | 986.8 | 141.1 | - | - | 0 | - |
| - | - | 1053 | 142.1 | - | - | 0 | - |
| - | - | 875.4 | 143.1 | - | - | 0 | - |
| - | - | 4.62E+04 | 144.1 | - | - | 0 | - |
| - | - | 2470 | 145.1 | - | - | 0 | - |
| - | - | 1899 | 147.1 | - | - | 0 | - |
| - | - | 567.5 | 149 | - | - | 0 | - |
| - | - | 1670 | 152.1 | - | - | 0 | - |
| - | - | 4068 | 153.1 | - | - | 0 | - |
| - | - | 2023 | 154 | - | - | 0 | - |
| - | - | 918.1 | 155.1 | - | - | 0 | - |
| - | - | 627.2 | 157.1 | - | - | 0 | - |
| - | - | 1071 | 157.1 | - | - | 0 | - |
| 7 | y | 8689 | 158.1 | 8.045E-05 | 0.5089 | +1 | 1 |
| - | - | 446.4 | 167 | - | - | 0 | - |
| - | - | 1.292E+04 | 169.1 | - | - | 0 | - |
| - | - | 4406 | 170.1 | - | - | 0 | - |
| - | - | 1227 | 170.1 | - | - | 0 | - |
| - | - | 414.7 | 170.3 | - | - | 0 | - |
| 2 | a | 7368 | 171.1 | 8.892E-05 | 0.5198 | +1 | 2 |
| - | - | 784.9 | 171.1 | - | - | 0 | - |
| - | - | 852.9 | 172.1 | - | - | 0 | - |
| - | - | 797.9 | 172.1 | - | - | 0 | - |
| - | - | 2326 | 173.1 | - | - | 0 | - |
| - | - | 2999 | 173.5 | - | - | 0 | - |
| 7 | y | 2.663E+04 | 175.1 | 0.0001274 | 0.7276 | +1 | 1 |
| - | - | 1352 | 176.1 | - | - | 0 | - |
| - | - | 1.621E+04 | 180.1 | - | - | 0 | - |
| - | - | 725.9 | 180.1 | - | - | 0 | - |
| - | - | 1354 | 181.1 | - | - | 0 | - |
| - | - | 840.2 | 181.1 | - | - | 0 | - |
| - | - | 834.3 | 182.1 | - | - | 0 | - |
| - | - | 724.3 | 183.1 | - | - | 0 | - |
| - | - | 563.7 | 185.1 | - | - | 0 | - |
| - | - | 1600 | 185.1 | - | - | 0 | - |
| - | - | 491.6 | 185.2 | - | - | 0 | - |
| - | - | 1.394E+04 | 187.1 | - | - | 0 | - |
| - | - | 1202 | 188.1 | - | - | 0 | - |
| 2 | a | 4.894E+04 | 189.1 | 9.857E-05 | 0.5213 | +1 | 2 |
| - | - | 3028 | 190.1 | - | - | 0 | - |
| - | - | 594.1 | 191.1 | - | - | 0 | - |
| - | - | 818 | 195.1 | - | - | 0 | - |
| - | - | 1.527E+04 | 197.1 | - | - | 0 | - |
| - | - | 2327 | 198.1 | - | - | 0 | - |
| - | - | 1431 | 198.1 | - | - | 0 | - |
| 2 | b | 1.785E+04 | 199.1 | 4.734E-05 | 0.2378 | +1 | 2 |
| - | - | 1574 | 200.1 | - | - | 0 | - |
| - | - | 2372 | 201.1 | - | - | 0 | - |
| - | - | 560.4 | 202.1 | - | - | 0 | - |
| - | - | 2347 | 202.1 | - | - | 0 | - |
| - | - | 1333 | 203.1 | - | - | 0 | - |
| - | - | 2325 | 203.1 | - | - | 0 | - |
| - | - | 695.2 | 207.1 | - | - | 0 | - |
| - | - | 515 | 209.1 | - | - | 0 | - |
| - | - | 1326 | 210.1 | - | - | 0 | - |
| - | - | 957.2 | 211.1 | - | - | 0 | - |
| - | - | 2996 | 215.1 | - | - | 0 | - |
| 2 | b | 2.83E+04 | 217.1 | 4.174E-05 | 0.1923 | +1 | 2 |
| - | - | 2061 | 218.1 | - | - | 0 | - |
| - | - | 742.3 | 219.1 | - | - | 0 | - |
| - | - | 1028 | 220.1 | - | - | 0 | - |
| - | - | 526.2 | 224.1 | - | - | 0 | - |
| - | - | 2556 | 225.1 | - | - | 0 | - |
| - | - | 833.1 | 226.2 | - | - | 0 | - |
| - | - | 2532 | 227.1 | - | - | 0 | - |
| - | - | 1158 | 228.1 | - | - | 0 | - |
| - | - | 2183 | 229 | - | - | 0 | - |
| - | - | 939.4 | 229 | - | - | 0 | - |
| - | - | 3153 | 229.6 | - | - | 0 | - |
| - | - | 625.1 | 230.1 | - | - | 0 | - |
| - | - | 595.3 | 230.1 | - | - | 0 | - |
| - | - | 3852 | 233.1 | - | - | 0 | - |
| - | - | 735.2 | 234.1 | - | - | 0 | - |
| - | - | 1236 | 238.2 | - | - | 0 | - |
| - | - | 669.1 | 239.1 | - | - | 0 | - |
| - | - | 1102 | 240.1 | - | - | 0 | - |
| - | - | 623.8 | 242.2 | - | - | 0 | - |
| - | - | 534.8 | 243.1 | - | - | 0 | - |
| - | - | 1506 | 244.1 | - | - | 0 | - |
| 6 | y | 2007 | 244.1 | 0.0002334 | 0.956 | +1 | 2 |
| - | - | 768.1 | 245.1 | - | - | 0 | - |
| 6 | y | 1.567E+04 | 245.1 | 4.973E-05 | 0.2029 | +1 | 2 |
| - | - | 1619 | 246.1 | - | - | 0 | - |
| - | - | 1095 | 249.1 | - | - | 0 | - |
| - | - | 873.4 | 251.1 | - | - | 0 | - |
| - | - | 6705 | 261.1 | - | - | 0 | - |
| 4 | y | 3127 | 261.6 | 0.002178 | 8.323 | +2 | 4 |
| 6 | y | 3.147E+04 | 262.2 | 7.115E-05 | 0.2714 | +1 | 2 |
| - | - | 2487 | 263.2 | - | - | 0 | - |
| - | - | 1140 | 266.2 | - | - | 0 | - |
| - | - | 1646 | 267.1 | - | - | 0 | - |
| - | - | 1521 | 271 | - | - | 0 | - |
| - | - | 897.9 | 272.1 | - | - | 0 | - |
| - | - | 838.5 | 277.1 | - | - | 0 | - |
| - | - | 1115 | 279 | - | - | 0 | - |
| - | - | 912.4 | 281 | - | - | 0 | - |
| 3 | a | 1.244E+04 | 284.2 | 8.224E-05 | 0.2894 | +1 | 3 |
| - | - | 1859 | 285.2 | - | - | 0 | - |
| - | - | 700.1 | 286.1 | - | - | 0 | - |
| - | - | 9710 | 286.2 | - | - | 0 | - |
| - | - | 3489 | 286.7 | - | - | 0 | - |
| - | - | 2241 | 293.9 | - | - | 0 | - |
| - | - | 1021 | 294.9 | - | - | 0 | - |
| - | - | 5834 | 297.1 | - | - | 0 | - |
| - | - | 625.4 | 298.1 | - | - | 0 | - |
| - | - | 1365 | 299.1 | - | - | 0 | - |
| - | - | 849.8 | 299.2 | - | - | 0 | - |
| 3 | a | 1174 | 302.2 | 0.0001563 | 0.5172 | +1 | 3 |
| - | - | 699.1 | 308.2 | - | - | 0 | - |
| - | - | 4726 | 311.9 | - | - | 0 | - |
| 3 | b | 2.109E+04 | 312.2 | 5.929E-05 | 0.1899 | +1 | 3 |
| - | - | 708.3 | 312.9 | - | - | 0 | - |
| - | - | 3308 | 313.2 | - | - | 0 | - |
| 3 | y | 2.585E+04 | 318.2 | 0.002473 | 7.774 | +2 | 5 |
| - | - | 6998 | 318.7 | - | - | 0 | - |
| - | - | 1454 | 319.2 | - | - | 0 | - |
| - | - | 1250 | 319.7 | - | - | 0 | - |
| - | - | 616.6 | 323.9 | - | - | 0 | - |
| 3 | b | 5959 | 330.2 | 0.0004809 | 1.457 | +1 | 3 |
| - | - | 928.8 | 332.7 | - | - | 0 | - |
| - | - | 1605 | 336.7 | - | - | 0 | - |
| - | - | 1136 | 340.2 | - | - | 0 | - |
| - | - | 1994 | 341.7 | - | - | 0 | - |
| - | - | 617.8 | 345.7 | - | - | 0 | - |
| - | - | 1018 | 346.1 | - | - | 0 | - |
| - | - | 1481 | 346.2 | - | - | 0 | - |
| - | - | 835.5 | 349.1 | - | - | 0 | - |
| - | - | 732.6 | 353.9 | - | - | 0 | - |
| - | - | 2829 | 354.9 | - | - | 0 | - |
| - | - | 785.3 | 355.7 | - | - | 0 | - |
| - | - | 1157 | 355.9 | - | - | 0 | - |
| 5 | y | 3027 | 357.2 | 0.0005114 | 1.431 | +1 | 3 |
| 5 | y | 2824 | 358.2 | 0.0005182 | 1.447 | +1 | 3 |
| - | - | 607.1 | 358.7 | - | - | 0 | - |
| 2 | y | 757 | 359.7 | 0.002624 | 7.294 | +2 | 6 |
| - | - | 2602 | 362.2 | - | - | 0 | - |
| - | - | 822.3 | 362.7 | - | - | 0 | - |
| - | - | 1047 | 364.7 | - | - | 0 | - |
| - | - | 689.5 | 367.7 | - | - | 0 | - |
| - | - | 771.2 | 368.2 | - | - | 0 | - |
| 2 | y | 1922 | 368.7 | 0.001766 | 4.791 | +2 | 6 |
| - | - | 773.1 | 371.9 | - | - | 0 | - |
| - | - | 2139 | 372.2 | - | - | 0 | - |
| - | - | 1471 | 372.9 | - | - | 0 | - |
| - | - | 801.8 | 373.7 | - | - | 0 | - |
| 5 | y | 5.435E+04 | 375.2 | 0.0003339 | 0.8897 | +1 | 3 |
| - | - | 1.086E+04 | 376.2 | - | - | 0 | - |
| - | - | 1339 | 377.2 | - | - | 0 | - |
| - | - | 7251 | 385.2 | - | - | 0 | - |
| - | - | 1653 | 385.7 | - | - | 0 | - |
| - | - | 636.5 | 386.2 | - | - | 0 | - |
| - | - | 8342 | 394.2 | - | - | 0 | - |
| - | - | 3461 | 394.7 | - | - | 0 | - |
| - | - | 1433 | 395.2 | - | - | 0 | - |
| - | - | 719.8 | 399.7 | - | - | 0 | - |
| - | - | 4428 | 402.2 | - | - | 0 | - |
| - | - | 1368 | 402.8 | - | - | 0 | - |
| - | - | 761.7 | 407.9 | - | - | 0 | - |
| - | - | 1268 | 408.2 | - | - | 0 | - |
| - | - | 791.4 | 413.2 | - | - | 0 | - |
| 0 | Precursor | 1.208E+04 | 417.2 | 0.002397 | 5.745 | +2 | -1 |
| - | - | 5736 | 417.7 | - | - | 0 | - |
| - | - | 1926 | 418.2 | - | - | 0 | - |
| - | - | 1023 | 421.6 | - | - | 0 | - |
| - | - | 2025 | 423.2 | - | - | 0 | - |
| - | - | 2375 | 425.2 | - | - | 0 | - |
| 0 | Precursor | 2132 | 426.2 | 0.0009293 | 2.18 | +2 | -1 |
| - | - | 805.4 | 430.2 | - | - | 0 | - |
| - | - | 1.097E+04 | 431.2 | - | - | 0 | - |
| - | - | 2742 | 432.3 | - | - | 0 | - |
| - | - | 2843 | 440.3 | - | - | 0 | - |
| - | - | 6092 | 441.2 | - | - | 0 | - |
| - | - | 1168 | 442.2 | - | - | 0 | - |
| - | - | 2585 | 444.3 | - | - | 0 | - |
| - | - | 6.395E+04 | 458.3 | - | - | 0 | - |
| 4 | b | 3792 | 459.2 | 0.005707 | 12.43 | +1 | 4 |
| - | - | 1.399E+04 | 459.3 | - | - | 0 | - |
| - | - | 1910 | 460.3 | - | - | 0 | - |
| - | - | 4713 | 476.3 | - | - | 0 | - |
| 4 | b | 4690 | 477.2 | 0.004451 | 9.327 | +1 | 4 |
| - | - | 1625 | 486.3 | - | - | 0 | - |
| - | - | 792.5 | 488.3 | - | - | 0 | - |
| - | - | 3360 | 500.8 | - | - | 0 | - |
| - | - | 4571 | 501.3 | - | - | 0 | - |
| 4 | y | 1827 | 504.3 | 0.004282 | 8.492 | +1 | 4 |
| - | - | 1340 | 507.2 | - | - | 0 | - |
| - | - | 944.4 | 515.3 | - | - | 0 | - |
| 4 | y | 6.658E+04 | 522.3 | 0.004307 | 8.247 | +1 | 4 |
| - | - | 1.507E+04 | 523.3 | - | - | 0 | - |
| - | - | 4328 | 524.3 | - | - | 0 | - |
| - | - | 6197 | 532.8 | - | - | 0 | - |
| - | - | 846.4 | 543.3 | - | - | 0 | - |
| - | - | 789.9 | 544.3 | - | - | 0 | - |
| - | - | 1469 | 553.3 | - | - | 0 | - |
| - | - | 1206 | 554.3 | - | - | 0 | - |
| - | - | 4.174E+04 | 571.4 | - | - | 0 | - |
| - | - | 1.455E+04 | 572.4 | - | - | 0 | - |
| - | - | 2156 | 573.4 | - | - | 0 | - |
| - | - | 1014 | 620.3 | - | - | 0 | - |
| - | - | 682.8 | 626.4 | - | - | 0 | - |
| 3 | y | 4.054E+04 | 635.3 | 0.003678 | 5.789 | +1 | 5 |
| - | - | 1.213E+04 | 636.4 | - | - | 0 | - |
| - | - | 4609 | 637.4 | - | - | 0 | - |
| - | - | 1147 | 638.4 | - | - | 0 | - |
| - | - | 866.3 | 664.4 | - | - | 0 | - |
| - | - | 3044 | 682.4 | - | - | 0 | - |
| - | - | 1673 | 683.4 | - | - | 0 | - |
| - | - | 671.3 | 684.4 | - | - | 0 | - |
| - | - | 1424 | 691.4 | - | - | 0 | - |
| 2 | y | 1210 | 718.4 | 0.006421 | 8.937 | +1 | 6 |
| 2 | y | 1161 | 736.4 | 0.006964 | 9.457 | +1 | 6 |
| - | - | 6208 | 746.4 | - | - | 0 | - |
| - | - | 2616 | 747.4 | - | - | 0 | - |
| - | - | 1406 | 748.4 | - | - | 0 | - |
| - | - | 2608 | 804.5 | - | - | 0 | - |
| - | - | 696.3 | 934.6 | - | - | 0 | - |
| - | - | 712.7 | 3080 | - | - | 0 | - |
| - | - | 800.9 | 3337 | - | - | 0 | - |
| - | - | 755.6 | 3425 | - | - | 0 | - |

m/z Charge Intensity FragmentType MassShift Position
120.0811767578125 0 674.50323
121.10116577148438 0 443.40912
123.08052825927734 0 363.18738
124.11221313476562 0 1282.8263
125.07088470458984 0 970.76855
125.10780334472656 0 384.40155
126.05516815185547 0 12671.541
126.64835357666016 0 406.65726
127.05068969726562 0 589.2221
127.05856323242188 0 469.96896
129.0660400390625 0 5438.4126
129.07102966308594 0 594.14215
129.07351684570312 0 402.41357
129.1024169921875 0 3349.7961
130.03196716308594 0 601.1678
130.050048828125 0 1230.0591
130.06126403808594 0 4110.706
130.08645629882812 0 1902.4834
130.09779357910156 0 945.30334
131.66221618652344 0 387.0614
136.039306640625 0 749.1437
136.07611083984375 0 635.9292
136.11248779296875 0 408.07208
139.05064392089844 0 469.02423
139.11167907714844 0 610.2471
139.76760864257812 0 433.79065
141.1023712158203 0 1097.5352
141.13864135742188 0 986.7876
142.12289428710938 0 1052.6857
143.11817932128906 0 875.3674
144.065673828125 0 46201.18
145.06910705566406 0 2470.432
147.11302185058594 0 1898.9042
148.95460510253906 0 567.4744
152.10720825195312 0 1669.6251
153.06594848632812 0 4068.4136
154.0499267578125 0 2023.0858
155.07032775878906 0 918.0731
157.09713745117188 0 627.20325
157.10888671875 0 1070.6074
158.0924835205078 0 8689.489 y Ammonia loss 6
167.02951049804688 0 446.39047
169.13363647460938 0 12922.911
170.09249877929688 0 4406.3066
170.1371307373047 0 1227.3661
170.3150177001953 0 414.7202
171.07650756835938 0 7368.08 a Water loss 1
171.1133575439453 0 784.90686
172.06072998046875 0 852.9252
172.0796661376953 0 797.92236
173.12860107421875 0 2326.0188
173.45033264160156 0 2998.747
175.11907958984375 0 26626.605 y 6
176.12246704101562 0 1351.6758
180.10202026367188 0 16206.447
180.1112823486328 0 725.9024
181.10531616210938 0 1354.1324
181.1337890625 0 840.22424
182.09254455566406 0 834.2557
183.1125030517578 0 724.3253
185.0918731689453 0 563.6606
185.13992309570312 0 1599.7141
185.16455078125 0 491.6436
187.1442108154297 0 13936.402
188.14767456054688 0 1202.4048
189.0870819091797 0 48942.38 a 1
190.09051513671875 0 3028.3435
191.09213256835938 0 594.06604
195.1129608154297 0 818.0383
197.12850952148438 0 15271.132
198.08734130859375 0 2327.3337
198.13180541992188 0 1430.6605
199.07138061523438 0 17846.467 b Water loss 1
200.07505798339844 0 1574.0399
201.12344360351562 0 2372.426
202.1091766357422 0 560.4216
202.11871337890625 0 2346.7788
203.10238647460938 0 1332.9313
203.12762451171875 0 2325.2961
207.11192321777344 0 695.16
209.07862854003906 0 514.99414
210.0875701904297 0 1326.369
211.10755920410156 0 957.2133
215.13893127441406 0 2995.8965
217.08193969726562 0 28302.14 b 1
218.0855712890625 0 2060.7668
219.08160400390625 0 742.3299
220.1295928955078 0 1027.8931
224.1398468017578 0 526.15173
225.12368774414062 0 2556.0154
226.15435791015625 0 833.11774
227.11402893066406 0 2531.6672
228.0980682373047 0 1157.7761
229.00125122070312 0 2182.8123
229.0123748779297 0 939.4119
229.63975524902344 0 3152.7563
230.12579345703125 0 625.0699
230.1400604248047 0 595.34283
233.1316680908203 0 3851.837
234.14561462402344 0 735.2186
238.15542602539062 0 1235.5352
239.13809204101562 0 669.0684
240.13433837890625 0 1101.5784
242.1501007080078 0 623.85
243.11669921875 0 534.7863
244.10018920898438 0 1506.4008
244.1401824951172 0 2006.5017 y Water loss 5
245.10311889648438 0 768.1188
245.12448120117188 0 15665.101 y Ammonia loss 5
246.1299285888672 0 1618.7375
249.12335205078125 0 1094.9989
251.10203552246094 0 873.36237
261.12664794921875 0 6705.099
261.6385803222656 0 3126.6367 y 3
262.1509094238281 0 31472.906 y 5
263.15399169921875 0 2486.9211
266.1502685546875 0 1139.7362
267.1338195800781 0 1646.0237
271.0274658203125 0 1520.6139
272.13580322265625 0 897.8782
277.1187438964844 0 838.51074
279.04791259765625 0 1114.7943
281.04364013671875 0 912.4239
284.160400390625 0 12441.379 a Water loss 2
285.1636657714844 0 1859.3682
286.1400146484375 0 700.08
286.18182373046875 0 9710.306
286.68310546875 0 3489.3147
293.9344787597656 0 2240.7627
294.9176330566406 0 1020.72675
297.0579833984375 0 5834.3247
298.0588684082031 0 625.35315
299.05426025390625 0 1365.1101
299.1716613769531 0 849.82104
302.17120361328125 0 1173.7655 a 2
308.15985107421875 0 699.13367
311.9450378417969 0 4725.665
312.15545654296875 0 21090.133 b Water loss 2
312.94403076171875 0 708.268
313.1588134765625 0 3308.2583
318.180908203125 0 25854.945 y 2
318.6822204589844 0 6998.1167
319.1815490722656 0 1453.8383
319.6854553222656 0 1250.1123
323.88238525390625 0 616.5868
330.16644287109375 0 5959.1685 b 2
332.6918640136719 0 928.80176
336.7059020996094 0 1604.7521
340.19805908203125 0 1136.4653
341.6972351074219 0 1993.9857
345.70263671875 0 617.7614
346.1426696777344 0 1018.18896
346.2090759277344 0 1481.4592
349.0752258300781 0 835.5188
353.9020080566406 0 732.5797
354.8849182128906 0 2828.538
355.6942138671875 0 785.33496
355.8833312988281 0 1156.6372
357.2239685058594 0 3026.6025 y Water loss 4
358.2079772949219 0 2824.2183 y Ammonia loss 4
358.6883239746094 0 607.1291
359.6996154785156 0 756.9973 y Water loss 1
362.2106018066406 0 2602.1433
362.7109375 0 822.34204
364.6902160644531 0 1047.0034
367.69561767578125 0 689.5386
368.1938781738281 0 771.2336
368.70404052734375 0 1921.5935 y 1
371.9116516113281 0 773.106
372.1589050292969 0 2138.9387
372.8957214355469 0 1471.122
373.6950378417969 0 801.77985
375.2347106933594 0 54350.44 y 4
376.23773193359375 0 10856.9795
377.2401428222656 0 1338.8091
385.2154235839844 0 7251.45
385.714111328125 0 1653.1938
386.22100830078125 0 636.5301
394.2164306640625 0 8342.194
394.7181091308594 0 3460.9165
395.2182312011719 0 1432.5963
399.69708251953125 0 719.824
402.2093200683594 0 4427.5664
402.75030517578125 0 1367.6534
407.8525085449219 0 761.70685
408.2087097167969 0 1268.4108
413.2077331542969 0 791.40564
417.2128601074219 0 12084.227 Precursor Water loss
417.7139892578125 0 5736.0303
418.21405029296875 0 1926.1338
421.564208984375 0 1023.10516
423.2348937988281 0 2025.2646
425.21478271484375 0 2375.2815
426.2166748046875 0 2132.3582 Precursor
430.23931884765625 0 805.3572
431.248291015625 0 10970.63
432.251953125 0 2741.6108
440.2611083984375 0 2843.0056
441.2452392578125 0 6092.475
442.24847412109375 0 1167.9634
444.26873779296875 0 2584.9385
458.2720031738281 0 63945.797
459.19158935546875 0 3791.5928 b Water loss 3
459.2749938964844 0 13985.022
460.27813720703125 0 1909.9143
476.2684020996094 0 4712.6255
477.2008972167969 0 4690.263 b 3
486.26708984375 0 1625.0007
488.26861572265625 0 792.5129
500.8105163574219 0 3360.0156
501.27783203125 0 4571.296
504.2592468261719 0 1827.2677 y Water loss 3
507.2466735839844 0 1339.862
515.2922973632812 0 944.41046
522.2698364257812 0 66576.45 y 3
523.2725219726562 0 15070.37
524.2702026367188 0 4327.6416
532.8095703125 0 6197.3164
543.3233032226562 0 846.3805
544.33056640625 0 789.89197
553.34375 0 1469.4447
554.3300170898438 0 1206.2798
571.355224609375 0 41742.746
572.3585205078125 0 14547.303
573.3602294921875 0 2156.0808
620.3275146484375 0 1014.4704
626.3951416015625 0 682.7704
635.353271484375 0 40535.973 y 2
636.3564453125 0 12125.246
637.3556518554688 0 4608.7344
638.3587646484375 0 1146.8765
664.3718872070312 0 866.2824
682.3873901367188 0 3044.3672
683.3910522460938 0 1673.1797
684.3963623046875 0 671.26746
691.4077758789062 0 1423.868
718.3931274414062 0 1209.5334 y Water loss 1
736.4042358398438 0 1160.5924 y 1
746.3848266601562 0 6208.2456
747.3870849609375 0 2616.4375
748.385986328125 0 1405.6914
804.4920654296875 0 2608.1335
934.5833740234375 0 696.31116
3079.57177734375 0 712.6948
3337.16552734375 0 800.87225
3424.82421875 0 755.61505

Spectrum Details

|  |  |
| --- | --- |
| Matched peaks? Matched peaks The total absolute number of peaks matched. Additionally in brackets the total fraction of peaks matched and the total number of peaks is shown. | 29 (11.79% of 246) |
| FDR? FDR The false discovery rate estimated for this peptide. It is calculated by matching all theoretical fragments with a non-integer shift with the raw peaks for this spectrum. This is done with 40 different shifts. The resulting percentage is the average number of annotated peaks over the number of annotated peaks with the correct spectrum. | 0.33% |
| Satellite FDR? Satellite FDR See the FDR for details on its calculation. This satellite ion specific FDR only contains the satellite ions (d/w) for I/L/J positions. | - |
| PSM Score? PSM Score The PSM Score as given by Hecklib to this annotated spectrum. It is shown with three significant figures. | 376 |

#### Reverse Lookup? Reverse Lookup All places where this read could be placed.

| Group | Segment | Template | Template Part | Read Part | Score | Unique |
| --- | --- | --- | --- | --- | --- | --- |
| Homo sapiens Heavy Chain | IGHC | IGHG1 | [131..138] | [0..7] | 56 | False |
| Homo sapiens Heavy Chain | IGHC | IGHG3 | [178..185] | [0..7] | 56 | False |
| Homo sapiens Heavy Chain | IGHC | IGHG2 | [127..134] | [0..7] | 56 | False |
| Homo sapiens Heavy Chain | IGHC | IGHG4 | [128..135] | [0..7] | 56 | False |

| Recombined | Template Part | Read Part | Score | Unique |
| --- | --- | --- | --- | --- |
| REC-0-1 | [254..261] | [0..7] | 56 | True |

#### Meta Information from Multiple reads

##### Number of combined reads

6

##### Intensity

1

##### TotalArea

0

##### Changes to the peptide sequence

DTJMJSR

I→JNo support for either Leucine or Isoleucine based on side chain ions (Position: 3)

J→ISupport for Isoleucine based on side chain ions (1 for I 0 for L) (Position: 3)

I→JNo support for either Leucine or Isoleucine based on side chain ions (Position: 5)

J→ISupport for Isoleucine based on side chain ions (1 for I 0 for L) (Position: 5)

L→JNo support for either Leucine or Isoleucine based on side chain ions (Position: 5)

L→JNo support for either Leucine or Isoleucine based on side chain ions (Position: 3)

#### Positional Score

Copy Data

##### Positional Score (TSV)

###### Preview

```
Loading example...
```

*Click on the button to copy the data to your clipboard.*

100123456

Label Value
"0" 0.612
"1" 0.617
"2" 0.66
"3" 0.66
"4" 0.662
"5" 0.662
"6" 0.667

#### Meta Information from PEAKS

##### Scan Identifier

F4:4447

##### Original sequence

D

T

L

M

+15.99

L

S

R

##### Posttranslational Modifications

Oxidation (M)

##### Source File

D:\separate\_stitch\_analyses\xle-disambiguation\raw\20210323\_F1\_UM1\_Peng0013\_SA\_F59\_ingel\_3ug\_tryp.raw

##### Fraction

4

##### Scan Feature

-

##### De Novo Score

98

##### ConfidenceScore

97

### m/z

426.2185

##### Mass

850.4219

##### Charge

2

##### Retention Time

23.82

##### Predicted Retention Time

23.12

##### Area

0

##### Parts Per Million

0.6

##### Fragmentation mode

HCD

##### Originating file

01 D:\separate\_stitch\_analyses\xle-disambiguation\20210325\_F59\_3ug\_DENOVO\_12.csv

#### Meta Information from PEAKS

##### Scan Identifier

F4:4337

##### Original sequence

D

T

L

M

+15.99

L

S

R

##### Posttranslational Modifications

Oxidation (M)

##### Source File

D:\separate\_stitch\_analyses\xle-disambiguation\raw\20210323\_F1\_UM1\_Peng0013\_SA\_F59\_ingel\_3ug\_tryp.raw

##### Fraction

4

##### Scan Feature

-

##### De Novo Score

98

##### ConfidenceScore

95

### m/z

426.2192

##### Mass

850.4219

##### Charge

2

##### Retention Time

23.26

##### Predicted Retention Time

23.12

##### Area

0

##### Parts Per Million

2.4

##### Fragmentation mode

ETHCD

##### Originating file

01 D:\separate\_stitch\_analyses\xle-disambiguation\20210325\_F59\_3ug\_DENOVO\_12.csv

#### Meta Information from PEAKS

##### Scan Identifier

F4:4286

##### Original sequence

D

T

L

M

+15.99

L

S

R

##### Posttranslational Modifications

Oxidation (M)

##### Source File

D:\separate\_stitch\_analyses\xle-disambiguation\raw\20210323\_F1\_UM1\_Peng0013\_SA\_F59\_ingel\_3ug\_tryp.raw

##### Fraction

4

##### Scan Feature

-

##### De Novo Score

98

##### ConfidenceScore

98

### m/z

426.2186

##### Mass

850.4219

##### Charge

2

##### Retention Time

22.98

##### Predicted Retention Time

-

##### Area

0

##### Parts Per Million

0.9

##### Fragmentation mode

HCD

##### Originating file

01 D:\separate\_stitch\_analyses\xle-disambiguation\20210325\_F59\_3ug\_DENOVO\_12.csv

#### Meta Information from PEAKS

##### Scan Identifier

F4:4554

##### Original sequence

D

T

L

M

+15.99

L

S

R

##### Posttranslational Modifications

Oxidation (M)

##### Source File

D:\separate\_stitch\_analyses\xle-disambiguation\raw\20210323\_F1\_UM1\_Peng0013\_SA\_F59\_ingel\_3ug\_tryp.raw

##### Fraction

4

##### Scan Feature

-

##### De Novo Score

97

##### ConfidenceScore

98

### m/z

426.2186

##### Mass

850.4219

##### Charge

2

##### Retention Time

24.39

##### Predicted Retention Time

23.12

##### Area

0

##### Parts Per Million

0.8

##### Fragmentation mode

HCD

##### Originating file

01 D:\separate\_stitch\_analyses\xle-disambiguation\20210325\_F59\_3ug\_DENOVO\_12.csv

#### Meta Information from PEAKS

##### Scan Identifier

F4:4616

##### Original sequence

D

T

L

M

+15.99

L

S

R

##### Posttranslational Modifications

Oxidation (M)

##### Source File

D:\separate\_stitch\_analyses\xle-disambiguation\raw\20210323\_F1\_UM1\_Peng0013\_SA\_F59\_ingel\_3ug\_tryp.raw

##### Fraction

4

##### Scan Feature

-

##### De Novo Score

96

##### ConfidenceScore

97

### m/z

426.2184

##### Mass

850.4219

##### Charge

2

##### Retention Time

24.72

##### Predicted Retention Time

23.12

##### Area

0

##### Parts Per Million

0.5

##### Fragmentation mode

HCD

##### Originating file

01 D:\separate\_stitch\_analyses\xle-disambiguation\20210325\_F59\_3ug\_DENOVO\_12.csv

#### Meta Information from PEAKS

##### Scan Identifier

F4:4676

##### Original sequence

D

T

L

M

+15.99

L

S

R

##### Posttranslational Modifications

Oxidation (M)

##### Source File

D:\separate\_stitch\_analyses\xle-disambiguation\raw\20210323\_F1\_UM1\_Peng0013\_SA\_F59\_ingel\_3ug\_tryp.raw

##### Fraction

4

##### Scan Feature

-

##### De Novo Score

95

##### ConfidenceScore

95

### m/z

426.2186

##### Mass

850.4219

##### Charge

2

##### Retention Time

25.04

##### Predicted Retention Time

23.12

##### Area

0

##### Parts Per Million

1

##### Fragmentation mode

HCD

##### Originating file

01 D:\separate\_stitch\_analyses\xle-disambiguation\20210325\_F59\_3ug\_DENOVO\_12.csv
