## Supplementary material for "A handle on mass coincidence errors in *de novo* sequencing of antibodies by bottom-up proteomics": Combined_079.html

Details Combined\_079 | Stitch OverviewUndefined

### Read Combined\_079

#### Sequence (length=16)

VSWNSGALTSGVHTWJ

#### Spectrum 9144? Spectrum 9144 The raw spectrum of this peptide as annotated by Hecklib. The fragments are coloured according to ion type (see legend). Any peaks with a star '\*' as text can be hovered over to see the full details, first the ion type second the mass shift type. By hovering over the amino acids in the peptide or ions in the legend the corresponding peaks are highlighted. By toggling the 'Unassigned' label you can turn the background (unassigned) peaks on or off in the plot. By updating the slider in the Ion legend you can update the spectrum to only show the top X% of the peaks with labels. The top X% means any peak that is within X% of the highest intensity. By dragging in the spectrum you can zoom in to a specific part of the spectrum and use 'Zoom Out' to get back to the original zoom level. The annotation of the spectrum is based on the given sequence in the peptides file and is done with different software so inconsistencies are likely. The peaks are annotated based on the given sequence, with 20 ppm tolerance.

Copy Data

##### Spectrum 9144 (TSV)

###### Preview

```
Loading example...
```

*Click on the button to copy the data to your clipboard.*

Mz MinMz MaxIntensity Max

WidthHeightPeptide font sizePeptide stroke widthSpectrum font sizeSpectrum stroke widthCompact peptide

Ion legend

wxyz

abcd

OtherUnassignedIon^Charge^_Position_Show for top:%

VSWNSGALTSGVHTWJ

05.68e+41.14e+51.70e+52.27e+5

Zoom Out

y^+1^_2_w^+2^_8_c^+1^_4_z^+1^_4_y^+1^_4_c^+1^_5_c^+1^_5_c^+1^_6_y^+1^_5_y^+2^_13_c^+2^_14_c^+1^_7_z^+1^_6_y^+1^_6_z^+2^_14_z^+2^_14_y^+2^_14_y^+2^_14_z^+2^_14_y^+2^_14_z^+1^_7_y^+1^_7_z^+1^_7_c^+2^_15_y^+2^_15_c^+1^_8_y^+2^_15_y^+1^_8_z^+1^_8_c^+1^_9_c^+1^_9_z^+1^_9_c^+1^_10_y^+1^_9_c^+1^_11_c^+1^_11_y^+1^_10_z^+1^_10_y^+1^_10_z^+1^_11_y^+1^_11_c^+1^_12_c^+1^_12_y^+1^_12_z^+1^_12_y^+1^_12_c^+1^_13_c^+1^_13_y^+1^_13_z^+1^_13_y^+1^_13_c^+1^_14_c^+1^_14_y^+1^_14_y^+1^_14_z^+1^_14_y^+1^_14_z^+1^_15_c^+1^_15_c^+1^_15_y^+1^_15_z^+1^_15_c^+1^_15_y^+1^_15_

0775155023253100

Fragment Matches Table

Show background peaks

| Position | Ion type | Intensity | mz Theoretical | mz Error (Th) | mz Error (ppm) | Charge | Series Number |
| --- | --- | --- | --- | --- | --- | --- | --- |
| - | - | 400.1 | 120.9 | - | - | 0 | - |
| - | - | 423.8 | 135.5 | - | - | 0 | - |
| - | - | 1066 | 148.9 | - | - | 0 | - |
| - | - | 2401 | 148.9 | - | - | 0 | - |
| - | - | 1689 | 149 | - | - | 0 | - |
| - | - | 1304 | 149 | - | - | 0 | - |
| - | - | 782.6 | 149 | - | - | 0 | - |
| - | - | 6566 | 159.1 | - | - | 0 | - |
| - | - | 1131 | 173.4 | - | - | 0 | - |
| - | - | 1175 | 173.5 | - | - | 0 | - |
| - | - | 3.099E+04 | 187.1 | - | - | 0 | - |
| - | - | 2526 | 188.1 | - | - | 0 | - |
| - | - | 495.9 | 231.3 | - | - | 0 | - |
| - | - | 815.4 | 246.1 | - | - | 0 | - |
| - | - | 531.2 | 250.3 | - | - | 0 | - |
| - | - | 610.6 | 256.1 | - | - | 0 | - |
| - | - | 1282 | 274.1 | - | - | 0 | - |
| - | - | 1058 | 284.1 | - | - | 0 | - |
| - | - | 3289 | 301.1 | - | - | 0 | - |
| 15 | y | 2502 | 334.2 | 0.005082 | 15.21 | +1 | 2 |
| - | - | 632.1 | 335.2 | - | - | 0 | - |
| - | - | 6186 | 355.2 | - | - | 0 | - |
| - | - | 1232 | 356.2 | - | - | 0 | - |
| - | - | 1105 | 373.2 | - | - | 0 | - |
| - | - | 2164 | 388.2 | - | - | 0 | - |
| - | - | 604.5 | 392.2 | - | - | 0 | - |
| - | - | 569 | 417 | - | - | 0 | - |
| 9 | w | 616.4 | 442.2 | 0.006398 | 14.47 | +2 | 8 |
| - | - | 808 | 457.2 | - | - | 0 | - |
| - | - | 830.7 | 459.2 | - | - | 0 | - |
| - | - | 718.7 | 469.2 | - | - | 0 | - |
| - | - | 533.5 | 476.7 | - | - | 0 | - |
| - | - | 894.7 | 482.2 | - | - | 0 | - |
| 4 | c | 4828 | 487.2 | 0.000418 | 0.8579 | +1 | 4 |
| - | - | 1095 | 488.2 | - | - | 0 | - |
| - | - | 653.9 | 524.3 | - | - | 0 | - |
| - | - | 751.7 | 542.3 | - | - | 0 | - |
| 13 | z | 2045 | 556.3 | 0.008108 | 14.58 | +1 | 4 |
| - | - | 1568 | 557.3 | - | - | 0 | - |
| - | - | 904.1 | 568.2 | - | - | 0 | - |
| 13 | y | 4826 | 572.3 | 0.005456 | 9.533 | +1 | 4 |
| 5 | c | 1414 | 573.3 | 0.008222 | 14.34 | +1 | 5 |
| 5 | c | 3613 | 574.3 | 0.0001584 | 0.2758 | +1 | 5 |
| - | - | 943.9 | 575.3 | - | - | 0 | - |
| - | - | 1881 | 585.2 | - | - | 0 | - |
| - | - | 923.5 | 603.3 | - | - | 0 | - |
| - | - | 729.5 | 611.3 | - | - | 0 | - |
| - | - | 731.7 | 612.3 | - | - | 0 | - |
| - | - | 1999 | 629.3 | - | - | 0 | - |
| 6 | c | 950.6 | 631.3 | 0.001408 | 2.23 | +1 | 6 |
| - | - | 667.2 | 667.3 | - | - | 0 | - |
| 12 | y | 692.1 | 671.3 | 0.001068 | 1.59 | +1 | 5 |
| - | - | 927.6 | 674.3 | - | - | 0 | - |
| 4 | y | 677.5 | 679.8 | 0.005908 | 8.69 | +2 | 13 |
| - | - | 914.4 | 680.3 | - | - | 0 | - |
| - | - | 3116 | 684.3 | - | - | 0 | - |
| - | - | 2459 | 685.3 | - | - | 0 | - |
| - | - | 883.3 | 690.3 | - | - | 0 | - |
| - | - | 832.5 | 691.3 | - | - | 0 | - |
| 14 | c | 1413 | 699.3 | 0.001877 | 2.684 | +2 | 14 |
| 7 | c | 2542 | 702.3 | 0.0001136 | 0.1618 | +1 | 7 |
| - | - | 796.1 | 703.3 | - | - | 0 | - |
| 11 | z | 1655 | 712.3 | 0.003638 | 5.107 | +1 | 6 |
| - | - | 6949 | 713.4 | - | - | 0 | - |
| - | - | 3267 | 714.4 | - | - | 0 | - |
| - | - | 1170 | 716.3 | - | - | 0 | - |
| 11 | y | 6376 | 728.4 | 0.005239 | 7.192 | +1 | 6 |
| - | - | 2752 | 729.4 | - | - | 0 | - |
| - | - | 3279 | 730.4 | - | - | 0 | - |
| - | - | 1071 | 731.4 | - | - | 0 | - |
| - | - | 1216 | 750.9 | - | - | 0 | - |
| - | - | 665.7 | 751.4 | - | - | 0 | - |
| - | - | 1073 | 754.9 | - | - | 0 | - |
| - | - | 1121 | 755.4 | - | - | 0 | - |
| 3 | z | 1325 | 755.9 | 0.001403 | 1.857 | +2 | 14 |
| 3 | z | 786.8 | 756.4 | 0.01324 | 17.51 | +2 | 14 |
| - | - | 2254 | 758.9 | - | - | 0 | - |
| - | - | 2185 | 759.4 | - | - | 0 | - |
| - | - | 799.5 | 759.9 | - | - | 0 | - |
| 3 | y | 3378 | 763.9 | 0.002417 | 3.165 | +2 | 14 |
| 3 | y | 5590 | 764.4 | 0.008151 | 10.66 | +2 | 14 |
| 3 | z | 4106 | 764.9 | 0.003689 | 4.824 | +2 | 14 |
| - | - | 1192 | 765.4 | - | - | 0 | - |
| - | - | 847.7 | 770.4 | - | - | 0 | - |
| 3 | y | 8613 | 772.9 | 0.002506 | 3.243 | +2 | 14 |
| - | - | 7920 | 773.4 | - | - | 0 | - |
| - | - | 4001 | 773.9 | - | - | 0 | - |
| - | - | 1660 | 774.4 | - | - | 0 | - |
| - | - | 793.7 | 780.4 | - | - | 0 | - |
| 10 | z | 1179 | 781.4 | 0.0004239 | 0.5425 | +1 | 7 |
| - | - | 726.2 | 788.4 | - | - | 0 | - |
| 10 | y | 2289 | 797.4 | 0.00233 | 2.922 | +1 | 7 |
| - | - | 3274 | 798.4 | - | - | 0 | - |
| 10 | z | 3873 | 799.4 | 0.005545 | 6.937 | +1 | 7 |
| 15 | c | 1.177E+04 | 800.4 | 0.01437 | 17.95 | +2 | 15 |
| - | - | 5963 | 801.4 | - | - | 0 | - |
| - | - | 1534 | 802.4 | - | - | 0 | - |
| 2 | y | 742.8 | 807.9 | 0.007091 | 8.777 | +2 | 15 |
| - | - | 628.7 | 814.4 | - | - | 0 | - |
| 8 | c | 1.169E+04 | 815.4 | 0.001011 | 1.239 | +1 | 8 |
| 2 | y | 6380 | 816.4 | 0.015 | 18.37 | +2 | 15 |
| - | - | 1969 | 816.9 | - | - | 0 | - |
| - | - | 2018 | 817.4 | - | - | 0 | - |
| - | - | 689.2 | 821.4 | - | - | 0 | - |
| - | - | 786.8 | 825.4 | - | - | 0 | - |
| - | - | 711.8 | 839.4 | - | - | 0 | - |
| - | - | 1753 | 843.4 | - | - | 0 | - |
| - | - | 747.2 | 844.4 | - | - | 0 | - |
| - | - | 1498 | 847.9 | - | - | 0 | - |
| - | - | 1997 | 848.4 | - | - | 0 | - |
| - | - | 2393 | 848.9 | - | - | 0 | - |
| - | - | 3788 | 856.9 | - | - | 0 | - |
| - | - | 5930 | 857.4 | - | - | 0 | - |
| - | - | 5454 | 857.9 | - | - | 0 | - |
| - | - | 2163 | 858.4 | - | - | 0 | - |
| - | - | 790.4 | 858.9 | - | - | 0 | - |
| - | - | 8752 | 865.9 | - | - | 0 | - |
| - | - | 1.272E+04 | 866.4 | - | - | 0 | - |
| - | - | 7744 | 866.9 | - | - | 0 | - |
| - | - | 3018 | 867.4 | - | - | 0 | - |
| - | - | 617.6 | 881.4 | - | - | 0 | - |
| 9 | y | 2390 | 898.4 | 0.004394 | 4.891 | +1 | 8 |
| - | - | 8247 | 899.4 | - | - | 0 | - |
| 9 | z | 5876 | 900.4 | 0.002972 | 3.3 | +1 | 8 |
| - | - | 8036 | 901.4 | - | - | 0 | - |
| - | - | 3389 | 902.4 | - | - | 0 | - |
| 9 | c | 1.636E+04 | 916.5 | 0.0002202 | 0.2402 | +1 | 9 |
| - | - | 8968 | 917.5 | - | - | 0 | - |
| - | - | 1873 | 918.5 | - | - | 0 | - |
| - | - | 768.7 | 932.5 | - | - | 0 | - |
| 9 | c | 2539 | 933.5 | 0.002772 | 2.969 | +1 | 9 |
| - | - | 1268 | 934.5 | - | - | 0 | - |
| - | - | 1287 | 971.5 | - | - | 0 | - |
| - | - | 784.7 | 1013 | - | - | 0 | - |
| 8 | z | 7676 | 1014 | 0.004906 | 4.841 | +1 | 9 |
| - | - | 9518 | 1015 | - | - | 0 | - |
| - | - | 4224 | 1016 | - | - | 0 | - |
| - | - | 1323 | 1017 | - | - | 0 | - |
| - | - | 1537 | 1020 | - | - | 0 | - |
| 10 | c | 3242 | 1021 | 0.001608 | 1.575 | +1 | 10 |
| - | - | 1917 | 1022 | - | - | 0 | - |
| - | - | 686.6 | 1023 | - | - | 0 | - |
| - | - | 1241 | 1028 | - | - | 0 | - |
| 8 | y | 1.251E+04 | 1030 | 0.004676 | 4.542 | +1 | 9 |
| - | - | 7567 | 1031 | - | - | 0 | - |
| - | - | 2736 | 1032 | - | - | 0 | - |
| - | - | 783.9 | 1036 | - | - | 0 | - |
| - | - | 664.3 | 1042 | - | - | 0 | - |
| - | - | 1529 | 1059 | - | - | 0 | - |
| 11 | c | 1447 | 1060 | 0.005071 | 4.786 | +1 | 11 |
| - | - | 3186 | 1077 | - | - | 0 | - |
| 11 | c | 2606 | 1078 | 0.001587 | 1.473 | +1 | 11 |
| - | - | 1547 | 1079 | - | - | 0 | - |
| 7 | y | 726 | 1083 | 0.008934 | 8.252 | +1 | 10 |
| 7 | z | 2.777E+04 | 1085 | 0.004719 | 4.351 | +1 | 10 |
| - | - | 3.078E+04 | 1086 | - | - | 0 | - |
| - | - | 1.488E+04 | 1087 | - | - | 0 | - |
| - | - | 4337 | 1088 | - | - | 0 | - |
| 7 | y | 6059 | 1101 | 0.004671 | 4.245 | +1 | 10 |
| - | - | 3727 | 1102 | - | - | 0 | - |
| - | - | 1351 | 1103 | - | - | 0 | - |
| - | - | 909.1 | 1133 | - | - | 0 | - |
| - | - | 1618 | 1134 | - | - | 0 | - |
| - | - | 725.2 | 1141 | - | - | 0 | - |
| 6 | z | 1.33E+04 | 1142 | 0.004129 | 3.617 | +1 | 11 |
| - | - | 1.247E+04 | 1143 | - | - | 0 | - |
| - | - | 6951 | 1144 | - | - | 0 | - |
| - | - | 2575 | 1145 | - | - | 0 | - |
| - | - | 651.6 | 1155 | - | - | 0 | - |
| - | - | 1224 | 1156 | - | - | 0 | - |
| - | - | 1544 | 1157 | - | - | 0 | - |
| 6 | y | 1.378E+04 | 1158 | 0.003349 | 2.893 | +1 | 11 |
| 12 | c | 9739 | 1159 | 0.006848 | 5.911 | +1 | 12 |
| - | - | 3203 | 1160 | - | - | 0 | - |
| - | - | 1636 | 1161 | - | - | 0 | - |
| - | - | 1897 | 1173 | - | - | 0 | - |
| - | - | 1023 | 1174 | - | - | 0 | - |
| - | - | 9649 | 1176 | - | - | 0 | - |
| 12 | c | 7491 | 1177 | 0.005182 | 4.404 | +1 | 12 |
| - | - | 2541 | 1178 | - | - | 0 | - |
| - | - | 972.9 | 1179 | - | - | 0 | - |
| - | - | 1419 | 1200 | - | - | 0 | - |
| - | - | 1160 | 1201 | - | - | 0 | - |
| 5 | y | 982.6 | 1227 | 0.0009024 | 0.7357 | +1 | 12 |
| - | - | 2366 | 1228 | - | - | 0 | - |
| 5 | z | 1.477E+04 | 1229 | 0.003961 | 3.224 | +1 | 12 |
| - | - | 1.626E+04 | 1230 | - | - | 0 | - |
| - | - | 8340 | 1231 | - | - | 0 | - |
| - | - | 2237 | 1232 | - | - | 0 | - |
| 5 | y | 1.152E+04 | 1245 | 0.004524 | 3.635 | +1 | 12 |
| - | - | 7723 | 1246 | - | - | 0 | - |
| - | - | 3545 | 1247 | - | - | 0 | - |
| - | - | 818.3 | 1253 | - | - | 0 | - |
| - | - | 4854 | 1270 | - | - | 0 | - |
| - | - | 5300 | 1271 | - | - | 0 | - |
| - | - | 2485 | 1272 | - | - | 0 | - |
| - | - | 1444 | 1273 | - | - | 0 | - |
| 13 | c | 1223 | 1297 | 0.0006792 | 0.5238 | +1 | 13 |
| - | - | 1168 | 1298 | - | - | 0 | - |
| - | - | 1.834E+04 | 1313 | - | - | 0 | - |
| - | - | 1046 | 1313 | - | - | 0 | - |
| 13 | c | 2.837E+04 | 1314 | 0.003913 | 2.979 | +1 | 13 |
| - | - | 1.704E+04 | 1315 | - | - | 0 | - |
| - | - | 6827 | 1316 | - | - | 0 | - |
| - | - | 1392 | 1317 | - | - | 0 | - |
| 4 | y | 1369 | 1342 | 0.00194 | 1.446 | +1 | 13 |
| 4 | z | 7838 | 1343 | 0.002537 | 1.89 | +1 | 13 |
| - | - | 6429 | 1344 | - | - | 0 | - |
| - | - | 3575 | 1345 | - | - | 0 | - |
| - | - | 664.1 | 1354 | - | - | 0 | - |
| 4 | y | 8577 | 1359 | 0.0005616 | 0.4133 | +1 | 13 |
| - | - | 7239 | 1360 | - | - | 0 | - |
| - | - | 3582 | 1361 | - | - | 0 | - |
| - | - | 880.5 | 1371 | - | - | 0 | - |
| 14 | c | 1655 | 1398 | 0.0007146 | 0.5113 | +1 | 14 |
| - | - | 1695 | 1399 | - | - | 0 | - |
| - | - | 6096 | 1414 | - | - | 0 | - |
| 14 | c | 2.584E+04 | 1415 | 0.003496 | 2.471 | +1 | 14 |
| - | - | 1.971E+04 | 1416 | - | - | 0 | - |
| - | - | 9094 | 1417 | - | - | 0 | - |
| - | - | 1502 | 1418 | - | - | 0 | - |
| - | - | 939.7 | 1447 | - | - | 0 | - |
| - | - | 737.7 | 1464 | - | - | 0 | - |
| - | - | 701.4 | 1501 | - | - | 0 | - |
| - | - | 916.9 | 1502 | - | - | 0 | - |
| - | - | 719.4 | 1517 | - | - | 0 | - |
| - | - | 1186 | 1518 | - | - | 0 | - |
| 3 | y | 2683 | 1527 | 0.0002702 | 0.177 | +1 | 14 |
| 3 | y | 3567 | 1528 | 0.01528 | 10 | +1 | 14 |
| 3 | z | 6974 | 1529 | 0.004157 | 2.719 | +1 | 14 |
| - | - | 5220 | 1530 | - | - | 0 | - |
| - | - | 2137 | 1531 | - | - | 0 | - |
| - | - | 768.6 | 1532 | - | - | 0 | - |
| 3 | y | 1.46E+04 | 1545 | 0.004476 | 2.898 | +1 | 14 |
| - | - | 1.513E+04 | 1546 | - | - | 0 | - |
| - | - | 6663 | 1547 | - | - | 0 | - |
| - | - | 2390 | 1548 | - | - | 0 | - |
| - | - | 1734 | 1560 | - | - | 0 | - |
| - | - | 1542 | 1573 | - | - | 0 | - |
| - | - | 958 | 1574 | - | - | 0 | - |
| - | - | 732.9 | 1575 | - | - | 0 | - |
| 2 | z | 1656 | 1598 | 0.006863 | 4.296 | +1 | 15 |
| 15 | c | 1794 | 1599 | 0.0233 | 14.58 | +1 | 15 |
| 15 | c | 760.6 | 1600 | 0.007807 | 4.88 | +1 | 15 |
| - | - | 1045 | 1601 | - | - | 0 | - |
| 2 | y | 1070 | 1615 | 0.01279 | 7.921 | +1 | 15 |
| 2 | z | 5.26E+04 | 1616 | 0.002158 | 1.336 | +1 | 15 |
| 15 | c | 5.556E+04 | 1617 | 0.01373 | 8.49 | +1 | 15 |
| - | - | 2.801E+04 | 1618 | - | - | 0 | - |
| - | - | 952.5 | 1618 | - | - | 0 | - |
| - | - | 8554 | 1619 | - | - | 0 | - |
| 2 | y | 1310 | 1632 | 0.01229 | 7.534 | +1 | 15 |
| - | - | 678.9 | 1643 | - | - | 0 | - |
| - | - | 2.079E+04 | 1659 | - | - | 0 | - |
| - | - | 1.83E+04 | 1660 | - | - | 0 | - |
| - | - | 1.236E+04 | 1661 | - | - | 0 | - |
| - | - | 3861 | 1662 | - | - | 0 | - |
| - | - | 1034 | 1670 | - | - | 0 | - |
| - | - | 1765 | 1673 | - | - | 0 | - |
| - | - | 1415 | 1674 | - | - | 0 | - |
| - | - | 1015 | 1675 | - | - | 0 | - |
| - | - | 1072 | 1676 | - | - | 0 | - |
| - | - | 2155 | 1686 | - | - | 0 | - |
| - | - | 2.458E+04 | 1687 | - | - | 0 | - |
| - | - | 2.575E+04 | 1688 | - | - | 0 | - |
| - | - | 1.555E+04 | 1689 | - | - | 0 | - |
| - | - | 5636 | 1690 | - | - | 0 | - |
| - | - | 1018 | 1698 | - | - | 0 | - |
| - | - | 2292 | 1704 | - | - | 0 | - |
| - | - | 3678 | 1705 | - | - | 0 | - |
| - | - | 1799 | 1706 | - | - | 0 | - |
| - | - | 3918 | 1714 | - | - | 0 | - |
| - | - | 3.116E+04 | 1715 | - | - | 0 | - |
| - | - | 3.363E+04 | 1716 | - | - | 0 | - |
| - | - | 1.947E+04 | 1717 | - | - | 0 | - |
| - | - | 6862 | 1718 | - | - | 0 | - |
| - | - | 1516 | 1719 | - | - | 0 | - |
| - | - | 2429 | 1730 | - | - | 0 | - |
| - | - | 8.399E+04 | 1731 | - | - | 0 | - |
| - | - | 2.25E+05 | 1732 | - | - | 0 | - |
| - | - | 1.929E+05 | 1733 | - | - | 0 | - |
| - | - | 9.803E+04 | 1734 | - | - | 0 | - |
| - | - | 3.006E+04 | 1735 | - | - | 0 | - |
| - | - | 3569 | 1736 | - | - | 0 | - |
| - | - | 707.3 | 2857 | - | - | 0 | - |
| - | - | 916.7 | 3069 | - | - | 0 | - |

m/z Charge Intensity FragmentType MassShift Position
120.87884521484375 0 400.12994
135.4641876220703 0 423.7762
148.93789672851562 0 1065.9425
148.94725036621094 0 2400.8064
148.95790100097656 0 1688.5863
148.96279907226562 0 1304.15
148.9676971435547 0 782.5658
159.091796875 0 6566.0103
173.43104553222656 0 1131.2808
173.45989990234375 0 1174.8192
187.10792541503906 0 30989.453
188.1115264892578 0 2526.0066
231.3017578125 0 495.92828
246.123779296875 0 815.36035
250.25445556640625 0 531.17804
256.10699462890625 0 610.5984
274.1184387207031 0 1281.8657
284.1029357910156 0 1057.6678
301.1296691894531 0 3288.981
334.1763000488281 0 2502.225 y 14
335.1816101074219 0 632.0912
355.17657470703125 0 6185.6553
356.1803894042969 0 1231.7217
373.1876525878906 0 1104.9131
388.1617431640625 0 2164.4097
392.2171630859375 0 604.53815
416.99920654296875 0 568.968
442.210205078125 0 616.4021 w 8
457.1845397949219 0 808.0349
459.2368469238281 0 830.6743
469.2203369140625 0 718.6508
476.6899108886719 0 533.4785
482.23828125 0 894.6896
487.2303771972656 0 4828.472 c Ammonia loss 3
488.2330627441406 0 1094.5608
524.25927734375 0 653.89844
542.27392578125 0 751.6655
556.2509765625 0 2045.0393 z 12
557.271484375 0 1568.0315
568.214111328125 0 904.0811
572.2832641601562 0 4825.8286 y 12
573.2861938476562 0 1413.5497 c Water loss 4
574.2621459960938 0 3613.3223 c Ammonia loss 4
575.265380859375 0 943.9329
585.2404174804688 0 1880.9347
603.2517700195312 0 923.455
611.2955932617188 0 729.4882
612.2825317382812 0 731.68524
629.3042602539062 0 1999.4219
631.2820434570312 0 950.63855 c Ammonia loss 5
667.2847290039062 0 667.22754
671.3472900390625 0 692.0711 y 11
674.3223876953125 0 927.602
679.842041015625 0 677.52905 y 3
680.3311157226562 0 914.4464
684.310302734375 0 3116.4795
685.30078125 0 2459.474
690.3395385742188 0 883.2723
691.3419799804688 0 832.5098
699.3421630859375 0 1412.7789 c Ammonia loss 13
702.3206787109375 0 2542.379 c Ammonia loss 6
703.3242797851562 0 796.14136
712.3526000976562 0 1654.8765 z 10
713.360595703125 0 6948.7227
714.3635864257812 0 3266.614
716.3370971679688 0 1170.2393
728.3729248046875 0 6375.589 y 10
729.3760375976562 0 2751.8853
730.3541870117188 0 3278.8826
731.3562622070312 0 1070.5431
750.8682861328125 0 1216.2162
751.3677368164062 0 665.7217
754.8735961914062 0 1072.6395
755.3615112304688 0 1120.9517
755.862548828125 0 1324.5768 z Water loss 2
756.3663940429688 0 786.8409 z Ammonia loss 2
758.8826904296875 0 2254.4968
759.3828125 0 2185.3306
759.8790283203125 0 799.46686
763.8729248046875 0 3378.1772 y Water loss 2
764.3706665039062 0 5590.375 y Ammonia loss 2
764.8701171875 0 4106.163 z 2
765.3698120117188 0 1192.4304
770.3923950195312 0 847.7342
772.8782958984375 0 8612.775 y 2
773.3798828125 0 7920.3647
773.8817749023438 0 4000.7817
774.379150390625 0 1660.2705
780.3748779296875 0 793.6941
781.370849609375 0 1178.6416 z Water loss 9
788.4141235351562 0 726.22925
797.3914794921875 0 2289.0466 y Water loss 9
798.3826904296875 0 3273.8784
799.3865356445312 0 3873.0332 z 9
800.39306640625 0 11772.605 c Ammonia loss 14
801.3961791992188 0 5962.6753
802.3986206054688 0 1534.314
807.8856201171875 0 742.7935 y Ammonia loss 1
814.3988647460938 0 628.7079
815.4056396484375 0 11687.191 c Ammonia loss 7
816.4067993164062 0 6380.348 y 1
816.8975830078125 0 1969.3604
817.4044189453125 0 2017.8171
821.4063720703125 0 689.2187
825.4283447265625 0 786.8349
839.3975830078125 0 711.785
843.4335327148438 0 1753.1362
844.4354248046875 0 747.1515
847.91650390625 0 1498.197
848.4130859375 0 1997.2344
848.916259765625 0 2392.615
856.922119140625 0 3787.942
857.4209594726562 0 5929.8164
857.9193725585938 0 5454.278
858.4199829101562 0 2163.3125
858.92041015625 0 790.43933
865.9287109375 0 8752.242
866.43017578125 0 12716.381
866.9309692382812 0 7744.0156
867.4315795898438 0 3017.7275
881.417236328125 0 617.60675
898.4324340820312 0 2389.7805 y Water loss 8
899.427978515625 0 8247.21
900.431640625 0 5875.7925 z 8
901.4395141601562 0 8036.131
902.4454345703125 0 3389.496
916.4520874023438 0 16355.727 c Ammonia loss 8
917.454345703125 0 8967.718
918.4581909179688 0 1873.1918
932.473876953125 0 768.6739
933.4816284179688 0 2538.7651 c 8
934.4789428710938 0 1268.1058
971.4912719726562 0 1286.7881
1012.5193481445312 0 784.697
1013.5176391601562 0 7675.761 z 7
1014.5216674804688 0 9518.303
1015.5261840820312 0 4223.822
1016.5277099609375 0 1322.907
1019.5109252929688 0 1536.8018
1020.50927734375 0 3241.71 c 9
1021.5135498046875 0 1916.873
1022.5060424804688 0 686.62164
1028.4873046875 0 1240.6133
1029.5361328125 0 12512.579 y 7
1030.5396728515625 0 7566.9004
1031.5413818359375 0 2736.1572
1035.528076171875 0 783.85443
1042.4908447265625 0 664.3306
1058.525390625 0 1528.9292
1059.52685546875 0 1447.063 c Water loss 10
1076.5240478515625 0 3186.3384
1077.53076171875 0 2605.8718 c 10
1078.534423828125 0 1547.2021
1082.549072265625 0 726.0412 y Water loss 6
1084.5545654296875 0 27768.62 z 6
1085.5595703125 0 30781.963
1086.5633544921875 0 14877.032
1087.5679931640625 0 4336.898
1100.5732421875 0 6058.7744 y 6
1101.5758056640625 0 3726.7258
1102.5780029296875 0 1351.0076
1132.586669921875 0 909.07806
1133.5875244140625 0 1617.7969
1140.57568359375 0 725.1899
1141.575439453125 0 13300.727 z 5
1142.580078125 0 12465.742
1143.584228515625 0 6950.735
1144.587890625 0 2575.2346
1154.564208984375 0 651.6409
1155.5770263671875 0 1224.0398
1156.5694580078125 0 1543.7305
1157.5933837890625 0 13782.944 y 5
1158.5970458984375 0 9739.364 c Water loss 11
1159.5985107421875 0 3202.801
1160.591064453125 0 1636.428
1172.568603515625 0 1896.7427
1173.5699462890625 0 1023.0162
1175.5926513671875 0 9648.874
1176.5955810546875 0 7490.9863 c 11
1177.603515625 0 2541.0845
1178.596923828125 0 972.93494
1199.5792236328125 0 1418.742
1200.5821533203125 0 1160.185
1226.610595703125 0 982.55304 y Water loss 4
1227.6085205078125 0 2366.4863
1228.6072998046875 0 14767.754 z 4
1229.61083984375 0 16257.038
1230.6165771484375 0 8340.159
1231.61669921875 0 2237.2024
1244.6265869140625 0 11515.954 y 4
1245.6298828125 0 7723.062
1246.625732421875 0 3544.5613
1253.335693359375 0 818.33844
1269.644775390625 0 4854.077
1270.6475830078125 0 5299.799
1271.648193359375 0 2484.6375
1272.6558837890625 0 1443.8296
1296.6324462890625 0 1223.2515 c Ammonia loss 12
1297.6295166015625 0 1167.5249
1312.6510009765625 0 18337.111
1312.8302001953125 0 1045.8763
1313.65576171875 0 28368.225 c 12
1314.6588134765625 0 17036.621
1315.661865234375 0 6826.5396
1316.6651611328125 0 1392.2056
1341.640380859375 0 1368.5872 y Ammonia loss 3
1342.6488037109375 0 7838.124 z 3
1343.6541748046875 0 6428.639
1344.65625 0 3575.4275
1353.68408203125 0 664.0638
1358.6644287109375 0 8577.1 y 3
1359.66796875 0 7238.624
1360.666259765625 0 3582.018
1370.679443359375 0 880.5257
1397.6815185546875 0 1654.695 c Ammonia loss 13
1398.68505859375 0 1694.5707
1413.698486328125 0 6095.795
1414.703857421875 0 25838.984 c 13
1415.7080078125 0 19710.43
1416.70947265625 0 9093.511
1417.7108154296875 0 1502.0085
1446.681884765625 0 939.6543
1463.8125 0 737.74927
1500.7491455078125 0 701.41187
1501.7462158203125 0 916.8705
1516.7567138671875 0 719.3877
1517.7562255859375 0 1185.8956
1526.7340087890625 0 2683.1736 y Water loss 2
1527.7330322265625 0 3566.843 y Ammonia loss 2
1528.729736328125 0 6973.5303 z 2
1529.732177734375 0 5219.7593
1530.736083984375 0 2136.6765
1531.743896484375 0 768.5809
1544.748779296875 0 14602.324 y 2
1545.74951171875 0 15126.257
1546.7550048828125 0 6662.8047
1547.7513427734375 0 2390.4595
1559.698486328125 0 1734.074
1572.7410888671875 0 1542.074
1573.748291015625 0 958.0313
1574.740234375 0 732.8722
1597.75390625 0 1656.2053 z Water loss 1
1598.7427978515625 0 1794.2789 c Water loss 14
1599.7423095703125 0 760.5803 c Ammonia loss 14
1600.768798828125 0 1045.3141
1614.7625732421875 0 1069.5558 y Ammonia loss 1
1615.759765625 0 52598.023 z 1
1616.762939453125 0 55561.9 c 14
1617.765380859375 0 28014.629
1618.015869140625 0 952.461
1618.77001953125 0 8553.798
1631.7640380859375 0 1309.783 y 1
1642.766845703125 0 678.8525
1658.8260498046875 0 20794.967
1659.830078125 0 18298.672
1660.83154296875 0 12356.152
1661.8343505859375 0 3860.9712
1669.835693359375 0 1034.026
1672.8206787109375 0 1764.9911
1673.80517578125 0 1415.1918
1674.8138427734375 0 1014.7079
1675.80224609375 0 1072.0616
1685.841064453125 0 2155.1626
1686.835205078125 0 24579.424
1687.841552734375 0 25751.049
1688.8441162109375 0 15552.549
1689.8441162109375 0 5636.424
1697.817138671875 0 1017.57825
1703.8612060546875 0 2292.4458
1704.8612060546875 0 3678.2712
1705.857421875 0 1799.3765
1713.8404541015625 0 3918.283
1714.831787109375 0 31157.629
1715.831787109375 0 33625.164
1716.8358154296875 0 19473.775
1717.836669921875 0 6862.1543
1718.8333740234375 0 1516.4672
1729.8446044921875 0 2429.1453
1730.8472900390625 0 83989.74
1731.8533935546875 0 224953.14
1732.857177734375 0 192890.11
1733.860595703125 0 98029.21
1734.862548828125 0 30056.537
1735.85986328125 0 3568.6282
2857.396484375 0 707.334
3069.26220703125 0 916.674

Spectrum Details

|  |  |
| --- | --- |
| Matched peaks? Matched peaks The total absolute number of peaks matched. Additionally in brackets the total fraction of peaks matched and the total number of peaks is shown. | 64 (22.38% of 286) |
| FDR? FDR The false discovery rate estimated for this peptide. It is calculated by matching all theoretical fragments with a non-integer shift with the raw peaks for this spectrum. This is done with 40 different shifts. The resulting percentage is the average number of annotated peaks over the number of annotated peaks with the correct spectrum. | 0.00% |
| Satellite FDR? Satellite FDR See the FDR for details on its calculation. This satellite ion specific FDR only contains the satellite ions (d/w) for I/L/J positions. | - |
| PSM Score? PSM Score The PSM Score as given by Hecklib to this annotated spectrum. It is shown with three significant figures. | 746 |

#### Spectrum 9105? Spectrum 9105 The raw spectrum of this peptide as annotated by Hecklib. The fragments are coloured according to ion type (see legend). Any peaks with a star '\*' as text can be hovered over to see the full details, first the ion type second the mass shift type. By hovering over the amino acids in the peptide or ions in the legend the corresponding peaks are highlighted. By toggling the 'Unassigned' label you can turn the background (unassigned) peaks on or off in the plot. By updating the slider in the Ion legend you can update the spectrum to only show the top X% of the peaks with labels. The top X% means any peak that is within X% of the highest intensity. By dragging in the spectrum you can zoom in to a specific part of the spectrum and use 'Zoom Out' to get back to the original zoom level. The annotation of the spectrum is based on the given sequence in the peptides file and is done with different software so inconsistencies are likely. The peaks are annotated based on the given sequence, with 20 ppm tolerance.

Copy Data

##### Spectrum 9105 (TSV)

###### Preview

```
Loading example...
```

*Click on the button to copy the data to your clipboard.*

Mz MinMz MaxIntensity Max

WidthHeightPeptide font sizePeptide stroke widthSpectrum font sizeSpectrum stroke widthCompact peptide

Ion legend

wxyz

abcd

OtherUnassignedIon^Charge^_Position_Show for top:%

VSWNSGALTSGVHTWJ

01.17e+42.33e+43.50e+44.66e+4

Zoom Out

c^+1^_2_z^+1^_2_y^+1^_3_y^+1^_3_c^+1^_4_c^+1^_4_z^+1^_4_y^+1^_4_c^+1^_5_c^+1^_5_c^+1^_5_c^+1^_6_w^+1^_5_c^+1^_6_z^+1^_5_c^+2^_13_c^+2^_14_c^+1^_7_c^+2^_14_z^+1^_6_c^+1^_7_y^+1^_6_z^+2^_14_y^+2^_14_y^+2^_14_z^+2^_14_y^+2^_14_z^+1^_7_c^+2^_15_z^+2^_15_c^+2^_15_c^+1^_8_c^+1^_8_y^+1^_8_z^+1^_8_c^+1^_9_c^+1^_9_w^+1^_9_z^+1^_9_c^+1^_10_c^+1^_11_z^+1^_10_y^+1^_10_z^+1^_11_y^+1^_11_c^+1^_12_z^+1^_12_y^+1^_12_c^+1^_13_z^+1^_13_c^+1^_14_z^+1^_14_y^+1^_14_z^+1^_15_c^+1^_15_

0690138120712761

Fragment Matches Table

Show background peaks

| Position | Ion type | Intensity | mz Theoretical | mz Error (Th) | mz Error (ppm) | Charge | Series Number |
| --- | --- | --- | --- | --- | --- | --- | --- |
| - | - | 650 | 120.1 | - | - | 0 | - |
| - | - | 351.7 | 122.6 | - | - | 0 | - |
| - | - | 355.9 | 126.6 | - | - | 0 | - |
| - | - | 347.6 | 129.9 | - | - | 0 | - |
| - | - | 2196 | 130.1 | - | - | 0 | - |
| - | - | 2125 | 133.1 | - | - | 0 | - |
| - | - | 1937 | 140.1 | - | - | 0 | - |
| - | - | 744.4 | 149 | - | - | 0 | - |
| - | - | 415.2 | 152.2 | - | - | 0 | - |
| - | - | 1004 | 153.1 | - | - | 0 | - |
| - | - | 984.9 | 159.1 | - | - | 0 | - |
| - | - | 529.1 | 173.1 | - | - | 0 | - |
| - | - | 460.1 | 174.1 | - | - | 0 | - |
| - | - | 1045 | 177.1 | - | - | 0 | - |
| - | - | 1027 | 182.1 | - | - | 0 | - |
| - | - | 922.4 | 186.1 | - | - | 0 | - |
| - | - | 1.774E+04 | 187.1 | - | - | 0 | - |
| - | - | 1542 | 188.1 | - | - | 0 | - |
| - | - | 682 | 194.1 | - | - | 0 | - |
| - | - | 990.4 | 195.1 | - | - | 0 | - |
| - | - | 1051 | 195.1 | - | - | 0 | - |
| - | - | 640.3 | 200.1 | - | - | 0 | - |
| - | - | 899.6 | 202.1 | - | - | 0 | - |
| 2 | c | 750.2 | 204.1 | 5.157E-05 | 0.2526 | +1 | 2 |
| - | - | 561.7 | 221.1 | - | - | 0 | - |
| - | - | 530.8 | 230.2 | - | - | 0 | - |
| - | - | 669.5 | 239.1 | - | - | 0 | - |
| - | - | 4209 | 239.1 | - | - | 0 | - |
| - | - | 492.5 | 239.2 | - | - | 0 | - |
| - | - | 559.2 | 240.2 | - | - | 0 | - |
| - | - | 524.4 | 248.6 | - | - | 0 | - |
| - | - | 527.8 | 253.1 | - | - | 0 | - |
| - | - | 957.3 | 256.2 | - | - | 0 | - |
| - | - | 3297 | 283.2 | - | - | 0 | - |
| - | - | 808 | 285.1 | - | - | 0 | - |
| - | - | 1032 | 296.2 | - | - | 0 | - |
| - | - | 2073 | 299.1 | - | - | 0 | - |
| - | - | 795.3 | 300.1 | - | - | 0 | - |
| - | - | 646.2 | 300.2 | - | - | 0 | - |
| - | - | 855.8 | 301.1 | - | - | 0 | - |
| - | - | 595.3 | 312.1 | - | - | 0 | - |
| 15 | z | 1517 | 318.2 | 0.004794 | 15.07 | +1 | 2 |
| - | - | 2439 | 327.2 | - | - | 0 | - |
| - | - | 733.3 | 329.2 | - | - | 0 | - |
| - | - | 1167 | 344.2 | - | - | 0 | - |
| - | - | 621.9 | 350.2 | - | - | 0 | - |
| - | - | 6637 | 355.1 | - | - | 0 | - |
| - | - | 769.2 | 355.2 | - | - | 0 | - |
| - | - | 1700 | 356.1 | - | - | 0 | - |
| - | - | 858.1 | 357.1 | - | - | 0 | - |
| - | - | 2090 | 371.2 | - | - | 0 | - |
| - | - | 916.4 | 373.2 | - | - | 0 | - |
| - | - | 828.9 | 388.3 | - | - | 0 | - |
| - | - | 843.4 | 415.3 | - | - | 0 | - |
| 14 | y | 698.8 | 417.2 | 0.004589 | 11 | +1 | 3 |
| - | - | 677.5 | 420.2 | - | - | 0 | - |
| 14 | y | 736 | 435.2 | 0.004858 | 11.16 | +1 | 3 |
| - | - | 2132 | 475 | - | - | 0 | - |
| - | - | 575.7 | 476.3 | - | - | 0 | - |
| - | - | 1395 | 479.2 | - | - | 0 | - |
| - | - | 627.9 | 482.2 | - | - | 0 | - |
| 4 | c | 1681 | 487.2 | 8.232E-05 | 0.169 | +1 | 4 |
| - | - | 660.3 | 489.2 | - | - | 0 | - |
| - | - | 960.9 | 499.3 | - | - | 0 | - |
| - | - | 651.1 | 503.1 | - | - | 0 | - |
| 4 | c | 2220 | 504.3 | 0.0002217 | 0.4396 | +1 | 4 |
| - | - | 2622 | 523.3 | - | - | 0 | - |
| - | - | 818.8 | 524.3 | - | - | 0 | - |
| - | - | 659.1 | 528.3 | - | - | 0 | - |
| 13 | z | 1081 | 556.3 | 0.002004 | 3.603 | +1 | 4 |
| - | - | 691.1 | 556.8 | - | - | 0 | - |
| - | - | 710.1 | 557.3 | - | - | 0 | - |
| - | - | 2673 | 567.3 | - | - | 0 | - |
| 13 | y | 1102 | 572.3 | 0.005334 | 9.32 | +1 | 4 |
| 5 | c | 794.8 | 573.3 | 0.004682 | 8.167 | +1 | 5 |
| 5 | c | 1438 | 574.3 | 0.0006467 | 1.126 | +1 | 5 |
| - | - | 842.4 | 578.3 | - | - | 0 | - |
| 5 | c | 1744 | 591.3 | 0.0002676 | 0.4527 | +1 | 5 |
| - | - | 1408 | 592.3 | - | - | 0 | - |
| - | - | 1771 | 611.3 | - | - | 0 | - |
| 6 | c | 789.1 | 631.3 | 0.0009725 | 1.541 | +1 | 6 |
| 12 | w | 1670 | 640.3 | 0.004265 | 6.662 | +1 | 5 |
| 6 | c | 7428 | 648.3 | 0.0002413 | 0.3722 | +1 | 6 |
| - | - | 2808 | 649.3 | - | - | 0 | - |
| 12 | z | 2007 | 655.3 | 0.0118 | 18 | +1 | 5 |
| 13 | c | 1197 | 657.3 | 0.001139 | 1.733 | +2 | 13 |
| - | - | 606.9 | 657.8 | - | - | 0 | - |
| 14 | c | 600.1 | 699.3 | 0.004135 | 5.913 | +2 | 14 |
| 7 | c | 1055 | 702.3 | 0.003165 | 4.507 | +1 | 7 |
| 14 | c | 1.042E+04 | 707.9 | 0.000137 | 0.1936 | +2 | 14 |
| - | - | 7404 | 708.4 | - | - | 0 | - |
| - | - | 2220 | 708.9 | - | - | 0 | - |
| - | - | 1444 | 709.4 | - | - | 0 | - |
| 11 | z | 1063 | 712.3 | 0.003455 | 4.85 | +1 | 6 |
| 7 | c | 2271 | 719.3 | 0.0002369 | 0.3293 | +1 | 7 |
| - | - | 875.7 | 720.4 | - | - | 0 | - |
| 11 | y | 1762 | 728.4 | 0.004262 | 5.852 | +1 | 6 |
| - | - | 1155 | 729.4 | - | - | 0 | - |
| - | - | 689.7 | 735.3 | - | - | 0 | - |
| - | - | 716.5 | 754.9 | - | - | 0 | - |
| 3 | z | 812.1 | 755.9 | 0.0006108 | 0.8081 | +2 | 14 |
| - | - | 803.2 | 758.9 | - | - | 0 | - |
| 3 | y | 3033 | 763.9 | 0.006555 | 8.581 | +2 | 14 |
| 3 | y | 2366 | 764.4 | 0.002719 | 3.557 | +2 | 14 |
| 3 | z | 2997 | 764.9 | 0.001193 | 1.56 | +2 | 14 |
| 3 | y | 1535 | 772.9 | 0.009952 | 12.88 | +2 | 14 |
| - | - | 2748 | 773.4 | - | - | 0 | - |
| - | - | 987.8 | 798.4 | - | - | 0 | - |
| 10 | z | 3798 | 799.4 | 0.003165 | 3.959 | +1 | 7 |
| 15 | c | 1537 | 800.4 | 0.01236 | 15.44 | +2 | 15 |
| - | - | 1889 | 801.4 | - | - | 0 | - |
| - | - | 811.4 | 807.4 | - | - | 0 | - |
| 2 | z | 1792 | 808.4 | 0.0004315 | 0.5338 | +2 | 15 |
| 15 | c | 1814 | 808.9 | 0.002811 | 3.475 | +2 | 15 |
| 8 | c | 3254 | 815.4 | 0.0006374 | 0.7817 | +1 | 8 |
| - | - | 592.1 | 816.1 | - | - | 0 | - |
| - | - | 1074 | 816.4 | - | - | 0 | - |
| - | - | 2636 | 829.9 | - | - | 0 | - |
| - | - | 2240 | 830.4 | - | - | 0 | - |
| - | - | 1259 | 830.9 | - | - | 0 | - |
| - | - | 808 | 831.4 | - | - | 0 | - |
| 8 | c | 3137 | 832.4 | 9.623E-05 | 0.1156 | +1 | 8 |
| - | - | 1651 | 833.4 | - | - | 0 | - |
| - | - | 766 | 834.4 | - | - | 0 | - |
| - | - | 1427 | 836.9 | - | - | 0 | - |
| - | - | 1387 | 837.4 | - | - | 0 | - |
| - | - | 870.9 | 844.4 | - | - | 0 | - |
| - | - | 996.3 | 844.9 | - | - | 0 | - |
| - | - | 602.8 | 845.4 | - | - | 0 | - |
| - | - | 655.1 | 856.3 | - | - | 0 | - |
| - | - | 9016 | 865.9 | - | - | 0 | - |
| - | - | 2.518E+04 | 866.4 | - | - | 0 | - |
| - | - | 1.672E+04 | 866.9 | - | - | 0 | - |
| - | - | 1.307E+04 | 867.4 | - | - | 0 | - |
| - | - | 5437 | 867.9 | - | - | 0 | - |
| - | - | 1082 | 868.4 | - | - | 0 | - |
| - | - | 834.1 | 868.9 | - | - | 0 | - |
| 9 | y | 1086 | 898.4 | 0.0119 | 13.25 | +1 | 8 |
| - | - | 1344 | 899.4 | - | - | 0 | - |
| 9 | z | 3870 | 900.4 | 0.004742 | 5.266 | +1 | 8 |
| - | - | 1508 | 901.4 | - | - | 0 | - |
| 9 | c | 1686 | 916.5 | 0.003564 | 3.889 | +1 | 9 |
| - | - | 1360 | 917.5 | - | - | 0 | - |
| 9 | c | 4400 | 933.5 | 9.689E-05 | 0.1038 | +1 | 9 |
| - | - | 2709 | 934.5 | - | - | 0 | - |
| - | - | 823 | 935.5 | - | - | 0 | - |
| 8 | w | 924.7 | 970.5 | 0.0003771 | 0.3885 | +1 | 9 |
| 8 | z | 1743 | 1014 | 0.007958 | 7.852 | +1 | 9 |
| - | - | 1104 | 1015 | - | - | 0 | - |
| 10 | c | 5957 | 1021 | 0.0002233 | 0.2188 | +1 | 10 |
| - | - | 3405 | 1022 | - | - | 0 | - |
| - | - | 898.1 | 1023 | - | - | 0 | - |
| - | - | 701.9 | 1028 | - | - | 0 | - |
| 11 | c | 5914 | 1078 | 0.0001219 | 0.1131 | +1 | 11 |
| - | - | 4073 | 1079 | - | - | 0 | - |
| - | - | 1443 | 1080 | - | - | 0 | - |
| 7 | z | 8779 | 1085 | 0.002888 | 2.663 | +1 | 10 |
| - | - | 4554 | 1086 | - | - | 0 | - |
| - | - | 1680 | 1087 | - | - | 0 | - |
| - | - | 792.1 | 1088 | - | - | 0 | - |
| - | - | 786.2 | 1089 | - | - | 0 | - |
| 7 | y | 2180 | 1101 | 0.005404 | 4.91 | +1 | 10 |
| - | - | 832.5 | 1102 | - | - | 0 | - |
| - | - | 4.618E+04 | 1137 | - | - | 0 | - |
| - | - | 2.827E+04 | 1138 | - | - | 0 | - |
| - | - | 1232 | 1138 | - | - | 0 | - |
| - | - | 752.5 | 1139 | - | - | 0 | - |
| - | - | 8950 | 1139 | - | - | 0 | - |
| - | - | 3471 | 1140 | - | - | 0 | - |
| 6 | z | 3975 | 1142 | 0.00303 | 2.655 | +1 | 11 |
| - | - | 2650 | 1143 | - | - | 0 | - |
| - | - | 1229 | 1144 | - | - | 0 | - |
| - | - | 852.2 | 1152 | - | - | 0 | - |
| - | - | 1242 | 1153 | - | - | 0 | - |
| - | - | 796.6 | 1158 | - | - | 0 | - |
| 6 | y | 973.7 | 1158 | 0.01324 | 11.44 | +1 | 11 |
| - | - | 1488 | 1159 | - | - | 0 | - |
| 12 | c | 2683 | 1177 | 0.0006777 | 0.576 | +1 | 12 |
| - | - | 2219 | 1178 | - | - | 0 | - |
| - | - | 1320 | 1179 | - | - | 0 | - |
| - | - | 642.6 | 1213 | - | - | 0 | - |
| - | - | 988.8 | 1228 | - | - | 0 | - |
| 5 | z | 3557 | 1229 | 0.0003114 | 0.2535 | +1 | 12 |
| - | - | 2748 | 1230 | - | - | 0 | - |
| - | - | 2035 | 1231 | - | - | 0 | - |
| 5 | y | 1666 | 1245 | 0.005501 | 4.42 | +1 | 12 |
| - | - | 933.8 | 1247 | - | - | 0 | - |
| - | - | 900.9 | 1313 | - | - | 0 | - |
| 13 | c | 1.169E+04 | 1314 | 0.002204 | 1.678 | +1 | 13 |
| - | - | 9019 | 1315 | - | - | 0 | - |
| - | - | 3710 | 1316 | - | - | 0 | - |
| - | - | 855.8 | 1317 | - | - | 0 | - |
| 4 | z | 1179 | 1343 | 0.01755 | 13.07 | +1 | 13 |
| - | - | 1039 | 1344 | - | - | 0 | - |
| - | - | 732 | 1414 | - | - | 0 | - |
| 14 | c | 8310 | 1415 | 0.003374 | 2.385 | +1 | 14 |
| - | - | 7044 | 1416 | - | - | 0 | - |
| - | - | 3964 | 1417 | - | - | 0 | - |
| - | - | 1261 | 1418 | - | - | 0 | - |
| - | - | 924.2 | 1487 | - | - | 0 | - |
| 3 | z | 3152 | 1529 | 0.001336 | 0.8739 | +1 | 14 |
| - | - | 2589 | 1530 | - | - | 0 | - |
| - | - | 1285 | 1531 | - | - | 0 | - |
| - | - | 721.3 | 1532 | - | - | 0 | - |
| 3 | y | 934.7 | 1545 | 0.009847 | 6.375 | +1 | 14 |
| - | - | 1157 | 1546 | - | - | 0 | - |
| - | - | 869.8 | 1547 | - | - | 0 | - |
| - | - | 609 | 1583 | - | - | 0 | - |
| 2 | z | 2076 | 1616 | 0.0001613 | 0.09982 | +1 | 15 |
| 15 | c | 6710 | 1617 | 0.01104 | 6.829 | +1 | 15 |
| - | - | 5822 | 1618 | - | - | 0 | - |
| - | - | 2438 | 1619 | - | - | 0 | - |
| - | - | 1363 | 1660 | - | - | 0 | - |
| - | - | 1468 | 1661 | - | - | 0 | - |
| - | - | 2601 | 1674 | - | - | 0 | - |
| - | - | 3388 | 1675 | - | - | 0 | - |
| - | - | 934.7 | 1676 | - | - | 0 | - |
| - | - | 1379 | 1688 | - | - | 0 | - |
| - | - | 770.7 | 1690 | - | - | 0 | - |
| - | - | 1795 | 1705 | - | - | 0 | - |
| - | - | 810.4 | 1707 | - | - | 0 | - |
| - | - | 9714 | 1716 | - | - | 0 | - |
| - | - | 8501 | 1717 | - | - | 0 | - |
| - | - | 5625 | 1718 | - | - | 0 | - |
| - | - | 2010 | 1719 | - | - | 0 | - |
| - | - | 1521 | 1731 | - | - | 0 | - |
| - | - | 3087 | 1732 | - | - | 0 | - |
| - | - | 1.197E+04 | 1733 | - | - | 0 | - |
| - | - | 1.015E+04 | 1734 | - | - | 0 | - |
| - | - | 6723 | 1735 | - | - | 0 | - |
| - | - | 2810 | 1736 | - | - | 0 | - |
| - | - | 662.9 | 2734 | - | - | 0 | - |

m/z Charge Intensity FragmentType MassShift Position
120.08120727539062 0 650.04974
122.58053588867188 0 351.73358
126.57894897460938 0 355.93695
129.8863067626953 0 347.64563
130.0653533935547 0 2196.2983
133.08619689941406 0 2124.6118
140.08209228515625 0 1937.3503
149.0452117919922 0 744.4491
152.1646270751953 0 415.20105
153.0550079345703 0 1003.756
159.09182739257812 0 984.9081
173.1196746826172 0 529.1305
174.0559844970703 0 460.1328
177.11215209960938 0 1044.5276
182.08154296875 0 1027.3925
186.10011291503906 0 922.3802
187.1079864501953 0 17738.383
188.11134338378906 0 1541.7795
194.12889099121094 0 681.97437
195.12261962890625 0 990.3664
195.13699340820312 0 1051.3867
200.107666015625 0 640.3167
202.12258911132812 0 899.60223
204.13421630859375 0 750.1667 c 1
221.08529663085938 0 561.747
230.15122985839844 0 530.78265
239.0938262939453 0 669.4965
239.14927673339844 0 4209.121
239.22036743164062 0 492.50223
240.15289306640625 0 559.16144
248.61915588378906 0 524.3588
253.1178436279297 0 527.76605
256.17523193359375 0 957.2931
283.1759338378906 0 3296.718
285.134033203125 0 808.0178
296.1713562011719 0 1032.1085
299.0622863769531 0 2072.8506
300.0620422363281 0 795.3454
300.2026672363281 0 646.17413
301.12994384765625 0 855.84283
312.0952453613281 0 595.2737
318.15728759765625 0 1517.1084 z 14
327.20208740234375 0 2438.6013
329.15869140625 0 733.28955
344.2281494140625 0 1166.7178
350.19207763671875 0 621.8766
355.0700378417969 0 6637.209
355.1772766113281 0 769.21045
356.0713806152344 0 1700.4957
357.0679016113281 0 858.1013
371.2281494140625 0 2090.2795
373.1856384277344 0 916.40753
388.2536926269531 0 828.926
415.25323486328125 0 843.42126
417.2129211425781 0 698.80664 y Water loss 13
420.21331787109375 0 677.53766
435.2237548828125 0 736.0444 y 13
475.0046081542969 0 2132.0244
476.304443359375 0 575.7411
479.2491149902344 0 1395.1355
482.2359313964844 0 627.8943
487.23004150390625 0 1680.9358 c Ammonia loss 3
489.2453918457031 0 660.325
499.2669372558594 0 960.8807
503.1092224121094 0 651.0524
504.25628662109375 0 2220.0586 c 3
523.2750244140625 0 2621.6914
524.2780151367188 0 818.84216
528.2727661132812 0 659.0562
556.257080078125 0 1080.5409 z 12
556.7716064453125 0 691.0884
557.2601928710938 0 710.14594
567.301513671875 0 2672.9797
572.2831420898438 0 1101.5352 y 12
573.2826538085938 0 794.8017 c Water loss 4
574.2626342773438 0 1438.0524 c Ammonia loss 4
578.3396606445312 0 842.4202
591.2882690429688 0 1744.3641 c 4
592.29150390625 0 1408.0836
611.3277587890625 0 1770.5999
631.284423828125 0 789.09204 c Ammonia loss 5
640.3082885742188 0 1670.0186 w 11
648.3102416992188 0 7427.958 c 5
649.3141479492188 0 2807.5264
655.3392944335938 0 2007.2363 z 11
657.3323364257812 0 1196.5874 c 12
657.8338012695312 0 606.8947
699.3399047851562 0 600.0815 c Ammonia loss 13
702.32373046875 0 1054.7856 c Ammonia loss 6
707.857177734375 0 10420.921 c 13
708.3588256835938 0 7403.668
708.8590698242188 0 2219.6006
709.3601684570312 0 1444.3477
712.3524169921875 0 1063.3223 z 10
719.3473510742188 0 2271.0176 c 6
720.3515014648438 0 875.6708
728.3719482421875 0 1762.0374 y 10
729.3762817382812 0 1154.6769
735.3458251953125 0 689.7102
754.8565063476562 0 716.51086
755.8605346679688 0 812.13776 z Water loss 2
758.8795776367188 0 803.185
763.8639526367188 0 3032.6013 y Water loss 2
764.365234375 0 2365.6885 y Ammonia loss 2
764.865234375 0 2997.3054 z 2
772.8857421875 0 1535.0317 y 2
773.3817749023438 0 2747.5713
798.3757934570312 0 987.7505
799.3841552734375 0 3798.4456 z 9
800.3910522460938 0 1537.294 c Ammonia loss 14
801.3944702148438 0 1888.862
807.3662719726562 0 811.43866
808.3828735351562 0 1791.9514 z 1
808.88916015625 0 1813.8916 c 14
815.4039916992188 0 3254.1028 c Ammonia loss 7
816.0848999023438 0 592.12683
816.4114379882812 0 1073.79
829.9190063476562 0 2636.1094
830.420654296875 0 2240.0476
830.9169311523438 0 1258.7969
831.4234619140625 0 807.9913
832.4312744140625 0 3136.7473 c 7
833.4314575195312 0 1651.398
834.4290161132812 0 766.03467
836.9166259765625 0 1427.4329
837.4183349609375 0 1387.467
844.4199829101562 0 870.94196
844.9315795898438 0 996.2662
845.4242553710938 0 602.8344
856.3373413085938 0 655.1249
865.9277954101562 0 9015.974
866.431640625 0 25179.227
866.932861328125 0 16722.348
867.43505859375 0 13068.726
867.9359741210938 0 5437.233
868.43798828125 0 1082.4019
868.9326171875 0 834.1178
898.4249267578125 0 1085.8539 y Water loss 8
899.4266357421875 0 1343.9238
900.4334106445312 0 3870.2957 z 8
901.433837890625 0 1507.9417
916.4558715820312 0 1685.5046 c Ammonia loss 8
917.4527587890625 0 1359.9216
933.478759765625 0 4400.1987 c 8
934.4815063476562 0 2709.116
935.4815673828125 0 823.0159
970.4575805664062 0 924.6862 w 7
1013.5206909179688 0 1743.338 z 7
1014.5210571289062 0 1103.9232
1020.5111083984375 0 5956.6016 c 9
1021.5101928710938 0 3405.1028
1022.5221557617188 0 898.0762
1028.4820556640625 0 701.87244
1077.532470703125 0 5913.7 c 10
1078.5350341796875 0 4072.8816
1079.533203125 0 1443.373
1084.552734375 0 8779.192 z 6
1085.5570068359375 0 4554.2246
1086.5596923828125 0 1679.7488
1087.6033935546875 0 792.08386
1088.6048583984375 0 786.2279
1100.573974609375 0 2180.0117 y 6
1101.5849609375 0 832.4882
1136.66259765625 0 46179.152
1137.6668701171875 0 28270.578
1137.8128662109375 0 1232.3834
1138.52734375 0 752.45404
1138.66845703125 0 8950.051
1139.67236328125 0 3470.6274
1141.5743408203125 0 3974.6008 z 5
1142.5787353515625 0 2649.531
1143.5723876953125 0 1229.0074
1151.6900634765625 0 852.22064
1152.683837890625 0 1242.372
1157.5089111328125 0 796.61926
1157.603271484375 0 973.7182 y 5
1158.511962890625 0 1488.4363
1176.6014404296875 0 2682.6348 c 11
1177.6036376953125 0 2218.5251
1178.6026611328125 0 1319.5555
1212.5867919921875 0 642.5851
1227.60107421875 0 988.81506
1228.60302734375 0 3556.89 z 4
1229.6138916015625 0 2748.4392
1230.6134033203125 0 2034.7537
1244.6275634765625 0 1666.0615 y 4
1246.623291015625 0 933.83636
1312.65185546875 0 900.90576
1313.657470703125 0 11694.008 c 12
1314.6605224609375 0 9019.084
1315.6629638671875 0 3709.9253
1316.6671142578125 0 855.82104
1342.663818359375 0 1178.9977 z 3
1343.6549072265625 0 1039.4094
1413.68212890625 0 732.03906
1414.7039794921875 0 8309.643 c 13
1415.7080078125 0 7044.261
1416.7115478515625 0 3964.1462
1417.71728515625 0 1261.2993
1486.6925048828125 0 924.22144
1528.7242431640625 0 3152.255 z 2
1529.730224609375 0 2588.517
1530.7264404296875 0 1285.4829
1531.743896484375 0 721.29346
1544.754150390625 0 934.7097 y 2
1545.7523193359375 0 1157.073
1546.738037109375 0 869.7728
1583.3736572265625 0 609.0234
1615.7574462890625 0 2076.0898 z 1
1616.765625 0 6710.0703 c 14
1617.7720947265625 0 5821.9644
1618.7685546875 0 2437.56
1659.834716796875 0 1363.1279
1660.8277587890625 0 1467.7612
1673.8277587890625 0 2600.8457
1674.825927734375 0 3388.2476
1675.8270263671875 0 934.72363
1687.85546875 0 1379.2125
1689.888427734375 0 770.6999
1704.8623046875 0 1795.2139
1706.876708984375 0 810.40173
1715.8375244140625 0 9713.792
1716.8397216796875 0 8500.941
1717.8404541015625 0 5625.464
1718.83984375 0 2010.2434
1730.851318359375 0 1521.3142
1731.8511962890625 0 3087.478
1732.861083984375 0 11974.408
1733.8621826171875 0 10150.539
1734.865966796875 0 6722.722
1735.87451171875 0 2810.3167
2733.791259765625 0 662.9025

Spectrum Details

|  |  |
| --- | --- |
| Matched peaks? Matched peaks The total absolute number of peaks matched. Additionally in brackets the total fraction of peaks matched and the total number of peaks is shown. | 55 (23.71% of 232) |
| FDR? FDR The false discovery rate estimated for this peptide. It is calculated by matching all theoretical fragments with a non-integer shift with the raw peaks for this spectrum. This is done with 40 different shifts. The resulting percentage is the average number of annotated peaks over the number of annotated peaks with the correct spectrum. | 0.69% |
| Satellite FDR? Satellite FDR See the FDR for details on its calculation. This satellite ion specific FDR only contains the satellite ions (d/w) for I/L/J positions. | 0.00% |
| PSM Score? PSM Score The PSM Score as given by Hecklib to this annotated spectrum. It is shown with three significant figures. | 553 |

#### Reverse Lookup? Reverse Lookup All places where this read could be placed.

| Group | Segment | Template | Template Part | Read Part | Score | Unique |
| --- | --- | --- | --- | --- | --- | --- |
| Homo sapiens Heavy Chain | IGHC | IGHG1 | [38..54] | [0..16] | 110 | False |
| Homo sapiens Heavy Chain | IGHC | IGHG3 | [38..54] | [0..16] | 110 | False |
| Homo sapiens Heavy Chain | IGHC | IGHG2 | [38..54] | [0..16] | 110 | False |
| Homo sapiens Heavy Chain | IGHC | IGHG4 | [38..54] | [0..16] | 110 | False |

| Recombined | Template Part | Read Part | Score | Unique |
| --- | --- | --- | --- | --- |
| REC-0-1 | [160..176] | [0..16] | 110 | True |

#### Meta Information from Multiple reads

##### Number of combined reads

2

##### Intensity

0.5188

##### TotalArea

1.02E+07

##### Changes to the peptide sequence

VSWNSGALTSGVHTWJ

J→LSupport for Leucine based on side chain ions (1 for L 0 for I) (Position: 8)

L→JNo support for either Leucine or Isoleucine based on side chain ions (Position: 16)

L→JNo support for either Leucine or Isoleucine based on side chain ions (Position: 8)

#### Positional Score

Copy Data

##### Positional Score (TSV)

###### Preview

```
Loading example...
```

*Click on the button to copy the data to your clipboard.*

000123456789101112131415

Label Value
"0" 0
"1" 0
"2" 0
"3" 0
"4" 0
"5" 0
"6" 0
"7" 0
"8" 0
"9" 0
"10" 0
"11" 0
"12" 0
"13" 0
"14" 0
"15" 0

#### Meta Information from PEAKS

##### Scan Identifier

F2:9144

##### Original sequence

V

S

W

N

S

G

A

L

T

S

G

V

H

T

W

+15.99

L

##### Posttranslational Modifications

Oxidation (HW)

##### Source File

D:\separate\_stitch\_analyses\xle-disambiguation\raw\20210323\_F1\_UM1\_Peng0013\_SA\_F59\_ingel\_3ug\_TL.raw

##### Fraction

2

##### Scan Feature

F2:16716

##### De Novo Score

98

##### ConfidenceScore

98

### m/z

865.9296

##### Mass

1729.8423

##### Charge

2

##### Retention Time

51.37

##### Predicted Retention Time

-

##### Area

9.363E+06

##### Parts Per Million

1.3

##### Fragmentation mode

ETHCD

##### Originating file

01 D:\separate\_stitch\_analyses\xle-disambiguation\20210325\_F59\_3ug\_DENOVO\_12.csv

#### Meta Information from PEAKS

##### Scan Identifier

F2:9105

##### Original sequence

V

S

W

N

S

G

A

L

T

S

G

V

H

T

W

+15.99

L

##### Posttranslational Modifications

Oxidation (HW)

##### Source File

D:\separate\_stitch\_analyses\xle-disambiguation\raw\20210323\_F1\_UM1\_Peng0013\_SA\_F59\_ingel\_3ug\_TL.raw

##### Fraction

2

##### Scan Feature

F2:7380

##### De Novo Score

98

##### ConfidenceScore

98

### m/z

577.6218

##### Mass

1729.8423

##### Charge

3

##### Retention Time

51.37

##### Predicted Retention Time

-

##### Area

8.358E+05

##### Parts Per Million

0.7

##### Fragmentation mode

ETHCD

##### Originating file

01 D:\separate\_stitch\_analyses\xle-disambiguation\20210325\_F59\_3ug\_DENOVO\_12.csv
