## Supplementary material for "A handle on mass coincidence errors in *de novo* sequencing of antibodies by bottom-up proteomics": Combined_080.html

Details Combined\_080 | Stitch OverviewUndefined

### Read Combined\_080

#### Sequence (length=12)

JJSDFYPGAVTV

#### Spectrum 11034? Spectrum 11034 The raw spectrum of this peptide as annotated by Hecklib. The fragments are coloured according to ion type (see legend). Any peaks with a star '\*' as text can be hovered over to see the full details, first the ion type second the mass shift type. By hovering over the amino acids in the peptide or ions in the legend the corresponding peaks are highlighted. By toggling the 'Unassigned' label you can turn the background (unassigned) peaks on or off in the plot. By updating the slider in the Ion legend you can update the spectrum to only show the top X% of the peaks with labels. The top X% means any peak that is within X% of the highest intensity. By dragging in the spectrum you can zoom in to a specific part of the spectrum and use 'Zoom Out' to get back to the original zoom level. The annotation of the spectrum is based on the given sequence in the peptides file and is done with different software so inconsistencies are likely. The peaks are annotated based on the given sequence, with 20 ppm tolerance.

Copy Data

##### Spectrum 11034 (TSV)

###### Preview

```
Loading example...
```

*Click on the button to copy the data to your clipboard.*

Mz MinMz MaxIntensity Max

WidthHeightPeptide font sizePeptide stroke widthSpectrum font sizeSpectrum stroke widthCompact peptide

Ion legend

wxyz

abcd

OtherUnassignedIon^Charge^_Position_Show for top:%

JJSDFYPGAVTV

04.47e+48.94e+41.34e+51.79e+5

Zoom Out

y^+1^_2_y^+1^_2_c^+1^_2_c^+1^_3_y^+1^_3_y^+1^_4_c^+1^_4_z^+1^_5_c^+1^_4_c^+2^_9_y^+1^_6_y^+1^_6_c^+1^_5_c^+2^_11_c^+1^_5_z^+1^_7_z^+1^_7_y^+1^_7_c^+1^_6_c^+1^_6_z^+1^_8_z^+1^_8_c^+1^_7_c^+1^_8_c^+1^_8_z^+1^_9_y^+1^_9_z^+1^_9_c^+1^_9_y^+1^_9_c^+1^_9_z^+1^_10_z^+1^_10_y^+1^_10_c^+1^_10_c^+1^_10_z^+1^_11_z^+1^_11_c^+1^_11_c^+1^_11_

0527105415822109

Fragment Matches Table

Show background peaks

| Position | Ion type | Intensity | mz Theoretical | mz Error (Th) | mz Error (ppm) | Charge | Series Number |
| --- | --- | --- | --- | --- | --- | --- | --- |
| - | - | 366 | 123.3 | - | - | 0 | - |
| - | - | 377.9 | 125.7 | - | - | 0 | - |
| - | - | 351.3 | 132.6 | - | - | 0 | - |
| - | - | 419 | 134.1 | - | - | 0 | - |
| - | - | 353.3 | 134.4 | - | - | 0 | - |
| - | - | 416.7 | 137.6 | - | - | 0 | - |
| - | - | 1085 | 149 | - | - | 0 | - |
| - | - | 2958 | 173.1 | - | - | 0 | - |
| - | - | 457.2 | 176.2 | - | - | 0 | - |
| - | - | 456.9 | 192.7 | - | - | 0 | - |
| - | - | 435.3 | 198.7 | - | - | 0 | - |
| - | - | 1.747E+04 | 199.2 | - | - | 0 | - |
| - | - | 447.9 | 199.9 | - | - | 0 | - |
| - | - | 1712 | 200.2 | - | - | 0 | - |
| 11 | y | 7580 | 201.1 | 0.0002121 | 1.055 | +1 | 2 |
| - | - | 617.6 | 202.1 | - | - | 0 | - |
| - | - | 482 | 203.5 | - | - | 0 | - |
| 11 | y | 4.786E+04 | 219.1 | 0.000298 | 1.36 | +1 | 2 |
| - | - | 524.7 | 219.9 | - | - | 0 | - |
| - | - | 3940 | 220.1 | - | - | 0 | - |
| - | - | 2399 | 226.1 | - | - | 0 | - |
| 2 | c | 1.267E+04 | 227.2 | 0.0003005 | 1.323 | +1 | 2 |
| - | - | 1404 | 228.2 | - | - | 0 | - |
| - | - | 526.2 | 229.6 | - | - | 0 | - |
| - | - | 483.7 | 278.2 | - | - | 0 | - |
| - | - | 3221 | 297.2 | - | - | 0 | - |
| - | - | 637.9 | 308 | - | - | 0 | - |
| - | - | 785.9 | 311.1 | - | - | 0 | - |
| 3 | c | 981.6 | 314.2 | 0.000514 | 1.636 | +1 | 3 |
| - | - | 575.6 | 316.2 | - | - | 0 | - |
| 10 | y | 1994 | 318.2 | 0.0001062 | 0.3336 | +1 | 3 |
| - | - | 8517 | 325.2 | - | - | 0 | - |
| - | - | 1571 | 326.2 | - | - | 0 | - |
| - | - | 499.3 | 326.6 | - | - | 0 | - |
| - | - | 839.6 | 329.2 | - | - | 0 | - |
| - | - | 573.4 | 331 | - | - | 0 | - |
| - | - | 754.1 | 335.2 | - | - | 0 | - |
| - | - | 593.9 | 339.2 | - | - | 0 | - |
| - | - | 2256 | 342.2 | - | - | 0 | - |
| - | - | 2950 | 350.1 | - | - | 0 | - |
| - | - | 608.3 | 351.1 | - | - | 0 | - |
| - | - | 1161 | 355.1 | - | - | 0 | - |
| 9 | y | 680.8 | 371.2 | 0.001572 | 4.235 | +1 | 4 |
| - | - | 781.3 | 389.2 | - | - | 0 | - |
| - | - | 3315 | 408.2 | - | - | 0 | - |
| - | - | 587.3 | 408.6 | - | - | 0 | - |
| - | - | 661.1 | 409.2 | - | - | 0 | - |
| - | - | 904.7 | 411.2 | - | - | 0 | - |
| - | - | 1302 | 424.2 | - | - | 0 | - |
| - | - | 783.2 | 426.2 | - | - | 0 | - |
| - | - | 6510 | 426.2 | - | - | 0 | - |
| - | - | 1597 | 427.2 | - | - | 0 | - |
| 4 | c | 2314 | 429.2 | 0.0006095 | 1.42 | +1 | 4 |
| 8 | z | 641.2 | 430.2 | 0.002272 | 5.28 | +1 | 5 |
| - | - | 620.8 | 434.1 | - | - | 0 | - |
| - | - | 829.1 | 435.2 | - | - | 0 | - |
| - | - | 1560 | 438.3 | - | - | 0 | - |
| 4 | c | 1215 | 446.3 | 0.0004574 | 1.025 | +1 | 4 |
| - | - | 697 | 447.7 | - | - | 0 | - |
| - | - | 676.3 | 460.3 | - | - | 0 | - |
| - | - | 662.4 | 461.2 | - | - | 0 | - |
| - | - | 3130 | 463.2 | - | - | 0 | - |
| - | - | 1043 | 464.2 | - | - | 0 | - |
| - | - | 802.5 | 470.2 | - | - | 0 | - |
| 9 | c | 3919 | 482.7 | 0.0006129 | 1.27 | +2 | 9 |
| - | - | 1886 | 483.2 | - | - | 0 | - |
| - | - | 1295 | 485.2 | - | - | 0 | - |
| - | - | 1566 | 488.3 | - | - | 0 | - |
| - | - | 1020 | 495.2 | - | - | 0 | - |
| - | - | 651.4 | 507.3 | - | - | 0 | - |
| - | - | 3407 | 513.2 | - | - | 0 | - |
| 7 | y | 3007 | 525.3 | 3.75E-05 | 0.07138 | +1 | 6 |
| - | - | 620.6 | 525.3 | - | - | 0 | - |
| - | - | 965.6 | 526.3 | - | - | 0 | - |
| - | - | 712.8 | 531.3 | - | - | 0 | - |
| - | - | 1470 | 537.3 | - | - | 0 | - |
| - | - | 1092 | 539.2 | - | - | 0 | - |
| - | - | 9302 | 542.3 | - | - | 0 | - |
| 7 | y | 1.012E+05 | 543.3 | 0.0003371 | 0.6204 | +1 | 6 |
| - | - | 2.63E+04 | 544.3 | - | - | 0 | - |
| - | - | 5285 | 545.3 | - | - | 0 | - |
| - | - | 1877 | 548.3 | - | - | 0 | - |
| - | - | 1806 | 551.4 | - | - | 0 | - |
| - | - | 578.2 | 553.3 | - | - | 0 | - |
| - | - | 1638 | 573.8 | - | - | 0 | - |
| - | - | 1498 | 574.3 | - | - | 0 | - |
| 5 | c | 3843 | 576.3 | 5.54E-05 | 0.09613 | +1 | 5 |
| - | - | 995 | 577.3 | - | - | 0 | - |
| - | - | 1216 | 578.3 | - | - | 0 | - |
| - | - | 749.2 | 579.3 | - | - | 0 | - |
| - | - | 687.1 | 581.3 | - | - | 0 | - |
| 11 | c | 786 | 582.8 | 0.0001823 | 0.3128 | +2 | 11 |
| - | - | 1158 | 583.3 | - | - | 0 | - |
| - | - | 1199 | 589.3 | - | - | 0 | - |
| - | - | 6374 | 592.3 | - | - | 0 | - |
| 5 | c | 2.932E+04 | 593.3 | 6.828E-06 | 0.01151 | +1 | 5 |
| - | - | 9940 | 594.3 | - | - | 0 | - |
| - | - | 1433 | 595.3 | - | - | 0 | - |
| - | - | 2105 | 598.3 | - | - | 0 | - |
| - | - | 1697 | 608.3 | - | - | 0 | - |
| - | - | 846.2 | 609.3 | - | - | 0 | - |
| - | - | 1.222E+04 | 626.3 | - | - | 0 | - |
| - | - | 678.3 | 626.3 | - | - | 0 | - |
| - | - | 3268 | 627.3 | - | - | 0 | - |
| - | - | 816.1 | 635.3 | - | - | 0 | - |
| - | - | 742.8 | 636.3 | - | - | 0 | - |
| - | - | 4584 | 637.3 | - | - | 0 | - |
| - | - | 1125 | 638.3 | - | - | 0 | - |
| - | - | 698.8 | 654.3 | - | - | 0 | - |
| - | - | 1072 | 670.3 | - | - | 0 | - |
| 6 | z | 2411 | 672.3 | 0.001842 | 2.74 | +1 | 7 |
| - | - | 1314 | 673.4 | - | - | 0 | - |
| - | - | 1.279E+04 | 688.3 | - | - | 0 | - |
| - | - | 4164 | 689.3 | - | - | 0 | - |
| 6 | z | 3543 | 690.4 | 0.002519 | 3.649 | +1 | 7 |
| - | - | 2670 | 691.4 | - | - | 0 | - |
| - | - | 798.1 | 692.4 | - | - | 0 | - |
| - | - | 1698 | 694.3 | - | - | 0 | - |
| - | - | 2.974E+04 | 705.4 | - | - | 0 | - |
| 6 | y | 3.017E+04 | 706.4 | 0.001773 | 2.51 | +1 | 7 |
| - | - | 9406 | 707.4 | - | - | 0 | - |
| - | - | 2170 | 708.4 | - | - | 0 | - |
| - | - | 3543 | 711.4 | - | - | 0 | - |
| - | - | 3690 | 712.4 | - | - | 0 | - |
| - | - | 7747 | 713.4 | - | - | 0 | - |
| - | - | 2208 | 714.4 | - | - | 0 | - |
| - | - | 2117 | 721.4 | - | - | 0 | - |
| - | - | 689.5 | 737.4 | - | - | 0 | - |
| - | - | 2057 | 738.3 | - | - | 0 | - |
| 6 | c | 1019 | 738.4 | 0.01272 | 17.22 | +1 | 6 |
| 6 | c | 2.22E+04 | 739.4 | 9.048E-05 | 0.1224 | +1 | 6 |
| - | - | 8701 | 740.4 | - | - | 0 | - |
| - | - | 2346 | 741.4 | - | - | 0 | - |
| - | - | 1501 | 752.4 | - | - | 0 | - |
| - | - | 710.7 | 765.4 | - | - | 0 | - |
| - | - | 6527 | 809.4 | - | - | 0 | - |
| - | - | 3791 | 810.4 | - | - | 0 | - |
| - | - | 849.8 | 811.4 | - | - | 0 | - |
| 5 | z | 3788 | 819.4 | 0.00147 | 1.794 | +1 | 8 |
| - | - | 1933 | 820.4 | - | - | 0 | - |
| - | - | 774 | 821.4 | - | - | 0 | - |
| - | - | 1205 | 822.4 | - | - | 0 | - |
| - | - | 850.7 | 836.4 | - | - | 0 | - |
| 5 | z | 2.053E+04 | 837.4 | 0.0003156 | 0.3769 | +1 | 8 |
| - | - | 1.469E+04 | 838.4 | - | - | 0 | - |
| - | - | 4517 | 839.4 | - | - | 0 | - |
| - | - | 865.1 | 840.4 | - | - | 0 | - |
| - | - | 1045 | 851.4 | - | - | 0 | - |
| - | - | 8066 | 852.4 | - | - | 0 | - |
| - | - | 691.3 | 853.4 | - | - | 0 | - |
| 7 | c | 1.478E+04 | 853.4 | 0.001767 | 2.07 | +1 | 7 |
| - | - | 5399 | 854.4 | - | - | 0 | - |
| - | - | 1530 | 855.5 | - | - | 0 | - |
| - | - | 1534 | 865.5 | - | - | 0 | - |
| - | - | 7207 | 866.5 | - | - | 0 | - |
| - | - | 3945 | 867.5 | - | - | 0 | - |
| 8 | c | 3864 | 893.4 | 0.001198 | 1.341 | +1 | 8 |
| - | - | 2660 | 894.4 | - | - | 0 | - |
| - | - | 1307 | 900.5 | - | - | 0 | - |
| - | - | 2754 | 908.5 | - | - | 0 | - |
| - | - | 3.472E+04 | 909.5 | - | - | 0 | - |
| 8 | c | 8.791E+04 | 910.5 | 0.001441 | 1.583 | +1 | 8 |
| - | - | 4.541E+04 | 911.5 | - | - | 0 | - |
| - | - | 1.167E+04 | 912.5 | - | - | 0 | - |
| - | - | 1322 | 913.5 | - | - | 0 | - |
| - | - | 1011 | 920.4 | - | - | 0 | - |
| - | - | 767.5 | 932.4 | - | - | 0 | - |
| 4 | z | 3988 | 934.4 | 0.0004569 | 0.489 | +1 | 9 |
| - | - | 2389 | 935.4 | - | - | 0 | - |
| - | - | 1222 | 936.5 | - | - | 0 | - |
| - | - | 2515 | 937.5 | - | - | 0 | - |
| - | - | 4144 | 938.4 | - | - | 0 | - |
| - | - | 2118 | 939.4 | - | - | 0 | - |
| - | - | 1391 | 946.5 | - | - | 0 | - |
| - | - | 2971 | 948.5 | - | - | 0 | - |
| - | - | 2188 | 949.5 | - | - | 0 | - |
| 4 | y | 3715 | 950.5 | 0.002607 | 2.743 | +1 | 9 |
| - | - | 2366 | 951.5 | - | - | 0 | - |
| 4 | z | 8068 | 952.5 | 0.001855 | 1.948 | +1 | 9 |
| - | - | 8443 | 953.5 | - | - | 0 | - |
| - | - | 3182 | 954.5 | - | - | 0 | - |
| - | - | 694.1 | 955.5 | - | - | 0 | - |
| 9 | c | 1.466E+04 | 964.5 | 0.0001404 | 0.1456 | +1 | 9 |
| - | - | 9514 | 965.5 | - | - | 0 | - |
| - | - | 2239 | 966.5 | - | - | 0 | - |
| - | - | 951.6 | 967.5 | - | - | 0 | - |
| 4 | y | 1174 | 968.5 | 0.003991 | 4.12 | +1 | 9 |
| - | - | 1082 | 969.5 | - | - | 0 | - |
| 9 | c | 4.87E+04 | 981.5 | 0.0005298 | 0.5398 | +1 | 9 |
| - | - | 2.809E+04 | 982.5 | - | - | 0 | - |
| - | - | 8455 | 983.5 | - | - | 0 | - |
| - | - | 807.9 | 984.5 | - | - | 0 | - |
| - | - | 904.5 | 1005 | - | - | 0 | - |
| 3 | z | 880.4 | 1021 | 0.001115 | 1.091 | +1 | 10 |
| - | - | 675.3 | 1035 | - | - | 0 | - |
| - | - | 954.7 | 1036 | - | - | 0 | - |
| - | - | 5061 | 1037 | - | - | 0 | - |
| - | - | 1502 | 1037 | - | - | 0 | - |
| - | - | 3549 | 1038 | - | - | 0 | - |
| - | - | 5276 | 1039 | - | - | 0 | - |
| 3 | z | 3481 | 1039 | 0.002891 | 2.781 | +1 | 10 |
| - | - | 2239 | 1040 | - | - | 0 | - |
| - | - | 5201 | 1040 | - | - | 0 | - |
| - | - | 697.5 | 1041 | - | - | 0 | - |
| - | - | 2538 | 1041 | - | - | 0 | - |
| - | - | 1880 | 1046 | - | - | 0 | - |
| - | - | 984.5 | 1047 | - | - | 0 | - |
| - | - | 1.333E+04 | 1048 | - | - | 0 | - |
| - | - | 8453 | 1049 | - | - | 0 | - |
| - | - | 3403 | 1050 | - | - | 0 | - |
| - | - | 802.6 | 1051 | - | - | 0 | - |
| - | - | 1890 | 1054 | - | - | 0 | - |
| 3 | y | 1.043E+04 | 1056 | 0.001961 | 1.858 | +1 | 10 |
| - | - | 3810 | 1057 | - | - | 0 | - |
| - | - | 908.9 | 1058 | - | - | 0 | - |
| 10 | c | 1.65E+04 | 1064 | 0.001074 | 1.01 | +1 | 10 |
| - | - | 9853 | 1065 | - | - | 0 | - |
| - | - | 5818 | 1066 | - | - | 0 | - |
| - | - | 2313 | 1067 | - | - | 0 | - |
| - | - | 755.4 | 1068 | - | - | 0 | - |
| 10 | c | 7.828E+04 | 1081 | 0.001012 | 0.9361 | +1 | 10 |
| - | - | 4.885E+04 | 1082 | - | - | 0 | - |
| - | - | 4.261E+04 | 1083 | - | - | 0 | - |
| - | - | 2.208E+04 | 1084 | - | - | 0 | - |
| - | - | 6259 | 1085 | - | - | 0 | - |
| - | - | 1.084E+04 | 1120 | - | - | 0 | - |
| - | - | 6201 | 1121 | - | - | 0 | - |
| - | - | 2468 | 1122 | - | - | 0 | - |
| 2 | z | 1164 | 1135 | 0.002468 | 2.175 | +1 | 11 |
| - | - | 1691 | 1138 | - | - | 0 | - |
| - | - | 1736 | 1139 | - | - | 0 | - |
| - | - | 768.8 | 1140 | - | - | 0 | - |
| - | - | 1060 | 1147 | - | - | 0 | - |
| - | - | 2418 | 1149 | - | - | 0 | - |
| - | - | 1362 | 1150 | - | - | 0 | - |
| 2 | z | 2.036E+04 | 1153 | 0.000895 | 0.7765 | +1 | 11 |
| - | - | 1.298E+04 | 1154 | - | - | 0 | - |
| - | - | 4747 | 1155 | - | - | 0 | - |
| - | - | 798.1 | 1156 | - | - | 0 | - |
| - | - | 1714 | 1164 | - | - | 0 | - |
| 11 | c | 992.7 | 1165 | 0.01665 | 14.29 | +1 | 11 |
| - | - | 1746 | 1165 | - | - | 0 | - |
| - | - | 1226 | 1166 | - | - | 0 | - |
| - | - | 9164 | 1181 | - | - | 0 | - |
| 11 | c | 2.877E+04 | 1182 | 0.002303 | 1.949 | +1 | 11 |
| - | - | 3109 | 1182 | - | - | 0 | - |
| - | - | 1.97E+04 | 1183 | - | - | 0 | - |
| - | - | 6504 | 1184 | - | - | 0 | - |
| - | - | 791 | 1185 | - | - | 0 | - |
| - | - | 752.7 | 1205 | - | - | 0 | - |
| - | - | 797.2 | 1210 | - | - | 0 | - |
| - | - | 667.3 | 1221 | - | - | 0 | - |
| - | - | 1.729E+04 | 1223 | - | - | 0 | - |
| - | - | 1.282E+04 | 1224 | - | - | 0 | - |
| - | - | 4238 | 1225 | - | - | 0 | - |
| - | - | 800.4 | 1226 | - | - | 0 | - |
| - | - | 795.8 | 1227 | - | - | 0 | - |
| - | - | 852.1 | 1230 | - | - | 0 | - |
| - | - | 1042 | 1237 | - | - | 0 | - |
| - | - | 1079 | 1238 | - | - | 0 | - |
| - | - | 1.008E+04 | 1239 | - | - | 0 | - |
| - | - | 904.4 | 1240 | - | - | 0 | - |
| - | - | 7580 | 1240 | - | - | 0 | - |
| - | - | 2984 | 1241 | - | - | 0 | - |
| - | - | 3242 | 1248 | - | - | 0 | - |
| - | - | 2463 | 1249 | - | - | 0 | - |
| - | - | 8923 | 1255 | - | - | 0 | - |
| - | - | 6479 | 1256 | - | - | 0 | - |
| - | - | 1944 | 1257 | - | - | 0 | - |
| - | - | 4663 | 1265 | - | - | 0 | - |
| - | - | 2.752E+04 | 1266 | - | - | 0 | - |
| - | - | 1.936E+04 | 1267 | - | - | 0 | - |
| - | - | 8077 | 1268 | - | - | 0 | - |
| - | - | 1756 | 1269 | - | - | 0 | - |
| - | - | 1203 | 1280 | - | - | 0 | - |
| - | - | 4206 | 1281 | - | - | 0 | - |
| - | - | 1246 | 1281 | - | - | 0 | - |
| - | - | 4.356E+04 | 1282 | - | - | 0 | - |
| - | - | 1.771E+05 | 1283 | - | - | 0 | - |
| - | - | 1.234E+05 | 1284 | - | - | 0 | - |
| - | - | 5.158E+04 | 1285 | - | - | 0 | - |
| - | - | 8551 | 1286 | - | - | 0 | - |
| - | - | 772.7 | 2088 | - | - | 0 | - |

m/z Charge Intensity FragmentType MassShift Position
123.33231353759766 0 365.9722
125.73748016357422 0 377.88763
132.56112670898438 0 351.2722
134.0703582763672 0 418.95847
134.4441680908203 0 353.26898
137.58364868164062 0 416.67953
148.95372009277344 0 1085.4149
173.12875366210938 0 2958.4905
176.18896484375 0 457.18222
192.650634765625 0 456.87634
198.66943359375 0 435.34634
199.1807403564453 0 17474.559
199.92637634277344 0 447.90536
200.184326171875 0 1712.0201
201.1235809326172 0 7579.7764 y Water loss 10
202.12713623046875 0 617.6095
203.47702026367188 0 482.0378
219.1342315673828 0 47856.59 y 10
219.9325714111328 0 524.68286
220.13755798339844 0 3940.2756
226.1187744140625 0 2399.294
227.1757049560547 0 12667.558 c Ammonia loss 1
228.1790313720703 0 1404.4885
229.5703125 0 526.15234
278.157470703125 0 483.7406
297.192626953125 0 3221.0415
307.9745788574219 0 637.92737
311.1397705078125 0 785.90485
314.20794677734375 0 981.6188 c Ammonia loss 2
316.15179443359375 0 575.6142
318.20245361328125 0 1993.859 y 9
325.1873779296875 0 8516.501
326.190185546875 0 1570.9441
326.56036376953125 0 499.2789
329.1823425292969 0 839.56537
330.953125 0 573.4004
335.1717834472656 0 754.06335
339.20233154296875 0 593.8501
342.2145690917969 0 2256.4333
350.1351318359375 0 2950.3926
351.1368713378906 0 608.33014
355.0702209472656 0 1160.964
371.23046875 0 680.81195 y Water loss 8
389.18353271484375 0 781.2533
408.2253723144531 0 3315.301
408.6200866699219 0 587.31415
409.2279052734375 0 661.14105
411.2239685058594 0 904.6956
424.22308349609375 0 1302.4203
426.1660461425781 0 783.2058
426.23486328125 0 6510.407
427.2399597167969 0 1597.3206
429.2349853515625 0 2313.5056 c Ammonia loss 3
430.23992919921875 0 641.17114 z 7
434.088623046875 0 620.79565
435.22430419921875 0 829.12634
438.2730407714844 0 1559.5695
446.2604675292969 0 1214.6063 c 3
447.7247009277344 0 697.0188
460.25506591796875 0 676.3113
461.20611572265625 0 662.3983
463.2187805175781 0 3130.187
464.222412109375 0 1042.9935
470.2169494628906 0 802.51666
482.74298095703125 0 3918.6272 c Ammonia loss 8
483.2447509765625 0 1886.1558
485.20404052734375 0 1295.3257
488.25189208984375 0 1565.9114
495.18658447265625 0 1020.4418
507.2945861816406 0 651.3957
513.1976928710938 0 3406.9885
525.3031616210938 0 3006.652 y Water loss 6
525.344970703125 0 620.62396
526.3085327148438 0 965.5762
531.28759765625 0 712.8451
537.3067016601562 0 1469.6898
539.24951171875 0 1091.5734
542.3062744140625 0 9302.042
543.3140258789062 0 101237.26 y 6
544.31689453125 0 26297.695
545.319580078125 0 5285.3745
548.307861328125 0 1877.3138
551.35546875 0 1806.295
553.259033203125 0 578.1855
573.7963256835938 0 1637.5776
574.2942504882812 0 1498.1138
576.302734375 0 3842.6006 c Ammonia loss 4
577.3074951171875 0 994.9909
578.3141479492188 0 1215.9
579.3182373046875 0 749.19696
581.2584838867188 0 687.13306
582.8002319335938 0 785.9816 c Ammonia loss 10
583.3036499023438 0 1157.8169
589.299072265625 0 1198.7345
592.3218994140625 0 6374.3066
593.329345703125 0 29323.434 c 4
594.3323974609375 0 9939.82
595.3342895507812 0 1433.2094
598.2880249023438 0 2105.2407
608.2710571289062 0 1697.3137
609.2749633789062 0 846.1936
626.2822875976562 0 12215.877
626.339599609375 0 678.2953
627.2843627929688 0 3268.4717
635.3212890625 0 816.1026
636.32275390625 0 742.75525
637.3341674804688 0 4583.7935
638.3382568359375 0 1124.9021
654.3428955078125 0 698.77905
670.3311767578125 0 1071.7799
672.3458862304688 0 2410.6555 z Water loss 5
673.3536987304688 0 1313.5142
688.3435668945312 0 12787.049
689.34619140625 0 4164.3545
690.3557739257812 0 3542.8674 z 5
691.36474609375 0 2670.05
692.367431640625 0 798.1286
694.3469848632812 0 1697.7007
705.368896484375 0 29738.262
706.375244140625 0 30165.344 y 5
707.3793334960938 0 9405.504
708.3829345703125 0 2169.692
711.371337890625 0 3543.4502
712.3767700195312 0 3690.14
713.385498046875 0 7747.4785
714.3897705078125 0 2208.1133
721.3548583984375 0 2117.0469
737.3734741210938 0 689.53937
738.3090209960938 0 2057.4368
738.369384765625 0 1018.5001 c Water loss 5
739.3660278320312 0 22199.414 c Ammonia loss 5
740.3697509765625 0 8700.542
741.3712768554688 0 2345.5276
752.3636474609375 0 1501.4796
765.3805541992188 0 710.70667
809.431640625 0 6526.504
810.4348754882812 0 3790.625
811.435546875 0 849.8279
819.4146728515625 0 3788.2244 z Water loss 4
820.4193115234375 0 1933.2211
821.35693359375 0 773.99774
822.3596801757812 0 1204.6962
836.3966064453125 0 850.6887
837.4263916015625 0 20531.13 z 4
838.4315795898438 0 14687.689
839.43115234375 0 4516.647
840.4395141601562 0 865.08356
851.3932495117188 0 1044.628
852.4370727539062 0 8065.9634
853.3506469726562 0 691.2537
853.4436645507812 0 14778.131 c 6
854.4481201171875 0 5398.75
855.452392578125 0 1529.8857
865.4730224609375 0 1533.652
866.4547119140625 0 7206.562
867.4583740234375 0 3944.939
893.4391479492188 0 3864.1182 c Ammonia loss 7
894.443359375 0 2660.0796
900.4812622070312 0 1306.8569
908.4630737304688 0 2754.2537
909.4590454101562 0 34721.56
910.4654541015625 0 87910.66 c 7
911.4688110351562 0 45411.63
912.4723510742188 0 11670.444
913.4755859375 0 1321.55
920.4160766601562 0 1011.4282
932.4461669921875 0 767.52924
934.4435424804688 0 3988.3125 z Water loss 3
935.44677734375 0 2388.946
936.4674072265625 0 1221.6434
937.4882202148438 0 2514.596
938.4251708984375 0 4143.745
939.4274291992188 0 2117.7578
946.47119140625 0 1390.5138
948.4595947265625 0 2971.0793
949.4619140625 0 2187.6328
950.4644165039062 0 3714.6743 y Water loss 3
951.4630126953125 0 2365.75
952.4555053710938 0 8067.563 z 3
953.4603881835938 0 8442.6875
954.4624633789062 0 3181.6538
955.4530029296875 0 694.0836
964.4776000976562 0 14655.207 c Ammonia loss 8
965.47900390625 0 9514.275
966.4823608398438 0 2239.1467
967.474365234375 0 951.60583
968.4683837890625 0 1173.9465 y 3
969.469970703125 0 1082.42
981.5034790039062 0 48696.824 c 8
982.5064086914062 0 28094.248
983.509033203125 0 8454.609
984.51611328125 0 807.93274
1005.470947265625 0 904.47174
1021.4739990234375 0 880.40643 z Water loss 2
1034.5184326171875 0 675.30365
1035.5645751953125 0 954.72186
1036.554443359375 0 5060.5747
1037.467529296875 0 1501.6489
1037.5618896484375 0 3548.7913
1038.5703125 0 5275.8696
1039.4827880859375 0 3481.1711 z 2
1039.580810546875 0 2238.6265
1040.4893798828125 0 5200.745
1040.583740234375 0 697.4646
1041.4940185546875 0 2538.468
1045.5333251953125 0 1879.8942
1046.531982421875 0 984.45953
1047.52783203125 0 13333.846
1048.5306396484375 0 8452.908
1049.534423828125 0 3402.8667
1050.5396728515625 0 802.57947
1054.4959716796875 0 1889.9647
1055.50244140625 0 10431.659 y 2
1056.507080078125 0 3810.311
1057.508544921875 0 908.9206
1063.5447998046875 0 16502.676 c Ammonia loss 9
1064.5477294921875 0 9853.1045
1065.5557861328125 0 5818.377
1066.559814453125 0 2312.881
1067.561767578125 0 755.41187
1080.5714111328125 0 78275.8 c 9
1081.57470703125 0 48852.27
1082.5841064453125 0 42608.715
1083.5894775390625 0 22080.334
1084.592529296875 0 6258.71
1119.595703125 0 10842.986
1120.597412109375 0 6200.684
1121.5966796875 0 2467.573
1134.5616455078125 0 1163.601 z Water loss 1
1137.607177734375 0 1691.1859
1138.6129150390625 0 1735.6196
1139.61865234375 0 768.8107
1146.582763671875 0 1060.2238
1148.5758056640625 0 2417.6672
1149.582763671875 0 1362.1571
1152.56884765625 0 20362.516 z 1
1153.572265625 0 12977.689
1154.572509765625 0 4747.2705
1155.58740234375 0 798.1166
1163.701416015625 0 1713.7751
1164.576904296875 0 992.67645 c Ammonia loss 10
1164.704345703125 0 1745.7227
1165.598388671875 0 1226.3175
1180.72412109375 0 9164.144
1181.6177978515625 0 28767.305 c 10
1181.733154296875 0 3109.3013
1182.6209716796875 0 19701.602
1183.625244140625 0 6504.387
1184.6209716796875 0 791.0287
1204.643798828125 0 752.7288
1209.59765625 0 797.193
1220.67041015625 0 667.30054
1222.65234375 0 17285.643
1223.6546630859375 0 12815.936
1224.657958984375 0 4237.505
1225.662841796875 0 800.438
1226.61181640625 0 795.7796
1229.635009765625 0 852.1011
1236.6676025390625 0 1041.82
1237.6795654296875 0 1078.8195
1238.68701171875 0 10078.218
1239.5479736328125 0 904.3593
1239.6907958984375 0 7579.6206
1240.6915283203125 0 2983.923
1247.6390380859375 0 3241.6306
1248.6455078125 0 2463.2712
1254.684326171875 0 8922.771
1255.68505859375 0 6479.1343
1256.6883544921875 0 1944.3462
1264.6632080078125 0 4662.605
1265.6539306640625 0 27523.117
1266.656982421875 0 19364.744
1267.6571044921875 0 8076.67
1268.6588134765625 0 1755.8467
1279.6552734375 0 1203.4303
1280.660400390625 0 4205.5884
1281.49755859375 0 1245.543
1281.6693115234375 0 43562.855
1282.677001953125 0 177061.64
1283.68017578125 0 123414.65
1284.68359375 0 51575.92
1285.68359375 0 8550.881
2087.91748046875 0 772.7013

Spectrum Details

|  |  |
| --- | --- |
| Matched peaks? Matched peaks The total absolute number of peaks matched. Additionally in brackets the total fraction of peaks matched and the total number of peaks is shown. | 40 (14.13% of 283) |
| FDR? FDR The false discovery rate estimated for this peptide. It is calculated by matching all theoretical fragments with a non-integer shift with the raw peaks for this spectrum. This is done with 40 different shifts. The resulting percentage is the average number of annotated peaks over the number of annotated peaks with the correct spectrum. | 1.31% |
| Satellite FDR? Satellite FDR See the FDR for details on its calculation. This satellite ion specific FDR only contains the satellite ions (d/w) for I/L/J positions. | ∞ |
| PSM Score? PSM Score The PSM Score as given by Hecklib to this annotated spectrum. It is shown with three significant figures. | 559 |

#### Spectrum 11187? Spectrum 11187 The raw spectrum of this peptide as annotated by Hecklib. The fragments are coloured according to ion type (see legend). Any peaks with a star '\*' as text can be hovered over to see the full details, first the ion type second the mass shift type. By hovering over the amino acids in the peptide or ions in the legend the corresponding peaks are highlighted. By toggling the 'Unassigned' label you can turn the background (unassigned) peaks on or off in the plot. By updating the slider in the Ion legend you can update the spectrum to only show the top X% of the peaks with labels. The top X% means any peak that is within X% of the highest intensity. By dragging in the spectrum you can zoom in to a specific part of the spectrum and use 'Zoom Out' to get back to the original zoom level. The annotation of the spectrum is based on the given sequence in the peptides file and is done with different software so inconsistencies are likely. The peaks are annotated based on the given sequence, with 20 ppm tolerance.

Copy Data

##### Spectrum 11187 (TSV)

###### Preview

```
Loading example...
```

*Click on the button to copy the data to your clipboard.*

Mz MinMz MaxIntensity Max

WidthHeightPeptide font sizePeptide stroke widthSpectrum font sizeSpectrum stroke widthCompact peptide

Ion legend

wxyz

abcd

OtherUnassignedIon^Charge^_Position_Show for top:%

JJSDFYPGAVTV

01.61e+43.21e+44.82e+46.42e+4

Zoom Out

y^+1^_2_y^+1^_2_c^+1^_2_c^+1^_3_y^+1^_3_c^+1^_4_c^+1^_4_c^+2^_8_c^+2^_9_y^+1^_6_y^+1^_6_c^+1^_5_c^+1^_5_z^+1^_7_z^+1^_7_y^+1^_7_c^+1^_6_z^+1^_8_c^+1^_7_z^+1^_8_c^+1^_7_c^+1^_8_c^+1^_8_z^+1^_9_y^+1^_9_z^+1^_9_c^+1^_9_c^+1^_9_z^+1^_10_y^+1^_10_c^+1^_10_c^+1^_10_z^+1^_11_c^+1^_11_c^+1^_11_

0595119017862381

Fragment Matches Table

Show background peaks

| Position | Ion type | Intensity | mz Theoretical | mz Error (Th) | mz Error (ppm) | Charge | Series Number |
| --- | --- | --- | --- | --- | --- | --- | --- |
| - | - | 422 | 127.7 | - | - | 0 | - |
| - | - | 704.9 | 148.9 | - | - | 0 | - |
| - | - | 486.3 | 159.4 | - | - | 0 | - |
| - | - | 430.9 | 167.1 | - | - | 0 | - |
| - | - | 491 | 170.6 | - | - | 0 | - |
| - | - | 573.8 | 173.1 | - | - | 0 | - |
| - | - | 707.9 | 173.5 | - | - | 0 | - |
| - | - | 487.1 | 190.3 | - | - | 0 | - |
| - | - | 507.3 | 191.6 | - | - | 0 | - |
| - | - | 7559 | 199.2 | - | - | 0 | - |
| 11 | y | 2307 | 201.1 | 1.534E-06 | 0.007626 | +1 | 2 |
| - | - | 794.9 | 219.1 | - | - | 0 | - |
| 11 | y | 1.512E+04 | 219.1 | 0.0001445 | 0.6593 | +1 | 2 |
| - | - | 1432 | 220.1 | - | - | 0 | - |
| - | - | 655.5 | 226.1 | - | - | 0 | - |
| 2 | c | 5208 | 227.2 | 1.063E-05 | 0.04677 | +1 | 2 |
| - | - | 593.7 | 239.1 | - | - | 0 | - |
| - | - | 516.6 | 279.9 | - | - | 0 | - |
| - | - | 619.7 | 297.2 | - | - | 0 | - |
| 3 | c | 649.7 | 314.2 | 0.00324 | 10.31 | +1 | 3 |
| 10 | y | 933.6 | 318.2 | 0.0001367 | 0.4296 | +1 | 3 |
| - | - | 3366 | 325.2 | - | - | 0 | - |
| - | - | 597.4 | 335.2 | - | - | 0 | - |
| - | - | 1237 | 350.1 | - | - | 0 | - |
| - | - | 856.5 | 355.1 | - | - | 0 | - |
| - | - | 502.6 | 362.1 | - | - | 0 | - |
| - | - | 509.5 | 392.8 | - | - | 0 | - |
| - | - | 1527 | 408.2 | - | - | 0 | - |
| - | - | 628.1 | 424.2 | - | - | 0 | - |
| - | - | 2071 | 426.2 | - | - | 0 | - |
| 4 | c | 1003 | 429.2 | 0.0001517 | 0.3535 | +1 | 4 |
| - | - | 558 | 433.1 | - | - | 0 | - |
| 4 | c | 696.3 | 446.3 | 0.001526 | 3.42 | +1 | 4 |
| 8 | c | 717.3 | 447.2 | 0.0007982 | 1.785 | +2 | 8 |
| - | - | 556.2 | 460.3 | - | - | 0 | - |
| - | - | 899.1 | 463.2 | - | - | 0 | - |
| 9 | c | 1160 | 482.7 | 0.001528 | 3.166 | +2 | 9 |
| - | - | 735.8 | 483.2 | - | - | 0 | - |
| - | - | 1302 | 513.2 | - | - | 0 | - |
| 7 | y | 1310 | 525.3 | 0.0001596 | 0.3038 | +1 | 6 |
| - | - | 2950 | 542.3 | - | - | 0 | - |
| 7 | y | 3.786E+04 | 543.3 | 0.0003343 | 0.6153 | +1 | 6 |
| - | - | 1.126E+04 | 544.3 | - | - | 0 | - |
| - | - | 1338 | 545.3 | - | - | 0 | - |
| - | - | 583.2 | 547.1 | - | - | 0 | - |
| - | - | 733.9 | 548.3 | - | - | 0 | - |
| - | - | 689.8 | 551.4 | - | - | 0 | - |
| - | - | 1031 | 573.8 | - | - | 0 | - |
| - | - | 696.7 | 574.8 | - | - | 0 | - |
| 5 | c | 1432 | 576.3 | 5.637E-06 | 0.009782 | +1 | 5 |
| - | - | 2333 | 592.3 | - | - | 0 | - |
| 5 | c | 1.25E+04 | 593.3 | 0.0006035 | 1.017 | +1 | 5 |
| - | - | 2912 | 594.3 | - | - | 0 | - |
| - | - | 631.3 | 595.3 | - | - | 0 | - |
| - | - | 924.8 | 598.3 | - | - | 0 | - |
| - | - | 622.7 | 624.3 | - | - | 0 | - |
| - | - | 3530 | 626.3 | - | - | 0 | - |
| - | - | 805 | 627.3 | - | - | 0 | - |
| - | - | 816.2 | 637.3 | - | - | 0 | - |
| - | - | 747.6 | 641.9 | - | - | 0 | - |
| 6 | z | 683.1 | 672.3 | 0.003285 | 4.885 | +1 | 7 |
| - | - | 934.6 | 673.4 | - | - | 0 | - |
| - | - | 4900 | 688.3 | - | - | 0 | - |
| - | - | 1365 | 689.3 | - | - | 0 | - |
| 6 | z | 1259 | 690.4 | 0.005205 | 7.539 | +1 | 7 |
| - | - | 1509 | 691.4 | - | - | 0 | - |
| - | - | 575.9 | 694.3 | - | - | 0 | - |
| - | - | 1.073E+04 | 705.4 | - | - | 0 | - |
| 6 | y | 1.133E+04 | 706.4 | 0.0022 | 3.115 | +1 | 7 |
| - | - | 4112 | 707.4 | - | - | 0 | - |
| - | - | 1390 | 711.4 | - | - | 0 | - |
| - | - | 1307 | 712.4 | - | - | 0 | - |
| - | - | 3408 | 713.4 | - | - | 0 | - |
| - | - | 1113 | 714.4 | - | - | 0 | - |
| - | - | 1166 | 738.3 | - | - | 0 | - |
| 6 | c | 8056 | 739.4 | 0.0005788 | 0.7828 | +1 | 6 |
| - | - | 3840 | 740.4 | - | - | 0 | - |
| - | - | 768.2 | 741.4 | - | - | 0 | - |
| - | - | 917.7 | 752.4 | - | - | 0 | - |
| - | - | 2864 | 809.4 | - | - | 0 | - |
| - | - | 1135 | 810.4 | - | - | 0 | - |
| 5 | z | 1039 | 819.4 | 0.00439 | 5.357 | +1 | 8 |
| - | - | 882.1 | 820.4 | - | - | 0 | - |
| - | - | 597.5 | 820.7 | - | - | 0 | - |
| 7 | c | 786 | 836.4 | 0.01092 | 13.06 | +1 | 7 |
| 5 | z | 8237 | 837.4 | 0.001109 | 1.324 | +1 | 8 |
| - | - | 5434 | 838.4 | - | - | 0 | - |
| - | - | 1522 | 839.4 | - | - | 0 | - |
| - | - | 820 | 851.4 | - | - | 0 | - |
| - | - | 3730 | 852.4 | - | - | 0 | - |
| 7 | c | 5299 | 853.4 | 0.002133 | 2.499 | +1 | 7 |
| - | - | 2376 | 854.4 | - | - | 0 | - |
| - | - | 753.5 | 855.4 | - | - | 0 | - |
| - | - | 726.8 | 865.5 | - | - | 0 | - |
| - | - | 1984 | 866.5 | - | - | 0 | - |
| - | - | 1101 | 867.5 | - | - | 0 | - |
| 8 | c | 1607 | 893.4 | 0.000511 | 0.572 | +1 | 8 |
| - | - | 1504 | 908.5 | - | - | 0 | - |
| - | - | 1.294E+04 | 909.5 | - | - | 0 | - |
| 8 | c | 3.389E+04 | 910.5 | 0.001624 | 1.784 | +1 | 8 |
| - | - | 1.802E+04 | 911.5 | - | - | 0 | - |
| - | - | 4139 | 912.5 | - | - | 0 | - |
| 4 | z | 680.7 | 934.4 | 0.003265 | 3.494 | +1 | 9 |
| - | - | 767.8 | 935.4 | - | - | 0 | - |
| - | - | 1439 | 938.4 | - | - | 0 | - |
| - | - | 1091 | 948.5 | - | - | 0 | - |
| 4 | y | 1488 | 950.5 | 0.002973 | 3.128 | +1 | 9 |
| - | - | 1004 | 951.5 | - | - | 0 | - |
| 4 | z | 3673 | 952.5 | 0.003381 | 3.55 | +1 | 9 |
| - | - | 2734 | 953.5 | - | - | 0 | - |
| - | - | 609.5 | 954.5 | - | - | 0 | - |
| - | - | 880.9 | 963.5 | - | - | 0 | - |
| 9 | c | 6388 | 964.5 | 0.00108 | 1.12 | +1 | 9 |
| - | - | 2444 | 965.5 | - | - | 0 | - |
| - | - | 996.5 | 966.5 | - | - | 0 | - |
| 9 | c | 1.823E+04 | 981.5 | 0.0004687 | 0.4776 | +1 | 9 |
| - | - | 1.071E+04 | 982.5 | - | - | 0 | - |
| - | - | 2395 | 983.5 | - | - | 0 | - |
| - | - | 2211 | 1037 | - | - | 0 | - |
| - | - | 903 | 1038 | - | - | 0 | - |
| - | - | 1845 | 1039 | - | - | 0 | - |
| 3 | z | 1548 | 1039 | 0.0003271 | 0.3147 | +1 | 10 |
| - | - | 731.6 | 1040 | - | - | 0 | - |
| - | - | 1696 | 1040 | - | - | 0 | - |
| - | - | 4334 | 1048 | - | - | 0 | - |
| - | - | 3700 | 1049 | - | - | 0 | - |
| - | - | 1496 | 1050 | - | - | 0 | - |
| - | - | 734.6 | 1050 | - | - | 0 | - |
| - | - | 1004 | 1054 | - | - | 0 | - |
| 3 | y | 3026 | 1056 | 0.003304 | 3.13 | +1 | 10 |
| - | - | 1526 | 1057 | - | - | 0 | - |
| 10 | c | 6398 | 1064 | 0.000269 | 0.2529 | +1 | 10 |
| - | - | 3548 | 1065 | - | - | 0 | - |
| - | - | 2209 | 1066 | - | - | 0 | - |
| - | - | 918.1 | 1067 | - | - | 0 | - |
| 10 | c | 2.821E+04 | 1081 | 0.001134 | 1.049 | +1 | 10 |
| - | - | 1.844E+04 | 1082 | - | - | 0 | - |
| - | - | 1.778E+04 | 1083 | - | - | 0 | - |
| - | - | 8979 | 1084 | - | - | 0 | - |
| - | - | 2599 | 1085 | - | - | 0 | - |
| - | - | 2768 | 1120 | - | - | 0 | - |
| - | - | 2258 | 1121 | - | - | 0 | - |
| 2 | z | 7295 | 1153 | 0.001139 | 0.9883 | +1 | 11 |
| - | - | 4800 | 1154 | - | - | 0 | - |
| - | - | 1741 | 1155 | - | - | 0 | - |
| 11 | c | 940.3 | 1165 | 0.009964 | 8.555 | +1 | 11 |
| 11 | c | 1.02E+04 | 1182 | 0.001327 | 1.123 | +1 | 11 |
| - | - | 7726 | 1183 | - | - | 0 | - |
| - | - | 2819 | 1184 | - | - | 0 | - |
| - | - | 7289 | 1223 | - | - | 0 | - |
| - | - | 5480 | 1224 | - | - | 0 | - |
| - | - | 2613 | 1225 | - | - | 0 | - |
| - | - | 4705 | 1239 | - | - | 0 | - |
| - | - | 3449 | 1240 | - | - | 0 | - |
| - | - | 873.1 | 1241 | - | - | 0 | - |
| - | - | 1596 | 1248 | - | - | 0 | - |
| - | - | 941.8 | 1249 | - | - | 0 | - |
| - | - | 767.7 | 1250 | - | - | 0 | - |
| - | - | 3915 | 1255 | - | - | 0 | - |
| - | - | 2400 | 1256 | - | - | 0 | - |
| - | - | 659.2 | 1257 | - | - | 0 | - |
| - | - | 838.7 | 1264 | - | - | 0 | - |
| - | - | 2138 | 1265 | - | - | 0 | - |
| - | - | 1.007E+04 | 1266 | - | - | 0 | - |
| - | - | 7319 | 1267 | - | - | 0 | - |
| - | - | 3374 | 1268 | - | - | 0 | - |
| - | - | 765.8 | 1280 | - | - | 0 | - |
| - | - | 1954 | 1281 | - | - | 0 | - |
| - | - | 1.567E+04 | 1282 | - | - | 0 | - |
| - | - | 6.358E+04 | 1283 | - | - | 0 | - |
| - | - | 4.803E+04 | 1284 | - | - | 0 | - |
| - | - | 1.758E+04 | 1285 | - | - | 0 | - |
| - | - | 2722 | 1286 | - | - | 0 | - |
| - | - | 651.5 | 1920 | - | - | 0 | - |
| - | - | 799.8 | 1945 | - | - | 0 | - |
| - | - | 691.8 | 1995 | - | - | 0 | - |
| - | - | 622.2 | 2357 | - | - | 0 | - |

m/z Charge Intensity FragmentType MassShift Position
127.74073028564453 0 422.02036
148.94760131835938 0 704.9037
159.35958862304688 0 486.2542
167.06187438964844 0 430.94815
170.5543212890625 0 490.97202
173.1282958984375 0 573.8297
173.45355224609375 0 707.91864
190.27842712402344 0 487.05157
191.5911407470703 0 507.2774
199.1802520751953 0 7559.031
201.1233673095703 0 2306.9062 y Water loss 10
219.12240600585938 0 794.9222
219.1337890625 0 15123.965 y 10
220.13743591308594 0 1431.5582
226.11843872070312 0 655.4851
227.1754150390625 0 5207.946 c Ammonia loss 1
239.09539794921875 0 593.6645
279.8812255859375 0 516.61194
297.19183349609375 0 619.68744
314.2041931152344 0 649.71027 c Ammonia loss 2
318.2024841308594 0 933.55524 y 9
325.1868896484375 0 3366.3926
335.17205810546875 0 597.44037
350.1337585449219 0 1236.762
355.0698547363281 0 856.52795
362.0887145996094 0 502.6145
392.8443298339844 0 509.4672
408.22467041015625 0 1526.604
424.2230224609375 0 628.1158
426.2347106933594 0 2071.404
429.2345275878906 0 1002.86346 c Ammonia loss 3
433.07476806640625 0 557.96484
446.262451171875 0 696.32825 c 3
447.224609375 0 717.2761 c Ammonia loss 7
460.2576904296875 0 556.2129
463.22052001953125 0 899.09906
482.743896484375 0 1160.0762 c Ammonia loss 8
483.2442932128906 0 735.77185
513.1978759765625 0 1302.3833
525.3032836914062 0 1309.8713 y Water loss 6
542.3045043945312 0 2950.42
543.3133544921875 0 37858.438 y 6
544.3164672851562 0 11261.79
545.31982421875 0 1337.563
547.1350708007812 0 583.17096
548.3107299804688 0 733.9488
551.3565063476562 0 689.76324
573.79541015625 0 1031.4482
574.7923583984375 0 696.7132
576.3027954101562 0 1431.81 c Ammonia loss 4
592.3204345703125 0 2333.1782
593.3287353515625 0 12503.425 c 4
594.33154296875 0 2911.6372
595.3357543945312 0 631.31665
598.2911376953125 0 924.75903
624.3043212890625 0 622.7196
626.282470703125 0 3530.3853
627.284423828125 0 804.9957
637.334716796875 0 816.22614
641.8543701171875 0 747.63934
672.3510131835938 0 683.1224 z Water loss 5
673.3513793945312 0 934.63806
688.3427734375 0 4900.4717
689.3455810546875 0 1365.3053
690.3530883789062 0 1258.7859 z 5
691.3649291992188 0 1508.7114
694.3428955078125 0 575.9469
705.3683471679688 0 10733.676
706.3748168945312 0 11328.854 y 5
707.378173828125 0 4111.73
711.3701171875 0 1390.4707
712.37841796875 0 1306.6293
713.3843383789062 0 3408.31
714.388916015625 0 1113.3322
738.3030395507812 0 1165.6971
739.3655395507812 0 8056.135 c Ammonia loss 5
740.3687744140625 0 3840.0034
741.3665771484375 0 768.23193
752.3579711914062 0 917.65985
809.4324340820312 0 2863.5457
810.4397583007812 0 1135.0123
819.4205322265625 0 1039.1195 z Water loss 4
820.4215698242188 0 882.0948
820.7330322265625 0 597.5292
836.407958984375 0 785.98865 c Ammonia loss 6
837.4255981445312 0 8237.415 z 4
838.4310913085938 0 5434.2554
839.434326171875 0 1522.4084
851.3889770507812 0 819.96924
852.4356079101562 0 3730.4485
853.4432983398438 0 5299.178 c 6
854.4471435546875 0 2376.2446
855.447021484375 0 753.48254
865.4685668945312 0 726.8395
866.4536743164062 0 1983.9867
867.4534912109375 0 1100.626
893.4408569335938 0 1606.8871 c Ammonia loss 7
908.4605102539062 0 1503.8722
909.4586181640625 0 12941.92
910.4652709960938 0 33891.754 c 7
911.4683227539062 0 18016.309
912.4705810546875 0 4139.313
934.4463500976562 0 680.69495 z Water loss 3
935.444091796875 0 767.81323
938.4251708984375 0 1438.5852
948.4639282226562 0 1090.5002
950.4647827148438 0 1488.1783 y Water loss 3
951.4623413085938 0 1003.5195
952.45703125 0 3673.1672 z 3
953.4592895507812 0 2733.963
954.4598999023438 0 609.5416
963.4740600585938 0 880.86316
964.4763793945312 0 6387.993 c Ammonia loss 8
965.4801025390625 0 2444.207
966.48193359375 0 996.52014
981.5035400390625 0 18233.086 c 8
982.5064697265625 0 10708.65
983.5074462890625 0 2394.833
1036.55712890625 0 2211.2305
1037.5692138671875 0 902.9588
1038.570068359375 0 1844.7949
1039.4853515625 0 1547.6589 z 2
1039.586669921875 0 731.57336
1040.49169921875 0 1695.5845
1047.5267333984375 0 4333.838
1048.5291748046875 0 3699.687
1049.5316162109375 0 1496.444
1049.6832275390625 0 734.6202
1054.4964599609375 0 1004.06995
1055.5010986328125 0 3026.3967 y 2
1056.5098876953125 0 1526.1486
1063.546142578125 0 6398.32 c Ammonia loss 9
1064.5487060546875 0 3548.334
1065.55810546875 0 2208.858
1066.560302734375 0 918.1428
1080.5712890625 0 28209.195 c 9
1081.574951171875 0 18438.508
1082.5838623046875 0 17779.666
1083.5899658203125 0 8979.309
1084.590087890625 0 2598.6016
1119.5955810546875 0 2768.3267
1120.595947265625 0 2257.594
1152.568603515625 0 7295.0903 z 1
1153.5728759765625 0 4800.3755
1154.57861328125 0 1741.37
1164.603515625 0 940.27527 c Ammonia loss 10
1181.6187744140625 0 10200.666 c 10
1182.6220703125 0 7726.2275
1183.625244140625 0 2818.942
1222.6551513671875 0 7289.316
1223.6551513671875 0 5480.3784
1224.659423828125 0 2612.991
1238.687255859375 0 4704.7036
1239.69091796875 0 3449.3206
1240.6890869140625 0 873.1485
1247.638916015625 0 1596.2322
1248.6322021484375 0 941.7946
1249.6494140625 0 767.74744
1254.6846923828125 0 3914.5874
1255.6875 0 2399.7593
1256.6878662109375 0 659.1803
1263.641845703125 0 838.73816
1264.6568603515625 0 2137.8777
1265.6536865234375 0 10071.137
1266.655517578125 0 7319.406
1267.65576171875 0 3373.652
1279.646484375 0 765.81744
1280.644775390625 0 1954.0951
1281.669189453125 0 15672.688
1282.6773681640625 0 63581.95
1283.680908203125 0 48029.51
1284.6844482421875 0 17577.37
1285.681396484375 0 2722.407
1920.0283203125 0 651.46564
1944.91015625 0 799.8234
1994.9542236328125 0 691.76135
2357.189697265625 0 622.1761

Spectrum Details

|  |  |
| --- | --- |
| Matched peaks? Matched peaks The total absolute number of peaks matched. Additionally in brackets the total fraction of peaks matched and the total number of peaks is shown. | 35 (19.77% of 177) |
| FDR? FDR The false discovery rate estimated for this peptide. It is calculated by matching all theoretical fragments with a non-integer shift with the raw peaks for this spectrum. This is done with 40 different shifts. The resulting percentage is the average number of annotated peaks over the number of annotated peaks with the correct spectrum. | 0.82% |
| Satellite FDR? Satellite FDR See the FDR for details on its calculation. This satellite ion specific FDR only contains the satellite ions (d/w) for I/L/J positions. | - |
| PSM Score? PSM Score The PSM Score as given by Hecklib to this annotated spectrum. It is shown with three significant figures. | 459 |

#### Spectrum 11086? Spectrum 11086 The raw spectrum of this peptide as annotated by Hecklib. The fragments are coloured according to ion type (see legend). Any peaks with a star '\*' as text can be hovered over to see the full details, first the ion type second the mass shift type. By hovering over the amino acids in the peptide or ions in the legend the corresponding peaks are highlighted. By toggling the 'Unassigned' label you can turn the background (unassigned) peaks on or off in the plot. By updating the slider in the Ion legend you can update the spectrum to only show the top X% of the peaks with labels. The top X% means any peak that is within X% of the highest intensity. By dragging in the spectrum you can zoom in to a specific part of the spectrum and use 'Zoom Out' to get back to the original zoom level. The annotation of the spectrum is based on the given sequence in the peptides file and is done with different software so inconsistencies are likely. The peaks are annotated based on the given sequence, with 20 ppm tolerance.

Copy Data

##### Spectrum 11086 (TSV)

###### Preview

```
Loading example...
```

*Click on the button to copy the data to your clipboard.*

Mz MinMz MaxIntensity Max

WidthHeightPeptide font sizePeptide stroke widthSpectrum font sizeSpectrum stroke widthCompact peptide

Ion legend

wxyz

abcd

OtherUnassignedIon^Charge^_Position_Show for top:%

JJSDFYPGAVTV

02.72e+45.45e+48.17e+41.09e+5

Zoom Out

y^+1^_2_y^+1^_2_c^+1^_2_y^+1^_3_c^+1^_4_c^+1^_4_c^+2^_9_y^+1^_6_y^+1^_6_c^+1^_5_c^+1^_5_z^+1^_7_z^+1^_7_y^+1^_7_c^+1^_6_z^+1^_8_z^+1^_8_c^+1^_7_c^+1^_8_c^+1^_8_c^+1^_8_z^+1^_9_y^+1^_9_z^+1^_9_c^+1^_9_y^+1^_9_c^+1^_9_z^+1^_10_y^+1^_10_c^+1^_10_c^+1^_10_z^+1^_11_c^+1^_11_c^+1^_11_

0742148322252966

Fragment Matches Table

Show background peaks

| Position | Ion type | Intensity | mz Theoretical | mz Error (Th) | mz Error (ppm) | Charge | Series Number |
| --- | --- | --- | --- | --- | --- | --- | --- |
| - | - | 387.1 | 122.4 | - | - | 0 | - |
| - | - | 313.4 | 126.1 | - | - | 0 | - |
| - | - | 382.8 | 132.3 | - | - | 0 | - |
| - | - | 404.7 | 135.7 | - | - | 0 | - |
| - | - | 458.6 | 146.3 | - | - | 0 | - |
| - | - | 475.7 | 147 | - | - | 0 | - |
| - | - | 425.7 | 159.1 | - | - | 0 | - |
| - | - | 463 | 163.6 | - | - | 0 | - |
| - | - | 432.6 | 165.8 | - | - | 0 | - |
| - | - | 1725 | 173.1 | - | - | 0 | - |
| - | - | 2605 | 173.5 | - | - | 0 | - |
| - | - | 757 | 177 | - | - | 0 | - |
| - | - | 627 | 183.1 | - | - | 0 | - |
| - | - | 437.2 | 195 | - | - | 0 | - |
| - | - | 1.017E+04 | 199.2 | - | - | 0 | - |
| - | - | 1441 | 200.2 | - | - | 0 | - |
| 11 | y | 5607 | 201.1 | 0.0001358 | 0.6752 | +1 | 2 |
| 11 | y | 2.867E+04 | 219.1 | 0.000237 | 1.082 | +1 | 2 |
| - | - | 2855 | 220.1 | - | - | 0 | - |
| - | - | 517.7 | 224.9 | - | - | 0 | - |
| - | - | 1859 | 226.1 | - | - | 0 | - |
| 2 | c | 7205 | 227.2 | 0.0002395 | 1.054 | +1 | 2 |
| - | - | 832.7 | 228.2 | - | - | 0 | - |
| - | - | 509.1 | 237.6 | - | - | 0 | - |
| - | - | 543.8 | 238.8 | - | - | 0 | - |
| - | - | 1660 | 297.2 | - | - | 0 | - |
| 10 | y | 1706 | 318.2 | 1.59E-05 | 0.04998 | +1 | 3 |
| - | - | 4708 | 325.2 | - | - | 0 | - |
| - | - | 1580 | 342.2 | - | - | 0 | - |
| - | - | 646.2 | 348.2 | - | - | 0 | - |
| - | - | 2810 | 350.1 | - | - | 0 | - |
| - | - | 1462 | 355.1 | - | - | 0 | - |
| - | - | 579.2 | 388.8 | - | - | 0 | - |
| - | - | 611.9 | 390.2 | - | - | 0 | - |
| - | - | 2170 | 408.2 | - | - | 0 | - |
| - | - | 1261 | 424.2 | - | - | 0 | - |
| - | - | 3421 | 426.2 | - | - | 0 | - |
| - | - | 1030 | 427.2 | - | - | 0 | - |
| 4 | c | 1176 | 429.2 | 0.0001212 | 0.2824 | +1 | 4 |
| - | - | 1328 | 438.3 | - | - | 0 | - |
| 4 | c | 800 | 446.3 | 0.0003659 | 0.8199 | +1 | 4 |
| - | - | 1900 | 463.2 | - | - | 0 | - |
| 9 | c | 2263 | 482.7 | 0.0001551 | 0.3214 | +2 | 9 |
| - | - | 1701 | 483.2 | - | - | 0 | - |
| - | - | 762.7 | 485.2 | - | - | 0 | - |
| - | - | 1209 | 488.3 | - | - | 0 | - |
| - | - | 735.2 | 495.2 | - | - | 0 | - |
| - | - | 1442 | 513.2 | - | - | 0 | - |
| - | - | 598.9 | 514.2 | - | - | 0 | - |
| 7 | y | 1877 | 525.3 | 0.001136 | 2.163 | +1 | 6 |
| - | - | 4864 | 542.3 | - | - | 0 | - |
| 7 | y | 6.005E+04 | 543.3 | 0.000215 | 0.3957 | +1 | 6 |
| - | - | 1.731E+04 | 544.3 | - | - | 0 | - |
| - | - | 2657 | 545.3 | - | - | 0 | - |
| - | - | 1007 | 548.3 | - | - | 0 | - |
| - | - | 932.3 | 551.4 | - | - | 0 | - |
| 5 | c | 2345 | 576.3 | 0.0007991 | 1.387 | +1 | 5 |
| - | - | 649.7 | 577.3 | - | - | 0 | - |
| - | - | 1117 | 589.3 | - | - | 0 | - |
| - | - | 4079 | 592.3 | - | - | 0 | - |
| 5 | c | 1.683E+04 | 593.3 | 6.828E-06 | 0.01151 | +1 | 5 |
| - | - | 6398 | 594.3 | - | - | 0 | - |
| - | - | 760.1 | 595.3 | - | - | 0 | - |
| - | - | 1023 | 598.3 | - | - | 0 | - |
| - | - | 1013 | 608.3 | - | - | 0 | - |
| - | - | 6591 | 626.3 | - | - | 0 | - |
| - | - | 2542 | 627.3 | - | - | 0 | - |
| - | - | 2031 | 637.3 | - | - | 0 | - |
| - | - | 714.6 | 654.3 | - | - | 0 | - |
| 6 | z | 1611 | 672.3 | 0.0002554 | 0.3799 | +1 | 7 |
| - | - | 5688 | 688.3 | - | - | 0 | - |
| - | - | 2513 | 689.3 | - | - | 0 | - |
| 6 | z | 2501 | 690.4 | 0.001055 | 1.527 | +1 | 7 |
| - | - | 1993 | 691.4 | - | - | 0 | - |
| - | - | 648.3 | 694.4 | - | - | 0 | - |
| - | - | 1.598E+04 | 705.4 | - | - | 0 | - |
| 6 | y | 1.697E+04 | 706.4 | 0.002445 | 3.461 | +1 | 7 |
| - | - | 6020 | 707.4 | - | - | 0 | - |
| - | - | 1184 | 708.4 | - | - | 0 | - |
| - | - | 1831 | 711.4 | - | - | 0 | - |
| - | - | 2267 | 712.4 | - | - | 0 | - |
| - | - | 3856 | 713.4 | - | - | 0 | - |
| - | - | 1185 | 714.4 | - | - | 0 | - |
| - | - | 1408 | 721.4 | - | - | 0 | - |
| - | - | 616.2 | 722.4 | - | - | 0 | - |
| - | - | 1041 | 738.3 | - | - | 0 | - |
| 6 | c | 1.292E+04 | 739.4 | 9.263E-05 | 0.1253 | +1 | 6 |
| - | - | 5736 | 740.4 | - | - | 0 | - |
| - | - | 1215 | 741.4 | - | - | 0 | - |
| - | - | 808.1 | 752.4 | - | - | 0 | - |
| - | - | 719.3 | 801.4 | - | - | 0 | - |
| - | - | 3395 | 809.4 | - | - | 0 | - |
| - | - | 1866 | 810.4 | - | - | 0 | - |
| - | - | 579.8 | 811.4 | - | - | 0 | - |
| 5 | z | 1268 | 819.4 | 0.0006762 | 0.8252 | +1 | 8 |
| - | - | 1503 | 820.4 | - | - | 0 | - |
| - | - | 912.5 | 822.4 | - | - | 0 | - |
| 5 | z | 1.188E+04 | 837.4 | 0.00117 | 1.397 | +1 | 8 |
| - | - | 8513 | 838.4 | - | - | 0 | - |
| - | - | 2885 | 839.4 | - | - | 0 | - |
| - | - | 1142 | 851.4 | - | - | 0 | - |
| - | - | 5298 | 852.4 | - | - | 0 | - |
| 7 | c | 8546 | 853.4 | 0.00195 | 2.285 | +1 | 7 |
| - | - | 3910 | 854.4 | - | - | 0 | - |
| - | - | 897.3 | 855.5 | - | - | 0 | - |
| - | - | 3919 | 866.5 | - | - | 0 | - |
| - | - | 2128 | 867.5 | - | - | 0 | - |
| - | - | 703.4 | 868.5 | - | - | 0 | - |
| 8 | c | 752.1 | 892.5 | 0.0001516 | 0.1699 | +1 | 8 |
| 8 | c | 2024 | 893.4 | 0.004189 | 4.688 | +1 | 8 |
| - | - | 1149 | 894.4 | - | - | 0 | - |
| - | - | 857.7 | 900.5 | - | - | 0 | - |
| - | - | 701.7 | 901.5 | - | - | 0 | - |
| - | - | 1986 | 908.5 | - | - | 0 | - |
| - | - | 1.874E+04 | 909.5 | - | - | 0 | - |
| 8 | c | 5.234E+04 | 910.5 | 0.001441 | 1.583 | +1 | 8 |
| - | - | 2.901E+04 | 911.5 | - | - | 0 | - |
| - | - | 5609 | 912.5 | - | - | 0 | - |
| 4 | z | 2938 | 934.4 | 0.00064 | 0.6849 | +1 | 9 |
| - | - | 1532 | 935.4 | - | - | 0 | - |
| - | - | 738.1 | 936.5 | - | - | 0 | - |
| - | - | 2019 | 937.5 | - | - | 0 | - |
| - | - | 2417 | 938.4 | - | - | 0 | - |
| - | - | 835.6 | 938.5 | - | - | 0 | - |
| - | - | 805.8 | 946.5 | - | - | 0 | - |
| - | - | 1015 | 948.5 | - | - | 0 | - |
| - | - | 1237 | 949.5 | - | - | 0 | - |
| 4 | y | 1670 | 950.5 | 0.001936 | 2.036 | +1 | 9 |
| - | - | 1098 | 951.5 | - | - | 0 | - |
| 4 | z | 4647 | 952.5 | 0.001184 | 1.243 | +1 | 9 |
| - | - | 6019 | 953.5 | - | - | 0 | - |
| - | - | 1994 | 954.5 | - | - | 0 | - |
| 9 | c | 9124 | 964.5 | 0.000592 | 0.6138 | +1 | 9 |
| - | - | 5348 | 965.5 | - | - | 0 | - |
| - | - | 1636 | 966.5 | - | - | 0 | - |
| - | - | 785.8 | 967.5 | - | - | 0 | - |
| 4 | y | 822 | 968.5 | 0.00454 | 4.688 | +1 | 9 |
| 9 | c | 2.716E+04 | 981.5 | 0.0005298 | 0.5398 | +1 | 9 |
| - | - | 1.577E+04 | 982.5 | - | - | 0 | - |
| - | - | 3880 | 983.5 | - | - | 0 | - |
| - | - | 778.4 | 1004 | - | - | 0 | - |
| - | - | 3564 | 1037 | - | - | 0 | - |
| - | - | 930.2 | 1037 | - | - | 0 | - |
| - | - | 1179 | 1038 | - | - | 0 | - |
| - | - | 2563 | 1039 | - | - | 0 | - |
| 3 | z | 1425 | 1039 | 0.006919 | 6.656 | +1 | 10 |
| - | - | 1870 | 1040 | - | - | 0 | - |
| - | - | 2693 | 1040 | - | - | 0 | - |
| - | - | 1207 | 1041 | - | - | 0 | - |
| - | - | 1497 | 1046 | - | - | 0 | - |
| - | - | 756.5 | 1047 | - | - | 0 | - |
| - | - | 8499 | 1048 | - | - | 0 | - |
| - | - | 6463 | 1049 | - | - | 0 | - |
| - | - | 2505 | 1050 | - | - | 0 | - |
| - | - | 739.7 | 1054 | - | - | 0 | - |
| 3 | y | 5558 | 1056 | 0.0009847 | 0.933 | +1 | 10 |
| - | - | 2772 | 1057 | - | - | 0 | - |
| - | - | 855.7 | 1058 | - | - | 0 | - |
| 10 | c | 9491 | 1064 | 0.0003414 | 0.321 | +1 | 10 |
| - | - | 6463 | 1065 | - | - | 0 | - |
| - | - | 4106 | 1066 | - | - | 0 | - |
| - | - | 1453 | 1067 | - | - | 0 | - |
| 10 | c | 4.402E+04 | 1081 | 0.001012 | 0.9361 | +1 | 10 |
| - | - | 2.979E+04 | 1082 | - | - | 0 | - |
| - | - | 2.643E+04 | 1083 | - | - | 0 | - |
| - | - | 1.354E+04 | 1084 | - | - | 0 | - |
| - | - | 3093 | 1085 | - | - | 0 | - |
| - | - | 5480 | 1120 | - | - | 0 | - |
| - | - | 3599 | 1121 | - | - | 0 | - |
| - | - | 1690 | 1122 | - | - | 0 | - |
| - | - | 886.7 | 1138 | - | - | 0 | - |
| - | - | 1111 | 1139 | - | - | 0 | - |
| - | - | 694.8 | 1147 | - | - | 0 | - |
| - | - | 2355 | 1149 | - | - | 0 | - |
| - | - | 668.8 | 1150 | - | - | 0 | - |
| 2 | z | 1.184E+04 | 1153 | 0.0007729 | 0.6706 | +1 | 11 |
| - | - | 7789 | 1154 | - | - | 0 | - |
| - | - | 3262 | 1155 | - | - | 0 | - |
| 11 | c | 1511 | 1165 | 0.004075 | 3.499 | +1 | 11 |
| - | - | 1394 | 1166 | - | - | 0 | - |
| 11 | c | 1.714E+04 | 1182 | 0.001815 | 1.536 | +1 | 11 |
| - | - | 1.271E+04 | 1183 | - | - | 0 | - |
| - | - | 3842 | 1184 | - | - | 0 | - |
| - | - | 774.3 | 1185 | - | - | 0 | - |
| - | - | 1.238E+04 | 1223 | - | - | 0 | - |
| - | - | 8548 | 1224 | - | - | 0 | - |
| - | - | 2919 | 1225 | - | - | 0 | - |
| - | - | 1309 | 1237 | - | - | 0 | - |
| - | - | 875.4 | 1238 | - | - | 0 | - |
| - | - | 6364 | 1239 | - | - | 0 | - |
| - | - | 3972 | 1240 | - | - | 0 | - |
| - | - | 1482 | 1241 | - | - | 0 | - |
| - | - | 2985 | 1248 | - | - | 0 | - |
| - | - | 1515 | 1249 | - | - | 0 | - |
| - | - | 5848 | 1255 | - | - | 0 | - |
| - | - | 4215 | 1256 | - | - | 0 | - |
| - | - | 1165 | 1257 | - | - | 0 | - |
| - | - | 2399 | 1265 | - | - | 0 | - |
| - | - | 1.424E+04 | 1266 | - | - | 0 | - |
| - | - | 1.499E+04 | 1267 | - | - | 0 | - |
| - | - | 3899 | 1268 | - | - | 0 | - |
| - | - | 1140 | 1280 | - | - | 0 | - |
| - | - | 2778 | 1281 | - | - | 0 | - |
| - | - | 2.327E+04 | 1282 | - | - | 0 | - |
| - | - | 1.078E+05 | 1283 | - | - | 0 | - |
| - | - | 7.728E+04 | 1284 | - | - | 0 | - |
| - | - | 2.886E+04 | 1285 | - | - | 0 | - |
| - | - | 4634 | 1286 | - | - | 0 | - |
| - | - | 672.5 | 1652 | - | - | 0 | - |
| - | - | 725.1 | 1798 | - | - | 0 | - |
| - | - | 783.5 | 1924 | - | - | 0 | - |
| - | - | 805.7 | 2937 | - | - | 0 | - |

m/z Charge Intensity FragmentType MassShift Position
122.40519714355469 0 387.14212
126.09398651123047 0 313.4181
132.33685302734375 0 382.7521
135.71783447265625 0 404.67142
146.2993927001953 0 458.6416
147.03688049316406 0 475.7247
159.08779907226562 0 425.69016
163.63409423828125 0 463.00128
165.83755493164062 0 432.6318
173.12864685058594 0 1725.2637
173.4514923095703 0 2604.9028
177.0399169921875 0 756.9868
183.11239624023438 0 627.0463
194.9919891357422 0 437.1624
199.1807098388672 0 10167.246
200.18392944335938 0 1440.6641
201.12350463867188 0 5607.158 y Water loss 10
219.13417053222656 0 28667.355 y 10
220.13754272460938 0 2855.1738
224.8908233642578 0 517.6752
226.11874389648438 0 1858.8488
227.17564392089844 0 7204.756 c Ammonia loss 1
228.1791534423828 0 832.69403
237.60203552246094 0 509.109
238.83172607421875 0 543.75336
297.1923828125 0 1660.1069
318.20233154296875 0 1706.2104 y 9
325.1875 0 4708.2925
342.2139892578125 0 1580.3992
348.2262268066406 0 646.1838
350.1344909667969 0 2810.0388
355.069580078125 0 1462.3722
388.7999267578125 0 579.2331
390.2401428222656 0 611.94025
408.2245788574219 0 2169.6821
424.22418212890625 0 1261.2732
426.2354431152344 0 3420.934
427.2375793457031 0 1029.8287
429.2344970703125 0 1175.8419 c Ammonia loss 3
438.2720947265625 0 1328.4044
446.26055908203125 0 799.9506 c 3
463.2196350097656 0 1899.5796
482.7425231933594 0 2262.6099 c Ammonia loss 8
483.2440185546875 0 1700.6467
485.2079162597656 0 762.6588
488.2509460449219 0 1209.3289
495.1915588378906 0 735.1973
513.1978759765625 0 1442.4009
514.2022705078125 0 598.8855
525.3042602539062 0 1876.667 y Water loss 6
542.3062133789062 0 4863.7354
543.3139038085938 0 60047.24 y 6
544.3168334960938 0 17311.46
545.3192749023438 0 2656.9705
548.3072509765625 0 1006.9748
551.3570556640625 0 932.3072
576.3035888671875 0 2345.1323 c Ammonia loss 4
577.308349609375 0 649.6929
589.2999877929688 0 1116.7659
592.3203735351562 0 4078.7615
593.329345703125 0 16825.238 c 4
594.3323974609375 0 6398.135
595.3380126953125 0 760.08685
598.283447265625 0 1023.13434
608.2702026367188 0 1013.1751
626.2822265625 0 6591.1924
627.2847290039062 0 2542.1177
637.3341064453125 0 2030.5496
654.3464965820312 0 714.5586
672.3474731445312 0 1611.0465 z Water loss 5
688.3433227539062 0 5688.403
689.3453369140625 0 2513.3755
690.3572387695312 0 2501.2017 z 5
691.36328125 0 1992.614
694.3510131835938 0 648.31177
705.3690185546875 0 15983.223
706.3745727539062 0 16974.074 y 5
707.3790893554688 0 6020.244
708.3820190429688 0 1184.3197
711.3729248046875 0 1830.7405
712.3773193359375 0 2267.4001
713.3858032226562 0 3856.4695
714.388671875 0 1185.1129
721.357177734375 0 1407.7839
722.3592529296875 0 616.2251
738.3040161132812 0 1041.2361
739.3662109375 0 12916.35 c Ammonia loss 5
740.3692016601562 0 5736.109
741.3709106445312 0 1215.0863
752.3601684570312 0 808.0773
801.41650390625 0 719.3208
809.4308471679688 0 3395.178
810.4351196289062 0 1865.8677
811.4346313476562 0 579.83594
819.4154663085938 0 1267.8599 z Water loss 4
820.4195556640625 0 1503.2921
822.3626098632812 0 912.4534
837.425537109375 0 11877.155 z 4
838.4315795898438 0 8513.362
839.4342041015625 0 2885.1348
851.391357421875 0 1142.4336
852.4380493164062 0 5298.482
853.4434814453125 0 8545.637 c 6
854.4453125 0 3909.661
855.4524536132812 0 897.29565
866.4551391601562 0 3918.854
867.4586181640625 0 2128.4717
868.468017578125 0 703.40594
892.4564819335938 0 752.1297 c Water loss 7
893.4361572265625 0 2024.4866 c Ammonia loss 7
894.4439697265625 0 1149.0016
900.4885864257812 0 857.68207
901.48291015625 0 701.69336
908.4613647460938 0 1985.8125
909.4593505859375 0 18737.926
910.4654541015625 0 52339.246 c 7
911.4688720703125 0 29013.46
912.4716796875 0 5608.837
934.4437255859375 0 2937.9348 z Water loss 3
935.4456176757812 0 1531.8693
936.46728515625 0 738.13385
937.4867553710938 0 2019.2744
938.4246826171875 0 2417.125
938.513916015625 0 835.60425
946.4714965820312 0 805.82294
948.4586181640625 0 1014.5216
949.4640502929688 0 1236.9656
950.4637451171875 0 1670.2 y Water loss 3
951.4640502929688 0 1098.4998
952.454833984375 0 4647.496 z 3
953.46044921875 0 6018.985
954.460693359375 0 1993.7769
964.4768676757812 0 9123.611 c Ammonia loss 8
965.4798583984375 0 5348.2803
966.482177734375 0 1635.9938
967.4682006835938 0 785.81775
968.4678344726562 0 822.0423 y 3
981.5034790039062 0 27157.967 c 8
982.5067138671875 0 15766.983
983.5098876953125 0 3880.1523
1004.4722290039062 0 778.4148
1036.5572509765625 0 3564.4558
1037.46875 0 930.2092
1037.5767822265625 0 1179.2423
1038.5711669921875 0 2562.8486
1039.478759765625 0 1425.2012 z 2
1039.5784912109375 0 1870.0251
1040.491943359375 0 2692.5762
1041.4959716796875 0 1206.9335
1045.5301513671875 0 1497.0325
1046.525146484375 0 756.5098
1047.527587890625 0 8499.263
1048.53173828125 0 6463.244
1049.53466796875 0 2504.5837
1054.4970703125 0 739.69794
1055.50341796875 0 5557.6323 y 2
1056.5072021484375 0 2771.5168
1057.51025390625 0 855.7232
1063.5455322265625 0 9491.238 c Ammonia loss 9
1064.5474853515625 0 6463.3027
1065.5548095703125 0 4106.376
1066.55908203125 0 1453.202
1080.5714111328125 0 44023.09 c 9
1081.5748291015625 0 29793.344
1082.5841064453125 0 26425.902
1083.5894775390625 0 13541.166
1084.59423828125 0 3093.3745
1119.59619140625 0 5479.5723
1120.5985107421875 0 3598.6533
1121.5943603515625 0 1690.1545
1137.6104736328125 0 886.7114
1138.6158447265625 0 1111.1606
1146.579345703125 0 694.7792
1148.5753173828125 0 2354.942
1149.5740966796875 0 668.8271
1152.5689697265625 0 11837.756 z 1
1153.5721435546875 0 7788.5864
1154.573486328125 0 3262.2627
1164.5894775390625 0 1511.1235 c Ammonia loss 10
1165.6011962890625 0 1394.3165
1181.6182861328125 0 17141.93 c 10
1182.6217041015625 0 12714.69
1183.6234130859375 0 3842.4875
1184.6231689453125 0 774.30237
1222.6534423828125 0 12380.607
1223.6556396484375 0 8547.867
1224.6585693359375 0 2918.6577
1236.6728515625 0 1309.3265
1237.6859130859375 0 875.41016
1238.6866455078125 0 6363.9424
1239.6920166015625 0 3972.4468
1240.6834716796875 0 1481.8966
1247.641845703125 0 2985.3438
1248.635986328125 0 1515.3615
1254.6837158203125 0 5848.4478
1255.686767578125 0 4214.78
1256.69091796875 0 1165.3105
1264.663818359375 0 2399.0378
1265.653564453125 0 14244.431
1266.6580810546875 0 14985.705
1267.656005859375 0 3898.9917
1279.6446533203125 0 1140.4637
1280.655517578125 0 2777.794
1281.6693115234375 0 23266.428
1282.6768798828125 0 107829.336
1283.680419921875 0 77275
1284.68359375 0 28862.658
1285.6854248046875 0 4634.0723
1651.637451171875 0 672.5024
1798.2120361328125 0 725.1444
1923.9058837890625 0 783.5488
2937.023193359375 0 805.65344

Spectrum Details

|  |  |
| --- | --- |
| Matched peaks? Matched peaks The total absolute number of peaks matched. Additionally in brackets the total fraction of peaks matched and the total number of peaks is shown. | 34 (16.04% of 212) |
| FDR? FDR The false discovery rate estimated for this peptide. It is calculated by matching all theoretical fragments with a non-integer shift with the raw peaks for this spectrum. This is done with 40 different shifts. The resulting percentage is the average number of annotated peaks over the number of annotated peaks with the correct spectrum. | 0.98% |
| Satellite FDR? Satellite FDR See the FDR for details on its calculation. This satellite ion specific FDR only contains the satellite ions (d/w) for I/L/J positions. | ∞ |
| PSM Score? PSM Score The PSM Score as given by Hecklib to this annotated spectrum. It is shown with three significant figures. | 459 |

#### Spectrum 11137? Spectrum 11137 The raw spectrum of this peptide as annotated by Hecklib. The fragments are coloured according to ion type (see legend). Any peaks with a star '\*' as text can be hovered over to see the full details, first the ion type second the mass shift type. By hovering over the amino acids in the peptide or ions in the legend the corresponding peaks are highlighted. By toggling the 'Unassigned' label you can turn the background (unassigned) peaks on or off in the plot. By updating the slider in the Ion legend you can update the spectrum to only show the top X% of the peaks with labels. The top X% means any peak that is within X% of the highest intensity. By dragging in the spectrum you can zoom in to a specific part of the spectrum and use 'Zoom Out' to get back to the original zoom level. The annotation of the spectrum is based on the given sequence in the peptides file and is done with different software so inconsistencies are likely. The peaks are annotated based on the given sequence, with 20 ppm tolerance.

Copy Data

##### Spectrum 11137 (TSV)

###### Preview

```
Loading example...
```

*Click on the button to copy the data to your clipboard.*

Mz MinMz MaxIntensity Max

WidthHeightPeptide font sizePeptide stroke widthSpectrum font sizeSpectrum stroke widthCompact peptide

Ion legend

wxyz

abcd

OtherUnassignedIon^Charge^_Position_Show for top:%

JJSDFYPGAVTV

01.97e+43.93e+45.90e+47.87e+4

Zoom Out

y^+1^_2_y^+1^_2_c^+1^_2_c^+1^_3_y^+1^_3_c^+1^_4_c^+2^_9_y^+1^_6_y^+1^_6_c^+1^_5_c^+1^_5_z^+1^_7_z^+1^_7_y^+1^_7_c^+1^_6_z^+1^_8_z^+1^_8_c^+1^_7_c^+1^_8_c^+1^_8_z^+1^_9_y^+1^_9_z^+1^_9_c^+1^_9_y^+1^_9_c^+1^_9_z^+1^_10_z^+1^_10_y^+1^_10_c^+1^_10_c^+1^_10_y^+1^_11_z^+1^_11_c^+1^_11_

0860172125813441

Fragment Matches Table

Show background peaks

| Position | Ion type | Intensity | mz Theoretical | mz Error (Th) | mz Error (ppm) | Charge | Series Number |
| --- | --- | --- | --- | --- | --- | --- | --- |
| - | - | 384.2 | 128.5 | - | - | 0 | - |
| - | - | 428.6 | 133.9 | - | - | 0 | - |
| - | - | 1179 | 173.1 | - | - | 0 | - |
| - | - | 527 | 173.4 | - | - | 0 | - |
| - | - | 622.4 | 173.5 | - | - | 0 | - |
| - | - | 472 | 173.5 | - | - | 0 | - |
| - | - | 540.2 | 177 | - | - | 0 | - |
| - | - | 8538 | 199.2 | - | - | 0 | - |
| - | - | 1599 | 200.2 | - | - | 0 | - |
| 11 | y | 3513 | 201.1 | 1.534E-06 | 0.007626 | +1 | 2 |
| 11 | y | 2.322E+04 | 219.1 | 3.864E-05 | 0.1763 | +1 | 2 |
| - | - | 2296 | 220.1 | - | - | 0 | - |
| - | - | 1146 | 226.1 | - | - | 0 | - |
| 2 | c | 6111 | 227.2 | 5.041E-05 | 0.2219 | +1 | 2 |
| - | - | 1077 | 297.2 | - | - | 0 | - |
| 3 | c | 915.2 | 314.2 | 0.0004834 | 1.539 | +1 | 3 |
| 10 | y | 1260 | 318.2 | 7.565E-05 | 0.2377 | +1 | 3 |
| - | - | 4057 | 325.2 | - | - | 0 | - |
| - | - | 588.5 | 326 | - | - | 0 | - |
| - | - | 536.5 | 339.2 | - | - | 0 | - |
| - | - | 1033 | 342.2 | - | - | 0 | - |
| - | - | 1677 | 350.1 | - | - | 0 | - |
| - | - | 943.7 | 355.1 | - | - | 0 | - |
| - | - | 501.9 | 397.3 | - | - | 0 | - |
| - | - | 1856 | 408.2 | - | - | 0 | - |
| - | - | 600.6 | 409.2 | - | - | 0 | - |
| - | - | 776.6 | 411.2 | - | - | 0 | - |
| - | - | 985.8 | 424.2 | - | - | 0 | - |
| - | - | 654.8 | 426.2 | - | - | 0 | - |
| - | - | 2581 | 426.2 | - | - | 0 | - |
| - | - | 581.3 | 427.2 | - | - | 0 | - |
| 4 | c | 894.8 | 429.2 | 9.241E-05 | 0.2153 | +1 | 4 |
| - | - | 1794 | 463.2 | - | - | 0 | - |
| 9 | c | 1988 | 482.7 | 0.0005519 | 1.143 | +2 | 9 |
| - | - | 860.5 | 483.2 | - | - | 0 | - |
| - | - | 756.4 | 488.3 | - | - | 0 | - |
| - | - | 531.3 | 495.6 | - | - | 0 | - |
| - | - | 1716 | 513.2 | - | - | 0 | - |
| 7 | y | 2294 | 525.3 | 0.0002066 | 0.3934 | +1 | 6 |
| - | - | 4345 | 542.3 | - | - | 0 | - |
| 7 | y | 4.795E+04 | 543.3 | 0.0001512 | 0.2783 | +1 | 6 |
| - | - | 1.299E+04 | 544.3 | - | - | 0 | - |
| - | - | 1637 | 545.3 | - | - | 0 | - |
| - | - | 711.6 | 548.3 | - | - | 0 | - |
| - | - | 842.7 | 551.4 | - | - | 0 | - |
| 5 | c | 1803 | 576.3 | 0.0001277 | 0.2216 | +1 | 5 |
| - | - | 771.6 | 577.3 | - | - | 0 | - |
| - | - | 695 | 589.3 | - | - | 0 | - |
| - | - | 2778 | 592.3 | - | - | 0 | - |
| 5 | c | 1.393E+04 | 593.3 | 0.0005425 | 0.9143 | +1 | 5 |
| - | - | 4081 | 594.3 | - | - | 0 | - |
| - | - | 757.9 | 595.3 | - | - | 0 | - |
| - | - | 735.6 | 608.3 | - | - | 0 | - |
| - | - | 4951 | 626.3 | - | - | 0 | - |
| - | - | 1929 | 627.3 | - | - | 0 | - |
| - | - | 1706 | 637.3 | - | - | 0 | - |
| - | - | 676.8 | 638.3 | - | - | 0 | - |
| - | - | 580.5 | 641.8 | - | - | 0 | - |
| - | - | 851 | 654.3 | - | - | 0 | - |
| - | - | 652.4 | 666.3 | - | - | 0 | - |
| 6 | z | 1163 | 672.3 | 0.0005606 | 0.8338 | +1 | 7 |
| - | - | 5688 | 688.3 | - | - | 0 | - |
| - | - | 2842 | 689.3 | - | - | 0 | - |
| 6 | z | 1682 | 690.4 | 0.003374 | 4.887 | +1 | 7 |
| - | - | 1104 | 691.4 | - | - | 0 | - |
| - | - | 1.3E+04 | 705.4 | - | - | 0 | - |
| 6 | y | 1.446E+04 | 706.4 | 0.002017 | 2.856 | +1 | 7 |
| - | - | 4359 | 707.4 | - | - | 0 | - |
| - | - | 1591 | 711.4 | - | - | 0 | - |
| - | - | 2271 | 712.4 | - | - | 0 | - |
| - | - | 2618 | 713.4 | - | - | 0 | - |
| - | - | 1145 | 714.4 | - | - | 0 | - |
| - | - | 1006 | 721.4 | - | - | 0 | - |
| - | - | 1240 | 738.3 | - | - | 0 | - |
| 6 | c | 1.131E+04 | 739.4 | 0.0006398 | 0.8653 | +1 | 6 |
| - | - | 4196 | 740.4 | - | - | 0 | - |
| - | - | 1008 | 741.4 | - | - | 0 | - |
| - | - | 3813 | 809.4 | - | - | 0 | - |
| - | - | 1914 | 810.4 | - | - | 0 | - |
| - | - | 611.7 | 811.4 | - | - | 0 | - |
| 5 | z | 1493 | 819.4 | 0.0007983 | 0.9742 | +1 | 8 |
| - | - | 1433 | 820.4 | - | - | 0 | - |
| - | - | 749 | 822.4 | - | - | 0 | - |
| 5 | z | 1.046E+04 | 837.4 | 0.0007428 | 0.8871 | +1 | 8 |
| - | - | 7199 | 838.4 | - | - | 0 | - |
| - | - | 1979 | 839.4 | - | - | 0 | - |
| - | - | 826.8 | 851.4 | - | - | 0 | - |
| - | - | 4500 | 852.4 | - | - | 0 | - |
| 7 | c | 8291 | 853.4 | 0.002865 | 3.357 | +1 | 7 |
| - | - | 2033 | 854.4 | - | - | 0 | - |
| - | - | 631.6 | 855.4 | - | - | 0 | - |
| - | - | 862 | 865.5 | - | - | 0 | - |
| - | - | 2842 | 866.5 | - | - | 0 | - |
| - | - | 1194 | 867.5 | - | - | 0 | - |
| 8 | c | 1481 | 893.4 | 0.002236 | 2.502 | +1 | 8 |
| - | - | 866.1 | 900.5 | - | - | 0 | - |
| - | - | 1801 | 908.5 | - | - | 0 | - |
| - | - | 1.651E+04 | 909.5 | - | - | 0 | - |
| 8 | c | 4.217E+04 | 910.5 | 0.002234 | 2.454 | +1 | 8 |
| - | - | 2.024E+04 | 911.5 | - | - | 0 | - |
| - | - | 4845 | 912.5 | - | - | 0 | - |
| 4 | z | 1362 | 934.4 | 0.002595 | 2.777 | +1 | 9 |
| - | - | 671.3 | 935.4 | - | - | 0 | - |
| - | - | 1696 | 937.5 | - | - | 0 | - |
| - | - | 1873 | 938.4 | - | - | 0 | - |
| - | - | 778.6 | 939.4 | - | - | 0 | - |
| - | - | 698.1 | 946.5 | - | - | 0 | - |
| - | - | 1195 | 948.5 | - | - | 0 | - |
| - | - | 900.2 | 949.5 | - | - | 0 | - |
| 4 | y | 1160 | 950.5 | 0.0004707 | 0.4952 | +1 | 9 |
| - | - | 1253 | 951.5 | - | - | 0 | - |
| 4 | z | 4151 | 952.5 | 0.0008786 | 0.9224 | +1 | 9 |
| - | - | 3703 | 953.5 | - | - | 0 | - |
| - | - | 975.9 | 954.5 | - | - | 0 | - |
| 9 | c | 7424 | 964.5 | 0.001446 | 1.5 | +1 | 9 |
| - | - | 3866 | 965.5 | - | - | 0 | - |
| - | - | 1068 | 966.5 | - | - | 0 | - |
| - | - | 1162 | 967.5 | - | - | 0 | - |
| 4 | y | 1482 | 968.5 | 0.001564 | 1.615 | +1 | 9 |
| 9 | c | 2.422E+04 | 981.5 | 0.001262 | 1.286 | +1 | 9 |
| - | - | 1.12E+04 | 982.5 | - | - | 0 | - |
| - | - | 4251 | 983.5 | - | - | 0 | - |
| 3 | z | 820.7 | 1021 | 0.002974 | 2.912 | +1 | 10 |
| - | - | 594.9 | 1036 | - | - | 0 | - |
| - | - | 2904 | 1037 | - | - | 0 | - |
| - | - | 1103 | 1038 | - | - | 0 | - |
| - | - | 3176 | 1039 | - | - | 0 | - |
| 3 | z | 1686 | 1039 | 0.0046 | 4.425 | +1 | 10 |
| - | - | 1365 | 1040 | - | - | 0 | - |
| - | - | 1870 | 1040 | - | - | 0 | - |
| - | - | 824.8 | 1041 | - | - | 0 | - |
| - | - | 645.6 | 1046 | - | - | 0 | - |
| - | - | 5744 | 1048 | - | - | 0 | - |
| - | - | 5463 | 1049 | - | - | 0 | - |
| - | - | 1476 | 1050 | - | - | 0 | - |
| - | - | 808.7 | 1054 | - | - | 0 | - |
| 3 | y | 4483 | 1056 | 0.002205 | 2.089 | +1 | 10 |
| - | - | 2140 | 1057 | - | - | 0 | - |
| - | - | 818.2 | 1058 | - | - | 0 | - |
| 10 | c | 7530 | 1064 | 0.0003414 | 0.321 | +1 | 10 |
| - | - | 5551 | 1065 | - | - | 0 | - |
| - | - | 3564 | 1066 | - | - | 0 | - |
| - | - | 736.9 | 1067 | - | - | 0 | - |
| 10 | c | 3.692E+04 | 1081 | 0.0015 | 1.388 | +1 | 10 |
| - | - | 2.374E+04 | 1082 | - | - | 0 | - |
| - | - | 1.898E+04 | 1083 | - | - | 0 | - |
| - | - | 9713 | 1084 | - | - | 0 | - |
| - | - | 2383 | 1085 | - | - | 0 | - |
| - | - | 3974 | 1120 | - | - | 0 | - |
| - | - | 2975 | 1121 | - | - | 0 | - |
| - | - | 1246 | 1122 | - | - | 0 | - |
| - | - | 925.9 | 1139 | - | - | 0 | - |
| - | - | 742.6 | 1150 | - | - | 0 | - |
| 2 | y | 1050 | 1151 | 0.009665 | 8.4 | +1 | 11 |
| 2 | z | 7938 | 1153 | 0.001139 | 0.9883 | +1 | 11 |
| - | - | 6413 | 1154 | - | - | 0 | - |
| - | - | 1848 | 1155 | - | - | 0 | - |
| - | - | 1002 | 1166 | - | - | 0 | - |
| 11 | c | 1.521E+04 | 1182 | 0.002914 | 2.466 | +1 | 11 |
| - | - | 9957 | 1183 | - | - | 0 | - |
| - | - | 2998 | 1184 | - | - | 0 | - |
| - | - | 897 | 1211 | - | - | 0 | - |
| - | - | 9422 | 1223 | - | - | 0 | - |
| - | - | 6861 | 1224 | - | - | 0 | - |
| - | - | 3533 | 1225 | - | - | 0 | - |
| - | - | 6274 | 1239 | - | - | 0 | - |
| - | - | 4174 | 1240 | - | - | 0 | - |
| - | - | 1527 | 1241 | - | - | 0 | - |
| - | - | 1478 | 1248 | - | - | 0 | - |
| - | - | 1037 | 1249 | - | - | 0 | - |
| - | - | 766.4 | 1250 | - | - | 0 | - |
| - | - | 3156 | 1255 | - | - | 0 | - |
| - | - | 2664 | 1256 | - | - | 0 | - |
| - | - | 1017 | 1257 | - | - | 0 | - |
| - | - | 1010 | 1264 | - | - | 0 | - |
| - | - | 2468 | 1265 | - | - | 0 | - |
| - | - | 1.218E+04 | 1266 | - | - | 0 | - |
| - | - | 1.105E+04 | 1267 | - | - | 0 | - |
| - | - | 4112 | 1268 | - | - | 0 | - |
| - | - | 768.9 | 1269 | - | - | 0 | - |
| - | - | 960.8 | 1280 | - | - | 0 | - |
| - | - | 2609 | 1281 | - | - | 0 | - |
| - | - | 1.773E+04 | 1282 | - | - | 0 | - |
| - | - | 7.788E+04 | 1283 | - | - | 0 | - |
| - | - | 5.995E+04 | 1284 | - | - | 0 | - |
| - | - | 2.381E+04 | 1285 | - | - | 0 | - |
| - | - | 3644 | 1286 | - | - | 0 | - |
| - | - | 935.3 | 1922 | - | - | 0 | - |
| - | - | 723.8 | 3100 | - | - | 0 | - |
| - | - | 710.6 | 3407 | - | - | 0 | - |

m/z Charge Intensity FragmentType MassShift Position
128.5386505126953 0 384.15656
133.93157958984375 0 428.5564
173.1288299560547 0 1178.9983
173.43850708007812 0 527.02637
173.45285034179688 0 622.4326
173.45889282226562 0 471.96075
177.0395965576172 0 540.19855
199.18057250976562 0 8538.198
200.183837890625 0 1599.3674
201.1233673095703 0 3512.7134 y Water loss 10
219.13397216796875 0 23224.545 y 10
220.13702392578125 0 2296.164
226.1190185546875 0 1146.2421
227.17535400390625 0 6111.074 c Ammonia loss 1
297.19183349609375 0 1076.6891
314.2079162597656 0 915.1783 c Ammonia loss 2
318.2024230957031 0 1260.435 y 9
325.18701171875 0 4057.2073
326.0419006347656 0 588.4816
339.2023620605469 0 536.54425
342.2129821777344 0 1032.696
350.1343078613281 0 1677.428
355.0696105957031 0 943.6976
397.3211975097656 0 501.87494
408.2233581542969 0 1856.4044
409.2261657714844 0 600.6042
411.2242126464844 0 776.61066
424.22174072265625 0 985.8463
426.16717529296875 0 654.78357
426.234375 0 2581.1287
427.23724365234375 0 581.34973
429.2342834472656 0 894.8361 c Ammonia loss 3
463.2191162109375 0 1794.3115
482.742919921875 0 1988.1361 c Ammonia loss 8
483.24407958984375 0 860.49713
488.2500915527344 0 756.38574
495.609619140625 0 531.2924
513.1978759765625 0 1715.6527
525.3029174804688 0 2293.873 y Water loss 6
542.3048095703125 0 4344.525
543.3135375976562 0 47945.348 y 6
544.3162841796875 0 12993.313
545.3196411132812 0 1637.3169
548.3080444335938 0 711.63635
551.3577270507812 0 842.74585
576.3029174804688 0 1802.6877 c Ammonia loss 4
577.3080444335938 0 771.60297
589.2988891601562 0 694.9955
592.3204345703125 0 2778.3083
593.3287963867188 0 13933.177 c 4
594.3316650390625 0 4080.764
595.33447265625 0 757.8577
608.2752075195312 0 735.56976
626.2816772460938 0 4950.7656
627.2850952148438 0 1928.8546
637.3345336914062 0 1706.2546
638.3451538085938 0 676.8372
641.8469848632812 0 580.54425
654.3425903320312 0 850.9777
666.34716796875 0 652.36176
672.34716796875 0 1163.2769 z Water loss 5
688.3425903320312 0 5687.902
689.3450317382812 0 2842.4087
690.3549194335938 0 1681.5599 z 5
691.3682861328125 0 1104.046
705.36865234375 0 12995.532
706.375 0 14460.257 y 5
707.379150390625 0 4358.9775
711.3705444335938 0 1590.9651
712.3754272460938 0 2270.6033
713.3826904296875 0 2618.0796
714.3892822265625 0 1145.3512
721.3541870117188 0 1005.63196
738.3052978515625 0 1239.8408
739.365478515625 0 11314.457 c Ammonia loss 5
740.3685913085938 0 4195.925
741.3742065429688 0 1007.57166
809.430908203125 0 3813.1199
810.4367065429688 0 1914.1643
811.4360961914062 0 611.7486
819.4153442382812 0 1493.1775 z Water loss 4
820.4160766601562 0 1433.4265
822.3638916015625 0 749.0311
837.4259643554688 0 10462.772 z 4
838.4307861328125 0 7198.701
839.4346923828125 0 1979.3862
851.3982543945312 0 826.82776
852.435546875 0 4500.419
853.4425659179688 0 8290.8545 c 6
854.4448852539062 0 2033.3037
855.4476928710938 0 631.6052
865.466064453125 0 861.9884
866.454833984375 0 2842.0337
867.45703125 0 1193.9946
893.4381103515625 0 1480.6943 c Ammonia loss 7
900.4833984375 0 866.072
908.4619140625 0 1800.5154
909.458251953125 0 16507.049
910.4646606445312 0 42168.01 c 7
911.4683227539062 0 20241.59
912.4716796875 0 4844.97
934.4404907226562 0 1361.8362 z Water loss 3
935.4434814453125 0 671.3083
937.486572265625 0 1696.2314
938.425537109375 0 1873.0454
939.4263305664062 0 778.6496
946.4771728515625 0 698.0808
948.4669189453125 0 1194.7834
949.4583129882812 0 900.1852
950.4622802734375 0 1159.8123 y Water loss 3
951.4551391601562 0 1253.2573
952.4545288085938 0 4150.884 z 3
953.4593505859375 0 3702.6206
954.4658813476562 0 975.86835
964.4760131835938 0 7423.7954 c Ammonia loss 8
965.4800415039062 0 3865.8672
966.478515625 0 1067.5052
967.4660034179688 0 1162.0923
968.4739379882812 0 1482.1643 y 3
981.5027465820312 0 24216.547 c 8
982.5059814453125 0 11198.956
983.5079345703125 0 4250.9346
1021.4780883789062 0 820.70544 z Water loss 2
1035.5390625 0 594.9103
1036.557861328125 0 2903.5354
1037.572998046875 0 1103.0079
1038.5714111328125 0 3175.8867
1039.4810791015625 0 1685.8285 z 2
1039.5799560546875 0 1365.2865
1040.490234375 0 1870.0651
1041.4837646484375 0 824.83746
1045.5361328125 0 645.581
1047.5267333984375 0 5743.5063
1048.528564453125 0 5462.869
1049.529541015625 0 1476.1306
1054.4869384765625 0 808.73615
1055.502197265625 0 4483.419 y 2
1056.5081787109375 0 2140.0933
1057.50537109375 0 818.24445
1063.5455322265625 0 7530.427 c Ammonia loss 9
1064.546630859375 0 5551.0176
1065.5565185546875 0 3563.95
1066.5626220703125 0 736.8524
1080.5709228515625 0 36921.29 c 9
1081.57421875 0 23743.818
1082.583251953125 0 18981.03
1083.588134765625 0 9712.519
1084.5963134765625 0 2383.051
1119.5946044921875 0 3973.5564
1120.597900390625 0 2974.9644
1121.5966796875 0 1246.3267
1138.6129150390625 0 925.875
1149.573486328125 0 742.6128
1150.5682373046875 0 1049.5319 y Water loss 1
1152.568603515625 0 7938.4155 z 1
1153.57080078125 0 6413.2476
1154.572021484375 0 1848.0374
1165.5906982421875 0 1001.5203
1181.6171875 0 15211.192 c 10
1182.62060546875 0 9956.759
1183.6221923828125 0 2998.3247
1210.593994140625 0 897.03864
1222.6527099609375 0 9422.334
1223.65380859375 0 6860.5107
1224.657958984375 0 3532.7693
1238.68505859375 0 6274.1943
1239.68994140625 0 4174.3247
1240.6851806640625 0 1527.4055
1247.646484375 0 1478.1431
1248.656005859375 0 1036.7263
1249.636474609375 0 766.4437
1254.6839599609375 0 3155.6272
1255.682373046875 0 2664.4155
1256.685791015625 0 1017.31805
1263.655029296875 0 1009.56573
1264.659423828125 0 2467.6719
1265.653564453125 0 12182.327
1266.65673828125 0 11046.163
1267.6566162109375 0 4112.138
1268.6463623046875 0 768.9353
1279.6556396484375 0 960.84076
1280.6585693359375 0 2609.2896
1281.6678466796875 0 17729.059
1282.6761474609375 0 77878.586
1283.6798095703125 0 59950.07
1284.6817626953125 0 23806.643
1285.684326171875 0 3644.1895
1921.954833984375 0 935.2758
3100.44140625 0 723.75256
3407.06005859375 0 710.6482

Spectrum Details

|  |  |
| --- | --- |
| Matched peaks? Matched peaks The total absolute number of peaks matched. Additionally in brackets the total fraction of peaks matched and the total number of peaks is shown. | 34 (17.89% of 190) |
| FDR? FDR The false discovery rate estimated for this peptide. It is calculated by matching all theoretical fragments with a non-integer shift with the raw peaks for this spectrum. This is done with 40 different shifts. The resulting percentage is the average number of annotated peaks over the number of annotated peaks with the correct spectrum. | 0.84% |
| Satellite FDR? Satellite FDR See the FDR for details on its calculation. This satellite ion specific FDR only contains the satellite ions (d/w) for I/L/J positions. | ∞ |
| PSM Score? PSM Score The PSM Score as given by Hecklib to this annotated spectrum. It is shown with three significant figures. | 459 |

#### Reverse Lookup? Reverse Lookup All places where this read could be placed.

| Group | Segment | Template | Template Part | Read Part | Score | Unique |
| --- | --- | --- | --- | --- | --- | --- |
| Homo sapiens Light Chain | IGLC | IGLC2 | [28..40] | [0..12] | 96 | False |
| Homo sapiens Light Chain | IGLC | IGLC3 | [26..38] | [0..12] | 96 | False |
| Homo sapiens Light Chain | IGLC | IGLC6 | [28..40] | [0..12] | 87 | False |
| Homo sapiens Light Chain | IGLC | IGLC7 | [28..40] | [0..12] | 87 | False |

| Recombined | Template Part | Read Part | Score | Unique |
| --- | --- | --- | --- | --- |
| REC-0-1\_002 | [139..151] | [0..12] | 96 | True |

#### Meta Information from Multiple reads

##### Number of combined reads

4

##### Intensity

0.8337

##### TotalArea

4.044E+08

##### Changes to the peptide sequence

JJSDFYPGAVTV

L→JNo support for either Leucine or Isoleucine based on side chain ions (Position: 2)

L→JNo support for either Leucine or Isoleucine based on side chain ions (Position: 1)

#### Positional Score

Copy Data

##### Positional Score (TSV)

###### Preview

```
Loading example...
```

*Click on the button to copy the data to your clipboard.*

1001234567891011

Label Value
"0" 0.5
"1" 0.49
"2" 0.473
"3" 0.485
"4" 0.495
"5" 0.485
"6" 0.465
"7" 0.473
"8" 0.487
"9" 0.495
"10" 0.495
"11" 0.49

#### Meta Information from PEAKS

##### Scan Identifier

F1:11034

##### Original sequence

L

L

S

D

F

Y

P

G

A

V

T

V

##### Posttranslational Modifications

##### Source File

D:\separate\_stitch\_analyses\xle-disambiguation\raw\20210323\_F1\_UM1\_Peng0013\_SA\_F59\_ingel\_3ug\_ELA.raw

##### Fraction

1

##### Scan Feature

F1:10281

##### De Novo Score

98

##### ConfidenceScore

98

### m/z

641.3415

##### Mass

1280.6653

##### Charge

2

##### Retention Time

60.86

##### Predicted Retention Time

-

##### Area

1.348E+08

##### Parts Per Million

2.5

##### Fragmentation mode

ETHCD

##### Originating file

01 D:\separate\_stitch\_analyses\xle-disambiguation\20210325\_F59\_3ug\_DENOVO\_12.csv

#### Meta Information from PEAKS

##### Scan Identifier

F1:11187

##### Original sequence

L

L

S

D

F

Y

P

G

A

V

T

V

##### Posttranslational Modifications

##### Source File

D:\separate\_stitch\_analyses\xle-disambiguation\raw\20210323\_F1\_UM1\_Peng0013\_SA\_F59\_ingel\_3ug\_ELA.raw

##### Fraction

1

##### Scan Feature

-

##### De Novo Score

98

##### ConfidenceScore

98

### m/z

641.341

##### Mass

1280.6653

##### Charge

2

##### Retention Time

62.28

##### Predicted Retention Time

-

##### Area

0

##### Parts Per Million

1.7

##### Fragmentation mode

ETHCD

##### Originating file

01 D:\separate\_stitch\_analyses\xle-disambiguation\20210325\_F59\_3ug\_DENOVO\_12.csv

#### Meta Information from PEAKS

##### Scan Identifier

F1:11086

##### Original sequence

L

L

S

D

F

Y

P

G

A

V

T

V

##### Posttranslational Modifications

##### Source File

D:\separate\_stitch\_analyses\xle-disambiguation\raw\20210323\_F1\_UM1\_Peng0013\_SA\_F59\_ingel\_3ug\_ELA.raw

##### Fraction

1

##### Scan Feature

F1:10281

##### De Novo Score

97

##### ConfidenceScore

97

### m/z

641.3415

##### Mass

1280.6653

##### Charge

2

##### Retention Time

60.86

##### Predicted Retention Time

-

##### Area

1.348E+08

##### Parts Per Million

2.5

##### Fragmentation mode

ETHCD

##### Originating file

01 D:\separate\_stitch\_analyses\xle-disambiguation\20210325\_F59\_3ug\_DENOVO\_12.csv

#### Meta Information from PEAKS

##### Scan Identifier

F1:11137

##### Original sequence

L

L

S

D

F

Y

P

G

A

V

T

V

##### Posttranslational Modifications

##### Source File

D:\separate\_stitch\_analyses\xle-disambiguation\raw\20210323\_F1\_UM1\_Peng0013\_SA\_F59\_ingel\_3ug\_ELA.raw

##### Fraction

1

##### Scan Feature

F1:10281

##### De Novo Score

96

##### ConfidenceScore

96

### m/z

641.3415

##### Mass

1280.6653

##### Charge

2

##### Retention Time

60.86

##### Predicted Retention Time

-

##### Area

1.348E+08

##### Parts Per Million

2.5

##### Fragmentation mode

ETHCD

##### Originating file

01 D:\separate\_stitch\_analyses\xle-disambiguation\20210325\_F59\_3ug\_DENOVO\_12.csv
