## Supplementary material for "A handle on mass coincidence errors in *de novo* sequencing of antibodies by bottom-up proteomics": Combined_081.html

Details Combined\_081 | Stitch OverviewUndefined

### Read Combined\_081

#### Sequence (length=7)

TVLHQDW

#### Spectrum 5429? Spectrum 5429 The raw spectrum of this peptide as annotated by Hecklib. The fragments are coloured according to ion type (see legend). Any peaks with a star '\*' as text can be hovered over to see the full details, first the ion type second the mass shift type. By hovering over the amino acids in the peptide or ions in the legend the corresponding peaks are highlighted. By toggling the 'Unassigned' label you can turn the background (unassigned) peaks on or off in the plot. By updating the slider in the Ion legend you can update the spectrum to only show the top X% of the peaks with labels. The top X% means any peak that is within X% of the highest intensity. By dragging in the spectrum you can zoom in to a specific part of the spectrum and use 'Zoom Out' to get back to the original zoom level. The annotation of the spectrum is based on the given sequence in the peptides file and is done with different software so inconsistencies are likely. The peaks are annotated based on the given sequence, with 20 ppm tolerance.

Copy Data

##### Spectrum 5429 (TSV)

###### Preview

```
Loading example...
```

*Click on the button to copy the data to your clipboard.*

Mz MinMz MaxIntensity Max

WidthHeightPeptide font sizePeptide stroke widthSpectrum font sizeSpectrum stroke widthCompact peptide

Ion legend

wxyz

abcd

OtherUnassignedIon^Charge^_Position_Show for top:%

TVLHQDW

02.27e+54.54e+56.81e+59.09e+5

Zoom Out

a^+1^_2_d^+1^_2_a^+1^_2_b^+1^_2_b^+1^_2_y^+1^_1_a^+1^_3_b^+1^_3_y^+1^_2_b^+1^_3_y^+1^_2_b^+2^_6_y^+2^_5_b^+2^_6_y^+2^_6_y^+1^_3_y^+1^_3_b^+1^_4_\*\*y^+1^_3_\*b^+1^_4_b^+1^_5_b^+1^_5_y^+1^_4_y^+1^_4_b^+1^_5_y^+1^_4_b^+1^_6_b^+1^_6_y^+1^_5_y^+1^_5_b^+1^_6_y^+1^_5_y^+1^_6_y^+1^_6_y^+1^_6_

0840167925193359

Fragment Matches Table

Show background peaks

| Position | Ion type | Intensity | mz Theoretical | mz Error (Th) | mz Error (ppm) | Charge | Series Number |
| --- | --- | --- | --- | --- | --- | --- | --- |
| - | - | 8368 | 120.1 | - | - | 0 | - |
| - | - | 2927 | 122.1 | - | - | 0 | - |
| - | - | 1565 | 123.1 | - | - | 0 | - |
| - | - | 887 | 124 | - | - | 0 | - |
| - | - | 780.2 | 125.1 | - | - | 0 | - |
| - | - | 1396 | 125.1 | - | - | 0 | - |
| - | - | 7025 | 126.1 | - | - | 0 | - |
| - | - | 763.2 | 126.1 | - | - | 0 | - |
| - | - | 609.3 | 127.1 | - | - | 0 | - |
| - | - | 2834 | 127.1 | - | - | 0 | - |
| - | - | 774.7 | 128.1 | - | - | 0 | - |
| - | - | 6.762E+04 | 128.1 | - | - | 0 | - |
| - | - | 3139 | 129.1 | - | - | 0 | - |
| - | - | 1.333E+05 | 129.1 | - | - | 0 | - |
| - | - | 4292 | 129.1 | - | - | 0 | - |
| - | - | 1254 | 130.1 | - | - | 0 | - |
| - | - | 4.394E+04 | 130.1 | - | - | 0 | - |
| - | - | 9274 | 130.1 | - | - | 0 | - |
| - | - | 829.8 | 130.1 | - | - | 0 | - |
| - | - | 822.2 | 130.1 | - | - | 0 | - |
| - | - | 3962 | 131.1 | - | - | 0 | - |
| - | - | 1184 | 131.1 | - | - | 0 | - |
| - | - | 1.805E+04 | 132.1 | - | - | 0 | - |
| - | - | 1311 | 132.1 | - | - | 0 | - |
| - | - | 9328 | 132.1 | - | - | 0 | - |
| - | - | 2060 | 133.1 | - | - | 0 | - |
| - | - | 1137 | 133.1 | - | - | 0 | - |
| - | - | 2068 | 136.1 | - | - | 0 | - |
| - | - | 529.1 | 136.9 | - | - | 0 | - |
| - | - | 9104 | 138.1 | - | - | 0 | - |
| - | - | 574.9 | 138.1 | - | - | 0 | - |
| - | - | 3672 | 138.1 | - | - | 0 | - |
| - | - | 1619 | 139.1 | - | - | 0 | - |
| - | - | 2077 | 139.1 | - | - | 0 | - |
| - | - | 881.5 | 140.1 | - | - | 0 | - |
| - | - | 586.7 | 140.2 | - | - | 0 | - |
| - | - | 3532 | 141.1 | - | - | 0 | - |
| - | - | 1897 | 142.1 | - | - | 0 | - |
| - | - | 803.1 | 142.1 | - | - | 0 | - |
| - | - | 2722 | 143 | - | - | 0 | - |
| - | - | 1918 | 144.1 | - | - | 0 | - |
| - | - | 5626 | 144.1 | - | - | 0 | - |
| - | - | 626.3 | 145.1 | - | - | 0 | - |
| - | - | 3.178E+04 | 146.1 | - | - | 0 | - |
| - | - | 2304 | 147.1 | - | - | 0 | - |
| - | - | 1056 | 148.8 | - | - | 0 | - |
| - | - | 771.3 | 148.9 | - | - | 0 | - |
| - | - | 1077 | 148.9 | - | - | 0 | - |
| - | - | 847.7 | 148.9 | - | - | 0 | - |
| - | - | 973.7 | 148.9 | - | - | 0 | - |
| - | - | 1810 | 148.9 | - | - | 0 | - |
| - | - | 1977 | 148.9 | - | - | 0 | - |
| - | - | 3502 | 148.9 | - | - | 0 | - |
| - | - | 5909 | 148.9 | - | - | 0 | - |
| - | - | 6030 | 149 | - | - | 0 | - |
| - | - | 3732 | 149 | - | - | 0 | - |
| - | - | 2153 | 149 | - | - | 0 | - |
| - | - | 1887 | 149 | - | - | 0 | - |
| - | - | 930.9 | 149 | - | - | 0 | - |
| - | - | 1069 | 149 | - | - | 0 | - |
| - | - | 1066 | 149 | - | - | 0 | - |
| - | - | 880.5 | 149 | - | - | 0 | - |
| - | - | 1545 | 149 | - | - | 0 | - |
| - | - | 1371 | 149 | - | - | 0 | - |
| - | - | 666.7 | 149.1 | - | - | 0 | - |
| - | - | 653.9 | 149.1 | - | - | 0 | - |
| - | - | 1233 | 150 | - | - | 0 | - |
| - | - | 1820 | 150.1 | - | - | 0 | - |
| - | - | 1050 | 151 | - | - | 0 | - |
| - | - | 747.8 | 151.1 | - | - | 0 | - |
| - | - | 713.5 | 153.1 | - | - | 0 | - |
| - | - | 3262 | 154.1 | - | - | 0 | - |
| - | - | 2933 | 155.1 | - | - | 0 | - |
| - | - | 4.01E+04 | 155.1 | - | - | 0 | - |
| 2 | a | 3.17E+04 | 155.1 | 0.0003508 | 2.262 | +1 | 2 |
| - | - | 2855 | 156.1 | - | - | 0 | - |
| - | - | 2531 | 156.1 | - | - | 0 | - |
| - | - | 1242 | 156.1 | - | - | 0 | - |
| - | - | 1568 | 156.1 | - | - | 0 | - |
| - | - | 8270 | 157.1 | - | - | 0 | - |
| - | - | 3679 | 157.1 | - | - | 0 | - |
| - | - | 1.474E+05 | 159.1 | - | - | 0 | - |
| 2 | d | 5846 | 159.1 | 0.0003092 | 1.944 | +1 | 2 |
| - | - | 1730 | 160.1 | - | - | 0 | - |
| - | - | 1346 | 160.1 | - | - | 0 | - |
| - | - | 1.364E+04 | 160.1 | - | - | 0 | - |
| - | - | 1413 | 165.1 | - | - | 0 | - |
| - | - | 6.238E+04 | 166.1 | - | - | 0 | - |
| - | - | 2476 | 166.1 | - | - | 0 | - |
| - | - | 2466 | 167.1 | - | - | 0 | - |
| - | - | 3055 | 167.1 | - | - | 0 | - |
| - | - | 2030 | 167.1 | - | - | 0 | - |
| - | - | 4730 | 167.1 | - | - | 0 | - |
| - | - | 753.4 | 168.1 | - | - | 0 | - |
| - | - | 1022 | 168.1 | - | - | 0 | - |
| - | - | 1866 | 169.1 | - | - | 0 | - |
| - | - | 1265 | 169.1 | - | - | 0 | - |
| - | - | 2378 | 169.1 | - | - | 0 | - |
| - | - | 1173 | 169.1 | - | - | 0 | - |
| - | - | 1445 | 169.1 | - | - | 0 | - |
| - | - | 7550 | 170.1 | - | - | 0 | - |
| - | - | 837.3 | 171.1 | - | - | 0 | - |
| - | - | 1366 | 171.1 | - | - | 0 | - |
| - | - | 1348 | 172.1 | - | - | 0 | - |
| 2 | a | 8.996E+05 | 173.1 | 0.0004825 | 2.787 | +1 | 2 |
| - | - | 7.149E+04 | 174.1 | - | - | 0 | - |
| - | - | 4739 | 175.1 | - | - | 0 | - |
| - | - | 3167 | 175.1 | - | - | 0 | - |
| - | - | 1640 | 176.1 | - | - | 0 | - |
| - | - | 3.156E+04 | 178.1 | - | - | 0 | - |
| - | - | 3261 | 179.1 | - | - | 0 | - |
| - | - | 1078 | 180.1 | - | - | 0 | - |
| - | - | 7406 | 181.1 | - | - | 0 | - |
| - | - | 1799 | 181.1 | - | - | 0 | - |
| - | - | 1531 | 183.1 | - | - | 0 | - |
| 2 | b | 1.222E+04 | 183.1 | 0.0003398 | 1.855 | +1 | 2 |
| - | - | 2770 | 183.1 | - | - | 0 | - |
| - | - | 883.1 | 185.1 | - | - | 0 | - |
| - | - | 1580 | 185.1 | - | - | 0 | - |
| - | - | 2822 | 185.1 | - | - | 0 | - |
| - | - | 4619 | 185.2 | - | - | 0 | - |
| - | - | 7465 | 187.1 | - | - | 0 | - |
| - | - | 1741 | 187.1 | - | - | 0 | - |
| - | - | 3226 | 187.1 | - | - | 0 | - |
| - | - | 2.437E+05 | 188.1 | - | - | 0 | - |
| - | - | 2.605E+04 | 189.1 | - | - | 0 | - |
| - | - | 3807 | 189.1 | - | - | 0 | - |
| - | - | 1672 | 190.1 | - | - | 0 | - |
| - | - | 6858 | 190.1 | - | - | 0 | - |
| - | - | 903.6 | 191.1 | - | - | 0 | - |
| - | - | 1859 | 194.1 | - | - | 0 | - |
| - | - | 738.1 | 195.1 | - | - | 0 | - |
| - | - | 5384 | 195.1 | - | - | 0 | - |
| - | - | 927.8 | 197.1 | - | - | 0 | - |
| - | - | 1.073E+04 | 198.1 | - | - | 0 | - |
| - | - | 5812 | 199.1 | - | - | 0 | - |
| - | - | 908.8 | 199.1 | - | - | 0 | - |
| - | - | 1501 | 199.1 | - | - | 0 | - |
| 2 | b | 1.644E+05 | 201.1 | 0.0003494 | 1.737 | +1 | 2 |
| - | - | 1.352E+04 | 202.1 | - | - | 0 | - |
| - | - | 7288 | 203.1 | - | - | 0 | - |
| 7 | y | 1.131E+05 | 205.1 | 0.0003801 | 1.853 | +1 | 1 |
| - | - | 2958 | 205.1 | - | - | 0 | - |
| - | - | 1.133E+04 | 206.1 | - | - | 0 | - |
| - | - | 3461 | 206.1 | - | - | 0 | - |
| - | - | 1747 | 207.1 | - | - | 0 | - |
| - | - | 5933 | 207.2 | - | - | 0 | - |
| - | - | 2521 | 208.1 | - | - | 0 | - |
| - | - | 1038 | 208.2 | - | - | 0 | - |
| - | - | 1342 | 209 | - | - | 0 | - |
| - | - | 1511 | 209.1 | - | - | 0 | - |
| - | - | 2158 | 209.1 | - | - | 0 | - |
| - | - | 2054 | 209.1 | - | - | 0 | - |
| - | - | 1670 | 211 | - | - | 0 | - |
| - | - | 1095 | 212.2 | - | - | 0 | - |
| - | - | 1434 | 213.2 | - | - | 0 | - |
| - | - | 3207 | 215.1 | - | - | 0 | - |
| - | - | 7562 | 216.1 | - | - | 0 | - |
| - | - | 6981 | 217.1 | - | - | 0 | - |
| - | - | 2731 | 220.1 | - | - | 0 | - |
| - | - | 973.7 | 221.1 | - | - | 0 | - |
| - | - | 5772 | 221.1 | - | - | 0 | - |
| - | - | 1631 | 221.1 | - | - | 0 | - |
| - | - | 6.027E+04 | 223.2 | - | - | 0 | - |
| - | - | 6966 | 224.2 | - | - | 0 | - |
| - | - | 2762 | 225 | - | - | 0 | - |
| - | - | 1288 | 225.2 | - | - | 0 | - |
| - | - | 819.1 | 226 | - | - | 0 | - |
| - | - | 2.463E+04 | 226.1 | - | - | 0 | - |
| - | - | 6964 | 226.1 | - | - | 0 | - |
| - | - | 6.978E+04 | 226.2 | - | - | 0 | - |
| - | - | 802.5 | 227 | - | - | 0 | - |
| - | - | 1.021E+04 | 227.1 | - | - | 0 | - |
| - | - | 1766 | 227.1 | - | - | 0 | - |
| - | - | 1543 | 227.1 | - | - | 0 | - |
| - | - | 1062 | 227.1 | - | - | 0 | - |
| - | - | 1.01E+04 | 227.2 | - | - | 0 | - |
| - | - | 747.5 | 228.2 | - | - | 0 | - |
| - | - | 855.7 | 230.1 | - | - | 0 | - |
| - | - | 1234 | 231.1 | - | - | 0 | - |
| - | - | 900 | 231.6 | - | - | 0 | - |
| - | - | 2542 | 232.1 | - | - | 0 | - |
| - | - | 1.654E+04 | 233.1 | - | - | 0 | - |
| - | - | 1.042E+04 | 233.2 | - | - | 0 | - |
| - | - | 3737 | 233.2 | - | - | 0 | - |
| - | - | 7.334E+04 | 234.1 | - | - | 0 | - |
| - | - | 1.086E+04 | 235.1 | - | - | 0 | - |
| - | - | 1562 | 235.2 | - | - | 0 | - |
| - | - | 1380 | 236.1 | - | - | 0 | - |
| - | - | 1745 | 238.1 | - | - | 0 | - |
| - | - | 1098 | 239.1 | - | - | 0 | - |
| - | - | 824.3 | 239.1 | - | - | 0 | - |
| - | - | 1090 | 240.1 | - | - | 0 | - |
| - | - | 959.9 | 241.1 | - | - | 0 | - |
| - | - | 2551 | 243.1 | - | - | 0 | - |
| - | - | 2336 | 243.1 | - | - | 0 | - |
| - | - | 3.798E+04 | 244.1 | - | - | 0 | - |
| - | - | 4059 | 244.1 | - | - | 0 | - |
| - | - | 1436 | 245.1 | - | - | 0 | - |
| - | - | 3672 | 245.1 | - | - | 0 | - |
| - | - | 1652 | 246.1 | - | - | 0 | - |
| - | - | 1.258E+04 | 248.1 | - | - | 0 | - |
| - | - | 974.2 | 248.5 | - | - | 0 | - |
| - | - | 2.613E+04 | 249.1 | - | - | 0 | - |
| - | - | 1352 | 249.1 | - | - | 0 | - |
| - | - | 1256 | 249.1 | - | - | 0 | - |
| - | - | 3849 | 249.1 | - | - | 0 | - |
| - | - | 3346 | 250.1 | - | - | 0 | - |
| - | - | 2473 | 250.2 | - | - | 0 | - |
| - | - | 826.2 | 250.9 | - | - | 0 | - |
| - | - | 2.911E+05 | 251.2 | - | - | 0 | - |
| - | - | 3.338E+04 | 252.2 | - | - | 0 | - |
| - | - | 1741 | 252.6 | - | - | 0 | - |
| - | - | 2655 | 253.2 | - | - | 0 | - |
| - | - | 865.6 | 253.5 | - | - | 0 | - |
| - | - | 2029 | 254.1 | - | - | 0 | - |
| - | - | 2195 | 254.1 | - | - | 0 | - |
| - | - | 3866 | 254.2 | - | - | 0 | - |
| - | - | 2329 | 256.1 | - | - | 0 | - |
| - | - | 1002 | 257.1 | - | - | 0 | - |
| - | - | 823 | 258.1 | - | - | 0 | - |
| - | - | 2004 | 260.1 | - | - | 0 | - |
| - | - | 1220 | 261.1 | - | - | 0 | - |
| - | - | 1996 | 261.1 | - | - | 0 | - |
| - | - | 1084 | 261.2 | - | - | 0 | - |
| - | - | 2675 | 262.1 | - | - | 0 | - |
| - | - | 1443 | 265.1 | - | - | 0 | - |
| - | - | 7.05E+04 | 266.1 | - | - | 0 | - |
| - | - | 7898 | 267.1 | - | - | 0 | - |
| - | - | 4273 | 268.2 | - | - | 0 | - |
| 3 | a | 986.4 | 268.2 | 0.0001339 | 0.4992 | +1 | 3 |
| - | - | 4171 | 270.1 | - | - | 0 | - |
| - | - | 3617 | 272.1 | - | - | 0 | - |
| - | - | 4460 | 272.1 | - | - | 0 | - |
| - | - | 966.2 | 273.1 | - | - | 0 | - |
| - | - | 5555 | 274.1 | - | - | 0 | - |
| - | - | 9650 | 274.1 | - | - | 0 | - |
| - | - | 932.7 | 274.7 | - | - | 0 | - |
| - | - | 1619 | 274.7 | - | - | 0 | - |
| - | - | 1415 | 275.1 | - | - | 0 | - |
| - | - | 1019 | 275.2 | - | - | 0 | - |
| - | - | 883.2 | 275.2 | - | - | 0 | - |
| - | - | 2527 | 276.1 | - | - | 0 | - |
| - | - | 1132 | 277.1 | - | - | 0 | - |
| - | - | 941.3 | 277.1 | - | - | 0 | - |
| - | - | 2216 | 279.1 | - | - | 0 | - |
| - | - | 1520 | 282.1 | - | - | 0 | - |
| - | - | 855.7 | 283.7 | - | - | 0 | - |
| - | - | 4289 | 284.1 | - | - | 0 | - |
| - | - | 756.8 | 284.1 | - | - | 0 | - |
| - | - | 2131 | 284.2 | - | - | 0 | - |
| - | - | 6882 | 285.1 | - | - | 0 | - |
| - | - | 1189 | 285.1 | - | - | 0 | - |
| - | - | 1384 | 287.2 | - | - | 0 | - |
| - | - | 2341 | 290.1 | - | - | 0 | - |
| - | - | 2036 | 293.2 | - | - | 0 | - |
| - | - | 5192 | 294.1 | - | - | 0 | - |
| - | - | 1479 | 294.2 | - | - | 0 | - |
| - | - | 997.8 | 295.1 | - | - | 0 | - |
| 3 | b | 1460 | 296.2 | 0.002076 | 7.009 | +1 | 3 |
| - | - | 899.4 | 297.2 | - | - | 0 | - |
| - | - | 1825 | 301.1 | - | - | 0 | - |
| 6 | y | 6679 | 302.1 | 0.0005423 | 1.795 | +1 | 2 |
| - | - | 1078 | 303.1 | - | - | 0 | - |
| - | - | 881.4 | 304.2 | - | - | 0 | - |
| - | - | 1351 | 305.2 | - | - | 0 | - |
| - | - | 1034 | 305.2 | - | - | 0 | - |
| - | - | 2848 | 306.2 | - | - | 0 | - |
| - | - | 1078 | 312.2 | - | - | 0 | - |
| 3 | b | 3453 | 314.2 | 5.619E-05 | 0.1788 | +1 | 3 |
| - | - | 1507 | 316.2 | - | - | 0 | - |
| - | - | 1294 | 317.2 | - | - | 0 | - |
| - | - | 1900 | 318.1 | - | - | 0 | - |
| - | - | 3006 | 319.2 | - | - | 0 | - |
| 6 | y | 3.104E+04 | 320.1 | 0.0005061 | 1.581 | +1 | 2 |
| - | - | 2016 | 320.9 | - | - | 0 | - |
| - | - | 5754 | 321.1 | - | - | 0 | - |
| - | - | 9556 | 323.2 | - | - | 0 | - |
| - | - | 1199 | 323.2 | - | - | 0 | - |
| - | - | 1.029E+04 | 323.2 | - | - | 0 | - |
| - | - | 1531 | 324.2 | - | - | 0 | - |
| - | - | 1837 | 324.2 | - | - | 0 | - |
| - | - | 1215 | 325.2 | - | - | 0 | - |
| - | - | 3975 | 326.7 | - | - | 0 | - |
| - | - | 4092 | 327.2 | - | - | 0 | - |
| - | - | 3229 | 330.2 | - | - | 0 | - |
| - | - | 8210 | 332.2 | - | - | 0 | - |
| - | - | 1147 | 332.7 | - | - | 0 | - |
| - | - | 2476 | 333.2 | - | - | 0 | - |
| - | - | 3303 | 334.2 | - | - | 0 | - |
| - | - | 1821 | 334.7 | - | - | 0 | - |
| 6 | b | 1650 | 338.7 | 0.0003993 | 1.179 | +2 | 6 |
| - | - | 1131 | 339.2 | - | - | 0 | - |
| 3 | y | 1314 | 341.2 | 0.004343 | 12.73 | +2 | 5 |
| - | - | 2.263E+04 | 341.2 | - | - | 0 | - |
| - | - | 3437 | 342.2 | - | - | 0 | - |
| - | - | 1133 | 343.2 | - | - | 0 | - |
| - | - | 7892 | 344.2 | - | - | 0 | - |
| - | - | 5589 | 345.1 | - | - | 0 | - |
| - | - | 4900 | 345.2 | - | - | 0 | - |
| - | - | 2.898E+04 | 346.1 | - | - | 0 | - |
| - | - | 1209 | 346.1 | - | - | 0 | - |
| - | - | 3704 | 347.1 | - | - | 0 | - |
| 6 | b | 5625 | 347.7 | 0.0005134 | 1.477 | +2 | 6 |
| - | - | 2362 | 348.2 | - | - | 0 | - |
| - | - | 2.57E+04 | 350.2 | - | - | 0 | - |
| - | - | 6745 | 351.2 | - | - | 0 | - |
| - | - | 2292 | 351.7 | - | - | 0 | - |
| - | - | 1334 | 352.2 | - | - | 0 | - |
| - | - | 1265 | 354.3 | - | - | 0 | - |
| - | - | 2011 | 355.2 | - | - | 0 | - |
| - | - | 1807 | 360 | - | - | 0 | - |
| - | - | 1408 | 360.7 | - | - | 0 | - |
| - | - | 6930 | 361 | - | - | 0 | - |
| - | - | 8910 | 361.2 | - | - | 0 | - |
| - | - | 7994 | 362 | - | - | 0 | - |
| - | - | 9391 | 362.2 | - | - | 0 | - |
| - | - | 5981 | 363 | - | - | 0 | - |
| - | - | 2.458E+04 | 363.1 | - | - | 0 | - |
| - | - | 3289 | 363.2 | - | - | 0 | - |
| - | - | 1668 | 364 | - | - | 0 | - |
| - | - | 1.428E+04 | 364.1 | - | - | 0 | - |
| - | - | 2227 | 364.1 | - | - | 0 | - |
| - | - | 2635 | 365.1 | - | - | 0 | - |
| - | - | 3159 | 367.1 | - | - | 0 | - |
| - | - | 3880 | 367.2 | - | - | 0 | - |
| - | - | 1731 | 367.7 | - | - | 0 | - |
| - | - | 1326 | 368.7 | - | - | 0 | - |
| - | - | 1338 | 369.2 | - | - | 0 | - |
| - | - | 1124 | 370.9 | - | - | 0 | - |
| - | - | 8042 | 373.2 | - | - | 0 | - |
| - | - | 6135 | 373.7 | - | - | 0 | - |
| - | - | 1819 | 374.2 | - | - | 0 | - |
| - | - | 3468 | 374.2 | - | - | 0 | - |
| - | - | 3705 | 376.2 | - | - | 0 | - |
| - | - | 868.8 | 377.2 | - | - | 0 | - |
| - | - | 3896 | 378.2 | - | - | 0 | - |
| - | - | 1.187E+05 | 379.2 | - | - | 0 | - |
| - | - | 2.145E+04 | 380.2 | - | - | 0 | - |
| - | - | 7.814E+04 | 381.2 | - | - | 0 | - |
| - | - | 2383 | 381.2 | - | - | 0 | - |
| - | - | 1.217E+04 | 382.2 | - | - | 0 | - |
| - | - | 1.405E+04 | 382.7 | - | - | 0 | - |
| - | - | 1.102E+04 | 383.2 | - | - | 0 | - |
| - | - | 2744 | 383.7 | - | - | 0 | - |
| - | - | 2084 | 384.2 | - | - | 0 | - |
| - | - | 4816 | 385.2 | - | - | 0 | - |
| - | - | 1019 | 386.2 | - | - | 0 | - |
| - | - | 1270 | 389.2 | - | - | 0 | - |
| - | - | 2469 | 390.7 | - | - | 0 | - |
| - | - | 3333 | 391.1 | - | - | 0 | - |
| - | - | 1877 | 391.2 | - | - | 0 | - |
| - | - | 1.539E+04 | 391.7 | - | - | 0 | - |
| - | - | 2217 | 392.1 | - | - | 0 | - |
| - | - | 8173 | 392.2 | - | - | 0 | - |
| - | - | 2986 | 392.7 | - | - | 0 | - |
| - | - | 4932 | 395.1 | - | - | 0 | - |
| - | - | 1198 | 398.2 | - | - | 0 | - |
| - | - | 4690 | 399.2 | - | - | 0 | - |
| 2 | y | 5086 | 399.2 | 1.317E-05 | 0.03299 | +2 | 6 |
| - | - | 3286 | 399.7 | - | - | 0 | - |
| - | - | 2.386E+04 | 399.7 | - | - | 0 | - |
| - | - | 1.514E+04 | 400.2 | - | - | 0 | - |
| - | - | 4633 | 400.7 | - | - | 0 | - |
| - | - | 1651 | 401.2 | - | - | 0 | - |
| - | - | 2855 | 402.2 | - | - | 0 | - |
| - | - | 1044 | 405.2 | - | - | 0 | - |
| - | - | 1846 | 405.3 | - | - | 0 | - |
| - | - | 1221 | 406.2 | - | - | 0 | - |
| - | - | 1674 | 407.2 | - | - | 0 | - |
| - | - | 7140 | 409.1 | - | - | 0 | - |
| - | - | 1294 | 409.2 | - | - | 0 | - |
| - | - | 1298 | 410.1 | - | - | 0 | - |
| - | - | 3728 | 412.2 | - | - | 0 | - |
| - | - | 7756 | 413.1 | - | - | 0 | - |
| - | - | 2433 | 414.1 | - | - | 0 | - |
| - | - | 3624 | 414.2 | - | - | 0 | - |
| - | - | 2052 | 414.9 | - | - | 0 | - |
| - | - | 1156 | 415.2 | - | - | 0 | - |
| - | - | 4691 | 417.7 | - | - | 0 | - |
| - | - | 7544 | 418.2 | - | - | 0 | - |
| - | - | 2862 | 418.7 | - | - | 0 | - |
| - | - | 1402 | 419.2 | - | - | 0 | - |
| - | - | 1290 | 422.3 | - | - | 0 | - |
| - | - | 1750 | 423.2 | - | - | 0 | - |
| - | - | 1.417E+04 | 423.3 | - | - | 0 | - |
| - | - | 1406 | 423.7 | - | - | 0 | - |
| - | - | 4007 | 424.2 | - | - | 0 | - |
| - | - | 1587 | 424.3 | - | - | 0 | - |
| - | - | 4212 | 425.3 | - | - | 0 | - |
| - | - | 2051 | 425.8 | - | - | 0 | - |
| - | - | 1257 | 426.3 | - | - | 0 | - |
| - | - | 2.295E+04 | 426.7 | - | - | 0 | - |
| - | - | 1138 | 426.8 | - | - | 0 | - |
| - | - | 1.024E+04 | 427.2 | - | - | 0 | - |
| - | - | 2884 | 427.7 | - | - | 0 | - |
| 5 | y | 7.096E+04 | 430.2 | 0.0006195 | 1.44 | +1 | 3 |
| - | - | 1084 | 430.8 | - | - | 0 | - |
| - | - | 1055 | 431.1 | - | - | 0 | - |
| 5 | y | 1.131E+04 | 431.2 | 0.0007605 | 1.764 | +1 | 3 |
| - | - | 8682 | 431.2 | - | - | 0 | - |
| - | - | 3476 | 431.7 | - | - | 0 | - |
| - | - | 2506 | 432.1 | - | - | 0 | - |
| - | - | 2937 | 432.2 | - | - | 0 | - |
| - | - | 2837 | 432.2 | - | - | 0 | - |
| - | - | 1376 | 432.7 | - | - | 0 | - |
| - | - | 1464 | 432.9 | - | - | 0 | - |
| - | - | 1207 | 433.1 | - | - | 0 | - |
| 4 | b | 4295 | 433.3 | 0.0003846 | 0.8876 | +1 | 4 |
| - | - | 1267 | 434.3 | - | - | 0 | - |
| - | - | 1.416E+04 | 439.3 | - | - | 0 | - |
| - | - | 1.048E+04 | 439.8 | - | - | 0 | - |
| - | - | 3811 | 440.3 | - | - | 0 | - |
| 0 | Precursor | 5907 | 440.7 | 1.096E-05 | 0.02487 | +2 | -1 |
| 0 | Precursor | 3322 | 441.2 | 0.004341 | 9.839 | +2 | -1 |
| - | - | 1070 | 441.7 | - | - | 0 | - |
| - | - | 1.093E+04 | 442.2 | - | - | 0 | - |
| - | - | 2609 | 443.2 | - | - | 0 | - |
| 5 | y | 6.516E+04 | 448.2 | 0.0006445 | 1.438 | +1 | 3 |
| - | - | 2.133E+05 | 448.3 | - | - | 0 | - |
| - | - | 1.518E+05 | 448.8 | - | - | 0 | - |
| - | - | 1.484E+04 | 449.2 | - | - | 0 | - |
| - | - | 5.804E+04 | 449.3 | - | - | 0 | - |
| 0 | Precursor | 1.201E+04 | 449.7 | 0.0008627 | 1.918 | +2 | -1 |
| - | - | 1.459E+04 | 449.8 | - | - | 0 | - |
| - | - | 2786 | 450.2 | - | - | 0 | - |
| - | - | 8524 | 450.2 | - | - | 0 | - |
| - | - | 3391 | 450.3 | - | - | 0 | - |
| - | - | 1357 | 450.7 | - | - | 0 | - |
| 4 | b | 2.462E+04 | 451.3 | 0.0004095 | 0.9074 | +1 | 4 |
| - | - | 2500 | 451.3 | - | - | 0 | - |
| - | - | 7009 | 452.3 | - | - | 0 | - |
| - | - | 1620 | 456.2 | - | - | 0 | - |
| - | - | 1837 | 458.3 | - | - | 0 | - |
| - | - | 4372 | 459.2 | - | - | 0 | - |
| - | - | 3674 | 460.2 | - | - | 0 | - |
| - | - | 1212 | 461.2 | - | - | 0 | - |
| - | - | 7245 | 468.3 | - | - | 0 | - |
| - | - | 1443 | 469.3 | - | - | 0 | - |
| - | - | 3137 | 476.2 | - | - | 0 | - |
| - | - | 9186 | 476.3 | - | - | 0 | - |
| - | - | 9251 | 477.2 | - | - | 0 | - |
| - | - | 2601 | 477.3 | - | - | 0 | - |
| - | - | 2874 | 478.2 | - | - | 0 | - |
| - | - | 9021 | 478.3 | - | - | 0 | - |
| - | - | 1980 | 479.3 | - | - | 0 | - |
| - | - | 1955 | 489.8 | - | - | 0 | - |
| - | - | 1.05E+05 | 494.2 | - | - | 0 | - |
| - | - | 2.584E+04 | 495.2 | - | - | 0 | - |
| - | - | 4602 | 496.2 | - | - | 0 | - |
| - | - | 3401 | 504.2 | - | - | 0 | - |
| - | - | 1.571E+04 | 508.3 | - | - | 0 | - |
| - | - | 3502 | 509.3 | - | - | 0 | - |
| - | - | 1631 | 511.3 | - | - | 0 | - |
| - | - | 5670 | 512.2 | - | - | 0 | - |
| - | - | 1025 | 522.2 | - | - | 0 | - |
| - | - | 948.5 | 523.2 | - | - | 0 | - |
| - | - | 2718 | 527.3 | - | - | 0 | - |
| - | - | 2469 | 532.2 | - | - | 0 | - |
| - | - | 1375 | 537.3 | - | - | 0 | - |
| - | - | 4259 | 539.2 | - | - | 0 | - |
| - | - | 981.7 | 540.2 | - | - | 0 | - |
| - | - | 1039 | 542.8 | - | - | 0 | - |
| - | - | 1592 | 547.3 | - | - | 0 | - |
| - | - | 2323 | 548.3 | - | - | 0 | - |
| - | - | 3791 | 548.4 | - | - | 0 | - |
| - | - | 1459 | 549.2 | - | - | 0 | - |
| - | - | 1849 | 549.4 | - | - | 0 | - |
| - | - | 2.619E+04 | 550.2 | - | - | 0 | - |
| - | - | 7146 | 551.2 | - | - | 0 | - |
| - | - | 1317 | 551.3 | - | - | 0 | - |
| - | - | 1025 | 552.2 | - | - | 0 | - |
| - | - | 1142 | 552.3 | - | - | 0 | - |
| - | - | 1161 | 555.3 | - | - | 0 | - |
| - | - | 1.006E+04 | 555.3 | - | - | 0 | - |
| - | - | 5049 | 556.3 | - | - | 0 | - |
| 5 | b | 4048 | 561.3 | 0.0008809 | 1.569 | +1 | 5 |
| 5 | b | 1049 | 562.3 | 0.005033 | 8.95 | +1 | 5 |
| - | - | 5142 | 566.3 | - | - | 0 | - |
| 4 | y | 8779 | 567.2 | 0.0005456 | 0.9618 | +1 | 4 |
| - | - | 1548 | 567.3 | - | - | 0 | - |
| 4 | y | 9109 | 568.2 | 0.001454 | 2.559 | +1 | 4 |
| - | - | 2607 | 569.2 | - | - | 0 | - |
| - | - | 3808 | 570.3 | - | - | 0 | - |
| - | - | 1169 | 571.3 | - | - | 0 | - |
| - | - | 6.887E+04 | 573.3 | - | - | 0 | - |
| - | - | 3.177E+04 | 574.3 | - | - | 0 | - |
| - | - | 4576 | 575.3 | - | - | 0 | - |
| - | - | 2641 | 576.3 | - | - | 0 | - |
| 5 | b | 1.486E+04 | 579.3 | 0.0003341 | 0.5768 | +1 | 5 |
| - | - | 4249 | 580.3 | - | - | 0 | - |
| - | - | 1081 | 581.3 | - | - | 0 | - |
| 4 | y | 8.904E+04 | 585.2 | 0.0004179 | 0.7141 | +1 | 4 |
| - | - | 2.398E+04 | 586.2 | - | - | 0 | - |
| - | - | 4465 | 587.2 | - | - | 0 | - |
| - | - | 1299 | 588.3 | - | - | 0 | - |
| - | - | 1.785E+04 | 593.3 | - | - | 0 | - |
| - | - | 6148 | 594.3 | - | - | 0 | - |
| - | - | 6456 | 595.2 | - | - | 0 | - |
| - | - | 1153 | 595.3 | - | - | 0 | - |
| - | - | 2421 | 596.2 | - | - | 0 | - |
| - | - | 1006 | 597.2 | - | - | 0 | - |
| - | - | 1402 | 598.3 | - | - | 0 | - |
| - | - | 1010 | 605.3 | - | - | 0 | - |
| - | - | 3510 | 607.3 | - | - | 0 | - |
| - | - | 1758 | 608.3 | - | - | 0 | - |
| - | - | 1536 | 610.3 | - | - | 0 | - |
| - | - | 1056 | 617.3 | - | - | 0 | - |
| - | - | 1629 | 622.4 | - | - | 0 | - |
| - | - | 1212 | 634.3 | - | - | 0 | - |
| - | - | 2451 | 635.3 | - | - | 0 | - |
| - | - | 2083 | 636.3 | - | - | 0 | - |
| - | - | 1130 | 637.3 | - | - | 0 | - |
| - | - | 1299 | 645.3 | - | - | 0 | - |
| - | - | 1190 | 645.4 | - | - | 0 | - |
| - | - | 2943 | 649.4 | - | - | 0 | - |
| - | - | 1080 | 650.4 | - | - | 0 | - |
| - | - | 1.151E+04 | 652.3 | - | - | 0 | - |
| - | - | 3987 | 653.3 | - | - | 0 | - |
| - | - | 1175 | 654.3 | - | - | 0 | - |
| - | - | 1457 | 659.3 | - | - | 0 | - |
| - | - | 1796 | 662.3 | - | - | 0 | - |
| - | - | 1.188E+04 | 663.3 | - | - | 0 | - |
| - | - | 1.112E+04 | 663.4 | - | - | 0 | - |
| - | - | 3838 | 664.3 | - | - | 0 | - |
| - | - | 4384 | 664.4 | - | - | 0 | - |
| - | - | 1564 | 665.4 | - | - | 0 | - |
| - | - | 1.11E+04 | 667.4 | - | - | 0 | - |
| - | - | 6192 | 668.4 | - | - | 0 | - |
| - | - | 1241 | 669.4 | - | - | 0 | - |
| 6 | b | 9631 | 676.3 | 0.0002361 | 0.3491 | +1 | 6 |
| 6 | b | 1.135E+04 | 677.3 | 0.00287 | 4.237 | +1 | 6 |
| - | - | 3759 | 678.3 | - | - | 0 | - |
| - | - | 1126 | 679.3 | - | - | 0 | - |
| 3 | y | 2.648E+04 | 680.3 | 0.0002218 | 0.3261 | +1 | 5 |
| 3 | y | 4.637E+04 | 681.3 | 0.00229 | 3.362 | +1 | 5 |
| - | - | 1.627E+04 | 682.3 | - | - | 0 | - |
| - | - | 3522 | 683.3 | - | - | 0 | - |
| - | - | 4064 | 683.4 | - | - | 0 | - |
| - | - | 1469 | 684.4 | - | - | 0 | - |
| - | - | 3506 | 685.4 | - | - | 0 | - |
| - | - | 1244 | 689.3 | - | - | 0 | - |
| - | - | 1071 | 690.3 | - | - | 0 | - |
| 6 | b | 9.738E+04 | 694.4 | 0.0001856 | 0.2672 | +1 | 6 |
| - | - | 3.661E+04 | 695.4 | - | - | 0 | - |
| - | - | 7416 | 696.4 | - | - | 0 | - |
| 3 | y | 3.062E+05 | 698.3 | 0.0002162 | 0.3096 | +1 | 5 |
| - | - | 1.093E+05 | 699.3 | - | - | 0 | - |
| - | - | 2.64E+04 | 700.3 | - | - | 0 | - |
| - | - | 1515 | 701.3 | - | - | 0 | - |
| - | - | 1.282E+04 | 701.4 | - | - | 0 | - |
| - | - | 6635 | 702.4 | - | - | 0 | - |
| - | - | 2119 | 703.4 | - | - | 0 | - |
| - | - | 1359 | 708.3 | - | - | 0 | - |
| - | - | 1173 | 709.3 | - | - | 0 | - |
| - | - | 1518 | 711.4 | - | - | 0 | - |
| - | - | 4460 | 712.4 | - | - | 0 | - |
| - | - | 1289 | 713.4 | - | - | 0 | - |
| - | - | 1133 | 721.4 | - | - | 0 | - |
| - | - | 1418 | 734.4 | - | - | 0 | - |
| - | - | 3167 | 751.4 | - | - | 0 | - |
| - | - | 1103 | 752.4 | - | - | 0 | - |
| - | - | 3024 | 764.4 | - | - | 0 | - |
| - | - | 1855 | 765.4 | - | - | 0 | - |
| - | - | 3354 | 769.4 | - | - | 0 | - |
| 2 | y | 2358 | 779.4 | 0.00151 | 1.937 | +1 | 6 |
| 2 | y | 7534 | 780.4 | 0.001931 | 2.474 | +1 | 6 |
| - | - | 6216 | 780.5 | - | - | 0 | - |
| - | - | 3372 | 781.4 | - | - | 0 | - |
| - | - | 3748 | 781.5 | - | - | 0 | - |
| - | - | 6420 | 782.4 | - | - | 0 | - |
| - | - | 2962 | 783.4 | - | - | 0 | - |
| - | - | 1584 | 784.4 | - | - | 0 | - |
| 2 | y | 3.404E+04 | 797.4 | 0.0003876 | 0.4861 | +1 | 6 |
| - | - | 1.462E+04 | 798.4 | - | - | 0 | - |
| - | - | 4.135E+04 | 798.5 | - | - | 0 | - |
| - | - | 3273 | 799.4 | - | - | 0 | - |
| - | - | 2.481E+04 | 799.5 | - | - | 0 | - |
| - | - | 9664 | 800.5 | - | - | 0 | - |
| - | - | 2043 | 801.5 | - | - | 0 | - |
| - | - | 1129 | 807.4 | - | - | 0 | - |
| - | - | 927.7 | 1357 | - | - | 0 | - |
| - | - | 1032 | 1565 | - | - | 0 | - |
| - | - | 1076 | 2354 | - | - | 0 | - |
| - | - | 1150 | 2805 | - | - | 0 | - |
| - | - | 1079 | 3085 | - | - | 0 | - |
| - | - | 1186 | 3325 | - | - | 0 | - |

m/z Charge Intensity FragmentType MassShift Position
120.08118438720703 0 8367.678
122.07164001464844 0 2927.3916
123.05574798583984 0 1565.082
124.01424407958984 0 887.0183
125.071533203125 0 780.15515
125.10770416259766 0 1396.3949
126.05531311035156 0 7025.3013
126.091552734375 0 763.16565
127.08682250976562 0 609.25745
127.12335205078125 0 2833.7207
128.0712127685547 0 774.68335
128.1073760986328 0 67621.805
129.06626892089844 0 3139.4487
129.10263061523438 0 133284.28
129.1111297607422 0 4291.566
130.05044555664062 0 1253.523
130.0655059814453 0 43936.125
130.1060028076172 0 9273.932
130.1109619140625 0 829.76715
130.11380004882812 0 822.24774
131.06874084472656 0 3962.2544
131.0821533203125 0 1183.6193
132.08114624023438 0 18053.969
132.08644104003906 0 1311.4482
132.1022491455078 0 9327.733
133.0845184326172 0 2059.9556
133.1054229736328 0 1136.5093
136.07601928710938 0 2067.9795
136.91156005859375 0 529.081
138.06663513183594 0 9104.218
138.0867156982422 0 574.91003
138.09173583984375 0 3672.2087
139.0507049560547 0 1619.3
139.08685302734375 0 2077.349
140.0825653076172 0 881.5351
140.18035888671875 0 586.6527
141.10264587402344 0 3531.5203
142.06544494628906 0 1896.5513
142.1228790283203 0 803.0691
143.04547119140625 0 2722.2473
144.06590270996094 0 1918.4335
144.08119201660156 0 5625.873
145.08383178710938 0 626.3231
146.0604248046875 0 31783.295
147.0637969970703 0 2304.1392
148.8195037841797 0 1055.6198
148.8623809814453 0 771.28107
148.8988800048828 0 1077.3567
148.90628051757812 0 847.7448
148.91334533691406 0 973.7118
148.92120361328125 0 1809.9027
148.92816162109375 0 1977.2057
148.9351348876953 0 3501.8982
148.94290161132812 0 5909.079
148.9596405029297 0 6030.28
148.9674072265625 0 3731.9902
148.97463989257812 0 2152.502
148.98162841796875 0 1886.9465
148.98928833007812 0 930.86646
148.99652099609375 0 1068.8387
149.00416564941406 0 1065.6636
149.01123046875 0 880.46045
149.02391052246094 0 1544.5933
149.04574584960938 0 1370.5889
149.05393981933594 0 666.72815
149.06149291992188 0 653.92596
150.02735900878906 0 1233.1844
150.06639099121094 0 1819.5223
151.04161071777344 0 1050.4615
151.12298583984375 0 747.7702
153.10299682617188 0 713.4792
154.0977783203125 0 3262.045
155.08160400390625 0 2932.7698
155.09312438964844 0 40103.676
155.1182403564453 0 31703.922 a Water loss 1
156.0770263671875 0 2855.3284
156.09629821777344 0 2531.2444
156.10243225097656 0 1242.2314
156.12130737304688 0 1568.0422
157.06114196777344 0 8270.308
157.0974884033203 0 3678.6353
159.0920867919922 0 147408.17
159.1131134033203 0 5846.333 d 1
160.075927734375 0 1729.7264
160.0887451171875 0 1345.7462
160.0955047607422 0 13639.509
165.07725524902344 0 1412.5435
166.06150817871094 0 62377.133
166.09765625 0 2475.7483
167.05613708496094 0 2466.4062
167.06492614746094 0 3054.7114
167.08168029785156 0 2030.1952
167.1183319091797 0 4729.836
168.05540466308594 0 753.4301
168.1220703125 0 1022.3177
169.05276489257812 0 1866.3151
169.0604705810547 0 1265.1372
169.07630920410156 0 2377.8755
169.09710693359375 0 1172.8048
169.13380432128906 0 1445.1741
170.0603790283203 0 7549.5166
171.063720703125 0 837.2791
171.0767059326172 0 1365.817
172.10818481445312 0 1348.2089
173.12893676757812 0 899638.2 a 1
174.13223266601562 0 71489.25
175.07183837890625 0 4738.5312
175.13414001464844 0 3166.8384
176.08206176757812 0 1639.8892
178.13421630859375 0 31561.686
179.1377716064453 0 3260.6965
180.0778045654297 0 1078.1714
181.0612030029297 0 7405.8574
181.13401794433594 0 1799.1995
183.0882568359375 0 1531.1271
183.11314392089844 0 12217.826 b Water loss 1
183.14971923828125 0 2770.1313
185.05679321289062 0 883.08936
185.0924530029297 0 1580.3162
185.1289825439453 0 2822.4612
185.16522216796875 0 4618.7827
187.0870361328125 0 7465.412
187.10780334472656 0 1741.3196
187.14454650878906 0 3226.4204
188.07102966308594 0 243731.39
189.07437133789062 0 26045.543
189.08726501464844 0 3806.7002
190.07688903808594 0 1672.0475
190.1342315673828 0 6858.386
191.13815307617188 0 903.55853
194.0925750732422 0 1859.1974
195.09295654296875 0 738.1469
195.11300659179688 0 5384.062
197.105712890625 0 927.83826
198.08773803710938 0 10733.34
199.07167053222656 0 5811.6406
199.0889434814453 0 908.8067
199.1078643798828 0 1501.404
201.12371826171875 0 164356.81 b 1
202.12709045410156 0 13521.217
203.06649780273438 0 7287.5605
205.0975341796875 0 113053.6 y 6
205.14508056640625 0 2957.789
206.10089111328125 0 11329.553
206.12936401367188 0 3460.8525
207.1033172607422 0 1747.1742
207.160888671875 0 5932.679
208.14427185058594 0 2521.4272
208.1641387939453 0 1037.5415
208.95321655273438 0 1342.0122
209.05532836914062 0 1510.7823
209.09230041503906 0 2158.4456
209.12892150878906 0 2053.6804
210.95018005371094 0 1670.1111
212.17623901367188 0 1094.8083
213.16012573242188 0 1433.9401
215.13946533203125 0 3206.6494
216.09820556640625 0 7561.9683
217.0821990966797 0 6980.852
220.11997985839844 0 2731.159
221.07032775878906 0 973.723
221.10369873046875 0 5771.6084
221.14073181152344 0 1631.4829
223.15574645996094 0 60273.617
224.15916442871094 0 6965.588
225.0434112548828 0 2762.2012
225.1710968017578 0 1287.8745
226.0436248779297 0 819.13153
226.08265686035156 0 24626.713
226.11904907226562 0 6964.161
226.15538024902344 0 69777.57
227.04086303710938 0 802.45105
227.06663513183594 0 10210.963
227.08631896972656 0 1765.7631
227.1021728515625 0 1542.9136
227.1227569580078 0 1061.8071
227.15869140625 0 10095.37
228.16183471679688 0 747.4579
230.10379028320312 0 855.71313
231.087646484375 0 1233.5963
231.63800048828125 0 900.03955
232.1405029296875 0 2541.9492
233.13987731933594 0 16537.566
233.1504364013672 0 10417.587
233.16566467285156 0 3737.3154
234.12417602539062 0 73338.88
235.1274871826172 0 10858.445
235.15576171875 0 1562.4089
236.13987731933594 0 1380.0747
238.12985229492188 0 1745.1442
239.08192443847656 0 1097.9258
239.117431640625 0 824.3008
240.13528442382812 0 1089.5133
241.09750366210938 0 959.87805
243.0765838623047 0 2551.039
243.10934448242188 0 2336.097
244.09320068359375 0 37978.293
244.1296844482422 0 4058.5903
245.07766723632812 0 1436.4164
245.09645080566406 0 3672.2883
246.13626098632812 0 1651.6893
248.11459350585938 0 12577.159
248.51113891601562 0 974.1584
249.09866333007812 0 26130.145
249.11489868164062 0 1351.5802
249.12046813964844 0 1255.852
249.1353302001953 0 3848.7627
250.10183715820312 0 3346.111
250.16671752929688 0 2472.8223
250.85211181640625 0 826.24164
251.15077209472656 0 291086.94
252.15394592285156 0 33378.05
252.6025848388672 0 1741.0518
253.15679931640625 0 2654.7615
253.5235137939453 0 865.5786
254.07762145996094 0 2029.4937
254.1142120361328 0 2194.6138
254.15020751953125 0 3866.393
256.10821533203125 0 2329.289
257.0918273925781 0 1001.926
258.14715576171875 0 822.98224
260.1029052734375 0 2004.247
261.1199951171875 0 1220.2727
261.13494873046875 0 1995.821
261.1602478027344 0 1083.762
262.1036071777344 0 2674.892
265.1410217285156 0 1443.2457
266.12506103515625 0 70503.98
267.1282043457031 0 7897.83
268.17706298828125 0 4272.641
268.20208740234375 0 986.3777 a Water loss 2
270.121826171875 0 4171.4414
272.0879821777344 0 3617.4575
272.1243591308594 0 4459.6533
273.0911865234375 0 966.195
274.1181335449219 0 5555.079
274.1303405761719 0 9650.327
274.6645202636719 0 932.7028
274.6816101074219 0 1619.4064
275.1357727050781 0 1414.5817
275.1501770019531 0 1018.58545
275.16644287109375 0 883.1799
276.1096496582031 0 2527.3672
277.0935363769531 0 1132.2402
277.1112976074219 0 941.31744
279.14508056640625 0 2215.8896
282.1451721191406 0 1519.5696
283.66961669921875 0 855.69775
284.1031799316406 0 4288.6143
284.13726806640625 0 756.75714
284.1605224609375 0 2131.0437
285.0873718261719 0 6882.1396
285.10498046875 0 1189.1958
287.1864318847656 0 1383.778
290.09893798828125 0 2341.3838
293.16107177734375 0 2036.2029
294.1199645996094 0 5191.502
294.21710205078125 0 1478.956
295.12359619140625 0 997.82416
296.1989440917969 0 1459.9294 b Water loss 2
297.15643310546875 0 899.3756
301.14044189453125 0 1825.0853
302.11407470703125 0 6678.9897 y Water loss 5
303.11614990234375 0 1077.9054
304.2139892578125 0 881.4333
305.1619873046875 0 1350.5739
305.19873046875 0 1033.8439
306.2290954589844 0 2848.0063
312.1564025878906 0 1078.1991
314.2074890136719 0 3453.0215 b 2
316.1783447265625 0 1507.043
317.1817932128906 0 1293.859
318.12103271484375 0 1899.5234
319.1521911621094 0 3006.4944
320.1246032714844 0 31036.059 y 5
320.92010498046875 0 2015.9152
321.1273498535156 0 5753.7275
323.1717224121094 0 9555.746
323.1899719238281 0 1198.6763
323.2083435058594 0 10292.808
324.17376708984375 0 1531.0492
324.2105407714844 0 1836.6351
325.15118408203125 0 1215.1982
326.66424560546875 0 3974.7163
327.1658935546875 0 4091.807
330.16656494140625 0 3229.151
332.2086181640625 0 8209.597
332.6983947753906 0 1147.0623
333.2107849121094 0 2475.8127
334.1916809082031 0 3303.384
334.6947937011719 0 1820.6658
338.67388916015625 0 1649.7974 b Water loss 5
339.1731872558594 0 1130.78
341.15753173828125 0 1313.5475 y Ammonia loss 2
341.18243408203125 0 22625.332
342.1842346191406 0 3436.7915
343.1613464355469 0 1133.0698
344.1722717285156 0 7891.7104
345.1307678222656 0 5589.492
345.1769714355469 0 4899.837
346.1150817871094 0 28982.223
346.1338195800781 0 1208.6667
347.1181945800781 0 3704.1033
347.6800842285156 0 5625.1743 b 5
348.1790771484375 0 2362.22
350.21905517578125 0 25698.46
351.2181701660156 0 6745.2017
351.7151794433594 0 2291.6829
352.220703125 0 1334.1448
354.250244140625 0 1264.5295
355.1606140136719 0 2010.8141
360.0286865234375 0 1807.2317
360.6960144042969 0 1408.1072
361.0267028808594 0 6930.1133
361.19866943359375 0 8910.413
362.0272521972656 0 7993.5376
362.1823425292969 0 9391.019
363.024169921875 0 5980.592
363.14141845703125 0 24582.719
363.18682861328125 0 3289.492
364.0242614746094 0 1668.1774
364.1253967285156 0 14284.296
364.14544677734375 0 2227.205
365.12823486328125 0 2635.2158
367.1405944824219 0 3159.226
367.24554443359375 0 3880.0884
367.6851501464844 0 1730.5403
368.7226257324219 0 1325.6646
369.2222595214844 0 1337.6519
370.88214111328125 0 1124.3572
373.1722717285156 0 8042.0986
373.7134094238281 0 6134.719
374.1758117675781 0 1819.0044
374.2156677246094 0 3468.039
376.1982727050781 0 3704.564
377.201904296875 0 868.7655
378.2138671875 0 3896.0125
379.20928955078125 0 118671.6
380.2121887207031 0 21454.658
381.1520690917969 0 78143.195
381.21368408203125 0 2383.318
382.15545654296875 0 12168.044
382.71905517578125 0 14053.818
383.2203674316406 0 11018.303
383.7215576171875 0 2744.218
384.1667785644531 0 2084.355
385.15142822265625 0 4816.47
386.15631103515625 0 1019.11005
389.1921691894531 0 1269.7352
390.7351989746094 0 2468.788
391.1363220214844 0 3333.1016
391.2359313964844 0 1877.0217
391.72442626953125 0 15391.353
392.1201477050781 0 2216.8857
392.2257385253906 0 8173.2524
392.7261962890625 0 2986.2065
395.1356201171875 0 4931.6123
398.177490234375 0 1197.6655
399.16314697265625 0 4690.351
399.20068359375 0 5086.329 y 1
399.70355224609375 0 3285.7139
399.7400207519531 0 23860.67
400.2416076660156 0 15144.893
400.7427062988281 0 4632.5356
401.2434387207031 0 1651.3154
402.17816162109375 0 2855.0679
405.22796630859375 0 1043.9875
405.25982666015625 0 1845.613
406.246337890625 0 1221.1584
407.20355224609375 0 1673.5774
409.1473388671875 0 7139.5635
409.2039489746094 0 1293.6821
410.1494140625 0 1297.8853
412.1623229980469 0 3727.9048
413.1462097167969 0 7755.6187
414.14776611328125 0 2432.6057
414.2351379394531 0 3624.091
414.8631286621094 0 2052.1377
415.2370910644531 0 1155.9664
417.7171936035156 0 4691.304
418.2101135253906 0 7543.904
418.7100830078125 0 2862.1455
419.211669921875 0 1401.5818
422.2520446777344 0 1290.272
423.2025451660156 0 1749.921
423.27203369140625 0 14169.604
423.7039794921875 0 1405.9128
424.2197570800781 0 4007.1013
424.2784423828125 0 1586.7495
425.2645263671875 0 4211.786
425.7657775878906 0 2051.3892
426.2586364746094 0 1256.9388
426.722412109375 0 22953.36
426.7527160644531 0 1137.6371
427.22210693359375 0 10244.963
427.7199401855469 0 2884.3105
430.1727294921875 0 70956.54 y Water loss 4
430.756103515625 0 1083.6642
431.087646484375 0 1055.4178
431.1553649902344 0 11310.779 y Ammonia loss 4
431.1786193847656 0 8682.072
431.7144775390625 0 3476.4382
432.0875244140625 0 2505.7742
432.15966796875 0 2937.1309
432.2135314941406 0 2837.066
432.7095947265625 0 1376.2706
432.8857727050781 0 1464.0886
433.0877380371094 0 1206.5695
433.25616455078125 0 4295.2725 b Water loss 3
434.2604064941406 0 1267.318
439.2613525390625 0 14162.5
439.76239013671875 0 10481.12
440.2623291015625 0 3810.5369
440.71923828125 0 5907.3726 Precursor Water loss
441.215576171875 0 3321.861 Precursor Ammonia loss
441.71612548828125 0 1070.0748
442.2301025390625 0 10930.5
443.23431396484375 0 2608.7605
448.1833190917969 0 65158.1 y 4
448.2667236328125 0 213289.77
448.7680358886719 0 151753.5
449.1861877441406 0 14837.575
449.26947021484375 0 58036.32
449.7253723144531 0 12007.199 Precursor
449.7702941894531 0 14591.321
450.188720703125 0 2785.9158
450.2270202636719 0 8523.544
450.2786865234375 0 3390.793
450.7269287109375 0 1356.7485
451.2667541503906 0 24615.78 b 3
451.3017883300781 0 2500.156
452.269775390625 0 7008.775
456.18389892578125 0 1620.3501
458.26019287109375 0 1836.6466
459.19927978515625 0 4371.8853
460.2428283691406 0 3673.553
461.24969482421875 0 1211.7572
468.2931823730469 0 7244.5713
469.27618408203125 0 1442.9202
476.2259826660156 0 3137.4758
476.2719421386719 0 9185.583
477.2099304199219 0 9250.728
477.2755126953125 0 2600.9158
478.21368408203125 0 2873.9949
478.27789306640625 0 9020.661
479.28094482421875 0 1979.6947
489.78790283203125 0 1954.5231
494.2362060546875 0 105010.05
495.238525390625 0 25838.838
496.24163818359375 0 4601.88
504.2003479003906 0 3401.214
508.2518005371094 0 15713.267
509.254638671875 0 3501.5005
511.2617492675781 0 1631.4224
512.2471313476562 0 5669.933
522.2066650390625 0 1025.4542
523.2402954101562 0 948.48004
527.3195190429688 0 2717.684
532.195556640625 0 2468.943
537.3023071289062 0 1374.9304
539.235595703125 0 4258.7065
540.2396240234375 0 981.7405
542.7996826171875 0 1039.0638
547.2861328125 0 1591.8381
548.3171997070312 0 2322.5845
548.3572387695312 0 3791.29
549.2194213867188 0 1458.5215
549.3601684570312 0 1849.1554
550.2049560546875 0 26193.758
551.2074584960938 0 7146.01
551.3291625976562 0 1317.1434
552.2108764648438 0 1025.3287
552.3131103515625 0 1141.8752
555.2677001953125 0 1161.0271
555.313720703125 0 10062.91
556.3173828125 0 5049.485
561.3134765625 0 4047.9985 b Water loss 4
562.3034057617188 0 1048.5107 b Ammonia loss 4
566.329833984375 0 5142.109
567.2315673828125 0 8778.738 y Water loss 3
567.3333740234375 0 1548.2595
568.2164916992188 0 9109.323 y Ammonia loss 3
569.2168579101562 0 2607.123
570.3251342773438 0 3808.3127
571.3262939453125 0 1169.0662
573.3244018554688 0 68874.46
574.3274536132812 0 31765.11
575.3302612304688 0 4575.937
576.2792358398438 0 2640.752
579.3252563476562 0 14863.863 b 4
580.3270874023438 0 4249.277
581.329833984375 0 1080.5966
585.2420043945312 0 89035.01 y 3
586.2448120117188 0 23975.902
587.2481079101562 0 4465.122
588.3375854492188 0 1298.8812
593.3047485351562 0 17847.148
594.3077392578125 0 6147.7163
595.2265014648438 0 6456.472
595.3104858398438 0 1153.394
596.221435546875 0 2421.367
597.2294921875 0 1005.6153
598.3148803710938 0 1402.1674
605.3364868164062 0 1009.77246
607.3206176757812 0 3509.949
608.32421875 0 1757.9019
610.3316040039062 0 1536.4276
617.2791748046875 0 1055.5292
622.36328125 0 1629.27
634.3095703125 0 1212.035
635.2979125976562 0 2450.739
636.3257446289062 0 2083.3076
637.3272094726562 0 1130.3639
645.2771606445312 0 1299.4368
645.3705444335938 0 1189.9049
649.3648071289062 0 2942.6226
650.3731079101562 0 1080.0647
652.320556640625 0 11514.624
653.3202514648438 0 3986.882
654.3255615234375 0 1174.8433
659.3172607421875 0 1457.4459
662.3056030273438 0 1795.539
663.2901611328125 0 11877.864
663.382568359375 0 11122.113
664.2913818359375 0 3838.0083
664.3851318359375 0 4384.1025
665.3897094726562 0 1564.057
667.3773803710938 0 11098.942
668.3803100585938 0 6192.0747
669.3846435546875 0 1240.5356
676.341064453125 0 9631.459 b Water loss 5
677.3281860351562 0 11354.886 b Ammonia loss 5
678.329833984375 0 3759.329
679.3301391601562 0 1126.1003
680.3153076171875 0 26484.445 y Water loss 2
681.3013916015625 0 46367.113 y Ammonia loss 2
682.3038330078125 0 16274.501
683.3062744140625 0 3521.5957
683.4081420898438 0 4063.83
684.409423828125 0 1468.6213
685.3911743164062 0 3505.6816
689.3115234375 0 1244.37
690.3123779296875 0 1070.9268
694.35205078125 0 97379.34 b 5
695.354736328125 0 36607.668
696.357177734375 0 7415.5312
698.3258666992188 0 306201.38 y 2
699.32861328125 0 109251.26
700.3309936523438 0 26397.443
701.3439331054688 0 1515.0615
701.4191284179688 0 12815.964
702.4223022460938 0 6635.187
703.4261474609375 0 2119.0815
708.3129272460938 0 1358.6443
709.296875 0 1172.5098
711.3840942382812 0 1517.9653
712.3624877929688 0 4460.446
713.3670654296875 0 1288.7192
721.3593139648438 0 1132.601
734.3635864257812 0 1418.3549
751.37890625 0 3166.8694
752.3923950195312 0 1103.0574
764.4307250976562 0 3023.7666
765.4352416992188 0 1854.9048
769.3818969726562 0 3353.7478
779.385009765625 0 2357.9768 y Water loss 1
780.3694458007812 0 7533.6685 y Ammonia loss 1
780.460693359375 0 6216.2183
781.3695678710938 0 3372.1655
781.46240234375 0 3748.148
782.4423217773438 0 6419.8794
783.4439086914062 0 2961.9119
784.4453735351562 0 1583.5063
797.3936767578125 0 34043.56 y 1
798.395751953125 0 14622.193
798.4722290039062 0 41349.176
799.3983764648438 0 3273.433
799.4749755859375 0 24806.559
800.4768676757812 0 9664.025
801.4774169921875 0 2043.459
807.3780517578125 0 1128.6047
1357.2733154296875 0 927.7292
1565.296630859375 0 1031.7548
2354.02490234375 0 1075.881
2805.151611328125 0 1149.5259
3084.649658203125 0 1078.9479
3325.41796875 0 1185.6544

Spectrum Details

|  |  |
| --- | --- |
| Matched peaks? Matched peaks The total absolute number of peaks matched. Additionally in brackets the total fraction of peaks matched and the total number of peaks is shown. | 38 (6.47% of 587) |
| FDR? FDR The false discovery rate estimated for this peptide. It is calculated by matching all theoretical fragments with a non-integer shift with the raw peaks for this spectrum. This is done with 40 different shifts. The resulting percentage is the average number of annotated peaks over the number of annotated peaks with the correct spectrum. | 4.32% |
| Satellite FDR? Satellite FDR See the FDR for details on its calculation. This satellite ion specific FDR only contains the satellite ions (d/w) for I/L/J positions. | - |
| PSM Score? PSM Score The PSM Score as given by Hecklib to this annotated spectrum. It is shown with three significant figures. | 554 |

#### Spectrum 5353? Spectrum 5353 The raw spectrum of this peptide as annotated by Hecklib. The fragments are coloured according to ion type (see legend). Any peaks with a star '\*' as text can be hovered over to see the full details, first the ion type second the mass shift type. By hovering over the amino acids in the peptide or ions in the legend the corresponding peaks are highlighted. By toggling the 'Unassigned' label you can turn the background (unassigned) peaks on or off in the plot. By updating the slider in the Ion legend you can update the spectrum to only show the top X% of the peaks with labels. The top X% means any peak that is within X% of the highest intensity. By dragging in the spectrum you can zoom in to a specific part of the spectrum and use 'Zoom Out' to get back to the original zoom level. The annotation of the spectrum is based on the given sequence in the peptides file and is done with different software so inconsistencies are likely. The peaks are annotated based on the given sequence, with 20 ppm tolerance.

Copy Data

##### Spectrum 5353 (TSV)

###### Preview

```
Loading example...
```

*Click on the button to copy the data to your clipboard.*

Mz MinMz MaxIntensity Max

WidthHeightPeptide font sizePeptide stroke widthSpectrum font sizeSpectrum stroke widthCompact peptide

Ion legend

wxyz

abcd

OtherUnassignedIon^Charge^_Position_Show for top:%

TVLHQDW

01.42e+62.85e+64.27e+65.70e+6

Zoom Out

y^+1^_1_y^+1^_2_c^+2^_6_y^+2^_6_y^+1^_3_y^+1^_3_z^+1^_3_y^+1^_3_c^+1^_4_c^+1^_4_z^+1^_4_y^+1^_4_y^+1^_4_z^+1^_4_c^+1^_5_c^+1^_5_y^+1^_4_c^+1^_5_w^+1^_5_y^+1^_5_y^+1^_5_z^+1^_5_c^+1^_6_y^+1^_5_c^+1^_6_w^+1^_6_y^+1^_6_y^+1^_6_z^+1^_6_y^+1^_6_

0514102915432057

Fragment Matches Table

Show background peaks

| Position | Ion type | Intensity | mz Theoretical | mz Error (Th) | mz Error (ppm) | Charge | Series Number |
| --- | --- | --- | --- | --- | --- | --- | --- |
| - | - | 4546 | 123.1 | - | - | 0 | - |
| - | - | 1.555E+04 | 128.1 | - | - | 0 | - |
| - | - | 1.687E+04 | 129.1 | - | - | 0 | - |
| - | - | 3.61E+04 | 130.1 | - | - | 0 | - |
| - | - | 6128 | 131.1 | - | - | 0 | - |
| - | - | 6311 | 132.1 | - | - | 0 | - |
| - | - | 4125 | 138.6 | - | - | 0 | - |
| - | - | 7729 | 146.1 | - | - | 0 | - |
| - | - | 4608 | 149 | - | - | 0 | - |
| - | - | 2.177E+05 | 155.1 | - | - | 0 | - |
| - | - | 3.259E+04 | 155.1 | - | - | 0 | - |
| - | - | 1.337E+04 | 156.1 | - | - | 0 | - |
| - | - | 1.893E+05 | 159.1 | - | - | 0 | - |
| - | - | 1.736E+04 | 160.1 | - | - | 0 | - |
| - | - | 8091 | 166.1 | - | - | 0 | - |
| - | - | 1.727E+06 | 173.1 | - | - | 0 | - |
| - | - | 1.475E+04 | 173.4 | - | - | 0 | - |
| - | - | 1.428E+05 | 174.1 | - | - | 0 | - |
| - | - | 5818 | 175.1 | - | - | 0 | - |
| - | - | 1.91E+04 | 183.1 | - | - | 0 | - |
| - | - | 8278 | 187.1 | - | - | 0 | - |
| - | - | 2.583E+05 | 188.1 | - | - | 0 | - |
| - | - | 2.965E+04 | 189.1 | - | - | 0 | - |
| - | - | 1.099E+04 | 194.1 | - | - | 0 | - |
| - | - | 4720 | 200.9 | - | - | 0 | - |
| - | - | 6.396E+05 | 201.1 | - | - | 0 | - |
| - | - | 5.837E+04 | 202.1 | - | - | 0 | - |
| 7 | y | 2.855E+05 | 205.1 | 0.0003496 | 1.704 | +1 | 1 |
| - | - | 8272 | 205.1 | - | - | 0 | - |
| - | - | 2.464E+04 | 206.1 | - | - | 0 | - |
| - | - | 9.331E+04 | 223.1 | - | - | 0 | - |
| - | - | 2.246E+04 | 223.2 | - | - | 0 | - |
| - | - | 8015 | 224.1 | - | - | 0 | - |
| - | - | 6485 | 233.1 | - | - | 0 | - |
| - | - | 2.731E+04 | 234.1 | - | - | 0 | - |
| - | - | 3.233E+04 | 244.1 | - | - | 0 | - |
| - | - | 6259 | 249.1 | - | - | 0 | - |
| - | - | 1.901E+04 | 250.1 | - | - | 0 | - |
| - | - | 1.382E+04 | 251.1 | - | - | 0 | - |
| - | - | 2.449E+05 | 251.2 | - | - | 0 | - |
| - | - | 2.585E+04 | 252.2 | - | - | 0 | - |
| - | - | 6913 | 260.1 | - | - | 0 | - |
| - | - | 4.23E+04 | 266.1 | - | - | 0 | - |
| - | - | 8385 | 267.1 | - | - | 0 | - |
| - | - | 1.624E+05 | 268.2 | - | - | 0 | - |
| - | - | 1.525E+04 | 269.2 | - | - | 0 | - |
| - | - | 1.358E+04 | 274.1 | - | - | 0 | - |
| - | - | 2.632E+04 | 283.2 | - | - | 0 | - |
| - | - | 6020 | 284.2 | - | - | 0 | - |
| - | - | 9920 | 285.1 | - | - | 0 | - |
| - | - | 7238 | 293.2 | - | - | 0 | - |
| - | - | 5902 | 301.9 | - | - | 0 | - |
| - | - | 4938 | 309.4 | - | - | 0 | - |
| - | - | 2.853E+04 | 314.2 | - | - | 0 | - |
| 6 | y | 7.314E+04 | 320.1 | 0.0005061 | 1.581 | +1 | 2 |
| - | - | 9481 | 321.1 | - | - | 0 | - |
| - | - | 1.57E+04 | 321.1 | - | - | 0 | - |
| - | - | 7867 | 322.1 | - | - | 0 | - |
| - | - | 1.274E+04 | 326.7 | - | - | 0 | - |
| - | - | 4878 | 335.5 | - | - | 0 | - |
| - | - | 6.848E+04 | 338.1 | - | - | 0 | - |
| - | - | 6956 | 338.7 | - | - | 0 | - |
| - | - | 1.063E+04 | 339.2 | - | - | 0 | - |
| - | - | 3.939E+04 | 339.2 | - | - | 0 | - |
| - | - | 8216 | 340.2 | - | - | 0 | - |
| 6 | c | 2.962E+04 | 347.7 | 0.0002387 | 0.6866 | +2 | 6 |
| - | - | 1.726E+04 | 350.2 | - | - | 0 | - |
| - | - | 5310 | 360 | - | - | 0 | - |
| - | - | 6731 | 360.7 | - | - | 0 | - |
| - | - | 5568 | 361 | - | - | 0 | - |
| - | - | 6302 | 361.2 | - | - | 0 | - |
| - | - | 7578 | 362 | - | - | 0 | - |
| - | - | 5348 | 362.2 | - | - | 0 | - |
| - | - | 5166 | 366.1 | - | - | 0 | - |
| - | - | 8793 | 367.2 | - | - | 0 | - |
| - | - | 6421 | 376.2 | - | - | 0 | - |
| - | - | 2.317E+04 | 378.2 | - | - | 0 | - |
| - | - | 1.935E+05 | 379.2 | - | - | 0 | - |
| - | - | 3.783E+04 | 380.2 | - | - | 0 | - |
| - | - | 8.235E+04 | 381.2 | - | - | 0 | - |
| - | - | 5847 | 381.2 | - | - | 0 | - |
| - | - | 1.065E+04 | 382.2 | - | - | 0 | - |
| - | - | 4.216E+04 | 383.2 | - | - | 0 | - |
| - | - | 7735 | 384.2 | - | - | 0 | - |
| - | - | 1.609E+04 | 396.2 | - | - | 0 | - |
| - | - | 6409 | 399.2 | - | - | 0 | - |
| 2 | y | 3.83E+04 | 399.2 | 0.0004709 | 1.18 | +2 | 6 |
| - | - | 1.262E+04 | 399.7 | - | - | 0 | - |
| - | - | 8397 | 400.2 | - | - | 0 | - |
| - | - | 5320 | 404.2 | - | - | 0 | - |
| - | - | 5749 | 405.3 | - | - | 0 | - |
| - | - | 6803 | 406.3 | - | - | 0 | - |
| - | - | 1.799E+04 | 408.3 | - | - | 0 | - |
| - | - | 6354 | 409.2 | - | - | 0 | - |
| - | - | 2.539E+04 | 409.2 | - | - | 0 | - |
| - | - | 5.45E+04 | 411.2 | - | - | 0 | - |
| - | - | 1.432E+04 | 412.2 | - | - | 0 | - |
| - | - | 1.212E+04 | 417.7 | - | - | 0 | - |
| - | - | 2.961E+04 | 418.2 | - | - | 0 | - |
| - | - | 1.503E+04 | 418.7 | - | - | 0 | - |
| - | - | 5344 | 419.3 | - | - | 0 | - |
| - | - | 6987 | 423.2 | - | - | 0 | - |
| - | - | 1.413E+05 | 423.3 | - | - | 0 | - |
| - | - | 1.007E+05 | 424.3 | - | - | 0 | - |
| - | - | 4.487E+04 | 425.3 | - | - | 0 | - |
| - | - | 6778 | 426.3 | - | - | 0 | - |
| - | - | 1.053E+05 | 426.7 | - | - | 0 | - |
| - | - | 4.862E+04 | 427.2 | - | - | 0 | - |
| - | - | 1.453E+04 | 427.7 | - | - | 0 | - |
| 5 | y | 9.564E+04 | 430.2 | 0.0004975 | 1.156 | +1 | 3 |
| 5 | y | 2.635E+04 | 431.2 | 0.0008264 | 1.917 | +1 | 3 |
| - | - | 1.441E+04 | 431.7 | - | - | 0 | - |
| 5 | z | 1.017E+04 | 432.2 | 0.003031 | 7.014 | +1 | 3 |
| - | - | 1.681E+04 | 432.2 | - | - | 0 | - |
| - | - | 1.15E+04 | 432.2 | - | - | 0 | - |
| - | - | 7008 | 432.7 | - | - | 0 | - |
| - | - | 1.009E+04 | 433.3 | - | - | 0 | - |
| - | - | 8019 | 437.3 | - | - | 0 | - |
| - | - | 5256 | 440.2 | - | - | 0 | - |
| - | - | 3.228E+04 | 440.7 | - | - | 0 | - |
| - | - | 1.967E+04 | 441.2 | - | - | 0 | - |
| - | - | 8079 | 441.7 | - | - | 0 | - |
| 5 | y | 1.948E+05 | 448.2 | 0.0007055 | 1.574 | +1 | 3 |
| - | - | 7.375E+04 | 448.3 | - | - | 0 | - |
| - | - | 5.693E+04 | 448.8 | - | - | 0 | - |
| - | - | 4.75E+04 | 449.2 | - | - | 0 | - |
| - | - | 1.447E+05 | 449.3 | - | - | 0 | - |
| - | - | 1.334E+05 | 449.7 | - | - | 0 | - |
| - | - | 4785 | 449.8 | - | - | 0 | - |
| - | - | 1.062E+04 | 450.2 | - | - | 0 | - |
| - | - | 5.444E+04 | 450.2 | - | - | 0 | - |
| 4 | c | 4.561E+04 | 450.3 | 0.001445 | 3.21 | +1 | 4 |
| - | - | 2.723E+04 | 450.7 | - | - | 0 | - |
| - | - | 3.835E+05 | 451.3 | - | - | 0 | - |
| - | - | 9.453E+04 | 452.3 | - | - | 0 | - |
| - | - | 6450 | 453.3 | - | - | 0 | - |
| - | - | 1.358E+05 | 467.3 | - | - | 0 | - |
| 4 | c | 4.325E+06 | 468.3 | 0.0007158 | 1.529 | +1 | 4 |
| - | - | 9.948E+05 | 469.3 | - | - | 0 | - |
| - | - | 1.369E+05 | 470.3 | - | - | 0 | - |
| - | - | 1.004E+04 | 477.2 | - | - | 0 | - |
| - | - | 1.672E+04 | 478.3 | - | - | 0 | - |
| - | - | 2.558E+05 | 494.2 | - | - | 0 | - |
| - | - | 9637 | 494.3 | - | - | 0 | - |
| - | - | 5.569E+04 | 495.2 | - | - | 0 | - |
| - | - | 1.531E+04 | 496.2 | - | - | 0 | - |
| - | - | 3.262E+04 | 498.2 | - | - | 0 | - |
| - | - | 1.415E+05 | 511.2 | - | - | 0 | - |
| - | - | 1.062E+04 | 511.3 | - | - | 0 | - |
| - | - | 4.193E+04 | 512.2 | - | - | 0 | - |
| - | - | 8325 | 512.2 | - | - | 0 | - |
| - | - | 7291 | 513.2 | - | - | 0 | - |
| - | - | 2.111E+04 | 525.2 | - | - | 0 | - |
| - | - | 1.272E+04 | 526.2 | - | - | 0 | - |
| - | - | 2.213E+04 | 536.3 | - | - | 0 | - |
| - | - | 6551 | 537.3 | - | - | 0 | - |
| - | - | 9788 | 539.2 | - | - | 0 | - |
| - | - | 1.305E+04 | 550.2 | - | - | 0 | - |
| 4 | z | 1.503E+04 | 551.2 | 0.0001046 | 0.1898 | +1 | 4 |
| - | - | 2.651E+04 | 551.3 | - | - | 0 | - |
| - | - | 5680 | 552.2 | - | - | 0 | - |
| - | - | 1.566E+04 | 552.3 | - | - | 0 | - |
| - | - | 6698 | 561.3 | - | - | 0 | - |
| - | - | 5917 | 564.4 | - | - | 0 | - |
| - | - | 6.842E+04 | 565.4 | - | - | 0 | - |
| - | - | 3.946E+04 | 566.4 | - | - | 0 | - |
| 4 | y | 8680 | 567.2 | 0.002811 | 4.956 | +1 | 4 |
| 4 | y | 1.46E+04 | 568.2 | 0.0005599 | 0.9853 | +1 | 4 |
| 4 | z | 2.141E+05 | 569.2 | 0.0002821 | 0.4956 | +1 | 4 |
| - | - | 9.315E+04 | 570.2 | - | - | 0 | - |
| - | - | 2.148E+04 | 571.2 | - | - | 0 | - |
| - | - | 6200 | 573.3 | - | - | 0 | - |
| - | - | 1.038E+04 | 577.3 | - | - | 0 | - |
| 5 | c | 7235 | 578.3 | 0.001734 | 2.999 | +1 | 5 |
| 5 | c | 1.744E+05 | 579.3 | 0.0002731 | 0.4714 | +1 | 5 |
| - | - | 6.423E+04 | 580.3 | - | - | 0 | - |
| - | - | 1.06E+04 | 581.3 | - | - | 0 | - |
| 4 | y | 2.664E+05 | 585.2 | 0.0004789 | 0.8183 | +1 | 4 |
| - | - | 6.424E+04 | 586.2 | - | - | 0 | - |
| - | - | 1.136E+04 | 587.2 | - | - | 0 | - |
| - | - | 3.307E+04 | 593.3 | - | - | 0 | - |
| - | - | 9265 | 594.3 | - | - | 0 | - |
| - | - | 1.16E+04 | 595.2 | - | - | 0 | - |
| 5 | c | 3.118E+06 | 596.4 | 0.0005184 | 0.8694 | +1 | 5 |
| - | - | 9.067E+05 | 597.4 | - | - | 0 | - |
| - | - | 1.782E+05 | 598.4 | - | - | 0 | - |
| - | - | 5599 | 599.4 | - | - | 0 | - |
| - | - | 2.006E+04 | 609.3 | - | - | 0 | - |
| - | - | 6419 | 612.2 | - | - | 0 | - |
| - | - | 9775 | 622.4 | - | - | 0 | - |
| - | - | 6016 | 624.3 | - | - | 0 | - |
| 3 | w | 2.591E+05 | 639.3 | 0.0004123 | 0.645 | +1 | 5 |
| - | - | 7.243E+04 | 640.3 | - | - | 0 | - |
| - | - | 1.492E+04 | 641.3 | - | - | 0 | - |
| - | - | 8.865E+04 | 651.3 | - | - | 0 | - |
| - | - | 3.934E+04 | 652.3 | - | - | 0 | - |
| - | - | 9371 | 653.3 | - | - | 0 | - |
| - | - | 2.591E+04 | 653.4 | - | - | 0 | - |
| - | - | 9254 | 654.4 | - | - | 0 | - |
| - | - | 9156 | 662.3 | - | - | 0 | - |
| - | - | 2.184E+04 | 663.3 | - | - | 0 | - |
| - | - | 1.047E+04 | 666.4 | - | - | 0 | - |
| - | - | 5.852E+04 | 667.4 | - | - | 0 | - |
| - | - | 2.591E+04 | 668.4 | - | - | 0 | - |
| - | - | 2.481E+04 | 669.3 | - | - | 0 | - |
| - | - | 1.057E+04 | 670.3 | - | - | 0 | - |
| - | - | 3.484E+04 | 676.3 | - | - | 0 | - |
| - | - | 3.891E+04 | 677.3 | - | - | 0 | - |
| - | - | 2.075E+04 | 678.3 | - | - | 0 | - |
| - | - | 2.202E+04 | 679.4 | - | - | 0 | - |
| 3 | y | 8.776E+04 | 680.3 | 0.0002829 | 0.4158 | +1 | 5 |
| - | - | 4.043E+04 | 680.4 | - | - | 0 | - |
| 3 | y | 1.518E+05 | 681.3 | 0.002656 | 3.899 | +1 | 5 |
| - | - | 2.356E+04 | 681.4 | - | - | 0 | - |
| 3 | z | 3.849E+05 | 682.3 | 0.0004689 | 0.6872 | +1 | 5 |
| - | - | 1.347E+05 | 683.3 | - | - | 0 | - |
| - | - | 3.203E+04 | 684.3 | - | - | 0 | - |
| - | - | 9998 | 686.4 | - | - | 0 | - |
| 6 | c | 3.535E+05 | 694.4 | 0.0001856 | 0.2672 | +1 | 6 |
| - | - | 1.335E+05 | 695.4 | - | - | 0 | - |
| - | - | 5.159E+04 | 696.4 | - | - | 0 | - |
| - | - | 1.222E+04 | 697.4 | - | - | 0 | - |
| 3 | y | 1.48E+06 | 698.3 | 0.0002773 | 0.397 | +1 | 5 |
| - | - | 5.078E+05 | 699.3 | - | - | 0 | - |
| - | - | 1.216E+05 | 700.3 | - | - | 0 | - |
| - | - | 6236 | 710.4 | - | - | 0 | - |
| 6 | c | 1.975E+06 | 711.4 | 0.0001257 | 0.1767 | +1 | 6 |
| - | - | 7.199E+05 | 712.4 | - | - | 0 | - |
| - | - | 1.09E+04 | 712.5 | - | - | 0 | - |
| - | - | 1.667E+05 | 713.4 | - | - | 0 | - |
| - | - | 7595 | 714.4 | - | - | 0 | - |
| - | - | 2.108E+04 | 722.4 | - | - | 0 | - |
| - | - | 9682 | 723.4 | - | - | 0 | - |
| - | - | 1.378E+05 | 725.3 | - | - | 0 | - |
| - | - | 4.61E+04 | 726.3 | - | - | 0 | - |
| - | - | 1.421E+04 | 727.3 | - | - | 0 | - |
| - | - | 5581 | 727.4 | - | - | 0 | - |
| - | - | 1.06E+04 | 737.4 | - | - | 0 | - |
| - | - | 1.277E+04 | 738.3 | - | - | 0 | - |
| - | - | 9437 | 748.4 | - | - | 0 | - |
| - | - | 9437 | 751.4 | - | - | 0 | - |
| - | - | 3.191E+04 | 753.4 | - | - | 0 | - |
| 2 | w | 4.924E+04 | 766.4 | 0.009097 | 11.87 | +1 | 6 |
| - | - | 1.334E+04 | 767.4 | - | - | 0 | - |
| - | - | 1.922E+05 | 768.4 | - | - | 0 | - |
| - | - | 8.063E+04 | 769.4 | - | - | 0 | - |
| - | - | 2.717E+04 | 770.4 | - | - | 0 | - |
| - | - | 9999 | 777.4 | - | - | 0 | - |
| 2 | y | 1.159E+04 | 779.4 | 0.0001989 | 0.2552 | +1 | 6 |
| 2 | y | 2.271E+04 | 780.4 | 0.003701 | 4.742 | +1 | 6 |
| 2 | z | 2.573E+05 | 781.4 | 0.0001572 | 0.2012 | +1 | 6 |
| - | - | 3.925E+04 | 781.5 | - | - | 0 | - |
| - | - | 9.155E+04 | 782.4 | - | - | 0 | - |
| - | - | 1.445E+04 | 782.5 | - | - | 0 | - |
| - | - | 2.749E+04 | 783.4 | - | - | 0 | - |
| - | - | 2.06E+04 | 794.4 | - | - | 0 | - |
| - | - | 8.535E+04 | 795.4 | - | - | 0 | - |
| - | - | 2.966E+04 | 796.4 | - | - | 0 | - |
| 2 | y | 1.621E+05 | 797.4 | 0.0002656 | 0.333 | +1 | 6 |
| - | - | 5.622E+04 | 798.4 | - | - | 0 | - |
| - | - | 1.677E+04 | 799.4 | - | - | 0 | - |
| - | - | 8232 | 807.4 | - | - | 0 | - |
| - | - | 6284 | 810.4 | - | - | 0 | - |
| - | - | 2.905E+04 | 811.4 | - | - | 0 | - |
| - | - | 1.343E+04 | 812.4 | - | - | 0 | - |
| - | - | 9266 | 817.4 | - | - | 0 | - |
| - | - | 1.757E+04 | 818.4 | - | - | 0 | - |
| - | - | 9966 | 820.4 | - | - | 0 | - |
| - | - | 7.683E+04 | 824.4 | - | - | 0 | - |
| - | - | 2.863E+04 | 825.4 | - | - | 0 | - |
| - | - | 1.882E+04 | 826.4 | - | - | 0 | - |
| - | - | 1.095E+04 | 827.4 | - | - | 0 | - |
| - | - | 1.207E+04 | 837.4 | - | - | 0 | - |
| - | - | 1.876E+05 | 838.4 | - | - | 0 | - |
| - | - | 7.226E+05 | 839.4 | - | - | 0 | - |
| - | - | 3.201E+05 | 840.4 | - | - | 0 | - |
| - | - | 7.005E+04 | 841.4 | - | - | 0 | - |
| - | - | 2.161E+04 | 853.4 | - | - | 0 | - |
| - | - | 3.182E+04 | 854.4 | - | - | 0 | - |
| - | - | 1.637E+04 | 855.4 | - | - | 0 | - |
| - | - | 8438 | 864.4 | - | - | 0 | - |
| - | - | 1.023E+05 | 865.4 | - | - | 0 | - |
| - | - | 4.89E+04 | 866.4 | - | - | 0 | - |
| - | - | 1.529E+05 | 867.4 | - | - | 0 | - |
| - | - | 7.799E+04 | 868.4 | - | - | 0 | - |
| - | - | 2.184E+04 | 869.4 | - | - | 0 | - |
| - | - | 1.07E+04 | 879.5 | - | - | 0 | - |
| - | - | 2.014E+04 | 880.4 | - | - | 0 | - |
| - | - | 6525 | 880.5 | - | - | 0 | - |
| - | - | 4.283E+04 | 881.4 | - | - | 0 | - |
| - | - | 3.078E+06 | 882.4 | - | - | 0 | - |
| - | - | 1.367E+06 | 883.4 | - | - | 0 | - |
| - | - | 4.195E+05 | 884.4 | - | - | 0 | - |
| - | - | 3817 | 884.5 | - | - | 0 | - |
| - | - | 3.086E+04 | 885.4 | - | - | 0 | - |
| - | - | 2.948E+04 | 895.5 | - | - | 0 | - |
| - | - | 1.27E+05 | 896.5 | - | - | 0 | - |
| - | - | 9.328E+04 | 897.5 | - | - | 0 | - |
| - | - | 2.003E+06 | 898.4 | - | - | 0 | - |
| - | - | 5.642E+06 | 899.4 | - | - | 0 | - |
| - | - | 2.449E+06 | 900.5 | - | - | 0 | - |
| - | - | 6.769E+05 | 901.5 | - | - | 0 | - |
| - | - | 5.595E+04 | 902.5 | - | - | 0 | - |
| - | - | 6233 | 911.7 | - | - | 0 | - |
| - | - | 7515 | 914.4 | - | - | 0 | - |
| - | - | 5506 | 1295 | - | - | 0 | - |
| - | - | 7079 | 1335 | - | - | 0 | - |
| - | - | 7954 | 2037 | - | - | 0 | - |

m/z Charge Intensity FragmentType MassShift Position
123.05580139160156 0 4545.919
128.10733032226562 0 15553.532
129.10255432128906 0 16871.115
130.06546020507812 0 36101.87
131.06878662109375 0 6127.603
132.08119201660156 0 6311.409
138.5928955078125 0 4125.0635
146.06044006347656 0 7729.373
149.0234832763672 0 4607.647
155.0930938720703 0 217686.6
155.11822509765625 0 32590.938
156.09652709960938 0 13366.894
159.09205627441406 0 189284.66
160.09527587890625 0 17359.572
166.0616912841797 0 8091.259
173.12887573242188 0 1726740
173.439208984375 0 14748.281
174.1322021484375 0 142787.9
175.1344757080078 0 5817.8003
183.11306762695312 0 19096.986
187.0869598388672 0 8278.048
188.07093811035156 0 258290.28
189.07432556152344 0 29646.79
194.09275817871094 0 10992.806
200.8511962890625 0 4719.569
201.12368774414062 0 639648.9
202.1271514892578 0 58366.24
205.09750366210938 0 285474.9 y 6
205.10861206054688 0 8271.769
206.10101318359375 0 24638.994
223.1193389892578 0 93313.89
223.1554718017578 0 22464.475
224.12611389160156 0 8014.9497
233.1405029296875 0 6484.834
234.12411499023438 0 27308.977
244.09303283691406 0 32329.72
249.0974884033203 0 6258.5645
250.1062774658203 0 19011.871
251.114013671875 0 13822.947
251.15072631835938 0 244925.02
252.15451049804688 0 25853.871
260.1033020019531 0 6913.3003
266.125 0 42301.004
267.10894775390625 0 8385.066
268.17706298828125 0 162402.1
269.1802978515625 0 15246.344
274.1185607910156 0 13579.699
283.15167236328125 0 26318.25
284.15472412109375 0 6020.253
285.0875244140625 0 9919.538
293.1617126464844 0 7237.729
301.9461364746094 0 5902.3022
309.3648986816406 0 4938.265
314.2083435058594 0 28530.67
320.1246032714844 0 73135.81 y 5
321.1256103515625 0 9480.543
321.1440734863281 0 15699.7705
322.1497497558594 0 7867.255
326.6634521484375 0 12736.802
335.457275390625 0 4877.8096
338.1463928222656 0 68480.21
338.6746520996094 0 6955.6387
339.1507568359375 0 10627.422
339.17779541015625 0 39388.207
340.17974853515625 0 8216.295
347.6798095703125 0 29621.973 c Ammonia loss 5
350.218994140625 0 17258.682
360.0306701660156 0 5310.29
360.696533203125 0 6730.5225
361.0264892578125 0 5567.518
361.198486328125 0 6301.628
362.02734375 0 7577.87
362.18194580078125 0 5348.1143
366.1405029296875 0 5166.033
367.2464904785156 0 8793.183
376.19781494140625 0 6420.628
378.2139587402344 0 23172.506
379.2092590332031 0 193505
380.21282958984375 0 37826.02
381.1520690917969 0 82348.5
381.215087890625 0 5846.542
382.1550598144531 0 10651.068
383.1676025390625 0 42163.336
384.1698913574219 0 7735.0273
396.2358093261719 0 16094.966
399.173828125 0 6409.152
399.2011413574219 0 38304.375 y 1
399.7035827636719 0 12621.477
400.2048034667969 0 8397.182
404.2084655761719 0 5320.2837
405.2620849609375 0 5749.2793
406.27069091796875 0 6802.97
408.2602233886719 0 17989.281
409.1505432128906 0 6353.913
409.18353271484375 0 25385.605
411.22296142578125 0 54495.97
412.2262878417969 0 14324.341
417.71826171875 0 12118.339
418.2092590332031 0 29614.416
418.7102966308594 0 15025.815
419.2530822753906 0 5344.026
423.2134704589844 0 6987.4087
423.27178955078125 0 141262.92
424.27862548828125 0 100726.73
425.28546142578125 0 44866.17
426.2872619628906 0 6778.321
426.7222595214844 0 105285.54
427.22332763671875 0 48616.902
427.7233581542969 0 14530.78
430.172607421875 0 95640.195 y Water loss 4
431.1569519042969 0 26348.027 y Ammonia loss 4
431.71466064453125 0 14412.926
432.1609191894531 0 10170.787 z 4
432.2117919921875 0 16808.309
432.248046875 0 11501.352
432.7056579589844 0 7007.604
433.2546691894531 0 10089.056
437.287353515625 0 8018.76
440.1658020019531 0 5255.855
440.71923828125 0 32276.562
441.21343994140625 0 19673.465
441.71820068359375 0 8079.1274
448.1833801269531 0 194812.78 y 4
448.2667236328125 0 73754.414
448.76763916015625 0 56932.766
449.1860046386719 0 47503.188
449.27447509765625 0 144667.75
449.7250061035156 0 133391.83
449.7572937011719 0 4785.4473
450.1907958984375 0 10615.51
450.2262268066406 0 54439.465
450.2808837890625 0 45608.34 c Water loss 3
450.72821044921875 0 27225.412
451.26702880859375 0 383460.6
452.2698059082031 0 94533.15
453.27215576171875 0 6449.933
467.2858581542969 0 135846.44
468.2936096191406 0 4324691 c 3
469.2963562011719 0 994762.9
470.2991638183594 0 136913.4
477.2073669433594 0 10039.586
478.2785339355469 0 16721.176
494.2362060546875 0 255797.69
494.3065185546875 0 9636.875
495.2391357421875 0 55694.97
496.2465515136719 0 15305.027
498.1853942871094 0 32616.338
511.1940612792969 0 141513.73
511.2611999511719 0 10616.021
512.1968383789062 0 41930.64
512.2422485351562 0 8325.178
513.2008056640625 0 7290.5396
525.2328491210938 0 21106.434
526.2359008789062 0 12718.675
536.3192749023438 0 22131.576
537.3247680664062 0 6550.5273
539.2356567382812 0 9787.804
550.2073364257812 0 13046.514
551.21240234375 0 15033.492 z Water loss 3
551.3300170898438 0 26514.484
552.22021484375 0 5679.848
552.3361206054688 0 15659.031
561.3154296875 0 6698.074
564.3712158203125 0 5916.937
565.3820190429688 0 68419.71
566.3853149414062 0 39456.37
567.2282104492188 0 8680.122 y Water loss 3
568.2144775390625 0 14604.535 y Ammonia loss 3
569.22314453125 0 214148.05 z 3
570.2283325195312 0 93153.234
571.231201171875 0 21479.078
573.3218994140625 0 6199.722
577.335205078125 0 10380.393
578.3391723632812 0 7235.4033 c Water loss 4
579.3251953125 0 174446.4 c Ammonia loss 4
580.3284301757812 0 64228.508
581.3287963867188 0 10600.598
585.2420654296875 0 266367.8 y 3
586.24462890625 0 64240.242
587.244873046875 0 11357.078
593.305908203125 0 33073.016
594.3057250976562 0 9264.896
595.2271118164062 0 11599.524
596.3519897460938 0 3117641.8 c 4
597.3545532226562 0 906719.5
598.3567504882812 0 178220.3
599.359619140625 0 5599.2217
609.3351440429688 0 20059.115
612.2322387695312 0 6419.3325
622.3682250976562 0 9775.165
624.271728515625 0 6016.308
639.2525634765625 0 259088.81 w 2
640.2549438476562 0 72425.74
641.2560424804688 0 14918.547
651.3466796875 0 88652.336
652.3223266601562 0 39340.95
653.31591796875 0 9370.959
653.3717651367188 0 25914.268
654.3775024414062 0 9253.627
662.3043823242188 0 9155.586
663.2904663085938 0 21839.604
666.358642578125 0 10473.394
667.3651123046875 0 58524.477
668.3667602539062 0 25913.93
669.2503662109375 0 24814.79
670.2508544921875 0 10570.027
676.3417358398438 0 34837.89
677.3281860351562 0 38908.234
678.3300170898438 0 20748.607
679.386962890625 0 22024.607
680.3153686523438 0 87759 y Water loss 2
680.4077758789062 0 40425.64
681.3017578125 0 151812.72 y Ammonia loss 2
681.4118041992188 0 23556.055
682.3064575195312 0 384884.88 z 2
683.3096313476562 0 134664.6
684.312744140625 0 32031.66
686.3995971679688 0 9997.517
694.35205078125 0 353546.06 c Ammonia loss 5
695.3546142578125 0 133465.17
696.36328125 0 51593.59
697.3705444335938 0 12218.188
698.325927734375 0 1480191.2 y 2
699.3287353515625 0 507763.97
700.3314208984375 0 121595.43
710.3693237304688 0 6236.4604
711.3785400390625 0 1974755.2 c 5
712.3809204101562 0 719871.25
712.4560546875 0 10899.468
713.38330078125 0 166655.44
714.3787231445312 0 7595.256
722.3842163085938 0 21077.389
723.381103515625 0 9681.503
725.3125610351562 0 137829.92
726.3154907226562 0 46103.953
727.3200073242188 0 14210.027
727.4228515625 0 5581.245
737.38671875 0 10598.466
738.3208618164062 0 12768.508
748.3583374023438 0 9436.733
751.3753662109375 0 9436.863
753.3661499023438 0 31909.895
766.3609619140625 0 49240.08 w 1
767.3666381835938 0 13338.668
768.3756103515625 0 192245.08
769.38037109375 0 80627.4
770.3819580078125 0 27171.82
777.4288330078125 0 9998.859
779.38330078125 0 11590.266 y Water loss 1
780.3712158203125 0 22711.914 y Ammonia loss 1
781.3751831054688 0 257310.64 z 1
781.4560546875 0 39245.11
782.37841796875 0 91551.67
782.4592895507812 0 14452.786
783.3807373046875 0 27491.012
794.4422607421875 0 20598.867
795.43994140625 0 85353.195
796.4407958984375 0 29664.234
797.393798828125 0 162083.9 y 1
798.39697265625 0 56221.867
799.3997802734375 0 16771.219
807.3756103515625 0 8232.441
810.4415283203125 0 6283.6274
811.3850708007812 0 29052.012
812.3904418945312 0 13429.847
817.40283203125 0 9266.288
818.4020385742188 0 17567.012
820.4237670898438 0 9966.482
824.3924560546875 0 76827.11
825.39599609375 0 28625.047
826.3627319335938 0 18816.615
827.3650512695312 0 10949.896
837.4469604492188 0 12070.043
838.43310546875 0 187615.77
839.4290771484375 0 722593.1
840.4315795898438 0 320141.34
841.4349365234375 0 70051.65
853.443603515625 0 21610.398
854.4356079101562 0 31816.828
855.4461059570312 0 16374.757
864.4083251953125 0 8438.377
865.419677734375 0 102349.86
866.422119140625 0 48897.605
867.4005126953125 0 152905.88
868.4033813476562 0 77986.15
869.4037475585938 0 21840.018
879.5087280273438 0 10701.083
880.3927001953125 0 20136.451
880.5044555664062 0 6525.3296
881.4344482421875 0 42825.004
882.4230346679688 0 3078338.8
883.425537109375 0 1366591.9
884.4280395507812 0 419522.6
884.5064697265625 0 3816.8555
885.4310302734375 0 30859.598
895.5242309570312 0 29479.67
896.531005859375 0 127029.125
897.5357055664062 0 93280.266
898.4415283203125 0 2003460.8
899.4483642578125 0 5642037
900.4515380859375 0 2448587
901.4547119140625 0 676936.6
902.45703125 0 55953.027
911.6732788085938 0 6233.1494
914.4098510742188 0 7515.3467
1294.5389404296875 0 5505.691
1335.3045654296875 0 7078.79
2036.83447265625 0 7954.2603

Spectrum Details

|  |  |
| --- | --- |
| Matched peaks? Matched peaks The total absolute number of peaks matched. Additionally in brackets the total fraction of peaks matched and the total number of peaks is shown. | 30 (9.74% of 308) |
| FDR? FDR The false discovery rate estimated for this peptide. It is calculated by matching all theoretical fragments with a non-integer shift with the raw peaks for this spectrum. This is done with 40 different shifts. The resulting percentage is the average number of annotated peaks over the number of annotated peaks with the correct spectrum. | 5.95% |
| Satellite FDR? Satellite FDR See the FDR for details on its calculation. This satellite ion specific FDR only contains the satellite ions (d/w) for I/L/J positions. | 2.38% |
| PSM Score? PSM Score The PSM Score as given by Hecklib to this annotated spectrum. It is shown with three significant figures. | 408 |

#### Spectrum 5494? Spectrum 5494 The raw spectrum of this peptide as annotated by Hecklib. The fragments are coloured according to ion type (see legend). Any peaks with a star '\*' as text can be hovered over to see the full details, first the ion type second the mass shift type. By hovering over the amino acids in the peptide or ions in the legend the corresponding peaks are highlighted. By toggling the 'Unassigned' label you can turn the background (unassigned) peaks on or off in the plot. By updating the slider in the Ion legend you can update the spectrum to only show the top X% of the peaks with labels. The top X% means any peak that is within X% of the highest intensity. By dragging in the spectrum you can zoom in to a specific part of the spectrum and use 'Zoom Out' to get back to the original zoom level. The annotation of the spectrum is based on the given sequence in the peptides file and is done with different software so inconsistencies are likely. The peaks are annotated based on the given sequence, with 20 ppm tolerance.

Copy Data

##### Spectrum 5494 (TSV)

###### Preview

```
Loading example...
```

*Click on the button to copy the data to your clipboard.*

Mz MinMz MaxIntensity Max

WidthHeightPeptide font sizePeptide stroke widthSpectrum font sizeSpectrum stroke widthCompact peptide

Ion legend

wxyz

abcd

OtherUnassignedIon^Charge^_Position_Show for top:%

TVLHQDW

01.40e+52.80e+54.20e+55.60e+5

Zoom Out

a^+1^_2_d^+1^_2_a^+1^_2_b^+1^_2_b^+1^_2_y^+1^_1_y^+2^_4_y^+1^_2_b^+1^_3_y^+1^_2_b^+2^_6_b^+2^_6_y^+2^_6_y^+1^_3_y^+1^_3_b^+1^_4_\*\*y^+1^_3_\*b^+1^_4_b^+1^_5_y^+1^_4_y^+1^_4_b^+1^_5_y^+1^_4_b^+1^_6_b^+1^_6_y^+1^_5_y^+1^_5_b^+1^_6_y^+1^_5_y^+1^_6_y^+1^_6_

0882176326453526

Fragment Matches Table

Show background peaks

| Position | Ion type | Intensity | mz Theoretical | mz Error (Th) | mz Error (ppm) | Charge | Series Number |
| --- | --- | --- | --- | --- | --- | --- | --- |
| - | - | 1.017E+04 | 120.1 | - | - | 0 | - |
| - | - | 786.8 | 120.5 | - | - | 0 | - |
| - | - | 1273 | 121.1 | - | - | 0 | - |
| - | - | 2274 | 122.1 | - | - | 0 | - |
| - | - | 1509 | 123.1 | - | - | 0 | - |
| - | - | 912.4 | 125.1 | - | - | 0 | - |
| - | - | 1.025E+04 | 126.1 | - | - | 0 | - |
| - | - | 1783 | 127.1 | - | - | 0 | - |
| - | - | 1023 | 128.1 | - | - | 0 | - |
| - | - | 4.288E+04 | 128.1 | - | - | 0 | - |
| - | - | 1748 | 129.1 | - | - | 0 | - |
| - | - | 1.619E+05 | 129.1 | - | - | 0 | - |
| - | - | 2.677E+04 | 130.1 | - | - | 0 | - |
| - | - | 1.011E+04 | 130.1 | - | - | 0 | - |
| - | - | 2535 | 131.1 | - | - | 0 | - |
| - | - | 1.196E+04 | 132.1 | - | - | 0 | - |
| - | - | 1.38E+04 | 132.1 | - | - | 0 | - |
| - | - | 870.4 | 132.1 | - | - | 0 | - |
| - | - | 993 | 133.1 | - | - | 0 | - |
| - | - | 1087 | 133.1 | - | - | 0 | - |
| - | - | 4096 | 136.1 | - | - | 0 | - |
| - | - | 817.4 | 136.3 | - | - | 0 | - |
| - | - | 4467 | 138.1 | - | - | 0 | - |
| - | - | 2691 | 138.1 | - | - | 0 | - |
| - | - | 2577 | 139.1 | - | - | 0 | - |
| - | - | 1112 | 140.1 | - | - | 0 | - |
| - | - | 959.5 | 140.1 | - | - | 0 | - |
| - | - | 2829 | 141.1 | - | - | 0 | - |
| - | - | 1174 | 142.1 | - | - | 0 | - |
| - | - | 1772 | 143 | - | - | 0 | - |
| - | - | 3748 | 144.1 | - | - | 0 | - |
| - | - | 2966 | 144.1 | - | - | 0 | - |
| - | - | 2.221E+04 | 146.1 | - | - | 0 | - |
| - | - | 2000 | 147.1 | - | - | 0 | - |
| - | - | 1596 | 149 | - | - | 0 | - |
| - | - | 2074 | 150.1 | - | - | 0 | - |
| - | - | 1264 | 151 | - | - | 0 | - |
| - | - | 1120 | 151.1 | - | - | 0 | - |
| - | - | 1341 | 154.1 | - | - | 0 | - |
| - | - | 2125 | 155.1 | - | - | 0 | - |
| - | - | 2.495E+04 | 155.1 | - | - | 0 | - |
| 2 | a | 1.782E+04 | 155.1 | 0.0002898 | 1.868 | +1 | 2 |
| - | - | 1791 | 156.1 | - | - | 0 | - |
| - | - | 1139 | 156.1 | - | - | 0 | - |
| - | - | 881.3 | 156.1 | - | - | 0 | - |
| - | - | 8827 | 157.1 | - | - | 0 | - |
| - | - | 4957 | 157.1 | - | - | 0 | - |
| - | - | 1134 | 158.1 | - | - | 0 | - |
| - | - | 1308 | 158.1 | - | - | 0 | - |
| - | - | 8.848E+04 | 159.1 | - | - | 0 | - |
| 2 | d | 7068 | 159.1 | 0.0003245 | 2.039 | +1 | 2 |
| - | - | 1.019E+04 | 160.1 | - | - | 0 | - |
| - | - | 1037 | 160.1 | - | - | 0 | - |
| - | - | 1418 | 165.1 | - | - | 0 | - |
| - | - | 1500 | 165.1 | - | - | 0 | - |
| - | - | 3.942E+04 | 166.1 | - | - | 0 | - |
| - | - | 1913 | 166.1 | - | - | 0 | - |
| - | - | 1702 | 166.1 | - | - | 0 | - |
| - | - | 1919 | 167.1 | - | - | 0 | - |
| - | - | 2933 | 167.1 | - | - | 0 | - |
| - | - | 4621 | 167.1 | - | - | 0 | - |
| - | - | 5301 | 167.1 | - | - | 0 | - |
| - | - | 1394 | 168.1 | - | - | 0 | - |
| - | - | 1505 | 169.1 | - | - | 0 | - |
| - | - | 1572 | 169.1 | - | - | 0 | - |
| - | - | 1121 | 170.1 | - | - | 0 | - |
| - | - | 4158 | 170.1 | - | - | 0 | - |
| - | - | 1306 | 171.1 | - | - | 0 | - |
| 2 | a | 5.544E+05 | 173.1 | 0.0004215 | 2.435 | +1 | 2 |
| - | - | 4292 | 173.4 | - | - | 0 | - |
| - | - | 4.354E+04 | 174.1 | - | - | 0 | - |
| - | - | 7237 | 175.1 | - | - | 0 | - |
| - | - | 1570 | 175.1 | - | - | 0 | - |
| - | - | 1117 | 175.1 | - | - | 0 | - |
| - | - | 1.884E+04 | 178.1 | - | - | 0 | - |
| - | - | 1475 | 179.1 | - | - | 0 | - |
| - | - | 1677 | 179.1 | - | - | 0 | - |
| - | - | 6136 | 181.1 | - | - | 0 | - |
| - | - | 1877 | 181.1 | - | - | 0 | - |
| 2 | b | 7908 | 183.1 | 0.0002177 | 1.189 | +1 | 2 |
| - | - | 3547 | 185.1 | - | - | 0 | - |
| - | - | 6017 | 185.1 | - | - | 0 | - |
| - | - | 3102 | 185.2 | - | - | 0 | - |
| - | - | 4979 | 187.1 | - | - | 0 | - |
| - | - | 2325 | 187.1 | - | - | 0 | - |
| - | - | 3174 | 187.1 | - | - | 0 | - |
| - | - | 1.52E+05 | 188.1 | - | - | 0 | - |
| - | - | 1.622E+04 | 189.1 | - | - | 0 | - |
| - | - | 6583 | 189.1 | - | - | 0 | - |
| - | - | 5062 | 190.1 | - | - | 0 | - |
| - | - | 1381 | 194.1 | - | - | 0 | - |
| - | - | 5979 | 195.1 | - | - | 0 | - |
| - | - | 1789 | 197.1 | - | - | 0 | - |
| - | - | 7353 | 198.1 | - | - | 0 | - |
| - | - | 5454 | 199.1 | - | - | 0 | - |
| - | - | 2966 | 199.1 | - | - | 0 | - |
| 2 | b | 9.586E+04 | 201.1 | 0.0002579 | 1.282 | +1 | 2 |
| - | - | 9379 | 202.1 | - | - | 0 | - |
| - | - | 9574 | 203.1 | - | - | 0 | - |
| 7 | y | 6.945E+04 | 205.1 | 0.0002733 | 1.332 | +1 | 1 |
| - | - | 2177 | 205.1 | - | - | 0 | - |
| - | - | 7071 | 206.1 | - | - | 0 | - |
| - | - | 2366 | 206.1 | - | - | 0 | - |
| - | - | 4435 | 207.2 | - | - | 0 | - |
| - | - | 2175 | 208.1 | - | - | 0 | - |
| - | - | 2167 | 209 | - | - | 0 | - |
| - | - | 1252 | 209.1 | - | - | 0 | - |
| - | - | 3622 | 209.1 | - | - | 0 | - |
| - | - | 2852 | 209.1 | - | - | 0 | - |
| - | - | 1256 | 211 | - | - | 0 | - |
| - | - | 3320 | 215.1 | - | - | 0 | - |
| - | - | 4095 | 216.1 | - | - | 0 | - |
| - | - | 8552 | 217.1 | - | - | 0 | - |
| - | - | 2021 | 220.1 | - | - | 0 | - |
| - | - | 3966 | 221.1 | - | - | 0 | - |
| - | - | 1694 | 221.1 | - | - | 0 | - |
| - | - | 3.737E+04 | 223.2 | - | - | 0 | - |
| - | - | 3307 | 224.2 | - | - | 0 | - |
| - | - | 1929 | 225 | - | - | 0 | - |
| - | - | 1137 | 225.8 | - | - | 0 | - |
| - | - | 1305 | 226 | - | - | 0 | - |
| - | - | 1.566E+04 | 226.1 | - | - | 0 | - |
| - | - | 1.115E+04 | 226.1 | - | - | 0 | - |
| - | - | 1.022E+05 | 226.2 | - | - | 0 | - |
| - | - | 7959 | 227.1 | - | - | 0 | - |
| - | - | 1556 | 227.1 | - | - | 0 | - |
| - | - | 1934 | 227.1 | - | - | 0 | - |
| - | - | 1.535E+04 | 227.2 | - | - | 0 | - |
| - | - | 2808 | 232.1 | - | - | 0 | - |
| - | - | 7484 | 233.1 | - | - | 0 | - |
| - | - | 1.649E+04 | 233.2 | - | - | 0 | - |
| - | - | 5531 | 233.2 | - | - | 0 | - |
| - | - | 4.764E+04 | 234.1 | - | - | 0 | - |
| - | - | 2112 | 234.2 | - | - | 0 | - |
| - | - | 6252 | 235.1 | - | - | 0 | - |
| - | - | 2283 | 236.1 | - | - | 0 | - |
| - | - | 1085 | 236.4 | - | - | 0 | - |
| - | - | 1576 | 238.1 | - | - | 0 | - |
| - | - | 2244 | 240.1 | - | - | 0 | - |
| - | - | 1989 | 243.1 | - | - | 0 | - |
| - | - | 1172 | 244.1 | - | - | 0 | - |
| - | - | 2.482E+04 | 244.1 | - | - | 0 | - |
| - | - | 8048 | 244.1 | - | - | 0 | - |
| - | - | 1640 | 245.1 | - | - | 0 | - |
| - | - | 2260 | 245.1 | - | - | 0 | - |
| - | - | 8294 | 248.1 | - | - | 0 | - |
| - | - | 1.782E+04 | 249.1 | - | - | 0 | - |
| - | - | 1750 | 249.1 | - | - | 0 | - |
| - | - | 1444 | 250.1 | - | - | 0 | - |
| - | - | 1.688E+05 | 251.2 | - | - | 0 | - |
| - | - | 2.097E+04 | 252.2 | - | - | 0 | - |
| - | - | 2063 | 252.2 | - | - | 0 | - |
| - | - | 1409 | 253.2 | - | - | 0 | - |
| - | - | 4969 | 254.1 | - | - | 0 | - |
| - | - | 4736 | 254.1 | - | - | 0 | - |
| - | - | 1438 | 261.1 | - | - | 0 | - |
| - | - | 1846 | 261.2 | - | - | 0 | - |
| - | - | 1225 | 265.2 | - | - | 0 | - |
| - | - | 4.18E+04 | 266.1 | - | - | 0 | - |
| - | - | 2188 | 266.2 | - | - | 0 | - |
| - | - | 3995 | 267.1 | - | - | 0 | - |
| - | - | 1487 | 268.2 | - | - | 0 | - |
| - | - | 4571 | 272.1 | - | - | 0 | - |
| - | - | 6320 | 272.1 | - | - | 0 | - |
| - | - | 1662 | 273.1 | - | - | 0 | - |
| - | - | 2415 | 274.1 | - | - | 0 | - |
| - | - | 8568 | 274.1 | - | - | 0 | - |
| - | - | 1691 | 276.1 | - | - | 0 | - |
| - | - | 1941 | 282.1 | - | - | 0 | - |
| - | - | 1631 | 282.2 | - | - | 0 | - |
| - | - | 2954 | 284.1 | - | - | 0 | - |
| 4 | y | 1346 | 284.1 | 0.002311 | 8.133 | +2 | 4 |
| - | - | 3950 | 284.2 | - | - | 0 | - |
| - | - | 3846 | 285.1 | - | - | 0 | - |
| - | - | 2346 | 290.1 | - | - | 0 | - |
| - | - | 2810 | 294.1 | - | - | 0 | - |
| - | - | 1178 | 294.2 | - | - | 0 | - |
| - | - | 1190 | 301.2 | - | - | 0 | - |
| 6 | y | 4614 | 302.1 | 0.0008474 | 2.805 | +1 | 2 |
| - | - | 3391 | 305.2 | - | - | 0 | - |
| - | - | 1556 | 305.2 | - | - | 0 | - |
| - | - | 1276 | 306.2 | - | - | 0 | - |
| - | - | 1541 | 308.1 | - | - | 0 | - |
| 3 | b | 1875 | 314.2 | 0.0001269 | 0.4039 | +1 | 3 |
| - | - | 1292 | 318.1 | - | - | 0 | - |
| - | - | 2034 | 319.2 | - | - | 0 | - |
| 6 | y | 2.1E+04 | 320.1 | 0.0004756 | 1.486 | +1 | 2 |
| - | - | 1590 | 320.2 | - | - | 0 | - |
| - | - | 2687 | 321.1 | - | - | 0 | - |
| - | - | 1.103E+04 | 323.2 | - | - | 0 | - |
| - | - | 1994 | 323.2 | - | - | 0 | - |
| - | - | 1.549E+04 | 323.2 | - | - | 0 | - |
| - | - | 2912 | 323.7 | - | - | 0 | - |
| - | - | 2503 | 324.2 | - | - | 0 | - |
| - | - | 3324 | 324.2 | - | - | 0 | - |
| - | - | 3526 | 327.2 | - | - | 0 | - |
| - | - | 6827 | 330.2 | - | - | 0 | - |
| - | - | 3474 | 332.2 | - | - | 0 | - |
| - | - | 3719 | 332.2 | - | - | 0 | - |
| - | - | 4753 | 334.2 | - | - | 0 | - |
| - | - | 2717 | 334.7 | - | - | 0 | - |
| - | - | 1647 | 336.1 | - | - | 0 | - |
| - | - | 2519 | 336.2 | - | - | 0 | - |
| 6 | b | 1302 | 338.7 | 0.000613 | 1.81 | +2 | 6 |
| - | - | 2.625E+04 | 341.2 | - | - | 0 | - |
| - | - | 1549 | 341.7 | - | - | 0 | - |
| - | - | 1194 | 342.2 | - | - | 0 | - |
| - | - | 6963 | 342.2 | - | - | 0 | - |
| - | - | 1684 | 342.2 | - | - | 0 | - |
| - | - | 1363 | 342.7 | - | - | 0 | - |
| - | - | 1710 | 343 | - | - | 0 | - |
| - | - | 3079 | 343.2 | - | - | 0 | - |
| - | - | 1406 | 343.2 | - | - | 0 | - |
| - | - | 1734 | 343.7 | - | - | 0 | - |
| - | - | 5426 | 344.2 | - | - | 0 | - |
| - | - | 1138 | 345 | - | - | 0 | - |
| - | - | 5095 | 345.1 | - | - | 0 | - |
| - | - | 5300 | 345.2 | - | - | 0 | - |
| - | - | 1.732E+04 | 346.1 | - | - | 0 | - |
| - | - | 1830 | 347.1 | - | - | 0 | - |
| 6 | b | 2493 | 347.7 | 3.593E-05 | 0.1033 | +2 | 6 |
| - | - | 3202 | 348.2 | - | - | 0 | - |
| - | - | 1.342E+04 | 350.2 | - | - | 0 | - |
| - | - | 4949 | 351.2 | - | - | 0 | - |
| - | - | 1316 | 351.7 | - | - | 0 | - |
| - | - | 1807 | 354.2 | - | - | 0 | - |
| - | - | 2305 | 355.2 | - | - | 0 | - |
| - | - | 2346 | 360 | - | - | 0 | - |
| - | - | 1716 | 360.7 | - | - | 0 | - |
| - | - | 6646 | 361 | - | - | 0 | - |
| - | - | 3070 | 361.2 | - | - | 0 | - |
| - | - | 3082 | 361.2 | - | - | 0 | - |
| - | - | 1.303E+04 | 362 | - | - | 0 | - |
| - | - | 6577 | 362.2 | - | - | 0 | - |
| - | - | 5921 | 363 | - | - | 0 | - |
| - | - | 1.428E+04 | 363.1 | - | - | 0 | - |
| - | - | 4427 | 363.2 | - | - | 0 | - |
| - | - | 1.055E+04 | 364.1 | - | - | 0 | - |
| - | - | 1526 | 364.1 | - | - | 0 | - |
| - | - | 1908 | 365.1 | - | - | 0 | - |
| - | - | 1254 | 365.2 | - | - | 0 | - |
| - | - | 1861 | 367.1 | - | - | 0 | - |
| - | - | 1393 | 367.2 | - | - | 0 | - |
| - | - | 2172 | 368.7 | - | - | 0 | - |
| - | - | 1393 | 369.7 | - | - | 0 | - |
| - | - | 1254 | 371.2 | - | - | 0 | - |
| - | - | 1.079E+04 | 373.2 | - | - | 0 | - |
| - | - | 9360 | 373.7 | - | - | 0 | - |
| - | - | 1342 | 374.2 | - | - | 0 | - |
| - | - | 4208 | 374.2 | - | - | 0 | - |
| - | - | 1386 | 376.7 | - | - | 0 | - |
| - | - | 1246 | 377.2 | - | - | 0 | - |
| - | - | 1854 | 378.2 | - | - | 0 | - |
| - | - | 6.894E+04 | 379.2 | - | - | 0 | - |
| - | - | 1.501E+04 | 380.2 | - | - | 0 | - |
| - | - | 4.867E+04 | 381.2 | - | - | 0 | - |
| - | - | 2132 | 381.2 | - | - | 0 | - |
| - | - | 7064 | 382.2 | - | - | 0 | - |
| - | - | 2.468E+04 | 382.7 | - | - | 0 | - |
| - | - | 1.753E+04 | 383.2 | - | - | 0 | - |
| - | - | 4832 | 383.7 | - | - | 0 | - |
| - | - | 3098 | 385.2 | - | - | 0 | - |
| - | - | 5492 | 390.7 | - | - | 0 | - |
| - | - | 1901 | 391.1 | - | - | 0 | - |
| - | - | 2936 | 391.2 | - | - | 0 | - |
| - | - | 1.579E+04 | 391.7 | - | - | 0 | - |
| - | - | 9413 | 392.2 | - | - | 0 | - |
| - | - | 3213 | 392.7 | - | - | 0 | - |
| - | - | 2351 | 395.1 | - | - | 0 | - |
| - | - | 1452 | 396.1 | - | - | 0 | - |
| - | - | 1789 | 398.2 | - | - | 0 | - |
| - | - | 1665 | 398.2 | - | - | 0 | - |
| - | - | 3354 | 399.2 | - | - | 0 | - |
| 2 | y | 7521 | 399.2 | 0.0004099 | 1.027 | +2 | 6 |
| - | - | 3.882E+04 | 399.7 | - | - | 0 | - |
| - | - | 2.373E+04 | 400.2 | - | - | 0 | - |
| - | - | 8995 | 400.7 | - | - | 0 | - |
| - | - | 1733 | 401.2 | - | - | 0 | - |
| - | - | 4652 | 409.1 | - | - | 0 | - |
| - | - | 2046 | 412.2 | - | - | 0 | - |
| - | - | 2211 | 413.1 | - | - | 0 | - |
| - | - | 4185 | 414.2 | - | - | 0 | - |
| - | - | 2453 | 417.7 | - | - | 0 | - |
| - | - | 3774 | 418.2 | - | - | 0 | - |
| - | - | 2011 | 418.7 | - | - | 0 | - |
| - | - | 1.109E+04 | 423.3 | - | - | 0 | - |
| - | - | 6912 | 424.2 | - | - | 0 | - |
| - | - | 4948 | 425.3 | - | - | 0 | - |
| - | - | 3831 | 425.8 | - | - | 0 | - |
| - | - | 1508 | 426.3 | - | - | 0 | - |
| - | - | 1.109E+04 | 426.7 | - | - | 0 | - |
| - | - | 6255 | 427.2 | - | - | 0 | - |
| 5 | y | 4.238E+04 | 430.2 | 0.0004059 | 0.9436 | +1 | 3 |
| - | - | 2128 | 430.3 | - | - | 0 | - |
| - | - | 2140 | 430.8 | - | - | 0 | - |
| - | - | 1819 | 431.1 | - | - | 0 | - |
| 5 | y | 8225 | 431.2 | 0.0009742 | 2.259 | +1 | 3 |
| - | - | 5839 | 431.2 | - | - | 0 | - |
| - | - | 1812 | 431.7 | - | - | 0 | - |
| - | - | 2175 | 432.2 | - | - | 0 | - |
| - | - | 1544 | 432.9 | - | - | 0 | - |
| - | - | 1261 | 433.1 | - | - | 0 | - |
| 4 | b | 3248 | 433.3 | 0.0009033 | 2.085 | +1 | 4 |
| - | - | 1230 | 437.3 | - | - | 0 | - |
| - | - | 2.143E+04 | 439.3 | - | - | 0 | - |
| - | - | 1.749E+04 | 439.8 | - | - | 0 | - |
| - | - | 5918 | 440.3 | - | - | 0 | - |
| 0 | Precursor | 3294 | 440.7 | 0.0004163 | 0.9446 | +2 | -1 |
| 0 | Precursor | 1523 | 441.2 | 0.003639 | 8.248 | +2 | -1 |
| - | - | 1.826E+04 | 442.2 | - | - | 0 | - |
| - | - | 5079 | 443.2 | - | - | 0 | - |
| 5 | y | 4.005E+04 | 448.2 | 0.0005834 | 1.302 | +1 | 3 |
| - | - | 2.899E+05 | 448.3 | - | - | 0 | - |
| - | - | 2.371E+05 | 448.8 | - | - | 0 | - |
| - | - | 6484 | 449.2 | - | - | 0 | - |
| - | - | 8.156E+04 | 449.3 | - | - | 0 | - |
| 0 | Precursor | 8612 | 449.7 | 0.0005575 | 1.24 | +2 | -1 |
| - | - | 1.591E+04 | 449.8 | - | - | 0 | - |
| - | - | 3712 | 450.2 | - | - | 0 | - |
| - | - | 1918 | 450.3 | - | - | 0 | - |
| 4 | b | 1.464E+04 | 451.3 | 0.0004705 | 1.043 | +1 | 4 |
| - | - | 3141 | 451.3 | - | - | 0 | - |
| - | - | 3589 | 452.3 | - | - | 0 | - |
| - | - | 3630 | 458.3 | - | - | 0 | - |
| - | - | 3010 | 459.2 | - | - | 0 | - |
| - | - | 7644 | 460.2 | - | - | 0 | - |
| - | - | 3382 | 461.2 | - | - | 0 | - |
| - | - | 3737 | 468.3 | - | - | 0 | - |
| - | - | 1730 | 469.3 | - | - | 0 | - |
| - | - | 1599 | 476.2 | - | - | 0 | - |
| - | - | 9873 | 476.3 | - | - | 0 | - |
| - | - | 5346 | 477.2 | - | - | 0 | - |
| - | - | 3220 | 477.3 | - | - | 0 | - |
| - | - | 1641 | 478.2 | - | - | 0 | - |
| - | - | 5562 | 478.3 | - | - | 0 | - |
| - | - | 6.622E+04 | 494.2 | - | - | 0 | - |
| - | - | 1.478E+04 | 495.2 | - | - | 0 | - |
| - | - | 1927 | 496.2 | - | - | 0 | - |
| - | - | 2926 | 497.3 | - | - | 0 | - |
| - | - | 2112 | 504.2 | - | - | 0 | - |
| - | - | 1.898E+04 | 508.3 | - | - | 0 | - |
| - | - | 5122 | 509.3 | - | - | 0 | - |
| - | - | 4270 | 512.2 | - | - | 0 | - |
| - | - | 1396 | 521.8 | - | - | 0 | - |
| - | - | 4466 | 527.3 | - | - | 0 | - |
| - | - | 1717 | 528.3 | - | - | 0 | - |
| - | - | 2205 | 537.3 | - | - | 0 | - |
| - | - | 2071 | 539.2 | - | - | 0 | - |
| - | - | 2029 | 542.8 | - | - | 0 | - |
| - | - | 1619 | 547.3 | - | - | 0 | - |
| - | - | 4828 | 548.3 | - | - | 0 | - |
| - | - | 5552 | 548.4 | - | - | 0 | - |
| - | - | 2024 | 549.3 | - | - | 0 | - |
| - | - | 3525 | 549.4 | - | - | 0 | - |
| - | - | 1.7E+04 | 550.2 | - | - | 0 | - |
| - | - | 4279 | 551.2 | - | - | 0 | - |
| - | - | 1.835E+04 | 555.3 | - | - | 0 | - |
| - | - | 7073 | 556.3 | - | - | 0 | - |
| 5 | b | 2232 | 561.3 | 3.458E-05 | 0.0616 | +1 | 5 |
| - | - | 6446 | 566.3 | - | - | 0 | - |
| 4 | y | 5005 | 567.2 | 0.0002404 | 0.4238 | +1 | 4 |
| - | - | 2992 | 567.3 | - | - | 0 | - |
| 4 | y | 4685 | 568.2 | 0.002187 | 3.848 | +1 | 4 |
| - | - | 1442 | 569.2 | - | - | 0 | - |
| - | - | 4207 | 570.3 | - | - | 0 | - |
| - | - | 2056 | 571.3 | - | - | 0 | - |
| - | - | 9.981E+04 | 573.3 | - | - | 0 | - |
| - | - | 3.895E+04 | 574.3 | - | - | 0 | - |
| - | - | 1.081E+04 | 575.3 | - | - | 0 | - |
| - | - | 1772 | 576.3 | - | - | 0 | - |
| 5 | b | 9314 | 579.3 | 0.0002121 | 0.3661 | +1 | 5 |
| - | - | 2851 | 580.3 | - | - | 0 | - |
| 4 | y | 5.094E+04 | 585.2 | 0.0001314 | 0.2246 | +1 | 4 |
| - | - | 1.307E+04 | 586.2 | - | - | 0 | - |
| - | - | 4610 | 587.2 | - | - | 0 | - |
| - | - | 1488 | 588.3 | - | - | 0 | - |
| - | - | 9272 | 593.3 | - | - | 0 | - |
| - | - | 2645 | 594.3 | - | - | 0 | - |
| - | - | 3010 | 595.2 | - | - | 0 | - |
| - | - | 1813 | 598.3 | - | - | 0 | - |
| - | - | 5577 | 607.3 | - | - | 0 | - |
| - | - | 2701 | 608.3 | - | - | 0 | - |
| - | - | 1665 | 622.4 | - | - | 0 | - |
| - | - | 1957 | 636.3 | - | - | 0 | - |
| - | - | 1483 | 639.4 | - | - | 0 | - |
| - | - | 1366 | 641.3 | - | - | 0 | - |
| - | - | 2078 | 645.4 | - | - | 0 | - |
| - | - | 1223 | 646.4 | - | - | 0 | - |
| - | - | 3564 | 649.4 | - | - | 0 | - |
| - | - | 1644 | 650.4 | - | - | 0 | - |
| - | - | 7751 | 652.3 | - | - | 0 | - |
| - | - | 3754 | 653.3 | - | - | 0 | - |
| - | - | 1546 | 662.3 | - | - | 0 | - |
| - | - | 8650 | 663.3 | - | - | 0 | - |
| - | - | 1.429E+04 | 663.4 | - | - | 0 | - |
| - | - | 2102 | 664.3 | - | - | 0 | - |
| - | - | 6568 | 664.4 | - | - | 0 | - |
| - | - | 2925 | 665.4 | - | - | 0 | - |
| - | - | 1.33E+04 | 667.4 | - | - | 0 | - |
| - | - | 8421 | 668.4 | - | - | 0 | - |
| - | - | 2509 | 669.4 | - | - | 0 | - |
| 6 | b | 6256 | 676.3 | 0.0004192 | 0.6198 | +1 | 6 |
| 6 | b | 6537 | 677.3 | 0.004091 | 6.039 | +1 | 6 |
| - | - | 2062 | 678.3 | - | - | 0 | - |
| 3 | y | 1.749E+04 | 680.3 | 3.872E-05 | 0.05691 | +1 | 5 |
| 3 | y | 2.728E+04 | 681.3 | 0.001802 | 2.645 | +1 | 5 |
| - | - | 8454 | 682.3 | - | - | 0 | - |
| - | - | 2090 | 683.3 | - | - | 0 | - |
| - | - | 3263 | 683.4 | - | - | 0 | - |
| - | - | 2831 | 684.4 | - | - | 0 | - |
| - | - | 3728 | 685.4 | - | - | 0 | - |
| - | - | 2295 | 686.4 | - | - | 0 | - |
| 6 | b | 6.676E+04 | 694.4 | 0.0001807 | 0.2602 | +1 | 6 |
| - | - | 2.206E+04 | 695.4 | - | - | 0 | - |
| - | - | 5434 | 696.4 | - | - | 0 | - |
| 3 | y | 1.811E+05 | 698.3 | 0.000211 | 0.3022 | +1 | 5 |
| - | - | 6.856E+04 | 699.3 | - | - | 0 | - |
| - | - | 1.235E+04 | 700.3 | - | - | 0 | - |
| - | - | 1.782E+04 | 701.4 | - | - | 0 | - |
| - | - | 9073 | 702.4 | - | - | 0 | - |
| - | - | 2441 | 703.4 | - | - | 0 | - |
| - | - | 1719 | 711.4 | - | - | 0 | - |
| - | - | 2896 | 712.4 | - | - | 0 | - |
| - | - | 1255 | 758.5 | - | - | 0 | - |
| - | - | 1259 | 758.5 | - | - | 0 | - |
| - | - | 1552 | 762.5 | - | - | 0 | - |
| - | - | 4839 | 764.4 | - | - | 0 | - |
| - | - | 2057 | 765.4 | - | - | 0 | - |
| - | - | 1481 | 776.3 | - | - | 0 | - |
| 2 | y | 4147 | 780.4 | 0.002114 | 2.708 | +1 | 6 |
| - | - | 9726 | 780.5 | - | - | 0 | - |
| - | - | 1300 | 781.4 | - | - | 0 | - |
| - | - | 7376 | 781.5 | - | - | 0 | - |
| - | - | 8756 | 782.4 | - | - | 0 | - |
| - | - | 4485 | 783.4 | - | - | 0 | - |
| 2 | y | 2.077E+04 | 797.4 | 0.0003876 | 0.4861 | +1 | 6 |
| - | - | 8064 | 798.4 | - | - | 0 | - |
| - | - | 5.475E+04 | 798.5 | - | - | 0 | - |
| - | - | 3.717E+04 | 799.5 | - | - | 0 | - |
| - | - | 1.381E+04 | 800.5 | - | - | 0 | - |
| - | - | 2985 | 801.5 | - | - | 0 | - |
| - | - | 1884 | 809.4 | - | - | 0 | - |
| - | - | 1283 | 1419 | - | - | 0 | - |
| - | - | 1510 | 2111 | - | - | 0 | - |
| - | - | 1381 | 2226 | - | - | 0 | - |
| - | - | 1429 | 2442 | - | - | 0 | - |
| - | - | 1374 | 3491 | - | - | 0 | - |

m/z Charge Intensity FragmentType MassShift Position
120.0811538696289 0 10165.345
120.47015380859375 0 786.80524
121.08470153808594 0 1273.0688
122.07158660888672 0 2274.008
123.05583190917969 0 1509.3256
125.10820770263672 0 912.41095
126.0552749633789 0 10246.729
127.1233139038086 0 1782.9839
128.0711212158203 0 1022.8263
128.1073455810547 0 42880.12
129.06622314453125 0 1747.8005
129.10260009765625 0 161929.34
130.0654754638672 0 26773.576
130.10598754882812 0 10113.642
131.06874084472656 0 2535.207
132.08111572265625 0 11956.7
132.10223388671875 0 13797.427
132.1150360107422 0 870.4114
133.08428955078125 0 993.0047
133.1058807373047 0 1087.0797
136.07594299316406 0 4096.117
136.31924438476562 0 817.3786
138.06654357910156 0 4466.9077
138.09164428710938 0 2691.1416
139.08721923828125 0 2576.8477
140.08216857910156 0 1111.519
140.10736083984375 0 959.4981
141.102294921875 0 2829.4724
142.06570434570312 0 1174.045
143.0450439453125 0 1771.6414
144.06585693359375 0 3748.449
144.08108520507812 0 2965.73
146.06033325195312 0 22206.678
147.0637664794922 0 1999.5887
149.04498291015625 0 1596.133
150.06661987304688 0 2073.5017
151.04168701171875 0 1264.1383
151.12362670898438 0 1120.4031
154.0979461669922 0 1340.7637
155.0819854736328 0 2124.9866
155.09303283691406 0 24946.492
155.11817932128906 0 17815.3 a Water loss 1
156.0769805908203 0 1790.6226
156.09640502929688 0 1138.5382
156.12158203125 0 881.29004
157.06106567382812 0 8826.821
157.0976104736328 0 4957.456
158.08444213867188 0 1133.9651
158.09202575683594 0 1307.7634
159.09202575683594 0 88479.164
159.11312866210938 0 7067.926 d 1
160.09530639648438 0 10185.819
160.11679077148438 0 1036.7949
165.10279846191406 0 1417.7914
165.1385955810547 0 1500.3529
166.06141662597656 0 39423.168
166.08665466308594 0 1912.742
166.0977325439453 0 1701.6456
167.05599975585938 0 1918.9603
167.06475830078125 0 2932.5945
167.08169555664062 0 4621.1406
167.1182403564453 0 5301.155
168.12171936035156 0 1393.5276
169.0974578857422 0 1504.6965
169.13392639160156 0 1571.6278
170.0528564453125 0 1121.2006
170.06044006347656 0 4158.4297
171.07623291015625 0 1306.3251
173.12887573242188 0 554405.25 a 1
173.4394073486328 0 4291.8296
174.13217163085938 0 43544.18
175.07162475585938 0 7236.74
175.0862579345703 0 1569.8026
175.13551330566406 0 1117.3339
178.13417053222656 0 18838.174
179.11830139160156 0 1475.2601
179.13734436035156 0 1677.2402
181.0611114501953 0 6136.3203
181.1337890625 0 1876.8134
183.11302185058594 0 7907.6987 b Water loss 1
185.09226989746094 0 3546.657
185.1287078857422 0 6016.6797
185.16505432128906 0 3101.9048
187.0869598388672 0 4979.258
187.10792541503906 0 2325.415
187.14459228515625 0 3174.4675
188.07093811035156 0 152017.8
189.07435607910156 0 16218.877
189.0872039794922 0 6583.122
190.13404846191406 0 5062.0225
194.0925750732422 0 1380.7599
195.1131134033203 0 5978.986
197.12869262695312 0 1788.6813
198.0874786376953 0 7352.6377
199.0715789794922 0 5453.66
199.10791015625 0 2965.6133
201.12362670898438 0 95857.49 b 1
202.12701416015625 0 9379.411
203.06654357910156 0 9574.128
205.09742736816406 0 69449.88 y 6
205.1448974609375 0 2176.51
206.1007537841797 0 7071.264
206.12991333007812 0 2366.0005
207.16065979003906 0 4435.2886
208.14466857910156 0 2175.2798
208.95404052734375 0 2167.382
209.0560760498047 0 1252.0272
209.09242248535156 0 3622.2312
209.12904357910156 0 2852.2642
210.95050048828125 0 1256.1594
215.13905334472656 0 3320.0952
216.09841918945312 0 4094.651
217.08224487304688 0 8551.589
220.11944580078125 0 2020.5073
221.10394287109375 0 3965.9153
221.13998413085938 0 1694.466
223.15565490722656 0 37373.227
224.15914916992188 0 3306.791
225.0436553955078 0 1929.483
225.80087280273438 0 1136.5164
226.04327392578125 0 1305.4594
226.08248901367188 0 15661.223
226.11895751953125 0 11154.816
226.15530395507812 0 102244.56
227.06663513183594 0 7959.4834
227.10272216796875 0 1555.7552
227.12283325195312 0 1933.843
227.15858459472656 0 15354.246
232.14073181152344 0 2807.7314
233.1395721435547 0 7484.0273
233.15017700195312 0 16487.305
233.16517639160156 0 5531.316
234.1240234375 0 47642.34
234.15367126464844 0 2111.5288
235.1273193359375 0 6252.064
236.13966369628906 0 2282.773
236.38087463378906 0 1084.9282
238.13031005859375 0 1576.4126
240.13523864746094 0 2243.562
243.0762176513672 0 1989.075
244.0808868408203 0 1172.0881
244.09312438964844 0 24817.146
244.12962341308594 0 8048.4077
245.0771026611328 0 1639.696
245.09710693359375 0 2260.401
248.11456298828125 0 8293.584
249.0985870361328 0 17819.805
249.13427734375 0 1750.4601
250.10047912597656 0 1444.1896
251.150634765625 0 168794.78
252.15394592285156 0 20965.486
252.16940307617188 0 2063.0466
253.15591430664062 0 1408.7645
254.11386108398438 0 4969.0947
254.14990234375 0 4736.0527
261.1195068359375 0 1437.8555
261.1598815917969 0 1846.2173
265.1536560058594 0 1224.832
266.1250305175781 0 41796.188
266.1503601074219 0 2187.5845
267.1280822753906 0 3995.2124
268.17718505859375 0 1487.1545
272.0877685546875 0 4570.922
272.1242370605469 0 6319.8706
273.1268005371094 0 1661.8157
274.1161804199219 0 2415.2205
274.1303405761719 0 8567.759
276.1094970703125 0 1691.0669
282.14581298828125 0 1941.0822
282.1811828613281 0 1631.2057
284.1026306152344 0 2953.9663
284.1214599609375 0 1345.6185 y Water loss 3
284.160888671875 0 3950.3708
285.08758544921875 0 3845.6238
290.0994567871094 0 2345.7747
294.1207580566406 0 2810.1401
294.21795654296875 0 1178.3484
301.19598388671875 0 1190.2952
302.1143798828125 0 4613.8843 y Water loss 5
305.1613464355469 0 3391.3767
305.1988830566406 0 1555.7782
306.2284851074219 0 1276.0751
308.1365966796875 0 1540.6842
314.2073059082031 0 1875.4188 b 2
318.1191101074219 0 1292.4257
319.15155029296875 0 2034.0049
320.12457275390625 0 21000.531 y 5
320.1949157714844 0 1590.0123
321.12750244140625 0 2687.3628
323.1717224121094 0 11027.33
323.1900329589844 0 1993.6948
323.20843505859375 0 15494.518
323.6920471191406 0 2912.3645
324.1744384765625 0 2502.9788
324.2120361328125 0 3323.8176
327.1660461425781 0 3525.5496
330.16607666015625 0 6826.5674
332.19390869140625 0 3474.0027
332.2099914550781 0 3718.6418
334.1925048828125 0 4753.2173
334.6936340332031 0 2716.5444
336.13043212890625 0 1647.2089
336.2393493652344 0 2518.8455
338.6736755371094 0 1302.4742 b Water loss 5
341.1822814941406 0 26252.422
341.7030944824219 0 1548.9413
342.1632385253906 0 1193.8396
342.1849060058594 0 6963.2817
342.206787109375 0 1684.4697
342.6993713378906 0 1362.6816
343.0182189941406 0 1710.2786
343.1629943847656 0 3078.9534
343.1987609863281 0 1405.77
343.6994323730469 0 1733.9283
344.172119140625 0 5426.369
344.9769592285156 0 1138.158
345.13067626953125 0 5094.5957
345.17694091796875 0 5299.6323
346.1148376464844 0 17317.203
347.119873046875 0 1829.5955
347.6795349121094 0 2492.5422 b 5
348.1807556152344 0 3201.538
350.21905517578125 0 13419.577
351.2146301269531 0 4949.078
351.7147521972656 0 1315.7013
354.24957275390625 0 1806.7498
355.16082763671875 0 2304.6501
360.02862548828125 0 2346.1704
360.703125 0 1716.0801
361.0267333984375 0 6645.545
361.180419921875 0 3070.1365
361.2001647949219 0 3081.9578
362.0265808105469 0 13027.09
362.18206787109375 0 6576.616
363.0248718261719 0 5921.0166
363.14111328125 0 14284.326
363.1869201660156 0 4427.26
364.12506103515625 0 10547.982
364.14642333984375 0 1526.3728
365.1282043457031 0 1908.2811
365.199951171875 0 1254.0593
367.1405944824219 0 1860.5464
367.24652099609375 0 1393.1477
368.7206726074219 0 2172.0325
369.72222900390625 0 1392.7158
371.1556701660156 0 1254.3177
373.1721496582031 0 10793.272
373.7138366699219 0 9360.195
374.1753845214844 0 1342.4608
374.2143249511719 0 4208.466
376.7370300292969 0 1385.6785
377.2368469238281 0 1245.9001
378.21484375 0 1853.5504
379.2090759277344 0 68938.65
380.2117004394531 0 15012.523
381.15203857421875 0 48674.58
381.21319580078125 0 2131.5974
382.1549377441406 0 7063.76
382.71893310546875 0 24681.002
383.2203369140625 0 17527.227
383.7223205566406 0 4831.5425
385.15179443359375 0 3097.6292
390.7343444824219 0 5491.5815
391.137451171875 0 1900.9557
391.23516845703125 0 2936.3657
391.7243957519531 0 15792.813
392.2259826660156 0 9413.46
392.7282409667969 0 3213.3655
395.1353454589844 0 2350.721
396.1380310058594 0 1451.8966
398.17816162109375 0 1788.5938
398.24114990234375 0 1664.5531
399.1625671386719 0 3354.1833
399.2010803222656 0 7521.3594 y 1
399.7398986816406 0 38820.727
400.2413024902344 0 23728.906
400.7430114746094 0 8995.2705
401.2444763183594 0 1733.4839
409.14691162109375 0 4651.6504
412.158447265625 0 2046.4474
413.14581298828125 0 2211.427
414.2349853515625 0 4184.5684
417.7170715332031 0 2453.226
418.21112060546875 0 3773.6423
418.7083740234375 0 2010.648
423.2719421386719 0 11089.104
424.2197265625 0 6911.718
425.26336669921875 0 4948.0513
425.7643127441406 0 3831.4194
426.2601623535156 0 1508.4095
426.7218017578125 0 11087.931
427.22247314453125 0 6254.645
430.1725158691406 0 42381.01 y Water loss 4
430.2583312988281 0 2128.0527
430.75567626953125 0 2139.5247
431.0873107910156 0 1818.6616
431.1551513671875 0 8224.703 y Ammonia loss 4
431.17864990234375 0 5839.0586
431.7142028808594 0 1811.8922
432.2090148925781 0 2174.7742
432.88543701171875 0 1543.6827
433.0839538574219 0 1260.8048
433.2566833496094 0 3247.7087 b Water loss 3
437.2904052734375 0 1229.7902
439.2607727050781 0 21433.787
439.7620849609375 0 17489.49
440.2626647949219 0 5918.343
440.71881103515625 0 3293.8105 Precursor Water loss
441.2148742675781 0 1522.7433 Precursor Ammonia loss
442.23016357421875 0 18263.027
443.2326354980469 0 5079.092
448.1832580566406 0 40045.35 y 4
448.2664489746094 0 289934.5
448.7678527832031 0 237134.03
449.1866760253906 0 6483.6157
449.26922607421875 0 81562.04
449.7250671386719 0 8611.533 Precursor
449.7704162597656 0 15911.012
450.2275390625 0 3712.4697
450.27618408203125 0 1917.8594
451.2668151855469 0 14640.871 b 3
451.30303955078125 0 3140.9524
452.26959228515625 0 3589.158
458.2607116699219 0 3629.5747
459.19854736328125 0 3009.9465
460.24176025390625 0 7644.218
461.2461242675781 0 3382.3345
468.2929992675781 0 3737.1296
469.2770690917969 0 1730.3428
476.22894287109375 0 1599.0568
476.2711486816406 0 9873.035
477.2098083496094 0 5345.891
477.27655029296875 0 3220.3176
478.2140197753906 0 1640.9197
478.2774963378906 0 5562.212
494.2359313964844 0 66221.32
495.2380065917969 0 14775.551
496.2415771484375 0 1927.3871
497.2710876464844 0 2926.054
504.19891357421875 0 2111.6643
508.2514343261719 0 18982.479
509.2563171386719 0 5122.3926
512.24658203125 0 4270.4033
521.7847900390625 0 1395.6722
527.3184814453125 0 4465.967
528.3214721679688 0 1717.0306
537.3036499023438 0 2204.9807
539.2362060546875 0 2070.6057
542.7969970703125 0 2028.7926
547.2843017578125 0 1619.2233
548.318603515625 0 4827.5215
548.3576049804688 0 5552.3647
549.3190307617188 0 2024.3667
549.3598022460938 0 3524.8462
550.2042846679688 0 17002.42
551.20751953125 0 4278.5547
555.3136596679688 0 18349.992
556.317138671875 0 7073.4453
561.3143920898438 0 2232.1125 b Water loss 4
566.330078125 0 6445.8867
567.2312622070312 0 5004.975 y Water loss 3
567.332275390625 0 2991.7188
568.2172241210938 0 4684.659 y Ammonia loss 3
569.2183227539062 0 1442.2161
570.3244018554688 0 4207.0312
571.3289794921875 0 2056.443
573.3240966796875 0 99808.8
574.3271484375 0 38951.07
575.3292846679688 0 10813.461
576.330810546875 0 1772.2103
579.3251342773438 0 9313.78 b 4
580.3250122070312 0 2850.881
585.241455078125 0 50935.14 y 3
586.2447509765625 0 13067.007
587.2467651367188 0 4609.699
588.3359985351562 0 1488.1085
593.3036499023438 0 9272.387
594.3076171875 0 2644.5015
595.2261352539062 0 3010.3914
598.3206787109375 0 1812.9076
607.3192138671875 0 5576.961
608.3235473632812 0 2701.3218
622.3700561523438 0 1664.6611
636.3291015625 0 1957.3716
639.3841552734375 0 1482.7307
641.3064575195312 0 1365.7076
645.37353515625 0 2077.528
646.3672485351562 0 1223.2239
649.3668212890625 0 3563.6934
650.3695678710938 0 1644.4362
652.3193969726562 0 7750.8657
653.3233642578125 0 3754.1345
662.3056030273438 0 1545.67
663.2890625 0 8649.557
663.3831176757812 0 14288.908
664.28955078125 0 2101.885
664.3855590820312 0 6568.3516
665.3900756835938 0 2924.6826
667.3771362304688 0 13298.31
668.3807373046875 0 8420.77
669.3811645507812 0 2508.6985
676.3408813476562 0 6255.927 b Water loss 5
677.3294067382812 0 6536.979 b Ammonia loss 5
678.3269653320312 0 2062.2993
680.3151245117188 0 17490.734 y Water loss 2
681.3009033203125 0 27277.996 y Ammonia loss 2
682.3029174804688 0 8453.987
683.3056030273438 0 2089.903
683.408447265625 0 3263.2014
684.4086303710938 0 2831.0828
685.3883056640625 0 3727.6245
686.391357421875 0 2294.9143
694.3516845703125 0 66759.09 b 5
695.3543701171875 0 22062.564
696.3583374023438 0 5434.415
698.325439453125 0 181090.83 y 2
699.3284912109375 0 68563.19
700.3318481445312 0 12349.858
701.4188842773438 0 17824.666
702.4215698242188 0 9073.447
703.425537109375 0 2440.682
711.3777465820312 0 1719.4458
712.3622436523438 0 2896.155
758.4508666992188 0 1255.1854
758.5311279296875 0 1259.3546
762.4532470703125 0 1552.1062
764.4315795898438 0 4839.0073
765.433837890625 0 2057.3535
776.2637939453125 0 1481.0847
780.36962890625 0 4146.568 y Ammonia loss 1
780.4612426757812 0 9726.286
781.380615234375 0 1299.6997
781.4633178710938 0 7376.381
782.441650390625 0 8755.907
783.4432983398438 0 4484.8086
797.3936767578125 0 20774.688 y 1
798.3955078125 0 8064.381
798.4716796875 0 54754.69
799.4740600585938 0 37171.625
800.4764404296875 0 13814.564
801.481201171875 0 2985.1917
809.3723754882812 0 1884.3733
1418.82958984375 0 1283.2095
2110.625 0 1510.0245
2225.983154296875 0 1380.603
2442.376708984375 0 1428.715
3491.30908203125 0 1374.0691

Spectrum Details

|  |  |
| --- | --- |
| Matched peaks? Matched peaks The total absolute number of peaks matched. Additionally in brackets the total fraction of peaks matched and the total number of peaks is shown. | 34 (7.61% of 447) |
| FDR? FDR The false discovery rate estimated for this peptide. It is calculated by matching all theoretical fragments with a non-integer shift with the raw peaks for this spectrum. This is done with 40 different shifts. The resulting percentage is the average number of annotated peaks over the number of annotated peaks with the correct spectrum. | 4.06% |
| Satellite FDR? Satellite FDR See the FDR for details on its calculation. This satellite ion specific FDR only contains the satellite ions (d/w) for I/L/J positions. | - |
| PSM Score? PSM Score The PSM Score as given by Hecklib to this annotated spectrum. It is shown with three significant figures. | 448 |

#### Spectrum 4081? Spectrum 4081 The raw spectrum of this peptide as annotated by Hecklib. The fragments are coloured according to ion type (see legend). Any peaks with a star '\*' as text can be hovered over to see the full details, first the ion type second the mass shift type. By hovering over the amino acids in the peptide or ions in the legend the corresponding peaks are highlighted. By toggling the 'Unassigned' label you can turn the background (unassigned) peaks on or off in the plot. By updating the slider in the Ion legend you can update the spectrum to only show the top X% of the peaks with labels. The top X% means any peak that is within X% of the highest intensity. By dragging in the spectrum you can zoom in to a specific part of the spectrum and use 'Zoom Out' to get back to the original zoom level. The annotation of the spectrum is based on the given sequence in the peptides file and is done with different software so inconsistencies are likely. The peaks are annotated based on the given sequence, with 20 ppm tolerance.

Copy Data

##### Spectrum 4081 (TSV)

###### Preview

```
Loading example...
```

*Click on the button to copy the data to your clipboard.*

Mz MinMz MaxIntensity Max

WidthHeightPeptide font sizePeptide stroke widthSpectrum font sizeSpectrum stroke widthCompact peptide

Ion legend

wxyz

abcd

OtherUnassignedIon^Charge^_Position_Show for top:%

TVLHQDW

01.01e+42.01e+43.02e+44.02e+4

Zoom Out

y^+1^_1_y^+1^_2_y^+2^_6_y^+1^_3_c^+1^_4_c^+1^_5_z^+1^_4_c^+1^_5_y^+1^_4_w^+1^_5_c^+1^_6_y^+1^_5_y^+1^_5_z^+1^_5_c^+1^_6_y^+1^_5_z^+1^_6_

0778155723353114

Fragment Matches Table

Show background peaks

| Position | Ion type | Intensity | mz Theoretical | mz Error (Th) | mz Error (ppm) | Charge | Series Number |
| --- | --- | --- | --- | --- | --- | --- | --- |
| - | - | 332.5 | 120.6 | - | - | 0 | - |
| - | - | 392.4 | 123.1 | - | - | 0 | - |
| - | - | 378.2 | 127.4 | - | - | 0 | - |
| - | - | 374.4 | 127.5 | - | - | 0 | - |
| - | - | 529.4 | 129.1 | - | - | 0 | - |
| - | - | 399.6 | 134.3 | - | - | 0 | - |
| - | - | 447.1 | 140.3 | - | - | 0 | - |
| - | - | 392.9 | 146.6 | - | - | 0 | - |
| - | - | 441.8 | 148.8 | - | - | 0 | - |
| - | - | 411 | 148.8 | - | - | 0 | - |
| - | - | 475.5 | 148.9 | - | - | 0 | - |
| - | - | 508.2 | 148.9 | - | - | 0 | - |
| - | - | 404.9 | 148.9 | - | - | 0 | - |
| - | - | 774.6 | 148.9 | - | - | 0 | - |
| - | - | 834.2 | 148.9 | - | - | 0 | - |
| - | - | 1289 | 148.9 | - | - | 0 | - |
| - | - | 1392 | 148.9 | - | - | 0 | - |
| - | - | 3242 | 148.9 | - | - | 0 | - |
| - | - | 4859 | 148.9 | - | - | 0 | - |
| - | - | 3721 | 149 | - | - | 0 | - |
| - | - | 1855 | 149 | - | - | 0 | - |
| - | - | 1382 | 149 | - | - | 0 | - |
| - | - | 936 | 149 | - | - | 0 | - |
| - | - | 790.4 | 149 | - | - | 0 | - |
| - | - | 747.7 | 149 | - | - | 0 | - |
| - | - | 461.6 | 149 | - | - | 0 | - |
| - | - | 433.4 | 149 | - | - | 0 | - |
| - | - | 3612 | 149 | - | - | 0 | - |
| - | - | 393.7 | 149.1 | - | - | 0 | - |
| - | - | 1862 | 155.1 | - | - | 0 | - |
| - | - | 1095 | 167.1 | - | - | 0 | - |
| - | - | 451.2 | 171.7 | - | - | 0 | - |
| - | - | 1.107E+04 | 173.1 | - | - | 0 | - |
| - | - | 1222 | 175.1 | - | - | 0 | - |
| - | - | 3970 | 201.1 | - | - | 0 | - |
| - | - | 1101 | 203.1 | - | - | 0 | - |
| - | - | 539 | 209.6 | - | - | 0 | - |
| 7 | y | 2040 | 221.1 | 0.003734 | 16.89 | +1 | 1 |
| - | - | 3003 | 221.1 | - | - | 0 | - |
| - | - | 562.8 | 223.1 | - | - | 0 | - |
| - | - | 3009 | 225 | - | - | 0 | - |
| - | - | 5260 | 239.1 | - | - | 0 | - |
| - | - | 1090 | 251.2 | - | - | 0 | - |
| - | - | 951.5 | 268.2 | - | - | 0 | - |
| - | - | 658 | 269.2 | - | - | 0 | - |
| - | - | 1258 | 281.1 | - | - | 0 | - |
| - | - | 4932 | 295.1 | - | - | 0 | - |
| - | - | 522.9 | 298.6 | - | - | 0 | - |
| - | - | 4469 | 299.1 | - | - | 0 | - |
| - | - | 710.7 | 313.1 | - | - | 0 | - |
| 6 | y | 636.9 | 336.1 | 0.004708 | 14.01 | +1 | 2 |
| - | - | 606.6 | 339.8 | - | - | 0 | - |
| - | - | 846.1 | 355.1 | - | - | 0 | - |
| - | - | 522 | 355.5 | - | - | 0 | - |
| - | - | 4719 | 369.1 | - | - | 0 | - |
| - | - | 658.4 | 379.2 | - | - | 0 | - |
| - | - | 680.4 | 381.2 | - | - | 0 | - |
| 2 | y | 713.5 | 407.2 | 0.003457 | 8.489 | +2 | 6 |
| - | - | 595.8 | 418 | - | - | 0 | - |
| - | - | 963.2 | 423.3 | - | - | 0 | - |
| - | - | 716.9 | 425.3 | - | - | 0 | - |
| - | - | 1145 | 448.7 | - | - | 0 | - |
| - | - | 721.7 | 449.3 | - | - | 0 | - |
| - | - | 2416 | 451.3 | - | - | 0 | - |
| - | - | 799.8 | 452.3 | - | - | 0 | - |
| - | - | 565.2 | 457.7 | - | - | 0 | - |
| - | - | 1132 | 458.3 | - | - | 0 | - |
| - | - | 1360 | 458.8 | - | - | 0 | - |
| 5 | y | 929.5 | 464.2 | 0.006036 | 13 | +1 | 3 |
| - | - | 1069 | 467.3 | - | - | 0 | - |
| - | - | 1095 | 468.3 | - | - | 0 | - |
| 4 | c | 2.188E+04 | 468.3 | 0.0003801 | 0.8118 | +1 | 4 |
| - | - | 3424 | 469.3 | - | - | 0 | - |
| - | - | 761 | 484.3 | - | - | 0 | - |
| - | - | 694 | 485.3 | - | - | 0 | - |
| - | - | 1424 | 494.2 | - | - | 0 | - |
| - | - | 749 | 499.3 | - | - | 0 | - |
| - | - | 2086 | 527.2 | - | - | 0 | - |
| - | - | 906.1 | 560.4 | - | - | 0 | - |
| - | - | 883.7 | 561.4 | - | - | 0 | - |
| 5 | c | 1328 | 579.3 | 0.0004562 | 0.7875 | +1 | 5 |
| 4 | z | 1392 | 585.2 | 0.005338 | 9.122 | +1 | 4 |
| - | - | 707.6 | 586.2 | - | - | 0 | - |
| 5 | c | 2.104E+04 | 596.4 | 0.0002743 | 0.46 | +1 | 5 |
| - | - | 6247 | 597.4 | - | - | 0 | - |
| - | - | 1032 | 598.4 | - | - | 0 | - |
| 4 | y | 1492 | 601.2 | 0.005718 | 9.511 | +1 | 4 |
| - | - | 631 | 630.4 | - | - | 0 | - |
| - | - | 2009 | 652.3 | - | - | 0 | - |
| 3 | w | 2672 | 655.2 | 0.005468 | 8.346 | +1 | 5 |
| - | - | 832.3 | 656.3 | - | - | 0 | - |
| - | - | 1068 | 656.4 | - | - | 0 | - |
| - | - | 712.9 | 666.4 | - | - | 0 | - |
| - | - | 585 | 667.4 | - | - | 0 | - |
| - | - | 2750 | 679.3 | - | - | 0 | - |
| - | - | 1333 | 680.3 | - | - | 0 | - |
| - | - | 993.2 | 688.2 | - | - | 0 | - |
| 6 | c | 2633 | 694.4 | 0.001284 | 1.849 | +1 | 6 |
| - | - | 933.4 | 695.4 | - | - | 0 | - |
| 3 | y | 825 | 696.3 | 0.001863 | 2.676 | +1 | 5 |
| - | - | 1167 | 696.4 | - | - | 0 | - |
| 3 | y | 1057 | 697.3 | 0.0107 | 15.35 | +1 | 5 |
| 3 | z | 2083 | 698.3 | 0.005503 | 7.88 | +1 | 5 |
| - | - | 883.7 | 699.4 | - | - | 0 | - |
| - | - | 2544 | 710.4 | - | - | 0 | - |
| 6 | c | 2.097E+04 | 711.4 | 0.0003015 | 0.4239 | +1 | 6 |
| - | - | 6262 | 712.4 | - | - | 0 | - |
| - | - | 1800 | 713.4 | - | - | 0 | - |
| 3 | y | 7697 | 714.3 | 0.00515 | 7.21 | +1 | 5 |
| - | - | 3499 | 715.3 | - | - | 0 | - |
| - | - | 785.3 | 716.3 | - | - | 0 | - |
| - | - | 4094 | 752.4 | - | - | 0 | - |
| - | - | 7084 | 753.4 | - | - | 0 | - |
| - | - | 2430 | 754.4 | - | - | 0 | - |
| - | - | 5496 | 768.4 | - | - | 0 | - |
| - | - | 2289 | 769.4 | - | - | 0 | - |
| - | - | 905.3 | 786.5 | - | - | 0 | - |
| 2 | z | 1644 | 797.4 | 0.007401 | 9.282 | +1 | 6 |
| - | - | 938 | 798.4 | - | - | 0 | - |
| - | - | 647.8 | 799.4 | - | - | 0 | - |
| - | - | 704.3 | 800.4 | - | - | 0 | - |
| - | - | 627.2 | 843.4 | - | - | 0 | - |
| - | - | 693.8 | 844.4 | - | - | 0 | - |
| - | - | 2901 | 853.4 | - | - | 0 | - |
| - | - | 1441 | 854.5 | - | - | 0 | - |
| - | - | 2120 | 855.4 | - | - | 0 | - |
| - | - | 919.7 | 856.4 | - | - | 0 | - |
| - | - | 2664 | 897.4 | - | - | 0 | - |
| - | - | 1.333E+04 | 898.4 | - | - | 0 | - |
| - | - | 6589 | 899.4 | - | - | 0 | - |
| - | - | 2553 | 900.4 | - | - | 0 | - |
| - | - | 980.7 | 901.4 | - | - | 0 | - |
| - | - | 1.551E+04 | 914.4 | - | - | 0 | - |
| - | - | 3.985E+04 | 915.4 | - | - | 0 | - |
| - | - | 1.848E+04 | 916.4 | - | - | 0 | - |
| - | - | 784.3 | 916.6 | - | - | 0 | - |
| - | - | 7671 | 917.5 | - | - | 0 | - |
| - | - | 1590 | 918.5 | - | - | 0 | - |
| - | - | 1063 | 1060 | - | - | 0 | - |
| - | - | 630.4 | 1167 | - | - | 0 | - |
| - | - | 641.2 | 1188 | - | - | 0 | - |
| - | - | 802.5 | 1353 | - | - | 0 | - |
| - | - | 991.2 | 1354 | - | - | 0 | - |
| - | - | 890.1 | 1369 | - | - | 0 | - |
| - | - | 998.2 | 1371 | - | - | 0 | - |
| - | - | 797.8 | 1376 | - | - | 0 | - |
| - | - | 1790 | 1376 | - | - | 0 | - |
| - | - | 690.7 | 2082 | - | - | 0 | - |
| - | - | 833.1 | 2831 | - | - | 0 | - |
| - | - | 745.7 | 3083 | - | - | 0 | - |

m/z Charge Intensity FragmentType MassShift Position
120.64678955078125 0 332.5434
123.12537384033203 0 392.35834
127.36466217041016 0 378.1849
127.54170227050781 0 374.43088
129.1026611328125 0 529.39264
134.29039001464844 0 399.57028
140.3062744140625 0 447.12155
146.64414978027344 0 392.93085
148.82284545898438 0 441.80597
148.8296661376953 0 411.04645
148.88717651367188 0 475.53275
148.894287109375 0 508.23737
148.9014892578125 0 404.90753
148.90859985351562 0 774.63385
148.91578674316406 0 834.20825
148.9232177734375 0 1289.1838
148.93043518066406 0 1392.2168
148.937744140625 0 3242.0898
148.94552612304688 0 4858.5225
148.96212768554688 0 3720.6553
148.96974182128906 0 1855.3486
148.9768829345703 0 1381.7322
148.98414611816406 0 936.01953
148.99160766601562 0 790.3692
148.99884033203125 0 747.67
149.0062255859375 0 461.6388
149.0208740234375 0 433.37177
149.04498291015625 0 3611.756
149.06361389160156 0 393.69873
155.09292602539062 0 1862.1315
167.05564880371094 0 1095.2103
171.7377166748047 0 451.17844
173.12867736816406 0 11071.708
175.08677673339844 0 1222.1659
201.12353515625 0 3970.376
203.0816650390625 0 1100.5244
209.6231231689453 0 539.03644
221.0834197998047 0 2039.74 y 6
221.0924530029297 0 3002.7913
223.11875915527344 0 562.80524
225.04318237304688 0 3008.9182
239.0950469970703 0 5260.385
251.15081787109375 0 1089.938
268.17724609375 0 951.48834
269.1816101074219 0 657.97253
281.0514221191406 0 1258.1271
295.1033630371094 0 4932.216
298.58642578125 0 522.91486
299.0618896484375 0 4468.6924
313.114501953125 0 710.7122
336.1188049316406 0 636.9293 y 5
339.8061828613281 0 606.5771
355.06982421875 0 846.08026
355.46429443359375 0 521.96906
369.1220703125 0 4719.4453
379.20941162109375 0 658.43384
381.1515197753906 0 680.4483
407.1991271972656 0 713.5055 y 1
418.0111083984375 0 595.7529
423.27130126953125 0 963.17596
425.2862548828125 0 716.85016
448.7349853515625 0 1144.942
449.27490234375 0 721.7497
451.2666931152344 0 2416.461
452.27301025390625 0 799.81525
457.7231750488281 0 565.15045
458.27838134765625 0 1132.2292
458.781982421875 0 1359.5452
464.1787109375 0 929.52515 y 4
467.2850646972656 0 1068.5165
468.25537109375 0 1095.2719
468.29327392578125 0 21884.533 c 3
469.29718017578125 0 3423.956
484.3011169433594 0 760.98834
485.3060607910156 0 694.0209
494.2368469238281 0 1424.1875
499.2619323730469 0 748.98663
527.1890869140625 0 2085.9639
560.3534545898438 0 906.1021
561.35693359375 0 883.71674
579.3253784179688 0 1327.5905 c Ammonia loss 4
585.2182006835938 0 1392.4948 z 3
586.2206420898438 0 707.6232
596.3517456054688 0 21044.812 c 4
597.3544921875 0 6246.9478
598.3605346679688 0 1032.4271
601.2373046875 0 1492.4811 y 3
630.382568359375 0 630.9665
652.3184814453125 0 2009.0471
655.2476196289062 0 2672.4045 w 2
656.25537109375 0 832.3304
656.3742065429688 0 1067.6324
666.3555908203125 0 712.9383
667.3707275390625 0 584.9679
679.3403930664062 0 2749.5752
680.3424072265625 0 1333.2578
688.1589965820312 0 993.18024
694.3531494140625 0 2632.5623 c Ammonia loss 5
695.3507080078125 0 933.4446
696.30322265625 0 824.9654 y Water loss 2
696.3678588867188 0 1166.8611
697.2998046875 0 1056.579 y Ammonia loss 2
698.3024291992188 0 2082.713 z 2
699.4378662109375 0 883.7378
710.3704833984375 0 2544.2622
711.3781127929688 0 20972.402 c 5
712.3809204101562 0 6261.9966
713.3843994140625 0 1800.4113
714.32080078125 0 7696.84 y 2
715.3233642578125 0 3499.4692
716.3232421875 0 785.3315
752.3803100585938 0 4093.7031
753.3689575195312 0 7083.59
754.3731689453125 0 2429.6235
768.3753051757812 0 5495.5576
769.3785400390625 0 2288.9211
786.4736328125 0 905.2967
797.3727416992188 0 1643.7153 z 1
798.37451171875 0 938.03595
799.3839111328125 0 647.8203
800.4411010742188 0 704.3012
843.4129638671875 0 627.2127
844.4234619140625 0 693.75165
853.4443359375 0 2900.6086
854.4505615234375 0 1441.2314
855.4285278320312 0 2119.535
856.4286499023438 0 919.7218
897.4344482421875 0 2664.1218
898.4195556640625 0 13327.459
899.4210205078125 0 6588.7656
900.4283447265625 0 2553.0823
901.4254150390625 0 980.6655
914.438232421875 0 15512.448
915.4459838867188 0 39845.41
916.4491577148438 0 18476.865
916.5664672851562 0 784.32825
917.4542236328125 0 7670.674
918.4522094726562 0 1590.42
1059.6031494140625 0 1063.444
1166.6719970703125 0 630.4058
1188.2010498046875 0 641.2337
1352.7779541015625 0 802.47473
1353.783203125 0 991.2273
1368.7847900390625 0 890.1014
1370.773193359375 0 998.1886
1375.7496337890625 0 797.79706
1376.3216552734375 0 1790.0732
2082.099609375 0 690.6888
2831.380126953125 0 833.05194
3082.968505859375 0 745.69434

Spectrum Details

|  |  |
| --- | --- |
| Matched peaks? Matched peaks The total absolute number of peaks matched. Additionally in brackets the total fraction of peaks matched and the total number of peaks is shown. | 17 (11.33% of 150) |
| FDR? FDR The false discovery rate estimated for this peptide. It is calculated by matching all theoretical fragments with a non-integer shift with the raw peaks for this spectrum. This is done with 40 different shifts. The resulting percentage is the average number of annotated peaks over the number of annotated peaks with the correct spectrum. | 1.82% |
| Satellite FDR? Satellite FDR See the FDR for details on its calculation. This satellite ion specific FDR only contains the satellite ions (d/w) for I/L/J positions. | 7.14% |
| PSM Score? PSM Score The PSM Score as given by Hecklib to this annotated spectrum. It is shown with three significant figures. | 191 |

#### Reverse Lookup? Reverse Lookup All places where this read could be placed.

| Group | Segment | Template | Template Part | Read Part | Score | Unique |
| --- | --- | --- | --- | --- | --- | --- |
| Homo sapiens Heavy Chain | IGHC | IGHG1 | [189..196] | [0..7] | 56 | False |
| Homo sapiens Heavy Chain | IGHC | IGHG3 | [236..243] | [0..7] | 56 | False |
| Homo sapiens Heavy Chain | IGHC | IGHG4 | [186..193] | [0..7] | 56 | False |

| Recombined | Template Part | Read Part | Score | Unique |
| --- | --- | --- | --- | --- |
| REC-0-1 | [314..321] | [0..7] | 56 | True |

#### Meta Information from Multiple reads

##### Number of combined reads

4

##### Intensity

0.7236

##### TotalArea

4.801E+08

##### Changes to the peptide sequence

TVLHQDW

J→LSupport for Leucine based on side chain ions (1 for L 0 for I) (Position: 3)

L→JNo support for either Leucine or Isoleucine based on side chain ions (Position: 3)

J→LSupport for Leucine based on side chain ions (1 for L 0 for I) (Position: 3)

L→JNo support for either Leucine or Isoleucine based on side chain ions (Position: 3)

#### Positional Score

Copy Data

##### Positional Score (TSV)

###### Preview

```
Loading example...
```

*Click on the button to copy the data to your clipboard.*

100123456

Label Value
"0" 0.47
"1" 0.473
"2" 0.49
"3" 0.477
"4" 0.48
"5" 0.497
"6" 0.5

#### Meta Information from PEAKS

##### Scan Identifier

F3:5429

##### Original sequence

T

V

L

H

Q

D

W

##### Posttranslational Modifications

##### Source File

D:\separate\_stitch\_analyses\xle-disambiguation\raw\20210323\_F1\_UM1\_Peng0013\_SA\_F59\_ingel\_3ug\_chymo.raw

##### Fraction

3

##### Scan Feature

F3:2304

##### De Novo Score

98

##### ConfidenceScore

98

### m/z

449.7257

##### Mass

897.4344

##### Charge

2

##### Retention Time

29.53

##### Predicted Retention Time

-

##### Area

1.59E+08

##### Parts Per Million

2.7

##### Fragmentation mode

HCD

##### Originating file

01 D:\separate\_stitch\_analyses\xle-disambiguation\20210325\_F59\_3ug\_DENOVO\_12.csv

#### Meta Information from PEAKS

##### Scan Identifier

F3:5353

##### Original sequence

T

V

L

H

Q

D

W

##### Posttranslational Modifications

##### Source File

D:\separate\_stitch\_analyses\xle-disambiguation\raw\20210323\_F1\_UM1\_Peng0013\_SA\_F59\_ingel\_3ug\_chymo.raw

##### Fraction

3

##### Scan Feature

F3:2304

##### De Novo Score

97

##### ConfidenceScore

97

### m/z

449.7257

##### Mass

897.4344

##### Charge

2

##### Retention Time

29.53

##### Predicted Retention Time

-

##### Area

1.59E+08

##### Parts Per Million

2.7

##### Fragmentation mode

ETHCD

##### Originating file

01 D:\separate\_stitch\_analyses\xle-disambiguation\20210325\_F59\_3ug\_DENOVO\_12.csv

#### Meta Information from PEAKS

##### Scan Identifier

F3:5494

##### Original sequence

T

V

L

H

Q

D

W

##### Posttranslational Modifications

##### Source File

D:\separate\_stitch\_analyses\xle-disambiguation\raw\20210323\_F1\_UM1\_Peng0013\_SA\_F59\_ingel\_3ug\_chymo.raw

##### Fraction

3

##### Scan Feature

F3:2304

##### De Novo Score

97

##### ConfidenceScore

97

### m/z

449.7257

##### Mass

897.4344

##### Charge

2

##### Retention Time

29.53

##### Predicted Retention Time

-

##### Area

1.59E+08

##### Parts Per Million

2.7

##### Fragmentation mode

HCD

##### Originating file

01 D:\separate\_stitch\_analyses\xle-disambiguation\20210325\_F59\_3ug\_DENOVO\_12.csv

#### Meta Information from PEAKS

##### Scan Identifier

F3:4081

##### Original sequence

T

V

L

H

Q

D

W

+15.99

##### Posttranslational Modifications

Oxidation (HW)

##### Source File

D:\separate\_stitch\_analyses\xle-disambiguation\raw\20210323\_F1\_UM1\_Peng0013\_SA\_F59\_ingel\_3ug\_chymo.raw

##### Fraction

3

##### Scan Feature

F3:2544

##### De Novo Score

95

##### ConfidenceScore

95

### m/z

457.7224

##### Mass

913.4294

##### Charge

2

##### Retention Time

22.31

##### Predicted Retention Time

-

##### Area

3.126E+06

##### Parts Per Million

1

##### Fragmentation mode

ETHCD

##### Originating file

01 D:\separate\_stitch\_analyses\xle-disambiguation\20210325\_F59\_3ug\_DENOVO\_12.csv
