## Supplementary material for "A handle on mass coincidence errors in *de novo* sequencing of antibodies by bottom-up proteomics": Combined_082.html

Details Combined\_082 | Stitch OverviewUndefined

### Read Combined\_082

#### Sequence (length=11)

TISRDWKNSJY

#### Spectrum 3575? Spectrum 3575 The raw spectrum of this peptide as annotated by Hecklib. The fragments are coloured according to ion type (see legend). Any peaks with a star '\*' as text can be hovered over to see the full details, first the ion type second the mass shift type. By hovering over the amino acids in the peptide or ions in the legend the corresponding peaks are highlighted. By toggling the 'Unassigned' label you can turn the background (unassigned) peaks on or off in the plot. By updating the slider in the Ion legend you can update the spectrum to only show the top X% of the peaks with labels. The top X% means any peak that is within X% of the highest intensity. By dragging in the spectrum you can zoom in to a specific part of the spectrum and use 'Zoom Out' to get back to the original zoom level. The annotation of the spectrum is based on the given sequence in the peptides file and is done with different software so inconsistencies are likely. The peaks are annotated based on the given sequence, with 20 ppm tolerance.

Copy Data

##### Spectrum 3575 (TSV)

###### Preview

```
Loading example...
```

*Click on the button to copy the data to your clipboard.*

Mz MinMz MaxIntensity Max

WidthHeightPeptide font sizePeptide stroke widthSpectrum font sizeSpectrum stroke widthCompact peptide

Ion legend

wxyz

abcd

OtherUnassignedIon^Charge^_Position_Show for top:%

TISRDWKNSJY

01.96e+53.91e+55.87e+57.83e+5

Zoom Out

y^+1^_1_c^+1^_2_y^+1^_2_z^+3^_7_c^+1^_3_z^+1^_3_y^+1^_3_c^+2^_7_c^+2^_7_z^+2^_7_c^+2^_7_c^+1^_4_c^+1^_4_z^+1^_4_y^+1^_4_c^+2^_8_c^+2^_8_c^+2^_8_y^+2^_8_z^+2^_8_y^+2^_8_c^+2^_9_c^+2^_9_c^+2^_9_z^+2^_9_z^+2^_9_c^+1^_5_y^+2^_9_y^+2^_9_z^+2^_9_y^+2^_9_c^+1^_5_c^+2^_10_c^+2^_10_z^+1^_5_c^+2^_10_w^+2^_10_y^+1^_5_y^+2^_10_y^+2^_10_z^+2^_10_y^+2^_10_c^+1^_6_c^+1^_6_c^+1^_6_y^+1^_6_z^+1^_6_y^+1^_6_w^+1^_7_c^+1^_7_c^+1^_7_c^+1^_7_y^+1^_7_z^+1^_7_y^+1^_7_c^+1^_8_c^+1^_8_c^+1^_8_z^+1^_8_y^+1^_8_y^+1^_8_z^+1^_8_y^+1^_8_c^+1^_9_c^+1^_9_z^+1^_9_z^+1^_9_y^+1^_9_z^+1^_9_y^+1^_9_c^+1^_10_c^+1^_10_z^+1^_10_

0779155723363114

Fragment Matches Table

Show background peaks

| Position | Ion type | Intensity | mz Theoretical | mz Error (Th) | mz Error (ppm) | Charge | Series Number |
| --- | --- | --- | --- | --- | --- | --- | --- |
| - | - | 1051 | 120.6 | - | - | 0 | - |
| - | - | 2017 | 123.1 | - | - | 0 | - |
| - | - | 1103 | 129 | - | - | 0 | - |
| - | - | 2511 | 129.1 | - | - | 0 | - |
| - | - | 1058 | 135.2 | - | - | 0 | - |
| - | - | 2.868E+04 | 136.1 | - | - | 0 | - |
| - | - | 3002 | 137.1 | - | - | 0 | - |
| - | - | 1236 | 140 | - | - | 0 | - |
| - | - | 2014 | 142.1 | - | - | 0 | - |
| - | - | 3356 | 143.1 | - | - | 0 | - |
| - | - | 1148 | 144.1 | - | - | 0 | - |
| - | - | 3851 | 149 | - | - | 0 | - |
| - | - | 1153 | 156.4 | - | - | 0 | - |
| - | - | 1505 | 158.1 | - | - | 0 | - |
| - | - | 1171 | 159.2 | - | - | 0 | - |
| - | - | 2.544E+04 | 165.1 | - | - | 0 | - |
| - | - | 2620 | 169.1 | - | - | 0 | - |
| - | - | 2142 | 173.1 | - | - | 0 | - |
| - | - | 3637 | 173.5 | - | - | 0 | - |
| 11 | y | 9.071E+04 | 182.1 | 0.0003275 | 1.799 | +1 | 1 |
| - | - | 6791 | 183.1 | - | - | 0 | - |
| - | - | 2517 | 186.1 | - | - | 0 | - |
| - | - | 2.018E+05 | 187.1 | - | - | 0 | - |
| - | - | 1.753E+04 | 188.1 | - | - | 0 | - |
| - | - | 1930 | 189.1 | - | - | 0 | - |
| - | - | 2789 | 200.1 | - | - | 0 | - |
| - | - | 6758 | 201.1 | - | - | 0 | - |
| - | - | 2195 | 202.1 | - | - | 0 | - |
| - | - | 1287 | 210 | - | - | 0 | - |
| - | - | 1639 | 213.1 | - | - | 0 | - |
| - | - | 4.953E+04 | 215.1 | - | - | 0 | - |
| - | - | 4783 | 216.1 | - | - | 0 | - |
| - | - | 1595 | 221.1 | - | - | 0 | - |
| - | - | 2045 | 225 | - | - | 0 | - |
| - | - | 1813 | 225.1 | - | - | 0 | - |
| - | - | 5668 | 229.1 | - | - | 0 | - |
| 2 | c | 5236 | 232.2 | 0.0005239 | 2.257 | +1 | 2 |
| - | - | 2721 | 239.1 | - | - | 0 | - |
| - | - | 2826 | 240.1 | - | - | 0 | - |
| - | - | 2222 | 244.1 | - | - | 0 | - |
| - | - | 1374 | 276.8 | - | - | 0 | - |
| - | - | 2000 | 281.1 | - | - | 0 | - |
| - | - | 8639 | 282.1 | - | - | 0 | - |
| - | - | 1890 | 283.1 | - | - | 0 | - |
| - | - | 1767 | 287.2 | - | - | 0 | - |
| 10 | y | 9572 | 295.2 | 0.0003852 | 1.305 | +1 | 2 |
| - | - | 1921 | 296.1 | - | - | 0 | - |
| - | - | 2863 | 297.2 | - | - | 0 | - |
| 5 | z | 3046 | 298.1 | 0.004034 | 13.53 | +3 | 7 |
| - | - | 1746 | 299.1 | - | - | 0 | - |
| - | - | 2445 | 299.2 | - | - | 0 | - |
| - | - | 3.268E+04 | 300.2 | - | - | 0 | - |
| - | - | 3146 | 301.1 | - | - | 0 | - |
| - | - | 3700 | 301.2 | - | - | 0 | - |
| - | - | 6914 | 315.2 | - | - | 0 | - |
| - | - | 1.562E+04 | 317.2 | - | - | 0 | - |
| - | - | 2257 | 318.2 | - | - | 0 | - |
| 3 | c | 1.902E+04 | 319.2 | 0.0005847 | 1.832 | +1 | 3 |
| - | - | 1577 | 320.2 | - | - | 0 | - |
| - | - | 1661 | 335.8 | - | - | 0 | - |
| - | - | 1.456E+04 | 341 | - | - | 0 | - |
| - | - | 2839 | 343.2 | - | - | 0 | - |
| - | - | 2299 | 355.1 | - | - | 0 | - |
| - | - | 4.989E+04 | 359 | - | - | 0 | - |
| 9 | z | 2054 | 366.2 | 0.001241 | 3.389 | +1 | 3 |
| - | - | 2061 | 367.2 | - | - | 0 | - |
| - | - | 3070 | 370.1 | - | - | 0 | - |
| - | - | 1790 | 371.1 | - | - | 0 | - |
| - | - | 3610 | 374.2 | - | - | 0 | - |
| - | - | 2417 | 379.9 | - | - | 0 | - |
| 9 | y | 6881 | 382.2 | 2.696E-05 | 0.07054 | +1 | 3 |
| - | - | 1.436E+04 | 385.2 | - | - | 0 | - |
| - | - | 2308 | 385.5 | - | - | 0 | - |
| - | - | 6908 | 385.9 | - | - | 0 | - |
| - | - | 6294 | 386.2 | - | - | 0 | - |
| - | - | 1436 | 386.5 | - | - | 0 | - |
| - | - | 2954 | 387.2 | - | - | 0 | - |
| - | - | 1.117E+04 | 391.6 | - | - | 0 | - |
| - | - | 1.018E+04 | 391.9 | - | - | 0 | - |
| - | - | 9783 | 392.2 | - | - | 0 | - |
| - | - | 2202 | 392.5 | - | - | 0 | - |
| - | - | 3287 | 394.7 | - | - | 0 | - |
| - | - | 4402 | 415.3 | - | - | 0 | - |
| - | - | 2377 | 419.2 | - | - | 0 | - |
| - | - | 1861 | 424.7 | - | - | 0 | - |
| - | - | 2372 | 426.2 | - | - | 0 | - |
| - | - | 1.275E+05 | 429.1 | - | - | 0 | - |
| - | - | 5710 | 429.8 | - | - | 0 | - |
| - | - | 4001 | 430.1 | - | - | 0 | - |
| - | - | 1465 | 430.2 | - | - | 0 | - |
| - | - | 6828 | 430.2 | - | - | 0 | - |
| - | - | 5335 | 430.3 | - | - | 0 | - |
| - | - | 2350 | 430.8 | - | - | 0 | - |
| - | - | 1709 | 431.2 | - | - | 0 | - |
| - | - | 3260 | 431.3 | - | - | 0 | - |
| - | - | 1616 | 437.2 | - | - | 0 | - |
| - | - | 4615 | 438.7 | - | - | 0 | - |
| - | - | 5508 | 439.2 | - | - | 0 | - |
| - | - | 1.084E+04 | 442.7 | - | - | 0 | - |
| - | - | 1.134E+04 | 443.2 | - | - | 0 | - |
| 7 | c | 5463 | 443.7 | 0.006498 | 14.64 | +2 | 7 |
| 7 | c | 5701 | 444.2 | 0.001586 | 3.57 | +2 | 7 |
| - | - | 6.572E+04 | 445.1 | - | - | 0 | - |
| - | - | 3106 | 446.1 | - | - | 0 | - |
| 5 | z | 9237 | 446.2 | 0.003437 | 7.702 | +2 | 7 |
| - | - | 3.378E+04 | 451.8 | - | - | 0 | - |
| - | - | 1.003E+05 | 452.3 | - | - | 0 | - |
| 7 | c | 6.345E+04 | 452.8 | 0.002106 | 4.652 | +2 | 7 |
| - | - | 2.248E+04 | 453.3 | - | - | 0 | - |
| - | - | 2804 | 453.8 | - | - | 0 | - |
| - | - | 1862 | 455.6 | - | - | 0 | - |
| 4 | c | 5404 | 458.3 | 0.0007655 | 1.67 | +1 | 4 |
| - | - | 3317 | 458.8 | - | - | 0 | - |
| - | - | 1693 | 461.2 | - | - | 0 | - |
| - | - | 2287 | 461.6 | - | - | 0 | - |
| - | - | 2044 | 461.9 | - | - | 0 | - |
| - | - | 2433 | 462.2 | - | - | 0 | - |
| - | - | 2216 | 465.8 | - | - | 0 | - |
| - | - | 3010 | 466.7 | - | - | 0 | - |
| - | - | 1931 | 467.7 | - | - | 0 | - |
| - | - | 3709 | 471.2 | - | - | 0 | - |
| - | - | 4845 | 472.3 | - | - | 0 | - |
| - | - | 2723 | 474.2 | - | - | 0 | - |
| - | - | 2682 | 474.3 | - | - | 0 | - |
| 4 | c | 4.275E+05 | 475.3 | 0.0009141 | 1.923 | +1 | 4 |
| - | - | 9.207E+04 | 476.3 | - | - | 0 | - |
| - | - | 1.385E+04 | 477.3 | - | - | 0 | - |
| - | - | 1967 | 478.3 | - | - | 0 | - |
| - | - | 2122 | 479.3 | - | - | 0 | - |
| - | - | 2.331E+04 | 479.8 | - | - | 0 | - |
| 8 | z | 4833 | 480.2 | 0.001006 | 2.096 | +1 | 4 |
| - | - | 1.825E+04 | 480.3 | - | - | 0 | - |
| - | - | 1.215E+04 | 480.8 | - | - | 0 | - |
| - | - | 2.202E+04 | 481.2 | - | - | 0 | - |
| - | - | 5965 | 482.2 | - | - | 0 | - |
| - | - | 1503 | 483.2 | - | - | 0 | - |
| - | - | 2522 | 484.8 | - | - | 0 | - |
| - | - | 4720 | 485.2 | - | - | 0 | - |
| - | - | 4213 | 485.7 | - | - | 0 | - |
| - | - | 4072 | 486.8 | - | - | 0 | - |
| - | - | 1.199E+04 | 487.3 | - | - | 0 | - |
| - | - | 6629 | 487.8 | - | - | 0 | - |
| - | - | 3950 | 488.3 | - | - | 0 | - |
| - | - | 2241 | 491.8 | - | - | 0 | - |
| - | - | 3884 | 492.3 | - | - | 0 | - |
| - | - | 6615 | 493.8 | - | - | 0 | - |
| - | - | 8324 | 494.3 | - | - | 0 | - |
| - | - | 2307 | 494.7 | - | - | 0 | - |
| - | - | 3119 | 495.2 | - | - | 0 | - |
| 8 | y | 1.453E+04 | 496.2 | 0.0006247 | 1.259 | +1 | 4 |
| - | - | 4204 | 497.2 | - | - | 0 | - |
| - | - | 2.978E+04 | 499.8 | - | - | 0 | - |
| - | - | 2.585E+04 | 500.3 | - | - | 0 | - |
| 8 | c | 2.398E+04 | 500.8 | 0.007423 | 14.82 | +2 | 8 |
| 8 | c | 1.725E+04 | 501.3 | 0.003551 | 7.084 | +2 | 8 |
| - | - | 7041 | 501.8 | - | - | 0 | - |
| - | - | 4.04E+04 | 502.3 | - | - | 0 | - |
| - | - | 4314 | 502.8 | - | - | 0 | - |
| - | - | 9474 | 503.3 | - | - | 0 | - |
| - | - | 4498 | 508.3 | - | - | 0 | - |
| - | - | 6.7E+04 | 508.8 | - | - | 0 | - |
| - | - | 1.215E+05 | 509.3 | - | - | 0 | - |
| 8 | c | 1.363E+05 | 509.8 | 0.004863 | 9.539 | +2 | 8 |
| - | - | 5.524E+04 | 510.3 | - | - | 0 | - |
| - | - | 2.041E+04 | 510.8 | - | - | 0 | - |
| - | - | 4661 | 511.3 | - | - | 0 | - |
| - | - | 2248 | 511.8 | - | - | 0 | - |
| - | - | 2845 | 521.3 | - | - | 0 | - |
| - | - | 2893 | 522.3 | - | - | 0 | - |
| - | - | 1952 | 522.8 | - | - | 0 | - |
| - | - | 1837 | 523.3 | - | - | 0 | - |
| - | - | 5807 | 526.3 | - | - | 0 | - |
| - | - | 5141 | 526.8 | - | - | 0 | - |
| - | - | 2089 | 527.3 | - | - | 0 | - |
| - | - | 7306 | 530.3 | - | - | 0 | - |
| - | - | 2462 | 530.8 | - | - | 0 | - |
| 4 | y | 9927 | 532.8 | 0.0051 | 9.573 | +2 | 8 |
| 4 | z | 1.302E+04 | 533.3 | 0.001742 | 3.267 | +2 | 8 |
| - | - | 6503 | 533.8 | - | - | 0 | - |
| - | - | 2279 | 534.3 | - | - | 0 | - |
| - | - | 6024 | 535.3 | - | - | 0 | - |
| - | - | 7214 | 535.8 | - | - | 0 | - |
| - | - | 4217 | 536.3 | - | - | 0 | - |
| - | - | 2013 | 536.8 | - | - | 0 | - |
| - | - | 1634 | 540.8 | - | - | 0 | - |
| 4 | y | 2014 | 541.3 | 0.001591 | 2.94 | +2 | 8 |
| - | - | 2591 | 541.8 | - | - | 0 | - |
| 9 | c | 1.127E+04 | 544.3 | 0.006439 | 11.83 | +2 | 9 |
| 9 | c | 1.788E+04 | 544.8 | 0.00278 | 5.104 | +2 | 9 |
| - | - | 4635 | 545.3 | - | - | 0 | - |
| - | - | 1.655E+04 | 545.3 | - | - | 0 | - |
| - | - | 2833 | 546.3 | - | - | 0 | - |
| - | - | 1.098E+04 | 546.3 | - | - | 0 | - |
| - | - | 1983 | 547.3 | - | - | 0 | - |
| - | - | 1.314E+04 | 548.3 | - | - | 0 | - |
| - | - | 1.285E+04 | 548.7 | - | - | 0 | - |
| - | - | 8721 | 549.2 | - | - | 0 | - |
| - | - | 2601 | 549.8 | - | - | 0 | - |
| - | - | 2639 | 550.8 | - | - | 0 | - |
| - | - | 1982 | 552.3 | - | - | 0 | - |
| - | - | 8.602E+04 | 552.8 | - | - | 0 | - |
| 9 | c | 1.128E+05 | 553.3 | 0.003726 | 6.734 | +2 | 9 |
| - | - | 6.082E+04 | 553.8 | - | - | 0 | - |
| - | - | 2.588E+04 | 554.3 | - | - | 0 | - |
| - | - | 1.124E+04 | 554.8 | - | - | 0 | - |
| - | - | 4162 | 555.3 | - | - | 0 | - |
| - | - | 2679 | 561.3 | - | - | 0 | - |
| - | - | 1798 | 563.3 | - | - | 0 | - |
| - | - | 1.295E+04 | 564.3 | - | - | 0 | - |
| - | - | 2606 | 565.3 | - | - | 0 | - |
| - | - | 8114 | 565.3 | - | - | 0 | - |
| - | - | 5234 | 565.8 | - | - | 0 | - |
| - | - | 8841 | 566.3 | - | - | 0 | - |
| - | - | 5720 | 566.8 | - | - | 0 | - |
| 3 | z | 2088 | 567.8 | 0.001015 | 1.787 | +2 | 9 |
| 3 | z | 5831 | 568.3 | 0.00504 | 8.869 | +2 | 9 |
| - | - | 5038 | 568.8 | - | - | 0 | - |
| - | - | 1981 | 569.3 | - | - | 0 | - |
| - | - | 2040 | 570.8 | - | - | 0 | - |
| 5 | c | 7186 | 573.3 | 0.00815 | 14.22 | +1 | 5 |
| - | - | 4237 | 574.3 | - | - | 0 | - |
| - | - | 2915 | 575.3 | - | - | 0 | - |
| 3 | y | 1.179E+04 | 575.8 | 0.00438 | 7.607 | +2 | 9 |
| 3 | y | 3.584E+04 | 576.3 | 0.004406 | 7.645 | +2 | 9 |
| 3 | z | 4.48E+04 | 576.8 | 0.001765 | 3.06 | +2 | 9 |
| - | - | 2.273E+04 | 577.3 | - | - | 0 | - |
| - | - | 8680 | 577.8 | - | - | 0 | - |
| - | - | 5671 | 578.3 | - | - | 0 | - |
| - | - | 5535 | 578.8 | - | - | 0 | - |
| - | - | 6553 | 579.3 | - | - | 0 | - |
| - | - | 4879 | 579.8 | - | - | 0 | - |
| - | - | 2395 | 580.3 | - | - | 0 | - |
| - | - | 4454 | 583.3 | - | - | 0 | - |
| - | - | 7876 | 583.8 | - | - | 0 | - |
| - | - | 3044 | 584.2 | - | - | 0 | - |
| - | - | 9.226E+04 | 584.3 | - | - | 0 | - |
| 3 | y | 1.289E+05 | 584.8 | 0.003681 | 6.294 | +2 | 9 |
| - | - | 6.833E+04 | 585.3 | - | - | 0 | - |
| - | - | 2.785E+04 | 585.8 | - | - | 0 | - |
| - | - | 4870 | 586.3 | - | - | 0 | - |
| - | - | 1.108E+04 | 586.8 | - | - | 0 | - |
| - | - | 1.88E+04 | 587.3 | - | - | 0 | - |
| - | - | 1.423E+04 | 587.8 | - | - | 0 | - |
| - | - | 5901 | 588.3 | - | - | 0 | - |
| - | - | 5465 | 588.3 | - | - | 0 | - |
| - | - | 9493 | 588.8 | - | - | 0 | - |
| - | - | 3530 | 589.3 | - | - | 0 | - |
| - | - | 4694 | 589.3 | - | - | 0 | - |
| 5 | c | 7.748E+05 | 590.3 | 0.0007045 | 1.193 | +1 | 5 |
| - | - | 2.343E+05 | 591.3 | - | - | 0 | - |
| - | - | 8026 | 591.8 | - | - | 0 | - |
| - | - | 3.93E+04 | 592.3 | - | - | 0 | - |
| - | - | 1.337E+04 | 592.8 | - | - | 0 | - |
| - | - | 2170 | 593.3 | - | - | 0 | - |
| - | - | 4725 | 593.3 | - | - | 0 | - |
| - | - | 2465 | 593.8 | - | - | 0 | - |
| - | - | 1929 | 598.3 | - | - | 0 | - |
| 10 | c | 1.257E+04 | 600.8 | 0.00715 | 11.9 | +2 | 10 |
| 10 | c | 2.227E+04 | 601.3 | 0.00221 | 3.675 | +2 | 10 |
| - | - | 1.252E+04 | 601.8 | - | - | 0 | - |
| - | - | 5864 | 602.3 | - | - | 0 | - |
| - | - | 2468 | 603.3 | - | - | 0 | - |
| - | - | 2471 | 607.3 | - | - | 0 | - |
| - | - | 5847 | 607.8 | - | - | 0 | - |
| 7 | z | 3.96E+04 | 608.3 | 0.001382 | 2.271 | +1 | 5 |
| - | - | 2414 | 608.8 | - | - | 0 | - |
| - | - | 2.165E+05 | 609.3 | - | - | 0 | - |
| 10 | c | 2.897E+05 | 609.8 | 0.003338 | 5.474 | +2 | 10 |
| - | - | 1.715E+05 | 610.3 | - | - | 0 | - |
| - | - | 6.505E+04 | 610.8 | - | - | 0 | - |
| - | - | 1.506E+04 | 611.3 | - | - | 0 | - |
| - | - | 2117 | 611.8 | - | - | 0 | - |
| - | - | 4878 | 616.3 | - | - | 0 | - |
| - | - | 1923 | 616.8 | - | - | 0 | - |
| - | - | 1.004E+04 | 618.3 | - | - | 0 | - |
| 2 | w | 1.439E+04 | 618.8 | 0.00275 | 4.444 | +2 | 10 |
| - | - | 8461 | 619.3 | - | - | 0 | - |
| - | - | 3960 | 619.8 | - | - | 0 | - |
| - | - | 1947 | 620.8 | - | - | 0 | - |
| - | - | 2270 | 621.3 | - | - | 0 | - |
| - | - | 1858 | 621.8 | - | - | 0 | - |
| 7 | y | 5.46E+04 | 624.3 | 0.0004798 | 0.7685 | +1 | 5 |
| - | - | 2.229E+04 | 625.3 | - | - | 0 | - |
| - | - | 3473 | 626.4 | - | - | 0 | - |
| - | - | 7047 | 630.3 | - | - | 0 | - |
| - | - | 1.239E+04 | 630.8 | - | - | 0 | - |
| - | - | 7980 | 631.3 | - | - | 0 | - |
| - | - | 3182 | 631.8 | - | - | 0 | - |
| 2 | y | 3222 | 632.3 | 0.007105 | 11.24 | +2 | 10 |
| 2 | y | 5626 | 632.8 | 0.006075 | 9.6 | +2 | 10 |
| 2 | z | 1.353E+04 | 633.3 | 0.001439 | 2.272 | +2 | 10 |
| - | - | 1.455E+04 | 633.8 | - | - | 0 | - |
| - | - | 5375 | 634.3 | - | - | 0 | - |
| - | - | 8069 | 636.3 | - | - | 0 | - |
| - | - | 3295 | 637.4 | - | - | 0 | - |
| - | - | 3843 | 637.8 | - | - | 0 | - |
| - | - | 7973 | 638.3 | - | - | 0 | - |
| - | - | 5472 | 638.8 | - | - | 0 | - |
| - | - | 3332 | 639.3 | - | - | 0 | - |
| - | - | 2923 | 639.8 | - | - | 0 | - |
| - | - | 4855 | 640.3 | - | - | 0 | - |
| - | - | 1.408E+04 | 640.8 | - | - | 0 | - |
| 2 | y | 2.31E+04 | 641.3 | 0.002805 | 4.374 | +2 | 10 |
| - | - | 1.226E+04 | 641.8 | - | - | 0 | - |
| - | - | 5499 | 642.3 | - | - | 0 | - |
| - | - | 2031 | 643.3 | - | - | 0 | - |
| - | - | 4195 | 644.3 | - | - | 0 | - |
| - | - | 2431 | 645.3 | - | - | 0 | - |
| - | - | 3994 | 646.3 | - | - | 0 | - |
| - | - | 1.018E+04 | 647.4 | - | - | 0 | - |
| - | - | 3.251E+04 | 647.8 | - | - | 0 | - |
| - | - | 1.822E+04 | 648.3 | - | - | 0 | - |
| - | - | 7351 | 648.9 | - | - | 0 | - |
| - | - | 2376 | 649.3 | - | - | 0 | - |
| - | - | 7647 | 652.8 | - | - | 0 | - |
| - | - | 7544 | 653.3 | - | - | 0 | - |
| - | - | 7384 | 653.8 | - | - | 0 | - |
| - | - | 3265 | 654.3 | - | - | 0 | - |
| - | - | 4688 | 654.8 | - | - | 0 | - |
| - | - | 1.601E+04 | 655.3 | - | - | 0 | - |
| - | - | 1.978E+04 | 655.8 | - | - | 0 | - |
| - | - | 1.192E+04 | 656.3 | - | - | 0 | - |
| - | - | 4372 | 656.8 | - | - | 0 | - |
| - | - | 2.231E+04 | 659.3 | - | - | 0 | - |
| - | - | 4.01E+04 | 660.3 | - | - | 0 | - |
| - | - | 2.973E+04 | 660.9 | - | - | 0 | - |
| - | - | 5.117E+04 | 661.3 | - | - | 0 | - |
| - | - | 6.224E+04 | 661.8 | - | - | 0 | - |
| - | - | 7.859E+04 | 662.3 | - | - | 0 | - |
| - | - | 5.867E+04 | 662.8 | - | - | 0 | - |
| - | - | 2.686E+04 | 663.3 | - | - | 0 | - |
| - | - | 5488 | 663.8 | - | - | 0 | - |
| - | - | 1.384E+04 | 668.8 | - | - | 0 | - |
| - | - | 3.977E+04 | 669.3 | - | - | 0 | - |
| - | - | 4.407E+04 | 669.8 | - | - | 0 | - |
| - | - | 4.062E+04 | 670.3 | - | - | 0 | - |
| - | - | 4.356E+04 | 670.8 | - | - | 0 | - |
| - | - | 2.916E+04 | 671.3 | - | - | 0 | - |
| - | - | 1.04E+04 | 671.8 | - | - | 0 | - |
| - | - | 7646 | 672.3 | - | - | 0 | - |
| - | - | 2162 | 673.3 | - | - | 0 | - |
| - | - | 4828 | 673.9 | - | - | 0 | - |
| - | - | 1.671E+04 | 674.3 | - | - | 0 | - |
| - | - | 3.267E+04 | 674.8 | - | - | 0 | - |
| - | - | 3.769E+04 | 675.3 | - | - | 0 | - |
| - | - | 3.007E+04 | 675.8 | - | - | 0 | - |
| - | - | 3.201E+04 | 676.3 | - | - | 0 | - |
| - | - | 1.6E+04 | 676.8 | - | - | 0 | - |
| - | - | 4915 | 677.3 | - | - | 0 | - |
| - | - | 2243 | 677.8 | - | - | 0 | - |
| - | - | 1.591E+05 | 679.4 | - | - | 0 | - |
| - | - | 7.089E+04 | 680.4 | - | - | 0 | - |
| - | - | 1.569E+04 | 681.4 | - | - | 0 | - |
| - | - | 3563 | 682.4 | - | - | 0 | - |
| - | - | 7723 | 682.9 | - | - | 0 | - |
| - | - | 1.477E+05 | 683.3 | - | - | 0 | - |
| - | - | 2.108E+05 | 683.8 | - | - | 0 | - |
| - | - | 1.258E+05 | 684.3 | - | - | 0 | - |
| - | - | 4.929E+04 | 684.8 | - | - | 0 | - |
| - | - | 1.459E+04 | 685.3 | - | - | 0 | - |
| - | - | 2122 | 685.8 | - | - | 0 | - |
| - | - | 4903 | 687.4 | - | - | 0 | - |
| - | - | 2782 | 688.3 | - | - | 0 | - |
| - | - | 1.201E+05 | 691.4 | - | - | 0 | - |
| - | - | 1.971E+05 | 691.9 | - | - | 0 | - |
| - | - | 1.222E+05 | 692.4 | - | - | 0 | - |
| - | - | 5.29E+04 | 692.9 | - | - | 0 | - |
| - | - | 1.83E+04 | 693.4 | - | - | 0 | - |
| - | - | 5309 | 693.9 | - | - | 0 | - |
| - | - | 6186 | 694.3 | - | - | 0 | - |
| - | - | 3.131E+04 | 695.4 | - | - | 0 | - |
| - | - | 1.54E+04 | 696.4 | - | - | 0 | - |
| - | - | 2299 | 697.4 | - | - | 0 | - |
| - | - | 1.455E+05 | 704.4 | - | - | 0 | - |
| - | - | 1.009E+05 | 705.4 | - | - | 0 | - |
| - | - | 2.926E+04 | 706.4 | - | - | 0 | - |
| - | - | 6324 | 707.4 | - | - | 0 | - |
| - | - | 3707 | 711.3 | - | - | 0 | - |
| - | - | 1.939E+04 | 731.4 | - | - | 0 | - |
| - | - | 1.932E+04 | 732.4 | - | - | 0 | - |
| - | - | 8053 | 733.4 | - | - | 0 | - |
| - | - | 8903 | 737.3 | - | - | 0 | - |
| - | - | 3180 | 738.3 | - | - | 0 | - |
| - | - | 5.623E+04 | 749.4 | - | - | 0 | - |
| - | - | 1.975E+05 | 750.4 | - | - | 0 | - |
| - | - | 7.293E+04 | 751.4 | - | - | 0 | - |
| - | - | 1.87E+04 | 752.4 | - | - | 0 | - |
| - | - | 2933 | 753.4 | - | - | 0 | - |
| 6 | c | 1.301E+04 | 758.4 | 0.01287 | 16.97 | +1 | 6 |
| 6 | c | 7109 | 759.4 | 0.005001 | 6.586 | +1 | 6 |
| - | - | 2638 | 760.4 | - | - | 0 | - |
| - | - | 1.34E+05 | 775.4 | - | - | 0 | - |
| 6 | c | 1.082E+05 | 776.4 | 0.00854 | 11 | +1 | 6 |
| - | - | 3.394E+04 | 777.4 | - | - | 0 | - |
| - | - | 7872 | 778.4 | - | - | 0 | - |
| - | - | 2704 | 782.4 | - | - | 0 | - |
| - | - | 5299 | 787.4 | - | - | 0 | - |
| - | - | 4696 | 788.4 | - | - | 0 | - |
| - | - | 2734 | 789.4 | - | - | 0 | - |
| 6 | y | 2.715E+05 | 793.4 | 0.009239 | 11.65 | +1 | 6 |
| 6 | z | 1.669E+05 | 794.4 | 0.003591 | 4.52 | +1 | 6 |
| - | - | 5.721E+04 | 795.4 | - | - | 0 | - |
| - | - | 1.303E+04 | 796.4 | - | - | 0 | - |
| - | - | 4407 | 797.4 | - | - | 0 | - |
| - | - | 3483 | 809.4 | - | - | 0 | - |
| 6 | y | 1.682E+04 | 810.4 | 0.01432 | 17.67 | +1 | 6 |
| - | - | 7645 | 811.4 | - | - | 0 | - |
| - | - | 2016 | 820.4 | - | - | 0 | - |
| - | - | 2310 | 821.4 | - | - | 0 | - |
| - | - | 3024 | 822.4 | - | - | 0 | - |
| - | - | 9571 | 832.4 | - | - | 0 | - |
| - | - | 4770 | 833.4 | - | - | 0 | - |
| - | - | 2333 | 843.5 | - | - | 0 | - |
| - | - | 1912 | 844.5 | - | - | 0 | - |
| - | - | 3306 | 853.4 | - | - | 0 | - |
| - | - | 4424 | 858.5 | - | - | 0 | - |
| - | - | 1.398E+04 | 859.5 | - | - | 0 | - |
| - | - | 1.796E+04 | 860.5 | - | - | 0 | - |
| - | - | 9480 | 861.5 | - | - | 0 | - |
| - | - | 3716 | 862.5 | - | - | 0 | - |
| 5 | w | 8.488E+04 | 864.4 | 0.008991 | 10.4 | +1 | 7 |
| - | - | 1.05E+05 | 865.4 | - | - | 0 | - |
| - | - | 3.505E+04 | 866.4 | - | - | 0 | - |
| - | - | 1.474E+04 | 867.4 | - | - | 0 | - |
| - | - | 2340 | 873.4 | - | - | 0 | - |
| - | - | 3410 | 875.4 | - | - | 0 | - |
| - | - | 5106 | 884.5 | - | - | 0 | - |
| - | - | 7839 | 885.5 | - | - | 0 | - |
| 7 | c | 1.801E+04 | 886.5 | 0.01451 | 16.37 | +1 | 7 |
| 7 | c | 1.1E+04 | 887.5 | 0.006458 | 7.277 | +1 | 7 |
| - | - | 6729 | 888.5 | - | - | 0 | - |
| - | - | 3300 | 889.5 | - | - | 0 | - |
| - | - | 1E+04 | 900.5 | - | - | 0 | - |
| - | - | 1.243E+04 | 901.5 | - | - | 0 | - |
| - | - | 4.012E+04 | 902.5 | - | - | 0 | - |
| - | - | 1.636E+05 | 903.5 | - | - | 0 | - |
| 7 | c | 1.507E+05 | 904.5 | 0.007556 | 8.354 | +1 | 7 |
| - | - | 5.087E+04 | 905.5 | - | - | 0 | - |
| - | - | 1.6E+04 | 906.5 | - | - | 0 | - |
| - | - | 2847 | 907.5 | - | - | 0 | - |
| 5 | y | 8.087E+04 | 908.4 | 0.008725 | 9.604 | +1 | 7 |
| 5 | z | 1.098E+05 | 909.4 | 0.009354 | 10.29 | +1 | 7 |
| - | - | 4.761E+04 | 910.4 | - | - | 0 | - |
| - | - | 1.372E+04 | 911.4 | - | - | 0 | - |
| - | - | 7111 | 921.5 | - | - | 0 | - |
| - | - | 2851 | 922.5 | - | - | 0 | - |
| - | - | 7162 | 924.4 | - | - | 0 | - |
| 5 | y | 6511 | 925.4 | 0.007571 | 8.181 | +1 | 7 |
| - | - | 3332 | 926.4 | - | - | 0 | - |
| - | - | 3997 | 969.5 | - | - | 0 | - |
| - | - | 2948 | 970.5 | - | - | 0 | - |
| - | - | 2509 | 973.5 | - | - | 0 | - |
| - | - | 4926 | 974.5 | - | - | 0 | - |
| - | - | 3429 | 975.5 | - | - | 0 | - |
| - | - | 6512 | 986.5 | - | - | 0 | - |
| - | - | 1.537E+04 | 987.5 | - | - | 0 | - |
| - | - | 1.282E+04 | 988.5 | - | - | 0 | - |
| - | - | 4600 | 989.5 | - | - | 0 | - |
| - | - | 5714 | 999.5 | - | - | 0 | - |
| 8 | c | 1.341E+04 | 1001 | 0.01007 | 10.07 | +1 | 8 |
| 8 | c | 1.17E+04 | 1002 | 0.002084 | 2.08 | +1 | 8 |
| - | - | 6899 | 1003 | - | - | 0 | - |
| - | - | 5406 | 1004 | - | - | 0 | - |
| - | - | 5068 | 1005 | - | - | 0 | - |
| - | - | 3440 | 1006 | - | - | 0 | - |
| - | - | 1.035E+04 | 1017 | - | - | 0 | - |
| - | - | 1.033E+05 | 1018 | - | - | 0 | - |
| 8 | c | 1.196E+05 | 1019 | 0.008552 | 8.396 | +1 | 8 |
| - | - | 5.416E+04 | 1020 | - | - | 0 | - |
| - | - | 2.038E+04 | 1021 | - | - | 0 | - |
| - | - | 9617 | 1022 | - | - | 0 | - |
| - | - | 2729 | 1023 | - | - | 0 | - |
| 4 | z | 2437 | 1048 | 0.006172 | 5.886 | +1 | 8 |
| - | - | 2185 | 1055 | - | - | 0 | - |
| - | - | 3992 | 1063 | - | - | 0 | - |
| 4 | y | 4342 | 1064 | 0.002347 | 2.207 | +1 | 8 |
| 4 | y | 3.516E+04 | 1065 | 0.008566 | 8.047 | +1 | 8 |
| 4 | z | 9.351E+04 | 1066 | 0.008415 | 7.897 | +1 | 8 |
| - | - | 6.463E+04 | 1067 | - | - | 0 | - |
| - | - | 2.369E+04 | 1068 | - | - | 0 | - |
| - | - | 6982 | 1069 | - | - | 0 | - |
| - | - | 3274 | 1081 | - | - | 0 | - |
| 4 | y | 4402 | 1082 | 0.01737 | 16.06 | +1 | 8 |
| 9 | c | 6507 | 1089 | 0.004767 | 4.38 | +1 | 9 |
| - | - | 9009 | 1090 | - | - | 0 | - |
| - | - | 6909 | 1091 | - | - | 0 | - |
| - | - | 2646 | 1092 | - | - | 0 | - |
| - | - | 2196 | 1094 | - | - | 0 | - |
| - | - | 2900 | 1095 | - | - | 0 | - |
| - | - | 3395 | 1096 | - | - | 0 | - |
| - | - | 4920 | 1097 | - | - | 0 | - |
| - | - | 2.408E+04 | 1105 | - | - | 0 | - |
| 9 | c | 4.289E+04 | 1106 | 0.007377 | 6.673 | +1 | 9 |
| - | - | 2.166E+04 | 1107 | - | - | 0 | - |
| - | - | 1E+04 | 1108 | - | - | 0 | - |
| - | - | 2757 | 1109 | - | - | 0 | - |
| - | - | 2919 | 1110 | - | - | 0 | - |
| 3 | z | 4489 | 1135 | 0.00479 | 4.222 | +1 | 9 |
| 3 | z | 4369 | 1136 | 0.01247 | 10.98 | +1 | 9 |
| - | - | 3031 | 1137 | - | - | 0 | - |
| 3 | y | 2.361E+04 | 1152 | 0.00852 | 7.398 | +1 | 9 |
| 3 | z | 6.507E+04 | 1153 | 0.002113 | 1.833 | +1 | 9 |
| - | - | 6.361E+04 | 1154 | - | - | 0 | - |
| - | - | 2.553E+04 | 1155 | - | - | 0 | - |
| - | - | 9922 | 1156 | - | - | 0 | - |
| - | - | 2414 | 1158 | - | - | 0 | - |
| - | - | 4015 | 1159 | - | - | 0 | - |
| - | - | 7136 | 1160 | - | - | 0 | - |
| - | - | 5107 | 1161 | - | - | 0 | - |
| 3 | y | 3869 | 1169 | 0.001794 | 1.535 | +1 | 9 |
| - | - | 4300 | 1170 | - | - | 0 | - |
| - | - | 6924 | 1174 | - | - | 0 | - |
| - | - | 1.659E+04 | 1175 | - | - | 0 | - |
| - | - | 2.074E+04 | 1176 | - | - | 0 | - |
| - | - | 1.32E+04 | 1177 | - | - | 0 | - |
| - | - | 4474 | 1178 | - | - | 0 | - |
| 10 | c | 1.551E+04 | 1202 | 0.008716 | 7.254 | +1 | 10 |
| - | - | 2.606E+04 | 1203 | - | - | 0 | - |
| - | - | 1.512E+04 | 1204 | - | - | 0 | - |
| - | - | 5154 | 1205 | - | - | 0 | - |
| - | - | 1.329E+04 | 1218 | - | - | 0 | - |
| 10 | c | 4.39E+04 | 1219 | 0.002574 | 2.112 | +1 | 10 |
| - | - | 3.99E+04 | 1220 | - | - | 0 | - |
| - | - | 1.643E+04 | 1221 | - | - | 0 | - |
| - | - | 4633 | 1222 | - | - | 0 | - |
| 2 | z | 1.02E+04 | 1266 | 0.007939 | 6.273 | +1 | 10 |
| - | - | 9982 | 1267 | - | - | 0 | - |
| - | - | 6010 | 1268 | - | - | 0 | - |
| - | - | 3933 | 1269 | - | - | 0 | - |
| - | - | 7199 | 1306 | - | - | 0 | - |
| - | - | 1.684E+04 | 1307 | - | - | 0 | - |
| - | - | 1.341E+04 | 1308 | - | - | 0 | - |
| - | - | 8571 | 1309 | - | - | 0 | - |
| - | - | 3812 | 1310 | - | - | 0 | - |
| - | - | 2797 | 1311 | - | - | 0 | - |
| - | - | 4636 | 1321 | - | - | 0 | - |
| - | - | 1.289E+04 | 1322 | - | - | 0 | - |
| - | - | 1.434E+04 | 1323 | - | - | 0 | - |
| - | - | 2.398E+04 | 1324 | - | - | 0 | - |
| - | - | 6.949E+04 | 1325 | - | - | 0 | - |
| - | - | 4.581E+04 | 1326 | - | - | 0 | - |
| - | - | 2.087E+04 | 1327 | - | - | 0 | - |
| - | - | 5622 | 1328 | - | - | 0 | - |
| - | - | 5363 | 1338 | - | - | 0 | - |
| - | - | 5.289E+04 | 1339 | - | - | 0 | - |
| - | - | 6.565E+04 | 1340 | - | - | 0 | - |
| - | - | 3.929E+04 | 1341 | - | - | 0 | - |
| - | - | 1.838E+04 | 1342 | - | - | 0 | - |
| - | - | 4984 | 1343 | - | - | 0 | - |
| - | - | 1.175E+04 | 1349 | - | - | 0 | - |
| - | - | 2.937E+04 | 1350 | - | - | 0 | - |
| - | - | 2.669E+04 | 1351 | - | - | 0 | - |
| - | - | 1.446E+04 | 1352 | - | - | 0 | - |
| - | - | 4473 | 1353 | - | - | 0 | - |
| - | - | 4917 | 1356 | - | - | 0 | - |
| - | - | 3009 | 1357 | - | - | 0 | - |
| - | - | 3.024E+04 | 1366 | - | - | 0 | - |
| - | - | 1.767E+05 | 1367 | - | - | 0 | - |
| - | - | 2.441E+05 | 1368 | - | - | 0 | - |
| - | - | 1.352E+05 | 1369 | - | - | 0 | - |
| - | - | 5.422E+04 | 1370 | - | - | 0 | - |
| - | - | 1.546E+04 | 1371 | - | - | 0 | - |
| - | - | 5185 | 1372 | - | - | 0 | - |
| - | - | 5196 | 1382 | - | - | 0 | - |
| - | - | 2.573E+04 | 1383 | - | - | 0 | - |
| - | - | 5.259E+04 | 1384 | - | - | 0 | - |
| - | - | 4.758E+04 | 1385 | - | - | 0 | - |
| - | - | 2.756E+04 | 1386 | - | - | 0 | - |
| - | - | 1.062E+04 | 1387 | - | - | 0 | - |
| - | - | 2314 | 3083 | - | - | 0 | - |

m/z Charge Intensity FragmentType MassShift Position
120.5738525390625 0 1050.8041
123.1045913696289 0 2017.4248
129.0089569091797 0 1102.7021
129.1023712158203 0 2510.9438
135.189208984375 0 1057.8927
136.07598876953125 0 28681.652
137.07913208007812 0 3001.5117
140.01486206054688 0 1236.3022
142.1226806640625 0 2013.8837
143.11819458007812 0 3356.4531
144.1219940185547 0 1148.1533
149.04501342773438 0 3850.9832
156.37342834472656 0 1153.4169
158.11741638183594 0 1505.263
159.23390197753906 0 1171.3567
165.05491638183594 0 25443.791
169.1339874267578 0 2620.472
173.1288299560547 0 2141.7334
173.45181274414062 0 3636.517
182.0814971923828 0 90711.97 y 10
183.08486938476562 0 6791.4937
186.11241149902344 0 2516.776
187.1444549560547 0 201804.92
188.14788818359375 0 17525.316
189.1489715576172 0 1929.9166
200.1396484375 0 2789.1772
201.1235809326172 0 6757.782
202.0824737548828 0 2194.9124
210.0427703857422 0 1287.4474
213.1467742919922 0 1639.3873
215.13929748535156 0 49530.44
216.14271545410156 0 4783.442
221.08444213867188 0 1594.9493
225.04322814941406 0 2044.6901
225.12432861328125 0 1812.8801
229.1297149658203 0 5667.703
232.1660919189453 0 5235.6016 c 1
239.0956573486328 0 2720.9792
240.09559631347656 0 2825.5676
244.1412811279297 0 2222.1511
276.8227233886719 0 1373.527
281.1368408203125 0 2000.2053
282.14520263671875 0 8638.954
283.12994384765625 0 1889.6987
287.1714172363281 0 1767.2898
295.1656188964844 0 9571.652 y 9
296.1032409667969 0 1921.0839
297.1567077636719 0 2862.887
298.14093017578125 0 3045.6333 z Ammonia loss 4
299.14715576171875 0 1745.9612
299.1728820800781 0 2445.0977
300.1557922363281 0 32678.082
301.1395568847656 0 3145.9358
301.15899658203125 0 3700.0754
315.16705322265625 0 6913.8745
317.1824645996094 0 15620.541
318.1854248046875 0 2256.5051
319.19818115234375 0 19016.004 c 2
320.2005615234375 0 1577.438
335.8025207519531 0 1660.5978
341.0184631347656 0 14562.822
343.2213439941406 0 2838.8806
355.06842041015625 0 2299.4207
359.02874755859375 0 49892.895
366.1797790527344 0 2054.3625 z 8
367.1860046386719 0 2061.4116
370.1229248046875 0 3070.1665
371.1199035644531 0 1789.8274
374.18060302734375 0 3610.3948
379.87530517578125 0 2417.2349
382.1972351074219 0 6880.559 y 8
385.2196044921875 0 14360.708
385.5472412109375 0 2307.6313
385.8768310546875 0 6907.984
386.2247009277344 0 6294.37
386.5384216308594 0 1436.0017
387.2344055175781 0 2954.0754
391.5513610839844 0 11170.557
391.8808288574219 0 10181.368
392.2139892578125 0 9782.577
392.5497131347656 0 2202.0366
394.706787109375 0 3286.8518
415.26690673828125 0 4401.596
419.21539306640625 0 2376.8672
424.73443603515625 0 1861.3513
426.23345947265625 0 2371.7615
429.0894470214844 0 127502.266
429.75518798828125 0 5710.098
430.0903625488281 0 4000.913
430.20440673828125 0 1465.0052
430.2442626953125 0 6827.9604
430.2784729003906 0 5335.436
430.7507629394531 0 2349.8188
431.2264404296875 0 1709.4299
431.28131103515625 0 3260.0762
437.2112121582031 0 1615.8209
438.73272705078125 0 4614.8164
439.2349548339844 0 5507.804
442.74615478515625 0 10843.653
443.2459716796875 0 11340.307
443.7418212890625 0 5462.5024 c Water loss 6
444.2419128417969 0 5700.59 c Ammonia loss 6
445.1208190917969 0 65715.945
446.1221923828125 0 3105.6765
446.213134765625 0 9237.128 z Water loss 4
451.75067138671875 0 33778.508
452.2539367675781 0 100296.016
452.7514953613281 0 63454.504 c 6
453.2511291503906 0 22478.998
453.7521057128906 0 2804.4766
455.5718078613281 0 1861.6115
458.2713928222656 0 5403.8467 c Ammonia loss 3
458.7580871582031 0 3316.6714
461.2297058105469 0 1693.2799
461.5699157714844 0 2287.2524
461.9034118652344 0 2044.1383
462.239013671875 0 2432.835
465.7674560546875 0 2215.7478
466.7303771972656 0 3009.9019
467.7265625 0 1931.2411
471.2492370605469 0 3709.076
472.2504577636719 0 4845.013
474.1943359375 0 2722.57
474.2930603027344 0 2681.8237
475.29962158203125 0 427472.06 c 3
476.30230712890625 0 92065.766
477.3052062988281 0 13854.897
478.3088684082031 0 1966.801
479.2583923339844 0 2121.7195
479.75738525390625 0 23312.887
480.220458984375 0 4833.0015 z 7
480.2552795410156 0 18253.615
480.754150390625 0 12149.62
481.2296447753906 0 22022.44
482.2327880859375 0 5964.6626
483.2323913574219 0 1503.0713
484.751220703125 0 2521.8088
485.24371337890625 0 4719.5073
485.7371826171875 0 4212.514
486.7679443359375 0 4072.079
487.2677307128906 0 11988.405
487.7650451660156 0 6629.2236
488.2666931152344 0 3950.4998
491.75714111328125 0 2241.4253
492.2528076171875 0 3884.2341
493.7548522949219 0 6614.511
494.25018310546875 0 8323.6455
494.7492370605469 0 2306.8167
495.2308044433594 0 3119.3398
496.2408142089844 0 14531.915 y 7
497.22564697265625 0 4204.044
499.7671813964844 0 29777.674
500.26336669921875 0 25847.3
500.7623596191406 0 23979.787 c Water loss 7
501.25823974609375 0 17254.115 c Ammonia loss 7
501.76275634765625 0 7040.5938
502.27447509765625 0 40400.008
502.76446533203125 0 4314.4404
503.27435302734375 0 9473.911
508.26422119140625 0 4497.567
508.77203369140625 0 66998.91
509.2725830078125 0 121501.55
509.77020263671875 0 136278.6 c 7
510.26934814453125 0 55240.938
510.76953125 0 20411.967
511.26824951171875 0 4661.3984
511.76580810546875 0 2247.9104
521.2738037109375 0 2845.4707
522.2737426757812 0 2893.2935
522.7762451171875 0 1952.4797
523.2667846679688 0 1836.6146
526.2559814453125 0 5806.7944
526.7532348632812 0 5140.709
527.25146484375 0 2088.5361
530.2931518554688 0 7305.955
530.7794189453125 0 2462.1072
532.7667236328125 0 9926.8125 y Ammonia loss 3
533.2637939453125 0 13021.5205 z 3
533.76220703125 0 6503.267
534.26171875 0 2278.7466
535.2748413085938 0 6023.9253
535.7697143554688 0 7214.205
536.2653198242188 0 4217.1035
536.7930297851562 0 2012.7396
540.7716064453125 0 1633.7706
541.2764892578125 0 2013.6311 y 3
541.7734985351562 0 2590.6936
544.2793579101562 0 11272.439 c Water loss 8
544.7750244140625 0 17878.182 c Ammonia loss 8
545.2689819335938 0 4635.0894
545.3052368164062 0 16552.266
546.259765625 0 2833.2322
546.3106689453125 0 10984.064
547.2575073242188 0 1983.4982
548.2515869140625 0 13142.577
548.7471313476562 0 12845.884
549.24755859375 0 8720.57
549.7539672851562 0 2600.6401
550.7944946289062 0 2638.8965
552.2527465820312 0 1981.9917
552.7919921875 0 86021.695
553.287353515625 0 112806.42 c 8
553.7860717773438 0 60823.656
554.2855834960938 0 25883.445
554.7827758789062 0 11235.988
555.2784423828125 0 4162.4795
561.2792358398438 0 2679.411
563.3206787109375 0 1798.4915
564.30419921875 0 12953.273
565.2621459960938 0 2605.626
565.3096923828125 0 8113.775
565.7993774414062 0 5234.0903
566.2955322265625 0 8840.735
566.7913208007812 0 5719.5244
567.7772827148438 0 2087.7834 z Water loss 2
568.2733154296875 0 5831.304 z Ammonia loss 2
568.777587890625 0 5038.2456
569.2765502929688 0 1980.9694
570.8023681640625 0 2039.6882
573.3072509765625 0 7186.4604 c Ammonia loss 4
574.3002319335938 0 4236.5776
575.2837524414062 0 2915.2903
575.78125 0 11794.317 y Water loss 2
576.2820434570312 0 35835.04 y Ammonia loss 2
576.77978515625 0 44801.133 z 2
577.2772216796875 0 22730.58
577.7781372070312 0 8679.559
578.315673828125 0 5670.535
578.809326171875 0 5535.2725
579.3098754882812 0 6553.1523
579.8114624023438 0 4879.1426
580.3126831054688 0 2395.4941
583.3035888671875 0 4453.601
583.7986450195312 0 7876.2383
584.237060546875 0 3044.0244
584.2919311523438 0 92264.13
584.7872314453125 0 128901.05 y 2
585.2864990234375 0 68325.32
585.7863159179688 0 27851.887
586.2874145507812 0 4870.171
586.8230590820312 0 11075.013
587.3201293945312 0 18802.922
587.8189086914062 0 14225.62
588.27392578125 0 5900.665
588.322265625 0 5465.0366
588.7721557617188 0 9493.327
589.2723388671875 0 3529.7031
589.3201293945312 0 4694.288
590.3263549804688 0 774828.94 c 4
591.3291015625 0 234303.03
591.8156127929688 0 8026.08
592.3284301757812 0 39296.684
592.8071899414062 0 13366.25
593.2959594726562 0 2169.6802
593.3336181640625 0 4724.521
593.8147583007812 0 2465.2742
598.2952880859375 0 1929.1027
600.8206787109375 0 12565.53 c Water loss 9
601.317626953125 0 22265.7 c Ammonia loss 9
601.818115234375 0 12518.257
602.3201293945312 0 5864.426
603.3201904296875 0 2467.955
607.3245239257812 0 2470.6548
607.8250732421875 0 5847.326
608.3178100585938 0 39599.617 z 6
608.8301391601562 0 2413.9768
609.3336181640625 0 216481.06
609.8297729492188 0 289707.2 c 9
610.3292236328125 0 171450.08
610.82861328125 0 65046.83
611.3287353515625 0 15058.89
611.8276977539062 0 2116.7268
616.34423828125 0 4877.789
616.824951171875 0 1923.2804
618.3045654296875 0 10039.675
618.80126953125 0 14388.312 w 1
619.3005981445312 0 8460.665
619.800048828125 0 3960.1067
620.828125 0 1947.4302
621.3291015625 0 2269.6404
621.8153686523438 0 1857.877
624.3356323242188 0 54601.65 y 6
625.3326416015625 0 22293.98
626.3512573242188 0 3472.6626
630.32763671875 0 7046.8184
630.8228149414062 0 12393.108
631.3255615234375 0 7980.3403
631.8251953125 0 3181.622
632.320556640625 0 3222.4119 y Water loss 1
632.8257446289062 0 5626.1484 y Ammonia loss 1
633.3221435546875 0 13533.614 z 1
633.8163452148438 0 14545.8955
634.3175048828125 0 5374.703
636.3489379882812 0 8068.702
637.3521728515625 0 3295.0396
637.8330078125 0 3842.6987
638.33349609375 0 7973.041
638.8369750976562 0 5472.054
639.3436889648438 0 3332.2942
639.82666015625 0 2923.2153
640.342529296875 0 4855.298
640.83349609375 0 14076.387
641.3301391601562 0 23102.447 y 1
641.828125 0 12264.551
642.3294067382812 0 5499.328
643.33251953125 0 2031.4951
644.3399047851562 0 4194.6567
645.3232421875 0 2430.752
646.3195190429688 0 3994.1743
647.35107421875 0 10182.7705
647.849853515625 0 32508.5
648.3495483398438 0 18217.955
648.8522338867188 0 7351.142
649.3388061523438 0 2376.2893
652.843994140625 0 7646.75
653.3381958007812 0 7544.428
653.839599609375 0 7384.236
654.3291625976562 0 3264.6235
654.8294677734375 0 4688.491
655.3193969726562 0 16008.615
655.8139038085938 0 19778.447
656.311279296875 0 11916.309
656.8154907226562 0 4372.3135
659.3478393554688 0 22313.234
660.3438720703125 0 40096.062
660.8504028320312 0 29733.688
661.3489990234375 0 51165.33
661.8477783203125 0 62243.043
662.3446655273438 0 78586.94
662.8436889648438 0 58672.117
663.3428344726562 0 26857.375
663.8460083007812 0 5487.535
668.8296508789062 0 13841.929
669.347412109375 0 39765.258
669.84716796875 0 44074.883
670.3458862304688 0 40617.56
670.843017578125 0 43562.582
671.341552734375 0 29158.033
671.841064453125 0 10400.355
672.3440551757812 0 7645.987
673.3353881835938 0 2161.51
673.851318359375 0 4827.5684
674.3446044921875 0 16705.1
674.8428955078125 0 32673.979
675.3424072265625 0 37691.316
675.8397827148438 0 30069.793
676.3360595703125 0 32010.814
676.8341674804688 0 16002.141
677.3330688476562 0 4915.1875
677.8363037109375 0 2243.0535
679.3543090820312 0 159145.94
680.3521728515625 0 70890.58
681.3529052734375 0 15690.561
682.3693237304688 0 3563.4897
682.8560180664062 0 7723.0303
683.3488159179688 0 147657.38
683.844970703125 0 210815.17
684.3440551757812 0 125767.47
684.8429565429688 0 49288.664
685.3441162109375 0 14593.024
685.8460083007812 0 2122.3872
687.3515625 0 4902.5337
688.3253784179688 0 2782.4062
691.3576049804688 0 120109.64
691.8538208007812 0 197094.58
692.3531494140625 0 122238.29
692.8522338867188 0 52896.484
693.3541870117188 0 18300.248
693.855224609375 0 5308.6943
694.3291625976562 0 6185.8975
695.3726196289062 0 31313.914
696.36865234375 0 15404.64
697.3670043945312 0 2299.1897
704.3692016601562 0 145535.2
705.3592529296875 0 100907.64
706.361328125 0 29257.88
707.3587646484375 0 6323.5234
711.3424682617188 0 3706.6223
731.392822265625 0 19393.768
732.3805541992188 0 19317.076
733.3828735351562 0 8053.1396
737.333984375 0 8902.609
738.3154296875 0 3179.7861
749.38525390625 0 56226.27
750.3900146484375 0 197480.2
751.3931884765625 0 72926.61
752.3950805664062 0 18703.342
753.3998413085938 0 2933.1733
758.3815307617188 0 13005.982 c Water loss 5
759.3734130859375 0 7109.4443 c Ammonia loss 5
760.3773803710938 0 2637.651
775.4061889648438 0 133966.66
776.3964233398438 0 108219.234 c 5
777.3973388671875 0 33943.01
778.3978271484375 0 7872.3164
782.3780517578125 0 2704.3877
787.3883666992188 0 5299.351
788.3899536132812 0 4695.6665
789.3905639648438 0 2734.488
793.3971557617188 0 271505.28 y Ammonia loss 5
794.3921508789062 0 166936.69 z 5
795.3876953125 0 57211.17
796.384765625 0 13026.66
797.3887939453125 0 4407.4146
809.4172973632812 0 3482.634
810.400146484375 0 16817.34 y 5
811.3958129882812 0 7644.951
820.4231567382812 0 2015.92
821.4338989257812 0 2310.0933
822.421142578125 0 3023.734
832.4434204101562 0 9570.711
833.4435424804688 0 4770.2646
843.474609375 0 2333.1033
844.4553833007812 0 1911.6423
853.353515625 0 3305.587
858.4965209960938 0 4423.529
859.4867553710938 0 13980.354
860.4835815429688 0 17964.428
861.4811401367188 0 9480.294
862.485595703125 0 3716.4058
864.4340209960938 0 84881.24 w 4
865.4232788085938 0 104960.93
866.4221801757812 0 35053.977
867.424072265625 0 14744.942
873.4255981445312 0 2340.3284
875.4165649414062 0 3409.8083
884.4815673828125 0 5105.946
885.47607421875 0 7838.7646
886.474853515625 0 18008.4 c Water loss 6
887.4669189453125 0 10998.401 c Ammonia loss 6
888.4788208007812 0 6729.0254
889.4844360351562 0 3300.4685
900.4763793945312 0 10000.855
901.470703125 0 12426.517
902.4906005859375 0 40123.69
903.4993286132812 0 163642.77
904.4923706054688 0 150711.6 c 6
905.4930419921875 0 50865.625
906.4937133789062 0 15998.723
907.494140625 0 2846.9812
908.423583984375 0 80868.445 y Ammonia loss 4
909.413330078125 0 109767.445 z 4
910.4122314453125 0 47612.07
911.413330078125 0 13718.782
921.4566040039062 0 7110.9766
922.4542236328125 0 2851.3765
924.4417724609375 0 7162.4116
925.433837890625 0 6511.0376 y 4
926.4330444335938 0 3332.2156
969.473876953125 0 3997.1653
970.4697265625 0 2947.5464
973.5266723632812 0 2508.9287
974.5225219726562 0 4926.2173
975.5281372070312 0 3429.2598
986.5029296875 0 6511.6396
987.4967651367188 0 15371.377
988.4933471679688 0 12821.236
989.4993896484375 0 4599.9873
999.5303955078125 0 5714.034
1000.522216796875 0 13408.878 c Water loss 7
1001.5142211914062 0 11702.1455 c Ammonia loss 7
1002.5191040039062 0 6898.5938
1003.5171508789062 0 5405.799
1004.5165405273438 0 5068.447
1005.5223999023438 0 3439.56
1016.532958984375 0 10347.569
1017.5419921875 0 103257.55
1018.5343017578125 0 119640.26 c 7
1019.53271484375 0 54156.15
1020.5298461914062 0 20380.299
1021.5311279296875 0 9617.38
1022.516845703125 0 2729.3381
1048.50341796875 0 2436.9236 z Ammonia loss 3
1054.5174560546875 0 2184.6414
1062.5595703125 0 3992.0906
1063.5343017578125 0 4341.8154 y Water loss 3
1064.5245361328125 0 35158.496 y Ammonia loss 3
1065.515380859375 0 93513.766 z 3
1066.5157470703125 0 64625.664
1067.5155029296875 0 23693.383
1068.525146484375 0 6981.939
1080.531982421875 0 3273.5742
1081.525146484375 0 4402.3193 y 3
1088.5531005859375 0 6507.091 c Ammonia loss 8
1089.5501708984375 0 9009.25
1090.5494384765625 0 6909.061
1091.546142578125 0 2645.781
1093.53076171875 0 2196.0862
1095.4949951171875 0 2899.6445
1096.48828125 0 3394.6975
1097.4937744140625 0 4919.6895
1104.5750732421875 0 24080.95
1105.5675048828125 0 42887.746 c 8
1106.5679931640625 0 21658.564
1107.56494140625 0 10002.059
1108.55126953125 0 2757.1614
1109.552490234375 0 2918.5315
1134.550048828125 0 4488.9854 z Water loss 2
1135.541748046875 0 4368.911 z Ammonia loss 2
1136.5379638671875 0 3031.1514
1151.5565185546875 0 23606.166 y Ammonia loss 2
1152.5537109375 0 65065.453 z 2
1153.552978515625 0 63607.363
1154.5533447265625 0 25525.662
1155.5540771484375 0 9921.771
1157.621337890625 0 2413.9187
1158.6202392578125 0 4015.022
1159.62646484375 0 7136.4443
1160.6251220703125 0 5106.9395
1168.57275390625 0 3868.8787 y 2
1169.5693359375 0 4300.4077
1173.6461181640625 0 6923.6455
1174.645263671875 0 16587.275
1175.6470947265625 0 20736.959
1176.643798828125 0 13196.1
1177.6427001953125 0 4474.4766
1201.64111328125 0 15513.48 c Ammonia loss 9
1202.632080078125 0 26057.68
1203.6280517578125 0 15120.004
1204.6395263671875 0 5154.3755
1217.6578369140625 0 13293.219
1218.6563720703125 0 43897.562 c 9
1219.6571044921875 0 39902.49
1220.658203125 0 16432.615
1221.6517333984375 0 4633.0127
1265.6478271484375 0 10196.715 z 1
1266.6376953125 0 9981.906
1267.6376953125 0 6010.1294
1268.6357421875 0 3933.324
1305.6717529296875 0 7199.306
1306.6517333984375 0 16842.33
1307.65234375 0 13407.75
1308.6492919921875 0 8570.699
1309.646484375 0 3811.5505
1310.6512451171875 0 2796.904
1320.6953125 0 4636.0347
1321.6893310546875 0 12891.789
1322.6893310546875 0 14337.265
1323.6810302734375 0 23976.758
1324.6778564453125 0 69490.875
1325.6761474609375 0 45811.844
1326.6749267578125 0 20874.826
1327.6715087890625 0 5622.1826
1337.69580078125 0 5363.2466
1338.699462890625 0 52892.43
1339.693359375 0 65654.05
1340.6923828125 0 39293.06
1341.68896484375 0 18383.898
1342.67626953125 0 4984.0117
1348.6849365234375 0 11750.197
1349.67333984375 0 29367.688
1350.6669921875 0 26692.24
1351.6636962890625 0 14456.822
1352.66064453125 0 4472.8374
1355.7174072265625 0 4917.0264
1356.7171630859375 0 3008.603
1365.6905517578125 0 30235.82
1366.6922607421875 0 176727.53
1367.6866455078125 0 244129.03
1368.685302734375 0 135223.5
1369.6846923828125 0 54216.676
1370.6866455078125 0 15463.151
1371.6954345703125 0 5184.5728
1381.7144775390625 0 5196.4043
1382.7054443359375 0 25734.13
1383.7093505859375 0 52587.77
1384.708740234375 0 47576.934
1385.7095947265625 0 27556.934
1386.7093505859375 0 10619.674
3083.314208984375 0 2313.5425

Spectrum Details

|  |  |
| --- | --- |
| Matched peaks? Matched peaks The total absolute number of peaks matched. Additionally in brackets the total fraction of peaks matched and the total number of peaks is shown. | 73 (12.81% of 570) |
| FDR? FDR The false discovery rate estimated for this peptide. It is calculated by matching all theoretical fragments with a non-integer shift with the raw peaks for this spectrum. This is done with 40 different shifts. The resulting percentage is the average number of annotated peaks over the number of annotated peaks with the correct spectrum. | 1.17% |
| Satellite FDR? Satellite FDR See the FDR for details on its calculation. This satellite ion specific FDR only contains the satellite ions (d/w) for I/L/J positions. | 0.00% |
| PSM Score? PSM Score The PSM Score as given by Hecklib to this annotated spectrum. It is shown with three significant figures. | 671 |

#### Spectrum 3718? Spectrum 3718 The raw spectrum of this peptide as annotated by Hecklib. The fragments are coloured according to ion type (see legend). Any peaks with a star '\*' as text can be hovered over to see the full details, first the ion type second the mass shift type. By hovering over the amino acids in the peptide or ions in the legend the corresponding peaks are highlighted. By toggling the 'Unassigned' label you can turn the background (unassigned) peaks on or off in the plot. By updating the slider in the Ion legend you can update the spectrum to only show the top X% of the peaks with labels. The top X% means any peak that is within X% of the highest intensity. By dragging in the spectrum you can zoom in to a specific part of the spectrum and use 'Zoom Out' to get back to the original zoom level. The annotation of the spectrum is based on the given sequence in the peptides file and is done with different software so inconsistencies are likely. The peaks are annotated based on the given sequence, with 20 ppm tolerance.

Copy Data

##### Spectrum 3718 (TSV)

###### Preview

```
Loading example...
```

*Click on the button to copy the data to your clipboard.*

Mz MinMz MaxIntensity Max

WidthHeightPeptide font sizePeptide stroke widthSpectrum font sizeSpectrum stroke widthCompact peptide

Ion legend

wxyz

abcd

OtherUnassignedIon^Charge^_Position_Show for top:%

TISRDWKNSJY

07.49e+41.50e+52.25e+52.99e+5

Zoom Out

y^+3^_4_y^+1^_1_c^+1^_2_y^+1^_2_z^+3^_7_c^+3^_7_c^+1^_3_y^+3^_8_y^+1^_3_z^+2^_6_c^+2^_7_c^+2^_7_z^+2^_7_c^+2^_7_c^+1^_4_z^+1^_4_y^+2^_7_c^+1^_4_z^+1^_4_y^+1^_4_c^+2^_8_c^+2^_8_c^+2^_8_y^+2^_8_z^+2^_8_y^+2^_8_c^+2^_9_c^+2^_9_w^+1^_5_c^+2^_9_z^+2^_9_w^+2^_9_c^+1^_5_y^+2^_9_y^+2^_9_z^+2^_9_y^+2^_9_c^+1^_5_c^+2^_10_c^+2^_10_z^+1^_5_c^+2^_10_w^+2^_10_y^+1^_5_w^+2^_10_y^+2^_10_z^+2^_10_y^+2^_10_c^+1^_6_c^+1^_6_c^+1^_6_y^+1^_6_z^+1^_6_y^+1^_6_w^+1^_7_c^+1^_7_c^+1^_7_z^+1^_7_c^+1^_7_y^+1^_7_z^+1^_7_y^+1^_7_c^+1^_8_c^+1^_8_c^+1^_8_z^+1^_8_y^+1^_8_y^+1^_8_z^+1^_8_y^+1^_8_c^+1^_9_c^+1^_9_z^+1^_9_z^+1^_9_y^+1^_9_z^+1^_9_y^+1^_9_c^+1^_10_c^+1^_10_y^+1^_10_z^+1^_10_y^+1^_10_

035070010501401

Fragment Matches Table

Show background peaks

| Position | Ion type | Intensity | mz Theoretical | mz Error (Th) | mz Error (ppm) | Charge | Series Number |
| --- | --- | --- | --- | --- | --- | --- | --- |
| - | - | 602 | 123 | - | - | 0 | - |
| - | - | 656.8 | 123.1 | - | - | 0 | - |
| - | - | 1280 | 129.1 | - | - | 0 | - |
| - | - | 681.2 | 131.1 | - | - | 0 | - |
| - | - | 1.396E+04 | 136.1 | - | - | 0 | - |
| - | - | 983.8 | 137.1 | - | - | 0 | - |
| - | - | 1141 | 142.1 | - | - | 0 | - |
| - | - | 415.5 | 142.6 | - | - | 0 | - |
| - | - | 1631 | 143.1 | - | - | 0 | - |
| - | - | 498.1 | 147 | - | - | 0 | - |
| - | - | 477.9 | 148.2 | - | - | 0 | - |
| - | - | 463.9 | 148.7 | - | - | 0 | - |
| - | - | 568.9 | 148.9 | - | - | 0 | - |
| - | - | 463.4 | 148.9 | - | - | 0 | - |
| - | - | 429.8 | 148.9 | - | - | 0 | - |
| - | - | 1059 | 148.9 | - | - | 0 | - |
| - | - | 1066 | 148.9 | - | - | 0 | - |
| - | - | 2062 | 148.9 | - | - | 0 | - |
| - | - | 3794 | 148.9 | - | - | 0 | - |
| - | - | 4694 | 149 | - | - | 0 | - |
| - | - | 2854 | 149 | - | - | 0 | - |
| - | - | 947.1 | 149 | - | - | 0 | - |
| - | - | 661.3 | 149 | - | - | 0 | - |
| - | - | 655.7 | 149 | - | - | 0 | - |
| - | - | 484.7 | 149 | - | - | 0 | - |
| - | - | 422.6 | 149 | - | - | 0 | - |
| - | - | 447.2 | 149 | - | - | 0 | - |
| - | - | 4524 | 149 | - | - | 0 | - |
| - | - | 714.9 | 149.1 | - | - | 0 | - |
| - | - | 1207 | 157.1 | - | - | 0 | - |
| - | - | 423.6 | 161 | - | - | 0 | - |
| - | - | 454.1 | 161.7 | - | - | 0 | - |
| - | - | 9897 | 165.1 | - | - | 0 | - |
| - | - | 820.1 | 166.1 | - | - | 0 | - |
| 8 | y | 618.3 | 166.1 | 0.00203 | 12.23 | +3 | 4 |
| - | - | 1148 | 167.1 | - | - | 0 | - |
| - | - | 430.6 | 168.1 | - | - | 0 | - |
| - | - | 721.2 | 169.1 | - | - | 0 | - |
| - | - | 1283 | 173.1 | - | - | 0 | - |
| - | - | 978.8 | 173.4 | - | - | 0 | - |
| - | - | 844.4 | 175.1 | - | - | 0 | - |
| - | - | 608.3 | 181.1 | - | - | 0 | - |
| 11 | y | 3.587E+04 | 182.1 | 0.0003428 | 1.883 | +1 | 1 |
| - | - | 3402 | 183.1 | - | - | 0 | - |
| - | - | 607.2 | 186.1 | - | - | 0 | - |
| - | - | 1179 | 186.1 | - | - | 0 | - |
| - | - | 469.6 | 186.9 | - | - | 0 | - |
| - | - | 617.3 | 187.1 | - | - | 0 | - |
| - | - | 8.538E+04 | 187.1 | - | - | 0 | - |
| - | - | 7672 | 188.1 | - | - | 0 | - |
| - | - | 733.9 | 196.1 | - | - | 0 | - |
| - | - | 1374 | 200.1 | - | - | 0 | - |
| - | - | 2705 | 201.1 | - | - | 0 | - |
| - | - | 612.1 | 201.1 | - | - | 0 | - |
| - | - | 868.8 | 201.1 | - | - | 0 | - |
| - | - | 1728 | 202.1 | - | - | 0 | - |
| - | - | 506.7 | 202.1 | - | - | 0 | - |
| - | - | 3313 | 203.1 | - | - | 0 | - |
| - | - | 716.2 | 203.1 | - | - | 0 | - |
| - | - | 1.94E+04 | 215.1 | - | - | 0 | - |
| - | - | 2365 | 216.1 | - | - | 0 | - |
| - | - | 2581 | 221.1 | - | - | 0 | - |
| - | - | 887 | 222.1 | - | - | 0 | - |
| - | - | 1055 | 223.1 | - | - | 0 | - |
| - | - | 1349 | 225 | - | - | 0 | - |
| - | - | 744 | 226 | - | - | 0 | - |
| - | - | 470.4 | 226.2 | - | - | 0 | - |
| - | - | 1472 | 229.1 | - | - | 0 | - |
| 2 | c | 2017 | 232.2 | 0.0005544 | 2.388 | +1 | 2 |
| - | - | 3540 | 239.1 | - | - | 0 | - |
| - | - | 1643 | 240.1 | - | - | 0 | - |
| - | - | 965.4 | 241.1 | - | - | 0 | - |
| - | - | 813.4 | 244.1 | - | - | 0 | - |
| - | - | 588.2 | 245.2 | - | - | 0 | - |
| - | - | 1206 | 260.1 | - | - | 0 | - |
| - | - | 574.6 | 267.5 | - | - | 0 | - |
| - | - | 750.5 | 282.1 | - | - | 0 | - |
| - | - | 3595 | 282.1 | - | - | 0 | - |
| - | - | 1019 | 283.1 | - | - | 0 | - |
| - | - | 719.8 | 287.2 | - | - | 0 | - |
| - | - | 786.5 | 292.1 | - | - | 0 | - |
| - | - | 1786 | 295.1 | - | - | 0 | - |
| 10 | y | 3528 | 295.2 | 0.0004768 | 1.615 | +1 | 2 |
| - | - | 2795 | 296.1 | - | - | 0 | - |
| - | - | 1714 | 297.1 | - | - | 0 | - |
| - | - | 1202 | 297.2 | - | - | 0 | - |
| 5 | z | 834.2 | 298.1 | 0.003485 | 11.69 | +3 | 7 |
| - | - | 1393 | 299.1 | - | - | 0 | - |
| - | - | 1561 | 299.1 | - | - | 0 | - |
| - | - | 1796 | 299.2 | - | - | 0 | - |
| - | - | 1694 | 300.1 | - | - | 0 | - |
| - | - | 1.439E+04 | 300.2 | - | - | 0 | - |
| - | - | 1100 | 300.2 | - | - | 0 | - |
| - | - | 1756 | 301.1 | - | - | 0 | - |
| - | - | 2046 | 301.2 | - | - | 0 | - |
| 7 | c | 1246 | 302.2 | 0.0003505 | 1.16 | +3 | 7 |
| - | - | 934 | 306.2 | - | - | 0 | - |
| - | - | 1074 | 307.2 | - | - | 0 | - |
| - | - | 3384 | 315.2 | - | - | 0 | - |
| - | - | 711.3 | 316.2 | - | - | 0 | - |
| - | - | 4284 | 317.1 | - | - | 0 | - |
| - | - | 6891 | 317.2 | - | - | 0 | - |
| - | - | 1637 | 318.2 | - | - | 0 | - |
| 3 | c | 6374 | 319.2 | 0.0005237 | 1.641 | +1 | 3 |
| - | - | 658.7 | 320.2 | - | - | 0 | - |
| - | - | 583.8 | 330.1 | - | - | 0 | - |
| - | - | 1.518E+04 | 341 | - | - | 0 | - |
| - | - | 660.3 | 342 | - | - | 0 | - |
| - | - | 571.6 | 343 | - | - | 0 | - |
| - | - | 1607 | 355.1 | - | - | 0 | - |
| - | - | 591 | 356.1 | - | - | 0 | - |
| - | - | 4.688E+04 | 359 | - | - | 0 | - |
| - | - | 680.2 | 359.2 | - | - | 0 | - |
| - | - | 1283 | 360 | - | - | 0 | - |
| 4 | y | 915.2 | 361.2 | 0.001487 | 4.116 | +3 | 8 |
| - | - | 2179 | 364.2 | - | - | 0 | - |
| - | - | 930.4 | 367.2 | - | - | 0 | - |
| - | - | 939.7 | 369.1 | - | - | 0 | - |
| - | - | 3178 | 370.1 | - | - | 0 | - |
| - | - | 2664 | 371.1 | - | - | 0 | - |
| - | - | 574 | 371.2 | - | - | 0 | - |
| - | - | 1042 | 372.7 | - | - | 0 | - |
| - | - | 1040 | 374.2 | - | - | 0 | - |
| - | - | 4123 | 379.2 | - | - | 0 | - |
| 9 | y | 3149 | 382.2 | 0.001041 | 2.724 | +1 | 3 |
| - | - | 5961 | 385.2 | - | - | 0 | - |
| - | - | 2179 | 385.5 | - | - | 0 | - |
| - | - | 2179 | 385.9 | - | - | 0 | - |
| - | - | 2878 | 386.2 | - | - | 0 | - |
| - | - | 1238 | 386.2 | - | - | 0 | - |
| - | - | 2036 | 388.2 | - | - | 0 | - |
| 6 | z | 640 | 389.2 | 0.001589 | 4.083 | +2 | 6 |
| - | - | 3364 | 391.6 | - | - | 0 | - |
| - | - | 3772 | 391.9 | - | - | 0 | - |
| - | - | 1270 | 392.2 | - | - | 0 | - |
| - | - | 745.9 | 394.2 | - | - | 0 | - |
| - | - | 1050 | 394.7 | - | - | 0 | - |
| - | - | 822.7 | 395.2 | - | - | 0 | - |
| - | - | 1084 | 400.3 | - | - | 0 | - |
| - | - | 2218 | 415.3 | - | - | 0 | - |
| - | - | 903.1 | 415.7 | - | - | 0 | - |
| - | - | 1852 | 416.2 | - | - | 0 | - |
| - | - | 854.6 | 425.3 | - | - | 0 | - |
| - | - | 850.1 | 428.7 | - | - | 0 | - |
| - | - | 1.138E+05 | 429.1 | - | - | 0 | - |
| - | - | 1167 | 429.8 | - | - | 0 | - |
| - | - | 4546 | 430.1 | - | - | 0 | - |
| - | - | 3375 | 430.2 | - | - | 0 | - |
| - | - | 1634 | 430.3 | - | - | 0 | - |
| - | - | 1710 | 431.3 | - | - | 0 | - |
| - | - | 2390 | 433.2 | - | - | 0 | - |
| - | - | 752.9 | 437.2 | - | - | 0 | - |
| - | - | 959.3 | 438.7 | - | - | 0 | - |
| - | - | 4323 | 442.7 | - | - | 0 | - |
| - | - | 1976 | 443.2 | - | - | 0 | - |
| - | - | 1540 | 443.3 | - | - | 0 | - |
| 7 | c | 2978 | 443.7 | 0.007017 | 15.81 | +2 | 7 |
| 7 | c | 1090 | 444.2 | 0.001525 | 3.433 | +2 | 7 |
| - | - | 763.8 | 444.7 | - | - | 0 | - |
| - | - | 6.161E+04 | 445.1 | - | - | 0 | - |
| - | - | 2324 | 446.1 | - | - | 0 | - |
| 5 | z | 4578 | 446.2 | 0.003345 | 7.497 | +2 | 7 |
| - | - | 751.4 | 447.1 | - | - | 0 | - |
| - | - | 787.6 | 447.2 | - | - | 0 | - |
| - | - | 1.47E+04 | 451.8 | - | - | 0 | - |
| - | - | 4.236E+04 | 452.3 | - | - | 0 | - |
| 7 | c | 2.953E+04 | 452.8 | 0.002564 | 5.663 | +2 | 7 |
| - | - | 9770 | 453.3 | - | - | 0 | - |
| - | - | 587.1 | 453.5 | - | - | 0 | - |
| - | - | 880.5 | 453.7 | - | - | 0 | - |
| - | - | 3162 | 453.8 | - | - | 0 | - |
| - | - | 831.1 | 454.3 | - | - | 0 | - |
| - | - | 922.7 | 455.6 | - | - | 0 | - |
| - | - | 1191 | 455.9 | - | - | 0 | - |
| - | - | 712.4 | 456.3 | - | - | 0 | - |
| 4 | c | 1871 | 458.3 | 0.00223 | 4.867 | +1 | 4 |
| - | - | 779.2 | 460.2 | - | - | 0 | - |
| - | - | 1197 | 461.2 | - | - | 0 | - |
| - | - | 1241 | 461.6 | - | - | 0 | - |
| - | - | 1083 | 461.9 | - | - | 0 | - |
| 8 | z | 910.3 | 462.2 | 0.002435 | 5.269 | +1 | 4 |
| - | - | 946.4 | 462.2 | - | - | 0 | - |
| 5 | y | 1240 | 463.2 | 0.005257 | 11.35 | +2 | 7 |
| - | - | 1183 | 465.3 | - | - | 0 | - |
| - | - | 1498 | 465.8 | - | - | 0 | - |
| - | - | 678.1 | 466.3 | - | - | 0 | - |
| - | - | 997.2 | 466.7 | - | - | 0 | - |
| - | - | 1012 | 470.8 | - | - | 0 | - |
| - | - | 1282 | 471.2 | - | - | 0 | - |
| - | - | 2068 | 472.2 | - | - | 0 | - |
| - | - | 1730 | 474.2 | - | - | 0 | - |
| - | - | 1002 | 474.3 | - | - | 0 | - |
| 4 | c | 1.746E+05 | 475.3 | 0.0009141 | 1.923 | +1 | 4 |
| - | - | 4.134E+04 | 476.3 | - | - | 0 | - |
| - | - | 5351 | 477.3 | - | - | 0 | - |
| - | - | 862.4 | 479.3 | - | - | 0 | - |
| - | - | 1.292E+04 | 479.8 | - | - | 0 | - |
| 8 | z | 1326 | 480.2 | 0.001403 | 2.922 | +1 | 4 |
| - | - | 1.077E+04 | 480.3 | - | - | 0 | - |
| - | - | 5346 | 480.8 | - | - | 0 | - |
| - | - | 1.14E+04 | 481.2 | - | - | 0 | - |
| - | - | 3155 | 482.2 | - | - | 0 | - |
| - | - | 751 | 483.2 | - | - | 0 | - |
| - | - | 833.8 | 484.7 | - | - | 0 | - |
| - | - | 3364 | 485.2 | - | - | 0 | - |
| - | - | 1555 | 485.7 | - | - | 0 | - |
| - | - | 887.5 | 486.2 | - | - | 0 | - |
| - | - | 643.4 | 486.3 | - | - | 0 | - |
| - | - | 2174 | 486.8 | - | - | 0 | - |
| - | - | 3565 | 487.3 | - | - | 0 | - |
| - | - | 2907 | 487.8 | - | - | 0 | - |
| - | - | 1826 | 488.3 | - | - | 0 | - |
| - | - | 805.9 | 490.2 | - | - | 0 | - |
| - | - | 931.5 | 491.8 | - | - | 0 | - |
| - | - | 2952 | 492.2 | - | - | 0 | - |
| - | - | 1906 | 493.2 | - | - | 0 | - |
| - | - | 1999 | 493.8 | - | - | 0 | - |
| - | - | 4071 | 494.3 | - | - | 0 | - |
| - | - | 1475 | 494.8 | - | - | 0 | - |
| - | - | 1274 | 495.2 | - | - | 0 | - |
| 8 | y | 7534 | 496.2 | 0.0008994 | 1.812 | +1 | 4 |
| - | - | 1027 | 497.2 | - | - | 0 | - |
| - | - | 1.352E+04 | 499.8 | - | - | 0 | - |
| - | - | 1.335E+04 | 500.3 | - | - | 0 | - |
| 8 | c | 1.032E+04 | 500.8 | 0.00547 | 10.92 | +2 | 8 |
| 8 | c | 5612 | 501.3 | 0.001659 | 3.309 | +2 | 8 |
| - | - | 2300 | 501.8 | - | - | 0 | - |
| - | - | 2.161E+04 | 502.3 | - | - | 0 | - |
| - | - | 1060 | 502.8 | - | - | 0 | - |
| - | - | 4679 | 503.3 | - | - | 0 | - |
| - | - | 770.1 | 504.3 | - | - | 0 | - |
| - | - | 1.299E+04 | 508.3 | - | - | 0 | - |
| - | - | 3.24E+04 | 508.8 | - | - | 0 | - |
| - | - | 5.564E+04 | 509.3 | - | - | 0 | - |
| 8 | c | 5.165E+04 | 509.8 | 0.003916 | 7.683 | +2 | 8 |
| - | - | 988.7 | 509.8 | - | - | 0 | - |
| - | - | 2.396E+04 | 510.3 | - | - | 0 | - |
| - | - | 5687 | 510.8 | - | - | 0 | - |
| - | - | 1588 | 511.3 | - | - | 0 | - |
| - | - | 835.2 | 511.8 | - | - | 0 | - |
| - | - | 782 | 518.2 | - | - | 0 | - |
| - | - | 994.3 | 521.3 | - | - | 0 | - |
| - | - | 895 | 522.3 | - | - | 0 | - |
| - | - | 1086 | 522.8 | - | - | 0 | - |
| - | - | 1540 | 523.3 | - | - | 0 | - |
| - | - | 1459 | 526.3 | - | - | 0 | - |
| - | - | 2576 | 526.8 | - | - | 0 | - |
| - | - | 814.6 | 527.3 | - | - | 0 | - |
| - | - | 696.4 | 529.3 | - | - | 0 | - |
| - | - | 2351 | 530.3 | - | - | 0 | - |
| - | - | 1828 | 530.8 | - | - | 0 | - |
| - | - | 965.9 | 531.3 | - | - | 0 | - |
| - | - | 773.4 | 532.3 | - | - | 0 | - |
| 4 | y | 5442 | 532.8 | 0.004734 | 8.886 | +2 | 8 |
| 4 | z | 4980 | 533.3 | 0.001803 | 3.381 | +2 | 8 |
| - | - | 2926 | 533.8 | - | - | 0 | - |
| - | - | 686.9 | 534.8 | - | - | 0 | - |
| - | - | 2057 | 535.3 | - | - | 0 | - |
| - | - | 719.6 | 535.3 | - | - | 0 | - |
| - | - | 1652 | 535.8 | - | - | 0 | - |
| - | - | 859.4 | 536.3 | - | - | 0 | - |
| - | - | 915.1 | 536.3 | - | - | 0 | - |
| - | - | 1814 | 540.8 | - | - | 0 | - |
| 4 | y | 1626 | 541.3 | 0.0002485 | 0.4591 | +2 | 8 |
| - | - | 822.3 | 541.8 | - | - | 0 | - |
| - | - | 696.5 | 543.3 | - | - | 0 | - |
| 9 | c | 7614 | 544.3 | 0.006317 | 11.61 | +2 | 9 |
| 9 | c | 5493 | 544.8 | 0.0008883 | 1.631 | +2 | 9 |
| - | - | 797.2 | 545.2 | - | - | 0 | - |
| - | - | 1877 | 545.3 | - | - | 0 | - |
| - | - | 4758 | 545.3 | - | - | 0 | - |
| - | - | 714.4 | 546.3 | - | - | 0 | - |
| - | - | 3646 | 546.3 | - | - | 0 | - |
| - | - | 5162 | 548.3 | - | - | 0 | - |
| - | - | 7756 | 548.7 | - | - | 0 | - |
| - | - | 3126 | 549.2 | - | - | 0 | - |
| - | - | 745 | 549.8 | - | - | 0 | - |
| 7 | w | 995.1 | 550.3 | 0.0005101 | 0.927 | +1 | 5 |
| - | - | 1460 | 550.8 | - | - | 0 | - |
| - | - | 4.035E+04 | 552.8 | - | - | 0 | - |
| 9 | c | 5.704E+04 | 553.3 | 0.004214 | 7.617 | +2 | 9 |
| - | - | 2.723E+04 | 553.8 | - | - | 0 | - |
| - | - | 1.169E+04 | 554.3 | - | - | 0 | - |
| - | - | 3746 | 554.8 | - | - | 0 | - |
| - | - | 1775 | 555.3 | - | - | 0 | - |
| - | - | 1097 | 562.3 | - | - | 0 | - |
| - | - | 762.2 | 563.3 | - | - | 0 | - |
| - | - | 5844 | 564.3 | - | - | 0 | - |
| - | - | 1181 | 565.3 | - | - | 0 | - |
| - | - | 2658 | 565.3 | - | - | 0 | - |
| - | - | 1916 | 565.8 | - | - | 0 | - |
| - | - | 4504 | 566.3 | - | - | 0 | - |
| - | - | 1651 | 566.8 | - | - | 0 | - |
| - | - | 1055 | 567.3 | - | - | 0 | - |
| 3 | z | 1884 | 567.8 | 0.003212 | 5.657 | +2 | 9 |
| 3 | w | 712.7 | 568.3 | 0.0007034 | 1.238 | +2 | 9 |
| - | - | 1284 | 568.8 | - | - | 0 | - |
| - | - | 961.4 | 569.3 | - | - | 0 | - |
| - | - | 700 | 569.8 | - | - | 0 | - |
| - | - | 762.3 | 570.8 | - | - | 0 | - |
| 5 | c | 3291 | 573.3 | 0.006929 | 12.09 | +1 | 5 |
| - | - | 974.3 | 574.3 | - | - | 0 | - |
| - | - | 887.9 | 574.8 | - | - | 0 | - |
| - | - | 1650 | 575.3 | - | - | 0 | - |
| 3 | y | 3048 | 575.8 | 0.004685 | 8.137 | +2 | 9 |
| 3 | y | 1.882E+04 | 576.3 | 0.004223 | 7.328 | +2 | 9 |
| 3 | z | 1.619E+04 | 576.8 | 0.001521 | 2.637 | +2 | 9 |
| - | - | 7147 | 577.3 | - | - | 0 | - |
| - | - | 1758 | 577.8 | - | - | 0 | - |
| - | - | 1887 | 577.8 | - | - | 0 | - |
| - | - | 3640 | 578.3 | - | - | 0 | - |
| - | - | 2995 | 578.8 | - | - | 0 | - |
| - | - | 2232 | 579.3 | - | - | 0 | - |
| - | - | 1203 | 579.8 | - | - | 0 | - |
| - | - | 1558 | 580.3 | - | - | 0 | - |
| - | - | 2326 | 583.3 | - | - | 0 | - |
| - | - | 2713 | 583.8 | - | - | 0 | - |
| - | - | 4.602E+04 | 584.3 | - | - | 0 | - |
| 3 | y | 6.994E+04 | 584.8 | 0.003376 | 5.772 | +2 | 9 |
| - | - | 3.482E+04 | 585.3 | - | - | 0 | - |
| - | - | 1.142E+04 | 585.8 | - | - | 0 | - |
| - | - | 4041 | 586.3 | - | - | 0 | - |
| - | - | 5417 | 586.8 | - | - | 0 | - |
| - | - | 7213 | 587.3 | - | - | 0 | - |
| - | - | 4965 | 587.8 | - | - | 0 | - |
| - | - | 2563 | 588.3 | - | - | 0 | - |
| - | - | 1747 | 588.3 | - | - | 0 | - |
| - | - | 2778 | 588.8 | - | - | 0 | - |
| - | - | 1118 | 588.8 | - | - | 0 | - |
| - | - | 1220 | 589.3 | - | - | 0 | - |
| - | - | 1417 | 589.3 | - | - | 0 | - |
| - | - | 994.4 | 589.8 | - | - | 0 | - |
| 5 | c | 2.965E+05 | 590.3 | 0.0007045 | 1.193 | +1 | 5 |
| - | - | 8.746E+04 | 591.3 | - | - | 0 | - |
| - | - | 1097 | 591.4 | - | - | 0 | - |
| - | - | 4479 | 591.8 | - | - | 0 | - |
| - | - | 1.686E+04 | 592.3 | - | - | 0 | - |
| - | - | 3917 | 592.8 | - | - | 0 | - |
| - | - | 2157 | 593.3 | - | - | 0 | - |
| - | - | 914.7 | 593.8 | - | - | 0 | - |
| - | - | 760.7 | 594.3 | - | - | 0 | - |
| - | - | 1480 | 598.3 | - | - | 0 | - |
| 10 | c | 7870 | 600.8 | 0.006845 | 11.39 | +2 | 10 |
| 10 | c | 8389 | 601.3 | 0.00221 | 3.675 | +2 | 10 |
| - | - | 5578 | 601.8 | - | - | 0 | - |
| - | - | 2568 | 602.3 | - | - | 0 | - |
| - | - | 877 | 602.8 | - | - | 0 | - |
| - | - | 757.6 | 603.3 | - | - | 0 | - |
| - | - | 6020 | 607.3 | - | - | 0 | - |
| - | - | 1561 | 607.8 | - | - | 0 | - |
| 7 | z | 2.048E+04 | 608.3 | 0.001931 | 3.174 | +1 | 5 |
| - | - | 1.02E+05 | 609.3 | - | - | 0 | - |
| 10 | c | 1.197E+05 | 609.8 | 0.002423 | 3.973 | +2 | 10 |
| - | - | 5.862E+04 | 610.3 | - | - | 0 | - |
| - | - | 2.229E+04 | 610.8 | - | - | 0 | - |
| - | - | 5810 | 611.3 | - | - | 0 | - |
| - | - | 1221 | 611.8 | - | - | 0 | - |
| - | - | 2270 | 616.3 | - | - | 0 | - |
| - | - | 1057 | 617.3 | - | - | 0 | - |
| - | - | 5166 | 618.3 | - | - | 0 | - |
| 2 | w | 9921 | 618.8 | 0.003544 | 5.726 | +2 | 10 |
| - | - | 3697 | 619.3 | - | - | 0 | - |
| - | - | 802 | 619.8 | - | - | 0 | - |
| - | - | 910.6 | 621.3 | - | - | 0 | - |
| - | - | 951.4 | 621.8 | - | - | 0 | - |
| - | - | 939.3 | 622.3 | - | - | 0 | - |
| 7 | y | 2.013E+04 | 624.3 | 0.0002356 | 0.3774 | +1 | 5 |
| - | - | 7124 | 625.3 | - | - | 0 | - |
| 2 | w | 829.9 | 625.8 | 0.008988 | 14.36 | +2 | 10 |
| - | - | 1114 | 626.3 | - | - | 0 | - |
| - | - | 2500 | 630.3 | - | - | 0 | - |
| - | - | 4504 | 630.8 | - | - | 0 | - |
| - | - | 2373 | 631.3 | - | - | 0 | - |
| - | - | 2018 | 631.8 | - | - | 0 | - |
| 2 | y | 763.4 | 632.3 | 0.001795 | 2.839 | +2 | 10 |
| - | - | 1978 | 632.8 | - | - | 0 | - |
| 2 | z | 5135 | 633.3 | 0.002293 | 3.621 | +2 | 10 |
| - | - | 4878 | 633.8 | - | - | 0 | - |
| - | - | 2079 | 634.3 | - | - | 0 | - |
| - | - | 1575 | 635.4 | - | - | 0 | - |
| - | - | 4603 | 636.3 | - | - | 0 | - |
| - | - | 935.1 | 637.3 | - | - | 0 | - |
| - | - | 969.7 | 637.8 | - | - | 0 | - |
| - | - | 2336 | 638.3 | - | - | 0 | - |
| - | - | 922.4 | 638.8 | - | - | 0 | - |
| - | - | 1677 | 639.3 | - | - | 0 | - |
| - | - | 2344 | 639.8 | - | - | 0 | - |
| - | - | 3273 | 640.3 | - | - | 0 | - |
| - | - | 6633 | 640.8 | - | - | 0 | - |
| 2 | y | 9324 | 641.3 | 0.003415 | 5.325 | +2 | 10 |
| - | - | 4302 | 641.8 | - | - | 0 | - |
| - | - | 2812 | 642.3 | - | - | 0 | - |
| - | - | 769.2 | 643.3 | - | - | 0 | - |
| - | - | 1255 | 644.3 | - | - | 0 | - |
| - | - | 737.9 | 645.3 | - | - | 0 | - |
| - | - | 977 | 646.3 | - | - | 0 | - |
| - | - | 8727 | 647.3 | - | - | 0 | - |
| - | - | 7712 | 647.9 | - | - | 0 | - |
| - | - | 3562 | 648.3 | - | - | 0 | - |
| - | - | 1007 | 648.9 | - | - | 0 | - |
| - | - | 903.3 | 652.3 | - | - | 0 | - |
| - | - | 1820 | 652.8 | - | - | 0 | - |
| - | - | 3937 | 653.3 | - | - | 0 | - |
| - | - | 2132 | 653.8 | - | - | 0 | - |
| - | - | 1517 | 654.4 | - | - | 0 | - |
| - | - | 1156 | 654.8 | - | - | 0 | - |
| - | - | 1036 | 654.9 | - | - | 0 | - |
| - | - | 9686 | 655.3 | - | - | 0 | - |
| - | - | 9116 | 655.8 | - | - | 0 | - |
| - | - | 6137 | 656.3 | - | - | 0 | - |
| - | - | 1709 | 656.8 | - | - | 0 | - |
| - | - | 927.7 | 657.3 | - | - | 0 | - |
| - | - | 1.164E+04 | 659.3 | - | - | 0 | - |
| - | - | 1.553E+04 | 660.3 | - | - | 0 | - |
| - | - | 1.164E+04 | 660.8 | - | - | 0 | - |
| - | - | 2.191E+04 | 661.4 | - | - | 0 | - |
| - | - | 2.421E+04 | 661.8 | - | - | 0 | - |
| - | - | 2.793E+04 | 662.3 | - | - | 0 | - |
| - | - | 1.899E+04 | 662.8 | - | - | 0 | - |
| - | - | 9298 | 663.3 | - | - | 0 | - |
| - | - | 2142 | 663.8 | - | - | 0 | - |
| - | - | 4840 | 668.8 | - | - | 0 | - |
| - | - | 2.106E+04 | 669.3 | - | - | 0 | - |
| - | - | 3.124E+04 | 669.8 | - | - | 0 | - |
| - | - | 2.518E+04 | 670.3 | - | - | 0 | - |
| - | - | 2.453E+04 | 670.8 | - | - | 0 | - |
| - | - | 1.275E+04 | 671.3 | - | - | 0 | - |
| - | - | 6019 | 671.8 | - | - | 0 | - |
| - | - | 4228 | 672.3 | - | - | 0 | - |
| - | - | 1196 | 673.9 | - | - | 0 | - |
| - | - | 7316 | 674.3 | - | - | 0 | - |
| - | - | 1.363E+04 | 674.8 | - | - | 0 | - |
| - | - | 1.193E+04 | 675.3 | - | - | 0 | - |
| - | - | 1.268E+04 | 675.8 | - | - | 0 | - |
| - | - | 1.098E+04 | 676.3 | - | - | 0 | - |
| - | - | 4623 | 676.8 | - | - | 0 | - |
| - | - | 1992 | 677.3 | - | - | 0 | - |
| - | - | 6.275E+04 | 679.4 | - | - | 0 | - |
| - | - | 2.67E+04 | 680.4 | - | - | 0 | - |
| - | - | 6845 | 681.4 | - | - | 0 | - |
| - | - | 1705 | 681.9 | - | - | 0 | - |
| - | - | 1708 | 682.4 | - | - | 0 | - |
| - | - | 4043 | 682.9 | - | - | 0 | - |
| - | - | 6.176E+04 | 683.3 | - | - | 0 | - |
| - | - | 9.598E+04 | 683.8 | - | - | 0 | - |
| - | - | 4.984E+04 | 684.3 | - | - | 0 | - |
| - | - | 2.022E+04 | 684.8 | - | - | 0 | - |
| - | - | 6010 | 685.3 | - | - | 0 | - |
| - | - | 1242 | 685.8 | - | - | 0 | - |
| - | - | 3069 | 687.4 | - | - | 0 | - |
| - | - | 1661 | 688.3 | - | - | 0 | - |
| - | - | 830.7 | 689.3 | - | - | 0 | - |
| - | - | 5988 | 690.4 | - | - | 0 | - |
| - | - | 5688 | 690.9 | - | - | 0 | - |
| - | - | 5.935E+04 | 691.4 | - | - | 0 | - |
| - | - | 9.424E+04 | 691.9 | - | - | 0 | - |
| - | - | 4.968E+04 | 692.4 | - | - | 0 | - |
| - | - | 2.273E+04 | 692.9 | - | - | 0 | - |
| - | - | 6686 | 693.4 | - | - | 0 | - |
| - | - | 1732 | 693.9 | - | - | 0 | - |
| - | - | 929.7 | 694.3 | - | - | 0 | - |
| - | - | 1.376E+04 | 695.4 | - | - | 0 | - |
| - | - | 5053 | 696.4 | - | - | 0 | - |
| - | - | 1223 | 697.4 | - | - | 0 | - |
| - | - | 6.051E+04 | 704.4 | - | - | 0 | - |
| - | - | 3.786E+04 | 705.4 | - | - | 0 | - |
| - | - | 9315 | 706.4 | - | - | 0 | - |
| - | - | 1633 | 707.4 | - | - | 0 | - |
| - | - | 1403 | 711.3 | - | - | 0 | - |
| - | - | 823.4 | 713.4 | - | - | 0 | - |
| - | - | 5621 | 721.4 | - | - | 0 | - |
| - | - | 2860 | 722.4 | - | - | 0 | - |
| - | - | 1376 | 727.3 | - | - | 0 | - |
| - | - | 1113 | 729.3 | - | - | 0 | - |
| - | - | 7022 | 731.4 | - | - | 0 | - |
| - | - | 6059 | 732.4 | - | - | 0 | - |
| - | - | 2193 | 733.4 | - | - | 0 | - |
| - | - | 1.041E+04 | 737.4 | - | - | 0 | - |
| - | - | 882.3 | 738.3 | - | - | 0 | - |
| - | - | 2862 | 738.4 | - | - | 0 | - |
| - | - | 1050 | 739.4 | - | - | 0 | - |
| - | - | 721.5 | 741.4 | - | - | 0 | - |
| - | - | 2.842E+04 | 749.4 | - | - | 0 | - |
| - | - | 2.158E+04 | 750.4 | - | - | 0 | - |
| - | - | 8319 | 751.4 | - | - | 0 | - |
| - | - | 1633 | 752.4 | - | - | 0 | - |
| 6 | c | 4070 | 758.4 | 0.01342 | 17.69 | +1 | 6 |
| 6 | c | 2380 | 759.4 | 0.002438 | 3.21 | +1 | 6 |
| - | - | 5.05E+04 | 775.4 | - | - | 0 | - |
| 6 | c | 4.6E+04 | 776.4 | 0.01007 | 12.96 | +1 | 6 |
| - | - | 1.457E+04 | 777.4 | - | - | 0 | - |
| - | - | 3426 | 778.4 | - | - | 0 | - |
| - | - | 915.8 | 786.4 | - | - | 0 | - |
| - | - | 1978 | 787.4 | - | - | 0 | - |
| - | - | 1156 | 788.4 | - | - | 0 | - |
| - | - | 971.9 | 789.4 | - | - | 0 | - |
| 6 | y | 1.327E+05 | 793.4 | 0.009239 | 11.65 | +1 | 6 |
| 6 | z | 1.109E+05 | 794.4 | 0.006886 | 8.669 | +1 | 6 |
| - | - | 3.888E+04 | 795.4 | - | - | 0 | - |
| - | - | 9718 | 796.4 | - | - | 0 | - |
| - | - | 1101 | 797.4 | - | - | 0 | - |
| - | - | 2414 | 809.4 | - | - | 0 | - |
| 6 | y | 4007 | 810.4 | 0.01292 | 15.94 | +1 | 6 |
| - | - | 2453 | 811.4 | - | - | 0 | - |
| - | - | 3823 | 821.4 | - | - | 0 | - |
| - | - | 1406 | 822.4 | - | - | 0 | - |
| - | - | 1096 | 825.4 | - | - | 0 | - |
| - | - | 1200 | 832.4 | - | - | 0 | - |
| - | - | 1596 | 847.4 | - | - | 0 | - |
| - | - | 1394 | 853.4 | - | - | 0 | - |
| - | - | 1626 | 855.4 | - | - | 0 | - |
| - | - | 1738 | 857.4 | - | - | 0 | - |
| - | - | 1665 | 858.5 | - | - | 0 | - |
| - | - | 5185 | 859.5 | - | - | 0 | - |
| - | - | 9503 | 860.5 | - | - | 0 | - |
| - | - | 4705 | 861.5 | - | - | 0 | - |
| - | - | 1458 | 862.5 | - | - | 0 | - |
| 5 | w | 4.108E+04 | 864.4 | 0.008197 | 9.483 | +1 | 7 |
| - | - | 3.893E+04 | 865.4 | - | - | 0 | - |
| - | - | 1.396E+04 | 866.4 | - | - | 0 | - |
| - | - | 3517 | 867.4 | - | - | 0 | - |
| - | - | 1102 | 873.4 | - | - | 0 | - |
| - | - | 1114 | 874.4 | - | - | 0 | - |
| - | - | 940.1 | 875.4 | - | - | 0 | - |
| - | - | 2175 | 879.4 | - | - | 0 | - |
| - | - | 1137 | 880.4 | - | - | 0 | - |
| - | - | 1400 | 881.4 | - | - | 0 | - |
| - | - | 2030 | 884.5 | - | - | 0 | - |
| - | - | 4266 | 885.5 | - | - | 0 | - |
| 7 | c | 9035 | 886.5 | 0.01469 | 16.57 | +1 | 7 |
| 7 | c | 5858 | 887.5 | 0.002674 | 3.013 | +1 | 7 |
| - | - | 1789 | 888.5 | - | - | 0 | - |
| - | - | 1029 | 889.5 | - | - | 0 | - |
| 5 | z | 1159 | 891.4 | 0.01366 | 15.33 | +1 | 7 |
| - | - | 1040 | 896.4 | - | - | 0 | - |
| - | - | 4758 | 900.5 | - | - | 0 | - |
| - | - | 3728 | 901.5 | - | - | 0 | - |
| - | - | 1.806E+04 | 902.5 | - | - | 0 | - |
| - | - | 7.171E+04 | 903.5 | - | - | 0 | - |
| 7 | c | 6.95E+04 | 904.5 | 0.007922 | 8.758 | +1 | 7 |
| - | - | 2.471E+04 | 905.5 | - | - | 0 | - |
| - | - | 6830 | 906.5 | - | - | 0 | - |
| - | - | 2.247E+04 | 907.4 | - | - | 0 | - |
| - | - | 512.2 | 907.5 | - | - | 0 | - |
| 5 | y | 4.589E+04 | 908.4 | 0.005246 | 5.774 | +1 | 7 |
| 5 | z | 5.343E+04 | 909.4 | 0.009782 | 10.76 | +1 | 7 |
| - | - | 2.159E+04 | 910.4 | - | - | 0 | - |
| - | - | 6562 | 911.4 | - | - | 0 | - |
| - | - | 1167 | 912.4 | - | - | 0 | - |
| - | - | 1675 | 920.5 | - | - | 0 | - |
| - | - | 907.8 | 921.5 | - | - | 0 | - |
| - | - | 5924 | 923.4 | - | - | 0 | - |
| - | - | 1.176E+04 | 924.4 | - | - | 0 | - |
| 5 | y | 1.054E+04 | 925.4 | 0.006777 | 7.323 | +1 | 7 |
| - | - | 2529 | 926.4 | - | - | 0 | - |
| - | - | 1101 | 935.6 | - | - | 0 | - |
| - | - | 1071 | 961.5 | - | - | 0 | - |
| - | - | 1061 | 968.5 | - | - | 0 | - |
| - | - | 2026 | 969.5 | - | - | 0 | - |
| - | - | 1220 | 973.5 | - | - | 0 | - |
| - | - | 1737 | 974.5 | - | - | 0 | - |
| - | - | 1474 | 975.5 | - | - | 0 | - |
| - | - | 4093 | 986.5 | - | - | 0 | - |
| - | - | 6744 | 987.5 | - | - | 0 | - |
| - | - | 4662 | 988.5 | - | - | 0 | - |
| - | - | 2048 | 989.5 | - | - | 0 | - |
| - | - | 2870 | 999.5 | - | - | 0 | - |
| 8 | c | 5194 | 1001 | 0.00995 | 9.945 | +1 | 8 |
| 8 | c | 5439 | 1002 | 0.0007408 | 0.7397 | +1 | 8 |
| - | - | 2285 | 1003 | - | - | 0 | - |
| - | - | 2630 | 1004 | - | - | 0 | - |
| - | - | 1808 | 1005 | - | - | 0 | - |
| - | - | 4744 | 1017 | - | - | 0 | - |
| - | - | 3.998E+04 | 1018 | - | - | 0 | - |
| 8 | c | 4.285E+04 | 1019 | 0.007454 | 7.318 | +1 | 8 |
| - | - | 1.757E+04 | 1020 | - | - | 0 | - |
| - | - | 6713 | 1021 | - | - | 0 | - |
| - | - | 2517 | 1022 | - | - | 0 | - |
| - | - | 977 | 1023 | - | - | 0 | - |
| - | - | 1114 | 1023 | - | - | 0 | - |
| - | - | 768.2 | 1026 | - | - | 0 | - |
| 4 | z | 1463 | 1048 | 0.01679 | 16.02 | +1 | 8 |
| - | - | 1274 | 1061 | - | - | 0 | - |
| - | - | 1071 | 1062 | - | - | 0 | - |
| - | - | 1191 | 1063 | - | - | 0 | - |
| 4 | y | 1560 | 1064 | 0.004788 | 4.502 | +1 | 8 |
| 4 | y | 1.554E+04 | 1065 | 0.008322 | 7.817 | +1 | 8 |
| 4 | z | 3.361E+04 | 1066 | 0.005119 | 4.804 | +1 | 8 |
| - | - | 2.328E+04 | 1067 | - | - | 0 | - |
| - | - | 7722 | 1068 | - | - | 0 | - |
| - | - | 2644 | 1069 | - | - | 0 | - |
| - | - | 1878 | 1081 | - | - | 0 | - |
| 4 | y | 2558 | 1082 | 0.01078 | 9.968 | +1 | 8 |
| 9 | c | 3695 | 1089 | 0.00611 | 5.613 | +1 | 9 |
| - | - | 5102 | 1090 | - | - | 0 | - |
| - | - | 3327 | 1091 | - | - | 0 | - |
| - | - | 1222 | 1092 | - | - | 0 | - |
| - | - | 824.6 | 1095 | - | - | 0 | - |
| - | - | 1468 | 1096 | - | - | 0 | - |
| - | - | 1902 | 1097 | - | - | 0 | - |
| - | - | 1203 | 1098 | - | - | 0 | - |
| - | - | 1.079E+04 | 1105 | - | - | 0 | - |
| 9 | c | 1.773E+04 | 1106 | 0.007499 | 6.783 | +1 | 9 |
| - | - | 9877 | 1107 | - | - | 0 | - |
| - | - | 3724 | 1108 | - | - | 0 | - |
| - | - | 1448 | 1109 | - | - | 0 | - |
| - | - | 982 | 1110 | - | - | 0 | - |
| - | - | 1469 | 1111 | - | - | 0 | - |
| - | - | 725 | 1126 | - | - | 0 | - |
| 3 | z | 1338 | 1135 | 0.00479 | 4.222 | +1 | 9 |
| 3 | z | 2387 | 1136 | 0.01565 | 13.78 | +1 | 9 |
| - | - | 1272 | 1137 | - | - | 0 | - |
| - | - | 791.8 | 1138 | - | - | 0 | - |
| 3 | y | 1.115E+04 | 1152 | 0.007299 | 6.338 | +1 | 9 |
| 3 | z | 2.602E+04 | 1153 | 0.0007702 | 0.6682 | +1 | 9 |
| - | - | 2.119E+04 | 1154 | - | - | 0 | - |
| - | - | 8917 | 1155 | - | - | 0 | - |
| - | - | 2473 | 1156 | - | - | 0 | - |
| - | - | 956 | 1158 | - | - | 0 | - |
| - | - | 774.6 | 1159 | - | - | 0 | - |
| - | - | 1909 | 1160 | - | - | 0 | - |
| - | - | 1168 | 1161 | - | - | 0 | - |
| - | - | 920.2 | 1168 | - | - | 0 | - |
| 3 | y | 2709 | 1169 | 0.006433 | 5.505 | +1 | 9 |
| - | - | 2084 | 1170 | - | - | 0 | - |
| - | - | 1125 | 1171 | - | - | 0 | - |
| - | - | 2334 | 1174 | - | - | 0 | - |
| - | - | 5685 | 1175 | - | - | 0 | - |
| - | - | 6468 | 1176 | - | - | 0 | - |
| - | - | 3735 | 1177 | - | - | 0 | - |
| - | - | 1111 | 1178 | - | - | 0 | - |
| 10 | c | 8270 | 1202 | 0.00774 | 6.441 | +1 | 10 |
| - | - | 1.123E+04 | 1203 | - | - | 0 | - |
| - | - | 4364 | 1204 | - | - | 0 | - |
| - | - | 1040 | 1205 | - | - | 0 | - |
| - | - | 5990 | 1218 | - | - | 0 | - |
| 10 | c | 1.858E+04 | 1219 | 0.001231 | 1.01 | +1 | 10 |
| - | - | 1.718E+04 | 1220 | - | - | 0 | - |
| - | - | 6839 | 1221 | - | - | 0 | - |
| - | - | 2171 | 1222 | - | - | 0 | - |
| 2 | y | 1149 | 1265 | 0.01515 | 11.98 | +1 | 10 |
| 2 | z | 3762 | 1266 | 0.00501 | 3.958 | +1 | 10 |
| - | - | 6445 | 1267 | - | - | 0 | - |
| - | - | 2908 | 1268 | - | - | 0 | - |
| - | - | 805.6 | 1269 | - | - | 0 | - |
| - | - | 976.5 | 1281 | - | - | 0 | - |
| 2 | y | 798.7 | 1282 | 0.002606 | 2.033 | +1 | 10 |
| - | - | 1329 | 1305 | - | - | 0 | - |
| - | - | 3591 | 1306 | - | - | 0 | - |
| - | - | 7268 | 1307 | - | - | 0 | - |
| - | - | 5157 | 1308 | - | - | 0 | - |
| - | - | 3814 | 1309 | - | - | 0 | - |
| - | - | 1777 | 1310 | - | - | 0 | - |
| - | - | 2277 | 1321 | - | - | 0 | - |
| - | - | 5871 | 1322 | - | - | 0 | - |
| - | - | 6180 | 1323 | - | - | 0 | - |
| - | - | 9234 | 1324 | - | - | 0 | - |
| - | - | 2.49E+04 | 1325 | - | - | 0 | - |
| - | - | 1.932E+04 | 1326 | - | - | 0 | - |
| - | - | 9184 | 1327 | - | - | 0 | - |
| - | - | 2928 | 1328 | - | - | 0 | - |
| - | - | 2882 | 1338 | - | - | 0 | - |
| - | - | 2.102E+04 | 1339 | - | - | 0 | - |
| - | - | 2.727E+04 | 1340 | - | - | 0 | - |
| - | - | 1.451E+04 | 1341 | - | - | 0 | - |
| - | - | 8149 | 1342 | - | - | 0 | - |
| - | - | 2159 | 1343 | - | - | 0 | - |
| - | - | 5912 | 1349 | - | - | 0 | - |
| - | - | 1.08E+04 | 1350 | - | - | 0 | - |
| - | - | 1.157E+04 | 1351 | - | - | 0 | - |
| - | - | 6414 | 1352 | - | - | 0 | - |
| - | - | 2173 | 1353 | - | - | 0 | - |
| - | - | 1795 | 1356 | - | - | 0 | - |
| - | - | 2269 | 1357 | - | - | 0 | - |
| - | - | 1182 | 1358 | - | - | 0 | - |
| - | - | 1275 | 1364 | - | - | 0 | - |
| - | - | 2533 | 1365 | - | - | 0 | - |
| - | - | 1.156E+04 | 1366 | - | - | 0 | - |
| - | - | 7.142E+04 | 1367 | - | - | 0 | - |
| - | - | 1.019E+05 | 1368 | - | - | 0 | - |
| - | - | 4.977E+04 | 1369 | - | - | 0 | - |
| - | - | 1.878E+04 | 1370 | - | - | 0 | - |
| - | - | 5588 | 1371 | - | - | 0 | - |
| - | - | 978.5 | 1372 | - | - | 0 | - |
| - | - | 996.4 | 1380 | - | - | 0 | - |
| - | - | 3235 | 1381 | - | - | 0 | - |
| - | - | 2931 | 1382 | - | - | 0 | - |
| - | - | 1.04E+04 | 1383 | - | - | 0 | - |
| - | - | 2.064E+04 | 1384 | - | - | 0 | - |
| - | - | 2.148E+04 | 1385 | - | - | 0 | - |
| - | - | 1.116E+04 | 1386 | - | - | 0 | - |
| - | - | 3767 | 1387 | - | - | 0 | - |

m/z Charge Intensity FragmentType MassShift Position
123.04434204101562 0 602.0403
123.10487365722656 0 656.84015
129.1026153564453 0 1279.735
131.11805725097656 0 681.1533
136.07601928710938 0 13959.224
137.07925415039062 0 983.78357
142.122802734375 0 1140.6002
142.63265991210938 0 415.47888
143.1181640625 0 1630.8527
147.0443572998047 0 498.05362
148.1647186279297 0 477.9334
148.66061401367188 0 463.88403
148.898681640625 0 568.9008
148.90562438964844 0 463.3644
148.91261291503906 0 429.81033
148.91983032226562 0 1059.2749
148.92779541015625 0 1066.1405
148.93418884277344 0 2061.9258
148.9419708251953 0 3793.9253
148.95855712890625 0 4693.5083
148.96627807617188 0 2854.1228
148.99539184570312 0 947.1402
149.00242614746094 0 661.3029
149.00958251953125 0 655.7129
149.0170135498047 0 484.69824
149.0243377685547 0 422.62473
149.0403289794922 0 447.18045
149.04510498046875 0 4524.086
149.05142211914062 0 714.903
157.0613250732422 0 1206.9542
161.01629638671875 0 423.59528
161.73629760742188 0 454.11432
165.05490112304688 0 9896.589
166.05810546875 0 820.11426
166.08694458007812 0 618.3105 y 7
167.0557098388672 0 1148.0378
168.10189819335938 0 430.57913
169.1339874267578 0 721.24963
173.12863159179688 0 1282.5911
173.44879150390625 0 978.81366
175.072021484375 0 844.37634
181.09732055664062 0 608.25195
182.08151245117188 0 35868.25 y 10
183.08489990234375 0 3401.5032
186.11314392089844 0 607.1857
186.12391662597656 0 1179.0338
186.8689727783203 0 469.61978
187.07269287109375 0 617.33057
187.14447021484375 0 85377.18
188.1478729248047 0 7671.6455
196.1207275390625 0 733.90137
200.13970947265625 0 1373.9048
201.12355041503906 0 2704.855
201.13328552246094 0 612.1006
201.14743041992188 0 868.82385
202.08236694335938 0 1728.2676
202.11904907226562 0 506.65323
203.06646728515625 0 3312.623
203.1393585205078 0 716.16907
215.1393280029297 0 19395.508
216.1424102783203 0 2364.5571
221.08460998535156 0 2581.1755
222.0846710205078 0 887.0323
223.06365966796875 0 1055.4766
225.04336547851562 0 1349.2993
226.0431365966797 0 744.02374
226.1676788330078 0 470.41608
229.12962341308594 0 1472.2021
232.16612243652344 0 2016.5452 c 1
239.0955047607422 0 3540.1155
240.096435546875 0 1642.6072
241.09259033203125 0 965.37646
244.1411590576172 0 813.41833
245.1602020263672 0 588.1953
260.12408447265625 0 1206.0199
267.5093078613281 0 574.55756
282.0507507324219 0 750.5315
282.14459228515625 0 3595.4023
283.14764404296875 0 1019.0217
287.1711730957031 0 719.8438
292.1309814453125 0 786.52545
295.1039123535156 0 1786.0458
295.16571044921875 0 3528.0422 y 9
296.1043701171875 0 2795.381
297.10137939453125 0 1713.8069
297.15594482421875 0 1202.4293
298.140380859375 0 834.15155 z Ammonia loss 4
299.0628356933594 0 1393.3656
299.1474304199219 0 1561.2006
299.17156982421875 0 1795.7379
300.062255859375 0 1694.1344
300.15582275390625 0 14389.128
300.1745300292969 0 1099.6171
301.05987548828125 0 1755.6482
301.15960693359375 0 2046.1329
302.171142578125 0 1245.7507 c 6
306.1700134277344 0 934.03284
307.1770935058594 0 1073.9827
315.16668701171875 0 3383.5227
316.15093994140625 0 711.34283
317.10980224609375 0 4283.6113
317.18243408203125 0 6890.6577
318.18536376953125 0 1636.667
319.1981201171875 0 6373.913 c 2
320.20123291015625 0 658.7401
330.1392822265625 0 583.80817
341.0184631347656 0 15177.899
342.0180969238281 0 660.3361
342.9969482421875 0 571.5643
355.0703430175781 0 1606.6744
356.0699157714844 0 590.9709
359.02880859375 0 46884.508
359.1669921875 0 680.2065
360.0285339355469 0 1283.1942
361.1842041015625 0 915.2316 y 3
364.1982116699219 0 2178.5671
367.1864013671875 0 930.40967
369.1214904785156 0 939.7418
370.122802734375 0 3178.3906
371.1207275390625 0 2664.1968
371.1564636230469 0 573.96106
372.697509765625 0 1042.158
374.1791076660156 0 1039.785
379.2093505859375 0 4123.093
382.19830322265625 0 3149.1367 y 8
385.21978759765625 0 5960.8813
385.54705810546875 0 2178.8765
385.87811279296875 0 2178.5774
386.2040710449219 0 2877.7444
386.2284240722656 0 1237.9988
388.18280029296875 0 2036.4225
389.1866455078125 0 640.01807 z Ammonia loss 5
391.5517578125 0 3363.9624
391.8830261230469 0 3772.2368
392.21466064453125 0 1270.0569
394.1911315917969 0 745.8644
394.7028503417969 0 1049.6787
395.205078125 0 822.7026
400.2566223144531 0 1083.8071
415.26690673828125 0 2217.717
415.7091369628906 0 903.14404
416.17840576171875 0 1852.482
425.2502746582031 0 854.63116
428.701171875 0 850.13776
429.0895080566406 0 113763.4
429.7554016113281 0 1166.849
430.09033203125 0 4546.249
430.2444763183594 0 3375.4531
430.27862548828125 0 1634.3601
431.2814636230469 0 1709.5106
433.2207336425781 0 2389.7708
437.2126770019531 0 752.8743
438.7330322265625 0 959.2528
442.74530029296875 0 4323.245
443.2377014160156 0 1975.7449
443.26312255859375 0 1539.905
443.7413024902344 0 2977.622 c Water loss 6
444.2418518066406 0 1090.1602 c Ammonia loss 6
444.6971435546875 0 763.7887
445.12078857421875 0 61610.875
446.1202087402344 0 2324.3225
446.2130432128906 0 4578.0566 z Water loss 4
447.1004943847656 0 751.4316
447.215576171875 0 787.5751
451.75054931640625 0 14701.088
452.2537841796875 0 42360.918
452.75103759765625 0 29534.46 c 6
453.2503967285156 0 9769.829
453.4760437011719 0 587.0705
453.7002258300781 0 880.5055
453.752197265625 0 3161.9958
454.25213623046875 0 831.05707
455.5665588378906 0 922.7425
455.9007568359375 0 1190.7699
456.26220703125 0 712.35144
458.2699279785156 0 1870.7595 c Ammonia loss 3
460.24041748046875 0 779.1801
461.2407531738281 0 1196.8003
461.5730285644531 0 1241.1422
461.9051208496094 0 1082.9478
462.2084655761719 0 910.299 z Water loss 7
462.2422790527344 0 946.36334
463.2190856933594 0 1239.767 y 4
465.2625732421875 0 1183.4469
465.76470947265625 0 1497.6145
466.2637023925781 0 678.1394
466.7329406738281 0 997.2024
470.7525329589844 0 1012.21576
471.2467956542969 0 1281.5507
472.2498474121094 0 2067.8828
474.1950378417969 0 1729.6294
474.2889709472656 0 1002.0095
475.29962158203125 0 174579.78 c 3
476.3021545410156 0 41342.223
477.30419921875 0 5350.564
479.25946044921875 0 862.3832
479.7585144042969 0 12918.223
480.2200622558594 0 1326.3005 z 7
480.2547912597656 0 10767.294
480.7552185058594 0 5346.149
481.2295837402344 0 11403.922
482.23272705078125 0 3154.825
483.2393493652344 0 751.0295
484.7482604980469 0 833.81366
485.24468994140625 0 3363.7034
485.7435302734375 0 1554.8475
486.2461853027344 0 887.54865
486.3037109375 0 643.40686
486.76953125 0 2173.8115
487.26824951171875 0 3565.244
487.7638244628906 0 2907.0278
488.2646484375 0 1826.4126
490.240234375 0 805.8666
491.7545471191406 0 931.45844
492.2356872558594 0 2952.308
493.2405700683594 0 1906.366
493.7532958984375 0 1999.0608
494.2506408691406 0 4071.1997
494.7515563964844 0 1475.247
495.226806640625 0 1274.1978
496.2410888671875 0 7534.229 y 7
497.2400817871094 0 1027.2646
499.7669982910156 0 13523.608
500.26373291015625 0 13354.35
500.7643127441406 0 10321.864 c Water loss 7
501.2601318359375 0 5611.957 c Ammonia loss 7
501.76654052734375 0 2300.163
502.2746887207031 0 21607.262
502.7620849609375 0 1060.3525
503.27447509765625 0 4679.3975
504.2750244140625 0 770.14575
508.25274658203125 0 12990.592
508.7721252441406 0 32402.227
509.27337646484375 0 55638.242
509.7711486816406 0 51652.64 c 7
509.8270568847656 0 988.6993
510.27069091796875 0 23960.469
510.7705078125 0 5686.9883
511.2701416015625 0 1588.4153
511.76727294921875 0 835.2417
518.2410278320312 0 781.9578
521.2716674804688 0 994.33203
522.2721557617188 0 894.9879
522.7744140625 0 1086.3488
523.2692260742188 0 1539.8049
526.2595825195312 0 1459.2681
526.7526245117188 0 2575.661
527.25146484375 0 814.63257
529.3125610351562 0 696.3863
530.2922973632812 0 2350.9644
530.7782592773438 0 1827.587
531.276611328125 0 965.88763
532.291748046875 0 773.41626
532.766357421875 0 5442.148 y Ammonia loss 3
533.2637329101562 0 4980.352 z 3
533.7611083984375 0 2925.553
534.79931640625 0 686.9377
535.2728271484375 0 2056.6824
535.3146362304688 0 719.6122
535.7703857421875 0 1651.8853
536.2603149414062 0 859.4271
536.3037719726562 0 915.14905
540.7745971679688 0 1814.2793
541.275146484375 0 1625.8691 y 3
541.7655029296875 0 822.282
543.282470703125 0 696.49023
544.2794799804688 0 7614.3955 c Water loss 8
544.7769165039062 0 5492.665 c Ammonia loss 8
545.2310180664062 0 797.1753
545.2683715820312 0 1877.3711
545.3048095703125 0 4758.2803
546.2655639648438 0 714.44684
546.3096923828125 0 3645.535
548.251220703125 0 5162.371
548.7467041015625 0 7755.576
549.2465209960938 0 3126.332
549.7532958984375 0 745.03125
550.250244140625 0 995.1388 w 6
550.7899780273438 0 1459.551
552.7918701171875 0 40351.855
553.286865234375 0 57038.45 c 8
553.787353515625 0 27225.781
554.286865234375 0 11685.002
554.78369140625 0 3745.9763
555.2814331054688 0 1774.6965
562.2852783203125 0 1096.8605
563.318603515625 0 762.2268
564.30419921875 0 5843.851
565.2615966796875 0 1181.4287
565.3096923828125 0 2657.7659
565.7993774414062 0 1915.6772
566.2965698242188 0 4504.233
566.7737426757812 0 1651.005
567.2970581054688 0 1055.1909
567.7794799804688 0 1884.4258 z Water loss 2
568.2808837890625 0 712.70056 w 2
568.7730102539062 0 1284.2952
569.3075561523438 0 961.4332
569.8018188476562 0 700.0487
570.8076171875 0 762.31964
573.3060302734375 0 3290.5896 c Ammonia loss 4
574.3004760742188 0 974.2623
574.789794921875 0 887.8949
575.2855224609375 0 1650.2725
575.7809448242188 0 3048.0413 y Water loss 2
576.2818603515625 0 18820.48 y Ammonia loss 2
576.780029296875 0 16191.721 z 2
577.2801513671875 0 7147.175
577.7781982421875 0 1758.1586
577.8170776367188 0 1887.1621
578.3125610351562 0 3639.6838
578.81298828125 0 2994.6013
579.3115234375 0 2231.896
579.8087158203125 0 1202.9152
580.3076171875 0 1557.6681
583.303466796875 0 2326.2163
583.7981567382812 0 2712.6997
584.291748046875 0 46021.992
584.7875366210938 0 69941.74 y 2
585.287841796875 0 34820.01
585.7877197265625 0 11415.344
586.287353515625 0 4040.9417
586.82275390625 0 5417.3237
587.3211669921875 0 7212.9556
587.8211669921875 0 4964.702
588.276123046875 0 2563.214
588.32080078125 0 1747.1145
588.771484375 0 2777.604
588.822265625 0 1117.7313
589.2741088867188 0 1220.0328
589.3204345703125 0 1417.4816
589.7737426757812 0 994.36127
590.3263549804688 0 296518.88 c 4
591.328857421875 0 87455.72
591.3859252929688 0 1096.7417
591.8154296875 0 4479.272
592.3275146484375 0 16859.895
592.8088989257812 0 3917.2493
593.3296508789062 0 2157.29
593.8057861328125 0 914.7249
594.2763061523438 0 760.6817
598.2830200195312 0 1479.9451
600.8209838867188 0 7870.251 c Water loss 9
601.317626953125 0 8389.106 c Ammonia loss 9
601.8204345703125 0 5578.2344
602.319580078125 0 2568.3523
602.7865600585938 0 876.9847
603.2860717773438 0 757.57056
607.3217163085938 0 6019.9307
607.8312377929688 0 1561.0085
608.318359375 0 20483.652 z 6
609.33349609375 0 101985.18
609.8306884765625 0 119725.234 c 9
610.3302001953125 0 58615.082
610.830322265625 0 22291.049
611.3294067382812 0 5809.942
611.8289184570312 0 1221.0522
616.3408203125 0 2269.7627
617.343994140625 0 1056.563
618.30517578125 0 5165.835
618.8004760742188 0 9920.89 w 1
619.3020629882812 0 3697.2292
619.7982177734375 0 801.9953
621.3253784179688 0 910.6441
621.8192138671875 0 951.4424
622.3102416992188 0 939.3078
624.3353881835938 0 20134.797 y 6
625.3348999023438 0 7124.0864
625.8028564453125 0 829.92554 w 1
626.339111328125 0 1113.5697
630.3276977539062 0 2500.488
630.823486328125 0 4504.2217
631.3214721679688 0 2372.5488
631.8312377929688 0 2018.1199
632.3258666992188 0 763.3525 y Water loss 1
632.83251953125 0 1978.4355
633.3212890625 0 5135.433 z 1
633.817626953125 0 4877.9277
634.3198852539062 0 2078.8298
635.3569946289062 0 1575.3397
636.3487548828125 0 4603.4067
637.3486328125 0 935.1075
637.8320922851562 0 969.739
638.3399047851562 0 2336.2788
638.8429565429688 0 922.35626
639.3408203125 0 1677.1735
639.8463134765625 0 2344.2964
640.345458984375 0 3272.8564
640.8347778320312 0 6632.9473
641.3295288085938 0 9323.7705 y 1
641.8284301757812 0 4302.3237
642.331298828125 0 2811.8057
643.3273315429688 0 769.18005
644.3383178710938 0 1255.0476
645.3414916992188 0 737.899
646.3218994140625 0 976.98254
647.3256225585938 0 8726.54
647.85107421875 0 7712.0117
648.3446655273438 0 3562.4492
648.8547973632812 0 1007.0706
652.3462524414062 0 903.25903
652.8428344726562 0 1820.4609
653.3395385742188 0 3936.9854
653.83984375 0 2131.581
654.3739013671875 0 1516.7947
654.8257446289062 0 1155.6688
654.8780517578125 0 1036.2239
655.3180541992188 0 9686.093
655.8131713867188 0 9116.465
656.3140869140625 0 6137.131
656.8158569335938 0 1708.7319
657.3162841796875 0 927.6574
659.3475952148438 0 11644.425
660.3471069335938 0 15526.151
660.8491821289062 0 11638.025
661.3505249023438 0 21908.799
661.8473510742188 0 24210.156
662.3447875976562 0 27930.88
662.8430786132812 0 18985.205
663.3447875976562 0 9297.712
663.8447875976562 0 2142.0164
668.829345703125 0 4839.9526
669.34765625 0 21055.898
669.8455200195312 0 31238.05
670.3463745117188 0 25176.682
670.8435668945312 0 24534.498
671.3432006835938 0 12750.78
671.8434448242188 0 6018.7466
672.343017578125 0 4228.2607
673.8514404296875 0 1195.863
674.3452758789062 0 7316.118
674.8427124023438 0 13631.875
675.3413696289062 0 11933.986
675.839111328125 0 12679.831
676.3355712890625 0 10977.638
676.83349609375 0 4622.9473
677.3349609375 0 1991.5795
679.3543090820312 0 62753.74
680.3540649414062 0 26703.975
681.3583374023438 0 6844.567
681.8748168945312 0 1704.711
682.3761596679688 0 1707.9799
682.8576049804688 0 4042.684
683.3486328125 0 61759.04
683.8450927734375 0 95980.52
684.3447875976562 0 49839.773
684.8447875976562 0 20219.266
685.3449096679688 0 6010.019
685.848388671875 0 1241.7526
687.3509521484375 0 3069.274
688.329345703125 0 1660.8308
689.3410034179688 0 830.68524
690.3820190429688 0 5987.762
690.8845825195312 0 5688.327
691.358154296875 0 59351.234
691.8538818359375 0 94235.08
692.3536376953125 0 49684.008
692.854248046875 0 22731.346
693.3543090820312 0 6686.327
693.857666015625 0 1732.4817
694.3289184570312 0 929.74536
695.3731689453125 0 13763.638
696.369873046875 0 5053.227
697.3674926757812 0 1222.9272
704.3690795898438 0 60514.66
705.3594360351562 0 37863.54
706.361083984375 0 9315.163
707.3658447265625 0 1632.879
711.3422241210938 0 1402.7524
713.39013671875 0 823.41644
721.364013671875 0 5620.9785
722.3636474609375 0 2859.9512
727.3167724609375 0 1375.6873
729.3324584960938 0 1113.383
731.3912963867188 0 7022.177
732.3818969726562 0 6059.024
733.3812866210938 0 2192.7153
737.3838500976562 0 10409.679
738.3194580078125 0 882.34753
738.3855590820312 0 2861.7927
739.3901977539062 0 1050.2638
741.3759155273438 0 721.4636
749.3845825195312 0 28417.057
750.3873291015625 0 21575.578
751.3897094726562 0 8319.271
752.3869018554688 0 1632.7162
758.3809814453125 0 4070.0188 c Water loss 5
759.3759765625 0 2379.5896 c Ammonia loss 5
775.4039306640625 0 50497.55
776.3948974609375 0 46004.24 c 5
777.3973999023438 0 14572.157
778.3969116210938 0 3425.887
786.3854370117188 0 915.77423
787.3873291015625 0 1978.341
788.387939453125 0 1156.4937
789.3836059570312 0 971.8869
793.3971557617188 0 132672.55 y Ammonia loss 5
794.3888549804688 0 110948.72 z 5
795.3890380859375 0 38880.18
796.39013671875 0 9718.286
797.3916015625 0 1100.8298
809.4140014648438 0 2414.2437
810.4015502929688 0 4007.4587 y 5
811.4022216796875 0 2453.021
821.4267578125 0 3822.6257
822.4283447265625 0 1406.0933
825.3800659179688 0 1095.9075
832.43994140625 0 1199.9503
847.3578491210938 0 1596.0361
853.3553466796875 0 1393.7008
855.4111938476562 0 1625.8135
857.4348754882812 0 1738.4718
858.4945678710938 0 1664.7069
859.486083984375 0 5184.596
860.481689453125 0 9502.596
861.4837646484375 0 4705.0605
862.4730224609375 0 1458.465
864.4332275390625 0 41078.574 w 4
865.4254150390625 0 38932.758
866.4252319335938 0 13962.632
867.4268188476562 0 3516.821
873.41845703125 0 1102.1569
874.4150390625 0 1114.4573
875.4111328125 0 940.1236
879.408203125 0 2175.3176
880.4118041992188 0 1136.6838
881.3968505859375 0 1400.469
884.4835205078125 0 2030.2863
885.4766845703125 0 4266.3467
886.4746704101562 0 9034.502 c Water loss 6
887.470703125 0 5858.1006 c Ammonia loss 6
888.474365234375 0 1788.7605
889.4818725585938 0 1029.2804
891.42578125 0 1159.2369 z Water loss 4
896.424072265625 0 1039.7838
900.4761352539062 0 4757.729
901.4740600585938 0 3727.852
902.4915161132812 0 18056.256
903.4981689453125 0 71708.19
904.4920043945312 0 69504.6 c 6
905.4921264648438 0 24712.584
906.494140625 0 6829.555
907.4031372070312 0 22470.51
907.48876953125 0 512.17725
908.4201049804688 0 45888.184 y Ammonia loss 4
909.4129028320312 0 53428.81 z 4
910.414306640625 0 21594.062
911.4164428710938 0 6561.946
912.4173583984375 0 1166.6819
920.4520263671875 0 1675.4113
921.45654296875 0 907.75574
923.4232788085938 0 5923.805
924.431884765625 0 11761.41
925.4346313476562 0 10541.623 y 4
926.4312744140625 0 2529.4023
935.5810546875 0 1100.9332
961.5118408203125 0 1071.3123
968.49609375 0 1061.3112
969.4824829101562 0 2025.5287
973.52880859375 0 1219.9094
974.5209350585938 0 1737.0454
975.52734375 0 1473.9324
986.5030517578125 0 4093.4646
987.4988403320312 0 6743.703
988.49951171875 0 4662.221
989.50439453125 0 2047.6488
999.5313110351562 0 2870.2173
1000.5223388671875 0 5194.306 c Water loss 7
1001.5155639648438 0 5438.6367 c Ammonia loss 7
1002.51806640625 0 2284.8374
1003.5194091796875 0 2630.12
1004.5203247070312 0 1808.1484
1016.534423828125 0 4744.0854
1017.541748046875 0 39984.22
1018.535400390625 0 42850.04 c 7
1019.535400390625 0 17567.055
1020.5371704101562 0 6713.033
1021.5335693359375 0 2517.1719
1022.5087280273438 0 976.9715
1023.4998779296875 0 1113.9559
1025.5223388671875 0 768.2119
1048.5140380859375 0 1462.6218 z Ammonia loss 3
1060.5560302734375 0 1274.3314
1061.5657958984375 0 1071.3757
1062.562255859375 0 1190.8026
1063.5367431640625 0 1560.252 y Water loss 3
1064.5242919921875 0 15539.494 y Ammonia loss 3
1065.5186767578125 0 33607.96 z 3
1066.5177001953125 0 23279.592
1067.51953125 0 7721.987
1068.5206298828125 0 2643.539
1080.5364990234375 0 1878.4055
1081.53173828125 0 2557.75 y 3
1088.554443359375 0 3695.457 c Ammonia loss 8
1089.5458984375 0 5102.1587
1090.548828125 0 3326.6763
1091.5489501953125 0 1221.5518
1095.4967041015625 0 824.6455
1096.4996337890625 0 1467.7197
1097.486572265625 0 1901.9987
1098.4908447265625 0 1203.449
1104.573974609375 0 10788.601
1105.5673828125 0 17729.52 c 8
1106.566650390625 0 9876.774
1107.5673828125 0 3723.9724
1108.5616455078125 0 1447.9269
1109.5308837890625 0 981.9607
1110.5372314453125 0 1469.3503
1125.5556640625 0 725.0366
1134.550048828125 0 1337.6064 z Water loss 2
1135.544921875 0 2387.4253 z Ammonia loss 2
1136.538330078125 0 1271.689
1137.547119140625 0 791.8051
1151.5552978515625 0 11146.849 y Ammonia loss 2
1152.5550537109375 0 26020.492 z 2
1153.5550537109375 0 21185.66
1154.5562744140625 0 8916.618
1155.5576171875 0 2472.5098
1157.63037109375 0 956.0312
1158.6199951171875 0 774.586
1159.6336669921875 0 1909.3384
1160.6182861328125 0 1168.3458
1167.5882568359375 0 920.2124
1168.568115234375 0 2708.9055 y 2
1169.5672607421875 0 2083.6047
1170.5673828125 0 1125.0663
1173.6451416015625 0 2333.6357
1174.6475830078125 0 5684.9014
1175.6458740234375 0 6468.054
1176.6435546875 0 3734.5916
1177.6412353515625 0 1111.2365
1201.64013671875 0 8269.778 c Ammonia loss 9
1202.633544921875 0 11228.87
1203.63330078125 0 4364.008
1204.639404296875 0 1040.413
1217.6588134765625 0 5989.9565
1218.65771484375 0 18576.375 c 9
1219.6572265625 0 17175.955
1220.6546630859375 0 6839.371
1221.6590576171875 0 2171.1443
1264.647216796875 0 1149.0764 y Ammonia loss 1
1265.6448974609375 0 3762.437 z 1
1266.6383056640625 0 6444.8555
1267.633544921875 0 2908.366
1268.6351318359375 0 805.592
1280.65478515625 0 976.464
1281.656005859375 0 798.7417 y 1
1304.6875 0 1329.4823
1305.662841796875 0 3591.4521
1306.6475830078125 0 7267.7217
1307.6492919921875 0 5156.5693
1308.6536865234375 0 3814.0542
1309.6422119140625 0 1777.4824
1320.6953125 0 2276.8345
1321.6953125 0 5871.2886
1322.6876220703125 0 6180.4253
1323.6785888671875 0 9234.261
1324.6728515625 0 24900.467
1325.672119140625 0 19320.387
1326.67236328125 0 9184.099
1327.675048828125 0 2927.5083
1337.6962890625 0 2882.4177
1338.6990966796875 0 21023.924
1339.6937255859375 0 27269.85
1340.69384765625 0 14512.045
1341.6884765625 0 8149.425
1342.6878662109375 0 2158.6333
1348.6844482421875 0 5911.596
1349.6724853515625 0 10803.977
1350.6656494140625 0 11567.374
1351.6685791015625 0 6414.313
1352.6611328125 0 2173.3735
1355.7203369140625 0 1794.9362
1356.7127685546875 0 2268.7046
1357.70947265625 0 1181.5549
1363.7327880859375 0 1274.946
1364.741943359375 0 2532.637
1365.69384765625 0 11560.355
1366.6920166015625 0 71421.38
1367.6868896484375 0 101885.195
1368.6873779296875 0 49769.055
1369.6881103515625 0 18781.387
1370.6915283203125 0 5588.411
1371.6942138671875 0 978.5345
1379.7498779296875 0 996.41046
1380.7607421875 0 3235.1892
1381.736572265625 0 2931.0403
1382.711181640625 0 10404.493
1383.712890625 0 20641.492
1384.710205078125 0 21478.062
1385.711669921875 0 11159.792
1386.7125244140625 0 3767.111

Spectrum Details

|  |  |
| --- | --- |
| Matched peaks? Matched peaks The total absolute number of peaks matched. Additionally in brackets the total fraction of peaks matched and the total number of peaks is shown. | 82 (11.85% of 692) |
| FDR? FDR The false discovery rate estimated for this peptide. It is calculated by matching all theoretical fragments with a non-integer shift with the raw peaks for this spectrum. This is done with 40 different shifts. The resulting percentage is the average number of annotated peaks over the number of annotated peaks with the correct spectrum. | 1.57% |
| Satellite FDR? Satellite FDR See the FDR for details on its calculation. This satellite ion specific FDR only contains the satellite ions (d/w) for I/L/J positions. | 0.00% |
| PSM Score? PSM Score The PSM Score as given by Hecklib to this annotated spectrum. It is shown with three significant figures. | 736 |

#### Reverse Lookup? Reverse Lookup All places where this read could be placed.

| Group | Segment | Template | Template Part | Read Part | Score | Unique |
| --- | --- | --- | --- | --- | --- | --- |
| Homo sapiens Heavy Chain | IGHV | IGHV3-9 | [68..80] | [0..11] | 67 | False |
| Homo sapiens Heavy Chain | IGHV | IGHV3-48 | [68..80] | [0..11] | 67 | False |
| Homo sapiens Heavy Chain | IGHV | IGHV3-21 | [68..80] | [0..11] | 67 | False |
| Homo sapiens Heavy Chain | IGHV | IGHV3-7 | [68..80] | [0..11] | 67 | False |
| Homo sapiens Heavy Chain | IGHV | IGHV3-43 | [68..80] | [0..11] | 67 | False |
| Homo sapiens Heavy Chain | IGHV | IGHV3-13 | [67..79] | [0..11] | 62 | False |
| Homo sapiens Heavy Chain | IGHV | IGHV3-11 | [68..80] | [0..11] | 67 | False |
| Homo sapiens Heavy Chain | IGHV | IGHV3-20 | [68..80] | [0..11] | 67 | False |
| Homo sapiens Heavy Chain | IGHV | IGHV3-72 | [70..82] | [0..11] | 67 | False |

| Recombined | Template Part | Read Part | Score | Unique |
| --- | --- | --- | --- | --- |
| REC-0-1 | [68..80] | [0..11] | 67 | True |

#### Meta Information from Multiple reads

##### Number of combined reads

2

##### Intensity

0.8581

##### TotalArea

5.34E+07

##### Changes to the peptide sequence

TISRDWKNSJY

L→JNo support for either Leucine or Isoleucine based on side chain ions (Position: 10)

L→ISupport for Isoleucine based on side chain ions (1 for I 0 for L) (Position: 2)

#### Positional Score

Copy Data

##### Positional Score (TSV)

###### Preview

```
Loading example...
```

*Click on the button to copy the data to your clipboard.*

00012345678910

Label Value
"0" 0
"1" 0
"2" 0
"3" 0
"4" 0
"5" 0
"6" 0
"7" 0
"8" 0
"9" 0
"10" 0

#### Meta Information from PEAKS

##### Scan Identifier

F3:3575

##### Original sequence

T

L

S

R

D

W

K

N

S

L

Y

##### Posttranslational Modifications

##### Source File

D:\separate\_stitch\_analyses\xle-disambiguation\raw\20210323\_F1\_UM1\_Peng0013\_SA\_F59\_ingel\_3ug\_chymo.raw

##### Fraction

3

##### Scan Feature

F3:2692

##### De Novo Score

98

##### ConfidenceScore

98

### m/z

461.5702

##### Mass

1381.699

##### Charge

3

##### Retention Time

19.23

##### Predicted Retention Time

-

##### Area

5.34E+07

##### Fragmentation mode

ETHCD

##### Originating file

01 D:\separate\_stitch\_analyses\xle-disambiguation\20210325\_F59\_3ug\_DENOVO\_12.csv

#### Meta Information from PEAKS

##### Scan Identifier

F3:3718

##### Original sequence

T

L

S

R

D

W

K

N

S

L

Y

##### Posttranslational Modifications

##### Source File

D:\separate\_stitch\_analyses\xle-disambiguation\raw\20210323\_F1\_UM1\_Peng0013\_SA\_F59\_ingel\_3ug\_chymo.raw

##### Fraction

3

##### Scan Feature

-

##### De Novo Score

97

##### ConfidenceScore

97

### m/z

461.572

##### Mass

1381.699

##### Charge

3

##### Retention Time

20.37

##### Predicted Retention Time

-

##### Area

0

##### Fragmentation mode

ETHCD

##### Originating file

01 D:\separate\_stitch\_analyses\xle-disambiguation\20210325\_F59\_3ug\_DENOVO\_12.csv
