## Supplementary material for "A handle on mass coincidence errors in *de novo* sequencing of antibodies by bottom-up proteomics": Combined_083.html

Details Combined\_083 | Stitch OverviewUndefined

### Read Combined\_083

#### Sequence (length=4)

VHYC

#### Spectrum 5790? Spectrum 5790 The raw spectrum of this peptide as annotated by Hecklib. The fragments are coloured according to ion type (see legend). Any peaks with a star '\*' as text can be hovered over to see the full details, first the ion type second the mass shift type. By hovering over the amino acids in the peptide or ions in the legend the corresponding peaks are highlighted. By toggling the 'Unassigned' label you can turn the background (unassigned) peaks on or off in the plot. By updating the slider in the Ion legend you can update the spectrum to only show the top X% of the peaks with labels. The top X% means any peak that is within X% of the highest intensity. By dragging in the spectrum you can zoom in to a specific part of the spectrum and use 'Zoom Out' to get back to the original zoom level. The annotation of the spectrum is based on the given sequence in the peptides file and is done with different software so inconsistencies are likely. The peaks are annotated based on the given sequence, with 20 ppm tolerance.

Copy Data

##### Spectrum 5790 (TSV)

###### Preview

```
Loading example...
```

*Click on the button to copy the data to your clipboard.*

Mz MinMz MaxIntensity Max

WidthHeightPeptide font sizePeptide stroke widthSpectrum font sizeSpectrum stroke widthCompact peptide

Ion legend

wxyz

abcd

OtherUnassignedIon^Charge^_Position_Show for top:%

VHYC

02.08e+44.15e+46.23e+48.30e+4

Zoom Out

c^+1^_2_y^+1^_2_c^+1^_3_

0775155123263102

Fragment Matches Table

Show background peaks

| Position | Ion type | Intensity | mz Theoretical | mz Error (Th) | mz Error (ppm) | Charge | Series Number |
| --- | --- | --- | --- | --- | --- | --- | --- |
| - | - | 532.4 | 120.1 | - | - | 0 | - |
| - | - | 345.1 | 122.6 | - | - | 0 | - |
| - | - | 366.2 | 128.8 | - | - | 0 | - |
| - | - | 484.9 | 131.1 | - | - | 0 | - |
| - | - | 924.3 | 133.1 | - | - | 0 | - |
| - | - | 3046 | 136.1 | - | - | 0 | - |
| - | - | 471.7 | 149 | - | - | 0 | - |
| - | - | 480.2 | 150.5 | - | - | 0 | - |
| - | - | 458.9 | 157.2 | - | - | 0 | - |
| - | - | 2209 | 166.1 | - | - | 0 | - |
| - | - | 479.9 | 166.7 | - | - | 0 | - |
| - | - | 2748 | 173.4 | - | - | 0 | - |
| - | - | 871.7 | 175.1 | - | - | 0 | - |
| - | - | 516.9 | 177.3 | - | - | 0 | - |
| - | - | 1177 | 180 | - | - | 0 | - |
| - | - | 548.5 | 187.9 | - | - | 0 | - |
| - | - | 473.5 | 216.4 | - | - | 0 | - |
| - | - | 677.4 | 219.1 | - | - | 0 | - |
| - | - | 4366 | 233.2 | - | - | 0 | - |
| - | - | 488.3 | 256.1 | - | - | 0 | - |
| - | - | 3479 | 261.2 | - | - | 0 | - |
| - | - | 6194 | 283.1 | - | - | 0 | - |
| - | - | 560.6 | 288.2 | - | - | 0 | - |
| - | - | 512.5 | 292.9 | - | - | 0 | - |
| - | - | 771.7 | 299.1 | - | - | 0 | - |
| - | - | 804.8 | 307.2 | - | - | 0 | - |
| 2 | c | 1.72E+04 | 311.1 | 0.004375 | 14.06 | +1 | 2 |
| - | - | 489.4 | 313.9 | - | - | 0 | - |
| - | - | 1229 | 327.1 | - | - | 0 | - |
| 3 | y | 3.738E+04 | 343.1 | 0.00459 | 13.38 | +1 | 2 |
| - | - | 618.6 | 352.2 | - | - | 0 | - |
| - | - | 3762 | 355.1 | - | - | 0 | - |
| - | - | 733.2 | 356.1 | - | - | 0 | - |
| - | - | 760.6 | 357.1 | - | - | 0 | - |
| - | - | 1163 | 359 | - | - | 0 | - |
| - | - | 564.4 | 383 | - | - | 0 | - |
| - | - | 653.1 | 421.1 | - | - | 0 | - |
| - | - | 1946 | 429.1 | - | - | 0 | - |
| - | - | 1689 | 429.2 | - | - | 0 | - |
| - | - | 1021 | 430.1 | - | - | 0 | - |
| - | - | 2360 | 439.3 | - | - | 0 | - |
| - | - | 936.3 | 439.8 | - | - | 0 | - |
| - | - | 608.6 | 448.4 | - | - | 0 | - |
| - | - | 710.9 | 473.3 | - | - | 0 | - |
| 3 | c | 6451 | 474.2 | 0.004553 | 9.602 | +1 | 3 |
| - | - | 2693 | 489.8 | - | - | 0 | - |
| - | - | 789.4 | 490.3 | - | - | 0 | - |
| - | - | 822.1 | 503.3 | - | - | 0 | - |
| - | - | 1664 | 506.2 | - | - | 0 | - |
| - | - | 969.1 | 512.8 | - | - | 0 | - |
| - | - | 6.285E+04 | 521.8 | - | - | 0 | - |
| - | - | 3.081E+04 | 522.3 | - | - | 0 | - |
| - | - | 1.359E+04 | 522.8 | - | - | 0 | - |
| - | - | 1842 | 523.3 | - | - | 0 | - |
| - | - | 1423 | 595.3 | - | - | 0 | - |
| - | - | 706.2 | 607.3 | - | - | 0 | - |
| - | - | 6101 | 608.3 | - | - | 0 | - |
| - | - | 1344 | 609.3 | - | - | 0 | - |
| - | - | 842.7 | 619.9 | - | - | 0 | - |
| - | - | 727.4 | 642.9 | - | - | 0 | - |
| - | - | 4093 | 650.4 | - | - | 0 | - |
| - | - | 3010 | 650.9 | - | - | 0 | - |
| - | - | 1253 | 651.4 | - | - | 0 | - |
| - | - | 2.469E+04 | 651.9 | - | - | 0 | - |
| - | - | 2.04E+04 | 652.4 | - | - | 0 | - |
| - | - | 1.039E+04 | 652.9 | - | - | 0 | - |
| - | - | 4495 | 653.2 | - | - | 0 | - |
| - | - | 3711 | 653.4 | - | - | 0 | - |
| - | - | 2.706E+04 | 696.4 | - | - | 0 | - |
| - | - | 3.581E+04 | 697.4 | - | - | 0 | - |
| - | - | 1.181E+04 | 698.4 | - | - | 0 | - |
| - | - | 2074 | 699.4 | - | - | 0 | - |
| - | - | 1178 | 720.4 | - | - | 0 | - |
| - | - | 2162 | 781.5 | - | - | 0 | - |
| - | - | 1530 | 782.5 | - | - | 0 | - |
| - | - | 2.179E+04 | 824.5 | - | - | 0 | - |
| - | - | 1.729E+04 | 825.5 | - | - | 0 | - |
| - | - | 6659 | 826.5 | - | - | 0 | - |
| - | - | 6980 | 832.4 | - | - | 0 | - |
| - | - | 3330 | 833.4 | - | - | 0 | - |
| - | - | 1163 | 877.5 | - | - | 0 | - |
| - | - | 884.2 | 878.5 | - | - | 0 | - |
| - | - | 704 | 897.5 | - | - | 0 | - |
| - | - | 1042 | 939.4 | - | - | 0 | - |
| - | - | 2.098E+04 | 939.6 | - | - | 0 | - |
| - | - | 2.357E+04 | 940.6 | - | - | 0 | - |
| - | - | 8693 | 941.6 | - | - | 0 | - |
| - | - | 1901 | 942.6 | - | - | 0 | - |
| - | - | 5329 | 945.5 | - | - | 0 | - |
| - | - | 2441 | 946.5 | - | - | 0 | - |
| - | - | 847.1 | 947.5 | - | - | 0 | - |
| - | - | 823.2 | 979.6 | - | - | 0 | - |
| - | - | 2665 | 1041 | - | - | 0 | - |
| - | - | 1.903E+04 | 1042 | - | - | 0 | - |
| - | - | 1.077E+04 | 1043 | - | - | 0 | - |
| - | - | 4836 | 1044 | - | - | 0 | - |
| - | - | 2667 | 1045 | - | - | 0 | - |
| - | - | 578.8 | 1071 | - | - | 0 | - |
| - | - | 583.8 | 1072 | - | - | 0 | - |
| - | - | 1088 | 1111 | - | - | 0 | - |
| - | - | 910.1 | 1112 | - | - | 0 | - |
| - | - | 725.9 | 1131 | - | - | 0 | - |
| - | - | 709.3 | 1138 | - | - | 0 | - |
| - | - | 3.055E+04 | 1155 | - | - | 0 | - |
| - | - | 2.228E+04 | 1156 | - | - | 0 | - |
| - | - | 8242 | 1157 | - | - | 0 | - |
| - | - | 718.6 | 1158 | - | - | 0 | - |
| - | - | 8911 | 1174 | - | - | 0 | - |
| - | - | 6267 | 1175 | - | - | 0 | - |
| - | - | 1866 | 1176 | - | - | 0 | - |
| - | - | 1754 | 1212 | - | - | 0 | - |
| - | - | 956.7 | 1213 | - | - | 0 | - |
| - | - | 993.8 | 1240 | - | - | 0 | - |
| - | - | 1119 | 1241 | - | - | 0 | - |
| - | - | 1898 | 1242 | - | - | 0 | - |
| - | - | 2434 | 1243 | - | - | 0 | - |
| - | - | 2804 | 1244 | - | - | 0 | - |
| - | - | 1442 | 1245 | - | - | 0 | - |
| - | - | 891.3 | 1246 | - | - | 0 | - |
| - | - | 690.5 | 1248 | - | - | 0 | - |
| - | - | 1675 | 1258 | - | - | 0 | - |
| - | - | 1790 | 1260 | - | - | 0 | - |
| - | - | 617.3 | 1269 | - | - | 0 | - |
| - | - | 877.9 | 1270 | - | - | 0 | - |
| - | - | 915.9 | 1271 | - | - | 0 | - |
| - | - | 874.1 | 1284 | - | - | 0 | - |
| - | - | 1850 | 1285 | - | - | 0 | - |
| - | - | 5283 | 1286 | - | - | 0 | - |
| - | - | 3.289E+04 | 1287 | - | - | 0 | - |
| - | - | 2.349E+04 | 1288 | - | - | 0 | - |
| - | - | 1.186E+04 | 1289 | - | - | 0 | - |
| - | - | 3262 | 1290 | - | - | 0 | - |
| - | - | 1345 | 1300 | - | - | 0 | - |
| - | - | 5213 | 1301 | - | - | 0 | - |
| - | - | 4285 | 1302 | - | - | 0 | - |
| - | - | 3.674E+04 | 1303 | - | - | 0 | - |
| - | - | 8.219E+04 | 1304 | - | - | 0 | - |
| - | - | 5.394E+04 | 1305 | - | - | 0 | - |
| - | - | 2.564E+04 | 1306 | - | - | 0 | - |
| - | - | 5266 | 1307 | - | - | 0 | - |
| - | - | 679 | 1378 | - | - | 0 | - |
| - | - | 619.4 | 1598 | - | - | 0 | - |
| - | - | 747.8 | 1625 | - | - | 0 | - |
| - | - | 704.2 | 1643 | - | - | 0 | - |
| - | - | 675 | 1892 | - | - | 0 | - |
| - | - | 708.5 | 2124 | - | - | 0 | - |
| - | - | 646.7 | 2531 | - | - | 0 | - |
| - | - | 685.9 | 2958 | - | - | 0 | - |
| - | - | 679.1 | 3071 | - | - | 0 | - |

m/z Charge Intensity FragmentType MassShift Position
120.08106994628906 0 532.40753
122.60386657714844 0 345.07803
128.7965545654297 0 366.22467
131.07022094726562 0 484.8897
133.08592224121094 0 924.3155
136.07571411132812 0 3046.1436
148.955078125 0 471.67252
150.50613403320312 0 480.24045
157.2384796142578 0 458.91025
166.0531463623047 0 2209.0576
166.72486877441406 0 479.8888
173.43942260742188 0 2748.1223
175.09683227539062 0 871.6926
177.30078125 0 516.9021
180.03248596191406 0 1177.3435
187.8978271484375 0 548.4625
216.36587524414062 0 473.508
219.12306213378906 0 677.36035
233.16464233398438 0 4366.248
256.1169128417969 0 488.30225
261.1595153808594 0 3479.401
283.1439514160156 0 6194.2427
288.1668701171875 0 560.5702
292.8728942871094 0 512.51086
299.06134033203125 0 771.6909
307.1752014160156 0 804.7743
311.13897705078125 0 17198.482 c Ammonia loss 1
313.8919677734375 0 489.35883
327.1336364746094 0 1229.3834
343.09576416015625 0 37381.7 y 2
352.1986389160156 0 618.5782
355.069580078125 0 3761.8901
356.0693359375 0 733.2152
357.0646057128906 0 760.58795
359.0281982421875 0 1162.895
383.026611328125 0 564.3881
421.0939025878906 0 653.1464
429.08782958984375 0 1946.0963
429.1803894042969 0 1689.0305
430.08880615234375 0 1021.22565
439.26031494140625 0 2360.0251
439.7621154785156 0 936.298
448.3795166015625 0 608.6334
473.25860595703125 0 710.8701
474.2024841308594 0 6450.5415 c Ammonia loss 2
489.7845764160156 0 2692.6228
490.2860107421875 0 789.3901
503.3050231933594 0 822.1435
506.1606140136719 0 1663.5264
512.7797241210938 0 969.07605
521.783447265625 0 62847.48
522.284912109375 0 30811.363
522.7848510742188 0 13585.759
523.2843627929688 0 1842.1586
595.318359375 0 1423.3391
607.287841796875 0 706.16833
608.295654296875 0 6100.785
609.2986450195312 0 1344.4218
619.8587036132812 0 842.658
642.85009765625 0 727.4183
650.3558349609375 0 4093.459
650.8534545898438 0 3009.7341
651.353759765625 0 1253.0985
651.859375 0 24691.492
652.3609619140625 0 20402.191
652.8601684570312 0 10389.811
653.2276000976562 0 4494.5776
653.3572387695312 0 3710.6602
696.4312133789062 0 27062.424
697.4378662109375 0 35807.89
698.4410400390625 0 11814.547
699.4447631835938 0 2074.0173
720.3585205078125 0 1178.3315
781.520263671875 0 2161.5723
782.5234375 0 1529.852
824.5257568359375 0 21789.99
825.5311279296875 0 17290.432
826.5343627929688 0 6659.0864
832.4351196289062 0 6980.043
833.4400634765625 0 3329.7437
877.509033203125 0 1162.5046
878.5089721679688 0 884.2073
897.5462646484375 0 704.04785
939.4430541992188 0 1041.5651
939.5526123046875 0 20979.453
940.5574951171875 0 23572.371
941.5615234375 0 8692.865
942.5640258789062 0 1900.9164
945.5078125 0 5329.1826
946.5074462890625 0 2441.292
947.5075073242188 0 847.0731
979.5643920898438 0 823.17316
1040.5987548828125 0 2665.1047
1041.6077880859375 0 19026.744
1042.5753173828125 0 10772.522
1043.5679931640625 0 4836.0254
1044.5634765625 0 2666.584
1070.5511474609375 0 578.7621
1071.5487060546875 0 583.758
1110.6689453125 0 1087.871
1111.6700439453125 0 910.12067
1130.6201171875 0 725.9137
1137.669921875 0 709.29926
1154.6917724609375 0 30547.1
1155.69482421875 0 22284.33
1156.696533203125 0 8241.962
1157.698974609375 0 718.64294
1173.6087646484375 0 8910.684
1174.6112060546875 0 6267.1826
1175.6107177734375 0 1866.146
1211.6903076171875 0 1753.5275
1212.687744140625 0 956.6646
1239.7120361328125 0 993.8349
1240.7220458984375 0 1119.0303
1241.7122802734375 0 1897.6294
1242.7078857421875 0 2434.2634
1243.69384765625 0 2804.467
1244.6861572265625 0 1441.9728
1245.6973876953125 0 891.27277
1247.663818359375 0 690.5011
1257.7091064453125 0 1674.9683
1259.728515625 0 1789.7034
1269.09130859375 0 617.32245
1269.5615234375 0 877.8676
1270.56005859375 0 915.9084
1283.68505859375 0 874.13934
1284.6702880859375 0 1849.6373
1285.702880859375 0 5282.5854
1286.6988525390625 0 32891.332
1287.7020263671875 0 23485.3
1288.7025146484375 0 11856.288
1289.7008056640625 0 3261.6204
1299.6951904296875 0 1344.8822
1300.698974609375 0 5213.499
1301.69873046875 0 4284.6367
1302.7098388671875 0 36739.348
1303.716064453125 0 82189.43
1304.7197265625 0 53936.977
1305.7186279296875 0 25639.678
1306.718994140625 0 5265.527
1378.2965087890625 0 678.96545
1598.218994140625 0 619.3573
1625.288330078125 0 747.7726
1643.055908203125 0 704.222
1892.0535888671875 0 674.98755
2123.98046875 0 708.47174
2531.4501953125 0 646.67096
2957.877197265625 0 685.8999
3070.833740234375 0 679.11194

Spectrum Details

|  |  |
| --- | --- |
| Matched peaks? Matched peaks The total absolute number of peaks matched. Additionally in brackets the total fraction of peaks matched and the total number of peaks is shown. | 3 (2.01% of 149) |
| FDR? FDR The false discovery rate estimated for this peptide. It is calculated by matching all theoretical fragments with a non-integer shift with the raw peaks for this spectrum. This is done with 40 different shifts. The resulting percentage is the average number of annotated peaks over the number of annotated peaks with the correct spectrum. | 0.79% |
| Satellite FDR? Satellite FDR See the FDR for details on its calculation. This satellite ion specific FDR only contains the satellite ions (d/w) for I/L/J positions. | - |
| PSM Score? PSM Score The PSM Score as given by Hecklib to this annotated spectrum. It is shown with three significant figures. | 36.9 |

#### Spectrum 5791? Spectrum 5791 The raw spectrum of this peptide as annotated by Hecklib. The fragments are coloured according to ion type (see legend). Any peaks with a star '\*' as text can be hovered over to see the full details, first the ion type second the mass shift type. By hovering over the amino acids in the peptide or ions in the legend the corresponding peaks are highlighted. By toggling the 'Unassigned' label you can turn the background (unassigned) peaks on or off in the plot. By updating the slider in the Ion legend you can update the spectrum to only show the top X% of the peaks with labels. The top X% means any peak that is within X% of the highest intensity. By dragging in the spectrum you can zoom in to a specific part of the spectrum and use 'Zoom Out' to get back to the original zoom level. The annotation of the spectrum is based on the given sequence in the peptides file and is done with different software so inconsistencies are likely. The peaks are annotated based on the given sequence, with 20 ppm tolerance.

Copy Data

##### Spectrum 5791 (TSV)

###### Preview

```
Loading example...
```

*Click on the button to copy the data to your clipboard.*

Mz MinMz MaxIntensity Max

WidthHeightPeptide font sizePeptide stroke widthSpectrum font sizeSpectrum stroke widthCompact peptide

Ion legend

wxyz

abcd

OtherUnassignedIon^Charge^_Position_Show for top:%

VHYC

01.17e+42.35e+43.52e+44.70e+4

Zoom Out

c^+1^_2_y^+1^_2_c^+1^_3_

0661132219832643

Fragment Matches Table

Show background peaks

| Position | Ion type | Intensity | mz Theoretical | mz Error (Th) | mz Error (ppm) | Charge | Series Number |
| --- | --- | --- | --- | --- | --- | --- | --- |
| - | - | 556 | 120.1 | - | - | 0 | - |
| - | - | 384.3 | 125.6 | - | - | 0 | - |
| - | - | 350.5 | 128.6 | - | - | 0 | - |
| - | - | 646.3 | 129.1 | - | - | 0 | - |
| - | - | 476.3 | 136.1 | - | - | 0 | - |
| - | - | 5468 | 136.1 | - | - | 0 | - |
| - | - | 474.4 | 137.1 | - | - | 0 | - |
| - | - | 376.3 | 140.1 | - | - | 0 | - |
| - | - | 539.8 | 148.9 | - | - | 0 | - |
| - | - | 622.3 | 148.9 | - | - | 0 | - |
| - | - | 731 | 148.9 | - | - | 0 | - |
| - | - | 846.9 | 148.9 | - | - | 0 | - |
| - | - | 902.5 | 148.9 | - | - | 0 | - |
| - | - | 1016 | 148.9 | - | - | 0 | - |
| - | - | 1358 | 148.9 | - | - | 0 | - |
| - | - | 3247 | 148.9 | - | - | 0 | - |
| - | - | 5459 | 148.9 | - | - | 0 | - |
| - | - | 3732 | 149 | - | - | 0 | - |
| - | - | 1599 | 149 | - | - | 0 | - |
| - | - | 1250 | 149 | - | - | 0 | - |
| - | - | 1084 | 149 | - | - | 0 | - |
| - | - | 656.5 | 149 | - | - | 0 | - |
| - | - | 526.8 | 149 | - | - | 0 | - |
| - | - | 1118 | 149 | - | - | 0 | - |
| - | - | 424 | 149 | - | - | 0 | - |
| - | - | 444.1 | 177.1 | - | - | 0 | - |
| - | - | 1800 | 180 | - | - | 0 | - |
| - | - | 457.9 | 188.7 | - | - | 0 | - |
| - | - | 479.4 | 215.4 | - | - | 0 | - |
| - | - | 597.8 | 225.1 | - | - | 0 | - |
| - | - | 855.8 | 233.2 | - | - | 0 | - |
| - | - | 531.5 | 239.1 | - | - | 0 | - |
| - | - | 539.5 | 274.1 | - | - | 0 | - |
| - | - | 9455 | 283.1 | - | - | 0 | - |
| - | - | 2142 | 284.1 | - | - | 0 | - |
| - | - | 533.9 | 287.5 | - | - | 0 | - |
| - | - | 782.9 | 299.1 | - | - | 0 | - |
| 2 | c | 2.348E+04 | 311.1 | 0.0041 | 13.18 | +1 | 2 |
| - | - | 4519 | 312.1 | - | - | 0 | - |
| - | - | 503.8 | 313.1 | - | - | 0 | - |
| - | - | 1562 | 327.1 | - | - | 0 | - |
| 3 | y | 4.649E+04 | 343.1 | 0.004987 | 14.53 | +1 | 2 |
| - | - | 8164 | 344.1 | - | - | 0 | - |
| - | - | 1558 | 345.1 | - | - | 0 | - |
| - | - | 1080 | 355.1 | - | - | 0 | - |
| - | - | 1225 | 356.1 | - | - | 0 | - |
| - | - | 626.8 | 381.2 | - | - | 0 | - |
| - | - | 610 | 415 | - | - | 0 | - |
| - | - | 1222 | 429.2 | - | - | 0 | - |
| - | - | 642.6 | 466.6 | - | - | 0 | - |
| 3 | c | 7502 | 474.2 | 0.003974 | 8.379 | +1 | 3 |
| - | - | 3405 | 475.2 | - | - | 0 | - |
| - | - | 1770 | 506.2 | - | - | 0 | - |
| - | - | 596.1 | 507.2 | - | - | 0 | - |
| - | - | 3062 | 522.3 | - | - | 0 | - |
| - | - | 8906 | 522.8 | - | - | 0 | - |
| - | - | 3519 | 523.3 | - | - | 0 | - |
| - | - | 619 | 523.8 | - | - | 0 | - |
| - | - | 580.5 | 599.9 | - | - | 0 | - |
| - | - | 858.3 | 609.3 | - | - | 0 | - |
| - | - | 1118 | 615.3 | - | - | 0 | - |
| - | - | 746.8 | 619.3 | - | - | 0 | - |
| - | - | 686 | 633.2 | - | - | 0 | - |
| - | - | 5689 | 652.9 | - | - | 0 | - |
| - | - | 3477 | 653.2 | - | - | 0 | - |
| - | - | 5678 | 653.4 | - | - | 0 | - |
| - | - | 4061 | 653.9 | - | - | 0 | - |
| - | - | 1748 | 654.2 | - | - | 0 | - |
| - | - | 7894 | 654.3 | - | - | 0 | - |
| - | - | 1457 | 654.4 | - | - | 0 | - |
| - | - | 6219 | 654.8 | - | - | 0 | - |
| - | - | 813 | 655.2 | - | - | 0 | - |
| - | - | 2669 | 655.3 | - | - | 0 | - |
| - | - | 1469 | 687.4 | - | - | 0 | - |
| - | - | 1746 | 696.4 | - | - | 0 | - |
| - | - | 1795 | 697.3 | - | - | 0 | - |
| - | - | 3758 | 697.4 | - | - | 0 | - |
| - | - | 698.4 | 698.3 | - | - | 0 | - |
| - | - | 4032 | 698.4 | - | - | 0 | - |
| - | - | 2050 | 699.4 | - | - | 0 | - |
| - | - | 665.4 | 700.4 | - | - | 0 | - |
| - | - | 1479 | 741.4 | - | - | 0 | - |
| - | - | 639.1 | 751.4 | - | - | 0 | - |
| - | - | 1218 | 824.5 | - | - | 0 | - |
| - | - | 2731 | 825.5 | - | - | 0 | - |
| - | - | 1715 | 825.5 | - | - | 0 | - |
| - | - | 1335 | 826.5 | - | - | 0 | - |
| - | - | 2702 | 826.5 | - | - | 0 | - |
| - | - | 1359 | 827.5 | - | - | 0 | - |
| - | - | 877.5 | 833.4 | - | - | 0 | - |
| - | - | 994.3 | 834.4 | - | - | 0 | - |
| - | - | 1435 | 835.4 | - | - | 0 | - |
| - | - | 788.6 | 867.4 | - | - | 0 | - |
| - | - | 818.6 | 868.4 | - | - | 0 | - |
| - | - | 709.6 | 871.4 | - | - | 0 | - |
| - | - | 1641 | 911.4 | - | - | 0 | - |
| - | - | 869 | 912.4 | - | - | 0 | - |
| - | - | 1596 | 927.4 | - | - | 0 | - |
| - | - | 641.5 | 935.9 | - | - | 0 | - |
| - | - | 818.9 | 939.6 | - | - | 0 | - |
| - | - | 3030 | 940.6 | - | - | 0 | - |
| - | - | 2619 | 941.6 | - | - | 0 | - |
| - | - | 2336 | 942.6 | - | - | 0 | - |
| - | - | 779.5 | 943.6 | - | - | 0 | - |
| - | - | 1006 | 947.5 | - | - | 0 | - |
| - | - | 1362 | 950.5 | - | - | 0 | - |
| - | - | 752.9 | 961.5 | - | - | 0 | - |
| - | - | 698.5 | 994.4 | - | - | 0 | - |
| - | - | 3146 | 1028 | - | - | 0 | - |
| - | - | 1257 | 1030 | - | - | 0 | - |
| - | - | 716.3 | 1039 | - | - | 0 | - |
| - | - | 1396 | 1043 | - | - | 0 | - |
| - | - | 2054 | 1044 | - | - | 0 | - |
| - | - | 1617 | 1045 | - | - | 0 | - |
| - | - | 555.1 | 1057 | - | - | 0 | - |
| - | - | 942.4 | 1102 | - | - | 0 | - |
| - | - | 3879 | 1111 | - | - | 0 | - |
| - | - | 2903 | 1112 | - | - | 0 | - |
| - | - | 771.7 | 1113 | - | - | 0 | - |
| - | - | 1096 | 1120 | - | - | 0 | - |
| - | - | 684.6 | 1121 | - | - | 0 | - |
| - | - | 1022 | 1143 | - | - | 0 | - |
| - | - | 1132 | 1155 | - | - | 0 | - |
| - | - | 2252 | 1156 | - | - | 0 | - |
| - | - | 4724 | 1157 | - | - | 0 | - |
| - | - | 1482 | 1158 | - | - | 0 | - |
| - | - | 1045 | 1176 | - | - | 0 | - |
| - | - | 1896 | 1177 | - | - | 0 | - |
| - | - | 821.6 | 1177 | - | - | 0 | - |
| - | - | 671.4 | 1178 | - | - | 0 | - |
| - | - | 702.3 | 1185 | - | - | 0 | - |
| - | - | 2525 | 1192 | - | - | 0 | - |
| - | - | 1893 | 1193 | - | - | 0 | - |
| - | - | 1382 | 1194 | - | - | 0 | - |
| - | - | 1464 | 1208 | - | - | 0 | - |
| - | - | 1508 | 1249 | - | - | 0 | - |
| - | - | 1543 | 1250 | - | - | 0 | - |
| - | - | 794 | 1251 | - | - | 0 | - |
| - | - | 715.1 | 1255 | - | - | 0 | - |
| - | - | 1552 | 1271 | - | - | 0 | - |
| - | - | 1123 | 1276 | - | - | 0 | - |
| - | - | 1264 | 1277 | - | - | 0 | - |
| - | - | 709.3 | 1278 | - | - | 0 | - |
| - | - | 888.8 | 1289 | - | - | 0 | - |
| - | - | 5165 | 1289 | - | - | 0 | - |
| - | - | 3955 | 1290 | - | - | 0 | - |
| - | - | 2378 | 1291 | - | - | 0 | - |
| - | - | 2522 | 1292 | - | - | 0 | - |
| - | - | 1003 | 1292 | - | - | 0 | - |
| - | - | 2607 | 1293 | - | - | 0 | - |
| - | - | 4570 | 1294 | - | - | 0 | - |
| - | - | 1343 | 1295 | - | - | 0 | - |
| - | - | 1257 | 1299 | - | - | 0 | - |
| - | - | 977.7 | 1300 | - | - | 0 | - |
| - | - | 987 | 1300 | - | - | 0 | - |
| - | - | 5884 | 1305 | - | - | 0 | - |
| - | - | 1126 | 1306 | - | - | 0 | - |
| - | - | 1.75E+04 | 1306 | - | - | 0 | - |
| - | - | 865.4 | 1307 | - | - | 0 | - |
| - | - | 1.438E+04 | 1307 | - | - | 0 | - |
| - | - | 2487 | 1308 | - | - | 0 | - |
| - | - | 5825 | 1308 | - | - | 0 | - |
| - | - | 1827 | 1308 | - | - | 0 | - |
| - | - | 1.31E+04 | 1309 | - | - | 0 | - |
| - | - | 1329 | 1309 | - | - | 0 | - |
| - | - | 1.046E+04 | 1310 | - | - | 0 | - |
| - | - | 6371 | 1311 | - | - | 0 | - |
| - | - | 1164 | 1312 | - | - | 0 | - |
| - | - | 678.1 | 1916 | - | - | 0 | - |
| - | - | 957.3 | 2286 | - | - | 0 | - |
| - | - | 652.2 | 2305 | - | - | 0 | - |
| - | - | 789.6 | 2598 | - | - | 0 | - |
| - | - | 1092 | 2599 | - | - | 0 | - |
| - | - | 844.8 | 2614 | - | - | 0 | - |
| - | - | 1335 | 2615 | - | - | 0 | - |
| - | - | 1689 | 2616 | - | - | 0 | - |
| - | - | 1416 | 2617 | - | - | 0 | - |

m/z Charge Intensity FragmentType MassShift Position
120.08090209960938 0 555.9658
125.61078643798828 0 384.2887
128.55352783203125 0 350.53693
129.1025848388672 0 646.2668
136.07095336914062 0 476.33838
136.07559204101562 0 5467.9175
137.0791473388672 0 474.36642
140.1125030517578 0 376.32132
148.87332153320312 0 539.7939
148.8801727294922 0 622.2896
148.9022979736328 0 730.98206
148.90919494628906 0 846.9265
148.91635131835938 0 902.51636
148.92349243164062 0 1015.951
148.93080139160156 0 1358.0842
148.93814086914062 0 3247.0635
148.9459228515625 0 5459.0635
148.96249389648438 0 3732.2378
148.9701385498047 0 1599.3103
148.977294921875 0 1250.3414
148.98443603515625 0 1083.6326
148.99172973632812 0 656.48694
148.99948120117188 0 526.77356
149.00648498535156 0 1117.6362
149.0129852294922 0 424.0266
177.13487243652344 0 444.08542
180.0322723388672 0 1800.3334
188.70362854003906 0 457.8803
215.366455078125 0 479.37845
225.068603515625 0 597.82776
233.16445922851562 0 855.78406
239.09457397460938 0 531.4788
274.100341796875 0 539.4671
283.1435852050781 0 9454.605
284.1470031738281 0 2142.4912
287.4753723144531 0 533.8897
299.13970947265625 0 782.8684
311.1387023925781 0 23483.56 c Ammonia loss 1
312.14208984375 0 4518.9624
313.1441650390625 0 503.79593
327.13385009765625 0 1562.2866
343.0953674316406 0 46487.395 y 2
344.0983581542969 0 8163.751
345.094482421875 0 1558.3964
355.0691223144531 0 1079.8959
356.0704345703125 0 1224.5294
381.1583251953125 0 626.78326
415.035400390625 0 610.0265
429.1802978515625 0 1221.6904
466.5512390136719 0 642.5986
474.201904296875 0 7502.1543 c Ammonia loss 2
475.2054138183594 0 3404.718
506.1590576171875 0 1769.8434
507.157958984375 0 596.12476
522.2846069335938 0 3062.0378
522.7845458984375 0 8905.543
523.2843627929688 0 3519.3142
523.787841796875 0 618.9534
599.8641357421875 0 580.54297
609.2993774414062 0 858.3279
615.3407592773438 0 1118.2551
619.30615234375 0 746.7771
633.22119140625 0 685.9836
652.8585205078125 0 5688.7705
653.2257080078125 0 3477.175
653.3605346679688 0 5678.411
653.86083984375 0 4060.987
654.227294921875 0 1748.488
654.2886352539062 0 7894.207
654.3529663085938 0 1456.7983
654.7905883789062 0 6218.686
655.2337036132812 0 813.01447
655.29248046875 0 2668.6704
687.3742065429688 0 1469.1705
696.4302978515625 0 1745.637
697.303955078125 0 1795.087
697.4365844726562 0 3757.9119
698.3160400390625 0 698.44543
698.4389038085938 0 4031.9375
699.4423828125 0 2049.9883
700.44482421875 0 665.3795
741.3880004882812 0 1479.1357
751.3594360351562 0 639.10767
824.5206909179688 0 1217.6855
825.4561157226562 0 2731.2217
825.5296630859375 0 1714.7312
826.4569702148438 0 1334.8928
826.5335083007812 0 2701.8838
827.5374145507812 0 1359.2607
833.4356079101562 0 877.4673
834.440673828125 0 994.31555
835.42333984375 0 1434.7354
867.4029541015625 0 788.58685
868.407470703125 0 818.5599
871.4391479492188 0 709.5657
911.4027709960938 0 1641.3845
912.399658203125 0 868.9611
927.4230346679688 0 1596.2504
935.9421997070312 0 641.4642
939.5563354492188 0 818.8989
940.556884765625 0 3030.0168
941.5615234375 0 2618.854
942.5663452148438 0 2335.8389
943.562744140625 0 779.5094
947.50732421875 0 1005.9443
950.4647827148438 0 1362.4305
961.4684448242188 0 752.8849
994.4288940429688 0 698.5022
1028.4964599609375 0 3146.1924
1029.5023193359375 0 1256.6115
1039.4913330078125 0 716.2915
1042.6031494140625 0 1396.0774
1043.60546875 0 2054.1516
1044.5635986328125 0 1616.8691
1057.2301025390625 0 555.1085
1101.5831298828125 0 942.43945
1110.531982421875 0 3878.8896
1111.53515625 0 2902.7437
1112.5386962890625 0 771.7447
1120.07666015625 0 1095.8602
1120.552978515625 0 684.5883
1142.53466796875 0 1021.5315
1154.6956787109375 0 1131.8083
1155.6934814453125 0 2251.869
1156.6976318359375 0 4724.259
1157.6932373046875 0 1481.6304
1175.614990234375 0 1044.881
1176.6123046875 0 1896.1595
1177.0931396484375 0 821.6497
1177.61181640625 0 671.3778
1185.3621826171875 0 702.2558
1191.5596923828125 0 2525.3167
1192.560546875 0 1893.0814
1193.56103515625 0 1381.5227
1207.579833984375 0 1463.5454
1248.559814453125 0 1508.1256
1249.5423583984375 0 1543.2473
1250.5433349609375 0 793.96173
1254.5654296875 0 715.1418
1270.5657958984375 0 1552.038
1275.593994140625 0 1123.1638
1276.6019287109375 0 1263.5583
1278.1114501953125 0 709.2796
1288.5911865234375 0 888.8171
1288.7000732421875 0 5164.9175
1289.6983642578125 0 3955.4243
1290.697998046875 0 2377.5554
1291.553955078125 0 2522.4844
1291.7142333984375 0 1002.65393
1292.56689453125 0 2607.07
1293.58837890625 0 4569.758
1294.591064453125 0 1343.0944
1299.138427734375 0 1257.4111
1299.651123046875 0 977.6742
1300.144775390625 0 987.00555
1304.7113037109375 0 5883.7915
1305.54638671875 0 1125.6522
1305.7169189453125 0 17501.916
1306.5609130859375 0 865.3893
1306.7174072265625 0 14379.388
1307.56982421875 0 2486.558
1307.7122802734375 0 5825.2534
1308.1527099609375 0 1826.5426
1308.5760498046875 0 13104.605
1308.6983642578125 0 1329.252
1309.582763671875 0 10457.061
1310.5933837890625 0 6371.18
1311.608642578125 0 1163.6687
1915.5550537109375 0 678.07794
2286.09375 0 957.2831
2304.691162109375 0 652.1579
2598.283935546875 0 789.6257
2599.28955078125 0 1092.0955
2614.32421875 0 844.84973
2615.28857421875 0 1334.5703
2616.30322265625 0 1689.3564
2617.294921875 0 1416.104

Spectrum Details

|  |  |
| --- | --- |
| Matched peaks? Matched peaks The total absolute number of peaks matched. Additionally in brackets the total fraction of peaks matched and the total number of peaks is shown. | 3 (1.69% of 177) |
| FDR? FDR The false discovery rate estimated for this peptide. It is calculated by matching all theoretical fragments with a non-integer shift with the raw peaks for this spectrum. This is done with 40 different shifts. The resulting percentage is the average number of annotated peaks over the number of annotated peaks with the correct spectrum. | 1.59% |
| Satellite FDR? Satellite FDR See the FDR for details on its calculation. This satellite ion specific FDR only contains the satellite ions (d/w) for I/L/J positions. | - |
| PSM Score? PSM Score The PSM Score as given by Hecklib to this annotated spectrum. It is shown with three significant figures. | 36.9 |

#### Reverse Lookup? Reverse Lookup All places where this read could be placed.

| Group | Segment | Template | Template Part | Read Part | Score | Unique |
| --- | --- | --- | --- | --- | --- | --- |
| Homo sapiens Heavy Chain | IGHV | IGHV3-48 | [92..96] | [0..4] | 23 | False |
| Homo sapiens Heavy Chain | IGHV | IGHV3-21 | [92..96] | [0..4] | 23 | False |
| Homo sapiens Heavy Chain | IGHV | IGHV3-7 | [92..96] | [0..4] | 23 | False |
| Homo sapiens Heavy Chain | IGHV | IGHV3-13 | [91..95] | [0..4] | 23 | False |
| Homo sapiens Heavy Chain | IGHV | IGHV3-11 | [92..96] | [0..4] | 23 | False |
| Homo sapiens Heavy Chain | IGHV | IGHV3-74 | [92..96] | [0..4] | 23 | False |
| Homo sapiens Heavy Chain | IGHV | IGHV3-23 | [92..96] | [0..4] | 23 | False |
| Homo sapiens Heavy Chain | IGHV | IGHV3-53 | [91..95] | [0..4] | 23 | False |
| Homo sapiens Heavy Chain | IGHV | IGHV3-66 | [91..95] | [0..4] | 23 | False |
| Homo sapiens Heavy Chain | IGHV | IGHV3-NL1 | [92..96] | [0..4] | 23 | False |
| Homo sapiens Heavy Chain | IGHV | IGHV3-30-5 | [92..96] | [0..4] | 23 | False |
| Homo sapiens Heavy Chain | IGHV | IGHV3-72 | [94..98] | [0..4] | 23 | False |
| Homo sapiens Heavy Chain | IGHV | IGHV3-30 | [92..96] | [0..4] | 23 | False |
| Homo sapiens Heavy Chain | IGHV | IGHV3-33 | [92..96] | [0..4] | 23 | False |
| Homo sapiens Heavy Chain | IGHV | IGHV3-64 | [92..96] | [0..4] | 23 | False |
| Homo sapiens Heavy Chain | IGHV | IGHV3-15 | [94..98] | [0..4] | 23 | False |
| Homo sapiens Heavy Chain | IGHV | IGHV3-49 | [94..98] | [0..4] | 23 | False |
| Homo sapiens Heavy Chain | IGHV | IGHV3-73 | [94..98] | [0..4] | 23 | False |
| Homo sapiens Heavy Chain | IGHV | IGHV1-18 | [92..96] | [0..4] | 23 | False |
| Homo sapiens Heavy Chain | IGHV | IGHV4-30-2 | [93..97] | [0..4] | 23 | False |
| Homo sapiens Heavy Chain | IGHV | IGHV4-31 | [93..97] | [0..4] | 23 | False |
| Homo sapiens Heavy Chain | IGHV | IGHV1-24 | [92..96] | [0..4] | 23 | False |
| Homo sapiens Heavy Chain | IGHV | IGHV4-39 | [93..97] | [0..4] | 23 | False |
| Homo sapiens Heavy Chain | IGHV | IGHV1-69 | [92..96] | [0..4] | 23 | False |
| Homo sapiens Heavy Chain | IGHV | IGHV1-2 | [92..96] | [0..4] | 23 | False |
| Homo sapiens Heavy Chain | IGHV | IGHV4-61 | [93..97] | [0..4] | 23 | False |
| Homo sapiens Heavy Chain | IGHV | IGHV1-69-2 | [92..96] | [0..4] | 23 | False |
| Homo sapiens Heavy Chain | IGHV | IGHV4-28 | [92..96] | [0..4] | 23 | False |
| Homo sapiens Heavy Chain | IGHV | IGHV4-30-4 | [93..97] | [0..4] | 23 | False |
| Homo sapiens Heavy Chain | IGHV | IGHV1-3 | [92..96] | [0..4] | 23 | False |
| Homo sapiens Heavy Chain | IGHV | IGHV7-4-1 | [92..96] | [0..4] | 23 | False |
| Homo sapiens Heavy Chain | IGHV | IGHV4-59 | [91..95] | [0..4] | 23 | False |
| Homo sapiens Heavy Chain | IGHV | IGHV4-34 | [91..95] | [0..4] | 23 | False |
| Homo sapiens Heavy Chain | IGHV | IGHV4-38-2 | [92..96] | [0..4] | 23 | False |
| Homo sapiens Heavy Chain | IGHV | IGHV1-46 | [92..96] | [0..4] | 23 | False |
| Homo sapiens Heavy Chain | IGHV | IGHV1-8 | [92..96] | [0..4] | 23 | False |
| Homo sapiens Heavy Chain | IGHV | IGHV6-1 | [95..99] | [0..4] | 23 | False |
| Homo sapiens Heavy Chain | IGHV | IGHV1-58 | [92..96] | [0..4] | 23 | False |
| Homo sapiens Light Chain | IGLV | IGKV2D-29 | [89..93] | [0..4] | 23 | False |
| Homo sapiens Light Chain | IGLV | IGKV2D-26 | [89..93] | [0..4] | 23 | False |
| Homo sapiens Light Chain | IGLV | IGKV2-30 | [89..93] | [0..4] | 23 | False |
| Homo sapiens Light Chain | IGLV | IGKV2D-30 | [89..93] | [0..4] | 23 | False |
| Homo sapiens Light Chain | IGLV | IGKV3-20 | [85..89] | [0..4] | 23 | False |
| Homo sapiens Light Chain | IGLV | IGKV4-1 | [90..94] | [0..4] | 23 | False |
| Homo sapiens Light Chain | IGLV | IGKV3D-7 | [85..89] | [0..4] | 23 | False |
| Homo sapiens Light Chain | IGLV | IGKV2-24 | [89..93] | [0..4] | 23 | False |
| Homo sapiens Light Chain | IGLV | IGKV2-40 | [90..94] | [0..4] | 23 | False |
| Homo sapiens Light Chain | IGLV | IGKV2-28 | [89..93] | [0..4] | 23 | False |
| Homo sapiens Light Chain | IGLV | IGKV3-15 | [84..88] | [0..4] | 23 | False |
| Homo sapiens Light Chain | IGLV | IGKV3D-20 | [85..89] | [0..4] | 23 | False |
| Homo sapiens Light Chain | IGLV | IGKV3-11 | [84..88] | [0..4] | 23 | False |
| Homo sapiens Light Chain | IGLV | IGKV3D-11 | [84..88] | [0..4] | 23 | False |
| Decoy | Decoy | THER | [335..339] | [0..4] | 23 | False |

| Recombined | Template Part | Read Part | Score | Unique |
| --- | --- | --- | --- | --- |
| THER | [335..339] | [0..4] | 23 | True |

#### Meta Information from Multiple reads

##### Number of combined reads

2

##### Intensity

0.4531

##### TotalArea

2.097E+06

#### Positional Score

Copy Data

##### Positional Score (TSV)

###### Preview

```
Loading example...
```

*Click on the button to copy the data to your clipboard.*

000123

Label Value
"0" 0
"1" 0
"2" 0
"3" 0

#### Meta Information from PEAKS

##### Scan Identifier

F2:5790

##### Original sequence

V

+58.01

H

+15.99

Y

C

+58.01

##### Posttranslational Modifications

Carboxymethyl (KW X@N-term); Oxidation (HW); Carboxymethyl

##### Source File

D:\separate\_stitch\_analyses\xle-disambiguation\raw\20210323\_F1\_UM1\_Peng0013\_SA\_F59\_ingel\_3ug\_TL.raw

##### Fraction

2

##### Scan Feature

F2:10545

##### De Novo Score

97

##### ConfidenceScore

97

### m/z

653.2285

##### Mass

652.2162

##### Charge

1

##### Retention Time

31.35

##### Predicted Retention Time

-

##### Area

1.048E+06

##### Parts Per Million

7.7

##### Fragmentation mode

ETHCD

##### Originating file

01 D:\separate\_stitch\_analyses\xle-disambiguation\20210325\_F59\_3ug\_DENOVO\_12.csv

#### Meta Information from PEAKS

##### Scan Identifier

F2:5791

##### Original sequence

V

+58.01

H

+15.99

Y

C

+58.01

##### Posttranslational Modifications

Carboxymethyl (KW X@N-term); Oxidation (HW); Carboxymethyl

##### Source File

D:\separate\_stitch\_analyses\xle-disambiguation\raw\20210323\_F1\_UM1\_Peng0013\_SA\_F59\_ingel\_3ug\_TL.raw

##### Fraction

2

##### Scan Feature

F2:10545

##### De Novo Score

97

##### ConfidenceScore

97

### m/z

653.2285

##### Mass

652.2162

##### Charge

1

##### Retention Time

31.35

##### Predicted Retention Time

-

##### Area

1.048E+06

##### Parts Per Million

7.7

##### Fragmentation mode

ETHCD

##### Originating file

01 D:\separate\_stitch\_analyses\xle-disambiguation\20210325\_F59\_3ug\_DENOVO\_12.csv
