## Supplementary material for "A handle on mass coincidence errors in *de novo* sequencing of antibodies by bottom-up proteomics": Combined_084.html

Details Combined\_084 | Stitch OverviewUndefined

### Read Combined\_084

#### Sequence (length=14)

VRQAPGRAJEWVSG

#### Spectrum 6352? Spectrum 6352 The raw spectrum of this peptide as annotated by Hecklib. The fragments are coloured according to ion type (see legend). Any peaks with a star '\*' as text can be hovered over to see the full details, first the ion type second the mass shift type. By hovering over the amino acids in the peptide or ions in the legend the corresponding peaks are highlighted. By toggling the 'Unassigned' label you can turn the background (unassigned) peaks on or off in the plot. By updating the slider in the Ion legend you can update the spectrum to only show the top X% of the peaks with labels. The top X% means any peak that is within X% of the highest intensity. By dragging in the spectrum you can zoom in to a specific part of the spectrum and use 'Zoom Out' to get back to the original zoom level. The annotation of the spectrum is based on the given sequence in the peptides file and is done with different software so inconsistencies are likely. The peaks are annotated based on the given sequence, with 20 ppm tolerance.

Copy Data

##### Spectrum 6352 (TSV)

###### Preview

```
Loading example...
```

*Click on the button to copy the data to your clipboard.*

Mz MinMz MaxIntensity Max

WidthHeightPeptide font sizePeptide stroke widthSpectrum font sizeSpectrum stroke widthCompact peptide

Ion legend

wxyz

abcd

OtherUnassignedIon^Charge^_Position_Show for top:%

VRQAPGRAJEWVSG

01.49e+52.99e+54.48e+55.98e+5

Zoom Out

y^+1^_2_y^+1^_2_y^+3^_5_y^+1^_3_c^+1^_2_y^+1^_3_c^+1^_2_c^+2^_7_c^+1^_3_c^+2^_7_c^+1^_3_c^+2^_8_c^+3^_11_c^+2^_8_y^+1^_4_c^+1^_4_c^+2^_9_c^+2^_9_c^+3^_13_c^+2^_10_c^+2^_10_c^+2^_10_z^+1^_5_c^+1^_5_c^+1^_6_c^+2^_11_c^+2^_11_c^+2^_11_c^+2^_12_c^+2^_12_c^+2^_12_z^+2^_13_y^+2^_13_c^+2^_13_c^+2^_13_z^+1^_7_y^+1^_7_c^+1^_7_c^+1^_8_z^+1^_8_y^+1^_8_z^+1^_9_c^+1^_9_y^+1^_9_w^+1^_10_y^+1^_10_c^+1^_10_y^+1^_11_y^+1^_11_z^+1^_11_y^+1^_11_y^+1^_12_z^+1^_12_y^+1^_12_c^+1^_11_w^+1^_13_c^+1^_12_c^+1^_12_z^+1^_13_y^+1^_13_c^+1^_13_

0777155323303107

Fragment Matches Table

Show background peaks

| Position | Ion type | Intensity | mz Theoretical | mz Error (Th) | mz Error (ppm) | Charge | Series Number |
| --- | --- | --- | --- | --- | --- | --- | --- |
| - | - | 670.7 | 121.9 | - | - | 0 | - |
| - | - | 693.4 | 126.1 | - | - | 0 | - |
| - | - | 653.5 | 126.8 | - | - | 0 | - |
| - | - | 707.5 | 128.1 | - | - | 0 | - |
| - | - | 732.5 | 128.4 | - | - | 0 | - |
| - | - | 1.539E+04 | 130.1 | - | - | 0 | - |
| - | - | 1871 | 131.1 | - | - | 0 | - |
| - | - | 1864 | 142.1 | - | - | 0 | - |
| 13 | y | 6591 | 145.1 | 0.0002971 | 2.048 | +1 | 2 |
| - | - | 2979 | 146.1 | - | - | 0 | - |
| - | - | 1462 | 147.1 | - | - | 0 | - |
| - | - | 2132 | 149 | - | - | 0 | - |
| - | - | 7034 | 157.1 | - | - | 0 | - |
| - | - | 1.142E+04 | 159.1 | - | - | 0 | - |
| - | - | 1357 | 159.1 | - | - | 0 | - |
| - | - | 758.2 | 162.6 | - | - | 0 | - |
| 13 | y | 1.614E+05 | 163.1 | 0.0003373 | 2.068 | +1 | 2 |
| - | - | 1211 | 164.1 | - | - | 0 | - |
| - | - | 7909 | 164.1 | - | - | 0 | - |
| - | - | 4495 | 173.5 | - | - | 0 | - |
| - | - | 3529 | 174.1 | - | - | 0 | - |
| - | - | 1247 | 184.1 | - | - | 0 | - |
| - | - | 4745 | 185.1 | - | - | 0 | - |
| 10 | y | 2196 | 187.1 | 0.00174 | 9.3 | +3 | 5 |
| - | - | 4304 | 187.1 | - | - | 0 | - |
| - | - | 1398 | 211.2 | - | - | 0 | - |
| - | - | 1353 | 213.1 | - | - | 0 | - |
| - | - | 2228 | 213.2 | - | - | 0 | - |
| - | - | 4005 | 214.2 | - | - | 0 | - |
| - | - | 1055 | 216.5 | - | - | 0 | - |
| - | - | 2501 | 228.2 | - | - | 0 | - |
| - | - | 1565 | 234.1 | - | - | 0 | - |
| - | - | 2358 | 238.2 | - | - | 0 | - |
| - | - | 3.64E+04 | 239.2 | - | - | 0 | - |
| - | - | 4420 | 240.2 | - | - | 0 | - |
| 12 | y | 2406 | 244.1 | 0.0007461 | 3.056 | +1 | 3 |
| - | - | 1200 | 246.1 | - | - | 0 | - |
| 2 | c | 1.814E+04 | 256.2 | 0.0003531 | 1.379 | +1 | 2 |
| - | - | 2409 | 257.2 | - | - | 0 | - |
| - | - | 1762 | 258.2 | - | - | 0 | - |
| 12 | y | 5561 | 262.1 | 0.0004811 | 1.835 | +1 | 3 |
| 2 | c | 4.694E+04 | 273.2 | 0.0002933 | 1.074 | +1 | 2 |
| - | - | 6677 | 274.2 | - | - | 0 | - |
| - | - | 2377 | 285.2 | - | - | 0 | - |
| - | - | 2148 | 286.2 | - | - | 0 | - |
| - | - | 1177 | 298.7 | - | - | 0 | - |
| - | - | 1.975E+04 | 299.2 | - | - | 0 | - |
| - | - | 2787 | 300.2 | - | - | 0 | - |
| - | - | 1355 | 302.6 | - | - | 0 | - |
| - | - | 5155 | 356.2 | - | - | 0 | - |
| - | - | 1915 | 357.2 | - | - | 0 | - |
| - | - | 1646 | 369.7 | - | - | 0 | - |
| - | - | 1823 | 370.3 | - | - | 0 | - |
| 7 | c | 2493 | 383.2 | 0.0009617 | 2.51 | +2 | 7 |
| 3 | c | 4412 | 384.2 | 0.0009187 | 2.391 | +1 | 3 |
| - | - | 3783 | 391.2 | - | - | 0 | - |
| 7 | c | 4.904E+04 | 391.7 | 0.0005961 | 1.522 | +2 | 7 |
| - | - | 1.839E+04 | 392.2 | - | - | 0 | - |
| - | - | 5215 | 392.7 | - | - | 0 | - |
| 3 | c | 2.326E+05 | 401.3 | 0.0008589 | 2.14 | +1 | 3 |
| - | - | 4.649E+04 | 402.3 | - | - | 0 | - |
| - | - | 6970 | 403.3 | - | - | 0 | - |
| - | - | 1302 | 405.3 | - | - | 0 | - |
| - | - | 2193 | 413.2 | - | - | 0 | - |
| - | - | 1686 | 413.3 | - | - | 0 | - |
| - | - | 1426 | 413.6 | - | - | 0 | - |
| - | - | 1114 | 416.2 | - | - | 0 | - |
| 8 | c | 2132 | 418.7 | 0.001936 | 4.624 | +2 | 8 |
| - | - | 1239 | 422.2 | - | - | 0 | - |
| 11 | c | 7561 | 422.2 | 0.0006228 | 1.475 | +3 | 11 |
| - | - | 8152 | 422.6 | - | - | 0 | - |
| - | - | 4103 | 422.9 | - | - | 0 | - |
| - | - | 2898 | 426.8 | - | - | 0 | - |
| 8 | c | 4.854E+04 | 427.3 | 0.0007465 | 1.747 | +2 | 8 |
| - | - | 2.535E+04 | 427.8 | - | - | 0 | - |
| - | - | 3491 | 428.3 | - | - | 0 | - |
| - | - | 3985 | 428.3 | - | - | 0 | - |
| - | - | 1432 | 428.8 | - | - | 0 | - |
| - | - | 6919 | 430.2 | - | - | 0 | - |
| - | - | 3233 | 430.9 | - | - | 0 | - |
| - | - | 2157 | 431.2 | - | - | 0 | - |
| - | - | 4536 | 431.3 | - | - | 0 | - |
| - | - | 2005 | 440.3 | - | - | 0 | - |
| - | - | 3288 | 445.9 | - | - | 0 | - |
| - | - | 2339 | 446.3 | - | - | 0 | - |
| - | - | 1355 | 446.6 | - | - | 0 | - |
| 11 | y | 4937 | 448.2 | 0.001124 | 2.508 | +1 | 4 |
| 4 | c | 3618 | 455.3 | 0.003969 | 8.717 | +1 | 4 |
| - | - | 1346 | 461.3 | - | - | 0 | - |
| - | - | 6106 | 461.8 | - | - | 0 | - |
| - | - | 4726 | 462.3 | - | - | 0 | - |
| - | - | 1559 | 469.3 | - | - | 0 | - |
| 9 | c | 1951 | 475.3 | 0.0003094 | 0.6509 | +2 | 9 |
| - | - | 2460 | 476.3 | - | - | 0 | - |
| - | - | 3.625E+04 | 483.3 | - | - | 0 | - |
| 9 | c | 2.126E+05 | 483.8 | 0.0007372 | 1.524 | +2 | 9 |
| 13 | c | 2.771E+04 | 484.3 | 0.0002042 | 0.4216 | +3 | 13 |
| - | - | 9.841E+04 | 484.3 | - | - | 0 | - |
| - | - | 2.304E+04 | 484.6 | - | - | 0 | - |
| - | - | 2.972E+04 | 484.8 | - | - | 0 | - |
| - | - | 7253 | 484.9 | - | - | 0 | - |
| - | - | 2036 | 485.3 | - | - | 0 | - |
| - | - | 5141 | 485.3 | - | - | 0 | - |
| - | - | 3.68E+04 | 496.8 | - | - | 0 | - |
| - | - | 1.674E+04 | 497.3 | - | - | 0 | - |
| - | - | 7500 | 497.8 | - | - | 0 | - |
| - | - | 1.19E+04 | 498.3 | - | - | 0 | - |
| - | - | 7953 | 498.8 | - | - | 0 | - |
| - | - | 2634 | 499.3 | - | - | 0 | - |
| - | - | 3874 | 499.3 | - | - | 0 | - |
| - | - | 1.551E+04 | 503.3 | - | - | 0 | - |
| - | - | 1.248E+04 | 503.6 | - | - | 0 | - |
| - | - | 7033 | 503.9 | - | - | 0 | - |
| - | - | 1.356E+05 | 509.3 | - | - | 0 | - |
| - | - | 6586 | 509.3 | - | - | 0 | - |
| - | - | 3.441E+04 | 510.3 | - | - | 0 | - |
| - | - | 1334 | 510.3 | - | - | 0 | - |
| - | - | 1618 | 518.3 | - | - | 0 | - |
| - | - | 2137 | 518.8 | - | - | 0 | - |
| - | - | 2004 | 519.8 | - | - | 0 | - |
| - | - | 4445 | 524.3 | - | - | 0 | - |
| - | - | 5374 | 525.3 | - | - | 0 | - |
| - | - | 1393 | 525.8 | - | - | 0 | - |
| - | - | 2.69E+04 | 526.3 | - | - | 0 | - |
| - | - | 1436 | 526.4 | - | - | 0 | - |
| - | - | 2.677E+04 | 526.8 | - | - | 0 | - |
| - | - | 1.259E+04 | 527.3 | - | - | 0 | - |
| - | - | 4632 | 527.8 | - | - | 0 | - |
| 10 | c | 1354 | 539.3 | 0.001147 | 2.127 | +2 | 10 |
| 10 | c | 5.509E+04 | 539.8 | 0.00123 | 2.278 | +2 | 10 |
| - | - | 3.271E+04 | 540.3 | - | - | 0 | - |
| - | - | 1.454E+04 | 540.8 | - | - | 0 | - |
| - | - | 4812 | 541.3 | - | - | 0 | - |
| - | - | 6.148E+04 | 547.8 | - | - | 0 | - |
| 10 | c | 2.56E+05 | 548.3 | 0.0006504 | 1.186 | +2 | 10 |
| - | - | 1.434E+05 | 548.8 | - | - | 0 | - |
| - | - | 2508 | 548.9 | - | - | 0 | - |
| - | - | 4.935E+04 | 549.3 | - | - | 0 | - |
| - | - | 8729 | 549.8 | - | - | 0 | - |
| - | - | 1169 | 559.3 | - | - | 0 | - |
| 10 | z | 2155 | 561.2 | 0.0007232 | 1.289 | +1 | 5 |
| - | - | 1535 | 562.3 | - | - | 0 | - |
| - | - | 1719 | 568.3 | - | - | 0 | - |
| 5 | c | 5.942E+04 | 569.4 | 0.0008555 | 1.503 | +1 | 5 |
| - | - | 1.676E+04 | 570.4 | - | - | 0 | - |
| - | - | 3957 | 571.4 | - | - | 0 | - |
| - | - | 5921 | 574.3 | - | - | 0 | - |
| - | - | 2931 | 574.8 | - | - | 0 | - |
| - | - | 2523 | 576.8 | - | - | 0 | - |
| - | - | 5053 | 582.4 | - | - | 0 | - |
| - | - | 3229 | 583.4 | - | - | 0 | - |
| - | - | 1660 | 597.4 | - | - | 0 | - |
| - | - | 2486 | 603.9 | - | - | 0 | - |
| - | - | 6355 | 604.4 | - | - | 0 | - |
| - | - | 3367 | 604.9 | - | - | 0 | - |
| - | - | 1506 | 605.3 | - | - | 0 | - |
| - | - | 2550 | 610.8 | - | - | 0 | - |
| - | - | 2.724E+04 | 611.3 | - | - | 0 | - |
| - | - | 1.998E+04 | 611.8 | - | - | 0 | - |
| - | - | 7238 | 612.4 | - | - | 0 | - |
| - | - | 2883 | 612.9 | - | - | 0 | - |
| - | - | 2234 | 613.3 | - | - | 0 | - |
| - | - | 2.399E+04 | 617.9 | - | - | 0 | - |
| - | - | 1.758E+04 | 618.4 | - | - | 0 | - |
| - | - | 4.334E+04 | 618.9 | - | - | 0 | - |
| - | - | 3.248E+04 | 619.4 | - | - | 0 | - |
| - | - | 2.318E+04 | 619.9 | - | - | 0 | - |
| - | - | 1.013E+04 | 620.4 | - | - | 0 | - |
| - | - | 4380 | 620.9 | - | - | 0 | - |
| - | - | 1502 | 624.4 | - | - | 0 | - |
| 6 | c | 3.402E+05 | 626.4 | 0.001242 | 1.983 | +1 | 6 |
| - | - | 1.153E+05 | 627.4 | - | - | 0 | - |
| - | - | 2.178E+04 | 628.4 | - | - | 0 | - |
| - | - | 3037 | 629.4 | - | - | 0 | - |
| - | - | 1329 | 631.8 | - | - | 0 | - |
| 11 | c | 2494 | 632.4 | 0.003694 | 5.842 | +2 | 11 |
| 11 | c | 1.866E+05 | 632.8 | 0.0009408 | 1.487 | +2 | 11 |
| - | - | 1.268E+05 | 633.4 | - | - | 0 | - |
| - | - | 5.839E+04 | 633.9 | - | - | 0 | - |
| - | - | 1.421E+04 | 634.4 | - | - | 0 | - |
| - | - | 3164 | 634.9 | - | - | 0 | - |
| - | - | 1427 | 638.4 | - | - | 0 | - |
| - | - | 1516 | 639.4 | - | - | 0 | - |
| - | - | 2154 | 639.9 | - | - | 0 | - |
| - | - | 1.261E+04 | 640.9 | - | - | 0 | - |
| 11 | c | 1.671E+05 | 641.4 | 0.0006668 | 1.04 | +2 | 11 |
| - | - | 1.208E+05 | 641.9 | - | - | 0 | - |
| - | - | 4.728E+04 | 642.4 | - | - | 0 | - |
| - | - | 9776 | 642.9 | - | - | 0 | - |
| - | - | 3163 | 643.4 | - | - | 0 | - |
| - | - | 1398 | 647.4 | - | - | 0 | - |
| - | - | 1471 | 660.4 | - | - | 0 | - |
| - | - | 5647 | 660.9 | - | - | 0 | - |
| - | - | 1.033E+04 | 661.4 | - | - | 0 | - |
| - | - | 6435 | 661.9 | - | - | 0 | - |
| - | - | 4211 | 662.4 | - | - | 0 | - |
| - | - | 1457 | 667.4 | - | - | 0 | - |
| - | - | 7.576E+04 | 668.4 | - | - | 0 | - |
| - | - | 9.237E+04 | 668.9 | - | - | 0 | - |
| - | - | 5.727E+04 | 669.4 | - | - | 0 | - |
| - | - | 3.332E+04 | 669.9 | - | - | 0 | - |
| - | - | 1.083E+04 | 670.4 | - | - | 0 | - |
| - | - | 1641 | 671.4 | - | - | 0 | - |
| - | - | 1701 | 673.4 | - | - | 0 | - |
| - | - | 1734 | 673.9 | - | - | 0 | - |
| - | - | 1842 | 676.4 | - | - | 0 | - |
| - | - | 6272 | 676.9 | - | - | 0 | - |
| - | - | 5574 | 677.4 | - | - | 0 | - |
| - | - | 1930 | 677.9 | - | - | 0 | - |
| - | - | 1768 | 678.4 | - | - | 0 | - |
| - | - | 2472 | 681.4 | - | - | 0 | - |
| 12 | c | 3998 | 681.9 | 0.009947 | 14.59 | +2 | 12 |
| 12 | c | 3.657E+05 | 682.4 | 0.0009746 | 1.428 | +2 | 12 |
| - | - | 2.729E+05 | 682.9 | - | - | 0 | - |
| - | - | 1.407E+05 | 683.4 | - | - | 0 | - |
| - | - | 4.096E+04 | 683.9 | - | - | 0 | - |
| - | - | 1.126E+04 | 684.4 | - | - | 0 | - |
| - | - | 2218 | 689.9 | - | - | 0 | - |
| - | - | 5466 | 690.4 | - | - | 0 | - |
| 12 | c | 2.222E+05 | 690.9 | 0.0006395 | 0.9256 | +2 | 12 |
| - | - | 1.786E+05 | 691.4 | - | - | 0 | - |
| - | - | 7.66E+04 | 691.9 | - | - | 0 | - |
| - | - | 2.34E+04 | 692.4 | - | - | 0 | - |
| - | - | 7103 | 692.9 | - | - | 0 | - |
| - | - | 3.566E+04 | 698.4 | - | - | 0 | - |
| - | - | 1.039E+04 | 698.9 | - | - | 0 | - |
| - | - | 7151 | 699.4 | - | - | 0 | - |
| - | - | 1827 | 699.9 | - | - | 0 | - |
| - | - | 1651 | 703.9 | - | - | 0 | - |
| - | - | 3499 | 704.4 | - | - | 0 | - |
| - | - | 4858 | 704.9 | - | - | 0 | - |
| - | - | 7929 | 705.4 | - | - | 0 | - |
| 2 | z | 3.926E+04 | 705.9 | 0.001285 | 1.82 | +2 | 13 |
| - | - | 2.579E+04 | 706.4 | - | - | 0 | - |
| - | - | 1.423E+04 | 706.9 | - | - | 0 | - |
| - | - | 3837 | 707.4 | - | - | 0 | - |
| - | - | 1493 | 707.9 | - | - | 0 | - |
| - | - | 3231 | 711.9 | - | - | 0 | - |
| - | - | 2.575E+04 | 712.4 | - | - | 0 | - |
| - | - | 2.211E+04 | 712.9 | - | - | 0 | - |
| - | - | 1.019E+04 | 713.4 | - | - | 0 | - |
| 2 | y | 5410 | 713.9 | 0.002665 | 3.733 | +2 | 13 |
| - | - | 5108 | 714.4 | - | - | 0 | - |
| - | - | 1456 | 716.9 | - | - | 0 | - |
| - | - | 1890 | 717.4 | - | - | 0 | - |
| - | - | 3194 | 717.9 | - | - | 0 | - |
| - | - | 1.091E+04 | 719.4 | - | - | 0 | - |
| - | - | 1.24E+04 | 719.9 | - | - | 0 | - |
| - | - | 8576 | 720.4 | - | - | 0 | - |
| - | - | 3245 | 720.9 | - | - | 0 | - |
| 13 | c | 1.704E+04 | 725.9 | 0.001806 | 2.488 | +2 | 13 |
| - | - | 2.232E+04 | 726.4 | - | - | 0 | - |
| - | - | 1.381E+04 | 726.9 | - | - | 0 | - |
| - | - | 1.265E+04 | 727.4 | - | - | 0 | - |
| - | - | 1.308E+04 | 727.9 | - | - | 0 | - |
| - | - | 1.464E+04 | 728.4 | - | - | 0 | - |
| - | - | 6033 | 728.9 | - | - | 0 | - |
| - | - | 2075 | 732.9 | - | - | 0 | - |
| - | - | 9606 | 733.4 | - | - | 0 | - |
| - | - | 6784 | 733.9 | - | - | 0 | - |
| 13 | c | 1.983E+05 | 734.4 | 0.0002503 | 0.3408 | +2 | 13 |
| - | - | 1.46E+05 | 734.9 | - | - | 0 | - |
| - | - | 8.115E+04 | 735.4 | - | - | 0 | - |
| - | - | 2.988E+04 | 735.9 | - | - | 0 | - |
| - | - | 1.987E+04 | 736.4 | - | - | 0 | - |
| - | - | 1.244E+04 | 736.9 | - | - | 0 | - |
| - | - | 3064 | 737.4 | - | - | 0 | - |
| - | - | 4136 | 738.5 | - | - | 0 | - |
| - | - | 2626 | 739.5 | - | - | 0 | - |
| - | - | 1872 | 740.5 | - | - | 0 | - |
| - | - | 8154 | 740.9 | - | - | 0 | - |
| - | - | 4.696E+04 | 741.4 | - | - | 0 | - |
| - | - | 3.856E+04 | 741.9 | - | - | 0 | - |
| - | - | 1.205E+05 | 742.4 | - | - | 0 | - |
| - | - | 8.566E+04 | 742.9 | - | - | 0 | - |
| - | - | 4.656E+04 | 743.4 | - | - | 0 | - |
| - | - | 1.178E+04 | 743.9 | - | - | 0 | - |
| - | - | 4283 | 744.4 | - | - | 0 | - |
| 8 | z | 5960 | 745.4 | 0.001616 | 2.168 | +1 | 7 |
| - | - | 2780 | 746.9 | - | - | 0 | - |
| - | - | 6284 | 747.4 | - | - | 0 | - |
| - | - | 3.002E+04 | 747.9 | - | - | 0 | - |
| - | - | 2.123E+04 | 748.4 | - | - | 0 | - |
| - | - | 1.419E+04 | 748.9 | - | - | 0 | - |
| - | - | 4289 | 749.4 | - | - | 0 | - |
| - | - | 5351 | 749.9 | - | - | 0 | - |
| - | - | 6127 | 750.4 | - | - | 0 | - |
| - | - | 4144 | 750.9 | - | - | 0 | - |
| - | - | 6323 | 754.4 | - | - | 0 | - |
| - | - | 3.146E+04 | 754.9 | - | - | 0 | - |
| - | - | 5.643E+05 | 755.4 | - | - | 0 | - |
| - | - | 4.481E+05 | 755.9 | - | - | 0 | - |
| - | - | 2.397E+05 | 756.4 | - | - | 0 | - |
| - | - | 6.337E+04 | 756.9 | - | - | 0 | - |
| - | - | 1940 | 757 | - | - | 0 | - |
| - | - | 1.731E+04 | 757.4 | - | - | 0 | - |
| 8 | y | 4446 | 761.4 | 0.001935 | 2.541 | +1 | 7 |
| - | - | 2797 | 762.4 | - | - | 0 | - |
| - | - | 1.712E+04 | 762.9 | - | - | 0 | - |
| - | - | 2.2E+05 | 763.4 | - | - | 0 | - |
| - | - | 5.918E+05 | 763.9 | - | - | 0 | - |
| - | - | 4.21E+05 | 764.4 | - | - | 0 | - |
| - | - | 2.07E+05 | 764.9 | - | - | 0 | - |
| - | - | 6.301E+04 | 765.4 | - | - | 0 | - |
| - | - | 2155 | 765.5 | - | - | 0 | - |
| - | - | 1.393E+04 | 765.9 | - | - | 0 | - |
| - | - | 2044 | 771.9 | - | - | 0 | - |
| 7 | c | 2.242E+04 | 782.5 | 0.0007173 | 0.9166 | +1 | 7 |
| - | - | 8973 | 783.5 | - | - | 0 | - |
| - | - | 1651 | 814.5 | - | - | 0 | - |
| - | - | 1799 | 852.5 | - | - | 0 | - |
| 8 | c | 1.087E+04 | 853.5 | 0.001323 | 1.55 | +1 | 8 |
| - | - | 4560 | 854.5 | - | - | 0 | - |
| - | - | 1612 | 855.5 | - | - | 0 | - |
| - | - | 2378 | 857.5 | - | - | 0 | - |
| - | - | 1847 | 886.5 | - | - | 0 | - |
| 7 | z | 2.915E+05 | 901.5 | 0.0009686 | 1.074 | +1 | 8 |
| - | - | 1.43E+05 | 902.5 | - | - | 0 | - |
| - | - | 4.374E+04 | 903.5 | - | - | 0 | - |
| - | - | 8236 | 904.5 | - | - | 0 | - |
| 7 | y | 1.521E+04 | 917.5 | 0.001471 | 1.603 | +1 | 8 |
| - | - | 7407 | 918.5 | - | - | 0 | - |
| - | - | 2459 | 919.5 | - | - | 0 | - |
| 6 | z | 4.357E+04 | 958.5 | 0.001783 | 1.86 | +1 | 9 |
| - | - | 2.604E+04 | 959.5 | - | - | 0 | - |
| - | - | 8334 | 960.5 | - | - | 0 | - |
| - | - | 3057 | 965.6 | - | - | 0 | - |
| 9 | c | 1.85E+04 | 966.6 | 0.000267 | 0.2763 | +1 | 9 |
| - | - | 1.038E+04 | 967.6 | - | - | 0 | - |
| - | - | 3664 | 968.6 | - | - | 0 | - |
| 6 | y | 8269 | 974.5 | 0.0006978 | 0.716 | +1 | 9 |
| - | - | 5551 | 975.5 | - | - | 0 | - |
| 5 | w | 3838 | 1029 | 0.007101 | 6.904 | +1 | 10 |
| - | - | 1855 | 1051 | - | - | 0 | - |
| - | - | 1.064E+04 | 1052 | - | - | 0 | - |
| - | - | 1.588E+04 | 1053 | - | - | 0 | - |
| - | - | 8112 | 1054 | - | - | 0 | - |
| - | - | 2323 | 1070 | - | - | 0 | - |
| - | - | 3243 | 1071 | - | - | 0 | - |
| 5 | y | 8041 | 1072 | 0.002628 | 2.452 | +1 | 10 |
| - | - | 4806 | 1073 | - | - | 0 | - |
| - | - | 2066 | 1080 | - | - | 0 | - |
| - | - | 4927 | 1081 | - | - | 0 | - |
| - | - | 4240 | 1082 | - | - | 0 | - |
| - | - | 1563 | 1083 | - | - | 0 | - |
| - | - | 7376 | 1095 | - | - | 0 | - |
| 10 | c | 9.166E+04 | 1096 | 0.0008868 | 0.8094 | +1 | 10 |
| - | - | 5.37E+04 | 1097 | - | - | 0 | - |
| - | - | 1307 | 1097 | - | - | 0 | - |
| - | - | 2.031E+04 | 1098 | - | - | 0 | - |
| - | - | 3945 | 1099 | - | - | 0 | - |
| - | - | 2521 | 1100 | - | - | 0 | - |
| - | - | 1713 | 1101 | - | - | 0 | - |
| - | - | 3821 | 1110 | - | - | 0 | - |
| - | - | 3250 | 1111 | - | - | 0 | - |
| - | - | 2022 | 1112 | - | - | 0 | - |
| 4 | y | 1747 | 1125 | 0.02002 | 17.8 | +1 | 11 |
| 4 | y | 2952 | 1126 | 0.003793 | 3.37 | +1 | 11 |
| 4 | z | 2.305E+05 | 1127 | 0.0005891 | 0.5229 | +1 | 11 |
| - | - | 1.422E+05 | 1128 | - | - | 0 | - |
| - | - | 5.238E+04 | 1129 | - | - | 0 | - |
| - | - | 1.45E+04 | 1130 | - | - | 0 | - |
| - | - | 3690 | 1131 | - | - | 0 | - |
| 4 | y | 2.479E+04 | 1143 | 0.001152 | 1.008 | +1 | 11 |
| - | - | 1.711E+04 | 1144 | - | - | 0 | - |
| - | - | 5524 | 1145 | - | - | 0 | - |
| - | - | 1665 | 1159 | - | - | 0 | - |
| - | - | 1710 | 1197 | - | - | 0 | - |
| - | - | 3111 | 1224 | - | - | 0 | - |
| - | - | 3604 | 1238 | - | - | 0 | - |
| - | - | 7425 | 1239 | - | - | 0 | - |
| - | - | 3512 | 1240 | - | - | 0 | - |
| 3 | y | 1845 | 1254 | 0.000862 | 0.6876 | +1 | 12 |
| 3 | z | 1.606E+05 | 1255 | 0.0001271 | 0.1013 | +1 | 12 |
| - | - | 1.168E+05 | 1256 | - | - | 0 | - |
| - | - | 5.274E+04 | 1257 | - | - | 0 | - |
| - | - | 1.348E+04 | 1258 | - | - | 0 | - |
| - | - | 2136 | 1259 | - | - | 0 | - |
| - | - | 8582 | 1266 | - | - | 0 | - |
| - | - | 8258 | 1267 | - | - | 0 | - |
| - | - | 3004 | 1268 | - | - | 0 | - |
| 3 | y | 1.315E+04 | 1271 | 0.001046 | 0.8234 | +1 | 12 |
| - | - | 9506 | 1272 | - | - | 0 | - |
| - | - | 4480 | 1273 | - | - | 0 | - |
| - | - | 2425 | 1281 | - | - | 0 | - |
| 11 | c | 3.631E+04 | 1282 | 0.0001791 | 0.1397 | +1 | 11 |
| - | - | 3.684E+04 | 1283 | - | - | 0 | - |
| - | - | 2.326E+04 | 1284 | - | - | 0 | - |
| - | - | 7691 | 1285 | - | - | 0 | - |
| - | - | 1743 | 1286 | - | - | 0 | - |
| - | - | 2468 | 1321 | - | - | 0 | - |
| - | - | 2413 | 1322 | - | - | 0 | - |
| - | - | 4077 | 1323 | - | - | 0 | - |
| - | - | 3582 | 1324 | - | - | 0 | - |
| 2 | w | 2004 | 1325 | 0.0001252 | 0.0945 | +1 | 13 |
| - | - | 3060 | 1326 | - | - | 0 | - |
| - | - | 1.507E+04 | 1337 | - | - | 0 | - |
| - | - | 3.379E+04 | 1338 | - | - | 0 | - |
| - | - | 2.489E+04 | 1339 | - | - | 0 | - |
| - | - | 9659 | 1340 | - | - | 0 | - |
| - | - | 1913 | 1341 | - | - | 0 | - |
| - | - | 1761 | 1353 | - | - | 0 | - |
| - | - | 1962 | 1354 | - | - | 0 | - |
| 12 | c | 3584 | 1364 | 0.001394 | 1.023 | +1 | 12 |
| - | - | 1.588E+04 | 1365 | - | - | 0 | - |
| - | - | 1.374E+04 | 1366 | - | - | 0 | - |
| - | - | 5927 | 1367 | - | - | 0 | - |
| - | - | 3188 | 1369 | - | - | 0 | - |
| - | - | 1691 | 1370 | - | - | 0 | - |
| - | - | 3744 | 1380 | - | - | 0 | - |
| 12 | c | 2.563E+04 | 1381 | 0.00121 | 0.8764 | +1 | 12 |
| - | - | 6.141E+04 | 1382 | - | - | 0 | - |
| - | - | 4.051E+04 | 1383 | - | - | 0 | - |
| - | - | 1.749E+04 | 1384 | - | - | 0 | - |
| - | - | 5775 | 1385 | - | - | 0 | - |
| - | - | 3901 | 1397 | - | - | 0 | - |
| - | - | 6710 | 1407 | - | - | 0 | - |
| - | - | 6503 | 1408 | - | - | 0 | - |
| - | - | 3541 | 1409 | - | - | 0 | - |
| - | - | 5413 | 1410 | - | - | 0 | - |
| 2 | z | 8950 | 1411 | 0.00301 | 2.134 | +1 | 13 |
| - | - | 3.799E+04 | 1412 | - | - | 0 | - |
| - | - | 2.545E+04 | 1413 | - | - | 0 | - |
| - | - | 1.088E+04 | 1414 | - | - | 0 | - |
| - | - | 4076 | 1415 | - | - | 0 | - |
| - | - | 4311 | 1424 | - | - | 0 | - |
| - | - | 1.859E+04 | 1425 | - | - | 0 | - |
| - | - | 1.677E+04 | 1426 | - | - | 0 | - |
| 2 | y | 1.108E+04 | 1427 | 0.007723 | 5.413 | +1 | 13 |
| - | - | 8177 | 1428 | - | - | 0 | - |
| - | - | 4614 | 1429 | - | - | 0 | - |
| - | - | 2418 | 1430 | - | - | 0 | - |
| - | - | 2807 | 1439 | - | - | 0 | - |
| - | - | 2344 | 1440 | - | - | 0 | - |
| - | - | 3164 | 1441 | - | - | 0 | - |
| - | - | 1.025E+04 | 1452 | - | - | 0 | - |
| - | - | 1.346E+04 | 1453 | - | - | 0 | - |
| - | - | 7838 | 1454 | - | - | 0 | - |
| - | - | 3606 | 1455 | - | - | 0 | - |
| - | - | 5318 | 1456 | - | - | 0 | - |
| - | - | 4822 | 1457 | - | - | 0 | - |
| - | - | 2273 | 1458 | - | - | 0 | - |
| - | - | 2658 | 1466 | - | - | 0 | - |
| - | - | 4956 | 1467 | - | - | 0 | - |
| 13 | c | 7.241E+04 | 1468 | 0.02884 | 19.65 | +1 | 13 |
| - | - | 1.223E+05 | 1469 | - | - | 0 | - |
| - | - | 8.097E+04 | 1470 | - | - | 0 | - |
| - | - | 3.291E+04 | 1471 | - | - | 0 | - |
| - | - | 9329 | 1472 | - | - | 0 | - |
| - | - | 3330 | 1473 | - | - | 0 | - |
| - | - | 7493 | 1482 | - | - | 0 | - |
| - | - | 3.245E+04 | 1483 | - | - | 0 | - |
| - | - | 3.17E+04 | 1484 | - | - | 0 | - |
| - | - | 2.373E+04 | 1485 | - | - | 0 | - |
| - | - | 1.139E+04 | 1486 | - | - | 0 | - |
| - | - | 4201 | 1487 | - | - | 0 | - |
| - | - | 3515 | 1493 | - | - | 0 | - |
| - | - | 2.79E+04 | 1494 | - | - | 0 | - |
| - | - | 2.597E+04 | 1495 | - | - | 0 | - |
| - | - | 1.268E+04 | 1496 | - | - | 0 | - |
| - | - | 4767 | 1497 | - | - | 0 | - |
| - | - | 2093 | 1498 | - | - | 0 | - |
| - | - | 3064 | 1499 | - | - | 0 | - |
| - | - | 2.08E+04 | 1500 | - | - | 0 | - |
| - | - | 1.566E+04 | 1501 | - | - | 0 | - |
| - | - | 7522 | 1502 | - | - | 0 | - |
| - | - | 1763 | 1503 | - | - | 0 | - |
| - | - | 2453 | 1509 | - | - | 0 | - |
| - | - | 4.713E+04 | 1510 | - | - | 0 | - |
| - | - | 3.014E+05 | 1511 | - | - | 0 | - |
| - | - | 2.352E+05 | 1512 | - | - | 0 | - |
| - | - | 1.22E+05 | 1513 | - | - | 0 | - |
| - | - | 3.576E+04 | 1514 | - | - | 0 | - |
| - | - | 1.216E+04 | 1515 | - | - | 0 | - |
| - | - | 1613 | 1525 | - | - | 0 | - |
| - | - | 2.278E+04 | 1526 | - | - | 0 | - |
| - | - | 9.86E+04 | 1527 | - | - | 0 | - |
| - | - | 4.047E+05 | 1528 | - | - | 0 | - |
| - | - | 3.071E+05 | 1529 | - | - | 0 | - |
| - | - | 1.544E+05 | 1530 | - | - | 0 | - |
| - | - | 5.037E+04 | 1531 | - | - | 0 | - |
| - | - | 1.657E+04 | 1532 | - | - | 0 | - |
| - | - | 1659 | 1985 | - | - | 0 | - |
| - | - | 1694 | 3076 | - | - | 0 | - |

m/z Charge Intensity FragmentType MassShift Position
121.9268798828125 0 670.7349
126.06670379638672 0 693.36523
126.7815933227539 0 653.50494
128.0823211669922 0 707.5352
128.37001037597656 0 732.54285
130.06544494628906 0 15391.295
131.0688018798828 0 1870.6681
142.09786987304688 0 1863.8629
145.06106567382812 0 6591.129 y Water loss 12
146.0928192138672 0 2979.189
147.11312866210938 0 1462.2716
148.95443725585938 0 2131.5483
157.10877990722656 0 7034.249
159.09205627441406 0 11424.732
159.11326599121094 0 1357.2672
162.59765625 0 758.19183
163.07167053222656 0 161380.6 y 12
164.0687255859375 0 1210.8398
164.07508850097656 0 7909.1523
173.45135498046875 0 4494.605
174.13522338867188 0 3529.3096
184.11227416992188 0 1246.5013
185.1037139892578 0 4744.8335
187.08680725097656 0 2195.63 y Water loss 9
187.10824584960938 0 4304.442
211.15542602539062 0 1398.45
213.13829040527344 0 1352.5671
213.1712646484375 0 2228.3687
214.1552734375 0 4005.1206
216.50758361816406 0 1055.1553
228.18212890625 0 2500.7246
234.1455078125 0 1565.4263
238.16639709472656 0 2358.2756
239.1506805419922 0 36401.613
240.15415954589844 0 4420.1147
244.1299285888672 0 2406.026 y Water loss 11
246.0684814453125 0 1199.7294
256.1771545410156 0 18141.55 c Ammonia loss 1
257.1804504394531 0 2408.6274
258.16094970703125 0 1761.8257
262.1402282714844 0 5560.7495 y 11
273.2036437988281 0 46937.105 c 1
274.2070617675781 0 6676.626
285.16876220703125 0 2376.9402
286.15545654296875 0 2148.4233
298.7378234863281 0 1176.9595
299.2196044921875 0 19753.738
300.2228698730469 0 2786.895
302.5639343261719 0 1354.6049
356.2407531738281 0 5154.5503
357.2478332519531 0 1914.5316
369.7334289550781 0 1645.9786
370.2565002441406 0 1823.2264
383.228515625 0 2492.708 c Ammonia loss 6
384.2362976074219 0 4412.1387 c Ammonia loss 2
391.23822021484375 0 3782.7583
391.7414245605469 0 49040.516 c 6
392.24267578125 0 18386.076
392.7435302734375 0 5215.4927
401.2627868652344 0 232646.11 c 2
402.26544189453125 0 46493.64
403.268310546875 0 6969.7803
405.2528076171875 0 1302.2628
413.23828125 0 2193.3005
413.27423095703125 0 1686.0536
413.5744934082031 0 1425.7988
416.2336120605469 0 1113.581
418.748046875 0 2131.8547 c Ammonia loss 7
422.18389892578125 0 1238.7444
422.23577880859375 0 7560.9 c Ammonia loss 10
422.56982421875 0 8152.3813
422.9043884277344 0 4103.451
426.7571716308594 0 2898.1113
427.2601318359375 0 48540.73 c 7
427.7619323730469 0 25349.25
428.260498046875 0 3491.2864
428.2865905761719 0 3984.6047
428.7647705078125 0 1432.4769
430.1857604980469 0 6918.604
430.9197082519531 0 3232.859
431.18975830078125 0 2157.2954
431.2533874511719 0 4535.611
440.2541198730469 0 2004.6832
445.92791748046875 0 3287.6973
446.2606201171875 0 2338.7322
446.591552734375 0 1355.0787
448.2201843261719 0 4937.076 y 10
455.2685241699219 0 3617.6123 c Ammonia loss 3
461.287109375 0 1345.7899
461.79559326171875 0 6106.3037
462.2974548339844 0 4726.263
469.2705078125 0 1559.0642
475.2884521484375 0 1950.863 c Ammonia loss 8
476.290283203125 0 2460.4536
483.2985534667969 0 36253.7
483.8021545410156 0 212575.86 c 8
484.2684326171875 0 27707.143 c Ammonia loss 12
484.30389404296875 0 98408.24
484.6040954589844 0 23036.887
484.8048400878906 0 29722.762
484.93731689453125 0 7252.982
485.27130126953125 0 2036.3746
485.3062744140625 0 5140.966
496.8101501464844 0 36799.97
497.3112487792969 0 16737.736
497.8131408691406 0 7500.3994
498.2851867675781 0 11898.104
498.7868347167969 0 7953.077
499.2872619628906 0 2634.4663
499.32342529296875 0 3873.777
503.27667236328125 0 15506.0625
503.6110534667969 0 12478.216
503.9459228515625 0 7032.7627
509.26153564453125 0 135629.02
509.3082275390625 0 6585.5483
510.2646789550781 0 34413.824
510.304931640625 0 1333.8463
518.3153076171875 0 1617.6603
518.8085327148438 0 2137.1357
519.7890014648438 0 2004.3468
524.3318481445312 0 4445.279
525.338134765625 0 5373.5977
525.8190307617188 0 1392.8014
526.31689453125 0 26904.428
526.35546875 0 1436.2233
526.8190307617188 0 26765.799
527.321044921875 0 12585.884
527.8232421875 0 4631.7534
539.3162841796875 0 1353.5768 c Water loss 9
539.8106689453125 0 55086.99 c Ammonia loss 9
540.3114624023438 0 32714.021
540.8145751953125 0 14543.506
541.3160400390625 0 4812.153
547.8197631835938 0 61480.8
548.3233642578125 0 255952.38 c 9
548.8245849609375 0 143386.53
548.8748168945312 0 2508.35
549.3262939453125 0 49348.168
549.827392578125 0 8728.718
559.3151245117188 0 1168.5834
561.24365234375 0 2154.7124 z 9
562.3147583007812 0 1534.5566
568.3214111328125 0 1719.0559
569.3526611328125 0 59418.168 c 4
570.355224609375 0 16763.34
571.359130859375 0 3956.6978
574.3211059570312 0 5921.072
574.819580078125 0 2930.7446
576.8342895507812 0 2522.5315
582.36083984375 0 5052.799
583.3548583984375 0 3229.2134
597.3512573242188 0 1659.9363
603.8584594726562 0 2485.5151
604.3603515625 0 6355.3037
604.8616943359375 0 3366.7866
605.343505859375 0 1505.8143
610.84521484375 0 2550.3125
611.347412109375 0 27235.352
611.8485717773438 0 19983.893
612.3507080078125 0 7237.6177
612.8512573242188 0 2882.5593
613.345947265625 0 2233.62
617.8548583984375 0 23987.898
618.3571166992188 0 17582.6
618.8536987304688 0 43338.547
619.3551025390625 0 32478.479
619.8572998046875 0 23175.086
620.3599243164062 0 10133.267
620.8582153320312 0 4380.348
624.3545532226562 0 1501.5043
626.37451171875 0 340162.12 c 5
627.3771362304688 0 115297
628.3792114257812 0 21779.637
629.3817749023438 0 3036.8445
631.8346557617188 0 1329.0352
632.3533935546875 0 2493.7168 c Water loss 10
632.8500366210938 0 186579.33 c Ammonia loss 10
633.3515625 0 126753.34
633.8529052734375 0 58386.24
634.3546142578125 0 14205.457
634.8514404296875 0 3163.629
638.3555908203125 0 1426.9438
639.381591796875 0 1516.0822
639.8844604492188 0 2153.7732
640.859375 0 12613.739
641.363037109375 0 167069.97 c 10
641.8646240234375 0 120818.72
642.365966796875 0 47277.207
642.8677368164062 0 9775.696
643.3694458007812 0 3162.6946
647.3864135742188 0 1397.6604
660.381591796875 0 1470.7228
660.880859375 0 5646.7163
661.3822631835938 0 10333.516
661.8846435546875 0 6435.407
662.3875122070312 0 4211.4077
667.370361328125 0 1456.6649
668.3869018554688 0 75763.78
668.8889770507812 0 92373.555
669.3914184570312 0 57267.46
669.8941040039062 0 33322.816
670.3970336914062 0 10830.886
671.3501586914062 0 1641.3339
673.3756713867188 0 1700.9105
673.8793334960938 0 1734.3422
676.3739013671875 0 1841.9331
676.8609008789062 0 6271.8
677.3633422851562 0 5574.0596
677.8558959960938 0 1929.667
678.351806640625 0 1767.989
681.3779907226562 0 2472.4604
681.88134765625 0 3998.2869 c Water loss 11
682.38427734375 0 365707.2 c Ammonia loss 11
682.8857421875 0 272875
683.387451171875 0 140654.78
683.888916015625 0 40959.76
684.3875732421875 0 11264.161
689.8895263671875 0 2218.1392
690.3899536132812 0 5465.6333
690.897216796875 0 222158.25 c 11
691.3984985351562 0 178574.62
691.8999633789062 0 76603.1
692.4015502929688 0 23404.082
692.9031372070312 0 7103.453
698.3826904296875 0 35662.22
698.8826904296875 0 10391.514
699.38623046875 0 7151.3784
699.8820190429688 0 1826.5957
703.9046630859375 0 1650.6747
704.401611328125 0 3499.1807
704.8966064453125 0 4858.0503
705.3897705078125 0 7928.887
705.873046875 0 39263.035 z 1
706.3748779296875 0 25794.17
706.8757934570312 0 14227.219
707.3787231445312 0 3837.055
707.885009765625 0 1493.4884
711.9048461914062 0 3230.545
712.4070434570312 0 25745.584
712.9080810546875 0 22111.701
713.4114379882812 0 10189.103
713.8837890625 0 5409.664 y 1
714.3822021484375 0 5107.8535
716.8993530273438 0 1455.9657
717.4000244140625 0 1889.7109
717.8954467773438 0 3194.0276
719.3950805664062 0 10911.155
719.891845703125 0 12400.973
720.3895263671875 0 8576.193
720.8905639648438 0 3244.6167
725.901123046875 0 17037.033 c Ammonia loss 12
726.3995361328125 0 22322.95
726.8994140625 0 13809.404
727.3883056640625 0 12645.137
727.8963012695312 0 13080.208
728.4042358398438 0 14642.956
728.9027099609375 0 6033.4834
732.917724609375 0 2075.4878
733.4124145507812 0 9605.669
733.9064331054688 0 6784.3013
734.412841796875 0 198254.98 c 12
734.9144287109375 0 145984.56
735.4159545898438 0 81147.39
735.9014892578125 0 29883.03
736.3927001953125 0 19867.926
736.8912963867188 0 12435.918
737.3920288085938 0 3063.9714
738.4593505859375 0 4135.9736
739.4717407226562 0 2625.9814
740.471435546875 0 1871.569
740.9151000976562 0 8153.8613
741.4102172851562 0 46964.652
741.9113159179688 0 38563.133
742.402099609375 0 120459.29
742.9024658203125 0 85663.46
743.4019165039062 0 46556.39
743.90380859375 0 11783.367
744.4067993164062 0 4283.1074
745.36572265625 0 5959.76 z 7
746.88916015625 0 2779.6987
747.3988037109375 0 6283.788
747.896240234375 0 30022.627
748.396484375 0 21231.695
748.9002075195312 0 14185.052
749.4036865234375 0 4288.8354
749.9197387695312 0 5351.0425
750.4244995117188 0 6126.972
750.9258422851562 0 4143.8433
754.4017944335938 0 6322.795
754.9115600585938 0 31461.035
755.4075927734375 0 564327.06
755.908935546875 0 448075.28
756.4098510742188 0 239676.16
756.9109497070312 0 63374.9
756.9916381835938 0 1939.8367
757.4142456054688 0 17310.867
761.384765625 0 4446.068 y 7
762.38916015625 0 2796.6858
762.91162109375 0 17124.887
763.4163818359375 0 220018.03
763.9194946289062 0 591840.2
764.4210205078125 0 421027.5
764.92236328125 0 206973.3
765.42333984375 0 63005.363
765.5059204101562 0 2155.005
765.9232177734375 0 13932.404
771.909912109375 0 2044.0938
782.47509765625 0 22415.867 c 6
783.4773559570312 0 8973.014
814.4501953125 0 1651.4906
852.5016479492188 0 1798.9854
853.5128173828125 0 10873.103 c 7
854.5137939453125 0 4559.8687
855.5177001953125 0 1612.1782
857.4769897460938 0 2378.0637
886.4610595703125 0 1846.6543
901.4661865234375 0 291520.62 z 6
902.4688720703125 0 142980.88
903.4715576171875 0 43739.805
904.4752807617188 0 8236.288
917.4854125976562 0 15205.757 y 6
918.4881591796875 0 7406.628
919.49072265625 0 2458.515
958.4884643554688 0 43570.62 z 5
959.4906616210938 0 26042.852
960.492919921875 0 8333.75
965.5897216796875 0 3056.6006
966.5958251953125 0 18500.78 c 8
967.600341796875 0 10384.315
968.6026611328125 0 3664.1113
974.506103515625 0 8269.266 y 5
975.50927734375 0 5551.216
1028.5230712890625 0 3837.5442 w 4
1050.6165771484375 0 1855.0186
1051.6256103515625 0 10637.105
1052.631103515625 0 15881.651
1053.630126953125 0 8111.553
1069.53515625 0 2322.812
1070.5479736328125 0 3242.591
1071.5555419921875 0 8041.4673 y 4
1072.564697265625 0 4805.6704
1079.6234130859375 0 2066.0134
1080.6258544921875 0 4926.7993
1081.62939453125 0 4240.1084
1082.6302490234375 0 1562.6792
1094.6287841796875 0 7375.605
1095.6390380859375 0 91656.97 c 9
1096.6416015625 0 53702.883
1096.7618408203125 0 1307.3236
1097.6455078125 0 20311.568
1098.649658203125 0 3944.7346
1099.5579833984375 0 2520.6162
1100.544921875 0 1713.2896
1109.601806640625 0 3821.336
1110.6033935546875 0 3249.6936
1111.594482421875 0 2022.4921
1124.564697265625 0 1746.6322 y Water loss 3
1125.56494140625 0 2951.6409 y Ammonia loss 3
1126.5771484375 0 230452.44 z 3
1127.5809326171875 0 142188.4
1128.5830078125 0 52381.355
1129.5853271484375 0 14496.731
1130.59130859375 0 3690.1184
1142.596435546875 0 24792.002 y 3
1143.598876953125 0 17113.814
1144.6011962890625 0 5523.5884
1158.562255859375 0 1664.5447
1196.6346435546875 0 1710.2837
1223.679931640625 0 3110.6846
1237.697265625 0 3604.0347
1238.705078125 0 7424.694
1239.716064453125 0 3512.1765
1253.628173828125 0 1845.4042 y Ammonia loss 2
1254.635009765625 0 160619.67 z 2
1255.637939453125 0 116845.125
1256.6409912109375 0 52743.74
1257.6427001953125 0 13480.911
1258.6427001953125 0 2136.446
1265.69921875 0 8582.123
1266.7015380859375 0 8257.579
1267.7027587890625 0 3004.3062
1270.6549072265625 0 13148.558 y 2
1271.6566162109375 0 9506.243
1272.6605224609375 0 4479.581
1280.716064453125 0 2425.496
1281.71728515625 0 36307.453 c 10
1282.7222900390625 0 36836.363
1283.7254638671875 0 23257.928
1284.727783203125 0 7691.222
1285.72900390625 0 1742.9222
1320.7403564453125 0 2468.3064
1321.7645263671875 0 2412.867
1322.742919921875 0 4077.2021
1323.7496337890625 0 3582.1897
1324.66455078125 0 2004.2048 w 1
1325.6796875 0 3059.5972
1336.76953125 0 15070.886
1337.7763671875 0 33792.016
1338.777587890625 0 24888.693
1339.778564453125 0 9659.33
1340.7841796875 0 1913.4219
1352.7337646484375 0 1760.5385
1353.7236328125 0 1961.9011
1363.7579345703125 0 3583.6511 c Ammonia loss 11
1364.76708984375 0 15882.632
1365.770751953125 0 13736.796
1366.7685546875 0 5927.3013
1368.712890625 0 3187.6624
1369.705810546875 0 1690.551
1379.7808837890625 0 3744.213
1380.78466796875 0 25633.299 c 11
1381.7913818359375 0 61406.17
1382.795166015625 0 40512.625
1383.797119140625 0 17485.807
1384.803466796875 0 5775.1147
1396.7552490234375 0 3901.3699
1406.8016357421875 0 6709.883
1407.792724609375 0 6503.2554
1408.8013916015625 0 3541.1584
1409.7711181640625 0 5413.111
1410.7392578125 0 8949.852 z 1
1411.743408203125 0 37987.277
1412.74609375 0 25446.803
1413.750732421875 0 10875.399
1414.7508544921875 0 4076.2405
1423.7838134765625 0 4311.4116
1424.8055419921875 0 18588.223
1425.7880859375 0 16771.508
1426.7626953125 0 11080.361 y 1
1427.7598876953125 0 8176.732
1428.7532958984375 0 4614.0713
1429.7474365234375 0 2417.886
1438.7362060546875 0 2807.115
1439.7548828125 0 2344.1948
1440.7869873046875 0 3164.181
1451.7919921875 0 10250.309
1452.7969970703125 0 13459.736
1453.8001708984375 0 7838.383
1454.79736328125 0 3606.4558
1455.7979736328125 0 5318.447
1456.796630859375 0 4821.8276
1457.7904052734375 0 2273.4617
1465.8060302734375 0 2657.6333
1466.7857666015625 0 4955.6987
1467.7890625 0 72410.266 c 12
1468.80029296875 0 122295.945
1469.8046875 0 80973.07
1470.8082275390625 0 32906.438
1471.8126220703125 0 9329.137
1472.8092041015625 0 3330.273
1481.8248291015625 0 7493.0986
1482.818603515625 0 32453.44
1483.8167724609375 0 31703.932
1484.8160400390625 0 23733.957
1485.8150634765625 0 11393.821
1486.8232421875 0 4201.472
1492.796875 0 3514.5874
1493.7861328125 0 27900.254
1494.78857421875 0 25972.59
1495.789306640625 0 12677.572
1496.7935791015625 0 4766.7056
1497.791259765625 0 2092.956
1498.8363037109375 0 3063.5334
1499.842529296875 0 20797.799
1500.8443603515625 0 15662.564
1501.850830078125 0 7521.9233
1502.854248046875 0 1763.2976
1508.7999267578125 0 2453.0703
1509.8094482421875 0 47132.992
1510.812255859375 0 301399.5
1511.8143310546875 0 235151.72
1512.8177490234375 0 122029.88
1513.81884765625 0 35757.105
1514.818115234375 0 12164.361
1524.8026123046875 0 1613.1458
1525.8223876953125 0 22779.492
1526.8297119140625 0 98596.766
1527.837646484375 0 404702.22
1528.8402099609375 0 307125.28
1529.8431396484375 0 154436.77
1530.844482421875 0 50369.773
1531.8426513671875 0 16565.787
1984.6434326171875 0 1659.1027
3076.1826171875 0 1694.1259

Spectrum Details

|  |  |
| --- | --- |
| Matched peaks? Matched peaks The total absolute number of peaks matched. Additionally in brackets the total fraction of peaks matched and the total number of peaks is shown. | 61 (12.60% of 484) |
| FDR? FDR The false discovery rate estimated for this peptide. It is calculated by matching all theoretical fragments with a non-integer shift with the raw peaks for this spectrum. This is done with 40 different shifts. The resulting percentage is the average number of annotated peaks over the number of annotated peaks with the correct spectrum. | 2.77% |
| Satellite FDR? Satellite FDR See the FDR for details on its calculation. This satellite ion specific FDR only contains the satellite ions (d/w) for I/L/J positions. | ∞ |
| PSM Score? PSM Score The PSM Score as given by Hecklib to this annotated spectrum. It is shown with three significant figures. | 488 |

#### Spectrum 6414? Spectrum 6414 The raw spectrum of this peptide as annotated by Hecklib. The fragments are coloured according to ion type (see legend). Any peaks with a star '\*' as text can be hovered over to see the full details, first the ion type second the mass shift type. By hovering over the amino acids in the peptide or ions in the legend the corresponding peaks are highlighted. By toggling the 'Unassigned' label you can turn the background (unassigned) peaks on or off in the plot. By updating the slider in the Ion legend you can update the spectrum to only show the top X% of the peaks with labels. The top X% means any peak that is within X% of the highest intensity. By dragging in the spectrum you can zoom in to a specific part of the spectrum and use 'Zoom Out' to get back to the original zoom level. The annotation of the spectrum is based on the given sequence in the peptides file and is done with different software so inconsistencies are likely. The peaks are annotated based on the given sequence, with 20 ppm tolerance.

Copy Data

##### Spectrum 6414 (TSV)

###### Preview

```
Loading example...
```

*Click on the button to copy the data to your clipboard.*

Mz MinMz MaxIntensity Max

WidthHeightPeptide font sizePeptide stroke widthSpectrum font sizeSpectrum stroke widthCompact peptide

Ion legend

wxyz

abcd

OtherUnassignedIon^Charge^_Position_Show for top:%

VRQAPGRAJEWVSG

02.64e+45.28e+47.92e+41.06e+5

Zoom Out

y^+1^_2_y^+1^_2_c^+1^_2_y^+1^_3_c^+1^_2_c^+1^_3_c^+2^_7_c^+1^_3_c^+3^_11_c^+2^_8_y^+1^_4_c^+1^_4_c^+2^_9_c^+3^_13_c^+2^_10_c^+2^_10_c^+1^_5_c^+1^_6_c^+2^_11_c^+2^_11_c^+2^_11_c^+2^_12_c^+2^_12_c^+2^_12_z^+2^_13_y^+2^_13_c^+2^_13_c^+2^_13_y^+1^_7_c^+1^_7_c^+1^_8_z^+1^_8_y^+1^_8_z^+1^_9_c^+1^_9_y^+1^_9_y^+1^_10_c^+1^_10_y^+1^_11_z^+1^_11_y^+1^_11_w^+1^_12_y^+1^_12_z^+1^_12_y^+1^_12_c^+1^_11_w^+1^_13_c^+1^_12_c^+1^_12_z^+1^_13_y^+1^_13_

0544108816322177

Fragment Matches Table

Show background peaks

| Position | Ion type | Intensity | mz Theoretical | mz Error (Th) | mz Error (ppm) | Charge | Series Number |
| --- | --- | --- | --- | --- | --- | --- | --- |
| - | - | 473.8 | 120.1 | - | - | 0 | - |
| - | - | 572 | 129.1 | - | - | 0 | - |
| - | - | 2367 | 130.1 | - | - | 0 | - |
| - | - | 686.9 | 142.1 | - | - | 0 | - |
| 13 | y | 1672 | 145.1 | 0.0002971 | 2.048 | +1 | 2 |
| - | - | 620.6 | 147.1 | - | - | 0 | - |
| - | - | 493.5 | 148.9 | - | - | 0 | - |
| - | - | 438.5 | 153 | - | - | 0 | - |
| - | - | 470.9 | 155.3 | - | - | 0 | - |
| - | - | 427.2 | 156.3 | - | - | 0 | - |
| - | - | 1731 | 157.1 | - | - | 0 | - |
| - | - | 457.3 | 157.3 | - | - | 0 | - |
| - | - | 1646 | 159.1 | - | - | 0 | - |
| - | - | 470.6 | 161.1 | - | - | 0 | - |
| 13 | y | 2.606E+04 | 163.1 | 0.0001999 | 1.226 | +1 | 2 |
| - | - | 1069 | 164.1 | - | - | 0 | - |
| - | - | 1718 | 173.4 | - | - | 0 | - |
| - | - | 1981 | 173.5 | - | - | 0 | - |
| - | - | 653.4 | 173.5 | - | - | 0 | - |
| - | - | 708.2 | 174.1 | - | - | 0 | - |
| - | - | 451.3 | 176.6 | - | - | 0 | - |
| - | - | 850 | 185.1 | - | - | 0 | - |
| - | - | 567.5 | 185.2 | - | - | 0 | - |
| - | - | 789.5 | 187.1 | - | - | 0 | - |
| - | - | 532.1 | 210.4 | - | - | 0 | - |
| - | - | 1202 | 214.2 | - | - | 0 | - |
| - | - | 488.1 | 214.2 | - | - | 0 | - |
| - | - | 820.2 | 228.2 | - | - | 0 | - |
| - | - | 392.2 | 239.1 | - | - | 0 | - |
| - | - | 7250 | 239.2 | - | - | 0 | - |
| - | - | 1648 | 240.2 | - | - | 0 | - |
| - | - | 549.3 | 252 | - | - | 0 | - |
| 2 | c | 4385 | 256.2 | 4.359E-05 | 0.1701 | +1 | 2 |
| 12 | y | 833.7 | 262.1 | 0.0001903 | 0.726 | +1 | 3 |
| 2 | c | 9815 | 273.2 | 0.0001102 | 0.4033 | +1 | 2 |
| - | - | 1623 | 274.2 | - | - | 0 | - |
| - | - | 4244 | 299.2 | - | - | 0 | - |
| - | - | 696.6 | 327.1 | - | - | 0 | - |
| - | - | 840.5 | 356.2 | - | - | 0 | - |
| - | - | 585.9 | 357.2 | - | - | 0 | - |
| 3 | c | 1567 | 384.2 | 0.0001558 | 0.4054 | +1 | 3 |
| - | - | 1240 | 391.2 | - | - | 0 | - |
| 7 | c | 1.153E+04 | 391.7 | 0.0001668 | 0.4258 | +2 | 7 |
| - | - | 3552 | 392.2 | - | - | 0 | - |
| - | - | 1343 | 392.7 | - | - | 0 | - |
| 3 | c | 4.732E+04 | 401.3 | 0.0002485 | 0.6193 | +1 | 3 |
| - | - | 1.079E+04 | 402.3 | - | - | 0 | - |
| - | - | 726.3 | 403.3 | - | - | 0 | - |
| 11 | c | 814.1 | 422.2 | 0.002118 | 5.017 | +3 | 11 |
| - | - | 962 | 422.9 | - | - | 0 | - |
| - | - | 928.8 | 426.8 | - | - | 0 | - |
| 8 | c | 8507 | 427.3 | 7.513E-05 | 0.1758 | +2 | 8 |
| - | - | 4224 | 427.8 | - | - | 0 | - |
| - | - | 1135 | 428.3 | - | - | 0 | - |
| - | - | 1003 | 428.3 | - | - | 0 | - |
| - | - | 977.6 | 429.1 | - | - | 0 | - |
| - | - | 2236 | 430.2 | - | - | 0 | - |
| - | - | 1286 | 431.3 | - | - | 0 | - |
| 11 | y | 1356 | 448.2 | 0.0006153 | 1.373 | +1 | 4 |
| 4 | c | 955.4 | 455.3 | 0.00522 | 11.47 | +1 | 4 |
| - | - | 1716 | 461.8 | - | - | 0 | - |
| - | - | 917.5 | 462.3 | - | - | 0 | - |
| - | - | 6994 | 483.3 | - | - | 0 | - |
| 9 | c | 3.76E+04 | 483.8 | 9.636E-05 | 0.1992 | +2 | 9 |
| 13 | c | 3165 | 484.3 | 0.0005399 | 1.115 | +3 | 13 |
| - | - | 2.021E+04 | 484.3 | - | - | 0 | - |
| - | - | 3389 | 484.6 | - | - | 0 | - |
| - | - | 8042 | 484.8 | - | - | 0 | - |
| - | - | 1017 | 484.9 | - | - | 0 | - |
| - | - | 863 | 485.3 | - | - | 0 | - |
| - | - | 927.2 | 485.3 | - | - | 0 | - |
| - | - | 5408 | 496.8 | - | - | 0 | - |
| - | - | 798.9 | 497.3 | - | - | 0 | - |
| - | - | 3347 | 497.3 | - | - | 0 | - |
| - | - | 1430 | 497.8 | - | - | 0 | - |
| - | - | 2140 | 498.3 | - | - | 0 | - |
| - | - | 830 | 498.8 | - | - | 0 | - |
| - | - | 2541 | 503.3 | - | - | 0 | - |
| - | - | 1037 | 503.6 | - | - | 0 | - |
| - | - | 885.9 | 503.9 | - | - | 0 | - |
| - | - | 607.7 | 508.2 | - | - | 0 | - |
| - | - | 1449 | 509.2 | - | - | 0 | - |
| - | - | 3.132E+04 | 509.3 | - | - | 0 | - |
| - | - | 9078 | 510.3 | - | - | 0 | - |
| - | - | 798.7 | 525.3 | - | - | 0 | - |
| - | - | 6421 | 526.3 | - | - | 0 | - |
| - | - | 5086 | 526.8 | - | - | 0 | - |
| - | - | 3184 | 527.3 | - | - | 0 | - |
| 10 | c | 8392 | 539.8 | 0.0006193 | 1.147 | +2 | 10 |
| - | - | 5624 | 540.3 | - | - | 0 | - |
| - | - | 2553 | 540.8 | - | - | 0 | - |
| - | - | 701 | 541.3 | - | - | 0 | - |
| - | - | 1.169E+04 | 547.8 | - | - | 0 | - |
| 10 | c | 4.693E+04 | 548.3 | 0.0003262 | 0.5948 | +2 | 10 |
| - | - | 2.628E+04 | 548.8 | - | - | 0 | - |
| - | - | 9809 | 549.3 | - | - | 0 | - |
| - | - | 2164 | 549.8 | - | - | 0 | - |
| - | - | 549.8 | 565.2 | - | - | 0 | - |
| 5 | c | 1.147E+04 | 569.4 | 9.892E-07 | 0.001737 | +1 | 5 |
| - | - | 3567 | 570.4 | - | - | 0 | - |
| - | - | 965.6 | 571.4 | - | - | 0 | - |
| - | - | 1102 | 574.3 | - | - | 0 | - |
| - | - | 1285 | 582.4 | - | - | 0 | - |
| - | - | 4995 | 611.3 | - | - | 0 | - |
| - | - | 3613 | 611.8 | - | - | 0 | - |
| - | - | 2192 | 612.3 | - | - | 0 | - |
| - | - | 3434 | 617.9 | - | - | 0 | - |
| - | - | 2141 | 618.4 | - | - | 0 | - |
| - | - | 6585 | 618.9 | - | - | 0 | - |
| - | - | 5171 | 619.4 | - | - | 0 | - |
| - | - | 5358 | 619.9 | - | - | 0 | - |
| - | - | 2116 | 620.4 | - | - | 0 | - |
| - | - | 725.8 | 624.3 | - | - | 0 | - |
| - | - | 805.2 | 624.8 | - | - | 0 | - |
| 6 | c | 6.695E+04 | 626.4 | 2.164E-05 | 0.03455 | +1 | 6 |
| - | - | 2.258E+04 | 627.4 | - | - | 0 | - |
| - | - | 4651 | 628.4 | - | - | 0 | - |
| 11 | c | 599 | 632.4 | 0.002291 | 3.622 | +2 | 11 |
| 11 | c | 2.655E+04 | 632.8 | 0.0002188 | 0.3458 | +2 | 11 |
| - | - | 1.866E+04 | 633.4 | - | - | 0 | - |
| - | - | 1.014E+04 | 633.9 | - | - | 0 | - |
| - | - | 2176 | 634.4 | - | - | 0 | - |
| - | - | 3009 | 640.9 | - | - | 0 | - |
| 11 | c | 2.626E+04 | 641.4 | 0.0004319 | 0.6733 | +2 | 11 |
| - | - | 2.12E+04 | 641.9 | - | - | 0 | - |
| - | - | 8916 | 642.4 | - | - | 0 | - |
| - | - | 2319 | 642.9 | - | - | 0 | - |
| - | - | 700.4 | 643.4 | - | - | 0 | - |
| - | - | 2049 | 660.9 | - | - | 0 | - |
| - | - | 1905 | 661.4 | - | - | 0 | - |
| - | - | 1956 | 661.9 | - | - | 0 | - |
| - | - | 1.101E+04 | 668.4 | - | - | 0 | - |
| - | - | 1.726E+04 | 668.9 | - | - | 0 | - |
| - | - | 1.133E+04 | 669.4 | - | - | 0 | - |
| - | - | 4960 | 669.9 | - | - | 0 | - |
| - | - | 2178 | 670.4 | - | - | 0 | - |
| - | - | 1133 | 670.9 | - | - | 0 | - |
| - | - | 1895 | 676.9 | - | - | 0 | - |
| - | - | 1349 | 677.4 | - | - | 0 | - |
| 12 | c | 1108 | 681.9 | 0.01044 | 15.3 | +2 | 12 |
| 12 | c | 5.275E+04 | 682.4 | 0.0004292 | 0.629 | +2 | 12 |
| - | - | 3.861E+04 | 682.9 | - | - | 0 | - |
| - | - | 1.928E+04 | 683.4 | - | - | 0 | - |
| - | - | 6902 | 683.9 | - | - | 0 | - |
| - | - | 1804 | 684.4 | - | - | 0 | - |
| - | - | 910.6 | 690.4 | - | - | 0 | - |
| 12 | c | 4.233E+04 | 690.9 | 0.0004591 | 0.6645 | +2 | 12 |
| - | - | 3.124E+04 | 691.4 | - | - | 0 | - |
| - | - | 1.475E+04 | 691.9 | - | - | 0 | - |
| - | - | 6571 | 692.4 | - | - | 0 | - |
| - | - | 1435 | 692.9 | - | - | 0 | - |
| - | - | 1.188E+04 | 698.4 | - | - | 0 | - |
| - | - | 3277 | 698.9 | - | - | 0 | - |
| - | - | 2125 | 699.4 | - | - | 0 | - |
| - | - | 670.3 | 703.9 | - | - | 0 | - |
| - | - | 703.7 | 705.4 | - | - | 0 | - |
| 2 | z | 6783 | 705.9 | 0.0009185 | 1.301 | +2 | 13 |
| - | - | 4448 | 706.4 | - | - | 0 | - |
| - | - | 1960 | 706.9 | - | - | 0 | - |
| - | - | 656.1 | 707.4 | - | - | 0 | - |
| - | - | 4492 | 712.4 | - | - | 0 | - |
| - | - | 3104 | 712.9 | - | - | 0 | - |
| - | - | 1751 | 713.4 | - | - | 0 | - |
| 2 | y | 1720 | 713.9 | 0.001913 | 2.679 | +2 | 13 |
| - | - | 1713 | 719.4 | - | - | 0 | - |
| - | - | 3848 | 719.9 | - | - | 0 | - |
| - | - | 2115 | 720.4 | - | - | 0 | - |
| - | - | 744.2 | 720.9 | - | - | 0 | - |
| 13 | c | 2695 | 725.9 | 0.001623 | 2.236 | +2 | 13 |
| - | - | 4153 | 726.4 | - | - | 0 | - |
| - | - | 3195 | 726.9 | - | - | 0 | - |
| - | - | 2255 | 727.4 | - | - | 0 | - |
| - | - | 2961 | 727.9 | - | - | 0 | - |
| - | - | 2330 | 728.4 | - | - | 0 | - |
| - | - | 1152 | 728.9 | - | - | 0 | - |
| - | - | 704.8 | 729.9 | - | - | 0 | - |
| - | - | 2381 | 733.4 | - | - | 0 | - |
| - | - | 1661 | 733.9 | - | - | 0 | - |
| 13 | c | 3.47E+04 | 734.4 | 0.001215 | 1.654 | +2 | 13 |
| - | - | 2.966E+04 | 734.9 | - | - | 0 | - |
| - | - | 1.428E+04 | 735.4 | - | - | 0 | - |
| - | - | 4273 | 735.9 | - | - | 0 | - |
| - | - | 3760 | 736.4 | - | - | 0 | - |
| - | - | 1395 | 736.9 | - | - | 0 | - |
| - | - | 1520 | 740.9 | - | - | 0 | - |
| - | - | 8688 | 741.4 | - | - | 0 | - |
| - | - | 7680 | 741.9 | - | - | 0 | - |
| - | - | 2.206E+04 | 742.4 | - | - | 0 | - |
| - | - | 1.53E+04 | 742.9 | - | - | 0 | - |
| - | - | 8625 | 743.4 | - | - | 0 | - |
| - | - | 1814 | 743.9 | - | - | 0 | - |
| - | - | 2254 | 747.4 | - | - | 0 | - |
| - | - | 4779 | 747.9 | - | - | 0 | - |
| - | - | 3871 | 748.4 | - | - | 0 | - |
| - | - | 3905 | 748.9 | - | - | 0 | - |
| - | - | 1095 | 749.4 | - | - | 0 | - |
| - | - | 768.9 | 750.4 | - | - | 0 | - |
| - | - | 777.3 | 750.9 | - | - | 0 | - |
| - | - | 1382 | 754.4 | - | - | 0 | - |
| - | - | 6587 | 754.9 | - | - | 0 | - |
| - | - | 1.008E+05 | 755.4 | - | - | 0 | - |
| - | - | 8.877E+04 | 755.9 | - | - | 0 | - |
| - | - | 4.435E+04 | 756.4 | - | - | 0 | - |
| - | - | 1.231E+04 | 756.9 | - | - | 0 | - |
| - | - | 4798 | 757.4 | - | - | 0 | - |
| 8 | y | 1518 | 761.4 | 0.001972 | 2.59 | +1 | 7 |
| - | - | 796.7 | 762.4 | - | - | 0 | - |
| - | - | 2088 | 762.9 | - | - | 0 | - |
| - | - | 4.406E+04 | 763.4 | - | - | 0 | - |
| - | - | 1.046E+05 | 763.9 | - | - | 0 | - |
| - | - | 7.889E+04 | 764.4 | - | - | 0 | - |
| - | - | 4.161E+04 | 764.9 | - | - | 0 | - |
| - | - | 1.29E+04 | 765.4 | - | - | 0 | - |
| - | - | 2431 | 765.9 | - | - | 0 | - |
| 7 | c | 4395 | 782.5 | 0.0004424 | 0.5654 | +1 | 7 |
| - | - | 1776 | 783.5 | - | - | 0 | - |
| - | - | 927.3 | 852.5 | - | - | 0 | - |
| 8 | c | 2396 | 853.5 | 0.0005689 | 0.6665 | +1 | 8 |
| 7 | z | 5.362E+04 | 901.5 | 0.001046 | 1.16 | +1 | 8 |
| - | - | 2.964E+04 | 902.5 | - | - | 0 | - |
| - | - | 423.2 | 902.5 | - | - | 0 | - |
| - | - | 715.6 | 902.6 | - | - | 0 | - |
| - | - | 8604 | 903.5 | - | - | 0 | - |
| - | - | 1997 | 904.5 | - | - | 0 | - |
| 7 | y | 2610 | 917.5 | 0.001398 | 1.524 | +1 | 8 |
| - | - | 1145 | 918.5 | - | - | 0 | - |
| 6 | z | 8859 | 958.5 | 0.0004146 | 0.4326 | +1 | 9 |
| - | - | 5698 | 959.5 | - | - | 0 | - |
| - | - | 1582 | 960.5 | - | - | 0 | - |
| 9 | c | 3792 | 966.6 | 0.003273 | 3.386 | +1 | 9 |
| - | - | 1731 | 967.6 | - | - | 0 | - |
| - | - | 917.6 | 968.6 | - | - | 0 | - |
| 6 | y | 1870 | 974.5 | 0.001072 | 1.1 | +1 | 9 |
| - | - | 832.6 | 975.5 | - | - | 0 | - |
| - | - | 610.6 | 1016 | - | - | 0 | - |
| - | - | 786.5 | 1018 | - | - | 0 | - |
| - | - | 854.8 | 1019 | - | - | 0 | - |
| - | - | 643.2 | 1020 | - | - | 0 | - |
| - | - | 2011 | 1052 | - | - | 0 | - |
| - | - | 3041 | 1053 | - | - | 0 | - |
| - | - | 1538 | 1054 | - | - | 0 | - |
| - | - | 695.9 | 1054 | - | - | 0 | - |
| 5 | y | 1832 | 1072 | 0.005313 | 4.958 | +1 | 10 |
| - | - | 794.6 | 1073 | - | - | 0 | - |
| - | - | 1534 | 1081 | - | - | 0 | - |
| - | - | 1943 | 1095 | - | - | 0 | - |
| 10 | c | 2.23E+04 | 1096 | 0.001921 | 1.753 | +1 | 10 |
| - | - | 1.129E+04 | 1097 | - | - | 0 | - |
| - | - | 3775 | 1098 | - | - | 0 | - |
| - | - | 1143 | 1099 | - | - | 0 | - |
| - | - | 1226 | 1110 | - | - | 0 | - |
| 4 | y | 874.2 | 1126 | 0.0001308 | 0.1162 | +1 | 11 |
| 4 | z | 4.519E+04 | 1127 | 0.001852 | 1.644 | +1 | 11 |
| - | - | 2.791E+04 | 1128 | - | - | 0 | - |
| - | - | 1.136E+04 | 1129 | - | - | 0 | - |
| - | - | 2584 | 1130 | - | - | 0 | - |
| - | - | 1129 | 1131 | - | - | 0 | - |
| 4 | y | 6520 | 1143 | 0.0006789 | 0.5942 | +1 | 11 |
| - | - | 3337 | 1144 | - | - | 0 | - |
| - | - | 1996 | 1145 | - | - | 0 | - |
| 3 | w | 691.2 | 1197 | 0.01024 | 8.558 | +1 | 12 |
| - | - | 1127 | 1239 | - | - | 0 | - |
| - | - | 775.1 | 1240 | - | - | 0 | - |
| - | - | 810.8 | 1249 | - | - | 0 | - |
| 3 | y | 887.6 | 1254 | 0.001839 | 1.467 | +1 | 12 |
| 3 | z | 3.121E+04 | 1255 | 0.002568 | 2.047 | +1 | 12 |
| - | - | 2.211E+04 | 1256 | - | - | 0 | - |
| - | - | 1.066E+04 | 1257 | - | - | 0 | - |
| - | - | 2813 | 1258 | - | - | 0 | - |
| - | - | 1462 | 1266 | - | - | 0 | - |
| - | - | 1156 | 1267 | - | - | 0 | - |
| - | - | 889.7 | 1268 | - | - | 0 | - |
| 3 | y | 3024 | 1271 | 0.002616 | 2.059 | +1 | 12 |
| - | - | 1941 | 1272 | - | - | 0 | - |
| - | - | 654.9 | 1273 | - | - | 0 | - |
| 11 | c | 7562 | 1282 | 0.003719 | 2.902 | +1 | 11 |
| - | - | 8268 | 1283 | - | - | 0 | - |
| - | - | 4050 | 1284 | - | - | 0 | - |
| - | - | 1992 | 1285 | - | - | 0 | - |
| - | - | 875.9 | 1321 | - | - | 0 | - |
| - | - | 1075 | 1323 | - | - | 0 | - |
| 2 | w | 686.9 | 1325 | 0.003659 | 2.762 | +1 | 13 |
| - | - | 3124 | 1337 | - | - | 0 | - |
| - | - | 7773 | 1338 | - | - | 0 | - |
| - | - | 5647 | 1339 | - | - | 0 | - |
| - | - | 2425 | 1340 | - | - | 0 | - |
| - | - | 644.9 | 1345 | - | - | 0 | - |
| - | - | 900.6 | 1354 | - | - | 0 | - |
| 12 | c | 1314 | 1364 | 0.005667 | 4.155 | +1 | 12 |
| - | - | 4779 | 1365 | - | - | 0 | - |
| - | - | 3088 | 1366 | - | - | 0 | - |
| - | - | 1199 | 1367 | - | - | 0 | - |
| - | - | 761.9 | 1370 | - | - | 0 | - |
| 12 | c | 5961 | 1381 | 0.003163 | 2.291 | +1 | 12 |
| - | - | 1.297E+04 | 1382 | - | - | 0 | - |
| - | - | 9606 | 1383 | - | - | 0 | - |
| - | - | 3369 | 1384 | - | - | 0 | - |
| - | - | 1116 | 1385 | - | - | 0 | - |
| - | - | 845.2 | 1395 | - | - | 0 | - |
| - | - | 1187 | 1397 | - | - | 0 | - |
| - | - | 789.7 | 1407 | - | - | 0 | - |
| - | - | 1708 | 1408 | - | - | 0 | - |
| - | - | 965.8 | 1409 | - | - | 0 | - |
| 2 | z | 2789 | 1411 | 0.004597 | 3.258 | +1 | 13 |
| - | - | 7567 | 1412 | - | - | 0 | - |
| - | - | 6853 | 1413 | - | - | 0 | - |
| - | - | 1906 | 1414 | - | - | 0 | - |
| - | - | 762.1 | 1415 | - | - | 0 | - |
| - | - | 760.3 | 1424 | - | - | 0 | - |
| - | - | 3982 | 1425 | - | - | 0 | - |
| - | - | 3682 | 1426 | - | - | 0 | - |
| 2 | y | 2310 | 1427 | 0.0002112 | 0.148 | +1 | 13 |
| - | - | 1811 | 1428 | - | - | 0 | - |
| - | - | 1101 | 1429 | - | - | 0 | - |
| - | - | 2693 | 1452 | - | - | 0 | - |
| - | - | 3081 | 1453 | - | - | 0 | - |
| - | - | 1655 | 1454 | - | - | 0 | - |
| - | - | 960.4 | 1455 | - | - | 0 | - |
| - | - | 1164 | 1456 | - | - | 0 | - |
| - | - | 761.3 | 1457 | - | - | 0 | - |
| - | - | 1429 | 1467 | - | - | 0 | - |
| - | - | 1.581E+04 | 1468 | - | - | 0 | - |
| - | - | 2.549E+04 | 1469 | - | - | 0 | - |
| - | - | 1.598E+04 | 1470 | - | - | 0 | - |
| - | - | 6994 | 1471 | - | - | 0 | - |
| - | - | 1754 | 1472 | - | - | 0 | - |
| - | - | 767.5 | 1473 | - | - | 0 | - |
| - | - | 1880 | 1482 | - | - | 0 | - |
| - | - | 7075 | 1483 | - | - | 0 | - |
| - | - | 6819 | 1484 | - | - | 0 | - |
| - | - | 5742 | 1485 | - | - | 0 | - |
| - | - | 2699 | 1486 | - | - | 0 | - |
| - | - | 7336 | 1494 | - | - | 0 | - |
| - | - | 5621 | 1495 | - | - | 0 | - |
| - | - | 3598 | 1496 | - | - | 0 | - |
| - | - | 1193 | 1499 | - | - | 0 | - |
| - | - | 4264 | 1500 | - | - | 0 | - |
| - | - | 4091 | 1501 | - | - | 0 | - |
| - | - | 1699 | 1502 | - | - | 0 | - |
| - | - | 1187 | 1509 | - | - | 0 | - |
| - | - | 1.289E+04 | 1510 | - | - | 0 | - |
| - | - | 7.094E+04 | 1511 | - | - | 0 | - |
| - | - | 5.382E+04 | 1512 | - | - | 0 | - |
| - | - | 2.662E+04 | 1513 | - | - | 0 | - |
| - | - | 8918 | 1514 | - | - | 0 | - |
| - | - | 3658 | 1515 | - | - | 0 | - |
| - | - | 5357 | 1526 | - | - | 0 | - |
| - | - | 2.368E+04 | 1527 | - | - | 0 | - |
| - | - | 8.841E+04 | 1528 | - | - | 0 | - |
| - | - | 6.851E+04 | 1529 | - | - | 0 | - |
| - | - | 3.355E+04 | 1530 | - | - | 0 | - |
| - | - | 1.049E+04 | 1531 | - | - | 0 | - |
| - | - | 5306 | 1532 | - | - | 0 | - |
| - | - | 766.3 | 2155 | - | - | 0 | - |

m/z Charge Intensity FragmentType MassShift Position
120.08096313476562 0 473.79593
129.1027374267578 0 572.03644
130.06533813476562 0 2366.5698
142.09788513183594 0 686.9267
145.06106567382812 0 1671.6493 y Water loss 12
147.1131134033203 0 620.619
148.945556640625 0 493.50613
153.03579711914062 0 438.54718
155.2716522216797 0 470.92126
156.299560546875 0 427.15817
157.10850524902344 0 1731.4425
157.2950897216797 0 457.2877
159.09165954589844 0 1645.7587
161.0721435546875 0 470.60913
163.071533203125 0 26058.604 y 12
164.07504272460938 0 1068.7062
173.4376983642578 0 1717.6567
173.45162963867188 0 1981.0331
173.45831298828125 0 653.4472
174.13504028320312 0 708.2174
176.62330627441406 0 451.3103
185.1038360595703 0 849.9563
185.1656036376953 0 567.456
187.10806274414062 0 789.53424
210.4208221435547 0 532.07196
214.15528869628906 0 1201.5559
214.24398803710938 0 488.09482
228.1823272705078 0 820.20764
239.13845825195312 0 392.2317
239.15048217773438 0 7249.7065
240.1535186767578 0 1647.9541
252.04612731933594 0 549.2871
256.1767578125 0 4385.0225 c Ammonia loss 1
262.1395568847656 0 833.65686 y 11
273.2034606933594 0 9814.537 c 1
274.20703125 0 1622.8988
299.21929931640625 0 4244.099
327.134033203125 0 696.61957
356.23980712890625 0 840.5225
357.24871826171875 0 585.892
384.23553466796875 0 1566.9229 c Ammonia loss 2
391.2378845214844 0 1240.4421
391.74066162109375 0 11531.705 c 6
392.242431640625 0 3552.285
392.74310302734375 0 1342.9972
401.2621765136719 0 47317.06 c 2
402.26495361328125 0 10786.263
403.26678466796875 0 726.2501
422.2372741699219 0 814.05994 c Ammonia loss 10
422.9046936035156 0 962.02875
426.7564697265625 0 928.7528
427.25946044921875 0 8506.648 c 7
427.7612609863281 0 4224.438
428.2596740722656 0 1135.2023
428.2886047363281 0 1003.21936
429.0901184082031 0 977.6135
430.1851806640625 0 2235.5125
431.2548522949219 0 1285.97
448.21844482421875 0 1356.185 y 10
455.26727294921875 0 955.37396 c Ammonia loss 3
461.7942810058594 0 1716.1569
462.29705810546875 0 917.4884
483.2977600097656 0 6994.319
483.801513671875 0 37603.582 c 8
484.2680969238281 0 3164.9893 c Ammonia loss 12
484.3031005859375 0 20213.02
484.6029052734375 0 3389.1287
484.80389404296875 0 8042.2734
484.9383239746094 0 1016.913
485.2710876464844 0 863.0385
485.30810546875 0 927.22705
496.8097229003906 0 5407.623
497.27301025390625 0 798.9336
497.3113098144531 0 3346.6921
497.81219482421875 0 1429.6553
498.285888671875 0 2140.0452
498.7862854003906 0 829.9654
503.2757263183594 0 2541.1904
503.6103515625 0 1037.352
503.9462890625 0 885.89435
508.2440490722656 0 607.6653
509.2155456542969 0 1448.6985
509.26092529296875 0 31316.182
510.263671875 0 9077.934
525.3350219726562 0 798.6778
526.3156127929688 0 6421.069
526.8172607421875 0 5085.609
527.3191528320312 0 3183.711
539.81005859375 0 8391.788 c Ammonia loss 9
540.31103515625 0 5624.458
540.8139038085938 0 2553.249
541.3129272460938 0 700.98846
547.8187866210938 0 11686.996
548.3223876953125 0 46930.69 c 9
548.8237915039062 0 26281.62
549.3250122070312 0 9808.836
549.82568359375 0 2164.3643
565.1993408203125 0 549.8101
569.351806640625 0 11470.312 c 4
570.354248046875 0 3566.5552
571.3588256835938 0 965.64996
574.3201904296875 0 1102.3754
582.3616333007812 0 1285.3752
611.3468627929688 0 4994.547
611.847900390625 0 3613.229
612.3494873046875 0 2192.2712
617.8539428710938 0 3433.791
618.35546875 0 2140.7063
618.8523559570312 0 6584.5386
619.352783203125 0 5171.417
619.85595703125 0 5357.508
620.35888671875 0 2115.8108
624.3482055664062 0 725.7542
624.8424682617188 0 805.17224
626.373291015625 0 66946.234 c 5
627.3760375976562 0 22581.1
628.3777465820312 0 4650.884
632.3547973632812 0 599.023 c Water loss 10
632.848876953125 0 26546.266 c Ammonia loss 10
633.3507080078125 0 18659.152
633.8517456054688 0 10140.535
634.3538208007812 0 2176.453
640.857666015625 0 3008.8665
641.3619384765625 0 26262.686 c 10
641.86376953125 0 21197.352
642.3651733398438 0 8915.967
642.8648681640625 0 2319.118
643.3700561523438 0 700.41736
660.8787231445312 0 2049.0735
661.3802490234375 0 1905.2845
661.8844604492188 0 1955.6718
668.3858642578125 0 11006.186
668.88818359375 0 17260.799
669.3901977539062 0 11333.467
669.8917846679688 0 4959.873
670.3947143554688 0 2177.5085
670.9049682617188 0 1133.381
676.8607788085938 0 1895.3523
677.3619995117188 0 1349.4138
681.880859375 0 1108.4905 c Water loss 11
682.3828735351562 0 52750.1 c Ammonia loss 11
682.884765625 0 38607.92
683.38623046875 0 19284.283
683.8869018554688 0 6901.701
684.3851928710938 0 1804.0228
690.3912963867188 0 910.5668
690.8961181640625 0 42330.34 c 11
691.3976440429688 0 31239.053
691.89892578125 0 14746.408
692.4017333984375 0 6571.081
692.8970336914062 0 1434.724
698.3814697265625 0 11882.22
698.8837280273438 0 3277.1138
699.3836059570312 0 2124.51
703.900390625 0 670.33624
705.3968505859375 0 703.701
705.8726806640625 0 6782.9443 z 1
706.371826171875 0 4448.218
706.8743286132812 0 1960.3964
707.3790893554688 0 656.1215
712.4053955078125 0 4491.961
712.9061889648438 0 3104.3337
713.408935546875 0 1751.1503
713.8792114257812 0 1719.6132 y 1
719.3947143554688 0 1712.6107
719.8915405273438 0 3847.7957
720.3914794921875 0 2115.4119
720.8798828125 0 744.2193
725.9009399414062 0 2694.6296 c Ammonia loss 12
726.397216796875 0 4153.1636
726.8994750976562 0 3195.4497
727.387939453125 0 2254.9763
727.8963012695312 0 2961.0393
728.4049072265625 0 2330.0593
728.9083251953125 0 1151.8816
729.9110717773438 0 704.7716
733.4091796875 0 2380.58
733.9149780273438 0 1661.0764
734.411376953125 0 34702.05 c 12
734.9131469726562 0 29655.953
735.41455078125 0 14282.395
735.89990234375 0 4272.8813
736.3902587890625 0 3760.2449
736.8917236328125 0 1394.6791
740.9152221679688 0 1520.4282
741.408447265625 0 8687.818
741.9103393554688 0 7680.153
742.4005737304688 0 22064.357
742.901611328125 0 15299.776
743.400634765625 0 8624.742
743.9029541015625 0 1813.5305
747.395751953125 0 2254.0386
747.8948974609375 0 4778.8027
748.3964233398438 0 3870.5676
748.8980102539062 0 3904.9739
749.3995971679688 0 1095.3033
750.4176635742188 0 768.9143
750.9244995117188 0 777.33484
754.3999633789062 0 1382.3649
754.9110717773438 0 6587.457
755.406005859375 0 100844.39
755.9070434570312 0 88766.61
756.4080200195312 0 44349.875
756.9094848632812 0 12313.47
757.4113159179688 0 4797.732
761.380859375 0 1517.7487 y 7
762.4005737304688 0 796.69635
762.9112548828125 0 2088.421
763.4144897460938 0 44057.426
763.9180908203125 0 104591.19
764.4196166992188 0 78890.98
764.9205322265625 0 41608.52
765.4215087890625 0 12903.91
765.9195556640625 0 2431.2231
782.4739379882812 0 4395.4185 c 6
783.4749145507812 0 1776.379
852.5095825195312 0 927.34705
853.5109252929688 0 2395.906 c 7
901.4641723632812 0 53621.008 z 6
902.4669799804688 0 29643.26
902.5452880859375 0 423.1578
902.5741577148438 0 715.5703
903.4698486328125 0 8604.189
904.4725341796875 0 1996.7836
917.4825439453125 0 2609.6514 y 6
918.48876953125 0 1144.9567
958.4862670898438 0 8858.844 z 5
959.4882202148438 0 5698.0874
960.4882202148438 0 1581.9657
966.59228515625 0 3791.7634 c 8
967.6013793945312 0 1730.555
968.5982055664062 0 917.643
974.5043334960938 0 1869.862 y 5
975.5079345703125 0 832.5963
1015.5164794921875 0 610.5825
1017.517578125 0 786.4956
1018.5413818359375 0 854.83563
1020.4800415039062 0 643.15656
1051.622802734375 0 2010.5662
1052.6268310546875 0 3040.783
1053.6295166015625 0 1537.7136
1054.03515625 0 695.9263
1071.5528564453125 0 1832.13 y 4
1072.553466796875 0 794.6144
1080.6239013671875 0 1534.0504
1094.6282958984375 0 1942.7286
1095.63623046875 0 22298.44 c 9
1096.6392822265625 0 11293.001
1097.6435546875 0 3774.858
1098.641845703125 0 1142.658
1109.595703125 0 1226.2041
1125.568603515625 0 874.2119 y Ammonia loss 3
1126.57470703125 0 45194.41 z 3
1127.5782470703125 0 27905.758
1128.5797119140625 0 11360.028
1129.5828857421875 0 2583.7493
1130.582275390625 0 1129.1721
1142.5946044921875 0 6519.9805 y 3
1143.5950927734375 0 3336.953
1144.603271484375 0 1995.8628
1196.6160888671875 0 691.2017 w 2
1238.703857421875 0 1126.8583
1239.708251953125 0 775.13873
1248.6640625 0 810.7847
1253.629150390625 0 887.56494 y Ammonia loss 2
1254.632568359375 0 31205.223 z 2
1255.63525390625 0 22105.316
1256.6380615234375 0 10663.257
1257.6356201171875 0 2812.6536
1265.6939697265625 0 1462.3013
1266.697265625 0 1156.1107
1267.700439453125 0 889.7068
1270.6512451171875 0 3024.2102 y 2
1271.6500244140625 0 1940.6348
1272.6636962890625 0 654.878
1281.7137451171875 0 7562.334 c 10
1282.7178955078125 0 8268.192
1283.721435546875 0 4050.0244
1284.72802734375 0 1991.9385
1320.7340087890625 0 875.86725
1322.750732421875 0 1075.0414
1324.6607666015625 0 686.9213 w 1
1336.771484375 0 3123.667
1337.77392578125 0 7773.136
1338.7733154296875 0 5646.849
1339.7777099609375 0 2424.7698
1344.95849609375 0 644.90857
1353.7254638671875 0 900.5699
1363.753662109375 0 1313.9421 c Ammonia loss 11
1364.7630615234375 0 4778.564
1365.7703857421875 0 3087.6064
1366.763671875 0 1198.984
1369.70263671875 0 761.9223
1380.78271484375 0 5961.022 c 11
1381.7877197265625 0 12965.193
1382.7913818359375 0 9606.157
1383.79296875 0 3369.3108
1384.79931640625 0 1116.0887
1394.69580078125 0 845.21857
1396.7669677734375 0 1187.2101
1406.78955078125 0 789.6726
1407.793701171875 0 1708.3385
1408.7943115234375 0 965.81647
1410.7408447265625 0 2789.0632 z 1
1411.740966796875 0 7566.8296
1412.7408447265625 0 6852.8535
1413.743408203125 0 1905.8448
1414.7318115234375 0 762.05884
1423.7738037109375 0 760.30035
1424.79931640625 0 3982.2925
1425.789306640625 0 3682.2312
1426.7547607421875 0 2309.6 y 1
1427.7640380859375 0 1810.5864
1428.7548828125 0 1101.1227
1451.7882080078125 0 2692.6047
1452.7977294921875 0 3081.327
1453.797607421875 0 1654.8113
1454.7957763671875 0 960.38275
1455.7965087890625 0 1164.1394
1456.7889404296875 0 761.3019
1466.79443359375 0 1428.6344
1467.78515625 0 15814.6875
1468.7957763671875 0 25494.406
1469.80126953125 0 15977.576
1470.8096923828125 0 6994.185
1471.80712890625 0 1754.4061
1472.800537109375 0 767.4871
1481.8189697265625 0 1880.2676
1482.8134765625 0 7075.4814
1483.81201171875 0 6818.661
1484.807373046875 0 5742.303
1485.8094482421875 0 2698.8335
1493.7821044921875 0 7335.6484
1494.78369140625 0 5621.118
1495.785888671875 0 3598.1333
1498.8392333984375 0 1192.9481
1499.837646484375 0 4263.869
1500.841064453125 0 4091.1792
1501.8472900390625 0 1698.8436
1508.7962646484375 0 1186.8866
1509.8060302734375 0 12892.736
1510.8076171875 0 70939.47
1511.810791015625 0 53818.19
1512.8138427734375 0 26619.104
1513.8155517578125 0 8918.035
1514.81591796875 0 3658.4465
1525.816650390625 0 5356.8394
1526.8250732421875 0 23680.74
1527.8328857421875 0 88410.984
1528.8365478515625 0 68505.24
1529.8382568359375 0 33547.066
1530.84033203125 0 10494.18
1531.83740234375 0 5306.145
2154.982421875 0 766.2571

Spectrum Details

|  |  |
| --- | --- |
| Matched peaks? Matched peaks The total absolute number of peaks matched. Additionally in brackets the total fraction of peaks matched and the total number of peaks is shown. | 51 (14.41% of 354) |
| FDR? FDR The false discovery rate estimated for this peptide. It is calculated by matching all theoretical fragments with a non-integer shift with the raw peaks for this spectrum. This is done with 40 different shifts. The resulting percentage is the average number of annotated peaks over the number of annotated peaks with the correct spectrum. | 2.33% |
| Satellite FDR? Satellite FDR See the FDR for details on its calculation. This satellite ion specific FDR only contains the satellite ions (d/w) for I/L/J positions. | ∞ |
| PSM Score? PSM Score The PSM Score as given by Hecklib to this annotated spectrum. It is shown with three significant figures. | 399 |

#### Spectrum 6466? Spectrum 6466 The raw spectrum of this peptide as annotated by Hecklib. The fragments are coloured according to ion type (see legend). Any peaks with a star '\*' as text can be hovered over to see the full details, first the ion type second the mass shift type. By hovering over the amino acids in the peptide or ions in the legend the corresponding peaks are highlighted. By toggling the 'Unassigned' label you can turn the background (unassigned) peaks on or off in the plot. By updating the slider in the Ion legend you can update the spectrum to only show the top X% of the peaks with labels. The top X% means any peak that is within X% of the highest intensity. By dragging in the spectrum you can zoom in to a specific part of the spectrum and use 'Zoom Out' to get back to the original zoom level. The annotation of the spectrum is based on the given sequence in the peptides file and is done with different software so inconsistencies are likely. The peaks are annotated based on the given sequence, with 20 ppm tolerance.

Copy Data

##### Spectrum 6466 (TSV)

###### Preview

```
Loading example...
```

*Click on the button to copy the data to your clipboard.*

Mz MinMz MaxIntensity Max

WidthHeightPeptide font sizePeptide stroke widthSpectrum font sizeSpectrum stroke widthCompact peptide

Ion legend

wxyz

abcd

OtherUnassignedIon^Charge^_Position_Show for top:%

VRQAPGRAJEWVSG

01.83e+43.67e+45.50e+47.34e+4

Zoom Out

y^+1^_2_y^+1^_2_c^+1^_2_y^+1^_3_c^+1^_2_c^+1^_3_c^+2^_7_c^+1^_3_c^+2^_8_c^+3^_12_z^+2^_9_c^+2^_9_c^+3^_13_c^+2^_10_c^+2^_10_z^+1^_5_c^+1^_5_c^+1^_6_c^+2^_11_c^+2^_11_c^+2^_12_c^+2^_12_z^+2^_13_c^+2^_13_c^+2^_13_c^+1^_7_c^+1^_8_z^+1^_8_y^+1^_8_z^+1^_9_c^+1^_9_y^+1^_9_y^+1^_10_c^+1^_10_z^+1^_11_y^+1^_11_z^+1^_12_y^+1^_12_c^+1^_11_c^+1^_12_z^+1^_13_y^+1^_13_

0783156623503133

Fragment Matches Table

Show background peaks

| Position | Ion type | Intensity | mz Theoretical | mz Error (Th) | mz Error (ppm) | Charge | Series Number |
| --- | --- | --- | --- | --- | --- | --- | --- |
| - | - | 777.1 | 120.1 | - | - | 0 | - |
| - | - | 1530 | 130.1 | - | - | 0 | - |
| - | - | 502 | 142.1 | - | - | 0 | - |
| 13 | y | 645.7 | 145.1 | 0.0005717 | 3.941 | +1 | 2 |
| - | - | 784.3 | 146.1 | - | - | 0 | - |
| - | - | 463.7 | 146.8 | - | - | 0 | - |
| - | - | 1222 | 157.1 | - | - | 0 | - |
| - | - | 755.6 | 159.1 | - | - | 0 | - |
| 13 | y | 1.853E+04 | 163.1 | 0.0001389 | 0.8517 | +1 | 2 |
| - | - | 824.1 | 164.1 | - | - | 0 | - |
| - | - | 903.8 | 173.4 | - | - | 0 | - |
| - | - | 1061 | 185.1 | - | - | 0 | - |
| - | - | 955.8 | 187.1 | - | - | 0 | - |
| - | - | 554.3 | 203.1 | - | - | 0 | - |
| - | - | 553.9 | 210.5 | - | - | 0 | - |
| - | - | 725.9 | 214.2 | - | - | 0 | - |
| - | - | 850.2 | 228.2 | - | - | 0 | - |
| - | - | 5669 | 239.2 | - | - | 0 | - |
| - | - | 607.1 | 240.2 | - | - | 0 | - |
| - | - | 599.8 | 256 | - | - | 0 | - |
| 2 | c | 2321 | 256.2 | 1.307E-05 | 0.05101 | +1 | 2 |
| - | - | 523.5 | 257.6 | - | - | 0 | - |
| 12 | y | 1035 | 262.1 | 3.772E-05 | 0.1439 | +1 | 3 |
| - | - | 622.3 | 271.1 | - | - | 0 | - |
| 2 | c | 7372 | 273.2 | 4.239E-05 | 0.1552 | +1 | 2 |
| - | - | 1048 | 274.2 | - | - | 0 | - |
| - | - | 2401 | 299.2 | - | - | 0 | - |
| - | - | 590.2 | 331.1 | - | - | 0 | - |
| 3 | c | 755.3 | 384.2 | 0.000363 | 0.9448 | +1 | 3 |
| 7 | c | 7740 | 391.7 | 0.0002889 | 0.7374 | +2 | 7 |
| - | - | 1886 | 392.2 | - | - | 0 | - |
| 3 | c | 3.093E+04 | 401.3 | 3.488E-05 | 0.08692 | +1 | 3 |
| - | - | 5929 | 402.3 | - | - | 0 | - |
| - | - | 693.5 | 403.3 | - | - | 0 | - |
| - | - | 904.7 | 412.9 | - | - | 0 | - |
| 8 | c | 7907 | 427.3 | 4.461E-05 | 0.1044 | +2 | 8 |
| - | - | 2256 | 427.8 | - | - | 0 | - |
| - | - | 1413 | 430.2 | - | - | 0 | - |
| 12 | c | 1452 | 455.3 | 0.001134 | 2.49 | +3 | 12 |
| - | - | 1020 | 461.8 | - | - | 0 | - |
| - | - | 673 | 462.3 | - | - | 0 | - |
| 6 | z | 543.6 | 479.7 | 0.0009152 | 1.908 | +2 | 9 |
| - | - | 4847 | 483.3 | - | - | 0 | - |
| 9 | c | 2.745E+04 | 483.8 | 0.0002699 | 0.5578 | +2 | 9 |
| 13 | c | 2148 | 484.3 | 0.000723 | 1.493 | +3 | 13 |
| - | - | 1.305E+04 | 484.3 | - | - | 0 | - |
| - | - | 581.3 | 484.6 | - | - | 0 | - |
| - | - | 4047 | 484.8 | - | - | 0 | - |
| - | - | 1206 | 484.9 | - | - | 0 | - |
| - | - | 545.2 | 490.1 | - | - | 0 | - |
| - | - | 4386 | 496.8 | - | - | 0 | - |
| - | - | 2163 | 497.3 | - | - | 0 | - |
| - | - | 637.7 | 497.8 | - | - | 0 | - |
| - | - | 581.2 | 498.7 | - | - | 0 | - |
| - | - | 1656 | 503.3 | - | - | 0 | - |
| - | - | 1383 | 503.6 | - | - | 0 | - |
| - | - | 823.1 | 509.2 | - | - | 0 | - |
| - | - | 6435 | 509.3 | - | - | 0 | - |
| - | - | 2089 | 510.3 | - | - | 0 | - |
| - | - | 717 | 518.3 | - | - | 0 | - |
| - | - | 775.2 | 524.3 | - | - | 0 | - |
| - | - | 3591 | 526.3 | - | - | 0 | - |
| - | - | 3433 | 526.8 | - | - | 0 | - |
| - | - | 1554 | 527.3 | - | - | 0 | - |
| 10 | c | 5309 | 539.8 | 0.0004793 | 0.888 | +2 | 10 |
| - | - | 3784 | 540.3 | - | - | 0 | - |
| - | - | 2097 | 540.8 | - | - | 0 | - |
| - | - | 741.3 | 541.3 | - | - | 0 | - |
| - | - | 8883 | 547.8 | - | - | 0 | - |
| 10 | c | 3.068E+04 | 548.3 | 0.0004482 | 0.8175 | +2 | 10 |
| - | - | 2.009E+04 | 548.8 | - | - | 0 | - |
| - | - | 6955 | 549.3 | - | - | 0 | - |
| - | - | 1471 | 549.8 | - | - | 0 | - |
| 10 | z | 852.3 | 561.2 | 0.001901 | 3.388 | +1 | 5 |
| 5 | c | 8326 | 569.4 | 0.0004263 | 0.7487 | +1 | 5 |
| - | - | 2737 | 570.4 | - | - | 0 | - |
| - | - | 728.6 | 574.8 | - | - | 0 | - |
| - | - | 983.8 | 582.4 | - | - | 0 | - |
| - | - | 595.2 | 603.9 | - | - | 0 | - |
| - | - | 4094 | 611.3 | - | - | 0 | - |
| - | - | 3043 | 611.8 | - | - | 0 | - |
| - | - | 1840 | 612.3 | - | - | 0 | - |
| - | - | 3258 | 617.9 | - | - | 0 | - |
| - | - | 1879 | 618.4 | - | - | 0 | - |
| - | - | 5436 | 618.9 | - | - | 0 | - |
| - | - | 4296 | 619.4 | - | - | 0 | - |
| - | - | 3353 | 619.9 | - | - | 0 | - |
| - | - | 1885 | 620.4 | - | - | 0 | - |
| 6 | c | 4.619E+04 | 626.4 | 3.939E-05 | 0.06289 | +1 | 6 |
| - | - | 1.599E+04 | 627.4 | - | - | 0 | - |
| - | - | 4413 | 628.4 | - | - | 0 | - |
| 11 | c | 2.004E+04 | 632.8 | 0.0002799 | 0.4422 | +2 | 11 |
| - | - | 1.737E+04 | 633.4 | - | - | 0 | - |
| - | - | 6974 | 633.9 | - | - | 0 | - |
| - | - | 1428 | 634.4 | - | - | 0 | - |
| - | - | 698.2 | 639.4 | - | - | 0 | - |
| - | - | 1377 | 640.9 | - | - | 0 | - |
| 11 | c | 1.712E+04 | 641.4 | 0.0007981 | 1.244 | +2 | 11 |
| - | - | 1.431E+04 | 641.9 | - | - | 0 | - |
| - | - | 6752 | 642.4 | - | - | 0 | - |
| - | - | 1077 | 642.9 | - | - | 0 | - |
| - | - | 615.8 | 652.4 | - | - | 0 | - |
| - | - | 868.2 | 661.4 | - | - | 0 | - |
| - | - | 1183 | 661.9 | - | - | 0 | - |
| - | - | 8928 | 668.4 | - | - | 0 | - |
| - | - | 1.226E+04 | 668.9 | - | - | 0 | - |
| - | - | 7352 | 669.4 | - | - | 0 | - |
| - | - | 3694 | 669.9 | - | - | 0 | - |
| - | - | 1739 | 670.4 | - | - | 0 | - |
| - | - | 1055 | 676.9 | - | - | 0 | - |
| - | - | 1221 | 677.4 | - | - | 0 | - |
| 12 | c | 3.542E+04 | 682.4 | 0.0006734 | 0.9868 | +2 | 12 |
| - | - | 2.612E+04 | 682.9 | - | - | 0 | - |
| - | - | 1.228E+04 | 683.4 | - | - | 0 | - |
| - | - | 3512 | 683.9 | - | - | 0 | - |
| - | - | 1022 | 684.4 | - | - | 0 | - |
| - | - | 1180 | 690.4 | - | - | 0 | - |
| 12 | c | 2.946E+04 | 690.9 | 0.001008 | 1.46 | +2 | 12 |
| - | - | 1.994E+04 | 691.4 | - | - | 0 | - |
| - | - | 1.117E+04 | 691.9 | - | - | 0 | - |
| - | - | 2484 | 692.4 | - | - | 0 | - |
| - | - | 1061 | 692.9 | - | - | 0 | - |
| - | - | 1.058E+04 | 698.4 | - | - | 0 | - |
| - | - | 1608 | 698.9 | - | - | 0 | - |
| - | - | 1025 | 699.4 | - | - | 0 | - |
| 2 | z | 4225 | 705.9 | 0.0009185 | 1.301 | +2 | 13 |
| - | - | 3518 | 706.4 | - | - | 0 | - |
| - | - | 1009 | 706.9 | - | - | 0 | - |
| - | - | 639.5 | 707.4 | - | - | 0 | - |
| - | - | 1628 | 712.4 | - | - | 0 | - |
| - | - | 1506 | 712.9 | - | - | 0 | - |
| - | - | 1167 | 713.4 | - | - | 0 | - |
| - | - | 932.4 | 714.4 | - | - | 0 | - |
| - | - | 660.7 | 718.4 | - | - | 0 | - |
| - | - | 1819 | 719.4 | - | - | 0 | - |
| - | - | 1614 | 719.9 | - | - | 0 | - |
| - | - | 1366 | 720.4 | - | - | 0 | - |
| 13 | c | 1447 | 725.9 | 0.001684 | 2.32 | +2 | 13 |
| - | - | 2672 | 726.4 | - | - | 0 | - |
| - | - | 2245 | 726.9 | - | - | 0 | - |
| - | - | 1819 | 727.9 | - | - | 0 | - |
| - | - | 2630 | 728.4 | - | - | 0 | - |
| - | - | 885 | 728.9 | - | - | 0 | - |
| - | - | 710 | 729.4 | - | - | 0 | - |
| - | - | 618.7 | 732.9 | - | - | 0 | - |
| - | - | 1000 | 733.4 | - | - | 0 | - |
| - | - | 794 | 733.9 | - | - | 0 | - |
| 13 | c | 2.431E+04 | 734.4 | 0.001581 | 2.152 | +2 | 13 |
| - | - | 2.109E+04 | 734.9 | - | - | 0 | - |
| - | - | 9025 | 735.4 | - | - | 0 | - |
| - | - | 4217 | 735.9 | - | - | 0 | - |
| - | - | 3250 | 736.4 | - | - | 0 | - |
| - | - | 1211 | 736.9 | - | - | 0 | - |
| - | - | 1184 | 740.9 | - | - | 0 | - |
| - | - | 5826 | 741.4 | - | - | 0 | - |
| - | - | 5349 | 741.9 | - | - | 0 | - |
| - | - | 1.471E+04 | 742.4 | - | - | 0 | - |
| - | - | 1.186E+04 | 742.9 | - | - | 0 | - |
| - | - | 5720 | 743.4 | - | - | 0 | - |
| - | - | 1973 | 743.9 | - | - | 0 | - |
| - | - | 829.4 | 744.4 | - | - | 0 | - |
| - | - | 1422 | 747.4 | - | - | 0 | - |
| - | - | 3217 | 747.9 | - | - | 0 | - |
| - | - | 3029 | 748.4 | - | - | 0 | - |
| - | - | 1507 | 748.9 | - | - | 0 | - |
| - | - | 812.2 | 749.9 | - | - | 0 | - |
| - | - | 744.7 | 754.4 | - | - | 0 | - |
| - | - | 3319 | 754.9 | - | - | 0 | - |
| - | - | 7.266E+04 | 755.4 | - | - | 0 | - |
| - | - | 6.19E+04 | 755.9 | - | - | 0 | - |
| - | - | 2.979E+04 | 756.4 | - | - | 0 | - |
| - | - | 9483 | 756.9 | - | - | 0 | - |
| - | - | 4243 | 757.4 | - | - | 0 | - |
| - | - | 2138 | 762.9 | - | - | 0 | - |
| - | - | 2.774E+04 | 763.4 | - | - | 0 | - |
| - | - | 7.095E+04 | 763.9 | - | - | 0 | - |
| - | - | 5.377E+04 | 764.4 | - | - | 0 | - |
| - | - | 2.542E+04 | 764.9 | - | - | 0 | - |
| - | - | 6838 | 765.4 | - | - | 0 | - |
| - | - | 1732 | 765.9 | - | - | 0 | - |
| 7 | c | 3598 | 782.5 | 0.0007476 | 0.9554 | +1 | 7 |
| - | - | 1065 | 783.5 | - | - | 0 | - |
| - | - | 907.9 | 817.5 | - | - | 0 | - |
| 8 | c | 1924 | 853.5 | 0.00179 | 2.097 | +1 | 8 |
| 7 | z | 3.763E+04 | 901.5 | 0.001473 | 1.634 | +1 | 8 |
| - | - | 1.969E+04 | 902.5 | - | - | 0 | - |
| - | - | 5622 | 903.5 | - | - | 0 | - |
| - | - | 1696 | 904.5 | - | - | 0 | - |
| 7 | y | 2315 | 917.5 | 0.0002384 | 0.2599 | +1 | 8 |
| - | - | 983.9 | 918.5 | - | - | 0 | - |
| 6 | z | 5751 | 958.5 | 0.001147 | 1.197 | +1 | 9 |
| - | - | 3667 | 959.5 | - | - | 0 | - |
| - | - | 1105 | 960.5 | - | - | 0 | - |
| 9 | c | 2772 | 966.6 | 0.003761 | 3.891 | +1 | 9 |
| - | - | 1225 | 967.6 | - | - | 0 | - |
| 6 | y | 1105 | 974.5 | 0.004429 | 4.545 | +1 | 9 |
| - | - | 1400 | 975.5 | - | - | 0 | - |
| - | - | 697.3 | 1003 | - | - | 0 | - |
| - | - | 749.5 | 1019 | - | - | 0 | - |
| - | - | 1211 | 1052 | - | - | 0 | - |
| - | - | 3234 | 1053 | - | - | 0 | - |
| - | - | 1722 | 1054 | - | - | 0 | - |
| 5 | y | 1273 | 1072 | 0.0069 | 6.439 | +1 | 10 |
| - | - | 1409 | 1095 | - | - | 0 | - |
| 10 | c | 1.297E+04 | 1096 | 0.002043 | 1.865 | +1 | 10 |
| - | - | 9387 | 1097 | - | - | 0 | - |
| - | - | 2698 | 1098 | - | - | 0 | - |
| - | - | 821.3 | 1110 | - | - | 0 | - |
| 4 | z | 2.98E+04 | 1127 | 0.002096 | 1.861 | +1 | 11 |
| - | - | 1.829E+04 | 1128 | - | - | 0 | - |
| - | - | 8296 | 1129 | - | - | 0 | - |
| - | - | 2091 | 1130 | - | - | 0 | - |
| - | - | 820.4 | 1131 | - | - | 0 | - |
| 4 | y | 3541 | 1143 | 0.004341 | 3.799 | +1 | 11 |
| - | - | 2397 | 1144 | - | - | 0 | - |
| - | - | 597.6 | 1224 | - | - | 0 | - |
| - | - | 631.3 | 1240 | - | - | 0 | - |
| 3 | z | 2.087E+04 | 1255 | 0.003423 | 2.728 | +1 | 12 |
| - | - | 1.408E+04 | 1256 | - | - | 0 | - |
| - | - | 6862 | 1257 | - | - | 0 | - |
| - | - | 1650 | 1258 | - | - | 0 | - |
| - | - | 1918 | 1266 | - | - | 0 | - |
| - | - | 1321 | 1267 | - | - | 0 | - |
| - | - | 688.7 | 1268 | - | - | 0 | - |
| 3 | y | 1974 | 1271 | 0.008719 | 6.862 | +1 | 12 |
| - | - | 1526 | 1272 | - | - | 0 | - |
| 11 | c | 5924 | 1282 | 0.002498 | 1.949 | +1 | 11 |
| - | - | 6228 | 1283 | - | - | 0 | - |
| - | - | 3994 | 1284 | - | - | 0 | - |
| - | - | 899 | 1285 | - | - | 0 | - |
| - | - | 823.3 | 1321 | - | - | 0 | - |
| - | - | 1614 | 1337 | - | - | 0 | - |
| - | - | 4851 | 1338 | - | - | 0 | - |
| - | - | 4571 | 1339 | - | - | 0 | - |
| - | - | 1287 | 1340 | - | - | 0 | - |
| - | - | 1183 | 1341 | - | - | 0 | - |
| - | - | 2498 | 1365 | - | - | 0 | - |
| - | - | 1978 | 1366 | - | - | 0 | - |
| - | - | 791.4 | 1367 | - | - | 0 | - |
| 12 | c | 5036 | 1381 | 0.002675 | 1.937 | +1 | 12 |
| - | - | 9897 | 1382 | - | - | 0 | - |
| - | - | 7173 | 1383 | - | - | 0 | - |
| - | - | 2718 | 1384 | - | - | 0 | - |
| - | - | 1085 | 1407 | - | - | 0 | - |
| - | - | 1141 | 1408 | - | - | 0 | - |
| - | - | 889.6 | 1410 | - | - | 0 | - |
| 2 | z | 1489 | 1411 | 0.004436 | 3.145 | +1 | 13 |
| - | - | 5610 | 1412 | - | - | 0 | - |
| - | - | 4992 | 1413 | - | - | 0 | - |
| - | - | 2075 | 1414 | - | - | 0 | - |
| - | - | 1208 | 1415 | - | - | 0 | - |
| - | - | 823.5 | 1424 | - | - | 0 | - |
| - | - | 2837 | 1425 | - | - | 0 | - |
| - | - | 2567 | 1426 | - | - | 0 | - |
| 2 | y | 1009 | 1427 | 0.005892 | 4.13 | +1 | 13 |
| - | - | 851.2 | 1428 | - | - | 0 | - |
| - | - | 2278 | 1452 | - | - | 0 | - |
| - | - | 2061 | 1453 | - | - | 0 | - |
| - | - | 1333 | 1454 | - | - | 0 | - |
| - | - | 812 | 1467 | - | - | 0 | - |
| - | - | 9893 | 1468 | - | - | 0 | - |
| - | - | 1.844E+04 | 1469 | - | - | 0 | - |
| - | - | 1.11E+04 | 1470 | - | - | 0 | - |
| - | - | 4721 | 1471 | - | - | 0 | - |
| - | - | 1906 | 1472 | - | - | 0 | - |
| - | - | 5127 | 1483 | - | - | 0 | - |
| - | - | 4416 | 1484 | - | - | 0 | - |
| - | - | 3799 | 1485 | - | - | 0 | - |
| - | - | 1756 | 1486 | - | - | 0 | - |
| - | - | 4788 | 1494 | - | - | 0 | - |
| - | - | 3845 | 1495 | - | - | 0 | - |
| - | - | 2375 | 1496 | - | - | 0 | - |
| - | - | 1035 | 1497 | - | - | 0 | - |
| - | - | 844.7 | 1499 | - | - | 0 | - |
| - | - | 3208 | 1500 | - | - | 0 | - |
| - | - | 3095 | 1501 | - | - | 0 | - |
| - | - | 1545 | 1502 | - | - | 0 | - |
| - | - | 6945 | 1510 | - | - | 0 | - |
| - | - | 4.903E+04 | 1511 | - | - | 0 | - |
| - | - | 3.821E+04 | 1512 | - | - | 0 | - |
| - | - | 1.876E+04 | 1513 | - | - | 0 | - |
| - | - | 6332 | 1514 | - | - | 0 | - |
| - | - | 3824 | 1515 | - | - | 0 | - |
| - | - | 3184 | 1526 | - | - | 0 | - |
| - | - | 1.471E+04 | 1527 | - | - | 0 | - |
| - | - | 6.299E+04 | 1528 | - | - | 0 | - |
| - | - | 5.203E+04 | 1529 | - | - | 0 | - |
| - | - | 2.545E+04 | 1530 | - | - | 0 | - |
| - | - | 7831 | 1531 | - | - | 0 | - |
| - | - | 4592 | 1532 | - | - | 0 | - |
| - | - | 725.6 | 3017 | - | - | 0 | - |
| - | - | 689.6 | 3069 | - | - | 0 | - |
| - | - | 688.5 | 3102 | - | - | 0 | - |

m/z Charge Intensity FragmentType MassShift Position
120.08078002929688 0 777.07837
130.06539916992188 0 1529.556
142.09799194335938 0 502.0091
145.06134033203125 0 645.6633 y Water loss 12
146.09283447265625 0 784.25726
146.76406860351562 0 463.67682
157.1088104248047 0 1222.2484
159.09178161621094 0 755.5621
163.07147216796875 0 18530.465 y 12
164.0754852294922 0 824.0724
173.43751525878906 0 903.82794
185.1037139892578 0 1061.3026
187.10751342773438 0 955.7707
203.0918426513672 0 554.2809
210.5455322265625 0 553.9315
214.15505981445312 0 725.8894
228.1816864013672 0 850.21155
239.15028381347656 0 5669.141
240.15350341796875 0 607.0532
255.97564697265625 0 599.8231
256.1767883300781 0 2320.7583 c Ammonia loss 1
257.599609375 0 523.4565
262.13970947265625 0 1034.7881 y 11
271.1322021484375 0 622.3372
273.20330810546875 0 7372.199 c 1
274.20721435546875 0 1048.2263
299.21917724609375 0 2401.3098
331.1397399902344 0 590.1805
384.2350158691406 0 755.25775 c Ammonia loss 2
391.74053955078125 0 7739.7915 c 6
392.2430725097656 0 1886.3864
401.261962890625 0 30926.572 c 2
402.2649230957031 0 5928.675
403.2666320800781 0 693.5405
412.9042663574219 0 904.70807
427.2594299316406 0 7907.484 c 7
427.7606201171875 0 2255.6677
430.18487548828125 0 1412.9703
455.25909423828125 0 1452.3934 c Ammonia loss 11
461.7942810058594 0 1019.98553
462.29583740234375 0 673.03
479.7478942871094 0 543.55286 z 5
483.2972106933594 0 4847.0894
483.8011474609375 0 27451.705 c 8
484.2679138183594 0 2147.5498 c Ammonia loss 12
484.30303955078125 0 13053.013
484.6040344238281 0 581.26544
484.8046569824219 0 4047.124
484.9356384277344 0 1206.2941
490.1033020019531 0 545.15967
496.8094177246094 0 4385.965
497.3102722167969 0 2162.8115
497.81207275390625 0 637.70026
498.7491455078125 0 581.24457
503.2738952636719 0 1656.3722
503.6109924316406 0 1382.8727
509.21697998046875 0 823.1195
509.2601318359375 0 6435.2886
510.2623596191406 0 2088.8762
518.3084716796875 0 716.9986
524.3280029296875 0 775.2158
526.31591796875 0 3590.7488
526.8173217773438 0 3432.7659
527.3190307617188 0 1553.5258
539.8089599609375 0 5309.145 c Ammonia loss 9
540.3107299804688 0 3783.919
540.8135375976562 0 2096.7192
541.3154907226562 0 741.3256
547.818359375 0 8882.822
548.322265625 0 30675.555 c 9
548.8235473632812 0 20091.377
549.3245849609375 0 6954.549
549.8267822265625 0 1470.7036
561.2410278320312 0 852.33374 z 9
569.3513793945312 0 8326.38 c 4
570.3544921875 0 2737.172
574.822998046875 0 728.607
582.362060546875 0 983.8197
603.8574829101562 0 595.2033
611.3457641601562 0 4093.645
611.8470458984375 0 3043.4478
612.3499755859375 0 1839.7621
617.8528442382812 0 3258.1675
618.3566284179688 0 1879.3014
618.8519287109375 0 5436.086
619.3543701171875 0 4296.01
619.8564453125 0 3352.837
620.3583984375 0 1884.8181
626.3732299804688 0 46193.387 c 5
627.375732421875 0 15988.197
628.3782958984375 0 4412.972
632.8488159179688 0 20035.512 c Ammonia loss 10
633.3505249023438 0 17371.576
633.8516845703125 0 6973.548
634.3513793945312 0 1427.7375
639.3804931640625 0 698.1818
640.8570556640625 0 1377.1685
641.361572265625 0 17117.316 c 10
641.8636474609375 0 14310.34
642.3645629882812 0 6752.103
642.8714599609375 0 1076.5321
652.3560791015625 0 615.82904
661.3853149414062 0 868.217
661.8834838867188 0 1183.2429
668.3851928710938 0 8927.582
668.8876342773438 0 12260.979
669.3900146484375 0 7352.3853
669.8934936523438 0 3694.3306
670.3939819335938 0 1738.5652
676.8562622070312 0 1054.9198
677.3549194335938 0 1221.4052
682.3826293945312 0 35417.848 c Ammonia loss 11
682.8843383789062 0 26120.227
683.3853149414062 0 12277.159
683.8865356445312 0 3512.165
684.3800659179688 0 1021.68274
690.3880615234375 0 1180.0436
690.8955688476562 0 29455.314 c 11
691.3972778320312 0 19942.201
691.898681640625 0 11170.645
692.4026489257812 0 2484.4348
692.903564453125 0 1061.1416
698.381591796875 0 10582.668
698.8832397460938 0 1607.6
699.384765625 0 1025.0659
705.8726806640625 0 4225.4863 z 1
706.3721313476562 0 3517.6187
706.8782958984375 0 1009.0697
707.373046875 0 639.53394
712.4052734375 0 1628.2023
712.9078979492188 0 1506.2991
713.4077758789062 0 1167.3375
714.3804321289062 0 932.38257
718.3984985351562 0 660.71344
719.3978881835938 0 1818.6552
719.8886108398438 0 1613.7562
720.3862915039062 0 1365.9562
725.9010009765625 0 1446.9998 c Ammonia loss 12
726.3947143554688 0 2672.4666
726.899658203125 0 2245.3523
727.899658203125 0 1818.9025
728.4041748046875 0 2629.762
728.9036865234375 0 884.95337
729.4007568359375 0 710.03094
732.9002075195312 0 618.73834
733.4147338867188 0 1000.2083
733.9110717773438 0 794.04364
734.4110107421875 0 24305.098 c 12
734.9133911132812 0 21088.227
735.4139404296875 0 9024.954
735.895263671875 0 4217.138
736.3905639648438 0 3250.0251
736.8926391601562 0 1210.8837
740.9099731445312 0 1183.9697
741.409912109375 0 5826.3496
741.9093017578125 0 5348.5835
742.4014282226562 0 14706.968
742.89990234375 0 11856.683
743.4010009765625 0 5719.5273
743.9066772460938 0 1972.8113
744.399658203125 0 829.4336
747.3971557617188 0 1421.595
747.8922729492188 0 3216.878
748.3930053710938 0 3029.1936
748.8985595703125 0 1507.2288
749.9210205078125 0 812.15875
754.3955078125 0 744.7086
754.9119873046875 0 3318.7588
755.4055786132812 0 72656.13
755.90673828125 0 61895.13
756.4077758789062 0 29789.871
756.9090576171875 0 9482.94
757.4088134765625 0 4243.136
762.9071044921875 0 2137.5588
763.4147338867188 0 27743.049
763.917724609375 0 70947.64
764.4193725585938 0 53768.074
764.9202880859375 0 25415.41
765.4204711914062 0 6838.0947
765.919189453125 0 1731.788
782.4736328125 0 3598.0066 c 6
783.4795532226562 0 1065.3616
817.4552612304688 0 907.94135
853.5097045898438 0 1924.2335 c 7
901.4637451171875 0 37627.31 z 6
902.4669189453125 0 19689.895
903.4696044921875 0 5621.839
904.4737548828125 0 1696.206
917.4837036132812 0 2314.6443 y 6
918.4863891601562 0 983.93616
958.4855346679688 0 5751.4004 z 5
959.4886474609375 0 3667.2605
960.48779296875 0 1105.2427
966.591796875 0 2771.641 c 8
967.596923828125 0 1224.5272
974.5009765625 0 1104.8041 y 5
975.50341796875 0 1399.8021
1003.4906005859375 0 697.31177
1018.508056640625 0 749.5151
1051.626220703125 0 1211.4786
1052.6312255859375 0 3233.624
1053.6300048828125 0 1722.2963
1071.55126953125 0 1273.0684 y 4
1094.624267578125 0 1409.4093
1095.6361083984375 0 12973.22 c 9
1096.6396484375 0 9387.472
1097.640869140625 0 2697.6958
1109.5850830078125 0 821.32043
1126.574462890625 0 29799.744 z 3
1127.577880859375 0 18294.338
1128.5789794921875 0 8296.164
1129.5814208984375 0 2090.691
1130.5821533203125 0 820.3978
1142.5909423828125 0 3540.9062 y 3
1143.59326171875 0 2397.2715
1223.677490234375 0 597.569
1239.705810546875 0 631.3028
1254.6317138671875 0 20870.049 z 2
1255.6357421875 0 14083.464
1256.636474609375 0 6862.0586
1257.640380859375 0 1649.8291
1265.6983642578125 0 1918.4764
1266.69921875 0 1321.3721
1267.71142578125 0 688.71906
1270.6451416015625 0 1973.7623 y 2
1271.6488037109375 0 1526.3564
1281.7149658203125 0 5923.7476 c 10
1282.718017578125 0 6228.097
1283.72314453125 0 3994.4856
1284.7183837890625 0 898.9864
1320.7469482421875 0 823.25684
1336.7652587890625 0 1614.0134
1337.775390625 0 4851.2812
1338.772705078125 0 4570.903
1339.77587890625 0 1287.0679
1340.7696533203125 0 1182.5052
1364.7596435546875 0 2498.1362
1365.764404296875 0 1977.6354
1366.765625 0 791.3555
1380.783203125 0 5036.133 c 11
1381.7874755859375 0 9896.803
1382.791259765625 0 7173.3794
1383.7940673828125 0 2717.5176
1406.7886962890625 0 1084.9979
1407.7864990234375 0 1140.5557
1409.7718505859375 0 889.6189
1410.7318115234375 0 1489.1555 z 1
1411.741455078125 0 5610.321
1412.7421875 0 4992.264
1413.7452392578125 0 2074.7317
1414.735595703125 0 1208.084
1423.7869873046875 0 823.50507
1424.798583984375 0 2836.6
1425.7904052734375 0 2566.8433
1426.7608642578125 0 1008.8015 y 1
1427.7572021484375 0 851.177
1451.7872314453125 0 2277.8425
1452.78662109375 0 2060.5435
1453.7994384765625 0 1333.3955
1466.7852783203125 0 812.0086
1467.7843017578125 0 9893.329
1468.798095703125 0 18440.725
1469.801025390625 0 11098.216
1470.8045654296875 0 4721.4966
1471.8193359375 0 1906.1241
1482.81201171875 0 5127.4126
1483.810546875 0 4415.8574
1484.8087158203125 0 3799.4053
1485.8060302734375 0 1755.8296
1493.781982421875 0 4787.942
1494.784423828125 0 3845.1165
1495.791015625 0 2374.5176
1496.7860107421875 0 1035.3174
1498.8509521484375 0 844.7456
1499.8382568359375 0 3207.926
1500.8419189453125 0 3094.9392
1501.8414306640625 0 1544.9459
1509.8046875 0 6944.6353
1510.806396484375 0 49034.996
1511.810302734375 0 38212.37
1512.8116455078125 0 18758.85
1513.8148193359375 0 6332.3525
1514.812255859375 0 3823.8225
1525.8192138671875 0 3184.1162
1526.82421875 0 14713.309
1527.8316650390625 0 62993.785
1528.835693359375 0 52031.43
1529.837158203125 0 25450.193
1530.83935546875 0 7830.84
1531.8326416015625 0 4591.67
3017.045654296875 0 725.6112
3069.453857421875 0 689.6216
3101.715087890625 0 688.4773

Spectrum Details

|  |  |
| --- | --- |
| Matched peaks? Matched peaks The total absolute number of peaks matched. Additionally in brackets the total fraction of peaks matched and the total number of peaks is shown. | 42 (14.33% of 293) |
| FDR? FDR The false discovery rate estimated for this peptide. It is calculated by matching all theoretical fragments with a non-integer shift with the raw peaks for this spectrum. This is done with 40 different shifts. The resulting percentage is the average number of annotated peaks over the number of annotated peaks with the correct spectrum. | 2.27% |
| Satellite FDR? Satellite FDR See the FDR for details on its calculation. This satellite ion specific FDR only contains the satellite ions (d/w) for I/L/J positions. | ∞ |
| PSM Score? PSM Score The PSM Score as given by Hecklib to this annotated spectrum. It is shown with three significant figures. | 314 |

#### Spectrum 6520? Spectrum 6520 The raw spectrum of this peptide as annotated by Hecklib. The fragments are coloured according to ion type (see legend). Any peaks with a star '\*' as text can be hovered over to see the full details, first the ion type second the mass shift type. By hovering over the amino acids in the peptide or ions in the legend the corresponding peaks are highlighted. By toggling the 'Unassigned' label you can turn the background (unassigned) peaks on or off in the plot. By updating the slider in the Ion legend you can update the spectrum to only show the top X% of the peaks with labels. The top X% means any peak that is within X% of the highest intensity. By dragging in the spectrum you can zoom in to a specific part of the spectrum and use 'Zoom Out' to get back to the original zoom level. The annotation of the spectrum is based on the given sequence in the peptides file and is done with different software so inconsistencies are likely. The peaks are annotated based on the given sequence, with 20 ppm tolerance.

Copy Data

##### Spectrum 6520 (TSV)

###### Preview

```
Loading example...
```

*Click on the button to copy the data to your clipboard.*

Mz MinMz MaxIntensity Max

WidthHeightPeptide font sizePeptide stroke widthSpectrum font sizeSpectrum stroke widthCompact peptide

Ion legend

wxyz

abcd

OtherUnassignedIon^Charge^_Position_Show for top:%

VRQAPGRAJEWVSG

01.38e+42.76e+44.14e+45.52e+4

Zoom Out

y^+1^_2_c^+1^_2_c^+1^_2_c^+1^_3_c^+2^_7_c^+1^_3_c^+2^_8_c^+3^_12_c^+2^_9_c^+3^_13_c^+2^_10_c^+2^_10_c^+1^_5_c^+1^_6_c^+2^_11_c^+2^_11_c^+2^_12_c^+2^_12_z^+2^_13_c^+2^_13_c^+2^_13_c^+1^_7_c^+1^_8_z^+1^_8_y^+1^_8_z^+1^_9_c^+1^_9_y^+1^_9_y^+1^_10_c^+1^_10_z^+1^_11_y^+1^_11_z^+1^_12_y^+1^_12_c^+1^_11_c^+1^_12_z^+1^_13_y^+1^_13_

0853170625593411

Fragment Matches Table

Show background peaks

| Position | Ion type | Intensity | mz Theoretical | mz Error (Th) | mz Error (ppm) | Charge | Series Number |
| --- | --- | --- | --- | --- | --- | --- | --- |
| - | - | 913.2 | 120.1 | - | - | 0 | - |
| - | - | 1556 | 130.1 | - | - | 0 | - |
| - | - | 1555 | 136.1 | - | - | 0 | - |
| - | - | 469.8 | 137 | - | - | 0 | - |
| - | - | 648.8 | 140.1 | - | - | 0 | - |
| - | - | 389.2 | 140.4 | - | - | 0 | - |
| - | - | 455.7 | 142.1 | - | - | 0 | - |
| - | - | 387.5 | 143.2 | - | - | 0 | - |
| - | - | 377.1 | 143.5 | - | - | 0 | - |
| - | - | 472.4 | 144.8 | - | - | 0 | - |
| - | - | 429 | 147 | - | - | 0 | - |
| - | - | 788.4 | 148.9 | - | - | 0 | - |
| - | - | 1081 | 157.1 | - | - | 0 | - |
| - | - | 1083 | 159.1 | - | - | 0 | - |
| 13 | y | 1.33E+04 | 163.1 | 7.473E-05 | 0.4583 | +1 | 2 |
| - | - | 1638 | 173.4 | - | - | 0 | - |
| - | - | 756.7 | 187.1 | - | - | 0 | - |
| - | - | 1352 | 219.1 | - | - | 0 | - |
| - | - | 468.4 | 222.6 | - | - | 0 | - |
| - | - | 506.7 | 233.1 | - | - | 0 | - |
| - | - | 526.9 | 235.1 | - | - | 0 | - |
| - | - | 3589 | 239.2 | - | - | 0 | - |
| - | - | 743.6 | 247.1 | - | - | 0 | - |
| - | - | 497.3 | 249.8 | - | - | 0 | - |
| 2 | c | 2139 | 256.2 | 0.0003182 | 1.242 | +1 | 2 |
| - | - | 654.1 | 263.1 | - | - | 0 | - |
| - | - | 474.1 | 266.4 | - | - | 0 | - |
| - | - | 474.2 | 266.4 | - | - | 0 | - |
| - | - | 594.9 | 270.6 | - | - | 0 | - |
| 2 | c | 4862 | 273.2 | 0.0003781 | 1.384 | +1 | 2 |
| - | - | 862.4 | 274.2 | - | - | 0 | - |
| - | - | 478.8 | 281.3 | - | - | 0 | - |
| - | - | 513.7 | 298.3 | - | - | 0 | - |
| - | - | 2116 | 299.2 | - | - | 0 | - |
| - | - | 558.4 | 346.2 | - | - | 0 | - |
| - | - | 470.7 | 348.7 | - | - | 0 | - |
| - | - | 472.9 | 361.1 | - | - | 0 | - |
| - | - | 576.8 | 362.2 | - | - | 0 | - |
| 3 | c | 632.6 | 384.2 | 0.0007356 | 1.914 | +1 | 3 |
| - | - | 889.7 | 391.2 | - | - | 0 | - |
| 7 | c | 5866 | 391.7 | 0.0003804 | 0.9711 | +2 | 7 |
| - | - | 2339 | 392.2 | - | - | 0 | - |
| - | - | 808 | 392.7 | - | - | 0 | - |
| - | - | 646.9 | 397.1 | - | - | 0 | - |
| - | - | 607.4 | 397.9 | - | - | 0 | - |
| 3 | c | 2.233E+04 | 401.3 | 0.0001482 | 0.3694 | +1 | 3 |
| - | - | 3790 | 402.3 | - | - | 0 | - |
| - | - | 1817 | 419.2 | - | - | 0 | - |
| - | - | 729.6 | 422.6 | - | - | 0 | - |
| - | - | 709.1 | 426.8 | - | - | 0 | - |
| 8 | c | 4361 | 427.3 | 0.0007794 | 1.824 | +2 | 8 |
| - | - | 1879 | 427.8 | - | - | 0 | - |
| - | - | 627.9 | 429.1 | - | - | 0 | - |
| - | - | 715.4 | 446.7 | - | - | 0 | - |
| 12 | c | 638.4 | 455.3 | 0.002651 | 5.822 | +3 | 12 |
| - | - | 543.3 | 462.3 | - | - | 0 | - |
| - | - | 1034 | 475.8 | - | - | 0 | - |
| - | - | 609.5 | 481.9 | - | - | 0 | - |
| - | - | 4715 | 483.3 | - | - | 0 | - |
| 9 | c | 2.107E+04 | 483.8 | 0.0004835 | 0.9993 | +2 | 9 |
| 13 | c | 2681 | 484.3 | 0.001059 | 2.186 | +3 | 13 |
| - | - | 9562 | 484.3 | - | - | 0 | - |
| - | - | 1398 | 484.6 | - | - | 0 | - |
| - | - | 2548 | 484.8 | - | - | 0 | - |
| - | - | 737.8 | 484.9 | - | - | 0 | - |
| - | - | 674.4 | 485.3 | - | - | 0 | - |
| - | - | 2161 | 496.8 | - | - | 0 | - |
| - | - | 1226 | 497.3 | - | - | 0 | - |
| - | - | 1209 | 497.8 | - | - | 0 | - |
| - | - | 1297 | 498.3 | - | - | 0 | - |
| - | - | 990.1 | 498.8 | - | - | 0 | - |
| - | - | 747.1 | 503.3 | - | - | 0 | - |
| - | - | 1118 | 503.6 | - | - | 0 | - |
| - | - | 1087 | 503.9 | - | - | 0 | - |
| - | - | 2915 | 509.3 | - | - | 0 | - |
| - | - | 602.4 | 509.8 | - | - | 0 | - |
| - | - | 1495 | 510.3 | - | - | 0 | - |
| - | - | 2816 | 526.3 | - | - | 0 | - |
| - | - | 2483 | 526.8 | - | - | 0 | - |
| - | - | 1139 | 527.3 | - | - | 0 | - |
| 10 | c | 4633 | 539.8 | 0.0007235 | 1.34 | +2 | 10 |
| - | - | 2607 | 540.3 | - | - | 0 | - |
| - | - | 5447 | 547.8 | - | - | 0 | - |
| 10 | c | 2.389E+04 | 548.3 | 0.0006313 | 1.151 | +2 | 10 |
| - | - | 1.552E+04 | 548.8 | - | - | 0 | - |
| - | - | 4023 | 549.3 | - | - | 0 | - |
| - | - | 842.5 | 566.3 | - | - | 0 | - |
| 5 | c | 5808 | 569.4 | 0.0003042 | 0.5343 | +1 | 5 |
| - | - | 1656 | 570.4 | - | - | 0 | - |
| - | - | 732.4 | 574.8 | - | - | 0 | - |
| - | - | 3166 | 611.3 | - | - | 0 | - |
| - | - | 2558 | 611.8 | - | - | 0 | - |
| - | - | 1301 | 612.3 | - | - | 0 | - |
| - | - | 618.9 | 612.9 | - | - | 0 | - |
| - | - | 2106 | 617.9 | - | - | 0 | - |
| - | - | 1555 | 618.4 | - | - | 0 | - |
| - | - | 4317 | 618.9 | - | - | 0 | - |
| - | - | 1320 | 619.4 | - | - | 0 | - |
| - | - | 2676 | 619.9 | - | - | 0 | - |
| - | - | 1160 | 620.4 | - | - | 0 | - |
| 6 | c | 3.639E+04 | 626.4 | 0.0005277 | 0.8424 | +1 | 6 |
| - | - | 9966 | 627.4 | - | - | 0 | - |
| - | - | 2531 | 628.4 | - | - | 0 | - |
| 11 | c | 1.503E+04 | 632.8 | 0.0009513 | 1.503 | +2 | 11 |
| - | - | 1.139E+04 | 633.4 | - | - | 0 | - |
| - | - | 4544 | 633.9 | - | - | 0 | - |
| - | - | 1593 | 634.4 | - | - | 0 | - |
| - | - | 878.2 | 640.9 | - | - | 0 | - |
| 11 | c | 1.683E+04 | 641.4 | 0.0009812 | 1.53 | +2 | 11 |
| - | - | 9003 | 641.9 | - | - | 0 | - |
| - | - | 5359 | 642.4 | - | - | 0 | - |
| - | - | 613.7 | 642.9 | - | - | 0 | - |
| - | - | 567.7 | 660.9 | - | - | 0 | - |
| - | - | 769.6 | 661.4 | - | - | 0 | - |
| - | - | 1003 | 661.9 | - | - | 0 | - |
| - | - | 1002 | 665.4 | - | - | 0 | - |
| - | - | 6639 | 668.4 | - | - | 0 | - |
| - | - | 9088 | 668.9 | - | - | 0 | - |
| - | - | 5948 | 669.4 | - | - | 0 | - |
| - | - | 3089 | 669.9 | - | - | 0 | - |
| - | - | 721.4 | 677.4 | - | - | 0 | - |
| 12 | c | 2.676E+04 | 682.4 | 0.0007954 | 1.166 | +2 | 12 |
| - | - | 1.946E+04 | 682.9 | - | - | 0 | - |
| - | - | 1.174E+04 | 683.4 | - | - | 0 | - |
| - | - | 3437 | 683.9 | - | - | 0 | - |
| - | - | 667 | 684.4 | - | - | 0 | - |
| - | - | 746.8 | 690.4 | - | - | 0 | - |
| 12 | c | 1.906E+04 | 690.9 | 0.001069 | 1.548 | +2 | 12 |
| - | - | 1.562E+04 | 691.4 | - | - | 0 | - |
| - | - | 9058 | 691.9 | - | - | 0 | - |
| - | - | 1747 | 692.4 | - | - | 0 | - |
| - | - | 1145 | 692.9 | - | - | 0 | - |
| - | - | 8606 | 698.4 | - | - | 0 | - |
| - | - | 1231 | 698.9 | - | - | 0 | - |
| - | - | 759.4 | 699.4 | - | - | 0 | - |
| - | - | 1358 | 705.4 | - | - | 0 | - |
| 2 | z | 3372 | 705.9 | 0.0007294 | 1.033 | +2 | 13 |
| - | - | 2594 | 706.4 | - | - | 0 | - |
| - | - | 1708 | 706.9 | - | - | 0 | - |
| - | - | 2012 | 712.4 | - | - | 0 | - |
| - | - | 2537 | 712.9 | - | - | 0 | - |
| - | - | 950 | 713.4 | - | - | 0 | - |
| - | - | 721.8 | 719.4 | - | - | 0 | - |
| - | - | 1571 | 719.9 | - | - | 0 | - |
| - | - | 761.3 | 720.4 | - | - | 0 | - |
| 13 | c | 1390 | 725.9 | 0.00149 | 2.052 | +2 | 13 |
| - | - | 2829 | 726.4 | - | - | 0 | - |
| - | - | 986.3 | 726.9 | - | - | 0 | - |
| - | - | 797.1 | 727.9 | - | - | 0 | - |
| - | - | 1216 | 728.4 | - | - | 0 | - |
| - | - | 613 | 732 | - | - | 0 | - |
| - | - | 665.7 | 733.4 | - | - | 0 | - |
| 13 | c | 1.866E+04 | 734.4 | 0.001825 | 2.485 | +2 | 13 |
| - | - | 1.287E+04 | 734.9 | - | - | 0 | - |
| - | - | 6065 | 735.4 | - | - | 0 | - |
| - | - | 2408 | 735.9 | - | - | 0 | - |
| - | - | 2534 | 736.4 | - | - | 0 | - |
| - | - | 767.7 | 737.4 | - | - | 0 | - |
| - | - | 4263 | 741.4 | - | - | 0 | - |
| - | - | 4292 | 741.9 | - | - | 0 | - |
| - | - | 9704 | 742.4 | - | - | 0 | - |
| - | - | 8703 | 742.9 | - | - | 0 | - |
| - | - | 4453 | 743.4 | - | - | 0 | - |
| - | - | 1362 | 743.9 | - | - | 0 | - |
| - | - | 1203 | 744.4 | - | - | 0 | - |
| - | - | 907.5 | 747.4 | - | - | 0 | - |
| - | - | 2399 | 747.9 | - | - | 0 | - |
| - | - | 3022 | 748.4 | - | - | 0 | - |
| - | - | 1151 | 748.9 | - | - | 0 | - |
| - | - | 838 | 750.4 | - | - | 0 | - |
| - | - | 752.2 | 754.4 | - | - | 0 | - |
| - | - | 1910 | 754.9 | - | - | 0 | - |
| - | - | 4.646E+04 | 755.4 | - | - | 0 | - |
| - | - | 3.603E+04 | 755.9 | - | - | 0 | - |
| - | - | 2.317E+04 | 756.4 | - | - | 0 | - |
| - | - | 5448 | 756.9 | - | - | 0 | - |
| - | - | 2294 | 757.4 | - | - | 0 | - |
| - | - | 1612 | 762.9 | - | - | 0 | - |
| - | - | 2.132E+04 | 763.4 | - | - | 0 | - |
| - | - | 5.461E+04 | 763.9 | - | - | 0 | - |
| - | - | 3.723E+04 | 764.4 | - | - | 0 | - |
| - | - | 2.08E+04 | 764.9 | - | - | 0 | - |
| - | - | 7533 | 765.4 | - | - | 0 | - |
| - | - | 2260 | 765.9 | - | - | 0 | - |
| - | - | 705.3 | 780.4 | - | - | 0 | - |
| 7 | c | 2226 | 782.5 | 0.0008086 | 1.033 | +1 | 7 |
| 8 | c | 1623 | 853.5 | 0.002705 | 3.169 | +1 | 8 |
| 7 | z | 2.737E+04 | 901.5 | 0.00129 | 1.431 | +1 | 8 |
| - | - | 1.524E+04 | 902.5 | - | - | 0 | - |
| - | - | 4511 | 903.5 | - | - | 0 | - |
| - | - | 1165 | 904.5 | - | - | 0 | - |
| - | - | 656 | 911.5 | - | - | 0 | - |
| 7 | y | 1328 | 917.5 | 0.004694 | 5.116 | +1 | 8 |
| - | - | 1062 | 926.5 | - | - | 0 | - |
| - | - | 752.1 | 927.5 | - | - | 0 | - |
| - | - | 1060 | 928.5 | - | - | 0 | - |
| - | - | 845.3 | 936.5 | - | - | 0 | - |
| - | - | 906.7 | 943.5 | - | - | 0 | - |
| - | - | 1547 | 944.5 | - | - | 0 | - |
| - | - | 2521 | 945.5 | - | - | 0 | - |
| - | - | 769 | 946.5 | - | - | 0 | - |
| 6 | z | 4005 | 958.5 | 0.001208 | 1.26 | +1 | 9 |
| - | - | 2552 | 959.5 | - | - | 0 | - |
| - | - | 1362 | 960.5 | - | - | 0 | - |
| - | - | 942.9 | 961.5 | - | - | 0 | - |
| - | - | 906.5 | 962.5 | - | - | 0 | - |
| 9 | c | 1575 | 966.6 | 0.004616 | 4.775 | +1 | 9 |
| - | - | 1391 | 967.6 | - | - | 0 | - |
| 6 | y | 728.2 | 974.5 | 0.0009502 | 0.9751 | +1 | 9 |
| - | - | 750.8 | 980.5 | - | - | 0 | - |
| - | - | 1020 | 1001 | - | - | 0 | - |
| - | - | 1067 | 1003 | - | - | 0 | - |
| - | - | 998.9 | 1004 | - | - | 0 | - |
| - | - | 1309 | 1017 | - | - | 0 | - |
| - | - | 1490 | 1019 | - | - | 0 | - |
| - | - | 930.2 | 1020 | - | - | 0 | - |
| - | - | 684 | 1020 | - | - | 0 | - |
| - | - | 1215 | 1052 | - | - | 0 | - |
| - | - | 1196 | 1053 | - | - | 0 | - |
| - | - | 1525 | 1054 | - | - | 0 | - |
| 5 | y | 1020 | 1072 | 0.004337 | 4.047 | +1 | 10 |
| 10 | c | 1.132E+04 | 1096 | 0.001555 | 1.419 | +1 | 10 |
| - | - | 6925 | 1097 | - | - | 0 | - |
| - | - | 2432 | 1098 | - | - | 0 | - |
| - | - | 875.7 | 1111 | - | - | 0 | - |
| 4 | z | 2.091E+04 | 1127 | 0.001608 | 1.427 | +1 | 11 |
| - | - | 1.432E+04 | 1128 | - | - | 0 | - |
| - | - | 4431 | 1129 | - | - | 0 | - |
| - | - | 1058 | 1130 | - | - | 0 | - |
| - | - | 725.7 | 1131 | - | - | 0 | - |
| 4 | y | 3361 | 1143 | 0.000923 | 0.8079 | +1 | 11 |
| - | - | 1911 | 1144 | - | - | 0 | - |
| - | - | 661.1 | 1187 | - | - | 0 | - |
| - | - | 1185 | 1239 | - | - | 0 | - |
| 3 | z | 1.463E+04 | 1255 | 0.002324 | 1.853 | +1 | 12 |
| - | - | 1.066E+04 | 1256 | - | - | 0 | - |
| - | - | 4449 | 1257 | - | - | 0 | - |
| - | - | 1142 | 1258 | - | - | 0 | - |
| - | - | 1433 | 1267 | - | - | 0 | - |
| 3 | y | 1432 | 1271 | 0.0001918 | 0.151 | +1 | 12 |
| 11 | c | 4165 | 1282 | 0.002743 | 2.14 | +1 | 11 |
| - | - | 5460 | 1283 | - | - | 0 | - |
| - | - | 2930 | 1284 | - | - | 0 | - |
| - | - | 977.8 | 1285 | - | - | 0 | - |
| - | - | 1641 | 1337 | - | - | 0 | - |
| - | - | 4345 | 1338 | - | - | 0 | - |
| - | - | 2736 | 1339 | - | - | 0 | - |
| - | - | 1596 | 1340 | - | - | 0 | - |
| - | - | 652 | 1354 | - | - | 0 | - |
| - | - | 1534 | 1365 | - | - | 0 | - |
| - | - | 1624 | 1366 | - | - | 0 | - |
| - | - | 789.9 | 1369 | - | - | 0 | - |
| 12 | c | 4056 | 1381 | 0.004018 | 2.91 | +1 | 12 |
| - | - | 6434 | 1382 | - | - | 0 | - |
| - | - | 4423 | 1383 | - | - | 0 | - |
| - | - | 1395 | 1384 | - | - | 0 | - |
| - | - | 768.8 | 1385 | - | - | 0 | - |
| - | - | 901.6 | 1408 | - | - | 0 | - |
| 2 | z | 922.4 | 1411 | 0.01424 | 10.09 | +1 | 13 |
| - | - | 4649 | 1412 | - | - | 0 | - |
| - | - | 2965 | 1413 | - | - | 0 | - |
| - | - | 1265 | 1414 | - | - | 0 | - |
| - | - | 2040 | 1425 | - | - | 0 | - |
| - | - | 1713 | 1426 | - | - | 0 | - |
| 2 | y | 2044 | 1427 | 0.004672 | 3.274 | +1 | 13 |
| - | - | 956.6 | 1428 | - | - | 0 | - |
| - | - | 1502 | 1452 | - | - | 0 | - |
| - | - | 888.4 | 1453 | - | - | 0 | - |
| - | - | 982.4 | 1467 | - | - | 0 | - |
| - | - | 7603 | 1468 | - | - | 0 | - |
| - | - | 1.262E+04 | 1469 | - | - | 0 | - |
| - | - | 9162 | 1470 | - | - | 0 | - |
| - | - | 3740 | 1471 | - | - | 0 | - |
| - | - | 1123 | 1472 | - | - | 0 | - |
| - | - | 2518 | 1483 | - | - | 0 | - |
| - | - | 3645 | 1484 | - | - | 0 | - |
| - | - | 3657 | 1485 | - | - | 0 | - |
| - | - | 1904 | 1486 | - | - | 0 | - |
| - | - | 3950 | 1494 | - | - | 0 | - |
| - | - | 2958 | 1495 | - | - | 0 | - |
| - | - | 2184 | 1496 | - | - | 0 | - |
| - | - | 2489 | 1500 | - | - | 0 | - |
| - | - | 2159 | 1501 | - | - | 0 | - |
| - | - | 5768 | 1510 | - | - | 0 | - |
| - | - | 3.824E+04 | 1511 | - | - | 0 | - |
| - | - | 2.758E+04 | 1512 | - | - | 0 | - |
| - | - | 1.439E+04 | 1513 | - | - | 0 | - |
| - | - | 5198 | 1514 | - | - | 0 | - |
| - | - | 2124 | 1515 | - | - | 0 | - |
| - | - | 2994 | 1526 | - | - | 0 | - |
| - | - | 1.284E+04 | 1527 | - | - | 0 | - |
| - | - | 4.814E+04 | 1528 | - | - | 0 | - |
| - | - | 3.549E+04 | 1529 | - | - | 0 | - |
| - | - | 1.874E+04 | 1530 | - | - | 0 | - |
| - | - | 5621 | 1531 | - | - | 0 | - |
| - | - | 3720 | 1532 | - | - | 0 | - |
| - | - | 745.3 | 2984 | - | - | 0 | - |
| - | - | 780.3 | 3378 | - | - | 0 | - |

m/z Charge Intensity FragmentType MassShift Position
120.08094024658203 0 913.22455
130.06515502929688 0 1556.1877
136.07577514648438 0 1554.9431
137.02395629882812 0 469.79114
140.08203125 0 648.7576
140.44439697265625 0 389.1975
142.0975341796875 0 455.74997
143.1647491455078 0 387.5408
143.50177001953125 0 377.08713
144.7917938232422 0 472.3589
147.04710388183594 0 429.02106
148.94703674316406 0 788.3905
157.1080780029297 0 1080.8828
159.09170532226562 0 1082.6046
163.07125854492188 0 13299.856 y 12
173.43861389160156 0 1637.951
187.10751342773438 0 756.726
219.14926147460938 0 1351.9475
222.6434326171875 0 468.4019
233.12796020507812 0 506.68707
235.14523315429688 0 526.9073
239.15020751953125 0 3588.6245
247.14402770996094 0 743.5973
249.81967163085938 0 497.31763
256.1764831542969 0 2139.2607 c Ammonia loss 1
263.1390075683594 0 654.0993
266.3793640136719 0 474.08386
266.39849853515625 0 474.23584
270.5878601074219 0 594.89874
273.2029724121094 0 4862.066 c 1
274.206787109375 0 862.36926
281.2919616699219 0 478.7718
298.3408508300781 0 513.6785
299.2186279296875 0 2116.3376
346.2091369628906 0 558.3812
348.652587890625 0 470.70975
361.1435546875 0 472.9233
362.20404052734375 0 576.7729
384.2361145019531 0 632.5628 c Ammonia loss 2
391.2351379394531 0 889.6977
391.7404479980469 0 5865.955 c 6
392.2418518066406 0 2338.9746
392.7410583496094 0 808.0159
397.1428527832031 0 646.89233
397.8848876953125 0 607.3577
401.26177978515625 0 22332.633 c 2
402.2638854980469 0 3790.1455
419.2249450683594 0 1816.9448
422.5702819824219 0 729.6488
426.7559509277344 0 709.14
427.25860595703125 0 4361.316 c 7
427.76025390625 0 1879.0692
429.08978271484375 0 627.93494
446.7204284667969 0 715.44073
455.25531005859375 0 638.41797 c Ammonia loss 11
462.2999267578125 0 543.32605
475.7867736816406 0 1033.928
481.9010009765625 0 609.4926
483.2974548339844 0 4715.406
483.8009338378906 0 21067.074 c 8
484.267578125 0 2680.6365 c Ammonia loss 12
484.3026123046875 0 9561.515
484.60296630859375 0 1398.1287
484.8032531738281 0 2548.0571
484.93853759765625 0 737.7572
485.3043212890625 0 674.3512
496.8088684082031 0 2161.2827
497.3099060058594 0 1225.9247
497.8134765625 0 1208.8386
498.2847595214844 0 1296.6561
498.7872009277344 0 990.13934
503.2754821777344 0 747.0906
503.6100158691406 0 1117.8717
503.9465637207031 0 1086.731
509.2615661621094 0 2915.4563
509.7654113769531 0 602.3531
510.2615661621094 0 1495.1068
526.315185546875 0 2815.7788
526.8167724609375 0 2483.04
527.3209228515625 0 1139.3795
539.8087158203125 0 4633.034 c Ammonia loss 9
540.3116455078125 0 2606.7053
547.8184814453125 0 5447.385
548.3220825195312 0 23886.605 c 9
548.8233642578125 0 15519.324
549.3246459960938 0 4023.1926
566.2937622070312 0 842.4664
569.3515014648438 0 5808.2104 c 4
570.3544311523438 0 1656.2257
574.8199462890625 0 732.4159
611.346435546875 0 3165.7498
611.8463745117188 0 2558.3494
612.347412109375 0 1300.8994
612.850341796875 0 618.8777
617.8516845703125 0 2105.8806
618.3555908203125 0 1555.027
618.8519287109375 0 4317.1064
619.3538818359375 0 1319.5337
619.8541870117188 0 2675.896
620.3578491210938 0 1160.1458
626.3727416992188 0 36394.805 c 5
627.3756103515625 0 9965.884
628.3777465820312 0 2530.9023
632.84814453125 0 15030.095 c Ammonia loss 10
633.3502197265625 0 11387.083
633.851318359375 0 4544.4233
634.3522338867188 0 1593.451
640.8594360351562 0 878.1931
641.3613891601562 0 16830.072 c 10
641.863037109375 0 9003.407
642.3644409179688 0 5359.217
642.8695068359375 0 613.7437
660.8724365234375 0 567.6732
661.3794555664062 0 769.61487
661.8834838867188 0 1003.37915
665.3616333007812 0 1001.6641
668.3855590820312 0 6638.859
668.8872680664062 0 9088.079
669.3892822265625 0 5947.552
669.8935546875 0 3088.549
677.3587036132812 0 721.3984
682.3825073242188 0 26760.346 c Ammonia loss 11
682.8840942382812 0 19456.84
683.3855590820312 0 11735.682
683.8880615234375 0 3436.8486
684.38623046875 0 667.0087
690.3863525390625 0 746.79016
690.8955078125 0 19057.854 c 11
691.39697265625 0 15624.601
691.8978881835938 0 9057.938
692.3999633789062 0 1746.8248
692.898193359375 0 1144.7925
698.3807983398438 0 8606.402
698.8839111328125 0 1231.2173
699.3875732421875 0 759.3903
705.3922729492188 0 1358.2289
705.8710327148438 0 3371.7913 z 1
706.3721923828125 0 2594.2368
706.8775634765625 0 1707.6144
712.4053344726562 0 2011.6879
712.906005859375 0 2537.409
713.414794921875 0 949.99255
719.3946533203125 0 721.7733
719.8878173828125 0 1571.2793
720.3868408203125 0 761.28625
725.8978271484375 0 1390.2151 c Ammonia loss 12
726.3959350585938 0 2829.2068
726.9027709960938 0 986.2676
727.8914184570312 0 797.05255
728.4081420898438 0 1215.8386
731.9818115234375 0 612.98926
733.3956909179688 0 665.6973
734.4107666015625 0 18664.492 c 12
734.9119262695312 0 12867.004
735.4125366210938 0 6064.915
735.8928833007812 0 2407.8157
736.3905639648438 0 2533.9368
737.392578125 0 767.7231
741.4094848632812 0 4262.5913
741.908447265625 0 4292.1494
742.3994750976562 0 9703.86
742.89990234375 0 8703.198
743.3995971679688 0 4453.3965
743.8941650390625 0 1362.4226
744.4000244140625 0 1203.1781
747.4036254882812 0 907.51794
747.89453125 0 2398.6707
748.396484375 0 3022.3906
748.8980712890625 0 1151.1982
750.42333984375 0 837.9735
754.3943481445312 0 752.22656
754.9110107421875 0 1909.9957
755.4056396484375 0 46464.11
755.9070434570312 0 36033.926
756.4076538085938 0 23172.137
756.9088134765625 0 5448.28
757.4088745117188 0 2294.3044
762.9110717773438 0 1611.8997
763.4137573242188 0 21321.748
763.91748046875 0 54614.516
764.4193725585938 0 37225.508
764.9205932617188 0 20802.29
765.4212036132812 0 7532.8896
765.91943359375 0 2260.0308
780.4285888671875 0 705.2958
782.4735717773438 0 2225.7195 c 6
853.5087890625 0 1622.8588 c 7
901.4639282226562 0 27372.197 z 6
902.4666748046875 0 15241.814
903.4696044921875 0 4511.319
904.469970703125 0 1165.1573
911.4976806640625 0 656.0407
917.479248046875 0 1327.5271 y 6
926.4896850585938 0 1061.7968
927.4882202148438 0 752.1012
928.4839477539062 0 1059.9595
936.466552734375 0 845.3448
943.4895629882812 0 906.7315
944.494873046875 0 1547.2449
945.5006103515625 0 2520.6675
946.5023193359375 0 769.0073
958.4854736328125 0 4004.8792 z 5
959.488525390625 0 2551.744
960.4898071289062 0 1361.5046
961.49609375 0 942.9265
962.498291015625 0 906.4953
966.5909423828125 0 1575.088 c 8
967.5938110351562 0 1391.1027
974.5044555664062 0 728.1986 y 5
980.4906616210938 0 750.76917
1000.5005493164062 0 1019.9233
1002.5076904296875 0 1066.5531
1003.506103515625 0 998.9441
1016.5159912109375 0 1308.9648
1018.5140991210938 0 1490.4575
1019.5155029296875 0 930.19354
1020.4873657226562 0 684.0482
1051.6300048828125 0 1215.4485
1052.634033203125 0 1195.5533
1053.6328125 0 1525.323
1071.5538330078125 0 1020.27765 y 4
1095.6365966796875 0 11322.913 c 9
1096.637451171875 0 6925.1333
1097.6414794921875 0 2431.6665
1110.60693359375 0 875.6769
1126.574951171875 0 20911.318 z 3
1127.5775146484375 0 14321.963
1128.580078125 0 4431.3335
1129.5831298828125 0 1057.7745
1130.5947265625 0 725.7042
1142.5943603515625 0 3360.9414 y 3
1143.59619140625 0 1911.4226
1187.3980712890625 0 661.1303
1238.707763671875 0 1184.9197
1254.6328125 0 14629.166 z 2
1255.634765625 0 10664.316
1256.637451171875 0 4449.245
1257.6444091796875 0 1141.5044
1266.700439453125 0 1432.7164
1270.654052734375 0 1432.4639 y 2
1281.7147216796875 0 4164.898 c 10
1282.716796875 0 5459.7563
1283.7181396484375 0 2930.4895
1284.7236328125 0 977.80145
1336.7659912109375 0 1641.4429
1337.77587890625 0 4345.0225
1338.7762451171875 0 2735.7808
1339.7840576171875 0 1596.4983
1353.7294921875 0 652.0092
1364.76806640625 0 1533.7352
1365.7677001953125 0 1623.9069
1368.719970703125 0 789.88306
1380.7818603515625 0 4056.4265 c 11
1381.7882080078125 0 6434.4297
1382.791015625 0 4422.9824
1383.7867431640625 0 1394.95
1384.80322265625 0 768.7504
1407.78857421875 0 901.5818
1410.75048828125 0 922.3651 z 1
1411.737548828125 0 4649.002
1412.739990234375 0 2965.2397
1413.7530517578125 0 1264.5343
1424.8031005859375 0 2040.2856
1425.7906494140625 0 1713.4547
1426.7596435546875 0 2044.2781 y 1
1427.7467041015625 0 956.5812
1451.7889404296875 0 1501.6008
1452.784423828125 0 888.38934
1466.777587890625 0 982.4217
1467.785888671875 0 7602.654
1468.796875 0 12624.559
1469.801025390625 0 9161.643
1470.8052978515625 0 3740.4866
1471.8046875 0 1122.7258
1482.8172607421875 0 2517.7263
1483.8128662109375 0 3645.2122
1484.807373046875 0 3656.994
1485.8096923828125 0 1904.3452
1493.77978515625 0 3950.0686
1494.7862548828125 0 2958.2437
1495.79443359375 0 2183.9001
1499.834716796875 0 2489.164
1500.8447265625 0 2159.0476
1509.8062744140625 0 5768.037
1510.807861328125 0 38240.547
1511.8111572265625 0 27581.412
1512.812744140625 0 14391.931
1513.81494140625 0 5197.9717
1514.8074951171875 0 2123.623
1525.8121337890625 0 2993.9019
1526.826171875 0 12837.178
1527.833251953125 0 48143.676
1528.8367919921875 0 35485.484
1529.8387451171875 0 18737.36
1530.8343505859375 0 5621.056
1531.8302001953125 0 3719.9285
2983.838623046875 0 745.30585
3377.6865234375 0 780.25226

Spectrum Details

|  |  |
| --- | --- |
| Matched peaks? Matched peaks The total absolute number of peaks matched. Additionally in brackets the total fraction of peaks matched and the total number of peaks is shown. | 38 (12.75% of 298) |
| FDR? FDR The false discovery rate estimated for this peptide. It is calculated by matching all theoretical fragments with a non-integer shift with the raw peaks for this spectrum. This is done with 40 different shifts. The resulting percentage is the average number of annotated peaks over the number of annotated peaks with the correct spectrum. | 2.44% |
| Satellite FDR? Satellite FDR See the FDR for details on its calculation. This satellite ion specific FDR only contains the satellite ions (d/w) for I/L/J positions. | ∞ |
| PSM Score? PSM Score The PSM Score as given by Hecklib to this annotated spectrum. It is shown with three significant figures. | 266 |

#### Spectrum 6576? Spectrum 6576 The raw spectrum of this peptide as annotated by Hecklib. The fragments are coloured according to ion type (see legend). Any peaks with a star '\*' as text can be hovered over to see the full details, first the ion type second the mass shift type. By hovering over the amino acids in the peptide or ions in the legend the corresponding peaks are highlighted. By toggling the 'Unassigned' label you can turn the background (unassigned) peaks on or off in the plot. By updating the slider in the Ion legend you can update the spectrum to only show the top X% of the peaks with labels. The top X% means any peak that is within X% of the highest intensity. By dragging in the spectrum you can zoom in to a specific part of the spectrum and use 'Zoom Out' to get back to the original zoom level. The annotation of the spectrum is based on the given sequence in the peptides file and is done with different software so inconsistencies are likely. The peaks are annotated based on the given sequence, with 20 ppm tolerance.

Copy Data

##### Spectrum 6576 (TSV)

###### Preview

```
Loading example...
```

*Click on the button to copy the data to your clipboard.*

Mz MinMz MaxIntensity Max

WidthHeightPeptide font sizePeptide stroke widthSpectrum font sizeSpectrum stroke widthCompact peptide

Ion legend

wxyz

abcd

OtherUnassignedIon^Charge^_Position_Show for top:%

VRQAPGRAJEWVSG

09.84e+31.97e+42.95e+43.94e+4

Zoom Out

y^+1^_2_y^+1^_2_c^+1^_2_c^+1^_2_z^+3^_11_c^+2^_7_c^+1^_3_c^+3^_11_c^+2^_8_c^+2^_9_c^+3^_13_c^+2^_10_c^+2^_10_c^+1^_5_c^+1^_6_c^+2^_11_c^+2^_11_c^+2^_11_c^+2^_12_c^+2^_12_z^+2^_13_c^+2^_13_c^+2^_13_c^+1^_7_c^+1^_8_z^+1^_8_y^+1^_8_z^+1^_9_c^+1^_9_y^+1^_9_y^+1^_10_c^+1^_10_z^+1^_11_y^+1^_11_z^+1^_12_y^+1^_12_c^+1^_11_c^+1^_12_

0778155623343112

Fragment Matches Table

Show background peaks

| Position | Ion type | Intensity | mz Theoretical | mz Error (Th) | mz Error (ppm) | Charge | Series Number |
| --- | --- | --- | --- | --- | --- | --- | --- |
| - | - | 587 | 120.1 | - | - | 0 | - |
| - | - | 6292 | 120.1 | - | - | 0 | - |
| - | - | 365.3 | 124.5 | - | - | 0 | - |
| - | - | 1429 | 130.1 | - | - | 0 | - |
| - | - | 445.9 | 130.6 | - | - | 0 | - |
| - | - | 369.1 | 135.8 | - | - | 0 | - |
| - | - | 1360 | 136.1 | - | - | 0 | - |
| 13 | y | 621 | 145.1 | 2.243E-05 | 0.1546 | +1 | 2 |
| - | - | 1252 | 146.1 | - | - | 0 | - |
| - | - | 541.2 | 156.8 | - | - | 0 | - |
| - | - | 554.6 | 157.1 | - | - | 0 | - |
| - | - | 1380 | 159.1 | - | - | 0 | - |
| - | - | 674.2 | 163.1 | - | - | 0 | - |
| 13 | y | 9668 | 163.1 | 0.0001205 | 0.739 | +1 | 2 |
| - | - | 1102 | 165.1 | - | - | 0 | - |
| - | - | 1988 | 173.5 | - | - | 0 | - |
| - | - | 576.3 | 174.1 | - | - | 0 | - |
| - | - | 513.1 | 174.1 | - | - | 0 | - |
| - | - | 2363 | 175.1 | - | - | 0 | - |
| - | - | 474.2 | 211.7 | - | - | 0 | - |
| - | - | 992.8 | 213.2 | - | - | 0 | - |
| - | - | 648.3 | 232 | - | - | 0 | - |
| - | - | 2917 | 239.2 | - | - | 0 | - |
| - | - | 502.3 | 256 | - | - | 0 | - |
| 2 | c | 2468 | 256.2 | 0.0004708 | 1.838 | +1 | 2 |
| - | - | 572.9 | 257.2 | - | - | 0 | - |
| - | - | 1107 | 273.1 | - | - | 0 | - |
| 2 | c | 4317 | 273.2 | 0.0003781 | 1.384 | +1 | 2 |
| - | - | 952.3 | 276.1 | - | - | 0 | - |
| - | - | 1251 | 276.1 | - | - | 0 | - |
| - | - | 2057 | 290.1 | - | - | 0 | - |
| - | - | 1046 | 291.1 | - | - | 0 | - |
| - | - | 1157 | 292.1 | - | - | 0 | - |
| - | - | 4269 | 293.1 | - | - | 0 | - |
| - | - | 599.6 | 299.1 | - | - | 0 | - |
| - | - | 1987 | 299.2 | - | - | 0 | - |
| - | - | 1692 | 306.1 | - | - | 0 | - |
| - | - | 597.8 | 308.1 | - | - | 0 | - |
| - | - | 606.9 | 320.1 | - | - | 0 | - |
| - | - | 675.6 | 325.1 | - | - | 0 | - |
| - | - | 536.8 | 329.1 | - | - | 0 | - |
| - | - | 1658 | 331.1 | - | - | 0 | - |
| - | - | 1130 | 336.1 | - | - | 0 | - |
| - | - | 847.7 | 345.1 | - | - | 0 | - |
| - | - | 1052 | 348.1 | - | - | 0 | - |
| - | - | 1758 | 363.1 | - | - | 0 | - |
| - | - | 747.5 | 364.1 | - | - | 0 | - |
| - | - | 612.9 | 365.1 | - | - | 0 | - |
| - | - | 766 | 366.1 | - | - | 0 | - |
| - | - | 1406 | 366.1 | - | - | 0 | - |
| - | - | 547.4 | 369.6 | - | - | 0 | - |
| - | - | 831.1 | 375.1 | - | - | 0 | - |
| 4 | z | 538.5 | 376.2 | 0.002425 | 6.447 | +3 | 11 |
| - | - | 698.4 | 391.2 | - | - | 0 | - |
| 7 | c | 5039 | 391.7 | 0.0004415 | 1.127 | +2 | 7 |
| - | - | 1550 | 392.2 | - | - | 0 | - |
| - | - | 2082 | 393.2 | - | - | 0 | - |
| 3 | c | 1.791E+04 | 401.3 | 0.0001787 | 0.4455 | +1 | 3 |
| - | - | 4260 | 402.3 | - | - | 0 | - |
| - | - | 890.4 | 403.1 | - | - | 0 | - |
| - | - | 769.9 | 404.1 | - | - | 0 | - |
| - | - | 710.3 | 419.2 | - | - | 0 | - |
| - | - | 5858 | 421.2 | - | - | 0 | - |
| 11 | c | 727.5 | 422.2 | 0.0001707 | 0.4042 | +3 | 11 |
| - | - | 907.5 | 422.9 | - | - | 0 | - |
| 8 | c | 4425 | 427.3 | 0.00023 | 0.5384 | +2 | 8 |
| - | - | 1433 | 427.8 | - | - | 0 | - |
| - | - | 1027 | 430.1 | - | - | 0 | - |
| - | - | 956.4 | 430.2 | - | - | 0 | - |
| - | - | 1609 | 437.2 | - | - | 0 | - |
| - | - | 1933 | 438.2 | - | - | 0 | - |
| - | - | 537.3 | 441.5 | - | - | 0 | - |
| - | - | 793.7 | 447.2 | - | - | 0 | - |
| - | - | 714.5 | 454.2 | - | - | 0 | - |
| - | - | 581 | 455.2 | - | - | 0 | - |
| - | - | 754.4 | 464.2 | - | - | 0 | - |
| - | - | 2689 | 465.2 | - | - | 0 | - |
| - | - | 848.1 | 466.2 | - | - | 0 | - |
| - | - | 606.2 | 481.7 | - | - | 0 | - |
| - | - | 3206 | 483.3 | - | - | 0 | - |
| 9 | c | 1.409E+04 | 483.8 | 0.0003614 | 0.747 | +2 | 9 |
| 13 | c | 1672 | 484.3 | 0.002127 | 4.392 | +3 | 13 |
| - | - | 7694 | 484.3 | - | - | 0 | - |
| - | - | 1703 | 484.6 | - | - | 0 | - |
| - | - | 2245 | 484.8 | - | - | 0 | - |
| - | - | 570.3 | 484.9 | - | - | 0 | - |
| - | - | 1049 | 492.2 | - | - | 0 | - |
| - | - | 2712 | 496.8 | - | - | 0 | - |
| - | - | 1796 | 497.3 | - | - | 0 | - |
| - | - | 666.5 | 503.3 | - | - | 0 | - |
| - | - | 933.9 | 503.6 | - | - | 0 | - |
| - | - | 1592 | 508.2 | - | - | 0 | - |
| - | - | 719.4 | 508.2 | - | - | 0 | - |
| - | - | 588.9 | 509.2 | - | - | 0 | - |
| - | - | 869.1 | 509.2 | - | - | 0 | - |
| - | - | 1830 | 509.3 | - | - | 0 | - |
| - | - | 4894 | 510.2 | - | - | 0 | - |
| - | - | 2160 | 526.3 | - | - | 0 | - |
| 10 | c | 2599 | 539.8 | 0.0006193 | 1.147 | +2 | 10 |
| - | - | 1956 | 540.3 | - | - | 0 | - |
| - | - | 1711 | 540.8 | - | - | 0 | - |
| - | - | 5044 | 547.8 | - | - | 0 | - |
| 10 | c | 1.839E+04 | 548.3 | 0.0003872 | 0.7062 | +2 | 10 |
| - | - | 1.071E+04 | 548.8 | - | - | 0 | - |
| - | - | 4122 | 549.3 | - | - | 0 | - |
| 5 | c | 3994 | 569.4 | 0.0003042 | 0.5343 | +1 | 5 |
| - | - | 1430 | 570.4 | - | - | 0 | - |
| - | - | 1099 | 604.4 | - | - | 0 | - |
| - | - | 649.6 | 605.3 | - | - | 0 | - |
| - | - | 3098 | 611.3 | - | - | 0 | - |
| - | - | 1673 | 611.8 | - | - | 0 | - |
| - | - | 1136 | 612.4 | - | - | 0 | - |
| - | - | 1813 | 617.9 | - | - | 0 | - |
| - | - | 1077 | 618.4 | - | - | 0 | - |
| - | - | 2624 | 618.9 | - | - | 0 | - |
| - | - | 3458 | 619.4 | - | - | 0 | - |
| - | - | 2868 | 619.9 | - | - | 0 | - |
| 6 | c | 2.761E+04 | 626.4 | 0.0002835 | 0.4527 | +1 | 6 |
| - | - | 8020 | 627.4 | - | - | 0 | - |
| - | - | 2085 | 628.4 | - | - | 0 | - |
| 11 | c | 585.7 | 632.4 | 0.01193 | 18.87 | +2 | 11 |
| 11 | c | 9178 | 632.8 | 0.0002799 | 0.4422 | +2 | 11 |
| - | - | 7687 | 633.4 | - | - | 0 | - |
| - | - | 3559 | 633.9 | - | - | 0 | - |
| - | - | 1389 | 634.4 | - | - | 0 | - |
| - | - | 1676 | 640.9 | - | - | 0 | - |
| 11 | c | 1.363E+04 | 641.4 | 0.0004319 | 0.6733 | +2 | 11 |
| - | - | 9722 | 641.9 | - | - | 0 | - |
| - | - | 3836 | 642.4 | - | - | 0 | - |
| - | - | 1258 | 642.9 | - | - | 0 | - |
| - | - | 1187 | 661.9 | - | - | 0 | - |
| - | - | 5139 | 668.4 | - | - | 0 | - |
| - | - | 6791 | 668.9 | - | - | 0 | - |
| - | - | 4515 | 669.4 | - | - | 0 | - |
| - | - | 1838 | 669.9 | - | - | 0 | - |
| - | - | 1327 | 670.4 | - | - | 0 | - |
| - | - | 795.6 | 676.9 | - | - | 0 | - |
| - | - | 825.3 | 677.4 | - | - | 0 | - |
| 12 | c | 2.071E+04 | 682.4 | 0.0006123 | 0.8973 | +2 | 12 |
| - | - | 1.367E+04 | 682.9 | - | - | 0 | - |
| - | - | 7321 | 683.4 | - | - | 0 | - |
| - | - | 2947 | 683.9 | - | - | 0 | - |
| - | - | 1593 | 684.4 | - | - | 0 | - |
| - | - | 686.5 | 690.4 | - | - | 0 | - |
| 12 | c | 1.556E+04 | 690.9 | 0.0007033 | 1.018 | +2 | 12 |
| - | - | 1.42E+04 | 691.4 | - | - | 0 | - |
| - | - | 6138 | 691.9 | - | - | 0 | - |
| - | - | 2550 | 692.4 | - | - | 0 | - |
| - | - | 1047 | 692.9 | - | - | 0 | - |
| - | - | 7226 | 698.4 | - | - | 0 | - |
| - | - | 968.8 | 698.9 | - | - | 0 | - |
| 2 | z | 1949 | 705.9 | 0.0006744 | 0.9554 | +2 | 13 |
| - | - | 1530 | 706.4 | - | - | 0 | - |
| - | - | 770.4 | 706.9 | - | - | 0 | - |
| - | - | 1333 | 712.4 | - | - | 0 | - |
| - | - | 1701 | 712.9 | - | - | 0 | - |
| - | - | 1076 | 719.4 | - | - | 0 | - |
| - | - | 1448 | 719.9 | - | - | 0 | - |
| - | - | 801.1 | 720.4 | - | - | 0 | - |
| 13 | c | 1059 | 725.9 | 0.001795 | 2.473 | +2 | 13 |
| - | - | 1757 | 726.4 | - | - | 0 | - |
| - | - | 681.5 | 726.9 | - | - | 0 | - |
| - | - | 773.6 | 727.9 | - | - | 0 | - |
| - | - | 929 | 733.9 | - | - | 0 | - |
| 13 | c | 1.547E+04 | 734.4 | 0.001031 | 1.404 | +2 | 13 |
| - | - | 9431 | 734.9 | - | - | 0 | - |
| - | - | 5705 | 735.4 | - | - | 0 | - |
| - | - | 1412 | 735.9 | - | - | 0 | - |
| - | - | 697.8 | 736.4 | - | - | 0 | - |
| - | - | 1207 | 740.9 | - | - | 0 | - |
| - | - | 2523 | 741.4 | - | - | 0 | - |
| - | - | 4684 | 741.9 | - | - | 0 | - |
| - | - | 6401 | 742.4 | - | - | 0 | - |
| - | - | 6869 | 742.9 | - | - | 0 | - |
| - | - | 4138 | 743.4 | - | - | 0 | - |
| - | - | 2912 | 747.9 | - | - | 0 | - |
| - | - | 2277 | 748.4 | - | - | 0 | - |
| - | - | 2801 | 754.9 | - | - | 0 | - |
| - | - | 3.768E+04 | 755.4 | - | - | 0 | - |
| - | - | 3.183E+04 | 755.9 | - | - | 0 | - |
| - | - | 1.684E+04 | 756.4 | - | - | 0 | - |
| - | - | 4759 | 756.9 | - | - | 0 | - |
| - | - | 2487 | 757.4 | - | - | 0 | - |
| - | - | 982 | 762.9 | - | - | 0 | - |
| - | - | 1.523E+04 | 763.4 | - | - | 0 | - |
| - | - | 3.856E+04 | 763.9 | - | - | 0 | - |
| - | - | 3.163E+04 | 764.4 | - | - | 0 | - |
| - | - | 1.37E+04 | 764.9 | - | - | 0 | - |
| - | - | 4776 | 765.4 | - | - | 0 | - |
| - | - | 2148 | 765.9 | - | - | 0 | - |
| 7 | c | 1640 | 782.5 | 0.0006562 | 0.8386 | +1 | 7 |
| 8 | c | 828.2 | 853.5 | 0.001018 | 1.193 | +1 | 8 |
| 7 | z | 2.317E+04 | 901.5 | 0.001046 | 1.16 | +1 | 8 |
| - | - | 1.129E+04 | 902.5 | - | - | 0 | - |
| - | - | 3419 | 903.5 | - | - | 0 | - |
| - | - | 693.2 | 904.5 | - | - | 0 | - |
| 7 | y | 1209 | 917.5 | 0.0008488 | 0.9251 | +1 | 8 |
| 6 | z | 2991 | 958.5 | 0.0002925 | 0.3052 | +1 | 9 |
| - | - | 1706 | 959.5 | - | - | 0 | - |
| 9 | c | 1385 | 966.6 | 0.001381 | 1.429 | +1 | 9 |
| - | - | 722.1 | 967.6 | - | - | 0 | - |
| 6 | y | 1363 | 974.5 | 0.0004619 | 0.474 | +1 | 9 |
| - | - | 767.1 | 1015 | - | - | 0 | - |
| - | - | 1263 | 1053 | - | - | 0 | - |
| 5 | y | 832 | 1072 | 0.004703 | 4.389 | +1 | 10 |
| - | - | 677.1 | 1082 | - | - | 0 | - |
| 10 | c | 9767 | 1096 | 0.000456 | 0.4162 | +1 | 10 |
| - | - | 4809 | 1097 | - | - | 0 | - |
| - | - | 1619 | 1098 | - | - | 0 | - |
| 4 | z | 1.747E+04 | 1127 | 0.0006316 | 0.5606 | +1 | 11 |
| - | - | 1.25E+04 | 1128 | - | - | 0 | - |
| - | - | 4920 | 1129 | - | - | 0 | - |
| - | - | 1145 | 1130 | - | - | 0 | - |
| 4 | y | 1493 | 1143 | 0.002388 | 2.09 | +1 | 11 |
| - | - | 1102 | 1144 | - | - | 0 | - |
| - | - | 687.7 | 1159 | - | - | 0 | - |
| - | - | 771.8 | 1240 | - | - | 0 | - |
| 3 | z | 1.282E+04 | 1255 | 0.002446 | 1.95 | +1 | 12 |
| - | - | 9168 | 1256 | - | - | 0 | - |
| - | - | 3779 | 1257 | - | - | 0 | - |
| - | - | 676.1 | 1258 | - | - | 0 | - |
| - | - | 814.1 | 1267 | - | - | 0 | - |
| 3 | y | 641.4 | 1271 | 0.01899 | 14.95 | +1 | 12 |
| - | - | 831.4 | 1272 | - | - | 0 | - |
| 11 | c | 2371 | 1282 | 0.001644 | 1.283 | +1 | 11 |
| - | - | 4454 | 1283 | - | - | 0 | - |
| - | - | 2290 | 1284 | - | - | 0 | - |
| - | - | 1040 | 1321 | - | - | 0 | - |
| - | - | 1082 | 1337 | - | - | 0 | - |
| - | - | 2857 | 1338 | - | - | 0 | - |
| - | - | 2300 | 1339 | - | - | 0 | - |
| - | - | 959.4 | 1365 | - | - | 0 | - |
| - | - | 1187 | 1366 | - | - | 0 | - |
| - | - | 920.5 | 1367 | - | - | 0 | - |
| 12 | c | 3054 | 1381 | 0.003774 | 2.733 | +1 | 12 |
| - | - | 5012 | 1382 | - | - | 0 | - |
| - | - | 3589 | 1383 | - | - | 0 | - |
| - | - | 1461 | 1384 | - | - | 0 | - |
| - | - | 786.3 | 1410 | - | - | 0 | - |
| - | - | 2900 | 1412 | - | - | 0 | - |
| - | - | 2495 | 1413 | - | - | 0 | - |
| - | - | 1854 | 1414 | - | - | 0 | - |
| - | - | 2048 | 1425 | - | - | 0 | - |
| - | - | 1683 | 1426 | - | - | 0 | - |
| - | - | 1095 | 1427 | - | - | 0 | - |
| - | - | 790.3 | 1452 | - | - | 0 | - |
| - | - | 1505 | 1453 | - | - | 0 | - |
| - | - | 5267 | 1468 | - | - | 0 | - |
| - | - | 1.027E+04 | 1469 | - | - | 0 | - |
| - | - | 7046 | 1470 | - | - | 0 | - |
| - | - | 2996 | 1471 | - | - | 0 | - |
| - | - | 1351 | 1472 | - | - | 0 | - |
| - | - | 3697 | 1483 | - | - | 0 | - |
| - | - | 2024 | 1484 | - | - | 0 | - |
| - | - | 2295 | 1485 | - | - | 0 | - |
| - | - | 1430 | 1486 | - | - | 0 | - |
| - | - | 2701 | 1494 | - | - | 0 | - |
| - | - | 1633 | 1495 | - | - | 0 | - |
| - | - | 901.4 | 1496 | - | - | 0 | - |
| - | - | 717.2 | 1499 | - | - | 0 | - |
| - | - | 2608 | 1500 | - | - | 0 | - |
| - | - | 1516 | 1501 | - | - | 0 | - |
| - | - | 877.5 | 1502 | - | - | 0 | - |
| - | - | 5336 | 1510 | - | - | 0 | - |
| - | - | 3.099E+04 | 1511 | - | - | 0 | - |
| - | - | 2.205E+04 | 1512 | - | - | 0 | - |
| - | - | 1.177E+04 | 1513 | - | - | 0 | - |
| - | - | 2632 | 1514 | - | - | 0 | - |
| - | - | 2245 | 1515 | - | - | 0 | - |
| - | - | 2422 | 1526 | - | - | 0 | - |
| - | - | 8477 | 1527 | - | - | 0 | - |
| - | - | 3.896E+04 | 1528 | - | - | 0 | - |
| - | - | 2.792E+04 | 1529 | - | - | 0 | - |
| - | - | 1.233E+04 | 1530 | - | - | 0 | - |
| - | - | 3862 | 1531 | - | - | 0 | - |
| - | - | 4064 | 1532 | - | - | 0 | - |
| - | - | 1026 | 3072 | - | - | 0 | - |
| - | - | 717.2 | 3081 | - | - | 0 | - |

m/z Charge Intensity FragmentType MassShift Position
120.07714080810547 0 586.98114
120.08076477050781 0 6292.1553
124.48892211914062 0 365.32242
130.0650634765625 0 1428.9762
130.58755493164062 0 445.93607
135.83961486816406 0 369.08905
136.0756072998047 0 1360.2015
145.060791015625 0 620.96014 y Water loss 12
146.05992126464844 0 1251.6339
156.84005737304688 0 541.18195
157.10801696777344 0 554.5803
159.09181213378906 0 1379.6682
163.06430053710938 0 674.2013
163.0712127685547 0 9667.947 y 12
165.10203552246094 0 1102.4506
173.4512176513672 0 1988.3132
174.05502319335938 0 576.27075
174.0879669189453 0 513.07513
175.08644104003906 0 2362.7651
211.706298828125 0 474.2453
213.15951538085938 0 992.8427
232.01144409179688 0 648.30457
239.1502685546875 0 2917.4954
255.9583740234375 0 502.32288
256.17633056640625 0 2467.8472 c Ammonia loss 1
257.1783752441406 0 572.92316
273.0863037109375 0 1106.6681
273.2029724121094 0 4317.0146 c 1
276.1104431152344 0 952.3023
276.1338806152344 0 1250.5823
290.1130676269531 0 2056.7375
291.1206970214844 0 1045.6326
292.1295471191406 0 1157.395
293.13671875 0 4269.414
299.13995361328125 0 599.6453
299.2185363769531 0 1987.4146
306.10858154296875 0 1692.4233
308.12298583984375 0 597.825
320.13946533203125 0 606.91644
325.1233825683594 0 675.5828
329.1231689453125 0 536.81165
331.1070556640625 0 1657.8345
336.133544921875 0 1129.5582
345.1184997558594 0 847.71716
348.13427734375 0 1052.1711
363.12939453125 0 1758.4413
364.1278991699219 0 747.47455
365.1379699707031 0 612.8728
366.1179504394531 0 765.9885
366.1448059082031 0 1406.2235
369.5783996582031 0 547.37585
375.144775390625 0 831.0587
376.199462890625 0 538.4876 z 3
391.23822021484375 0 698.39777
391.7403869628906 0 5039.3403 c 6
392.24102783203125 0 1550.0719
393.1561279296875 0 2082.093
401.2617492675781 0 17910.314 c 2
402.2642517089844 0 4259.9937
403.140380859375 0 890.3527
404.12457275390625 0 769.92865
419.168701171875 0 710.3114
421.1503601074219 0 5857.802
422.2349853515625 0 727.53925 c Ammonia loss 10
422.9041748046875 0 907.51636
427.2591552734375 0 4424.5996 c 7
427.76123046875 0 1432.9906
430.0952453613281 0 1027.4287
430.1839294433594 0 956.35767
437.1812744140625 0 1608.7979
438.1891784667969 0 1932.5894
441.5286560058594 0 537.2995
447.16314697265625 0 793.65094
454.1837463378906 0 714.45123
455.1901550292969 0 581.04376
464.1914978027344 0 754.419
465.1715087890625 0 2689.208
466.1731872558594 0 848.0548
481.6947326660156 0 606.21454
483.2972106933594 0 3205.8845
483.8010559082031 0 14089.163 c 8
484.2665100097656 0 1671.771 c Ammonia loss 12
484.302734375 0 7693.922
484.6023254394531 0 1702.5372
484.80517578125 0 2244.6228
484.94134521484375 0 570.32074
492.1871337890625 0 1049.3691
496.8091125488281 0 2712.306
497.3114929199219 0 1795.5028
503.279541015625 0 666.54474
503.6100769042969 0 933.9408
508.1766662597656 0 1591.5443
508.21368408203125 0 719.35956
509.1638488769531 0 588.8919
509.220458984375 0 869.1267
509.26275634765625 0 1830.0946
510.1986999511719 0 4893.673
526.3163452148438 0 2160.3003
539.81005859375 0 2599.0967 c Ammonia loss 9
540.311279296875 0 1956.039
540.8125 0 1711.4407
547.8182983398438 0 5043.82
548.3223266601562 0 18390.727 c 9
548.8234252929688 0 10710.589
549.3244018554688 0 4121.97
569.3515014648438 0 3994.0098 c 4
570.353759765625 0 1429.903
604.3616333007812 0 1099.2008
605.3378295898438 0 649.597
611.3445434570312 0 3098.3247
611.8465576171875 0 1672.8044
612.3501586914062 0 1135.5579
617.8563232421875 0 1813.2809
618.3550415039062 0 1076.7216
618.8516235351562 0 2624.101
619.353271484375 0 3458.1782
619.8560180664062 0 2867.6353
626.3729858398438 0 27607.186 c 5
627.3758544921875 0 8019.589
628.3792724609375 0 2085.3008
632.3690185546875 0 585.71924 c Water loss 10
632.8488159179688 0 9177.66 c Ammonia loss 10
633.3507080078125 0 7687.301
633.8508911132812 0 3558.7417
634.3557739257812 0 1388.6876
640.8569946289062 0 1676.0485
641.3619384765625 0 13633.688 c 10
641.8638305664062 0 9721.587
642.3641967773438 0 3835.9185
642.8655395507812 0 1257.7709
661.8776245117188 0 1186.5712
668.3861083984375 0 5139.27
668.8873901367188 0 6791.3496
669.389404296875 0 4514.9214
669.8939208984375 0 1837.5712
670.3914794921875 0 1327.1089
676.8572998046875 0 795.5796
677.363525390625 0 825.26447
682.3826904296875 0 20713.205 c Ammonia loss 11
682.8841552734375 0 13668.489
683.3856201171875 0 7321.0547
683.8884887695312 0 2946.7275
684.3827514648438 0 1593.0736
690.3917236328125 0 686.5022
690.8958740234375 0 15556.201 c 11
691.3970947265625 0 14197.959
691.89892578125 0 6138.345
692.4017944335938 0 2550.1155
692.8988037109375 0 1046.9585
698.38134765625 0 7226.2603
698.8797607421875 0 968.78064
705.8724365234375 0 1949.3992 z 1
706.373291015625 0 1529.6207
706.874755859375 0 770.3863
712.4058227539062 0 1333.1522
712.9092407226562 0 1700.7076
719.3931884765625 0 1075.9086
719.8925170898438 0 1447.7681
720.4002685546875 0 801.13696
725.8975219726562 0 1058.5277 c Ammonia loss 12
726.3980712890625 0 1756.6577
726.8934326171875 0 681.4533
727.9049682617188 0 773.5553
733.9130859375 0 928.9561
734.4115600585938 0 15468.1875 c 12
734.9130859375 0 9430.566
735.4129638671875 0 5705.3213
735.8880615234375 0 1412.1896
736.3901977539062 0 697.76495
740.9130249023438 0 1206.7665
741.4083251953125 0 2523.0098
741.9100952148438 0 4684.4434
742.399658203125 0 6401.2095
742.9003295898438 0 6868.5127
743.4031372070312 0 4137.508
747.892578125 0 2912.0063
748.3967895507812 0 2277.4624
754.910400390625 0 2801.296
755.4058837890625 0 37678.973
755.906982421875 0 31830.795
756.4081420898438 0 16836.143
756.908447265625 0 4758.758
757.4075927734375 0 2486.5117
762.9119873046875 0 982.014
763.4147338867188 0 15234.771
763.917724609375 0 38556.78
764.4193115234375 0 31629.506
764.9205322265625 0 13700.252
765.4200439453125 0 4776.2085
765.9190673828125 0 2147.8193
782.4750366210938 0 1640.0096 c 6
853.5125122070312 0 828.23486 c 7
901.4641723632812 0 23167.54 z 6
902.4669799804688 0 11285.26
903.4674682617188 0 3419.1167
904.4732055664062 0 693.179
917.4830932617188 0 1208.626 y 6
958.4863891601562 0 2990.604 z 5
959.4810791015625 0 1705.6198
966.5941772460938 0 1385.4069 c 8
967.5963134765625 0 722.1444
974.5049438476562 0 1362.5946 y 5
1014.5269775390625 0 767.0909
1052.6331787109375 0 1263.3549
1071.553466796875 0 832.0461 y 4
1081.63720703125 0 677.141
1095.6376953125 0 9767.0205 c 9
1096.640625 0 4808.938
1097.64111328125 0 1619.0024
1126.575927734375 0 17467.76 z 3
1127.5792236328125 0 12495.754
1128.5810546875 0 4919.9316
1129.590576171875 0 1144.8757
1142.5928955078125 0 1493.085 y 3
1143.5999755859375 0 1101.6302
1158.5809326171875 0 687.6719
1239.6995849609375 0 771.7696
1254.6326904296875 0 12816.426 z 2
1255.6348876953125 0 9167.533
1256.638427734375 0 3778.8906
1257.646484375 0 676.14844
1266.6976318359375 0 814.0889
1270.6728515625 0 641.4427 y 2
1271.658203125 0 831.4179
1281.7158203125 0 2370.737 c 10
1282.7197265625 0 4453.857
1283.7242431640625 0 2289.518
1320.7530517578125 0 1040.0204
1336.77587890625 0 1082.3967
1337.7734375 0 2856.747
1338.7706298828125 0 2300.17
1364.7701416015625 0 959.4384
1365.769287109375 0 1187.1311
1366.76416015625 0 920.51605
1380.7821044921875 0 3053.5254 c 11
1381.788818359375 0 5012.3657
1382.79541015625 0 3589.4214
1383.7945556640625 0 1460.584
1409.7662353515625 0 786.347
1411.73876953125 0 2899.726
1412.74560546875 0 2495.453
1413.7506103515625 0 1853.7963
1424.8021240234375 0 2048.1995
1425.7900390625 0 1683.3644
1426.7913818359375 0 1094.7898
1451.7880859375 0 790.3415
1452.7899169921875 0 1505.3248
1467.78515625 0 5266.9805
1468.797607421875 0 10269.335
1469.804931640625 0 7045.7104
1470.806884765625 0 2995.744
1471.804443359375 0 1350.8308
1482.8193359375 0 3696.9465
1483.81787109375 0 2024.3464
1484.81298828125 0 2295.072
1485.8157958984375 0 1430.2518
1493.7843017578125 0 2701.0842
1494.7784423828125 0 1633.1743
1495.789794921875 0 901.3644
1498.8392333984375 0 717.22144
1499.8380126953125 0 2608.3083
1500.8406982421875 0 1515.5345
1501.852294921875 0 877.4894
1509.8106689453125 0 5335.531
1510.81005859375 0 30991.96
1511.8123779296875 0 22047.121
1512.81494140625 0 11773.611
1513.8177490234375 0 2631.799
1514.808349609375 0 2244.5862
1525.822509765625 0 2422.135
1526.8277587890625 0 8476.702
1527.835205078125 0 38962.55
1528.837890625 0 27918.168
1529.8402099609375 0 12330.69
1530.839599609375 0 3861.7112
1531.8289794921875 0 4064.444
3071.603271484375 0 1026.4425
3081.39990234375 0 717.236

Spectrum Details

|  |  |
| --- | --- |
| Matched peaks? Matched peaks The total absolute number of peaks matched. Additionally in brackets the total fraction of peaks matched and the total number of peaks is shown. | 38 (13.67% of 278) |
| FDR? FDR The false discovery rate estimated for this peptide. It is calculated by matching all theoretical fragments with a non-integer shift with the raw peaks for this spectrum. This is done with 40 different shifts. The resulting percentage is the average number of annotated peaks over the number of annotated peaks with the correct spectrum. | 3.07% |
| Satellite FDR? Satellite FDR See the FDR for details on its calculation. This satellite ion specific FDR only contains the satellite ions (d/w) for I/L/J positions. | ∞ |
| PSM Score? PSM Score The PSM Score as given by Hecklib to this annotated spectrum. It is shown with three significant figures. | 236 |

#### Reverse Lookup? Reverse Lookup All places where this read could be placed.

| Group | Segment | Template | Template Part | Read Part | Score | Unique |
| --- | --- | --- | --- | --- | --- | --- |
| Homo sapiens Heavy Chain | IGHV | IGHV3-9 | [36..50] | [0..14] | 94 | False |
| Homo sapiens Heavy Chain | IGHV | IGHV3-48 | [36..50] | [0..14] | 85 | False |
| Homo sapiens Heavy Chain | IGHV | IGHV3-21 | [36..50] | [0..14] | 85 | False |
| Homo sapiens Heavy Chain | IGHV | IGHV3-43 | [36..50] | [0..14] | 85 | False |
| Homo sapiens Heavy Chain | IGHV | IGHV3-20 | [36..50] | [0..14] | 94 | False |
| Homo sapiens Heavy Chain | IGHV | IGHV3-23 | [36..50] | [0..14] | 85 | False |
| Homo sapiens Heavy Chain | IGHV | IGHV3-53 | [36..50] | [0..14] | 85 | False |
| Homo sapiens Heavy Chain | IGHV | IGHV3-66 | [36..50] | [0..14] | 85 | False |
| Homo sapiens Heavy Chain | IGHV | IGHV3-NL1 | [36..50] | [0..14] | 85 | False |

| Recombined | Template Part | Read Part | Score | Unique |
| --- | --- | --- | --- | --- |
| REC-0-1 | [36..50] | [0..14] | 112 | True |

#### Meta Information from Multiple reads

##### Number of combined reads

5

##### Intensity

1

##### TotalArea

0

##### Changes to the peptide sequence

VRQAPGRAJEWVSG

L→JNo support for either Leucine or Isoleucine based on side chain ions (Position: 9)

#### Positional Score

Copy Data

##### Positional Score (TSV)

###### Preview

```
Loading example...
```

*Click on the button to copy the data to your clipboard.*

10012345678910111213

Label Value
"0" 0.598
"1" 0.588
"2" 0.574
"3" 0.536
"4" 0.566
"5" 0.596
"6" 0.584
"7" 0.568
"8" 0.574
"9" 0.584
"10" 0.57
"11" 0.588
"12" 0.592
"13" 0.586

#### Meta Information from PEAKS

##### Scan Identifier

F2:6352

##### Original sequence

V

R

Q

A

P

G

R

A

L

E

W

V

S

G

##### Posttranslational Modifications

##### Source File

D:\separate\_stitch\_analyses\xle-disambiguation\raw\20210323\_F1\_UM1\_Peng0013\_SA\_F59\_ingel\_3ug\_TL.raw

##### Fraction

2

##### Scan Feature

-

##### De Novo Score

98

##### ConfidenceScore

98

### m/z

509.2805

##### Mass

1524.8162

##### Charge

3

##### Retention Time

34.96

##### Predicted Retention Time

-

##### Area

0

##### Parts Per Million

2.2

##### Fragmentation mode

ETHCD

##### Originating file

01 D:\separate\_stitch\_analyses\xle-disambiguation\20210325\_F59\_3ug\_DENOVO\_12.csv

#### Meta Information from PEAKS

##### Scan Identifier

F2:6414

##### Original sequence

V

R

Q

A

P

G

R

A

L

E

W

V

S

G

##### Posttranslational Modifications

##### Source File

D:\separate\_stitch\_analyses\xle-disambiguation\raw\20210323\_F1\_UM1\_Peng0013\_SA\_F59\_ingel\_3ug\_TL.raw

##### Fraction

2

##### Scan Feature

-

##### De Novo Score

97

##### ConfidenceScore

97

### m/z

509.2802

##### Mass

1524.8162

##### Charge

3

##### Retention Time

35.32

##### Predicted Retention Time

-

##### Area

0

##### Parts Per Million

1.6

##### Fragmentation mode

ETHCD

##### Originating file

01 D:\separate\_stitch\_analyses\xle-disambiguation\20210325\_F59\_3ug\_DENOVO\_12.csv

#### Meta Information from PEAKS

##### Scan Identifier

F2:6466

##### Original sequence

V

R

Q

A

P

G

R

A

L

E

W

V

S

G

##### Posttranslational Modifications

##### Source File

D:\separate\_stitch\_analyses\xle-disambiguation\raw\20210323\_F1\_UM1\_Peng0013\_SA\_F59\_ingel\_3ug\_TL.raw

##### Fraction

2

##### Scan Feature

-

##### De Novo Score

97

##### ConfidenceScore

97

### m/z

509.2802

##### Mass

1524.8162

##### Charge

3

##### Retention Time

35.63

##### Predicted Retention Time

-

##### Area

0

##### Parts Per Million

1.7

##### Fragmentation mode

ETHCD

##### Originating file

01 D:\separate\_stitch\_analyses\xle-disambiguation\20210325\_F59\_3ug\_DENOVO\_12.csv

#### Meta Information from PEAKS

##### Scan Identifier

F2:6520

##### Original sequence

V

R

Q

A

P

G

R

A

L

E

W

V

S

G

##### Posttranslational Modifications

##### Source File

D:\separate\_stitch\_analyses\xle-disambiguation\raw\20210323\_F1\_UM1\_Peng0013\_SA\_F59\_ingel\_3ug\_TL.raw

##### Fraction

2

##### Scan Feature

-

##### De Novo Score

96

##### ConfidenceScore

96

### m/z

509.2799

##### Mass

1524.8162

##### Charge

3

##### Retention Time

35.96

##### Predicted Retention Time

-

##### Area

0

##### Parts Per Million

1.2

##### Fragmentation mode

ETHCD

##### Originating file

01 D:\separate\_stitch\_analyses\xle-disambiguation\20210325\_F59\_3ug\_DENOVO\_12.csv

#### Meta Information from PEAKS

##### Scan Identifier

F2:6576

##### Original sequence

V

R

Q

A

P

G

R

A

L

E

W

V

S

G

##### Posttranslational Modifications

##### Source File

D:\separate\_stitch\_analyses\xle-disambiguation\raw\20210323\_F1\_UM1\_Peng0013\_SA\_F59\_ingel\_3ug\_TL.raw

##### Fraction

2

##### Scan Feature

-

##### De Novo Score

95

##### ConfidenceScore

95

### m/z

509.2801

##### Mass

1524.8162

##### Charge

3

##### Retention Time

36.29

##### Predicted Retention Time

-

##### Area

0

##### Parts Per Million

1.6

##### Fragmentation mode

ETHCD

##### Originating file

01 D:\separate\_stitch\_analyses\xle-disambiguation\20210325\_F59\_3ug\_DENOVO\_12.csv
