## Supplementary material for "A handle on mass coincidence errors in *de novo* sequencing of antibodies by bottom-up proteomics": Combined_085.html

Details Combined\_085 | Stitch OverviewUndefined

### Read Combined\_085

#### Sequence (length=8)

SJSJSPGK

#### Spectrum 4021? Spectrum 4021 The raw spectrum of this peptide as annotated by Hecklib. The fragments are coloured according to ion type (see legend). Any peaks with a star '\*' as text can be hovered over to see the full details, first the ion type second the mass shift type. By hovering over the amino acids in the peptide or ions in the legend the corresponding peaks are highlighted. By toggling the 'Unassigned' label you can turn the background (unassigned) peaks on or off in the plot. By updating the slider in the Ion legend you can update the spectrum to only show the top X% of the peaks with labels. The top X% means any peak that is within X% of the highest intensity. By dragging in the spectrum you can zoom in to a specific part of the spectrum and use 'Zoom Out' to get back to the original zoom level. The annotation of the spectrum is based on the given sequence in the peptides file and is done with different software so inconsistencies are likely. The peaks are annotated based on the given sequence, with 20 ppm tolerance.

Copy Data

##### Spectrum 4021 (TSV)

###### Preview

```
Loading example...
```

*Click on the button to copy the data to your clipboard.*

Mz MinMz MaxIntensity Max

WidthHeightPeptide font sizePeptide stroke widthSpectrum font sizeSpectrum stroke widthCompact peptide

Ion legend

wxyz

abcd

OtherUnassignedIon^Charge^_Position_Show for top:%

SJSJSPGK

03.76e+57.52e+51.13e+61.50e+6

Zoom Out

y^+1^_1_d^+1^_2_y^+1^_1_a^+1^_2_a^+1^_2_b^+1^_2_y^+2^_4_y^+1^_2_y^+2^_4_b^+1^_2_y^+1^_2_a^+1^_3_d^+1^_3_a^+1^_3_b^+1^_3_b^+2^_6_y^+2^_6_b^+1^_3_b^+2^_6_y^+2^_6_y^+1^_3_y^+2^_7_y^+2^_7_y^+1^_4_b^+1^_4_\*y^+1^_4_\*b^+1^_4_b^+1^_5_y^+1^_5_b^+1^_5_y^+1^_5_b^+1^_6_y^+1^_6_y^+1^_6_b^+1^_7_y^+1^_7_y^+1^_7_

0570114017102280

Fragment Matches Table

Show background peaks

| Position | Ion type | Intensity | mz Theoretical | mz Error (Th) | mz Error (ppm) | Charge | Series Number |
| --- | --- | --- | --- | --- | --- | --- | --- |
| - | - | 499.3 | 123.1 | - | - | 0 | - |
| - | - | 469.1 | 124.2 | - | - | 0 | - |
| - | - | 1437 | 125.1 | - | - | 0 | - |
| - | - | 1144 | 126.1 | - | - | 0 | - |
| - | - | 5093 | 126.1 | - | - | 0 | - |
| - | - | 1.456E+04 | 127.1 | - | - | 0 | - |
| - | - | 1051 | 127.1 | - | - | 0 | - |
| - | - | 377 | 127.5 | - | - | 0 | - |
| - | - | 1274 | 128.1 | - | - | 0 | - |
| - | - | 1247 | 128.1 | - | - | 0 | - |
| - | - | 6.934E+04 | 128.1 | - | - | 0 | - |
| - | - | 1408 | 129.1 | - | - | 0 | - |
| - | - | 9.612E+04 | 129.1 | - | - | 0 | - |
| - | - | 4805 | 129.1 | - | - | 0 | - |
| 8 | y | 5.458E+04 | 130.1 | 0.0003843 | 2.955 | +1 | 1 |
| - | - | 5278 | 130.1 | - | - | 0 | - |
| 2 | d | 430.6 | 131.1 | 0.002496 | 19.04 | +1 | 2 |
| - | - | 2385 | 131.1 | - | - | 0 | - |
| - | - | 919.5 | 137.1 | - | - | 0 | - |
| - | - | 390.6 | 137.1 | - | - | 0 | - |
| - | - | 1685 | 138.1 | - | - | 0 | - |
| - | - | 4281 | 139.1 | - | - | 0 | - |
| - | - | 461.8 | 139.7 | - | - | 0 | - |
| - | - | 5020 | 141.1 | - | - | 0 | - |
| - | - | 629 | 142.1 | - | - | 0 | - |
| - | - | 3230 | 143.1 | - | - | 0 | - |
| - | - | 439.8 | 144 | - | - | 0 | - |
| - | - | 2907 | 144.1 | - | - | 0 | - |
| 8 | y | 9.987E+04 | 147.1 | 0.0004161 | 2.828 | +1 | 1 |
| - | - | 456.8 | 147.5 | - | - | 0 | - |
| - | - | 877.6 | 148.1 | - | - | 0 | - |
| - | - | 6411 | 148.1 | - | - | 0 | - |
| - | - | 586.9 | 148.9 | - | - | 0 | - |
| - | - | 597.8 | 148.9 | - | - | 0 | - |
| - | - | 687 | 148.9 | - | - | 0 | - |
| - | - | 823.4 | 148.9 | - | - | 0 | - |
| - | - | 1084 | 148.9 | - | - | 0 | - |
| - | - | 1167 | 148.9 | - | - | 0 | - |
| - | - | 1592 | 148.9 | - | - | 0 | - |
| - | - | 3360 | 148.9 | - | - | 0 | - |
| - | - | 5814 | 149 | - | - | 0 | - |
| - | - | 3635 | 149 | - | - | 0 | - |
| - | - | 1442 | 149 | - | - | 0 | - |
| - | - | 1210 | 149 | - | - | 0 | - |
| - | - | 872.6 | 149 | - | - | 0 | - |
| - | - | 918.7 | 149 | - | - | 0 | - |
| - | - | 621.8 | 149 | - | - | 0 | - |
| - | - | 532.1 | 149 | - | - | 0 | - |
| - | - | 458.7 | 149 | - | - | 0 | - |
| - | - | 3690 | 149 | - | - | 0 | - |
| - | - | 462.1 | 149 | - | - | 0 | - |
| - | - | 439.9 | 149.1 | - | - | 0 | - |
| - | - | 431.2 | 149.1 | - | - | 0 | - |
| - | - | 412.9 | 149.2 | - | - | 0 | - |
| - | - | 495.2 | 149.4 | - | - | 0 | - |
| - | - | 3165 | 150 | - | - | 0 | - |
| - | - | 1526 | 151 | - | - | 0 | - |
| - | - | 1203 | 151.1 | - | - | 0 | - |
| - | - | 836.4 | 153.1 | - | - | 0 | - |
| - | - | 852.3 | 155 | - | - | 0 | - |
| - | - | 3.494E+04 | 155.1 | - | - | 0 | - |
| 2 | a | 1.242E+04 | 155.1 | 0.0003661 | 2.36 | +1 | 2 |
| - | - | 1785 | 156.1 | - | - | 0 | - |
| - | - | 4060 | 156.1 | - | - | 0 | - |
| - | - | 942.1 | 156.1 | - | - | 0 | - |
| - | - | 5.234E+04 | 157.1 | - | - | 0 | - |
| - | - | 708.7 | 158.1 | - | - | 0 | - |
| - | - | 3095 | 158.1 | - | - | 0 | - |
| - | - | 799.2 | 159.1 | - | - | 0 | - |
| - | - | 622.6 | 165.1 | - | - | 0 | - |
| - | - | 5933 | 167.1 | - | - | 0 | - |
| - | - | 986 | 167.1 | - | - | 0 | - |
| - | - | 1120 | 168.1 | - | - | 0 | - |
| - | - | 5402 | 169.1 | - | - | 0 | - |
| - | - | 589.7 | 169.1 | - | - | 0 | - |
| - | - | 2027 | 171.1 | - | - | 0 | - |
| - | - | 783.7 | 171.1 | - | - | 0 | - |
| - | - | 466.7 | 171.1 | - | - | 0 | - |
| - | - | 5074 | 172.1 | - | - | 0 | - |
| - | - | 2320 | 173.1 | - | - | 0 | - |
| - | - | 2.509E+04 | 173.1 | - | - | 0 | - |
| 2 | a | 1.489E+06 | 173.1 | 0.0004673 | 2.699 | +1 | 2 |
| - | - | 734.5 | 173.4 | - | - | 0 | - |
| - | - | 514.2 | 173.8 | - | - | 0 | - |
| - | - | 2096 | 174.1 | - | - | 0 | - |
| - | - | 1.261E+05 | 174.1 | - | - | 0 | - |
| - | - | 1344 | 175.1 | - | - | 0 | - |
| - | - | 6699 | 175.1 | - | - | 0 | - |
| - | - | 914.7 | 178.1 | - | - | 0 | - |
| - | - | 2672 | 179.1 | - | - | 0 | - |
| - | - | 889.6 | 181.1 | - | - | 0 | - |
| - | - | 461.1 | 181.1 | - | - | 0 | - |
| 2 | b | 1.197E+04 | 183.1 | 0.000355 | 1.939 | +1 | 2 |
| - | - | 661.4 | 183.1 | - | - | 0 | - |
| - | - | 1429 | 184.1 | - | - | 0 | - |
| - | - | 823.3 | 185.1 | - | - | 0 | - |
| - | - | 3.493E+04 | 185.1 | - | - | 0 | - |
| - | - | 883.2 | 185.1 | - | - | 0 | - |
| - | - | 3.259E+04 | 185.2 | - | - | 0 | - |
| 5 | y | 651.5 | 185.6 | 5.209E-05 | 0.2807 | +2 | 4 |
| - | - | 2476 | 186.1 | - | - | 0 | - |
| - | - | 5.147E+04 | 186.1 | - | - | 0 | - |
| - | - | 2301 | 186.2 | - | - | 0 | - |
| - | - | 763.3 | 187.1 | - | - | 0 | - |
| 7 | y | 1.296E+04 | 187.1 | 0.0003745 | 2.001 | +1 | 2 |
| - | - | 3809 | 187.1 | - | - | 0 | - |
| - | - | 2101 | 187.1 | - | - | 0 | - |
| - | - | 1250 | 188.1 | - | - | 0 | - |
| - | - | 794.5 | 189.1 | - | - | 0 | - |
| - | - | 4471 | 189.1 | - | - | 0 | - |
| - | - | 2729 | 193.1 | - | - | 0 | - |
| - | - | 1566 | 194.1 | - | - | 0 | - |
| 5 | y | 732.2 | 194.6 | 0.0004944 | 2.541 | +2 | 4 |
| - | - | 1629 | 195.1 | - | - | 0 | - |
| - | - | 985.3 | 195.1 | - | - | 0 | - |
| - | - | 2562 | 196.1 | - | - | 0 | - |
| - | - | 814.2 | 197.1 | - | - | 0 | - |
| - | - | 1573 | 197.1 | - | - | 0 | - |
| - | - | 837 | 199.1 | - | - | 0 | - |
| - | - | 792.8 | 200.1 | - | - | 0 | - |
| 2 | b | 2.988E+05 | 201.1 | 0.0003952 | 1.965 | +1 | 2 |
| - | - | 2.794E+04 | 202.1 | - | - | 0 | - |
| - | - | 5737 | 203.1 | - | - | 0 | - |
| - | - | 2041 | 203.1 | - | - | 0 | - |
| 7 | y | 2.277E+05 | 204.1 | 0.0004062 | 1.99 | +1 | 2 |
| - | - | 1.833E+04 | 205.1 | - | - | 0 | - |
| - | - | 3598 | 206.1 | - | - | 0 | - |
| - | - | 1173 | 206.1 | - | - | 0 | - |
| - | - | 3282 | 207.1 | - | - | 0 | - |
| - | - | 605.3 | 210.1 | - | - | 0 | - |
| - | - | 3336 | 211.1 | - | - | 0 | - |
| - | - | 1064 | 213.1 | - | - | 0 | - |
| - | - | 896.8 | 213.1 | - | - | 0 | - |
| - | - | 6067 | 213.2 | - | - | 0 | - |
| - | - | 4480 | 214.1 | - | - | 0 | - |
| - | - | 580.7 | 215.1 | - | - | 0 | - |
| - | - | 4534 | 215.1 | - | - | 0 | - |
| - | - | 578.6 | 216.1 | - | - | 0 | - |
| - | - | 7461 | 217.1 | - | - | 0 | - |
| - | - | 580.1 | 222.9 | - | - | 0 | - |
| - | - | 2104 | 223.1 | - | - | 0 | - |
| - | - | 2.154E+04 | 224.1 | - | - | 0 | - |
| - | - | 2636 | 224.1 | - | - | 0 | - |
| - | - | 566.5 | 224.5 | - | - | 0 | - |
| - | - | 2631 | 225.1 | - | - | 0 | - |
| - | - | 1028 | 225.1 | - | - | 0 | - |
| - | - | 737.8 | 229.1 | - | - | 0 | - |
| - | - | 2065 | 230.2 | - | - | 0 | - |
| - | - | 1131 | 235.1 | - | - | 0 | - |
| - | - | 4634 | 240.1 | - | - | 0 | - |
| - | - | 1102 | 241.1 | - | - | 0 | - |
| - | - | 1020 | 241.2 | - | - | 0 | - |
| - | - | 2.332E+04 | 242.1 | - | - | 0 | - |
| 3 | a | 5176 | 242.1 | 0.0005032 | 2.078 | +1 | 3 |
| - | - | 3019 | 243.1 | - | - | 0 | - |
| - | - | 5163 | 243.1 | - | - | 0 | - |
| - | - | 623.1 | 244.1 | - | - | 0 | - |
| 3 | d | 663.5 | 244.2 | 0.0008749 | 3.583 | +1 | 3 |
| - | - | 728.8 | 245.1 | - | - | 0 | - |
| - | - | 1.201E+04 | 248.2 | - | - | 0 | - |
| - | - | 1046 | 249.2 | - | - | 0 | - |
| - | - | 1875 | 251.1 | - | - | 0 | - |
| - | - | 3144 | 252.1 | - | - | 0 | - |
| - | - | 823.1 | 252.2 | - | - | 0 | - |
| - | - | 1150 | 253.1 | - | - | 0 | - |
| - | - | 1069 | 254.1 | - | - | 0 | - |
| - | - | 1466 | 258.1 | - | - | 0 | - |
| - | - | 1080 | 259.1 | - | - | 0 | - |
| - | - | 1.144E+04 | 260.1 | - | - | 0 | - |
| 3 | a | 3254 | 260.2 | 0.000406 | 1.561 | +1 | 3 |
| - | - | 762.9 | 261.2 | - | - | 0 | - |
| - | - | 1431 | 262.2 | - | - | 0 | - |
| - | - | 663.8 | 266.1 | - | - | 0 | - |
| - | - | 753.6 | 268.1 | - | - | 0 | - |
| - | - | 1816 | 268.2 | - | - | 0 | - |
| - | - | 1054 | 269.2 | - | - | 0 | - |
| - | - | 581.6 | 269.2 | - | - | 0 | - |
| 3 | b | 4.799E+04 | 270.1 | 0.0003701 | 1.37 | +1 | 3 |
| - | - | 7110 | 271.1 | - | - | 0 | - |
| - | - | 1151 | 272.2 | - | - | 0 | - |
| - | - | 945.2 | 276.7 | - | - | 0 | - |
| - | - | 572.9 | 280.1 | - | - | 0 | - |
| - | - | 1745 | 280.2 | - | - | 0 | - |
| - | - | 5534 | 283.1 | - | - | 0 | - |
| - | - | 5.236E+04 | 283.2 | - | - | 0 | - |
| - | - | 1016 | 284.1 | - | - | 0 | - |
| 6 | b | 3225 | 284.2 | 0.0006502 | 2.288 | +2 | 6 |
| - | - | 7284 | 284.2 | - | - | 0 | - |
| 3 | y | 2.852E+04 | 285.7 | 0.0003887 | 1.361 | +2 | 6 |
| - | - | 7777 | 286.2 | - | - | 0 | - |
| - | - | 1869 | 286.7 | - | - | 0 | - |
| 3 | b | 5.361E+04 | 288.2 | 0.0005171 | 1.794 | +1 | 3 |
| - | - | 7120 | 289.2 | - | - | 0 | - |
| - | - | 813.5 | 290.2 | - | - | 0 | - |
| - | - | 836.5 | 291.2 | - | - | 0 | - |
| 6 | b | 620.8 | 293.2 | 0.005334 | 18.19 | +2 | 6 |
| - | - | 551 | 294.1 | - | - | 0 | - |
| 3 | y | 4500 | 294.7 | 0.000691 | 2.345 | +2 | 6 |
| - | - | 2058 | 295.2 | - | - | 0 | - |
| - | - | 3847 | 296.2 | - | - | 0 | - |
| - | - | 2375 | 298.1 | - | - | 0 | - |
| - | - | 932.5 | 298.2 | - | - | 0 | - |
| - | - | 657.4 | 299.2 | - | - | 0 | - |
| - | - | 1453 | 299.7 | - | - | 0 | - |
| - | - | 562.5 | 300.2 | - | - | 0 | - |
| 6 | y | 7.074E+05 | 301.2 | 0.0006209 | 2.061 | +1 | 3 |
| - | - | 9.875E+04 | 302.2 | - | - | 0 | - |
| - | - | 1.311E+04 | 303.2 | - | - | 0 | - |
| - | - | 4864 | 308.2 | - | - | 0 | - |
| - | - | 728.7 | 309.2 | - | - | 0 | - |
| - | - | 734.2 | 309.2 | - | - | 0 | - |
| - | - | 617.5 | 311.1 | - | - | 0 | - |
| - | - | 1179 | 311.2 | - | - | 0 | - |
| - | - | 2815 | 312.2 | - | - | 0 | - |
| - | - | 2058 | 313.2 | - | - | 0 | - |
| - | - | 2705 | 316.2 | - | - | 0 | - |
| - | - | 913.8 | 321.2 | - | - | 0 | - |
| - | - | 856.3 | 322.2 | - | - | 0 | - |
| - | - | 2475 | 323.2 | - | - | 0 | - |
| - | - | 2764 | 325.2 | - | - | 0 | - |
| - | - | 1627 | 325.2 | - | - | 0 | - |
| - | - | 573.7 | 325.4 | - | - | 0 | - |
| - | - | 2638 | 326.2 | - | - | 0 | - |
| - | - | 784.7 | 326.2 | - | - | 0 | - |
| - | - | 1826 | 328.2 | - | - | 0 | - |
| - | - | 844.1 | 328.7 | - | - | 0 | - |
| - | - | 788.8 | 329.2 | - | - | 0 | - |
| - | - | 5288 | 330.2 | - | - | 0 | - |
| - | - | 829.2 | 334.2 | - | - | 0 | - |
| - | - | 7081 | 335.2 | - | - | 0 | - |
| - | - | 1459 | 336.2 | - | - | 0 | - |
| - | - | 692.9 | 336.7 | - | - | 0 | - |
| - | - | 1088 | 337.2 | - | - | 0 | - |
| - | - | 795.7 | 337.2 | - | - | 0 | - |
| - | - | 1085 | 338.2 | - | - | 0 | - |
| - | - | 1236 | 339.2 | - | - | 0 | - |
| - | - | 780.1 | 339.2 | - | - | 0 | - |
| - | - | 1648 | 340.2 | - | - | 0 | - |
| 2 | y | 9044 | 342.2 | 0.000593 | 1.733 | +2 | 7 |
| - | - | 3896 | 342.7 | - | - | 0 | - |
| - | - | 5878 | 343.2 | - | - | 0 | - |
| - | - | 601.1 | 343.2 | - | - | 0 | - |
| - | - | 657.4 | 343.2 | - | - | 0 | - |
| - | - | 1016 | 343.2 | - | - | 0 | - |
| - | - | 3174 | 343.7 | - | - | 0 | - |
| - | - | 558.9 | 344.2 | - | - | 0 | - |
| - | - | 1066 | 349.2 | - | - | 0 | - |
| 2 | y | 1.086E+04 | 351.2 | 0.0004681 | 1.333 | +2 | 7 |
| - | - | 3814 | 351.7 | - | - | 0 | - |
| - | - | 1.076E+04 | 352.2 | - | - | 0 | - |
| - | - | 8420 | 353.2 | - | - | 0 | - |
| - | - | 1192 | 353.2 | - | - | 0 | - |
| - | - | 1167 | 354.2 | - | - | 0 | - |
| - | - | 3170 | 355.2 | - | - | 0 | - |
| - | - | 1.542E+04 | 355.2 | - | - | 0 | - |
| - | - | 1015 | 355.3 | - | - | 0 | - |
| - | - | 3767 | 356.2 | - | - | 0 | - |
| - | - | 2231 | 356.2 | - | - | 0 | - |
| - | - | 1941 | 356.7 | - | - | 0 | - |
| - | - | 1598 | 357.2 | - | - | 0 | - |
| - | - | 6279 | 358.2 | - | - | 0 | - |
| - | - | 1589 | 358.9 | - | - | 0 | - |
| - | - | 999.5 | 359.2 | - | - | 0 | - |
| - | - | 6780 | 361.2 | - | - | 0 | - |
| - | - | 1321 | 362.2 | - | - | 0 | - |
| - | - | 1359 | 365.2 | - | - | 0 | - |
| - | - | 4953 | 367.2 | - | - | 0 | - |
| - | - | 1781 | 368.2 | - | - | 0 | - |
| 5 | y | 3.926E+04 | 370.2 | 0.0003974 | 1.073 | +1 | 4 |
| - | - | 7743 | 371.2 | - | - | 0 | - |
| - | - | 1063 | 372.2 | - | - | 0 | - |
| - | - | 1391 | 373.2 | - | - | 0 | - |
| - | - | 761.5 | 374.2 | - | - | 0 | - |
| - | - | 816.6 | 375.2 | - | - | 0 | - |
| - | - | 1039 | 375.2 | - | - | 0 | - |
| - | - | 735.6 | 375.9 | - | - | 0 | - |
| - | - | 3970 | 376.7 | - | - | 0 | - |
| - | - | 1828 | 376.8 | - | - | 0 | - |
| - | - | 768.7 | 377.2 | - | - | 0 | - |
| - | - | 1176 | 380.2 | - | - | 0 | - |
| 4 | b | 8943 | 383.2 | 0.0006567 | 1.714 | +1 | 4 |
| - | - | 1778 | 384.2 | - | - | 0 | - |
| - | - | 3424 | 385.2 | - | - | 0 | - |
| 0 | Precursor | 5.403E+04 | 385.7 | 0.0003869 | 1.003 | +2 | -1 |
| - | - | 2.369E+04 | 386.2 | - | - | 0 | - |
| - | - | 6490 | 386.7 | - | - | 0 | - |
| 5 | y | 2.543E+05 | 388.2 | 0.0005749 | 1.481 | +1 | 4 |
| - | - | 4.654E+04 | 389.2 | - | - | 0 | - |
| - | - | 6110 | 390.2 | - | - | 0 | - |
| - | - | 1675 | 393.2 | - | - | 0 | - |
| - | - | 1474 | 393.3 | - | - | 0 | - |
| - | - | 891.5 | 394.2 | - | - | 0 | - |
| - | - | 2819 | 394.2 | - | - | 0 | - |
| - | - | 844.6 | 394.2 | - | - | 0 | - |
| 0 | Precursor | 1513 | 394.7 | 0.00163 | 4.13 | +2 | -1 |
| - | - | 662.3 | 395.2 | - | - | 0 | - |
| - | - | 1182 | 396.2 | - | - | 0 | - |
| - | - | 4495 | 398.2 | - | - | 0 | - |
| - | - | 1010 | 399.2 | - | - | 0 | - |
| 4 | b | 2036 | 401.2 | 0.0004069 | 1.014 | +1 | 4 |
| - | - | 3954 | 406.2 | - | - | 0 | - |
| - | - | 1042 | 411.2 | - | - | 0 | - |
| - | - | 1919 | 413.2 | - | - | 0 | - |
| - | - | 4159 | 424.2 | - | - | 0 | - |
| - | - | 1393 | 424.3 | - | - | 0 | - |
| - | - | 3152 | 431.2 | - | - | 0 | - |
| - | - | 1723 | 441.2 | - | - | 0 | - |
| - | - | 5866 | 442.2 | - | - | 0 | - |
| - | - | 2314 | 442.3 | - | - | 0 | - |
| - | - | 952.9 | 443.2 | - | - | 0 | - |
| - | - | 2095 | 452.3 | - | - | 0 | - |
| - | - | 1495 | 456.3 | - | - | 0 | - |
| - | - | 644.3 | 458.3 | - | - | 0 | - |
| - | - | 1712 | 460.2 | - | - | 0 | - |
| - | - | 1952 | 460.3 | - | - | 0 | - |
| - | - | 595.6 | 461.2 | - | - | 0 | - |
| - | - | 918.4 | 461.3 | - | - | 0 | - |
| - | - | 1609 | 465.3 | - | - | 0 | - |
| 5 | b | 4927 | 470.3 | 0.0005802 | 1.234 | +1 | 5 |
| - | - | 1987 | 471.3 | - | - | 0 | - |
| - | - | 2662 | 476.3 | - | - | 0 | - |
| - | - | 1037 | 481.3 | - | - | 0 | - |
| 4 | y | 1.226E+04 | 483.3 | 0.0005924 | 1.226 | +1 | 5 |
| - | - | 2968 | 484.3 | - | - | 0 | - |
| 5 | b | 2041 | 488.3 | 0.0002694 | 0.5517 | +1 | 5 |
| - | - | 1337 | 493.3 | - | - | 0 | - |
| 4 | y | 6.497E+04 | 501.3 | 0.0005258 | 1.049 | +1 | 5 |
| - | - | 1.771E+04 | 502.3 | - | - | 0 | - |
| - | - | 3522 | 503.3 | - | - | 0 | - |
| - | - | 1594 | 511.3 | - | - | 0 | - |
| - | - | 865 | 517.3 | - | - | 0 | - |
| - | - | 821.8 | 533.3 | - | - | 0 | - |
| - | - | 844.2 | 534.3 | - | - | 0 | - |
| - | - | 3377 | 535.3 | - | - | 0 | - |
| - | - | 1032 | 536.3 | - | - | 0 | - |
| - | - | 1283 | 540.3 | - | - | 0 | - |
| - | - | 782.2 | 550.3 | - | - | 0 | - |
| - | - | 6451 | 552.3 | - | - | 0 | - |
| - | - | 2118 | 553.3 | - | - | 0 | - |
| - | - | 3584 | 558.3 | - | - | 0 | - |
| - | - | 1631 | 559.3 | - | - | 0 | - |
| - | - | 924.3 | 562.3 | - | - | 0 | - |
| 6 | b | 733.8 | 567.3 | 0.002226 | 3.924 | +1 | 6 |
| - | - | 760.9 | 568.3 | - | - | 0 | - |
| - | - | 1682 | 569.3 | - | - | 0 | - |
| 3 | y | 4.383E+04 | 570.3 | 0.0004854 | 0.8511 | +1 | 6 |
| - | - | 1.349E+04 | 571.3 | - | - | 0 | - |
| - | - | 2145 | 572.3 | - | - | 0 | - |
| - | - | 931.8 | 573.4 | - | - | 0 | - |
| - | - | 4.263E+04 | 577.3 | - | - | 0 | - |
| - | - | 1.121E+04 | 578.3 | - | - | 0 | - |
| - | - | 698.7 | 579.3 | - | - | 0 | - |
| - | - | 3800 | 580.3 | - | - | 0 | - |
| - | - | 991.7 | 581.3 | - | - | 0 | - |
| 3 | y | 7.733E+05 | 588.3 | 0.0006629 | 1.127 | +1 | 6 |
| - | - | 2.343E+05 | 589.3 | - | - | 0 | - |
| - | - | 5.109E+04 | 590.3 | - | - | 0 | - |
| - | - | 3026 | 591.3 | - | - | 0 | - |
| - | - | 1.933E+04 | 598.3 | - | - | 0 | - |
| - | - | 5680 | 599.3 | - | - | 0 | - |
| - | - | 828.6 | 600.3 | - | - | 0 | - |
| - | - | 942.3 | 606.3 | - | - | 0 | - |
| 7 | b | 1244 | 624.3 | 0.003671 | 5.879 | +1 | 7 |
| - | - | 2159 | 650.1 | - | - | 0 | - |
| - | - | 1990 | 650.2 | - | - | 0 | - |
| - | - | 1040 | 655.3 | - | - | 0 | - |
| - | - | 2574 | 658.4 | - | - | 0 | - |
| - | - | 1245 | 659.4 | - | - | 0 | - |
| - | - | 8280 | 667.3 | - | - | 0 | - |
| - | - | 2706 | 668.3 | - | - | 0 | - |
| - | - | 777.4 | 669.3 | - | - | 0 | - |
| - | - | 1.274E+04 | 676.4 | - | - | 0 | - |
| - | - | 4591 | 677.4 | - | - | 0 | - |
| 2 | y | 4099 | 683.4 | 0.001321 | 1.933 | +1 | 7 |
| - | - | 1396 | 684.4 | - | - | 0 | - |
| - | - | 2226 | 685.4 | - | - | 0 | - |
| - | - | 699.6 | 685.4 | - | - | 0 | - |
| - | - | 1397 | 693.4 | - | - | 0 | - |
| 2 | y | 1.212E+04 | 701.4 | 0.0006374 | 0.9087 | +1 | 7 |
| - | - | 5607 | 702.4 | - | - | 0 | - |
| - | - | 933.6 | 703.4 | - | - | 0 | - |
| - | - | 2761 | 711.4 | - | - | 0 | - |
| - | - | 742.4 | 712.4 | - | - | 0 | - |
| - | - | 912 | 827.1 | - | - | 0 | - |
| - | - | 676.2 | 863.6 | - | - | 0 | - |
| - | - | 667 | 906.1 | - | - | 0 | - |
| - | - | 672.7 | 1386 | - | - | 0 | - |
| - | - | 663.1 | 2257 | - | - | 0 | - |

m/z Charge Intensity FragmentType MassShift Position
123.0918197631836 0 499.29297
124.2302474975586 0 469.10782
125.10787963867188 0 1437.3625
126.05545043945312 0 1143.5208
126.09173583984375 0 5093.012
127.08699035644531 0 14559.813
127.12330627441406 0 1050.6558
127.5369644165039 0 377.03748
128.07093811035156 0 1273.5386
128.09027099609375 0 1247.4226
128.1073760986328 0 69338.07
129.06614685058594 0 1408.3087
129.10264587402344 0 96116.78
129.1109161376953 0 4805.4175
130.08663940429688 0 54578.055 y Ammonia loss 7
130.10597229003906 0 5277.9727
131.08399963378906 0 430.56357 d 1
131.0899658203125 0 2384.7156
137.0717010498047 0 919.46216
137.094482421875 0 390.58972
138.091796875 0 1685.3718
139.08689880371094 0 4280.534
139.6983184814453 0 461.79504
141.10260009765625 0 5019.877
142.10597229003906 0 629.04065
143.11830139160156 0 3230.4246
143.98883056640625 0 439.7638
144.06597900390625 0 2906.9639
147.11322021484375 0 99869.47 y 7
147.4835662841797 0 456.83163
148.10995483398438 0 877.6265
148.11659240722656 0 6411.2915
148.87506103515625 0 586.9353
148.8899688720703 0 597.75964
148.90367126464844 0 686.9983
148.9105987548828 0 823.4333
148.91839599609375 0 1084.493
148.92572021484375 0 1166.7524
148.93251037597656 0 1592.2957
148.9401092529297 0 3359.7598
148.95643615722656 0 5813.9106
148.96412658691406 0 3634.815
148.9717254638672 0 1442.3585
148.97866821289062 0 1210.3402
148.9860076904297 0 872.61554
148.99273681640625 0 918.7076
149.000244140625 0 621.82635
149.00762939453125 0 532.06
149.01577758789062 0 458.67496
149.02366638183594 0 3690.2334
149.0359344482422 0 462.0698
149.07135009765625 0 439.87317
149.1079864501953 0 431.19327
149.15078735351562 0 412.86542
149.4384002685547 0 495.21878
150.02713012695312 0 3165.255
151.0286865234375 0 1526.057
151.0867156982422 0 1202.9562
153.0658721923828 0 836.44775
155.0453338623047 0 852.2623
155.08187866210938 0 34938.574
155.11825561523438 0 12424.85 a Water loss 1
156.08543395996094 0 1784.6912
156.1023406982422 0 4060.4314
156.12161254882812 0 942.137
157.09756469726562 0 52337.65
158.09454345703125 0 708.6743
158.10089111328125 0 3094.57
159.1131591796875 0 799.2072
165.06639099121094 0 622.5577
167.08181762695312 0 5933.4375
167.1180877685547 0 985.9725
168.11375427246094 0 1119.5546
169.0975799560547 0 5401.6997
169.13414001464844 0 589.67926
171.07716369628906 0 2027.3574
171.11279296875 0 783.6757
171.14883422851562 0 466.65198
172.10853576660156 0 5073.586
173.05636596679688 0 2319.6401
173.0931396484375 0 25093.822
173.12892150878906 0 1488628.6 a 1
173.4401092529297 0 734.517
173.75730895996094 0 514.1667
174.09591674804688 0 2096.3513
174.1322021484375 0 126078.23
175.11915588378906 0 1344.3638
175.13389587402344 0 6699.0054
178.09744262695312 0 914.70715
179.0817413330078 0 2672.3726
181.09747314453125 0 889.6299
181.13328552246094 0 461.0622
183.1131591796875 0 11969.677 b Water loss 1
183.1488494873047 0 661.40027
184.11651611328125 0 1428.5034
185.05648803710938 0 823.2878
185.0924530029297 0 34932.914
185.12884521484375 0 883.2046
185.16525268554688 0 32588.156
185.6078338623047 0 651.46094 y Water loss 4
186.09576416015625 0 2476.354
186.12408447265625 0 51474.22
186.1685333251953 0 2300.6982
187.0985870361328 0 763.28406
187.10809326171875 0 12956.085 y Ammonia loss 6
187.1276092529297 0 3809.012
187.1444854736328 0 2101.151
188.11141967773438 0 1249.7631
189.06661987304688 0 794.50073
189.08734130859375 0 4470.6255
193.09747314453125 0 2728.5027
194.0926055908203 0 1566.3854
194.61366271972656 0 732.2348 y 4
195.07630920410156 0 1629.478
195.1131134033203 0 985.2839
196.10862731933594 0 2561.585
197.09193420410156 0 814.2238
197.1289825439453 0 1572.926
199.0714874267578 0 836.9919
200.1396026611328 0 792.8256
201.12376403808594 0 298824.38 b 1
202.1271514892578 0 27942.53
203.0664825439453 0 5737.0957
203.12899780273438 0 2040.9167
204.13467407226562 0 227654.72 y 6
205.13796997070312 0 18325.66
206.09266662597656 0 3598.1436
206.13880920410156 0 1172.8027
207.07672119140625 0 3281.8013
210.12364196777344 0 605.2905
211.1082305908203 0 3335.6902
213.08782958984375 0 1064.1566
213.1242218017578 0 896.8386
213.16012573242188 0 6066.6504
214.11911010742188 0 4479.999
215.1223602294922 0 580.68536
215.1393585205078 0 4533.984
216.13394165039062 0 578.6217
217.08226013183594 0 7461.4624
222.8984832763672 0 580.0552
223.10772705078125 0 2103.9434
224.1033935546875 0 21543.715
224.13949584960938 0 2636.4297
224.45703125 0 566.5377
225.10704040527344 0 2630.894
225.12362670898438 0 1027.98
229.11880493164062 0 737.75867
230.15036010742188 0 2065.0278
235.10794067382812 0 1131.4459
240.13462829589844 0 4634.302
241.1186065673828 0 1102.0425
241.19163513183594 0 1019.8546
242.11395263671875 0 23321.354
242.15042114257812 0 5175.513 a Water loss 2
243.11727905273438 0 3018.5815
243.13427734375 0 5163.353
244.1378173828125 0 623.12604
244.16644287109375 0 663.4819 d 2
245.1245880126953 0 728.84125
248.1609649658203 0 12009.93
249.16432189941406 0 1045.8488
251.13966369628906 0 1875.267
252.13458251953125 0 3143.7412
252.17019653320312 0 823.0838
253.1182403564453 0 1149.9417
254.1143341064453 0 1068.5771
258.1450500488281 0 1465.6287
259.1406555175781 0 1079.6893
260.12445068359375 0 11441.397
260.160888671875 0 3253.9216 a 2
261.1653747558594 0 762.92883
262.15130615234375 0 1431.0562
266.1498107910156 0 663.8176
268.1296081542969 0 753.5913
268.2022705078125 0 1816.1204
269.18701171875 0 1054.3645
269.205078125 0 581.6147
270.14520263671875 0 47985.133 b Water loss 2
271.14898681640625 0 7110.326
272.15020751953125 0 1150.5848
276.660888671875 0 945.18463
280.1288757324219 0 572.8898
280.166015625 0 1744.8036
283.14031982421875 0 5533.66
283.17694091796875 0 52361.574
284.1436767578125 0 1015.6702
284.1611328125 0 3225.2537 b Water loss 5
284.1801452636719 0 7284.393
285.66632080078125 0 28520.129 y Water loss 2
286.167724609375 0 7777.0317
286.6694641113281 0 1868.7239
288.1559143066406 0 53613.438 b 2
289.15924072265625 0 7119.586
290.1614074707031 0 813.54376
291.1671142578125 0 836.5022
293.1604309082031 0 620.759 b 5
294.1460876464844 0 550.96985
294.6719055175781 0 4499.5557 y 2
295.17340087890625 0 2058.2117
296.1975402832031 0 3847.0105
298.140380859375 0 2374.642
298.1788330078125 0 932.5033
299.17333984375 0 657.41534
299.663818359375 0 1452.9839
300.1662902832031 0 562.5273
301.1876525878906 0 707373.94 y 5
302.1905822753906 0 98752.945
303.19293212890625 0 13114.406
308.16082763671875 0 4864.1294
309.16436767578125 0 728.66046
309.2041320800781 0 734.19794
311.1346130371094 0 617.477
311.17156982421875 0 1179.1906
312.1561584472656 0 2814.7988
313.1872253417969 0 2057.6724
316.1506042480469 0 2704.7822
321.156494140625 0 913.77026
322.1755676269531 0 856.29144
323.171875 0 2474.5674
325.1877136230469 0 2764.4275
325.2242736816406 0 1627.0851
325.36334228515625 0 573.6716
326.1716613769531 0 2637.8337
326.2077331542969 0 784.73926
328.1748352050781 0 1825.9473
328.675048828125 0 844.07385
329.1820373535156 0 788.8213
330.166748046875 0 5287.7144
334.17431640625 0 829.2057
335.1717224121094 0 7080.7334
336.174560546875 0 1459.0612
336.7060852050781 0 692.8956
337.1871337890625 0 1088.0612
337.2245178222656 0 795.676
338.2080383300781 0 1085.0687
339.16650390625 0 1236.3591
339.20196533203125 0 780.1348
340.198974609375 0 1648.447
342.20855712890625 0 9043.938 y Water loss 1
342.70977783203125 0 3895.8628
343.1800842285156 0 5878.2476
343.2029113769531 0 601.1142
343.2127380371094 0 657.3923
343.2332763671875 0 1015.91504
343.6811218261719 0 3173.8826
344.18408203125 0 558.90515
349.1873779296875 0 1066.1189
351.2137145996094 0 10859.197 y 1
351.715087890625 0 3813.5867
352.19842529296875 0 10761.551
353.1821594238281 0 8420.262
353.202392578125 0 1192.0891
354.18505859375 0 1166.6903
355.19781494140625 0 3170.0105
355.23443603515625 0 15422.142
355.2582092285156 0 1015.0417
356.2055358886719 0 3766.8535
356.2374267578125 0 2230.7473
356.7079162597656 0 1941.0945
357.1763610839844 0 1597.63
358.2088317871094 0 6278.531
358.88946533203125 0 1589.2286
359.2107849121094 0 999.5327
361.2447814941406 0 6779.6
362.2470703125 0 1320.6282
365.2187805175781 0 1359.2897
367.1980895996094 0 4952.5566
368.1998291015625 0 1780.8508
370.2088928222656 0 39259.27 y Water loss 4
371.2121276855469 0 7742.831
372.2152404785156 0 1063.0175
373.24493408203125 0 1390.6771
374.1759338378906 0 761.54865
375.1861877441406 0 816.62067
375.23553466796875 0 1039.3135
375.8588562011719 0 735.5599
376.71868896484375 0 3969.8625
376.8439025878906 0 1828.2206
377.2198486328125 0 768.70233
380.19427490234375 0 1176.307
383.22955322265625 0 8942.529 b Water loss 3
384.23223876953125 0 1777.947
385.2093811035156 0 3424.477
385.724365234375 0 54028.285 Precursor Water loss
386.2259521484375 0 23686.906
386.7274475097656 0 6490.04
388.2196350097656 0 254262.55 y 4
389.22247314453125 0 46541.74
390.2246398925781 0 6109.818
393.24664306640625 0 1674.9489
393.29779052734375 0 1474.4161
394.18231201171875 0 891.517
394.2178649902344 0 2818.5884
394.2466735839844 0 844.5912
394.7276306152344 0 1512.949 Precursor
395.1925048828125 0 662.2889
396.2258605957031 0 1182.0046
398.2037353515625 0 4495.3433
399.2057189941406 0 1010.3516
401.2398681640625 0 2036.3015 b 3
406.2087707519531 0 3953.703
411.2245178222656 0 1042.309
413.20306396484375 0 1919.2968
424.21929931640625 0 4158.801
424.25469970703125 0 1392.9022
431.2137451171875 0 3151.8184
441.24591064453125 0 1723.2842
442.2299499511719 0 5866.498
442.2665100097656 0 2313.9487
443.2303771972656 0 952.8836
452.25152587890625 0 2095.395
456.2569274902344 0 1494.5225
458.26544189453125 0 644.3307
460.240966796875 0 1711.9807
460.2867431640625 0 1951.7773
461.246826171875 0 595.6262
461.2893981933594 0 918.36865
465.2830810546875 0 1608.9498
470.2615051269531 0 4927.2876 b Water loss 4
471.2640686035156 0 1986.6788
476.2722473144531 0 2662.0383
481.27716064453125 0 1036.6456
483.29315185546875 0 12263.761 y Water loss 3
484.2962646484375 0 2967.9592
488.2717590332031 0 2040.9991 b 4
493.2763977050781 0 1336.8351
501.30364990234375 0 64970.11 y 3
502.30645751953125 0 17714.006
503.30877685546875 0 3522.2305
511.28619384765625 0 1594.3337
517.2784423828125 0 864.9952
533.2911376953125 0 821.83954
534.3005981445312 0 844.1602
535.2877197265625 0 3377.2346
536.2903442382812 0 1032.3252
540.3132934570312 0 1282.9705
550.298095703125 0 782.17633
552.3145141601562 0 6450.918
553.31201171875 0 2117.7234
558.3244018554688 0 3583.8982
559.314453125 0 1631.168
562.2977905273438 0 924.33075
567.3114624023438 0 733.77405 b Water loss 5
568.3114013671875 0 760.8855
569.3406982421875 0 1681.7856
570.3250732421875 0 43825.676 y Water loss 2
571.3275756835938 0 13490.366
572.3304443359375 0 2144.9807
573.3709716796875 0 931.83124
577.3195190429688 0 42630.73
578.3225708007812 0 11210.284
579.3256225585938 0 698.74457
580.3092651367188 0 3800.117
581.3154907226562 0 991.6519
588.3358154296875 0 773320.1 y 2
589.3385009765625 0 234270.95
590.3408813476562 0 51085.566
591.343994140625 0 3025.738
598.3197631835938 0 19329.021
599.32275390625 0 5679.789
600.3226318359375 0 828.56433
606.3255615234375 0 942.27966
624.3314819335938 0 1243.5358 b Water loss 6
650.0861206054688 0 2158.7588
650.1591796875 0 1989.5646
655.3412475585938 0 1039.8651
658.3768310546875 0 2573.7913
659.3797607421875 0 1244.7465
667.3411254882812 0 8279.987
668.3439331054688 0 2706.3337
669.3472900390625 0 777.40216
676.3878173828125 0 12735.938
677.39111328125 0 4591.187
683.4099731445312 0 4099.2886 y Water loss 1
684.4136352539062 0 1395.7112
685.351318359375 0 2225.6055
685.4129638671875 0 699.6155
693.39453125 0 1397.0286
701.4185791015625 0 12116.609 y 1
702.4223022460938 0 5606.5146
703.4237670898438 0 933.60065
711.4042358398438 0 2760.625
712.4054565429688 0 742.35376
827.0673217773438 0 911.9986
863.6317749023438 0 676.20685
906.1305541992188 0 666.9559
1385.509521484375 0 672.7124
2257.444091796875 0 663.0551

Spectrum Details

|  |  |
| --- | --- |
| Matched peaks? Matched peaks The total absolute number of peaks matched. Additionally in brackets the total fraction of peaks matched and the total number of peaks is shown. | 39 (10.05% of 388) |
| FDR? FDR The false discovery rate estimated for this peptide. It is calculated by matching all theoretical fragments with a non-integer shift with the raw peaks for this spectrum. This is done with 40 different shifts. The resulting percentage is the average number of annotated peaks over the number of annotated peaks with the correct spectrum. | 0.24% |
| Satellite FDR? Satellite FDR See the FDR for details on its calculation. This satellite ion specific FDR only contains the satellite ions (d/w) for I/L/J positions. | 0.00% |
| PSM Score? PSM Score The PSM Score as given by Hecklib to this annotated spectrum. It is shown with three significant figures. | 479 |

#### Spectrum 4071? Spectrum 4071 The raw spectrum of this peptide as annotated by Hecklib. The fragments are coloured according to ion type (see legend). Any peaks with a star '\*' as text can be hovered over to see the full details, first the ion type second the mass shift type. By hovering over the amino acids in the peptide or ions in the legend the corresponding peaks are highlighted. By toggling the 'Unassigned' label you can turn the background (unassigned) peaks on or off in the plot. By updating the slider in the Ion legend you can update the spectrum to only show the top X% of the peaks with labels. The top X% means any peak that is within X% of the highest intensity. By dragging in the spectrum you can zoom in to a specific part of the spectrum and use 'Zoom Out' to get back to the original zoom level. The annotation of the spectrum is based on the given sequence in the peptides file and is done with different software so inconsistencies are likely. The peaks are annotated based on the given sequence, with 20 ppm tolerance.

Copy Data

##### Spectrum 4071 (TSV)

###### Preview

```
Loading example...
```

*Click on the button to copy the data to your clipboard.*

Mz MinMz MaxIntensity Max

WidthHeightPeptide font sizePeptide stroke widthSpectrum font sizeSpectrum stroke widthCompact peptide

Ion legend

wxyz

abcd

OtherUnassignedIon^Charge^_Position_Show for top:%

SJSJSPGK

01.06e+52.11e+53.17e+54.22e+5

Zoom Out

y^+1^_1_y^+1^_1_a^+1^_2_a^+1^_2_b^+1^_2_y^+1^_2_b^+1^_2_y^+1^_2_a^+1^_3_a^+1^_3_b^+1^_3_b^+2^_6_y^+2^_6_b^+1^_3_y^+2^_6_y^+1^_3_y^+2^_7_y^+2^_7_y^+1^_4_b^+1^_4_\*y^+1^_4_b^+1^_5_y^+1^_5_y^+1^_5_y^+1^_6_y^+1^_6_b^+1^_7_y^+1^_7_

0807161324203227

Fragment Matches Table

Show background peaks

| Position | Ion type | Intensity | mz Theoretical | mz Error (Th) | mz Error (ppm) | Charge | Series Number |
| --- | --- | --- | --- | --- | --- | --- | --- |
| - | - | 403.2 | 120.6 | - | - | 0 | - |
| - | - | 409.5 | 125.1 | - | - | 0 | - |
| - | - | 703.5 | 125.1 | - | - | 0 | - |
| - | - | 717.1 | 126.1 | - | - | 0 | - |
| - | - | 1560 | 126.1 | - | - | 0 | - |
| - | - | 4172 | 127.1 | - | - | 0 | - |
| - | - | 373.7 | 127.9 | - | - | 0 | - |
| - | - | 965.6 | 128.1 | - | - | 0 | - |
| - | - | 1.817E+04 | 128.1 | - | - | 0 | - |
| - | - | 2.538E+04 | 129.1 | - | - | 0 | - |
| - | - | 1096 | 129.1 | - | - | 0 | - |
| 8 | y | 1.53E+04 | 130.1 | 0.0002165 | 1.664 | +1 | 1 |
| - | - | 1665 | 130.1 | - | - | 0 | - |
| - | - | 441.3 | 131.3 | - | - | 0 | - |
| - | - | 364.7 | 135.8 | - | - | 0 | - |
| - | - | 481.5 | 137.7 | - | - | 0 | - |
| - | - | 998.4 | 138.1 | - | - | 0 | - |
| - | - | 1251 | 139.1 | - | - | 0 | - |
| - | - | 378.5 | 139.9 | - | - | 0 | - |
| - | - | 2516 | 141.1 | - | - | 0 | - |
| - | - | 1181 | 143.1 | - | - | 0 | - |
| - | - | 2184 | 144.1 | - | - | 0 | - |
| 8 | y | 2.746E+04 | 147.1 | 0.0001872 | 1.272 | +1 | 1 |
| - | - | 1492 | 148.1 | - | - | 0 | - |
| - | - | 499 | 148.9 | - | - | 0 | - |
| - | - | 498.3 | 148.9 | - | - | 0 | - |
| - | - | 610.6 | 148.9 | - | - | 0 | - |
| - | - | 662.8 | 148.9 | - | - | 0 | - |
| - | - | 523.5 | 148.9 | - | - | 0 | - |
| - | - | 871.8 | 148.9 | - | - | 0 | - |
| - | - | 1168 | 148.9 | - | - | 0 | - |
| - | - | 1140 | 148.9 | - | - | 0 | - |
| - | - | 2381 | 148.9 | - | - | 0 | - |
| - | - | 3675 | 148.9 | - | - | 0 | - |
| - | - | 4503 | 149 | - | - | 0 | - |
| - | - | 2820 | 149 | - | - | 0 | - |
| - | - | 1398 | 149 | - | - | 0 | - |
| - | - | 886.1 | 149 | - | - | 0 | - |
| - | - | 842.7 | 149 | - | - | 0 | - |
| - | - | 848.9 | 149 | - | - | 0 | - |
| - | - | 709 | 149 | - | - | 0 | - |
| - | - | 755.7 | 149 | - | - | 0 | - |
| - | - | 5629 | 149 | - | - | 0 | - |
| - | - | 467.1 | 149 | - | - | 0 | - |
| - | - | 5215 | 150 | - | - | 0 | - |
| - | - | 2011 | 151 | - | - | 0 | - |
| - | - | 7623 | 155.1 | - | - | 0 | - |
| 2 | a | 3756 | 155.1 | 0.0001372 | 0.8845 | +1 | 2 |
| - | - | 623.1 | 156.1 | - | - | 0 | - |
| - | - | 903.1 | 156.1 | - | - | 0 | - |
| - | - | 1.397E+04 | 157.1 | - | - | 0 | - |
| - | - | 1039 | 158.1 | - | - | 0 | - |
| - | - | 922.7 | 167.1 | - | - | 0 | - |
| - | - | 964.6 | 167.1 | - | - | 0 | - |
| - | - | 625.9 | 168 | - | - | 0 | - |
| - | - | 1292 | 169.1 | - | - | 0 | - |
| - | - | 503.5 | 171.1 | - | - | 0 | - |
| - | - | 731.6 | 171.1 | - | - | 0 | - |
| - | - | 828.5 | 172.1 | - | - | 0 | - |
| - | - | 894.1 | 173.1 | - | - | 0 | - |
| - | - | 8278 | 173.1 | - | - | 0 | - |
| 2 | a | 4.181E+05 | 173.1 | 0.0002537 | 1.465 | +1 | 2 |
| - | - | 459.8 | 173.6 | - | - | 0 | - |
| - | - | 3.272E+04 | 174.1 | - | - | 0 | - |
| - | - | 1262 | 175.1 | - | - | 0 | - |
| - | - | 2231 | 175.1 | - | - | 0 | - |
| - | - | 735 | 179.1 | - | - | 0 | - |
| - | - | 472.4 | 179.3 | - | - | 0 | - |
| - | - | 524.3 | 183.1 | - | - | 0 | - |
| 2 | b | 3064 | 183.1 | 4.985E-05 | 0.2722 | +1 | 2 |
| - | - | 538.5 | 183.1 | - | - | 0 | - |
| - | - | 1.07E+04 | 185.1 | - | - | 0 | - |
| - | - | 4378 | 185.2 | - | - | 0 | - |
| - | - | 1075 | 186.1 | - | - | 0 | - |
| - | - | 1.577E+04 | 186.1 | - | - | 0 | - |
| 7 | y | 3537 | 187.1 | 6.931E-05 | 0.3704 | +1 | 2 |
| - | - | 1291 | 187.1 | - | - | 0 | - |
| - | - | 1173 | 187.1 | - | - | 0 | - |
| - | - | 2480 | 189.1 | - | - | 0 | - |
| - | - | 1019 | 193.1 | - | - | 0 | - |
| - | - | 589.7 | 194.1 | - | - | 0 | - |
| - | - | 753.1 | 195.1 | - | - | 0 | - |
| - | - | 809.2 | 197.1 | - | - | 0 | - |
| - | - | 669.5 | 199.1 | - | - | 0 | - |
| 2 | b | 8.293E+04 | 201.1 | 4.424E-05 | 0.22 | +1 | 2 |
| - | - | 8016 | 202.1 | - | - | 0 | - |
| - | - | 1635 | 203.1 | - | - | 0 | - |
| 7 | y | 6.317E+04 | 204.1 | 5.524E-05 | 0.2706 | +1 | 2 |
| - | - | 5063 | 205.1 | - | - | 0 | - |
| - | - | 956.1 | 206.1 | - | - | 0 | - |
| - | - | 668 | 206.1 | - | - | 0 | - |
| - | - | 1104 | 207.1 | - | - | 0 | - |
| - | - | 1682 | 211.1 | - | - | 0 | - |
| - | - | 1388 | 213.2 | - | - | 0 | - |
| - | - | 1037 | 214.1 | - | - | 0 | - |
| - | - | 1199 | 215.1 | - | - | 0 | - |
| - | - | 3187 | 217.1 | - | - | 0 | - |
| - | - | 823.4 | 223.1 | - | - | 0 | - |
| - | - | 6388 | 224.1 | - | - | 0 | - |
| - | - | 708.7 | 224.1 | - | - | 0 | - |
| - | - | 555.7 | 225 | - | - | 0 | - |
| - | - | 609.7 | 229 | - | - | 0 | - |
| - | - | 587.9 | 230.2 | - | - | 0 | - |
| - | - | 1169 | 240.1 | - | - | 0 | - |
| - | - | 572.5 | 241.1 | - | - | 0 | - |
| - | - | 6244 | 242.1 | - | - | 0 | - |
| 3 | a | 977.4 | 242.1 | 0.0004879 | 2.015 | +1 | 3 |
| - | - | 1015 | 243.1 | - | - | 0 | - |
| - | - | 1152 | 243.1 | - | - | 0 | - |
| - | - | 575.3 | 245.1 | - | - | 0 | - |
| - | - | 2348 | 248.2 | - | - | 0 | - |
| - | - | 596.2 | 251.1 | - | - | 0 | - |
| - | - | 742.9 | 252.1 | - | - | 0 | - |
| - | - | 2697 | 260.1 | - | - | 0 | - |
| 3 | a | 1416 | 260.2 | 0.0001619 | 0.6223 | +1 | 3 |
| - | - | 853 | 262.2 | - | - | 0 | - |
| - | - | 719.9 | 268.2 | - | - | 0 | - |
| 3 | b | 1.163E+04 | 270.1 | 0.0002403 | 0.8895 | +1 | 3 |
| - | - | 1675 | 271.1 | - | - | 0 | - |
| - | - | 1298 | 283.1 | - | - | 0 | - |
| - | - | 1.481E+04 | 283.2 | - | - | 0 | - |
| 6 | b | 1135 | 284.2 | 0.0002229 | 0.7845 | +2 | 6 |
| - | - | 1393 | 284.2 | - | - | 0 | - |
| 3 | y | 6866 | 285.7 | 3.86E-05 | 0.1351 | +2 | 6 |
| - | - | 1500 | 286.2 | - | - | 0 | - |
| - | - | 896.2 | 286.2 | - | - | 0 | - |
| 3 | b | 1.465E+04 | 288.2 | 0.0001849 | 0.6415 | +1 | 3 |
| - | - | 1997 | 289.2 | - | - | 0 | - |
| 3 | y | 1335 | 294.7 | 0.0003858 | 1.309 | +2 | 6 |
| - | - | 945.5 | 296.2 | - | - | 0 | - |
| - | - | 503 | 296.9 | - | - | 0 | - |
| - | - | 789 | 298.1 | - | - | 0 | - |
| - | - | 609.5 | 299.7 | - | - | 0 | - |
| 6 | y | 1.796E+05 | 301.2 | 4.102E-05 | 0.1362 | +1 | 3 |
| - | - | 2.516E+04 | 302.2 | - | - | 0 | - |
| - | - | 2333 | 303.2 | - | - | 0 | - |
| - | - | 744 | 308.2 | - | - | 0 | - |
| - | - | 1064 | 309.2 | - | - | 0 | - |
| - | - | 777.1 | 312.2 | - | - | 0 | - |
| - | - | 917.6 | 326.2 | - | - | 0 | - |
| - | - | 1133 | 330.2 | - | - | 0 | - |
| - | - | 862.3 | 333.2 | - | - | 0 | - |
| - | - | 2195 | 335.2 | - | - | 0 | - |
| - | - | 1145 | 336.7 | - | - | 0 | - |
| 2 | y | 2285 | 342.2 | 0.0002005 | 0.5858 | +2 | 7 |
| - | - | 759.7 | 342.7 | - | - | 0 | - |
| - | - | 1465 | 343.2 | - | - | 0 | - |
| 2 | y | 2092 | 351.2 | 7.139E-05 | 0.2033 | +2 | 7 |
| - | - | 2036 | 352.2 | - | - | 0 | - |
| - | - | 1907 | 353.2 | - | - | 0 | - |
| - | - | 785 | 355.2 | - | - | 0 | - |
| - | - | 3866 | 355.2 | - | - | 0 | - |
| - | - | 652.2 | 356.2 | - | - | 0 | - |
| - | - | 571.7 | 357.2 | - | - | 0 | - |
| - | - | 1378 | 358.2 | - | - | 0 | - |
| - | - | 1759 | 358.9 | - | - | 0 | - |
| - | - | 796 | 359.9 | - | - | 0 | - |
| - | - | 960.7 | 361.2 | - | - | 0 | - |
| - | - | 802.1 | 365.9 | - | - | 0 | - |
| - | - | 1043 | 367.2 | - | - | 0 | - |
| - | - | 767.1 | 368.2 | - | - | 0 | - |
| 5 | y | 1.018E+04 | 370.2 | 0.0004876 | 1.317 | +1 | 4 |
| - | - | 1415 | 371.2 | - | - | 0 | - |
| - | - | 1378 | 375.2 | - | - | 0 | - |
| - | - | 585.7 | 375.9 | - | - | 0 | - |
| - | - | 1009 | 376.7 | - | - | 0 | - |
| - | - | 1300 | 376.8 | - | - | 0 | - |
| - | - | 500.6 | 377.8 | - | - | 0 | - |
| 4 | b | 1483 | 383.2 | 0.001754 | 4.577 | +1 | 4 |
| - | - | 875.7 | 385.2 | - | - | 0 | - |
| 0 | Precursor | 1.354E+04 | 385.7 | 0.0005592 | 1.45 | +2 | -1 |
| - | - | 6418 | 386.2 | - | - | 0 | - |
| - | - | 1552 | 386.7 | - | - | 0 | - |
| 5 | y | 6.588E+04 | 388.2 | 0.0004932 | 1.271 | +1 | 4 |
| - | - | 1.211E+04 | 389.2 | - | - | 0 | - |
| - | - | 1556 | 390.2 | - | - | 0 | - |
| - | - | 726.1 | 393.2 | - | - | 0 | - |
| - | - | 1412 | 393.3 | - | - | 0 | - |
| - | - | 797.8 | 393.9 | - | - | 0 | - |
| - | - | 2036 | 394.2 | - | - | 0 | - |
| - | - | 751.3 | 394.2 | - | - | 0 | - |
| - | - | 1005 | 394.3 | - | - | 0 | - |
| - | - | 1567 | 394.7 | - | - | 0 | - |
| - | - | 1632 | 398.2 | - | - | 0 | - |
| - | - | 1074 | 406.2 | - | - | 0 | - |
| - | - | 1040 | 424.2 | - | - | 0 | - |
| - | - | 1770 | 441.2 | - | - | 0 | - |
| - | - | 1713 | 442.2 | - | - | 0 | - |
| - | - | 613.7 | 442.3 | - | - | 0 | - |
| - | - | 948.4 | 456.3 | - | - | 0 | - |
| - | - | 632 | 457.3 | - | - | 0 | - |
| - | - | 2262 | 460.3 | - | - | 0 | - |
| - | - | 669 | 461.3 | - | - | 0 | - |
| 5 | b | 1721 | 470.3 | 0.0003659 | 0.778 | +1 | 5 |
| - | - | 571.7 | 471.3 | - | - | 0 | - |
| 4 | y | 2887 | 483.3 | 0.0008114 | 1.679 | +1 | 5 |
| 4 | y | 1.689E+04 | 501.3 | 0.0004508 | 0.8992 | +1 | 5 |
| - | - | 4857 | 502.3 | - | - | 0 | - |
| - | - | 1108 | 552.3 | - | - | 0 | - |
| - | - | 873.7 | 558.3 | - | - | 0 | - |
| - | - | 652.7 | 569.3 | - | - | 0 | - |
| 3 | y | 9306 | 570.3 | 0.001102 | 1.931 | +1 | 6 |
| - | - | 2853 | 571.3 | - | - | 0 | - |
| - | - | 1076 | 573.4 | - | - | 0 | - |
| - | - | 5397 | 577.3 | - | - | 0 | - |
| - | - | 1021 | 578.3 | - | - | 0 | - |
| 3 | y | 1.904E+05 | 588.3 | 0.001168 | 1.986 | +1 | 6 |
| - | - | 6.059E+04 | 589.3 | - | - | 0 | - |
| - | - | 1.254E+04 | 590.3 | - | - | 0 | - |
| - | - | 672.7 | 591.3 | - | - | 0 | - |
| - | - | 4748 | 598.3 | - | - | 0 | - |
| - | - | 1319 | 599.3 | - | - | 0 | - |
| 7 | b | 792.4 | 624.3 | 0.000863 | 1.382 | +1 | 7 |
| - | - | 855.5 | 658.4 | - | - | 0 | - |
| - | - | 1842 | 667.3 | - | - | 0 | - |
| - | - | 966.3 | 668.3 | - | - | 0 | - |
| - | - | 983.3 | 674.4 | - | - | 0 | - |
| - | - | 1335 | 676.4 | - | - | 0 | - |
| 2 | y | 3492 | 701.4 | 0.001187 | 1.692 | +1 | 7 |
| - | - | 639.5 | 1072 | - | - | 0 | - |
| - | - | 748.2 | 1436 | - | - | 0 | - |
| - | - | 585 | 1500 | - | - | 0 | - |
| - | - | 739.3 | 2173 | - | - | 0 | - |
| - | - | 608.8 | 2775 | - | - | 0 | - |
| - | - | 587.1 | 3027 | - | - | 0 | - |
| - | - | 710.4 | 3195 | - | - | 0 | - |

m/z Charge Intensity FragmentType MassShift Position
120.57836151123047 0 403.1666
125.07132720947266 0 409.50305
125.10774230957031 0 703.4597
126.05537414550781 0 717.136
126.09159088134766 0 1559.5801
127.08677673339844 0 4172.0474
127.90579223632812 0 373.68765
128.101806640625 0 965.57544
128.10719299316406 0 18172.346
129.1024169921875 0 25378.938
129.11058044433594 0 1095.5016
130.0864715576172 0 15299.106 y Ammonia loss 7
130.10577392578125 0 1664.815
131.33323669433594 0 441.32135
135.75767517089844 0 364.69183
137.73428344726562 0 481.47372
138.09181213378906 0 998.3552
139.0867919921875 0 1250.8433
139.85801696777344 0 378.46802
141.10244750976562 0 2516.4563
143.1179656982422 0 1180.951
144.06565856933594 0 2184.1416
147.1129913330078 0 27462.602 y 7
148.11630249023438 0 1492.1783
148.87890625 0 498.9996
148.88571166992188 0 498.32492
148.8921356201172 0 610.5879
148.899658203125 0 662.8499
148.90638732910156 0 523.45844
148.91372680664062 0 871.7865
148.9207763671875 0 1168.4913
148.9281005859375 0 1139.5995
148.93484497070312 0 2381.4065
148.94244384765625 0 3675.1794
148.95858764648438 0 4502.8022
148.96620178222656 0 2820.0225
148.97340393066406 0 1398.0354
148.9803466796875 0 886.055
148.9876251220703 0 842.7003
148.99459838867188 0 848.9191
149.00157165527344 0 708.962
149.00845336914062 0 755.671
149.02342224121094 0 5628.638
149.04385375976562 0 467.08008
150.02687072753906 0 5214.6694
151.02882385253906 0 2010.8389
155.08164978027344 0 7623.077
155.11802673339844 0 3755.7344 a Water loss 1
156.0850830078125 0 623.14856
156.1021728515625 0 903.1003
157.09730529785156 0 13974.9795
158.10064697265625 0 1038.5181
167.0813446044922 0 922.74066
167.1183624267578 0 964.5807
168.03785705566406 0 625.94116
169.0972137451172 0 1291.8412
171.07664489746094 0 503.4876
171.1132049560547 0 731.62494
172.10809326171875 0 828.5114
173.05587768554688 0 894.0852
173.09214782714844 0 8278.37
173.1287078857422 0 418144.56 a 1
173.576904296875 0 459.77548
174.13194274902344 0 32723.021
175.11892700195312 0 1261.8369
175.13392639160156 0 2230.545
179.08172607421875 0 734.97687
179.31137084960938 0 472.44052
183.06784057617188 0 524.32275
183.11285400390625 0 3064.3975 b Water loss 1
183.13262939453125 0 538.49615
185.09222412109375 0 10704.112
185.1649627685547 0 4378.443
186.09571838378906 0 1074.6057
186.12380981445312 0 15768.373
187.1077880859375 0 3536.5667 y Ammonia loss 6
187.1274871826172 0 1290.5289
187.1442108154297 0 1173.2461
189.08712768554688 0 2480.085
193.09715270996094 0 1018.64325
194.0921630859375 0 589.66516
195.0762176513672 0 753.08496
197.12843322753906 0 809.1886
199.07106018066406 0 669.5466
201.1234130859375 0 82934.2 b 1
202.1267547607422 0 8015.893
203.06613159179688 0 1635.0991
204.1343231201172 0 63165.324 y 6
205.1375732421875 0 5062.926
206.09217834472656 0 956.13916
206.14007568359375 0 668.00586
207.07656860351562 0 1103.9316
211.1074981689453 0 1682.2941
213.15951538085938 0 1388.2158
214.1187744140625 0 1037.3447
215.13900756835938 0 1199.3682
217.0819091796875 0 3187.2612
223.1077880859375 0 823.3722
224.1029510498047 0 6388.2915
224.13966369628906 0 708.7253
225.00721740722656 0 555.6829
228.96876525878906 0 609.7231
230.15078735351562 0 587.9273
240.13421630859375 0 1169.1394
241.11953735351562 0 572.5186
242.11361694335938 0 6244.3647
242.15040588378906 0 977.3862 a Water loss 2
243.11642456054688 0 1014.9558
243.1334686279297 0 1151.7274
245.12396240234375 0 575.3377
248.1603240966797 0 2347.8438
251.13931274414062 0 596.17413
252.13453674316406 0 742.90643
260.1239318847656 0 2697.1455
260.16064453125 0 1416.2731 a 2
262.15032958984375 0 853.01666
268.2015686035156 0 719.89813
270.14459228515625 0 11631.939 b Water loss 2
271.1485595703125 0 1675.0503
283.1395263671875 0 1297.6416
283.1763610839844 0 14807.407
284.16070556640625 0 1135.3463 b Water loss 5
284.1796569824219 0 1392.7019
285.6658935546875 0 6865.5117 y Water loss 2
286.1673278808594 0 1500.4629
286.18292236328125 0 896.2008
288.15521240234375 0 14649.976 b 2
289.15814208984375 0 1996.5681
294.6716003417969 0 1335.0323 y 2
296.19647216796875 0 945.4523
296.9256896972656 0 503.0246
298.1394348144531 0 789.0463
299.6633605957031 0 609.52734
301.18707275390625 0 179619.28 y 5
302.1900329589844 0 25158.209
303.19232177734375 0 2333.3818
308.1609802246094 0 744.00183
309.2033996582031 0 1064.0034
312.15545654296875 0 777.1452
326.17181396484375 0 917.6084
330.1661682128906 0 1132.615
333.1911926269531 0 862.2521
335.1708984375 0 2195.2278
336.70477294921875 0 1145.2367
342.207763671875 0 2284.765 y Water loss 1
342.7081298828125 0 759.73267
343.1791076660156 0 1464.9878
351.21331787109375 0 2091.6816 y 1
352.1976013183594 0 2035.9728
353.1810607910156 0 1906.903
355.1953430175781 0 784.9622
355.2333679199219 0 3865.8843
356.2057800292969 0 652.2324
357.17462158203125 0 571.711
358.2080993652344 0 1377.7062
358.88824462890625 0 1759.3345
359.8873291015625 0 795.99054
361.2448425292969 0 960.71375
365.89617919921875 0 802.0664
367.1973571777344 0 1043.4252
368.1933898925781 0 767.1376
370.2080078125 0 10180.525 y Water loss 4
371.21026611328125 0 1414.9425
375.23388671875 0 1377.7063
375.883544921875 0 585.69354
376.71881103515625 0 1008.89514
376.843017578125 0 1299.6808
377.8469543457031 0 500.61377
383.2271423339844 0 1483.2721 b Water loss 3
385.2110290527344 0 875.6655
385.7234191894531 0 13537.579 Precursor Water loss
386.2250671386719 0 6418.1787
386.7249755859375 0 1551.5806
388.21856689453125 0 65878.88 y 4
389.2214660644531 0 12105.713
390.22454833984375 0 1555.6727
393.2462463378906 0 726.09674
393.29681396484375 0 1412.0394
393.87103271484375 0 797.8297
394.21795654296875 0 2036.2339
394.24542236328125 0 751.2967
394.3003845214844 0 1005.4055
394.7205505371094 0 1567.0482
398.2016906738281 0 1631.8794
406.207763671875 0 1074.2374
424.217529296875 0 1039.9167
441.2441711425781 0 1769.7325
442.2295837402344 0 1713.4678
442.2667541503906 0 613.66254
456.2554626464844 0 948.35187
457.2613525390625 0 632.0253
460.2873840332031 0 2261.8843
461.2895202636719 0 669.04944
470.26055908203125 0 1720.6239 b Water loss 4
471.26617431640625 0 571.67084
483.291748046875 0 2886.7715 y Water loss 3
501.30267333984375 0 16892.238 y 3
502.3055725097656 0 4856.647
552.3134765625 0 1107.997
558.3258056640625 0 873.68884
569.3397216796875 0 652.6922
570.323486328125 0 9305.604 y Water loss 2
571.3250732421875 0 2853.088
573.3701171875 0 1076.3273
577.3173828125 0 5397.199
578.3202514648438 0 1020.76654
588.333984375 0 190364 y 2
589.3367919921875 0 60588.664
590.3389282226562 0 12536.457
591.3421630859375 0 672.7305
598.3182983398438 0 4748.2026
599.322021484375 0 1319.4321
624.3342895507812 0 792.384 b Water loss 6
658.3749389648438 0 855.54065
667.3392333984375 0 1841.7452
668.3389282226562 0 966.3058
674.3978881835938 0 983.29236
676.3876953125 0 1334.5764
701.4180297851562 0 3492.378 y 1
1072.1065673828125 0 639.4995
1436.4774169921875 0 748.1933
1499.5516357421875 0 585.0283
2172.546630859375 0 739.32745
2774.839599609375 0 608.82574
3027.38720703125 0 587.1096
3194.580322265625 0 710.44495

Spectrum Details

|  |  |
| --- | --- |
| Matched peaks? Matched peaks The total absolute number of peaks matched. Additionally in brackets the total fraction of peaks matched and the total number of peaks is shown. | 29 (12.83% of 226) |
| FDR? FDR The false discovery rate estimated for this peptide. It is calculated by matching all theoretical fragments with a non-integer shift with the raw peaks for this spectrum. This is done with 40 different shifts. The resulting percentage is the average number of annotated peaks over the number of annotated peaks with the correct spectrum. | 0.33% |
| Satellite FDR? Satellite FDR See the FDR for details on its calculation. This satellite ion specific FDR only contains the satellite ions (d/w) for I/L/J positions. | - |
| PSM Score? PSM Score The PSM Score as given by Hecklib to this annotated spectrum. It is shown with three significant figures. | 385 |

#### Reverse Lookup? Reverse Lookup All places where this read could be placed.

| Group | Segment | Template | Template Part | Read Part | Score | Unique |
| --- | --- | --- | --- | --- | --- | --- |
| Homo sapiens Heavy Chain | IGHC | IGHG1 | [322..330] | [0..8] | 64 | False |
| Homo sapiens Heavy Chain | IGHC | IGHG3 | [369..377] | [0..8] | 64 | False |
| Homo sapiens Heavy Chain | IGHC | IGHG2 | [318..326] | [0..8] | 64 | False |

| Recombined | Template Part | Read Part | Score | Unique |
| --- | --- | --- | --- | --- |
| REC-0-1 | [447..455] | [0..8] | 55 | True |

#### Meta Information from Multiple reads

##### Number of combined reads

2

##### Intensity

0.5801

##### TotalArea

1.397E+07

##### Changes to the peptide sequence

SJSJSPGK

L→JNo support for either Leucine or Isoleucine based on side chain ions (Position: 4)

L→JEqual support for both Leucine and Isoleucine based on side chain ions (1 ions for both) (Position: 2)

#### Positional Score

Copy Data

##### Positional Score (TSV)

###### Preview

```
Loading example...
```

*Click on the button to copy the data to your clipboard.*

0001234567

Label Value
"0" 0
"1" 0
"2" 0
"3" 0
"4" 0
"5" 0
"6" 0
"7" 0

#### Meta Information from PEAKS

##### Scan Identifier

F4:4021

##### Original sequence

S

L

S

L

S

P

G

K

##### Posttranslational Modifications

##### Source File

D:\separate\_stitch\_analyses\xle-disambiguation\raw\20210323\_F1\_UM1\_Peng0013\_SA\_F59\_ingel\_3ug\_tryp.raw

##### Fraction

4

##### Scan Feature

F4:610

##### De Novo Score

98

##### ConfidenceScore

98

### m/z

394.7301

##### Mass

787.4439

##### Charge

2

##### Retention Time

21.59

##### Predicted Retention Time

18.36

##### Area

6.983E+06

##### Parts Per Million

2.2

##### Fragmentation mode

HCD

##### Originating file

01 D:\separate\_stitch\_analyses\xle-disambiguation\20210325\_F59\_3ug\_DENOVO\_12.csv

#### Meta Information from PEAKS

##### Scan Identifier

F4:4071

##### Original sequence

S

L

S

L

S

P

G

K

##### Posttranslational Modifications

##### Source File

D:\separate\_stitch\_analyses\xle-disambiguation\raw\20210323\_F1\_UM1\_Peng0013\_SA\_F59\_ingel\_3ug\_tryp.raw

##### Fraction

4

##### Scan Feature

F4:610

##### De Novo Score

97

##### ConfidenceScore

98

### m/z

394.7301

##### Mass

787.4439

##### Charge

2

##### Retention Time

21.59

##### Predicted Retention Time

18.36

##### Area

6.983E+06

##### Parts Per Million

2.2

##### Fragmentation mode

HCD

##### Originating file

01 D:\separate\_stitch\_analyses\xle-disambiguation\20210325\_F59\_3ug\_DENOVO\_12.csv
