## Supplementary material for "A handle on mass coincidence errors in *de novo* sequencing of antibodies by bottom-up proteomics": Combined_086.html

Details Combined\_086 | Stitch OverviewUndefined

### Read Combined\_086

#### Sequence (length=9)

AQYEDJAQK

#### Spectrum 3073? Spectrum 3073 The raw spectrum of this peptide as annotated by Hecklib. The fragments are coloured according to ion type (see legend). Any peaks with a star '\*' as text can be hovered over to see the full details, first the ion type second the mass shift type. By hovering over the amino acids in the peptide or ions in the legend the corresponding peaks are highlighted. By toggling the 'Unassigned' label you can turn the background (unassigned) peaks on or off in the plot. By updating the slider in the Ion legend you can update the spectrum to only show the top X% of the peaks with labels. The top X% means any peak that is within X% of the highest intensity. By dragging in the spectrum you can zoom in to a specific part of the spectrum and use 'Zoom Out' to get back to the original zoom level. The annotation of the spectrum is based on the given sequence in the peptides file and is done with different software so inconsistencies are likely. The peaks are annotated based on the given sequence, with 20 ppm tolerance.

Copy Data

##### Spectrum 3073 (TSV)

###### Preview

```
Loading example...
```

*Click on the button to copy the data to your clipboard.*

Mz MinMz MaxIntensity Max

WidthHeightPeptide font sizePeptide stroke widthSpectrum font sizeSpectrum stroke widthCompact peptide

Ion legend

wxyz

abcd

OtherUnassignedIon^Charge^_Position_Show for top:%

AQYEDJAQK

03.41e+46.83e+41.02e+51.37e+5

Zoom Out

y^+1^_1_y^+1^_1_c^+1^_2_y^+1^_2_z^+1^_2_y^+1^_2_z^+1^_3_y^+1^_3_c^+1^_3_w^+1^_4_y^+2^_7_z^+1^_4_y^+1^_4_y^+2^_8_y^+2^_8_z^+2^_8_c^+1^_4_y^+2^_8_z^+1^_5_y^+1^_5_c^+1^_5_w^+1^_6_y^+1^_6_z^+1^_6_y^+1^_6_c^+1^_6_c^+1^_6_c^+1^_7_z^+1^_7_y^+1^_7_c^+1^_8_w^+1^_8_c^+1^_8_y^+1^_8_z^+1^_8_y^+1^_8_

0847169425413388

Fragment Matches Table

Show background peaks

| Position | Ion type | Intensity | mz Theoretical | mz Error (Th) | mz Error (ppm) | Charge | Series Number |
| --- | --- | --- | --- | --- | --- | --- | --- |
| - | - | 351.8 | 120.6 | - | - | 0 | - |
| - | - | 349.5 | 120.8 | - | - | 0 | - |
| - | - | 401 | 129 | - | - | 0 | - |
| - | - | 851.9 | 129.1 | - | - | 0 | - |
| 9 | y | 789 | 130.1 | 0.0002318 | 1.782 | +1 | 1 |
| - | - | 387 | 131.2 | - | - | 0 | - |
| - | - | 416.2 | 136.1 | - | - | 0 | - |
| - | - | 434.1 | 138.8 | - | - | 0 | - |
| - | - | 406 | 139.5 | - | - | 0 | - |
| - | - | 375.4 | 139.7 | - | - | 0 | - |
| 9 | y | 2771 | 147.1 | 0.0002635 | 1.791 | +1 | 1 |
| - | - | 405.8 | 148.5 | - | - | 0 | - |
| - | - | 2315 | 149 | - | - | 0 | - |
| - | - | 427.1 | 154.4 | - | - | 0 | - |
| - | - | 1184 | 155.1 | - | - | 0 | - |
| - | - | 467.9 | 156.6 | - | - | 0 | - |
| - | - | 1369 | 166.1 | - | - | 0 | - |
| - | - | 585.5 | 167.1 | - | - | 0 | - |
| - | - | 3126 | 172.1 | - | - | 0 | - |
| - | - | 1.953E+04 | 183.1 | - | - | 0 | - |
| - | - | 1017 | 184.1 | - | - | 0 | - |
| 2 | c | 5.221E+04 | 200.1 | 0.0001511 | 0.7553 | +1 | 2 |
| - | - | 4596 | 201.1 | - | - | 0 | - |
| - | - | 507.9 | 211.7 | - | - | 0 | - |
| - | - | 8406 | 221.1 | - | - | 0 | - |
| - | - | 1160 | 222.1 | - | - | 0 | - |
| - | - | 3802 | 225 | - | - | 0 | - |
| - | - | 1.15E+04 | 239.1 | - | - | 0 | - |
| - | - | 1090 | 240.1 | - | - | 0 | - |
| - | - | 592.9 | 247.1 | - | - | 0 | - |
| - | - | 1004 | 257.2 | - | - | 0 | - |
| 8 | y | 1354 | 258.1 | 3.438E-05 | 0.1332 | +1 | 2 |
| 8 | z | 794.6 | 259.2 | 0.0001439 | 0.5553 | +1 | 2 |
| - | - | 829.1 | 272.2 | - | - | 0 | - |
| - | - | 585.6 | 273.6 | - | - | 0 | - |
| - | - | 658.7 | 275.1 | - | - | 0 | - |
| 8 | y | 3756 | 275.2 | 0.0002797 | 1.016 | +1 | 2 |
| - | - | 1028 | 281.1 | - | - | 0 | - |
| - | - | 593.4 | 288.8 | - | - | 0 | - |
| - | - | 850.8 | 293.1 | - | - | 0 | - |
| - | - | 6441 | 295.1 | - | - | 0 | - |
| - | - | 1211 | 296.1 | - | - | 0 | - |
| - | - | 3616 | 299.1 | - | - | 0 | - |
| - | - | 952.8 | 313.1 | - | - | 0 | - |
| - | - | 683.6 | 318.1 | - | - | 0 | - |
| 7 | z | 1.133E+04 | 330.2 | 0.0003226 | 0.977 | +1 | 3 |
| - | - | 8561 | 331.2 | - | - | 0 | - |
| - | - | 991.9 | 332.2 | - | - | 0 | - |
| - | - | 836.9 | 346.1 | - | - | 0 | - |
| 7 | y | 6848 | 346.2 | 3.115E-05 | 0.08999 | +1 | 3 |
| - | - | 1289 | 347.2 | - | - | 0 | - |
| - | - | 1329 | 355.1 | - | - | 0 | - |
| 3 | c | 3833 | 363.2 | 5.503E-05 | 0.1515 | +1 | 3 |
| - | - | 705.9 | 364.2 | - | - | 0 | - |
| - | - | 6472 | 369.1 | - | - | 0 | - |
| - | - | 1671 | 370.1 | - | - | 0 | - |
| - | - | 824.4 | 408.1 | - | - | 0 | - |
| 6 | w | 6263 | 414.2 | 0.0003362 | 0.8116 | +1 | 4 |
| - | - | 1028 | 415.2 | - | - | 0 | - |
| 3 | y | 2890 | 433.7 | 0.001149 | 2.648 | +2 | 7 |
| - | - | 842 | 434.2 | - | - | 0 | - |
| 6 | z | 3.516E+04 | 443.3 | 0.0004871 | 1.099 | +1 | 4 |
| - | - | 2.437E+04 | 444.3 | - | - | 0 | - |
| - | - | 4637 | 445.3 | - | - | 0 | - |
| - | - | 552.7 | 446.3 | - | - | 0 | - |
| - | - | 1143 | 458.2 | - | - | 0 | - |
| 6 | y | 6082 | 459.3 | 0.0002872 | 0.6254 | +1 | 4 |
| - | - | 1495 | 460.3 | - | - | 0 | - |
| - | - | 3785 | 485.3 | - | - | 0 | - |
| - | - | 876.3 | 486.3 | - | - | 0 | - |
| 2 | y | 5848 | 488.7 | 0.0003354 | 0.6864 | +2 | 8 |
| 2 | y | 3739 | 489.2 | 0.009396 | 19.21 | +2 | 8 |
| 2 | z | 1426 | 489.7 | 0.008474 | 17.3 | +2 | 8 |
| 4 | c | 928.5 | 492.2 | 0.0005833 | 1.185 | +1 | 4 |
| 2 | y | 2031 | 497.7 | 0.0003937 | 0.7909 | +2 | 8 |
| - | - | 1305 | 498.2 | - | - | 0 | - |
| - | - | 599.1 | 498.7 | - | - | 0 | - |
| - | - | 1.254E+04 | 514.3 | - | - | 0 | - |
| - | - | 8546 | 515.3 | - | - | 0 | - |
| - | - | 3913 | 515.7 | - | - | 0 | - |
| - | - | 3910 | 516.2 | - | - | 0 | - |
| - | - | 1967 | 516.3 | - | - | 0 | - |
| - | - | 983.3 | 516.7 | - | - | 0 | - |
| - | - | 678.5 | 519.2 | - | - | 0 | - |
| - | - | 5248 | 524.8 | - | - | 0 | - |
| - | - | 2839 | 525.3 | - | - | 0 | - |
| - | - | 949.5 | 525.8 | - | - | 0 | - |
| - | - | 860.3 | 529.3 | - | - | 0 | - |
| - | - | 4466 | 533.3 | - | - | 0 | - |
| - | - | 2974 | 533.8 | - | - | 0 | - |
| - | - | 1497 | 534.3 | - | - | 0 | - |
| 5 | z | 2.043E+04 | 558.3 | 0.0001249 | 0.2238 | +1 | 5 |
| - | - | 8.187E+04 | 559.3 | - | - | 0 | - |
| - | - | 2.58E+04 | 560.3 | - | - | 0 | - |
| - | - | 3563 | 561.3 | - | - | 0 | - |
| 5 | y | 1.069E+04 | 574.3 | 0.0002607 | 0.454 | +1 | 5 |
| - | - | 2618 | 575.3 | - | - | 0 | - |
| - | - | 918.1 | 592.3 | - | - | 0 | - |
| 5 | c | 1154 | 607.2 | 0.0004809 | 0.7919 | +1 | 5 |
| 4 | w | 1.162E+04 | 628.3 | 0.0004993 | 0.7946 | +1 | 6 |
| - | - | 4075 | 629.3 | - | - | 0 | - |
| - | - | 770.7 | 630.3 | - | - | 0 | - |
| - | - | 843.5 | 632.3 | - | - | 0 | - |
| - | - | 2418 | 643.4 | - | - | 0 | - |
| - | - | 712.8 | 644.4 | - | - | 0 | - |
| - | - | 644.1 | 682.4 | - | - | 0 | - |
| 4 | y | 1180 | 685.4 | 0.002046 | 2.985 | +1 | 6 |
| - | - | 780.3 | 686.4 | - | - | 0 | - |
| 4 | z | 2.41E+04 | 687.3 | 0.0001708 | 0.2485 | +1 | 6 |
| - | - | 2.54E+04 | 688.3 | - | - | 0 | - |
| - | - | 8943 | 689.4 | - | - | 0 | - |
| - | - | 1433 | 690.4 | - | - | 0 | - |
| - | - | 624.7 | 702.3 | - | - | 0 | - |
| - | - | 897.9 | 703.3 | - | - | 0 | - |
| 4 | y | 1.783E+04 | 703.4 | 0.0004012 | 0.5704 | +1 | 6 |
| - | - | 5453 | 704.4 | - | - | 0 | - |
| - | - | 1338 | 705.4 | - | - | 0 | - |
| 6 | c | 2702 | 720.3 | 0.00121 | 1.679 | +1 | 6 |
| - | - | 1091 | 721.3 | - | - | 0 | - |
| - | - | 1.658E+04 | 736.3 | - | - | 0 | - |
| 6 | c | 1.121E+04 | 737.3 | 0.002207 | 2.993 | +1 | 6 |
| - | - | 3418 | 738.3 | - | - | 0 | - |
| - | - | 750.1 | 739.3 | - | - | 0 | - |
| - | - | 2951 | 764.4 | - | - | 0 | - |
| - | - | 3561 | 765.4 | - | - | 0 | - |
| - | - | 1033 | 766.4 | - | - | 0 | - |
| - | - | 1441 | 778.4 | - | - | 0 | - |
| - | - | 2026 | 779.4 | - | - | 0 | - |
| - | - | 1062 | 780.4 | - | - | 0 | - |
| - | - | 3722 | 791.4 | - | - | 0 | - |
| - | - | 2132 | 792.4 | - | - | 0 | - |
| - | - | 5489 | 794.3 | - | - | 0 | - |
| - | - | 3277 | 795.3 | - | - | 0 | - |
| - | - | 1147 | 806.4 | - | - | 0 | - |
| - | - | 1.013E+04 | 807.4 | - | - | 0 | - |
| 7 | c | 1.486E+04 | 808.4 | 0.001479 | 1.83 | +1 | 7 |
| - | - | 5959 | 809.4 | - | - | 0 | - |
| - | - | 835.5 | 810.4 | - | - | 0 | - |
| - | - | 1064 | 834.4 | - | - | 0 | - |
| 3 | z | 5.415E+04 | 850.4 | 0.000328 | 0.3857 | +1 | 7 |
| - | - | 2.65E+04 | 851.4 | - | - | 0 | - |
| - | - | 6523 | 852.4 | - | - | 0 | - |
| 3 | y | 8.492E+04 | 866.4 | 0.0006194 | 0.7149 | +1 | 7 |
| - | - | 4.335E+04 | 867.4 | - | - | 0 | - |
| - | - | 783.7 | 867.5 | - | - | 0 | - |
| - | - | 1.191E+04 | 868.4 | - | - | 0 | - |
| - | - | 899.6 | 869.4 | - | - | 0 | - |
| - | - | 4615 | 892.4 | - | - | 0 | - |
| - | - | 2519 | 893.4 | - | - | 0 | - |
| - | - | 933.2 | 894.4 | - | - | 0 | - |
| 8 | c | 4672 | 919.4 | 0.002015 | 2.191 | +1 | 8 |
| 2 | w | 5237 | 920.4 | 0.009292 | 10.1 | +1 | 8 |
| - | - | 2829 | 921.4 | - | - | 0 | - |
| - | - | 2748 | 935.4 | - | - | 0 | - |
| 8 | c | 1.352E+05 | 936.4 | 0.0006085 | 0.6498 | +1 | 8 |
| - | - | 7.075E+04 | 937.4 | - | - | 0 | - |
| - | - | 1.811E+04 | 938.4 | - | - | 0 | - |
| - | - | 1673 | 939.4 | - | - | 0 | - |
| - | - | 8167 | 950.5 | - | - | 0 | - |
| - | - | 4618 | 951.5 | - | - | 0 | - |
| - | - | 1202 | 952.5 | - | - | 0 | - |
| 2 | y | 6118 | 977.5 | 0.00141 | 1.442 | +1 | 8 |
| 2 | z | 2.895E+04 | 978.5 | 0.0003117 | 0.3186 | +1 | 8 |
| - | - | 1.5E+04 | 979.5 | - | - | 0 | - |
| - | - | 5042 | 980.5 | - | - | 0 | - |
| - | - | 707.5 | 991.5 | - | - | 0 | - |
| - | - | 2.932E+04 | 993.5 | - | - | 0 | - |
| 2 | y | 1.493E+04 | 994.5 | 0.005913 | 5.946 | +1 | 8 |
| - | - | 6081 | 995.5 | - | - | 0 | - |
| - | - | 1098 | 996.5 | - | - | 0 | - |
| - | - | 664.9 | 1006 | - | - | 0 | - |
| - | - | 2.182E+04 | 1007 | - | - | 0 | - |
| - | - | 1.379E+04 | 1008 | - | - | 0 | - |
| - | - | 3128 | 1009 | - | - | 0 | - |
| - | - | 7127 | 1021 | - | - | 0 | - |
| - | - | 4.341E+04 | 1022 | - | - | 0 | - |
| - | - | 2.502E+04 | 1023 | - | - | 0 | - |
| - | - | 8465 | 1024 | - | - | 0 | - |
| - | - | 2736 | 1039 | - | - | 0 | - |
| - | - | 2287 | 1040 | - | - | 0 | - |
| - | - | 781 | 1041 | - | - | 0 | - |
| - | - | 1141 | 1049 | - | - | 0 | - |
| - | - | 1.252E+05 | 1050 | - | - | 0 | - |
| - | - | 7.497E+04 | 1051 | - | - | 0 | - |
| - | - | 2.339E+04 | 1052 | - | - | 0 | - |
| - | - | 2389 | 1053 | - | - | 0 | - |
| - | - | 7.871E+04 | 1066 | - | - | 0 | - |
| - | - | 1.062E+05 | 1067 | - | - | 0 | - |
| - | - | 4.864E+04 | 1068 | - | - | 0 | - |
| - | - | 1.313E+04 | 1069 | - | - | 0 | - |
| - | - | 1344 | 1070 | - | - | 0 | - |
| - | - | 1184 | 1081 | - | - | 0 | - |
| - | - | 707.3 | 1114 | - | - | 0 | - |
| - | - | 617.7 | 1227 | - | - | 0 | - |
| - | - | 907.9 | 3014 | - | - | 0 | - |
| - | - | 759.3 | 3077 | - | - | 0 | - |
| - | - | 662.5 | 3354 | - | - | 0 | - |

m/z Charge Intensity FragmentType MassShift Position
120.55233764648438 0 351.7893
120.79913330078125 0 349.49417
128.99656677246094 0 401.0451
129.10247802734375 0 851.91364
130.08648681640625 0 789.00714 y Ammonia loss 8
131.2035675048828 0 387.0328
136.07608032226562 0 416.1826
138.78582763671875 0 434.10516
139.45782470703125 0 405.98737
139.70384216308594 0 375.43793
147.11306762695312 0 2770.7263 y 8
148.4741668701172 0 405.84344
149.04519653320312 0 2315.0276
154.42031860351562 0 427.0983
155.0818634033203 0 1183.9598
156.63043212890625 0 467.9184
166.05032348632812 0 1368.6396
167.05557250976562 0 585.5174
172.10829162597656 0 3126.005
183.0766143798828 0 19530.342
184.08033752441406 0 1017.1462
200.10311889648438 0 52210.16 c Ammonia loss 1
201.10655212402344 0 4595.9624
211.70565795898438 0 507.88672
221.08460998535156 0 8406.333
222.0851287841797 0 1160.1879
225.04307556152344 0 3801.8357
239.09521484375 0 11504.675
240.0965576171875 0 1090.2296
247.10794067382812 0 592.86444
257.16070556640625 0 1004.0419
258.1448669433594 0 1353.5967 y Ammonia loss 7
259.1528015136719 0 794.60645 z 7
272.1604309082031 0 829.0607
273.6229248046875 0 585.55597
275.1027526855469 0 658.73956
275.1716613769531 0 3755.7483 y 7
281.0508117675781 0 1027.5016
288.8396301269531 0 593.3939
293.1130676269531 0 850.8355
295.1033935546875 0 6440.7
296.10491943359375 0 1211.2065
299.0621337890625 0 3616.359
313.1134033203125 0 952.7635
318.145751953125 0 683.62317
330.1900939941406 0 11332.858 z 6
331.1972961425781 0 8561.288
332.20001220703125 0 991.87714
346.1396179199219 0 836.8701
346.2085266113281 0 6848.2207 y 6
347.2115478515625 0 1288.9591
355.07037353515625 0 1328.6117
363.1663513183594 0 3833.0625 c Ammonia loss 2
364.1716613769531 0 705.8878
369.1220397949219 0 6471.8677
370.1233825683594 0 1671.2313
408.1409606933594 0 824.35205
414.23504638671875 0 6262.7207 w 5
415.2366638183594 0 1028.177
433.7174987792969 0 2890.2742 y 2
434.21746826171875 0 841.9726
443.2743225097656 0 35164.527 z 5
444.2804260253906 0 24365.363
445.2834777832031 0 4636.6274
446.28924560546875 0 552.7181
458.2476501464844 0 1142.7712
459.2928466796875 0 6082.044 y 5
460.29534912109375 0 1494.5989
485.272705078125 0 3784.543
486.2768249511719 0 876.31165
488.7406921386719 0 5847.8076 y Water loss 1
489.24176025390625 0 3739.4626 y Ammonia loss 1
489.7447509765625 0 1426.4786 z 1
492.20947265625 0 928.49536 c Ammonia loss 3
497.74603271484375 0 2030.8723 y 1
498.2463684082031 0 1305.439
498.7487487792969 0 599.12054
514.31103515625 0 12540.696
515.316650390625 0 8545.563
515.7454223632812 0 3912.8982
516.24169921875 0 3910.1606
516.3206176757812 0 1967.3379
516.7398071289062 0 983.3176
519.169677734375 0 678.5335
524.7514038085938 0 5248.4805
525.2533569335938 0 2839.4905
525.7554321289062 0 949.46747
529.2626953125 0 860.2552
533.264404296875 0 4465.611
533.7655029296875 0 2973.8398
534.2662353515625 0 1496.5226
558.3009033203125 0 20434.768 z 4
559.3082885742188 0 81872.07
560.3113403320312 0 25804.209
561.3135375976562 0 3563.2427
574.3197631835938 0 10688.6875 y 4
575.3223266601562 0 2617.5283
592.2630615234375 0 918.08527
607.2353515625 0 1153.7467 c Ammonia loss 4
628.33056640625 0 11619.29 w 3
629.3328857421875 0 4074.7715
630.3342895507812 0 770.73346
632.25439453125 0 843.47034
643.3533325195312 0 2418.0762
644.3570556640625 0 712.80237
682.3728637695312 0 644.1188
685.3535766601562 0 1180.245 y Water loss 3
686.3504028320312 0 780.3291
687.3432006835938 0 24100.408 z 3
688.3494873046875 0 25402.29
689.3529663085938 0 8943.023
690.3585815429688 0 1432.9587
702.3107299804688 0 624.72076
703.2926025390625 0 897.856
703.3616943359375 0 17829.076 y 3
704.3644409179688 0 5453.3203
705.3642578125 0 1337.9694
720.3211059570312 0 2702.3916 c Ammonia loss 5
721.321533203125 0 1091.0344
736.3386840820312 0 16577.6
737.34423828125 0 11212.366 c 5
738.3470458984375 0 3417.7666
739.3486938476562 0 750.1115
764.3695068359375 0 2951.2065
765.3758544921875 0 3561.2913
766.38330078125 0 1033.0255
778.3858642578125 0 1440.8212
779.372314453125 0 2026.2367
780.3720092773438 0 1062.3748
791.3923950195312 0 3722.2205
792.3880004882812 0 2132.1304
794.3429565429688 0 5488.994
795.3468627929688 0 3277.3796
806.4205932617188 0 1146.7343
807.3763427734375 0 10129.414
808.382080078125 0 14863.723 c 6
809.3856811523438 0 5959.1567
810.3900146484375 0 835.4934
834.4013061523438 0 1063.7896
850.4063720703125 0 54148.28 z 2
851.40966796875 0 26499.607
852.4129028320312 0 6523.0728
866.4248046875 0 84919.89 y 2
867.4276733398438 0 43347.97
867.5289306640625 0 783.739
868.4304809570312 0 11905.599
869.4327392578125 0 899.62726
892.42529296875 0 4614.978
893.4281616210938 0 2519.325
894.4304809570312 0 933.17975
919.4176025390625 0 4671.9146 c Ammonia loss 7
920.4266967773438 0 5236.815 w 1
921.434326171875 0 2829.1487
935.4354248046875 0 2747.6726
936.4415283203125 0 135229.95 c 7
937.4443359375 0 70748.086
938.4467163085938 0 18108.885
939.4473876953125 0 1672.7173
950.4685668945312 0 8167.228
951.4729614257812 0 4618.0586
952.4716796875 0 1201.5825
977.4588623046875 0 6117.576 y Ammonia loss 1
978.4649658203125 0 28945.941 z 1
979.4680786132812 0 14995.3545
980.47021484375 0 5041.7056
991.4694213867188 0 707.4981
993.4750366210938 0 29323.426
994.4780883789062 0 14932.338 y 1
995.4833374023438 0 6080.975
996.4923095703125 0 1097.612
1005.5111083984375 0 664.8615
1006.5069580078125 0 21817.05
1007.5103149414062 0 13794.7
1008.5126342773438 0 3127.8413
1020.5216674804688 0 7126.558
1021.5075073242188 0 43412.27
1022.510498046875 0 25018.758
1023.513671875 0 8465.074
1038.5303955078125 0 2736.396
1039.533935546875 0 2286.7153
1040.541748046875 0 780.9707
1048.5137939453125 0 1141.0786
1049.50146484375 0 125189.92
1050.504638671875 0 74972.695
1051.5074462890625 0 23393.867
1052.508544921875 0 2389.3284
1065.520263671875 0 78712.07
1066.5252685546875 0 106172.37
1067.529052734375 0 48635.64
1068.5322265625 0 13127.656
1069.5396728515625 0 1343.6605
1081.4921875 0 1184.1942
1113.74609375 0 707.3002
1226.788330078125 0 617.6611
3013.605712890625 0 907.88544
3076.909423828125 0 759.31934
3354.1767578125 0 662.506

Spectrum Details

|  |  |
| --- | --- |
| Matched peaks? Matched peaks The total absolute number of peaks matched. Additionally in brackets the total fraction of peaks matched and the total number of peaks is shown. | 36 (18.27% of 197) |
| FDR? FDR The false discovery rate estimated for this peptide. It is calculated by matching all theoretical fragments with a non-integer shift with the raw peaks for this spectrum. This is done with 40 different shifts. The resulting percentage is the average number of annotated peaks over the number of annotated peaks with the correct spectrum. | 1.19% |
| Satellite FDR? Satellite FDR See the FDR for details on its calculation. This satellite ion specific FDR only contains the satellite ions (d/w) for I/L/J positions. | 2.38% |
| PSM Score? PSM Score The PSM Score as given by Hecklib to this annotated spectrum. It is shown with three significant figures. | 394 |

#### Spectrum 3127? Spectrum 3127 The raw spectrum of this peptide as annotated by Hecklib. The fragments are coloured according to ion type (see legend). Any peaks with a star '\*' as text can be hovered over to see the full details, first the ion type second the mass shift type. By hovering over the amino acids in the peptide or ions in the legend the corresponding peaks are highlighted. By toggling the 'Unassigned' label you can turn the background (unassigned) peaks on or off in the plot. By updating the slider in the Ion legend you can update the spectrum to only show the top X% of the peaks with labels. The top X% means any peak that is within X% of the highest intensity. By dragging in the spectrum you can zoom in to a specific part of the spectrum and use 'Zoom Out' to get back to the original zoom level. The annotation of the spectrum is based on the given sequence in the peptides file and is done with different software so inconsistencies are likely. The peaks are annotated based on the given sequence, with 20 ppm tolerance.

Copy Data

##### Spectrum 3127 (TSV)

###### Preview

```
Loading example...
```

*Click on the button to copy the data to your clipboard.*

Mz MinMz MaxIntensity Max

WidthHeightPeptide font sizePeptide stroke widthSpectrum font sizeSpectrum stroke widthCompact peptide

Ion legend

wxyz

abcd

OtherUnassignedIon^Charge^_Position_Show for top:%

AQYEDJAQK

01.12e+42.23e+43.35e+44.46e+4

Zoom Out

y^+1^_1_y^+1^_1_a^+1^_2_a^+1^_2_b^+1^_2_b^+1^_2_y^+1^_2_y^+1^_2_y^+1^_3_y^+1^_3_b^+1^_3_y^+2^_7_y^+1^_4_y^+2^_8_y^+2^_8_b^+1^_4_y^+2^_8_\*\*y^+1^_5_b^+1^_5_b^+1^_5_y^+1^_6_b^+1^_6_y^+1^_6_b^+1^_6_y^+1^_7_y^+1^_7_b^+1^_8_y^+1^_8_

047394614191892

Fragment Matches Table

Show background peaks

| Position | Ion type | Intensity | mz Theoretical | mz Error (Th) | mz Error (ppm) | Charge | Series Number |
| --- | --- | --- | --- | --- | --- | --- | --- |
| - | - | 396.2 | 120.1 | - | - | 0 | - |
| - | - | 355.8 | 122.5 | - | - | 0 | - |
| - | - | 1276 | 129.1 | - | - | 0 | - |
| - | - | 4571 | 129.1 | - | - | 0 | - |
| 9 | y | 2770 | 130.1 | 0.0001402 | 1.078 | +1 | 1 |
| - | - | 1.169E+04 | 136.1 | - | - | 0 | - |
| - | - | 392.7 | 136.5 | - | - | 0 | - |
| - | - | 498.3 | 137.1 | - | - | 0 | - |
| - | - | 584.9 | 138.1 | - | - | 0 | - |
| 9 | y | 4798 | 147.1 | 0.0002177 | 1.48 | +1 | 1 |
| - | - | 714.2 | 148.9 | - | - | 0 | - |
| - | - | 5683 | 149 | - | - | 0 | - |
| - | - | 386.4 | 151.7 | - | - | 0 | - |
| - | - | 447 | 154.7 | - | - | 0 | - |
| 2 | a | 2422 | 155.1 | 0.0001305 | 0.8414 | +1 | 2 |
| - | - | 453 | 162.3 | - | - | 0 | - |
| - | - | 9158 | 166.1 | - | - | 0 | - |
| - | - | 1.024E+04 | 167.1 | - | - | 0 | - |
| - | - | 764.2 | 168.1 | - | - | 0 | - |
| 2 | a | 1582 | 172.1 | 0.0001622 | 0.9424 | +1 | 2 |
| - | - | 516.9 | 173.1 | - | - | 0 | - |
| - | - | 711.7 | 173.4 | - | - | 0 | - |
| 2 | b | 2.2E+04 | 183.1 | 0.0001652 | 0.9024 | +1 | 2 |
| - | - | 1491 | 184.1 | - | - | 0 | - |
| - | - | 552.9 | 194 | - | - | 0 | - |
| 2 | b | 3.297E+04 | 200.1 | 0.0001359 | 0.6791 | +1 | 2 |
| - | - | 571.2 | 201.1 | - | - | 0 | - |
| - | - | 2966 | 201.1 | - | - | 0 | - |
| - | - | 3409 | 201.1 | - | - | 0 | - |
| - | - | 882.1 | 202.1 | - | - | 0 | - |
| - | - | 682.7 | 207.1 | - | - | 0 | - |
| - | - | 515.4 | 217 | - | - | 0 | - |
| - | - | 1.273E+04 | 221.1 | - | - | 0 | - |
| - | - | 1748 | 222.1 | - | - | 0 | - |
| - | - | 588.8 | 223.1 | - | - | 0 | - |
| - | - | 1.932E+04 | 225 | - | - | 0 | - |
| - | - | 2039 | 226 | - | - | 0 | - |
| - | - | 2263 | 227 | - | - | 0 | - |
| - | - | 1309 | 227.1 | - | - | 0 | - |
| - | - | 1898 | 229.1 | - | - | 0 | - |
| - | - | 706.3 | 235.1 | - | - | 0 | - |
| - | - | 548.6 | 235.9 | - | - | 0 | - |
| - | - | 3.207E+04 | 239.1 | - | - | 0 | - |
| - | - | 1205 | 239.2 | - | - | 0 | - |
| - | - | 4126 | 240.1 | - | - | 0 | - |
| - | - | 2267 | 240.1 | - | - | 0 | - |
| - | - | 3000 | 245.1 | - | - | 0 | - |
| - | - | 623.8 | 246.1 | - | - | 0 | - |
| - | - | 4020 | 247.1 | - | - | 0 | - |
| - | - | 783.2 | 248.1 | - | - | 0 | - |
| - | - | 3934 | 257.2 | - | - | 0 | - |
| 8 | y | 7291 | 258.1 | 9.541E-05 | 0.3696 | +1 | 2 |
| - | - | 3634 | 265.1 | - | - | 0 | - |
| - | - | 1103 | 274.1 | - | - | 0 | - |
| - | - | 2364 | 275.1 | - | - | 0 | - |
| 8 | y | 4666 | 275.2 | 5.599E-05 | 0.2035 | +1 | 2 |
| - | - | 646 | 276.2 | - | - | 0 | - |
| - | - | 533.6 | 279.5 | - | - | 0 | - |
| - | - | 1216 | 281.1 | - | - | 0 | - |
| - | - | 991 | 285 | - | - | 0 | - |
| - | - | 3860 | 293.1 | - | - | 0 | - |
| - | - | 1123 | 294.1 | - | - | 0 | - |
| - | - | 5266 | 295.1 | - | - | 0 | - |
| - | - | 1491 | 296.1 | - | - | 0 | - |
| - | - | 1.291E+04 | 299.1 | - | - | 0 | - |
| - | - | 2238 | 300.1 | - | - | 0 | - |
| - | - | 1137 | 300.2 | - | - | 0 | - |
| - | - | 512.9 | 301.1 | - | - | 0 | - |
| - | - | 637.2 | 309 | - | - | 0 | - |
| - | - | 1997 | 311.2 | - | - | 0 | - |
| - | - | 1088 | 312.2 | - | - | 0 | - |
| - | - | 1771 | 313.1 | - | - | 0 | - |
| - | - | 646.3 | 314.1 | - | - | 0 | - |
| 7 | y | 667.6 | 329.2 | 0.0006708 | 2.038 | +1 | 3 |
| - | - | 552 | 330.2 | - | - | 0 | - |
| - | - | 1468 | 340.2 | - | - | 0 | - |
| 7 | y | 9251 | 346.2 | 0.0002753 | 0.7952 | +1 | 3 |
| - | - | 1331 | 347.2 | - | - | 0 | - |
| - | - | 1828 | 355.1 | - | - | 0 | - |
| - | - | 2182 | 358.2 | - | - | 0 | - |
| 3 | b | 2557 | 363.2 | 0.0004027 | 1.109 | +1 | 3 |
| - | - | 5710 | 369.1 | - | - | 0 | - |
| - | - | 1517 | 370.1 | - | - | 0 | - |
| - | - | 514.3 | 385.7 | - | - | 0 | - |
| - | - | 1511 | 404.1 | - | - | 0 | - |
| - | - | 8402 | 408.1 | - | - | 0 | - |
| - | - | 2054 | 409.1 | - | - | 0 | - |
| - | - | 551.6 | 409.4 | - | - | 0 | - |
| - | - | 640.6 | 416.8 | - | - | 0 | - |
| - | - | 1302 | 429.2 | - | - | 0 | - |
| 3 | y | 703.3 | 433.7 | 0.0004994 | 1.152 | +2 | 7 |
| - | - | 658.5 | 442.1 | - | - | 0 | - |
| 6 | y | 6015 | 459.3 | 0.0005924 | 1.29 | +1 | 4 |
| - | - | 1162 | 460.3 | - | - | 0 | - |
| - | - | 929 | 479.2 | - | - | 0 | - |
| - | - | 725.4 | 480.2 | - | - | 0 | - |
| 2 | y | 4099 | 488.7 | 0.0003049 | 0.6239 | +2 | 8 |
| 2 | y | 1799 | 489.2 | 0.009762 | 19.95 | +2 | 8 |
| 4 | b | 711.1 | 492.2 | 0.002933 | 5.959 | +1 | 4 |
| - | - | 1444 | 493.2 | - | - | 0 | - |
| 2 | y | 1192 | 497.7 | 0.001559 | 3.133 | +2 | 8 |
| - | - | 1029 | 515.7 | - | - | 0 | - |
| - | - | 1103 | 516.2 | - | - | 0 | - |
| - | - | 713.6 | 516.7 | - | - | 0 | - |
| - | - | 1030 | 518.2 | - | - | 0 | - |
| - | - | 1492 | 519.2 | - | - | 0 | - |
| - | - | 645.8 | 520.2 | - | - | 0 | - |
| 0 | Precursor | 2458 | 524.8 | 6.689E-05 | 0.1275 | +2 | -1 |
| - | - | 1663 | 525.3 | - | - | 0 | - |
| 0 | Precursor | 3134 | 533.3 | 0.001318 | 2.471 | +2 | -1 |
| - | - | 1324 | 533.8 | - | - | 0 | - |
| - | - | 656.9 | 557.3 | - | - | 0 | - |
| 5 | y | 7663 | 574.3 | 0.0005659 | 0.9853 | +1 | 5 |
| - | - | 1385 | 575.3 | - | - | 0 | - |
| 5 | b | 627.6 | 589.2 | 0.0004403 | 0.7472 | +1 | 5 |
| - | - | 1304 | 592.3 | - | - | 0 | - |
| 5 | b | 1187 | 607.2 | 0.001946 | 3.204 | +1 | 5 |
| 4 | y | 1135 | 685.4 | 0.0006398 | 0.9335 | +1 | 6 |
| 6 | b | 810 | 703.3 | 0.00487 | 6.925 | +1 | 6 |
| 4 | y | 9130 | 703.4 | 0.0003312 | 0.4709 | +1 | 6 |
| - | - | 3052 | 704.4 | - | - | 0 | - |
| - | - | 1085 | 705.4 | - | - | 0 | - |
| - | - | 600.4 | 718.7 | - | - | 0 | - |
| 6 | b | 924.2 | 720.3 | 0.001454 | 2.018 | +1 | 6 |
| - | - | 688.2 | 721.3 | - | - | 0 | - |
| - | - | 652.6 | 830.4 | - | - | 0 | - |
| 3 | y | 830.5 | 849.4 | 0.002818 | 3.317 | +1 | 7 |
| 3 | y | 4.419E+04 | 866.4 | 0.0001311 | 0.1514 | +1 | 7 |
| - | - | 2.08E+04 | 867.4 | - | - | 0 | - |
| - | - | 6218 | 868.4 | - | - | 0 | - |
| - | - | 724.5 | 869.4 | - | - | 0 | - |
| 8 | b | 710.3 | 919.4 | 0.00232 | 2.523 | +1 | 8 |
| 2 | y | 5469 | 977.5 | 0.0003112 | 0.3183 | +1 | 8 |
| - | - | 2592 | 978.5 | - | - | 0 | - |
| - | - | 693.1 | 979.5 | - | - | 0 | - |
| - | - | 607.7 | 1575 | - | - | 0 | - |
| - | - | 609.1 | 1696 | - | - | 0 | - |
| - | - | 653.5 | 1797 | - | - | 0 | - |
| - | - | 585.6 | 1873 | - | - | 0 | - |

m/z Charge Intensity FragmentType MassShift Position
120.08121490478516 0 396.23328
122.5324935913086 0 355.83044
129.06617736816406 0 1275.5623
129.10247802734375 0 4570.6694
130.08639526367188 0 2770.307 y Ammonia loss 8
136.07589721679688 0 11685.4375
136.45155334472656 0 392.67093
137.079345703125 0 498.31256
138.05528259277344 0 584.9316
147.11302185058594 0 4797.5483 y 8
148.94805908203125 0 714.2495
149.04507446289062 0 5683.224
151.681884765625 0 386.36496
154.70098876953125 0 447.00775
155.08163452148438 0 2422.2014 a Ammonia loss 1
162.32235717773438 0 453.01163
166.05006408691406 0 9157.847
167.0554962158203 0 10235.526
168.0542755126953 0 764.17346
172.10821533203125 0 1581.5587 a 1
173.12864685058594 0 516.85614
173.4383544921875 0 711.65063
183.0765838623047 0 22004.191 b Ammonia loss 1
184.07994079589844 0 1490.8284
194.01954650878906 0 552.86914
200.1031036376953 0 32971.406 b 1
201.08859252929688 0 571.2352
201.1065673828125 0 2966.0735
201.1235809326172 0 3408.9045
202.0862579345703 0 882.07635
207.11265563964844 0 682.70526
217.0305633544922 0 515.35657
221.0845489501953 0 12725.733
222.08506774902344 0 1748.3608
223.10830688476562 0 588.83453
225.04310607910156 0 19316.416
226.04415893554688 0 2038.7886
227.02215576171875 0 2262.535
227.06663513183594 0 1309.1943
229.11849975585938 0 1897.8108
235.10752868652344 0 706.3182
235.9456024169922 0 548.6393
239.09518432617188 0 32065.943
239.15025329589844 0 1204.87
240.09605407714844 0 4125.8613
240.13436889648438 0 2267.4534
245.07708740234375 0 2999.7908
246.12420654296875 0 623.84436
247.10797119140625 0 4020.2886
248.11175537109375 0 783.1854
257.1607360839844 0 3934.4585
258.1449279785156 0 7290.933 y Ammonia loss 7
265.1184997558594 0 3633.653
274.1181945800781 0 1103.3466
275.10296630859375 0 2364.2202
275.17132568359375 0 4665.8423 y 7
276.1741638183594 0 645.9995
279.5225830078125 0 533.6347
281.05133056640625 0 1216.298
285.0101623535156 0 990.9823
293.1134033203125 0 3859.731
294.1163330078125 0 1123.1646
295.103515625 0 5266.402
296.103759765625 0 1491.2163
299.0619201660156 0 12914.347
300.0626525878906 0 2237.76
300.1553955078125 0 1137.1693
301.12054443359375 0 512.9228
309.0196533203125 0 637.2117
311.1716003417969 0 1996.7556
312.156005859375 0 1087.5385
313.11376953125 0 1770.6847
314.11553955078125 0 646.31793
329.1826171875 0 667.5995 y Ammonia loss 6
330.1667785644531 0 552.03876
340.15081787109375 0 1467.5215
346.2087707519531 0 9251.17 y 6
347.2120056152344 0 1330.914
355.0693664550781 0 1828.4518
358.1610412597656 0 2181.6377
363.1658935546875 0 2557.1526 b 2
369.1217956542969 0 5709.5093
370.1227111816406 0 1517.1016
385.7403259277344 0 514.31805
404.1452331542969 0 1511.1423
408.1407470703125 0 8401.71
409.1435546875 0 2053.8838
409.4364929199219 0 551.64435
416.7725524902344 0 640.5813
429.1986083984375 0 1301.9917
433.7158508300781 0 703.3171 y 2
442.0972595214844 0 658.4507
459.29315185546875 0 6015.0728 y 5
460.2952575683594 0 1162.0067
479.178466796875 0 928.97266
480.1787414550781 0 725.43176
488.74066162109375 0 4098.859 y Water loss 1
489.24212646484375 0 1799.4 y Ammonia loss 1
492.2118225097656 0 711.0962 b 3
493.2296142578125 0 1444.4507
497.74407958984375 0 1192.1466 y 1
515.7471923828125 0 1029.2107
516.2400512695312 0 1102.8008
516.74658203125 0 713.5828
518.1875610351562 0 1029.9685
519.1736450195312 0 1492.2877
520.1753540039062 0 645.82526
524.7508544921875 0 2458.012 Precursor Ammonia loss
525.253173828125 0 1663.3962
533.2628784179688 0 3134.2046 Precursor
533.7681884765625 0 1324.2451
557.25390625 0 656.9077
574.320068359375 0 7663.248 y 4
575.3211669921875 0 1385.2102
589.2257080078125 0 627.561 b Water loss 4
592.26025390625 0 1304.4005
607.23388671875 0 1186.8783 b 4
685.3508911132812 0 1135.0228 y Water loss 3
703.2982177734375 0 810.0242 b Ammonia loss 5
703.3624267578125 0 9130.199 y 3
704.3644409179688 0 3052.439
705.3674926757812 0 1085.2006
718.7261352539062 0 600.3998
720.3213500976562 0 924.22565 b 5
721.3193969726562 0 688.1828
830.3958740234375 0 652.6166
849.3960571289062 0 830.5468 y Ammonia loss 2
866.42529296875 0 44188.09 y 2
867.427978515625 0 20804.404
868.4301147460938 0 6218.0796
869.4378662109375 0 724.46216
919.4179077148438 0 710.2677 b 7
977.457763671875 0 5469.269 y Ammonia loss 1
978.4597778320312 0 2592.1465
979.4613647460938 0 693.082
1575.221923828125 0 607.74927
1696.153076171875 0 609.11914
1796.78466796875 0 653.4989
1873.4166259765625 0 585.5676

Spectrum Details

|  |  |
| --- | --- |
| Matched peaks? Matched peaks The total absolute number of peaks matched. Additionally in brackets the total fraction of peaks matched and the total number of peaks is shown. | 30 (21.58% of 139) |
| FDR? FDR The false discovery rate estimated for this peptide. It is calculated by matching all theoretical fragments with a non-integer shift with the raw peaks for this spectrum. This is done with 40 different shifts. The resulting percentage is the average number of annotated peaks over the number of annotated peaks with the correct spectrum. | 0.32% |
| Satellite FDR? Satellite FDR See the FDR for details on its calculation. This satellite ion specific FDR only contains the satellite ions (d/w) for I/L/J positions. | - |
| PSM Score? PSM Score The PSM Score as given by Hecklib to this annotated spectrum. It is shown with three significant figures. | 352 |

#### Reverse Lookup? Reverse Lookup All places where this read could be placed.

| Group | Segment | Template | Template Part | Read Part | Score | Unique |
| --- | --- | --- | --- | --- | --- | --- |
| Homo sapiens Light Chain | IGLV | IGKV1-33 | [78..86] | [0..9] | 36 | False |
| Homo sapiens Light Chain | IGLV | IGLV3-19 | [76..85] | [0..9] | 36 | False |
| Homo sapiens Light Chain | IGLV | IGLV3-27 | [76..85] | [0..9] | 36 | False |
| Homo sapiens Light Chain | IGLV | IGLV7-46 | [79..88] | [0..9] | 36 | False |
| Homo sapiens Light Chain | IGLV | IGLV3-10 | [76..85] | [0..9] | 36 | False |
| Decoy | Decoy | THER | [148..156] | [0..9] | 36 | False |
| Decoy | Decoy | K1C20 | [245..252] | [0..9] | 36 | False |

| Recombined | Template Part | Read Part | Score | Unique |
| --- | --- | --- | --- | --- |
| K1C20 | [245..252] | [0..9] | 36 | False |
| THER | [148..156] | [0..9] | 36 | False |

#### Meta Information from Multiple reads

##### Number of combined reads

2

##### Intensity

0.7807

##### TotalArea

5.278E+06

##### Changes to the peptide sequence

AQYEDJAQK

I→JNo support for either Leucine or Isoleucine based on side chain ions (Position: 6)

L→ISupport for Isoleucine based on side chain ions (1 for I 0 for L) (Position: 6)

#### Positional Score

Copy Data

##### Positional Score (TSV)

###### Preview

```
Loading example...
```

*Click on the button to copy the data to your clipboard.*

00012345678

Label Value
"0" 0
"1" 0
"2" 0
"3" 0
"4" 0
"5" 0
"6" 0
"7" 0
"8" 0

#### Meta Information from PEAKS

##### Scan Identifier

F4:3073

##### Original sequence

A

Q

Y

E

D

L

A

Q

K

##### Posttranslational Modifications

##### Source File

D:\separate\_stitch\_analyses\xle-disambiguation\raw\20210323\_F1\_UM1\_Peng0013\_SA\_F59\_ingel\_3ug\_tryp.raw

##### Fraction

4

##### Scan Feature

F4:3976

##### De Novo Score

98

##### ConfidenceScore

98

### m/z

533.2651

##### Mass

1064.5139

##### Charge

2

##### Retention Time

16.39

##### Predicted Retention Time

18.20

##### Area

5.278E+06

##### Parts Per Million

1.7

##### Fragmentation mode

ETHCD

##### Originating file

01 D:\separate\_stitch\_analyses\xle-disambiguation\20210325\_F59\_3ug\_DENOVO\_12.csv

#### Meta Information from PEAKS

##### Scan Identifier

F4:3127

##### Original sequence

A

Q

Y

E

D

L

A

Q

K

##### Posttranslational Modifications

##### Source File

D:\separate\_stitch\_analyses\xle-disambiguation\raw\20210323\_F1\_UM1\_Peng0013\_SA\_F59\_ingel\_3ug\_tryp.raw

##### Fraction

4

##### Scan Feature

-

##### De Novo Score

97

##### ConfidenceScore

96

### m/z

533.2651

##### Mass

1064.5139

##### Charge

2

##### Retention Time

16.63

##### Predicted Retention Time

18.20

##### Area

0

##### Parts Per Million

1.6

##### Fragmentation mode

HCD

##### Originating file

01 D:\separate\_stitch\_analyses\xle-disambiguation\20210325\_F59\_3ug\_DENOVO\_12.csv
