## Supplementary material for "A handle on mass coincidence errors in *de novo* sequencing of antibodies by bottom-up proteomics": Combined_087.html

Details Combined\_087 | Stitch OverviewUndefined

### Read Combined\_087

#### Sequence (length=8)

ALPPAIEK

#### Spectrum 4576? Spectrum 4576 The raw spectrum of this peptide as annotated by Hecklib. The fragments are coloured according to ion type (see legend). Any peaks with a star '\*' as text can be hovered over to see the full details, first the ion type second the mass shift type. By hovering over the amino acids in the peptide or ions in the legend the corresponding peaks are highlighted. By toggling the 'Unassigned' label you can turn the background (unassigned) peaks on or off in the plot. By updating the slider in the Ion legend you can update the spectrum to only show the top X% of the peaks with labels. The top X% means any peak that is within X% of the highest intensity. By dragging in the spectrum you can zoom in to a specific part of the spectrum and use 'Zoom Out' to get back to the original zoom level. The annotation of the spectrum is based on the given sequence in the peptides file and is done with different software so inconsistencies are likely. The peaks are annotated based on the given sequence, with 20 ppm tolerance.

Copy Data

##### Spectrum 4576 (TSV)

###### Preview

```
Loading example...
```

*Click on the button to copy the data to your clipboard.*

Mz MinMz MaxIntensity Max

WidthHeightPeptide font sizePeptide stroke widthSpectrum font sizeSpectrum stroke widthCompact peptide

Ion legend

wxyz

abcd

OtherUnassignedIon^Charge^_Position_Show for top:%

ALPPAIEK

02.55e+55.09e+57.64e+51.02e+6

Zoom Out

y^+1^_1_y^+1^_1_w^+1^_2_y^+1^_2_z^+1^_2_y^+1^_2_c^+1^_3_y^+2^_6_y^+2^_6_w^+1^_3_w^+1^_3_y^+1^_3_z^+1^_3_y^+2^_7_y^+2^_7_y^+1^_3_c^+1^_4_y^+1^_4_z^+1^_4_c^+1^_5_w^+1^_5_y^+1^_5_z^+1^_5_y^+1^_5_c^+1^_6_w^+1^_6_y^+1^_6_y^+1^_6_w^+1^_7_c^+1^_7_z^+1^_7_

0563112516882251

Fragment Matches Table

Show background peaks

| Position | Ion type | Intensity | mz Theoretical | mz Error (Th) | mz Error (ppm) | Charge | Series Number |
| --- | --- | --- | --- | --- | --- | --- | --- |
| - | - | 362.6 | 120.9 | - | - | 0 | - |
| - | - | 421.2 | 123.6 | - | - | 0 | - |
| - | - | 353.3 | 125.5 | - | - | 0 | - |
| - | - | 5083 | 129.1 | - | - | 0 | - |
| 8 | y | 3316 | 130.1 | 0.0003081 | 2.368 | +1 | 1 |
| - | - | 566.4 | 130.1 | - | - | 0 | - |
| - | - | 455.7 | 133.1 | - | - | 0 | - |
| - | - | 2.998E+04 | 141.1 | - | - | 0 | - |
| - | - | 1745 | 142.1 | - | - | 0 | - |
| - | - | 827.3 | 143.1 | - | - | 0 | - |
| 8 | y | 1.412E+04 | 147.1 | 0.0003398 | 2.31 | +1 | 1 |
| - | - | 499.4 | 148.1 | - | - | 0 | - |
| - | - | 1.87E+04 | 149 | - | - | 0 | - |
| - | - | 1626 | 150 | - | - | 0 | - |
| - | - | 3.112E+05 | 157.1 | - | - | 0 | - |
| - | - | 848.4 | 158.1 | - | - | 0 | - |
| - | - | 2.569E+04 | 158.1 | - | - | 0 | - |
| - | - | 599.6 | 159.1 | - | - | 0 | - |
| - | - | 851.8 | 159.1 | - | - | 0 | - |
| - | - | 3006 | 167 | - | - | 0 | - |
| - | - | 6.623E+04 | 169.1 | - | - | 0 | - |
| - | - | 4442 | 170.1 | - | - | 0 | - |
| - | - | 861.7 | 171.1 | - | - | 0 | - |
| - | - | 451 | 172.2 | - | - | 0 | - |
| - | - | 4484 | 173.4 | - | - | 0 | - |
| - | - | 4786 | 183.1 | - | - | 0 | - |
| - | - | 3824 | 184.1 | - | - | 0 | - |
| - | - | 1.37E+05 | 185.1 | - | - | 0 | - |
| - | - | 1.143E+04 | 186.1 | - | - | 0 | - |
| - | - | 506.9 | 194.3 | - | - | 0 | - |
| - | - | 570 | 197.1 | - | - | 0 | - |
| - | - | 1072 | 198.1 | - | - | 0 | - |
| 7 | w | 1366 | 201.1 | 0.0001816 | 0.9028 | +1 | 2 |
| - | - | 467.5 | 207.3 | - | - | 0 | - |
| - | - | 5891 | 211.1 | - | - | 0 | - |
| - | - | 480.8 | 219.8 | - | - | 0 | - |
| - | - | 2.452E+04 | 228.2 | - | - | 0 | - |
| - | - | 2253 | 229.2 | - | - | 0 | - |
| - | - | 3089 | 240.1 | - | - | 0 | - |
| - | - | 9285 | 243.7 | - | - | 0 | - |
| - | - | 1777 | 244.2 | - | - | 0 | - |
| - | - | 535.1 | 246.1 | - | - | 0 | - |
| - | - | 580.2 | 254.2 | - | - | 0 | - |
| 7 | y | 5221 | 258.1 | 0.0002175 | 0.8425 | +1 | 2 |
| 7 | z | 3413 | 260.1 | 0.0003507 | 1.348 | +1 | 2 |
| - | - | 816.7 | 261.1 | - | - | 0 | - |
| - | - | 6839 | 266.1 | - | - | 0 | - |
| - | - | 3377 | 266.2 | - | - | 0 | - |
| - | - | 823.1 | 267.2 | - | - | 0 | - |
| - | - | 1376 | 267.2 | - | - | 0 | - |
| - | - | 389 | 276.1 | - | - | 0 | - |
| 7 | y | 1.62E+04 | 276.2 | 0.0003339 | 1.209 | +1 | 2 |
| - | - | 1681 | 277.2 | - | - | 0 | - |
| - | - | 4105 | 282.2 | - | - | 0 | - |
| - | - | 1.998E+04 | 283.2 | - | - | 0 | - |
| - | - | 2935 | 284.2 | - | - | 0 | - |
| - | - | 1053 | 293.2 | - | - | 0 | - |
| 3 | c | 725.4 | 299.2 | 0.001553 | 5.19 | +1 | 3 |
| - | - | 1086 | 312.2 | - | - | 0 | - |
| 3 | y | 1255 | 318.7 | 0.0001374 | 0.4312 | +2 | 6 |
| - | - | 1248 | 325.2 | - | - | 0 | - |
| - | - | 2635 | 325.2 | - | - | 0 | - |
| - | - | 1419 | 327.2 | - | - | 0 | - |
| 3 | y | 6.952E+05 | 327.7 | 0.0006229 | 1.901 | +2 | 6 |
| - | - | 2.35E+05 | 328.2 | - | - | 0 | - |
| - | - | 5.148E+04 | 328.7 | - | - | 0 | - |
| - | - | 3618 | 329.2 | - | - | 0 | - |
| - | - | 2.02E+04 | 340.2 | - | - | 0 | - |
| - | - | 2746 | 341.2 | - | - | 0 | - |
| 6 | w | 1.728E+04 | 344.2 | 0.0004559 | 1.324 | +1 | 3 |
| - | - | 667.9 | 344.2 | - | - | 0 | - |
| - | - | 2499 | 345.2 | - | - | 0 | - |
| - | - | 6876 | 351.2 | - | - | 0 | - |
| - | - | 4821 | 352.2 | - | - | 0 | - |
| - | - | 1483 | 352.2 | - | - | 0 | - |
| - | - | 1.638E+04 | 353.2 | - | - | 0 | - |
| - | - | 741.4 | 353.3 | - | - | 0 | - |
| - | - | 2299 | 354.2 | - | - | 0 | - |
| - | - | 2.23E+04 | 357.2 | - | - | 0 | - |
| 6 | w | 4983 | 358.2 | 0.0002101 | 0.5865 | +1 | 3 |
| - | - | 3506 | 358.2 | - | - | 0 | - |
| - | - | 1100 | 359.2 | - | - | 0 | - |
| - | - | 706.9 | 368.2 | - | - | 0 | - |
| - | - | 4157 | 368.3 | - | - | 0 | - |
| 6 | y | 561.6 | 372.2 | 0.0008025 | 2.156 | +1 | 3 |
| 6 | z | 4.118E+04 | 373.2 | 0.0003322 | 0.89 | +1 | 3 |
| - | - | 2.476E+04 | 374.2 | - | - | 0 | - |
| 2 | y | 4390 | 375.2 | 0.0004822 | 1.285 | +2 | 7 |
| - | - | 2.115E+04 | 379.2 | - | - | 0 | - |
| - | - | 3514 | 380.2 | - | - | 0 | - |
| 2 | y | 5797 | 384.2 | 0.0005831 | 1.518 | +2 | 7 |
| - | - | 1864 | 384.7 | - | - | 0 | - |
| - | - | 649.1 | 388.2 | - | - | 0 | - |
| 6 | y | 5097 | 389.2 | 0.000529 | 1.359 | +1 | 3 |
| - | - | 769.8 | 395.2 | - | - | 0 | - |
| 4 | c | 1.707E+05 | 396.3 | 0.0005774 | 1.457 | +1 | 4 |
| - | - | 3.418E+04 | 397.3 | - | - | 0 | - |
| - | - | 690.1 | 398.2 | - | - | 0 | - |
| - | - | 4557 | 398.3 | - | - | 0 | - |
| - | - | 1010 | 399.2 | - | - | 0 | - |
| - | - | 1513 | 410.2 | - | - | 0 | - |
| - | - | 1860 | 411.2 | - | - | 0 | - |
| - | - | 860.5 | 412.2 | - | - | 0 | - |
| - | - | 1242 | 412.3 | - | - | 0 | - |
| - | - | 2020 | 413.2 | - | - | 0 | - |
| - | - | 1855 | 415.2 | - | - | 0 | - |
| - | - | 690.7 | 418.7 | - | - | 0 | - |
| 5 | y | 1.13E+04 | 443.3 | 0.000737 | 1.663 | +1 | 4 |
| 5 | z | 3473 | 444.3 | 0.002236 | 5.032 | +1 | 4 |
| - | - | 932.4 | 445.3 | - | - | 0 | - |
| - | - | 1725 | 450.3 | - | - | 0 | - |
| - | - | 677.3 | 451.3 | - | - | 0 | - |
| - | - | 913.7 | 454.3 | - | - | 0 | - |
| - | - | 1707 | 456.3 | - | - | 0 | - |
| - | - | 1432 | 462.3 | - | - | 0 | - |
| - | - | 5.952E+04 | 466.3 | - | - | 0 | - |
| 5 | c | 4.629E+04 | 467.3 | 0.0006783 | 1.451 | +1 | 5 |
| - | - | 6876 | 468.3 | - | - | 0 | - |
| - | - | 5742 | 469.3 | - | - | 0 | - |
| - | - | 1662 | 470.2 | - | - | 0 | - |
| - | - | 1387 | 470.3 | - | - | 0 | - |
| - | - | 3573 | 471.3 | - | - | 0 | - |
| - | - | 3165 | 480.3 | - | - | 0 | - |
| - | - | 778.2 | 481.3 | - | - | 0 | - |
| - | - | 5027 | 482.3 | - | - | 0 | - |
| - | - | 1246 | 483.3 | - | - | 0 | - |
| - | - | 3.029E+04 | 484.3 | - | - | 0 | - |
| - | - | 1850 | 485.2 | - | - | 0 | - |
| - | - | 1.525E+04 | 485.3 | - | - | 0 | - |
| - | - | 2.288E+05 | 486.3 | - | - | 0 | - |
| - | - | 5.802E+04 | 487.3 | - | - | 0 | - |
| - | - | 1.028E+04 | 488.3 | - | - | 0 | - |
| - | - | 669.1 | 489.3 | - | - | 0 | - |
| - | - | 1223 | 490.3 | - | - | 0 | - |
| - | - | 3.042E+04 | 508.3 | - | - | 0 | - |
| - | - | 8352 | 509.3 | - | - | 0 | - |
| - | - | 2305 | 509.3 | - | - | 0 | - |
| - | - | 1436 | 510.3 | - | - | 0 | - |
| - | - | 9019 | 512.3 | - | - | 0 | - |
| - | - | 2522 | 513.3 | - | - | 0 | - |
| 4 | w | 3773 | 514.3 | 0.0002134 | 0.415 | +1 | 5 |
| - | - | 986 | 515.3 | - | - | 0 | - |
| - | - | 7891 | 523.3 | - | - | 0 | - |
| - | - | 2120 | 524.3 | - | - | 0 | - |
| - | - | 8.312E+04 | 525.3 | - | - | 0 | - |
| - | - | 2.116E+04 | 526.3 | - | - | 0 | - |
| - | - | 4408 | 527.3 | - | - | 0 | - |
| - | - | 4295 | 535.4 | - | - | 0 | - |
| - | - | 4632 | 536.4 | - | - | 0 | - |
| - | - | 7687 | 537.4 | - | - | 0 | - |
| - | - | 1941 | 538.4 | - | - | 0 | - |
| 4 | y | 2341 | 539.3 | 0.0001955 | 0.3626 | +1 | 5 |
| - | - | 555.7 | 539.4 | - | - | 0 | - |
| - | - | 961.6 | 540.3 | - | - | 0 | - |
| 4 | z | 1.013E+05 | 541.3 | 0.0004814 | 0.8893 | +1 | 5 |
| - | - | 3.011E+04 | 542.3 | - | - | 0 | - |
| - | - | 5044 | 543.3 | - | - | 0 | - |
| 4 | y | 4.503E+04 | 557.3 | 0.000251 | 0.4503 | +1 | 5 |
| - | - | 1.323E+04 | 558.3 | - | - | 0 | - |
| - | - | 2582 | 559.3 | - | - | 0 | - |
| - | - | 2066 | 562.4 | - | - | 0 | - |
| - | - | 1.421E+04 | 563.4 | - | - | 0 | - |
| - | - | 3817 | 564.4 | - | - | 0 | - |
| - | - | 937.5 | 565.4 | - | - | 0 | - |
| - | - | 1764 | 570.4 | - | - | 0 | - |
| - | - | 1.189E+05 | 579.4 | - | - | 0 | - |
| 6 | c | 1.875E+05 | 580.4 | 0.0005442 | 0.9377 | +1 | 6 |
| - | - | 5.314E+04 | 581.4 | - | - | 0 | - |
| - | - | 9360 | 582.4 | - | - | 0 | - |
| - | - | 756.6 | 584.3 | - | - | 0 | - |
| - | - | 790.7 | 585.3 | - | - | 0 | - |
| - | - | 927.6 | 597.3 | - | - | 0 | - |
| - | - | 1798 | 599.3 | - | - | 0 | - |
| - | - | 1.414E+04 | 606.4 | - | - | 0 | - |
| - | - | 5006 | 607.4 | - | - | 0 | - |
| - | - | 798.4 | 608.4 | - | - | 0 | - |
| - | - | 1100 | 609.3 | - | - | 0 | - |
| 3 | w | 5071 | 611.3 | 0.0006726 | 1.1 | +1 | 6 |
| - | - | 3184 | 612.3 | - | - | 0 | - |
| - | - | 1095 | 613.3 | - | - | 0 | - |
| - | - | 1035 | 623.4 | - | - | 0 | - |
| 3 | y | 7063 | 636.4 | 0.0003492 | 0.5487 | +1 | 6 |
| - | - | 3232 | 637.4 | - | - | 0 | - |
| - | - | 1128 | 638.4 | - | - | 0 | - |
| - | - | 9041 | 652.4 | - | - | 0 | - |
| - | - | 1913 | 653.4 | - | - | 0 | - |
| 3 | y | 5.896E+05 | 654.4 | 0.0005877 | 0.8981 | +1 | 6 |
| - | - | 2.196E+05 | 655.4 | - | - | 0 | - |
| - | - | 1288 | 655.5 | - | - | 0 | - |
| - | - | 5.009E+04 | 656.4 | - | - | 0 | - |
| - | - | 3434 | 657.4 | - | - | 0 | - |
| - | - | 4566 | 665.4 | - | - | 0 | - |
| - | - | 1929 | 666.4 | - | - | 0 | - |
| - | - | 1106 | 670.4 | - | - | 0 | - |
| - | - | 2804 | 671.4 | - | - | 0 | - |
| - | - | 5033 | 679.4 | - | - | 0 | - |
| - | - | 1542 | 680.4 | - | - | 0 | - |
| - | - | 759.8 | 681.4 | - | - | 0 | - |
| - | - | 3103 | 682.4 | - | - | 0 | - |
| - | - | 3967 | 683.4 | - | - | 0 | - |
| - | - | 1021 | 684.4 | - | - | 0 | - |
| - | - | 1.727E+04 | 692.4 | - | - | 0 | - |
| - | - | 6320 | 693.4 | - | - | 0 | - |
| - | - | 2412 | 694.4 | - | - | 0 | - |
| - | - | 5847 | 695.4 | - | - | 0 | - |
| - | - | 2564 | 696.4 | - | - | 0 | - |
| 2 | w | 6.66E+04 | 708.4 | 0.00339 | 4.785 | +1 | 7 |
| 7 | c | 1.008E+06 | 709.4 | 0.0003807 | 0.5367 | +1 | 7 |
| - | - | 3.841E+05 | 710.4 | - | - | 0 | - |
| - | - | 9.493E+04 | 711.4 | - | - | 0 | - |
| - | - | 7381 | 712.4 | - | - | 0 | - |
| - | - | 1044 | 726.4 | - | - | 0 | - |
| - | - | 932.1 | 742.5 | - | - | 0 | - |
| - | - | 8382 | 750.5 | - | - | 0 | - |
| 2 | z | 5.882E+04 | 751.4 | 0.0006164 | 0.8203 | +1 | 7 |
| - | - | 2.199E+04 | 752.5 | - | - | 0 | - |
| - | - | 5433 | 753.5 | - | - | 0 | - |
| - | - | 729.5 | 754.5 | - | - | 0 | - |
| - | - | 1569 | 763.5 | - | - | 0 | - |
| - | - | 3188 | 764.4 | - | - | 0 | - |
| - | - | 1781 | 765.4 | - | - | 0 | - |
| - | - | 4.685E+05 | 766.4 | - | - | 0 | - |
| - | - | 1.806E+05 | 767.4 | - | - | 0 | - |
| - | - | 5.035E+04 | 768.4 | - | - | 0 | - |
| - | - | 5216 | 769.4 | - | - | 0 | - |
| - | - | 1080 | 779.4 | - | - | 0 | - |
| - | - | 2677 | 780.5 | - | - | 0 | - |
| - | - | 1634 | 781.5 | - | - | 0 | - |
| - | - | 5.569E+04 | 783.4 | - | - | 0 | - |
| - | - | 2.283E+04 | 784.5 | - | - | 0 | - |
| - | - | 6825 | 785.5 | - | - | 0 | - |
| - | - | 997.3 | 786.5 | - | - | 0 | - |
| - | - | 2663 | 793.5 | - | - | 0 | - |
| - | - | 1849 | 794.5 | - | - | 0 | - |
| - | - | 4272 | 796.5 | - | - | 0 | - |
| - | - | 1122 | 797.5 | - | - | 0 | - |
| - | - | 831 | 799.5 | - | - | 0 | - |
| - | - | 2565 | 810.5 | - | - | 0 | - |
| - | - | 3626 | 821.5 | - | - | 0 | - |
| - | - | 3.199E+05 | 822.5 | - | - | 0 | - |
| - | - | 1.424E+05 | 823.5 | - | - | 0 | - |
| - | - | 4.46E+04 | 824.5 | - | - | 0 | - |
| - | - | 7196 | 825.5 | - | - | 0 | - |
| - | - | 3.489E+05 | 838.5 | - | - | 0 | - |
| - | - | 1.008E+06 | 839.5 | - | - | 0 | - |
| - | - | 4.18E+05 | 840.5 | - | - | 0 | - |
| - | - | 1.195E+05 | 841.5 | - | - | 0 | - |
| - | - | 1.376E+04 | 842.5 | - | - | 0 | - |
| - | - | 1613 | 854.5 | - | - | 0 | - |
| - | - | 1099 | 855.5 | - | - | 0 | - |
| - | - | 1504 | 871.5 | - | - | 0 | - |
| - | - | 710.9 | 2228 | - | - | 0 | - |

m/z Charge Intensity FragmentType MassShift Position
120.9053955078125 0 362.6404
123.62415313720703 0 421.2179
125.50856018066406 0 353.26392
129.1025848388672 0 5082.767
130.08656311035156 0 3315.7703 y Ammonia loss 7
130.1059112548828 0 566.418
133.1356201171875 0 455.71692
141.1025848388672 0 29984.047
142.10592651367188 0 1745.2745
143.11834716796875 0 827.3372
147.11314392089844 0 14122.582 y 7
148.11685180664062 0 499.38547
149.02366638183594 0 18695.453
150.0269317626953 0 1625.9973
157.1339111328125 0 311210.06
158.1167449951172 0 848.4003
158.13720703125 0 25693.373
159.12442016601562 0 599.56464
159.14036560058594 0 851.8066
167.0342559814453 0 3006.248
169.09750366210938 0 66231.03
170.1009521484375 0 4442.435
171.11293029785156 0 861.73206
172.1613006591797 0 451.00854
173.44041442871094 0 4483.583
183.14962768554688 0 4785.682
184.12107849121094 0 3824.1416
185.12881469726562 0 137008.02
186.13217163085938 0 11425.732
194.33718872070312 0 506.88925
197.09275817871094 0 569.9807
198.0767059326172 0 1071.7452
201.12355041503906 0 1366.1929 w 6
207.3159942626953 0 467.48148
211.14427185058594 0 5891.2603
219.78684997558594 0 480.78
228.1710205078125 0 24521.016
229.1745147705078 0 2252.9385
240.13453674316406 0 3088.8428
243.650146484375 0 9284.706
244.1512908935547 0 1777.3718
246.13351440429688 0 535.10657
254.1505889892578 0 580.1608
258.1450500488281 0 5221.473 y Water loss 6
260.13702392578125 0 3412.717 z 6
261.1426086425781 0 816.6865
266.1499328613281 0 6839.39
266.1634521484375 0 3377.4385
267.15277099609375 0 823.101
267.169189453125 0 1375.7657
276.1411437988281 0 388.99918
276.1557312011719 0 16196.432 y 6
277.15875244140625 0 1681.4517
282.1816101074219 0 4104.7856
283.177001953125 0 19983.953
284.1798095703125 0 2935.1602
293.17547607421875 0 1053.4258
299.2093200683594 0 725.408 c 2
312.1918029785156 0 1086.3451
318.6895446777344 0 1254.9603 y Water loss 2
325.2000427246094 0 1248.0999
325.2244567871094 0 2634.906
327.209228515625 0 1419.2406
327.6953125 0 695168.3 y 2
328.19659423828125 0 235027.11
328.6978759765625 0 51476.516
329.1996765136719 0 3618.4797
340.1871643066406 0 20195.725
341.1903076171875 0 2746.1409
344.18206787109375 0 17283.895 w 5
344.2060852050781 0 667.8833
345.18426513671875 0 2499.4412
351.2393798828125 0 6875.6514
352.2112731933594 0 4821.431
352.24273681640625 0 1482.8156
353.2184753417969 0 16380.953
353.2535095214844 0 741.3562
354.22088623046875 0 2298.707
357.213623046875 0 22302.922
358.1970520019531 0 4982.9146 w 5
358.21759033203125 0 3505.818
359.2000732421875 0 1099.6796
368.2174072265625 0 706.9389
368.25396728515625 0 4156.861
372.2137145996094 0 561.5817 y Ammonia loss 5
373.2210693359375 0 41175.637 z 5
374.2275085449219 0 24762.684
375.23095703125 0 4389.581 y Water loss 1
379.2343444824219 0 21153.152
380.2378234863281 0 3514.1716
384.2373046875 0 5796.885 y 1
384.7384948730469 0 1864.1538
388.2341003417969 0 649.1233
389.239990234375 0 5096.624 y 5
395.20654296875 0 769.84436
396.2611083984375 0 170650.17 c 3
397.2638244628906 0 34184.047
398.216552734375 0 690.10767
398.2666015625 0 4557.1577
399.2218017578125 0 1010.1451
410.22833251953125 0 1513.1
411.22491455078125 0 1860.1517
412.227294921875 0 860.50397
412.26153564453125 0 1242.4
413.2272033691406 0 2019.7205
415.2196350097656 0 1854.6273
418.6782531738281 0 690.7202
443.2507629394531 0 11296.944 y Ammonia loss 4
444.255615234375 0 3472.5203 z 4
445.25592041015625 0 932.35626
450.27215576171875 0 1725.2166
451.2711181640625 0 677.2997
454.2774658203125 0 913.73895
456.2942199707031 0 1706.8176
462.2707214355469 0 1431.9366
466.2904052734375 0 59517.805
467.2969665527344 0 46287.883 c 4
468.29998779296875 0 6875.6646
469.28900146484375 0 5741.9863
470.23681640625 0 1662.2415
470.2935791015625 0 1386.8308
471.3039855957031 0 3573.3757
480.28216552734375 0 3165.1995
481.2846374511719 0 778.2478
482.2976379394531 0 5026.627
483.30218505859375 0 1246.0363
484.27728271484375 0 30294.215
485.2479248046875 0 1850.0782
485.2828674316406 0 15251.511
486.2927551269531 0 228765.84
487.2958068847656 0 58021.51
488.29864501953125 0 10284.62
489.2976379394531 0 669.12286
490.2669677734375 0 1223.454
508.2770690917969 0 30423.023
509.2798767089844 0 8351.565
509.3453369140625 0 2304.9263
510.28216552734375 0 1436.0417
512.2719116210938 0 9019.42
513.2755126953125 0 2522.27
514.2869262695312 0 3773.028 w 3
515.2918701171875 0 986.03265
523.3111572265625 0 7890.8804
524.3113403320312 0 2120.0725
525.3035888671875 0 83117.73
526.3046875 0 21156.656
527.3048095703125 0 4407.6953
535.3604125976562 0 4294.537
536.3663330078125 0 4631.5996
537.3753051757812 0 7687.172
538.3776245117188 0 1940.9287
539.3189697265625 0 2341.4426 y Water loss 3
539.371826171875 0 555.68463
540.320068359375 0 961.5569
541.3110961914062 0 101328.65 z 3
542.3140258789062 0 30111.46
543.3168334960938 0 5043.571
557.32958984375 0 45025.305 y 3
558.332763671875 0 13232.087
559.3348388671875 0 2582.0208
562.3507690429688 0 2066.1326
563.355224609375 0 14213.216
564.358154296875 0 3817.143
565.3673706054688 0 937.4679
570.372802734375 0 1763.8108
579.3743286132812 0 118907.54
580.3811645507812 0 187504.9 c 5
581.3843994140625 0 53143.586
582.3872680664062 0 9360.419
584.31005859375 0 756.5644
585.3228759765625 0 790.6683
597.3226928710938 0 927.6072
599.3361206054688 0 1797.6295
606.3976440429688 0 14141.723
607.400634765625 0 5006.0645
608.4053955078125 0 798.3762
609.3495483398438 0 1100.1971
611.340576171875 0 5071.384 w 2
612.345703125 0 3184.14
613.3489379882812 0 1095.0486
623.36083984375 0 1035.4397
636.3718872070312 0 7062.985 y Water loss 2
637.3759765625 0 3231.9014
638.3859252929688 0 1127.7609
652.3560791015625 0 9041.076
653.3626708984375 0 1913.2014
654.3826904296875 0 589604.3 y 2
655.3860473632812 0 219581.73
655.5149536132812 0 1288.3182
656.388916015625 0 50094.008
657.3924560546875 0 3433.6128
665.41064453125 0 4565.613
666.41357421875 0 1928.8289
670.376953125 0 1105.7654
671.4197998046875 0 2804.4956
679.4257202148438 0 5033.413
680.4300537109375 0 1542.1108
681.3663330078125 0 759.8101
682.3784790039062 0 3102.6577
683.384033203125 0 3967.2075
684.3882446289062 0 1021.3758
692.3984985351562 0 17269.889
693.4003295898438 0 6320.359
694.4029541015625 0 2412.3555
695.3873291015625 0 5847.375
696.385498046875 0 2564.4014
708.3960571289062 0 66601.55 w 1
709.4246826171875 0 1008269.1 c 6
710.4276733398438 0 384070.06
711.4300537109375 0 94932.42
712.4334716796875 0 7380.6724
726.4408569335938 0 1044.3312
742.4638061523438 0 932.09515
750.463134765625 0 8382.262
751.4480590820312 0 58815.316 z 1
752.451171875 0 21992.072
753.4542236328125 0 5433.3
754.4617309570312 0 729.48737
763.46533203125 0 1569.4004
764.4296264648438 0 3188.1914
765.4312133789062 0 1780.8245
766.4222412109375 0 468477.3
767.425048828125 0 180616.78
768.4276733398438 0 50347.316
769.4349365234375 0 5215.8354
779.4259033203125 0 1079.7754
780.4963989257812 0 2677.3071
781.4984741210938 0 1633.645
783.4486694335938 0 55692.566
784.45166015625 0 22830.348
785.4549560546875 0 6825.232
786.4721069335938 0 997.27466
793.5067749023438 0 2663.3782
794.5075073242188 0 1848.8281
796.458740234375 0 4272.4194
797.4559326171875 0 1122.0599
799.5149536132812 0 831.0357
810.4720458984375 0 2564.5525
821.4969482421875 0 3626.2954
822.484619140625 0 319884.3
823.48779296875 0 142443.06
824.4900512695312 0 44603.133
825.5005493164062 0 7195.5947
838.50341796875 0 348923.72
839.5106811523438 0 1008408.44
840.5136108398438 0 418047.75
841.5167846679688 0 119513.42
842.5234375 0 13755.095
854.4755249023438 0 1612.9119
855.4746704101562 0 1098.7377
871.49658203125 0 1504.1
2228.2939453125 0 710.9431

Spectrum Details

|  |  |
| --- | --- |
| Matched peaks? Matched peaks The total absolute number of peaks matched. Additionally in brackets the total fraction of peaks matched and the total number of peaks is shown. | 31 (12.30% of 252) |
| FDR? FDR The false discovery rate estimated for this peptide. It is calculated by matching all theoretical fragments with a non-integer shift with the raw peaks for this spectrum. This is done with 40 different shifts. The resulting percentage is the average number of annotated peaks over the number of annotated peaks with the correct spectrum. | 0.08% |
| Satellite FDR? Satellite FDR See the FDR for details on its calculation. This satellite ion specific FDR only contains the satellite ions (d/w) for I/L/J positions. | 0.00% |
| PSM Score? PSM Score The PSM Score as given by Hecklib to this annotated spectrum. It is shown with three significant figures. | 290 |

#### Spectrum 4522? Spectrum 4522 The raw spectrum of this peptide as annotated by Hecklib. The fragments are coloured according to ion type (see legend). Any peaks with a star '\*' as text can be hovered over to see the full details, first the ion type second the mass shift type. By hovering over the amino acids in the peptide or ions in the legend the corresponding peaks are highlighted. By toggling the 'Unassigned' label you can turn the background (unassigned) peaks on or off in the plot. By updating the slider in the Ion legend you can update the spectrum to only show the top X% of the peaks with labels. The top X% means any peak that is within X% of the highest intensity. By dragging in the spectrum you can zoom in to a specific part of the spectrum and use 'Zoom Out' to get back to the original zoom level. The annotation of the spectrum is based on the given sequence in the peptides file and is done with different software so inconsistencies are likely. The peaks are annotated based on the given sequence, with 20 ppm tolerance.

Copy Data

##### Spectrum 4522 (TSV)

###### Preview

```
Loading example...
```

*Click on the button to copy the data to your clipboard.*

Mz MinMz MaxIntensity Max

WidthHeightPeptide font sizePeptide stroke widthSpectrum font sizeSpectrum stroke widthCompact peptide

Ion legend

wxyz

abcd

OtherUnassignedIon^Charge^_Position_Show for top:%

ALPPAIEK

04.96e+59.93e+51.49e+61.99e+6

Zoom Out

y^+1^_1_y^+1^_1_w^+1^_2_y^+1^_2_z^+1^_2_y^+1^_2_y^+2^_6_y^+2^_6_w^+1^_3_w^+1^_3_z^+1^_3_y^+2^_7_y^+2^_7_y^+1^_3_c^+1^_4_y^+1^_4_z^+1^_4_c^+1^_5_w^+1^_5_y^+1^_5_z^+1^_5_y^+1^_5_c^+1^_6_w^+1^_6_y^+1^_6_y^+1^_6_w^+1^_7_c^+1^_7_z^+1^_7_

0220440660880

Fragment Matches Table

Show background peaks

| Position | Ion type | Intensity | mz Theoretical | mz Error (Th) | mz Error (ppm) | Charge | Series Number |
| --- | --- | --- | --- | --- | --- | --- | --- |
| - | - | 697 | 126.3 | - | - | 0 | - |
| - | - | 1319 | 129.1 | - | - | 0 | - |
| - | - | 1.317E+04 | 129.1 | - | - | 0 | - |
| - | - | 816.7 | 129.1 | - | - | 0 | - |
| 8 | y | 6244 | 130.1 | 0.0004606 | 3.541 | +1 | 1 |
| - | - | 761.3 | 135.5 | - | - | 0 | - |
| - | - | 1005 | 137.2 | - | - | 0 | - |
| - | - | 722.1 | 137.6 | - | - | 0 | - |
| - | - | 851.1 | 139.2 | - | - | 0 | - |
| - | - | 5.684E+04 | 141.1 | - | - | 0 | - |
| - | - | 3849 | 142.1 | - | - | 0 | - |
| 8 | y | 2.816E+04 | 147.1 | 0.0004466 | 3.036 | +1 | 1 |
| - | - | 1241 | 148.1 | - | - | 0 | - |
| - | - | 1.943E+04 | 149 | - | - | 0 | - |
| - | - | 1480 | 150 | - | - | 0 | - |
| - | - | 5.683E+05 | 157.1 | - | - | 0 | - |
| - | - | 3459 | 158.1 | - | - | 0 | - |
| - | - | 4.701E+04 | 158.1 | - | - | 0 | - |
| - | - | 1240 | 159.1 | - | - | 0 | - |
| - | - | 889.4 | 163.4 | - | - | 0 | - |
| - | - | 3756 | 167 | - | - | 0 | - |
| - | - | 1282 | 168.1 | - | - | 0 | - |
| - | - | 1.193E+05 | 169.1 | - | - | 0 | - |
| - | - | 9980 | 170.1 | - | - | 0 | - |
| - | - | 1026 | 173.4 | - | - | 0 | - |
| - | - | 2714 | 173.5 | - | - | 0 | - |
| - | - | 1003 | 173.5 | - | - | 0 | - |
| - | - | 7205 | 183.1 | - | - | 0 | - |
| - | - | 8668 | 184.1 | - | - | 0 | - |
| - | - | 2.441E+05 | 185.1 | - | - | 0 | - |
| - | - | 934.4 | 185.8 | - | - | 0 | - |
| - | - | 1934 | 186.1 | - | - | 0 | - |
| - | - | 2.4E+04 | 186.1 | - | - | 0 | - |
| - | - | 908.6 | 187.6 | - | - | 0 | - |
| - | - | 1274 | 197.1 | - | - | 0 | - |
| - | - | 1358 | 198.1 | - | - | 0 | - |
| 7 | w | 3131 | 201.1 | 0.0007004 | 3.482 | +1 | 2 |
| - | - | 1.265E+04 | 211.1 | - | - | 0 | - |
| - | - | 1614 | 212.1 | - | - | 0 | - |
| - | - | 5.051E+04 | 228.2 | - | - | 0 | - |
| - | - | 5405 | 229.2 | - | - | 0 | - |
| - | - | 6029 | 240.1 | - | - | 0 | - |
| - | - | 2.035E+04 | 243.7 | - | - | 0 | - |
| - | - | 2833 | 244.2 | - | - | 0 | - |
| - | - | 1294 | 244.7 | - | - | 0 | - |
| - | - | 1120 | 251.3 | - | - | 0 | - |
| 7 | y | 1.205E+04 | 258.1 | 0.0004921 | 1.906 | +1 | 2 |
| 7 | z | 1.011E+04 | 260.1 | 0.0004118 | 1.583 | +1 | 2 |
| - | - | 2021 | 261.1 | - | - | 0 | - |
| - | - | 1.031E+04 | 266.2 | - | - | 0 | - |
| - | - | 6712 | 266.2 | - | - | 0 | - |
| - | - | 1108 | 267.2 | - | - | 0 | - |
| - | - | 3062 | 267.2 | - | - | 0 | - |
| 7 | y | 2.989E+04 | 276.2 | 0.0006391 | 2.314 | +1 | 2 |
| - | - | 2911 | 277.2 | - | - | 0 | - |
| - | - | 9890 | 282.2 | - | - | 0 | - |
| - | - | 4.206E+04 | 283.2 | - | - | 0 | - |
| - | - | 5049 | 284.2 | - | - | 0 | - |
| - | - | 1402 | 293.2 | - | - | 0 | - |
| - | - | 1218 | 298.1 | - | - | 0 | - |
| - | - | 1391 | 300.2 | - | - | 0 | - |
| - | - | 1394 | 314.2 | - | - | 0 | - |
| - | - | 1857 | 318.2 | - | - | 0 | - |
| 3 | y | 1597 | 318.7 | 0.0006562 | 2.059 | +2 | 6 |
| - | - | 2818 | 325.2 | - | - | 0 | - |
| - | - | 1133 | 326.2 | - | - | 0 | - |
| 3 | y | 1.268E+06 | 327.7 | 0.0008976 | 2.739 | +2 | 6 |
| - | - | 4.404E+05 | 328.2 | - | - | 0 | - |
| - | - | 9.907E+04 | 328.7 | - | - | 0 | - |
| - | - | 7953 | 329.2 | - | - | 0 | - |
| - | - | 4.063E+04 | 340.2 | - | - | 0 | - |
| - | - | 5837 | 341.2 | - | - | 0 | - |
| 6 | w | 3.435E+04 | 344.2 | 0.0008831 | 2.566 | +1 | 3 |
| - | - | 5538 | 345.2 | - | - | 0 | - |
| - | - | 1249 | 347.5 | - | - | 0 | - |
| - | - | 1.423E+04 | 351.2 | - | - | 0 | - |
| - | - | 8823 | 352.2 | - | - | 0 | - |
| - | - | 2203 | 352.2 | - | - | 0 | - |
| - | - | 3.235E+04 | 353.2 | - | - | 0 | - |
| - | - | 5661 | 354.2 | - | - | 0 | - |
| - | - | 4.718E+04 | 357.2 | - | - | 0 | - |
| 6 | w | 1.07E+04 | 358.2 | 6.459E-05 | 0.1803 | +1 | 3 |
| - | - | 6230 | 358.2 | - | - | 0 | - |
| - | - | 2587 | 359.2 | - | - | 0 | - |
| - | - | 1506 | 368.2 | - | - | 0 | - |
| 6 | z | 7.67E+04 | 373.2 | 0.0006373 | 1.708 | +1 | 3 |
| - | - | 4.672E+04 | 374.2 | - | - | 0 | - |
| 2 | y | 7243 | 375.2 | 0.0002075 | 0.5531 | +2 | 7 |
| - | - | 4.337E+04 | 379.2 | - | - | 0 | - |
| - | - | 6685 | 380.2 | - | - | 0 | - |
| - | - | 2936 | 383.2 | - | - | 0 | - |
| 2 | y | 1.38E+04 | 384.2 | 0.0006747 | 1.756 | +2 | 7 |
| - | - | 1959 | 384.7 | - | - | 0 | - |
| 6 | y | 1.092E+04 | 389.2 | 0.0007121 | 1.829 | +1 | 3 |
| 4 | c | 3.106E+05 | 396.3 | 0.0009131 | 2.304 | +1 | 4 |
| - | - | 7.014E+04 | 397.3 | - | - | 0 | - |
| - | - | 6690 | 398.3 | - | - | 0 | - |
| - | - | 3268 | 410.2 | - | - | 0 | - |
| - | - | 3417 | 411.2 | - | - | 0 | - |
| - | - | 3329 | 413.2 | - | - | 0 | - |
| - | - | 1413 | 413.3 | - | - | 0 | - |
| - | - | 3521 | 415.2 | - | - | 0 | - |
| - | - | 6229 | 418.7 | - | - | 0 | - |
| - | - | 3773 | 419.2 | - | - | 0 | - |
| 5 | y | 2.254E+04 | 443.3 | 0.0009201 | 2.076 | +1 | 4 |
| 5 | z | 6081 | 444.3 | 0.00309 | 6.956 | +1 | 4 |
| - | - | 4150 | 450.3 | - | - | 0 | - |
| - | - | 2461 | 462.3 | - | - | 0 | - |
| - | - | 1.143E+05 | 466.3 | - | - | 0 | - |
| 5 | c | 8.349E+04 | 467.3 | 0.0002815 | 0.6025 | +1 | 5 |
| - | - | 1.122E+04 | 468.3 | - | - | 0 | - |
| - | - | 9718 | 469.3 | - | - | 0 | - |
| - | - | 3007 | 470.2 | - | - | 0 | - |
| - | - | 2453 | 470.3 | - | - | 0 | - |
| - | - | 7896 | 480.3 | - | - | 0 | - |
| - | - | 9265 | 482.3 | - | - | 0 | - |
| - | - | 2570 | 483.3 | - | - | 0 | - |
| - | - | 6.073E+04 | 484.3 | - | - | 0 | - |
| - | - | 4768 | 485.2 | - | - | 0 | - |
| - | - | 2.638E+04 | 485.3 | - | - | 0 | - |
| - | - | 4.277E+05 | 486.3 | - | - | 0 | - |
| - | - | 1.013E+05 | 487.3 | - | - | 0 | - |
| - | - | 1.736E+04 | 488.3 | - | - | 0 | - |
| - | - | 1958 | 489.3 | - | - | 0 | - |
| - | - | 1950 | 490.3 | - | - | 0 | - |
| - | - | 6.092E+04 | 508.3 | - | - | 0 | - |
| - | - | 1.463E+04 | 509.3 | - | - | 0 | - |
| - | - | 5152 | 509.3 | - | - | 0 | - |
| - | - | 2840 | 510.3 | - | - | 0 | - |
| - | - | 1.783E+04 | 512.3 | - | - | 0 | - |
| - | - | 5282 | 513.3 | - | - | 0 | - |
| 4 | w | 6254 | 514.3 | 0.0002138 | 0.4157 | +1 | 5 |
| - | - | 2208 | 515.3 | - | - | 0 | - |
| - | - | 1.342E+04 | 523.3 | - | - | 0 | - |
| - | - | 5330 | 524.3 | - | - | 0 | - |
| - | - | 1.664E+05 | 525.3 | - | - | 0 | - |
| - | - | 3.96E+04 | 526.3 | - | - | 0 | - |
| - | - | 7863 | 527.3 | - | - | 0 | - |
| - | - | 8409 | 535.4 | - | - | 0 | - |
| - | - | 1.112E+04 | 536.4 | - | - | 0 | - |
| - | - | 1.517E+04 | 537.4 | - | - | 0 | - |
| - | - | 3177 | 538.4 | - | - | 0 | - |
| 4 | y | 3284 | 539.3 | 0.00105 | 1.947 | +1 | 5 |
| - | - | 1679 | 540.3 | - | - | 0 | - |
| 4 | z | 2.004E+05 | 541.3 | 0.001031 | 1.904 | +1 | 5 |
| - | - | 5.849E+04 | 542.3 | - | - | 0 | - |
| - | - | 1.253E+04 | 543.3 | - | - | 0 | - |
| 4 | y | 8.42E+04 | 557.3 | 0.0009224 | 1.655 | +1 | 5 |
| - | - | 1811 | 557.4 | - | - | 0 | - |
| - | - | 2.189E+04 | 558.3 | - | - | 0 | - |
| - | - | 3953 | 559.3 | - | - | 0 | - |
| - | - | 3099 | 562.3 | - | - | 0 | - |
| - | - | 2.599E+04 | 563.4 | - | - | 0 | - |
| - | - | 7799 | 564.4 | - | - | 0 | - |
| - | - | 1935 | 565.4 | - | - | 0 | - |
| - | - | 2474 | 573.3 | - | - | 0 | - |
| - | - | 2.341E+05 | 579.4 | - | - | 0 | - |
| 6 | c | 3.681E+05 | 580.4 | 6.611E-05 | 0.1139 | +1 | 6 |
| - | - | 1.008E+05 | 581.4 | - | - | 0 | - |
| - | - | 1471 | 582.3 | - | - | 0 | - |
| - | - | 1.906E+04 | 582.4 | - | - | 0 | - |
| - | - | 2057 | 583.3 | - | - | 0 | - |
| - | - | 2.306E+04 | 606.4 | - | - | 0 | - |
| - | - | 9341 | 607.4 | - | - | 0 | - |
| - | - | 1454 | 608.4 | - | - | 0 | - |
| - | - | 2185 | 609.4 | - | - | 0 | - |
| 3 | w | 9053 | 611.3 | 0.001405 | 2.298 | +1 | 6 |
| - | - | 7151 | 612.3 | - | - | 0 | - |
| - | - | 2413 | 622.3 | - | - | 0 | - |
| - | - | 1763 | 623.4 | - | - | 0 | - |
| 3 | y | 1.617E+04 | 636.4 | 0.0005323 | 0.8364 | +1 | 6 |
| - | - | 5190 | 637.4 | - | - | 0 | - |
| - | - | 1433 | 639.3 | - | - | 0 | - |
| - | - | 1.598E+04 | 652.4 | - | - | 0 | - |
| - | - | 6589 | 653.4 | - | - | 0 | - |
| 3 | y | 1.104E+06 | 654.4 | 0.001137 | 1.738 | +1 | 6 |
| - | - | 3.972E+05 | 655.4 | - | - | 0 | - |
| - | - | 9.622E+04 | 656.4 | - | - | 0 | - |
| - | - | 8635 | 657.4 | - | - | 0 | - |
| - | - | 9157 | 665.4 | - | - | 0 | - |
| - | - | 4236 | 666.4 | - | - | 0 | - |
| - | - | 1.259E+04 | 679.4 | - | - | 0 | - |
| - | - | 3134 | 680.4 | - | - | 0 | - |
| - | - | 1654 | 681.4 | - | - | 0 | - |
| - | - | 7748 | 682.4 | - | - | 0 | - |
| - | - | 8896 | 683.4 | - | - | 0 | - |
| - | - | 2100 | 684.4 | - | - | 0 | - |
| - | - | 3.594E+04 | 692.4 | - | - | 0 | - |
| - | - | 1.151E+04 | 693.4 | - | - | 0 | - |
| - | - | 5407 | 694.4 | - | - | 0 | - |
| - | - | 1.265E+04 | 695.4 | - | - | 0 | - |
| - | - | 5497 | 696.4 | - | - | 0 | - |
| 2 | w | 1.299E+05 | 708.4 | 0.004183 | 5.905 | +1 | 7 |
| 7 | c | 1.961E+06 | 709.4 | 0.001052 | 1.483 | +1 | 7 |
| - | - | 7.283E+05 | 710.4 | - | - | 0 | - |
| - | - | 4699 | 710.6 | - | - | 0 | - |
| - | - | 1.802E+05 | 711.4 | - | - | 0 | - |
| - | - | 1.42E+04 | 712.4 | - | - | 0 | - |
| - | - | 1468 | 737.4 | - | - | 0 | - |
| - | - | 1.608E+04 | 750.5 | - | - | 0 | - |
| 2 | z | 1.133E+05 | 751.4 | 0.00141 | 1.876 | +1 | 7 |
| - | - | 4.146E+04 | 752.5 | - | - | 0 | - |
| - | - | 1.28E+04 | 753.5 | - | - | 0 | - |
| - | - | 3708 | 760.4 | - | - | 0 | - |
| - | - | 1993 | 760.9 | - | - | 0 | - |
| - | - | 1830 | 763.5 | - | - | 0 | - |
| - | - | 6685 | 764.4 | - | - | 0 | - |
| - | - | 1841 | 765.4 | - | - | 0 | - |
| - | - | 9.004E+05 | 766.4 | - | - | 0 | - |
| - | - | 3.403E+05 | 767.4 | - | - | 0 | - |
| - | - | 9.894E+04 | 768.4 | - | - | 0 | - |
| - | - | 8633 | 769.4 | - | - | 0 | - |
| - | - | 3174 | 779.3 | - | - | 0 | - |
| - | - | 3236 | 779.4 | - | - | 0 | - |
| - | - | 1861 | 780.3 | - | - | 0 | - |
| - | - | 4772 | 780.5 | - | - | 0 | - |
| - | - | 1853 | 781.5 | - | - | 0 | - |
| - | - | 1627 | 782.4 | - | - | 0 | - |
| - | - | 1.111E+05 | 783.4 | - | - | 0 | - |
| - | - | 4.646E+04 | 784.5 | - | - | 0 | - |
| - | - | 1.197E+04 | 785.5 | - | - | 0 | - |
| - | - | 3746 | 793.5 | - | - | 0 | - |
| - | - | 1916 | 794.5 | - | - | 0 | - |
| - | - | 2412 | 795.5 | - | - | 0 | - |
| - | - | 6651 | 796.5 | - | - | 0 | - |
| - | - | 2127 | 797.5 | - | - | 0 | - |
| - | - | 4096 | 810.5 | - | - | 0 | - |
| - | - | 2012 | 811.5 | - | - | 0 | - |
| - | - | 1499 | 820.4 | - | - | 0 | - |
| - | - | 5048 | 821.5 | - | - | 0 | - |
| - | - | 6.163E+05 | 822.5 | - | - | 0 | - |
| - | - | 2.667E+05 | 823.5 | - | - | 0 | - |
| - | - | 8.316E+04 | 824.5 | - | - | 0 | - |
| - | - | 7750 | 825.5 | - | - | 0 | - |
| - | - | 2988 | 831.4 | - | - | 0 | - |
| - | - | 2740 | 831.9 | - | - | 0 | - |
| - | - | 2368 | 832.4 | - | - | 0 | - |
| - | - | 6031 | 837.4 | - | - | 0 | - |
| - | - | 6.73E+05 | 838.5 | - | - | 0 | - |
| - | - | 1.966E+06 | 839.5 | - | - | 0 | - |
| - | - | 8.067E+05 | 840.5 | - | - | 0 | - |
| - | - | 2.243E+05 | 841.5 | - | - | 0 | - |
| - | - | 1.988E+04 | 842.5 | - | - | 0 | - |
| - | - | 4014 | 854.5 | - | - | 0 | - |
| - | - | 1560 | 855.5 | - | - | 0 | - |
| - | - | 2017 | 871.5 | - | - | 0 | - |

m/z Charge Intensity FragmentType MassShift Position
126.3384017944336 0 697.0138
129.09767150878906 0 1318.8217
129.10272216796875 0 13168.751
129.13946533203125 0 816.6779
130.0867156982422 0 6244.064 y Ammonia loss 7
135.46009826660156 0 761.32416
137.2176971435547 0 1005.3858
137.55181884765625 0 722.0998
139.23861694335938 0 851.087
141.1027069091797 0 56839.047
142.10610961914062 0 3848.7412
147.11325073242188 0 28164.775 y 7
148.11685180664062 0 1241.3699
149.02377319335938 0 19426.684
150.0269317626953 0 1480.1073
157.13406372070312 0 568332.4
158.13092041015625 0 3459.3662
158.13735961914062 0 47005.84
159.14039611816406 0 1240.2499
163.38311767578125 0 889.44995
167.034423828125 0 3755.6028
168.1022491455078 0 1281.6277
169.09765625 0 119271.46
170.1009979248047 0 9979.569
173.4312744140625 0 1026.178
173.45286560058594 0 2713.538
173.45730590820312 0 1003.37714
183.14979553222656 0 7205.1475
184.12130737304688 0 8668.11
185.1289825439453 0 244136
185.76266479492188 0 934.4326
186.12399291992188 0 1933.7386
186.1324005126953 0 24002.035
187.60736083984375 0 908.59106
197.0922393798828 0 1274.0662
198.07717895507812 0 1358.0061
201.1240692138672 0 3130.6753 w 6
211.14462280273438 0 12652.737
212.14833068847656 0 1614.4854
228.17123413085938 0 50512.812
229.17471313476562 0 5404.5645
240.1349639892578 0 6028.532
243.650390625 0 20350.76
244.1520538330078 0 2833.1892
244.654296875 0 1293.5266
251.28817749023438 0 1120.483
258.14532470703125 0 12045.534 y Water loss 6
260.1370849609375 0 10109.924 z 6
261.1439514160156 0 2020.7784
266.1501770019531 0 10310.463
266.1634216308594 0 6711.756
267.1535949707031 0 1107.6868
267.169677734375 0 3061.922
276.1560363769531 0 29892.262 y 6
277.15863037109375 0 2910.7317
282.181884765625 0 9889.722
283.17718505859375 0 42055.99
284.18035888671875 0 5048.681
293.1755676269531 0 1401.7008
298.13897705078125 0 1218.0568
300.1919860839844 0 1390.6908
314.2099609375 0 1393.5775
318.1634216308594 0 1857.4169
318.6900634765625 0 1596.6079 y Water loss 2
325.22454833984375 0 2817.8472
326.2294616699219 0 1132.5696
327.6955871582031 0 1268367.5 y 2
328.197021484375 0 440424.47
328.6981201171875 0 99073.46
329.19927978515625 0 7953.245
340.1874694824219 0 40628.605
341.1905517578125 0 5837.309
344.1824951171875 0 34347.51 w 5
345.1856689453125 0 5538.038
347.46246337890625 0 1248.9302
351.2395935058594 0 14226.275
352.21124267578125 0 8822.52
352.24285888671875 0 2202.6943
353.21893310546875 0 32351.006
354.2224426269531 0 5660.7188
357.21392822265625 0 47177.83
358.19732666015625 0 10695.168 w 5
358.21832275390625 0 6229.6733
359.2010192871094 0 2587.2754
368.21722412109375 0 1505.6229
373.22137451171875 0 76700.625 z 5
374.2282409667969 0 46721.535
375.2312316894531 0 7242.7075 y Water loss 1
379.2347106933594 0 43370.35
380.237548828125 0 6684.909
383.1590270996094 0 2936.426
384.2373962402344 0 13803.301 y 1
384.73944091796875 0 1959.1586
389.24017333984375 0 10918.105 y 5
396.2614440917969 0 310568.53 c 3
397.264404296875 0 70141.98
398.26641845703125 0 6690.042
410.22845458984375 0 3267.7634
411.2254333496094 0 3417.2983
413.2276306152344 0 3328.851
413.2789001464844 0 1412.6141
415.2198791503906 0 3521.2595
418.67864990234375 0 6229.3936
419.17913818359375 0 3772.6023
443.2509460449219 0 22536.54 y Ammonia loss 4
444.2547607421875 0 6080.944 z 4
450.2716979980469 0 4149.6367
462.27203369140625 0 2461.0173
466.29083251953125 0 114272.37
467.29736328125 0 83493.01 c 4
468.29931640625 0 11221.776
469.29058837890625 0 9717.811
470.2367248535156 0 3007.491
470.29461669921875 0 2453.0457
480.2828674316406 0 7895.777
482.2976379394531 0 9264.508
483.30230712890625 0 2569.5928
484.2777099609375 0 60725.8
485.24786376953125 0 4767.788
485.2832336425781 0 26380.537
486.2931823730469 0 427745.75
487.29608154296875 0 101251.34
488.29840087890625 0 17357.768
489.2962646484375 0 1957.9597
490.2669677734375 0 1950.4216
508.2775573730469 0 60915.344
509.2804870605469 0 14632.922
509.344970703125 0 5151.871
510.283203125 0 2840.4504
512.2722778320312 0 17832.123
513.27685546875 0 5281.812
514.287353515625 0 6253.6284 w 3
515.2926635742188 0 2207.9182
523.3115844726562 0 13423.682
524.3134155273438 0 5329.9673
525.3040771484375 0 166427.83
526.304931640625 0 39597.18
527.30712890625 0 7862.9434
535.360595703125 0 8409.173
536.3677978515625 0 11118.4375
537.3757934570312 0 15166.046
538.3798828125 0 3177.397
539.31982421875 0 3283.7595 y Water loss 3
540.3267822265625 0 1679.2842
541.3116455078125 0 200381.02 z 3
542.314697265625 0 58487.72
543.3176879882812 0 12529.356
557.3302612304688 0 84202.38 y 3
557.38134765625 0 1811.1941
558.3328247070312 0 21889.799
559.3338623046875 0 3952.8022
562.3483276367188 0 3099.4849
563.355712890625 0 25985.195
564.358154296875 0 7799.2827
565.367431640625 0 1934.6361
573.303955078125 0 2473.9019
579.374755859375 0 234065.52
580.3817749023438 0 368119.12 c 5
581.385009765625 0 100785.875
582.304443359375 0 1471.0405
582.38818359375 0 19062.055
583.3096923828125 0 2057.2349
606.3983764648438 0 23058.844
607.40087890625 0 9340.9375
608.4056396484375 0 1453.8083
609.3512573242188 0 2185.1685
611.34130859375 0 9052.873 w 2
612.3467407226562 0 7150.882
622.2665405273438 0 2413.2522
623.3607177734375 0 1763.1123
636.3720703125 0 16174.795 y Water loss 2
637.3770751953125 0 5190.006
639.3221435546875 0 1432.5062
652.3565063476562 0 15977.354
653.35986328125 0 6589.194
654.3832397460938 0 1103672.9 y 2
655.3866577148438 0 397243.44
656.3894653320312 0 96217.03
657.3935546875 0 8634.703
665.4111938476562 0 9156.93
666.4161987304688 0 4235.769
679.4255981445312 0 12594.213
680.4317626953125 0 3133.797
681.3656616210938 0 1653.6115
682.377685546875 0 7748.429
683.384033203125 0 8895.83
684.3887329101562 0 2099.95
692.3992919921875 0 35940.496
693.4025268554688 0 11513.395
694.4057006835938 0 5406.884
695.3876953125 0 12651.234
696.3876342773438 0 5496.833
708.3968505859375 0 129911.18 w 1
709.4253540039062 0 1961462 c 6
710.4283447265625 0 728287.75
710.5728149414062 0 4698.649
711.4307250976562 0 180151.31
712.433837890625 0 14202.724
737.4495239257812 0 1467.725
750.4642333984375 0 16083.919
751.4488525390625 0 113347.15 z 1
752.4515380859375 0 41457.08
753.4545288085938 0 12804.113
760.3800048828125 0 3707.8916
760.8873291015625 0 1992.8818
763.4674072265625 0 1830.4656
764.4310913085938 0 6685.1333
765.4283447265625 0 1841.4045
766.4229736328125 0 900357
767.4257202148438 0 340251.94
768.4287109375 0 98944.46
769.4307861328125 0 8633.328
779.3263549804688 0 3174.3823
779.4290771484375 0 3236.1785
780.3236083984375 0 1860.8386
780.4985961914062 0 4771.8955
781.4989013671875 0 1853.3689
782.4341430664062 0 1626.9694
783.4495849609375 0 111072.38
784.4522705078125 0 46464.56
785.4549560546875 0 11967.15
793.5049438476562 0 3745.6929
794.5126342773438 0 1916.2267
795.5166625976562 0 2411.864
796.459716796875 0 6651.168
797.4512329101562 0 2126.5505
810.47216796875 0 4095.8005
811.46728515625 0 2012.2854
820.4200439453125 0 1498.8555
821.49951171875 0 5048.2705
822.485595703125 0 616260.1
823.48876953125 0 266716.56
824.4910278320312 0 83164.914
825.4916381835938 0 7750.144
831.4202880859375 0 2987.9084
831.9232788085938 0 2739.9065
832.4142456054688 0 2368.4072
837.3536376953125 0 6030.641
838.5042114257812 0 673013.94
839.5116577148438 0 1965507.8
840.5143432617188 0 806720.5
841.5173950195312 0 224348.56
842.5189208984375 0 19884.582
854.4752807617188 0 4014.2097
855.4747924804688 0 1559.6959
871.4956665039062 0 2017.1573

Spectrum Details

|  |  |
| --- | --- |
| Matched peaks? Matched peaks The total absolute number of peaks matched. Additionally in brackets the total fraction of peaks matched and the total number of peaks is shown. | 29 (11.79% of 246) |
| FDR? FDR The false discovery rate estimated for this peptide. It is calculated by matching all theoretical fragments with a non-integer shift with the raw peaks for this spectrum. This is done with 40 different shifts. The resulting percentage is the average number of annotated peaks over the number of annotated peaks with the correct spectrum. | 0.33% |
| Satellite FDR? Satellite FDR See the FDR for details on its calculation. This satellite ion specific FDR only contains the satellite ions (d/w) for I/L/J positions. | 0.00% |
| PSM Score? PSM Score The PSM Score as given by Hecklib to this annotated spectrum. It is shown with three significant figures. | 253 |

#### Spectrum 4468? Spectrum 4468 The raw spectrum of this peptide as annotated by Hecklib. The fragments are coloured according to ion type (see legend). Any peaks with a star '\*' as text can be hovered over to see the full details, first the ion type second the mass shift type. By hovering over the amino acids in the peptide or ions in the legend the corresponding peaks are highlighted. By toggling the 'Unassigned' label you can turn the background (unassigned) peaks on or off in the plot. By updating the slider in the Ion legend you can update the spectrum to only show the top X% of the peaks with labels. The top X% means any peak that is within X% of the highest intensity. By dragging in the spectrum you can zoom in to a specific part of the spectrum and use 'Zoom Out' to get back to the original zoom level. The annotation of the spectrum is based on the given sequence in the peptides file and is done with different software so inconsistencies are likely. The peaks are annotated based on the given sequence, with 20 ppm tolerance.

Copy Data

##### Spectrum 4468 (TSV)

###### Preview

```
Loading example...
```

*Click on the button to copy the data to your clipboard.*

Mz MinMz MaxIntensity Max

WidthHeightPeptide font sizePeptide stroke widthSpectrum font sizeSpectrum stroke widthCompact peptide

Ion legend

wxyz

abcd

OtherUnassignedIon^Charge^_Position_Show for top:%

ALPPAIEK

01.49e+62.98e+64.48e+65.97e+6

Zoom Out

y^+1^_1_y^+1^_1_w^+1^_2_y^+1^_2_z^+1^_2_y^+1^_2_y^+2^_5_y^+2^_6_y^+2^_6_w^+1^_3_w^+1^_3_z^+1^_3_y^+2^_7_y^+2^_7_y^+1^_3_c^+1^_4_y^+1^_4_z^+1^_4_c^+1^_5_w^+1^_5_y^+1^_5_z^+1^_5_y^+1^_5_c^+1^_6_w^+1^_6_y^+1^_6_y^+1^_6_w^+1^_7_c^+1^_7_z^+1^_7_

0220440660880

Fragment Matches Table

Show background peaks

| Position | Ion type | Intensity | mz Theoretical | mz Error (Th) | mz Error (ppm) | Charge | Series Number |
| --- | --- | --- | --- | --- | --- | --- | --- |
| - | - | 1872 | 124.1 | - | - | 0 | - |
| - | - | 2457 | 124.4 | - | - | 0 | - |
| - | - | 1990 | 128.2 | - | - | 0 | - |
| - | - | 3.91E+04 | 129.1 | - | - | 0 | - |
| 8 | y | 1.824E+04 | 130.1 | 0.0004454 | 3.424 | +1 | 1 |
| - | - | 1.778E+05 | 141.1 | - | - | 0 | - |
| - | - | 1.253E+04 | 142.1 | - | - | 0 | - |
| 8 | y | 8.527E+04 | 147.1 | 0.0005686 | 3.865 | +1 | 1 |
| - | - | 3594 | 148.1 | - | - | 0 | - |
| - | - | 3121 | 148.9 | - | - | 0 | - |
| - | - | 2775 | 148.9 | - | - | 0 | - |
| - | - | 4633 | 148.9 | - | - | 0 | - |
| - | - | 4325 | 148.9 | - | - | 0 | - |
| - | - | 3970 | 148.9 | - | - | 0 | - |
| - | - | 1.105E+04 | 148.9 | - | - | 0 | - |
| - | - | 1.337E+04 | 148.9 | - | - | 0 | - |
| - | - | 2.747E+04 | 149 | - | - | 0 | - |
| - | - | 1.645E+04 | 149 | - | - | 0 | - |
| - | - | 7867 | 149 | - | - | 0 | - |
| - | - | 4728 | 149 | - | - | 0 | - |
| - | - | 4778 | 149 | - | - | 0 | - |
| - | - | 3289 | 149 | - | - | 0 | - |
| - | - | 3686 | 149 | - | - | 0 | - |
| - | - | 1.72E+04 | 149 | - | - | 0 | - |
| - | - | 2457 | 152.3 | - | - | 0 | - |
| - | - | 2343 | 152.8 | - | - | 0 | - |
| - | - | 1916 | 154.9 | - | - | 0 | - |
| - | - | 1.927E+06 | 157.1 | - | - | 0 | - |
| - | - | 8967 | 158.1 | - | - | 0 | - |
| - | - | 1.582E+05 | 158.1 | - | - | 0 | - |
| - | - | 2728 | 158.7 | - | - | 0 | - |
| - | - | 3542 | 159.1 | - | - | 0 | - |
| - | - | 2468 | 161 | - | - | 0 | - |
| - | - | 3469 | 168.1 | - | - | 0 | - |
| - | - | 3191 | 168.1 | - | - | 0 | - |
| - | - | 4.037E+05 | 169.1 | - | - | 0 | - |
| - | - | 3980 | 170.1 | - | - | 0 | - |
| - | - | 3.594E+04 | 170.1 | - | - | 0 | - |
| - | - | 2780 | 173.5 | - | - | 0 | - |
| - | - | 2.487E+04 | 183.1 | - | - | 0 | - |
| - | - | 2.029E+04 | 184.1 | - | - | 0 | - |
| - | - | 2449 | 184.2 | - | - | 0 | - |
| - | - | 2557 | 184.6 | - | - | 0 | - |
| - | - | 8.609E+05 | 185.1 | - | - | 0 | - |
| - | - | 7.863E+04 | 186.1 | - | - | 0 | - |
| - | - | 2520 | 187.1 | - | - | 0 | - |
| - | - | 5478 | 198.1 | - | - | 0 | - |
| 7 | w | 7911 | 201.1 | 0.0006546 | 3.255 | +1 | 2 |
| - | - | 3.628E+04 | 211.1 | - | - | 0 | - |
| - | - | 2822 | 211.2 | - | - | 0 | - |
| - | - | 4308 | 212.1 | - | - | 0 | - |
| - | - | 1.381E+05 | 228.2 | - | - | 0 | - |
| - | - | 1.424E+04 | 229.2 | - | - | 0 | - |
| - | - | 1.737E+04 | 240.1 | - | - | 0 | - |
| - | - | 6.182E+04 | 243.7 | - | - | 0 | - |
| - | - | 8608 | 244.2 | - | - | 0 | - |
| - | - | 5568 | 254.2 | - | - | 0 | - |
| 7 | y | 3.817E+04 | 258.1 | 0.0007058 | 2.734 | +1 | 2 |
| - | - | 4834 | 259.1 | - | - | 0 | - |
| 7 | z | 2.437E+04 | 260.1 | 0.000778 | 2.991 | +1 | 2 |
| - | - | 5385 | 261.1 | - | - | 0 | - |
| - | - | 3.817E+04 | 266.2 | - | - | 0 | - |
| - | - | 1.951E+04 | 266.2 | - | - | 0 | - |
| - | - | 4579 | 267.2 | - | - | 0 | - |
| - | - | 8334 | 267.2 | - | - | 0 | - |
| 7 | y | 1.042E+05 | 276.2 | 0.0007917 | 2.867 | +1 | 2 |
| - | - | 1.139E+04 | 277.2 | - | - | 0 | - |
| 4 | y | 5954 | 279.2 | 0.0009123 | 3.268 | +2 | 5 |
| - | - | 2.604E+04 | 282.2 | - | - | 0 | - |
| - | - | 1.175E+05 | 283.2 | - | - | 0 | - |
| - | - | 1.446E+04 | 284.2 | - | - | 0 | - |
| - | - | 5164 | 304.2 | - | - | 0 | - |
| - | - | 4279 | 305.2 | - | - | 0 | - |
| - | - | 5116 | 309.7 | - | - | 0 | - |
| - | - | 4705 | 312.2 | - | - | 0 | - |
| - | - | 2714 | 314.2 | - | - | 0 | - |
| 3 | y | 7558 | 318.7 | 7.619E-05 | 0.2391 | +2 | 6 |
| - | - | 5654 | 325.2 | - | - | 0 | - |
| - | - | 4269 | 327.2 | - | - | 0 | - |
| 3 | y | 4.309E+06 | 327.7 | 0.001111 | 3.391 | +2 | 6 |
| - | - | 1.499E+06 | 328.2 | - | - | 0 | - |
| - | - | 3.492E+05 | 328.7 | - | - | 0 | - |
| - | - | 1.694E+04 | 329.2 | - | - | 0 | - |
| - | - | 1.28E+05 | 340.2 | - | - | 0 | - |
| - | - | 2.223E+04 | 341.2 | - | - | 0 | - |
| 6 | w | 8.695E+04 | 344.2 | 0.001097 | 3.187 | +1 | 3 |
| - | - | 1.344E+04 | 345.2 | - | - | 0 | - |
| - | - | 4.787E+04 | 351.2 | - | - | 0 | - |
| - | - | 1.835E+04 | 352.2 | - | - | 0 | - |
| - | - | 5724 | 352.2 | - | - | 0 | - |
| - | - | 9.974E+04 | 353.2 | - | - | 0 | - |
| - | - | 3991 | 353.3 | - | - | 0 | - |
| - | - | 1.877E+04 | 354.2 | - | - | 0 | - |
| - | - | 1.245E+05 | 357.2 | - | - | 0 | - |
| 6 | w | 2.951E+04 | 358.2 | 0.0006139 | 1.714 | +1 | 3 |
| - | - | 1.74E+04 | 358.2 | - | - | 0 | - |
| - | - | 5704 | 359.2 | - | - | 0 | - |
| - | - | 3184 | 368.2 | - | - | 0 | - |
| 6 | z | 2.093E+05 | 373.2 | 0.0009425 | 2.525 | +1 | 3 |
| - | - | 1.328E+05 | 374.2 | - | - | 0 | - |
| 2 | y | 2.655E+04 | 375.2 | 0.0001587 | 0.4228 | +2 | 7 |
| - | - | 1.323E+05 | 379.2 | - | - | 0 | - |
| - | - | 2.298E+04 | 380.2 | - | - | 0 | - |
| - | - | 3383 | 381.2 | - | - | 0 | - |
| 2 | y | 4.082E+04 | 384.2 | 0.0008578 | 2.232 | +2 | 7 |
| - | - | 1.388E+04 | 384.7 | - | - | 0 | - |
| 6 | y | 2.944E+04 | 389.2 | 0.0007731 | 1.986 | +1 | 3 |
| - | - | 5267 | 390.2 | - | - | 0 | - |
| - | - | 2537 | 391.2 | - | - | 0 | - |
| - | - | 3934 | 395.2 | - | - | 0 | - |
| 4 | c | 9.429E+05 | 396.3 | 0.001188 | 2.997 | +1 | 4 |
| - | - | 1.907E+05 | 397.3 | - | - | 0 | - |
| - | - | 2.494E+04 | 398.3 | - | - | 0 | - |
| - | - | 3901 | 399.2 | - | - | 0 | - |
| - | - | 3092 | 400.2 | - | - | 0 | - |
| - | - | 1.091E+04 | 410.2 | - | - | 0 | - |
| - | - | 1.72E+04 | 411.2 | - | - | 0 | - |
| - | - | 1.107E+04 | 413.2 | - | - | 0 | - |
| - | - | 1.502E+04 | 415.2 | - | - | 0 | - |
| - | - | 3269 | 416.2 | - | - | 0 | - |
| - | - | 4343 | 422.3 | - | - | 0 | - |
| - | - | 3634 | 423.3 | - | - | 0 | - |
| - | - | 4514 | 429.2 | - | - | 0 | - |
| 5 | y | 6.799E+04 | 443.3 | 0.001317 | 2.971 | +1 | 4 |
| 5 | z | 1.43E+04 | 444.3 | 0.001534 | 3.453 | +1 | 4 |
| - | - | 3533 | 445.3 | - | - | 0 | - |
| - | - | 1.033E+04 | 450.3 | - | - | 0 | - |
| - | - | 9061 | 462.3 | - | - | 0 | - |
| - | - | 3.299E+05 | 466.3 | - | - | 0 | - |
| 5 | c | 2.612E+05 | 467.3 | 8.468E-05 | 0.1812 | +1 | 5 |
| - | - | 4.08E+04 | 468.3 | - | - | 0 | - |
| - | - | 3.006E+04 | 469.3 | - | - | 0 | - |
| - | - | 7801 | 470.2 | - | - | 0 | - |
| - | - | 8697 | 470.3 | - | - | 0 | - |
| - | - | 2.765E+04 | 480.3 | - | - | 0 | - |
| - | - | 7379 | 481.3 | - | - | 0 | - |
| - | - | 3.224E+04 | 482.3 | - | - | 0 | - |
| - | - | 7397 | 483.3 | - | - | 0 | - |
| - | - | 1.935E+05 | 484.3 | - | - | 0 | - |
| - | - | 1.354E+04 | 485.2 | - | - | 0 | - |
| - | - | 8.171E+04 | 485.3 | - | - | 0 | - |
| - | - | 1.467E+06 | 486.3 | - | - | 0 | - |
| - | - | 3.311E+05 | 487.3 | - | - | 0 | - |
| - | - | 6.125E+04 | 488.3 | - | - | 0 | - |
| - | - | 4285 | 490.3 | - | - | 0 | - |
| - | - | 2.019E+05 | 508.3 | - | - | 0 | - |
| - | - | 5.233E+04 | 509.3 | - | - | 0 | - |
| - | - | 1.33E+04 | 509.3 | - | - | 0 | - |
| - | - | 1.231E+04 | 510.3 | - | - | 0 | - |
| - | - | 4.938E+04 | 512.3 | - | - | 0 | - |
| - | - | 1.663E+04 | 513.3 | - | - | 0 | - |
| 4 | w | 1.546E+04 | 514.3 | 0.0002134 | 0.415 | +1 | 5 |
| - | - | 5322 | 515.3 | - | - | 0 | - |
| - | - | 3745 | 518.4 | - | - | 0 | - |
| - | - | 5.312E+04 | 523.3 | - | - | 0 | - |
| - | - | 1.2E+04 | 524.3 | - | - | 0 | - |
| - | - | 5.071E+05 | 525.3 | - | - | 0 | - |
| - | - | 1.203E+05 | 526.3 | - | - | 0 | - |
| - | - | 1.452E+04 | 527.3 | - | - | 0 | - |
| - | - | 2.215E+04 | 535.4 | - | - | 0 | - |
| - | - | 2.845E+04 | 536.4 | - | - | 0 | - |
| - | - | 3.876E+04 | 537.4 | - | - | 0 | - |
| - | - | 1.343E+04 | 538.4 | - | - | 0 | - |
| 4 | y | 1.175E+04 | 539.3 | 0.0003786 | 0.7021 | +1 | 5 |
| - | - | 3726 | 540.3 | - | - | 0 | - |
| 4 | z | 5.842E+05 | 541.3 | 0.001458 | 2.693 | +1 | 5 |
| - | - | 1.594E+05 | 542.3 | - | - | 0 | - |
| - | - | 2.951E+04 | 543.3 | - | - | 0 | - |
| 4 | y | 2.898E+05 | 557.3 | 0.001472 | 2.641 | +1 | 5 |
| - | - | 8.035E+04 | 558.3 | - | - | 0 | - |
| - | - | 1.66E+04 | 559.3 | - | - | 0 | - |
| - | - | 1.137E+04 | 562.3 | - | - | 0 | - |
| - | - | 6.505E+04 | 563.4 | - | - | 0 | - |
| - | - | 2.086E+04 | 564.4 | - | - | 0 | - |
| - | - | 5026 | 565.4 | - | - | 0 | - |
| - | - | 4809 | 573.3 | - | - | 0 | - |
| - | - | 6.801E+05 | 579.4 | - | - | 0 | - |
| 6 | c | 1.087E+06 | 580.4 | 0.0004934 | 0.85 | +1 | 6 |
| - | - | 3.13E+05 | 581.4 | - | - | 0 | - |
| - | - | 5959 | 582.3 | - | - | 0 | - |
| - | - | 5.396E+04 | 582.4 | - | - | 0 | - |
| - | - | 3969 | 583.3 | - | - | 0 | - |
| - | - | 4979 | 583.4 | - | - | 0 | - |
| - | - | 3999 | 585.3 | - | - | 0 | - |
| - | - | 5696 | 597.3 | - | - | 0 | - |
| - | - | 7.212E+04 | 606.4 | - | - | 0 | - |
| - | - | 1.951E+04 | 607.4 | - | - | 0 | - |
| - | - | 4915 | 608.4 | - | - | 0 | - |
| - | - | 9353 | 609.3 | - | - | 0 | - |
| 3 | w | 3.235E+04 | 611.3 | 0.001039 | 1.699 | +1 | 6 |
| - | - | 1.576E+04 | 612.3 | - | - | 0 | - |
| - | - | 6318 | 613.3 | - | - | 0 | - |
| - | - | 5882 | 623.4 | - | - | 0 | - |
| 3 | y | 4.802E+04 | 636.4 | 0.001326 | 2.083 | +1 | 6 |
| - | - | 1.71E+04 | 637.4 | - | - | 0 | - |
| - | - | 4.971E+04 | 652.4 | - | - | 0 | - |
| - | - | 1.591E+04 | 653.4 | - | - | 0 | - |
| 3 | y | 3.839E+06 | 654.4 | 0.001747 | 2.67 | +1 | 6 |
| - | - | 1.35E+06 | 655.4 | - | - | 0 | - |
| - | - | 3.167E+05 | 656.4 | - | - | 0 | - |
| - | - | 2.201E+04 | 657.4 | - | - | 0 | - |
| - | - | 2.195E+04 | 665.4 | - | - | 0 | - |
| - | - | 9470 | 666.4 | - | - | 0 | - |
| - | - | 2.124E+04 | 679.4 | - | - | 0 | - |
| - | - | 8394 | 680.4 | - | - | 0 | - |
| - | - | 2.132E+04 | 682.4 | - | - | 0 | - |
| - | - | 2.773E+04 | 683.4 | - | - | 0 | - |
| - | - | 7869 | 684.4 | - | - | 0 | - |
| - | - | 4039 | 685.4 | - | - | 0 | - |
| - | - | 1.029E+05 | 692.4 | - | - | 0 | - |
| - | - | 4.204E+04 | 693.4 | - | - | 0 | - |
| - | - | 2.076E+04 | 694.4 | - | - | 0 | - |
| - | - | 4.371E+04 | 695.4 | - | - | 0 | - |
| - | - | 1.031E+04 | 696.4 | - | - | 0 | - |
| 2 | w | 4.057E+05 | 708.4 | 0.004671 | 6.594 | +1 | 7 |
| 7 | c | 5.908E+06 | 709.4 | 0.001723 | 2.429 | +1 | 7 |
| - | - | 2.214E+06 | 710.4 | - | - | 0 | - |
| - | - | 1.262E+04 | 710.6 | - | - | 0 | - |
| - | - | 5.335E+05 | 711.4 | - | - | 0 | - |
| - | - | 3.358E+04 | 712.4 | - | - | 0 | - |
| - | - | 4813 | 737.4 | - | - | 0 | - |
| - | - | 3.991E+04 | 750.5 | - | - | 0 | - |
| 2 | z | 3.253E+05 | 751.4 | 0.00202 | 2.688 | +1 | 7 |
| - | - | 1.317E+05 | 752.5 | - | - | 0 | - |
| - | - | 3.654E+04 | 753.5 | - | - | 0 | - |
| - | - | 9433 | 763.5 | - | - | 0 | - |
| - | - | 2.028E+04 | 764.4 | - | - | 0 | - |
| - | - | 6554 | 765.4 | - | - | 0 | - |
| - | - | 2.691E+06 | 766.4 | - | - | 0 | - |
| - | - | 1.068E+06 | 767.4 | - | - | 0 | - |
| - | - | 2.612E+05 | 768.4 | - | - | 0 | - |
| - | - | 2.207E+04 | 769.4 | - | - | 0 | - |
| - | - | 4041 | 769.5 | - | - | 0 | - |
| - | - | 1.309E+04 | 779.4 | - | - | 0 | - |
| - | - | 1.16E+04 | 780.5 | - | - | 0 | - |
| - | - | 6532 | 781.4 | - | - | 0 | - |
| - | - | 3.229E+05 | 783.5 | - | - | 0 | - |
| - | - | 1.283E+05 | 784.5 | - | - | 0 | - |
| - | - | 3.411E+04 | 785.5 | - | - | 0 | - |
| - | - | 2.025E+04 | 793.5 | - | - | 0 | - |
| - | - | 1.179E+04 | 794.5 | - | - | 0 | - |
| - | - | 2E+04 | 796.5 | - | - | 0 | - |
| - | - | 7134 | 797.5 | - | - | 0 | - |
| - | - | 9601 | 810.5 | - | - | 0 | - |
| - | - | 1.832E+04 | 821.5 | - | - | 0 | - |
| - | - | 1.836E+06 | 822.5 | - | - | 0 | - |
| - | - | 7.964E+05 | 823.5 | - | - | 0 | - |
| - | - | 2.375E+05 | 824.5 | - | - | 0 | - |
| - | - | 2.292E+04 | 825.5 | - | - | 0 | - |
| - | - | 1.97E+06 | 838.5 | - | - | 0 | - |
| - | - | 5.814E+06 | 839.5 | - | - | 0 | - |
| - | - | 2.414E+06 | 840.5 | - | - | 0 | - |
| - | - | 6.493E+05 | 841.5 | - | - | 0 | - |
| - | - | 5.592E+04 | 842.5 | - | - | 0 | - |
| - | - | 1.19E+04 | 854.5 | - | - | 0 | - |
| - | - | 9178 | 871.5 | - | - | 0 | - |

m/z Charge Intensity FragmentType MassShift Position
124.13031005859375 0 1872.1733
124.42993927001953 0 2457.0657
128.20065307617188 0 1989.9904
129.102783203125 0 39097.35
130.08670043945312 0 18243.488 y Ammonia loss 7
141.10276794433594 0 177831.27
142.106201171875 0 12526.265
147.11337280273438 0 85266.445 y 7
148.11695861816406 0 3594.1086
148.90750122070312 0 3121.4077
148.91317749023438 0 2775.1006
148.91915893554688 0 4632.9014
148.92498779296875 0 4325.2256
148.9304656982422 0 3969.7522
148.93624877929688 0 11054.857
148.94259643554688 0 13368.839
148.95596313476562 0 27471.418
148.96163940429688 0 16453.252
148.96836853027344 0 7867.489
148.97373962402344 0 4727.824
148.97955322265625 0 4777.577
148.99114990234375 0 3289.2253
149.00277709960938 0 3686.131
149.0240478515625 0 17196.56
152.28807067871094 0 2457.4158
152.79296875 0 2343.4155
154.8961639404297 0 1916.4092
157.13412475585938 0 1927282.8
158.13148498535156 0 8966.562
158.137451171875 0 158155.19
158.6644744873047 0 2727.922
159.13975524902344 0 3541.5725
160.99855041503906 0 2468.3843
168.0899658203125 0 3469.475
168.10238647460938 0 3190.837
169.0977325439453 0 403703.8
170.09373474121094 0 3979.8447
170.10108947753906 0 35936.363
173.51551818847656 0 2779.576
183.14987182617188 0 24865.23
184.12132263183594 0 20290.656
184.15365600585938 0 2449.2695
184.587158203125 0 2557.4207
185.1291046142578 0 860925.06
186.1324920654297 0 78629.555
187.13497924804688 0 2519.9915
198.0767364501953 0 5478.1445
201.1240234375 0 7911.0923 w 6
211.1446990966797 0 36278.17
211.19590759277344 0 2822.415
212.1479034423828 0 4307.7124
228.17138671875 0 138056.86
229.1747283935547 0 14244.287
240.13511657714844 0 17365.152
243.65049743652344 0 61815.875
244.15187072753906 0 8608.302
254.18743896484375 0 5568.093
258.1455383300781 0 38172.066 y Water loss 6
259.14910888671875 0 4833.803
260.137451171875 0 24373.855 z 6
261.1438293457031 0 5385.3716
266.150390625 0 38171.016
266.1634826660156 0 19505.818
267.1533508300781 0 4578.7266
267.170166015625 0 8334.086
276.15618896484375 0 104192.23 y 6
277.15887451171875 0 11385.974
279.1692199707031 0 5953.9937 y 3
282.18212890625 0 26037.188
283.1773986816406 0 117456.734
284.18023681640625 0 14455.505
304.1513977050781 0 5163.7544
305.19488525390625 0 4279.0635
309.68505859375 0 5115.7876
312.192138671875 0 4705.201
314.20648193359375 0 2713.8218
318.6893310546875 0 7557.866 y Water loss 2
325.2250671386719 0 5654.099
327.18798828125 0 4268.591
327.69580078125 0 4308859 y 2
328.1972351074219 0 1499138.2
328.6983642578125 0 349215.72
329.1998291015625 0 16944.97
340.187744140625 0 127972.84
341.1911315917969 0 22232.344
344.1827087402344 0 86952.875 w 5
345.18597412109375 0 13441.013
351.239990234375 0 47873.63
352.2116394042969 0 18350.107
352.2430419921875 0 5723.6167
353.21917724609375 0 99738.17
353.2552185058594 0 3991.4353
354.2222900390625 0 18774.53
357.2142028808594 0 124492.586
358.1978759765625 0 29513.414 w 5
358.2180480957031 0 17402.938
359.2005920410156 0 5704.1426
368.21783447265625 0 3184.027
373.2216796875 0 209274.61 z 5
374.2283935546875 0 132800.77
375.2315979003906 0 26546.83 y Water loss 1
379.23504638671875 0 132290.69
380.2381896972656 0 22981.387
381.2406005859375 0 3382.568
384.2375793457031 0 40816.367 y 1
384.7394104003906 0 13883.902
389.240234375 0 29439.715 y 5
390.24261474609375 0 5266.5244
391.2155456542969 0 2536.5654
395.20849609375 0 3933.624
396.26171875 0 942949.6 c 3
397.2647705078125 0 190733.58
398.2671203613281 0 24943.285
399.2240295410156 0 3901.295
400.2200622558594 0 3091.8044
410.2278747558594 0 10912.388
411.22552490234375 0 17202.955
413.2298889160156 0 11069.108
415.22088623046875 0 15017.518
416.2247009277344 0 3269.0405
422.2772521972656 0 4342.971
423.28302001953125 0 3633.9216
429.23553466796875 0 4514.0903
443.2513427734375 0 67989.63 y Ammonia loss 4
444.2563171386719 0 14299.823 z 4
445.2581787109375 0 3532.503
450.2721862792969 0 10328.777
462.2732849121094 0 9061.443
466.2912902832031 0 329937.72
467.2977294921875 0 261158.34 c 4
468.2999572753906 0 40795.96
469.2900695800781 0 30062.76
470.23828125 0 7801.362
470.29449462890625 0 8697.152
480.2832946777344 0 27652.613
481.28375244140625 0 7378.6704
482.2984313964844 0 32235.133
483.30255126953125 0 7396.6514
484.2781066894531 0 193523.62
485.2479553222656 0 13541.135
485.2839050292969 0 81711.445
486.29364013671875 0 1467285.9
487.29656982421875 0 331092.2
488.2989807128906 0 61247.582
490.2680969238281 0 4285.198
508.2779235839844 0 201948.55
509.28118896484375 0 52326.516
509.3462219238281 0 13297.487
510.2831726074219 0 12307.592
512.27294921875 0 49377.047
513.2764282226562 0 16633.68
514.2869262695312 0 15460.842 w 3
515.2933959960938 0 5321.9463
518.3570556640625 0 3744.7336
523.3119506835938 0 53115.08
524.3138427734375 0 12001.827
525.304443359375 0 507051.25
526.304931640625 0 120281.22
527.3030395507812 0 14523.111
535.3612670898438 0 22152.299
536.3692016601562 0 28448.838
537.376220703125 0 38757.48
538.3792114257812 0 13427.2705
539.3191528320312 0 11749.692 y Water loss 3
540.324462890625 0 3726.0225
541.3120727539062 0 584208.75 z 3
542.3150024414062 0 159421.17
543.3180541992188 0 29514.38
557.330810546875 0 289773.88 y 3
558.3336181640625 0 80352.21
559.33544921875 0 16599.465
562.3492431640625 0 11365.1875
563.3562622070312 0 65053.66
564.3594970703125 0 20857.371
565.3644409179688 0 5025.939
573.3031616210938 0 4809.364
579.3753051757812 0 680052
580.3822021484375 0 1087129 c 5
581.385498046875 0 313035.25
582.3056030273438 0 5959.436
582.387939453125 0 53960.016
583.3116455078125 0 3968.6025
583.3705444335938 0 4979.488
585.3192749023438 0 3998.8086
597.3240966796875 0 5696.471
606.3986206054688 0 72122.445
607.4020385742188 0 19505.824
608.4060668945312 0 4915.46
609.3488159179688 0 9353.368
611.3409423828125 0 32345.955 w 2
612.3460693359375 0 15755.792
613.3491821289062 0 6318.0186
623.3627319335938 0 5882.2817
636.3728637695312 0 48015.93 y Water loss 2
637.3753662109375 0 17101.994
652.3567504882812 0 49706.688
653.3639526367188 0 15912.854
654.3838500976562 0 3839341.2 y 2
655.3871459960938 0 1349694
656.389892578125 0 316741
657.39404296875 0 22011.055
665.4114379882812 0 21945.914
666.4154663085938 0 9469.69
679.4277954101562 0 21243.195
680.4302368164062 0 8393.816
682.3779296875 0 21322.875
683.3854370117188 0 27733.09
684.39013671875 0 7868.539
685.386474609375 0 4039.2834
692.4000244140625 0 102941.48
693.4026489257812 0 42036.43
694.4052734375 0 20755.938
695.38818359375 0 43711.137
696.3890380859375 0 10310.233
708.3973388671875 0 405662.62 w 1
709.426025390625 0 5907790.5 c 6
710.4290771484375 0 2214471.2
710.5758666992188 0 12615.533
711.4313354492188 0 533513.56
712.4345092773438 0 33577.094
737.4482421875 0 4813.468
750.4650268554688 0 39905.703
751.449462890625 0 325327.84 z 1
752.4524536132812 0 131672.86
753.4547729492188 0 36543.645
763.4710083007812 0 9432.534
764.4317016601562 0 20282.334
765.4312744140625 0 6553.7866
766.4237060546875 0 2690813
767.4265747070312 0 1068168
768.429443359375 0 261222.55
769.4332275390625 0 22066.66
769.5025634765625 0 4040.5398
779.4304809570312 0 13086.3955
780.5003051757812 0 11599.514
781.4361572265625 0 6532.045
783.4502563476562 0 322939.94
784.4530639648438 0 128290.96
785.4561767578125 0 34106.71
793.5070190429688 0 20245.416
794.5120239257812 0 11788.801
796.4608764648438 0 20004.25
797.4605712890625 0 7134.145
810.477294921875 0 9600.659
821.5007934570312 0 18316.389
822.4866943359375 0 1836166
823.4896240234375 0 796417.6
824.4920654296875 0 237543.98
825.4937744140625 0 22924.09
838.5050659179688 0 1970384.9
839.5127563476562 0 5813569
840.5151977539062 0 2413801.8
841.5180053710938 0 649308
842.5203857421875 0 55924.445
854.4755859375 0 11896.927
871.4996337890625 0 9178.416

Spectrum Details

|  |  |
| --- | --- |
| Matched peaks? Matched peaks The total absolute number of peaks matched. Additionally in brackets the total fraction of peaks matched and the total number of peaks is shown. | 30 (11.72% of 256) |
| FDR? FDR The false discovery rate estimated for this peptide. It is calculated by matching all theoretical fragments with a non-integer shift with the raw peaks for this spectrum. This is done with 40 different shifts. The resulting percentage is the average number of annotated peaks over the number of annotated peaks with the correct spectrum. | 0.08% |
| Satellite FDR? Satellite FDR See the FDR for details on its calculation. This satellite ion specific FDR only contains the satellite ions (d/w) for I/L/J positions. | 0.00% |
| PSM Score? PSM Score The PSM Score as given by Hecklib to this annotated spectrum. It is shown with three significant figures. | 253 |

#### Reverse Lookup? Reverse Lookup All places where this read could be placed.

| Group | Segment | Template | Template Part | Read Part | Score | Unique |
| --- | --- | --- | --- | --- | --- | --- |
| Homo sapiens Heavy Chain | IGHC | IGHG1 | [209..217] | [0..8] | 52 | False |
| Homo sapiens Heavy Chain | IGHC | IGHG3 | [256..264] | [0..8] | 52 | False |

| Recombined | Template Part | Read Part | Score | Unique |
| --- | --- | --- | --- | --- |
| REC-0-1 | [334..342] | [0..8] | 52 | True |

#### Meta Information from Multiple reads

##### Number of combined reads

3

##### Intensity

0.9187

##### TotalArea

3.291E+09

##### Changes to the peptide sequence

ALPPAIEK

L→ISupport for Isoleucine based on side chain ions (2 for I 0 for L) (Position: 6)

#### Positional Score

Copy Data

##### Positional Score (TSV)

###### Preview

```
Loading example...
```

*Click on the button to copy the data to your clipboard.*

1001234567

Label Value
"0" 0.323
"1" 0.307
"2" 0.31
"3" 0.317
"4" 0.327
"5" 0.33
"6" 0.333
"7" 0.33

#### Meta Information from PEAKS

##### Scan Identifier

F4:4576

##### Original sequence

A

L

P

P

A

L

E

K

##### Posttranslational Modifications

##### Source File

D:\separate\_stitch\_analyses\xle-disambiguation\raw\20210323\_F1\_UM1\_Peng0013\_SA\_F59\_ingel\_3ug\_tryp.raw

##### Fraction

4

##### Scan Feature

F4:1103

##### De Novo Score

98

##### ConfidenceScore

95

### m/z

419.756

##### Mass

837.496

##### Charge

2

##### Retention Time

23.45

##### Predicted Retention Time

24.61

##### Area

1.097E+09

##### Parts Per Million

1.8

##### Fragmentation mode

ETHCD

##### Originating file

01 D:\separate\_stitch\_analyses\xle-disambiguation\20210325\_F59\_3ug\_DENOVO\_12.csv

#### Meta Information from PEAKS

##### Scan Identifier

F4:4522

##### Original sequence

A

L

P

P

A

L

E

K

##### Posttranslational Modifications

##### Source File

D:\separate\_stitch\_analyses\xle-disambiguation\raw\20210323\_F1\_UM1\_Peng0013\_SA\_F59\_ingel\_3ug\_tryp.raw

##### Fraction

4

##### Scan Feature

F4:1103

##### De Novo Score

97

##### ConfidenceScore

95

### m/z

419.756

##### Mass

837.496

##### Charge

2

##### Retention Time

23.45

##### Predicted Retention Time

24.61

##### Area

1.097E+09

##### Parts Per Million

1.8

##### Fragmentation mode

ETHCD

##### Originating file

01 D:\separate\_stitch\_analyses\xle-disambiguation\20210325\_F59\_3ug\_DENOVO\_12.csv

#### Meta Information from PEAKS

##### Scan Identifier

F4:4468

##### Original sequence

A

L

P

P

A

L

E

K

##### Posttranslational Modifications

##### Source File

D:\separate\_stitch\_analyses\xle-disambiguation\raw\20210323\_F1\_UM1\_Peng0013\_SA\_F59\_ingel\_3ug\_tryp.raw

##### Fraction

4

##### Scan Feature

F4:1103

##### De Novo Score

97

##### ConfidenceScore

96

### m/z

419.756

##### Mass

837.496

##### Charge

2

##### Retention Time

23.45

##### Predicted Retention Time

24.61

##### Area

1.097E+09

##### Parts Per Million

1.8

##### Fragmentation mode

ETHCD

##### Originating file

01 D:\separate\_stitch\_analyses\xle-disambiguation\20210325\_F59\_3ug\_DENOVO\_12.csv
