## Supplementary material for "A handle on mass coincidence errors in *de novo* sequencing of antibodies by bottom-up proteomics": Combined_088.html

Details Combined\_088 | Stitch OverviewUndefined

### Read Combined\_088

#### Sequence (length=14)

TVJHQDMSJDGKEY

#### Spectrum 5825? Spectrum 5825 The raw spectrum of this peptide as annotated by Hecklib. The fragments are coloured according to ion type (see legend). Any peaks with a star '\*' as text can be hovered over to see the full details, first the ion type second the mass shift type. By hovering over the amino acids in the peptide or ions in the legend the corresponding peaks are highlighted. By toggling the 'Unassigned' label you can turn the background (unassigned) peaks on or off in the plot. By updating the slider in the Ion legend you can update the spectrum to only show the top X% of the peaks with labels. The top X% means any peak that is within X% of the highest intensity. By dragging in the spectrum you can zoom in to a specific part of the spectrum and use 'Zoom Out' to get back to the original zoom level. The annotation of the spectrum is based on the given sequence in the peptides file and is done with different software so inconsistencies are likely. The peaks are annotated based on the given sequence, with 20 ppm tolerance.

Copy Data

##### Spectrum 5825 (TSV)

###### Preview

```
Loading example...
```

*Click on the button to copy the data to your clipboard.*

Mz MinMz MaxIntensity Max

WidthHeightPeptide font sizePeptide stroke widthSpectrum font sizeSpectrum stroke widthCompact peptide

Ion legend

wxyz

abcd

OtherUnassignedIon^Charge^_Position_Show for top:%

TVJHQDMSJDGKEY

04.79e+59.59e+51.44e+61.92e+6

Zoom Out

a^+1^_2_y^+2^_2_y^+3^_4_a^+1^_2_y^+1^_1_b^+1^_2_b^+1^_2_y^+2^_3_b^+3^_6_b^+2^_4_y^+3^_7_b^+2^_5_y^+1^_2_b^+1^_3_y^+2^_5_y^+2^_5_y^+1^_2_b^+1^_3_b^+2^_6_b^+2^_6_b^+2^_6_y^+2^_6_y^+2^_6_b^+2^_7_y^+2^_7_y^+1^_3_y^+1^_3_b^+1^_4_y^+1^_3_b^+2^_8_b^+2^_8_b^+1^_4_b^+2^_8_y^+2^_8_y^+1^_4_b^+3^_13_y^+1^_4_b^+2^_9_b^+2^_9_y^+3^_13_b^+2^_9_\*\*\*b^+1^_5_b^+1^_5_b^+2^_10_b^+1^_5_y^+2^_10_y^+2^_10_y^+1^_5_y^+1^_5_y^+1^_5_y^+2^_11_y^+2^_11_b^+2^_12_y^+2^_11_b^+1^_6_b^+1^_6_b^+1^_6_y^+1^_6_y^+2^_12_y^+2^_12_y^+2^_12_b^+2^_13_b^+2^_13_y^+1^_6_b^+2^_13_y^+2^_13_y^+2^_13_y^+2^_13_y^+1^_7_b^+1^_7_b^+1^_7_\*y^+1^_7_b^+1^_7_b^+1^_8_b^+1^_8_b^+1^_8_y^+1^_8_y^+1^_8_y^+1^_8_b^+1^_9_b^+1^_9_b^+1^_9_y^+1^_9_y^+1^_9_b^+1^_10_b^+1^_10_y^+1^_10_y^+1^_10_b^+1^_11_b^+1^_11_y^+1^_10_b^+1^_11_y^+1^_11_

0779155823363115

Fragment Matches Table

Show background peaks

| Position | Ion type | Intensity | mz Theoretical | mz Error (Th) | mz Error (ppm) | Charge | Series Number |
| --- | --- | --- | --- | --- | --- | --- | --- |
| - | - | 1.859E+04 | 120 | - | - | 0 | - |
| - | - | 803.8 | 120.1 | - | - | 0 | - |
| - | - | 1189 | 120.1 | - | - | 0 | - |
| - | - | 1.218E+04 | 120.1 | - | - | 0 | - |
| - | - | 678.7 | 120.4 | - | - | 0 | - |
| - | - | 1366 | 121 | - | - | 0 | - |
| - | - | 1868 | 121 | - | - | 0 | - |
| - | - | 789.2 | 121.1 | - | - | 0 | - |
| - | - | 6417 | 122.1 | - | - | 0 | - |
| - | - | 1.056E+04 | 123 | - | - | 0 | - |
| - | - | 2390 | 123.1 | - | - | 0 | - |
| - | - | 1169 | 124.1 | - | - | 0 | - |
| - | - | 3459 | 125.1 | - | - | 0 | - |
| - | - | 3211 | 126.1 | - | - | 0 | - |
| - | - | 2.269E+04 | 127.1 | - | - | 0 | - |
| - | - | 2301 | 127.1 | - | - | 0 | - |
| - | - | 2435 | 127.1 | - | - | 0 | - |
| - | - | 1223 | 128 | - | - | 0 | - |
| - | - | 819.3 | 128.1 | - | - | 0 | - |
| - | - | 1.207E+05 | 128.1 | - | - | 0 | - |
| - | - | 5009 | 129.1 | - | - | 0 | - |
| - | - | 8.885E+05 | 129.1 | - | - | 0 | - |
| - | - | 897 | 130 | - | - | 0 | - |
| - | - | 1485 | 130.1 | - | - | 0 | - |
| - | - | 3369 | 130.1 | - | - | 0 | - |
| - | - | 1968 | 130.1 | - | - | 0 | - |
| - | - | 6142 | 130.1 | - | - | 0 | - |
| - | - | 5.898E+04 | 130.1 | - | - | 0 | - |
| - | - | 977 | 131.1 | - | - | 0 | - |
| - | - | 3334 | 132 | - | - | 0 | - |
| - | - | 984.2 | 132.1 | - | - | 0 | - |
| - | - | 6638 | 133.1 | - | - | 0 | - |
| - | - | 1.494E+05 | 136.1 | - | - | 0 | - |
| - | - | 1577 | 137.1 | - | - | 0 | - |
| - | - | 1.168E+04 | 137.1 | - | - | 0 | - |
| - | - | 1.352E+04 | 138.1 | - | - | 0 | - |
| - | - | 5836 | 138.1 | - | - | 0 | - |
| - | - | 944.7 | 139.1 | - | - | 0 | - |
| - | - | 9165 | 139.1 | - | - | 0 | - |
| - | - | 1.53E+04 | 141.1 | - | - | 0 | - |
| - | - | 1534 | 142.1 | - | - | 0 | - |
| - | - | 2757 | 143 | - | - | 0 | - |
| - | - | 1185 | 144.1 | - | - | 0 | - |
| - | - | 1.135E+04 | 145.1 | - | - | 0 | - |
| - | - | 2.338E+05 | 146.1 | - | - | 0 | - |
| - | - | 7857 | 147 | - | - | 0 | - |
| - | - | 2.375E+04 | 147.1 | - | - | 0 | - |
| - | - | 8.124E+04 | 148 | - | - | 0 | - |
| - | - | 1565 | 148.1 | - | - | 0 | - |
| - | - | 839.7 | 148.9 | - | - | 0 | - |
| - | - | 6698 | 149 | - | - | 0 | - |
| - | - | 4996 | 150.1 | - | - | 0 | - |
| - | - | 1025 | 150.1 | - | - | 0 | - |
| - | - | 1.883E+04 | 151.1 | - | - | 0 | - |
| - | - | 1045 | 152.1 | - | - | 0 | - |
| - | - | 1777 | 152.1 | - | - | 0 | - |
| - | - | 1929 | 153.1 | - | - | 0 | - |
| - | - | 1667 | 154.1 | - | - | 0 | - |
| - | - | 1478 | 154.1 | - | - | 0 | - |
| - | - | 2671 | 155.1 | - | - | 0 | - |
| - | - | 6.294E+04 | 155.1 | - | - | 0 | - |
| 2 | a | 4.961E+04 | 155.1 | 0.0005187 | 3.344 | +1 | 2 |
| - | - | 7502 | 156 | - | - | 0 | - |
| 13 | y | 1492 | 156.1 | 0.0005052 | 3.237 | +2 | 2 |
| - | - | 4477 | 156.1 | - | - | 0 | - |
| - | - | 1429 | 156.1 | - | - | 0 | - |
| - | - | 5233 | 156.1 | - | - | 0 | - |
| - | - | 1698 | 156.1 | - | - | 0 | - |
| - | - | 4481 | 156.1 | - | - | 0 | - |
| - | - | 3389 | 157.1 | - | - | 0 | - |
| - | - | 4175 | 158.1 | - | - | 0 | - |
| - | - | 1341 | 160 | - | - | 0 | - |
| - | - | 765.3 | 161.2 | - | - | 0 | - |
| - | - | 1.145E+04 | 163.1 | - | - | 0 | - |
| - | - | 1697 | 164.1 | - | - | 0 | - |
| - | - | 1218 | 164.1 | - | - | 0 | - |
| - | - | 897.6 | 164.1 | - | - | 0 | - |
| - | - | 7.996E+04 | 165.1 | - | - | 0 | - |
| - | - | 3371 | 165.1 | - | - | 0 | - |
| - | - | 1574 | 165.1 | - | - | 0 | - |
| - | - | 1.359E+05 | 166.1 | - | - | 0 | - |
| 11 | y | 6796 | 166.1 | 0.001954 | 11.77 | +3 | 4 |
| - | - | 7848 | 166.1 | - | - | 0 | - |
| - | - | 1262 | 167.1 | - | - | 0 | - |
| - | - | 1.233E+04 | 167.1 | - | - | 0 | - |
| - | - | 850 | 167.1 | - | - | 0 | - |
| - | - | 855.3 | 167.1 | - | - | 0 | - |
| - | - | 4081 | 167.1 | - | - | 0 | - |
| - | - | 9098 | 168.1 | - | - | 0 | - |
| - | - | 6639 | 168.1 | - | - | 0 | - |
| - | - | 873.5 | 168.1 | - | - | 0 | - |
| - | - | 1.911E+04 | 169.1 | - | - | 0 | - |
| - | - | 838.1 | 169.1 | - | - | 0 | - |
| - | - | 998.7 | 169.1 | - | - | 0 | - |
| - | - | 1394 | 170.1 | - | - | 0 | - |
| - | - | 7228 | 172.1 | - | - | 0 | - |
| - | - | 1.22E+04 | 173.1 | - | - | 0 | - |
| - | - | 2.388E+04 | 173.1 | - | - | 0 | - |
| 2 | a | 1.898E+06 | 173.1 | 0.0006199 | 3.58 | +1 | 2 |
| - | - | 4.026E+05 | 174.1 | - | - | 0 | - |
| - | - | 1.631E+05 | 174.1 | - | - | 0 | - |
| - | - | 4.658E+04 | 175.1 | - | - | 0 | - |
| - | - | 978.1 | 175.1 | - | - | 0 | - |
| - | - | 1.063E+04 | 175.1 | - | - | 0 | - |
| - | - | 1786 | 176.1 | - | - | 0 | - |
| - | - | 2231 | 176.1 | - | - | 0 | - |
| - | - | 2950 | 177.1 | - | - | 0 | - |
| - | - | 5.943E+04 | 178.1 | - | - | 0 | - |
| - | - | 1316 | 179.1 | - | - | 0 | - |
| - | - | 4885 | 179.1 | - | - | 0 | - |
| - | - | 7399 | 181.1 | - | - | 0 | - |
| 14 | y | 1.631E+05 | 182.1 | 0.0006327 | 3.475 | +1 | 1 |
| - | - | 1039 | 183.1 | - | - | 0 | - |
| - | - | 1.543E+04 | 183.1 | - | - | 0 | - |
| 2 | b | 2.855E+04 | 183.1 | 0.0006297 | 3.439 | +1 | 2 |
| - | - | 1756 | 183.1 | - | - | 0 | - |
| - | - | 4093 | 184 | - | - | 0 | - |
| - | - | 917.2 | 184.1 | - | - | 0 | - |
| - | - | 2227 | 184.1 | - | - | 0 | - |
| - | - | 5043 | 185.1 | - | - | 0 | - |
| - | - | 2294 | 185.1 | - | - | 0 | - |
| - | - | 1.349E+04 | 185.2 | - | - | 0 | - |
| - | - | 1096 | 186.1 | - | - | 0 | - |
| - | - | 1.919E+05 | 186.1 | - | - | 0 | - |
| - | - | 990.3 | 187.1 | - | - | 0 | - |
| - | - | 1.522E+04 | 187.1 | - | - | 0 | - |
| - | - | 1834 | 187.1 | - | - | 0 | - |
| - | - | 1135 | 188.1 | - | - | 0 | - |
| - | - | 1969 | 189.1 | - | - | 0 | - |
| - | - | 1.726E+04 | 190.1 | - | - | 0 | - |
| - | - | 1.162E+04 | 190.1 | - | - | 0 | - |
| - | - | 2.393E+04 | 191.1 | - | - | 0 | - |
| - | - | 1982 | 191.1 | - | - | 0 | - |
| - | - | 1629 | 192.1 | - | - | 0 | - |
| - | - | 2962 | 192.1 | - | - | 0 | - |
| - | - | 2690 | 193.1 | - | - | 0 | - |
| - | - | 928.4 | 193.1 | - | - | 0 | - |
| - | - | 6367 | 194.1 | - | - | 0 | - |
| - | - | 6.434E+04 | 195.1 | - | - | 0 | - |
| - | - | 908.7 | 196.1 | - | - | 0 | - |
| - | - | 3836 | 196.1 | - | - | 0 | - |
| - | - | 6900 | 196.1 | - | - | 0 | - |
| - | - | 4090 | 197.1 | - | - | 0 | - |
| - | - | 2321 | 197.2 | - | - | 0 | - |
| - | - | 8219 | 198.1 | - | - | 0 | - |
| - | - | 6682 | 199.1 | - | - | 0 | - |
| - | - | 2840 | 201.1 | - | - | 0 | - |
| 2 | b | 4.665E+05 | 201.1 | 0.0005783 | 2.875 | +1 | 2 |
| - | - | 1.402E+05 | 202.1 | - | - | 0 | - |
| - | - | 3889 | 202.1 | - | - | 0 | - |
| - | - | 4.371E+04 | 202.1 | - | - | 0 | - |
| - | - | 1.618E+04 | 203.1 | - | - | 0 | - |
| - | - | 892.1 | 203.1 | - | - | 0 | - |
| - | - | 3398 | 203.1 | - | - | 0 | - |
| - | - | 1481 | 203.2 | - | - | 0 | - |
| - | - | 1702 | 204.1 | - | - | 0 | - |
| - | - | 1293 | 204.1 | - | - | 0 | - |
| - | - | 1865 | 204.1 | - | - | 0 | - |
| - | - | 2.4E+04 | 205.1 | - | - | 0 | - |
| - | - | 2063 | 205.1 | - | - | 0 | - |
| - | - | 5958 | 205.1 | - | - | 0 | - |
| - | - | 2567 | 206.1 | - | - | 0 | - |
| - | - | 7145 | 206.1 | - | - | 0 | - |
| - | - | 1097 | 206.1 | - | - | 0 | - |
| - | - | 1073 | 207.1 | - | - | 0 | - |
| - | - | 1.255E+04 | 207.2 | - | - | 0 | - |
| - | - | 1573 | 208.1 | - | - | 0 | - |
| - | - | 1943 | 208.2 | - | - | 0 | - |
| - | - | 2926 | 209.1 | - | - | 0 | - |
| - | - | 1660 | 209.1 | - | - | 0 | - |
| - | - | 1016 | 209.1 | - | - | 0 | - |
| - | - | 7154 | 210.1 | - | - | 0 | - |
| 12 | y | 4486 | 211.1 | 0.0007407 | 3.509 | +2 | 3 |
| - | - | 9095 | 212.1 | - | - | 0 | - |
| - | - | 2185 | 213.1 | - | - | 0 | - |
| - | - | 5179 | 213.1 | - | - | 0 | - |
| - | - | 1657 | 213.1 | - | - | 0 | - |
| - | - | 2.545E+04 | 213.1 | - | - | 0 | - |
| - | - | 1805 | 213.1 | - | - | 0 | - |
| - | - | 4752 | 213.2 | - | - | 0 | - |
| - | - | 2061 | 214.1 | - | - | 0 | - |
| - | - | 2349 | 215.1 | - | - | 0 | - |
| - | - | 4786 | 216.1 | - | - | 0 | - |
| - | - | 2761 | 219.1 | - | - | 0 | - |
| - | - | 3181 | 220.1 | - | - | 0 | - |
| - | - | 2041 | 221.1 | - | - | 0 | - |
| - | - | 9168 | 221.1 | - | - | 0 | - |
| - | - | 4600 | 221.1 | - | - | 0 | - |
| - | - | 4275 | 221.1 | - | - | 0 | - |
| - | - | 1.187E+04 | 222.1 | - | - | 0 | - |
| - | - | 1231 | 222.1 | - | - | 0 | - |
| - | - | 1.507E+04 | 223.1 | - | - | 0 | - |
| - | - | 1.423E+05 | 223.2 | - | - | 0 | - |
| - | - | 1665 | 224.1 | - | - | 0 | - |
| - | - | 1.504E+04 | 224.2 | - | - | 0 | - |
| - | - | 2846 | 225.1 | - | - | 0 | - |
| - | - | 1510 | 225.1 | - | - | 0 | - |
| - | - | 2403 | 225.2 | - | - | 0 | - |
| - | - | 2.457E+04 | 226.1 | - | - | 0 | - |
| 6 | b | 1297 | 226.1 | 0.000889 | 3.932 | +3 | 6 |
| 4 | b | 2140 | 226.1 | 0.0003507 | 1.551 | +2 | 4 |
| - | - | 1512 | 226.6 | - | - | 0 | - |
| - | - | 3.601E+04 | 227.1 | - | - | 0 | - |
| - | - | 2509 | 227.1 | - | - | 0 | - |
| - | - | 3337 | 228.1 | - | - | 0 | - |
| - | - | 6559 | 229.1 | - | - | 0 | - |
| - | - | 3753 | 230.2 | - | - | 0 | - |
| - | - | 2117 | 230.2 | - | - | 0 | - |
| - | - | 1275 | 231.1 | - | - | 0 | - |
| - | - | 736.5 | 233.1 | - | - | 0 | - |
| - | - | 2.596E+04 | 233.1 | - | - | 0 | - |
| - | - | 1.672E+05 | 234.1 | - | - | 0 | - |
| - | - | 2612 | 234.6 | - | - | 0 | - |
| - | - | 2.173E+04 | 235.1 | - | - | 0 | - |
| - | - | 1767 | 235.1 | - | - | 0 | - |
| - | - | 3183 | 235.2 | - | - | 0 | - |
| - | - | 1280 | 236.1 | - | - | 0 | - |
| - | - | 1402 | 236.1 | - | - | 0 | - |
| - | - | 4266 | 237.1 | - | - | 0 | - |
| - | - | 4668 | 237.1 | - | - | 0 | - |
| - | - | 987.5 | 238.1 | - | - | 0 | - |
| - | - | 8306 | 238.1 | - | - | 0 | - |
| - | - | 3188 | 238.1 | - | - | 0 | - |
| - | - | 3071 | 239.1 | - | - | 0 | - |
| - | - | 1818 | 239.1 | - | - | 0 | - |
| - | - | 5.231E+04 | 240.1 | - | - | 0 | - |
| - | - | 6457 | 241.1 | - | - | 0 | - |
| - | - | 4.509E+04 | 241.1 | - | - | 0 | - |
| - | - | 6307 | 241.2 | - | - | 0 | - |
| - | - | 8279 | 242.1 | - | - | 0 | - |
| - | - | 3.497E+04 | 243.1 | - | - | 0 | - |
| - | - | 4.1E+04 | 244.1 | - | - | 0 | - |
| - | - | 3092 | 244.1 | - | - | 0 | - |
| - | - | 4536 | 245.1 | - | - | 0 | - |
| - | - | 1041 | 246.1 | - | - | 0 | - |
| - | - | 7.819E+04 | 247.1 | - | - | 0 | - |
| - | - | 4387 | 247.1 | - | - | 0 | - |
| - | - | 1607 | 247.2 | - | - | 0 | - |
| - | - | 980.1 | 247.6 | - | - | 0 | - |
| - | - | 2.914E+04 | 248.1 | - | - | 0 | - |
| - | - | 1016 | 248.2 | - | - | 0 | - |
| - | - | 5.209E+04 | 249.1 | - | - | 0 | - |
| - | - | 1651 | 249.1 | - | - | 0 | - |
| - | - | 1601 | 249.1 | - | - | 0 | - |
| - | - | 9399 | 249.1 | - | - | 0 | - |
| - | - | 4888 | 250.1 | - | - | 0 | - |
| - | - | 2057 | 250.1 | - | - | 0 | - |
| - | - | 2722 | 250.2 | - | - | 0 | - |
| - | - | 5.505E+05 | 251.2 | - | - | 0 | - |
| - | - | 7.075E+04 | 252.2 | - | - | 0 | - |
| - | - | 2576 | 252.2 | - | - | 0 | - |
| - | - | 1602 | 253.1 | - | - | 0 | - |
| - | - | 5585 | 253.2 | - | - | 0 | - |
| - | - | 1.191E+04 | 255.1 | - | - | 0 | - |
| - | - | 1726 | 256.1 | - | - | 0 | - |
| - | - | 1946 | 256.1 | - | - | 0 | - |
| - | - | 1093 | 257.1 | - | - | 0 | - |
| - | - | 7798 | 257.1 | - | - | 0 | - |
| - | - | 1.899E+04 | 258.1 | - | - | 0 | - |
| - | - | 802.9 | 258.2 | - | - | 0 | - |
| - | - | 2853 | 259.1 | - | - | 0 | - |
| - | - | 3251 | 259.1 | - | - | 0 | - |
| - | - | 2.715E+04 | 261.1 | - | - | 0 | - |
| - | - | 5161 | 261.1 | - | - | 0 | - |
| - | - | 1273 | 261.2 | - | - | 0 | - |
| - | - | 6706 | 262.1 | - | - | 0 | - |
| - | - | 994.4 | 262.1 | - | - | 0 | - |
| - | - | 1087 | 263.1 | - | - | 0 | - |
| - | - | 888.1 | 263.1 | - | - | 0 | - |
| 8 | y | 4.346E+04 | 265.1 | 0.0009949 | 3.752 | +3 | 7 |
| - | - | 1322 | 265.6 | - | - | 0 | - |
| - | - | 1.269E+05 | 266.1 | - | - | 0 | - |
| - | - | 1.641E+04 | 267.1 | - | - | 0 | - |
| - | - | 5287 | 268.1 | - | - | 0 | - |
| - | - | 3175 | 268.1 | - | - | 0 | - |
| - | - | 7513 | 268.2 | - | - | 0 | - |
| - | - | 4.628E+04 | 269.1 | - | - | 0 | - |
| - | - | 2433 | 269.2 | - | - | 0 | - |
| - | - | 6061 | 269.2 | - | - | 0 | - |
| - | - | 7657 | 270.1 | - | - | 0 | - |
| - | - | 2054 | 270.1 | - | - | 0 | - |
| - | - | 1236 | 271.1 | - | - | 0 | - |
| - | - | 1322 | 271.1 | - | - | 0 | - |
| - | - | 1.255E+04 | 274.1 | - | - | 0 | - |
| - | - | 6548 | 275.1 | - | - | 0 | - |
| - | - | 1287 | 275.1 | - | - | 0 | - |
| - | - | 1946 | 275.2 | - | - | 0 | - |
| - | - | 5526 | 276.1 | - | - | 0 | - |
| - | - | 1.013E+04 | 276.2 | - | - | 0 | - |
| - | - | 1556 | 276.2 | - | - | 0 | - |
| - | - | 991.9 | 277.1 | - | - | 0 | - |
| - | - | 1318 | 278.2 | - | - | 0 | - |
| - | - | 8450 | 279.1 | - | - | 0 | - |
| - | - | 8502 | 280.1 | - | - | 0 | - |
| - | - | 1515 | 280.1 | - | - | 0 | - |
| - | - | 2062 | 282.2 | - | - | 0 | - |
| - | - | 2.989E+05 | 283.1 | - | - | 0 | - |
| - | - | 2453 | 283.6 | - | - | 0 | - |
| - | - | 4897 | 284.1 | - | - | 0 | - |
| - | - | 4.076E+04 | 284.1 | - | - | 0 | - |
| - | - | 965.2 | 285.1 | - | - | 0 | - |
| - | - | 1989 | 285.1 | - | - | 0 | - |
| - | - | 1264 | 285.2 | - | - | 0 | - |
| - | - | 1.643E+04 | 286.1 | - | - | 0 | - |
| - | - | 2322 | 286.2 | - | - | 0 | - |
| - | - | 2.349E+04 | 287.1 | - | - | 0 | - |
| - | - | 8544 | 287.2 | - | - | 0 | - |
| - | - | 1806 | 287.2 | - | - | 0 | - |
| - | - | 2289 | 288.1 | - | - | 0 | - |
| - | - | 1671 | 288.1 | - | - | 0 | - |
| - | - | 3780 | 288.1 | - | - | 0 | - |
| 5 | b | 1594 | 290.2 | 0.0001142 | 0.3936 | +2 | 5 |
| - | - | 2010 | 291.2 | - | - | 0 | - |
| 13 | y | 2.032E+05 | 293.1 | 0.0009987 | 3.407 | +1 | 2 |
| - | - | 4790 | 293.2 | - | - | 0 | - |
| - | - | 3.952E+04 | 294.1 | - | - | 0 | - |
| - | - | 985.1 | 294.2 | - | - | 0 | - |
| - | - | 3996 | 295.1 | - | - | 0 | - |
| - | - | 2762 | 296.1 | - | - | 0 | - |
| - | - | 1890 | 296.1 | - | - | 0 | - |
| - | - | 6996 | 296.1 | - | - | 0 | - |
| 3 | b | 5545 | 296.2 | 0.0007637 | 2.578 | +1 | 3 |
| 10 | y | 3251 | 297.1 | 0.002081 | 7.002 | +2 | 5 |
| - | - | 5.41E+04 | 297.2 | - | - | 0 | - |
| - | - | 1582 | 297.2 | - | - | 0 | - |
| - | - | 1135 | 297.6 | - | - | 0 | - |
| - | - | 1220 | 297.7 | - | - | 0 | - |
| - | - | 5479 | 298.1 | - | - | 0 | - |
| - | - | 7389 | 298.2 | - | - | 0 | - |
| - | - | 2768 | 300.2 | - | - | 0 | - |
| - | - | 1.669E+05 | 301.2 | - | - | 0 | - |
| - | - | 3567 | 302.1 | - | - | 0 | - |
| - | - | 2.281E+04 | 302.2 | - | - | 0 | - |
| - | - | 2006 | 303.2 | - | - | 0 | - |
| - | - | 1.896E+04 | 304.2 | - | - | 0 | - |
| - | - | 1220 | 304.2 | - | - | 0 | - |
| - | - | 2632 | 305.2 | - | - | 0 | - |
| - | - | 8714 | 306.1 | - | - | 0 | - |
| 10 | y | 5.649E+04 | 306.1 | 0.001071 | 3.497 | +2 | 5 |
| - | - | 5038 | 306.2 | - | - | 0 | - |
| - | - | 1.844E+04 | 306.6 | - | - | 0 | - |
| - | - | 4824 | 307.1 | - | - | 0 | - |
| 13 | y | 3.465E+04 | 311.1 | 0.001115 | 3.584 | +1 | 2 |
| - | - | 4950 | 312.1 | - | - | 0 | - |
| - | - | 1118 | 313.1 | - | - | 0 | - |
| - | - | 2888 | 314.2 | - | - | 0 | - |
| 3 | b | 1.498E+04 | 314.2 | 0.001033 | 3.287 | +1 | 3 |
| - | - | 9163 | 315.1 | - | - | 0 | - |
| - | - | 9.975E+04 | 315.2 | - | - | 0 | - |
| - | - | 3191 | 315.2 | - | - | 0 | - |
| - | - | 2807 | 316.1 | - | - | 0 | - |
| - | - | 1675 | 316.1 | - | - | 0 | - |
| - | - | 1.495E+04 | 316.2 | - | - | 0 | - |
| - | - | 1259 | 317.1 | - | - | 0 | - |
| - | - | 1426 | 317.2 | - | - | 0 | - |
| - | - | 2640 | 318.1 | - | - | 0 | - |
| - | - | 2692 | 319.2 | - | - | 0 | - |
| - | - | 2338 | 319.2 | - | - | 0 | - |
| - | - | 1651 | 320.2 | - | - | 0 | - |
| - | - | 2371 | 321.2 | - | - | 0 | - |
| - | - | 1570 | 322.2 | - | - | 0 | - |
| - | - | 1935 | 324.1 | - | - | 0 | - |
| - | - | 1109 | 324.1 | - | - | 0 | - |
| - | - | 1365 | 327.1 | - | - | 0 | - |
| - | - | 1294 | 330.7 | - | - | 0 | - |
| - | - | 5989 | 332.2 | - | - | 0 | - |
| - | - | 1.106E+04 | 332.2 | - | - | 0 | - |
| - | - | 2.891E+04 | 333.2 | - | - | 0 | - |
| - | - | 2246 | 333.2 | - | - | 0 | - |
| - | - | 1021 | 333.7 | - | - | 0 | - |
| - | - | 2.748E+04 | 334.1 | - | - | 0 | - |
| - | - | 4854 | 334.2 | - | - | 0 | - |
| - | - | 5835 | 335.1 | - | - | 0 | - |
| - | - | 1201 | 335.1 | - | - | 0 | - |
| - | - | 2771 | 338.2 | - | - | 0 | - |
| - | - | 4413 | 338.6 | - | - | 0 | - |
| 6 | b | 3139 | 338.7 | 0.001523 | 4.498 | +2 | 6 |
| - | - | 2011 | 339.1 | - | - | 0 | - |
| 6 | b | 2555 | 339.2 | 0.00213 | 6.281 | +2 | 6 |
| - | - | 1192 | 342.7 | - | - | 0 | - |
| - | - | 1646 | 343.2 | - | - | 0 | - |
| - | - | 9358 | 344.2 | - | - | 0 | - |
| - | - | 1.024E+04 | 345.1 | - | - | 0 | - |
| - | - | 3.017E+04 | 346.1 | - | - | 0 | - |
| - | - | 2353 | 346.1 | - | - | 0 | - |
| - | - | 5174 | 347.1 | - | - | 0 | - |
| - | - | 2729 | 347.2 | - | - | 0 | - |
| 6 | b | 1.367E+04 | 347.7 | 0.001307 | 3.759 | +2 | 6 |
| - | - | 5417 | 348.2 | - | - | 0 | - |
| - | - | 1447 | 348.7 | - | - | 0 | - |
| - | - | 3240 | 349.2 | - | - | 0 | - |
| - | - | 4.395E+04 | 350.2 | - | - | 0 | - |
| - | - | 8379 | 351.2 | - | - | 0 | - |
| - | - | 1096 | 352.2 | - | - | 0 | - |
| - | - | 1258 | 352.7 | - | - | 0 | - |
| 9 | y | 4515 | 353.7 | 0.001034 | 2.923 | +2 | 6 |
| - | - | 1211 | 354.2 | - | - | 0 | - |
| - | - | 1.46E+04 | 356.2 | - | - | 0 | - |
| - | - | 3951 | 356.7 | - | - | 0 | - |
| - | - | 5239 | 356.7 | - | - | 0 | - |
| - | - | 1388 | 357.2 | - | - | 0 | - |
| - | - | 1632 | 360.2 | - | - | 0 | - |
| - | - | 1.125E+04 | 361.2 | - | - | 0 | - |
| - | - | 1.007E+04 | 362.2 | - | - | 0 | - |
| 9 | y | 4.065E+04 | 362.7 | 0.001153 | 3.179 | +2 | 6 |
| - | - | 7.301E+04 | 363.1 | - | - | 0 | - |
| - | - | 1.662E+04 | 363.2 | - | - | 0 | - |
| - | - | 4227 | 363.7 | - | - | 0 | - |
| - | - | 5672 | 364.1 | - | - | 0 | - |
| - | - | 1.169E+04 | 364.1 | - | - | 0 | - |
| - | - | 1344 | 366.1 | - | - | 0 | - |
| - | - | 7979 | 366.2 | - | - | 0 | - |
| - | - | 6102 | 367.2 | - | - | 0 | - |
| - | - | 5834 | 367.2 | - | - | 0 | - |
| - | - | 1983 | 369.2 | - | - | 0 | - |
| - | - | 1972 | 373.2 | - | - | 0 | - |
| - | - | 2447 | 376.2 | - | - | 0 | - |
| - | - | 1488 | 376.2 | - | - | 0 | - |
| - | - | 2242 | 377.1 | - | - | 0 | - |
| - | - | 1149 | 377.3 | - | - | 0 | - |
| - | - | 2.072E+04 | 378.2 | - | - | 0 | - |
| - | - | 1.228E+05 | 379.2 | - | - | 0 | - |
| - | - | 8006 | 380.2 | - | - | 0 | - |
| - | - | 2.585E+04 | 380.2 | - | - | 0 | - |
| - | - | 9.262E+04 | 381.2 | - | - | 0 | - |
| - | - | 2540 | 381.2 | - | - | 0 | - |
| - | - | 1.584E+04 | 382.2 | - | - | 0 | - |
| - | - | 1966 | 384.2 | - | - | 0 | - |
| - | - | 1.765E+04 | 384.2 | - | - | 0 | - |
| - | - | 9376 | 385.2 | - | - | 0 | - |
| - | - | 1622 | 385.2 | - | - | 0 | - |
| - | - | 3339 | 386.2 | - | - | 0 | - |
| - | - | 1389 | 387.2 | - | - | 0 | - |
| - | - | 3178 | 389.1 | - | - | 0 | - |
| - | - | 2270 | 389.2 | - | - | 0 | - |
| - | - | 2029 | 390.2 | - | - | 0 | - |
| - | - | 1594 | 390.2 | - | - | 0 | - |
| - | - | 4567 | 391.1 | - | - | 0 | - |
| - | - | 5418 | 392.2 | - | - | 0 | - |
| - | - | 1934 | 392.7 | - | - | 0 | - |
| - | - | 1585 | 393.2 | - | - | 0 | - |
| - | - | 2.183E+04 | 394.2 | - | - | 0 | - |
| - | - | 9932 | 395.2 | - | - | 0 | - |
| - | - | 3131 | 395.2 | - | - | 0 | - |
| - | - | 3599 | 395.7 | - | - | 0 | - |
| - | - | 3107 | 396.2 | - | - | 0 | - |
| - | - | 2181 | 396.2 | - | - | 0 | - |
| - | - | 1.935E+04 | 396.2 | - | - | 0 | - |
| - | - | 4340 | 397.2 | - | - | 0 | - |
| - | - | 8.397E+04 | 398.2 | - | - | 0 | - |
| - | - | 1378 | 398.7 | - | - | 0 | - |
| - | - | 5519 | 399.1 | - | - | 0 | - |
| - | - | 1.425E+04 | 399.2 | - | - | 0 | - |
| - | - | 1680 | 400.1 | - | - | 0 | - |
| - | - | 1624 | 400.2 | - | - | 0 | - |
| - | - | 3611 | 401.2 | - | - | 0 | - |
| - | - | 2.641E+04 | 402.2 | - | - | 0 | - |
| - | - | 2247 | 402.7 | - | - | 0 | - |
| - | - | 1.154E+04 | 403.2 | - | - | 0 | - |
| - | - | 2092 | 404.2 | - | - | 0 | - |
| 7 | b | 1210 | 404.7 | 0.004257 | 10.52 | +2 | 7 |
| - | - | 9654 | 405.3 | - | - | 0 | - |
| 8 | y | 6136 | 406.2 | 0.004089 | 10.07 | +2 | 7 |
| - | - | 3044 | 406.2 | - | - | 0 | - |
| - | - | 4102 | 406.2 | - | - | 0 | - |
| - | - | 5382 | 406.7 | - | - | 0 | - |
| - | - | 1336 | 407.2 | - | - | 0 | - |
| - | - | 1945 | 407.2 | - | - | 0 | - |
| - | - | 2113 | 407.2 | - | - | 0 | - |
| - | - | 2170 | 407.7 | - | - | 0 | - |
| - | - | 3839 | 408.2 | - | - | 0 | - |
| - | - | 1818 | 409.1 | - | - | 0 | - |
| - | - | 8977 | 409.1 | - | - | 0 | - |
| - | - | 1380 | 409.2 | - | - | 0 | - |
| - | - | 1486 | 410.2 | - | - | 0 | - |
| - | - | 1159 | 410.2 | - | - | 0 | - |
| - | - | 3957 | 411.2 | - | - | 0 | - |
| - | - | 1943 | 411.2 | - | - | 0 | - |
| - | - | 1.208E+05 | 412.2 | - | - | 0 | - |
| - | - | 1.984E+04 | 413.2 | - | - | 0 | - |
| - | - | 1.438E+04 | 413.2 | - | - | 0 | - |
| - | - | 1432 | 413.7 | - | - | 0 | - |
| - | - | 6219 | 414.2 | - | - | 0 | - |
| - | - | 3.233E+04 | 414.2 | - | - | 0 | - |
| - | - | 1060 | 415.2 | - | - | 0 | - |
| - | - | 8153 | 415.2 | - | - | 0 | - |
| - | - | 1.644E+04 | 416.2 | - | - | 0 | - |
| - | - | 1443 | 416.2 | - | - | 0 | - |
| - | - | 2.651E+04 | 417.1 | - | - | 0 | - |
| - | - | 5666 | 418.1 | - | - | 0 | - |
| - | - | 4095 | 419.2 | - | - | 0 | - |
| - | - | 1373 | 419.7 | - | - | 0 | - |
| 12 | y | 5682 | 421.2 | 0.00241 | 5.722 | +1 | 3 |
| 12 | y | 2837 | 422.2 | 0.000908 | 2.151 | +1 | 3 |
| - | - | 1176 | 422.3 | - | - | 0 | - |
| - | - | 1346 | 423.2 | - | - | 0 | - |
| - | - | 5.071E+04 | 423.3 | - | - | 0 | - |
| - | - | 2806 | 424.2 | - | - | 0 | - |
| - | - | 1.161E+04 | 424.3 | - | - | 0 | - |
| - | - | 2064 | 424.7 | - | - | 0 | - |
| - | - | 5001 | 425.2 | - | - | 0 | - |
| - | - | 1172 | 425.3 | - | - | 0 | - |
| - | - | 2027 | 425.7 | - | - | 0 | - |
| - | - | 1372 | 426.2 | - | - | 0 | - |
| - | - | 5911 | 427.1 | - | - | 0 | - |
| - | - | 1531 | 428.1 | - | - | 0 | - |
| - | - | 1142 | 428.2 | - | - | 0 | - |
| - | - | 1730 | 429.2 | - | - | 0 | - |
| - | - | 5892 | 429.2 | - | - | 0 | - |
| - | - | 3.33E+05 | 430.2 | - | - | 0 | - |
| - | - | 7.285E+04 | 431.2 | - | - | 0 | - |
| - | - | 1.289E+04 | 432.2 | - | - | 0 | - |
| - | - | 3116 | 433.2 | - | - | 0 | - |
| 4 | b | 1.096E+04 | 433.3 | 0.001514 | 3.494 | +1 | 4 |
| - | - | 1.278E+04 | 433.7 | - | - | 0 | - |
| - | - | 2.333E+04 | 434.2 | - | - | 0 | - |
| - | - | 1.24E+04 | 434.2 | - | - | 0 | - |
| - | - | 2297 | 434.3 | - | - | 0 | - |
| - | - | 3744 | 434.7 | - | - | 0 | - |
| - | - | 3837 | 435.2 | - | - | 0 | - |
| - | - | 3662 | 438.7 | - | - | 0 | - |
| - | - | 2087 | 438.7 | - | - | 0 | - |
| 12 | y | 8.777E+04 | 439.2 | 0.00152 | 3.46 | +1 | 3 |
| - | - | 2811 | 439.7 | - | - | 0 | - |
| - | - | 3897 | 440.2 | - | - | 0 | - |
| - | - | 2.271E+04 | 440.2 | - | - | 0 | - |
| - | - | 2993 | 440.7 | - | - | 0 | - |
| - | - | 4220 | 441.2 | - | - | 0 | - |
| - | - | 6.657E+04 | 442.7 | - | - | 0 | - |
| - | - | 3.264E+04 | 443.2 | - | - | 0 | - |
| - | - | 8400 | 443.7 | - | - | 0 | - |
| - | - | 2.734E+04 | 444.2 | - | - | 0 | - |
| - | - | 1993 | 444.2 | - | - | 0 | - |
| - | - | 1993 | 444.2 | - | - | 0 | - |
| - | - | 5.717E+04 | 445.1 | - | - | 0 | - |
| - | - | 1.465E+04 | 446.1 | - | - | 0 | - |
| - | - | 1694 | 446.2 | - | - | 0 | - |
| - | - | 1531 | 447.1 | - | - | 0 | - |
| - | - | 1014 | 447.2 | - | - | 0 | - |
| - | - | 8136 | 447.2 | - | - | 0 | - |
| - | - | 1634 | 447.3 | - | - | 0 | - |
| 8 | b | 2.705E+04 | 447.7 | 0.0001568 | 0.3503 | +2 | 8 |
| 8 | b | 3.976E+04 | 448.2 | 0.004143 | 9.243 | +2 | 8 |
| - | - | 7653 | 448.7 | - | - | 0 | - |
| - | - | 2300 | 448.7 | - | - | 0 | - |
| - | - | 7071 | 449.2 | - | - | 0 | - |
| - | - | 1515 | 449.2 | - | - | 0 | - |
| - | - | 5365 | 450.2 | - | - | 0 | - |
| - | - | 5034 | 450.3 | - | - | 0 | - |
| 4 | b | 9.567E+04 | 451.3 | 0.001691 | 3.748 | +1 | 4 |
| - | - | 3642 | 452.2 | - | - | 0 | - |
| - | - | 2.373E+04 | 452.3 | - | - | 0 | - |
| - | - | 3612 | 453.3 | - | - | 0 | - |
| - | - | 1.169E+04 | 454.2 | - | - | 0 | - |
| - | - | 3988 | 454.7 | - | - | 0 | - |
| 8 | b | 1.805E+05 | 456.7 | 9.859E-05 | 0.2159 | +2 | 8 |
| - | - | 9.383E+04 | 457.2 | - | - | 0 | - |
| - | - | 2.933E+04 | 457.7 | - | - | 0 | - |
| - | - | 1647 | 458.2 | - | - | 0 | - |
| - | - | 6428 | 458.2 | - | - | 0 | - |
| - | - | 6445 | 459.2 | - | - | 0 | - |
| - | - | 1653 | 460.2 | - | - | 0 | - |
| - | - | 9573 | 461.2 | - | - | 0 | - |
| - | - | 2001 | 461.3 | - | - | 0 | - |
| - | - | 6.182E+04 | 462.2 | - | - | 0 | - |
| - | - | 2345 | 462.2 | - | - | 0 | - |
| - | - | 1804 | 462.2 | - | - | 0 | - |
| - | - | 1.52E+04 | 463.2 | - | - | 0 | - |
| - | - | 1756 | 463.7 | - | - | 0 | - |
| - | - | 2050 | 464.2 | - | - | 0 | - |
| - | - | 1830 | 464.2 | - | - | 0 | - |
| - | - | 1684 | 467.3 | - | - | 0 | - |
| - | - | 1581 | 467.7 | - | - | 0 | - |
| - | - | 1713 | 468.3 | - | - | 0 | - |
| - | - | 2.516E+04 | 468.3 | - | - | 0 | - |
| - | - | 1492 | 468.7 | - | - | 0 | - |
| - | - | 1542 | 469.2 | - | - | 0 | - |
| - | - | 6774 | 469.3 | - | - | 0 | - |
| 7 | y | 3279 | 471.7 | 0.0007408 | 1.57 | +2 | 8 |
| - | - | 1043 | 475.2 | - | - | 0 | - |
| - | - | 9732 | 476.2 | - | - | 0 | - |
| - | - | 6158 | 476.3 | - | - | 0 | - |
| - | - | 4013 | 476.8 | - | - | 0 | - |
| - | - | 1.022E+04 | 477.2 | - | - | 0 | - |
| - | - | 2083 | 477.2 | - | - | 0 | - |
| 11 | y | 2.1E+04 | 478.2 | 0.001302 | 2.722 | +1 | 4 |
| - | - | 1.295E+04 | 478.3 | - | - | 0 | - |
| - | - | 1380 | 479.2 | - | - | 0 | - |
| - | - | 3556 | 479.2 | - | - | 0 | - |
| - | - | 1180 | 479.3 | - | - | 0 | - |
| - | - | 1200 | 479.7 | - | - | 0 | - |
| - | - | 5358 | 481.2 | - | - | 0 | - |
| - | - | 3294 | 481.7 | - | - | 0 | - |
| - | - | 2904 | 482.2 | - | - | 0 | - |
| - | - | 2477 | 484.2 | - | - | 0 | - |
| - | - | 1585 | 484.7 | - | - | 0 | - |
| - | - | 1800 | 485.2 | - | - | 0 | - |
| - | - | 1462 | 485.3 | - | - | 0 | - |
| 13 | b | 3036 | 485.6 | 0.001484 | 3.056 | +3 | 13 |
| - | - | 2149 | 485.8 | - | - | 0 | - |
| - | - | 1251 | 486.2 | - | - | 0 | - |
| - | - | 3872 | 486.2 | - | - | 0 | - |
| - | - | 2626 | 486.7 | - | - | 0 | - |
| - | - | 1414 | 487.2 | - | - | 0 | - |
| - | - | 2193 | 488.7 | - | - | 0 | - |
| - | - | 2.064E+04 | 490.3 | - | - | 0 | - |
| - | - | 1.682E+04 | 490.8 | - | - | 0 | - |
| - | - | 8458 | 491.3 | - | - | 0 | - |
| - | - | 2862 | 491.8 | - | - | 0 | - |
| - | - | 6015 | 493.3 | - | - | 0 | - |
| - | - | 1531 | 493.7 | - | - | 0 | - |
| - | - | 1.167E+05 | 494.2 | - | - | 0 | - |
| - | - | 4.68E+04 | 495.2 | - | - | 0 | - |
| - | - | 9471 | 495.7 | - | - | 0 | - |
| 11 | y | 2.621E+05 | 496.2 | 0.001662 | 3.35 | +1 | 4 |
| - | - | 2173 | 496.9 | - | - | 0 | - |
| - | - | 7.009E+04 | 497.2 | - | - | 0 | - |
| - | - | 3377 | 497.7 | - | - | 0 | - |
| - | - | 1.206E+04 | 498.2 | - | - | 0 | - |
| - | - | 5.468E+04 | 499.3 | - | - | 0 | - |
| - | - | 2713 | 499.6 | - | - | 0 | - |
| - | - | 1836 | 499.7 | - | - | 0 | - |
| - | - | 3.598E+04 | 499.8 | - | - | 0 | - |
| - | - | 2874 | 499.9 | - | - | 0 | - |
| - | - | 8395 | 500.3 | - | - | 0 | - |
| - | - | 1727 | 500.8 | - | - | 0 | - |
| - | - | 1901 | 501.2 | - | - | 0 | - |
| - | - | 2840 | 502.2 | - | - | 0 | - |
| - | - | 1561 | 503.2 | - | - | 0 | - |
| - | - | 3013 | 504.2 | - | - | 0 | - |
| 9 | b | 1.1E+04 | 504.3 | 0.0001085 | 0.2153 | +2 | 9 |
| 9 | b | 5843 | 504.7 | 0.008009 | 15.87 | +2 | 9 |
| - | - | 2667 | 505.3 | - | - | 0 | - |
| - | - | 1528 | 505.9 | - | - | 0 | - |
| 2 | y | 2165 | 506.2 | 0.003224 | 6.369 | +3 | 13 |
| - | - | 2587 | 506.3 | - | - | 0 | - |
| - | - | 1272 | 506.9 | - | - | 0 | - |
| - | - | 3849 | 507.3 | - | - | 0 | - |
| - | - | 5.7E+04 | 511.3 | - | - | 0 | - |
| - | - | 1416 | 511.7 | - | - | 0 | - |
| - | - | 3339 | 512.2 | - | - | 0 | - |
| - | - | 1.29E+04 | 512.3 | - | - | 0 | - |
| - | - | 2709 | 512.6 | - | - | 0 | - |
| - | - | 1426 | 513.2 | - | - | 0 | - |
| 9 | b | 2.514E+04 | 513.3 | 0.0002583 | 0.5033 | +2 | 9 |
| - | - | 1.052E+04 | 513.8 | - | - | 0 | - |
| - | - | 3904 | 514.3 | - | - | 0 | - |
| - | - | 3365 | 515.3 | - | - | 0 | - |
| - | - | 2205 | 516.2 | - | - | 0 | - |
| - | - | 1902 | 518.6 | - | - | 0 | - |
| - | - | 1274 | 519.3 | - | - | 0 | - |
| - | - | 1242 | 520.3 | - | - | 0 | - |
| - | - | 2608 | 521.2 | - | - | 0 | - |
| - | - | 1976 | 522.2 | - | - | 0 | - |
| - | - | 3177 | 524.6 | - | - | 0 | - |
| - | - | 3322 | 524.9 | - | - | 0 | - |
| - | - | 3.051E+04 | 525.3 | - | - | 0 | - |
| - | - | 7497 | 526.3 | - | - | 0 | - |
| - | - | 3110 | 527.3 | - | - | 0 | - |
| - | - | 1621 | 527.9 | - | - | 0 | - |
| - | - | 3023 | 528.2 | - | - | 0 | - |
| - | - | 2242 | 528.2 | - | - | 0 | - |
| - | - | 1.544E+04 | 529.2 | - | - | 0 | - |
| - | - | 2154 | 529.7 | - | - | 0 | - |
| - | - | 1.319E+04 | 530.2 | - | - | 0 | - |
| - | - | 6299 | 530.6 | - | - | 0 | - |
| - | - | 5699 | 530.9 | - | - | 0 | - |
| - | - | 2672 | 531.2 | - | - | 0 | - |
| - | - | 1769 | 531.3 | - | - | 0 | - |
| - | - | 6119 | 533.9 | - | - | 0 | - |
| - | - | 6436 | 534.3 | - | - | 0 | - |
| - | - | 2446 | 534.3 | - | - | 0 | - |
| - | - | 3732 | 534.6 | - | - | 0 | - |
| - | - | 1510 | 534.9 | - | - | 0 | - |
| - | - | 1794 | 535.3 | - | - | 0 | - |
| - | - | 1.694E+04 | 536.2 | - | - | 0 | - |
| - | - | 3513 | 536.6 | - | - | 0 | - |
| - | - | 4416 | 537.2 | - | - | 0 | - |
| - | - | 4078 | 539.2 | - | - | 0 | - |
| - | - | 1436 | 539.6 | - | - | 0 | - |
| 0 | Precursor | 1.716E+04 | 539.9 | 0.000404 | 0.7482 | +3 | -1 |
| - | - | 1765 | 540.2 | - | - | 0 | - |
| 0 | Precursor | 1.84E+04 | 540.3 | 0.007238 | 13.4 | +3 | -1 |
| - | - | 8368 | 540.6 | - | - | 0 | - |
| - | - | 3124 | 540.9 | - | - | 0 | - |
| - | - | 1343 | 543.1 | - | - | 0 | - |
| - | - | 3.558E+04 | 543.3 | - | - | 0 | - |
| - | - | 1.246E+04 | 544.3 | - | - | 0 | - |
| - | - | 4360 | 544.7 | - | - | 0 | - |
| - | - | 2812 | 545.3 | - | - | 0 | - |
| - | - | 2142 | 545.8 | - | - | 0 | - |
| 0 | Precursor | 3.575E+04 | 545.9 | 0.0005445 | 0.9975 | +3 | -1 |
| - | - | 2036 | 546.2 | - | - | 0 | - |
| - | - | 2.494E+04 | 546.3 | - | - | 0 | - |
| - | - | 2.067E+04 | 546.6 | - | - | 0 | - |
| - | - | 2254 | 546.8 | - | - | 0 | - |
| - | - | 7432 | 546.9 | - | - | 0 | - |
| - | - | 2819 | 547.1 | - | - | 0 | - |
| - | - | 1.167E+04 | 547.3 | - | - | 0 | - |
| - | - | 1683 | 547.3 | - | - | 0 | - |
| - | - | 1806 | 547.8 | - | - | 0 | - |
| - | - | 5293 | 548.2 | - | - | 0 | - |
| - | - | 2059 | 549.2 | - | - | 0 | - |
| - | - | 2619 | 551.3 | - | - | 0 | - |
| - | - | 5087 | 551.3 | - | - | 0 | - |
| - | - | 1225 | 553.2 | - | - | 0 | - |
| - | - | 9047 | 553.7 | - | - | 0 | - |
| - | - | 3088 | 554.2 | - | - | 0 | - |
| - | - | 7650 | 554.3 | - | - | 0 | - |
| - | - | 2893 | 554.8 | - | - | 0 | - |
| - | - | 3316 | 555.3 | - | - | 0 | - |
| - | - | 1292 | 555.8 | - | - | 0 | - |
| - | - | 1.361E+04 | 557.2 | - | - | 0 | - |
| - | - | 5853 | 557.8 | - | - | 0 | - |
| - | - | 1.152E+04 | 558.2 | - | - | 0 | - |
| - | - | 1719 | 559.2 | - | - | 0 | - |
| 5 | b | 7577 | 561.3 | 0.0008891 | 1.584 | +1 | 5 |
| - | - | 2.174E+04 | 562.2 | - | - | 0 | - |
| 5 | b | 4694 | 562.3 | 0.003751 | 6.671 | +1 | 5 |
| - | - | 1.084E+04 | 562.7 | - | - | 0 | - |
| - | - | 1756 | 563.2 | - | - | 0 | - |
| - | - | 6577 | 563.3 | - | - | 0 | - |
| - | - | 6981 | 563.8 | - | - | 0 | - |
| - | - | 1.459E+04 | 564.2 | - | - | 0 | - |
| - | - | 2255 | 564.3 | - | - | 0 | - |
| - | - | 3955 | 565.2 | - | - | 0 | - |
| - | - | 2.743E+04 | 565.3 | - | - | 0 | - |
| - | - | 2096 | 565.3 | - | - | 0 | - |
| - | - | 8089 | 566.3 | - | - | 0 | - |
| - | - | 1627 | 567.3 | - | - | 0 | - |
| 10 | b | 5172 | 570.8 | 0.001983 | 3.474 | +2 | 10 |
| - | - | 2.813E+04 | 571.2 | - | - | 0 | - |
| - | - | 2.753E+04 | 571.2 | - | - | 0 | - |
| - | - | 2.432E+04 | 571.8 | - | - | 0 | - |
| - | - | 9228 | 572.2 | - | - | 0 | - |
| - | - | 1381 | 572.8 | - | - | 0 | - |
| - | - | 2925 | 573.2 | - | - | 0 | - |
| - | - | 2.24E+04 | 575.2 | - | - | 0 | - |
| - | - | 1663 | 575.3 | - | - | 0 | - |
| - | - | 8876 | 576.2 | - | - | 0 | - |
| - | - | 2901 | 577.2 | - | - | 0 | - |
| - | - | 1568 | 577.8 | - | - | 0 | - |
| 5 | b | 6.51E+04 | 579.3 | 0.001677 | 2.895 | +1 | 5 |
| - | - | 3337 | 580.3 | - | - | 0 | - |
| - | - | 2.293E+04 | 580.3 | - | - | 0 | - |
| - | - | 2418 | 580.8 | - | - | 0 | - |
| - | - | 8336 | 581.2 | - | - | 0 | - |
| - | - | 3215 | 581.3 | - | - | 0 | - |
| - | - | 4486 | 581.3 | - | - | 0 | - |
| - | - | 7060 | 582.2 | - | - | 0 | - |
| - | - | 1697 | 583.2 | - | - | 0 | - |
| 5 | y | 9065 | 584.3 | 0.0001419 | 0.2428 | +2 | 10 |
| 5 | y | 3659 | 584.7 | 0.007219 | 12.34 | +2 | 10 |
| - | - | 3298 | 585.3 | - | - | 0 | - |
| - | - | 2603 | 592.3 | - | - | 0 | - |
| - | - | 3070 | 592.3 | - | - | 0 | - |
| 10 | y | 9.169E+04 | 593.3 | 0.001611 | 2.715 | +1 | 5 |
| - | - | 1.878E+04 | 593.3 | - | - | 0 | - |
| - | - | 4184 | 593.8 | - | - | 0 | - |
| 10 | y | 2.788E+04 | 594.2 | 0.01167 | 19.65 | +1 | 5 |
| - | - | 4176 | 594.3 | - | - | 0 | - |
| - | - | 7502 | 595.3 | - | - | 0 | - |
| - | - | 1357 | 595.3 | - | - | 0 | - |
| - | - | 1513 | 596.3 | - | - | 0 | - |
| - | - | 2633 | 596.4 | - | - | 0 | - |
| - | - | 2289 | 596.8 | - | - | 0 | - |
| - | - | 7.961E+04 | 599.2 | - | - | 0 | - |
| - | - | 2.43E+04 | 600.2 | - | - | 0 | - |
| - | - | 4687 | 601.2 | - | - | 0 | - |
| - | - | 2759 | 601.3 | - | - | 0 | - |
| - | - | 1152 | 603.8 | - | - | 0 | - |
| - | - | 4764 | 604.8 | - | - | 0 | - |
| - | - | 3851 | 605.3 | - | - | 0 | - |
| - | - | 1582 | 605.3 | - | - | 0 | - |
| - | - | 2618 | 605.8 | - | - | 0 | - |
| - | - | 1488 | 606.3 | - | - | 0 | - |
| - | - | 1331 | 609.2 | - | - | 0 | - |
| - | - | 4721 | 609.8 | - | - | 0 | - |
| - | - | 2065 | 610.2 | - | - | 0 | - |
| - | - | 1503 | 610.2 | - | - | 0 | - |
| - | - | 1.987E+04 | 610.3 | - | - | 0 | - |
| - | - | 1.073E+04 | 610.3 | - | - | 0 | - |
| - | - | 5884 | 610.8 | - | - | 0 | - |
| 10 | y | 8.364E+05 | 611.3 | 0.002032 | 3.325 | +1 | 5 |
| - | - | 2.639E+05 | 612.3 | - | - | 0 | - |
| - | - | 5.594E+04 | 613.3 | - | - | 0 | - |
| - | - | 1.45E+04 | 613.8 | - | - | 0 | - |
| - | - | 1.032E+04 | 614.3 | - | - | 0 | - |
| - | - | 1981 | 614.8 | - | - | 0 | - |
| - | - | 2174 | 615.3 | - | - | 0 | - |
| - | - | 2.863E+04 | 618.8 | - | - | 0 | - |
| - | - | 1120 | 619.2 | - | - | 0 | - |
| - | - | 2.33E+04 | 619.3 | - | - | 0 | - |
| - | - | 1.148E+04 | 619.8 | - | - | 0 | - |
| - | - | 3102 | 620.3 | - | - | 0 | - |
| - | - | 4421 | 621.3 | - | - | 0 | - |
| - | - | 1454 | 622.3 | - | - | 0 | - |
| - | - | 1581 | 622.4 | - | - | 0 | - |
| - | - | 1390 | 626.3 | - | - | 0 | - |
| - | - | 7632 | 627.2 | - | - | 0 | - |
| - | - | 2229 | 627.3 | - | - | 0 | - |
| - | - | 7.863E+04 | 627.8 | - | - | 0 | - |
| - | - | 5.942E+04 | 628.3 | - | - | 0 | - |
| - | - | 2.471E+04 | 628.8 | - | - | 0 | - |
| - | - | 5942 | 629.3 | - | - | 0 | - |
| - | - | 1838 | 629.8 | - | - | 0 | - |
| - | - | 1822 | 631.3 | - | - | 0 | - |
| - | - | 2972 | 632.3 | - | - | 0 | - |
| - | - | 2353 | 635.3 | - | - | 0 | - |
| - | - | 2788 | 635.8 | - | - | 0 | - |
| - | - | 8289 | 636.8 | - | - | 0 | - |
| - | - | 4026 | 637.3 | - | - | 0 | - |
| - | - | 2153 | 637.8 | - | - | 0 | - |
| - | - | 1800 | 638.3 | - | - | 0 | - |
| - | - | 5267 | 638.8 | - | - | 0 | - |
| - | - | 4144 | 639.3 | - | - | 0 | - |
| - | - | 1847 | 639.8 | - | - | 0 | - |
| - | - | 1369 | 640.3 | - | - | 0 | - |
| - | - | 1.128E+04 | 644.3 | - | - | 0 | - |
| - | - | 3074 | 644.6 | - | - | 0 | - |
| - | - | 4903 | 644.8 | - | - | 0 | - |
| - | - | 2953 | 645 | - | - | 0 | - |
| - | - | 3190 | 645.3 | - | - | 0 | - |
| - | - | 3595 | 647.8 | - | - | 0 | - |
| - | - | 4863 | 648.3 | - | - | 0 | - |
| - | - | 1975 | 648.3 | - | - | 0 | - |
| - | - | 5203 | 648.8 | - | - | 0 | - |
| - | - | 1.124E+04 | 649.3 | - | - | 0 | - |
| - | - | 1404 | 649.3 | - | - | 0 | - |
| - | - | 2183 | 649.8 | - | - | 0 | - |
| - | - | 2266 | 650.3 | - | - | 0 | - |
| - | - | 1881 | 650.3 | - | - | 0 | - |
| 4 | y | 2.085E+04 | 652.8 | 0.000349 | 0.5347 | +2 | 11 |
| 4 | y | 1.665E+04 | 653.3 | 0.008891 | 13.61 | +2 | 11 |
| - | - | 4979 | 653.8 | - | - | 0 | - |
| 12 | b | 2466 | 654.8 | 0.01051 | 16.05 | +2 | 12 |
| - | - | 3446 | 655.3 | - | - | 0 | - |
| - | - | 2681 | 657.8 | - | - | 0 | - |
| - | - | 1884 | 658.3 | - | - | 0 | - |
| - | - | 1902 | 659.3 | - | - | 0 | - |
| - | - | 3930 | 659.3 | - | - | 0 | - |
| - | - | 1408 | 660.3 | - | - | 0 | - |
| 4 | y | 1.19E+05 | 661.8 | 0.0003157 | 0.477 | +2 | 11 |
| - | - | 8.667E+04 | 662.3 | - | - | 0 | - |
| - | - | 3.396E+04 | 662.8 | - | - | 0 | - |
| - | - | 8755 | 663.3 | - | - | 0 | - |
| - | - | 1922 | 663.8 | - | - | 0 | - |
| - | - | 1517 | 664.3 | - | - | 0 | - |
| - | - | 1.03E+04 | 666.3 | - | - | 0 | - |
| - | - | 3594 | 666.8 | - | - | 0 | - |
| - | - | 1.774E+04 | 667.3 | - | - | 0 | - |
| - | - | 2220 | 667.8 | - | - | 0 | - |
| - | - | 6180 | 668.3 | - | - | 0 | - |
| - | - | 2672 | 668.8 | - | - | 0 | - |
| - | - | 3011 | 669.3 | - | - | 0 | - |
| - | - | 1392 | 669.8 | - | - | 0 | - |
| - | - | 1753 | 670.3 | - | - | 0 | - |
| - | - | 1.603E+04 | 672.3 | - | - | 0 | - |
| - | - | 6358 | 673.3 | - | - | 0 | - |
| - | - | 1769 | 675.4 | - | - | 0 | - |
| - | - | 2127 | 676.3 | - | - | 0 | - |
| 6 | b | 2.09E+04 | 676.3 | 0.002999 | 4.434 | +1 | 6 |
| - | - | 5851 | 677.3 | - | - | 0 | - |
| 6 | b | 1.969E+04 | 677.3 | 0.007509 | 11.09 | +1 | 6 |
| - | - | 7775 | 677.8 | - | - | 0 | - |
| - | - | 8858 | 678.3 | - | - | 0 | - |
| - | - | 2425 | 679.3 | - | - | 0 | - |
| - | - | 1546 | 682.8 | - | - | 0 | - |
| - | - | 1.069E+05 | 684.3 | - | - | 0 | - |
| - | - | 4.153E+04 | 685.3 | - | - | 0 | - |
| - | - | 1.267E+04 | 686.3 | - | - | 0 | - |
| - | - | 5295 | 686.8 | - | - | 0 | - |
| - | - | 4700 | 687.3 | - | - | 0 | - |
| - | - | 1687 | 687.8 | - | - | 0 | - |
| - | - | 2533 | 688.3 | - | - | 0 | - |
| - | - | 1467 | 689.3 | - | - | 0 | - |
| - | - | 8263 | 690.3 | - | - | 0 | - |
| - | - | 2702 | 691.3 | - | - | 0 | - |
| - | - | 3282 | 691.8 | - | - | 0 | - |
| - | - | 2903 | 692.3 | - | - | 0 | - |
| - | - | 1297 | 693.3 | - | - | 0 | - |
| - | - | 2.42E+04 | 693.4 | - | - | 0 | - |
| - | - | 1.042E+04 | 694.3 | - | - | 0 | - |
| 6 | b | 1.216E+05 | 694.4 | 0.003115 | 4.487 | +1 | 6 |
| - | - | 1.232E+04 | 695.3 | - | - | 0 | - |
| - | - | 4.485E+04 | 695.4 | - | - | 0 | - |
| - | - | 1.411E+04 | 695.8 | - | - | 0 | - |
| - | - | 2827 | 696.3 | - | - | 0 | - |
| - | - | 1.113E+04 | 696.4 | - | - | 0 | - |
| - | - | 1700 | 696.8 | - | - | 0 | - |
| - | - | 1740 | 697.3 | - | - | 0 | - |
| - | - | 9805 | 700.3 | - | - | 0 | - |
| - | - | 2.647E+04 | 700.8 | - | - | 0 | - |
| - | - | 1.455E+04 | 701.3 | - | - | 0 | - |
| - | - | 1.005E+04 | 701.8 | - | - | 0 | - |
| - | - | 1473 | 702.3 | - | - | 0 | - |
| - | - | 1.715E+04 | 704.3 | - | - | 0 | - |
| - | - | 1.169E+04 | 704.8 | - | - | 0 | - |
| - | - | 7160 | 705.3 | - | - | 0 | - |
| - | - | 2794 | 705.8 | - | - | 0 | - |
| 9 | y | 2.14E+04 | 706.3 | 0.002142 | 3.032 | +1 | 6 |
| - | - | 8310 | 707.3 | - | - | 0 | - |
| - | - | 1809 | 708.3 | - | - | 0 | - |
| - | - | 2155 | 708.8 | - | - | 0 | - |
| 3 | y | 9.802E+04 | 709.3 | 0.0002482 | 0.3499 | +2 | 12 |
| 3 | y | 1.042E+05 | 709.8 | 0.007447 | 10.49 | +2 | 12 |
| - | - | 6.112E+04 | 710.3 | - | - | 0 | - |
| - | - | 2.282E+04 | 710.8 | - | - | 0 | - |
| - | - | 1.815E+05 | 711.4 | - | - | 0 | - |
| - | - | 2.017E+05 | 712.3 | - | - | 0 | - |
| - | - | 6.796E+04 | 712.4 | - | - | 0 | - |
| - | - | 8.386E+04 | 713.3 | - | - | 0 | - |
| - | - | 1.699E+04 | 713.4 | - | - | 0 | - |
| - | - | 6921 | 713.9 | - | - | 0 | - |
| - | - | 1.768E+04 | 714.3 | - | - | 0 | - |
| - | - | 2459 | 714.9 | - | - | 0 | - |
| - | - | 1780 | 715.3 | - | - | 0 | - |
| - | - | 1262 | 716.3 | - | - | 0 | - |
| - | - | 1522 | 717.3 | - | - | 0 | - |
| - | - | 1.585E+04 | 717.8 | - | - | 0 | - |
| 3 | y | 6.39E+05 | 718.3 | 0.0007642 | 1.064 | +2 | 12 |
| 13 | b | 5.27E+05 | 718.8 | 0.01347 | 18.73 | +2 | 13 |
| 13 | b | 2.546E+05 | 719.3 | 0.004253 | 5.913 | +2 | 13 |
| - | - | 7.158E+04 | 719.8 | - | - | 0 | - |
| - | - | 1.548E+04 | 720.3 | - | - | 0 | - |
| - | - | 6367 | 721.4 | - | - | 0 | - |
| - | - | 2122 | 722.4 | - | - | 0 | - |
| - | - | 1731 | 723.3 | - | - | 0 | - |
| - | - | 2158 | 723.4 | - | - | 0 | - |
| - | - | 2625 | 723.8 | - | - | 0 | - |
| 9 | y | 1.651E+05 | 724.4 | 0.002197 | 3.033 | +1 | 6 |
| - | - | 6.931E+04 | 725.4 | - | - | 0 | - |
| - | - | 1.57E+04 | 726.4 | - | - | 0 | - |
| - | - | 4447 | 727.4 | - | - | 0 | - |
| 13 | b | 3.692E+04 | 727.9 | 0.0005386 | 0.74 | +2 | 13 |
| - | - | 3.231E+04 | 728.4 | - | - | 0 | - |
| - | - | 1.234E+04 | 728.9 | - | - | 0 | - |
| - | - | 2877 | 729.3 | - | - | 0 | - |
| - | - | 6238 | 729.3 | - | - | 0 | - |
| - | - | 4150 | 730.3 | - | - | 0 | - |
| - | - | 1802 | 733.4 | - | - | 0 | - |
| - | - | 1649 | 734.3 | - | - | 0 | - |
| - | - | 4885 | 736.9 | - | - | 0 | - |
| - | - | 7103 | 737.4 | - | - | 0 | - |
| - | - | 4608 | 738.4 | - | - | 0 | - |
| - | - | 8119 | 740.3 | - | - | 0 | - |
| - | - | 2246 | 741.3 | - | - | 0 | - |
| - | - | 2186 | 743.3 | - | - | 0 | - |
| - | - | 9025 | 744.3 | - | - | 0 | - |
| - | - | 3431 | 745.3 | - | - | 0 | - |
| - | - | 2144 | 746.3 | - | - | 0 | - |
| - | - | 4005 | 747.3 | - | - | 0 | - |
| - | - | 1957 | 750.3 | - | - | 0 | - |
| - | - | 1366 | 751.3 | - | - | 0 | - |
| - | - | 2032 | 757.4 | - | - | 0 | - |
| 2 | y | 9845 | 758.9 | 8.426E-05 | 0.111 | +2 | 13 |
| 2 | y | 1.119E+04 | 759.4 | 0.005344 | 7.038 | +2 | 13 |
| - | - | 6526 | 759.9 | - | - | 0 | - |
| - | - | 4083 | 760.4 | - | - | 0 | - |
| - | - | 5987 | 761.4 | - | - | 0 | - |
| - | - | 4021 | 762.4 | - | - | 0 | - |
| - | - | 2260 | 764.3 | - | - | 0 | - |
| - | - | 2059 | 766.4 | - | - | 0 | - |
| - | - | 3067 | 767.4 | - | - | 0 | - |
| 2 | y | 5.75E+04 | 767.9 | 0.0007369 | 0.9597 | +2 | 13 |
| - | - | 4.814E+04 | 768.4 | - | - | 0 | - |
| - | - | 2.278E+04 | 768.9 | - | - | 0 | - |
| - | - | 7441 | 769.4 | - | - | 0 | - |
| - | - | 1806 | 769.9 | - | - | 0 | - |
| - | - | 2378 | 773.4 | - | - | 0 | - |
| - | - | 1482 | 775.4 | - | - | 0 | - |
| - | - | 1.078E+04 | 779.4 | - | - | 0 | - |
| - | - | 8181 | 780.4 | - | - | 0 | - |
| - | - | 1698 | 781.3 | - | - | 0 | - |
| - | - | 2691 | 781.4 | - | - | 0 | - |
| - | - | 1804 | 782.3 | - | - | 0 | - |
| - | - | 1.022E+04 | 783.4 | - | - | 0 | - |
| - | - | 6141 | 784.4 | - | - | 0 | - |
| - | - | 1851 | 785.4 | - | - | 0 | - |
| - | - | 2192 | 789.4 | - | - | 0 | - |
| - | - | 2825 | 790.4 | - | - | 0 | - |
| - | - | 2165 | 791.4 | - | - | 0 | - |
| - | - | 5602 | 792.3 | - | - | 0 | - |
| - | - | 1819 | 793.3 | - | - | 0 | - |
| 8 | y | 4966 | 794.4 | 0.003954 | 4.977 | +1 | 7 |
| - | - | 8508 | 795.3 | - | - | 0 | - |
| - | - | 4094 | 796.3 | - | - | 0 | - |
| - | - | 2.708E+04 | 797.4 | - | - | 0 | - |
| - | - | 1.397E+04 | 798.4 | - | - | 0 | - |
| - | - | 3768 | 799.3 | - | - | 0 | - |
| - | - | 2934 | 799.4 | - | - | 0 | - |
| - | - | 3051 | 803.4 | - | - | 0 | - |
| 7 | b | 4718 | 807.4 | 0.002024 | 2.507 | +1 | 7 |
| 7 | b | 7916 | 808.4 | 0.00157 | 1.942 | +1 | 7 |
| 0 | Precursor | 3052 | 809.4 | 0.001585 | 1.958 | +2 | -1 |
| - | - | 2984 | 810.3 | - | - | 0 | - |
| 8 | y | 2.649E+04 | 811.4 | 0.007798 | 9.61 | +1 | 7 |
| - | - | 1.22E+04 | 812.4 | - | - | 0 | - |
| - | - | 8533 | 813.4 | - | - | 0 | - |
| - | - | 2245 | 814.4 | - | - | 0 | - |
| - | - | 1813 | 822.3 | - | - | 0 | - |
| - | - | 2172 | 824.5 | - | - | 0 | - |
| 7 | b | 7.161E+04 | 825.4 | 0.001725 | 2.09 | +1 | 7 |
| - | - | 3.464E+04 | 826.4 | - | - | 0 | - |
| - | - | 2.11E+04 | 827.3 | - | - | 0 | - |
| - | - | 6228 | 827.4 | - | - | 0 | - |
| - | - | 9137 | 828.3 | - | - | 0 | - |
| - | - | 2125 | 828.4 | - | - | 0 | - |
| - | - | 1.316E+04 | 839.4 | - | - | 0 | - |
| - | - | 8137 | 840.4 | - | - | 0 | - |
| - | - | 4111 | 841.4 | - | - | 0 | - |
| - | - | 4555 | 843.4 | - | - | 0 | - |
| - | - | 1735 | 844.4 | - | - | 0 | - |
| - | - | 1824 | 848.4 | - | - | 0 | - |
| - | - | 1994 | 849.3 | - | - | 0 | - |
| - | - | 1891 | 849.4 | - | - | 0 | - |
| - | - | 1990 | 850.4 | - | - | 0 | - |
| - | - | 1477 | 851.4 | - | - | 0 | - |
| - | - | 2025 | 853.4 | - | - | 0 | - |
| - | - | 3452 | 856.4 | - | - | 0 | - |
| - | - | 7589 | 857.4 | - | - | 0 | - |
| - | - | 1.041E+04 | 858.4 | - | - | 0 | - |
| - | - | 6090 | 859.4 | - | - | 0 | - |
| - | - | 1882 | 860.4 | - | - | 0 | - |
| - | - | 4520 | 866.3 | - | - | 0 | - |
| - | - | 5869 | 866.4 | - | - | 0 | - |
| - | - | 7751 | 867.4 | - | - | 0 | - |
| - | - | 7527 | 868.4 | - | - | 0 | - |
| - | - | 2644 | 869.4 | - | - | 0 | - |
| - | - | 4374 | 875.4 | - | - | 0 | - |
| - | - | 1.524E+04 | 876.4 | - | - | 0 | - |
| - | - | 1.442E+04 | 877.4 | - | - | 0 | - |
| - | - | 4181 | 878.4 | - | - | 0 | - |
| - | - | 2004 | 883.4 | - | - | 0 | - |
| - | - | 1.347E+04 | 884.4 | - | - | 0 | - |
| - | - | 5.156E+04 | 884.4 | - | - | 0 | - |
| - | - | 6665 | 885.4 | - | - | 0 | - |
| - | - | 2.722E+04 | 885.4 | - | - | 0 | - |
| - | - | 1914 | 886.3 | - | - | 0 | - |
| - | - | 7036 | 886.4 | - | - | 0 | - |
| 8 | b | 4.425E+04 | 894.4 | 0.001948 | 2.178 | +1 | 8 |
| 8 | b | 3.639E+04 | 895.4 | 0.007566 | 8.45 | +1 | 8 |
| - | - | 1.523E+04 | 896.4 | - | - | 0 | - |
| - | - | 4674 | 897.4 | - | - | 0 | - |
| - | - | 4165 | 901.4 | - | - | 0 | - |
| - | - | 2122 | 902.4 | - | - | 0 | - |
| - | - | 2565 | 906.4 | - | - | 0 | - |
| - | - | 1803 | 907.4 | - | - | 0 | - |
| - | - | 5836 | 908.4 | - | - | 0 | - |
| - | - | 2820 | 909.4 | - | - | 0 | - |
| 8 | b | 3.831E+05 | 912.4 | 0.001344 | 1.473 | +1 | 8 |
| - | - | 2.02E+05 | 913.4 | - | - | 0 | - |
| - | - | 6.006E+04 | 914.4 | - | - | 0 | - |
| - | - | 1.182E+04 | 915.4 | - | - | 0 | - |
| - | - | 1802 | 916.4 | - | - | 0 | - |
| - | - | 5325 | 922.4 | - | - | 0 | - |
| - | - | 6827 | 923.4 | - | - | 0 | - |
| 7 | y | 4915 | 924.4 | 0.01068 | 11.55 | +1 | 8 |
| - | - | 5605 | 924.5 | - | - | 0 | - |
| 7 | y | 1.58E+04 | 925.4 | 0.0003615 | 0.3907 | +1 | 8 |
| - | - | 8223 | 926.4 | - | - | 0 | - |
| - | - | 2081 | 927.4 | - | - | 0 | - |
| - | - | 2081 | 939.4 | - | - | 0 | - |
| - | - | 5.214E+04 | 940.4 | - | - | 0 | - |
| - | - | 2.746E+04 | 941.4 | - | - | 0 | - |
| 7 | y | 3.909E+04 | 942.4 | 0.001285 | 1.364 | +1 | 8 |
| - | - | 1.81E+04 | 943.4 | - | - | 0 | - |
| - | - | 5071 | 944.4 | - | - | 0 | - |
| - | - | 1960 | 950.4 | - | - | 0 | - |
| - | - | 1767 | 951.4 | - | - | 0 | - |
| - | - | 2313 | 952.4 | - | - | 0 | - |
| - | - | 1627 | 958.4 | - | - | 0 | - |
| - | - | 5166 | 959.4 | - | - | 0 | - |
| - | - | 3063 | 960.4 | - | - | 0 | - |
| - | - | 2146 | 962.4 | - | - | 0 | - |
| - | - | 6843 | 968.4 | - | - | 0 | - |
| - | - | 1.079E+04 | 969.4 | - | - | 0 | - |
| - | - | 6054 | 970.4 | - | - | 0 | - |
| - | - | 3488 | 976.4 | - | - | 0 | - |
| - | - | 1942 | 977.4 | - | - | 0 | - |
| - | - | 1338 | 978.4 | - | - | 0 | - |
| - | - | 3798 | 979.4 | - | - | 0 | - |
| - | - | 7443 | 980.4 | - | - | 0 | - |
| - | - | 3179 | 981.4 | - | - | 0 | - |
| - | - | 1573 | 982.4 | - | - | 0 | - |
| - | - | 1.022E+04 | 986.4 | - | - | 0 | - |
| - | - | 2.261E+04 | 987.4 | - | - | 0 | - |
| - | - | 1.04E+04 | 988.4 | - | - | 0 | - |
| - | - | 3409 | 989.4 | - | - | 0 | - |
| - | - | 1253 | 990.4 | - | - | 0 | - |
| - | - | 3879 | 994.4 | - | - | 0 | - |
| - | - | 1774 | 995.4 | - | - | 0 | - |
| - | - | 4196 | 996.5 | - | - | 0 | - |
| - | - | 3.799E+04 | 997.4 | - | - | 0 | - |
| - | - | 1.746E+04 | 998.4 | - | - | 0 | - |
| - | - | 2046 | 998.5 | - | - | 0 | - |
| - | - | 5701 | 999.4 | - | - | 0 | - |
| - | - | 1623 | 1000 | - | - | 0 | - |
| - | - | 1.423E+04 | 1004 | - | - | 0 | - |
| - | - | 8233 | 1005 | - | - | 0 | - |
| - | - | 1597 | 1006 | - | - | 0 | - |
| 9 | b | 4849 | 1007 | 0.001296 | 1.286 | +1 | 9 |
| 9 | b | 3279 | 1008 | 0.01158 | 11.48 | +1 | 9 |
| - | - | 1716 | 1009 | - | - | 0 | - |
| - | - | 7676 | 1012 | - | - | 0 | - |
| - | - | 4940 | 1013 | - | - | 0 | - |
| - | - | 6195 | 1014 | - | - | 0 | - |
| - | - | 3176 | 1015 | - | - | 0 | - |
| - | - | 3201 | 1022 | - | - | 0 | - |
| - | - | 1991 | 1023 | - | - | 0 | - |
| 9 | b | 2.925E+04 | 1026 | 0.001973 | 1.923 | +1 | 9 |
| - | - | 1.792E+04 | 1027 | - | - | 0 | - |
| - | - | 5826 | 1028 | - | - | 0 | - |
| - | - | 2508 | 1029 | - | - | 0 | - |
| 6 | y | 6985 | 1039 | 0.01401 | 13.48 | +1 | 9 |
| - | - | 4406 | 1040 | - | - | 0 | - |
| - | - | 1609 | 1041 | - | - | 0 | - |
| - | - | 1802 | 1044 | - | - | 0 | - |
| 6 | y | 3.296E+04 | 1057 | 0.00119 | 1.125 | +1 | 9 |
| - | - | 1.694E+04 | 1058 | - | - | 0 | - |
| - | - | 5754 | 1059 | - | - | 0 | - |
| - | - | 1461 | 1070 | - | - | 0 | - |
| - | - | 1224 | 1079 | - | - | 0 | - |
| - | - | 1501 | 1090 | - | - | 0 | - |
| - | - | 5288 | 1097 | - | - | 0 | - |
| - | - | 2837 | 1098 | - | - | 0 | - |
| - | - | 1316 | 1105 | - | - | 0 | - |
| - | - | 5377 | 1108 | - | - | 0 | - |
| - | - | 4702 | 1109 | - | - | 0 | - |
| - | - | 2758 | 1122 | - | - | 0 | - |
| 10 | b | 5298 | 1124 | 0.01726 | 15.36 | +1 | 10 |
| - | - | 2800 | 1124 | - | - | 0 | - |
| - | - | 2.084E+04 | 1126 | - | - | 0 | - |
| - | - | 1.142E+04 | 1127 | - | - | 0 | - |
| - | - | 5066 | 1128 | - | - | 0 | - |
| - | - | 2498 | 1140 | - | - | 0 | - |
| 10 | b | 6257 | 1141 | 0.01683 | 14.76 | +1 | 10 |
| - | - | 5675 | 1142 | - | - | 0 | - |
| - | - | 1523 | 1143 | - | - | 0 | - |
| - | - | 5623 | 1149 | - | - | 0 | - |
| - | - | 9185 | 1150 | - | - | 0 | - |
| - | - | 5633 | 1151 | - | - | 0 | - |
| - | - | 2345 | 1152 | - | - | 0 | - |
| - | - | 2045 | 1158 | - | - | 0 | - |
| 5 | y | 2.512E+04 | 1167 | 0.001351 | 1.157 | +1 | 10 |
| 5 | y | 5.075E+04 | 1168 | 0.003769 | 3.226 | +1 | 10 |
| - | - | 2.54E+04 | 1169 | - | - | 0 | - |
| - | - | 1.141E+04 | 1170 | - | - | 0 | - |
| - | - | 3328 | 1171 | - | - | 0 | - |
| 11 | b | 2195 | 1180 | 0.0002245 | 0.1903 | +1 | 11 |
| 11 | b | 1345 | 1181 | 0.01267 | 10.73 | +1 | 11 |
| - | - | 1770 | 1185 | - | - | 0 | - |
| 5 | y | 7.015E+04 | 1186 | 0.00154 | 1.299 | +1 | 10 |
| - | - | 4.758E+04 | 1187 | - | - | 0 | - |
| - | - | 1.99E+04 | 1188 | - | - | 0 | - |
| - | - | 4676 | 1189 | - | - | 0 | - |
| 11 | b | 1.029E+04 | 1198 | 0.001551 | 1.295 | +1 | 11 |
| - | - | 5701 | 1199 | - | - | 0 | - |
| - | - | 2047 | 1200 | - | - | 0 | - |
| - | - | 1832 | 1215 | - | - | 0 | - |
| - | - | 1967 | 1225 | - | - | 0 | - |
| - | - | 1644 | 1237 | - | - | 0 | - |
| - | - | 2694 | 1255 | - | - | 0 | - |
| 4 | y | 5062 | 1323 | 0.002224 | 1.682 | +1 | 11 |
| - | - | 2594 | 1324 | - | - | 0 | - |
| - | - | 1280 | 1823 | - | - | 0 | - |
| - | - | 1383 | 3084 | - | - | 0 | - |

m/z Charge Intensity FragmentType MassShift Position
120.04490661621094 0 18594.05
120.0522689819336 0 803.81323
120.05630493164062 0 1188.9734
120.08123779296875 0 12179.846
120.437744140625 0 678.6839
121.0400619506836 0 1365.6902
121.04826354980469 0 1867.9576
121.08460235595703 0 789.21893
122.07178497314453 0 6416.941
123.0445327758789 0 10563.824
123.055908203125 0 2389.7312
124.076416015625 0 1169.4941
125.10785675048828 0 3458.853
126.09187316894531 0 3211.24
127.05071258544922 0 22685.762
127.0873031616211 0 2300.6177
127.12350463867188 0 2434.7603
128.03477478027344 0 1222.924
128.053955078125 0 819.31274
128.1074981689453 0 120722.01
129.06637573242188 0 5008.5664
129.10276794433594 0 888479.3
130.0289306640625 0 897.0385
130.05030822753906 0 1485.192
130.065673828125 0 3369.3042
130.08688354492188 0 1968.2653
130.0999298095703 0 6141.803
130.1060791015625 0 58977.285
131.1078338623047 0 976.9711
132.04501342773438 0 3333.5706
132.10198974609375 0 984.18304
133.06124877929688 0 6637.5127
136.07618713378906 0 149379.53
137.07403564453125 0 1577.0602
137.07955932617188 0 11680.789
138.06671142578125 0 13516.709
138.0918731689453 0 5835.9175
139.06997680664062 0 944.72406
139.08706665039062 0 9165.464
141.102783203125 0 15303.303
142.12310791015625 0 1533.9348
143.0455322265625 0 2756.7432
144.07720947265625 0 1185.4315
145.06126403808594 0 11352.268
146.0605926513672 0 233830.72
147.04458618164062 0 7857.1455
147.06390380859375 0 23750.705
148.03985595703125 0 81240.77
148.0609893798828 0 1564.7657
148.9466552734375 0 839.6659
149.04327392578125 0 6698.4473
150.06663513183594 0 4996.141
150.09190368652344 0 1025.425
151.08714294433594 0 18832.96
152.0712127685547 0 1044.8966
152.0904998779297 0 1776.6205
153.10287475585938 0 1929.4962
154.06199645996094 0 1666.7106
154.09812927246094 0 1477.7566
155.08181762695312 0 2670.9756
155.09327697753906 0 62935.47
155.118408203125 0 49606.062 a Water loss 1
156.044921875 0 7501.624
156.06602478027344 0 1492.0267 y 12
156.07733154296875 0 4476.7515
156.0906219482422 0 1429.3601
156.09664916992188 0 5232.933
156.10211181640625 0 1698.3641
156.121826171875 0 4481.0386
157.0977325439453 0 3389.0093
158.06057739257812 0 4175.335
160.0397186279297 0 1340.8674
161.24281311035156 0 765.3331
163.08714294433594 0 11451.989
164.0710906982422 0 1697.4607
164.0834197998047 0 1217.528
164.091064453125 0 897.6324
165.05517578125 0 79958.35
165.07752990722656 0 3371.4473
165.10264587402344 0 1574.4144
166.0615997314453 0 135931.03
166.0868682861328 0 6796.255 y 10
166.0979766845703 0 7847.585
167.05874633789062 0 1262.2041
167.06512451171875 0 12330.394
167.081787109375 0 850.00604
167.1012420654297 0 855.259
167.11839294433594 0 4081.3516
168.10240173339844 0 9098.074
168.11383056640625 0 6638.5425
168.12026977539062 0 873.46967
169.0977325439453 0 19106.86
169.10586547851562 0 838.064
169.11659240722656 0 998.68964
170.10108947753906 0 1394.4468
172.072265625 0 7228.4307
173.05642700195312 0 12195.991
173.07183837890625 0 23882.818
173.1290740966797 0 1898264 a 1
174.05548095703125 0 402557.78
174.13232421875 0 163072.47
175.05880737304688 0 46580.91
175.0870819091797 0 978.0995
175.13418579101562 0 10626.096
176.0617218017578 0 1786.4528
176.08242797851562 0 2230.6897
177.10260009765625 0 2949.6917
178.1344451904297 0 59428.395
179.1182861328125 0 1316.0769
179.13780212402344 0 4884.8486
181.06149291992188 0 7399.1987
182.08180236816406 0 163107.1 y 13
183.05641174316406 0 1038.8114
183.08554077148438 0 15432.733
183.11343383789062 0 28551.18 b Water loss 1
183.14944458007812 0 1755.9994
184.0397186279297 0 4092.7573
184.08677673339844 0 917.18787
184.1170654296875 0 2226.5955
185.07150268554688 0 5042.8027
185.12908935546875 0 2293.917
185.1654510498047 0 13489.525
186.05593872070312 0 1095.9171
186.1243133544922 0 191873.12
187.1082000732422 0 990.3151
187.1277313232422 0 15215.7705
187.1443634033203 0 1833.54
188.07118225097656 0 1135.4097
189.06643676757812 0 1968.9879
190.0828094482422 0 17255.582
190.1345672607422 0 11621.002
191.08216857910156 0 23931.252
191.13800048828125 0 1982.0804
192.06642150878906 0 1628.9818
192.08584594726562 0 2961.8655
193.09744262695312 0 2689.6394
193.107421875 0 928.4399
194.0929718017578 0 6367.351
195.11341857910156 0 64336.26
196.09962463378906 0 908.73505
196.1085968017578 0 3835.9475
196.11680603027344 0 6900.039
197.12899780273438 0 4089.532
197.16519165039062 0 2321.3726
198.08798217773438 0 8219.359
199.0717315673828 0 6681.5586
201.06675720214844 0 2840.1968
201.1239471435547 0 466451.3 b 1
202.0504608154297 0 140228.73
202.0867919921875 0 3888.8022
202.12721252441406 0 43709.67
203.0539093017578 0 16177.5625
203.0916748046875 0 892.0965
203.12925720214844 0 3397.6575
203.15133666992188 0 1481.3518
204.05703735351562 0 1702.4841
204.07766723632812 0 1292.8417
204.1345672607422 0 1864.615
205.09774780273438 0 23998.666
205.1088104248047 0 2063.4888
205.14532470703125 0 5957.984
206.10093688964844 0 2566.8
206.1293487548828 0 7145.0073
206.14840698242188 0 1097.0371
207.1331329345703 0 1073.0565
207.1609344482422 0 12551.028
208.1092071533203 0 1573.4281
208.16400146484375 0 1943.425
209.05599975585938 0 2925.56
209.09263610839844 0 1659.7482
209.12890625 0 1015.99506
210.12432861328125 0 7153.6743
211.10845947265625 0 4485.59 y Water loss 11
212.13999938964844 0 9095.175
213.05181884765625 0 2184.65
213.06668090820312 0 5178.863
213.11260986328125 0 1656.853
213.1240692138672 0 25452.51
213.14102172851562 0 1804.8368
213.16038513183594 0 4752.184
214.1273651123047 0 2061.1523
215.13961791992188 0 2349.0251
216.0985870361328 0 4786.0303
219.0770721435547 0 2761.2915
220.1202850341797 0 3180.6658
221.0932159423828 0 2041.1532
221.10379028320312 0 9168.351
221.12905883789062 0 4600.2207
221.14068603515625 0 4274.6333
222.12440490722656 0 11865.567
222.1352996826172 0 1231.2592
223.10838317871094 0 15071.762
223.15606689453125 0 142274.62
224.11148071289062 0 1664.8799
224.1593475341797 0 15035.641
225.0989227294922 0 2845.6738
225.1233367919922 0 1510.1141
225.1611785888672 0 2403.1155
226.08299255371094 0 24569.295
226.1195068359375 0 1297.4756 b Water loss 5
226.1371612548828 0 2140.0522 b 3
226.63967895507812 0 1512.0858
227.06695556640625 0 36012.39
227.08668518066406 0 2509.495
228.06993103027344 0 3337.4836
229.11875915527344 0 6558.6445
230.150634765625 0 3752.5469
230.18679809570312 0 2117.076
231.08827209472656 0 1274.9263
233.1284942626953 0 736.5415
233.14044189453125 0 25958.48
234.1244659423828 0 167212.48
234.6499481201172 0 2612.2437
235.12777709960938 0 21733.637
235.14308166503906 0 1767.4948
235.15640258789062 0 3182.5837
236.10414123535156 0 1279.9916
236.13002014160156 0 1402.1631
237.08779907226562 0 4265.5845
237.13536071777344 0 4668.2876
238.0996856689453 0 987.50775
238.11920166015625 0 8306.062
238.13116455078125 0 3188.1077
239.10342407226562 0 3071.275
239.11508178710938 0 1818.3018
240.13499450683594 0 52306.598
241.11911010742188 0 6457.0596
241.13461303710938 0 45087.113
241.191650390625 0 6307.2583
242.13775634765625 0 8278.558
243.1095428466797 0 34965.312
244.0936279296875 0 41000.68
244.11279296875 0 3091.619
245.0969696044922 0 4535.9126
246.1350555419922 0 1041.1613
247.1085968017578 0 78193.414
247.1448516845703 0 4387.4443
247.1567840576172 0 1607.3232
247.57264709472656 0 980.0848
248.11407470703125 0 29143.688
248.1592559814453 0 1016.45197
249.09906005859375 0 52088.984
249.11480712890625 0 1650.6062
249.1209259033203 0 1601.277
249.1355438232422 0 9398.582
250.10205078125 0 4888.391
250.1173553466797 0 2057.252
250.16708374023438 0 2721.6482
251.15109252929688 0 550453
252.1542510986328 0 70754.89
252.17095947265625 0 2576.3513
253.11927795410156 0 1601.5219
253.15635681152344 0 5585.1226
255.1459197998047 0 11909.111
256.1304626464844 0 1726.3953
256.1487121582031 0 1945.7618
257.0927734375 0 1092.8035
257.11383056640625 0 7797.908
258.1455383300781 0 18989.99
258.1607971191406 0 802.9436
259.1293640136719 0 2852.6997
259.14599609375 0 3251.189
261.12005615234375 0 27150.32
261.1355285644531 0 5161.075
261.1618957519531 0 1273.2013
262.1206359863281 0 6705.729
262.1374816894531 0 994.4436
263.1024475097656 0 1086.5574
263.1224060058594 0 888.1321
265.13006591796875 0 43464.074 y Water loss 7
265.6265563964844 0 1322.2222
266.12554931640625 0 126906.99
267.1286926269531 0 16412.258
268.1300048828125 0 5286.5312
268.14556884765625 0 3175.3528
268.1775207519531 0 7512.596
269.12921142578125 0 46279.492
269.1615295410156 0 2432.9717
269.1862487792969 0 6061.006
270.13226318359375 0 7656.9434
270.1462097167969 0 2053.909
271.1041259765625 0 1236.4526
271.1460266113281 0 1322.4796
274.13043212890625 0 12545.902
275.1033630371094 0 6548.3086
275.1353454589844 0 1286.6317
275.1506042480469 0 1946.0463
276.1097717285156 0 5526.119
276.15618896484375 0 10130.385
276.1708068847656 0 1556.3398
277.1114196777344 0 991.9297
278.1615295410156 0 1317.9207
279.14605712890625 0 8450.475
280.1300964355469 0 8501.68
280.1462097167969 0 1515.2147
282.1578369140625 0 2061.7954
283.140869140625 0 298882.28
283.637451171875 0 2452.6245
284.1250915527344 0 4897.1416
284.14410400390625 0 40759.926
285.127685546875 0 965.2272
285.1432189941406 0 1988.7526
285.1589050292969 0 1263.549
286.140625 0 16426.477
286.15679931640625 0 2321.648
287.14019775390625 0 23487.305
287.172119140625 0 8543.7
287.1880187988281 0 1805.5923
288.0986633300781 0 2289.0955
288.12677001953125 0 1670.6105
288.1436767578125 0 3780.0369
290.1659851074219 0 1594.183 b 4
291.2187194824219 0 2010.3777
293.11419677734375 0 203239.16 y Water loss 12
293.1625061035156 0 4790.1816
294.1181335449219 0 39521.86
294.15814208984375 0 985.1184
295.1201171875 0 3995.603
296.0886535644531 0 2761.658
296.1254577636719 0 1890.3718
296.14031982421875 0 6995.8667
296.1976318359375 0 5545.258 b Water loss 2
297.1340026855469 0 3250.971 y Water loss 9
297.15673828125 0 54098.39
297.1988830566406 0 1582.3196
297.6352233886719 0 1135.4294
297.6579895019531 0 1220.2437
298.14019775390625 0 5478.8154
298.1596984863281 0 7388.81
300.167236328125 0 2768.3916
301.1516418457031 0 166933.47
302.12579345703125 0 3566.9326
302.1548156738281 0 22810.24
303.1672668457031 0 2005.7249
304.1666564941406 0 18957.982
304.2148132324219 0 1219.516
305.1689758300781 0 2631.574
306.1089172363281 0 8714.002
306.1382751464844 0 56488.754 y 9
306.2300720214844 0 5037.948
306.63946533203125 0 18441.219
307.1405944824219 0 4823.7915
311.1248779296875 0 34649.69 y 12
312.1284484863281 0 4950.5
313.130615234375 0 1118.2555
314.15118408203125 0 2887.7659
314.2084655761719 0 14984.481 b 2
315.1346130371094 0 9162.795
315.1673889160156 0 99749.77
315.2111511230469 0 3191.2292
316.0939636230469 0 2806.7974
316.1377868652344 0 1675.465
316.17071533203125 0 14952.064
317.0784912109375 0 1259.4781
317.17437744140625 0 1426.1759
318.120361328125 0 2640.3672
319.1514587402344 0 2692.168
319.21429443359375 0 2337.6577
320.1726989746094 0 1650.8699
321.1571960449219 0 2371.371
322.2239685058594 0 1569.8499
324.08270263671875 0 1935.3724
324.1191101074219 0 1109.2316
327.13006591796875 0 1365.0796
330.667236328125 0 1293.5941
332.161376953125 0 5989.212
332.2093505859375 0 11055.4375
333.1778564453125 0 28911.14
333.2127380371094 0 2245.7747
333.656005859375 0 1021.1831
334.1045227050781 0 27478.852
334.1844787597656 0 4854.0596
335.1073913574219 0 5834.927
335.146240234375 0 1200.6365
338.18365478515625 0 2770.7017
338.64691162109375 0 4413.293
338.6758117675781 0 3138.5425 b Water loss 5
339.1477966308594 0 2011.1411
339.1684265136719 0 2555.3489 b Ammonia loss 5
342.658935546875 0 1191.6635
343.16143798828125 0 1645.5653
344.1729736328125 0 9357.905
345.1317138671875 0 10235.181
346.1156005859375 0 30172.248
346.1375427246094 0 2352.6072
347.1185607910156 0 5174.01
347.186767578125 0 2729.443
347.6808776855469 0 13667.64 b 5
348.1822814941406 0 5417.077
348.6824951171875 0 1446.5352
349.15191650390625 0 3239.5486
350.2197570800781 0 43951.582
351.22198486328125 0 8378.648
352.22442626953125 0 1096.0619
352.67108154296875 0 1257.572
353.67498779296875 0 4515.486 y Water loss 8
354.17547607421875 0 1211.3529
356.19390869140625 0 14595.028
356.6573486328125 0 3951.1272
356.69525146484375 0 5238.7686
357.1984558105469 0 1388.3129
360.2038879394531 0 1632.4867
361.19879150390625 0 11246.472
362.183349609375 0 10065.891
362.6803894042969 0 40650.137 y 8
363.1421813964844 0 73005.76
363.1824645996094 0 16620.809
363.6833190917969 0 4226.9233
364.1252746582031 0 5672.2993
364.145751953125 0 11692.127
366.1466979980469 0 1343.9285
366.17803955078125 0 7978.768
367.1619567871094 0 6101.636
367.2462158203125 0 5833.879
369.20269775390625 0 1982.9181
373.1896057128906 0 1971.9901
376.1619567871094 0 2446.8499
376.19964599609375 0 1488.3267
377.14605712890625 0 2242.128
377.26812744140625 0 1148.5652
378.21478271484375 0 20722.084
379.2101745605469 0 122775.56
380.16864013671875 0 8005.6777
380.2131042480469 0 25845.324
381.15283203125 0 92615.81
381.2154235839844 0 2539.9585
382.1559143066406 0 15841.17
384.1594543457031 0 1966.4967
384.188720703125 0 17653.527
385.1725158691406 0 9376.045
385.1968994140625 0 1621.8925
386.1748352050781 0 3339.0574
387.18048095703125 0 1389.1792
389.1463317871094 0 3177.6963
389.1932678222656 0 2270.4897
390.1516418457031 0 2028.8184
390.1954345703125 0 1593.9128
391.1370544433594 0 4566.9834
392.1942443847656 0 5418.083
392.69317626953125 0 1934.184
393.2159118652344 0 1585.444
394.17340087890625 0 21832.031
395.1574401855469 0 9932.076
395.18292236328125 0 3130.682
395.6896667480469 0 3599.009
396.1600646972656 0 3106.917
396.18994140625 0 2180.881
396.22576904296875 0 19349.129
397.2268981933594 0 4339.648
398.1794128417969 0 83965.17
398.6764221191406 0 1377.7802
399.1310119628906 0 5519.2554
399.1819763183594 0 14249.088
400.13494873046875 0 1680.1681
400.1839294433594 0 1624.2623
401.18353271484375 0 3611.2249
402.199462890625 0 26411.602
402.685302734375 0 2247.0217
403.2010192871094 0 11536.958
404.2052917480469 0 2092.2554
404.6907958984375 0 1209.8171 b Ammonia loss 6
405.2623596191406 0 9653.6455
406.191162109375 0 6135.5933 y 7
406.2145690917969 0 3044.24
406.2466125488281 0 4102.2734
406.69287109375 0 5382.499
407.1798400878906 0 1336.1904
407.2073059082031 0 1945.2947
407.2423400878906 0 2113.1606
407.6819763183594 0 2169.767
408.16400146484375 0 3839.2458
409.1175231933594 0 1817.7906
409.1480712890625 0 8976.645
409.1737976074219 0 1379.5508
410.150146484375 0 1485.8196
410.18353271484375 0 1159.3778
411.167724609375 0 3956.7588
411.2005310058594 0 1943.3899
412.1840515136719 0 120785.84
413.16729736328125 0 19836.857
413.1896057128906 0 14381.145
413.7015075683594 0 1432.1793
414.171630859375 0 6219.348
414.236083984375 0 32326.508
415.1725158691406 0 1059.634
415.2392578125 0 8152.513
416.15753173828125 0 16438.684
416.1864013671875 0 1443.2908
417.1419372558594 0 26509.48
418.1451416015625 0 5665.757
419.1943664550781 0 4094.941
419.71539306640625 0 1373.4255
421.2105712890625 0 5682.1685 y Water loss 11
422.1930847167969 0 2836.849 y Ammonia loss 11
422.25152587890625 0 1175.8104
423.16876220703125 0 1346.4196
423.27288818359375 0 50714.348
424.2200012207031 0 2806.2197
424.2755126953125 0 11606.356
424.7084045410156 0 2063.711
425.2010192871094 0 5000.67
425.2770080566406 0 1171.8115
425.69921875 0 2026.8315
426.2002258300781 0 1371.8403
427.12640380859375 0 5910.9937
428.1309814453125 0 1530.756
428.1923828125 0 1141.8997
429.1810607910156 0 1729.5615
429.2112121582031 0 5891.579
430.19476318359375 0 332959.22
431.19775390625 0 72845.125
432.2000427246094 0 12885.567
433.20867919921875 0 3116.1538
433.2572937011719 0 10957.668 b Water loss 3
433.71258544921875 0 12782.181
434.1685791015625 0 23327.604
434.2098693847656 0 12397.315
434.26092529296875 0 2297.1038
434.7110595703125 0 3743.8228
435.1711730957031 0 3837.4116
438.7041931152344 0 3662.284
438.7342529296875 0 2086.5322
439.2202453613281 0 87774.414 y 11
439.6990966796875 0 2811.1304
440.1977233886719 0 3897.263
440.223876953125 0 22709.375
440.7007141113281 0 2993.288
441.22601318359375 0 4219.5005
442.7181396484375 0 66572.055
443.21978759765625 0 32642.014
443.7214660644531 0 8400.322
444.1528625488281 0 27341.904
444.18267822265625 0 1992.6443
444.22308349609375 0 1992.8491
445.13702392578125 0 57173.152
446.1401672363281 0 14652.623
446.2052307128906 0 1694.2821
447.1404724121094 0 1530.8992
447.1576843261719 0 1013.9414
447.1889343261719 0 8136.115
447.27056884765625 0 1634.2897
447.71038818359375 0 27053.217 b Water loss 7
448.2066955566406 0 39759.14 b Ammonia loss 7
448.70758056640625 0 7652.894
448.73797607421875 0 2299.8533
449.2079162597656 0 7070.868
449.2402648925781 0 1515.1195
450.2360534667969 0 5364.6465
450.28448486328125 0 5033.7754
451.2680358886719 0 95674.75 b 3
452.2148742675781 0 3642.0488
452.27081298828125 0 23727.746
453.2728576660156 0 3611.7246
454.1973571777344 0 11692.065
454.69879150390625 0 3988.0671
456.7157287597656 0 180499.45 b 7
457.2170715332031 0 93827.125
457.71832275390625 0 29329.23
458.1871337890625 0 1647.2047
458.2201232910156 0 6427.808
459.1991882324219 0 6444.768
460.2192687988281 0 1653.0164
461.20477294921875 0 9573.133
461.2508850097656 0 2000.7062
462.1634521484375 0 61817.426
462.20361328125 0 2345.4597
462.24810791015625 0 1804.1465
463.1665954589844 0 15196.708
463.7253723144531 0 1755.5181
464.169677734375 0 2050.0134
464.2204284667969 0 1830.2413
467.25201416015625 0 1683.9442
467.74566650390625 0 1580.969
468.2542724609375 0 1712.9364
468.2944030761719 0 25160.863
468.73724365234375 0 1492.1974
469.17236328125 0 1542.4258
469.29681396484375 0 6774.4585
471.7147521972656 0 3278.968 y 6
475.2290954589844 0 1042.8323
476.2251281738281 0 9732.231
476.2586364746094 0 6158.0757
476.756103515625 0 4012.7349
477.21124267578125 0 10216.433
477.24749755859375 0 2082.9368
478.2309265136719 0 21002.264 y Water loss 10
478.27880859375 0 12946.548
479.1910095214844 0 1380.2341
479.2327880859375 0 3555.51
479.2762756347656 0 1179.6345
479.7159729003906 0 1199.7517
481.2489929199219 0 5357.795
481.7475891113281 0 3293.76
482.2463684082031 0 2904.0908
484.2220153808594 0 2477.2515
484.70947265625 0 1584.5421
485.2256164550781 0 1800.0043
485.2640075683594 0 1462.302
485.5711364746094 0 3036.423 b 12
485.763427734375 0 2148.6243
486.2044677734375 0 1250.944
486.24127197265625 0 3871.7783
486.74310302734375 0 2625.6973
487.1893005371094 0 1414.4703
488.7191162109375 0 2192.8276
490.2550048828125 0 20644.096
490.75177001953125 0 16817.803
491.250732421875 0 8457.899
491.75262451171875 0 2862.2683
493.2528381347656 0 6015.4614
493.71575927734375 0 1531.1862
494.237548828125 0 116735.5
495.2428894042969 0 46802.863
495.7472229003906 0 9470.858
496.2418518066406 0 262064.8 y 10
496.912109375 0 2172.587
497.244873046875 0 70092.56
497.724853515625 0 3376.7944
498.24774169921875 0 12057.629
499.2604064941406 0 54684.668
499.5738525390625 0 2712.8699
499.7197570800781 0 1836.1805
499.7616882324219 0 35978.49
499.9059143066406 0 2874.1936
500.2630310058594 0 8394.732
500.7642517089844 0 1727.2848
501.2475891113281 0 1900.76
502.2301940917969 0 2840.1414
503.23480224609375 0 1561.0823
504.2153015136719 0 3013.0486
504.252685546875 0 11000.135 b Water loss 8
504.7525939941406 0 5842.5415 b Ammonia loss 8
505.2527160644531 0 2666.5251
505.9148254394531 0 1527.6918
506.2383117675781 0 2164.718 y Water loss 1
506.2752380371094 0 2587.0515
506.906982421875 0 1271.9749
507.25823974609375 0 3848.6326
511.26397705078125 0 57004
511.7184143066406 0 1416.1207
512.2159423828125 0 3339.123
512.2666015625 0 12897.636
512.5813598632812 0 2709.034
513.2192993164062 0 1426.1484
513.2581176757812 0 25138.89 b 8
513.759033203125 0 10522.936
514.2607421875 0 3904.3564
515.2833862304688 0 3365.4722
516.2099609375 0 2205.0576
518.5829467773438 0 1901.6764
519.328125 0 1274.069
520.2516479492188 0 1241.5629
521.246337890625 0 2607.6238
522.1974487304688 0 1976.0687
524.5890502929688 0 3177.3735
524.9219360351562 0 3321.9275
525.2680053710938 0 30508.398
526.2711181640625 0 7496.833
527.2667846679688 0 3109.9167
527.9171752929688 0 1620.7109
528.1736450195312 0 3023.338
528.2498779296875 0 2242.1584
529.2413940429688 0 15435.176
529.727783203125 0 2153.6665
530.227783203125 0 13189.552
530.5926513671875 0 6298.789
530.9262084960938 0 5699.483
531.2261352539062 0 2671.8508
531.2643432617188 0 1768.9506
533.9216918945312 0 6118.9756
534.2544555664062 0 6436.252
534.3056640625 0 2445.5676
534.5885009765625 0 3732.4956
534.916259765625 0 1509.9995
535.255859375 0 1793.6768
536.190673828125 0 16940.502
536.5968627929688 0 3513.0496
537.1917724609375 0 4415.727
539.2271728515625 0 4077.8735
539.59765625 0 1436.4764
539.9244995117188 0 17156.133 Precursor Water loss
540.2144165039062 0 1765.0266
540.2593383789062 0 18396.65 Precursor Ammonia loss
540.5921020507812 0 8368.042
540.9262084960938 0 3123.6558
543.0615234375 0 1342.9716
543.2789306640625 0 35580.457
544.28271484375 0 12462.425
544.7269897460938 0 4359.759
545.2777709960938 0 2811.719
545.755859375 0 2142.3516
545.9281616210938 0 35747.258 Precursor
546.1866455078125 0 2036.3805
546.2625732421875 0 24936.623
546.5967407226562 0 20669.922
546.81787109375 0 2254.4988
546.9305419921875 0 7432.298
547.0673217773438 0 2819.0044
547.253662109375 0 11671.449
547.3017578125 0 1682.7936
547.77001953125 0 1805.8108
548.2453002929688 0 5293.135
549.2498779296875 0 2059.4087
551.2598876953125 0 2619.4783
551.3306884765625 0 5087.278
553.2462158203125 0 1225.484
553.7335815429688 0 9047.265
554.1990356445312 0 3087.865
554.266845703125 0 7649.5425
554.7672729492188 0 2893.2268
555.2698364257812 0 3316.4514
555.7669677734375 0 1292.3798
557.2442626953125 0 13607.406
557.7549438476562 0 5852.99
558.2230834960938 0 11521.697
559.2257690429688 0 1718.848
561.3152465820312 0 7577.2246 b Water loss 4
562.2455444335938 0 21737.494
562.3021240234375 0 4693.811 b Ammonia loss 4
562.7464599609375 0 10838.059
563.2001953125 0 1755.7375
563.2657470703125 0 6576.597
563.7714233398438 0 6981.125
564.1857299804688 0 14585.003
564.2745361328125 0 2255.4502
565.18603515625 0 3954.5684
565.26318359375 0 27425.938
565.312255859375 0 2095.5981
566.26513671875 0 8088.7397
567.2677612304688 0 1627.3474
570.7693481445312 0 5172.4224 b 9
571.23046875 0 28128.543
571.247314453125 0 27532.904
571.7520141601562 0 24317.473
572.2340087890625 0 9228.209
572.7542724609375 0 1381.3263
573.2326049804688 0 2925.0967
575.2479248046875 0 22403.307
575.2943725585938 0 1663.321
576.2422485351562 0 8876.279
577.238037109375 0 2900.9724
577.845458984375 0 1567.891
579.3265991210938 0 65102.81 b 4
580.2568969726562 0 3337.306
580.3294067382812 0 22927.865
580.7559204101562 0 2418.488
581.2118530273438 0 8335.828
581.2640380859375 0 3214.8542
581.3316650390625 0 4486.0425
582.2003173828125 0 7059.59
583.1976928710938 0 1696.5355
584.2531127929688 0 9065.378 y Water loss 4
584.752197265625 0 3658.86 y Ammonia loss 4
585.2540893554688 0 3298.1736
592.2674560546875 0 2602.9368
592.3221435546875 0 3069.9421
593.2581787109375 0 91689.95 y Water loss 9
593.3071899414062 0 18777.69
593.7605590820312 0 4183.6377
594.2522583007812 0 27883.94 y Ammonia loss 9
594.3115234375 0 4176.3115
595.2565307617188 0 7501.5728
595.3095703125 0 1357.4902
596.285888671875 0 1513.2898
596.354736328125 0 2632.9136
596.7772216796875 0 2289.2039
599.222900390625 0 79610.96
600.2256469726562 0 24296.86
601.22705078125 0 4686.908
601.2704467773438 0 2758.897
603.8035278320312 0 1152.2893
604.7910766601562 0 4763.7715
605.2865600585938 0 3851.009
605.3428344726562 0 1582.2551
605.7888793945312 0 2618.173
606.2891235351562 0 1487.7776
609.2051391601562 0 1331.0859
609.7846069335938 0 4721.4478
610.1990966796875 0 2064.9243
610.2224731445312 0 1502.5155
610.2818603515625 0 19871.672
610.3334350585938 0 10731.477
610.7789306640625 0 5884.1353
611.2691650390625 0 836368 y 9
612.2718505859375 0 263877.2
613.2742919921875 0 55942.492
613.7955322265625 0 14498.353
614.2896728515625 0 10319.173
614.800048828125 0 1980.7416
615.2896728515625 0 2174.1497
618.7879028320312 0 28629.45
619.2329711914062 0 1119.579
619.2866821289062 0 23304.352
619.7868041992188 0 11475.107
620.2854614257812 0 3102.1855
621.2575073242188 0 4420.937
622.2583618164062 0 1453.6622
622.372314453125 0 1581.1528
626.256591796875 0 1389.6595
627.2193603515625 0 7632.015
627.3010864257812 0 2229.4165
627.793212890625 0 78629.664
628.2941284179688 0 59424.6
628.7958374023438 0 24709.19
629.2969360351562 0 5942.025
629.7940673828125 0 1837.6583
631.2658081054688 0 1822.0488
632.3089599609375 0 2972.4124
635.265380859375 0 2352.9648
635.7649536132812 0 2788.0889
636.7987060546875 0 8289.371
637.2979125976562 0 4025.5354
637.7996215820312 0 2153.0854
638.3064575195312 0 1800.032
638.785888671875 0 5266.6177
639.2857666015625 0 4144.383
639.7910766601562 0 1846.5953
640.2817993164062 0 1368.7052
644.2708740234375 0 11276.66
644.6338500976562 0 3074.0862
644.7733154296875 0 4903.234
644.9694213867188 0 2952.5605
645.2623291015625 0 3189.6226
647.7891235351562 0 3595.1313
648.291259765625 0 4862.8945
648.3482666015625 0 1974.7
648.7990112304688 0 5203.0356
649.2774047851562 0 11241.857
649.3421630859375 0 1403.8015
649.8006591796875 0 2183.1626
650.2763061523438 0 2266.0068
650.335205078125 0 1880.8802
652.7827758789062 0 20847.74 y Water loss 3
653.2833251953125 0 16651.322 y Ammonia loss 3
653.7823486328125 0 4979.3296
654.8267822265625 0 2465.7686 b Ammonia loss 11
655.3247680664062 0 3446.1633
657.7997436523438 0 2681.3345
658.3206787109375 0 1883.7587
659.2616577148438 0 1902.1626
659.32177734375 0 3930.1702
660.3240966796875 0 1408.385
661.7880249023438 0 118974.5 y 3
662.2894287109375 0 86670.086
662.790771484375 0 33959.82
663.2913818359375 0 8755.216
663.836669921875 0 1922.224
664.3334350585938 0 1517.0402
666.3010864257812 0 10302.426
666.7819213867188 0 3594.3564
667.287353515625 0 17737.299
667.7789306640625 0 2219.66
668.2919921875 0 6179.5273
668.8235473632812 0 2671.708
669.3216552734375 0 3010.591
669.8274536132812 0 1391.8903
670.3243408203125 0 1752.868
672.2650146484375 0 16031.025
673.2645874023438 0 6357.9253
675.3623657226562 0 1768.5753
676.2839965820312 0 2126.5898
676.3442993164062 0 20900.71 b Water loss 5
677.2685546875 0 5851.3496
677.3328247070312 0 19687.826 b Ammonia loss 5
677.8268432617188 0 7774.5405
678.336181640625 0 8857.813
679.3468017578125 0 2425.0076
682.8125610351562 0 1546.3229
684.3119506835938 0 106938.64
685.3150024414062 0 41528.57
686.319091796875 0 12666.446
686.8224487304688 0 5294.7114
687.324462890625 0 4700.352
687.82666015625 0 1687.4175
688.3298950195312 0 2532.728
689.326171875 0 1467.1248
690.2760009765625 0 8263.273
691.2776489257812 0 2702.477
691.8098754882812 0 3282.2717
692.3014526367188 0 2902.6428
693.3079833984375 0 1297.0455
693.370361328125 0 24195.668
694.2947387695312 0 10422.019
694.35498046875 0 121637.98 b 5
695.2816162109375 0 12316.419
695.3548583984375 0 44853.047
695.8291625976562 0 14112.422
696.2859497070312 0 2827.3313
696.3535766601562 0 11127.215
696.8265991210938 0 1700.1791
697.3375854492188 0 1740.1552
700.3207397460938 0 9805.299
700.8148803710938 0 26466.375
701.31396484375 0 14548.135
701.8154907226562 0 10050.06
702.317626953125 0 1473.0977
704.3335571289062 0 17148.584
704.8357543945312 0 11692.131
705.3389282226562 0 7160.2456
705.8427124023438 0 2794.3062
706.3427734375 0 21402.584 y Water loss 8
707.343505859375 0 8309.831
708.3489990234375 0 1809.3806
708.8306274414062 0 2155.1511
709.32470703125 0 98017.76 y Water loss 2
709.8239135742188 0 104189.46 y Ammonia loss 2
710.324462890625 0 61116.902
710.8251953125 0 22821.922
711.3804931640625 0 181508.5
712.30712890625 0 201682.58
712.3831787109375 0 67957.43
713.3094482421875 0 83855.51
713.3856201171875 0 16990.672
713.85498046875 0 6920.548
714.31201171875 0 17679.354
714.8558959960938 0 2459.0764
715.3046875 0 1779.5059
716.3233642578125 0 1261.7828
717.3242797851562 0 1521.9009
717.8375854492188 0 15847.106
718.3305053710938 0 638975.6 y 2
718.8320922851562 0 526953.3 b Water loss 12
719.3333129882812 0 254574.78 b Ammonia loss 12
719.8348999023438 0 71576.164
720.3359375 0 15484.388
721.3662719726562 0 6366.6436
722.3648071289062 0 2121.5698
723.3111572265625 0 1731.4238
723.3750610351562 0 2158.0923
723.8200073242188 0 2625.0366
724.3533935546875 0 165084.75 y 8
725.3558959960938 0 69313.7
726.3593139648438 0 15703.62
727.360107421875 0 4446.8267
727.8513793945312 0 36915.69 b 12
728.3527221679688 0 32310.885
728.8543701171875 0 12340.948
729.2786865234375 0 2877.4143
729.3427734375 0 6238.453
730.3264770507812 0 4149.761
733.3530883789062 0 1802.048
734.3390502929688 0 1648.7793
736.8566284179688 0 4884.5874
737.395751953125 0 7102.506
738.3959350585938 0 4607.9546
740.302734375 0 8119.315
741.304931640625 0 2245.8645
743.3406372070312 0 2186.2505
744.3234252929688 0 9025.183
745.3250122070312 0 3430.7236
746.3037109375 0 2143.7217
747.2994384765625 0 4004.5303
750.3455200195312 0 1956.5232
751.348876953125 0 1365.7728
757.358154296875 0 2031.6206
758.8585815429688 0 9844.709 y Water loss 1
759.3560180664062 0 11191.172 y Ammonia loss 1
759.8588256835938 0 6526.4478
760.3602294921875 0 4082.9905
761.3512573242188 0 5986.901
762.3547973632812 0 4020.516
764.3115234375 0 2260.284
766.3607788085938 0 2058.7126
767.3598022460938 0 3066.75
767.8646850585938 0 57500.715 y 1
768.3657836914062 0 48138.695
768.8665771484375 0 22775.521
769.3668212890625 0 7441.0747
769.872314453125 0 1806.2836
773.3580322265625 0 2378.3672
775.3692626953125 0 1481.6669
779.3838500976562 0 10778.497
780.3748779296875 0 8180.8086
781.31884765625 0 1697.866
781.3876953125 0 2691.1755
782.3217163085938 0 1804.0927
783.381103515625 0 10217.387
784.3794555664062 0 6141.1855
785.3673706054688 0 1850.7194
789.3739013671875 0 2192.2693
790.3587036132812 0 2824.5913
791.3699951171875 0 2165.038
792.2975463867188 0 5601.9233
793.2970581054688 0 1818.877
794.3527221679688 0 4965.751 y Ammonia loss 7
795.3455810546875 0 8508.247
796.33984375 0 4093.6707
797.3961181640625 0 27077.35
798.3988647460938 0 13969.873
799.3397827148438 0 3767.7458
799.4043579101562 0 2933.7349
803.3621215820312 0 3051.4736
807.3797607421875 0 4718.103 b Water loss 6
808.3673706054688 0 7916.2515 b Ammonia loss 6
809.3809204101562 0 3051.7986 Precursor Water loss
810.3184814453125 0 2983.8936
811.3754272460938 0 26490.053 y 7
812.3768310546875 0 12195.107
813.3598022460938 0 8533.428
814.362548828125 0 2245.0486
822.33935546875 0 1813.4961
824.4661254882812 0 2172.349
825.390625 0 71605.62 b 6
826.3932495117188 0 34639.082
827.3333129882812 0 21095.215
827.4005126953125 0 6228.0083
828.3367919921875 0 9136.659
828.4117431640625 0 2124.6245
839.3700561523438 0 13160.33
840.3682250976562 0 8136.892
841.3668212890625 0 4110.837
843.4014282226562 0 4554.766
844.4072875976562 0 1734.6681
848.3895874023438 0 1824.423
849.3150024414062 0 1993.7408
849.3974609375 0 1891.0586
850.3994140625 0 1990.2986
851.4059448242188 0 1476.9028
853.387939453125 0 2025.4106
856.3604125976562 0 3451.6821
857.3761596679688 0 7588.5767
858.3685302734375 0 10409.053
859.36865234375 0 6090.43
860.3704833984375 0 1882.3448
866.3425903320312 0 4520.47
866.4193115234375 0 5869.107
867.4083862304688 0 7750.817
868.4044189453125 0 7526.7124
869.4052124023438 0 2644.4875
875.3903198242188 0 4373.7036
876.3792114257812 0 15237.196
877.3843383789062 0 14420.617
878.3865356445312 0 4181.3296
883.3749389648438 0 2003.8269
884.3508911132812 0 13471.499
884.4283447265625 0 51561.07
885.353515625 0 6664.507
885.4317626953125 0 27219.35
886.3493041992188 0 1913.8708
886.43310546875 0 7035.712
894.411865234375 0 44246.387 b Water loss 7
895.4053955078125 0 36388.926 b Ammonia loss 7
896.4053955078125 0 15226.037
897.4038696289062 0 4673.808
901.3820190429688 0 4164.687
902.385498046875 0 2122.014
906.3881225585938 0 2565.0872
907.3782958984375 0 1803.037
908.421875 0 5835.818
909.426025390625 0 2819.9573
912.4230346679688 0 383062.03 b 7
913.4254760742188 0 202034.4
914.4281616210938 0 60059.387
915.4296264648438 0 11816.458
916.430419921875 0 1801.6266
922.4063110351562 0 5325.1846
923.3988647460938 0 6826.7754
924.4024658203125 0 4915.369 y Water loss 6
924.4597778320312 0 5605.2256
925.3975219726562 0 15795.166 y Ammonia loss 6
926.4059448242188 0 8223.019
927.4039306640625 0 2081.4824
939.4303588867188 0 2081.4465
940.4178466796875 0 52141.2
941.4197998046875 0 27464.232
942.4224243164062 0 39093.53 y 6
943.4248046875 0 18095.092
944.4284057617188 0 5071.26
950.4033203125 0 1960.4602
951.4028930664062 0 1767.1199
952.4210815429688 0 2313.0168
958.4329833984375 0 1627.1462
959.414794921875 0 5166.217
960.4175415039062 0 3062.939
962.3999633789062 0 2145.7854
968.4137573242188 0 6842.5713
969.4237060546875 0 10787.267
970.4115600585938 0 6054.1685
976.4415893554688 0 3488.112
977.4273071289062 0 1942.206
978.4265747070312 0 1337.5131
979.4282836914062 0 3797.5872
980.42041015625 0 7443.4937
981.4265747070312 0 3179.3115
982.4180297851562 0 1572.6802
986.4249877929688 0 10222.662
987.411865234375 0 22606.61
988.4119873046875 0 10398.433
989.4154663085938 0 3409.455
990.4122924804688 0 1252.9274
994.4385375976562 0 3879.009
995.4364013671875 0 1774.4167
996.451904296875 0 4195.6177
997.4395751953125 0 37988.816
998.4423217773438 0 17464.451
998.53857421875 0 2045.9778
999.4456787109375 0 5700.7573
1000.4464721679688 0 1623.0735
1004.43408203125 0 14225.831
1005.4328002929688 0 8233.203
1006.4328002929688 0 1596.6448
1007.49658203125 0 4848.8853 b Water loss 8
1008.4934692382812 0 3278.5308 b Ammonia loss 8
1009.48876953125 0 1715.6527
1012.4523315429688 0 7676.342
1013.4515991210938 0 4939.7812
1014.4631958007812 0 6195.2803
1015.4632568359375 0 3176.4607
1022.4387817382812 0 3201.207
1023.45263671875 0 1991.2507
1025.5064697265625 0 29249.322 b 8
1026.5091552734375 0 17916.76
1027.5093994140625 0 5825.547
1028.5120849609375 0 2507.817
1039.4541015625 0 6984.764 y Water loss 5
1040.464111328125 0 4405.505
1041.448486328125 0 1608.9514
1043.52197265625 0 1802.239
1057.449462890625 0 32956.363 y 5
1058.452880859375 0 16939.1
1059.45556640625 0 5753.8066
1069.5172119140625 0 1461.1576
1079.4949951171875 0 1223.8552
1090.4906005859375 0 1501.2612
1096.5106201171875 0 5288.2236
1097.5146484375 0 2837.0315
1105.47265625 0 1316.3724
1107.52099609375 0 5377.3037
1108.516845703125 0 4701.733
1122.490234375 0 2757.6099
1123.4915771484375 0 5297.554 b Ammonia loss 9
1124.4910888671875 0 2799.9138
1125.532958984375 0 20839.459
1126.5360107421875 0 11417.837
1127.5421142578125 0 5066.468
1139.5040283203125 0 2497.5476
1140.5185546875 0 6257.004 b 9
1141.5108642578125 0 5674.659
1142.5054931640625 0 1522.7834
1149.48583984375 0 5622.898
1150.4757080078125 0 9184.717
1151.474853515625 0 5633.4854
1152.4840087890625 0 2344.8743
1157.513916015625 0 2044.7966
1167.497314453125 0 25115.188 y Water loss 4
1168.4864501953125 0 50754.484 y Ammonia loss 4
1169.4886474609375 0 25395.854
1170.490234375 0 11407.725
1171.4908447265625 0 3327.5027
1179.5465087890625 0 2194.6084 b Water loss 10
1180.54296875 0 1345.4004 b Ammonia loss 10
1184.5294189453125 0 1769.8611
1185.5076904296875 0 70147.72 y 4
1186.5107421875 0 47578.88
1187.5133056640625 0 19902.756
1188.517333984375 0 4675.9756
1197.5552978515625 0 10286.809 b 10
1198.557373046875 0 5700.914
1199.567138671875 0 2047.474
1214.5848388671875 0 1831.7068
1224.59228515625 0 1966.5585
1236.569580078125 0 1643.5321
1254.5792236328125 0 2693.7266
1322.56591796875 0 5061.8687 y 3
1323.570556640625 0 2593.6982
1823.2110595703125 0 1279.6033
3084.389892578125 0 1383.0034

Spectrum Details

|  |  |
| --- | --- |
| Matched peaks? Matched peaks The total absolute number of peaks matched. Additionally in brackets the total fraction of peaks matched and the total number of peaks is shown. | 97 (8.31% of 1167) |
| FDR? FDR The false discovery rate estimated for this peptide. It is calculated by matching all theoretical fragments with a non-integer shift with the raw peaks for this spectrum. This is done with 40 different shifts. The resulting percentage is the average number of annotated peaks over the number of annotated peaks with the correct spectrum. | 0.96% |
| Satellite FDR? Satellite FDR See the FDR for details on its calculation. This satellite ion specific FDR only contains the satellite ions (d/w) for I/L/J positions. | - |
| PSM Score? PSM Score The PSM Score as given by Hecklib to this annotated spectrum. It is shown with three significant figures. | 845 |

#### Spectrum 6077? Spectrum 6077 The raw spectrum of this peptide as annotated by Hecklib. The fragments are coloured according to ion type (see legend). Any peaks with a star '\*' as text can be hovered over to see the full details, first the ion type second the mass shift type. By hovering over the amino acids in the peptide or ions in the legend the corresponding peaks are highlighted. By toggling the 'Unassigned' label you can turn the background (unassigned) peaks on or off in the plot. By updating the slider in the Ion legend you can update the spectrum to only show the top X% of the peaks with labels. The top X% means any peak that is within X% of the highest intensity. By dragging in the spectrum you can zoom in to a specific part of the spectrum and use 'Zoom Out' to get back to the original zoom level. The annotation of the spectrum is based on the given sequence in the peptides file and is done with different software so inconsistencies are likely. The peaks are annotated based on the given sequence, with 20 ppm tolerance.

Copy Data

##### Spectrum 6077 (TSV)

###### Preview

```
Loading example...
```

*Click on the button to copy the data to your clipboard.*

Mz MinMz MaxIntensity Max

WidthHeightPeptide font sizePeptide stroke widthSpectrum font sizeSpectrum stroke widthCompact peptide

Ion legend

wxyz

abcd

OtherUnassignedIon^Charge^_Position_Show for top:%

TVJHQDMSJDGKEY

07.92e+41.58e+52.38e+53.17e+5

Zoom Out

a^+1^_2_d^+1^_2_y^+3^_4_a^+1^_2_y^+1^_1_b^+1^_2_b^+1^_2_y^+2^_3_b^+3^_6_y^+3^_7_b^+3^_7_b^+2^_5_y^+1^_2_b^+1^_3_y^+2^_5_y^+1^_2_b^+1^_3_b^+2^_6_b^+2^_6_y^+2^_6_y^+2^_6_y^+2^_7_y^+1^_3_b^+1^_4_y^+1^_3_b^+2^_8_b^+2^_8_b^+1^_4_b^+2^_8_y^+1^_4_y^+1^_4_b^+2^_9_b^+2^_9_b^+2^_9_\*\*\*b^+1^_5_b^+2^_10_b^+1^_5_y^+2^_10_y^+2^_10_y^+1^_5_y^+1^_5_y^+2^_11_y^+2^_11_b^+2^_12_b^+2^_12_y^+2^_11_b^+1^_6_b^+1^_6_b^+1^_6_y^+1^_6_y^+2^_12_y^+2^_12_y^+2^_12_b^+2^_13_y^+1^_6_b^+2^_13_y^+2^_13_y^+2^_13_y^+2^_13_b^+1^_7_y^+1^_7_b^+1^_7_b^+1^_8_b^+1^_8_b^+1^_8_y^+1^_8_y^+1^_8_b^+1^_9_b^+1^_9_b^+1^_9_y^+1^_9_b^+1^_10_b^+1^_10_y^+1^_10_y^+1^_10_y^+1^_10_b^+1^_11_

0800160024003200

Fragment Matches Table

Show background peaks

| Position | Ion type | Intensity | mz Theoretical | mz Error (Th) | mz Error (ppm) | Charge | Series Number |
| --- | --- | --- | --- | --- | --- | --- | --- |
| - | - | 2500 | 120 | - | - | 0 | - |
| - | - | 2.579E+04 | 120.1 | - | - | 0 | - |
| - | - | 1668 | 121.1 | - | - | 0 | - |
| - | - | 1199 | 122.1 | - | - | 0 | - |
| - | - | 1916 | 123 | - | - | 0 | - |
| - | - | 1734 | 127.1 | - | - | 0 | - |
| - | - | 854.4 | 127.1 | - | - | 0 | - |
| - | - | 793.8 | 127.1 | - | - | 0 | - |
| - | - | 965.8 | 128.1 | - | - | 0 | - |
| - | - | 1.949E+04 | 128.1 | - | - | 0 | - |
| - | - | 1398 | 129.1 | - | - | 0 | - |
| - | - | 1.426E+05 | 129.1 | - | - | 0 | - |
| - | - | 683.3 | 130.1 | - | - | 0 | - |
| - | - | 542 | 130.1 | - | - | 0 | - |
| - | - | 3606 | 130.1 | - | - | 0 | - |
| - | - | 916 | 130.1 | - | - | 0 | - |
| - | - | 7751 | 130.1 | - | - | 0 | - |
| - | - | 491.6 | 131 | - | - | 0 | - |
| - | - | 448.6 | 131.1 | - | - | 0 | - |
| - | - | 729.3 | 132.1 | - | - | 0 | - |
| - | - | 601.6 | 132.1 | - | - | 0 | - |
| - | - | 1483 | 133.1 | - | - | 0 | - |
| - | - | 3.42E+04 | 136.1 | - | - | 0 | - |
| - | - | 2468 | 137.1 | - | - | 0 | - |
| - | - | 2236 | 138.1 | - | - | 0 | - |
| - | - | 1201 | 138.1 | - | - | 0 | - |
| - | - | 1404 | 139.1 | - | - | 0 | - |
| - | - | 3403 | 141.1 | - | - | 0 | - |
| - | - | 620.7 | 142.1 | - | - | 0 | - |
| - | - | 555.5 | 143 | - | - | 0 | - |
| - | - | 1205 | 143.1 | - | - | 0 | - |
| - | - | 1728 | 145.1 | - | - | 0 | - |
| - | - | 3.543E+04 | 146.1 | - | - | 0 | - |
| - | - | 1364 | 147 | - | - | 0 | - |
| - | - | 2813 | 147.1 | - | - | 0 | - |
| - | - | 1.072E+04 | 148 | - | - | 0 | - |
| - | - | 994.3 | 149 | - | - | 0 | - |
| - | - | 3186 | 151.1 | - | - | 0 | - |
| - | - | 664.3 | 153.1 | - | - | 0 | - |
| - | - | 808.3 | 155.1 | - | - | 0 | - |
| - | - | 9726 | 155.1 | - | - | 0 | - |
| 2 | a | 8983 | 155.1 | 0.0002898 | 1.868 | +1 | 2 |
| - | - | 812.7 | 156 | - | - | 0 | - |
| - | - | 599.3 | 156.1 | - | - | 0 | - |
| - | - | 680.9 | 156.1 | - | - | 0 | - |
| - | - | 1124 | 156.1 | - | - | 0 | - |
| - | - | 1400 | 157.1 | - | - | 0 | - |
| - | - | 468 | 157.1 | - | - | 0 | - |
| - | - | 818.9 | 158.1 | - | - | 0 | - |
| - | - | 571.1 | 159.1 | - | - | 0 | - |
| - | - | 4805 | 159.1 | - | - | 0 | - |
| 2 | d | 804 | 159.1 | 0.0001567 | 0.9846 | +1 | 2 |
| - | - | 473.4 | 160.1 | - | - | 0 | - |
| - | - | 1127 | 163.1 | - | - | 0 | - |
| - | - | 1.649E+04 | 165.1 | - | - | 0 | - |
| - | - | 500.9 | 165.1 | - | - | 0 | - |
| - | - | 584 | 165.1 | - | - | 0 | - |
| - | - | 2.395E+04 | 166.1 | - | - | 0 | - |
| 11 | y | 1.625E+04 | 166.1 | 0.001603 | 9.653 | +3 | 4 |
| - | - | 866.6 | 166.1 | - | - | 0 | - |
| - | - | 924.2 | 166.1 | - | - | 0 | - |
| - | - | 1169 | 167.1 | - | - | 0 | - |
| - | - | 665.1 | 167.1 | - | - | 0 | - |
| - | - | 2008 | 167.1 | - | - | 0 | - |
| - | - | 497.8 | 167.1 | - | - | 0 | - |
| - | - | 2234 | 168.1 | - | - | 0 | - |
| - | - | 1529 | 168.1 | - | - | 0 | - |
| - | - | 582.4 | 169.1 | - | - | 0 | - |
| - | - | 700.6 | 169.1 | - | - | 0 | - |
| - | - | 4752 | 169.1 | - | - | 0 | - |
| - | - | 555.6 | 169.1 | - | - | 0 | - |
| - | - | 478.2 | 170.1 | - | - | 0 | - |
| - | - | 1331 | 171.1 | - | - | 0 | - |
| - | - | 1294 | 172.1 | - | - | 0 | - |
| - | - | 1495 | 173.1 | - | - | 0 | - |
| - | - | 3648 | 173.1 | - | - | 0 | - |
| 2 | a | 3.136E+05 | 173.1 | 0.0003452 | 1.994 | +1 | 2 |
| - | - | 5.977E+04 | 174.1 | - | - | 0 | - |
| - | - | 617.6 | 174.1 | - | - | 0 | - |
| - | - | 2.777E+04 | 174.1 | - | - | 0 | - |
| - | - | 6431 | 175.1 | - | - | 0 | - |
| - | - | 642.9 | 175.1 | - | - | 0 | - |
| - | - | 1641 | 175.1 | - | - | 0 | - |
| - | - | 724.9 | 177.1 | - | - | 0 | - |
| - | - | 1.085E+04 | 178.1 | - | - | 0 | - |
| - | - | 1057 | 179.1 | - | - | 0 | - |
| - | - | 1613 | 181.1 | - | - | 0 | - |
| 14 | y | 3.186E+04 | 182.1 | 0.0002665 | 1.464 | +1 | 1 |
| - | - | 2174 | 183.1 | - | - | 0 | - |
| 2 | b | 5859 | 183.1 | 0.0002482 | 1.356 | +1 | 2 |
| - | - | 1208 | 183.1 | - | - | 0 | - |
| - | - | 754.9 | 184 | - | - | 0 | - |
| - | - | 664.8 | 184.1 | - | - | 0 | - |
| - | - | 778.5 | 185.1 | - | - | 0 | - |
| - | - | 649.4 | 185.1 | - | - | 0 | - |
| - | - | 2136 | 185.2 | - | - | 0 | - |
| - | - | 4.743E+04 | 186.1 | - | - | 0 | - |
| - | - | 532.9 | 187.1 | - | - | 0 | - |
| - | - | 3767 | 187.1 | - | - | 0 | - |
| - | - | 1363 | 187.1 | - | - | 0 | - |
| - | - | 3529 | 188.1 | - | - | 0 | - |
| - | - | 1654 | 189.1 | - | - | 0 | - |
| - | - | 1278 | 190.1 | - | - | 0 | - |
| - | - | 1277 | 190.1 | - | - | 0 | - |
| - | - | 4721 | 191.1 | - | - | 0 | - |
| - | - | 932.1 | 193.1 | - | - | 0 | - |
| - | - | 1479 | 194.1 | - | - | 0 | - |
| - | - | 1.104E+04 | 195.1 | - | - | 0 | - |
| - | - | 4002 | 196.1 | - | - | 0 | - |
| - | - | 1255 | 196.1 | - | - | 0 | - |
| - | - | 3138 | 197.1 | - | - | 0 | - |
| - | - | 1112 | 198.1 | - | - | 0 | - |
| - | - | 686.8 | 198.1 | - | - | 0 | - |
| - | - | 940.2 | 199.1 | - | - | 0 | - |
| - | - | 713.1 | 201.1 | - | - | 0 | - |
| 2 | b | 7.407E+04 | 201.1 | 0.0001968 | 0.9787 | +1 | 2 |
| - | - | 1.853E+04 | 202.1 | - | - | 0 | - |
| - | - | 891.3 | 202.1 | - | - | 0 | - |
| - | - | 7396 | 202.1 | - | - | 0 | - |
| - | - | 2246 | 203.1 | - | - | 0 | - |
| - | - | 545.1 | 203.5 | - | - | 0 | - |
| - | - | 6040 | 205.1 | - | - | 0 | - |
| - | - | 1141 | 206.1 | - | - | 0 | - |
| - | - | 1591 | 207.2 | - | - | 0 | - |
| - | - | 877.7 | 208.1 | - | - | 0 | - |
| - | - | 994.1 | 209.1 | - | - | 0 | - |
| - | - | 530.5 | 209.1 | - | - | 0 | - |
| 12 | y | 956.1 | 211.1 | 0.0003592 | 1.702 | +2 | 3 |
| - | - | 874.1 | 211.1 | - | - | 0 | - |
| - | - | 1445 | 212.1 | - | - | 0 | - |
| - | - | 5025 | 213.1 | - | - | 0 | - |
| - | - | 712.8 | 213.2 | - | - | 0 | - |
| - | - | 943.1 | 215.1 | - | - | 0 | - |
| - | - | 521 | 216.1 | - | - | 0 | - |
| - | - | 807.9 | 216.1 | - | - | 0 | - |
| - | - | 683.1 | 216.1 | - | - | 0 | - |
| - | - | 1026 | 216.1 | - | - | 0 | - |
| - | - | 618.1 | 217.1 | - | - | 0 | - |
| - | - | 510.1 | 217.5 | - | - | 0 | - |
| - | - | 655.9 | 219.1 | - | - | 0 | - |
| - | - | 707.1 | 220.1 | - | - | 0 | - |
| - | - | 752.6 | 221.1 | - | - | 0 | - |
| - | - | 1853 | 221.1 | - | - | 0 | - |
| - | - | 849 | 221.1 | - | - | 0 | - |
| - | - | 1823 | 222.1 | - | - | 0 | - |
| - | - | 3793 | 223.1 | - | - | 0 | - |
| - | - | 2.193E+04 | 223.2 | - | - | 0 | - |
| - | - | 1246 | 224.1 | - | - | 0 | - |
| - | - | 2379 | 224.2 | - | - | 0 | - |
| - | - | 1607 | 225.1 | - | - | 0 | - |
| - | - | 3915 | 226.1 | - | - | 0 | - |
| 6 | b | 1474 | 226.1 | 0.0002634 | 1.165 | +3 | 6 |
| - | - | 833.7 | 226.2 | - | - | 0 | - |
| - | - | 5633 | 227.1 | - | - | 0 | - |
| - | - | 3365 | 229.1 | - | - | 0 | - |
| - | - | 526.6 | 231.7 | - | - | 0 | - |
| - | - | 3592 | 233.1 | - | - | 0 | - |
| - | - | 2438 | 233.2 | - | - | 0 | - |
| - | - | 2.839E+04 | 234.1 | - | - | 0 | - |
| - | - | 3046 | 235.1 | - | - | 0 | - |
| - | - | 973.4 | 235.1 | - | - | 0 | - |
| - | - | 617.7 | 235.2 | - | - | 0 | - |
| - | - | 1938 | 237.1 | - | - | 0 | - |
| - | - | 1055 | 237.1 | - | - | 0 | - |
| - | - | 1218 | 238.1 | - | - | 0 | - |
| - | - | 1062 | 238.1 | - | - | 0 | - |
| - | - | 650 | 239.1 | - | - | 0 | - |
| - | - | 691.8 | 239.2 | - | - | 0 | - |
| - | - | 8243 | 240.1 | - | - | 0 | - |
| - | - | 641.9 | 241.1 | - | - | 0 | - |
| - | - | 6352 | 241.1 | - | - | 0 | - |
| - | - | 1039 | 241.2 | - | - | 0 | - |
| - | - | 1221 | 242.1 | - | - | 0 | - |
| - | - | 1303 | 242.2 | - | - | 0 | - |
| - | - | 5410 | 243.1 | - | - | 0 | - |
| - | - | 503.1 | 243.3 | - | - | 0 | - |
| - | - | 5936 | 244.1 | - | - | 0 | - |
| - | - | 606.1 | 244.1 | - | - | 0 | - |
| - | - | 924.9 | 245.1 | - | - | 0 | - |
| - | - | 1.362E+04 | 247.1 | - | - | 0 | - |
| - | - | 573 | 247.1 | - | - | 0 | - |
| - | - | 3378 | 248.1 | - | - | 0 | - |
| - | - | 6889 | 249.1 | - | - | 0 | - |
| - | - | 1351 | 249.1 | - | - | 0 | - |
| - | - | 938.9 | 250.1 | - | - | 0 | - |
| - | - | 674.7 | 250.1 | - | - | 0 | - |
| - | - | 8.972E+04 | 251.2 | - | - | 0 | - |
| - | - | 1003 | 252.1 | - | - | 0 | - |
| - | - | 982.9 | 252.1 | - | - | 0 | - |
| - | - | 1.145E+04 | 252.2 | - | - | 0 | - |
| - | - | 781.6 | 253.2 | - | - | 0 | - |
| - | - | 1498 | 255.1 | - | - | 0 | - |
| - | - | 600.2 | 256.1 | - | - | 0 | - |
| - | - | 640.5 | 256.6 | - | - | 0 | - |
| - | - | 1634 | 257.1 | - | - | 0 | - |
| - | - | 3269 | 258.1 | - | - | 0 | - |
| - | - | 3764 | 261.1 | - | - | 0 | - |
| - | - | 1041 | 261.1 | - | - | 0 | - |
| - | - | 758.2 | 261.2 | - | - | 0 | - |
| - | - | 3043 | 262.1 | - | - | 0 | - |
| 8 | y | 5140 | 265.1 | 0.0003845 | 1.45 | +3 | 7 |
| - | - | 1.823E+04 | 266.1 | - | - | 0 | - |
| - | - | 2325 | 267.1 | - | - | 0 | - |
| - | - | 723.3 | 268.1 | - | - | 0 | - |
| - | - | 1073 | 268.2 | - | - | 0 | - |
| - | - | 1013 | 269.1 | - | - | 0 | - |
| - | - | 6925 | 269.1 | - | - | 0 | - |
| - | - | 926.7 | 269.2 | - | - | 0 | - |
| - | - | 1097 | 269.2 | - | - | 0 | - |
| 7 | b | 1228 | 270.1 | 0.004044 | 14.97 | +3 | 7 |
| - | - | 1216 | 270.1 | - | - | 0 | - |
| - | - | 653.5 | 272.1 | - | - | 0 | - |
| - | - | 2487 | 273.1 | - | - | 0 | - |
| - | - | 1690 | 274.1 | - | - | 0 | - |
| - | - | 1278 | 275.1 | - | - | 0 | - |
| - | - | 859.1 | 276.1 | - | - | 0 | - |
| - | - | 1019 | 279.1 | - | - | 0 | - |
| - | - | 2132 | 280.1 | - | - | 0 | - |
| - | - | 877.3 | 282.2 | - | - | 0 | - |
| - | - | 3.566E+04 | 283.1 | - | - | 0 | - |
| - | - | 990.4 | 284.1 | - | - | 0 | - |
| - | - | 572.5 | 284.1 | - | - | 0 | - |
| - | - | 4189 | 284.1 | - | - | 0 | - |
| - | - | 3495 | 286.1 | - | - | 0 | - |
| - | - | 3603 | 287.1 | - | - | 0 | - |
| - | - | 1278 | 287.2 | - | - | 0 | - |
| 5 | b | 918.5 | 290.2 | 3.837E-05 | 0.1322 | +2 | 5 |
| 13 | y | 3.755E+04 | 293.1 | 0.0001747 | 0.5961 | +1 | 2 |
| - | - | 6502 | 294.1 | - | - | 0 | - |
| - | - | 604.5 | 295.1 | - | - | 0 | - |
| - | - | 571.2 | 296.1 | - | - | 0 | - |
| 3 | b | 601.4 | 296.2 | 0.0008247 | 2.784 | +1 | 3 |
| - | - | 714.8 | 297.1 | - | - | 0 | - |
| - | - | 8689 | 297.2 | - | - | 0 | - |
| - | - | 645.1 | 298.1 | - | - | 0 | - |
| - | - | 1218 | 298.2 | - | - | 0 | - |
| - | - | 683.1 | 300.2 | - | - | 0 | - |
| - | - | 1011 | 301.1 | - | - | 0 | - |
| - | - | 1.864E+04 | 301.2 | - | - | 0 | - |
| - | - | 1008 | 301.2 | - | - | 0 | - |
| - | - | 2704 | 302.2 | - | - | 0 | - |
| - | - | 3169 | 304.2 | - | - | 0 | - |
| - | - | 675.6 | 305.2 | - | - | 0 | - |
| - | - | 1421 | 306.1 | - | - | 0 | - |
| 10 | y | 3605 | 306.1 | 0.0005213 | 1.703 | +2 | 5 |
| - | - | 905.3 | 306.6 | - | - | 0 | - |
| 13 | y | 5447 | 311.1 | 0.0007489 | 2.407 | +1 | 2 |
| - | - | 969 | 312.1 | - | - | 0 | - |
| 3 | b | 2298 | 314.2 | 2.567E-05 | 0.0817 | +1 | 3 |
| - | - | 633.2 | 315.1 | - | - | 0 | - |
| - | - | 4226 | 315.1 | - | - | 0 | - |
| - | - | 1.614E+04 | 315.2 | - | - | 0 | - |
| - | - | 673 | 316.1 | - | - | 0 | - |
| - | - | 2305 | 316.2 | - | - | 0 | - |
| - | - | 599 | 321.1 | - | - | 0 | - |
| - | - | 628.1 | 323.2 | - | - | 0 | - |
| - | - | 1114 | 325.2 | - | - | 0 | - |
| - | - | 1102 | 332.2 | - | - | 0 | - |
| - | - | 1670 | 332.2 | - | - | 0 | - |
| - | - | 2250 | 333.2 | - | - | 0 | - |
| - | - | 5117 | 334.1 | - | - | 0 | - |
| - | - | 995 | 335.1 | - | - | 0 | - |
| 6 | b | 715.4 | 338.7 | 0.001462 | 4.318 | +2 | 6 |
| - | - | 689.1 | 339.1 | - | - | 0 | - |
| - | - | 690.7 | 342.4 | - | - | 0 | - |
| - | - | 645.5 | 343.1 | - | - | 0 | - |
| - | - | 1247 | 344.2 | - | - | 0 | - |
| - | - | 2029 | 345.1 | - | - | 0 | - |
| - | - | 3827 | 346.1 | - | - | 0 | - |
| - | - | 679.8 | 347.1 | - | - | 0 | - |
| 6 | b | 1849 | 347.7 | 0.000158 | 0.4544 | +2 | 6 |
| - | - | 6285 | 350.2 | - | - | 0 | - |
| - | - | 1174 | 351.2 | - | - | 0 | - |
| 9 | y | 941.8 | 353.7 | 6.499E-05 | 0.1837 | +2 | 6 |
| - | - | 680.6 | 354.2 | - | - | 0 | - |
| - | - | 1535 | 356.2 | - | - | 0 | - |
| - | - | 651.5 | 356.7 | - | - | 0 | - |
| - | - | 1973 | 361.2 | - | - | 0 | - |
| - | - | 1652 | 362.2 | - | - | 0 | - |
| 9 | y | 4146 | 362.7 | 9.831E-05 | 0.2711 | +2 | 6 |
| - | - | 1.098E+04 | 363.1 | - | - | 0 | - |
| - | - | 1439 | 363.2 | - | - | 0 | - |
| - | - | 717.7 | 363.7 | - | - | 0 | - |
| - | - | 1676 | 364.1 | - | - | 0 | - |
| - | - | 817.1 | 366.2 | - | - | 0 | - |
| - | - | 1209 | 367.2 | - | - | 0 | - |
| - | - | 604.2 | 367.9 | - | - | 0 | - |
| - | - | 669.6 | 376.2 | - | - | 0 | - |
| - | - | 2770 | 378.2 | - | - | 0 | - |
| - | - | 2.045E+04 | 379.2 | - | - | 0 | - |
| - | - | 990.8 | 379.2 | - | - | 0 | - |
| - | - | 1220 | 380.2 | - | - | 0 | - |
| - | - | 3332 | 380.2 | - | - | 0 | - |
| - | - | 1.558E+04 | 381.2 | - | - | 0 | - |
| - | - | 1903 | 382.2 | - | - | 0 | - |
| - | - | 784.7 | 384.2 | - | - | 0 | - |
| - | - | 1550 | 384.2 | - | - | 0 | - |
| - | - | 1867 | 385.2 | - | - | 0 | - |
| - | - | 962.1 | 392.2 | - | - | 0 | - |
| - | - | 850.4 | 392.7 | - | - | 0 | - |
| - | - | 3798 | 394.2 | - | - | 0 | - |
| - | - | 1109 | 395.2 | - | - | 0 | - |
| - | - | 5034 | 396.2 | - | - | 0 | - |
| - | - | 779.2 | 397.2 | - | - | 0 | - |
| - | - | 1.154E+04 | 398.2 | - | - | 0 | - |
| - | - | 927.5 | 399.1 | - | - | 0 | - |
| - | - | 1744 | 399.2 | - | - | 0 | - |
| - | - | 2435 | 402.2 | - | - | 0 | - |
| - | - | 775.1 | 403.2 | - | - | 0 | - |
| - | - | 1197 | 405.3 | - | - | 0 | - |
| 8 | y | 913.5 | 406.2 | 0.006499 | 16 | +2 | 7 |
| - | - | 681.2 | 406.2 | - | - | 0 | - |
| - | - | 788.6 | 407.2 | - | - | 0 | - |
| - | - | 921.3 | 408.2 | - | - | 0 | - |
| - | - | 1203 | 409.1 | - | - | 0 | - |
| - | - | 711.7 | 411.2 | - | - | 0 | - |
| - | - | 984 | 412.2 | - | - | 0 | - |
| - | - | 1.403E+04 | 412.2 | - | - | 0 | - |
| - | - | 2275 | 413.2 | - | - | 0 | - |
| - | - | 3027 | 414.2 | - | - | 0 | - |
| - | - | 1235 | 415.2 | - | - | 0 | - |
| - | - | 2203 | 416.2 | - | - | 0 | - |
| - | - | 5301 | 417.1 | - | - | 0 | - |
| - | - | 942.7 | 418.1 | - | - | 0 | - |
| 12 | y | 719.9 | 421.2 | 0.004546 | 10.79 | +1 | 3 |
| - | - | 7678 | 423.3 | - | - | 0 | - |
| - | - | 1415 | 424.3 | - | - | 0 | - |
| - | - | 907.1 | 427.1 | - | - | 0 | - |
| - | - | 794.3 | 429.2 | - | - | 0 | - |
| - | - | 1318 | 429.2 | - | - | 0 | - |
| - | - | 3.859E+04 | 430.2 | - | - | 0 | - |
| - | - | 8232 | 431.2 | - | - | 0 | - |
| - | - | 807.8 | 432.2 | - | - | 0 | - |
| - | - | 627.5 | 433.2 | - | - | 0 | - |
| 4 | b | 707.8 | 433.3 | 0.0001709 | 0.3945 | +1 | 4 |
| - | - | 2695 | 433.7 | - | - | 0 | - |
| - | - | 3170 | 434.2 | - | - | 0 | - |
| - | - | 1169 | 434.2 | - | - | 0 | - |
| 12 | y | 1.149E+04 | 439.2 | 8.524E-05 | 0.1941 | +1 | 3 |
| - | - | 727.1 | 439.7 | - | - | 0 | - |
| - | - | 2725 | 440.2 | - | - | 0 | - |
| - | - | 882.5 | 441.2 | - | - | 0 | - |
| - | - | 638.3 | 442.2 | - | - | 0 | - |
| - | - | 8423 | 442.7 | - | - | 0 | - |
| - | - | 2242 | 443.2 | - | - | 0 | - |
| - | - | 1439 | 443.7 | - | - | 0 | - |
| - | - | 3767 | 444.2 | - | - | 0 | - |
| - | - | 9928 | 445.1 | - | - | 0 | - |
| - | - | 2456 | 446.1 | - | - | 0 | - |
| - | - | 2108 | 447.2 | - | - | 0 | - |
| 8 | b | 5116 | 447.7 | 0.0007672 | 1.714 | +2 | 8 |
| 8 | b | 4588 | 448.2 | 0.003044 | 6.792 | +2 | 8 |
| - | - | 811.6 | 448.7 | - | - | 0 | - |
| - | - | 1219 | 449.2 | - | - | 0 | - |
| - | - | 821.3 | 450.2 | - | - | 0 | - |
| - | - | 734.9 | 451.2 | - | - | 0 | - |
| 4 | b | 1.305E+04 | 451.3 | 0.0001653 | 0.3664 | +1 | 4 |
| - | - | 2563 | 452.3 | - | - | 0 | - |
| - | - | 1957 | 454.2 | - | - | 0 | - |
| - | - | 1109 | 454.7 | - | - | 0 | - |
| 8 | b | 2.317E+04 | 456.7 | 0.001594 | 3.49 | +2 | 8 |
| - | - | 1.115E+04 | 457.2 | - | - | 0 | - |
| - | - | 998.4 | 457.6 | - | - | 0 | - |
| - | - | 4372 | 457.7 | - | - | 0 | - |
| - | - | 893.6 | 458.2 | - | - | 0 | - |
| - | - | 683.8 | 459.2 | - | - | 0 | - |
| - | - | 1215 | 461.2 | - | - | 0 | - |
| - | - | 7908 | 462.2 | - | - | 0 | - |
| - | - | 2469 | 463.2 | - | - | 0 | - |
| - | - | 628.3 | 466.2 | - | - | 0 | - |
| - | - | 583 | 468.2 | - | - | 0 | - |
| - | - | 4036 | 468.3 | - | - | 0 | - |
| - | - | 980.6 | 469.3 | - | - | 0 | - |
| - | - | 990.9 | 474.8 | - | - | 0 | - |
| - | - | 1446 | 475.2 | - | - | 0 | - |
| - | - | 910.5 | 475.9 | - | - | 0 | - |
| - | - | 1256 | 476.2 | - | - | 0 | - |
| - | - | 1412 | 477.2 | - | - | 0 | - |
| - | - | 966.3 | 477.8 | - | - | 0 | - |
| 11 | y | 3895 | 478.2 | 0.000743 | 1.554 | +1 | 4 |
| - | - | 1149 | 478.3 | - | - | 0 | - |
| - | - | 754.7 | 480.2 | - | - | 0 | - |
| - | - | 1492 | 481.2 | - | - | 0 | - |
| - | - | 633.5 | 481.7 | - | - | 0 | - |
| - | - | 2525 | 490.3 | - | - | 0 | - |
| - | - | 2094 | 490.8 | - | - | 0 | - |
| - | - | 1128 | 491.3 | - | - | 0 | - |
| - | - | 953.5 | 492.8 | - | - | 0 | - |
| - | - | 836 | 493.3 | - | - | 0 | - |
| - | - | 686.3 | 493.8 | - | - | 0 | - |
| - | - | 1.85E+04 | 494.2 | - | - | 0 | - |
| - | - | 6703 | 495.2 | - | - | 0 | - |
| - | - | 1524 | 495.7 | - | - | 0 | - |
| 11 | y | 4.756E+04 | 496.2 | 0.0002585 | 0.5209 | +1 | 4 |
| - | - | 1.324E+04 | 497.2 | - | - | 0 | - |
| - | - | 1073 | 497.7 | - | - | 0 | - |
| - | - | 1975 | 498.2 | - | - | 0 | - |
| - | - | 7120 | 499.3 | - | - | 0 | - |
| - | - | 4174 | 499.8 | - | - | 0 | - |
| - | - | 928.2 | 500.3 | - | - | 0 | - |
| 9 | b | 1699 | 504.3 | 0.001021 | 2.024 | +2 | 9 |
| 9 | b | 1174 | 504.7 | 0.006911 | 13.69 | +2 | 9 |
| - | - | 879.7 | 506.3 | - | - | 0 | - |
| - | - | 7096 | 511.3 | - | - | 0 | - |
| - | - | 649.7 | 512.2 | - | - | 0 | - |
| - | - | 1790 | 512.3 | - | - | 0 | - |
| - | - | 671.2 | 512.6 | - | - | 0 | - |
| 9 | b | 3468 | 513.3 | 0.0007793 | 1.518 | +2 | 9 |
| - | - | 1962 | 513.8 | - | - | 0 | - |
| - | - | 2326 | 525.3 | - | - | 0 | - |
| - | - | 1037 | 526.3 | - | - | 0 | - |
| - | - | 1505 | 529.2 | - | - | 0 | - |
| - | - | 2528 | 530.2 | - | - | 0 | - |
| - | - | 725.7 | 530.6 | - | - | 0 | - |
| - | - | 825 | 533.6 | - | - | 0 | - |
| - | - | 1771 | 533.9 | - | - | 0 | - |
| - | - | 815.1 | 534.3 | - | - | 0 | - |
| - | - | 3047 | 534.3 | - | - | 0 | - |
| - | - | 1170 | 534.6 | - | - | 0 | - |
| - | - | 829.4 | 535.3 | - | - | 0 | - |
| - | - | 1579 | 536.2 | - | - | 0 | - |
| 0 | Precursor | 2021 | 539.9 | 0.001244 | 2.304 | +3 | -1 |
| 0 | Precursor | 2272 | 540.3 | 0.00559 | 10.35 | +3 | -1 |
| - | - | 1636 | 540.6 | - | - | 0 | - |
| - | - | 686.2 | 542.2 | - | - | 0 | - |
| - | - | 8113 | 543.3 | - | - | 0 | - |
| - | - | 2271 | 544.3 | - | - | 0 | - |
| - | - | 1020 | 544.7 | - | - | 0 | - |
| - | - | 990.3 | 545.2 | - | - | 0 | - |
| - | - | 687.3 | 545.8 | - | - | 0 | - |
| 0 | Precursor | 5595 | 545.9 | 0.001348 | 2.468 | +3 | -1 |
| - | - | 4631 | 546.3 | - | - | 0 | - |
| - | - | 2239 | 546.6 | - | - | 0 | - |
| - | - | 2693 | 546.8 | - | - | 0 | - |
| - | - | 3061 | 547.1 | - | - | 0 | - |
| - | - | 1232 | 547.3 | - | - | 0 | - |
| - | - | 2049 | 547.3 | - | - | 0 | - |
| - | - | 784.1 | 548.7 | - | - | 0 | - |
| - | - | 1606 | 553.7 | - | - | 0 | - |
| - | - | 3242 | 554.3 | - | - | 0 | - |
| - | - | 1495 | 554.8 | - | - | 0 | - |
| - | - | 1427 | 557.2 | - | - | 0 | - |
| - | - | 1562 | 558.2 | - | - | 0 | - |
| 5 | b | 2182 | 561.3 | 9.561E-05 | 0.1703 | +1 | 5 |
| - | - | 4484 | 562.2 | - | - | 0 | - |
| - | - | 612 | 562.3 | - | - | 0 | - |
| - | - | 2177 | 562.7 | - | - | 0 | - |
| - | - | 954.1 | 563.3 | - | - | 0 | - |
| - | - | 1722 | 563.8 | - | - | 0 | - |
| - | - | 1971 | 564.2 | - | - | 0 | - |
| - | - | 1697 | 565.3 | - | - | 0 | - |
| 10 | b | 1441 | 570.8 | 0.0007028 | 1.231 | +2 | 10 |
| - | - | 3173 | 571.2 | - | - | 0 | - |
| - | - | 2775 | 571.2 | - | - | 0 | - |
| - | - | 2741 | 571.8 | - | - | 0 | - |
| - | - | 887.5 | 572.2 | - | - | 0 | - |
| - | - | 822.6 | 574.3 | - | - | 0 | - |
| - | - | 3815 | 575.2 | - | - | 0 | - |
| - | - | 920.8 | 576.2 | - | - | 0 | - |
| 5 | b | 9329 | 579.3 | 0.0004593 | 0.7928 | +1 | 5 |
| - | - | 1067 | 580.3 | - | - | 0 | - |
| - | - | 2139 | 580.3 | - | - | 0 | - |
| - | - | 873.3 | 580.8 | - | - | 0 | - |
| - | - | 1441 | 581.2 | - | - | 0 | - |
| - | - | 891.7 | 581.3 | - | - | 0 | - |
| - | - | 818.4 | 582.2 | - | - | 0 | - |
| 5 | y | 1091 | 584.3 | 0.001506 | 2.578 | +2 | 10 |
| 5 | y | 901.3 | 584.7 | 0.008378 | 14.33 | +2 | 10 |
| - | - | 1225 | 584.8 | - | - | 0 | - |
| 10 | y | 1.015E+04 | 593.3 | 0.0004643 | 0.7827 | +1 | 5 |
| - | - | 1744 | 593.3 | - | - | 0 | - |
| - | - | 1185 | 593.8 | - | - | 0 | - |
| - | - | 2412 | 594.3 | - | - | 0 | - |
| - | - | 725.2 | 597.3 | - | - | 0 | - |
| - | - | 765.8 | 597.8 | - | - | 0 | - |
| - | - | 1.061E+04 | 599.2 | - | - | 0 | - |
| - | - | 3067 | 600.2 | - | - | 0 | - |
| - | - | 812.3 | 600.3 | - | - | 0 | - |
| - | - | 923 | 601.2 | - | - | 0 | - |
| - | - | 746.4 | 604.8 | - | - | 0 | - |
| - | - | 748.7 | 609.8 | - | - | 0 | - |
| - | - | 5965 | 610.3 | - | - | 0 | - |
| - | - | 1815 | 610.3 | - | - | 0 | - |
| - | - | 1381 | 610.8 | - | - | 0 | - |
| 10 | y | 8.519E+04 | 611.3 | 0.0002258 | 0.3694 | +1 | 5 |
| - | - | 2.737E+04 | 612.3 | - | - | 0 | - |
| - | - | 5850 | 613.3 | - | - | 0 | - |
| - | - | 2696 | 613.8 | - | - | 0 | - |
| - | - | 1935 | 614.3 | - | - | 0 | - |
| - | - | 6052 | 618.8 | - | - | 0 | - |
| - | - | 4701 | 619.3 | - | - | 0 | - |
| - | - | 2444 | 619.8 | - | - | 0 | - |
| - | - | 885.8 | 620.3 | - | - | 0 | - |
| - | - | 1628 | 621.3 | - | - | 0 | - |
| - | - | 727.4 | 627.2 | - | - | 0 | - |
| - | - | 1.737E+04 | 627.8 | - | - | 0 | - |
| - | - | 1026 | 628.2 | - | - | 0 | - |
| - | - | 1.228E+04 | 628.3 | - | - | 0 | - |
| - | - | 5589 | 628.8 | - | - | 0 | - |
| - | - | 1797 | 629.3 | - | - | 0 | - |
| - | - | 1028 | 636.8 | - | - | 0 | - |
| - | - | 1429 | 637.3 | - | - | 0 | - |
| - | - | 1923 | 644.3 | - | - | 0 | - |
| - | - | 1254 | 644.8 | - | - | 0 | - |
| - | - | 768.4 | 645 | - | - | 0 | - |
| - | - | 1044 | 645.3 | - | - | 0 | - |
| - | - | 1748 | 649.3 | - | - | 0 | - |
| - | - | 1885 | 649.3 | - | - | 0 | - |
| 4 | y | 3530 | 652.8 | 0.002153 | 3.299 | +2 | 11 |
| 4 | y | 2610 | 653.3 | 0.009318 | 14.26 | +2 | 11 |
| 12 | b | 1536 | 654.3 | 0.00694 | 10.61 | +2 | 12 |
| 12 | b | 1391 | 654.8 | 0.004897 | 7.479 | +2 | 12 |
| - | - | 822.3 | 655.3 | - | - | 0 | - |
| - | - | 1714 | 659.3 | - | - | 0 | - |
| - | - | 763.1 | 660.3 | - | - | 0 | - |
| - | - | 1717 | 661.3 | - | - | 0 | - |
| 4 | y | 1.806E+04 | 661.8 | 0.001515 | 2.29 | +2 | 11 |
| - | - | 1.794E+04 | 662.3 | - | - | 0 | - |
| - | - | 5914 | 662.8 | - | - | 0 | - |
| - | - | 1423 | 663.3 | - | - | 0 | - |
| - | - | 1099 | 666.3 | - | - | 0 | - |
| - | - | 962.5 | 666.8 | - | - | 0 | - |
| - | - | 1726 | 666.8 | - | - | 0 | - |
| - | - | 1528 | 667.3 | - | - | 0 | - |
| - | - | 1127 | 668.3 | - | - | 0 | - |
| - | - | 1206 | 668.8 | - | - | 0 | - |
| - | - | 1160 | 669.3 | - | - | 0 | - |
| - | - | 3159 | 672.3 | - | - | 0 | - |
| - | - | 791.7 | 672.3 | - | - | 0 | - |
| - | - | 2033 | 673.3 | - | - | 0 | - |
| - | - | 712.7 | 673.3 | - | - | 0 | - |
| 6 | b | 3503 | 676.3 | 0.002311 | 3.417 | +1 | 6 |
| - | - | 3621 | 676.8 | - | - | 0 | - |
| - | - | 964.6 | 677.3 | - | - | 0 | - |
| 6 | b | 6355 | 677.3 | 0.001161 | 1.714 | +1 | 6 |
| - | - | 2498 | 677.8 | - | - | 0 | - |
| - | - | 1646 | 678.3 | - | - | 0 | - |
| - | - | 1.477E+04 | 684.3 | - | - | 0 | - |
| - | - | 1.027E+04 | 685.3 | - | - | 0 | - |
| - | - | 5348 | 685.8 | - | - | 0 | - |
| - | - | 4843 | 686.3 | - | - | 0 | - |
| - | - | 2318 | 686.8 | - | - | 0 | - |
| - | - | 875.2 | 687.3 | - | - | 0 | - |
| - | - | 2428 | 687.9 | - | - | 0 | - |
| - | - | 1816 | 688.4 | - | - | 0 | - |
| - | - | 1101 | 688.9 | - | - | 0 | - |
| - | - | 2085 | 689.8 | - | - | 0 | - |
| - | - | 3142 | 690.3 | - | - | 0 | - |
| - | - | 788.2 | 690.3 | - | - | 0 | - |
| - | - | 1165 | 691.3 | - | - | 0 | - |
| - | - | 1088 | 691.8 | - | - | 0 | - |
| - | - | 3321 | 693.4 | - | - | 0 | - |
| - | - | 2453 | 694.3 | - | - | 0 | - |
| 6 | b | 1.943E+04 | 694.4 | 0.0006128 | 0.8825 | +1 | 6 |
| - | - | 1746 | 695.3 | - | - | 0 | - |
| - | - | 6920 | 695.4 | - | - | 0 | - |
| - | - | 1666 | 695.8 | - | - | 0 | - |
| - | - | 1657 | 696.4 | - | - | 0 | - |
| - | - | 2824 | 700.3 | - | - | 0 | - |
| - | - | 4189 | 700.8 | - | - | 0 | - |
| - | - | 3109 | 701.3 | - | - | 0 | - |
| - | - | 1187 | 701.8 | - | - | 0 | - |
| - | - | 1029 | 703.4 | - | - | 0 | - |
| - | - | 2562 | 704.3 | - | - | 0 | - |
| - | - | 2373 | 704.8 | - | - | 0 | - |
| - | - | 1273 | 705.3 | - | - | 0 | - |
| - | - | 755.8 | 705.8 | - | - | 0 | - |
| 9 | y | 3042 | 706.3 | 6.64E-05 | 0.09401 | +1 | 6 |
| - | - | 1602 | 707.3 | - | - | 0 | - |
| - | - | 943.8 | 708.8 | - | - | 0 | - |
| 3 | y | 1.425E+04 | 709.3 | 0.00201 | 2.834 | +2 | 12 |
| 3 | y | 1.839E+04 | 709.8 | 0.005067 | 7.138 | +2 | 12 |
| - | - | 1.159E+04 | 710.3 | - | - | 0 | - |
| - | - | 2807 | 710.8 | - | - | 0 | - |
| - | - | 2.632E+04 | 711.4 | - | - | 0 | - |
| - | - | 2.615E+04 | 712.3 | - | - | 0 | - |
| - | - | 8791 | 712.4 | - | - | 0 | - |
| - | - | 1.087E+04 | 713.3 | - | - | 0 | - |
| - | - | 2105 | 713.4 | - | - | 0 | - |
| - | - | 974 | 713.8 | - | - | 0 | - |
| - | - | 1623 | 714.3 | - | - | 0 | - |
| - | - | 972.3 | 717.3 | - | - | 0 | - |
| - | - | 6528 | 717.8 | - | - | 0 | - |
| 3 | y | 1.147E+05 | 718.3 | 0.002288 | 3.185 | +2 | 12 |
| - | - | 9.494E+04 | 718.8 | - | - | 0 | - |
| 13 | b | 4.437E+04 | 719.3 | 0.006939 | 9.646 | +2 | 13 |
| - | - | 1.341E+04 | 719.8 | - | - | 0 | - |
| - | - | 3070 | 720.3 | - | - | 0 | - |
| - | - | 2185 | 720.9 | - | - | 0 | - |
| - | - | 2522 | 721.4 | - | - | 0 | - |
| - | - | 901.3 | 721.9 | - | - | 0 | - |
| - | - | 924.5 | 723.4 | - | - | 0 | - |
| 9 | y | 2.224E+04 | 724.4 | 0.0007327 | 1.011 | +1 | 6 |
| - | - | 9067 | 725.4 | - | - | 0 | - |
| - | - | 1295 | 725.9 | - | - | 0 | - |
| - | - | 5077 | 726.4 | - | - | 0 | - |
| - | - | 1903 | 726.9 | - | - | 0 | - |
| - | - | 2760 | 727.4 | - | - | 0 | - |
| 13 | b | 9043 | 727.9 | 0.001476 | 2.027 | +2 | 13 |
| - | - | 7065 | 728.4 | - | - | 0 | - |
| - | - | 2530 | 728.8 | - | - | 0 | - |
| - | - | 786.9 | 729.3 | - | - | 0 | - |
| - | - | 685.6 | 734.3 | - | - | 0 | - |
| - | - | 5197 | 734.9 | - | - | 0 | - |
| - | - | 5705 | 735.4 | - | - | 0 | - |
| - | - | 2637 | 735.9 | - | - | 0 | - |
| - | - | 825.1 | 736.4 | - | - | 0 | - |
| - | - | 2355 | 736.9 | - | - | 0 | - |
| - | - | 1612 | 737.4 | - | - | 0 | - |
| - | - | 897.5 | 740.3 | - | - | 0 | - |
| - | - | 2280 | 744.3 | - | - | 0 | - |
| - | - | 790.5 | 751.3 | - | - | 0 | - |
| 2 | y | 2016 | 758.9 | 0.002526 | 3.328 | +2 | 13 |
| 2 | y | 2094 | 759.4 | 0.00272 | 3.582 | +2 | 13 |
| - | - | 1553 | 761.3 | - | - | 0 | - |
| - | - | 1188 | 762.3 | - | - | 0 | - |
| 2 | y | 1.148E+04 | 767.9 | 0.002803 | 3.651 | +2 | 13 |
| - | - | 8908 | 768.4 | - | - | 0 | - |
| - | - | 4679 | 768.9 | - | - | 0 | - |
| - | - | 840.6 | 777.4 | - | - | 0 | - |
| - | - | 1249 | 779.4 | - | - | 0 | - |
| - | - | 925.5 | 780.4 | - | - | 0 | - |
| - | - | 1157 | 782.3 | - | - | 0 | - |
| - | - | 2177 | 783.4 | - | - | 0 | - |
| - | - | 1566 | 784.4 | - | - | 0 | - |
| - | - | 1320 | 792.3 | - | - | 0 | - |
| - | - | 764.9 | 793.3 | - | - | 0 | - |
| - | - | 3794 | 797.4 | - | - | 0 | - |
| - | - | 2451 | 798.4 | - | - | 0 | - |
| - | - | 990.3 | 799.3 | - | - | 0 | - |
| 7 | b | 811 | 807.4 | 0.01423 | 17.63 | +1 | 7 |
| 8 | y | 3195 | 811.4 | 0.008835 | 10.89 | +1 | 7 |
| - | - | 1934 | 812.4 | - | - | 0 | - |
| - | - | 954 | 813.4 | - | - | 0 | - |
| - | - | 714.7 | 819.4 | - | - | 0 | - |
| 7 | b | 7960 | 825.4 | 0.0038 | 4.604 | +1 | 7 |
| - | - | 4226 | 826.4 | - | - | 0 | - |
| - | - | 6291 | 827.3 | - | - | 0 | - |
| - | - | 3243 | 828.3 | - | - | 0 | - |
| - | - | 739.4 | 829.3 | - | - | 0 | - |
| - | - | 704.4 | 837.9 | - | - | 0 | - |
| - | - | 2280 | 839.4 | - | - | 0 | - |
| - | - | 1744 | 840.4 | - | - | 0 | - |
| - | - | 868.1 | 841.4 | - | - | 0 | - |
| - | - | 2030 | 857.4 | - | - | 0 | - |
| - | - | 2511 | 858.4 | - | - | 0 | - |
| - | - | 950.1 | 859.4 | - | - | 0 | - |
| - | - | 708.4 | 860.4 | - | - | 0 | - |
| - | - | 1226 | 866.4 | - | - | 0 | - |
| - | - | 930.6 | 867.4 | - | - | 0 | - |
| - | - | 1208 | 875.4 | - | - | 0 | - |
| - | - | 2965 | 876.4 | - | - | 0 | - |
| - | - | 2434 | 877.4 | - | - | 0 | - |
| - | - | 862.3 | 878.4 | - | - | 0 | - |
| - | - | 948.3 | 879.4 | - | - | 0 | - |
| - | - | 2155 | 884.3 | - | - | 0 | - |
| - | - | 5429 | 884.4 | - | - | 0 | - |
| - | - | 1507 | 885.3 | - | - | 0 | - |
| - | - | 3513 | 885.4 | - | - | 0 | - |
| - | - | 878.8 | 886.4 | - | - | 0 | - |
| 8 | b | 6324 | 894.4 | 0.004817 | 5.386 | +1 | 8 |
| 8 | b | 5949 | 895.4 | 0.004942 | 5.519 | +1 | 8 |
| - | - | 1404 | 896.4 | - | - | 0 | - |
| - | - | 1002 | 897.4 | - | - | 0 | - |
| - | - | 1129 | 901.4 | - | - | 0 | - |
| - | - | 906.4 | 905.4 | - | - | 0 | - |
| 8 | b | 5.215E+04 | 912.4 | 0.00525 | 5.754 | +1 | 8 |
| - | - | 2.589E+04 | 913.4 | - | - | 0 | - |
| - | - | 8722 | 914.4 | - | - | 0 | - |
| - | - | 1353 | 915.4 | - | - | 0 | - |
| - | - | 2112 | 922.4 | - | - | 0 | - |
| - | - | 2214 | 923.4 | - | - | 0 | - |
| - | - | 1456 | 924.4 | - | - | 0 | - |
| 7 | y | 2975 | 925.4 | 0.001958 | 2.116 | +1 | 8 |
| - | - | 1083 | 926.4 | - | - | 0 | - |
| - | - | 1.55E+04 | 940.4 | - | - | 0 | - |
| - | - | 8453 | 941.4 | - | - | 0 | - |
| 7 | y | 6259 | 942.4 | 0.005253 | 5.573 | +1 | 8 |
| - | - | 4131 | 943.4 | - | - | 0 | - |
| - | - | 881.1 | 944.4 | - | - | 0 | - |
| - | - | 1099 | 968.4 | - | - | 0 | - |
| - | - | 3435 | 969.4 | - | - | 0 | - |
| - | - | 1487 | 970.4 | - | - | 0 | - |
| - | - | 1195 | 979.4 | - | - | 0 | - |
| - | - | 1717 | 980.4 | - | - | 0 | - |
| - | - | 716 | 985.5 | - | - | 0 | - |
| - | - | 2337 | 986.4 | - | - | 0 | - |
| - | - | 4677 | 987.4 | - | - | 0 | - |
| - | - | 2805 | 988.4 | - | - | 0 | - |
| - | - | 1061 | 994.4 | - | - | 0 | - |
| - | - | 1484 | 996.4 | - | - | 0 | - |
| - | - | 5774 | 997.4 | - | - | 0 | - |
| - | - | 3648 | 998.4 | - | - | 0 | - |
| - | - | 1370 | 999.4 | - | - | 0 | - |
| - | - | 4000 | 1004 | - | - | 0 | - |
| - | - | 1400 | 1005 | - | - | 0 | - |
| 9 | b | 971 | 1007 | 0.005141 | 5.102 | +1 | 9 |
| 9 | b | 803.2 | 1008 | 0.01206 | 11.96 | +1 | 9 |
| - | - | 2539 | 1012 | - | - | 0 | - |
| - | - | 952.1 | 1013 | - | - | 0 | - |
| - | - | 1957 | 1014 | - | - | 0 | - |
| - | - | 863 | 1015 | - | - | 0 | - |
| - | - | 1135 | 1022 | - | - | 0 | - |
| 9 | b | 3821 | 1026 | 0.004536 | 4.423 | +1 | 9 |
| - | - | 3165 | 1027 | - | - | 0 | - |
| - | - | 919.4 | 1028 | - | - | 0 | - |
| - | - | 1592 | 1039 | - | - | 0 | - |
| - | - | 1259 | 1040 | - | - | 0 | - |
| 6 | y | 5726 | 1057 | 0.004974 | 4.704 | +1 | 9 |
| - | - | 3991 | 1058 | - | - | 0 | - |
| - | - | 1185 | 1059 | - | - | 0 | - |
| - | - | 1218 | 1070 | - | - | 0 | - |
| - | - | 1406 | 1071 | - | - | 0 | - |
| - | - | 1051 | 1098 | - | - | 0 | - |
| - | - | 1545 | 1108 | - | - | 0 | - |
| - | - | 820.7 | 1109 | - | - | 0 | - |
| 10 | b | 1220 | 1124 | 0.02019 | 17.97 | +1 | 10 |
| - | - | 5020 | 1126 | - | - | 0 | - |
| - | - | 3655 | 1127 | - | - | 0 | - |
| - | - | 1013 | 1128 | - | - | 0 | - |
| 10 | b | 1956 | 1141 | 0.00914 | 8.014 | +1 | 10 |
| - | - | 1154 | 1142 | - | - | 0 | - |
| - | - | 1157 | 1149 | - | - | 0 | - |
| - | - | 1002 | 1150 | - | - | 0 | - |
| - | - | 876 | 1151 | - | - | 0 | - |
| 5 | y | 3590 | 1167 | 0.005257 | 4.503 | +1 | 10 |
| 5 | y | 8285 | 1168 | 0.0008695 | 0.7441 | +1 | 10 |
| - | - | 5245 | 1169 | - | - | 0 | - |
| - | - | 1905 | 1170 | - | - | 0 | - |
| - | - | 714.7 | 1176 | - | - | 0 | - |
| - | - | 1256 | 1185 | - | - | 0 | - |
| 5 | y | 1.284E+04 | 1186 | 0.007033 | 5.932 | +1 | 10 |
| - | - | 7658 | 1187 | - | - | 0 | - |
| - | - | 2205 | 1188 | - | - | 0 | - |
| - | - | 1069 | 1189 | - | - | 0 | - |
| 11 | b | 1338 | 1198 | 0.004359 | 3.64 | +1 | 11 |
| - | - | 866.3 | 1199 | - | - | 0 | - |
| - | - | 1277 | 1200 | - | - | 0 | - |
| - | - | 769.2 | 1216 | - | - | 0 | - |
| - | - | 780.4 | 3169 | - | - | 0 | - |

m/z Charge Intensity FragmentType MassShift Position
120.04472351074219 0 2500.3308
120.08106994628906 0 25788.758
121.08430480957031 0 1668.4723
122.07170104980469 0 1199.4453
123.04434204101562 0 1915.5227
127.05056762695312 0 1734.441
127.08678436279297 0 854.3582
127.12318420410156 0 793.78564
128.10194396972656 0 965.7861
128.10728454589844 0 19491.557
129.0659942626953 0 1397.8312
129.10255432128906 0 142556.92
130.05050659179688 0 683.30804
130.0608367919922 0 542.00244
130.06541442871094 0 3605.8047
130.1002197265625 0 915.9689
130.1059112548828 0 7750.774
131.049560546875 0 491.57245
131.06915283203125 0 448.5688
132.08087158203125 0 729.33826
132.10223388671875 0 601.56067
133.0610809326172 0 1483.4152
136.0759735107422 0 34204
137.0792236328125 0 2467.9915
138.06643676757812 0 2236.4885
138.09181213378906 0 1201.4031
139.0869598388672 0 1403.6274
141.1025390625 0 3403.1587
142.06158447265625 0 620.6851
143.04554748535156 0 555.54156
143.08135986328125 0 1205.334
145.06097412109375 0 1728.2029
146.06033325195312 0 35429.477
147.0445098876953 0 1363.9043
147.0635223388672 0 2812.7761
148.03956604003906 0 10715.102
149.0429229736328 0 994.27136
151.08680725097656 0 3185.6528
153.10260009765625 0 664.29474
155.08181762695312 0 808.3115
155.0929718017578 0 9726.21
155.11817932128906 0 8982.787 a Water loss 1
156.04458618164062 0 812.6702
156.0772247314453 0 599.3022
156.10275268554688 0 680.9355
156.1212921142578 0 1124.4785
157.0975799560547 0 1400.0046
157.134033203125 0 467.97742
158.06008911132812 0 818.9049
159.07691955566406 0 571.06573
159.09188842773438 0 4805.438
159.1129608154297 0 803.99554 d 1
160.095703125 0 473.42975
163.08692932128906 0 1127.3351
165.0548553466797 0 16494.578
165.07693481445312 0 500.88757
165.10252380371094 0 584.01074
166.06129455566406 0 23947.955
166.08651733398438 0 16246.371 y 10
166.09518432617188 0 866.57916
166.097900390625 0 924.16156
167.0647735595703 0 1168.5011
167.0821533203125 0 665.073
167.09017944335938 0 2007.5533
167.1184539794922 0 497.84592
168.10211181640625 0 2233.6777
168.11331176757812 0 1529.3267
169.06118774414062 0 582.40094
169.0764617919922 0 700.58276
169.09739685058594 0 4752.433
169.13333129882812 0 555.58606
170.10145568847656 0 478.15317
171.076904296875 0 1331.0043
172.07188415527344 0 1294.1659
173.0561065673828 0 1494.5758
173.07130432128906 0 3647.901
173.12879943847656 0 313643.22 a 1
174.0552215576172 0 59765.88
174.0733184814453 0 617.55493
174.13209533691406 0 27771.049
175.05857849121094 0 6430.858
175.0870361328125 0 642.8988
175.1346435546875 0 1641.1553
177.10311889648438 0 724.87775
178.1340789794922 0 10851.53
179.1374053955078 0 1057.2416
181.06080627441406 0 1613.3136
182.08143615722656 0 31862.387 y 13
183.08502197265625 0 2173.6616
183.11305236816406 0 5859.482 b Water loss 1
183.1495361328125 0 1207.9175
184.03948974609375 0 754.8985
184.11639404296875 0 664.80646
185.07110595703125 0 778.5336
185.09207153320312 0 649.41504
185.16526794433594 0 2136.3838
186.1239471435547 0 47428.957
187.1077880859375 0 532.91046
187.1273651123047 0 3767.0288
187.14407348632812 0 1363.1525
188.07080078125 0 3528.9124
189.0658721923828 0 1654.3152
190.08242797851562 0 1278.0978
190.13409423828125 0 1276.8291
191.08168029785156 0 4721.1465
193.0966796875 0 932.0968
194.09210205078125 0 1478.8638
195.11305236816406 0 11041.337
196.10824584960938 0 4001.6543
196.11697387695312 0 1254.9127
197.128662109375 0 3137.5781
198.08755493164062 0 1111.5332
198.1230926513672 0 686.8226
199.07159423828125 0 940.23724
201.06617736816406 0 713.0983
201.12356567382812 0 74068.19 b 1
202.05007934570312 0 18525.879
202.0860137939453 0 891.3163
202.12684631347656 0 7395.862
203.0533905029297 0 2246.3816
203.45236206054688 0 545.1153
205.09727478027344 0 6039.689
206.12911987304688 0 1141.4407
207.16053771972656 0 1590.6558
208.10800170898438 0 877.6919
209.09243774414062 0 994.1131
209.12908935546875 0 530.53345
211.1080780029297 0 956.10645 y Water loss 11
211.14402770996094 0 874.1252
212.13929748535156 0 1444.5486
213.12367248535156 0 5024.7515
213.16026306152344 0 712.81323
215.1390380859375 0 943.1336
216.06552124023438 0 521.0117
216.09857177734375 0 807.9403
216.11256408691406 0 683.0551
216.13409423828125 0 1025.8566
217.08245849609375 0 618.1135
217.53384399414062 0 510.10788
219.07647705078125 0 655.8759
220.10804748535156 0 707.0807
221.0924530029297 0 752.6203
221.103515625 0 1853.4425
221.12899780273438 0 848.9506
222.12387084960938 0 1823.2904
223.1078643798828 0 3793.3813
223.15557861328125 0 21925.918
224.13983154296875 0 1245.5791
224.15887451171875 0 2379.1611
225.1236572265625 0 1607.2539
226.0824737548828 0 3914.671
226.11888122558594 0 1473.8313 b Water loss 5
226.1558837890625 0 833.7314
227.06658935546875 0 5633.45
229.1185302734375 0 3365.3228
231.74444580078125 0 526.5732
233.13973999023438 0 3591.771
233.1647491455078 0 2438.3716
234.1239471435547 0 28387.693
235.12730407714844 0 3045.856
235.14315795898438 0 973.4359
235.15553283691406 0 617.72687
237.0870819091797 0 1938.0342
237.13446044921875 0 1054.8057
238.11863708496094 0 1218.0037
238.1304168701172 0 1062.1704
239.10182189941406 0 649.9888
239.15121459960938 0 691.8303
240.1344451904297 0 8243.14
241.11952209472656 0 641.8849
241.1341094970703 0 6351.6577
241.1914520263672 0 1038.9923
242.1369171142578 0 1220.7704
242.1501007080078 0 1303.0043
243.10894775390625 0 5409.869
243.34934997558594 0 503.138
244.09300231933594 0 5935.629
244.10755920410156 0 606.08997
245.09634399414062 0 924.8795
247.10791015625 0 13616.303
247.14453125 0 573.0038
248.11337280273438 0 3378.3528
249.09848022460938 0 6889.0396
249.13507080078125 0 1350.7657
250.10247802734375 0 938.87274
250.11813354492188 0 674.74194
251.15049743652344 0 89718.14
252.10858154296875 0 1003.32
252.13426208496094 0 982.9049
252.1536865234375 0 11450.452
253.15577697753906 0 781.5723
255.1457061767578 0 1497.9365
256.10882568359375 0 600.1824
256.6111755371094 0 640.5213
257.113037109375 0 1634.4429
258.1448059082031 0 3269.2815
261.11968994140625 0 3764.4136
261.134033203125 0 1040.5459
261.1588134765625 0 758.1665
262.1192321777344 0 3043.2454
265.12945556640625 0 5140.0674 y Water loss 7
266.1249694824219 0 18232.94
267.12725830078125 0 2325.1404
268.1300048828125 0 723.2753
268.1770324707031 0 1072.9249
269.1131286621094 0 1012.62067
269.1285705566406 0 6924.593
269.160888671875 0 926.6623
269.1854248046875 0 1097.4067
270.1308288574219 0 1228.2401 b Ammonia loss 6
270.14532470703125 0 1215.7637
272.1233215332031 0 653.5288
273.1340026855469 0 2487.345
274.1300354003906 0 1690.2461
275.1024169921875 0 1278.3173
276.108642578125 0 859.1118
279.1466979980469 0 1019.3401
280.1291809082031 0 2131.8118
282.15667724609375 0 877.3105
283.1402587890625 0 35663.402
284.1036682128906 0 990.39716
284.1249694824219 0 572.527
284.1430969238281 0 4188.6895
286.13983154296875 0 3494.6772
287.1391296386719 0 3602.526
287.169921875 0 1278.4209
290.1661376953125 0 918.5081 b 4
293.1133728027344 0 37552.344 y Water loss 12
294.1172180175781 0 6502.0537
295.1192321777344 0 604.47644
296.1401672363281 0 571.17975
296.19769287109375 0 601.43384 b Water loss 2
297.1257019042969 0 714.7641
297.156005859375 0 8688.806
298.1407775878906 0 645.1093
298.15838623046875 0 1218.2686
300.1675109863281 0 683.1304
301.1317443847656 0 1011.15424
301.1509094238281 0 18636.275
301.17071533203125 0 1007.5228
302.1541748046875 0 2703.9473
304.165771484375 0 3169.2324
305.1671447753906 0 675.633
306.1089782714844 0 1420.6587
306.1377258300781 0 3605.3003 y 9
306.63818359375 0 905.2541
311.12451171875 0 5447.334 y 12
312.1274719238281 0 969.0136
314.20745849609375 0 2297.5796 b 2
315.11273193359375 0 633.23193
315.1341552734375 0 4225.704
315.16656494140625 0 16135.793
316.1352233886719 0 673.0473
316.17059326171875 0 2304.9468
321.08154296875 0 598.9541
323.2076721191406 0 628.1231
325.1504211425781 0 1114.2168
332.1611022949219 0 1101.8093
332.2085266113281 0 1670.448
333.17657470703125 0 2250.3022
334.1036682128906 0 5117.4014
335.1069030761719 0 995.01465
338.6757507324219 0 715.37524 b Water loss 5
339.14703369140625 0 689.14154
342.4300231933594 0 690.69104
343.1278076171875 0 645.4692
344.171142578125 0 1247.2092
345.1314697265625 0 2029.0314
346.1150817871094 0 3826.9548
347.11578369140625 0 679.8242
347.6794128417969 0 1848.5787 b 5
350.2187805175781 0 6285.2446
351.2215576171875 0 1174.4636
353.67388916015625 0 941.8088 y Water loss 8
354.1736145019531 0 680.5888
356.1922607421875 0 1535.424
356.6545104980469 0 651.5032
361.1983642578125 0 1973.0046
362.1831359863281 0 1651.912
362.67913818359375 0 4146.413 y 8
363.1409606933594 0 10975.679
363.183349609375 0 1439.0475
363.68157958984375 0 717.6796
364.1447448730469 0 1675.8041
366.1761779785156 0 817.0913
367.24468994140625 0 1209.4991
367.9445495605469 0 604.2348
376.16021728515625 0 669.55164
378.2132568359375 0 2769.5906
379.20904541015625 0 20453.084
379.2402648925781 0 990.7977
380.16790771484375 0 1220.2024
380.2112121582031 0 3331.7876
381.1518859863281 0 15579.294
382.1539306640625 0 1902.7498
384.15655517578125 0 784.7284
384.1866455078125 0 1550.0399
385.1709899902344 0 1867.0421
392.19512939453125 0 962.0584
392.6955261230469 0 850.4306
394.1724853515625 0 3797.9722
395.15576171875 0 1109.2955
396.2245788574219 0 5034.2334
397.2289123535156 0 779.17865
398.17822265625 0 11535.949
399.1317443847656 0 927.4934
399.1814270019531 0 1744.419
402.19818115234375 0 2434.9478
403.19757080078125 0 775.1344
405.2622985839844 0 1196.7839
406.1887512207031 0 913.46185 y 7
406.21466064453125 0 681.22565
407.239990234375 0 788.63885
408.1631164550781 0 921.30817
409.14697265625 0 1203.0361
411.1649169921875 0 711.7166
412.15179443359375 0 983.9922
412.1829528808594 0 14029.224
413.1871337890625 0 2275.1843
414.2349853515625 0 3027.3904
415.2373046875 0 1234.5201
416.156494140625 0 2202.7114
417.14080810546875 0 5301.084
418.14617919921875 0 942.6603
421.21270751953125 0 719.94086 y Water loss 11
423.2715759277344 0 7678.4214
424.2739562988281 0 1414.8873
427.1248779296875 0 907.0717
429.17913818359375 0 794.29736
429.21044921875 0 1317.6003
430.19342041015625 0 38585.67
431.1964111328125 0 8232.181
432.1972961425781 0 807.82733
433.2061462402344 0 627.47266
433.2559509277344 0 707.7581 b Water loss 3
433.7112731933594 0 2694.5432
434.16717529296875 0 3169.5327
434.2081298828125 0 1168.7955
439.21881103515625 0 11490.802 y 11
439.6947937011719 0 727.0653
440.22222900390625 0 2725.1492
441.2239990234375 0 882.50256
442.22027587890625 0 638.30286
442.7168884277344 0 8423.416
443.21771240234375 0 2242.4412
443.72088623046875 0 1439.3553
444.1512451171875 0 3766.739
445.1357116699219 0 9927.639
446.13836669921875 0 2456.0537
447.1874084472656 0 2108.2175
447.70977783203125 0 5116.3066 b Water loss 7
448.2055969238281 0 4587.5996 b Ammonia loss 7
448.7371826171875 0 811.59216
449.2066345214844 0 1219.0392
450.23583984375 0 821.2631
451.2356262207031 0 734.9353
451.2665100097656 0 13045.007 b 3
452.26947021484375 0 2563.2036
454.1963806152344 0 1957.3789
454.6980285644531 0 1109.1229
456.7142333984375 0 23167.771 b 7
457.2162780761719 0 11153.366
457.55889892578125 0 998.37744
457.7175598144531 0 4372.2085
458.22015380859375 0 893.5959
459.2001037597656 0 683.839
461.2024841308594 0 1214.8276
462.16192626953125 0 7908.4087
463.1660461425781 0 2468.7441
466.2422790527344 0 628.2872
468.24737548828125 0 583.00977
468.2923583984375 0 4035.6138
469.29229736328125 0 980.63214
474.77978515625 0 990.8537
475.2397155761719 0 1445.6526
475.910888671875 0 910.51917
476.2265930175781 0 1255.5759
477.2088928222656 0 1411.5994
477.7518310546875 0 966.2879
478.2288818359375 0 3894.6926 y Water loss 10
478.27520751953125 0 1149.2834
480.214111328125 0 754.70154
481.24981689453125 0 1492.4614
481.7413635253906 0 633.51886
490.2535705566406 0 2524.6892
490.7509765625 0 2093.8489
491.2500305175781 0 1127.6031
492.8431701660156 0 953.4538
493.2524719238281 0 835.98047
493.84283447265625 0 686.3272
494.2358093261719 0 18497.508
495.2411193847656 0 6703.377
495.74273681640625 0 1523.7266
496.2404479980469 0 47564.99 y 10
497.24310302734375 0 13237.593
497.725341796875 0 1073.4795
498.24578857421875 0 1974.9268
499.2583923339844 0 7119.931
499.7611999511719 0 4174.4395
500.26123046875 0 928.1781
504.2515563964844 0 1698.9465 b Water loss 8
504.7514953613281 0 1173.7006 b Ammonia loss 8
506.2698974609375 0 879.7344
511.2623596191406 0 7096.1636
512.2117919921875 0 649.70593
512.2639770507812 0 1789.8171
512.577880859375 0 671.2225
513.257080078125 0 3467.8904 b 8
513.7603149414062 0 1961.779
525.2664794921875 0 2325.639
526.2659912109375 0 1037.0707
529.2396240234375 0 1504.8282
530.2252197265625 0 2528.4148
530.5883178710938 0 725.6584
533.58251953125 0 824.9902
533.923583984375 0 1770.9993
534.2591552734375 0 815.08997
534.3038330078125 0 3047.2869
534.5899658203125 0 1169.6617
535.30517578125 0 829.37506
536.1883544921875 0 1579.3605
539.9228515625 0 2021.2876 Precursor Water loss
540.2576904296875 0 2272.4448 Precursor Ammonia loss
540.5908203125 0 1635.6455
542.24853515625 0 686.22577
543.2772827148438 0 8113.186
544.2803955078125 0 2270.5256
544.7265625 0 1019.863
545.1837768554688 0 990.34467
545.750732421875 0 687.25073
545.92626953125 0 5595.0015 Precursor
546.2606811523438 0 4631.066
546.5940551757812 0 2238.9421
546.8159790039062 0 2693.324
547.0664672851562 0 3061.1338
547.2522583007812 0 1231.9125
547.3146362304688 0 2048.6985
548.7374267578125 0 784.13086
553.7335815429688 0 1605.9165
554.2639770507812 0 3241.6196
554.7664184570312 0 1495.4424
557.239013671875 0 1427.2163
558.2210083007812 0 1561.6848
561.314453125 0 2181.6157 b Water loss 4
562.24365234375 0 4484.373
562.3179321289062 0 611.95667
562.7437133789062 0 2177.4211
563.2639770507812 0 954.14905
563.7714233398438 0 1721.8568
564.1835327148438 0 1971.4683
565.2603759765625 0 1697.4202
570.7720336914062 0 1440.7256 b 9
571.22998046875 0 3172.939
571.247802734375 0 2775.093
571.7526245117188 0 2740.8086
572.224609375 0 887.5442
574.2605590820312 0 822.6332
575.2459716796875 0 3814.781
576.2421264648438 0 920.83844
579.324462890625 0 9328.959 b 4
580.2555541992188 0 1066.8425
580.328369140625 0 2138.7576
580.7581787109375 0 873.3001
581.212646484375 0 1441.2804
581.3294677734375 0 891.7267
582.2015380859375 0 818.44556
584.25146484375 0 1091.4062 y Water loss 4
584.7533569335938 0 901.2509 y Ammonia loss 4
584.8027954101562 0 1225.4663
593.256103515625 0 10149.773 y Water loss 9
593.3047485351562 0 1743.9124
593.7576904296875 0 1185.367
594.2568969726562 0 2412.111
597.3296508789062 0 725.1671
597.8338012695312 0 765.807
599.22021484375 0 10608.19
600.222412109375 0 3066.8184
600.30029296875 0 812.25867
601.2234497070312 0 923.0289
604.7882690429688 0 746.3887
609.7786865234375 0 748.73395
610.2805786132812 0 5964.83
610.3322143554688 0 1815.483
610.7733154296875 0 1380.6342
611.2669067382812 0 85188.86 y 9
612.2698364257812 0 27369.652
613.273193359375 0 5849.844
613.7943725585938 0 2695.7114
614.2915649414062 0 1934.8015
618.785400390625 0 6051.808
619.2855834960938 0 4700.8047
619.7882080078125 0 2444.2908
620.2907104492188 0 885.7551
621.2540283203125 0 1628.0076
627.2110595703125 0 727.44806
627.791015625 0 17369.785
628.2355346679688 0 1025.6685
628.2922973632812 0 12281.197
628.794921875 0 5589.3735
629.292236328125 0 1797.148
636.796630859375 0 1028.0507
637.3009643554688 0 1429.3506
644.2708740234375 0 1923.0803
644.7683715820312 0 1254.2577
644.9624633789062 0 768.39764
645.3145751953125 0 1044.4749
649.2706909179688 0 1748.3892
649.3319702148438 0 1884.9952
652.7802734375 0 3530.1262 y Water loss 3
653.2837524414062 0 2609.8188 y Ammonia loss 3
654.3173217773438 0 1536.1216 b Water loss 11
654.8211669921875 0 1391.3921 b Ammonia loss 11
655.3203735351562 0 822.2635
659.3140258789062 0 1713.6344
660.3201293945312 0 763.0533
661.295166015625 0 1716.7657
661.7861938476562 0 18062.91 y 3
662.2868041992188 0 17941.064
662.7885131835938 0 5914.061
663.2857666015625 0 1422.6229
666.2933349609375 0 1098.8953
666.7788696289062 0 962.4641
666.8349609375 0 1726.4519
667.2838134765625 0 1528.0676
668.313720703125 0 1126.8008
668.8164672851562 0 1206.0001
669.3106689453125 0 1159.754
672.2623901367188 0 3158.6838
672.3233642578125 0 791.703
673.2569580078125 0 2032.5676
673.3275756835938 0 712.7274
676.3389892578125 0 3502.9885 b Water loss 5
676.823974609375 0 3620.7588
677.2635498046875 0 964.62103
677.3264770507812 0 6354.965 b Ammonia loss 5
677.825927734375 0 2497.9346
678.3358764648438 0 1646.0309
684.3096923828125 0 14773.202
685.3253784179688 0 10269.415
685.8344116210938 0 5348.0386
686.330810546875 0 4842.584
686.8316650390625 0 2317.9873
687.3347778320312 0 875.1676
687.8646240234375 0 2428.0493
688.366455078125 0 1815.7795
688.8675537109375 0 1100.7224
689.8457641601562 0 2085.2273
690.2725830078125 0 3141.657
690.3326416015625 0 788.2282
691.2767333984375 0 1164.6638
691.8043823242188 0 1087.7794
693.3677368164062 0 3321.006
694.2927856445312 0 2452.7617
694.3524780273438 0 19431.809 b 5
695.2833862304688 0 1745.8906
695.3521728515625 0 6920.496
695.8275756835938 0 1666.0739
696.3541870117188 0 1657.1218
700.3171997070312 0 2824.013
700.8104248046875 0 4188.887
701.3118896484375 0 3109.3892
701.8153686523438 0 1186.9417
703.3533325195312 0 1029.3286
704.3309936523438 0 2561.748
704.8383178710938 0 2372.649
705.3408203125 0 1273.324
705.8391723632812 0 755.7698
706.3406982421875 0 3042.0251 y Water loss 8
707.3388671875 0 1602.0787
708.8206176757812 0 943.81616
709.3224487304688 0 14252.451 y Water loss 2
709.821533203125 0 18387.656 y Ammonia loss 2
710.3219604492188 0 11587.235
710.8234252929688 0 2807.2756
711.3777465820312 0 26320.932
712.3043212890625 0 26148.45
712.3790893554688 0 8791.08
713.3068237304688 0 10874.658
713.3763427734375 0 2104.6802
713.8479614257812 0 974.0247
714.3089599609375 0 1623.357
717.3312377929688 0 972.30237
717.8346557617188 0 6527.6167
718.3274536132812 0 114724.82 y 2
718.8294067382812 0 94937.766
719.3306274414062 0 44367.1 b Ammonia loss 12
719.8328857421875 0 13409.132
720.3345947265625 0 3069.934
720.87109375 0 2184.793
721.3681030273438 0 2522.4243
721.8709106445312 0 901.2752
723.3702392578125 0 924.4709
724.3504638671875 0 22236.371 y 8
725.35302734375 0 9067.141
725.862060546875 0 1295.189
726.3571166992188 0 5076.9272
726.8601684570312 0 1902.7826
727.358154296875 0 2760.1875
727.849365234375 0 9042.815 b 12
728.350830078125 0 7065.24
728.8497924804688 0 2529.8113
729.3335571289062 0 786.89246
734.3370971679688 0 685.6368
734.866455078125 0 5196.9736
735.3678588867188 0 5704.684
735.8704833984375 0 2637.2288
736.3644409179688 0 825.0558
736.8637084960938 0 2354.8652
737.3953247070312 0 1612.2439
740.2960815429688 0 897.4629
744.3211669921875 0 2279.888
751.3384399414062 0 790.5405
758.8561401367188 0 2015.8104 y Water loss 1
759.3533935546875 0 2093.6934 y Ammonia loss 1
761.3488159179688 0 1552.8378
762.3442993164062 0 1187.5245
767.8611450195312 0 11479.813 y 1
768.3626708984375 0 8908.016
768.8642578125 0 4678.9385
777.4210205078125 0 840.64984
779.37841796875 0 1248.9607
780.3796997070312 0 925.4984
782.3115234375 0 1157.2667
783.3753051757812 0 2176.5308
784.3779907226562 0 1565.7386
792.29443359375 0 1319.5477
793.2935791015625 0 764.943
797.3934936523438 0 3794.4856
798.3992919921875 0 2450.7815
799.3335571289062 0 990.27356
807.3675537109375 0 811.0237 b Water loss 6
811.3743896484375 0 3195.0244 y 7
812.3755493164062 0 1933.5642
813.359130859375 0 953.9803
819.396728515625 0 714.65216
825.3885498046875 0 7959.767 b 6
826.3904418945312 0 4226.2993
827.3302612304688 0 6290.8545
828.3346557617188 0 3242.7231
829.3324584960938 0 739.3746
837.8761596679688 0 704.426
839.3692626953125 0 2279.99
840.3672485351562 0 1743.9728
841.3612060546875 0 868.1104
857.3847045898438 0 2030.1229
858.3675537109375 0 2511.163
859.3729858398438 0 950.0999
860.373046875 0 708.3819
866.4182739257812 0 1226.0995
867.4157104492188 0 930.5647
875.3878173828125 0 1208.2605
876.3770141601562 0 2964.8213
877.3797607421875 0 2433.6416
878.3851318359375 0 862.29456
879.3914184570312 0 948.2541
884.3482666015625 0 2154.6084
884.4262084960938 0 5428.691
885.3451538085938 0 1507.0463
885.42724609375 0 3513.09
886.4342041015625 0 878.8059
894.4089965820312 0 6323.5176 b Water loss 7
895.4027709960938 0 5949.08 b Ammonia loss 7
896.39501953125 0 1403.6924
897.4054565429688 0 1001.9787
901.382568359375 0 1128.6781
905.3760986328125 0 906.4182
912.4191284179688 0 52145.3 b 7
913.421630859375 0 25889.387
914.4246826171875 0 8722.123
915.4267578125 0 1352.6343
922.4039306640625 0 2112.2842
923.3959350585938 0 2213.8787
924.394287109375 0 1455.5948
925.3952026367188 0 2975.0398 y Ammonia loss 6
926.3857421875 0 1083.4174
940.414306640625 0 15498.669
941.4166870117188 0 8453.198
942.41845703125 0 6258.758 y 6
943.4222412109375 0 4131.151
944.42919921875 0 881.07043
968.41064453125 0 1098.7734
969.400634765625 0 3434.73
970.39990234375 0 1487.3077
979.4219970703125 0 1195.2478
980.4168090820312 0 1717.1416
985.4896850585938 0 715.9802
986.420166015625 0 2337.1826
987.4076538085938 0 4677.2466
988.4086303710938 0 2804.703
994.4404296875 0 1060.8469
996.44921875 0 1483.8054
997.4363403320312 0 5774.1606
998.439208984375 0 3647.7856
999.4368286132812 0 1370.4438
1004.4309692382812 0 4000.3025
1005.4284057617188 0 1400.2112
1007.4927368164062 0 970.9969 b Water loss 8
1008.4939575195312 0 803.2169 b Ammonia loss 8
1012.4464721679688 0 2539.2703
1013.4601440429688 0 952.1118
1014.45703125 0 1957.0842
1015.470703125 0 862.9594
1022.4459228515625 0 1135.122
1025.50390625 0 3821.1038 b 8
1026.5057373046875 0 3165.417
1027.5084228515625 0 919.44635
1039.462158203125 0 1591.9897
1040.469970703125 0 1259.3798
1057.4456787109375 0 5725.977 y 5
1058.4486083984375 0 3991.15
1059.452392578125 0 1184.6493
1069.533203125 0 1217.8805
1070.5361328125 0 1405.5348
1097.5205078125 0 1050.7422
1107.5162353515625 0 1544.6072
1108.519287109375 0 820.74493
1123.4886474609375 0 1219.7341 b Ammonia loss 9
1125.5283203125 0 5020.49
1126.5323486328125 0 3654.5918
1127.5279541015625 0 1013.27795
1140.5262451171875 0 1956.0625 b 9
1141.5244140625 0 1153.8303
1149.4853515625 0 1157.3611
1150.4639892578125 0 1001.89307
1151.485595703125 0 876.0148
1167.493408203125 0 3590.3716 y Water loss 4
1168.4818115234375 0 8285.112 y Ammonia loss 4
1169.4847412109375 0 5245.204
1170.4857177734375 0 1905.064
1176.225830078125 0 714.6525
1184.515380859375 0 1255.9246
1185.502197265625 0 12835.044 y 4
1186.50537109375 0 7658.123
1187.508544921875 0 2204.7505
1188.5146484375 0 1068.7098
1197.552490234375 0 1337.5118 b 10
1198.54931640625 0 866.2554
1199.5819091796875 0 1276.9744
1215.5943603515625 0 769.16266
3168.72900390625 0 780.3617

Spectrum Details

|  |  |
| --- | --- |
| Matched peaks? Matched peaks The total absolute number of peaks matched. Additionally in brackets the total fraction of peaks matched and the total number of peaks is shown. | 80 (10.81% of 740) |
| FDR? FDR The false discovery rate estimated for this peptide. It is calculated by matching all theoretical fragments with a non-integer shift with the raw peaks for this spectrum. This is done with 40 different shifts. The resulting percentage is the average number of annotated peaks over the number of annotated peaks with the correct spectrum. | 0.80% |
| Satellite FDR? Satellite FDR See the FDR for details on its calculation. This satellite ion specific FDR only contains the satellite ions (d/w) for I/L/J positions. | - |
| PSM Score? PSM Score The PSM Score as given by Hecklib to this annotated spectrum. It is shown with three significant figures. | 625 |

#### Spectrum 6141? Spectrum 6141 The raw spectrum of this peptide as annotated by Hecklib. The fragments are coloured according to ion type (see legend). Any peaks with a star '\*' as text can be hovered over to see the full details, first the ion type second the mass shift type. By hovering over the amino acids in the peptide or ions in the legend the corresponding peaks are highlighted. By toggling the 'Unassigned' label you can turn the background (unassigned) peaks on or off in the plot. By updating the slider in the Ion legend you can update the spectrum to only show the top X% of the peaks with labels. The top X% means any peak that is within X% of the highest intensity. By dragging in the spectrum you can zoom in to a specific part of the spectrum and use 'Zoom Out' to get back to the original zoom level. The annotation of the spectrum is based on the given sequence in the peptides file and is done with different software so inconsistencies are likely. The peaks are annotated based on the given sequence, with 20 ppm tolerance.

Copy Data

##### Spectrum 6141 (TSV)

###### Preview

```
Loading example...
```

*Click on the button to copy the data to your clipboard.*

Mz MinMz MaxIntensity Max

WidthHeightPeptide font sizePeptide stroke widthSpectrum font sizeSpectrum stroke widthCompact peptide

Ion legend

wxyz

abcd

OtherUnassignedIon^Charge^_Position_Show for top:%

TVJHQDMSJDGKEY

02.96e+45.93e+48.89e+41.19e+5

Zoom Out

y^+3^_3_a^+1^_2_d^+1^_2_y^+3^_4_a^+1^_2_y^+1^_1_b^+1^_2_b^+1^_2_b^+3^_6_y^+3^_7_b^+3^_7_y^+1^_2_b^+1^_3_b^+3^_8_y^+2^_5_y^+1^_2_b^+1^_3_y^+2^_6_b^+1^_4_y^+1^_3_b^+2^_8_b^+2^_8_b^+1^_4_b^+2^_8_y^+1^_4_y^+1^_4_b^+2^_9_b^+2^_9_\*\*\*b^+1^_5_b^+2^_10_b^+1^_5_y^+2^_10_y^+1^_5_y^+1^_5_y^+2^_11_y^+2^_11_b^+2^_12_y^+2^_11_b^+1^_6_b^+1^_6_b^+1^_6_y^+1^_6_y^+2^_12_y^+2^_12_y^+2^_12_b^+2^_13_y^+1^_6_b^+2^_13_y^+2^_13_y^+1^_7_b^+1^_7_b^+1^_8_b^+1^_8_b^+1^_8_y^+1^_8_y^+1^_8_b^+1^_9_y^+1^_9_b^+1^_10_y^+1^_10_y^+1^_10_y^+1^_10_

0778155723353114

Fragment Matches Table

Show background peaks

| Position | Ion type | Intensity | mz Theoretical | mz Error (Th) | mz Error (ppm) | Charge | Series Number |
| --- | --- | --- | --- | --- | --- | --- | --- |
| - | - | 691.2 | 120 | - | - | 0 | - |
| - | - | 1.709E+04 | 120.1 | - | - | 0 | - |
| - | - | 1024 | 120.1 | - | - | 0 | - |
| - | - | 1023 | 121.1 | - | - | 0 | - |
| - | - | 521.1 | 122.1 | - | - | 0 | - |
| - | - | 720.6 | 123 | - | - | 0 | - |
| - | - | 413.5 | 126.1 | - | - | 0 | - |
| - | - | 747.3 | 127.1 | - | - | 0 | - |
| - | - | 409.4 | 127.1 | - | - | 0 | - |
| - | - | 7684 | 128.1 | - | - | 0 | - |
| - | - | 779.4 | 129.1 | - | - | 0 | - |
| - | - | 5.51E+04 | 129.1 | - | - | 0 | - |
| - | - | 1124 | 130.1 | - | - | 0 | - |
| - | - | 547.5 | 130.1 | - | - | 0 | - |
| - | - | 3314 | 130.1 | - | - | 0 | - |
| - | - | 504.5 | 132.1 | - | - | 0 | - |
| - | - | 1359 | 132.1 | - | - | 0 | - |
| - | - | 1.475E+04 | 136.1 | - | - | 0 | - |
| - | - | 906.3 | 137.1 | - | - | 0 | - |
| - | - | 957.5 | 138.1 | - | - | 0 | - |
| - | - | 625.9 | 139.1 | - | - | 0 | - |
| - | - | 2751 | 141.1 | - | - | 0 | - |
| - | - | 641.7 | 143.1 | - | - | 0 | - |
| - | - | 399.5 | 144.1 | - | - | 0 | - |
| - | - | 532.1 | 145.1 | - | - | 0 | - |
| - | - | 1.2E+04 | 146.1 | - | - | 0 | - |
| - | - | 1521 | 147.1 | - | - | 0 | - |
| 12 | y | 447.4 | 147.1 | 0.0009163 | 6.23 | +3 | 3 |
| - | - | 3252 | 148 | - | - | 0 | - |
| - | - | 472.8 | 148.9 | - | - | 0 | - |
| - | - | 1532 | 151.1 | - | - | 0 | - |
| - | - | 672 | 155.1 | - | - | 0 | - |
| - | - | 4099 | 155.1 | - | - | 0 | - |
| 2 | a | 3010 | 155.1 | 0.0002745 | 1.77 | +1 | 2 |
| - | - | 434.1 | 156 | - | - | 0 | - |
| - | - | 536 | 157.1 | - | - | 0 | - |
| - | - | 697.5 | 157.1 | - | - | 0 | - |
| - | - | 936.1 | 157.1 | - | - | 0 | - |
| - | - | 1830 | 159.1 | - | - | 0 | - |
| 2 | d | 760.5 | 159.1 | 8.036E-05 | 0.5051 | +1 | 2 |
| - | - | 6809 | 165.1 | - | - | 0 | - |
| - | - | 6709 | 166.1 | - | - | 0 | - |
| 11 | y | 7698 | 166.1 | 0.001557 | 9.377 | +3 | 4 |
| - | - | 549.9 | 167.1 | - | - | 0 | - |
| - | - | 792.8 | 167.1 | - | - | 0 | - |
| - | - | 2281 | 169.1 | - | - | 0 | - |
| - | - | 742.1 | 170.1 | - | - | 0 | - |
| - | - | 7487 | 171.1 | - | - | 0 | - |
| - | - | 1034 | 172.1 | - | - | 0 | - |
| - | - | 682.2 | 172.2 | - | - | 0 | - |
| - | - | 620.1 | 173.1 | - | - | 0 | - |
| - | - | 2342 | 173.1 | - | - | 0 | - |
| 2 | a | 1.173E+05 | 173.1 | 0.0002689 | 1.553 | +1 | 2 |
| - | - | 1.972E+04 | 174.1 | - | - | 0 | - |
| - | - | 637.5 | 174.1 | - | - | 0 | - |
| - | - | 9080 | 174.1 | - | - | 0 | - |
| - | - | 2255 | 175.1 | - | - | 0 | - |
| - | - | 876.6 | 175.1 | - | - | 0 | - |
| - | - | 4045 | 178.1 | - | - | 0 | - |
| - | - | 889.4 | 181.1 | - | - | 0 | - |
| 14 | y | 1.485E+04 | 182.1 | 0.0002054 | 1.128 | +1 | 1 |
| - | - | 1402 | 183.1 | - | - | 0 | - |
| 2 | b | 2195 | 183.1 | 0.000355 | 1.939 | +1 | 2 |
| - | - | 1737 | 183.1 | - | - | 0 | - |
| - | - | 1410 | 185.1 | - | - | 0 | - |
| - | - | 1.541E+04 | 186.1 | - | - | 0 | - |
| - | - | 965 | 186.1 | - | - | 0 | - |
| - | - | 743.8 | 187.1 | - | - | 0 | - |
| - | - | 1572 | 187.1 | - | - | 0 | - |
| - | - | 2106 | 187.1 | - | - | 0 | - |
| - | - | 3341 | 188.1 | - | - | 0 | - |
| - | - | 1096 | 189.1 | - | - | 0 | - |
| - | - | 689.5 | 190.1 | - | - | 0 | - |
| - | - | 783.9 | 190.1 | - | - | 0 | - |
| - | - | 1573 | 191.1 | - | - | 0 | - |
| - | - | 552.1 | 193.1 | - | - | 0 | - |
| - | - | 4727 | 195.1 | - | - | 0 | - |
| - | - | 2113 | 196.1 | - | - | 0 | - |
| - | - | 654.9 | 197.1 | - | - | 0 | - |
| - | - | 941.2 | 198.1 | - | - | 0 | - |
| - | - | 5099 | 199.1 | - | - | 0 | - |
| - | - | 643.4 | 200.1 | - | - | 0 | - |
| - | - | 790.8 | 201.1 | - | - | 0 | - |
| 2 | b | 2.761E+04 | 201.1 | 0.0001511 | 0.7511 | +1 | 2 |
| - | - | 5939 | 202.1 | - | - | 0 | - |
| - | - | 2098 | 202.1 | - | - | 0 | - |
| - | - | 766.5 | 203.1 | - | - | 0 | - |
| - | - | 3056 | 205.1 | - | - | 0 | - |
| - | - | 614.6 | 206.1 | - | - | 0 | - |
| - | - | 779.9 | 207.2 | - | - | 0 | - |
| - | - | 514.9 | 208.1 | - | - | 0 | - |
| - | - | 1217 | 211.1 | - | - | 0 | - |
| - | - | 1271 | 212.1 | - | - | 0 | - |
| - | - | 1916 | 213.1 | - | - | 0 | - |
| - | - | 1452 | 215.1 | - | - | 0 | - |
| - | - | 683.3 | 216.1 | - | - | 0 | - |
| - | - | 679.9 | 221.1 | - | - | 0 | - |
| - | - | 893 | 221.1 | - | - | 0 | - |
| - | - | 2514 | 223.1 | - | - | 0 | - |
| - | - | 7711 | 223.2 | - | - | 0 | - |
| - | - | 577 | 224.1 | - | - | 0 | - |
| - | - | 859.8 | 224.2 | - | - | 0 | - |
| - | - | 1157 | 226.1 | - | - | 0 | - |
| 6 | b | 781.1 | 226.1 | 0.0002176 | 0.9625 | +3 | 6 |
| - | - | 1735 | 226.2 | - | - | 0 | - |
| - | - | 2089 | 227.1 | - | - | 0 | - |
| - | - | 577.6 | 227.1 | - | - | 0 | - |
| - | - | 1126 | 229.1 | - | - | 0 | - |
| - | - | 1629 | 229.2 | - | - | 0 | - |
| - | - | 763.8 | 230.2 | - | - | 0 | - |
| - | - | 1541 | 233.1 | - | - | 0 | - |
| - | - | 5330 | 233.2 | - | - | 0 | - |
| - | - | 9624 | 234.1 | - | - | 0 | - |
| - | - | 903.9 | 234.2 | - | - | 0 | - |
| - | - | 573.6 | 235.1 | - | - | 0 | - |
| - | - | 1262 | 235.1 | - | - | 0 | - |
| - | - | 576 | 237.1 | - | - | 0 | - |
| - | - | 666.4 | 238.1 | - | - | 0 | - |
| - | - | 3413 | 240.1 | - | - | 0 | - |
| - | - | 746.9 | 241.1 | - | - | 0 | - |
| - | - | 1386 | 241.1 | - | - | 0 | - |
| - | - | 1343 | 243.1 | - | - | 0 | - |
| - | - | 2932 | 244.1 | - | - | 0 | - |
| - | - | 492.6 | 244.1 | - | - | 0 | - |
| - | - | 531.4 | 244.1 | - | - | 0 | - |
| - | - | 5251 | 247.1 | - | - | 0 | - |
| - | - | 1423 | 248.1 | - | - | 0 | - |
| - | - | 2510 | 249.1 | - | - | 0 | - |
| - | - | 3.013E+04 | 251.2 | - | - | 0 | - |
| - | - | 4155 | 252.2 | - | - | 0 | - |
| - | - | 735.8 | 255.1 | - | - | 0 | - |
| - | - | 760.8 | 257.1 | - | - | 0 | - |
| - | - | 1084 | 258.1 | - | - | 0 | - |
| - | - | 652.4 | 259.1 | - | - | 0 | - |
| - | - | 712.2 | 260.2 | - | - | 0 | - |
| - | - | 1423 | 261.1 | - | - | 0 | - |
| - | - | 1400 | 261.2 | - | - | 0 | - |
| - | - | 1459 | 262.1 | - | - | 0 | - |
| 8 | y | 1594 | 265.1 | 0.000293 | 1.105 | +3 | 7 |
| - | - | 6786 | 266.1 | - | - | 0 | - |
| - | - | 1239 | 267.1 | - | - | 0 | - |
| - | - | 1152 | 269.1 | - | - | 0 | - |
| - | - | 2420 | 269.1 | - | - | 0 | - |
| 7 | b | 868.8 | 270.1 | 0.004411 | 16.33 | +3 | 7 |
| - | - | 675.7 | 279.1 | - | - | 0 | - |
| - | - | 567.3 | 279.5 | - | - | 0 | - |
| - | - | 565.1 | 280.1 | - | - | 0 | - |
| - | - | 628.7 | 282.2 | - | - | 0 | - |
| - | - | 1.389E+04 | 283.1 | - | - | 0 | - |
| - | - | 2035 | 284.1 | - | - | 0 | - |
| - | - | 972.9 | 286.1 | - | - | 0 | - |
| - | - | 551.3 | 286.2 | - | - | 0 | - |
| - | - | 1525 | 287.1 | - | - | 0 | - |
| - | - | 595 | 290.7 | - | - | 0 | - |
| 13 | y | 1.255E+04 | 293.1 | 8.316E-05 | 0.2837 | +1 | 2 |
| - | - | 2073 | 294.1 | - | - | 0 | - |
| - | - | 635.1 | 295.1 | - | - | 0 | - |
| 3 | b | 695.3 | 296.2 | 0.0003059 | 1.033 | +1 | 3 |
| - | - | 3757 | 297.2 | - | - | 0 | - |
| 8 | b | 569 | 299.1 | 0.001303 | 4.355 | +3 | 8 |
| - | - | 643.1 | 300.2 | - | - | 0 | - |
| - | - | 7314 | 301.2 | - | - | 0 | - |
| - | - | 1087 | 302.2 | - | - | 0 | - |
| - | - | 1602 | 304.2 | - | - | 0 | - |
| 10 | y | 1344 | 306.1 | 9.408E-05 | 0.3073 | +2 | 5 |
| - | - | 571.9 | 309.2 | - | - | 0 | - |
| 13 | y | 1566 | 311.1 | 0.0006269 | 2.015 | +1 | 2 |
| 3 | b | 715.3 | 314.2 | 0.0001477 | 0.4702 | +1 | 3 |
| - | - | 755.1 | 315.1 | - | - | 0 | - |
| - | - | 5179 | 315.2 | - | - | 0 | - |
| - | - | 720.1 | 316.2 | - | - | 0 | - |
| - | - | 630.6 | 321.7 | - | - | 0 | - |
| - | - | 720.2 | 323.2 | - | - | 0 | - |
| - | - | 988.8 | 332.2 | - | - | 0 | - |
| - | - | 1840 | 334.1 | - | - | 0 | - |
| - | - | 1612 | 340.2 | - | - | 0 | - |
| - | - | 879.2 | 345.1 | - | - | 0 | - |
| - | - | 1464 | 346.1 | - | - | 0 | - |
| - | - | 2817 | 350.2 | - | - | 0 | - |
| - | - | 671 | 351.2 | - | - | 0 | - |
| - | - | 730.2 | 362.2 | - | - | 0 | - |
| 9 | y | 1318 | 362.7 | 0.0002509 | 0.6918 | +2 | 6 |
| - | - | 3637 | 363.1 | - | - | 0 | - |
| - | - | 906.7 | 363.7 | - | - | 0 | - |
| - | - | 926.8 | 364.1 | - | - | 0 | - |
| - | - | 1462 | 378.2 | - | - | 0 | - |
| - | - | 6826 | 379.2 | - | - | 0 | - |
| - | - | 1676 | 380.2 | - | - | 0 | - |
| - | - | 5717 | 381.2 | - | - | 0 | - |
| - | - | 846.1 | 382.2 | - | - | 0 | - |
| - | - | 631.7 | 385.2 | - | - | 0 | - |
| - | - | 1021 | 394.2 | - | - | 0 | - |
| - | - | 1531 | 396.2 | - | - | 0 | - |
| - | - | 732.6 | 397.2 | - | - | 0 | - |
| - | - | 3367 | 398.2 | - | - | 0 | - |
| - | - | 660.2 | 399.2 | - | - | 0 | - |
| - | - | 856.5 | 402.2 | - | - | 0 | - |
| - | - | 671.7 | 406.7 | - | - | 0 | - |
| - | - | 768.5 | 409.1 | - | - | 0 | - |
| - | - | 697.8 | 411.2 | - | - | 0 | - |
| - | - | 5593 | 412.2 | - | - | 0 | - |
| - | - | 619.1 | 413.2 | - | - | 0 | - |
| - | - | 1441 | 414.2 | - | - | 0 | - |
| - | - | 986.5 | 416.2 | - | - | 0 | - |
| - | - | 1701 | 417.1 | - | - | 0 | - |
| - | - | 1743 | 420.2 | - | - | 0 | - |
| - | - | 667.2 | 420.7 | - | - | 0 | - |
| - | - | 3320 | 423.3 | - | - | 0 | - |
| - | - | 736.5 | 424.3 | - | - | 0 | - |
| - | - | 1.505E+04 | 429.2 | - | - | 0 | - |
| - | - | 3925 | 429.7 | - | - | 0 | - |
| - | - | 1.292E+04 | 430.2 | - | - | 0 | - |
| - | - | 1107 | 430.2 | - | - | 0 | - |
| - | - | 2712 | 431.2 | - | - | 0 | - |
| - | - | 811.7 | 432.2 | - | - | 0 | - |
| 4 | b | 861.7 | 433.3 | 0.0008056 | 1.859 | +1 | 4 |
| - | - | 633.6 | 433.7 | - | - | 0 | - |
| - | - | 760.6 | 434.2 | - | - | 0 | - |
| 12 | y | 4077 | 439.2 | 0.0003294 | 0.7499 | +1 | 3 |
| - | - | 2681 | 439.3 | - | - | 0 | - |
| - | - | 1271 | 439.8 | - | - | 0 | - |
| - | - | 1600 | 440.2 | - | - | 0 | - |
| - | - | 550 | 440.3 | - | - | 0 | - |
| - | - | 3246 | 442.7 | - | - | 0 | - |
| - | - | 1319 | 443.2 | - | - | 0 | - |
| - | - | 1197 | 444.2 | - | - | 0 | - |
| - | - | 3435 | 445.1 | - | - | 0 | - |
| - | - | 1029 | 446.1 | - | - | 0 | - |
| 8 | b | 1918 | 447.7 | 9.579E-05 | 0.214 | +2 | 8 |
| 8 | b | 2085 | 448.2 | 0.005424 | 12.1 | +2 | 8 |
| 4 | b | 5256 | 451.3 | 0.0001399 | 0.3099 | +1 | 4 |
| - | - | 1289 | 452.3 | - | - | 0 | - |
| 8 | b | 7176 | 456.7 | 0.001808 | 3.958 | +2 | 8 |
| - | - | 3800 | 457.2 | - | - | 0 | - |
| - | - | 743 | 457.7 | - | - | 0 | - |
| - | - | 899.7 | 460.2 | - | - | 0 | - |
| - | - | 3029 | 462.2 | - | - | 0 | - |
| - | - | 882.6 | 468.3 | - | - | 0 | - |
| - | - | 641.9 | 474.8 | - | - | 0 | - |
| - | - | 689.7 | 474.8 | - | - | 0 | - |
| 11 | y | 965.8 | 478.2 | 0.0003251 | 0.6798 | +1 | 4 |
| - | - | 818.5 | 478.3 | - | - | 0 | - |
| - | - | 1248 | 490.3 | - | - | 0 | - |
| - | - | 1075 | 490.8 | - | - | 0 | - |
| - | - | 2134 | 492.8 | - | - | 0 | - |
| - | - | 750.6 | 493.7 | - | - | 0 | - |
| - | - | 965.5 | 493.8 | - | - | 0 | - |
| - | - | 8120 | 494.2 | - | - | 0 | - |
| - | - | 1529 | 495.2 | - | - | 0 | - |
| - | - | 720.3 | 495.7 | - | - | 0 | - |
| 11 | y | 1.534E+04 | 496.2 | 4.486E-05 | 0.0904 | +1 | 4 |
| - | - | 4081 | 497.2 | - | - | 0 | - |
| - | - | 835.9 | 498.2 | - | - | 0 | - |
| - | - | 3019 | 499.3 | - | - | 0 | - |
| - | - | 1444 | 499.8 | - | - | 0 | - |
| 9 | b | 766.9 | 504.3 | 0.003065 | 6.079 | +2 | 9 |
| - | - | 584.2 | 505.9 | - | - | 0 | - |
| - | - | 1673 | 510.3 | - | - | 0 | - |
| - | - | 2363 | 511.3 | - | - | 0 | - |
| 9 | b | 1116 | 513.3 | 7.523E-05 | 0.1466 | +2 | 9 |
| - | - | 1435 | 513.8 | - | - | 0 | - |
| - | - | 1619 | 525.3 | - | - | 0 | - |
| - | - | 881.3 | 530.2 | - | - | 0 | - |
| - | - | 827.1 | 530.9 | - | - | 0 | - |
| - | - | 2985 | 532.3 | - | - | 0 | - |
| - | - | 734 | 533.3 | - | - | 0 | - |
| - | - | 1470 | 534.3 | - | - | 0 | - |
| - | - | 893.5 | 536.2 | - | - | 0 | - |
| - | - | 902 | 538.3 | - | - | 0 | - |
| 0 | Precursor | 707.9 | 539.9 | 0.002159 | 4 | +3 | -1 |
| 0 | Precursor | 1124 | 540.3 | 0.009252 | 17.13 | +3 | -1 |
| - | - | 1061 | 540.6 | - | - | 0 | - |
| - | - | 909.3 | 542.3 | - | - | 0 | - |
| - | - | 2681 | 543.3 | - | - | 0 | - |
| - | - | 1015 | 544.3 | - | - | 0 | - |
| - | - | 729.8 | 545.1 | - | - | 0 | - |
| 0 | Precursor | 1739 | 545.9 | 0.0007372 | 1.35 | +3 | -1 |
| - | - | 2649 | 546.3 | - | - | 0 | - |
| - | - | 1460 | 546.3 | - | - | 0 | - |
| - | - | 1949 | 546.6 | - | - | 0 | - |
| - | - | 1213 | 546.8 | - | - | 0 | - |
| - | - | 2619 | 547.1 | - | - | 0 | - |
| - | - | 1155 | 547.2 | - | - | 0 | - |
| - | - | 909.3 | 547.6 | - | - | 0 | - |
| - | - | 1660 | 553.8 | - | - | 0 | - |
| 5 | b | 850.1 | 561.3 | 0.0004618 | 0.8228 | +1 | 5 |
| - | - | 1449 | 562.2 | - | - | 0 | - |
| - | - | 1566 | 562.7 | - | - | 0 | - |
| - | - | 799.8 | 563.8 | - | - | 0 | - |
| 10 | b | 928.8 | 570.8 | 0.002959 | 5.185 | +2 | 10 |
| - | - | 1651 | 571.2 | - | - | 0 | - |
| - | - | 1408 | 571.7 | - | - | 0 | - |
| - | - | 716.4 | 573.3 | - | - | 0 | - |
| - | - | 970 | 575.2 | - | - | 0 | - |
| 5 | b | 2850 | 579.3 | 0.001985 | 3.427 | +1 | 5 |
| - | - | 667 | 580.3 | - | - | 0 | - |
| - | - | 1329 | 580.3 | - | - | 0 | - |
| - | - | 832.2 | 580.8 | - | - | 0 | - |
| - | - | 727.9 | 581.2 | - | - | 0 | - |
| 5 | y | 831.1 | 584.3 | 0.001201 | 2.055 | +2 | 10 |
| - | - | 843.1 | 588.3 | - | - | 0 | - |
| 10 | y | 4041 | 593.3 | 0.0009526 | 1.606 | +1 | 5 |
| - | - | 1580 | 593.3 | - | - | 0 | - |
| - | - | 1052 | 594.3 | - | - | 0 | - |
| - | - | 957.1 | 597.3 | - | - | 0 | - |
| - | - | 4399 | 599.2 | - | - | 0 | - |
| - | - | 866.6 | 600.2 | - | - | 0 | - |
| - | - | 5608 | 610.3 | - | - | 0 | - |
| - | - | 733.9 | 610.3 | - | - | 0 | - |
| 10 | y | 3.146E+04 | 611.3 | 0.0001037 | 0.1697 | +1 | 5 |
| - | - | 8682 | 612.3 | - | - | 0 | - |
| - | - | 923.7 | 613.1 | - | - | 0 | - |
| - | - | 2174 | 613.3 | - | - | 0 | - |
| - | - | 796.3 | 613.8 | - | - | 0 | - |
| - | - | 1139 | 618.8 | - | - | 0 | - |
| - | - | 998.1 | 619.3 | - | - | 0 | - |
| - | - | 620.5 | 619.8 | - | - | 0 | - |
| - | - | 1007 | 621.3 | - | - | 0 | - |
| - | - | 4771 | 627.8 | - | - | 0 | - |
| - | - | 3367 | 628.3 | - | - | 0 | - |
| - | - | 635.3 | 644.3 | - | - | 0 | - |
| 4 | y | 1491 | 652.8 | 0.0007496 | 1.148 | +2 | 11 |
| 4 | y | 1114 | 653.3 | 0.008341 | 12.77 | +2 | 11 |
| - | - | 678.6 | 653.8 | - | - | 0 | - |
| 12 | b | 801.9 | 654.3 | 0.0002873 | 0.439 | +2 | 12 |
| - | - | 1608 | 656.8 | - | - | 0 | - |
| - | - | 760.4 | 661.3 | - | - | 0 | - |
| 4 | y | 6962 | 661.8 | 0.001515 | 2.29 | +2 | 11 |
| - | - | 3381 | 662.3 | - | - | 0 | - |
| - | - | 2415 | 662.8 | - | - | 0 | - |
| - | - | 952.4 | 669.3 | - | - | 0 | - |
| - | - | 749.1 | 672.8 | - | - | 0 | - |
| 6 | b | 1343 | 676.3 | 0.0007854 | 1.161 | +1 | 6 |
| - | - | 695.5 | 677.3 | - | - | 0 | - |
| 6 | b | 2195 | 677.3 | 0.002748 | 4.057 | +1 | 6 |
| - | - | 861.1 | 678.3 | - | - | 0 | - |
| - | - | 1.134E+04 | 679.4 | - | - | 0 | - |
| - | - | 3083 | 680.4 | - | - | 0 | - |
| - | - | 785 | 683.3 | - | - | 0 | - |
| - | - | 4689 | 684.3 | - | - | 0 | - |
| - | - | 2396 | 685.3 | - | - | 0 | - |
| - | - | 1516 | 685.8 | - | - | 0 | - |
| - | - | 663.8 | 686.3 | - | - | 0 | - |
| - | - | 775.4 | 686.8 | - | - | 0 | - |
| - | - | 758.4 | 687.3 | - | - | 0 | - |
| - | - | 3071 | 687.9 | - | - | 0 | - |
| - | - | 2270 | 688.4 | - | - | 0 | - |
| - | - | 1017 | 688.9 | - | - | 0 | - |
| - | - | 908.1 | 689.4 | - | - | 0 | - |
| - | - | 676.8 | 690.3 | - | - | 0 | - |
| - | - | 645.8 | 692.4 | - | - | 0 | - |
| - | - | 951.5 | 693.4 | - | - | 0 | - |
| - | - | 1397 | 694.3 | - | - | 0 | - |
| 6 | b | 7292 | 694.4 | 0.0004297 | 0.6188 | +1 | 6 |
| - | - | 2159 | 695.4 | - | - | 0 | - |
| - | - | 766.4 | 695.8 | - | - | 0 | - |
| - | - | 904.6 | 700.3 | - | - | 0 | - |
| - | - | 916.3 | 700.8 | - | - | 0 | - |
| - | - | 1086 | 701.3 | - | - | 0 | - |
| - | - | 964.5 | 704.8 | - | - | 0 | - |
| 9 | y | 1040 | 706.3 | 0.004949 | 7.007 | +1 | 6 |
| - | - | 1121 | 708.8 | - | - | 0 | - |
| 3 | y | 6283 | 709.3 | 0.002376 | 3.35 | +2 | 12 |
| 3 | y | 8414 | 709.8 | 0.005555 | 7.826 | +2 | 12 |
| - | - | 3985 | 710.3 | - | - | 0 | - |
| - | - | 849.1 | 710.8 | - | - | 0 | - |
| - | - | 8381 | 711.4 | - | - | 0 | - |
| - | - | 8689 | 712.3 | - | - | 0 | - |
| - | - | 3415 | 712.4 | - | - | 0 | - |
| - | - | 3540 | 713.3 | - | - | 0 | - |
| - | - | 1022 | 713.4 | - | - | 0 | - |
| - | - | 4752 | 717.8 | - | - | 0 | - |
| 3 | y | 3.326E+04 | 718.3 | 0.001982 | 2.76 | +2 | 12 |
| - | - | 2.625E+04 | 718.8 | - | - | 0 | - |
| 13 | b | 1.184E+04 | 719.3 | 0.007488 | 10.41 | +2 | 13 |
| - | - | 3910 | 719.8 | - | - | 0 | - |
| - | - | 692.3 | 720.3 | - | - | 0 | - |
| 9 | y | 8535 | 724.4 | 0.0007937 | 1.096 | +1 | 6 |
| - | - | 2970 | 725.4 | - | - | 0 | - |
| - | - | 1573 | 726.4 | - | - | 0 | - |
| - | - | 1143 | 727.4 | - | - | 0 | - |
| 13 | b | 2044 | 727.9 | 0.002736 | 3.759 | +2 | 13 |
| - | - | 1541 | 728.4 | - | - | 0 | - |
| - | - | 1686 | 728.9 | - | - | 0 | - |
| - | - | 1217 | 735.4 | - | - | 0 | - |
| - | - | 1268 | 735.9 | - | - | 0 | - |
| - | - | 1543 | 736.4 | - | - | 0 | - |
| - | - | 704.9 | 744.3 | - | - | 0 | - |
| - | - | 731.4 | 746.4 | - | - | 0 | - |
| - | - | 776.5 | 749.9 | - | - | 0 | - |
| - | - | 823.1 | 750.7 | - | - | 0 | - |
| - | - | 654.3 | 751.3 | - | - | 0 | - |
| 2 | y | 3033 | 767.9 | 0.001094 | 1.425 | +2 | 13 |
| - | - | 1533 | 768.4 | - | - | 0 | - |
| - | - | 1286 | 768.9 | - | - | 0 | - |
| - | - | 745.5 | 779.3 | - | - | 0 | - |
| - | - | 756.7 | 782.3 | - | - | 0 | - |
| - | - | 917.9 | 783.4 | - | - | 0 | - |
| - | - | 1424 | 797.4 | - | - | 0 | - |
| - | - | 644.9 | 805.4 | - | - | 0 | - |
| 8 | y | 1246 | 811.4 | 0.01152 | 14.2 | +1 | 7 |
| 7 | b | 2398 | 825.4 | 0.00734 | 8.893 | +1 | 7 |
| - | - | 780.7 | 826.4 | - | - | 0 | - |
| - | - | 1876 | 827.3 | - | - | 0 | - |
| - | - | 783.7 | 839.4 | - | - | 0 | - |
| - | - | 664.5 | 857.4 | - | - | 0 | - |
| - | - | 731.1 | 866.3 | - | - | 0 | - |
| - | - | 857.5 | 876.4 | - | - | 0 | - |
| - | - | 972 | 877.5 | - | - | 0 | - |
| - | - | 2660 | 884.4 | - | - | 0 | - |
| - | - | 1164 | 885.4 | - | - | 0 | - |
| 8 | b | 2649 | 894.4 | 0.004634 | 5.181 | +1 | 8 |
| 8 | b | 2202 | 895.4 | 0.007933 | 8.859 | +1 | 8 |
| - | - | 1041 | 896.4 | - | - | 0 | - |
| 8 | b | 1.772E+04 | 912.4 | 0.006104 | 6.69 | +1 | 8 |
| - | - | 9256 | 913.4 | - | - | 0 | - |
| - | - | 2487 | 914.4 | - | - | 0 | - |
| - | - | 806.3 | 914.9 | - | - | 0 | - |
| - | - | 736.3 | 922.4 | - | - | 0 | - |
| - | - | 893.1 | 923.4 | - | - | 0 | - |
| 7 | y | 863.4 | 924.4 | 0.01184 | 12.81 | +1 | 8 |
| - | - | 4169 | 940.4 | - | - | 0 | - |
| - | - | 1918 | 941.4 | - | - | 0 | - |
| 7 | y | 1987 | 942.4 | 0.001407 | 1.493 | +1 | 8 |
| - | - | 1208 | 943.4 | - | - | 0 | - |
| - | - | 991.4 | 986.4 | - | - | 0 | - |
| - | - | 1468 | 987.4 | - | - | 0 | - |
| - | - | 2155 | 997.4 | - | - | 0 | - |
| - | - | 1154 | 1004 | - | - | 0 | - |
| - | - | 666.9 | 1009 | - | - | 0 | - |
| - | - | 764 | 1020 | - | - | 0 | - |
| 9 | b | 2021 | 1026 | 0.006977 | 6.804 | +1 | 9 |
| - | - | 834.4 | 1027 | - | - | 0 | - |
| 6 | y | 1445 | 1057 | 0.003753 | 3.549 | +1 | 9 |
| - | - | 1038 | 1058 | - | - | 0 | - |
| 10 | b | 898.4 | 1141 | 0.01231 | 10.8 | +1 | 10 |
| - | - | 843.6 | 1150 | - | - | 0 | - |
| 5 | y | 1687 | 1167 | 0.008797 | 7.535 | +1 | 10 |
| 5 | y | 1970 | 1168 | 0.003433 | 2.938 | +1 | 10 |
| - | - | 1776 | 1169 | - | - | 0 | - |
| 5 | y | 4200 | 1186 | 0.007887 | 6.653 | +1 | 10 |
| - | - | 3288 | 1187 | - | - | 0 | - |
| - | - | 1232 | 1188 | - | - | 0 | - |
| - | - | 748.1 | 1926 | - | - | 0 | - |
| - | - | 626.7 | 1945 | - | - | 0 | - |
| - | - | 729.4 | 2570 | - | - | 0 | - |
| - | - | 786.2 | 3083 | - | - | 0 | - |

m/z Charge Intensity FragmentType MassShift Position
120.0445785522461 0 691.15906
120.08103942871094 0 17087.855
120.08594512939453 0 1024.3119
121.08452606201172 0 1022.7406
122.07150268554688 0 521.09827
123.04450988769531 0 720.56555
126.0917739868164 0 413.4815
127.05028533935547 0 747.31226
127.08718872070312 0 409.41956
128.1072540283203 0 7684.355
129.06619262695312 0 779.3526
129.10250854492188 0 55098.863
130.06521606445312 0 1123.9982
130.1001739501953 0 547.48285
130.10585021972656 0 3313.6555
132.08139038085938 0 504.5358
132.1022186279297 0 1359.1279
136.07591247558594 0 14754.091
137.0794219970703 0 906.2546
138.0662384033203 0 957.5016
139.08685302734375 0 625.9489
141.1025390625 0 2751.2546
143.0818634033203 0 641.6856
144.0803985595703 0 399.47318
145.06101989746094 0 532.06104
146.06027221679688 0 11996.243
147.0638427734375 0 1520.9532
147.07684326171875 0 447.438 y 11
148.0394744873047 0 3252.0803
148.94642639160156 0 472.80756
151.08680725097656 0 1532.125
155.0815887451172 0 671.97504
155.09295654296875 0 4098.954
155.1181640625 0 3010.2502 a Water loss 1
156.04400634765625 0 434.0831
157.06056213378906 0 535.99023
157.09738159179688 0 697.5145
157.13381958007812 0 936.1282
159.09188842773438 0 1830.2501
159.11288452148438 0 760.4891 d 1
165.05482482910156 0 6808.883
166.06121826171875 0 6709.4004
166.0864715576172 0 7697.52 y 10
167.0646514892578 0 549.8917
167.09014892578125 0 792.83984
169.0972442626953 0 2280.5117
170.1000213623047 0 742.053
171.1492919921875 0 7486.695
172.0719757080078 0 1034.0543
172.1525115966797 0 682.16064
173.0560302734375 0 620.08875
173.07130432128906 0 2341.7207
173.12872314453125 0 117346.67 a 1
174.05517578125 0 19718.023
174.1240234375 0 637.53986
174.13206481933594 0 9079.915
175.0584716796875 0 2255.188
175.0865478515625 0 876.5834
178.13414001464844 0 4044.9268
181.06100463867188 0 889.3617
182.0813751220703 0 14846.467 y 13
183.0847625732422 0 1401.8463
183.1131591796875 0 2194.8901 b Water loss 1
183.14927673339844 0 1737.356
185.12893676757812 0 1410.1823
186.1239013671875 0 15407.202
186.13302612304688 0 964.9568
187.10775756835938 0 743.84564
187.12730407714844 0 1572.0431
187.14462280273438 0 2106.1135
188.0707550048828 0 3341.3074
189.06597900390625 0 1096.0316
190.08270263671875 0 689.4851
190.13404846191406 0 783.9355
191.08168029785156 0 1572.5651
193.09771728515625 0 552.14264
195.11314392089844 0 4726.5645
196.10818481445312 0 2113.131
197.12925720214844 0 654.90265
198.0878448486328 0 941.21747
199.14425659179688 0 5098.5645
200.14761352539062 0 643.4476
201.06614685058594 0 790.8453
201.12351989746094 0 27606.293 b 1
202.05007934570312 0 5938.9097
202.12696838378906 0 2097.915
203.0537872314453 0 766.5483
205.0974578857422 0 3055.794
206.12811279296875 0 614.5597
207.1605987548828 0 779.86456
208.09524536132812 0 514.91174
211.14431762695312 0 1217.0654
212.13934326171875 0 1270.8455
213.12351989746094 0 1916.415
215.13966369628906 0 1452.4271
216.1334686279297 0 683.25903
221.10333251953125 0 679.9155
221.12818908691406 0 893.0026
223.1077880859375 0 2514.3374
223.15550231933594 0 7710.589
224.10350036621094 0 577.0306
224.15858459472656 0 859.8156
226.0826873779297 0 1156.9464
226.11883544921875 0 781.12384 b Water loss 5
226.1548309326172 0 1734.9271
227.06625366210938 0 2089.2656
227.10203552246094 0 577.6233
229.11854553222656 0 1126.4442
229.15496826171875 0 1629.1747
230.15061950683594 0 763.7625
233.13986206054688 0 1541.1111
233.16519165039062 0 5329.8296
234.12391662597656 0 9624.452
234.16842651367188 0 903.9182
235.06033325195312 0 573.5949
235.12744140625 0 1262.2731
237.08665466308594 0 576.0291
238.11846923828125 0 666.3769
240.1344757080078 0 3412.625
241.11892700195312 0 746.8867
241.13348388671875 0 1386.2195
243.10928344726562 0 1343.0627
244.09303283691406 0 2931.883
244.10743713378906 0 492.63037
244.12960815429688 0 531.39105
247.10784912109375 0 5251.489
248.11380004882812 0 1422.604
249.09825134277344 0 2510.1675
251.150390625 0 30131.654
252.15380859375 0 4155.3623
255.14622497558594 0 735.8172
257.1136779785156 0 760.8472
258.1448059082031 0 1083.9707
259.142822265625 0 652.35565
260.1966552734375 0 712.15326
261.1197814941406 0 1422.6063
261.1601867675781 0 1400.2776
262.1192626953125 0 1458.9087
265.1293640136719 0 1593.6682 y Water loss 7
266.124755859375 0 6786.362
267.1287536621094 0 1239.1412
269.1132507324219 0 1152.4884
269.1286315917969 0 2420.4292
270.1311950683594 0 868.8383 b Ammonia loss 6
279.1455993652344 0 675.7433
279.489990234375 0 567.2825
280.1282958984375 0 565.1363
282.1564025878906 0 628.73663
283.1402282714844 0 13892.478
284.1426696777344 0 2034.8873
286.1399230957031 0 972.9165
286.1587219238281 0 551.2791
287.1387634277344 0 1524.8867
290.70733642578125 0 594.97015
293.11328125 0 12549.024 y Water loss 12
294.1170349121094 0 2073.2742
295.1194763183594 0 635.0802
296.1971740722656 0 695.28973 b Water loss 2
297.15594482421875 0 3756.535
299.1387634277344 0 569.0215 b Ammonia loss 7
300.1673583984375 0 643.1098
301.1508483886719 0 7313.5234
302.15509033203125 0 1086.9956
304.1658020019531 0 1602.3129
306.1372985839844 0 1343.905 y 9
309.2049560546875 0 571.8864
311.1243896484375 0 1566.0521 y 12
314.20758056640625 0 715.2514 b 2
315.1339111328125 0 755.12
315.1664733886719 0 5179.13
316.1684875488281 0 720.1252
321.70782470703125 0 630.5738
323.2074279785156 0 720.24615
332.2087097167969 0 988.8139
334.1035461425781 0 1840.4641
340.1907653808594 0 1612.4335
345.1297912597656 0 879.1816
346.1141357421875 0 1463.7645
350.2190246582031 0 2816.5562
351.21990966796875 0 671.0385
362.18255615234375 0 730.2499
362.6789855957031 0 1317.6981 y 8
363.14056396484375 0 3637.1743
363.6794738769531 0 906.6516
364.1448059082031 0 926.75885
378.2143859863281 0 1462.1942
379.20867919921875 0 6826.169
380.2106018066406 0 1675.7626
381.15179443359375 0 5717.25
382.15582275390625 0 846.0874
385.1688232421875 0 631.74805
394.1726989746094 0 1020.95526
396.2244873046875 0 1530.6483
397.2283020019531 0 732.5588
398.1783447265625 0 3367.4155
399.1827087402344 0 660.2094
402.20037841796875 0 856.5343
406.691162109375 0 671.67926
409.1465148925781 0 768.4945
411.1966857910156 0 697.80334
412.1827392578125 0 5592.5195
413.16259765625 0 619.0748
414.2351989746094 0 1441.4182
416.15643310546875 0 986.47754
417.1408386230469 0 1700.9675
420.22265625 0 1743.2344
420.7180480957031 0 667.1999
423.27154541015625 0 3319.911
424.272705078125 0 736.4971
429.2264404296875 0 15047.461
429.7279357910156 0 3924.6736
430.1933898925781 0 12920.673
430.2236328125 0 1106.9381
431.1960754394531 0 2712.3826
432.20025634765625 0 811.68317
433.2549743652344 0 861.6852 b Water loss 3
433.7107238769531 0 633.55865
434.2123718261719 0 760.55994
439.21905517578125 0 4077.2085 y 11
439.2613525390625 0 2681.1768
439.7626953125 0 1270.8208
440.22161865234375 0 1600.4015
440.2628479003906 0 550.02673
442.7161560058594 0 3246.3079
443.2162170410156 0 1319.4738
444.1524353027344 0 1196.6302
445.1353759765625 0 3435.1184
446.1405029296875 0 1029.3124
447.71044921875 0 1918.1658 b Water loss 7
448.2079772949219 0 2084.5764 b Ammonia loss 7
451.2662048339844 0 5256.1904 b 3
452.26849365234375 0 1289.411
456.7140197753906 0 7176.083 b 7
457.2148742675781 0 3799.76
457.7167053222656 0 742.95013
460.2331237792969 0 899.6564
462.1619567871094 0 3028.8408
468.29345703125 0 882.58203
474.7792663574219 0 641.94324
474.8325500488281 0 689.7214
478.2299499511719 0 965.7803 y Water loss 10
478.2752990722656 0 818.5049
490.2541198730469 0 1248.2465
490.7533264160156 0 1075.3828
492.8417053222656 0 2134.4158
493.7135009765625 0 750.603
493.84130859375 0 965.4691
494.2359924316406 0 8119.8154
495.2407531738281 0 1529.3999
495.7474365234375 0 720.2523
496.240234375 0 15338.542 y 10
497.2430725097656 0 4080.8748
498.2481689453125 0 835.9171
499.2596740722656 0 3019.2756
499.75872802734375 0 1443.5541
504.24951171875 0 766.8556 b Water loss 8
505.9088439941406 0 584.19653
510.2962341308594 0 1673.4314
511.26220703125 0 2363.0203
513.2579345703125 0 1116.4982 b 8
513.7579345703125 0 1434.8165
525.2660522460938 0 1618.7303
530.2222900390625 0 881.2937
530.9234619140625 0 827.06055
532.307861328125 0 2984.8616
533.3111572265625 0 733.989
534.3021850585938 0 1470.2699
536.1861572265625 0 893.5041
538.2606201171875 0 901.96246
539.9219360351562 0 707.93463 Precursor Water loss
540.2613525390625 0 1123.7267 Precursor Ammonia loss
540.5890502929688 0 1060.6117
542.2954711914062 0 909.3046
543.2777099609375 0 2681.2732
544.2779541015625 0 1014.9769
545.1063842773438 0 729.75354
545.9268798828125 0 1738.8865 Precursor
546.2589721679688 0 2649.3503
546.3046875 0 1460.2483
546.64404296875 0 1948.8132
546.8120727539062 0 1212.7424
547.0648193359375 0 2619.4458
547.246826171875 0 1155.2667
547.5640869140625 0 909.3324
553.8134765625 0 1660.2874
561.3148193359375 0 850.06714 b Water loss 4
562.2440795898438 0 1449.04
562.7440185546875 0 1566.2511
563.77001953125 0 799.8087
570.7683715820312 0 928.8203 b 9
571.2484130859375 0 1650.8584
571.7484130859375 0 1408.1428
573.3226318359375 0 716.3904
575.2445678710938 0 970.0166
579.3229370117188 0 2849.6458 b 4
580.2593383789062 0 667.01556
580.3264770507812 0 1328.8893
580.7561645507812 0 832.1997
581.2093505859375 0 727.87695
584.2517700195312 0 831.14453 y Water loss 4
588.3273315429688 0 843.1246
593.255615234375 0 4040.5066 y Water loss 9
593.3049926757812 0 1580.1755
594.2549438476562 0 1051.9493
597.3287353515625 0 957.107
599.2200927734375 0 4398.837
600.2230224609375 0 866.6301
610.2821655273438 0 5607.869
610.3296508789062 0 733.88434
611.2670288085938 0 31464.664 y 9
612.27001953125 0 8682.066
613.1012573242188 0 923.67224
613.2743530273438 0 2173.7861
613.7935180664062 0 796.2572
618.786376953125 0 1138.5193
619.2852172851562 0 998.05133
619.7791748046875 0 620.51245
621.25439453125 0 1006.5022
627.7904663085938 0 4771.367
628.292236328125 0 3367.3936
644.273681640625 0 635.2841
652.7816772460938 0 1490.8186 y Water loss 3
653.2827758789062 0 1114.0703 y Ammonia loss 3
653.8348999023438 0 678.6377
654.323974609375 0 801.9056 b Water loss 11
656.8215942382812 0 1608.1232
661.294677734375 0 760.3654
661.7861938476562 0 6962.1265 y 3
662.2876586914062 0 3381.3618
662.7871704101562 0 2414.7092
669.3206787109375 0 952.4345
672.83642578125 0 749.1226
676.3405151367188 0 1343.2131 b Water loss 5
677.2625732421875 0 695.51
677.3280639648438 0 2195.4707 b Ammonia loss 5
678.3261108398438 0 861.1136
679.3763427734375 0 11341.013
680.3795776367188 0 3083.454
683.3250122070312 0 785.0259
684.30859375 0 4688.9507
685.3120727539062 0 2396.3655
685.83740234375 0 1516.015
686.3408203125 0 663.8125
686.8440551757812 0 775.408
687.3317260742188 0 758.4234
687.864501953125 0 3071.4785
688.367431640625 0 2270.466
688.8689575195312 0 1017.40234
689.3753051757812 0 908.119
690.3453369140625 0 676.78265
692.3739624023438 0 645.7998
693.3677368164062 0 951.49176
694.2921142578125 0 1397.1509
694.352294921875 0 7292.449 b 5
695.3536376953125 0 2158.5964
695.8291015625 0 766.44946
700.3129272460938 0 904.566
700.8129272460938 0 916.2613
701.3128662109375 0 1085.7935
704.8372802734375 0 964.47864
706.3455810546875 0 1039.6149 y Water loss 8
708.8292236328125 0 1120.9639
709.3220825195312 0 6282.761 y Water loss 2
709.822021484375 0 8414.181 y Ammonia loss 2
710.3209228515625 0 3984.7458
710.8233642578125 0 849.08746
711.3768920898438 0 8381.426
712.3040771484375 0 8689.028
712.379638671875 0 3415.0193
713.3067016601562 0 3540.4722
713.3794555664062 0 1021.6586
717.8341064453125 0 4752.427
718.3277587890625 0 33264.918 y 2
718.8292236328125 0 26245.479
719.330078125 0 11843.959 b Ammonia loss 12
719.83349609375 0 3909.8328
720.3345336914062 0 692.27856
724.3504028320312 0 8534.623 y 8
725.3533935546875 0 2970.3108
726.3535766601562 0 1573.1376
727.3536376953125 0 1143.2969
727.8535766601562 0 2044.1471 b 12
728.35107421875 0 1541.3113
728.8504028320312 0 1685.9109
735.3739013671875 0 1217.2596
735.885986328125 0 1268.0835
736.3804321289062 0 1543.461
744.3241577148438 0 704.87585
746.4157104492188 0 731.4164
749.851318359375 0 776.52716
750.67138671875 0 823.063
751.338623046875 0 654.28406
767.8628540039062 0 3033.4082 y 1
768.366455078125 0 1533.389
768.864013671875 0 1285.525
779.2827758789062 0 745.50037
782.3126831054688 0 756.6957
783.37158203125 0 917.9345
797.3855590820312 0 1424.032
805.439453125 0 644.85693
811.3717041015625 0 1245.5386 y 7
825.385009765625 0 2398.1282 b 6
826.3900146484375 0 780.6627
827.3286743164062 0 1875.8792
839.3623657226562 0 783.6636
857.3740844726562 0 664.45264
866.344482421875 0 731.06146
876.3685913085938 0 857.48175
877.5137329101562 0 972.0466
884.4268188476562 0 2660.0515
885.4277954101562 0 1164.3839
894.4091796875 0 2649.3535 b Water loss 7
895.40576171875 0 2201.6272 b Ammonia loss 7
896.4072875976562 0 1041.4038
912.4182739257812 0 17720.92 b 7
913.4213256835938 0 9256.166
914.4251708984375 0 2487.083
914.9462280273438 0 806.33453
922.4099731445312 0 736.30975
923.3936157226562 0 893.0808
924.4013061523438 0 863.42236 y Water loss 6
940.4147338867188 0 4169.0713
941.41796875 0 1917.8802
942.4223022460938 0 1986.6749 y 6
943.425537109375 0 1208.1921
986.4171752929688 0 991.398
987.4064331054688 0 1468.006
997.4376831054688 0 2154.5615
1004.432373046875 0 1154.2794
1009.314697265625 0 666.9494
1019.5816040039062 0 763.9586
1025.50146484375 0 2020.9708 b 8
1026.503173828125 0 834.4151
1057.4468994140625 0 1444.9741 y 5
1058.4554443359375 0 1038.0117
1140.5230712890625 0 898.3812 b 9
1150.4742431640625 0 843.595
1167.4898681640625 0 1687.4075 y Water loss 4
1168.479248046875 0 1969.9988 y Ammonia loss 4
1169.4781494140625 0 1776.2933
1185.5013427734375 0 4200.3975 y 4
1186.507568359375 0 3287.511
1187.5072021484375 0 1232.3912
1926.4412841796875 0 748.1316
1944.72509765625 0 626.72314
2569.806884765625 0 729.3972
3083.055908203125 0 786.15735

Spectrum Details

|  |  |
| --- | --- |
| Matched peaks? Matched peaks The total absolute number of peaks matched. Additionally in brackets the total fraction of peaks matched and the total number of peaks is shown. | 65 (14.54% of 447) |
| FDR? FDR The false discovery rate estimated for this peptide. It is calculated by matching all theoretical fragments with a non-integer shift with the raw peaks for this spectrum. This is done with 40 different shifts. The resulting percentage is the average number of annotated peaks over the number of annotated peaks with the correct spectrum. | 0.51% |
| Satellite FDR? Satellite FDR See the FDR for details on its calculation. This satellite ion specific FDR only contains the satellite ions (d/w) for I/L/J positions. | ∞ |
| PSM Score? PSM Score The PSM Score as given by Hecklib to this annotated spectrum. It is shown with three significant figures. | 504 |

#### Reverse Lookup? Reverse Lookup All places where this read could be placed.

| Group | Segment | Template | Template Part | Read Part | Score | Unique |
| --- | --- | --- | --- | --- | --- | --- |
| Homo sapiens Heavy Chain | IGHC | IGHG1 | [189..202] | [0..14] | 78 | False |
| Homo sapiens Heavy Chain | IGHC | IGHG3 | [236..249] | [0..14] | 78 | False |
| Homo sapiens Heavy Chain | IGHC | IGHG4 | [186..199] | [0..14] | 78 | False |

| Recombined | Template Part | Read Part | Score | Unique |
| --- | --- | --- | --- | --- |
| REC-0-1 | [314..327] | [0..14] | 83 | True |

#### Meta Information from Multiple reads

##### Number of combined reads

3

##### Intensity

0.7255

##### TotalArea

1.838E+08

##### Changes to the peptide sequence

TVJHQDMSJDGKEY

L→JNo support for either Leucine or Isoleucine based on side chain ions (Position: 9)

L→JNo support for either Leucine or Isoleucine based on side chain ions (Position: 3)

#### Positional Score

Copy Data

##### Positional Score (TSV)

###### Preview

```
Loading example...
```

*Click on the button to copy the data to your clipboard.*

10012345678910111213

Label Value
"0" 0.327
"1" 0.33
"2" 0.333
"3" 0.33
"4" 0.323
"5" 0.327
"6" 0.28
"7" 0.307
"8" 0.333
"9" 0.33
"10" 0.32
"11" 0.317
"12" 0.327
"13" 0.333

#### Meta Information from PEAKS

##### Scan Identifier

F3:5825

##### Original sequence

T

V

L

H

Q

D

M

S

L

D

G

K

E

Y

##### Posttranslational Modifications

##### Source File

D:\separate\_stitch\_analyses\xle-disambiguation\raw\20210323\_F1\_UM1\_Peng0013\_SA\_F59\_ingel\_3ug\_chymo.raw

##### Fraction

3

##### Scan Feature

F3:6132

##### De Novo Score

98

##### ConfidenceScore

98

### m/z

545.9277

##### Mass

1634.761

##### Charge

3

##### Retention Time

32.22

##### Predicted Retention Time

-

##### Area

6.127E+07

##### Parts Per Million

0.1

##### Fragmentation mode

HCD

##### Originating file

01 D:\separate\_stitch\_analyses\xle-disambiguation\20210325\_F59\_3ug\_DENOVO\_12.csv

#### Meta Information from PEAKS

##### Scan Identifier

F3:6077

##### Original sequence

T

V

L

H

Q

D

M

S

L

D

G

K

E

Y

##### Posttranslational Modifications

##### Source File

D:\separate\_stitch\_analyses\xle-disambiguation\raw\20210323\_F1\_UM1\_Peng0013\_SA\_F59\_ingel\_3ug\_chymo.raw

##### Fraction

3

##### Scan Feature

F3:6132

##### De Novo Score

97

##### ConfidenceScore

97

### m/z

545.9277

##### Mass

1634.761

##### Charge

3

##### Retention Time

32.22

##### Predicted Retention Time

-

##### Area

6.127E+07

##### Parts Per Million

0.1

##### Fragmentation mode

HCD

##### Originating file

01 D:\separate\_stitch\_analyses\xle-disambiguation\20210325\_F59\_3ug\_DENOVO\_12.csv

#### Meta Information from PEAKS

##### Scan Identifier

F3:6141

##### Original sequence

T

V

L

H

Q

D

M

S

L

D

G

K

E

Y

##### Posttranslational Modifications

##### Source File

D:\separate\_stitch\_analyses\xle-disambiguation\raw\20210323\_F1\_UM1\_Peng0013\_SA\_F59\_ingel\_3ug\_chymo.raw

##### Fraction

3

##### Scan Feature

F3:6132

##### De Novo Score

96

##### ConfidenceScore

96

### m/z

545.9277

##### Mass

1634.761

##### Charge

3

##### Retention Time

32.22

##### Predicted Retention Time

-

##### Area

6.127E+07

##### Parts Per Million

0.1

##### Fragmentation mode

HCD

##### Originating file

01 D:\separate\_stitch\_analyses\xle-disambiguation\20210325\_F59\_3ug\_DENOVO\_12.csv
