## Supplementary material for "A handle on mass coincidence errors in *de novo* sequencing of antibodies by bottom-up proteomics": Combined_089.html

Details Combined\_089 | Stitch OverviewUndefined

### Read Combined\_089

#### Sequence (length=12)

GFTFDDYAMHWV

#### Spectrum 11056? Spectrum 11056 The raw spectrum of this peptide as annotated by Hecklib. The fragments are coloured according to ion type (see legend). Any peaks with a star '\*' as text can be hovered over to see the full details, first the ion type second the mass shift type. By hovering over the amino acids in the peptide or ions in the legend the corresponding peaks are highlighted. By toggling the 'Unassigned' label you can turn the background (unassigned) peaks on or off in the plot. By updating the slider in the Ion legend you can update the spectrum to only show the top X% of the peaks with labels. The top X% means any peak that is within X% of the highest intensity. By dragging in the spectrum you can zoom in to a specific part of the spectrum and use 'Zoom Out' to get back to the original zoom level. The annotation of the spectrum is based on the given sequence in the peptides file and is done with different software so inconsistencies are likely. The peaks are annotated based on the given sequence, with 20 ppm tolerance.

Copy Data

##### Spectrum 11056 (TSV)

###### Preview

```
Loading example...
```

*Click on the button to copy the data to your clipboard.*

Mz MinMz MaxIntensity Max

WidthHeightPeptide font sizePeptide stroke widthSpectrum font sizeSpectrum stroke widthCompact peptide

Ion legend

wxyz

abcd

OtherUnassignedIon^Charge^_Position_Show for top:%

GFTFDDYAMHWV

06.21e+41.24e+51.86e+52.49e+5

Zoom Out

a^+1^_2_a^+1^_2_b^+1^_2_b^+1^_2_a^+1^_3_a^+1^_3_a^+1^_3_b^+1^_3_b^+1^_3_y^+1^_2_b^+1^_3_y^+2^_6_b^+1^_4_y^+1^_3_b^+1^_4_y^+2^_7_y^+2^_8_b^+1^_5_b^+1^_5_y^+1^_4_y^+2^_9_b^+2^_10_y^+2^_9_y^+2^_10_y^+2^_10_y^+1^_5_b^+1^_6_b^+1^_6_b^+2^_11_b^+2^_11_b^+2^_11_y^+2^_11_y^+2^_11_\*\*y^+1^_6_b^+1^_7_b^+1^_7_b^+1^_8_b^+1^_8_b^+1^_8_y^+1^_7_y^+1^_7_y^+1^_8_b^+1^_9_y^+1^_8_b^+1^_9_y^+1^_9_b^+1^_10_y^+1^_9_y^+1^_10_y^+1^_10_b^+1^_11_

0520104015592079

Fragment Matches Table

Show background peaks

| Position | Ion type | Intensity | mz Theoretical | mz Error (Th) | mz Error (ppm) | Charge | Series Number |
| --- | --- | --- | --- | --- | --- | --- | --- |
| - | - | 2.285E+05 | 120.1 | - | - | 0 | - |
| - | - | 869.5 | 121.1 | - | - | 0 | - |
| - | - | 1.764E+04 | 121.1 | - | - | 0 | - |
| - | - | 1147 | 122.1 | - | - | 0 | - |
| - | - | 536.7 | 122.1 | - | - | 0 | - |
| - | - | 2053 | 127.1 | - | - | 0 | - |
| - | - | 662.7 | 127.1 | - | - | 0 | - |
| - | - | 368.2 | 128 | - | - | 0 | - |
| - | - | 852.4 | 129.1 | - | - | 0 | - |
| - | - | 1744 | 129.1 | - | - | 0 | - |
| - | - | 890.3 | 130.1 | - | - | 0 | - |
| - | - | 1.618E+04 | 130.1 | - | - | 0 | - |
| - | - | 939.2 | 131 | - | - | 0 | - |
| - | - | 1578 | 131.1 | - | - | 0 | - |
| - | - | 5453 | 132.1 | - | - | 0 | - |
| - | - | 710.8 | 133.1 | - | - | 0 | - |
| - | - | 411.5 | 133.1 | - | - | 0 | - |
| - | - | 3.266E+04 | 136.1 | - | - | 0 | - |
| - | - | 1924 | 136.1 | - | - | 0 | - |
| - | - | 2544 | 137.1 | - | - | 0 | - |
| - | - | 3411 | 138.1 | - | - | 0 | - |
| - | - | 1871 | 138.1 | - | - | 0 | - |
| - | - | 441.4 | 139.1 | - | - | 0 | - |
| - | - | 1011 | 141.1 | - | - | 0 | - |
| - | - | 452.9 | 143 | - | - | 0 | - |
| - | - | 454.5 | 143.1 | - | - | 0 | - |
| - | - | 3892 | 144.1 | - | - | 0 | - |
| - | - | 717 | 146.1 | - | - | 0 | - |
| - | - | 2409 | 146.1 | - | - | 0 | - |
| - | - | 1770 | 148.1 | - | - | 0 | - |
| - | - | 1480 | 148.1 | - | - | 0 | - |
| - | - | 702.1 | 148.9 | - | - | 0 | - |
| - | - | 1037 | 155.1 | - | - | 0 | - |
| - | - | 2421 | 155.1 | - | - | 0 | - |
| - | - | 2218 | 156.1 | - | - | 0 | - |
| - | - | 458.3 | 157.1 | - | - | 0 | - |
| - | - | 996.1 | 158.1 | - | - | 0 | - |
| - | - | 5768 | 158.1 | - | - | 0 | - |
| - | - | 3.721E+04 | 159.1 | - | - | 0 | - |
| 2 | a | 2676 | 160.1 | 0.0002984 | 1.864 | +1 | 2 |
| - | - | 588.3 | 160.1 | - | - | 0 | - |
| - | - | 2931 | 160.1 | - | - | 0 | - |
| - | - | 2325 | 160.1 | - | - | 0 | - |
| - | - | 675.7 | 164.1 | - | - | 0 | - |
| - | - | 4158 | 165.1 | - | - | 0 | - |
| - | - | 587.5 | 165.1 | - | - | 0 | - |
| - | - | 955.2 | 166.1 | - | - | 0 | - |
| - | - | 1781 | 166.1 | - | - | 0 | - |
| - | - | 1034 | 166.1 | - | - | 0 | - |
| - | - | 617.2 | 166.1 | - | - | 0 | - |
| - | - | 1943 | 167.1 | - | - | 0 | - |
| - | - | 3555 | 169.1 | - | - | 0 | - |
| - | - | 1640 | 169.1 | - | - | 0 | - |
| - | - | 2727 | 170.1 | - | - | 0 | - |
| - | - | 2687 | 171.1 | - | - | 0 | - |
| - | - | 1157 | 171.1 | - | - | 0 | - |
| - | - | 750.9 | 171.1 | - | - | 0 | - |
| - | - | 672.1 | 173.1 | - | - | 0 | - |
| - | - | 539.8 | 173.1 | - | - | 0 | - |
| - | - | 4645 | 173.5 | - | - | 0 | - |
| - | - | 859.2 | 174.1 | - | - | 0 | - |
| - | - | 1109 | 175.1 | - | - | 0 | - |
| - | - | 532.1 | 175.1 | - | - | 0 | - |
| - | - | 1875 | 175.1 | - | - | 0 | - |
| - | - | 3099 | 176.1 | - | - | 0 | - |
| - | - | 1.646E+04 | 176.1 | - | - | 0 | - |
| 2 | a | 2.461E+05 | 177.1 | 0.000498 | 2.812 | +1 | 2 |
| - | - | 622.1 | 178.1 | - | - | 0 | - |
| - | - | 1399 | 178.1 | - | - | 0 | - |
| - | - | 2.512E+04 | 178.1 | - | - | 0 | - |
| - | - | 880.7 | 179.1 | - | - | 0 | - |
| - | - | 618.9 | 179.1 | - | - | 0 | - |
| - | - | 2333 | 180.1 | - | - | 0 | - |
| - | - | 1430 | 185.1 | - | - | 0 | - |
| - | - | 700.4 | 185.1 | - | - | 0 | - |
| - | - | 1.187E+04 | 186.1 | - | - | 0 | - |
| - | - | 3227 | 187.1 | - | - | 0 | - |
| - | - | 713.9 | 187.1 | - | - | 0 | - |
| - | - | 858.4 | 187.1 | - | - | 0 | - |
| - | - | 1205 | 187.1 | - | - | 0 | - |
| 2 | b | 578.2 | 188.1 | 2.574E-06 | 0.01369 | +1 | 2 |
| - | - | 1788 | 191.1 | - | - | 0 | - |
| - | - | 2659 | 191.1 | - | - | 0 | - |
| - | - | 1648 | 193.1 | - | - | 0 | - |
| - | - | 371.6 | 193.1 | - | - | 0 | - |
| - | - | 1.085E+04 | 193.1 | - | - | 0 | - |
| - | - | 583.3 | 194.1 | - | - | 0 | - |
| - | - | 1173 | 194.1 | - | - | 0 | - |
| - | - | 3410 | 195.1 | - | - | 0 | - |
| - | - | 529.2 | 197.6 | - | - | 0 | - |
| - | - | 550.5 | 199.1 | - | - | 0 | - |
| - | - | 797.4 | 201.1 | - | - | 0 | - |
| - | - | 1107 | 202.1 | - | - | 0 | - |
| - | - | 1678 | 203.1 | - | - | 0 | - |
| - | - | 2093 | 203.1 | - | - | 0 | - |
| - | - | 1.782E+04 | 203.1 | - | - | 0 | - |
| - | - | 5806 | 204.1 | - | - | 0 | - |
| - | - | 737.1 | 204.1 | - | - | 0 | - |
| - | - | 1321 | 204.1 | - | - | 0 | - |
| 2 | b | 1.162E+05 | 205.1 | 0.0004716 | 2.3 | +1 | 2 |
| - | - | 1.353E+04 | 206.1 | - | - | 0 | - |
| - | - | 779 | 207.1 | - | - | 0 | - |
| - | - | 2151 | 207.1 | - | - | 0 | - |
| - | - | 516.6 | 208.1 | - | - | 0 | - |
| - | - | 1843 | 208.1 | - | - | 0 | - |
| - | - | 507.4 | 209.9 | - | - | 0 | - |
| - | - | 3084 | 212.1 | - | - | 0 | - |
| - | - | 558.8 | 213.1 | - | - | 0 | - |
| - | - | 791.1 | 214.1 | - | - | 0 | - |
| - | - | 3021 | 215.1 | - | - | 0 | - |
| - | - | 5305 | 217.1 | - | - | 0 | - |
| - | - | 2174 | 219.1 | - | - | 0 | - |
| - | - | 920.9 | 219.1 | - | - | 0 | - |
| - | - | 588.4 | 219.1 | - | - | 0 | - |
| - | - | 942.9 | 220.1 | - | - | 0 | - |
| - | - | 2.961E+04 | 221.1 | - | - | 0 | - |
| - | - | 1.405E+05 | 221.1 | - | - | 0 | - |
| - | - | 2532 | 222.1 | - | - | 0 | - |
| - | - | 1.383E+04 | 222.1 | - | - | 0 | - |
| - | - | 670.3 | 223.1 | - | - | 0 | - |
| - | - | 723.5 | 225.1 | - | - | 0 | - |
| - | - | 2556 | 228.1 | - | - | 0 | - |
| - | - | 968.5 | 229.1 | - | - | 0 | - |
| - | - | 624 | 230.1 | - | - | 0 | - |
| - | - | 5351 | 231.1 | - | - | 0 | - |
| - | - | 2.149E+04 | 231.1 | - | - | 0 | - |
| - | - | 1093 | 231.1 | - | - | 0 | - |
| - | - | 729.7 | 231.1 | - | - | 0 | - |
| - | - | 2568 | 232.1 | - | - | 0 | - |
| - | - | 688 | 233.1 | - | - | 0 | - |
| - | - | 7487 | 233.1 | - | - | 0 | - |
| - | - | 942.8 | 234.1 | - | - | 0 | - |
| - | - | 1.334E+04 | 235.1 | - | - | 0 | - |
| - | - | 1769 | 236.1 | - | - | 0 | - |
| - | - | 792.6 | 238.1 | - | - | 0 | - |
| - | - | 803.4 | 241.1 | - | - | 0 | - |
| - | - | 1090 | 243.1 | - | - | 0 | - |
| - | - | 1557 | 244.1 | - | - | 0 | - |
| - | - | 1049 | 245.1 | - | - | 0 | - |
| - | - | 1290 | 246.1 | - | - | 0 | - |
| - | - | 1504 | 248.1 | - | - | 0 | - |
| - | - | 711.8 | 248.2 | - | - | 0 | - |
| - | - | 4.702E+04 | 249.1 | - | - | 0 | - |
| - | - | 5709 | 250.1 | - | - | 0 | - |
| - | - | 1.037E+04 | 251.1 | - | - | 0 | - |
| - | - | 1762 | 251.1 | - | - | 0 | - |
| - | - | 969 | 252.1 | - | - | 0 | - |
| - | - | 633 | 252.1 | - | - | 0 | - |
| - | - | 540.9 | 253 | - | - | 0 | - |
| - | - | 795.8 | 256.1 | - | - | 0 | - |
| - | - | 8684 | 257.1 | - | - | 0 | - |
| - | - | 1061 | 258.1 | - | - | 0 | - |
| - | - | 814.8 | 258.2 | - | - | 0 | - |
| - | - | 2239 | 259.1 | - | - | 0 | - |
| - | - | 2231 | 259.1 | - | - | 0 | - |
| 3 | a | 2.396E+04 | 260.1 | 0.0004478 | 1.721 | +1 | 3 |
| - | - | 890.6 | 261.1 | - | - | 0 | - |
| - | - | 1146 | 261.1 | - | - | 0 | - |
| 3 | a | 1.746E+04 | 261.1 | 0.0004715 | 1.806 | +1 | 3 |
| - | - | 2706 | 261.1 | - | - | 0 | - |
| - | - | 6662 | 262.1 | - | - | 0 | - |
| - | - | 731 | 262.1 | - | - | 0 | - |
| - | - | 779.7 | 263.1 | - | - | 0 | - |
| - | - | 684.6 | 263.1 | - | - | 0 | - |
| - | - | 1.419E+04 | 263.1 | - | - | 0 | - |
| - | - | 2066 | 264.1 | - | - | 0 | - |
| - | - | 1758 | 264.1 | - | - | 0 | - |
| - | - | 1316 | 267.1 | - | - | 0 | - |
| - | - | 1050 | 268.1 | - | - | 0 | - |
| - | - | 665 | 268.1 | - | - | 0 | - |
| - | - | 730 | 270.1 | - | - | 0 | - |
| - | - | 1246 | 271.1 | - | - | 0 | - |
| - | - | 658.7 | 273.1 | - | - | 0 | - |
| - | - | 6356 | 274.1 | - | - | 0 | - |
| - | - | 756.4 | 275.1 | - | - | 0 | - |
| - | - | 1192 | 275.1 | - | - | 0 | - |
| - | - | 2249 | 276.1 | - | - | 0 | - |
| - | - | 697.9 | 277.1 | - | - | 0 | - |
| 3 | a | 3794 | 278.1 | 0.0004422 | 1.59 | +1 | 3 |
| - | - | 9257 | 279.1 | - | - | 0 | - |
| - | - | 1939 | 279.1 | - | - | 0 | - |
| - | - | 3937 | 279.2 | - | - | 0 | - |
| - | - | 858.6 | 280.1 | - | - | 0 | - |
| - | - | 707 | 283.2 | - | - | 0 | - |
| - | - | 1.747E+04 | 285.1 | - | - | 0 | - |
| - | - | 2192 | 286.1 | - | - | 0 | - |
| - | - | 615.7 | 286.1 | - | - | 0 | - |
| - | - | 610 | 286.2 | - | - | 0 | - |
| - | - | 624.5 | 286.3 | - | - | 0 | - |
| - | - | 1004 | 287.1 | - | - | 0 | - |
| - | - | 3.268E+04 | 287.1 | - | - | 0 | - |
| 3 | b | 7.348E+04 | 288.1 | 0.0008334 | 2.893 | +1 | 3 |
| 3 | b | 1297 | 289.1 | 0.0004299 | 1.487 | +1 | 3 |
| - | - | 1.212E+04 | 289.1 | - | - | 0 | - |
| - | - | 1013 | 290.1 | - | - | 0 | - |
| - | - | 992.6 | 290.1 | - | - | 0 | - |
| - | - | 1.367E+04 | 292.1 | - | - | 0 | - |
| - | - | 1160 | 292.2 | - | - | 0 | - |
| - | - | 1235 | 293.1 | - | - | 0 | - |
| - | - | 1584 | 295.1 | - | - | 0 | - |
| - | - | 3.29E+04 | 296.2 | - | - | 0 | - |
| - | - | 4434 | 297.2 | - | - | 0 | - |
| - | - | 808.9 | 299.2 | - | - | 0 | - |
| - | - | 690.6 | 302.1 | - | - | 0 | - |
| - | - | 792.8 | 302.1 | - | - | 0 | - |
| - | - | 577 | 303.1 | - | - | 0 | - |
| 11 | y | 2.336E+04 | 304.2 | 0.0007528 | 2.475 | +1 | 2 |
| - | - | 733.1 | 305.1 | - | - | 0 | - |
| - | - | 4160 | 305.2 | - | - | 0 | - |
| 3 | b | 9.553E+04 | 306.1 | 0.0007363 | 2.405 | +1 | 3 |
| - | - | 1187 | 307.1 | - | - | 0 | - |
| - | - | 1.592E+04 | 307.1 | - | - | 0 | - |
| - | - | 1835 | 308.1 | - | - | 0 | - |
| - | - | 884.2 | 313.1 | - | - | 0 | - |
| - | - | 1227 | 314.1 | - | - | 0 | - |
| - | - | 1178 | 314.2 | - | - | 0 | - |
| - | - | 3609 | 318.1 | - | - | 0 | - |
| - | - | 989.3 | 320.1 | - | - | 0 | - |
| - | - | 2293 | 323.1 | - | - | 0 | - |
| - | - | 1220 | 323.1 | - | - | 0 | - |
| - | - | 1440 | 323.2 | - | - | 0 | - |
| - | - | 3.884E+04 | 324.1 | - | - | 0 | - |
| - | - | 5595 | 325.1 | - | - | 0 | - |
| - | - | 886.8 | 327.1 | - | - | 0 | - |
| - | - | 1946 | 328.1 | - | - | 0 | - |
| - | - | 1201 | 328.1 | - | - | 0 | - |
| - | - | 833 | 329.1 | - | - | 0 | - |
| - | - | 1492 | 332.1 | - | - | 0 | - |
| - | - | 1697 | 333.1 | - | - | 0 | - |
| - | - | 916.2 | 333.2 | - | - | 0 | - |
| - | - | 1684 | 334.1 | - | - | 0 | - |
| - | - | 3060 | 338.1 | - | - | 0 | - |
| - | - | 700.5 | 339.1 | - | - | 0 | - |
| - | - | 1564 | 339.2 | - | - | 0 | - |
| - | - | 5695 | 342.2 | - | - | 0 | - |
| - | - | 958.4 | 343.2 | - | - | 0 | - |
| - | - | 5935 | 346.1 | - | - | 0 | - |
| - | - | 1463 | 347.1 | - | - | 0 | - |
| - | - | 2656 | 348.1 | - | - | 0 | - |
| - | - | 8444 | 350.1 | - | - | 0 | - |
| - | - | 3508 | 350.2 | - | - | 0 | - |
| - | - | 1112 | 351.1 | - | - | 0 | - |
| - | - | 738.6 | 351.2 | - | - | 0 | - |
| - | - | 1316 | 352.1 | - | - | 0 | - |
| - | - | 663.8 | 352.2 | - | - | 0 | - |
| - | - | 885.2 | 355.1 | - | - | 0 | - |
| - | - | 1.049E+04 | 356.1 | - | - | 0 | - |
| - | - | 926.9 | 357.1 | - | - | 0 | - |
| - | - | 942.9 | 358.1 | - | - | 0 | - |
| - | - | 1661 | 360.1 | - | - | 0 | - |
| - | - | 835.1 | 360.1 | - | - | 0 | - |
| - | - | 736.6 | 361.2 | - | - | 0 | - |
| - | - | 1749 | 362.2 | - | - | 0 | - |
| - | - | 2.42E+04 | 364.2 | - | - | 0 | - |
| - | - | 5365 | 365.2 | - | - | 0 | - |
| - | - | 2718 | 366.1 | - | - | 0 | - |
| - | - | 719.8 | 367.2 | - | - | 0 | - |
| - | - | 1134 | 371.1 | - | - | 0 | - |
| - | - | 728.5 | 372.1 | - | - | 0 | - |
| - | - | 672.9 | 373.2 | - | - | 0 | - |
| - | - | 1177 | 374.1 | - | - | 0 | - |
| - | - | 537.2 | 375.1 | - | - | 0 | - |
| - | - | 2645 | 376.1 | - | - | 0 | - |
| - | - | 1102 | 378.1 | - | - | 0 | - |
| - | - | 2.194E+04 | 378.1 | - | - | 0 | - |
| - | - | 1772 | 378.2 | - | - | 0 | - |
| - | - | 3558 | 379.1 | - | - | 0 | - |
| - | - | 1859 | 381.2 | - | - | 0 | - |
| - | - | 3348 | 382.1 | - | - | 0 | - |
| - | - | 940.6 | 384.1 | - | - | 0 | - |
| - | - | 959.9 | 385.2 | - | - | 0 | - |
| - | - | 709.1 | 389.1 | - | - | 0 | - |
| - | - | 1025 | 390.2 | - | - | 0 | - |
| - | - | 5182 | 390.2 | - | - | 0 | - |
| - | - | 1053 | 391.2 | - | - | 0 | - |
| - | - | 1.6E+04 | 394.1 | - | - | 0 | - |
| - | - | 2767 | 395.1 | - | - | 0 | - |
| - | - | 774.4 | 395.2 | - | - | 0 | - |
| - | - | 772.6 | 395.2 | - | - | 0 | - |
| - | - | 703.7 | 399.2 | - | - | 0 | - |
| - | - | 3627 | 403.2 | - | - | 0 | - |
| - | - | 942.6 | 404.1 | - | - | 0 | - |
| - | - | 1535 | 404.1 | - | - | 0 | - |
| - | - | 2272 | 405.2 | - | - | 0 | - |
| - | - | 605.8 | 406.2 | - | - | 0 | - |
| - | - | 6400 | 407.2 | - | - | 0 | - |
| - | - | 2.614E+04 | 407.2 | - | - | 0 | - |
| - | - | 1188 | 408.2 | - | - | 0 | - |
| - | - | 5683 | 408.2 | - | - | 0 | - |
| - | - | 2179 | 409.2 | - | - | 0 | - |
| - | - | 1145 | 410.1 | - | - | 0 | - |
| - | - | 849.1 | 410.2 | - | - | 0 | - |
| 7 | y | 1016 | 411.7 | 0.003381 | 8.213 | +2 | 6 |
| - | - | 1001 | 416.1 | - | - | 0 | - |
| - | - | 785 | 419.2 | - | - | 0 | - |
| - | - | 1808 | 421.2 | - | - | 0 | - |
| - | - | 761.2 | 422.2 | - | - | 0 | - |
| - | - | 3763 | 423.2 | - | - | 0 | - |
| - | - | 967 | 424.2 | - | - | 0 | - |
| - | - | 1157 | 425.2 | - | - | 0 | - |
| - | - | 4091 | 425.2 | - | - | 0 | - |
| - | - | 1125 | 426.2 | - | - | 0 | - |
| - | - | 3984 | 427.2 | - | - | 0 | - |
| - | - | 688 | 428.2 | - | - | 0 | - |
| - | - | 1132 | 431.2 | - | - | 0 | - |
| - | - | 2534 | 433.2 | - | - | 0 | - |
| - | - | 806.5 | 433.2 | - | - | 0 | - |
| - | - | 643.3 | 434.2 | - | - | 0 | - |
| - | - | 512.7 | 435.2 | - | - | 0 | - |
| 4 | b | 2.173E+04 | 435.2 | 0.001237 | 2.842 | +1 | 4 |
| - | - | 5144 | 436.2 | - | - | 0 | - |
| 10 | y | 6.906E+04 | 441.2 | 0.001106 | 2.507 | +1 | 3 |
| - | - | 1623 | 442.2 | - | - | 0 | - |
| - | - | 1.639E+04 | 442.2 | - | - | 0 | - |
| - | - | 629.2 | 443.2 | - | - | 0 | - |
| - | - | 1562 | 443.2 | - | - | 0 | - |
| - | - | 2425 | 443.2 | - | - | 0 | - |
| - | - | 1660 | 447.2 | - | - | 0 | - |
| - | - | 1399 | 449.2 | - | - | 0 | - |
| - | - | 2263 | 450.2 | - | - | 0 | - |
| - | - | 589.7 | 451.1 | - | - | 0 | - |
| - | - | 1113 | 451.2 | - | - | 0 | - |
| 4 | b | 3897 | 453.2 | 0.001689 | 3.726 | +1 | 4 |
| - | - | 996.9 | 454.2 | - | - | 0 | - |
| - | - | 2231 | 455.2 | - | - | 0 | - |
| - | - | 880.2 | 456.2 | - | - | 0 | - |
| - | - | 6302 | 461.2 | - | - | 0 | - |
| - | - | 1094 | 462.2 | - | - | 0 | - |
| - | - | 1.197E+04 | 465.2 | - | - | 0 | - |
| - | - | 2645 | 466.2 | - | - | 0 | - |
| 6 | y | 697.9 | 469.2 | 0.005169 | 11.02 | +2 | 7 |
| - | - | 1843 | 470.2 | - | - | 0 | - |
| - | - | 8833 | 471.2 | - | - | 0 | - |
| - | - | 2600 | 472.2 | - | - | 0 | - |
| - | - | 712.5 | 472.2 | - | - | 0 | - |
| - | - | 620.9 | 473.2 | - | - | 0 | - |
| - | - | 770.3 | 476.2 | - | - | 0 | - |
| - | - | 1202 | 477.2 | - | - | 0 | - |
| - | - | 2797 | 478.2 | - | - | 0 | - |
| - | - | 1.391E+04 | 479.2 | - | - | 0 | - |
| - | - | 3392 | 480.2 | - | - | 0 | - |
| - | - | 946.9 | 483.2 | - | - | 0 | - |
| - | - | 2708 | 486.2 | - | - | 0 | - |
| - | - | 646.5 | 488.2 | - | - | 0 | - |
| - | - | 1401 | 489.2 | - | - | 0 | - |
| - | - | 1630 | 489.2 | - | - | 0 | - |
| - | - | 899.9 | 490.2 | - | - | 0 | - |
| - | - | 681.4 | 490.2 | - | - | 0 | - |
| - | - | 4928 | 491.2 | - | - | 0 | - |
| - | - | 828.5 | 492.2 | - | - | 0 | - |
| - | - | 2270 | 493.2 | - | - | 0 | - |
| - | - | 1735 | 495.2 | - | - | 0 | - |
| - | - | 948.5 | 496.2 | - | - | 0 | - |
| - | - | 3124 | 497.2 | - | - | 0 | - |
| - | - | 2468 | 500.2 | - | - | 0 | - |
| - | - | 2202 | 504.2 | - | - | 0 | - |
| - | - | 2580 | 507.2 | - | - | 0 | - |
| - | - | 1813 | 508.2 | - | - | 0 | - |
| - | - | 1983 | 511.2 | - | - | 0 | - |
| - | - | 3424 | 513.2 | - | - | 0 | - |
| - | - | 6626 | 514.2 | - | - | 0 | - |
| - | - | 1415 | 515.2 | - | - | 0 | - |
| - | - | 2185 | 518.2 | - | - | 0 | - |
| - | - | 5066 | 519.2 | - | - | 0 | - |
| - | - | 1782 | 520.2 | - | - | 0 | - |
| - | - | 1563 | 521.2 | - | - | 0 | - |
| - | - | 2170 | 522.2 | - | - | 0 | - |
| - | - | 728.3 | 523.2 | - | - | 0 | - |
| - | - | 8574 | 524.3 | - | - | 0 | - |
| - | - | 3055 | 525.3 | - | - | 0 | - |
| 5 | y | 1272 | 526.7 | 0.002409 | 4.573 | +2 | 8 |
| - | - | 982 | 527.2 | - | - | 0 | - |
| - | - | 2206 | 527.7 | - | - | 0 | - |
| - | - | 1124 | 528.2 | - | - | 0 | - |
| - | - | 825.2 | 528.7 | - | - | 0 | - |
| - | - | 1241 | 532.2 | - | - | 0 | - |
| - | - | 802.6 | 533.2 | - | - | 0 | - |
| - | - | 833 | 535.2 | - | - | 0 | - |
| - | - | 1388 | 540.2 | - | - | 0 | - |
| - | - | 2056 | 541.2 | - | - | 0 | - |
| - | - | 1228 | 541.7 | - | - | 0 | - |
| - | - | 8818 | 542.2 | - | - | 0 | - |
| - | - | 2068 | 543.2 | - | - | 0 | - |
| - | - | 4257 | 548.2 | - | - | 0 | - |
| - | - | 1476 | 549.2 | - | - | 0 | - |
| 5 | b | 1.223E+04 | 550.2 | 0.0005998 | 1.09 | +1 | 5 |
| - | - | 3749 | 551.2 | - | - | 0 | - |
| - | - | 1190 | 553.3 | - | - | 0 | - |
| - | - | 1675 | 557.2 | - | - | 0 | - |
| - | - | 1132 | 560.2 | - | - | 0 | - |
| - | - | 898.2 | 560.3 | - | - | 0 | - |
| - | - | 1235 | 566.2 | - | - | 0 | - |
| - | - | 903.4 | 567.2 | - | - | 0 | - |
| 5 | b | 8486 | 568.2 | 0.001937 | 3.409 | +1 | 5 |
| - | - | 3797 | 569.2 | - | - | 0 | - |
| - | - | 1786 | 569.7 | - | - | 0 | - |
| - | - | 5818 | 570.2 | - | - | 0 | - |
| - | - | 1798 | 571.2 | - | - | 0 | - |
| - | - | 8823 | 578.2 | - | - | 0 | - |
| - | - | 5103 | 578.7 | - | - | 0 | - |
| - | - | 2201 | 579.2 | - | - | 0 | - |
| - | - | 2225 | 583.2 | - | - | 0 | - |
| - | - | 1276 | 583.7 | - | - | 0 | - |
| - | - | 782.7 | 584.2 | - | - | 0 | - |
| 9 | y | 2.845E+04 | 588.3 | 0.006205 | 10.55 | +1 | 4 |
| - | - | 991.1 | 589.2 | - | - | 0 | - |
| - | - | 9576 | 589.3 | - | - | 0 | - |
| - | - | 1498 | 590.3 | - | - | 0 | - |
| 4 | y | 1470 | 591.2 | 0.00272 | 4.6 | +2 | 9 |
| - | - | 1215 | 591.7 | - | - | 0 | - |
| 10 | b | 6418 | 592.2 | 0.005214 | 8.804 | +2 | 10 |
| - | - | 5058 | 592.7 | - | - | 0 | - |
| - | - | 2554 | 593.2 | - | - | 0 | - |
| - | - | 1.051E+04 | 595.3 | - | - | 0 | - |
| - | - | 2174 | 596.2 | - | - | 0 | - |
| - | - | 3348 | 596.3 | - | - | 0 | - |
| - | - | 1330 | 597.2 | - | - | 0 | - |
| - | - | 1116 | 598.2 | - | - | 0 | - |
| 4 | y | 1.491E+04 | 600.2 | 0.003724 | 6.204 | +2 | 9 |
| - | - | 1.05E+04 | 600.7 | - | - | 0 | - |
| - | - | 4324 | 601.2 | - | - | 0 | - |
| - | - | 1934 | 601.7 | - | - | 0 | - |
| - | - | 991.8 | 605.2 | - | - | 0 | - |
| - | - | 5023 | 606.2 | - | - | 0 | - |
| - | - | 1865 | 607.2 | - | - | 0 | - |
| - | - | 2177 | 608.2 | - | - | 0 | - |
| - | - | 1965 | 609.8 | - | - | 0 | - |
| - | - | 1602 | 610.3 | - | - | 0 | - |
| - | - | 3849 | 612.2 | - | - | 0 | - |
| - | - | 965.8 | 613.2 | - | - | 0 | - |
| - | - | 3383 | 613.3 | - | - | 0 | - |
| - | - | 2645 | 614.2 | - | - | 0 | - |
| - | - | 2225 | 618.8 | - | - | 0 | - |
| - | - | 2764 | 619.3 | - | - | 0 | - |
| - | - | 928.6 | 620.2 | - | - | 0 | - |
| - | - | 638 | 621.2 | - | - | 0 | - |
| - | - | 4754 | 624.2 | - | - | 0 | - |
| - | - | 769.2 | 624.8 | - | - | 0 | - |
| - | - | 1041 | 625.2 | - | - | 0 | - |
| - | - | 925.8 | 626.2 | - | - | 0 | - |
| - | - | 1759 | 628.3 | - | - | 0 | - |
| - | - | 1078 | 631.2 | - | - | 0 | - |
| - | - | 1342 | 632.8 | - | - | 0 | - |
| - | - | 543.9 | 633.2 | - | - | 0 | - |
| - | - | 7881 | 634.2 | - | - | 0 | - |
| - | - | 2141 | 635.2 | - | - | 0 | - |
| - | - | 2478 | 637.3 | - | - | 0 | - |
| - | - | 1002 | 638.3 | - | - | 0 | - |
| - | - | 1243 | 639.3 | - | - | 0 | - |
| - | - | 822.3 | 639.8 | - | - | 0 | - |
| - | - | 1346 | 640.2 | - | - | 0 | - |
| - | - | 657.9 | 641.2 | - | - | 0 | - |
| - | - | 6484 | 641.3 | - | - | 0 | - |
| 3 | y | 1.639E+04 | 641.8 | 0.003905 | 6.085 | +2 | 10 |
| - | - | 1.126E+04 | 642.3 | - | - | 0 | - |
| - | - | 4163 | 642.8 | - | - | 0 | - |
| - | - | 770.4 | 643.3 | - | - | 0 | - |
| - | - | 4199 | 647.2 | - | - | 0 | - |
| - | - | 1633 | 648.2 | - | - | 0 | - |
| - | - | 677.6 | 649.2 | - | - | 0 | - |
| - | - | 779 | 650.2 | - | - | 0 | - |
| 3 | y | 3.228E+04 | 650.8 | 0.003811 | 5.856 | +2 | 10 |
| - | - | 2.506E+04 | 651.3 | - | - | 0 | - |
| - | - | 1.079E+04 | 651.8 | - | - | 0 | - |
| - | - | 2368 | 652.3 | - | - | 0 | - |
| - | - | 1477 | 653.3 | - | - | 0 | - |
| - | - | 788.8 | 655.3 | - | - | 0 | - |
| - | - | 5999 | 657.3 | - | - | 0 | - |
| - | - | 2456 | 658.3 | - | - | 0 | - |
| 8 | y | 5.443E+04 | 659.3 | 0.006262 | 9.497 | +1 | 5 |
| - | - | 1.678E+04 | 660.3 | - | - | 0 | - |
| - | - | 4009 | 661.3 | - | - | 0 | - |
| - | - | 1274 | 662.3 | - | - | 0 | - |
| - | - | 776 | 662.8 | - | - | 0 | - |
| - | - | 855 | 663.3 | - | - | 0 | - |
| 6 | b | 1.072E+04 | 665.3 | 0.001123 | 1.687 | +1 | 6 |
| - | - | 4096 | 666.3 | - | - | 0 | - |
| - | - | 1999 | 667.3 | - | - | 0 | - |
| - | - | 951 | 668.3 | - | - | 0 | - |
| - | - | 930.8 | 669.3 | - | - | 0 | - |
| - | - | 1772 | 670.3 | - | - | 0 | - |
| - | - | 9284 | 671.3 | - | - | 0 | - |
| - | - | 6860 | 671.8 | - | - | 0 | - |
| - | - | 3180 | 672.3 | - | - | 0 | - |
| - | - | 694.4 | 675.3 | - | - | 0 | - |
| - | - | 2312 | 676.3 | - | - | 0 | - |
| - | - | 4113 | 677.3 | - | - | 0 | - |
| - | - | 2236 | 678.3 | - | - | 0 | - |
| - | - | 8389 | 680.3 | - | - | 0 | - |
| - | - | 7279 | 680.8 | - | - | 0 | - |
| - | - | 3174 | 681.3 | - | - | 0 | - |
| - | - | 971.6 | 682.3 | - | - | 0 | - |
| 6 | b | 4297 | 683.3 | 0.0001404 | 0.2055 | +1 | 6 |
| - | - | 2685 | 684.3 | - | - | 0 | - |
| - | - | 823.7 | 684.8 | - | - | 0 | - |
| 11 | b | 1.469E+04 | 685.3 | 0.002521 | 3.679 | +2 | 11 |
| 11 | b | 5224 | 685.8 | 0.01212 | 17.68 | +2 | 11 |
| - | - | 5847 | 686.3 | - | - | 0 | - |
| - | - | 737.1 | 686.8 | - | - | 0 | - |
| - | - | 933.4 | 687.3 | - | - | 0 | - |
| 11 | b | 3111 | 694.3 | 0.002939 | 4.233 | +2 | 11 |
| - | - | 1790 | 694.8 | - | - | 0 | - |
| - | - | 6028 | 695.3 | - | - | 0 | - |
| - | - | 1809 | 696.3 | - | - | 0 | - |
| - | - | 652.1 | 699.3 | - | - | 0 | - |
| - | - | 743.7 | 700.3 | - | - | 0 | - |
| - | - | 1036 | 703.3 | - | - | 0 | - |
| - | - | 658.9 | 704.3 | - | - | 0 | - |
| - | - | 7752 | 705.3 | - | - | 0 | - |
| - | - | 3018 | 706.3 | - | - | 0 | - |
| - | - | 1004 | 707.3 | - | - | 0 | - |
| - | - | 7843 | 711.8 | - | - | 0 | - |
| - | - | 5384 | 712.3 | - | - | 0 | - |
| - | - | 2243 | 712.8 | - | - | 0 | - |
| - | - | 2315 | 713.3 | - | - | 0 | - |
| 2 | y | 1943 | 715.3 | 0.0005819 | 0.8136 | +2 | 11 |
| - | - | 1348 | 715.8 | - | - | 0 | - |
| - | - | 1534 | 716.3 | - | - | 0 | - |
| - | - | 2633 | 720.8 | - | - | 0 | - |
| - | - | 9162 | 721.3 | - | - | 0 | - |
| - | - | 1460 | 721.3 | - | - | 0 | - |
| - | - | 2284 | 721.8 | - | - | 0 | - |
| - | - | 3483 | 722.3 | - | - | 0 | - |
| - | - | 860.7 | 723.3 | - | - | 0 | - |
| 2 | y | 1825 | 724.3 | 0.002502 | 3.454 | +2 | 11 |
| - | - | 2841 | 724.8 | - | - | 0 | - |
| - | - | 973.9 | 725.3 | - | - | 0 | - |
| - | - | 922 | 726.3 | - | - | 0 | - |
| - | - | 2111 | 728.3 | - | - | 0 | - |
| - | - | 857.5 | 729.3 | - | - | 0 | - |
| - | - | 1482 | 729.8 | - | - | 0 | - |
| - | - | 2113 | 730.3 | - | - | 0 | - |
| - | - | 673.6 | 734.3 | - | - | 0 | - |
| - | - | 4399 | 734.8 | - | - | 0 | - |
| - | - | 5013 | 735.3 | - | - | 0 | - |
| - | - | 2506 | 735.8 | - | - | 0 | - |
| - | - | 1791 | 736.3 | - | - | 0 | - |
| - | - | 1078 | 737.3 | - | - | 0 | - |
| 0 | Precursor | 4.288E+04 | 743.8 | 0.003888 | 5.227 | +2 | -1 |
| - | - | 4.084E+04 | 744.3 | - | - | 0 | - |
| - | - | 2.127E+04 | 744.8 | - | - | 0 | - |
| - | - | 4393 | 745.3 | - | - | 0 | - |
| - | - | 1.356E+04 | 749.3 | - | - | 0 | - |
| - | - | 5386 | 750.3 | - | - | 0 | - |
| - | - | 1384 | 751.3 | - | - | 0 | - |
| - | - | 923.6 | 752.3 | - | - | 0 | - |
| 0 | Precursor | 1.309E+04 | 752.8 | 0.00355 | 4.715 | +2 | -1 |
| - | - | 1.015E+04 | 753.3 | - | - | 0 | - |
| - | - | 5125 | 753.8 | - | - | 0 | - |
| - | - | 1443 | 754.3 | - | - | 0 | - |
| - | - | 3184 | 756.3 | - | - | 0 | - |
| - | - | 1460 | 757.3 | - | - | 0 | - |
| - | - | 9846 | 758.4 | - | - | 0 | - |
| - | - | 1440 | 759.3 | - | - | 0 | - |
| - | - | 5187 | 759.4 | - | - | 0 | - |
| - | - | 936 | 760.3 | - | - | 0 | - |
| - | - | 968.3 | 778.3 | - | - | 0 | - |
| - | - | 1202 | 782.3 | - | - | 0 | - |
| - | - | 920.1 | 783.3 | - | - | 0 | - |
| - | - | 660.2 | 784.3 | - | - | 0 | - |
| - | - | 789.6 | 786.3 | - | - | 0 | - |
| - | - | 696.8 | 787.3 | - | - | 0 | - |
| - | - | 4498 | 792.3 | - | - | 0 | - |
| - | - | 1838 | 793.3 | - | - | 0 | - |
| - | - | 901.2 | 796.3 | - | - | 0 | - |
| - | - | 711.1 | 797.3 | - | - | 0 | - |
| - | - | 722.1 | 798.3 | - | - | 0 | - |
| - | - | 2003 | 800.3 | - | - | 0 | - |
| - | - | 1661 | 801.3 | - | - | 0 | - |
| - | - | 5166 | 804.3 | - | - | 0 | - |
| - | - | 2271 | 805.3 | - | - | 0 | - |
| - | - | 1848 | 810.3 | - | - | 0 | - |
| - | - | 1174 | 811.3 | - | - | 0 | - |
| - | - | 836.5 | 817.3 | - | - | 0 | - |
| - | - | 2267 | 818.3 | - | - | 0 | - |
| - | - | 778.4 | 819.3 | - | - | 0 | - |
| - | - | 8191 | 820.3 | - | - | 0 | - |
| - | - | 3218 | 821.3 | - | - | 0 | - |
| 7 | y | 3.732E+04 | 822.4 | 0.006288 | 7.646 | +1 | 6 |
| - | - | 1.847E+04 | 823.4 | - | - | 0 | - |
| - | - | 5546 | 824.4 | - | - | 0 | - |
| - | - | 849.3 | 827.3 | - | - | 0 | - |
| 7 | b | 8891 | 828.3 | 0.002491 | 3.008 | +1 | 7 |
| - | - | 3206 | 829.3 | - | - | 0 | - |
| - | - | 816.6 | 830.3 | - | - | 0 | - |
| - | - | 8652 | 832.3 | - | - | 0 | - |
| - | - | 2809 | 833.3 | - | - | 0 | - |
| - | - | 1719 | 842.3 | - | - | 0 | - |
| - | - | 4397 | 843.3 | - | - | 0 | - |
| - | - | 1926 | 844.3 | - | - | 0 | - |
| - | - | 898.8 | 845.4 | - | - | 0 | - |
| 7 | b | 5698 | 846.3 | 0.002486 | 2.937 | +1 | 7 |
| - | - | 2595 | 847.3 | - | - | 0 | - |
| - | - | 758.6 | 850.3 | - | - | 0 | - |
| - | - | 850.8 | 860.3 | - | - | 0 | - |
| - | - | 911.7 | 861.3 | - | - | 0 | - |
| - | - | 4387 | 868.3 | - | - | 0 | - |
| - | - | 1755 | 869.3 | - | - | 0 | - |
| - | - | 8176 | 871.3 | - | - | 0 | - |
| - | - | 4537 | 872.3 | - | - | 0 | - |
| - | - | 1.138E+04 | 873.4 | - | - | 0 | - |
| - | - | 6254 | 874.4 | - | - | 0 | - |
| - | - | 1405 | 878.3 | - | - | 0 | - |
| - | - | 1799 | 881.3 | - | - | 0 | - |
| - | - | 1241 | 882.3 | - | - | 0 | - |
| - | - | 2218 | 889.4 | - | - | 0 | - |
| - | - | 8822 | 896.3 | - | - | 0 | - |
| - | - | 4144 | 897.3 | - | - | 0 | - |
| - | - | 873.7 | 898.3 | - | - | 0 | - |
| 8 | b | 6189 | 899.4 | 0.001664 | 1.85 | +1 | 8 |
| 8 | b | 3048 | 900.3 | 0.01749 | 19.43 | +1 | 8 |
| - | - | 1921 | 905.4 | - | - | 0 | - |
| - | - | 9398 | 907.3 | - | - | 0 | - |
| - | - | 4619 | 908.3 | - | - | 0 | - |
| - | - | 1475 | 909.3 | - | - | 0 | - |
| - | - | 1781 | 915.4 | - | - | 0 | - |
| 8 | b | 3806 | 917.4 | 0.002768 | 3.017 | +1 | 8 |
| - | - | 929.5 | 917.5 | - | - | 0 | - |
| - | - | 1748 | 918.4 | - | - | 0 | - |
| 6 | y | 2122 | 919.4 | 0.003276 | 3.563 | +1 | 7 |
| - | - | 1191 | 920.4 | - | - | 0 | - |
| - | - | 2371 | 933.4 | - | - | 0 | - |
| - | - | 2217 | 934.4 | - | - | 0 | - |
| - | - | 1.57E+04 | 935.3 | - | - | 0 | - |
| - | - | 8368 | 936.3 | - | - | 0 | - |
| 6 | y | 5.038E+04 | 937.4 | 0.00559 | 5.963 | +1 | 7 |
| - | - | 2.538E+04 | 938.4 | - | - | 0 | - |
| - | - | 8360 | 939.4 | - | - | 0 | - |
| - | - | 1036 | 940.4 | - | - | 0 | - |
| - | - | 1959 | 953.3 | - | - | 0 | - |
| - | - | 702.8 | 954.4 | - | - | 0 | - |
| - | - | 905.6 | 961.4 | - | - | 0 | - |
| - | - | 869.5 | 964.4 | - | - | 0 | - |
| - | - | 1923 | 969.4 | - | - | 0 | - |
| - | - | 3521 | 970.4 | - | - | 0 | - |
| - | - | 2339 | 971.4 | - | - | 0 | - |
| - | - | 822 | 972.4 | - | - | 0 | - |
| - | - | 1796 | 979.4 | - | - | 0 | - |
| - | - | 1107 | 980.4 | - | - | 0 | - |
| - | - | 2026 | 982.4 | - | - | 0 | - |
| - | - | 1253 | 983.4 | - | - | 0 | - |
| - | - | 2.926E+04 | 988.4 | - | - | 0 | - |
| - | - | 1.417E+04 | 989.4 | - | - | 0 | - |
| - | - | 7191 | 990.4 | - | - | 0 | - |
| - | - | 2315 | 991.4 | - | - | 0 | - |
| - | - | 5883 | 997.4 | - | - | 0 | - |
| - | - | 2502 | 998.4 | - | - | 0 | - |
| - | - | 1290 | 1000 | - | - | 0 | - |
| - | - | 1021 | 1006 | - | - | 0 | - |
| - | - | 5648 | 1018 | - | - | 0 | - |
| - | - | 4076 | 1019 | - | - | 0 | - |
| - | - | 1164 | 1020 | - | - | 0 | - |
| 5 | y | 5607 | 1034 | 0.006789 | 6.564 | +1 | 8 |
| - | - | 4295 | 1035 | - | - | 0 | - |
| - | - | 2807 | 1036 | - | - | 0 | - |
| - | - | 1329 | 1037 | - | - | 0 | - |
| 9 | b | 2918 | 1046 | 0.007647 | 7.308 | +1 | 9 |
| - | - | 3192 | 1047 | - | - | 0 | - |
| 5 | y | 1.212E+05 | 1052 | 0.006479 | 6.156 | +1 | 8 |
| - | - | 6.92E+04 | 1053 | - | - | 0 | - |
| - | - | 3.002E+04 | 1054 | - | - | 0 | - |
| - | - | 6358 | 1055 | - | - | 0 | - |
| - | - | 1152 | 1056 | - | - | 0 | - |
| - | - | 982.6 | 1062 | - | - | 0 | - |
| 9 | b | 2906 | 1064 | 0.006115 | 5.745 | +1 | 9 |
| - | - | 1976 | 1065 | - | - | 0 | - |
| - | - | 8758 | 1082 | - | - | 0 | - |
| - | - | 4989 | 1083 | - | - | 0 | - |
| - | - | 1786 | 1084 | - | - | 0 | - |
| - | - | 2352 | 1091 | - | - | 0 | - |
| - | - | 1436 | 1092 | - | - | 0 | - |
| - | - | 986.5 | 1101 | - | - | 0 | - |
| - | - | 2235 | 1117 | - | - | 0 | - |
| - | - | 1004 | 1118 | - | - | 0 | - |
| - | - | 3732 | 1119 | - | - | 0 | - |
| - | - | 2562 | 1120 | - | - | 0 | - |
| - | - | 921 | 1121 | - | - | 0 | - |
| - | - | 1.726E+04 | 1135 | - | - | 0 | - |
| - | - | 1.063E+04 | 1136 | - | - | 0 | - |
| - | - | 5386 | 1137 | - | - | 0 | - |
| - | - | 1188 | 1138 | - | - | 0 | - |
| - | - | 906.7 | 1153 | - | - | 0 | - |
| - | - | 865.5 | 1154 | - | - | 0 | - |
| - | - | 3807 | 1155 | - | - | 0 | - |
| - | - | 2360 | 1156 | - | - | 0 | - |
| - | - | 1484 | 1157 | - | - | 0 | - |
| - | - | 1500 | 1165 | - | - | 0 | - |
| - | - | 1358 | 1173 | - | - | 0 | - |
| 4 | y | 2750 | 1181 | 0.008566 | 7.25 | +1 | 9 |
| - | - | 2335 | 1182 | - | - | 0 | - |
| 10 | b | 1.017E+04 | 1183 | 0.007695 | 6.502 | +1 | 10 |
| - | - | 6988 | 1184 | - | - | 0 | - |
| - | - | 2039 | 1185 | - | - | 0 | - |
| 4 | y | 6.935E+04 | 1199 | 0.005936 | 4.949 | +1 | 9 |
| - | - | 4.665E+04 | 1200 | - | - | 0 | - |
| - | - | 2.693E+04 | 1201 | - | - | 0 | - |
| - | - | 5774 | 1202 | - | - | 0 | - |
| - | - | 1421 | 1203 | - | - | 0 | - |
| - | - | 2012 | 1219 | - | - | 0 | - |
| - | - | 1626 | 1220 | - | - | 0 | - |
| - | - | 8286 | 1237 | - | - | 0 | - |
| - | - | 6265 | 1238 | - | - | 0 | - |
| - | - | 2633 | 1239 | - | - | 0 | - |
| - | - | 846 | 1258 | - | - | 0 | - |
| 3 | y | 3468 | 1283 | 0.005809 | 4.53 | +1 | 10 |
| - | - | 2819 | 1284 | - | - | 0 | - |
| - | - | 943.4 | 1285 | - | - | 0 | - |
| 3 | y | 3.487E+04 | 1301 | 0.005499 | 4.228 | +1 | 10 |
| - | - | 2.863E+04 | 1302 | - | - | 0 | - |
| - | - | 1.157E+04 | 1303 | - | - | 0 | - |
| - | - | 1840 | 1304 | - | - | 0 | - |
| - | - | 881.1 | 1311 | - | - | 0 | - |
| - | - | 954.2 | 1312 | - | - | 0 | - |
| - | - | 784.5 | 1340 | - | - | 0 | - |
| - | - | 1008 | 1359 | - | - | 0 | - |
| 11 | b | 799 | 1388 | 0.006318 | 4.554 | +1 | 11 |
| - | - | 651.6 | 2059 | - | - | 0 | - |

m/z Charge Intensity FragmentType MassShift Position
120.0811996459961 0 228508.86
121.07916259765625 0 869.52423
121.08451843261719 0 17636.174
122.07173156738281 0 1146.5847
122.08769989013672 0 536.68506
127.05055236816406 0 2053.3767
127.08683776855469 0 662.7044
128.04559326171875 0 368.24866
129.0663604736328 0 852.41656
129.10267639160156 0 1743.6624
130.05032348632812 0 890.2903
130.0655059814453 0 16182.215
131.04525756835938 0 939.205
131.06887817382812 0 1578.4137
132.08116149902344 0 5453.271
133.06100463867188 0 710.84705
133.0856475830078 0 411.51318
136.07608032226562 0 32663.94
136.0872344970703 0 1924.1555
137.07945251464844 0 2543.7542
138.0553436279297 0 3410.522
138.0665740966797 0 1870.5408
139.0701446533203 0 441.42505
141.06643676757812 0 1010.7477
143.04493713378906 0 452.8882
143.11886596679688 0 454.4828
144.0812225341797 0 3891.665
146.0606689453125 0 717.04816
146.09681701660156 0 2409.335
148.07606506347656 0 1770.3341
148.08761596679688 0 1480.0897
148.94662475585938 0 702.0711
155.08229064941406 0 1036.7546
155.09335327148438 0 2421.3372
156.07728576660156 0 2217.9248
157.0977325439453 0 458.28696
158.0843963623047 0 996.09424
158.09681701660156 0 5768.2495
159.0920867919922 0 37207.38
160.07598876953125 0 2675.5955 a Ammonia loss 1
160.08860778808594 0 588.3213
160.09548950195312 0 2931.2368
160.1124267578125 0 2324.8376
164.08200073242188 0 675.7299
165.07754516601562 0 4158.492
165.10252380371094 0 587.47925
166.05355834960938 0 955.15137
166.06153869628906 0 1780.509
166.08604431152344 0 1033.6826
166.09780883789062 0 617.18445
167.0930938720703 0 1943.032
169.06105041503906 0 3555.434
169.07615661621094 0 1639.508
170.0605926513672 0 2726.9795
171.06817626953125 0 2687.151
171.07630920410156 0 1157.0802
171.0919952392578 0 750.85956
173.09259033203125 0 672.1113
173.12901306152344 0 539.83887
173.4502716064453 0 4644.6963
174.0663604736328 0 859.16473
175.08717346191406 0 1109.3945
175.09759521484375 0 532.08606
175.1231231689453 0 1875.2361
176.08230590820312 0 3099.2805
176.107421875 0 16459.148
177.1027374267578 0 246106.98 a 1
178.06219482421875 0 622.0582
178.0996551513672 0 1399.063
178.10604858398438 0 25118.357
179.09304809570312 0 880.6788
179.1089630126953 0 618.8979
180.07720947265625 0 2332.9905
185.0562286376953 0 1430.0347
185.07106018066406 0 700.35834
186.09176635742188 0 11866.832
187.0715789794922 0 3226.5457
187.087158203125 0 713.88586
187.09548950195312 0 858.4421
187.1443328857422 0 1205.2279
188.0706024169922 0 578.1875 b Ammonia loss 1
191.09304809570312 0 1787.6964
191.11830139160156 0 2659.3262
193.0725555419922 0 1648.4723
193.0989227294922 0 371.61517
193.1088104248047 0 10854.557
194.09327697753906 0 583.2829
194.1122589111328 0 1172.656
195.08816528320312 0 3410.4473
197.60238647460938 0 529.15393
199.08607482910156 0 550.4789
201.1241912841797 0 797.3696
202.0530548095703 0 1107.3768
203.06654357910156 0 1677.7744
203.09335327148438 0 2093.1714
203.11831665039062 0 17822.42
204.07711791992188 0 5805.585
204.10105895996094 0 737.10034
204.12159729003906 0 1320.9626
205.09762573242188 0 116203.74 b 1
206.10093688964844 0 13527.186
207.10345458984375 0 778.97064
207.11334228515625 0 2151.2249
208.0607452392578 0 516.59796
208.07192993164062 0 1843.2141
209.89537048339844 0 507.39795
212.1184539794922 0 3083.9011
213.12245178222656 0 558.77673
214.08641052246094 0 791.1026
215.13934326171875 0 3021.3298
217.0975341796875 0 5304.8135
219.08004760742188 0 2173.6863
219.11328125 0 920.9015
219.1342010498047 0 588.4347
220.11959838867188 0 942.9095
221.10372924804688 0 29613.795
221.12901306152344 0 140458.67
222.10751342773438 0 2532.3462
222.1321563720703 0 13830.912
223.1338653564453 0 670.31134
225.10150146484375 0 723.4871
228.11361694335938 0 2556.411
229.10855102539062 0 968.45215
230.07730102539062 0 624.0323
231.0618133544922 0 5351.1743
231.11329650878906 0 21494.924
231.1261444091797 0 1092.5635
231.1488800048828 0 729.7148
232.11660766601562 0 2568.0317
233.0926513671875 0 687.9506
233.12896728515625 0 7486.663
234.13330078125 0 942.7645
235.10824584960938 0 13335.547
236.11053466796875 0 1769.1741
238.13099670410156 0 792.6307
241.0960693359375 0 803.4139
243.11312866210938 0 1090.0056
244.10882568359375 0 1556.6859
245.09274291992188 0 1049.3197
246.13514709472656 0 1289.9418
248.13951110839844 0 1504.4125
248.15211486816406 0 711.83014
249.1239013671875 0 47015.633
250.12718200683594 0 5708.7793
251.10311889648438 0 10370.667
251.1293487548828 0 1761.9408
252.10702514648438 0 968.95966
252.13320922851562 0 632.99554
253.046630859375 0 540.85443
256.1074523925781 0 795.77106
257.10699462890625 0 8684.123
258.1097106933594 0 1060.8484
258.16070556640625 0 814.8319
259.10736083984375 0 2239.4268
259.1441345214844 0 2230.6729
260.1398010253906 0 23957.357 a Water loss 2
261.08807373046875 0 890.6059
261.109375 0 1146.2029
261.12384033203125 0 17462.496 a Ammonia loss 2
261.1426086425781 0 2705.786
262.1192321777344 0 6661.916
262.1432189941406 0 730.9651
263.0704345703125 0 779.69073
263.08697509765625 0 684.6459
263.1029357910156 0 14191.048
264.1073303222656 0 2066.1619
264.1453552246094 0 1758.4363
267.09149169921875 0 1316.4135
268.07537841796875 0 1050.3518
268.14154052734375 0 665.031
270.1235656738281 0 729.9696
271.10699462890625 0 1246.0308
273.1227111816406 0 658.72394
274.1303405761719 0 6355.9336
275.1148376464844 0 756.3535
275.13525390625 0 1192.2981
276.1459045410156 0 2249.272
277.11822509765625 0 697.8749
278.1503601074219 0 3794.3538 a 2
279.09808349609375 0 9257.175
279.124267578125 0 1938.8016
279.1502990722656 0 3937.2605
280.1011962890625 0 858.62335
283.1528015136719 0 706.9618
285.10223388671875 0 17465.326
286.1050720214844 0 2191.787
286.1327209472656 0 615.66095
286.15545654296875 0 610.0004
286.2646789550781 0 624.5422
287.1014099121094 0 1004.21545
287.13970947265625 0 32676.1
288.1351013183594 0 73476.41 b Water loss 2
289.11871337890625 0 1297.2285 b Ammonia loss 2
289.1380615234375 0 12115.458
290.12554931640625 0 1013.313
290.1412048339844 0 992.5909
292.14111328125 0 13672.051
292.1672058105469 0 1159.7368
293.1440734863281 0 1234.7323
295.1444091796875 0 1583.9377
296.1512145996094 0 32903.457
297.1542663574219 0 4433.553
299.1506652832031 0 808.9034
302.11260986328125 0 690.56464
302.1292724609375 0 792.7576
303.1134948730469 0 576.9798
304.16632080078125 0 23356.81 y 10
305.11407470703125 0 733.1235
305.1687927246094 0 4159.827
306.14556884765625 0 95527.29 b 2
307.11962890625 0 1187.1509
307.14825439453125 0 15920.045
308.14996337890625 0 1834.911
313.0977478027344 0 884.2158
314.0992736816406 0 1227.4327
314.15087890625 0 1177.6907
318.1452941894531 0 3609.296
320.12255859375 0 989.34937
323.0992126464844 0 2292.5745
323.1393127441406 0 1219.9199
323.1759338378906 0 1439.6445
324.14630126953125 0 38843.074
325.1492919921875 0 5594.8755
327.1462097167969 0 886.7917
328.1298828125 0 1946.4442
328.1472473144531 0 1201.1077
329.13336181640625 0 832.9537
332.125 0 1492.079
333.1079406738281 0 1696.7065
333.1616516113281 0 916.216
334.1315002441406 0 1684.1757
338.128662109375 0 3060.2925
339.1319885253906 0 700.4808
339.17169189453125 0 1563.9677
342.15667724609375 0 5695.3267
343.1607666015625 0 958.4225
346.1406555175781 0 5934.995
347.14300537109375 0 1463.1995
348.1199951171875 0 2656.4963
350.1352233886719 0 8443.895
350.1871643066406 0 3507.529
351.1383972167969 0 1111.7732
351.1910400390625 0 738.5702
352.1418151855469 0 1315.5112
352.1661071777344 0 663.80927
355.1402282714844 0 885.17975
356.139404296875 0 10491.079
357.1390075683594 0 926.9209
358.13958740234375 0 942.911
360.1195983886719 0 1661.2938
360.1446838378906 0 835.0566
361.1772155761719 0 736.616
362.1612854003906 0 1748.9865
364.15093994140625 0 24199.836
365.15386962890625 0 5364.8633
366.1297302246094 0 2718.077
367.16455078125 0 719.79456
371.1382141113281 0 1133.556
372.11737060546875 0 728.53735
373.1676940917969 0 672.8989
374.13348388671875 0 1176.5164
375.1407165527344 0 537.16516
376.1147155761719 0 2644.826
378.10064697265625 0 1102.4496
378.1302795410156 0 21938.994
378.1824645996094 0 1771.6398
379.1332702636719 0 3557.8926
381.1925964355469 0 1858.6934
382.1435546875 0 3348.295
384.133544921875 0 940.5625
385.1515197753906 0 959.94794
389.1467590332031 0 709.0626
390.15716552734375 0 1024.9691
390.1820068359375 0 5181.9614
391.1830139160156 0 1052.9126
394.12548828125 0 16003.152
395.1283264160156 0 2767.149
395.154541015625 0 774.363
395.22198486328125 0 772.57556
399.1707458496094 0 703.71844
403.1626281738281 0 3626.6682
404.11309814453125 0 942.6078
404.1451721191406 0 1535.0215
405.20513916015625 0 2272.0881
406.17913818359375 0 605.80975
407.18218994140625 0 6400.066
407.2089538574219 0 26140.91
408.1860656738281 0 1188.2483
408.2120056152344 0 5682.628
409.18927001953125 0 2178.501
410.1382751464844 0 1144.9475
410.1883544921875 0 849.0571
411.6847229003906 0 1015.7504 y 6
416.1471862792969 0 1001.3911
419.1564636230469 0 785.04004
421.17291259765625 0 1807.5034
422.1729431152344 0 761.21985
423.21466064453125 0 3762.9827
424.214599609375 0 966.9807
425.19134521484375 0 1157.0248
425.2196960449219 0 4091.2712
426.2231750488281 0 1125.283
427.20928955078125 0 3983.6816
428.2132568359375 0 687.97546
431.15740966796875 0 1132.2919
433.1728210449219 0 2533.7502
433.224609375 0 806.54675
434.1744384765625 0 643.27185
435.17742919921875 0 512.7251
435.20391845703125 0 21727.602 b Water loss 3
436.20697021484375 0 5144.025
441.2255859375 0 69064.02 y 9
442.1718444824219 0 1623.0106
442.22833251953125 0 16386.94
443.1564636230469 0 629.16766
443.1871337890625 0 1562.4294
443.23077392578125 0 2424.857
447.1520080566406 0 1660.2855
449.1675109863281 0 1399.2446
450.22686767578125 0 2263.061
451.1455078125 0 589.7095
451.2044372558594 0 1112.5691
453.2149353027344 0 3897.0554 b 3
454.2203063964844 0 996.8865
455.2054443359375 0 2230.6763
456.2077331542969 0 880.21185
461.16754150390625 0 6301.5723
462.17010498046875 0 1094.407
465.1628112792969 0 11967.569
466.16522216796875 0 2645.4238
469.1999816894531 0 697.93256 y 5
470.16705322265625 0 1842.7468
471.1818542480469 0 8832.987
472.1844787597656 0 2599.5964
472.22430419921875 0 712.52997
473.21527099609375 0 620.9401
476.18511962890625 0 770.30756
477.1713562011719 0 1201.8712
478.220458984375 0 2797.1543
479.1786804199219 0 13906.212
480.18157958984375 0 3391.699
483.1922607421875 0 946.938
486.16357421875 0 2708.07
488.21380615234375 0 646.45905
489.1628723144531 0 1401.0256
489.2261962890625 0 1630.2279
490.1947326660156 0 899.8858
490.2306213378906 0 681.3945
491.207763671875 0 4927.6064
492.20758056640625 0 828.51373
493.2096862792969 0 2269.7363
495.1868591308594 0 1735.4048
496.17230224609375 0 948.51166
497.1720886230469 0 3124.0393
500.18084716796875 0 2468.1052
504.2249755859375 0 2202.0928
507.2371520996094 0 2580.0735
508.24053955078125 0 1813.0458
511.2202453613281 0 1982.5792
513.19873046875 0 3423.7737
514.2234497070312 0 6625.609
515.2279052734375 0 1414.9323
518.1896362304688 0 2185.4104
519.2015991210938 0 5066.0986
520.2061157226562 0 1781.6486
521.2055053710938 0 1562.9832
522.2371826171875 0 2169.951
523.184326171875 0 728.3142
524.2628173828125 0 8574.163
525.26513671875 0 3055.0613
526.710693359375 0 1272.0374 y 4
527.2129516601562 0 981.9989
527.7099609375 0 2205.5435
528.2102661132812 0 1123.9194
528.7130126953125 0 825.1548
532.2168579101562 0 1241.3635
533.20703125 0 802.6332
535.1821899414062 0 833.03876
540.2452392578125 0 1387.5892
541.1947021484375 0 2055.7788
541.70751953125 0 1227.5903
542.2197875976562 0 8817.895
543.2218017578125 0 2067.6384
548.2008666992188 0 4256.986
549.2044677734375 0 1476.379
550.230224609375 0 12232.343 b Water loss 4
551.2318725585938 0 3749.097
553.2642822265625 0 1190.1023
557.2007446289062 0 1675.1964
560.22412109375 0 1131.6625
560.2670288085938 0 898.23004
566.2256469726562 0 1235.2468
567.2091064453125 0 903.38794
568.2421264648438 0 8485.977 b 4
569.2355346679688 0 3796.9841
569.7296752929688 0 1786.1073
570.234619140625 0 5818.027
571.233154296875 0 1798.1495
578.2327880859375 0 8822.554
578.7340087890625 0 5102.681
579.23388671875 0 2201.1436
583.2268676757812 0 2224.5654
583.7252807617188 0 1275.662
584.2332153320312 0 782.69293
588.2611694335938 0 28445.295 y 8
589.2119140625 0 991.0748
589.2639770507812 0 9576.413
590.2647094726562 0 1497.8202
591.2399291992188 0 1470.119 y Water loss 3
591.740966796875 0 1215.3055
592.2320556640625 0 6417.803 b Water loss 9
592.7316284179688 0 5058.458
593.232421875 0 2554.1716
595.3001098632812 0 10505.285
596.2368774414062 0 2173.9934
596.3021240234375 0 3347.802
597.22119140625 0 1330.2988
598.2256469726562 0 1115.5763
600.2462158203125 0 14905.227 y 3
600.7476806640625 0 10504.773
601.2451782226562 0 4324.1123
601.74267578125 0 1934.2686
605.2347412109375 0 991.7631
606.2343139648438 0 5022.8457
607.2365112304688 0 1865.1047
608.238037109375 0 2177.0374
609.7646484375 0 1964.5529
610.2662963867188 0 1601.9753
612.1981201171875 0 3848.9175
613.1994018554688 0 965.78644
613.29052734375 0 3382.7407
614.245849609375 0 2644.593
618.7687377929688 0 2224.893
619.2619018554688 0 2764.3733
620.2417602539062 0 928.59326
621.2318115234375 0 637.9502
624.2312622070312 0 4753.517
624.754150390625 0 769.18756
625.235107421875 0 1040.8169
626.24267578125 0 925.7717
628.2679443359375 0 1758.5443
631.2473754882812 0 1078.3191
632.7573852539062 0 1341.8872
633.2391967773438 0 543.9435
634.2304077148438 0 7880.862
635.2322998046875 0 2141.0286
637.2630615234375 0 2478.385
638.2636108398438 0 1001.8543
639.27001953125 0 1242.5902
639.7728881835938 0 822.2839
640.2354125976562 0 1346.0438
641.2408447265625 0 657.9033
641.2853393554688 0 6483.9287
641.7649536132812 0 16385.072 y Water loss 2
642.2669067382812 0 11260.761
642.7647094726562 0 4162.5874
643.2567138671875 0 770.3885
647.247802734375 0 4199.017
648.2457275390625 0 1632.9652
649.2426147460938 0 677.5821
650.24560546875 0 779.0253
650.7701416015625 0 32277.102 y 2
651.2714233398438 0 25057.39
651.7717895507812 0 10793.48
652.2711791992188 0 2368.0574
653.2665405273438 0 1476.8721
655.2733154296875 0 788.756
657.26416015625 0 5998.7896
658.267822265625 0 2455.5667
659.29833984375 0 54429.332 y 7
660.3009643554688 0 16775.566
661.296875 0 4009.4785
662.271728515625 0 1273.9739
662.7680053710938 0 775.9674
663.2505493164062 0 855.01843
665.2576904296875 0 10724.361 b Water loss 5
666.26123046875 0 4095.954
667.260986328125 0 1999.0227
668.2554931640625 0 950.9894
669.2755126953125 0 930.77954
670.27490234375 0 1771.6216
671.2728881835938 0 9283.828
671.7747802734375 0 6859.567
672.2745971679688 0 3179.928
675.270263671875 0 694.4117
676.2694091796875 0 2312.0374
677.2847290039062 0 4113.098
678.2890625 0 2236.2073
680.2781372070312 0 8389.48
680.779296875 0 7278.5166
681.260498046875 0 3174.057
682.2529296875 0 971.57837
683.2672729492188 0 4297.1963 b 5
684.27197265625 0 2685.2825
684.7726440429688 0 823.66833
685.2639770507812 0 14694.837 b Water loss 10
685.7706298828125 0 5223.944 b Ammonia loss 10
686.26513671875 0 5847.357
686.7714233398438 0 737.10126
687.266845703125 0 933.35455
694.2747192382812 0 3111.4202 b 10
694.7767333984375 0 1789.7502
695.2692260742188 0 6027.884
696.2710571289062 0 1809.0168
699.2603759765625 0 652.092
700.2667236328125 0 743.70374
703.2980346679688 0 1035.7021
704.2730102539062 0 658.8748
705.2818603515625 0 7752.0317
706.2841796875 0 3017.7358
707.2921142578125 0 1003.9612
711.810302734375 0 7842.586
712.3118286132812 0 5383.899
712.8145141601562 0 2243.1987
713.2830200195312 0 2315.116
715.2958374023438 0 1943.0238 y Water loss 1
715.8030395507812 0 1347.5515
716.2982177734375 0 1534.3739
720.81201171875 0 2633.001
721.2619018554688 0 9162.389
721.3170776367188 0 1460.3433
721.8058471679688 0 2283.8809
722.2656860351562 0 3483.1323
723.2933349609375 0 860.69434
724.3030395507812 0 1824.5452 y 1
724.8046875 0 2840.5474
725.30517578125 0 973.8709
726.2940063476562 0 922.039
728.3174438476562 0 2110.6824
729.3167724609375 0 857.48083
729.8075561523438 0 1481.9662
730.3130493164062 0 2113.1685
734.2839965820312 0 673.5734
734.8051147460938 0 4399.043
735.3016357421875 0 5013.2515
735.8047485351562 0 2506.1746
736.300048828125 0 1791.1909
737.3004150390625 0 1077.6716
743.8098754882812 0 42877.715 Precursor Water loss
744.3112182617188 0 40837.805
744.8121337890625 0 21269.766
745.3121337890625 0 4393.2095
749.25732421875 0 13564.805
750.2610473632812 0 5385.6357
751.2573852539062 0 1383.6602
752.2952270507812 0 923.64813
752.8148193359375 0 13088.992 Precursor
753.3154907226562 0 10146.313
753.8165283203125 0 5125.0347
754.3129272460938 0 1442.744
756.3104248046875 0 3184.4873
757.312255859375 0 1460.0067
758.3633422851562 0 9846.427
759.2633056640625 0 1439.9547
759.365234375 0 5186.5327
760.2713012695312 0 935.96045
778.3056640625 0 968.29535
782.3121337890625 0 1201.6877
783.3060913085938 0 920.06085
784.3026733398438 0 660.1543
786.31787109375 0 789.5982
787.305908203125 0 696.8026
792.3150634765625 0 4498.1597
793.316162109375 0 1838.1492
796.3145751953125 0 901.15375
797.3148193359375 0 711.1437
798.3209838867188 0 722.11334
800.3240966796875 0 2003.3256
801.3175659179688 0 1660.9465
804.33447265625 0 5166.1333
805.3375854492188 0 2270.9185
810.3115234375 0 1847.541
811.3062744140625 0 1173.6016
817.3255615234375 0 836.4886
818.3369140625 0 2266.943
819.33544921875 0 778.42804
820.3106689453125 0 8190.775
821.3140258789062 0 3217.9846
822.3616943359375 0 37315.074 y 6
823.3642578125 0 18472.123
824.3641357421875 0 5546.361
827.3460083007812 0 849.33887
828.3223876953125 0 8891.406 b Water loss 6
829.3248291015625 0 3206.109
830.3258666992188 0 816.6244
832.328125 0 8651.66
833.3284301757812 0 2808.8835
842.310546875 0 1719.1295
843.3417358398438 0 4396.5625
844.34912109375 0 1925.7756
845.356689453125 0 898.84686
846.3329467773438 0 5698.3213 b 6
847.3347778320312 0 2595.3115
850.325439453125 0 758.6158
860.3148803710938 0 850.8293
861.3153076171875 0 911.7346
868.3308715820312 0 4386.874
869.3333129882812 0 1754.7484
871.340087890625 0 8175.577
872.3430786132812 0 4536.799
873.3890380859375 0 11375.602
874.3932495117188 0 6253.619
878.31591796875 0 1404.5277
881.3485107421875 0 1798.7826
882.3453979492188 0 1241.2526
889.350830078125 0 2218.4578
896.3253173828125 0 8822.228
897.33056640625 0 4143.751
898.330078125 0 873.72815
899.3553466796875 0 6188.8735 b Water loss 7
900.3585205078125 0 3047.9338 b Ammonia loss 7
905.3759155273438 0 1920.5454
907.3412475585938 0 9397.517
908.344970703125 0 4618.607
909.3444213867188 0 1475.3274
915.3604736328125 0 1780.5974
917.3648071289062 0 3805.9639 b 7
917.4706420898438 0 929.5147
918.3724365234375 0 1747.65
919.3750610351562 0 2121.707 y Water loss 5
920.3829345703125 0 1190.571
933.3737182617188 0 2371.1345
934.3787841796875 0 2216.9358
935.3370361328125 0 15697.821
936.3392944335938 0 8367.644
937.387939453125 0 50378.13 y 5
938.3912963867188 0 25375.252
939.391357421875 0 8360.377
940.3873291015625 0 1035.8911
953.3482055664062 0 1958.501
954.3673706054688 0 702.7936
961.4033813476562 0 905.64294
964.3883056640625 0 869.4898
969.3784790039062 0 1923.1189
970.4011840820312 0 3521.3215
971.3966674804688 0 2338.931
972.401123046875 0 822.0201
979.362548828125 0 1795.771
980.3709106445312 0 1107.3348
982.3948974609375 0 2025.9802
983.3994140625 0 1252.5519
988.417724609375 0 29255.727
989.420166015625 0 14165.124
990.4187622070312 0 7190.7104
991.4119873046875 0 2315.1294
997.3729248046875 0 5882.533
998.3770751953125 0 2501.8992
1000.395751953125 0 1289.75
1006.4105834960938 0 1020.9628
1018.4058837890625 0 5647.556
1019.4071044921875 0 4075.8862
1020.4058837890625 0 1164.3715
1034.405517578125 0 5607.256 y Water loss 4
1035.4046630859375 0 4294.7856
1036.406005859375 0 2806.865
1037.39599609375 0 1329.0388
1046.3951416015625 0 2918.4844 b Water loss 8
1047.399658203125 0 3192.3828
1052.415771484375 0 121181.125 y 4
1053.4185791015625 0 69203.2
1054.4169921875 0 30020.88
1055.41796875 0 6357.9663
1056.40869140625 0 1151.9375
1062.395751953125 0 982.5907
1064.4041748046875 0 2905.992 b 8
1065.406982421875 0 1976.3333
1082.4049072265625 0 8757.543
1083.4083251953125 0 4989.3315
1084.4080810546875 0 1786.356
1091.4583740234375 0 2352.2808
1092.4600830078125 0 1436.2253
1101.434814453125 0 986.5417
1117.476318359375 0 2235.266
1118.482421875 0 1004.0147
1119.45458984375 0 3731.934
1120.4573974609375 0 2561.551
1121.4547119140625 0 921.01794
1135.485107421875 0 17257.822
1136.4886474609375 0 10628.515
1137.482177734375 0 5386.4595
1138.4801025390625 0 1187.8214
1153.47607421875 0 906.7102
1154.47021484375 0 865.45276
1155.461181640625 0 3806.7195
1156.4605712890625 0 2360.3296
1157.4639892578125 0 1484.4741
1165.4482421875 0 1499.9445
1173.4658203125 0 1357.8921
1181.4757080078125 0 2749.801 y Water loss 3
1182.4764404296875 0 2334.8118
1183.4541015625 0 10172.07 b Water loss 9
1184.4560546875 0 6988.4067
1185.45654296875 0 2039.2803
1199.483642578125 0 69350.9 y 3
1200.4864501953125 0 46651.66
1201.4827880859375 0 26928.209
1202.474853515625 0 5773.6636
1203.47314453125 0 1421.3212
1218.50732421875 0 2012.145
1219.5045166015625 0 1626.1649
1236.531982421875 0 8286.304
1237.5347900390625 0 6265.2046
1238.5308837890625 0 2633.2136
1258.4921875 0 846.0295
1282.5206298828125 0 3467.7634 y Water loss 2
1283.5247802734375 0 2818.693
1284.5294189453125 0 943.3682
1300.5308837890625 0 34872.887 y 2
1301.5343017578125 0 28632.162
1302.5340576171875 0 11571.54
1303.5350341796875 0 1840.0315
1310.5225830078125 0 881.13916
1311.5286865234375 0 954.19745
1339.546630859375 0 784.5161
1358.552001953125 0 1007.66034
1387.5426025390625 0 799.0398 b 10
2058.510009765625 0 651.6457

Spectrum Details

|  |  |
| --- | --- |
| Matched peaks? Matched peaks The total absolute number of peaks matched. Additionally in brackets the total fraction of peaks matched and the total number of peaks is shown. | 53 (7.39% of 717) |
| FDR? FDR The false discovery rate estimated for this peptide. It is calculated by matching all theoretical fragments with a non-integer shift with the raw peaks for this spectrum. This is done with 40 different shifts. The resulting percentage is the average number of annotated peaks over the number of annotated peaks with the correct spectrum. | 0.18% |
| Satellite FDR? Satellite FDR See the FDR for details on its calculation. This satellite ion specific FDR only contains the satellite ions (d/w) for I/L/J positions. | - |
| PSM Score? PSM Score The PSM Score as given by Hecklib to this annotated spectrum. It is shown with three significant figures. | 638 |

#### Spectrum 11109? Spectrum 11109 The raw spectrum of this peptide as annotated by Hecklib. The fragments are coloured according to ion type (see legend). Any peaks with a star '\*' as text can be hovered over to see the full details, first the ion type second the mass shift type. By hovering over the amino acids in the peptide or ions in the legend the corresponding peaks are highlighted. By toggling the 'Unassigned' label you can turn the background (unassigned) peaks on or off in the plot. By updating the slider in the Ion legend you can update the spectrum to only show the top X% of the peaks with labels. The top X% means any peak that is within X% of the highest intensity. By dragging in the spectrum you can zoom in to a specific part of the spectrum and use 'Zoom Out' to get back to the original zoom level. The annotation of the spectrum is based on the given sequence in the peptides file and is done with different software so inconsistencies are likely. The peaks are annotated based on the given sequence, with 20 ppm tolerance.

Copy Data

##### Spectrum 11109 (TSV)

###### Preview

```
Loading example...
```

*Click on the button to copy the data to your clipboard.*

Mz MinMz MaxIntensity Max

WidthHeightPeptide font sizePeptide stroke widthSpectrum font sizeSpectrum stroke widthCompact peptide

Ion legend

wxyz

abcd

OtherUnassignedIon^Charge^_Position_Show for top:%

GFTFDDYAMHWV

01.07e+42.15e+43.22e+44.29e+4

Zoom Out

a^+1^_2_a^+1^_2_b^+1^_2_a^+1^_3_a^+1^_3_a^+1^_3_b^+1^_3_y^+1^_2_b^+1^_3_b^+1^_4_y^+1^_3_b^+1^_5_b^+1^_5_y^+1^_4_y^+2^_9_y^+2^_10_y^+2^_10_y^+1^_5_b^+1^_6_b^+2^_11_b^+2^_11_y^+2^_11_\*\*y^+1^_6_b^+1^_7_y^+1^_7_y^+1^_8_y^+1^_9_b^+1^_10_y^+1^_9_b^+1^_10_y^+1^_10_

0542108516272170

Fragment Matches Table

Show background peaks

| Position | Ion type | Intensity | mz Theoretical | mz Error (Th) | mz Error (ppm) | Charge | Series Number |
| --- | --- | --- | --- | --- | --- | --- | --- |
| - | - | 4.248E+04 | 120.1 | - | - | 0 | - |
| - | - | 3388 | 121.1 | - | - | 0 | - |
| - | - | 422.5 | 125.8 | - | - | 0 | - |
| - | - | 839.5 | 129.1 | - | - | 0 | - |
| - | - | 2961 | 129.1 | - | - | 0 | - |
| - | - | 539.5 | 130.1 | - | - | 0 | - |
| - | - | 3039 | 130.1 | - | - | 0 | - |
| - | - | 414.9 | 130.1 | - | - | 0 | - |
| - | - | 548.6 | 131 | - | - | 0 | - |
| - | - | 954.5 | 132.1 | - | - | 0 | - |
| - | - | 738.3 | 133.1 | - | - | 0 | - |
| - | - | 1656 | 133.1 | - | - | 0 | - |
| - | - | 8428 | 136.1 | - | - | 0 | - |
| - | - | 909.6 | 138.1 | - | - | 0 | - |
| - | - | 596.2 | 138.1 | - | - | 0 | - |
| - | - | 471.3 | 140.5 | - | - | 0 | - |
| - | - | 507.2 | 143.1 | - | - | 0 | - |
| - | - | 420.1 | 145.3 | - | - | 0 | - |
| - | - | 522.5 | 146.1 | - | - | 0 | - |
| - | - | 430.8 | 147.1 | - | - | 0 | - |
| - | - | 400.9 | 148.1 | - | - | 0 | - |
| - | - | 1068 | 149 | - | - | 0 | - |
| - | - | 453.8 | 150.1 | - | - | 0 | - |
| - | - | 431.9 | 151.8 | - | - | 0 | - |
| - | - | 592.9 | 152.1 | - | - | 0 | - |
| - | - | 701 | 153.1 | - | - | 0 | - |
| - | - | 429.3 | 154.1 | - | - | 0 | - |
| - | - | 463.7 | 154.6 | - | - | 0 | - |
| - | - | 1580 | 155.1 | - | - | 0 | - |
| - | - | 629.9 | 155.1 | - | - | 0 | - |
| - | - | 1714 | 155.1 | - | - | 0 | - |
| - | - | 937.8 | 156.1 | - | - | 0 | - |
| - | - | 477.3 | 158.1 | - | - | 0 | - |
| - | - | 620.8 | 158.1 | - | - | 0 | - |
| - | - | 6492 | 159.1 | - | - | 0 | - |
| 2 | a | 585.7 | 160.1 | 0.0001306 | 0.8156 | +1 | 2 |
| - | - | 1050 | 160.1 | - | - | 0 | - |
| - | - | 657.6 | 165.1 | - | - | 0 | - |
| - | - | 507.5 | 166.1 | - | - | 0 | - |
| - | - | 439.2 | 167.1 | - | - | 0 | - |
| - | - | 548 | 169.1 | - | - | 0 | - |
| - | - | 471 | 171.1 | - | - | 0 | - |
| - | - | 466.2 | 173.1 | - | - | 0 | - |
| - | - | 1745 | 173.1 | - | - | 0 | - |
| - | - | 2841 | 176.1 | - | - | 0 | - |
| 2 | a | 3.856E+04 | 177.1 | 0.0002538 | 1.433 | +1 | 2 |
| - | - | 3534 | 178.1 | - | - | 0 | - |
| - | - | 591.9 | 181.1 | - | - | 0 | - |
| - | - | 1001 | 183.1 | - | - | 0 | - |
| - | - | 559.6 | 184.2 | - | - | 0 | - |
| - | - | 546.3 | 185.1 | - | - | 0 | - |
| - | - | 2214 | 186.1 | - | - | 0 | - |
| - | - | 554.5 | 187.1 | - | - | 0 | - |
| - | - | 2497 | 187.1 | - | - | 0 | - |
| - | - | 572.9 | 188.1 | - | - | 0 | - |
| - | - | 1982 | 193.1 | - | - | 0 | - |
| - | - | 1007 | 195.1 | - | - | 0 | - |
| - | - | 502.2 | 197.1 | - | - | 0 | - |
| - | - | 669.7 | 199.1 | - | - | 0 | - |
| - | - | 591.8 | 199.2 | - | - | 0 | - |
| - | - | 1917 | 201.1 | - | - | 0 | - |
| - | - | 535.9 | 203.1 | - | - | 0 | - |
| - | - | 3009 | 203.1 | - | - | 0 | - |
| - | - | 703.4 | 204.1 | - | - | 0 | - |
| 2 | b | 1.761E+04 | 205.1 | 0.0001817 | 0.886 | +1 | 2 |
| - | - | 1148 | 205.1 | - | - | 0 | - |
| - | - | 2401 | 206.1 | - | - | 0 | - |
| - | - | 739.5 | 207.1 | - | - | 0 | - |
| - | - | 548.1 | 212.1 | - | - | 0 | - |
| - | - | 559.9 | 215.1 | - | - | 0 | - |
| - | - | 5444 | 215.1 | - | - | 0 | - |
| - | - | 804.1 | 216.1 | - | - | 0 | - |
| - | - | 1119 | 217.1 | - | - | 0 | - |
| - | - | 481.6 | 217.9 | - | - | 0 | - |
| - | - | 5207 | 219.1 | - | - | 0 | - |
| - | - | 5000 | 221.1 | - | - | 0 | - |
| - | - | 2.367E+04 | 221.1 | - | - | 0 | - |
| - | - | 2762 | 222.1 | - | - | 0 | - |
| - | - | 1823 | 226.1 | - | - | 0 | - |
| - | - | 704.3 | 231.1 | - | - | 0 | - |
| - | - | 3654 | 231.1 | - | - | 0 | - |
| - | - | 805.8 | 233.1 | - | - | 0 | - |
| - | - | 547.1 | 234.1 | - | - | 0 | - |
| - | - | 2152 | 235.1 | - | - | 0 | - |
| - | - | 478.4 | 236.5 | - | - | 0 | - |
| - | - | 615.8 | 238.1 | - | - | 0 | - |
| - | - | 495.1 | 241.1 | - | - | 0 | - |
| - | - | 8325 | 249.1 | - | - | 0 | - |
| - | - | 1687 | 250.1 | - | - | 0 | - |
| - | - | 1954 | 251.1 | - | - | 0 | - |
| - | - | 865.5 | 257.1 | - | - | 0 | - |
| 3 | a | 3980 | 260.1 | 0.0001121 | 0.4308 | +1 | 3 |
| 3 | a | 2567 | 261.1 | 0.0001968 | 0.7538 | +1 | 3 |
| - | - | 713.8 | 261.1 | - | - | 0 | - |
| - | - | 846 | 262.1 | - | - | 0 | - |
| - | - | 3176 | 263.1 | - | - | 0 | - |
| - | - | 1124 | 274.1 | - | - | 0 | - |
| 3 | a | 829.4 | 278.1 | 0.0006253 | 2.248 | +1 | 3 |
| - | - | 1914 | 279.1 | - | - | 0 | - |
| - | - | 1050 | 279.1 | - | - | 0 | - |
| - | - | 575.7 | 283.1 | - | - | 0 | - |
| - | - | 1214 | 284.2 | - | - | 0 | - |
| - | - | 2702 | 285.1 | - | - | 0 | - |
| - | - | 4988 | 287.1 | - | - | 0 | - |
| 3 | b | 1.18E+04 | 288.1 | 0.0002841 | 0.9861 | +1 | 3 |
| - | - | 1608 | 289.1 | - | - | 0 | - |
| - | - | 2242 | 292.1 | - | - | 0 | - |
| - | - | 5592 | 296.2 | - | - | 0 | - |
| - | - | 1229 | 297.2 | - | - | 0 | - |
| - | - | 1270 | 297.2 | - | - | 0 | - |
| 11 | y | 4365 | 304.2 | 0.0006307 | 2.074 | +1 | 2 |
| - | - | 864.9 | 305.2 | - | - | 0 | - |
| 3 | b | 1.529E+04 | 306.1 | 0.0003701 | 1.209 | +1 | 3 |
| - | - | 2372 | 307.1 | - | - | 0 | - |
| - | - | 870.1 | 314.2 | - | - | 0 | - |
| - | - | 5812 | 324.1 | - | - | 0 | - |
| - | - | 913.3 | 325.2 | - | - | 0 | - |
| - | - | 888.2 | 325.2 | - | - | 0 | - |
| - | - | 584.5 | 328.2 | - | - | 0 | - |
| - | - | 652.5 | 342.2 | - | - | 0 | - |
| - | - | 582.8 | 346.1 | - | - | 0 | - |
| - | - | 2008 | 350.1 | - | - | 0 | - |
| - | - | 510.7 | 350.2 | - | - | 0 | - |
| - | - | 2366 | 356.1 | - | - | 0 | - |
| - | - | 546.1 | 362.2 | - | - | 0 | - |
| - | - | 4325 | 364.2 | - | - | 0 | - |
| - | - | 683.1 | 365.2 | - | - | 0 | - |
| - | - | 688.3 | 366.1 | - | - | 0 | - |
| - | - | 964.7 | 376.1 | - | - | 0 | - |
| - | - | 2980 | 378.1 | - | - | 0 | - |
| - | - | 755.4 | 379.1 | - | - | 0 | - |
| - | - | 573.9 | 379.2 | - | - | 0 | - |
| - | - | 967.8 | 382.1 | - | - | 0 | - |
| - | - | 669.4 | 390.2 | - | - | 0 | - |
| - | - | 2165 | 394.1 | - | - | 0 | - |
| - | - | 737.7 | 403.2 | - | - | 0 | - |
| - | - | 827.2 | 407.2 | - | - | 0 | - |
| - | - | 4374 | 407.2 | - | - | 0 | - |
| - | - | 890 | 424.3 | - | - | 0 | - |
| - | - | 847 | 425.2 | - | - | 0 | - |
| - | - | 593 | 426.2 | - | - | 0 | - |
| 4 | b | 2707 | 435.2 | 0.0002296 | 0.5275 | +1 | 4 |
| - | - | 795.9 | 436.2 | - | - | 0 | - |
| 10 | y | 1.001E+04 | 441.2 | 0.0005262 | 1.193 | +1 | 3 |
| - | - | 2229 | 442.2 | - | - | 0 | - |
| - | - | 524.8 | 444.5 | - | - | 0 | - |
| - | - | 800.9 | 449.2 | - | - | 0 | - |
| - | - | 1320 | 461.2 | - | - | 0 | - |
| - | - | 2342 | 465.2 | - | - | 0 | - |
| - | - | 1758 | 471.2 | - | - | 0 | - |
| - | - | 681.2 | 472.2 | - | - | 0 | - |
| - | - | 2662 | 479.2 | - | - | 0 | - |
| - | - | 817.4 | 480.2 | - | - | 0 | - |
| - | - | 758 | 497.2 | - | - | 0 | - |
| - | - | 675.3 | 513.2 | - | - | 0 | - |
| - | - | 993 | 514.2 | - | - | 0 | - |
| - | - | 647.7 | 519.2 | - | - | 0 | - |
| - | - | 1817 | 524.3 | - | - | 0 | - |
| - | - | 627.4 | 525.3 | - | - | 0 | - |
| - | - | 687.8 | 527.7 | - | - | 0 | - |
| - | - | 796.7 | 529.7 | - | - | 0 | - |
| - | - | 666.4 | 541.2 | - | - | 0 | - |
| - | - | 1497 | 542.2 | - | - | 0 | - |
| - | - | 725.9 | 543.2 | - | - | 0 | - |
| - | - | 1977 | 543.3 | - | - | 0 | - |
| - | - | 574.3 | 544.3 | - | - | 0 | - |
| 5 | b | 1942 | 550.2 | 0.0005998 | 1.09 | +1 | 5 |
| 5 | b | 1546 | 568.2 | 0.0008096 | 1.425 | +1 | 5 |
| - | - | 631.2 | 570.2 | - | - | 0 | - |
| - | - | 1371 | 578.2 | - | - | 0 | - |
| - | - | 913.4 | 578.7 | - | - | 0 | - |
| 9 | y | 4943 | 588.3 | 0.00474 | 8.058 | +1 | 4 |
| - | - | 1272 | 589.3 | - | - | 0 | - |
| - | - | 1394 | 595.3 | - | - | 0 | - |
| 4 | y | 3546 | 600.2 | 0.00171 | 2.849 | +2 | 9 |
| - | - | 1994 | 600.7 | - | - | 0 | - |
| - | - | 1556 | 606.2 | - | - | 0 | - |
| - | - | 1053 | 624.2 | - | - | 0 | - |
| - | - | 650.7 | 625.4 | - | - | 0 | - |
| - | - | 799 | 634.2 | - | - | 0 | - |
| - | - | 1356 | 641.3 | - | - | 0 | - |
| 3 | y | 2739 | 641.8 | 0.00244 | 3.802 | +2 | 10 |
| - | - | 2245 | 642.3 | - | - | 0 | - |
| - | - | 739.6 | 642.8 | - | - | 0 | - |
| - | - | 794 | 647.2 | - | - | 0 | - |
| 3 | y | 4435 | 650.8 | 0.002468 | 3.792 | +2 | 10 |
| - | - | 3913 | 651.3 | - | - | 0 | - |
| - | - | 1366 | 651.8 | - | - | 0 | - |
| - | - | 1104 | 657.3 | - | - | 0 | - |
| 8 | y | 6413 | 659.3 | 0.004858 | 7.368 | +1 | 5 |
| - | - | 2786 | 660.3 | - | - | 0 | - |
| 6 | b | 1383 | 665.3 | 0.001197 | 1.799 | +1 | 6 |
| - | - | 2030 | 671.3 | - | - | 0 | - |
| - | - | 944.3 | 671.8 | - | - | 0 | - |
| - | - | 1391 | 680.3 | - | - | 0 | - |
| - | - | 973.4 | 680.8 | - | - | 0 | - |
| 11 | b | 2582 | 685.3 | 0.005145 | 7.509 | +2 | 11 |
| 11 | b | 897.1 | 685.8 | 0.0128 | 18.66 | +2 | 11 |
| - | - | 880.4 | 686.3 | - | - | 0 | - |
| - | - | 650.2 | 689.8 | - | - | 0 | - |
| - | - | 688.8 | 694.8 | - | - | 0 | - |
| - | - | 1077 | 705.3 | - | - | 0 | - |
| - | - | 891.3 | 711.8 | - | - | 0 | - |
| - | - | 1028 | 712.3 | - | - | 0 | - |
| - | - | 1145 | 721.3 | - | - | 0 | - |
| 2 | y | 699.7 | 724.3 | 0.004395 | 6.068 | +2 | 11 |
| - | - | 1167 | 734.8 | - | - | 0 | - |
| - | - | 707 | 735.3 | - | - | 0 | - |
| - | - | 643 | 735.8 | - | - | 0 | - |
| 0 | Precursor | 6125 | 743.8 | 0.003156 | 4.243 | +2 | -1 |
| - | - | 4820 | 744.3 | - | - | 0 | - |
| - | - | 2858 | 744.8 | - | - | 0 | - |
| - | - | 3009 | 749.3 | - | - | 0 | - |
| 0 | Precursor | 1624 | 752.8 | 0.003428 | 4.553 | +2 | -1 |
| - | - | 1038 | 753.3 | - | - | 0 | - |
| - | - | 794.4 | 753.8 | - | - | 0 | - |
| - | - | 895.8 | 756.3 | - | - | 0 | - |
| - | - | 1583 | 758.4 | - | - | 0 | - |
| - | - | 846.9 | 798.3 | - | - | 0 | - |
| - | - | 627.5 | 801.3 | - | - | 0 | - |
| - | - | 1541 | 820.3 | - | - | 0 | - |
| 7 | y | 6058 | 822.4 | 0.005494 | 6.681 | +1 | 6 |
| - | - | 2840 | 823.4 | - | - | 0 | - |
| 7 | b | 885.8 | 828.3 | 0.001515 | 1.829 | +1 | 7 |
| - | - | 1129 | 832.3 | - | - | 0 | - |
| - | - | 821.3 | 843.3 | - | - | 0 | - |
| - | - | 1064 | 871.3 | - | - | 0 | - |
| - | - | 2325 | 873.4 | - | - | 0 | - |
| - | - | 962.3 | 874.4 | - | - | 0 | - |
| - | - | 942.5 | 896.3 | - | - | 0 | - |
| - | - | 855.7 | 897.3 | - | - | 0 | - |
| - | - | 2018 | 907.3 | - | - | 0 | - |
| - | - | 2556 | 935.3 | - | - | 0 | - |
| - | - | 1267 | 936.3 | - | - | 0 | - |
| 6 | y | 8180 | 937.4 | 0.004125 | 4.4 | +1 | 7 |
| - | - | 4842 | 938.4 | - | - | 0 | - |
| - | - | 1925 | 939.4 | - | - | 0 | - |
| - | - | 724.8 | 955 | - | - | 0 | - |
| - | - | 1663 | 961.4 | - | - | 0 | - |
| - | - | 1227 | 962.4 | - | - | 0 | - |
| - | - | 709.4 | 983.4 | - | - | 0 | - |
| - | - | 4523 | 988.4 | - | - | 0 | - |
| - | - | 2679 | 989.4 | - | - | 0 | - |
| - | - | 1225 | 990.4 | - | - | 0 | - |
| - | - | 644 | 991.4 | - | - | 0 | - |
| - | - | 677.4 | 997.4 | - | - | 0 | - |
| - | - | 696.8 | 1011 | - | - | 0 | - |
| - | - | 767.2 | 1035 | - | - | 0 | - |
| 5 | y | 1.744E+04 | 1052 | 0.003793 | 3.604 | +1 | 8 |
| - | - | 1.022E+04 | 1053 | - | - | 0 | - |
| - | - | 4637 | 1054 | - | - | 0 | - |
| - | - | 1149 | 1055 | - | - | 0 | - |
| - | - | 1083 | 1082 | - | - | 0 | - |
| - | - | 704.4 | 1091 | - | - | 0 | - |
| - | - | 2514 | 1135 | - | - | 0 | - |
| - | - | 1878 | 1136 | - | - | 0 | - |
| - | - | 1024 | 1137 | - | - | 0 | - |
| 4 | y | 828.9 | 1181 | 0.007223 | 6.114 | +1 | 9 |
| 10 | b | 1433 | 1183 | 0.003789 | 3.201 | +1 | 10 |
| - | - | 730.6 | 1184 | - | - | 0 | - |
| - | - | 840.7 | 1186 | - | - | 0 | - |
| 4 | y | 1.113E+04 | 1199 | 0.003372 | 2.811 | +1 | 9 |
| - | - | 7634 | 1200 | - | - | 0 | - |
| 10 | b | 3671 | 1201 | 0.02081 | 17.32 | +1 | 10 |
| - | - | 1145 | 1202 | - | - | 0 | - |
| - | - | 1009 | 1237 | - | - | 0 | - |
| - | - | 838.7 | 1238 | - | - | 0 | - |
| - | - | 1251 | 1286 | - | - | 0 | - |
| - | - | 904.6 | 1287 | - | - | 0 | - |
| 3 | y | 5164 | 1301 | 0.002813 | 2.163 | +1 | 10 |
| - | - | 2619 | 1302 | - | - | 0 | - |
| - | - | 1606 | 1303 | - | - | 0 | - |
| - | - | 659.9 | 2148 | - | - | 0 | - |

m/z Charge Intensity FragmentType MassShift Position
120.0810317993164 0 42484.605
121.08441162109375 0 3388.0615
125.82366943359375 0 422.46555
129.0661163330078 0 839.45526
129.1024169921875 0 2961.069
130.06089782714844 0 539.5314
130.06546020507812 0 3038.8132
130.0868682861328 0 414.9269
131.04498291015625 0 548.6106
132.0811309814453 0 954.47064
133.06100463867188 0 738.34564
133.08616638183594 0 1656.458
136.075927734375 0 8427.805
138.0551300048828 0 909.631
138.0659637451172 0 596.1775
140.45382690429688 0 471.31818
143.1182861328125 0 507.17853
145.29559326171875 0 420.13995
146.0967254638672 0 522.50354
147.07672119140625 0 430.83386
148.07687377929688 0 400.93796
148.9536590576172 0 1068.0536
150.06607055664062 0 453.77774
151.79385375976562 0 431.9177
152.07102966308594 0 592.93317
153.0776824951172 0 701.0494
154.06109619140625 0 429.2835
154.6063232421875 0 463.7102
155.0817413330078 0 1579.5431
155.09317016601562 0 629.89655
155.1182098388672 0 1713.5507
156.0768280029297 0 937.7722
158.0556640625 0 477.2952
158.09730529785156 0 620.7914
159.0919647216797 0 6491.672
160.07582092285156 0 585.7191 a Ammonia loss 1
160.0952606201172 0 1050.3065
165.077880859375 0 657.62646
166.08628845214844 0 507.54575
167.08123779296875 0 439.1898
169.09652709960938 0 548.02234
171.06802368164062 0 471.0156
173.0712890625 0 466.23828
173.12841796875 0 1745.1494
176.10704040527344 0 2840.567
177.1024932861328 0 38558.81 a 1
178.10580444335938 0 3534.0496
181.0977783203125 0 591.88745
183.11289978027344 0 1000.6846
184.16024780273438 0 559.6195
185.09201049804688 0 546.3346
186.09153747558594 0 2214.4043
187.09548950195312 0 554.4869
187.14443969726562 0 2497.3154
188.14797973632812 0 572.88617
193.10867309570312 0 1982.2357
195.0879364013672 0 1006.897
197.1291961669922 0 502.16788
199.10862731933594 0 669.72296
199.1802215576172 0 591.76886
201.1233673095703 0 1916.6393
203.06626892089844 0 535.8686
203.11802673339844 0 3008.8198
204.07667541503906 0 703.40497
205.0973358154297 0 17613.729 b 1
205.10824584960938 0 1147.7144
206.1005859375 0 2401.269
207.11328125 0 739.45447
212.117919921875 0 548.1041
215.07728576660156 0 559.90784
215.13916015625 0 5443.6294
216.142822265625 0 804.07855
217.097412109375 0 1119.3789
217.9287109375 0 481.59897
219.1341552734375 0 5207.48
221.10353088378906 0 4999.739
221.1287078857422 0 23666.496
222.13198852539062 0 2761.6897
226.1188507080078 0 1823.2563
231.06088256835938 0 704.3388
231.1129150390625 0 3653.6682
233.1287384033203 0 805.76587
234.1334686279297 0 547.1439
235.1082305908203 0 2151.5837
236.54150390625 0 478.36453
238.12945556640625 0 615.7987
241.0638885498047 0 495.1378
249.12359619140625 0 8325.179
250.12716674804688 0 1687.0575
251.1029510498047 0 1954.1677
257.1058654785156 0 865.54474
260.13946533203125 0 3979.8848 a Water loss 2
261.1235656738281 0 2567.4507 a Ammonia loss 2
261.140869140625 0 713.82776
262.11865234375 0 845.98303
263.10260009765625 0 3176.2803
274.13037109375 0 1124.4121
278.1505432128906 0 829.4392 a 2
279.0980529785156 0 1914.0901
279.1495666503906 0 1050.0482
283.14300537109375 0 575.67474
284.16021728515625 0 1213.9254
285.10174560546875 0 2701.969
287.1391296386719 0 4988.176
288.1345520019531 0 11797.49 b Water loss 2
289.137939453125 0 1607.8982
292.1407470703125 0 2241.5105
296.1510009765625 0 5591.6187
297.1536865234375 0 1229.4408
297.19232177734375 0 1270.0992
304.16619873046875 0 4365.203 y 10
305.1683349609375 0 864.929
306.14520263671875 0 15294.009 b 2
307.147705078125 0 2371.595
314.207275390625 0 870.1379
324.14605712890625 0 5811.6543
325.1500549316406 0 913.30054
325.1880798339844 0 888.1727
328.16839599609375 0 584.45483
342.1567077636719 0 652.5017
346.1408386230469 0 582.83417
350.1349182128906 0 2008.3215
350.1875915527344 0 510.68546
356.1383056640625 0 2365.593
362.15765380859375 0 546.0996
364.15069580078125 0 4325.064
365.1536865234375 0 683.1239
366.1298522949219 0 688.328
376.1130065917969 0 964.68134
378.1297607421875 0 2980.4893
379.1325378417969 0 755.4042
379.1867370605469 0 573.8747
382.1429748535156 0 967.761
390.1816711425781 0 669.41785
394.12567138671875 0 2165.3875
403.16156005859375 0 737.73676
407.18218994140625 0 827.1835
407.2083740234375 0 4374.3276
424.2705078125 0 889.9561
425.2180480957031 0 846.95483
426.2292175292969 0 592.9761
435.2029113769531 0 2706.5315 b Water loss 3
436.20819091796875 0 795.9153
441.2250061035156 0 10006.432 y 9
442.2287292480469 0 2229.364
444.53192138671875 0 524.75836
449.1894226074219 0 800.901
461.1666564941406 0 1320.4222
465.16204833984375 0 2341.9272
471.18060302734375 0 1757.5953
472.1840515136719 0 681.2145
479.1781921386719 0 2661.71
480.183349609375 0 817.4123
497.1702880859375 0 758.046
513.1986083984375 0 675.3456
514.2252197265625 0 992.9621
519.2000732421875 0 647.7137
524.261962890625 0 1816.8726
525.308837890625 0 627.4325
527.7078857421875 0 687.8344
529.66845703125 0 796.7473
541.1942138671875 0 666.361
542.2185668945312 0 1497.0905
543.2338256835938 0 725.8816
543.3148193359375 0 1977.035
544.315185546875 0 574.3199
550.230224609375 0 1941.9137 b Water loss 4
568.2393798828125 0 1545.5659 b 4
570.2352905273438 0 631.24475
578.2324829101562 0 1370.7274
578.7329711914062 0 913.3764
588.2597045898438 0 4943.0493 y 8
589.2625732421875 0 1272.2751
595.2970581054688 0 1393.5682
600.2442016601562 0 3546.2576 y 3
600.7462768554688 0 1994.4814
606.233154296875 0 1555.9368
624.2276611328125 0 1052.5791
625.3908081054688 0 650.6937
634.229248046875 0 798.9657
641.2843627929688 0 1356.4489
641.7634887695312 0 2738.5923 y Water loss 2
642.26318359375 0 2244.5874
642.7655029296875 0 739.6067
647.2440185546875 0 793.9868
650.768798828125 0 4435.094 y 2
651.2686157226562 0 3913.4792
651.7727661132812 0 1366.1029
657.2630615234375 0 1104.1431
659.2969360351562 0 6413.071 y 7
660.3009033203125 0 2785.7065
665.25537109375 0 1383.4617 b Water loss 5
671.2716064453125 0 2030.2891
671.7703247070312 0 944.32104
680.2760620117188 0 1390.7255
680.7770385742188 0 973.44775
685.2613525390625 0 2582.2102 b Water loss 10
685.7713012695312 0 897.0655 b Ammonia loss 10
686.2679443359375 0 880.38965
689.8052368164062 0 650.21545
694.773193359375 0 688.77893
705.2821044921875 0 1077.2292
711.8079833984375 0 891.2841
712.3104858398438 0 1028.398
721.2583618164062 0 1144.6362
724.296142578125 0 699.7258 y 1
734.8001098632812 0 1166.9889
735.2982177734375 0 706.9757
735.8035278320312 0 642.966
743.8091430664062 0 6125.4517 Precursor Water loss
744.30908203125 0 4820.111
744.8103637695312 0 2858.3198
749.2555541992188 0 3008.7722
752.814697265625 0 1624.0632 Precursor
753.3140258789062 0 1037.9478
753.821533203125 0 794.414
756.3165893554688 0 895.81067
758.3637084960938 0 1582.7588
798.313720703125 0 846.872
801.3167724609375 0 627.4907
820.3114624023438 0 1540.7278
822.3609008789062 0 6058.0103 y 6
823.3645629882812 0 2840.2803
828.3214111328125 0 885.7694 b Water loss 6
832.3286743164062 0 1129.4756
843.3455810546875 0 821.32196
871.3411865234375 0 1063.651
873.38818359375 0 2324.7686
874.394287109375 0 962.33295
896.317138671875 0 942.45544
897.328369140625 0 855.7112
907.34130859375 0 2017.6068
935.3358764648438 0 2556.205
936.3390502929688 0 1267.2106
937.386474609375 0 8179.838 y 5
938.3902587890625 0 4841.821
939.39013671875 0 1925.2101
955.0137329101562 0 724.753
961.38330078125 0 1663.0988
962.3822631835938 0 1226.5322
983.3999633789062 0 709.36456
988.416015625 0 4522.913
989.4178466796875 0 2679.1162
990.4179077148438 0 1224.5428
991.41552734375 0 644.0182
997.374267578125 0 677.4444
1010.9706420898438 0 696.7722
1035.4072265625 0 767.19604
1052.4130859375 0 17435.865 y 4
1053.4168701171875 0 10216.478
1054.415771484375 0 4637.3027
1055.412353515625 0 1149.341
1082.4019775390625 0 1082.9037
1091.4456787109375 0 704.43384
1135.4857177734375 0 2514.254
1136.48388671875 0 1877.9236
1137.4759521484375 0 1023.52295
1181.474365234375 0 828.9091 y Water loss 3
1183.4501953125 0 1433.3583 b Water loss 9
1184.46533203125 0 730.5926
1186.4947509765625 0 840.74695
1199.4810791015625 0 11132.686 y 3
1200.4837646484375 0 7633.6216
1201.477783203125 0 3670.5698 b 9
1202.46533203125 0 1145.0239
1236.533935546875 0 1009.45984
1237.5352783203125 0 838.73083
1285.5557861328125 0 1251.0994
1286.554443359375 0 904.64874
1300.5281982421875 0 5164.0537 y 2
1301.528076171875 0 2619.1458
1302.5274658203125 0 1606.2565
2148.29345703125 0 659.9066

Spectrum Details

|  |  |
| --- | --- |
| Matched peaks? Matched peaks The total absolute number of peaks matched. Additionally in brackets the total fraction of peaks matched and the total number of peaks is shown. | 33 (12.09% of 273) |
| FDR? FDR The false discovery rate estimated for this peptide. It is calculated by matching all theoretical fragments with a non-integer shift with the raw peaks for this spectrum. This is done with 40 different shifts. The resulting percentage is the average number of annotated peaks over the number of annotated peaks with the correct spectrum. | 0.00% |
| Satellite FDR? Satellite FDR See the FDR for details on its calculation. This satellite ion specific FDR only contains the satellite ions (d/w) for I/L/J positions. | - |
| PSM Score? PSM Score The PSM Score as given by Hecklib to this annotated spectrum. It is shown with three significant figures. | 354 |

#### Spectrum 11000? Spectrum 11000 The raw spectrum of this peptide as annotated by Hecklib. The fragments are coloured according to ion type (see legend). Any peaks with a star '\*' as text can be hovered over to see the full details, first the ion type second the mass shift type. By hovering over the amino acids in the peptide or ions in the legend the corresponding peaks are highlighted. By toggling the 'Unassigned' label you can turn the background (unassigned) peaks on or off in the plot. By updating the slider in the Ion legend you can update the spectrum to only show the top X% of the peaks with labels. The top X% means any peak that is within X% of the highest intensity. By dragging in the spectrum you can zoom in to a specific part of the spectrum and use 'Zoom Out' to get back to the original zoom level. The annotation of the spectrum is based on the given sequence in the peptides file and is done with different software so inconsistencies are likely. The peaks are annotated based on the given sequence, with 20 ppm tolerance.

Copy Data

##### Spectrum 11000 (TSV)

###### Preview

```
Loading example...
```

*Click on the button to copy the data to your clipboard.*

Mz MinMz MaxIntensity Max

WidthHeightPeptide font sizePeptide stroke widthSpectrum font sizeSpectrum stroke widthCompact peptide

Ion legend

wxyz

abcd

OtherUnassignedIon^Charge^_Position_Show for top:%

GFTFDDYAMHWV

01.86e+43.72e+45.57e+47.43e+4

Zoom Out

c^+1^_2_c^+1^_3_y^+1^_3_y^+1^_4_y^+2^_9_y^+2^_10_z^+1^_5_y^+2^_10_y^+1^_5_z^+1^_6_y^+1^_6_z^+1^_7_y^+1^_7_z^+1^_8_y^+1^_8_c^+1^_9_z^+1^_9_y^+1^_9_c^+1^_10_c^+1^_10_z^+1^_10_y^+1^_10_c^+1^_11_z^+1^_11_y^+1^_11_

038376611491531

Fragment Matches Table

Show background peaks

| Position | Ion type | Intensity | mz Theoretical | mz Error (Th) | mz Error (ppm) | Charge | Series Number |
| --- | --- | --- | --- | --- | --- | --- | --- |
| - | - | 1509 | 120.1 | - | - | 0 | - |
| - | - | 366.5 | 122.1 | - | - | 0 | - |
| - | - | 378.6 | 123 | - | - | 0 | - |
| - | - | 455.6 | 123.8 | - | - | 0 | - |
| - | - | 376.7 | 127.9 | - | - | 0 | - |
| - | - | 501.1 | 133.1 | - | - | 0 | - |
| - | - | 521.9 | 159.1 | - | - | 0 | - |
| - | - | 4102 | 177.1 | - | - | 0 | - |
| 2 | c | 3556 | 205.1 | 0.0001207 | 0.5884 | +1 | 2 |
| - | - | 2426 | 215.1 | - | - | 0 | - |
| - | - | 1933 | 221.1 | - | - | 0 | - |
| - | - | 460.3 | 236.5 | - | - | 0 | - |
| - | - | 503.5 | 245.6 | - | - | 0 | - |
| - | - | 2254 | 249.1 | - | - | 0 | - |
| - | - | 657.3 | 260.1 | - | - | 0 | - |
| - | - | 869.3 | 261.1 | - | - | 0 | - |
| - | - | 574 | 266.7 | - | - | 0 | - |
| - | - | 2109 | 288.1 | - | - | 0 | - |
| - | - | 582.3 | 298.2 | - | - | 0 | - |
| 3 | c | 4876 | 306.1 | 0.0001564 | 0.511 | +1 | 3 |
| - | - | 2536 | 314.2 | - | - | 0 | - |
| - | - | 1037 | 315.2 | - | - | 0 | - |
| - | - | 1688 | 330.2 | - | - | 0 | - |
| - | - | 1405 | 364.1 | - | - | 0 | - |
| - | - | 491 | 371.7 | - | - | 0 | - |
| - | - | 993.8 | 383.2 | - | - | 0 | - |
| - | - | 856.1 | 397.2 | - | - | 0 | - |
| - | - | 1744 | 407.2 | - | - | 0 | - |
| - | - | 569.9 | 429.3 | - | - | 0 | - |
| - | - | 1314 | 435.2 | - | - | 0 | - |
| 10 | y | 3059 | 441.2 | 0.001045 | 2.368 | +1 | 3 |
| - | - | 628.2 | 442.2 | - | - | 0 | - |
| - | - | 496 | 446.3 | - | - | 0 | - |
| - | - | 526 | 454.8 | - | - | 0 | - |
| - | - | 667.9 | 457.2 | - | - | 0 | - |
| - | - | 1035 | 494.3 | - | - | 0 | - |
| - | - | 1042 | 496.3 | - | - | 0 | - |
| - | - | 928.7 | 512.3 | - | - | 0 | - |
| - | - | 743.6 | 513.3 | - | - | 0 | - |
| - | - | 565.6 | 542.2 | - | - | 0 | - |
| - | - | 654.3 | 550.2 | - | - | 0 | - |
| - | - | 2055 | 573.2 | - | - | 0 | - |
| - | - | 606.1 | 583.7 | - | - | 0 | - |
| 9 | y | 2442 | 588.3 | 0.004862 | 8.265 | +1 | 4 |
| 4 | y | 1130 | 600.2 | 0.002687 | 4.476 | +2 | 9 |
| - | - | 2064 | 625.4 | - | - | 0 | - |
| - | - | 1357 | 626.4 | - | - | 0 | - |
| - | - | 662.7 | 630.8 | - | - | 0 | - |
| 3 | y | 1110 | 641.8 | 0.003844 | 5.99 | +2 | 10 |
| - | - | 791.4 | 642.3 | - | - | 0 | - |
| 8 | z | 1421 | 643.3 | 0.003257 | 5.063 | +1 | 5 |
| - | - | 2216 | 643.4 | - | - | 0 | - |
| - | - | 2819 | 644.3 | - | - | 0 | - |
| - | - | 869 | 644.4 | - | - | 0 | - |
| - | - | 684.2 | 645.3 | - | - | 0 | - |
| 3 | y | 1877 | 650.8 | 0.001919 | 2.948 | +2 | 10 |
| - | - | 1710 | 651.3 | - | - | 0 | - |
| - | - | 634.9 | 651.8 | - | - | 0 | - |
| 8 | y | 2877 | 659.3 | 0.004919 | 7.461 | +1 | 5 |
| - | - | 1271 | 660.3 | - | - | 0 | - |
| - | - | 784.6 | 665.3 | - | - | 0 | - |
| - | - | 845.5 | 680.8 | - | - | 0 | - |
| - | - | 712.5 | 711.8 | - | - | 0 | - |
| - | - | 610.9 | 721.3 | - | - | 0 | - |
| - | - | 4459 | 743.8 | - | - | 0 | - |
| - | - | 2121 | 744.3 | - | - | 0 | - |
| - | - | 960.8 | 744.8 | - | - | 0 | - |
| - | - | 1195 | 751.4 | - | - | 0 | - |
| - | - | 973.9 | 752.4 | - | - | 0 | - |
| - | - | 864.2 | 752.8 | - | - | 0 | - |
| - | - | 563.1 | 753.4 | - | - | 0 | - |
| 7 | z | 8260 | 806.3 | 0.005786 | 7.175 | +1 | 6 |
| - | - | 4968 | 807.3 | - | - | 0 | - |
| - | - | 1978 | 808.3 | - | - | 0 | - |
| 7 | y | 2777 | 822.4 | 0.005006 | 6.087 | +1 | 6 |
| - | - | 1260 | 823.4 | - | - | 0 | - |
| - | - | 4452 | 877.4 | - | - | 0 | - |
| - | - | 2914 | 878.4 | - | - | 0 | - |
| 6 | z | 1.484E+04 | 921.4 | 0.004477 | 4.859 | +1 | 7 |
| - | - | 1.521E+04 | 922.4 | - | - | 0 | - |
| - | - | 6091 | 923.4 | - | - | 0 | - |
| - | - | 971.3 | 924.4 | - | - | 0 | - |
| 6 | y | 5333 | 937.4 | 0.00443 | 4.726 | +1 | 7 |
| - | - | 2759 | 938.4 | - | - | 0 | - |
| - | - | 958.5 | 988.4 | - | - | 0 | - |
| - | - | 2351 | 992.4 | - | - | 0 | - |
| - | - | 1753 | 993.4 | - | - | 0 | - |
| - | - | 801.4 | 994.4 | - | - | 0 | - |
| 5 | z | 4730 | 1036 | 0.004695 | 4.53 | +1 | 8 |
| - | - | 5820 | 1037 | - | - | 0 | - |
| - | - | 4187 | 1038 | - | - | 0 | - |
| - | - | 1521 | 1039 | - | - | 0 | - |
| 5 | y | 7585 | 1052 | 0.004892 | 4.648 | +1 | 8 |
| - | - | 4274 | 1053 | - | - | 0 | - |
| - | - | 1559 | 1054 | - | - | 0 | - |
| - | - | 5477 | 1080 | - | - | 0 | - |
| 9 | c | 1.312E+04 | 1081 | 0.003614 | 3.342 | +1 | 9 |
| - | - | 7707 | 1082 | - | - | 0 | - |
| - | - | 2904 | 1083 | - | - | 0 | - |
| - | - | 678.3 | 1111 | - | - | 0 | - |
| - | - | 1630 | 1174 | - | - | 0 | - |
| - | - | 1003 | 1175 | - | - | 0 | - |
| 4 | z | 3385 | 1183 | 0.002077 | 1.755 | +1 | 9 |
| - | - | 2513 | 1184 | - | - | 0 | - |
| - | - | 1394 | 1185 | - | - | 0 | - |
| 4 | y | 5261 | 1199 | 0.003372 | 2.811 | +1 | 9 |
| 10 | c | 3571 | 1200 | 0.01288 | 10.73 | +1 | 10 |
| - | - | 1717 | 1201 | - | - | 0 | - |
| - | - | 874.2 | 1202 | - | - | 0 | - |
| 10 | c | 5690 | 1218 | 0.004639 | 3.807 | +1 | 10 |
| - | - | 3329 | 1219 | - | - | 0 | - |
| - | - | 1518 | 1220 | - | - | 0 | - |
| - | - | 1054 | 1241 | - | - | 0 | - |
| - | - | 687.8 | 1242 | - | - | 0 | - |
| - | - | 748.1 | 1283 | - | - | 0 | - |
| 3 | z | 1.058E+04 | 1285 | 0.004447 | 3.462 | +1 | 10 |
| - | - | 9146 | 1286 | - | - | 0 | - |
| - | - | 3026 | 1287 | - | - | 0 | - |
| - | - | 780.2 | 1288 | - | - | 0 | - |
| 3 | y | 5305 | 1301 | 0.005376 | 4.134 | +1 | 10 |
| - | - | 5742 | 1302 | - | - | 0 | - |
| - | - | 2552 | 1303 | - | - | 0 | - |
| - | - | 660.1 | 1315 | - | - | 0 | - |
| - | - | 807.8 | 1359 | - | - | 0 | - |
| - | - | 1092 | 1361 | - | - | 0 | - |
| - | - | 1075 | 1375 | - | - | 0 | - |
| - | - | 780.8 | 1376 | - | - | 0 | - |
| 11 | c | 1.506E+04 | 1405 | 0.00284 | 2.022 | +1 | 11 |
| - | - | 1.25E+04 | 1406 | - | - | 0 | - |
| - | - | 5680 | 1407 | - | - | 0 | - |
| - | - | 905.7 | 1408 | - | - | 0 | - |
| 2 | z | 3426 | 1432 | 0.005491 | 3.836 | +1 | 11 |
| - | - | 3760 | 1433 | - | - | 0 | - |
| - | - | 1509 | 1434 | - | - | 0 | - |
| - | - | 827.1 | 1435 | - | - | 0 | - |
| - | - | 1114 | 1442 | - | - | 0 | - |
| - | - | 1780 | 1444 | - | - | 0 | - |
| - | - | 2313 | 1445 | - | - | 0 | - |
| - | - | 7617 | 1446 | - | - | 0 | - |
| - | - | 7197 | 1447 | - | - | 0 | - |
| 2 | y | 2763 | 1448 | 0.01533 | 10.59 | +1 | 11 |
| - | - | 770.3 | 1449 | - | - | 0 | - |
| - | - | 765.6 | 1460 | - | - | 0 | - |
| - | - | 1118 | 1461 | - | - | 0 | - |
| - | - | 851.3 | 1478 | - | - | 0 | - |
| - | - | 3674 | 1488 | - | - | 0 | - |
| - | - | 2.289E+04 | 1489 | - | - | 0 | - |
| - | - | 2.156E+04 | 1490 | - | - | 0 | - |
| - | - | 1.201E+04 | 1491 | - | - | 0 | - |
| - | - | 3078 | 1492 | - | - | 0 | - |
| - | - | 2.396E+04 | 1505 | - | - | 0 | - |
| - | - | 7.358E+04 | 1506 | - | - | 0 | - |
| - | - | 6E+04 | 1507 | - | - | 0 | - |
| - | - | 2.736E+04 | 1508 | - | - | 0 | - |
| - | - | 5429 | 1509 | - | - | 0 | - |
| - | - | 638.7 | 1516 | - | - | 0 | - |

m/z Charge Intensity FragmentType MassShift Position
120.08096313476562 0 1509.4695
122.05021667480469 0 366.4756
123.02568054199219 0 378.55927
123.84135437011719 0 455.60812
127.90550994873047 0 376.69437
133.0858154296875 0 501.1442
159.0921630859375 0 521.8781
177.10232543945312 0 4102.1533
205.09727478027344 0 3556.044 c Ammonia loss 1
215.13905334472656 0 2425.53
221.128662109375 0 1933.4194
236.48483276367188 0 460.2543
245.62051391601562 0 503.5028
249.12353515625 0 2253.5535
260.13916015625 0 657.2876
261.1238098144531 0 869.32294
266.68017578125 0 574.02344
288.1342468261719 0 2108.6394
298.1773681640625 0 582.2659
306.1449890136719 0 4876.4697 c Ammonia loss 2
314.2076416015625 0 2535.5046
315.21246337890625 0 1037.3582
330.2027893066406 0 1688.3131
364.1498107910156 0 1405.0454
371.7429504394531 0 490.9775
383.2295837402344 0 993.765
397.242919921875 0 856.1085
407.20745849609375 0 1743.731
429.2717590332031 0 569.92596
435.2036437988281 0 1314.3694
441.22552490234375 0 3058.7183 y 9
442.228271484375 0 628.215
446.34844970703125 0 496.0046
454.84185791015625 0 526.0016
457.1727600097656 0 667.9139
494.2962341308594 0 1034.9012
496.31231689453125 0 1042.4398
512.3101806640625 0 928.6919
513.3113403320312 0 743.57574
542.2092895507812 0 565.5952
550.2243041992188 0 654.31134
573.2476196289062 0 2055.4746
583.718017578125 0 606.1326
588.2598266601562 0 2442.2927 y 8
600.2451782226562 0 1130.154 y 3
625.3914794921875 0 2063.873
626.3959350585938 0 1356.5547
630.8403930664062 0 662.7153
641.764892578125 0 1110.3628 y Water loss 2
642.2620239257812 0 791.3625
643.276611328125 0 1421.0348 z 7
643.402587890625 0 2215.8076
644.2841186523438 0 2819.0437
644.4011840820312 0 869.0275
645.2884521484375 0 684.17084
650.7682495117188 0 1876.5022 y 2
651.2677612304688 0 1709.8508
651.766845703125 0 634.8712
659.2969970703125 0 2876.6777 y 7
660.3016967773438 0 1270.8954
665.2535400390625 0 784.57056
680.779541015625 0 845.46985
711.8033447265625 0 712.4553
721.3092651367188 0 610.94055
743.808349609375 0 4458.8257
744.3084106445312 0 2121.4656
744.8155517578125 0 960.8254
751.3870239257812 0 1195.3987
752.3950805664062 0 973.9222
752.8160400390625 0 864.178
753.3970947265625 0 563.1219
806.3424682617188 0 8260.383 z 6
807.3468627929688 0 4967.94
808.3479614257812 0 1978.4143
822.3604125976562 0 2776.7356 y 6
823.3675537109375 0 1259.7167
877.3798828125 0 4452.322
878.3839721679688 0 2914.4448
921.3681030273438 0 14838.824 z 5
922.373779296875 0 15205.133
923.3760375976562 0 6091.077
924.3780517578125 0 971.3496
937.3867797851562 0 5333.1123 y 5
938.3905029296875 0 2759.072
988.4136962890625 0 958.47363
992.4058837890625 0 2350.5166
993.4083251953125 0 1753.2576
994.4013061523438 0 801.37476
1036.395263671875 0 4729.852 z 4
1037.4031982421875 0 5820.253
1038.41015625 0 4186.7114
1039.4110107421875 0 1520.9128
1052.4141845703125 0 7584.659 y 4
1053.4188232421875 0 4273.894
1054.4173583984375 0 1558.8574
1080.4215087890625 0 5477.135
1081.42822265625 0 13116.204 c 8
1082.4310302734375 0 7707.06
1083.4298095703125 0 2903.9495
1110.70556640625 0 678.2581
1174.4761962890625 0 1629.5061
1175.477294921875 0 1002.78107
1183.4610595703125 0 3384.9395 z 3
1184.4652099609375 0 2513.445
1185.46923828125 0 1393.5918
1199.4810791015625 0 5261.4214 y 3
1200.48583984375 0 3570.7437 c Water loss 9
1201.4810791015625 0 1717.1289
1202.491943359375 0 874.2032
1218.4881591796875 0 5689.9336 c 9
1219.490478515625 0 3328.7803
1220.4906005859375 0 1517.9963
1240.52685546875 0 1053.7205
1241.53271484375 0 687.8323
1283.4991455078125 0 748.13885
1284.5111083984375 0 10583.158 z 2
1285.513671875 0 9145.872
1286.51416015625 0 3025.7502
1287.5146484375 0 780.1767
1300.53076171875 0 5305.1665 y 2
1301.530517578125 0 5741.5513
1302.533935546875 0 2551.6665
1314.5675048828125 0 660.1078
1358.5526123046875 0 807.757
1360.5579833984375 0 1091.9501
1374.5540771484375 0 1075.1097
1375.5606689453125 0 780.82336
1404.565673828125 0 15060.756 c 10
1405.569580078125 0 12502.297
1406.5694580078125 0 5679.787
1407.5740966796875 0 905.72754
1431.58056640625 0 3426.3152 z 1
1432.5797119140625 0 3759.6782
1433.5904541015625 0 1508.8187
1434.568115234375 0 827.10944
1441.624755859375 0 1114.4069
1443.6251220703125 0 1780.0046
1444.6298828125 0 2313.1313
1445.6064453125 0 7617.43
1446.608642578125 0 7196.723
1447.609130859375 0 2762.7566 y 1
1448.602783203125 0 770.25806
1459.6182861328125 0 765.5887
1460.630859375 0 1118.4271
1477.6181640625 0 851.262
1487.616943359375 0 3673.8364
1488.60498046875 0 22890.166
1489.607177734375 0 21563.764
1490.608642578125 0 12005.851
1491.610595703125 0 3078.165
1504.6180419921875 0 23963.346
1505.6246337890625 0 73578.8
1506.62841796875 0 59995.156
1507.6309814453125 0 27361.238
1508.6353759765625 0 5428.5747
1516.3017578125 0 638.66486

Spectrum Details

|  |  |
| --- | --- |
| Matched peaks? Matched peaks The total absolute number of peaks matched. Additionally in brackets the total fraction of peaks matched and the total number of peaks is shown. | 25 (16.03% of 156) |
| FDR? FDR The false discovery rate estimated for this peptide. It is calculated by matching all theoretical fragments with a non-integer shift with the raw peaks for this spectrum. This is done with 40 different shifts. The resulting percentage is the average number of annotated peaks over the number of annotated peaks with the correct spectrum. | 0.38% |
| Satellite FDR? Satellite FDR See the FDR for details on its calculation. This satellite ion specific FDR only contains the satellite ions (d/w) for I/L/J positions. | - |
| PSM Score? PSM Score The PSM Score as given by Hecklib to this annotated spectrum. It is shown with three significant figures. | 278 |

#### Reverse Lookup? Reverse Lookup All places where this read could be placed.

| Group | Segment | Template | Template Part | Read Part | Score | Unique |
| --- | --- | --- | --- | --- | --- | --- |
| Homo sapiens Heavy Chain | IGHV | IGHV3-9 | [25..37] | [0..12] | 96 | False |
| Homo sapiens Heavy Chain | IGHV | IGHV3-43 | [25..37] | [0..12] | 87 | False |

| Recombined | Template Part | Read Part | Score | Unique |
| --- | --- | --- | --- | --- |
| REC-0-1 | [25..37] | [0..12] | 96 | True |

#### Meta Information from Multiple reads

##### Number of combined reads

3

##### Intensity

0.616

##### TotalArea

3.583E+07

#### Positional Score

Copy Data

##### Positional Score (TSV)

###### Preview

```
Loading example...
```

*Click on the button to copy the data to your clipboard.*

1001234567891011

Label Value
"0" 0.333
"1" 0.333
"2" 0.333
"3" 0.33
"4" 0.33
"5" 0.33
"6" 0.32
"7" 0.283
"8" 0.287
"9" 0.323
"10" 0.323
"11" 0.33

#### Meta Information from PEAKS

##### Scan Identifier

F1:11056

##### Original sequence

G

F

T

F

D

D

Y

A

M

+15.99

H

W

V

##### Posttranslational Modifications

Oxidation (M)

##### Source File

D:\separate\_stitch\_analyses\xle-disambiguation\raw\20210323\_F1\_UM1\_Peng0013\_SA\_F59\_ingel\_3ug\_ELA.raw

##### Fraction

1

##### Scan Feature

F1:14918

##### De Novo Score

98

##### ConfidenceScore

98

### m/z

752.8145

##### Mass

1503.6128

##### Charge

2

##### Retention Time

61.5

##### Predicted Retention Time

-

##### Area

1.194E+07

##### Parts Per Million

1

##### Fragmentation mode

HCD

##### Originating file

01 D:\separate\_stitch\_analyses\xle-disambiguation\20210325\_F59\_3ug\_DENOVO\_12.csv

#### Meta Information from PEAKS

##### Scan Identifier

F1:11109

##### Original sequence

G

F

T

F

D

D

Y

A

M

+15.99

H

W

V

##### Posttranslational Modifications

Oxidation (M)

##### Source File

D:\separate\_stitch\_analyses\xle-disambiguation\raw\20210323\_F1\_UM1\_Peng0013\_SA\_F59\_ingel\_3ug\_ELA.raw

##### Fraction

1

##### Scan Feature

F1:14918

##### De Novo Score

97

##### ConfidenceScore

97

### m/z

752.8145

##### Mass

1503.6128

##### Charge

2

##### Retention Time

61.5

##### Predicted Retention Time

-

##### Area

1.194E+07

##### Parts Per Million

1

##### Fragmentation mode

HCD

##### Originating file

01 D:\separate\_stitch\_analyses\xle-disambiguation\20210325\_F59\_3ug\_DENOVO\_12.csv

#### Meta Information from PEAKS

##### Scan Identifier

F1:11000

##### Original sequence

G

F

T

F

D

D

Y

A

M

+15.99

H

W

V

##### Posttranslational Modifications

Oxidation (M)

##### Source File

D:\separate\_stitch\_analyses\xle-disambiguation\raw\20210323\_F1\_UM1\_Peng0013\_SA\_F59\_ingel\_3ug\_ELA.raw

##### Fraction

1

##### Scan Feature

F1:14918

##### De Novo Score

96

##### ConfidenceScore

96

### m/z

752.8145

##### Mass

1503.6128

##### Charge

2

##### Retention Time

61.5

##### Predicted Retention Time

-

##### Area

1.194E+07

##### Parts Per Million

1

##### Fragmentation mode

ETHCD

##### Originating file

01 D:\separate\_stitch\_analyses\xle-disambiguation\20210325\_F59\_3ug\_DENOVO\_12.csv
