## Supplementary material for "A handle on mass coincidence errors in *de novo* sequencing of antibodies by bottom-up proteomics": Combined_090.html

Details Combined\_090 | Stitch OverviewUndefined

### Read Combined\_090

#### Sequence (length=13)

CSSVMHEALHNHY

#### Spectrum 4496? Spectrum 4496 The raw spectrum of this peptide as annotated by Hecklib. The fragments are coloured according to ion type (see legend). Any peaks with a star '\*' as text can be hovered over to see the full details, first the ion type second the mass shift type. By hovering over the amino acids in the peptide or ions in the legend the corresponding peaks are highlighted. By toggling the 'Unassigned' label you can turn the background (unassigned) peaks on or off in the plot. By updating the slider in the Ion legend you can update the spectrum to only show the top X% of the peaks with labels. The top X% means any peak that is within X% of the highest intensity. By dragging in the spectrum you can zoom in to a specific part of the spectrum and use 'Zoom Out' to get back to the original zoom level. The annotation of the spectrum is based on the given sequence in the peptides file and is done with different software so inconsistencies are likely. The peaks are annotated based on the given sequence, with 20 ppm tolerance.

Copy Data

##### Spectrum 4496 (TSV)

###### Preview

```
Loading example...
```

*Click on the button to copy the data to your clipboard.*

Mz MinMz MaxIntensity Max

WidthHeightPeptide font sizePeptide stroke widthSpectrum font sizeSpectrum stroke widthCompact peptide

Ion legend

wxyz

abcd

OtherUnassignedIon^Charge^_Position_Show for top:%

CSSVMHEALHNHY

04.18e+58.36e+51.25e+61.67e+6

Zoom Out

y^+1^_1_z^+3^_5_z^+3^_6_z^+2^_4_y^+2^_4_z^+1^_2_y^+1^_2_y^+2^_5_c^+1^_3_w^+1^_3_y^+2^_6_y^+3^_9_c^+2^_7_y^+3^_10_y^+1^_3_z^+3^_11_z^+3^_11_y^+3^_11_y^+3^_11_z^+3^_11_y^+2^_7_y^+3^_11_c^+1^_4_z^+3^_12_z^+3^_12_c^+3^_12_y^+3^_12_z^+2^_8_y^+2^_8_y^+2^_8_z^+2^_8_c^+2^_9_y^+2^_8_c^+2^_9_w^+2^_9_y^+1^_4_z^+1^_4_z^+2^_9_y^+2^_9_y^+2^_9_z^+2^_9_y^+1^_4_y^+2^_9_c^+2^_10_c^+1^_5_c^+2^_10_z^+2^_10_z^+2^_10_w^+2^_10_y^+2^_10_y^+2^_10_z^+2^_10_w^+1^_5_y^+2^_10_c^+2^_11_c^+2^_11_c^+2^_11_z^+2^_11_z^+2^_11_y^+2^_11_y^+2^_11_z^+2^_11_y^+1^_5_z^+1^_5_y^+2^_11_y^+1^_5_z^+2^_12_w^+2^_12_c^+2^_12_y^+2^_12_z^+2^_12_c^+2^_12_y^+2^_12_c^+1^_6_z^+1^_6_y^+1^_6_w^+1^_7_c^+1^_7_y^+1^_7_y^+1^_7_z^+1^_7_y^+1^_7_c^+1^_8_y^+1^_8_z^+1^_8_y^+1^_8_c^+1^_9_w^+1^_9_y^+1^_9_z^+1^_9_y^+1^_9_c^+1^_10_c^+1^_10_z^+1^_10_y^+1^_10_c^+1^_11_c^+1^_11_w^+1^_11_z^+1^_11_y^+1^_11_w^+1^_12_c^+1^_12_y^+1^_12_

040280412061608

Fragment Matches Table

Show background peaks

| Position | Ion type | Intensity | mz Theoretical | mz Error (Th) | mz Error (ppm) | Charge | Series Number |
| --- | --- | --- | --- | --- | --- | --- | --- |
| - | - | 759.4 | 120.1 | - | - | 0 | - |
| - | - | 602.9 | 120.1 | - | - | 0 | - |
| - | - | 9149 | 120.1 | - | - | 0 | - |
| - | - | 751.9 | 121.1 | - | - | 0 | - |
| - | - | 1.607E+04 | 123.1 | - | - | 0 | - |
| - | - | 632.9 | 124.1 | - | - | 0 | - |
| - | - | 459.6 | 124.8 | - | - | 0 | - |
| - | - | 5456 | 129.1 | - | - | 0 | - |
| - | - | 1097 | 131.1 | - | - | 0 | - |
| - | - | 586.3 | 134 | - | - | 0 | - |
| - | - | 1.079E+04 | 134 | - | - | 0 | - |
| - | - | 2.562E+04 | 136.1 | - | - | 0 | - |
| - | - | 2399 | 137.1 | - | - | 0 | - |
| - | - | 1412 | 138.1 | - | - | 0 | - |
| - | - | 6462 | 140.1 | - | - | 0 | - |
| - | - | 454.3 | 141.1 | - | - | 0 | - |
| - | - | 696.9 | 141.1 | - | - | 0 | - |
| - | - | 430.7 | 141.9 | - | - | 0 | - |
| - | - | 1168 | 145.1 | - | - | 0 | - |
| - | - | 993.9 | 148.9 | - | - | 0 | - |
| - | - | 799.8 | 155.1 | - | - | 0 | - |
| - | - | 1649 | 156.1 | - | - | 0 | - |
| - | - | 5215 | 157.1 | - | - | 0 | - |
| - | - | 4623 | 158 | - | - | 0 | - |
| - | - | 8455 | 159.1 | - | - | 0 | - |
| - | - | 725.5 | 160.1 | - | - | 0 | - |
| - | - | 2789 | 161 | - | - | 0 | - |
| - | - | 1479 | 164 | - | - | 0 | - |
| - | - | 1849 | 164.1 | - | - | 0 | - |
| - | - | 4719 | 165.1 | - | - | 0 | - |
| - | - | 3.009E+04 | 166.1 | - | - | 0 | - |
| - | - | 2276 | 167.1 | - | - | 0 | - |
| - | - | 990.1 | 168 | - | - | 0 | - |
| - | - | 1337 | 173.1 | - | - | 0 | - |
| - | - | 1163 | 173.5 | - | - | 0 | - |
| - | - | 1161 | 174.1 | - | - | 0 | - |
| - | - | 5295 | 176 | - | - | 0 | - |
| - | - | 2271 | 177.1 | - | - | 0 | - |
| 13 | y | 2.575E+04 | 182.1 | 0.0006022 | 3.307 | +1 | 1 |
| - | - | 1250 | 182.1 | - | - | 0 | - |
| - | - | 2009 | 183.1 | - | - | 0 | - |
| - | - | 954.4 | 185 | - | - | 0 | - |
| - | - | 1.486E+04 | 186 | - | - | 0 | - |
| - | - | 947.3 | 187 | - | - | 0 | - |
| - | - | 8136 | 187.1 | - | - | 0 | - |
| - | - | 571.4 | 188 | - | - | 0 | - |
| - | - | 836.5 | 188.1 | - | - | 0 | - |
| - | - | 1.275E+04 | 192.1 | - | - | 0 | - |
| - | - | 702.3 | 195.1 | - | - | 0 | - |
| - | - | 579.4 | 196.1 | - | - | 0 | - |
| - | - | 502.1 | 196.2 | - | - | 0 | - |
| - | - | 2379 | 197.1 | - | - | 0 | - |
| - | - | 728.5 | 198.1 | - | - | 0 | - |
| - | - | 741.4 | 198.5 | - | - | 0 | - |
| - | - | 819.4 | 198.5 | - | - | 0 | - |
| - | - | 545.4 | 198.7 | - | - | 0 | - |
| - | - | 2208 | 199.1 | - | - | 0 | - |
| - | - | 930.7 | 201.1 | - | - | 0 | - |
| - | - | 5287 | 203 | - | - | 0 | - |
| - | - | 891.1 | 203.1 | - | - | 0 | - |
| - | - | 834.3 | 205.1 | - | - | 0 | - |
| - | - | 981.9 | 208.1 | - | - | 0 | - |
| - | - | 1.044E+04 | 209.1 | - | - | 0 | - |
| - | - | 2835 | 210.1 | - | - | 0 | - |
| - | - | 473.9 | 213.5 | - | - | 0 | - |
| - | - | 542.2 | 214 | - | - | 0 | - |
| - | - | 490.2 | 214.8 | - | - | 0 | - |
| - | - | 2399 | 216.1 | - | - | 0 | - |
| - | - | 446.7 | 219.1 | - | - | 0 | - |
| - | - | 3.616E+05 | 221.1 | - | - | 0 | - |
| - | - | 2.699E+04 | 222.1 | - | - | 0 | - |
| - | - | 1.322E+04 | 223.1 | - | - | 0 | - |
| - | - | 1881 | 223.1 | - | - | 0 | - |
| 9 | z | 3.29E+04 | 223.1 | 0.001051 | 4.71 | +3 | 5 |
| - | - | 857.3 | 224.1 | - | - | 0 | - |
| - | - | 5456 | 224.1 | - | - | 0 | - |
| - | - | 853.3 | 225.1 | - | - | 0 | - |
| - | - | 6356 | 226.1 | - | - | 0 | - |
| - | - | 569.8 | 227.1 | - | - | 0 | - |
| - | - | 2912 | 231 | - | - | 0 | - |
| - | - | 755.1 | 232 | - | - | 0 | - |
| - | - | 1385 | 235.1 | - | - | 0 | - |
| - | - | 3882 | 236.1 | - | - | 0 | - |
| - | - | 3895 | 237.1 | - | - | 0 | - |
| 8 | z | 1698 | 241.1 | 0.002015 | 8.359 | +3 | 6 |
| - | - | 3.735E+04 | 244.1 | - | - | 0 | - |
| - | - | 914.8 | 244.2 | - | - | 0 | - |
| - | - | 3178 | 245.1 | - | - | 0 | - |
| - | - | 1.877E+05 | 249.1 | - | - | 0 | - |
| - | - | 1.722E+04 | 250.1 | - | - | 0 | - |
| - | - | 7067 | 251.1 | - | - | 0 | - |
| - | - | 730.3 | 252.1 | - | - | 0 | - |
| - | - | 6836 | 252.1 | - | - | 0 | - |
| - | - | 1159 | 253.1 | - | - | 0 | - |
| - | - | 2773 | 254.1 | - | - | 0 | - |
| - | - | 502.7 | 254.8 | - | - | 0 | - |
| - | - | 6483 | 258.1 | - | - | 0 | - |
| - | - | 937.6 | 259.1 | - | - | 0 | - |
| - | - | 1613 | 267.1 | - | - | 0 | - |
| 10 | z | 7555 | 269.1 | 0.005368 | 19.95 | +2 | 4 |
| - | - | 1087 | 270.1 | - | - | 0 | - |
| - | - | 1378 | 270.2 | - | - | 0 | - |
| - | - | 625.4 | 270.6 | - | - | 0 | - |
| - | - | 860.6 | 273.1 | - | - | 0 | - |
| - | - | 1919 | 278.1 | - | - | 0 | - |
| - | - | 978.4 | 278.7 | - | - | 0 | - |
| 10 | y | 1.332E+04 | 285.6 | 0.0008286 | 2.901 | +2 | 4 |
| - | - | 4064 | 286.1 | - | - | 0 | - |
| - | - | 670.2 | 286.2 | - | - | 0 | - |
| - | - | 1149 | 292.1 | - | - | 0 | - |
| - | - | 1293 | 301.1 | - | - | 0 | - |
| 12 | z | 2949 | 303.1 | 0.0007434 | 2.452 | +1 | 2 |
| - | - | 604 | 304.1 | - | - | 0 | - |
| - | - | 1105 | 308.1 | - | - | 0 | - |
| - | - | 1484 | 310.1 | - | - | 0 | - |
| - | - | 1.399E+04 | 310.1 | - | - | 0 | - |
| - | - | 928.3 | 311.1 | - | - | 0 | - |
| - | - | 1622 | 311.1 | - | - | 0 | - |
| - | - | 653 | 313.7 | - | - | 0 | - |
| - | - | 1213 | 317.1 | - | - | 0 | - |
| - | - | 1.915E+04 | 318.1 | - | - | 0 | - |
| - | - | 1950 | 318.1 | - | - | 0 | - |
| - | - | 3004 | 319.1 | - | - | 0 | - |
| 12 | y | 2.72E+04 | 319.1 | 0.001032 | 3.233 | +1 | 2 |
| - | - | 4529 | 320.1 | - | - | 0 | - |
| - | - | 1304 | 325.2 | - | - | 0 | - |
| - | - | 1368 | 329.7 | - | - | 0 | - |
| - | - | 2501 | 330.1 | - | - | 0 | - |
| - | - | 2.56E+04 | 336.1 | - | - | 0 | - |
| - | - | 2199 | 337.1 | - | - | 0 | - |
| 9 | y | 3987 | 342.2 | 0.001002 | 2.93 | +2 | 5 |
| - | - | 1925 | 342.7 | - | - | 0 | - |
| - | - | 731.2 | 343.1 | - | - | 0 | - |
| - | - | 892.1 | 343.2 | - | - | 0 | - |
| - | - | 2662 | 346.2 | - | - | 0 | - |
| - | - | 1647 | 346.2 | - | - | 0 | - |
| - | - | 2496 | 348.1 | - | - | 0 | - |
| - | - | 1063 | 348.2 | - | - | 0 | - |
| - | - | 1649 | 352.1 | - | - | 0 | - |
| 3 | c | 5649 | 353.1 | 0.003389 | 9.597 | +1 | 3 |
| - | - | 774.5 | 354.1 | - | - | 0 | - |
| - | - | 1546 | 355.1 | - | - | 0 | - |
| - | - | 1401 | 357.2 | - | - | 0 | - |
| - | - | 2188 | 357.2 | - | - | 0 | - |
| - | - | 1681 | 364.2 | - | - | 0 | - |
| - | - | 2235 | 368.2 | - | - | 0 | - |
| - | - | 1118 | 371.2 | - | - | 0 | - |
| - | - | 2309 | 372.2 | - | - | 0 | - |
| 11 | w | 2E+04 | 373.2 | 0.0009651 | 2.586 | +1 | 3 |
| - | - | 1892 | 373.7 | - | - | 0 | - |
| - | - | 3629 | 374.2 | - | - | 0 | - |
| - | - | 772 | 374.2 | - | - | 0 | - |
| - | - | 5607 | 374.3 | - | - | 0 | - |
| - | - | 959.1 | 375.2 | - | - | 0 | - |
| - | - | 948.9 | 375.3 | - | - | 0 | - |
| - | - | 2314 | 376.7 | - | - | 0 | - |
| 8 | y | 6912 | 377.7 | 0.0009697 | 2.567 | +2 | 6 |
| - | - | 3443 | 378.2 | - | - | 0 | - |
| - | - | 999.3 | 378.7 | - | - | 0 | - |
| - | - | 1864 | 381.1 | - | - | 0 | - |
| - | - | 1937 | 381.2 | - | - | 0 | - |
| 5 | y | 2380 | 384.5 | 0.000533 | 1.386 | +3 | 9 |
| - | - | 921 | 385.7 | - | - | 0 | - |
| - | - | 4475 | 389.2 | - | - | 0 | - |
| - | - | 2826 | 392.7 | - | - | 0 | - |
| - | - | 904.3 | 393.2 | - | - | 0 | - |
| - | - | 895.8 | 393.2 | - | - | 0 | - |
| - | - | 631.2 | 397.7 | - | - | 0 | - |
| - | - | 3310 | 398.2 | - | - | 0 | - |
| - | - | 1607 | 399.1 | - | - | 0 | - |
| - | - | 1392 | 400.3 | - | - | 0 | - |
| - | - | 637.1 | 401.3 | - | - | 0 | - |
| - | - | 1546 | 401.7 | - | - | 0 | - |
| - | - | 1122 | 406.2 | - | - | 0 | - |
| - | - | 5695 | 407.2 | - | - | 0 | - |
| - | - | 997.8 | 408.2 | - | - | 0 | - |
| - | - | 1114 | 409.2 | - | - | 0 | - |
| - | - | 6244 | 412.2 | - | - | 0 | - |
| - | - | 677 | 412.5 | - | - | 0 | - |
| - | - | 1391 | 413.2 | - | - | 0 | - |
| - | - | 863.5 | 415.2 | - | - | 0 | - |
| 7 | c | 3116 | 416.2 | 0.00401 | 9.636 | +2 | 7 |
| - | - | 4076 | 416.7 | - | - | 0 | - |
| - | - | 1.788E+04 | 417.1 | - | - | 0 | - |
| 4 | y | 2108 | 417.5 | 0.001441 | 3.45 | +3 | 10 |
| - | - | 805.3 | 417.7 | - | - | 0 | - |
| - | - | 1461 | 417.9 | - | - | 0 | - |
| - | - | 4018 | 418.1 | - | - | 0 | - |
| - | - | 1361 | 419.1 | - | - | 0 | - |
| - | - | 2834 | 420.2 | - | - | 0 | - |
| - | - | 1256 | 420.7 | - | - | 0 | - |
| - | - | 3595 | 421.2 | - | - | 0 | - |
| - | - | 852.9 | 422.2 | - | - | 0 | - |
| - | - | 1423 | 422.9 | - | - | 0 | - |
| - | - | 941.4 | 423.2 | - | - | 0 | - |
| - | - | 1863 | 423.2 | - | - | 0 | - |
| - | - | 3385 | 427.2 | - | - | 0 | - |
| - | - | 1743 | 429.2 | - | - | 0 | - |
| 11 | y | 2.386E+04 | 433.2 | 0.001744 | 4.027 | +1 | 3 |
| - | - | 1971 | 433.7 | - | - | 0 | - |
| - | - | 5013 | 434.2 | - | - | 0 | - |
| - | - | 2977 | 434.9 | - | - | 0 | - |
| - | - | 1.318E+04 | 435.2 | - | - | 0 | - |
| 3 | z | 1715 | 435.2 | 0.003572 | 8.208 | +3 | 11 |
| 3 | z | 1017 | 435.5 | 0.007412 | 17.02 | +3 | 11 |
| - | - | 3196 | 436.2 | - | - | 0 | - |
| - | - | 744 | 437.2 | - | - | 0 | - |
| - | - | 838.2 | 437.2 | - | - | 0 | - |
| - | - | 751.7 | 437.7 | - | - | 0 | - |
| - | - | 4429 | 438.2 | - | - | 0 | - |
| - | - | 1962 | 438.7 | - | - | 0 | - |
| - | - | 1285 | 439.2 | - | - | 0 | - |
| 3 | y | 3.699E+04 | 440.5 | 0.001488 | 3.378 | +3 | 11 |
| 3 | y | 2.952E+04 | 440.9 | 0.007742 | 17.56 | +3 | 11 |
| 3 | z | 1.746E+04 | 441.2 | 0.005418 | 12.28 | +3 | 11 |
| - | - | 4800 | 441.5 | - | - | 0 | - |
| 7 | y | 8212 | 442.2 | 0.001432 | 3.239 | +2 | 7 |
| - | - | 3316 | 442.7 | - | - | 0 | - |
| - | - | 2223 | 443.2 | - | - | 0 | - |
| - | - | 1113 | 443.2 | - | - | 0 | - |
| 3 | y | 2.034E+04 | 446.5 | 0.00169 | 3.784 | +3 | 11 |
| - | - | 1.719E+04 | 446.9 | - | - | 0 | - |
| - | - | 5644 | 447.2 | - | - | 0 | - |
| - | - | 2439 | 447.5 | - | - | 0 | - |
| - | - | 2782 | 448.2 | - | - | 0 | - |
| - | - | 772.7 | 449.9 | - | - | 0 | - |
| - | - | 1074 | 451.2 | - | - | 0 | - |
| 4 | c | 7616 | 452.2 | 0.00738 | 16.32 | +1 | 4 |
| - | - | 1724 | 452.7 | - | - | 0 | - |
| - | - | 1645 | 453.2 | - | - | 0 | - |
| - | - | 831.2 | 453.5 | - | - | 0 | - |
| - | - | 8898 | 455.2 | - | - | 0 | - |
| - | - | 1645 | 456.2 | - | - | 0 | - |
| - | - | 1056 | 457.9 | - | - | 0 | - |
| - | - | 874.3 | 458.2 | - | - | 0 | - |
| - | - | 4141 | 459.2 | - | - | 0 | - |
| - | - | 3710 | 459.5 | - | - | 0 | - |
| - | - | 2427 | 459.7 | - | - | 0 | - |
| - | - | 2705 | 459.9 | - | - | 0 | - |
| - | - | 1320 | 460.2 | - | - | 0 | - |
| - | - | 1137 | 460.2 | - | - | 0 | - |
| - | - | 903.9 | 461.7 | - | - | 0 | - |
| - | - | 689 | 463.2 | - | - | 0 | - |
| - | - | 3675 | 463.5 | - | - | 0 | - |
| - | - | 3086 | 463.9 | - | - | 0 | - |
| 2 | z | 2147 | 464.2 | 0.006443 | 13.88 | +3 | 12 |
| 2 | z | 1001 | 464.5 | 0.008516 | 18.33 | +3 | 12 |
| - | - | 795 | 464.9 | - | - | 0 | - |
| - | - | 1652 | 468.2 | - | - | 0 | - |
| - | - | 872.2 | 468.7 | - | - | 0 | - |
| 12 | c | 6708 | 468.9 | 0.0004751 | 1.013 | +3 | 12 |
| - | - | 5650 | 469.2 | - | - | 0 | - |
| - | - | 4320 | 469.5 | - | - | 0 | - |
| 2 | y | 851.4 | 469.9 | 0.008183 | 17.42 | +3 | 12 |
| - | - | 2243 | 471.7 | - | - | 0 | - |
| - | - | 1678 | 472.2 | - | - | 0 | - |
| - | - | 785.3 | 472.9 | - | - | 0 | - |
| - | - | 2789 | 474.9 | - | - | 0 | - |
| - | - | 8040 | 475.2 | - | - | 0 | - |
| - | - | 782.7 | 475.5 | - | - | 0 | - |
| - | - | 3664 | 475.7 | - | - | 0 | - |
| - | - | 927.9 | 475.9 | - | - | 0 | - |
| - | - | 1698 | 476.2 | - | - | 0 | - |
| - | - | 807.8 | 476.7 | - | - | 0 | - |
| - | - | 1260 | 477.2 | - | - | 0 | - |
| - | - | 949.5 | 480.9 | - | - | 0 | - |
| - | - | 753.7 | 481.2 | - | - | 0 | - |
| - | - | 6215 | 484.2 | - | - | 0 | - |
| - | - | 2699 | 484.7 | - | - | 0 | - |
| - | - | 901.5 | 485.2 | - | - | 0 | - |
| - | - | 5435 | 485.7 | - | - | 0 | - |
| - | - | 3167 | 486.2 | - | - | 0 | - |
| - | - | 1805 | 486.6 | - | - | 0 | - |
| - | - | 1089 | 486.7 | - | - | 0 | - |
| - | - | 4735 | 486.9 | - | - | 0 | - |
| - | - | 3343 | 487.2 | - | - | 0 | - |
| - | - | 1308 | 487.6 | - | - | 0 | - |
| - | - | 1125 | 488.2 | - | - | 0 | - |
| - | - | 3524 | 490.2 | - | - | 0 | - |
| - | - | 826.8 | 491.2 | - | - | 0 | - |
| - | - | 4.393E+04 | 492.6 | - | - | 0 | - |
| - | - | 4.397E+04 | 492.9 | - | - | 0 | - |
| - | - | 2.049E+04 | 493.2 | - | - | 0 | - |
| - | - | 6823 | 493.6 | - | - | 0 | - |
| - | - | 1387 | 493.9 | - | - | 0 | - |
| 6 | z | 9688 | 494.2 | 0.003312 | 6.702 | +2 | 8 |
| - | - | 9294 | 494.5 | - | - | 0 | - |
| - | - | 4842 | 494.7 | - | - | 0 | - |
| - | - | 5654 | 494.9 | - | - | 0 | - |
| - | - | 4044 | 495.2 | - | - | 0 | - |
| - | - | 1285 | 495.5 | - | - | 0 | - |
| - | - | 1301 | 495.7 | - | - | 0 | - |
| - | - | 1313 | 497.2 | - | - | 0 | - |
| - | - | 3567 | 498.6 | - | - | 0 | - |
| - | - | 3161 | 498.9 | - | - | 0 | - |
| - | - | 2205 | 499.2 | - | - | 0 | - |
| - | - | 1008 | 499.7 | - | - | 0 | - |
| - | - | 3.849E+04 | 500.2 | - | - | 0 | - |
| - | - | 3.026E+04 | 500.5 | - | - | 0 | - |
| - | - | 1.687E+04 | 500.9 | - | - | 0 | - |
| - | - | 4881 | 501.2 | - | - | 0 | - |
| - | - | 1114 | 501.5 | - | - | 0 | - |
| 6 | y | 1225 | 501.7 | 0.001764 | 3.516 | +2 | 8 |
| 6 | y | 4748 | 502.2 | 0.001364 | 2.716 | +2 | 8 |
| 6 | z | 1667 | 502.7 | 0.001419 | 2.822 | +2 | 8 |
| - | - | 839.9 | 502.9 | - | - | 0 | - |
| - | - | 771.6 | 503.2 | - | - | 0 | - |
| - | - | 722.2 | 504.2 | - | - | 0 | - |
| - | - | 937.8 | 505.2 | - | - | 0 | - |
| - | - | 1163 | 505.9 | - | - | 0 | - |
| - | - | 1043 | 507.5 | - | - | 0 | - |
| - | - | 1091 | 507.9 | - | - | 0 | - |
| 9 | c | 915.2 | 508.2 | 0.008935 | 17.58 | +2 | 9 |
| - | - | 1064 | 508.7 | - | - | 0 | - |
| - | - | 5600 | 510.2 | - | - | 0 | - |
| 6 | y | 6.553E+04 | 510.7 | 0.001792 | 3.509 | +2 | 8 |
| - | - | 3.545E+04 | 511.2 | - | - | 0 | - |
| - | - | 9249 | 511.5 | - | - | 0 | - |
| - | - | 9762 | 511.7 | - | - | 0 | - |
| - | - | 6623 | 511.9 | - | - | 0 | - |
| - | - | 1610 | 512.2 | - | - | 0 | - |
| - | - | 2190 | 512.2 | - | - | 0 | - |
| - | - | 2746 | 513.9 | - | - | 0 | - |
| - | - | 1699 | 514.2 | - | - | 0 | - |
| - | - | 3593 | 514.6 | - | - | 0 | - |
| 9 | c | 1647 | 517.2 | 0.009456 | 18.28 | +2 | 9 |
| - | - | 7392 | 517.5 | - | - | 0 | - |
| - | - | 6417 | 517.9 | - | - | 0 | - |
| - | - | 3269 | 518.2 | - | - | 0 | - |
| - | - | 908.8 | 518.5 | - | - | 0 | - |
| - | - | 1895 | 519.2 | - | - | 0 | - |
| - | - | 1034 | 521.2 | - | - | 0 | - |
| - | - | 1762 | 521.3 | - | - | 0 | - |
| - | - | 1127 | 521.8 | - | - | 0 | - |
| - | - | 2245 | 522.3 | - | - | 0 | - |
| - | - | 3.414E+04 | 523.2 | - | - | 0 | - |
| - | - | 4.513E+04 | 523.6 | - | - | 0 | - |
| - | - | 2.929E+04 | 523.9 | - | - | 0 | - |
| - | - | 1.406E+04 | 524.2 | - | - | 0 | - |
| - | - | 5622 | 524.6 | - | - | 0 | - |
| - | - | 1533 | 524.9 | - | - | 0 | - |
| - | - | 2486 | 528.3 | - | - | 0 | - |
| - | - | 5.207E+05 | 529.2 | - | - | 0 | - |
| - | - | 4.387E+05 | 529.6 | - | - | 0 | - |
| - | - | 2.73E+05 | 529.9 | - | - | 0 | - |
| - | - | 1.059E+05 | 530.2 | - | - | 0 | - |
| - | - | 3.165E+04 | 530.6 | - | - | 0 | - |
| - | - | 1137 | 531.2 | - | - | 0 | - |
| - | - | 917 | 531.7 | - | - | 0 | - |
| - | - | 1323 | 535.2 | - | - | 0 | - |
| - | - | 4447 | 535.3 | - | - | 0 | - |
| - | - | 1787 | 535.8 | - | - | 0 | - |
| - | - | 770.3 | 536.2 | - | - | 0 | - |
| - | - | 2129 | 536.3 | - | - | 0 | - |
| 5 | w | 5836 | 537.7 | 0.001026 | 1.908 | +2 | 9 |
| - | - | 1648 | 538.2 | - | - | 0 | - |
| - | - | 4781 | 538.2 | - | - | 0 | - |
| - | - | 2609 | 538.7 | - | - | 0 | - |
| - | - | 1699 | 539.2 | - | - | 0 | - |
| - | - | 2.118E+04 | 540.2 | - | - | 0 | - |
| - | - | 4833 | 541.2 | - | - | 0 | - |
| - | - | 2403 | 542.2 | - | - | 0 | - |
| - | - | 1780 | 544.3 | - | - | 0 | - |
| - | - | 1702 | 544.8 | - | - | 0 | - |
| - | - | 1470 | 545.3 | - | - | 0 | - |
| - | - | 949.8 | 545.8 | - | - | 0 | - |
| - | - | 938.8 | 546.2 | - | - | 0 | - |
| - | - | 2972 | 548.2 | - | - | 0 | - |
| - | - | 978.5 | 549.2 | - | - | 0 | - |
| - | - | 1382 | 552.2 | - | - | 0 | - |
| - | - | 1013 | 552.7 | - | - | 0 | - |
| 10 | y | 4963 | 553.2 | 0.002951 | 5.334 | +1 | 4 |
| 10 | z | 6520 | 554.2 | 0.0007412 | 1.337 | +1 | 4 |
| - | - | 2270 | 555.2 | - | - | 0 | - |
| - | - | 2026 | 555.3 | - | - | 0 | - |
| - | - | 2399 | 556.3 | - | - | 0 | - |
| - | - | 2369 | 557.3 | - | - | 0 | - |
| - | - | 2033 | 557.8 | - | - | 0 | - |
| - | - | 1880 | 558.7 | - | - | 0 | - |
| 5 | z | 1136 | 559.2 | 0.0001439 | 0.2573 | +2 | 9 |
| - | - | 1087 | 561.3 | - | - | 0 | - |
| - | - | 984.3 | 562.3 | - | - | 0 | - |
| - | - | 2499 | 563.2 | - | - | 0 | - |
| - | - | 1557 | 563.3 | - | - | 0 | - |
| - | - | 1234 | 563.7 | - | - | 0 | - |
| - | - | 745.1 | 564.2 | - | - | 0 | - |
| - | - | 1033 | 564.2 | - | - | 0 | - |
| - | - | 1.589E+04 | 564.8 | - | - | 0 | - |
| - | - | 1.27E+04 | 565.3 | - | - | 0 | - |
| - | - | 6108 | 565.8 | - | - | 0 | - |
| - | - | 4859 | 566.2 | - | - | 0 | - |
| - | - | 1711 | 566.3 | - | - | 0 | - |
| - | - | 1262 | 567.2 | - | - | 0 | - |
| 5 | y | 3669 | 567.3 | 0.00502 | 8.85 | +2 | 9 |
| 5 | y | 1.074E+04 | 567.7 | 0.003613 | 6.364 | +2 | 9 |
| 5 | z | 8573 | 568.2 | 0.001349 | 2.373 | +2 | 9 |
| - | - | 3840 | 568.7 | - | - | 0 | - |
| - | - | 1874 | 569.2 | - | - | 0 | - |
| - | - | 1494 | 569.8 | - | - | 0 | - |
| 10 | y | 5.03E+04 | 570.2 | 0.002342 | 4.107 | +1 | 4 |
| - | - | 799 | 570.8 | - | - | 0 | - |
| - | - | 1.758E+04 | 571.2 | - | - | 0 | - |
| - | - | 3733 | 572.2 | - | - | 0 | - |
| - | - | 1027 | 575.8 | - | - | 0 | - |
| 5 | y | 1.176E+05 | 576.3 | 0.001813 | 3.147 | +2 | 9 |
| 10 | c | 7.397E+04 | 576.8 | 0.007203 | 12.49 | +2 | 10 |
| - | - | 1026 | 576.8 | - | - | 0 | - |
| - | - | 3.328E+04 | 577.3 | - | - | 0 | - |
| - | - | 8774 | 577.8 | - | - | 0 | - |
| - | - | 2293 | 578.3 | - | - | 0 | - |
| - | - | 2.784E+04 | 578.8 | - | - | 0 | - |
| - | - | 1.868E+04 | 579.3 | - | - | 0 | - |
| - | - | 8375 | 579.8 | - | - | 0 | - |
| - | - | 3051 | 580.3 | - | - | 0 | - |
| - | - | 1370 | 580.8 | - | - | 0 | - |
| - | - | 762.1 | 581.3 | - | - | 0 | - |
| - | - | 1017 | 581.8 | - | - | 0 | - |
| - | - | 2032 | 582.3 | - | - | 0 | - |
| 5 | c | 9923 | 583.2 | 0.002882 | 4.941 | +1 | 5 |
| - | - | 1.001E+04 | 584.2 | - | - | 0 | - |
| - | - | 2330 | 585.3 | - | - | 0 | - |
| 10 | c | 826.9 | 585.8 | 0.004794 | 8.184 | +2 | 10 |
| - | - | 881.6 | 586.3 | - | - | 0 | - |
| - | - | 5138 | 587.3 | - | - | 0 | - |
| - | - | 1.75E+04 | 587.8 | - | - | 0 | - |
| - | - | 9825 | 588.3 | - | - | 0 | - |
| - | - | 4738 | 588.8 | - | - | 0 | - |
| - | - | 1151 | 589.7 | - | - | 0 | - |
| - | - | 1760 | 590.3 | - | - | 0 | - |
| - | - | 1304 | 590.8 | - | - | 0 | - |
| - | - | 1464 | 594.8 | - | - | 0 | - |
| - | - | 944.4 | 597.2 | - | - | 0 | - |
| - | - | 1039 | 602.8 | - | - | 0 | - |
| - | - | 1265 | 603.3 | - | - | 0 | - |
| - | - | 1267 | 606.3 | - | - | 0 | - |
| - | - | 2884 | 608.3 | - | - | 0 | - |
| 4 | z | 2998 | 608.8 | 0.003308 | 5.434 | +2 | 10 |
| 4 | z | 1484 | 609.3 | 0.01215 | 19.95 | +2 | 10 |
| 4 | w | 2591 | 610.3 | 0.002073 | 3.396 | +2 | 10 |
| - | - | 1039 | 610.8 | - | - | 0 | - |
| - | - | 1112 | 611.3 | - | - | 0 | - |
| - | - | 1500 | 616.2 | - | - | 0 | - |
| 4 | y | 2439 | 616.8 | 0.002613 | 4.236 | +2 | 10 |
| 4 | y | 7844 | 617.3 | 0.002121 | 3.436 | +2 | 10 |
| 4 | z | 2.27E+04 | 617.8 | 0.001688 | 2.732 | +2 | 10 |
| - | - | 1.833E+04 | 618.3 | - | - | 0 | - |
| - | - | 8102 | 618.8 | - | - | 0 | - |
| - | - | 1956 | 619.3 | - | - | 0 | - |
| - | - | 7663 | 620.3 | - | - | 0 | - |
| - | - | 4460 | 620.8 | - | - | 0 | - |
| - | - | 2149 | 621.3 | - | - | 0 | - |
| - | - | 771.8 | 622.3 | - | - | 0 | - |
| 9 | w | 3.799E+04 | 624.3 | 0.002153 | 3.449 | +1 | 5 |
| - | - | 4974 | 624.8 | - | - | 0 | - |
| - | - | 9696 | 625.3 | - | - | 0 | - |
| - | - | 7219 | 625.3 | - | - | 0 | - |
| 4 | y | 1.056E+05 | 625.8 | 0.001969 | 3.147 | +2 | 10 |
| - | - | 7.283E+04 | 626.3 | - | - | 0 | - |
| - | - | 3.364E+04 | 626.8 | - | - | 0 | - |
| - | - | 7604 | 627.3 | - | - | 0 | - |
| - | - | 2710 | 627.8 | - | - | 0 | - |
| - | - | 1.062E+05 | 630.8 | - | - | 0 | - |
| - | - | 7.666E+04 | 631.3 | - | - | 0 | - |
| - | - | 3.541E+04 | 631.8 | - | - | 0 | - |
| - | - | 9409 | 632.3 | - | - | 0 | - |
| - | - | 2838 | 632.8 | - | - | 0 | - |
| - | - | 3813 | 633.3 | - | - | 0 | - |
| 11 | c | 3193 | 633.8 | 0.003275 | 5.167 | +2 | 11 |
| 11 | c | 5786 | 634.3 | 0.0002618 | 0.4127 | +2 | 11 |
| - | - | 2821 | 634.8 | - | - | 0 | - |
| - | - | 2320 | 635.3 | - | - | 0 | - |
| - | - | 1500 | 637.3 | - | - | 0 | - |
| - | - | 1789 | 637.8 | - | - | 0 | - |
| - | - | 4311 | 638.3 | - | - | 0 | - |
| - | - | 3974 | 638.8 | - | - | 0 | - |
| - | - | 1937 | 639.3 | - | - | 0 | - |
| - | - | 1323 | 640.3 | - | - | 0 | - |
| 11 | c | 1311 | 642.8 | 0.008411 | 13.08 | +2 | 11 |
| - | - | 1092 | 643.3 | - | - | 0 | - |
| - | - | 805.9 | 643.4 | - | - | 0 | - |
| - | - | 762.9 | 643.8 | - | - | 0 | - |
| - | - | 3371 | 645.3 | - | - | 0 | - |
| - | - | 2016 | 645.8 | - | - | 0 | - |
| - | - | 4607 | 646.3 | - | - | 0 | - |
| - | - | 3289 | 646.8 | - | - | 0 | - |
| - | - | 3021 | 647.3 | - | - | 0 | - |
| - | - | 1233 | 647.8 | - | - | 0 | - |
| - | - | 1314 | 648.3 | - | - | 0 | - |
| - | - | 987.1 | 648.8 | - | - | 0 | - |
| - | - | 6553 | 651.8 | - | - | 0 | - |
| 3 | z | 5159 | 652.3 | 0.0006214 | 0.9527 | +2 | 11 |
| 3 | z | 1.124E+04 | 652.8 | 0.01231 | 18.86 | +2 | 11 |
| - | - | 6125 | 653.3 | - | - | 0 | - |
| - | - | 4410 | 653.8 | - | - | 0 | - |
| - | - | 2661 | 654.3 | - | - | 0 | - |
| - | - | 2100 | 654.8 | - | - | 0 | - |
| - | - | 8867 | 655.3 | - | - | 0 | - |
| - | - | 2756 | 656.3 | - | - | 0 | - |
| - | - | 909.1 | 657.3 | - | - | 0 | - |
| - | - | 4837 | 658.3 | - | - | 0 | - |
| - | - | 4664 | 659.3 | - | - | 0 | - |
| - | - | 5792 | 659.8 | - | - | 0 | - |
| 3 | y | 2.53E+04 | 660.3 | 0.001064 | 1.611 | +2 | 11 |
| 3 | y | 5.233E+04 | 660.8 | 0.004845 | 7.332 | +2 | 11 |
| 3 | z | 9.507E+04 | 661.3 | 0.002092 | 3.163 | +2 | 11 |
| - | - | 6.916E+04 | 661.8 | - | - | 0 | - |
| - | - | 3.381E+04 | 662.3 | - | - | 0 | - |
| - | - | 1.042E+04 | 662.8 | - | - | 0 | - |
| - | - | 1502 | 663.3 | - | - | 0 | - |
| 9 | y | 2343 | 666.3 | 0.001157 | 1.736 | +1 | 5 |
| 9 | z | 7329 | 667.3 | 0.001047 | 1.57 | +1 | 5 |
| - | - | 2734 | 668.3 | - | - | 0 | - |
| - | - | 1650 | 668.8 | - | - | 0 | - |
| 3 | y | 5.738E+05 | 669.3 | 0.002373 | 3.546 | +2 | 11 |
| - | - | 4.225E+05 | 669.8 | - | - | 0 | - |
| - | - | 2.029E+05 | 670.3 | - | - | 0 | - |
| - | - | 6.822E+04 | 670.8 | - | - | 0 | - |
| - | - | 1.837E+04 | 671.3 | - | - | 0 | - |
| - | - | 4264 | 674.3 | - | - | 0 | - |
| - | - | 4268 | 674.8 | - | - | 0 | - |
| - | - | 2556 | 675.3 | - | - | 0 | - |
| - | - | 3340 | 675.3 | - | - | 0 | - |
| - | - | 1426 | 676.3 | - | - | 0 | - |
| - | - | 6968 | 676.3 | - | - | 0 | - |
| - | - | 1003 | 676.8 | - | - | 0 | - |
| - | - | 2130 | 677.3 | - | - | 0 | - |
| - | - | 1291 | 678.3 | - | - | 0 | - |
| - | - | 1709 | 679.8 | - | - | 0 | - |
| - | - | 2999 | 680.3 | - | - | 0 | - |
| - | - | 1505 | 680.8 | - | - | 0 | - |
| - | - | 2.286E+04 | 682.3 | - | - | 0 | - |
| 9 | y | 1.826E+04 | 683.3 | 0.002689 | 3.936 | +1 | 5 |
| - | - | 7064 | 684.3 | - | - | 0 | - |
| - | - | 1318 | 685.3 | - | - | 0 | - |
| - | - | 1033 | 687.8 | - | - | 0 | - |
| - | - | 1229 | 688.3 | - | - | 0 | - |
| - | - | 1.838E+04 | 688.8 | - | - | 0 | - |
| - | - | 1.951E+04 | 689.3 | - | - | 0 | - |
| - | - | 1.245E+04 | 689.8 | - | - | 0 | - |
| - | - | 5217 | 690.3 | - | - | 0 | - |
| - | - | 1756 | 690.8 | - | - | 0 | - |
| - | - | 2553 | 691.4 | - | - | 0 | - |
| - | - | 1191 | 691.9 | - | - | 0 | - |
| - | - | 4065 | 692.3 | - | - | 0 | - |
| - | - | 1989 | 693.3 | - | - | 0 | - |
| - | - | 1085 | 693.8 | - | - | 0 | - |
| - | - | 1341 | 694.3 | - | - | 0 | - |
| - | - | 1668 | 694.3 | - | - | 0 | - |
| - | - | 3922 | 694.8 | - | - | 0 | - |
| - | - | 6861 | 695.3 | - | - | 0 | - |
| 2 | z | 1401 | 695.8 | 0.0009425 | 1.355 | +2 | 12 |
| 2 | w | 1.126E+05 | 696.3 | 0.002157 | 3.098 | +2 | 12 |
| - | - | 9.1E+04 | 696.8 | - | - | 0 | - |
| - | - | 4.216E+04 | 697.3 | - | - | 0 | - |
| - | - | 1.336E+04 | 697.8 | - | - | 0 | - |
| - | - | 4175 | 698.3 | - | - | 0 | - |
| - | - | 894.4 | 699.9 | - | - | 0 | - |
| - | - | 1157 | 702.3 | - | - | 0 | - |
| 12 | c | 2.783E+04 | 702.8 | 0.0005299 | 0.7541 | +2 | 12 |
| - | - | 2.114E+04 | 703.3 | - | - | 0 | - |
| - | - | 1.116E+04 | 703.8 | - | - | 0 | - |
| 2 | y | 6187 | 704.3 | 0.009094 | 12.91 | +2 | 12 |
| 2 | z | 2306 | 704.8 | 0.009223 | 13.09 | +2 | 12 |
| - | - | 940.1 | 705.3 | - | - | 0 | - |
| - | - | 1825 | 705.4 | - | - | 0 | - |
| - | - | 1625 | 708.9 | - | - | 0 | - |
| - | - | 1196 | 710.8 | - | - | 0 | - |
| 12 | c | 2.613E+05 | 711.3 | 0.0001949 | 0.2739 | +2 | 12 |
| - | - | 2.103E+05 | 711.8 | - | - | 0 | - |
| - | - | 1.31E+05 | 712.3 | - | - | 0 | - |
| 2 | y | 4.285E+04 | 712.8 | 0.01327 | 18.62 | +2 | 12 |
| - | - | 1.343E+04 | 713.3 | - | - | 0 | - |
| - | - | 4690 | 713.4 | - | - | 0 | - |
| - | - | 2777 | 714.4 | - | - | 0 | - |
| - | - | 1317 | 717.3 | - | - | 0 | - |
| - | - | 1785 | 717.8 | - | - | 0 | - |
| - | - | 1273 | 718.3 | - | - | 0 | - |
| - | - | 1354 | 718.8 | - | - | 0 | - |
| - | - | 1022 | 719.3 | - | - | 0 | - |
| 6 | c | 1.127E+05 | 720.3 | 0.002101 | 2.917 | +1 | 6 |
| - | - | 4.239E+04 | 721.3 | - | - | 0 | - |
| - | - | 1.766E+04 | 722.3 | - | - | 0 | - |
| - | - | 2895 | 723.3 | - | - | 0 | - |
| - | - | 4332 | 724.3 | - | - | 0 | - |
| - | - | 2965 | 724.8 | - | - | 0 | - |
| - | - | 2287 | 725.3 | - | - | 0 | - |
| - | - | 1748 | 725.8 | - | - | 0 | - |
| - | - | 1210 | 726.3 | - | - | 0 | - |
| - | - | 2715 | 726.8 | - | - | 0 | - |
| - | - | 3851 | 727.3 | - | - | 0 | - |
| - | - | 1808 | 727.8 | - | - | 0 | - |
| - | - | 1185 | 731.3 | - | - | 0 | - |
| - | - | 2172 | 731.8 | - | - | 0 | - |
| - | - | 2885 | 732.3 | - | - | 0 | - |
| - | - | 2256 | 732.8 | - | - | 0 | - |
| - | - | 3440 | 733.3 | - | - | 0 | - |
| - | - | 2956 | 733.8 | - | - | 0 | - |
| - | - | 2333 | 734.3 | - | - | 0 | - |
| - | - | 994.3 | 734.8 | - | - | 0 | - |
| - | - | 1317 | 735.8 | - | - | 0 | - |
| 8 | z | 2.272E+04 | 738.3 | 0.002244 | 3.039 | +1 | 6 |
| - | - | 995.5 | 738.8 | - | - | 0 | - |
| - | - | 1.006E+04 | 739.3 | - | - | 0 | - |
| - | - | 3.111E+04 | 739.8 | - | - | 0 | - |
| - | - | 3.433E+04 | 740.3 | - | - | 0 | - |
| - | - | 2.092E+04 | 740.8 | - | - | 0 | - |
| - | - | 1.539E+04 | 741.3 | - | - | 0 | - |
| - | - | 3.488E+04 | 741.8 | - | - | 0 | - |
| - | - | 2.57E+04 | 742.3 | - | - | 0 | - |
| - | - | 1.401E+04 | 742.8 | - | - | 0 | - |
| - | - | 3784 | 743.3 | - | - | 0 | - |
| - | - | 1125 | 743.8 | - | - | 0 | - |
| - | - | 835.8 | 745.3 | - | - | 0 | - |
| - | - | 8126 | 745.3 | - | - | 0 | - |
| - | - | 1612 | 746.3 | - | - | 0 | - |
| - | - | 4654 | 746.3 | - | - | 0 | - |
| - | - | 5691 | 747.3 | - | - | 0 | - |
| - | - | 5486 | 747.8 | - | - | 0 | - |
| - | - | 9189 | 748.3 | - | - | 0 | - |
| - | - | 6581 | 748.8 | - | - | 0 | - |
| - | - | 5890 | 749.3 | - | - | 0 | - |
| - | - | 9.589E+04 | 749.8 | - | - | 0 | - |
| - | - | 7.358E+04 | 750.3 | - | - | 0 | - |
| - | - | 3.984E+04 | 750.8 | - | - | 0 | - |
| - | - | 1.551E+04 | 751.3 | - | - | 0 | - |
| - | - | 4684 | 751.8 | - | - | 0 | - |
| - | - | 7395 | 752.8 | - | - | 0 | - |
| - | - | 4854 | 753.3 | - | - | 0 | - |
| - | - | 3683 | 753.8 | - | - | 0 | - |
| 8 | y | 2.96E+04 | 754.4 | 0.002929 | 3.883 | +1 | 6 |
| - | - | 2.502E+04 | 754.8 | - | - | 0 | - |
| - | - | 1.829E+04 | 755.3 | - | - | 0 | - |
| - | - | 7218 | 755.4 | - | - | 0 | - |
| - | - | 1.78E+04 | 755.8 | - | - | 0 | - |
| - | - | 7547 | 756.3 | - | - | 0 | - |
| - | - | 1577 | 756.4 | - | - | 0 | - |
| - | - | 5952 | 756.8 | - | - | 0 | - |
| - | - | 2265 | 757.3 | - | - | 0 | - |
| - | - | 1264 | 757.8 | - | - | 0 | - |
| - | - | 2456 | 761.8 | - | - | 0 | - |
| - | - | 3774 | 762.3 | - | - | 0 | - |
| - | - | 1123 | 762.8 | - | - | 0 | - |
| - | - | 8938 | 763.3 | - | - | 0 | - |
| - | - | 1.484E+04 | 763.8 | - | - | 0 | - |
| - | - | 2.046E+05 | 764.3 | - | - | 0 | - |
| - | - | 1.66E+05 | 764.8 | - | - | 0 | - |
| - | - | 9.725E+04 | 765.3 | - | - | 0 | - |
| - | - | 3.715E+04 | 765.8 | - | - | 0 | - |
| - | - | 1.202E+04 | 766.3 | - | - | 0 | - |
| - | - | 7767 | 768.4 | - | - | 0 | - |
| - | - | 3110 | 769.4 | - | - | 0 | - |
| - | - | 8897 | 770.4 | - | - | 0 | - |
| - | - | 4674 | 770.8 | - | - | 0 | - |
| - | - | 4.322E+04 | 771.3 | - | - | 0 | - |
| - | - | 3.688E+04 | 771.8 | - | - | 0 | - |
| - | - | 2.184E+04 | 772.3 | - | - | 0 | - |
| - | - | 8425 | 772.8 | - | - | 0 | - |
| - | - | 1295 | 773.3 | - | - | 0 | - |
| - | - | 2167 | 777.8 | - | - | 0 | - |
| - | - | 1402 | 778.3 | - | - | 0 | - |
| - | - | 1263 | 783.4 | - | - | 0 | - |
| - | - | 1504 | 783.9 | - | - | 0 | - |
| - | - | 2693 | 784.8 | - | - | 0 | - |
| - | - | 1.524E+04 | 785.3 | - | - | 0 | - |
| - | - | 1.325E+04 | 785.8 | - | - | 0 | - |
| - | - | 1.039E+04 | 786.3 | - | - | 0 | - |
| - | - | 7801 | 786.8 | - | - | 0 | - |
| - | - | 2460 | 787.3 | - | - | 0 | - |
| - | - | 1222 | 791.9 | - | - | 0 | - |
| - | - | 3.22E+05 | 793.3 | - | - | 0 | - |
| - | - | 1.655E+06 | 793.8 | - | - | 0 | - |
| - | - | 1.328E+06 | 794.3 | - | - | 0 | - |
| - | - | 7.765E+05 | 794.8 | - | - | 0 | - |
| - | - | 2.904E+05 | 795.3 | - | - | 0 | - |
| - | - | 9.531E+04 | 795.8 | - | - | 0 | - |
| - | - | 1678 | 796.3 | - | - | 0 | - |
| - | - | 9358 | 802.4 | - | - | 0 | - |
| - | - | 4584 | 803.4 | - | - | 0 | - |
| - | - | 2563 | 805.3 | - | - | 0 | - |
| 7 | w | 3.278E+04 | 808.4 | 0.002374 | 2.937 | +1 | 7 |
| - | - | 1.38E+04 | 809.4 | - | - | 0 | - |
| - | - | 4009 | 810.4 | - | - | 0 | - |
| - | - | 1231 | 811.4 | - | - | 0 | - |
| - | - | 1311 | 816.3 | - | - | 0 | - |
| - | - | 6940 | 820.4 | - | - | 0 | - |
| - | - | 3245 | 821.4 | - | - | 0 | - |
| - | - | 1472 | 822.4 | - | - | 0 | - |
| - | - | 1163 | 823.4 | - | - | 0 | - |
| - | - | 1341 | 824.4 | - | - | 0 | - |
| - | - | 5330 | 832.3 | - | - | 0 | - |
| - | - | 1917 | 833.3 | - | - | 0 | - |
| - | - | 7252 | 833.4 | - | - | 0 | - |
| - | - | 2847 | 834.4 | - | - | 0 | - |
| - | - | 1229 | 835.4 | - | - | 0 | - |
| - | - | 1262 | 839.4 | - | - | 0 | - |
| 7 | c | 5.523E+04 | 849.3 | 0.001909 | 2.247 | +1 | 7 |
| - | - | 2.312E+04 | 850.3 | - | - | 0 | - |
| - | - | 1182 | 850.4 | - | - | 0 | - |
| - | - | 1.04E+04 | 851.3 | - | - | 0 | - |
| - | - | 2814 | 852.3 | - | - | 0 | - |
| 7 | y | 2740 | 865.4 | 0.0004418 | 0.5105 | +1 | 7 |
| 7 | y | 2639 | 866.4 | 0.01124 | 12.97 | +1 | 7 |
| 7 | z | 7.14E+04 | 867.4 | 0.002681 | 3.091 | +1 | 7 |
| - | - | 3.88E+04 | 868.4 | - | - | 0 | - |
| - | - | 1.093E+04 | 869.4 | - | - | 0 | - |
| - | - | 1916 | 870.4 | - | - | 0 | - |
| - | - | 2743 | 876.3 | - | - | 0 | - |
| - | - | 1361 | 877.4 | - | - | 0 | - |
| 7 | y | 3.9E+04 | 883.4 | 0.002511 | 2.843 | +1 | 7 |
| - | - | 1.742E+04 | 884.4 | - | - | 0 | - |
| - | - | 5553 | 885.4 | - | - | 0 | - |
| - | - | 4380 | 890.4 | - | - | 0 | - |
| - | - | 2749 | 891.4 | - | - | 0 | - |
| - | - | 1192 | 892.4 | - | - | 0 | - |
| - | - | 1.117E+04 | 903.3 | - | - | 0 | - |
| - | - | 4985 | 904.3 | - | - | 0 | - |
| - | - | 1106 | 904.4 | - | - | 0 | - |
| - | - | 2331 | 905.3 | - | - | 0 | - |
| - | - | 5950 | 905.4 | - | - | 0 | - |
| - | - | 2970 | 906.4 | - | - | 0 | - |
| - | - | 5381 | 907.4 | - | - | 0 | - |
| - | - | 2808 | 908.4 | - | - | 0 | - |
| 8 | c | 5.154E+04 | 920.4 | 0.001791 | 1.946 | +1 | 8 |
| - | - | 2.053E+04 | 921.4 | - | - | 0 | - |
| - | - | 861 | 921.5 | - | - | 0 | - |
| - | - | 1.099E+04 | 922.4 | - | - | 0 | - |
| - | - | 1034 | 922.5 | - | - | 0 | - |
| - | - | 2854 | 923.4 | - | - | 0 | - |
| - | - | 1032 | 929.4 | - | - | 0 | - |
| - | - | 1620 | 930.4 | - | - | 0 | - |
| - | - | 1187 | 932.4 | - | - | 0 | - |
| - | - | 7483 | 935.5 | - | - | 0 | - |
| - | - | 5176 | 936.5 | - | - | 0 | - |
| - | - | 2068 | 937.5 | - | - | 0 | - |
| - | - | 1123 | 948.4 | - | - | 0 | - |
| - | - | 2078 | 971.4 | - | - | 0 | - |
| - | - | 1062 | 972.4 | - | - | 0 | - |
| - | - | 1004 | 975.5 | - | - | 0 | - |
| - | - | 1335 | 976.5 | - | - | 0 | - |
| - | - | 1309 | 985.4 | - | - | 0 | - |
| - | - | 1.442E+04 | 989.4 | - | - | 0 | - |
| - | - | 7723 | 990.4 | - | - | 0 | - |
| - | - | 4155 | 991.4 | - | - | 0 | - |
| - | - | 1688 | 992.4 | - | - | 0 | - |
| 6 | y | 1675 | 1003 | 0.007746 | 7.72 | +1 | 8 |
| 6 | z | 1.445E+04 | 1004 | 0.003278 | 3.264 | +1 | 8 |
| - | - | 1.137E+04 | 1005 | - | - | 0 | - |
| - | - | 5654 | 1006 | - | - | 0 | - |
| - | - | 1460 | 1007 | - | - | 0 | - |
| - | - | 6349 | 1016 | - | - | 0 | - |
| - | - | 3939 | 1017 | - | - | 0 | - |
| - | - | 2165 | 1018 | - | - | 0 | - |
| - | - | 2.409E+04 | 1019 | - | - | 0 | - |
| 6 | y | 4.826E+04 | 1020 | 0.002621 | 2.568 | +1 | 8 |
| - | - | 2.439E+04 | 1021 | - | - | 0 | - |
| - | - | 5883 | 1022 | - | - | 0 | - |
| - | - | 951 | 1023 | - | - | 0 | - |
| 9 | c | 2.096E+04 | 1033 | 0.001505 | 1.456 | +1 | 9 |
| - | - | 1.045E+04 | 1034 | - | - | 0 | - |
| - | - | 5907 | 1035 | - | - | 0 | - |
| - | - | 1787 | 1036 | - | - | 0 | - |
| - | - | 8507 | 1039 | - | - | 0 | - |
| - | - | 5977 | 1040 | - | - | 0 | - |
| - | - | 1612 | 1041 | - | - | 0 | - |
| - | - | 2907 | 1055 | - | - | 0 | - |
| - | - | 5749 | 1056 | - | - | 0 | - |
| - | - | 3307 | 1057 | - | - | 0 | - |
| - | - | 1468 | 1058 | - | - | 0 | - |
| 5 | w | 2074 | 1074 | 0.005056 | 4.706 | +1 | 9 |
| - | - | 1651 | 1075 | - | - | 0 | - |
| - | - | 1508 | 1125 | - | - | 0 | - |
| - | - | 3091 | 1126 | - | - | 0 | - |
| - | - | 1888 | 1127 | - | - | 0 | - |
| - | - | 909.7 | 1128 | - | - | 0 | - |
| 5 | y | 929.4 | 1134 | 0.004981 | 4.391 | +1 | 9 |
| 5 | z | 3508 | 1135 | 0.001917 | 1.688 | +1 | 9 |
| - | - | 4591 | 1136 | - | - | 0 | - |
| - | - | 1720 | 1137 | - | - | 0 | - |
| 5 | y | 6057 | 1152 | 0.003213 | 2.79 | +1 | 9 |
| 10 | c | 3802 | 1152 | 0.01692 | 14.68 | +1 | 10 |
| - | - | 2423 | 1154 | - | - | 0 | - |
| - | - | 1451 | 1154 | - | - | 0 | - |
| - | - | 987.8 | 1158 | - | - | 0 | - |
| - | - | 1291 | 1159 | - | - | 0 | - |
| 10 | c | 6.17E+04 | 1171 | 0.001456 | 1.244 | +1 | 10 |
| - | - | 3.911E+04 | 1172 | - | - | 0 | - |
| - | - | 1.899E+04 | 1173 | - | - | 0 | - |
| - | - | 6346 | 1174 | - | - | 0 | - |
| - | - | 1587 | 1175 | - | - | 0 | - |
| - | - | 1565 | 1180 | - | - | 0 | - |
| - | - | 1234 | 1181 | - | - | 0 | - |
| 4 | z | 3599 | 1235 | 0.001618 | 1.311 | +1 | 10 |
| - | - | 5215 | 1236 | - | - | 0 | - |
| - | - | 3321 | 1237 | - | - | 0 | - |
| - | - | 1145 | 1238 | - | - | 0 | - |
| - | - | 915.2 | 1240 | - | - | 0 | - |
| - | - | 5457 | 1241 | - | - | 0 | - |
| - | - | 4306 | 1242 | - | - | 0 | - |
| - | - | 2161 | 1243 | - | - | 0 | - |
| - | - | 957.1 | 1244 | - | - | 0 | - |
| 4 | y | 2682 | 1251 | 0.005722 | 4.575 | +1 | 10 |
| - | - | 1236 | 1252 | - | - | 0 | - |
| - | - | 928.6 | 1253 | - | - | 0 | - |
| 11 | c | 2384 | 1268 | 0.003186 | 2.514 | +1 | 11 |
| - | - | 1372 | 1269 | - | - | 0 | - |
| - | - | 1414 | 1270 | - | - | 0 | - |
| 11 | c | 4.594E+04 | 1285 | 0.002636 | 2.052 | +1 | 11 |
| - | - | 3.244E+04 | 1286 | - | - | 0 | - |
| - | - | 1.638E+04 | 1287 | - | - | 0 | - |
| - | - | 5896 | 1288 | - | - | 0 | - |
| - | - | 1254 | 1289 | - | - | 0 | - |
| 3 | w | 1752 | 1305 | 0.00712 | 5.457 | +1 | 11 |
| - | - | 1020 | 1306 | - | - | 0 | - |
| 3 | z | 3615 | 1322 | 0.002183 | 1.652 | +1 | 11 |
| - | - | 2.295E+04 | 1323 | - | - | 0 | - |
| - | - | 1.456E+04 | 1324 | - | - | 0 | - |
| - | - | 8374 | 1325 | - | - | 0 | - |
| - | - | 2423 | 1326 | - | - | 0 | - |
| 3 | y | 1113 | 1338 | 0.00653 | 4.882 | +1 | 11 |
| - | - | 971.1 | 1339 | - | - | 0 | - |
| - | - | 3513 | 1364 | - | - | 0 | - |
| - | - | 2860 | 1365 | - | - | 0 | - |
| - | - | 1344 | 1366 | - | - | 0 | - |
| - | - | 1599 | 1378 | - | - | 0 | - |
| - | - | 2622 | 1379 | - | - | 0 | - |
| - | - | 1688 | 1380 | - | - | 0 | - |
| 2 | w | 3066 | 1392 | 0.004632 | 3.329 | +1 | 12 |
| - | - | 9922 | 1393 | - | - | 0 | - |
| - | - | 7114 | 1394 | - | - | 0 | - |
| - | - | 3254 | 1395 | - | - | 0 | - |
| 12 | c | 2664 | 1422 | 0.002344 | 1.649 | +1 | 12 |
| - | - | 1.044E+04 | 1423 | - | - | 0 | - |
| - | - | 7246 | 1424 | - | - | 0 | - |
| 2 | y | 4534 | 1425 | 0.02062 | 14.47 | +1 | 12 |
| - | - | 997.3 | 1426 | - | - | 0 | - |
| - | - | 990.9 | 1480 | - | - | 0 | - |
| - | - | 1497 | 1512 | - | - | 0 | - |
| - | - | 1811 | 1513 | - | - | 0 | - |
| - | - | 1.185E+04 | 1528 | - | - | 0 | - |
| - | - | 2.129E+04 | 1529 | - | - | 0 | - |
| - | - | 1.685E+04 | 1530 | - | - | 0 | - |
| - | - | 8092 | 1531 | - | - | 0 | - |
| - | - | 3495 | 1532 | - | - | 0 | - |
| - | - | 1005 | 1542 | - | - | 0 | - |
| - | - | 4774 | 1543 | - | - | 0 | - |
| - | - | 3364 | 1544 | - | - | 0 | - |
| - | - | 2399 | 1545 | - | - | 0 | - |
| - | - | 3431 | 1560 | - | - | 0 | - |
| - | - | 1717 | 1561 | - | - | 0 | - |
| - | - | 1038 | 1562 | - | - | 0 | - |
| - | - | 2335 | 1570 | - | - | 0 | - |
| - | - | 9201 | 1571 | - | - | 0 | - |
| - | - | 7905 | 1572 | - | - | 0 | - |
| - | - | 4649 | 1573 | - | - | 0 | - |
| - | - | 1772 | 1574 | - | - | 0 | - |
| - | - | 835 | 1575 | - | - | 0 | - |
| - | - | 2986 | 1586 | - | - | 0 | - |
| - | - | 1.986E+04 | 1587 | - | - | 0 | - |
| - | - | 7.797E+04 | 1588 | - | - | 0 | - |
| - | - | 5.969E+04 | 1589 | - | - | 0 | - |
| - | - | 3.45E+04 | 1590 | - | - | 0 | - |
| - | - | 1.135E+04 | 1591 | - | - | 0 | - |
| - | - | 3193 | 1592 | - | - | 0 | - |

m/z Charge Intensity FragmentType MassShift Position
120.05757904052734 0 759.40967
120.0770034790039 0 602.884
120.08123016357422 0 9149.081
121.08465576171875 0 751.8665
123.05575561523438 0 16071.781
124.05912780761719 0 632.94836
124.80400085449219 0 459.5841
129.0663299560547 0 5455.52
131.08212280273438 0 1096.8334
134.0221405029297 0 586.2516
134.0275421142578 0 10788.126
136.076171875 0 25621.332
137.07955932617188 0 2398.9763
138.06671142578125 0 1412.2781
140.08236694335938 0 6462.2183
141.0852508544922 0 454.343
141.10279846191406 0 696.89557
141.89886474609375 0 430.6555
145.06121826171875 0 1168.4219
148.94827270507812 0 993.93646
155.09347534179688 0 799.8203
156.07736206054688 0 1649.1483
157.06124877929688 0 5214.6743
158.02755737304688 0 4623.002
159.1133270263672 0 8455.114
160.11695861816406 0 725.46967
161.0384063720703 0 2789.4119
164.04727172851562 0 1478.9891
164.0824432373047 0 1848.5752
165.05517578125 0 4719.451
166.08680725097656 0 30093.086
167.0901641845703 0 2275.715
168.01206970214844 0 990.06775
173.1291961669922 0 1337.2432
173.4520721435547 0 1162.5073
174.1354217529297 0 1161.3546
176.0381622314453 0 5294.585
177.1028594970703 0 2270.8794
182.08177185058594 0 25752.55 y 12
182.091064453125 0 1250.1683
183.08514404296875 0 2009.3949
185.03831481933594 0 954.4239
186.02255249023438 0 14859.685
187.0257568359375 0 947.3055
187.10829162597656 0 8135.7344
188.01806640625 0 571.3636
188.11155700683594 0 836.52765
192.0773468017578 0 12752.623
195.06410217285156 0 702.33545
196.0718231201172 0 579.443
196.24810791015625 0 502.0777
197.080078125 0 2378.956
198.088134765625 0 728.46625
198.5165557861328 0 741.4052
198.52584838867188 0 819.3666
198.65066528320312 0 545.3749
199.0718536376953 0 2207.6013
201.12440490722656 0 930.6802
203.04910278320312 0 5287.456
203.1219940185547 0 891.0644
205.0974884033203 0 834.2624
208.09580993652344 0 981.94403
209.10391235351562 0 10443.322
210.11077880859375 0 2835.431
213.49864196777344 0 473.90222
214.01707458496094 0 542.15906
214.75538635253906 0 490.2308
216.0987548828125 0 2399.2954
219.10638427734375 0 446.67618
221.0597381591797 0 361554.53
222.0628204345703 0 26989.8
223.05551147460938 0 13223.552
223.0644989013672 0 1881.0167
223.1083221435547 0 32895.098 z 8
224.05921936035156 0 857.26587
224.11253356933594 0 5455.609
225.1142578125 0 853.31104
226.08282470703125 0 6356.4395
227.08734130859375 0 569.827
231.04420471191406 0 2911.6345
232.04937744140625 0 755.13995
235.08375549316406 0 1385.4149
236.09103393554688 0 3881.7751
237.09861755371094 0 3894.6646
241.11280822753906 0 1698.1133 z Ammonia loss 7
244.09359741210938 0 37346.64
244.21319580078125 0 914.7633
245.0967559814453 0 3178.4243
249.05479431152344 0 187679.17
250.05783081054688 0 17224.69
251.05055236816406 0 7067.2607
252.0801239013672 0 730.29333
252.10997009277344 0 6836.1177
253.11380004882812 0 1159.0505
254.12571716308594 0 2772.7454
254.7978057861328 0 502.72238
258.12432861328125 0 6482.726
259.1282958984375 0 937.641
267.1092834472656 0 1612.9427
269.1073303222656 0 7555.467 z Ammonia loss 9
270.10784912109375 0 1086.6664
270.19281005859375 0 1378.241
270.56353759765625 0 625.4147
273.134521484375 0 860.63245
278.1488037109375 0 1919.3271
278.6505432128906 0 978.39014
285.62542724609375 0 13316.686 y 9
286.12701416015625 0 4064.314
286.2122802734375 0 670.1878
292.1294250488281 0 1149.198
301.1296081542969 0 1293.4578
303.1221008300781 0 2949.4426 z 11
304.12518310546875 0 603.9798
308.0917053222656 0 1105.3915
310.1076354980469 0 1484.1901
310.1408386230469 0 13992.27
311.1231994628906 0 928.2682
311.14373779296875 0 1621.7721
313.66656494140625 0 652.98395
317.1257019042969 0 1213.0504
318.0766296386719 0 19149.83
318.1498107910156 0 1950.1456
319.07989501953125 0 3004.2495
319.14111328125 0 27201.893 y 11
320.14434814453125 0 4528.8804
325.15130615234375 0 1304.0073
329.6544494628906 0 1368.3684
330.11273193359375 0 2500.6873
336.0872497558594 0 25595.283
337.0900573730469 0 2198.5618
342.1676330566406 0 3986.5078 y 8
342.6688537597656 0 1925.3763
343.1260986328125 0 731.2477
343.1666564941406 0 892.081
346.1632385253906 0 2661.923
346.1815490722656 0 1646.7914
348.1229248046875 0 2496.3818
348.17889404296875 0 1062.7855
352.1318054199219 0 1649.2899
353.1136779785156 0 5648.667 c 2
354.11322021484375 0 774.5313
355.1412048339844 0 1545.9211
357.18267822265625 0 1400.8442
357.2256774902344 0 2187.8696
364.1651611328125 0 1681.4912
368.17608642578125 0 2235.2563
371.1590576171875 0 1118.4802
372.1675109863281 0 2308.5156
373.151611328125 0 20000.81 w 10
373.6792907714844 0 1892.217
374.15521240234375 0 3629.3164
374.1783447265625 0 772.0036
374.2521667480469 0 5606.6475
375.1561279296875 0 959.1001
375.255615234375 0 948.9313
376.68988037109375 0 2313.6587
377.6861572265625 0 6912.2734 y 7
378.1872253417969 0 3443.3228
378.6871032714844 0 999.3152
381.1205139160156 0 1863.644
381.17828369140625 0 1937.4008
384.507080078125 0 2379.8757 y 4
385.6938171386719 0 921.0409
389.15008544921875 0 4475.3994
392.6753845214844 0 2826.133
393.1775817871094 0 904.30444
393.2259826660156 0 895.75287
397.6710205078125 0 631.1981
398.1501770019531 0 3310.4153
399.1322937011719 0 1607.081
400.26953125 0 1392.185
401.2706604003906 0 637.0936
401.6819152832031 0 1546.0223
406.20281982421875 0 1121.9493
407.1607360839844 0 5695.499
408.16351318359375 0 997.8237
409.1758117675781 0 1113.5304
412.2028503417969 0 6243.509
412.52423095703125 0 676.9624
413.20574951171875 0 1391.3005
415.1748046875 0 863.5009
416.1580810546875 0 3116.2588 c Water loss 6
416.65325927734375 0 4076.2883
417.1456604003906 0 17883.816
417.5307922363281 0 2107.5073 y 3
417.6945495605469 0 805.3146
417.86566162109375 0 1461.4589
418.1487731933594 0 4018.1936
419.14447021484375 0 1361.1842
420.2000427246094 0 2833.6475
420.7016906738281 0 1256.3776
421.15850830078125 0 3595.1594
422.16180419921875 0 852.9359
422.8671875 0 1423.0121
423.1539611816406 0 941.38715
423.1957092285156 0 1863.3871
427.2139587402344 0 3385.065
429.2068786621094 0 1743.1381
433.18475341796875 0 23864.03 y 10
433.70361328125 0 1971.3042
434.18780517578125 0 5013.1084
434.8625793457031 0 2977.4424
435.15594482421875 0 13184.031
435.193359375 0 1715.3801 z Water loss 2
435.5323486328125 0 1016.6214 z Ammonia loss 2
436.1587219238281 0 3195.5032
437.15362548828125 0 743.9559
437.2027282714844 0 838.1639
437.705078125 0 751.73804
438.17413330078125 0 4429.1035
438.67547607421875 0 1961.6855
439.17437744140625 0 1284.8053
440.5379943847656 0 36989.805 y Water loss 2
440.87225341796875 0 29518.635 y Ammonia loss 2
441.20587158203125 0 17464.037 z 2
441.5390625 0 4799.779
442.2079162597656 0 8211.795 y 6
442.7093200683594 0 3315.955
443.16790771484375 0 2222.8071
443.210205078125 0 1112.5122
446.5417175292969 0 20342.17 y 2
446.87579345703125 0 17192.818
447.21002197265625 0 5644.4385
447.5450134277344 0 2439.2898
448.1829833984375 0 2781.6035
449.8724365234375 0 772.72217
451.23223876953125 0 1073.8832
452.1781005859375 0 7615.9365 c 3
452.6724548339844 0 1723.6924
453.18365478515625 0 1644.8895
453.5301513671875 0 831.24536
455.20867919921875 0 8898.273
456.21185302734375 0 1645.3597
457.87017822265625 0 1055.8066
458.2207946777344 0 874.26666
459.22900390625 0 4140.6934
459.53466796875 0 3710.1672
459.7301025390625 0 2427.033
459.8688659667969 0 2705.1206
460.1713562011719 0 1320.3969
460.20135498046875 0 1137.4952
461.7195129394531 0 903.85364
463.1932678222656 0 688.98114
463.5445556640625 0 3674.6436
463.8786926269531 0 3086.0925
464.21405029296875 0 2147.0537 z Water loss 1
464.54412841796875 0 1001.3354 z Ammonia loss 1
464.87548828125 0 795.0433
468.23297119140625 0 1651.7007
468.7359313964844 0 872.1763
468.8653259277344 0 6707.9463 c Ammonia loss 11
469.19940185546875 0 5650.4175
469.5335998535156 0 4319.9863
469.86700439453125 0 851.37213 y Ammonia loss 1
471.7232666015625 0 2243.1199
472.2251281738281 0 1678.2838
472.8733215332031 0 785.27435
474.8689880371094 0 2788.5222
475.2137451171875 0 8040.4683
475.53570556640625 0 782.6664
475.7165222167969 0 3663.9497
475.87176513671875 0 927.89056
476.216552734375 0 1697.9706
476.71612548828125 0 807.82983
477.2115173339844 0 1259.8064
480.87921142578125 0 949.5357
481.20733642578125 0 753.7357
484.2206115722656 0 6215.0806
484.721923828125 0 2698.7563
485.2236022949219 0 901.4915
485.7210388183594 0 5435.44
486.2213439941406 0 3166.5674
486.5527038574219 0 1805.3556
486.7249450683594 0 1088.9644
486.882568359375 0 4735.4424
487.2154541015625 0 3342.8457
487.554443359375 0 1307.7312
488.2098693847656 0 1125.4645
490.1976623535156 0 3524.1423
491.2009582519531 0 826.78253
492.5559997558594 0 43932.082
492.890380859375 0 43969.137
493.2242126464844 0 20488.957
493.55743408203125 0 6822.9893
493.89276123046875 0 1387.4467
494.2099914550781 0 9688.406 z Ammonia loss 5
494.5417175292969 0 9294.018
494.720947265625 0 4841.5835
494.87567138671875 0 5654.4053
495.2132873535156 0 4043.9194
495.54156494140625 0 1285.4231
495.7258605957031 0 1301.2751
497.21917724609375 0 1312.6194
498.5605773925781 0 3566.7874
498.8913269042969 0 3160.7224
499.2264709472656 0 2205.4216
499.70855712890625 0 1007.78656
500.21319580078125 0 38491.168
500.5475769042969 0 30258.38
500.8809509277344 0 16869.416
501.21417236328125 0 4881.0244
501.5431213378906 0 1113.7524
501.732421875 0 1224.8887 y Water loss 5
502.2240295410156 0 4748.335 y Ammonia loss 5
502.7279968261719 0 1666.9198 z 5
502.8786315917969 0 839.87164
503.22021484375 0 771.6079
504.2109680175781 0 722.18964
505.2084045410156 0 937.7583
505.8687744140625 0 1163.1987
507.5434875488281 0 1043.3822
507.8774108886719 0 1091.3322
508.2137451171875 0 915.23486 c Water loss 8
508.7137451171875 0 1064.0476
510.21136474609375 0 5599.517
510.73773193359375 0 65529.066 y 5
511.2390441894531 0 35445.555
511.5450134277344 0 9249.13
511.7406005859375 0 9762.138
511.87738037109375 0 6623.47
512.205078125 0 1610.3464
512.2412109375 0 2190.4731
513.888916015625 0 2746.1616
514.2211303710938 0 1699.4243
514.5570068359375 0 3592.5586
517.218505859375 0 1647.4834 c 8
517.5469360351562 0 7391.883
517.8795776367188 0 6416.9688
518.212646484375 0 3268.5786
518.5469360351562 0 908.76135
519.2403564453125 0 1895.072
521.2015991210938 0 1033.5944
521.2584228515625 0 1761.8232
521.7583618164062 0 1126.7732
522.2691650390625 0 2244.804
523.2206420898438 0 34136.13
523.5530395507812 0 45127.035
523.886474609375 0 29292.9
524.2188720703125 0 14057.404
524.551513671875 0 5621.933
524.8832397460938 0 1532.5322
528.279296875 0 2485.649
529.2239990234375 0 520673.78
529.5582275390625 0 438650.22
529.8917236328125 0 272982.28
530.2251586914062 0 105858.06
530.5589599609375 0 31645.777
531.2400512695312 0 1137.1064
531.7406005859375 0 917.0034
535.2071533203125 0 1323.3259
535.25439453125 0 4447.0464
535.7537841796875 0 1787.0117
536.2118530273438 0 770.3229
536.256103515625 0 2129.4158
537.7422485351562 0 5836.359 w 4
538.1998291015625 0 1647.5947
538.24462890625 0 4780.54
538.7454833984375 0 2608.9492
539.2093505859375 0 1699.1189
540.2171630859375 0 21179.037
541.2190551757812 0 4832.8696
542.2146606445312 0 2402.6724
544.2605590820312 0 1780.4797
544.7615356445312 0 1701.8672
545.2501831054688 0 1469.596
545.750732421875 0 949.7739
546.2479858398438 0 938.7549
548.185546875 0 2971.7612
549.190185546875 0 978.5383
552.2333984375 0 1382.2583
552.7449951171875 0 1013.21606
553.2183227539062 0 4963.3896 y Ammonia loss 9
554.2239379882812 0 6519.581 z 9
555.2291259765625 0 2269.955
555.2901000976562 0 2025.9619
556.259521484375 0 2398.9797
557.2672119140625 0 2368.592
557.7696533203125 0 2033.491
558.7401123046875 0 1879.6753
559.2413940429688 0 1136.4205 z Water loss 4
561.26953125 0 1086.8347
562.2921142578125 0 984.3444
563.2459106445312 0 2499.042
563.29736328125 0 1556.6267
563.7484741210938 0 1233.9067
564.19970703125 0 745.09406
564.249267578125 0 1033.3647
564.7733154296875 0 15891.565
565.2747192382812 0 12698.531
565.7755737304688 0 6107.856
566.19580078125 0 4858.68
566.2755737304688 0 1711.2776
567.2020874023438 0 1261.748
567.2559204101562 0 3668.821 y Water loss 4
567.7465209960938 0 10735.147 y Ammonia loss 4
568.2481689453125 0 8572.577 z 4
568.748046875 0 3840.1008
569.2492065429688 0 1874.1389
569.7670288085938 0 1494.0342
570.2442626953125 0 50295.773 y 9
570.764404296875 0 799.0457
571.2470092773438 0 17579.021
572.2492065429688 0 3732.6323
575.7584838867188 0 1027.1458
576.2579956054688 0 117622.84 y 4
576.7593383789062 0 73970.63 c Water loss 9
576.8131713867188 0 1025.7208
577.25830078125 0 33284.965
577.7576904296875 0 8774.401
578.2572631835938 0 2292.857
578.7711181640625 0 27844.457
579.2713623046875 0 18682.15
579.7722778320312 0 8374.858
580.27392578125 0 3051.2676
580.77294921875 0 1370.0223
581.267578125 0 762.1174
581.7748413085938 0 1017.1544
582.2730102539062 0 2032.2924
583.2230834960938 0 9923.135 c 4
584.2494506835938 0 10011.84
585.2552490234375 0 2329.6353
585.7526245117188 0 826.9291 c 9
586.2552490234375 0 881.6033
587.2775268554688 0 5137.9487
587.7774047851562 0 17499.572
588.2789306640625 0 9824.926
588.7779541015625 0 4738.0293
589.7492065429688 0 1151.4021
590.25146484375 0 1760.0927
590.750244140625 0 1303.8248
594.787841796875 0 1464.2705
597.2421264648438 0 944.42267
602.7882690429688 0 1039.2949
603.3002319335938 0 1265.1464
606.27685546875 0 1267.3882
608.2762451171875 0 2884.354
608.779052734375 0 2998.2976 z Water loss 3
609.2799072265625 0 1483.8182 z Ammonia loss 3
610.2713623046875 0 2591.317 w 3
610.77197265625 0 1039.1493
611.2711181640625 0 1112.4702
616.223388671875 0 1499.9216
616.7877197265625 0 2439.131 y Water loss 3
617.2792358398438 0 7844.456 y Ammonia loss 3
617.78271484375 0 22700.494 z 3
618.283935546875 0 18326.613
618.7847900390625 0 8102.0386
619.2828369140625 0 1955.8092
620.2680053710938 0 7662.515
620.7695922851562 0 4459.872
621.2683715820312 0 2148.5413
622.2715454101562 0 771.80054
624.254638671875 0 37986.01 w 8
624.7908935546875 0 4973.696
625.2562255859375 0 9695.809
625.2933349609375 0 7218.892
625.7923583984375 0 105643 y 3
626.2935791015625 0 72832.5
626.79443359375 0 33635.016
627.2941284179688 0 7603.887
627.7955322265625 0 2709.6687
630.7935180664062 0 106199.61
631.2947387695312 0 76656.41
631.7957153320312 0 35406.12
632.2968139648438 0 9409.375
632.7976684570312 0 2837.6797
633.267578125 0 3812.5017
633.7703247070312 0 3193.0935 c Water loss 10
634.265869140625 0 5786.2524 c Ammonia loss 10
634.767578125 0 2820.705
635.2682495117188 0 2320.228
637.2799682617188 0 1499.5283
637.7974243164062 0 1788.9679
638.302490234375 0 4310.531
638.802978515625 0 3974.3403
639.29833984375 0 1937.2484
640.2747192382812 0 1323.1764
642.7872924804688 0 1311.0166 c 10
643.285400390625 0 1092.434
643.3517456054688 0 805.9372
643.8064575195312 0 762.9478
645.2828979492188 0 3371.009
645.784423828125 0 2016.1688
646.3038940429688 0 4607.1196
646.8056030273438 0 3289.2156
647.3052368164062 0 3020.6113
647.7898559570312 0 1232.5255
648.2879028320312 0 1314.3896
648.7882080078125 0 987.0797
651.7924194335938 0 6553.052
652.2911376953125 0 5159.359 z Water loss 2
652.7960815429688 0 11244.513 z Ammonia loss 2
653.2981567382812 0 6125.119
653.7920532226562 0 4410.0303
654.2957153320312 0 2661.1555
654.7914428710938 0 2100.1519
655.2892456054688 0 8866.685
656.2921142578125 0 2756.3752
657.2962036132812 0 909.0816
658.3004760742188 0 4836.87
659.2791137695312 0 4663.8604
659.7876586914062 0 5791.903
660.3021850585938 0 25303.219 y Water loss 2
660.7979736328125 0 52326.395 y Ammonia loss 2
661.2991333007812 0 95069.56 z 2
661.8005981445312 0 69155.24
662.301025390625 0 33806.87
662.8023681640625 0 10423.238
663.30224609375 0 1502.401
666.2982788085938 0 2343.1636 y Ammonia loss 8
667.3062133789062 0 7328.8423 z 8
668.30517578125 0 2734.4922
668.7900390625 0 1649.999
669.3087768554688 0 573849.9 y 2
669.8101806640625 0 422512.03
670.3106689453125 0 202863.27
670.8114013671875 0 68221.77
671.3118286132812 0 18366.467
674.299560546875 0 4263.598
674.8026123046875 0 4268.2275
675.2592163085938 0 2556.3967
675.3106689453125 0 3339.8623
676.2575073242188 0 1425.9182
676.3115844726562 0 6968.0396
676.77734375 0 1003.0315
677.3130493164062 0 2129.978
678.3104248046875 0 1291.1967
679.7940063476562 0 1708.8275
680.2896728515625 0 2998.7676
680.7947387695312 0 1504.7981
682.2842407226562 0 22858.49
683.3286743164062 0 18263.047 y 8
684.33203125 0 7063.8853
685.333740234375 0 1318.4899
687.8055419921875 0 1033.0967
688.3016967773438 0 1228.6604
688.7979736328125 0 18376.346
689.300048828125 0 19514.463
689.7996826171875 0 12451.89
690.3015747070312 0 5217.1265
690.803955078125 0 1756.0424
691.3510131835938 0 2552.674
691.8601684570312 0 1191.1907
692.3425903320312 0 4064.665
693.3387451171875 0 1989.445
693.7909545898438 0 1085.471
694.290283203125 0 1340.5199
694.3484497070312 0 1668.3606
694.8131713867188 0 3921.6418
695.2955322265625 0 6860.645
695.8087158203125 0 1401.3885 z Water loss 1
696.3138427734375 0 112647.23 w 1
696.8152465820312 0 90998.77
697.316162109375 0 42156.473
697.8167724609375 0 13363.72
698.316162109375 0 4175.412
699.85546875 0 894.4192
702.3366088867188 0 1156.824
702.7955932617188 0 27832.291 c Ammonia loss 11
703.296875 0 21141.818
703.7979125976562 0 11159.151
704.300048828125 0 6186.7095 y Ammonia loss 1
704.8038330078125 0 2305.5325 z 1
705.2559204101562 0 940.07855
705.4066162109375 0 1825.1875
708.8606567382812 0 1625.3597
710.8157348632812 0 1196.0084
711.3085327148438 0 261281.72 c 11
711.80908203125 0 210311.36
712.3091430664062 0 131033.91
712.8091430664062 0 42848.87 y 1
713.3084106445312 0 13428.09
713.36279296875 0 4690.4653
714.3626098632812 0 2776.7986
717.3353271484375 0 1317.4729
717.8160400390625 0 1784.9366
718.3049926757812 0 1272.8386
718.8167114257812 0 1353.9984
719.3209228515625 0 1021.51105
720.2827758789062 0 112662.086 c 5
721.2857666015625 0 42385.18
722.283203125 0 17662.762
723.2865600585938 0 2895.2139
724.3414306640625 0 4332.004
724.8417358398438 0 2965.0283
725.334228515625 0 2286.8347
725.8336181640625 0 1748.2274
726.337158203125 0 1209.5951
726.8165893554688 0 2715.0999
727.3152465820312 0 3851.3044
727.8139038085938 0 1807.998
731.315673828125 0 1185.2993
731.8148803710938 0 2171.619
732.3054809570312 0 2884.527
732.8054809570312 0 2256.2957
733.3134155273438 0 3440.2415
733.8197021484375 0 2956.2678
734.3152465820312 0 2333.35
734.81787109375 0 994.3368
735.796630859375 0 1317.1755
738.3466186523438 0 22722.832 z 7
738.8256225585938 0 995.45624
739.3489990234375 0 10059.438
739.8295288085938 0 31113.277
740.33056640625 0 34325.777
740.8273315429688 0 20924.607
741.3157958984375 0 15393.899
741.8177490234375 0 34884.41
742.3182983398438 0 25700.932
742.8201904296875 0 14009.483
743.3199462890625 0 3784.3801
743.8213500976562 0 1125.4755
745.2661743164062 0 835.75757
745.3319091796875 0 8125.961
746.2875366210938 0 1612.409
746.3473510742188 0 4654.3926
747.3382568359375 0 5690.8687
747.8373413085938 0 5485.762
748.3174438476562 0 9189.017
748.81787109375 0 6580.536
749.3151245117188 0 5889.692
749.816162109375 0 95890.89
750.3178100585938 0 73582.75
750.8180541992188 0 39835.945
751.318115234375 0 15505.555
751.8182983398438 0 4684.385
752.8087158203125 0 7395.149
753.31103515625 0 4853.8896
753.8109741210938 0 3682.7173
754.3660278320312 0 29600.37 y 7
754.8170776367188 0 25020.723
755.318115234375 0 18287.867
755.3760986328125 0 7217.774
755.8186645507812 0 17802.098
756.3186645507812 0 7546.6724
756.3736572265625 0 1576.6189
756.81689453125 0 5952.4727
757.3172607421875 0 2264.8818
757.8119506835938 0 1264.4531
761.8265991210938 0 2456.1077
762.3272705078125 0 3774.429
762.8298950195312 0 1122.9696
763.3385620117188 0 8938.132
763.825439453125 0 14840.279
764.3292846679688 0 204649.06
764.830322265625 0 166002.12
765.3307495117188 0 97252.4
765.8311767578125 0 37149.68
766.33154296875 0 12015.429
768.3734741210938 0 7766.5547
769.3789672851562 0 3109.762
770.3814086914062 0 8896.879
770.8319091796875 0 4674.068
771.3273315429688 0 43222.812
771.82763671875 0 36884.383
772.3276977539062 0 21837.99
772.8278198242188 0 8425.198
773.32861328125 0 1295.1063
777.816162109375 0 2167.4097
778.3223266601562 0 1401.9196
783.3983154296875 0 1263.1854
783.9013061523438 0 1504.0966
784.8235473632812 0 2693.3904
785.3258666992188 0 15237.761
785.8274536132812 0 13247.89
786.3251342773438 0 10393.969
786.824951171875 0 7800.88
787.3255004882812 0 2460.39
791.9049682617188 0 1222.1743
793.3331298828125 0 321956.66
793.8364868164062 0 1655478.8
794.3374633789062 0 1327937.1
794.8380737304688 0 776493.5
795.3381958007812 0 290437.03
795.8388061523438 0 95311.69
796.338623046875 0 1677.9832
802.353759765625 0 9358.3
803.3564453125 0 4583.816
805.3107299804688 0 2563.168
808.3760375976562 0 32776.58 w 6
809.3792114257812 0 13799.364
810.3800048828125 0 4009.486
811.4010009765625 0 1231.0792
816.2998657226562 0 1311.1141
820.3655395507812 0 6940.267
821.36767578125 0 3244.9146
822.3665161132812 0 1472.4757
823.3880004882812 0 1163.1024
824.3827514648438 0 1340.6194
832.2987670898438 0 5330.44
833.2979736328125 0 1917.1719
833.3833618164062 0 7251.5654
834.385009765625 0 2847.309
835.3787231445312 0 1228.5873
839.3924560546875 0 1261.6971
849.3255615234375 0 55232.996 c 6
850.3280029296875 0 23120.596
850.417236328125 0 1182.3801
851.3260498046875 0 10402.795
852.3265380859375 0 2814.1562
865.3955688476562 0 2740.3545 y Water loss 6
866.390380859375 0 2639.3599 y Ammonia loss 6
867.3896484375 0 71402.89 z 6
868.3927612304688 0 38802.457
869.394775390625 0 10933.942
870.3968505859375 0 1916.4175
876.3495483398438 0 2742.5356
877.4422607421875 0 1360.8977
883.408203125 0 38995.445 y 6
884.41162109375 0 17418.857
885.4135131835938 0 5552.988
890.4069213867188 0 4380.3823
891.4072265625 0 2749.0063
892.4153442382812 0 1191.5757
903.3360595703125 0 11166.682
904.3390502929688 0 4984.8726
904.4412841796875 0 1105.5233
905.3361206054688 0 2330.8008
905.4330444335938 0 5950.198
906.4373779296875 0 2970.147
907.43408203125 0 5381.48
908.4341430664062 0 2808.164
920.36279296875 0 51536.734 c 7
921.3652954101562 0 20530.727
921.4564208984375 0 861.0006
922.3646240234375 0 10988.434
922.4623413085938 0 1034.3488
923.3673706054688 0 2853.9707
929.3811645507812 0 1032.3843
930.40966796875 0 1619.7628
932.4370727539062 0 1187.4637
935.4611206054688 0 7483.415
936.4645385742188 0 5176.3096
937.4652099609375 0 2068.0542
948.3848876953125 0 1123.3615
971.4246826171875 0 2077.8735
972.427490234375 0 1062.497
975.4542846679688 0 1003.9562
976.4747314453125 0 1335.4142
985.44482421875 0 1308.5692
989.4337158203125 0 14419.279
990.4374389648438 0 7723.324
991.4376831054688 0 4154.6484
992.4415283203125 0 1687.6974
1003.44580078125 0 1675.4669 y Ammonia loss 5
1004.4491577148438 0 14453.512 z 5
1005.4525146484375 0 11373.768
1006.4534912109375 0 5653.6626
1007.4646606445312 0 1460.2349
1016.4214477539062 0 6348.511
1017.4244995117188 0 3938.8735
1018.4212036132812 0 2164.5676
1019.4638671875 0 24086.787
1020.4672241210938 0 48255.46 y 5
1021.4708862304688 0 24392.11
1022.4718627929688 0 5883.3486
1023.4795532226562 0 951.03314
1033.4471435546875 0 20963.572 c 8
1034.44970703125 0 10452.739
1035.4512939453125 0 5907.023
1036.4534912109375 0 1786.7614
1039.4678955078125 0 8506.814
1040.4747314453125 0 5977.3906
1041.46826171875 0 1612.2123
1055.4881591796875 0 2907.2996
1056.4920654296875 0 5749.239
1057.494873046875 0 3307.2188
1058.4964599609375 0 1468.0175
1074.480224609375 0 2074.111 w 4
1075.4808349609375 0 1650.5164
1125.485595703125 0 1507.9594
1126.49169921875 0 3090.87
1127.4954833984375 0 1888.4155
1128.49951171875 0 909.68646
1134.4835205078125 0 929.41614 y Ammonia loss 4
1135.48828125 0 3507.8462 z 4
1136.4942626953125 0 4591.1743
1137.494384765625 0 1720.2156
1151.50830078125 0 6057.437 y 4
1152.513916015625 0 3801.6829 c Water loss 9
1153.505126953125 0 2423.474
1154.49267578125 0 1451.3356
1157.5421142578125 0 987.8431
1158.556396484375 0 1290.7278
1170.506103515625 0 61701.56 c 9
1171.509033203125 0 39106.863
1172.5086669921875 0 18991.793
1173.5113525390625 0 6346.186
1174.5399169921875 0 1587.3013
1180.4918212890625 0 1565.3131
1181.4990234375 0 1233.8533
1234.556396484375 0 3598.8247 z 3
1235.5621337890625 0 5215.4854
1236.565185546875 0 3321.3596
1237.5635986328125 0 1145.2456
1239.5413818359375 0 915.2109
1240.53369140625 0 5457.411
1241.5389404296875 0 4306.305
1242.54345703125 0 2160.7512
1243.5352783203125 0 957.13477
1250.5792236328125 0 2681.8496 y 3
1251.57080078125 0 1235.7592
1252.5791015625 0 928.58246
1267.520751953125 0 2383.776 c Ammonia loss 10
1268.53271484375 0 1372.2795
1269.5206298828125 0 1414.2745
1284.5478515625 0 45939.9 c 10
1285.55078125 0 32443.96
1286.5506591796875 0 16379.827
1287.5516357421875 0 5896.2007
1288.5513916015625 0 1254.2449
1304.5911865234375 0 1752.0485 w 2
1305.5902099609375 0 1020.2592
1321.5889892578125 0 3614.639 z 2
1322.595947265625 0 22952.072
1323.5980224609375 0 14555.718
1324.601318359375 0 8373.992
1325.6041259765625 0 2423.0017
1337.612060546875 0 1112.9502 y 2
1338.616943359375 0 971.1452
1363.5989990234375 0 3512.6853
1364.6055908203125 0 2860.1265
1365.6107177734375 0 1344.2264
1377.5968017578125 0 1599.081
1378.598388671875 0 2622.1953
1379.5982666015625 0 1687.5326
1391.6207275390625 0 3066.2312 w 1
1392.6251220703125 0 9922.169
1393.6280517578125 0 7114.1123
1394.6298828125 0 3254.1016
1421.6070556640625 0 2664.1238 c 11
1422.61279296875 0 10444.591
1423.614990234375 0 7245.517
1424.616943359375 0 4534.165 y 1
1425.61279296875 0 997.2548
1479.6502685546875 0 990.88727
1511.627197265625 0 1496.6024
1512.62841796875 0 1810.5775
1527.649169921875 0 11853.785
1528.653076171875 0 21286.07
1529.6549072265625 0 16853.99
1530.65625 0 8091.77
1531.6519775390625 0 3495.2422
1541.660400390625 0 1005.00885
1542.6513671875 0 4774.4443
1543.6527099609375 0 3364.3608
1544.6549072265625 0 2399.231
1559.6763916015625 0 3430.6333
1560.6767578125 0 1717.4493
1561.668212890625 0 1037.6595
1569.6422119140625 0 2334.5422
1570.6446533203125 0 9200.779
1571.645751953125 0 7905.0913
1572.6451416015625 0 4648.7104
1573.6490478515625 0 1771.9409
1574.648193359375 0 835.01337
1585.649169921875 0 2986.3801
1586.659423828125 0 19859.514
1587.668212890625 0 77974.75
1588.67041015625 0 59690.97
1589.6719970703125 0 34495.676
1590.6732177734375 0 11346.826
1591.6734619140625 0 3193.342

Spectrum Details

|  |  |
| --- | --- |
| Matched peaks? Matched peaks The total absolute number of peaks matched. Additionally in brackets the total fraction of peaks matched and the total number of peaks is shown. | 103 (11.94% of 863) |
| FDR? FDR The false discovery rate estimated for this peptide. It is calculated by matching all theoretical fragments with a non-integer shift with the raw peaks for this spectrum. This is done with 40 different shifts. The resulting percentage is the average number of annotated peaks over the number of annotated peaks with the correct spectrum. | 0.39% |
| Satellite FDR? Satellite FDR See the FDR for details on its calculation. This satellite ion specific FDR only contains the satellite ions (d/w) for I/L/J positions. | 4.76% |
| PSM Score? PSM Score The PSM Score as given by Hecklib to this annotated spectrum. It is shown with three significant figures. | 571 |

#### Spectrum 2851? Spectrum 2851 The raw spectrum of this peptide as annotated by Hecklib. The fragments are coloured according to ion type (see legend). Any peaks with a star '\*' as text can be hovered over to see the full details, first the ion type second the mass shift type. By hovering over the amino acids in the peptide or ions in the legend the corresponding peaks are highlighted. By toggling the 'Unassigned' label you can turn the background (unassigned) peaks on or off in the plot. By updating the slider in the Ion legend you can update the spectrum to only show the top X% of the peaks with labels. The top X% means any peak that is within X% of the highest intensity. By dragging in the spectrum you can zoom in to a specific part of the spectrum and use 'Zoom Out' to get back to the original zoom level. The annotation of the spectrum is based on the given sequence in the peptides file and is done with different software so inconsistencies are likely. The peaks are annotated based on the given sequence, with 20 ppm tolerance.

Copy Data

##### Spectrum 2851 (TSV)

###### Preview

```
Loading example...
```

*Click on the button to copy the data to your clipboard.*

Mz MinMz MaxIntensity Max

WidthHeightPeptide font sizePeptide stroke widthSpectrum font sizeSpectrum stroke widthCompact peptide

Ion legend

wxyz

abcd

OtherUnassignedIon^Charge^_Position_Show for top:%

CSSVMHEALHNHY

09.66e+51.93e+62.90e+63.86e+6

Zoom Out

y^+1^_1_y^+1^_2_w^+1^_3_y^+2^_6_z^+1^_3_y^+1^_3_y^+3^_11_z^+3^_11_y^+3^_11_c^+1^_4_c^+3^_12_y^+2^_8_y^+1^_4_y^+2^_9_c^+2^_10_c^+1^_5_w^+1^_5_z^+2^_10_y^+2^_10_c^+2^_11_c^+2^_11_z^+2^_11_z^+1^_5_y^+2^_11_y^+2^_11_z^+2^_11_y^+2^_11_y^+1^_5_z^+2^_12_c^+2^_12_y^+2^_12_c^+2^_12_y^+2^_12_z^+1^_6_c^+1^_6_z^+1^_6_y^+1^_6_w^+1^_7_c^+1^_7_z^+1^_7_y^+1^_7_c^+1^_8_z^+1^_8_y^+1^_8_c^+1^_9_z^+1^_9_c^+1^_10_z^+1^_10_y^+1^_10_c^+1^_11_w^+1^_11_z^+1^_11_y^+1^_11_z^+1^_12_y^+1^_12_c^+1^_12_y^+1^_12_

0666133219982664

Fragment Matches Table

Show background peaks

| Position | Ion type | Intensity | mz Theoretical | mz Error (Th) | mz Error (ppm) | Charge | Series Number |
| --- | --- | --- | --- | --- | --- | --- | --- |
| - | - | 5.884E+04 | 123.1 | - | - | 0 | - |
| - | - | 8506 | 134 | - | - | 0 | - |
| - | - | 2.496E+04 | 136.1 | - | - | 0 | - |
| - | - | 2.376E+04 | 140.1 | - | - | 0 | - |
| - | - | 8758 | 164 | - | - | 0 | - |
| - | - | 1.326E+04 | 173.4 | - | - | 0 | - |
| 13 | y | 2.752E+04 | 182.1 | 0.0002981 | 1.637 | +1 | 1 |
| - | - | 3.814E+04 | 192.1 | - | - | 0 | - |
| - | - | 3.309E+04 | 209.1 | - | - | 0 | - |
| - | - | 2.221E+05 | 221.1 | - | - | 0 | - |
| - | - | 1.198E+04 | 222.1 | - | - | 0 | - |
| - | - | 8634 | 232.7 | - | - | 0 | - |
| - | - | 1.367E+04 | 237.1 | - | - | 0 | - |
| - | - | 9096 | 237.4 | - | - | 0 | - |
| - | - | 1.664E+04 | 239.1 | - | - | 0 | - |
| - | - | 1.82E+04 | 244.1 | - | - | 0 | - |
| - | - | 9.614E+04 | 249.1 | - | - | 0 | - |
| - | - | 1.19E+04 | 250.1 | - | - | 0 | - |
| - | - | 8501 | 254.1 | - | - | 0 | - |
| - | - | 7961 | 258 | - | - | 0 | - |
| - | - | 2.134E+04 | 258.1 | - | - | 0 | - |
| - | - | 7597 | 268.1 | - | - | 0 | - |
| - | - | 7891 | 311.1 | - | - | 0 | - |
| 12 | y | 2.545E+04 | 319.1 | 0.0009824 | 3.078 | +1 | 2 |
| - | - | 1.263E+04 | 336.1 | - | - | 0 | - |
| - | - | 1.483E+04 | 372.2 | - | - | 0 | - |
| 11 | w | 6.869E+04 | 373.2 | 0.0007134 | 1.912 | +1 | 3 |
| 8 | y | 1.079E+04 | 377.7 | 0.002113 | 5.594 | +2 | 6 |
| - | - | 9213 | 393.2 | - | - | 0 | - |
| - | - | 2.285E+04 | 407.2 | - | - | 0 | - |
| - | - | 8362 | 408.5 | - | - | 0 | - |
| 11 | z | 2.579E+04 | 417.2 | 5.224E-05 | 0.1252 | +1 | 3 |
| - | - | 1.33E+04 | 417.3 | - | - | 0 | - |
| - | - | 1.07E+04 | 418.2 | - | - | 0 | - |
| - | - | 2.702E+04 | 428.2 | - | - | 0 | - |
| 11 | y | 2.367E+04 | 433.2 | 0.001063 | 2.454 | +1 | 3 |
| - | - | 2.325E+04 | 435.2 | - | - | 0 | - |
| 3 | y | 2.39E+04 | 445.9 | 0.00236 | 5.292 | +3 | 11 |
| 3 | z | 1.162E+04 | 446.5 | 0.001438 | 3.22 | +3 | 11 |
| 3 | y | 2.3E+04 | 451.9 | 0.0006082 | 1.346 | +3 | 11 |
| 4 | c | 1.331E+04 | 452.2 | 0.007838 | 17.33 | +1 | 4 |
| - | - | 2.094E+04 | 452.2 | - | - | 0 | - |
| 12 | c | 1.35E+04 | 474.2 | 0.0007632 | 1.609 | +3 | 12 |
| - | - | 1.453E+04 | 476.6 | - | - | 0 | - |
| - | - | 2.369E+04 | 497.9 | - | - | 0 | - |
| - | - | 2.455E+04 | 498.2 | - | - | 0 | - |
| - | - | 1.27E+04 | 498.6 | - | - | 0 | - |
| - | - | 2.539E+04 | 505.5 | - | - | 0 | - |
| - | - | 2.53E+04 | 505.9 | - | - | 0 | - |
| - | - | 2.594E+04 | 510.2 | - | - | 0 | - |
| 6 | y | 1.779E+04 | 510.7 | 0.0003443 | 0.6741 | +2 | 8 |
| - | - | 3.935E+04 | 513.2 | - | - | 0 | - |
| - | - | 2.534E+04 | 513.6 | - | - | 0 | - |
| - | - | 1.254E+04 | 513.9 | - | - | 0 | - |
| - | - | 9148 | 514.2 | - | - | 0 | - |
| - | - | 1.726E+04 | 522.3 | - | - | 0 | - |
| - | - | 2.44E+04 | 528.5 | - | - | 0 | - |
| - | - | 3.14E+04 | 528.9 | - | - | 0 | - |
| - | - | 1.029E+04 | 532.7 | - | - | 0 | - |
| - | - | 3.941E+05 | 534.6 | - | - | 0 | - |
| - | - | 2.868E+05 | 534.9 | - | - | 0 | - |
| - | - | 2.169E+05 | 535.2 | - | - | 0 | - |
| - | - | 8.951E+04 | 535.6 | - | - | 0 | - |
| - | - | 2.441E+04 | 535.9 | - | - | 0 | - |
| - | - | 1.204E+04 | 544.2 | - | - | 0 | - |
| - | - | 2.504E+04 | 545.3 | - | - | 0 | - |
| - | - | 1.134E+04 | 545.8 | - | - | 0 | - |
| - | - | 2.205E+04 | 555.2 | - | - | 0 | - |
| - | - | 1.095E+04 | 556.2 | - | - | 0 | - |
| 10 | y | 9.348E+04 | 570.2 | 0.0005878 | 1.031 | +1 | 4 |
| - | - | 2.325E+04 | 571.2 | - | - | 0 | - |
| - | - | 1.199E+04 | 572.8 | - | - | 0 | - |
| - | - | 1.149E+04 | 583.3 | - | - | 0 | - |
| 5 | y | 4.1E+04 | 584.3 | 0.002663 | 4.558 | +2 | 9 |
| 10 | c | 3.773E+04 | 584.7 | 0.008174 | 13.98 | +2 | 10 |
| - | - | 1.869E+04 | 586.8 | - | - | 0 | - |
| - | - | 1.565E+04 | 595.8 | - | - | 0 | - |
| 5 | c | 7.104E+04 | 599.2 | 0.0002061 | 0.3439 | +1 | 5 |
| - | - | 1.705E+04 | 600.2 | - | - | 0 | - |
| 9 | w | 1.281E+05 | 624.3 | 0.0005934 | 0.9507 | +1 | 5 |
| - | - | 1.13E+04 | 624.3 | - | - | 0 | - |
| - | - | 5.766E+04 | 625.3 | - | - | 0 | - |
| 4 | z | 4.105E+04 | 625.8 | 0.002354 | 3.762 | +2 | 10 |
| - | - | 3.337E+04 | 626.3 | - | - | 0 | - |
| - | - | 1.06E+04 | 626.8 | - | - | 0 | - |
| - | - | 1.796E+04 | 628.3 | - | - | 0 | - |
| - | - | 1.997E+04 | 628.8 | - | - | 0 | - |
| - | - | 2.767E+04 | 633.3 | - | - | 0 | - |
| 4 | y | 7.826E+04 | 633.8 | 0.001232 | 1.944 | +2 | 10 |
| - | - | 6.871E+04 | 634.3 | - | - | 0 | - |
| - | - | 1.566E+04 | 634.8 | - | - | 0 | - |
| - | - | 1.241E+04 | 637.3 | - | - | 0 | - |
| - | - | 1.163E+04 | 637.8 | - | - | 0 | - |
| - | - | 2.464E+04 | 638.3 | - | - | 0 | - |
| - | - | 1.376E+04 | 638.8 | - | - | 0 | - |
| - | - | 1.829E+04 | 639.8 | - | - | 0 | - |
| - | - | 2.082E+04 | 640.3 | - | - | 0 | - |
| - | - | 3.616E+04 | 641.3 | - | - | 0 | - |
| 11 | c | 1.65E+04 | 641.8 | 0.003707 | 5.776 | +2 | 11 |
| 11 | c | 1.38E+04 | 642.3 | 0.005811 | 9.048 | +2 | 11 |
| - | - | 4.62E+04 | 645.3 | - | - | 0 | - |
| - | - | 5.286E+04 | 645.8 | - | - | 0 | - |
| 3 | z | 5.378E+04 | 660.8 | 0.01292 | 19.55 | +2 | 11 |
| - | - | 4.391E+04 | 661.3 | - | - | 0 | - |
| - | - | 1.481E+04 | 661.8 | - | - | 0 | - |
| 9 | z | 2.44E+04 | 667.3 | 0.002126 | 3.187 | +1 | 5 |
| 3 | y | 2.813E+04 | 668.3 | 0.0003266 | 0.4887 | +2 | 11 |
| 3 | y | 5.693E+04 | 668.8 | 0.005328 | 7.967 | +2 | 11 |
| 3 | z | 1.782E+05 | 669.3 | 0.001904 | 2.845 | +2 | 11 |
| - | - | 1.359E+05 | 669.8 | - | - | 0 | - |
| - | - | 6.149E+04 | 670.3 | - | - | 0 | - |
| - | - | 1.777E+04 | 670.8 | - | - | 0 | - |
| - | - | 2.583E+04 | 674.3 | - | - | 0 | - |
| 3 | y | 3.244E+05 | 677.3 | 0.00188 | 2.776 | +2 | 11 |
| - | - | 2.712E+05 | 677.8 | - | - | 0 | - |
| - | - | 1.185E+05 | 678.3 | - | - | 0 | - |
| - | - | 3.252E+04 | 678.8 | - | - | 0 | - |
| - | - | 8.665E+04 | 682.3 | - | - | 0 | - |
| 9 | y | 2.32E+04 | 683.3 | 0.003788 | 5.544 | +1 | 5 |
| - | - | 1.233E+04 | 684.3 | - | - | 0 | - |
| - | - | 1.715E+04 | 687.8 | - | - | 0 | - |
| - | - | 1.278E+04 | 689.3 | - | - | 0 | - |
| - | - | 2.625E+04 | 695.3 | - | - | 0 | - |
| - | - | 1.06E+04 | 696.3 | - | - | 0 | - |
| - | - | 1.007E+04 | 696.8 | - | - | 0 | - |
| - | - | 2.393E+04 | 697.3 | - | - | 0 | - |
| - | - | 1.318E+04 | 697.8 | - | - | 0 | - |
| 2 | z | 1.537E+05 | 704.3 | 0.01393 | 19.79 | +2 | 12 |
| - | - | 1.391E+05 | 704.8 | - | - | 0 | - |
| - | - | 4.56E+04 | 705.3 | - | - | 0 | - |
| - | - | 1.321E+04 | 705.8 | - | - | 0 | - |
| 12 | c | 3.083E+04 | 710.8 | 0.00138 | 1.941 | +2 | 12 |
| - | - | 2.032E+04 | 711.3 | - | - | 0 | - |
| - | - | 1.689E+04 | 711.8 | - | - | 0 | - |
| 2 | y | 1.375E+04 | 712.3 | 0.004949 | 6.948 | +2 | 12 |
| - | - | 3.027E+04 | 718.9 | - | - | 0 | - |
| 12 | c | 6.937E+05 | 719.3 | 0.0004814 | 0.6693 | +2 | 12 |
| - | - | 4.979E+05 | 719.8 | - | - | 0 | - |
| - | - | 2.855E+05 | 720.3 | - | - | 0 | - |
| 2 | y | 1.072E+05 | 720.8 | 0.01249 | 17.32 | +2 | 12 |
| 8 | z | 2.342E+04 | 721.3 | 0.01112 | 15.42 | +1 | 6 |
| - | - | 4.209E+04 | 724.3 | - | - | 0 | - |
| - | - | 4.387E+04 | 724.8 | - | - | 0 | - |
| - | - | 1.429E+04 | 725.3 | - | - | 0 | - |
| - | - | 1.137E+04 | 726.3 | - | - | 0 | - |
| - | - | 1.338E+04 | 730.3 | - | - | 0 | - |
| 6 | c | 6.877E+05 | 736.3 | 9.696E-05 | 0.1317 | +1 | 6 |
| - | - | 2.611E+05 | 737.3 | - | - | 0 | - |
| - | - | 1.192E+05 | 738.3 | - | - | 0 | - |
| 8 | z | 6.828E+04 | 738.3 | 0.0001363 | 0.1846 | +1 | 6 |
| - | - | 1.583E+04 | 739.3 | - | - | 0 | - |
| - | - | 4.724E+04 | 739.3 | - | - | 0 | - |
| - | - | 1.695E+04 | 739.8 | - | - | 0 | - |
| - | - | 2.711E+04 | 740.3 | - | - | 0 | - |
| - | - | 1.473E+04 | 740.8 | - | - | 0 | - |
| - | - | 2.524E+04 | 747.3 | - | - | 0 | - |
| - | - | 5.375E+04 | 747.8 | - | - | 0 | - |
| - | - | 3.504E+04 | 748.3 | - | - | 0 | - |
| - | - | 2.959E+04 | 748.8 | - | - | 0 | - |
| - | - | 1.253E+04 | 749.3 | - | - | 0 | - |
| - | - | 4.793E+04 | 749.8 | - | - | 0 | - |
| - | - | 4.658E+04 | 750.3 | - | - | 0 | - |
| - | - | 2.347E+04 | 750.8 | - | - | 0 | - |
| 8 | y | 2.909E+04 | 754.4 | 6.056E-05 | 0.08028 | +1 | 6 |
| - | - | 1.19E+04 | 755.4 | - | - | 0 | - |
| - | - | 1.423E+04 | 755.8 | - | - | 0 | - |
| - | - | 3.637E+04 | 757.8 | - | - | 0 | - |
| - | - | 3.074E+04 | 758.3 | - | - | 0 | - |
| - | - | 2.496E+04 | 758.8 | - | - | 0 | - |
| - | - | 1.836E+04 | 760.8 | - | - | 0 | - |
| - | - | 2.012E+04 | 761.3 | - | - | 0 | - |
| - | - | 1.295E+04 | 762.3 | - | - | 0 | - |
| - | - | 3.352E+04 | 762.8 | - | - | 0 | - |
| - | - | 3.323E+04 | 763.3 | - | - | 0 | - |
| - | - | 4.456E+04 | 763.8 | - | - | 0 | - |
| - | - | 3.469E+04 | 764.3 | - | - | 0 | - |
| - | - | 1.867E+04 | 764.8 | - | - | 0 | - |
| - | - | 1.464E+04 | 770.3 | - | - | 0 | - |
| - | - | 1.966E+04 | 770.8 | - | - | 0 | - |
| - | - | 1.564E+04 | 771.3 | - | - | 0 | - |
| - | - | 4.25E+04 | 771.8 | - | - | 0 | - |
| - | - | 7.122E+05 | 772.3 | - | - | 0 | - |
| - | - | 5.337E+05 | 772.8 | - | - | 0 | - |
| - | - | 2.825E+05 | 773.3 | - | - | 0 | - |
| - | - | 1.227E+05 | 773.8 | - | - | 0 | - |
| - | - | 3.919E+04 | 774.3 | - | - | 0 | - |
| - | - | 1.118E+04 | 778.8 | - | - | 0 | - |
| - | - | 1.215E+04 | 778.9 | - | - | 0 | - |
| - | - | 5.986E+04 | 779.3 | - | - | 0 | - |
| - | - | 8.374E+04 | 779.8 | - | - | 0 | - |
| - | - | 4.776E+04 | 780.3 | - | - | 0 | - |
| - | - | 1.874E+04 | 785.3 | - | - | 0 | - |
| - | - | 1.328E+04 | 786.3 | - | - | 0 | - |
| - | - | 3.844E+04 | 792.8 | - | - | 0 | - |
| - | - | 2.397E+05 | 793.3 | - | - | 0 | - |
| - | - | 2.412E+05 | 793.8 | - | - | 0 | - |
| - | - | 1.468E+05 | 794.3 | - | - | 0 | - |
| - | - | 6.007E+04 | 794.8 | - | - | 0 | - |
| - | - | 1.854E+04 | 795.3 | - | - | 0 | - |
| - | - | 2.915E+04 | 800.8 | - | - | 0 | - |
| - | - | 7.15E+05 | 801.3 | - | - | 0 | - |
| - | - | 3.826E+06 | 801.8 | - | - | 0 | - |
| - | - | 3.168E+06 | 802.3 | - | - | 0 | - |
| - | - | 1.839E+06 | 802.8 | - | - | 0 | - |
| - | - | 6.306E+05 | 803.3 | - | - | 0 | - |
| - | - | 2.261E+05 | 803.8 | - | - | 0 | - |
| 7 | w | 2.298E+05 | 808.4 | 0.0003113 | 0.385 | +1 | 7 |
| - | - | 9.06E+04 | 809.4 | - | - | 0 | - |
| - | - | 2.701E+04 | 810.4 | - | - | 0 | - |
| 7 | c | 2.252E+05 | 865.3 | 0.0002096 | 0.2422 | +1 | 7 |
| - | - | 9.605E+04 | 866.3 | - | - | 0 | - |
| 7 | z | 4.009E+05 | 867.4 | 0.000432 | 0.498 | +1 | 7 |
| - | - | 2.09E+05 | 868.4 | - | - | 0 | - |
| - | - | 6.551E+04 | 869.4 | - | - | 0 | - |
| 7 | y | 8.383E+04 | 883.4 | 0.0002352 | 0.2662 | +1 | 7 |
| - | - | 3.658E+04 | 884.4 | - | - | 0 | - |
| 8 | c | 1.981E+05 | 936.4 | 0.0007633 | 0.8152 | +1 | 8 |
| - | - | 9.383E+04 | 937.4 | - | - | 0 | - |
| - | - | 3.385E+04 | 938.4 | - | - | 0 | - |
| - | - | 1.196E+04 | 967 | - | - | 0 | - |
| 6 | z | 1.074E+05 | 1004 | 0.001849 | 1.841 | +1 | 8 |
| - | - | 1.16E+05 | 1005 | - | - | 0 | - |
| - | - | 4.066E+04 | 1006 | - | - | 0 | - |
| - | - | 1.422E+04 | 1007 | - | - | 0 | - |
| 6 | y | 1.303E+05 | 1020 | 0.000126 | 0.1235 | +1 | 8 |
| - | - | 8.247E+04 | 1021 | - | - | 0 | - |
| - | - | 3.028E+04 | 1022 | - | - | 0 | - |
| 9 | c | 7.647E+04 | 1049 | 0.0007819 | 0.7451 | +1 | 9 |
| - | - | 4.096E+04 | 1050 | - | - | 0 | - |
| - | - | 2.345E+04 | 1051 | - | - | 0 | - |
| - | - | 1.28E+04 | 1089 | - | - | 0 | - |
| - | - | 1.122E+04 | 1130 | - | - | 0 | - |
| 5 | z | 1.576E+04 | 1151 | 0.01375 | 11.94 | +1 | 9 |
| - | - | 2.835E+04 | 1152 | - | - | 0 | - |
| - | - | 2.318E+04 | 1153 | - | - | 0 | - |
| - | - | 4.246E+04 | 1174 | - | - | 0 | - |
| - | - | 2.056E+04 | 1175 | - | - | 0 | - |
| 10 | c | 2.428E+05 | 1186 | 0.0006117 | 0.5156 | +1 | 10 |
| - | - | 1.712E+04 | 1187 | - | - | 0 | - |
| - | - | 1.383E+05 | 1187 | - | - | 0 | - |
| - | - | 1.591E+04 | 1188 | - | - | 0 | - |
| - | - | 7.034E+04 | 1188 | - | - | 0 | - |
| - | - | 2.566E+04 | 1190 | - | - | 0 | - |
| - | - | 1.569E+04 | 1244 | - | - | 0 | - |
| 4 | z | 2.332E+04 | 1251 | 0.008811 | 7.046 | +1 | 10 |
| - | - | 3.353E+04 | 1252 | - | - | 0 | - |
| - | - | 2.598E+04 | 1253 | - | - | 0 | - |
| - | - | 1.875E+04 | 1254 | - | - | 0 | - |
| - | - | 1.382E+04 | 1257 | - | - | 0 | - |
| - | - | 1.164E+04 | 1258 | - | - | 0 | - |
| 4 | y | 1.47E+04 | 1267 | 0.003637 | 2.871 | +1 | 10 |
| 11 | c | 2.192E+05 | 1301 | 0.0002841 | 0.2184 | +1 | 11 |
| - | - | 1.334E+05 | 1302 | - | - | 0 | - |
| - | - | 9.02E+04 | 1303 | - | - | 0 | - |
| - | - | 2.603E+04 | 1304 | - | - | 0 | - |
| 3 | w | 1.483E+04 | 1321 | 0.005401 | 4.09 | +1 | 11 |
| - | - | 3.064E+04 | 1322 | - | - | 0 | - |
| 3 | z | 8.887E+04 | 1338 | 0.002905 | 2.172 | +1 | 11 |
| - | - | 3.332E+05 | 1339 | - | - | 0 | - |
| - | - | 2.011E+04 | 1339 | - | - | 0 | - |
| - | - | 2.713E+05 | 1340 | - | - | 0 | - |
| - | - | 1.085E+05 | 1341 | - | - | 0 | - |
| - | - | 3.662E+04 | 1342 | - | - | 0 | - |
| - | - | 1.224E+04 | 1343 | - | - | 0 | - |
| 3 | y | 2.449E+04 | 1354 | 0.005055 | 3.735 | +1 | 11 |
| - | - | 4.365E+04 | 1355 | - | - | 0 | - |
| - | - | 1.93E+04 | 1356 | - | - | 0 | - |
| - | - | 4.505E+04 | 1380 | - | - | 0 | - |
| - | - | 3.525E+04 | 1381 | - | - | 0 | - |
| - | - | 1.824E+04 | 1382 | - | - | 0 | - |
| - | - | 1.876E+04 | 1394 | - | - | 0 | - |
| - | - | 4.051E+04 | 1395 | - | - | 0 | - |
| - | - | 1.705E+04 | 1396 | - | - | 0 | - |
| - | - | 1.912E+04 | 1397 | - | - | 0 | - |
| 2 | z | 5.862E+04 | 1408 | 0.02794 | 19.85 | +1 | 12 |
| - | - | 9.382E+04 | 1409 | - | - | 0 | - |
| - | - | 7.726E+04 | 1410 | - | - | 0 | - |
| - | - | 3.835E+04 | 1411 | - | - | 0 | - |
| - | - | 3.013E+04 | 1422 | - | - | 0 | - |
| 2 | y | 1.137E+04 | 1423 | 0.02154 | 15.14 | +1 | 12 |
| 12 | c | 5.751E+04 | 1438 | 0.002353 | 1.637 | +1 | 12 |
| - | - | 1.671E+05 | 1439 | - | - | 0 | - |
| - | - | 1.311E+05 | 1440 | - | - | 0 | - |
| 2 | y | 8.515E+04 | 1441 | 0.01953 | 13.55 | +1 | 12 |
| - | - | 2.855E+04 | 1442 | - | - | 0 | - |
| - | - | 2.846E+04 | 1449 | - | - | 0 | - |
| - | - | 2.793E+04 | 1450 | - | - | 0 | - |
| - | - | 1.683E+04 | 1500 | - | - | 0 | - |
| - | - | 1.421E+04 | 1501 | - | - | 0 | - |
| - | - | 1.287E+04 | 1502 | - | - | 0 | - |
| - | - | 1.513E+04 | 1522 | - | - | 0 | - |
| - | - | 1.123E+04 | 1523 | - | - | 0 | - |
| - | - | 3.273E+04 | 1527 | - | - | 0 | - |
| - | - | 4.295E+04 | 1528 | - | - | 0 | - |
| - | - | 4.89E+04 | 1529 | - | - | 0 | - |
| - | - | 2.041E+04 | 1530 | - | - | 0 | - |
| - | - | 3.849E+04 | 1542 | - | - | 0 | - |
| - | - | 3.784E+04 | 1543 | - | - | 0 | - |
| - | - | 2.207E+05 | 1544 | - | - | 0 | - |
| - | - | 4.744E+05 | 1545 | - | - | 0 | - |
| - | - | 3.733E+05 | 1546 | - | - | 0 | - |
| - | - | 2.119E+05 | 1547 | - | - | 0 | - |
| - | - | 7.129E+04 | 1548 | - | - | 0 | - |
| - | - | 1.76E+04 | 1549 | - | - | 0 | - |
| - | - | 1.611E+04 | 1558 | - | - | 0 | - |
| - | - | 7.937E+04 | 1559 | - | - | 0 | - |
| - | - | 7.011E+04 | 1560 | - | - | 0 | - |
| - | - | 3.528E+04 | 1561 | - | - | 0 | - |
| - | - | 2.679E+04 | 1569 | - | - | 0 | - |
| - | - | 1.983E+04 | 1570 | - | - | 0 | - |
| - | - | 3.574E+04 | 1576 | - | - | 0 | - |
| - | - | 4.421E+04 | 1577 | - | - | 0 | - |
| - | - | 1.636E+04 | 1578 | - | - | 0 | - |
| - | - | 1.171E+04 | 1579 | - | - | 0 | - |
| - | - | 9.827E+04 | 1586 | - | - | 0 | - |
| - | - | 2.454E+05 | 1587 | - | - | 0 | - |
| - | - | 1.878E+05 | 1588 | - | - | 0 | - |
| - | - | 1.335E+05 | 1589 | - | - | 0 | - |
| - | - | 4.94E+04 | 1590 | - | - | 0 | - |
| - | - | 5.81E+04 | 1602 | - | - | 0 | - |
| - | - | 3.896E+05 | 1603 | - | - | 0 | - |
| - | - | 2.133E+04 | 1603 | - | - | 0 | - |
| - | - | 1.474E+06 | 1604 | - | - | 0 | - |
| - | - | 1.174E+06 | 1605 | - | - | 0 | - |
| - | - | 6.808E+05 | 1606 | - | - | 0 | - |
| - | - | 2.621E+05 | 1607 | - | - | 0 | - |
| - | - | 7.319E+04 | 1608 | - | - | 0 | - |
| - | - | 1.09E+04 | 2637 | - | - | 0 | - |

m/z Charge Intensity FragmentType MassShift Position
123.05501556396484 0 58840.64
134.0269317626953 0 8505.6
136.0753173828125 0 24960.678
140.0816650390625 0 23755.533
164.04722595214844 0 8757.838
173.43833923339844 0 13263.132
182.08087158203125 0 27521.785 y 12
192.07647705078125 0 38135.934
209.10256958007812 0 33087.504
221.05865478515625 0 222106.7
222.06227111816406 0 11983.109
232.689208984375 0 8634.103
237.09771728515625 0 13667.268
237.41041564941406 0 9096.145
239.09478759765625 0 16636.209
244.0928192138672 0 18196.377
249.05337524414062 0 96137.99
250.05743408203125 0 11896.593
254.1241912841797 0 8501.458
258.0421142578125 0 7961.104
258.1230163574219 0 21338.283
268.0606689453125 0 7596.7334
311.1224365234375 0 7891.4624
319.13909912109375 0 25446.885 y 11
336.0863037109375 0 12625.165
372.16461181640625 0 14832.346
373.1499328613281 0 68690.66 w 10
377.6830749511719 0 10785.8955 y 7
393.15899658203125 0 9213.464
407.1591491699219 0 22850.943
408.50103759765625 0 8362.197
417.1643371582031 0 25794.059 z 10
417.2820739746094 0 13303.3955
418.1702880859375 0 10701.029
428.1954650878906 0 27021.518
433.18194580078125 0 23670.375 y 10
435.15362548828125 0 23246.139
445.8688659667969 0 23902.852 y Water loss 2
446.5318908691406 0 11624.892 z 2
451.8706359863281 0 23002.475 y 2
452.1776428222656 0 13309.367 c 3
452.2081604003906 0 20939.984
474.1950378417969 0 13495.1045 c Ammonia loss 11
476.5511474609375 0 14525.11
497.8862609863281 0 23685.873
498.22015380859375 0 24551.873
498.5543212890625 0 12697.333
505.54193115234375 0 25392.041
505.8753967285156 0 25301.062
510.2083740234375 0 25935.064
510.735595703125 0 17790.98 y 5
513.22021484375 0 39354.867
513.5550537109375 0 25344.424
513.8875732421875 0 12542.929
514.2199096679688 0 9147.589
522.2694091796875 0 17260.154
528.5498657226562 0 24395.006
528.8838500976562 0 31404.262
532.6652221679688 0 10287.955
534.553466796875 0 394090.97
534.8873291015625 0 286829.53
535.2210083007812 0 216888.7
535.5547485351562 0 89506.65
535.890380859375 0 24411.441
544.2428588867188 0 12038.32
545.250244140625 0 25039.922
545.7528076171875 0 11335.088
555.2037353515625 0 22046.13
556.209228515625 0 10945.606
570.2413330078125 0 93482.98 y 9
571.2449951171875 0 23253.045
572.7715454101562 0 11991.707
583.2673950195312 0 11494.284
584.2538452148438 0 40995.625 y 4
584.7553100585938 0 37732.133 c Water loss 9
586.7686767578125 0 18691.732
595.7764282226562 0 15648.074
599.2157592773438 0 71038.125 c 4
600.2194213867188 0 17053.305
624.2518920898438 0 128144.59 w 8
624.3005981445312 0 11299.649
625.2562866210938 0 57658.652
625.7783813476562 0 41045.055 z 3
626.277587890625 0 33372.344
626.7805786132812 0 10602.513
628.2611083984375 0 17961.531
628.7648315429688 0 19973.59
633.2825317382812 0 27670.785
633.78662109375 0 78257.164 y 3
634.287353515625 0 68709.945
634.7869262695312 0 15658.916
637.2939453125 0 12407.353
637.7954711914062 0 11626.462
638.2988891601562 0 24637.969
638.802001953125 0 13761.701
639.7914428710938 0 18292.21
640.292724609375 0 20822.389
641.2636108398438 0 36159.168
641.764892578125 0 16502.742 c Water loss 10
642.2664184570312 0 13802.52 c Ammonia loss 10
645.3041381835938 0 46200.22
645.804443359375 0 52857.074
660.7916870117188 0 53782.81 z Ammonia loss 2
661.2927856445312 0 43914.645
661.7855224609375 0 14814.673
667.3093872070312 0 24396.04 z 8
668.2964477539062 0 28132.143 y Water loss 2
668.79345703125 0 56928.055 y Ammonia loss 2
669.2939453125 0 178236.36 z 2
669.7958984375 0 135911.31
670.2972412109375 0 61487.695
670.79541015625 0 17768.512
674.2509155273438 0 25832.898
677.3032836914062 0 324394.4 y 2
677.804931640625 0 271174.47
678.3055419921875 0 118528.51
678.8045043945312 0 32517.646
682.28173828125 0 86646.95
683.3297729492188 0 23203.045 y 8
684.3305053710938 0 12331.913
687.806884765625 0 17151.535
689.2932739257812 0 12778.214
695.290771484375 0 26249.033
696.2908935546875 0 10598.269
696.7958374023438 0 10068.112
697.29541015625 0 23931.84
697.7979125976562 0 13183.779
704.3087158203125 0 153719.58 z Ammonia loss 1
704.8098754882812 0 139147.77
705.310791015625 0 45595.363
705.809814453125 0 13213.533
710.7914428710938 0 30830.113 c Ammonia loss 11
711.2899169921875 0 20322.025
711.7933959960938 0 16889.582
712.2991943359375 0 13746.841 y Ammonia loss 1
718.922119140625 0 30265.824
719.3028564453125 0 693675.1 c 11
719.8041381835938 0 497863.06
720.3040771484375 0 285541.22
720.804931640625 0 107155.836 y 1
721.3067016601562 0 23422.555 z Ammonia loss 7
724.3382568359375 0 42093.59
724.8407592773438 0 43872.312
725.3426513671875 0 14287.517
726.3079223632812 0 11366.498
730.3006591796875 0 13379.768
736.2747802734375 0 687689.7 c 5
737.2772827148438 0 261064.38
738.2741088867188 0 119209.77
738.34423828125 0 68282.85 z 7
739.2777709960938 0 15833.708
739.3482666015625 0 47235.043
739.802490234375 0 16950.35
740.33154296875 0 27109.824
740.8167114257812 0 14726.793
747.3310546875 0 25244.564
747.82861328125 0 53747.44
748.326416015625 0 35039.184
748.830810546875 0 29587.756
749.3092041015625 0 12530.556
749.8139038085938 0 47925.87
750.3155517578125 0 46576.152
750.815185546875 0 23474.258
754.3631591796875 0 29090.758 y 7
755.3655395507812 0 11899.485
755.8203125 0 14230.688
757.8116455078125 0 36368.293
758.3138427734375 0 30741.633
758.8128662109375 0 24958.312
760.80322265625 0 18360.531
761.3046875 0 20118.06
762.3075561523438 0 12945.607
762.8134155273438 0 33519.613
763.3167724609375 0 33231.242
763.810302734375 0 44564.83
764.3089599609375 0 34691.34
764.8111572265625 0 18667.285
770.326416015625 0 14637.53
770.82666015625 0 19662.465
771.3362426757812 0 15642.473
771.822021484375 0 42495.53
772.3228149414062 0 712233.5
772.8245239257812 0 533658.75
773.32470703125 0 282456.38
773.8245239257812 0 122668.5
774.3258056640625 0 39191.473
778.8267211914062 0 11177.906
778.940673828125 0 12148.378
779.320556640625 0 59857.906
779.8214721679688 0 83737.43
780.3206787109375 0 47758.76
785.3150024414062 0 18740.52
786.3128662109375 0 13275.473
792.8231201171875 0 38435.66
793.327880859375 0 239731.16
793.8289184570312 0 241164.1
794.328369140625 0 146758.66
794.8256225585938 0 60071.395
795.3257446289062 0 18535.354
800.8417358398438 0 29154.78
801.3272094726562 0 715031.6
801.8301391601562 0 3826366.8
802.3318481445312 0 3167580
802.8323364257812 0 1838618.4
803.3324584960938 0 630551.3
803.8331298828125 0 226118.97
808.3733520507812 0 229845.38 w 6
809.3757934570312 0 90596.664
810.377685546875 0 27009.027
865.3172607421875 0 225246.19 c 6
866.3207397460938 0 96053.836
867.3865356445312 0 400922.78 z 6
868.3893432617188 0 209032.06
869.391357421875 0 65505.227
883.4054565429688 0 83829.75 y 6
884.409423828125 0 36583.465
936.3538208007812 0 198084.7 c 7
937.3572998046875 0 93827.1
938.3577880859375 0 33852.016
967.00244140625 0 11962.091
1004.4440307617188 0 107395.08 z 5
1005.4401245117188 0 115969.055
1006.44140625 0 40656.02
1007.4407958984375 0 14220.657
1020.4644775390625 0 130304.93 y 5
1021.466796875 0 82474.375
1022.4708862304688 0 30278.652
1049.4378662109375 0 76469.03 c 8
1050.4432373046875 0 40959.586
1051.4346923828125 0 23447.535
1089.491455078125 0 12798.839
1129.516357421875 0 11219.522
1151.4901123046875 0 15761.861 z 4
1152.4859619140625 0 28353.875
1153.4891357421875 0 23177.955
1173.5355224609375 0 42456.45
1174.533447265625 0 20558.746
1186.4969482421875 0 242801.25 c 9
1186.6358642578125 0 17117.627
1187.4998779296875 0 138287.1
1187.630615234375 0 15906.372
1188.496337890625 0 70336.33
1189.510498046875 0 25660.758
1243.5560302734375 0 15692.835
1250.5535888671875 0 23315.82 z 3
1251.5567626953125 0 33534.297
1252.5555419921875 0 25983.12
1253.5648193359375 0 18753.898
1256.52099609375 0 13816.748
1257.521484375 0 11636.075
1266.567138671875 0 14695.27 y 3
1300.540771484375 0 219169.78 c 10
1301.5400390625 0 133404.45
1302.54296875 0 90202.086
1303.54736328125 0 26028.2
1320.5794677734375 0 14831.214 w 2
1321.5870361328125 0 30636.838
1337.5797119140625 0 88873.02 z 2
1338.587890625 0 333154.16
1338.77001953125 0 20111.975
1339.5904541015625 0 271261.06
1340.59423828125 0 108477.57
1341.5989990234375 0 36621.062
1342.598388671875 0 12242.471
1353.6005859375 0 24487.395 y 2
1354.607421875 0 43651.805
1355.6103515625 0 19295.727
1379.591552734375 0 45047.586
1380.58837890625 0 35245.152
1381.601806640625 0 18241.559
1393.583251953125 0 18755.691
1394.596435546875 0 40512.414
1395.583740234375 0 17046.252
1396.587890625 0 19123.264
1407.6102294921875 0 58615.92 z Ammonia loss 1
1408.614990234375 0 93820.76
1409.6168212890625 0 77262.76
1410.62109375 0 38349.16
1421.592041015625 0 30129.154
1422.595458984375 0 11372.995 y Water loss 1
1437.5970458984375 0 57511.074 c 11
1438.60498046875 0 167098.98
1439.6083984375 0 131106.27
1440.6080322265625 0 85149.4 y 1
1441.60498046875 0 28545.963
1448.6715087890625 0 28456.156
1449.6854248046875 0 27925.887
1499.6307373046875 0 16833.75
1500.613037109375 0 14214.724
1501.628662109375 0 12869.776
1521.5732421875 0 15127.62
1522.60888671875 0 11225.861
1526.6356201171875 0 32725.717
1527.6356201171875 0 42951.17
1528.63330078125 0 48903.633
1529.62548828125 0 20410.277
1541.6611328125 0 38490.367
1542.647216796875 0 37841.71
1543.64013671875 0 220733.84
1544.6439208984375 0 474363.7
1545.6473388671875 0 373314.3
1546.648193359375 0 211912.98
1547.6505126953125 0 71288.91
1548.6575927734375 0 17602.2
1557.62646484375 0 16111.017
1558.6434326171875 0 79372.39
1559.6400146484375 0 70105.09
1560.6455078125 0 35277.17
1568.624755859375 0 26785.166
1569.630126953125 0 19830.951
1575.660888671875 0 35737.37
1576.6666259765625 0 44212.008
1577.661376953125 0 16362.663
1578.6546630859375 0 11711.79
1585.64794921875 0 98272.64
1586.6451416015625 0 245433.47
1587.647705078125 0 187818.75
1588.64697265625 0 133528.02
1589.6529541015625 0 49404.965
1601.65087890625 0 58101.82
1602.6531982421875 0 389627.44
1602.8746337890625 0 21328.732
1603.660400390625 0 1474229.8
1604.6629638671875 0 1173551.6
1605.6639404296875 0 680844.1
1606.6644287109375 0 262058.56
1607.66650390625 0 73190.75
2637.438232421875 0 10904.63

Spectrum Details

|  |  |
| --- | --- |
| Matched peaks? Matched peaks The total absolute number of peaks matched. Additionally in brackets the total fraction of peaks matched and the total number of peaks is shown. | 57 (17.38% of 328) |
| FDR? FDR The false discovery rate estimated for this peptide. It is calculated by matching all theoretical fragments with a non-integer shift with the raw peaks for this spectrum. This is done with 40 different shifts. The resulting percentage is the average number of annotated peaks over the number of annotated peaks with the correct spectrum. | 0.42% |
| Satellite FDR? Satellite FDR See the FDR for details on its calculation. This satellite ion specific FDR only contains the satellite ions (d/w) for I/L/J positions. | 0.00% |
| PSM Score? PSM Score The PSM Score as given by Hecklib to this annotated spectrum. It is shown with three significant figures. | 401 |

#### Reverse Lookup? Reverse Lookup All places where this read could be placed.

| Group | Segment | Template | Template Part | Read Part | Score | Unique |
| --- | --- | --- | --- | --- | --- | --- |
| Homo sapiens Heavy Chain | IGHC | IGHG1 | [306..319] | [0..13] | 92 | False |
| Homo sapiens Heavy Chain | IGHC | IGHG2 | [302..315] | [0..13] | 92 | False |
| Homo sapiens Heavy Chain | IGHC | IGHG4 | [303..316] | [0..13] | 92 | False |

| Recombined | Template Part | Read Part | Score | Unique |
| --- | --- | --- | --- | --- |
| REC-0-1 | [431..444] | [0..13] | 92 | True |

#### Meta Information from Multiple reads

##### Number of combined reads

2

##### Intensity

0.7075

##### TotalArea

1.855E+08

#### Positional Score

Copy Data

##### Positional Score (TSV)

###### Preview

```
Loading example...
```

*Click on the button to copy the data to your clipboard.*

000123456789101112

Label Value
"0" 0
"1" 0
"2" 0
"3" 0
"4" 0
"5" 0
"6" 0
"7" 0
"8" 0
"9" 0
"10" 0
"11" 0
"12" 0

#### Meta Information from PEAKS

##### Scan Identifier

F3:4496

##### Original sequence

C

+58.01

S

S

V

M

H

E

A

L

H

N

H

Y

##### Posttranslational Modifications

Carboxymethyl

##### Source File

D:\separate\_stitch\_analyses\xle-disambiguation\raw\20210323\_F1\_UM1\_Peng0013\_SA\_F59\_ingel\_3ug\_chymo.raw

##### Fraction

3

##### Scan Feature

F3:5269

##### De Novo Score

98

##### ConfidenceScore

98

### m/z

529.2233

##### Mass

1584.645

##### Charge

3

##### Retention Time

24.77

##### Predicted Retention Time

-

##### Area

1.269E+07

##### Parts Per Million

1.9

##### Fragmentation mode

ETHCD

##### Originating file

01 D:\separate\_stitch\_analyses\xle-disambiguation\20210325\_F59\_3ug\_DENOVO\_12.csv

#### Meta Information from PEAKS

##### Scan Identifier

F3:2851

##### Original sequence

C

+58.01

S

S

V

M

+15.99

H

E

A

L

H

N

H

Y

##### Posttranslational Modifications

Carboxymethyl; Oxidation (M)

##### Source File

D:\separate\_stitch\_analyses\xle-disambiguation\raw\20210323\_F1\_UM1\_Peng0013\_SA\_F59\_ingel\_3ug\_chymo.raw

##### Fraction

3

##### Scan Feature

F3:5548

##### De Novo Score

97

##### ConfidenceScore

97

### m/z

534.5554

##### Mass

1600.6399

##### Charge

3

##### Retention Time

15.16

##### Predicted Retention Time

-

##### Area

1.728E+08

##### Parts Per Million

2.8

##### Fragmentation mode

ETHCD

##### Originating file

01 D:\separate\_stitch\_analyses\xle-disambiguation\20210325\_F59\_3ug\_DENOVO\_12.csv
