## Supplementary material for "A handle on mass coincidence errors in *de novo* sequencing of antibodies by bottom-up proteomics": Combined_091.html

Details Combined\_091 | Stitch OverviewUndefined

### Read Combined\_091

#### Sequence (length=8)

AIPAPJEK

#### Spectrum 4860? Spectrum 4860 The raw spectrum of this peptide as annotated by Hecklib. The fragments are coloured according to ion type (see legend). Any peaks with a star '\*' as text can be hovered over to see the full details, first the ion type second the mass shift type. By hovering over the amino acids in the peptide or ions in the legend the corresponding peaks are highlighted. By toggling the 'Unassigned' label you can turn the background (unassigned) peaks on or off in the plot. By updating the slider in the Ion legend you can update the spectrum to only show the top X% of the peaks with labels. The top X% means any peak that is within X% of the highest intensity. By dragging in the spectrum you can zoom in to a specific part of the spectrum and use 'Zoom Out' to get back to the original zoom level. The annotation of the spectrum is based on the given sequence in the peptides file and is done with different software so inconsistencies are likely. The peaks are annotated based on the given sequence, with 20 ppm tolerance.

Copy Data

##### Spectrum 4860 (TSV)

###### Preview

```
Loading example...
```

*Click on the button to copy the data to your clipboard.*

Mz MinMz MaxIntensity Max

WidthHeightPeptide font sizePeptide stroke widthSpectrum font sizeSpectrum stroke widthCompact peptide

Ion legend

wxyz

abcd

OtherUnassignedIon^Charge^_Position_Show for top:%

AIPAPJEK

08.00e+41.60e+52.40e+53.20e+5

Zoom Out

d^+1^_2_y^+1^_1_y^+1^_1_a^+1^_2_b^+1^_2_y^+2^_4_a^+1^_3_y^+1^_2_y^+1^_2_y^+1^_2_y^+2^_5_b^+1^_3_y^+2^_6_b^+1^_4_y^+1^_3_y^+2^_7_y^+2^_7_y^+1^_3_y^+1^_4_y^+1^_4_y^+1^_5_y^+1^_5_y^+1^_6_y^+1^_6_

0778155523333111

Fragment Matches Table

Show background peaks

| Position | Ion type | Intensity | mz Theoretical | mz Error (Th) | mz Error (ppm) | Charge | Series Number |
| --- | --- | --- | --- | --- | --- | --- | --- |
| - | - | 1313 | 120.1 | - | - | 0 | - |
| - | - | 350.6 | 121.4 | - | - | 0 | - |
| - | - | 383.4 | 121.9 | - | - | 0 | - |
| - | - | 342.8 | 122.1 | - | - | 0 | - |
| - | - | 507 | 123.1 | - | - | 0 | - |
| - | - | 388.7 | 124.1 | - | - | 0 | - |
| - | - | 365.4 | 125.1 | - | - | 0 | - |
| - | - | 1431 | 126.1 | - | - | 0 | - |
| - | - | 1183 | 129 | - | - | 0 | - |
| - | - | 434.5 | 129.1 | - | - | 0 | - |
| 2 | d | 4.266E+04 | 129.1 | 0.0002843 | 2.202 | +1 | 2 |
| 8 | y | 2.276E+04 | 130.1 | 0.0003081 | 2.368 | +1 | 1 |
| - | - | 2641 | 130.1 | - | - | 0 | - |
| - | - | 1521 | 131.1 | - | - | 0 | - |
| - | - | 515.1 | 136.1 | - | - | 0 | - |
| - | - | 717.6 | 139.1 | - | - | 0 | - |
| - | - | 1.193E+05 | 141.1 | - | - | 0 | - |
| - | - | 851.9 | 142.1 | - | - | 0 | - |
| - | - | 9206 | 142.1 | - | - | 0 | - |
| - | - | 1747 | 144.1 | - | - | 0 | - |
| 8 | y | 3.512E+04 | 147.1 | 0.000294 | 1.998 | +1 | 1 |
| - | - | 448.5 | 148.1 | - | - | 0 | - |
| - | - | 517.6 | 148.1 | - | - | 0 | - |
| - | - | 1702 | 148.1 | - | - | 0 | - |
| - | - | 6.186E+04 | 149 | - | - | 0 | - |
| - | - | 4561 | 150 | - | - | 0 | - |
| - | - | 553.9 | 151.1 | - | - | 0 | - |
| - | - | 502.6 | 157 | - | - | 0 | - |
| - | - | 2879 | 157.1 | - | - | 0 | - |
| 2 | a | 2.938E+05 | 157.1 | 0.000341 | 2.17 | +1 | 2 |
| - | - | 660.9 | 158.1 | - | - | 0 | - |
| - | - | 1633 | 158.1 | - | - | 0 | - |
| - | - | 2.445E+04 | 158.1 | - | - | 0 | - |
| - | - | 850.5 | 159.1 | - | - | 0 | - |
| - | - | 439.1 | 162.1 | - | - | 0 | - |
| - | - | 3828 | 167 | - | - | 0 | - |
| - | - | 496.1 | 168.1 | - | - | 0 | - |
| - | - | 1.234E+05 | 169.1 | - | - | 0 | - |
| - | - | 459.1 | 169.7 | - | - | 0 | - |
| - | - | 572.9 | 170.1 | - | - | 0 | - |
| - | - | 9632 | 170.1 | - | - | 0 | - |
| - | - | 480.9 | 171.1 | - | - | 0 | - |
| - | - | 451.7 | 171.1 | - | - | 0 | - |
| - | - | 456.1 | 171.5 | - | - | 0 | - |
| - | - | 455.3 | 171.6 | - | - | 0 | - |
| - | - | 2201 | 173.5 | - | - | 0 | - |
| - | - | 1206 | 175.1 | - | - | 0 | - |
| - | - | 932.8 | 176.1 | - | - | 0 | - |
| - | - | 590.5 | 176.6 | - | - | 0 | - |
| - | - | 431.9 | 178.2 | - | - | 0 | - |
| - | - | 432.5 | 181 | - | - | 0 | - |
| - | - | 1693 | 183.1 | - | - | 0 | - |
| - | - | 1.153E+05 | 183.1 | - | - | 0 | - |
| - | - | 1.149E+04 | 184.2 | - | - | 0 | - |
| - | - | 917.9 | 185.1 | - | - | 0 | - |
| 2 | b | 6.864E+04 | 185.1 | 0.0002537 | 1.37 | +1 | 2 |
| - | - | 6609 | 186.1 | - | - | 0 | - |
| - | - | 615 | 187.1 | - | - | 0 | - |
| - | - | 1100 | 189.1 | - | - | 0 | - |
| - | - | 4151 | 195.1 | - | - | 0 | - |
| - | - | 3051 | 197.1 | - | - | 0 | - |
| - | - | 897.4 | 197.1 | - | - | 0 | - |
| - | - | 617.6 | 201.1 | - | - | 0 | - |
| - | - | 504.1 | 205.6 | - | - | 0 | - |
| - | - | 6.13E+04 | 211.1 | - | - | 0 | - |
| - | - | 6499 | 212.1 | - | - | 0 | - |
| - | - | 568.6 | 213.1 | - | - | 0 | - |
| - | - | 1690 | 213.1 | - | - | 0 | - |
| - | - | 706.7 | 213.1 | - | - | 0 | - |
| - | - | 5599 | 215.1 | - | - | 0 | - |
| - | - | 718.2 | 216.1 | - | - | 0 | - |
| - | - | 492.3 | 217 | - | - | 0 | - |
| - | - | 1181 | 217.1 | - | - | 0 | - |
| - | - | 1128 | 217.1 | - | - | 0 | - |
| - | - | 5379 | 222.1 | - | - | 0 | - |
| - | - | 592.6 | 223.1 | - | - | 0 | - |
| - | - | 585.3 | 223.1 | - | - | 0 | - |
| - | - | 652.9 | 226.2 | - | - | 0 | - |
| - | - | 770.4 | 227.1 | - | - | 0 | - |
| - | - | 1175 | 236.2 | - | - | 0 | - |
| - | - | 831.1 | 237.1 | - | - | 0 | - |
| - | - | 2.527E+04 | 240.1 | - | - | 0 | - |
| - | - | 3227 | 241.1 | - | - | 0 | - |
| - | - | 2682 | 241.1 | - | - | 0 | - |
| - | - | 6422 | 243.1 | - | - | 0 | - |
| 5 | y | 1.868E+04 | 243.6 | 0.0002126 | 0.8726 | +2 | 4 |
| - | - | 808.6 | 244.1 | - | - | 0 | - |
| - | - | 4725 | 244.2 | - | - | 0 | - |
| - | - | 2539 | 244.2 | - | - | 0 | - |
| - | - | 662 | 244.7 | - | - | 0 | - |
| - | - | 1084 | 245.1 | - | - | 0 | - |
| - | - | 1548 | 253.1 | - | - | 0 | - |
| 3 | a | 1929 | 254.2 | 0.0004336 | 1.706 | +1 | 3 |
| 7 | y | 8.517E+04 | 258.1 | 0.0001259 | 0.4878 | +1 | 2 |
| 7 | y | 1410 | 259.1 | 8.862E-05 | 0.342 | +1 | 2 |
| - | - | 9732 | 259.1 | - | - | 0 | - |
| - | - | 512.1 | 259.8 | - | - | 0 | - |
| - | - | 921.1 | 260.1 | - | - | 0 | - |
| - | - | 1620 | 261.1 | - | - | 0 | - |
| - | - | 822.4 | 262.2 | - | - | 0 | - |
| - | - | 7609 | 264.2 | - | - | 0 | - |
| - | - | 1357 | 265.2 | - | - | 0 | - |
| - | - | 2.404E+04 | 266.2 | - | - | 0 | - |
| - | - | 3109 | 267.2 | - | - | 0 | - |
| 7 | y | 7.191E+04 | 276.2 | 0.0001203 | 0.4357 | +1 | 2 |
| - | - | 8153 | 277.2 | - | - | 0 | - |
| - | - | 893.2 | 278.2 | - | - | 0 | - |
| 4 | y | 618 | 279.2 | 0.0006376 | 2.284 | +2 | 5 |
| - | - | 777.7 | 280.1 | - | - | 0 | - |
| - | - | 497.4 | 281.6 | - | - | 0 | - |
| 3 | b | 6984 | 282.2 | 5.634E-05 | 0.1997 | +1 | 3 |
| - | - | 701.2 | 283.2 | - | - | 0 | - |
| - | - | 538 | 284.2 | - | - | 0 | - |
| - | - | 874.5 | 286.1 | - | - | 0 | - |
| - | - | 698.5 | 294.2 | - | - | 0 | - |
| - | - | 547.3 | 296.2 | - | - | 0 | - |
| - | - | 810.8 | 298.1 | - | - | 0 | - |
| - | - | 536.5 | 312.2 | - | - | 0 | - |
| - | - | 5100 | 312.2 | - | - | 0 | - |
| - | - | 2733 | 322.2 | - | - | 0 | - |
| 3 | y | 3.169E+05 | 327.7 | 0.0003483 | 1.063 | +2 | 6 |
| - | - | 1.092E+05 | 328.2 | - | - | 0 | - |
| - | - | 2.537E+04 | 328.7 | - | - | 0 | - |
| - | - | 1585 | 329.2 | - | - | 0 | - |
| - | - | 6.514E+04 | 340.2 | - | - | 0 | - |
| - | - | 1.225E+04 | 341.2 | - | - | 0 | - |
| - | - | 686.5 | 342.2 | - | - | 0 | - |
| - | - | 1257 | 349.2 | - | - | 0 | - |
| - | - | 1.374E+04 | 351.2 | - | - | 0 | - |
| - | - | 3077 | 352.2 | - | - | 0 | - |
| 4 | b | 1372 | 353.2 | 0.0001922 | 0.5442 | +1 | 4 |
| - | - | 1533 | 357.2 | - | - | 0 | - |
| - | - | 1.417E+04 | 358.2 | - | - | 0 | - |
| - | - | 2412 | 359.2 | - | - | 0 | - |
| 6 | y | 705.8 | 371.2 | 0.0007166 | 1.93 | +1 | 3 |
| 2 | y | 2046 | 375.2 | 0.004095 | 10.91 | +2 | 7 |
| - | - | 745.2 | 377.2 | - | - | 0 | - |
| - | - | 1.952E+04 | 379.2 | - | - | 0 | - |
| - | - | 4162 | 380.2 | - | - | 0 | - |
| - | - | 829.2 | 381.2 | - | - | 0 | - |
| - | - | 2769 | 383.2 | - | - | 0 | - |
| 2 | y | 2921 | 384.2 | 0.0003629 | 0.9446 | +2 | 7 |
| - | - | 758.5 | 384.7 | - | - | 0 | - |
| 6 | y | 2.195E+04 | 389.2 | 4.072E-05 | 0.1046 | +1 | 3 |
| - | - | 3850 | 390.2 | - | - | 0 | - |
| - | - | 932.1 | 393.2 | - | - | 0 | - |
| - | - | 823.6 | 395.2 | - | - | 0 | - |
| - | - | 4213 | 399.2 | - | - | 0 | - |
| - | - | 1829 | 400.2 | - | - | 0 | - |
| - | - | 1.301E+04 | 411.2 | - | - | 0 | - |
| - | - | 2707 | 412.2 | - | - | 0 | - |
| - | - | 921.5 | 419.8 | - | - | 0 | - |
| - | - | 802.7 | 420.9 | - | - | 0 | - |
| - | - | 1675 | 429.2 | - | - | 0 | - |
| - | - | 1436 | 437.2 | - | - | 0 | - |
| - | - | 1639 | 462.3 | - | - | 0 | - |
| 5 | y | 1.035E+04 | 468.3 | 1.367E-05 | 0.0292 | +1 | 4 |
| - | - | 3058 | 469.3 | - | - | 0 | - |
| - | - | 7373 | 480.3 | - | - | 0 | - |
| - | - | 2021 | 481.3 | - | - | 0 | - |
| 5 | y | 2.694E+05 | 486.3 | 7.228E-05 | 0.1486 | +1 | 4 |
| - | - | 6.364E+04 | 487.3 | - | - | 0 | - |
| - | - | 1.079E+04 | 488.3 | - | - | 0 | - |
| - | - | 1766 | 490.3 | - | - | 0 | - |
| - | - | 3585 | 506.3 | - | - | 0 | - |
| - | - | 1756 | 507.3 | - | - | 0 | - |
| - | - | 2.339E+04 | 508.3 | - | - | 0 | - |
| - | - | 6977 | 509.3 | - | - | 0 | - |
| - | - | 1332 | 510.3 | - | - | 0 | - |
| - | - | 1022 | 525.3 | - | - | 0 | - |
| - | - | 5623 | 526.3 | - | - | 0 | - |
| - | - | 1944 | 527.3 | - | - | 0 | - |
| 4 | y | 7190 | 539.3 | 1.243E-05 | 0.02305 | +1 | 5 |
| - | - | 1450 | 540.3 | - | - | 0 | - |
| - | - | 1495 | 542.3 | - | - | 0 | - |
| 4 | y | 6.864E+04 | 557.3 | 0.0006035 | 1.083 | +1 | 5 |
| - | - | 1.859E+04 | 558.3 | - | - | 0 | - |
| - | - | 3143 | 559.3 | - | - | 0 | - |
| - | - | 1870 | 619.4 | - | - | 0 | - |
| - | - | 904.7 | 620.4 | - | - | 0 | - |
| 3 | y | 5781 | 636.4 | 0.0003222 | 0.5063 | +1 | 6 |
| - | - | 2105 | 637.4 | - | - | 0 | - |
| 3 | y | 2.655E+05 | 654.4 | 0.000694 | 1.061 | +1 | 6 |
| - | - | 9.3E+04 | 655.4 | - | - | 0 | - |
| - | - | 2.113E+04 | 656.4 | - | - | 0 | - |
| - | - | 1147 | 657.4 | - | - | 0 | - |
| - | - | 1737 | 752.4 | - | - | 0 | - |
| - | - | 744.2 | 753.4 | - | - | 0 | - |
| - | - | 615.5 | 803 | - | - | 0 | - |
| - | - | 637.2 | 1802 | - | - | 0 | - |
| - | - | 663.7 | 1964 | - | - | 0 | - |
| - | - | 754.3 | 3080 | - | - | 0 | - |

m/z Charge Intensity FragmentType MassShift Position
120.0810317993164 0 1313.0583
121.37792205810547 0 350.63742
121.93342590332031 0 383.43082
122.13141632080078 0 342.75586
123.09208679199219 0 506.96112
124.14273071289062 0 388.72318
125.05992126464844 0 365.4218
126.05543518066406 0 1430.5419
129.0187225341797 0 1183.3605
129.06619262695312 0 434.53375
129.10252380371094 0 42660.66 d 1
130.08656311035156 0 22761.576 y Ammonia loss 7
130.1059112548828 0 2641.237
131.08995056152344 0 1520.5271
136.0762939453125 0 515.1138
139.08677673339844 0 717.56305
141.10256958007812 0 119263.5
142.09976196289062 0 851.89844
142.10586547851562 0 9206.322
144.0656280517578 0 1746.5715
147.11309814453125 0 35122.938 y 7
148.06134033203125 0 448.51227
148.11007690429688 0 517.63635
148.11636352539062 0 1702.2281
149.02362060546875 0 61861.406
150.0269012451172 0 4561.324
151.08697509765625 0 553.9449
157.0135040283203 0 502.61676
157.0974884033203 0 2879.3313
157.13388061523438 0 293787.5 a 1
158.09298706054688 0 660.9405
158.1308135986328 0 1633.3236
158.1371612548828 0 24447.549
159.14024353027344 0 850.4641
162.14678955078125 0 439.13623
167.0342254638672 0 3827.939
168.10194396972656 0 496.1252
169.09747314453125 0 123379.164
169.73509216308594 0 459.078
170.09368896484375 0 572.9474
170.10079956054688 0 9631.723
171.1040496826172 0 480.90976
171.13771057128906 0 451.73175
171.4636688232422 0 456.0725
171.64886474609375 0 455.31073
173.45095825195312 0 2200.6096
175.1190185546875 0 1206.3016
176.12306213378906 0 932.764
176.62513732910156 0 590.46155
178.17840576171875 0 431.87503
180.9729461669922 0 432.4543
183.11294555664062 0 1693.3182
183.1494903564453 0 115266.19
184.1528778076172 0 11492.116
185.0811309814453 0 917.9251
185.1287078857422 0 68643.414 b 1
186.13206481933594 0 6608.815
187.13394165039062 0 615.04175
189.08692932128906 0 1100.2806
195.11306762695312 0 4150.523
197.09226989746094 0 3051.4468
197.12924194335938 0 897.3698
201.1232147216797 0 617.59656
205.5668182373047 0 504.07983
211.1443328857422 0 61300.19
212.14776611328125 0 6498.58
213.07662963867188 0 568.5661
213.1236114501953 0 1689.8704
213.1493682861328 0 706.6862
215.13922119140625 0 5598.6147
216.14321899414062 0 718.23206
217.0338897705078 0 492.26144
217.08192443847656 0 1181.3384
217.13694763183594 0 1128.0844
222.12380981445312 0 5378.909
223.10829162597656 0 592.5775
223.12771606445312 0 585.3041
226.15457153320312 0 652.9463
227.10311889648438 0 770.365
236.17567443847656 0 1175.1156
237.09136962890625 0 831.09674
240.13449096679688 0 25266.576
241.11846923828125 0 3226.627
241.137939453125 0 2682.2363
243.13421630859375 0 6422.4277
243.64996337890625 0 18682.398 y 4
244.13888549804688 0 808.5903
244.1515655517578 0 4725.331
244.16580200195312 0 2539.3052
244.65383911132812 0 661.95917
245.12460327148438 0 1084.2762
253.1183624267578 0 1547.5581
254.18673706054688 0 1928.5759 a 2
258.14495849609375 0 85167.62 y Water loss 6
259.1289367675781 0 1409.9384 y Ammonia loss 6
259.14849853515625 0 9731.711
259.7874755859375 0 512.091
260.149169921875 0 921.13446
261.1451721191406 0 1620.37
262.15069580078125 0 822.4423
264.17071533203125 0 7608.985
265.1737365722656 0 1357.0021
266.1500244140625 0 24040.912
267.1535339355469 0 3108.8748
276.155517578125 0 71906.72 y 6
277.15875244140625 0 8153.202
278.16143798828125 0 893.2044
279.1689453125 0 617.95325 y 3
280.1299743652344 0 777.7331
281.5781555175781 0 497.3975
282.1812744140625 0 6984.2183 b 2
283.183837890625 0 701.1533
284.1603698730469 0 537.9519
286.1402893066406 0 874.4873
294.18011474609375 0 698.54846
296.1970520019531 0 547.28046
298.1392822265625 0 810.7787
312.1553955078125 0 536.49945
312.1919860839844 0 5100.492
322.176025390625 0 2732.5908
327.6950378417969 0 316853.4 y 2
328.1964111328125 0 109195.15
328.69775390625 0 25365.074
329.19915771484375 0 1585.3123
340.1868591308594 0 65138.3
341.1901550292969 0 12253.168
342.1904296875 0 686.47473
349.18670654296875 0 1256.7468
351.23907470703125 0 13740.781
352.2419128417969 0 3076.906
353.2181396484375 0 1372.3312 b 3
357.2137451171875 0 1533.4677
358.1969909667969 0 14168.842
359.1997985839844 0 2412.144
371.2281799316406 0 705.8383 y Water loss 5
375.23553466796875 0 2046.0615 y Water loss 1
377.18304443359375 0 745.1903
379.2339172363281 0 19522.66
380.2367248535156 0 4162.3545
381.2136535644531 0 829.21735
383.22857666015625 0 2769.1304
384.2363586425781 0 2921.3943 y 1
384.73907470703125 0 758.45856
389.239501953125 0 21946.45 y 5
390.24298095703125 0 3850.0354
393.2138671875 0 932.1246
395.19122314453125 0 823.55646
399.22393798828125 0 4212.6943
400.2266845703125 0 1828.725
411.2236328125 0 13011.208
412.22674560546875 0 2706.651
419.7660827636719 0 921.4981
420.8605041503906 0 802.6731
429.2366943359375 0 1675.2158
437.2394714355469 0 1436.3397
462.2702331542969 0 1639.3429
468.2816467285156 0 10354.548 y Water loss 4
469.2840270996094 0 3058.1052
480.2814025878906 0 7373.442
481.2852478027344 0 2020.7885
486.29229736328125 0 269357.7 y 4
487.29522705078125 0 63636.008
488.2974548339844 0 10794.395
490.266357421875 0 1766.4681
506.2750549316406 0 3585.1125
507.2771301269531 0 1755.755
508.276611328125 0 23386.28
509.2793273925781 0 6977.064
510.283447265625 0 1331.6396
525.3034057617188 0 1022.49567
526.2865600585938 0 5623.3623
527.289306640625 0 1944.2299
539.3187866210938 0 7189.879 y Water loss 3
540.3223266601562 0 1450.0264
542.329345703125 0 1494.7614
557.3287353515625 0 68643.664 y 3
558.3319702148438 0 18587.305
559.3348999023438 0 3143.1785
619.3587036132812 0 1870.2268
620.36279296875 0 904.65765
636.3712158203125 0 5781.382 y Water loss 2
637.3721313476562 0 2105.0815
654.3814086914062 0 265533.03 y 2
655.3842163085938 0 93002.4
656.38720703125 0 21134.977
657.3892822265625 0 1146.7649
752.4270629882812 0 1737.0032
753.43505859375 0 744.24457
802.9746704101562 0 615.4743
1802.4609375 0 637.2411
1963.757080078125 0 663.66376
3079.86865234375 0 754.32574

Spectrum Details

|  |  |
| --- | --- |
| Matched peaks? Matched peaks The total absolute number of peaks matched. Additionally in brackets the total fraction of peaks matched and the total number of peaks is shown. | 24 (12.50% of 192) |
| FDR? FDR The false discovery rate estimated for this peptide. It is calculated by matching all theoretical fragments with a non-integer shift with the raw peaks for this spectrum. This is done with 40 different shifts. The resulting percentage is the average number of annotated peaks over the number of annotated peaks with the correct spectrum. | 0.30% |
| Satellite FDR? Satellite FDR See the FDR for details on its calculation. This satellite ion specific FDR only contains the satellite ions (d/w) for I/L/J positions. | 0.00% |
| PSM Score? PSM Score The PSM Score as given by Hecklib to this annotated spectrum. It is shown with three significant figures. | 274 |

#### Spectrum 5744? Spectrum 5744 The raw spectrum of this peptide as annotated by Hecklib. The fragments are coloured according to ion type (see legend). Any peaks with a star '\*' as text can be hovered over to see the full details, first the ion type second the mass shift type. By hovering over the amino acids in the peptide or ions in the legend the corresponding peaks are highlighted. By toggling the 'Unassigned' label you can turn the background (unassigned) peaks on or off in the plot. By updating the slider in the Ion legend you can update the spectrum to only show the top X% of the peaks with labels. The top X% means any peak that is within X% of the highest intensity. By dragging in the spectrum you can zoom in to a specific part of the spectrum and use 'Zoom Out' to get back to the original zoom level. The annotation of the spectrum is based on the given sequence in the peptides file and is done with different software so inconsistencies are likely. The peaks are annotated based on the given sequence, with 20 ppm tolerance.

Copy Data

##### Spectrum 5744 (TSV)

###### Preview

```
Loading example...
```

*Click on the button to copy the data to your clipboard.*

Mz MinMz MaxIntensity Max

WidthHeightPeptide font sizePeptide stroke widthSpectrum font sizeSpectrum stroke widthCompact peptide

Ion legend

wxyz

abcd

OtherUnassignedIon^Charge^_Position_Show for top:%

AIPAPJEK

02.22e+44.43e+46.65e+48.86e+4

Zoom Out

d^+1^_2_y^+1^_1_y^+1^_1_a^+1^_2_b^+1^_2_y^+2^_4_y^+1^_2_y^+1^_2_b^+1^_3_y^+2^_6_y^+2^_7_y^+1^_3_y^+1^_4_y^+1^_4_y^+1^_5_y^+1^_5_y^+1^_6_y^+1^_6_

0665133019962661

Fragment Matches Table

Show background peaks

| Position | Ion type | Intensity | mz Theoretical | mz Error (Th) | mz Error (ppm) | Charge | Series Number |
| --- | --- | --- | --- | --- | --- | --- | --- |
| - | - | 707.2 | 120.1 | - | - | 0 | - |
| - | - | 5688 | 120.1 | - | - | 0 | - |
| - | - | 837 | 121 | - | - | 0 | - |
| - | - | 405.6 | 127.2 | - | - | 0 | - |
| - | - | 1001 | 129 | - | - | 0 | - |
| - | - | 734.4 | 129.1 | - | - | 0 | - |
| 2 | d | 1.249E+04 | 129.1 | 0.0002691 | 2.084 | +1 | 2 |
| - | - | 855.9 | 129.1 | - | - | 0 | - |
| 8 | y | 5221 | 130.1 | 0.0002928 | 2.251 | +1 | 1 |
| - | - | 460.4 | 130.1 | - | - | 0 | - |
| - | - | 796.9 | 131.1 | - | - | 0 | - |
| - | - | 1055 | 136.1 | - | - | 0 | - |
| - | - | 3.005E+04 | 141.1 | - | - | 0 | - |
| - | - | 2443 | 142.1 | - | - | 0 | - |
| 8 | y | 8839 | 147.1 | 0.0002787 | 1.895 | +1 | 1 |
| - | - | 474.3 | 148.1 | - | - | 0 | - |
| - | - | 8.776E+04 | 149 | - | - | 0 | - |
| - | - | 6986 | 150 | - | - | 0 | - |
| - | - | 1327 | 157 | - | - | 0 | - |
| - | - | 673.4 | 157.1 | - | - | 0 | - |
| 2 | a | 7.093E+04 | 157.1 | 0.0002647 | 1.685 | +1 | 2 |
| - | - | 5197 | 158.1 | - | - | 0 | - |
| - | - | 4850 | 167 | - | - | 0 | - |
| - | - | 2.742E+04 | 169.1 | - | - | 0 | - |
| - | - | 1956 | 170.1 | - | - | 0 | - |
| - | - | 678.4 | 173.1 | - | - | 0 | - |
| - | - | 517.7 | 179 | - | - | 0 | - |
| - | - | 2.84E+04 | 183.1 | - | - | 0 | - |
| - | - | 3281 | 184.2 | - | - | 0 | - |
| - | - | 1217 | 185.1 | - | - | 0 | - |
| 2 | b | 1.611E+04 | 185.1 | 0.0001926 | 1.041 | +1 | 2 |
| - | - | 1071 | 186.1 | - | - | 0 | - |
| - | - | 1408 | 191.1 | - | - | 0 | - |
| - | - | 1136 | 195.1 | - | - | 0 | - |
| - | - | 884.3 | 197.1 | - | - | 0 | - |
| - | - | 1.17E+04 | 211.1 | - | - | 0 | - |
| - | - | 1685 | 212.1 | - | - | 0 | - |
| - | - | 1338 | 215.1 | - | - | 0 | - |
| - | - | 1070 | 222.1 | - | - | 0 | - |
| - | - | 964.9 | 226.2 | - | - | 0 | - |
| - | - | 512.1 | 227.2 | - | - | 0 | - |
| - | - | 2655 | 233.2 | - | - | 0 | - |
| - | - | 801.1 | 237.1 | - | - | 0 | - |
| - | - | 6419 | 240.1 | - | - | 0 | - |
| - | - | 599.8 | 241.1 | - | - | 0 | - |
| - | - | 578.2 | 241.1 | - | - | 0 | - |
| - | - | 1818 | 243.1 | - | - | 0 | - |
| 5 | y | 4584 | 243.6 | 0.0002279 | 0.9352 | +2 | 4 |
| - | - | 837 | 244.2 | - | - | 0 | - |
| - | - | 661.4 | 253.1 | - | - | 0 | - |
| 7 | y | 1.958E+04 | 258.1 | 0.0001259 | 0.4878 | +1 | 2 |
| - | - | 2257 | 259.1 | - | - | 0 | - |
| - | - | 1015 | 262.1 | - | - | 0 | - |
| - | - | 1476 | 264.2 | - | - | 0 | - |
| - | - | 5598 | 266.2 | - | - | 0 | - |
| - | - | 616.2 | 267.2 | - | - | 0 | - |
| - | - | 572.3 | 270.2 | - | - | 0 | - |
| 7 | y | 1.85E+04 | 276.2 | 0.0001203 | 0.4357 | +1 | 2 |
| - | - | 2613 | 277.2 | - | - | 0 | - |
| 3 | b | 1823 | 282.2 | 0.0001784 | 0.6323 | +1 | 3 |
| - | - | 1290 | 293.2 | - | - | 0 | - |
| - | - | 1196 | 312.2 | - | - | 0 | - |
| - | - | 584.8 | 313.5 | - | - | 0 | - |
| - | - | 861.4 | 323.2 | - | - | 0 | - |
| 3 | y | 7.543E+04 | 327.7 | 0.0002872 | 0.8765 | +2 | 6 |
| - | - | 2.632E+04 | 328.2 | - | - | 0 | - |
| - | - | 6559 | 328.7 | - | - | 0 | - |
| - | - | 1.498E+04 | 340.2 | - | - | 0 | - |
| - | - | 3086 | 341.2 | - | - | 0 | - |
| - | - | 2586 | 351.2 | - | - | 0 | - |
| - | - | 1156 | 356.9 | - | - | 0 | - |
| - | - | 2823 | 358.2 | - | - | 0 | - |
| - | - | 5263 | 379.2 | - | - | 0 | - |
| - | - | 785.4 | 380.2 | - | - | 0 | - |
| - | - | 853.8 | 383.8 | - | - | 0 | - |
| 2 | y | 1127 | 384.2 | 0.001095 | 2.851 | +2 | 7 |
| - | - | 710.1 | 384.9 | - | - | 0 | - |
| 6 | y | 5406 | 389.2 | 0.0003154 | 0.8102 | +1 | 3 |
| - | - | 676.8 | 390.2 | - | - | 0 | - |
| - | - | 729.6 | 399.2 | - | - | 0 | - |
| - | - | 989.2 | 401.8 | - | - | 0 | - |
| - | - | 2518 | 411.2 | - | - | 0 | - |
| - | - | 725.1 | 418.2 | - | - | 0 | - |
| - | - | 2191 | 420.9 | - | - | 0 | - |
| - | - | 1117 | 424.3 | - | - | 0 | - |
| - | - | 716 | 429.2 | - | - | 0 | - |
| - | - | 2311 | 438.3 | - | - | 0 | - |
| - | - | 1093 | 438.8 | - | - | 0 | - |
| - | - | 610.1 | 444.2 | - | - | 0 | - |
| - | - | 690.5 | 462.3 | - | - | 0 | - |
| 5 | y | 2448 | 468.3 | 4.736E-05 | 0.1011 | +1 | 4 |
| - | - | 878.8 | 469.3 | - | - | 0 | - |
| - | - | 1348 | 480.3 | - | - | 0 | - |
| - | - | 622.5 | 480.8 | - | - | 0 | - |
| 5 | y | 5.828E+04 | 486.3 | 7.228E-05 | 0.1486 | +1 | 4 |
| - | - | 1.4E+04 | 487.3 | - | - | 0 | - |
| - | - | 2642 | 488.3 | - | - | 0 | - |
| - | - | 1379 | 497.8 | - | - | 0 | - |
| - | - | 1687 | 498.3 | - | - | 0 | - |
| - | - | 5939 | 508.3 | - | - | 0 | - |
| - | - | 1663 | 509.3 | - | - | 0 | - |
| - | - | 624.7 | 510.3 | - | - | 0 | - |
| - | - | 1271 | 526.3 | - | - | 0 | - |
| 4 | y | 1261 | 539.3 | 0.000989 | 1.834 | +1 | 5 |
| - | - | 802 | 553.3 | - | - | 0 | - |
| 4 | y | 1.361E+04 | 557.3 | 0.0004204 | 0.7543 | +1 | 5 |
| - | - | 3935 | 558.3 | - | - | 0 | - |
| - | - | 1328 | 559.3 | - | - | 0 | - |
| - | - | 534.7 | 560.6 | - | - | 0 | - |
| - | - | 1368 | 620.3 | - | - | 0 | - |
| - | - | 527.1 | 622.8 | - | - | 0 | - |
| 3 | y | 1216 | 636.4 | 0.002607 | 4.097 | +1 | 6 |
| 3 | y | 5.74E+04 | 654.4 | 0.0005109 | 0.7808 | +1 | 6 |
| - | - | 2.153E+04 | 655.4 | - | - | 0 | - |
| - | - | 4895 | 656.4 | - | - | 0 | - |
| - | - | 922.9 | 762.4 | - | - | 0 | - |
| - | - | 635.4 | 2635 | - | - | 0 | - |

m/z Charge Intensity FragmentType MassShift Position
120.06597900390625 0 707.1878
120.08100891113281 0 5687.572
121.02875518798828 0 836.95483
127.2130126953125 0 405.62842
129.01846313476562 0 1001.1167
129.0660400390625 0 734.36597
129.10250854492188 0 12492.603 d 1
129.10765075683594 0 855.9031
130.0865478515625 0 5221.094 y Ammonia loss 7
130.10589599609375 0 460.44772
131.0901336669922 0 796.92126
136.07606506347656 0 1054.9834
141.10252380371094 0 30052.93
142.1058807373047 0 2442.993
147.1130828857422 0 8838.596 y 7
148.1162109375 0 474.25186
149.0236053466797 0 87757.92
150.02696228027344 0 6986.4
157.01329040527344 0 1326.7826
157.09735107421875 0 673.3988
157.13380432128906 0 70934.48 a 1
158.13714599609375 0 5197.363
167.03404235839844 0 4850.1143
169.097412109375 0 27418.455
170.1007537841797 0 1956.1403
173.12806701660156 0 678.4169
179.00314331054688 0 517.7438
183.1494598388672 0 28401.375
184.1527862548828 0 3280.6619
185.08079528808594 0 1216.9672
185.12864685058594 0 16105.492 b 1
186.13241577148438 0 1071.4131
191.11825561523438 0 1407.5448
195.1128692626953 0 1136.3799
197.091552734375 0 884.2697
211.14431762695312 0 11700.899
212.14791870117188 0 1685.1538
215.1387939453125 0 1338.1495
222.12301635742188 0 1070.306
226.15513610839844 0 964.893
227.15907287597656 0 512.07153
233.1649932861328 0 2655.0674
237.0913848876953 0 801.0634
240.1344451904297 0 6418.8096
241.11672973632812 0 599.8309
241.13795471191406 0 578.2194
243.13426208496094 0 1818.4163
243.6499786376953 0 4583.9717 y 4
244.1515655517578 0 837.0053
253.11880493164062 0 661.3784
258.14495849609375 0 19579.373 y Water loss 6
259.14886474609375 0 2257.122
262.0504150390625 0 1014.79095
264.1705627441406 0 1476.3984
266.1500244140625 0 5598.4214
267.152587890625 0 616.1905
270.15576171875 0 572.2699
276.155517578125 0 18497.473 y 6
277.1590270996094 0 2613.3748
282.181396484375 0 1823.1044 b 2
293.1746520996094 0 1289.742
312.19219970703125 0 1196.2218
313.4596252441406 0 584.7908
323.208251953125 0 861.4121
327.6949768066406 0 75434.93 y 2
328.196533203125 0 26317.617
328.69793701171875 0 6558.603
340.1869201660156 0 14982.3545
341.1903381347656 0 3085.9524
351.2391052246094 0 2585.9648
356.8935241699219 0 1156.115
358.1973876953125 0 2823.2722
379.2339782714844 0 5262.512
380.2377624511719 0 785.37463
383.80694580078125 0 853.7553
384.2356262207031 0 1127.138 y 1
384.88604736328125 0 710.12616
389.2397766113281 0 5406.1284 y 5
390.2433776855469 0 676.79016
399.2242736816406 0 729.55634
401.8142395019531 0 989.1587
411.2237548828125 0 2518.295
418.2322998046875 0 725.0685
420.8577575683594 0 2190.6736
424.27484130859375 0 1117.2625
429.2332763671875 0 716.0004
438.27166748046875 0 2310.5603
438.77386474609375 0 1093.333
444.21923828125 0 610.1113
462.2743225097656 0 690.4766
468.2817077636719 0 2448.375 y Water loss 4
469.2851257324219 0 878.8477
480.2820129394531 0 1347.7695
480.7505187988281 0 622.54205
486.29229736328125 0 58276.64 y 4
487.29522705078125 0 13997.009
488.2983703613281 0 2642.3845
497.8016357421875 0 1379.4666
498.3028564453125 0 1687.325
508.2766418457031 0 5939.1216
509.2803955078125 0 1663.087
510.2957458496094 0 624.7131
526.2879028320312 0 1270.8545
539.3197631835938 0 1260.9619 y Water loss 3
553.3377075195312 0 801.9975
557.3289184570312 0 13611.425 y 3
558.3322143554688 0 3934.637
559.3333129882812 0 1328.1696
560.561279296875 0 534.74493
620.276123046875 0 1368.0796
622.7647705078125 0 527.09644
636.3741455078125 0 1215.9436 y Water loss 2
654.381591796875 0 57403.168 y 2
655.3843383789062 0 21534.863
656.3873901367188 0 4895.266
762.449462890625 0 922.93634
2634.529541015625 0 635.38696

Spectrum Details

|  |  |
| --- | --- |
| Matched peaks? Matched peaks The total absolute number of peaks matched. Additionally in brackets the total fraction of peaks matched and the total number of peaks is shown. | 18 (15.38% of 117) |
| FDR? FDR The false discovery rate estimated for this peptide. It is calculated by matching all theoretical fragments with a non-integer shift with the raw peaks for this spectrum. This is done with 40 different shifts. The resulting percentage is the average number of annotated peaks over the number of annotated peaks with the correct spectrum. | 0.53% |
| Satellite FDR? Satellite FDR See the FDR for details on its calculation. This satellite ion specific FDR only contains the satellite ions (d/w) for I/L/J positions. | 0.00% |
| PSM Score? PSM Score The PSM Score as given by Hecklib to this annotated spectrum. It is shown with three significant figures. | 194 |

#### Spectrum 5570? Spectrum 5570 The raw spectrum of this peptide as annotated by Hecklib. The fragments are coloured according to ion type (see legend). Any peaks with a star '\*' as text can be hovered over to see the full details, first the ion type second the mass shift type. By hovering over the amino acids in the peptide or ions in the legend the corresponding peaks are highlighted. By toggling the 'Unassigned' label you can turn the background (unassigned) peaks on or off in the plot. By updating the slider in the Ion legend you can update the spectrum to only show the top X% of the peaks with labels. The top X% means any peak that is within X% of the highest intensity. By dragging in the spectrum you can zoom in to a specific part of the spectrum and use 'Zoom Out' to get back to the original zoom level. The annotation of the spectrum is based on the given sequence in the peptides file and is done with different software so inconsistencies are likely. The peaks are annotated based on the given sequence, with 20 ppm tolerance.

Copy Data

##### Spectrum 5570 (TSV)

###### Preview

```
Loading example...
```

*Click on the button to copy the data to your clipboard.*

Mz MinMz MaxIntensity Max

WidthHeightPeptide font sizePeptide stroke widthSpectrum font sizeSpectrum stroke widthCompact peptide

Ion legend

wxyz

abcd

OtherUnassignedIon^Charge^_Position_Show for top:%

AIPAPJEK

02.34e+44.67e+47.01e+49.34e+4

Zoom Out

d^+1^_2_y^+1^_1_y^+1^_1_a^+1^_2_b^+1^_2_y^+2^_4_y^+1^_2_y^+1^_2_y^+1^_2_b^+1^_3_y^+2^_6_y^+1^_3_y^+2^_7_y^+1^_3_y^+1^_4_y^+1^_4_y^+1^_5_y^+1^_5_y^+1^_6_y^+1^_6_

0755151022653020

Fragment Matches Table

Show background peaks

| Position | Ion type | Intensity | mz Theoretical | mz Error (Th) | mz Error (ppm) | Charge | Series Number |
| --- | --- | --- | --- | --- | --- | --- | --- |
| - | - | 993.4 | 120.1 | - | - | 0 | - |
| - | - | 7540 | 120.1 | - | - | 0 | - |
| - | - | 1100 | 121.1 | - | - | 0 | - |
| - | - | 407.1 | 127 | - | - | 0 | - |
| - | - | 1938 | 129 | - | - | 0 | - |
| 2 | d | 1.37E+04 | 129.1 | 0.0002691 | 2.084 | +1 | 2 |
| 8 | y | 6547 | 130.1 | 0.0002775 | 2.133 | +1 | 1 |
| - | - | 1521 | 130.1 | - | - | 0 | - |
| - | - | 389.7 | 131.6 | - | - | 0 | - |
| - | - | 1324 | 133.1 | - | - | 0 | - |
| - | - | 730.8 | 134.1 | - | - | 0 | - |
| - | - | 854.9 | 136.1 | - | - | 0 | - |
| - | - | 3.339E+04 | 141.1 | - | - | 0 | - |
| - | - | 709.5 | 142.1 | - | - | 0 | - |
| - | - | 2581 | 142.1 | - | - | 0 | - |
| - | - | 1395 | 145.1 | - | - | 0 | - |
| - | - | 1650 | 147 | - | - | 0 | - |
| 8 | y | 1.023E+04 | 147.1 | 0.0002635 | 1.791 | +1 | 1 |
| - | - | 604.9 | 148.1 | - | - | 0 | - |
| - | - | 654.5 | 148.9 | - | - | 0 | - |
| - | - | 8.088E+04 | 149 | - | - | 0 | - |
| - | - | 6423 | 150 | - | - | 0 | - |
| - | - | 662.1 | 157 | - | - | 0 | - |
| - | - | 1200 | 157.1 | - | - | 0 | - |
| 2 | a | 8.531E+04 | 157.1 | 0.00028 | 1.782 | +1 | 2 |
| - | - | 506.8 | 158.1 | - | - | 0 | - |
| - | - | 7730 | 158.1 | - | - | 0 | - |
| - | - | 1338 | 159.1 | - | - | 0 | - |
| - | - | 460.6 | 159.6 | - | - | 0 | - |
| - | - | 3688 | 167 | - | - | 0 | - |
| - | - | 3.405E+04 | 169.1 | - | - | 0 | - |
| - | - | 2861 | 170.1 | - | - | 0 | - |
| - | - | 647.1 | 175.1 | - | - | 0 | - |
| - | - | 463.6 | 178.2 | - | - | 0 | - |
| - | - | 2.956E+04 | 183.1 | - | - | 0 | - |
| - | - | 2711 | 184.2 | - | - | 0 | - |
| - | - | 1433 | 185.1 | - | - | 0 | - |
| 2 | b | 2.097E+04 | 185.1 | 0.0002384 | 1.288 | +1 | 2 |
| - | - | 1735 | 186.1 | - | - | 0 | - |
| - | - | 3255 | 191.1 | - | - | 0 | - |
| - | - | 416.9 | 193.8 | - | - | 0 | - |
| - | - | 900.3 | 195.1 | - | - | 0 | - |
| - | - | 628.5 | 197.1 | - | - | 0 | - |
| - | - | 487.5 | 204.6 | - | - | 0 | - |
| - | - | 523.4 | 209.5 | - | - | 0 | - |
| - | - | 1.663E+04 | 211.1 | - | - | 0 | - |
| - | - | 1374 | 212.1 | - | - | 0 | - |
| - | - | 563.7 | 213.1 | - | - | 0 | - |
| - | - | 470.1 | 214.9 | - | - | 0 | - |
| - | - | 1757 | 215.1 | - | - | 0 | - |
| - | - | 597.9 | 219.1 | - | - | 0 | - |
| - | - | 467.4 | 219.4 | - | - | 0 | - |
| - | - | 1622 | 222.1 | - | - | 0 | - |
| - | - | 488.4 | 223.9 | - | - | 0 | - |
| - | - | 532.3 | 225.1 | - | - | 0 | - |
| - | - | 1282 | 226.2 | - | - | 0 | - |
| - | - | 2364 | 233.2 | - | - | 0 | - |
| - | - | 848.4 | 237.1 | - | - | 0 | - |
| - | - | 8107 | 240.1 | - | - | 0 | - |
| - | - | 994.7 | 241.1 | - | - | 0 | - |
| - | - | 753.5 | 241.1 | - | - | 0 | - |
| - | - | 2511 | 243.1 | - | - | 0 | - |
| 5 | y | 6467 | 243.6 | 0.0003042 | 1.248 | +2 | 4 |
| - | - | 1995 | 244.2 | - | - | 0 | - |
| - | - | 580.7 | 244.2 | - | - | 0 | - |
| 7 | y | 2.324E+04 | 258.1 | 9.541E-05 | 0.3696 | +1 | 2 |
| 7 | y | 561.6 | 259.1 | 0.001706 | 6.584 | +1 | 2 |
| - | - | 2583 | 259.1 | - | - | 0 | - |
| - | - | 583.8 | 263.1 | - | - | 0 | - |
| - | - | 2155 | 264.2 | - | - | 0 | - |
| - | - | 6052 | 266.2 | - | - | 0 | - |
| - | - | 531.2 | 267 | - | - | 0 | - |
| - | - | 822.5 | 267.2 | - | - | 0 | - |
| - | - | 628.1 | 269 | - | - | 0 | - |
| - | - | 809 | 273.1 | - | - | 0 | - |
| 7 | y | 1.966E+04 | 276.2 | 8.981E-05 | 0.3252 | +1 | 2 |
| - | - | 1990 | 277.2 | - | - | 0 | - |
| 3 | b | 1844 | 282.2 | 4.695E-06 | 0.01664 | +1 | 3 |
| - | - | 484.6 | 286.2 | - | - | 0 | - |
| - | - | 1108 | 293.1 | - | - | 0 | - |
| - | - | 1255 | 293.2 | - | - | 0 | - |
| - | - | 1274 | 312.2 | - | - | 0 | - |
| - | - | 945.9 | 322.2 | - | - | 0 | - |
| - | - | 794 | 323.2 | - | - | 0 | - |
| 3 | y | 9.252E+04 | 327.7 | 0.0003177 | 0.9696 | +2 | 6 |
| - | - | 3.059E+04 | 328.2 | - | - | 0 | - |
| - | - | 6040 | 328.7 | - | - | 0 | - |
| - | - | 1.711E+04 | 340.2 | - | - | 0 | - |
| - | - | 3013 | 341.2 | - | - | 0 | - |
| - | - | 583.9 | 342.2 | - | - | 0 | - |
| - | - | 3636 | 351.2 | - | - | 0 | - |
| - | - | 731.6 | 352.2 | - | - | 0 | - |
| - | - | 1431 | 356.9 | - | - | 0 | - |
| - | - | 3642 | 358.2 | - | - | 0 | - |
| - | - | 752.1 | 359 | - | - | 0 | - |
| - | - | 1172 | 365.2 | - | - | 0 | - |
| 6 | y | 644.9 | 371.2 | 4.523E-05 | 0.1218 | +1 | 3 |
| - | - | 1395 | 373.2 | - | - | 0 | - |
| - | - | 5307 | 379.2 | - | - | 0 | - |
| 2 | y | 1206 | 384.2 | 0.0006441 | 1.676 | +2 | 7 |
| - | - | 602.7 | 387.8 | - | - | 0 | - |
| 6 | y | 6111 | 389.2 | 0.0001933 | 0.4966 | +1 | 3 |
| - | - | 634.8 | 389.9 | - | - | 0 | - |
| - | - | 1409 | 390.2 | - | - | 0 | - |
| - | - | 1080 | 399.2 | - | - | 0 | - |
| - | - | 611.5 | 402.8 | - | - | 0 | - |
| - | - | 1351 | 403 | - | - | 0 | - |
| - | - | 2809 | 411.2 | - | - | 0 | - |
| - | - | 739.7 | 412.2 | - | - | 0 | - |
| - | - | 934.9 | 418.9 | - | - | 0 | - |
| - | - | 1840 | 420.9 | - | - | 0 | - |
| - | - | 4065 | 421.2 | - | - | 0 | - |
| - | - | 784.4 | 429.2 | - | - | 0 | - |
| - | - | 1136 | 438.3 | - | - | 0 | - |
| - | - | 950.5 | 438.8 | - | - | 0 | - |
| 5 | y | 3387 | 468.3 | 0.0001084 | 0.2315 | +1 | 4 |
| - | - | 1753 | 480.3 | - | - | 0 | - |
| 5 | y | 7.458E+04 | 486.3 | 4.176E-05 | 0.08587 | +1 | 4 |
| - | - | 1.797E+04 | 487.3 | - | - | 0 | - |
| - | - | 3452 | 488.3 | - | - | 0 | - |
| - | - | 961.9 | 497.8 | - | - | 0 | - |
| - | - | 692.4 | 498.3 | - | - | 0 | - |
| - | - | 622.7 | 498.8 | - | - | 0 | - |
| - | - | 1567 | 505.3 | - | - | 0 | - |
| - | - | 6033 | 508.3 | - | - | 0 | - |
| - | - | 2094 | 509.3 | - | - | 0 | - |
| - | - | 1987 | 526.3 | - | - | 0 | - |
| 4 | y | 1877 | 539.3 | 0.0005007 | 0.9284 | +1 | 5 |
| 4 | y | 1.599E+04 | 557.3 | 0.0005425 | 0.9734 | +1 | 5 |
| - | - | 5766 | 558.3 | - | - | 0 | - |
| - | - | 1306 | 559.3 | - | - | 0 | - |
| - | - | 566.5 | 587.3 | - | - | 0 | - |
| - | - | 3975 | 620.3 | - | - | 0 | - |
| - | - | 929.7 | 621.3 | - | - | 0 | - |
| 3 | y | 1355 | 636.4 | 0.0008715 | 1.37 | +1 | 6 |
| 3 | y | 7.66E+04 | 654.4 | 0.000633 | 0.9673 | +1 | 6 |
| - | - | 2.685E+04 | 655.4 | - | - | 0 | - |
| - | - | 6058 | 656.4 | - | - | 0 | - |
| - | - | 696 | 875.5 | - | - | 0 | - |
| - | - | 585.8 | 1017 | - | - | 0 | - |
| - | - | 756.1 | 2369 | - | - | 0 | - |
| - | - | 579.8 | 2990 | - | - | 0 | - |

m/z Charge Intensity FragmentType MassShift Position
120.06597137451172 0 993.3826
120.08104705810547 0 7539.596
121.0843276977539 0 1099.5955
126.98592376708984 0 407.05762
129.01853942871094 0 1937.9906
129.10250854492188 0 13703.037 d 1
130.08653259277344 0 6546.7812 y Ammonia loss 7
130.1059112548828 0 1520.9155
131.59466552734375 0 389.69968
133.08619689941406 0 1324.1892
134.08935546875 0 730.84753
136.0759735107422 0 854.928
141.10252380371094 0 33386.758
142.0999298095703 0 709.4884
142.1058349609375 0 2580.7979
145.06842041015625 0 1394.7842
147.04759216308594 0 1649.9984
147.11306762695312 0 10231.359 y 7
148.11679077148438 0 604.91473
148.94683837890625 0 654.531
149.02362060546875 0 80875.62
150.02687072753906 0 6423.2925
157.01370239257812 0 662.11414
157.0973358154297 0 1199.7633
157.13381958007812 0 85309.914 a 1
158.13047790527344 0 506.82935
158.1371612548828 0 7730.373
159.09190368652344 0 1338.264
159.64451599121094 0 460.5996
167.0341033935547 0 3687.7095
169.09739685058594 0 34047.625
170.10073852539062 0 2860.945
175.11929321289062 0 647.12103
178.24485778808594 0 463.62274
183.14944458007812 0 29562.623
184.15298461914062 0 2710.6367
185.08106994628906 0 1433.0371
185.12869262695312 0 20967.984 b 1
186.13211059570312 0 1735.4712
191.11814880371094 0 3255.162
193.83786010742188 0 416.91708
195.1125946044922 0 900.30115
197.09274291992188 0 628.50055
204.60479736328125 0 487.5202
209.484375 0 523.3546
211.14434814453125 0 16632.008
212.14772033691406 0 1373.7217
213.1232452392578 0 563.6612
214.87158203125 0 470.133
215.13929748535156 0 1757.3473
219.0814208984375 0 597.9325
219.36734008789062 0 467.36533
222.1238250732422 0 1621.8007
223.9439239501953 0 488.39603
225.12310791015625 0 532.2938
226.15538024902344 0 1281.9329
233.16485595703125 0 2364.0798
237.09080505371094 0 848.35913
240.13446044921875 0 8107.4805
241.1187744140625 0 994.69586
241.1378936767578 0 753.47546
243.1342315673828 0 2511.2131
243.65005493164062 0 6466.841 y 4
244.15147399902344 0 1995.2688
244.1647186279297 0 580.6583
258.1449279785156 0 23236.926 y Water loss 6
259.13055419921875 0 561.58997 y Ammonia loss 6
259.1487121582031 0 2583.235
263.1039733886719 0 583.8466
264.17108154296875 0 2154.9292
266.1500244140625 0 6051.985
267.0320129394531 0 531.1508
267.15289306640625 0 822.49146
269.0086364746094 0 628.14667
273.0931396484375 0 809.01776
276.1554870605469 0 19659.562 y 6
277.1585388183594 0 1989.5713
282.18121337890625 0 1844.4404 b 2
286.2314453125 0 484.5521
293.0997009277344 0 1108.4498
293.17486572265625 0 1255.1759
312.192138671875 0 1274.3422
322.1759033203125 0 945.94635
323.2081604003906 0 794.0036
327.69500732421875 0 92521.5 y 2
328.1963806152344 0 30592.615
328.6978759765625 0 6040.388
340.18695068359375 0 17109.4
341.1899108886719 0 3012.6023
342.1942138671875 0 583.8752
351.2388916015625 0 3636.3752
352.24176025390625 0 731.6144
356.8923034667969 0 1431.4413
358.19708251953125 0 3642.2546
359.0147399902344 0 752.1168
365.20013427734375 0 1172.3384
371.2288513183594 0 644.9355 y Water loss 5
373.23175048828125 0 1395.149
379.23388671875 0 5307.4927
384.23736572265625 0 1206.0946 y 1
387.79071044921875 0 602.7278
389.2396545410156 0 6110.9067 y 5
389.8570861816406 0 634.82385
390.2421875 0 1408.5654
399.2229919433594 0 1079.9327
402.8219299316406 0 611.4641
403.0039978027344 0 1350.7031
411.2245788574219 0 2808.7336
412.2254943847656 0 739.70593
418.8631591796875 0 934.88336
420.8583068847656 0 1840.3726
421.2043151855469 0 4064.5535
429.23455810546875 0 784.4142
438.2715148925781 0 1135.5599
438.76873779296875 0 950.5129
468.2817687988281 0 3386.9785 y Water loss 4
480.2804870605469 0 1753.4705
486.2922668457031 0 74578.12 y 4
487.29522705078125 0 17967.504
488.2975769042969 0 3451.9365
497.8013916015625 0 961.8799
498.3016052246094 0 692.40643
498.806396484375 0 622.69336
505.2525329589844 0 1567.3441
508.2764587402344 0 6033.213
509.2789611816406 0 2094.131
526.28564453125 0 1986.5029
539.3192749023438 0 1876.9272 y Water loss 3
557.3287963867188 0 15994.378 y 3
558.3322143554688 0 5765.877
559.3336791992188 0 1305.8943
587.2801513671875 0 566.5424
620.2781982421875 0 3975.0022
621.281982421875 0 929.696
636.3706665039062 0 1354.5216 y Water loss 2
654.3814697265625 0 76602.664 y 2
655.3845825195312 0 26848.66
656.3870239257812 0 6057.872
875.5339965820312 0 695.98584
1016.778564453125 0 585.8379
2369.11181640625 0 756.108
2990.08056640625 0 579.81494

Spectrum Details

|  |  |
| --- | --- |
| Matched peaks? Matched peaks The total absolute number of peaks matched. Additionally in brackets the total fraction of peaks matched and the total number of peaks is shown. | 20 (14.08% of 142) |
| FDR? FDR The false discovery rate estimated for this peptide. It is calculated by matching all theoretical fragments with a non-integer shift with the raw peaks for this spectrum. This is done with 40 different shifts. The resulting percentage is the average number of annotated peaks over the number of annotated peaks with the correct spectrum. | 0.36% |
| Satellite FDR? Satellite FDR See the FDR for details on its calculation. This satellite ion specific FDR only contains the satellite ions (d/w) for I/L/J positions. | 0.00% |
| PSM Score? PSM Score The PSM Score as given by Hecklib to this annotated spectrum. It is shown with three significant figures. | 233 |

#### Spectrum 6013? Spectrum 6013 The raw spectrum of this peptide as annotated by Hecklib. The fragments are coloured according to ion type (see legend). Any peaks with a star '\*' as text can be hovered over to see the full details, first the ion type second the mass shift type. By hovering over the amino acids in the peptide or ions in the legend the corresponding peaks are highlighted. By toggling the 'Unassigned' label you can turn the background (unassigned) peaks on or off in the plot. By updating the slider in the Ion legend you can update the spectrum to only show the top X% of the peaks with labels. The top X% means any peak that is within X% of the highest intensity. By dragging in the spectrum you can zoom in to a specific part of the spectrum and use 'Zoom Out' to get back to the original zoom level. The annotation of the spectrum is based on the given sequence in the peptides file and is done with different software so inconsistencies are likely. The peaks are annotated based on the given sequence, with 20 ppm tolerance.

Copy Data

##### Spectrum 6013 (TSV)

###### Preview

```
Loading example...
```

*Click on the button to copy the data to your clipboard.*

Mz MinMz MaxIntensity Max

WidthHeightPeptide font sizePeptide stroke widthSpectrum font sizeSpectrum stroke widthCompact peptide

Ion legend

wxyz

abcd

OtherUnassignedIon^Charge^_Position_Show for top:%

AIPAPJEK

02.24e+44.49e+46.73e+48.98e+4

Zoom Out

d^+1^_2_y^+1^_1_y^+1^_1_a^+1^_2_b^+1^_2_y^+2^_4_y^+1^_2_y^+1^_2_y^+1^_2_b^+1^_3_y^+2^_6_y^+2^_7_y^+1^_3_y^+1^_4_y^+1^_4_y^+1^_5_y^+1^_5_y^+1^_6_y^+1^_6_

0864172825923455

Fragment Matches Table

Show background peaks

| Position | Ion type | Intensity | mz Theoretical | mz Error (Th) | mz Error (ppm) | Charge | Series Number |
| --- | --- | --- | --- | --- | --- | --- | --- |
| - | - | 4622 | 120.1 | - | - | 0 | - |
| - | - | 1.179E+04 | 120.1 | - | - | 0 | - |
| - | - | 934 | 121 | - | - | 0 | - |
| - | - | 572.1 | 121.1 | - | - | 0 | - |
| - | - | 986.3 | 126.1 | - | - | 0 | - |
| - | - | 534.2 | 127.1 | - | - | 0 | - |
| - | - | 1144 | 129 | - | - | 0 | - |
| 2 | d | 1.182E+04 | 129.1 | 0.0003301 | 2.557 | +1 | 2 |
| - | - | 863.1 | 129.1 | - | - | 0 | - |
| 8 | y | 3587 | 130.1 | 0.0003538 | 2.72 | +1 | 1 |
| - | - | 1130 | 130.1 | - | - | 0 | - |
| - | - | 470.4 | 135.9 | - | - | 0 | - |
| - | - | 1121 | 136.1 | - | - | 0 | - |
| - | - | 1143 | 141.1 | - | - | 0 | - |
| - | - | 2.04E+04 | 141.1 | - | - | 0 | - |
| - | - | 1404 | 142.1 | - | - | 0 | - |
| 8 | y | 6014 | 147.1 | 0.0003092 | 2.102 | +1 | 1 |
| - | - | 532.1 | 147.8 | - | - | 0 | - |
| - | - | 532.1 | 148.9 | - | - | 0 | - |
| - | - | 8.886E+04 | 149 | - | - | 0 | - |
| - | - | 7263 | 150 | - | - | 0 | - |
| - | - | 1027 | 157 | - | - | 0 | - |
| - | - | 710 | 157.1 | - | - | 0 | - |
| 2 | a | 5.213E+04 | 157.1 | 0.0003258 | 2.073 | +1 | 2 |
| - | - | 4684 | 158.1 | - | - | 0 | - |
| - | - | 3670 | 167 | - | - | 0 | - |
| - | - | 508.8 | 167.1 | - | - | 0 | - |
| - | - | 2.003E+04 | 169.1 | - | - | 0 | - |
| - | - | 1618 | 170.1 | - | - | 0 | - |
| - | - | 497.6 | 171.1 | - | - | 0 | - |
| - | - | 550.9 | 175.1 | - | - | 0 | - |
| - | - | 1.932E+04 | 183.1 | - | - | 0 | - |
| - | - | 1670 | 184.2 | - | - | 0 | - |
| - | - | 1211 | 185.1 | - | - | 0 | - |
| 2 | b | 1.181E+04 | 185.1 | 0.0002689 | 1.453 | +1 | 2 |
| - | - | 1227 | 186.1 | - | - | 0 | - |
| - | - | 894.6 | 191.1 | - | - | 0 | - |
| - | - | 1348 | 195.1 | - | - | 0 | - |
| - | - | 755.7 | 197.1 | - | - | 0 | - |
| - | - | 647 | 197.1 | - | - | 0 | - |
| - | - | 9516 | 211.1 | - | - | 0 | - |
| - | - | 644.6 | 212.1 | - | - | 0 | - |
| - | - | 577.8 | 212.2 | - | - | 0 | - |
| - | - | 494.1 | 212.3 | - | - | 0 | - |
| - | - | 885.8 | 215.1 | - | - | 0 | - |
| - | - | 577 | 217.1 | - | - | 0 | - |
| - | - | 1527 | 217.1 | - | - | 0 | - |
| - | - | 829.5 | 222.1 | - | - | 0 | - |
| - | - | 602.8 | 226.1 | - | - | 0 | - |
| - | - | 3933 | 226.2 | - | - | 0 | - |
| - | - | 572.3 | 227.2 | - | - | 0 | - |
| - | - | 7047 | 233.2 | - | - | 0 | - |
| - | - | 1619 | 234.2 | - | - | 0 | - |
| - | - | 3561 | 240.1 | - | - | 0 | - |
| - | - | 561.9 | 241.1 | - | - | 0 | - |
| - | - | 947.2 | 243.1 | - | - | 0 | - |
| 5 | y | 3238 | 243.6 | 0.0001058 | 0.4342 | +2 | 4 |
| - | - | 555.3 | 244.7 | - | - | 0 | - |
| - | - | 526.8 | 247.1 | - | - | 0 | - |
| - | - | 546 | 250.3 | - | - | 0 | - |
| 7 | y | 1.326E+04 | 258.1 | 0.0001564 | 0.606 | +1 | 2 |
| 7 | y | 636.1 | 259.1 | 0.0009126 | 3.522 | +1 | 2 |
| - | - | 1637 | 259.2 | - | - | 0 | - |
| - | - | 1715 | 261.2 | - | - | 0 | - |
| - | - | 527 | 262 | - | - | 0 | - |
| - | - | 647.7 | 264.2 | - | - | 0 | - |
| - | - | 4114 | 266.2 | - | - | 0 | - |
| 7 | y | 1.128E+04 | 276.2 | 0.0001508 | 0.5462 | +1 | 2 |
| - | - | 1517 | 277.2 | - | - | 0 | - |
| - | - | 563.9 | 277.2 | - | - | 0 | - |
| 3 | b | 1009 | 282.2 | 0.001033 | 3.66 | +1 | 3 |
| - | - | 543.6 | 292 | - | - | 0 | - |
| - | - | 1175 | 293.1 | - | - | 0 | - |
| - | - | 1257 | 293.2 | - | - | 0 | - |
| - | - | 596.4 | 303.9 | - | - | 0 | - |
| - | - | 946.3 | 307 | - | - | 0 | - |
| - | - | 1112 | 312.2 | - | - | 0 | - |
| - | - | 619.5 | 323.2 | - | - | 0 | - |
| - | - | 1591 | 323.2 | - | - | 0 | - |
| 3 | y | 5.311E+04 | 327.7 | 0.0004093 | 1.249 | +2 | 6 |
| - | - | 1.986E+04 | 328.2 | - | - | 0 | - |
| - | - | 3618 | 328.7 | - | - | 0 | - |
| - | - | 939.2 | 340.2 | - | - | 0 | - |
| - | - | 1.017E+04 | 340.2 | - | - | 0 | - |
| - | - | 2205 | 341.2 | - | - | 0 | - |
| - | - | 2463 | 351.2 | - | - | 0 | - |
| - | - | 667.2 | 356.9 | - | - | 0 | - |
| - | - | 1420 | 358.1 | - | - | 0 | - |
| - | - | 2103 | 358.2 | - | - | 0 | - |
| - | - | 514.6 | 359.6 | - | - | 0 | - |
| - | - | 524.8 | 361.1 | - | - | 0 | - |
| - | - | 544.4 | 367.1 | - | - | 0 | - |
| - | - | 632.8 | 367.2 | - | - | 0 | - |
| - | - | 864.7 | 367.7 | - | - | 0 | - |
| - | - | 654.4 | 371.2 | - | - | 0 | - |
| - | - | 3598 | 379.2 | - | - | 0 | - |
| - | - | 747.7 | 380.2 | - | - | 0 | - |
| - | - | 1462 | 381.7 | - | - | 0 | - |
| - | - | 1221 | 382.2 | - | - | 0 | - |
| 2 | y | 961.1 | 384.2 | 0.0005221 | 1.359 | +2 | 7 |
| - | - | 973.6 | 388.7 | - | - | 0 | - |
| 6 | y | 4468 | 389.2 | 8.135E-05 | 0.209 | +1 | 3 |
| - | - | 1004 | 401.8 | - | - | 0 | - |
| - | - | 994.9 | 402.1 | - | - | 0 | - |
| - | - | 1977 | 411.2 | - | - | 0 | - |
| - | - | 642.6 | 418.9 | - | - | 0 | - |
| - | - | 878.9 | 419 | - | - | 0 | - |
| - | - | 808.1 | 420.2 | - | - | 0 | - |
| - | - | 1448 | 420.9 | - | - | 0 | - |
| - | - | 3267 | 424.3 | - | - | 0 | - |
| - | - | 1988 | 424.8 | - | - | 0 | - |
| - | - | 5832 | 438.3 | - | - | 0 | - |
| - | - | 2732 | 438.8 | - | - | 0 | - |
| - | - | 705 | 439.3 | - | - | 0 | - |
| - | - | 921.1 | 440.3 | - | - | 0 | - |
| 5 | y | 2320 | 468.3 | 0.001115 | 2.382 | +1 | 4 |
| - | - | 752 | 480.3 | - | - | 0 | - |
| 5 | y | 4.365E+04 | 486.3 | 0.0002249 | 0.4624 | +1 | 4 |
| - | - | 1.058E+04 | 487.3 | - | - | 0 | - |
| - | - | 2166 | 488.3 | - | - | 0 | - |
| - | - | 4049 | 497.8 | - | - | 0 | - |
| - | - | 3570 | 498.3 | - | - | 0 | - |
| - | - | 1117 | 498.8 | - | - | 0 | - |
| - | - | 4674 | 508.3 | - | - | 0 | - |
| - | - | 709.2 | 509.3 | - | - | 0 | - |
| - | - | 1287 | 525.3 | - | - | 0 | - |
| - | - | 848.8 | 526.3 | - | - | 0 | - |
| 4 | y | 1198 | 539.3 | 0.001269 | 2.354 | +1 | 5 |
| - | - | 2418 | 553.3 | - | - | 0 | - |
| 4 | y | 1.048E+04 | 557.3 | 0.0001152 | 0.2068 | +1 | 5 |
| - | - | 2864 | 558.3 | - | - | 0 | - |
| - | - | 723.1 | 620.3 | - | - | 0 | - |
| 3 | y | 1192 | 636.4 | 0.002153 | 3.384 | +1 | 6 |
| 3 | y | 4.304E+04 | 654.4 | 0.0003889 | 0.5942 | +1 | 6 |
| - | - | 1.418E+04 | 655.4 | - | - | 0 | - |
| - | - | 2489 | 656.4 | - | - | 0 | - |
| - | - | 646.7 | 664.4 | - | - | 0 | - |
| - | - | 1190 | 665.4 | - | - | 0 | - |
| - | - | 714.9 | 666.4 | - | - | 0 | - |
| - | - | 1249 | 672.4 | - | - | 0 | - |
| - | - | 697.7 | 673.4 | - | - | 0 | - |
| - | - | 600.9 | 750.5 | - | - | 0 | - |
| - | - | 2067 | 762.5 | - | - | 0 | - |
| - | - | 1066 | 763.5 | - | - | 0 | - |
| - | - | 2470 | 778.5 | - | - | 0 | - |
| - | - | 1561 | 779.5 | - | - | 0 | - |
| - | - | 1468 | 875.5 | - | - | 0 | - |
| - | - | 943.7 | 897.5 | - | - | 0 | - |
| - | - | 788.4 | 898.5 | - | - | 0 | - |
| - | - | 685.4 | 1012 | - | - | 0 | - |
| - | - | 700.6 | 1099 | - | - | 0 | - |
| - | - | 715.6 | 1497 | - | - | 0 | - |
| - | - | 622.7 | 2508 | - | - | 0 | - |
| - | - | 820.6 | 2862 | - | - | 0 | - |
| - | - | 721.9 | 3421 | - | - | 0 | - |

m/z Charge Intensity FragmentType MassShift Position
120.06586456298828 0 4622.4517
120.08110046386719 0 11794.998
121.02870178222656 0 934.03375
121.0844497680664 0 572.1366
126.05532836914062 0 986.28644
127.08692169189453 0 534.1868
129.01876831054688 0 1144.3228
129.10256958007812 0 11817.063 d 1
129.10765075683594 0 863.1496
130.08660888671875 0 3586.9434 y Ammonia loss 7
130.1057586669922 0 1129.5299
135.90968322753906 0 470.37457
136.0759735107422 0 1121.1239
141.09628295898438 0 1143.314
141.10256958007812 0 20395.752
142.10614013671875 0 1403.621
147.1131134033203 0 6014.2617 y 7
147.83892822265625 0 532.14343
148.94677734375 0 532.0929
149.02369689941406 0 88863.54
150.02699279785156 0 7263.2803
157.0135955810547 0 1027.4656
157.09780883789062 0 710.024
157.1338653564453 0 52133.016 a 1
158.1372833251953 0 4683.617
167.03416442871094 0 3670.3884
167.1182861328125 0 508.79407
169.0974578857422 0 20034.576
170.10092163085938 0 1618.4198
171.1385955810547 0 497.6476
175.11953735351562 0 550.89
183.14950561523438 0 19320.322
184.1531524658203 0 1670.4381
185.08140563964844 0 1211.2527
185.12872314453125 0 11805.785 b 1
186.13204956054688 0 1227.4706
191.1179656982422 0 894.5944
195.1128387451172 0 1347.7207
197.09286499023438 0 755.67523
197.12799072265625 0 647.0276
211.1444549560547 0 9516.44
212.14764404296875 0 644.591
212.2455596923828 0 577.785
212.25660705566406 0 494.06494
215.13894653320312 0 885.7701
217.09756469726562 0 577.03986
217.13356018066406 0 1526.8445
222.12454223632812 0 829.45953
226.11892700195312 0 602.7742
226.15521240234375 0 3933.4663
227.1598358154297 0 572.3442
233.165283203125 0 7046.9644
234.16830444335938 0 1618.9551
240.13453674316406 0 3560.716
241.11878967285156 0 561.94904
243.13441467285156 0 947.17163
243.6498565673828 0 3238.1243 y 4
244.65264892578125 0 555.3436
247.09063720703125 0 526.7832
250.33877563476562 0 546.02905
258.1449890136719 0 13262.204 y Water loss 6
259.1297607421875 0 636.1287 y Ammonia loss 6
259.1500549316406 0 1637.1934
261.1600036621094 0 1715.3196
262.04986572265625 0 526.95416
264.17095947265625 0 647.70526
266.150146484375 0 4113.667
276.1555480957031 0 11280.408 y 6
277.158447265625 0 1517.0566
277.2027282714844 0 563.9133
282.1822509765625 0 1008.884 b 2
292.0409851074219 0 543.58203
293.1007995605469 0 1175.0211
293.17498779296875 0 1257.2401
303.90130615234375 0 596.371
306.9866027832031 0 946.2805
312.1922302246094 0 1111.7158
323.1723327636719 0 619.51917
323.20831298828125 0 1590.8121
327.6950988769531 0 53114.824 y 2
328.1964416503906 0 19857.004
328.6979064941406 0 3618.2537
340.1709289550781 0 939.21185
340.18719482421875 0 10170.759
341.1859436035156 0 2204.9114
351.23907470703125 0 2462.5615
356.8927001953125 0 667.21014
358.0821228027344 0 1419.7859
358.1966247558594 0 2103.4387
359.6048889160156 0 514.5867
361.1180725097656 0 524.8109
367.13775634765625 0 544.38153
367.2331848144531 0 632.79285
367.73028564453125 0 864.69055
371.1910705566406 0 654.3927
379.2336120605469 0 3598.4639
380.2373352050781 0 747.74634
381.7298889160156 0 1461.7246
382.2317199707031 0 1220.5065
384.23724365234375 0 961.09515 y 1
388.7353515625 0 973.5541
389.2393798828125 0 4467.527 y 5
401.8184814453125 0 1003.753
402.1109619140625 0 994.9377
411.2236633300781 0 1977.1298
418.8648376464844 0 642.6002
418.97442626953125 0 878.9111
420.22100830078125 0 808.10986
420.8582458496094 0 1448.3315
424.2745056152344 0 3266.9727
424.7755432128906 0 1987.5355
438.271484375 0 5832.433
438.7724609375 0 2732.0283
439.27294921875 0 704.96936
440.25189208984375 0 921.1447
468.28277587890625 0 2319.755 y Water loss 4
480.28216552734375 0 752.04974
486.2924499511719 0 43653.02 y 4
487.2953186035156 0 10576.911
488.29827880859375 0 2165.9653
497.80029296875 0 4049.123
498.3022766113281 0 3570.2646
498.8026428222656 0 1116.9059
508.27679443359375 0 4673.9697
509.281494140625 0 709.1773
525.3403930664062 0 1287.3501
526.285400390625 0 848.8322
539.3175048828125 0 1197.9945 y Water loss 3
553.3343505859375 0 2417.6904
557.3292236328125 0 10482.093 y 3
558.3319091796875 0 2864.006
620.2788696289062 0 723.05707
636.369384765625 0 1191.751 y Water loss 2
654.3817138671875 0 43035.867 y 2
655.3847045898438 0 14175.883
656.3887939453125 0 2489.2454
664.3876953125 0 646.72644
665.3993530273438 0 1189.9736
666.4011840820312 0 714.92413
672.3923950195312 0 1249.2579
673.3980712890625 0 697.7486
750.4855346679688 0 600.88916
762.4500122070312 0 2067.2915
763.450927734375 0 1066.3563
778.4797973632812 0 2469.515
779.4827880859375 0 1560.8484
875.5347900390625 0 1467.8087
897.5377807617188 0 943.68756
898.5413208007812 0 788.4178
1012.0990600585938 0 685.4134
1098.5126953125 0 700.63544
1496.6041259765625 0 715.56256
2508.080810546875 0 622.6603
2862.341552734375 0 820.6397
3421.214599609375 0 721.9106

Spectrum Details

|  |  |
| --- | --- |
| Matched peaks? Matched peaks The total absolute number of peaks matched. Additionally in brackets the total fraction of peaks matched and the total number of peaks is shown. | 19 (12.26% of 155) |
| FDR? FDR The false discovery rate estimated for this peptide. It is calculated by matching all theoretical fragments with a non-integer shift with the raw peaks for this spectrum. This is done with 40 different shifts. The resulting percentage is the average number of annotated peaks over the number of annotated peaks with the correct spectrum. | 0.88% |
| Satellite FDR? Satellite FDR See the FDR for details on its calculation. This satellite ion specific FDR only contains the satellite ions (d/w) for I/L/J positions. | 0.00% |
| PSM Score? PSM Score The PSM Score as given by Hecklib to this annotated spectrum. It is shown with three significant figures. | 213 |

#### Spectrum 4798? Spectrum 4798 The raw spectrum of this peptide as annotated by Hecklib. The fragments are coloured according to ion type (see legend). Any peaks with a star '\*' as text can be hovered over to see the full details, first the ion type second the mass shift type. By hovering over the amino acids in the peptide or ions in the legend the corresponding peaks are highlighted. By toggling the 'Unassigned' label you can turn the background (unassigned) peaks on or off in the plot. By updating the slider in the Ion legend you can update the spectrum to only show the top X% of the peaks with labels. The top X% means any peak that is within X% of the highest intensity. By dragging in the spectrum you can zoom in to a specific part of the spectrum and use 'Zoom Out' to get back to the original zoom level. The annotation of the spectrum is based on the given sequence in the peptides file and is done with different software so inconsistencies are likely. The peaks are annotated based on the given sequence, with 20 ppm tolerance.

Copy Data

##### Spectrum 4798 (TSV)

###### Preview

```
Loading example...
```

*Click on the button to copy the data to your clipboard.*

Mz MinMz MaxIntensity Max

WidthHeightPeptide font sizePeptide stroke widthSpectrum font sizeSpectrum stroke widthCompact peptide

Ion legend

wxyz

abcd

OtherUnassignedIon^Charge^_Position_Show for top:%

AIPAPJEK

08.12e+41.62e+52.43e+53.25e+5

Zoom Out

y^+1^_1_c^+2^_4_w^+1^_2_y^+2^_4_y^+1^_2_z^+1^_2_z^+2^_5_y^+1^_2_y^+2^_6_w^+1^_3_w^+1^_3_z^+1^_3_y^+2^_7_y^+2^_7_y^+1^_3_w^+1^_4_c^+1^_5_y^+1^_4_z^+1^_5_y^+1^_5_c^+1^_6_w^+1^_6_z^+1^_6_y^+1^_6_y^+1^_6_w^+1^_7_c^+1^_7_z^+1^_7_

0879175826373516

Fragment Matches Table

Show background peaks

| Position | Ion type | Intensity | mz Theoretical | mz Error (Th) | mz Error (ppm) | Charge | Series Number |
| --- | --- | --- | --- | --- | --- | --- | --- |
| - | - | 416.1 | 123.7 | - | - | 0 | - |
| - | - | 682.3 | 127.1 | - | - | 0 | - |
| - | - | 1491 | 129.1 | - | - | 0 | - |
| - | - | 425.6 | 135.2 | - | - | 0 | - |
| - | - | 389.8 | 140.6 | - | - | 0 | - |
| - | - | 4849 | 141.1 | - | - | 0 | - |
| 8 | y | 3392 | 147.1 | 0.000233 | 1.583 | +1 | 1 |
| - | - | 468.5 | 148.9 | - | - | 0 | - |
| - | - | 2.222E+04 | 149 | - | - | 0 | - |
| - | - | 1907 | 150 | - | - | 0 | - |
| - | - | 472.1 | 155.1 | - | - | 0 | - |
| - | - | 476.3 | 157 | - | - | 0 | - |
| - | - | 6.025E+04 | 157.1 | - | - | 0 | - |
| - | - | 4407 | 158.1 | - | - | 0 | - |
| - | - | 576.4 | 159.1 | - | - | 0 | - |
| - | - | 1194 | 160.1 | - | - | 0 | - |
| - | - | 3265 | 167 | - | - | 0 | - |
| - | - | 1.384E+04 | 169.1 | - | - | 0 | - |
| - | - | 939 | 170.1 | - | - | 0 | - |
| - | - | 420.1 | 170.3 | - | - | 0 | - |
| - | - | 443.7 | 170.6 | - | - | 0 | - |
| - | - | 1167 | 173.4 | - | - | 0 | - |
| - | - | 428.3 | 175.4 | - | - | 0 | - |
| - | - | 556 | 183.1 | - | - | 0 | - |
| - | - | 684.6 | 183.1 | - | - | 0 | - |
| - | - | 1240 | 184.1 | - | - | 0 | - |
| - | - | 703.8 | 185.1 | - | - | 0 | - |
| - | - | 1112 | 185.1 | - | - | 0 | - |
| - | - | 2.345E+04 | 185.1 | - | - | 0 | - |
| 4 | c | 446.9 | 185.6 | 0.0008744 | 4.711 | +2 | 4 |
| - | - | 2088 | 186.1 | - | - | 0 | - |
| - | - | 834 | 189.1 | - | - | 0 | - |
| - | - | 562 | 197.1 | - | - | 0 | - |
| - | - | 482.1 | 199.1 | - | - | 0 | - |
| 7 | w | 687.2 | 201.1 | 0.0001816 | 0.9028 | +1 | 2 |
| - | - | 1722 | 211.1 | - | - | 0 | - |
| - | - | 488.1 | 216.7 | - | - | 0 | - |
| - | - | 9689 | 228.2 | - | - | 0 | - |
| - | - | 596.6 | 231.7 | - | - | 0 | - |
| 5 | y | 1713 | 243.6 | 0.0004568 | 1.875 | +2 | 4 |
| - | - | 1546 | 244.2 | - | - | 0 | - |
| - | - | 1714 | 247.1 | - | - | 0 | - |
| 7 | y | 1329 | 258.1 | 0.0005837 | 2.261 | +1 | 2 |
| - | - | 718.2 | 259.2 | - | - | 0 | - |
| 7 | z | 1112 | 260.1 | 0.0007479 | 2.875 | +1 | 2 |
| 4 | z | 917.2 | 262.2 | 0.002845 | 10.85 | +2 | 5 |
| - | - | 1442 | 266.1 | - | - | 0 | - |
| - | - | 1027 | 266.2 | - | - | 0 | - |
| - | - | 589.8 | 267.2 | - | - | 0 | - |
| 7 | y | 2541 | 276.2 | 0.0001814 | 0.6567 | +1 | 2 |
| - | - | 710.9 | 277.2 | - | - | 0 | - |
| - | - | 1477 | 282.2 | - | - | 0 | - |
| - | - | 7416 | 283.2 | - | - | 0 | - |
| - | - | 911.8 | 284.2 | - | - | 0 | - |
| - | - | 1312 | 293.2 | - | - | 0 | - |
| - | - | 7930 | 298.2 | - | - | 0 | - |
| - | - | 1397 | 299.2 | - | - | 0 | - |
| 3 | y | 1.14E+05 | 327.7 | 0.0004703 | 1.435 | +2 | 6 |
| - | - | 4.093E+04 | 328.2 | - | - | 0 | - |
| - | - | 8940 | 328.7 | - | - | 0 | - |
| - | - | 3411 | 340.2 | - | - | 0 | - |
| 6 | w | 5015 | 344.2 | 0.0004864 | 1.413 | +1 | 3 |
| - | - | 571.3 | 345.2 | - | - | 0 | - |
| - | - | 1254 | 351.2 | - | - | 0 | - |
| - | - | 1169 | 352.2 | - | - | 0 | - |
| - | - | 5366 | 353.2 | - | - | 0 | - |
| - | - | 944 | 354.2 | - | - | 0 | - |
| - | - | 999.5 | 356.2 | - | - | 0 | - |
| - | - | 9139 | 357.2 | - | - | 0 | - |
| 6 | w | 828.2 | 358.2 | 0.002255 | 6.295 | +1 | 3 |
| - | - | 1084 | 358.2 | - | - | 0 | - |
| 6 | z | 1.195E+04 | 373.2 | 0.0001796 | 0.4812 | +1 | 3 |
| - | - | 7513 | 374.2 | - | - | 0 | - |
| 2 | y | 2462 | 375.2 | 0.001624 | 4.327 | +2 | 7 |
| - | - | 4795 | 379.2 | - | - | 0 | - |
| - | - | 877.1 | 380.2 | - | - | 0 | - |
| 2 | y | 621 | 384.2 | 0.0006071 | 1.58 | +2 | 7 |
| 6 | y | 870.8 | 389.2 | 0.001048 | 2.692 | +1 | 3 |
| - | - | 4.971E+04 | 396.3 | - | - | 0 | - |
| - | - | 815 | 397.2 | - | - | 0 | - |
| - | - | 9797 | 397.3 | - | - | 0 | - |
| - | - | 1466 | 398.3 | - | - | 0 | - |
| - | - | 582.5 | 411.2 | - | - | 0 | - |
| - | - | 1059 | 414.2 | - | - | 0 | - |
| - | - | 714.2 | 415.2 | - | - | 0 | - |
| - | - | 1119 | 419.3 | - | - | 0 | - |
| - | - | 3085 | 429.2 | - | - | 0 | - |
| - | - | 890.4 | 430.2 | - | - | 0 | - |
| 5 | w | 4290 | 443.3 | 0.0005539 | 1.25 | +1 | 4 |
| - | - | 1363 | 444.3 | - | - | 0 | - |
| - | - | 599.7 | 444.7 | - | - | 0 | - |
| - | - | 1.819E+04 | 466.3 | - | - | 0 | - |
| 5 | c | 1.492E+04 | 467.3 | 0.0008003 | 1.713 | +1 | 5 |
| - | - | 2444 | 468.3 | - | - | 0 | - |
| - | - | 1484 | 469.3 | - | - | 0 | - |
| - | - | 623 | 480.3 | - | - | 0 | - |
| - | - | 1654 | 482.3 | - | - | 0 | - |
| - | - | 4890 | 483.3 | - | - | 0 | - |
| - | - | 9978 | 484.3 | - | - | 0 | - |
| - | - | 947.9 | 485.2 | - | - | 0 | - |
| - | - | 3870 | 485.3 | - | - | 0 | - |
| 5 | y | 4.646E+04 | 486.3 | 0.0003164 | 0.6507 | +1 | 4 |
| - | - | 1.141E+04 | 487.3 | - | - | 0 | - |
| - | - | 1677 | 488.3 | - | - | 0 | - |
| - | - | 2330 | 506.3 | - | - | 0 | - |
| - | - | 640.1 | 507.3 | - | - | 0 | - |
| - | - | 6731 | 508.3 | - | - | 0 | - |
| - | - | 1938 | 509.3 | - | - | 0 | - |
| - | - | 864.8 | 509.3 | - | - | 0 | - |
| - | - | 2435 | 512.3 | - | - | 0 | - |
| - | - | 813.4 | 513.3 | - | - | 0 | - |
| - | - | 1595 | 514.3 | - | - | 0 | - |
| - | - | 1797 | 523.3 | - | - | 0 | - |
| - | - | 3.026E+04 | 525.3 | - | - | 0 | - |
| - | - | 6593 | 526.3 | - | - | 0 | - |
| - | - | 1469 | 527.3 | - | - | 0 | - |
| - | - | 1326 | 535.4 | - | - | 0 | - |
| - | - | 1636 | 536.4 | - | - | 0 | - |
| - | - | 2254 | 537.4 | - | - | 0 | - |
| 4 | z | 3.029E+04 | 541.3 | 0.0001762 | 0.3255 | +1 | 5 |
| - | - | 9574 | 542.3 | - | - | 0 | - |
| - | - | 1807 | 543.3 | - | - | 0 | - |
| 4 | y | 6918 | 557.3 | 0.0006782 | 1.217 | +1 | 5 |
| - | - | 2197 | 558.3 | - | - | 0 | - |
| - | - | 2216 | 560.3 | - | - | 0 | - |
| - | - | 1051 | 561.3 | - | - | 0 | - |
| - | - | 3718 | 563.4 | - | - | 0 | - |
| - | - | 1678 | 564.4 | - | - | 0 | - |
| - | - | 3.576E+04 | 579.4 | - | - | 0 | - |
| 6 | c | 5.948E+04 | 580.4 | 0.0007274 | 1.253 | +1 | 6 |
| - | - | 1.501E+04 | 581.4 | - | - | 0 | - |
| - | - | 3003 | 582.4 | - | - | 0 | - |
| - | - | 4648 | 606.4 | - | - | 0 | - |
| - | - | 1703 | 607.4 | - | - | 0 | - |
| 3 | w | 1056 | 611.3 | 0.002015 | 3.297 | +1 | 6 |
| - | - | 1230 | 612.3 | - | - | 0 | - |
| - | - | 2096 | 619.4 | - | - | 0 | - |
| 3 | z | 1405 | 620.4 | 0.0065 | 10.48 | +1 | 6 |
| 3 | y | 1570 | 636.4 | 0.0009326 | 1.465 | +1 | 6 |
| - | - | 2115 | 652.4 | - | - | 0 | - |
| 3 | y | 1.15E+05 | 654.4 | 0.0001605 | 0.2452 | +1 | 6 |
| - | - | 4.131E+04 | 655.4 | - | - | 0 | - |
| - | - | 9789 | 656.4 | - | - | 0 | - |
| - | - | 967.4 | 657.4 | - | - | 0 | - |
| - | - | 741 | 665.4 | - | - | 0 | - |
| - | - | 1211 | 679.4 | - | - | 0 | - |
| - | - | 1309 | 682.4 | - | - | 0 | - |
| - | - | 2251 | 683.4 | - | - | 0 | - |
| - | - | 5019 | 692.4 | - | - | 0 | - |
| - | - | 2522 | 693.4 | - | - | 0 | - |
| - | - | 874.1 | 694.4 | - | - | 0 | - |
| - | - | 1993 | 695.4 | - | - | 0 | - |
| - | - | 813.3 | 705.4 | - | - | 0 | - |
| 2 | w | 2.162E+04 | 708.4 | 0.003085 | 4.354 | +1 | 7 |
| 7 | c | 3.214E+05 | 709.4 | 4.652E-05 | 0.06558 | +1 | 7 |
| - | - | 1.186E+05 | 710.4 | - | - | 0 | - |
| - | - | 3.095E+04 | 711.4 | - | - | 0 | - |
| - | - | 2444 | 712.4 | - | - | 0 | - |
| - | - | 716.8 | 724.4 | - | - | 0 | - |
| - | - | 1054 | 750.5 | - | - | 0 | - |
| 2 | z | 1.664E+04 | 751.4 | 0.0004944 | 0.6579 | +1 | 7 |
| - | - | 8475 | 752.4 | - | - | 0 | - |
| - | - | 2547 | 753.4 | - | - | 0 | - |
| - | - | 912.3 | 758.4 | - | - | 0 | - |
| - | - | 1219 | 764.4 | - | - | 0 | - |
| - | - | 1216 | 765.4 | - | - | 0 | - |
| - | - | 1.45E+05 | 766.4 | - | - | 0 | - |
| - | - | 5.675E+04 | 767.4 | - | - | 0 | - |
| - | - | 1.501E+04 | 768.4 | - | - | 0 | - |
| - | - | 998.7 | 769.4 | - | - | 0 | - |
| - | - | 2264 | 775.4 | - | - | 0 | - |
| - | - | 2219 | 776.4 | - | - | 0 | - |
| - | - | 2793 | 778.4 | - | - | 0 | - |
| - | - | 985.8 | 779.4 | - | - | 0 | - |
| - | - | 1.707E+04 | 783.4 | - | - | 0 | - |
| - | - | 7476 | 784.5 | - | - | 0 | - |
| - | - | 2144 | 785.5 | - | - | 0 | - |
| - | - | 1271 | 793.4 | - | - | 0 | - |
| - | - | 1393 | 793.5 | - | - | 0 | - |
| - | - | 1092 | 796.5 | - | - | 0 | - |
| - | - | 2896 | 819.4 | - | - | 0 | - |
| - | - | 2055 | 820.4 | - | - | 0 | - |
| - | - | 771.7 | 821.4 | - | - | 0 | - |
| - | - | 9123 | 821.5 | - | - | 0 | - |
| - | - | 1.006E+05 | 822.5 | - | - | 0 | - |
| - | - | 4.146E+04 | 823.5 | - | - | 0 | - |
| - | - | 1.272E+04 | 824.5 | - | - | 0 | - |
| - | - | 1118 | 825.5 | - | - | 0 | - |
| - | - | 1987 | 835.4 | - | - | 0 | - |
| - | - | 4308 | 836.4 | - | - | 0 | - |
| - | - | 2312 | 837.4 | - | - | 0 | - |
| - | - | 2253 | 837.5 | - | - | 0 | - |
| - | - | 1.112E+05 | 838.5 | - | - | 0 | - |
| - | - | 3.016E+05 | 839.5 | - | - | 0 | - |
| - | - | 1.28E+05 | 840.5 | - | - | 0 | - |
| - | - | 3.585E+04 | 841.5 | - | - | 0 | - |
| - | - | 3726 | 842.5 | - | - | 0 | - |
| - | - | 595.9 | 876 | - | - | 0 | - |
| - | - | 607.5 | 1559 | - | - | 0 | - |
| - | - | 665.1 | 2856 | - | - | 0 | - |
| - | - | 738 | 3078 | - | - | 0 | - |
| - | - | 961.8 | 3078 | - | - | 0 | - |
| - | - | 731.1 | 3482 | - | - | 0 | - |

m/z Charge Intensity FragmentType MassShift Position
123.72128295898438 0 416.05127
127.14825439453125 0 682.28125
129.10252380371094 0 1490.5437
135.18557739257812 0 425.63846
140.56246948242188 0 389.7561
141.10250854492188 0 4849.2983
147.113037109375 0 3392.416 y 7
148.94679260253906 0 468.4605
149.02359008789062 0 22219.107
150.02688598632812 0 1906.6986
155.1470947265625 0 472.05396
157.01336669921875 0 476.33765
157.13380432128906 0 60253.656
158.1371612548828 0 4406.6323
159.10018920898438 0 576.3919
160.10830688476562 0 1194.3722
167.03411865234375 0 3264.8328
169.097412109375 0 13835.252
170.1007537841797 0 938.9528
170.32814025878906 0 420.1445
170.62673950195312 0 443.67984
173.4386749267578 0 1166.7595
175.3697967529297 0 428.27957
183.1124267578125 0 555.9599
183.1495819091797 0 684.57196
184.12071228027344 0 1240.34
185.08116149902344 0 703.7524
185.1191864013672 0 1112.4238
185.1287078857422 0 23451.965
185.626953125 0 446.87653 c 3
186.13226318359375 0 2087.5063
189.08750915527344 0 834.0005
197.12855529785156 0 562.0098
199.0718231201172 0 482.08063
201.12355041503906 0 687.19275 w 6
211.1439208984375 0 1721.602
216.70458984375 0 488.10205
228.1708984375 0 9688.759
231.69305419921875 0 596.6383
243.65020751953125 0 1712.6394 y 4
244.1659393310547 0 1545.6927
247.14004516601562 0 1714.1437
258.1454162597656 0 1328.6775 y Water loss 6
259.1531677246094 0 718.1732
260.13592529296875 0 1111.6921 z 6
262.15081787109375 0 917.16235 z Water loss 3
266.1495666503906 0 1441.6671
266.1634216308594 0 1026.8492
267.1689453125 0 589.7783
276.15557861328125 0 2540.6262 y 6
277.1582946777344 0 710.85657
282.1815185546875 0 1477.2059
283.1767578125 0 7415.5244
284.1795654296875 0 911.78937
293.17529296875 0 1312.0969
298.1764831542969 0 7929.875
299.1796569824219 0 1396.6082
327.6951599121094 0 114044.72 y 2
328.196533203125 0 40926.18
328.69793701171875 0 8940.468
340.1866760253906 0 3410.8374
344.1820983886719 0 5014.7056 w 5
345.18597412109375 0 571.25836
351.23870849609375 0 1254.1619
352.2110900878906 0 1169.0172
353.21832275390625 0 5366.1343
354.22015380859375 0 943.961
356.20526123046875 0 999.45935
357.21319580078125 0 9138.836
358.19500732421875 0 828.16925 w 5
358.21783447265625 0 1084.4532
373.2209167480469 0 11952.555 z 5
374.2273254394531 0 7513.2563
375.2330627441406 0 2462.205 y Water loss 1
379.2344055175781 0 4795.1064
380.2365417480469 0 877.12256
384.2361145019531 0 620.9506 y 1
389.2405090332031 0 870.7885 y 5
396.2607727050781 0 49712.324
397.20916748046875 0 815.03284
397.263916015625 0 9797.292
398.26593017578125 0 1466.398
411.221435546875 0 582.5426
414.2350158691406 0 1058.8395
415.219970703125 0 714.16345
419.2644348144531 0 1119.4572
429.24505615234375 0 3085.0027
430.2480773925781 0 890.3722
443.2505798339844 0 4290.1934 w 4
444.25396728515625 0 1363.3586
444.7087707519531 0 599.7365
466.2901611328125 0 18186.162
467.2968444824219 0 14923.642 c 4
468.29937744140625 0 2444.2544
469.2885437011719 0 1483.6547
480.28033447265625 0 623.01733
482.2978515625 0 1653.8015
483.2558288574219 0 4889.7046
484.2763977050781 0 9977.843
485.2464599609375 0 947.8589
485.2822265625 0 3869.6667
486.29254150390625 0 46456.195 y 4
487.2953796386719 0 11409.951
488.2969055175781 0 1676.7507
506.27569580078125 0 2329.7388
507.2762756347656 0 640.1174
508.27679443359375 0 6730.5767
509.28021240234375 0 1938.0579
509.3454895019531 0 864.7982
512.2728271484375 0 2435.4243
513.2767333984375 0 813.3689
514.2881469726562 0 1595.3138
523.3111572265625 0 1796.9137
525.3032836914062 0 30260.43
526.3053588867188 0 6592.9443
527.3082275390625 0 1469.1506
535.3604125976562 0 1326.2025
536.3661499023438 0 1635.8081
537.3748168945312 0 2253.8677
541.310791015625 0 30288.203 z 3
542.3154296875 0 9573.521
543.3212890625 0 1806.508
557.3300170898438 0 6918.0415 y 3
558.3321533203125 0 2197.0476
560.2853393554688 0 2216.0994
561.2860107421875 0 1051.442
563.3554077148438 0 3718.086
564.3595581054688 0 1678.3547
579.3741455078125 0 35764.4
580.3809814453125 0 59480.332 c 5
581.38427734375 0 15005.261
582.3864135742188 0 3003.2146
606.3980712890625 0 4648.213
607.4014892578125 0 1702.77
611.3419189453125 0 1056.0698 w 2
612.3436889648438 0 1230.2273
619.359130859375 0 2095.8728
620.3593139648438 0 1405.2854 z Water loss 2
636.37060546875 0 1569.7429 y Water loss 2
652.3569946289062 0 2115.462
654.3822631835938 0 115015.04 y 2
655.3856811523438 0 41308.832
656.38916015625 0 9789.279
657.3909301757812 0 967.4291
665.4122924804688 0 740.98517
679.424560546875 0 1210.844
682.3760986328125 0 1308.7037
683.3820190429688 0 2250.9885
692.3986206054688 0 5018.576
693.4026489257812 0 2522.487
694.4005737304688 0 874.05817
695.3864135742188 0 1992.9974
705.37548828125 0 813.2547
708.395751953125 0 21618.734 w 1
709.4242553710938 0 321431.56 c 6
710.4271850585938 0 118584.62
711.429931640625 0 30950.559
712.432373046875 0 2444.2637
724.4332885742188 0 716.84485
750.461181640625 0 1054.4878
751.4479370117188 0 16638.305 z 1
752.4437255859375 0 8475.063
753.444091796875 0 2547.3284
758.4027099609375 0 912.30536
764.42529296875 0 1219.1025
765.4329223632812 0 1216.2327
766.421875 0 144981.78
767.4247436523438 0 56753.812
768.4276123046875 0 15006.848
769.4315795898438 0 998.7102
775.4263305664062 0 2264.4187
776.4259033203125 0 2218.548
778.4425659179688 0 2793.2717
779.4330444335938 0 985.8335
783.4481811523438 0 17069.95
784.4517822265625 0 7475.751
785.4523315429688 0 2143.6438
793.4104614257812 0 1271.2935
793.5068969726562 0 1393.369
796.4586181640625 0 1092.1511
819.4153442382812 0 2895.778
820.4168090820312 0 2055.142
821.4183349609375 0 771.7461
821.49951171875 0 9123.423
822.4847412109375 0 100610.71
823.4878540039062 0 41459.043
824.4901733398438 0 12723.773
825.4904174804688 0 1118.2383
835.4320678710938 0 1986.785
836.4431762695312 0 4307.9233
837.4443969726562 0 2311.594
837.5204467773438 0 2252.9082
838.5043334960938 0 111205.01
839.51025390625 0 301641.97
840.5133666992188 0 128038.81
841.5164184570312 0 35846.496
842.5200805664062 0 3725.51
875.9890747070312 0 595.8815
1558.742431640625 0 607.53827
2856.30224609375 0 665.1485
3077.814208984375 0 738.00073
3078.41748046875 0 961.77515
3481.588623046875 0 731.1038

Spectrum Details

|  |  |
| --- | --- |
| Matched peaks? Matched peaks The total absolute number of peaks matched. Additionally in brackets the total fraction of peaks matched and the total number of peaks is shown. | 28 (13.79% of 203) |
| FDR? FDR The false discovery rate estimated for this peptide. It is calculated by matching all theoretical fragments with a non-integer shift with the raw peaks for this spectrum. This is done with 40 different shifts. The resulting percentage is the average number of annotated peaks over the number of annotated peaks with the correct spectrum. | 0.17% |
| Satellite FDR? Satellite FDR See the FDR for details on its calculation. This satellite ion specific FDR only contains the satellite ions (d/w) for I/L/J positions. | 0.00% |
| PSM Score? PSM Score The PSM Score as given by Hecklib to this annotated spectrum. It is shown with three significant figures. | 200 |

#### Spectrum 4630? Spectrum 4630 The raw spectrum of this peptide as annotated by Hecklib. The fragments are coloured according to ion type (see legend). Any peaks with a star '\*' as text can be hovered over to see the full details, first the ion type second the mass shift type. By hovering over the amino acids in the peptide or ions in the legend the corresponding peaks are highlighted. By toggling the 'Unassigned' label you can turn the background (unassigned) peaks on or off in the plot. By updating the slider in the Ion legend you can update the spectrum to only show the top X% of the peaks with labels. The top X% means any peak that is within X% of the highest intensity. By dragging in the spectrum you can zoom in to a specific part of the spectrum and use 'Zoom Out' to get back to the original zoom level. The annotation of the spectrum is based on the given sequence in the peptides file and is done with different software so inconsistencies are likely. The peaks are annotated based on the given sequence, with 20 ppm tolerance.

Copy Data

##### Spectrum 4630 (TSV)

###### Preview

```
Loading example...
```

*Click on the button to copy the data to your clipboard.*

Mz MinMz MaxIntensity Max

WidthHeightPeptide font sizePeptide stroke widthSpectrum font sizeSpectrum stroke widthCompact peptide

Ion legend

wxyz

abcd

OtherUnassignedIon^Charge^_Position_Show for top:%

AIPAPJEK

02.15e+54.31e+56.46e+58.62e+5

Zoom Out

d^+1^_2_y^+1^_1_d^+1^_2_y^+1^_1_a^+1^_2_b^+1^_2_y^+2^_3_b^+2^_5_d^+1^_3_y^+2^_4_a^+1^_3_y^+1^_2_y^+1^_2_y^+1^_2_y^+2^_5_b^+1^_3_y^+2^_6_y^+2^_6_b^+1^_4_y^+1^_3_y^+2^_7_y^+1^_3_b^+1^_5_y^+1^_4_y^+1^_4_y^+1^_5_y^+1^_5_y^+1^_6_y^+1^_6_y^+1^_7_

0867173326003467

Fragment Matches Table

Show background peaks

| Position | Ion type | Intensity | mz Theoretical | mz Error (Th) | mz Error (ppm) | Charge | Series Number |
| --- | --- | --- | --- | --- | --- | --- | --- |
| - | - | 337.6 | 120.3 | - | - | 0 | - |
| - | - | 393.4 | 121 | - | - | 0 | - |
| - | - | 483.2 | 123.1 | - | - | 0 | - |
| - | - | 1024 | 126.1 | - | - | 0 | - |
| - | - | 505.6 | 128.1 | - | - | 0 | - |
| - | - | 783.5 | 129 | - | - | 0 | - |
| - | - | 1526 | 129.1 | - | - | 0 | - |
| 2 | d | 1.056E+05 | 129.1 | 0.0004827 | 3.739 | +1 | 2 |
| - | - | 582.2 | 130.1 | - | - | 0 | - |
| 8 | y | 5.761E+04 | 130.1 | 0.0004454 | 3.424 | +1 | 1 |
| - | - | 6415 | 130.1 | - | - | 0 | - |
| - | - | 562.3 | 131.1 | - | - | 0 | - |
| - | - | 3539 | 131.1 | - | - | 0 | - |
| - | - | 1984 | 139.1 | - | - | 0 | - |
| - | - | 2.965E+05 | 141.1 | - | - | 0 | - |
| - | - | 541.8 | 142.1 | - | - | 0 | - |
| - | - | 2393 | 142.1 | - | - | 0 | - |
| - | - | 2.292E+04 | 142.1 | - | - | 0 | - |
| - | - | 992.6 | 143.1 | - | - | 0 | - |
| 2 | d | 527.9 | 143.1 | 0.000595 | 4.157 | +1 | 2 |
| - | - | 439.5 | 143.9 | - | - | 0 | - |
| 8 | y | 8.622E+04 | 147.1 | 0.0004771 | 3.243 | +1 | 1 |
| - | - | 447.6 | 148.1 | - | - | 0 | - |
| - | - | 708.8 | 148.1 | - | - | 0 | - |
| - | - | 4977 | 148.1 | - | - | 0 | - |
| - | - | 454.9 | 148.8 | - | - | 0 | - |
| - | - | 571 | 148.9 | - | - | 0 | - |
| - | - | 986.4 | 148.9 | - | - | 0 | - |
| - | - | 873 | 148.9 | - | - | 0 | - |
| - | - | 868.8 | 148.9 | - | - | 0 | - |
| - | - | 1361 | 148.9 | - | - | 0 | - |
| - | - | 2292 | 148.9 | - | - | 0 | - |
| - | - | 4002 | 148.9 | - | - | 0 | - |
| - | - | 4291 | 149 | - | - | 0 | - |
| - | - | 2491 | 149 | - | - | 0 | - |
| - | - | 1444 | 149 | - | - | 0 | - |
| - | - | 1379 | 149 | - | - | 0 | - |
| - | - | 922.9 | 149 | - | - | 0 | - |
| - | - | 842.3 | 149 | - | - | 0 | - |
| - | - | 4.048E+04 | 149 | - | - | 0 | - |
| - | - | 446.2 | 149.1 | - | - | 0 | - |
| - | - | 2448 | 150 | - | - | 0 | - |
| - | - | 1028 | 151.1 | - | - | 0 | - |
| - | - | 486.3 | 151.5 | - | - | 0 | - |
| - | - | 610.2 | 152.1 | - | - | 0 | - |
| - | - | 721.5 | 156.1 | - | - | 0 | - |
| - | - | 8571 | 157.1 | - | - | 0 | - |
| 2 | a | 7.601E+05 | 157.1 | 0.0005394 | 3.433 | +1 | 2 |
| - | - | 4250 | 158.1 | - | - | 0 | - |
| - | - | 5.9E+04 | 158.1 | - | - | 0 | - |
| - | - | 2174 | 159.1 | - | - | 0 | - |
| - | - | 542.3 | 163.5 | - | - | 0 | - |
| - | - | 876.8 | 165.1 | - | - | 0 | - |
| - | - | 1549 | 167 | - | - | 0 | - |
| - | - | 2052 | 167.1 | - | - | 0 | - |
| - | - | 1253 | 168.1 | - | - | 0 | - |
| - | - | 3.105E+05 | 169.1 | - | - | 0 | - |
| - | - | 1311 | 170.1 | - | - | 0 | - |
| - | - | 2.362E+04 | 170.1 | - | - | 0 | - |
| - | - | 1080 | 171.1 | - | - | 0 | - |
| - | - | 1035 | 173.1 | - | - | 0 | - |
| - | - | 4547 | 176.1 | - | - | 0 | - |
| - | - | 662.3 | 176.6 | - | - | 0 | - |
| - | - | 722.6 | 183.1 | - | - | 0 | - |
| - | - | 2.947E+05 | 183.1 | - | - | 0 | - |
| - | - | 2.931E+04 | 184.2 | - | - | 0 | - |
| 2 | b | 1.775E+05 | 185.1 | 0.0004978 | 2.689 | +1 | 2 |
| - | - | 1.663E+04 | 186.1 | - | - | 0 | - |
| - | - | 1031 | 187.1 | - | - | 0 | - |
| - | - | 1292 | 193.1 | - | - | 0 | - |
| - | - | 9502 | 195.1 | - | - | 0 | - |
| 6 | y | 627.9 | 195.1 | 0.0001694 | 0.8681 | +2 | 3 |
| - | - | 1207 | 196.1 | - | - | 0 | - |
| - | - | 9000 | 197.1 | - | - | 0 | - |
| - | - | 718.3 | 197.1 | - | - | 0 | - |
| - | - | 571 | 199.1 | - | - | 0 | - |
| - | - | 1164 | 201.1 | - | - | 0 | - |
| - | - | 957.7 | 209.2 | - | - | 0 | - |
| - | - | 1.565E+05 | 211.1 | - | - | 0 | - |
| - | - | 512 | 211.7 | - | - | 0 | - |
| - | - | 1.7E+04 | 212.1 | - | - | 0 | - |
| - | - | 3817 | 213.1 | - | - | 0 | - |
| - | - | 692.1 | 214.1 | - | - | 0 | - |
| - | - | 785.7 | 215.1 | - | - | 0 | - |
| - | - | 1.554E+04 | 215.1 | - | - | 0 | - |
| - | - | 1395 | 216.1 | - | - | 0 | - |
| - | - | 1.202E+04 | 222.1 | - | - | 0 | - |
| - | - | 1373 | 223.1 | - | - | 0 | - |
| - | - | 1469 | 223.1 | - | - | 0 | - |
| - | - | 1694 | 225.1 | - | - | 0 | - |
| 5 | b | 1064 | 225.6 | 8.696E-05 | 0.3854 | +2 | 5 |
| - | - | 2016 | 227.1 | - | - | 0 | - |
| 3 | d | 1317 | 228.2 | 0.000657 | 2.88 | +1 | 3 |
| - | - | 1333 | 229.2 | - | - | 0 | - |
| - | - | 2607 | 236.2 | - | - | 0 | - |
| - | - | 554.6 | 237.2 | - | - | 0 | - |
| - | - | 1482 | 238.2 | - | - | 0 | - |
| - | - | 7.116E+04 | 240.1 | - | - | 0 | - |
| - | - | 9514 | 241.1 | - | - | 0 | - |
| - | - | 7207 | 241.1 | - | - | 0 | - |
| - | - | 558 | 242.1 | - | - | 0 | - |
| - | - | 1.742E+04 | 243.1 | - | - | 0 | - |
| 5 | y | 4.757E+04 | 243.6 | 0.0005636 | 2.313 | +2 | 4 |
| - | - | 1798 | 244.1 | - | - | 0 | - |
| - | - | 1.041E+04 | 244.2 | - | - | 0 | - |
| - | - | 992.6 | 244.2 | - | - | 0 | - |
| - | - | 1993 | 244.7 | - | - | 0 | - |
| - | - | 521.3 | 250.6 | - | - | 0 | - |
| - | - | 1865 | 252.1 | - | - | 0 | - |
| - | - | 4590 | 253.1 | - | - | 0 | - |
| - | - | 1102 | 254.1 | - | - | 0 | - |
| - | - | 721.2 | 254.2 | - | - | 0 | - |
| 3 | a | 5958 | 254.2 | 0.0005709 | 2.246 | +1 | 3 |
| - | - | 591.7 | 255.2 | - | - | 0 | - |
| 7 | y | 2.172E+05 | 258.1 | 0.0004921 | 1.906 | +1 | 2 |
| 7 | y | 4735 | 259.1 | 0.0005159 | 1.991 | +1 | 2 |
| - | - | 2.304E+04 | 259.1 | - | - | 0 | - |
| - | - | 744.2 | 260.1 | - | - | 0 | - |
| - | - | 2517 | 260.2 | - | - | 0 | - |
| - | - | 5442 | 261.1 | - | - | 0 | - |
| - | - | 743.2 | 262.2 | - | - | 0 | - |
| - | - | 1.943E+04 | 264.2 | - | - | 0 | - |
| - | - | 2429 | 265.2 | - | - | 0 | - |
| - | - | 6.184E+04 | 266.2 | - | - | 0 | - |
| - | - | 8999 | 267.2 | - | - | 0 | - |
| - | - | 834.6 | 268.1 | - | - | 0 | - |
| - | - | 590.1 | 268.2 | - | - | 0 | - |
| 7 | y | 1.91E+05 | 276.2 | 0.0005476 | 1.983 | +1 | 2 |
| - | - | 2.222E+04 | 277.2 | - | - | 0 | - |
| - | - | 2625 | 278.2 | - | - | 0 | - |
| 4 | y | 1955 | 279.2 | 0.0006376 | 2.284 | +2 | 5 |
| - | - | 1831 | 280.1 | - | - | 0 | - |
| 3 | b | 1.953E+04 | 282.2 | 0.0006057 | 2.146 | +1 | 3 |
| - | - | 1733 | 283.2 | - | - | 0 | - |
| - | - | 3050 | 286.1 | - | - | 0 | - |
| - | - | 607.4 | 293.1 | - | - | 0 | - |
| - | - | 630.9 | 293.5 | - | - | 0 | - |
| - | - | 493.3 | 296.2 | - | - | 0 | - |
| - | - | 3716 | 298.1 | - | - | 0 | - |
| - | - | 1055 | 299.1 | - | - | 0 | - |
| - | - | 1107 | 305.2 | - | - | 0 | - |
| - | - | 496.1 | 307.2 | - | - | 0 | - |
| - | - | 1756 | 308.2 | - | - | 0 | - |
| - | - | 1.141E+04 | 312.2 | - | - | 0 | - |
| - | - | 1755 | 313.2 | - | - | 0 | - |
| - | - | 560.9 | 313.7 | - | - | 0 | - |
| - | - | 602.4 | 314.2 | - | - | 0 | - |
| 3 | y | 2251 | 318.7 | 0.0006257 | 1.963 | +2 | 6 |
| - | - | 569.6 | 319.2 | - | - | 0 | - |
| - | - | 6548 | 322.2 | - | - | 0 | - |
| - | - | 785.5 | 323.2 | - | - | 0 | - |
| - | - | 771 | 325.2 | - | - | 0 | - |
| 3 | y | 8.531E+05 | 327.7 | 0.000806 | 2.46 | +2 | 6 |
| - | - | 2.915E+05 | 328.2 | - | - | 0 | - |
| - | - | 6.461E+04 | 328.7 | - | - | 0 | - |
| - | - | 1076 | 328.7 | - | - | 0 | - |
| - | - | 678.2 | 328.7 | - | - | 0 | - |
| - | - | 3702 | 329.2 | - | - | 0 | - |
| - | - | 576.7 | 335.2 | - | - | 0 | - |
| - | - | 629.6 | 339.2 | - | - | 0 | - |
| - | - | 1.779E+05 | 340.2 | - | - | 0 | - |
| - | - | 3.017E+04 | 341.2 | - | - | 0 | - |
| - | - | 3498 | 342.2 | - | - | 0 | - |
| - | - | 1561 | 349.2 | - | - | 0 | - |
| - | - | 3.564E+04 | 351.2 | - | - | 0 | - |
| - | - | 6081 | 352.2 | - | - | 0 | - |
| 4 | b | 4818 | 353.2 | 0.0004487 | 1.27 | +1 | 4 |
| - | - | 1091 | 353.2 | - | - | 0 | - |
| - | - | 736.9 | 354.2 | - | - | 0 | - |
| - | - | 3403 | 357.2 | - | - | 0 | - |
| - | - | 3.482E+04 | 358.2 | - | - | 0 | - |
| - | - | 6785 | 359.2 | - | - | 0 | - |
| - | - | 747.2 | 360.2 | - | - | 0 | - |
| - | - | 1011 | 361.2 | - | - | 0 | - |
| - | - | 1600 | 366.2 | - | - | 0 | - |
| 6 | y | 2411 | 371.2 | 0.001298 | 3.495 | +1 | 3 |
| - | - | 1861 | 377.2 | - | - | 0 | - |
| - | - | 5.288E+04 | 379.2 | - | - | 0 | - |
| - | - | 1.09E+04 | 380.2 | - | - | 0 | - |
| - | - | 1752 | 381.2 | - | - | 0 | - |
| - | - | 1466 | 381.2 | - | - | 0 | - |
| - | - | 7054 | 383.2 | - | - | 0 | - |
| 2 | y | 8498 | 384.2 | 6.431E-05 | 0.1674 | +2 | 7 |
| - | - | 2864 | 384.7 | - | - | 0 | - |
| 6 | y | 5.915E+04 | 389.2 | 0.0007426 | 1.908 | +1 | 3 |
| - | - | 9996 | 390.2 | - | - | 0 | - |
| - | - | 1622 | 391.2 | - | - | 0 | - |
| - | - | 1502 | 393.2 | - | - | 0 | - |
| - | - | 1490 | 395.2 | - | - | 0 | - |
| - | - | 1.125E+04 | 399.2 | - | - | 0 | - |
| - | - | 2593 | 400.2 | - | - | 0 | - |
| - | - | 587.6 | 401.2 | - | - | 0 | - |
| - | - | 665.6 | 402.2 | - | - | 0 | - |
| - | - | 3.271E+04 | 411.2 | - | - | 0 | - |
| - | - | 6174 | 412.2 | - | - | 0 | - |
| - | - | 4430 | 429.2 | - | - | 0 | - |
| - | - | 1114 | 430.2 | - | - | 0 | - |
| - | - | 2712 | 437.2 | - | - | 0 | - |
| - | - | 838.5 | 438.2 | - | - | 0 | - |
| 5 | b | 1159 | 450.3 | 0.000999 | 2.219 | +1 | 5 |
| - | - | 4582 | 462.3 | - | - | 0 | - |
| - | - | 728.8 | 463.3 | - | - | 0 | - |
| 5 | y | 2.804E+04 | 468.3 | 0.0009324 | 1.991 | +1 | 4 |
| - | - | 7455 | 469.3 | - | - | 0 | - |
| - | - | 890.3 | 470.3 | - | - | 0 | - |
| - | - | 1623 | 471.3 | - | - | 0 | - |
| - | - | 1.539E+04 | 480.3 | - | - | 0 | - |
| - | - | 5032 | 481.3 | - | - | 0 | - |
| 5 | y | 7.248E+05 | 486.3 | 0.001049 | 2.157 | +1 | 4 |
| - | - | 1.745E+05 | 487.3 | - | - | 0 | - |
| - | - | 2.812E+04 | 488.3 | - | - | 0 | - |
| - | - | 1580 | 489.3 | - | - | 0 | - |
| - | - | 2211 | 490.3 | - | - | 0 | - |
| - | - | 917.8 | 491.3 | - | - | 0 | - |
| - | - | 6.631E+04 | 508.3 | - | - | 0 | - |
| - | - | 1.92E+04 | 509.3 | - | - | 0 | - |
| - | - | 4198 | 510.3 | - | - | 0 | - |
| - | - | 823.2 | 525.3 | - | - | 0 | - |
| - | - | 1.602E+04 | 526.3 | - | - | 0 | - |
| - | - | 3340 | 527.3 | - | - | 0 | - |
| 4 | y | 1.752E+04 | 539.3 | 0.0003786 | 0.7021 | +1 | 5 |
| - | - | 4216 | 540.3 | - | - | 0 | - |
| - | - | 741.1 | 541.3 | - | - | 0 | - |
| 4 | y | 1.734E+05 | 557.3 | 0.0007393 | 1.326 | +1 | 5 |
| - | - | 4.67E+04 | 558.3 | - | - | 0 | - |
| - | - | 1.089E+04 | 559.3 | - | - | 0 | - |
| - | - | 1224 | 567.3 | - | - | 0 | - |
| - | - | 1008 | 585.3 | - | - | 0 | - |
| - | - | 690.6 | 609.4 | - | - | 0 | - |
| 3 | y | 1.456E+04 | 636.4 | 0.0007764 | 1.22 | +1 | 6 |
| - | - | 4637 | 637.4 | - | - | 0 | - |
| - | - | 1583 | 638.4 | - | - | 0 | - |
| 3 | y | 7.262E+05 | 654.4 | 0.0009539 | 1.458 | +1 | 6 |
| - | - | 2.51E+05 | 655.4 | - | - | 0 | - |
| - | - | 5.667E+04 | 656.4 | - | - | 0 | - |
| - | - | 3607 | 657.4 | - | - | 0 | - |
| 2 | y | 1354 | 767.5 | 0.00533 | 6.945 | +1 | 7 |
| - | - | 676.5 | 1056 | - | - | 0 | - |
| - | - | 710.5 | 3432 | - | - | 0 | - |

m/z Charge Intensity FragmentType MassShift Position
120.34355926513672 0 337.60855
121.02906036376953 0 393.3967
123.09197235107422 0 483.1666
126.05545043945312 0 1023.57355
128.07164001464844 0 505.57278
129.0186309814453 0 783.465
129.06626892089844 0 1525.5028
129.10272216796875 0 105604.19 d 1
130.0500030517578 0 582.2211
130.08670043945312 0 57612.49 y Ammonia loss 7
130.1060333251953 0 6415.2456
131.0847625732422 0 562.256
131.09005737304688 0 3539.2314
139.0869598388672 0 1983.692
141.1027374267578 0 296494.28
142.08680725097656 0 541.79
142.0998077392578 0 2392.9502
142.10604858398438 0 22923.129
143.10728454589844 0 992.5816
143.1184844970703 0 527.9366 d 1
143.87161254882812 0 439.4725
147.11328125 0 86219.68 y 7
148.0605926513672 0 447.64807
148.11029052734375 0 708.7552
148.11671447753906 0 4977.3823
148.84788513183594 0 454.85617
148.8990020751953 0 570.9828
148.90626525878906 0 986.39734
148.9133758544922 0 872.9989
148.92074584960938 0 868.809
148.927490234375 0 1360.655
148.9346466064453 0 2292.2104
148.9425811767578 0 4002.3406
148.95913696289062 0 4290.826
148.9669189453125 0 2490.527
148.97412109375 0 1444.1941
148.98146057128906 0 1379.1508
148.9888458251953 0 922.86414
148.99586486816406 0 842.2796
149.02378845214844 0 40484.688
149.05337524414062 0 446.18658
150.027099609375 0 2448.0894
151.08741760253906 0 1027.7817
151.45962524414062 0 486.27615
152.07144165039062 0 610.2119
156.06585693359375 0 721.5118
157.0977020263672 0 8570.898
157.1340789794922 0 760134 a 1
158.1309356689453 0 4249.854
158.13734436035156 0 59001.32
159.13980102539062 0 2173.9207
163.54722595214844 0 542.3485
165.13934326171875 0 876.8475
167.0344696044922 0 1548.8784
167.1181640625 0 2051.8528
168.10232543945312 0 1252.8486
169.09767150878906 0 310517.56
170.09498596191406 0 1311.4646
170.1010284423828 0 23623.271
171.1035614013672 0 1079.6938
173.12887573242188 0 1034.7622
176.12367248535156 0 4547.3745
176.62509155273438 0 662.33746
183.07679748535156 0 722.63806
183.1497344970703 0 294721.62
184.15310668945312 0 29308.744
185.1289520263672 0 177541.39 b 1
186.13230895996094 0 16630.082
187.1346435546875 0 1031.0966
193.13458251953125 0 1292.0173
195.11326599121094 0 9501.703
195.12319946289062 0 627.8653 y 5
196.1160125732422 0 1206.7059
197.0925750732422 0 9000.209
197.12864685058594 0 718.3245
199.1081085205078 0 570.9672
201.08746337890625 0 1164.3253
209.16578674316406 0 957.6915
211.14463806152344 0 156469.77
211.7115478515625 0 512.04956
212.1479949951172 0 17002.812
213.1239776611328 0 3817.1267
214.12738037109375 0 692.0896
215.12789916992188 0 785.68585
215.13951110839844 0 15544.742
216.1427001953125 0 1395.3613
222.1243133544922 0 12020.542
223.10842895507812 0 1373.1277
223.12765502929688 0 1469.3949
225.1238555908203 0 1694.3522
225.63909912109375 0 1064.4058 b 4
227.10296630859375 0 2015.6595
228.1713104248047 0 1317.4779 d 2
229.15545654296875 0 1333.1144
236.17637634277344 0 2607.2517
237.1593780517578 0 554.56775
238.15528869628906 0 1481.8033
240.13487243652344 0 71162.42
241.11883544921875 0 9514.399
241.1380157470703 0 7207.295
242.12290954589844 0 557.9559
243.134521484375 0 17421.348
243.6503143310547 0 47569.7 y 4
244.1376953125 0 1798.47
244.15196228027344 0 10405.265
244.1654510498047 0 992.5509
244.65338134765625 0 1992.8362
250.6225128173828 0 521.28326
252.13519287109375 0 1864.6908
253.11883544921875 0 4590.4146
254.12213134765625 0 1102.4712
254.1630859375 0 721.2012
254.18687438964844 0 5957.925 a 2
255.24513244628906 0 591.7309
258.14532470703125 0 217170.28 y Water loss 6
259.1293640136719 0 4735.252 y Ammonia loss 6
259.148681640625 0 23035.598
260.13336181640625 0 744.2141
260.15081787109375 0 2516.6887
261.14495849609375 0 5442.462
262.1500244140625 0 743.1606
264.1711120605469 0 19429.127
265.1741638183594 0 2429.32
266.150390625 0 61843.043
267.1536560058594 0 8998.857
268.1302490234375 0 834.6435
268.1576843261719 0 590.0845
276.15594482421875 0 190981.17 y 6
277.1591796875 0 22220.375
278.1609802246094 0 2624.6826
279.1689453125 0 1955.1077 y 3
280.1294250488281 0 1831.4258
282.18182373046875 0 19531.404 b 2
283.1840515136719 0 1732.7728
286.1406555175781 0 3049.6204
293.0984802246094 0 607.4237
293.4749755859375 0 630.909
296.1997375488281 0 493.3154
298.1405029296875 0 3715.7046
299.1435852050781 0 1055.472
305.19512939453125 0 1107.2711
307.21234130859375 0 496.12643
308.1974792480469 0 1756.0508
312.19244384765625 0 11413.166
313.1956481933594 0 1755.1333
313.7159118652344 0 560.9049
314.1706237792969 0 602.41705
318.6900329589844 0 2251.4995 y Water loss 2
319.1924133300781 0 569.63403
322.1769714355469 0 6547.535
323.179931640625 0 785.48083
325.2250671386719 0 770.95557
327.69549560546875 0 853071.94 y 2
328.19696044921875 0 291547.38
328.6981201171875 0 64605.375
328.7217712402344 0 1076.4855
328.7449035644531 0 678.2123
329.1993408203125 0 3702.4395
335.20941162109375 0 576.6685
339.2030944824219 0 629.63416
340.1875 0 177854.7
341.1905212402344 0 30171.314
342.1927490234375 0 3498.4175
349.1874084472656 0 1561.3477
351.2396545410156 0 35643.703
352.24267578125 0 6081.3525
353.2187805175781 0 4817.915 b 3
353.2432861328125 0 1090.7122
354.2231750488281 0 736.9028
357.2133483886719 0 3403.1943
358.1978454589844 0 34815.734
359.2011413574219 0 6785.294
360.2026062011719 0 747.2283
361.2238464355469 0 1011.067
366.20269775390625 0 1599.5834
371.2301940917969 0 2411.3225 y Water loss 5
377.1826477050781 0 1861.1973
379.234619140625 0 52875.992
380.2376403808594 0 10900.85
381.2132873535156 0 1752.3331
381.2406311035156 0 1466.1725
383.22943115234375 0 7053.8164
384.2367858886719 0 8498.319 y 1
384.7388916015625 0 2864.214
389.2402038574219 0 59146.934 y 5
390.2432556152344 0 9995.786
391.24542236328125 0 1621.9031
393.21514892578125 0 1502.2675
395.192626953125 0 1490.0804
399.2245788574219 0 11249.971
400.2267761230469 0 2593.425
401.23114013671875 0 587.60956
402.16802978515625 0 665.56726
411.2247314453125 0 32711.793
412.22735595703125 0 6174.0073
429.23529052734375 0 4430.2124
430.2379455566406 0 1113.5327
437.2401123046875 0 2712.0454
438.24224853515625 0 838.48303
450.2720947265625 0 1159.4834 b 4
462.2722473144531 0 4582.14
463.27960205078125 0 728.8039
468.2825927734375 0 28038.07 y Water loss 4
469.28533935546875 0 7455.0474
470.2918701171875 0 890.3225
471.3056640625 0 1623.2671
480.2825622558594 0 15392.035
481.2859191894531 0 5031.737
486.29327392578125 0 724764.4 y 4
487.2961120605469 0 174549.45
488.2986755371094 0 28117.053
489.3004455566406 0 1579.8141
490.26666259765625 0 2211.2644
491.2713623046875 0 917.7969
508.2773742675781 0 66305.31
509.28057861328125 0 19201.107
510.28350830078125 0 4197.6675
525.3019409179688 0 823.23724
526.2879028320312 0 16016.778
527.2905883789062 0 3340.3865
539.3191528320312 0 17524.04 y Water loss 3
540.32177734375 0 4216.319
541.320068359375 0 741.0901
557.330078125 0 173359.42 y 3
558.3328247070312 0 46704.67
559.3353881835938 0 10894.971
567.315185546875 0 1224.0144
585.324951171875 0 1008.06104
609.3870239257812 0 690.5843
636.372314453125 0 14558.719 y Water loss 2
637.3739013671875 0 4637.181
638.3795776367188 0 1583.4602
654.383056640625 0 726231.25 y 2
655.3860473632812 0 250985.48
656.3883666992188 0 56673.01
657.3905029296875 0 3606.7517
767.4714965820312 0 1354.4768 y 1
1056.36376953125 0 676.5369
3432.48779296875 0 710.4796

Spectrum Details

|  |  |
| --- | --- |
| Matched peaks? Matched peaks The total absolute number of peaks matched. Additionally in brackets the total fraction of peaks matched and the total number of peaks is shown. | 30 (12.55% of 239) |
| FDR? FDR The false discovery rate estimated for this peptide. It is calculated by matching all theoretical fragments with a non-integer shift with the raw peaks for this spectrum. This is done with 40 different shifts. The resulting percentage is the average number of annotated peaks over the number of annotated peaks with the correct spectrum. | 0.24% |
| Satellite FDR? Satellite FDR See the FDR for details on its calculation. This satellite ion specific FDR only contains the satellite ions (d/w) for I/L/J positions. | 0.00% |
| PSM Score? PSM Score The PSM Score as given by Hecklib to this annotated spectrum. It is shown with three significant figures. | 317 |

#### Reverse Lookup? Reverse Lookup All places where this read could be placed.

| Group | Segment | Template | Template Part | Read Part | Score | Unique |
| --- | --- | --- | --- | --- | --- | --- |
| Homo sapiens Heavy Chain | IGHC | IGHG1 | [209..217] | [0..8] | 64 | False |
| Homo sapiens Heavy Chain | IGHC | IGHG3 | [256..264] | [0..8] | 64 | False |

| Recombined | Template Part | Read Part | Score | Unique |
| --- | --- | --- | --- | --- |
| REC-0-1 | [334..342] | [0..8] | 64 | True |

#### Meta Information from Multiple reads

##### Number of combined reads

6

##### Intensity

1

##### TotalArea

0

##### Changes to the peptide sequence

AIPAPJEK

I→JNo support for either Leucine or Isoleucine based on side chain ions (Position: 6)

L→ISupport for Isoleucine based on side chain ions (2 for I 0 for L) (Position: 2)

J→ISupport for Isoleucine based on side chain ions (2 for I 0 for L) (Position: 6)

I→LSupport for Leucine based on side chain ions (1 for L 0 for I) (Position: 2)

L→JNo support for either Leucine or Isoleucine based on side chain ions (Position: 6)

L→ISupport for Isoleucine based on side chain ions (1 for I 0 for L) (Position: 2)

#### Positional Score

Copy Data

##### Positional Score (TSV)

###### Preview

```
Loading example...
```

*Click on the button to copy the data to your clipboard.*

1001234567

Label Value
"0" 0.608
"1" 0.623
"2" 0.65
"3" 0.635
"4" 0.633
"5" 0.652
"6" 0.662
"7" 0.66

#### Meta Information from PEAKS

##### Scan Identifier

F4:4860

##### Original sequence

A

L

P

A

P

L

E

K

##### Posttranslational Modifications

##### Source File

D:\separate\_stitch\_analyses\xle-disambiguation\raw\20210323\_F1\_UM1\_Peng0013\_SA\_F59\_ingel\_3ug\_tryp.raw

##### Fraction

4

##### Scan Feature

-

##### De Novo Score

98

##### ConfidenceScore

97

### m/z

419.7557

##### Mass

837.496

##### Charge

2

##### Retention Time

25.99

##### Predicted Retention Time

26.59

##### Area

0

##### Parts Per Million

1.1

##### Fragmentation mode

HCD

##### Originating file

01 D:\separate\_stitch\_analyses\xle-disambiguation\20210325\_F59\_3ug\_DENOVO\_12.csv

#### Meta Information from PEAKS

##### Scan Identifier

F4:5744

##### Original sequence

A

L

P

A

P

L

E

K

##### Posttranslational Modifications

##### Source File

D:\separate\_stitch\_analyses\xle-disambiguation\raw\20210323\_F1\_UM1\_Peng0013\_SA\_F59\_ingel\_3ug\_tryp.raw

##### Fraction

4

##### Scan Feature

-

##### De Novo Score

97

##### ConfidenceScore

96

### m/z

419.7559

##### Mass

837.496

##### Charge

2

##### Retention Time

30.75

##### Predicted Retention Time

26.59

##### Area

0

##### Parts Per Million

1.6

##### Fragmentation mode

HCD

##### Originating file

01 D:\separate\_stitch\_analyses\xle-disambiguation\20210325\_F59\_3ug\_DENOVO\_12.csv

#### Meta Information from PEAKS

##### Scan Identifier

F4:5570

##### Original sequence

A

L

P

A

P

L

E

K

##### Posttranslational Modifications

##### Source File

D:\separate\_stitch\_analyses\xle-disambiguation\raw\20210323\_F1\_UM1\_Peng0013\_SA\_F59\_ingel\_3ug\_tryp.raw

##### Fraction

4

##### Scan Feature

-

##### De Novo Score

97

##### ConfidenceScore

96

### m/z

419.7559

##### Mass

837.496

##### Charge

2

##### Retention Time

29.84

##### Predicted Retention Time

26.59

##### Area

0

##### Parts Per Million

1.5

##### Fragmentation mode

HCD

##### Originating file

01 D:\separate\_stitch\_analyses\xle-disambiguation\20210325\_F59\_3ug\_DENOVO\_12.csv

#### Meta Information from PEAKS

##### Scan Identifier

F4:6013

##### Original sequence

A

L

P

A

P

L

E

K

##### Posttranslational Modifications

##### Source File

D:\separate\_stitch\_analyses\xle-disambiguation\raw\20210323\_F1\_UM1\_Peng0013\_SA\_F59\_ingel\_3ug\_tryp.raw

##### Fraction

4

##### Scan Feature

-

##### De Novo Score

97

##### ConfidenceScore

95

### m/z

419.7559

##### Mass

837.496

##### Charge

2

##### Retention Time

32.15

##### Predicted Retention Time

26.59

##### Area

0

##### Parts Per Million

1.5

##### Fragmentation mode

HCD

##### Originating file

01 D:\separate\_stitch\_analyses\xle-disambiguation\20210325\_F59\_3ug\_DENOVO\_12.csv

#### Meta Information from PEAKS

##### Scan Identifier

F4:4798

##### Original sequence

A

L

P

A

P

L

E

K

##### Posttranslational Modifications

##### Source File

D:\separate\_stitch\_analyses\xle-disambiguation\raw\20210323\_F1\_UM1\_Peng0013\_SA\_F59\_ingel\_3ug\_tryp.raw

##### Fraction

4

##### Scan Feature

-

##### De Novo Score

97

##### ConfidenceScore

95

### m/z

419.7562

##### Mass

837.496

##### Charge

2

##### Retention Time

25.67

##### Predicted Retention Time

26.59

##### Area

0

##### Parts Per Million

2.2

##### Fragmentation mode

ETHCD

##### Originating file

01 D:\separate\_stitch\_analyses\xle-disambiguation\20210325\_F59\_3ug\_DENOVO\_12.csv

#### Meta Information from PEAKS

##### Scan Identifier

F4:4630

##### Original sequence

A

L

P

A

P

L

E

K

##### Posttranslational Modifications

##### Source File

D:\separate\_stitch\_analyses\xle-disambiguation\raw\20210323\_F1\_UM1\_Peng0013\_SA\_F59\_ingel\_3ug\_tryp.raw

##### Fraction

4

##### Scan Feature

-

##### De Novo Score

96

##### ConfidenceScore

97

### m/z

419.7558

##### Mass

837.496

##### Charge

2

##### Retention Time

24.8

##### Predicted Retention Time

26.59

##### Area

0

##### Parts Per Million

1.2

##### Fragmentation mode

HCD

##### Originating file

01 D:\separate\_stitch\_analyses\xle-disambiguation\20210325\_F59\_3ug\_DENOVO\_12.csv
