## Supplementary material for "A handle on mass coincidence errors in *de novo* sequencing of antibodies by bottom-up proteomics": Combined_092.html

Details Combined\_092 | Stitch OverviewUndefined

### Read Combined\_092

#### Sequence (length=9)

VVFGGGTKJ

#### Spectrum 6223? Spectrum 6223 The raw spectrum of this peptide as annotated by Hecklib. The fragments are coloured according to ion type (see legend). Any peaks with a star '\*' as text can be hovered over to see the full details, first the ion type second the mass shift type. By hovering over the amino acids in the peptide or ions in the legend the corresponding peaks are highlighted. By toggling the 'Unassigned' label you can turn the background (unassigned) peaks on or off in the plot. By updating the slider in the Ion legend you can update the spectrum to only show the top X% of the peaks with labels. The top X% means any peak that is within X% of the highest intensity. By dragging in the spectrum you can zoom in to a specific part of the spectrum and use 'Zoom Out' to get back to the original zoom level. The annotation of the spectrum is based on the given sequence in the peptides file and is done with different software so inconsistencies are likely. The peaks are annotated based on the given sequence, with 20 ppm tolerance.

Copy Data

##### Spectrum 6223 (TSV)

###### Preview

```
Loading example...
```

*Click on the button to copy the data to your clipboard.*

Mz MinMz MaxIntensity Max

WidthHeightPeptide font sizePeptide stroke widthSpectrum font sizeSpectrum stroke widthCompact peptide

Ion legend

wxyz

abcd

OtherUnassignedIon^Charge^_Position_Show for top:%

VVFGGGTKJ

05.69e+41.14e+51.71e+52.28e+5

Zoom Out

y^+1^_1_a^+1^_2_b^+1^_2_y^+1^_2_y^+1^_2_y^+2^_7_y^+2^_7_y^+1^_3_b^+1^_3_y^+1^_3_b^+2^_8_y^+2^_8_y^+1^_4_b^+1^_4_y^+1^_4_y^+1^_5_y^+1^_5_y^+1^_6_y^+1^_6_b^+1^_7_y^+1^_7_y^+1^_7_b^+1^_8_y^+1^_8_y^+1^_8_

0243485728971

Fragment Matches Table

Show background peaks

| Position | Ion type | Intensity | mz Theoretical | mz Error (Th) | mz Error (ppm) | Charge | Series Number |
| --- | --- | --- | --- | --- | --- | --- | --- |
| - | - | 1.099E+05 | 120.1 | - | - | 0 | - |
| - | - | 7924 | 121.1 | - | - | 0 | - |
| - | - | 373.6 | 123.2 | - | - | 0 | - |
| - | - | 449.9 | 125.1 | - | - | 0 | - |
| - | - | 640.3 | 126.1 | - | - | 0 | - |
| - | - | 584.5 | 126.1 | - | - | 0 | - |
| - | - | 1661 | 127.1 | - | - | 0 | - |
| - | - | 945.2 | 127.1 | - | - | 0 | - |
| - | - | 1128 | 128.1 | - | - | 0 | - |
| - | - | 673.5 | 129.1 | - | - | 0 | - |
| - | - | 1.064E+05 | 129.1 | - | - | 0 | - |
| - | - | 5393 | 129.1 | - | - | 0 | - |
| - | - | 478.2 | 129.9 | - | - | 0 | - |
| - | - | 6816 | 130.1 | - | - | 0 | - |
| - | - | 1264 | 130.1 | - | - | 0 | - |
| - | - | 1126 | 130.1 | - | - | 0 | - |
| - | - | 6005 | 130.1 | - | - | 0 | - |
| - | - | 3858 | 131.1 | - | - | 0 | - |
| - | - | 657.8 | 132 | - | - | 0 | - |
| - | - | 1760 | 132.1 | - | - | 0 | - |
| 9 | y | 4431 | 132.1 | 0.0002067 | 1.565 | +1 | 1 |
| - | - | 412.6 | 133.7 | - | - | 0 | - |
| - | - | 3124 | 136.1 | - | - | 0 | - |
| - | - | 458.5 | 138.1 | - | - | 0 | - |
| - | - | 830.5 | 139.1 | - | - | 0 | - |
| - | - | 1067 | 140.1 | - | - | 0 | - |
| - | - | 441.3 | 140.8 | - | - | 0 | - |
| - | - | 2767 | 141.1 | - | - | 0 | - |
| - | - | 759.3 | 141.1 | - | - | 0 | - |
| - | - | 487.7 | 142.1 | - | - | 0 | - |
| - | - | 654.8 | 143.1 | - | - | 0 | - |
| - | - | 2834 | 146.1 | - | - | 0 | - |
| - | - | 4714 | 146.2 | - | - | 0 | - |
| - | - | 7.81E+04 | 147 | - | - | 0 | - |
| - | - | 4578 | 147.1 | - | - | 0 | - |
| - | - | 484.9 | 147.2 | - | - | 0 | - |
| - | - | 7438 | 148 | - | - | 0 | - |
| - | - | 425.8 | 148.1 | - | - | 0 | - |
| - | - | 590.7 | 149 | - | - | 0 | - |
| - | - | 440.7 | 150 | - | - | 0 | - |
| - | - | 574.3 | 151.1 | - | - | 0 | - |
| - | - | 494.4 | 152.1 | - | - | 0 | - |
| - | - | 553.9 | 153.1 | - | - | 0 | - |
| - | - | 515.7 | 153.1 | - | - | 0 | - |
| - | - | 2448 | 153.1 | - | - | 0 | - |
| - | - | 1117 | 154.1 | - | - | 0 | - |
| - | - | 662.1 | 154.1 | - | - | 0 | - |
| - | - | 949.9 | 155.1 | - | - | 0 | - |
| - | - | 5007 | 155.1 | - | - | 0 | - |
| - | - | 808.4 | 156.1 | - | - | 0 | - |
| - | - | 544.7 | 156.1 | - | - | 0 | - |
| - | - | 810.4 | 156.1 | - | - | 0 | - |
| - | - | 1085 | 157.1 | - | - | 0 | - |
| - | - | 951.5 | 157.1 | - | - | 0 | - |
| - | - | 1354 | 158.1 | - | - | 0 | - |
| - | - | 799.5 | 158.1 | - | - | 0 | - |
| - | - | 3423 | 159.1 | - | - | 0 | - |
| - | - | 860.4 | 159.1 | - | - | 0 | - |
| - | - | 1931 | 159.1 | - | - | 0 | - |
| - | - | 710.2 | 162.1 | - | - | 0 | - |
| - | - | 2141 | 166.1 | - | - | 0 | - |
| - | - | 448.9 | 167.1 | - | - | 0 | - |
| - | - | 1290 | 167.1 | - | - | 0 | - |
| - | - | 676.3 | 169.1 | - | - | 0 | - |
| - | - | 640.1 | 171.1 | - | - | 0 | - |
| 2 | a | 2.254E+05 | 171.1 | 0.0002397 | 1.4 | +1 | 2 |
| - | - | 4409 | 172.1 | - | - | 0 | - |
| - | - | 557.1 | 172.1 | - | - | 0 | - |
| - | - | 2422 | 172.1 | - | - | 0 | - |
| - | - | 1.981E+04 | 172.2 | - | - | 0 | - |
| - | - | 847.4 | 173.2 | - | - | 0 | - |
| - | - | 1529 | 173.5 | - | - | 0 | - |
| - | - | 1014 | 174.1 | - | - | 0 | - |
| - | - | 444.3 | 175 | - | - | 0 | - |
| - | - | 476.6 | 175 | - | - | 0 | - |
| - | - | 2970 | 175.1 | - | - | 0 | - |
| - | - | 6979 | 176.1 | - | - | 0 | - |
| - | - | 7272 | 177.1 | - | - | 0 | - |
| - | - | 1003 | 177.1 | - | - | 0 | - |
| - | - | 631.2 | 178.1 | - | - | 0 | - |
| - | - | 476.2 | 180 | - | - | 0 | - |
| - | - | 620.2 | 180.3 | - | - | 0 | - |
| - | - | 476.9 | 181.1 | - | - | 0 | - |
| - | - | 1064 | 182.1 | - | - | 0 | - |
| - | - | 713.5 | 183.1 | - | - | 0 | - |
| - | - | 1416 | 185.1 | - | - | 0 | - |
| - | - | 1503 | 186.1 | - | - | 0 | - |
| - | - | 1517 | 187.1 | - | - | 0 | - |
| - | - | 712.1 | 188.1 | - | - | 0 | - |
| - | - | 1198 | 188.1 | - | - | 0 | - |
| - | - | 647 | 194.1 | - | - | 0 | - |
| - | - | 1354 | 195.1 | - | - | 0 | - |
| - | - | 615.8 | 196.1 | - | - | 0 | - |
| - | - | 1141 | 197.2 | - | - | 0 | - |
| - | - | 4670 | 198.1 | - | - | 0 | - |
| 2 | b | 8.516E+04 | 199.1 | 7.601E-05 | 0.3817 | +1 | 2 |
| - | - | 7886 | 200.1 | - | - | 0 | - |
| - | - | 519.4 | 201.1 | - | - | 0 | - |
| - | - | 628.3 | 201.1 | - | - | 0 | - |
| - | - | 488.1 | 203.1 | - | - | 0 | - |
| - | - | 668.3 | 203.1 | - | - | 0 | - |
| - | - | 2557 | 203.1 | - | - | 0 | - |
| - | - | 7.721E+04 | 204.1 | - | - | 0 | - |
| - | - | 1.366E+04 | 205.1 | - | - | 0 | - |
| - | - | 1.108E+04 | 205.1 | - | - | 0 | - |
| - | - | 1802 | 206.1 | - | - | 0 | - |
| - | - | 1130 | 208.1 | - | - | 0 | - |
| - | - | 1018 | 209.1 | - | - | 0 | - |
| - | - | 538.3 | 210.1 | - | - | 0 | - |
| - | - | 692.1 | 211.1 | - | - | 0 | - |
| - | - | 1.2E+04 | 212.1 | - | - | 0 | - |
| - | - | 1683 | 213.1 | - | - | 0 | - |
| - | - | 586.3 | 214.1 | - | - | 0 | - |
| - | - | 950.2 | 215.1 | - | - | 0 | - |
| - | - | 2872 | 216.1 | - | - | 0 | - |
| - | - | 3198 | 217.1 | - | - | 0 | - |
| - | - | 1.388E+04 | 218.1 | - | - | 0 | - |
| - | - | 2686 | 219.1 | - | - | 0 | - |
| - | - | 7636 | 219.1 | - | - | 0 | - |
| - | - | 775.1 | 220.2 | - | - | 0 | - |
| - | - | 4507 | 221.1 | - | - | 0 | - |
| - | - | 922 | 225.1 | - | - | 0 | - |
| - | - | 1929 | 226.2 | - | - | 0 | - |
| - | - | 561.6 | 227.1 | - | - | 0 | - |
| - | - | 5125 | 227.1 | - | - | 0 | - |
| - | - | 2461 | 228.1 | - | - | 0 | - |
| - | - | 585.9 | 229.1 | - | - | 0 | - |
| - | - | 1.958E+04 | 230.2 | - | - | 0 | - |
| - | - | 677.6 | 231.1 | - | - | 0 | - |
| - | - | 1452 | 231.2 | - | - | 0 | - |
| - | - | 1219 | 233.1 | - | - | 0 | - |
| - | - | 3797 | 233.2 | - | - | 0 | - |
| - | - | 642.1 | 234.1 | - | - | 0 | - |
| - | - | 845.4 | 234.2 | - | - | 0 | - |
| - | - | 1444 | 235.1 | - | - | 0 | - |
| - | - | 1480 | 237.1 | - | - | 0 | - |
| - | - | 1121 | 239.2 | - | - | 0 | - |
| - | - | 848.5 | 242.2 | - | - | 0 | - |
| - | - | 2668 | 243.1 | - | - | 0 | - |
| 8 | y | 7709 | 243.2 | 4.326E-07 | 0.001779 | +1 | 2 |
| - | - | 609.5 | 244.2 | - | - | 0 | - |
| - | - | 585.1 | 245.1 | - | - | 0 | - |
| - | - | 2688 | 245.1 | - | - | 0 | - |
| - | - | 661.1 | 246.1 | - | - | 0 | - |
| - | - | 5743 | 247.1 | - | - | 0 | - |
| - | - | 838.7 | 248.1 | - | - | 0 | - |
| - | - | 1190 | 248.2 | - | - | 0 | - |
| - | - | 967.6 | 249.2 | - | - | 0 | - |
| - | - | 1.079E+04 | 255.1 | - | - | 0 | - |
| - | - | 1105 | 256.1 | - | - | 0 | - |
| 8 | y | 3.471E+04 | 260.2 | 2.976E-05 | 0.1144 | +1 | 2 |
| - | - | 2754 | 261.1 | - | - | 0 | - |
| - | - | 1039 | 261.2 | - | - | 0 | - |
| - | - | 4477 | 261.2 | - | - | 0 | - |
| - | - | 4631 | 262.1 | - | - | 0 | - |
| - | - | 706.6 | 263.1 | - | - | 0 | - |
| - | - | 2357 | 263.1 | - | - | 0 | - |
| - | - | 9888 | 269.2 | - | - | 0 | - |
| - | - | 1161 | 270.2 | - | - | 0 | - |
| - | - | 747.4 | 270.2 | - | - | 0 | - |
| - | - | 649.8 | 271.1 | - | - | 0 | - |
| - | - | 616.2 | 272.1 | - | - | 0 | - |
| - | - | 8693 | 273.1 | - | - | 0 | - |
| - | - | 7822 | 274.1 | - | - | 0 | - |
| - | - | 3546 | 274.2 | - | - | 0 | - |
| - | - | 1112 | 275.1 | - | - | 0 | - |
| - | - | 1.807E+04 | 275.2 | - | - | 0 | - |
| - | - | 3721 | 276.2 | - | - | 0 | - |
| - | - | 844.2 | 277.1 | - | - | 0 | - |
| - | - | 923.8 | 277.2 | - | - | 0 | - |
| - | - | 1219 | 282.2 | - | - | 0 | - |
| - | - | 2162 | 287.2 | - | - | 0 | - |
| - | - | 1071 | 288.1 | - | - | 0 | - |
| - | - | 2981 | 291.1 | - | - | 0 | - |
| - | - | 1935 | 292.1 | - | - | 0 | - |
| - | - | 8.176E+04 | 292.2 | - | - | 0 | - |
| - | - | 1.778E+04 | 293.2 | - | - | 0 | - |
| - | - | 1505 | 294.2 | - | - | 0 | - |
| - | - | 998.8 | 300.1 | - | - | 0 | - |
| - | - | 3960 | 301.2 | - | - | 0 | - |
| - | - | 1354 | 304.2 | - | - | 0 | - |
| - | - | 1819 | 317.2 | - | - | 0 | - |
| - | - | 1195 | 318.1 | - | - | 0 | - |
| - | - | 938.6 | 318.2 | - | - | 0 | - |
| - | - | 4272 | 319.1 | - | - | 0 | - |
| - | - | 3661 | 322.2 | - | - | 0 | - |
| - | - | 9086 | 326.2 | - | - | 0 | - |
| - | - | 807.9 | 327.2 | - | - | 0 | - |
| - | - | 604.4 | 327.8 | - | - | 0 | - |
| 3 | y | 1222 | 331.2 | 0.0008345 | 2.52 | +2 | 7 |
| - | - | 6335 | 333.1 | - | - | 0 | - |
| - | - | 1371 | 334.1 | - | - | 0 | - |
| - | - | 1573 | 339.2 | - | - | 0 | - |
| 3 | y | 1.912E+04 | 340.2 | 9.925E-05 | 0.2918 | +2 | 7 |
| - | - | 7267 | 340.7 | - | - | 0 | - |
| - | - | 1292 | 341.2 | - | - | 0 | - |
| 7 | y | 1431 | 343.2 | 0.001064 | 3.101 | +1 | 3 |
| - | - | 2479 | 344.2 | - | - | 0 | - |
| - | - | 711.5 | 345.2 | - | - | 0 | - |
| 3 | b | 3407 | 346.2 | 0.0002379 | 0.6872 | +1 | 3 |
| - | - | 2778 | 348.2 | - | - | 0 | - |
| - | - | 3849 | 350.1 | - | - | 0 | - |
| - | - | 868 | 356.2 | - | - | 0 | - |
| - | - | 3306 | 357.2 | - | - | 0 | - |
| - | - | 905.4 | 358.2 | - | - | 0 | - |
| - | - | 744.4 | 361.2 | - | - | 0 | - |
| 7 | y | 6653 | 361.2 | 0.0002534 | 0.7015 | +1 | 3 |
| - | - | 1563 | 362.2 | - | - | 0 | - |
| - | - | 1208 | 362.2 | - | - | 0 | - |
| 8 | b | 1821 | 365.2 | 0.007207 | 19.73 | +2 | 8 |
| - | - | 4815 | 374.2 | - | - | 0 | - |
| - | - | 2629 | 375.2 | - | - | 0 | - |
| - | - | 892 | 375.2 | - | - | 0 | - |
| - | - | 1987 | 376.2 | - | - | 0 | - |
| - | - | 7616 | 383.2 | - | - | 0 | - |
| - | - | 790.1 | 384.2 | - | - | 0 | - |
| - | - | 1126 | 384.2 | - | - | 0 | - |
| - | - | 1214 | 385.1 | - | - | 0 | - |
| - | - | 557.4 | 388.7 | - | - | 0 | - |
| 2 | y | 1712 | 389.7 | 0.0007825 | 2.008 | +2 | 8 |
| - | - | 6781 | 392.2 | - | - | 0 | - |
| - | - | 941.6 | 393.2 | - | - | 0 | - |
| 6 | y | 1188 | 400.3 | 0.00136 | 3.397 | +1 | 4 |
| - | - | 4384 | 401.2 | - | - | 0 | - |
| - | - | 1.717E+04 | 402.2 | - | - | 0 | - |
| - | - | 572.6 | 403 | - | - | 0 | - |
| - | - | 3335 | 403.2 | - | - | 0 | - |
| 4 | b | 1035 | 403.2 | 0.002537 | 6.291 | +1 | 4 |
| - | - | 776.4 | 405.2 | - | - | 0 | - |
| 6 | y | 5438 | 418.3 | 0.000164 | 0.392 | +1 | 4 |
| - | - | 3234 | 419.2 | - | - | 0 | - |
| - | - | 1218 | 419.3 | - | - | 0 | - |
| - | - | 9894 | 420.2 | - | - | 0 | - |
| - | - | 2997 | 420.2 | - | - | 0 | - |
| - | - | 3.51E+04 | 421.2 | - | - | 0 | - |
| - | - | 953.2 | 421.8 | - | - | 0 | - |
| - | - | 913.8 | 422.2 | - | - | 0 | - |
| - | - | 1.337E+04 | 422.2 | - | - | 0 | - |
| - | - | 1954 | 423.2 | - | - | 0 | - |
| - | - | 758.9 | 437.8 | - | - | 0 | - |
| - | - | 1.743E+05 | 438.2 | - | - | 0 | - |
| - | - | 6.787E+04 | 439.2 | - | - | 0 | - |
| - | - | 848.4 | 439.8 | - | - | 0 | - |
| - | - | 7479 | 439.8 | - | - | 0 | - |
| - | - | 469.6 | 440.2 | - | - | 0 | - |
| - | - | 1.125E+04 | 440.2 | - | - | 0 | - |
| 5 | y | 2023 | 457.3 | 6.817E-05 | 0.1491 | +1 | 5 |
| - | - | 702.7 | 470.3 | - | - | 0 | - |
| - | - | 1343 | 473.3 | - | - | 0 | - |
| 5 | y | 2.129E+04 | 475.3 | 5.951E-05 | 0.1252 | +1 | 5 |
| - | - | 899.7 | 475.8 | - | - | 0 | - |
| - | - | 4820 | 476.3 | - | - | 0 | - |
| - | - | 988.7 | 477.3 | - | - | 0 | - |
| - | - | 751.7 | 486.3 | - | - | 0 | - |
| - | - | 2307 | 496.3 | - | - | 0 | - |
| - | - | 848.8 | 497.3 | - | - | 0 | - |
| - | - | 870.6 | 501.2 | - | - | 0 | - |
| - | - | 2277 | 512.3 | - | - | 0 | - |
| 4 | y | 4503 | 514.3 | 9.429E-05 | 0.1833 | +1 | 6 |
| - | - | 923.9 | 515.3 | - | - | 0 | - |
| - | - | 1810 | 519.8 | - | - | 0 | - |
| - | - | 1881 | 521.8 | - | - | 0 | - |
| - | - | 964.4 | 522.3 | - | - | 0 | - |
| - | - | 1032 | 525.8 | - | - | 0 | - |
| - | - | 722.8 | 526.3 | - | - | 0 | - |
| - | - | 618.3 | 529.3 | - | - | 0 | - |
| - | - | 1697 | 529.8 | - | - | 0 | - |
| - | - | 8751 | 530.3 | - | - | 0 | - |
| - | - | 3020 | 531.3 | - | - | 0 | - |
| 4 | y | 7.256E+04 | 532.3 | 0.0004661 | 0.8756 | +1 | 6 |
| - | - | 1.666E+04 | 533.3 | - | - | 0 | - |
| - | - | 2366 | 534.3 | - | - | 0 | - |
| - | - | 1387 | 539.8 | - | - | 0 | - |
| - | - | 998.2 | 540.3 | - | - | 0 | - |
| - | - | 638.5 | 540.8 | - | - | 0 | - |
| - | - | 1.533E+04 | 548.3 | - | - | 0 | - |
| - | - | 3514 | 549.3 | - | - | 0 | - |
| - | - | 979.4 | 564.3 | - | - | 0 | - |
| - | - | 9124 | 566.3 | - | - | 0 | - |
| - | - | 2435 | 567.3 | - | - | 0 | - |
| - | - | 654.3 | 572.8 | - | - | 0 | - |
| - | - | 927.5 | 586.8 | - | - | 0 | - |
| 7 | b | 737.8 | 600.3 | 0.005374 | 8.952 | +1 | 7 |
| - | - | 977.5 | 608.3 | - | - | 0 | - |
| - | - | 1009 | 617.3 | - | - | 0 | - |
| - | - | 1612 | 633.4 | - | - | 0 | - |
| - | - | 1067 | 635.4 | - | - | 0 | - |
| - | - | 4979 | 643.4 | - | - | 0 | - |
| - | - | 1614 | 644.4 | - | - | 0 | - |
| - | - | 1139 | 647.4 | - | - | 0 | - |
| 3 | y | 1.136E+04 | 661.4 | 0.000515 | 0.7788 | +1 | 7 |
| - | - | 4145 | 662.4 | - | - | 0 | - |
| - | - | 1036 | 663.4 | - | - | 0 | - |
| - | - | 856.5 | 677.4 | - | - | 0 | - |
| 3 | y | 2.172E+05 | 679.4 | 0.0008258 | 1.216 | +1 | 7 |
| - | - | 7.102E+04 | 680.4 | - | - | 0 | - |
| - | - | 1.657E+04 | 681.4 | - | - | 0 | - |
| - | - | 1505 | 682.4 | - | - | 0 | - |
| - | - | 772.7 | 720.4 | - | - | 0 | - |
| 8 | b | 793.5 | 746.4 | 0.001103 | 1.478 | +1 | 8 |
| 2 | y | 847.4 | 760.4 | 2.027E-05 | 0.02665 | +1 | 8 |
| - | - | 631.5 | 777.3 | - | - | 0 | - |
| 2 | y | 2.097E+04 | 778.4 | 0.001735 | 2.229 | +1 | 8 |
| - | - | 9075 | 779.4 | - | - | 0 | - |
| - | - | 3326 | 780.5 | - | - | 0 | - |
| - | - | 717.1 | 961 | - | - | 0 | - |

m/z Charge Intensity FragmentType MassShift Position
120.08101654052734 0 109914.48
121.08432006835938 0 7924.1636
123.16796112060547 0 373.55133
125.07096099853516 0 449.8619
126.05536651611328 0 640.2825
126.10273742675781 0 584.4976
127.05049896240234 0 1660.9288
127.0867919921875 0 945.2387
128.1071319580078 0 1128.2184
129.06619262695312 0 673.5082
129.1024627685547 0 106390.555
129.1388397216797 0 5392.8506
129.9318389892578 0 478.2046
130.0653533935547 0 6815.8706
130.08670043945312 0 1263.8768
130.10025024414062 0 1125.6322
130.10580444335938 0 6004.927
131.0816650390625 0 3857.6047
132.0449676513672 0 657.7625
132.08103942871094 0 1760.4642
132.10211181640625 0 4430.5625 y 8
133.71420288085938 0 412.5879
136.0758514404297 0 3124.3733
138.0662078857422 0 458.49176
139.08694458007812 0 830.4997
140.08192443847656 0 1066.7332
140.8058624267578 0 441.3212
141.0659942626953 0 2766.7468
141.1022186279297 0 759.2873
142.0978240966797 0 487.68237
143.08148193359375 0 654.7934
146.06024169921875 0 2834.3728
146.16539001464844 0 4713.75
147.0442657470703 0 78100.555
147.11302185058594 0 4577.638
147.1691436767578 0 484.90372
148.04762268066406 0 7437.9224
148.0751953125 0 425.76495
148.955322265625 0 590.71387
149.96949768066406 0 440.70737
151.08657836914062 0 574.3095
152.08187866210938 0 494.39865
153.05487060546875 0 553.9259
153.06617736816406 0 515.68024
153.0772705078125 0 2447.5427
154.0613555908203 0 1117.4583
154.09779357910156 0 662.09705
155.08148193359375 0 949.8917
155.11802673339844 0 5006.9595
156.07684326171875 0 808.39087
156.11410522460938 0 544.7473
156.12132263183594 0 810.4298
157.06106567382812 0 1085.2678
157.0765838623047 0 951.46844
158.06040954589844 0 1353.881
158.0923309326172 0 799.52386
159.07662963867188 0 3422.7817
159.09193420410156 0 860.4059
159.11288452148438 0 1930.8333
162.0556182861328 0 710.16693
166.0533447265625 0 2140.727
167.0814666748047 0 448.8918
167.11817932128906 0 1289.6052
169.13392639160156 0 676.27216
171.07704162597656 0 640.0762
171.14942932128906 0 225447.7 a 1
172.07179260253906 0 4408.7744
172.1078643798828 0 557.09155
172.14527893066406 0 2421.533
172.15274047851562 0 19807.684
173.1553955078125 0 847.4406
173.45176696777344 0 1528.5803
174.0550537109375 0 1013.6333
174.991455078125 0 444.2865
175.03021240234375 0 476.60165
175.08663940429688 0 2969.9568
176.1071319580078 0 6978.8984
177.1023712158203 0 7271.968
177.11053466796875 0 1003.4233
178.10643005371094 0 631.1576
179.95870971679688 0 476.22418
180.3291778564453 0 620.1756
181.07205200195312 0 476.88815
182.0817108154297 0 1064.0674
183.11282348632812 0 713.45215
185.12850952148438 0 1416.1857
186.1239471435547 0 1503.4185
187.10792541503906 0 1516.7463
188.0709991455078 0 712.0826
188.1031036376953 0 1198.0648
194.12855529785156 0 646.98584
195.11300659179688 0 1353.8214
196.10862731933594 0 615.8488
197.1652374267578 0 1140.6643
198.08740234375 0 4669.597
199.14418029785156 0 85157.26 b 1
200.14761352539062 0 7885.731
201.10084533691406 0 519.43176
201.14840698242188 0 628.293
203.0805206298828 0 488.0705
203.0917510986328 0 668.31683
203.1179962158203 0 2557.4604
204.10195922851562 0 77212.1
205.0969696044922 0 13656.971
205.10572814941406 0 11076.7
206.10060119628906 0 1801.7424
208.10818481445312 0 1130.4602
209.10299682617188 0 1018.4646
210.08702087402344 0 538.31683
211.1442413330078 0 692.0932
212.13946533203125 0 12004.7705
213.14280700683594 0 1682.5598
214.1202392578125 0 586.2821
215.1156005859375 0 950.1688
216.09803771972656 0 2871.7632
217.0972900390625 0 3198.0408
218.1176300048828 0 13879.046
219.12120056152344 0 2685.636
219.14927673339844 0 7636.382
220.15280151367188 0 775.115
221.12855529785156 0 4506.8623
225.13514709472656 0 921.96484
226.1551513671875 0 1929.2296
227.08212280273438 0 561.647
227.11392211914062 0 5124.9146
228.09774780273438 0 2460.7979
229.0941619873047 0 585.89703
230.15000915527344 0 19578.686
231.11264038085938 0 677.5905
231.15277099609375 0 1452.0552
233.12860107421875 0 1218.6373
233.16493225097656 0 3796.7656
234.1234893798828 0 642.1174
234.16932678222656 0 845.4423
235.1442108154297 0 1443.7144
237.09832763671875 0 1480.3179
239.15040588378906 0 1120.9869
242.186767578125 0 848.5119
243.11253356933594 0 2667.8794
243.17031860351562 0 7709.4316 y Ammonia loss 7
244.1738739013672 0 609.5471
245.11178588867188 0 585.065
245.1248779296875 0 2687.7168
246.1239776611328 0 661.1452
247.14418029785156 0 5743.4277
248.14793395996094 0 838.74524
248.16099548339844 0 1189.5841
249.15956115722656 0 967.619
255.1087646484375 0 10792.857
256.11151123046875 0 1105.3348
260.19683837890625 0 34707.633 y 7
261.12335205078125 0 2753.589
261.159423828125 0 1038.6343
261.20013427734375 0 4477.202
262.11859130859375 0 4631.1323
263.12457275390625 0 706.649
263.1388854980469 0 2357.116
269.1607360839844 0 9887.568
270.1647644042969 0 1161.113
270.1807861328125 0 747.3877
271.10748291015625 0 649.83075
272.13494873046875 0 616.164
273.1193542480469 0 8692.811
274.1188049316406 0 7821.9336
274.1910705566406 0 3545.9636
275.123779296875 0 1112.4078
275.1753845214844 0 18070.543
276.17840576171875 0 3720.845
277.13311767578125 0 844.1894
277.1815185546875 0 923.84924
282.1556701660156 0 1218.7003
287.171875 0 2161.792
288.13427734375 0 1071.0159
291.14532470703125 0 2981.2297
292.128662109375 0 1935.4058
292.2019958496094 0 81757.89
293.2051696777344 0 17781.26
294.2076416015625 0 1505.0908
300.1351318359375 0 998.81354
301.1907958984375 0 3960.1956
304.165771484375 0 1354.0553
317.1861572265625 0 1818.584
318.1459655761719 0 1194.722
318.18267822265625 0 938.62604
319.13970947265625 0 4272.028
322.1550598144531 0 3661.0073
326.1824035644531 0 9085.904
327.18536376953125 0 807.854
327.82769775390625 0 604.35315
331.1878662109375 0 1221.6124 y Water loss 2
333.1234130859375 0 6335.1685
334.1266174316406 0 1371.2853
339.1772766113281 0 1573.4863
340.1924133300781 0 19115.477 y 2
340.69384765625 0 7267.4043
341.19671630859375 0 1292.0552
343.23504638671875 0 1431.2023 y Water loss 6
344.19293212890625 0 2478.9014
345.154052734375 0 711.48267
346.2122802734375 0 3407.145 b 2
348.16607666015625 0 2778.28
350.1498718261719 0 3848.77
356.1730041503906 0 867.99036
357.1549987792969 0 3305.789
358.1570129394531 0 905.40704
361.1875 0 744.3966
361.2442932128906 0 6653.399 y 6
362.2037353515625 0 1562.9789
362.24835205078125 0 1207.7001
365.19293212890625 0 1821.2214 b Ammonia loss 7
374.18218994140625 0 4814.8945
375.1663818359375 0 2628.5823
375.2400817871094 0 892.0372
376.16180419921875 0 1986.5967
383.2035217285156 0 7616.3643
384.16473388671875 0 790.09033
384.2066345214844 0 1126.3116
385.14947509765625 0 1213.9095
388.6844482421875 0 557.43146
389.7257385253906 0 1711.8671 y 1
392.1925354003906 0 6780.5396
393.19268798828125 0 941.64655
400.2568054199219 0 1188.1561 y Water loss 5
401.214111328125 0 4384.2085
402.17706298828125 0 17172.51
402.9637756347656 0 572.6317
403.1800231933594 0 3335.2805
403.2314453125 0 1035.3057 b 3
405.1750183105469 0 776.41205
418.26617431640625 0 5438.048 y 5
419.2249450683594 0 3234.4238
419.2681579589844 0 1217.5463
420.187744140625 0 9893.956
420.2283935546875 0 2996.9734
421.2122497558594 0 35104.57
421.833251953125 0 953.19293
422.1867370605469 0 913.8296
422.21539306640625 0 13369.697
423.2180480957031 0 1953.8944
437.84832763671875 0 758.937
438.23876953125 0 174295.61
439.241943359375 0 67866.664
439.7629089355469 0 848.4442
439.8437194824219 0 7479.192
440.21160888671875 0 469.63242
440.2446594238281 0 11250.642
457.2769775390625 0 2023.4286 y Water loss 4
470.2730407714844 0 702.7346
473.2518310546875 0 1343.1172
475.28741455078125 0 21291.502 y 4
475.7881164550781 0 899.70325
476.2908630371094 0 4820.324
477.29217529296875 0 988.69666
486.3018798828125 0 751.7222
496.28790283203125 0 2306.8325
497.273681640625 0 848.8083
501.2447509765625 0 870.5602
512.26025390625 0 2276.9436
514.2982788085938 0 4503.0977 y Water loss 3
515.3009643554688 0 923.9443
519.7687377929688 0 1810.1931
521.7831420898438 0 1881.27
522.2808837890625 0 964.35706
525.8092041015625 0 1032.1067
526.3133544921875 0 722.7778
529.2747192382812 0 618.338
529.7793579101562 0 1696.6646
530.2726440429688 0 8750.583
531.2743530273438 0 3020.1018
532.3084716796875 0 72559.375 y 3
533.3111572265625 0 16656.36
534.3131713867188 0 2366.2178
539.8079833984375 0 1386.7817
540.3107299804688 0 998.23785
540.8151245117188 0 638.48065
548.2822265625 0 15327.9
549.284912109375 0 3514.0828
564.2617797851562 0 979.3678
566.2922973632812 0 9124.233
567.2952270507812 0 2435.3564
572.7635498046875 0 654.34674
586.766357421875 0 927.4561
600.3193969726562 0 737.8463 b Water loss 6
608.2781372070312 0 977.5014
617.3419799804688 0 1009.2742
633.36865234375 0 1612.0385
635.3568115234375 0 1067.3921
643.3548583984375 0 4978.729
644.3532104492188 0 1613.7484
647.3517456054688 0 1139.4069
661.3662719726562 0 11364.346 y Water loss 2
662.3677978515625 0 4144.9634
663.3758544921875 0 1036.4014
677.3593139648438 0 856.4881
679.3765258789062 0 217195.56 y 2
680.3792114257812 0 71024.14
681.3819580078125 0 16574.09
682.385498046875 0 1505.4274
720.3556518554688 0 772.7116
746.420654296875 0 793.51697 b 7
760.4351806640625 0 847.4215 y Water loss 1
777.3461303710938 0 631.4988
778.4440307617188 0 20974.242 y 1
779.4468383789062 0 9075.08
780.4509887695312 0 3326.4243
961.0185546875 0 717.1312

Spectrum Details

|  |  |
| --- | --- |
| Matched peaks? Matched peaks The total absolute number of peaks matched. Additionally in brackets the total fraction of peaks matched and the total number of peaks is shown. | 25 (8.17% of 306) |
| FDR? FDR The false discovery rate estimated for this peptide. It is calculated by matching all theoretical fragments with a non-integer shift with the raw peaks for this spectrum. This is done with 40 different shifts. The resulting percentage is the average number of annotated peaks over the number of annotated peaks with the correct spectrum. | 0.67% |
| Satellite FDR? Satellite FDR See the FDR for details on its calculation. This satellite ion specific FDR only contains the satellite ions (d/w) for I/L/J positions. | - |
| PSM Score? PSM Score The PSM Score as given by Hecklib to this annotated spectrum. It is shown with three significant figures. | 323 |

#### Spectrum 6556? Spectrum 6556 The raw spectrum of this peptide as annotated by Hecklib. The fragments are coloured according to ion type (see legend). Any peaks with a star '\*' as text can be hovered over to see the full details, first the ion type second the mass shift type. By hovering over the amino acids in the peptide or ions in the legend the corresponding peaks are highlighted. By toggling the 'Unassigned' label you can turn the background (unassigned) peaks on or off in the plot. By updating the slider in the Ion legend you can update the spectrum to only show the top X% of the peaks with labels. The top X% means any peak that is within X% of the highest intensity. By dragging in the spectrum you can zoom in to a specific part of the spectrum and use 'Zoom Out' to get back to the original zoom level. The annotation of the spectrum is based on the given sequence in the peptides file and is done with different software so inconsistencies are likely. The peaks are annotated based on the given sequence, with 20 ppm tolerance.

Copy Data

##### Spectrum 6556 (TSV)

###### Preview

```
Loading example...
```

*Click on the button to copy the data to your clipboard.*

Mz MinMz MaxIntensity Max

WidthHeightPeptide font sizePeptide stroke widthSpectrum font sizeSpectrum stroke widthCompact peptide

Ion legend

wxyz

abcd

OtherUnassignedIon^Charge^_Position_Show for top:%

VVFGGGTKJ

04.47e+48.93e+41.34e+51.79e+5

Zoom Out

y^+1^_1_d^+1^_2_a^+1^_2_b^+1^_2_y^+2^_4_y^+1^_2_y^+1^_2_y^+2^_7_y^+1^_3_b^+1^_3_y^+1^_3_b^+2^_8_b^+2^_8_y^+2^_8_y^+1^_4_b^+1^_4_y^+1^_4_\*\*y^+1^_5_y^+1^_5_y^+1^_6_y^+1^_6_b^+1^_7_y^+1^_7_y^+1^_7_b^+1^_8_y^+1^_8_y^+1^_8_\*

0778155623343112

Fragment Matches Table

Show background peaks

| Position | Ion type | Intensity | mz Theoretical | mz Error (Th) | mz Error (ppm) | Charge | Series Number |
| --- | --- | --- | --- | --- | --- | --- | --- |
| - | - | 9.097E+04 | 120.1 | - | - | 0 | - |
| - | - | 405.8 | 120.5 | - | - | 0 | - |
| - | - | 7246 | 121.1 | - | - | 0 | - |
| - | - | 732.4 | 123 | - | - | 0 | - |
| - | - | 928.7 | 124.1 | - | - | 0 | - |
| - | - | 445.6 | 125.1 | - | - | 0 | - |
| - | - | 764.9 | 126.1 | - | - | 0 | - |
| - | - | 1352 | 127.1 | - | - | 0 | - |
| - | - | 617.7 | 127.1 | - | - | 0 | - |
| - | - | 667.6 | 128.1 | - | - | 0 | - |
| - | - | 1062 | 129.1 | - | - | 0 | - |
| - | - | 9.067E+04 | 129.1 | - | - | 0 | - |
| - | - | 425.1 | 130.1 | - | - | 0 | - |
| - | - | 1147 | 130.1 | - | - | 0 | - |
| - | - | 1478 | 130.1 | - | - | 0 | - |
| - | - | 737.4 | 130.1 | - | - | 0 | - |
| - | - | 5188 | 130.1 | - | - | 0 | - |
| - | - | 4487 | 131.1 | - | - | 0 | - |
| - | - | 803.7 | 131.1 | - | - | 0 | - |
| - | - | 452.7 | 132 | - | - | 0 | - |
| - | - | 1482 | 132.1 | - | - | 0 | - |
| 9 | y | 5296 | 132.1 | 0.000344 | 2.604 | +1 | 1 |
| - | - | 530.9 | 133.1 | - | - | 0 | - |
| - | - | 459 | 134.1 | - | - | 0 | - |
| - | - | 7611 | 136.1 | - | - | 0 | - |
| - | - | 470 | 136.1 | - | - | 0 | - |
| - | - | 405.1 | 138.4 | - | - | 0 | - |
| - | - | 703.4 | 139.1 | - | - | 0 | - |
| - | - | 1639 | 141.1 | - | - | 0 | - |
| - | - | 1274 | 141.1 | - | - | 0 | - |
| - | - | 561.1 | 144.1 | - | - | 0 | - |
| - | - | 3938 | 146.1 | - | - | 0 | - |
| - | - | 1001 | 147 | - | - | 0 | - |
| - | - | 4292 | 147.1 | - | - | 0 | - |
| - | - | 422.7 | 148 | - | - | 0 | - |
| - | - | 552.1 | 148 | - | - | 0 | - |
| - | - | 417.1 | 148.9 | - | - | 0 | - |
| - | - | 474.7 | 148.9 | - | - | 0 | - |
| - | - | 593.4 | 148.9 | - | - | 0 | - |
| - | - | 781.5 | 148.9 | - | - | 0 | - |
| - | - | 1025 | 148.9 | - | - | 0 | - |
| - | - | 1042 | 148.9 | - | - | 0 | - |
| - | - | 1237 | 148.9 | - | - | 0 | - |
| - | - | 2587 | 148.9 | - | - | 0 | - |
| - | - | 4092 | 148.9 | - | - | 0 | - |
| - | - | 4156 | 149 | - | - | 0 | - |
| - | - | 2084 | 149 | - | - | 0 | - |
| - | - | 1395 | 149 | - | - | 0 | - |
| - | - | 1349 | 149 | - | - | 0 | - |
| - | - | 716.3 | 149 | - | - | 0 | - |
| - | - | 513.7 | 149 | - | - | 0 | - |
| - | - | 477.8 | 149 | - | - | 0 | - |
| - | - | 587.6 | 149 | - | - | 0 | - |
| - | - | 421.2 | 149.8 | - | - | 0 | - |
| - | - | 666.1 | 152.1 | - | - | 0 | - |
| - | - | 1512 | 155.1 | - | - | 0 | - |
| - | - | 2546 | 155.1 | - | - | 0 | - |
| - | - | 630.9 | 157.1 | - | - | 0 | - |
| 2 | d | 601.2 | 157.1 | 0.0004326 | 2.753 | +1 | 2 |
| - | - | 489.2 | 158.1 | - | - | 0 | - |
| - | - | 3491 | 159.1 | - | - | 0 | - |
| - | - | 1339 | 159.1 | - | - | 0 | - |
| - | - | 3927 | 165.1 | - | - | 0 | - |
| - | - | 1132 | 166.1 | - | - | 0 | - |
| - | - | 715.5 | 167.1 | - | - | 0 | - |
| - | - | 1271 | 167.1 | - | - | 0 | - |
| - | - | 486 | 168 | - | - | 0 | - |
| - | - | 739.3 | 169.1 | - | - | 0 | - |
| - | - | 1230 | 169.1 | - | - | 0 | - |
| - | - | 695.8 | 171.1 | - | - | 0 | - |
| 2 | a | 1.769E+05 | 171.1 | 0.0004228 | 2.47 | +1 | 2 |
| - | - | 3661 | 172.1 | - | - | 0 | - |
| - | - | 1221 | 172.1 | - | - | 0 | - |
| - | - | 1.467E+04 | 172.2 | - | - | 0 | - |
| - | - | 6581 | 173.1 | - | - | 0 | - |
| - | - | 722 | 173.2 | - | - | 0 | - |
| - | - | 2666 | 174.1 | - | - | 0 | - |
| - | - | 455.2 | 174.1 | - | - | 0 | - |
| - | - | 2281 | 175.1 | - | - | 0 | - |
| - | - | 4800 | 176.1 | - | - | 0 | - |
| - | - | 6295 | 177.1 | - | - | 0 | - |
| - | - | 1126 | 177.1 | - | - | 0 | - |
| - | - | 1185 | 178.1 | - | - | 0 | - |
| - | - | 536 | 179.8 | - | - | 0 | - |
| - | - | 837.6 | 181.1 | - | - | 0 | - |
| - | - | 4230 | 182.1 | - | - | 0 | - |
| - | - | 1582 | 183.1 | - | - | 0 | - |
| - | - | 1610 | 185.1 | - | - | 0 | - |
| - | - | 1871 | 186.1 | - | - | 0 | - |
| - | - | 1609 | 187.1 | - | - | 0 | - |
| - | - | 1374 | 188.1 | - | - | 0 | - |
| - | - | 1454 | 188.1 | - | - | 0 | - |
| - | - | 556.6 | 189.1 | - | - | 0 | - |
| - | - | 602.2 | 191.1 | - | - | 0 | - |
| - | - | 1041 | 192.1 | - | - | 0 | - |
| - | - | 1561 | 195.1 | - | - | 0 | - |
| - | - | 996.4 | 197.1 | - | - | 0 | - |
| - | - | 1358 | 197.2 | - | - | 0 | - |
| - | - | 3841 | 198.1 | - | - | 0 | - |
| - | - | 1118 | 199.1 | - | - | 0 | - |
| 2 | b | 6.951E+04 | 199.1 | 0.0003354 | 1.684 | +1 | 2 |
| - | - | 6642 | 200.1 | - | - | 0 | - |
| - | - | 801.9 | 200.2 | - | - | 0 | - |
| 6 | y | 1874 | 201.1 | 0.0002884 | 1.434 | +2 | 4 |
| - | - | 569.8 | 201.1 | - | - | 0 | - |
| - | - | 924.1 | 202.1 | - | - | 0 | - |
| - | - | 633.3 | 203.1 | - | - | 0 | - |
| - | - | 1130 | 203.1 | - | - | 0 | - |
| - | - | 1.193E+04 | 205.1 | - | - | 0 | - |
| - | - | 895.8 | 206.1 | - | - | 0 | - |
| - | - | 1692 | 208.1 | - | - | 0 | - |
| - | - | 489.3 | 209.1 | - | - | 0 | - |
| - | - | 861.3 | 209.1 | - | - | 0 | - |
| - | - | 501.4 | 209.3 | - | - | 0 | - |
| - | - | 1871 | 211.1 | - | - | 0 | - |
| - | - | 9716 | 212.1 | - | - | 0 | - |
| - | - | 719 | 213.1 | - | - | 0 | - |
| - | - | 691 | 215.1 | - | - | 0 | - |
| - | - | 873.8 | 215.1 | - | - | 0 | - |
| - | - | 3126 | 216.1 | - | - | 0 | - |
| - | - | 1595 | 217.1 | - | - | 0 | - |
| - | - | 1112 | 217.1 | - | - | 0 | - |
| - | - | 6001 | 219.1 | - | - | 0 | - |
| - | - | 499.9 | 221.1 | - | - | 0 | - |
| - | - | 1458 | 221.1 | - | - | 0 | - |
| - | - | 1364 | 221.1 | - | - | 0 | - |
| - | - | 482.3 | 221.1 | - | - | 0 | - |
| - | - | 797.3 | 225.1 | - | - | 0 | - |
| - | - | 893.4 | 226.1 | - | - | 0 | - |
| - | - | 4448 | 226.2 | - | - | 0 | - |
| - | - | 517.4 | 227.1 | - | - | 0 | - |
| - | - | 1427 | 227.1 | - | - | 0 | - |
| - | - | 3903 | 227.1 | - | - | 0 | - |
| - | - | 1394 | 228.1 | - | - | 0 | - |
| - | - | 658.5 | 229.1 | - | - | 0 | - |
| - | - | 480 | 229.5 | - | - | 0 | - |
| - | - | 1.641E+04 | 230.2 | - | - | 0 | - |
| - | - | 562.5 | 231.1 | - | - | 0 | - |
| - | - | 1958 | 231.2 | - | - | 0 | - |
| - | - | 952.9 | 233.1 | - | - | 0 | - |
| - | - | 842.7 | 233.2 | - | - | 0 | - |
| - | - | 9640 | 233.2 | - | - | 0 | - |
| - | - | 641.9 | 234.2 | - | - | 0 | - |
| - | - | 600.4 | 237.1 | - | - | 0 | - |
| - | - | 566 | 239.1 | - | - | 0 | - |
| - | - | 1199 | 239.2 | - | - | 0 | - |
| - | - | 4153 | 242.2 | - | - | 0 | - |
| - | - | 2020 | 243.1 | - | - | 0 | - |
| 8 | y | 7519 | 243.2 | 0.0004268 | 1.755 | +1 | 2 |
| - | - | 895.1 | 244.1 | - | - | 0 | - |
| - | - | 647 | 244.1 | - | - | 0 | - |
| - | - | 2118 | 245.1 | - | - | 0 | - |
| - | - | 725.2 | 246.1 | - | - | 0 | - |
| - | - | 1492 | 247.1 | - | - | 0 | - |
| - | - | 1003 | 248.2 | - | - | 0 | - |
| - | - | 670.6 | 251.1 | - | - | 0 | - |
| - | - | 865.6 | 254.1 | - | - | 0 | - |
| - | - | 8729 | 255.1 | - | - | 0 | - |
| - | - | 584.5 | 255.2 | - | - | 0 | - |
| - | - | 524.3 | 256.1 | - | - | 0 | - |
| - | - | 2125 | 259.1 | - | - | 0 | - |
| 8 | y | 2.65E+04 | 260.2 | 0.0002754 | 1.058 | +1 | 2 |
| - | - | 1887 | 261.1 | - | - | 0 | - |
| - | - | 1554 | 261.2 | - | - | 0 | - |
| - | - | 3273 | 261.2 | - | - | 0 | - |
| - | - | 4511 | 262.1 | - | - | 0 | - |
| - | - | 655.8 | 262.2 | - | - | 0 | - |
| - | - | 888.5 | 263.1 | - | - | 0 | - |
| - | - | 8502 | 269.2 | - | - | 0 | - |
| - | - | 881.4 | 270.2 | - | - | 0 | - |
| - | - | 8724 | 273.1 | - | - | 0 | - |
| - | - | 1017 | 273.1 | - | - | 0 | - |
| - | - | 2065 | 274.1 | - | - | 0 | - |
| - | - | 737.5 | 276.2 | - | - | 0 | - |
| - | - | 1040 | 282.2 | - | - | 0 | - |
| - | - | 2032 | 287.2 | - | - | 0 | - |
| - | - | 623.6 | 288.2 | - | - | 0 | - |
| - | - | 2396 | 291.1 | - | - | 0 | - |
| - | - | 1449 | 300.1 | - | - | 0 | - |
| - | - | 2448 | 301.2 | - | - | 0 | - |
| - | - | 915.2 | 304.2 | - | - | 0 | - |
| - | - | 679.2 | 305.2 | - | - | 0 | - |
| - | - | 804.4 | 318.1 | - | - | 0 | - |
| - | - | 4066 | 319.1 | - | - | 0 | - |
| - | - | 1127 | 320.1 | - | - | 0 | - |
| - | - | 692.3 | 321.2 | - | - | 0 | - |
| - | - | 1234 | 323.2 | - | - | 0 | - |
| - | - | 1917 | 323.2 | - | - | 0 | - |
| - | - | 8008 | 326.2 | - | - | 0 | - |
| - | - | 634.6 | 327.2 | - | - | 0 | - |
| - | - | 818.1 | 331.1 | - | - | 0 | - |
| - | - | 596.9 | 332.1 | - | - | 0 | - |
| - | - | 1082 | 339.2 | - | - | 0 | - |
| - | - | 1005 | 339.2 | - | - | 0 | - |
| 3 | y | 1.334E+04 | 340.2 | 0.0005265 | 1.548 | +2 | 7 |
| - | - | 6848 | 340.7 | - | - | 0 | - |
| - | - | 999.1 | 341.2 | - | - | 0 | - |
| 7 | y | 1377 | 343.2 | 0.0006067 | 1.768 | +1 | 3 |
| - | - | 1728 | 344.2 | - | - | 0 | - |
| - | - | 647.2 | 345.2 | - | - | 0 | - |
| 3 | b | 2389 | 346.2 | 0.0005555 | 1.605 | +1 | 3 |
| - | - | 2431 | 348.2 | - | - | 0 | - |
| - | - | 1458 | 350.2 | - | - | 0 | - |
| - | - | 955.3 | 350.7 | - | - | 0 | - |
| - | - | 1813 | 355.2 | - | - | 0 | - |
| - | - | 3695 | 357.2 | - | - | 0 | - |
| 7 | y | 4130 | 361.2 | 0.0003264 | 0.9036 | +1 | 3 |
| - | - | 1222 | 362.2 | - | - | 0 | - |
| - | - | 1447 | 362.2 | - | - | 0 | - |
| 8 | b | 710.6 | 364.7 | 0.001463 | 4.013 | +2 | 8 |
| 8 | b | 703.6 | 365.2 | 0.004979 | 13.63 | +2 | 8 |
| - | - | 3727 | 374.2 | - | - | 0 | - |
| - | - | 1986 | 375.2 | - | - | 0 | - |
| - | - | 1111 | 376.7 | - | - | 0 | - |
| - | - | 823.8 | 382.7 | - | - | 0 | - |
| - | - | 5368 | 383.2 | - | - | 0 | - |
| - | - | 871.1 | 383.2 | - | - | 0 | - |
| 2 | y | 870.7 | 389.7 | 0.000752 | 1.93 | +2 | 8 |
| - | - | 796.8 | 390.7 | - | - | 0 | - |
| - | - | 4762 | 392.2 | - | - | 0 | - |
| - | - | 706.9 | 393.2 | - | - | 0 | - |
| 6 | y | 981.3 | 400.3 | 0.0006884 | 1.72 | +1 | 4 |
| - | - | 3469 | 401.2 | - | - | 0 | - |
| - | - | 1.413E+04 | 402.2 | - | - | 0 | - |
| - | - | 2598 | 403.2 | - | - | 0 | - |
| 4 | b | 1188 | 403.2 | 0.001316 | 3.263 | +1 | 4 |
| - | - | 1785 | 418.2 | - | - | 0 | - |
| 6 | y | 4204 | 418.3 | 0.0004692 | 1.122 | +1 | 4 |
| - | - | 2828 | 419.2 | - | - | 0 | - |
| - | - | 901.1 | 419.3 | - | - | 0 | - |
| - | - | 6524 | 420.2 | - | - | 0 | - |
| - | - | 1042 | 421.2 | - | - | 0 | - |
| - | - | 3288 | 425.3 | - | - | 0 | - |
| 0 | Precursor | 1908 | 430.3 | 0.001329 | 3.089 | +2 | -1 |
| - | - | 787.6 | 437.9 | - | - | 0 | - |
| - | - | 1828 | 438.2 | - | - | 0 | - |
| - | - | 686 | 438.3 | - | - | 0 | - |
| - | - | 2296 | 438.3 | - | - | 0 | - |
| - | - | 2016 | 439.2 | - | - | 0 | - |
| 0 | Precursor | 9337 | 439.3 | 0.001357 | 3.089 | +2 | -1 |
| - | - | 3068 | 439.8 | - | - | 0 | - |
| - | - | 6767 | 439.8 | - | - | 0 | - |
| - | - | 1141 | 440.3 | - | - | 0 | - |
| - | - | 2798 | 448.3 | - | - | 0 | - |
| - | - | 1413 | 448.8 | - | - | 0 | - |
| - | - | 879.4 | 451.3 | - | - | 0 | - |
| 5 | y | 1658 | 457.3 | 6.817E-05 | 0.1491 | +1 | 5 |
| - | - | 687 | 466.8 | - | - | 0 | - |
| - | - | 2470 | 469.3 | - | - | 0 | - |
| 5 | y | 1.82E+04 | 475.3 | 0.0007645 | 1.608 | +1 | 5 |
| - | - | 3666 | 476.3 | - | - | 0 | - |
| - | - | 932.5 | 481.3 | - | - | 0 | - |
| - | - | 714 | 486.3 | - | - | 0 | - |
| - | - | 1098 | 489.3 | - | - | 0 | - |
| - | - | 972.4 | 496.3 | - | - | 0 | - |
| - | - | 760.6 | 497.3 | - | - | 0 | - |
| - | - | 842.2 | 501.2 | - | - | 0 | - |
| - | - | 1505 | 512.3 | - | - | 0 | - |
| 4 | y | 3905 | 514.3 | 3.325E-05 | 0.06466 | +1 | 6 |
| - | - | 738.4 | 519.3 | - | - | 0 | - |
| - | - | 635 | 521.8 | - | - | 0 | - |
| - | - | 861 | 526.3 | - | - | 0 | - |
| - | - | 1391 | 527.3 | - | - | 0 | - |
| - | - | 9989 | 529.8 | - | - | 0 | - |
| - | - | 4978 | 530.3 | - | - | 0 | - |
| - | - | 1451 | 530.3 | - | - | 0 | - |
| - | - | 1750 | 531.3 | - | - | 0 | - |
| 4 | y | 5.26E+04 | 532.3 | 0.0002663 | 0.5003 | +1 | 6 |
| - | - | 1.321E+04 | 533.3 | - | - | 0 | - |
| - | - | 2405 | 534.3 | - | - | 0 | - |
| - | - | 636 | 537.3 | - | - | 0 | - |
| - | - | 2136 | 539.8 | - | - | 0 | - |
| - | - | 673.3 | 540.3 | - | - | 0 | - |
| - | - | 9595 | 548.3 | - | - | 0 | - |
| - | - | 1392 | 548.3 | - | - | 0 | - |
| - | - | 944.8 | 548.8 | - | - | 0 | - |
| - | - | 2979 | 549.3 | - | - | 0 | - |
| - | - | 2353 | 555.3 | - | - | 0 | - |
| - | - | 6475 | 566.3 | - | - | 0 | - |
| - | - | 1507 | 567.3 | - | - | 0 | - |
| - | - | 1962 | 573.3 | - | - | 0 | - |
| - | - | 1175 | 598.3 | - | - | 0 | - |
| 7 | b | 640.2 | 600.3 | 0.0006741 | 1.123 | +1 | 7 |
| - | - | 590.8 | 604 | - | - | 0 | - |
| - | - | 2774 | 616.3 | - | - | 0 | - |
| - | - | 1164 | 617.3 | - | - | 0 | - |
| - | - | 1196 | 633.4 | - | - | 0 | - |
| - | - | 3819 | 643.4 | - | - | 0 | - |
| - | - | 1031 | 644.3 | - | - | 0 | - |
| - | - | 711.8 | 647.4 | - | - | 0 | - |
| 3 | y | 8391 | 661.4 | 0.0002784 | 0.421 | +1 | 7 |
| - | - | 2549 | 662.4 | - | - | 0 | - |
| - | - | 965.7 | 663.4 | - | - | 0 | - |
| - | - | 747 | 665.4 | - | - | 0 | - |
| 3 | y | 1.685E+05 | 679.4 | 0.0003338 | 0.4914 | +1 | 7 |
| - | - | 6.015E+04 | 680.4 | - | - | 0 | - |
| - | - | 1.194E+04 | 681.4 | - | - | 0 | - |
| - | - | 749.5 | 682.4 | - | - | 0 | - |
| - | - | 587.2 | 697.4 | - | - | 0 | - |
| - | - | 1284 | 699.4 | - | - | 0 | - |
| - | - | 1704 | 736.4 | - | - | 0 | - |
| - | - | 800.5 | 737.4 | - | - | 0 | - |
| 8 | b | 934.3 | 746.4 | 0.001164 | 1.56 | +1 | 8 |
| 2 | y | 704.4 | 760.4 | 0.00529 | 6.956 | +1 | 8 |
| - | - | 1609 | 764.4 | - | - | 0 | - |
| 2 | y | 1.691E+04 | 778.4 | 3.516E-05 | 0.04517 | +1 | 8 |
| - | - | 6979 | 779.4 | - | - | 0 | - |
| - | - | 4152 | 780.5 | - | - | 0 | - |
| - | - | 801.7 | 781.5 | - | - | 0 | - |
| - | - | 983.9 | 798.5 | - | - | 0 | - |
| - | - | 643.2 | 826.5 | - | - | 0 | - |
| 0 | Precursor | 2293 | 877.5 | 0.004292 | 4.891 | +1 | -1 |
| - | - | 642 | 878.5 | - | - | 0 | - |
| - | - | 1194 | 895.5 | - | - | 0 | - |
| - | - | 626.8 | 896.5 | - | - | 0 | - |
| - | - | 550.4 | 921 | - | - | 0 | - |
| - | - | 1371 | 961.5 | - | - | 0 | - |
| - | - | 695 | 1072 | - | - | 0 | - |
| - | - | 623.4 | 1712 | - | - | 0 | - |
| - | - | 648.4 | 2007 | - | - | 0 | - |
| - | - | 762.8 | 3081 | - | - | 0 | - |

m/z Charge Intensity FragmentType MassShift Position
120.0811538696289 0 90973.21
120.45833587646484 0 405.77866
121.08446502685547 0 7246.407
123.04436492919922 0 732.4093
124.11259460449219 0 928.7195
125.07150268554688 0 445.55356
126.05550384521484 0 764.87946
127.05062103271484 0 1352.3447
127.08676147460938 0 617.69293
128.1071319580078 0 667.5595
129.06607055664062 0 1062.3038
129.10260009765625 0 90665.84
130.0508270263672 0 425.11975
130.06541442871094 0 1146.5514
130.0867462158203 0 1477.6118
130.1003875732422 0 737.4353
130.10598754882812 0 5187.705
131.08189392089844 0 4486.579
131.11810302734375 0 803.7284
132.04473876953125 0 452.70453
132.0811309814453 0 1481.7128
132.1022491455078 0 5295.6797 y 8
133.06077575683594 0 530.86945
134.06085205078125 0 459.00195
136.07603454589844 0 7610.8735
136.12655639648438 0 469.96143
138.3702850341797 0 405.05127
139.08689880371094 0 703.37134
141.06588745117188 0 1639.1079
141.10240173339844 0 1274.4968
144.07687377929688 0 561.09534
146.06039428710938 0 3938.394
147.04432678222656 0 1001.4983
147.11318969726562 0 4292.078
148.01498413085938 0 422.65744
148.03982543945312 0 552.11926
148.87831115722656 0 417.0602
148.89263916015625 0 474.67398
148.89976501464844 0 593.39
148.90713500976562 0 781.533
148.91448974609375 0 1024.773
148.92178344726562 0 1042.1941
148.92904663085938 0 1236.7974
148.93618774414062 0 2587.2656
148.94400024414062 0 4091.8696
148.96070861816406 0 4156.0405
148.96847534179688 0 2083.797
148.9757080078125 0 1394.9934
148.98300170898438 0 1348.7006
148.99034118652344 0 716.2667
149.0044403076172 0 513.67377
149.01171875 0 477.78745
149.04832458496094 0 587.5656
149.81390380859375 0 421.1802
152.1437530517578 0 666.14276
155.0816192626953 0 1512.4357
155.1182098388672 0 2546.1626
157.09742736816406 0 630.9085
157.13397216796875 0 601.1719 d 1
158.0603485107422 0 489.1795
159.07672119140625 0 3490.703
159.09188842773438 0 1338.5842
165.05503845214844 0 3927.2615
166.0867919921875 0 1131.612
167.08189392089844 0 715.5397
167.1183319091797 0 1270.7225
168.04576110839844 0 486.04617
169.0972137451172 0 739.2559
169.13394165039062 0 1230.4177
171.07708740234375 0 695.8076
171.1496124267578 0 176912.69 a 1
172.0720977783203 0 3661.1157
172.1453094482422 0 1221.4897
172.15292358398438 0 14673.801
173.1288604736328 0 6581.359
173.15538024902344 0 721.98865
174.05545043945312 0 2666.4946
174.13133239746094 0 455.24353
175.0868682861328 0 2280.6042
176.107421875 0 4800.3374
177.10252380371094 0 6294.8296
177.1105194091797 0 1126.494
178.1062469482422 0 1184.5424
179.81236267089844 0 535.96967
181.0970916748047 0 837.5837
182.08157348632812 0 4229.9346
183.14984130859375 0 1582.0787
185.12872314453125 0 1610.2999
186.12405395507812 0 1871.4594
187.1443328857422 0 1609.0775
188.07102966308594 0 1373.6136
188.10317993164062 0 1454.4037
189.10263061523438 0 556.565
191.118408203125 0 602.2467
192.0658721923828 0 1040.9645
195.11318969726562 0 1560.8295
197.12887573242188 0 996.4264
197.16485595703125 0 1358.074
198.0875244140625 0 3841.449
199.10842895507812 0 1118.3623
199.14443969726562 0 69509.64 b 1
200.14788818359375 0 6642.128
200.1644287109375 0 801.9
201.1236572265625 0 1874.1111 y Ammonia loss 5
201.14988708496094 0 569.83496
202.05015563964844 0 924.1044
203.06622314453125 0 633.2936
203.08180236816406 0 1130.2632
205.09747314453125 0 11930.226
206.1009521484375 0 895.8174
208.10858154296875 0 1691.6533
209.0929718017578 0 489.29742
209.1037139892578 0 861.25836
209.316650390625 0 501.4006
211.14451599121094 0 1871.406
212.13973999023438 0 9715.696
213.12399291992188 0 719.0441
215.11700439453125 0 690.9973
215.1389923095703 0 873.83905
216.0982666015625 0 3125.826
217.09771728515625 0 1595.0798
217.1339874267578 0 1111.5511
219.1495819091797 0 6000.72
221.05929565429688 0 499.9399
221.09286499023438 0 1458.1135
221.1284942626953 0 1363.5526
221.14007568359375 0 482.33334
225.12364196777344 0 797.26404
226.11895751953125 0 893.38824
226.15533447265625 0 4447.652
227.0842742919922 0 517.44574
227.10279846191406 0 1427.2407
227.11416625976562 0 3902.9382
228.09825134277344 0 1394.1141
229.0929412841797 0 658.4564
229.51510620117188 0 479.99033
230.15029907226562 0 16410.824
231.11338806152344 0 562.46356
231.15365600585938 0 1957.8506
233.1287841796875 0 952.9266
233.15162658691406 0 842.7409
233.16522216796875 0 9639.686
234.16888427734375 0 641.9468
237.0987548828125 0 600.3979
239.1386260986328 0 566.0307
239.1503448486328 0 1199.1764
242.15042114257812 0 4153.2725
243.1134796142578 0 2020.1528
243.17074584960938 0 7518.7524 y Ammonia loss 7
244.1077423095703 0 895.0918
244.129638671875 0 647.0313
245.1255645751953 0 2118.1423
246.12692260742188 0 725.1534
247.14437866210938 0 1491.6367
248.16061401367188 0 1003.35455
251.14988708496094 0 670.6398
254.1130828857422 0 865.61926
255.1092071533203 0 8729.0625
255.16940307617188 0 584.47504
256.11004638671875 0 524.3295
259.1181335449219 0 2124.5122
260.1971435546875 0 26495.27 y 7
261.12335205078125 0 1886.9415
261.1598205566406 0 1553.9706
261.200439453125 0 3273.058
262.1190185546875 0 4510.6533
262.2029113769531 0 655.8395
263.1221923828125 0 888.538
269.1610412597656 0 8501.515
270.16497802734375 0 881.36835
273.1195983886719 0 8723.751
273.13543701171875 0 1016.7414
274.1204528808594 0 2064.5742
276.1689453125 0 737.504
282.1562805175781 0 1039.7527
287.1709899902344 0 2031.9137
288.1734924316406 0 623.5944
291.1454162597656 0 2395.7036
300.1339416503906 0 1448.9917
301.1913757324219 0 2448.3774
304.1654052734375 0 915.21497
305.1831970214844 0 679.16675
318.14544677734375 0 804.425
319.1412658691406 0 4066.3735
320.1433410644531 0 1127.0175
321.17706298828125 0 692.2775
323.1716613769531 0 1233.5295
323.2088928222656 0 1917.387
326.182861328125 0 8008.436
327.18621826171875 0 634.5899
331.14080810546875 0 818.098
332.12530517578125 0 596.9194
339.1787109375 0 1082.2579
339.2015686035156 0 1005.1084
340.1928405761719 0 13340.589 y 2
340.6941223144531 0 6847.892
341.1809387207031 0 999.14
343.2345886230469 0 1377.1782 y Water loss 6
344.1931457519531 0 1727.9972
345.1552429199219 0 647.17926
346.21307373046875 0 2388.5342 b 2
348.1670837402344 0 2430.9048
350.2054443359375 0 1458.2781
350.7060546875 0 955.3488
355.16119384765625 0 1812.7565
357.15576171875 0 3694.854
361.244873046875 0 4129.573 y 6
362.2032775878906 0 1222.4167
362.248046875 0 1447.2418
364.7095947265625 0 710.6192 b Water loss 7
365.1951599121094 0 703.59045 b Ammonia loss 7
374.1825866699219 0 3726.8938
375.1661376953125 0 1986.3553
376.73797607421875 0 1110.977
382.7187805175781 0 823.8493
383.2038879394531 0 5368.1636
383.2275695800781 0 871.0731
389.72576904296875 0 870.6735 y 1
390.7343444824219 0 796.8154
392.1929626464844 0 4761.5303
393.1952209472656 0 706.9344
400.2561340332031 0 981.3437 y Water loss 5
401.214599609375 0 3468.9387
402.1778259277344 0 14128.517
403.1805114746094 0 2597.5042
403.232666015625 0 1187.622 b 3
418.2306823730469 0 1785.379
418.2664794921875 0 4204.4175 y 5
419.2264099121094 0 2827.7173
419.2685241699219 0 901.13416
420.1885681152344 0 6524.136
421.1922912597656 0 1042.0474
425.2637023925781 0 3288.2734
430.25677490234375 0 1907.9187 Precursor Water loss
437.8511047363281 0 787.5969
438.24053955078125 0 1827.5947
438.2734069824219 0 686.04456
438.3065490722656 0 2295.6096
439.22918701171875 0 2015.9564
439.2620849609375 0 9337.468 Precursor
439.7635192871094 0 3068.1702
439.84381103515625 0 6767.2153
440.265380859375 0 1140.5812
448.2665710449219 0 2798.4607
448.7687072753906 0 1412.7526
451.2669372558594 0 879.39264
457.2769775390625 0 1658.2029 y Water loss 4
466.7764587402344 0 687.0224
469.27752685546875 0 2470.023
475.2882385253906 0 18202.176 y 4
476.2911071777344 0 3665.83
481.27239990234375 0 932.5233
486.3057861328125 0 714.00055
489.265380859375 0 1098.2571
496.2905578613281 0 972.44464
497.2712097167969 0 760.5529
501.2454528808594 0 842.1778
512.2623291015625 0 1505.3972
514.29833984375 0 3905.1404 y Water loss 3
519.2548217773438 0 738.41516
521.797119140625 0 635.0171
526.3118286132812 0 861.02075
527.3186645507812 0 1391.1232
529.7982177734375 0 9988.882
530.2722778320312 0 4977.8193
530.3075561523438 0 1450.8112
531.2745971679688 0 1749.8386
532.3092041015625 0 52600.39 y 3
533.312255859375 0 13209.522
534.3129272460938 0 2405.34
537.3011474609375 0 636.007
539.8101196289062 0 2136.187
540.312255859375 0 673.34357
548.2827758789062 0 9595.312
548.3232421875 0 1391.5964
548.8248291015625 0 944.8474
549.2842407226562 0 2978.7131
555.3143920898438 0 2353.1516
566.2935180664062 0 6475.019
567.2925415039062 0 1506.86
573.3246459960938 0 1962.4346
598.334228515625 0 1175.1183
600.314697265625 0 640.184 b Water loss 6
604.0259399414062 0 590.7781
616.345703125 0 2774.377
617.3458862304688 0 1163.6055
633.3700561523438 0 1195.9244
643.3560791015625 0 3819.1372
644.3483276367188 0 1030.5635
647.3501586914062 0 711.7678
661.3670654296875 0 8391.242 y Water loss 2
662.3692016601562 0 2548.6912
663.3784790039062 0 965.6504
665.3568115234375 0 746.9552
679.377685546875 0 168465.39 y 2
680.380859375 0 60151.223
681.3837280273438 0 11939.105
682.3860473632812 0 749.4888
697.4014892578125 0 587.2146
699.4017333984375 0 1283.9321
736.3869018554688 0 1704.0366
737.3921508789062 0 800.4851
746.4207153320312 0 934.2881 b 7
760.4404907226562 0 704.4469 y Water loss 1
764.4287719726562 0 1609.061
778.44580078125 0 16911.332 y 1
779.4486694335938 0 6979.271
780.4593505859375 0 4152.0522
781.4620361328125 0 801.72504
798.4744873046875 0 983.88074
826.4803466796875 0 643.1704
877.5098876953125 0 2293.3577 Precursor
878.5114135742188 0 641.98254
895.525634765625 0 1194.1885
896.5303344726562 0 626.8226
921.024658203125 0 550.35376
961.5320434570312 0 1370.5258
1072.3197021484375 0 695.0394
1711.594970703125 0 623.42596
2006.857666015625 0 648.3778
3080.9052734375 0 762.8469

Spectrum Details

|  |  |
| --- | --- |
| Matched peaks? Matched peaks The total absolute number of peaks matched. Additionally in brackets the total fraction of peaks matched and the total number of peaks is shown. | 30 (9.35% of 321) |
| FDR? FDR The false discovery rate estimated for this peptide. It is calculated by matching all theoretical fragments with a non-integer shift with the raw peaks for this spectrum. This is done with 40 different shifts. The resulting percentage is the average number of annotated peaks over the number of annotated peaks with the correct spectrum. | 0.40% |
| Satellite FDR? Satellite FDR See the FDR for details on its calculation. This satellite ion specific FDR only contains the satellite ions (d/w) for I/L/J positions. | - |
| PSM Score? PSM Score The PSM Score as given by Hecklib to this annotated spectrum. It is shown with three significant figures. | 345 |

#### Spectrum 6150? Spectrum 6150 The raw spectrum of this peptide as annotated by Hecklib. The fragments are coloured according to ion type (see legend). Any peaks with a star '\*' as text can be hovered over to see the full details, first the ion type second the mass shift type. By hovering over the amino acids in the peptide or ions in the legend the corresponding peaks are highlighted. By toggling the 'Unassigned' label you can turn the background (unassigned) peaks on or off in the plot. By updating the slider in the Ion legend you can update the spectrum to only show the top X% of the peaks with labels. The top X% means any peak that is within X% of the highest intensity. By dragging in the spectrum you can zoom in to a specific part of the spectrum and use 'Zoom Out' to get back to the original zoom level. The annotation of the spectrum is based on the given sequence in the peptides file and is done with different software so inconsistencies are likely. The peaks are annotated based on the given sequence, with 20 ppm tolerance.

Copy Data

##### Spectrum 6150 (TSV)

###### Preview

```
Loading example...
```

*Click on the button to copy the data to your clipboard.*

Mz MinMz MaxIntensity Max

WidthHeightPeptide font sizePeptide stroke widthSpectrum font sizeSpectrum stroke widthCompact peptide

Ion legend

wxyz

abcd

OtherUnassignedIon^Charge^_Position_Show for top:%

VVFGGGTKJ

02.88e+65.77e+68.65e+61.15e+7

Zoom Out

y^+1^_1_y^+1^_2_z^+1^_2_y^+1^_2_y^+2^_7_y^+2^_7_y^+1^_3_y^+1^_3_z^+1^_3_y^+1^_3_c^+1^_3_c^+2^_8_y^+2^_8_z^+1^_4_y^+1^_4_c^+1^_4_z^+1^_5_y^+1^_5_z^+1^_5_y^+1^_5_c^+1^_5_z^+1^_6_y^+1^_6_z^+1^_6_y^+1^_6_c^+1^_6_c^+1^_7_c^+1^_7_z^+1^_7_y^+1^_7_z^+1^_7_y^+1^_7_c^+1^_8_y^+1^_8_y^+1^_8_z^+1^_8_c^+1^_8_y^+1^_8_

0240481721962

Fragment Matches Table

Show background peaks

| Position | Ion type | Intensity | mz Theoretical | mz Error (Th) | mz Error (ppm) | Charge | Series Number |
| --- | --- | --- | --- | --- | --- | --- | --- |
| - | - | 4.922E+05 | 120.1 | - | - | 0 | - |
| - | - | 3.138E+04 | 121.1 | - | - | 0 | - |
| - | - | 5448 | 122 | - | - | 0 | - |
| - | - | 6467 | 122.1 | - | - | 0 | - |
| - | - | 1.891E+04 | 128.1 | - | - | 0 | - |
| - | - | 5.38E+05 | 129.1 | - | - | 0 | - |
| - | - | 3.402E+04 | 130.1 | - | - | 0 | - |
| 9 | y | 1.446E+05 | 132.1 | 0.0004814 | 3.644 | +1 | 1 |
| - | - | 7708 | 133.1 | - | - | 0 | - |
| - | - | 6579 | 134.8 | - | - | 0 | - |
| - | - | 9155 | 143.2 | - | - | 0 | - |
| - | - | 4.946E+06 | 171.1 | - | - | 0 | - |
| - | - | 4.658E+05 | 172.2 | - | - | 0 | - |
| - | - | 1.311E+04 | 173.2 | - | - | 0 | - |
| - | - | 4.435E+04 | 177.1 | - | - | 0 | - |
| - | - | 1.461E+04 | 188.1 | - | - | 0 | - |
| - | - | 8036 | 191 | - | - | 0 | - |
| - | - | 3.775E+04 | 198.1 | - | - | 0 | - |
| - | - | 3.333E+06 | 199.1 | - | - | 0 | - |
| - | - | 3.398E+05 | 200.1 | - | - | 0 | - |
| - | - | 9683 | 201.1 | - | - | 0 | - |
| - | - | 1.6E+04 | 201.2 | - | - | 0 | - |
| - | - | 1.946E+05 | 205.1 | - | - | 0 | - |
| - | - | 2.025E+04 | 206.1 | - | - | 0 | - |
| - | - | 2.191E+04 | 212.1 | - | - | 0 | - |
| - | - | 3.078E+04 | 212.1 | - | - | 0 | - |
| - | - | 1.315E+04 | 216.1 | - | - | 0 | - |
| - | - | 8.365E+04 | 219.1 | - | - | 0 | - |
| - | - | 1.498E+05 | 230.2 | - | - | 0 | - |
| - | - | 1.773E+04 | 231.2 | - | - | 0 | - |
| 8 | y | 1.457E+04 | 243.2 | 0.001434 | 5.897 | +1 | 2 |
| 8 | z | 1.166E+04 | 244.2 | 0.0008568 | 3.509 | +1 | 2 |
| - | - | 1.481E+04 | 245.1 | - | - | 0 | - |
| - | - | 1.745E+04 | 245.2 | - | - | 0 | - |
| - | - | 9.923E+04 | 247.1 | - | - | 0 | - |
| - | - | 1.79E+04 | 248.1 | - | - | 0 | - |
| - | - | 1.248E+04 | 254.1 | - | - | 0 | - |
| - | - | 2.261E+04 | 255.1 | - | - | 0 | - |
| - | - | 2.565E+04 | 256.2 | - | - | 0 | - |
| - | - | 8061 | 257.1 | - | - | 0 | - |
| - | - | 8538 | 257.7 | - | - | 0 | - |
| - | - | 4.034E+04 | 258.1 | - | - | 0 | - |
| - | - | 1.439E+04 | 259.2 | - | - | 0 | - |
| 8 | y | 4.317E+05 | 260.2 | 0.0005501 | 2.114 | +1 | 2 |
| - | - | 4.156E+04 | 261.2 | - | - | 0 | - |
| - | - | 2.715E+04 | 262.1 | - | - | 0 | - |
| - | - | 1.761E+04 | 269.2 | - | - | 0 | - |
| - | - | 1.23E+04 | 272.1 | - | - | 0 | - |
| - | - | 1.029E+04 | 272.2 | - | - | 0 | - |
| - | - | 8.748E+04 | 273.1 | - | - | 0 | - |
| - | - | 9505 | 273.1 | - | - | 0 | - |
| - | - | 4.958E+04 | 279.1 | - | - | 0 | - |
| - | - | 1.312E+04 | 283.7 | - | - | 0 | - |
| - | - | 2.452E+04 | 287.1 | - | - | 0 | - |
| - | - | 3.855E+04 | 287.2 | - | - | 0 | - |
| - | - | 2.663E+04 | 290.1 | - | - | 0 | - |
| - | - | 1.212E+04 | 300.2 | - | - | 0 | - |
| - | - | 1.068E+05 | 301.2 | - | - | 0 | - |
| - | - | 3.229E+04 | 302.2 | - | - | 0 | - |
| - | - | 2.998E+04 | 304.2 | - | - | 0 | - |
| - | - | 1.194E+05 | 311.2 | - | - | 0 | - |
| - | - | 2.14E+04 | 312.2 | - | - | 0 | - |
| - | - | 1.092E+04 | 318.2 | - | - | 0 | - |
| - | - | 1.848E+04 | 319.1 | - | - | 0 | - |
| - | - | 1.895E+04 | 322.2 | - | - | 0 | - |
| - | - | 7.932E+04 | 326.2 | - | - | 0 | - |
| - | - | 1.181E+04 | 327.2 | - | - | 0 | - |
| - | - | 1.169E+04 | 328.2 | - | - | 0 | - |
| - | - | 3.321E+04 | 329.2 | - | - | 0 | - |
| 3 | y | 2.091E+04 | 331.2 | 0.001262 | 3.81 | +2 | 7 |
| - | - | 1.573E+04 | 332.2 | - | - | 0 | - |
| - | - | 1.231E+04 | 335.2 | - | - | 0 | - |
| - | - | 1.807E+05 | 336.2 | - | - | 0 | - |
| - | - | 1.739E+04 | 337.2 | - | - | 0 | - |
| 3 | y | 1.073E+06 | 340.2 | 0.001045 | 3.073 | +2 | 7 |
| - | - | 3.659E+05 | 340.7 | - | - | 0 | - |
| - | - | 8.35E+04 | 341.2 | - | - | 0 | - |
| - | - | 3.834E+05 | 343.2 | - | - | 0 | - |
| 7 | y | 1.525E+04 | 343.2 | 0.0005835 | 1.7 | +1 | 3 |
| 7 | y | 8.924E+05 | 344.2 | 0.0006915 | 2.009 | +1 | 3 |
| 7 | z | 1.389E+05 | 345.2 | 0.003838 | 11.12 | +1 | 3 |
| - | - | 1.671E+04 | 346.2 | - | - | 0 | - |
| - | - | 1.507E+05 | 346.2 | - | - | 0 | - |
| - | - | 3.467E+05 | 346.2 | - | - | 0 | - |
| - | - | 3.439E+04 | 347.2 | - | - | 0 | - |
| - | - | 2.337E+04 | 347.2 | - | - | 0 | - |
| - | - | 4.89E+04 | 347.2 | - | - | 0 | - |
| - | - | 1.055E+04 | 357.2 | - | - | 0 | - |
| - | - | 5.044E+04 | 358.2 | - | - | 0 | - |
| - | - | 1.509E+04 | 359.2 | - | - | 0 | - |
| - | - | 1.543E+04 | 361.2 | - | - | 0 | - |
| 7 | y | 1.205E+05 | 361.2 | 0.0008452 | 2.34 | +1 | 3 |
| - | - | 1.256E+04 | 362.2 | - | - | 0 | - |
| 3 | c | 2.658E+04 | 363.2 | 0.001106 | 3.045 | +1 | 3 |
| - | - | 3.786E+04 | 368.2 | - | - | 0 | - |
| - | - | 2.963E+05 | 369.2 | - | - | 0 | - |
| - | - | 5.044E+04 | 370.2 | - | - | 0 | - |
| - | - | 1.742E+04 | 371.1 | - | - | 0 | - |
| - | - | 1.951E+04 | 372.2 | - | - | 0 | - |
| 8 | c | 1.455E+04 | 373.7 | 0.001644 | 4.398 | +2 | 8 |
| - | - | 2.99E+04 | 374.2 | - | - | 0 | - |
| - | - | 1.26E+04 | 375.2 | - | - | 0 | - |
| - | - | 1.755E+04 | 375.2 | - | - | 0 | - |
| - | - | 1.69E+04 | 376.2 | - | - | 0 | - |
| - | - | 9394 | 376.4 | - | - | 0 | - |
| - | - | 5.668E+04 | 383.2 | - | - | 0 | - |
| - | - | 4.698E+04 | 385.2 | - | - | 0 | - |
| - | - | 1.177E+04 | 385.2 | - | - | 0 | - |
| - | - | 4.027E+04 | 386.2 | - | - | 0 | - |
| - | - | 1.179E+04 | 388.2 | - | - | 0 | - |
| - | - | 9514 | 389.2 | - | - | 0 | - |
| 2 | y | 9.361E+04 | 389.7 | 0.0009875 | 2.534 | +2 | 8 |
| - | - | 3.651E+04 | 390.2 | - | - | 0 | - |
| - | - | 1.618E+04 | 390.7 | - | - | 0 | - |
| - | - | 6.768E+04 | 392.2 | - | - | 0 | - |
| - | - | 1.204E+04 | 393.2 | - | - | 0 | - |
| - | - | 1.229E+04 | 394.7 | - | - | 0 | - |
| - | - | 7.273E+04 | 401.2 | - | - | 0 | - |
| - | - | 2.327E+05 | 402.2 | - | - | 0 | - |
| - | - | 1.281E+04 | 402.2 | - | - | 0 | - |
| - | - | 1.455E+04 | 402.2 | - | - | 0 | - |
| 6 | z | 7.39E+04 | 402.2 | 0.001005 | 2.498 | +1 | 4 |
| - | - | 1.182E+06 | 403.2 | - | - | 0 | - |
| - | - | 3.938E+06 | 403.3 | - | - | 0 | - |
| - | - | 2.159E+05 | 404.2 | - | - | 0 | - |
| - | - | 7.489E+05 | 404.3 | - | - | 0 | - |
| - | - | 1.383E+04 | 405.2 | - | - | 0 | - |
| - | - | 3.169E+04 | 405.2 | - | - | 0 | - |
| - | - | 9.064E+04 | 405.3 | - | - | 0 | - |
| - | - | 2.361E+05 | 416.2 | - | - | 0 | - |
| - | - | 1.392E+04 | 416.2 | - | - | 0 | - |
| - | - | 3.834E+04 | 417.2 | - | - | 0 | - |
| - | - | 7.635E+04 | 417.3 | - | - | 0 | - |
| 6 | y | 2.721E+05 | 418.3 | 0.001232 | 2.946 | +1 | 4 |
| - | - | 1.766E+04 | 419.2 | - | - | 0 | - |
| - | - | 5.03E+04 | 419.3 | - | - | 0 | - |
| - | - | 1.791E+05 | 420.2 | - | - | 0 | - |
| 4 | c | 3.502E+04 | 420.3 | 0.002714 | 6.457 | +1 | 4 |
| - | - | 4.712E+04 | 421.2 | - | - | 0 | - |
| - | - | 9046 | 422.2 | - | - | 0 | - |
| - | - | 1.668E+04 | 424.3 | - | - | 0 | - |
| - | - | 9.047E+04 | 429.2 | - | - | 0 | - |
| - | - | 3.191E+04 | 430.3 | - | - | 0 | - |
| - | - | 8540 | 436 | - | - | 0 | - |
| - | - | 1.306E+05 | 437.2 | - | - | 0 | - |
| - | - | 1.327E+04 | 438.2 | - | - | 0 | - |
| - | - | 1.03E+05 | 439.3 | - | - | 0 | - |
| - | - | 5.696E+04 | 439.8 | - | - | 0 | - |
| 5 | z | 1.165E+04 | 441.3 | 0.00141 | 3.196 | +1 | 5 |
| - | - | 4.071E+04 | 442.3 | - | - | 0 | - |
| - | - | 1.085E+04 | 443.3 | - | - | 0 | - |
| - | - | 6.066E+04 | 445.2 | - | - | 0 | - |
| 5 | y | 1.315E+04 | 457.3 | 0.0008474 | 1.853 | +1 | 5 |
| 5 | z | 4.782E+05 | 459.3 | 0.001422 | 3.097 | +1 | 5 |
| - | - | 5.535E+06 | 460.3 | - | - | 0 | - |
| - | - | 1.239E+06 | 461.3 | - | - | 0 | - |
| - | - | 1.755E+05 | 462.3 | - | - | 0 | - |
| - | - | 1.501E+04 | 463.3 | - | - | 0 | - |
| - | - | 1.385E+04 | 471.3 | - | - | 0 | - |
| - | - | 2.401E+04 | 472.3 | - | - | 0 | - |
| - | - | 4.654E+05 | 473.2 | - | - | 0 | - |
| - | - | 9.639E+04 | 474.2 | - | - | 0 | - |
| - | - | 2.831E+05 | 474.3 | - | - | 0 | - |
| 5 | y | 7.534E+05 | 475.3 | 0.0011 | 2.315 | +1 | 5 |
| - | - | 1.04E+06 | 476.3 | - | - | 0 | - |
| 5 | c | 5.802E+05 | 477.3 | 0.0005616 | 1.177 | +1 | 5 |
| - | - | 1.057E+05 | 478.3 | - | - | 0 | - |
| - | - | 3.588E+04 | 490.3 | - | - | 0 | - |
| - | - | 3.061E+04 | 491.3 | - | - | 0 | - |
| - | - | 1.471E+04 | 496.3 | - | - | 0 | - |
| 4 | z | 3.548E+04 | 498.3 | 0.0005939 | 1.192 | +1 | 6 |
| - | - | 4.01E+04 | 499.3 | - | - | 0 | - |
| - | - | 1.082E+04 | 500.3 | - | - | 0 | - |
| - | - | 1.977E+04 | 501.2 | - | - | 0 | - |
| - | - | 5.638E+04 | 505.3 | - | - | 0 | - |
| - | - | 2.465E+04 | 506.3 | - | - | 0 | - |
| - | - | 1.952E+04 | 512.3 | - | - | 0 | - |
| 4 | y | 5.181E+04 | 514.3 | 0.001615 | 3.14 | +1 | 6 |
| 4 | z | 4.101E+05 | 516.3 | 0.001351 | 2.617 | +1 | 6 |
| - | - | 3.639E+06 | 517.3 | - | - | 0 | - |
| - | - | 8.826E+05 | 518.3 | - | - | 0 | - |
| - | - | 2.441E+04 | 519.3 | - | - | 0 | - |
| - | - | 1.429E+05 | 519.3 | - | - | 0 | - |
| - | - | 1.598E+05 | 530.3 | - | - | 0 | - |
| - | - | 5.407E+05 | 531.3 | - | - | 0 | - |
| 4 | y | 3.346E+06 | 532.3 | 0.001121 | 2.106 | +1 | 6 |
| - | - | 1.8E+06 | 533.3 | - | - | 0 | - |
| 6 | c | 7.928E+05 | 534.3 | 0.0002358 | 0.4414 | +1 | 6 |
| - | - | 1.776E+05 | 535.3 | - | - | 0 | - |
| - | - | 2.218E+04 | 536.3 | - | - | 0 | - |
| - | - | 4.153E+05 | 548.3 | - | - | 0 | - |
| - | - | 1.178E+05 | 549.3 | - | - | 0 | - |
| - | - | 2.85E+04 | 550.3 | - | - | 0 | - |
| - | - | 1.919E+04 | 558.3 | - | - | 0 | - |
| - | - | 1.404E+04 | 559.3 | - | - | 0 | - |
| - | - | 1.111E+04 | 563.3 | - | - | 0 | - |
| - | - | 1.118E+05 | 566.3 | - | - | 0 | - |
| - | - | 2.392E+04 | 567.3 | - | - | 0 | - |
| - | - | 3.844E+04 | 573.3 | - | - | 0 | - |
| - | - | 7.726E+04 | 576.3 | - | - | 0 | - |
| - | - | 1.732E+04 | 577.3 | - | - | 0 | - |
| - | - | 2.362E+04 | 590.3 | - | - | 0 | - |
| - | - | 2.065E+04 | 591.3 | - | - | 0 | - |
| - | - | 8.651E+04 | 592.3 | - | - | 0 | - |
| - | - | 1.516E+04 | 592.3 | - | - | 0 | - |
| - | - | 2.246E+04 | 593.3 | - | - | 0 | - |
| - | - | 6.761E+04 | 600.3 | - | - | 0 | - |
| - | - | 2.438E+04 | 601.3 | - | - | 0 | - |
| - | - | 1.015E+04 | 602.3 | - | - | 0 | - |
| - | - | 1.113E+04 | 604.3 | - | - | 0 | - |
| - | - | 1.42E+05 | 607.3 | - | - | 0 | - |
| - | - | 4.126E+04 | 608.3 | - | - | 0 | - |
| - | - | 1.26E+04 | 609.3 | - | - | 0 | - |
| - | - | 2.622E+04 | 616.3 | - | - | 0 | - |
| 7 | c | 1.609E+05 | 617.3 | 0.001103 | 1.786 | +1 | 7 |
| - | - | 2.635E+05 | 618.3 | - | - | 0 | - |
| - | - | 7.453E+04 | 619.3 | - | - | 0 | - |
| - | - | 1.174E+05 | 619.4 | - | - | 0 | - |
| - | - | 1.732E+05 | 620.3 | - | - | 0 | - |
| - | - | 5.243E+04 | 620.4 | - | - | 0 | - |
| - | - | 5.942E+04 | 621.3 | - | - | 0 | - |
| - | - | 1.71E+04 | 621.4 | - | - | 0 | - |
| - | - | 1.844E+04 | 622.3 | - | - | 0 | - |
| - | - | 1.47E+04 | 633.4 | - | - | 0 | - |
| - | - | 4.336E+05 | 634.3 | - | - | 0 | - |
| 7 | c | 5.084E+06 | 635.4 | 0.001402 | 2.207 | +1 | 7 |
| - | - | 1.595E+06 | 636.4 | - | - | 0 | - |
| - | - | 3.236E+05 | 637.4 | - | - | 0 | - |
| - | - | 6.695E+04 | 643.4 | - | - | 0 | - |
| - | - | 3.566E+04 | 644.4 | - | - | 0 | - |
| 3 | z | 2.105E+05 | 645.3 | 0.001546 | 2.396 | +1 | 7 |
| - | - | 6.729E+04 | 646.4 | - | - | 0 | - |
| - | - | 4.209E+04 | 647.4 | - | - | 0 | - |
| - | - | 1.108E+04 | 648.4 | - | - | 0 | - |
| 3 | y | 2.588E+05 | 661.4 | 0.001194 | 1.805 | +1 | 7 |
| - | - | 8.904E+04 | 662.4 | - | - | 0 | - |
| - | - | 2.115E+04 | 663.3 | - | - | 0 | - |
| 3 | z | 2.555E+06 | 663.4 | 0.001541 | 2.323 | +1 | 7 |
| - | - | 8.844E+05 | 664.4 | - | - | 0 | - |
| - | - | 2.045E+05 | 665.4 | - | - | 0 | - |
| 3 | y | 9.067E+06 | 679.4 | 0.001616 | 2.378 | +1 | 7 |
| - | - | 3.098E+06 | 680.4 | - | - | 0 | - |
| - | - | 6.619E+05 | 681.4 | - | - | 0 | - |
| - | - | 3.876E+04 | 682.4 | - | - | 0 | - |
| - | - | 1.118E+05 | 706.4 | - | - | 0 | - |
| - | - | 4.298E+04 | 707.4 | - | - | 0 | - |
| - | - | 7.326E+04 | 719.4 | - | - | 0 | - |
| - | - | 2.489E+04 | 720.4 | - | - | 0 | - |
| - | - | 1.904E+04 | 728.4 | - | - | 0 | - |
| - | - | 3.54E+04 | 729.4 | - | - | 0 | - |
| - | - | 1.036E+04 | 731.4 | - | - | 0 | - |
| 8 | c | 1.826E+05 | 746.4 | 0.001592 | 2.132 | +1 | 8 |
| - | - | 6.818E+04 | 747.4 | - | - | 0 | - |
| - | - | 2.464E+04 | 748.4 | - | - | 0 | - |
| 2 | y | 1.105E+04 | 760.4 | 0.001445 | 1.9 | +1 | 8 |
| 2 | y | 1.207E+04 | 761.4 | 0.01133 | 14.87 | +1 | 8 |
| 2 | z | 8.66E+05 | 762.4 | 0.001608 | 2.11 | +1 | 8 |
| 8 | c | 1.142E+07 | 763.4 | 0.001349 | 1.767 | +1 | 8 |
| - | - | 4.59E+06 | 764.5 | - | - | 0 | - |
| - | - | 1.096E+06 | 765.5 | - | - | 0 | - |
| - | - | 6.903E+04 | 766.5 | - | - | 0 | - |
| 2 | y | 7.679E+05 | 778.4 | 0.001561 | 2.005 | +1 | 8 |
| - | - | 2.977E+05 | 779.5 | - | - | 0 | - |
| - | - | 8.615E+04 | 780.5 | - | - | 0 | - |
| - | - | 1.138E+04 | 782.5 | - | - | 0 | - |
| - | - | 1.269E+04 | 788.4 | - | - | 0 | - |
| - | - | 3.126E+04 | 790.4 | - | - | 0 | - |
| - | - | 6.242E+04 | 805.4 | - | - | 0 | - |
| - | - | 2.869E+04 | 806.4 | - | - | 0 | - |
| - | - | 1.416E+04 | 817.5 | - | - | 0 | - |
| - | - | 8.408E+04 | 818.5 | - | - | 0 | - |
| - | - | 4.019E+04 | 819.4 | - | - | 0 | - |
| - | - | 2.783E+04 | 819.5 | - | - | 0 | - |
| - | - | 1.304E+04 | 820.5 | - | - | 0 | - |
| - | - | 2.27E+04 | 822.5 | - | - | 0 | - |
| - | - | 6.232E+04 | 832.5 | - | - | 0 | - |
| - | - | 1.98E+04 | 833.5 | - | - | 0 | - |
| - | - | 1.784E+04 | 834.5 | - | - | 0 | - |
| - | - | 2.674E+04 | 843.5 | - | - | 0 | - |
| - | - | 2.058E+04 | 844.5 | - | - | 0 | - |
| - | - | 2.409E+05 | 846.5 | - | - | 0 | - |
| - | - | 1.175E+05 | 847.5 | - | - | 0 | - |
| - | - | 2.849E+04 | 848.5 | - | - | 0 | - |
| - | - | 4.059E+04 | 850.5 | - | - | 0 | - |
| - | - | 1.55E+04 | 851.5 | - | - | 0 | - |
| - | - | 2.57E+04 | 860.5 | - | - | 0 | - |
| - | - | 9.23E+06 | 861.5 | - | - | 0 | - |
| - | - | 4.282E+06 | 862.5 | - | - | 0 | - |
| - | - | 1.273E+06 | 863.5 | - | - | 0 | - |
| - | - | 9.347E+04 | 864.5 | - | - | 0 | - |
| - | - | 4.214E+06 | 877.5 | - | - | 0 | - |
| - | - | 5.965E+06 | 878.5 | - | - | 0 | - |
| - | - | 2.457E+06 | 879.5 | - | - | 0 | - |
| - | - | 5.889E+05 | 880.5 | - | - | 0 | - |
| - | - | 4.185E+04 | 881.5 | - | - | 0 | - |
| - | - | 2.231E+04 | 893.5 | - | - | 0 | - |
| - | - | 1.556E+04 | 894.5 | - | - | 0 | - |
| - | - | 1.336E+04 | 952.3 | - | - | 0 | - |

m/z Charge Intensity FragmentType MassShift Position
120.08126068115234 0 492163
121.08456420898438 0 31380.836
121.9585189819336 0 5448.225
122.10327911376953 0 6467.137
128.09494018554688 0 18907.148
129.10272216796875 0 538008.2
130.10614013671875 0 34020.3
132.10238647460938 0 144635.22 y 8
133.10568237304688 0 7708.173
134.7915802001953 0 6578.54
143.1548614501953 0 9155.233
171.14979553222656 0 4945997.5
172.15309143066406 0 465809.53
173.15621948242188 0 13106.196
177.1028594970703 0 44346.207
188.11544799804688 0 14607.476
190.99110412597656 0 8036.2427
198.13673400878906 0 37746.93
199.1446990966797 0 3332885.8
200.1480712890625 0 339803.38
201.08767700195312 0 9683.205
201.1503448486328 0 15996.521
205.09768676757812 0 194639.47
206.10089111328125 0 20254.139
212.1151123046875 0 21905.873
212.14002990722656 0 30778.018
216.0988311767578 0 13151.309
219.14984130859375 0 83652.71
230.150634765625 0 149842.3
231.1537322998047 0 17730.607
243.1717529296875 0 14572.931 y Ammonia loss 7
244.1790008544922 0 11658.666 z 7
245.12506103515625 0 14806.157
245.1867218017578 0 17447.959
247.1446533203125 0 99233.21
248.14840698242188 0 17895.455
254.14967346191406 0 12481.336
255.10983276367188 0 22612.445
256.1662902832031 0 25650.285
257.1236267089844 0 8060.884
257.7110290527344 0 8538.145
258.1092529296875 0 40343.64
259.18963623046875 0 14391.99
260.1974182128906 0 431748.28 y 7
261.200927734375 0 41557.062
262.1190490722656 0 27146.758
269.1611633300781 0 17606.2
272.13653564453125 0 12298.11
272.16107177734375 0 10289.199
273.1198425292969 0 87475.305
273.13330078125 0 9505.402
279.14630126953125 0 49582.227
283.6503601074219 0 13116.677
287.14874267578125 0 24523.688
287.1723327636719 0 38546.086
290.14654541015625 0 26629.041
300.156494140625 0 12115.202
301.19171142578125 0 106812.43
302.1947021484375 0 32285.363
304.1661682128906 0 29981.977
311.1722412109375 0 119365.29
312.1756896972656 0 21398.559
318.2187194824219 0 10924.747
319.1404113769531 0 18482.795
322.1823425292969 0 18952.754
326.1834716796875 0 79315.32
327.1846008300781 0 11813.659
328.1753845214844 0 11687.857
329.1826171875 0 33209.926
331.18829345703125 0 20908.232 y Water loss 2
332.16217041015625 0 15734.131
335.15985107421875 0 12306.539
336.1676025390625 0 180736.81
337.1710205078125 0 17392.914
340.193359375 0 1073054.1 y 2
340.69476318359375 0 365924.7
341.1954650878906 0 83499.984
343.2112121582031 0 383448.2
343.2333984375 0 15247.88 y Water loss 6
344.21868896484375 0 892354.3 y Ammonia loss 6
345.22198486328125 0 138858.3 z 6
346.1869812011719 0 16709.297
346.2131652832031 0 150675.86
346.2347717285156 0 346655.12
347.19317626953125 0 34386.984
347.2159118652344 0 23367.86
347.2379150390625 0 48899.64
357.1566467285156 0 10554.941
358.20953369140625 0 50435.805
359.21533203125 0 15089.851
361.1879577636719 0 15428.05
361.2453918457031 0 120543.86 y 6
362.2489929199219 0 12559.692
363.24017333984375 0 26581.303 c 2
368.19378662109375 0 37863.49
369.21392822265625 0 296284.4
370.21728515625 0 50439.785
371.1440734863281 0 17423.152
372.22357177734375 0 19508.31
373.7150573730469 0 14546.276 c Ammonia loss 7
374.1830749511719 0 29903.99
375.1658935546875 0 12603.988
375.2377014160156 0 17548.729
376.16326904296875 0 16898.316
376.4389343261719 0 9394.389
383.2043762207031 0 56679.242
385.19635009765625 0 46976.723
385.244384765625 0 11768.569
386.2032470703125 0 40266.246
388.1966857910156 0 11786.93
389.15374755859375 0 9514.355
389.7275085449219 0 93605.586 y 1
390.22906494140625 0 36511.125
390.72955322265625 0 16184.123
392.1939697265625 0 67682.77
393.1976623535156 0 12040.104
394.7192687988281 0 12289.689
401.2154235839844 0 72729.46
402.1783447265625 0 232725.31
402.210205078125 0 12809.888
402.2214050292969 0 14553.35
402.248291015625 0 73895.33 z 5
403.2073669433594 0 1181574.1
403.2562255859375 0 3937956
404.2103576660156 0 215864.72
404.259521484375 0 748906.9
405.1775207519531 0 13831.215
405.2127380371094 0 31686.66
405.26165771484375 0 90641.64
416.2151794433594 0 236094.78
416.2445068359375 0 13922.059
417.2182312011719 0 38338.17
417.25909423828125 0 76352.74
418.2672424316406 0 272124.38 y 5
419.2272644042969 0 17660.564
419.2696228027344 0 50298.984
420.18896484375 0 179116.16
420.26324462890625 0 35024.668 c 3
421.19219970703125 0 47118.41
422.20465087890625 0 9046.162
424.2580261230469 0 16682.977
429.24700927734375 0 90470.53
430.25201416015625 0 31906.941
436.0419006347656 0 8540.459
437.2159423828125 0 130584.26
438.2177734375 0 13274.402
439.2625732421875 0 102983.13
439.763671875 0 56963.027
441.25677490234375 0 11648.141 z Water loss 4
442.2674560546875 0 40711.1
443.2688903808594 0 10848.992
445.2176818847656 0 60659.16
457.27606201171875 0 13152.61 y Water loss 4
459.2701721191406 0 478169.34 z 4
460.2780456542969 0 5534781
461.28094482421875 0 1239492.5
462.283203125 0 175518.78
463.2878112792969 0 15006.452
471.2948303222656 0 13854.946
472.3017578125 0 24006.58
473.2369689941406 0 465365.56
474.2398986816406 0 96392.06
474.2808532714844 0 283135.97
475.28857421875 0 753364.8 y 4
476.2763977050781 0 1039865.7
477.28143310546875 0 580225 c 4
478.2846984863281 0 105666.625
490.29095458984375 0 35883.73
491.29925537109375 0 30607.975
496.28857421875 0 14711.39
498.2802429199219 0 35479.707 z Water loss 3
499.2882080078125 0 40099.973
500.29034423828125 0 10817.0625
501.2450256347656 0 19774.957
505.2784118652344 0 56380.883
506.28082275390625 0 24651.191
512.2622680664062 0 19523.87
514.2999877929688 0 51813.7 y Water loss 3
516.2915649414062 0 410068.22 z 3
517.2991943359375 0 3639155
518.3023071289062 0 882567
519.25537109375 0 24407.21
519.3048095703125 0 142897.23
530.2736206054688 0 159760.94
531.3024291992188 0 540703.4
532.31005859375 0 3345939.8 y 3
533.3009033203125 0 1799741.2
534.30322265625 0 792819.5 c 5
535.3051147460938 0 177624.47
536.3092041015625 0 22183.686
548.2840576171875 0 415253.56
549.28662109375 0 117775.18
550.2890014648438 0 28501.764
558.2890014648438 0 19191.322
559.293701171875 0 14040.332
563.3074951171875 0 11106.185
566.2940673828125 0 111830.69
567.2991333007812 0 23920.379
573.3267211914062 0 38444.934
576.3162841796875 0 77261.44
577.3194580078125 0 17318.31
590.33056640625 0 23615.932
591.33740234375 0 20647.947
592.2864990234375 0 86511.52
592.3394775390625 0 15158.052
593.2897338867188 0 22463.766
600.3154296875 0 67609.98
601.3196411132812 0 24382.037
602.3236694335938 0 10152.849
604.2832641601562 0 11130.059
607.2973022460938 0 142021.64
608.3007202148438 0 41259.434
609.2996826171875 0 12598.726
616.3331298828125 0 26219.986
617.3416748046875 0 160882.14 c Water loss 6
618.3278198242188 0 263538.34
619.327392578125 0 74529.195
619.371337890625 0 117393.445
620.3053588867188 0 173215.25
620.3641357421875 0 52433.83
621.3078002929688 0 59417.008
621.36474609375 0 17095.502
622.3076171875 0 18441.012
633.3704223632812 0 14699.355
634.3447875976562 0 433601.66
635.3525390625 0 5083802.5 c 6
636.35546875 0 1594529.4
637.357666015625 0 323649.4
643.3585815429688 0 66946.75
644.353271484375 0 35655.305
645.349609375 0 210476.12 z Water loss 2
646.3525390625 0 67290.09
647.3531494140625 0 42086.582
648.3572387695312 0 11076.406
661.3679809570312 0 258773.38 y Water loss 2
662.3689575195312 0 89043.7
663.3101196289062 0 21146.87
663.3601684570312 0 2555223.5 z 2
664.3632202148438 0 884399.06
665.3656616210938 0 204548.58
679.3789672851562 0 9067130 y 2
680.3818969726562 0 3097550.5
681.38427734375 0 661897.1
682.3849487304688 0 38756.887
706.3657836914062 0 111832.85
707.369873046875 0 42975.41
719.43359375 0 73261.04
720.4365844726562 0 24885.232
728.4112548828125 0 19040.52
729.4197387695312 0 35395.516
731.4204711914062 0 10355.36
746.421142578125 0 182605.03 c Ammonia loss 7
747.4190063476562 0 68175.84
748.4172973632812 0 24640.49
760.4366455078125 0 11052.816 y Water loss 1
761.4305419921875 0 12071.743 y Ammonia loss 1
762.4286499023438 0 866021.8 z 1
763.4474487304688 0 11424741 c 7
764.450439453125 0 4589994.5
765.4530029296875 0 1096233.4
766.4540405273438 0 69026.76
778.4473266601562 0 767926.4 y 1
779.450439453125 0 297654.75
780.45263671875 0 86151.055
782.4706420898438 0 11384.059
788.4322509765625 0 12690.69
790.421875 0 31261.049
805.4336547851562 0 62423.695
806.4407958984375 0 28690.686
817.4894409179688 0 14164.107
818.4902954101562 0 84076.125
819.4428100585938 0 40193.613
819.5018310546875 0 27829.87
820.4985961914062 0 13044.127
822.4600830078125 0 22702.516
832.5170288085938 0 62322.887
833.5167236328125 0 19802.738
834.5186157226562 0 17838.363
843.4839477539062 0 26741.574
844.4843139648438 0 20582.65
846.4742431640625 0 240917
847.4766845703125 0 117453.22
848.4810791015625 0 28490.91
850.529052734375 0 40592.164
851.5376586914062 0 15497.674
860.5120239257812 0 25702.107
861.4974975585938 0 9230081
862.5001831054688 0 4282441.5
863.5027465820312 0 1272866.6
864.5054931640625 0 93474.21
877.5159301757812 0 4214387
878.522216796875 0 5964529.5
879.5249633789062 0 2456683.2
880.52880859375 0 588919.8
881.5311279296875 0 41849.383
893.4871215820312 0 22313.568
894.4964599609375 0 15562.377
952.3272094726562 0 13363.658

Spectrum Details

|  |  |
| --- | --- |
| Matched peaks? Matched peaks The total absolute number of peaks matched. Additionally in brackets the total fraction of peaks matched and the total number of peaks is shown. | 38 (12.75% of 298) |
| FDR? FDR The false discovery rate estimated for this peptide. It is calculated by matching all theoretical fragments with a non-integer shift with the raw peaks for this spectrum. This is done with 40 different shifts. The resulting percentage is the average number of annotated peaks over the number of annotated peaks with the correct spectrum. | 0.06% |
| Satellite FDR? Satellite FDR See the FDR for details on its calculation. This satellite ion specific FDR only contains the satellite ions (d/w) for I/L/J positions. | - |
| PSM Score? PSM Score The PSM Score as given by Hecklib to this annotated spectrum. It is shown with three significant figures. | 544 |

#### Spectrum 6481? Spectrum 6481 The raw spectrum of this peptide as annotated by Hecklib. The fragments are coloured according to ion type (see legend). Any peaks with a star '\*' as text can be hovered over to see the full details, first the ion type second the mass shift type. By hovering over the amino acids in the peptide or ions in the legend the corresponding peaks are highlighted. By toggling the 'Unassigned' label you can turn the background (unassigned) peaks on or off in the plot. By updating the slider in the Ion legend you can update the spectrum to only show the top X% of the peaks with labels. The top X% means any peak that is within X% of the highest intensity. By dragging in the spectrum you can zoom in to a specific part of the spectrum and use 'Zoom Out' to get back to the original zoom level. The annotation of the spectrum is based on the given sequence in the peptides file and is done with different software so inconsistencies are likely. The peaks are annotated based on the given sequence, with 20 ppm tolerance.

Copy Data

##### Spectrum 6481 (TSV)

###### Preview

```
Loading example...
```

*Click on the button to copy the data to your clipboard.*

Mz MinMz MaxIntensity Max

WidthHeightPeptide font sizePeptide stroke widthSpectrum font sizeSpectrum stroke widthCompact peptide

Ion legend

wxyz

abcd

OtherUnassignedIon^Charge^_Position_Show for top:%

VVFGGGTKJ

05.59e+41.12e+51.68e+52.24e+5

Zoom Out

y^+1^_1_d^+1^_2_a^+1^_2_b^+1^_2_y^+2^_4_y^+1^_2_b^+2^_6_y^+1^_2_y^+2^_7_b^+1^_3_y^+1^_3_b^+2^_8_y^+2^_8_y^+1^_4_b^+1^_4_y^+1^_4_\*\*y^+1^_5_y^+1^_5_y^+1^_6_y^+1^_6_y^+1^_7_y^+1^_7_b^+1^_8_y^+1^_8_\*

0867173426013468

Fragment Matches Table

Show background peaks

| Position | Ion type | Intensity | mz Theoretical | mz Error (Th) | mz Error (ppm) | Charge | Series Number |
| --- | --- | --- | --- | --- | --- | --- | --- |
| - | - | 1.134E+05 | 120.1 | - | - | 0 | - |
| - | - | 611.9 | 121.1 | - | - | 0 | - |
| - | - | 9705 | 121.1 | - | - | 0 | - |
| - | - | 1116 | 123 | - | - | 0 | - |
| - | - | 399.5 | 124.1 | - | - | 0 | - |
| - | - | 992 | 126.1 | - | - | 0 | - |
| - | - | 431.4 | 126.1 | - | - | 0 | - |
| - | - | 1404 | 127.1 | - | - | 0 | - |
| - | - | 916.5 | 127.1 | - | - | 0 | - |
| - | - | 972.6 | 128.1 | - | - | 0 | - |
| - | - | 635.4 | 129.1 | - | - | 0 | - |
| - | - | 1.108E+05 | 129.1 | - | - | 0 | - |
| - | - | 519.1 | 130.1 | - | - | 0 | - |
| - | - | 1297 | 130.1 | - | - | 0 | - |
| - | - | 2089 | 130.1 | - | - | 0 | - |
| - | - | 1027 | 130.1 | - | - | 0 | - |
| - | - | 6631 | 130.1 | - | - | 0 | - |
| - | - | 6065 | 131.1 | - | - | 0 | - |
| - | - | 1549 | 132.1 | - | - | 0 | - |
| - | - | 688.5 | 132.1 | - | - | 0 | - |
| 9 | y | 7591 | 132.1 | 0.0002372 | 1.796 | +1 | 1 |
| - | - | 1116 | 133.1 | - | - | 0 | - |
| - | - | 1.019E+04 | 136.1 | - | - | 0 | - |
| - | - | 855.6 | 139.1 | - | - | 0 | - |
| - | - | 1933 | 141.1 | - | - | 0 | - |
| - | - | 1135 | 141.1 | - | - | 0 | - |
| - | - | 453.8 | 142.1 | - | - | 0 | - |
| - | - | 446.7 | 143.1 | - | - | 0 | - |
| - | - | 3284 | 146.1 | - | - | 0 | - |
| - | - | 1095 | 147 | - | - | 0 | - |
| - | - | 627 | 147.1 | - | - | 0 | - |
| - | - | 4881 | 147.1 | - | - | 0 | - |
| - | - | 591 | 148 | - | - | 0 | - |
| - | - | 650.4 | 149 | - | - | 0 | - |
| - | - | 858.9 | 152.1 | - | - | 0 | - |
| - | - | 591.3 | 153.1 | - | - | 0 | - |
| - | - | 602 | 153.1 | - | - | 0 | - |
| - | - | 615.1 | 154.1 | - | - | 0 | - |
| - | - | 1387 | 155.1 | - | - | 0 | - |
| - | - | 3468 | 155.1 | - | - | 0 | - |
| 2 | d | 633.8 | 157.1 | 0.0003258 | 2.073 | +1 | 2 |
| - | - | 1357 | 158.1 | - | - | 0 | - |
| - | - | 862.4 | 158.1 | - | - | 0 | - |
| - | - | 4298 | 159.1 | - | - | 0 | - |
| - | - | 1410 | 159.1 | - | - | 0 | - |
| - | - | 635.2 | 160.1 | - | - | 0 | - |
| - | - | 621.6 | 165 | - | - | 0 | - |
| - | - | 6755 | 165.1 | - | - | 0 | - |
| - | - | 1385 | 166.1 | - | - | 0 | - |
| - | - | 479.8 | 166.2 | - | - | 0 | - |
| - | - | 592.3 | 167.1 | - | - | 0 | - |
| - | - | 1179 | 167.1 | - | - | 0 | - |
| - | - | 908.6 | 169.1 | - | - | 0 | - |
| - | - | 679.2 | 169.1 | - | - | 0 | - |
| - | - | 875.9 | 171.1 | - | - | 0 | - |
| 2 | a | 2.216E+05 | 171.1 | 0.0003007 | 1.757 | +1 | 2 |
| - | - | 612.2 | 172.1 | - | - | 0 | - |
| - | - | 3998 | 172.1 | - | - | 0 | - |
| - | - | 1141 | 172.1 | - | - | 0 | - |
| - | - | 1.948E+04 | 172.2 | - | - | 0 | - |
| - | - | 6894 | 173.1 | - | - | 0 | - |
| - | - | 781.1 | 173.2 | - | - | 0 | - |
| - | - | 1134 | 173.4 | - | - | 0 | - |
| - | - | 2761 | 174.1 | - | - | 0 | - |
| - | - | 2554 | 175.1 | - | - | 0 | - |
| - | - | 5158 | 176.1 | - | - | 0 | - |
| - | - | 6998 | 177.1 | - | - | 0 | - |
| - | - | 819 | 177.1 | - | - | 0 | - |
| - | - | 7565 | 182.1 | - | - | 0 | - |
| - | - | 1210 | 183.1 | - | - | 0 | - |
| - | - | 1829 | 183.1 | - | - | 0 | - |
| - | - | 648.7 | 185.1 | - | - | 0 | - |
| - | - | 1677 | 185.1 | - | - | 0 | - |
| - | - | 1176 | 186.1 | - | - | 0 | - |
| - | - | 3436 | 187.1 | - | - | 0 | - |
| - | - | 3216 | 188.1 | - | - | 0 | - |
| - | - | 871.3 | 188.1 | - | - | 0 | - |
| - | - | 586 | 189.1 | - | - | 0 | - |
| - | - | 602.5 | 189.1 | - | - | 0 | - |
| - | - | 558.9 | 191.1 | - | - | 0 | - |
| - | - | 586.9 | 191.1 | - | - | 0 | - |
| - | - | 654.2 | 192.1 | - | - | 0 | - |
| - | - | 792.1 | 194.1 | - | - | 0 | - |
| - | - | 2377 | 195.1 | - | - | 0 | - |
| - | - | 1153 | 197.2 | - | - | 0 | - |
| - | - | 4028 | 198.1 | - | - | 0 | - |
| - | - | 1355 | 199.1 | - | - | 0 | - |
| 2 | b | 8.241E+04 | 199.1 | 0.0001523 | 0.7648 | +1 | 2 |
| - | - | 9370 | 200.1 | - | - | 0 | - |
| 6 | y | 2682 | 201.1 | 5.95E-05 | 0.2958 | +2 | 4 |
| - | - | 580.1 | 201.2 | - | - | 0 | - |
| - | - | 863.6 | 202.1 | - | - | 0 | - |
| - | - | 1168 | 203.1 | - | - | 0 | - |
| - | - | 866.7 | 203.1 | - | - | 0 | - |
| - | - | 1.581E+04 | 205.1 | - | - | 0 | - |
| - | - | 1705 | 206.1 | - | - | 0 | - |
| - | - | 1710 | 208.1 | - | - | 0 | - |
| - | - | 574.1 | 209.1 | - | - | 0 | - |
| - | - | 1117 | 209.1 | - | - | 0 | - |
| - | - | 545.6 | 209.6 | - | - | 0 | - |
| - | - | 676 | 210.1 | - | - | 0 | - |
| - | - | 1090 | 211.1 | - | - | 0 | - |
| - | - | 986.1 | 212.1 | - | - | 0 | - |
| - | - | 1.123E+04 | 212.1 | - | - | 0 | - |
| - | - | 689 | 213.1 | - | - | 0 | - |
| - | - | 1214 | 213.1 | - | - | 0 | - |
| - | - | 1099 | 215.1 | - | - | 0 | - |
| - | - | 793.7 | 215.1 | - | - | 0 | - |
| - | - | 2528 | 216.1 | - | - | 0 | - |
| - | - | 743.5 | 217.1 | - | - | 0 | - |
| - | - | 2900 | 217.1 | - | - | 0 | - |
| - | - | 2302 | 217.1 | - | - | 0 | - |
| - | - | 7229 | 219.1 | - | - | 0 | - |
| - | - | 1108 | 220.2 | - | - | 0 | - |
| - | - | 1569 | 221.1 | - | - | 0 | - |
| - | - | 953.1 | 221.1 | - | - | 0 | - |
| - | - | 606.4 | 222.1 | - | - | 0 | - |
| - | - | 624.3 | 224.1 | - | - | 0 | - |
| - | - | 630.5 | 225.1 | - | - | 0 | - |
| - | - | 778.7 | 225.2 | - | - | 0 | - |
| - | - | 892.3 | 226.1 | - | - | 0 | - |
| - | - | 7290 | 226.2 | - | - | 0 | - |
| - | - | 1319 | 227.1 | - | - | 0 | - |
| - | - | 3520 | 227.1 | - | - | 0 | - |
| - | - | 555.3 | 227.1 | - | - | 0 | - |
| - | - | 1488 | 228.1 | - | - | 0 | - |
| - | - | 678.8 | 228.1 | - | - | 0 | - |
| - | - | 796.5 | 229.1 | - | - | 0 | - |
| - | - | 616.5 | 229.1 | - | - | 0 | - |
| - | - | 1.901E+04 | 230.2 | - | - | 0 | - |
| - | - | 1740 | 231.1 | - | - | 0 | - |
| - | - | 2225 | 231.2 | - | - | 0 | - |
| - | - | 1291 | 233.1 | - | - | 0 | - |
| - | - | 991.6 | 233.2 | - | - | 0 | - |
| - | - | 1.446E+04 | 233.2 | - | - | 0 | - |
| - | - | 1459 | 234.1 | - | - | 0 | - |
| - | - | 1506 | 234.2 | - | - | 0 | - |
| - | - | 959.8 | 237.1 | - | - | 0 | - |
| - | - | 859.2 | 239.2 | - | - | 0 | - |
| - | - | 662.8 | 240.1 | - | - | 0 | - |
| - | - | 679.5 | 241.2 | - | - | 0 | - |
| - | - | 935 | 242.2 | - | - | 0 | - |
| - | - | 1819 | 243.1 | - | - | 0 | - |
| 8 | y | 7630 | 243.2 | 0.0002284 | 0.9395 | +1 | 2 |
| - | - | 942.1 | 244.1 | - | - | 0 | - |
| - | - | 916.1 | 244.1 | - | - | 0 | - |
| - | - | 712 | 244.2 | - | - | 0 | - |
| - | - | 2772 | 245.1 | - | - | 0 | - |
| - | - | 2726 | 247.1 | - | - | 0 | - |
| - | - | 1688 | 248.2 | - | - | 0 | - |
| - | - | 938.3 | 251.2 | - | - | 0 | - |
| - | - | 701.8 | 254.1 | - | - | 0 | - |
| - | - | 1.03E+04 | 255.1 | - | - | 0 | - |
| - | - | 1488 | 256.1 | - | - | 0 | - |
| 6 | b | 607.8 | 259.1 | 0.001004 | 3.874 | +2 | 6 |
| 8 | y | 3.301E+04 | 260.2 | 7.587E-07 | 0.002916 | +1 | 2 |
| - | - | 531.4 | 260.3 | - | - | 0 | - |
| - | - | 2766 | 261.1 | - | - | 0 | - |
| - | - | 2964 | 261.2 | - | - | 0 | - |
| - | - | 3664 | 261.2 | - | - | 0 | - |
| - | - | 5147 | 262.1 | - | - | 0 | - |
| - | - | 9839 | 269.2 | - | - | 0 | - |
| - | - | 1113 | 270.2 | - | - | 0 | - |
| - | - | 653.5 | 272.1 | - | - | 0 | - |
| - | - | 9781 | 273.1 | - | - | 0 | - |
| - | - | 3457 | 274.1 | - | - | 0 | - |
| - | - | 948.3 | 282.2 | - | - | 0 | - |
| - | - | 2099 | 287.2 | - | - | 0 | - |
| - | - | 999.4 | 288.1 | - | - | 0 | - |
| - | - | 2789 | 291.1 | - | - | 0 | - |
| - | - | 754.3 | 292.1 | - | - | 0 | - |
| - | - | 1373 | 300.1 | - | - | 0 | - |
| - | - | 3292 | 301.2 | - | - | 0 | - |
| - | - | 844 | 302.2 | - | - | 0 | - |
| - | - | 1306 | 304.2 | - | - | 0 | - |
| - | - | 978.9 | 309.2 | - | - | 0 | - |
| - | - | 1504 | 318.1 | - | - | 0 | - |
| - | - | 4657 | 319.1 | - | - | 0 | - |
| - | - | 869.1 | 320.1 | - | - | 0 | - |
| - | - | 1106 | 321.2 | - | - | 0 | - |
| - | - | 1497 | 323.2 | - | - | 0 | - |
| - | - | 2684 | 323.2 | - | - | 0 | - |
| - | - | 7434 | 326.2 | - | - | 0 | - |
| - | - | 932.7 | 327.2 | - | - | 0 | - |
| - | - | 696.7 | 328.1 | - | - | 0 | - |
| - | - | 659.2 | 330.2 | - | - | 0 | - |
| - | - | 885.1 | 331.1 | - | - | 0 | - |
| - | - | 906.6 | 333.2 | - | - | 0 | - |
| 3 | y | 1.934E+04 | 340.2 | 0.0002518 | 0.7403 | +2 | 7 |
| - | - | 6944 | 340.7 | - | - | 0 | - |
| - | - | 1116 | 341.2 | - | - | 0 | - |
| - | - | 815 | 341.2 | - | - | 0 | - |
| - | - | 770.6 | 342.2 | - | - | 0 | - |
| - | - | 2352 | 344.2 | - | - | 0 | - |
| 3 | b | 3054 | 346.2 | 0.0001893 | 0.5468 | +1 | 3 |
| - | - | 2230 | 348.2 | - | - | 0 | - |
| - | - | 764.6 | 350.2 | - | - | 0 | - |
| - | - | 652.8 | 353.9 | - | - | 0 | - |
| - | - | 1167 | 355.2 | - | - | 0 | - |
| - | - | 708.6 | 356.2 | - | - | 0 | - |
| - | - | 2420 | 357.2 | - | - | 0 | - |
| - | - | 598.7 | 358.2 | - | - | 0 | - |
| - | - | 728.1 | 360.6 | - | - | 0 | - |
| - | - | 1013 | 361.2 | - | - | 0 | - |
| 7 | y | 5804 | 361.2 | 0.0001313 | 0.3635 | +1 | 3 |
| - | - | 1392 | 362.2 | - | - | 0 | - |
| - | - | 819.1 | 362.2 | - | - | 0 | - |
| - | - | 2211 | 365.2 | - | - | 0 | - |
| - | - | 849.3 | 366.2 | - | - | 0 | - |
| - | - | 729.2 | 373.2 | - | - | 0 | - |
| 8 | b | 934.1 | 373.7 | 0.0002704 | 0.7236 | +2 | 8 |
| - | - | 4457 | 374.2 | - | - | 0 | - |
| - | - | 2437 | 375.2 | - | - | 0 | - |
| - | - | 999.8 | 376.2 | - | - | 0 | - |
| - | - | 675.6 | 382.2 | - | - | 0 | - |
| - | - | 960.7 | 382.7 | - | - | 0 | - |
| - | - | 7409 | 383.2 | - | - | 0 | - |
| - | - | 873.9 | 383.2 | - | - | 0 | - |
| - | - | 695.6 | 383.7 | - | - | 0 | - |
| - | - | 986.9 | 384.2 | - | - | 0 | - |
| - | - | 815.6 | 385.2 | - | - | 0 | - |
| 2 | y | 1634 | 389.7 | 0.001018 | 2.612 | +2 | 8 |
| - | - | 1385 | 390.7 | - | - | 0 | - |
| - | - | 6941 | 392.2 | - | - | 0 | - |
| - | - | 1167 | 393.2 | - | - | 0 | - |
| 6 | y | 1216 | 400.3 | 0.0005968 | 1.491 | +1 | 4 |
| - | - | 4369 | 401.2 | - | - | 0 | - |
| - | - | 1.587E+04 | 402.2 | - | - | 0 | - |
| - | - | 1098 | 402.2 | - | - | 0 | - |
| - | - | 726.2 | 403 | - | - | 0 | - |
| - | - | 3595 | 403.2 | - | - | 0 | - |
| 4 | b | 1343 | 403.2 | 0.0003931 | 0.9748 | +1 | 4 |
| - | - | 1352 | 403.8 | - | - | 0 | - |
| - | - | 2988 | 418.2 | - | - | 0 | - |
| 6 | y | 5315 | 418.3 | 0.0003243 | 0.7753 | +1 | 4 |
| - | - | 2643 | 419.2 | - | - | 0 | - |
| - | - | 1261 | 419.3 | - | - | 0 | - |
| - | - | 9541 | 420.2 | - | - | 0 | - |
| - | - | 977.3 | 420.2 | - | - | 0 | - |
| - | - | 2094 | 421.2 | - | - | 0 | - |
| - | - | 704.6 | 421.8 | - | - | 0 | - |
| - | - | 739.9 | 424.2 | - | - | 0 | - |
| - | - | 2879 | 425.3 | - | - | 0 | - |
| - | - | 1383 | 425.8 | - | - | 0 | - |
| 0 | Precursor | 1982 | 430.3 | 0.0005323 | 1.237 | +2 | -1 |
| - | - | 583.2 | 431.5 | - | - | 0 | - |
| - | - | 3269 | 438.2 | - | - | 0 | - |
| - | - | 1931 | 439.2 | - | - | 0 | - |
| 0 | Precursor | 1.124E+04 | 439.3 | 0.0009297 | 2.117 | +2 | -1 |
| - | - | 2528 | 439.8 | - | - | 0 | - |
| - | - | 8110 | 439.8 | - | - | 0 | - |
| - | - | 945.4 | 440.2 | - | - | 0 | - |
| - | - | 1017 | 442.2 | - | - | 0 | - |
| - | - | 2550 | 448.3 | - | - | 0 | - |
| - | - | 1194 | 448.8 | - | - | 0 | - |
| - | - | 658.7 | 451.3 | - | - | 0 | - |
| 5 | y | 2779 | 457.3 | 0.0003428 | 0.7497 | +1 | 5 |
| - | - | 699.5 | 461.3 | - | - | 0 | - |
| - | - | 1750 | 469.3 | - | - | 0 | - |
| - | - | 1138 | 470.3 | - | - | 0 | - |
| - | - | 960 | 473.3 | - | - | 0 | - |
| 5 | y | 2.198E+04 | 475.3 | 0.0002151 | 0.4527 | +1 | 5 |
| - | - | 830.2 | 475.8 | - | - | 0 | - |
| - | - | 4356 | 476.3 | - | - | 0 | - |
| - | - | 2319 | 481.3 | - | - | 0 | - |
| - | - | 731.9 | 486.2 | - | - | 0 | - |
| - | - | 2075 | 489.3 | - | - | 0 | - |
| - | - | 632.7 | 490.3 | - | - | 0 | - |
| - | - | 2268 | 496.3 | - | - | 0 | - |
| - | - | 941.9 | 501.2 | - | - | 0 | - |
| - | - | 757.3 | 509.3 | - | - | 0 | - |
| - | - | 2145 | 512.3 | - | - | 0 | - |
| 4 | y | 5067 | 514.3 | 0.0003995 | 0.7767 | +1 | 6 |
| - | - | 1147 | 515.3 | - | - | 0 | - |
| - | - | 716.9 | 520.8 | - | - | 0 | - |
| - | - | 1379 | 525.8 | - | - | 0 | - |
| - | - | 2270 | 527.3 | - | - | 0 | - |
| - | - | 1.701E+04 | 529.8 | - | - | 0 | - |
| - | - | 6037 | 530.3 | - | - | 0 | - |
| - | - | 4755 | 530.3 | - | - | 0 | - |
| - | - | 2028 | 531.3 | - | - | 0 | - |
| 4 | y | 6.615E+04 | 532.3 | 0.000344 | 0.6463 | +1 | 6 |
| - | - | 1.516E+04 | 533.3 | - | - | 0 | - |
| - | - | 2897 | 534.3 | - | - | 0 | - |
| - | - | 1129 | 537.3 | - | - | 0 | - |
| - | - | 1847 | 539.8 | - | - | 0 | - |
| - | - | 1154 | 540.3 | - | - | 0 | - |
| - | - | 849.5 | 540.8 | - | - | 0 | - |
| - | - | 1.28E+04 | 548.3 | - | - | 0 | - |
| - | - | 1383 | 548.3 | - | - | 0 | - |
| - | - | 1184 | 548.8 | - | - | 0 | - |
| - | - | 3335 | 549.3 | - | - | 0 | - |
| - | - | 1136 | 550.3 | - | - | 0 | - |
| - | - | 3245 | 555.3 | - | - | 0 | - |
| - | - | 687.4 | 556.3 | - | - | 0 | - |
| - | - | 8461 | 566.3 | - | - | 0 | - |
| - | - | 3122 | 567.3 | - | - | 0 | - |
| - | - | 1755 | 573.3 | - | - | 0 | - |
| - | - | 828.9 | 588.4 | - | - | 0 | - |
| - | - | 1120 | 597.3 | - | - | 0 | - |
| - | - | 2897 | 598.3 | - | - | 0 | - |
| - | - | 946.8 | 599.3 | - | - | 0 | - |
| - | - | 3395 | 616.3 | - | - | 0 | - |
| - | - | 1968 | 617.3 | - | - | 0 | - |
| - | - | 888 | 618.3 | - | - | 0 | - |
| - | - | 696.4 | 625.3 | - | - | 0 | - |
| - | - | 1337 | 633.4 | - | - | 0 | - |
| - | - | 4241 | 643.4 | - | - | 0 | - |
| - | - | 2486 | 644.3 | - | - | 0 | - |
| - | - | 868.1 | 645.4 | - | - | 0 | - |
| - | - | 1096 | 647.4 | - | - | 0 | - |
| - | - | 880.4 | 649.4 | - | - | 0 | - |
| 3 | y | 1.114E+04 | 661.4 | 0.0005761 | 0.871 | +1 | 7 |
| - | - | 3609 | 662.4 | - | - | 0 | - |
| - | - | 1591 | 663.4 | - | - | 0 | - |
| - | - | 1443 | 667.4 | - | - | 0 | - |
| 3 | y | 2.02E+05 | 679.4 | 0.0004596 | 0.6765 | +1 | 7 |
| - | - | 7.007E+04 | 680.4 | - | - | 0 | - |
| - | - | 1.67E+04 | 681.4 | - | - | 0 | - |
| - | - | 917.4 | 681.5 | - | - | 0 | - |
| - | - | 1171 | 682.4 | - | - | 0 | - |
| - | - | 911.3 | 697.4 | - | - | 0 | - |
| - | - | 2217 | 699.4 | - | - | 0 | - |
| - | - | 715.3 | 700.4 | - | - | 0 | - |
| - | - | 754.8 | 701.4 | - | - | 0 | - |
| - | - | 587.3 | 733.4 | - | - | 0 | - |
| - | - | 3795 | 736.4 | - | - | 0 | - |
| - | - | 1261 | 737.4 | - | - | 0 | - |
| 8 | b | 1712 | 746.4 | 0.001948 | 2.61 | +1 | 8 |
| - | - | 610.3 | 747.4 | - | - | 0 | - |
| - | - | 2558 | 764.4 | - | - | 0 | - |
| - | - | 760.9 | 765.4 | - | - | 0 | - |
| 2 | y | 2.008E+04 | 778.4 | 0.001247 | 1.601 | +1 | 8 |
| - | - | 8290 | 779.4 | - | - | 0 | - |
| - | - | 571.8 | 779.5 | - | - | 0 | - |
| - | - | 5801 | 780.5 | - | - | 0 | - |
| - | - | 1445 | 781.5 | - | - | 0 | - |
| - | - | 793.8 | 798.5 | - | - | 0 | - |
| 0 | Precursor | 3296 | 877.5 | 0.0006297 | 0.7176 | +1 | -1 |
| - | - | 936.2 | 878.5 | - | - | 0 | - |
| - | - | 990.8 | 895.5 | - | - | 0 | - |
| - | - | 582.1 | 913.8 | - | - | 0 | - |
| - | - | 3613 | 961.5 | - | - | 0 | - |
| - | - | 815.3 | 962.5 | - | - | 0 | - |
| - | - | 639 | 1273 | - | - | 0 | - |
| - | - | 594 | 1614 | - | - | 0 | - |
| - | - | 685.6 | 3021 | - | - | 0 | - |
| - | - | 682.8 | 3081 | - | - | 0 | - |
| - | - | 716.2 | 3434 | - | - | 0 | - |

m/z Charge Intensity FragmentType MassShift Position
120.0810546875 0 113397.984
121.07923889160156 0 611.8573
121.08436584472656 0 9704.785
123.04419708251953 0 1115.581
124.11238098144531 0 399.50995
126.0551528930664 0 992.03613
126.0915298461914 0 431.40408
127.05042266845703 0 1403.7395
127.08695220947266 0 916.4929
128.1073760986328 0 972.6269
129.06629943847656 0 635.35376
129.10250854492188 0 110818.12
130.05044555664062 0 519.0908
130.06541442871094 0 1297.4081
130.0865020751953 0 2088.755
130.1001739501953 0 1027.3436
130.10581970214844 0 6631.055
131.0817108154297 0 6064.7163
132.08106994628906 0 1549.3541
132.09716796875 0 688.5462
132.10214233398438 0 7590.6387 y 8
133.1055908203125 0 1116.2144
136.07591247558594 0 10186.939
139.086669921875 0 855.5511
141.066162109375 0 1933.2393
141.10264587402344 0 1134.8201
142.1227264404297 0 453.78918
143.11854553222656 0 446.66818
146.0602569580078 0 3284.267
147.0442657470703 0 1094.5256
147.06382751464844 0 627.0293
147.113037109375 0 4881.081
148.03929138183594 0 591.01794
148.955078125 0 650.41534
152.14373779296875 0 858.9247
153.06591796875 0 591.2923
153.10269165039062 0 601.9534
154.06097412109375 0 615.1034
155.08155822753906 0 1386.6382
155.11813354492188 0 3468.1912
157.1338653564453 0 633.7956 d 1
158.06031799316406 0 1356.9773
158.09263610839844 0 862.3695
159.07664489746094 0 4298.0723
159.09194946289062 0 1409.7172
160.07589721679688 0 635.23206
165.04827880859375 0 621.64594
165.0547332763672 0 6755.194
166.0863494873047 0 1384.9332
166.1628875732422 0 479.82886
167.08152770996094 0 592.3217
167.11795043945312 0 1178.9146
169.09716796875 0 908.579
169.13417053222656 0 679.1991
171.0767822265625 0 875.9456
171.1494903564453 0 221581.94 a 1
172.06422424316406 0 612.21985
172.07180786132812 0 3997.8806
172.1465301513672 0 1141.4192
172.15281677246094 0 19479.38
173.12852478027344 0 6893.785
173.15591430664062 0 781.081
173.4394989013672 0 1134.2252
174.05523681640625 0 2761.151
175.08676147460938 0 2553.8657
176.10714721679688 0 5157.645
177.10247802734375 0 6998.065
177.11045837402344 0 818.9831
182.08135986328125 0 7565.173
183.11294555664062 0 1209.6455
183.14938354492188 0 1828.637
185.07061767578125 0 648.73413
185.12875366210938 0 1677.0458
186.1239013671875 0 1176.2148
187.14431762695312 0 3435.8416
188.07081604003906 0 3215.6138
188.1031036376953 0 871.34973
189.07376098632812 0 585.99646
189.10260009765625 0 602.54
191.08152770996094 0 558.93774
191.1184844970703 0 586.92096
192.0654754638672 0 654.23126
194.12905883789062 0 792.11237
195.11305236816406 0 2376.6655
197.1650848388672 0 1153.0458
198.08743286132812 0 4028.4116
199.10768127441406 0 1354.9471
199.14425659179688 0 82412.65 b 1
200.1476593017578 0 9370.129
201.12342834472656 0 2681.5894 y Ammonia loss 5
201.15077209472656 0 580.0666
202.05032348632812 0 863.62885
203.08203125 0 1168.1576
203.1182861328125 0 866.6887
205.0972900390625 0 15810.911
206.10061645507812 0 1705.4089
208.10836791992188 0 1710.2656
209.09278869628906 0 574.06824
209.1028594970703 0 1116.6367
209.61863708496094 0 545.5842
210.08702087402344 0 675.97986
211.1442413330078 0 1090.0875
212.12863159179688 0 986.0637
212.13949584960938 0 11233.623
213.1240692138672 0 689.0008
213.14300537109375 0 1214.051
215.11785888671875 0 1099.3191
215.13906860351562 0 793.6785
216.09786987304688 0 2527.8015
217.08303833007812 0 743.52966
217.09744262695312 0 2899.998
217.1337127685547 0 2301.6917
219.1493682861328 0 7229.055
220.1537322998047 0 1107.9211
221.09239196777344 0 1568.6838
221.12823486328125 0 953.09357
222.1232147216797 0 606.3626
224.10203552246094 0 624.2545
225.1342315673828 0 630.5432
225.1596221923828 0 778.7354
226.11878967285156 0 892.3071
226.1551055908203 0 7290.287
227.10287475585938 0 1319.473
227.11399841308594 0 3520.197
227.12478637695312 0 555.33185
228.09780883789062 0 1488.1273
228.1179962158203 0 678.78326
229.09349060058594 0 796.4911
229.1053466796875 0 616.45715
230.15013122558594 0 19007.041
231.1132354736328 0 1740.2128
231.153076171875 0 2224.9617
233.1282958984375 0 1291.0388
233.1515350341797 0 991.6469
233.16493225097656 0 14463.417
234.1449737548828 0 1458.9907
234.16864013671875 0 1506.2603
237.0983428955078 0 959.8274
239.1501007080078 0 859.1556
240.13499450683594 0 662.7958
241.15524291992188 0 679.54083
242.18701171875 0 935.0282
243.1133270263672 0 1818.8433
243.17054748535156 0 7629.924 y Ammonia loss 7
244.10824584960938 0 942.0901
244.129150390625 0 916.132
244.17393493652344 0 712.01086
245.12513732910156 0 2772.309
247.14419555664062 0 2725.5657
248.1610565185547 0 1687.8225
251.150390625 0 938.29297
254.1143035888672 0 701.7504
255.10888671875 0 10302.446
256.1109619140625 0 1487.7
259.1430969238281 0 607.83105 b 5
260.1968688964844 0 33010.9 y 7
260.3064270019531 0 531.438
261.1233215332031 0 2766.3066
261.1596984863281 0 2963.6914
261.2004089355469 0 3664.3926
262.1185607910156 0 5147.1685
269.1608581542969 0 9839.4
270.16485595703125 0 1113.2943
272.1235656738281 0 653.52435
273.1194763183594 0 9781.46
274.12005615234375 0 3456.566
282.15606689453125 0 948.28033
287.1720275878906 0 2098.8533
288.1351623535156 0 999.39276
291.14508056640625 0 2789.1333
292.1497497558594 0 754.32227
300.1341247558594 0 1372.7737
301.19146728515625 0 3292.135
302.1727600097656 0 844.0036
304.1661071777344 0 1305.6844
309.2041015625 0 978.8612
318.14495849609375 0 1503.7263
319.1405029296875 0 4657.143
320.14324951171875 0 869.1473
321.1776123046875 0 1106.1377
323.1712341308594 0 1496.9536
323.2074890136719 0 2683.5344
326.18243408203125 0 7434.142
327.1849365234375 0 932.72437
328.1278381347656 0 696.7303
330.1636657714844 0 659.18994
331.1400146484375 0 885.10065
333.19293212890625 0 906.5693
340.19256591796875 0 19339.04 y 2
340.69384765625 0 6944.355
341.1793212890625 0 1115.874
341.1987609863281 0 814.9902
342.1838073730469 0 770.6106
344.1927795410156 0 2352.4102
346.21270751953125 0 3053.8445 b 2
348.16693115234375 0 2230.454
350.2047119140625 0 764.5541
353.8694763183594 0 652.8119
355.1609802246094 0 1166.9365
356.172119140625 0 708.605
357.1557922363281 0 2419.6611
358.15679931640625 0 598.7441
360.56170654296875 0 728.0959
361.1863708496094 0 1013.2116
361.2444152832031 0 5804.1436 y 6
362.2031555175781 0 1391.9982
362.2453918457031 0 819.14
365.1925048828125 0 2211.1572
366.17694091796875 0 849.33246
373.1702880859375 0 729.20935
373.71368408203125 0 934.1035 b 7
374.18231201171875 0 4456.651
375.16552734375 0 2437.4778
376.1660461425781 0 999.83466
382.2440185546875 0 675.59674
382.7178039550781 0 960.7234
383.2037048339844 0 7409.234
383.2273864746094 0 873.8531
383.7278747558594 0 695.56036
384.2088928222656 0 986.9179
385.1513977050781 0 815.6417
389.7275390625 0 1633.9568 y 1
390.7330322265625 0 1384.6921
392.19281005859375 0 6940.9185
393.1935729980469 0 1166.9368
400.25604248046875 0 1216.0706 y Water loss 5
401.2145690917969 0 4369.398
402.1773376464844 0 15869.988
402.2172546386719 0 1098.0082
402.9640197753906 0 726.1982
403.18072509765625 0 3595.319
403.234375 0 1342.9727 b 3
403.8143615722656 0 1352.2268
418.2293395996094 0 2987.6077
418.26568603515625 0 5315.299 y 5
419.22705078125 0 2643.0737
419.2701416015625 0 1261.2192
420.1878967285156 0 9541.465
420.2235107421875 0 977.3286
421.19085693359375 0 2093.8752
421.8345031738281 0 704.6182
424.2174072265625 0 739.8641
425.2635498046875 0 2878.8938
425.76531982421875 0 1383.2693
430.2549133300781 0 1981.8524 Precursor Water loss
431.54010009765625 0 583.1795
438.2133483886719 0 3269.398
439.22601318359375 0 1931.0249
439.26165771484375 0 11244.762 Precursor
439.76312255859375 0 2527.6416
439.8435974121094 0 8110.457
440.2286682128906 0 945.42957
442.2287292480469 0 1017.3998
448.266357421875 0 2550.4346
448.7685852050781 0 1193.9956
451.2655334472656 0 658.7096
457.2772521972656 0 2778.516 y Water loss 4
461.291748046875 0 699.47943
469.2765197753906 0 1749.8964
470.2790222167969 0 1138.1727
473.25140380859375 0 960.03723
475.2876892089844 0 21978 y 4
475.7912292480469 0 830.1548
476.2906188964844 0 4355.7925
481.27117919921875 0 2318.6736
486.2467346191406 0 731.9078
489.2653503417969 0 2074.8394
490.271728515625 0 632.6653
496.2886962890625 0 2268.1572
501.2440185546875 0 941.881
509.3117980957031 0 757.3206
512.2622680664062 0 2145.2937
514.2979736328125 0 5067.4736 y Water loss 3
515.3009643554688 0 1147.0049
520.7922973632812 0 716.8567
525.810546875 0 1379.4906
527.3184204101562 0 2269.691
529.7976684570312 0 17013.22
530.2700805664062 0 6036.6436
530.3023071289062 0 4755.027
531.2744750976562 0 2027.8383
532.30859375 0 66154.48 y 3
533.3114624023438 0 15160.879
534.3130493164062 0 2897.338
537.3029174804688 0 1128.5829
539.8110961914062 0 1847.3152
540.3101806640625 0 1154.1951
540.7998657226562 0 849.4518
548.2822265625 0 12795.53
548.32373046875 0 1383.0248
548.8229370117188 0 1183.5425
549.2848510742188 0 3334.7402
550.2865600585938 0 1136.0444
555.3126220703125 0 3244.9204
556.3160400390625 0 687.37286
566.2928466796875 0 8460.803
567.2957763671875 0 3121.7612
573.3226318359375 0 1754.8016
588.353271484375 0 828.90784
597.3291625976562 0 1119.7196
598.337158203125 0 2897.0781
599.3338012695312 0 946.7639
616.3446044921875 0 3394.8474
617.3453369140625 0 1967.789
618.3391723632812 0 888.0159
625.3478393554688 0 696.3844
633.3721313476562 0 1336.6703
643.3550415039062 0 4240.9194
644.349609375 0 2485.5105
645.3562622070312 0 868.0678
647.3518676757812 0 1095.7205
649.3674926757812 0 880.4127
661.3662109375 0 11141.193 y Water loss 2
662.3689575195312 0 3609.1387
663.3770141601562 0 1591.1392
667.3765258789062 0 1442.5664
679.3768920898438 0 201986.48 y 2
680.3800048828125 0 70071.6
681.3827514648438 0 16704.229
681.45458984375 0 917.40283
682.3868408203125 0 1171.0731
697.4063110351562 0 911.34424
699.4036865234375 0 2217.4778
700.4109497070312 0 715.3218
701.4130249023438 0 754.8272
733.3687744140625 0 587.30615
736.3870239257812 0 3794.764
737.3868408203125 0 1260.9868
746.4176025390625 0 1712.1711 b 7
747.41796875 0 610.26624
764.4292602539062 0 2557.5872
765.4354858398438 0 760.9057
778.4445190429688 0 20081.701 y 1
779.4473876953125 0 8289.949
779.5303344726562 0 571.7561
780.4578247070312 0 5801.4756
781.460205078125 0 1444.775
798.4672241210938 0 793.78534
877.5135498046875 0 3295.531 Precursor
878.5164794921875 0 936.2034
895.5281982421875 0 990.819
913.8070068359375 0 582.1305
961.5338745117188 0 3612.6643
962.5404663085938 0 815.33325
1272.5028076171875 0 639.0087
1613.665771484375 0 594.00635
3021.288818359375 0 685.571
3080.653076171875 0 682.80786
3434.021240234375 0 716.1863

Spectrum Details

|  |  |
| --- | --- |
| Matched peaks? Matched peaks The total absolute number of peaks matched. Additionally in brackets the total fraction of peaks matched and the total number of peaks is shown. | 27 (7.74% of 349) |
| FDR? FDR The false discovery rate estimated for this peptide. It is calculated by matching all theoretical fragments with a non-integer shift with the raw peaks for this spectrum. This is done with 40 different shifts. The resulting percentage is the average number of annotated peaks over the number of annotated peaks with the correct spectrum. | 0.71% |
| Satellite FDR? Satellite FDR See the FDR for details on its calculation. This satellite ion specific FDR only contains the satellite ions (d/w) for I/L/J positions. | - |
| PSM Score? PSM Score The PSM Score as given by Hecklib to this annotated spectrum. It is shown with three significant figures. | 281 |

#### Spectrum 6753? Spectrum 6753 The raw spectrum of this peptide as annotated by Hecklib. The fragments are coloured according to ion type (see legend). Any peaks with a star '\*' as text can be hovered over to see the full details, first the ion type second the mass shift type. By hovering over the amino acids in the peptide or ions in the legend the corresponding peaks are highlighted. By toggling the 'Unassigned' label you can turn the background (unassigned) peaks on or off in the plot. By updating the slider in the Ion legend you can update the spectrum to only show the top X% of the peaks with labels. The top X% means any peak that is within X% of the highest intensity. By dragging in the spectrum you can zoom in to a specific part of the spectrum and use 'Zoom Out' to get back to the original zoom level. The annotation of the spectrum is based on the given sequence in the peptides file and is done with different software so inconsistencies are likely. The peaks are annotated based on the given sequence, with 20 ppm tolerance.

Copy Data

##### Spectrum 6753 (TSV)

###### Preview

```
Loading example...
```

*Click on the button to copy the data to your clipboard.*

Mz MinMz MaxIntensity Max

WidthHeightPeptide font sizePeptide stroke widthSpectrum font sizeSpectrum stroke widthCompact peptide

Ion legend

wxyz

abcd

OtherUnassignedIon^Charge^_Position_Show for top:%

VVFGGGTKJ

02.77e+45.53e+48.30e+41.11e+5

Zoom Out

y^+1^_1_a^+1^_2_b^+1^_2_y^+2^_4_y^+1^_2_y^+1^_2_y^+2^_7_y^+1^_3_b^+1^_3_y^+1^_3_y^+2^_8_y^+1^_4_\*y^+1^_5_y^+1^_5_y^+1^_6_y^+1^_6_y^+1^_7_y^+1^_7_b^+1^_8_y^+1^_8_

0727145321802906

Fragment Matches Table

Show background peaks

| Position | Ion type | Intensity | mz Theoretical | mz Error (Th) | mz Error (ppm) | Charge | Series Number |
| --- | --- | --- | --- | --- | --- | --- | --- |
| - | - | 5.394E+04 | 120.1 | - | - | 0 | - |
| - | - | 479.4 | 121.1 | - | - | 0 | - |
| - | - | 3966 | 121.1 | - | - | 0 | - |
| - | - | 400.2 | 127 | - | - | 0 | - |
| - | - | 939.9 | 127.1 | - | - | 0 | - |
| - | - | 437.7 | 127.1 | - | - | 0 | - |
| - | - | 731.3 | 129.1 | - | - | 0 | - |
| - | - | 5.76E+04 | 129.1 | - | - | 0 | - |
| - | - | 1501 | 130.1 | - | - | 0 | - |
| - | - | 754.8 | 130.1 | - | - | 0 | - |
| - | - | 921.3 | 130.1 | - | - | 0 | - |
| - | - | 3756 | 130.1 | - | - | 0 | - |
| - | - | 2010 | 131.1 | - | - | 0 | - |
| - | - | 872.8 | 132.1 | - | - | 0 | - |
| 9 | y | 4407 | 132.1 | 0.0003135 | 2.373 | +1 | 1 |
| - | - | 438.6 | 133.1 | - | - | 0 | - |
| - | - | 1.813E+04 | 136.1 | - | - | 0 | - |
| - | - | 837 | 137.1 | - | - | 0 | - |
| - | - | 855 | 139.1 | - | - | 0 | - |
| - | - | 380.7 | 139.1 | - | - | 0 | - |
| - | - | 1480 | 141.1 | - | - | 0 | - |
| - | - | 876.1 | 141.1 | - | - | 0 | - |
| - | - | 479.4 | 145.3 | - | - | 0 | - |
| - | - | 3108 | 146.1 | - | - | 0 | - |
| - | - | 861.7 | 147 | - | - | 0 | - |
| - | - | 1750 | 147.1 | - | - | 0 | - |
| - | - | 1066 | 148.9 | - | - | 0 | - |
| - | - | 444.5 | 154.1 | - | - | 0 | - |
| - | - | 1432 | 155.1 | - | - | 0 | - |
| - | - | 1688 | 155.1 | - | - | 0 | - |
| - | - | 607.1 | 156.1 | - | - | 0 | - |
| - | - | 739.2 | 157.1 | - | - | 0 | - |
| - | - | 719.7 | 158.1 | - | - | 0 | - |
| - | - | 1851 | 159.1 | - | - | 0 | - |
| - | - | 1079 | 159.1 | - | - | 0 | - |
| - | - | 444.7 | 162.1 | - | - | 0 | - |
| - | - | 1189 | 166.1 | - | - | 0 | - |
| - | - | 739.1 | 167.1 | - | - | 0 | - |
| - | - | 444.8 | 169.1 | - | - | 0 | - |
| - | - | 834.9 | 171.1 | - | - | 0 | - |
| 2 | a | 1.095E+05 | 171.1 | 0.0003312 | 1.935 | +1 | 2 |
| - | - | 2092 | 172.1 | - | - | 0 | - |
| - | - | 552.8 | 172.1 | - | - | 0 | - |
| - | - | 1.008E+04 | 172.2 | - | - | 0 | - |
| - | - | 4888 | 173.1 | - | - | 0 | - |
| - | - | 589.1 | 173.2 | - | - | 0 | - |
| - | - | 2427 | 173.5 | - | - | 0 | - |
| - | - | 1903 | 174.1 | - | - | 0 | - |
| - | - | 1198 | 175.1 | - | - | 0 | - |
| - | - | 3280 | 176.1 | - | - | 0 | - |
| - | - | 3836 | 177.1 | - | - | 0 | - |
| - | - | 712.2 | 177.1 | - | - | 0 | - |
| - | - | 846.1 | 181.1 | - | - | 0 | - |
| - | - | 1346 | 182.1 | - | - | 0 | - |
| - | - | 539.5 | 183.1 | - | - | 0 | - |
| - | - | 420.3 | 183.1 | - | - | 0 | - |
| - | - | 2206 | 183.1 | - | - | 0 | - |
| - | - | 530.2 | 185.1 | - | - | 0 | - |
| - | - | 1035 | 185.1 | - | - | 0 | - |
| - | - | 607.6 | 186.1 | - | - | 0 | - |
| - | - | 707.8 | 187.1 | - | - | 0 | - |
| - | - | 929.6 | 188.1 | - | - | 0 | - |
| - | - | 686.6 | 189.1 | - | - | 0 | - |
| - | - | 758.8 | 192.1 | - | - | 0 | - |
| - | - | 646.2 | 195.1 | - | - | 0 | - |
| - | - | 614.3 | 196.1 | - | - | 0 | - |
| - | - | 480.8 | 197.1 | - | - | 0 | - |
| - | - | 652.7 | 197.2 | - | - | 0 | - |
| - | - | 2390 | 198.1 | - | - | 0 | - |
| - | - | 812 | 198.1 | - | - | 0 | - |
| - | - | 1355 | 199.1 | - | - | 0 | - |
| 2 | b | 3.926E+04 | 199.1 | 0.0002439 | 1.225 | +1 | 2 |
| - | - | 3802 | 200.1 | - | - | 0 | - |
| 6 | y | 2732 | 201.1 | 0.0001816 | 0.9028 | +2 | 4 |
| - | - | 873.7 | 202.1 | - | - | 0 | - |
| - | - | 1043 | 203.1 | - | - | 0 | - |
| - | - | 745.2 | 203.1 | - | - | 0 | - |
| - | - | 8060 | 205.1 | - | - | 0 | - |
| - | - | 727.6 | 206.1 | - | - | 0 | - |
| - | - | 811.4 | 208.1 | - | - | 0 | - |
| - | - | 516.9 | 209.1 | - | - | 0 | - |
| - | - | 3567 | 211.1 | - | - | 0 | - |
| - | - | 5491 | 212.1 | - | - | 0 | - |
| - | - | 584.3 | 212.1 | - | - | 0 | - |
| - | - | 560.5 | 212.9 | - | - | 0 | - |
| - | - | 551.4 | 213.1 | - | - | 0 | - |
| - | - | 626.4 | 215.1 | - | - | 0 | - |
| - | - | 774.1 | 215.1 | - | - | 0 | - |
| - | - | 1676 | 216.1 | - | - | 0 | - |
| - | - | 518.3 | 217.1 | - | - | 0 | - |
| - | - | 1327 | 217.1 | - | - | 0 | - |
| - | - | 2827 | 217.1 | - | - | 0 | - |
| - | - | 3569 | 219.1 | - | - | 0 | - |
| - | - | 1812 | 221.1 | - | - | 0 | - |
| - | - | 510.8 | 221.1 | - | - | 0 | - |
| - | - | 704.6 | 221.1 | - | - | 0 | - |
| - | - | 1733 | 226.1 | - | - | 0 | - |
| - | - | 4069 | 226.2 | - | - | 0 | - |
| - | - | 896.1 | 227.1 | - | - | 0 | - |
| - | - | 1560 | 227.1 | - | - | 0 | - |
| - | - | 677.6 | 228.1 | - | - | 0 | - |
| - | - | 9858 | 230.2 | - | - | 0 | - |
| - | - | 656.4 | 230.2 | - | - | 0 | - |
| - | - | 896.8 | 231.1 | - | - | 0 | - |
| - | - | 672.6 | 231.2 | - | - | 0 | - |
| - | - | 1203 | 233.1 | - | - | 0 | - |
| - | - | 592 | 233.2 | - | - | 0 | - |
| - | - | 3866 | 233.2 | - | - | 0 | - |
| - | - | 674.7 | 234.2 | - | - | 0 | - |
| - | - | 583.4 | 237.1 | - | - | 0 | - |
| - | - | 856.7 | 239.1 | - | - | 0 | - |
| - | - | 562.3 | 242.1 | - | - | 0 | - |
| - | - | 598.9 | 243.1 | - | - | 0 | - |
| 8 | y | 2988 | 243.2 | 0.0006252 | 2.571 | +1 | 2 |
| - | - | 931 | 244.1 | - | - | 0 | - |
| - | - | 679 | 244.2 | - | - | 0 | - |
| - | - | 2343 | 245.1 | - | - | 0 | - |
| - | - | 1497 | 247.1 | - | - | 0 | - |
| - | - | 4694 | 249.2 | - | - | 0 | - |
| - | - | 519.8 | 250.2 | - | - | 0 | - |
| - | - | 5990 | 255.1 | - | - | 0 | - |
| - | - | 811.4 | 256.2 | - | - | 0 | - |
| 8 | y | 1.487E+04 | 260.2 | 9.231E-05 | 0.3548 | +1 | 2 |
| - | - | 958.6 | 261.1 | - | - | 0 | - |
| - | - | 842.7 | 261.2 | - | - | 0 | - |
| - | - | 1604 | 261.2 | - | - | 0 | - |
| - | - | 3233 | 262.1 | - | - | 0 | - |
| - | - | 5367 | 269.2 | - | - | 0 | - |
| - | - | 585 | 270.2 | - | - | 0 | - |
| - | - | 5122 | 273.1 | - | - | 0 | - |
| - | - | 768.8 | 274.1 | - | - | 0 | - |
| - | - | 735.3 | 276.2 | - | - | 0 | - |
| - | - | 634.1 | 277.2 | - | - | 0 | - |
| - | - | 4621 | 279.1 | - | - | 0 | - |
| - | - | 789.2 | 280.1 | - | - | 0 | - |
| - | - | 600.4 | 282.2 | - | - | 0 | - |
| - | - | 963 | 287.2 | - | - | 0 | - |
| - | - | 1371 | 291.1 | - | - | 0 | - |
| - | - | 604.6 | 296.1 | - | - | 0 | - |
| - | - | 607.1 | 300.1 | - | - | 0 | - |
| - | - | 2036 | 301.2 | - | - | 0 | - |
| - | - | 737 | 304.2 | - | - | 0 | - |
| - | - | 544.6 | 314.8 | - | - | 0 | - |
| - | - | 961.2 | 318.1 | - | - | 0 | - |
| - | - | 3228 | 319.1 | - | - | 0 | - |
| - | - | 869.6 | 323.2 | - | - | 0 | - |
| - | - | 1348 | 323.2 | - | - | 0 | - |
| - | - | 4474 | 326.2 | - | - | 0 | - |
| - | - | 570.8 | 329.4 | - | - | 0 | - |
| - | - | 658.9 | 337.2 | - | - | 0 | - |
| - | - | 1148 | 339.2 | - | - | 0 | - |
| 3 | y | 9022 | 340.2 | 0.000496 | 1.458 | +2 | 7 |
| - | - | 2889 | 340.7 | - | - | 0 | - |
| - | - | 1500 | 341.2 | - | - | 0 | - |
| 7 | y | 786.1 | 343.2 | 0.0004541 | 1.323 | +1 | 3 |
| - | - | 953.8 | 344.2 | - | - | 0 | - |
| 3 | b | 1394 | 346.2 | 0.0003295 | 0.9517 | +1 | 3 |
| - | - | 561.7 | 347.2 | - | - | 0 | - |
| - | - | 936.5 | 350.2 | - | - | 0 | - |
| - | - | 1433 | 355.2 | - | - | 0 | - |
| - | - | 1892 | 357.2 | - | - | 0 | - |
| 7 | y | 3338 | 361.2 | 0.0003144 | 0.8704 | +1 | 3 |
| - | - | 961.3 | 362.2 | - | - | 0 | - |
| - | - | 598.9 | 362.2 | - | - | 0 | - |
| - | - | 564.1 | 364 | - | - | 0 | - |
| - | - | 872.9 | 365.2 | - | - | 0 | - |
| - | - | 553.9 | 371.5 | - | - | 0 | - |
| - | - | 2731 | 374.2 | - | - | 0 | - |
| - | - | 2354 | 374.2 | - | - | 0 | - |
| - | - | 1035 | 375.2 | - | - | 0 | - |
| - | - | 924.8 | 382.7 | - | - | 0 | - |
| - | - | 2580 | 383.2 | - | - | 0 | - |
| - | - | 1011 | 384.2 | - | - | 0 | - |
| 2 | y | 1137 | 389.7 | 0.0006299 | 1.616 | +2 | 8 |
| - | - | 3192 | 392.2 | - | - | 0 | - |
| - | - | 2656 | 401.2 | - | - | 0 | - |
| - | - | 7002 | 402.2 | - | - | 0 | - |
| - | - | 1162 | 403.2 | - | - | 0 | - |
| - | - | 915.5 | 418.2 | - | - | 0 | - |
| 6 | y | 3068 | 418.3 | 0.0006523 | 1.559 | +1 | 4 |
| - | - | 1436 | 419.2 | - | - | 0 | - |
| - | - | 4776 | 420.2 | - | - | 0 | - |
| - | - | 816.7 | 420.2 | - | - | 0 | - |
| - | - | 819.1 | 421.2 | - | - | 0 | - |
| - | - | 567.4 | 423.3 | - | - | 0 | - |
| - | - | 1025 | 425.3 | - | - | 0 | - |
| - | - | 869.9 | 437.8 | - | - | 0 | - |
| - | - | 1656 | 439.2 | - | - | 0 | - |
| 0 | Precursor | 2086 | 439.3 | 0.001479 | 3.367 | +2 | -1 |
| - | - | 956.2 | 439.8 | - | - | 0 | - |
| - | - | 7678 | 439.8 | - | - | 0 | - |
| - | - | 7347 | 448.3 | - | - | 0 | - |
| - | - | 1242 | 448.8 | - | - | 0 | - |
| - | - | 709.4 | 455.3 | - | - | 0 | - |
| 5 | y | 1272 | 457.3 | 0.001075 | 2.351 | +1 | 5 |
| - | - | 1311 | 461.3 | - | - | 0 | - |
| - | - | 1298 | 469.3 | - | - | 0 | - |
| 5 | y | 1.094E+04 | 475.3 | 0.0004593 | 0.9663 | +1 | 5 |
| - | - | 2363 | 476.3 | - | - | 0 | - |
| - | - | 593.2 | 488.3 | - | - | 0 | - |
| - | - | 871.4 | 489.3 | - | - | 0 | - |
| - | - | 916.3 | 496.3 | - | - | 0 | - |
| - | - | 1830 | 512.3 | - | - | 0 | - |
| 4 | y | 2089 | 514.3 | 0.0005161 | 1.003 | +1 | 6 |
| - | - | 946.2 | 515.3 | - | - | 0 | - |
| - | - | 3093 | 521.8 | - | - | 0 | - |
| - | - | 849.4 | 522.3 | - | - | 0 | - |
| - | - | 919.2 | 525.8 | - | - | 0 | - |
| - | - | 1799 | 529.8 | - | - | 0 | - |
| - | - | 4502 | 530.3 | - | - | 0 | - |
| - | - | 854.6 | 531.3 | - | - | 0 | - |
| 4 | y | 3.098E+04 | 532.3 | 0.0001609 | 0.3023 | +1 | 6 |
| - | - | 7860 | 533.3 | - | - | 0 | - |
| - | - | 1488 | 534.3 | - | - | 0 | - |
| - | - | 2158 | 539.8 | - | - | 0 | - |
| - | - | 1996 | 540.3 | - | - | 0 | - |
| - | - | 7267 | 548.3 | - | - | 0 | - |
| - | - | 882.5 | 548.3 | - | - | 0 | - |
| - | - | 983.5 | 548.8 | - | - | 0 | - |
| - | - | 1677 | 549.3 | - | - | 0 | - |
| - | - | 991 | 555.3 | - | - | 0 | - |
| - | - | 3708 | 566.3 | - | - | 0 | - |
| - | - | 1306 | 567.3 | - | - | 0 | - |
| - | - | 4733 | 573.3 | - | - | 0 | - |
| - | - | 641.2 | 574.3 | - | - | 0 | - |
| - | - | 1712 | 599.3 | - | - | 0 | - |
| - | - | 1395 | 616.3 | - | - | 0 | - |
| - | - | 649.9 | 617.3 | - | - | 0 | - |
| - | - | 670.5 | 626.4 | - | - | 0 | - |
| - | - | 1646 | 643.4 | - | - | 0 | - |
| - | - | 1018 | 644.4 | - | - | 0 | - |
| 3 | y | 4886 | 661.4 | 0.000454 | 0.6865 | +1 | 7 |
| - | - | 2096 | 662.4 | - | - | 0 | - |
| - | - | 708.8 | 667.4 | - | - | 0 | - |
| 3 | y | 1.029E+05 | 679.4 | 0.0004596 | 0.6765 | +1 | 7 |
| - | - | 3.432E+04 | 680.4 | - | - | 0 | - |
| - | - | 943.5 | 680.4 | - | - | 0 | - |
| - | - | 7525 | 681.4 | - | - | 0 | - |
| - | - | 869.1 | 699.4 | - | - | 0 | - |
| 8 | b | 618.1 | 746.4 | 0.002315 | 3.101 | +1 | 8 |
| - | - | 893.6 | 764.4 | - | - | 0 | - |
| 2 | y | 1.057E+04 | 778.4 | 0.001308 | 1.68 | +1 | 8 |
| - | - | 3791 | 779.4 | - | - | 0 | - |
| - | - | 1728 | 780.5 | - | - | 0 | - |
| - | - | 3864 | 798.5 | - | - | 0 | - |
| - | - | 941.4 | 799.5 | - | - | 0 | - |
| - | - | 540.5 | 821.7 | - | - | 0 | - |
| - | - | 644.8 | 824.4 | - | - | 0 | - |
| - | - | 1713 | 895.5 | - | - | 0 | - |
| - | - | 1045 | 896.5 | - | - | 0 | - |
| - | - | 644 | 1043 | - | - | 0 | - |
| - | - | 564.9 | 1130 | - | - | 0 | - |
| - | - | 661 | 1216 | - | - | 0 | - |
| - | - | 604.1 | 1280 | - | - | 0 | - |
| - | - | 551.6 | 1369 | - | - | 0 | - |
| - | - | 645.3 | 1629 | - | - | 0 | - |
| - | - | 707.2 | 1837 | - | - | 0 | - |
| - | - | 657.3 | 2877 | - | - | 0 | - |

m/z Charge Intensity FragmentType MassShift Position
120.08109283447266 0 53939.754
121.06539916992188 0 479.36063
121.08438873291016 0 3966.1042
127.03959655761719 0 400.1916
127.05034637451172 0 939.9261
127.0751953125 0 437.73773
129.06619262695312 0 731.30634
129.1025390625 0 57595.527
130.0653839111328 0 1500.6461
130.086669921875 0 754.7913
130.10006713867188 0 921.29987
130.10592651367188 0 3756.257
131.08181762695312 0 2010.3622
132.08116149902344 0 872.7904
132.1022186279297 0 4406.851 y 8
133.105224609375 0 438.6407
136.07601928710938 0 18127.396
137.07957458496094 0 836.9764
139.08702087402344 0 854.9999
139.0941162109375 0 380.7479
141.06631469726562 0 1479.7742
141.1026153564453 0 876.07355
145.26173400878906 0 479.43756
146.0603485107422 0 3107.7046
147.0442657470703 0 861.7395
147.11317443847656 0 1749.776
148.94717407226562 0 1065.98
154.06163024902344 0 444.46927
155.0816650390625 0 1432.0035
155.11830139160156 0 1688.4227
156.0771484375 0 607.0788
157.07647705078125 0 739.2099
158.06044006347656 0 719.70825
159.0766143798828 0 1851.0903
159.0919189453125 0 1078.9163
162.05511474609375 0 444.70996
166.08663940429688 0 1188.8291
167.11781311035156 0 739.0604
169.10826110839844 0 444.75055
171.07601928710938 0 834.9144
171.14952087402344 0 109547.68 a 1
172.0720977783203 0 2092.1392
172.14601135253906 0 552.84827
172.1529083251953 0 10075.364
173.1287078857422 0 4887.6577
173.15505981445312 0 589.12933
173.4516143798828 0 2426.5757
174.05506896972656 0 1902.7002
175.08673095703125 0 1197.8599
176.10728454589844 0 3280.188
177.10240173339844 0 3835.6924
177.110595703125 0 712.1525
181.09725952148438 0 846.06854
182.08148193359375 0 1345.7739
183.11355590820312 0 539.4894
183.14077758789062 0 420.316
183.14952087402344 0 2206.4126
185.0924530029297 0 530.1675
185.12893676757812 0 1035.0172
186.1236114501953 0 607.5619
187.1445770263672 0 707.7773
188.0708770751953 0 929.573
189.06605529785156 0 686.6044
192.0662078857422 0 758.81335
195.11288452148438 0 646.24585
196.10816955566406 0 614.3069
197.12774658203125 0 480.84216
197.1652069091797 0 652.6546
198.0876007080078 0 2390.0474
198.12391662597656 0 811.9667
199.1079864501953 0 1354.9645
199.14434814453125 0 39263.33 b 1
200.1477508544922 0 3802.108
201.12355041503906 0 2731.5269 y Ammonia loss 5
202.05010986328125 0 873.65906
203.08139038085938 0 1043.019
203.11865234375 0 745.23914
205.09742736816406 0 8059.9995
206.1011199951172 0 727.63776
208.10812377929688 0 811.4466
209.0926971435547 0 516.8958
211.14451599121094 0 3567.3982
212.13958740234375 0 5490.6953
212.1480712890625 0 584.33453
212.947509765625 0 560.5287
213.12356567382812 0 551.40375
215.11647033691406 0 626.36536
215.1397705078125 0 774.14374
216.09800720214844 0 1675.9968
217.08177185058594 0 518.27893
217.09762573242188 0 1326.9034
217.1337127685547 0 2826.9136
219.1494903564453 0 3568.5408
221.0917205810547 0 1811.975
221.10342407226562 0 510.7904
221.1288604736328 0 704.5502
226.1187286376953 0 1733.2277
226.15505981445312 0 4069.0635
227.1024169921875 0 896.09686
227.1138458251953 0 1560.1603
228.09812927246094 0 677.5991
230.1502685546875 0 9857.984
230.16311645507812 0 656.42035
231.1127471923828 0 896.76935
231.15426635742188 0 672.5794
233.12852478027344 0 1202.8856
233.1533203125 0 591.9942
233.16510009765625 0 3866.2527
234.16795349121094 0 674.7283
237.09873962402344 0 583.4433
239.14993286132812 0 856.6503
242.1492462158203 0 562.34045
243.11302185058594 0 598.93274
243.1709442138672 0 2987.9094 y Ammonia loss 7
244.12924194335938 0 931.0305
244.1735382080078 0 678.96344
245.1261749267578 0 2342.5215
247.14414978027344 0 1496.8098
249.1598663330078 0 4693.947
250.16302490234375 0 519.7827
255.1089324951172 0 5990.316
256.1778869628906 0 811.364
260.19696044921875 0 14873.246 y 7
261.1231689453125 0 958.5841
261.1599426269531 0 842.664
261.2001037597656 0 1603.6503
262.1187438964844 0 3233.4507
269.1609802246094 0 5367.134
270.1821594238281 0 584.9517
273.1195068359375 0 5121.968
274.1184387207031 0 768.80035
276.1673889160156 0 735.25287
277.1544189453125 0 634.0643
279.1343078613281 0 4620.8823
280.13653564453125 0 789.2268
282.1558532714844 0 600.4044
287.1710205078125 0 962.98694
291.14593505859375 0 1371.071
296.13470458984375 0 604.6081
300.135009765625 0 607.0754
301.19134521484375 0 2035.8442
304.1669921875 0 737.0126
314.76922607421875 0 544.5587
318.1461181640625 0 961.1847
319.14031982421875 0 3228.082
323.17218017578125 0 869.5862
323.2078857421875 0 1347.6821
326.18255615234375 0 4473.6943
329.4281921386719 0 570.7901
337.1517639160156 0 658.88196
339.1778869628906 0 1147.9668
340.19281005859375 0 9022.268 y 2
340.694580078125 0 2889.1675
341.18194580078125 0 1500.2091
343.23443603515625 0 786.09827 y Water loss 6
344.19244384765625 0 953.8177
346.2121887207031 0 1393.6959 b 2
347.2161865234375 0 561.74945
350.20587158203125 0 936.48236
355.1612854003906 0 1433.1444
357.1553039550781 0 1891.7482
361.2442321777344 0 3337.9177 y 6
362.2045593261719 0 961.327
362.2486572265625 0 598.9008
364.0281066894531 0 564.13605
365.1924133300781 0 872.9438
371.5202941894531 0 553.894
374.18157958984375 0 2731.1592
374.2075500488281 0 2354.2354
375.16583251953125 0 1034.5515
382.7176208496094 0 924.7839
383.20391845703125 0 2580.0527
384.2057189941406 0 1010.6231
389.72589111328125 0 1136.8701 y 1
392.1935119628906 0 3192.0474
401.21435546875 0 2655.7012
402.17730712890625 0 7001.559
403.1798095703125 0 1161.8234
418.2338562011719 0 915.5229
418.26666259765625 0 3068.3315 y 5
419.2245178222656 0 1436.3118
420.18780517578125 0 4776.009
420.2197570800781 0 816.6983
421.1912841796875 0 819.0507
423.2626647949219 0 567.408
425.2647399902344 0 1025.1666
437.8481750488281 0 869.9298
439.2300720214844 0 1656.0378
439.26220703125 0 2086.0076 Precursor
439.76123046875 0 956.18915
439.84405517578125 0 7678.3345
448.26605224609375 0 7346.8555
448.76837158203125 0 1242.3743
455.2711486816406 0 709.3942
457.2779846191406 0 1271.7662 y Water loss 4
461.2895812988281 0 1311.2977
469.2776184082031 0 1298.2777
475.2879333496094 0 10939.655 y 4
476.28912353515625 0 2362.9785
488.2830810546875 0 593.1756
489.2649230957031 0 871.37317
496.2841491699219 0 916.2735
512.2626342773438 0 1830.3324
514.2988891601562 0 2089.0488 y Water loss 3
515.3004760742188 0 946.16864
521.7996215820312 0 3093.412
522.301025390625 0 849.37305
525.8101196289062 0 919.20734
529.7969360351562 0 1798.5366
530.2721557617188 0 4502.111
531.2739868164062 0 854.61005
532.3087768554688 0 30984.465 y 3
533.3115844726562 0 7860.2954
534.3126831054688 0 1487.5375
539.8106079101562 0 2158.0066
540.3093872070312 0 1996.0969
548.2823486328125 0 7266.5044
548.3235473632812 0 882.52765
548.8219604492188 0 983.4561
549.2849731445312 0 1677.4858
555.313232421875 0 991.034
566.2928466796875 0 3707.8591
567.2950439453125 0 1306.1847
573.3237915039062 0 4733.136
574.3284301757812 0 641.18463
599.3176879882812 0 1712.0636
616.3441772460938 0 1394.546
617.3447265625 0 649.91943
626.377197265625 0 670.4969
643.3534545898438 0 1646.2076
644.3546142578125 0 1018.32776
661.3663330078125 0 4885.674 y Water loss 2
662.3685302734375 0 2096.0708
667.3787841796875 0 708.78186
679.3768920898438 0 102908.93 y 2
680.3797607421875 0 34320.74
680.4490356445312 0 943.5369
681.3826904296875 0 7525.1323
699.4030151367188 0 869.0673
746.417236328125 0 618.0943 b 7
764.4292602539062 0 893.5998
778.4444580078125 0 10571.109 y 1
779.4473266601562 0 3791.1917
780.4552612304688 0 1728.178
798.4708251953125 0 3864.1323
799.475830078125 0 941.3981
821.743896484375 0 540.499
824.4447021484375 0 644.79956
895.527099609375 0 1713.2632
896.5245361328125 0 1044.5643
1042.815673828125 0 644.00037
1129.7490234375 0 564.9354
1215.7568359375 0 661.0249
1279.865478515625 0 604.1058
1368.585693359375 0 551.6337
1628.7083740234375 0 645.2731
1837.3699951171875 0 707.21014
2877.34423828125 0 657.33636

Spectrum Details

|  |  |
| --- | --- |
| Matched peaks? Matched peaks The total absolute number of peaks matched. Additionally in brackets the total fraction of peaks matched and the total number of peaks is shown. | 21 (8.14% of 258) |
| FDR? FDR The false discovery rate estimated for this peptide. It is calculated by matching all theoretical fragments with a non-integer shift with the raw peaks for this spectrum. This is done with 40 different shifts. The resulting percentage is the average number of annotated peaks over the number of annotated peaks with the correct spectrum. | 0.68% |
| Satellite FDR? Satellite FDR See the FDR for details on its calculation. This satellite ion specific FDR only contains the satellite ions (d/w) for I/L/J positions. | - |
| PSM Score? PSM Score The PSM Score as given by Hecklib to this annotated spectrum. It is shown with three significant figures. | 241 |

#### Spectrum 6940? Spectrum 6940 The raw spectrum of this peptide as annotated by Hecklib. The fragments are coloured according to ion type (see legend). Any peaks with a star '\*' as text can be hovered over to see the full details, first the ion type second the mass shift type. By hovering over the amino acids in the peptide or ions in the legend the corresponding peaks are highlighted. By toggling the 'Unassigned' label you can turn the background (unassigned) peaks on or off in the plot. By updating the slider in the Ion legend you can update the spectrum to only show the top X% of the peaks with labels. The top X% means any peak that is within X% of the highest intensity. By dragging in the spectrum you can zoom in to a specific part of the spectrum and use 'Zoom Out' to get back to the original zoom level. The annotation of the spectrum is based on the given sequence in the peptides file and is done with different software so inconsistencies are likely. The peaks are annotated based on the given sequence, with 20 ppm tolerance.

Copy Data

##### Spectrum 6940 (TSV)

###### Preview

```
Loading example...
```

*Click on the button to copy the data to your clipboard.*

Mz MinMz MaxIntensity Max

WidthHeightPeptide font sizePeptide stroke widthSpectrum font sizeSpectrum stroke widthCompact peptide

Ion legend

wxyz

abcd

OtherUnassignedIon^Charge^_Position_Show for top:%

VVFGGGTKJ

02.11e+44.21e+46.32e+48.43e+4

Zoom Out

y^+1^_1_a^+1^_2_b^+1^_2_y^+2^_4_y^+1^_2_b^+2^_6_y^+1^_2_y^+2^_7_b^+1^_3_y^+1^_3_b^+2^_8_y^+1^_4_\*y^+1^_5_y^+1^_5_y^+1^_6_y^+1^_6_b^+1^_7_y^+1^_7_y^+1^_7_b^+1^_8_y^+1^_8_\*

0778155723353113

Fragment Matches Table

Show background peaks

| Position | Ion type | Intensity | mz Theoretical | mz Error (Th) | mz Error (ppm) | Charge | Series Number |
| --- | --- | --- | --- | --- | --- | --- | --- |
| - | - | 402.5 | 120 | - | - | 0 | - |
| - | - | 5.978E+04 | 120.1 | - | - | 0 | - |
| - | - | 481.7 | 121.1 | - | - | 0 | - |
| - | - | 3316 | 121.1 | - | - | 0 | - |
| - | - | 330.2 | 122.3 | - | - | 0 | - |
| - | - | 424.8 | 124.1 | - | - | 0 | - |
| - | - | 1479 | 126.1 | - | - | 0 | - |
| - | - | 756 | 127.1 | - | - | 0 | - |
| - | - | 626.7 | 129.1 | - | - | 0 | - |
| - | - | 6.36E+04 | 129.1 | - | - | 0 | - |
| - | - | 903.2 | 130.1 | - | - | 0 | - |
| - | - | 1085 | 130.1 | - | - | 0 | - |
| - | - | 653.7 | 130.1 | - | - | 0 | - |
| - | - | 3007 | 130.1 | - | - | 0 | - |
| - | - | 1440 | 131.1 | - | - | 0 | - |
| - | - | 629.7 | 132.1 | - | - | 0 | - |
| 9 | y | 4809 | 132.1 | 0.0002982 | 2.258 | +1 | 1 |
| - | - | 740.3 | 133.1 | - | - | 0 | - |
| - | - | 372.2 | 133.3 | - | - | 0 | - |
| - | - | 420.9 | 134.3 | - | - | 0 | - |
| - | - | 469.7 | 135 | - | - | 0 | - |
| - | - | 361.2 | 135.8 | - | - | 0 | - |
| - | - | 8.345E+04 | 136.1 | - | - | 0 | - |
| - | - | 700.9 | 137.1 | - | - | 0 | - |
| - | - | 4564 | 137.1 | - | - | 0 | - |
| - | - | 716.5 | 139.1 | - | - | 0 | - |
| - | - | 375.6 | 139.8 | - | - | 0 | - |
| - | - | 785.7 | 141.1 | - | - | 0 | - |
| - | - | 476.9 | 141.1 | - | - | 0 | - |
| - | - | 404.2 | 141.2 | - | - | 0 | - |
| - | - | 695.3 | 144.1 | - | - | 0 | - |
| - | - | 2981 | 146.1 | - | - | 0 | - |
| - | - | 5023 | 147 | - | - | 0 | - |
| - | - | 2479 | 147.1 | - | - | 0 | - |
| - | - | 597.7 | 148 | - | - | 0 | - |
| - | - | 1041 | 149 | - | - | 0 | - |
| - | - | 739.7 | 155.1 | - | - | 0 | - |
| - | - | 916.8 | 155.1 | - | - | 0 | - |
| - | - | 421.9 | 157.1 | - | - | 0 | - |
| - | - | 647.4 | 158.1 | - | - | 0 | - |
| - | - | 1374 | 159.1 | - | - | 0 | - |
| - | - | 1379 | 159.1 | - | - | 0 | - |
| - | - | 466.7 | 163 | - | - | 0 | - |
| - | - | 428.2 | 163.2 | - | - | 0 | - |
| - | - | 458.4 | 163.2 | - | - | 0 | - |
| - | - | 403.2 | 163.3 | - | - | 0 | - |
| - | - | 744.6 | 165.1 | - | - | 0 | - |
| - | - | 689 | 165.1 | - | - | 0 | - |
| - | - | 858 | 166.1 | - | - | 0 | - |
| - | - | 1404 | 167.1 | - | - | 0 | - |
| 2 | a | 8.258E+04 | 171.1 | 0.0003312 | 1.935 | +1 | 2 |
| - | - | 1599 | 172.1 | - | - | 0 | - |
| - | - | 772.2 | 172.1 | - | - | 0 | - |
| - | - | 727.9 | 172.1 | - | - | 0 | - |
| - | - | 7319 | 172.2 | - | - | 0 | - |
| - | - | 3920 | 173.1 | - | - | 0 | - |
| - | - | 1984 | 174.1 | - | - | 0 | - |
| - | - | 658.7 | 175.1 | - | - | 0 | - |
| - | - | 2538 | 176.1 | - | - | 0 | - |
| - | - | 460.1 | 176.6 | - | - | 0 | - |
| - | - | 3036 | 177.1 | - | - | 0 | - |
| - | - | 705.1 | 177.1 | - | - | 0 | - |
| - | - | 1250 | 182.1 | - | - | 0 | - |
| - | - | 1691 | 183.1 | - | - | 0 | - |
| - | - | 886.7 | 185.1 | - | - | 0 | - |
| - | - | 773.9 | 186.1 | - | - | 0 | - |
| - | - | 1067 | 187.1 | - | - | 0 | - |
| - | - | 977.4 | 188.1 | - | - | 0 | - |
| - | - | 964.5 | 189.1 | - | - | 0 | - |
| - | - | 552.9 | 191.1 | - | - | 0 | - |
| - | - | 1353 | 192.1 | - | - | 0 | - |
| - | - | 1381 | 195.1 | - | - | 0 | - |
| - | - | 608.4 | 197.1 | - | - | 0 | - |
| - | - | 508.2 | 197.2 | - | - | 0 | - |
| - | - | 1175 | 198.1 | - | - | 0 | - |
| - | - | 1134 | 199.1 | - | - | 0 | - |
| 2 | b | 2.877E+04 | 199.1 | 0.0002744 | 1.378 | +1 | 2 |
| - | - | 2775 | 200.1 | - | - | 0 | - |
| 6 | y | 1291 | 201.1 | 5.95E-05 | 0.2958 | +2 | 4 |
| - | - | 630.8 | 202.1 | - | - | 0 | - |
| - | - | 574.2 | 203.1 | - | - | 0 | - |
| - | - | 753.8 | 203.1 | - | - | 0 | - |
| - | - | 548.3 | 204.1 | - | - | 0 | - |
| - | - | 804.4 | 204.1 | - | - | 0 | - |
| - | - | 5083 | 205.1 | - | - | 0 | - |
| - | - | 467.2 | 205.9 | - | - | 0 | - |
| - | - | 757.2 | 208.1 | - | - | 0 | - |
| - | - | 1186 | 209.1 | - | - | 0 | - |
| - | - | 1316 | 211.1 | - | - | 0 | - |
| - | - | 5564 | 212.1 | - | - | 0 | - |
| - | - | 759.3 | 215.1 | - | - | 0 | - |
| - | - | 1395 | 216.1 | - | - | 0 | - |
| - | - | 1349 | 217.1 | - | - | 0 | - |
| - | - | 952.6 | 217.1 | - | - | 0 | - |
| - | - | 1.058E+04 | 217.1 | - | - | 0 | - |
| - | - | 743.5 | 218.1 | - | - | 0 | - |
| - | - | 2578 | 219.1 | - | - | 0 | - |
| - | - | 1209 | 221.1 | - | - | 0 | - |
| - | - | 1049 | 221.1 | - | - | 0 | - |
| - | - | 536.4 | 221.9 | - | - | 0 | - |
| - | - | 2284 | 226.1 | - | - | 0 | - |
| - | - | 1190 | 226.1 | - | - | 0 | - |
| - | - | 2.05E+04 | 226.2 | - | - | 0 | - |
| - | - | 1190 | 227.1 | - | - | 0 | - |
| - | - | 1220 | 227.1 | - | - | 0 | - |
| - | - | 1338 | 227.2 | - | - | 0 | - |
| - | - | 487.6 | 228.3 | - | - | 0 | - |
| - | - | 1509 | 229.1 | - | - | 0 | - |
| - | - | 1775 | 229.2 | - | - | 0 | - |
| - | - | 6994 | 230.2 | - | - | 0 | - |
| - | - | 731.2 | 231.1 | - | - | 0 | - |
| - | - | 824.4 | 231.2 | - | - | 0 | - |
| - | - | 718.1 | 233.1 | - | - | 0 | - |
| - | - | 1568 | 233.2 | - | - | 0 | - |
| - | - | 9498 | 233.2 | - | - | 0 | - |
| - | - | 972.5 | 234.2 | - | - | 0 | - |
| - | - | 949.5 | 239.2 | - | - | 0 | - |
| - | - | 957.4 | 240.1 | - | - | 0 | - |
| - | - | 583.9 | 243.1 | - | - | 0 | - |
| 8 | y | 1925 | 243.2 | 4.621E-05 | 0.19 | +1 | 2 |
| - | - | 1836 | 244.1 | - | - | 0 | - |
| - | - | 5731 | 245.1 | - | - | 0 | - |
| - | - | 1019 | 247.1 | - | - | 0 | - |
| - | - | 2.847E+04 | 249.2 | - | - | 0 | - |
| - | - | 2507 | 250.2 | - | - | 0 | - |
| - | - | 1178 | 254.1 | - | - | 0 | - |
| - | - | 752.6 | 254.1 | - | - | 0 | - |
| - | - | 3578 | 255.1 | - | - | 0 | - |
| - | - | 770.8 | 256.2 | - | - | 0 | - |
| 6 | b | 1106 | 259.1 | 0.002652 | 10.23 | +2 | 6 |
| 8 | y | 1.18E+04 | 260.2 | 0.0002144 | 0.8239 | +1 | 2 |
| - | - | 1676 | 261.1 | - | - | 0 | - |
| - | - | 2670 | 261.2 | - | - | 0 | - |
| - | - | 1672 | 261.2 | - | - | 0 | - |
| - | - | 2583 | 262.1 | - | - | 0 | - |
| - | - | 3241 | 269.2 | - | - | 0 | - |
| - | - | 1117 | 272.1 | - | - | 0 | - |
| - | - | 4223 | 273.1 | - | - | 0 | - |
| - | - | 952.9 | 274.1 | - | - | 0 | - |
| - | - | 6779 | 277.2 | - | - | 0 | - |
| - | - | 703.7 | 278.2 | - | - | 0 | - |
| - | - | 1431 | 287.2 | - | - | 0 | - |
| - | - | 852.8 | 291.1 | - | - | 0 | - |
| - | - | 1152 | 301.2 | - | - | 0 | - |
| - | - | 623.8 | 302.2 | - | - | 0 | - |
| - | - | 655.7 | 302.2 | - | - | 0 | - |
| - | - | 653 | 309.2 | - | - | 0 | - |
| - | - | 2099 | 319.1 | - | - | 0 | - |
| - | - | 2121 | 323.2 | - | - | 0 | - |
| - | - | 7126 | 323.2 | - | - | 0 | - |
| - | - | 1924 | 326.2 | - | - | 0 | - |
| - | - | 745 | 330.2 | - | - | 0 | - |
| 3 | y | 6500 | 340.2 | 0.0006181 | 1.817 | +2 | 7 |
| - | - | 2427 | 340.7 | - | - | 0 | - |
| - | - | 6490 | 341.2 | - | - | 0 | - |
| - | - | 590 | 341.2 | - | - | 0 | - |
| 3 | b | 1190 | 346.2 | 0.0002379 | 0.6872 | +1 | 3 |
| - | - | 850.6 | 348.2 | - | - | 0 | - |
| - | - | 705.8 | 351.2 | - | - | 0 | - |
| - | - | 1058 | 355.2 | - | - | 0 | - |
| - | - | 1927 | 357.2 | - | - | 0 | - |
| 7 | y | 2103 | 361.2 | 9.259E-06 | 0.02563 | +1 | 3 |
| - | - | 605.9 | 363.9 | - | - | 0 | - |
| - | - | 876.4 | 365.2 | - | - | 0 | - |
| - | - | 614.3 | 370.5 | - | - | 0 | - |
| - | - | 2259 | 373.2 | - | - | 0 | - |
| 8 | b | 1574 | 373.7 | 0.0008503 | 2.275 | +2 | 8 |
| - | - | 1451 | 374.2 | - | - | 0 | - |
| - | - | 587.9 | 374.2 | - | - | 0 | - |
| - | - | 980.9 | 375.2 | - | - | 0 | - |
| - | - | 890.6 | 379.2 | - | - | 0 | - |
| - | - | 3990 | 382.7 | - | - | 0 | - |
| - | - | 2133 | 383.2 | - | - | 0 | - |
| - | - | 785.4 | 383.2 | - | - | 0 | - |
| - | - | 737.8 | 384.2 | - | - | 0 | - |
| - | - | 1766 | 391.7 | - | - | 0 | - |
| - | - | 2654 | 392.2 | - | - | 0 | - |
| - | - | 4760 | 399.7 | - | - | 0 | - |
| - | - | 850.8 | 400.2 | - | - | 0 | - |
| - | - | 1800 | 401.2 | - | - | 0 | - |
| - | - | 5664 | 402.2 | - | - | 0 | - |
| - | - | 741.6 | 403 | - | - | 0 | - |
| - | - | 1024 | 403.2 | - | - | 0 | - |
| - | - | 901.7 | 414.2 | - | - | 0 | - |
| - | - | 937.8 | 418.2 | - | - | 0 | - |
| 6 | y | 2307 | 418.3 | 0.0007133 | 1.705 | +1 | 4 |
| - | - | 1412 | 419.2 | - | - | 0 | - |
| - | - | 2506 | 420.2 | - | - | 0 | - |
| - | - | 1051 | 421.2 | - | - | 0 | - |
| - | - | 1876 | 424.2 | - | - | 0 | - |
| - | - | 1877 | 425.3 | - | - | 0 | - |
| - | - | 822.9 | 437.8 | - | - | 0 | - |
| - | - | 892.8 | 439.1 | - | - | 0 | - |
| - | - | 708.9 | 439.2 | - | - | 0 | - |
| - | - | 1400 | 439.2 | - | - | 0 | - |
| 0 | Precursor | 3880 | 439.3 | 0.0009297 | 2.117 | +2 | -1 |
| - | - | 1305 | 439.8 | - | - | 0 | - |
| - | - | 7109 | 439.8 | - | - | 0 | - |
| - | - | 621.5 | 440.6 | - | - | 0 | - |
| - | - | 3281 | 442.2 | - | - | 0 | - |
| - | - | 4.109E+04 | 448.3 | - | - | 0 | - |
| - | - | 1.302E+04 | 448.8 | - | - | 0 | - |
| - | - | 737.9 | 451.3 | - | - | 0 | - |
| 5 | y | 656.1 | 458.3 | 0.0007626 | 1.664 | +1 | 5 |
| - | - | 1741 | 460.2 | - | - | 0 | - |
| - | - | 873.9 | 469.3 | - | - | 0 | - |
| - | - | 1579 | 470.3 | - | - | 0 | - |
| 5 | y | 7675 | 475.3 | 0.0005814 | 1.223 | +1 | 5 |
| - | - | 1474 | 476.3 | - | - | 0 | - |
| - | - | 892.6 | 476.3 | - | - | 0 | - |
| - | - | 730.7 | 496.3 | - | - | 0 | - |
| - | - | 1499 | 512.8 | - | - | 0 | - |
| 4 | y | 1810 | 514.3 | 2.778E-05 | 0.05402 | +1 | 6 |
| - | - | 1.096E+04 | 521.8 | - | - | 0 | - |
| - | - | 4425 | 522.3 | - | - | 0 | - |
| - | - | 1538 | 525.8 | - | - | 0 | - |
| - | - | 844.8 | 527.3 | - | - | 0 | - |
| - | - | 1618 | 529.8 | - | - | 0 | - |
| - | - | 3291 | 530.3 | - | - | 0 | - |
| - | - | 943.9 | 531.3 | - | - | 0 | - |
| 4 | y | 2.363E+04 | 532.3 | 8.321E-05 | 0.1563 | +1 | 6 |
| - | - | 5664 | 533.3 | - | - | 0 | - |
| - | - | 1185 | 534.3 | - | - | 0 | - |
| - | - | 1585 | 539.8 | - | - | 0 | - |
| - | - | 1432 | 540.3 | - | - | 0 | - |
| - | - | 5423 | 548.3 | - | - | 0 | - |
| - | - | 1334 | 548.3 | - | - | 0 | - |
| - | - | 866.5 | 548.4 | - | - | 0 | - |
| - | - | 904.9 | 548.8 | - | - | 0 | - |
| - | - | 1417 | 549.3 | - | - | 0 | - |
| - | - | 3411 | 555.3 | - | - | 0 | - |
| - | - | 2576 | 566.3 | - | - | 0 | - |
| - | - | 1099 | 566.3 | - | - | 0 | - |
| - | - | 821.6 | 570.3 | - | - | 0 | - |
| - | - | 2.279E+04 | 573.3 | - | - | 0 | - |
| - | - | 4641 | 574.3 | - | - | 0 | - |
| - | - | 1484 | 578.3 | - | - | 0 | - |
| - | - | 1156 | 578.8 | - | - | 0 | - |
| - | - | 641.7 | 580.2 | - | - | 0 | - |
| 7 | b | 625.5 | 600.3 | 0.00043 | 0.7162 | +1 | 7 |
| - | - | 661.5 | 615.4 | - | - | 0 | - |
| - | - | 665.2 | 617.3 | - | - | 0 | - |
| - | - | 574 | 626.7 | - | - | 0 | - |
| - | - | 1386 | 643.4 | - | - | 0 | - |
| - | - | 981.5 | 644.4 | - | - | 0 | - |
| - | - | 725.8 | 649.4 | - | - | 0 | - |
| 3 | y | 4584 | 661.4 | 3.428E-05 | 0.05183 | +1 | 7 |
| - | - | 1690 | 662.4 | - | - | 0 | - |
| - | - | 3258 | 663.4 | - | - | 0 | - |
| - | - | 711.7 | 664.4 | - | - | 0 | - |
| - | - | 2762 | 667.4 | - | - | 0 | - |
| - | - | 755.7 | 668.4 | - | - | 0 | - |
| 3 | y | 7.691E+04 | 679.4 | 0.0001507 | 0.2219 | +1 | 7 |
| - | - | 2.636E+04 | 680.4 | - | - | 0 | - |
| - | - | 5606 | 681.4 | - | - | 0 | - |
| - | - | 749.2 | 699.4 | - | - | 0 | - |
| - | - | 3202 | 701.4 | - | - | 0 | - |
| - | - | 1095 | 702.4 | - | - | 0 | - |
| - | - | 579.4 | 718.9 | - | - | 0 | - |
| 8 | b | 1004 | 746.4 | 0.000371 | 0.4971 | +1 | 8 |
| - | - | 1968 | 764.4 | - | - | 0 | - |
| - | - | 585.1 | 768.8 | - | - | 0 | - |
| 2 | y | 8105 | 778.4 | 0.0002183 | 0.2804 | +1 | 8 |
| - | - | 2595 | 779.4 | - | - | 0 | - |
| - | - | 4354 | 780.5 | - | - | 0 | - |
| - | - | 790 | 782.4 | - | - | 0 | - |
| - | - | 2.095E+04 | 798.5 | - | - | 0 | - |
| - | - | 6418 | 799.5 | - | - | 0 | - |
| 0 | Precursor | 752.2 | 877.5 | 0.0002025 | 0.2307 | +1 | -1 |
| - | - | 9164 | 895.5 | - | - | 0 | - |
| - | - | 3406 | 896.5 | - | - | 0 | - |
| - | - | 720 | 1576 | - | - | 0 | - |
| - | - | 731 | 2031 | - | - | 0 | - |
| - | - | 733.8 | 3038 | - | - | 0 | - |
| - | - | 1105 | 3082 | - | - | 0 | - |
| - | - | 1206 | 3082 | - | - | 0 | - |

m/z Charge Intensity FragmentType MassShift Position
120.04487609863281 0 402.53183
120.0810775756836 0 59782.848
121.0794677734375 0 481.71768
121.08443450927734 0 3316.0254
122.30280303955078 0 330.228
124.11260986328125 0 424.7957
126.05528259277344 0 1479.0591
127.0509033203125 0 755.97626
129.0662384033203 0 626.72626
129.1025390625 0 63595.26
130.06532287597656 0 903.1834
130.08645629882812 0 1084.5634
130.10018920898438 0 653.7116
130.10597229003906 0 3007.3733
131.08177185058594 0 1440.2874
132.08116149902344 0 629.70245
132.10220336914062 0 4809.1187 y 8
133.0611114501953 0 740.29443
133.29440307617188 0 372.21674
134.33416748046875 0 420.88876
134.99356079101562 0 469.6596
135.8223419189453 0 361.17902
136.07601928710938 0 83451.945
137.07327270507812 0 700.8821
137.079345703125 0 4563.5415
139.08712768554688 0 716.4629
139.80123901367188 0 375.57227
141.0659637451172 0 785.713
141.10275268554688 0 476.90356
141.15438842773438 0 404.24463
144.06527709960938 0 695.3127
146.06028747558594 0 2980.8333
147.04429626464844 0 5022.7524
147.1131591796875 0 2479.123
148.03929138183594 0 597.6709
148.95411682128906 0 1041.1095
155.081787109375 0 739.7365
155.11819458007812 0 916.8301
157.08544921875 0 421.87604
158.06027221679688 0 647.39496
159.07664489746094 0 1374.3569
159.0918731689453 0 1379.1492
163.04029846191406 0 466.67627
163.15695190429688 0 428.16272
163.2217559814453 0 458.38123
163.2596435546875 0 403.21713
165.05484008789062 0 744.57135
165.10293579101562 0 689.0146
166.086669921875 0 857.9803
167.11805725097656 0 1403.5895
171.14952087402344 0 82578.08 a 1
172.07188415527344 0 1598.8376
172.10806274414062 0 772.234
172.1452178955078 0 727.8766
172.15286254882812 0 7318.95
173.1287078857422 0 3920.18
174.05523681640625 0 1984.3374
175.08712768554688 0 658.68365
176.10739135742188 0 2538.3452
176.55628967285156 0 460.09705
177.10244750976562 0 3035.862
177.11012268066406 0 705.07117
182.08090209960938 0 1250.0538
183.14942932128906 0 1690.9342
185.1280975341797 0 886.7024
186.1237335205078 0 773.9095
187.14454650878906 0 1067.4263
188.07119750976562 0 977.44543
189.08741760253906 0 964.4538
191.08132934570312 0 552.8731
192.06561279296875 0 1352.7922
195.11314392089844 0 1380.5178
197.12889099121094 0 608.42004
197.1646728515625 0 508.16196
198.0877227783203 0 1174.6318
199.1081085205078 0 1134.2238
199.14437866210938 0 28772.445 b 1
200.14772033691406 0 2775.0618
201.12342834472656 0 1291.4738 y Ammonia loss 5
202.05001831054688 0 630.7598
203.08236694335938 0 574.1893
203.11846923828125 0 753.8226
204.06568908691406 0 548.34546
204.1389923095703 0 804.40247
205.0974578857422 0 5083.3467
205.91099548339844 0 467.19708
208.10775756835938 0 757.21765
209.09217834472656 0 1186.4984
211.14456176757812 0 1316.2125
212.13973999023438 0 5564.344
215.1392364501953 0 759.2918
216.09840393066406 0 1394.6812
217.08265686035156 0 1349.0635
217.09747314453125 0 952.63226
217.13377380371094 0 10580.699
218.1372833251953 0 743.48206
219.1494598388672 0 2577.618
221.0925750732422 0 1208.5675
221.1290283203125 0 1048.7295
221.91973876953125 0 536.4214
226.11892700195312 0 2283.828
226.142578125 0 1190.3258
226.15528869628906 0 20503.902
227.10272216796875 0 1189.67
227.11407470703125 0 1219.5576
227.15914916992188 0 1337.5205
228.3062286376953 0 487.58896
229.09747314453125 0 1508.5151
229.15518188476562 0 1774.5177
230.150146484375 0 6993.87
231.1126708984375 0 731.2072
231.15261840820312 0 824.38995
233.12857055664062 0 718.11194
233.15023803710938 0 1568.0309
233.16506958007812 0 9497.876
234.16848754882812 0 972.4804
239.15023803710938 0 949.5024
240.13462829589844 0 957.356
243.1143798828125 0 583.8573
243.17027282714844 0 1924.6606 y Ammonia loss 7
244.12965393066406 0 1836.2928
245.12818908691406 0 5730.5005
247.1440887451172 0 1018.8851
249.1600341796875 0 28471.076
250.1634521484375 0 2507.4304
254.11404418945312 0 1178.4292
254.14967346191406 0 752.5721
255.1089630126953 0 3578.5
256.1776123046875 0 770.78357
259.1447448730469 0 1105.6278 b 5
260.19708251953125 0 11798.968 y 7
261.12384033203125 0 1675.8708
261.1600036621094 0 2669.555
261.200439453125 0 1672.4021
262.118896484375 0 2582.9338
269.1609191894531 0 3240.506
272.1240539550781 0 1117.163
273.1195983886719 0 4223.353
274.12103271484375 0 952.8995
277.1548767089844 0 6779.08
278.1590576171875 0 703.6871
287.1673889160156 0 1431.1337
291.1448669433594 0 852.8262
301.19158935546875 0 1151.9935
302.1735534667969 0 623.796
302.1943054199219 0 655.6799
309.20440673828125 0 652.99457
319.1407775878906 0 2099.4639
323.1719055175781 0 2121.375
323.2081604003906 0 7126.421
326.18280029296875 0 1924.3271
330.16650390625 0 745.03265
340.19293212890625 0 6500.345 y 2
340.6942443847656 0 2426.7168
341.18267822265625 0 6489.695
341.2020568847656 0 590.0488
346.2122802734375 0 1189.7386 b 2
348.16864013671875 0 850.6006
351.2101135253906 0 705.82196
355.16156005859375 0 1057.827
357.1563720703125 0 1927.4213
361.2445373535156 0 2102.6128 y 6
363.9011535644531 0 605.9054
365.1919860839844 0 876.4359
370.53509521484375 0 614.25116
373.1723937988281 0 2258.9778
373.7142639160156 0 1573.7273 b 7
374.18182373046875 0 1451.0994
374.21087646484375 0 587.94
375.1658630371094 0 980.8778
379.20245361328125 0 890.63794
382.7193298339844 0 3990.0461
383.20330810546875 0 2133.4941
383.2275695800781 0 785.4143
384.20654296875 0 737.822
391.7239990234375 0 1766.0178
392.1932067871094 0 2653.93
399.74029541015625 0 4760.093
400.24188232421875 0 850.8255
401.2154235839844 0 1800.4888
402.1776428222656 0 5663.846
402.9627380371094 0 741.55963
403.1811828613281 0 1023.91644
414.23492431640625 0 901.6592
418.23248291015625 0 937.7803
418.2667236328125 0 2307.326 y 5
419.22576904296875 0 1412.0177
420.18768310546875 0 2505.5842
421.1889343261719 0 1050.6383
424.2215576171875 0 1875.6776
425.2632141113281 0 1877.3646
437.849609375 0 822.9334
439.1419372558594 0 892.7655
439.1723327636719 0 708.85724
439.22943115234375 0 1400.335
439.26165771484375 0 3880.313 Precursor
439.761962890625 0 1304.5016
439.8441162109375 0 7108.6064
440.5880126953125 0 621.51776
442.2303771972656 0 3280.614
448.2665100097656 0 41091.418
448.76788330078125 0 13021.411
451.2674255371094 0 737.92114
458.2601623535156 0 656.0773 y Ammonia loss 4
460.24127197265625 0 1741.1028
469.2762145996094 0 873.9119
470.2754211425781 0 1579.0804
475.2880554199219 0 7674.9155 y 4
476.26983642578125 0 1473.8689
476.2973327636719 0 892.58734
496.2862548828125 0 730.66046
512.7958984375 0 1499.3064
514.2984008789062 0 1809.6478 y Water loss 3
521.8007202148438 0 10955.796
522.3013305664062 0 4425.2065
525.8118286132812 0 1537.949
527.3226318359375 0 844.8047
529.7981567382812 0 1617.5197
530.272216796875 0 3291.2017
531.2763671875 0 943.8981
532.3090209960938 0 23627.549 y 3
533.3123779296875 0 5663.8843
534.3153686523438 0 1185.0426
539.8109130859375 0 1585.0072
540.309814453125 0 1431.931
548.2824096679688 0 5423.2227
548.3223876953125 0 1334.2397
548.3602905273438 0 866.50366
548.8241577148438 0 904.9097
549.2852172851562 0 1416.7936
555.3130493164062 0 3410.5251
566.2921142578125 0 2575.9883
566.334716796875 0 1098.8528
570.3264770507812 0 821.6081
573.3243408203125 0 22791.562
574.327392578125 0 4641.373
578.3427124023438 0 1484.1727
578.8438110351562 0 1156.435
580.2318115234375 0 641.67596
600.314453125 0 625.4614 b Water loss 6
615.35986328125 0 661.5
617.3462524414062 0 665.23553
626.7388916015625 0 574.0399
643.3556518554688 0 1385.51
644.3500366210938 0 981.46313
649.36669921875 0 725.7625
661.3668212890625 0 4584.339 y Water loss 2
662.3692016601562 0 1690.3055
663.38232421875 0 3258.1125
664.3860473632812 0 711.7429
667.3772583007812 0 2762.2761
668.3829956054688 0 755.6891
679.3775024414062 0 76914.414 y 2
680.3804321289062 0 26362.834
681.38330078125 0 5605.805
699.4037475585938 0 749.234
701.418212890625 0 3202.111
702.4227905273438 0 1094.8954
718.9349365234375 0 579.43567
746.419921875 0 1004.19965 b 7
764.4290771484375 0 1967.9526
768.7815551757812 0 585.06024
778.4459838867188 0 8105.2354 y 1
779.4476318359375 0 2595.3254
780.4591674804688 0 4354.0596
782.4390258789062 0 789.9533
798.4721069335938 0 20951.615
799.474365234375 0 6418.4424
877.5139770507812 0 752.2497 Precursor
895.5242919921875 0 9163.729
896.5277099609375 0 3406.0134
1575.9708251953125 0 720.0395
2030.9129638671875 0 731.0294
3037.6015625 0 733.8493
3081.80615234375 0 1104.6674
3082.469482421875 0 1206.2214

Spectrum Details

|  |  |
| --- | --- |
| Matched peaks? Matched peaks The total absolute number of peaks matched. Additionally in brackets the total fraction of peaks matched and the total number of peaks is shown. | 23 (8.33% of 276) |
| FDR? FDR The false discovery rate estimated for this peptide. It is calculated by matching all theoretical fragments with a non-integer shift with the raw peaks for this spectrum. This is done with 40 different shifts. The resulting percentage is the average number of annotated peaks over the number of annotated peaks with the correct spectrum. | 0.41% |
| Satellite FDR? Satellite FDR See the FDR for details on its calculation. This satellite ion specific FDR only contains the satellite ions (d/w) for I/L/J positions. | - |
| PSM Score? PSM Score The PSM Score as given by Hecklib to this annotated spectrum. It is shown with three significant figures. | 261 |

#### Spectrum 7725? Spectrum 7725 The raw spectrum of this peptide as annotated by Hecklib. The fragments are coloured according to ion type (see legend). Any peaks with a star '\*' as text can be hovered over to see the full details, first the ion type second the mass shift type. By hovering over the amino acids in the peptide or ions in the legend the corresponding peaks are highlighted. By toggling the 'Unassigned' label you can turn the background (unassigned) peaks on or off in the plot. By updating the slider in the Ion legend you can update the spectrum to only show the top X% of the peaks with labels. The top X% means any peak that is within X% of the highest intensity. By dragging in the spectrum you can zoom in to a specific part of the spectrum and use 'Zoom Out' to get back to the original zoom level. The annotation of the spectrum is based on the given sequence in the peptides file and is done with different software so inconsistencies are likely. The peaks are annotated based on the given sequence, with 20 ppm tolerance.

Copy Data

##### Spectrum 7725 (TSV)

###### Preview

```
Loading example...
```

*Click on the button to copy the data to your clipboard.*

Mz MinMz MaxIntensity Max

WidthHeightPeptide font sizePeptide stroke widthSpectrum font sizeSpectrum stroke widthCompact peptide

Ion legend

wxyz

abcd

OtherUnassignedIon^Charge^_Position_Show for top:%

VVFGGGTKJ

01.09e+42.18e+43.27e+44.36e+4

Zoom Out

y^+1^_1_a^+1^_2_b^+1^_2_y^+1^_2_y^+1^_2_y^+2^_7_y^+1^_3_b^+2^_8_y^+1^_4_\*y^+1^_5_y^+1^_6_y^+1^_7_y^+1^_7_y^+1^_8_

0778155623343111

Fragment Matches Table

Show background peaks

| Position | Ion type | Intensity | mz Theoretical | mz Error (Th) | mz Error (ppm) | Charge | Series Number |
| --- | --- | --- | --- | --- | --- | --- | --- |
| - | - | 2.553E+04 | 120.1 | - | - | 0 | - |
| - | - | 2043 | 121.1 | - | - | 0 | - |
| - | - | 459.4 | 123.1 | - | - | 0 | - |
| - | - | 500.4 | 127.1 | - | - | 0 | - |
| - | - | 487 | 129.1 | - | - | 0 | - |
| - | - | 1127 | 129.1 | - | - | 0 | - |
| - | - | 2.303E+04 | 129.1 | - | - | 0 | - |
| - | - | 1863 | 130.1 | - | - | 0 | - |
| - | - | 472.4 | 130.1 | - | - | 0 | - |
| - | - | 1140 | 130.1 | - | - | 0 | - |
| - | - | 826.7 | 131.1 | - | - | 0 | - |
| 9 | y | 2019 | 132.1 | 9.989E-05 | 0.7561 | +1 | 1 |
| - | - | 548.9 | 133.1 | - | - | 0 | - |
| - | - | 518.8 | 135.1 | - | - | 0 | - |
| - | - | 7801 | 136.1 | - | - | 0 | - |
| - | - | 439.7 | 139.1 | - | - | 0 | - |
| - | - | 536.8 | 141.1 | - | - | 0 | - |
| - | - | 562.2 | 141.1 | - | - | 0 | - |
| - | - | 2797 | 146.1 | - | - | 0 | - |
| - | - | 695.2 | 147.1 | - | - | 0 | - |
| - | - | 781.4 | 148 | - | - | 0 | - |
| - | - | 525.8 | 148.1 | - | - | 0 | - |
| - | - | 934.4 | 149 | - | - | 0 | - |
| - | - | 491.9 | 151.1 | - | - | 0 | - |
| - | - | 459.9 | 153.1 | - | - | 0 | - |
| - | - | 441.3 | 155.1 | - | - | 0 | - |
| - | - | 568.7 | 155.1 | - | - | 0 | - |
| - | - | 625.8 | 155.1 | - | - | 0 | - |
| - | - | 698.4 | 155.1 | - | - | 0 | - |
| - | - | 566.5 | 156.1 | - | - | 0 | - |
| - | - | 808.1 | 158.1 | - | - | 0 | - |
| - | - | 445.5 | 158.1 | - | - | 0 | - |
| - | - | 1574 | 159.1 | - | - | 0 | - |
| - | - | 470 | 160.1 | - | - | 0 | - |
| - | - | 676.5 | 165.1 | - | - | 0 | - |
| - | - | 3067 | 166.1 | - | - | 0 | - |
| - | - | 1048 | 170.1 | - | - | 0 | - |
| - | - | 477.3 | 170.7 | - | - | 0 | - |
| 2 | a | 4.317E+04 | 171.1 | 0.0001786 | 1.044 | +1 | 2 |
| - | - | 3594 | 172.2 | - | - | 0 | - |
| - | - | 473.9 | 173.1 | - | - | 0 | - |
| - | - | 1318 | 173.1 | - | - | 0 | - |
| - | - | 1194 | 173.4 | - | - | 0 | - |
| - | - | 2009 | 174.1 | - | - | 0 | - |
| - | - | 1467 | 175.1 | - | - | 0 | - |
| - | - | 1233 | 176.1 | - | - | 0 | - |
| - | - | 1227 | 177.1 | - | - | 0 | - |
| - | - | 742.5 | 182.1 | - | - | 0 | - |
| - | - | 1301 | 183.1 | - | - | 0 | - |
| - | - | 535.3 | 185.1 | - | - | 0 | - |
| - | - | 1384 | 188.1 | - | - | 0 | - |
| - | - | 1007 | 192.1 | - | - | 0 | - |
| - | - | 525 | 195.1 | - | - | 0 | - |
| - | - | 640.9 | 198.1 | - | - | 0 | - |
| 2 | b | 1.605E+04 | 199.1 | 0.000137 | 0.6882 | +1 | 2 |
| - | - | 1335 | 200.1 | - | - | 0 | - |
| - | - | 3335 | 205.1 | - | - | 0 | - |
| - | - | 690.2 | 207.1 | - | - | 0 | - |
| - | - | 1662 | 212.1 | - | - | 0 | - |
| - | - | 547.5 | 212.9 | - | - | 0 | - |
| - | - | 638 | 216.1 | - | - | 0 | - |
| - | - | 1330 | 217.1 | - | - | 0 | - |
| - | - | 1453 | 219.1 | - | - | 0 | - |
| - | - | 659.7 | 221.1 | - | - | 0 | - |
| - | - | 1528 | 226.2 | - | - | 0 | - |
| - | - | 913.9 | 227.1 | - | - | 0 | - |
| - | - | 483.7 | 228.4 | - | - | 0 | - |
| - | - | 4317 | 230.2 | - | - | 0 | - |
| - | - | 1301 | 233.2 | - | - | 0 | - |
| - | - | 552.9 | 234.1 | - | - | 0 | - |
| - | - | 709.7 | 237.1 | - | - | 0 | - |
| - | - | 797.7 | 239.2 | - | - | 0 | - |
| 8 | y | 1395 | 243.2 | 0.0002132 | 0.8767 | +1 | 2 |
| - | - | 677.4 | 244.1 | - | - | 0 | - |
| - | - | 1013 | 245.1 | - | - | 0 | - |
| - | - | 1371 | 249.2 | - | - | 0 | - |
| - | - | 2006 | 255.1 | - | - | 0 | - |
| 8 | y | 6311 | 260.2 | 6.028E-05 | 0.2317 | +1 | 2 |
| - | - | 500.8 | 260.4 | - | - | 0 | - |
| - | - | 741.1 | 261.2 | - | - | 0 | - |
| - | - | 1036 | 262.1 | - | - | 0 | - |
| - | - | 557.6 | 262.2 | - | - | 0 | - |
| - | - | 598.4 | 263.5 | - | - | 0 | - |
| - | - | 499.7 | 266.3 | - | - | 0 | - |
| - | - | 1915 | 269.2 | - | - | 0 | - |
| - | - | 1819 | 273.1 | - | - | 0 | - |
| - | - | 913.6 | 276.2 | - | - | 0 | - |
| - | - | 732.3 | 277.2 | - | - | 0 | - |
| - | - | 618.9 | 287.2 | - | - | 0 | - |
| - | - | 1366 | 309.2 | - | - | 0 | - |
| - | - | 615 | 312.7 | - | - | 0 | - |
| - | - | 1011 | 326.2 | - | - | 0 | - |
| 3 | y | 3635 | 340.2 | 0.0001908 | 0.5609 | +2 | 7 |
| - | - | 1470 | 340.7 | - | - | 0 | - |
| - | - | 713.3 | 348.2 | - | - | 0 | - |
| 7 | y | 1111 | 361.2 | 0.00115 | 3.185 | +1 | 3 |
| 8 | b | 770.3 | 365.2 | 0.006841 | 18.73 | +2 | 8 |
| - | - | 850.3 | 374.2 | - | - | 0 | - |
| - | - | 1173 | 383.2 | - | - | 0 | - |
| - | - | 773.8 | 383.2 | - | - | 0 | - |
| - | - | 1611 | 392.2 | - | - | 0 | - |
| - | - | 716.6 | 401.2 | - | - | 0 | - |
| - | - | 3042 | 402.2 | - | - | 0 | - |
| - | - | 662.6 | 418.2 | - | - | 0 | - |
| 6 | y | 1090 | 418.3 | 0.001087 | 2.599 | +1 | 4 |
| - | - | 575.7 | 420 | - | - | 0 | - |
| - | - | 1558 | 420.2 | - | - | 0 | - |
| - | - | 1151 | 421.8 | - | - | 0 | - |
| - | - | 1152 | 437.8 | - | - | 0 | - |
| - | - | 1098 | 439.2 | - | - | 0 | - |
| 0 | Precursor | 873.7 | 439.3 | 0.004165 | 9.481 | +2 | -1 |
| - | - | 7497 | 439.8 | - | - | 0 | - |
| - | - | 722 | 440.3 | - | - | 0 | - |
| - | - | 2542 | 448.3 | - | - | 0 | - |
| - | - | 777.7 | 448.8 | - | - | 0 | - |
| - | - | 697.7 | 466.8 | - | - | 0 | - |
| 5 | y | 4512 | 475.3 | 6.256E-05 | 0.1316 | +1 | 5 |
| - | - | 648.3 | 476.3 | - | - | 0 | - |
| - | - | 699.3 | 496.3 | - | - | 0 | - |
| - | - | 675.8 | 499.3 | - | - | 0 | - |
| - | - | 654.2 | 521.8 | - | - | 0 | - |
| - | - | 1616 | 525.8 | - | - | 0 | - |
| - | - | 1826 | 530.3 | - | - | 0 | - |
| 4 | y | 1.183E+04 | 532.3 | 0.0004661 | 0.8756 | +1 | 6 |
| - | - | 2237 | 533.3 | - | - | 0 | - |
| - | - | 830.3 | 539.8 | - | - | 0 | - |
| - | - | 1021 | 540.3 | - | - | 0 | - |
| - | - | 2585 | 548.3 | - | - | 0 | - |
| - | - | 839.1 | 548.3 | - | - | 0 | - |
| - | - | 1122 | 548.8 | - | - | 0 | - |
| - | - | 1478 | 566.3 | - | - | 0 | - |
| - | - | 1667 | 573.3 | - | - | 0 | - |
| - | - | 887.7 | 643.4 | - | - | 0 | - |
| 3 | y | 1756 | 661.4 | 0.002041 | 3.086 | +1 | 7 |
| 3 | y | 3.772E+04 | 679.4 | 0.0008869 | 1.305 | +1 | 7 |
| - | - | 1.247E+04 | 680.4 | - | - | 0 | - |
| - | - | 2942 | 681.4 | - | - | 0 | - |
| 2 | y | 3387 | 778.4 | 0.0009414 | 1.209 | +1 | 8 |
| - | - | 1468 | 779.4 | - | - | 0 | - |
| - | - | 1832 | 798.5 | - | - | 0 | - |
| - | - | 1173 | 3081 | - | - | 0 | - |

m/z Charge Intensity FragmentType MassShift Position
120.08100128173828 0 25531.932
121.08431243896484 0 2042.577
123.08068084716797 0 459.41663
127.07559204101562 0 500.36472
129.06597900390625 0 487.00845
129.0970001220703 0 1126.9185
129.1024627685547 0 23030.523
130.0654296875 0 1863.1814
130.08644104003906 0 472.40472
130.10574340820312 0 1139.9277
131.08160400390625 0 826.6583
132.1020050048828 0 2019.4456 y 8
133.0607452392578 0 548.93585
135.08091735839844 0 518.7536
136.07589721679688 0 7800.795
139.07546997070312 0 439.6732
141.06591796875 0 536.83417
141.1025390625 0 562.1905
146.06024169921875 0 2797.3364
147.11260986328125 0 695.24695
148.03944396972656 0 781.43164
148.0963897705078 0 525.81146
148.95445251464844 0 934.3696
151.07467651367188 0 491.93802
153.0661163330078 0 459.86896
155.07064819335938 0 441.2542
155.0811767578125 0 568.71497
155.1072540283203 0 625.76044
155.11805725097656 0 698.449
156.07667541503906 0 566.4605
158.06033325195312 0 808.0519
158.1219482421875 0 445.53128
159.0919647216797 0 1574.3455
160.0752716064453 0 469.9768
165.05490112304688 0 676.5152
166.0865936279297 0 3066.6465
170.060302734375 0 1048.2896
170.71543884277344 0 477.28296
171.1493682861328 0 43171.67 a 1
172.1527862548828 0 3593.8933
173.07139587402344 0 473.89822
173.1289520263672 0 1318.4701
173.43882751464844 0 1194.3898
174.0551300048828 0 2009.0249
175.08688354492188 0 1466.9026
176.1071014404297 0 1233.1785
177.1024169921875 0 1227.4302
182.08103942871094 0 742.46716
183.14915466308594 0 1300.5042
185.05506896972656 0 535.2819
188.07066345214844 0 1384.2844
192.06582641601562 0 1007.21356
195.1136474609375 0 524.98755
198.08773803710938 0 640.927
199.1442413330078 0 16053.133 b 1
200.1477813720703 0 1335.2874
205.09715270996094 0 3334.687
207.11289978027344 0 690.22546
212.13905334472656 0 1661.5769
212.9468994140625 0 547.5481
216.09832763671875 0 638.0042
217.1339874267578 0 1329.8784
219.1494140625 0 1452.9412
221.09292602539062 0 659.7274
226.1553955078125 0 1527.764
227.11412048339844 0 913.8557
228.39183044433594 0 483.66785
230.15003967285156 0 4317.291
233.16505432128906 0 1300.5869
234.12332153320312 0 552.9327
237.08734130859375 0 709.7221
239.15081787109375 0 797.65515
243.1705322265625 0 1395.1951 y Ammonia loss 7
244.12939453125 0 677.3605
245.12606811523438 0 1013.163
249.15945434570312 0 1371.2803
255.10899353027344 0 2005.6085
260.1968078613281 0 6310.947 y 7
260.38848876953125 0 500.75674
261.1998596191406 0 741.0911
262.1190185546875 0 1036.0233
262.20220947265625 0 557.6301
263.51177978515625 0 598.3707
266.3135986328125 0 499.6891
269.1607360839844 0 1914.5327
273.1191101074219 0 1818.9271
276.16656494140625 0 913.6398
277.1554870605469 0 732.255
287.1697998046875 0 618.91864
309.2040710449219 0 1365.551
312.7352294921875 0 615.03925
326.1820068359375 0 1011.35876
340.1925048828125 0 3635.4304 y 2
340.69427490234375 0 1470.156
348.1660461425781 0 713.3026
361.2456970214844 0 1110.5668 y 6
365.19329833984375 0 770.27386 b Ammonia loss 7
374.1810302734375 0 850.2537
383.202880859375 0 1172.7278
383.2280578613281 0 773.8255
392.1929931640625 0 1610.8668
401.2149658203125 0 716.58185
402.1770324707031 0 3042.1812
418.2304382324219 0 662.5771
418.2649230957031 0 1089.7664 y 5
419.9967346191406 0 575.6942
420.1882629394531 0 1557.8455
421.8323974609375 0 1151.4431
437.84893798828125 0 1152.0189
439.2340087890625 0 1098.1372
439.264892578125 0 873.7153 Precursor
439.8439636230469 0 7497.425
440.31146240234375 0 721.99805
448.2648620605469 0 2542.2603
448.7666015625 0 777.69073
466.7748107910156 0 697.70996
475.28753662109375 0 4512.073 y 4
476.2910461425781 0 648.2968
496.2921447753906 0 699.3104
499.2980651855469 0 675.7641
521.7986450195312 0 654.20386
525.8121948242188 0 1616.3759
530.27197265625 0 1826.1134
532.3084716796875 0 11828.287 y 3
533.3112182617188 0 2237.4478
539.8082885742188 0 830.30786
540.3112182617188 0 1020.8874
548.2815551757812 0 2584.5527
548.32080078125 0 839.11694
548.8244018554688 0 1121.8262
566.2921142578125 0 1477.9956
573.3236694335938 0 1666.5817
643.3545532226562 0 887.66064
661.36474609375 0 1755.9753 y Water loss 2
679.37646484375 0 37716.633 y 2
680.3792724609375 0 12473.921
681.3818969726562 0 2941.5386
778.44482421875 0 3386.548 y 1
779.4461669921875 0 1468.3477
798.47119140625 0 1831.9513
3080.595703125 0 1172.5946

Spectrum Details

|  |  |
| --- | --- |
| Matched peaks? Matched peaks The total absolute number of peaks matched. Additionally in brackets the total fraction of peaks matched and the total number of peaks is shown. | 15 (10.64% of 141) |
| FDR? FDR The false discovery rate estimated for this peptide. It is calculated by matching all theoretical fragments with a non-integer shift with the raw peaks for this spectrum. This is done with 40 different shifts. The resulting percentage is the average number of annotated peaks over the number of annotated peaks with the correct spectrum. | 0.48% |
| Satellite FDR? Satellite FDR See the FDR for details on its calculation. This satellite ion specific FDR only contains the satellite ions (d/w) for I/L/J positions. | - |
| PSM Score? PSM Score The PSM Score as given by Hecklib to this annotated spectrum. It is shown with three significant figures. | 150 |

#### Spectrum 8222? Spectrum 8222 The raw spectrum of this peptide as annotated by Hecklib. The fragments are coloured according to ion type (see legend). Any peaks with a star '\*' as text can be hovered over to see the full details, first the ion type second the mass shift type. By hovering over the amino acids in the peptide or ions in the legend the corresponding peaks are highlighted. By toggling the 'Unassigned' label you can turn the background (unassigned) peaks on or off in the plot. By updating the slider in the Ion legend you can update the spectrum to only show the top X% of the peaks with labels. The top X% means any peak that is within X% of the highest intensity. By dragging in the spectrum you can zoom in to a specific part of the spectrum and use 'Zoom Out' to get back to the original zoom level. The annotation of the spectrum is based on the given sequence in the peptides file and is done with different software so inconsistencies are likely. The peaks are annotated based on the given sequence, with 20 ppm tolerance.

Copy Data

##### Spectrum 8222 (TSV)

###### Preview

```
Loading example...
```

*Click on the button to copy the data to your clipboard.*

Mz MinMz MaxIntensity Max

WidthHeightPeptide font sizePeptide stroke widthSpectrum font sizeSpectrum stroke widthCompact peptide

Ion legend

wxyz

abcd

OtherUnassignedIon^Charge^_Position_Show for top:%

VVFGGGTKJ

08.85e+31.77e+42.65e+43.54e+4

Zoom Out

y^+1^_1_a^+1^_2_b^+1^_2_y^+2^_4_y^+1^_2_y^+1^_2_y^+2^_7_y^+1^_3_y^+1^_4_\*y^+1^_5_y^+1^_6_y^+1^_7_y^+1^_7_y^+1^_8_

0777155523323109

Fragment Matches Table

Show background peaks

| Position | Ion type | Intensity | mz Theoretical | mz Error (Th) | mz Error (ppm) | Charge | Series Number |
| --- | --- | --- | --- | --- | --- | --- | --- |
| - | - | 2.067E+04 | 120.1 | - | - | 0 | - |
| - | - | 479.8 | 121.1 | - | - | 0 | - |
| - | - | 1733 | 121.1 | - | - | 0 | - |
| - | - | 515.1 | 123 | - | - | 0 | - |
| - | - | 535.9 | 127.1 | - | - | 0 | - |
| - | - | 2.16E+04 | 129.1 | - | - | 0 | - |
| - | - | 564.4 | 130.1 | - | - | 0 | - |
| - | - | 733.5 | 130.1 | - | - | 0 | - |
| - | - | 1092 | 130.1 | - | - | 0 | - |
| - | - | 487.2 | 131.1 | - | - | 0 | - |
| - | - | 364.1 | 131.3 | - | - | 0 | - |
| 9 | y | 1510 | 132.1 | 0.0001914 | 1.449 | +1 | 1 |
| - | - | 922.4 | 133.1 | - | - | 0 | - |
| - | - | 633.4 | 133.1 | - | - | 0 | - |
| - | - | 5550 | 136.1 | - | - | 0 | - |
| - | - | 467.7 | 139.1 | - | - | 0 | - |
| - | - | 2243 | 146.1 | - | - | 0 | - |
| - | - | 1299 | 147.1 | - | - | 0 | - |
| - | - | 508.5 | 151.1 | - | - | 0 | - |
| - | - | 533 | 155.1 | - | - | 0 | - |
| - | - | 632.2 | 155.1 | - | - | 0 | - |
| - | - | 918.8 | 159.1 | - | - | 0 | - |
| - | - | 1032 | 159.1 | - | - | 0 | - |
| - | - | 544.3 | 161.1 | - | - | 0 | - |
| - | - | 717.3 | 165.1 | - | - | 0 | - |
| - | - | 511.7 | 165.1 | - | - | 0 | - |
| - | - | 680.8 | 166.1 | - | - | 0 | - |
| - | - | 597.2 | 167.1 | - | - | 0 | - |
| 2 | a | 3.505E+04 | 171.1 | 0.0002397 | 1.4 | +1 | 2 |
| - | - | 706.7 | 172.1 | - | - | 0 | - |
| - | - | 3906 | 172.2 | - | - | 0 | - |
| - | - | 1833 | 173.1 | - | - | 0 | - |
| - | - | 1787 | 173.4 | - | - | 0 | - |
| - | - | 1577 | 174.1 | - | - | 0 | - |
| - | - | 974.6 | 176.1 | - | - | 0 | - |
| - | - | 1762 | 177.1 | - | - | 0 | - |
| - | - | 592 | 182.1 | - | - | 0 | - |
| - | - | 526.5 | 182.5 | - | - | 0 | - |
| - | - | 603.6 | 183.1 | - | - | 0 | - |
| - | - | 931 | 188.1 | - | - | 0 | - |
| - | - | 536.8 | 191.1 | - | - | 0 | - |
| - | - | 511.9 | 198.1 | - | - | 0 | - |
| 2 | b | 1.263E+04 | 199.1 | 0.0001523 | 0.7648 | +1 | 2 |
| - | - | 1393 | 200.1 | - | - | 0 | - |
| 6 | y | 915.7 | 201.1 | 0.0003952 | 1.965 | +2 | 4 |
| - | - | 509.1 | 202 | - | - | 0 | - |
| - | - | 2614 | 205.1 | - | - | 0 | - |
| - | - | 660.5 | 209.1 | - | - | 0 | - |
| - | - | 480.4 | 209.1 | - | - | 0 | - |
| - | - | 2028 | 212.1 | - | - | 0 | - |
| - | - | 577.6 | 213.1 | - | - | 0 | - |
| - | - | 838.8 | 216.1 | - | - | 0 | - |
| - | - | 631 | 217.1 | - | - | 0 | - |
| - | - | 1769 | 219.1 | - | - | 0 | - |
| - | - | 840.2 | 221.1 | - | - | 0 | - |
| - | - | 867.9 | 221.1 | - | - | 0 | - |
| - | - | 1351 | 226.2 | - | - | 0 | - |
| - | - | 797.1 | 227.1 | - | - | 0 | - |
| - | - | 628.5 | 227.2 | - | - | 0 | - |
| - | - | 2491 | 230.2 | - | - | 0 | - |
| - | - | 1243 | 233.2 | - | - | 0 | - |
| - | - | 852 | 239.1 | - | - | 0 | - |
| - | - | 1026 | 239.2 | - | - | 0 | - |
| 8 | y | 1093 | 243.2 | 1.483E-05 | 0.06097 | +1 | 2 |
| - | - | 774.4 | 249.1 | - | - | 0 | - |
| - | - | 1227 | 249.2 | - | - | 0 | - |
| - | - | 540.6 | 251.1 | - | - | 0 | - |
| - | - | 1754 | 255.1 | - | - | 0 | - |
| - | - | 570.3 | 256.9 | - | - | 0 | - |
| 8 | y | 5981 | 260.2 | 3.128E-05 | 0.1202 | +1 | 2 |
| - | - | 1005 | 262.1 | - | - | 0 | - |
| - | - | 4852 | 267.1 | - | - | 0 | - |
| - | - | 711.6 | 268.1 | - | - | 0 | - |
| - | - | 2059 | 269.2 | - | - | 0 | - |
| - | - | 1885 | 273.1 | - | - | 0 | - |
| - | - | 771 | 274.1 | - | - | 0 | - |
| - | - | 682.1 | 276.2 | - | - | 0 | - |
| - | - | 1154 | 293.1 | - | - | 0 | - |
| - | - | 695.4 | 301.2 | - | - | 0 | - |
| - | - | 1420 | 309.2 | - | - | 0 | - |
| - | - | 759 | 321.2 | - | - | 0 | - |
| - | - | 1219 | 326.2 | - | - | 0 | - |
| 3 | y | 3063 | 340.2 | 0.0001754 | 0.5156 | +2 | 7 |
| - | - | 1126 | 340.7 | - | - | 0 | - |
| - | - | 659.8 | 348.2 | - | - | 0 | - |
| - | - | 622.6 | 356.4 | - | - | 0 | - |
| 7 | y | 970.6 | 361.2 | 0.0005281 | 1.462 | +1 | 3 |
| - | - | 572.2 | 362.2 | - | - | 0 | - |
| - | - | 787.3 | 365.1 | - | - | 0 | - |
| - | - | 623.7 | 374.2 | - | - | 0 | - |
| - | - | 1528 | 383.2 | - | - | 0 | - |
| - | - | 1272 | 392.2 | - | - | 0 | - |
| - | - | 1883 | 402.2 | - | - | 0 | - |
| - | - | 742.2 | 403.2 | - | - | 0 | - |
| - | - | 1213 | 409.1 | - | - | 0 | - |
| 6 | y | 798.4 | 418.3 | 0.001087 | 2.599 | +1 | 4 |
| - | - | 1334 | 420.2 | - | - | 0 | - |
| - | - | 726.7 | 421.8 | - | - | 0 | - |
| - | - | 819.9 | 438.1 | - | - | 0 | - |
| - | - | 1118 | 439.2 | - | - | 0 | - |
| - | - | 937.1 | 439.2 | - | - | 0 | - |
| - | - | 5544 | 439.2 | - | - | 0 | - |
| 0 | Precursor | 921.5 | 439.3 | 0.004744 | 10.8 | +2 | -1 |
| - | - | 1.046E+04 | 439.8 | - | - | 0 | - |
| - | - | 715.8 | 440.2 | - | - | 0 | - |
| - | - | 1418 | 440.2 | - | - | 0 | - |
| - | - | 2732 | 440.3 | - | - | 0 | - |
| - | - | 4324 | 440.8 | - | - | 0 | - |
| - | - | 755.3 | 441.2 | - | - | 0 | - |
| - | - | 907.4 | 441.2 | - | - | 0 | - |
| - | - | 1291 | 441.3 | - | - | 0 | - |
| - | - | 2233 | 448.3 | - | - | 0 | - |
| - | - | 664.2 | 469.3 | - | - | 0 | - |
| 5 | y | 3173 | 475.3 | 0.0003952 | 0.8315 | +1 | 5 |
| - | - | 809.8 | 476.3 | - | - | 0 | - |
| - | - | 609.7 | 489.3 | - | - | 0 | - |
| - | - | 597.9 | 498.5 | - | - | 0 | - |
| - | - | 815.4 | 530.3 | - | - | 0 | - |
| 4 | y | 1.059E+04 | 532.3 | 0.000222 | 0.417 | +1 | 6 |
| - | - | 3439 | 533.3 | - | - | 0 | - |
| - | - | 1676 | 539.8 | - | - | 0 | - |
| - | - | 913 | 540.3 | - | - | 0 | - |
| - | - | 1535 | 548.3 | - | - | 0 | - |
| - | - | 1051 | 548.8 | - | - | 0 | - |
| - | - | 1365 | 566.3 | - | - | 0 | - |
| - | - | 597.1 | 642.1 | - | - | 0 | - |
| - | - | 935.4 | 643.4 | - | - | 0 | - |
| 3 | y | 1913 | 661.4 | 0.0006371 | 0.9633 | +1 | 7 |
| 3 | y | 3.088E+04 | 679.4 | 0.0007038 | 1.036 | +1 | 7 |
| - | - | 1.191E+04 | 680.4 | - | - | 0 | - |
| - | - | 2207 | 681.4 | - | - | 0 | - |
| 2 | y | 2851 | 778.4 | 0.002345 | 3.013 | +1 | 8 |
| - | - | 1434 | 779.4 | - | - | 0 | - |
| - | - | 754 | 896.5 | - | - | 0 | - |
| - | - | 597.6 | 1414 | - | - | 0 | - |
| - | - | 694.3 | 1484 | - | - | 0 | - |
| - | - | 578.4 | 1560 | - | - | 0 | - |
| - | - | 684.6 | 2008 | - | - | 0 | - |
| - | - | 705.6 | 2197 | - | - | 0 | - |
| - | - | 705.1 | 2497 | - | - | 0 | - |
| - | - | 717.1 | 2568 | - | - | 0 | - |
| - | - | 632.5 | 3079 | - | - | 0 | - |

m/z Charge Intensity FragmentType MassShift Position
120.08103942871094 0 20667.91
121.06507110595703 0 479.84848
121.08436584472656 0 1732.8693
123.04415893554688 0 515.1108
127.07552337646484 0 535.8973
129.1024932861328 0 21603.5
130.0655517578125 0 564.3537
130.08636474609375 0 733.5477
130.10586547851562 0 1092.1151
131.0814666748047 0 487.21194
131.26805114746094 0 364.11237
132.1020965576172 0 1509.7487 y 8
133.06076049804688 0 922.3566
133.0859375 0 633.3958
136.07595825195312 0 5550.417
139.0757293701172 0 467.71558
146.06031799316406 0 2242.9985
147.11302185058594 0 1298.6578
151.0757598876953 0 508.45886
155.0820770263672 0 532.96625
155.11805725097656 0 632.23883
159.07666015625 0 918.81415
159.0919647216797 0 1031.5521
161.06016540527344 0 544.2779
165.05516052246094 0 717.2793
165.09141540527344 0 511.72314
166.0865020751953 0 680.7552
167.0820770263672 0 597.22943
171.14942932128906 0 35047.855 a 1
172.07156372070312 0 706.67426
172.1526641845703 0 3905.6975
173.1285858154297 0 1833.3773
173.43919372558594 0 1786.8044
174.05538940429688 0 1577.0366
176.10684204101562 0 974.5991
177.1024932861328 0 1762.139
182.081298828125 0 591.9757
182.49008178710938 0 526.51373
183.14962768554688 0 603.5904
188.07073974609375 0 931.0162
191.117431640625 0 536.79877
198.0880126953125 0 511.92493
199.14425659179688 0 12634.814 b 1
200.1478729248047 0 1393.0271
201.12376403808594 0 915.7131 y Ammonia loss 5
202.04998779296875 0 509.1182
205.09738159179688 0 2613.9417
209.07965087890625 0 660.50494
209.09121704101562 0 480.44888
212.13975524902344 0 2027.5107
213.1116485595703 0 577.61395
216.09849548339844 0 838.8379
217.09799194335938 0 630.9626
219.14947509765625 0 1769.3562
221.08094787597656 0 840.20746
221.09181213378906 0 867.87305
226.15499877929688 0 1350.5737
227.11399841308594 0 797.08203
227.17543029785156 0 628.46533
230.15003967285156 0 2491.086
233.1651611328125 0 1243.2616
239.09068298339844 0 852.001
239.15011596679688 0 1026.162
243.1703338623047 0 1093.3436 y Ammonia loss 7
249.07566833496094 0 774.4362
249.1597900390625 0 1226.5281
251.0890655517578 0 540.6499
255.10885620117188 0 1753.7644
256.8667907714844 0 570.2951
260.1968994140625 0 5981.357 y 7
262.11859130859375 0 1005.1149
267.0861511230469 0 4852.0386
268.09063720703125 0 711.565
269.1601257324219 0 2058.8274
273.1194152832031 0 1885.0743
274.1190490722656 0 770.9728
276.1669921875 0 682.0996
293.1358947753906 0 1153.5929
301.1921081542969 0 695.40594
309.20355224609375 0 1419.6228
321.1705627441406 0 759.0245
326.18133544921875 0 1219.28
340.192138671875 0 3062.6936 y 2
340.6941833496094 0 1126.0807
348.1654052734375 0 659.7587
356.39581298828125 0 622.61035
361.2440185546875 0 970.5504 y 6
362.20635986328125 0 572.1619
365.1234436035156 0 787.3241
374.1825866699219 0 623.69354
383.20391845703125 0 1527.766
392.1936950683594 0 1271.851
402.1766052246094 0 1882.7504
403.1812438964844 0 742.17847
409.1124267578125 0 1213.3176
418.2649230957031 0 798.35974 y 5
420.18707275390625 0 1333.5317
421.8345031738281 0 726.71277
438.14947509765625 0 819.947
439.1510009765625 0 1117.9364
439.1800842285156 0 937.13416
439.212890625 0 5544.1333
439.2654724121094 0 921.50476 Precursor
439.84375 0 10456.128
440.1842041015625 0 715.83167
440.2154541015625 0 1418.4658
440.3115539550781 0 2732.1125
440.8442077636719 0 4324.0483
441.18975830078125 0 755.2582
441.225341796875 0 907.3724
441.29840087890625 0 1291.0052
448.2655029296875 0 2232.985
469.274169921875 0 664.22504
475.2870788574219 0 3172.9976 y 4
476.2908630371094 0 809.7774
489.2685546875 0 609.7283
498.4608154296875 0 597.94324
530.2713623046875 0 815.36957
532.3087158203125 0 10586.264 y 3
533.3119506835938 0 3438.7126
539.8104248046875 0 1676.3506
540.3112182617188 0 912.99426
548.2826538085938 0 1534.6516
548.8237915039062 0 1051.1775
566.2937622070312 0 1364.7966
642.1019897460938 0 597.1029
643.3531494140625 0 935.40985
661.3661499023438 0 1912.6641 y Water loss 2
679.3766479492188 0 30875.402 y 2
680.3794555664062 0 11905.369
681.3834838867188 0 2206.7378
778.4434204101562 0 2851.499 y 1
779.4490966796875 0 1433.8708
896.531005859375 0 753.9597
1414.4852294921875 0 597.5525
1484.1416015625 0 694.30695
1559.7554931640625 0 578.4103
2007.595947265625 0 684.60034
2197.111328125 0 705.6249
2496.86328125 0 705.1317
2568.295654296875 0 717.06726
3078.570556640625 0 632.5465

Spectrum Details

|  |  |
| --- | --- |
| Matched peaks? Matched peaks The total absolute number of peaks matched. Additionally in brackets the total fraction of peaks matched and the total number of peaks is shown. | 15 (10.56% of 142) |
| FDR? FDR The false discovery rate estimated for this peptide. It is calculated by matching all theoretical fragments with a non-integer shift with the raw peaks for this spectrum. This is done with 40 different shifts. The resulting percentage is the average number of annotated peaks over the number of annotated peaks with the correct spectrum. | 1.43% |
| Satellite FDR? Satellite FDR See the FDR for details on its calculation. This satellite ion specific FDR only contains the satellite ions (d/w) for I/L/J positions. | - |
| PSM Score? PSM Score The PSM Score as given by Hecklib to this annotated spectrum. It is shown with three significant figures. | 150 |

#### Reverse Lookup? Reverse Lookup All places where this read could be placed.

| Group | Segment | Template | Template Part | Read Part | Score | Unique |
| --- | --- | --- | --- | --- | --- | --- |
| Homo sapiens Light Chain | IGLJ | IGLJ2 | [0..9] | [0..9] | 72 | True |

| Recombined | Template Part | Read Part | Score | Unique |
| --- | --- | --- | --- | --- |
| REC-0-1\_002 | [99..108] | [0..9] | 72 | True |

#### Meta Information from Multiple reads

##### Number of combined reads

8

##### Intensity

0.7014

##### TotalArea

2.67E+08

##### Changes to the peptide sequence

VVFGGGTKJ

L→JNo support for either Leucine or Isoleucine based on side chain ions (Position: 9)

#### Positional Score

Copy Data

##### Positional Score (TSV)

###### Preview

```
Loading example...
```

*Click on the button to copy the data to your clipboard.*

10012345678

Label Value
"0" 0.748
"1" 0.75
"2" 0.746
"3" 0.739
"4" 0.709
"5" 0.689
"6" 0.711
"7" 0.714
"8" 0.695

#### Meta Information from PEAKS

##### Scan Identifier

F2:6223

##### Original sequence

V

V

F

G

G

G

T

K

L

##### Posttranslational Modifications

##### Source File

D:\separate\_stitch\_analyses\xle-disambiguation\raw\20210323\_F1\_UM1\_Peng0013\_SA\_F59\_ingel\_3ug\_TL.raw

##### Fraction

2

##### Scan Feature

F2:1858

##### De Novo Score

98

##### ConfidenceScore

98

### m/z

439.2608

##### Mass

876.5069

##### Charge

2

##### Retention Time

34.19

##### Predicted Retention Time

-

##### Area

8.424E+05

##### Parts Per Million

0.2

##### Fragmentation mode

HCD

##### Originating file

01 D:\separate\_stitch\_analyses\xle-disambiguation\20210325\_F59\_3ug\_DENOVO\_12.csv

#### Meta Information from PEAKS

##### Scan Identifier

F3:6556

##### Original sequence

V

V

F

G

G

G

T

K

L

##### Posttranslational Modifications

##### Source File

D:\separate\_stitch\_analyses\xle-disambiguation\raw\20210323\_F1\_UM1\_Peng0013\_SA\_F59\_ingel\_3ug\_chymo.raw

##### Fraction

3

##### Scan Feature

-

##### De Novo Score

97

##### ConfidenceScore

97

### m/z

439.2606

##### Mass

876.5069

##### Charge

2

##### Retention Time

36.38

##### Predicted Retention Time

-

##### Area

0

##### Fragmentation mode

HCD

##### Originating file

01 D:\separate\_stitch\_analyses\xle-disambiguation\20210325\_F59\_3ug\_DENOVO\_12.csv

#### Meta Information from PEAKS

##### Scan Identifier

F3:6150

##### Original sequence

V

V

F

G

G

G

T

K

L

##### Posttranslational Modifications

##### Source File

D:\separate\_stitch\_analyses\xle-disambiguation\raw\20210323\_F1\_UM1\_Peng0013\_SA\_F59\_ingel\_3ug\_chymo.raw

##### Fraction

3

##### Scan Feature

F3:1931

##### De Novo Score

97

##### ConfidenceScore

97

### m/z

439.2618

##### Mass

876.5069

##### Charge

2

##### Retention Time

34.09

##### Predicted Retention Time

-

##### Area

2.626E+08

##### Parts Per Million

2.4

##### Fragmentation mode

ETHCD

##### Originating file

01 D:\separate\_stitch\_analyses\xle-disambiguation\20210325\_F59\_3ug\_DENOVO\_12.csv

#### Meta Information from PEAKS

##### Scan Identifier

F3:6481

##### Original sequence

V

V

F

G

G

G

T

K

L

##### Posttranslational Modifications

##### Source File

D:\separate\_stitch\_analyses\xle-disambiguation\raw\20210323\_F1\_UM1\_Peng0013\_SA\_F59\_ingel\_3ug\_chymo.raw

##### Fraction

3

##### Scan Feature

-

##### De Novo Score

96

##### ConfidenceScore

96

### m/z

439.2615

##### Mass

876.5069

##### Charge

2

##### Retention Time

35.96

##### Predicted Retention Time

-

##### Area

0

##### Parts Per Million

1.7

##### Fragmentation mode

HCD

##### Originating file

01 D:\separate\_stitch\_analyses\xle-disambiguation\20210325\_F59\_3ug\_DENOVO\_12.csv

#### Meta Information from PEAKS

##### Scan Identifier

F3:6753

##### Original sequence

V

V

F

G

G

G

T

K

L

##### Posttranslational Modifications

##### Source File

D:\separate\_stitch\_analyses\xle-disambiguation\raw\20210323\_F1\_UM1\_Peng0013\_SA\_F59\_ingel\_3ug\_chymo.raw

##### Fraction

3

##### Scan Feature

-

##### De Novo Score

96

##### ConfidenceScore

96

### m/z

439.2612

##### Mass

876.5069

##### Charge

2

##### Retention Time

37.53

##### Predicted Retention Time

-

##### Area

0

##### Parts Per Million

1.2

##### Fragmentation mode

HCD

##### Originating file

01 D:\separate\_stitch\_analyses\xle-disambiguation\20210325\_F59\_3ug\_DENOVO\_12.csv

#### Meta Information from PEAKS

##### Scan Identifier

F3:6940

##### Original sequence

V

V

F

G

G

G

T

K

L

##### Posttranslational Modifications

##### Source File

D:\separate\_stitch\_analyses\xle-disambiguation\raw\20210323\_F1\_UM1\_Peng0013\_SA\_F59\_ingel\_3ug\_chymo.raw

##### Fraction

3

##### Scan Feature

F3:1927

##### De Novo Score

96

##### ConfidenceScore

96

### m/z

439.2614

##### Mass

876.5069

##### Charge

2

##### Retention Time

39.45

##### Predicted Retention Time

-

##### Area

2.272E+06

##### Parts Per Million

1.4

##### Fragmentation mode

HCD

##### Originating file

01 D:\separate\_stitch\_analyses\xle-disambiguation\20210325\_F59\_3ug\_DENOVO\_12.csv

#### Meta Information from PEAKS

##### Scan Identifier

F3:7725

##### Original sequence

V

V

F

G

G

G

T

K

L

##### Posttranslational Modifications

##### Source File

D:\separate\_stitch\_analyses\xle-disambiguation\raw\20210323\_F1\_UM1\_Peng0013\_SA\_F59\_ingel\_3ug\_chymo.raw

##### Fraction

3

##### Scan Feature

F3:1924

##### De Novo Score

95

##### ConfidenceScore

95

### m/z

439.261

##### Mass

876.5069

##### Charge

2

##### Retention Time

43.25

##### Predicted Retention Time

-

##### Area

6.956E+05

##### Parts Per Million

0.7

##### Fragmentation mode

HCD

##### Originating file

01 D:\separate\_stitch\_analyses\xle-disambiguation\20210325\_F59\_3ug\_DENOVO\_12.csv

#### Meta Information from PEAKS

##### Scan Identifier

F3:8222

##### Original sequence

V

V

F

G

G

G

T

K

L

##### Posttranslational Modifications

##### Source File

D:\separate\_stitch\_analyses\xle-disambiguation\raw\20210323\_F1\_UM1\_Peng0013\_SA\_F59\_ingel\_3ug\_chymo.raw

##### Fraction

3

##### Scan Feature

F3:1934

##### De Novo Score

95

##### ConfidenceScore

95

### m/z

439.262

##### Mass

876.5069

##### Charge

2

##### Retention Time

45.32

##### Predicted Retention Time

-

##### Area

6.348E+05

##### Parts Per Million

3

##### Fragmentation mode

HCD

##### Originating file

01 D:\separate\_stitch\_analyses\xle-disambiguation\20210325\_F59\_3ug\_DENOVO\_12.csv
