## Supplementary material for "A handle on mass coincidence errors in *de novo* sequencing of antibodies by bottom-up proteomics": Combined_093.html

Details Combined\_093 | Stitch OverviewUndefined

### Read Combined\_093

#### Sequence (length=7)

MTJDDFR

#### Spectrum 4722? Spectrum 4722 The raw spectrum of this peptide as annotated by Hecklib. The fragments are coloured according to ion type (see legend). Any peaks with a star '\*' as text can be hovered over to see the full details, first the ion type second the mass shift type. By hovering over the amino acids in the peptide or ions in the legend the corresponding peaks are highlighted. By toggling the 'Unassigned' label you can turn the background (unassigned) peaks on or off in the plot. By updating the slider in the Ion legend you can update the spectrum to only show the top X% of the peaks with labels. The top X% means any peak that is within X% of the highest intensity. By dragging in the spectrum you can zoom in to a specific part of the spectrum and use 'Zoom Out' to get back to the original zoom level. The annotation of the spectrum is based on the given sequence in the peptides file and is done with different software so inconsistencies are likely. The peaks are annotated based on the given sequence, with 20 ppm tolerance.

Copy Data

##### Spectrum 4722 (TSV)

###### Preview

```
Loading example...
```

*Click on the button to copy the data to your clipboard.*

Mz MinMz MaxIntensity Max

WidthHeightPeptide font sizePeptide stroke widthSpectrum font sizeSpectrum stroke widthCompact peptide

Ion legend

wxyz

abcd

OtherUnassignedIon^Charge^_Position_Show for top:%

MTJDDFR

04.37e+48.75e+41.31e+51.75e+5

Zoom Out

y^+1^_1_y^+1^_1_b^+1^_2_y^+1^_2_a^+1^_3_y^+1^_2_y^+2^_5_b^+1^_3_b^+1^_3_y^+2^_6_y^+1^_3_y^+1^_3_y^+1^_3_\*\*b^+1^_4_b^+1^_4_y^+1^_4_y^+1^_4_y^+1^_4_b^+1^_5_b^+1^_5_y^+1^_5_y^+1^_5_y^+1^_5_y^+1^_6_y^+1^_6_

0761152222823043

Fragment Matches Table

Show background peaks

| Position | Ion type | Intensity | mz Theoretical | mz Error (Th) | mz Error (ppm) | Charge | Series Number |
| --- | --- | --- | --- | --- | --- | --- | --- |
| - | - | 2.616E+04 | 120.1 | - | - | 0 | - |
| - | - | 2218 | 121.1 | - | - | 0 | - |
| - | - | 402.5 | 121.7 | - | - | 0 | - |
| - | - | 1207 | 122.1 | - | - | 0 | - |
| - | - | 620.8 | 124.1 | - | - | 0 | - |
| - | - | 1529 | 124.1 | - | - | 0 | - |
| - | - | 1679 | 125.1 | - | - | 0 | - |
| - | - | 735.3 | 126.1 | - | - | 0 | - |
| - | - | 1910 | 126.1 | - | - | 0 | - |
| - | - | 437.2 | 127.1 | - | - | 0 | - |
| - | - | 959.9 | 128.1 | - | - | 0 | - |
| - | - | 475.8 | 128.1 | - | - | 0 | - |
| - | - | 380.8 | 128.5 | - | - | 0 | - |
| - | - | 8957 | 129.1 | - | - | 0 | - |
| - | - | 7770 | 129.1 | - | - | 0 | - |
| - | - | 948.5 | 130 | - | - | 0 | - |
| - | - | 462.2 | 130.1 | - | - | 0 | - |
| - | - | 3359 | 130.1 | - | - | 0 | - |
| - | - | 1596 | 130.1 | - | - | 0 | - |
| - | - | 1120 | 130.1 | - | - | 0 | - |
| - | - | 572.2 | 133.1 | - | - | 0 | - |
| - | - | 1549 | 136.1 | - | - | 0 | - |
| - | - | 1166 | 139.1 | - | - | 0 | - |
| - | - | 3601 | 139.1 | - | - | 0 | - |
| - | - | 461 | 139.8 | - | - | 0 | - |
| - | - | 2657 | 140.1 | - | - | 0 | - |
| - | - | 480 | 140.1 | - | - | 0 | - |
| - | - | 410.4 | 140.2 | - | - | 0 | - |
| - | - | 1709 | 141.1 | - | - | 0 | - |
| - | - | 729.1 | 141.1 | - | - | 0 | - |
| - | - | 4072 | 141.1 | - | - | 0 | - |
| - | - | 5429 | 142.1 | - | - | 0 | - |
| - | - | 1249 | 143.1 | - | - | 0 | - |
| - | - | 1492 | 143.1 | - | - | 0 | - |
| - | - | 469.1 | 146.4 | - | - | 0 | - |
| - | - | 6435 | 147.1 | - | - | 0 | - |
| - | - | 2737 | 148 | - | - | 0 | - |
| - | - | 497.2 | 148.1 | - | - | 0 | - |
| - | - | 891.1 | 148.9 | - | - | 0 | - |
| - | - | 588 | 149 | - | - | 0 | - |
| - | - | 435.9 | 149 | - | - | 0 | - |
| - | - | 3022 | 152.1 | - | - | 0 | - |
| - | - | 505 | 153.1 | - | - | 0 | - |
| - | - | 531.9 | 155.1 | - | - | 0 | - |
| - | - | 830.9 | 155.1 | - | - | 0 | - |
| - | - | 2139 | 157.1 | - | - | 0 | - |
| - | - | 1.821E+04 | 157.1 | - | - | 0 | - |
| - | - | 2283 | 157.1 | - | - | 0 | - |
| 7 | y | 1.433E+04 | 158.1 | 4.161E-06 | 0.02632 | +1 | 1 |
| - | - | 1405 | 158.1 | - | - | 0 | - |
| - | - | 552.8 | 159.1 | - | - | 0 | - |
| - | - | 875.4 | 165.1 | - | - | 0 | - |
| - | - | 2541 | 166.1 | - | - | 0 | - |
| - | - | 2595 | 167.1 | - | - | 0 | - |
| - | - | 718.2 | 167.1 | - | - | 0 | - |
| - | - | 834.1 | 169.1 | - | - | 0 | - |
| - | - | 1.821E+04 | 169.1 | - | - | 0 | - |
| - | - | 1439 | 170.1 | - | - | 0 | - |
| - | - | 443.6 | 171.1 | - | - | 0 | - |
| - | - | 2043 | 171.1 | - | - | 0 | - |
| - | - | 1.241E+04 | 171.1 | - | - | 0 | - |
| - | - | 1252 | 172.2 | - | - | 0 | - |
| - | - | 5051 | 173.1 | - | - | 0 | - |
| - | - | 977 | 173.4 | - | - | 0 | - |
| - | - | 1418 | 175.1 | - | - | 0 | - |
| 7 | y | 3.399E+04 | 175.1 | 9.907E-06 | 0.05657 | +1 | 1 |
| - | - | 1987 | 176.1 | - | - | 0 | - |
| - | - | 1109 | 177.1 | - | - | 0 | - |
| - | - | 876.5 | 179.1 | - | - | 0 | - |
| - | - | 1351 | 180.1 | - | - | 0 | - |
| - | - | 3990 | 181.1 | - | - | 0 | - |
| - | - | 1072 | 181.1 | - | - | 0 | - |
| - | - | 539.6 | 182.1 | - | - | 0 | - |
| - | - | 2403 | 183.1 | - | - | 0 | - |
| - | - | 555.1 | 183.1 | - | - | 0 | - |
| - | - | 528.2 | 183.5 | - | - | 0 | - |
| - | - | 583.3 | 183.5 | - | - | 0 | - |
| - | - | 582.8 | 185 | - | - | 0 | - |
| - | - | 1487 | 185.1 | - | - | 0 | - |
| - | - | 7693 | 185.1 | - | - | 0 | - |
| - | - | 1073 | 185.1 | - | - | 0 | - |
| - | - | 3330 | 185.2 | - | - | 0 | - |
| - | - | 1207 | 186.1 | - | - | 0 | - |
| - | - | 1.752E+04 | 187.1 | - | - | 0 | - |
| - | - | 814.5 | 188.1 | - | - | 0 | - |
| - | - | 812.5 | 188.1 | - | - | 0 | - |
| - | - | 1533 | 189.1 | - | - | 0 | - |
| - | - | 1.226E+04 | 190.1 | - | - | 0 | - |
| - | - | 834.1 | 191.1 | - | - | 0 | - |
| - | - | 760.7 | 195.1 | - | - | 0 | - |
| - | - | 3.484E+04 | 197.1 | - | - | 0 | - |
| - | - | 2563 | 198.1 | - | - | 0 | - |
| - | - | 615.7 | 199.1 | - | - | 0 | - |
| - | - | 580.1 | 200.1 | - | - | 0 | - |
| - | - | 4100 | 201.1 | - | - | 0 | - |
| - | - | 1236 | 203.1 | - | - | 0 | - |
| - | - | 2624 | 203.1 | - | - | 0 | - |
| - | - | 1423 | 204.1 | - | - | 0 | - |
| - | - | 2326 | 205.1 | - | - | 0 | - |
| - | - | 1269 | 207.1 | - | - | 0 | - |
| - | - | 1486 | 208.1 | - | - | 0 | - |
| - | - | 4846 | 209.1 | - | - | 0 | - |
| - | - | 954.6 | 209.1 | - | - | 0 | - |
| - | - | 600.7 | 209.2 | - | - | 0 | - |
| - | - | 585.4 | 210.1 | - | - | 0 | - |
| - | - | 3028 | 211.1 | - | - | 0 | - |
| - | - | 3412 | 213.2 | - | - | 0 | - |
| - | - | 757.8 | 214.1 | - | - | 0 | - |
| - | - | 1515 | 215 | - | - | 0 | - |
| - | - | 1.108E+04 | 215.1 | - | - | 0 | - |
| - | - | 608.9 | 217.1 | - | - | 0 | - |
| - | - | 481 | 217.1 | - | - | 0 | - |
| - | - | 879.6 | 217.1 | - | - | 0 | - |
| - | - | 1.757E+04 | 221.1 | - | - | 0 | - |
| - | - | 1470 | 222.1 | - | - | 0 | - |
| - | - | 621.6 | 222.2 | - | - | 0 | - |
| - | - | 922.4 | 223.1 | - | - | 0 | - |
| - | - | 660.7 | 223.1 | - | - | 0 | - |
| - | - | 5657 | 224.1 | - | - | 0 | - |
| - | - | 1.13E+04 | 225.1 | - | - | 0 | - |
| - | - | 1329 | 226.1 | - | - | 0 | - |
| - | - | 819.9 | 226.1 | - | - | 0 | - |
| - | - | 6963 | 226.2 | - | - | 0 | - |
| - | - | 2128 | 227.1 | - | - | 0 | - |
| - | - | 1407 | 227.2 | - | - | 0 | - |
| - | - | 3570 | 229.1 | - | - | 0 | - |
| - | - | 572 | 230.1 | - | - | 0 | - |
| - | - | 1997 | 231.1 | - | - | 0 | - |
| - | - | 4269 | 231.1 | - | - | 0 | - |
| - | - | 1311 | 232.1 | - | - | 0 | - |
| - | - | 603.7 | 233.1 | - | - | 0 | - |
| - | - | 8681 | 235.1 | - | - | 0 | - |
| - | - | 920.5 | 236.1 | - | - | 0 | - |
| - | - | 1265 | 237.2 | - | - | 0 | - |
| - | - | 668.7 | 241 | - | - | 0 | - |
| - | - | 693.6 | 242.1 | - | - | 0 | - |
| - | - | 564.7 | 242.2 | - | - | 0 | - |
| - | - | 955.8 | 243.1 | - | - | 0 | - |
| - | - | 3.752E+04 | 244.2 | - | - | 0 | - |
| - | - | 773.1 | 245.1 | - | - | 0 | - |
| - | - | 739.9 | 245.1 | - | - | 0 | - |
| - | - | 4044 | 245.2 | - | - | 0 | - |
| - | - | 781 | 246.2 | - | - | 0 | - |
| 2 | b | 2.63E+04 | 249.1 | 0.00477 | 19.15 | +1 | 2 |
| - | - | 2636 | 250.1 | - | - | 0 | - |
| - | - | 1023 | 251.1 | - | - | 0 | - |
| - | - | 3956 | 251.1 | - | - | 0 | - |
| - | - | 2520 | 252.1 | - | - | 0 | - |
| - | - | 1398 | 252.2 | - | - | 0 | - |
| - | - | 948.9 | 253.1 | - | - | 0 | - |
| - | - | 1683 | 254.1 | - | - | 0 | - |
| - | - | 1396 | 254.1 | - | - | 0 | - |
| - | - | 571.9 | 255.1 | - | - | 0 | - |
| - | - | 617.4 | 255.2 | - | - | 0 | - |
| - | - | 797.3 | 260.1 | - | - | 0 | - |
| - | - | 2152 | 262.2 | - | - | 0 | - |
| - | - | 753 | 263.1 | - | - | 0 | - |
| - | - | 1585 | 263.1 | - | - | 0 | - |
| - | - | 698.5 | 266.1 | - | - | 0 | - |
| - | - | 1.333E+04 | 268.2 | - | - | 0 | - |
| - | - | 809.4 | 269.1 | - | - | 0 | - |
| - | - | 1206 | 270.1 | - | - | 0 | - |
| - | - | 1923 | 270.2 | - | - | 0 | - |
| - | - | 643.7 | 271.1 | - | - | 0 | - |
| - | - | 613.8 | 273.1 | - | - | 0 | - |
| - | - | 658.7 | 273.1 | - | - | 0 | - |
| - | - | 3356 | 280.2 | - | - | 0 | - |
| - | - | 4986 | 286.2 | - | - | 0 | - |
| - | - | 2451 | 287.2 | - | - | 0 | - |
| - | - | 925.8 | 287.2 | - | - | 0 | - |
| - | - | 1213 | 288.1 | - | - | 0 | - |
| - | - | 751.1 | 288.2 | - | - | 0 | - |
| - | - | 985.5 | 290.6 | - | - | 0 | - |
| - | - | 562.9 | 294.1 | - | - | 0 | - |
| - | - | 603.4 | 296.1 | - | - | 0 | - |
| - | - | 1540 | 296.2 | - | - | 0 | - |
| - | - | 8118 | 296.2 | - | - | 0 | - |
| - | - | 612.4 | 298.1 | - | - | 0 | - |
| - | - | 597.7 | 298.1 | - | - | 0 | - |
| - | - | 1775 | 298.2 | - | - | 0 | - |
| - | - | 1672 | 299.1 | - | - | 0 | - |
| - | - | 3632 | 299.1 | - | - | 0 | - |
| - | - | 900.4 | 299.2 | - | - | 0 | - |
| - | - | 741 | 299.6 | - | - | 0 | - |
| - | - | 1785 | 300.1 | - | - | 0 | - |
| - | - | 1241 | 300.1 | - | - | 0 | - |
| - | - | 1722 | 301.1 | - | - | 0 | - |
| - | - | 977.3 | 302.1 | - | - | 0 | - |
| - | - | 1660 | 304.2 | - | - | 0 | - |
| - | - | 1657 | 304.6 | - | - | 0 | - |
| 6 | y | 1.569E+04 | 305.2 | 8.09E-05 | 0.2651 | +1 | 2 |
| - | - | 1226 | 305.6 | - | - | 0 | - |
| - | - | 2443 | 306.2 | - | - | 0 | - |
| - | - | 575.1 | 306.6 | - | - | 0 | - |
| - | - | 637.6 | 310.1 | - | - | 0 | - |
| - | - | 1302 | 312.2 | - | - | 0 | - |
| - | - | 2470 | 313.1 | - | - | 0 | - |
| - | - | 2069 | 313.6 | - | - | 0 | - |
| - | - | 2236 | 314.2 | - | - | 0 | - |
| 3 | a | 2805 | 316.2 | 0.004677 | 14.79 | +1 | 3 |
| - | - | 2207 | 318.1 | - | - | 0 | - |
| 6 | y | 1.733E+05 | 322.2 | 1.185E-05 | 0.03677 | +1 | 2 |
| - | - | 2.905E+04 | 323.2 | - | - | 0 | - |
| - | - | 866.9 | 324.2 | - | - | 0 | - |
| - | - | 3930 | 324.2 | - | - | 0 | - |
| - | - | 930.4 | 325.2 | - | - | 0 | - |
| - | - | 848.5 | 326.1 | - | - | 0 | - |
| - | - | 1926 | 327.1 | - | - | 0 | - |
| - | - | 3219 | 328.1 | - | - | 0 | - |
| - | - | 699.2 | 332.1 | - | - | 0 | - |
| - | - | 851.5 | 332.2 | - | - | 0 | - |
| 3 | y | 5643 | 333.2 | 5.503E-05 | 0.1652 | +2 | 5 |
| - | - | 1710 | 333.7 | - | - | 0 | - |
| - | - | 540.6 | 334 | - | - | 0 | - |
| - | - | 750.1 | 335.1 | - | - | 0 | - |
| - | - | 1211 | 338.2 | - | - | 0 | - |
| - | - | 3323 | 340.1 | - | - | 0 | - |
| - | - | 1138 | 340.7 | - | - | 0 | - |
| - | - | 1696 | 341.1 | - | - | 0 | - |
| - | - | 630.6 | 341.7 | - | - | 0 | - |
| - | - | 1133 | 342.2 | - | - | 0 | - |
| - | - | 2606 | 344.1 | - | - | 0 | - |
| - | - | 3133 | 344.1 | - | - | 0 | - |
| 3 | b | 1973 | 344.2 | 0.005673 | 16.48 | +1 | 3 |
| - | - | 1570 | 346.2 | - | - | 0 | - |
| - | - | 813.8 | 346.7 | - | - | 0 | - |
| - | - | 850.8 | 347.2 | - | - | 0 | - |
| - | - | 2120 | 349.7 | - | - | 0 | - |
| - | - | 942.2 | 351.7 | - | - | 0 | - |
| - | - | 665.9 | 353.1 | - | - | 0 | - |
| - | - | 1162 | 353.6 | - | - | 0 | - |
| - | - | 784.5 | 354.1 | - | - | 0 | - |
| - | - | 903.2 | 354.1 | - | - | 0 | - |
| - | - | 1209 | 354.2 | - | - | 0 | - |
| - | - | 835.9 | 355.1 | - | - | 0 | - |
| - | - | 1765 | 355.2 | - | - | 0 | - |
| - | - | 3897 | 356.1 | - | - | 0 | - |
| - | - | 820.2 | 356.1 | - | - | 0 | - |
| - | - | 742.4 | 357.1 | - | - | 0 | - |
| - | - | 1305 | 360.7 | - | - | 0 | - |
| - | - | 687.9 | 361.7 | - | - | 0 | - |
| - | - | 1.447E+04 | 362.2 | - | - | 0 | - |
| 3 | b | 4232 | 362.2 | 0.005118 | 14.13 | +1 | 3 |
| - | - | 8086 | 362.7 | - | - | 0 | - |
| - | - | 6533 | 363.1 | - | - | 0 | - |
| - | - | 2576 | 363.7 | - | - | 0 | - |
| - | - | 1921 | 364.2 | - | - | 0 | - |
| - | - | 1426 | 365.7 | - | - | 0 | - |
| - | - | 2283 | 366.7 | - | - | 0 | - |
| - | - | 2203 | 367.2 | - | - | 0 | - |
| - | - | 757.9 | 367.2 | - | - | 0 | - |
| - | - | 1605 | 368.1 | - | - | 0 | - |
| - | - | 2020 | 369.1 | - | - | 0 | - |
| - | - | 1048 | 371.7 | - | - | 0 | - |
| 2 | y | 5769 | 374.7 | 0.0006491 | 1.732 | +2 | 6 |
| - | - | 684.7 | 374.8 | - | - | 0 | - |
| - | - | 3619 | 375.2 | - | - | 0 | - |
| - | - | 1025 | 375.7 | - | - | 0 | - |
| - | - | 1068 | 378.1 | - | - | 0 | - |
| - | - | 711.1 | 379.2 | - | - | 0 | - |
| - | - | 1057 | 383.1 | - | - | 0 | - |
| - | - | 1117 | 384.1 | - | - | 0 | - |
| - | - | 2381 | 384.7 | - | - | 0 | - |
| - | - | 3133 | 388.7 | - | - | 0 | - |
| - | - | 1116 | 389.7 | - | - | 0 | - |
| - | - | 735.7 | 393.2 | - | - | 0 | - |
| - | - | 644.2 | 393.7 | - | - | 0 | - |
| - | - | 1030 | 397.2 | - | - | 0 | - |
| - | - | 863.7 | 399.3 | - | - | 0 | - |
| - | - | 777.4 | 401.1 | - | - | 0 | - |
| - | - | 7058 | 402.2 | - | - | 0 | - |
| - | - | 3237 | 402.7 | - | - | 0 | - |
| - | - | 832.8 | 403.2 | - | - | 0 | - |
| - | - | 665.3 | 409.2 | - | - | 0 | - |
| - | - | 991.4 | 409.7 | - | - | 0 | - |
| - | - | 771.2 | 411.2 | - | - | 0 | - |
| - | - | 821.4 | 412.1 | - | - | 0 | - |
| - | - | 3531 | 412.2 | - | - | 0 | - |
| - | - | 653.7 | 413.1 | - | - | 0 | - |
| - | - | 816.5 | 413.2 | - | - | 0 | - |
| - | - | 1.001E+04 | 416.2 | - | - | 0 | - |
| - | - | 3666 | 416.7 | - | - | 0 | - |
| - | - | 1898 | 417.2 | - | - | 0 | - |
| - | - | 821 | 417.7 | - | - | 0 | - |
| - | - | 1939 | 418.2 | - | - | 0 | - |
| 5 | y | 1337 | 419.2 | 0.001199 | 2.861 | +1 | 3 |
| 5 | y | 2407 | 420.2 | 0.0004114 | 0.979 | +1 | 3 |
| - | - | 682.7 | 421.2 | - | - | 0 | - |
| - | - | 1.504E+04 | 425.2 | - | - | 0 | - |
| - | - | 1.162E+04 | 425.7 | - | - | 0 | - |
| - | - | 989 | 426.2 | - | - | 0 | - |
| - | - | 5238 | 426.2 | - | - | 0 | - |
| - | - | 1976 | 426.7 | - | - | 0 | - |
| - | - | 1095 | 427.3 | - | - | 0 | - |
| - | - | 665.3 | 429.2 | - | - | 0 | - |
| - | - | 2564 | 433.7 | - | - | 0 | - |
| - | - | 2168 | 434.2 | - | - | 0 | - |
| - | - | 2.461E+04 | 434.7 | - | - | 0 | - |
| - | - | 9690 | 435.2 | - | - | 0 | - |
| - | - | 2527 | 435.7 | - | - | 0 | - |
| - | - | 1.045E+04 | 436.2 | - | - | 0 | - |
| - | - | 1081 | 436.7 | - | - | 0 | - |
| 5 | y | 3.513E+04 | 437.2 | 0.0007776 | 1.779 | +1 | 3 |
| - | - | 7172 | 438.2 | - | - | 0 | - |
| - | - | 1705 | 439.7 | - | - | 0 | - |
| - | - | 1102 | 439.8 | - | - | 0 | - |
| - | - | 1462 | 440.2 | - | - | 0 | - |
| - | - | 3563 | 440.3 | - | - | 0 | - |
| - | - | 1.223E+04 | 447.2 | - | - | 0 | - |
| - | - | 8935 | 447.7 | - | - | 0 | - |
| 0 | Precursor | 1.621E+04 | 448.2 | 0.003091 | 6.895 | +2 | -1 |
| - | - | 3652 | 448.2 | - | - | 0 | - |
| - | - | 8059 | 448.7 | - | - | 0 | - |
| - | - | 2614 | 449.2 | - | - | 0 | - |
| - | - | 811.8 | 449.2 | - | - | 0 | - |
| - | - | 1352 | 449.3 | - | - | 0 | - |
| - | - | 851.2 | 449.7 | - | - | 0 | - |
| - | - | 1485 | 452.1 | - | - | 0 | - |
| - | - | 1.253E+04 | 453.2 | - | - | 0 | - |
| - | - | 927.2 | 454.2 | - | - | 0 | - |
| - | - | 2056 | 455.2 | - | - | 0 | - |
| - | - | 827.6 | 456.2 | - | - | 0 | - |
| - | - | 2627 | 456.2 | - | - | 0 | - |
| - | - | 7136 | 456.7 | - | - | 0 | - |
| 0 | Precursor | 7814 | 457.2 | 0.003698 | 8.088 | +2 | -1 |
| - | - | 3125 | 457.7 | - | - | 0 | - |
| - | - | 5361 | 458.2 | - | - | 0 | - |
| 4 | b | 854.7 | 459.2 | 0.004639 | 10.1 | +1 | 4 |
| - | - | 750.1 | 463.2 | - | - | 0 | - |
| - | - | 1860 | 465.2 | - | - | 0 | - |
| - | - | 1846 | 471.3 | - | - | 0 | - |
| 4 | b | 1090 | 477.2 | 0.004634 | 9.71 | +1 | 4 |
| - | - | 1.435E+04 | 480.1 | - | - | 0 | - |
| - | - | 2730 | 481.1 | - | - | 0 | - |
| - | - | 6872 | 482.1 | - | - | 0 | - |
| - | - | 1269 | 483.1 | - | - | 0 | - |
| - | - | 949.6 | 485.2 | - | - | 0 | - |
| - | - | 977.7 | 486.2 | - | - | 0 | - |
| - | - | 733.2 | 487.2 | - | - | 0 | - |
| - | - | 1360 | 497.2 | - | - | 0 | - |
| - | - | 2831 | 510.2 | - | - | 0 | - |
| - | - | 1181 | 511.2 | - | - | 0 | - |
| - | - | 2950 | 512.2 | - | - | 0 | - |
| - | - | 958.3 | 513.2 | - | - | 0 | - |
| - | - | 5666 | 514.2 | - | - | 0 | - |
| - | - | 975.6 | 515.2 | - | - | 0 | - |
| - | - | 1190 | 517.2 | - | - | 0 | - |
| - | - | 2085 | 528.2 | - | - | 0 | - |
| 4 | y | 2245 | 534.2 | 0.0002797 | 0.5236 | +1 | 4 |
| 4 | y | 2080 | 535.2 | 0.0005069 | 0.9471 | +1 | 4 |
| - | - | 876.8 | 539.2 | - | - | 0 | - |
| 4 | y | 6.846E+04 | 552.2 | 0.0004685 | 0.8483 | +1 | 4 |
| - | - | 1.808E+04 | 553.2 | - | - | 0 | - |
| - | - | 3075 | 554.2 | - | - | 0 | - |
| - | - | 2975 | 562.2 | - | - | 0 | - |
| - | - | 1221 | 563.2 | - | - | 0 | - |
| - | - | 1.67E+04 | 566.3 | - | - | 0 | - |
| 5 | b | 4098 | 574.2 | 0.003789 | 6.598 | +1 | 5 |
| - | - | 1040 | 575.2 | - | - | 0 | - |
| - | - | 2073 | 584.3 | - | - | 0 | - |
| - | - | 640 | 586.3 | - | - | 0 | - |
| 5 | b | 2595 | 592.2 | 0.004394 | 7.419 | +1 | 5 |
| - | - | 611.4 | 598.2 | - | - | 0 | - |
| - | - | 1038 | 599.2 | - | - | 0 | - |
| - | - | 661.1 | 607.3 | - | - | 0 | - |
| - | - | 1523 | 608.2 | - | - | 0 | - |
| - | - | 563.7 | 621.9 | - | - | 0 | - |
| - | - | 785.7 | 622.3 | - | - | 0 | - |
| - | - | 860.9 | 624.3 | - | - | 0 | - |
| - | - | 1.149E+04 | 625.3 | - | - | 0 | - |
| - | - | 3527 | 626.3 | - | - | 0 | - |
| 3 | y | 996 | 647.3 | 0.0007256 | 1.121 | +1 | 5 |
| 3 | y | 1700 | 648.3 | 0.001526 | 2.354 | +1 | 5 |
| - | - | 1539 | 650.3 | - | - | 0 | - |
| - | - | 1237 | 651.3 | - | - | 0 | - |
| - | - | 820 | 652.3 | - | - | 0 | - |
| 3 | y | 7.683E+04 | 665.3 | 0.0005481 | 0.8238 | +1 | 5 |
| - | - | 1.646E+04 | 665.4 | - | - | 0 | - |
| - | - | 2.51E+04 | 666.3 | - | - | 0 | - |
| - | - | 1238 | 666.4 | - | - | 0 | - |
| - | - | 5053 | 667.3 | - | - | 0 | - |
| - | - | 1831 | 675.3 | - | - | 0 | - |
| - | - | 733.3 | 676.3 | - | - | 0 | - |
| - | - | 2860 | 683.4 | - | - | 0 | - |
| - | - | 2252 | 691.3 | - | - | 0 | - |
| - | - | 840.9 | 692.3 | - | - | 0 | - |
| - | - | 1036 | 698.3 | - | - | 0 | - |
| - | - | 1314 | 722.3 | - | - | 0 | - |
| - | - | 8527 | 723.3 | - | - | 0 | - |
| - | - | 3221 | 724.3 | - | - | 0 | - |
| - | - | 5026 | 725.3 | - | - | 0 | - |
| - | - | 1974 | 726.3 | - | - | 0 | - |
| - | - | 1221 | 732.3 | - | - | 0 | - |
| - | - | 797.6 | 733.3 | - | - | 0 | - |
| - | - | 1366 | 740.3 | - | - | 0 | - |
| - | - | 1004 | 741.3 | - | - | 0 | - |
| 2 | y | 3698 | 748.4 | 0.001651 | 2.206 | +1 | 6 |
| - | - | 1466 | 749.4 | - | - | 0 | - |
| - | - | 632 | 750.4 | - | - | 0 | - |
| - | - | 1092 | 755.3 | - | - | 0 | - |
| - | - | 1004 | 758.3 | - | - | 0 | - |
| 2 | y | 2.28E+04 | 766.4 | 0.001352 | 1.764 | +1 | 6 |
| - | - | 4260 | 766.4 | - | - | 0 | - |
| - | - | 7083 | 767.4 | - | - | 0 | - |
| - | - | 2526 | 768.4 | - | - | 0 | - |
| - | - | 3.013E+04 | 776.4 | - | - | 0 | - |
| - | - | 1.064E+04 | 777.4 | - | - | 0 | - |
| - | - | 3261 | 778.4 | - | - | 0 | - |
| - | - | 2017 | 784.5 | - | - | 0 | - |
| - | - | 1079 | 794.4 | - | - | 0 | - |
| - | - | 5646 | 849.4 | - | - | 0 | - |
| - | - | 1049 | 850.5 | - | - | 0 | - |
| - | - | 2536 | 877.5 | - | - | 0 | - |
| - | - | 609.7 | 1152 | - | - | 0 | - |
| - | - | 686.5 | 3013 | - | - | 0 | - |

m/z Charge Intensity FragmentType MassShift Position
120.08088684082031 0 26159.674
121.08416748046875 0 2217.8594
121.74068450927734 0 402.50095
122.06013488769531 0 1207.261
124.07585144042969 0 620.82635
124.11219024658203 0 1528.8563
125.1074447631836 0 1679.063
126.09146118164062 0 735.3183
126.1278305053711 0 1910.0596
127.08738708496094 0 437.22183
128.0706787109375 0 959.87396
128.10699462890625 0 475.8014
128.5409698486328 0 380.8007
129.06594848632812 0 8957.467
129.1023406982422 0 7769.6367
130.0323028564453 0 948.5169
130.0504913330078 0 462.21365
130.0863800048828 0 3359.0342
130.09768676757812 0 1596.4506
130.10569763183594 0 1119.8126
133.09727478027344 0 572.2149
136.07568359375 0 1549.0997
139.05032348632812 0 1165.5381
139.086669921875 0 3601.1465
139.79330444335938 0 461.0497
140.0706329345703 0 2657.067
140.0819549560547 0 479.98962
140.24887084960938 0 410.35468
141.06591796875 0 1708.7046
141.10223388671875 0 729.0995
141.13864135742188 0 4072.435
142.12274169921875 0 5429.142
143.0815887451172 0 1248.7778
143.11793518066406 0 1491.5703
146.44947814941406 0 469.14838
147.1128692626953 0 6435.407
148.04275512695312 0 2737.053
148.11572265625 0 497.2368
148.94737243652344 0 891.0607
149.02658081054688 0 588.0476
149.04531860351562 0 435.91144
152.1071319580078 0 3021.978
153.10226440429688 0 504.97507
155.08114624023438 0 531.90295
155.1180877685547 0 830.91437
157.060791015625 0 2139.3613
157.09715270996094 0 18207.467
157.1084442138672 0 2283.4888
158.0924072265625 0 14330.141 y Ammonia loss 6
158.10023498535156 0 1404.9043
159.07672119140625 0 552.7888
165.10279846191406 0 875.3941
166.05323791503906 0 2540.7324
167.0814666748047 0 2595.0132
167.1180877685547 0 718.2317
169.06076049804688 0 834.0651
169.133544921875 0 18214.332
170.13690185546875 0 1438.7347
171.07763671875 0 443.5975
171.11276245117188 0 2042.5132
171.14920043945312 0 12408.959
172.1526641845703 0 1252.0261
173.12841796875 0 5050.7095
173.43907165527344 0 976.9778
175.08631896972656 0 1418.1017
175.1189422607422 0 33992.957 y 6
176.1223602294922 0 1986.5034
177.06918334960938 0 1108.7307
179.1177520751953 0 876.4978
180.11334228515625 0 1350.5985
181.09722900390625 0 3989.8418
181.13340759277344 0 1072.0068
182.08042907714844 0 539.5721
183.1129150390625 0 2403.3044
183.1497802734375 0 555.09
183.46017456054688 0 528.1879
183.4694061279297 0 583.3353
184.9847869873047 0 582.7665
185.0553436279297 0 1487.3263
185.09202575683594 0 7692.8135
185.1017608642578 0 1073.3699
185.1647491455078 0 3330.0344
186.08753967285156 0 1207.2286
187.14404296875 0 17519.756
188.13926696777344 0 814.5307
188.14785766601562 0 812.46906
189.12301635742188 0 1532.7622
190.09742736816406 0 12260.298
191.100830078125 0 834.1241
195.1126251220703 0 760.6561
197.12840270996094 0 34843.273
198.13185119628906 0 2562.765
199.14393615722656 0 615.74646
200.0701141357422 0 580.1369
201.12335205078125 0 4100.126
203.06605529785156 0 1235.5231
203.08477783203125 0 2624.124
204.06893920898438 0 1423.2244
205.0640869140625 0 2326.1887
207.11233520507812 0 1269.4504
208.10794067382812 0 1486.396
209.09194946289062 0 4846.444
209.12847900390625 0 954.57135
209.1649932861328 0 600.69604
210.08670043945312 0 585.39856
211.14425659179688 0 3027.5654
213.15966796875 0 3412.0854
214.0529327392578 0 757.7637
215.0099334716797 0 1514.8693
215.13897705078125 0 11081.532
217.0967254638672 0 608.9434
217.12118530273438 0 480.9544
217.13316345214844 0 879.6291
221.09539794921875 0 17566.938
222.09869384765625 0 1469.7424
222.16006469726562 0 621.55444
223.0916290283203 0 922.3995
223.14407348632812 0 660.668
224.13925170898438 0 5657.4966
225.1232452392578 0 11297.081
226.11798095703125 0 1329.4015
226.12838745117188 0 819.9272
226.1549530029297 0 6963.0684
227.13870239257812 0 2127.7532
227.1581268310547 0 1407.1957
229.11810302734375 0 3570.4844
230.12225341796875 0 572.022
231.0609130859375 0 1996.564
231.07972717285156 0 4268.505
232.06381225585938 0 1311.2383
233.1286163330078 0 603.68536
235.1076202392578 0 8680.519
236.13941955566406 0 920.5327
237.16018676757812 0 1264.6691
241.04763793945312 0 668.6819
242.12828063964844 0 693.6152
242.15089416503906 0 564.72107
243.0638427734375 0 955.84344
244.16554260253906 0 37520.11
245.0576171875 0 773.11536
245.128173828125 0 739.866
245.16885375976562 0 4043.6445
246.170654296875 0 780.98364
249.0902099609375 0 26304.3 b 1
250.09335327148438 0 2635.5356
251.08570861816406 0 1022.9892
251.13885498046875 0 3956.3755
252.13426208496094 0 2519.8662
252.17080688476562 0 1398.4634
253.1181640625 0 948.9406
254.0792236328125 0 1683.2502
254.14971923828125 0 1395.6652
255.07708740234375 0 571.8723
255.15345764160156 0 617.3519
260.13934326171875 0 797.25287
262.1549377441406 0 2152.4475
263.1031799316406 0 753.01526
263.1387939453125 0 1584.7604
266.1497497558594 0 698.50757
268.1653137207031 0 13332.719
269.0789794921875 0 809.4062
270.1449279785156 0 1206.1135
270.18084716796875 0 1923.002
271.0752258300781 0 643.71454
273.0870361328125 0 613.8073
273.1208190917969 0 658.6768
280.1656494140625 0 3356.311
286.17596435546875 0 4985.5728
287.15032958984375 0 2451.0771
287.20770263671875 0 925.7723
288.1343994140625 0 1212.6696
288.15234375 0 751.1303
290.6243896484375 0 985.4659
294.1442565917969 0 562.9215
296.1060485839844 0 603.36584
296.1605529785156 0 1540.1083
296.19677734375 0 8118.273
298.1059875488281 0 612.43506
298.141357421875 0 597.7239
298.1762390136719 0 1774.8359
299.1041259765625 0 1672.0942
299.13763427734375 0 3632.3494
299.17156982421875 0 900.40204
299.6380615234375 0 741.026
300.08502197265625 0 1784.548
300.1389465332031 0 1241.3024
301.0842590332031 0 1721.5912
302.0805969238281 0 977.2611
304.1769104003906 0 1659.7054
304.620361328125 0 1656.9724
305.1607360839844 0 15685.601 y Ammonia loss 5
305.61859130859375 0 1225.5088
306.16363525390625 0 2442.828
306.64263916015625 0 575.1059
310.0686950683594 0 637.6408
312.1560363769531 0 1302.4841
313.1349182128906 0 2469.9258
313.63623046875 0 2069.324
314.207763671875 0 2236.0742
316.168701171875 0 2805.145 a Water loss 2
318.1488342285156 0 2207.0176
322.1873779296875 0 173253.77 y 5
323.1904296875 0 29052.508
324.1714172363281 0 866.8525
324.1927795410156 0 3929.6907
325.1869201660156 0 930.35565
326.1004333496094 0 848.5384
327.09759521484375 0 1926.411
328.0950012207031 0 3218.5586
332.12396240234375 0 699.2039
332.1716613769531 0 851.4566
333.1663513183594 0 5643.028 y 2
333.6679382324219 0 1709.5977
334.0006408691406 0 540.5636
335.1480407714844 0 750.11
338.15814208984375 0 1210.5067
340.1498107910156 0 3322.912
340.6554870605469 0 1138.466
341.14923095703125 0 1695.6873
341.6517028808594 0 630.5964
342.2097473144531 0 1132.6176
344.1245422363281 0 2605.9106
344.1454772949219 0 3133.0579
344.16461181640625 0 1972.8435 b Water loss 2
346.1737060546875 0 1569.6842
346.67474365234375 0 813.8353
347.17041015625 0 850.7603
349.6611328125 0 2120.2554
351.6824951171875 0 942.164
353.146728515625 0 665.86414
353.6391296386719 0 1161.5547
354.09552001953125 0 784.4618
354.1444091796875 0 903.1811
354.2134704589844 0 1208.5603
355.0910339355469 0 835.89166
355.1969299316406 0 1764.6189
356.0888366699219 0 3897.326
356.1260986328125 0 820.2484
357.0882568359375 0 742.4052
360.6868896484375 0 1304.8534
361.66302490234375 0 687.8879
362.1517333984375 0 14466.312
362.17462158203125 0 4231.751 b 2
362.65313720703125 0 8085.8296
363.1498718261719 0 6533.132
363.6506042480469 0 2575.6528
364.1500549316406 0 1921.2394
365.6797790527344 0 1425.6411
366.6684265136719 0 2283.2603
367.1618347167969 0 2202.6184
367.231689453125 0 757.8614
368.14593505859375 0 1604.9839
369.1436767578125 0 2020.1622
371.6610107421875 0 1047.6067
374.6842041015625 0 5768.854 y Water loss 1
374.7825927734375 0 684.65326
375.1863098144531 0 3619.4385
375.6873474121094 0 1024.5447
378.12896728515625 0 1067.7961
379.1979675292969 0 711.08527
383.1369934082031 0 1056.9999
384.1207275390625 0 1117.3696
384.6684875488281 0 2381.2092
388.68255615234375 0 3133.2964
389.6885681152344 0 1116.4976
393.1786193847656 0 735.6966
393.67333984375 0 644.16864
397.2081298828125 0 1030.3718
399.296875 0 863.7104
401.146484375 0 777.449
402.18634033203125 0 7057.9946
402.6886291503906 0 3237.017
403.1911315917969 0 832.8128
409.2287292480469 0 665.3292
409.72064208984375 0 991.3936
411.19189453125 0 771.1907
412.1152038574219 0 821.3957
412.18804931640625 0 3531.4614
413.1140441894531 0 653.7461
413.1827087402344 0 816.5099
416.20330810546875 0 10007.162
416.705078125 0 3666.4443
417.2027893066406 0 1898.4811
417.6952819824219 0 820.9887
418.2347412109375 0 1939.0616
419.2025451660156 0 1336.7804 y Water loss 4
420.18817138671875 0 2406.6982 y Ammonia loss 4
421.18939208984375 0 682.7003
425.2084045410156 0 15039.469
425.7093505859375 0 11623.092
426.16644287109375 0 988.97327
426.206787109375 0 5237.552
426.705078125 0 1975.544
427.2914123535156 0 1094.7053
429.2444152832031 0 665.2784
433.716552734375 0 2563.9119
434.2157897949219 0 2167.979
434.71429443359375 0 24608.213
435.2154235839844 0 9690.087
435.71661376953125 0 2527.0054
436.2193908691406 0 10447.391
436.72021484375 0 1080.9794
437.2135314941406 0 35126.21 y 4
438.216064453125 0 7172.0366
439.6911315917969 0 1704.623
439.8436584472656 0 1102.4901
440.1876220703125 0 1462.3317
440.26861572265625 0 3563.22
447.22308349609375 0 12226.945
447.7237548828125 0 8935.379
448.20318603515625 0 16209.868 Precursor Water loss
448.2271423339844 0 3651.6948
448.7037658691406 0 8058.6865
449.20306396484375 0 2614.3943
449.2298278808594 0 811.7591
449.2744140625 0 1351.6263
449.72039794921875 0 851.2076
452.14404296875 0 1484.5172
453.2454528808594 0 12533.97
454.1617736816406 0 927.21564
455.15643310546875 0 2055.719
456.1575622558594 0 827.615
456.22735595703125 0 2627.434
456.72802734375 0 7135.8726
457.2090759277344 0 7814.3994 Precursor
457.7094421386719 0 3125.28
458.2325134277344 0 5360.7007
459.1905212402344 0 854.6976 b Water loss 3
463.21844482421875 0 750.0981
465.20147705078125 0 1860.463
471.25531005859375 0 1846.1147
477.2010803222656 0 1089.9652 b 3
480.138671875 0 14354.32
481.1405334472656 0 2729.884
482.1341857910156 0 6872.1406
483.1354064941406 0 1268.748
485.165771484375 0 949.5885
486.16375732421875 0 977.71216
487.16046142578125 0 733.1765
497.201904296875 0 1360.0033
510.22003173828125 0 2830.578
511.2220153808594 0 1180.7764
512.17919921875 0 2949.5186
513.1799926757812 0 958.2687
514.2305908203125 0 5665.558
515.232421875 0 975.6225
517.204833984375 0 1189.9491
528.2294921875 0 2084.8599
534.2304077148438 0 2245.0386 y Water loss 3
535.2152099609375 0 2080.007 y Ammonia loss 3
539.2496337890625 0 876.7838
552.2407836914062 0 68463.22 y 3
553.2437744140625 0 18077.736
554.2459106445312 0 3075.3477
562.2242431640625 0 2975.2341
563.227294921875 0 1220.8279
566.3291015625 0 16700.754
574.2166137695312 0 4097.902 b Water loss 4
575.21826171875 0 1039.7191
584.3392944335938 0 2072.7205
586.294189453125 0 640.0257
592.227783203125 0 2594.5664 b 4
598.1553955078125 0 611.3804
599.244873046875 0 1037.72
607.251953125 0 661.1291
608.2346801757812 0 1522.8445
621.8744506835938 0 563.7168
622.2850341796875 0 785.6513
624.2677001953125 0 860.9483
625.2623901367188 0 11485.2295
626.264892578125 0 3526.9202
647.3140258789062 0 995.96967 y Water loss 2
648.30029296875 0 1700.0104 y Ammonia loss 2
650.2788696289062 0 1538.5266
651.27978515625 0 1236.5304
652.2784423828125 0 819.9617
665.3247680664062 0 76828.76 y 2
665.3961791992188 0 16462.945
666.3270874023438 0 25099.955
666.3937377929688 0 1238.024
667.329833984375 0 5052.536
675.308349609375 0 1831.3717
676.3109741210938 0 733.3345
683.4075317382812 0 2860.063
691.302978515625 0 2251.996
692.3081665039062 0 840.85034
698.3165893554688 0 1036.4414
722.3462524414062 0 1313.5327
723.2965698242188 0 8526.517
724.2998657226562 0 3221.012
725.2930908203125 0 5026.493
726.2925415039062 0 1974.4314
732.3253173828125 0 1221.2655
733.3323974609375 0 797.58673
740.3247680664062 0 1366.1954
741.3265380859375 0 1004.0475
748.3607788085938 0 3698.3018 y Water loss 1
749.3626098632812 0 1466.3695
750.355224609375 0 632.0318
755.33203125 0 1091.9242
758.3424682617188 0 1003.98517
766.3716430664062 0 22795.867 y 1
766.4451904296875 0 4260.38
767.3740234375 0 7083.334
768.375244140625 0 2526.044
776.3562622070312 0 30133.756
777.359130859375 0 10641.084
778.3619995117188 0 3260.7825
784.4556274414062 0 2016.6671
794.4383544921875 0 1078.8533
849.4494018554688 0 5645.9976
850.4533081054688 0 1048.571
877.5101928710938 0 2536.2942
1152.443603515625 0 609.7231
3012.87744140625 0 686.54144

Spectrum Details

|  |  |
| --- | --- |
| Matched peaks? Matched peaks The total absolute number of peaks matched. Additionally in brackets the total fraction of peaks matched and the total number of peaks is shown. | 27 (6.51% of 415) |
| FDR? FDR The false discovery rate estimated for this peptide. It is calculated by matching all theoretical fragments with a non-integer shift with the raw peaks for this spectrum. This is done with 40 different shifts. The resulting percentage is the average number of annotated peaks over the number of annotated peaks with the correct spectrum. | 0.88% |
| Satellite FDR? Satellite FDR See the FDR for details on its calculation. This satellite ion specific FDR only contains the satellite ions (d/w) for I/L/J positions. | - |
| PSM Score? PSM Score The PSM Score as given by Hecklib to this annotated spectrum. It is shown with three significant figures. | 381 |

#### Spectrum 4667? Spectrum 4667 The raw spectrum of this peptide as annotated by Hecklib. The fragments are coloured according to ion type (see legend). Any peaks with a star '\*' as text can be hovered over to see the full details, first the ion type second the mass shift type. By hovering over the amino acids in the peptide or ions in the legend the corresponding peaks are highlighted. By toggling the 'Unassigned' label you can turn the background (unassigned) peaks on or off in the plot. By updating the slider in the Ion legend you can update the spectrum to only show the top X% of the peaks with labels. The top X% means any peak that is within X% of the highest intensity. By dragging in the spectrum you can zoom in to a specific part of the spectrum and use 'Zoom Out' to get back to the original zoom level. The annotation of the spectrum is based on the given sequence in the peptides file and is done with different software so inconsistencies are likely. The peaks are annotated based on the given sequence, with 20 ppm tolerance.

Copy Data

##### Spectrum 4667 (TSV)

###### Preview

```
Loading example...
```

*Click on the button to copy the data to your clipboard.*

Mz MinMz MaxIntensity Max

WidthHeightPeptide font sizePeptide stroke widthSpectrum font sizeSpectrum stroke widthCompact peptide

Ion legend

wxyz

abcd

OtherUnassignedIon^Charge^_Position_Show for top:%

MTJDDFR

06.22e+41.24e+51.87e+52.49e+5

Zoom Out

y^+1^_1_y^+1^_1_b^+1^_2_y^+1^_2_a^+1^_3_y^+1^_2_y^+2^_5_a^+1^_3_b^+1^_3_b^+1^_3_y^+2^_6_y^+1^_3_y^+1^_3_y^+1^_3_\*\*\*b^+1^_4_b^+1^_4_y^+1^_4_y^+1^_4_y^+1^_4_b^+1^_5_b^+1^_5_y^+1^_5_y^+1^_5_y^+1^_5_y^+1^_6_y^+1^_6_

0524104915732097

Fragment Matches Table

Show background peaks

| Position | Ion type | Intensity | mz Theoretical | mz Error (Th) | mz Error (ppm) | Charge | Series Number |
| --- | --- | --- | --- | --- | --- | --- | --- |
| - | - | 3.984E+04 | 120.1 | - | - | 0 | - |
| - | - | 476.3 | 121 | - | - | 0 | - |
| - | - | 2994 | 121.1 | - | - | 0 | - |
| - | - | 2154 | 122.1 | - | - | 0 | - |
| - | - | 766.5 | 123.1 | - | - | 0 | - |
| - | - | 799.6 | 124 | - | - | 0 | - |
| - | - | 2094 | 124.1 | - | - | 0 | - |
| - | - | 448 | 126.1 | - | - | 0 | - |
| - | - | 2233 | 126.1 | - | - | 0 | - |
| - | - | 355 | 128.1 | - | - | 0 | - |
| - | - | 1.267E+04 | 129.1 | - | - | 0 | - |
| - | - | 611.4 | 129.1 | - | - | 0 | - |
| - | - | 1318 | 130 | - | - | 0 | - |
| - | - | 589.8 | 130.1 | - | - | 0 | - |
| - | - | 1708 | 130.1 | - | - | 0 | - |
| - | - | 395.2 | 131.2 | - | - | 0 | - |
| - | - | 1448 | 133.1 | - | - | 0 | - |
| - | - | 1652 | 136.1 | - | - | 0 | - |
| - | - | 1128 | 139.1 | - | - | 0 | - |
| - | - | 4523 | 139.1 | - | - | 0 | - |
| - | - | 4579 | 140.1 | - | - | 0 | - |
| - | - | 719 | 140.1 | - | - | 0 | - |
| - | - | 2556 | 141.1 | - | - | 0 | - |
| - | - | 865.7 | 141.1 | - | - | 0 | - |
| - | - | 7938 | 141.1 | - | - | 0 | - |
| - | - | 695.2 | 142.1 | - | - | 0 | - |
| - | - | 621.6 | 142.1 | - | - | 0 | - |
| - | - | 896.6 | 143.1 | - | - | 0 | - |
| - | - | 667.1 | 144.1 | - | - | 0 | - |
| - | - | 3937 | 148 | - | - | 0 | - |
| - | - | 468.4 | 149 | - | - | 0 | - |
| - | - | 538.2 | 150.1 | - | - | 0 | - |
| - | - | 538.1 | 150.1 | - | - | 0 | - |
| - | - | 1360 | 152.1 | - | - | 0 | - |
| - | - | 399 | 153.4 | - | - | 0 | - |
| - | - | 1605 | 157.1 | - | - | 0 | - |
| - | - | 2.257E+04 | 157.1 | - | - | 0 | - |
| - | - | 2282 | 157.1 | - | - | 0 | - |
| 7 | y | 2.01E+04 | 158.1 | 9.571E-05 | 0.6054 | +1 | 1 |
| - | - | 1460 | 158.1 | - | - | 0 | - |
| - | - | 624.5 | 159.1 | - | - | 0 | - |
| - | - | 824.1 | 159.1 | - | - | 0 | - |
| - | - | 1122 | 165.1 | - | - | 0 | - |
| - | - | 4058 | 166.1 | - | - | 0 | - |
| - | - | 511.5 | 167.1 | - | - | 0 | - |
| - | - | 3350 | 167.1 | - | - | 0 | - |
| - | - | 2.292E+04 | 169.1 | - | - | 0 | - |
| - | - | 436.4 | 170.1 | - | - | 0 | - |
| - | - | 2117 | 170.1 | - | - | 0 | - |
| - | - | 6385 | 173.4 | - | - | 0 | - |
| - | - | 1378 | 175.1 | - | - | 0 | - |
| 7 | y | 5.143E+04 | 175.1 | 0.0001122 | 0.6405 | +1 | 1 |
| - | - | 2779 | 176.1 | - | - | 0 | - |
| - | - | 1332 | 177.1 | - | - | 0 | - |
| - | - | 810.4 | 179.1 | - | - | 0 | - |
| - | - | 2194 | 180.1 | - | - | 0 | - |
| - | - | 4847 | 181.1 | - | - | 0 | - |
| - | - | 1468 | 183.1 | - | - | 0 | - |
| - | - | 2120 | 185.1 | - | - | 0 | - |
| - | - | 8997 | 185.1 | - | - | 0 | - |
| - | - | 1385 | 185.1 | - | - | 0 | - |
| - | - | 8892 | 187.1 | - | - | 0 | - |
| - | - | 511.4 | 188 | - | - | 0 | - |
| - | - | 491.7 | 189.1 | - | - | 0 | - |
| - | - | 1.785E+04 | 190.1 | - | - | 0 | - |
| - | - | 1219 | 191.1 | - | - | 0 | - |
| - | - | 577.1 | 191.3 | - | - | 0 | - |
| - | - | 1158 | 196.1 | - | - | 0 | - |
| - | - | 3.901E+04 | 197.1 | - | - | 0 | - |
| - | - | 4119 | 198.1 | - | - | 0 | - |
| - | - | 805.2 | 199.1 | - | - | 0 | - |
| - | - | 2419 | 201.1 | - | - | 0 | - |
| - | - | 2149 | 203.1 | - | - | 0 | - |
| - | - | 4872 | 203.1 | - | - | 0 | - |
| - | - | 1975 | 204.1 | - | - | 0 | - |
| - | - | 3001 | 205.1 | - | - | 0 | - |
| - | - | 1138 | 207.1 | - | - | 0 | - |
| - | - | 910.6 | 208.1 | - | - | 0 | - |
| - | - | 6251 | 209.1 | - | - | 0 | - |
| - | - | 1009 | 209.1 | - | - | 0 | - |
| - | - | 856.5 | 214.1 | - | - | 0 | - |
| - | - | 2861 | 215.1 | - | - | 0 | - |
| - | - | 1164 | 217.1 | - | - | 0 | - |
| - | - | 2.561E+04 | 221.1 | - | - | 0 | - |
| - | - | 2466 | 222.1 | - | - | 0 | - |
| - | - | 1119 | 223.1 | - | - | 0 | - |
| - | - | 897.2 | 223.1 | - | - | 0 | - |
| - | - | 6593 | 224.1 | - | - | 0 | - |
| - | - | 1.648E+04 | 225.1 | - | - | 0 | - |
| - | - | 888.7 | 225.1 | - | - | 0 | - |
| - | - | 1526 | 226.1 | - | - | 0 | - |
| - | - | 815.4 | 226.1 | - | - | 0 | - |
| - | - | 609.3 | 227.1 | - | - | 0 | - |
| - | - | 3763 | 229.1 | - | - | 0 | - |
| - | - | 777.7 | 229.1 | - | - | 0 | - |
| - | - | 4006 | 231.1 | - | - | 0 | - |
| - | - | 5011 | 231.1 | - | - | 0 | - |
| - | - | 850.5 | 231.1 | - | - | 0 | - |
| - | - | 937.3 | 232.1 | - | - | 0 | - |
| - | - | 739.7 | 232.1 | - | - | 0 | - |
| - | - | 1048 | 233.1 | - | - | 0 | - |
| - | - | 1.186E+04 | 235.1 | - | - | 0 | - |
| - | - | 657.8 | 235.1 | - | - | 0 | - |
| - | - | 1198 | 236.1 | - | - | 0 | - |
| - | - | 1092 | 236.1 | - | - | 0 | - |
| - | - | 1130 | 242.1 | - | - | 0 | - |
| - | - | 769.7 | 245.1 | - | - | 0 | - |
| - | - | 1423 | 245.1 | - | - | 0 | - |
| - | - | 666.1 | 246.1 | - | - | 0 | - |
| 2 | b | 3.489E+04 | 249.1 | 0.004938 | 19.83 | +1 | 2 |
| - | - | 768.3 | 249.2 | - | - | 0 | - |
| - | - | 2556 | 250.1 | - | - | 0 | - |
| - | - | 1101 | 251.1 | - | - | 0 | - |
| - | - | 6636 | 251.1 | - | - | 0 | - |
| - | - | 4455 | 252.1 | - | - | 0 | - |
| - | - | 1394 | 252.2 | - | - | 0 | - |
| - | - | 1948 | 253.1 | - | - | 0 | - |
| - | - | 2987 | 254.1 | - | - | 0 | - |
| - | - | 1188 | 260.1 | - | - | 0 | - |
| - | - | 3614 | 262.2 | - | - | 0 | - |
| - | - | 1881 | 263.1 | - | - | 0 | - |
| - | - | 3170 | 263.1 | - | - | 0 | - |
| - | - | 689.8 | 270.1 | - | - | 0 | - |
| - | - | 1022 | 270.2 | - | - | 0 | - |
| - | - | 682.4 | 279.1 | - | - | 0 | - |
| - | - | 3828 | 280.2 | - | - | 0 | - |
| - | - | 4438 | 287.2 | - | - | 0 | - |
| - | - | 1903 | 288.1 | - | - | 0 | - |
| - | - | 1088 | 288.2 | - | - | 0 | - |
| - | - | 876.1 | 294.1 | - | - | 0 | - |
| - | - | 732.8 | 297.2 | - | - | 0 | - |
| - | - | 1911 | 298.2 | - | - | 0 | - |
| - | - | 632.8 | 304.1 | - | - | 0 | - |
| - | - | 2615 | 304.2 | - | - | 0 | - |
| 6 | y | 2.293E+04 | 305.2 | 8.09E-05 | 0.2651 | +1 | 2 |
| - | - | 3964 | 306.2 | - | - | 0 | - |
| - | - | 826.3 | 309.2 | - | - | 0 | - |
| - | - | 2295 | 312.2 | - | - | 0 | - |
| - | - | 952.7 | 314.1 | - | - | 0 | - |
| - | - | 607.8 | 315.1 | - | - | 0 | - |
| 3 | a | 4793 | 316.2 | 0.004891 | 15.47 | +1 | 3 |
| - | - | 3899 | 318.1 | - | - | 0 | - |
| - | - | 5700 | 322.1 | - | - | 0 | - |
| 6 | y | 2.465E+05 | 322.2 | 0.0001034 | 0.3209 | +1 | 2 |
| - | - | 4.118E+04 | 323.2 | - | - | 0 | - |
| - | - | 3690 | 324.2 | - | - | 0 | - |
| - | - | 760.6 | 328.2 | - | - | 0 | - |
| - | - | 756.5 | 332.1 | - | - | 0 | - |
| - | - | 1753 | 332.2 | - | - | 0 | - |
| 3 | y | 7542 | 333.2 | 0.0002076 | 0.6232 | +2 | 5 |
| - | - | 2946 | 333.7 | - | - | 0 | - |
| 3 | a | 898.7 | 334.2 | 0.002378 | 7.117 | +1 | 3 |
| - | - | 1169 | 338.2 | - | - | 0 | - |
| - | - | 672.9 | 338.7 | - | - | 0 | - |
| - | - | 4203 | 340.2 | - | - | 0 | - |
| - | - | 622.3 | 341.2 | - | - | 0 | - |
| - | - | 4610 | 344.1 | - | - | 0 | - |
| 3 | b | 3928 | 344.2 | 0.005429 | 15.77 | +1 | 3 |
| - | - | 1715 | 346.2 | - | - | 0 | - |
| - | - | 993.4 | 346.7 | - | - | 0 | - |
| - | - | 768.7 | 349.2 | - | - | 0 | - |
| - | - | 822.3 | 351.7 | - | - | 0 | - |
| - | - | 637.2 | 360.1 | - | - | 0 | - |
| - | - | 2527 | 360.7 | - | - | 0 | - |
| - | - | 1799 | 361.2 | - | - | 0 | - |
| 3 | b | 6755 | 362.2 | 0.004599 | 12.7 | +1 | 3 |
| - | - | 1623 | 363.2 | - | - | 0 | - |
| - | - | 1316 | 365.7 | - | - | 0 | - |
| - | - | 777.1 | 366.2 | - | - | 0 | - |
| - | - | 2590 | 366.7 | - | - | 0 | - |
| - | - | 2824 | 367.2 | - | - | 0 | - |
| - | - | 653.5 | 367.7 | - | - | 0 | - |
| - | - | 1942 | 368.1 | - | - | 0 | - |
| - | - | 1775 | 371.7 | - | - | 0 | - |
| 2 | y | 1.136E+04 | 374.7 | 0.0004355 | 1.162 | +2 | 6 |
| - | - | 5632 | 375.2 | - | - | 0 | - |
| - | - | 777.4 | 375.7 | - | - | 0 | - |
| - | - | 659.6 | 377.2 | - | - | 0 | - |
| - | - | 1010 | 379.7 | - | - | 0 | - |
| - | - | 1484 | 380.2 | - | - | 0 | - |
| - | - | 1898 | 384.7 | - | - | 0 | - |
| - | - | 736.2 | 385.2 | - | - | 0 | - |
| - | - | 839.3 | 385.7 | - | - | 0 | - |
| - | - | 4216 | 388.7 | - | - | 0 | - |
| - | - | 1756 | 389.2 | - | - | 0 | - |
| - | - | 1131 | 393.7 | - | - | 0 | - |
| - | - | 1105 | 395.2 | - | - | 0 | - |
| - | - | 1.174E+04 | 402.2 | - | - | 0 | - |
| - | - | 4695 | 402.7 | - | - | 0 | - |
| - | - | 669.7 | 403.1 | - | - | 0 | - |
| - | - | 849.2 | 403.2 | - | - | 0 | - |
| - | - | 1.34E+04 | 416.2 | - | - | 0 | - |
| - | - | 5889 | 416.7 | - | - | 0 | - |
| - | - | 1121 | 417.2 | - | - | 0 | - |
| 5 | y | 1308 | 419.2 | 0.0005889 | 1.405 | +1 | 3 |
| 5 | y | 4007 | 420.2 | 0.000199 | 0.4735 | +1 | 3 |
| - | - | 844.5 | 421.2 | - | - | 0 | - |
| - | - | 2.296E+04 | 425.2 | - | - | 0 | - |
| - | - | 7623 | 425.7 | - | - | 0 | - |
| - | - | 2364 | 426.2 | - | - | 0 | - |
| - | - | 4449 | 426.7 | - | - | 0 | - |
| - | - | 4368 | 427.2 | - | - | 0 | - |
| - | - | 773.7 | 427.7 | - | - | 0 | - |
| 5 | y | 4.661E+04 | 437.2 | 0.0001673 | 0.3826 | +1 | 3 |
| - | - | 9807 | 438.2 | - | - | 0 | - |
| - | - | 668 | 439.2 | - | - | 0 | - |
| - | - | 1405 | 439.2 | - | - | 0 | - |
| - | - | 2757 | 439.7 | - | - | 0 | - |
| - | - | 814.4 | 439.8 | - | - | 0 | - |
| - | - | 2025 | 440.2 | - | - | 0 | - |
| - | - | 1424 | 445.2 | - | - | 0 | - |
| 0 | Precursor | 3.695E+04 | 448.2 | 0.002206 | 4.921 | +2 | -1 |
| 0 | Precursor | 1.891E+04 | 448.7 | 0.008092 | 18.03 | +2 | -1 |
| - | - | 9760 | 449.2 | - | - | 0 | - |
| - | - | 1390 | 449.7 | - | - | 0 | - |
| - | - | 4227 | 455.2 | - | - | 0 | - |
| - | - | 888.4 | 456.2 | - | - | 0 | - |
| - | - | 432.3 | 457.2 | - | - | 0 | - |
| 0 | Precursor | 1.199E+04 | 457.2 | 0.002538 | 5.552 | +2 | -1 |
| - | - | 6062 | 457.7 | - | - | 0 | - |
| - | - | 2197 | 458.2 | - | - | 0 | - |
| - | - | 1071 | 458.3 | - | - | 0 | - |
| 4 | b | 1602 | 459.2 | 0.005799 | 12.63 | +1 | 4 |
| - | - | 1317 | 463.2 | - | - | 0 | - |
| 4 | b | 1302 | 477.2 | 0.003474 | 7.28 | +1 | 4 |
| - | - | 850.3 | 478.2 | - | - | 0 | - |
| - | - | 1520 | 482.2 | - | - | 0 | - |
| - | - | 4154 | 510.2 | - | - | 0 | - |
| - | - | 678.5 | 511.2 | - | - | 0 | - |
| - | - | 881.1 | 516.2 | - | - | 0 | - |
| - | - | 1250 | 517.2 | - | - | 0 | - |
| - | - | 3622 | 528.2 | - | - | 0 | - |
| - | - | 1141 | 529.2 | - | - | 0 | - |
| - | - | 2967 | 531.2 | - | - | 0 | - |
| 4 | y | 5196 | 534.2 | 0.000768 | 1.438 | +1 | 4 |
| 4 | y | 3103 | 535.2 | 0.0009952 | 1.859 | +1 | 4 |
| 4 | y | 1.051E+05 | 552.2 | 0.0005905 | 1.069 | +1 | 4 |
| - | - | 2.806E+04 | 553.2 | - | - | 0 | - |
| - | - | 5333 | 554.2 | - | - | 0 | - |
| - | - | 602.5 | 557.2 | - | - | 0 | - |
| - | - | 4878 | 562.2 | - | - | 0 | - |
| - | - | 1010 | 563.2 | - | - | 0 | - |
| - | - | 663.4 | 566.2 | - | - | 0 | - |
| 5 | b | 6690 | 574.2 | 0.003789 | 6.598 | +1 | 5 |
| - | - | 2338 | 575.2 | - | - | 0 | - |
| - | - | 1029 | 576.2 | - | - | 0 | - |
| 5 | b | 5544 | 592.2 | 0.003722 | 6.285 | +1 | 5 |
| - | - | 1495 | 593.2 | - | - | 0 | - |
| - | - | 569.6 | 594.2 | - | - | 0 | - |
| - | - | 761.3 | 630.3 | - | - | 0 | - |
| - | - | 787.2 | 631.3 | - | - | 0 | - |
| 3 | y | 1350 | 647.3 | 0.000312 | 0.482 | +1 | 5 |
| 3 | y | 1632 | 648.3 | 0.0007324 | 1.13 | +1 | 5 |
| - | - | 1151 | 649.3 | - | - | 0 | - |
| - | - | 907.8 | 654.4 | - | - | 0 | - |
| 3 | y | 9.513E+04 | 665.3 | 0.001036 | 1.558 | +1 | 5 |
| - | - | 3.169E+04 | 666.3 | - | - | 0 | - |
| - | - | 7012 | 667.3 | - | - | 0 | - |
| - | - | 2901 | 675.3 | - | - | 0 | - |
| - | - | 994.1 | 676.3 | - | - | 0 | - |
| - | - | 2481 | 691.3 | - | - | 0 | - |
| - | - | 670.7 | 692.3 | - | - | 0 | - |
| - | - | 1482 | 698.3 | - | - | 0 | - |
| - | - | 604.7 | 700.7 | - | - | 0 | - |
| - | - | 1304 | 722.3 | - | - | 0 | - |
| - | - | 2075 | 732.3 | - | - | 0 | - |
| - | - | 793.6 | 733.3 | - | - | 0 | - |
| 2 | y | 5346 | 748.4 | 0.003177 | 4.245 | +1 | 6 |
| - | - | 3006 | 749.4 | - | - | 0 | - |
| - | - | 594.7 | 750.4 | - | - | 0 | - |
| - | - | 1166 | 758.3 | - | - | 0 | - |
| - | - | 660.9 | 759.4 | - | - | 0 | - |
| - | - | 718.4 | 766.2 | - | - | 0 | - |
| 2 | y | 3.231E+04 | 766.4 | 0.001657 | 2.162 | +1 | 6 |
| - | - | 1.165E+04 | 767.4 | - | - | 0 | - |
| - | - | 3336 | 768.4 | - | - | 0 | - |
| - | - | 4.132E+04 | 776.4 | - | - | 0 | - |
| - | - | 1.708E+04 | 777.4 | - | - | 0 | - |
| - | - | 5030 | 778.4 | - | - | 0 | - |
| - | - | 620.5 | 1127 | - | - | 0 | - |
| - | - | 635.9 | 1339 | - | - | 0 | - |
| - | - | 666.2 | 1683 | - | - | 0 | - |
| - | - | 613.3 | 1785 | - | - | 0 | - |
| - | - | 592.4 | 1835 | - | - | 0 | - |
| - | - | 744.1 | 1923 | - | - | 0 | - |
| - | - | 717.4 | 2077 | - | - | 0 | - |

m/z Charge Intensity FragmentType MassShift Position
120.08097076416016 0 39839.977
121.03195190429688 0 476.25885
121.08440399169922 0 2993.6355
122.0602035522461 0 2153.8013
123.0552978515625 0 766.465
124.0394515991211 0 799.60583
124.11221313476562 0 2094.4817
126.10324096679688 0 448.03714
126.12787628173828 0 2232.8914
128.07000732421875 0 355.04486
129.0660400390625 0 12666.905
129.10247802734375 0 611.3983
130.03236389160156 0 1317.5745
130.05027770996094 0 589.84216
130.09771728515625 0 1707.6676
131.17127990722656 0 395.22577
133.09735107421875 0 1448.3324
136.0757598876953 0 1652.0226
139.05032348632812 0 1127.5739
139.0867919921875 0 4523.11
140.0707550048828 0 4579.065
140.0816192626953 0 718.9527
141.0659637451172 0 2556.2175
141.1022491455078 0 865.6991
141.13876342773438 0 7938.0845
142.12255859375 0 695.17285
142.1422576904297 0 621.6084
143.11846923828125 0 896.5767
144.10137939453125 0 667.0751
148.0428009033203 0 3937.2832
149.04684448242188 0 468.4266
150.0667724609375 0 538.2298
150.09164428710938 0 538.1055
152.10723876953125 0 1359.9305
153.3931427001953 0 398.96387
157.0609130859375 0 1605.0336
157.09725952148438 0 22566.26
157.1086883544922 0 2282.3418
158.09249877929688 0 20101.768 y Ammonia loss 6
158.1002960205078 0 1459.69
159.07664489746094 0 624.5013
159.09622192382812 0 824.1273
165.10240173339844 0 1122.3765
166.05323791503906 0 4057.6506
167.05665588378906 0 511.45956
167.08157348632812 0 3349.7812
169.13368225097656 0 22921.887
170.11622619628906 0 436.41754
170.1370849609375 0 2117.0781
173.4405517578125 0 6385.452
175.0867919921875 0 1378.3346
175.1190643310547 0 51433.996 y 6
176.12254333496094 0 2779.146
177.06961059570312 0 1332.4463
179.1180877685547 0 810.37225
180.1133270263672 0 2194.3872
181.09719848632812 0 4846.863
183.11280822753906 0 1468.459
185.05584716796875 0 2120.453
185.09217834472656 0 8997.382
185.1029815673828 0 1385.2264
187.14422607421875 0 8892.388
188.03697204589844 0 511.41714
189.10214233398438 0 491.73727
190.0975799560547 0 17845.18
191.10110473632812 0 1219.2087
191.27249145507812 0 577.13525
196.14492797851562 0 1157.5441
197.12850952148438 0 39013.895
198.13177490234375 0 4118.6855
199.1080780029297 0 805.197
201.12319946289062 0 2418.6138
203.0665283203125 0 2149.0498
203.0849151611328 0 4872.0776
204.0690155029297 0 1974.6552
205.06423950195312 0 3000.8794
207.11288452148438 0 1137.831
208.14385986328125 0 910.5696
209.09207153320312 0 6250.6846
209.12841796875 0 1009.213
214.05307006835938 0 856.45325
215.13890075683594 0 2860.8843
217.0972900390625 0 1163.9193
221.09544372558594 0 25607.03
222.09902954101562 0 2465.6648
223.090576171875 0 1119.0103
223.14430236816406 0 897.2308
224.13943481445312 0 6592.7437
225.12339782714844 0 16477.951
225.142578125 0 888.72906
226.11795043945312 0 1526.3033
226.1284637451172 0 815.4256
227.13926696777344 0 609.29785
229.11819458007812 0 3762.817
229.12973022460938 0 777.67786
231.06117248535156 0 4006.4333
231.0798797607422 0 5011.301
231.13290405273438 0 850.50714
232.06419372558594 0 937.3196
232.0834503173828 0 739.74805
233.12847900390625 0 1047.9597
235.1077117919922 0 11857.66
235.14364624023438 0 657.841
236.11056518554688 0 1198.1154
236.13868713378906 0 1091.8391
242.14956665039062 0 1130.3901
245.09274291992188 0 769.65125
245.12847900390625 0 1422.7493
246.07540893554688 0 666.10443
249.0903778076172 0 34888.438 b 1
249.15943908691406 0 768.29065
250.09361267089844 0 2555.505
251.0854034423828 0 1101.289
251.1389617919922 0 6636.314
252.1342010498047 0 4455.0923
252.17112731933594 0 1393.5377
253.11814880371094 0 1948.2391
254.14991760253906 0 2986.502
260.1392822265625 0 1188.1886
262.1547546386719 0 3614.0813
263.1025085449219 0 1881.1483
263.1387634277344 0 3169.5515
270.1442565917969 0 689.795
270.1814270019531 0 1022.0121
279.1343994140625 0 682.4308
280.1654968261719 0 3828.3757
287.1502380371094 0 4438.3994
288.1337585449219 0 1902.5243
288.1524658203125 0 1087.767
294.1450500488281 0 876.13654
297.2017517089844 0 732.8117
298.1758117675781 0 1911.4294
304.14422607421875 0 632.8131
304.177001953125 0 2615.3037
305.1607360839844 0 22930.316 y Ammonia loss 5
306.1641540527344 0 3964.159
309.2041320800781 0 826.31934
312.1564025878906 0 2294.5542
314.1139831542969 0 952.7019
315.0959777832031 0 607.83105
316.1689147949219 0 4792.62 a Water loss 2
318.148193359375 0 3899.4434
322.14105224609375 0 5699.6245
322.1874694824219 0 246523.05 y 5
323.1904296875 0 41184.99
324.1930847167969 0 3689.732
328.1950988769531 0 760.6037
332.123291015625 0 756.4758
332.17205810546875 0 1753.127
333.16650390625 0 7542.1743 y 2
333.66729736328125 0 2945.674
334.1722106933594 0 898.67236 a 2
338.1593322753906 0 1168.9576
338.6583251953125 0 672.8742
340.1501159667969 0 4202.8154
341.1536865234375 0 622.2758
344.1445007324219 0 4609.532
344.16436767578125 0 3928.4404 b Water loss 2
346.1736755371094 0 1715.4021
346.6761169433594 0 993.39636
349.1514892578125 0 768.7018
351.6803283691406 0 822.32837
360.11932373046875 0 637.20416
360.6871337890625 0 2527.1091
361.18865966796875 0 1798.7739
362.1741027832031 0 6754.5293 b 2
363.1770935058594 0 1622.9485
365.67889404296875 0 1316.2888
366.1801452636719 0 777.0954
366.6684265136719 0 2590.0952
367.161865234375 0 2824.308
367.6634216308594 0 653.51184
368.14495849609375 0 1941.682
371.6615295410156 0 1775.4138
374.6844177246094 0 11363.644 y Water loss 1
375.1858825683594 0 5632.204
375.6865234375 0 777.4401
377.193603515625 0 659.59845
379.6773376464844 0 1010.0486
380.1749267578125 0 1484.1321
384.6692199707031 0 1897.9578
385.1714172363281 0 736.15906
385.67620849609375 0 839.33276
388.681396484375 0 4216
389.183837890625 0 1756.2275
393.6768493652344 0 1131.0018
395.1922302246094 0 1105.2607
402.1866455078125 0 11740.432
402.68841552734375 0 4695.1035
403.0928039550781 0 669.655
403.18890380859375 0 849.2438
416.2032775878906 0 13397.345
416.70428466796875 0 5888.7505
417.2047424316406 0 1121.0864
419.2031555175781 0 1308.1423 y Water loss 4
420.18756103515625 0 4006.6929 y Ammonia loss 4
421.19183349609375 0 844.457
425.20849609375 0 22956.002
425.7096252441406 0 7623.26
426.212158203125 0 2363.6921
426.6814270019531 0 4448.565
427.1828918457031 0 4367.993
427.68060302734375 0 773.7462
437.2141418457031 0 46608.875 y 4
438.21685791015625 0 9807.243
439.19244384765625 0 668.0409
439.2217712402344 0 1404.7577
439.6913757324219 0 2756.6746
439.8434753417969 0 814.351
440.192626953125 0 2025.1893
445.1922302246094 0 1423.7472
448.2023010253906 0 36953.844 Precursor Water loss
448.7001953125 0 18908.713 Precursor Ammonia loss
449.2012634277344 0 9759.563
449.6998596191406 0 1389.5935
455.17657470703125 0 4226.6855
456.1799011230469 0 888.359
457.17449951171875 0 432.3014
457.2079162597656 0 11988.071 Precursor
457.709228515625 0 6061.797
458.2092590332031 0 2197.4456
458.29449462890625 0 1070.8301
459.1916809082031 0 1602.0447 b Water loss 3
463.21746826171875 0 1316.5312
477.1999206542969 0 1301.561 b 3
478.2035217285156 0 850.34406
482.1876525878906 0 1520.3806
510.21978759765625 0 4154.3594
511.2225036621094 0 678.53796
516.2208251953125 0 881.0504
517.2006225585938 0 1250.4642
528.2288818359375 0 3622.3513
529.231689453125 0 1140.8434
531.1749877929688 0 2967.2812
534.2299194335938 0 5196.2085 y Water loss 3
535.2156982421875 0 3102.979 y Ammonia loss 3
552.2406616210938 0 105085.96 y 3
553.2435302734375 0 28062.434
554.2444458007812 0 5332.94
557.1968383789062 0 602.53436
562.224609375 0 4877.705
563.2271118164062 0 1010.01056
566.22265625 0 663.37524
574.2166137695312 0 6689.9023 b Water loss 4
575.2138061523438 0 2337.6077
576.2109985351562 0 1029.341
592.2271118164062 0 5544.198 b 4
593.2286987304688 0 1495.3965
594.230712890625 0 569.62494
630.2889404296875 0 761.32855
631.271484375 0 787.15814
647.3150634765625 0 1349.9441 y Water loss 2
648.2980346679688 0 1632.2665 y Ammonia loss 2
649.3019409179688 0 1150.8795
654.3842163085938 0 907.8398
665.3242797851562 0 95126.65 y 2
666.3270263671875 0 31692.562
667.32958984375 0 7012.1763
675.306640625 0 2901.1855
676.3124389648438 0 994.0603
691.3023071289062 0 2481.0059
692.3065185546875 0 670.6625
698.2778930664062 0 1482.2241
700.6790771484375 0 604.6787
722.3435668945312 0 1303.7129
732.330322265625 0 2075.2498
733.343994140625 0 793.6029
748.3592529296875 0 5345.7603 y Water loss 1
749.3614501953125 0 3006.1897
750.363037109375 0 594.6862
758.3436889648438 0 1165.5164
759.3505249023438 0 660.8718
766.2489624023438 0 718.36884
766.371337890625 0 32312.414 y 1
767.3743896484375 0 11648.957
768.3777465820312 0 3336.0107
776.3555908203125 0 41324.105
777.3585815429688 0 17078.834
778.3607788085938 0 5029.8086
1126.6474609375 0 620.4717
1338.946533203125 0 635.89886
1682.573486328125 0 666.21674
1785.451416015625 0 613.2816
1835.432861328125 0 592.42975
1923.0875244140625 0 744.13873
2076.71923828125 0 717.3637

Spectrum Details

|  |  |
| --- | --- |
| Matched peaks? Matched peaks The total absolute number of peaks matched. Additionally in brackets the total fraction of peaks matched and the total number of peaks is shown. | 29 (10.14% of 286) |
| FDR? FDR The false discovery rate estimated for this peptide. It is calculated by matching all theoretical fragments with a non-integer shift with the raw peaks for this spectrum. This is done with 40 different shifts. The resulting percentage is the average number of annotated peaks over the number of annotated peaks with the correct spectrum. | 0.08% |
| Satellite FDR? Satellite FDR See the FDR for details on its calculation. This satellite ion specific FDR only contains the satellite ions (d/w) for I/L/J positions. | - |
| PSM Score? PSM Score The PSM Score as given by Hecklib to this annotated spectrum. It is shown with three significant figures. | 405 |

#### Reverse Lookup? Reverse Lookup All places where this read could be placed.

| Group | Segment | Template | Template Part | Read Part | Score | Unique |
| --- | --- | --- | --- | --- | --- | --- |
| Homo sapiens Heavy Chain | IGHC | IGHD | [307..314] | [0..7] | 29 | False |
| Decoy | Decoy | ASPN | [138..144] | [0..7] | 29 | False |

| Recombined | Template Part | Read Part | Score | Unique |
| --- | --- | --- | --- | --- |
| ASPN | [138..144] | [0..7] | 29 | True |

#### Meta Information from Multiple reads

##### Number of combined reads

2

##### Intensity

0.48

##### TotalArea

3.148E+06

##### Changes to the peptide sequence

MTJDDFR

L→JNo support for either Leucine or Isoleucine based on side chain ions (Position: 3)

#### Positional Score

Copy Data

##### Positional Score (TSV)

###### Preview

```
Loading example...
```

*Click on the button to copy the data to your clipboard.*

000123456

Label Value
"0" 0
"1" 0
"2" 0
"3" 0
"4" 0
"5" 0
"6" 0

#### Meta Information from PEAKS

##### Scan Identifier

F4:4722

##### Original sequence

M

+15.99

T

L

D

D

F

R

##### Posttranslational Modifications

Oxidation (M)

##### Source File

D:\separate\_stitch\_analyses\xle-disambiguation\raw\20210323\_F1\_UM1\_Peng0013\_SA\_F59\_ingel\_3ug\_tryp.raw

##### Fraction

4

##### Scan Feature

F4:2296

##### De Novo Score

97

##### ConfidenceScore

98

### m/z

457.2081

##### Mass

912.4011

##### Charge

2

##### Retention Time

25.26

##### Predicted Retention Time

26.83

##### Area

1.432E+06

##### Parts Per Million

0.6

##### Fragmentation mode

HCD

##### Originating file

01 D:\separate\_stitch\_analyses\xle-disambiguation\20210325\_F59\_3ug\_DENOVO\_12.csv

#### Meta Information from PEAKS

##### Scan Identifier

F4:4667

##### Original sequence

M

+15.99

T

L

D

D

F

R

##### Posttranslational Modifications

Oxidation (M)

##### Source File

D:\separate\_stitch\_analyses\xle-disambiguation\raw\20210323\_F1\_UM1\_Peng0013\_SA\_F59\_ingel\_3ug\_tryp.raw

##### Fraction

4

##### Scan Feature

F4:2295

##### De Novo Score

97

##### ConfidenceScore

99

### m/z

457.208

##### Mass

912.4011

##### Charge

2

##### Retention Time

25.05

##### Predicted Retention Time

26.83

##### Area

1.716E+06

##### Parts Per Million

0.4

##### Fragmentation mode

HCD

##### Originating file

01 D:\separate\_stitch\_analyses\xle-disambiguation\20210325\_F59\_3ug\_DENOVO\_12.csv
