## Supplementary material for "A handle on mass coincidence errors in *de novo* sequencing of antibodies by bottom-up proteomics": Combined_094.html

Details Combined\_094 | Stitch OverviewUndefined

### Read Combined\_094

#### Sequence (length=11)

VRQAPGRAJEW

#### Spectrum 5723? Spectrum 5723 The raw spectrum of this peptide as annotated by Hecklib. The fragments are coloured according to ion type (see legend). Any peaks with a star '\*' as text can be hovered over to see the full details, first the ion type second the mass shift type. By hovering over the amino acids in the peptide or ions in the legend the corresponding peaks are highlighted. By toggling the 'Unassigned' label you can turn the background (unassigned) peaks on or off in the plot. By updating the slider in the Ion legend you can update the spectrum to only show the top X% of the peaks with labels. The top X% means any peak that is within X% of the highest intensity. By dragging in the spectrum you can zoom in to a specific part of the spectrum and use 'Zoom Out' to get back to the original zoom level. The annotation of the spectrum is based on the given sequence in the peptides file and is done with different software so inconsistencies are likely. The peaks are annotated based on the given sequence, with 20 ppm tolerance.

Copy Data

##### Spectrum 5723 (TSV)

###### Preview

```
Loading example...
```

*Click on the button to copy the data to your clipboard.*

Mz MinMz MaxIntensity Max

WidthHeightPeptide font sizePeptide stroke widthSpectrum font sizeSpectrum stroke widthCompact peptide

Ion legend

wxyz

abcd

OtherUnassignedIon^Charge^_Position_Show for top:%

VRQAPGRAJEW

04.34e+58.68e+51.30e+61.74e+6

Zoom Out

z^+1^_1_c^+2^_3_y^+1^_1_z^+2^_3_c^+2^_4_c^+1^_2_c^+1^_2_c^+2^_5_c^+3^_8_c^+2^_6_y^+1^_2_c^+3^_9_y^+1^_2_c^+3^_10_c^+2^_7_z^+3^_10_c^+1^_3_c^+2^_7_c^+1^_3_y^+2^_7_y^+2^_7_c^+2^_8_c^+2^_8_c^+1^_4_c^+2^_9_c^+2^_9_z^+1^_4_c^+2^_10_c^+2^_10_c^+2^_10_c^+1^_5_y^+2^_10_z^+2^_10_y^+2^_10_c^+1^_6_c^+1^_6_y^+1^_5_z^+1^_5_y^+1^_5_y^+1^_6_z^+1^_6_y^+1^_6_c^+1^_7_w^+1^_7_y^+1^_7_z^+1^_7_y^+1^_7_c^+1^_8_y^+1^_8_z^+1^_8_y^+1^_8_w^+1^_9_c^+1^_9_z^+1^_9_y^+1^_9_c^+1^_10_w^+1^_10_c^+1^_10_y^+1^_10_z^+1^_10_y^+1^_10_

0831166124923323

Fragment Matches Table

Show background peaks

| Position | Ion type | Intensity | mz Theoretical | mz Error (Th) | mz Error (ppm) | Charge | Series Number |
| --- | --- | --- | --- | --- | --- | --- | --- |
| - | - | 398.6 | 120.7 | - | - | 0 | - |
| - | - | 769.6 | 125.1 | - | - | 0 | - |
| - | - | 534.1 | 125.9 | - | - | 0 | - |
| - | - | 522.5 | 129.1 | - | - | 0 | - |
| - | - | 1840 | 130.1 | - | - | 0 | - |
| - | - | 5.748E+04 | 130.1 | - | - | 0 | - |
| - | - | 4915 | 131.1 | - | - | 0 | - |
| - | - | 1.011E+04 | 132.1 | - | - | 0 | - |
| - | - | 1509 | 133.1 | - | - | 0 | - |
| - | - | 605.3 | 134.1 | - | - | 0 | - |
| - | - | 416.7 | 134.7 | - | - | 0 | - |
| - | - | 473.3 | 135 | - | - | 0 | - |
| - | - | 414.2 | 140.1 | - | - | 0 | - |
| - | - | 3399 | 142.1 | - | - | 0 | - |
| - | - | 1160 | 143.1 | - | - | 0 | - |
| - | - | 4098 | 144.1 | - | - | 0 | - |
| - | - | 698.9 | 145.1 | - | - | 0 | - |
| - | - | 2.582E+04 | 146.1 | - | - | 0 | - |
| - | - | 2358 | 147.1 | - | - | 0 | - |
| - | - | 406.6 | 147.4 | - | - | 0 | - |
| - | - | 1494 | 153.1 | - | - | 0 | - |
| - | - | 792.8 | 157.1 | - | - | 0 | - |
| - | - | 4996 | 157.1 | - | - | 0 | - |
| - | - | 1029 | 157.1 | - | - | 0 | - |
| - | - | 457.4 | 157.6 | - | - | 0 | - |
| - | - | 1.734E+05 | 159.1 | - | - | 0 | - |
| - | - | 1218 | 160.1 | - | - | 0 | - |
| - | - | 1.94E+04 | 160.1 | - | - | 0 | - |
| - | - | 1120 | 161.1 | - | - | 0 | - |
| - | - | 509.1 | 162.1 | - | - | 0 | - |
| - | - | 1542 | 169.1 | - | - | 0 | - |
| - | - | 708.5 | 169.1 | - | - | 0 | - |
| - | - | 5340 | 170.1 | - | - | 0 | - |
| - | - | 560.6 | 171.1 | - | - | 0 | - |
| - | - | 797.6 | 171.1 | - | - | 0 | - |
| - | - | 611.3 | 171.1 | - | - | 0 | - |
| - | - | 969.6 | 173.4 | - | - | 0 | - |
| - | - | 1474 | 174.1 | - | - | 0 | - |
| - | - | 2395 | 174.1 | - | - | 0 | - |
| - | - | 745.5 | 175.1 | - | - | 0 | - |
| - | - | 489.8 | 175.4 | - | - | 0 | - |
| - | - | 5126 | 178.6 | - | - | 0 | - |
| - | - | 505.7 | 179.1 | - | - | 0 | - |
| - | - | 3830 | 185.1 | - | - | 0 | - |
| - | - | 2177 | 185.1 | - | - | 0 | - |
| - | - | 592.7 | 185.2 | - | - | 0 | - |
| - | - | 1447 | 187.1 | - | - | 0 | - |
| - | - | 2.961E+04 | 187.1 | - | - | 0 | - |
| - | - | 5.72E+05 | 188.1 | - | - | 0 | - |
| 11 | z | 6.71E+04 | 189.1 | 0.003738 | 19.77 | +1 | 1 |
| - | - | 1317 | 190.1 | - | - | 0 | - |
| - | - | 4519 | 190.1 | - | - | 0 | - |
| - | - | 467.1 | 191 | - | - | 0 | - |
| 3 | c | 1.712E+04 | 192.6 | 0.0007731 | 4.014 | +2 | 3 |
| - | - | 3007 | 193.1 | - | - | 0 | - |
| - | - | 1373 | 195.2 | - | - | 0 | - |
| - | - | 1064 | 196.1 | - | - | 0 | - |
| - | - | 1715 | 197.1 | - | - | 0 | - |
| - | - | 1442 | 197.1 | - | - | 0 | - |
| - | - | 536.3 | 199.1 | - | - | 0 | - |
| - | - | 823.7 | 200.1 | - | - | 0 | - |
| - | - | 870.2 | 203.1 | - | - | 0 | - |
| - | - | 493.3 | 203.1 | - | - | 0 | - |
| - | - | 528.3 | 203.9 | - | - | 0 | - |
| 11 | y | 3.546E+05 | 205.1 | 0.0007768 | 3.788 | +1 | 1 |
| - | - | 4013 | 205.6 | - | - | 0 | - |
| - | - | 4.222E+04 | 206.1 | - | - | 0 | - |
| - | - | 1081 | 206.1 | - | - | 0 | - |
| 9 | z | 3152 | 207.1 | 0.002906 | 14.03 | +2 | 3 |
| - | - | 6305 | 209 | - | - | 0 | - |
| - | - | 921 | 213.2 | - | - | 0 | - |
| - | - | 6895 | 213.2 | - | - | 0 | - |
| - | - | 5.981E+04 | 214.1 | - | - | 0 | - |
| - | - | 4856 | 214.2 | - | - | 0 | - |
| - | - | 1.251E+04 | 214.6 | - | - | 0 | - |
| - | - | 885.6 | 215.1 | - | - | 0 | - |
| - | - | 697 | 215.2 | - | - | 0 | - |
| - | - | 1394 | 219.6 | - | - | 0 | - |
| 4 | c | 1.849E+04 | 228.1 | 0.000893 | 3.914 | +2 | 4 |
| - | - | 3031 | 228.2 | - | - | 0 | - |
| - | - | 4658 | 228.6 | - | - | 0 | - |
| - | - | 506 | 228.9 | - | - | 0 | - |
| - | - | 1414 | 233.2 | - | - | 0 | - |
| - | - | 3682 | 234.2 | - | - | 0 | - |
| - | - | 917.4 | 234.7 | - | - | 0 | - |
| - | - | 1549 | 238.2 | - | - | 0 | - |
| - | - | 3.69E+04 | 239.2 | - | - | 0 | - |
| - | - | 3957 | 240.2 | - | - | 0 | - |
| - | - | 6464 | 242.2 | - | - | 0 | - |
| - | - | 3116 | 243.1 | - | - | 0 | - |
| - | - | 574.1 | 243.1 | - | - | 0 | - |
| - | - | 1481 | 243.2 | - | - | 0 | - |
| - | - | 791.6 | 244.1 | - | - | 0 | - |
| - | - | 1276 | 248.2 | - | - | 0 | - |
| - | - | 2134 | 256.1 | - | - | 0 | - |
| 2 | c | 2.801E+04 | 256.2 | 0.000933 | 3.642 | +1 | 2 |
| - | - | 2888 | 257.2 | - | - | 0 | - |
| - | - | 561.1 | 258.8 | - | - | 0 | - |
| - | - | 494.6 | 261.2 | - | - | 0 | - |
| - | - | 3042 | 262.7 | - | - | 0 | - |
| - | - | 543.2 | 262.7 | - | - | 0 | - |
| - | - | 673.6 | 263.2 | - | - | 0 | - |
| - | - | 537.5 | 263.7 | - | - | 0 | - |
| - | - | 497.6 | 268.9 | - | - | 0 | - |
| - | - | 2.109E+04 | 270.1 | - | - | 0 | - |
| - | - | 465.9 | 271.1 | - | - | 0 | - |
| - | - | 3102 | 271.1 | - | - | 0 | - |
| 2 | c | 5.417E+04 | 273.2 | 0.0009647 | 3.531 | +1 | 2 |
| - | - | 6080 | 274.2 | - | - | 0 | - |
| 5 | c | 5334 | 276.7 | 0.001183 | 4.278 | +2 | 5 |
| - | - | 1780 | 277.2 | - | - | 0 | - |
| 8 | c | 1245 | 279.5 | 0.0006251 | 2.236 | +3 | 8 |
| - | - | 4952 | 280.1 | - | - | 0 | - |
| - | - | 1073 | 281.1 | - | - | 0 | - |
| - | - | 759 | 291.7 | - | - | 0 | - |
| - | - | 1385 | 294.2 | - | - | 0 | - |
| - | - | 1.197E+04 | 298.1 | - | - | 0 | - |
| - | - | 1698 | 298.2 | - | - | 0 | - |
| - | - | 4718 | 298.7 | - | - | 0 | - |
| - | - | 2353 | 299.1 | - | - | 0 | - |
| - | - | 1481 | 299.2 | - | - | 0 | - |
| - | - | 1.706E+04 | 299.2 | - | - | 0 | - |
| - | - | 2371 | 300.2 | - | - | 0 | - |
| - | - | 627.8 | 302.2 | - | - | 0 | - |
| 6 | c | 1231 | 305.2 | 0.000736 | 2.412 | +2 | 6 |
| - | - | 5914 | 307.9 | - | - | 0 | - |
| - | - | 4154 | 308.2 | - | - | 0 | - |
| - | - | 880.1 | 308.5 | - | - | 0 | - |
| - | - | 1345 | 311.2 | - | - | 0 | - |
| - | - | 1110 | 312.2 | - | - | 0 | - |
| - | - | 1.017E+04 | 312.7 | - | - | 0 | - |
| - | - | 792.8 | 312.7 | - | - | 0 | - |
| - | - | 1478 | 313.2 | - | - | 0 | - |
| - | - | 3588 | 313.2 | - | - | 0 | - |
| - | - | 630.7 | 313.7 | - | - | 0 | - |
| - | - | 1463 | 314.2 | - | - | 0 | - |
| - | - | 837.5 | 314.2 | - | - | 0 | - |
| 10 | y | 8.162E+04 | 316.1 | 0.001341 | 4.243 | +1 | 2 |
| - | - | 1.516E+04 | 317.1 | - | - | 0 | - |
| 9 | c | 2440 | 317.2 | 0.001097 | 3.458 | +3 | 9 |
| - | - | 2268 | 318.1 | - | - | 0 | - |
| - | - | 5358 | 318.9 | - | - | 0 | - |
| 10 | y | 7907 | 334.1 | 0.001397 | 4.18 | +1 | 2 |
| - | - | 1461 | 335.1 | - | - | 0 | - |
| - | - | 708.2 | 339.2 | - | - | 0 | - |
| - | - | 1774 | 341 | - | - | 0 | - |
| - | - | 2153 | 341.2 | - | - | 0 | - |
| - | - | 738 | 342.2 | - | - | 0 | - |
| - | - | 1461 | 347.7 | - | - | 0 | - |
| - | - | 712.8 | 350.2 | - | - | 0 | - |
| - | - | 1812 | 350.9 | - | - | 0 | - |
| - | - | 752.8 | 351.2 | - | - | 0 | - |
| - | - | 5781 | 356.2 | - | - | 0 | - |
| - | - | 2975 | 357.2 | - | - | 0 | - |
| - | - | 5811 | 359 | - | - | 0 | - |
| 10 | c | 1686 | 360.2 | 0.0008459 | 2.348 | +3 | 10 |
| - | - | 739.2 | 360.5 | - | - | 0 | - |
| - | - | 787.7 | 361.2 | - | - | 0 | - |
| - | - | 5235 | 361.7 | - | - | 0 | - |
| - | - | 2913 | 362.2 | - | - | 0 | - |
| - | - | 600.3 | 362.7 | - | - | 0 | - |
| - | - | 639.4 | 366.2 | - | - | 0 | - |
| - | - | 1142 | 367.2 | - | - | 0 | - |
| - | - | 5030 | 367.2 | - | - | 0 | - |
| - | - | 1588 | 368.2 | - | - | 0 | - |
| - | - | 4652 | 369.2 | - | - | 0 | - |
| - | - | 1.989E+04 | 369.7 | - | - | 0 | - |
| - | - | 7930 | 370.2 | - | - | 0 | - |
| - | - | 2244 | 370.3 | - | - | 0 | - |
| - | - | 1903 | 370.7 | - | - | 0 | - |
| - | - | 766.9 | 371.3 | - | - | 0 | - |
| - | - | 2971 | 374.7 | - | - | 0 | - |
| - | - | 802.4 | 375.2 | - | - | 0 | - |
| - | - | 684.2 | 375.7 | - | - | 0 | - |
| - | - | 1062 | 382.2 | - | - | 0 | - |
| - | - | 1413 | 382.7 | - | - | 0 | - |
| 7 | c | 3.055E+04 | 383.2 | 0.001358 | 3.545 | +2 | 7 |
| - | - | 1.286E+04 | 383.7 | - | - | 0 | - |
| 2 | z | 6208 | 384.2 | 0.001 | 2.604 | +3 | 10 |
| 3 | c | 6.26E+04 | 384.2 | 0.001163 | 3.026 | +1 | 3 |
| - | - | 854.2 | 384.7 | - | - | 0 | - |
| - | - | 1400 | 385.2 | - | - | 0 | - |
| - | - | 1.288E+04 | 385.2 | - | - | 0 | - |
| - | - | 1233 | 385.3 | - | - | 0 | - |
| - | - | 1682 | 386.2 | - | - | 0 | - |
| - | - | 1019 | 388.7 | - | - | 0 | - |
| - | - | 952.6 | 390.7 | - | - | 0 | - |
| - | - | 9.379E+04 | 391.2 | - | - | 0 | - |
| 7 | c | 2.943E+05 | 391.7 | 0.001206 | 3.08 | +2 | 7 |
| - | - | 1.221E+05 | 392.2 | - | - | 0 | - |
| - | - | 2.77E+04 | 392.7 | - | - | 0 | - |
| - | - | 2925 | 393.2 | - | - | 0 | - |
| - | - | 1344 | 396.2 | - | - | 0 | - |
| - | - | 2668 | 396.7 | - | - | 0 | - |
| - | - | 1153 | 397.2 | - | - | 0 | - |
| - | - | 1045 | 397.7 | - | - | 0 | - |
| - | - | 606.8 | 400.9 | - | - | 0 | - |
| 3 | c | 1.708E+05 | 401.3 | 0.001561 | 3.89 | +1 | 3 |
| - | - | 1557 | 401.6 | - | - | 0 | - |
| - | - | 1506 | 401.9 | - | - | 0 | - |
| - | - | 3.281E+04 | 402.3 | - | - | 0 | - |
| - | - | 4367 | 403.3 | - | - | 0 | - |
| - | - | 1.532E+04 | 404.8 | - | - | 0 | - |
| - | - | 7677 | 405.3 | - | - | 0 | - |
| 5 | y | 2.119E+04 | 405.7 | 0.001591 | 3.921 | +2 | 7 |
| - | - | 2256 | 405.8 | - | - | 0 | - |
| - | - | 1.071E+04 | 406.2 | - | - | 0 | - |
| - | - | 2956 | 406.7 | - | - | 0 | - |
| - | - | 4.878E+04 | 406.9 | - | - | 0 | - |
| - | - | 3.408E+04 | 407.2 | - | - | 0 | - |
| - | - | 1.436E+04 | 407.6 | - | - | 0 | - |
| - | - | 2048 | 407.9 | - | - | 0 | - |
| - | - | 1.049E+04 | 410.2 | - | - | 0 | - |
| - | - | 4536 | 410.7 | - | - | 0 | - |
| - | - | 1430 | 411.2 | - | - | 0 | - |
| - | - | 595.6 | 412.2 | - | - | 0 | - |
| - | - | 4.773E+04 | 412.9 | - | - | 0 | - |
| - | - | 3.457E+04 | 413.2 | - | - | 0 | - |
| - | - | 2016 | 413.3 | - | - | 0 | - |
| - | - | 1.132E+04 | 413.6 | - | - | 0 | - |
| - | - | 3261 | 413.9 | - | - | 0 | - |
| 5 | y | 2.554E+04 | 414.7 | 0.001588 | 3.829 | +2 | 7 |
| - | - | 1.19E+04 | 415.2 | - | - | 0 | - |
| - | - | 3501 | 415.7 | - | - | 0 | - |
| - | - | 7158 | 416.2 | - | - | 0 | - |
| - | - | 3669 | 416.6 | - | - | 0 | - |
| - | - | 890.3 | 416.9 | - | - | 0 | - |
| - | - | 897.9 | 417.7 | - | - | 0 | - |
| - | - | 6420 | 418.3 | - | - | 0 | - |
| 8 | c | 4.754E+04 | 418.7 | 0.001997 | 4.769 | +2 | 8 |
| - | - | 2.102E+04 | 419.2 | - | - | 0 | - |
| - | - | 6497 | 419.8 | - | - | 0 | - |
| - | - | 1160 | 420.3 | - | - | 0 | - |
| - | - | 1.488E+04 | 422.2 | - | - | 0 | - |
| - | - | 1.124E+04 | 422.6 | - | - | 0 | - |
| - | - | 5699 | 422.9 | - | - | 0 | - |
| - | - | 978.3 | 423.2 | - | - | 0 | - |
| - | - | 2037 | 426.3 | - | - | 0 | - |
| - | - | 3.497E+04 | 426.8 | - | - | 0 | - |
| 8 | c | 2.428E+05 | 427.3 | 0.001723 | 4.033 | +2 | 8 |
| - | - | 773.9 | 427.6 | - | - | 0 | - |
| - | - | 1.124E+05 | 427.8 | - | - | 0 | - |
| - | - | 2.171E+04 | 428.3 | - | - | 0 | - |
| - | - | 1518 | 428.6 | - | - | 0 | - |
| - | - | 6191 | 428.8 | - | - | 0 | - |
| - | - | 2767 | 428.9 | - | - | 0 | - |
| - | - | 5196 | 429.1 | - | - | 0 | - |
| - | - | 731.3 | 429.2 | - | - | 0 | - |
| - | - | 1563 | 429.3 | - | - | 0 | - |
| - | - | 3998 | 438.2 | - | - | 0 | - |
| - | - | 617.9 | 439.3 | - | - | 0 | - |
| - | - | 678.5 | 439.8 | - | - | 0 | - |
| - | - | 2348 | 440.3 | - | - | 0 | - |
| - | - | 788.7 | 440.8 | - | - | 0 | - |
| - | - | 4006 | 452.8 | - | - | 0 | - |
| - | - | 1674 | 453.3 | - | - | 0 | - |
| - | - | 1948 | 453.8 | - | - | 0 | - |
| - | - | 719.7 | 454.3 | - | - | 0 | - |
| 4 | c | 6.957E+04 | 455.3 | 0.00186 | 4.086 | +1 | 4 |
| - | - | 917.4 | 455.8 | - | - | 0 | - |
| - | - | 1.57E+04 | 456.3 | - | - | 0 | - |
| - | - | 2074 | 457.3 | - | - | 0 | - |
| - | - | 1.145E+05 | 461.3 | - | - | 0 | - |
| - | - | 6.089E+04 | 461.8 | - | - | 0 | - |
| - | - | 1.874E+04 | 462.3 | - | - | 0 | - |
| - | - | 6232 | 462.8 | - | - | 0 | - |
| - | - | 6831 | 466.8 | - | - | 0 | - |
| - | - | 2864 | 467.3 | - | - | 0 | - |
| - | - | 2863 | 467.3 | - | - | 0 | - |
| - | - | 1573 | 467.8 | - | - | 0 | - |
| - | - | 846.4 | 469.3 | - | - | 0 | - |
| - | - | 785.6 | 470.3 | - | - | 0 | - |
| - | - | 1346 | 472.3 | - | - | 0 | - |
| - | - | 1175 | 473.3 | - | - | 0 | - |
| - | - | 1.125E+04 | 474.8 | - | - | 0 | - |
| 9 | c | 3.027E+05 | 475.3 | 0.00211 | 4.439 | +2 | 9 |
| - | - | 1.596E+05 | 475.8 | - | - | 0 | - |
| - | - | 4.762E+04 | 476.3 | - | - | 0 | - |
| - | - | 9511 | 476.8 | - | - | 0 | - |
| - | - | 1893 | 481.3 | - | - | 0 | - |
| - | - | 1376 | 481.8 | - | - | 0 | - |
| - | - | 1931 | 482.3 | - | - | 0 | - |
| - | - | 996.6 | 482.8 | - | - | 0 | - |
| - | - | 9458 | 483.3 | - | - | 0 | - |
| 9 | c | 1.208E+05 | 483.8 | 0.001836 | 3.795 | +2 | 9 |
| - | - | 6.326E+04 | 484.3 | - | - | 0 | - |
| - | - | 1.945E+04 | 484.8 | - | - | 0 | - |
| - | - | 4385 | 485.3 | - | - | 0 | - |
| - | - | 1452 | 486.3 | - | - | 0 | - |
| - | - | 1246 | 489.8 | - | - | 0 | - |
| - | - | 1062 | 490.3 | - | - | 0 | - |
| - | - | 1180 | 490.8 | - | - | 0 | - |
| - | - | 4310 | 495.3 | - | - | 0 | - |
| - | - | 988.3 | 496.3 | - | - | 0 | - |
| - | - | 6.835E+04 | 496.8 | - | - | 0 | - |
| - | - | 4.068E+04 | 497.3 | - | - | 0 | - |
| - | - | 1.259E+04 | 497.8 | - | - | 0 | - |
| - | - | 1.284E+04 | 498.3 | - | - | 0 | - |
| - | - | 3008 | 498.3 | - | - | 0 | - |
| - | - | 9022 | 498.8 | - | - | 0 | - |
| - | - | 2029 | 499.3 | - | - | 0 | - |
| - | - | 4209 | 499.3 | - | - | 0 | - |
| - | - | 803.8 | 500.3 | - | - | 0 | - |
| 8 | z | 789.1 | 502.2 | 0.000166 | 0.3305 | +1 | 4 |
| - | - | 886 | 503.8 | - | - | 0 | - |
| - | - | 877.2 | 504.3 | - | - | 0 | - |
| - | - | 5177 | 504.8 | - | - | 0 | - |
| - | - | 2506 | 505.3 | - | - | 0 | - |
| - | - | 1389 | 505.8 | - | - | 0 | - |
| - | - | 928.7 | 508.8 | - | - | 0 | - |
| - | - | 1149 | 509.3 | - | - | 0 | - |
| - | - | 1739 | 510.3 | - | - | 0 | - |
| - | - | 2798 | 511.3 | - | - | 0 | - |
| - | - | 3013 | 512.8 | - | - | 0 | - |
| - | - | 1465 | 513.3 | - | - | 0 | - |
| - | - | 1016 | 516.8 | - | - | 0 | - |
| - | - | 1602 | 517.3 | - | - | 0 | - |
| - | - | 1.4E+04 | 517.8 | - | - | 0 | - |
| - | - | 939 | 518.3 | - | - | 0 | - |
| - | - | 8017 | 518.3 | - | - | 0 | - |
| - | - | 5255 | 518.8 | - | - | 0 | - |
| - | - | 1241 | 519.3 | - | - | 0 | - |
| - | - | 907.7 | 519.8 | - | - | 0 | - |
| - | - | 864.3 | 521.3 | - | - | 0 | - |
| - | - | 2333 | 522.3 | - | - | 0 | - |
| - | - | 2545 | 522.8 | - | - | 0 | - |
| - | - | 784 | 523.3 | - | - | 0 | - |
| - | - | 2769 | 524.3 | - | - | 0 | - |
| - | - | 1668 | 524.8 | - | - | 0 | - |
| - | - | 4630 | 525.3 | - | - | 0 | - |
| - | - | 2.99E+05 | 525.8 | - | - | 0 | - |
| - | - | 1.87E+05 | 526.3 | - | - | 0 | - |
| - | - | 5.874E+04 | 526.8 | - | - | 0 | - |
| - | - | 1.383E+04 | 527.3 | - | - | 0 | - |
| - | - | 4050 | 527.8 | - | - | 0 | - |
| - | - | 1463 | 528.3 | - | - | 0 | - |
| - | - | 2037 | 529.3 | - | - | 0 | - |
| - | - | 7243 | 530.8 | - | - | 0 | - |
| - | - | 1.647E+04 | 531.3 | - | - | 0 | - |
| - | - | 7426 | 531.8 | - | - | 0 | - |
| - | - | 4842 | 532.3 | - | - | 0 | - |
| - | - | 2728 | 532.8 | - | - | 0 | - |
| - | - | 1208 | 533.3 | - | - | 0 | - |
| - | - | 1051 | 533.8 | - | - | 0 | - |
| - | - | 7797 | 538.8 | - | - | 0 | - |
| 10 | c | 1.107E+04 | 539.3 | 0.001758 | 3.259 | +2 | 10 |
| 10 | c | 1.719E+06 | 539.8 | 0.002328 | 4.313 | +2 | 10 |
| - | - | 9.904E+05 | 540.3 | - | - | 0 | - |
| - | - | 3.366E+05 | 540.8 | - | - | 0 | - |
| - | - | 8.187E+04 | 541.3 | - | - | 0 | - |
| - | - | 1.139E+04 | 541.8 | - | - | 0 | - |
| - | - | 6146 | 542.3 | - | - | 0 | - |
| - | - | 1727 | 543.3 | - | - | 0 | - |
| - | - | 655 | 546.8 | - | - | 0 | - |
| - | - | 7999 | 547.3 | - | - | 0 | - |
| - | - | 7982 | 547.8 | - | - | 0 | - |
| 10 | c | 6.001E+05 | 548.3 | 0.001749 | 3.19 | +2 | 10 |
| - | - | 3.575E+05 | 548.8 | - | - | 0 | - |
| - | - | 1.14E+05 | 549.3 | - | - | 0 | - |
| - | - | 2.811E+04 | 549.8 | - | - | 0 | - |
| - | - | 5661 | 550.3 | - | - | 0 | - |
| - | - | 756.8 | 553.8 | - | - | 0 | - |
| - | - | 1712 | 554.3 | - | - | 0 | - |
| - | - | 1647 | 554.8 | - | - | 0 | - |
| - | - | 2.633E+04 | 555.3 | - | - | 0 | - |
| - | - | 2700 | 555.3 | - | - | 0 | - |
| - | - | 1.919E+04 | 555.8 | - | - | 0 | - |
| - | - | 5620 | 556.3 | - | - | 0 | - |
| - | - | 614 | 556.8 | - | - | 0 | - |
| - | - | 896.2 | 556.8 | - | - | 0 | - |
| - | - | 1103 | 558.3 | - | - | 0 | - |
| - | - | 2457 | 559.3 | - | - | 0 | - |
| - | - | 2447 | 559.8 | - | - | 0 | - |
| - | - | 911.1 | 560.3 | - | - | 0 | - |
| - | - | 2.088E+04 | 561.3 | - | - | 0 | - |
| - | - | 1.697E+04 | 561.8 | - | - | 0 | - |
| - | - | 5282 | 562.3 | - | - | 0 | - |
| - | - | 2491 | 562.8 | - | - | 0 | - |
| - | - | 1416 | 562.8 | - | - | 0 | - |
| - | - | 2592 | 563.3 | - | - | 0 | - |
| - | - | 758.8 | 563.8 | - | - | 0 | - |
| - | - | 2.082E+04 | 568.3 | - | - | 0 | - |
| - | - | 1.902E+04 | 568.8 | - | - | 0 | - |
| 5 | c | 6.118E+04 | 569.4 | 0.002015 | 3.539 | +1 | 5 |
| - | - | 1864 | 569.8 | - | - | 0 | - |
| - | - | 1041 | 570.3 | - | - | 0 | - |
| - | - | 1.698E+04 | 570.4 | - | - | 0 | - |
| - | - | 970.5 | 571.3 | - | - | 0 | - |
| - | - | 2986 | 571.4 | - | - | 0 | - |
| - | - | 658 | 574.3 | - | - | 0 | - |
| - | - | 6565 | 575.8 | - | - | 0 | - |
| - | - | 8350 | 576.3 | - | - | 0 | - |
| - | - | 3.973E+04 | 576.8 | - | - | 0 | - |
| - | - | 7.673E+04 | 577.3 | - | - | 0 | - |
| - | - | 4.257E+04 | 577.8 | - | - | 0 | - |
| - | - | 1.517E+04 | 578.3 | - | - | 0 | - |
| - | - | 3646 | 578.8 | - | - | 0 | - |
| - | - | 7011 | 582.4 | - | - | 0 | - |
| - | - | 4699 | 582.8 | - | - | 0 | - |
| 2 | y | 3065 | 583.3 | 0.009818 | 16.83 | +2 | 10 |
| - | - | 4120 | 583.8 | - | - | 0 | - |
| 2 | z | 2.967E+04 | 584.3 | 0.002301 | 3.938 | +2 | 10 |
| - | - | 4387 | 584.4 | - | - | 0 | - |
| - | - | 2.006E+04 | 584.8 | - | - | 0 | - |
| - | - | 5813 | 585.3 | - | - | 0 | - |
| - | - | 2261 | 585.8 | - | - | 0 | - |
| - | - | 1531 | 586.3 | - | - | 0 | - |
| - | - | 792.4 | 589.8 | - | - | 0 | - |
| - | - | 2340 | 590.3 | - | - | 0 | - |
| - | - | 2226 | 590.9 | - | - | 0 | - |
| - | - | 1632 | 591.3 | - | - | 0 | - |
| - | - | 1616 | 591.4 | - | - | 0 | - |
| - | - | 1589 | 591.8 | - | - | 0 | - |
| 2 | y | 5079 | 592.3 | 0.002155 | 3.639 | +2 | 10 |
| - | - | 3490 | 592.8 | - | - | 0 | - |
| - | - | 1590 | 593.3 | - | - | 0 | - |
| - | - | 8810 | 596.4 | - | - | 0 | - |
| - | - | 2298 | 597.4 | - | - | 0 | - |
| - | - | 4605 | 597.8 | - | - | 0 | - |
| - | - | 8048 | 598.3 | - | - | 0 | - |
| - | - | 6902 | 598.8 | - | - | 0 | - |
| - | - | 5168 | 599.3 | - | - | 0 | - |
| - | - | 1925 | 600.3 | - | - | 0 | - |
| - | - | 4721 | 602.3 | - | - | 0 | - |
| - | - | 2002 | 603.3 | - | - | 0 | - |
| - | - | 919 | 603.3 | - | - | 0 | - |
| - | - | 1263 | 603.8 | - | - | 0 | - |
| - | - | 1.177E+04 | 604.3 | - | - | 0 | - |
| - | - | 2.663E+04 | 604.8 | - | - | 0 | - |
| - | - | 1.836E+04 | 605.3 | - | - | 0 | - |
| - | - | 9081 | 605.8 | - | - | 0 | - |
| - | - | 4293 | 606.3 | - | - | 0 | - |
| - | - | 4228 | 606.3 | - | - | 0 | - |
| - | - | 4920 | 606.8 | - | - | 0 | - |
| - | - | 1594 | 607.3 | - | - | 0 | - |
| 6 | c | 2044 | 609.3 | 0.000997 | 1.636 | +1 | 6 |
| - | - | 1498 | 610.8 | - | - | 0 | - |
| - | - | 8916 | 611.4 | - | - | 0 | - |
| - | - | 4.718E+04 | 611.9 | - | - | 0 | - |
| - | - | 3.328E+04 | 612.4 | - | - | 0 | - |
| - | - | 1903 | 612.8 | - | - | 0 | - |
| - | - | 1.258E+04 | 612.9 | - | - | 0 | - |
| - | - | 8778 | 613.4 | - | - | 0 | - |
| - | - | 2681 | 613.8 | - | - | 0 | - |
| - | - | 1.703E+04 | 614.4 | - | - | 0 | - |
| - | - | 2142 | 614.8 | - | - | 0 | - |
| - | - | 9519 | 615.3 | - | - | 0 | - |
| - | - | 6415 | 615.4 | - | - | 0 | - |
| - | - | 3162 | 616.3 | - | - | 0 | - |
| - | - | 1303 | 616.4 | - | - | 0 | - |
| - | - | 871 | 617.3 | - | - | 0 | - |
| - | - | 1893 | 617.8 | - | - | 0 | - |
| - | - | 1.011E+04 | 619.4 | - | - | 0 | - |
| - | - | 3.611E+04 | 619.9 | - | - | 0 | - |
| - | - | 4.453E+04 | 620.4 | - | - | 0 | - |
| - | - | 8.899E+04 | 620.8 | - | - | 0 | - |
| - | - | 6.611E+04 | 621.3 | - | - | 0 | - |
| - | - | 2.398E+04 | 621.8 | - | - | 0 | - |
| - | - | 7329 | 622.3 | - | - | 0 | - |
| - | - | 1513 | 622.8 | - | - | 0 | - |
| - | - | 2.725E+04 | 624.3 | - | - | 0 | - |
| - | - | 1.266E+04 | 625.4 | - | - | 0 | - |
| - | - | 6162 | 625.8 | - | - | 0 | - |
| 6 | c | 2.27E+05 | 626.4 | 0.002524 | 4.03 | +1 | 6 |
| - | - | 3.188E+04 | 626.8 | - | - | 0 | - |
| - | - | 7.333E+04 | 627.4 | - | - | 0 | - |
| - | - | 3840 | 627.8 | - | - | 0 | - |
| - | - | 1.501E+04 | 628.4 | - | - | 0 | - |
| - | - | 2641 | 629.4 | - | - | 0 | - |
| - | - | 3634 | 632.8 | - | - | 0 | - |
| - | - | 1.304E+04 | 633.3 | - | - | 0 | - |
| - | - | 4.654E+05 | 633.8 | - | - | 0 | - |
| - | - | 3.346E+05 | 634.3 | - | - | 0 | - |
| - | - | 1.287E+05 | 634.9 | - | - | 0 | - |
| - | - | 4.133E+04 | 635.4 | - | - | 0 | - |
| - | - | 6848 | 635.9 | - | - | 0 | - |
| - | - | 1060 | 640.3 | - | - | 0 | - |
| - | - | 1565 | 640.8 | - | - | 0 | - |
| - | - | 7182 | 641.4 | - | - | 0 | - |
| - | - | 2.409E+05 | 641.9 | - | - | 0 | - |
| - | - | 3.118E+05 | 642.4 | - | - | 0 | - |
| - | - | 1.612E+05 | 642.9 | - | - | 0 | - |
| - | - | 5.742E+04 | 643.4 | - | - | 0 | - |
| - | - | 1.558E+04 | 643.9 | - | - | 0 | - |
| - | - | 3264 | 644.4 | - | - | 0 | - |
| - | - | 1677 | 652.4 | - | - | 0 | - |
| - | - | 1396 | 653.4 | - | - | 0 | - |
| 7 | y | 2687 | 656.4 | 0.0008847 | 1.348 | +1 | 5 |
| - | - | 1424 | 657.4 | - | - | 0 | - |
| 7 | z | 1.346E+05 | 658.3 | 0.002452 | 3.725 | +1 | 5 |
| - | - | 5.453E+04 | 659.3 | - | - | 0 | - |
| - | - | 1.078E+04 | 660.4 | - | - | 0 | - |
| - | - | 1889 | 661.4 | - | - | 0 | - |
| - | - | 1056 | 666.4 | - | - | 0 | - |
| - | - | 993.4 | 668.4 | - | - | 0 | - |
| - | - | 8115 | 671.4 | - | - | 0 | - |
| - | - | 2577 | 672.4 | - | - | 0 | - |
| - | - | 1001 | 673.4 | - | - | 0 | - |
| 7 | y | 1.161E+04 | 674.4 | 0.002222 | 3.295 | +1 | 5 |
| - | - | 3896 | 675.4 | - | - | 0 | - |
| - | - | 985.2 | 682.4 | - | - | 0 | - |
| - | - | 991.3 | 683.4 | - | - | 0 | - |
| - | - | 2470 | 694.4 | - | - | 0 | - |
| - | - | 5452 | 695.4 | - | - | 0 | - |
| - | - | 3000 | 696.4 | - | - | 0 | - |
| - | - | 850.8 | 699.4 | - | - | 0 | - |
| 6 | y | 1.381E+04 | 713.4 | 0.002675 | 3.75 | +1 | 6 |
| - | - | 6271 | 714.4 | - | - | 0 | - |
| 6 | z | 3.173E+04 | 715.4 | 0.002839 | 3.969 | +1 | 6 |
| - | - | 1.469E+04 | 716.4 | - | - | 0 | - |
| - | - | 4127 | 717.4 | - | - | 0 | - |
| - | - | 1212 | 723.4 | - | - | 0 | - |
| 6 | y | 2.93E+04 | 731.4 | 0.002792 | 3.817 | +1 | 6 |
| - | - | 1.256E+04 | 732.4 | - | - | 0 | - |
| - | - | 2726 | 733.4 | - | - | 0 | - |
| - | - | 1443 | 738.5 | - | - | 0 | - |
| - | - | 914 | 739.5 | - | - | 0 | - |
| - | - | 2656 | 748.4 | - | - | 0 | - |
| - | - | 694.8 | 758.4 | - | - | 0 | - |
| - | - | 1752 | 764.4 | - | - | 0 | - |
| - | - | 1444 | 781.5 | - | - | 0 | - |
| 7 | c | 1.053E+04 | 782.5 | 0.002426 | 3.101 | +1 | 7 |
| - | - | 3598 | 783.5 | - | - | 0 | - |
| - | - | 855.3 | 784.5 | - | - | 0 | - |
| 5 | w | 7147 | 785.4 | 0.003336 | 4.247 | +1 | 7 |
| - | - | 4076 | 786.4 | - | - | 0 | - |
| - | - | 1096 | 787.4 | - | - | 0 | - |
| - | - | 952.2 | 791.5 | - | - | 0 | - |
| - | - | 1602 | 792.4 | - | - | 0 | - |
| - | - | 951 | 793.4 | - | - | 0 | - |
| - | - | 4121 | 795.5 | - | - | 0 | - |
| - | - | 1926 | 796.5 | - | - | 0 | - |
| - | - | 2150 | 806.4 | - | - | 0 | - |
| - | - | 1433 | 807.4 | - | - | 0 | - |
| 5 | y | 2.651E+04 | 810.4 | 0.002585 | 3.189 | +1 | 7 |
| - | - | 1.714E+04 | 811.4 | - | - | 0 | - |
| 5 | z | 5119 | 812.4 | 0.01447 | 17.81 | +1 | 7 |
| - | - | 9334 | 819.5 | - | - | 0 | - |
| - | - | 4589 | 820.5 | - | - | 0 | - |
| - | - | 1159 | 821.5 | - | - | 0 | - |
| - | - | 1.446E+04 | 823.4 | - | - | 0 | - |
| - | - | 7231 | 824.4 | - | - | 0 | - |
| - | - | 3055 | 825.5 | - | - | 0 | - |
| - | - | 5540 | 826.4 | - | - | 0 | - |
| - | - | 3181 | 827.4 | - | - | 0 | - |
| 5 | y | 5.371E+04 | 828.4 | 0.002335 | 2.819 | +1 | 7 |
| - | - | 2.514E+04 | 829.4 | - | - | 0 | - |
| - | - | 8029 | 830.4 | - | - | 0 | - |
| - | - | 1706 | 831.4 | - | - | 0 | - |
| - | - | 3212 | 839.5 | - | - | 0 | - |
| - | - | 1978 | 840.5 | - | - | 0 | - |
| - | - | 769.3 | 852.5 | - | - | 0 | - |
| 8 | c | 7872 | 853.5 | 0.002727 | 3.195 | +1 | 8 |
| - | - | 2923 | 854.5 | - | - | 0 | - |
| - | - | 1193 | 855.5 | - | - | 0 | - |
| - | - | 1740 | 856.4 | - | - | 0 | - |
| - | - | 878.6 | 857.4 | - | - | 0 | - |
| - | - | 1435 | 881.4 | - | - | 0 | - |
| 4 | y | 1206 | 882.4 | 0.004466 | 5.061 | +1 | 8 |
| 4 | z | 1.466E+05 | 883.5 | 0.002439 | 2.761 | +1 | 8 |
| - | - | 7.33E+04 | 884.5 | - | - | 0 | - |
| - | - | 1027 | 884.6 | - | - | 0 | - |
| - | - | 2.215E+04 | 885.5 | - | - | 0 | - |
| - | - | 5054 | 886.5 | - | - | 0 | - |
| - | - | 877.7 | 887.5 | - | - | 0 | - |
| - | - | 1094 | 898.5 | - | - | 0 | - |
| - | - | 1144 | 899.4 | - | - | 0 | - |
| 4 | y | 2.185E+04 | 899.5 | 0.002209 | 2.455 | +1 | 8 |
| - | - | 1.144E+04 | 900.5 | - | - | 0 | - |
| - | - | 2984 | 901.5 | - | - | 0 | - |
| - | - | 774.4 | 915.5 | - | - | 0 | - |
| - | - | 766.1 | 923.6 | - | - | 0 | - |
| - | - | 3134 | 932.5 | - | - | 0 | - |
| - | - | 2551 | 933.5 | - | - | 0 | - |
| - | - | 1991 | 939.5 | - | - | 0 | - |
| - | - | 911.4 | 950.6 | - | - | 0 | - |
| - | - | 860.1 | 951.6 | - | - | 0 | - |
| 3 | w | 5578 | 953.5 | 0.003546 | 3.719 | +1 | 9 |
| - | - | 2172 | 954.5 | - | - | 0 | - |
| - | - | 1133 | 955.5 | - | - | 0 | - |
| - | - | 963.9 | 962.5 | - | - | 0 | - |
| - | - | 939.2 | 963.5 | - | - | 0 | - |
| - | - | 982.9 | 965.6 | - | - | 0 | - |
| 9 | c | 9821 | 966.6 | 0.001732 | 1.792 | +1 | 9 |
| - | - | 969.8 | 967.5 | - | - | 0 | - |
| - | - | 3862 | 967.6 | - | - | 0 | - |
| - | - | 1355 | 968.6 | - | - | 0 | - |
| - | - | 1262 | 980.6 | - | - | 0 | - |
| - | - | 882.9 | 995.6 | - | - | 0 | - |
| - | - | 718.5 | 1011 | - | - | 0 | - |
| 3 | z | 1.072E+05 | 1012 | 0.00276 | 2.729 | +1 | 9 |
| - | - | 6.083E+04 | 1013 | - | - | 0 | - |
| - | - | 2.001E+04 | 1014 | - | - | 0 | - |
| - | - | 4111 | 1015 | - | - | 0 | - |
| - | - | 858.8 | 1016 | - | - | 0 | - |
| - | - | 1410 | 1025 | - | - | 0 | - |
| - | - | 1114 | 1026 | - | - | 0 | - |
| 3 | y | 2.499E+04 | 1028 | 0.002835 | 2.759 | +1 | 9 |
| - | - | 1058 | 1028 | - | - | 0 | - |
| - | - | 1.445E+04 | 1029 | - | - | 0 | - |
| - | - | 5014 | 1030 | - | - | 0 | - |
| - | - | 1269 | 1031 | - | - | 0 | - |
| - | - | 676.2 | 1036 | - | - | 0 | - |
| - | - | 1743 | 1038 | - | - | 0 | - |
| - | - | 8172 | 1052 | - | - | 0 | - |
| - | - | 1.018E+04 | 1053 | - | - | 0 | - |
| - | - | 5756 | 1054 | - | - | 0 | - |
| - | - | 2770 | 1055 | - | - | 0 | - |
| - | - | 1035 | 1067 | - | - | 0 | - |
| - | - | 1168 | 1069 | - | - | 0 | - |
| - | - | 997.5 | 1071 | - | - | 0 | - |
| 10 | c | 1334 | 1079 | 0.001923 | 1.783 | +1 | 10 |
| - | - | 8522 | 1080 | - | - | 0 | - |
| - | - | 5185 | 1081 | - | - | 0 | - |
| 2 | w | 9793 | 1082 | 0.002524 | 2.334 | +1 | 10 |
| - | - | 8322 | 1083 | - | - | 0 | - |
| - | - | 3309 | 1084 | - | - | 0 | - |
| - | - | 985.6 | 1085 | - | - | 0 | - |
| 10 | c | 5704 | 1096 | 0.001497 | 1.366 | +1 | 10 |
| - | - | 1.397E+04 | 1097 | - | - | 0 | - |
| - | - | 8105 | 1098 | - | - | 0 | - |
| - | - | 2694 | 1099 | - | - | 0 | - |
| - | - | 699.3 | 1100 | - | - | 0 | - |
| - | - | 1967 | 1111 | - | - | 0 | - |
| - | - | 1047 | 1112 | - | - | 0 | - |
| - | - | 1032 | 1124 | - | - | 0 | - |
| - | - | 653.4 | 1126 | - | - | 0 | - |
| - | - | 743.6 | 1137 | - | - | 0 | - |
| - | - | 839.2 | 1138 | - | - | 0 | - |
| - | - | 1072 | 1141 | - | - | 0 | - |
| - | - | 1816 | 1153 | - | - | 0 | - |
| - | - | 3639 | 1154 | - | - | 0 | - |
| - | - | 1932 | 1155 | - | - | 0 | - |
| - | - | 1100 | 1156 | - | - | 0 | - |
| 2 | y | 780.9 | 1167 | 0.01763 | 15.11 | +1 | 10 |
| 2 | z | 4379 | 1168 | 0.006065 | 5.195 | +1 | 10 |
| - | - | 1.251E+04 | 1169 | - | - | 0 | - |
| - | - | 7496 | 1170 | - | - | 0 | - |
| - | - | 2125 | 1171 | - | - | 0 | - |
| - | - | 9766 | 1181 | - | - | 0 | - |
| - | - | 7382 | 1182 | - | - | 0 | - |
| - | - | 5243 | 1183 | - | - | 0 | - |
| 2 | y | 2743 | 1184 | 0.006845 | 5.783 | +1 | 10 |
| - | - | 1228 | 1185 | - | - | 0 | - |
| - | - | 2127 | 1195 | - | - | 0 | - |
| - | - | 7292 | 1196 | - | - | 0 | - |
| - | - | 5393 | 1197 | - | - | 0 | - |
| - | - | 2906 | 1198 | - | - | 0 | - |
| - | - | 813.9 | 1199 | - | - | 0 | - |
| - | - | 925.2 | 1209 | - | - | 0 | - |
| - | - | 1245 | 1210 | - | - | 0 | - |
| - | - | 844.8 | 1223 | - | - | 0 | - |
| - | - | 2281 | 1224 | - | - | 0 | - |
| - | - | 6.865E+04 | 1225 | - | - | 0 | - |
| - | - | 5.185E+04 | 1226 | - | - | 0 | - |
| - | - | 1.874E+04 | 1227 | - | - | 0 | - |
| - | - | 5336 | 1228 | - | - | 0 | - |
| - | - | 2004 | 1229 | - | - | 0 | - |
| - | - | 705.9 | 1239 | - | - | 0 | - |
| - | - | 3483 | 1240 | - | - | 0 | - |
| - | - | 3781 | 1241 | - | - | 0 | - |
| - | - | 2683 | 1242 | - | - | 0 | - |
| - | - | 1617 | 1243 | - | - | 0 | - |
| - | - | 1168 | 1244 | - | - | 0 | - |
| - | - | 4110 | 1251 | - | - | 0 | - |
| - | - | 3705 | 1252 | - | - | 0 | - |
| - | - | 1235 | 1253 | - | - | 0 | - |
| - | - | 4705 | 1267 | - | - | 0 | - |
| - | - | 3.871E+04 | 1268 | - | - | 0 | - |
| - | - | 2.853E+04 | 1269 | - | - | 0 | - |
| - | - | 8973 | 1270 | - | - | 0 | - |
| - | - | 3350 | 1271 | - | - | 0 | - |
| - | - | 1608 | 1283 | - | - | 0 | - |
| - | - | 9968 | 1284 | - | - | 0 | - |
| - | - | 3.239E+04 | 1285 | - | - | 0 | - |
| - | - | 2.157E+04 | 1286 | - | - | 0 | - |
| - | - | 855.7 | 1286 | - | - | 0 | - |
| - | - | 7646 | 1287 | - | - | 0 | - |
| - | - | 2741 | 1288 | - | - | 0 | - |
| - | - | 665.7 | 2558 | - | - | 0 | - |
| - | - | 1063 | 3071 | - | - | 0 | - |
| - | - | 593.2 | 3290 | - | - | 0 | - |

m/z Charge Intensity FragmentType MassShift Position
120.67501831054688 0 398.60208
125.07147216796875 0 769.6249
125.93717193603516 0 534.07526
129.06597900390625 0 522.5304
130.0506134033203 0 1839.5149
130.0657501220703 0 57480.086
131.06915283203125 0 4915.2783
132.0813751220703 0 10113.624
133.08558654785156 0 1508.8235
134.06068420410156 0 605.26587
134.6582489013672 0 416.67902
135.01290893554688 0 473.27393
140.08216857910156 0 414.17722
142.0980987548828 0 3399.027
143.0736846923828 0 1160.0226
144.08143615722656 0 4098.0786
145.0851287841797 0 698.93884
146.06069946289062 0 25817.969
147.06423950195312 0 2358.061
147.4088592529297 0 406.6124
153.06642150878906 0 1493.6356
157.0775604248047 0 792.82965
157.10903930664062 0 4996.0986
157.13417053222656 0 1028.879
157.5950469970703 0 457.39502
159.09234619140625 0 173356
160.08932495117188 0 1217.9734
160.09571838378906 0 19401.771
161.09922790527344 0 1119.5083
162.05584716796875 0 509.0575
169.0766143798828 0 1542.1644
169.13433837890625 0 708.4645
170.06069946289062 0 5340.172
171.0645294189453 0 560.55457
171.09219360351562 0 797.6173
171.1250457763672 0 611.2882
173.43943786621094 0 969.63403
174.1033935546875 0 1474.4882
174.135498046875 0 2395.247
175.08734130859375 0 745.486
175.420654296875 0 489.81982
178.62474060058594 0 5125.651
179.12677001953125 0 505.67715
185.10400390625 0 3829.9243
185.12925720214844 0 2177.376
185.16470336914062 0 592.6772
187.06349182128906 0 1447.0627
187.08740234375 0 29613.488
188.07142639160156 0 571952.06
189.07469177246094 0 67096.79 z 10
190.05078125 0 1317.4824
190.07728576660156 0 4518.8906
190.997802734375 0 467.1373
192.62210083007812 0 17124.006 c Ammonia loss 2
193.123291015625 0 3007.3267
195.16102600097656 0 1373.0647
196.1454315185547 0 1064.3767
197.12925720214844 0 1714.5593
197.14064025878906 0 1441.7695
199.07212829589844 0 536.26227
200.10360717773438 0 823.715
203.08226013183594 0 870.23706
203.12872314453125 0 493.26108
203.8613739013672 0 528.2679
205.09793090820312 0 354618.56 y 10
205.62979125976562 0 4012.7725
206.10128784179688 0 42222.188
206.13111877441406 0 1080.7383
207.1038055419922 0 3152.0547 z Water loss 8
208.9539337158203 0 6304.975
213.16070556640625 0 920.99805
213.1717987060547 0 6895.4766
214.1432342529297 0 59812.78
214.15570068359375 0 4856.3115
214.64474487304688 0 12506.589
215.1473846435547 0 885.5696
215.15869140625 0 696.97174
219.6274871826172 0 1394.3689
228.14077758789062 0 18489.35 c Ammonia loss 3
228.18272399902344 0 3031.0552
228.64236450195312 0 4658.0693
228.94659423828125 0 506.0148
233.1655731201172 0 1414.1498
234.15902709960938 0 3682.1877
234.66046142578125 0 917.3959
238.1675567626953 0 1549.4845
239.15121459960938 0 36903.402
240.1544189453125 0 3957.2175
242.16220092773438 0 6463.998
243.1348114013672 0 3115.7615
243.1466827392578 0 574.066
243.1673126220703 0 1481.081
244.1380615234375 0 791.63275
248.15625 0 1275.6249
256.1210632324219 0 2134.3774
256.177734375 0 28010.428 c Ammonia loss 1
257.1803894042969 0 2888.162
258.8445739746094 0 561.0939
261.1592102050781 0 494.5661
262.6698303222656 0 3042.449
262.68328857421875 0 543.1729
263.171142578125 0 673.64984
263.6719055175781 0 537.5037
268.94866943359375 0 497.57895
270.1246032714844 0 21093.354
271.11181640625 0 465.87354
271.1280517578125 0 3101.6873
273.2043151855469 0 54165.19 c 1
274.20703125 0 6079.741
276.6674499511719 0 5334.305 c Ammonia loss 4
277.1689758300781 0 1780.2318
279.5004577636719 0 1245.1055 c Ammonia loss 7
280.1091613769531 0 4952.0503
281.11151123046875 0 1073.0558
291.6837158203125 0 758.9548
294.15728759765625 0 1384.7521
298.1197814941406 0 11971.542
298.2250671386719 0 1697.6451
298.6805114746094 0 4718.1978
299.12335205078125 0 2352.9321
299.18231201171875 0 1480.6201
299.22021484375 0 17059.588
300.2230529785156 0 2370.7595
302.18829345703125 0 627.8257
305.177734375 0 1230.541 c Ammonia loss 5
307.86419677734375 0 5913.8877
308.1983337402344 0 4154.352
308.5324401855469 0 880.0626
311.1834411621094 0 1345.2219
312.1908264160156 0 1110.0732
312.6778564453125 0 10167.676
312.697509765625 0 792.7642
313.16229248046875 0 1478.4272
313.18011474609375 0 3587.8965
313.6824645996094 0 630.6764
314.1719665527344 0 1463.498
314.2081604003906 0 837.4625
316.1305236816406 0 81622.88 y Water loss 9
317.1336975097656 0 15157.108
317.19561767578125 0 2439.7407 c Ammonia loss 8
318.1359558105469 0 2267.9917
318.92352294921875 0 5358.0337
334.1411437988281 0 7906.864 y 9
335.1439514160156 0 1460.9832
339.21636962890625 0 708.20404
341.0184020996094 0 1774.2327
341.2297668457031 0 2152.633
342.23870849609375 0 738.01495
347.7286071777344 0 1460.8746
350.1835632324219 0 712.7671
350.87823486328125 0 1811.5497
351.21405029296875 0 752.8393
356.241943359375 0 5781.2153
357.2474670410156 0 2974.9976
359.0296325683594 0 5811.374
360.2095642089844 0 1685.6191 c Ammonia loss 9
360.54473876953125 0 739.1892
361.2223205566406 0 787.71625
361.7257995605469 0 5234.8887
362.226318359375 0 2912.5605
362.71929931640625 0 600.27527
366.2268981933594 0 639.4025
367.173095703125 0 1141.7777
367.21002197265625 0 5029.595
368.2139587402344 0 1588.1683
369.23150634765625 0 4652.3296
369.7350158691406 0 19885.904
370.23663330078125 0 7930.166
370.2587890625 0 2243.996
370.7388000488281 0 1902.6422
371.2618103027344 0 766.8556
374.7162780761719 0 2970.7104
375.2156677246094 0 802.3759
375.7325134277344 0 684.1599
382.219970703125 0 1061.8837
382.71661376953125 0 1412.5929
383.2289123535156 0 30552.166 c Ammonia loss 6
383.729736328125 0 12864.729
384.2017822265625 0 6208.188 z Ammonia loss 1
384.2365417480469 0 62603.49 c Ammonia loss 2
384.7324523925781 0 854.1608
385.20361328125 0 1399.77
385.23968505859375 0 12879.846
385.2825927734375 0 1233.231
386.2422790527344 0 1681.5085
388.7323913574219 0 1019.2942
390.7350769042969 0 952.64966
391.2383728027344 0 93791.65
391.7420349121094 0 294262.22 c 6
392.24334716796875 0 122055.414
392.74481201171875 0 27695.615
393.24639892578125 0 2924.5996
396.23712158203125 0 1343.8231
396.71270751953125 0 2667.515
397.2129821777344 0 1153.3119
397.74481201171875 0 1045.3646
400.899658203125 0 606.7841
401.26348876953125 0 170807 c 2
401.5600280761719 0 1557.3939
401.89520263671875 0 1505.576
402.26617431640625 0 32807.953
403.2686767578125 0 4367.0117
404.7505187988281 0 15315.146
405.2523193359375 0 7677.077
405.71807861328125 0 21190.553 y Water loss 4
405.7543029785156 0 2256.0386
406.21966552734375 0 10708.891
406.7210388183594 0 2955.9849
406.9015808105469 0 48775.824
407.23577880859375 0 34084.98
407.5699462890625 0 14357.703
407.90301513671875 0 2047.5725
410.2342834472656 0 10487.743
410.73590087890625 0 4535.9185
411.2365417480469 0 1430.3412
412.2304992675781 0 595.6257
412.9051818847656 0 47730.72
413.2394714355469 0 34571.914
413.27435302734375 0 2015.6982
413.5734558105469 0 11321.159
413.9085388183594 0 3261.2336
414.7233581542969 0 25537.404 y 4
415.22479248046875 0 11902.859
415.7259521484375 0 3501.1812
416.232666015625 0 7158.2114
416.5668640136719 0 3669.2026
416.8995361328125 0 890.2519
417.7370300292969 0 897.898
418.2554016113281 0 6420.349
418.74810791015625 0 47543.594 c Ammonia loss 7
419.2492370605469 0 21022.521
419.75018310546875 0 6497.316
420.2522277832031 0 1160.4017
422.2366638183594 0 14884.539
422.5707092285156 0 11238.675
422.9046936035156 0 5699.4243
423.23797607421875 0 978.344
426.2531433105469 0 2036.5465
426.7572021484375 0 34971.24
427.2611083984375 0 242806.03 c 7
427.5669250488281 0 773.9131
427.76226806640625 0 112433.984
428.2637634277344 0 21712.133
428.5752258300781 0 1517.8329
428.7648620605469 0 6191.2515
428.9031982421875 0 2767.073
429.0907897949219 0 5196.187
429.2439880371094 0 731.31793
429.2894287109375 0 1562.8689
438.2479248046875 0 3998.3809
439.25067138671875 0 617.88684
439.764404296875 0 678.45734
440.26708984375 0 2347.716
440.7677917480469 0 788.71826
452.77734375 0 4006.4253
453.2884826660156 0 1674.2667
453.78668212890625 0 1947.9521
454.2648010253906 0 719.6879
455.27435302734375 0 69567.68 c Ammonia loss 3
455.76995849609375 0 917.3751
456.27703857421875 0 15698.906
457.279296875 0 2074.1953
461.29266357421875 0 114510.32
461.79412841796875 0 60891.65
462.29547119140625 0 18741.53
462.7969665527344 0 6232.47
466.7772216796875 0 6830.52
467.2792053222656 0 2864.4077
467.3096618652344 0 2863.0789
467.7829284667969 0 1573.2324
469.2746887207031 0 846.4008
470.2805480957031 0 785.6339
472.2881164550781 0 1345.9963
473.2860107421875 0 1175.1318
474.7976989746094 0 11254.594
475.2902526855469 0 302712.44 c Ammonia loss 8
475.79156494140625 0 159606.98
476.2925109863281 0 47618.957
476.7937316894531 0 9511.255
481.2549133300781 0 1892.9565
481.7615051269531 0 1376.3931
482.2655029296875 0 1931.3152
482.79986572265625 0 996.60443
483.2994384765625 0 9457.539
483.8032531738281 0 120804.375 c 8
484.3043212890625 0 63255.695
484.8056335449219 0 19453.85
485.3062744140625 0 4384.6157
486.2713623046875 0 1451.7119
489.8056335449219 0 1246.2042
490.2760009765625 0 1061.9823
490.781494140625 0 1180.0632
495.3059387207031 0 4309.8535
496.31060791015625 0 988.3397
496.81109619140625 0 68352.49
497.31231689453125 0 40684.785
497.8135681152344 0 12589.442
498.2861633300781 0 12837.798
498.3182678222656 0 3008.2568
498.7879333496094 0 9022.046
499.289306640625 0 2029.0984
499.32354736328125 0 4209.0366
500.3282470703125 0 803.8194
502.2420349121094 0 789.1126 z 7
503.7725524902344 0 886.0174
504.30804443359375 0 877.2236
504.7904968261719 0 5177.3984
505.2923889160156 0 2505.6309
505.79608154296875 0 1388.9163
508.81536865234375 0 928.6893
509.3099365234375 0 1148.698
510.286865234375 0 1739.0513
511.281494140625 0 2797.5132
512.7857055664062 0 3013.387
513.2880859375 0 1465.3103
516.8120727539062 0 1016.27106
517.3050537109375 0 1602.3699
517.815185546875 0 14002.916
518.2752075195312 0 939.02673
518.3153076171875 0 8016.909
518.8118896484375 0 5254.6216
519.3079223632812 0 1241.113
519.7881469726562 0 907.6975
521.2620849609375 0 864.2607
522.2930908203125 0 2332.8088
522.788330078125 0 2545.2473
523.2908935546875 0 784.03705
524.3321533203125 0 2769.1682
524.8060913085938 0 1668.0499
525.3390502929688 0 4630.1465
525.8140258789062 0 299036.97
526.3153686523438 0 186967.34
526.8167724609375 0 58738.65
527.3168334960938 0 13830.556
527.8211669921875 0 4050.4602
528.3016357421875 0 1463.3334
529.2890014648438 0 2037.4868
530.8059692382812 0 7242.587
531.3011474609375 0 16468.133
531.8012084960938 0 7426.0254
532.2994995117188 0 4842.0996
532.8059692382812 0 2727.9424
533.3109741210938 0 1208.102
533.8080444335938 0 1050.5455
538.8045043945312 0 7796.957
539.315673828125 0 11066.294 c Water loss 9
539.811767578125 0 1719348.9 c Ammonia loss 9
540.3131103515625 0 990441.3
540.8140869140625 0 336612.22
541.3152465820312 0 81866.766
541.8159790039062 0 11386.158
542.3345336914062 0 6146.376
543.338623046875 0 1726.9856
546.8259887695312 0 655.00146
547.3172607421875 0 7999.169
547.81591796875 0 7981.573
548.324462890625 0 600089.4 c 9
548.8260498046875 0 357479.6
549.3270874023438 0 114044.42
549.8284301757812 0 28111.482
550.3297119140625 0 5661.324
553.8045043945312 0 756.7578
554.3212280273438 0 1711.9421
554.8094482421875 0 1647.1213
555.2981567382812 0 26333.203
555.3384399414062 0 2699.986
555.7993774414062 0 19187.143
556.3012084960938 0 5620.4395
556.7905883789062 0 614.0033
556.8235473632812 0 896.18634
558.2903442382812 0 1102.8196
559.3147583007812 0 2456.8396
559.8163452148438 0 2447.1418
560.3150634765625 0 911.0576
561.33251953125 0 20879.434
561.8340454101562 0 16966.746
562.3322143554688 0 5281.674
562.7852783203125 0 2491.1082
562.8276977539062 0 1416.2579
563.2882080078125 0 2591.8257
563.78955078125 0 758.79364
568.3223266601562 0 20821.682
568.8232421875 0 19020.371
569.3538208007812 0 61175.56 c 4
569.810546875 0 1863.6893
570.30712890625 0 1041.3772
570.3566284179688 0 16976.625
571.3060913085938 0 970.50183
571.3585815429688 0 2985.5234
574.3252563476562 0 658.0291
575.8370971679688 0 6565.4395
576.33935546875 0 8349.526
576.8233032226562 0 39730.527
577.3268432617188 0 76730.92
577.828369140625 0 42571.016
578.3291015625 0 15171.31
578.828369140625 0 3646.4897
582.353271484375 0 7011.3765
582.8438720703125 0 4698.957
583.32470703125 0 3064.5737 y Water loss 1
583.8251342773438 0 4119.755
584.3131103515625 0 29673.367 z 1
584.353515625 0 4386.5747
584.81494140625 0 20058.492
585.31591796875 0 5813.1646
585.8161010742188 0 2260.7456
586.3231811523438 0 1531.379
589.843017578125 0 792.38525
590.3409423828125 0 2340.1548
590.855712890625 0 2225.6648
591.3170776367188 0 1631.9658
591.360595703125 0 1615.9531
591.8184814453125 0 1589.4545
592.3223266601562 0 5078.9775 y 1
592.8243408203125 0 3490.458
593.3261108398438 0 1590.4852
596.3528442382812 0 8810.447
597.3560180664062 0 2297.5085
597.8377075195312 0 4605.25
598.3286743164062 0 8048.2705
598.82861328125 0 6902.2266
599.3334350585938 0 5167.584
600.3366088867188 0 1925.3519
602.2832641601562 0 4721.4634
603.2852172851562 0 2001.5449
603.3358154296875 0 919.0267
603.8404541015625 0 1262.8872
604.340576171875 0 11769.898
604.8348999023438 0 26627.865
605.3351440429688 0 18360.445
605.827880859375 0 9080.758
606.3221435546875 0 4293.158
606.3425903320312 0 4227.906
606.8417358398438 0 4920.322
607.3262329101562 0 1593.5345
609.3477172851562 0 2044.353 c Ammonia loss 5
610.8455200195312 0 1498.058
611.3557739257812 0 8915.936
611.8530883789062 0 47175.938
612.3530883789062 0 33280.05
612.81689453125 0 1903.1318
612.854248046875 0 12584.557
613.35107421875 0 8777.684
613.8428955078125 0 2681.261
614.3544921875 0 17025.908
614.8330078125 0 2141.6565
615.2911987304688 0 9519.107
615.3578491210938 0 6414.659
616.2941284179688 0 3162.3042
616.3555297851562 0 1302.9032
617.2987060546875 0 871.0305
617.8221435546875 0 1893.4867
619.356689453125 0 10111.287
619.8529663085938 0 36114.77
620.3596801757812 0 44530.027
620.8482666015625 0 88987.83
621.348876953125 0 66105.37
621.8495483398438 0 23976.34
622.3489990234375 0 7328.624
622.849609375 0 1512.807
624.3491821289062 0 27249.166
625.3502197265625 0 12657.836
625.83740234375 0 6161.5264
626.3757934570312 0 226951.95 c 5
626.8372802734375 0 31875.572
627.3783569335938 0 73325.695
627.8418579101562 0 3840.4473
628.3802490234375 0 15011.197
629.3815307617188 0 2640.769
632.841796875 0 3633.7576
633.3475341796875 0 13035.994
633.8475341796875 0 465398.78
634.3486938476562 0 334628.78
634.8500366210938 0 128688.06
635.3512573242188 0 41334.895
635.8516235351562 0 6847.551
640.3470458984375 0 1060.1735
640.8467407226562 0 1564.9857
641.3529663085938 0 7182.3716
641.8565673828125 0 240922.8
642.3590698242188 0 311846.4
642.860595703125 0 161172.05
643.3617553710938 0 57421.867
643.8623046875 0 15583.961
644.360107421875 0 3264.352
652.3911743164062 0 1677.3865
653.3951416015625 0 1396.1295
656.3523559570312 0 2686.9224 y Water loss 6
657.3548583984375 0 1423.7911
658.3457641601562 0 134628.48 z 6
659.3487548828125 0 54532.348
660.3512573242188 0 10777.632
661.3540649414062 0 1889.4705
666.407958984375 0 1055.749
668.3922119140625 0 993.3637
671.3772583007812 0 8115.229
672.3778686523438 0 2577.145
673.38232421875 0 1000.51306
674.3642578125 0 11614.324 y 6
675.3662109375 0 3895.6436
682.388427734375 0 985.16534
683.395751953125 0 991.25916
694.4033203125 0 2469.7778
695.3853759765625 0 5451.977
696.3916015625 0 2999.8694
699.3817749023438 0 850.8041
713.3756103515625 0 13809.533 y Water loss 5
714.37841796875 0 6271.087
715.3676147460938 0 31734.545 z 5
716.3701171875 0 14694.18
717.3726196289062 0 4127.0117
723.4290161132812 0 1211.9192
731.3862915039062 0 29299.092 y 5
732.388916015625 0 12557.682
733.388916015625 0 2725.8477
738.4638671875 0 1443.0037
739.4596557617188 0 914.0005
748.4235229492188 0 2656.1484
758.3746337890625 0 694.82605
764.4212036132812 0 1752.2665
781.4656982421875 0 1444.1985
782.476806640625 0 10526.191 c 6
783.4791870117188 0 3597.7751
784.4908447265625 0 855.3079
785.3973999023438 0 7146.6235 w 4
786.403076171875 0 4075.8875
787.4048461914062 0 1095.5251
791.4658813476562 0 952.2467
792.416748046875 0 1602.2242
793.41357421875 0 951.00714
795.45068359375 0 4120.9688
796.4525756835938 0 1925.5287
806.4185180664062 0 2150.467
807.42333984375 0 1433.1842
810.4282836914062 0 26505.686 y Water loss 4
811.4298095703125 0 17139.701
812.4320068359375 0 5118.8594 z 4
819.4612426757812 0 9334.13
820.4642944335938 0 4588.5303
821.4638061523438 0 1158.507
823.4449462890625 0 14462.264
824.4480590820312 0 7231.343
825.4532470703125 0 3055.415
826.424560546875 0 5540.4385
827.4116821289062 0 3181.4475
828.4385986328125 0 53711.51 y 4
829.4414672851562 0 25139.695
830.44384765625 0 8028.9526
831.44482421875 0 1705.5325
839.4692993164062 0 3212.2236
840.46923828125 0 1978.0857
852.5132446289062 0 769.26746
853.5142211914062 0 7872.018 c 7
854.5169677734375 0 2923.339
855.5238647460938 0 1193.028
856.43212890625 0 1740.3611
857.4308471679688 0 878.5645
881.4444580078125 0 1434.994
882.4512939453125 0 1205.8038 y Ammonia loss 3
883.4570922851562 0 146588.45 z 3
884.4598999023438 0 73299.93
884.5650634765625 0 1027.138
885.4622802734375 0 22152.56
886.464599609375 0 5054.4517
887.458251953125 0 877.6554
898.4716796875 0 1093.8153
899.3740844726562 0 1143.786
899.4755859375 0 21847.383 y 3
900.4781494140625 0 11439.862
901.4801025390625 0 2983.782
915.4518432617188 0 774.4409
923.5809936523438 0 766.05115
932.5420532226562 0 3133.799
933.5493774414062 0 2550.5403
939.4935913085938 0 1991.4562
950.5814819335938 0 911.3685
951.5869750976562 0 860.11957
953.4874877929688 0 5578.025 w 2
954.4910278320312 0 2172.2395
955.4765625 0 1133.2964
962.5230712890625 0 963.87714
963.5234985351562 0 939.21594
965.553955078125 0 982.8526
966.5972900390625 0 9821.047 c 8
967.5084838867188 0 969.76666
967.6025390625 0 3862.2834
968.6065063476562 0 1354.5791
980.5534057617188 0 1261.7197
995.5662231445312 0 882.9211
1010.6138305664062 0 718.4595
1011.5159912109375 0 107244.766 z 2
1012.5193481445312 0 60830.734
1013.5209350585938 0 20012.91
1014.5228881835938 0 4110.8013
1015.5162353515625 0 858.79584
1024.5245361328125 0 1409.9594
1025.522216796875 0 1113.9036
1027.5347900390625 0 24986.354 y 2
1028.4202880859375 0 1057.9369
1028.537109375 0 14450.37
1029.53955078125 0 5013.8184
1030.5460205078125 0 1269.0394
1035.6201171875 0 676.2323
1037.5997314453125 0 1742.5399
1051.6265869140625 0 8171.635
1052.631103515625 0 10182.363
1053.631591796875 0 5756.463
1054.631103515625 0 2770.1782
1066.5701904296875 0 1034.5121
1068.5472412109375 0 1167.502
1070.5340576171875 0 997.4683
1078.613525390625 0 1334.1864 c Ammonia loss 9
1079.622314453125 0 8522.436
1080.6260986328125 0 5184.944
1081.5450439453125 0 9793.3 w 1
1082.5494384765625 0 8322.231
1083.5516357421875 0 3309.3462
1084.5567626953125 0 985.5966
1095.6396484375 0 5704.1826 c 9
1096.64697265625 0 13967.735
1097.6494140625 0 8105.199
1098.65380859375 0 2693.8506
1099.6580810546875 0 699.2529
1110.6141357421875 0 1967.2255
1111.6151123046875 0 1046.9692
1123.624267578125 0 1031.7198
1125.6153564453125 0 653.4138
1136.6328125 0 743.5627
1137.6279296875 0 839.21265
1140.6343994140625 0 1071.961
1152.630126953125 0 1815.6793
1153.6346435546875 0 3639.243
1154.6353759765625 0 1932.0105
1155.645263671875 0 1099.6572
1166.6241455078125 0 780.9075 y Ammonia loss 1
1167.6082763671875 0 4378.6055 z 1
1168.6207275390625 0 12507.392
1169.6260986328125 0 7495.804
1170.6279296875 0 2125.3435
1180.6181640625 0 9766.195
1181.621337890625 0 7381.7544
1182.6204833984375 0 5242.6997
1183.626220703125 0 2742.9856 y 1
1184.6328125 0 1227.8727
1194.6248779296875 0 2127.1018
1195.625244140625 0 7292.207
1196.62841796875 0 5392.508
1197.6322021484375 0 2906.2593
1198.647216796875 0 813.86444
1208.643798828125 0 925.15234
1209.6383056640625 0 1244.9962
1222.69775390625 0 844.8065
1223.695556640625 0 2281.0938
1224.6632080078125 0 68645.52
1225.665771484375 0 51851.637
1226.668701171875 0 18737.629
1227.6717529296875 0 5336.0327
1228.6700439453125 0 2003.5253
1238.6998291015625 0 705.88617
1239.698486328125 0 3482.8682
1240.705078125 0 3780.7083
1241.697998046875 0 2682.925
1242.6953125 0 1616.869
1243.7010498046875 0 1167.8055
1250.6678466796875 0 4109.5283
1251.6717529296875 0 3704.9229
1252.6776123046875 0 1234.9208
1266.691162109375 0 4704.841
1267.6923828125 0 38705.74
1268.695556640625 0 28525.94
1269.6968994140625 0 8972.66
1270.7001953125 0 3350.0562
1282.6981201171875 0 1607.7671
1283.709716796875 0 9967.516
1284.717529296875 0 32386.992
1285.7200927734375 0 21567.898
1285.8973388671875 0 855.6948
1286.72314453125 0 7645.7153
1287.7254638671875 0 2740.7546
2558.17529296875 0 665.6943
3071.175537109375 0 1062.96
3289.736328125 0 593.22327

Spectrum Details

|  |  |
| --- | --- |
| Matched peaks? Matched peaks The total absolute number of peaks matched. Additionally in brackets the total fraction of peaks matched and the total number of peaks is shown. | 61 (8.94% of 682) |
| FDR? FDR The false discovery rate estimated for this peptide. It is calculated by matching all theoretical fragments with a non-integer shift with the raw peaks for this spectrum. This is done with 40 different shifts. The resulting percentage is the average number of annotated peaks over the number of annotated peaks with the correct spectrum. | 1.72% |
| Satellite FDR? Satellite FDR See the FDR for details on its calculation. This satellite ion specific FDR only contains the satellite ions (d/w) for I/L/J positions. | ∞ |
| PSM Score? PSM Score The PSM Score as given by Hecklib to this annotated spectrum. It is shown with three significant figures. | 504 |

#### Spectrum 5776? Spectrum 5776 The raw spectrum of this peptide as annotated by Hecklib. The fragments are coloured according to ion type (see legend). Any peaks with a star '\*' as text can be hovered over to see the full details, first the ion type second the mass shift type. By hovering over the amino acids in the peptide or ions in the legend the corresponding peaks are highlighted. By toggling the 'Unassigned' label you can turn the background (unassigned) peaks on or off in the plot. By updating the slider in the Ion legend you can update the spectrum to only show the top X% of the peaks with labels. The top X% means any peak that is within X% of the highest intensity. By dragging in the spectrum you can zoom in to a specific part of the spectrum and use 'Zoom Out' to get back to the original zoom level. The annotation of the spectrum is based on the given sequence in the peptides file and is done with different software so inconsistencies are likely. The peaks are annotated based on the given sequence, with 20 ppm tolerance.

Copy Data

##### Spectrum 5776 (TSV)

###### Preview

```
Loading example...
```

*Click on the button to copy the data to your clipboard.*

Mz MinMz MaxIntensity Max

WidthHeightPeptide font sizePeptide stroke widthSpectrum font sizeSpectrum stroke widthCompact peptide

Ion legend

wxyz

abcd

OtherUnassignedIon^Charge^_Position_Show for top:%

VRQAPGRAJEW

03.48e+46.97e+41.05e+51.39e+5

Zoom Out

c^+2^_3_y^+1^_1_c^+2^_4_c^+1^_2_c^+1^_2_y^+1^_2_c^+2^_7_c^+1^_3_c^+2^_7_c^+1^_3_y^+2^_7_y^+2^_7_c^+2^_8_c^+2^_8_y^+1^_3_c^+1^_4_c^+2^_9_c^+2^_9_c^+2^_10_c^+2^_10_c^+1^_5_z^+2^_10_y^+2^_10_c^+1^_6_z^+1^_5_y^+1^_5_z^+1^_6_y^+1^_6_c^+1^_7_w^+1^_7_y^+1^_7_y^+1^_7_c^+1^_8_z^+1^_8_y^+1^_8_w^+1^_9_c^+1^_9_y^+1^_9_z^+1^_9_y^+1^_9_w^+1^_10_c^+1^_10_z^+1^_10_y^+1^_10_

03256519761302

Fragment Matches Table

Show background peaks

| Position | Ion type | Intensity | mz Theoretical | mz Error (Th) | mz Error (ppm) | Charge | Series Number |
| --- | --- | --- | --- | --- | --- | --- | --- |
| - | - | 423 | 120.1 | - | - | 0 | - |
| - | - | 1918 | 120.1 | - | - | 0 | - |
| - | - | 409.8 | 125.1 | - | - | 0 | - |
| - | - | 377.9 | 127.1 | - | - | 0 | - |
| - | - | 438 | 129.1 | - | - | 0 | - |
| - | - | 3416 | 130.1 | - | - | 0 | - |
| - | - | 580.4 | 131.1 | - | - | 0 | - |
| - | - | 383.2 | 132 | - | - | 0 | - |
| - | - | 782.1 | 132.1 | - | - | 0 | - |
| - | - | 1480 | 133.1 | - | - | 0 | - |
| - | - | 401.1 | 134.3 | - | - | 0 | - |
| - | - | 554.9 | 142.1 | - | - | 0 | - |
| - | - | 3083 | 142.1 | - | - | 0 | - |
| - | - | 648.1 | 144.1 | - | - | 0 | - |
| - | - | 4057 | 146.1 | - | - | 0 | - |
| - | - | 425 | 148.3 | - | - | 0 | - |
| - | - | 1236 | 149 | - | - | 0 | - |
| - | - | 420.8 | 154.4 | - | - | 0 | - |
| - | - | 2309 | 157.1 | - | - | 0 | - |
| - | - | 9071 | 159.1 | - | - | 0 | - |
| - | - | 976.5 | 160.1 | - | - | 0 | - |
| - | - | 1069 | 165.1 | - | - | 0 | - |
| - | - | 515.9 | 166 | - | - | 0 | - |
| - | - | 484.6 | 170.1 | - | - | 0 | - |
| - | - | 464.8 | 171.1 | - | - | 0 | - |
| - | - | 2118 | 173.5 | - | - | 0 | - |
| - | - | 1031 | 174.1 | - | - | 0 | - |
| - | - | 516.6 | 174.1 | - | - | 0 | - |
| - | - | 731.3 | 174.1 | - | - | 0 | - |
| - | - | 604.6 | 177.1 | - | - | 0 | - |
| - | - | 754.8 | 185.1 | - | - | 0 | - |
| - | - | 1538 | 185.1 | - | - | 0 | - |
| - | - | 1489 | 187.1 | - | - | 0 | - |
| - | - | 2.494E+04 | 188.1 | - | - | 0 | - |
| - | - | 3052 | 189.1 | - | - | 0 | - |
| 3 | c | 1035 | 192.6 | 0.0005137 | 2.667 | +2 | 3 |
| - | - | 817.4 | 201.1 | - | - | 0 | - |
| - | - | 636.4 | 203.1 | - | - | 0 | - |
| - | - | 1634 | 203.1 | - | - | 0 | - |
| 11 | y | 1.417E+04 | 205.1 | 0.0001817 | 0.886 | +1 | 1 |
| - | - | 1385 | 206.1 | - | - | 0 | - |
| - | - | 2.031E+04 | 209 | - | - | 0 | - |
| - | - | 630.2 | 211.2 | - | - | 0 | - |
| - | - | 762.1 | 212.1 | - | - | 0 | - |
| - | - | 2247 | 213.2 | - | - | 0 | - |
| - | - | 2631 | 214.1 | - | - | 0 | - |
| - | - | 1419 | 214.2 | - | - | 0 | - |
| - | - | 727.9 | 214.6 | - | - | 0 | - |
| - | - | 950.6 | 215.1 | - | - | 0 | - |
| 4 | c | 817.8 | 228.1 | 0.0002216 | 0.9714 | +2 | 4 |
| - | - | 946.1 | 228.2 | - | - | 0 | - |
| - | - | 561.9 | 228.6 | - | - | 0 | - |
| - | - | 1028 | 238.2 | - | - | 0 | - |
| - | - | 503 | 239.1 | - | - | 0 | - |
| - | - | 9448 | 239.2 | - | - | 0 | - |
| - | - | 839.2 | 240.2 | - | - | 0 | - |
| - | - | 1498 | 242.2 | - | - | 0 | - |
| - | - | 2210 | 245.1 | - | - | 0 | - |
| - | - | 609.1 | 246.1 | - | - | 0 | - |
| 2 | c | 4836 | 256.2 | 0.000109 | 0.4255 | +1 | 2 |
| - | - | 651 | 257.2 | - | - | 0 | - |
| 2 | c | 2.063E+04 | 273.2 | 0.0001407 | 0.515 | +1 | 2 |
| - | - | 2017 | 274.2 | - | - | 0 | - |
| - | - | 541.7 | 280.1 | - | - | 0 | - |
| - | - | 646.9 | 290 | - | - | 0 | - |
| - | - | 555.5 | 296.2 | - | - | 0 | - |
| - | - | 666.4 | 298.2 | - | - | 0 | - |
| - | - | 597.3 | 298.2 | - | - | 0 | - |
| - | - | 5268 | 299.2 | - | - | 0 | - |
| - | - | 738.9 | 300.2 | - | - | 0 | - |
| - | - | 524.8 | 314.9 | - | - | 0 | - |
| 10 | y | 3588 | 316.1 | 0.0002763 | 0.8739 | +1 | 2 |
| - | - | 1.439E+04 | 318.9 | - | - | 0 | - |
| - | - | 630.2 | 327.2 | - | - | 0 | - |
| - | - | 4596 | 338.1 | - | - | 0 | - |
| - | - | 1131 | 339.2 | - | - | 0 | - |
| - | - | 3869 | 341 | - | - | 0 | - |
| - | - | 1068 | 356.2 | - | - | 0 | - |
| - | - | 1109 | 357.2 | - | - | 0 | - |
| - | - | 1.372E+04 | 359 | - | - | 0 | - |
| - | - | 1101 | 361.7 | - | - | 0 | - |
| - | - | 647.7 | 362.2 | - | - | 0 | - |
| - | - | 513.5 | 364.7 | - | - | 0 | - |
| - | - | 563.7 | 369.2 | - | - | 0 | - |
| - | - | 4491 | 369.7 | - | - | 0 | - |
| - | - | 2357 | 370.2 | - | - | 0 | - |
| - | - | 664.5 | 370.3 | - | - | 0 | - |
| 7 | c | 2970 | 383.2 | 4.622E-05 | 0.1206 | +2 | 7 |
| - | - | 647.5 | 383.7 | - | - | 0 | - |
| 3 | c | 3928 | 384.2 | 0.0003325 | 0.8654 | +1 | 3 |
| - | - | 2.465E+04 | 391.2 | - | - | 0 | - |
| 7 | c | 7.363E+04 | 391.7 | 0.0001973 | 0.5037 | +2 | 7 |
| - | - | 3.069E+04 | 392.2 | - | - | 0 | - |
| - | - | 6261 | 392.7 | - | - | 0 | - |
| - | - | 961.3 | 393.2 | - | - | 0 | - |
| 3 | c | 6.521E+04 | 401.3 | 0.0003095 | 0.7714 | +1 | 3 |
| - | - | 1.336E+04 | 402.3 | - | - | 0 | - |
| - | - | 1468 | 403.3 | - | - | 0 | - |
| - | - | 1122 | 404.7 | - | - | 0 | - |
| 5 | y | 1239 | 405.7 | 0.0006144 | 1.514 | +2 | 7 |
| - | - | 1481 | 406.9 | - | - | 0 | - |
| - | - | 1552 | 407.2 | - | - | 0 | - |
| - | - | 1304 | 409.2 | - | - | 0 | - |
| - | - | 855.4 | 410.2 | - | - | 0 | - |
| - | - | 1646 | 412.9 | - | - | 0 | - |
| - | - | 1046 | 413.2 | - | - | 0 | - |
| - | - | 951.8 | 413.6 | - | - | 0 | - |
| 5 | y | 924.4 | 414.7 | 0.000459 | 1.107 | +2 | 7 |
| 8 | c | 3709 | 418.7 | 0.0007764 | 1.854 | +2 | 8 |
| - | - | 1752 | 419.2 | - | - | 0 | - |
| - | - | 8713 | 426.8 | - | - | 0 | - |
| 8 | c | 6.072E+04 | 427.3 | 0.0003193 | 0.7472 | +2 | 8 |
| - | - | 2.414E+04 | 427.8 | - | - | 0 | - |
| - | - | 1711 | 428.2 | - | - | 0 | - |
| - | - | 6271 | 428.3 | - | - | 0 | - |
| - | - | 2944 | 428.9 | - | - | 0 | - |
| - | - | 1.546E+04 | 429.1 | - | - | 0 | - |
| 9 | y | 1204 | 429.2 | 0.001821 | 4.242 | +1 | 3 |
| - | - | 664.7 | 429.2 | - | - | 0 | - |
| 4 | c | 3607 | 455.3 | 0.0001208 | 0.2654 | +1 | 4 |
| - | - | 841.2 | 456.3 | - | - | 0 | - |
| - | - | 6293 | 461.3 | - | - | 0 | - |
| - | - | 4257 | 461.8 | - | - | 0 | - |
| - | - | 1527 | 462.3 | - | - | 0 | - |
| - | - | 1305 | 474.8 | - | - | 0 | - |
| 9 | c | 1.293E+04 | 475.3 | 0.0004315 | 0.9078 | +2 | 9 |
| - | - | 7894 | 475.8 | - | - | 0 | - |
| - | - | 1728 | 476.3 | - | - | 0 | - |
| - | - | 1413 | 476.8 | - | - | 0 | - |
| - | - | 703.4 | 481.8 | - | - | 0 | - |
| - | - | 581.1 | 482.8 | - | - | 0 | - |
| - | - | 2117 | 483.3 | - | - | 0 | - |
| 9 | c | 3.064E+04 | 483.8 | 0.0001879 | 0.3884 | +2 | 9 |
| - | - | 1.819E+04 | 484.3 | - | - | 0 | - |
| - | - | 4207 | 484.8 | - | - | 0 | - |
| - | - | 702.9 | 486.3 | - | - | 0 | - |
| - | - | 1.721E+04 | 496.8 | - | - | 0 | - |
| - | - | 8511 | 497.3 | - | - | 0 | - |
| - | - | 2170 | 497.8 | - | - | 0 | - |
| - | - | 1856 | 498.3 | - | - | 0 | - |
| - | - | 1006 | 498.3 | - | - | 0 | - |
| - | - | 1154 | 498.8 | - | - | 0 | - |
| - | - | 911.4 | 499.3 | - | - | 0 | - |
| - | - | 1980 | 504.8 | - | - | 0 | - |
| - | - | 766.4 | 519.8 | - | - | 0 | - |
| - | - | 1293 | 524.3 | - | - | 0 | - |
| - | - | 1722 | 525.3 | - | - | 0 | - |
| - | - | 1.757E+04 | 525.8 | - | - | 0 | - |
| - | - | 1.037E+04 | 526.3 | - | - | 0 | - |
| - | - | 3538 | 526.8 | - | - | 0 | - |
| - | - | 948.8 | 527.3 | - | - | 0 | - |
| - | - | 705.5 | 530.8 | - | - | 0 | - |
| 10 | c | 7.644E+04 | 539.8 | 0.0003141 | 0.5819 | +2 | 10 |
| - | - | 4.097E+04 | 540.3 | - | - | 0 | - |
| - | - | 1.416E+04 | 540.8 | - | - | 0 | - |
| - | - | 3529 | 541.3 | - | - | 0 | - |
| - | - | 796.7 | 541.8 | - | - | 0 | - |
| - | - | 2683 | 542.3 | - | - | 0 | - |
| - | - | 708.6 | 543.3 | - | - | 0 | - |
| - | - | 960.9 | 547.3 | - | - | 0 | - |
| - | - | 983.8 | 547.8 | - | - | 0 | - |
| 10 | c | 1.38E+05 | 548.3 | 0.0002232 | 0.407 | +2 | 10 |
| - | - | 8.065E+04 | 548.8 | - | - | 0 | - |
| - | - | 2.95E+04 | 549.3 | - | - | 0 | - |
| - | - | 6908 | 549.8 | - | - | 0 | - |
| - | - | 1155 | 550.3 | - | - | 0 | - |
| - | - | 8055 | 555.3 | - | - | 0 | - |
| - | - | 1255 | 555.3 | - | - | 0 | - |
| - | - | 4424 | 555.8 | - | - | 0 | - |
| - | - | 989.9 | 556.3 | - | - | 0 | - |
| - | - | 5902 | 561.3 | - | - | 0 | - |
| - | - | 794.8 | 561.8 | - | - | 0 | - |
| - | - | 4354 | 561.8 | - | - | 0 | - |
| - | - | 1641 | 562.3 | - | - | 0 | - |
| - | - | 1186 | 562.8 | - | - | 0 | - |
| - | - | 1597 | 568.3 | - | - | 0 | - |
| 5 | c | 2.269E+04 | 569.4 | 6.005E-05 | 0.1055 | +1 | 5 |
| - | - | 8269 | 570.4 | - | - | 0 | - |
| - | - | 860.8 | 571.3 | - | - | 0 | - |
| - | - | 1604 | 571.4 | - | - | 0 | - |
| - | - | 2430 | 575.8 | - | - | 0 | - |
| - | - | 1123 | 576.3 | - | - | 0 | - |
| - | - | 1.034E+04 | 576.8 | - | - | 0 | - |
| - | - | 8982 | 577.3 | - | - | 0 | - |
| - | - | 2942 | 577.8 | - | - | 0 | - |
| - | - | 733.9 | 578.3 | - | - | 0 | - |
| - | - | 2775 | 582.4 | - | - | 0 | - |
| - | - | 831 | 582.8 | - | - | 0 | - |
| - | - | 1413 | 583.8 | - | - | 0 | - |
| 2 | z | 5047 | 584.3 | 0.001569 | 2.685 | +2 | 10 |
| - | - | 4590 | 584.8 | - | - | 0 | - |
| - | - | 1344 | 584.9 | - | - | 0 | - |
| - | - | 1676 | 585.3 | - | - | 0 | - |
| - | - | 863.6 | 586.3 | - | - | 0 | - |
| - | - | 833.7 | 590.3 | - | - | 0 | - |
| - | - | 813.4 | 590.9 | - | - | 0 | - |
| - | - | 1009 | 591.4 | - | - | 0 | - |
| - | - | 751.1 | 591.8 | - | - | 0 | - |
| 2 | y | 607.4 | 592.3 | 0.01143 | 19.3 | +2 | 10 |
| - | - | 1172 | 597.8 | - | - | 0 | - |
| - | - | 3080 | 598.3 | - | - | 0 | - |
| - | - | 1959 | 598.8 | - | - | 0 | - |
| - | - | 2427 | 599.3 | - | - | 0 | - |
| - | - | 1029 | 600.3 | - | - | 0 | - |
| - | - | 1512 | 602.3 | - | - | 0 | - |
| - | - | 649.7 | 603.3 | - | - | 0 | - |
| - | - | 670.7 | 603.3 | - | - | 0 | - |
| - | - | 2562 | 604.3 | - | - | 0 | - |
| - | - | 5505 | 604.8 | - | - | 0 | - |
| - | - | 4823 | 605.3 | - | - | 0 | - |
| - | - | 2341 | 605.8 | - | - | 0 | - |
| - | - | 1060 | 606.3 | - | - | 0 | - |
| - | - | 733.2 | 606.8 | - | - | 0 | - |
| - | - | 2248 | 611.4 | - | - | 0 | - |
| - | - | 1.082E+04 | 611.9 | - | - | 0 | - |
| - | - | 7313 | 612.4 | - | - | 0 | - |
| - | - | 3068 | 612.9 | - | - | 0 | - |
| - | - | 2083 | 613.3 | - | - | 0 | - |
| - | - | 765.8 | 613.8 | - | - | 0 | - |
| - | - | 6872 | 614.4 | - | - | 0 | - |
| - | - | 1433 | 614.8 | - | - | 0 | - |
| - | - | 4829 | 615.3 | - | - | 0 | - |
| - | - | 2506 | 615.4 | - | - | 0 | - |
| - | - | 1432 | 616.3 | - | - | 0 | - |
| - | - | 1327 | 619.4 | - | - | 0 | - |
| - | - | 6656 | 619.8 | - | - | 0 | - |
| - | - | 1.115E+04 | 620.4 | - | - | 0 | - |
| - | - | 1.955E+04 | 620.8 | - | - | 0 | - |
| - | - | 1.407E+04 | 621.3 | - | - | 0 | - |
| - | - | 6047 | 621.8 | - | - | 0 | - |
| - | - | 2290 | 622.3 | - | - | 0 | - |
| - | - | 1076 | 624.4 | - | - | 0 | - |
| - | - | 4865 | 625.4 | - | - | 0 | - |
| - | - | 1408 | 625.8 | - | - | 0 | - |
| 6 | c | 9.969E+04 | 626.4 | 0.0002658 | 0.4243 | +1 | 6 |
| - | - | 7680 | 626.8 | - | - | 0 | - |
| - | - | 3.382E+04 | 627.4 | - | - | 0 | - |
| - | - | 1092 | 627.4 | - | - | 0 | - |
| - | - | 1027 | 627.8 | - | - | 0 | - |
| - | - | 6407 | 628.4 | - | - | 0 | - |
| - | - | 1286 | 632.8 | - | - | 0 | - |
| - | - | 2840 | 633.3 | - | - | 0 | - |
| - | - | 1.084E+05 | 633.8 | - | - | 0 | - |
| - | - | 7.971E+04 | 634.3 | - | - | 0 | - |
| - | - | 3.129E+04 | 634.8 | - | - | 0 | - |
| - | - | 8965 | 635.3 | - | - | 0 | - |
| - | - | 1911 | 635.9 | - | - | 0 | - |
| - | - | 673.2 | 640.8 | - | - | 0 | - |
| - | - | 2139 | 641.4 | - | - | 0 | - |
| - | - | 4.94E+04 | 641.9 | - | - | 0 | - |
| - | - | 7.075E+04 | 642.4 | - | - | 0 | - |
| - | - | 3.718E+04 | 642.9 | - | - | 0 | - |
| - | - | 1.374E+04 | 643.4 | - | - | 0 | - |
| - | - | 4608 | 643.9 | - | - | 0 | - |
| - | - | 1395 | 644.4 | - | - | 0 | - |
| - | - | 833 | 652.4 | - | - | 0 | - |
| 7 | z | 5.483E+04 | 658.3 | 1.084E-05 | 0.01646 | +1 | 5 |
| - | - | 1.914E+04 | 659.3 | - | - | 0 | - |
| - | - | 4131 | 660.3 | - | - | 0 | - |
| - | - | 3499 | 671.4 | - | - | 0 | - |
| - | - | 1156 | 672.4 | - | - | 0 | - |
| 7 | y | 3315 | 674.4 | 0.0002077 | 0.3079 | +1 | 5 |
| - | - | 817.7 | 675.4 | - | - | 0 | - |
| - | - | 742.5 | 696.4 | - | - | 0 | - |
| 6 | z | 1.5E+04 | 715.4 | 2.955E-05 | 0.0413 | +1 | 6 |
| - | - | 5368 | 716.4 | - | - | 0 | - |
| - | - | 1144 | 717.4 | - | - | 0 | - |
| 6 | y | 3116 | 731.4 | 0.001876 | 2.565 | +1 | 6 |
| - | - | 1235 | 732.4 | - | - | 0 | - |
| - | - | 762.3 | 738.5 | - | - | 0 | - |
| - | - | 1053 | 740.5 | - | - | 0 | - |
| - | - | 573.3 | 755.2 | - | - | 0 | - |
| - | - | 870.8 | 780.4 | - | - | 0 | - |
| - | - | 1968 | 781.5 | - | - | 0 | - |
| 7 | c | 2.068E+04 | 782.5 | 0.0004424 | 0.5654 | +1 | 7 |
| - | - | 9425 | 783.5 | - | - | 0 | - |
| - | - | 2703 | 784.5 | - | - | 0 | - |
| 5 | w | 1625 | 785.4 | 0.001932 | 2.46 | +1 | 7 |
| - | - | 876.5 | 786.4 | - | - | 0 | - |
| - | - | 915.7 | 787.4 | - | - | 0 | - |
| 5 | y | 1162 | 810.4 | 0.003336 | 4.116 | +1 | 7 |
| - | - | 1919 | 811.4 | - | - | 0 | - |
| - | - | 915.8 | 819.5 | - | - | 0 | - |
| - | - | 3548 | 824.4 | - | - | 0 | - |
| - | - | 2146 | 825.5 | - | - | 0 | - |
| - | - | 1298 | 825.5 | - | - | 0 | - |
| - | - | 743.9 | 826.4 | - | - | 0 | - |
| - | - | 1033 | 827.4 | - | - | 0 | - |
| 5 | y | 4420 | 828.4 | 0.0007777 | 0.9388 | +1 | 7 |
| - | - | 2307 | 829.4 | - | - | 0 | - |
| - | - | 1122 | 830.4 | - | - | 0 | - |
| - | - | 1389 | 839.5 | - | - | 0 | - |
| - | - | 802.2 | 840.5 | - | - | 0 | - |
| - | - | 1396 | 852.5 | - | - | 0 | - |
| 8 | c | 1.57E+04 | 853.5 | 0.0005079 | 0.595 | +1 | 8 |
| - | - | 877.7 | 854.4 | - | - | 0 | - |
| - | - | 7338 | 854.5 | - | - | 0 | - |
| - | - | 1693 | 855.5 | - | - | 0 | - |
| - | - | 865.9 | 856.4 | - | - | 0 | - |
| - | - | 740.2 | 858.4 | - | - | 0 | - |
| 4 | z | 5.605E+04 | 883.5 | 0.0009179 | 1.039 | +1 | 8 |
| - | - | 2.843E+04 | 884.5 | - | - | 0 | - |
| - | - | 7856 | 885.5 | - | - | 0 | - |
| - | - | 1894 | 886.5 | - | - | 0 | - |
| 4 | y | 8205 | 899.5 | 0.0007211 | 0.8017 | +1 | 8 |
| - | - | 3858 | 900.5 | - | - | 0 | - |
| - | - | 718 | 901.5 | - | - | 0 | - |
| - | - | 1141 | 915.4 | - | - | 0 | - |
| - | - | 817.9 | 924.6 | - | - | 0 | - |
| - | - | 930.5 | 950.6 | - | - | 0 | - |
| - | - | 1383 | 951.6 | - | - | 0 | - |
| - | - | 976 | 952.6 | - | - | 0 | - |
| 3 | w | 1485 | 953.5 | 0.002569 | 2.695 | +1 | 9 |
| - | - | 705.9 | 965.6 | - | - | 0 | - |
| 9 | c | 1.793E+04 | 966.6 | 0.001076 | 1.113 | +1 | 9 |
| - | - | 679.7 | 967.5 | - | - | 0 | - |
| - | - | 8821 | 967.6 | - | - | 0 | - |
| - | - | 2484 | 968.6 | - | - | 0 | - |
| - | - | 1915 | 980.6 | - | - | 0 | - |
| - | - | 1342 | 981.6 | - | - | 0 | - |
| - | - | 813.3 | 1010 | - | - | 0 | - |
| 3 | y | 780.9 | 1011 | 0.002964 | 2.934 | +1 | 9 |
| - | - | 682.4 | 1011 | - | - | 0 | - |
| 3 | z | 4.662E+04 | 1012 | 0.001085 | 1.072 | +1 | 9 |
| - | - | 2.744E+04 | 1013 | - | - | 0 | - |
| - | - | 469.4 | 1013 | - | - | 0 | - |
| - | - | 820.3 | 1013 | - | - | 0 | - |
| - | - | 9603 | 1014 | - | - | 0 | - |
| - | - | 2487 | 1015 | - | - | 0 | - |
| - | - | 775.1 | 1016 | - | - | 0 | - |
| - | - | 803.7 | 1025 | - | - | 0 | - |
| 3 | y | 1.185E+04 | 1028 | 0.0007049 | 0.686 | +1 | 9 |
| - | - | 6982 | 1029 | - | - | 0 | - |
| - | - | 2339 | 1030 | - | - | 0 | - |
| - | - | 855.6 | 1031 | - | - | 0 | - |
| - | - | 1297 | 1036 | - | - | 0 | - |
| - | - | 836 | 1037 | - | - | 0 | - |
| - | - | 2629 | 1038 | - | - | 0 | - |
| - | - | 1637 | 1039 | - | - | 0 | - |
| - | - | 628.3 | 1040 | - | - | 0 | - |
| - | - | 6117 | 1052 | - | - | 0 | - |
| - | - | 1.527E+04 | 1053 | - | - | 0 | - |
| - | - | 9861 | 1054 | - | - | 0 | - |
| - | - | 3121 | 1055 | - | - | 0 | - |
| - | - | 1.149E+04 | 1080 | - | - | 0 | - |
| - | - | 8144 | 1081 | - | - | 0 | - |
| 2 | w | 975.1 | 1082 | 0.01774 | 16.4 | +1 | 10 |
| - | - | 2111 | 1082 | - | - | 0 | - |
| - | - | 1967 | 1083 | - | - | 0 | - |
| - | - | 1073 | 1084 | - | - | 0 | - |
| 10 | c | 1.116E+04 | 1096 | 0.001066 | 0.9732 | +1 | 10 |
| - | - | 3.188E+04 | 1097 | - | - | 0 | - |
| - | - | 1.658E+04 | 1098 | - | - | 0 | - |
| - | - | 4617 | 1099 | - | - | 0 | - |
| - | - | 850 | 1100 | - | - | 0 | - |
| - | - | 619.2 | 1106 | - | - | 0 | - |
| - | - | 1888 | 1154 | - | - | 0 | - |
| - | - | 1735 | 1155 | - | - | 0 | - |
| 2 | z | 2792 | 1168 | 0.003502 | 2.999 | +1 | 10 |
| - | - | 1.885E+04 | 1169 | - | - | 0 | - |
| - | - | 1.274E+04 | 1170 | - | - | 0 | - |
| - | - | 4205 | 1171 | - | - | 0 | - |
| - | - | 1467 | 1172 | - | - | 0 | - |
| - | - | 3089 | 1181 | - | - | 0 | - |
| - | - | 2148 | 1182 | - | - | 0 | - |
| - | - | 2224 | 1183 | - | - | 0 | - |
| 2 | y | 1786 | 1184 | 0.01441 | 12.18 | +1 | 10 |
| - | - | 1010 | 1185 | - | - | 0 | - |
| - | - | 757.6 | 1186 | - | - | 0 | - |
| - | - | 1579 | 1196 | - | - | 0 | - |
| - | - | 843.4 | 1197 | - | - | 0 | - |
| - | - | 761 | 1198 | - | - | 0 | - |
| - | - | 973.8 | 1207 | - | - | 0 | - |
| - | - | 676.6 | 1208 | - | - | 0 | - |
| - | - | 844.2 | 1222 | - | - | 0 | - |
| - | - | 2696 | 1223 | - | - | 0 | - |
| - | - | 5256 | 1224 | - | - | 0 | - |
| - | - | 3.283E+04 | 1225 | - | - | 0 | - |
| - | - | 2.314E+04 | 1226 | - | - | 0 | - |
| - | - | 9797 | 1227 | - | - | 0 | - |
| - | - | 3570 | 1228 | - | - | 0 | - |
| - | - | 1730 | 1229 | - | - | 0 | - |
| - | - | 2625 | 1239 | - | - | 0 | - |
| - | - | 9543 | 1240 | - | - | 0 | - |
| - | - | 1.007E+04 | 1241 | - | - | 0 | - |
| - | - | 7042 | 1242 | - | - | 0 | - |
| - | - | 3435 | 1243 | - | - | 0 | - |
| - | - | 1449 | 1244 | - | - | 0 | - |
| - | - | 1170 | 1250 | - | - | 0 | - |
| - | - | 1.017E+04 | 1251 | - | - | 0 | - |
| - | - | 8600 | 1252 | - | - | 0 | - |
| - | - | 3065 | 1253 | - | - | 0 | - |
| - | - | 1134 | 1254 | - | - | 0 | - |
| - | - | 1959 | 1257 | - | - | 0 | - |
| - | - | 1963 | 1258 | - | - | 0 | - |
| - | - | 1518 | 1266 | - | - | 0 | - |
| - | - | 1.237E+04 | 1267 | - | - | 0 | - |
| - | - | 9.679E+04 | 1268 | - | - | 0 | - |
| - | - | 6.564E+04 | 1269 | - | - | 0 | - |
| - | - | 2.699E+04 | 1270 | - | - | 0 | - |
| - | - | 7034 | 1271 | - | - | 0 | - |
| - | - | 1782 | 1272 | - | - | 0 | - |
| - | - | 928.6 | 1282 | - | - | 0 | - |
| - | - | 5179 | 1283 | - | - | 0 | - |
| - | - | 2.233E+04 | 1284 | - | - | 0 | - |
| - | - | 8.178E+04 | 1285 | - | - | 0 | - |
| - | - | 5.486E+04 | 1286 | - | - | 0 | - |
| - | - | 2.211E+04 | 1287 | - | - | 0 | - |
| - | - | 6807 | 1288 | - | - | 0 | - |
| - | - | 1061 | 1289 | - | - | 0 | - |

m/z Charge Intensity FragmentType MassShift Position
120.06690216064453 0 422.9981
120.08111572265625 0 1918.3389
125.0714111328125 0 409.84033
127.1117935180664 0 377.94586
129.1025848388672 0 438.04855
130.06539916992188 0 3415.5542
131.07022094726562 0 580.3765
132.04188537597656 0 383.24963
132.08145141601562 0 782.07385
133.08612060546875 0 1480.4019
134.33242797851562 0 401.10284
142.06137084960938 0 554.87476
142.0977325439453 0 3082.6436
144.08123779296875 0 648.1389
146.06028747558594 0 4056.7097
148.2520751953125 0 424.9662
149.04518127441406 0 1235.9642
154.42532348632812 0 420.75684
157.1086883544922 0 2309.356
159.09182739257812 0 9071.466
160.09510803222656 0 976.46814
165.1024932861328 0 1069.4922
165.98464965820312 0 515.8522
170.11572265625 0 484.55972
171.092041015625 0 464.7966
173.450927734375 0 2118.0664
174.05519104003906 0 1031.0732
174.10276794433594 0 516.60223
174.13568115234375 0 731.33563
177.11219787597656 0 604.5586
185.10308837890625 0 754.7779
185.11770629882812 0 1538.4374
187.086669921875 0 1489.3193
188.07081604003906 0 24935.6
189.07432556152344 0 3051.6887
192.62184143066406 0 1035.3485 c Ammonia loss 2
201.12411499023438 0 817.4369
203.10255432128906 0 636.35754
203.12811279296875 0 1634.0492
205.0973358154297 0 14170.9795 y 10
206.10110473632812 0 1384.7156
208.95335388183594 0 20314.518
211.1551055908203 0 630.22345
212.13938903808594 0 762.072
213.17117309570312 0 2247.3074
214.1428985595703 0 2630.5383
214.15530395507812 0 1419.3573
214.64483642578125 0 727.8942
215.13916015625 0 950.5948
228.14010620117188 0 817.8356 c Ammonia loss 3
228.18173217773438 0 946.08057
228.6402587890625 0 561.92413
238.16702270507812 0 1027.6182
239.05531311035156 0 503.00058
239.15045166015625 0 9448.007
240.15443420410156 0 839.1558
242.16168212890625 0 1497.5029
245.09217834472656 0 2209.722
246.09512329101562 0 609.1227
256.1769104003906 0 4836.483 c Ammonia loss 1
257.1809387207031 0 651.01764
273.2034912109375 0 20632.229 c 1
274.2065734863281 0 2017.3558
280.1298828125 0 541.678
290.0459899902344 0 646.87585
296.1969299316406 0 555.53595
298.1775207519531 0 666.4474
298.22454833984375 0 597.2876
299.21942138671875 0 5268.458
300.2219543457031 0 738.939
314.9366760253906 0 524.7705
316.12890625 0 3587.8018 y Water loss 9
318.9225769042969 0 14394.432
327.2026062011719 0 630.2098
338.1498718261719 0 4595.718
339.1539306640625 0 1131.2097
341.01824951171875 0 3868.6926
356.2403869628906 0 1068.4617
357.2472229003906 0 1108.8391
359.028564453125 0 13715.053
361.7226867675781 0 1100.7328
362.2251281738281 0 647.73096
364.7184143066406 0 513.4659
369.2283020019531 0 563.7144
369.7342224121094 0 4490.718
370.2351989746094 0 2356.8667
370.25921630859375 0 664.51483
383.22760009765625 0 2970.0654 c Ammonia loss 6
383.7295837402344 0 647.4707
384.23504638671875 0 3927.7126 c Ammonia loss 2
391.2370300292969 0 24645.182
391.7406311035156 0 73626.02 c 6
392.2423095703125 0 30687.422
392.7438659667969 0 6260.889
393.2434997558594 0 961.28864
401.2622375488281 0 65208.03 c 2
402.2650146484375 0 13358.886
403.2671813964844 0 1468.368
404.74951171875 0 1122.1609
405.71710205078125 0 1238.9578 y Water loss 4
406.8998107910156 0 1480.5055
407.2345886230469 0 1551.5872
409.1878662109375 0 1304.0206
410.192626953125 0 855.38086
412.9032897949219 0 1646.4742
413.2386474609375 0 1045.6544
413.5727233886719 0 951.8251
414.72222900390625 0 924.38116 y 4
418.74688720703125 0 3709.356 c Ammonia loss 7
419.2489318847656 0 1751.6882
426.7555236816406 0 8713.374
427.25970458984375 0 60718.926 c 7
427.7607421875 0 24137.549
428.15167236328125 0 1710.764
428.2622985839844 0 6271.2886
428.8920593261719 0 2944.335
429.0890197753906 0 15462.107
429.21142578125 0 1204.0077 y Water loss 8
429.2482604980469 0 664.7379
455.2726135253906 0 3606.5784 c Ammonia loss 3
456.2745666503906 0 841.17377
461.2912902832031 0 6292.8267
461.7934265136719 0 4257.405
462.29510498046875 0 1527.0435
474.79644775390625 0 1304.5485
475.28857421875 0 12934.767 c Ammonia loss 8
475.78997802734375 0 7894.264
476.2893371582031 0 1728.3618
476.79180908203125 0 1413.4548
481.7610168457031 0 703.4385
482.7939147949219 0 581.1455
483.2983703613281 0 2117.234
483.8016052246094 0 30638.068 c 8
484.30291748046875 0 18193.314
484.80426025390625 0 4206.603
486.2680969238281 0 702.8598
496.80950927734375 0 17213.195
497.310791015625 0 8510.798
497.8114318847656 0 2170.4504
498.2841796875 0 1855.6448
498.317626953125 0 1005.90216
498.7887878417969 0 1154.3639
499.3228454589844 0 911.39233
504.78936767578125 0 1980.4528
519.7860717773438 0 766.3877
524.333984375 0 1293.4105
525.33740234375 0 1721.6117
525.8123779296875 0 17574.277
526.31396484375 0 10370.739
526.8153076171875 0 3537.924
527.318603515625 0 948.7593
530.802490234375 0 705.4585
539.8097534179688 0 76437.19 c Ammonia loss 9
540.3106689453125 0 40967.137
540.812255859375 0 14156.282
541.3142700195312 0 3529.3855
541.8167724609375 0 796.7217
542.3320922851562 0 2683.2908
543.33837890625 0 708.57196
547.3142700195312 0 960.93933
547.8141479492188 0 983.81744
548.3229370117188 0 137996.52 c 9
548.82421875 0 80650.766
549.325439453125 0 29495.94
549.8262939453125 0 6908.4863
550.3272705078125 0 1155.079
555.2962036132812 0 8055.069
555.3406982421875 0 1254.799
555.7979125976562 0 4423.8027
556.2943725585938 0 989.94684
561.3303833007812 0 5902.228
561.788330078125 0 794.7736
561.8329467773438 0 4354.3916
562.330810546875 0 1640.8539
562.7824096679688 0 1186.2775
568.3463134765625 0 1597.3287
569.3517456054688 0 22685.16 c 4
570.3548583984375 0 8268.8545
571.3002319335938 0 860.8434
571.3562622070312 0 1603.7941
575.836669921875 0 2429.991
576.3384399414062 0 1123.3917
576.8222045898438 0 10336.044
577.3240356445312 0 8982.361
577.824462890625 0 2942.3213
578.3235473632812 0 733.9252
582.3528442382812 0 2775.415
582.8447875976562 0 830.99066
583.8273315429688 0 1412.6901
584.3123779296875 0 5046.985 z 1
584.8114013671875 0 4589.731
584.85546875 0 1344.4116
585.314697265625 0 1676.2086
586.319580078125 0 863.6035
590.3431396484375 0 833.7004
590.8529663085938 0 813.392
591.3587036132812 0 1009.4933
591.8123168945312 0 751.05066
592.3316040039062 0 607.3992 y 1
597.832275390625 0 1171.8711
598.3262329101562 0 3080.2231
598.8298950195312 0 1959.0104
599.3333740234375 0 2426.6396
600.3335571289062 0 1028.6089
602.2814331054688 0 1511.8982
603.2846069335938 0 649.7265
603.3419189453125 0 670.66693
604.3381958007812 0 2561.6912
604.8323364257812 0 5504.5464
605.333740234375 0 4823.464
605.8239135742188 0 2341.1316
606.3440551757812 0 1060.4376
606.8392333984375 0 733.15656
611.354736328125 0 2247.7637
611.8501586914062 0 10822.577
612.3525390625 0 7313.0747
612.8523559570312 0 3067.8127
613.3424682617188 0 2083.496
613.8471069335938 0 765.8404
614.3526000976562 0 6871.73
614.8284301757812 0 1432.5701
615.2885131835938 0 4829.4155
615.3568725585938 0 2506.3413
616.2928466796875 0 1431.9851
619.357177734375 0 1326.5171
619.8496704101562 0 6656.169
620.35888671875 0 11153.533
620.8460083007812 0 19548.312
621.3469848632812 0 14070.86
621.8467407226562 0 6047.2207
622.3472290039062 0 2290.4058
624.3505249023438 0 1076.0645
625.3648681640625 0 4864.7017
625.8369750976562 0 1407.5327
626.37353515625 0 99692.37 c 5
626.8348388671875 0 7679.6655
627.3764038085938 0 33816.715
627.4367065429688 0 1092.1628
627.8382568359375 0 1027.3702
628.3779296875 0 6407.459
632.8397827148438 0 1285.9343
633.3442993164062 0 2840.1252
633.8450927734375 0 108367.445
634.346435546875 0 79705.57
634.8474731445312 0 31294.67
635.3489990234375 0 8965.32
635.8509521484375 0 1910.9583
640.8474731445312 0 673.2117
641.3502197265625 0 2138.9514
641.8543090820312 0 49403.176
642.356689453125 0 70751.266
642.8587646484375 0 37183.152
643.359619140625 0 13737.692
643.8599853515625 0 4608.1343
644.3551635742188 0 1394.6648
652.3851318359375 0 832.9978
658.3433227539062 0 54831.707 z 6
659.34619140625 0 19140.766
660.348388671875 0 4131.1426
671.3754272460938 0 3499.4246
672.3773803710938 0 1155.8444
674.3622436523438 0 3314.8994 y 6
675.363525390625 0 817.73627
696.3930053710938 0 742.5033
715.36474609375 0 15002.848 z 5
716.366943359375 0 5367.8706
717.3676147460938 0 1144.201
731.3853759765625 0 3116.1064 y 5
732.3880615234375 0 1234.6912
738.4593505859375 0 762.3242
740.4595947265625 0 1053.0977
755.1654052734375 0 573.3035
780.4400024414062 0 870.8048
781.464599609375 0 1968.2013
782.4739379882812 0 20677.625 c 6
783.47607421875 0 9425.015
784.4808349609375 0 2703.2217
785.39599609375 0 1624.733 w 4
786.3997802734375 0 876.5087
787.3997192382812 0 915.687
810.42236328125 0 1161.8458 y Water loss 4
811.4282836914062 0 1919.1688
819.4577026367188 0 915.8011
824.44775390625 0 3547.6052
825.4534301757812 0 2145.8418
825.5335083007812 0 1298.0205
826.4382934570312 0 743.9138
827.4049072265625 0 1033.4514
828.4354858398438 0 4419.9346 y 4
829.4393310546875 0 2307.2126
830.4425048828125 0 1121.9211
839.4619140625 0 1388.9967
840.4707641601562 0 802.1691
852.5034790039062 0 1396.0125
853.510986328125 0 15702.396 c 7
854.43701171875 0 877.6832
854.5147094726562 0 7337.8823
855.5192260742188 0 1693.3604
856.4348754882812 0 865.9236
858.4281616210938 0 740.2041
883.4537353515625 0 56045.656 z 3
884.4569702148438 0 28429.299
885.4595336914062 0 7856.0513
886.4630126953125 0 1893.5793
899.47265625 0 8204.638 y 3
900.475830078125 0 3857.954
901.476318359375 0 717.9789
915.444580078125 0 1141.4557
924.57763671875 0 817.8796
950.5732421875 0 930.51843
951.5830688476562 0 1383.249
952.5868530273438 0 975.9729
953.4865112304688 0 1484.5206 w 2
965.5875244140625 0 705.8953
966.594482421875 0 17934.906 c 8
967.5127563476562 0 679.715
967.5975341796875 0 8820.725
968.5999755859375 0 2483.6282
980.551513671875 0 1914.9244
981.5507202148438 0 1341.5841
1009.6027221679688 0 813.2958
1010.50244140625 0 780.8691 y Ammonia loss 2
1010.6083984375 0 682.39923
1011.5121459960938 0 46619.137 z 2
1012.514892578125 0 27443.453
1012.6150512695312 0 469.4123
1012.6410522460938 0 820.291
1013.516845703125 0 9603.113
1014.515869140625 0 2487.0254
1015.5213623046875 0 775.14
1024.6258544921875 0 803.71826
1027.53125 0 11849.926 y 2
1028.534912109375 0 6981.555
1029.537109375 0 2338.8718
1030.5372314453125 0 855.62744
1035.603759765625 0 1296.8236
1036.608642578125 0 835.9545
1037.5975341796875 0 2628.9956
1038.5963134765625 0 1636.7057
1039.609130859375 0 628.274
1051.62353515625 0 6117.429
1052.6270751953125 0 15266.735
1053.6259765625 0 9861.232
1054.6287841796875 0 3121.091
1079.6181640625 0 11491.364
1080.62158203125 0 8143.825
1081.5247802734375 0 975.12524 w 1
1081.6231689453125 0 2110.7727
1082.546630859375 0 1967.4387
1083.547607421875 0 1073.0012
1095.6370849609375 0 11161.574 c 9
1096.6436767578125 0 31883.379
1097.6466064453125 0 16577.736
1098.648681640625 0 4616.978
1099.65234375 0 850.0254
1106.0472412109375 0 619.243
1153.634033203125 0 1887.8533
1154.6328125 0 1734.955
1167.61083984375 0 2792.1462 z 1
1168.6199951171875 0 18846.428
1169.623046875 0 12741.516
1170.62841796875 0 4204.642
1171.6259765625 0 1467.2858
1180.6243896484375 0 3089.0508
1181.6259765625 0 2147.8813
1182.6097412109375 0 2223.8474
1183.61865234375 0 1786.0184 y 1
1184.630615234375 0 1009.9132
1185.6314697265625 0 757.6419
1195.62353515625 0 1578.8357
1196.6180419921875 0 843.438
1197.6351318359375 0 761.0133
1206.6697998046875 0 973.7651
1207.675537109375 0 676.61566
1221.691162109375 0 844.23474
1222.7037353515625 0 2696.159
1223.6951904296875 0 5255.8145
1224.6607666015625 0 32826.793
1225.6632080078125 0 23140.463
1226.6656494140625 0 9796.561
1227.6680908203125 0 3569.851
1228.6632080078125 0 1730.4427
1238.6998291015625 0 2625.0608
1239.69580078125 0 9543.106
1240.6973876953125 0 10066.806
1241.6964111328125 0 7041.6973
1242.6986083984375 0 3435.3948
1243.69482421875 0 1448.7292
1249.6800537109375 0 1170.1422
1250.6614990234375 0 10171.325
1251.6651611328125 0 8599.814
1252.663818359375 0 3065.4878
1253.6728515625 0 1133.8125
1256.7237548828125 0 1959.0768
1257.715576171875 0 1963.2606
1265.685302734375 0 1517.9615
1266.6861572265625 0 12368.487
1267.68798828125 0 96787.18
1268.6900634765625 0 65644.266
1269.6937255859375 0 26989.97
1270.6944580078125 0 7034.21
1271.69775390625 0 1782.3842
1281.6905517578125 0 928.6477
1282.698974609375 0 5179.4067
1283.7049560546875 0 22325.547
1284.71337890625 0 81781.164
1285.7156982421875 0 54857.668
1286.7188720703125 0 22112.693
1287.720947265625 0 6806.906
1288.715087890625 0 1060.6786

Spectrum Details

|  |  |
| --- | --- |
| Matched peaks? Matched peaks The total absolute number of peaks matched. Additionally in brackets the total fraction of peaks matched and the total number of peaks is shown. | 44 (10.73% of 410) |
| FDR? FDR The false discovery rate estimated for this peptide. It is calculated by matching all theoretical fragments with a non-integer shift with the raw peaks for this spectrum. This is done with 40 different shifts. The resulting percentage is the average number of annotated peaks over the number of annotated peaks with the correct spectrum. | 2.06% |
| Satellite FDR? Satellite FDR See the FDR for details on its calculation. This satellite ion specific FDR only contains the satellite ions (d/w) for I/L/J positions. | ∞ |
| PSM Score? PSM Score The PSM Score as given by Hecklib to this annotated spectrum. It is shown with three significant figures. | 351 |

#### Spectrum 5752? Spectrum 5752 The raw spectrum of this peptide as annotated by Hecklib. The fragments are coloured according to ion type (see legend). Any peaks with a star '\*' as text can be hovered over to see the full details, first the ion type second the mass shift type. By hovering over the amino acids in the peptide or ions in the legend the corresponding peaks are highlighted. By toggling the 'Unassigned' label you can turn the background (unassigned) peaks on or off in the plot. By updating the slider in the Ion legend you can update the spectrum to only show the top X% of the peaks with labels. The top X% means any peak that is within X% of the highest intensity. By dragging in the spectrum you can zoom in to a specific part of the spectrum and use 'Zoom Out' to get back to the original zoom level. The annotation of the spectrum is based on the given sequence in the peptides file and is done with different software so inconsistencies are likely. The peaks are annotated based on the given sequence, with 20 ppm tolerance.

Copy Data

##### Spectrum 5752 (TSV)

###### Preview

```
Loading example...
```

*Click on the button to copy the data to your clipboard.*

Mz MinMz MaxIntensity Max

WidthHeightPeptide font sizePeptide stroke widthSpectrum font sizeSpectrum stroke widthCompact peptide

Ion legend

wxyz

abcd

OtherUnassignedIon^Charge^_Position_Show for top:%

VRQAPGRAJEW

09.95e+41.99e+52.98e+53.98e+5

Zoom Out

c^+2^_3_y^+1^_1_c^+2^_4_c^+1^_2_c^+1^_2_c^+2^_5_y^+1^_2_y^+1^_2_c^+2^_7_z^+3^_10_c^+1^_3_c^+2^_7_c^+1^_3_y^+2^_7_y^+2^_7_c^+2^_8_c^+2^_8_c^+1^_4_c^+2^_9_c^+2^_9_y^+1^_4_c^+2^_10_c^+2^_10_c^+2^_10_c^+1^_5_z^+2^_10_y^+2^_10_c^+1^_6_y^+1^_5_z^+1^_5_y^+1^_5_y^+1^_6_z^+1^_6_y^+1^_6_c^+1^_7_w^+1^_7_y^+1^_7_z^+1^_7_y^+1^_7_c^+1^_8_z^+1^_8_y^+1^_8_w^+1^_9_c^+1^_9_z^+1^_9_y^+1^_9_c^+1^_10_w^+1^_10_c^+1^_10_z^+1^_10_y^+1^_10_

0847169425423389

Fragment Matches Table

Show background peaks

| Position | Ion type | Intensity | mz Theoretical | mz Error (Th) | mz Error (ppm) | Charge | Series Number |
| --- | --- | --- | --- | --- | --- | --- | --- |
| - | - | 368.5 | 122.4 | - | - | 0 | - |
| - | - | 361.1 | 122.8 | - | - | 0 | - |
| - | - | 380.3 | 126.1 | - | - | 0 | - |
| - | - | 332.1 | 128 | - | - | 0 | - |
| - | - | 649.4 | 129.1 | - | - | 0 | - |
| - | - | 390.9 | 129.3 | - | - | 0 | - |
| - | - | 7998 | 130.1 | - | - | 0 | - |
| - | - | 723.8 | 131.1 | - | - | 0 | - |
| - | - | 1760 | 132.1 | - | - | 0 | - |
| - | - | 1331 | 133.1 | - | - | 0 | - |
| - | - | 370.5 | 135.5 | - | - | 0 | - |
| - | - | 512 | 142.1 | - | - | 0 | - |
| - | - | 1646 | 142.1 | - | - | 0 | - |
| - | - | 410.7 | 143.9 | - | - | 0 | - |
| - | - | 1702 | 144.1 | - | - | 0 | - |
| - | - | 8057 | 146.1 | - | - | 0 | - |
| - | - | 947.4 | 147.1 | - | - | 0 | - |
| - | - | 424.1 | 147.4 | - | - | 0 | - |
| - | - | 977.6 | 149 | - | - | 0 | - |
| - | - | 1385 | 149 | - | - | 0 | - |
| - | - | 416.5 | 154.5 | - | - | 0 | - |
| - | - | 3776 | 157.1 | - | - | 0 | - |
| - | - | 2.621E+04 | 159.1 | - | - | 0 | - |
| - | - | 531.9 | 159.2 | - | - | 0 | - |
| - | - | 2698 | 160.1 | - | - | 0 | - |
| - | - | 447 | 161.1 | - | - | 0 | - |
| - | - | 844.3 | 166.1 | - | - | 0 | - |
| - | - | 1515 | 170.1 | - | - | 0 | - |
| - | - | 491.5 | 171.1 | - | - | 0 | - |
| - | - | 1867 | 173.5 | - | - | 0 | - |
| - | - | 633.9 | 173.5 | - | - | 0 | - |
| - | - | 1163 | 174.1 | - | - | 0 | - |
| - | - | 1230 | 174.1 | - | - | 0 | - |
| - | - | 1975 | 185.1 | - | - | 0 | - |
| - | - | 530.4 | 185.1 | - | - | 0 | - |
| - | - | 748.3 | 185.1 | - | - | 0 | - |
| - | - | 5264 | 187.1 | - | - | 0 | - |
| - | - | 8.798E+04 | 188.1 | - | - | 0 | - |
| - | - | 1.007E+04 | 189.1 | - | - | 0 | - |
| - | - | 742.3 | 190.1 | - | - | 0 | - |
| 3 | c | 2265 | 192.6 | 0.0005137 | 2.667 | +2 | 3 |
| - | - | 578.9 | 196.1 | - | - | 0 | - |
| - | - | 517.8 | 196.4 | - | - | 0 | - |
| - | - | 1751 | 197.1 | - | - | 0 | - |
| - | - | 504.6 | 198.1 | - | - | 0 | - |
| - | - | 1245 | 203.1 | - | - | 0 | - |
| - | - | 509.7 | 204.7 | - | - | 0 | - |
| 11 | y | 5.286E+04 | 205.1 | 0.0004259 | 2.076 | +1 | 1 |
| - | - | 569.1 | 205.6 | - | - | 0 | - |
| - | - | 6665 | 206.1 | - | - | 0 | - |
| - | - | 2.302E+04 | 209 | - | - | 0 | - |
| - | - | 494.4 | 212.4 | - | - | 0 | - |
| - | - | 5231 | 213.2 | - | - | 0 | - |
| - | - | 7963 | 214.1 | - | - | 0 | - |
| - | - | 3574 | 214.2 | - | - | 0 | - |
| - | - | 2252 | 214.6 | - | - | 0 | - |
| - | - | 760.5 | 215.2 | - | - | 0 | - |
| 4 | c | 2697 | 228.1 | 0.0006794 | 2.978 | +2 | 4 |
| - | - | 2115 | 228.2 | - | - | 0 | - |
| - | - | 778.1 | 228.6 | - | - | 0 | - |
| - | - | 1931 | 238.2 | - | - | 0 | - |
| - | - | 2.256E+04 | 239.2 | - | - | 0 | - |
| - | - | 3047 | 240.2 | - | - | 0 | - |
| - | - | 672.7 | 240.9 | - | - | 0 | - |
| - | - | 4579 | 242.2 | - | - | 0 | - |
| - | - | 571.5 | 243.2 | - | - | 0 | - |
| - | - | 533.8 | 252.2 | - | - | 0 | - |
| - | - | 1567 | 256.1 | - | - | 0 | - |
| 2 | c | 1.226E+04 | 256.2 | 0.0004142 | 1.617 | +1 | 2 |
| - | - | 1923 | 257.2 | - | - | 0 | - |
| - | - | 3390 | 270.1 | - | - | 0 | - |
| - | - | 704.6 | 271.1 | - | - | 0 | - |
| 2 | c | 4.838E+04 | 273.2 | 0.0003849 | 1.409 | +1 | 2 |
| - | - | 6548 | 274.2 | - | - | 0 | - |
| - | - | 713.1 | 275.2 | - | - | 0 | - |
| 5 | c | 636.1 | 276.7 | 0.0003424 | 1.238 | +2 | 5 |
| - | - | 699.7 | 280.1 | - | - | 0 | - |
| - | - | 1664 | 298.1 | - | - | 0 | - |
| - | - | 979 | 298.2 | - | - | 0 | - |
| - | - | 1.27E+04 | 299.2 | - | - | 0 | - |
| - | - | 2276 | 300.2 | - | - | 0 | - |
| - | - | 799.1 | 307.9 | - | - | 0 | - |
| - | - | 678.5 | 312.2 | - | - | 0 | - |
| - | - | 1859 | 312.7 | - | - | 0 | - |
| 10 | y | 1.369E+04 | 316.1 | 0.0006393 | 2.022 | +1 | 2 |
| - | - | 1.421E+04 | 318.9 | - | - | 0 | - |
| - | - | 565.7 | 324.4 | - | - | 0 | - |
| - | - | 577.9 | 325 | - | - | 0 | - |
| 10 | y | 2043 | 334.1 | 0.0009083 | 2.718 | +1 | 2 |
| - | - | 5198 | 341 | - | - | 0 | - |
| - | - | 978.3 | 341.2 | - | - | 0 | - |
| - | - | 582.4 | 350.2 | - | - | 0 | - |
| - | - | 2804 | 356.2 | - | - | 0 | - |
| - | - | 2760 | 357.2 | - | - | 0 | - |
| - | - | 1.654E+04 | 359 | - | - | 0 | - |
| - | - | 557.8 | 361.2 | - | - | 0 | - |
| - | - | 3326 | 361.7 | - | - | 0 | - |
| - | - | 1720 | 362.2 | - | - | 0 | - |
| - | - | 1479 | 367.2 | - | - | 0 | - |
| - | - | 1389 | 369.2 | - | - | 0 | - |
| - | - | 1.111E+04 | 369.7 | - | - | 0 | - |
| - | - | 1029 | 370.2 | - | - | 0 | - |
| - | - | 6739 | 370.2 | - | - | 0 | - |
| - | - | 921.9 | 370.3 | - | - | 0 | - |
| - | - | 1230 | 370.7 | - | - | 0 | - |
| 7 | c | 4676 | 383.2 | 4.622E-05 | 0.1206 | +2 | 7 |
| - | - | 1850 | 383.7 | - | - | 0 | - |
| 2 | z | 918.7 | 384.2 | 0.005334 | 13.88 | +3 | 10 |
| 3 | c | 1.013E+04 | 384.2 | 0.0003999 | 1.041 | +1 | 3 |
| - | - | 2165 | 385.2 | - | - | 0 | - |
| - | - | 740.8 | 385.3 | - | - | 0 | - |
| - | - | 6.294E+04 | 391.2 | - | - | 0 | - |
| 7 | c | 2.034E+05 | 391.7 | 0.000352 | 0.8985 | +2 | 7 |
| - | - | 8.213E+04 | 392.2 | - | - | 0 | - |
| - | - | 1.84E+04 | 392.7 | - | - | 0 | - |
| - | - | 2231 | 393.2 | - | - | 0 | - |
| 3 | c | 1.63E+05 | 401.3 | 0.0008894 | 2.216 | +1 | 3 |
| - | - | 892.5 | 401.6 | - | - | 0 | - |
| - | - | 3.075E+04 | 402.3 | - | - | 0 | - |
| - | - | 3967 | 403.3 | - | - | 0 | - |
| - | - | 3212 | 404.7 | - | - | 0 | - |
| - | - | 2404 | 405.3 | - | - | 0 | - |
| 5 | y | 4461 | 405.7 | 0.0007365 | 1.815 | +2 | 7 |
| - | - | 2523 | 406.2 | - | - | 0 | - |
| - | - | 6491 | 406.9 | - | - | 0 | - |
| - | - | 5388 | 407.2 | - | - | 0 | - |
| - | - | 2731 | 407.6 | - | - | 0 | - |
| - | - | 608.7 | 407.9 | - | - | 0 | - |
| - | - | 952.4 | 410.2 | - | - | 0 | - |
| - | - | 7079 | 412.9 | - | - | 0 | - |
| - | - | 2704 | 413.2 | - | - | 0 | - |
| - | - | 1124 | 413.3 | - | - | 0 | - |
| - | - | 1094 | 413.6 | - | - | 0 | - |
| 5 | y | 3777 | 414.7 | 0.001039 | 2.505 | +2 | 7 |
| - | - | 1535 | 415.2 | - | - | 0 | - |
| 8 | c | 7074 | 418.7 | 0.001326 | 3.166 | +2 | 8 |
| - | - | 2372 | 419.2 | - | - | 0 | - |
| - | - | 817.5 | 419.7 | - | - | 0 | - |
| - | - | 2722 | 422.2 | - | - | 0 | - |
| - | - | 995.9 | 422.6 | - | - | 0 | - |
| - | - | 2.872E+04 | 426.8 | - | - | 0 | - |
| 8 | c | 1.597E+05 | 427.3 | 0.000777 | 1.819 | +2 | 8 |
| - | - | 7.062E+04 | 427.8 | - | - | 0 | - |
| - | - | 608.1 | 428.1 | - | - | 0 | - |
| - | - | 1.684E+04 | 428.3 | - | - | 0 | - |
| - | - | 2458 | 428.8 | - | - | 0 | - |
| - | - | 1719 | 428.9 | - | - | 0 | - |
| - | - | 1.637E+04 | 429.1 | - | - | 0 | - |
| - | - | 667.7 | 429.2 | - | - | 0 | - |
| - | - | 1015 | 429.3 | - | - | 0 | - |
| - | - | 1522 | 453.8 | - | - | 0 | - |
| - | - | 1623 | 454.3 | - | - | 0 | - |
| 4 | c | 1.151E+04 | 455.3 | 0.0006701 | 1.472 | +1 | 4 |
| - | - | 2931 | 456.3 | - | - | 0 | - |
| - | - | 1.927E+04 | 461.3 | - | - | 0 | - |
| - | - | 1.418E+04 | 461.8 | - | - | 0 | - |
| - | - | 5825 | 462.3 | - | - | 0 | - |
| - | - | 1587 | 462.8 | - | - | 0 | - |
| - | - | 1069 | 466.8 | - | - | 0 | - |
| - | - | 786.7 | 467.3 | - | - | 0 | - |
| - | - | 863.2 | 467.3 | - | - | 0 | - |
| - | - | 833.7 | 468.8 | - | - | 0 | - |
| - | - | 1120 | 469.3 | - | - | 0 | - |
| - | - | 2153 | 472.3 | - | - | 0 | - |
| - | - | 2158 | 474.8 | - | - | 0 | - |
| 9 | c | 4.727E+04 | 475.3 | 0.0009808 | 2.064 | +2 | 9 |
| - | - | 2.425E+04 | 475.8 | - | - | 0 | - |
| - | - | 7673 | 476.3 | - | - | 0 | - |
| - | - | 1213 | 476.8 | - | - | 0 | - |
| - | - | 1058 | 483.3 | - | - | 0 | - |
| - | - | 7878 | 483.3 | - | - | 0 | - |
| 9 | c | 7.845E+04 | 483.8 | 0.0007372 | 1.524 | +2 | 9 |
| - | - | 4.521E+04 | 484.3 | - | - | 0 | - |
| - | - | 1.288E+04 | 484.8 | - | - | 0 | - |
| - | - | 2756 | 485.3 | - | - | 0 | - |
| - | - | 1788 | 486.3 | - | - | 0 | - |
| - | - | 858.1 | 490.8 | - | - | 0 | - |
| - | - | 729.8 | 495.3 | - | - | 0 | - |
| - | - | 4.601E+04 | 496.8 | - | - | 0 | - |
| - | - | 2.442E+04 | 497.3 | - | - | 0 | - |
| - | - | 9329 | 497.8 | - | - | 0 | - |
| - | - | 8329 | 498.3 | - | - | 0 | - |
| - | - | 2408 | 498.3 | - | - | 0 | - |
| - | - | 5062 | 498.8 | - | - | 0 | - |
| - | - | 1399 | 499.3 | - | - | 0 | - |
| - | - | 3334 | 499.3 | - | - | 0 | - |
| - | - | 987.8 | 500.3 | - | - | 0 | - |
| - | - | 4024 | 504.8 | - | - | 0 | - |
| - | - | 1874 | 505.3 | - | - | 0 | - |
| - | - | 630.1 | 510.3 | - | - | 0 | - |
| - | - | 960.4 | 517.3 | - | - | 0 | - |
| - | - | 2304 | 517.8 | - | - | 0 | - |
| 8 | y | 1062 | 518.3 | 6.07E-05 | 0.1171 | +1 | 4 |
| - | - | 3640 | 518.3 | - | - | 0 | - |
| - | - | 1802 | 518.8 | - | - | 0 | - |
| - | - | 942.2 | 519.3 | - | - | 0 | - |
| - | - | 646.8 | 524.3 | - | - | 0 | - |
| - | - | 2441 | 524.3 | - | - | 0 | - |
| - | - | 4144 | 525.3 | - | - | 0 | - |
| - | - | 4.286E+04 | 525.8 | - | - | 0 | - |
| - | - | 2.975E+04 | 526.3 | - | - | 0 | - |
| - | - | 1.204E+04 | 526.8 | - | - | 0 | - |
| - | - | 3767 | 527.3 | - | - | 0 | - |
| - | - | 1492 | 529.3 | - | - | 0 | - |
| - | - | 1012 | 530.8 | - | - | 0 | - |
| - | - | 2954 | 531.3 | - | - | 0 | - |
| - | - | 2320 | 531.8 | - | - | 0 | - |
| - | - | 1220 | 532.3 | - | - | 0 | - |
| - | - | 760.6 | 532.8 | - | - | 0 | - |
| - | - | 744.8 | 533.3 | - | - | 0 | - |
| 10 | c | 2125 | 539.3 | 0.0001954 | 0.3624 | +2 | 10 |
| 10 | c | 2.723E+05 | 539.8 | 0.0009855 | 1.826 | +2 | 10 |
| - | - | 1.509E+05 | 540.3 | - | - | 0 | - |
| - | - | 5.092E+04 | 540.8 | - | - | 0 | - |
| - | - | 1.487E+04 | 541.3 | - | - | 0 | - |
| - | - | 2627 | 541.8 | - | - | 0 | - |
| - | - | 7179 | 542.3 | - | - | 0 | - |
| - | - | 1507 | 543.3 | - | - | 0 | - |
| - | - | 634.4 | 545.2 | - | - | 0 | - |
| - | - | 2177 | 547.8 | - | - | 0 | - |
| 10 | c | 3.939E+05 | 548.3 | 0.001017 | 1.854 | +2 | 10 |
| - | - | 2.256E+05 | 548.8 | - | - | 0 | - |
| - | - | 1263 | 548.9 | - | - | 0 | - |
| - | - | 7.49E+04 | 549.3 | - | - | 0 | - |
| - | - | 1.788E+04 | 549.8 | - | - | 0 | - |
| - | - | 2766 | 550.3 | - | - | 0 | - |
| - | - | 840.1 | 553.8 | - | - | 0 | - |
| - | - | 1138 | 554.3 | - | - | 0 | - |
| - | - | 1800 | 554.8 | - | - | 0 | - |
| - | - | 1.797E+04 | 555.3 | - | - | 0 | - |
| - | - | 3028 | 555.3 | - | - | 0 | - |
| - | - | 1.123E+04 | 555.8 | - | - | 0 | - |
| - | - | 4561 | 556.3 | - | - | 0 | - |
| - | - | 1758 | 558.3 | - | - | 0 | - |
| - | - | 1.371E+04 | 561.3 | - | - | 0 | - |
| - | - | 804.2 | 561.8 | - | - | 0 | - |
| - | - | 9205 | 561.8 | - | - | 0 | - |
| - | - | 3711 | 562.3 | - | - | 0 | - |
| - | - | 2147 | 562.8 | - | - | 0 | - |
| - | - | 1185 | 563.3 | - | - | 0 | - |
| - | - | 788.1 | 567.3 | - | - | 0 | - |
| - | - | 3582 | 568.3 | - | - | 0 | - |
| - | - | 2665 | 568.8 | - | - | 0 | - |
| 5 | c | 5.963E+04 | 569.4 | 0.0007944 | 1.395 | +1 | 5 |
| - | - | 1037 | 569.8 | - | - | 0 | - |
| - | - | 443.4 | 570.3 | - | - | 0 | - |
| - | - | 1.913E+04 | 570.4 | - | - | 0 | - |
| - | - | 3957 | 571.4 | - | - | 0 | - |
| - | - | 6880 | 575.8 | - | - | 0 | - |
| - | - | 3580 | 576.3 | - | - | 0 | - |
| - | - | 2.338E+04 | 576.8 | - | - | 0 | - |
| - | - | 1.985E+04 | 577.3 | - | - | 0 | - |
| - | - | 7749 | 577.8 | - | - | 0 | - |
| - | - | 2317 | 578.3 | - | - | 0 | - |
| - | - | 985.5 | 578.8 | - | - | 0 | - |
| - | - | 5973 | 582.4 | - | - | 0 | - |
| - | - | 2652 | 582.8 | - | - | 0 | - |
| - | - | 2178 | 583.4 | - | - | 0 | - |
| - | - | 1800 | 583.8 | - | - | 0 | - |
| 2 | z | 1.937E+04 | 584.3 | 0.001569 | 2.685 | +2 | 10 |
| - | - | 1.208E+04 | 584.8 | - | - | 0 | - |
| - | - | 4784 | 585.3 | - | - | 0 | - |
| - | - | 1538 | 585.8 | - | - | 0 | - |
| - | - | 1742 | 586.3 | - | - | 0 | - |
| - | - | 586.8 | 590.8 | - | - | 0 | - |
| - | - | 986.8 | 590.9 | - | - | 0 | - |
| - | - | 967.7 | 591.3 | - | - | 0 | - |
| - | - | 1407 | 591.8 | - | - | 0 | - |
| 2 | y | 2256 | 592.3 | 0.003498 | 5.906 | +2 | 10 |
| - | - | 1468 | 592.8 | - | - | 0 | - |
| - | - | 811.5 | 593.3 | - | - | 0 | - |
| - | - | 2129 | 596.4 | - | - | 0 | - |
| - | - | 930.3 | 597.4 | - | - | 0 | - |
| - | - | 3117 | 597.8 | - | - | 0 | - |
| - | - | 6365 | 598.3 | - | - | 0 | - |
| - | - | 4601 | 598.8 | - | - | 0 | - |
| - | - | 5976 | 599.3 | - | - | 0 | - |
| - | - | 1000 | 600.3 | - | - | 0 | - |
| - | - | 5576 | 602.3 | - | - | 0 | - |
| - | - | 1308 | 603.3 | - | - | 0 | - |
| - | - | 931 | 603.3 | - | - | 0 | - |
| - | - | 6538 | 604.3 | - | - | 0 | - |
| - | - | 1.591E+04 | 604.8 | - | - | 0 | - |
| - | - | 1.265E+04 | 605.3 | - | - | 0 | - |
| - | - | 7314 | 605.8 | - | - | 0 | - |
| - | - | 3115 | 606.3 | - | - | 0 | - |
| - | - | 2956 | 606.8 | - | - | 0 | - |
| - | - | 5970 | 611.4 | - | - | 0 | - |
| - | - | 3.281E+04 | 611.9 | - | - | 0 | - |
| - | - | 1.721E+04 | 612.4 | - | - | 0 | - |
| - | - | 1259 | 612.8 | - | - | 0 | - |
| - | - | 1.011E+04 | 612.9 | - | - | 0 | - |
| - | - | 4573 | 613.3 | - | - | 0 | - |
| - | - | 1795 | 613.8 | - | - | 0 | - |
| - | - | 1.836E+04 | 614.4 | - | - | 0 | - |
| - | - | 1969 | 614.8 | - | - | 0 | - |
| - | - | 9173 | 615.3 | - | - | 0 | - |
| - | - | 5406 | 615.4 | - | - | 0 | - |
| - | - | 3353 | 616.3 | - | - | 0 | - |
| - | - | 1447 | 616.4 | - | - | 0 | - |
| - | - | 716.1 | 617.3 | - | - | 0 | - |
| - | - | 3358 | 619.4 | - | - | 0 | - |
| - | - | 2.123E+04 | 619.8 | - | - | 0 | - |
| - | - | 2.64E+04 | 620.4 | - | - | 0 | - |
| - | - | 6.008E+04 | 620.8 | - | - | 0 | - |
| - | - | 3.935E+04 | 621.3 | - | - | 0 | - |
| - | - | 1.452E+04 | 621.8 | - | - | 0 | - |
| - | - | 4406 | 622.3 | - | - | 0 | - |
| - | - | 4108 | 624.3 | - | - | 0 | - |
| - | - | 8726 | 625.4 | - | - | 0 | - |
| - | - | 4884 | 625.8 | - | - | 0 | - |
| 6 | c | 2.399E+05 | 626.4 | 0.001303 | 2.081 | +1 | 6 |
| - | - | 2.365E+04 | 626.8 | - | - | 0 | - |
| - | - | 7.854E+04 | 627.4 | - | - | 0 | - |
| - | - | 1852 | 627.8 | - | - | 0 | - |
| - | - | 1.592E+04 | 628.4 | - | - | 0 | - |
| - | - | 2276 | 629.4 | - | - | 0 | - |
| - | - | 6438 | 633.3 | - | - | 0 | - |
| - | - | 3.192E+05 | 633.8 | - | - | 0 | - |
| - | - | 2.204E+05 | 634.3 | - | - | 0 | - |
| - | - | 8.768E+04 | 634.8 | - | - | 0 | - |
| - | - | 2.434E+04 | 635.3 | - | - | 0 | - |
| - | - | 4674 | 635.9 | - | - | 0 | - |
| - | - | 1.432E+05 | 641.9 | - | - | 0 | - |
| - | - | 1.896E+05 | 642.4 | - | - | 0 | - |
| - | - | 9.801E+04 | 642.9 | - | - | 0 | - |
| - | - | 3.567E+04 | 643.4 | - | - | 0 | - |
| - | - | 9976 | 643.9 | - | - | 0 | - |
| - | - | 2685 | 644.4 | - | - | 0 | - |
| - | - | 1471 | 652.4 | - | - | 0 | - |
| - | - | 866.9 | 653.4 | - | - | 0 | - |
| 7 | y | 1014 | 656.4 | 0.001617 | 2.464 | +1 | 5 |
| 7 | z | 1.434E+05 | 658.3 | 0.0009264 | 1.407 | +1 | 5 |
| - | - | 5.184E+04 | 659.3 | - | - | 0 | - |
| - | - | 1.197E+04 | 660.3 | - | - | 0 | - |
| - | - | 1591 | 661.4 | - | - | 0 | - |
| - | - | 759.6 | 667.4 | - | - | 0 | - |
| - | - | 9571 | 671.4 | - | - | 0 | - |
| - | - | 878.7 | 672.3 | - | - | 0 | - |
| - | - | 3250 | 672.4 | - | - | 0 | - |
| 7 | y | 8843 | 674.4 | 0.000818 | 1.213 | +1 | 5 |
| - | - | 3009 | 675.4 | - | - | 0 | - |
| - | - | 1886 | 696.4 | - | - | 0 | - |
| - | - | 1172 | 697.4 | - | - | 0 | - |
| - | - | 674.4 | 704.4 | - | - | 0 | - |
| 6 | y | 1861 | 713.4 | 0.000417 | 0.5846 | +1 | 6 |
| 6 | z | 3.399E+04 | 715.4 | 0.0007639 | 1.068 | +1 | 6 |
| - | - | 1.388E+04 | 716.4 | - | - | 0 | - |
| - | - | 3087 | 717.4 | - | - | 0 | - |
| 6 | y | 9221 | 731.4 | 0.001327 | 1.814 | +1 | 6 |
| - | - | 4803 | 732.4 | - | - | 0 | - |
| - | - | 1366 | 733.4 | - | - | 0 | - |
| - | - | 890.1 | 738.5 | - | - | 0 | - |
| - | - | 2063 | 739.5 | - | - | 0 | - |
| - | - | 2422 | 740.5 | - | - | 0 | - |
| - | - | 2106 | 767.5 | - | - | 0 | - |
| - | - | 906.1 | 780.4 | - | - | 0 | - |
| - | - | 3776 | 781.5 | - | - | 0 | - |
| 7 | c | 4.282E+04 | 782.5 | 0.0008393 | 1.073 | +1 | 7 |
| - | - | 1.65E+04 | 783.5 | - | - | 0 | - |
| - | - | 4439 | 784.5 | - | - | 0 | - |
| 5 | w | 3758 | 785.4 | 0.001321 | 1.682 | +1 | 7 |
| - | - | 1000 | 785.5 | - | - | 0 | - |
| - | - | 3146 | 786.4 | - | - | 0 | - |
| 5 | y | 3966 | 810.4 | 0.001547 | 1.909 | +1 | 7 |
| - | - | 5445 | 811.4 | - | - | 0 | - |
| - | - | 795.2 | 811.5 | - | - | 0 | - |
| 5 | z | 1814 | 812.4 | 0.01428 | 17.58 | +1 | 7 |
| - | - | 2033 | 819.5 | - | - | 0 | - |
| - | - | 841.2 | 820.5 | - | - | 0 | - |
| - | - | 2490 | 823.4 | - | - | 0 | - |
| - | - | 6972 | 824.4 | - | - | 0 | - |
| - | - | 4775 | 825.5 | - | - | 0 | - |
| - | - | 2368 | 826.4 | - | - | 0 | - |
| - | - | 3036 | 827.4 | - | - | 0 | - |
| 5 | y | 1.197E+04 | 828.4 | 0.0001063 | 0.1284 | +1 | 7 |
| - | - | 6336 | 829.4 | - | - | 0 | - |
| - | - | 1571 | 830.4 | - | - | 0 | - |
| - | - | 3245 | 839.5 | - | - | 0 | - |
| - | - | 2691 | 840.5 | - | - | 0 | - |
| - | - | 770.2 | 841.5 | - | - | 0 | - |
| - | - | 889.1 | 844.5 | - | - | 0 | - |
| - | - | 685.5 | 850 | - | - | 0 | - |
| - | - | 1152 | 851.5 | - | - | 0 | - |
| - | - | 3470 | 852.5 | - | - | 0 | - |
| - | - | 781.4 | 853 | - | - | 0 | - |
| 8 | c | 3.154E+04 | 853.5 | 0.0002856 | 0.3346 | +1 | 8 |
| - | - | 1.475E+04 | 854.5 | - | - | 0 | - |
| - | - | 4570 | 855.5 | - | - | 0 | - |
| - | - | 964.4 | 856.5 | - | - | 0 | - |
| 4 | z | 1.545E+05 | 883.5 | 0.0008521 | 0.9645 | +1 | 8 |
| - | - | 7.901E+04 | 884.5 | - | - | 0 | - |
| - | - | 2.181E+04 | 885.5 | - | - | 0 | - |
| - | - | 4626 | 886.5 | - | - | 0 | - |
| 4 | y | 2.056E+04 | 899.5 | 0.0006217 | 0.6912 | +1 | 8 |
| - | - | 9753 | 900.5 | - | - | 0 | - |
| - | - | 3513 | 901.5 | - | - | 0 | - |
| - | - | 1139 | 908.5 | - | - | 0 | - |
| - | - | 1440 | 915.4 | - | - | 0 | - |
| - | - | 1150 | 916.5 | - | - | 0 | - |
| - | - | 948.7 | 922.6 | - | - | 0 | - |
| - | - | 1098 | 923.6 | - | - | 0 | - |
| - | - | 2656 | 924.6 | - | - | 0 | - |
| - | - | 964.8 | 925.6 | - | - | 0 | - |
| - | - | 791.7 | 936.5 | - | - | 0 | - |
| - | - | 1405 | 939.5 | - | - | 0 | - |
| - | - | 860.9 | 940.5 | - | - | 0 | - |
| - | - | 2305 | 950.6 | - | - | 0 | - |
| - | - | 3134 | 951.6 | - | - | 0 | - |
| - | - | 773 | 952.6 | - | - | 0 | - |
| 3 | w | 5173 | 953.5 | 0.002203 | 2.31 | +1 | 9 |
| - | - | 2277 | 954.5 | - | - | 0 | - |
| - | - | 632.4 | 955.5 | - | - | 0 | - |
| - | - | 1165 | 963.5 | - | - | 0 | - |
| - | - | 1576 | 965.6 | - | - | 0 | - |
| 9 | c | 3.705E+04 | 966.6 | 0.0003281 | 0.3394 | +1 | 9 |
| - | - | 1.972E+04 | 967.6 | - | - | 0 | - |
| - | - | 871 | 968.5 | - | - | 0 | - |
| - | - | 5264 | 968.6 | - | - | 0 | - |
| - | - | 1533 | 969.6 | - | - | 0 | - |
| - | - | 4491 | 980.6 | - | - | 0 | - |
| - | - | 3112 | 981.6 | - | - | 0 | - |
| - | - | 854.8 | 995.6 | - | - | 0 | - |
| - | - | 818.4 | 1008 | - | - | 0 | - |
| - | - | 1839 | 1010 | - | - | 0 | - |
| 3 | z | 1.103E+05 | 1012 | 0.0005021 | 0.4964 | +1 | 9 |
| - | - | 6.461E+04 | 1013 | - | - | 0 | - |
| - | - | 2.252E+04 | 1014 | - | - | 0 | - |
| - | - | 4691 | 1015 | - | - | 0 | - |
| - | - | 1688 | 1025 | - | - | 0 | - |
| - | - | 1112 | 1025 | - | - | 0 | - |
| 3 | y | 3.05E+04 | 1028 | 0.0006379 | 0.6208 | +1 | 9 |
| - | - | 1.615E+04 | 1029 | - | - | 0 | - |
| - | - | 5343 | 1030 | - | - | 0 | - |
| - | - | 994.5 | 1031 | - | - | 0 | - |
| - | - | 1458 | 1035 | - | - | 0 | - |
| - | - | 2639 | 1036 | - | - | 0 | - |
| - | - | 1535 | 1037 | - | - | 0 | - |
| - | - | 5224 | 1038 | - | - | 0 | - |
| - | - | 3224 | 1039 | - | - | 0 | - |
| - | - | 825.6 | 1040 | - | - | 0 | - |
| - | - | 1.36E+04 | 1052 | - | - | 0 | - |
| - | - | 3.291E+04 | 1053 | - | - | 0 | - |
| - | - | 1.847E+04 | 1054 | - | - | 0 | - |
| - | - | 6701 | 1055 | - | - | 0 | - |
| - | - | 938.9 | 1056 | - | - | 0 | - |
| - | - | 879 | 1063 | - | - | 0 | - |
| 10 | c | 2435 | 1079 | 0.001679 | 1.557 | +1 | 10 |
| - | - | 2.476E+04 | 1080 | - | - | 0 | - |
| - | - | 1.361E+04 | 1081 | - | - | 0 | - |
| 2 | w | 6416 | 1082 | 0.0001612 | 0.149 | +1 | 10 |
| - | - | 3623 | 1082 | - | - | 0 | - |
| - | - | 5441 | 1083 | - | - | 0 | - |
| - | - | 2379 | 1084 | - | - | 0 | - |
| 10 | c | 2.485E+04 | 1096 | 0.0002765 | 0.2523 | +1 | 10 |
| - | - | 5.87E+04 | 1097 | - | - | 0 | - |
| - | - | 3.204E+04 | 1098 | - | - | 0 | - |
| - | - | 9653 | 1099 | - | - | 0 | - |
| - | - | 2311 | 1100 | - | - | 0 | - |
| - | - | 755.5 | 1111 | - | - | 0 | - |
| - | - | 817.3 | 1112 | - | - | 0 | - |
| - | - | 871.4 | 1124 | - | - | 0 | - |
| - | - | 1141 | 1125 | - | - | 0 | - |
| - | - | 1002 | 1126 | - | - | 0 | - |
| - | - | 1010 | 1127 | - | - | 0 | - |
| - | - | 2006 | 1153 | - | - | 0 | - |
| - | - | 4541 | 1154 | - | - | 0 | - |
| - | - | 1801 | 1155 | - | - | 0 | - |
| 2 | z | 9346 | 1168 | 0.002403 | 2.058 | +1 | 10 |
| - | - | 4.666E+04 | 1169 | - | - | 0 | - |
| - | - | 2.866E+04 | 1170 | - | - | 0 | - |
| - | - | 8726 | 1171 | - | - | 0 | - |
| - | - | 2741 | 1172 | - | - | 0 | - |
| - | - | 8184 | 1181 | - | - | 0 | - |
| - | - | 5913 | 1182 | - | - | 0 | - |
| - | - | 5272 | 1183 | - | - | 0 | - |
| 2 | y | 3922 | 1184 | 0.00477 | 4.03 | +1 | 10 |
| - | - | 1614 | 1185 | - | - | 0 | - |
| - | - | 924.5 | 1187 | - | - | 0 | - |
| - | - | 1497 | 1195 | - | - | 0 | - |
| - | - | 6023 | 1196 | - | - | 0 | - |
| - | - | 4114 | 1197 | - | - | 0 | - |
| - | - | 2064 | 1198 | - | - | 0 | - |
| - | - | 1039 | 1199 | - | - | 0 | - |
| - | - | 2032 | 1207 | - | - | 0 | - |
| - | - | 789.3 | 1208 | - | - | 0 | - |
| - | - | 1238 | 1209 | - | - | 0 | - |
| - | - | 835.3 | 1210 | - | - | 0 | - |
| - | - | 1064 | 1211 | - | - | 0 | - |
| - | - | 2091 | 1212 | - | - | 0 | - |
| - | - | 1970 | 1213 | - | - | 0 | - |
| - | - | 4146 | 1223 | - | - | 0 | - |
| - | - | 1.075E+04 | 1224 | - | - | 0 | - |
| - | - | 8.013E+04 | 1225 | - | - | 0 | - |
| - | - | 6.098E+04 | 1226 | - | - | 0 | - |
| - | - | 2.642E+04 | 1227 | - | - | 0 | - |
| - | - | 9042 | 1228 | - | - | 0 | - |
| - | - | 3242 | 1229 | - | - | 0 | - |
| - | - | 3638 | 1239 | - | - | 0 | - |
| - | - | 1.445E+04 | 1240 | - | - | 0 | - |
| - | - | 1.637E+04 | 1241 | - | - | 0 | - |
| - | - | 1.611E+04 | 1242 | - | - | 0 | - |
| - | - | 7465 | 1243 | - | - | 0 | - |
| - | - | 2305 | 1244 | - | - | 0 | - |
| - | - | 2983 | 1250 | - | - | 0 | - |
| - | - | 2.325E+04 | 1251 | - | - | 0 | - |
| - | - | 1.593E+04 | 1252 | - | - | 0 | - |
| - | - | 5749 | 1253 | - | - | 0 | - |
| - | - | 2160 | 1254 | - | - | 0 | - |
| - | - | 927.3 | 1256 | - | - | 0 | - |
| - | - | 4777 | 1257 | - | - | 0 | - |
| - | - | 4213 | 1258 | - | - | 0 | - |
| - | - | 1075 | 1259 | - | - | 0 | - |
| - | - | 2.555E+04 | 1267 | - | - | 0 | - |
| - | - | 1.96E+05 | 1268 | - | - | 0 | - |
| - | - | 1.374E+05 | 1269 | - | - | 0 | - |
| - | - | 5.39E+04 | 1270 | - | - | 0 | - |
| - | - | 1.295E+04 | 1271 | - | - | 0 | - |
| - | - | 3145 | 1272 | - | - | 0 | - |
| - | - | 7990 | 1283 | - | - | 0 | - |
| - | - | 4.146E+04 | 1284 | - | - | 0 | - |
| - | - | 1.705E+05 | 1285 | - | - | 0 | - |
| - | - | 1.053E+05 | 1286 | - | - | 0 | - |
| - | - | 4.332E+04 | 1287 | - | - | 0 | - |
| - | - | 1.239E+04 | 1288 | - | - | 0 | - |
| - | - | 3409 | 1289 | - | - | 0 | - |
| - | - | 702.9 | 1720 | - | - | 0 | - |
| - | - | 597.3 | 1895 | - | - | 0 | - |
| - | - | 691.3 | 2913 | - | - | 0 | - |
| - | - | 677.7 | 3242 | - | - | 0 | - |
| - | - | 666.1 | 3326 | - | - | 0 | - |
| - | - | 806.8 | 3355 | - | - | 0 | - |

m/z Charge Intensity FragmentType MassShift Position
122.35016632080078 0 368.45328
122.81430053710938 0 361.10086
126.07386779785156 0 380.30093
128.00990295410156 0 332.1108
129.1027374267578 0 649.3564
129.3446044921875 0 390.9359
130.0654754638672 0 7997.8584
131.06890869140625 0 723.7848
132.0812530517578 0 1759.9878
133.0863800048828 0 1330.7803
135.53497314453125 0 370.48
142.06602478027344 0 512.0346
142.09779357910156 0 1646.3228
143.91326904296875 0 410.67337
144.0812530517578 0 1702.149
146.06045532226562 0 8056.9375
147.0640869140625 0 947.4123
147.42221069335938 0 424.08597
148.95477294921875 0 977.64276
149.04510498046875 0 1385.0784
154.5377655029297 0 416.45587
157.1088104248047 0 3775.7522
159.0920867919922 0 26208.201
159.2250518798828 0 531.88995
160.095458984375 0 2697.937
161.09898376464844 0 446.9933
166.0867156982422 0 844.28015
170.06039428710938 0 1514.8817
171.0921173095703 0 491.5411
173.4518585205078 0 1866.864
173.46131896972656 0 633.85846
174.10287475585938 0 1163.0001
174.134765625 0 1230.4003
185.10379028320312 0 1974.5629
185.11737060546875 0 530.35876
185.14002990722656 0 748.2911
187.08700561523438 0 5263.515
188.07102966308594 0 87982.59
189.07444763183594 0 10065.158
190.07737731933594 0 742.2675
192.62184143066406 0 2265.0051 c Ammonia loss 2
196.14407348632812 0 578.86847
196.37449645996094 0 517.77966
197.1400909423828 0 1750.9851
198.14317321777344 0 504.6308
203.12843322753906 0 1245.176
204.66836547851562 0 509.66235
205.0975799560547 0 52859.05 y 10
205.62887573242188 0 569.0808
206.1010284423828 0 6664.9277
208.95355224609375 0 23020.705
212.42263793945312 0 494.39297
213.1714324951172 0 5231.0864
214.142822265625 0 7962.6016
214.15542602539062 0 3573.5608
214.6440887451172 0 2251.9946
215.15818786621094 0 760.516
228.14056396484375 0 2696.683 c Ammonia loss 3
228.1823272705078 0 2115.2737
228.6417236328125 0 778.06433
238.16665649414062 0 1930.8401
239.1507568359375 0 22564.213
240.1542205810547 0 3047.403
240.9071502685547 0 672.7339
242.16162109375 0 4578.8135
243.16810607910156 0 571.4777
252.24559020996094 0 533.8284
256.121337890625 0 1567.3665
256.1772155761719 0 12258.554 c Ammonia loss 1
257.18072509765625 0 1922.7876
270.1241760253906 0 3389.7654
271.1265563964844 0 704.6273
273.2037353515625 0 48377.91 c 1
274.20721435546875 0 6547.8203
275.2113037109375 0 713.13446
276.6659240722656 0 636.0958 c Ammonia loss 4
280.1095275878906 0 699.6558
298.11956787109375 0 1664.1398
298.2237243652344 0 979.0052
299.21966552734375 0 12696.429
300.2225036621094 0 2275.5356
307.8637390136719 0 799.07245
312.1908264160156 0 678.5059
312.6773986816406 0 1858.9176
316.12982177734375 0 13694.185 y Water loss 9
318.92279052734375 0 14207.082
324.4042053222656 0 565.7362
324.9857482910156 0 577.8664
334.1406555175781 0 2043.0682 y 9
341.0185241699219 0 5198.475
341.2311706542969 0 978.26184
350.1817321777344 0 582.3777
356.24102783203125 0 2803.7402
357.2478942871094 0 2759.7073
359.02886962890625 0 16541.748
361.2202453613281 0 557.8026
361.7247619628906 0 3325.5017
362.2256164550781 0 1719.5647
367.209228515625 0 1479.3899
369.23150634765625 0 1389.2361
369.7344970703125 0 11111.074
370.2124938964844 0 1029.0391
370.23651123046875 0 6739.0386
370.25836181640625 0 921.90717
370.73876953125 0 1229.7834
383.22760009765625 0 4675.9634 c Ammonia loss 6
383.7295227050781 0 1849.6564
384.20611572265625 0 918.7323 z Ammonia loss 1
384.23577880859375 0 10127.821 c Ammonia loss 2
385.2395324707031 0 2165.412
385.27862548828125 0 740.81915
391.2375793457031 0 62941.46
391.7411804199219 0 203412.56 c 6
392.24273681640625 0 82126.99
392.7437438964844 0 18402.336
393.2455749511719 0 2230.8938
401.2628173828125 0 163041.55 c 2
401.5614013671875 0 892.48737
402.2656555175781 0 30748.695
403.2686767578125 0 3967.45
404.7498779296875 0 3212.087
405.25146484375 0 2404.0933
405.71722412109375 0 4461.4146 y Water loss 4
406.21954345703125 0 2523.3325
406.900390625 0 6490.523
407.2347106933594 0 5387.537
407.5697326660156 0 2730.9626
407.90350341796875 0 608.7165
410.2339782714844 0 952.40314
412.90435791015625 0 7078.787
413.2388916015625 0 2703.9814
413.27197265625 0 1124.0082
413.57379150390625 0 1093.5095
414.7228088378906 0 3777.0156 y 4
415.2247314453125 0 1534.779
418.7474365234375 0 7073.595 c Ammonia loss 7
419.24871826171875 0 2372.063
419.749755859375 0 817.4659
422.23602294921875 0 2721.572
422.56915283203125 0 995.92737
426.7563781738281 0 28722.936
427.2601623535156 0 159718.34 c 7
427.7615051269531 0 70617.11
428.1487121582031 0 608.13226
428.2630310058594 0 16842.707
428.7644348144531 0 2458.213
428.89215087890625 0 1719.1644
429.0897521972656 0 16365.976
429.15472412109375 0 667.7346
429.29058837890625 0 1015.1748
453.78521728515625 0 1522.3352
454.2876892089844 0 1622.7666
455.2731628417969 0 11508.081 c Ammonia loss 3
456.276123046875 0 2931.331
461.2919006347656 0 19272.102
461.7936706542969 0 14184.82
462.29498291015625 0 5824.725
462.79888916015625 0 1587.1641
466.7738342285156 0 1069.1537
467.2759704589844 0 786.7273
467.30987548828125 0 863.24243
468.77679443359375 0 833.7047
469.2772521972656 0 1120.0916
472.2889709472656 0 2153.4966
474.7970275878906 0 2158.4832
475.28912353515625 0 47272.793 c Ammonia loss 8
475.7904968261719 0 24254.842
476.2918701171875 0 7673.4834
476.7939758300781 0 1213.0105
483.26055908203125 0 1058.1201
483.298583984375 0 7877.859
483.8021545410156 0 78452.09 c 8
484.3037109375 0 45210.277
484.8054504394531 0 12880.086
485.30419921875 0 2755.8762
486.27239990234375 0 1788.3995
490.7997741699219 0 858.12366
495.3039245605469 0 729.82635
496.8102111816406 0 46012.652
497.3113708496094 0 24422.15
497.8124084472656 0 9328.55
498.2850646972656 0 8328.568
498.3168640136719 0 2408.2212
498.7868347167969 0 5062.4263
499.2889099121094 0 1399.2626
499.3229675292969 0 3334.0571
500.3272399902344 0 987.7515
504.78973388671875 0 4024.44
505.29266357421875 0 1873.6663
510.2814636230469 0 630.1068
517.3003540039062 0 960.3538
517.8129272460938 0 2303.7383
518.2608642578125 0 1062.439 y 7
518.30908203125 0 3640.4812
518.8072509765625 0 1802.4695
519.3096313476562 0 942.1713
524.2904052734375 0 646.7663
524.3306274414062 0 2440.8782
525.3386840820312 0 4143.5093
525.8128662109375 0 42861.93
526.3145751953125 0 29748.47
526.8167724609375 0 12035.556
527.3162841796875 0 3766.899
529.2874755859375 0 1491.8038
530.8045654296875 0 1011.8632
531.2990112304688 0 2953.788
531.7982177734375 0 2320.2112
532.2981567382812 0 1220.4583
532.8099365234375 0 760.5653
533.30615234375 0 744.8294
539.317626953125 0 2124.9675 c Water loss 9
539.8104248046875 0 272260.94 c Ammonia loss 9
540.3118896484375 0 150925.17
540.81298828125 0 50922.457
541.3145141601562 0 14868.237
541.81640625 0 2626.8972
542.3326416015625 0 7178.6694
543.3365478515625 0 1507.4127
545.1884765625 0 634.38715
547.8132934570312 0 2177.0078
548.32373046875 0 393944.2 c 9
548.8248291015625 0 225598.02
548.9248046875 0 1263.4299
549.326171875 0 74902.43
549.8276977539062 0 17875.139
550.3285522460938 0 2766.4048
553.8007202148438 0 840.1381
554.329833984375 0 1137.7065
554.8095703125 0 1799.8787
555.2970581054688 0 17966.078
555.3414306640625 0 3028.449
555.7984619140625 0 11231.263
556.3006591796875 0 4560.5015
558.2933959960938 0 1758.3899
561.331298828125 0 13712.721
561.7850341796875 0 804.22974
561.83349609375 0 9205.078
562.3256225585938 0 3710.9883
562.7848510742188 0 2147.3071
563.2852783203125 0 1184.8346
567.3430786132812 0 788.06305
568.3456420898438 0 3581.5388
568.8226928710938 0 2664.8005
569.3526000976562 0 59628.805 c 4
569.8058471679688 0 1037.4243
570.3101196289062 0 443.39685
570.355224609375 0 19134.479
571.3582763671875 0 3956.8882
575.8378295898438 0 6880.24
576.3370971679688 0 3579.9023
576.8226928710938 0 23382.729
577.3245849609375 0 19854.404
577.8268432617188 0 7748.573
578.3236083984375 0 2317.4543
578.8234252929688 0 985.5314
582.3551025390625 0 5972.901
582.842041015625 0 2651.7297
583.3536987304688 0 2178.2139
583.8263549804688 0 1799.6718
584.3123779296875 0 19370.865 z 1
584.8136596679688 0 12079.785
585.31494140625 0 4783.738
585.8179931640625 0 1538.3649
586.3197631835938 0 1742.2129
590.827880859375 0 586.82184
590.8563232421875 0 986.79443
591.3136596679688 0 967.6823
591.812744140625 0 1406.7638
592.3236694335938 0 2256.4673 y 1
592.8226318359375 0 1468.2777
593.3257446289062 0 811.53265
596.352783203125 0 2128.9473
597.3568725585938 0 930.2769
597.8372192382812 0 3117.4302
598.3281860351562 0 6364.6333
598.8284301757812 0 4600.6704
599.33203125 0 5975.8076
600.3375854492188 0 1000.41583
602.2813720703125 0 5575.8384
603.284912109375 0 1307.6447
603.3399047851562 0 931.02527
604.3389892578125 0 6537.8193
604.83349609375 0 15909.078
605.3330078125 0 12652.791
605.8258666992188 0 7314.133
606.3404541015625 0 3114.933
606.8416137695312 0 2956.0938
611.3554077148438 0 5970.3027
611.8509521484375 0 32813.39
612.3525390625 0 17209.785
612.8084716796875 0 1259.068
612.8533935546875 0 10114.532
613.3489990234375 0 4573.0464
613.8469848632812 0 1795.1566
614.3533325195312 0 18363.373
614.8285522460938 0 1969.272
615.2900390625 0 9173.079
615.3565673828125 0 5405.57
616.2926025390625 0 3352.6318
616.3551025390625 0 1447.4312
617.2943115234375 0 716.10675
619.3551635742188 0 3357.6997
619.8492431640625 0 21230.643
620.3585815429688 0 26398.92
620.8461303710938 0 60078.336
621.3468017578125 0 39348.56
621.8475952148438 0 14523.613
622.3491821289062 0 4406.25
624.34814453125 0 4108.272
625.3650512695312 0 8725.82
625.8368530273438 0 4883.53
626.3745727539062 0 239854.78 c 5
626.835693359375 0 23654.209
627.3771362304688 0 78543.8
627.8397216796875 0 1851.8479
628.3793334960938 0 15922.675
629.3810424804688 0 2275.5266
633.3496704101562 0 6437.555
633.84619140625 0 319218.88
634.347412109375 0 220435.27
634.8487548828125 0 87682.05
635.3499145507812 0 24340.426
635.8509521484375 0 4673.738
641.8554077148438 0 143234.14
642.3577880859375 0 189603.5
642.8595581054688 0 98013.28
643.3607177734375 0 35672.055
643.8617553710938 0 9975.608
644.3599853515625 0 2684.8354
652.3855590820312 0 1471.0844
653.3984985351562 0 866.8981
656.3530883789062 0 1013.8273 y Water loss 6
658.34423828125 0 143393.52 z 6
659.3472900390625 0 51843.805
660.3499145507812 0 11966.76
661.3530883789062 0 1590.8267
667.3871459960938 0 759.5558
671.3760375976562 0 9571.235
672.3141479492188 0 878.7019
672.37841796875 0 3250.207
674.3628540039062 0 8842.903 y 6
675.3652954101562 0 3008.9065
696.3942260742188 0 1886.4015
697.3967895507812 0 1172.2606
704.3919677734375 0 674.3751
713.3733520507812 0 1860.5334 y Water loss 5
715.3655395507812 0 33987.637 z 5
716.3689575195312 0 13880.383
717.37158203125 0 3087.2854
731.3848266601562 0 9221.492 y 5
732.3866577148438 0 4802.786
733.3932495117188 0 1365.676
738.4625244140625 0 890.06635
739.462890625 0 2062.9133
740.4550170898438 0 2422.1914
767.4592895507812 0 2106.452
780.4487915039062 0 906.1102
781.4633178710938 0 3776.16
782.4752197265625 0 42822.457 c 6
783.4775390625 0 16498.164
784.4805908203125 0 4439.254
785.3953857421875 0 3758.0583 w 4
785.4796142578125 0 1000.21796
786.4004516601562 0 3146.0938
810.42724609375 0 3966.1829 y Water loss 4
811.427734375 0 5444.554
811.4957885742188 0 795.2149
812.4318237304688 0 1814.0259 z 4
819.4589233398438 0 2033.4569
820.460693359375 0 841.21576
823.4439086914062 0 2490.3928
824.44970703125 0 6972.085
825.4544677734375 0 4774.772
826.4336547851562 0 2367.7305
827.4021606445312 0 3035.9504
828.4361572265625 0 11971.508 y 4
829.439453125 0 6336.308
830.4426879882812 0 1570.6841
839.4661865234375 0 3245.1404
840.4689331054688 0 2691.1577
841.4736328125 0 770.15314
844.465087890625 0 889.06537
849.9970092773438 0 685.45135
851.501708984375 0 1152.0743
852.5050659179688 0 3470.1533
852.983154296875 0 781.37524
853.5117797851562 0 31538.31 c 7
854.5147705078125 0 14750.834
855.5213623046875 0 4570.498
856.53125 0 964.4011
883.4555053710938 0 154513.11 z 3
884.4584350585938 0 79010.95
885.4609985351562 0 21805.082
886.4641723632812 0 4625.887
899.4739990234375 0 20558.154 y 3
900.475830078125 0 9753.234
901.4783325195312 0 3513.3923
908.548095703125 0 1138.6613
915.4442749023438 0 1440.3844
916.4530639648438 0 1150.236
922.5846557617188 0 948.67944
923.5795288085938 0 1097.8427
924.5747680664062 0 2656.445
925.5811767578125 0 964.7539
936.54052734375 0 791.67664
939.4918823242188 0 1404.5914
940.4959716796875 0 860.87146
950.574951171875 0 2304.784
951.5843505859375 0 3134.216
952.5911865234375 0 773.0059
953.4861450195312 0 5173.4277 w 2
954.4866943359375 0 2277.2666
955.47314453125 0 632.43567
963.5272216796875 0 1165.2528
965.5863037109375 0 1575.5017
966.5958862304688 0 37047.24 c 8
967.59912109375 0 19721.045
968.506103515625 0 870.9592
968.6033935546875 0 5264.1587
969.61279296875 0 1533.4822
980.5523071289062 0 4491.4585
981.5531005859375 0 3112.365
995.556884765625 0 854.8195
1007.5948486328125 0 818.44867
1009.6021728515625 0 1838.7405
1011.5137329101562 0 110253.055 z 2
1012.5166015625 0 64614.49
1013.5194091796875 0 22522.883
1014.5227661132812 0 4690.8286
1024.5191650390625 0 1688.0371
1024.6287841796875 0 1112.1493
1027.5325927734375 0 30500.846 y 2
1028.5360107421875 0 16145.31
1029.5377197265625 0 5342.694
1030.551025390625 0 994.4662
1034.620361328125 0 1457.6156
1035.605712890625 0 2639.109
1036.6094970703125 0 1534.992
1037.598388671875 0 5223.554
1038.5985107421875 0 3224.2896
1039.602294921875 0 825.64246
1051.6260986328125 0 13601.327
1052.62939453125 0 32910.32
1053.627685546875 0 18469.605
1054.6300048828125 0 6701.291
1055.6396484375 0 938.9481
1062.5904541015625 0 878.99023
1078.61328125 0 2435.285 c Ammonia loss 9
1079.6190185546875 0 24760.484
1080.622802734375 0 13608.968
1081.5423583984375 0 6416.2124 w 1
1081.6376953125 0 3622.5608
1082.5478515625 0 5441.48
1083.552978515625 0 2379.1072
1095.638427734375 0 24851.656 c 9
1096.6453857421875 0 58702.527
1097.648193359375 0 32044.629
1098.6514892578125 0 9653.471
1099.655029296875 0 2311.4973
1110.620849609375 0 755.522
1111.6109619140625 0 817.2999
1123.6358642578125 0 871.4218
1124.6175537109375 0 1140.607
1125.592529296875 0 1001.856
1126.598388671875 0 1010.084
1152.6273193359375 0 2005.6019
1153.6334228515625 0 4541.1533
1154.63037109375 0 1801.1714
1167.6119384765625 0 9345.763 z 1
1168.6214599609375 0 46657.883
1169.625 0 28657.234
1170.62744140625 0 8725.6875
1171.6309814453125 0 2740.615
1180.6202392578125 0 8183.814
1181.6221923828125 0 5913.0996
1182.621826171875 0 5271.922
1183.6282958984375 0 3921.704 y 1
1184.6297607421875 0 1613.6932
1186.6273193359375 0 924.53217
1194.618896484375 0 1496.5592
1195.6275634765625 0 6022.638
1196.631103515625 0 4114.454
1197.633056640625 0 2063.5193
1198.6500244140625 0 1038.9432
1206.677734375 0 2031.845
1207.6800537109375 0 789.29944
1208.6588134765625 0 1237.9669
1209.6500244140625 0 835.2897
1210.649658203125 0 1064.3206
1211.67041015625 0 2091.324
1212.6683349609375 0 1970.0583
1222.7061767578125 0 4145.8677
1223.6998291015625 0 10752.04
1224.6622314453125 0 80134.25
1225.6649169921875 0 60979.42
1226.6673583984375 0 26417.102
1227.6693115234375 0 9041.966
1228.6708984375 0 3241.7673
1238.70068359375 0 3638.4548
1239.695556640625 0 14450.15
1240.69873046875 0 16365.397
1241.6964111328125 0 16113.913
1242.69482421875 0 7465.193
1243.6942138671875 0 2304.5725
1249.680908203125 0 2982.9824
1250.6650390625 0 23246.062
1251.6668701171875 0 15928.335
1252.6695556640625 0 5749.31
1253.6717529296875 0 2160.1023
1255.717529296875 0 927.33813
1256.72021484375 0 4777.134
1257.723388671875 0 4213.262
1258.7149658203125 0 1074.5476
1266.68896484375 0 25547.842
1267.6903076171875 0 195978.1
1268.6932373046875 0 137355.05
1269.6954345703125 0 53900.72
1270.6986083984375 0 12949.037
1271.6966552734375 0 3144.9976
1282.700927734375 0 7989.886
1283.707763671875 0 41462.598
1284.7156982421875 0 170512.1
1285.7181396484375 0 105317.27
1286.7213134765625 0 43319.625
1287.723388671875 0 12385.679
1288.7208251953125 0 3408.7578
1719.9827880859375 0 702.8969
1895.2852783203125 0 597.27893
2913.396728515625 0 691.2789
3241.941162109375 0 677.6775
3325.681396484375 0 666.0987
3355.33154296875 0 806.777

Spectrum Details

|  |  |
| --- | --- |
| Matched peaks? Matched peaks The total absolute number of peaks matched. Additionally in brackets the total fraction of peaks matched and the total number of peaks is shown. | 51 (9.59% of 532) |
| FDR? FDR The false discovery rate estimated for this peptide. It is calculated by matching all theoretical fragments with a non-integer shift with the raw peaks for this spectrum. This is done with 40 different shifts. The resulting percentage is the average number of annotated peaks over the number of annotated peaks with the correct spectrum. | 2.71% |
| Satellite FDR? Satellite FDR See the FDR for details on its calculation. This satellite ion specific FDR only contains the satellite ions (d/w) for I/L/J positions. | ∞ |
| PSM Score? PSM Score The PSM Score as given by Hecklib to this annotated spectrum. It is shown with three significant figures. | 426 |

#### Spectrum 5690? Spectrum 5690 The raw spectrum of this peptide as annotated by Hecklib. The fragments are coloured according to ion type (see legend). Any peaks with a star '\*' as text can be hovered over to see the full details, first the ion type second the mass shift type. By hovering over the amino acids in the peptide or ions in the legend the corresponding peaks are highlighted. By toggling the 'Unassigned' label you can turn the background (unassigned) peaks on or off in the plot. By updating the slider in the Ion legend you can update the spectrum to only show the top X% of the peaks with labels. The top X% means any peak that is within X% of the highest intensity. By dragging in the spectrum you can zoom in to a specific part of the spectrum and use 'Zoom Out' to get back to the original zoom level. The annotation of the spectrum is based on the given sequence in the peptides file and is done with different software so inconsistencies are likely. The peaks are annotated based on the given sequence, with 20 ppm tolerance.

Copy Data

##### Spectrum 5690 (TSV)

###### Preview

```
Loading example...
```

*Click on the button to copy the data to your clipboard.*

Mz MinMz MaxIntensity Max

WidthHeightPeptide font sizePeptide stroke widthSpectrum font sizeSpectrum stroke widthCompact peptide

Ion legend

wxyz

abcd

OtherUnassignedIon^Charge^_Position_Show for top:%

VRQAPGRAJEW

03.17e+56.35e+59.52e+51.27e+6

Zoom Out

c^+2^_3_y^+1^_1_c^+2^_4_c^+1^_2_c^+1^_2_c^+2^_5_y^+1^_2_y^+1^_2_c^+2^_7_z^+3^_10_c^+1^_3_c^+2^_7_c^+1^_3_y^+2^_7_y^+2^_7_c^+2^_8_c^+2^_8_c^+1^_4_c^+2^_9_c^+2^_9_y^+1^_4_c^+2^_10_c^+2^_10_c^+2^_10_c^+1^_5_z^+2^_10_y^+2^_10_c^+1^_6_y^+1^_5_z^+1^_5_y^+1^_5_y^+1^_6_z^+1^_6_y^+1^_6_c^+1^_7_w^+1^_7_y^+1^_7_z^+1^_7_y^+1^_7_c^+1^_8_z^+1^_8_y^+1^_8_w^+1^_9_c^+1^_9_y^+1^_9_z^+1^_9_y^+1^_9_c^+1^_10_w^+1^_10_c^+1^_10_z^+1^_10_y^+1^_10_

0870174026093479

Fragment Matches Table

Show background peaks

| Position | Ion type | Intensity | mz Theoretical | mz Error (Th) | mz Error (ppm) | Charge | Series Number |
| --- | --- | --- | --- | --- | --- | --- | --- |
| - | - | 1198 | 123.8 | - | - | 0 | - |
| - | - | 2012 | 130.1 | - | - | 0 | - |
| - | - | 2.731E+04 | 130.1 | - | - | 0 | - |
| - | - | 1581 | 131.1 | - | - | 0 | - |
| - | - | 7562 | 132.1 | - | - | 0 | - |
| - | - | 1.323E+04 | 133.1 | - | - | 0 | - |
| - | - | 3190 | 142.1 | - | - | 0 | - |
| - | - | 1241 | 142.1 | - | - | 0 | - |
| - | - | 5515 | 144.1 | - | - | 0 | - |
| - | - | 1439 | 145.1 | - | - | 0 | - |
| - | - | 2.634E+04 | 146.1 | - | - | 0 | - |
| - | - | 2565 | 147.1 | - | - | 0 | - |
| - | - | 1.462E+04 | 157.1 | - | - | 0 | - |
| - | - | 9.627E+04 | 159.1 | - | - | 0 | - |
| - | - | 1677 | 160.1 | - | - | 0 | - |
| - | - | 9766 | 160.1 | - | - | 0 | - |
| - | - | 1674 | 166.1 | - | - | 0 | - |
| - | - | 1448 | 169.1 | - | - | 0 | - |
| - | - | 4574 | 170.1 | - | - | 0 | - |
| - | - | 2148 | 171.1 | - | - | 0 | - |
| - | - | 2835 | 171.1 | - | - | 0 | - |
| - | - | 1427 | 171.5 | - | - | 0 | - |
| - | - | 1358 | 172.1 | - | - | 0 | - |
| - | - | 3607 | 173.5 | - | - | 0 | - |
| - | - | 2930 | 174.1 | - | - | 0 | - |
| - | - | 4727 | 174.1 | - | - | 0 | - |
| - | - | 8330 | 177.1 | - | - | 0 | - |
| - | - | 1701 | 178.6 | - | - | 0 | - |
| - | - | 7340 | 185.1 | - | - | 0 | - |
| - | - | 3468 | 185.1 | - | - | 0 | - |
| - | - | 1.632E+04 | 187.1 | - | - | 0 | - |
| - | - | 3.182E+05 | 188.1 | - | - | 0 | - |
| - | - | 3.701E+04 | 189.1 | - | - | 0 | - |
| - | - | 1390 | 190.7 | - | - | 0 | - |
| 3 | c | 1.019E+04 | 192.6 | 0.0002849 | 1.479 | +2 | 3 |
| - | - | 2110 | 195.2 | - | - | 0 | - |
| - | - | 2507 | 196.1 | - | - | 0 | - |
| - | - | 3509 | 197.1 | - | - | 0 | - |
| - | - | 3479 | 197.1 | - | - | 0 | - |
| - | - | 1361 | 199.1 | - | - | 0 | - |
| - | - | 1633 | 203.1 | - | - | 0 | - |
| 11 | y | 2.006E+05 | 205.1 | 0.0003801 | 1.853 | +1 | 1 |
| - | - | 1486 | 205.6 | - | - | 0 | - |
| - | - | 2.232E+04 | 206.1 | - | - | 0 | - |
| - | - | 2.926E+04 | 209 | - | - | 0 | - |
| - | - | 1.937E+04 | 213.2 | - | - | 0 | - |
| - | - | 3.464E+04 | 214.1 | - | - | 0 | - |
| - | - | 1.297E+04 | 214.2 | - | - | 0 | - |
| - | - | 1655 | 214.2 | - | - | 0 | - |
| - | - | 5634 | 214.6 | - | - | 0 | - |
| - | - | 1457 | 215.2 | - | - | 0 | - |
| - | - | 1398 | 218.9 | - | - | 0 | - |
| 4 | c | 1.228E+04 | 228.1 | 0.0004047 | 1.774 | +2 | 4 |
| - | - | 8131 | 228.2 | - | - | 0 | - |
| - | - | 1902 | 228.6 | - | - | 0 | - |
| - | - | 1579 | 235.2 | - | - | 0 | - |
| - | - | 4693 | 238.2 | - | - | 0 | - |
| - | - | 7.09E+04 | 239.2 | - | - | 0 | - |
| - | - | 8751 | 240.2 | - | - | 0 | - |
| - | - | 1.339E+04 | 242.2 | - | - | 0 | - |
| - | - | 2620 | 243.2 | - | - | 0 | - |
| - | - | 2281 | 248.2 | - | - | 0 | - |
| - | - | 1513 | 249 | - | - | 0 | - |
| - | - | 4640 | 256.1 | - | - | 0 | - |
| 2 | c | 4.695E+04 | 256.2 | 0.0003531 | 1.379 | +1 | 2 |
| - | - | 5365 | 257.2 | - | - | 0 | - |
| - | - | 1.258E+04 | 270.1 | - | - | 0 | - |
| - | - | 3211 | 271.1 | - | - | 0 | - |
| 2 | c | 1.721E+05 | 273.2 | 0.0003238 | 1.185 | +1 | 2 |
| - | - | 1.768E+04 | 274.2 | - | - | 0 | - |
| 5 | c | 2798 | 276.7 | 0.0003595 | 1.299 | +2 | 5 |
| - | - | 2378 | 279.2 | - | - | 0 | - |
| - | - | 2191 | 280.1 | - | - | 0 | - |
| - | - | 6549 | 298.1 | - | - | 0 | - |
| - | - | 5241 | 298.2 | - | - | 0 | - |
| - | - | 3441 | 298.7 | - | - | 0 | - |
| - | - | 4.477E+04 | 299.2 | - | - | 0 | - |
| - | - | 3475 | 300.2 | - | - | 0 | - |
| - | - | 6426 | 307.9 | - | - | 0 | - |
| - | - | 2691 | 308.2 | - | - | 0 | - |
| - | - | 3741 | 312.2 | - | - | 0 | - |
| - | - | 5310 | 312.7 | - | - | 0 | - |
| - | - | 2620 | 313.2 | - | - | 0 | - |
| 10 | y | 4.729E+04 | 316.1 | 0.0006088 | 1.926 | +1 | 2 |
| - | - | 1.894E+04 | 318.9 | - | - | 0 | - |
| 10 | y | 4605 | 334.1 | 0.0007252 | 2.17 | +1 | 2 |
| - | - | 4485 | 341 | - | - | 0 | - |
| - | - | 2712 | 341.2 | - | - | 0 | - |
| - | - | 2903 | 347.7 | - | - | 0 | - |
| - | - | 1847 | 348.2 | - | - | 0 | - |
| - | - | 1.044E+04 | 356.2 | - | - | 0 | - |
| - | - | 7845 | 357.2 | - | - | 0 | - |
| - | - | 1.927E+04 | 359 | - | - | 0 | - |
| - | - | 1.143E+04 | 361.7 | - | - | 0 | - |
| - | - | 7661 | 362.2 | - | - | 0 | - |
| - | - | 1845 | 362.7 | - | - | 0 | - |
| - | - | 4283 | 367.2 | - | - | 0 | - |
| - | - | 5472 | 369.2 | - | - | 0 | - |
| - | - | 3.089E+04 | 369.7 | - | - | 0 | - |
| - | - | 1701 | 369.8 | - | - | 0 | - |
| - | - | 1.674E+04 | 370.2 | - | - | 0 | - |
| - | - | 3786 | 370.3 | - | - | 0 | - |
| - | - | 5317 | 370.7 | - | - | 0 | - |
| 7 | c | 1.777E+04 | 383.2 | 0.0002598 | 0.678 | +2 | 7 |
| - | - | 6000 | 383.7 | - | - | 0 | - |
| 2 | z | 3202 | 384.2 | 0.002557 | 6.655 | +3 | 10 |
| 3 | c | 3.51E+04 | 384.2 | 0.0005525 | 1.438 | +1 | 3 |
| - | - | 6857 | 385.2 | - | - | 0 | - |
| - | - | 3744 | 385.3 | - | - | 0 | - |
| - | - | 2.142E+05 | 391.2 | - | - | 0 | - |
| 7 | c | 6.842E+05 | 391.7 | 0.0002604 | 0.6648 | +2 | 7 |
| - | - | 2.564E+05 | 392.2 | - | - | 0 | - |
| - | - | 6.285E+04 | 392.7 | - | - | 0 | - |
| - | - | 8045 | 393.2 | - | - | 0 | - |
| - | - | 2413 | 396.7 | - | - | 0 | - |
| 3 | c | 5.351E+05 | 401.3 | 0.0007368 | 1.836 | +1 | 3 |
| - | - | 1.015E+05 | 402.3 | - | - | 0 | - |
| - | - | 1.206E+04 | 403.3 | - | - | 0 | - |
| - | - | 8454 | 404.7 | - | - | 0 | - |
| - | - | 7208 | 405.3 | - | - | 0 | - |
| 5 | y | 1.308E+04 | 405.7 | 0.0009501 | 2.342 | +2 | 7 |
| - | - | 2143 | 405.8 | - | - | 0 | - |
| - | - | 4498 | 406.2 | - | - | 0 | - |
| - | - | 2298 | 406.7 | - | - | 0 | - |
| - | - | 2.44E+04 | 406.9 | - | - | 0 | - |
| - | - | 1.872E+04 | 407.2 | - | - | 0 | - |
| - | - | 6173 | 407.6 | - | - | 0 | - |
| - | - | 6973 | 410.2 | - | - | 0 | - |
| - | - | 2.924E+04 | 412.9 | - | - | 0 | - |
| - | - | 1.192E+04 | 413.2 | - | - | 0 | - |
| - | - | 3027 | 413.3 | - | - | 0 | - |
| - | - | 2427 | 413.6 | - | - | 0 | - |
| - | - | 1699 | 413.9 | - | - | 0 | - |
| 5 | y | 1.385E+04 | 414.7 | 0.0011 | 2.652 | +2 | 7 |
| - | - | 5281 | 415.2 | - | - | 0 | - |
| - | - | 2720 | 416.2 | - | - | 0 | - |
| - | - | 2157 | 416.6 | - | - | 0 | - |
| - | - | 2137 | 418.3 | - | - | 0 | - |
| 8 | c | 2.861E+04 | 418.7 | 0.001143 | 2.729 | +2 | 8 |
| - | - | 9335 | 419.2 | - | - | 0 | - |
| - | - | 9362 | 422.2 | - | - | 0 | - |
| - | - | 5502 | 422.6 | - | - | 0 | - |
| - | - | 2929 | 422.9 | - | - | 0 | - |
| - | - | 7.912E+04 | 426.8 | - | - | 0 | - |
| 8 | c | 5.261E+05 | 427.3 | 0.000655 | 1.533 | +2 | 8 |
| - | - | 2.513E+05 | 427.8 | - | - | 0 | - |
| - | - | 6.097E+04 | 428.3 | - | - | 0 | - |
| - | - | 1.162E+04 | 428.8 | - | - | 0 | - |
| - | - | 3880 | 428.9 | - | - | 0 | - |
| - | - | 1.598E+04 | 429.1 | - | - | 0 | - |
| - | - | 2918 | 429.3 | - | - | 0 | - |
| - | - | 1724 | 438.2 | - | - | 0 | - |
| - | - | 2923 | 440.3 | - | - | 0 | - |
| - | - | 1754 | 453.3 | - | - | 0 | - |
| - | - | 2575 | 453.8 | - | - | 0 | - |
| 4 | c | 3.718E+04 | 455.3 | 0.0007617 | 1.673 | +1 | 4 |
| - | - | 1.05E+04 | 456.3 | - | - | 0 | - |
| - | - | 6.506E+04 | 461.3 | - | - | 0 | - |
| - | - | 4.749E+04 | 461.8 | - | - | 0 | - |
| - | - | 1.721E+04 | 462.3 | - | - | 0 | - |
| - | - | 2506 | 462.8 | - | - | 0 | - |
| - | - | 2985 | 466.8 | - | - | 0 | - |
| - | - | 3294 | 468.8 | - | - | 0 | - |
| - | - | 5184 | 472.3 | - | - | 0 | - |
| - | - | 2245 | 473.3 | - | - | 0 | - |
| - | - | 7737 | 474.8 | - | - | 0 | - |
| 9 | c | 1.787E+05 | 475.3 | 0.0008587 | 1.807 | +2 | 9 |
| - | - | 8.976E+04 | 475.8 | - | - | 0 | - |
| - | - | 2.442E+04 | 476.3 | - | - | 0 | - |
| - | - | 6031 | 476.8 | - | - | 0 | - |
| - | - | 2256 | 482.8 | - | - | 0 | - |
| - | - | 2.637E+04 | 483.3 | - | - | 0 | - |
| 9 | c | 2.831E+05 | 483.8 | 0.0006457 | 1.335 | +2 | 9 |
| - | - | 1.319E+05 | 484.3 | - | - | 0 | - |
| - | - | 4.13E+04 | 484.8 | - | - | 0 | - |
| - | - | 7811 | 485.3 | - | - | 0 | - |
| - | - | 5928 | 486.3 | - | - | 0 | - |
| - | - | 2501 | 489.3 | - | - | 0 | - |
| - | - | 2672 | 490.8 | - | - | 0 | - |
| - | - | 2647 | 495.3 | - | - | 0 | - |
| - | - | 1.47E+05 | 496.8 | - | - | 0 | - |
| - | - | 8.924E+04 | 497.3 | - | - | 0 | - |
| - | - | 3.024E+04 | 497.8 | - | - | 0 | - |
| - | - | 2.463E+04 | 498.3 | - | - | 0 | - |
| - | - | 6075 | 498.3 | - | - | 0 | - |
| - | - | 1.485E+04 | 498.8 | - | - | 0 | - |
| - | - | 4631 | 499.3 | - | - | 0 | - |
| - | - | 8684 | 499.3 | - | - | 0 | - |
| - | - | 2396 | 500.3 | - | - | 0 | - |
| - | - | 1.045E+04 | 504.8 | - | - | 0 | - |
| - | - | 2708 | 505.3 | - | - | 0 | - |
| - | - | 2342 | 505.8 | - | - | 0 | - |
| - | - | 2750 | 511.3 | - | - | 0 | - |
| - | - | 9703 | 517.8 | - | - | 0 | - |
| 8 | y | 2858 | 518.3 | 0.001953 | 3.769 | +1 | 4 |
| - | - | 1.056E+04 | 518.3 | - | - | 0 | - |
| - | - | 6332 | 518.8 | - | - | 0 | - |
| - | - | 2348 | 520.8 | - | - | 0 | - |
| - | - | 6628 | 524.3 | - | - | 0 | - |
| - | - | 1.503E+04 | 525.3 | - | - | 0 | - |
| - | - | 1.659E+05 | 525.8 | - | - | 0 | - |
| - | - | 1.12E+05 | 526.3 | - | - | 0 | - |
| - | - | 3.845E+04 | 526.8 | - | - | 0 | - |
| - | - | 1.313E+04 | 527.3 | - | - | 0 | - |
| - | - | 2651 | 527.8 | - | - | 0 | - |
| - | - | 2292 | 528.3 | - | - | 0 | - |
| - | - | 4687 | 529.3 | - | - | 0 | - |
| - | - | 5884 | 530.8 | - | - | 0 | - |
| - | - | 5394 | 531.3 | - | - | 0 | - |
| - | - | 6248 | 531.8 | - | - | 0 | - |
| - | - | 3315 | 532.3 | - | - | 0 | - |
| - | - | 2344 | 532.8 | - | - | 0 | - |
| 10 | c | 8104 | 539.3 | 0.0002565 | 0.4755 | +2 | 10 |
| 10 | c | 9.876E+05 | 539.8 | 0.0008634 | 1.599 | +2 | 10 |
| - | - | 5.586E+05 | 540.3 | - | - | 0 | - |
| - | - | 3691 | 540.4 | - | - | 0 | - |
| - | - | 1.96E+05 | 540.8 | - | - | 0 | - |
| - | - | 4.681E+04 | 541.3 | - | - | 0 | - |
| - | - | 8948 | 541.8 | - | - | 0 | - |
| - | - | 1499 | 542.3 | - | - | 0 | - |
| - | - | 1.616E+04 | 542.3 | - | - | 0 | - |
| - | - | 1811 | 543.3 | - | - | 0 | - |
| - | - | 6126 | 543.3 | - | - | 0 | - |
| - | - | 2608 | 547.3 | - | - | 0 | - |
| - | - | 1.271E+04 | 547.8 | - | - | 0 | - |
| 10 | c | 1.257E+06 | 548.3 | 0.0007725 | 1.409 | +2 | 10 |
| - | - | 7.61E+05 | 548.8 | - | - | 0 | - |
| - | - | 2.604E+05 | 549.3 | - | - | 0 | - |
| - | - | 5.314E+04 | 549.8 | - | - | 0 | - |
| - | - | 1.078E+04 | 550.3 | - | - | 0 | - |
| - | - | 3818 | 553.8 | - | - | 0 | - |
| - | - | 1889 | 554.3 | - | - | 0 | - |
| - | - | 3256 | 554.8 | - | - | 0 | - |
| - | - | 6.189E+04 | 555.3 | - | - | 0 | - |
| - | - | 1.087E+04 | 555.3 | - | - | 0 | - |
| - | - | 3.703E+04 | 555.8 | - | - | 0 | - |
| - | - | 1.254E+04 | 556.3 | - | - | 0 | - |
| - | - | 3579 | 558.3 | - | - | 0 | - |
| - | - | 3103 | 559.3 | - | - | 0 | - |
| - | - | 3.96E+04 | 561.3 | - | - | 0 | - |
| - | - | 3.56E+04 | 561.8 | - | - | 0 | - |
| - | - | 8155 | 562.3 | - | - | 0 | - |
| - | - | 7955 | 562.8 | - | - | 0 | - |
| - | - | 4553 | 562.8 | - | - | 0 | - |
| - | - | 2712 | 563.3 | - | - | 0 | - |
| - | - | 2520 | 567.3 | - | - | 0 | - |
| - | - | 9351 | 568.3 | - | - | 0 | - |
| - | - | 6410 | 568.8 | - | - | 0 | - |
| 5 | c | 1.919E+05 | 569.4 | 0.0004893 | 0.8593 | +1 | 5 |
| - | - | 2133 | 569.8 | - | - | 0 | - |
| - | - | 1664 | 570.3 | - | - | 0 | - |
| - | - | 5.975E+04 | 570.4 | - | - | 0 | - |
| - | - | 1.143E+04 | 571.4 | - | - | 0 | - |
| - | - | 2.289E+04 | 575.8 | - | - | 0 | - |
| - | - | 1.601E+04 | 576.3 | - | - | 0 | - |
| - | - | 6.21E+04 | 576.8 | - | - | 0 | - |
| - | - | 6.93E+04 | 577.3 | - | - | 0 | - |
| - | - | 3.009E+04 | 577.8 | - | - | 0 | - |
| - | - | 1.082E+04 | 578.3 | - | - | 0 | - |
| - | - | 3411 | 578.8 | - | - | 0 | - |
| - | - | 2.097E+04 | 582.4 | - | - | 0 | - |
| - | - | 9779 | 582.8 | - | - | 0 | - |
| - | - | 5905 | 583.4 | - | - | 0 | - |
| - | - | 7704 | 583.8 | - | - | 0 | - |
| 2 | z | 6.25E+04 | 584.3 | 0.00163 | 2.789 | +2 | 10 |
| - | - | 3.85E+04 | 584.8 | - | - | 0 | - |
| - | - | 1.364E+04 | 585.3 | - | - | 0 | - |
| - | - | 3243 | 585.4 | - | - | 0 | - |
| - | - | 4231 | 585.8 | - | - | 0 | - |
| - | - | 4065 | 586.3 | - | - | 0 | - |
| - | - | 2198 | 587.3 | - | - | 0 | - |
| - | - | 3290 | 590.3 | - | - | 0 | - |
| - | - | 6537 | 590.9 | - | - | 0 | - |
| - | - | 4819 | 591.3 | - | - | 0 | - |
| - | - | 3928 | 591.8 | - | - | 0 | - |
| 2 | y | 7392 | 592.3 | 0.001911 | 3.227 | +2 | 10 |
| - | - | 4655 | 592.8 | - | - | 0 | - |
| - | - | 5882 | 596.4 | - | - | 0 | - |
| - | - | 2346 | 597.4 | - | - | 0 | - |
| - | - | 7533 | 597.8 | - | - | 0 | - |
| - | - | 1.951E+04 | 598.3 | - | - | 0 | - |
| - | - | 1.467E+04 | 598.8 | - | - | 0 | - |
| - | - | 1.51E+04 | 599.3 | - | - | 0 | - |
| - | - | 2447 | 599.8 | - | - | 0 | - |
| - | - | 3308 | 600.3 | - | - | 0 | - |
| - | - | 1.453E+04 | 602.3 | - | - | 0 | - |
| - | - | 4205 | 603.3 | - | - | 0 | - |
| - | - | 2.677E+04 | 604.3 | - | - | 0 | - |
| - | - | 5.914E+04 | 604.8 | - | - | 0 | - |
| - | - | 3.466E+04 | 605.3 | - | - | 0 | - |
| - | - | 2.621E+04 | 605.8 | - | - | 0 | - |
| - | - | 1.134E+04 | 606.3 | - | - | 0 | - |
| - | - | 6483 | 606.3 | - | - | 0 | - |
| - | - | 8371 | 606.8 | - | - | 0 | - |
| - | - | 4831 | 607.3 | - | - | 0 | - |
| - | - | 3252 | 607.8 | - | - | 0 | - |
| - | - | 1.718E+04 | 611.4 | - | - | 0 | - |
| - | - | 1.109E+05 | 611.9 | - | - | 0 | - |
| - | - | 2707 | 612.3 | - | - | 0 | - |
| - | - | 6.796E+04 | 612.4 | - | - | 0 | - |
| - | - | 7582 | 612.8 | - | - | 0 | - |
| - | - | 2.871E+04 | 612.9 | - | - | 0 | - |
| - | - | 1.719E+04 | 613.3 | - | - | 0 | - |
| - | - | 4637 | 613.8 | - | - | 0 | - |
| - | - | 5.24E+04 | 614.4 | - | - | 0 | - |
| - | - | 7510 | 614.8 | - | - | 0 | - |
| - | - | 3.099E+04 | 615.3 | - | - | 0 | - |
| - | - | 1.738E+04 | 615.4 | - | - | 0 | - |
| - | - | 9854 | 616.3 | - | - | 0 | - |
| - | - | 4709 | 616.4 | - | - | 0 | - |
| - | - | 3212 | 617.8 | - | - | 0 | - |
| - | - | 9613 | 619.4 | - | - | 0 | - |
| - | - | 7.086E+04 | 619.8 | - | - | 0 | - |
| - | - | 9.03E+04 | 620.4 | - | - | 0 | - |
| - | - | 2.027E+05 | 620.8 | - | - | 0 | - |
| - | - | 1.383E+05 | 621.3 | - | - | 0 | - |
| - | - | 5.378E+04 | 621.8 | - | - | 0 | - |
| - | - | 1.203E+04 | 622.3 | - | - | 0 | - |
| - | - | 3289 | 622.8 | - | - | 0 | - |
| - | - | 1.586E+04 | 624.3 | - | - | 0 | - |
| - | - | 2.63E+04 | 625.4 | - | - | 0 | - |
| - | - | 1.926E+04 | 625.8 | - | - | 0 | - |
| 6 | c | 7.435E+05 | 626.4 | 0.00112 | 1.789 | +1 | 6 |
| - | - | 7.871E+04 | 626.8 | - | - | 0 | - |
| - | - | 2.351E+05 | 627.4 | - | - | 0 | - |
| - | - | 8902 | 627.8 | - | - | 0 | - |
| - | - | 4.604E+04 | 628.4 | - | - | 0 | - |
| - | - | 4670 | 629.4 | - | - | 0 | - |
| - | - | 1.934E+04 | 633.3 | - | - | 0 | - |
| - | - | 1.039E+06 | 633.8 | - | - | 0 | - |
| - | - | 7.531E+05 | 634.3 | - | - | 0 | - |
| - | - | 3.009E+05 | 634.8 | - | - | 0 | - |
| - | - | 9.1E+04 | 635.3 | - | - | 0 | - |
| - | - | 1.788E+04 | 635.8 | - | - | 0 | - |
| - | - | 4.917E+05 | 641.9 | - | - | 0 | - |
| - | - | 6.123E+05 | 642.4 | - | - | 0 | - |
| - | - | 3.317E+05 | 642.9 | - | - | 0 | - |
| - | - | 1.118E+05 | 643.4 | - | - | 0 | - |
| - | - | 3.205E+04 | 643.9 | - | - | 0 | - |
| - | - | 3117 | 644.4 | - | - | 0 | - |
| - | - | 5053 | 652.4 | - | - | 0 | - |
| - | - | 2459 | 653.4 | - | - | 0 | - |
| 7 | y | 3136 | 656.4 | 0.003387 | 5.16 | +1 | 5 |
| 7 | z | 4.478E+05 | 658.3 | 0.0006822 | 1.036 | +1 | 5 |
| - | - | 1.555E+05 | 659.3 | - | - | 0 | - |
| - | - | 3.513E+04 | 660.3 | - | - | 0 | - |
| - | - | 5666 | 661.4 | - | - | 0 | - |
| - | - | 3.058E+04 | 671.4 | - | - | 0 | - |
| - | - | 3071 | 672.3 | - | - | 0 | - |
| - | - | 1.115E+04 | 672.4 | - | - | 0 | - |
| - | - | 2005 | 673.4 | - | - | 0 | - |
| 7 | y | 2.473E+04 | 674.4 | 0.0005128 | 0.7605 | +1 | 5 |
| - | - | 9849 | 675.4 | - | - | 0 | - |
| - | - | 2802 | 676.4 | - | - | 0 | - |
| - | - | 2591 | 682.4 | - | - | 0 | - |
| - | - | 2999 | 695.4 | - | - | 0 | - |
| - | - | 3588 | 696.4 | - | - | 0 | - |
| - | - | 1951 | 697.4 | - | - | 0 | - |
| 6 | y | 8737 | 713.4 | 0.001638 | 2.296 | +1 | 6 |
| - | - | 3311 | 714.4 | - | - | 0 | - |
| 6 | z | 1.025E+05 | 715.4 | 0.000886 | 1.239 | +1 | 6 |
| - | - | 4.226E+04 | 716.4 | - | - | 0 | - |
| - | - | 9872 | 717.4 | - | - | 0 | - |
| - | - | 3417 | 723.4 | - | - | 0 | - |
| - | - | 2276 | 724.4 | - | - | 0 | - |
| 6 | y | 3.407E+04 | 731.4 | 0.0002894 | 0.3956 | +1 | 6 |
| - | - | 1.616E+04 | 732.4 | - | - | 0 | - |
| - | - | 3526 | 733.4 | - | - | 0 | - |
| - | - | 2290 | 738.5 | - | - | 0 | - |
| - | - | 3277 | 739.5 | - | - | 0 | - |
| - | - | 5255 | 740.5 | - | - | 0 | - |
| - | - | 2960 | 741.5 | - | - | 0 | - |
| - | - | 4239 | 767.5 | - | - | 0 | - |
| - | - | 2640 | 768.5 | - | - | 0 | - |
| - | - | 6440 | 780.4 | - | - | 0 | - |
| - | - | 1.131E+04 | 781.5 | - | - | 0 | - |
| 7 | c | 1.188E+05 | 782.5 | 0.0004731 | 0.6046 | +1 | 7 |
| - | - | 4.527E+04 | 783.5 | - | - | 0 | - |
| - | - | 9551 | 784.5 | - | - | 0 | - |
| 5 | w | 1.51E+04 | 785.4 | 0.001504 | 1.916 | +1 | 7 |
| - | - | 3059 | 785.5 | - | - | 0 | - |
| - | - | 1.066E+04 | 786.4 | - | - | 0 | - |
| - | - | 4301 | 787.4 | - | - | 0 | - |
| - | - | 2858 | 795.4 | - | - | 0 | - |
| 5 | y | 1.364E+04 | 810.4 | 0.0002044 | 0.2523 | +1 | 7 |
| - | - | 1.938E+04 | 811.4 | - | - | 0 | - |
| 5 | z | 5554 | 812.4 | 0.008669 | 10.67 | +1 | 7 |
| - | - | 6807 | 819.5 | - | - | 0 | - |
| - | - | 7062 | 823.4 | - | - | 0 | - |
| - | - | 2.062E+04 | 824.5 | - | - | 0 | - |
| - | - | 1.56E+04 | 825.5 | - | - | 0 | - |
| - | - | 1.045E+04 | 826.4 | - | - | 0 | - |
| - | - | 8471 | 827.4 | - | - | 0 | - |
| 5 | y | 4.943E+04 | 828.4 | 0.0001378 | 0.1663 | +1 | 7 |
| - | - | 1.935E+04 | 829.4 | - | - | 0 | - |
| - | - | 5657 | 830.4 | - | - | 0 | - |
| - | - | 2214 | 837.5 | - | - | 0 | - |
| - | - | 2356 | 838.5 | - | - | 0 | - |
| - | - | 1.217E+04 | 839.5 | - | - | 0 | - |
| - | - | 7478 | 840.5 | - | - | 0 | - |
| - | - | 3979 | 851.5 | - | - | 0 | - |
| - | - | 1.069E+04 | 852.5 | - | - | 0 | - |
| 8 | c | 8.986E+04 | 853.5 | 8.062E-05 | 0.09445 | +1 | 8 |
| - | - | 4.25E+04 | 854.5 | - | - | 0 | - |
| - | - | 1.154E+04 | 855.5 | - | - | 0 | - |
| - | - | 5639 | 856.4 | - | - | 0 | - |
| - | - | 2724 | 856.5 | - | - | 0 | - |
| - | - | 2237 | 857 | - | - | 0 | - |
| 4 | z | 4.781E+05 | 883.5 | 0.0001807 | 0.2045 | +1 | 8 |
| - | - | 2.38E+05 | 884.5 | - | - | 0 | - |
| - | - | 6.96E+04 | 885.5 | - | - | 0 | - |
| - | - | 1.297E+04 | 886.5 | - | - | 0 | - |
| 4 | y | 7.313E+04 | 899.5 | 0.0003775 | 0.4197 | +1 | 8 |
| - | - | 2.991E+04 | 900.5 | - | - | 0 | - |
| - | - | 7017 | 901.5 | - | - | 0 | - |
| - | - | 4005 | 908.6 | - | - | 0 | - |
| - | - | 3814 | 915.4 | - | - | 0 | - |
| - | - | 6412 | 923.6 | - | - | 0 | - |
| - | - | 6732 | 924.6 | - | - | 0 | - |
| - | - | 2174 | 933.6 | - | - | 0 | - |
| - | - | 4810 | 939.5 | - | - | 0 | - |
| - | - | 3415 | 940.5 | - | - | 0 | - |
| - | - | 1942 | 942.4 | - | - | 0 | - |
| - | - | 6010 | 950.6 | - | - | 0 | - |
| - | - | 6300 | 951.6 | - | - | 0 | - |
| - | - | 4221 | 952.6 | - | - | 0 | - |
| 3 | w | 1.005E+04 | 953.5 | 0.002447 | 2.567 | +1 | 9 |
| - | - | 7441 | 954.5 | - | - | 0 | - |
| - | - | 3323 | 955.5 | - | - | 0 | - |
| - | - | 3812 | 963.5 | - | - | 0 | - |
| - | - | 5565 | 965.6 | - | - | 0 | - |
| 9 | c | 1.059E+05 | 966.6 | 0.000145 | 0.15 | +1 | 9 |
| - | - | 2577 | 967.5 | - | - | 0 | - |
| - | - | 5.115E+04 | 967.6 | - | - | 0 | - |
| - | - | 1.603E+04 | 968.6 | - | - | 0 | - |
| - | - | 3605 | 969.6 | - | - | 0 | - |
| - | - | 1.12E+04 | 980.5 | - | - | 0 | - |
| - | - | 6957 | 981.6 | - | - | 0 | - |
| - | - | 2599 | 982.6 | - | - | 0 | - |
| - | - | 2663 | 1008 | - | - | 0 | - |
| - | - | 2572 | 1009 | - | - | 0 | - |
| - | - | 5753 | 1010 | - | - | 0 | - |
| 3 | y | 2322 | 1011 | 0.003514 | 3.477 | +1 | 9 |
| - | - | 3226 | 1011 | - | - | 0 | - |
| 3 | z | 3.702E+05 | 1012 | 7.487E-05 | 0.07402 | +1 | 9 |
| - | - | 2.032E+05 | 1013 | - | - | 0 | - |
| - | - | 6.844E+04 | 1014 | - | - | 0 | - |
| - | - | 1.787E+04 | 1015 | - | - | 0 | - |
| - | - | 2301 | 1016 | - | - | 0 | - |
| - | - | 3866 | 1025 | - | - | 0 | - |
| - | - | 3202 | 1025 | - | - | 0 | - |
| 3 | y | 9.106E+04 | 1028 | 9.451E-05 | 0.09198 | +1 | 9 |
| - | - | 4.781E+04 | 1029 | - | - | 0 | - |
| - | - | 1.877E+04 | 1030 | - | - | 0 | - |
| - | - | 3421 | 1031 | - | - | 0 | - |
| - | - | 4935 | 1035 | - | - | 0 | - |
| - | - | 6182 | 1036 | - | - | 0 | - |
| - | - | 4298 | 1037 | - | - | 0 | - |
| - | - | 1.824E+04 | 1038 | - | - | 0 | - |
| - | - | 7680 | 1039 | - | - | 0 | - |
| - | - | 4393 | 1040 | - | - | 0 | - |
| - | - | 2549 | 1051 | - | - | 0 | - |
| - | - | 3.863E+04 | 1052 | - | - | 0 | - |
| - | - | 8.444E+04 | 1053 | - | - | 0 | - |
| - | - | 5.791E+04 | 1054 | - | - | 0 | - |
| - | - | 2.103E+04 | 1055 | - | - | 0 | - |
| - | - | 3675 | 1056 | - | - | 0 | - |
| - | - | 2634 | 1063 | - | - | 0 | - |
| 10 | c | 5114 | 1079 | 0.003448 | 3.197 | +1 | 10 |
| - | - | 7.239E+04 | 1080 | - | - | 0 | - |
| - | - | 4.645E+04 | 1081 | - | - | 0 | - |
| 2 | w | 1.945E+04 | 1082 | 0.0004053 | 0.3747 | +1 | 10 |
| - | - | 9277 | 1082 | - | - | 0 | - |
| - | - | 1.43E+04 | 1083 | - | - | 0 | - |
| - | - | 5110 | 1084 | - | - | 0 | - |
| 10 | c | 7.173E+04 | 1096 | 3.232E-05 | 0.0295 | +1 | 10 |
| - | - | 1.787E+05 | 1097 | - | - | 0 | - |
| - | - | 9.664E+04 | 1098 | - | - | 0 | - |
| - | - | 2.969E+04 | 1099 | - | - | 0 | - |
| - | - | 6661 | 1100 | - | - | 0 | - |
| - | - | 4908 | 1111 | - | - | 0 | - |
| - | - | 4349 | 1112 | - | - | 0 | - |
| - | - | 3167 | 1124 | - | - | 0 | - |
| - | - | 3775 | 1125 | - | - | 0 | - |
| - | - | 2750 | 1126 | - | - | 0 | - |
| - | - | 3015 | 1141 | - | - | 0 | - |
| - | - | 2825 | 1142 | - | - | 0 | - |
| - | - | 5201 | 1152 | - | - | 0 | - |
| - | - | 6532 | 1153 | - | - | 0 | - |
| - | - | 1.122E+04 | 1154 | - | - | 0 | - |
| - | - | 7431 | 1155 | - | - | 0 | - |
| 2 | z | 2.536E+04 | 1168 | 0.002647 | 2.267 | +1 | 10 |
| - | - | 1.244E+05 | 1169 | - | - | 0 | - |
| - | - | 7.995E+04 | 1170 | - | - | 0 | - |
| - | - | 2.831E+04 | 1171 | - | - | 0 | - |
| - | - | 7505 | 1172 | - | - | 0 | - |
| - | - | 2.332E+04 | 1181 | - | - | 0 | - |
| - | - | 1.611E+04 | 1182 | - | - | 0 | - |
| - | - | 1.413E+04 | 1183 | - | - | 0 | - |
| 2 | y | 1.493E+04 | 1184 | 0.008066 | 6.815 | +1 | 10 |
| - | - | 9674 | 1185 | - | - | 0 | - |
| - | - | 4343 | 1186 | - | - | 0 | - |
| - | - | 3472 | 1195 | - | - | 0 | - |
| - | - | 1.991E+04 | 1196 | - | - | 0 | - |
| - | - | 1.38E+04 | 1197 | - | - | 0 | - |
| - | - | 6157 | 1198 | - | - | 0 | - |
| - | - | 2030 | 1199 | - | - | 0 | - |
| - | - | 5800 | 1207 | - | - | 0 | - |
| - | - | 3736 | 1208 | - | - | 0 | - |
| - | - | 2756 | 1209 | - | - | 0 | - |
| - | - | 4091 | 1210 | - | - | 0 | - |
| - | - | 2880 | 1212 | - | - | 0 | - |
| - | - | 5753 | 1213 | - | - | 0 | - |
| - | - | 4610 | 1214 | - | - | 0 | - |
| - | - | 2698 | 1222 | - | - | 0 | - |
| - | - | 1.515E+04 | 1223 | - | - | 0 | - |
| - | - | 3.238E+04 | 1224 | - | - | 0 | - |
| - | - | 2.399E+05 | 1225 | - | - | 0 | - |
| - | - | 1.851E+05 | 1226 | - | - | 0 | - |
| - | - | 7.951E+04 | 1227 | - | - | 0 | - |
| - | - | 2.344E+04 | 1228 | - | - | 0 | - |
| - | - | 5172 | 1229 | - | - | 0 | - |
| - | - | 2477 | 1238 | - | - | 0 | - |
| - | - | 1.177E+04 | 1239 | - | - | 0 | - |
| - | - | 4.527E+04 | 1240 | - | - | 0 | - |
| - | - | 5.011E+04 | 1241 | - | - | 0 | - |
| - | - | 3.752E+04 | 1242 | - | - | 0 | - |
| - | - | 1.856E+04 | 1243 | - | - | 0 | - |
| - | - | 6653 | 1244 | - | - | 0 | - |
| - | - | 5086 | 1250 | - | - | 0 | - |
| - | - | 6.469E+04 | 1251 | - | - | 0 | - |
| - | - | 4.307E+04 | 1252 | - | - | 0 | - |
| - | - | 1.558E+04 | 1253 | - | - | 0 | - |
| - | - | 5988 | 1254 | - | - | 0 | - |
| - | - | 4478 | 1256 | - | - | 0 | - |
| - | - | 1.76E+04 | 1257 | - | - | 0 | - |
| - | - | 5700 | 1258 | - | - | 0 | - |
| - | - | 5798 | 1259 | - | - | 0 | - |
| - | - | 6.468E+04 | 1267 | - | - | 0 | - |
| - | - | 5.322E+05 | 1268 | - | - | 0 | - |
| - | - | 3.706E+05 | 1269 | - | - | 0 | - |
| - | - | 1.407E+05 | 1270 | - | - | 0 | - |
| - | - | 3.775E+04 | 1271 | - | - | 0 | - |
| - | - | 8567 | 1272 | - | - | 0 | - |
| - | - | 1.615E+04 | 1283 | - | - | 0 | - |
| - | - | 1.184E+05 | 1284 | - | - | 0 | - |
| - | - | 4.728E+05 | 1285 | - | - | 0 | - |
| - | - | 3.177E+05 | 1286 | - | - | 0 | - |
| - | - | 4317 | 1286 | - | - | 0 | - |
| - | - | 1.206E+05 | 1287 | - | - | 0 | - |
| - | - | 2.929E+04 | 1288 | - | - | 0 | - |
| - | - | 8765 | 1289 | - | - | 0 | - |
| - | - | 2005 | 1819 | - | - | 0 | - |
| - | - | 1963 | 1876 | - | - | 0 | - |
| - | - | 2151 | 1916 | - | - | 0 | - |
| - | - | 2063 | 2491 | - | - | 0 | - |
| - | - | 2442 | 2731 | - | - | 0 | - |
| - | - | 2227 | 3197 | - | - | 0 | - |
| - | - | 2105 | 3445 | - | - | 0 | - |

m/z Charge Intensity FragmentType MassShift Position
123.84258270263672 0 1197.8959
130.0606231689453 0 2011.9478
130.06549072265625 0 27313.752
131.0686492919922 0 1581.4567
132.0811309814453 0 7562.444
133.08633422851562 0 13230.644
142.09823608398438 0 3189.8923
142.1037139892578 0 1240.5062
144.08106994628906 0 5514.5713
145.08551025390625 0 1439.1211
146.0603790283203 0 26344.986
147.06410217285156 0 2565.3538
157.1087188720703 0 14624.53
159.09207153320312 0 96265.07
160.08859252929688 0 1677.378
160.09544372558594 0 9765.537
166.08673095703125 0 1673.8761
169.13380432128906 0 1447.7239
170.06031799316406 0 4573.6953
171.0919189453125 0 2147.8113
171.12440490722656 0 2835.288
171.47161865234375 0 1427.2942
172.07960510253906 0 1357.5474
173.4517059326172 0 3607.2378
174.1030731201172 0 2930.1335
174.1352996826172 0 4727.2393
177.11241149902344 0 8329.731
178.625 0 1700.9431
185.103759765625 0 7340.0522
185.13999938964844 0 3468.0005
187.08697509765625 0 16319.101
188.07098388671875 0 318171.16
189.07432556152344 0 37008.684
190.6595458984375 0 1390.065
192.62161254882812 0 10187.334 c Ammonia loss 2
195.1611328125 0 2110.478
196.14451599121094 0 2506.7979
197.12876892089844 0 3509.2935
197.1393585205078 0 3478.881
199.0897979736328 0 1360.5076
203.1278076171875 0 1632.646
205.0975341796875 0 200608.7 y 10
205.62855529785156 0 1486.4119
206.1009063720703 0 22322.893
208.95361328125 0 29261.223
213.17135620117188 0 19365.861
214.14285278320312 0 34644.723
214.15536499023438 0 12974.986
214.17442321777344 0 1655.1031
214.6443634033203 0 5634.285
215.1580352783203 0 1457.4369
218.868408203125 0 1397.9426
228.14028930664062 0 12282.756 c Ammonia loss 3
228.18223571777344 0 8130.543
228.64149475097656 0 1902.0045
235.15370178222656 0 1579.0452
238.16651916503906 0 4692.565
239.15066528320312 0 70904.13
240.15426635742188 0 8751.097
242.16148376464844 0 13394.236
243.16639709472656 0 2619.6313
248.1565399169922 0 2280.6204
248.97364807128906 0 1512.639
256.12054443359375 0 4640.265
256.1771545410156 0 46945.312 c Ammonia loss 1
257.1800842285156 0 5364.6274
270.1242370605469 0 12580.115
271.1278991699219 0 3210.8298
273.20367431640625 0 172120.67 c 1
274.206787109375 0 17681.357
276.6666259765625 0 2798.0674 c Ammonia loss 4
279.1808166503906 0 2378.3264
280.10888671875 0 2190.7952
298.1192626953125 0 6548.8823
298.223876953125 0 5240.735
298.6798400878906 0 3440.6047
299.2196350097656 0 44769.203
300.2216796875 0 3474.7266
307.8631896972656 0 6426.3677
308.19775390625 0 2690.6887
312.19085693359375 0 3741.073
312.6771545410156 0 5310.1694
313.235107421875 0 2619.987
316.1297912597656 0 47292.098 y Water loss 9
318.9227294921875 0 18939.371
334.1404724121094 0 4604.9243 y 9
341.019287109375 0 4484.7354
341.2301330566406 0 2711.729
347.7277526855469 0 2902.8748
348.2287292480469 0 1846.7798
356.24090576171875 0 10438.955
357.2473449707031 0 7844.583
359.02880859375 0 19267.875
361.7254943847656 0 11427.632
362.2257385253906 0 7661.1245
362.7249450683594 0 1845.2556
367.20916748046875 0 4282.503
369.22991943359375 0 5471.5176
369.7342224121094 0 30892.445
369.7608337402344 0 1700.693
370.2363586425781 0 16735.076
370.25860595703125 0 3786.3853
370.738037109375 0 5316.9194
383.2278137207031 0 17773.393 c Ammonia loss 6
383.730224609375 0 6000.165
384.2033386230469 0 3202.212 z Ammonia loss 1
384.2359313964844 0 35097.18 c Ammonia loss 2
385.239013671875 0 6857.4644
385.27984619140625 0 3744.1348
391.23748779296875 0 214159.08
391.7410888671875 0 684163.56 c 6
392.24249267578125 0 256355.66
392.7436218261719 0 62848.867
393.2448425292969 0 8045.0874
396.7120666503906 0 2412.6172
401.2626647949219 0 535088.3 c 2
402.2651672363281 0 101451.55
403.2674865722656 0 12059.787
404.7493896484375 0 8454.328
405.2520446777344 0 7208.0093
405.7174377441406 0 13079.086 y Water loss 4
405.7511291503906 0 2143.4246
406.2188415527344 0 4497.6387
406.721435546875 0 2298.3167
406.9003601074219 0 24402.623
407.234619140625 0 18720.732
407.5679931640625 0 6173.0938
410.2336120605469 0 6972.8296
412.9042663574219 0 29242.033
413.23822021484375 0 11916.041
413.27294921875 0 3027.14
413.5731506347656 0 2426.8987
413.9057922363281 0 1698.7191
414.7228698730469 0 13853.21 y 4
415.2243347167969 0 5281.446
416.2305603027344 0 2719.518
416.5673522949219 0 2156.934
418.2532653808594 0 2137.3765
418.74725341796875 0 28605.877 c Ammonia loss 7
419.2486267089844 0 9335.367
422.2352600097656 0 9361.552
422.5699462890625 0 5501.8833
422.90509033203125 0 2928.7764
426.75634765625 0 79115.734
427.2600402832031 0 526094.1 c 7
427.76141357421875 0 251321.77
428.2628173828125 0 60967.33
428.76361083984375 0 11617.732
428.890380859375 0 3879.7078
429.0895690917969 0 15978.045
429.2679138183594 0 2917.501
438.2440490722656 0 1724.2656
440.2688903808594 0 2923.3042
453.28863525390625 0 1754.1642
453.78582763671875 0 2574.9226
455.27325439453125 0 37181.22 c Ammonia loss 3
456.2757873535156 0 10495.628
461.29156494140625 0 65061.258
461.7936706542969 0 47494.008
462.29608154296875 0 17209.43
462.80224609375 0 2505.8442
466.77587890625 0 2985.2725
468.7758483886719 0 3294.434
472.2890930175781 0 5183.9053
473.28509521484375 0 2245.343
474.7967529296875 0 7737.06
475.28900146484375 0 178674.52 c Ammonia loss 8
475.7903137207031 0 89758.44
476.2916564941406 0 24424.389
476.7928466796875 0 6030.946
482.7963562011719 0 2256.4858
483.2984924316406 0 26367.174
483.80206298828125 0 283073.25 c 8
484.3033142089844 0 131887.03
484.8048400878906 0 41303.15
485.3052978515625 0 7810.527
486.2714538574219 0 5928.397
489.30230712890625 0 2501.125
490.78155517578125 0 2671.7063
495.3034362792969 0 2646.824
496.8099060058594 0 146971.23
497.3111877441406 0 89242.734
497.8124084472656 0 30239.025
498.2853088378906 0 24632.863
498.3167724609375 0 6074.66
498.78619384765625 0 14847.259
499.2881164550781 0 4631.1606
499.3231506347656 0 8684.5
500.32635498046875 0 2396.0742
504.7904357910156 0 10448.385
505.291748046875 0 2707.8594
505.79296875 0 2341.562
511.3117980957031 0 2749.8223
517.8140869140625 0 9703.083
518.2628784179688 0 2858.2478 y 7
518.30810546875 0 10558.035
518.8082275390625 0 6332.014
520.7640991210938 0 2347.6606
524.3320922851562 0 6628.0723
525.3377075195312 0 15034.522
525.8126831054688 0 165871.89
526.314208984375 0 111958.12
526.8153076171875 0 38447.547
527.3165893554688 0 13127.302
527.8219604492188 0 2650.7083
528.2982788085938 0 2292.3835
529.285888671875 0 4687.348
530.8058471679688 0 5884.1104
531.3006591796875 0 5394.0684
531.800048828125 0 6247.871
532.2942504882812 0 3314.9846
532.8060913085938 0 2343.8801
539.3176879882812 0 8103.743 c Water loss 9
539.810302734375 0 987567.56 c Ammonia loss 9
540.3116455078125 0 558616.44
540.4088134765625 0 3691.013
540.8129272460938 0 196048.4
541.314208984375 0 46811.496
541.81689453125 0 8948.18
542.298828125 0 1498.6085
542.3325805664062 0 16157.108
543.2928466796875 0 1811.3962
543.337158203125 0 6125.5234
547.3142700195312 0 2608.085
547.8120727539062 0 12709.992
548.323486328125 0 1257123.2 c 9
548.82470703125 0 760954.25
549.3261108398438 0 260429.12
549.826904296875 0 53141.92
550.3294067382812 0 10784.962
553.8027954101562 0 3817.788
554.3248291015625 0 1888.9075
554.80419921875 0 3256.4504
555.2966918945312 0 61890.7
555.3409423828125 0 10865.311
555.7980346679688 0 37029.74
556.2980346679688 0 12537.744
558.2906494140625 0 3579.3594
559.3180541992188 0 3103.4553
561.3308715820312 0 39597.58
561.8331909179688 0 35599.348
562.3307495117188 0 8154.858
562.7844848632812 0 7955.1377
562.8289794921875 0 4552.8604
563.2840576171875 0 2711.932
567.3461303710938 0 2520.22
568.34619140625 0 9350.885
568.8209228515625 0 6409.605
569.352294921875 0 191865.77 c 4
569.8069458007812 0 2132.6267
570.3065795898438 0 1664.0885
570.3552856445312 0 59745.62
571.3576049804688 0 11433.202
575.8384399414062 0 22893.234
576.3385009765625 0 16005.034
576.8225708007812 0 62095.203
577.3246459960938 0 69302.98
577.8265991210938 0 30088.021
578.3246459960938 0 10817.13
578.8233642578125 0 3410.9507
582.3549194335938 0 20968.26
582.8411254882812 0 9779.195
583.35693359375 0 5905.1987
583.8292846679688 0 7704.3154
584.3124389648438 0 62500.98 z 1
584.8134765625 0 38495.09
585.3145751953125 0 13635.448
585.3594360351562 0 3242.797
585.815185546875 0 4230.8647
586.3206176757812 0 4065.2083
587.32470703125 0 2198.3293
590.3374633789062 0 3290.2463
590.8541870117188 0 6537.1875
591.3157958984375 0 4819.121
591.8157958984375 0 3927.9207
592.3220825195312 0 7391.545 y 1
592.8240356445312 0 4655.4453
596.3517456054688 0 5881.931
597.3561401367188 0 2345.866
597.8394165039062 0 7532.7124
598.3265380859375 0 19507.273
598.8259887695312 0 14672.288
599.3323974609375 0 15098.528
599.8302612304688 0 2446.8962
600.3300170898438 0 3307.9148
602.2811889648438 0 14527.757
603.2846069335938 0 4205.0537
604.3394165039062 0 26765.879
604.833740234375 0 59141.13
605.3327026367188 0 34662.707
605.828369140625 0 26213.895
606.3182373046875 0 11337.449
606.3490600585938 0 6482.98
606.8450317382812 0 8370.876
607.3438110351562 0 4830.922
607.8475341796875 0 3251.9602
611.3555297851562 0 17183.793
611.8510131835938 0 110916.13
612.3049926757812 0 2707.3188
612.352783203125 0 67958.1
612.8131103515625 0 7582.46
612.8532104492188 0 28708.65
613.347412109375 0 17188.766
613.8484497070312 0 4637.391
614.3526000976562 0 52402.848
614.8297119140625 0 7510.448
615.2894897460938 0 30990.48
615.3551635742188 0 17376.318
616.2928466796875 0 9853.747
616.3580932617188 0 4708.5005
617.8234252929688 0 3212.0083
619.35498046875 0 9612.682
619.8496704101562 0 70863.67
620.3572387695312 0 90301.86
620.846435546875 0 202727.2
621.3465576171875 0 138294.88
621.8468627929688 0 53784.586
622.3485107421875 0 12032.127
622.8495483398438 0 3288.8362
624.347412109375 0 15861.824
625.3638916015625 0 26295.385
625.8363647460938 0 19261.613
626.3743896484375 0 743471.9 c 5
626.8360595703125 0 78705.89
627.3770141601562 0 235106.83
627.8399658203125 0 8902.298
628.3787841796875 0 46042.477
629.3807373046875 0 4669.7417
633.3482666015625 0 19340.805
633.8460083007812 0 1039222.25
634.3472900390625 0 753120.1
634.8484497070312 0 300883.84
635.3494873046875 0 90998.84
635.8494262695312 0 17877.363
641.8551635742188 0 491745.25
642.3574829101562 0 612304.4
642.8594970703125 0 331744.1
643.3602905273438 0 111828.95
643.862060546875 0 32051.994
644.3594360351562 0 3116.8625
652.3849487304688 0 5052.6177
653.3964233398438 0 2459.2424
656.3548583984375 0 3136.461 y Water loss 6
658.343994140625 0 447777.97 z 6
659.3468017578125 0 155506.1
660.349853515625 0 35133.79
661.351806640625 0 5665.8696
671.3756103515625 0 30581.75
672.317626953125 0 3070.6746
672.37841796875 0 11150.838
673.3795166015625 0 2004.9515
674.362548828125 0 24733.666 y 6
675.36474609375 0 9849.295
676.3715209960938 0 2801.8408
682.3890991210938 0 2591.0227
695.392333984375 0 2999.475
696.3916625976562 0 3588.4653
697.3993530273438 0 1951.2533
713.3745727539062 0 8737.115 y Water loss 5
714.3768310546875 0 3311.4165
715.3656616210938 0 102505.26 z 5
716.3682250976562 0 42261.94
717.3699951171875 0 9871.602
723.43408203125 0 3417.4158
724.4324340820312 0 2275.972
731.3837890625 0 34066.902 y 5
732.3873291015625 0 16161.741
733.389404296875 0 3525.576
738.4605712890625 0 2289.6184
739.4661254882812 0 3276.6353
740.456787109375 0 5254.9834
741.4533081054688 0 2959.6401
767.4561767578125 0 4238.9194
768.459228515625 0 2640.0479
780.4398803710938 0 6440.4126
781.4649658203125 0 11305.066
782.474853515625 0 118771.63 c 6
783.477294921875 0 45265.46
784.4815063476562 0 9550.862
785.3955688476562 0 15099.416 w 4
785.4796142578125 0 3058.9487
786.3984375 0 10662.354
787.4058837890625 0 4301.401
795.4482421875 0 2857.8086
810.4259033203125 0 13635.128 y Water loss 4
811.4295654296875 0 19378.54
812.4262084960938 0 5554.157 z 4
819.4581909179688 0 6807.4985
823.4425048828125 0 7062.4365
824.4505615234375 0 20615.129
825.4548950195312 0 15596.489
826.4236450195312 0 10450.039
827.408203125 0 8471.191
828.4364013671875 0 49430.465 y 4
829.439697265625 0 19348.023
830.4413452148438 0 5657.435
837.4881591796875 0 2214.2246
838.4916381835938 0 2356.103
839.4635620117188 0 12169.9
840.4698486328125 0 7478.319
851.5076293945312 0 3978.7822
852.5032348632812 0 10686.207
853.5114135742188 0 89856.984 c 7
854.5144653320312 0 42504.066
855.51806640625 0 11536.556
856.429443359375 0 5639.354
856.5187377929688 0 2724.3125
856.9888916015625 0 2237.1135
883.454833984375 0 478149.5 z 3
884.4581298828125 0 238016.48
885.4607543945312 0 69603.24
886.4637451171875 0 12970.546
899.4737548828125 0 73131.44 y 3
900.47607421875 0 29914.488
901.4775390625 0 7016.5493
908.5552978515625 0 4004.9102
915.446044921875 0 3813.7869
923.5737915039062 0 6412.4404
924.5742797851562 0 6731.7876
933.5581665039062 0 2173.9194
939.4905395507812 0 4810.0474
940.4946899414062 0 3415.3633
942.4484252929688 0 1942.4777
950.5760498046875 0 6010.309
951.5813598632812 0 6299.5576
952.5849609375 0 4221.431
953.4863891601562 0 10045.349 w 2
954.4883422851562 0 7441.4473
955.47119140625 0 3323.3455
963.5247192382812 0 3811.7478
965.5780639648438 0 5564.8315
966.595703125 0 105939.54 c 8
967.498291015625 0 2577.0527
967.5980834960938 0 51148.105
968.602783203125 0 16034.545
969.6119995117188 0 3605.4329
980.5491943359375 0 11200.868
981.5557861328125 0 6957.4854
982.558837890625 0 2599.4219
1007.6024169921875 0 2662.9226
1008.5985107421875 0 2571.981
1009.60302734375 0 5753.216
1010.5018920898438 0 2322.3418 y Ammonia loss 2
1010.6104736328125 0 3225.5752
1011.5133056640625 0 370219.5 z 2
1012.5161743164062 0 203227.73
1013.5191040039062 0 68438.766
1014.5223388671875 0 17872.174
1015.5213623046875 0 2301.002
1024.5206298828125 0 3865.8223
1024.63134765625 0 3201.6565
1027.5318603515625 0 91058.73 y 2
1028.534912109375 0 47810.39
1029.5394287109375 0 18766.643
1030.5389404296875 0 3420.9187
1034.6177978515625 0 4934.729
1035.60986328125 0 6181.989
1036.610595703125 0 4298.0195
1037.5970458984375 0 18238.393
1038.5989990234375 0 7680.021
1039.60400390625 0 4392.6587
1050.6278076171875 0 2549.415
1051.6243896484375 0 38626.434
1052.6278076171875 0 84435.766
1053.6280517578125 0 57906.47
1054.62890625 0 21026.709
1055.6328125 0 3675.4827
1062.59716796875 0 2634.28
1078.608154296875 0 5114.202 c Ammonia loss 9
1079.619140625 0 72388.99
1080.622314453125 0 46450.79
1081.5421142578125 0 19448.781 w 1
1081.6378173828125 0 9277.454
1082.54638671875 0 14301.107
1083.5479736328125 0 5110.1055
1095.63818359375 0 71726.25 c 9
1096.64501953125 0 178719.25
1097.6475830078125 0 96640.125
1098.6495361328125 0 29688.492
1099.6524658203125 0 6661.065
1110.6063232421875 0 4908.384
1111.60107421875 0 4349.3467
1123.6278076171875 0 3166.7715
1124.6219482421875 0 3774.8198
1125.60205078125 0 2750.3901
1140.6334228515625 0 3015.4944
1141.6378173828125 0 2825.4653
1151.617919921875 0 5201.0107
1152.6217041015625 0 6531.6504
1153.625 0 11215.45
1154.6285400390625 0 7431.3774
1167.6116943359375 0 25358.385 z 1
1168.62060546875 0 124395.01
1169.6239013671875 0 79946.6
1170.627685546875 0 28305.146
1171.6307373046875 0 7505.1416
1180.6165771484375 0 23320.736
1181.62255859375 0 16112.639
1182.620849609375 0 14128.716
1183.625 0 14929.552 y 1
1184.6309814453125 0 9674.487
1185.6317138671875 0 4343.1514
1194.6297607421875 0 3471.6538
1195.623779296875 0 19911.54
1196.6297607421875 0 13803.757
1197.6336669921875 0 6157.4146
1198.646484375 0 2030.1597
1206.6744384765625 0 5799.53
1207.6832275390625 0 3735.8406
1208.6597900390625 0 2755.818
1209.6331787109375 0 4091.1304
1211.6549072265625 0 2879.7634
1212.6759033203125 0 5753.245
1213.6851806640625 0 4609.618
1221.6861572265625 0 2698.4312
1222.70703125 0 15153.672
1223.69873046875 0 32379.11
1224.6619873046875 0 239883.67
1225.663818359375 0 185101.92
1226.6663818359375 0 79511.67
1227.6673583984375 0 23442.893
1228.67236328125 0 5171.8315
1237.7054443359375 0 2476.7463
1238.7010498046875 0 11768.305
1239.6954345703125 0 45270.445
1240.6981201171875 0 50113.242
1241.69482421875 0 37520.316
1242.6932373046875 0 18556.434
1243.693115234375 0 6653.488
1249.684814453125 0 5085.6377
1250.663818359375 0 64687.07
1251.6668701171875 0 43066.957
1252.6697998046875 0 15575.437
1253.662109375 0 5988.3936
1255.712646484375 0 4477.8657
1256.716552734375 0 17602.26
1257.7274169921875 0 5699.7124
1258.72119140625 0 5797.926
1266.6890869140625 0 64683.082
1267.689453125 0 532220.7
1268.6920166015625 0 370628.88
1269.6947021484375 0 140740.39
1270.6966552734375 0 37753.863
1271.6956787109375 0 8567.415
1282.700439453125 0 16148.47
1283.70703125 0 118381.984
1284.71484375 0 472829.88
1285.7174072265625 0 317669.8
1285.8468017578125 0 4317.1113
1286.720947265625 0 120588.875
1287.7225341796875 0 29293.166
1288.7218017578125 0 8765.421
1818.972900390625 0 2005.0298
1876.4677734375 0 1962.8079
1916.426513671875 0 2151.2688
2491.115966796875 0 2062.539
2731.48046875 0 2441.7131
3197.045654296875 0 2226.6367
3444.630615234375 0 2105.079

Spectrum Details

|  |  |
| --- | --- |
| Matched peaks? Matched peaks The total absolute number of peaks matched. Additionally in brackets the total fraction of peaks matched and the total number of peaks is shown. | 52 (9.30% of 559) |
| FDR? FDR The false discovery rate estimated for this peptide. It is calculated by matching all theoretical fragments with a non-integer shift with the raw peaks for this spectrum. This is done with 40 different shifts. The resulting percentage is the average number of annotated peaks over the number of annotated peaks with the correct spectrum. | 2.52% |
| Satellite FDR? Satellite FDR See the FDR for details on its calculation. This satellite ion specific FDR only contains the satellite ions (d/w) for I/L/J positions. | ∞ |
| PSM Score? PSM Score The PSM Score as given by Hecklib to this annotated spectrum. It is shown with three significant figures. | 445 |

#### Spectrum 4540? Spectrum 4540 The raw spectrum of this peptide as annotated by Hecklib. The fragments are coloured according to ion type (see legend). Any peaks with a star '\*' as text can be hovered over to see the full details, first the ion type second the mass shift type. By hovering over the amino acids in the peptide or ions in the legend the corresponding peaks are highlighted. By toggling the 'Unassigned' label you can turn the background (unassigned) peaks on or off in the plot. By updating the slider in the Ion legend you can update the spectrum to only show the top X% of the peaks with labels. The top X% means any peak that is within X% of the highest intensity. By dragging in the spectrum you can zoom in to a specific part of the spectrum and use 'Zoom Out' to get back to the original zoom level. The annotation of the spectrum is based on the given sequence in the peptides file and is done with different software so inconsistencies are likely. The peaks are annotated based on the given sequence, with 20 ppm tolerance.

Copy Data

##### Spectrum 4540 (TSV)

###### Preview

```
Loading example...
```

*Click on the button to copy the data to your clipboard.*

Mz MinMz MaxIntensity Max

WidthHeightPeptide font sizePeptide stroke widthSpectrum font sizeSpectrum stroke widthCompact peptide

Ion legend

wxyz

abcd

OtherUnassignedIon^Charge^_Position_Show for top:%

VRQAPGRAJEW

02.82e+45.64e+48.46e+41.13e+5

Zoom Out

c^+2^_3_c^+2^_4_c^+1^_2_c^+1^_2_y^+1^_2_y^+1^_2_c^+2^_7_c^+1^_3_c^+2^_7_c^+1^_3_y^+2^_7_c^+2^_8_y^+2^_7_c^+2^_8_c^+1^_4_c^+2^_9_c^+2^_9_y^+1^_4_c^+2^_10_c^+2^_10_c^+2^_10_c^+1^_5_z^+2^_10_y^+2^_10_c^+1^_6_z^+1^_5_y^+1^_5_z^+1^_6_y^+1^_6_c^+1^_7_y^+1^_7_y^+1^_7_y^+1^_7_c^+1^_8_y^+1^_8_z^+1^_8_y^+1^_8_c^+1^_9_w^+1^_9_y^+1^_9_z^+1^_9_y^+1^_9_c^+1^_10_w^+1^_10_z^+1^_10_y^+1^_10_

0778155723353114

Fragment Matches Table

Show background peaks

| Position | Ion type | Intensity | mz Theoretical | mz Error (Th) | mz Error (ppm) | Charge | Series Number |
| --- | --- | --- | --- | --- | --- | --- | --- |
| - | - | 949.4 | 130.1 | - | - | 0 | - |
| - | - | 796.1 | 142.1 | - | - | 0 | - |
| - | - | 503 | 146.1 | - | - | 0 | - |
| - | - | 476.5 | 149 | - | - | 0 | - |
| - | - | 702.6 | 157.1 | - | - | 0 | - |
| - | - | 1634 | 157.1 | - | - | 0 | - |
| - | - | 1125 | 158.1 | - | - | 0 | - |
| - | - | 494.2 | 165.9 | - | - | 0 | - |
| - | - | 1165 | 166.1 | - | - | 0 | - |
| - | - | 482.7 | 168.9 | - | - | 0 | - |
| - | - | 3340 | 173.4 | - | - | 0 | - |
| - | - | 868.3 | 174.1 | - | - | 0 | - |
| - | - | 594.6 | 174.1 | - | - | 0 | - |
| - | - | 924.9 | 175.1 | - | - | 0 | - |
| - | - | 1.624E+04 | 175.1 | - | - | 0 | - |
| - | - | 1793 | 176.1 | - | - | 0 | - |
| - | - | 668.5 | 185.1 | - | - | 0 | - |
| 3 | c | 640.2 | 192.6 | 0.0003459 | 1.796 | +2 | 3 |
| - | - | 483 | 195.1 | - | - | 0 | - |
| - | - | 547.4 | 197.1 | - | - | 0 | - |
| - | - | 538.9 | 197.8 | - | - | 0 | - |
| - | - | 502.4 | 203.1 | - | - | 0 | - |
| - | - | 525.4 | 207.3 | - | - | 0 | - |
| - | - | 488.2 | 207.8 | - | - | 0 | - |
| - | - | 1.905E+04 | 209 | - | - | 0 | - |
| - | - | 461.6 | 210.3 | - | - | 0 | - |
| - | - | 3.029E+04 | 211 | - | - | 0 | - |
| - | - | 1176 | 212 | - | - | 0 | - |
| - | - | 476.3 | 212.3 | - | - | 0 | - |
| - | - | 5240 | 212.9 | - | - | 0 | - |
| - | - | 1797 | 213.2 | - | - | 0 | - |
| - | - | 2350 | 214.1 | - | - | 0 | - |
| - | - | 1314 | 214.2 | - | - | 0 | - |
| - | - | 2.482E+04 | 221.1 | - | - | 0 | - |
| - | - | 2859 | 222.1 | - | - | 0 | - |
| - | - | 525.2 | 225.3 | - | - | 0 | - |
| 4 | c | 941.4 | 228.1 | 0.0002514 | 1.102 | +2 | 4 |
| - | - | 719.5 | 228.2 | - | - | 0 | - |
| - | - | 9406 | 239.2 | - | - | 0 | - |
| - | - | 1552 | 240.2 | - | - | 0 | - |
| - | - | 1573 | 242.2 | - | - | 0 | - |
| - | - | 553.6 | 249.5 | - | - | 0 | - |
| 2 | c | 4630 | 256.2 | 0.0003837 | 1.498 | +1 | 2 |
| 2 | c | 1.886E+04 | 273.2 | 0.0002017 | 0.7384 | +1 | 2 |
| - | - | 2108 | 274.2 | - | - | 0 | - |
| - | - | 992.6 | 286.1 | - | - | 0 | - |
| - | - | 5417 | 299.2 | - | - | 0 | - |
| - | - | 2175 | 314.1 | - | - | 0 | - |
| - | - | 1693 | 318.9 | - | - | 0 | - |
| - | - | 1.742E+04 | 320.9 | - | - | 0 | - |
| - | - | 537.6 | 321.8 | - | - | 0 | - |
| - | - | 1305 | 321.9 | - | - | 0 | - |
| - | - | 1.8E+04 | 322.9 | - | - | 0 | - |
| - | - | 697.6 | 323.9 | - | - | 0 | - |
| - | - | 1871 | 324.9 | - | - | 0 | - |
| 10 | y | 3239 | 332.1 | 0.005146 | 15.49 | +1 | 2 |
| - | - | 820.6 | 333.1 | - | - | 0 | - |
| - | - | 616.1 | 337.1 | - | - | 0 | - |
| - | - | 796 | 341.2 | - | - | 0 | - |
| - | - | 591.3 | 346.2 | - | - | 0 | - |
| 10 | y | 938.4 | 350.1 | 0.005018 | 14.33 | +1 | 2 |
| - | - | 753.7 | 356.2 | - | - | 0 | - |
| - | - | 1092 | 357.2 | - | - | 0 | - |
| - | - | 549.3 | 363 | - | - | 0 | - |
| - | - | 1569 | 369.2 | - | - | 0 | - |
| - | - | 1415 | 369.7 | - | - | 0 | - |
| - | - | 3609 | 370.2 | - | - | 0 | - |
| - | - | 830.6 | 370.7 | - | - | 0 | - |
| 7 | c | 1638 | 383.2 | 0.0003514 | 0.9169 | +2 | 7 |
| - | - | 863.1 | 383.7 | - | - | 0 | - |
| 3 | c | 3117 | 384.2 | 0.0002168 | 0.5642 | +1 | 3 |
| - | - | 972.1 | 385.2 | - | - | 0 | - |
| - | - | 690 | 390.7 | - | - | 0 | - |
| - | - | 630.2 | 391.2 | - | - | 0 | - |
| - | - | 1.524E+04 | 391.2 | - | - | 0 | - |
| 7 | c | 6.157E+04 | 391.7 | 0.0001058 | 0.27 | +2 | 7 |
| - | - | 2.297E+04 | 392.2 | - | - | 0 | - |
| - | - | 4813 | 392.7 | - | - | 0 | - |
| - | - | 825.8 | 393.2 | - | - | 0 | - |
| - | - | 962.7 | 393.7 | - | - | 0 | - |
| 3 | c | 5.825E+04 | 401.3 | 0.0003401 | 0.8475 | +1 | 3 |
| - | - | 1.12E+04 | 402.3 | - | - | 0 | - |
| - | - | 1597 | 402.7 | - | - | 0 | - |
| - | - | 1085 | 403.3 | - | - | 0 | - |
| - | - | 1648 | 404.8 | - | - | 0 | - |
| - | - | 701 | 405.3 | - | - | 0 | - |
| 5 | y | 1888 | 413.7 | 0.002624 | 6.342 | +2 | 7 |
| - | - | 542.3 | 414.2 | - | - | 0 | - |
| - | - | 702.9 | 415 | - | - | 0 | - |
| - | - | 1426 | 415.3 | - | - | 0 | - |
| - | - | 1156 | 416 | - | - | 0 | - |
| - | - | 1351 | 417 | - | - | 0 | - |
| 8 | c | 1989 | 418.7 | 0.0004713 | 1.125 | +2 | 8 |
| - | - | 799.7 | 419.2 | - | - | 0 | - |
| 5 | y | 811.3 | 422.7 | 0.004086 | 9.665 | +2 | 7 |
| - | - | 667 | 424.7 | - | - | 0 | - |
| - | - | 9271 | 426.8 | - | - | 0 | - |
| 8 | c | 4.58E+04 | 427.3 | 0.0002582 | 0.6044 | +2 | 8 |
| - | - | 624.6 | 427.6 | - | - | 0 | - |
| - | - | 2.146E+04 | 427.8 | - | - | 0 | - |
| - | - | 657.6 | 427.9 | - | - | 0 | - |
| - | - | 4066 | 428.3 | - | - | 0 | - |
| - | - | 712.5 | 428.3 | - | - | 0 | - |
| - | - | 839.5 | 428.8 | - | - | 0 | - |
| - | - | 977.4 | 432.1 | - | - | 0 | - |
| - | - | 1304 | 432.2 | - | - | 0 | - |
| - | - | 4252 | 432.9 | - | - | 0 | - |
| - | - | 607.3 | 433.9 | - | - | 0 | - |
| - | - | 1770 | 434.9 | - | - | 0 | - |
| - | - | 1573 | 440.3 | - | - | 0 | - |
| - | - | 859.5 | 440.8 | - | - | 0 | - |
| 4 | c | 4009 | 455.3 | 9.28E-05 | 0.2038 | +1 | 4 |
| - | - | 1438 | 456.3 | - | - | 0 | - |
| - | - | 6371 | 461.3 | - | - | 0 | - |
| - | - | 1.253E+04 | 461.8 | - | - | 0 | - |
| - | - | 5057 | 462.3 | - | - | 0 | - |
| - | - | 1810 | 462.8 | - | - | 0 | - |
| 9 | c | 1.457E+04 | 475.3 | 0.0006756 | 1.421 | +2 | 9 |
| - | - | 6261 | 475.8 | - | - | 0 | - |
| - | - | 2623 | 476.3 | - | - | 0 | - |
| - | - | 2403 | 483.3 | - | - | 0 | - |
| 9 | c | 2.65E+04 | 483.8 | 0.0004015 | 0.83 | +2 | 9 |
| - | - | 1.295E+04 | 484.3 | - | - | 0 | - |
| - | - | 3588 | 484.8 | - | - | 0 | - |
| - | - | 4792 | 496.8 | - | - | 0 | - |
| - | - | 2042 | 497.3 | - | - | 0 | - |
| - | - | 1179 | 497.8 | - | - | 0 | - |
| - | - | 767.9 | 518.3 | - | - | 0 | - |
| - | - | 885.5 | 518.8 | - | - | 0 | - |
| - | - | 4007 | 519.8 | - | - | 0 | - |
| - | - | 1820 | 520.3 | - | - | 0 | - |
| - | - | 1042 | 524.3 | - | - | 0 | - |
| - | - | 1736 | 525.3 | - | - | 0 | - |
| - | - | 1.252E+04 | 525.8 | - | - | 0 | - |
| - | - | 6392 | 526.3 | - | - | 0 | - |
| - | - | 4257 | 526.8 | - | - | 0 | - |
| - | - | 1481 | 527.3 | - | - | 0 | - |
| - | - | 1.844E+04 | 529.3 | - | - | 0 | - |
| - | - | 4917 | 530.3 | - | - | 0 | - |
| - | - | 1441 | 530.8 | - | - | 0 | - |
| - | - | 1627 | 531.3 | - | - | 0 | - |
| - | - | 901.2 | 531.8 | - | - | 0 | - |
| - | - | 692.5 | 532.3 | - | - | 0 | - |
| - | - | 1414 | 533.3 | - | - | 0 | - |
| - | - | 1118 | 533.8 | - | - | 0 | - |
| 8 | y | 626.5 | 534.3 | 0.007254 | 13.58 | +1 | 4 |
| 10 | c | 1428 | 539.3 | 0.002002 | 3.712 | +2 | 10 |
| 10 | c | 6.646E+04 | 539.8 | 0.0003751 | 0.695 | +2 | 10 |
| - | - | 3.332E+04 | 540.3 | - | - | 0 | - |
| - | - | 1.336E+04 | 540.8 | - | - | 0 | - |
| - | - | 5523 | 541.3 | - | - | 0 | - |
| - | - | 1063 | 541.8 | - | - | 0 | - |
| - | - | 2521 | 542.3 | - | - | 0 | - |
| - | - | 765.1 | 547.8 | - | - | 0 | - |
| 10 | c | 8.801E+04 | 548.3 | 0.0004063 | 0.7409 | +2 | 10 |
| - | - | 5.126E+04 | 548.8 | - | - | 0 | - |
| - | - | 1.665E+04 | 549.3 | - | - | 0 | - |
| - | - | 622.3 | 549.6 | - | - | 0 | - |
| - | - | 3711 | 549.8 | - | - | 0 | - |
| - | - | 675.8 | 550.3 | - | - | 0 | - |
| - | - | 1279 | 553.3 | - | - | 0 | - |
| - | - | 2414 | 553.8 | - | - | 0 | - |
| - | - | 1140 | 554.3 | - | - | 0 | - |
| - | - | 929.1 | 554.8 | - | - | 0 | - |
| - | - | 3657 | 555.3 | - | - | 0 | - |
| - | - | 2775 | 555.8 | - | - | 0 | - |
| - | - | 741.5 | 556.3 | - | - | 0 | - |
| - | - | 2245 | 556.8 | - | - | 0 | - |
| - | - | 1284 | 557.3 | - | - | 0 | - |
| - | - | 3014 | 561.3 | - | - | 0 | - |
| - | - | 854.4 | 561.8 | - | - | 0 | - |
| - | - | 1532 | 562.3 | - | - | 0 | - |
| - | - | 6314 | 563.3 | - | - | 0 | - |
| - | - | 3755 | 563.8 | - | - | 0 | - |
| - | - | 1736 | 564.3 | - | - | 0 | - |
| - | - | 4586 | 567.8 | - | - | 0 | - |
| - | - | 2628 | 568.3 | - | - | 0 | - |
| - | - | 1.755E+04 | 568.8 | - | - | 0 | - |
| - | - | 3.702E+04 | 569.3 | - | - | 0 | - |
| 5 | c | 1.383E+04 | 569.4 | 0.003358 | 5.898 | +1 | 5 |
| - | - | 2.705E+04 | 569.8 | - | - | 0 | - |
| - | - | 8501 | 570.3 | - | - | 0 | - |
| - | - | 2937 | 570.4 | - | - | 0 | - |
| - | - | 947.3 | 570.8 | - | - | 0 | - |
| - | - | 1526 | 570.8 | - | - | 0 | - |
| - | - | 1186 | 571.4 | - | - | 0 | - |
| - | - | 3467 | 575.8 | - | - | 0 | - |
| - | - | 2763 | 576.3 | - | - | 0 | - |
| - | - | 7724 | 576.8 | - | - | 0 | - |
| - | - | 4420 | 577.3 | - | - | 0 | - |
| - | - | 1.978E+04 | 577.8 | - | - | 0 | - |
| - | - | 1.101E+04 | 578.3 | - | - | 0 | - |
| - | - | 668.9 | 578.6 | - | - | 0 | - |
| - | - | 4967 | 578.8 | - | - | 0 | - |
| - | - | 2376 | 582.4 | - | - | 0 | - |
| - | - | 4693 | 583.3 | - | - | 0 | - |
| - | - | 3059 | 583.8 | - | - | 0 | - |
| - | - | 2814 | 584.3 | - | - | 0 | - |
| - | - | 871.9 | 585.3 | - | - | 0 | - |
| - | - | 1.09E+04 | 586.3 | - | - | 0 | - |
| - | - | 2954 | 587.3 | - | - | 0 | - |
| - | - | 4277 | 589.8 | - | - | 0 | - |
| - | - | 2262 | 590.3 | - | - | 0 | - |
| - | - | 741 | 590.8 | - | - | 0 | - |
| - | - | 746.9 | 591.3 | - | - | 0 | - |
| 2 | z | 4773 | 592.3 | 0.002968 | 5.01 | +2 | 10 |
| - | - | 3441 | 592.8 | - | - | 0 | - |
| - | - | 1807 | 593.3 | - | - | 0 | - |
| - | - | 911.2 | 597.8 | - | - | 0 | - |
| - | - | 1225 | 598.8 | - | - | 0 | - |
| - | - | 1522 | 599.3 | - | - | 0 | - |
| 2 | y | 1341 | 600.3 | 0.002151 | 3.582 | +2 | 10 |
| - | - | 1397 | 606.3 | - | - | 0 | - |
| - | - | 865.3 | 606.8 | - | - | 0 | - |
| - | - | 712.7 | 611.3 | - | - | 0 | - |
| - | - | 1023 | 612.3 | - | - | 0 | - |
| - | - | 4899 | 612.8 | - | - | 0 | - |
| - | - | 3414 | 613.3 | - | - | 0 | - |
| - | - | 2090 | 613.8 | - | - | 0 | - |
| - | - | 724.1 | 614.3 | - | - | 0 | - |
| - | - | 1047 | 618.3 | - | - | 0 | - |
| - | - | 2.216E+04 | 619.4 | - | - | 0 | - |
| - | - | 1.364E+04 | 619.9 | - | - | 0 | - |
| - | - | 1456 | 620.3 | - | - | 0 | - |
| - | - | 5323 | 620.4 | - | - | 0 | - |
| - | - | 2096 | 620.8 | - | - | 0 | - |
| - | - | 1545 | 620.9 | - | - | 0 | - |
| - | - | 1145 | 622.3 | - | - | 0 | - |
| - | - | 996.3 | 624.3 | - | - | 0 | - |
| - | - | 2979 | 625.4 | - | - | 0 | - |
| 6 | c | 7.688E+04 | 626.4 | 0.0002658 | 0.4243 | +1 | 6 |
| - | - | 2.46E+04 | 627.4 | - | - | 0 | - |
| - | - | 4090 | 627.8 | - | - | 0 | - |
| - | - | 2664 | 628.3 | - | - | 0 | - |
| - | - | 2861 | 628.4 | - | - | 0 | - |
| - | - | 1.778E+04 | 628.8 | - | - | 0 | - |
| - | - | 1.038E+04 | 629.3 | - | - | 0 | - |
| - | - | 4694 | 629.8 | - | - | 0 | - |
| - | - | 756.4 | 630.3 | - | - | 0 | - |
| - | - | 2628 | 631.3 | - | - | 0 | - |
| - | - | 1993 | 633.3 | - | - | 0 | - |
| - | - | 2921 | 633.8 | - | - | 0 | - |
| - | - | 8974 | 634.3 | - | - | 0 | - |
| - | - | 6076 | 634.8 | - | - | 0 | - |
| - | - | 2226 | 635.3 | - | - | 0 | - |
| - | - | 694.6 | 635.8 | - | - | 0 | - |
| - | - | 3011 | 640.8 | - | - | 0 | - |
| - | - | 2733 | 641.3 | - | - | 0 | - |
| - | - | 8.504E+04 | 641.8 | - | - | 0 | - |
| - | - | 6.292E+04 | 642.3 | - | - | 0 | - |
| - | - | 2.365E+04 | 642.8 | - | - | 0 | - |
| - | - | 6827 | 643.3 | - | - | 0 | - |
| - | - | 933.1 | 648.8 | - | - | 0 | - |
| - | - | 2617 | 649.3 | - | - | 0 | - |
| - | - | 4.706E+04 | 649.9 | - | - | 0 | - |
| - | - | 8.142E+04 | 650.4 | - | - | 0 | - |
| - | - | 3.917E+04 | 650.9 | - | - | 0 | - |
| - | - | 1.328E+04 | 651.4 | - | - | 0 | - |
| - | - | 4041 | 651.9 | - | - | 0 | - |
| - | - | 992 | 672.3 | - | - | 0 | - |
| 7 | z | 3.986E+04 | 674.3 | 0.005006 | 7.424 | +1 | 5 |
| - | - | 1.415E+04 | 675.3 | - | - | 0 | - |
| - | - | 3764 | 676.3 | - | - | 0 | - |
| - | - | 840.4 | 677.3 | - | - | 0 | - |
| 7 | y | 4136 | 690.4 | 0.005264 | 7.625 | +1 | 5 |
| - | - | 1392 | 691.4 | - | - | 0 | - |
| - | - | 2110 | 700.4 | - | - | 0 | - |
| - | - | 954.2 | 704.3 | - | - | 0 | - |
| - | - | 984.9 | 718.4 | - | - | 0 | - |
| - | - | 887.6 | 730.4 | - | - | 0 | - |
| 6 | z | 8627 | 731.4 | 0.004843 | 6.623 | +1 | 6 |
| - | - | 3591 | 732.4 | - | - | 0 | - |
| - | - | 613.7 | 740.4 | - | - | 0 | - |
| 6 | y | 3124 | 747.4 | 0.0062 | 8.296 | +1 | 6 |
| - | - | 1869 | 748.4 | - | - | 0 | - |
| - | - | 4759 | 754.4 | - | - | 0 | - |
| - | - | 1731 | 755.4 | - | - | 0 | - |
| - | - | 781.3 | 756.4 | - | - | 0 | - |
| - | - | 621.6 | 763.9 | - | - | 0 | - |
| - | - | 1296 | 781.5 | - | - | 0 | - |
| 7 | c | 1.22E+04 | 782.5 | 0.0005645 | 0.7214 | +1 | 7 |
| - | - | 5050 | 783.5 | - | - | 0 | - |
| - | - | 3680 | 824.4 | - | - | 0 | - |
| - | - | 2081 | 825.5 | - | - | 0 | - |
| 5 | y | 1797 | 826.4 | 0.007824 | 9.467 | +1 | 7 |
| 5 | y | 1167 | 827.4 | 0.01655 | 20 | +1 | 7 |
| - | - | 1097 | 842.4 | - | - | 0 | - |
| 5 | y | 5878 | 844.4 | 0.003607 | 4.272 | +1 | 7 |
| - | - | 1817 | 845.4 | - | - | 0 | - |
| - | - | 960.3 | 846.4 | - | - | 0 | - |
| - | - | 874.1 | 848.4 | - | - | 0 | - |
| - | - | 857.4 | 852.5 | - | - | 0 | - |
| 8 | c | 9982 | 853.5 | 0.001362 | 1.596 | +1 | 8 |
| - | - | 3392 | 854.5 | - | - | 0 | - |
| - | - | 1117 | 855.5 | - | - | 0 | - |
| - | - | 854.4 | 864.4 | - | - | 0 | - |
| - | - | 690.5 | 872.4 | - | - | 0 | - |
| - | - | 1432 | 882.5 | - | - | 0 | - |
| - | - | 815.7 | 895.5 | - | - | 0 | - |
| - | - | 1450 | 897.4 | - | - | 0 | - |
| 4 | y | 1625 | 898.4 | 0.0007931 | 0.8827 | +1 | 8 |
| 4 | z | 4.996E+04 | 899.4 | 0.004077 | 4.533 | +1 | 8 |
| - | - | 2.729E+04 | 900.5 | - | - | 0 | - |
| - | - | 836.3 | 900.6 | - | - | 0 | - |
| - | - | 7061 | 901.5 | - | - | 0 | - |
| - | - | 2278 | 902.5 | - | - | 0 | - |
| 4 | y | 6733 | 915.5 | 0.003908 | 4.269 | +1 | 8 |
| - | - | 3705 | 916.5 | - | - | 0 | - |
| - | - | 849.6 | 951.6 | - | - | 0 | - |
| - | - | 833.1 | 966.5 | - | - | 0 | - |
| 9 | c | 9461 | 966.6 | 0.0005264 | 0.5446 | +1 | 9 |
| - | - | 6963 | 967.6 | - | - | 0 | - |
| - | - | 1139 | 968.6 | - | - | 0 | - |
| 3 | w | 701.1 | 969.5 | 0.005192 | 5.356 | +1 | 9 |
| - | - | 2009 | 980.5 | - | - | 0 | - |
| - | - | 1054 | 981.5 | - | - | 0 | - |
| 3 | y | 782.5 | 1026 | 0.0002886 | 0.2811 | +1 | 9 |
| 3 | z | 3.935E+04 | 1028 | 0.004704 | 4.578 | +1 | 9 |
| - | - | 2.062E+04 | 1029 | - | - | 0 | - |
| - | - | 7608 | 1030 | - | - | 0 | - |
| - | - | 1153 | 1031 | - | - | 0 | - |
| - | - | 1709 | 1038 | - | - | 0 | - |
| - | - | 804.4 | 1039 | - | - | 0 | - |
| 3 | y | 4931 | 1044 | 0.004412 | 4.228 | +1 | 9 |
| - | - | 2657 | 1045 | - | - | 0 | - |
| - | - | 969.8 | 1046 | - | - | 0 | - |
| - | - | 1574 | 1052 | - | - | 0 | - |
| - | - | 5318 | 1053 | - | - | 0 | - |
| - | - | 3394 | 1054 | - | - | 0 | - |
| - | - | 1159 | 1055 | - | - | 0 | - |
| - | - | 7665 | 1080 | - | - | 0 | - |
| - | - | 5329 | 1081 | - | - | 0 | - |
| - | - | 3271 | 1082 | - | - | 0 | - |
| - | - | 1373 | 1083 | - | - | 0 | - |
| 10 | c | 9502 | 1096 | 0.006804 | 6.21 | +1 | 10 |
| - | - | 2.219E+04 | 1097 | - | - | 0 | - |
| 2 | w | 1564 | 1098 | 0.0004394 | 0.4004 | +1 | 10 |
| - | - | 1.008E+04 | 1098 | - | - | 0 | - |
| - | - | 1215 | 1099 | - | - | 0 | - |
| - | - | 3110 | 1099 | - | - | 0 | - |
| - | - | 939.4 | 1123 | - | - | 0 | - |
| - | - | 1385 | 1136 | - | - | 0 | - |
| - | - | 858.9 | 1137 | - | - | 0 | - |
| - | - | 809.9 | 1138 | - | - | 0 | - |
| - | - | 1064 | 1139 | - | - | 0 | - |
| - | - | 1182 | 1152 | - | - | 0 | - |
| - | - | 935.3 | 1168 | - | - | 0 | - |
| - | - | 1160 | 1169 | - | - | 0 | - |
| - | - | 1015 | 1170 | - | - | 0 | - |
| 2 | z | 2260 | 1184 | 0.002226 | 1.88 | +1 | 10 |
| - | - | 2.083E+04 | 1185 | - | - | 0 | - |
| - | - | 1.133E+04 | 1186 | - | - | 0 | - |
| - | - | 4235 | 1187 | - | - | 0 | - |
| - | - | 1860 | 1188 | - | - | 0 | - |
| - | - | 2295 | 1197 | - | - | 0 | - |
| - | - | 1535 | 1198 | - | - | 0 | - |
| - | - | 3481 | 1199 | - | - | 0 | - |
| 2 | y | 2399 | 1200 | 0.008198 | 6.834 | +1 | 10 |
| - | - | 1295 | 1201 | - | - | 0 | - |
| - | - | 2048 | 1212 | - | - | 0 | - |
| - | - | 1719 | 1213 | - | - | 0 | - |
| - | - | 929.3 | 1222 | - | - | 0 | - |
| - | - | 1253 | 1225 | - | - | 0 | - |
| - | - | 2466 | 1238 | - | - | 0 | - |
| - | - | 2.244E+04 | 1239 | - | - | 0 | - |
| - | - | 1.264E+04 | 1240 | - | - | 0 | - |
| - | - | 2.341E+04 | 1241 | - | - | 0 | - |
| - | - | 2.201E+04 | 1242 | - | - | 0 | - |
| - | - | 1.005E+04 | 1243 | - | - | 0 | - |
| - | - | 3549 | 1244 | - | - | 0 | - |
| - | - | 1508 | 1255 | - | - | 0 | - |
| - | - | 7537 | 1256 | - | - | 0 | - |
| - | - | 7324 | 1257 | - | - | 0 | - |
| - | - | 7105 | 1258 | - | - | 0 | - |
| - | - | 2528 | 1259 | - | - | 0 | - |
| - | - | 1089 | 1260 | - | - | 0 | - |
| - | - | 2725 | 1266 | - | - | 0 | - |
| - | - | 7699 | 1267 | - | - | 0 | - |
| - | - | 4978 | 1268 | - | - | 0 | - |
| - | - | 2641 | 1269 | - | - | 0 | - |
| - | - | 744.4 | 1271 | - | - | 0 | - |
| - | - | 3421 | 1273 | - | - | 0 | - |
| - | - | 1130 | 1274 | - | - | 0 | - |
| - | - | 797.6 | 1275 | - | - | 0 | - |
| - | - | 3337 | 1282 | - | - | 0 | - |
| - | - | 3.054E+04 | 1283 | - | - | 0 | - |
| - | - | 1.116E+05 | 1284 | - | - | 0 | - |
| - | - | 6.768E+04 | 1285 | - | - | 0 | - |
| - | - | 2.806E+04 | 1286 | - | - | 0 | - |
| - | - | 6983 | 1287 | - | - | 0 | - |
| - | - | 1819 | 1288 | - | - | 0 | - |
| - | - | 5420 | 1299 | - | - | 0 | - |
| - | - | 2.229E+04 | 1300 | - | - | 0 | - |
| - | - | 9.432E+04 | 1301 | - | - | 0 | - |
| - | - | 6.358E+04 | 1302 | - | - | 0 | - |
| - | - | 2.279E+04 | 1303 | - | - | 0 | - |
| - | - | 6293 | 1304 | - | - | 0 | - |
| - | - | 1182 | 1305 | - | - | 0 | - |
| - | - | 974.8 | 3083 | - | - | 0 | - |

m/z Charge Intensity FragmentType MassShift Position
130.06533813476562 0 949.44336
142.09774780273438 0 796.1132
146.06036376953125 0 503.04968
149.02442932128906 0 476.4677
157.07589721679688 0 702.56744
157.108642578125 0 1633.8718
158.06024169921875 0 1124.6456
165.93600463867188 0 494.2373
166.08631896972656 0 1164.596
168.9337158203125 0 482.71945
173.44046020507812 0 3339.8958
174.05516052246094 0 868.30927
174.13504028320312 0 594.6368
175.07867431640625 0 924.8586
175.08682250976562 0 16242.466
176.09005737304688 0 1793.4918
185.10379028320312 0 668.51013
192.62167358398438 0 640.19916 c Ammonia loss 2
195.06509399414062 0 482.9972
197.1297149658203 0 547.35156
197.7725372314453 0 538.9056
203.08184814453125 0 502.37692
207.28659057617188 0 525.3827
207.78065490722656 0 488.154
208.95335388183594 0 19048.127
210.33212280273438 0 461.62814
210.95046997070312 0 30287.572
211.9537811279297 0 1175.9939
212.2747039794922 0 476.25394
212.94744873046875 0 5239.7617
213.17124938964844 0 1797.2089
214.14266967773438 0 2350.2522
214.1552276611328 0 1314.3629
221.09231567382812 0 24816.648
222.09573364257812 0 2858.9292
225.28627014160156 0 525.17145
228.13963317871094 0 941.40466 c Ammonia loss 3
228.1822052001953 0 719.48596
239.1504669189453 0 9406.116
240.15365600585938 0 1551.554
242.1614532470703 0 1572.5364
249.53079223632812 0 553.5712
256.17718505859375 0 4629.607 c Ammonia loss 1
273.20355224609375 0 18863.846 c 1
274.2060546875 0 2107.6873
286.1181640625 0 992.5876
299.2195739746094 0 5417.184
314.1141052246094 0 2174.8152
318.9228210449219 0 1692.8499
320.9195251464844 0 17420.955
321.794921875 0 537.5945
321.9224853515625 0 1304.9033
322.9168701171875 0 17997.113
323.9208984375 0 697.6005
324.913330078125 0 1870.7935
332.12432861328125 0 3238.559 y Water loss 9
333.1269836425781 0 820.5713
337.05072021484375 0 616.09515
341.228515625 0 796.0404
346.2438659667969 0 591.258
350.134765625 0 938.4029 y 9
356.2411804199219 0 753.66504
357.2484436035156 0 1092.3785
363.02398681640625 0 549.2514
369.2305603027344 0 1568.5426
369.734130859375 0 1415.3965
370.2372741699219 0 3608.6785
370.7387390136719 0 830.60645
383.2279052734375 0 1637.518 c Ammonia loss 6
383.7283935546875 0 863.1281
384.235595703125 0 3116.5457 c Ammonia loss 2
385.23773193359375 0 972.11646
390.7099304199219 0 690.0294
391.208251953125 0 630.2274
391.2372131347656 0 15240.195
391.74072265625 0 61570.297 c 6
392.24237060546875 0 22973.467
392.74310302734375 0 4813.415
393.2448425292969 0 825.83093
393.6900329589844 0 962.6519
401.26226806640625 0 58245.79 c 2
402.264892578125 0 11196.727
402.6951904296875 0 1597.058
403.26904296875 0 1085.4735
404.75067138671875 0 1648.1831
405.2523498535156 0 701.0084
413.714111328125 0 1888.1118 y Water loss 4
414.2153625488281 0 542.316
415.03887939453125 0 702.8793
415.25384521484375 0 1425.8014
416.03912353515625 0 1156.2784
417.03472900390625 0 1350.976
418.74658203125 0 1989.41 c Ammonia loss 7
419.24603271484375 0 799.73755
422.7208557128906 0 811.3237 y 4
424.7084655761719 0 666.9662
426.7559814453125 0 9271.048
427.2596435546875 0 45801.3 c 7
427.5692443847656 0 624.6434
427.7611083984375 0 21459.719
427.902099609375 0 657.56244
428.2627868652344 0 4065.8884
428.2918395996094 0 712.5301
428.7611389160156 0 839.4885
432.08685302734375 0 977.39624
432.2038879394531 0 1303.5717
432.8866882324219 0 4252.0996
433.8879699707031 0 607.2914
434.88128662109375 0 1770.1725
440.26715087890625 0 1573.191
440.7694091796875 0 859.4664
455.27239990234375 0 4009.4275 c Ammonia loss 3
456.2762145996094 0 1437.6398
461.2906188964844 0 6371.0195
461.7945251464844 0 12528.172
462.2962951660156 0 5056.819
462.7979736328125 0 1809.8438
475.288818359375 0 14567.665 c Ammonia loss 8
475.7899475097656 0 6260.7856
476.2916564941406 0 2623.139
483.2980651855469 0 2403.4626
483.80181884765625 0 26500.838 c 8
484.3033752441406 0 12954.32
484.80487060546875 0 3588.383
496.8095397949219 0 4791.8047
497.3091735839844 0 2042.0435
497.8129577636719 0 1178.5507
518.3115844726562 0 767.93427
518.8078002929688 0 885.54663
519.7818603515625 0 4007.208
520.2831420898438 0 1819.5242
524.3310546875 0 1041.5747
525.3380737304688 0 1736.2001
525.8121948242188 0 12517.081
526.3147583007812 0 6391.7046
526.8170776367188 0 4256.9727
527.3211669921875 0 1481.3584
529.28564453125 0 18442.963
530.2886962890625 0 4917.2554
530.8046875 0 1440.8518
531.2974853515625 0 1627.3195
531.8034057617188 0 901.2105
532.299560546875 0 692.5151
533.3064575195312 0 1414.4513
533.8094482421875 0 1118.1738
534.2581787109375 0 626.5234 y 7
539.3154296875 0 1428.0457 c Water loss 9
539.809814453125 0 66455.74 c Ammonia loss 9
540.3111572265625 0 33324.133
540.8131103515625 0 13361.491
541.3156127929688 0 5522.7183
541.81982421875 0 1063.4807
542.2928466796875 0 2521.2175
547.8140869140625 0 765.12585
548.3231201171875 0 88012.11 c 9
548.8241577148438 0 51255.98
549.3252563476562 0 16646.055
549.594482421875 0 622.252
549.8267211914062 0 3710.799
550.3269653320312 0 675.78046
553.3300170898438 0 1278.7378
553.8275146484375 0 2414.2039
554.3279418945312 0 1139.8602
554.82861328125 0 929.0818
555.3179931640625 0 3657.3147
555.8165283203125 0 2775.388
556.318359375 0 741.5231
556.7904052734375 0 2244.8682
557.2924194335938 0 1283.6161
561.3324584960938 0 3014.0408
561.8350219726562 0 854.35284
562.3282470703125 0 1531.5714
563.2940063476562 0 6314.432
563.794677734375 0 3755.1792
564.2974853515625 0 1736.4266
567.8168334960938 0 4585.832
568.3184204101562 0 2628.0376
568.8235473632812 0 17546.605
569.3171997070312 0 37024.117
569.3551635742188 0 13827.716 c 4
569.8185424804688 0 27048.38
570.3189086914062 0 8501.063
570.3557739257812 0 2937.4624
570.7777099609375 0 947.2992
570.822021484375 0 1525.6615
571.3592529296875 0 1186.1743
575.8203735351562 0 3466.6333
576.3218994140625 0 2762.875
576.8221435546875 0 7723.812
577.3225708007812 0 4419.9272
577.8291015625 0 19776.133
578.33056640625 0 11014.922
578.5523071289062 0 668.85315
578.8323364257812 0 4967.299
582.3600463867188 0 2375.759
583.3448486328125 0 4692.7446
583.8458251953125 0 3058.6467
584.3394165039062 0 2813.5852
585.3016967773438 0 871.9318
586.30712890625 0 10896.912
587.3099975585938 0 2953.7039
589.8499145507812 0 4276.8975
590.3485107421875 0 2262.4868
590.8474731445312 0 741.0175
591.3255615234375 0 746.8656
592.3087768554688 0 4772.5195 z 1
592.8095092773438 0 3440.9167
593.3125610351562 0 1807.1957
597.8291625976562 0 911.1508
598.8314208984375 0 1225.2871
599.3150634765625 0 1522.3099
600.3173217773438 0 1341.4177 y 1
606.3226318359375 0 1396.9095
606.8155517578125 0 865.3448
611.3493041992188 0 712.6778
612.3316650390625 0 1022.822
612.8290405273438 0 4898.6313
613.3306884765625 0 3413.85
613.8214111328125 0 2090.1235
614.3475341796875 0 724.0755
618.2772216796875 0 1046.8086
619.355712890625 0 22158.7
619.8568115234375 0 13644.798
620.3140258789062 0 1456.2656
620.3583374023438 0 5322.9487
620.81591796875 0 2096.4902
620.8609008789062 0 1544.5537
622.3242797851562 0 1145.2772
624.3484497070312 0 996.3378
625.3641967773438 0 2979.4236
626.37353515625 0 76882.15 c 5
627.3760986328125 0 24597.418
627.8445434570312 0 4089.9268
628.3385009765625 0 2663.9814
628.3812866210938 0 2861.055
628.84130859375 0 17778.406
629.34228515625 0 10382.738
629.8433227539062 0 4694.231
630.3440551757812 0 756.42584
631.2847290039062 0 2628.269
633.3302001953125 0 1993.2084
633.8328857421875 0 2921.3337
634.3314819335938 0 8973.925
634.8320922851562 0 6075.738
635.33349609375 0 2226.119
635.8441162109375 0 694.5806
640.8363037109375 0 3011.4558
641.345458984375 0 2733.358
641.8425903320312 0 85037.664
642.3441162109375 0 62922.367
642.8455200195312 0 23653.447
643.3470458984375 0 6826.7593
648.837646484375 0 933.1058
649.3486938476562 0 2617.1887
649.8519287109375 0 47063.24
650.354736328125 0 81424.04
650.8563232421875 0 39170.406
651.3575439453125 0 13277.365
651.8571166992188 0 4040.775
672.3256225585938 0 992.0185
674.3383178710938 0 39859.035 z 6
675.3412475585938 0 14147.422
676.3433227539062 0 3764.147
677.3432006835938 0 840.43536
690.3572998046875 0 4136.1865 y 6
691.358642578125 0 1391.6943
700.3613891601562 0 2110.194
704.2942504882812 0 954.2297
718.41015625 0 984.8766
730.3707885742188 0 887.58405
731.359619140625 0 8626.881 z 5
732.363525390625 0 3591.0784
740.4464721679688 0 613.71576
747.3796997070312 0 3123.614 y 5
748.3819580078125 0 1868.7715
754.3973999023438 0 4758.8076
755.3974609375 0 1731.2151
756.40283203125 0 781.2954
763.8765869140625 0 621.63104
781.4649658203125 0 1296.3737
782.4738159179688 0 12203.802 c 6
783.475830078125 0 5049.913
824.4496459960938 0 3679.5664
825.4537963867188 0 2080.8557
826.4235229492188 0 1797.1763 y Water loss 4
827.416259765625 0 1166.7643 y Ammonia loss 4
842.4156494140625 0 1097.4749
844.4298706054688 0 5878.3354 y 4
845.4331665039062 0 1817.1353
846.4370727539062 0 960.2947
848.4089965820312 0 874.0741
852.5077514648438 0 857.4224
853.5101318359375 0 9981.958 c 7
854.513671875 0 3391.73
855.51904296875 0 1117.302
864.41845703125 0 854.4372
872.4234619140625 0 690.50494
882.4579467773438 0 1431.938
895.4647827148438 0 815.666
897.4335327148438 0 1450.2877
898.43603515625 0 1625.1074 y Ammonia loss 3
899.44873046875 0 49961.996 z 3
900.4520874023438 0 27288.637
900.5548706054688 0 836.3109
901.4556274414062 0 7061.2075
902.4570922851562 0 2277.619
915.46728515625 0 6733.3643 y 3
916.469970703125 0 3704.9739
951.5765380859375 0 849.573
966.4969482421875 0 833.07117
966.5950317382812 0 9461.192 c 8
967.5977172851562 0 6963.4565
968.6041259765625 0 1139.2538
969.46875 0 701.112 w 2
980.5488891601562 0 2009.4775
981.5468139648438 0 1053.7877
1026.4951171875 0 782.5182 y Ammonia loss 2
1027.5079345703125 0 39350.668 z 2
1028.51025390625 0 20624.076
1029.513427734375 0 7607.7065
1030.51123046875 0 1152.7151
1037.5994873046875 0 1708.6057
1038.60986328125 0 804.3705
1043.5263671875 0 4931.2144 y 2
1044.5308837890625 0 2657.477
1045.5286865234375 0 969.76807
1051.6187744140625 0 1573.7435
1052.62744140625 0 5318.1914
1053.6259765625 0 3394.1511
1054.6298828125 0 1158.9705
1079.61865234375 0 7665.06
1080.6231689453125 0 5329.007
1081.6278076171875 0 3270.8586
1082.6297607421875 0 1372.8033
1095.63134765625 0 9501.927 c 9
1096.6429443359375 0 22189.207
1097.532958984375 0 1564.2473 w 1
1097.6461181640625 0 10080.276
1098.5362548828125 0 1214.9631
1098.6553955078125 0 3109.7644
1122.6343994140625 0 939.43134
1135.63427734375 0 1385.0106
1136.6392822265625 0 858.94745
1137.6446533203125 0 809.94653
1138.6395263671875 0 1064.1145
1151.632568359375 0 1181.8931
1167.653076171875 0 935.2892
1168.6766357421875 0 1159.9529
1169.6314697265625 0 1014.89575
1183.6065673828125 0 2260.2668 z 1
1184.6151123046875 0 20828.256
1185.61767578125 0 11330.452
1186.6185302734375 0 4235.442
1187.625 0 1860.0413
1196.6064453125 0 2295.3916
1197.612060546875 0 1534.6508
1198.6077880859375 0 3480.5005
1199.6148681640625 0 2399.2476 y 1
1200.6201171875 0 1294.7417
1211.6181640625 0 2047.8229
1212.622802734375 0 1718.7549
1221.689208984375 0 929.2616
1224.6441650390625 0 1252.644
1237.696533203125 0 2465.5586
1238.708740234375 0 22444.205
1239.7099609375 0 12638.988
1240.6585693359375 0 23411.945
1241.6600341796875 0 22007.469
1242.663330078125 0 10054.837
1243.664306640625 0 3548.5435
1254.685302734375 0 1508.4385
1255.6871337890625 0 7536.534
1256.685791015625 0 7323.9917
1257.6827392578125 0 7104.9165
1258.6822509765625 0 2528.343
1259.6873779296875 0 1088.5154
1265.66943359375 0 2724.5159
1266.6597900390625 0 7699.106
1267.660400390625 0 4977.924
1268.6617431640625 0 2640.854
1270.6878662109375 0 744.42474
1272.7099609375 0 3421.4705
1273.71240234375 0 1129.6903
1274.7139892578125 0 797.55237
1281.6783447265625 0 3336.6274
1282.6922607421875 0 30541.271
1283.685302734375 0 111648.83
1284.687744140625 0 67679.08
1285.68896484375 0 28060.922
1286.690673828125 0 6982.9756
1287.689453125 0 1819.3978
1298.69677734375 0 5419.741
1299.701171875 0 22288.535
1300.708984375 0 94320.3
1301.7120361328125 0 63584.21
1302.7138671875 0 22792.992
1303.712890625 0 6292.9126
1304.7196044921875 0 1181.6509
3083.029541015625 0 974.77234

Spectrum Details

|  |  |
| --- | --- |
| Matched peaks? Matched peaks The total absolute number of peaks matched. Additionally in brackets the total fraction of peaks matched and the total number of peaks is shown. | 46 (11.53% of 399) |
| FDR? FDR The false discovery rate estimated for this peptide. It is calculated by matching all theoretical fragments with a non-integer shift with the raw peaks for this spectrum. This is done with 40 different shifts. The resulting percentage is the average number of annotated peaks over the number of annotated peaks with the correct spectrum. | 2.17% |
| Satellite FDR? Satellite FDR See the FDR for details on its calculation. This satellite ion specific FDR only contains the satellite ions (d/w) for I/L/J positions. | ∞ |
| PSM Score? PSM Score The PSM Score as given by Hecklib to this annotated spectrum. It is shown with three significant figures. | 388 |

#### Spectrum 5805? Spectrum 5805 The raw spectrum of this peptide as annotated by Hecklib. The fragments are coloured according to ion type (see legend). Any peaks with a star '\*' as text can be hovered over to see the full details, first the ion type second the mass shift type. By hovering over the amino acids in the peptide or ions in the legend the corresponding peaks are highlighted. By toggling the 'Unassigned' label you can turn the background (unassigned) peaks on or off in the plot. By updating the slider in the Ion legend you can update the spectrum to only show the top X% of the peaks with labels. The top X% means any peak that is within X% of the highest intensity. By dragging in the spectrum you can zoom in to a specific part of the spectrum and use 'Zoom Out' to get back to the original zoom level. The annotation of the spectrum is based on the given sequence in the peptides file and is done with different software so inconsistencies are likely. The peaks are annotated based on the given sequence, with 20 ppm tolerance.

Copy Data

##### Spectrum 5805 (TSV)

###### Preview

```
Loading example...
```

*Click on the button to copy the data to your clipboard.*

Mz MinMz MaxIntensity Max

WidthHeightPeptide font sizePeptide stroke widthSpectrum font sizeSpectrum stroke widthCompact peptide

Ion legend

wxyz

abcd

OtherUnassignedIon^Charge^_Position_Show for top:%

VRQAPGRAJEW

09.41e+41.88e+52.82e+53.77e+5

Zoom Out

c^+2^_3_y^+1^_1_c^+2^_4_c^+1^_2_c^+1^_2_c^+2^_5_y^+1^_2_y^+1^_2_c^+2^_7_c^+1^_3_c^+2^_7_c^+1^_3_y^+2^_7_y^+2^_7_c^+2^_8_c^+2^_8_c^+1^_4_c^+2^_9_c^+2^_9_y^+1^_4_c^+2^_10_c^+2^_10_c^+2^_10_c^+1^_5_z^+2^_10_y^+2^_10_c^+1^_6_y^+1^_5_z^+1^_5_y^+1^_5_y^+1^_6_z^+1^_6_y^+1^_6_c^+1^_7_w^+1^_7_y^+1^_7_z^+1^_7_y^+1^_7_c^+1^_8_y^+1^_8_z^+1^_8_y^+1^_8_w^+1^_9_c^+1^_9_z^+1^_9_y^+1^_9_c^+1^_10_w^+1^_10_c^+1^_10_y^+1^_10_z^+1^_10_y^+1^_10_

0778155723353114

Fragment Matches Table

Show background peaks

| Position | Ion type | Intensity | mz Theoretical | mz Error (Th) | mz Error (ppm) | Charge | Series Number |
| --- | --- | --- | --- | --- | --- | --- | --- |
| - | - | 6926 | 130.1 | - | - | 0 | - |
| - | - | 606.2 | 131.1 | - | - | 0 | - |
| - | - | 1929 | 132.1 | - | - | 0 | - |
| - | - | 1319 | 142.1 | - | - | 0 | - |
| - | - | 1338 | 144.1 | - | - | 0 | - |
| - | - | 7274 | 146.1 | - | - | 0 | - |
| - | - | 472.7 | 148.5 | - | - | 0 | - |
| - | - | 1877 | 149 | - | - | 0 | - |
| - | - | 4429 | 157.1 | - | - | 0 | - |
| - | - | 480.4 | 159.1 | - | - | 0 | - |
| - | - | 2.286E+04 | 159.1 | - | - | 0 | - |
| - | - | 1925 | 160.1 | - | - | 0 | - |
| - | - | 459.9 | 165 | - | - | 0 | - |
| - | - | 708.4 | 166.1 | - | - | 0 | - |
| - | - | 421.9 | 168.6 | - | - | 0 | - |
| - | - | 565.3 | 169.1 | - | - | 0 | - |
| - | - | 803.4 | 169.1 | - | - | 0 | - |
| - | - | 1443 | 170.1 | - | - | 0 | - |
| - | - | 736.4 | 171.1 | - | - | 0 | - |
| - | - | 1324 | 173.5 | - | - | 0 | - |
| - | - | 554.6 | 174.1 | - | - | 0 | - |
| - | - | 1654 | 174.1 | - | - | 0 | - |
| - | - | 480 | 185 | - | - | 0 | - |
| - | - | 1961 | 185.1 | - | - | 0 | - |
| - | - | 1228 | 185.1 | - | - | 0 | - |
| - | - | 714.9 | 185.1 | - | - | 0 | - |
| - | - | 3957 | 187.1 | - | - | 0 | - |
| - | - | 7.083E+04 | 188.1 | - | - | 0 | - |
| - | - | 7636 | 189.1 | - | - | 0 | - |
| - | - | 728.6 | 190.1 | - | - | 0 | - |
| 3 | c | 1216 | 192.6 | 0.0002187 | 1.135 | +2 | 3 |
| - | - | 761.6 | 195.2 | - | - | 0 | - |
| - | - | 482.3 | 195.7 | - | - | 0 | - |
| - | - | 851.8 | 197.1 | - | - | 0 | - |
| - | - | 1184 | 197.1 | - | - | 0 | - |
| - | - | 498 | 200.2 | - | - | 0 | - |
| - | - | 1223 | 203.1 | - | - | 0 | - |
| 11 | y | 4.437E+04 | 205.1 | 0.0001665 | 0.8116 | +1 | 1 |
| - | - | 558.1 | 205.6 | - | - | 0 | - |
| - | - | 4821 | 206.1 | - | - | 0 | - |
| - | - | 494.6 | 208.9 | - | - | 0 | - |
| - | - | 2.213E+04 | 209 | - | - | 0 | - |
| - | - | 4353 | 213.2 | - | - | 0 | - |
| - | - | 7520 | 214.1 | - | - | 0 | - |
| - | - | 4288 | 214.2 | - | - | 0 | - |
| - | - | 1586 | 214.6 | - | - | 0 | - |
| - | - | 753.6 | 215.1 | - | - | 0 | - |
| - | - | 557.4 | 225 | - | - | 0 | - |
| 4 | c | 1212 | 228.1 | 0.000404 | 1.771 | +2 | 4 |
| - | - | 1910 | 228.2 | - | - | 0 | - |
| - | - | 525.7 | 234.2 | - | - | 0 | - |
| - | - | 1628 | 238.2 | - | - | 0 | - |
| - | - | 2.134E+04 | 239.2 | - | - | 0 | - |
| - | - | 1043 | 239.2 | - | - | 0 | - |
| - | - | 609.8 | 239.2 | - | - | 0 | - |
| - | - | 2388 | 240.2 | - | - | 0 | - |
| - | - | 4576 | 242.2 | - | - | 0 | - |
| - | - | 587.6 | 243.2 | - | - | 0 | - |
| - | - | 1155 | 256.1 | - | - | 0 | - |
| 2 | c | 1.151E+04 | 256.2 | 0.000109 | 0.4255 | +1 | 2 |
| - | - | 986.3 | 257.2 | - | - | 0 | - |
| - | - | 499.4 | 260.6 | - | - | 0 | - |
| - | - | 2853 | 270.1 | - | - | 0 | - |
| 2 | c | 4.733E+04 | 273.2 | 0.0001102 | 0.4033 | +1 | 2 |
| - | - | 6472 | 274.2 | - | - | 0 | - |
| 5 | c | 671.2 | 276.7 | 0.001153 | 4.167 | +2 | 5 |
| - | - | 697.4 | 280.1 | - | - | 0 | - |
| - | - | 545.7 | 298 | - | - | 0 | - |
| - | - | 938.2 | 298.1 | - | - | 0 | - |
| - | - | 1512 | 298.2 | - | - | 0 | - |
| - | - | 1.256E+04 | 299.2 | - | - | 0 | - |
| - | - | 1487 | 300.2 | - | - | 0 | - |
| - | - | 824.7 | 307.9 | - | - | 0 | - |
| - | - | 673.8 | 314.2 | - | - | 0 | - |
| 10 | y | 1.107E+04 | 316.1 | 0.0003036 | 0.9603 | +1 | 2 |
| - | - | 1.672E+04 | 318.9 | - | - | 0 | - |
| 10 | y | 1296 | 334.1 | 3.772E-05 | 0.1129 | +1 | 2 |
| - | - | 734 | 339.2 | - | - | 0 | - |
| - | - | 4721 | 341 | - | - | 0 | - |
| - | - | 950.6 | 341.2 | - | - | 0 | - |
| - | - | 607.5 | 344.5 | - | - | 0 | - |
| - | - | 767.6 | 355.7 | - | - | 0 | - |
| - | - | 2295 | 356.2 | - | - | 0 | - |
| - | - | 2151 | 357.2 | - | - | 0 | - |
| - | - | 1.621E+04 | 359 | - | - | 0 | - |
| - | - | 3198 | 361.7 | - | - | 0 | - |
| - | - | 2050 | 362.2 | - | - | 0 | - |
| - | - | 1061 | 367.2 | - | - | 0 | - |
| - | - | 1906 | 369.2 | - | - | 0 | - |
| - | - | 8920 | 369.7 | - | - | 0 | - |
| - | - | 4551 | 370.2 | - | - | 0 | - |
| - | - | 1000 | 370.3 | - | - | 0 | - |
| - | - | 978.3 | 370.7 | - | - | 0 | - |
| - | - | 647.4 | 374.7 | - | - | 0 | - |
| 7 | c | 3694 | 383.2 | 0.0002895 | 0.7554 | +2 | 7 |
| - | - | 1871 | 383.7 | - | - | 0 | - |
| 3 | c | 7767 | 384.2 | 2.735E-05 | 0.07117 | +1 | 3 |
| - | - | 2192 | 385.2 | - | - | 0 | - |
| - | - | 872.7 | 385.3 | - | - | 0 | - |
| - | - | 1201 | 390.7 | - | - | 0 | - |
| - | - | 5.76E+04 | 391.2 | - | - | 0 | - |
| 7 | c | 1.863E+05 | 391.7 | 0.0001363 | 0.3479 | +2 | 7 |
| - | - | 7.349E+04 | 392.2 | - | - | 0 | - |
| - | - | 1.465E+04 | 392.7 | - | - | 0 | - |
| - | - | 2029 | 393.2 | - | - | 0 | - |
| 3 | c | 1.494E+05 | 401.3 | 0.000279 | 0.6954 | +1 | 3 |
| - | - | 2.974E+04 | 402.3 | - | - | 0 | - |
| - | - | 3179 | 403.3 | - | - | 0 | - |
| - | - | 1632 | 404.7 | - | - | 0 | - |
| - | - | 1959 | 405.3 | - | - | 0 | - |
| 5 | y | 3466 | 405.7 | 0.0006449 | 1.59 | +2 | 7 |
| - | - | 755.3 | 405.8 | - | - | 0 | - |
| - | - | 1357 | 406.2 | - | - | 0 | - |
| - | - | 4693 | 406.9 | - | - | 0 | - |
| - | - | 4492 | 407.2 | - | - | 0 | - |
| - | - | 2478 | 407.6 | - | - | 0 | - |
| - | - | 537.3 | 407.9 | - | - | 0 | - |
| - | - | 640.1 | 409.2 | - | - | 0 | - |
| - | - | 1710 | 410.2 | - | - | 0 | - |
| - | - | 6252 | 412.9 | - | - | 0 | - |
| - | - | 4925 | 413.2 | - | - | 0 | - |
| - | - | 791.7 | 413.3 | - | - | 0 | - |
| - | - | 2173 | 413.6 | - | - | 0 | - |
| 5 | y | 2391 | 414.7 | 0.0007337 | 1.769 | +2 | 7 |
| - | - | 1107 | 415.2 | - | - | 0 | - |
| - | - | 765.1 | 418.3 | - | - | 0 | - |
| 8 | c | 6059 | 418.7 | 0.0006849 | 1.636 | +2 | 8 |
| - | - | 2861 | 419.2 | - | - | 0 | - |
| - | - | 934.7 | 422.2 | - | - | 0 | - |
| - | - | 1621 | 422.6 | - | - | 0 | - |
| - | - | 2.192E+04 | 426.8 | - | - | 0 | - |
| 8 | c | 1.378E+05 | 427.3 | 0.0003193 | 0.7472 | +2 | 8 |
| - | - | 6.237E+04 | 427.8 | - | - | 0 | - |
| - | - | 1157 | 427.8 | - | - | 0 | - |
| - | - | 652.9 | 428.2 | - | - | 0 | - |
| - | - | 1.53E+04 | 428.3 | - | - | 0 | - |
| - | - | 1872 | 428.8 | - | - | 0 | - |
| - | - | 1788 | 428.9 | - | - | 0 | - |
| - | - | 1.682E+04 | 429.1 | - | - | 0 | - |
| - | - | 1770 | 440.3 | - | - | 0 | - |
| - | - | 822.4 | 440.8 | - | - | 0 | - |
| - | - | 964.3 | 453.8 | - | - | 0 | - |
| - | - | 802.6 | 454.8 | - | - | 0 | - |
| 4 | c | 8423 | 455.3 | 0.0004565 | 1.003 | +1 | 4 |
| - | - | 2550 | 456.3 | - | - | 0 | - |
| - | - | 1.562E+04 | 461.3 | - | - | 0 | - |
| - | - | 1.148E+04 | 461.8 | - | - | 0 | - |
| - | - | 5131 | 462.3 | - | - | 0 | - |
| - | - | 1020 | 462.8 | - | - | 0 | - |
| - | - | 892.1 | 468.8 | - | - | 0 | - |
| - | - | 1401 | 472.3 | - | - | 0 | - |
| - | - | 2698 | 474.8 | - | - | 0 | - |
| 9 | c | 3.845E+04 | 475.3 | 0.0004925 | 1.036 | +2 | 9 |
| - | - | 1.924E+04 | 475.8 | - | - | 0 | - |
| - | - | 5828 | 476.3 | - | - | 0 | - |
| - | - | 1881 | 476.8 | - | - | 0 | - |
| - | - | 5106 | 483.3 | - | - | 0 | - |
| 9 | c | 7.239E+04 | 483.8 | 0.0002184 | 0.4515 | +2 | 9 |
| - | - | 3.609E+04 | 484.3 | - | - | 0 | - |
| - | - | 1.029E+04 | 484.8 | - | - | 0 | - |
| - | - | 2591 | 485.3 | - | - | 0 | - |
| - | - | 661.4 | 485.8 | - | - | 0 | - |
| - | - | 1014 | 486.3 | - | - | 0 | - |
| - | - | 4.275E+04 | 496.8 | - | - | 0 | - |
| - | - | 2.399E+04 | 497.3 | - | - | 0 | - |
| - | - | 5603 | 497.8 | - | - | 0 | - |
| - | - | 7150 | 498.3 | - | - | 0 | - |
| - | - | 1980 | 498.3 | - | - | 0 | - |
| - | - | 4452 | 498.8 | - | - | 0 | - |
| - | - | 1028 | 499.3 | - | - | 0 | - |
| - | - | 3223 | 499.3 | - | - | 0 | - |
| - | - | 625.3 | 499.8 | - | - | 0 | - |
| - | - | 1885 | 504.8 | - | - | 0 | - |
| - | - | 995.7 | 505.3 | - | - | 0 | - |
| - | - | 1966 | 517.8 | - | - | 0 | - |
| 8 | y | 1053 | 518.3 | 0.002747 | 5.3 | +1 | 4 |
| - | - | 2639 | 518.3 | - | - | 0 | - |
| - | - | 2515 | 518.8 | - | - | 0 | - |
| - | - | 854.2 | 519.8 | - | - | 0 | - |
| - | - | 1622 | 524.3 | - | - | 0 | - |
| - | - | 3932 | 525.3 | - | - | 0 | - |
| - | - | 4.045E+04 | 525.8 | - | - | 0 | - |
| - | - | 2.578E+04 | 526.3 | - | - | 0 | - |
| - | - | 1.034E+04 | 526.8 | - | - | 0 | - |
| - | - | 3062 | 527.3 | - | - | 0 | - |
| - | - | 991.5 | 529.3 | - | - | 0 | - |
| - | - | 799 | 530.8 | - | - | 0 | - |
| - | - | 1480 | 531.3 | - | - | 0 | - |
| - | - | 1201 | 531.8 | - | - | 0 | - |
| - | - | 966 | 532.8 | - | - | 0 | - |
| - | - | 938.7 | 533.8 | - | - | 0 | - |
| 10 | c | 2186 | 539.3 | 0.0008422 | 1.562 | +2 | 10 |
| 10 | c | 2.056E+05 | 539.8 | 0.0003751 | 0.695 | +2 | 10 |
| - | - | 1.187E+05 | 540.3 | - | - | 0 | - |
| - | - | 4.137E+04 | 540.8 | - | - | 0 | - |
| - | - | 1.005E+04 | 541.3 | - | - | 0 | - |
| - | - | 1021 | 541.8 | - | - | 0 | - |
| - | - | 5615 | 542.3 | - | - | 0 | - |
| - | - | 1715 | 543.3 | - | - | 0 | - |
| - | - | 2140 | 547.8 | - | - | 0 | - |
| 10 | c | 3.728E+05 | 548.3 | 0.0004063 | 0.7409 | +2 | 10 |
| - | - | 2.063E+05 | 548.8 | - | - | 0 | - |
| - | - | 6.686E+04 | 549.3 | - | - | 0 | - |
| - | - | 1.886E+04 | 549.8 | - | - | 0 | - |
| - | - | 2645 | 550.3 | - | - | 0 | - |
| - | - | 985.8 | 553.8 | - | - | 0 | - |
| - | - | 1822 | 554.8 | - | - | 0 | - |
| - | - | 1.526E+04 | 555.3 | - | - | 0 | - |
| - | - | 2541 | 555.3 | - | - | 0 | - |
| - | - | 8621 | 555.8 | - | - | 0 | - |
| - | - | 3328 | 556.3 | - | - | 0 | - |
| - | - | 747.6 | 556.3 | - | - | 0 | - |
| - | - | 1278 | 558.3 | - | - | 0 | - |
| - | - | 1.094E+04 | 561.3 | - | - | 0 | - |
| - | - | 9861 | 561.8 | - | - | 0 | - |
| - | - | 3429 | 562.3 | - | - | 0 | - |
| - | - | 1685 | 562.8 | - | - | 0 | - |
| - | - | 1217 | 562.8 | - | - | 0 | - |
| - | - | 1038 | 567.3 | - | - | 0 | - |
| - | - | 1681 | 568.3 | - | - | 0 | - |
| - | - | 2326 | 568.3 | - | - | 0 | - |
| - | - | 1573 | 568.8 | - | - | 0 | - |
| 5 | c | 5.567E+04 | 569.4 | 6.005E-05 | 0.1055 | +1 | 5 |
| - | - | 661.3 | 570.3 | - | - | 0 | - |
| - | - | 1.747E+04 | 570.4 | - | - | 0 | - |
| - | - | 691.4 | 571.3 | - | - | 0 | - |
| - | - | 2386 | 571.4 | - | - | 0 | - |
| - | - | 5250 | 575.8 | - | - | 0 | - |
| - | - | 3553 | 576.3 | - | - | 0 | - |
| - | - | 2.299E+04 | 576.8 | - | - | 0 | - |
| - | - | 1.763E+04 | 577.3 | - | - | 0 | - |
| - | - | 7917 | 577.8 | - | - | 0 | - |
| - | - | 2352 | 578.3 | - | - | 0 | - |
| - | - | 4323 | 582.4 | - | - | 0 | - |
| - | - | 3013 | 582.8 | - | - | 0 | - |
| - | - | 2233 | 583.4 | - | - | 0 | - |
| - | - | 2812 | 583.8 | - | - | 0 | - |
| 2 | z | 1.917E+04 | 584.3 | 0.000226 | 0.3868 | +2 | 10 |
| - | - | 1.034E+04 | 584.8 | - | - | 0 | - |
| - | - | 3669 | 585.3 | - | - | 0 | - |
| - | - | 780.9 | 585.4 | - | - | 0 | - |
| - | - | 1081 | 585.8 | - | - | 0 | - |
| - | - | 1349 | 586.3 | - | - | 0 | - |
| - | - | 851.1 | 587.3 | - | - | 0 | - |
| - | - | 1792 | 590.9 | - | - | 0 | - |
| - | - | 1214 | 591.3 | - | - | 0 | - |
| - | - | 1112 | 591.8 | - | - | 0 | - |
| 2 | y | 1728 | 592.3 | 0.004292 | 7.246 | +2 | 10 |
| - | - | 1758 | 596.4 | - | - | 0 | - |
| - | - | 2770 | 597.8 | - | - | 0 | - |
| - | - | 6850 | 598.3 | - | - | 0 | - |
| - | - | 3915 | 598.8 | - | - | 0 | - |
| - | - | 5160 | 599.3 | - | - | 0 | - |
| - | - | 856.8 | 600.3 | - | - | 0 | - |
| - | - | 4848 | 602.3 | - | - | 0 | - |
| - | - | 1183 | 603.3 | - | - | 0 | - |
| - | - | 756.3 | 603.3 | - | - | 0 | - |
| - | - | 5564 | 604.3 | - | - | 0 | - |
| - | - | 1.376E+04 | 604.8 | - | - | 0 | - |
| - | - | 9897 | 605.3 | - | - | 0 | - |
| - | - | 5144 | 605.8 | - | - | 0 | - |
| - | - | 2062 | 606.3 | - | - | 0 | - |
| - | - | 2664 | 606.3 | - | - | 0 | - |
| - | - | 1723 | 606.8 | - | - | 0 | - |
| - | - | 746.3 | 607.4 | - | - | 0 | - |
| - | - | 5449 | 611.4 | - | - | 0 | - |
| - | - | 2.912E+04 | 611.9 | - | - | 0 | - |
| - | - | 1.608E+04 | 612.4 | - | - | 0 | - |
| - | - | 7137 | 612.9 | - | - | 0 | - |
| - | - | 905.6 | 613.3 | - | - | 0 | - |
| - | - | 4003 | 613.4 | - | - | 0 | - |
| - | - | 1533 | 613.8 | - | - | 0 | - |
| - | - | 1.539E+04 | 614.4 | - | - | 0 | - |
| - | - | 2128 | 614.8 | - | - | 0 | - |
| - | - | 7626 | 615.3 | - | - | 0 | - |
| - | - | 5937 | 615.4 | - | - | 0 | - |
| - | - | 2679 | 616.3 | - | - | 0 | - |
| - | - | 872.2 | 616.4 | - | - | 0 | - |
| - | - | 1293 | 617.8 | - | - | 0 | - |
| - | - | 4152 | 619.4 | - | - | 0 | - |
| - | - | 1.723E+04 | 619.8 | - | - | 0 | - |
| - | - | 2.544E+04 | 620.4 | - | - | 0 | - |
| - | - | 5.49E+04 | 620.8 | - | - | 0 | - |
| - | - | 3.916E+04 | 621.3 | - | - | 0 | - |
| - | - | 1.264E+04 | 621.8 | - | - | 0 | - |
| - | - | 3651 | 622.3 | - | - | 0 | - |
| - | - | 3136 | 624.3 | - | - | 0 | - |
| - | - | 837.2 | 624.8 | - | - | 0 | - |
| - | - | 7949 | 625.4 | - | - | 0 | - |
| - | - | 4857 | 625.8 | - | - | 0 | - |
| 6 | c | 2.256E+05 | 626.4 | 0.000571 | 0.9115 | +1 | 6 |
| - | - | 1.779E+04 | 626.8 | - | - | 0 | - |
| - | - | 6.689E+04 | 627.4 | - | - | 0 | - |
| - | - | 2626 | 627.8 | - | - | 0 | - |
| - | - | 1.258E+04 | 628.4 | - | - | 0 | - |
| - | - | 2149 | 629.4 | - | - | 0 | - |
| - | - | 4779 | 633.3 | - | - | 0 | - |
| - | - | 2.785E+05 | 633.8 | - | - | 0 | - |
| - | - | 1.921E+05 | 634.3 | - | - | 0 | - |
| - | - | 7.015E+04 | 634.8 | - | - | 0 | - |
| - | - | 2.103E+04 | 635.3 | - | - | 0 | - |
| - | - | 4713 | 635.8 | - | - | 0 | - |
| - | - | 1.341E+05 | 641.9 | - | - | 0 | - |
| - | - | 1.619E+05 | 642.4 | - | - | 0 | - |
| - | - | 8.095E+04 | 642.9 | - | - | 0 | - |
| - | - | 2.875E+04 | 643.4 | - | - | 0 | - |
| - | - | 7494 | 643.9 | - | - | 0 | - |
| - | - | 1282 | 644.4 | - | - | 0 | - |
| - | - | 2045 | 652.4 | - | - | 0 | - |
| 7 | y | 1148 | 656.4 | 0.001861 | 2.836 | +1 | 5 |
| 7 | z | 1.256E+05 | 658.3 | 7.187E-05 | 0.1092 | +1 | 5 |
| - | - | 4.708E+04 | 659.3 | - | - | 0 | - |
| - | - | 9996 | 660.3 | - | - | 0 | - |
| - | - | 1585 | 661.4 | - | - | 0 | - |
| - | - | 8060 | 671.4 | - | - | 0 | - |
| - | - | 2957 | 672.4 | - | - | 0 | - |
| 7 | y | 7441 | 674.4 | 0.0001466 | 0.2174 | +1 | 5 |
| - | - | 2415 | 675.4 | - | - | 0 | - |
| - | - | 778.7 | 676.4 | - | - | 0 | - |
| - | - | 962.8 | 695.4 | - | - | 0 | - |
| - | - | 1789 | 696.4 | - | - | 0 | - |
| - | - | 917.9 | 697.4 | - | - | 0 | - |
| 6 | y | 1435 | 713.4 | 0.00121 | 1.697 | +1 | 6 |
| - | - | 717.2 | 714.4 | - | - | 0 | - |
| 6 | z | 3.259E+04 | 715.4 | 3.149E-05 | 0.04402 | +1 | 6 |
| - | - | 1.107E+04 | 716.4 | - | - | 0 | - |
| - | - | 2714 | 717.4 | - | - | 0 | - |
| - | - | 1159 | 723.4 | - | - | 0 | - |
| 6 | y | 9174 | 731.4 | 0.0005945 | 0.8129 | +1 | 6 |
| - | - | 3007 | 732.4 | - | - | 0 | - |
| - | - | 887.6 | 733.4 | - | - | 0 | - |
| - | - | 1111 | 738.5 | - | - | 0 | - |
| - | - | 1669 | 739.5 | - | - | 0 | - |
| - | - | 2411 | 740.5 | - | - | 0 | - |
| - | - | 776.5 | 741.5 | - | - | 0 | - |
| - | - | 1621 | 767.5 | - | - | 0 | - |
| - | - | 1023 | 780.4 | - | - | 0 | - |
| - | - | 4429 | 781.5 | - | - | 0 | - |
| 7 | c | 3.868E+04 | 782.5 | 0.0003203 | 0.4094 | +1 | 7 |
| - | - | 1.55E+04 | 783.5 | - | - | 0 | - |
| - | - | 4260 | 784.5 | - | - | 0 | - |
| 5 | w | 4077 | 785.4 | 0.0004058 | 0.5167 | +1 | 7 |
| - | - | 1062 | 785.5 | - | - | 0 | - |
| - | - | 1863 | 786.4 | - | - | 0 | - |
| 5 | y | 2784 | 810.4 | 0.00126 | 1.555 | +1 | 7 |
| - | - | 5554 | 811.4 | - | - | 0 | - |
| 5 | z | 1757 | 812.4 | 0.008913 | 10.97 | +1 | 7 |
| - | - | 1024 | 819.5 | - | - | 0 | - |
| - | - | 1818 | 823.4 | - | - | 0 | - |
| - | - | 7802 | 824.4 | - | - | 0 | - |
| - | - | 4632 | 825.5 | - | - | 0 | - |
| - | - | 2769 | 826.4 | - | - | 0 | - |
| - | - | 2306 | 827.4 | - | - | 0 | - |
| 5 | y | 9585 | 828.4 | 0.001083 | 1.307 | +1 | 7 |
| - | - | 4645 | 829.4 | - | - | 0 | - |
| - | - | 2034 | 830.4 | - | - | 0 | - |
| - | - | 1068 | 837.5 | - | - | 0 | - |
| - | - | 714.2 | 838.5 | - | - | 0 | - |
| - | - | 3084 | 839.5 | - | - | 0 | - |
| - | - | 2702 | 840.5 | - | - | 0 | - |
| - | - | 1193 | 851.5 | - | - | 0 | - |
| - | - | 2907 | 852.5 | - | - | 0 | - |
| 8 | c | 3.232E+04 | 853.5 | 0.0005689 | 0.6665 | +1 | 8 |
| - | - | 1.219E+04 | 854.5 | - | - | 0 | - |
| - | - | 3729 | 855.5 | - | - | 0 | - |
| - | - | 1685 | 856.4 | - | - | 0 | - |
| - | - | 990.6 | 857.4 | - | - | 0 | - |
| 4 | y | 969 | 882.4 | 0.002207 | 2.501 | +1 | 8 |
| 4 | z | 1.395E+05 | 883.5 | 0.0006738 | 0.7627 | +1 | 8 |
| - | - | 6.568E+04 | 884.5 | - | - | 0 | - |
| - | - | 2.12E+04 | 885.5 | - | - | 0 | - |
| - | - | 4359 | 886.5 | - | - | 0 | - |
| 4 | y | 1.676E+04 | 899.5 | 0.0009652 | 1.073 | +1 | 8 |
| - | - | 9824 | 900.5 | - | - | 0 | - |
| - | - | 2665 | 901.5 | - | - | 0 | - |
| - | - | 1186 | 908.6 | - | - | 0 | - |
| - | - | 1027 | 915.4 | - | - | 0 | - |
| - | - | 942.2 | 922.6 | - | - | 0 | - |
| - | - | 2418 | 923.6 | - | - | 0 | - |
| - | - | 2022 | 924.6 | - | - | 0 | - |
| - | - | 1011 | 936.5 | - | - | 0 | - |
| - | - | 1221 | 939.5 | - | - | 0 | - |
| - | - | 1369 | 940.5 | - | - | 0 | - |
| - | - | 2173 | 950.6 | - | - | 0 | - |
| - | - | 2371 | 951.6 | - | - | 0 | - |
| 3 | w | 3894 | 953.5 | 0.0001164 | 0.122 | +1 | 9 |
| - | - | 1960 | 954.5 | - | - | 0 | - |
| - | - | 783.2 | 955.5 | - | - | 0 | - |
| - | - | 780.7 | 956.5 | - | - | 0 | - |
| - | - | 962.2 | 963.5 | - | - | 0 | - |
| - | - | 2121 | 965.6 | - | - | 0 | - |
| 9 | c | 3.261E+04 | 966.6 | 0.001076 | 1.113 | +1 | 9 |
| - | - | 1.911E+04 | 967.6 | - | - | 0 | - |
| - | - | 5910 | 968.6 | - | - | 0 | - |
| - | - | 779 | 969.6 | - | - | 0 | - |
| - | - | 4136 | 980.5 | - | - | 0 | - |
| - | - | 2010 | 981.6 | - | - | 0 | - |
| - | - | 760.4 | 1008 | - | - | 0 | - |
| - | - | 686.8 | 1009 | - | - | 0 | - |
| - | - | 1550 | 1010 | - | - | 0 | - |
| 3 | z | 1.026E+05 | 1012 | 0.0008407 | 0.8311 | +1 | 9 |
| - | - | 5.966E+04 | 1013 | - | - | 0 | - |
| - | - | 908.7 | 1013 | - | - | 0 | - |
| - | - | 1.901E+04 | 1014 | - | - | 0 | - |
| - | - | 4117 | 1015 | - | - | 0 | - |
| - | - | 1556 | 1025 | - | - | 0 | - |
| 3 | y | 2.597E+04 | 1028 | 0.001437 | 1.399 | +1 | 9 |
| - | - | 1.307E+04 | 1029 | - | - | 0 | - |
| - | - | 4726 | 1030 | - | - | 0 | - |
| - | - | 1711 | 1035 | - | - | 0 | - |
| - | - | 2380 | 1036 | - | - | 0 | - |
| - | - | 743.8 | 1037 | - | - | 0 | - |
| - | - | 5998 | 1038 | - | - | 0 | - |
| - | - | 2993 | 1039 | - | - | 0 | - |
| - | - | 1.228E+04 | 1052 | - | - | 0 | - |
| - | - | 2.935E+04 | 1053 | - | - | 0 | - |
| - | - | 1.742E+04 | 1054 | - | - | 0 | - |
| - | - | 6380 | 1055 | - | - | 0 | - |
| - | - | 1656 | 1056 | - | - | 0 | - |
| - | - | 865.6 | 1063 | - | - | 0 | - |
| - | - | 970.2 | 1064 | - | - | 0 | - |
| 10 | c | 1535 | 1079 | 0.004913 | 4.555 | +1 | 10 |
| - | - | 2.212E+04 | 1080 | - | - | 0 | - |
| - | - | 1.386E+04 | 1081 | - | - | 0 | - |
| 2 | w | 3667 | 1082 | 0.003335 | 3.084 | +1 | 10 |
| - | - | 2604 | 1082 | - | - | 0 | - |
| - | - | 4752 | 1083 | - | - | 0 | - |
| - | - | 2202 | 1084 | - | - | 0 | - |
| 10 | c | 2.128E+04 | 1096 | 0.001433 | 1.307 | +1 | 10 |
| - | - | 5.888E+04 | 1097 | - | - | 0 | - |
| - | - | 3.01E+04 | 1098 | - | - | 0 | - |
| - | - | 9410 | 1099 | - | - | 0 | - |
| - | - | 3316 | 1100 | - | - | 0 | - |
| - | - | 757.8 | 1111 | - | - | 0 | - |
| - | - | 761 | 1112 | - | - | 0 | - |
| - | - | 823.2 | 1124 | - | - | 0 | - |
| - | - | 938.5 | 1125 | - | - | 0 | - |
| - | - | 1720 | 1126 | - | - | 0 | - |
| - | - | 653.6 | 1137 | - | - | 0 | - |
| - | - | 834.5 | 1140 | - | - | 0 | - |
| - | - | 1101 | 1141 | - | - | 0 | - |
| - | - | 1142 | 1152 | - | - | 0 | - |
| - | - | 1859 | 1153 | - | - | 0 | - |
| - | - | 3749 | 1154 | - | - | 0 | - |
| - | - | 1909 | 1155 | - | - | 0 | - |
| - | - | 1143 | 1156 | - | - | 0 | - |
| 2 | y | 971.3 | 1167 | 0.02105 | 18.04 | +1 | 10 |
| 2 | z | 7228 | 1168 | 0.003258 | 2.79 | +1 | 10 |
| - | - | 4.205E+04 | 1169 | - | - | 0 | - |
| - | - | 2.894E+04 | 1170 | - | - | 0 | - |
| - | - | 9588 | 1171 | - | - | 0 | - |
| - | - | 3212 | 1172 | - | - | 0 | - |
| - | - | 6787 | 1181 | - | - | 0 | - |
| - | - | 5313 | 1182 | - | - | 0 | - |
| - | - | 4941 | 1183 | - | - | 0 | - |
| 2 | y | 3533 | 1184 | 0.007333 | 6.196 | +1 | 10 |
| - | - | 3064 | 1185 | - | - | 0 | - |
| - | - | 745.5 | 1186 | - | - | 0 | - |
| - | - | 989.3 | 1195 | - | - | 0 | - |
| - | - | 3919 | 1196 | - | - | 0 | - |
| - | - | 3715 | 1197 | - | - | 0 | - |
| - | - | 1907 | 1198 | - | - | 0 | - |
| - | - | 1037 | 1199 | - | - | 0 | - |
| - | - | 1854 | 1207 | - | - | 0 | - |
| - | - | 934.6 | 1208 | - | - | 0 | - |
| - | - | 1344 | 1209 | - | - | 0 | - |
| - | - | 1247 | 1212 | - | - | 0 | - |
| - | - | 985.4 | 1213 | - | - | 0 | - |
| - | - | 1164 | 1214 | - | - | 0 | - |
| - | - | 4183 | 1223 | - | - | 0 | - |
| - | - | 9956 | 1224 | - | - | 0 | - |
| - | - | 7.136E+04 | 1225 | - | - | 0 | - |
| - | - | 5.469E+04 | 1226 | - | - | 0 | - |
| - | - | 2.202E+04 | 1227 | - | - | 0 | - |
| - | - | 7977 | 1228 | - | - | 0 | - |
| - | - | 1966 | 1229 | - | - | 0 | - |
| - | - | 3646 | 1239 | - | - | 0 | - |
| - | - | 1.547E+04 | 1240 | - | - | 0 | - |
| - | - | 1.726E+04 | 1241 | - | - | 0 | - |
| - | - | 1.407E+04 | 1242 | - | - | 0 | - |
| - | - | 5926 | 1243 | - | - | 0 | - |
| - | - | 1927 | 1244 | - | - | 0 | - |
| - | - | 2359 | 1250 | - | - | 0 | - |
| - | - | 2.226E+04 | 1251 | - | - | 0 | - |
| - | - | 1.594E+04 | 1252 | - | - | 0 | - |
| - | - | 5954 | 1253 | - | - | 0 | - |
| - | - | 1490 | 1254 | - | - | 0 | - |
| - | - | 1332 | 1256 | - | - | 0 | - |
| - | - | 4912 | 1257 | - | - | 0 | - |
| - | - | 2590 | 1258 | - | - | 0 | - |
| - | - | 2.334E+04 | 1267 | - | - | 0 | - |
| - | - | 1.871E+05 | 1268 | - | - | 0 | - |
| - | - | 1.24E+05 | 1269 | - | - | 0 | - |
| - | - | 4.771E+04 | 1270 | - | - | 0 | - |
| - | - | 1.435E+04 | 1271 | - | - | 0 | - |
| - | - | 4490 | 1272 | - | - | 0 | - |
| - | - | 7945 | 1283 | - | - | 0 | - |
| - | - | 3.947E+04 | 1284 | - | - | 0 | - |
| - | - | 1.586E+05 | 1285 | - | - | 0 | - |
| - | - | 1.066E+05 | 1286 | - | - | 0 | - |
| - | - | 3.95E+04 | 1287 | - | - | 0 | - |
| - | - | 1.215E+04 | 1288 | - | - | 0 | - |
| - | - | 3646 | 1289 | - | - | 0 | - |
| - | - | 732.4 | 2110 | - | - | 0 | - |
| - | - | 688.1 | 2207 | - | - | 0 | - |
| - | - | 1060 | 3083 | - | - | 0 | - |

m/z Charge Intensity FragmentType MassShift Position
130.0653839111328 0 6926.3804
131.0687713623047 0 606.1674
132.08103942871094 0 1928.7974
142.0975799560547 0 1318.9221
144.08096313476562 0 1337.8636
146.06028747558594 0 7273.813
148.50469970703125 0 472.66336
149.0450897216797 0 1877.2844
157.10862731933594 0 4429.068
159.06439208984375 0 480.3639
159.0918426513672 0 22857.037
160.0951690673828 0 1924.8907
164.95582580566406 0 459.88712
166.0865478515625 0 708.40405
168.55938720703125 0 421.91467
169.07669067382812 0 565.2961
169.1334686279297 0 803.43835
170.06040954589844 0 1443.3522
171.1241912841797 0 736.4174
173.45176696777344 0 1323.6674
174.10275268554688 0 554.6158
174.13504028320312 0 1654.3805
184.97427368164062 0 480.01794
185.10342407226562 0 1960.5673
185.11727905273438 0 1227.7849
185.1401824951172 0 714.87244
187.0868377685547 0 3957.262
188.07083129882812 0 70830.51
189.0741729736328 0 7636.1514
190.0770263671875 0 728.6438
192.62110900878906 0 1216.1265 c Ammonia loss 2
195.1606903076172 0 761.62854
195.65457153320312 0 482.2914
197.12864685058594 0 851.75336
197.1395721435547 0 1183.6575
200.2338409423828 0 498.02054
203.1279296875 0 1223.1882
205.09732055664062 0 44367.402 y 10
205.6291046142578 0 558.1091
206.1006622314453 0 4820.862
208.8605194091797 0 494.64072
208.9533233642578 0 22131.068
213.1712188720703 0 4352.655
214.1425323486328 0 7519.7773
214.15518188476562 0 4287.762
214.64398193359375 0 1585.6963
215.13845825195312 0 753.61273
225.043212890625 0 557.38184
228.1394805908203 0 1212.2133 c Ammonia loss 3
228.18218994140625 0 1910.4562
234.15757751464844 0 525.7221
238.16653442382812 0 1627.9677
239.15045166015625 0 21338.16
239.16432189941406 0 1042.6782
239.17965698242188 0 609.78906
240.1540069580078 0 2387.7402
242.161376953125 0 4575.671
243.1671600341797 0 587.55347
256.12091064453125 0 1155.1573
256.1769104003906 0 11513.207 c Ammonia loss 1
257.1802673339844 0 986.26715
260.55322265625 0 499.37656
270.1238098144531 0 2852.5474
273.2034606933594 0 47330.27 c 1
274.2066650390625 0 6471.57
276.66741943359375 0 671.1539 c Ammonia loss 4
280.1084289550781 0 697.4108
298.02099609375 0 545.7131
298.1191711425781 0 938.167
298.2239074707031 0 1512.3506
299.2192687988281 0 12560.81
300.2221374511719 0 1487.4021
307.86431884765625 0 824.7422
314.20599365234375 0 673.7677
316.1294860839844 0 11069.778 y Water loss 9
318.9224853515625 0 16724.701
334.13970947265625 0 1296.4574 y 9
339.21466064453125 0 734.0144
341.0182189941406 0 4720.659
341.2296447753906 0 950.5624
344.5381774902344 0 607.4745
355.7169494628906 0 767.5924
356.24017333984375 0 2295.4788
357.2464904785156 0 2150.6785
359.0284118652344 0 16205.294
361.7242736816406 0 3197.605
362.2245178222656 0 2050.2734
367.2090759277344 0 1060.523
369.2298278808594 0 1905.8303
369.734130859375 0 8919.863
370.2359619140625 0 4551.036
370.2554016113281 0 1000.3156
370.7388916015625 0 978.33813
374.7154235839844 0 647.3577
383.2272644042969 0 3694.3748 c Ammonia loss 6
383.72979736328125 0 1870.5336
384.2353515625 0 7767.4165 c Ammonia loss 2
385.2383728027344 0 2191.6743
385.27960205078125 0 872.6682
390.73419189453125 0 1201.1035
391.2370910644531 0 57599.832
391.7406921386719 0 186325.08 c 6
392.2421875 0 73486.69
392.7434387207031 0 14648.118
393.245361328125 0 2029.1818
401.26220703125 0 149430.62 c 2
402.2649841308594 0 29744.87
403.26708984375 0 3179.208
404.7491455078125 0 1632.422
405.2519836425781 0 1958.7494
405.7171325683594 0 3466.2974 y Water loss 4
405.7527160644531 0 755.2617
406.2171630859375 0 1357.3358
406.9002380371094 0 4693.26
407.2349548339844 0 4492.0684
407.5686950683594 0 2478.1824
407.90234375 0 537.3279
409.2014465332031 0 640.14685
410.2332763671875 0 1709.8324
412.9036865234375 0 6251.7446
413.23773193359375 0 4924.503
413.2690734863281 0 791.7259
413.57147216796875 0 2173.4548
414.7225036621094 0 2391.1455 y 4
415.22589111328125 0 1107.4376
418.2540588378906 0 765.07733
418.7467956542969 0 6059.247 c Ammonia loss 7
419.24725341796875 0 2861.49
422.2342834472656 0 934.7341
422.57049560546875 0 1620.6914
426.7557067871094 0 21919.627
427.25970458984375 0 137815.28 c 7
427.76092529296875 0 62369.875
427.796142578125 0 1156.9445
428.20758056640625 0 652.87555
428.2623291015625 0 15303.403
428.7635498046875 0 1872.3333
428.89276123046875 0 1787.6973
429.0890808105469 0 16815.484
440.26776123046875 0 1769.8662
440.7733459472656 0 822.41504
453.78570556640625 0 964.314
454.7838439941406 0 802.5546
455.27294921875 0 8423.375 c Ammonia loss 3
456.2759094238281 0 2550.4392
461.29107666015625 0 15621.211
461.7930908203125 0 11481.708
462.294921875 0 5130.8115
462.7970886230469 0 1020.0933
468.7710266113281 0 892.10956
472.2887268066406 0 1400.5417
474.7958679199219 0 2698.0107
475.28863525390625 0 38452.395 c Ammonia loss 8
475.7900390625 0 19241.373
476.29058837890625 0 5828.2007
476.7926025390625 0 1880.858
483.29833984375 0 5105.7646
483.8016357421875 0 72392.484 c 8
484.30291748046875 0 36092.91
484.804443359375 0 10293.303
485.3050231933594 0 2590.853
485.80682373046875 0 661.44946
486.2716064453125 0 1013.7497
496.80950927734375 0 42749.547
497.3109130859375 0 23994.45
497.8117370605469 0 5603.33
498.2840270996094 0 7150.1416
498.3166198730469 0 1980.0675
498.78558349609375 0 4452.2456
499.28521728515625 0 1027.789
499.32244873046875 0 3222.6675
499.7838439941406 0 625.327
504.7883605957031 0 1885.0154
505.2892761230469 0 995.7355
517.8123779296875 0 1965.9401
518.263671875 0 1052.6754 y 7
518.310302734375 0 2639.2576
518.8071899414062 0 2515.2368
519.7892456054688 0 854.22784
524.331298828125 0 1621.8971
525.337646484375 0 3931.7036
525.812255859375 0 40447.39
526.313720703125 0 25779.838
526.8148193359375 0 10342.681
527.3161010742188 0 3062.031
529.2822875976562 0 991.51764
530.8046264648438 0 798.9705
531.2980346679688 0 1479.7563
531.8005981445312 0 1201.4541
532.809326171875 0 965.953
533.8021850585938 0 938.69495
539.3165893554688 0 2185.5232 c Water loss 9
539.809814453125 0 205579.89 c Ammonia loss 9
540.3109741210938 0 118672.81
540.8125 0 41366.785
541.3140258789062 0 10048.499
541.8189086914062 0 1020.61786
542.3323364257812 0 5615.413
543.3372802734375 0 1714.6814
547.8108520507812 0 2140.1008
548.3231201171875 0 372811.4 c 9
548.8240966796875 0 206346.31
549.3253173828125 0 66861.64
549.8263549804688 0 18860.23
550.3269653320312 0 2644.5317
553.8037109375 0 985.7623
554.808837890625 0 1821.5996
555.296630859375 0 15255.668
555.3392944335938 0 2541.1484
555.7982177734375 0 8621.32
556.2991943359375 0 3327.89
556.3446044921875 0 747.6251
558.2910766601562 0 1277.6918
561.3302001953125 0 10936.499
561.83251953125 0 9861.404
562.322021484375 0 3429.0713
562.7823486328125 0 1685.3948
562.8338012695312 0 1216.7664
567.3441772460938 0 1038.2181
568.3135375976562 0 1681.0106
568.3465576171875 0 2326.1418
568.8212890625 0 1573.4985
569.3517456054688 0 55667.42 c 4
570.3076782226562 0 661.28375
570.3545532226562 0 17468.172
571.3070678710938 0 691.3603
571.3580322265625 0 2386.0483
575.8357543945312 0 5250.346
576.3349609375 0 3552.99
576.8218383789062 0 22985.486
577.3236083984375 0 17632.973
577.8262329101562 0 7917.3374
578.3251953125 0 2351.6357
582.353515625 0 4323.33
582.8441772460938 0 3013.1887
583.3603515625 0 2233.3176
583.8231201171875 0 2812.3928
584.31103515625 0 19167.613 z 1
584.8133544921875 0 10337.036
585.3140258789062 0 3669.1775
585.361328125 0 780.88336
585.8157348632812 0 1080.9023
586.3209228515625 0 1349.1863
587.3265380859375 0 851.1076
590.85205078125 0 1792.0085
591.311767578125 0 1213.7073
591.8162841796875 0 1112.0471
592.324462890625 0 1728.1584 y 1
596.35107421875 0 1758.4219
597.8352661132812 0 2769.7073
598.3291015625 0 6849.604
598.8298950195312 0 3914.898
599.330810546875 0 5159.943
600.3350219726562 0 856.811
602.2816162109375 0 4847.94
603.2853393554688 0 1182.511
603.3339233398438 0 756.3451
604.3381958007812 0 5563.6416
604.8327026367188 0 13759.474
605.3324584960938 0 9897.3545
605.8248291015625 0 5144.336
606.3147583007812 0 2062.1855
606.3453979492188 0 2664.2676
606.840087890625 0 1723.2854
607.35009765625 0 746.3486
611.3551025390625 0 5449.3965
611.8502807617188 0 29123.133
612.3518676757812 0 16084.278
612.8521118164062 0 7137.2764
613.3141479492188 0 905.60315
613.3521728515625 0 4002.8442
613.8456420898438 0 1533.3398
614.3529663085938 0 15387.357
614.828125 0 2127.8088
615.2890625 0 7625.8086
615.35595703125 0 5937.311
616.2911987304688 0 2678.9805
616.35400390625 0 872.2173
617.8192138671875 0 1292.7822
619.3557739257812 0 4152.0728
619.8492431640625 0 17226.756
620.3574829101562 0 25444.484
620.8451538085938 0 54897.875
621.345703125 0 39155.5
621.8468627929688 0 12643.605
622.3482666015625 0 3650.697
624.346923828125 0 3135.7217
624.844482421875 0 837.20966
625.3645629882812 0 7948.946
625.8357543945312 0 4857.467
626.3738403320312 0 225642.7 c 5
626.8353271484375 0 17790.31
627.3764038085938 0 66892.73
627.8397216796875 0 2626.083
628.3782958984375 0 12578.118
629.3804931640625 0 2148.5637
633.3457641601562 0 4778.8896
633.8453369140625 0 278463.22
634.3465576171875 0 192054.12
634.8477172851562 0 70152.695
635.3493041992188 0 21032.633
635.8492431640625 0 4713.0454
641.8545532226562 0 134131.77
642.3568725585938 0 161867.64
642.8587036132812 0 80948.23
643.3596801757812 0 28751.754
643.8606567382812 0 7494.492
644.3604125976562 0 1282.2312
652.383056640625 0 2044.6604
656.3533325195312 0 1147.7599 y Water loss 6
658.3433837890625 0 125597.56 z 6
659.34619140625 0 47075.625
660.3492431640625 0 9995.877
661.3501586914062 0 1584.7002
671.3746948242188 0 8060.1895
672.3778076171875 0 2957.1917
674.3621826171875 0 7440.5874 y 6
675.3643798828125 0 2415.0051
676.3698120117188 0 778.6532
695.4006958007812 0 962.84625
696.398193359375 0 1788.5612
697.4019165039062 0 917.89764
713.3741455078125 0 1434.5901 y Water loss 5
714.3804321289062 0 717.2138
715.3648071289062 0 32585.53 z 5
716.3678588867188 0 11073.333
717.3697509765625 0 2713.6738
723.4301147460938 0 1159.4167
731.3840942382812 0 9173.896 y 5
732.3880615234375 0 3006.7742
733.3880615234375 0 887.5954
738.4575805664062 0 1111.4517
739.4588623046875 0 1668.8947
740.452880859375 0 2410.9272
741.4607543945312 0 776.4937
767.4609375 0 1620.982
780.43603515625 0 1022.7091
781.46484375 0 4429.1523
782.4740600585938 0 38677.566 c 6
783.4767456054688 0 15497.973
784.4795532226562 0 4260.462
785.3944702148438 0 4076.5994 w 4
785.475341796875 0 1061.6885
786.400634765625 0 1862.8223
810.4244384765625 0 2783.8774 y Water loss 4
811.427978515625 0 5553.5625
812.4264526367188 0 1756.9905 z 4
819.4552612304688 0 1023.50604
823.4421997070312 0 1817.5004
824.4489135742188 0 7801.9243
825.4544067382812 0 4631.8286
826.4227905273438 0 2768.5852
827.4030151367188 0 2305.6155
828.4351806640625 0 9585.345 y 4
829.437255859375 0 4644.6606
830.4429931640625 0 2033.81
837.4915771484375 0 1067.611
838.5043334960938 0 714.1814
839.469970703125 0 3084.428
840.468994140625 0 2701.7327
851.5050659179688 0 1193.3069
852.5032348632812 0 2907.349
853.5109252929688 0 32324.777 c 7
854.513427734375 0 12188.092
855.5181884765625 0 3729.2324
856.4303588867188 0 1684.9286
857.4365234375 0 990.55524
882.4490356445312 0 968.97003 y Ammonia loss 3
883.4539794921875 0 139491.84 z 3
884.4573364257812 0 65681.586
885.459716796875 0 21196.732
886.4642944335938 0 4359.3345
899.472412109375 0 16755.617 y 3
900.4754638671875 0 9824.098
901.4774169921875 0 2665.3933
908.5618896484375 0 1186.2721
915.4491577148438 0 1026.8622
922.57763671875 0 942.18207
923.5726318359375 0 2417.665
924.5759887695312 0 2022.3397
936.5348510742188 0 1010.53705
939.4862670898438 0 1221.1306
940.4927368164062 0 1369.067
950.5758666992188 0 2172.6199
951.5824584960938 0 2371.2869
953.4838256835938 0 3894.391 w 2
954.4854736328125 0 1959.8241
955.473388671875 0 783.2473
956.4529418945312 0 780.6788
963.525146484375 0 962.2435
965.5787963867188 0 2120.5298
966.594482421875 0 32606.805 c 8
967.5979614257812 0 19106.965
968.6006469726562 0 5910.1787
969.609375 0 778.98566
980.549560546875 0 4136.443
981.5503540039062 0 2009.5712
1007.5928955078125 0 760.36755
1008.6092529296875 0 686.78705
1009.6055297851562 0 1550.1322
1011.5123901367188 0 102555.75 z 2
1012.5151977539062 0 59657.96
1012.6434326171875 0 908.6996
1013.517822265625 0 19007.617
1014.520751953125 0 4117.3433
1024.5185546875 0 1556.476
1027.530517578125 0 25967.36 y 2
1028.534423828125 0 13065.398
1029.5372314453125 0 4725.775
1034.6190185546875 0 1710.7048
1035.6109619140625 0 2379.9163
1036.6168212890625 0 743.75995
1037.59619140625 0 5997.8315
1038.5977783203125 0 2992.6565
1051.6234130859375 0 12278.853
1052.628173828125 0 29347.018
1053.6265869140625 0 17415.941
1054.6297607421875 0 6379.917
1055.63427734375 0 1656.2627
1062.596435546875 0 865.56036
1063.59716796875 0 970.161
1078.606689453125 0 1535.1215 c Ammonia loss 9
1079.6180419921875 0 22118.697
1080.6212158203125 0 13864.533
1081.5391845703125 0 3667.1113 w 1
1081.6331787109375 0 2603.7656
1082.54736328125 0 4752.285
1083.5518798828125 0 2201.7925
1095.63671875 0 21277.07 c 9
1096.6439208984375 0 58877.54
1097.646484375 0 30104.531
1098.649658203125 0 9409.526
1099.65283203125 0 3316.1184
1110.609619140625 0 757.8045
1111.60693359375 0 761.0104
1123.6092529296875 0 823.23706
1124.6295166015625 0 938.4539
1125.602783203125 0 1719.5813
1136.6260986328125 0 653.6148
1139.6148681640625 0 834.4501
1140.6314697265625 0 1101.3083
1151.614501953125 0 1141.856
1152.6087646484375 0 1859.4498
1153.6240234375 0 3748.8232
1154.635009765625 0 1909.2814
1155.623779296875 0 1143.1539
1166.6275634765625 0 971.34595 y Ammonia loss 1
1167.611083984375 0 7227.9946 z 1
1168.61962890625 0 42045.812
1169.623046875 0 28941.195
1170.6265869140625 0 9587.827
1171.6258544921875 0 3212.009
1180.617919921875 0 6787.3193
1181.6209716796875 0 5313.34
1182.6171875 0 4941.001
1183.625732421875 0 3533.0818 y 1
1184.63037109375 0 3063.861
1185.6336669921875 0 745.4717
1194.62255859375 0 989.26416
1195.6229248046875 0 3918.941
1196.6309814453125 0 3715.0222
1197.637451171875 0 1906.7577
1198.649658203125 0 1036.8296
1206.6756591796875 0 1854.3638
1207.6800537109375 0 934.5742
1208.652587890625 0 1344.4532
1211.6533203125 0 1247.0736
1212.6937255859375 0 985.412
1213.6817626953125 0 1164.0721
1222.70361328125 0 4183.082
1223.69873046875 0 9955.594
1224.6610107421875 0 71360.48
1225.6629638671875 0 54692.316
1226.666259765625 0 22024.693
1227.6673583984375 0 7977.1025
1228.6697998046875 0 1965.5104
1238.7003173828125 0 3645.76
1239.69287109375 0 15471.297
1240.6976318359375 0 17259.76
1241.69580078125 0 14072.048
1242.6956787109375 0 5925.6157
1243.6990966796875 0 1926.6704
1249.6796875 0 2359.2332
1250.662109375 0 22261.572
1251.664794921875 0 15939.266
1252.6695556640625 0 5954.2134
1253.6685791015625 0 1490.3002
1255.7161865234375 0 1331.509
1256.717529296875 0 4911.895
1257.721923828125 0 2590.114
1266.688232421875 0 23344.455
1267.688232421875 0 187055.36
1268.6907958984375 0 123964.734
1269.69384765625 0 47706.023
1270.6954345703125 0 14351.269
1271.69677734375 0 4489.8257
1282.6982421875 0 7945.315
1283.705810546875 0 39467.22
1284.7139892578125 0 158614.02
1285.716552734375 0 106583.96
1286.7198486328125 0 39502.824
1287.72265625 0 12153.189
1288.7181396484375 0 3645.9788
2110.315673828125 0 732.43756
2206.671875 0 688.0835
3083.003662109375 0 1059.8783

Spectrum Details

|  |  |
| --- | --- |
| Matched peaks? Matched peaks The total absolute number of peaks matched. Additionally in brackets the total fraction of peaks matched and the total number of peaks is shown. | 52 (10.28% of 506) |
| FDR? FDR The false discovery rate estimated for this peptide. It is calculated by matching all theoretical fragments with a non-integer shift with the raw peaks for this spectrum. This is done with 40 different shifts. The resulting percentage is the average number of annotated peaks over the number of annotated peaks with the correct spectrum. | 2.43% |
| Satellite FDR? Satellite FDR See the FDR for details on its calculation. This satellite ion specific FDR only contains the satellite ions (d/w) for I/L/J positions. | ∞ |
| PSM Score? PSM Score The PSM Score as given by Hecklib to this annotated spectrum. It is shown with three significant figures. | 464 |

#### Spectrum 5860? Spectrum 5860 The raw spectrum of this peptide as annotated by Hecklib. The fragments are coloured according to ion type (see legend). Any peaks with a star '\*' as text can be hovered over to see the full details, first the ion type second the mass shift type. By hovering over the amino acids in the peptide or ions in the legend the corresponding peaks are highlighted. By toggling the 'Unassigned' label you can turn the background (unassigned) peaks on or off in the plot. By updating the slider in the Ion legend you can update the spectrum to only show the top X% of the peaks with labels. The top X% means any peak that is within X% of the highest intensity. By dragging in the spectrum you can zoom in to a specific part of the spectrum and use 'Zoom Out' to get back to the original zoom level. The annotation of the spectrum is based on the given sequence in the peptides file and is done with different software so inconsistencies are likely. The peaks are annotated based on the given sequence, with 20 ppm tolerance.

Copy Data

##### Spectrum 5860 (TSV)

###### Preview

```
Loading example...
```

*Click on the button to copy the data to your clipboard.*

Mz MinMz MaxIntensity Max

WidthHeightPeptide font sizePeptide stroke widthSpectrum font sizeSpectrum stroke widthCompact peptide

Ion legend

wxyz

abcd

OtherUnassignedIon^Charge^_Position_Show for top:%

VRQAPGRAJEW

07.65e+41.53e+52.29e+53.06e+5

Zoom Out

c^+2^_3_y^+1^_1_c^+2^_4_c^+1^_2_c^+1^_2_y^+1^_2_c^+2^_7_c^+1^_3_c^+2^_7_c^+1^_3_y^+2^_7_y^+2^_7_c^+2^_8_c^+2^_8_c^+1^_4_c^+2^_9_c^+2^_9_y^+1^_4_c^+2^_10_c^+2^_10_c^+2^_10_c^+1^_5_z^+2^_10_y^+2^_10_c^+1^_6_z^+1^_5_y^+1^_5_y^+1^_6_z^+1^_6_y^+1^_6_c^+1^_7_w^+1^_7_y^+1^_7_z^+1^_7_y^+1^_7_c^+1^_8_z^+1^_8_y^+1^_8_w^+1^_9_c^+1^_9_z^+1^_9_y^+1^_9_c^+1^_10_w^+1^_10_c^+1^_10_z^+1^_10_y^+1^_10_

0593118617782371

Fragment Matches Table

Show background peaks

| Position | Ion type | Intensity | mz Theoretical | mz Error (Th) | mz Error (ppm) | Charge | Series Number |
| --- | --- | --- | --- | --- | --- | --- | --- |
| - | - | 5446 | 130.1 | - | - | 0 | - |
| - | - | 551 | 131.1 | - | - | 0 | - |
| - | - | 1151 | 132.1 | - | - | 0 | - |
| - | - | 1114 | 142.1 | - | - | 0 | - |
| - | - | 773.1 | 144.1 | - | - | 0 | - |
| - | - | 7711 | 146.1 | - | - | 0 | - |
| - | - | 1578 | 149 | - | - | 0 | - |
| - | - | 2947 | 157.1 | - | - | 0 | - |
| - | - | 1.698E+04 | 159.1 | - | - | 0 | - |
| - | - | 2180 | 160.1 | - | - | 0 | - |
| - | - | 667 | 166.1 | - | - | 0 | - |
| - | - | 660.5 | 170.1 | - | - | 0 | - |
| - | - | 458.7 | 171.1 | - | - | 0 | - |
| - | - | 915.8 | 173.4 | - | - | 0 | - |
| - | - | 545.8 | 174.1 | - | - | 0 | - |
| - | - | 1090 | 174.1 | - | - | 0 | - |
| - | - | 1803 | 185.1 | - | - | 0 | - |
| - | - | 1064 | 185.1 | - | - | 0 | - |
| - | - | 586.3 | 185.1 | - | - | 0 | - |
| - | - | 2894 | 187.1 | - | - | 0 | - |
| - | - | 6.073E+04 | 188.1 | - | - | 0 | - |
| - | - | 6543 | 189.1 | - | - | 0 | - |
| - | - | 500.9 | 192 | - | - | 0 | - |
| 3 | c | 1093 | 192.6 | 0.0003459 | 1.796 | +2 | 3 |
| - | - | 540.9 | 193.1 | - | - | 0 | - |
| - | - | 478.9 | 194.9 | - | - | 0 | - |
| - | - | 757.6 | 196.1 | - | - | 0 | - |
| - | - | 1070 | 197.1 | - | - | 0 | - |
| - | - | 1399 | 203.1 | - | - | 0 | - |
| 11 | y | 3.381E+04 | 205.1 | 0.0003038 | 1.481 | +1 | 1 |
| - | - | 4076 | 206.1 | - | - | 0 | - |
| - | - | 2.337E+04 | 209 | - | - | 0 | - |
| - | - | 4283 | 213.2 | - | - | 0 | - |
| - | - | 5441 | 214.1 | - | - | 0 | - |
| - | - | 3149 | 214.2 | - | - | 0 | - |
| - | - | 1136 | 214.6 | - | - | 0 | - |
| - | - | 934 | 215.1 | - | - | 0 | - |
| 4 | c | 1092 | 228.1 | 0.0003437 | 1.507 | +2 | 4 |
| - | - | 1600 | 228.2 | - | - | 0 | - |
| - | - | 560 | 235.3 | - | - | 0 | - |
| - | - | 1206 | 238.2 | - | - | 0 | - |
| - | - | 1.781E+04 | 239.2 | - | - | 0 | - |
| - | - | 1670 | 240.2 | - | - | 0 | - |
| - | - | 3592 | 242.2 | - | - | 0 | - |
| - | - | 529.5 | 243.2 | - | - | 0 | - |
| - | - | 1459 | 256.1 | - | - | 0 | - |
| 2 | c | 1.07E+04 | 256.2 | 0.0003531 | 1.379 | +1 | 2 |
| - | - | 2442 | 270.1 | - | - | 0 | - |
| 2 | c | 3.846E+04 | 273.2 | 0.0002933 | 1.074 | +1 | 2 |
| - | - | 5503 | 274.2 | - | - | 0 | - |
| - | - | 1489 | 298.1 | - | - | 0 | - |
| - | - | 1231 | 298.2 | - | - | 0 | - |
| - | - | 1.061E+04 | 299.2 | - | - | 0 | - |
| - | - | 1828 | 300.2 | - | - | 0 | - |
| - | - | 819.3 | 307.9 | - | - | 0 | - |
| - | - | 814.3 | 312.2 | - | - | 0 | - |
| - | - | 1642 | 312.7 | - | - | 0 | - |
| 10 | y | 7417 | 316.1 | 0.0004257 | 1.346 | +1 | 2 |
| - | - | 1.657E+04 | 318.9 | - | - | 0 | - |
| - | - | 731.8 | 339.2 | - | - | 0 | - |
| - | - | 6144 | 341 | - | - | 0 | - |
| - | - | 1054 | 341.2 | - | - | 0 | - |
| - | - | 2728 | 356.2 | - | - | 0 | - |
| - | - | 1912 | 357.2 | - | - | 0 | - |
| - | - | 1.878E+04 | 359 | - | - | 0 | - |
| - | - | 2526 | 361.7 | - | - | 0 | - |
| - | - | 1390 | 362.2 | - | - | 0 | - |
| - | - | 758.7 | 362.7 | - | - | 0 | - |
| - | - | 1439 | 367.2 | - | - | 0 | - |
| - | - | 1454 | 369.2 | - | - | 0 | - |
| - | - | 7683 | 369.7 | - | - | 0 | - |
| - | - | 4229 | 370.2 | - | - | 0 | - |
| - | - | 859.9 | 370.3 | - | - | 0 | - |
| - | - | 1138 | 370.7 | - | - | 0 | - |
| - | - | 779.6 | 375.9 | - | - | 0 | - |
| 7 | c | 2186 | 383.2 | 0.0002284 | 0.5961 | +2 | 7 |
| - | - | 2006 | 383.7 | - | - | 0 | - |
| 3 | c | 6468 | 384.2 | 9.472E-05 | 0.2465 | +1 | 3 |
| - | - | 1476 | 385.2 | - | - | 0 | - |
| - | - | 802.4 | 385.3 | - | - | 0 | - |
| - | - | 5.558E+04 | 391.2 | - | - | 0 | - |
| 7 | c | 1.569E+05 | 391.7 | 0.0001384 | 0.3532 | +2 | 7 |
| - | - | 6.133E+04 | 392.2 | - | - | 0 | - |
| - | - | 1.196E+04 | 392.7 | - | - | 0 | - |
| - | - | 1188 | 393.2 | - | - | 0 | - |
| - | - | 602.1 | 397.7 | - | - | 0 | - |
| 3 | c | 1.265E+05 | 401.3 | 0.0005842 | 1.456 | +1 | 3 |
| - | - | 2.43E+04 | 402.3 | - | - | 0 | - |
| - | - | 4000 | 403.3 | - | - | 0 | - |
| - | - | 2760 | 404.7 | - | - | 0 | - |
| - | - | 1534 | 405.3 | - | - | 0 | - |
| 5 | y | 2007 | 405.7 | 0.0003397 | 0.8374 | +2 | 7 |
| - | - | 1091 | 405.8 | - | - | 0 | - |
| - | - | 925.8 | 406.2 | - | - | 0 | - |
| - | - | 5036 | 406.9 | - | - | 0 | - |
| - | - | 3441 | 407.2 | - | - | 0 | - |
| - | - | 1013 | 407.6 | - | - | 0 | - |
| - | - | 1101 | 410.2 | - | - | 0 | - |
| - | - | 4671 | 412.9 | - | - | 0 | - |
| - | - | 1959 | 413.2 | - | - | 0 | - |
| - | - | 982.4 | 413.3 | - | - | 0 | - |
| 5 | y | 2643 | 414.7 | 0.0003369 | 0.8124 | +2 | 7 |
| - | - | 642.4 | 415.2 | - | - | 0 | - |
| - | - | 697.3 | 418.3 | - | - | 0 | - |
| 8 | c | 4587 | 418.7 | 0.001082 | 2.583 | +2 | 8 |
| - | - | 2126 | 419.2 | - | - | 0 | - |
| - | - | 1074 | 419.7 | - | - | 0 | - |
| - | - | 1790 | 422.2 | - | - | 0 | - |
| - | - | 1128 | 422.6 | - | - | 0 | - |
| - | - | 1.886E+04 | 426.8 | - | - | 0 | - |
| 8 | c | 1.223E+05 | 427.3 | 0.0005939 | 1.39 | +2 | 8 |
| - | - | 5.191E+04 | 427.8 | - | - | 0 | - |
| - | - | 1.405E+04 | 428.3 | - | - | 0 | - |
| - | - | 1972 | 428.8 | - | - | 0 | - |
| - | - | 2747 | 428.9 | - | - | 0 | - |
| - | - | 1.724E+04 | 429.1 | - | - | 0 | - |
| - | - | 1376 | 440.3 | - | - | 0 | - |
| - | - | 717.5 | 440.8 | - | - | 0 | - |
| - | - | 1244 | 453.8 | - | - | 0 | - |
| 4 | c | 9696 | 455.3 | 5.979E-05 | 0.1313 | +1 | 4 |
| - | - | 1641 | 456.3 | - | - | 0 | - |
| - | - | 1.469E+04 | 461.3 | - | - | 0 | - |
| - | - | 9969 | 461.8 | - | - | 0 | - |
| - | - | 3085 | 462.3 | - | - | 0 | - |
| - | - | 960.4 | 467.3 | - | - | 0 | - |
| - | - | 712.3 | 468.8 | - | - | 0 | - |
| - | - | 702.8 | 469.3 | - | - | 0 | - |
| - | - | 1125 | 472.3 | - | - | 0 | - |
| - | - | 2053 | 474.8 | - | - | 0 | - |
| 9 | c | 3.47E+04 | 475.3 | 0.0007061 | 1.486 | +2 | 9 |
| - | - | 1.282E+04 | 475.8 | - | - | 0 | - |
| - | - | 5974 | 476.3 | - | - | 0 | - |
| - | - | 4669 | 483.3 | - | - | 0 | - |
| 9 | c | 6.28E+04 | 483.8 | 0.0004931 | 1.019 | +2 | 9 |
| - | - | 3.525E+04 | 484.3 | - | - | 0 | - |
| - | - | 1.041E+04 | 484.8 | - | - | 0 | - |
| - | - | 1875 | 485.3 | - | - | 0 | - |
| - | - | 576.5 | 496.3 | - | - | 0 | - |
| - | - | 3.399E+04 | 496.8 | - | - | 0 | - |
| - | - | 2.13E+04 | 497.3 | - | - | 0 | - |
| - | - | 6812 | 497.8 | - | - | 0 | - |
| - | - | 6277 | 498.3 | - | - | 0 | - |
| - | - | 1955 | 498.3 | - | - | 0 | - |
| - | - | 4346 | 498.8 | - | - | 0 | - |
| - | - | 1103 | 499.3 | - | - | 0 | - |
| - | - | 2300 | 499.3 | - | - | 0 | - |
| - | - | 3437 | 504.8 | - | - | 0 | - |
| - | - | 954.3 | 505.3 | - | - | 0 | - |
| - | - | 805.7 | 512.3 | - | - | 0 | - |
| - | - | 2315 | 517.8 | - | - | 0 | - |
| 8 | y | 1283 | 518.3 | 0.0009762 | 1.884 | +1 | 4 |
| - | - | 985.6 | 518.3 | - | - | 0 | - |
| - | - | 2113 | 518.8 | - | - | 0 | - |
| - | - | 1804 | 524.3 | - | - | 0 | - |
| - | - | 3763 | 525.3 | - | - | 0 | - |
| - | - | 3.026E+04 | 525.8 | - | - | 0 | - |
| - | - | 2.386E+04 | 526.3 | - | - | 0 | - |
| - | - | 8774 | 526.8 | - | - | 0 | - |
| - | - | 2792 | 527.3 | - | - | 0 | - |
| - | - | 795.7 | 529.3 | - | - | 0 | - |
| - | - | 2034 | 531.3 | - | - | 0 | - |
| - | - | 1596 | 531.8 | - | - | 0 | - |
| - | - | 617.2 | 532.3 | - | - | 0 | - |
| - | - | 846 | 532.8 | - | - | 0 | - |
| 10 | c | 1005 | 539.3 | 0.004138 | 7.673 | +2 | 10 |
| 10 | c | 1.678E+05 | 539.8 | 0.0006803 | 1.26 | +2 | 10 |
| - | - | 9.736E+04 | 540.3 | - | - | 0 | - |
| - | - | 3.493E+04 | 540.8 | - | - | 0 | - |
| - | - | 8894 | 541.3 | - | - | 0 | - |
| - | - | 1725 | 541.8 | - | - | 0 | - |
| - | - | 798.6 | 542.3 | - | - | 0 | - |
| - | - | 4480 | 542.3 | - | - | 0 | - |
| - | - | 1072 | 543.3 | - | - | 0 | - |
| - | - | 709.1 | 547.3 | - | - | 0 | - |
| - | - | 2107 | 547.8 | - | - | 0 | - |
| 10 | c | 3.028E+05 | 548.3 | 0.0007114 | 1.297 | +2 | 10 |
| - | - | 1.719E+05 | 548.8 | - | - | 0 | - |
| - | - | 5.635E+04 | 549.3 | - | - | 0 | - |
| - | - | 769.6 | 549.8 | - | - | 0 | - |
| - | - | 1.458E+04 | 549.8 | - | - | 0 | - |
| - | - | 2356 | 550.3 | - | - | 0 | - |
| - | - | 1.398E+04 | 555.3 | - | - | 0 | - |
| - | - | 2338 | 555.3 | - | - | 0 | - |
| - | - | 8538 | 555.8 | - | - | 0 | - |
| - | - | 2319 | 556.3 | - | - | 0 | - |
| - | - | 1230 | 558.3 | - | - | 0 | - |
| - | - | 1.081E+04 | 561.3 | - | - | 0 | - |
| - | - | 7202 | 561.8 | - | - | 0 | - |
| - | - | 1655 | 562.3 | - | - | 0 | - |
| - | - | 960.1 | 562.8 | - | - | 0 | - |
| - | - | 918.5 | 562.8 | - | - | 0 | - |
| - | - | 866.7 | 563.3 | - | - | 0 | - |
| - | - | 542.5 | 567.3 | - | - | 0 | - |
| - | - | 3195 | 568.3 | - | - | 0 | - |
| - | - | 622.4 | 568.6 | - | - | 0 | - |
| - | - | 1631 | 568.8 | - | - | 0 | - |
| 5 | c | 4.809E+04 | 569.4 | 0.0003062 | 0.5377 | +1 | 5 |
| - | - | 792 | 570.3 | - | - | 0 | - |
| - | - | 1.419E+04 | 570.4 | - | - | 0 | - |
| - | - | 1001 | 571.3 | - | - | 0 | - |
| - | - | 3111 | 571.4 | - | - | 0 | - |
| - | - | 4204 | 575.8 | - | - | 0 | - |
| - | - | 4036 | 576.3 | - | - | 0 | - |
| - | - | 1.751E+04 | 576.8 | - | - | 0 | - |
| - | - | 1.592E+04 | 577.3 | - | - | 0 | - |
| - | - | 6811 | 577.8 | - | - | 0 | - |
| - | - | 1529 | 578.3 | - | - | 0 | - |
| - | - | 5211 | 582.4 | - | - | 0 | - |
| - | - | 1476 | 582.8 | - | - | 0 | - |
| - | - | 1557 | 583.3 | - | - | 0 | - |
| - | - | 1114 | 583.8 | - | - | 0 | - |
| 2 | z | 1.614E+04 | 584.3 | 0.001019 | 1.745 | +2 | 10 |
| - | - | 8948 | 584.8 | - | - | 0 | - |
| - | - | 954.4 | 584.9 | - | - | 0 | - |
| - | - | 2511 | 585.3 | - | - | 0 | - |
| - | - | 796.2 | 585.8 | - | - | 0 | - |
| - | - | 1438 | 586.3 | - | - | 0 | - |
| - | - | 718.1 | 590.3 | - | - | 0 | - |
| - | - | 1039 | 590.9 | - | - | 0 | - |
| - | - | 1005 | 591.3 | - | - | 0 | - |
| - | - | 769.1 | 591.8 | - | - | 0 | - |
| 2 | y | 1364 | 592.3 | 0.0001638 | 0.2766 | +2 | 10 |
| - | - | 606.3 | 593.3 | - | - | 0 | - |
| - | - | 1143 | 596.4 | - | - | 0 | - |
| - | - | 2138 | 597.8 | - | - | 0 | - |
| - | - | 5444 | 598.3 | - | - | 0 | - |
| - | - | 3041 | 598.8 | - | - | 0 | - |
| - | - | 4352 | 599.3 | - | - | 0 | - |
| - | - | 1371 | 600.3 | - | - | 0 | - |
| - | - | 3599 | 602.3 | - | - | 0 | - |
| - | - | 745.7 | 603.3 | - | - | 0 | - |
| - | - | 4969 | 604.3 | - | - | 0 | - |
| - | - | 1.254E+04 | 604.8 | - | - | 0 | - |
| - | - | 9411 | 605.3 | - | - | 0 | - |
| - | - | 4579 | 605.8 | - | - | 0 | - |
| - | - | 2160 | 606.3 | - | - | 0 | - |
| - | - | 1770 | 606.8 | - | - | 0 | - |
| - | - | 847.1 | 607.4 | - | - | 0 | - |
| - | - | 3711 | 611.4 | - | - | 0 | - |
| - | - | 2.387E+04 | 611.9 | - | - | 0 | - |
| - | - | 1.392E+04 | 612.4 | - | - | 0 | - |
| - | - | 5422 | 612.9 | - | - | 0 | - |
| - | - | 3508 | 613.4 | - | - | 0 | - |
| - | - | 1555 | 613.8 | - | - | 0 | - |
| - | - | 1.317E+04 | 614.4 | - | - | 0 | - |
| - | - | 2170 | 614.8 | - | - | 0 | - |
| - | - | 8188 | 615.3 | - | - | 0 | - |
| - | - | 3567 | 615.4 | - | - | 0 | - |
| - | - | 2542 | 616.3 | - | - | 0 | - |
| - | - | 1044 | 616.4 | - | - | 0 | - |
| - | - | 2593 | 619.4 | - | - | 0 | - |
| - | - | 1.416E+04 | 619.8 | - | - | 0 | - |
| - | - | 1.944E+04 | 620.4 | - | - | 0 | - |
| - | - | 4.798E+04 | 620.8 | - | - | 0 | - |
| - | - | 3.034E+04 | 621.3 | - | - | 0 | - |
| - | - | 1.261E+04 | 621.8 | - | - | 0 | - |
| - | - | 3579 | 622.3 | - | - | 0 | - |
| - | - | 724.5 | 622.9 | - | - | 0 | - |
| - | - | 2904 | 624.3 | - | - | 0 | - |
| - | - | 1046 | 624.8 | - | - | 0 | - |
| - | - | 7154 | 625.4 | - | - | 0 | - |
| - | - | 3651 | 625.8 | - | - | 0 | - |
| 6 | c | 1.891E+05 | 626.4 | 0.0008761 | 1.399 | +1 | 6 |
| - | - | 1.635E+04 | 626.8 | - | - | 0 | - |
| - | - | 5.995E+04 | 627.4 | - | - | 0 | - |
| - | - | 843 | 627.8 | - | - | 0 | - |
| - | - | 628.2 | 628.3 | - | - | 0 | - |
| - | - | 1.187E+04 | 628.4 | - | - | 0 | - |
| - | - | 1320 | 629.4 | - | - | 0 | - |
| - | - | 3728 | 633.4 | - | - | 0 | - |
| - | - | 2.309E+05 | 633.8 | - | - | 0 | - |
| - | - | 1.58E+05 | 634.3 | - | - | 0 | - |
| - | - | 6.481E+04 | 634.8 | - | - | 0 | - |
| - | - | 1.845E+04 | 635.3 | - | - | 0 | - |
| - | - | 3869 | 635.8 | - | - | 0 | - |
| - | - | 1.087E+05 | 641.9 | - | - | 0 | - |
| - | - | 1.367E+05 | 642.4 | - | - | 0 | - |
| - | - | 7.244E+04 | 642.9 | - | - | 0 | - |
| - | - | 2.727E+04 | 643.4 | - | - | 0 | - |
| - | - | 6072 | 643.9 | - | - | 0 | - |
| - | - | 1394 | 644.4 | - | - | 0 | - |
| - | - | 1533 | 652.4 | - | - | 0 | - |
| 7 | z | 1.086E+05 | 658.3 | 0.0004381 | 0.6654 | +1 | 5 |
| - | - | 4.404E+04 | 659.3 | - | - | 0 | - |
| - | - | 8274 | 660.3 | - | - | 0 | - |
| - | - | 1004 | 661.4 | - | - | 0 | - |
| - | - | 6981 | 671.4 | - | - | 0 | - |
| - | - | 1795 | 672.4 | - | - | 0 | - |
| 7 | y | 6913 | 674.4 | 0.0002077 | 0.3079 | +1 | 5 |
| - | - | 2271 | 675.4 | - | - | 0 | - |
| - | - | 627 | 682.4 | - | - | 0 | - |
| - | - | 1190 | 695.4 | - | - | 0 | - |
| - | - | 996.9 | 696.4 | - | - | 0 | - |
| 6 | y | 1627 | 713.4 | 0.00176 | 2.467 | +1 | 6 |
| 6 | z | 2.568E+04 | 715.4 | 0.0005198 | 0.7266 | +1 | 6 |
| - | - | 1.019E+04 | 716.4 | - | - | 0 | - |
| - | - | 1982 | 717.4 | - | - | 0 | - |
| - | - | 1181 | 723.4 | - | - | 0 | - |
| 6 | y | 6433 | 731.4 | 0.0004725 | 0.646 | +1 | 6 |
| - | - | 2814 | 732.4 | - | - | 0 | - |
| - | - | 835.1 | 738.5 | - | - | 0 | - |
| - | - | 2158 | 739.5 | - | - | 0 | - |
| - | - | 1352 | 740.5 | - | - | 0 | - |
| - | - | 1652 | 767.5 | - | - | 0 | - |
| - | - | 1412 | 780.4 | - | - | 0 | - |
| - | - | 3556 | 781.5 | - | - | 0 | - |
| 7 | c | 3.575E+04 | 782.5 | 0.000351 | 0.4486 | +1 | 7 |
| - | - | 1.696E+04 | 783.5 | - | - | 0 | - |
| - | - | 3540 | 784.5 | - | - | 0 | - |
| 5 | w | 2705 | 785.4 | 0.0005707 | 0.7267 | +1 | 7 |
| - | - | 907.3 | 785.5 | - | - | 0 | - |
| - | - | 2197 | 786.4 | - | - | 0 | - |
| - | - | 883.3 | 787.4 | - | - | 0 | - |
| - | - | 659.5 | 796.4 | - | - | 0 | - |
| 5 | y | 2766 | 810.4 | 0.001242 | 1.533 | +1 | 7 |
| - | - | 4546 | 811.4 | - | - | 0 | - |
| - | - | 630.8 | 811.5 | - | - | 0 | - |
| 5 | z | 1187 | 812.4 | 0.009279 | 11.42 | +1 | 7 |
| - | - | 961.8 | 819.5 | - | - | 0 | - |
| - | - | 1093 | 823.4 | - | - | 0 | - |
| - | - | 5420 | 824.5 | - | - | 0 | - |
| - | - | 5225 | 825.5 | - | - | 0 | - |
| - | - | 2580 | 826.4 | - | - | 0 | - |
| - | - | 1214 | 827.4 | - | - | 0 | - |
| 5 | y | 9711 | 828.4 | 0.0001674 | 0.202 | +1 | 7 |
| - | - | 4343 | 829.4 | - | - | 0 | - |
| - | - | 992.3 | 830.4 | - | - | 0 | - |
| - | - | 1556 | 838.5 | - | - | 0 | - |
| - | - | 2565 | 839.5 | - | - | 0 | - |
| - | - | 1450 | 840.5 | - | - | 0 | - |
| - | - | 1294 | 851.5 | - | - | 0 | - |
| - | - | 2660 | 852.5 | - | - | 0 | - |
| - | - | 944.4 | 853 | - | - | 0 | - |
| 8 | c | 2.799E+04 | 853.5 | 0.0002637 | 0.309 | +1 | 8 |
| - | - | 1.238E+04 | 854.5 | - | - | 0 | - |
| - | - | 3652 | 855.5 | - | - | 0 | - |
| - | - | 970.2 | 856.4 | - | - | 0 | - |
| 4 | z | 1.152E+05 | 883.5 | 6.344E-05 | 0.07181 | +1 | 8 |
| - | - | 5.996E+04 | 884.5 | - | - | 0 | - |
| - | - | 1.718E+04 | 885.5 | - | - | 0 | - |
| - | - | 4195 | 886.5 | - | - | 0 | - |
| 4 | y | 1.485E+04 | 899.5 | 0.0003549 | 0.3946 | +1 | 8 |
| - | - | 6755 | 900.5 | - | - | 0 | - |
| - | - | 2720 | 901.5 | - | - | 0 | - |
| - | - | 1147 | 908.6 | - | - | 0 | - |
| - | - | 944.1 | 915.4 | - | - | 0 | - |
| - | - | 722 | 922.6 | - | - | 0 | - |
| - | - | 1193 | 923.6 | - | - | 0 | - |
| - | - | 1695 | 924.6 | - | - | 0 | - |
| - | - | 630 | 936.5 | - | - | 0 | - |
| - | - | 949.9 | 939.5 | - | - | 0 | - |
| - | - | 2554 | 950.6 | - | - | 0 | - |
| - | - | 1869 | 951.6 | - | - | 0 | - |
| - | - | 948.2 | 952.6 | - | - | 0 | - |
| 3 | w | 2332 | 953.5 | 0.001898 | 1.99 | +1 | 9 |
| - | - | 2057 | 954.5 | - | - | 0 | - |
| - | - | 1523 | 955.5 | - | - | 0 | - |
| - | - | 1320 | 963.5 | - | - | 0 | - |
| - | - | 1118 | 965.6 | - | - | 0 | - |
| 9 | c | 3.058E+04 | 966.6 | 0.0005264 | 0.5446 | +1 | 9 |
| - | - | 1.704E+04 | 967.6 | - | - | 0 | - |
| - | - | 4293 | 968.6 | - | - | 0 | - |
| - | - | 736.5 | 969.6 | - | - | 0 | - |
| - | - | 2765 | 980.6 | - | - | 0 | - |
| - | - | 2039 | 981.6 | - | - | 0 | - |
| - | - | 684.8 | 992.6 | - | - | 0 | - |
| - | - | 721.5 | 1009 | - | - | 0 | - |
| - | - | 1122 | 1010 | - | - | 0 | - |
| 3 | z | 9.089E+04 | 1012 | 0.0001693 | 0.1673 | +1 | 9 |
| - | - | 4.941E+04 | 1013 | - | - | 0 | - |
| - | - | 1.79E+04 | 1014 | - | - | 0 | - |
| - | - | 3193 | 1015 | - | - | 0 | - |
| - | - | 980 | 1016 | - | - | 0 | - |
| 3 | y | 2.144E+04 | 1028 | 0.0002166 | 0.2108 | +1 | 9 |
| - | - | 1.13E+04 | 1029 | - | - | 0 | - |
| - | - | 4115 | 1030 | - | - | 0 | - |
| - | - | 920.9 | 1035 | - | - | 0 | - |
| - | - | 2933 | 1036 | - | - | 0 | - |
| - | - | 1042 | 1037 | - | - | 0 | - |
| - | - | 5801 | 1038 | - | - | 0 | - |
| - | - | 2654 | 1039 | - | - | 0 | - |
| - | - | 1.077E+04 | 1052 | - | - | 0 | - |
| - | - | 2.803E+04 | 1053 | - | - | 0 | - |
| - | - | 1.76E+04 | 1054 | - | - | 0 | - |
| - | - | 5678 | 1055 | - | - | 0 | - |
| - | - | 1406 | 1056 | - | - | 0 | - |
| - | - | 768.7 | 1064 | - | - | 0 | - |
| 10 | c | 1067 | 1079 | 0.001313 | 1.217 | +1 | 10 |
| - | - | 2.266E+04 | 1080 | - | - | 0 | - |
| - | - | 1.304E+04 | 1081 | - | - | 0 | - |
| 2 | w | 4381 | 1082 | 0.001016 | 0.9391 | +1 | 10 |
| - | - | 2558 | 1082 | - | - | 0 | - |
| - | - | 2991 | 1083 | - | - | 0 | - |
| 10 | c | 1.986E+04 | 1096 | 0.0007001 | 0.639 | +1 | 10 |
| - | - | 5.44E+04 | 1097 | - | - | 0 | - |
| - | - | 2.852E+04 | 1098 | - | - | 0 | - |
| - | - | 8428 | 1099 | - | - | 0 | - |
| - | - | 1332 | 1100 | - | - | 0 | - |
| - | - | 867.7 | 1111 | - | - | 0 | - |
| - | - | 970.5 | 1126 | - | - | 0 | - |
| - | - | 1009 | 1127 | - | - | 0 | - |
| - | - | 818.5 | 1141 | - | - | 0 | - |
| - | - | 978.7 | 1152 | - | - | 0 | - |
| - | - | 1945 | 1153 | - | - | 0 | - |
| - | - | 3240 | 1154 | - | - | 0 | - |
| - | - | 1621 | 1155 | - | - | 0 | - |
| 2 | z | 6303 | 1168 | 0.001793 | 1.536 | +1 | 10 |
| - | - | 3.633E+04 | 1169 | - | - | 0 | - |
| - | - | 2.485E+04 | 1170 | - | - | 0 | - |
| - | - | 8571 | 1171 | - | - | 0 | - |
| - | - | 1825 | 1172 | - | - | 0 | - |
| - | - | 999.5 | 1173 | - | - | 0 | - |
| - | - | 5156 | 1181 | - | - | 0 | - |
| - | - | 3722 | 1182 | - | - | 0 | - |
| - | - | 4269 | 1183 | - | - | 0 | - |
| 2 | y | 3354 | 1184 | 0.005869 | 4.958 | +1 | 10 |
| - | - | 1958 | 1185 | - | - | 0 | - |
| - | - | 863.2 | 1186 | - | - | 0 | - |
| - | - | 4193 | 1196 | - | - | 0 | - |
| - | - | 2684 | 1197 | - | - | 0 | - |
| - | - | 1726 | 1198 | - | - | 0 | - |
| - | - | 1656 | 1207 | - | - | 0 | - |
| - | - | 798.5 | 1209 | - | - | 0 | - |
| - | - | 1464 | 1210 | - | - | 0 | - |
| - | - | 1366 | 1212 | - | - | 0 | - |
| - | - | 1174 | 1213 | - | - | 0 | - |
| - | - | 4003 | 1223 | - | - | 0 | - |
| - | - | 9922 | 1224 | - | - | 0 | - |
| - | - | 6.115E+04 | 1225 | - | - | 0 | - |
| - | - | 4.82E+04 | 1226 | - | - | 0 | - |
| - | - | 2.114E+04 | 1227 | - | - | 0 | - |
| - | - | 5928 | 1228 | - | - | 0 | - |
| - | - | 1871 | 1229 | - | - | 0 | - |
| - | - | 3167 | 1239 | - | - | 0 | - |
| - | - | 1.518E+04 | 1240 | - | - | 0 | - |
| - | - | 1.47E+04 | 1241 | - | - | 0 | - |
| - | - | 1.28E+04 | 1242 | - | - | 0 | - |
| - | - | 4906 | 1243 | - | - | 0 | - |
| - | - | 1898 | 1244 | - | - | 0 | - |
| - | - | 867.7 | 1245 | - | - | 0 | - |
| - | - | 1870 | 1250 | - | - | 0 | - |
| - | - | 2.068E+04 | 1251 | - | - | 0 | - |
| - | - | 1.266E+04 | 1252 | - | - | 0 | - |
| - | - | 871.8 | 1252 | - | - | 0 | - |
| - | - | 6687 | 1253 | - | - | 0 | - |
| - | - | 1623 | 1254 | - | - | 0 | - |
| - | - | 925.3 | 1256 | - | - | 0 | - |
| - | - | 5384 | 1257 | - | - | 0 | - |
| - | - | 2460 | 1258 | - | - | 0 | - |
| - | - | 1291 | 1259 | - | - | 0 | - |
| - | - | 868 | 1266 | - | - | 0 | - |
| - | - | 2.08E+04 | 1267 | - | - | 0 | - |
| - | - | 1.684E+05 | 1268 | - | - | 0 | - |
| - | - | 1.157E+05 | 1269 | - | - | 0 | - |
| - | - | 4.292E+04 | 1270 | - | - | 0 | - |
| - | - | 1.251E+04 | 1271 | - | - | 0 | - |
| - | - | 3870 | 1272 | - | - | 0 | - |
| - | - | 6290 | 1283 | - | - | 0 | - |
| - | - | 3.615E+04 | 1284 | - | - | 0 | - |
| - | - | 1.509E+05 | 1285 | - | - | 0 | - |
| - | - | 9.753E+04 | 1286 | - | - | 0 | - |
| - | - | 3.816E+04 | 1287 | - | - | 0 | - |
| - | - | 1.141E+04 | 1288 | - | - | 0 | - |
| - | - | 3408 | 1289 | - | - | 0 | - |
| - | - | 739.7 | 2348 | - | - | 0 | - |

m/z Charge Intensity FragmentType MassShift Position
130.06546020507812 0 5446.4233
131.06942749023438 0 550.9958
132.08111572265625 0 1151.3806
142.09791564941406 0 1113.7517
144.08108520507812 0 773.11237
146.0603790283203 0 7710.8467
149.04531860351562 0 1578.2217
157.1087188720703 0 2946.7195
159.09197998046875 0 16977.332
160.0954132080078 0 2180.1794
166.0866241455078 0 666.9928
170.06069946289062 0 660.50244
171.12440490722656 0 458.66254
173.43861389160156 0 915.8302
174.1029510498047 0 545.76324
174.13516235351562 0 1089.848
185.10366821289062 0 1802.6681
185.1175994873047 0 1064.3337
185.13951110839844 0 586.2614
187.08700561523438 0 2893.7014
188.07093811035156 0 60730.59
189.0743408203125 0 6542.642
191.96656799316406 0 500.90833
192.62167358398438 0 1093.1265 c Ammonia loss 2
193.12237548828125 0 540.85156
194.92173767089844 0 478.86975
196.14422607421875 0 757.57007
197.1399688720703 0 1069.9697
203.1280975341797 0 1399.2137
205.0974578857422 0 33812.344 y 10
206.10093688964844 0 4076.3694
208.95347595214844 0 23365.475
213.17135620117188 0 4283.4756
214.14279174804688 0 5441.4883
214.1552734375 0 3149.0754
214.64419555664062 0 1135.6997
215.1381072998047 0 933.9864
228.14022827148438 0 1092.1018 c Ammonia loss 3
228.1822509765625 0 1600.0739
235.2523956298828 0 560.0083
238.16648864746094 0 1206.3984
239.15066528320312 0 17811.002
240.15414428710938 0 1669.5674
242.161376953125 0 3591.5244
243.168212890625 0 529.50446
256.120849609375 0 1459.4641
256.1771545410156 0 10704.886 c Ammonia loss 1
270.1239013671875 0 2442.2783
273.2036437988281 0 38464.82 c 1
274.206787109375 0 5503.487
298.11907958984375 0 1489.306
298.22412109375 0 1231.3134
299.21954345703125 0 10614.262
300.22314453125 0 1828.1904
307.8643493652344 0 819.28735
312.1903381347656 0 814.34894
312.6776428222656 0 1641.6356
316.1296081542969 0 7417.2026 y Water loss 9
318.92266845703125 0 16570.254
339.215087890625 0 731.83594
341.01849365234375 0 6144.3115
341.2289733886719 0 1054.3145
356.2410583496094 0 2728.37
357.24761962890625 0 1912.1033
359.0285949707031 0 18782.932
361.7250671386719 0 2525.9736
362.2253723144531 0 1390.3077
362.72528076171875 0 758.74915
367.20880126953125 0 1438.9229
369.2303771972656 0 1454.4128
369.73406982421875 0 7682.878
370.23577880859375 0 4229.3438
370.2564697265625 0 859.8624
370.7403564453125 0 1137.5325
375.86199951171875 0 779.5564
383.2273254394531 0 2185.9568 c Ammonia loss 6
383.728759765625 0 2005.8499
384.2354736328125 0 6468.0894 c Ammonia loss 2
385.2392578125 0 1476.3922
385.27911376953125 0 802.37427
391.2372741699219 0 55579.543
391.740966796875 0 156860.72 c 6
392.2425231933594 0 61333.75
392.74359130859375 0 11957.316
393.2444152832031 0 1188.2225
397.7464294433594 0 602.1458
401.26251220703125 0 126483.086 c 2
402.265380859375 0 24299.453
403.267333984375 0 3999.6426
404.7499694824219 0 2759.7837
405.2520446777344 0 1533.7395
405.7168273925781 0 2007.1709 y Water loss 4
405.75225830078125 0 1090.7458
406.2173767089844 0 925.81494
406.9008483886719 0 5035.6084
407.2344970703125 0 3441.302
407.5691833496094 0 1013.3551
410.2330627441406 0 1101.0212
412.90460205078125 0 4671.26
413.2386474609375 0 1958.5973
413.2728576660156 0 982.39575
414.72210693359375 0 2643.28 y 4
415.2231140136719 0 642.43365
418.25445556640625 0 697.3462
418.7471923828125 0 4586.9395 c Ammonia loss 7
419.247802734375 0 2125.593
419.7485046386719 0 1074.1432
422.23565673828125 0 1790.4755
422.568603515625 0 1128.1384
426.75616455078125 0 18861.334
427.2599792480469 0 122307.555 c 7
427.7611389160156 0 51912.383
428.2628479003906 0 14052.685
428.76385498046875 0 1972.1128
428.8915710449219 0 2746.5933
429.08941650390625 0 17239.906
440.2677307128906 0 1375.837
440.7688903808594 0 717.4648
453.78759765625 0 1243.6925
455.2725524902344 0 9695.551 c Ammonia loss 3
456.276611328125 0 1641.0585
461.2911376953125 0 14692.454
461.7936096191406 0 9968.55
462.2950439453125 0 3084.909
467.3075256347656 0 960.4374
468.7740478515625 0 712.331
469.2717590332031 0 702.76843
472.2888488769531 0 1125.132
474.7966003417969 0 2053.1646
475.2888488769531 0 34701.094 c Ammonia loss 8
475.78997802734375 0 12819.182
476.29144287109375 0 5973.661
483.2984924316406 0 4668.9707
483.8019104003906 0 62798.914 c 8
484.30340576171875 0 35245.305
484.80474853515625 0 10405.254
485.3045349121094 0 1875.2585
496.3118591308594 0 576.4933
496.8099060058594 0 33987.816
497.31121826171875 0 21296.314
497.8121337890625 0 6811.9756
498.28460693359375 0 6276.9043
498.3159484863281 0 1954.8745
498.78564453125 0 4345.691
499.2873229980469 0 1103.0348
499.3233642578125 0 2300.4438
504.7903137207031 0 3437.06
505.2893981933594 0 954.3157
512.2974243164062 0 805.72363
517.8135375976562 0 2314.9192
518.2599487304688 0 1283.0248 y 7
518.3110961914062 0 985.5595
518.8090209960938 0 2112.7175
524.33154296875 0 1804.2548
525.337158203125 0 3763.4707
525.8123779296875 0 30255.535
526.3141479492188 0 23857.418
526.8158569335938 0 8773.964
527.3170166015625 0 2792.4426
529.281005859375 0 795.66675
531.2988891601562 0 2034.2482
531.801025390625 0 1595.9421
532.2941284179688 0 617.1938
532.8064575195312 0 845.98785
539.3132934570312 0 1005.29315 c Water loss 9
539.8101196289062 0 167783.33 c Ammonia loss 9
540.3114624023438 0 97356.984
540.81298828125 0 34932.387
541.3142700195312 0 8894.387
541.8176879882812 0 1725.2482
542.2921752929688 0 798.6118
542.332275390625 0 4480.151
543.3368530273438 0 1072.0684
547.3219604492188 0 709.0731
547.810546875 0 2107.3345
548.3234252929688 0 302835.94 c 9
548.8244018554688 0 171943.61
549.3258666992188 0 56354.926
549.785400390625 0 769.63794
549.8270874023438 0 14581.607
550.3291625976562 0 2356.4478
555.2965087890625 0 13981.047
555.3409423828125 0 2337.5374
555.797607421875 0 8537.954
556.29931640625 0 2318.87
558.2892456054688 0 1230.2375
561.330810546875 0 10811.024
561.832275390625 0 7202.2783
562.3348999023438 0 1654.8578
562.7792358398438 0 960.1395
562.8268432617188 0 918.5467
563.2815551757812 0 866.7293
567.34814453125 0 542.4608
568.3464965820312 0 3195.157
568.6487426757812 0 622.4178
568.8228759765625 0 1631.2317
569.3521118164062 0 48093.246 c 4
570.3074951171875 0 792.03125
570.35498046875 0 14187.234
571.3048095703125 0 1000.777
571.3572387695312 0 3111.2585
575.8372192382812 0 4203.7227
576.3381958007812 0 4036.1626
576.8223876953125 0 17507.768
577.323974609375 0 15921.62
577.8253784179688 0 6811.1353
578.3265991210938 0 1529.4034
582.3556518554688 0 5210.5986
582.8406372070312 0 1475.9149
583.340087890625 0 1556.8676
583.8191528320312 0 1113.5194
584.3118286132812 0 16143.665 z 1
584.812744140625 0 8948.059
584.8543090820312 0 954.40717
585.3167724609375 0 2511.292
585.8190307617188 0 796.2428
586.3212890625 0 1437.6385
590.3270874023438 0 718.0917
590.8577880859375 0 1039.0756
591.3167114257812 0 1005.25555
591.814208984375 0 769.1366
592.3200073242188 0 1364.301 y 1
593.3226928710938 0 606.2582
596.3516235351562 0 1143.0765
597.8385620117188 0 2138.4214
598.326416015625 0 5443.9775
598.8291625976562 0 3040.759
599.3314819335938 0 4352.1187
600.3351440429688 0 1371.4337
602.2811279296875 0 3599.4917
603.3349609375 0 745.67114
604.3395385742188 0 4969.1875
604.8330078125 0 12544.104
605.3331909179688 0 9411.214
605.8253784179688 0 4578.7256
606.3352661132812 0 2160.464
606.843505859375 0 1769.6371
607.3512573242188 0 847.11536
611.3550415039062 0 3710.881
611.8507080078125 0 23867.31
612.3530883789062 0 13916.112
612.8521728515625 0 5421.8804
613.3508911132812 0 3507.6904
613.8480224609375 0 1555.494
614.3533935546875 0 13167.105
614.8275756835938 0 2170.2393
615.2891235351562 0 8188.1353
615.356201171875 0 3567.322
616.2926025390625 0 2542.0332
616.3544921875 0 1044.2694
619.3560180664062 0 2593.0222
619.8493041992188 0 14155.56
620.3583374023438 0 19438.719
620.845947265625 0 47975.348
621.3462524414062 0 30336.969
621.84716796875 0 12612.737
622.3477172851562 0 3578.6606
622.8505859375 0 724.5118
624.34716796875 0 2904.0435
624.8359375 0 1045.8717
625.3650512695312 0 7154.397
625.8356323242188 0 3650.7053
626.3741455078125 0 189069.86 c 5
626.8353271484375 0 16353.863
627.376708984375 0 59950.37
627.8469848632812 0 842.99963
628.3241577148438 0 628.1934
628.3788452148438 0 11865.516
629.3812255859375 0 1319.5941
633.3502807617188 0 3727.8323
633.8456420898438 0 230865.97
634.3469848632812 0 158010.19
634.8483276367188 0 64805.668
635.3496704101562 0 18446.102
635.8499755859375 0 3868.622
641.8549194335938 0 108686.62
642.3572998046875 0 136686.52
642.8592529296875 0 72438.07
643.3602294921875 0 27270.201
643.8626708984375 0 6071.975
644.3598022460938 0 1394.4996
652.38623046875 0 1533.4283
658.34375 0 108614.016 z 6
659.3468017578125 0 44038.016
660.3497314453125 0 8273.531
661.3506469726562 0 1004.36707
671.3753662109375 0 6980.9546
672.3803100585938 0 1795.1887
674.3622436523438 0 6912.757 y 6
675.3646240234375 0 2271.1816
682.3872680664062 0 627.03687
695.3981323242188 0 1190.4728
696.39453125 0 996.9474
713.3746948242188 0 1626.5544 y Water loss 5
715.3652954101562 0 25679.127 z 5
716.367919921875 0 10190.423
717.3719482421875 0 1981.9846
723.4285278320312 0 1181.2941
731.3839721679688 0 6432.8867 y 5
732.3890380859375 0 2814.023
738.4610595703125 0 835.0558
739.4608154296875 0 2158.3467
740.455078125 0 1351.5422
767.4625854492188 0 1652.0223
780.43896484375 0 1412.0543
781.4660034179688 0 3555.7244
782.4747314453125 0 35754.19 c 6
783.4768676757812 0 16960.818
784.4808349609375 0 3539.9517
785.3934936523438 0 2705.2869 w 4
785.4744873046875 0 907.3357
786.401123046875 0 2196.9988
787.3993530273438 0 883.25146
796.44970703125 0 659.5108
810.4269409179688 0 2765.577 y Water loss 4
811.4282836914062 0 4546.2095
811.5055541992188 0 630.77374
812.4268188476562 0 1186.7921 z 4
819.4617919921875 0 961.8128
823.4447631835938 0 1092.9814
824.4507446289062 0 5420.105
825.4557495117188 0 5224.7866
826.4271240234375 0 2580.1956
827.4057006835938 0 1214.2544
828.4360961914062 0 9711.306 y 4
829.4393920898438 0 4343.1816
830.436767578125 0 992.34143
838.498779296875 0 1556.1759
839.4664306640625 0 2565.1382
840.4736938476562 0 1450.2225
851.503173828125 0 1294.4747
852.5047607421875 0 2659.702
852.9813842773438 0 944.36163
853.51123046875 0 27988.47 c 7
854.5140991210938 0 12381.401
855.51611328125 0 3652.3726
856.4310913085938 0 970.24866
883.45458984375 0 115161.69 z 3
884.4576416015625 0 59957.246
885.4600830078125 0 17180.336
886.4620971679688 0 4194.592
899.4730224609375 0 14850.226 y 3
900.4755859375 0 6755.2515
901.4781494140625 0 2719.8489
908.5576171875 0 1146.8999
915.4497680664062 0 944.13605
922.5830078125 0 721.97815
923.574951171875 0 1192.9547
924.5731201171875 0 1695.4368
936.537353515625 0 629.98505
939.4896850585938 0 949.91284
950.5767211914062 0 2553.628
951.5800170898438 0 1868.59
952.5903930664062 0 948.2265
953.48583984375 0 2332.4458 w 2
954.487548828125 0 2057.1108
955.464111328125 0 1522.9783
963.5245361328125 0 1319.8948
965.5758666992188 0 1118.01
966.5950317382812 0 30581.76 c 8
967.59912109375 0 17039.088
968.6021118164062 0 4292.975
969.609375 0 736.49835
980.5509033203125 0 2765.0527
981.5546875 0 2039.3768
992.6091918945312 0 684.7963
1008.5999755859375 0 721.5334
1009.6007080078125 0 1121.894
1011.5130615234375 0 90892.5 z 2
1012.5157470703125 0 49410.137
1013.5183715820312 0 17898.143
1014.5195922851562 0 3192.6553
1015.5125122070312 0 980.04626
1027.53173828125 0 21442.713 y 2
1028.5355224609375 0 11299.057
1029.537109375 0 4115.2305
1034.6229248046875 0 920.91547
1035.6090087890625 0 2932.6416
1036.6119384765625 0 1041.8512
1037.59716796875 0 5801.226
1038.60400390625 0 2653.6582
1051.62353515625 0 10774.482
1052.6278076171875 0 28026.055
1053.6278076171875 0 17598.041
1054.6246337890625 0 5678.293
1055.6318359375 0 1406.2134
1063.593994140625 0 768.72284
1078.6129150390625 0 1067.1965 c Ammonia loss 9
1079.6187744140625 0 22659.947
1080.6224365234375 0 13035.36
1081.54150390625 0 4380.8086 w 1
1081.6361083984375 0 2557.584
1082.546875 0 2991.1453
1095.637451171875 0 19864.201 c 9
1096.64453125 0 54398.664
1097.6475830078125 0 28521.96
1098.6505126953125 0 8428.436
1099.650634765625 0 1331.5371
1110.607177734375 0 867.6973
1125.6064453125 0 970.52954
1126.6005859375 0 1009.264
1140.627685546875 0 818.4773
1151.614013671875 0 978.6688
1152.6258544921875 0 1945.3271
1153.629638671875 0 3239.8035
1154.6312255859375 0 1621.1914
1167.612548828125 0 6302.699 z 1
1168.620849609375 0 36333.4
1169.623291015625 0 24850.547
1170.6265869140625 0 8571.033
1171.627197265625 0 1825.2789
1172.6285400390625 0 999.4556
1180.61962890625 0 5156.468
1181.6226806640625 0 3721.907
1182.6231689453125 0 4269.2637
1183.627197265625 0 3353.5974 y 1
1184.635498046875 0 1957.974
1185.6427001953125 0 863.1637
1195.622314453125 0 4192.735
1196.630859375 0 2683.7173
1197.6422119140625 0 1725.5071
1206.673095703125 0 1655.731
1208.644775390625 0 798.5249
1209.6534423828125 0 1463.8772
1211.6658935546875 0 1365.7705
1212.6710205078125 0 1173.8976
1222.7060546875 0 4002.7344
1223.6982421875 0 9921.576
1224.662109375 0 61152.684
1225.663818359375 0 48203.355
1226.665771484375 0 21143.926
1227.6688232421875 0 5927.6943
1228.672607421875 0 1870.6052
1238.695068359375 0 3167.1658
1239.6944580078125 0 15178.202
1240.7005615234375 0 14701.491
1241.695556640625 0 12799.798
1242.6962890625 0 4906.3467
1243.7012939453125 0 1897.7559
1244.689697265625 0 867.7116
1249.679443359375 0 1870.2762
1250.663330078125 0 20677.453
1251.6650390625 0 12658.648
1251.832275390625 0 871.786
1252.66796875 0 6687.0083
1253.663330078125 0 1623.3215
1255.710693359375 0 925.25275
1256.719970703125 0 5383.8237
1257.721923828125 0 2460.1582
1258.716552734375 0 1291.189
1265.7042236328125 0 867.97705
1266.6883544921875 0 20803.434
1267.6890869140625 0 168351.12
1268.6917724609375 0 115653.33
1269.6942138671875 0 42922.24
1270.6971435546875 0 12507.052
1271.6922607421875 0 3870.2441
1282.7000732421875 0 6290.414
1283.7061767578125 0 36150.555
1284.7147216796875 0 150933.31
1285.71728515625 0 97534.03
1286.720458984375 0 38162.406
1287.722412109375 0 11412.479
1288.71923828125 0 3408.4084
2347.80810546875 0 739.6513

Spectrum Details

|  |  |
| --- | --- |
| Matched peaks? Matched peaks The total absolute number of peaks matched. Additionally in brackets the total fraction of peaks matched and the total number of peaks is shown. | 47 (10.11% of 465) |
| FDR? FDR The false discovery rate estimated for this peptide. It is calculated by matching all theoretical fragments with a non-integer shift with the raw peaks for this spectrum. This is done with 40 different shifts. The resulting percentage is the average number of annotated peaks over the number of annotated peaks with the correct spectrum. | 2.38% |
| Satellite FDR? Satellite FDR See the FDR for details on its calculation. This satellite ion specific FDR only contains the satellite ions (d/w) for I/L/J positions. | ∞ |
| PSM Score? PSM Score The PSM Score as given by Hecklib to this annotated spectrum. It is shown with three significant figures. | 388 |

#### Spectrum 6378? Spectrum 6378 The raw spectrum of this peptide as annotated by Hecklib. The fragments are coloured according to ion type (see legend). Any peaks with a star '\*' as text can be hovered over to see the full details, first the ion type second the mass shift type. By hovering over the amino acids in the peptide or ions in the legend the corresponding peaks are highlighted. By toggling the 'Unassigned' label you can turn the background (unassigned) peaks on or off in the plot. By updating the slider in the Ion legend you can update the spectrum to only show the top X% of the peaks with labels. The top X% means any peak that is within X% of the highest intensity. By dragging in the spectrum you can zoom in to a specific part of the spectrum and use 'Zoom Out' to get back to the original zoom level. The annotation of the spectrum is based on the given sequence in the peptides file and is done with different software so inconsistencies are likely. The peaks are annotated based on the given sequence, with 20 ppm tolerance.

Copy Data

##### Spectrum 6378 (TSV)

###### Preview

```
Loading example...
```

*Click on the button to copy the data to your clipboard.*

Mz MinMz MaxIntensity Max

WidthHeightPeptide font sizePeptide stroke widthSpectrum font sizeSpectrum stroke widthCompact peptide

Ion legend

wxyz

abcd

OtherUnassignedIon^Charge^_Position_Show for top:%

VRQAPGRAJEW

07.01e+41.40e+52.10e+52.80e+5

Zoom Out

y^+1^_1_c^+1^_2_c^+1^_2_y^+1^_2_c^+2^_7_c^+1^_3_c^+2^_7_c^+1^_3_y^+2^_7_c^+2^_8_c^+2^_8_c^+1^_4_c^+2^_9_c^+2^_9_c^+2^_10_c^+2^_10_c^+1^_5_z^+2^_10_c^+1^_6_z^+1^_5_y^+1^_5_z^+1^_6_y^+1^_6_c^+1^_7_w^+1^_7_z^+1^_7_y^+1^_7_c^+1^_8_z^+1^_8_y^+1^_8_w^+1^_9_c^+1^_9_z^+1^_9_y^+1^_9_w^+1^_10_c^+1^_10_z^+1^_10_y^+1^_10_

0839167825173356

Fragment Matches Table

Show background peaks

| Position | Ion type | Intensity | mz Theoretical | mz Error (Th) | mz Error (ppm) | Charge | Series Number |
| --- | --- | --- | --- | --- | --- | --- | --- |
| - | - | 856 | 120.1 | - | - | 0 | - |
| - | - | 1437 | 130.1 | - | - | 0 | - |
| - | - | 374.1 | 131 | - | - | 0 | - |
| - | - | 644.8 | 133.1 | - | - | 0 | - |
| - | - | 406.8 | 138 | - | - | 0 | - |
| - | - | 463.9 | 142.7 | - | - | 0 | - |
| - | - | 2518 | 146.1 | - | - | 0 | - |
| - | - | 2307 | 149 | - | - | 0 | - |
| - | - | 525.6 | 153.1 | - | - | 0 | - |
| - | - | 1323 | 157.1 | - | - | 0 | - |
| - | - | 922.3 | 157.1 | - | - | 0 | - |
| - | - | 4097 | 159.1 | - | - | 0 | - |
| - | - | 450.9 | 160.1 | - | - | 0 | - |
| - | - | 595.2 | 166.1 | - | - | 0 | - |
| - | - | 465.3 | 166.8 | - | - | 0 | - |
| - | - | 8992 | 173.1 | - | - | 0 | - |
| - | - | 1567 | 173.4 | - | - | 0 | - |
| - | - | 541.8 | 183.1 | - | - | 0 | - |
| - | - | 615.3 | 185.1 | - | - | 0 | - |
| - | - | 1268 | 185.1 | - | - | 0 | - |
| - | - | 986.2 | 185.1 | - | - | 0 | - |
| - | - | 1.423E+04 | 188.1 | - | - | 0 | - |
| - | - | 2259 | 189.1 | - | - | 0 | - |
| - | - | 7819 | 201.1 | - | - | 0 | - |
| - | - | 1464 | 203.1 | - | - | 0 | - |
| 11 | y | 8967 | 205.1 | 1.664E-05 | 0.08114 | +1 | 1 |
| - | - | 674.6 | 206.1 | - | - | 0 | - |
| - | - | 2.98E+04 | 209 | - | - | 0 | - |
| - | - | 2605 | 212.1 | - | - | 0 | - |
| - | - | 1280 | 213.2 | - | - | 0 | - |
| - | - | 1332 | 214.1 | - | - | 0 | - |
| - | - | 950.7 | 214.2 | - | - | 0 | - |
| - | - | 715.6 | 223.1 | - | - | 0 | - |
| - | - | 2.776E+05 | 229.2 | - | - | 0 | - |
| - | - | 652.1 | 231 | - | - | 0 | - |
| - | - | 1507 | 233.2 | - | - | 0 | - |
| - | - | 541.8 | 234.1 | - | - | 0 | - |
| - | - | 507.2 | 238.2 | - | - | 0 | - |
| - | - | 5791 | 239.2 | - | - | 0 | - |
| - | - | 1782 | 242.2 | - | - | 0 | - |
| 2 | c | 3744 | 256.2 | 4.359E-05 | 0.1701 | +1 | 2 |
| - | - | 1824 | 260.2 | - | - | 0 | - |
| - | - | 776.8 | 261.2 | - | - | 0 | - |
| - | - | 674.4 | 270.1 | - | - | 0 | - |
| 2 | c | 1.195E+04 | 273.2 | 0.0001339 | 0.4903 | +1 | 2 |
| - | - | 2075 | 274.2 | - | - | 0 | - |
| - | - | 655.9 | 290 | - | - | 0 | - |
| - | - | 3603 | 298.2 | - | - | 0 | - |
| - | - | 3568 | 299.2 | - | - | 0 | - |
| - | - | 609.5 | 300.2 | - | - | 0 | - |
| - | - | 1028 | 312.7 | - | - | 0 | - |
| - | - | 774.7 | 315.2 | - | - | 0 | - |
| 10 | y | 1416 | 316.1 | 9.315E-05 | 0.2946 | +1 | 2 |
| - | - | 2.008E+04 | 318.9 | - | - | 0 | - |
| - | - | 7243 | 341 | - | - | 0 | - |
| - | - | 873.9 | 342.2 | - | - | 0 | - |
| - | - | 1378 | 350.1 | - | - | 0 | - |
| - | - | 749.6 | 351.1 | - | - | 0 | - |
| - | - | 643.6 | 356.2 | - | - | 0 | - |
| - | - | 918.1 | 357.2 | - | - | 0 | - |
| - | - | 596.9 | 358.3 | - | - | 0 | - |
| - | - | 2.582E+04 | 359 | - | - | 0 | - |
| - | - | 748.4 | 361.7 | - | - | 0 | - |
| - | - | 2977 | 369.7 | - | - | 0 | - |
| - | - | 1078 | 370.2 | - | - | 0 | - |
| - | - | 611.8 | 371.2 | - | - | 0 | - |
| - | - | 516.8 | 375.8 | - | - | 0 | - |
| - | - | 719.4 | 375.9 | - | - | 0 | - |
| 7 | c | 1092 | 383.2 | 0.0005031 | 1.313 | +2 | 7 |
| 3 | c | 1845 | 384.2 | 0.000241 | 0.6271 | +1 | 3 |
| - | - | 657.9 | 385.2 | - | - | 0 | - |
| - | - | 1.276E+04 | 391.2 | - | - | 0 | - |
| 7 | c | 4.592E+04 | 391.7 | 0.0005635 | 1.439 | +2 | 7 |
| - | - | 1035 | 391.9 | - | - | 0 | - |
| - | - | 1.711E+04 | 392.2 | - | - | 0 | - |
| - | - | 4944 | 392.7 | - | - | 0 | - |
| 3 | c | 3.965E+04 | 401.3 | 0.0001482 | 0.3694 | +1 | 3 |
| - | - | 8424 | 402.3 | - | - | 0 | - |
| - | - | 833.5 | 403.3 | - | - | 0 | - |
| 5 | y | 772.5 | 405.7 | 0.001705 | 4.202 | +2 | 7 |
| - | - | 2011 | 406.9 | - | - | 0 | - |
| - | - | 809.2 | 407.2 | - | - | 0 | - |
| - | - | 995.3 | 412.8 | - | - | 0 | - |
| - | - | 1158 | 412.9 | - | - | 0 | - |
| - | - | 2844 | 413.3 | - | - | 0 | - |
| - | - | 610.8 | 413.6 | - | - | 0 | - |
| 8 | c | 1375 | 418.7 | 0.0001391 | 0.3322 | +2 | 8 |
| - | - | 635.1 | 419.2 | - | - | 0 | - |
| - | - | 4836 | 426.8 | - | - | 0 | - |
| - | - | 775.3 | 427.2 | - | - | 0 | - |
| 8 | c | 3.467E+04 | 427.3 | 1.409E-05 | 0.03298 | +2 | 8 |
| - | - | 1.439E+04 | 427.8 | - | - | 0 | - |
| - | - | 536.5 | 428.1 | - | - | 0 | - |
| - | - | 4246 | 428.3 | - | - | 0 | - |
| - | - | 501.6 | 428.3 | - | - | 0 | - |
| - | - | 2324 | 428.9 | - | - | 0 | - |
| - | - | 2.466E+04 | 429.1 | - | - | 0 | - |
| - | - | 4432 | 429.3 | - | - | 0 | - |
| - | - | 1172 | 439.3 | - | - | 0 | - |
| - | - | 1392 | 447.8 | - | - | 0 | - |
| - | - | 629.1 | 453.6 | - | - | 0 | - |
| 4 | c | 2214 | 455.3 | 0.0008252 | 1.813 | +1 | 4 |
| - | - | 3567 | 461.3 | - | - | 0 | - |
| - | - | 2138 | 461.8 | - | - | 0 | - |
| - | - | 1196 | 462.3 | - | - | 0 | - |
| - | - | 1145 | 464.2 | - | - | 0 | - |
| - | - | 670.1 | 470.3 | - | - | 0 | - |
| - | - | 2040 | 470.8 | - | - | 0 | - |
| 9 | c | 7871 | 475.3 | 5.682E-05 | 0.1196 | +2 | 9 |
| - | - | 4113 | 475.8 | - | - | 0 | - |
| - | - | 1629 | 476.3 | - | - | 0 | - |
| - | - | 1780 | 483.3 | - | - | 0 | - |
| 9 | c | 1.883E+04 | 483.8 | 0.0002393 | 0.4947 | +2 | 9 |
| - | - | 8809 | 484.3 | - | - | 0 | - |
| - | - | 3006 | 484.8 | - | - | 0 | - |
| - | - | 1.009E+04 | 496.8 | - | - | 0 | - |
| - | - | 6745 | 497.3 | - | - | 0 | - |
| - | - | 1549 | 497.8 | - | - | 0 | - |
| - | - | 2263 | 498.3 | - | - | 0 | - |
| - | - | 642.8 | 499.3 | - | - | 0 | - |
| - | - | 690.4 | 504.8 | - | - | 0 | - |
| - | - | 4669 | 513.8 | - | - | 0 | - |
| - | - | 548 | 514.8 | - | - | 0 | - |
| - | - | 2860 | 514.8 | - | - | 0 | - |
| - | - | 836 | 518.8 | - | - | 0 | - |
| - | - | 516.6 | 520.8 | - | - | 0 | - |
| - | - | 1114 | 525.3 | - | - | 0 | - |
| - | - | 9121 | 525.8 | - | - | 0 | - |
| - | - | 6548 | 526.3 | - | - | 0 | - |
| - | - | 2904 | 526.8 | - | - | 0 | - |
| 10 | c | 4.513E+04 | 539.8 | 0.0004793 | 0.888 | +2 | 10 |
| - | - | 2.536E+04 | 540.3 | - | - | 0 | - |
| - | - | 9180 | 540.8 | - | - | 0 | - |
| - | - | 3733 | 541.3 | - | - | 0 | - |
| - | - | 1716 | 542.3 | - | - | 0 | - |
| 10 | c | 7.903E+04 | 548.3 | 0.0003262 | 0.5948 | +2 | 10 |
| - | - | 4.336E+04 | 548.8 | - | - | 0 | - |
| - | - | 1.702E+04 | 549.3 | - | - | 0 | - |
| - | - | 5011 | 549.8 | - | - | 0 | - |
| - | - | 3964 | 555.3 | - | - | 0 | - |
| - | - | 916.1 | 555.3 | - | - | 0 | - |
| - | - | 2880 | 555.8 | - | - | 0 | - |
| - | - | 2976 | 561.3 | - | - | 0 | - |
| - | - | 3124 | 561.8 | - | - | 0 | - |
| - | - | 969.9 | 565.4 | - | - | 0 | - |
| - | - | 1454 | 568.3 | - | - | 0 | - |
| 5 | c | 1.545E+04 | 569.4 | 0.0007925 | 1.392 | +1 | 5 |
| - | - | 4437 | 570.4 | - | - | 0 | - |
| - | - | 841.7 | 571.4 | - | - | 0 | - |
| - | - | 1778 | 575.8 | - | - | 0 | - |
| - | - | 693.6 | 576.3 | - | - | 0 | - |
| - | - | 6822 | 576.8 | - | - | 0 | - |
| - | - | 4156 | 577.3 | - | - | 0 | - |
| - | - | 5351 | 577.8 | - | - | 0 | - |
| - | - | 648.3 | 578.8 | - | - | 0 | - |
| - | - | 2200 | 579.3 | - | - | 0 | - |
| - | - | 988.5 | 582.4 | - | - | 0 | - |
| 2 | z | 3826 | 584.3 | 0.0004454 | 0.7622 | +2 | 10 |
| - | - | 996.5 | 584.4 | - | - | 0 | - |
| - | - | 2309 | 584.8 | - | - | 0 | - |
| - | - | 1535 | 585.3 | - | - | 0 | - |
| - | - | 614.5 | 591.3 | - | - | 0 | - |
| - | - | 812.3 | 598.3 | - | - | 0 | - |
| - | - | 678.7 | 598.8 | - | - | 0 | - |
| - | - | 1177 | 599.3 | - | - | 0 | - |
| - | - | 1569 | 602.3 | - | - | 0 | - |
| - | - | 1774 | 604.3 | - | - | 0 | - |
| - | - | 2951 | 604.8 | - | - | 0 | - |
| - | - | 1726 | 605.3 | - | - | 0 | - |
| - | - | 1695 | 605.8 | - | - | 0 | - |
| - | - | 788.6 | 606.8 | - | - | 0 | - |
| - | - | 895.6 | 607.4 | - | - | 0 | - |
| - | - | 1365 | 611.4 | - | - | 0 | - |
| - | - | 7076 | 611.8 | - | - | 0 | - |
| - | - | 3991 | 612.4 | - | - | 0 | - |
| - | - | 1505 | 612.9 | - | - | 0 | - |
| - | - | 1133 | 613.3 | - | - | 0 | - |
| - | - | 829.4 | 613.9 | - | - | 0 | - |
| - | - | 6264 | 614.3 | - | - | 0 | - |
| - | - | 3733 | 614.4 | - | - | 0 | - |
| - | - | 2254 | 615.3 | - | - | 0 | - |
| - | - | 1882 | 615.4 | - | - | 0 | - |
| - | - | 882.9 | 616.3 | - | - | 0 | - |
| - | - | 645.9 | 618.1 | - | - | 0 | - |
| - | - | 967.6 | 619.4 | - | - | 0 | - |
| - | - | 4789 | 619.8 | - | - | 0 | - |
| - | - | 7490 | 620.4 | - | - | 0 | - |
| - | - | 1.277E+04 | 620.8 | - | - | 0 | - |
| - | - | 9522 | 621.3 | - | - | 0 | - |
| - | - | 2469 | 621.8 | - | - | 0 | - |
| - | - | 816.6 | 622.3 | - | - | 0 | - |
| - | - | 687.4 | 624.3 | - | - | 0 | - |
| - | - | 2929 | 625.4 | - | - | 0 | - |
| - | - | 1444 | 625.8 | - | - | 0 | - |
| 6 | c | 5.885E+04 | 626.4 | 0.0004056 | 0.6475 | +1 | 6 |
| - | - | 4649 | 626.8 | - | - | 0 | - |
| - | - | 2.004E+04 | 627.4 | - | - | 0 | - |
| - | - | 3313 | 628.4 | - | - | 0 | - |
| - | - | 942.4 | 633.3 | - | - | 0 | - |
| - | - | 6.429E+04 | 633.8 | - | - | 0 | - |
| - | - | 4.457E+04 | 634.3 | - | - | 0 | - |
| - | - | 1.66E+04 | 634.8 | - | - | 0 | - |
| - | - | 5107 | 635.3 | - | - | 0 | - |
| - | - | 2399 | 635.8 | - | - | 0 | - |
| - | - | 695.7 | 641.4 | - | - | 0 | - |
| - | - | 3.16E+04 | 641.9 | - | - | 0 | - |
| - | - | 3.729E+04 | 642.4 | - | - | 0 | - |
| - | - | 2.036E+04 | 642.9 | - | - | 0 | - |
| - | - | 7223 | 643.4 | - | - | 0 | - |
| - | - | 1.612E+04 | 643.9 | - | - | 0 | - |
| - | - | 6424 | 644.4 | - | - | 0 | - |
| 7 | z | 3.341E+04 | 658.3 | 0.0007826 | 1.189 | +1 | 5 |
| - | - | 1.179E+04 | 659.3 | - | - | 0 | - |
| - | - | 2828 | 660.3 | - | - | 0 | - |
| - | - | 2684 | 671.4 | - | - | 0 | - |
| - | - | 927.9 | 672.4 | - | - | 0 | - |
| 7 | y | 2223 | 674.4 | 0.002112 | 3.131 | +1 | 5 |
| - | - | 826.9 | 675.4 | - | - | 0 | - |
| - | - | 824.8 | 680.4 | - | - | 0 | - |
| - | - | 4083 | 697.4 | - | - | 0 | - |
| - | - | 592.9 | 704.4 | - | - | 0 | - |
| 6 | z | 8276 | 715.4 | 0.0005178 | 0.7239 | +1 | 6 |
| - | - | 3590 | 716.4 | - | - | 0 | - |
| 6 | y | 2154 | 731.4 | 0.002274 | 3.109 | +1 | 6 |
| - | - | 1258 | 732.4 | - | - | 0 | - |
| - | - | 1588 | 781.5 | - | - | 0 | - |
| 7 | c | 1.431E+04 | 782.5 | 0.0009307 | 1.189 | +1 | 7 |
| - | - | 6191 | 783.5 | - | - | 0 | - |
| - | - | 1479 | 784.5 | - | - | 0 | - |
| 5 | w | 1413 | 785.4 | 0.001669 | 2.126 | +1 | 7 |
| - | - | 939.8 | 785.5 | - | - | 0 | - |
| 5 | z | 738 | 812.4 | 0.006228 | 7.666 | +1 | 7 |
| - | - | 4051 | 816.4 | - | - | 0 | - |
| - | - | 1589 | 824.4 | - | - | 0 | - |
| - | - | 4013 | 824.5 | - | - | 0 | - |
| - | - | 1349 | 825.5 | - | - | 0 | - |
| - | - | 1.031E+04 | 825.5 | - | - | 0 | - |
| 5 | y | 2543 | 828.4 | 0.001937 | 2.339 | +1 | 7 |
| - | - | 1263 | 829.4 | - | - | 0 | - |
| - | - | 718.9 | 845.5 | - | - | 0 | - |
| - | - | 1148 | 852.5 | - | - | 0 | - |
| 8 | c | 1.059E+04 | 853.5 | 0.002034 | 2.383 | +1 | 8 |
| - | - | 5681 | 854.5 | - | - | 0 | - |
| - | - | 1718 | 855.5 | - | - | 0 | - |
| - | - | 912.7 | 856.4 | - | - | 0 | - |
| 4 | z | 3.717E+04 | 883.5 | 0.001711 | 1.937 | +1 | 8 |
| - | - | 1.933E+04 | 884.5 | - | - | 0 | - |
| - | - | 5000 | 885.5 | - | - | 0 | - |
| - | - | 1308 | 886.5 | - | - | 0 | - |
| - | - | 734.8 | 887.5 | - | - | 0 | - |
| 4 | y | 5215 | 899.5 | 0.00243 | 2.702 | +1 | 8 |
| - | - | 2428 | 900.5 | - | - | 0 | - |
| - | - | 875.5 | 901.5 | - | - | 0 | - |
| - | - | 719.4 | 923.6 | - | - | 0 | - |
| - | - | 4682 | 940.6 | - | - | 0 | - |
| 3 | w | 1255 | 953.5 | 0.0006046 | 0.6341 | +1 | 9 |
| 9 | c | 1.046E+04 | 966.6 | 0.001686 | 1.744 | +1 | 9 |
| - | - | 5238 | 967.6 | - | - | 0 | - |
| - | - | 1496 | 968.6 | - | - | 0 | - |
| - | - | 1881 | 980.5 | - | - | 0 | - |
| 3 | z | 2.538E+04 | 1012 | 0.001939 | 1.917 | +1 | 9 |
| - | - | 1.625E+04 | 1013 | - | - | 0 | - |
| - | - | 4201 | 1014 | - | - | 0 | - |
| - | - | 1632 | 1015 | - | - | 0 | - |
| - | - | 807.2 | 1025 | - | - | 0 | - |
| 3 | y | 6551 | 1028 | 0.001559 | 1.518 | +1 | 9 |
| - | - | 3331 | 1029 | - | - | 0 | - |
| - | - | 1096 | 1030 | - | - | 0 | - |
| - | - | 1474 | 1036 | - | - | 0 | - |
| - | - | 2301 | 1038 | - | - | 0 | - |
| - | - | 811.5 | 1039 | - | - | 0 | - |
| - | - | 931.7 | 1042 | - | - | 0 | - |
| - | - | 3090 | 1052 | - | - | 0 | - |
| - | - | 8700 | 1053 | - | - | 0 | - |
| - | - | 6680 | 1054 | - | - | 0 | - |
| - | - | 2072 | 1055 | - | - | 0 | - |
| - | - | 7270 | 1080 | - | - | 0 | - |
| - | - | 4177 | 1081 | - | - | 0 | - |
| 2 | w | 807.7 | 1082 | 0.0142 | 13.13 | +1 | 10 |
| - | - | 843 | 1083 | - | - | 0 | - |
| 10 | c | 7116 | 1096 | 0.002287 | 2.087 | +1 | 10 |
| - | - | 2.074E+04 | 1097 | - | - | 0 | - |
| - | - | 1.068E+04 | 1098 | - | - | 0 | - |
| - | - | 3122 | 1099 | - | - | 0 | - |
| - | - | 979.2 | 1100 | - | - | 0 | - |
| - | - | 1604 | 1112 | - | - | 0 | - |
| - | - | 1616 | 1154 | - | - | 0 | - |
| - | - | 1182 | 1155 | - | - | 0 | - |
| - | - | 1808 | 1156 | - | - | 0 | - |
| - | - | 763.7 | 1158 | - | - | 0 | - |
| - | - | 2547 | 1159 | - | - | 0 | - |
| 2 | z | 1773 | 1168 | 0.004479 | 3.836 | +1 | 10 |
| - | - | 1.387E+04 | 1169 | - | - | 0 | - |
| - | - | 9428 | 1170 | - | - | 0 | - |
| - | - | 2913 | 1171 | - | - | 0 | - |
| - | - | 947.8 | 1181 | - | - | 0 | - |
| - | - | 897 | 1182 | - | - | 0 | - |
| - | - | 1133 | 1183 | - | - | 0 | - |
| 2 | y | 1381 | 1184 | 0.007089 | 5.989 | +1 | 10 |
| - | - | 1009 | 1196 | - | - | 0 | - |
| - | - | 742.6 | 1212 | - | - | 0 | - |
| - | - | 2035 | 1223 | - | - | 0 | - |
| - | - | 3129 | 1224 | - | - | 0 | - |
| - | - | 1.76E+04 | 1225 | - | - | 0 | - |
| - | - | 1.46E+04 | 1226 | - | - | 0 | - |
| - | - | 7705 | 1227 | - | - | 0 | - |
| - | - | 2437 | 1228 | - | - | 0 | - |
| - | - | 2644 | 1229 | - | - | 0 | - |
| - | - | 876.9 | 1239 | - | - | 0 | - |
| - | - | 4986 | 1240 | - | - | 0 | - |
| - | - | 5171 | 1241 | - | - | 0 | - |
| - | - | 4824 | 1242 | - | - | 0 | - |
| - | - | 2778 | 1243 | - | - | 0 | - |
| - | - | 1214 | 1244 | - | - | 0 | - |
| - | - | 7926 | 1251 | - | - | 0 | - |
| - | - | 5704 | 1252 | - | - | 0 | - |
| - | - | 2750 | 1253 | - | - | 0 | - |
| - | - | 1674 | 1257 | - | - | 0 | - |
| - | - | 7263 | 1267 | - | - | 0 | - |
| - | - | 6.257E+04 | 1268 | - | - | 0 | - |
| - | - | 4.267E+04 | 1269 | - | - | 0 | - |
| - | - | 1214 | 1269 | - | - | 0 | - |
| - | - | 1.773E+04 | 1270 | - | - | 0 | - |
| - | - | 5580 | 1271 | - | - | 0 | - |
| - | - | 7981 | 1272 | - | - | 0 | - |
| - | - | 2411 | 1283 | - | - | 0 | - |
| - | - | 1.359E+04 | 1284 | - | - | 0 | - |
| - | - | 5.309E+04 | 1285 | - | - | 0 | - |
| - | - | 3.441E+04 | 1286 | - | - | 0 | - |
| - | - | 526.1 | 1286 | - | - | 0 | - |
| - | - | 837.6 | 1286 | - | - | 0 | - |
| - | - | 1.448E+04 | 1287 | - | - | 0 | - |
| - | - | 6517 | 1288 | - | - | 0 | - |
| - | - | 1.009E+04 | 1289 | - | - | 0 | - |
| - | - | 641.7 | 1449 | - | - | 0 | - |
| - | - | 746.9 | 2332 | - | - | 0 | - |
| - | - | 1169 | 3084 | - | - | 0 | - |
| - | - | 895.6 | 3323 | - | - | 0 | - |

m/z Charge Intensity FragmentType MassShift Position
120.0811767578125 0 855.9635
130.06541442871094 0 1437.489
130.96176147460938 0 374.09036
133.08607482910156 0 644.75995
138.03370666503906 0 406.7858
142.7002410888672 0 463.92767
146.06019592285156 0 2517.7808
149.04495239257812 0 2307.4773
153.0717010498047 0 525.5959
157.1084747314453 0 1323.4937
157.13340759277344 0 922.2538
159.09178161621094 0 4096.807
160.09608459472656 0 450.90976
166.0869140625 0 595.21606
166.76109313964844 0 465.25473
173.1284942626953 0 8992.329
173.4398651123047 0 1566.7875
183.1498260498047 0 541.8208
185.1033172607422 0 615.33777
185.11756896972656 0 1268.3131
185.1283721923828 0 986.19824
188.07064819335938 0 14226.072
189.07412719726562 0 2259.3267
201.1233367919922 0 7818.7217
203.12779235839844 0 1464.2794
205.09713745117188 0 8967.4 y 10
206.10032653808594 0 674.62836
208.95318603515625 0 29804.812
212.13937377929688 0 2604.6233
213.17091369628906 0 1280.2827
214.14273071289062 0 1331.9502
214.15460205078125 0 950.652
223.10740661621094 0 715.6085
229.15476989746094 0 277552.8
230.9986572265625 0 652.148
233.1640625 0 1506.7214
234.11087036132812 0 541.77844
238.16494750976562 0 507.2013
239.15025329589844 0 5790.9116
242.161376953125 0 1782.3013
256.1767578125 0 3744.3743 c Ammonia loss 1
260.15155029296875 0 1824.0469
261.1601867675781 0 776.77356
270.123291015625 0 674.377
273.2032165527344 0 11952.251 c 1
274.2061767578125 0 2074.541
290.04302978515625 0 655.9058
298.176025390625 0 3602.7056
299.2187805175781 0 3568.428
300.2233581542969 0 609.4962
312.6766357421875 0 1027.9342
315.1567077636719 0 774.7425
316.12908935546875 0 1416.3835 y Water loss 9
318.9222717285156 0 20084.16
341.0178527832031 0 7243.429
342.2376403808594 0 873.925
350.074951171875 0 1377.9126
351.07525634765625 0 749.64624
356.2400207519531 0 643.59814
357.24786376953125 0 918.06647
358.2535095214844 0 596.92804
359.02813720703125 0 25824.668
361.7244567871094 0 748.3789
369.7336120605469 0 2976.676
370.2356262207031 0 1078.0604
371.2403869628906 0 611.7996
375.7548522949219 0 516.76715
375.8616943359375 0 719.44604
383.22705078125 0 1091.6266 c Ammonia loss 6
384.2351379394531 0 1845.0532 c Ammonia loss 2
385.2393493652344 0 657.91943
391.236572265625 0 12755.399
391.7402648925781 0 45922.78 c 6
391.855224609375 0 1035.1365
392.2418212890625 0 17107.602
392.743408203125 0 4943.8765
401.26177978515625 0 39654.992 c 2
402.2645263671875 0 8424.113
403.2662658691406 0 833.45074
405.71478271484375 0 772.48676 y Water loss 4
406.900390625 0 2010.9531
407.23284912109375 0 809.2399
412.76678466796875 0 995.2982
412.90399169921875 0 1158.1187
413.2716979980469 0 2843.5586
413.57122802734375 0 610.76764
418.7459716796875 0 1375.2712 c Ammonia loss 7
419.2486572265625 0 635.09143
426.7552185058594 0 4836.2026
427.1565246582031 0 775.3455
427.2593994140625 0 34672.6 c 7
427.76031494140625 0 14387.553
428.1484375 0 536.5234
428.2613525390625 0 4246.4414
428.2877197265625 0 501.55624
428.89080810546875 0 2323.765
429.0885925292969 0 24656.902
429.2706604003906 0 4431.9395
439.2601318359375 0 1171.7666
447.77325439453125 0 1391.647
453.6052551269531 0 629.1054
455.27166748046875 0 2213.9482 c Ammonia loss 3
461.2908935546875 0 3567.4934
461.7926940917969 0 2138.3596
462.2976989746094 0 1196.4618
464.2048034667969 0 1145.4286
470.2787170410156 0 670.06165
470.7851257324219 0 2040.3071
475.2880859375 0 7871.0796 c Ammonia loss 8
475.7896728515625 0 4113.239
476.29107666015625 0 1628.8741
483.2978820800781 0 1780.1735
483.8011779785156 0 18832.32 c 8
484.3025817871094 0 8808.551
484.8043212890625 0 3006.4834
496.8089294433594 0 10089.768
497.3102111816406 0 6745.0537
497.8099060058594 0 1549.3804
498.2842102050781 0 2263.0818
499.3201599121094 0 642.8234
504.7871398925781 0 690.3789
513.7857666015625 0 4669.177
514.754150390625 0 548.03436
514.793701171875 0 2860.167
518.8063354492188 0 835.9598
520.7698974609375 0 516.5693
525.3377075195312 0 1114.0428
525.8116455078125 0 9121.3
526.3133544921875 0 6548.264
526.8150634765625 0 2904.0774
539.8089599609375 0 45134.2 c Ammonia loss 9
540.3102416992188 0 25359.223
540.8117065429688 0 9179.681
541.3131103515625 0 3733.2485
542.33056640625 0 1716.1738
548.3223876953125 0 79033.74 c 9
548.8236083984375 0 43358.707
549.3247680664062 0 17023.33
549.8248291015625 0 5011.17
555.2955932617188 0 3963.5056
555.3389282226562 0 916.1397
555.797607421875 0 2880.4863
561.32958984375 0 2975.756
561.832763671875 0 3123.83
565.3809814453125 0 969.8521
568.3457641601562 0 1454.1814
569.3510131835938 0 15446.271 c 4
570.3539428710938 0 4437.1313
571.3577270507812 0 841.68384
575.83642578125 0 1778.0106
576.3342895507812 0 693.6337
576.8216552734375 0 6821.882
577.322509765625 0 4156.3755
577.8469848632812 0 5350.8525
578.8126220703125 0 648.2921
579.3111572265625 0 2199.8398
582.3561401367188 0 988.4633
584.3103637695312 0 3825.9133 z 1
584.3570556640625 0 996.48737
584.8112182617188 0 2308.6323
585.31787109375 0 1535.3132
591.2952880859375 0 614.51324
598.32568359375 0 812.29333
598.8363037109375 0 678.7227
599.3345336914062 0 1177.3856
602.2781982421875 0 1569.1152
604.3386840820312 0 1773.5007
604.8320922851562 0 2951.4539
605.3350830078125 0 1725.9629
605.816650390625 0 1694.7905
606.846923828125 0 788.5572
607.3543090820312 0 895.59827
611.35400390625 0 1365.4861
611.8496704101562 0 7076.4404
612.3526611328125 0 3991.249
612.8512573242188 0 1505.2998
613.3428344726562 0 1133.3245
613.8554077148438 0 829.3856
614.3263549804688 0 6263.9985
614.3538208007812 0 3733.0254
615.2888793945312 0 2253.6172
615.3546142578125 0 1882.0986
616.2905883789062 0 882.87115
618.0547485351562 0 645.8653
619.3513793945312 0 967.5536
619.84716796875 0 4788.821
620.356201171875 0 7489.768
620.8446044921875 0 12768.037
621.3448486328125 0 9522.388
621.8463745117188 0 2469.252
622.348388671875 0 816.64966
624.34375 0 687.3951
625.3640747070312 0 2928.7256
625.8302001953125 0 1443.7576
626.3728637695312 0 58845.406 c 5
626.834228515625 0 4648.8984
627.3756713867188 0 20036.883
628.3778686523438 0 3312.549
633.348876953125 0 942.40424
633.8442993164062 0 64292.582
634.3458251953125 0 44574.098
634.8472900390625 0 16604.96
635.3484497070312 0 5106.845
635.8479614257812 0 2398.6396
641.39306640625 0 695.6933
641.8539428710938 0 31600.06
642.3562622070312 0 37285.97
642.8579711914062 0 20356.842
643.3599243164062 0 7223.4478
643.8614501953125 0 16122.344
644.36572265625 0 6424.4233
658.342529296875 0 33405.438 z 6
659.3455810546875 0 11786.088
660.348388671875 0 2828.2466
671.3735961914062 0 2683.9595
672.3778076171875 0 927.9052
674.3599243164062 0 2222.7686 y 6
675.36572265625 0 826.8521
680.408203125 0 824.79895
697.438720703125 0 4082.7754
704.390869140625 0 592.8882
715.3642578125 0 8275.853 z 5
716.3659057617188 0 3590.3386
731.3812255859375 0 2154.2275 y 5
732.38232421875 0 1258.0891
781.4646606445312 0 1587.6609
782.4734497070312 0 14313.67 c 6
783.4758911132812 0 6191.3677
784.4788208007812 0 1478.6025
785.3923950195312 0 1413.4507 w 4
785.4754028320312 0 939.7855
812.4237670898438 0 737.98914 z 4
816.4395141601562 0 4051.2192
824.446533203125 0 1588.5271
824.5245971679688 0 4012.6924
825.451416015625 0 1348.774
825.5340576171875 0 10309.159
828.434326171875 0 2543.4495 y 4
829.4371337890625 0 1262.5286
845.45703125 0 718.94305
852.5077514648438 0 1147.68
853.5094604492188 0 10589.304 c 7
854.5133666992188 0 5681.24
855.5174560546875 0 1718.3748
856.4364624023438 0 912.686
883.4529418945312 0 37166.63 z 3
884.455810546875 0 19326.18
885.4588623046875 0 5000.3833
886.4622192382812 0 1308.4227
887.4511108398438 0 734.79895
899.470947265625 0 5214.629 y 3
900.4761352539062 0 2427.7761
901.4720458984375 0 875.5001
923.5736083984375 0 719.4278
940.5608520507812 0 4682.23
953.4833374023438 0 1254.5316 w 2
966.5938720703125 0 10460.036 c 8
967.5962524414062 0 5237.7354
968.5989379882812 0 1495.6344
980.5468139648438 0 1880.5991
1011.5112915039062 0 25380.053 z 2
1012.51416015625 0 16251.551
1013.5181274414062 0 4201.41
1014.52001953125 0 1632.081
1024.6337890625 0 807.20734
1027.5303955078125 0 6551.075 y 2
1028.542724609375 0 3331.3262
1029.5380859375 0 1096.2844
1035.604736328125 0 1473.8262
1037.5970458984375 0 2300.5293
1038.5994873046875 0 811.4537
1041.609619140625 0 931.74884
1051.6217041015625 0 3089.6675
1052.626708984375 0 8699.836
1053.626708984375 0 6679.6133
1054.6258544921875 0 2071.5327
1079.61767578125 0 7269.7427
1080.6182861328125 0 4177.339
1081.5283203125 0 807.6516 w 1
1082.5467529296875 0 843.028
1095.6358642578125 0 7116.277 c 9
1096.642333984375 0 20737.99
1097.644775390625 0 10677.032
1098.65087890625 0 3121.9146
1099.654296875 0 979.2049
1111.6824951171875 0 1603.5133
1153.6356201171875 0 1615.5167
1154.693359375 0 1182.0278
1155.68896484375 0 1807.7073
1157.6102294921875 0 763.69934
1158.6185302734375 0 2546.8804
1167.60986328125 0 1772.892 z 1
1168.618896484375 0 13866.753
1169.621337890625 0 9428.229
1170.625732421875 0 2912.505
1180.6187744140625 0 947.8441
1181.603759765625 0 897.0112
1182.6251220703125 0 1132.8463
1183.6259765625 0 1381.1357 y 1
1195.6220703125 0 1009.2818
1211.65771484375 0 742.6324
1222.706787109375 0 2034.91
1223.6947021484375 0 3129.3184
1224.6600341796875 0 17601.943
1225.6622314453125 0 14595.672
1226.665283203125 0 7705.4497
1227.6654052734375 0 2436.8599
1228.6949462890625 0 2644.225
1238.7015380859375 0 876.85876
1239.695556640625 0 4986.092
1240.692626953125 0 5170.7163
1241.6942138671875 0 4824.459
1242.6962890625 0 2777.8481
1243.6829833984375 0 1214.2756
1250.6595458984375 0 7926.2476
1251.660888671875 0 5703.5347
1252.669677734375 0 2750.256
1256.7158203125 0 1674.1914
1266.6861572265625 0 7262.863
1267.6865234375 0 62566.85
1268.6890869140625 0 42672.684
1268.8673095703125 0 1214.2974
1269.6923828125 0 17726.475
1270.6954345703125 0 5580.4995
1271.7003173828125 0 7980.9014
1282.7015380859375 0 2411.299
1283.7052001953125 0 13587.919
1284.7119140625 0 53085.582
1285.715087890625 0 34412.637
1285.8641357421875 0 526.0796
1285.896728515625 0 837.5771
1286.7196044921875 0 14477.052
1287.718994140625 0 6517.1357
1288.7266845703125 0 10088.442
1449.0528564453125 0 641.71655
2332.375 0 746.8849
3084.143798828125 0 1168.9198
3323.182861328125 0 895.6066

Spectrum Details

|  |  |
| --- | --- |
| Matched peaks? Matched peaks The total absolute number of peaks matched. Additionally in brackets the total fraction of peaks matched and the total number of peaks is shown. | 38 (11.24% of 338) |
| FDR? FDR The false discovery rate estimated for this peptide. It is calculated by matching all theoretical fragments with a non-integer shift with the raw peaks for this spectrum. This is done with 40 different shifts. The resulting percentage is the average number of annotated peaks over the number of annotated peaks with the correct spectrum. | 2.38% |
| Satellite FDR? Satellite FDR See the FDR for details on its calculation. This satellite ion specific FDR only contains the satellite ions (d/w) for I/L/J positions. | ∞ |
| PSM Score? PSM Score The PSM Score as given by Hecklib to this annotated spectrum. It is shown with three significant figures. | 316 |

#### Spectrum 7046? Spectrum 7046 The raw spectrum of this peptide as annotated by Hecklib. The fragments are coloured according to ion type (see legend). Any peaks with a star '\*' as text can be hovered over to see the full details, first the ion type second the mass shift type. By hovering over the amino acids in the peptide or ions in the legend the corresponding peaks are highlighted. By toggling the 'Unassigned' label you can turn the background (unassigned) peaks on or off in the plot. By updating the slider in the Ion legend you can update the spectrum to only show the top X% of the peaks with labels. The top X% means any peak that is within X% of the highest intensity. By dragging in the spectrum you can zoom in to a specific part of the spectrum and use 'Zoom Out' to get back to the original zoom level. The annotation of the spectrum is based on the given sequence in the peptides file and is done with different software so inconsistencies are likely. The peaks are annotated based on the given sequence, with 20 ppm tolerance.

Copy Data

##### Spectrum 7046 (TSV)

###### Preview

```
Loading example...
```

*Click on the button to copy the data to your clipboard.*

Mz MinMz MaxIntensity Max

WidthHeightPeptide font sizePeptide stroke widthSpectrum font sizeSpectrum stroke widthCompact peptide

Ion legend

wxyz

abcd

OtherUnassignedIon^Charge^_Position_Show for top:%

VRQAPGRAJEW

01.15e+42.30e+43.45e+44.60e+4

Zoom Out

y^+1^_1_c^+1^_2_c^+1^_2_y^+1^_2_c^+2^_7_c^+1^_3_c^+2^_7_c^+1^_3_c^+2^_8_c^+2^_8_c^+1^_4_c^+2^_9_c^+2^_9_c^+2^_10_c^+2^_10_c^+1^_5_z^+2^_10_c^+1^_6_z^+1^_5_y^+1^_5_z^+1^_6_y^+1^_6_c^+1^_7_y^+1^_7_c^+1^_8_z^+1^_8_y^+1^_8_c^+1^_9_z^+1^_9_y^+1^_9_c^+1^_10_

0665133119962661

Fragment Matches Table

Show background peaks

| Position | Ion type | Intensity | mz Theoretical | mz Error (Th) | mz Error (ppm) | Charge | Series Number |
| --- | --- | --- | --- | --- | --- | --- | --- |
| - | - | 487.9 | 120.1 | - | - | 0 | - |
| - | - | 646.8 | 130.1 | - | - | 0 | - |
| - | - | 528.3 | 144.1 | - | - | 0 | - |
| - | - | 993.3 | 146.1 | - | - | 0 | - |
| - | - | 917.4 | 149 | - | - | 0 | - |
| - | - | 2258 | 149 | - | - | 0 | - |
| - | - | 604.6 | 157.1 | - | - | 0 | - |
| - | - | 2526 | 159.1 | - | - | 0 | - |
| - | - | 669.2 | 167.1 | - | - | 0 | - |
| - | - | 724 | 173 | - | - | 0 | - |
| - | - | 490.2 | 173.1 | - | - | 0 | - |
| - | - | 766.2 | 173.5 | - | - | 0 | - |
| - | - | 485.3 | 181.5 | - | - | 0 | - |
| - | - | 524.8 | 184.9 | - | - | 0 | - |
| - | - | 1455 | 185.1 | - | - | 0 | - |
| - | - | 578.3 | 187.1 | - | - | 0 | - |
| - | - | 8505 | 188.1 | - | - | 0 | - |
| - | - | 704.7 | 189.1 | - | - | 0 | - |
| - | - | 435.1 | 202.2 | - | - | 0 | - |
| - | - | 1243 | 203.1 | - | - | 0 | - |
| 11 | y | 5294 | 205.1 | 2.913E-05 | 0.1421 | +1 | 1 |
| - | - | 3.632E+04 | 209 | - | - | 0 | - |
| - | - | 758.7 | 212.1 | - | - | 0 | - |
| - | - | 1051 | 214.1 | - | - | 0 | - |
| - | - | 607.5 | 217.1 | - | - | 0 | - |
| - | - | 1219 | 219.1 | - | - | 0 | - |
| - | - | 517.2 | 219.7 | - | - | 0 | - |
| - | - | 4511 | 229.2 | - | - | 0 | - |
| - | - | 653.7 | 233.2 | - | - | 0 | - |
| - | - | 3366 | 239.2 | - | - | 0 | - |
| - | - | 811 | 242.2 | - | - | 0 | - |
| 2 | c | 2200 | 256.2 | 0.0001657 | 0.6466 | +1 | 2 |
| - | - | 800.7 | 263.1 | - | - | 0 | - |
| 2 | c | 6948 | 273.2 | 4.916E-05 | 0.1799 | +1 | 2 |
| - | - | 630.6 | 274.2 | - | - | 0 | - |
| - | - | 1636 | 299.2 | - | - | 0 | - |
| 10 | y | 696.4 | 316.1 | 0.0009445 | 2.988 | +1 | 2 |
| - | - | 526.9 | 316.7 | - | - | 0 | - |
| - | - | 2.431E+04 | 318.9 | - | - | 0 | - |
| - | - | 6788 | 341 | - | - | 0 | - |
| - | - | 699.4 | 343 | - | - | 0 | - |
| - | - | 2.65E+04 | 359 | - | - | 0 | - |
| - | - | 1855 | 370.2 | - | - | 0 | - |
| 7 | c | 990.3 | 383.2 | 0.0006566 | 1.713 | +2 | 7 |
| 3 | c | 1059 | 384.2 | 0.001071 | 2.788 | +1 | 3 |
| - | - | 7223 | 391.2 | - | - | 0 | - |
| 7 | c | 2.665E+04 | 391.7 | 0.0004415 | 1.127 | +2 | 7 |
| - | - | 684.1 | 391.9 | - | - | 0 | - |
| - | - | 9801 | 392.2 | - | - | 0 | - |
| - | - | 2111 | 392.7 | - | - | 0 | - |
| - | - | 921 | 393.2 | - | - | 0 | - |
| - | - | 597.2 | 393.2 | - | - | 0 | - |
| 3 | c | 2.18E+04 | 401.3 | 0.0001264 | 0.3151 | +1 | 3 |
| - | - | 3795 | 402.3 | - | - | 0 | - |
| - | - | 633.3 | 406.9 | - | - | 0 | - |
| - | - | 636.4 | 413.3 | - | - | 0 | - |
| 8 | c | 1066 | 418.7 | 0.0002271 | 0.5424 | +2 | 8 |
| - | - | 621.6 | 419.2 | - | - | 0 | - |
| - | - | 2337 | 426.8 | - | - | 0 | - |
| - | - | 698.2 | 427.2 | - | - | 0 | - |
| - | - | 1131 | 427.2 | - | - | 0 | - |
| 8 | c | 1.667E+04 | 427.3 | 1.409E-05 | 0.03298 | +2 | 8 |
| - | - | 9128 | 427.8 | - | - | 0 | - |
| - | - | 1868 | 428.1 | - | - | 0 | - |
| - | - | 1775 | 428.3 | - | - | 0 | - |
| - | - | 3593 | 428.9 | - | - | 0 | - |
| - | - | 2.451E+04 | 429.1 | - | - | 0 | - |
| - | - | 810.3 | 429.2 | - | - | 0 | - |
| - | - | 1100 | 439.3 | - | - | 0 | - |
| 4 | c | 1236 | 455.3 | 0.0002429 | 0.5335 | +1 | 4 |
| - | - | 1663 | 461.3 | - | - | 0 | - |
| - | - | 1021 | 461.8 | - | - | 0 | - |
| 9 | c | 3705 | 475.3 | 0.000462 | 0.972 | +2 | 9 |
| - | - | 1729 | 475.8 | - | - | 0 | - |
| - | - | 880.4 | 476.3 | - | - | 0 | - |
| 9 | c | 9984 | 483.8 | 0.000371 | 0.7669 | +2 | 9 |
| - | - | 5422 | 484.3 | - | - | 0 | - |
| - | - | 1616 | 484.8 | - | - | 0 | - |
| - | - | 6144 | 496.8 | - | - | 0 | - |
| - | - | 3039 | 497.3 | - | - | 0 | - |
| - | - | 921.3 | 497.8 | - | - | 0 | - |
| - | - | 978.3 | 513.8 | - | - | 0 | - |
| - | - | 564 | 521.8 | - | - | 0 | - |
| - | - | 703.3 | 525.3 | - | - | 0 | - |
| - | - | 4575 | 525.8 | - | - | 0 | - |
| - | - | 3006 | 526.3 | - | - | 0 | - |
| - | - | 1639 | 526.8 | - | - | 0 | - |
| - | - | 651.4 | 528.3 | - | - | 0 | - |
| 10 | c | 2.45E+04 | 539.8 | 8.932E-06 | 0.01655 | +2 | 10 |
| - | - | 1.362E+04 | 540.3 | - | - | 0 | - |
| - | - | 4066 | 540.8 | - | - | 0 | - |
| - | - | 1422 | 541.3 | - | - | 0 | - |
| - | - | 1071 | 542.3 | - | - | 0 | - |
| 10 | c | 4.55E+04 | 548.3 | 0.0001011 | 0.1843 | +2 | 10 |
| - | - | 2.835E+04 | 548.8 | - | - | 0 | - |
| - | - | 690 | 549.3 | - | - | 0 | - |
| - | - | 9085 | 549.3 | - | - | 0 | - |
| - | - | 2404 | 549.8 | - | - | 0 | - |
| - | - | 3203 | 555.3 | - | - | 0 | - |
| - | - | 2201 | 555.8 | - | - | 0 | - |
| - | - | 600.2 | 556.3 | - | - | 0 | - |
| - | - | 2061 | 561.3 | - | - | 0 | - |
| - | - | 1147 | 561.8 | - | - | 0 | - |
| - | - | 816.6 | 568.3 | - | - | 0 | - |
| 5 | c | 9040 | 569.4 | 0.0001841 | 0.3233 | +1 | 5 |
| - | - | 2557 | 570.4 | - | - | 0 | - |
| - | - | 4908 | 576.8 | - | - | 0 | - |
| - | - | 2453 | 577.3 | - | - | 0 | - |
| - | - | 2292 | 577.8 | - | - | 0 | - |
| - | - | 833.9 | 582.4 | - | - | 0 | - |
| 2 | z | 2757 | 584.3 | 1.812E-05 | 0.03101 | +2 | 10 |
| - | - | 1212 | 584.8 | - | - | 0 | - |
| - | - | 580.8 | 585.8 | - | - | 0 | - |
| - | - | 1604 | 604.3 | - | - | 0 | - |
| - | - | 1891 | 604.8 | - | - | 0 | - |
| - | - | 850.8 | 605.3 | - | - | 0 | - |
| - | - | 819.2 | 605.8 | - | - | 0 | - |
| - | - | 4029 | 611.9 | - | - | 0 | - |
| - | - | 1018 | 612.4 | - | - | 0 | - |
| - | - | 1748 | 614.3 | - | - | 0 | - |
| - | - | 2013 | 614.3 | - | - | 0 | - |
| - | - | 1159 | 615.3 | - | - | 0 | - |
| - | - | 677.8 | 615.4 | - | - | 0 | - |
| - | - | 2333 | 619.8 | - | - | 0 | - |
| - | - | 3659 | 620.4 | - | - | 0 | - |
| - | - | 6854 | 620.8 | - | - | 0 | - |
| - | - | 5097 | 621.3 | - | - | 0 | - |
| - | - | 2409 | 621.8 | - | - | 0 | - |
| - | - | 1737 | 625.4 | - | - | 0 | - |
| - | - | 1005 | 625.8 | - | - | 0 | - |
| 6 | c | 3.108E+04 | 626.4 | 0.0002658 | 0.4243 | +1 | 6 |
| - | - | 3047 | 626.8 | - | - | 0 | - |
| - | - | 1.032E+04 | 627.4 | - | - | 0 | - |
| - | - | 2587 | 628.4 | - | - | 0 | - |
| - | - | 759.1 | 629.4 | - | - | 0 | - |
| - | - | 4.071E+04 | 633.8 | - | - | 0 | - |
| - | - | 2.287E+04 | 634.3 | - | - | 0 | - |
| - | - | 9299 | 634.8 | - | - | 0 | - |
| - | - | 3188 | 635.3 | - | - | 0 | - |
| - | - | 709.7 | 635.8 | - | - | 0 | - |
| - | - | 1.73E+04 | 641.9 | - | - | 0 | - |
| - | - | 1.908E+04 | 642.4 | - | - | 0 | - |
| - | - | 1.08E+04 | 642.9 | - | - | 0 | - |
| - | - | 3931 | 643.4 | - | - | 0 | - |
| - | - | 5645 | 643.9 | - | - | 0 | - |
| - | - | 2909 | 644.4 | - | - | 0 | - |
| 7 | z | 1.801E+04 | 658.3 | 0.0001723 | 0.2617 | +1 | 5 |
| - | - | 6710 | 659.3 | - | - | 0 | - |
| - | - | 1338 | 660.4 | - | - | 0 | - |
| 7 | y | 1223 | 674.4 | 0.000952 | 1.412 | +1 | 5 |
| - | - | 1665 | 697.4 | - | - | 0 | - |
| 6 | z | 5148 | 715.4 | 0.0001516 | 0.2119 | +1 | 6 |
| - | - | 1683 | 716.4 | - | - | 0 | - |
| 6 | y | 913.6 | 731.4 | 0.002946 | 4.027 | +1 | 6 |
| - | - | 855.1 | 732.4 | - | - | 0 | - |
| - | - | 1394 | 781.5 | - | - | 0 | - |
| 7 | c | 8825 | 782.5 | 0.0001983 | 0.2534 | +1 | 7 |
| - | - | 3320 | 783.5 | - | - | 0 | - |
| - | - | 988.4 | 816.4 | - | - | 0 | - |
| - | - | 1097 | 824.5 | - | - | 0 | - |
| - | - | 1751 | 824.5 | - | - | 0 | - |
| - | - | 3506 | 825.5 | - | - | 0 | - |
| 5 | y | 1136 | 828.4 | 0.003739 | 4.513 | +1 | 7 |
| 8 | c | 5346 | 853.5 | 0.001606 | 1.882 | +1 | 8 |
| - | - | 5578 | 854.5 | - | - | 0 | - |
| - | - | 1996 | 855.5 | - | - | 0 | - |
| 4 | z | 1.894E+04 | 883.5 | 0.0004907 | 0.5554 | +1 | 8 |
| - | - | 9751 | 884.5 | - | - | 0 | - |
| - | - | 3030 | 885.5 | - | - | 0 | - |
| 4 | y | 2077 | 899.5 | 0.000538 | 0.5981 | +1 | 8 |
| - | - | 1513 | 900.5 | - | - | 0 | - |
| - | - | 1522 | 940.6 | - | - | 0 | - |
| 9 | c | 6438 | 966.6 | 0.001259 | 1.302 | +1 | 9 |
| - | - | 3383 | 967.6 | - | - | 0 | - |
| - | - | 699.8 | 981.6 | - | - | 0 | - |
| 3 | z | 1.447E+04 | 1012 | 0.0009017 | 0.8914 | +1 | 9 |
| - | - | 8306 | 1013 | - | - | 0 | - |
| - | - | 3499 | 1014 | - | - | 0 | - |
| 3 | y | 3500 | 1028 | 0.0004607 | 0.4484 | +1 | 9 |
| - | - | 1753 | 1029 | - | - | 0 | - |
| - | - | 1219 | 1038 | - | - | 0 | - |
| - | - | 1063 | 1039 | - | - | 0 | - |
| - | - | 2008 | 1052 | - | - | 0 | - |
| - | - | 5457 | 1053 | - | - | 0 | - |
| - | - | 2920 | 1054 | - | - | 0 | - |
| - | - | 782 | 1055 | - | - | 0 | - |
| - | - | 3404 | 1080 | - | - | 0 | - |
| - | - | 2399 | 1081 | - | - | 0 | - |
| 10 | c | 3500 | 1096 | 0.001066 | 0.9732 | +1 | 10 |
| - | - | 1.084E+04 | 1097 | - | - | 0 | - |
| - | - | 5046 | 1098 | - | - | 0 | - |
| - | - | 1369 | 1099 | - | - | 0 | - |
| - | - | 966.6 | 1154 | - | - | 0 | - |
| - | - | 969.6 | 1156 | - | - | 0 | - |
| - | - | 1153 | 1159 | - | - | 0 | - |
| - | - | 6284 | 1169 | - | - | 0 | - |
| - | - | 4668 | 1170 | - | - | 0 | - |
| - | - | 2127 | 1171 | - | - | 0 | - |
| - | - | 694 | 1181 | - | - | 0 | - |
| - | - | 1456 | 1224 | - | - | 0 | - |
| - | - | 9801 | 1225 | - | - | 0 | - |
| - | - | 8757 | 1226 | - | - | 0 | - |
| - | - | 3710 | 1227 | - | - | 0 | - |
| - | - | 793 | 1228 | - | - | 0 | - |
| - | - | 927.8 | 1239 | - | - | 0 | - |
| - | - | 2540 | 1240 | - | - | 0 | - |
| - | - | 3235 | 1241 | - | - | 0 | - |
| - | - | 2515 | 1242 | - | - | 0 | - |
| - | - | 1055 | 1243 | - | - | 0 | - |
| - | - | 4191 | 1251 | - | - | 0 | - |
| - | - | 3019 | 1252 | - | - | 0 | - |
| - | - | 1040 | 1253 | - | - | 0 | - |
| - | - | 622.2 | 1257 | - | - | 0 | - |
| - | - | 5114 | 1267 | - | - | 0 | - |
| - | - | 3.292E+04 | 1268 | - | - | 0 | - |
| - | - | 1043 | 1269 | - | - | 0 | - |
| - | - | 2.147E+04 | 1269 | - | - | 0 | - |
| - | - | 1.069E+04 | 1270 | - | - | 0 | - |
| - | - | 2577 | 1271 | - | - | 0 | - |
| - | - | 3375 | 1272 | - | - | 0 | - |
| - | - | 1662 | 1283 | - | - | 0 | - |
| - | - | 7474 | 1284 | - | - | 0 | - |
| - | - | 2.951E+04 | 1285 | - | - | 0 | - |
| - | - | 1.935E+04 | 1286 | - | - | 0 | - |
| - | - | 7631 | 1287 | - | - | 0 | - |
| - | - | 4181 | 1288 | - | - | 0 | - |
| - | - | 3745 | 1289 | - | - | 0 | - |
| - | - | 699.2 | 2635 | - | - | 0 | - |

m/z Charge Intensity FragmentType MassShift Position
120.0810317993164 0 487.8814
130.06533813476562 0 646.8041
144.08099365234375 0 528.3498
146.0602264404297 0 993.33466
148.95480346679688 0 917.37683
149.0448760986328 0 2257.5876
157.10894775390625 0 604.6033
159.09188842773438 0 2525.8264
167.05548095703125 0 669.22296
172.9772186279297 0 724.0239
173.1284942626953 0 490.20532
173.45211791992188 0 766.16205
181.49166870117188 0 485.31177
184.91343688964844 0 524.76245
185.11773681640625 0 1455.0623
187.0869903564453 0 578.32733
188.0707244873047 0 8505.365
189.0740203857422 0 704.7174
202.18246459960938 0 435.0636
203.12786865234375 0 1243.1704
205.09718322753906 0 5294.4727 y 10
208.9532470703125 0 36322.496
212.1393280029297 0 758.67834
214.1420440673828 0 1051.2261
217.1366424560547 0 607.4691
219.13388061523438 0 1218.6681
219.6771697998047 0 517.1812
229.1548309326172 0 4510.539
233.16546630859375 0 653.74066
239.15042114257812 0 3366.2185
242.16183471679688 0 811.02966
256.1766357421875 0 2200.4675 c Ammonia loss 1
263.1429138183594 0 800.6621
273.2033996582031 0 6948.2227 c 1
274.20635986328125 0 630.6342
299.2185363769531 0 1635.5043
316.130126953125 0 696.44824 y Water loss 9
316.7297668457031 0 526.9339
318.9223937988281 0 24307.592
341.0181884765625 0 6788.1245
342.9986267089844 0 699.38776
359.0283508300781 0 26502.947
370.23541259765625 0 1854.6631
383.22821044921875 0 990.29095 c Ammonia loss 6
384.2364501953125 0 1059.4242 c Ammonia loss 2
391.23687744140625 0 7223.1562
391.7403869628906 0 26647.363 c 6
391.855224609375 0 684.0845
392.24212646484375 0 9801.157
392.74200439453125 0 2111.1077
393.17742919921875 0 920.9888
393.21038818359375 0 597.2077
401.2620544433594 0 21797.87 c 2
402.2644958496094 0 3794.8843
406.9000549316406 0 633.2872
413.2745056152344 0 636.3672
418.746337890625 0 1065.993 c Ammonia loss 7
419.2127380371094 0 621.6367
426.7555847167969 0 2337.165
427.1598205566406 0 698.1565
427.20770263671875 0 1130.7277
427.2593994140625 0 16673.463 c 7
427.76080322265625 0 9127.958
428.1485900878906 0 1867.5815
428.2621765136719 0 1775.1174
428.8919372558594 0 3593.2502
429.0890808105469 0 24511.979
429.17626953125 0 810.30164
439.2601623535156 0 1100.1111
455.2727355957031 0 1235.719 c Ammonia loss 3
461.2908630371094 0 1663.244
461.7948913574219 0 1020.52545
475.2886047363281 0 3705.3223 c Ammonia loss 8
475.7901916503906 0 1728.9696
476.2901306152344 0 880.44415
483.8017883300781 0 9983.67 c 8
484.302734375 0 5421.5864
484.804931640625 0 1615.6871
496.80908203125 0 6144.2734
497.31085205078125 0 3039.4287
497.81201171875 0 921.3281
513.78662109375 0 978.3474
521.8104248046875 0 563.9696
525.3385620117188 0 703.3387
525.811767578125 0 4574.68
526.3135375976562 0 3006.1863
526.8148193359375 0 1638.6466
528.28662109375 0 651.40497
539.8094482421875 0 24501.982 c Ammonia loss 9
540.3111572265625 0 13618.689
540.8126220703125 0 4066.2278
541.3142700195312 0 1421.79
542.3322143554688 0 1070.8055
548.3228149414062 0 45499.51 c 9
548.8240356445312 0 28354.316
549.2860717773438 0 690.03174
549.3255004882812 0 9085.003
549.826416015625 0 2403.9995
555.2972412109375 0 3203.062
555.797119140625 0 2200.608
556.301025390625 0 600.1883
561.3302001953125 0 2061.3457
561.8322143554688 0 1147.4902
568.345947265625 0 816.5712
569.3519897460938 0 9039.882 c 4
570.3545532226562 0 2557.2886
576.8201293945312 0 4907.8384
577.32373046875 0 2453.3557
577.8490600585938 0 2292.2764
582.3604125976562 0 833.9415
584.310791015625 0 2756.505 z 1
584.8101196289062 0 1211.8689
585.808349609375 0 580.77747
604.33837890625 0 1603.5038
604.8331298828125 0 1890.5477
605.3385620117188 0 850.8248
605.8281860351562 0 819.1899
611.8506469726562 0 4029.274
612.354248046875 0 1018.4972
614.328369140625 0 1748.0857
614.3496704101562 0 2012.6113
615.2885131835938 0 1159.2604
615.3570556640625 0 677.8494
619.8489990234375 0 2333.278
620.3567504882812 0 3658.6057
620.8464965820312 0 6854.4536
621.346923828125 0 5097.105
621.8475341796875 0 2408.7983
625.3655395507812 0 1736.6772
625.8358764648438 0 1004.7046
626.37353515625 0 31084.625 c 5
626.8350830078125 0 3046.6567
627.3760375976562 0 10322.449
628.3783569335938 0 2586.5803
629.3804321289062 0 759.14343
633.8450927734375 0 40707.105
634.3464965820312 0 22874.32
634.84814453125 0 9298.535
635.3486328125 0 3187.7466
635.847900390625 0 709.6563
641.8544311523438 0 17297.758
642.3567504882812 0 19077.26
642.8587646484375 0 10799.045
643.3599243164062 0 3931.2405
643.8618774414062 0 5645.2354
644.3618774414062 0 2908.7622
658.3431396484375 0 18014.195 z 6
659.3458251953125 0 6709.556
660.3504028320312 0 1338.2959
674.361083984375 0 1222.8346 y 6
697.4371337890625 0 1665.3502
715.3646240234375 0 5148.381 z 5
716.3658447265625 0 1682.9457
731.3805541992188 0 913.5846 y 5
732.392822265625 0 855.11035
781.46533203125 0 1394.284
782.4741821289062 0 8825.254 c 6
783.4766845703125 0 3320.105
816.4421997070312 0 988.42053
824.4508056640625 0 1097.2743
824.530517578125 0 1750.6177
825.5340576171875 0 3505.652
828.4400024414062 0 1136.0767 y 4
853.5098876953125 0 5346.2935 c 7
854.5247192382812 0 5578.364
855.5296020507812 0 1996.3811
883.4541625976562 0 18936.104 z 3
884.45703125 0 9750.655
885.4592895507812 0 3030.13
899.4728393554688 0 2077.4185 y 3
900.4717407226562 0 1513.3715
940.562744140625 0 1521.9431
966.5942993164062 0 6438.1963 c 8
967.5960693359375 0 3382.7095
981.5519409179688 0 699.78754
1011.5123291015625 0 14470.433 z 2
1012.5150756835938 0 8306.472
1013.5172119140625 0 3498.503
1027.531494140625 0 3500.3572 y 2
1028.5303955078125 0 1752.882
1037.5966796875 0 1219.1925
1038.591064453125 0 1062.5693
1051.6229248046875 0 2008.0197
1052.62744140625 0 5456.8955
1053.62646484375 0 2919.9927
1054.6253662109375 0 781.98175
1079.6162109375 0 3404.428
1080.622314453125 0 2398.6492
1095.6370849609375 0 3499.5889 c 9
1096.6441650390625 0 10836.557
1097.647705078125 0 5045.8496
1098.6527099609375 0 1368.7527
1153.6263427734375 0 966.5629
1155.705322265625 0 969.63745
1158.6241455078125 0 1152.6246
1168.61962890625 0 6284.0537
1169.6248779296875 0 4668.4053
1170.6256103515625 0 2126.7495
1180.6063232421875 0 693.988
1223.702880859375 0 1455.9186
1224.6612548828125 0 9800.548
1225.6634521484375 0 8756.695
1226.6669921875 0 3709.9463
1227.676513671875 0 792.9847
1238.701416015625 0 927.82227
1239.6943359375 0 2540.263
1240.698486328125 0 3235.247
1241.6951904296875 0 2515.0776
1242.7052001953125 0 1054.929
1250.66064453125 0 4191.1914
1251.6646728515625 0 3019.047
1252.6656494140625 0 1039.707
1256.7086181640625 0 622.1986
1266.69140625 0 5113.6074
1267.6888427734375 0 32915.473
1268.521728515625 0 1042.5652
1268.6905517578125 0 21468.488
1269.6943359375 0 10686.361
1270.697021484375 0 2576.9885
1271.7027587890625 0 3374.9287
1282.69287109375 0 1662.2178
1283.70458984375 0 7474.1406
1284.7144775390625 0 29508.465
1285.7164306640625 0 19345.506
1286.7200927734375 0 7631.2974
1287.720458984375 0 4181.2583
1288.7276611328125 0 3745.1375
2634.945068359375 0 699.1855

Spectrum Details

|  |  |
| --- | --- |
| Matched peaks? Matched peaks The total absolute number of peaks matched. Additionally in brackets the total fraction of peaks matched and the total number of peaks is shown. | 31 (13.60% of 228) |
| FDR? FDR The false discovery rate estimated for this peptide. It is calculated by matching all theoretical fragments with a non-integer shift with the raw peaks for this spectrum. This is done with 40 different shifts. The resulting percentage is the average number of annotated peaks over the number of annotated peaks with the correct spectrum. | 1.61% |
| Satellite FDR? Satellite FDR See the FDR for details on its calculation. This satellite ion specific FDR only contains the satellite ions (d/w) for I/L/J positions. | ∞ |
| PSM Score? PSM Score The PSM Score as given by Hecklib to this annotated spectrum. It is shown with three significant figures. | 264 |

#### Reverse Lookup? Reverse Lookup All places where this read could be placed.

| Group | Segment | Template | Template Part | Read Part | Score | Unique |
| --- | --- | --- | --- | --- | --- | --- |
| Homo sapiens Heavy Chain | IGHV | IGHV1-45 | [36..47] | [0..11] | 79 | True |

| Recombined | Template Part | Read Part | Score | Unique |
| --- | --- | --- | --- | --- |
| REC-0-1 | [36..47] | [0..11] | 88 | True |

#### Meta Information from Multiple reads

##### Number of combined reads

9

##### Intensity

0.8417

##### TotalArea

3.42E+09

##### Changes to the peptide sequence

VRQAPGRAJEW

L→JNo support for either Leucine or Isoleucine based on side chain ions (Position: 9)

#### Positional Score

Copy Data

##### Positional Score (TSV)

###### Preview

```
Loading example...
```

*Click on the button to copy the data to your clipboard.*

10012345678910

Label Value
"0" 0.713
"1" 0.706
"2" 0.728
"3" 0.726
"4" 0.748
"5" 0.77
"6" 0.766
"7" 0.757
"8" 0.768
"9" 0.777
"10" 0.776

#### Meta Information from PEAKS

##### Scan Identifier

F2:5723

##### Original sequence

V

R

Q

A

P

G

R

A

L

E

W

##### Posttranslational Modifications

##### Source File

D:\separate\_stitch\_analyses\xle-disambiguation\raw\20210323\_F1\_UM1\_Peng0013\_SA\_F59\_ingel\_3ug\_TL.raw

##### Fraction

2

##### Scan Feature

F2:1495

##### De Novo Score

99

##### ConfidenceScore

99

### m/z

428.2395

##### Mass

1281.6941

##### Charge

3

##### Retention Time

31.25

##### Predicted Retention Time

-

##### Area

2.74E+07

##### Parts Per Million

2

##### Fragmentation mode

ETHCD

##### Originating file

01 D:\separate\_stitch\_analyses\xle-disambiguation\20210325\_F59\_3ug\_DENOVO\_12.csv

#### Meta Information from PEAKS

##### Scan Identifier

F2:5776

##### Original sequence

V

R

Q

A

P

G

R

A

L

E

W

##### Posttranslational Modifications

##### Source File

D:\separate\_stitch\_analyses\xle-disambiguation\raw\20210323\_F1\_UM1\_Peng0013\_SA\_F59\_ingel\_3ug\_TL.raw

##### Fraction

2

##### Scan Feature

F2:1495

##### De Novo Score

97

##### ConfidenceScore

97

### m/z

428.2395

##### Mass

1281.6941

##### Charge

3

##### Retention Time

31.25

##### Predicted Retention Time

-

##### Area

2.74E+07

##### Parts Per Million

2

##### Fragmentation mode

ETHCD

##### Originating file

01 D:\separate\_stitch\_analyses\xle-disambiguation\20210325\_F59\_3ug\_DENOVO\_12.csv

#### Meta Information from PEAKS

##### Scan Identifier

F3:5752

##### Original sequence

V

R

Q

A

P

G

R

A

L

E

W

##### Posttranslational Modifications

##### Source File

D:\separate\_stitch\_analyses\xle-disambiguation\raw\20210323\_F1\_UM1\_Peng0013\_SA\_F59\_ingel\_3ug\_chymo.raw

##### Fraction

3

##### Scan Feature

F3:1539

##### De Novo Score

97

##### ConfidenceScore

97

### m/z

428.2397

##### Mass

1281.6941

##### Charge

3

##### Retention Time

30.93

##### Predicted Retention Time

-

##### Area

8.377E+08

##### Parts Per Million

2.4

##### Fragmentation mode

ETHCD

##### Originating file

01 D:\separate\_stitch\_analyses\xle-disambiguation\20210325\_F59\_3ug\_DENOVO\_12.csv

#### Meta Information from PEAKS

##### Scan Identifier

F3:5690

##### Original sequence

V

R

Q

A

P

G

R

A

L

E

W

##### Posttranslational Modifications

##### Source File

D:\separate\_stitch\_analyses\xle-disambiguation\raw\20210323\_F1\_UM1\_Peng0013\_SA\_F59\_ingel\_3ug\_chymo.raw

##### Fraction

3

##### Scan Feature

F3:1539

##### De Novo Score

96

##### ConfidenceScore

96

### m/z

428.2397

##### Mass

1281.6941

##### Charge

3

##### Retention Time

30.93

##### Predicted Retention Time

-

##### Area

8.377E+08

##### Parts Per Million

2.4

##### Fragmentation mode

ETHCD

##### Originating file

01 D:\separate\_stitch\_analyses\xle-disambiguation\20210325\_F59\_3ug\_DENOVO\_12.csv

#### Meta Information from PEAKS

##### Scan Identifier

F3:4540

##### Original sequence

V

R

Q

A

P

G

R

A

L

E

W

+15.99

##### Posttranslational Modifications

Oxidation (HW)

##### Source File

D:\separate\_stitch\_analyses\xle-disambiguation\raw\20210323\_F1\_UM1\_Peng0013\_SA\_F59\_ingel\_3ug\_chymo.raw

##### Fraction

3

##### Scan Feature

F3:1749

##### De Novo Score

96

##### ConfidenceScore

96

### m/z

433.5705

##### Mass

1297.689

##### Charge

3

##### Retention Time

25.09

##### Predicted Retention Time

-

##### Area

1.46E+07

##### Parts Per Million

0.5

##### Fragmentation mode

ETHCD

##### Originating file

01 D:\separate\_stitch\_analyses\xle-disambiguation\20210325\_F59\_3ug\_DENOVO\_12.csv

#### Meta Information from PEAKS

##### Scan Identifier

F3:5805

##### Original sequence

V

R

Q

A

P

G

R

A

L

E

W

##### Posttranslational Modifications

##### Source File

D:\separate\_stitch\_analyses\xle-disambiguation\raw\20210323\_F1\_UM1\_Peng0013\_SA\_F59\_ingel\_3ug\_chymo.raw

##### Fraction

3

##### Scan Feature

F3:1539

##### De Novo Score

96

##### ConfidenceScore

96

### m/z

428.2397

##### Mass

1281.6941

##### Charge

3

##### Retention Time

30.93

##### Predicted Retention Time

-

##### Area

8.377E+08

##### Parts Per Million

2.4

##### Fragmentation mode

ETHCD

##### Originating file

01 D:\separate\_stitch\_analyses\xle-disambiguation\20210325\_F59\_3ug\_DENOVO\_12.csv

#### Meta Information from PEAKS

##### Scan Identifier

F3:5860

##### Original sequence

V

R

Q

A

P

G

R

A

L

E

W

##### Posttranslational Modifications

##### Source File

D:\separate\_stitch\_analyses\xle-disambiguation\raw\20210323\_F1\_UM1\_Peng0013\_SA\_F59\_ingel\_3ug\_chymo.raw

##### Fraction

3

##### Scan Feature

F3:1539

##### De Novo Score

95

##### ConfidenceScore

95

### m/z

428.2397

##### Mass

1281.6941

##### Charge

3

##### Retention Time

30.93

##### Predicted Retention Time

-

##### Area

8.377E+08

##### Parts Per Million

2.4

##### Fragmentation mode

ETHCD

##### Originating file

01 D:\separate\_stitch\_analyses\xle-disambiguation\20210325\_F59\_3ug\_DENOVO\_12.csv

#### Meta Information from PEAKS

##### Scan Identifier

F3:6378

##### Original sequence

V

R

Q

A

P

G

R

A

L

E

W

##### Posttranslational Modifications

##### Source File

D:\separate\_stitch\_analyses\xle-disambiguation\raw\20210323\_F1\_UM1\_Peng0013\_SA\_F59\_ingel\_3ug\_chymo.raw

##### Fraction

3

##### Scan Feature

-

##### De Novo Score

95

##### ConfidenceScore

95

### m/z

428.2392

##### Mass

1281.6941

##### Charge

3

##### Retention Time

35.39

##### Predicted Retention Time

-

##### Area

0

##### Parts Per Million

1.3

##### Fragmentation mode

ETHCD

##### Originating file

01 D:\separate\_stitch\_analyses\xle-disambiguation\20210325\_F59\_3ug\_DENOVO\_12.csv

#### Meta Information from PEAKS

##### Scan Identifier

F3:7046

##### Original sequence

V

R

Q

A

P

G

R

A

L

E

W

##### Posttranslational Modifications

##### Source File

D:\separate\_stitch\_analyses\xle-disambiguation\raw\20210323\_F1\_UM1\_Peng0013\_SA\_F59\_ingel\_3ug\_chymo.raw

##### Fraction

3

##### Scan Feature

-

##### De Novo Score

95

##### ConfidenceScore

95

### m/z

428.2395

##### Mass

1281.6941

##### Charge

3

##### Retention Time

39.16

##### Predicted Retention Time

-

##### Area

0

##### Parts Per Million

2.1

##### Fragmentation mode

ETHCD

##### Originating file

01 D:\separate\_stitch\_analyses\xle-disambiguation\20210325\_F59\_3ug\_DENOVO\_12.csv
