## Supplementary material for "A handle on mass coincidence errors in *de novo* sequencing of antibodies by bottom-up proteomics": Combined_095.html

Details Combined\_095 | Stitch OverviewUndefined

### Read Combined\_095

#### Sequence (length=13)

FYYCAKDVRPYYD

#### Spectrum 6983? Spectrum 6983 The raw spectrum of this peptide as annotated by Hecklib. The fragments are coloured according to ion type (see legend). Any peaks with a star '\*' as text can be hovered over to see the full details, first the ion type second the mass shift type. By hovering over the amino acids in the peptide or ions in the legend the corresponding peaks are highlighted. By toggling the 'Unassigned' label you can turn the background (unassigned) peaks on or off in the plot. By updating the slider in the Ion legend you can update the spectrum to only show the top X% of the peaks with labels. The top X% means any peak that is within X% of the highest intensity. By dragging in the spectrum you can zoom in to a specific part of the spectrum and use 'Zoom Out' to get back to the original zoom level. The annotation of the spectrum is based on the given sequence in the peptides file and is done with different software so inconsistencies are likely. The peaks are annotated based on the given sequence, with 20 ppm tolerance.

Copy Data

##### Spectrum 6983 (TSV)

###### Preview

```
Loading example...
```

*Click on the button to copy the data to your clipboard.*

Mz MinMz MaxIntensity Max

WidthHeightPeptide font sizePeptide stroke widthSpectrum font sizeSpectrum stroke widthCompact peptide

Ion legend

wxyz

abcd

OtherUnassignedIon^Charge^_Position_Show for top:%

FYYCAKDVRPYYD

08.18e+41.64e+52.45e+53.27e+5

Zoom Out

y^+1^_1_y^+1^_2_a^+1^_2_y^+1^_2_b^+1^_2_b^+3^_9_y^+3^_10_y^+1^_3_b^+1^_3_b^+2^_7_b^+3^_11_y^+2^_8_y^+2^_8_y^+2^_8_y^+1^_4_b^+3^_12_y^+2^_9_y^+2^_9_y^+1^_4_y^+2^_9_\*\*\*b^+2^_9_b^+1^_4_y^+2^_10_y^+2^_10_y^+2^_10_b^+2^_10_y^+1^_5_y^+1^_5_b^+1^_5_y^+1^_5_y^+2^_11_y^+2^_11_y^+2^_11_b^+2^_11_y^+1^_6_y^+2^_12_y^+2^_12_y^+2^_12_y^+1^_6_b^+2^_12_y^+1^_7_y^+1^_7_y^+1^_7_b^+1^_7_y^+1^_8_y^+1^_8_y^+1^_8_y^+1^_9_y^+1^_9_y^+1^_9_b^+1^_9_y^+1^_10_y^+1^_10_y^+1^_10_

0543108616292172

Fragment Matches Table

Show background peaks

| Position | Ion type | Intensity | mz Theoretical | mz Error (Th) | mz Error (ppm) | Charge | Series Number |
| --- | --- | --- | --- | --- | --- | --- | --- |
| - | - | 3.022E+05 | 120.1 | - | - | 0 | - |
| - | - | 743.4 | 121.1 | - | - | 0 | - |
| - | - | 862.4 | 121.1 | - | - | 0 | - |
| - | - | 2.642E+04 | 121.1 | - | - | 0 | - |
| - | - | 920.5 | 122.1 | - | - | 0 | - |
| - | - | 464.3 | 125.1 | - | - | 0 | - |
| - | - | 375.8 | 126.2 | - | - | 0 | - |
| - | - | 442.7 | 128.2 | - | - | 0 | - |
| - | - | 5.466E+04 | 129.1 | - | - | 0 | - |
| - | - | 1757 | 130.1 | - | - | 0 | - |
| - | - | 3246 | 130.1 | - | - | 0 | - |
| - | - | 659.9 | 131 | - | - | 0 | - |
| - | - | 1517 | 132.1 | - | - | 0 | - |
| - | - | 504.3 | 133.1 | - | - | 0 | - |
| - | - | 650 | 134 | - | - | 0 | - |
| - | - | 9918 | 134 | - | - | 0 | - |
| 13 | y | 3.479E+04 | 134 | 0.000275 | 2.052 | +1 | 1 |
| - | - | 1487 | 135 | - | - | 0 | - |
| - | - | 464.3 | 135.4 | - | - | 0 | - |
| - | - | 457.2 | 135.5 | - | - | 0 | - |
| - | - | 3.239E+05 | 136.1 | - | - | 0 | - |
| - | - | 1171 | 137.1 | - | - | 0 | - |
| - | - | 2.889E+04 | 137.1 | - | - | 0 | - |
| - | - | 511.4 | 138.1 | - | - | 0 | - |
| - | - | 1140 | 138.1 | - | - | 0 | - |
| - | - | 764.3 | 139.1 | - | - | 0 | - |
| - | - | 920.4 | 140.1 | - | - | 0 | - |
| - | - | 994.6 | 141.1 | - | - | 0 | - |
| - | - | 2862 | 143.1 | - | - | 0 | - |
| - | - | 1945 | 146.1 | - | - | 0 | - |
| - | - | 2305 | 147 | - | - | 0 | - |
| - | - | 597 | 152.1 | - | - | 0 | - |
| - | - | 1178 | 155.1 | - | - | 0 | - |
| - | - | 2426 | 157.1 | - | - | 0 | - |
| - | - | 591.4 | 158.1 | - | - | 0 | - |
| - | - | 832.2 | 158.1 | - | - | 0 | - |
| - | - | 5594 | 159.1 | - | - | 0 | - |
| - | - | 2359 | 159.1 | - | - | 0 | - |
| - | - | 844.6 | 160.1 | - | - | 0 | - |
| - | - | 1095 | 160.1 | - | - | 0 | - |
| - | - | 638.3 | 162 | - | - | 0 | - |
| - | - | 522.3 | 162.1 | - | - | 0 | - |
| - | - | 1160 | 164.1 | - | - | 0 | - |
| - | - | 574.9 | 165.1 | - | - | 0 | - |
| - | - | 594.6 | 166.1 | - | - | 0 | - |
| - | - | 504.5 | 167.6 | - | - | 0 | - |
| - | - | 1085 | 169.1 | - | - | 0 | - |
| - | - | 948.9 | 170.1 | - | - | 0 | - |
| - | - | 542.8 | 171.1 | - | - | 0 | - |
| - | - | 583.2 | 171.1 | - | - | 0 | - |
| - | - | 2799 | 171.1 | - | - | 0 | - |
| - | - | 781.9 | 173.1 | - | - | 0 | - |
| - | - | 561.1 | 173.1 | - | - | 0 | - |
| - | - | 556.9 | 173.1 | - | - | 0 | - |
| - | - | 3146 | 173.5 | - | - | 0 | - |
| - | - | 1745 | 174.1 | - | - | 0 | - |
| - | - | 807.7 | 174.1 | - | - | 0 | - |
| - | - | 1.049E+04 | 175.1 | - | - | 0 | - |
| - | - | 473.9 | 175.9 | - | - | 0 | - |
| - | - | 492.9 | 176 | - | - | 0 | - |
| - | - | 1792 | 176.1 | - | - | 0 | - |
| - | - | 630.8 | 180.1 | - | - | 0 | - |
| - | - | 678.8 | 181.1 | - | - | 0 | - |
| - | - | 1211 | 182.1 | - | - | 0 | - |
| - | - | 1002 | 183.1 | - | - | 0 | - |
| - | - | 1411 | 185.1 | - | - | 0 | - |
| - | - | 2001 | 185.1 | - | - | 0 | - |
| - | - | 935.3 | 186.1 | - | - | 0 | - |
| - | - | 878.6 | 187.1 | - | - | 0 | - |
| - | - | 892.4 | 187.1 | - | - | 0 | - |
| - | - | 7433 | 187.1 | - | - | 0 | - |
| - | - | 539.7 | 187.5 | - | - | 0 | - |
| - | - | 1340 | 189.1 | - | - | 0 | - |
| - | - | 4393 | 191.1 | - | - | 0 | - |
| - | - | 742.1 | 192 | - | - | 0 | - |
| - | - | 588.7 | 192.1 | - | - | 0 | - |
| - | - | 1094 | 192.1 | - | - | 0 | - |
| - | - | 484.5 | 192.2 | - | - | 0 | - |
| - | - | 604.2 | 195.1 | - | - | 0 | - |
| - | - | 1498 | 197.1 | - | - | 0 | - |
| - | - | 1716 | 199.1 | - | - | 0 | - |
| - | - | 1.306E+04 | 200.1 | - | - | 0 | - |
| - | - | 1966 | 201.1 | - | - | 0 | - |
| - | - | 1256 | 201.1 | - | - | 0 | - |
| - | - | 836.9 | 201.1 | - | - | 0 | - |
| - | - | 908.6 | 202 | - | - | 0 | - |
| - | - | 1173 | 202.1 | - | - | 0 | - |
| - | - | 1052 | 204.1 | - | - | 0 | - |
| - | - | 8836 | 205.1 | - | - | 0 | - |
| - | - | 770.8 | 206.1 | - | - | 0 | - |
| - | - | 980.1 | 209.1 | - | - | 0 | - |
| - | - | 758.5 | 210.1 | - | - | 0 | - |
| - | - | 717.9 | 214.2 | - | - | 0 | - |
| - | - | 843.5 | 215 | - | - | 0 | - |
| - | - | 2014 | 215.1 | - | - | 0 | - |
| - | - | 1069 | 215.1 | - | - | 0 | - |
| - | - | 1352 | 216.1 | - | - | 0 | - |
| - | - | 601 | 220.1 | - | - | 0 | - |
| - | - | 626.6 | 222.1 | - | - | 0 | - |
| - | - | 3653 | 226.1 | - | - | 0 | - |
| - | - | 830 | 227.1 | - | - | 0 | - |
| - | - | 566 | 228.1 | - | - | 0 | - |
| - | - | 2959 | 229.1 | - | - | 0 | - |
| - | - | 528.8 | 231.1 | - | - | 0 | - |
| - | - | 2823 | 233.1 | - | - | 0 | - |
| - | - | 1.142E+04 | 233.1 | - | - | 0 | - |
| - | - | 2.074E+04 | 233.1 | - | - | 0 | - |
| - | - | 1096 | 233.1 | - | - | 0 | - |
| - | - | 1633 | 234.1 | - | - | 0 | - |
| - | - | 923.9 | 234.1 | - | - | 0 | - |
| - | - | 2719 | 234.1 | - | - | 0 | - |
| - | - | 779 | 237.1 | - | - | 0 | - |
| - | - | 2902 | 237.1 | - | - | 0 | - |
| - | - | 1200 | 238.1 | - | - | 0 | - |
| - | - | 1013 | 238.1 | - | - | 0 | - |
| - | - | 2.109E+04 | 239.2 | - | - | 0 | - |
| - | - | 2158 | 240.2 | - | - | 0 | - |
| - | - | 2595 | 244.1 | - | - | 0 | - |
| - | - | 625.1 | 246.1 | - | - | 0 | - |
| - | - | 1043 | 247.1 | - | - | 0 | - |
| - | - | 480.6 | 247.4 | - | - | 0 | - |
| - | - | 585.3 | 250.2 | - | - | 0 | - |
| - | - | 5843 | 252.1 | - | - | 0 | - |
| - | - | 590 | 253.1 | - | - | 0 | - |
| - | - | 2241 | 254.1 | - | - | 0 | - |
| - | - | 1270 | 254.2 | - | - | 0 | - |
| - | - | 9023 | 256.2 | - | - | 0 | - |
| - | - | 948.7 | 257.2 | - | - | 0 | - |
| - | - | 1.674E+04 | 261.1 | - | - | 0 | - |
| - | - | 680.6 | 262.1 | - | - | 0 | - |
| - | - | 537.8 | 262.1 | - | - | 0 | - |
| - | - | 2424 | 262.1 | - | - | 0 | - |
| - | - | 829.9 | 266.1 | - | - | 0 | - |
| - | - | 576.4 | 270.1 | - | - | 0 | - |
| - | - | 2886 | 270.2 | - | - | 0 | - |
| - | - | 557.1 | 271 | - | - | 0 | - |
| - | - | 1847 | 272.1 | - | - | 0 | - |
| - | - | 781.3 | 274.1 | - | - | 0 | - |
| - | - | 734.7 | 274.2 | - | - | 0 | - |
| - | - | 858 | 276.1 | - | - | 0 | - |
| - | - | 645.3 | 278.2 | - | - | 0 | - |
| 12 | y | 710.1 | 279.1 | 0.0003829 | 1.372 | +1 | 2 |
| - | - | 1736 | 280.1 | - | - | 0 | - |
| - | - | 644.6 | 280.1 | - | - | 0 | - |
| - | - | 1235 | 282.2 | - | - | 0 | - |
| 2 | a | 1.572E+05 | 283.1 | 0.0003659 | 1.292 | +1 | 2 |
| - | - | 481.7 | 283.9 | - | - | 0 | - |
| - | - | 2.563E+04 | 284.1 | - | - | 0 | - |
| - | - | 2479 | 285.2 | - | - | 0 | - |
| - | - | 803 | 286.1 | - | - | 0 | - |
| - | - | 3.477E+04 | 297.1 | - | - | 0 | - |
| 12 | y | 1.79E+04 | 297.1 | 0.0006825 | 2.297 | +1 | 2 |
| - | - | 1444 | 297.2 | - | - | 0 | - |
| - | - | 5972 | 298.1 | - | - | 0 | - |
| - | - | 2668 | 298.1 | - | - | 0 | - |
| - | - | 2082 | 299.1 | - | - | 0 | - |
| - | - | 4.533E+04 | 299.1 | - | - | 0 | - |
| - | - | 6413 | 300.1 | - | - | 0 | - |
| - | - | 1413 | 300.2 | - | - | 0 | - |
| - | - | 698.8 | 301.1 | - | - | 0 | - |
| - | - | 7888 | 304.1 | - | - | 0 | - |
| - | - | 1403 | 305.1 | - | - | 0 | - |
| - | - | 1173 | 306.1 | - | - | 0 | - |
| - | - | 592.4 | 309.2 | - | - | 0 | - |
| 2 | b | 3.628E+04 | 311.1 | 0.0004159 | 1.337 | +1 | 2 |
| - | - | 6146 | 312.1 | - | - | 0 | - |
| - | - | 997.7 | 313.1 | - | - | 0 | - |
| - | - | 3560 | 315.2 | - | - | 0 | - |
| - | - | 2066 | 318.2 | - | - | 0 | - |
| - | - | 1334 | 319.2 | - | - | 0 | - |
| - | - | 1.744E+04 | 325.1 | - | - | 0 | - |
| - | - | 2701 | 326.1 | - | - | 0 | - |
| - | - | 691.9 | 326.2 | - | - | 0 | - |
| - | - | 1.658E+04 | 327.1 | - | - | 0 | - |
| - | - | 2921 | 328.1 | - | - | 0 | - |
| - | - | 740.3 | 336.2 | - | - | 0 | - |
| - | - | 775.9 | 337.2 | - | - | 0 | - |
| - | - | 713.4 | 340.2 | - | - | 0 | - |
| - | - | 716.8 | 342.2 | - | - | 0 | - |
| - | - | 869.5 | 343.1 | - | - | 0 | - |
| - | - | 1005 | 343.2 | - | - | 0 | - |
| - | - | 699.9 | 343.2 | - | - | 0 | - |
| - | - | 2098 | 350.1 | - | - | 0 | - |
| - | - | 763.1 | 353.2 | - | - | 0 | - |
| - | - | 610.9 | 353.8 | - | - | 0 | - |
| - | - | 2219 | 354.2 | - | - | 0 | - |
| - | - | 1142 | 355.2 | - | - | 0 | - |
| - | - | 2701 | 361.2 | - | - | 0 | - |
| - | - | 817.2 | 364.2 | - | - | 0 | - |
| - | - | 879.6 | 365.1 | - | - | 0 | - |
| - | - | 660.7 | 366.1 | - | - | 0 | - |
| - | - | 1181 | 366.1 | - | - | 0 | - |
| - | - | 515 | 366.2 | - | - | 0 | - |
| - | - | 2368 | 370.3 | - | - | 0 | - |
| - | - | 511.6 | 371.2 | - | - | 0 | - |
| - | - | 5790 | 371.2 | - | - | 0 | - |
| - | - | 951.3 | 371.2 | - | - | 0 | - |
| - | - | 573 | 372.2 | - | - | 0 | - |
| - | - | 1597 | 372.2 | - | - | 0 | - |
| - | - | 3491 | 382.1 | - | - | 0 | - |
| - | - | 1722 | 383.1 | - | - | 0 | - |
| - | - | 743.1 | 383.1 | - | - | 0 | - |
| - | - | 733.2 | 383.2 | - | - | 0 | - |
| - | - | 961.6 | 386.2 | - | - | 0 | - |
| - | - | 1514 | 387.2 | - | - | 0 | - |
| - | - | 587.5 | 389.2 | - | - | 0 | - |
| - | - | 4057 | 396.1 | - | - | 0 | - |
| - | - | 3586 | 396.2 | - | - | 0 | - |
| 9 | b | 1437 | 396.2 | 0.006474 | 16.34 | +3 | 9 |
| - | - | 1253 | 397.1 | - | - | 0 | - |
| - | - | 3023 | 399.2 | - | - | 0 | - |
| - | - | 7108 | 400.2 | - | - | 0 | - |
| - | - | 797.8 | 400.2 | - | - | 0 | - |
| - | - | 1769 | 401.2 | - | - | 0 | - |
| - | - | 725.9 | 401.2 | - | - | 0 | - |
| - | - | 849.8 | 405.2 | - | - | 0 | - |
| - | - | 941 | 414.2 | - | - | 0 | - |
| - | - | 554.2 | 415.1 | - | - | 0 | - |
| - | - | 2318 | 417.2 | - | - | 0 | - |
| 4 | y | 3684 | 424.2 | 0.001525 | 3.596 | +3 | 10 |
| - | - | 927.7 | 425.2 | - | - | 0 | - |
| - | - | 996.2 | 429.2 | - | - | 0 | - |
| - | - | 798.8 | 439.2 | - | - | 0 | - |
| - | - | 540.9 | 443.2 | - | - | 0 | - |
| - | - | 1334 | 447.7 | - | - | 0 | - |
| - | - | 1362 | 447.7 | - | - | 0 | - |
| - | - | 812.2 | 448.2 | - | - | 0 | - |
| - | - | 588.3 | 456.2 | - | - | 0 | - |
| - | - | 2750 | 458.2 | - | - | 0 | - |
| - | - | 803.4 | 459.2 | - | - | 0 | - |
| 11 | y | 1188 | 460.2 | 0.0005863 | 1.274 | +1 | 3 |
| - | - | 676.9 | 461.2 | - | - | 0 | - |
| - | - | 2278 | 467.2 | - | - | 0 | - |
| - | - | 666.4 | 471.3 | - | - | 0 | - |
| - | - | 819.1 | 473.2 | - | - | 0 | - |
| 3 | b | 4546 | 474.2 | 0.0005029 | 1.061 | +1 | 3 |
| 7 | b | 948.3 | 475.2 | 0.006373 | 13.41 | +2 | 7 |
| - | - | 7433 | 476.2 | - | - | 0 | - |
| - | - | 1561 | 477.2 | - | - | 0 | - |
| - | - | 1038 | 479.6 | - | - | 0 | - |
| - | - | 844.9 | 479.9 | - | - | 0 | - |
| - | - | 2038 | 482.2 | - | - | 0 | - |
| - | - | 2236 | 482.3 | - | - | 0 | - |
| - | - | 722.1 | 482.7 | - | - | 0 | - |
| - | - | 944.5 | 483.3 | - | - | 0 | - |
| - | - | 997.7 | 483.8 | - | - | 0 | - |
| - | - | 1480 | 487.2 | - | - | 0 | - |
| - | - | 6425 | 488.1 | - | - | 0 | - |
| - | - | 919.8 | 488.2 | - | - | 0 | - |
| - | - | 1.117E+04 | 488.3 | - | - | 0 | - |
| 11 | b | 1853 | 488.9 | 0.000355 | 0.7261 | +3 | 11 |
| - | - | 1613 | 489.2 | - | - | 0 | - |
| - | - | 1716 | 489.2 | - | - | 0 | - |
| - | - | 3131 | 489.3 | - | - | 0 | - |
| - | - | 1229 | 489.6 | - | - | 0 | - |
| - | - | 1662 | 496.2 | - | - | 0 | - |
| - | - | 1024 | 496.7 | - | - | 0 | - |
| - | - | 1523 | 497.3 | - | - | 0 | - |
| - | - | 2867 | 499.3 | - | - | 0 | - |
| - | - | 7745 | 516.3 | - | - | 0 | - |
| - | - | 2165 | 517.3 | - | - | 0 | - |
| 6 | y | 612.3 | 519.3 | 0.003305 | 6.365 | +2 | 8 |
| 6 | y | 981.4 | 519.7 | 0.003431 | 6.601 | +2 | 8 |
| - | - | 1775 | 524.2 | - | - | 0 | - |
| 6 | y | 6634 | 528.3 | 0.0006897 | 1.306 | +2 | 8 |
| - | - | 1008 | 528.3 | - | - | 0 | - |
| - | - | 3932 | 528.8 | - | - | 0 | - |
| - | - | 702.4 | 529.3 | - | - | 0 | - |
| - | - | 551.1 | 531.2 | - | - | 0 | - |
| - | - | 2822 | 533.3 | - | - | 0 | - |
| - | - | 4525 | 533.9 | - | - | 0 | - |
| - | - | 3704 | 534.2 | - | - | 0 | - |
| - | - | 922.4 | 535.3 | - | - | 0 | - |
| - | - | 679.2 | 538.9 | - | - | 0 | - |
| 10 | y | 1326 | 539.2 | 0.003374 | 6.258 | +1 | 4 |
| 12 | b | 677.1 | 543.2 | 0.003144 | 5.787 | +3 | 12 |
| - | - | 886.6 | 543.6 | - | - | 0 | - |
| - | - | 1188 | 543.9 | - | - | 0 | - |
| - | - | 670.3 | 546.3 | - | - | 0 | - |
| - | - | 1452 | 547.3 | - | - | 0 | - |
| - | - | 1042 | 552.3 | - | - | 0 | - |
| - | - | 946.5 | 553.3 | - | - | 0 | - |
| 5 | y | 928.9 | 554.8 | 0.006542 | 11.79 | +2 | 9 |
| 5 | y | 1509 | 555.3 | 0.0003202 | 0.5766 | +2 | 9 |
| 10 | y | 1.496E+04 | 557.2 | 0.0005874 | 1.054 | +1 | 4 |
| - | - | 5859 | 558.2 | - | - | 0 | - |
| - | - | 981.5 | 559.2 | - | - | 0 | - |
| - | - | 860.1 | 559.2 | - | - | 0 | - |
| 5 | y | 1.652E+04 | 563.8 | 0.003341 | 5.926 | +2 | 9 |
| - | - | 8660 | 564.3 | - | - | 0 | - |
| - | - | 4425 | 564.8 | - | - | 0 | - |
| - | - | 1478 | 565.3 | - | - | 0 | - |
| - | - | 643.9 | 569.3 | - | - | 0 | - |
| - | - | 3643 | 570.3 | - | - | 0 | - |
| - | - | 1084 | 571.3 | - | - | 0 | - |
| - | - | 2480 | 575.2 | - | - | 0 | - |
| - | - | 6255 | 577.8 | - | - | 0 | - |
| - | - | 3550 | 578.3 | - | - | 0 | - |
| - | - | 2729 | 578.8 | - | - | 0 | - |
| - | - | 1047 | 579.3 | - | - | 0 | - |
| - | - | 1398 | 580.3 | - | - | 0 | - |
| - | - | 1037 | 581.3 | - | - | 0 | - |
| 0 | Precursor | 1422 | 581.6 | 0.0006705 | 1.153 | +3 | -1 |
| 0 | Precursor | 1243 | 581.9 | 0.006407 | 11.01 | +3 | -1 |
| - | - | 875.7 | 582.3 | - | - | 0 | - |
| - | - | 2423 | 586.3 | - | - | 0 | - |
| - | - | 1158 | 586.4 | - | - | 0 | - |
| - | - | 1036 | 587.3 | - | - | 0 | - |
| - | - | 1658 | 587.3 | - | - | 0 | - |
| 0 | Precursor | 1.126E+04 | 587.6 | 0.001323 | 2.252 | +3 | -1 |
| - | - | 1.453E+04 | 587.9 | - | - | 0 | - |
| - | - | 9574 | 588.3 | - | - | 0 | - |
| - | - | 1464 | 588.3 | - | - | 0 | - |
| - | - | 2833 | 588.6 | - | - | 0 | - |
| - | - | 1231 | 588.9 | - | - | 0 | - |
| - | - | 1013 | 594.2 | - | - | 0 | - |
| - | - | 791.8 | 596.4 | - | - | 0 | - |
| - | - | 1232 | 598.3 | - | - | 0 | - |
| - | - | 799.3 | 599.3 | - | - | 0 | - |
| 9 | b | 4931 | 602.8 | 0.001333 | 2.211 | +2 | 9 |
| - | - | 2308 | 603.3 | - | - | 0 | - |
| - | - | 4023 | 603.3 | - | - | 0 | - |
| - | - | 1647 | 603.8 | - | - | 0 | - |
| - | - | 1440 | 604.3 | - | - | 0 | - |
| - | - | 1999 | 621.2 | - | - | 0 | - |
| - | - | 742.4 | 622.2 | - | - | 0 | - |
| - | - | 2218 | 631.3 | - | - | 0 | - |
| 4 | b | 1053 | 635.2 | 0.001552 | 2.443 | +1 | 4 |
| 4 | y | 2977 | 635.3 | 0.0008758 | 1.379 | +2 | 10 |
| 4 | y | 1857 | 635.8 | 0.005957 | 9.369 | +2 | 10 |
| - | - | 778.7 | 636.2 | - | - | 0 | - |
| - | - | 816.5 | 636.3 | - | - | 0 | - |
| - | - | 1.175E+04 | 639.2 | - | - | 0 | - |
| - | - | 3476 | 640.2 | - | - | 0 | - |
| - | - | 1318 | 641.2 | - | - | 0 | - |
| 4 | y | 2.541E+04 | 644.3 | 0.001642 | 2.548 | +2 | 10 |
| - | - | 2.145E+04 | 644.8 | - | - | 0 | - |
| - | - | 1.946E+04 | 645.3 | - | - | 0 | - |
| - | - | 1.136E+04 | 645.8 | - | - | 0 | - |
| - | - | 4068 | 646.3 | - | - | 0 | - |
| - | - | 1214 | 646.8 | - | - | 0 | - |
| - | - | 929.6 | 648.3 | - | - | 0 | - |
| 10 | b | 1257 | 651.3 | 0.001653 | 2.538 | +2 | 10 |
| - | - | 8676 | 651.4 | - | - | 0 | - |
| - | - | 802.5 | 652.3 | - | - | 0 | - |
| - | - | 3034 | 652.4 | - | - | 0 | - |
| - | - | 1112 | 653.4 | - | - | 0 | - |
| - | - | 796.9 | 655.3 | - | - | 0 | - |
| - | - | 1.064E+04 | 659.3 | - | - | 0 | - |
| - | - | 8353 | 659.8 | - | - | 0 | - |
| - | - | 4479 | 660.3 | - | - | 0 | - |
| - | - | 919.1 | 660.8 | - | - | 0 | - |
| - | - | 1725 | 668.3 | - | - | 0 | - |
| - | - | 900 | 668.8 | - | - | 0 | - |
| - | - | 4960 | 679.4 | - | - | 0 | - |
| - | - | 1968 | 680.4 | - | - | 0 | - |
| 9 | y | 1383 | 695.3 | 6.789E-05 | 0.09764 | +1 | 5 |
| 9 | y | 6265 | 696.3 | 0.001953 | 2.805 | +1 | 5 |
| - | - | 2593 | 697.3 | - | - | 0 | - |
| - | - | 3190 | 697.4 | - | - | 0 | - |
| - | - | 1966 | 698.4 | - | - | 0 | - |
| - | - | 714.1 | 699.4 | - | - | 0 | - |
| 5 | b | 764.3 | 706.3 | 0.003397 | 4.809 | +1 | 5 |
| - | - | 1138 | 707.4 | - | - | 0 | - |
| - | - | 1505 | 708.3 | - | - | 0 | - |
| - | - | 781.8 | 708.8 | - | - | 0 | - |
| - | - | 1207 | 710.3 | - | - | 0 | - |
| 9 | y | 1.546E+04 | 713.3 | 0.0007947 | 1.114 | +1 | 5 |
| - | - | 7217 | 714.3 | - | - | 0 | - |
| - | - | 849.8 | 714.4 | - | - | 0 | - |
| - | - | 1922 | 715.3 | - | - | 0 | - |
| 3 | y | 6199 | 716.8 | 0.002633 | 3.673 | +2 | 11 |
| 3 | y | 4990 | 717.3 | 0.007312 | 10.19 | +2 | 11 |
| - | - | 3267 | 717.8 | - | - | 0 | - |
| - | - | 1186 | 718.3 | - | - | 0 | - |
| - | - | 7596 | 718.8 | - | - | 0 | - |
| - | - | 5936 | 719.3 | - | - | 0 | - |
| - | - | 3376 | 719.8 | - | - | 0 | - |
| 3 | y | 1.75E+05 | 725.8 | 0.00169 | 2.328 | +2 | 11 |
| - | - | 1.39E+05 | 726.3 | - | - | 0 | - |
| - | - | 7.643E+04 | 726.8 | - | - | 0 | - |
| - | - | 2.623E+04 | 727.3 | - | - | 0 | - |
| - | - | 8115 | 727.8 | - | - | 0 | - |
| - | - | 3875 | 731.4 | - | - | 0 | - |
| - | - | 2226 | 731.4 | - | - | 0 | - |
| - | - | 2485 | 732.4 | - | - | 0 | - |
| 11 | b | 2.411E+04 | 732.8 | 0.001945 | 2.654 | +2 | 11 |
| - | - | 1.922E+04 | 733.3 | - | - | 0 | - |
| - | - | 1.139E+04 | 733.8 | - | - | 0 | - |
| - | - | 4660 | 734.3 | - | - | 0 | - |
| - | - | 2644 | 738.3 | - | - | 0 | - |
| - | - | 936.8 | 739.3 | - | - | 0 | - |
| - | - | 2688 | 740.8 | - | - | 0 | - |
| - | - | 3781 | 741.3 | - | - | 0 | - |
| - | - | 1635 | 741.8 | - | - | 0 | - |
| - | - | 756 | 742.3 | - | - | 0 | - |
| - | - | 704.2 | 742.4 | - | - | 0 | - |
| - | - | 1048 | 749.8 | - | - | 0 | - |
| - | - | 1159 | 750.3 | - | - | 0 | - |
| - | - | 698.3 | 753.9 | - | - | 0 | - |
| - | - | 1124 | 756.9 | - | - | 0 | - |
| - | - | 1001 | 757.3 | - | - | 0 | - |
| - | - | 2140 | 759.4 | - | - | 0 | - |
| - | - | 878.7 | 760.4 | - | - | 0 | - |
| - | - | 3992 | 766.4 | - | - | 0 | - |
| - | - | 2277 | 767.4 | - | - | 0 | - |
| 8 | y | 6967 | 794.4 | 0.0009899 | 1.246 | +1 | 6 |
| - | - | 2944 | 795.4 | - | - | 0 | - |
| 2 | y | 2990 | 798.3 | 0.001216 | 1.523 | +2 | 12 |
| 2 | y | 2818 | 798.8 | 0.008851 | 11.08 | +2 | 12 |
| - | - | 2222 | 799.3 | - | - | 0 | - |
| - | - | 7774 | 800.4 | - | - | 0 | - |
| - | - | 8939 | 800.9 | - | - | 0 | - |
| - | - | 3621 | 801.4 | - | - | 0 | - |
| - | - | 1051 | 801.9 | - | - | 0 | - |
| - | - | 4301 | 802.3 | - | - | 0 | - |
| - | - | 2356 | 802.5 | - | - | 0 | - |
| - | - | 1934 | 803.3 | - | - | 0 | - |
| - | - | 1220 | 803.5 | - | - | 0 | - |
| 2 | y | 7.268E+04 | 807.4 | 0.001738 | 2.152 | +2 | 12 |
| - | - | 7.358E+04 | 807.9 | - | - | 0 | - |
| - | - | 4.01E+04 | 808.4 | - | - | 0 | - |
| - | - | 1.325E+04 | 808.9 | - | - | 0 | - |
| - | - | 2796 | 809.4 | - | - | 0 | - |
| 8 | y | 6.486E+04 | 812.4 | 0.000435 | 0.5354 | +1 | 6 |
| - | - | 2.749E+04 | 813.4 | - | - | 0 | - |
| 12 | b | 1.558E+04 | 814.4 | 0.004111 | 5.047 | +2 | 12 |
| - | - | 1.927E+04 | 814.9 | - | - | 0 | - |
| - | - | 9457 | 815.4 | - | - | 0 | - |
| - | - | 3555 | 815.9 | - | - | 0 | - |
| - | - | 753 | 816.4 | - | - | 0 | - |
| - | - | 898.1 | 822.4 | - | - | 0 | - |
| - | - | 813.4 | 828.4 | - | - | 0 | - |
| - | - | 3866 | 830.5 | - | - | 0 | - |
| - | - | 1592 | 831.5 | - | - | 0 | - |
| - | - | 743.9 | 832.5 | - | - | 0 | - |
| - | - | 733.7 | 866.4 | - | - | 0 | - |
| - | - | 1756 | 877.4 | - | - | 0 | - |
| - | - | 737.9 | 878.4 | - | - | 0 | - |
| - | - | 6264 | 894.4 | - | - | 0 | - |
| - | - | 1621 | 894.5 | - | - | 0 | - |
| - | - | 3978 | 895.4 | - | - | 0 | - |
| - | - | 1067 | 896.4 | - | - | 0 | - |
| - | - | 1053 | 901.4 | - | - | 0 | - |
| - | - | 858 | 908.5 | - | - | 0 | - |
| 7 | y | 2522 | 909.4 | 0.0005625 | 0.6185 | +1 | 7 |
| 7 | y | 2797 | 910.4 | 0.009196 | 10.1 | +1 | 7 |
| - | - | 4169 | 922.5 | - | - | 0 | - |
| - | - | 1750 | 923.5 | - | - | 0 | - |
| 7 | y | 5.301E+04 | 927.4 | 0.0005305 | 0.5721 | +1 | 7 |
| - | - | 2.706E+04 | 928.4 | - | - | 0 | - |
| - | - | 8233 | 929.4 | - | - | 0 | - |
| - | - | 2063 | 930.4 | - | - | 0 | - |
| - | - | 1567 | 940.5 | - | - | 0 | - |
| 7 | b | 4991 | 949.4 | 0.003904 | 4.112 | +1 | 7 |
| - | - | 2652 | 950.4 | - | - | 0 | - |
| - | - | 1325 | 951.4 | - | - | 0 | - |
| - | - | 4437 | 963.5 | - | - | 0 | - |
| - | - | 2173 | 964.5 | - | - | 0 | - |
| - | - | 2307 | 965.5 | - | - | 0 | - |
| - | - | 1282 | 966.5 | - | - | 0 | - |
| - | - | 709.4 | 976.5 | - | - | 0 | - |
| - | - | 6071 | 991.5 | - | - | 0 | - |
| - | - | 3777 | 992.5 | - | - | 0 | - |
| - | - | 4601 | 993.5 | - | - | 0 | - |
| - | - | 1239 | 994.5 | - | - | 0 | - |
| - | - | 1020 | 995.5 | - | - | 0 | - |
| - | - | 1781 | 1012 | - | - | 0 | - |
| - | - | 670.3 | 1013 | - | - | 0 | - |
| 6 | y | 708.8 | 1038 | 0.0007989 | 0.7701 | +1 | 8 |
| 6 | y | 1103 | 1038 | 0.01104 | 10.63 | +1 | 8 |
| 6 | y | 1.812E+04 | 1056 | 0.0003551 | 0.3364 | +1 | 8 |
| - | - | 1.148E+04 | 1057 | - | - | 0 | - |
| - | - | 3954 | 1058 | - | - | 0 | - |
| - | - | 2364 | 1058 | - | - | 0 | - |
| - | - | 877.8 | 1059 | - | - | 0 | - |
| - | - | 687.7 | 1067 | - | - | 0 | - |
| 5 | y | 881.9 | 1109 | 0.007029 | 6.341 | +1 | 9 |
| 5 | y | 1343 | 1110 | 0.007369 | 6.641 | +1 | 9 |
| - | - | 906.5 | 1111 | - | - | 0 | - |
| 5 | y | 2.828E+04 | 1127 | 0.003556 | 3.156 | +1 | 9 |
| - | - | 2.003E+04 | 1128 | - | - | 0 | - |
| - | - | 6112 | 1129 | - | - | 0 | - |
| - | - | 1247 | 1130 | - | - | 0 | - |
| - | - | 1050 | 1138 | - | - | 0 | - |
| - | - | 1.206E+04 | 1155 | - | - | 0 | - |
| - | - | 8071 | 1156 | - | - | 0 | - |
| - | - | 3013 | 1157 | - | - | 0 | - |
| - | - | 1190 | 1158 | - | - | 0 | - |
| - | - | 839.2 | 1173 | - | - | 0 | - |
| - | - | 879.3 | 1174 | - | - | 0 | - |
| - | - | 1018 | 1196 | - | - | 0 | - |
| - | - | 911.1 | 1197 | - | - | 0 | - |
| 9 | b | 1015 | 1205 | 0.0198 | 16.44 | +1 | 9 |
| - | - | 618.2 | 1257 | - | - | 0 | - |
| 4 | y | 1497 | 1270 | 0.004729 | 3.725 | +1 | 10 |
| 4 | y | 1152 | 1271 | 0.003687 | 2.902 | +1 | 10 |
| - | - | 897.3 | 1272 | - | - | 0 | - |
| 4 | y | 1.658E+04 | 1288 | 0.00504 | 3.914 | +1 | 10 |
| - | - | 1.083E+04 | 1289 | - | - | 0 | - |
| - | - | 6167 | 1290 | - | - | 0 | - |
| - | - | 2302 | 1291 | - | - | 0 | - |
| - | - | 839 | 1292 | - | - | 0 | - |
| - | - | 4483 | 1318 | - | - | 0 | - |
| - | - | 3789 | 1319 | - | - | 0 | - |
| - | - | 1494 | 1320 | - | - | 0 | - |
| - | - | 1230 | 1452 | - | - | 0 | - |
| - | - | 698.3 | 1482 | - | - | 0 | - |
| - | - | 629.1 | 2150 | - | - | 0 | - |

m/z Charge Intensity FragmentType MassShift Position
120.08112335205078 0 302234.44
121.06513977050781 0 743.43823
121.07926177978516 0 862.4497
121.08438873291016 0 26420.58
122.08748626708984 0 920.46027
125.07137298583984 0 464.2884
126.22425079345703 0 375.80118
128.23023986816406 0 442.68073
129.1024932861328 0 54661.207
130.06536865234375 0 1757.1477
130.10592651367188 0 3246.0725
131.0495147705078 0 659.93066
132.10202026367188 0 1516.9236
133.08621215820312 0 504.3183
134.0220184326172 0 650.02795
134.0272979736328 0 9917.903
134.04505920410156 0 34790.367 y 12
135.04849243164062 0 1487.1191
135.4372100830078 0 464.2545
135.52285766601562 0 457.2396
136.07603454589844 0 323905.97
137.0737762451172 0 1170.5925
137.07928466796875 0 28890.234
138.06646728515625 0 511.43585
138.0823211669922 0 1139.7272
139.08700561523438 0 764.2939
140.08233642578125 0 920.4065
141.1028289794922 0 994.56085
143.1181640625 0 2862.4907
146.06036376953125 0 1944.8751
147.0442657470703 0 2305.0208
152.07064819335938 0 596.99036
155.1182403564453 0 1178.2234
157.10867309570312 0 2426.051
158.0600128173828 0 591.3754
158.09286499023438 0 832.2495
159.09181213378906 0 5593.7563
159.1128692626953 0 2358.9395
160.07574462890625 0 844.5878
160.0955352783203 0 1094.7723
162.02256774902344 0 638.31464
162.06704711914062 0 522.3401
164.0706787109375 0 1159.5089
165.05543518066406 0 574.8625
166.0868682861328 0 594.5534
167.6372528076172 0 504.49374
169.09751892089844 0 1085.4487
170.06036376953125 0 948.8623
171.06826782226562 0 542.80554
171.076416015625 0 583.15485
171.11305236816406 0 2798.8176
173.0711669921875 0 781.92554
173.09242248535156 0 561.12573
173.1295928955078 0 556.8782
173.45086669921875 0 3146.1423
174.05516052246094 0 1745.0807
174.07925415039062 0 807.72314
175.0868682861328 0 10491.675
175.90191650390625 0 473.93744
176.02133178710938 0 492.94016
176.0902099609375 0 1792.168
180.10205078125 0 630.77057
181.0978240966797 0 678.7972
182.0814666748047 0 1211.3368
183.11306762695312 0 1002.2725
185.07115173339844 0 1410.8945
185.10391235351562 0 2000.8497
186.07913208007812 0 935.32416
187.05349731445312 0 878.6168
187.08685302734375 0 892.43835
187.10804748535156 0 7432.6763
187.5230712890625 0 539.6915
189.12359619140625 0 1340.1329
191.08177185058594 0 4392.8975
192.04800415039062 0 742.07336
192.08570861816406 0 588.6963
192.11361694335938 0 1093.9777
192.15394592285156 0 484.54645
195.11338806152344 0 604.2362
197.12818908691406 0 1498.1443
199.10792541503906 0 1715.8838
200.13958740234375 0 13062.64
201.06594848632812 0 1965.717
201.12344360351562 0 1255.9595
201.1433868408203 0 836.94775
202.04986572265625 0 908.63385
202.12310791015625 0 1172.8245
204.1344451904297 0 1052.2445
205.06439208984375 0 8835.715
206.06723022460938 0 770.7513
209.0924530029297 0 980.1282
210.12379455566406 0 758.4554
214.1552276611328 0 717.875
215.0487060546875 0 843.5433
215.1029052734375 0 2014.3275
215.1390380859375 0 1068.5555
216.06558227539062 0 1351.8644
220.10787963867188 0 601.0221
222.12413024902344 0 626.632
226.11892700195312 0 3653.413
227.10269165039062 0 829.98627
228.13377380371094 0 565.9858
229.11843872070312 0 2958.7847
231.11386108398438 0 528.8023
233.059326171875 0 2823.3484
233.0923309326172 0 11417.562
233.1287384033203 0 20740.498
233.14883422851562 0 1096.2189
234.05908203125 0 1633.3927
234.0963134765625 0 923.88873
234.1322021484375 0 2718.5747
237.08714294433594 0 779.02313
237.13487243652344 0 2901.6365
238.09471130371094 0 1200.3447
238.12306213378906 0 1012.5099
239.15057373046875 0 21090.814
240.1538543701172 0 2158.5
244.12924194335938 0 2595.2715
246.12437438964844 0 625.10266
247.14405822753906 0 1043.3204
247.44732666015625 0 480.5846
250.15306091308594 0 585.3001
252.06918334960938 0 5843.01
253.07290649414062 0 590.0481
254.1164093017578 0 2241.2864
254.16148376464844 0 1270.3472
256.17706298828125 0 9023.156
257.18035888671875 0 948.69446
261.1235656738281 0 16735.857
262.0534973144531 0 680.56683
262.0718994140625 0 537.7556
262.12725830078125 0 2423.9575
266.1177673339844 0 829.8922
270.08831787109375 0 576.3852
270.1812438964844 0 2885.9038
271.01776123046875 0 557.0804
272.12457275390625 0 1847.1022
274.11895751953125 0 781.27795
274.1887512207031 0 734.6585
276.1346130371094 0 858.0408
278.1501159667969 0 645.2822
279.0979309082031 0 710.1031 y Water loss 11
280.08184814453125 0 1735.879
280.1307067871094 0 644.61273
282.1568603515625 0 1235.015
283.14447021484375 0 157213.88 a 1
283.9369812011719 0 481.6916
284.1477355957031 0 25626.074
285.1506652832031 0 2479.2449
286.1392517089844 0 803.0119
297.0906982421875 0 34765.2
297.1087951660156 0 17903.604 y 11
297.1567687988281 0 1444.0355
298.09368896484375 0 5972.1323
298.1117248535156 0 2667.7998
299.0876159667969 0 2082.4885
299.1394348144531 0 45334.23
300.1424560546875 0 6413.4473
300.1569519042969 0 1412.554
301.1452331542969 0 698.7578
304.12969970703125 0 7887.567
305.1329650878906 0 1403.3312
306.10943603515625 0 1172.6188
309.1601257324219 0 592.42596
311.1394348144531 0 36278.133 b 1
312.1427001953125 0 6146.496
313.1478271484375 0 997.6605
315.16748046875 0 3559.6116
318.16650390625 0 2066.021
319.17718505859375 0 1334.1808
325.0857849121094 0 17443.174
326.0889892578125 0 2701.469
326.1846923828125 0 691.8926
327.13446044921875 0 16579.406
328.1378479003906 0 2921.3867
336.20458984375 0 740.31903
337.1549072265625 0 775.86926
340.1998596191406 0 713.4065
342.2163391113281 0 716.75305
343.1424865722656 0 869.4579
343.1632080078125 0 1004.8353
343.19549560546875 0 699.9041
350.1268615722656 0 2097.7607
353.1613464355469 0 763.0971
353.7716369628906 0 610.90466
354.177001953125 0 2218.7642
355.17730712890625 0 1141.9323
361.15496826171875 0 2701.418
364.1988525390625 0 817.18744
365.11358642578125 0 879.58575
366.11663818359375 0 660.69946
366.1445617675781 0 1180.9014
366.1743469238281 0 515.01263
370.2559814453125 0 2367.862
371.1746520996094 0 511.63443
371.2034606933594 0 5789.591
371.2284240722656 0 951.30756
372.1790466308594 0 573.03204
372.20556640625 0 1597.2148
382.1402587890625 0 3491.4753
383.1242980957031 0 1722.4862
383.14691162109375 0 743.0786
383.20379638671875 0 733.17334
386.24041748046875 0 961.5804
387.1669921875 0 1513.6313
389.2290954589844 0 587.45013
396.122802734375 0 4057.2815
396.1557312011719 0 3585.5796
396.191162109375 0 1436.7429 b Water loss 8
397.1253356933594 0 1253.2233
399.2232971191406 0 3023.0906
400.1507873535156 0 7108.3813
400.2266845703125 0 797.84247
401.15374755859375 0 1769.2373
401.1816101074219 0 725.9188
405.1759033203125 0 849.7891
414.2349548339844 0 941.0242
415.13037109375 0 554.1835
417.23052978515625 0 2318.0083
424.18817138671875 0 3684.479 y Ammonia loss 3
425.1914978027344 0 927.6737
429.18023681640625 0 996.2302
439.2006530761719 0 798.7879
443.1702575683594 0 540.9322
447.7107849121094 0 1334.1345
447.74609375 0 1361.5249
448.2131042480469 0 812.2143
456.2470703125 0 588.31555
458.1711730957031 0 2749.6404
459.1513366699219 0 803.39777
460.1720275878906 0 1188.1342 y 10
461.17572021484375 0 676.94696
467.19384765625 0 2277.8164
471.272216796875 0 666.4048
473.2333068847656 0 819.14246
474.2028503417969 0 4546.4634 b 2
475.2002868652344 0 948.26013 b 6
476.1817321777344 0 7433.336
477.1838073730469 0 1560.5789
479.5611572265625 0 1037.8187
479.8957824707031 0 844.89716
482.2392578125 0 2037.6289
482.2737731933594 0 2236.4434
482.7425231933594 0 722.0551
483.2672119140625 0 944.5081
483.7657165527344 0 997.6698
487.1807556152344 0 1479.8826
488.1490173339844 0 6424.8394
488.1859436035156 0 919.7553
488.2989501953125 0 11172.245
488.8932189941406 0 1853.2379 b 10
489.15301513671875 0 1613.4244
489.22784423828125 0 1716.1566
489.3011474609375 0 3130.7927
489.5616455078125 0 1229.4452
496.2393798828125 0 1662.0363
496.7389221191406 0 1023.5232
497.2634582519531 0 1523.2196
499.2998962402344 0 2866.6985
516.293701171875 0 7744.7427
517.2977905273438 0 2165.4736
519.2528686523438 0 612.3257 y Water loss 5
519.7447509765625 0 981.44556 y Ammonia loss 5
524.216796875 0 1774.9913
528.2621459960938 0 6633.875 y 5
528.3099365234375 0 1007.99756
528.763671875 0 3931.8662
529.2684936523438 0 702.38947
531.1946411132812 0 551.0838
533.3198852539062 0 2821.508
533.9150390625 0 4524.657
534.249755859375 0 3704.3704
535.2659912109375 0 922.4256
538.9070434570312 0 679.18353
539.2102661132812 0 1326.155 y Water loss 9
543.244873046875 0 677.0634 b 11
543.5836181640625 0 886.6088
543.9160766601562 0 1187.8081
546.2575073242188 0 670.33215
547.2562255859375 0 1451.6636
552.2921142578125 0 1041.8772
553.308349609375 0 946.4649
554.7681884765625 0 928.8852 y Water loss 4
555.2664184570312 0 1508.5352 y Ammonia loss 4
557.2247924804688 0 14961.463 y 9
558.2286987304688 0 5858.6177
559.1846923828125 0 981.4544
559.2294921875 0 860.13086
563.7766723632812 0 16518.783 y 4
564.27783203125 0 8660.084
564.7765502929688 0 4424.6006
565.2811889648438 0 1477.6359
569.2612915039062 0 643.8672
570.3361206054688 0 3643.451
571.3394165039062 0 1083.8284
575.248779296875 0 2479.8804
577.7691040039062 0 6255.2627
578.2717895507812 0 3550.397
578.7721557617188 0 2728.8677
579.2697143554688 0 1046.953
580.2890014648438 0 1398.2798
581.28564453125 0 1037.1593
581.5896606445312 0 1421.9908 Precursor Water loss
581.9247436523438 0 1243.02 Precursor Ammonia loss
582.2573852539062 0 875.69073
586.2525634765625 0 2423.2803
586.3817749023438 0 1158.3386
587.2548217773438 0 1036.2207
587.340576171875 0 1658.0942
587.592529296875 0 11262.093 Precursor
587.927001953125 0 14534.784
588.260986328125 0 9573.8955
588.3400268554688 0 1464.2102
588.5950927734375 0 2832.9688
588.931640625 0 1231.3152
594.2217407226562 0 1012.57306
596.352783203125 0 791.75006
598.2938232421875 0 1232.1766
599.3056640625 0 799.32574
602.77734375 0 4930.9326 b 8
603.2766723632812 0 2307.788
603.325927734375 0 4022.7207
603.7772827148438 0 1647.2532
604.327392578125 0 1439.8531
621.233642578125 0 1998.6537
622.2342529296875 0 742.4221
631.3198852539062 0 2218.0557
635.2230834960938 0 1052.6019 b 3
635.283447265625 0 2977.3696 y Water loss 3
635.7822875976562 0 1857.037 y Ammonia loss 3
636.2174072265625 0 778.7302
636.2764282226562 0 816.5119
639.2447509765625 0 11747.414
640.2472534179688 0 3475.859
641.2463989257812 0 1317.6193
644.2879638671875 0 25412.18 y 3
644.7894287109375 0 21451.512
645.2972412109375 0 19460.438
645.8016357421875 0 11360.041
646.3057250976562 0 4068.1128
646.806640625 0 1214.0885
648.3475341796875 0 929.5986
651.3034057617188 0 1256.7595 b 9
651.3617553710938 0 8676.492
652.3099975585938 0 802.53094
652.3649291992188 0 3034.121
653.3692016601562 0 1111.9767
655.2763061523438 0 796.91394
659.3006591796875 0 10635.644
659.8030395507812 0 8353.38
660.3037719726562 0 4478.71
660.8033447265625 0 919.12775
668.3054809570312 0 1724.5172
668.8026733398438 0 900.0211
679.357177734375 0 4959.8193
680.3589477539062 0 1968.4513
695.3148193359375 0 1383.4156 y Water loss 8
696.3007202148438 0 6265.4346 y Ammonia loss 8
697.3032836914062 0 2593.1108
697.3678588867188 0 3190.4932
698.3699340820312 0 1965.8607
699.3698120117188 0 714.07776
706.2552490234375 0 764.26855 b 4
707.3663940429688 0 1137.8049
708.3037719726562 0 1504.5391
708.8009643554688 0 781.8407
710.3220825195312 0 1207.2732
713.3261108398438 0 15461.385 y 8
714.328369140625 0 7217.161
714.3919067382812 0 849.82983
715.3296508789062 0 1922.425
716.8133544921875 0 6198.607 y Water loss 2
717.3153076171875 0 4990.3003 y Ammonia loss 2
717.8170166015625 0 3266.6838
718.314697265625 0 1186.329
718.8367919921875 0 7595.8545
719.33837890625 0 5935.7637
719.8386840820312 0 3375.9062
725.819580078125 0 175037.69 y 2
726.3208618164062 0 138973.25
726.822265625 0 76434.54
727.3238525390625 0 26233.744
727.8259887695312 0 8115.2285
731.3507080078125 0 3874.7092
731.4202270507812 0 2225.689
732.3525390625 0 2484.695
732.8347778320312 0 24114.344 b 10
733.336669921875 0 19218.463
733.8373413085938 0 11388.108
734.3387451171875 0 4660.433
738.3136596679688 0 2643.8076
739.3128662109375 0 936.8239
740.8324584960938 0 2687.5417
741.3366088867188 0 3780.658
741.8372802734375 0 1634.6232
742.3302001953125 0 756.0404
742.3961181640625 0 704.22986
749.8389892578125 0 1048.0695
750.344482421875 0 1159.2355
753.8509521484375 0 698.2797
756.8966064453125 0 1123.9011
757.2898559570312 0 1001.4216
759.4149780273438 0 2139.8628
760.42041015625 0 878.71967
766.3878784179688 0 3992.295
767.3897094726562 0 2277.4275
794.3841552734375 0 6966.588 y Water loss 7
795.3847045898438 0 2943.937
798.346435546875 0 2990.2656 y Water loss 1
798.8485107421875 0 2818.4001 y Ammonia loss 1
799.3475341796875 0 2221.7534
800.3690185546875 0 7774.255
800.8698120117188 0 8939.041
801.37255859375 0 3621.0505
801.8704223632812 0 1051.2092
802.3082275390625 0 4301.234
802.4569091796875 0 2355.5442
803.3084106445312 0 1933.6652
803.462646484375 0 1219.6392
807.3511962890625 0 72682.586 y 1
807.8524780273438 0 73580.54
808.3534545898438 0 40096.97
808.854248046875 0 13246.567
809.354736328125 0 2795.6836
812.3941650390625 0 64862.4 y 7
813.3976440429688 0 27485.613
814.3724975585938 0 15576.762 b 11
814.8681640625 0 19272.68
815.3701171875 0 9457.423
815.8701782226562 0 3554.775
816.37060546875 0 752.95013
822.379638671875 0 898.07947
828.404296875 0 813.40186
830.450927734375 0 3865.9338
831.4564208984375 0 1592.1792
832.4552001953125 0 743.93805
866.4107055664062 0 733.6619
877.3876342773438 0 1755.701
878.3853149414062 0 737.9237
894.413330078125 0 6264.0654
894.4905395507812 0 1620.5977
895.4158935546875 0 3978.3057
896.4139404296875 0 1067.3273
901.3746948242188 0 1053.0995
908.4860229492188 0 857.97845
909.4095458984375 0 2522.2195 y Water loss 6
910.4033203125 0 2796.8176 y Ammonia loss 6
922.4808349609375 0 4168.843
923.4774169921875 0 1749.9302
927.4212036132812 0 53009.99 y 6
928.4241943359375 0 27064.773
929.4268798828125 0 8233.015
930.4295654296875 0 2063.227
940.4834594726562 0 1566.9067
949.3766479492188 0 4991.272 b 6
950.3796997070312 0 2652.3806
951.3805541992188 0 1324.5295
963.4710693359375 0 4436.581
964.473388671875 0 2172.9302
965.5143432617188 0 2307.1316
966.5185546875 0 1281.5145
976.4632568359375 0 709.3851
991.466796875 0 6070.7437
992.4683837890625 0 3777.0503
993.510498046875 0 4600.9673
994.5193481445312 0 1239.369
995.51806640625 0 1019.894
1011.5223999023438 0 1781.1292
1012.5338745117188 0 670.3385
1037.5042724609375 0 708.79706 y Water loss 5
1038.5001220703125 0 1103.3098 y Ammonia loss 5
1055.5159912109375 0 18121.111 y 5
1056.518798828125 0 11480.478
1057.501220703125 0 3954.1807
1058.489501953125 0 2364.2349
1059.480712890625 0 877.76465
1067.4208984375 0 687.6811
1108.53515625 0 881.8753 y Water loss 4
1109.5335693359375 0 1343.3583 y Ammonia loss 4
1110.5318603515625 0 906.45215
1126.5491943359375 0 28275.86 y 4
1127.552001953125 0 20026.344
1128.5540771484375 0 6111.945
1129.55517578125 0 1246.6633
1137.503173828125 0 1049.6051
1154.5299072265625 0 12064.872
1155.5325927734375 0 8071.097
1156.5330810546875 0 3013.2944
1157.53857421875 0 1189.997
1172.5438232421875 0 839.17914
1173.547607421875 0 879.2604
1195.57373046875 0 1017.53174
1196.5728759765625 0 911.0989
1204.5302734375 0 1015.02966 b 8
1256.5643310546875 0 618.1585
1269.556640625 0 1497.1367 y Water loss 3
1270.549072265625 0 1151.9459 y Ammonia loss 3
1271.5479736328125 0 897.2678
1287.56689453125 0 16581.703 y 3
1288.568359375 0 10825.683
1289.57568359375 0 6166.857
1290.585693359375 0 2301.8792
1291.5743408203125 0 838.9638
1317.5927734375 0 4482.5967
1318.597900390625 0 3789.225
1319.6082763671875 0 1494.4731
1451.6343994140625 0 1230.1378
1481.6412353515625 0 698.258
2150.4970703125 0 629.07935

Spectrum Details

|  |  |
| --- | --- |
| Matched peaks? Matched peaks The total absolute number of peaks matched. Additionally in brackets the total fraction of peaks matched and the total number of peaks is shown. | 57 (11.20% of 509) |
| FDR? FDR The false discovery rate estimated for this peptide. It is calculated by matching all theoretical fragments with a non-integer shift with the raw peaks for this spectrum. This is done with 40 different shifts. The resulting percentage is the average number of annotated peaks over the number of annotated peaks with the correct spectrum. | 1.59% |
| Satellite FDR? Satellite FDR See the FDR for details on its calculation. This satellite ion specific FDR only contains the satellite ions (d/w) for I/L/J positions. | - |
| PSM Score? PSM Score The PSM Score as given by Hecklib to this annotated spectrum. It is shown with three significant figures. | 428 |

#### Spectrum 7041? Spectrum 7041 The raw spectrum of this peptide as annotated by Hecklib. The fragments are coloured according to ion type (see legend). Any peaks with a star '\*' as text can be hovered over to see the full details, first the ion type second the mass shift type. By hovering over the amino acids in the peptide or ions in the legend the corresponding peaks are highlighted. By toggling the 'Unassigned' label you can turn the background (unassigned) peaks on or off in the plot. By updating the slider in the Ion legend you can update the spectrum to only show the top X% of the peaks with labels. The top X% means any peak that is within X% of the highest intensity. By dragging in the spectrum you can zoom in to a specific part of the spectrum and use 'Zoom Out' to get back to the original zoom level. The annotation of the spectrum is based on the given sequence in the peptides file and is done with different software so inconsistencies are likely. The peaks are annotated based on the given sequence, with 20 ppm tolerance.

Copy Data

##### Spectrum 7041 (TSV)

###### Preview

```
Loading example...
```

*Click on the button to copy the data to your clipboard.*

Mz MinMz MaxIntensity Max

WidthHeightPeptide font sizePeptide stroke widthSpectrum font sizeSpectrum stroke widthCompact peptide

Ion legend

wxyz

abcd

OtherUnassignedIon^Charge^_Position_Show for top:%

FYYCAKDVRPYYD

04.28e+48.56e+41.28e+51.71e+5

Zoom Out

y^+1^_1_a^+1^_2_y^+1^_2_b^+1^_2_b^+3^_9_y^+3^_10_y^+1^_3_b^+1^_3_b^+3^_11_y^+2^_8_y^+1^_4_y^+1^_4_y^+2^_9_\*b^+2^_9_b^+1^_4_y^+2^_10_y^+2^_10_y^+1^_5_y^+1^_5_y^+1^_5_y^+2^_11_y^+2^_11_y^+2^_11_b^+2^_11_y^+1^_6_y^+2^_12_y^+2^_12_y^+2^_12_y^+1^_6_b^+2^_12_y^+1^_7_y^+1^_7_b^+1^_7_y^+1^_8_y^+1^_9_y^+1^_9_y^+1^_9_y^+1^_10_y^+1^_10_y^+1^_10_

0502100315052007

Fragment Matches Table

Show background peaks

| Position | Ion type | Intensity | mz Theoretical | mz Error (Th) | mz Error (ppm) | Charge | Series Number |
| --- | --- | --- | --- | --- | --- | --- | --- |
| - | - | 1.633E+05 | 120.1 | - | - | 0 | - |
| - | - | 1.295E+04 | 121.1 | - | - | 0 | - |
| - | - | 584.7 | 122.1 | - | - | 0 | - |
| - | - | 840 | 125.1 | - | - | 0 | - |
| - | - | 3.261E+04 | 129.1 | - | - | 0 | - |
| - | - | 1566 | 130.1 | - | - | 0 | - |
| - | - | 643.2 | 131 | - | - | 0 | - |
| - | - | 466.1 | 131.1 | - | - | 0 | - |
| - | - | 4783 | 133.1 | - | - | 0 | - |
| - | - | 4804 | 134 | - | - | 0 | - |
| - | - | 1018 | 134 | - | - | 0 | - |
| 13 | y | 1.779E+04 | 134 | 0.0001987 | 1.483 | +1 | 1 |
| - | - | 1033 | 135 | - | - | 0 | - |
| - | - | 1.696E+05 | 136.1 | - | - | 0 | - |
| - | - | 1067 | 137.1 | - | - | 0 | - |
| - | - | 1.322E+04 | 137.1 | - | - | 0 | - |
| - | - | 502.5 | 138.1 | - | - | 0 | - |
| - | - | 495.7 | 139.1 | - | - | 0 | - |
| - | - | 610 | 143.1 | - | - | 0 | - |
| - | - | 1221 | 147 | - | - | 0 | - |
| - | - | 669.8 | 152.1 | - | - | 0 | - |
| - | - | 380.8 | 153.1 | - | - | 0 | - |
| - | - | 2364 | 155.1 | - | - | 0 | - |
| - | - | 596.9 | 155.1 | - | - | 0 | - |
| - | - | 916.9 | 155.1 | - | - | 0 | - |
| - | - | 1043 | 157.1 | - | - | 0 | - |
| - | - | 536.4 | 159.1 | - | - | 0 | - |
| - | - | 471.4 | 160.1 | - | - | 0 | - |
| - | - | 709.1 | 162 | - | - | 0 | - |
| - | - | 1010 | 164.1 | - | - | 0 | - |
| - | - | 1049 | 165.1 | - | - | 0 | - |
| - | - | 1986 | 171.1 | - | - | 0 | - |
| - | - | 447.4 | 174.1 | - | - | 0 | - |
| - | - | 710.6 | 175.1 | - | - | 0 | - |
| - | - | 845.6 | 175.1 | - | - | 0 | - |
| - | - | 1894 | 177.1 | - | - | 0 | - |
| - | - | 804.4 | 181.1 | - | - | 0 | - |
| - | - | 902 | 183.1 | - | - | 0 | - |
| - | - | 1517 | 185.1 | - | - | 0 | - |
| - | - | 481.8 | 185.6 | - | - | 0 | - |
| - | - | 4180 | 187.1 | - | - | 0 | - |
| - | - | 743.9 | 187.1 | - | - | 0 | - |
| - | - | 497.8 | 191.1 | - | - | 0 | - |
| - | - | 684.1 | 195.1 | - | - | 0 | - |
| - | - | 741.8 | 195.1 | - | - | 0 | - |
| - | - | 900.8 | 197.1 | - | - | 0 | - |
| - | - | 717.1 | 199.1 | - | - | 0 | - |
| - | - | 902 | 199.1 | - | - | 0 | - |
| - | - | 869.7 | 199.1 | - | - | 0 | - |
| - | - | 522.9 | 199.2 | - | - | 0 | - |
| - | - | 6972 | 200.1 | - | - | 0 | - |
| - | - | 708.1 | 201.1 | - | - | 0 | - |
| - | - | 752.7 | 201.1 | - | - | 0 | - |
| - | - | 883.1 | 202.1 | - | - | 0 | - |
| - | - | 5382 | 205.1 | - | - | 0 | - |
| - | - | 1023 | 209.1 | - | - | 0 | - |
| - | - | 1581 | 215.1 | - | - | 0 | - |
| - | - | 561.4 | 215.1 | - | - | 0 | - |
| - | - | 659.9 | 221.1 | - | - | 0 | - |
| - | - | 660.7 | 224.1 | - | - | 0 | - |
| - | - | 1268 | 226.1 | - | - | 0 | - |
| - | - | 1207 | 227.1 | - | - | 0 | - |
| - | - | 956.8 | 233.1 | - | - | 0 | - |
| - | - | 5412 | 233.1 | - | - | 0 | - |
| - | - | 1.219E+04 | 233.1 | - | - | 0 | - |
| - | - | 524.5 | 233.2 | - | - | 0 | - |
| - | - | 989.5 | 234.1 | - | - | 0 | - |
| - | - | 827.2 | 234.1 | - | - | 0 | - |
| - | - | 1216 | 234.1 | - | - | 0 | - |
| - | - | 637.3 | 235.1 | - | - | 0 | - |
| - | - | 867.1 | 237.1 | - | - | 0 | - |
| - | - | 1806 | 237.1 | - | - | 0 | - |
| - | - | 1.292E+04 | 239.2 | - | - | 0 | - |
| - | - | 1222 | 240.2 | - | - | 0 | - |
| - | - | 1579 | 244.1 | - | - | 0 | - |
| - | - | 3155 | 252.1 | - | - | 0 | - |
| - | - | 924.8 | 254.1 | - | - | 0 | - |
| - | - | 4885 | 256.2 | - | - | 0 | - |
| - | - | 901.2 | 257.2 | - | - | 0 | - |
| - | - | 7374 | 261.1 | - | - | 0 | - |
| - | - | 571.1 | 262.1 | - | - | 0 | - |
| - | - | 996.1 | 262.1 | - | - | 0 | - |
| - | - | 1510 | 272.1 | - | - | 0 | - |
| - | - | 585.5 | 276.8 | - | - | 0 | - |
| - | - | 889.2 | 280.1 | - | - | 0 | - |
| - | - | 709.4 | 282.2 | - | - | 0 | - |
| 2 | a | 7.699E+04 | 283.1 | 0.0001828 | 0.6457 | +1 | 2 |
| - | - | 1.48E+04 | 284.1 | - | - | 0 | - |
| - | - | 1642 | 285.2 | - | - | 0 | - |
| - | - | 769.3 | 287.1 | - | - | 0 | - |
| - | - | 1.829E+04 | 297.1 | - | - | 0 | - |
| 12 | y | 9054 | 297.1 | 0.0004078 | 1.373 | +1 | 2 |
| - | - | 816.9 | 297.2 | - | - | 0 | - |
| - | - | 3218 | 298.1 | - | - | 0 | - |
| - | - | 1377 | 298.1 | - | - | 0 | - |
| - | - | 2.332E+04 | 299.1 | - | - | 0 | - |
| - | - | 4295 | 300.1 | - | - | 0 | - |
| - | - | 4986 | 304.1 | - | - | 0 | - |
| - | - | 779.8 | 305.1 | - | - | 0 | - |
| 2 | b | 1.864E+04 | 311.1 | 0.0003549 | 1.141 | +1 | 2 |
| - | - | 3958 | 312.1 | - | - | 0 | - |
| - | - | 1492 | 315.2 | - | - | 0 | - |
| - | - | 8839 | 325.1 | - | - | 0 | - |
| - | - | 1346 | 326.1 | - | - | 0 | - |
| - | - | 8642 | 327.1 | - | - | 0 | - |
| - | - | 1879 | 328.1 | - | - | 0 | - |
| - | - | 1819 | 333.2 | - | - | 0 | - |
| - | - | 732.3 | 343.1 | - | - | 0 | - |
| - | - | 813.2 | 343.2 | - | - | 0 | - |
| - | - | 641.8 | 343.2 | - | - | 0 | - |
| - | - | 697.2 | 349.2 | - | - | 0 | - |
| - | - | 1079 | 354.2 | - | - | 0 | - |
| - | - | 906.8 | 361.2 | - | - | 0 | - |
| - | - | 1874 | 370.3 | - | - | 0 | - |
| - | - | 3654 | 371.2 | - | - | 0 | - |
| - | - | 650.9 | 372.2 | - | - | 0 | - |
| - | - | 1045 | 377.2 | - | - | 0 | - |
| - | - | 593.5 | 382.1 | - | - | 0 | - |
| - | - | 660.2 | 389.2 | - | - | 0 | - |
| - | - | 565.5 | 391.1 | - | - | 0 | - |
| - | - | 2191 | 396.1 | - | - | 0 | - |
| - | - | 1894 | 396.2 | - | - | 0 | - |
| 9 | b | 755.3 | 396.2 | 0.00742 | 18.73 | +3 | 9 |
| - | - | 585.3 | 401.2 | - | - | 0 | - |
| 4 | y | 1919 | 424.2 | 0.0008845 | 2.085 | +3 | 10 |
| - | - | 1768 | 447.7 | - | - | 0 | - |
| - | - | 679.1 | 447.7 | - | - | 0 | - |
| - | - | 609.2 | 448.2 | - | - | 0 | - |
| - | - | 1796 | 458.2 | - | - | 0 | - |
| 11 | y | 785.4 | 460.2 | 0.003333 | 7.243 | +1 | 3 |
| - | - | 1586 | 467.2 | - | - | 0 | - |
| 3 | b | 1869 | 474.2 | 0.0007165 | 1.511 | +1 | 3 |
| - | - | 4528 | 476.2 | - | - | 0 | - |
| - | - | 888.8 | 482.2 | - | - | 0 | - |
| - | - | 856.8 | 482.7 | - | - | 0 | - |
| - | - | 850.4 | 483.3 | - | - | 0 | - |
| - | - | 922.3 | 483.8 | - | - | 0 | - |
| - | - | 2892 | 488.1 | - | - | 0 | - |
| - | - | 6130 | 488.3 | - | - | 0 | - |
| 11 | b | 972.6 | 488.9 | 0.003529 | 7.218 | +3 | 11 |
| - | - | 881.1 | 489.2 | - | - | 0 | - |
| - | - | 1927 | 489.3 | - | - | 0 | - |
| - | - | 778 | 490.3 | - | - | 0 | - |
| - | - | 882.6 | 496.2 | - | - | 0 | - |
| - | - | 1020 | 497.3 | - | - | 0 | - |
| - | - | 1477 | 499.3 | - | - | 0 | - |
| - | - | 3845 | 516.3 | - | - | 0 | - |
| - | - | 1316 | 517.3 | - | - | 0 | - |
| - | - | 1285 | 524.2 | - | - | 0 | - |
| 6 | y | 3570 | 528.3 | 7.937E-05 | 0.1502 | +2 | 8 |
| - | - | 2078 | 528.8 | - | - | 0 | - |
| - | - | 1276 | 533.3 | - | - | 0 | - |
| - | - | 2525 | 533.9 | - | - | 0 | - |
| - | - | 1959 | 534.2 | - | - | 0 | - |
| - | - | 1342 | 534.6 | - | - | 0 | - |
| - | - | 1083 | 535.3 | - | - | 0 | - |
| 10 | y | 1142 | 539.2 | 0.0002878 | 0.5338 | +1 | 4 |
| - | - | 746.1 | 539.6 | - | - | 0 | - |
| - | - | 6265 | 543.3 | - | - | 0 | - |
| - | - | 695.2 | 553.3 | - | - | 0 | - |
| 10 | y | 7257 | 557.2 | 3.806E-05 | 0.06831 | +1 | 4 |
| - | - | 1946 | 558.2 | - | - | 0 | - |
| 5 | y | 8256 | 563.8 | 0.003951 | 7.008 | +2 | 9 |
| - | - | 5854 | 564.3 | - | - | 0 | - |
| - | - | 1652 | 564.8 | - | - | 0 | - |
| - | - | 640 | 568 | - | - | 0 | - |
| - | - | 655.5 | 569.3 | - | - | 0 | - |
| - | - | 2022 | 570.3 | - | - | 0 | - |
| - | - | 8283 | 571.3 | - | - | 0 | - |
| - | - | 815.3 | 571.3 | - | - | 0 | - |
| - | - | 1097 | 575.2 | - | - | 0 | - |
| - | - | 3337 | 577.8 | - | - | 0 | - |
| - | - | 3031 | 578.3 | - | - | 0 | - |
| - | - | 1370 | 578.8 | - | - | 0 | - |
| - | - | 729.2 | 581.3 | - | - | 0 | - |
| - | - | 622.4 | 582.3 | - | - | 0 | - |
| 0 | Precursor | 5683 | 587.6 | 0.001995 | 3.395 | +3 | -1 |
| - | - | 7605 | 587.9 | - | - | 0 | - |
| - | - | 4546 | 588.3 | - | - | 0 | - |
| - | - | 4848 | 588.3 | - | - | 0 | - |
| - | - | 1211 | 588.6 | - | - | 0 | - |
| 9 | b | 2654 | 602.8 | 0.00292 | 4.844 | +2 | 9 |
| - | - | 1644 | 603.3 | - | - | 0 | - |
| - | - | 1401 | 603.3 | - | - | 0 | - |
| - | - | 705.2 | 603.8 | - | - | 0 | - |
| - | - | 696.5 | 604.3 | - | - | 0 | - |
| - | - | 993.2 | 621.2 | - | - | 0 | - |
| - | - | 676.3 | 631.3 | - | - | 0 | - |
| 4 | b | 702.3 | 635.2 | 0.002345 | 3.692 | +1 | 4 |
| 4 | y | 1368 | 635.3 | 0.001425 | 2.243 | +2 | 10 |
| - | - | 4391 | 639.2 | - | - | 0 | - |
| - | - | 1567 | 640.2 | - | - | 0 | - |
| 4 | y | 1.19E+04 | 644.3 | 0.002069 | 3.211 | +2 | 10 |
| - | - | 1.108E+04 | 644.8 | - | - | 0 | - |
| - | - | 1.124E+04 | 645.3 | - | - | 0 | - |
| - | - | 5031 | 645.8 | - | - | 0 | - |
| - | - | 3477 | 646.3 | - | - | 0 | - |
| - | - | 679.1 | 646.8 | - | - | 0 | - |
| - | - | 4241 | 651.4 | - | - | 0 | - |
| - | - | 1623 | 652.4 | - | - | 0 | - |
| - | - | 6570 | 659.3 | - | - | 0 | - |
| - | - | 4864 | 659.8 | - | - | 0 | - |
| - | - | 3064 | 660.3 | - | - | 0 | - |
| - | - | 1178 | 660.8 | - | - | 0 | - |
| - | - | 929 | 668.3 | - | - | 0 | - |
| - | - | 776.4 | 668.8 | - | - | 0 | - |
| - | - | 2357 | 679.4 | - | - | 0 | - |
| - | - | 956.8 | 679.8 | - | - | 0 | - |
| - | - | 1157 | 680.4 | - | - | 0 | - |
| 9 | y | 909.8 | 695.3 | 0.001885 | 2.711 | +1 | 5 |
| 9 | y | 3209 | 696.3 | 0.0002442 | 0.3507 | +1 | 5 |
| - | - | 940.7 | 697.3 | - | - | 0 | - |
| - | - | 2446 | 697.4 | - | - | 0 | - |
| - | - | 944.5 | 698.4 | - | - | 0 | - |
| - | - | 1560 | 700.8 | - | - | 0 | - |
| - | - | 1009 | 707.4 | - | - | 0 | - |
| 9 | y | 8980 | 713.3 | 6.229E-05 | 0.08732 | +1 | 5 |
| - | - | 3955 | 714.3 | - | - | 0 | - |
| 3 | y | 3136 | 716.8 | 0.002022 | 2.821 | +2 | 11 |
| 3 | y | 3767 | 717.3 | 0.006946 | 9.684 | +2 | 11 |
| - | - | 1906 | 717.8 | - | - | 0 | - |
| - | - | 3144 | 718.8 | - | - | 0 | - |
| - | - | 2734 | 719.3 | - | - | 0 | - |
| - | - | 1322 | 719.8 | - | - | 0 | - |
| 3 | y | 8.44E+04 | 725.8 | 0.002544 | 3.505 | +2 | 11 |
| - | - | 7.319E+04 | 726.3 | - | - | 0 | - |
| - | - | 3.86E+04 | 726.8 | - | - | 0 | - |
| - | - | 1.154E+04 | 727.3 | - | - | 0 | - |
| - | - | 3461 | 727.8 | - | - | 0 | - |
| - | - | 1998 | 731.4 | - | - | 0 | - |
| - | - | 767.6 | 731.4 | - | - | 0 | - |
| - | - | 753.8 | 732.3 | - | - | 0 | - |
| 11 | b | 1.338E+04 | 732.8 | 0.002555 | 3.487 | +2 | 11 |
| - | - | 1.264E+04 | 733.3 | - | - | 0 | - |
| - | - | 5136 | 733.8 | - | - | 0 | - |
| - | - | 2123 | 734.3 | - | - | 0 | - |
| - | - | 1085 | 734.8 | - | - | 0 | - |
| - | - | 1482 | 738.3 | - | - | 0 | - |
| - | - | 775.2 | 739.3 | - | - | 0 | - |
| - | - | 1664 | 740.8 | - | - | 0 | - |
| - | - | 2605 | 741.3 | - | - | 0 | - |
| - | - | 1006 | 741.8 | - | - | 0 | - |
| - | - | 1309 | 756.9 | - | - | 0 | - |
| - | - | 1126 | 759.4 | - | - | 0 | - |
| - | - | 1979 | 766.4 | - | - | 0 | - |
| - | - | 942.1 | 767.4 | - | - | 0 | - |
| 8 | y | 3942 | 794.4 | 7.439E-05 | 0.09364 | +1 | 6 |
| - | - | 1937 | 795.4 | - | - | 0 | - |
| 2 | y | 683.7 | 798.3 | 0.004512 | 5.652 | +2 | 12 |
| 2 | y | 837.3 | 798.8 | 0.006776 | 8.482 | +2 | 12 |
| - | - | 1236 | 799.3 | - | - | 0 | - |
| - | - | 5332 | 800.4 | - | - | 0 | - |
| - | - | 4516 | 800.9 | - | - | 0 | - |
| - | - | 2306 | 801.4 | - | - | 0 | - |
| - | - | 880.4 | 801.9 | - | - | 0 | - |
| - | - | 2445 | 802.3 | - | - | 0 | - |
| - | - | 2033 | 802.5 | - | - | 0 | - |
| - | - | 1506 | 803.3 | - | - | 0 | - |
| - | - | 679.7 | 806.8 | - | - | 0 | - |
| 2 | y | 3.778E+04 | 807.4 | 0.002592 | 3.211 | +2 | 12 |
| - | - | 3.741E+04 | 807.9 | - | - | 0 | - |
| - | - | 1.996E+04 | 808.4 | - | - | 0 | - |
| - | - | 8558 | 808.9 | - | - | 0 | - |
| - | - | 1194 | 809.4 | - | - | 0 | - |
| 8 | y | 3.178E+04 | 812.4 | 0.0001143 | 0.1407 | +1 | 6 |
| - | - | 1.489E+04 | 813.4 | - | - | 0 | - |
| 12 | b | 9111 | 814.4 | 0.003012 | 3.698 | +2 | 12 |
| - | - | 6514 | 814.9 | - | - | 0 | - |
| - | - | 4504 | 815.4 | - | - | 0 | - |
| - | - | 1164 | 815.9 | - | - | 0 | - |
| - | - | 2334 | 830.5 | - | - | 0 | - |
| - | - | 913.5 | 877.4 | - | - | 0 | - |
| - | - | 2445 | 894.4 | - | - | 0 | - |
| - | - | 1909 | 895.4 | - | - | 0 | - |
| - | - | 715.5 | 901.4 | - | - | 0 | - |
| - | - | 660.1 | 908 | - | - | 0 | - |
| 7 | y | 954.2 | 910.4 | 0.00114 | 1.252 | +1 | 7 |
| - | - | 1638 | 922.5 | - | - | 0 | - |
| - | - | 891.7 | 923.5 | - | - | 0 | - |
| 7 | y | 2.697E+04 | 927.4 | 0.000385 | 0.4151 | +1 | 7 |
| - | - | 1.312E+04 | 928.4 | - | - | 0 | - |
| - | - | 5197 | 929.4 | - | - | 0 | - |
| 7 | b | 2237 | 949.4 | 0.005491 | 5.783 | +1 | 7 |
| - | - | 1577 | 950.4 | - | - | 0 | - |
| - | - | 1887 | 963.5 | - | - | 0 | - |
| - | - | 1302 | 964.5 | - | - | 0 | - |
| - | - | 1001 | 965.5 | - | - | 0 | - |
| - | - | 2942 | 991.5 | - | - | 0 | - |
| - | - | 2445 | 992.5 | - | - | 0 | - |
| - | - | 2377 | 993.5 | - | - | 0 | - |
| - | - | 1715 | 994.5 | - | - | 0 | - |
| - | - | 823.6 | 1012 | - | - | 0 | - |
| 6 | y | 9786 | 1056 | 0.0006214 | 0.5888 | +1 | 8 |
| - | - | 4775 | 1057 | - | - | 0 | - |
| - | - | 2197 | 1058 | - | - | 0 | - |
| - | - | 959.4 | 1058 | - | - | 0 | - |
| 5 | y | 853.8 | 1109 | 0.008128 | 7.332 | +1 | 9 |
| 5 | y | 805.3 | 1110 | 0.006303 | 5.681 | +1 | 9 |
| 5 | y | 1.392E+04 | 1127 | 0.004898 | 4.348 | +1 | 9 |
| - | - | 1.019E+04 | 1128 | - | - | 0 | - |
| - | - | 3017 | 1129 | - | - | 0 | - |
| - | - | 1172 | 1130 | - | - | 0 | - |
| - | - | 600.1 | 1138 | - | - | 0 | - |
| - | - | 6897 | 1155 | - | - | 0 | - |
| - | - | 4131 | 1156 | - | - | 0 | - |
| - | - | 1443 | 1157 | - | - | 0 | - |
| - | - | 758 | 1174 | - | - | 0 | - |
| 4 | y | 735.8 | 1270 | 0.003997 | 3.148 | +1 | 10 |
| 4 | y | 744.8 | 1271 | 0.001001 | 0.7882 | +1 | 10 |
| 4 | y | 7793 | 1288 | 0.007847 | 6.095 | +1 | 10 |
| - | - | 5453 | 1289 | - | - | 0 | - |
| - | - | 4341 | 1290 | - | - | 0 | - |
| - | - | 1375 | 1291 | - | - | 0 | - |
| - | - | 693.7 | 1300 | - | - | 0 | - |
| - | - | 3248 | 1318 | - | - | 0 | - |
| - | - | 2126 | 1319 | - | - | 0 | - |
| - | - | 1002 | 1320 | - | - | 0 | - |
| - | - | 1015 | 1452 | - | - | 0 | - |
| - | - | 755.3 | 1987 | - | - | 0 | - |

m/z Charge Intensity FragmentType MassShift Position
120.08102416992188 0 163288.86
121.08430480957031 0 12951.072
122.08780670166016 0 584.66235
125.06005096435547 0 839.9832
129.1024169921875 0 32609.045
130.10572814941406 0 1565.5726
131.04945373535156 0 643.2202
131.07070922851562 0 466.10995
133.08612060546875 0 4782.5605
134.02719116210938 0 4803.79
134.0395965576172 0 1017.54266
134.04498291015625 0 17790.15 y 12
135.04837036132812 0 1033.2892
136.07594299316406 0 169577.67
137.07339477539062 0 1066.8157
137.07923889160156 0 13219.176
138.06597900390625 0 502.52444
139.0868682861328 0 495.74554
143.1182403564453 0 610.0152
147.04408264160156 0 1220.9558
152.0709686279297 0 669.76917
153.0765380859375 0 380.8091
155.07041931152344 0 2363.7156
155.08189392089844 0 596.8693
155.11831665039062 0 916.8863
157.10855102539062 0 1043.0624
159.092041015625 0 536.44104
160.0756072998047 0 471.3575
162.02239990234375 0 709.05707
164.07078552246094 0 1010.36755
165.1026153564453 0 1049.2698
171.1494903564453 0 1986.3777
174.0555877685547 0 447.40048
175.08682250976562 0 710.60315
175.09645080566406 0 845.6431
177.11227416992188 0 1893.7385
181.0970916748047 0 804.3747
183.11294555664062 0 902.04156
185.103759765625 0 1516.8011
185.60543823242188 0 481.84592
187.10789489746094 0 4180.2812
187.1444091796875 0 743.93274
191.0819854736328 0 497.75604
195.1129913330078 0 684.12866
195.12258911132812 0 741.79193
197.12887573242188 0 900.7725
199.09689331054688 0 717.0939
199.1072998046875 0 901.9591
199.1443634033203 0 869.6984
199.18077087402344 0 522.9165
200.1395263671875 0 6972.4775
201.12301635742188 0 708.14355
201.1428680419922 0 752.66
202.1232147216797 0 883.103
205.06414794921875 0 5382.067
209.09193420410156 0 1022.61053
215.1024169921875 0 1580.9692
215.138671875 0 561.4362
221.138427734375 0 659.9118
224.13925170898438 0 660.69135
226.11875915527344 0 1268.1864
227.10340881347656 0 1207.305
233.0594024658203 0 956.84076
233.09217834472656 0 5412.27
233.12852478027344 0 12189.702
233.1642608642578 0 524.49414
234.05873107910156 0 989.4966
234.09561157226562 0 827.18115
234.13238525390625 0 1216.0507
235.13125610351562 0 637.34644
237.06979370117188 0 867.06885
237.13485717773438 0 1806.4552
239.15037536621094 0 12922.095
240.1539764404297 0 1222.1846
244.12940979003906 0 1578.7458
252.06903076171875 0 3155.0037
254.1157989501953 0 924.83765
256.177001953125 0 4884.8174
257.179931640625 0 901.24
261.12322998046875 0 7373.63
262.07037353515625 0 571.06256
262.1273193359375 0 996.1439
272.124267578125 0 1510.348
276.7696228027344 0 585.5259
280.0815124511719 0 889.217
282.1568603515625 0 709.4216
283.144287109375 0 76991.55 a 1
284.1476135253906 0 14798.062
285.1501159667969 0 1642.4904
287.1479797363281 0 769.27795
297.0904846191406 0 18293.98
297.1085205078125 0 9054.449 y 11
297.1553649902344 0 816.90265
298.0935974121094 0 3218.3716
298.1123046875 0 1376.6973
299.1392517089844 0 23322.994
300.1427307128906 0 4294.554
304.1294250488281 0 4986.098
305.1329345703125 0 779.80457
311.1393737792969 0 18643.523 b 1
312.1424255371094 0 3957.621
315.16632080078125 0 1492.3616
325.08563232421875 0 8838.62
326.0897216796875 0 1346.0791
327.1343994140625 0 8642.453
328.1373291015625 0 1879.3945
333.1552429199219 0 1819.0571
343.14056396484375 0 732.2592
343.1618347167969 0 813.2463
343.1982727050781 0 641.77924
349.18487548828125 0 697.1843
354.17767333984375 0 1079.4364
361.15509033203125 0 906.7589
370.257080078125 0 1873.8381
371.2034606933594 0 3653.9912
372.20269775390625 0 650.85486
377.18182373046875 0 1044.9602
382.1387939453125 0 593.5329
389.23004150390625 0 660.1628
391.0614318847656 0 565.5331
396.1220397949219 0 2190.56
396.155029296875 0 1894.2595
396.1921081542969 0 755.3441 b Water loss 8
401.18389892578125 0 585.28705
424.1875305175781 0 1918.576 y Ammonia loss 3
447.71044921875 0 1768.1343
447.7439880371094 0 679.13855
448.21240234375 0 609.1712
458.1706237792969 0 1796.1952
460.1747741699219 0 785.4298 y 10
467.19189453125 0 1586.3945
474.20306396484375 0 1869.3196 b 2
476.1817321777344 0 4528.013
482.2389831542969 0 888.8424
482.739501953125 0 856.8412
483.2635192871094 0 850.3759
483.7653503417969 0 922.28143
488.1495361328125 0 2891.7297
488.29876708984375 0 6130.0674
488.8900451660156 0 972.5815 b 10
489.1545715332031 0 881.076
489.3021240234375 0 1927.2455
490.305908203125 0 778.00616
496.2362976074219 0 882.56793
497.25921630859375 0 1020.3794
499.3002014160156 0 1477.0818
516.293701171875 0 3845.0723
517.2960205078125 0 1315.9889
524.2186889648438 0 1285.1846
528.2615356445312 0 3570.252 y 5
528.7623291015625 0 2077.649
533.3192138671875 0 1276.1687
533.9146118164062 0 2524.5007
534.24951171875 0 1959.444
534.5827026367188 0 1341.7487
535.2625122070312 0 1083.3014
539.2139282226562 0 1142.0149 y Water loss 9
539.5734252929688 0 746.08527
543.30126953125 0 6264.572
553.3023071289062 0 695.1545
557.2242431640625 0 7256.8896 y 9
558.22900390625 0 1945.98
563.7760620117188 0 8256.263 y 4
564.2774047851562 0 5854.0137
564.7743530273438 0 1651.7109
567.9707641601562 0 640.0315
569.2667236328125 0 655.45624
570.3355712890625 0 2022.1627
571.295654296875 0 8283.319
571.3429565429688 0 815.256
575.2498168945312 0 1097.2198
577.7695922851562 0 3337.27
578.2705078125 0 3031.1843
578.77294921875 0 1370.4115
581.2692260742188 0 729.2149
582.284912109375 0 622.4412
587.5918579101562 0 5682.737 Precursor
587.9264526367188 0 7604.7827
588.2604370117188 0 4546.479
588.3226318359375 0 4847.7227
588.5941162109375 0 1211.1431
602.7757568359375 0 2653.6538 b 8
603.2767333984375 0 1644.1887
603.3267822265625 0 1401.0565
603.7794799804688 0 705.21484
604.326904296875 0 696.4708
621.2364501953125 0 993.159
631.3194580078125 0 676.2754
635.223876953125 0 702.32916 b 3
635.2828979492188 0 1367.5557 y Water loss 3
639.243896484375 0 4391.4634
640.248046875 0 1567.0831
644.2875366210938 0 11902.196 y 3
644.78857421875 0 11081.993
645.2973022460938 0 11243.814
645.8018798828125 0 5030.9434
646.3048095703125 0 3477.4075
646.8065795898438 0 679.0589
651.361328125 0 4241.1646
652.366455078125 0 1623.4651
659.3002319335938 0 6569.5195
659.8018798828125 0 4864.2954
660.3046875 0 3064.3235
660.8051147460938 0 1177.9998
668.3089599609375 0 929.01715
668.8081665039062 0 776.41724
679.3570556640625 0 2356.63
679.8258056640625 0 956.8229
680.3642578125 0 1157.4481
695.3128662109375 0 909.81537 y Water loss 8
696.2990112304688 0 3208.7297 y Ammonia loss 8
697.3047485351562 0 940.6572
697.3676147460938 0 2445.9753
698.3681640625 0 944.5166
700.8387451171875 0 1560.3279
707.3641967773438 0 1009.3231
713.3253784179688 0 8979.943 y 8
714.328369140625 0 3955.1736
716.81396484375 0 3135.8728 y Water loss 2
717.31494140625 0 3766.6626 y Ammonia loss 2
717.814697265625 0 1906.3524
718.8363647460938 0 3143.7668
719.3373413085938 0 2734.098
719.839111328125 0 1321.908
725.8187255859375 0 84402.52 y 2
726.3201904296875 0 73188.16
726.8215942382812 0 38599.023
727.3225708007812 0 11541.73
727.8258056640625 0 3461.433
731.3502807617188 0 1997.7109
731.4193115234375 0 767.5884
732.349609375 0 753.77606
732.8341674804688 0 13382.132 b 10
733.3361206054688 0 12641.665
733.8360595703125 0 5135.9585
734.3359985351562 0 2122.5813
734.8397827148438 0 1084.6959
738.3134765625 0 1482.0166
739.3134765625 0 775.2112
740.8304443359375 0 1663.8386
741.3359985351562 0 2604.6072
741.8381958007812 0 1006.18616
756.8981323242188 0 1309.398
759.4149780273438 0 1126.1003
766.3844604492188 0 1978.9988
767.3963012695312 0 942.09283
794.3832397460938 0 3941.5842 y Water loss 7
795.386474609375 0 1936.5396
798.3431396484375 0 683.7147 y Water loss 1
798.846435546875 0 837.2751 y Ammonia loss 1
799.3423461914062 0 1236.1228
800.3671875 0 5332.1587
800.8697509765625 0 4516.309
801.3720703125 0 2305.5198
801.8753662109375 0 880.3526
802.3067016601562 0 2444.8015
802.4525146484375 0 2033.2322
803.311767578125 0 1505.5696
806.8482666015625 0 679.69165
807.350341796875 0 37784.676 y 1
807.851806640625 0 37405.992
808.3527221679688 0 19957.637
808.8538818359375 0 8557.604
809.3549194335938 0 1194.2478
812.3936157226562 0 31775.674 y 7
813.3965454101562 0 14894.637
814.3713989257812 0 9111.353 b 11
814.8673095703125 0 6513.6084
815.3704833984375 0 4504.015
815.8665161132812 0 1164.2838
830.4501953125 0 2334.02
877.3896484375 0 913.5447
894.4149780273438 0 2445.4302
895.4146728515625 0 1908.9092
901.3694458007812 0 715.4785
907.9503173828125 0 660.12115
910.395263671875 0 954.2366 y Ammonia loss 6
922.4776611328125 0 1637.9132
923.4832153320312 0 891.7484
927.4202880859375 0 26965.316 y 6
928.4234619140625 0 13121.355
929.4247436523438 0 5196.673
949.3750610351562 0 2236.8896 b 6
950.3809814453125 0 1576.6055
963.4732666015625 0 1886.9055
964.4752197265625 0 1301.6377
965.5132446289062 0 1000.72424
991.4662475585938 0 2941.639
992.4678955078125 0 2445.0183
993.50927734375 0 2377.2935
994.5093383789062 0 1715.2712
1011.5270385742188 0 823.58997
1055.5150146484375 0 9785.746 y 5
1056.517333984375 0 4775.1885
1057.50341796875 0 2197.079
1058.489013671875 0 959.42096
1108.5340576171875 0 853.788 y Water loss 4
1109.5198974609375 0 805.2642 y Ammonia loss 4
1126.5478515625 0 13916.485 y 4
1127.549560546875 0 10187.59
1128.5552978515625 0 3016.8647
1129.553955078125 0 1171.8475
1137.5106201171875 0 600.067
1154.5294189453125 0 6896.721
1155.533203125 0 4131.2773
1156.5364990234375 0 1442.8329
1173.544677734375 0 758.0211
1269.557373046875 0 735.76227 y Water loss 3
1270.54638671875 0 744.77826 y Ammonia loss 3
1287.5640869140625 0 7793.1694 y 3
1288.5667724609375 0 5452.8955
1289.571533203125 0 4341.319
1290.581298828125 0 1375.0863
1300.31103515625 0 693.7143
1317.58740234375 0 3248.3716
1318.5941162109375 0 2126.3713
1319.58056640625 0 1002.44226
1451.640869140625 0 1014.50385
1986.94482421875 0 755.3089

Spectrum Details

|  |  |
| --- | --- |
| Matched peaks? Matched peaks The total absolute number of peaks matched. Additionally in brackets the total fraction of peaks matched and the total number of peaks is shown. | 41 (12.85% of 319) |
| FDR? FDR The false discovery rate estimated for this peptide. It is calculated by matching all theoretical fragments with a non-integer shift with the raw peaks for this spectrum. This is done with 40 different shifts. The resulting percentage is the average number of annotated peaks over the number of annotated peaks with the correct spectrum. | 1.39% |
| Satellite FDR? Satellite FDR See the FDR for details on its calculation. This satellite ion specific FDR only contains the satellite ions (d/w) for I/L/J positions. | - |
| PSM Score? PSM Score The PSM Score as given by Hecklib to this annotated spectrum. It is shown with three significant figures. | 314 |

#### Reverse Lookup? Reverse Lookup All places where this read could be placed.

| Group | Segment | Template | Template Part | Read Part | Score | Unique |
| --- | --- | --- | --- | --- | --- | --- |
| Homo sapiens Heavy Chain | IGHV | IGHV3-9 | [93..105] | [0..13] | 53 | False |
| Homo sapiens Heavy Chain | IGHV | IGHV3-43 | [93..105] | [0..13] | 53 | False |

| Recombined | Template Part | Read Part | Score | Unique |
| --- | --- | --- | --- | --- |
| REC-0-1 | [92..105] | [0..13] | 104 | True |

#### Meta Information from Multiple reads

##### Number of combined reads

2

##### Intensity

0.7008

##### TotalArea

8.471E+07

#### Positional Score

Copy Data

##### Positional Score (TSV)

###### Preview

```
Loading example...
```

*Click on the button to copy the data to your clipboard.*

000123456789101112

Label Value
"0" 0
"1" 0
"2" 0
"3" 0
"4" 0
"5" 0
"6" 0
"7" 0
"8" 0
"9" 0
"10" 0
"11" 0
"12" 0

#### Meta Information from PEAKS

##### Scan Identifier

F2:6983

##### Original sequence

F

Y

Y

C

+58.01

A

K

D

V

R

P

Y

Y

D

##### Posttranslational Modifications

Carboxymethyl

##### Source File

D:\separate\_stitch\_analyses\xle-disambiguation\raw\20210323\_F1\_UM1\_Peng0013\_SA\_F59\_ingel\_3ug\_TL.raw

##### Fraction

2

##### Scan Feature

F2:7719

##### De Novo Score

98

##### ConfidenceScore

98

### m/z

587.5934

##### Mass

1759.7551

##### Charge

3

##### Retention Time

38.48

##### Predicted Retention Time

-

##### Area

4.236E+07

##### Parts Per Million

1.8

##### Fragmentation mode

HCD

##### Originating file

01 D:\separate\_stitch\_analyses\xle-disambiguation\20210325\_F59\_3ug\_DENOVO\_12.csv

#### Meta Information from PEAKS

##### Scan Identifier

F2:7041

##### Original sequence

F

Y

Y

C

+58.01

A

K

D

V

R

P

Y

Y

D

##### Posttranslational Modifications

Carboxymethyl

##### Source File

D:\separate\_stitch\_analyses\xle-disambiguation\raw\20210323\_F1\_UM1\_Peng0013\_SA\_F59\_ingel\_3ug\_TL.raw

##### Fraction

2

##### Scan Feature

F2:7719

##### De Novo Score

97

##### ConfidenceScore

97

### m/z

587.5934

##### Mass

1759.7551

##### Charge

3

##### Retention Time

38.48

##### Predicted Retention Time

-

##### Area

4.236E+07

##### Parts Per Million

1.8

##### Fragmentation mode

HCD

##### Originating file

01 D:\separate\_stitch\_analyses\xle-disambiguation\20210325\_F59\_3ug\_DENOVO\_12.csv
