## Supplementary material for "A handle on mass coincidence errors in *de novo* sequencing of antibodies by bottom-up proteomics": Combined_096.html

Details Combined\_096 | Stitch OverviewUndefined

### Read Combined\_096

#### Sequence (length=17)

TLPPSREEMTKNQVSTJ

#### Spectrum 3817? Spectrum 3817 The raw spectrum of this peptide as annotated by Hecklib. The fragments are coloured according to ion type (see legend). Any peaks with a star '\*' as text can be hovered over to see the full details, first the ion type second the mass shift type. By hovering over the amino acids in the peptide or ions in the legend the corresponding peaks are highlighted. By toggling the 'Unassigned' label you can turn the background (unassigned) peaks on or off in the plot. By updating the slider in the Ion legend you can update the spectrum to only show the top X% of the peaks with labels. The top X% means any peak that is within X% of the highest intensity. By dragging in the spectrum you can zoom in to a specific part of the spectrum and use 'Zoom Out' to get back to the original zoom level. The annotation of the spectrum is based on the given sequence in the peptides file and is done with different software so inconsistencies are likely. The peaks are annotated based on the given sequence, with 20 ppm tolerance.

Copy Data

##### Spectrum 3817 (TSV)

###### Preview

```
Loading example...
```

*Click on the button to copy the data to your clipboard.*

Mz MinMz MaxIntensity Max

WidthHeightPeptide font sizePeptide stroke widthSpectrum font sizeSpectrum stroke widthCompact peptide

Ion legend

wxyz

abcd

OtherUnassignedIon^Charge^_Position_Show for top:%

TLPPSREEMTKNQVSTJ

01.30e+52.59e+53.89e+55.19e+5

Zoom Out

y^+2^_4_y^+1^_2_y^+1^_2_y^+1^_3_y^+1^_3_c^+1^_4_w^+1^_5_c^+1^_5_z^+1^_5_y^+3^_14_y^+1^_5_z^+3^_15_c^+3^_15_y^+3^_15_z^+3^_15_c^+3^_15_y^+3^_15_c^+2^_10_c^+2^_11_y^+1^_6_z^+1^_6_c^+2^_11_y^+1^_6_c^+1^_6_c^+2^_12_c^+2^_12_c^+2^_12_z^+2^_12_y^+2^_12_y^+2^_13_z^+2^_13_c^+2^_13_c^+2^_13_y^+2^_13_y^+1^_7_c^+2^_13_y^+1^_7_c^+1^_7_y^+2^_14_c^+2^_14_c^+2^_14_y^+2^_14_c^+2^_14_w^+2^_15_z^+2^_15_z^+2^_15_z^+1^_8_w^+1^_8_c^+2^_15_y^+2^_15_z^+2^_15_w^+1^_8_c^+2^_15_y^+2^_15_y^+1^_8_z^+1^_8_y^+1^_8_w^+2^_16_z^+2^_16_c^+2^_16_c^+2^_16_y^+2^_16_y^+2^_16_z^+2^_16_c^+2^_16_y^+2^_16_c^+1^_8_z^+1^_9_z^+1^_9_y^+1^_9_c^+1^_9_w^+1^_10_z^+1^_10_z^+1^_10_y^+1^_10_c^+1^_10_w^+1^_11_z^+1^_11_c^+1^_11_y^+1^_11_c^+1^_11_c^+1^_12_y^+1^_12_z^+1^_12_y^+1^_12_z^+1^_13_c^+1^_13_y^+1^_13_c^+1^_13_c^+1^_14_c^+1^_14_y^+1^_14_c^+1^_14_c^+1^_15_z^+1^_15_c^+1^_15_y^+1^_15_y^+1^_16_z^+1^_16_c^+1^_16_y^+1^_16_

0880175926393519

Fragment Matches Table

Show background peaks

| Position | Ion type | Intensity | mz Theoretical | mz Error (Th) | mz Error (ppm) | Charge | Series Number |
| --- | --- | --- | --- | --- | --- | --- | --- |
| - | - | 1061 | 120.1 | - | - | 0 | - |
| - | - | 1.57E+04 | 120.1 | - | - | 0 | - |
| - | - | 1512 | 143.1 | - | - | 0 | - |
| - | - | 998.1 | 168.4 | - | - | 0 | - |
| - | - | 1389 | 169.1 | - | - | 0 | - |
| - | - | 1335 | 173.1 | - | - | 0 | - |
| - | - | 3048 | 173.5 | - | - | 0 | - |
| - | - | 1437 | 185.1 | - | - | 0 | - |
| - | - | 1.554E+05 | 187.1 | - | - | 0 | - |
| - | - | 1268 | 188.1 | - | - | 0 | - |
| - | - | 1.665E+04 | 188.1 | - | - | 0 | - |
| - | - | 1228 | 192.9 | - | - | 0 | - |
| 14 | y | 4514 | 201.1 | 0.0002121 | 1.055 | +2 | 4 |
| - | - | 1066 | 201.1 | - | - | 0 | - |
| - | - | 5217 | 212.1 | - | - | 0 | - |
| 16 | y | 1.019E+05 | 215.1 | 0.0002786 | 1.295 | +1 | 2 |
| - | - | 8884 | 216.1 | - | - | 0 | - |
| 16 | y | 1.036E+04 | 233.1 | 0.000334 | 1.433 | +1 | 2 |
| - | - | 1295 | 239.4 | - | - | 0 | - |
| - | - | 2188 | 256.2 | - | - | 0 | - |
| - | - | 1.854E+04 | 299.2 | - | - | 0 | - |
| - | - | 3107 | 300.2 | - | - | 0 | - |
| 15 | y | 3241 | 302.2 | 0.0007056 | 2.335 | +1 | 3 |
| - | - | 4825 | 312.2 | - | - | 0 | - |
| - | - | 1701 | 313.2 | - | - | 0 | - |
| 15 | y | 6218 | 320.2 | 0.0009441 | 2.949 | +1 | 3 |
| - | - | 1533 | 326.2 | - | - | 0 | - |
| - | - | 1439 | 327.2 | - | - | 0 | - |
| - | - | 2075 | 369.2 | - | - | 0 | - |
| - | - | 2982 | 382.3 | - | - | 0 | - |
| - | - | 1436 | 383.3 | - | - | 0 | - |
| - | - | 6792 | 407.3 | - | - | 0 | - |
| - | - | 1563 | 413.2 | - | - | 0 | - |
| - | - | 3716 | 425.3 | - | - | 0 | - |
| 4 | c | 7047 | 426.3 | 0.0008931 | 2.095 | +1 | 4 |
| - | - | 1957 | 427.3 | - | - | 0 | - |
| - | - | 1451 | 428.3 | - | - | 0 | - |
| - | - | 1348 | 449.7 | - | - | 0 | - |
| - | - | 2311 | 451.3 | - | - | 0 | - |
| - | - | 9359 | 470.3 | - | - | 0 | - |
| - | - | 2529 | 471.3 | - | - | 0 | - |
| - | - | 2336 | 471.9 | - | - | 0 | - |
| - | - | 1681 | 472.7 | - | - | 0 | - |
| 13 | w | 1762 | 473.3 | 6.003E-05 | 0.1268 | +1 | 5 |
| - | - | 1264 | 494.2 | - | - | 0 | - |
| - | - | 4341 | 494.3 | - | - | 0 | - |
| - | - | 1915 | 495.2 | - | - | 0 | - |
| - | - | 2553 | 499.9 | - | - | 0 | - |
| - | - | 6535 | 500.9 | - | - | 0 | - |
| - | - | 5246 | 501.2 | - | - | 0 | - |
| - | - | 1652 | 501.3 | - | - | 0 | - |
| - | - | 1677 | 501.6 | - | - | 0 | - |
| - | - | 1904 | 508.3 | - | - | 0 | - |
| - | - | 2058 | 509.3 | - | - | 0 | - |
| - | - | 3008 | 512.3 | - | - | 0 | - |
| 5 | c | 9.937E+04 | 513.3 | 0.0005868 | 1.143 | +1 | 5 |
| - | - | 2.69E+04 | 514.3 | - | - | 0 | - |
| - | - | 4984 | 515.3 | - | - | 0 | - |
| - | - | 2436 | 523.3 | - | - | 0 | - |
| - | - | 2587 | 523.6 | - | - | 0 | - |
| - | - | 2839 | 526.3 | - | - | 0 | - |
| - | - | 2401 | 526.9 | - | - | 0 | - |
| - | - | 2681 | 527.3 | - | - | 0 | - |
| - | - | 1587 | 528.3 | - | - | 0 | - |
| - | - | 7575 | 529.3 | - | - | 0 | - |
| - | - | 2457 | 529.3 | - | - | 0 | - |
| - | - | 6603 | 529.6 | - | - | 0 | - |
| - | - | 3894 | 529.9 | - | - | 0 | - |
| 13 | z | 9807 | 531.3 | 0.0007701 | 1.449 | +1 | 5 |
| - | - | 9607 | 532.3 | - | - | 0 | - |
| - | - | 1.326E+04 | 532.6 | - | - | 0 | - |
| - | - | 1.047E+04 | 532.9 | - | - | 0 | - |
| - | - | 5831 | 533.3 | - | - | 0 | - |
| - | - | 1961 | 533.3 | - | - | 0 | - |
| - | - | 2601 | 533.6 | - | - | 0 | - |
| - | - | 2.328E+04 | 538.6 | - | - | 0 | - |
| - | - | 2.714E+04 | 538.9 | - | - | 0 | - |
| - | - | 1.366E+04 | 539.3 | - | - | 0 | - |
| - | - | 6089 | 539.6 | - | - | 0 | - |
| 4 | y | 1763 | 539.9 | 0.006635 | 12.29 | +3 | 14 |
| - | - | 1746 | 544.3 | - | - | 0 | - |
| - | - | 1386 | 544.9 | - | - | 0 | - |
| 13 | y | 5372 | 547.3 | 7.072E-05 | 0.1292 | +1 | 5 |
| - | - | 5947 | 557 | - | - | 0 | - |
| - | - | 7515 | 557.3 | - | - | 0 | - |
| - | - | 7379 | 557.6 | - | - | 0 | - |
| - | - | 2841 | 558 | - | - | 0 | - |
| - | - | 1643 | 563.3 | - | - | 0 | - |
| - | - | 1935 | 563.6 | - | - | 0 | - |
| - | - | 3474 | 566.3 | - | - | 0 | - |
| - | - | 3467 | 566.6 | - | - | 0 | - |
| 3 | z | 2309 | 566.9 | 0.004505 | 7.946 | +3 | 15 |
| - | - | 3772 | 570.3 | - | - | 0 | - |
| - | - | 1.025E+04 | 571.3 | - | - | 0 | - |
| 15 | c | 2.812E+04 | 572.3 | 0.001967 | 3.436 | +3 | 15 |
| 3 | y | 2.332E+04 | 572.6 | 0.007885 | 13.77 | +3 | 15 |
| 3 | z | 1.219E+04 | 572.9 | 0.004341 | 7.576 | +3 | 15 |
| - | - | 9986 | 573.3 | - | - | 0 | - |
| - | - | 4353 | 573.3 | - | - | 0 | - |
| - | - | 2316 | 573.6 | - | - | 0 | - |
| - | - | 1667 | 573.9 | - | - | 0 | - |
| - | - | 1484 | 574.3 | - | - | 0 | - |
| 15 | c | 3581 | 578 | 0.001804 | 3.122 | +3 | 15 |
| 3 | y | 5.136E+05 | 578.3 | 0.002595 | 4.488 | +3 | 15 |
| - | - | 4.533E+05 | 578.6 | - | - | 0 | - |
| - | - | 2.786E+05 | 579 | - | - | 0 | - |
| 10 | c | 1.007E+05 | 579.3 | 0.01009 | 17.42 | +2 | 10 |
| - | - | 2.487E+04 | 579.6 | - | - | 0 | - |
| - | - | 1470 | 607.3 | - | - | 0 | - |
| - | - | 2954 | 607.4 | - | - | 0 | - |
| - | - | 2120 | 615.3 | - | - | 0 | - |
| - | - | 1.13E+04 | 625.4 | - | - | 0 | - |
| - | - | 6801 | 626.4 | - | - | 0 | - |
| - | - | 1582 | 627.4 | - | - | 0 | - |
| - | - | 2647 | 628.3 | - | - | 0 | - |
| - | - | 3558 | 628.4 | - | - | 0 | - |
| - | - | 6878 | 643.7 | - | - | 0 | - |
| 11 | c | 1455 | 643.8 | 0.003131 | 4.863 | +2 | 11 |
| - | - | 3787 | 644 | - | - | 0 | - |
| 12 | y | 6431 | 644.3 | 0.007294 | 11.32 | +1 | 6 |
| 12 | z | 5668 | 645.3 | 0.001361 | 2.108 | +1 | 6 |
| - | - | 1.005E+04 | 646.3 | - | - | 0 | - |
| - | - | 2605 | 647.3 | - | - | 0 | - |
| - | - | 2535 | 649.8 | - | - | 0 | - |
| - | - | 2659 | 650.3 | - | - | 0 | - |
| - | - | 3383 | 650.8 | - | - | 0 | - |
| 11 | c | 1.771E+04 | 652.3 | 0.002979 | 4.566 | +2 | 11 |
| - | - | 9093 | 652.8 | - | - | 0 | - |
| - | - | 4046 | 653.3 | - | - | 0 | - |
| - | - | 1917 | 657.8 | - | - | 0 | - |
| 12 | y | 1.943E+04 | 661.4 | 0.000764 | 1.155 | +1 | 6 |
| - | - | 6756 | 662.4 | - | - | 0 | - |
| 6 | c | 1.566E+05 | 669.4 | 0.0007331 | 1.095 | +1 | 6 |
| - | - | 6.072E+04 | 670.4 | - | - | 0 | - |
| - | - | 1.092E+04 | 671.4 | - | - | 0 | - |
| - | - | 4773 | 672.4 | - | - | 0 | - |
| - | - | 2906 | 686.8 | - | - | 0 | - |
| - | - | 2292 | 687.4 | - | - | 0 | - |
| - | - | 3612 | 695.4 | - | - | 0 | - |
| - | - | 1676 | 696.4 | - | - | 0 | - |
| - | - | 4906 | 699.8 | - | - | 0 | - |
| 12 | c | 5728 | 700.4 | 0.001141 | 1.629 | +2 | 12 |
| 12 | c | 2542 | 700.8 | 0.008339 | 11.9 | +2 | 12 |
| - | - | 1.387E+04 | 707.3 | - | - | 0 | - |
| - | - | 1.268E+04 | 707.8 | - | - | 0 | - |
| - | - | 6024 | 708.4 | - | - | 0 | - |
| - | - | 1.873E+04 | 708.9 | - | - | 0 | - |
| 12 | c | 1.097E+05 | 709.4 | 0.002694 | 3.798 | +2 | 12 |
| - | - | 8.531E+04 | 709.9 | - | - | 0 | - |
| - | - | 4.298E+04 | 710.4 | - | - | 0 | - |
| - | - | 1.425E+04 | 710.9 | - | - | 0 | - |
| - | - | 3464 | 711.4 | - | - | 0 | - |
| - | - | 2135 | 711.4 | - | - | 0 | - |
| 6 | z | 8139 | 718.3 | 0.004739 | 6.597 | +2 | 12 |
| - | - | 1.016E+04 | 718.9 | - | - | 0 | - |
| - | - | 4008 | 719.4 | - | - | 0 | - |
| - | - | 2255 | 719.9 | - | - | 0 | - |
| - | - | 1901 | 720.4 | - | - | 0 | - |
| - | - | 3638 | 722.4 | - | - | 0 | - |
| - | - | 4378 | 722.9 | - | - | 0 | - |
| - | - | 2198 | 725.4 | - | - | 0 | - |
| - | - | 5146 | 725.9 | - | - | 0 | - |
| 6 | y | 2.176E+04 | 726.4 | 0.003006 | 4.139 | +2 | 12 |
| - | - | 1.554E+04 | 726.9 | - | - | 0 | - |
| - | - | 7724 | 727.4 | - | - | 0 | - |
| - | - | 2.045E+04 | 729.4 | - | - | 0 | - |
| - | - | 7426 | 730.4 | - | - | 0 | - |
| - | - | 3226 | 741.9 | - | - | 0 | - |
| - | - | 1912 | 742.4 | - | - | 0 | - |
| - | - | 1.525E+04 | 750.9 | - | - | 0 | - |
| - | - | 1.489E+04 | 751.4 | - | - | 0 | - |
| - | - | 6749 | 751.9 | - | - | 0 | - |
| - | - | 2358 | 752.4 | - | - | 0 | - |
| - | - | 2773 | 754.4 | - | - | 0 | - |
| - | - | 2040 | 755.4 | - | - | 0 | - |
| - | - | 1762 | 757.9 | - | - | 0 | - |
| - | - | 1719 | 759.4 | - | - | 0 | - |
| 5 | y | 2543 | 761.4 | 0.00576 | 7.565 | +2 | 13 |
| 5 | z | 1.406E+04 | 761.9 | 0.003251 | 4.267 | +2 | 13 |
| - | - | 2.153E+04 | 762.4 | - | - | 0 | - |
| - | - | 9984 | 762.9 | - | - | 0 | - |
| - | - | 3708 | 763.4 | - | - | 0 | - |
| - | - | 2081 | 763.9 | - | - | 0 | - |
| 13 | c | 2499 | 764.4 | 0.001881 | 2.461 | +2 | 13 |
| 13 | c | 2719 | 764.9 | 0.007615 | 9.956 | +2 | 13 |
| - | - | 4423 | 765.4 | - | - | 0 | - |
| - | - | 2728 | 765.9 | - | - | 0 | - |
| - | - | 2924 | 766.4 | - | - | 0 | - |
| - | - | 4417 | 769.4 | - | - | 0 | - |
| 5 | y | 1.373E+04 | 769.9 | 0.001946 | 2.527 | +2 | 13 |
| - | - | 1.361E+04 | 770.4 | - | - | 0 | - |
| - | - | 5360 | 770.9 | - | - | 0 | - |
| 11 | y | 2800 | 771.4 | 0.005538 | 7.179 | +1 | 7 |
| - | - | 8177 | 772.9 | - | - | 0 | - |
| 13 | c | 1.468E+05 | 773.4 | 0.004716 | 6.098 | +2 | 13 |
| - | - | 2.344E+04 | 773.5 | - | - | 0 | - |
| - | - | 1.61E+05 | 773.9 | - | - | 0 | - |
| - | - | 5.868E+04 | 774.4 | - | - | 0 | - |
| - | - | 1.093E+04 | 774.5 | - | - | 0 | - |
| - | - | 2.41E+04 | 774.9 | - | - | 0 | - |
| - | - | 3578 | 775.4 | - | - | 0 | - |
| - | - | 6194 | 775.4 | - | - | 0 | - |
| - | - | 3668 | 779.4 | - | - | 0 | - |
| - | - | 2148 | 786.4 | - | - | 0 | - |
| - | - | 2923 | 786.9 | - | - | 0 | - |
| - | - | 1835 | 787.4 | - | - | 0 | - |
| 11 | y | 2.147E+04 | 789.4 | 0.001321 | 1.673 | +1 | 7 |
| - | - | 8706 | 790.4 | - | - | 0 | - |
| - | - | 2361 | 791.9 | - | - | 0 | - |
| - | - | 2093 | 792.4 | - | - | 0 | - |
| - | - | 9216 | 792.9 | - | - | 0 | - |
| - | - | 1.107E+04 | 793.4 | - | - | 0 | - |
| - | - | 7731 | 793.9 | - | - | 0 | - |
| - | - | 7405 | 794.4 | - | - | 0 | - |
| - | - | 2865 | 794.9 | - | - | 0 | - |
| - | - | 3361 | 797.4 | - | - | 0 | - |
| 7 | c | 2.345E+05 | 798.4 | 0.001231 | 1.542 | +1 | 7 |
| - | - | 6977 | 798.9 | - | - | 0 | - |
| - | - | 1.055E+05 | 799.5 | - | - | 0 | - |
| - | - | 2.441E+04 | 800.5 | - | - | 0 | - |
| - | - | 1.712E+04 | 800.9 | - | - | 0 | - |
| - | - | 7913 | 801.4 | - | - | 0 | - |
| - | - | 7719 | 801.9 | - | - | 0 | - |
| - | - | 3827 | 802.4 | - | - | 0 | - |
| - | - | 2072 | 803.4 | - | - | 0 | - |
| - | - | 2318 | 805.4 | - | - | 0 | - |
| - | - | 1862 | 806.4 | - | - | 0 | - |
| - | - | 3.579E+04 | 807.4 | - | - | 0 | - |
| - | - | 3.521E+04 | 807.9 | - | - | 0 | - |
| - | - | 1.923E+04 | 808.4 | - | - | 0 | - |
| - | - | 5414 | 808.9 | - | - | 0 | - |
| 4 | y | 2470 | 809.9 | 0.007271 | 8.978 | +2 | 14 |
| - | - | 2000 | 812.5 | - | - | 0 | - |
| - | - | 4.937E+04 | 813.4 | - | - | 0 | - |
| 14 | c | 4.604E+04 | 813.9 | 0.001076 | 1.322 | +2 | 14 |
| 14 | c | 3.277E+04 | 814.4 | 0.007527 | 9.242 | +2 | 14 |
| - | - | 2477 | 814.5 | - | - | 0 | - |
| - | - | 1.514E+04 | 814.9 | - | - | 0 | - |
| - | - | 9663 | 815.4 | - | - | 0 | - |
| - | - | 4252 | 816.4 | - | - | 0 | - |
| - | - | 3447 | 817.9 | - | - | 0 | - |
| 4 | y | 8.07E+04 | 818.4 | 0.003884 | 4.746 | +2 | 14 |
| - | - | 6.389E+04 | 818.9 | - | - | 0 | - |
| - | - | 3.142E+04 | 819.4 | - | - | 0 | - |
| - | - | 1.105E+04 | 819.9 | - | - | 0 | - |
| - | - | 2703 | 820.4 | - | - | 0 | - |
| - | - | 7.672E+04 | 822.4 | - | - | 0 | - |
| 14 | c | 1.083E+05 | 822.9 | 0.001698 | 2.064 | +2 | 14 |
| - | - | 8.434E+04 | 823.4 | - | - | 0 | - |
| - | - | 4.456E+04 | 823.9 | - | - | 0 | - |
| - | - | 1.156E+04 | 824.4 | - | - | 0 | - |
| - | - | 3960 | 824.9 | - | - | 0 | - |
| - | - | 4366 | 825.5 | - | - | 0 | - |
| - | - | 6751 | 830.5 | - | - | 0 | - |
| - | - | 3891 | 831.5 | - | - | 0 | - |
| - | - | 1.149E+04 | 834.9 | - | - | 0 | - |
| - | - | 1.792E+04 | 835.4 | - | - | 0 | - |
| - | - | 9194 | 835.9 | - | - | 0 | - |
| - | - | 6322 | 836.4 | - | - | 0 | - |
| - | - | 4704 | 836.9 | - | - | 0 | - |
| - | - | 1833 | 843.9 | - | - | 0 | - |
| - | - | 1867 | 844.4 | - | - | 0 | - |
| - | - | 7686 | 844.9 | - | - | 0 | - |
| 3 | w | 9372 | 845.4 | 0.01081 | 12.79 | +2 | 15 |
| - | - | 7853 | 845.9 | - | - | 0 | - |
| - | - | 3470 | 846.4 | - | - | 0 | - |
| - | - | 2469 | 848.9 | - | - | 0 | - |
| - | - | 5401 | 849.4 | - | - | 0 | - |
| 3 | z | 2718 | 849.9 | 0.01333 | 15.68 | +2 | 15 |
| 3 | z | 2742 | 850.4 | 0.006792 | 7.986 | +2 | 15 |
| 10 | z | 2925 | 856.5 | 0.0001432 | 0.1672 | +1 | 8 |
| - | - | 1981 | 856.9 | - | - | 0 | - |
| 10 | w | 1.377E+04 | 857.5 | 0.0002355 | 0.2746 | +1 | 8 |
| 15 | c | 1.884E+04 | 857.9 | 0.004452 | 5.189 | +2 | 15 |
| 3 | y | 1.394E+04 | 858.4 | 0.01342 | 15.63 | +2 | 15 |
| 3 | z | 1.249E+04 | 858.9 | 0.009325 | 10.86 | +2 | 15 |
| 10 | w | 2.252E+04 | 859.5 | 0.001717 | 1.998 | +1 | 8 |
| - | - | 2931 | 859.9 | - | - | 0 | - |
| - | - | 1.065E+04 | 860.5 | - | - | 0 | - |
| - | - | 1979 | 861.5 | - | - | 0 | - |
| - | - | 3354 | 864.9 | - | - | 0 | - |
| - | - | 2889 | 865.4 | - | - | 0 | - |
| - | - | 1.531E+04 | 865.9 | - | - | 0 | - |
| 15 | c | 2.297E+05 | 866.4 | 0.00314 | 3.624 | +2 | 15 |
| 3 | y | 5.002E+05 | 866.9 | 0.006433 | 7.42 | +2 | 15 |
| - | - | 3.842E+05 | 867.4 | - | - | 0 | - |
| - | - | 2.113E+05 | 867.9 | - | - | 0 | - |
| - | - | 7.309E+04 | 868.4 | - | - | 0 | - |
| - | - | 1.386E+04 | 868.9 | - | - | 0 | - |
| - | - | 3855 | 872.5 | - | - | 0 | - |
| - | - | 3232 | 872.9 | - | - | 0 | - |
| 10 | y | 2.033E+04 | 873.5 | 0.002347 | 2.687 | +1 | 8 |
| 10 | z | 1.324E+05 | 874.5 | 0.000748 | 0.8554 | +1 | 8 |
| - | - | 6.279E+04 | 875.5 | - | - | 0 | - |
| - | - | 2.148E+04 | 876.5 | - | - | 0 | - |
| - | - | 4989 | 877.5 | - | - | 0 | - |
| - | - | 9616 | 879.4 | - | - | 0 | - |
| - | - | 9096 | 880 | - | - | 0 | - |
| - | - | 5512 | 880.5 | - | - | 0 | - |
| - | - | 3924 | 880.9 | - | - | 0 | - |
| - | - | 5572 | 881.4 | - | - | 0 | - |
| - | - | 3304 | 881.9 | - | - | 0 | - |
| - | - | 4511 | 883.5 | - | - | 0 | - |
| - | - | 4645 | 884.5 | - | - | 0 | - |
| - | - | 5349 | 887.4 | - | - | 0 | - |
| - | - | 5211 | 887.9 | - | - | 0 | - |
| - | - | 2127 | 888.4 | - | - | 0 | - |
| - | - | 3598 | 889.5 | - | - | 0 | - |
| 10 | y | 3.874E+04 | 890.5 | 0.0005176 | 0.5812 | +1 | 8 |
| - | - | 1.887E+04 | 891.5 | - | - | 0 | - |
| - | - | 5058 | 892 | - | - | 0 | - |
| - | - | 5550 | 892.5 | - | - | 0 | - |
| - | - | 5734 | 893 | - | - | 0 | - |
| - | - | 7473 | 893.5 | - | - | 0 | - |
| 2 | w | 8.607E+04 | 893.9 | 0.003897 | 4.359 | +2 | 16 |
| - | - | 8.71E+04 | 894.4 | - | - | 0 | - |
| - | - | 5.599E+04 | 894.9 | - | - | 0 | - |
| - | - | 1.863E+04 | 895.4 | - | - | 0 | - |
| - | - | 5109 | 895.9 | - | - | 0 | - |
| - | - | 7084 | 896.4 | - | - | 0 | - |
| - | - | 3089 | 896.5 | - | - | 0 | - |
| - | - | 2795 | 897.4 | - | - | 0 | - |
| - | - | 2222 | 897.5 | - | - | 0 | - |
| - | - | 2060 | 898.5 | - | - | 0 | - |
| - | - | 2250 | 900.5 | - | - | 0 | - |
| - | - | 1.255E+04 | 901 | - | - | 0 | - |
| - | - | 1.579E+04 | 901.5 | - | - | 0 | - |
| - | - | 1.2E+04 | 902 | - | - | 0 | - |
| - | - | 7998 | 902.5 | - | - | 0 | - |
| - | - | 3485 | 903 | - | - | 0 | - |
| - | - | 2612 | 903.5 | - | - | 0 | - |
| - | - | 1908 | 904.5 | - | - | 0 | - |
| - | - | 2351 | 906 | - | - | 0 | - |
| 2 | z | 2825 | 906.5 | 0.01213 | 13.38 | +2 | 16 |
| - | - | 6402 | 907 | - | - | 0 | - |
| - | - | 6965 | 907.5 | - | - | 0 | - |
| 16 | c | 4342 | 908 | 0.01687 | 18.58 | +2 | 16 |
| 16 | c | 3671 | 908.4 | 0.006858 | 7.549 | +2 | 16 |
| - | - | 2430 | 908.9 | - | - | 0 | - |
| 2 | y | 1.436E+04 | 914.5 | 0.004473 | 4.892 | +2 | 16 |
| 2 | y | 1.245E+04 | 915 | 0.01045 | 11.42 | +2 | 16 |
| 2 | z | 5.285E+04 | 915.5 | 0.0066 | 7.209 | +2 | 16 |
| - | - | 5.499E+04 | 916 | - | - | 0 | - |
| - | - | 3.241E+04 | 916.5 | - | - | 0 | - |
| 16 | c | 1.682E+04 | 917 | 0.003227 | 3.519 | +2 | 16 |
| - | - | 6631 | 917.5 | - | - | 0 | - |
| - | - | 2933 | 917.9 | - | - | 0 | - |
| - | - | 2119 | 920 | - | - | 0 | - |
| - | - | 8563 | 920.5 | - | - | 0 | - |
| - | - | 2350 | 921 | - | - | 0 | - |
| - | - | 3699 | 921.5 | - | - | 0 | - |
| - | - | 7075 | 922.5 | - | - | 0 | - |
| - | - | 2.079E+05 | 923 | - | - | 0 | - |
| 2 | y | 1.967E+05 | 923.5 | 0.01207 | 13.07 | +2 | 16 |
| - | - | 3268 | 923.6 | - | - | 0 | - |
| - | - | 1.444E+05 | 924 | - | - | 0 | - |
| - | - | 6.025E+04 | 924.5 | - | - | 0 | - |
| - | - | 1.945E+04 | 925 | - | - | 0 | - |
| - | - | 2069 | 925.5 | - | - | 0 | - |
| - | - | 6096 | 926.5 | - | - | 0 | - |
| 8 | c | 2.306E+05 | 927.5 | 0.0009961 | 1.074 | +1 | 8 |
| - | - | 2470 | 928 | - | - | 0 | - |
| - | - | 1.191E+05 | 928.5 | - | - | 0 | - |
| - | - | 7482 | 929 | - | - | 0 | - |
| - | - | 3.821E+04 | 929.5 | - | - | 0 | - |
| - | - | 1.217E+04 | 930 | - | - | 0 | - |
| - | - | 1.677E+04 | 930.5 | - | - | 0 | - |
| - | - | 8215 | 931 | - | - | 0 | - |
| - | - | 8668 | 931.5 | - | - | 0 | - |
| - | - | 1885 | 933.5 | - | - | 0 | - |
| - | - | 2946 | 934.5 | - | - | 0 | - |
| - | - | 6615 | 935 | - | - | 0 | - |
| - | - | 5152 | 935.5 | - | - | 0 | - |
| - | - | 2904 | 936 | - | - | 0 | - |
| - | - | 8458 | 936.5 | - | - | 0 | - |
| - | - | 9571 | 937 | - | - | 0 | - |
| - | - | 1.482E+04 | 937.5 | - | - | 0 | - |
| - | - | 8.509E+04 | 938 | - | - | 0 | - |
| - | - | 7.714E+04 | 938.5 | - | - | 0 | - |
| - | - | 5.524E+04 | 939 | - | - | 0 | - |
| - | - | 2.535E+04 | 939.5 | - | - | 0 | - |
| - | - | 1.015E+04 | 940 | - | - | 0 | - |
| - | - | 2136 | 940.5 | - | - | 0 | - |
| - | - | 3809 | 941.5 | - | - | 0 | - |
| - | - | 2.224E+04 | 942.5 | - | - | 0 | - |
| - | - | 2.778E+04 | 943 | - | - | 0 | - |
| - | - | 2.993E+04 | 943.5 | - | - | 0 | - |
| - | - | 2.641E+04 | 944 | - | - | 0 | - |
| - | - | 1.711E+04 | 944.5 | - | - | 0 | - |
| - | - | 4.296E+04 | 945 | - | - | 0 | - |
| - | - | 4.556E+04 | 945.5 | - | - | 0 | - |
| - | - | 2.79E+04 | 946 | - | - | 0 | - |
| - | - | 3.018E+04 | 946.5 | - | - | 0 | - |
| - | - | 2.559E+04 | 947 | - | - | 0 | - |
| - | - | 1.613E+04 | 947.5 | - | - | 0 | - |
| - | - | 6647 | 948 | - | - | 0 | - |
| - | - | 2063 | 950.5 | - | - | 0 | - |
| - | - | 6693 | 951.5 | - | - | 0 | - |
| - | - | 5.055E+04 | 952 | - | - | 0 | - |
| - | - | 6.333E+04 | 952.5 | - | - | 0 | - |
| - | - | 4.46E+04 | 953 | - | - | 0 | - |
| - | - | 2.962E+04 | 953.5 | - | - | 0 | - |
| - | - | 1.127E+04 | 954 | - | - | 0 | - |
| - | - | 4012 | 954.5 | - | - | 0 | - |
| - | - | 1.878E+04 | 956.5 | - | - | 0 | - |
| - | - | 3.829E+04 | 957 | - | - | 0 | - |
| - | - | 3.375E+04 | 957.5 | - | - | 0 | - |
| - | - | 2.598E+04 | 958 | - | - | 0 | - |
| - | - | 3.352E+04 | 958.5 | - | - | 0 | - |
| - | - | 5634 | 959 | - | - | 0 | - |
| - | - | 1.089E+05 | 959.5 | - | - | 0 | - |
| - | - | 5.308E+04 | 960.5 | - | - | 0 | - |
| - | - | 1.341E+04 | 961.5 | - | - | 0 | - |
| - | - | 7.218E+04 | 965.5 | - | - | 0 | - |
| - | - | 3.2E+05 | 966 | - | - | 0 | - |
| - | - | 3.152E+05 | 966.5 | - | - | 0 | - |
| - | - | 2.189E+05 | 967 | - | - | 0 | - |
| - | - | 8.648E+04 | 967.5 | - | - | 0 | - |
| - | - | 2.816E+04 | 968 | - | - | 0 | - |
| - | - | 1.018E+04 | 968.5 | - | - | 0 | - |
| - | - | 5722 | 969.5 | - | - | 0 | - |
| - | - | 2817 | 970.5 | - | - | 0 | - |
| - | - | 3841 | 973.5 | - | - | 0 | - |
| - | - | 2.069E+05 | 974 | - | - | 0 | - |
| - | - | 3.827E+05 | 974.5 | - | - | 0 | - |
| - | - | 3.01E+05 | 975 | - | - | 0 | - |
| - | - | 3269 | 975.1 | - | - | 0 | - |
| - | - | 1.61E+05 | 975.5 | - | - | 0 | - |
| - | - | 5.736E+04 | 976 | - | - | 0 | - |
| - | - | 1.482E+04 | 976.5 | - | - | 0 | - |
| - | - | 2460 | 990.5 | - | - | 0 | - |
| 9 | z | 2545 | 1003 | 0.001458 | 1.453 | +1 | 9 |
| - | - | 4467 | 1006 | - | - | 0 | - |
| - | - | 2285 | 1007 | - | - | 0 | - |
| - | - | 3306 | 1013 | - | - | 0 | - |
| - | - | 3569 | 1013 | - | - | 0 | - |
| - | - | 2228 | 1015 | - | - | 0 | - |
| - | - | 2644 | 1015 | - | - | 0 | - |
| 9 | z | 2.76E+04 | 1022 | 0.006884 | 6.74 | +1 | 9 |
| - | - | 2.197E+04 | 1023 | - | - | 0 | - |
| - | - | 1.003E+04 | 1024 | - | - | 0 | - |
| - | - | 3277 | 1025 | - | - | 0 | - |
| - | - | 2.756E+04 | 1031 | - | - | 0 | - |
| - | - | 2.252E+04 | 1032 | - | - | 0 | - |
| - | - | 8917 | 1033 | - | - | 0 | - |
| - | - | 4173 | 1034 | - | - | 0 | - |
| 9 | y | 1.503E+04 | 1038 | 0.005616 | 5.413 | +1 | 9 |
| - | - | 7692 | 1039 | - | - | 0 | - |
| - | - | 2416 | 1040 | - | - | 0 | - |
| - | - | 1.091E+04 | 1050 | - | - | 0 | - |
| - | - | 9220 | 1051 | - | - | 0 | - |
| - | - | 4345 | 1052 | - | - | 0 | - |
| - | - | 2473 | 1056 | - | - | 0 | - |
| - | - | 2454 | 1059 | - | - | 0 | - |
| - | - | 4552 | 1074 | - | - | 0 | - |
| 9 | c | 2.53E+05 | 1075 | 0.005973 | 5.559 | +1 | 9 |
| - | - | 1.349E+05 | 1076 | - | - | 0 | - |
| - | - | 5.821E+04 | 1077 | - | - | 0 | - |
| - | - | 1.586E+04 | 1078 | - | - | 0 | - |
| - | - | 1775 | 1079 | - | - | 0 | - |
| - | - | 3441 | 1080 | - | - | 0 | - |
| - | - | 8279 | 1088 | - | - | 0 | - |
| - | - | 5508 | 1089 | - | - | 0 | - |
| 8 | w | 3.415E+04 | 1092 | 0.00616 | 5.644 | +1 | 10 |
| - | - | 1.751E+04 | 1093 | - | - | 0 | - |
| - | - | 9671 | 1094 | - | - | 0 | - |
| - | - | 5819 | 1095 | - | - | 0 | - |
| - | - | 3128 | 1095 | - | - | 0 | - |
| - | - | 2443 | 1107 | - | - | 0 | - |
| - | - | 5220 | 1114 | - | - | 0 | - |
| - | - | 3708 | 1115 | - | - | 0 | - |
| - | - | 2498 | 1116 | - | - | 0 | - |
| - | - | 6092 | 1132 | - | - | 0 | - |
| 8 | z | 6847 | 1133 | 0.02259 | 19.94 | +1 | 10 |
| - | - | 3224 | 1134 | - | - | 0 | - |
| - | - | 4433 | 1135 | - | - | 0 | - |
| - | - | 3090 | 1136 | - | - | 0 | - |
| - | - | 2903 | 1150 | - | - | 0 | - |
| 8 | z | 1.136E+05 | 1151 | 0.005673 | 4.931 | +1 | 10 |
| - | - | 6.926E+04 | 1152 | - | - | 0 | - |
| - | - | 3.472E+04 | 1153 | - | - | 0 | - |
| - | - | 8195 | 1154 | - | - | 0 | - |
| 8 | y | 8111 | 1167 | 0.007335 | 6.288 | +1 | 10 |
| - | - | 4998 | 1168 | - | - | 0 | - |
| 10 | c | 1.198E+05 | 1176 | 0.005292 | 4.501 | +1 | 10 |
| - | - | 7.986E+04 | 1177 | - | - | 0 | - |
| - | - | 3.13E+04 | 1178 | - | - | 0 | - |
| - | - | 1.161E+04 | 1179 | - | - | 0 | - |
| - | - | 2614 | 1180 | - | - | 0 | - |
| - | - | 2599 | 1208 | - | - | 0 | - |
| - | - | 3425 | 1209 | - | - | 0 | - |
| - | - | 1895 | 1217 | - | - | 0 | - |
| 7 | w | 1.636E+04 | 1221 | 0.007146 | 5.855 | +1 | 11 |
| - | - | 1.464E+04 | 1222 | - | - | 0 | - |
| - | - | 6980 | 1223 | - | - | 0 | - |
| - | - | 2154 | 1225 | - | - | 0 | - |
| - | - | 1880 | 1237 | - | - | 0 | - |
| - | - | 8170 | 1260 | - | - | 0 | - |
| - | - | 6212 | 1261 | - | - | 0 | - |
| - | - | 2208 | 1262 | - | - | 0 | - |
| 7 | z | 8.338E+04 | 1280 | 0.005439 | 4.25 | +1 | 11 |
| - | - | 5.898E+04 | 1281 | - | - | 0 | - |
| - | - | 3.014E+04 | 1282 | - | - | 0 | - |
| - | - | 8038 | 1283 | - | - | 0 | - |
| - | - | 2775 | 1285 | - | - | 0 | - |
| 11 | c | 1861 | 1286 | 0.001226 | 0.9535 | +1 | 11 |
| - | - | 2943 | 1295 | - | - | 0 | - |
| 7 | y | 2.539E+04 | 1296 | 0.006124 | 4.726 | +1 | 11 |
| - | - | 1.939E+04 | 1297 | - | - | 0 | - |
| - | - | 6379 | 1298 | - | - | 0 | - |
| - | - | 2157 | 1299 | - | - | 0 | - |
| - | - | 8509 | 1303 | - | - | 0 | - |
| 11 | c | 2.436E+04 | 1304 | 0.00359 | 2.754 | +1 | 11 |
| - | - | 1.739E+04 | 1305 | - | - | 0 | - |
| - | - | 8071 | 1306 | - | - | 0 | - |
| - | - | 2460 | 1334 | - | - | 0 | - |
| - | - | 6800 | 1335 | - | - | 0 | - |
| - | - | 3339 | 1336 | - | - | 0 | - |
| - | - | 2098 | 1364 | - | - | 0 | - |
| - | - | 2436 | 1365 | - | - | 0 | - |
| - | - | 2505 | 1366 | - | - | 0 | - |
| - | - | 2746 | 1374 | - | - | 0 | - |
| - | - | 2860 | 1375 | - | - | 0 | - |
| - | - | 1949 | 1393 | - | - | 0 | - |
| - | - | 5635 | 1417 | - | - | 0 | - |
| 12 | c | 4.646E+04 | 1418 | 0.0018 | 1.27 | +1 | 12 |
| - | - | 3.319E+04 | 1419 | - | - | 0 | - |
| - | - | 1.575E+04 | 1420 | - | - | 0 | - |
| - | - | 9029 | 1421 | - | - | 0 | - |
| - | - | 3997 | 1422 | - | - | 0 | - |
| 6 | y | 2941 | 1435 | 0.01713 | 11.94 | +1 | 12 |
| 6 | z | 6.594E+04 | 1436 | 0.004547 | 3.167 | +1 | 12 |
| - | - | 9.937E+04 | 1437 | - | - | 0 | - |
| - | - | 6.001E+04 | 1438 | - | - | 0 | - |
| - | - | 2.602E+04 | 1439 | - | - | 0 | - |
| - | - | 6409 | 1440 | - | - | 0 | - |
| - | - | 8569 | 1451 | - | - | 0 | - |
| 6 | y | 3.465E+04 | 1452 | 0.00389 | 2.679 | +1 | 12 |
| - | - | 2.37E+04 | 1453 | - | - | 0 | - |
| - | - | 1.237E+04 | 1454 | - | - | 0 | - |
| - | - | 4571 | 1455 | - | - | 0 | - |
| - | - | 3487 | 1503 | - | - | 0 | - |
| 5 | z | 4774 | 1523 | 0.002792 | 1.834 | +1 | 13 |
| - | - | 2.558E+04 | 1524 | - | - | 0 | - |
| - | - | 1.994E+04 | 1525 | - | - | 0 | - |
| - | - | 9004 | 1526 | - | - | 0 | - |
| - | - | 3819 | 1527 | - | - | 0 | - |
| 13 | c | 2867 | 1529 | 0.01933 | 12.65 | +1 | 13 |
| 5 | y | 7747 | 1539 | 0.002623 | 1.705 | +1 | 13 |
| - | - | 5079 | 1540 | - | - | 0 | - |
| - | - | 4400 | 1541 | - | - | 0 | - |
| - | - | 2796 | 1545 | - | - | 0 | - |
| 13 | c | 5.66E+04 | 1546 | 0.003892 | 2.518 | +1 | 13 |
| - | - | 4.91E+04 | 1547 | - | - | 0 | - |
| - | - | 2.392E+04 | 1548 | - | - | 0 | - |
| - | - | 9010 | 1549 | - | - | 0 | - |
| - | - | 2174 | 1601 | - | - | 0 | - |
| - | - | 5803 | 1602 | - | - | 0 | - |
| - | - | 5327 | 1603 | - | - | 0 | - |
| - | - | 2538 | 1604 | - | - | 0 | - |
| - | - | 6620 | 1615 | - | - | 0 | - |
| - | - | 4085 | 1616 | - | - | 0 | - |
| - | - | 1967 | 1617 | - | - | 0 | - |
| 14 | c | 4016 | 1627 | 0.008669 | 5.329 | +1 | 14 |
| 14 | c | 5968 | 1628 | 0.01476 | 9.068 | +1 | 14 |
| - | - | 4508 | 1629 | - | - | 0 | - |
| 4 | y | 1.376E+04 | 1636 | 0.001129 | 0.6899 | +1 | 14 |
| - | - | 1.515E+04 | 1637 | - | - | 0 | - |
| - | - | 8509 | 1638 | - | - | 0 | - |
| - | - | 5008 | 1639 | - | - | 0 | - |
| - | - | 8987 | 1644 | - | - | 0 | - |
| 14 | c | 6.208E+04 | 1645 | 0.004326 | 2.63 | +1 | 14 |
| - | - | 5.361E+04 | 1646 | - | - | 0 | - |
| - | - | 3.201E+04 | 1647 | - | - | 0 | - |
| - | - | 1.101E+04 | 1648 | - | - | 0 | - |
| - | - | 3492 | 1649 | - | - | 0 | - |
| - | - | 2664 | 1673 | - | - | 0 | - |
| - | - | 2239 | 1675 | - | - | 0 | - |
| - | - | 2619 | 1688 | - | - | 0 | - |
| - | - | 6131 | 1689 | - | - | 0 | - |
| - | - | 5483 | 1690 | - | - | 0 | - |
| - | - | 2206 | 1691 | - | - | 0 | - |
| 15 | c | 1886 | 1715 | 0.02778 | 16.2 | +1 | 15 |
| - | - | 4610 | 1716 | - | - | 0 | - |
| 3 | z | 5885 | 1717 | 0.01995 | 11.62 | +1 | 15 |
| - | - | 2603 | 1728 | - | - | 0 | - |
| - | - | 7251 | 1729 | - | - | 0 | - |
| - | - | 4378 | 1730 | - | - | 0 | - |
| - | - | 1711 | 1731 | - | - | 0 | - |
| 15 | c | 3.615E+04 | 1732 | 0.005012 | 2.894 | +1 | 15 |
| 3 | y | 6.029E+04 | 1733 | 0.02661 | 15.36 | +1 | 15 |
| - | - | 4.512E+04 | 1734 | - | - | 0 | - |
| - | - | 2.432E+04 | 1735 | - | - | 0 | - |
| - | - | 8749 | 1736 | - | - | 0 | - |
| - | - | 2282 | 1737 | - | - | 0 | - |
| - | - | 2047 | 1785 | - | - | 0 | - |
| - | - | 4808 | 1786 | - | - | 0 | - |
| - | - | 2878 | 1787 | - | - | 0 | - |
| - | - | 4165 | 1788 | - | - | 0 | - |
| - | - | 4424 | 1801 | - | - | 0 | - |
| - | - | 9424 | 1802 | - | - | 0 | - |
| - | - | 1.336E+04 | 1803 | - | - | 0 | - |
| - | - | 7863 | 1804 | - | - | 0 | - |
| - | - | 3584 | 1805 | - | - | 0 | - |
| - | - | 1888 | 1820 | - | - | 0 | - |
| 2 | y | 7990 | 1829 | 0.01548 | 8.466 | +1 | 16 |
| 2 | z | 1.532E+04 | 1830 | 0.01486 | 8.121 | +1 | 16 |
| - | - | 5.558E+04 | 1831 | - | - | 0 | - |
| - | - | 5.238E+04 | 1832 | - | - | 0 | - |
| 16 | c | 3.268E+04 | 1833 | 0.01727 | 9.422 | +1 | 16 |
| - | - | 1.41E+04 | 1834 | - | - | 0 | - |
| - | - | 5508 | 1835 | - | - | 0 | - |
| - | - | 1.84E+04 | 1845 | - | - | 0 | - |
| 2 | y | 4.517E+04 | 1846 | 0.02397 | 12.98 | +1 | 16 |
| - | - | 3.803E+04 | 1847 | - | - | 0 | - |
| - | - | 2.427E+04 | 1848 | - | - | 0 | - |
| - | - | 1.082E+04 | 1849 | - | - | 0 | - |
| - | - | 5063 | 1850 | - | - | 0 | - |
| - | - | 5008 | 1871 | - | - | 0 | - |
| - | - | 3026 | 1872 | - | - | 0 | - |
| - | - | 3427 | 1873 | - | - | 0 | - |
| - | - | 2329 | 1874 | - | - | 0 | - |
| - | - | 2884 | 1876 | - | - | 0 | - |
| - | - | 4037 | 1885 | - | - | 0 | - |
| - | - | 9372 | 1886 | - | - | 0 | - |
| - | - | 1.109E+04 | 1887 | - | - | 0 | - |
| - | - | 8113 | 1888 | - | - | 0 | - |
| - | - | 9168 | 1889 | - | - | 0 | - |
| - | - | 2.158E+04 | 1890 | - | - | 0 | - |
| - | - | 1.714E+04 | 1891 | - | - | 0 | - |
| - | - | 9383 | 1892 | - | - | 0 | - |
| - | - | 3900 | 1893 | - | - | 0 | - |
| - | - | 2862 | 1902 | - | - | 0 | - |
| - | - | 1.396E+04 | 1903 | - | - | 0 | - |
| - | - | 4.119E+04 | 1904 | - | - | 0 | - |
| - | - | 3.572E+04 | 1905 | - | - | 0 | - |
| - | - | 2.382E+04 | 1906 | - | - | 0 | - |
| - | - | 9140 | 1907 | - | - | 0 | - |
| - | - | 3664 | 1908 | - | - | 0 | - |
| - | - | 2244 | 1913 | - | - | 0 | - |
| - | - | 9239 | 1914 | - | - | 0 | - |
| - | - | 1.354E+04 | 1915 | - | - | 0 | - |
| - | - | 9642 | 1916 | - | - | 0 | - |
| - | - | 7347 | 1917 | - | - | 0 | - |
| - | - | 1886 | 1918 | - | - | 0 | - |
| - | - | 7558 | 1920 | - | - | 0 | - |
| - | - | 2.895E+04 | 1921 | - | - | 0 | - |
| - | - | 2.763E+04 | 1922 | - | - | 0 | - |
| - | - | 1.798E+04 | 1923 | - | - | 0 | - |
| - | - | 7297 | 1924 | - | - | 0 | - |
| - | - | 2196 | 1930 | - | - | 0 | - |
| - | - | 4.047E+04 | 1931 | - | - | 0 | - |
| - | - | 1.317E+05 | 1932 | - | - | 0 | - |
| - | - | 1.235E+05 | 1933 | - | - | 0 | - |
| - | - | 8.135E+04 | 1934 | - | - | 0 | - |
| - | - | 3.468E+04 | 1935 | - | - | 0 | - |
| - | - | 9479 | 1936 | - | - | 0 | - |
| - | - | 1.743E+04 | 1947 | - | - | 0 | - |
| - | - | 9.963E+04 | 1948 | - | - | 0 | - |
| - | - | 4.012E+05 | 1949 | - | - | 0 | - |
| - | - | 3.749E+05 | 1950 | - | - | 0 | - |
| - | - | 2.231E+05 | 1951 | - | - | 0 | - |
| - | - | 9.09E+04 | 1952 | - | - | 0 | - |
| - | - | 2.876E+04 | 1953 | - | - | 0 | - |
| - | - | 1719 | 3083 | - | - | 0 | - |
| - | - | 1899 | 3484 | - | - | 0 | - |

m/z Charge Intensity FragmentType MassShift Position
120.06156158447266 0 1060.6595
120.0657730102539 0 15698.042
143.1180877685547 0 1512.2297
168.38775634765625 0 998.14435
169.13377380371094 0 1388.8689
173.1283721923828 0 1335.3822
173.45291137695312 0 3048.1096
185.12892150878906 0 1437.0466
187.1444091796875 0 155377
188.1400146484375 0 1268.1268
188.14772033691406 0 16654.078
192.8508758544922 0 1228.0968
201.1235809326172 0 4513.778 y Water loss 13
201.1343231201172 0 1065.7544
212.13963317871094 0 5217.254
215.13929748535156 0 101853.6 y Water loss 15
216.1426544189453 0 8884.124
233.14991760253906 0 10357.536 y 15
239.41883850097656 0 1294.5116
256.16607666015625 0 2187.715
299.17181396484375 0 18544.916
300.175048828125 0 3106.5903
302.1717529296875 0 3241.3923 y Water loss 14
312.1920471191406 0 4825.1606
313.1958312988281 0 1701.4177
320.18255615234375 0 6217.769 y 14
326.15966796875 0 1532.8531
327.1647033691406 0 1439.3547
369.1900634765625 0 2075.3909
382.2584228515625 0 2981.6323
383.2638854980469 0 1436.328
407.2528076171875 0 6791.525
413.23956298828125 0 1562.7109
425.26446533203125 0 3715.5
426.27020263671875 0 7047.0337 c 3
427.2720031738281 0 1957.0942
428.2747802734375 0 1451.1031
449.7096252441406 0 1347.6134
451.2783508300781 0 2310.6575
470.29840087890625 0 9358.816
471.30120849609375 0 2529.4275
471.8973388671875 0 2336.2373
472.71112060546875 0 1681.1033
473.2606506347656 0 1762.0421 w 12
494.24029541015625 0 1264.3926
494.28582763671875 0 4340.9233
495.23980712890625 0 1915.2169
499.908935546875 0 2552.6912
500.909423828125 0 6535.273
501.24456787109375 0 5246.0854
501.3085021972656 0 1651.9417
501.57867431640625 0 1677.1104
508.2646789550781 0 1904.1351
509.2727355957031 0 2057.9133
512.2965087890625 0 3008.4683
513.3037109375 0 99366.37 c 4
514.3067626953125 0 26901
515.3094482421875 0 4983.9463
523.269287109375 0 2435.7988
523.6026611328125 0 2587.4219
526.2758178710938 0 2839.2651
526.9234008789062 0 2401.0635
527.2840576171875 0 2681.1826
528.3206176757812 0 1586.7416
529.27197265625 0 7575.493
529.3179931640625 0 2456.7322
529.6065673828125 0 6602.872
529.9390869140625 0 3894.2922
531.2906494140625 0 9807.355 z 12
532.2970581054688 0 9607.31
532.5997314453125 0 13259.649
532.931884765625 0 10474.232
533.2675170898438 0 5830.7427
533.3067016601562 0 1961.484
533.6001586914062 0 2600.6672
538.6038208007812 0 23275.066
538.9380493164062 0 27140.111
539.2723388671875 0 13655.131
539.6069946289062 0 6089.232
539.9398803710938 0 1763.4584 y Water loss 3
544.3086547851562 0 1746.3582
544.9481201171875 0 1386.1333
547.3085327148438 0 5371.8486 y 12
556.95751953125 0 5947.4624
557.2919311523438 0 7515.016
557.6243896484375 0 7379.209
557.9586791992188 0 2840.6113
563.2809448242188 0 1643.2804
563.6209716796875 0 1934.5574
566.2833251953125 0 3473.688
566.61669921875 0 3466.7158
566.9490966796875 0 2308.8826 z Water loss 2
570.31201171875 0 3771.6511
571.319580078125 0 10246.715
572.2861328125 0 28117.137 c Ammonia loss 14
572.6200561523438 0 23321.074 y Ammonia loss 2
572.9524536132812 0 12193.387 z 2
573.2853393554688 0 9985.891
573.3362426757812 0 4353.469
573.6180419921875 0 2315.756
573.9473266601562 0 1667.1832
574.2872924804688 0 1484.0118
577.9614868164062 0 3581.243 c 14
578.290283203125 0 513593.22 y 2
578.6243896484375 0 453304.34
578.9581298828125 0 278587.88
579.292236328125 0 100713.54 c Water loss 9
579.6258544921875 0 24871.064
607.3300170898438 0 1469.9073
607.3822021484375 0 2954.049
615.3472290039062 0 2120.2844
625.3912353515625 0 11303.595
626.3948974609375 0 6800.8027
627.3939819335938 0 1581.5768
628.334228515625 0 2646.8047
628.3936767578125 0 3557.973
643.6624755859375 0 6878.4414
643.8247680664062 0 1455.4661 c Ammonia loss 10
643.9979248046875 0 3786.927
644.332275390625 0 6430.809 y Ammonia loss 11
645.3341674804688 0 5667.582 z 11
646.3402709960938 0 10054.152
647.3458862304688 0 2605.2148
649.8460083007812 0 2535.035
650.345458984375 0 2659.3835
650.8417358398438 0 3383.4783
652.337890625 0 17706.225 c 10
652.8399047851562 0 9092.699
653.340576171875 0 4045.7708
657.8118286132812 0 1917.0144
661.352294921875 0 19431.549 y 11
662.3555908203125 0 6755.888
669.4049682617188 0 156593.03 c 5
670.4075927734375 0 60715.695
671.4104614257812 0 10924.425
672.4083251953125 0 4772.83
686.849853515625 0 2905.5608
687.3525390625 0 2292.2466
695.4205322265625 0 3612.2617
696.4196166992188 0 1676.3348
699.8487548828125 0 4906.4507
700.3522338867188 0 5728.2085 c Water loss 11
700.8514404296875 0 2542.0588 c Ammonia loss 11
707.34423828125 0 13867.443
707.8452758789062 0 12675.572
708.3505249023438 0 6023.774
708.854736328125 0 18727.486
709.3590698242188 0 109747.78 c 11
709.86083984375 0 85314.336
710.3616333007812 0 42984.58
710.8619384765625 0 14252.545
711.361083984375 0 3463.7847
711.4277954101562 0 2135.2427
718.3544921875 0 8138.7676 z 5
718.8554077148438 0 10159.179
719.3572387695312 0 4007.9917
719.853515625 0 2254.586
720.3673706054688 0 1901.0974
722.3711547851562 0 3638.455
722.8679809570312 0 4378.354
725.3604125976562 0 2198.4302
725.8622436523438 0 5145.9507
726.3621215820312 0 21762.2 y 5
726.8636474609375 0 15536.961
727.3618774414062 0 7724.2153
729.426513671875 0 20446.102
730.4285888671875 0 7425.7827
741.8582153320312 0 3225.5227
742.3576049804688 0 1912.1858
750.8600463867188 0 15247.223
751.3654174804688 0 14894.446
751.8680419921875 0 6748.9473
752.3723754882812 0 2358.0173
754.4348754882812 0 2772.7444
755.4337768554688 0 2039.9158
757.8812866210938 0 1761.641
759.3746948242188 0 1718.6671
761.3676147460938 0 2542.7488 y Ammonia loss 4
761.8690185546875 0 14056.696 z 4
762.372314453125 0 21532.527
762.872314453125 0 9983.524
763.3770141601562 0 3708.3774
763.8753662109375 0 2081.0107
764.3822631835938 0 2498.9907 c Water loss 12
764.8800048828125 0 2719.2458 c Ammonia loss 12
765.3922729492188 0 4422.7725
765.8842163085938 0 2728.1755
766.38623046875 0 2924.4187
769.3759155273438 0 4416.808
769.8770751953125 0 13725.901 y 4
770.3804931640625 0 13607.834
770.8826904296875 0 5360.378
771.4414672851562 0 2799.7256 y Water loss 10
772.8868408203125 0 8176.5083
773.390380859375 0 146829.7 c 12
773.4511108398438 0 23437.688
773.8902587890625 0 160969.97
774.391845703125 0 58680.668
774.4560546875 0 10930.764
774.8895874023438 0 24104.293
775.3903198242188 0 3578.267
775.4431762695312 0 6193.809
779.4261474609375 0 3668.4902
786.3964233398438 0 2148.2761
786.8991088867188 0 2923.4956
787.39404296875 0 1835.464
789.4478149414062 0 21473.566 y 10
790.4498291015625 0 8706.001
791.91455078125 0 2361.0024
792.4100341796875 0 2092.644
792.908935546875 0 9215.824
793.4097290039062 0 11072.358
793.9111938476562 0 7730.719
794.4057006835938 0 7404.9297
794.8967895507812 0 2864.9521
797.4411010742188 0 3360.885
798.4480590820312 0 234470.77 c 6
798.8958740234375 0 6976.6484
799.4505004882812 0 105451.74
800.4517822265625 0 24413.08
800.9154052734375 0 17123.521
801.41943359375 0 7913.402
801.9180908203125 0 7719.437
802.4226684570312 0 3826.7898
803.4000854492188 0 2072.3032
805.4006958007812 0 2317.807
806.396484375 0 1862.0688
807.40185546875 0 35786.54
807.9030151367188 0 35209.3
808.4046630859375 0 19227.502
808.9055786132812 0 5413.5054
809.8955078125 0 2470.285 y Ammonia loss 3
812.4766845703125 0 1999.6561
813.4137573242188 0 49367.12
813.9135131835938 0 46035.906 c Water loss 13
814.4141235351562 0 32773.418 c Ammonia loss 13
814.4951171875 0 2476.6108
814.914306640625 0 15142.271
815.4158325195312 0 9662.781
816.4149169921875 0 4251.901
817.9041137695312 0 3447.1748
818.4053955078125 0 80703.85 y 3
818.9065551757812 0 63892.137
819.4080200195312 0 31421.707
819.9088134765625 0 11049.077
820.411865234375 0 2703.03
822.4190063476562 0 76717.164
822.9215698242188 0 108338.516 c 13
823.4228515625 0 84335.21
823.9246826171875 0 44556.984
824.4332275390625 0 11561.946
824.9303588867188 0 3959.9443
825.4622802734375 0 4365.867
830.4649047851562 0 6751.153
831.4655151367188 0 3891.3428
834.9327392578125 0 11493.862
835.4320068359375 0 17924.234
835.9318237304688 0 9194.292
836.4306640625 0 6322.447
836.9281616210938 0 4703.726
843.9365844726562 0 1833.3778
844.432373046875 0 1866.6688
844.9302978515625 0 7686.238
845.4176025390625 0 9371.83 w 2
845.9130249023438 0 7852.5024
846.4156494140625 0 3469.5322
848.9188842773438 0 2468.8782
849.41748046875 0 5401.0303
849.9265747070312 0 2717.9277 z Water loss 2
850.4120483398438 0 2741.874 z Ammonia loss 2
856.4650268554688 0 2924.8438 z Water loss 9
856.9332275390625 0 1980.5116
857.4724731445312 0 13774.166 w 9
857.9270629882812 0 18840.035 c Ammonia loss 14
858.4280395507812 0 13935.595 y Ammonia loss 2
858.9278564453125 0 12491.254 z 2
859.4502563476562 0 22517.64 w 9
859.9248657226562 0 2931.08
860.45556640625 0 10646.143
861.459228515625 0 1978.5356
864.9490966796875 0 3353.6074
865.447509765625 0 2889.1628
865.9375610351562 0 15306.293
866.4390258789062 0 229663.8 c 14
866.934326171875 0 500152.66 y 2
867.4348754882812 0 384236.9
867.9349365234375 0 211259.67
868.4357299804688 0 73093.19
868.93603515625 0 13864.426
872.4600219726562 0 3854.7124
872.941650390625 0 3231.8699
873.469970703125 0 20327.258 y Ammonia loss 9
874.4761962890625 0 132394.98 z 9
875.4794921875 0 62792.316
876.4821166992188 0 21481.719
877.4837036132812 0 4988.509
879.4488525390625 0 9616.336
879.952392578125 0 9096.16
880.4513549804688 0 5512.474
880.9425048828125 0 3924.4949
881.4329833984375 0 5572.069
881.9318237304688 0 3304.2341
883.4766845703125 0 4511.2886
884.480712890625 0 4645.491
887.4349975585938 0 5348.684
887.9373779296875 0 5210.9
888.4259643554688 0 2126.7405
889.487548828125 0 3598.1846
890.4946899414062 0 38742.3 y 9
891.4974975585938 0 18871.607
891.9666748046875 0 5057.6724
892.4807739257812 0 5550.1836
892.9680786132812 0 5733.6006
893.4646606445312 0 7473.4316
893.9370727539062 0 86072.91 w 1
894.4380493164062 0 87096.586
894.9392700195312 0 55994.79
895.4403686523438 0 18626.947
895.9461059570312 0 5108.7124
896.4110717773438 0 7083.8857
896.5095825195312 0 3089.269
897.4147338867188 0 2795.0164
897.5106811523438 0 2222.3042
898.4570922851562 0 2060.1736
900.4708862304688 0 2250.3672
900.9729614257812 0 12551.82
901.4764404296875 0 15785.85
901.9771118164062 0 11997.763
902.4741821289062 0 7997.975
902.9811401367188 0 3485.458
903.4660034179688 0 2611.7349
904.463623046875 0 1907.9128
905.96435546875 0 2351.4663
906.4674072265625 0 2824.7344 z Water loss 1
906.968994140625 0 6401.8
907.4686279296875 0 6964.777
907.9713134765625 0 4341.899 c Water loss 15
908.4533081054688 0 3670.8079 c Ammonia loss 15
908.9423828125 0 2429.6216
914.4691162109375 0 14355.531 y Water loss 1
914.9671020507812 0 12450.153 y Ammonia loss 1
915.4671630859375 0 52845.21 z 1
915.9669189453125 0 54987.75
916.4705200195312 0 32409.816
916.9629516601562 0 16819.672 c 15
917.4581909179688 0 6631.11
917.9465942382812 0 2933.246
919.9857177734375 0 2119.0405
920.4898681640625 0 8562.796
920.9968872070312 0 2349.9946
921.489013671875 0 3698.694
922.4796752929688 0 7074.603
922.9812622070312 0 207923.05
923.4819946289062 0 196746.98 y 1
923.5916137695312 0 3267.5208
923.9830932617188 0 144443.11
924.4835205078125 0 60252.18
924.9837646484375 0 19448.61
925.4857788085938 0 2069.079
926.4833984375 0 6095.541
927.4904174804688 0 230647.92 c 7
927.9724731445312 0 2469.6045
928.4929809570312 0 119107.766
928.976318359375 0 7482.152
929.4940185546875 0 38214.965
929.9703979492188 0 12170.034
930.4827880859375 0 16774.775
930.9730834960938 0 8214.915
931.4860229492188 0 8667.778
933.4984130859375 0 1885.2377
934.4910888671875 0 2946.4392
934.9898681640625 0 6615.314
935.4895629882812 0 5151.815
935.9835205078125 0 2904.3867
936.4813232421875 0 8457.749
936.9813232421875 0 9571.045
937.4700317382812 0 14821.202
937.9603271484375 0 85087.06
938.4622192382812 0 77138.92
938.9655151367188 0 55237.76
939.4681396484375 0 25353.021
939.9691772460938 0 10150.907
940.4950561523438 0 2135.5635
941.516357421875 0 3808.6802
942.49658203125 0 22240.193
942.9959106445312 0 27778.947
943.4901733398438 0 29929.508
943.9869995117188 0 26412.043
944.48046875 0 17112.125
944.9811401367188 0 42963.21
945.4828491210938 0 45563.89
945.9843139648438 0 27895.084
946.4751586914062 0 30183.059
946.97314453125 0 25585.168
947.4728393554688 0 16132.366
947.9732055664062 0 6646.581
950.5064697265625 0 2062.627
951.4973754882812 0 6693.065
951.9911499023438 0 50554.01
952.492919921875 0 63329.21
952.990966796875 0 44600.945
953.49072265625 0 29624.816
953.9895629882812 0 11267.829
954.4974975585938 0 4012.4026
956.4901733398438 0 18777.602
956.9871215820312 0 38292.33
957.4891357421875 0 33753.793
957.9840087890625 0 25980.748
958.5155029296875 0 33517.836
958.980712890625 0 5634.4194
959.5280151367188 0 108900.79
960.5308837890625 0 53078.9
961.53369140625 0 13409.5625
965.4946899414062 0 72184.49
965.9915771484375 0 320018.28
966.491943359375 0 315217.66
966.9924926757812 0 218935.33
967.4935302734375 0 86479.79
967.9942626953125 0 28161.244
968.5260009765625 0 10182.351
969.5326538085938 0 5721.732
970.539306640625 0 2817.2341
973.501220703125 0 3840.5002
973.9977416992188 0 206927.84
974.4996948242188 0 382712
975.0009155273438 0 301036.72
975.1223754882812 0 3268.542
975.5023193359375 0 161012.83
976.0028076171875 0 57361.6
976.5031127929688 0 14823.007
990.4896850585938 0 2459.5508
1003.496826171875 0 2545.0364 z Water loss 8
1005.5136108398438 0 4466.667
1006.5133056640625 0 2284.9097
1012.506591796875 0 3306.3916
1013.4938354492188 0 3568.8936
1014.5008544921875 0 2228.2097
1015.4965209960938 0 2643.7893
1021.5128173828125 0 27601.443 z 8
1022.51708984375 0 21970.47
1023.517578125 0 10025.826
1024.52392578125 0 3277.3965
1030.51171875 0 27558.111
1031.515625 0 22516.682
1032.5189208984375 0 8916.746
1033.519287109375 0 4172.643
1037.5302734375 0 15031.351 y 8
1038.531982421875 0 7691.909
1039.533203125 0 2416.3894
1049.531005859375 0 10908.493
1050.5328369140625 0 9220.014
1051.532470703125 0 4344.6694
1055.5091552734375 0 2473.2993
1058.5040283203125 0 2454.4844
1073.5211181640625 0 4551.9004
1074.52587890625 0 253030.7 c 8
1075.528564453125 0 134907.9
1076.5303955078125 0 58211.22
1077.529541015625 0 15863.0625
1078.53076171875 0 1774.6987
1079.515869140625 0 3440.8452
1087.567626953125 0 8278.918
1088.564453125 0 5507.72
1091.5413818359375 0 34148.57 w 7
1092.5435791015625 0 17508.695
1093.5458984375 0 9670.822
1094.501953125 0 5819.0337
1095.4986572265625 0 3128.203
1106.54052734375 0 2442.769
1113.549072265625 0 5220.2705
1114.5479736328125 0 3708.264
1115.5401611328125 0 2497.6584
1131.5546875 0 6092.352
1132.560546875 0 6846.983 z Water loss 7
1133.56396484375 0 3223.8398
1134.5631103515625 0 4433.022
1135.554443359375 0 3090.3342
1149.571533203125 0 2902.9363
1150.55419921875 0 113594.03 z 7
1151.55810546875 0 69262.52
1152.5589599609375 0 34724.89
1153.56005859375 0 8195.033
1166.5745849609375 0 8111.3174 y 7
1167.577392578125 0 4997.589
1175.5728759765625 0 119764.51 c 9
1176.5755615234375 0 79858.6
1177.576416015625 0 31295.172
1178.5780029296875 0 11614.985
1179.573486328125 0 2614.2688
1207.57080078125 0 2598.8464
1208.5670166015625 0 3425.1523
1216.63916015625 0 1894.7346
1220.5849609375 0 16356.856 w 6
1221.58447265625 0 14640.538
1222.5899658203125 0 6980.018
1224.53955078125 0 2153.676
1236.5926513671875 0 1880.0665
1259.6536865234375 0 8170.3696
1260.657958984375 0 6211.7915
1261.656494140625 0 2208.2954
1279.5965576171875 0 83379.875 z 6
1280.59912109375 0 58979.47
1281.6015625 0 30142.926
1282.5997314453125 0 8038.006
1284.638916015625 0 2775.3528
1285.6507568359375 0 1861.4758 c Water loss 10
1294.6151123046875 0 2942.6465
1295.615966796875 0 25385.646 y 6
1296.6207275390625 0 19389.295
1297.6207275390625 0 6379.373
1298.625 0 2156.7593
1302.6627197265625 0 8508.679
1303.6661376953125 0 24356.752 c 10
1304.669189453125 0 17386.604
1305.6737060546875 0 8070.5317
1333.644775390625 0 2460.205
1334.6673583984375 0 6800.3477
1335.6732177734375 0 3339.4058
1363.671142578125 0 2098.1511
1364.659912109375 0 2436.2715
1365.66259765625 0 2505.4622
1373.7015380859375 0 2745.7554
1374.706787109375 0 2860.002
1392.6773681640625 0 1949.3801
1416.7071533203125 0 5635.2393
1417.707275390625 0 46460.73 c 11
1418.7115478515625 0 33192.734
1419.7115478515625 0 15749.84
1420.7060546875 0 9028.793
1421.6961669921875 0 3997.2104
1434.7015380859375 0 2941.05 y Ammonia loss 5
1435.69677734375 0 65938.94 z 5
1436.7021484375 0 99370.06
1437.7039794921875 0 60006.93
1438.7066650390625 0 26017.49
1439.7076416015625 0 6408.853
1450.7099609375 0 8568.702
1451.71484375 0 34650.22 y 5
1452.717041015625 0 23698.984
1453.718994140625 0 12367.901
1454.730712890625 0 4571.2407
1502.75634765625 0 3487.2402
1522.72705078125 0 4773.5605 z 4
1523.7353515625 0 25579.668
1524.7386474609375 0 19944.184
1525.739990234375 0 9003.512
1526.7440185546875 0 3819.3376
1528.7568359375 0 2866.6968 c Ammonia loss 12
1538.74560546875 0 7747.1553 y 4
1539.7484130859375 0 5079.167
1540.7490234375 0 4400.029
1544.7578125 0 2796.0334
1545.7679443359375 0 56601.688 c 12
1546.7708740234375 0 49097.52
1547.7716064453125 0 23924.486
1548.7755126953125 0 9010.148
1600.822021484375 0 2173.9531
1601.8294677734375 0 5803.154
1602.8199462890625 0 5326.646
1603.83349609375 0 2538.331
1614.8087158203125 0 6620.2217
1615.8089599609375 0 4085.3823
1616.8140869140625 0 1967.1719
1626.813232421875 0 4015.6365 c Water loss 13
1627.8206787109375 0 5967.7334 c Ammonia loss 13
1628.82568359375 0 4507.926
1635.796875 0 13755.203 y 3
1636.8011474609375 0 15151.553
1637.8055419921875 0 8509.368
1638.8070068359375 0 5007.9824
1643.8294677734375 0 8986.955
1644.8367919921875 0 62082.695 c 13
1645.8402099609375 0 53609.273
1646.841064453125 0 32013.697
1647.84423828125 0 11007.061
1648.8472900390625 0 3491.6572
1672.860107421875 0 2664.137
1674.8280029296875 0 2238.5657
1687.8695068359375 0 2618.8823
1688.8677978515625 0 6131.0283
1689.8612060546875 0 5482.7363
1690.8612060546875 0 2206.4263
1714.86572265625 0 1886.4376 c Ammonia loss 14
1715.8660888671875 0 4610.433
1716.8497314453125 0 5884.9424 z 2
1727.8896484375 0 2602.9739
1728.8935546875 0 7250.5605
1729.8961181640625 0 4378.4023
1730.84033203125 0 1711.193
1731.8695068359375 0 36151.67 c 14
1732.8751220703125 0 60288.086 y 2
1733.8743896484375 0 45117.45
1734.8775634765625 0 24322.18
1735.87744140625 0 8748.736
1736.8927001953125 0 2281.6738
1784.911865234375 0 2046.967
1785.91015625 0 4807.7905
1786.907470703125 0 2878.27
1787.9197998046875 0 4165.156
1800.9334716796875 0 4424.032
1801.9422607421875 0 9423.879
1802.947998046875 0 13364.538
1803.9481201171875 0 7863.1187
1804.944091796875 0 3584.2717
1819.9658203125 0 1888.1765
1828.9215087890625 0 7990.3853 y Ammonia loss 1
1829.9287109375 0 15320.561 z 1
1830.92919921875 0 55576.383
1831.928955078125 0 52384.883
1832.929443359375 0 32680.877 c 15
1833.9287109375 0 14104.711
1834.9354248046875 0 5508.3237
1844.95166015625 0 18399.67
1845.95654296875 0 45174.38 y 1
1846.9610595703125 0 38033.645
1847.9654541015625 0 24270.854
1848.9669189453125 0 10817.047
1849.9462890625 0 5062.9004
1870.9339599609375 0 5008.39
1871.937255859375 0 3025.9114
1872.9193115234375 0 3426.9524
1873.9232177734375 0 2329.0447
1875.9468994140625 0 2884.2688
1884.9954833984375 0 4036.7578
1885.989013671875 0 9372.292
1886.977783203125 0 11087.608
1887.9669189453125 0 8113.4233
1888.9620361328125 0 9167.924
1889.9644775390625 0 21582.33
1890.962646484375 0 17144.875
1891.962890625 0 9382.984
1892.9691162109375 0 3899.5142
1901.9903564453125 0 2861.5637
1902.9847412109375 0 13957.722
1903.9827880859375 0 41193.81
1904.9825439453125 0 35717.637
1905.9849853515625 0 23824.13
1906.987060546875 0 9140.3545
1907.9921875 0 3664.4285
1912.946533203125 0 2243.889
1913.9664306640625 0 9239.04
1914.9671630859375 0 13539.243
1915.9622802734375 0 9642.051
1916.9556884765625 0 7347.3315
1917.9371337890625 0 1886.2792
1920.0050048828125 0 7558.453
1921.00537109375 0 28947.307
1922.007568359375 0 27625.668
1923.009033203125 0 17979.271
1924.011962890625 0 7297.177
1929.9786376953125 0 2195.8335
1930.9776611328125 0 40471.86
1931.9757080078125 0 131723.45
1932.9783935546875 0 123474.01
1933.9788818359375 0 81351.17
1934.980224609375 0 34679.312
1935.984130859375 0 9479.478
1946.984130859375 0 17431.143
1947.9910888671875 0 99627.96
1948.99658203125 0 401233.62
1950.00048828125 0 374886.78
1951.0023193359375 0 223121.36
1952.0030517578125 0 90902.555
1953.0076904296875 0 28755.727
3082.86767578125 0 1718.6299
3483.71240234375 0 1898.8467

Spectrum Details

|  |  |
| --- | --- |
| Matched peaks? Matched peaks The total absolute number of peaks matched. Additionally in brackets the total fraction of peaks matched and the total number of peaks is shown. | 101 (15.17% of 666) |
| FDR? FDR The false discovery rate estimated for this peptide. It is calculated by matching all theoretical fragments with a non-integer shift with the raw peaks for this spectrum. This is done with 40 different shifts. The resulting percentage is the average number of annotated peaks over the number of annotated peaks with the correct spectrum. | 0.80% |
| Satellite FDR? Satellite FDR See the FDR for details on its calculation. This satellite ion specific FDR only contains the satellite ions (d/w) for I/L/J positions. | 11.90% |
| PSM Score? PSM Score The PSM Score as given by Hecklib to this annotated spectrum. It is shown with three significant figures. | 681 |

#### Spectrum 4052? Spectrum 4052 The raw spectrum of this peptide as annotated by Hecklib. The fragments are coloured according to ion type (see legend). Any peaks with a star '\*' as text can be hovered over to see the full details, first the ion type second the mass shift type. By hovering over the amino acids in the peptide or ions in the legend the corresponding peaks are highlighted. By toggling the 'Unassigned' label you can turn the background (unassigned) peaks on or off in the plot. By updating the slider in the Ion legend you can update the spectrum to only show the top X% of the peaks with labels. The top X% means any peak that is within X% of the highest intensity. By dragging in the spectrum you can zoom in to a specific part of the spectrum and use 'Zoom Out' to get back to the original zoom level. The annotation of the spectrum is based on the given sequence in the peptides file and is done with different software so inconsistencies are likely. The peaks are annotated based on the given sequence, with 20 ppm tolerance.

Copy Data

##### Spectrum 4052 (TSV)

###### Preview

```
Loading example...
```

*Click on the button to copy the data to your clipboard.*

Mz MinMz MaxIntensity Max

WidthHeightPeptide font sizePeptide stroke widthSpectrum font sizeSpectrum stroke widthCompact peptide

Ion legend

wxyz

abcd

OtherUnassignedIon^Charge^_Position_Show for top:%

TLPPSREEMTKNQVSTJ

09.20e+41.84e+52.76e+53.68e+5

Zoom Out

y^+2^_4_y^+1^_2_y^+1^_2_y^+1^_3_y^+1^_3_c^+1^_4_c^+1^_5_z^+1^_5_y^+3^_14_y^+1^_5_c^+3^_15_y^+3^_15_z^+3^_15_y^+3^_15_c^+2^_10_y^+1^_6_z^+1^_6_c^+2^_11_y^+1^_6_c^+1^_6_c^+2^_12_c^+2^_12_c^+2^_12_z^+2^_12_y^+2^_12_z^+2^_13_c^+2^_13_c^+2^_13_y^+2^_13_y^+1^_7_c^+2^_13_c^+1^_7_y^+1^_7_c^+1^_7_y^+2^_14_c^+2^_14_c^+2^_14_y^+2^_14_c^+2^_14_w^+2^_15_z^+2^_15_z^+1^_8_w^+1^_8_c^+2^_15_y^+2^_15_z^+2^_15_w^+1^_8_c^+2^_15_y^+2^_15_y^+1^_8_z^+1^_8_y^+1^_8_w^+2^_16_z^+2^_16_z^+2^_16_c^+2^_16_c^+2^_16_y^+2^_16_y^+2^_16_z^+2^_16_c^+2^_16_y^+2^_16_c^+1^_8_z^+1^_9_z^+1^_9_z^+1^_9_y^+1^_9_c^+1^_9_w^+1^_10_z^+1^_10_y^+1^_10_c^+1^_10_w^+1^_11_z^+1^_11_c^+1^_11_y^+1^_11_c^+1^_11_c^+1^_12_y^+1^_12_z^+1^_12_y^+1^_12_z^+1^_13_y^+1^_13_c^+1^_13_c^+1^_14_y^+1^_14_c^+1^_14_z^+1^_15_c^+1^_15_y^+1^_15_y^+1^_16_z^+1^_16_c^+1^_16_y^+1^_16_

0637127419112548

Fragment Matches Table

Show background peaks

| Position | Ion type | Intensity | mz Theoretical | mz Error (Th) | mz Error (ppm) | Charge | Series Number |
| --- | --- | --- | --- | --- | --- | --- | --- |
| - | - | 1.09E+04 | 120.1 | - | - | 0 | - |
| - | - | 994.6 | 139.2 | - | - | 0 | - |
| - | - | 920.5 | 140.1 | - | - | 0 | - |
| - | - | 1013 | 141.2 | - | - | 0 | - |
| - | - | 1193 | 143.1 | - | - | 0 | - |
| - | - | 1149 | 145.8 | - | - | 0 | - |
| - | - | 1062 | 147.3 | - | - | 0 | - |
| - | - | 1540 | 148.9 | - | - | 0 | - |
| - | - | 1180 | 151.9 | - | - | 0 | - |
| - | - | 1639 | 173.1 | - | - | 0 | - |
| - | - | 9.546E+04 | 187.1 | - | - | 0 | - |
| - | - | 7653 | 188.1 | - | - | 0 | - |
| 14 | y | 3984 | 201.1 | 0.0001968 | 0.9787 | +2 | 4 |
| - | - | 4783 | 212.1 | - | - | 0 | - |
| 16 | y | 6.922E+04 | 215.1 | 0.0002786 | 1.295 | +1 | 2 |
| - | - | 7897 | 216.1 | - | - | 0 | - |
| 16 | y | 6663 | 233.1 | 0.0002577 | 1.105 | +1 | 2 |
| - | - | 1808 | 256.2 | - | - | 0 | - |
| - | - | 1419 | 298.6 | - | - | 0 | - |
| - | - | 1.156E+04 | 299.2 | - | - | 0 | - |
| - | - | 2326 | 300.2 | - | - | 0 | - |
| 15 | y | 2606 | 302.2 | 0.0006141 | 2.032 | +1 | 3 |
| - | - | 2657 | 312.2 | - | - | 0 | - |
| 15 | y | 2987 | 320.2 | 0.0002728 | 0.8519 | +1 | 3 |
| - | - | 1609 | 344.2 | - | - | 0 | - |
| - | - | 1313 | 377 | - | - | 0 | - |
| - | - | 1466 | 378.4 | - | - | 0 | - |
| - | - | 1746 | 382.3 | - | - | 0 | - |
| - | - | 1634 | 383.3 | - | - | 0 | - |
| - | - | 1483 | 401.4 | - | - | 0 | - |
| - | - | 3627 | 407.3 | - | - | 0 | - |
| - | - | 1497 | 412.2 | - | - | 0 | - |
| - | - | 1641 | 413.2 | - | - | 0 | - |
| - | - | 1351 | 421.4 | - | - | 0 | - |
| - | - | 2321 | 425.3 | - | - | 0 | - |
| 4 | c | 4543 | 426.3 | 0.0007854 | 1.842 | +1 | 4 |
| - | - | 1343 | 427.3 | - | - | 0 | - |
| - | - | 2161 | 451.3 | - | - | 0 | - |
| - | - | 1456 | 462.6 | - | - | 0 | - |
| - | - | 1445 | 463.2 | - | - | 0 | - |
| - | - | 6464 | 470.3 | - | - | 0 | - |
| - | - | 1663 | 472.2 | - | - | 0 | - |
| - | - | 1682 | 472.3 | - | - | 0 | - |
| - | - | 2458 | 493.9 | - | - | 0 | - |
| - | - | 4458 | 494.3 | - | - | 0 | - |
| - | - | 1619 | 494.9 | - | - | 0 | - |
| - | - | 1618 | 499.9 | - | - | 0 | - |
| - | - | 2575 | 500.9 | - | - | 0 | - |
| - | - | 1500 | 501.2 | - | - | 0 | - |
| - | - | 1839 | 501.6 | - | - | 0 | - |
| - | - | 1959 | 509.3 | - | - | 0 | - |
| 5 | c | 7.688E+04 | 513.3 | 0.0006478 | 1.262 | +1 | 5 |
| - | - | 2.247E+04 | 514.3 | - | - | 0 | - |
| - | - | 4063 | 515.3 | - | - | 0 | - |
| - | - | 2275 | 523.3 | - | - | 0 | - |
| - | - | 2500 | 523.6 | - | - | 0 | - |
| - | - | 2676 | 526.3 | - | - | 0 | - |
| - | - | 1995 | 527.3 | - | - | 0 | - |
| - | - | 1428 | 528.3 | - | - | 0 | - |
| - | - | 6352 | 529.3 | - | - | 0 | - |
| - | - | 1502 | 529.3 | - | - | 0 | - |
| - | - | 5670 | 529.6 | - | - | 0 | - |
| 13 | z | 9718 | 531.3 | 0.001258 | 2.368 | +1 | 5 |
| - | - | 7629 | 532.3 | - | - | 0 | - |
| - | - | 8076 | 532.6 | - | - | 0 | - |
| - | - | 1.221E+04 | 532.9 | - | - | 0 | - |
| - | - | 6768 | 533.3 | - | - | 0 | - |
| - | - | 1729 | 533.6 | - | - | 0 | - |
| - | - | 1.829E+04 | 538.6 | - | - | 0 | - |
| - | - | 1.495E+04 | 538.9 | - | - | 0 | - |
| - | - | 5785 | 539.3 | - | - | 0 | - |
| - | - | 6904 | 539.6 | - | - | 0 | - |
| 4 | y | 2131 | 539.9 | 0.008528 | 15.79 | +3 | 14 |
| 13 | y | 4870 | 547.3 | 0.001394 | 2.547 | +1 | 5 |
| - | - | 2712 | 557 | - | - | 0 | - |
| - | - | 2433 | 557.3 | - | - | 0 | - |
| - | - | 4796 | 557.6 | - | - | 0 | - |
| - | - | 1815 | 563.3 | - | - | 0 | - |
| - | - | 2545 | 566.6 | - | - | 0 | - |
| - | - | 1250 | 567.3 | - | - | 0 | - |
| - | - | 2255 | 570.3 | - | - | 0 | - |
| - | - | 5895 | 571.3 | - | - | 0 | - |
| 15 | c | 1.476E+04 | 572.3 | 0.002394 | 4.183 | +3 | 15 |
| 3 | y | 1.521E+04 | 572.6 | 0.007335 | 12.81 | +3 | 15 |
| 3 | z | 1.242E+04 | 572.9 | 0.004341 | 7.576 | +3 | 15 |
| - | - | 2790 | 573.3 | - | - | 0 | - |
| - | - | 3209 | 573.3 | - | - | 0 | - |
| - | - | 2750 | 573.6 | - | - | 0 | - |
| 3 | y | 3.452E+05 | 578.3 | 0.002534 | 4.383 | +3 | 15 |
| - | - | 3.263E+05 | 578.6 | - | - | 0 | - |
| - | - | 1.775E+05 | 579 | - | - | 0 | - |
| 10 | c | 7.076E+04 | 579.3 | 0.009234 | 15.94 | +2 | 10 |
| - | - | 1.812E+04 | 579.6 | - | - | 0 | - |
| - | - | 2784 | 607.4 | - | - | 0 | - |
| - | - | 1491 | 612.3 | - | - | 0 | - |
| - | - | 1640 | 615.3 | - | - | 0 | - |
| - | - | 7194 | 625.4 | - | - | 0 | - |
| - | - | 5601 | 626.4 | - | - | 0 | - |
| - | - | 2212 | 643.7 | - | - | 0 | - |
| - | - | 1701 | 644 | - | - | 0 | - |
| 12 | y | 2368 | 644.3 | 0.00827 | 12.84 | +1 | 6 |
| 12 | z | 4395 | 645.3 | 0.001264 | 1.958 | +1 | 6 |
| - | - | 6666 | 646.3 | - | - | 0 | - |
| - | - | 3316 | 647.3 | - | - | 0 | - |
| - | - | 2012 | 649 | - | - | 0 | - |
| - | - | 1807 | 650.3 | - | - | 0 | - |
| - | - | 1618 | 651.8 | - | - | 0 | - |
| 11 | c | 1.423E+04 | 652.3 | 0.002796 | 4.285 | +2 | 11 |
| - | - | 1.025E+04 | 652.8 | - | - | 0 | - |
| - | - | 4344 | 653.3 | - | - | 0 | - |
| 12 | y | 1.181E+04 | 661.4 | 0.001008 | 1.524 | +1 | 6 |
| - | - | 2775 | 662.4 | - | - | 0 | - |
| 6 | c | 1.329E+05 | 669.4 | 0.0009772 | 1.46 | +1 | 6 |
| - | - | 5.288E+04 | 670.4 | - | - | 0 | - |
| - | - | 1.036E+04 | 671.4 | - | - | 0 | - |
| - | - | 4364 | 672.4 | - | - | 0 | - |
| - | - | 1813 | 680.8 | - | - | 0 | - |
| - | - | 3124 | 687.8 | - | - | 0 | - |
| - | - | 4577 | 695.4 | - | - | 0 | - |
| - | - | 2507 | 699.8 | - | - | 0 | - |
| 12 | c | 3953 | 700.4 | 0.0006295 | 0.8988 | +2 | 12 |
| 12 | c | 1791 | 700.8 | 0.01029 | 14.69 | +2 | 12 |
| - | - | 8606 | 707.3 | - | - | 0 | - |
| - | - | 8397 | 707.8 | - | - | 0 | - |
| - | - | 6096 | 708.3 | - | - | 0 | - |
| - | - | 1.393E+04 | 708.9 | - | - | 0 | - |
| 12 | c | 8.696E+04 | 709.4 | 0.002694 | 3.798 | +2 | 12 |
| - | - | 6.606E+04 | 709.9 | - | - | 0 | - |
| - | - | 3.539E+04 | 710.4 | - | - | 0 | - |
| - | - | 9049 | 710.9 | - | - | 0 | - |
| - | - | 1886 | 711.4 | - | - | 0 | - |
| - | - | 1814 | 716.4 | - | - | 0 | - |
| 6 | z | 7746 | 718.3 | 0.005288 | 7.362 | +2 | 12 |
| - | - | 6360 | 718.9 | - | - | 0 | - |
| - | - | 4811 | 719.4 | - | - | 0 | - |
| - | - | 4895 | 722.4 | - | - | 0 | - |
| - | - | 4704 | 722.9 | - | - | 0 | - |
| - | - | 3865 | 723.4 | - | - | 0 | - |
| - | - | 3670 | 725.9 | - | - | 0 | - |
| 6 | y | 1.724E+04 | 726.4 | 0.003067 | 4.223 | +2 | 12 |
| - | - | 9411 | 726.9 | - | - | 0 | - |
| - | - | 3955 | 727.4 | - | - | 0 | - |
| - | - | 2004 | 727.9 | - | - | 0 | - |
| - | - | 1.557E+04 | 729.4 | - | - | 0 | - |
| - | - | 5899 | 730.4 | - | - | 0 | - |
| - | - | 2795 | 741.9 | - | - | 0 | - |
| - | - | 2189 | 742.4 | - | - | 0 | - |
| - | - | 1948 | 742.9 | - | - | 0 | - |
| - | - | 1.384E+04 | 750.9 | - | - | 0 | - |
| - | - | 1.301E+04 | 751.4 | - | - | 0 | - |
| - | - | 7132 | 751.9 | - | - | 0 | - |
| - | - | 2687 | 752.4 | - | - | 0 | - |
| - | - | 2849 | 754.4 | - | - | 0 | - |
| - | - | 1687 | 755.4 | - | - | 0 | - |
| 5 | z | 7374 | 761.9 | 0.004411 | 5.789 | +2 | 13 |
| - | - | 1.516E+04 | 762.4 | - | - | 0 | - |
| - | - | 7278 | 762.9 | - | - | 0 | - |
| - | - | 3369 | 763.4 | - | - | 0 | - |
| 13 | c | 2144 | 764.4 | 0.003834 | 5.016 | +2 | 13 |
| 13 | c | 2076 | 764.9 | 0.01006 | 13.15 | +2 | 13 |
| - | - | 6524 | 765.4 | - | - | 0 | - |
| 5 | y | 9593 | 769.9 | 0.002434 | 3.161 | +2 | 13 |
| - | - | 6058 | 770.4 | - | - | 0 | - |
| - | - | 4110 | 770.9 | - | - | 0 | - |
| 11 | y | 1528 | 771.4 | 0.007979 | 10.34 | +1 | 7 |
| - | - | 7733 | 772.9 | - | - | 0 | - |
| 13 | c | 1.049E+05 | 773.4 | 0.004961 | 6.414 | +2 | 13 |
| - | - | 2.312E+04 | 773.5 | - | - | 0 | - |
| - | - | 1.052E+05 | 773.9 | - | - | 0 | - |
| - | - | 4.422E+04 | 774.4 | - | - | 0 | - |
| - | - | 7620 | 774.5 | - | - | 0 | - |
| - | - | 1.736E+04 | 774.9 | - | - | 0 | - |
| - | - | 3498 | 775.4 | - | - | 0 | - |
| - | - | 4397 | 775.4 | - | - | 0 | - |
| - | - | 1707 | 775.9 | - | - | 0 | - |
| - | - | 2727 | 779.4 | - | - | 0 | - |
| 7 | c | 1923 | 780.4 | 0.004989 | 6.393 | +1 | 7 |
| - | - | 2574 | 786.4 | - | - | 0 | - |
| - | - | 1641 | 787.4 | - | - | 0 | - |
| 11 | y | 1.907E+04 | 789.4 | 0.001443 | 1.828 | +1 | 7 |
| - | - | 6099 | 790.4 | - | - | 0 | - |
| - | - | 2558 | 791.9 | - | - | 0 | - |
| - | - | 2577 | 792.4 | - | - | 0 | - |
| - | - | 4393 | 792.9 | - | - | 0 | - |
| - | - | 7375 | 793.4 | - | - | 0 | - |
| - | - | 7916 | 793.9 | - | - | 0 | - |
| - | - | 2881 | 794.4 | - | - | 0 | - |
| - | - | 2142 | 794.9 | - | - | 0 | - |
| - | - | 2072 | 795.4 | - | - | 0 | - |
| - | - | 2464 | 797.4 | - | - | 0 | - |
| 7 | c | 1.798E+05 | 798.4 | 0.00117 | 1.465 | +1 | 7 |
| - | - | 3510 | 798.9 | - | - | 0 | - |
| - | - | 7.366E+04 | 799.5 | - | - | 0 | - |
| - | - | 2460 | 799.9 | - | - | 0 | - |
| - | - | 1.609E+04 | 800.5 | - | - | 0 | - |
| - | - | 9007 | 800.9 | - | - | 0 | - |
| - | - | 6591 | 801.4 | - | - | 0 | - |
| - | - | 7365 | 801.9 | - | - | 0 | - |
| - | - | 2027 | 806.4 | - | - | 0 | - |
| - | - | 2.744E+04 | 807.4 | - | - | 0 | - |
| - | - | 1.83E+04 | 807.9 | - | - | 0 | - |
| - | - | 1.329E+04 | 808.4 | - | - | 0 | - |
| - | - | 5054 | 808.9 | - | - | 0 | - |
| 4 | y | 2619 | 809.9 | 0.005257 | 6.491 | +2 | 14 |
| - | - | 3.039E+04 | 813.4 | - | - | 0 | - |
| 14 | c | 3.626E+04 | 813.9 | 0.001137 | 1.397 | +2 | 14 |
| 14 | c | 2.261E+04 | 814.4 | 0.007527 | 9.242 | +2 | 14 |
| - | - | 1.399E+04 | 814.9 | - | - | 0 | - |
| - | - | 9337 | 815.4 | - | - | 0 | - |
| - | - | 2453 | 815.9 | - | - | 0 | - |
| - | - | 1929 | 817.4 | - | - | 0 | - |
| - | - | 4141 | 817.9 | - | - | 0 | - |
| 4 | y | 4.738E+04 | 818.4 | 0.004311 | 5.268 | +2 | 14 |
| - | - | 4.188E+04 | 818.9 | - | - | 0 | - |
| - | - | 2.681E+04 | 819.4 | - | - | 0 | - |
| - | - | 1.029E+04 | 819.9 | - | - | 0 | - |
| - | - | 2611 | 820.4 | - | - | 0 | - |
| - | - | 4.991E+04 | 822.4 | - | - | 0 | - |
| 14 | c | 7.551E+04 | 822.9 | 0.001759 | 2.138 | +2 | 14 |
| - | - | 6.716E+04 | 823.4 | - | - | 0 | - |
| - | - | 3.295E+04 | 823.9 | - | - | 0 | - |
| - | - | 8867 | 824.4 | - | - | 0 | - |
| - | - | 3242 | 825.5 | - | - | 0 | - |
| - | - | 7790 | 830.5 | - | - | 0 | - |
| - | - | 4778 | 831.5 | - | - | 0 | - |
| - | - | 8847 | 834.9 | - | - | 0 | - |
| - | - | 7594 | 835.4 | - | - | 0 | - |
| - | - | 1.139E+04 | 835.9 | - | - | 0 | - |
| - | - | 4988 | 836.4 | - | - | 0 | - |
| - | - | 3468 | 836.9 | - | - | 0 | - |
| - | - | 1833 | 844.4 | - | - | 0 | - |
| - | - | 2998 | 844.9 | - | - | 0 | - |
| 3 | w | 5283 | 845.4 | 0.007086 | 8.381 | +2 | 15 |
| - | - | 5561 | 845.9 | - | - | 0 | - |
| - | - | 2553 | 846.4 | - | - | 0 | - |
| - | - | 2016 | 846.9 | - | - | 0 | - |
| - | - | 2502 | 849.4 | - | - | 0 | - |
| 3 | z | 2992 | 850.4 | 0.0132 | 15.52 | +2 | 15 |
| 10 | z | 2236 | 856.5 | 0.005533 | 6.46 | +1 | 8 |
| 10 | w | 1.024E+04 | 857.5 | 0.002189 | 2.552 | +1 | 8 |
| 15 | c | 1.452E+04 | 857.9 | 0.004147 | 4.834 | +2 | 15 |
| 3 | y | 1.051E+04 | 858.4 | 0.01452 | 16.91 | +2 | 15 |
| 3 | z | 7559 | 858.9 | 0.007799 | 9.08 | +2 | 15 |
| 10 | w | 2.016E+04 | 859.5 | 0.001168 | 1.358 | +1 | 8 |
| - | - | 2296 | 859.9 | - | - | 0 | - |
| - | - | 6507 | 860.5 | - | - | 0 | - |
| - | - | 1930 | 861.5 | - | - | 0 | - |
| - | - | 1864 | 864.9 | - | - | 0 | - |
| - | - | 2595 | 865.5 | - | - | 0 | - |
| - | - | 1.09E+04 | 865.9 | - | - | 0 | - |
| 15 | c | 1.643E+05 | 866.4 | 0.002896 | 3.343 | +2 | 15 |
| 3 | y | 3.628E+05 | 866.9 | 0.006799 | 7.843 | +2 | 15 |
| - | - | 2.881E+05 | 867.4 | - | - | 0 | - |
| - | - | 1.351E+05 | 867.9 | - | - | 0 | - |
| - | - | 6.111E+04 | 868.4 | - | - | 0 | - |
| - | - | 1.506E+04 | 868.9 | - | - | 0 | - |
| - | - | 3632 | 872.5 | - | - | 0 | - |
| 10 | y | 1.516E+04 | 873.5 | 0.0003381 | 0.3871 | +1 | 8 |
| 10 | z | 9.314E+04 | 874.5 | 0.0005649 | 0.646 | +1 | 8 |
| - | - | 4.84E+04 | 875.5 | - | - | 0 | - |
| - | - | 1.33E+04 | 876.5 | - | - | 0 | - |
| - | - | 3578 | 877.5 | - | - | 0 | - |
| - | - | 6541 | 879.5 | - | - | 0 | - |
| - | - | 8601 | 880 | - | - | 0 | - |
| - | - | 3461 | 880.9 | - | - | 0 | - |
| - | - | 4013 | 881.4 | - | - | 0 | - |
| - | - | 2578 | 881.9 | - | - | 0 | - |
| - | - | 4176 | 883.5 | - | - | 0 | - |
| - | - | 2849 | 884.5 | - | - | 0 | - |
| - | - | 3097 | 887.4 | - | - | 0 | - |
| - | - | 4269 | 887.9 | - | - | 0 | - |
| - | - | 1870 | 888.4 | - | - | 0 | - |
| - | - | 1775 | 889.5 | - | - | 0 | - |
| 10 | y | 2.802E+04 | 890.5 | 2.929E-05 | 0.03289 | +1 | 8 |
| - | - | 1.376E+04 | 891.5 | - | - | 0 | - |
| - | - | 5058 | 892 | - | - | 0 | - |
| - | - | 8094 | 892.5 | - | - | 0 | - |
| - | - | 3524 | 893 | - | - | 0 | - |
| - | - | 5854 | 893.5 | - | - | 0 | - |
| 2 | w | 6.355E+04 | 893.9 | 0.003348 | 3.745 | +2 | 16 |
| - | - | 6.264E+04 | 894.4 | - | - | 0 | - |
| - | - | 3.366E+04 | 894.9 | - | - | 0 | - |
| - | - | 1.683E+04 | 895.4 | - | - | 0 | - |
| - | - | 4299 | 895.9 | - | - | 0 | - |
| - | - | 2234 | 898.5 | - | - | 0 | - |
| - | - | 7449 | 901 | - | - | 0 | - |
| - | - | 1.113E+04 | 901.5 | - | - | 0 | - |
| - | - | 8741 | 902 | - | - | 0 | - |
| - | - | 5359 | 902.5 | - | - | 0 | - |
| - | - | 2557 | 903 | - | - | 0 | - |
| 2 | z | 2846 | 906.5 | 0.01097 | 12.1 | +2 | 16 |
| 2 | z | 3490 | 906.9 | 0.01121 | 12.36 | +2 | 16 |
| - | - | 3139 | 907.5 | - | - | 0 | - |
| 16 | c | 3582 | 908 | 0.00149 | 1.641 | +2 | 16 |
| 16 | c | 3290 | 908.4 | 0.01791 | 19.71 | +2 | 16 |
| 2 | y | 8424 | 914.5 | 0.005389 | 5.893 | +2 | 16 |
| 2 | y | 5203 | 915 | 0.01051 | 11.49 | +2 | 16 |
| 2 | z | 3.557E+04 | 915.5 | 0.005379 | 5.876 | +2 | 16 |
| - | - | 4.285E+04 | 916 | - | - | 0 | - |
| - | - | 2.316E+04 | 916.5 | - | - | 0 | - |
| 16 | c | 1.114E+04 | 917 | 0.005607 | 6.115 | +2 | 16 |
| - | - | 5263 | 917.5 | - | - | 0 | - |
| - | - | 2333 | 918 | - | - | 0 | - |
| - | - | 6025 | 920.5 | - | - | 0 | - |
| - | - | 4029 | 921 | - | - | 0 | - |
| - | - | 1879 | 921.5 | - | - | 0 | - |
| - | - | 3349 | 922.5 | - | - | 0 | - |
| - | - | 1.443E+05 | 923 | - | - | 0 | - |
| 2 | y | 1.613E+05 | 923.5 | 0.01183 | 12.81 | +2 | 16 |
| - | - | 1.102E+05 | 924 | - | - | 0 | - |
| - | - | 5.032E+04 | 924.5 | - | - | 0 | - |
| - | - | 1.962E+04 | 925 | - | - | 0 | - |
| - | - | 5264 | 926.5 | - | - | 0 | - |
| 8 | c | 1.792E+05 | 927.5 | 0.001057 | 1.14 | +1 | 8 |
| - | - | 9.016E+04 | 928.5 | - | - | 0 | - |
| - | - | 3778 | 929 | - | - | 0 | - |
| - | - | 2.636E+04 | 929.5 | - | - | 0 | - |
| - | - | 9564 | 930 | - | - | 0 | - |
| - | - | 1.188E+04 | 930.5 | - | - | 0 | - |
| - | - | 6044 | 931 | - | - | 0 | - |
| - | - | 6926 | 931.5 | - | - | 0 | - |
| - | - | 2037 | 934 | - | - | 0 | - |
| - | - | 2056 | 934.5 | - | - | 0 | - |
| - | - | 3999 | 935 | - | - | 0 | - |
| - | - | 4850 | 935.5 | - | - | 0 | - |
| - | - | 4674 | 936 | - | - | 0 | - |
| - | - | 5253 | 936.5 | - | - | 0 | - |
| - | - | 7461 | 937 | - | - | 0 | - |
| - | - | 1.043E+04 | 937.5 | - | - | 0 | - |
| - | - | 6.008E+04 | 938 | - | - | 0 | - |
| - | - | 6.506E+04 | 938.5 | - | - | 0 | - |
| - | - | 3.765E+04 | 939 | - | - | 0 | - |
| - | - | 1.653E+04 | 939.5 | - | - | 0 | - |
| - | - | 5838 | 940 | - | - | 0 | - |
| - | - | 2236 | 941.5 | - | - | 0 | - |
| - | - | 1.572E+04 | 942.5 | - | - | 0 | - |
| - | - | 1.778E+04 | 943 | - | - | 0 | - |
| - | - | 1.78E+04 | 943.5 | - | - | 0 | - |
| - | - | 1.645E+04 | 944 | - | - | 0 | - |
| - | - | 1.506E+04 | 944.5 | - | - | 0 | - |
| - | - | 3.245E+04 | 945 | - | - | 0 | - |
| - | - | 3.358E+04 | 945.5 | - | - | 0 | - |
| - | - | 2.228E+04 | 946 | - | - | 0 | - |
| - | - | 3E+04 | 946.5 | - | - | 0 | - |
| - | - | 2.261E+04 | 947 | - | - | 0 | - |
| - | - | 1.027E+04 | 947.5 | - | - | 0 | - |
| - | - | 6535 | 948 | - | - | 0 | - |
| - | - | 3345 | 948.5 | - | - | 0 | - |
| - | - | 2637 | 951.5 | - | - | 0 | - |
| - | - | 3.583E+04 | 952 | - | - | 0 | - |
| - | - | 4.108E+04 | 952.5 | - | - | 0 | - |
| - | - | 3.386E+04 | 953 | - | - | 0 | - |
| - | - | 2.44E+04 | 953.5 | - | - | 0 | - |
| - | - | 9414 | 954 | - | - | 0 | - |
| - | - | 4916 | 954.5 | - | - | 0 | - |
| - | - | 1.454E+04 | 956.5 | - | - | 0 | - |
| - | - | 2.522E+04 | 957 | - | - | 0 | - |
| - | - | 2.636E+04 | 957.5 | - | - | 0 | - |
| - | - | 1.903E+04 | 958 | - | - | 0 | - |
| - | - | 2.409E+04 | 958.5 | - | - | 0 | - |
| - | - | 3880 | 959 | - | - | 0 | - |
| - | - | 7.821E+04 | 959.5 | - | - | 0 | - |
| - | - | 1791 | 960 | - | - | 0 | - |
| - | - | 4.313E+04 | 960.5 | - | - | 0 | - |
| - | - | 1.284E+04 | 961.5 | - | - | 0 | - |
| - | - | 2179 | 962.5 | - | - | 0 | - |
| - | - | 5.367E+04 | 965.5 | - | - | 0 | - |
| - | - | 2.373E+05 | 966 | - | - | 0 | - |
| - | - | 2.34E+05 | 966.5 | - | - | 0 | - |
| - | - | 1.505E+05 | 967 | - | - | 0 | - |
| - | - | 7.284E+04 | 967.5 | - | - | 0 | - |
| - | - | 2.36E+04 | 968 | - | - | 0 | - |
| - | - | 9432 | 968.5 | - | - | 0 | - |
| - | - | 4963 | 969.5 | - | - | 0 | - |
| - | - | 1.572E+05 | 974 | - | - | 0 | - |
| - | - | 2.661E+05 | 974.5 | - | - | 0 | - |
| - | - | 2.226E+05 | 975 | - | - | 0 | - |
| - | - | 1.241E+05 | 975.5 | - | - | 0 | - |
| - | - | 4.626E+04 | 976 | - | - | 0 | - |
| - | - | 1.11E+04 | 976.5 | - | - | 0 | - |
| 9 | z | 1858 | 1003 | 0.009148 | 9.117 | +1 | 9 |
| 9 | z | 1602 | 1004 | 0.01921 | 19.13 | +1 | 9 |
| - | - | 2826 | 1006 | - | - | 0 | - |
| - | - | 2574 | 1007 | - | - | 0 | - |
| - | - | 1988 | 1012 | - | - | 0 | - |
| - | - | 2497 | 1015 | - | - | 0 | - |
| 9 | z | 1.877E+04 | 1022 | 0.005969 | 5.843 | +1 | 9 |
| - | - | 1.943E+04 | 1023 | - | - | 0 | - |
| - | - | 7556 | 1024 | - | - | 0 | - |
| - | - | 3634 | 1025 | - | - | 0 | - |
| - | - | 2.137E+04 | 1031 | - | - | 0 | - |
| - | - | 1.544E+04 | 1032 | - | - | 0 | - |
| - | - | 6907 | 1033 | - | - | 0 | - |
| - | - | 2472 | 1034 | - | - | 0 | - |
| 9 | y | 9801 | 1038 | 0.002321 | 2.237 | +1 | 9 |
| - | - | 6411 | 1039 | - | - | 0 | - |
| - | - | 1832 | 1040 | - | - | 0 | - |
| - | - | 7989 | 1050 | - | - | 0 | - |
| - | - | 6218 | 1051 | - | - | 0 | - |
| - | - | 2747 | 1060 | - | - | 0 | - |
| - | - | 3862 | 1074 | - | - | 0 | - |
| 9 | c | 2.001E+05 | 1075 | 0.005973 | 5.559 | +1 | 9 |
| - | - | 1.055E+05 | 1076 | - | - | 0 | - |
| - | - | 4.542E+04 | 1077 | - | - | 0 | - |
| - | - | 1975 | 1077 | - | - | 0 | - |
| - | - | 1.109E+04 | 1078 | - | - | 0 | - |
| - | - | 6509 | 1088 | - | - | 0 | - |
| - | - | 4832 | 1089 | - | - | 0 | - |
| 8 | w | 2.084E+04 | 1092 | 0.006648 | 6.091 | +1 | 10 |
| - | - | 1.734E+04 | 1093 | - | - | 0 | - |
| - | - | 6121 | 1094 | - | - | 0 | - |
| - | - | 4701 | 1095 | - | - | 0 | - |
| - | - | 3356 | 1096 | - | - | 0 | - |
| - | - | 2634 | 1107 | - | - | 0 | - |
| - | - | 3715 | 1114 | - | - | 0 | - |
| - | - | 2773 | 1115 | - | - | 0 | - |
| - | - | 4316 | 1132 | - | - | 0 | - |
| - | - | 6783 | 1133 | - | - | 0 | - |
| - | - | 2565 | 1134 | - | - | 0 | - |
| - | - | 4454 | 1135 | - | - | 0 | - |
| - | - | 2955 | 1136 | - | - | 0 | - |
| 8 | z | 8.656E+04 | 1151 | 0.005551 | 4.825 | +1 | 10 |
| - | - | 5.703E+04 | 1152 | - | - | 0 | - |
| - | - | 2.628E+04 | 1153 | - | - | 0 | - |
| - | - | 7650 | 1154 | - | - | 0 | - |
| 8 | y | 6117 | 1167 | 0.004405 | 3.776 | +1 | 10 |
| - | - | 3706 | 1168 | - | - | 0 | - |
| 10 | c | 9.78E+04 | 1176 | 0.005414 | 4.605 | +1 | 10 |
| - | - | 6.29E+04 | 1177 | - | - | 0 | - |
| - | - | 2.243E+04 | 1178 | - | - | 0 | - |
| - | - | 9187 | 1179 | - | - | 0 | - |
| - | - | 2887 | 1180 | - | - | 0 | - |
| - | - | 3193 | 1209 | - | - | 0 | - |
| 7 | w | 1.476E+04 | 1221 | 0.005803 | 4.755 | +1 | 11 |
| - | - | 1.347E+04 | 1222 | - | - | 0 | - |
| - | - | 4281 | 1223 | - | - | 0 | - |
| - | - | 2606 | 1224 | - | - | 0 | - |
| - | - | 2263 | 1225 | - | - | 0 | - |
| - | - | 5947 | 1260 | - | - | 0 | - |
| - | - | 5331 | 1261 | - | - | 0 | - |
| - | - | 2461 | 1262 | - | - | 0 | - |
| 7 | z | 6.67E+04 | 1280 | 0.005072 | 3.964 | +1 | 11 |
| - | - | 4.667E+04 | 1281 | - | - | 0 | - |
| - | - | 2.34E+04 | 1282 | - | - | 0 | - |
| - | - | 6690 | 1283 | - | - | 0 | - |
| 11 | c | 2230 | 1286 | 0.003423 | 2.663 | +1 | 11 |
| 7 | y | 2.231E+04 | 1296 | 0.005269 | 4.067 | +1 | 11 |
| - | - | 1.486E+04 | 1297 | - | - | 0 | - |
| - | - | 5226 | 1298 | - | - | 0 | - |
| - | - | 2782 | 1299 | - | - | 0 | - |
| - | - | 6330 | 1303 | - | - | 0 | - |
| 11 | c | 2.201E+04 | 1304 | 0.004079 | 3.129 | +1 | 11 |
| - | - | 1.601E+04 | 1305 | - | - | 0 | - |
| - | - | 4894 | 1306 | - | - | 0 | - |
| - | - | 5022 | 1335 | - | - | 0 | - |
| - | - | 2810 | 1336 | - | - | 0 | - |
| - | - | 3810 | 1374 | - | - | 0 | - |
| - | - | 2383 | 1375 | - | - | 0 | - |
| - | - | 2219 | 1377 | - | - | 0 | - |
| - | - | 3769 | 1417 | - | - | 0 | - |
| 12 | c | 3.606E+04 | 1418 | 0.002533 | 1.787 | +1 | 12 |
| - | - | 2.653E+04 | 1419 | - | - | 0 | - |
| - | - | 1.299E+04 | 1420 | - | - | 0 | - |
| - | - | 7871 | 1421 | - | - | 0 | - |
| - | - | 2367 | 1422 | - | - | 0 | - |
| 6 | y | 1856 | 1435 | 0.001055 | 0.7357 | +1 | 12 |
| 6 | z | 4.929E+04 | 1436 | 0.004791 | 3.337 | +1 | 12 |
| - | - | 7.618E+04 | 1437 | - | - | 0 | - |
| - | - | 5.53E+04 | 1438 | - | - | 0 | - |
| - | - | 2.546E+04 | 1439 | - | - | 0 | - |
| - | - | 7348 | 1440 | - | - | 0 | - |
| - | - | 5888 | 1451 | - | - | 0 | - |
| 6 | y | 2.622E+04 | 1452 | 0.002913 | 2.007 | +1 | 12 |
| - | - | 2.048E+04 | 1453 | - | - | 0 | - |
| - | - | 1.181E+04 | 1454 | - | - | 0 | - |
| - | - | 3317 | 1455 | - | - | 0 | - |
| - | - | 4873 | 1503 | - | - | 0 | - |
| 5 | z | 4772 | 1523 | 0.004745 | 3.116 | +1 | 13 |
| - | - | 2.3E+04 | 1524 | - | - | 0 | - |
| - | - | 1.504E+04 | 1525 | - | - | 0 | - |
| - | - | 7927 | 1526 | - | - | 0 | - |
| 5 | y | 7601 | 1539 | 6.269E-05 | 0.04074 | +1 | 13 |
| - | - | 5259 | 1540 | - | - | 0 | - |
| - | - | 2314 | 1541 | - | - | 0 | - |
| - | - | 3243 | 1545 | - | - | 0 | - |
| 13 | c | 4.773E+04 | 1546 | 0.003892 | 2.518 | +1 | 13 |
| - | - | 4.439E+04 | 1547 | - | - | 0 | - |
| - | - | 2.297E+04 | 1548 | - | - | 0 | - |
| - | - | 1.069E+04 | 1549 | - | - | 0 | - |
| - | - | 3032 | 1550 | - | - | 0 | - |
| - | - | 3554 | 1602 | - | - | 0 | - |
| - | - | 4469 | 1603 | - | - | 0 | - |
| - | - | 3540 | 1604 | - | - | 0 | - |
| - | - | 4848 | 1615 | - | - | 0 | - |
| - | - | 3955 | 1616 | - | - | 0 | - |
| 14 | c | 5746 | 1628 | 0.01818 | 11.17 | +1 | 14 |
| - | - | 4274 | 1629 | - | - | 0 | - |
| - | - | 3973 | 1630 | - | - | 0 | - |
| 4 | y | 1.398E+04 | 1636 | 0.00357 | 2.182 | +1 | 14 |
| - | - | 1.265E+04 | 1637 | - | - | 0 | - |
| - | - | 8898 | 1638 | - | - | 0 | - |
| - | - | 3307 | 1639 | - | - | 0 | - |
| - | - | 9400 | 1644 | - | - | 0 | - |
| 14 | c | 6.172E+04 | 1645 | 0.004204 | 2.556 | +1 | 14 |
| - | - | 5.652E+04 | 1646 | - | - | 0 | - |
| - | - | 3.019E+04 | 1647 | - | - | 0 | - |
| - | - | 1.276E+04 | 1648 | - | - | 0 | - |
| - | - | 4614 | 1649 | - | - | 0 | - |
| - | - | 2965 | 1671 | - | - | 0 | - |
| - | - | 1815 | 1672 | - | - | 0 | - |
| - | - | 2150 | 1688 | - | - | 0 | - |
| - | - | 5912 | 1689 | - | - | 0 | - |
| - | - | 4711 | 1690 | - | - | 0 | - |
| - | - | 1892 | 1691 | - | - | 0 | - |
| - | - | 4630 | 1716 | - | - | 0 | - |
| 3 | z | 3530 | 1717 | 0.01592 | 9.271 | +1 | 15 |
| - | - | 1910 | 1719 | - | - | 0 | - |
| - | - | 2643 | 1728 | - | - | 0 | - |
| - | - | 6054 | 1729 | - | - | 0 | - |
| - | - | 5109 | 1730 | - | - | 0 | - |
| - | - | 3198 | 1731 | - | - | 0 | - |
| 15 | c | 3.113E+04 | 1732 | 0.005867 | 3.387 | +1 | 15 |
| 3 | y | 5.133E+04 | 1733 | 0.02441 | 14.09 | +1 | 15 |
| - | - | 4.364E+04 | 1734 | - | - | 0 | - |
| - | - | 2.174E+04 | 1735 | - | - | 0 | - |
| - | - | 9140 | 1736 | - | - | 0 | - |
| - | - | 3583 | 1786 | - | - | 0 | - |
| - | - | 3633 | 1787 | - | - | 0 | - |
| - | - | 2287 | 1788 | - | - | 0 | - |
| - | - | 2872 | 1801 | - | - | 0 | - |
| - | - | 1.262E+04 | 1802 | - | - | 0 | - |
| - | - | 1.474E+04 | 1803 | - | - | 0 | - |
| - | - | 1.038E+04 | 1804 | - | - | 0 | - |
| - | - | 4721 | 1805 | - | - | 0 | - |
| 2 | y | 6734 | 1829 | 0.02305 | 12.6 | +1 | 16 |
| 2 | z | 1.542E+04 | 1830 | 0.01266 | 6.92 | +1 | 16 |
| - | - | 5.18E+04 | 1831 | - | - | 0 | - |
| - | - | 5.359E+04 | 1832 | - | - | 0 | - |
| 16 | c | 2.745E+04 | 1833 | 0.01556 | 8.49 | +1 | 16 |
| - | - | 1.582E+04 | 1834 | - | - | 0 | - |
| - | - | 5583 | 1835 | - | - | 0 | - |
| - | - | 1.545E+04 | 1845 | - | - | 0 | - |
| 2 | y | 4.115E+04 | 1846 | 0.02446 | 13.25 | +1 | 16 |
| - | - | 4.196E+04 | 1847 | - | - | 0 | - |
| - | - | 2.76E+04 | 1848 | - | - | 0 | - |
| - | - | 1.009E+04 | 1849 | - | - | 0 | - |
| - | - | 3950 | 1850 | - | - | 0 | - |
| - | - | 2249 | 1851 | - | - | 0 | - |
| - | - | 3216 | 1871 | - | - | 0 | - |
| - | - | 2047 | 1872 | - | - | 0 | - |
| - | - | 2764 | 1873 | - | - | 0 | - |
| - | - | 1781 | 1875 | - | - | 0 | - |
| - | - | 1731 | 1877 | - | - | 0 | - |
| - | - | 4672 | 1885 | - | - | 0 | - |
| - | - | 7536 | 1886 | - | - | 0 | - |
| - | - | 1.107E+04 | 1887 | - | - | 0 | - |
| - | - | 8352 | 1888 | - | - | 0 | - |
| - | - | 7590 | 1889 | - | - | 0 | - |
| - | - | 1.714E+04 | 1890 | - | - | 0 | - |
| - | - | 1.548E+04 | 1891 | - | - | 0 | - |
| - | - | 9950 | 1892 | - | - | 0 | - |
| - | - | 5020 | 1893 | - | - | 0 | - |
| - | - | 1.229E+04 | 1903 | - | - | 0 | - |
| - | - | 3.777E+04 | 1904 | - | - | 0 | - |
| - | - | 3.226E+04 | 1905 | - | - | 0 | - |
| - | - | 2.277E+04 | 1906 | - | - | 0 | - |
| - | - | 1.047E+04 | 1907 | - | - | 0 | - |
| - | - | 3216 | 1908 | - | - | 0 | - |
| - | - | 6503 | 1914 | - | - | 0 | - |
| - | - | 1.29E+04 | 1915 | - | - | 0 | - |
| - | - | 7338 | 1916 | - | - | 0 | - |
| - | - | 3753 | 1917 | - | - | 0 | - |
| - | - | 2874 | 1918 | - | - | 0 | - |
| - | - | 4729 | 1920 | - | - | 0 | - |
| - | - | 2.983E+04 | 1921 | - | - | 0 | - |
| - | - | 2.669E+04 | 1922 | - | - | 0 | - |
| - | - | 1.666E+04 | 1923 | - | - | 0 | - |
| - | - | 7279 | 1924 | - | - | 0 | - |
| - | - | 3049 | 1925 | - | - | 0 | - |
| - | - | 3495 | 1930 | - | - | 0 | - |
| - | - | 3.597E+04 | 1931 | - | - | 0 | - |
| - | - | 1.301E+05 | 1932 | - | - | 0 | - |
| - | - | 1.31E+05 | 1933 | - | - | 0 | - |
| - | - | 7.699E+04 | 1934 | - | - | 0 | - |
| - | - | 3.188E+04 | 1935 | - | - | 0 | - |
| - | - | 1.03E+04 | 1936 | - | - | 0 | - |
| - | - | 1.627E+04 | 1947 | - | - | 0 | - |
| - | - | 8.667E+04 | 1948 | - | - | 0 | - |
| - | - | 3.642E+05 | 1949 | - | - | 0 | - |
| - | - | 3.499E+05 | 1950 | - | - | 0 | - |
| - | - | 2.268E+05 | 1951 | - | - | 0 | - |
| - | - | 9.735E+04 | 1952 | - | - | 0 | - |
| - | - | 3.136E+04 | 1953 | - | - | 0 | - |
| - | - | 1857 | 2523 | - | - | 0 | - |

m/z Charge Intensity FragmentType MassShift Position
120.06574249267578 0 10899.234
139.19180297851562 0 994.6061
140.12977600097656 0 920.4734
141.1693878173828 0 1013.046
143.11819458007812 0 1192.7388
145.79598999023438 0 1149.3345
147.32431030273438 0 1061.791
148.9476318359375 0 1540.1034
151.85096740722656 0 1179.517
173.1289825439453 0 1638.6254
187.1444091796875 0 95464.23
188.14776611328125 0 7653.1914
201.12356567382812 0 3984.479 y Water loss 13
212.13995361328125 0 4783.177
215.13929748535156 0 69224.336 y Water loss 15
216.1426544189453 0 7896.619
233.14984130859375 0 6663.234 y 15
256.1661071777344 0 1808.4307
298.58599853515625 0 1419.085
299.1719970703125 0 11558.099
300.1748046875 0 2326.23
302.1716613769531 0 2605.6655 y Water loss 14
312.1924133300781 0 2657.0503
320.181884765625 0 2986.9915 y 14
344.1947326660156 0 1609.0353
376.9773864746094 0 1313.3026
378.3929443359375 0 1465.8015
382.2582702636719 0 1746.2493
383.2620544433594 0 1633.9047
401.44366455078125 0 1483.3807
407.25335693359375 0 3627.1125
412.2302551269531 0 1496.7755
413.2376708984375 0 1641.1646
421.3855895996094 0 1350.5427
425.26531982421875 0 2320.901
426.2718811035156 0 4542.6274 c 3
427.26971435546875 0 1342.5299
451.27947998046875 0 2160.9224
462.5653076171875 0 1455.8501
463.22955322265625 0 1444.8237
470.2978515625 0 6463.712
472.2412414550781 0 1662.8115
472.2880859375 0 1681.9122
493.90179443359375 0 2458.1309
494.2856140136719 0 4458.445
494.9115905761719 0 1618.5114
499.9115905761719 0 1618.1176
500.9089660644531 0 2575.1038
501.24365234375 0 1499.7458
501.575927734375 0 1838.7451
509.2737121582031 0 1958.7382
513.3037719726562 0 76880.484 c 4
514.3067016601562 0 22472.477
515.3087768554688 0 4062.6418
523.2695922851562 0 2275.42
523.6004028320312 0 2500.3938
526.2720336914062 0 2675.7053
527.2843017578125 0 1995.19
528.3136596679688 0 1427.9187
529.2723388671875 0 6351.8354
529.317626953125 0 1502.4629
529.6058349609375 0 5670.4844
531.2911376953125 0 9718.238 z 12
532.2969970703125 0 7629.1543
532.60009765625 0 8076.1147
532.9331665039062 0 12214.04
533.266845703125 0 6767.9897
533.59765625 0 1729.3119
538.6038818359375 0 18291.543
538.938720703125 0 14949.234
539.2711791992188 0 5785.4946
539.6048583984375 0 6904.408
539.9417724609375 0 2131.031 y Water loss 3
547.3099975585938 0 4869.841 y 12
556.9545288085938 0 2712.4873
557.293212890625 0 2433.1736
557.6221313476562 0 4795.8237
563.2854614257812 0 1814.8295
566.6139526367188 0 2544.776
567.284912109375 0 1249.8015
570.3124389648438 0 2255.3518
571.3189697265625 0 5894.6763
572.2865600585938 0 14756.455 c Ammonia loss 14
572.6195068359375 0 15212.314 y Ammonia loss 2
572.9524536132812 0 12422.444 z 2
573.2868041992188 0 2790.1658
573.3356323242188 0 3209.2224
573.6160278320312 0 2750.2708
578.2902221679688 0 345179.53 y 2
578.6240234375 0 326271.75
578.9579467773438 0 177477.2
579.2913818359375 0 70762.805 c Water loss 9
579.62548828125 0 18118.252
607.3815307617188 0 2784.0547
612.3358154296875 0 1490.6146
615.345703125 0 1640.0194
625.391357421875 0 7194.4404
626.3955688476562 0 5600.5522
643.6636352539062 0 2211.6873
643.99609375 0 1700.8308
644.333251953125 0 2368.4143 y Ammonia loss 11
645.33154296875 0 4394.9834 z 11
646.3384399414062 0 6666.1934
647.3421020507812 0 3315.6116
649.0169677734375 0 2012.141
650.34423828125 0 1806.8624
651.8374633789062 0 1617.8049
652.3377075195312 0 14233.678 c 10
652.8401489257812 0 10245.451
653.3383178710938 0 4344.0845
661.3525390625 0 11807.806 y 11
662.3527221679688 0 2775.0422
669.4052124023438 0 132899.67 c 5
670.4078979492188 0 52880.6
671.4093627929688 0 10364.369
672.4060668945312 0 4364.4346
680.8302001953125 0 1812.8102
687.8494262695312 0 3123.6965
695.4188842773438 0 4576.589
699.8497314453125 0 2507.3142
700.3504638671875 0 3952.9514 c Water loss 11
700.8533935546875 0 1791.4108 c Ammonia loss 11
707.3438110351562 0 8606.086
707.8460083007812 0 8396.779
708.3446655273438 0 6095.7437
708.8541259765625 0 13934.124
709.3590698242188 0 86959.484 c 11
709.8607177734375 0 66055.54
710.361328125 0 35390.918
710.861328125 0 9049.308
711.4292602539062 0 1885.7874
716.3795166015625 0 1813.8749
718.3550415039062 0 7746.1753 z 5
718.8543090820312 0 6359.5815
719.3566284179688 0 4810.979
722.3658447265625 0 4894.691
722.868896484375 0 4703.6245
723.3723754882812 0 3865.021
725.8580322265625 0 3669.539
726.3621826171875 0 17242.998 y 5
726.8629150390625 0 9410.809
727.362548828125 0 3954.813
727.8618774414062 0 2004.3264
729.4268188476562 0 15574.549
730.4288330078125 0 5899.1714
741.8555297851562 0 2794.6428
742.3583984375 0 2189.0427
742.860107421875 0 1947.814
750.8599853515625 0 13841.72
751.3673095703125 0 13006.489
751.8671875 0 7132.1533
752.3606567382812 0 2687.4548
754.4331665039062 0 2849.31
755.4337158203125 0 1687.1439
761.8701782226562 0 7374.3047 z 4
762.3709716796875 0 15160.3
762.8736572265625 0 7277.8447
763.37060546875 0 3369.1904
764.3842163085938 0 2144.4355 c Water loss 12
764.8824462890625 0 2076.4062 c Ammonia loss 12
765.3901977539062 0 6524.4697
769.8775634765625 0 9592.775 y 4
770.3798828125 0 6058.154
770.8773803710938 0 4109.674
771.4439086914062 0 1527.698 y Water loss 10
772.8842163085938 0 7733.26
773.390625 0 104901.5 c 12
773.4509887695312 0 23123.268
773.8897094726562 0 105228.37
774.3917846679688 0 44221.17
774.4519653320312 0 7620.4897
774.8908081054688 0 17357.748
775.3931884765625 0 3497.9998
775.4415893554688 0 4397.0513
775.896240234375 0 1706.9901
779.4301147460938 0 2726.741
780.4312744140625 0 1922.6298 c Water loss 6
786.3904418945312 0 2574.1953
787.40234375 0 1640.5386
789.4479370117188 0 19068.07 y 10
790.4494018554688 0 6099.377
791.914306640625 0 2558.4666
792.4060668945312 0 2576.6445
792.9063720703125 0 4393.2104
793.408935546875 0 7374.5195
793.91015625 0 7916.4067
794.408447265625 0 2881.3376
794.8932495117188 0 2142.4143
795.40234375 0 2072.0337
797.4424438476562 0 2463.6934
798.447998046875 0 179783.03 c 6
798.8978271484375 0 3510.3755
799.450439453125 0 73662.1
799.9005126953125 0 2459.723
800.4524536132812 0 16090.729
800.91455078125 0 9006.556
801.4175415039062 0 6591.324
801.91650390625 0 7364.8223
806.401611328125 0 2027.3793
807.4024047851562 0 27436.863
807.9029541015625 0 18295.326
808.4025268554688 0 13291.808
808.9036865234375 0 5053.8423
809.8934936523438 0 2619.146 y Ammonia loss 3
813.4139404296875 0 30387.586
813.9134521484375 0 36261.61 c Water loss 13
814.4141235351562 0 22612.053 c Ammonia loss 13
814.9127807617188 0 13993.994
815.4163208007812 0 9337.358
815.9154052734375 0 2452.5305
817.3952026367188 0 1929.0039
817.9034423828125 0 4140.598
818.4058227539062 0 47378.395 y 3
818.9059448242188 0 41880.47
819.4068603515625 0 26809.89
819.9083251953125 0 10293.817
820.41064453125 0 2610.667
822.4191284179688 0 49909.043
822.921630859375 0 75507.61 c 13
823.4224853515625 0 67164.57
823.9237060546875 0 32952.56
824.4293212890625 0 8867.057
825.4633178710938 0 3241.922
830.4630737304688 0 7790.0703
831.4669799804688 0 4777.935
834.934814453125 0 8847.216
835.4313354492188 0 7593.6343
835.9319458007812 0 11389.724
836.4295043945312 0 4987.714
836.9326171875 0 3467.9053
844.4385375976562 0 1833.1573
844.9297485351562 0 2998.1382
845.4138793945312 0 5283.16 w 2
845.9137573242188 0 5561.2144
846.4107055664062 0 2553.3264
846.909912109375 0 2015.9237
849.4210205078125 0 2502.3818
850.41845703125 0 2991.636 z Ammonia loss 2
856.4593505859375 0 2235.6968 z Water loss 9
857.4705200195312 0 10238.04 w 9
857.9267578125 0 14523.749 c Ammonia loss 14
858.4291381835938 0 10509.752 y Ammonia loss 2
858.9263305664062 0 7559.262 z 2
859.4508056640625 0 20160.008 w 9
859.9246826171875 0 2296.3967
860.4552612304688 0 6507.2773
861.4609375 0 1929.7323
864.9334106445312 0 1864.252
865.451416015625 0 2594.9749
865.93603515625 0 10901.53
866.4387817382812 0 164303.17 c 14
866.9346923828125 0 362826.62 y 2
867.4347534179688 0 288069.06
867.9342041015625 0 135098.62
868.4349365234375 0 61107.273
868.934326171875 0 15055.61
872.460205078125 0 3632.0242
873.46728515625 0 15160.162 y Ammonia loss 9
874.4760131835938 0 93138.77 z 9
875.478515625 0 48402.094
876.4806518554688 0 13298.755
877.4857788085938 0 3577.8408
879.4530029296875 0 6540.5796
879.9500732421875 0 8601.3545
880.9432983398438 0 3461.101
881.4247436523438 0 4013.1384
881.9390869140625 0 2578.0376
883.4763793945312 0 4176.475
884.4754638671875 0 2849.2268
887.4271850585938 0 3097.4421
887.9347534179688 0 4269.4
888.4243774414062 0 1870.0153
889.4907836914062 0 1775.2529
890.4942016601562 0 28019.035 y 9
891.4963989257812 0 13758.732
891.9653930664062 0 5058.432
892.483154296875 0 8093.633
892.9749145507812 0 3523.9465
893.4636840820312 0 5853.799
893.9365234375 0 63553.285 w 1
894.4375610351562 0 62643.63
894.9380493164062 0 33662.4
895.4380493164062 0 16826.781
895.93896484375 0 4299.2627
898.4531860351562 0 2234.433
900.9730224609375 0 7449.375
901.4732666015625 0 11131.046
901.9752197265625 0 8741.043
902.4786376953125 0 5358.5522
902.9567260742188 0 2557.1304
906.4662475585938 0 2846.1892 z Water loss 1
906.95849609375 0 3489.6987 z Ammonia loss 1
907.475341796875 0 3139.4678
907.9559326171875 0 3581.9983 c Water loss 15
908.46435546875 0 3290.2747 c Ammonia loss 15
914.4700317382812 0 8424.235 y Water loss 1
914.9671630859375 0 5203.2583 y Ammonia loss 1
915.4659423828125 0 35565.93 z 1
915.9658813476562 0 42850.29
916.4668579101562 0 23156.5
916.96533203125 0 11143.716 c 15
917.4635620117188 0 5262.9893
917.9500122070312 0 2332.7832
920.4931640625 0 6024.808
920.9924926757812 0 4029.0298
921.4927978515625 0 1878.8546
922.4778442382812 0 3348.733
922.9812622070312 0 144319.98
923.4817504882812 0 161264.47 y 1
923.983154296875 0 110176.25
924.4830932617188 0 50322.402
924.98095703125 0 19622.326
926.4813232421875 0 5263.9795
927.490478515625 0 179151.92 c 7
928.4932861328125 0 90164.2
928.9757690429688 0 3777.784
929.494140625 0 26358.586
929.9727783203125 0 9564.36
930.4827880859375 0 11875.889
930.96826171875 0 6043.752
931.4869995117188 0 6925.8687
933.9948120117188 0 2036.9891
934.5028076171875 0 2056.0645
934.982421875 0 3999.379
935.4893188476562 0 4849.7207
935.9807739257812 0 4674.2
936.4790649414062 0 5252.6196
936.9807739257812 0 7461.0903
937.4694213867188 0 10433.413
937.959228515625 0 60084.223
938.4617309570312 0 65064.824
938.9632568359375 0 37650.773
939.468994140625 0 16525.107
939.96875 0 5837.767
941.5088500976562 0 2235.9421
942.4990234375 0 15716.737
942.995361328125 0 17775.754
943.4893798828125 0 17802.45
943.9860229492188 0 16445.2
944.4781494140625 0 15061.667
944.9799194335938 0 32450.58
945.482666015625 0 33577.23
945.9832153320312 0 22277.014
946.47412109375 0 30001.623
946.9718627929688 0 22611.84
947.4730834960938 0 10268.242
947.9699096679688 0 6535.1606
948.473876953125 0 3344.599
951.493896484375 0 2636.8865
951.9910888671875 0 35826.87
952.49267578125 0 41081.09
952.991455078125 0 33859.902
953.4929809570312 0 24403.102
953.9872436523438 0 9413.666
954.4892578125 0 4916.3813
956.4902954101562 0 14539.26
956.9869384765625 0 25223.531
957.489013671875 0 26356.186
957.982177734375 0 19027.31
958.5119018554688 0 24092.979
958.9800415039062 0 3879.6375
959.5277099609375 0 78212.125
959.9878540039062 0 1790.9962
960.5303955078125 0 43127.223
961.5328979492188 0 12842.676
962.5272827148438 0 2179.492
965.4947509765625 0 53673.543
965.991455078125 0 237267.8
966.4912109375 0 234004.45
966.9917602539062 0 150539.4
967.4918823242188 0 72839.67
967.9927978515625 0 23599.137
968.5278930664062 0 9431.552
969.5267944335938 0 4962.8535
973.9972534179688 0 157206.08
974.4988403320312 0 266137.2
975.0004272460938 0 222618.83
975.5013427734375 0 124122.14
976.001953125 0 46258.727
976.5042114257812 0 11095.796
1003.5045166015625 0 1858.1528 z Water loss 8
1004.4985961914062 0 1602.2052 z Ammonia loss 8
1005.50927734375 0 2826.1858
1006.5146484375 0 2573.5642
1012.4992065429688 0 1988.2506
1015.4979858398438 0 2497.1033
1021.5119018554688 0 18769.73 z 8
1022.5144653320312 0 19433.848
1023.5159301757812 0 7556.1274
1024.5177001953125 0 3634.1404
1030.512451171875 0 21372.375
1031.5162353515625 0 15437.225
1032.517822265625 0 6907.09
1033.51904296875 0 2472.4546
1037.5269775390625 0 9800.816 y 8
1038.5306396484375 0 6411.03
1039.5404052734375 0 1832.2377
1049.5284423828125 0 7988.746
1050.532958984375 0 6217.776
1059.5103759765625 0 2746.7402
1073.520263671875 0 3862.0803
1074.52587890625 0 200061.52 c 8
1075.5286865234375 0 105526.91
1076.5299072265625 0 45416.023
1076.662353515625 0 1975.0758
1077.5306396484375 0 11086.215
1087.5650634765625 0 6509.1123
1088.5657958984375 0 4832.0825
1091.5418701171875 0 20835.264 w 7
1092.542724609375 0 17343.002
1093.5418701171875 0 6120.763
1094.5096435546875 0 4700.9126
1095.5029296875 0 3355.5168
1106.5435791015625 0 2633.5806
1113.5499267578125 0 3714.738
1114.5538330078125 0 2773.4526
1131.559814453125 0 4316.1475
1132.564453125 0 6783.08
1133.5589599609375 0 2564.9246
1134.55908203125 0 4454.302
1135.5521240234375 0 2955.3257
1150.5540771484375 0 86558.19 z 7
1151.55615234375 0 57032.414
1152.5560302734375 0 26278.645
1153.5550537109375 0 7649.587
1166.5716552734375 0 6116.9814 y 7
1167.57421875 0 3706.4465
1175.572998046875 0 97803.4 c 9
1176.57568359375 0 62897.7
1177.5762939453125 0 22428.836
1178.579833984375 0 9186.51
1179.57763671875 0 2887.0994
1208.5653076171875 0 3193.0444
1220.5836181640625 0 14759.037 w 6
1221.5859375 0 13467.206
1222.576904296875 0 4281.0586
1223.543701171875 0 2606.123
1224.540283203125 0 2263.317
1259.6583251953125 0 5947.367
1260.6505126953125 0 5331.0635
1261.6688232421875 0 2461.36
1279.59619140625 0 66702.37 z 6
1280.5977783203125 0 46667.688
1281.598388671875 0 23398.428
1282.60205078125 0 6689.962
1285.6485595703125 0 2229.6826 c Water loss 10
1295.6151123046875 0 22312.955 y 6
1296.6158447265625 0 14855.88
1297.6177978515625 0 5226.1924
1298.61865234375 0 2782.1113
1302.6622314453125 0 6330.414
1303.6666259765625 0 22008.432 c 10
1304.6680908203125 0 16009.618
1305.6693115234375 0 4893.623
1334.669189453125 0 5022.1094
1335.6768798828125 0 2809.586
1373.6947021484375 0 3809.7625
1374.698974609375 0 2383.1624
1376.6990966796875 0 2218.921
1416.7001953125 0 3769.0972
1417.7080078125 0 36063.61 c 11
1418.7109375 0 26526.018
1419.7113037109375 0 12988.1
1420.706298828125 0 7871.228
1421.6976318359375 0 2367.1243
1434.683349609375 0 1856.0212 y Ammonia loss 5
1435.697021484375 0 49290.23 z 5
1436.701904296875 0 76178.36
1437.703857421875 0 55304.832
1438.7049560546875 0 25461.883
1439.707763671875 0 7348.464
1450.7100830078125 0 5887.805
1451.7138671875 0 26215.25 y 5
1452.714111328125 0 20480.934
1453.7197265625 0 11812.931
1454.7198486328125 0 3317.3503
1502.75634765625 0 4873.121
1522.72900390625 0 4772.167 z 4
1523.7342529296875 0 23002.56
1524.73876953125 0 15038.708
1525.739501953125 0 7927.1177
1538.742919921875 0 7601.0693 y 4
1539.7520751953125 0 5258.8145
1540.7501220703125 0 2314.464
1544.765625 0 3243.2236
1545.7679443359375 0 47729.805 c 12
1546.7703857421875 0 44392.656
1547.7705078125 0 22965.557
1548.7735595703125 0 10694.317
1549.7703857421875 0 3032.0703
1601.82568359375 0 3553.5847
1602.8380126953125 0 4468.926
1603.8333740234375 0 3539.7722
1614.8021240234375 0 4847.573
1615.813232421875 0 3955.4712
1627.8240966796875 0 5745.747 c Ammonia loss 13
1628.8167724609375 0 4274.462
1629.8131103515625 0 3972.8335
1635.79931640625 0 13982.046 y 3
1636.80029296875 0 12650.79
1637.8043212890625 0 8898.307
1638.8128662109375 0 3306.7485
1643.8343505859375 0 9400.425
1644.836669921875 0 61716.875 c 13
1645.8389892578125 0 56515.07
1646.84326171875 0 30186.107
1647.8441162109375 0 12757.188
1648.849365234375 0 4613.547
1670.857666015625 0 2965.1116
1671.863525390625 0 1814.6116
1687.8431396484375 0 2149.5867
1688.8626708984375 0 5912.0605
1689.862548828125 0 4710.568
1690.8714599609375 0 1892.1803
1715.859130859375 0 4630.043
1716.845703125 0 3530.4958 z 2
1718.870849609375 0 1910.332
1727.875 0 2643.3457
1728.8914794921875 0 6054.4653
1729.89599609375 0 5108.745
1730.8675537109375 0 3198.343
1731.870361328125 0 31125.902 c 14
1732.8729248046875 0 51329.555 y 2
1733.87451171875 0 43640.375
1734.87451171875 0 21740.035
1735.8746337890625 0 9140.038
1785.911376953125 0 3582.9956
1786.9183349609375 0 3633.2122
1787.9312744140625 0 2287.3105
1800.9302978515625 0 2871.6125
1801.947021484375 0 12615.097
1802.9495849609375 0 14736.4795
1803.9451904296875 0 10379.389
1804.94287109375 0 4720.5386
1828.9290771484375 0 6734.397 y Ammonia loss 1
1829.926513671875 0 15421.049 z 1
1830.92919921875 0 51795.395
1831.927734375 0 53592.105
1832.927734375 0 27453.912 c 15
1833.9227294921875 0 15824.515
1834.93359375 0 5583.0083
1844.9534912109375 0 15453.165
1845.95703125 0 41147.08 y 1
1846.9620361328125 0 41964.91
1847.9627685546875 0 27599.635
1848.9637451171875 0 10086.761
1849.9461669921875 0 3950.2148
1850.92431640625 0 2248.9358
1870.9439697265625 0 3215.961
1871.934326171875 0 2047.3268
1872.9515380859375 0 2763.6658
1874.911865234375 0 1781.0321
1877 0 1730.5065
1884.992919921875 0 4672.1533
1886.00634765625 0 7536.163
1886.974609375 0 11071.512
1887.966552734375 0 8351.681
1888.9669189453125 0 7590.131
1889.959228515625 0 17140.244
1890.9677734375 0 15484.925
1891.966064453125 0 9949.977
1892.9600830078125 0 5019.749
1902.9827880859375 0 12286.151
1903.982421875 0 37767.402
1904.9853515625 0 32262.863
1905.982421875 0 22767.098
1906.9844970703125 0 10468.496
1907.994140625 0 3216.4692
1913.966552734375 0 6502.8647
1914.9593505859375 0 12899.481
1915.9635009765625 0 7337.5156
1916.9671630859375 0 3753.3855
1917.95849609375 0 2873.848
1919.9896240234375 0 4729.301
1921.00537109375 0 29834.502
1922.0068359375 0 26691.668
1923.0067138671875 0 16664.623
1924.005859375 0 7278.792
1925.0206298828125 0 3048.7876
1929.9735107421875 0 3494.9924
1930.97900390625 0 35967.98
1931.975341796875 0 130075.9
1932.9769287109375 0 130959.586
1933.978759765625 0 76987.46
1934.979248046875 0 31875.041
1935.9769287109375 0 10300.2
1946.984619140625 0 16272.998
1947.9898681640625 0 86668.19
1948.9959716796875 0 364244.9
1949.999755859375 0 349882.56
1951.00146484375 0 226787.73
1952.0020751953125 0 97346.13
1953.0040283203125 0 31357.893
2522.538330078125 0 1857.1827

Spectrum Details

|  |  |
| --- | --- |
| Matched peaks? Matched peaks The total absolute number of peaks matched. Additionally in brackets the total fraction of peaks matched and the total number of peaks is shown. | 94 (15.82% of 594) |
| FDR? FDR The false discovery rate estimated for this peptide. It is calculated by matching all theoretical fragments with a non-integer shift with the raw peaks for this spectrum. This is done with 40 different shifts. The resulting percentage is the average number of annotated peaks over the number of annotated peaks with the correct spectrum. | 0.68% |
| Satellite FDR? Satellite FDR See the FDR for details on its calculation. This satellite ion specific FDR only contains the satellite ions (d/w) for I/L/J positions. | 11.90% |
| PSM Score? PSM Score The PSM Score as given by Hecklib to this annotated spectrum. It is shown with three significant figures. | 645 |

#### Spectrum 4280? Spectrum 4280 The raw spectrum of this peptide as annotated by Hecklib. The fragments are coloured according to ion type (see legend). Any peaks with a star '\*' as text can be hovered over to see the full details, first the ion type second the mass shift type. By hovering over the amino acids in the peptide or ions in the legend the corresponding peaks are highlighted. By toggling the 'Unassigned' label you can turn the background (unassigned) peaks on or off in the plot. By updating the slider in the Ion legend you can update the spectrum to only show the top X% of the peaks with labels. The top X% means any peak that is within X% of the highest intensity. By dragging in the spectrum you can zoom in to a specific part of the spectrum and use 'Zoom Out' to get back to the original zoom level. The annotation of the spectrum is based on the given sequence in the peptides file and is done with different software so inconsistencies are likely. The peaks are annotated based on the given sequence, with 20 ppm tolerance.

Copy Data

##### Spectrum 4280 (TSV)

###### Preview

```
Loading example...
```

*Click on the button to copy the data to your clipboard.*

Mz MinMz MaxIntensity Max

WidthHeightPeptide font sizePeptide stroke widthSpectrum font sizeSpectrum stroke widthCompact peptide

Ion legend

wxyz

abcd

OtherUnassignedIon^Charge^_Position_Show for top:%

TLPPSREEMTKNQVSTJ

03.08e+46.17e+49.25e+41.23e+5

Zoom Out

y^+2^_4_y^+1^_2_y^+1^_2_y^+1^_3_y^+1^_3_c^+1^_4_c^+1^_5_z^+1^_5_y^+1^_5_z^+3^_15_c^+3^_15_y^+3^_15_z^+3^_15_y^+3^_15_c^+2^_10_c^+2^_11_z^+1^_6_c^+2^_11_y^+1^_6_c^+1^_6_c^+2^_12_c^+2^_12_c^+2^_12_z^+2^_12_y^+2^_12_z^+1^_7_y^+2^_13_z^+2^_13_y^+2^_13_c^+2^_13_y^+1^_7_w^+2^_14_c^+1^_7_c^+2^_14_c^+2^_14_y^+2^_14_c^+2^_14_w^+2^_15_w^+1^_8_c^+2^_15_y^+2^_15_z^+2^_15_w^+1^_8_c^+2^_15_y^+2^_15_y^+1^_8_z^+1^_8_y^+1^_8_w^+2^_16_z^+2^_16_c^+2^_16_y^+2^_16_y^+2^_16_z^+2^_16_c^+2^_16_y^+2^_16_c^+1^_8_z^+1^_9_z^+1^_9_y^+1^_9_c^+1^_9_w^+1^_10_z^+1^_10_y^+1^_10_c^+1^_10_w^+1^_11_z^+1^_11_c^+1^_11_c^+1^_11_y^+1^_11_c^+1^_11_c^+1^_12_z^+1^_12_y^+1^_12_z^+1^_13_y^+1^_13_c^+1^_13_c^+1^_14_c^+1^_14_y^+1^_14_c^+1^_14_y^+1^_15_z^+1^_15_c^+1^_15_y^+1^_15_y^+1^_16_z^+1^_16_c^+1^_16_y^+1^_16_

0778155623333111

Fragment Matches Table

Show background peaks

| Position | Ion type | Intensity | mz Theoretical | mz Error (Th) | mz Error (ppm) | Charge | Series Number |
| --- | --- | --- | --- | --- | --- | --- | --- |
| - | - | 3450 | 120.1 | - | - | 0 | - |
| - | - | 344 | 121.1 | - | - | 0 | - |
| - | - | 382.5 | 130.9 | - | - | 0 | - |
| - | - | 417.9 | 133 | - | - | 0 | - |
| - | - | 396.8 | 137 | - | - | 0 | - |
| - | - | 414.7 | 141.2 | - | - | 0 | - |
| - | - | 446.1 | 149 | - | - | 0 | - |
| - | - | 456.1 | 170.9 | - | - | 0 | - |
| - | - | 1917 | 173.4 | - | - | 0 | - |
| - | - | 3.404E+04 | 187.1 | - | - | 0 | - |
| - | - | 3012 | 188.1 | - | - | 0 | - |
| 14 | y | 1648 | 201.1 | 0.0006241 | 3.103 | +2 | 4 |
| - | - | 1301 | 212.1 | - | - | 0 | - |
| 16 | y | 2.261E+04 | 215.1 | 0.0002938 | 1.366 | +1 | 2 |
| - | - | 1468 | 216.1 | - | - | 0 | - |
| - | - | 575.9 | 228 | - | - | 0 | - |
| - | - | 554.9 | 228.4 | - | - | 0 | - |
| 16 | y | 1629 | 233.1 | 0.0001814 | 0.7781 | +1 | 2 |
| - | - | 582 | 292.9 | - | - | 0 | - |
| - | - | 3000 | 299.2 | - | - | 0 | - |
| - | - | 712.2 | 300.2 | - | - | 0 | - |
| 15 | y | 703.5 | 302.2 | 0.0008887 | 2.941 | +1 | 3 |
| - | - | 959.1 | 312.2 | - | - | 0 | - |
| 15 | y | 1079 | 320.2 | 0.000185 | 0.5778 | +1 | 3 |
| - | - | 591 | 328.2 | - | - | 0 | - |
| - | - | 493.4 | 329.9 | - | - | 0 | - |
| - | - | 642.6 | 374.4 | - | - | 0 | - |
| - | - | 657.7 | 382.3 | - | - | 0 | - |
| - | - | 1318 | 407.3 | - | - | 0 | - |
| - | - | 623 | 412.2 | - | - | 0 | - |
| 4 | c | 976.1 | 426.3 | 0.001426 | 3.346 | +1 | 4 |
| - | - | 1793 | 429.1 | - | - | 0 | - |
| - | - | 1364 | 470.3 | - | - | 0 | - |
| - | - | 994 | 471.9 | - | - | 0 | - |
| - | - | 577.8 | 479.5 | - | - | 0 | - |
| - | - | 951.1 | 494.3 | - | - | 0 | - |
| - | - | 799.4 | 495.2 | - | - | 0 | - |
| - | - | 703.9 | 500.2 | - | - | 0 | - |
| - | - | 1840 | 500.9 | - | - | 0 | - |
| - | - | 1242 | 501.2 | - | - | 0 | - |
| - | - | 662.7 | 501.6 | - | - | 0 | - |
| - | - | 790.3 | 512.3 | - | - | 0 | - |
| 5 | c | 2.33E+04 | 513.3 | 0.0007089 | 1.381 | +1 | 5 |
| - | - | 6506 | 514.3 | - | - | 0 | - |
| - | - | 1247 | 515.3 | - | - | 0 | - |
| - | - | 1076 | 523.6 | - | - | 0 | - |
| - | - | 1158 | 527.3 | - | - | 0 | - |
| - | - | 1569 | 529.3 | - | - | 0 | - |
| - | - | 875.3 | 529.3 | - | - | 0 | - |
| - | - | 1513 | 529.6 | - | - | 0 | - |
| - | - | 1119 | 529.9 | - | - | 0 | - |
| 13 | z | 2454 | 531.3 | 0.0005869 | 1.105 | +1 | 5 |
| - | - | 1335 | 532.3 | - | - | 0 | - |
| - | - | 2002 | 532.6 | - | - | 0 | - |
| - | - | 1873 | 532.9 | - | - | 0 | - |
| - | - | 1907 | 533.3 | - | - | 0 | - |
| - | - | 651 | 533.3 | - | - | 0 | - |
| - | - | 1038 | 533.6 | - | - | 0 | - |
| - | - | 5616 | 538.6 | - | - | 0 | - |
| - | - | 5185 | 538.9 | - | - | 0 | - |
| - | - | 3086 | 539.3 | - | - | 0 | - |
| - | - | 1087 | 539.6 | - | - | 0 | - |
| 13 | y | 692.9 | 547.3 | 0.0009669 | 1.767 | +1 | 5 |
| - | - | 1644 | 557 | - | - | 0 | - |
| - | - | 1493 | 557.3 | - | - | 0 | - |
| - | - | 1360 | 557.6 | - | - | 0 | - |
| - | - | 1037 | 558 | - | - | 0 | - |
| - | - | 763.9 | 558.3 | - | - | 0 | - |
| - | - | 572.8 | 563.6 | - | - | 0 | - |
| - | - | 648.2 | 564.2 | - | - | 0 | - |
| 3 | z | 1098 | 566.9 | 0.001759 | 3.102 | +3 | 15 |
| - | - | 2927 | 571.3 | - | - | 0 | - |
| 15 | c | 4434 | 572.3 | 0.00276 | 4.823 | +3 | 15 |
| 3 | y | 5016 | 572.6 | 0.007519 | 13.13 | +3 | 15 |
| 3 | z | 3850 | 572.9 | 0.003547 | 6.191 | +3 | 15 |
| - | - | 1129 | 573.3 | - | - | 0 | - |
| - | - | 801.2 | 573.3 | - | - | 0 | - |
| - | - | 895.4 | 573.6 | - | - | 0 | - |
| 3 | y | 9.515E+04 | 578.3 | 0.002534 | 4.383 | +3 | 15 |
| - | - | 9.254E+04 | 578.6 | - | - | 0 | - |
| - | - | 5.618E+04 | 579 | - | - | 0 | - |
| 10 | c | 2.524E+04 | 579.3 | 0.009051 | 15.62 | +2 | 10 |
| - | - | 7000 | 579.6 | - | - | 0 | - |
| - | - | 2024 | 580 | - | - | 0 | - |
| - | - | 1142 | 607.4 | - | - | 0 | - |
| - | - | 761.4 | 608.4 | - | - | 0 | - |
| - | - | 3090 | 625.4 | - | - | 0 | - |
| - | - | 1331 | 626.4 | - | - | 0 | - |
| 11 | c | 770.4 | 643.3 | 0.00466 | 7.243 | +2 | 11 |
| - | - | 640.6 | 644 | - | - | 0 | - |
| 12 | z | 1417 | 645.3 | 0.001361 | 2.108 | +1 | 6 |
| - | - | 2077 | 646.3 | - | - | 0 | - |
| - | - | 1819 | 649.9 | - | - | 0 | - |
| - | - | 1041 | 650.4 | - | - | 0 | - |
| - | - | 2788 | 651.4 | - | - | 0 | - |
| 11 | c | 3597 | 652.3 | 0.00304 | 4.66 | +2 | 11 |
| - | - | 4025 | 652.8 | - | - | 0 | - |
| - | - | 1477 | 653.3 | - | - | 0 | - |
| 12 | y | 3561 | 661.4 | 0.000703 | 1.063 | +1 | 6 |
| - | - | 1464 | 662.3 | - | - | 0 | - |
| 6 | c | 3.946E+04 | 669.4 | 0.001038 | 1.551 | +1 | 6 |
| - | - | 1.43E+04 | 670.4 | - | - | 0 | - |
| - | - | 2988 | 671.4 | - | - | 0 | - |
| - | - | 1088 | 672.4 | - | - | 0 | - |
| - | - | 610.5 | 673.4 | - | - | 0 | - |
| - | - | 1025 | 695.4 | - | - | 0 | - |
| 12 | c | 876.2 | 700.4 | 0.0006295 | 0.8988 | +2 | 12 |
| 12 | c | 815.8 | 700.8 | 0.008034 | 11.46 | +2 | 12 |
| - | - | 3628 | 707.3 | - | - | 0 | - |
| - | - | 2797 | 707.8 | - | - | 0 | - |
| - | - | 1673 | 708.3 | - | - | 0 | - |
| - | - | 3137 | 708.9 | - | - | 0 | - |
| 12 | c | 2.476E+04 | 709.4 | 0.002633 | 3.712 | +2 | 12 |
| - | - | 1.595E+04 | 709.9 | - | - | 0 | - |
| - | - | 9396 | 710.4 | - | - | 0 | - |
| - | - | 4551 | 710.9 | - | - | 0 | - |
| - | - | 1030 | 711.4 | - | - | 0 | - |
| - | - | 619.1 | 715.4 | - | - | 0 | - |
| 6 | z | 1497 | 718.3 | 0.006692 | 9.316 | +2 | 12 |
| - | - | 2903 | 718.9 | - | - | 0 | - |
| - | - | 927.9 | 719.4 | - | - | 0 | - |
| - | - | 969.4 | 719.9 | - | - | 0 | - |
| - | - | 1713 | 722.4 | - | - | 0 | - |
| - | - | 708.8 | 723.4 | - | - | 0 | - |
| - | - | 826.5 | 725.9 | - | - | 0 | - |
| 6 | y | 4741 | 726.4 | 0.001908 | 2.626 | +2 | 12 |
| - | - | 4018 | 726.9 | - | - | 0 | - |
| - | - | 1791 | 727.4 | - | - | 0 | - |
| - | - | 668.6 | 728.4 | - | - | 0 | - |
| - | - | 4819 | 729.4 | - | - | 0 | - |
| - | - | 2268 | 730.4 | - | - | 0 | - |
| - | - | 807.2 | 742.9 | - | - | 0 | - |
| - | - | 2733 | 750.9 | - | - | 0 | - |
| - | - | 3571 | 751.4 | - | - | 0 | - |
| - | - | 2180 | 751.9 | - | - | 0 | - |
| - | - | 1063 | 752.4 | - | - | 0 | - |
| 11 | z | 850.2 | 755.4 | 0.007539 | 9.979 | +1 | 7 |
| - | - | 809.1 | 758.9 | - | - | 0 | - |
| - | - | 640.7 | 759.4 | - | - | 0 | - |
| 5 | y | 684.3 | 761.4 | 0.0006938 | 0.9113 | +2 | 13 |
| 5 | z | 2003 | 761.9 | 0.004838 | 6.35 | +2 | 13 |
| - | - | 4391 | 762.4 | - | - | 0 | - |
| - | - | 2639 | 762.9 | - | - | 0 | - |
| - | - | 1636 | 763.4 | - | - | 0 | - |
| - | - | 865.6 | 765.4 | - | - | 0 | - |
| 5 | y | 2156 | 769.9 | 0.002861 | 3.716 | +2 | 13 |
| - | - | 1661 | 770.4 | - | - | 0 | - |
| - | - | 853.7 | 770.9 | - | - | 0 | - |
| - | - | 2068 | 772.9 | - | - | 0 | - |
| 13 | c | 3.05E+04 | 773.4 | 0.004961 | 6.414 | +2 | 13 |
| - | - | 6264 | 773.5 | - | - | 0 | - |
| - | - | 3.155E+04 | 773.9 | - | - | 0 | - |
| - | - | 1.223E+04 | 774.4 | - | - | 0 | - |
| - | - | 2836 | 774.4 | - | - | 0 | - |
| - | - | 6467 | 774.9 | - | - | 0 | - |
| - | - | 1807 | 775.4 | - | - | 0 | - |
| - | - | 722.3 | 779.4 | - | - | 0 | - |
| 11 | y | 5007 | 789.4 | 0.0005276 | 0.6683 | +1 | 7 |
| - | - | 1781 | 790.4 | - | - | 0 | - |
| - | - | 1418 | 791.9 | - | - | 0 | - |
| - | - | 847.6 | 792.9 | - | - | 0 | - |
| - | - | 3266 | 793.4 | - | - | 0 | - |
| - | - | 2660 | 793.9 | - | - | 0 | - |
| - | - | 1867 | 794.4 | - | - | 0 | - |
| - | - | 787.9 | 794.9 | - | - | 0 | - |
| - | - | 683 | 796.2 | - | - | 0 | - |
| 4 | w | 678.3 | 796.9 | 0.007406 | 9.293 | +2 | 14 |
| 7 | c | 5.681E+04 | 798.4 | 0.001109 | 1.389 | +1 | 7 |
| - | - | 2.454E+04 | 799.5 | - | - | 0 | - |
| - | - | 877.5 | 799.9 | - | - | 0 | - |
| - | - | 6188 | 800.5 | - | - | 0 | - |
| - | - | 2046 | 800.9 | - | - | 0 | - |
| - | - | 3241 | 801.4 | - | - | 0 | - |
| - | - | 2028 | 801.9 | - | - | 0 | - |
| - | - | 7417 | 807.4 | - | - | 0 | - |
| - | - | 8511 | 807.9 | - | - | 0 | - |
| - | - | 3938 | 808.4 | - | - | 0 | - |
| - | - | 2455 | 808.9 | - | - | 0 | - |
| - | - | 7818 | 813.4 | - | - | 0 | - |
| 14 | c | 1.221E+04 | 813.9 | 0.0004655 | 0.5719 | +2 | 14 |
| 14 | c | 7532 | 814.4 | 0.006245 | 7.668 | +2 | 14 |
| - | - | 2785 | 814.9 | - | - | 0 | - |
| - | - | 2380 | 815.4 | - | - | 0 | - |
| - | - | 687.8 | 817.4 | - | - | 0 | - |
| - | - | 700.4 | 817.9 | - | - | 0 | - |
| 4 | y | 1.842E+04 | 818.4 | 0.003884 | 4.746 | +2 | 14 |
| - | - | 1.398E+04 | 818.9 | - | - | 0 | - |
| - | - | 5794 | 819.4 | - | - | 0 | - |
| - | - | 3411 | 819.9 | - | - | 0 | - |
| - | - | 1.505E+04 | 822.4 | - | - | 0 | - |
| 14 | c | 2.562E+04 | 822.9 | 0.001332 | 1.619 | +2 | 14 |
| - | - | 2.219E+04 | 823.4 | - | - | 0 | - |
| - | - | 1.034E+04 | 823.9 | - | - | 0 | - |
| - | - | 4152 | 824.4 | - | - | 0 | - |
| - | - | 1448 | 824.9 | - | - | 0 | - |
| - | - | 775.5 | 825.5 | - | - | 0 | - |
| - | - | 1691 | 830.5 | - | - | 0 | - |
| - | - | 1890 | 834.9 | - | - | 0 | - |
| - | - | 3589 | 835.4 | - | - | 0 | - |
| - | - | 2250 | 835.9 | - | - | 0 | - |
| - | - | 1949 | 836.4 | - | - | 0 | - |
| - | - | 665.5 | 836.9 | - | - | 0 | - |
| - | - | 735.3 | 843.4 | - | - | 0 | - |
| - | - | 676.2 | 844.4 | - | - | 0 | - |
| 3 | w | 1259 | 845.4 | 0.006231 | 7.371 | +2 | 15 |
| - | - | 1529 | 845.9 | - | - | 0 | - |
| - | - | 1350 | 846.4 | - | - | 0 | - |
| - | - | 1022 | 849.4 | - | - | 0 | - |
| 10 | w | 3538 | 857.5 | 0.001334 | 1.556 | +1 | 8 |
| 15 | c | 4181 | 857.9 | 0.003475 | 4.051 | +2 | 15 |
| 3 | y | 5282 | 858.4 | 0.01568 | 18.27 | +2 | 15 |
| 3 | z | 2298 | 858.9 | 0.005175 | 6.025 | +2 | 15 |
| 10 | w | 5398 | 859.5 | 0.004219 | 4.909 | +1 | 8 |
| - | - | 750.4 | 859.9 | - | - | 0 | - |
| - | - | 2539 | 860.5 | - | - | 0 | - |
| - | - | 953.5 | 864.9 | - | - | 0 | - |
| - | - | 2535 | 865.9 | - | - | 0 | - |
| 15 | c | 4.342E+04 | 866.4 | 0.003323 | 3.836 | +2 | 15 |
| 3 | y | 1.042E+05 | 866.9 | 0.006555 | 7.561 | +2 | 15 |
| - | - | 8.287E+04 | 867.4 | - | - | 0 | - |
| - | - | 5.178E+04 | 867.9 | - | - | 0 | - |
| - | - | 2.027E+04 | 868.4 | - | - | 0 | - |
| - | - | 5222 | 868.9 | - | - | 0 | - |
| - | - | 778.4 | 869.4 | - | - | 0 | - |
| 10 | y | 5000 | 873.5 | 0.0003943 | 0.4514 | +1 | 8 |
| 10 | z | 2.98E+04 | 874.5 | 0.0003207 | 0.3668 | +1 | 8 |
| - | - | 1.688E+04 | 875.5 | - | - | 0 | - |
| - | - | 5660 | 876.5 | - | - | 0 | - |
| - | - | 1906 | 879.4 | - | - | 0 | - |
| - | - | 1639 | 880 | - | - | 0 | - |
| - | - | 974.3 | 880.4 | - | - | 0 | - |
| - | - | 1007 | 880.9 | - | - | 0 | - |
| - | - | 1013 | 881.4 | - | - | 0 | - |
| - | - | 927.6 | 881.9 | - | - | 0 | - |
| - | - | 1532 | 883.5 | - | - | 0 | - |
| - | - | 1208 | 884.5 | - | - | 0 | - |
| - | - | 737.9 | 887.4 | - | - | 0 | - |
| 10 | y | 9377 | 890.5 | 3.175E-05 | 0.03565 | +1 | 8 |
| - | - | 5632 | 891.5 | - | - | 0 | - |
| - | - | 2222 | 892 | - | - | 0 | - |
| - | - | 2134 | 892.5 | - | - | 0 | - |
| - | - | 1011 | 893 | - | - | 0 | - |
| 2 | w | 1.609E+04 | 893.9 | 0.003897 | 4.359 | +2 | 16 |
| - | - | 1.784E+04 | 894.4 | - | - | 0 | - |
| - | - | 1.137E+04 | 894.9 | - | - | 0 | - |
| - | - | 5286 | 895.4 | - | - | 0 | - |
| - | - | 1669 | 895.9 | - | - | 0 | - |
| - | - | 1976 | 901 | - | - | 0 | - |
| - | - | 3819 | 901.5 | - | - | 0 | - |
| - | - | 2835 | 902 | - | - | 0 | - |
| - | - | 1158 | 902.5 | - | - | 0 | - |
| - | - | 1095 | 903 | - | - | 0 | - |
| - | - | 805.9 | 906 | - | - | 0 | - |
| 2 | z | 941 | 906.5 | 0.01091 | 12.03 | +2 | 16 |
| - | - | 1449 | 907 | - | - | 0 | - |
| - | - | 1490 | 907.5 | - | - | 0 | - |
| 16 | c | 709.2 | 908 | 0.005213 | 5.742 | +2 | 16 |
| - | - | 1213 | 909.5 | - | - | 0 | - |
| 2 | y | 2386 | 914.5 | 0.00667 | 7.294 | +2 | 16 |
| 2 | y | 2207 | 915 | 0.0121 | 13.22 | +2 | 16 |
| 2 | z | 1.036E+04 | 915.5 | 0.006417 | 7.009 | +2 | 16 |
| - | - | 1.431E+04 | 916 | - | - | 0 | - |
| - | - | 7129 | 916.5 | - | - | 0 | - |
| 16 | c | 3286 | 917 | 0.002006 | 2.188 | +2 | 16 |
| - | - | 1779 | 917.5 | - | - | 0 | - |
| - | - | 725.4 | 917.9 | - | - | 0 | - |
| - | - | 1901 | 920.5 | - | - | 0 | - |
| - | - | 1101 | 921 | - | - | 0 | - |
| - | - | 1100 | 921.5 | - | - | 0 | - |
| - | - | 1127 | 922.5 | - | - | 0 | - |
| - | - | 4.924E+04 | 923 | - | - | 0 | - |
| 2 | y | 5.461E+04 | 923.5 | 0.01128 | 12.21 | +2 | 16 |
| - | - | 3.052E+04 | 924 | - | - | 0 | - |
| - | - | 1.766E+04 | 924.5 | - | - | 0 | - |
| - | - | 5286 | 925 | - | - | 0 | - |
| - | - | 1528 | 925.5 | - | - | 0 | - |
| - | - | 1524 | 926.5 | - | - | 0 | - |
| 8 | c | 5.868E+04 | 927.5 | 0.001179 | 1.271 | +1 | 8 |
| - | - | 2.784E+04 | 928.5 | - | - | 0 | - |
| - | - | 1981 | 929 | - | - | 0 | - |
| - | - | 1.024E+04 | 929.5 | - | - | 0 | - |
| - | - | 3458 | 930 | - | - | 0 | - |
| - | - | 3946 | 930.5 | - | - | 0 | - |
| - | - | 2934 | 931 | - | - | 0 | - |
| - | - | 2047 | 931.5 | - | - | 0 | - |
| - | - | 759.1 | 934 | - | - | 0 | - |
| - | - | 945.9 | 934.5 | - | - | 0 | - |
| - | - | 1074 | 935 | - | - | 0 | - |
| - | - | 1270 | 935.5 | - | - | 0 | - |
| - | - | 1207 | 936.5 | - | - | 0 | - |
| - | - | 2251 | 937 | - | - | 0 | - |
| - | - | 2732 | 937.5 | - | - | 0 | - |
| - | - | 1.614E+04 | 938 | - | - | 0 | - |
| - | - | 1.735E+04 | 938.5 | - | - | 0 | - |
| - | - | 1.138E+04 | 939 | - | - | 0 | - |
| - | - | 7574 | 939.5 | - | - | 0 | - |
| - | - | 2376 | 940 | - | - | 0 | - |
| - | - | 1038 | 940.5 | - | - | 0 | - |
| - | - | 3455 | 942.5 | - | - | 0 | - |
| - | - | 7155 | 943 | - | - | 0 | - |
| - | - | 6001 | 943.5 | - | - | 0 | - |
| - | - | 5466 | 944 | - | - | 0 | - |
| - | - | 4238 | 944.5 | - | - | 0 | - |
| - | - | 1.116E+04 | 945 | - | - | 0 | - |
| - | - | 1E+04 | 945.5 | - | - | 0 | - |
| - | - | 7777 | 946 | - | - | 0 | - |
| - | - | 8378 | 946.5 | - | - | 0 | - |
| - | - | 7641 | 947 | - | - | 0 | - |
| - | - | 4507 | 947.5 | - | - | 0 | - |
| - | - | 1386 | 948 | - | - | 0 | - |
| - | - | 797.2 | 949.5 | - | - | 0 | - |
| - | - | 1720 | 951.5 | - | - | 0 | - |
| - | - | 9324 | 952 | - | - | 0 | - |
| - | - | 1.287E+04 | 952.5 | - | - | 0 | - |
| - | - | 1.037E+04 | 953 | - | - | 0 | - |
| - | - | 8228 | 953.5 | - | - | 0 | - |
| - | - | 3174 | 954 | - | - | 0 | - |
| - | - | 2084 | 954.5 | - | - | 0 | - |
| - | - | 1665 | 955.4 | - | - | 0 | - |
| - | - | 3502 | 956.5 | - | - | 0 | - |
| - | - | 8564 | 957 | - | - | 0 | - |
| - | - | 1.031E+04 | 957.5 | - | - | 0 | - |
| - | - | 5086 | 958 | - | - | 0 | - |
| - | - | 8323 | 958.5 | - | - | 0 | - |
| - | - | 2.433E+04 | 959.5 | - | - | 0 | - |
| - | - | 1.257E+04 | 960.5 | - | - | 0 | - |
| - | - | 4064 | 961.5 | - | - | 0 | - |
| - | - | 776.8 | 962.5 | - | - | 0 | - |
| - | - | 1.356E+04 | 965.5 | - | - | 0 | - |
| - | - | 6.794E+04 | 966 | - | - | 0 | - |
| - | - | 6.936E+04 | 966.5 | - | - | 0 | - |
| - | - | 4.807E+04 | 967 | - | - | 0 | - |
| - | - | 2.139E+04 | 967.5 | - | - | 0 | - |
| - | - | 9404 | 968 | - | - | 0 | - |
| - | - | 3989 | 968.5 | - | - | 0 | - |
| - | - | 1744 | 969.5 | - | - | 0 | - |
| - | - | 1073 | 973.5 | - | - | 0 | - |
| - | - | 654.8 | 973.9 | - | - | 0 | - |
| - | - | 4.704E+04 | 974 | - | - | 0 | - |
| - | - | 8.497E+04 | 974.5 | - | - | 0 | - |
| - | - | 6.919E+04 | 975 | - | - | 0 | - |
| - | - | 4.168E+04 | 975.5 | - | - | 0 | - |
| - | - | 1.775E+04 | 976 | - | - | 0 | - |
| - | - | 6823 | 976.5 | - | - | 0 | - |
| - | - | 958.1 | 977 | - | - | 0 | - |
| - | - | 1475 | 992.4 | - | - | 0 | - |
| 9 | z | 943 | 1003 | 0.01086 | 10.82 | +1 | 9 |
| - | - | 1685 | 1006 | - | - | 0 | - |
| - | - | 860.3 | 1013 | - | - | 0 | - |
| 9 | z | 6092 | 1022 | 0.005237 | 5.126 | +1 | 9 |
| - | - | 5974 | 1023 | - | - | 0 | - |
| - | - | 2442 | 1024 | - | - | 0 | - |
| - | - | 1574 | 1025 | - | - | 0 | - |
| - | - | 6508 | 1031 | - | - | 0 | - |
| - | - | 4695 | 1032 | - | - | 0 | - |
| - | - | 3634 | 1033 | - | - | 0 | - |
| - | - | 768.9 | 1034 | - | - | 0 | - |
| 9 | y | 2692 | 1038 | 0.007081 | 6.825 | +1 | 9 |
| - | - | 2105 | 1039 | - | - | 0 | - |
| - | - | 913.4 | 1040 | - | - | 0 | - |
| - | - | 3434 | 1050 | - | - | 0 | - |
| - | - | 1722 | 1051 | - | - | 0 | - |
| - | - | 868.5 | 1056 | - | - | 0 | - |
| - | - | 995.1 | 1074 | - | - | 0 | - |
| 9 | c | 5.681E+04 | 1075 | 0.005851 | 5.445 | +1 | 9 |
| - | - | 3.67E+04 | 1076 | - | - | 0 | - |
| - | - | 1.449E+04 | 1077 | - | - | 0 | - |
| - | - | 4906 | 1078 | - | - | 0 | - |
| - | - | 805.6 | 1079 | - | - | 0 | - |
| - | - | 864.5 | 1080 | - | - | 0 | - |
| - | - | 1019 | 1082 | - | - | 0 | - |
| - | - | 926.3 | 1082 | - | - | 0 | - |
| - | - | 1722 | 1088 | - | - | 0 | - |
| - | - | 1540 | 1089 | - | - | 0 | - |
| 8 | w | 7736 | 1092 | 0.004939 | 4.525 | +1 | 10 |
| - | - | 3797 | 1093 | - | - | 0 | - |
| - | - | 2773 | 1094 | - | - | 0 | - |
| - | - | 1188 | 1095 | - | - | 0 | - |
| - | - | 1151 | 1095 | - | - | 0 | - |
| - | - | 1019 | 1096 | - | - | 0 | - |
| - | - | 1859 | 1114 | - | - | 0 | - |
| - | - | 1045 | 1115 | - | - | 0 | - |
| - | - | 744.5 | 1126 | - | - | 0 | - |
| - | - | 1363 | 1132 | - | - | 0 | - |
| - | - | 1399 | 1133 | - | - | 0 | - |
| - | - | 734.8 | 1134 | - | - | 0 | - |
| - | - | 1479 | 1135 | - | - | 0 | - |
| 8 | z | 2.335E+04 | 1151 | 0.005795 | 5.037 | +1 | 10 |
| - | - | 1.905E+04 | 1152 | - | - | 0 | - |
| - | - | 7611 | 1153 | - | - | 0 | - |
| - | - | 3320 | 1154 | - | - | 0 | - |
| - | - | 887.1 | 1155 | - | - | 0 | - |
| 8 | y | 1333 | 1167 | 0.001109 | 0.9509 | +1 | 10 |
| - | - | 1538 | 1168 | - | - | 0 | - |
| 10 | c | 3.074E+04 | 1176 | 0.004925 | 4.19 | +1 | 10 |
| - | - | 1.883E+04 | 1177 | - | - | 0 | - |
| - | - | 7734 | 1178 | - | - | 0 | - |
| - | - | 2642 | 1179 | - | - | 0 | - |
| - | - | 1586 | 1180 | - | - | 0 | - |
| 7 | w | 4434 | 1221 | 0.001287 | 1.054 | +1 | 11 |
| - | - | 3069 | 1222 | - | - | 0 | - |
| - | - | 1672 | 1223 | - | - | 0 | - |
| - | - | 890.2 | 1224 | - | - | 0 | - |
| - | - | 741.6 | 1242 | - | - | 0 | - |
| - | - | 1620 | 1260 | - | - | 0 | - |
| - | - | 2093 | 1261 | - | - | 0 | - |
| - | - | 902.2 | 1262 | - | - | 0 | - |
| 7 | z | 2.054E+04 | 1280 | 0.005194 | 4.059 | +1 | 11 |
| - | - | 1.44E+04 | 1281 | - | - | 0 | - |
| - | - | 6482 | 1282 | - | - | 0 | - |
| - | - | 1924 | 1283 | - | - | 0 | - |
| 11 | c | 1093 | 1286 | 0.0008597 | 0.6687 | +1 | 11 |
| 11 | c | 863.5 | 1287 | 0.01012 | 7.865 | +1 | 11 |
| 7 | y | 6255 | 1296 | 0.004781 | 3.69 | +1 | 11 |
| - | - | 5835 | 1297 | - | - | 0 | - |
| - | - | 3411 | 1298 | - | - | 0 | - |
| - | - | 957.2 | 1299 | - | - | 0 | - |
| - | - | 2845 | 1303 | - | - | 0 | - |
| 11 | c | 6928 | 1304 | 0.003346 | 2.567 | +1 | 11 |
| - | - | 4141 | 1305 | - | - | 0 | - |
| - | - | 2329 | 1306 | - | - | 0 | - |
| - | - | 749.7 | 1307 | - | - | 0 | - |
| - | - | 936.9 | 1318 | - | - | 0 | - |
| - | - | 1717 | 1335 | - | - | 0 | - |
| - | - | 1123 | 1336 | - | - | 0 | - |
| - | - | 920.5 | 1374 | - | - | 0 | - |
| - | - | 1010 | 1377 | - | - | 0 | - |
| - | - | 1837 | 1417 | - | - | 0 | - |
| 12 | c | 1.151E+04 | 1418 | 0.003143 | 2.217 | +1 | 12 |
| - | - | 9415 | 1419 | - | - | 0 | - |
| - | - | 3552 | 1420 | - | - | 0 | - |
| - | - | 2227 | 1421 | - | - | 0 | - |
| - | - | 893.3 | 1423 | - | - | 0 | - |
| 6 | z | 1.398E+04 | 1436 | 0.004303 | 2.997 | +1 | 12 |
| - | - | 2.744E+04 | 1437 | - | - | 0 | - |
| - | - | 1.562E+04 | 1438 | - | - | 0 | - |
| - | - | 7204 | 1439 | - | - | 0 | - |
| - | - | 2926 | 1440 | - | - | 0 | - |
| - | - | 1066 | 1441 | - | - | 0 | - |
| - | - | 2441 | 1451 | - | - | 0 | - |
| 6 | y | 9912 | 1452 | 0.002058 | 1.418 | +1 | 12 |
| - | - | 5603 | 1453 | - | - | 0 | - |
| - | - | 3976 | 1454 | - | - | 0 | - |
| - | - | 1705 | 1455 | - | - | 0 | - |
| - | - | 1001 | 1504 | - | - | 0 | - |
| 5 | z | 2225 | 1523 | 0.000595 | 0.3907 | +1 | 13 |
| - | - | 6197 | 1524 | - | - | 0 | - |
| - | - | 5594 | 1525 | - | - | 0 | - |
| - | - | 2266 | 1526 | - | - | 0 | - |
| - | - | 1480 | 1527 | - | - | 0 | - |
| 5 | y | 2307 | 1539 | 0.002501 | 1.625 | +1 | 13 |
| - | - | 1109 | 1540 | - | - | 0 | - |
| - | - | 731.1 | 1545 | - | - | 0 | - |
| 13 | c | 1.566E+04 | 1546 | 0.002793 | 1.807 | +1 | 13 |
| - | - | 1.395E+04 | 1547 | - | - | 0 | - |
| - | - | 9137 | 1548 | - | - | 0 | - |
| - | - | 3899 | 1549 | - | - | 0 | - |
| - | - | 1606 | 1602 | - | - | 0 | - |
| - | - | 1659 | 1603 | - | - | 0 | - |
| - | - | 1382 | 1615 | - | - | 0 | - |
| - | - | 953.8 | 1616 | - | - | 0 | - |
| - | - | 757.7 | 1622 | - | - | 0 | - |
| 14 | c | 1055 | 1627 | 0.009153 | 5.626 | +1 | 14 |
| 14 | c | 2067 | 1628 | 0.01427 | 8.768 | +1 | 14 |
| - | - | 1624 | 1629 | - | - | 0 | - |
| - | - | 1437 | 1630 | - | - | 0 | - |
| - | - | 1012 | 1631 | - | - | 0 | - |
| 4 | y | 4661 | 1636 | 0.001129 | 0.6899 | +1 | 14 |
| - | - | 3820 | 1637 | - | - | 0 | - |
| - | - | 3910 | 1638 | - | - | 0 | - |
| - | - | 2036 | 1639 | - | - | 0 | - |
| - | - | 2728 | 1644 | - | - | 0 | - |
| 14 | c | 2.015E+04 | 1645 | 0.004204 | 2.556 | +1 | 14 |
| - | - | 1.847E+04 | 1646 | - | - | 0 | - |
| - | - | 1.081E+04 | 1647 | - | - | 0 | - |
| - | - | 5144 | 1648 | - | - | 0 | - |
| - | - | 1941 | 1649 | - | - | 0 | - |
| - | - | 1051 | 1689 | - | - | 0 | - |
| - | - | 1983 | 1690 | - | - | 0 | - |
| 3 | y | 1641 | 1716 | 0.02655 | 15.47 | +1 | 15 |
| 3 | z | 1893 | 1717 | 0.03191 | 18.59 | +1 | 15 |
| - | - | 962.8 | 1718 | - | - | 0 | - |
| - | - | 884.3 | 1728 | - | - | 0 | - |
| - | - | 1583 | 1729 | - | - | 0 | - |
| - | - | 1920 | 1730 | - | - | 0 | - |
| - | - | 957.7 | 1731 | - | - | 0 | - |
| 15 | c | 8494 | 1732 | 0.006599 | 3.81 | +1 | 15 |
| 3 | y | 1.745E+04 | 1733 | 0.02417 | 13.95 | +1 | 15 |
| - | - | 1.528E+04 | 1734 | - | - | 0 | - |
| - | - | 8604 | 1735 | - | - | 0 | - |
| - | - | 3252 | 1736 | - | - | 0 | - |
| - | - | 829.7 | 1787 | - | - | 0 | - |
| - | - | 947.4 | 1788 | - | - | 0 | - |
| - | - | 836.6 | 1801 | - | - | 0 | - |
| - | - | 3665 | 1802 | - | - | 0 | - |
| - | - | 4491 | 1803 | - | - | 0 | - |
| - | - | 3028 | 1804 | - | - | 0 | - |
| - | - | 1511 | 1805 | - | - | 0 | - |
| 2 | y | 1647 | 1829 | 0.02232 | 12.2 | +1 | 16 |
| 2 | z | 4852 | 1830 | 0.01413 | 7.721 | +1 | 16 |
| - | - | 1.793E+04 | 1831 | - | - | 0 | - |
| - | - | 1.549E+04 | 1832 | - | - | 0 | - |
| 16 | c | 9920 | 1833 | 0.01519 | 8.29 | +1 | 16 |
| - | - | 4386 | 1834 | - | - | 0 | - |
| - | - | 1574 | 1835 | - | - | 0 | - |
| - | - | 4678 | 1845 | - | - | 0 | - |
| 2 | y | 1.219E+04 | 1846 | 0.02568 | 13.91 | +1 | 16 |
| - | - | 1.238E+04 | 1847 | - | - | 0 | - |
| - | - | 7792 | 1848 | - | - | 0 | - |
| - | - | 4251 | 1849 | - | - | 0 | - |
| - | - | 1394 | 1850 | - | - | 0 | - |
| - | - | 1340 | 1871 | - | - | 0 | - |
| - | - | 1063 | 1872 | - | - | 0 | - |
| - | - | 898.9 | 1873 | - | - | 0 | - |
| - | - | 918.4 | 1875 | - | - | 0 | - |
| - | - | 762.6 | 1877 | - | - | 0 | - |
| - | - | 1988 | 1885 | - | - | 0 | - |
| - | - | 870.5 | 1886 | - | - | 0 | - |
| - | - | 3654 | 1887 | - | - | 0 | - |
| - | - | 2650 | 1888 | - | - | 0 | - |
| - | - | 3108 | 1889 | - | - | 0 | - |
| - | - | 7226 | 1890 | - | - | 0 | - |
| - | - | 5348 | 1891 | - | - | 0 | - |
| - | - | 2300 | 1892 | - | - | 0 | - |
| - | - | 1641 | 1893 | - | - | 0 | - |
| - | - | 971.4 | 1902 | - | - | 0 | - |
| - | - | 3663 | 1903 | - | - | 0 | - |
| - | - | 1.171E+04 | 1904 | - | - | 0 | - |
| - | - | 1.196E+04 | 1905 | - | - | 0 | - |
| - | - | 8134 | 1906 | - | - | 0 | - |
| - | - | 3508 | 1907 | - | - | 0 | - |
| - | - | 1332 | 1908 | - | - | 0 | - |
| - | - | 2701 | 1914 | - | - | 0 | - |
| - | - | 4286 | 1915 | - | - | 0 | - |
| - | - | 2763 | 1916 | - | - | 0 | - |
| - | - | 2734 | 1917 | - | - | 0 | - |
| - | - | 860.4 | 1918 | - | - | 0 | - |
| - | - | 2347 | 1920 | - | - | 0 | - |
| - | - | 9149 | 1921 | - | - | 0 | - |
| - | - | 8750 | 1922 | - | - | 0 | - |
| - | - | 5565 | 1923 | - | - | 0 | - |
| - | - | 2264 | 1924 | - | - | 0 | - |
| - | - | 1052 | 1930 | - | - | 0 | - |
| - | - | 1.126E+04 | 1931 | - | - | 0 | - |
| - | - | 3.969E+04 | 1932 | - | - | 0 | - |
| - | - | 4.15E+04 | 1933 | - | - | 0 | - |
| - | - | 2.449E+04 | 1934 | - | - | 0 | - |
| - | - | 1.303E+04 | 1935 | - | - | 0 | - |
| - | - | 4028 | 1936 | - | - | 0 | - |
| - | - | 1173 | 1937 | - | - | 0 | - |
| - | - | 6043 | 1947 | - | - | 0 | - |
| - | - | 2.496E+04 | 1948 | - | - | 0 | - |
| - | - | 1.221E+05 | 1949 | - | - | 0 | - |
| - | - | 1.213E+05 | 1950 | - | - | 0 | - |
| - | - | 8.017E+04 | 1951 | - | - | 0 | - |
| - | - | 3.609E+04 | 1952 | - | - | 0 | - |
| - | - | 1.241E+04 | 1953 | - | - | 0 | - |
| - | - | 4038 | 1954 | - | - | 0 | - |
| - | - | 1018 | 3080 | - | - | 0 | - |

m/z Charge Intensity FragmentType MassShift Position
120.06581115722656 0 3449.856
121.06837463378906 0 343.9661
130.90467834472656 0 382.46304
132.98416137695312 0 417.86526
136.95968627929688 0 396.80505
141.22879028320312 0 414.68253
148.95513916015625 0 446.1221
170.92190551757812 0 456.09854
173.4391326904297 0 1916.8376
187.14447021484375 0 34038.844
188.14781188964844 0 3011.8953
201.12399291992188 0 1647.5219 y Water loss 13
212.13954162597656 0 1300.8916
215.13931274414062 0 22607.21 y Water loss 15
216.14260864257812 0 1467.8481
228.017578125 0 575.8598
228.3984832763672 0 554.8667
233.14976501464844 0 1628.574 y 15
292.9458923339844 0 581.95746
299.171630859375 0 2999.7847
300.17413330078125 0 712.19037
302.17193603515625 0 703.52094 y Water loss 14
312.1927185058594 0 959.0877
320.1814270019531 0 1079.3424 y 14
328.19976806640625 0 591.0399
329.8642272949219 0 493.42056
374.41485595703125 0 642.62933
382.25836181640625 0 657.6737
407.2537536621094 0 1318.3174
412.23095703125 0 622.99603
426.27252197265625 0 976.1289 c 3
429.08935546875 0 1792.9384
470.2972412109375 0 1363.7761
471.89788818359375 0 993.9947
479.496337890625 0 577.8305
494.2852478027344 0 951.0711
495.2345275878906 0 799.36896
500.24432373046875 0 703.88153
500.91009521484375 0 1839.7698
501.24261474609375 0 1241.7776
501.57977294921875 0 662.6589
512.294189453125 0 790.317
513.3038330078125 0 23304.508 c 4
514.3071899414062 0 6506.433
515.30908203125 0 1246.7715
523.5986938476562 0 1076.3123
527.2835083007812 0 1157.9515
529.2720947265625 0 1568.834
529.3191528320312 0 875.3383
529.604248046875 0 1512.7148
529.9425048828125 0 1119.137
531.2904663085938 0 2453.9707 z 12
532.2965698242188 0 1335.2487
532.6013793945312 0 2001.5629
532.9332275390625 0 1872.8654
533.2655639648438 0 1906.6309
533.3056030273438 0 651.0238
533.5999755859375 0 1037.902
538.6039428710938 0 5615.736
538.9376220703125 0 5185.0234
539.271728515625 0 3085.704
539.6041870117188 0 1086.5983
547.3095703125 0 692.86444 y 12
556.9569702148438 0 1644.0046
557.288818359375 0 1492.6764
557.62451171875 0 1359.6417
557.953857421875 0 1037.3463
558.2933959960938 0 763.9293
563.6114501953125 0 572.8347
564.24267578125 0 648.2273
566.9463500976562 0 1098.152 z Water loss 2
571.3186645507812 0 2926.8394
572.2869262695312 0 4434.2974 c Ammonia loss 14
572.6196899414062 0 5015.5396 y Ammonia loss 2
572.95166015625 0 3849.5889 z 2
573.2864990234375 0 1128.6067
573.3355712890625 0 801.2308
573.6119384765625 0 895.3731
578.2902221679688 0 95147.22 y 2
578.6240234375 0 92540.305
578.9577026367188 0 56176.234
579.2911987304688 0 25243.068 c Water loss 9
579.6248168945312 0 6999.616
579.960693359375 0 2023.6522
607.3823852539062 0 1142.332
608.3858032226562 0 761.3621
625.3916015625 0 3090.003
626.3969116210938 0 1330.7701
643.3342895507812 0 770.4448 c Water loss 10
643.9926147460938 0 640.6112
645.3341674804688 0 1417.1885 z 11
646.3406372070312 0 2077.2585
649.8511962890625 0 1818.8292
650.3549194335938 0 1040.9585
651.3585205078125 0 2788.3005
652.3379516601562 0 3596.944 c 10
652.8389282226562 0 4025.2898
653.3400268554688 0 1476.6064
661.3522338867188 0 3561.3894 y 11
662.345947265625 0 1463.9518
669.4052734375 0 39460.773 c 5
670.4080200195312 0 14298.548
671.4102172851562 0 2987.6409
672.40869140625 0 1087.7235
673.3978271484375 0 610.50684
695.42041015625 0 1024.8679
700.3504638671875 0 876.23425 c Water loss 11
700.8511352539062 0 815.7718 c Ammonia loss 11
707.343994140625 0 3628.2212
707.84423828125 0 2797.281
708.3449096679688 0 1673.377
708.8536987304688 0 3137.4353
709.3590087890625 0 24760.87 c 11
709.8601684570312 0 15948.665
710.3602905273438 0 9396.331
710.8607788085938 0 4551.179
711.3643188476562 0 1030.2542
715.3975219726562 0 619.11584
718.3564453125 0 1496.9513 z 5
718.8539428710938 0 2902.7427
719.3570556640625 0 927.9354
719.856201171875 0 969.43396
722.366455078125 0 1712.795
723.37646484375 0 708.8187
725.8580932617188 0 826.4646
726.3610229492188 0 4741.1377 y 5
726.8627319335938 0 4018.0288
727.3614501953125 0 1791.4272
728.4319458007812 0 668.6113
729.4253540039062 0 4819.3726
730.4263916015625 0 2268.3052
742.8624267578125 0 807.2284
750.8623657226562 0 2732.9324
751.3650512695312 0 3570.9204
751.8675537109375 0 2180.0923
752.371826171875 0 1063.2202
755.4247436523438 0 850.1528 z Water loss 10
758.8853149414062 0 809.10187
759.3868408203125 0 640.70575
761.362548828125 0 684.26904 y Ammonia loss 4
761.87060546875 0 2002.8702 z 4
762.3714599609375 0 4390.9717
762.8701782226562 0 2639.269
763.3731689453125 0 1636.1764
765.3949584960938 0 865.61957
769.8779907226562 0 2156.354 y 4
770.3804321289062 0 1661.4536
770.8824462890625 0 853.67633
772.886474609375 0 2068.1082
773.390625 0 30498.686 c 12
773.4520874023438 0 6263.9653
773.8899536132812 0 31553.91
774.390869140625 0 12225.195
774.44873046875 0 2835.7893
774.8912963867188 0 6467.4214
775.3969116210938 0 1806.659
779.4298706054688 0 722.2665
789.447021484375 0 5007.3013 y 10
790.4476318359375 0 1781.3059
791.9093627929688 0 1417.9604
792.9056396484375 0 847.6316
793.4110717773438 0 3265.9949
793.9093627929688 0 2659.5374
794.4093627929688 0 1867.1447
794.8936767578125 0 787.85266
796.2047119140625 0 683.00104
796.8878173828125 0 678.3287 w 3
798.4479370117188 0 56806.93 c 6
799.4506225585938 0 24536.16
799.8931274414062 0 877.45465
800.453369140625 0 6187.9995
800.9153442382812 0 2045.6337
801.4180908203125 0 3240.6082
801.9161987304688 0 2028.2003
807.4024658203125 0 7417.0938
807.9033813476562 0 8510.849
808.4025268554688 0 3937.5745
808.9019775390625 0 2454.7393
813.414306640625 0 7817.5454
813.9141235351562 0 12206.6875 c Water loss 13
814.412841796875 0 7531.582 c Ammonia loss 13
814.9127807617188 0 2784.7031
815.41162109375 0 2380.5
817.4036254882812 0 687.7902
817.9068603515625 0 700.37054
818.4053955078125 0 18421.312 y 3
818.9059448242188 0 13979.475
819.40771484375 0 5793.8965
819.9076538085938 0 3411.0693
822.4190673828125 0 15053.184
822.9212036132812 0 25618.09 c 13
823.4219360351562 0 22185.797
823.9224243164062 0 10344.616
824.4255981445312 0 4152.3057
824.923828125 0 1447.8069
825.4622192382812 0 775.5113
830.4652709960938 0 1690.8074
834.9339599609375 0 1889.6705
835.4306640625 0 3589.3804
835.9317016601562 0 2249.6711
836.4328002929688 0 1949.4705
836.9259033203125 0 665.5252
843.447509765625 0 735.30176
844.426025390625 0 676.19275
845.4130249023438 0 1258.6956 w 2
845.91796875 0 1528.9583
846.4160766601562 0 1349.786
849.4146118164062 0 1021.7401
857.4713745117188 0 3537.8303 w 9
857.9260864257812 0 4181.3047 c Ammonia loss 14
858.4302978515625 0 5282.254 y Ammonia loss 2
858.9237060546875 0 2297.722 z 2
859.44775390625 0 5397.8413 w 9
859.9345703125 0 750.4251
860.453369140625 0 2539.026
864.94140625 0 953.45483
865.9368896484375 0 2535.1687
866.439208984375 0 43420.082 c 14
866.9344482421875 0 104173.586 y 2
867.4345703125 0 82874.445
867.933837890625 0 51783.16
868.4345092773438 0 20271.404
868.9349975585938 0 5221.7466
869.4324340820312 0 778.4083
873.468017578125 0 4999.6426 y Ammonia loss 9
874.4757690429688 0 29801.572 z 9
875.4772338867188 0 16883.916
876.4794921875 0 5659.596
879.4494018554688 0 1906.3727
879.9503784179688 0 1638.5724
880.4450073242188 0 974.3405
880.9380493164062 0 1006.53735
881.433349609375 0 1012.6639
881.9251708984375 0 927.6024
883.4761962890625 0 1532.1233
884.475341796875 0 1207.6768
887.4266357421875 0 737.91296
890.494140625 0 9377.006 y 9
891.4967041015625 0 5632.064
891.9700317382812 0 2222.4219
892.4913940429688 0 2133.9497
892.9728393554688 0 1011.0358
893.9370727539062 0 16092.319 w 1
894.4374389648438 0 17843.963
894.9385375976562 0 11365.03
895.4375610351562 0 5285.595
895.9449462890625 0 1669.4974
900.9754638671875 0 1976.1152
901.4775390625 0 3819.083
901.976318359375 0 2834.6533
902.4769897460938 0 1157.5792
902.9755859375 0 1094.8359
905.9614868164062 0 805.9343
906.4661865234375 0 941.03345 z Water loss 1
906.9718017578125 0 1448.9867
907.470703125 0 1489.8572
907.9596557617188 0 709.1567 c Water loss 15
909.4566040039062 0 1213.0461
914.4713134765625 0 2386.0698 y Water loss 1
914.96875 0 2207.3647 y Ammonia loss 1
915.4669799804688 0 10357.654 z 1
915.9661865234375 0 14309.332
916.4692993164062 0 7129.313
916.9617309570312 0 3286.1907 c 15
917.4600219726562 0 1778.9459
917.9462280273438 0 725.43146
920.4910888671875 0 1901.0117
920.9931640625 0 1101.2329
921.4898681640625 0 1099.7057
922.4868774414062 0 1126.6757
922.9811401367188 0 49240.207
923.481201171875 0 54606.16 y 1
923.9820556640625 0 30519.396
924.4827270507812 0 17661.785
924.9817504882812 0 5286.13
925.47509765625 0 1527.7925
926.4822998046875 0 1523.9187
927.4906005859375 0 58677.477 c 7
928.4932250976562 0 27838.043
928.97265625 0 1981.0518
929.492919921875 0 10242.351
929.9742431640625 0 3458.0203
930.4833374023438 0 3945.5264
930.9732666015625 0 2934.2974
931.4842529296875 0 2047.3479
933.9912109375 0 759.0599
934.4959106445312 0 945.91907
934.9832763671875 0 1073.6174
935.4944458007812 0 1270.4165
936.4756469726562 0 1206.9791
936.9806518554688 0 2250.6865
937.4705200195312 0 2732.2646
937.9588012695312 0 16139.781
938.4615478515625 0 17353.67
938.9647827148438 0 11380.014
939.4652709960938 0 7573.893
939.9691162109375 0 2375.6277
940.484375 0 1037.9524
942.4985961914062 0 3454.6821
942.99462890625 0 7154.72
943.4915161132812 0 6001.363
943.98876953125 0 5465.893
944.4844360351562 0 4238.3086
944.982177734375 0 11155.348
945.4827270507812 0 10000.312
945.9832763671875 0 7776.8013
946.4760131835938 0 8378.108
946.9722290039062 0 7640.9087
947.4712524414062 0 4507.343
947.974609375 0 1386.3744
949.4618530273438 0 797.1958
951.49609375 0 1720.2565
951.9923706054688 0 9324.23
952.4918823242188 0 12874.893
952.9896240234375 0 10370.398
953.4915771484375 0 8227.745
953.9857788085938 0 3173.7483
954.4879150390625 0 2083.8142
955.4273681640625 0 1665.2783
956.4917602539062 0 3501.8335
956.987060546875 0 8564.082
957.489013671875 0 10305.111
957.9855346679688 0 5086.077
958.513671875 0 8323.495
959.5275268554688 0 24327.9
960.5285034179688 0 12572.479
961.53125 0 4064.4358
962.5291748046875 0 776.7675
965.49658203125 0 13559.524
965.9908447265625 0 67937.69
966.4907836914062 0 69357.26
966.9916381835938 0 48070.29
967.49169921875 0 21394.965
967.9906005859375 0 9404.145
968.5186767578125 0 3988.8418
969.5313720703125 0 1744.0946
973.5087890625 0 1072.8082
973.8775024414062 0 654.84705
973.9974975585938 0 47044.74
974.4988403320312 0 84965.13
975.0000610351562 0 69190.9
975.5006103515625 0 41683.434
976.0005493164062 0 17751.484
976.5011596679688 0 6822.947
976.9969482421875 0 958.064
992.4169311523438 0 1475.1206
1003.5062255859375 0 943.0302 z Water loss 8
1005.51416015625 0 1684.919
1013.4981079101562 0 860.25226
1021.5111694335938 0 6092.219 z 8
1022.5161743164062 0 5974.14
1023.51513671875 0 2441.658
1024.517333984375 0 1574.4753
1030.5133056640625 0 6508.0366
1031.5159912109375 0 4695.4897
1032.51806640625 0 3633.997
1033.5234375 0 768.90546
1037.53173828125 0 2691.9082 y 8
1038.5302734375 0 2105.4077
1039.5294189453125 0 913.3579
1049.528564453125 0 3434.155
1050.5267333984375 0 1722.1377
1055.50439453125 0 868.516
1073.518310546875 0 995.06793
1074.5257568359375 0 56805.44 c 8
1075.5289306640625 0 36702.535
1076.529541015625 0 14486.085
1077.5301513671875 0 4905.9565
1078.5333251953125 0 805.55225
1079.5174560546875 0 864.4737
1081.8031005859375 0 1018.5354
1082.1439208984375 0 926.29816
1087.5640869140625 0 1721.8994
1088.5662841796875 0 1540.2441
1091.5401611328125 0 7735.951 w 7
1092.541015625 0 3797.2058
1093.540283203125 0 2773.4548
1094.50537109375 0 1187.8519
1095.490966796875 0 1150.882
1096.4873046875 0 1018.5489
1113.554443359375 0 1858.8015
1114.5438232421875 0 1044.641
1125.995849609375 0 744.5463
1131.560791015625 0 1363.4692
1132.5667724609375 0 1399.4966
1133.569580078125 0 734.7813
1134.552978515625 0 1478.6409
1150.5543212890625 0 23351.514 z 7
1151.5552978515625 0 19047.6
1152.553466796875 0 7610.9805
1153.5596923828125 0 3320.178
1154.560302734375 0 887.0734
1166.568359375 0 1333.3003 y 7
1167.578125 0 1538.381
1175.572509765625 0 30736.828 c 9
1176.5762939453125 0 18825.621
1177.5760498046875 0 7734.023
1178.5806884765625 0 2641.6345
1179.571533203125 0 1586.4221
1220.5791015625 0 4433.862 w 6
1221.5811767578125 0 3069.2944
1222.5819091796875 0 1671.9926
1223.5533447265625 0 890.1553
1241.6612548828125 0 741.6063
1259.657958984375 0 1620.3481
1260.6593017578125 0 2092.9895
1261.66357421875 0 902.2119
1279.5963134765625 0 20541.135 z 6
1280.5970458984375 0 14400.606
1281.59619140625 0 6481.771
1282.5999755859375 0 1924.0419
1285.651123046875 0 1092.7019 c Water loss 10
1286.6461181640625 0 863.5395 c Ammonia loss 10
1295.6146240234375 0 6255.3726 y 6
1296.615234375 0 5834.584
1297.615234375 0 3411.1335
1298.6224365234375 0 957.15436
1302.654541015625 0 2844.5713
1303.6658935546875 0 6928.085 c 10
1304.6683349609375 0 4140.6147
1305.6688232421875 0 2328.9849
1306.6602783203125 0 749.6662
1317.630126953125 0 936.88824
1334.6734619140625 0 1716.5226
1335.6690673828125 0 1123.1322
1373.69677734375 0 920.4665
1376.6806640625 0 1009.75977
1416.7054443359375 0 1836.5424
1417.7086181640625 0 11514.264 c 11
1418.7098388671875 0 9414.573
1419.712890625 0 3552.2417
1420.70947265625 0 2227.3481
1422.7135009765625 0 893.3245
1435.696533203125 0 13979.83 z 5
1436.701171875 0 27440.666
1437.70263671875 0 15615.551
1438.705810546875 0 7204.0537
1439.706787109375 0 2925.9148
1440.7039794921875 0 1065.6202
1450.7081298828125 0 2441.283
1451.7130126953125 0 9912.1045 y 5
1452.7142333984375 0 5603.483
1453.7169189453125 0 3976.2896
1454.7122802734375 0 1704.9845
1503.766845703125 0 1000.8698
1522.724853515625 0 2224.8782 z 4
1523.7332763671875 0 6197.1533
1524.738037109375 0 5593.67
1525.7410888671875 0 2266.452
1526.74609375 0 1480.3097
1538.7454833984375 0 2306.8967 y 4
1539.74072265625 0 1109.4543
1544.761474609375 0 731.1078
1545.766845703125 0 15655.069 c 12
1546.7689208984375 0 13953.416
1547.770263671875 0 9136.728
1548.775634765625 0 3899.409
1601.8287353515625 0 1605.6532
1602.8253173828125 0 1659.1602
1614.779052734375 0 1381.9584
1615.8011474609375 0 953.77136
1622.202392578125 0 757.6716
1626.8310546875 0 1054.6522 c Water loss 13
1627.8201904296875 0 2066.5706 c Ammonia loss 13
1628.81640625 0 1624.4991
1629.813720703125 0 1436.7073
1630.805419921875 0 1011.6021
1635.796875 0 4660.5396 y 3
1636.801513671875 0 3819.9834
1637.8040771484375 0 3910.0264
1638.8040771484375 0 2036.1641
1643.83056640625 0 2727.5715
1644.836669921875 0 20150.412 c 13
1645.8392333984375 0 18466.832
1646.8399658203125 0 10808.122
1647.8408203125 0 5143.8076
1648.8367919921875 0 1941.1274
1688.85791015625 0 1050.7947
1689.856201171875 0 1983.0599
1715.8485107421875 0 1641.2341 y Ammonia loss 2
1716.8616943359375 0 1892.8451 z 2
1717.85302734375 0 962.769
1727.8912353515625 0 884.27716
1728.8870849609375 0 1583.4185
1729.8966064453125 0 1920.4014
1730.895751953125 0 957.6713
1731.87109375 0 8493.809 c 14
1732.8726806640625 0 17452.953 y 2
1733.873779296875 0 15275.798
1734.8719482421875 0 8604.403
1735.8753662109375 0 3252.2024
1786.9112548828125 0 829.6798
1787.9080810546875 0 947.4065
1800.925048828125 0 836.5746
1801.943359375 0 3664.6208
1802.947021484375 0 4490.8286
1803.9412841796875 0 3028.1782
1804.9373779296875 0 1511.488
1828.9283447265625 0 1647.4574 y Ammonia loss 1
1829.927978515625 0 4852.4683 z 1
1830.927490234375 0 17926.4
1831.927490234375 0 15486.643
1832.9273681640625 0 9920.452 c 15
1833.9217529296875 0 4386.1885
1834.91796875 0 1574.0547
1844.9556884765625 0 4678.026
1845.958251953125 0 12193.63 y 1
1846.9608154296875 0 12382.052
1847.9642333984375 0 7792.3203
1848.96337890625 0 4251.1973
1849.932373046875 0 1394.4884
1870.934326171875 0 1339.6932
1871.9271240234375 0 1063.1611
1872.9266357421875 0 898.9017
1874.9283447265625 0 918.3947
1876.9854736328125 0 762.64514
1884.9951171875 0 1987.6556
1886.0128173828125 0 870.5219
1886.9791259765625 0 3654.0647
1887.9765625 0 2650.16
1888.963623046875 0 3108.182
1889.96044921875 0 7226.206
1890.9617919921875 0 5348.439
1891.963623046875 0 2299.778
1892.959716796875 0 1641.1577
1901.9649658203125 0 971.4416
1902.985107421875 0 3663.4468
1903.9813232421875 0 11713.147
1904.9810791015625 0 11961.228
1905.98046875 0 8133.5938
1906.9884033203125 0 3508.0588
1907.9774169921875 0 1332.0227
1913.9613037109375 0 2701.0688
1914.958984375 0 4285.8
1915.947998046875 0 2763.1562
1916.9644775390625 0 2734.4507
1917.9490966796875 0 860.4104
1919.9876708984375 0 2347.1567
1921.00341796875 0 9149.461
1922.0067138671875 0 8749.757
1923.0037841796875 0 5565.2144
1924.010986328125 0 2264.2578
1929.998046875 0 1051.6102
1930.9783935546875 0 11256.527
1931.9747314453125 0 39689.05
1932.9755859375 0 41495.004
1933.9774169921875 0 24486.79
1934.9775390625 0 13027.3955
1935.975341796875 0 4027.679
1936.989013671875 0 1172.8932
1946.9835205078125 0 6042.9644
1947.9898681640625 0 24955.969
1948.9959716796875 0 122103.836
1949.9984130859375 0 121305.33
1950.999755859375 0 80170.46
1952.0001220703125 0 36090.184
1953.000244140625 0 12413.3125
1953.9986572265625 0 4038.1003
3080.40771484375 0 1018.13074

Spectrum Details

|  |  |
| --- | --- |
| Matched peaks? Matched peaks The total absolute number of peaks matched. Additionally in brackets the total fraction of peaks matched and the total number of peaks is shown. | 89 (15.92% of 559) |
| FDR? FDR The false discovery rate estimated for this peptide. It is calculated by matching all theoretical fragments with a non-integer shift with the raw peaks for this spectrum. This is done with 40 different shifts. The resulting percentage is the average number of annotated peaks over the number of annotated peaks with the correct spectrum. | 0.70% |
| Satellite FDR? Satellite FDR See the FDR for details on its calculation. This satellite ion specific FDR only contains the satellite ions (d/w) for I/L/J positions. | 11.90% |
| PSM Score? PSM Score The PSM Score as given by Hecklib to this annotated spectrum. It is shown with three significant figures. | 627 |

#### Spectrum 3951? Spectrum 3951 The raw spectrum of this peptide as annotated by Hecklib. The fragments are coloured according to ion type (see legend). Any peaks with a star '\*' as text can be hovered over to see the full details, first the ion type second the mass shift type. By hovering over the amino acids in the peptide or ions in the legend the corresponding peaks are highlighted. By toggling the 'Unassigned' label you can turn the background (unassigned) peaks on or off in the plot. By updating the slider in the Ion legend you can update the spectrum to only show the top X% of the peaks with labels. The top X% means any peak that is within X% of the highest intensity. By dragging in the spectrum you can zoom in to a specific part of the spectrum and use 'Zoom Out' to get back to the original zoom level. The annotation of the spectrum is based on the given sequence in the peptides file and is done with different software so inconsistencies are likely. The peaks are annotated based on the given sequence, with 20 ppm tolerance.

Copy Data

##### Spectrum 3951 (TSV)

###### Preview

```
Loading example...
```

*Click on the button to copy the data to your clipboard.*

Mz MinMz MaxIntensity Max

WidthHeightPeptide font sizePeptide stroke widthSpectrum font sizeSpectrum stroke widthCompact peptide

Ion legend

wxyz

abcd

OtherUnassignedIon^Charge^_Position_Show for top:%

TLPPSREEMTKNQVSTJ

01.25e+52.50e+53.74e+54.99e+5

Zoom Out

y^+2^_4_y^+1^_2_y^+1^_2_y^+1^_3_y^+1^_3_c^+1^_4_c^+1^_5_c^+1^_5_z^+1^_5_y^+1^_5_z^+3^_15_c^+3^_15_y^+3^_15_z^+3^_15_c^+3^_15_y^+3^_15_c^+2^_10_y^+1^_6_z^+1^_6_c^+2^_11_y^+1^_6_c^+1^_6_c^+2^_12_c^+2^_12_c^+2^_12_z^+2^_12_y^+2^_12_y^+2^_13_z^+2^_13_y^+2^_13_c^+2^_13_y^+1^_7_c^+1^_7_y^+2^_14_y^+2^_14_c^+2^_14_c^+2^_14_y^+2^_14_c^+2^_14_w^+2^_15_z^+1^_8_w^+1^_8_c^+2^_15_y^+2^_15_z^+2^_15_w^+1^_8_c^+2^_15_y^+2^_15_y^+1^_8_y^+1^_8_z^+1^_8_y^+1^_8_w^+2^_16_z^+2^_16_c^+2^_16_y^+2^_16_y^+2^_16_z^+2^_16_c^+2^_16_y^+2^_16_c^+1^_8_z^+1^_9_z^+1^_9_y^+1^_9_c^+1^_9_w^+1^_10_z^+1^_10_z^+1^_10_y^+1^_10_c^+1^_10_w^+1^_11_z^+1^_11_c^+1^_11_c^+1^_11_y^+1^_11_c^+1^_11_c^+1^_12_z^+1^_12_y^+1^_12_z^+1^_13_y^+1^_13_c^+1^_13_y^+1^_14_c^+1^_14_c^+1^_14_y^+1^_14_c^+1^_14_z^+1^_15_c^+1^_15_y^+1^_15_y^+1^_16_z^+1^_16_c^+1^_16_y^+1^_16_

0779155823373115

Fragment Matches Table

Show background peaks

| Position | Ion type | Intensity | mz Theoretical | mz Error (Th) | mz Error (ppm) | Charge | Series Number |
| --- | --- | --- | --- | --- | --- | --- | --- |
| - | - | 1317 | 120.1 | - | - | 0 | - |
| - | - | 1.695E+04 | 120.1 | - | - | 0 | - |
| - | - | 1313 | 148.2 | - | - | 0 | - |
| - | - | 2080 | 149 | - | - | 0 | - |
| - | - | 1623 | 167.7 | - | - | 0 | - |
| - | - | 1482 | 183.3 | - | - | 0 | - |
| - | - | 1.384E+05 | 187.1 | - | - | 0 | - |
| - | - | 1.054E+04 | 188.1 | - | - | 0 | - |
| - | - | 1473 | 197 | - | - | 0 | - |
| 14 | y | 4511 | 201.1 | 0.0001511 | 0.7511 | +2 | 4 |
| - | - | 5211 | 212.1 | - | - | 0 | - |
| 16 | y | 9.314E+04 | 215.1 | 0.0002633 | 1.224 | +1 | 2 |
| - | - | 1.073E+04 | 216.1 | - | - | 0 | - |
| 16 | y | 7161 | 233.1 | 0.0003187 | 1.367 | +1 | 2 |
| - | - | 3409 | 256.2 | - | - | 0 | - |
| - | - | 1571 | 276 | - | - | 0 | - |
| - | - | 1.742E+04 | 299.2 | - | - | 0 | - |
| - | - | 3697 | 300.2 | - | - | 0 | - |
| 15 | y | 2131 | 302.2 | 3.423E-05 | 0.1133 | +1 | 3 |
| - | - | 3607 | 312.2 | - | - | 0 | - |
| 15 | y | 4429 | 320.2 | 0.0004864 | 1.519 | +1 | 3 |
| - | - | 2318 | 326.2 | - | - | 0 | - |
| - | - | 2289 | 369.2 | - | - | 0 | - |
| - | - | 1932 | 382.3 | - | - | 0 | - |
| - | - | 5583 | 407.3 | - | - | 0 | - |
| - | - | 3525 | 425.3 | - | - | 0 | - |
| 4 | c | 5959 | 426.3 | 0.0001912 | 0.4485 | +1 | 4 |
| - | - | 2334 | 427.3 | - | - | 0 | - |
| - | - | 1982 | 428.3 | - | - | 0 | - |
| - | - | 1981 | 466.8 | - | - | 0 | - |
| - | - | 8683 | 470.3 | - | - | 0 | - |
| - | - | 2302 | 471.3 | - | - | 0 | - |
| - | - | 2264 | 471.9 | - | - | 0 | - |
| - | - | 3161 | 472.3 | - | - | 0 | - |
| - | - | 4829 | 494.3 | - | - | 0 | - |
| 5 | c | 1923 | 495.3 | 0.005572 | 11.25 | +1 | 5 |
| - | - | 3171 | 499.9 | - | - | 0 | - |
| - | - | 4620 | 500.9 | - | - | 0 | - |
| - | - | 3813 | 501.2 | - | - | 0 | - |
| - | - | 2011 | 501.6 | - | - | 0 | - |
| - | - | 1825 | 501.9 | - | - | 0 | - |
| - | - | 3097 | 512.3 | - | - | 0 | - |
| 5 | c | 9.348E+04 | 513.3 | 0.0004647 | 0.9054 | +1 | 5 |
| - | - | 2.83E+04 | 514.3 | - | - | 0 | - |
| - | - | 4947 | 515.3 | - | - | 0 | - |
| - | - | 2817 | 517.6 | - | - | 0 | - |
| - | - | 2188 | 523.3 | - | - | 0 | - |
| - | - | 2795 | 523.6 | - | - | 0 | - |
| - | - | 2946 | 526.3 | - | - | 0 | - |
| - | - | 3532 | 527.3 | - | - | 0 | - |
| - | - | 6607 | 529.3 | - | - | 0 | - |
| - | - | 8507 | 529.6 | - | - | 0 | - |
| - | - | 4525 | 529.9 | - | - | 0 | - |
| 13 | z | 8642 | 531.3 | 0.0002065 | 0.3887 | +1 | 5 |
| - | - | 1.056E+04 | 532.3 | - | - | 0 | - |
| - | - | 9155 | 532.6 | - | - | 0 | - |
| - | - | 1.381E+04 | 532.9 | - | - | 0 | - |
| - | - | 6509 | 533.3 | - | - | 0 | - |
| - | - | 5136 | 533.6 | - | - | 0 | - |
| - | - | 3.072E+04 | 538.6 | - | - | 0 | - |
| - | - | 3.228E+04 | 538.9 | - | - | 0 | - |
| - | - | 1.226E+04 | 539.3 | - | - | 0 | - |
| - | - | 6604 | 539.6 | - | - | 0 | - |
| - | - | 1816 | 544.3 | - | - | 0 | - |
| 13 | y | 5597 | 547.3 | 0.002249 | 4.109 | +1 | 5 |
| - | - | 5231 | 557 | - | - | 0 | - |
| - | - | 6935 | 557.3 | - | - | 0 | - |
| - | - | 6733 | 557.6 | - | - | 0 | - |
| - | - | 3285 | 558 | - | - | 0 | - |
| - | - | 2295 | 558.3 | - | - | 0 | - |
| - | - | 1894 | 561.1 | - | - | 0 | - |
| - | - | 3268 | 566.3 | - | - | 0 | - |
| - | - | 3922 | 566.6 | - | - | 0 | - |
| 3 | z | 1852 | 567.3 | 0.01103 | 19.45 | +3 | 15 |
| - | - | 2507 | 569.3 | - | - | 0 | - |
| - | - | 3297 | 570.3 | - | - | 0 | - |
| - | - | 7330 | 571.3 | - | - | 0 | - |
| 15 | c | 1.868E+04 | 572.3 | 0.002577 | 4.503 | +3 | 15 |
| 3 | y | 2.068E+04 | 572.6 | 0.007213 | 12.6 | +3 | 15 |
| 3 | z | 1.576E+04 | 572.9 | 0.00605 | 10.56 | +3 | 15 |
| - | - | 5151 | 573.3 | - | - | 0 | - |
| - | - | 4811 | 573.3 | - | - | 0 | - |
| - | - | 3410 | 573.6 | - | - | 0 | - |
| - | - | 1884 | 573.9 | - | - | 0 | - |
| 15 | c | 2760 | 578 | 0.001621 | 2.805 | +3 | 15 |
| 3 | y | 4.648E+05 | 578.3 | 0.002473 | 4.277 | +3 | 15 |
| - | - | 4.553E+05 | 578.6 | - | - | 0 | - |
| - | - | 2.629E+05 | 579 | - | - | 0 | - |
| 10 | c | 1.041E+05 | 579.3 | 0.009478 | 16.36 | +2 | 10 |
| - | - | 3.061E+04 | 579.6 | - | - | 0 | - |
| - | - | 1.059E+04 | 580 | - | - | 0 | - |
| - | - | 2338 | 607.4 | - | - | 0 | - |
| - | - | 3220 | 615.3 | - | - | 0 | - |
| - | - | 9499 | 625.4 | - | - | 0 | - |
| - | - | 6087 | 626.4 | - | - | 0 | - |
| - | - | 3786 | 643.7 | - | - | 0 | - |
| - | - | 6844 | 644 | - | - | 0 | - |
| 12 | y | 1915 | 644.3 | 0.00943 | 14.64 | +1 | 6 |
| 12 | z | 5876 | 645.3 | 0.0009587 | 1.486 | +1 | 6 |
| - | - | 8506 | 646.3 | - | - | 0 | - |
| - | - | 3840 | 647.3 | - | - | 0 | - |
| - | - | 1855 | 649.3 | - | - | 0 | - |
| 11 | c | 1.601E+04 | 652.3 | 0.002612 | 4.005 | +2 | 11 |
| - | - | 1.455E+04 | 652.8 | - | - | 0 | - |
| - | - | 6835 | 653.3 | - | - | 0 | - |
| - | - | 4411 | 657.8 | - | - | 0 | - |
| 12 | y | 1.799E+04 | 661.4 | 0.001252 | 1.894 | +1 | 6 |
| - | - | 5192 | 662.4 | - | - | 0 | - |
| 6 | c | 1.595E+05 | 669.4 | 0.0007941 | 1.186 | +1 | 6 |
| - | - | 5.529E+04 | 670.4 | - | - | 0 | - |
| - | - | 1.356E+04 | 671.4 | - | - | 0 | - |
| - | - | 5055 | 672.4 | - | - | 0 | - |
| - | - | 2438 | 686.9 | - | - | 0 | - |
| - | - | 3659 | 687.4 | - | - | 0 | - |
| - | - | 5888 | 699.9 | - | - | 0 | - |
| 12 | c | 8557 | 700.4 | 0.002705 | 3.862 | +2 | 12 |
| 12 | c | 3049 | 700.8 | 0.005471 | 7.806 | +2 | 12 |
| - | - | 1.38E+04 | 707.3 | - | - | 0 | - |
| - | - | 1.182E+04 | 707.8 | - | - | 0 | - |
| - | - | 3934 | 708.3 | - | - | 0 | - |
| - | - | 2.644E+04 | 708.9 | - | - | 0 | - |
| 12 | c | 1.046E+05 | 709.4 | 0.002572 | 3.626 | +2 | 12 |
| - | - | 8.59E+04 | 709.9 | - | - | 0 | - |
| - | - | 4.378E+04 | 710.4 | - | - | 0 | - |
| - | - | 1.479E+04 | 710.9 | - | - | 0 | - |
| - | - | 3682 | 711.4 | - | - | 0 | - |
| - | - | 2771 | 711.4 | - | - | 0 | - |
| 6 | z | 7390 | 718.3 | 0.00596 | 8.296 | +2 | 12 |
| - | - | 5786 | 718.9 | - | - | 0 | - |
| - | - | 5731 | 719.4 | - | - | 0 | - |
| - | - | 2622 | 719.9 | - | - | 0 | - |
| - | - | 2534 | 720.4 | - | - | 0 | - |
| - | - | 8376 | 722.4 | - | - | 0 | - |
| - | - | 4779 | 722.9 | - | - | 0 | - |
| - | - | 5574 | 725.9 | - | - | 0 | - |
| 6 | y | 1.942E+04 | 726.4 | 0.002274 | 3.13 | +2 | 12 |
| - | - | 1.114E+04 | 726.9 | - | - | 0 | - |
| - | - | 9614 | 727.4 | - | - | 0 | - |
| - | - | 1.97E+04 | 729.4 | - | - | 0 | - |
| - | - | 1.014E+04 | 730.4 | - | - | 0 | - |
| - | - | 3409 | 748.9 | - | - | 0 | - |
| - | - | 2627 | 749.4 | - | - | 0 | - |
| - | - | 1.36E+04 | 750.9 | - | - | 0 | - |
| - | - | 1.7E+04 | 751.4 | - | - | 0 | - |
| - | - | 6259 | 751.9 | - | - | 0 | - |
| - | - | 3190 | 752.4 | - | - | 0 | - |
| - | - | 3173 | 753.9 | - | - | 0 | - |
| - | - | 3865 | 754.4 | - | - | 0 | - |
| - | - | 3174 | 755.4 | - | - | 0 | - |
| - | - | 2594 | 756.4 | - | - | 0 | - |
| - | - | 2122 | 757.9 | - | - | 0 | - |
| - | - | 2576 | 758.9 | - | - | 0 | - |
| 5 | y | 2543 | 761.4 | 0.0046 | 6.042 | +2 | 13 |
| 5 | z | 1.593E+04 | 761.9 | 0.003556 | 4.668 | +2 | 13 |
| - | - | 1.547E+04 | 762.4 | - | - | 0 | - |
| - | - | 1.133E+04 | 762.9 | - | - | 0 | - |
| - | - | 4162 | 763.4 | - | - | 0 | - |
| - | - | 3735 | 765.4 | - | - | 0 | - |
| - | - | 3878 | 765.9 | - | - | 0 | - |
| - | - | 2129 | 766.9 | - | - | 0 | - |
| 5 | y | 1.353E+04 | 769.9 | 0.002068 | 2.686 | +2 | 13 |
| - | - | 7796 | 770.4 | - | - | 0 | - |
| - | - | 4918 | 770.9 | - | - | 0 | - |
| - | - | 2457 | 772.4 | - | - | 0 | - |
| - | - | 7806 | 772.9 | - | - | 0 | - |
| 13 | c | 1.411E+05 | 773.4 | 0.004716 | 6.098 | +2 | 13 |
| - | - | 2.899E+04 | 773.5 | - | - | 0 | - |
| - | - | 1.395E+05 | 773.9 | - | - | 0 | - |
| - | - | 6.257E+04 | 774.4 | - | - | 0 | - |
| - | - | 8989 | 774.5 | - | - | 0 | - |
| - | - | 3.096E+04 | 774.9 | - | - | 0 | - |
| - | - | 9223 | 775.4 | - | - | 0 | - |
| - | - | 2111 | 779.4 | - | - | 0 | - |
| - | - | 2693 | 784.4 | - | - | 0 | - |
| - | - | 2265 | 786.4 | - | - | 0 | - |
| 11 | y | 2.29E+04 | 789.4 | 0.0001614 | 0.2044 | +1 | 7 |
| - | - | 7299 | 790.4 | - | - | 0 | - |
| - | - | 3192 | 791.9 | - | - | 0 | - |
| - | - | 2879 | 792.4 | - | - | 0 | - |
| - | - | 5432 | 792.9 | - | - | 0 | - |
| - | - | 1.445E+04 | 793.4 | - | - | 0 | - |
| - | - | 1.034E+04 | 793.9 | - | - | 0 | - |
| - | - | 5223 | 794.4 | - | - | 0 | - |
| - | - | 3343 | 794.9 | - | - | 0 | - |
| - | - | 4877 | 797.4 | - | - | 0 | - |
| 7 | c | 2.491E+05 | 798.4 | 0.0009867 | 1.236 | +1 | 7 |
| - | - | 3381 | 798.9 | - | - | 0 | - |
| - | - | 1.061E+05 | 799.5 | - | - | 0 | - |
| - | - | 2.62E+04 | 800.5 | - | - | 0 | - |
| - | - | 1.224E+04 | 800.9 | - | - | 0 | - |
| - | - | 1.117E+04 | 801.4 | - | - | 0 | - |
| - | - | 8533 | 801.9 | - | - | 0 | - |
| - | - | 4833 | 802.4 | - | - | 0 | - |
| - | - | 5531 | 803.4 | - | - | 0 | - |
| - | - | 2452 | 805.4 | - | - | 0 | - |
| - | - | 3.587E+04 | 807.4 | - | - | 0 | - |
| - | - | 3.185E+04 | 807.9 | - | - | 0 | - |
| - | - | 1.752E+04 | 808.4 | - | - | 0 | - |
| - | - | 9830 | 808.9 | - | - | 0 | - |
| 4 | y | 3843 | 809.4 | 0.006847 | 8.459 | +2 | 14 |
| 4 | y | 3531 | 809.9 | 0.008736 | 10.79 | +2 | 14 |
| - | - | 2730 | 812.5 | - | - | 0 | - |
| - | - | 4.61E+04 | 813.4 | - | - | 0 | - |
| 14 | c | 4.327E+04 | 813.9 | 0.0005265 | 0.6469 | +2 | 14 |
| 14 | c | 3.357E+04 | 814.4 | 0.006123 | 7.518 | +2 | 14 |
| - | - | 3180 | 814.5 | - | - | 0 | - |
| - | - | 1.51E+04 | 814.9 | - | - | 0 | - |
| - | - | 1.169E+04 | 815.4 | - | - | 0 | - |
| - | - | 4555 | 815.9 | - | - | 0 | - |
| - | - | 2232 | 816.4 | - | - | 0 | - |
| - | - | 2670 | 817.4 | - | - | 0 | - |
| - | - | 2846 | 817.9 | - | - | 0 | - |
| 4 | y | 7.179E+04 | 818.4 | 0.004433 | 5.417 | +2 | 14 |
| - | - | 6.531E+04 | 818.9 | - | - | 0 | - |
| - | - | 2.924E+04 | 819.4 | - | - | 0 | - |
| - | - | 1.332E+04 | 819.9 | - | - | 0 | - |
| - | - | 3795 | 820.4 | - | - | 0 | - |
| - | - | 7.939E+04 | 822.4 | - | - | 0 | - |
| 14 | c | 1.011E+05 | 822.9 | 0.001454 | 1.767 | +2 | 14 |
| - | - | 8.965E+04 | 823.4 | - | - | 0 | - |
| - | - | 4.357E+04 | 823.9 | - | - | 0 | - |
| - | - | 1.478E+04 | 824.4 | - | - | 0 | - |
| - | - | 4877 | 824.9 | - | - | 0 | - |
| - | - | 4412 | 825.5 | - | - | 0 | - |
| - | - | 7453 | 830.5 | - | - | 0 | - |
| - | - | 5174 | 831.5 | - | - | 0 | - |
| - | - | 2702 | 832.4 | - | - | 0 | - |
| - | - | 3624 | 832.9 | - | - | 0 | - |
| - | - | 1.366E+04 | 834.9 | - | - | 0 | - |
| - | - | 1.714E+04 | 835.4 | - | - | 0 | - |
| - | - | 7406 | 835.9 | - | - | 0 | - |
| - | - | 6941 | 836.4 | - | - | 0 | - |
| - | - | 4741 | 836.9 | - | - | 0 | - |
| - | - | 3373 | 843.9 | - | - | 0 | - |
| - | - | 2963 | 844.4 | - | - | 0 | - |
| - | - | 4839 | 844.9 | - | - | 0 | - |
| 3 | w | 9581 | 845.4 | 0.01533 | 18.13 | +2 | 15 |
| - | - | 9269 | 845.9 | - | - | 0 | - |
| - | - | 3577 | 846.4 | - | - | 0 | - |
| - | - | 5183 | 849.4 | - | - | 0 | - |
| 10 | z | 4662 | 856.5 | 0.0004484 | 0.5236 | +1 | 8 |
| - | - | 2510 | 856.9 | - | - | 0 | - |
| 10 | w | 1.275E+04 | 857.5 | 0.0001134 | 0.1323 | +1 | 8 |
| 15 | c | 1.476E+04 | 857.9 | 0.005123 | 5.972 | +2 | 15 |
| 3 | y | 1.556E+04 | 858.4 | 0.01397 | 16.27 | +2 | 15 |
| 3 | z | 1.192E+04 | 858.9 | 0.008532 | 9.933 | +2 | 15 |
| 10 | w | 2.533E+04 | 859.5 | 0.002266 | 2.637 | +1 | 8 |
| - | - | 5065 | 859.9 | - | - | 0 | - |
| - | - | 1.024E+04 | 860.5 | - | - | 0 | - |
| - | - | 6555 | 865.4 | - | - | 0 | - |
| - | - | 1.902E+04 | 865.9 | - | - | 0 | - |
| 15 | c | 2.135E+05 | 866.4 | 0.002957 | 3.413 | +2 | 15 |
| 3 | y | 4.858E+05 | 866.9 | 0.006677 | 7.702 | +2 | 15 |
| - | - | 3.813E+05 | 867.4 | - | - | 0 | - |
| - | - | 2.053E+05 | 867.9 | - | - | 0 | - |
| - | - | 2961 | 868 | - | - | 0 | - |
| - | - | 7.645E+04 | 868.4 | - | - | 0 | - |
| - | - | 2.296E+04 | 868.9 | - | - | 0 | - |
| - | - | 5346 | 869.4 | - | - | 0 | - |
| 10 | y | 4533 | 872.5 | 0.01467 | 16.82 | +1 | 8 |
| - | - | 3086 | 872.9 | - | - | 0 | - |
| 10 | y | 2.317E+04 | 873.5 | 0.001315 | 1.505 | +1 | 8 |
| 10 | z | 1.268E+05 | 874.5 | 0.0003818 | 0.4366 | +1 | 8 |
| - | - | 6.921E+04 | 875.5 | - | - | 0 | - |
| - | - | 2.183E+04 | 876.5 | - | - | 0 | - |
| - | - | 5361 | 877.5 | - | - | 0 | - |
| - | - | 1.067E+04 | 879.5 | - | - | 0 | - |
| - | - | 1.269E+04 | 880 | - | - | 0 | - |
| - | - | 4541 | 880.4 | - | - | 0 | - |
| - | - | 3520 | 880.9 | - | - | 0 | - |
| - | - | 6416 | 881.4 | - | - | 0 | - |
| - | - | 2622 | 882.4 | - | - | 0 | - |
| - | - | 4883 | 883.5 | - | - | 0 | - |
| - | - | 3098 | 884.5 | - | - | 0 | - |
| - | - | 7116 | 887.4 | - | - | 0 | - |
| - | - | 5720 | 887.9 | - | - | 0 | - |
| - | - | 3447 | 888.5 | - | - | 0 | - |
| - | - | 3547 | 889.5 | - | - | 0 | - |
| 10 | y | 4.504E+04 | 890.5 | 0.0002734 | 0.3071 | +1 | 8 |
| - | - | 1.728E+04 | 891.5 | - | - | 0 | - |
| - | - | 7110 | 892 | - | - | 0 | - |
| - | - | 9625 | 892.5 | - | - | 0 | - |
| - | - | 5250 | 893 | - | - | 0 | - |
| - | - | 3960 | 893.5 | - | - | 0 | - |
| 2 | w | 8.395E+04 | 893.9 | 0.003409 | 3.813 | +2 | 16 |
| - | - | 8.231E+04 | 894.4 | - | - | 0 | - |
| - | - | 4.506E+04 | 894.9 | - | - | 0 | - |
| - | - | 2.715E+04 | 895.4 | - | - | 0 | - |
| - | - | 5388 | 895.9 | - | - | 0 | - |
| - | - | 8978 | 901 | - | - | 0 | - |
| - | - | 1.509E+04 | 901.5 | - | - | 0 | - |
| - | - | 1.417E+04 | 902 | - | - | 0 | - |
| - | - | 1.105E+04 | 902.5 | - | - | 0 | - |
| - | - | 2720 | 903 | - | - | 0 | - |
| - | - | 4171 | 903.5 | - | - | 0 | - |
| 2 | z | 5055 | 906.5 | 0.0167 | 18.43 | +2 | 16 |
| - | - | 6196 | 907 | - | - | 0 | - |
| - | - | 5732 | 907.5 | - | - | 0 | - |
| 16 | c | 2478 | 908.4 | 0.01205 | 13.26 | +2 | 16 |
| - | - | 2591 | 909.5 | - | - | 0 | - |
| - | - | 2272 | 913.5 | - | - | 0 | - |
| - | - | 3098 | 914 | - | - | 0 | - |
| 2 | y | 1.184E+04 | 914.5 | 0.009539 | 10.43 | +2 | 16 |
| 2 | y | 1.544E+04 | 915 | 0.008986 | 9.822 | +2 | 16 |
| 2 | z | 4.84E+04 | 915.5 | 0.006966 | 7.609 | +2 | 16 |
| - | - | 4.646E+04 | 916 | - | - | 0 | - |
| - | - | 3.253E+04 | 916.5 | - | - | 0 | - |
| 16 | c | 1.837E+04 | 917 | 0.0028 | 3.053 | +2 | 16 |
| - | - | 8548 | 917.5 | - | - | 0 | - |
| - | - | 7853 | 920.5 | - | - | 0 | - |
| - | - | 3857 | 921 | - | - | 0 | - |
| - | - | 4604 | 921.5 | - | - | 0 | - |
| - | - | 7630 | 922.5 | - | - | 0 | - |
| - | - | 1.978E+05 | 923 | - | - | 0 | - |
| 2 | y | 1.912E+05 | 923.5 | 0.01146 | 12.41 | +2 | 16 |
| - | - | 1.41E+05 | 924 | - | - | 0 | - |
| - | - | 7.292E+04 | 924.5 | - | - | 0 | - |
| - | - | 2.778E+04 | 925 | - | - | 0 | - |
| - | - | 8175 | 925.5 | - | - | 0 | - |
| - | - | 7400 | 926.5 | - | - | 0 | - |
| 8 | c | 2.413E+05 | 927.5 | 0.000752 | 0.8108 | +1 | 8 |
| - | - | 1.248E+05 | 928.5 | - | - | 0 | - |
| - | - | 8024 | 929 | - | - | 0 | - |
| - | - | 4.157E+04 | 929.5 | - | - | 0 | - |
| - | - | 1.356E+04 | 930 | - | - | 0 | - |
| - | - | 2.123E+04 | 930.5 | - | - | 0 | - |
| - | - | 6054 | 931 | - | - | 0 | - |
| - | - | 7408 | 931.5 | - | - | 0 | - |
| - | - | 3510 | 932 | - | - | 0 | - |
| - | - | 2489 | 932.5 | - | - | 0 | - |
| - | - | 4623 | 934.5 | - | - | 0 | - |
| - | - | 7182 | 935 | - | - | 0 | - |
| - | - | 5623 | 935.5 | - | - | 0 | - |
| - | - | 4813 | 936 | - | - | 0 | - |
| - | - | 7490 | 936.5 | - | - | 0 | - |
| - | - | 6272 | 937 | - | - | 0 | - |
| - | - | 1.427E+04 | 937.5 | - | - | 0 | - |
| - | - | 8.273E+04 | 938 | - | - | 0 | - |
| - | - | 8.393E+04 | 938.5 | - | - | 0 | - |
| - | - | 5.928E+04 | 939 | - | - | 0 | - |
| - | - | 2.788E+04 | 939.5 | - | - | 0 | - |
| - | - | 9269 | 940 | - | - | 0 | - |
| - | - | 3387 | 940.5 | - | - | 0 | - |
| - | - | 5585 | 941.5 | - | - | 0 | - |
| - | - | 2.212E+04 | 942.5 | - | - | 0 | - |
| - | - | 2.705E+04 | 943 | - | - | 0 | - |
| - | - | 3.133E+04 | 943.5 | - | - | 0 | - |
| - | - | 2.288E+04 | 944 | - | - | 0 | - |
| - | - | 2.013E+04 | 944.5 | - | - | 0 | - |
| - | - | 4.869E+04 | 945 | - | - | 0 | - |
| - | - | 4.4E+04 | 945.5 | - | - | 0 | - |
| - | - | 2.435E+04 | 946 | - | - | 0 | - |
| - | - | 3.743E+04 | 946.5 | - | - | 0 | - |
| - | - | 3.528E+04 | 947 | - | - | 0 | - |
| - | - | 1.284E+04 | 947.5 | - | - | 0 | - |
| - | - | 6574 | 948 | - | - | 0 | - |
| - | - | 3824 | 948.5 | - | - | 0 | - |
| - | - | 1.158E+04 | 951.5 | - | - | 0 | - |
| - | - | 4.393E+04 | 952 | - | - | 0 | - |
| - | - | 4.898E+04 | 952.5 | - | - | 0 | - |
| - | - | 5.051E+04 | 953 | - | - | 0 | - |
| - | - | 3.201E+04 | 953.5 | - | - | 0 | - |
| - | - | 1.376E+04 | 954 | - | - | 0 | - |
| - | - | 7238 | 954.5 | - | - | 0 | - |
| - | - | 1.815E+04 | 956.5 | - | - | 0 | - |
| - | - | 3.937E+04 | 957 | - | - | 0 | - |
| - | - | 4.306E+04 | 957.5 | - | - | 0 | - |
| - | - | 2.457E+04 | 958 | - | - | 0 | - |
| - | - | 3.464E+04 | 958.5 | - | - | 0 | - |
| - | - | 8934 | 959 | - | - | 0 | - |
| - | - | 9.839E+04 | 959.5 | - | - | 0 | - |
| - | - | 2359 | 960 | - | - | 0 | - |
| - | - | 5.527E+04 | 960.5 | - | - | 0 | - |
| - | - | 1.57E+04 | 961.5 | - | - | 0 | - |
| - | - | 4890 | 962.5 | - | - | 0 | - |
| - | - | 6.116E+04 | 965.5 | - | - | 0 | - |
| - | - | 3.028E+05 | 966 | - | - | 0 | - |
| - | - | 2.93E+05 | 966.5 | - | - | 0 | - |
| - | - | 1.862E+05 | 967 | - | - | 0 | - |
| - | - | 1.025E+05 | 967.5 | - | - | 0 | - |
| - | - | 2.989E+04 | 968 | - | - | 0 | - |
| - | - | 1.365E+04 | 968.5 | - | - | 0 | - |
| - | - | 7937 | 969.5 | - | - | 0 | - |
| - | - | 2.197E+05 | 974 | - | - | 0 | - |
| - | - | 3.702E+05 | 974.5 | - | - | 0 | - |
| - | - | 3.015E+05 | 975 | - | - | 0 | - |
| - | - | 1.78E+05 | 975.5 | - | - | 0 | - |
| - | - | 7.758E+04 | 976 | - | - | 0 | - |
| - | - | 2.096E+04 | 976.5 | - | - | 0 | - |
| - | - | 4786 | 977 | - | - | 0 | - |
| 9 | z | 3057 | 1003 | 0.007256 | 7.231 | +1 | 9 |
| - | - | 3011 | 1005 | - | - | 0 | - |
| - | - | 3266 | 1006 | - | - | 0 | - |
| - | - | 4307 | 1007 | - | - | 0 | - |
| - | - | 6504 | 1013 | - | - | 0 | - |
| - | - | 2917 | 1013 | - | - | 0 | - |
| - | - | 4565 | 1016 | - | - | 0 | - |
| - | - | 3293 | 1017 | - | - | 0 | - |
| 9 | z | 2.835E+04 | 1022 | 0.00603 | 5.903 | +1 | 9 |
| - | - | 2.541E+04 | 1023 | - | - | 0 | - |
| - | - | 1.001E+04 | 1024 | - | - | 0 | - |
| - | - | 2.606E+04 | 1031 | - | - | 0 | - |
| - | - | 1.998E+04 | 1032 | - | - | 0 | - |
| - | - | 1.101E+04 | 1033 | - | - | 0 | - |
| - | - | 4320 | 1034 | - | - | 0 | - |
| 9 | y | 1.504E+04 | 1038 | 0.006593 | 6.355 | +1 | 9 |
| - | - | 1.083E+04 | 1039 | - | - | 0 | - |
| - | - | 1.489E+04 | 1050 | - | - | 0 | - |
| - | - | 6269 | 1051 | - | - | 0 | - |
| - | - | 3525 | 1056 | - | - | 0 | - |
| - | - | 5039 | 1074 | - | - | 0 | - |
| 9 | c | 2.56E+05 | 1075 | 0.005851 | 5.445 | +1 | 9 |
| - | - | 1.495E+05 | 1076 | - | - | 0 | - |
| - | - | 5.795E+04 | 1077 | - | - | 0 | - |
| - | - | 2.14E+04 | 1078 | - | - | 0 | - |
| - | - | 4931 | 1079 | - | - | 0 | - |
| - | - | 2185 | 1080 | - | - | 0 | - |
| - | - | 6588 | 1088 | - | - | 0 | - |
| - | - | 4409 | 1089 | - | - | 0 | - |
| 8 | w | 3.348E+04 | 1092 | 0.006771 | 6.203 | +1 | 10 |
| - | - | 2.273E+04 | 1093 | - | - | 0 | - |
| - | - | 9918 | 1094 | - | - | 0 | - |
| - | - | 6537 | 1095 | - | - | 0 | - |
| - | - | 2481 | 1095 | - | - | 0 | - |
| - | - | 6293 | 1114 | - | - | 0 | - |
| - | - | 3190 | 1115 | - | - | 0 | - |
| - | - | 5836 | 1132 | - | - | 0 | - |
| 8 | z | 9066 | 1133 | 0.01441 | 12.72 | +1 | 10 |
| - | - | 2719 | 1134 | - | - | 0 | - |
| - | - | 4826 | 1135 | - | - | 0 | - |
| - | - | 2366 | 1150 | - | - | 0 | - |
| 8 | z | 1.05E+05 | 1151 | 0.005429 | 4.719 | +1 | 10 |
| - | - | 7.677E+04 | 1152 | - | - | 0 | - |
| - | - | 3.83E+04 | 1153 | - | - | 0 | - |
| - | - | 1.147E+04 | 1154 | - | - | 0 | - |
| - | - | 2374 | 1155 | - | - | 0 | - |
| 8 | y | 7618 | 1167 | 0.00587 | 5.032 | +1 | 10 |
| - | - | 4930 | 1168 | - | - | 0 | - |
| 10 | c | 1.205E+05 | 1176 | 0.005414 | 4.605 | +1 | 10 |
| - | - | 7.404E+04 | 1177 | - | - | 0 | - |
| - | - | 3.59E+04 | 1178 | - | - | 0 | - |
| - | - | 1.605E+04 | 1179 | - | - | 0 | - |
| - | - | 4281 | 1180 | - | - | 0 | - |
| - | - | 2711 | 1208 | - | - | 0 | - |
| - | - | 3886 | 1209 | - | - | 0 | - |
| 7 | w | 2.027E+04 | 1221 | 0.002019 | 1.654 | +1 | 11 |
| - | - | 1.432E+04 | 1222 | - | - | 0 | - |
| - | - | 8258 | 1223 | - | - | 0 | - |
| - | - | 3207 | 1224 | - | - | 0 | - |
| - | - | 8130 | 1260 | - | - | 0 | - |
| - | - | 8031 | 1261 | - | - | 0 | - |
| - | - | 2568 | 1262 | - | - | 0 | - |
| 7 | z | 8.39E+04 | 1280 | 0.00434 | 3.392 | +1 | 11 |
| - | - | 5.761E+04 | 1281 | - | - | 0 | - |
| - | - | 3.11E+04 | 1282 | - | - | 0 | - |
| - | - | 8314 | 1283 | - | - | 0 | - |
| - | - | 2782 | 1285 | - | - | 0 | - |
| 11 | c | 3380 | 1286 | 0.008916 | 6.935 | +1 | 11 |
| 11 | c | 2839 | 1287 | 0.01488 | 11.57 | +1 | 11 |
| - | - | 2309 | 1295 | - | - | 0 | - |
| 7 | y | 2.729E+04 | 1296 | 0.005635 | 4.35 | +1 | 11 |
| - | - | 1.715E+04 | 1297 | - | - | 0 | - |
| - | - | 9531 | 1298 | - | - | 0 | - |
| - | - | 5114 | 1299 | - | - | 0 | - |
| - | - | 1.023E+04 | 1303 | - | - | 0 | - |
| 11 | c | 2.356E+04 | 1304 | 0.002003 | 1.537 | +1 | 11 |
| - | - | 1.66E+04 | 1305 | - | - | 0 | - |
| - | - | 6551 | 1306 | - | - | 0 | - |
| - | - | 3128 | 1307 | - | - | 0 | - |
| - | - | 2709 | 1334 | - | - | 0 | - |
| - | - | 6555 | 1335 | - | - | 0 | - |
| - | - | 2746 | 1362 | - | - | 0 | - |
| - | - | 3605 | 1374 | - | - | 0 | - |
| - | - | 3224 | 1375 | - | - | 0 | - |
| - | - | 3382 | 1377 | - | - | 0 | - |
| - | - | 3012 | 1380 | - | - | 0 | - |
| - | - | 6251 | 1417 | - | - | 0 | - |
| 12 | c | 4.724E+04 | 1418 | 0.002411 | 1.701 | +1 | 12 |
| - | - | 4.196E+04 | 1419 | - | - | 0 | - |
| - | - | 1.672E+04 | 1420 | - | - | 0 | - |
| - | - | 8052 | 1421 | - | - | 0 | - |
| - | - | 4451 | 1422 | - | - | 0 | - |
| - | - | 2433 | 1423 | - | - | 0 | - |
| 6 | z | 6.303E+04 | 1436 | 0.004547 | 3.167 | +1 | 12 |
| - | - | 1.103E+05 | 1437 | - | - | 0 | - |
| - | - | 6.652E+04 | 1438 | - | - | 0 | - |
| - | - | 3.425E+04 | 1439 | - | - | 0 | - |
| - | - | 9010 | 1440 | - | - | 0 | - |
| - | - | 8014 | 1451 | - | - | 0 | - |
| 6 | y | 3.849E+04 | 1452 | 0.002181 | 1.502 | +1 | 12 |
| - | - | 2.473E+04 | 1453 | - | - | 0 | - |
| - | - | 1.26E+04 | 1454 | - | - | 0 | - |
| - | - | 5919 | 1455 | - | - | 0 | - |
| - | - | 6044 | 1503 | - | - | 0 | - |
| 5 | z | 5560 | 1523 | 0.005234 | 3.437 | +1 | 13 |
| - | - | 2.693E+04 | 1524 | - | - | 0 | - |
| - | - | 2.392E+04 | 1525 | - | - | 0 | - |
| - | - | 1.164E+04 | 1526 | - | - | 0 | - |
| - | - | 5703 | 1527 | - | - | 0 | - |
| - | - | 2696 | 1530 | - | - | 0 | - |
| 5 | y | 7581 | 1539 | 0.004332 | 2.815 | +1 | 13 |
| - | - | 7726 | 1540 | - | - | 0 | - |
| - | - | 2969 | 1545 | - | - | 0 | - |
| 13 | c | 6.892E+04 | 1546 | 0.003037 | 1.965 | +1 | 13 |
| - | - | 5.484E+04 | 1547 | - | - | 0 | - |
| - | - | 2.944E+04 | 1548 | - | - | 0 | - |
| - | - | 1.061E+04 | 1549 | - | - | 0 | - |
| - | - | 4600 | 1550 | - | - | 0 | - |
| - | - | 5231 | 1602 | - | - | 0 | - |
| - | - | 2494 | 1604 | - | - | 0 | - |
| - | - | 6269 | 1615 | - | - | 0 | - |
| - | - | 5520 | 1616 | - | - | 0 | - |
| 4 | y | 2685 | 1618 | 0.02463 | 15.23 | +1 | 14 |
| 14 | c | 2961 | 1627 | 0.0008567 | 0.5266 | +1 | 14 |
| 14 | c | 7083 | 1628 | 0.0133 | 8.168 | +1 | 14 |
| - | - | 6529 | 1629 | - | - | 0 | - |
| - | - | 5012 | 1630 | - | - | 0 | - |
| 4 | y | 2.144E+04 | 1636 | 0.00357 | 2.182 | +1 | 14 |
| - | - | 1.935E+04 | 1637 | - | - | 0 | - |
| - | - | 1.178E+04 | 1638 | - | - | 0 | - |
| - | - | 5040 | 1639 | - | - | 0 | - |
| - | - | 9183 | 1644 | - | - | 0 | - |
| 14 | c | 7.581E+04 | 1645 | 0.002617 | 1.591 | +1 | 14 |
| - | - | 6.678E+04 | 1646 | - | - | 0 | - |
| - | - | 3.928E+04 | 1647 | - | - | 0 | - |
| - | - | 1.629E+04 | 1648 | - | - | 0 | - |
| - | - | 6923 | 1649 | - | - | 0 | - |
| - | - | 2487 | 1650 | - | - | 0 | - |
| - | - | 3356 | 1671 | - | - | 0 | - |
| - | - | 3315 | 1672 | - | - | 0 | - |
| - | - | 2864 | 1675 | - | - | 0 | - |
| - | - | 2572 | 1688 | - | - | 0 | - |
| - | - | 6613 | 1689 | - | - | 0 | - |
| - | - | 5234 | 1690 | - | - | 0 | - |
| - | - | 5333 | 1716 | - | - | 0 | - |
| 3 | z | 6063 | 1717 | 0.02507 | 14.6 | +1 | 15 |
| - | - | 4308 | 1718 | - | - | 0 | - |
| - | - | 7913 | 1729 | - | - | 0 | - |
| - | - | 6991 | 1730 | - | - | 0 | - |
| - | - | 4696 | 1731 | - | - | 0 | - |
| 15 | c | 5.133E+04 | 1732 | 0.004402 | 2.542 | +1 | 15 |
| 3 | y | 6.834E+04 | 1733 | 0.02564 | 14.79 | +1 | 15 |
| - | - | 4.747E+04 | 1734 | - | - | 0 | - |
| - | - | 2.912E+04 | 1735 | - | - | 0 | - |
| - | - | 1.239E+04 | 1736 | - | - | 0 | - |
| - | - | 4631 | 1737 | - | - | 0 | - |
| - | - | 3009 | 1786 | - | - | 0 | - |
| - | - | 4491 | 1787 | - | - | 0 | - |
| - | - | 3023 | 1788 | - | - | 0 | - |
| - | - | 4304 | 1801 | - | - | 0 | - |
| - | - | 1.745E+04 | 1802 | - | - | 0 | - |
| - | - | 1.647E+04 | 1803 | - | - | 0 | - |
| - | - | 9177 | 1804 | - | - | 0 | - |
| - | - | 6143 | 1805 | - | - | 0 | - |
| - | - | 3055 | 1817 | - | - | 0 | - |
| 2 | y | 7234 | 1829 | 0.02342 | 12.8 | +1 | 16 |
| 2 | z | 2.148E+04 | 1830 | 0.01132 | 6.186 | +1 | 16 |
| - | - | 7.785E+04 | 1831 | - | - | 0 | - |
| - | - | 6.963E+04 | 1832 | - | - | 0 | - |
| 16 | c | 4.399E+04 | 1833 | 0.01214 | 6.625 | +1 | 16 |
| - | - | 1.944E+04 | 1834 | - | - | 0 | - |
| - | - | 6319 | 1835 | - | - | 0 | - |
| - | - | 2.097E+04 | 1845 | - | - | 0 | - |
| 2 | y | 4.881E+04 | 1846 | 0.02641 | 14.31 | +1 | 16 |
| - | - | 5.2E+04 | 1847 | - | - | 0 | - |
| - | - | 3.443E+04 | 1848 | - | - | 0 | - |
| - | - | 1.767E+04 | 1849 | - | - | 0 | - |
| - | - | 7751 | 1850 | - | - | 0 | - |
| - | - | 3032 | 1851 | - | - | 0 | - |
| - | - | 4964 | 1872 | - | - | 0 | - |
| - | - | 3618 | 1873 | - | - | 0 | - |
| - | - | 2915 | 1878 | - | - | 0 | - |
| - | - | 6850 | 1885 | - | - | 0 | - |
| - | - | 8371 | 1886 | - | - | 0 | - |
| - | - | 1.255E+04 | 1887 | - | - | 0 | - |
| - | - | 1.255E+04 | 1888 | - | - | 0 | - |
| - | - | 1.16E+04 | 1889 | - | - | 0 | - |
| - | - | 2.454E+04 | 1890 | - | - | 0 | - |
| - | - | 2.303E+04 | 1891 | - | - | 0 | - |
| - | - | 1.132E+04 | 1892 | - | - | 0 | - |
| - | - | 6842 | 1893 | - | - | 0 | - |
| - | - | 1.738E+04 | 1903 | - | - | 0 | - |
| - | - | 5.529E+04 | 1904 | - | - | 0 | - |
| - | - | 4.682E+04 | 1905 | - | - | 0 | - |
| - | - | 2.808E+04 | 1906 | - | - | 0 | - |
| - | - | 1.24E+04 | 1907 | - | - | 0 | - |
| - | - | 4642 | 1908 | - | - | 0 | - |
| - | - | 1.221E+04 | 1914 | - | - | 0 | - |
| - | - | 1.649E+04 | 1915 | - | - | 0 | - |
| - | - | 1.524E+04 | 1916 | - | - | 0 | - |
| - | - | 6062 | 1917 | - | - | 0 | - |
| - | - | 3011 | 1918 | - | - | 0 | - |
| - | - | 9057 | 1920 | - | - | 0 | - |
| - | - | 3.921E+04 | 1921 | - | - | 0 | - |
| - | - | 3.728E+04 | 1922 | - | - | 0 | - |
| - | - | 2.201E+04 | 1923 | - | - | 0 | - |
| - | - | 9844 | 1924 | - | - | 0 | - |
| - | - | 4769 | 1925 | - | - | 0 | - |
| - | - | 4425 | 1930 | - | - | 0 | - |
| - | - | 5.508E+04 | 1931 | - | - | 0 | - |
| - | - | 1.767E+05 | 1932 | - | - | 0 | - |
| - | - | 1.689E+05 | 1933 | - | - | 0 | - |
| - | - | 1.066E+05 | 1934 | - | - | 0 | - |
| - | - | 4.466E+04 | 1935 | - | - | 0 | - |
| - | - | 1.728E+04 | 1936 | - | - | 0 | - |
| - | - | 5892 | 1937 | - | - | 0 | - |
| - | - | 2.083E+04 | 1947 | - | - | 0 | - |
| - | - | 1.09E+05 | 1948 | - | - | 0 | - |
| - | - | 4.942E+05 | 1949 | - | - | 0 | - |
| - | - | 4.696E+05 | 1950 | - | - | 0 | - |
| - | - | 2.97E+05 | 1951 | - | - | 0 | - |
| - | - | 1.378E+05 | 1952 | - | - | 0 | - |
| - | - | 4.606E+04 | 1953 | - | - | 0 | - |
| - | - | 1.401E+04 | 1954 | - | - | 0 | - |
| - | - | 2861 | 2152 | - | - | 0 | - |
| - | - | 2208 | 3084 | - | - | 0 | - |
| - | - | 2336 | 3085 | - | - | 0 | - |

m/z Charge Intensity FragmentType MassShift Position
120.06224060058594 0 1317.3342
120.0657730102539 0 16945.455
148.15476989746094 0 1313.1896
148.95465087890625 0 2079.7207
167.69308471679688 0 1623.258
183.25787353515625 0 1481.7021
187.14439392089844 0 138428.77
188.14772033691406 0 10539.407
197.04884338378906 0 1473.1488
201.12351989746094 0 4511.198 y Water loss 13
212.1397247314453 0 5210.7266
215.1392822265625 0 93136.78 y Water loss 15
216.14247131347656 0 10734.094
233.14990234375 0 7160.8687 y 15
256.16571044921875 0 3408.5598
276.0448913574219 0 1570.7991
299.171875 0 17422.291
300.1758728027344 0 3697.025
302.17108154296875 0 2130.5798 y Water loss 14
312.191650390625 0 3606.5483
320.1820983886719 0 4429.1025 y 14
326.1598205566406 0 2317.8252
369.1893005371094 0 2288.8628
382.2575988769531 0 1932.113
407.2532958984375 0 5583.308
425.2630615234375 0 3524.5518
426.2709045410156 0 5958.8813 c 3
427.2713928222656 0 2334.2944
428.2736511230469 0 1981.6381
466.8216552734375 0 1980.6085
470.2976379394531 0 8682.56
471.3027648925781 0 2302.4346
471.9010009765625 0 2263.8267
472.2889709472656 0 3160.982
494.2856140136719 0 4828.5996
495.2869873046875 0 1922.857 c Water loss 4
499.907958984375 0 3170.7014
500.90985107421875 0 4619.841
501.2451171875 0 3812.9517
501.57696533203125 0 2011.204
501.9144287109375 0 1824.74
512.2965087890625 0 3096.802
513.3035888671875 0 93476.81 c 4
514.3064575195312 0 28297.334
515.30810546875 0 4946.98
517.5961303710938 0 2816.7546
523.2676391601562 0 2187.9622
523.5995483398438 0 2794.8865
526.27685546875 0 2945.6997
527.2820434570312 0 3531.692
529.2725219726562 0 6606.763
529.6065063476562 0 8507.2295
529.9371948242188 0 4524.574
531.2896728515625 0 8642.352 z 12
532.2971801757812 0 10559.961
532.6007080078125 0 9155.0205
532.9330444335938 0 13811.606
533.2662353515625 0 6509.095
533.5995483398438 0 5136.2876
538.6040649414062 0 30721.719
538.9379272460938 0 32281.797
539.2723388671875 0 12255.257
539.6067504882812 0 6603.662
544.2741088867188 0 1816.3397
547.3108520507812 0 5597.4204 y 12
556.9575805664062 0 5230.9233
557.2908325195312 0 6935.456
557.6231689453125 0 6732.664
557.9595336914062 0 3284.5854
558.2924194335938 0 2295.445
561.1382446289062 0 1894.1481
566.282470703125 0 3267.8887
566.6148071289062 0 3922.0232
567.2836303710938 0 1851.9148 z Ammonia loss 2
569.3286743164062 0 2506.637
570.3109741210938 0 3297.2654
571.3186645507812 0 7330.1753
572.2867431640625 0 18675.62 c Ammonia loss 14
572.619384765625 0 20675.594 y Ammonia loss 2
572.9541625976562 0 15762.723 z 2
573.283935546875 0 5151.25
573.3351440429688 0 4811.45
573.6216430664062 0 3409.8894
573.9487915039062 0 1883.862
577.9613037109375 0 2760.3738 c 14
578.2901611328125 0 464821.06 y 2
578.6241455078125 0 455280.56
578.9580688476562 0 262908.1
579.2916259765625 0 104055.39 c Water loss 9
579.6253662109375 0 30607.885
579.9589233398438 0 10585.377
607.3829956054688 0 2337.8186
615.3474731445312 0 3220.0012
625.3914184570312 0 9498.696
626.3956298828125 0 6086.5454
643.6649780273438 0 3786.1243
643.99755859375 0 6844.3213
644.3344116210938 0 1915.2247 y Ammonia loss 11
645.3318481445312 0 5875.896 z 11
646.3392333984375 0 8505.954
647.3404541015625 0 3840.1562
649.3217163085938 0 1854.727
652.3375244140625 0 16005.984 c 10
652.8402099609375 0 14549.073
653.3397216796875 0 6834.6597
657.8074340820312 0 4410.599
661.352783203125 0 17990.033 y 11
662.3524780273438 0 5192.43
669.405029296875 0 159469.22 c 5
670.4075927734375 0 55286.25
671.4102172851562 0 13563.334
672.408935546875 0 5054.744
686.8511962890625 0 2437.6108
687.350830078125 0 3659.497
699.8513793945312 0 5888.4194
700.348388671875 0 8556.699 c Water loss 11
700.8485717773438 0 3048.9287 c Ammonia loss 11
707.3436889648438 0 13800.857
707.8451538085938 0 11821.016
708.3432006835938 0 3933.6025
708.8543090820312 0 26439.383
709.3589477539062 0 104627 c 11
709.8602294921875 0 85895.03
710.3609008789062 0 43777.773
710.8619384765625 0 14788.468
711.362060546875 0 3682.0596
711.4314575195312 0 2770.6
718.355712890625 0 7390.353 z 5
718.854736328125 0 5785.8584
719.3563842773438 0 5731.4805
719.8548583984375 0 2622.1453
720.3622436523438 0 2533.6128
722.3660278320312 0 8376.181
722.8656616210938 0 4778.5107
725.857666015625 0 5573.9336
726.3613891601562 0 19423.184 y 5
726.86328125 0 11136.965
727.3629150390625 0 9613.826
729.4225463867188 0 19702.252
730.4332275390625 0 10140.5625
748.8639526367188 0 3408.6765
749.3652954101562 0 2626.9927
750.8612670898438 0 13604.9795
751.3643798828125 0 17004.234
751.8671264648438 0 6259.2974
752.3626708984375 0 3189.7944
753.8682250976562 0 3173.445
754.4373168945312 0 3865.0754
755.4337768554688 0 3173.7495
756.4185791015625 0 2594.0566
757.893798828125 0 2121.8494
758.8792114257812 0 2576.044
761.366455078125 0 2542.9126 y Ammonia loss 4
761.8693237304688 0 15933.892 z 4
762.37060546875 0 15469.904
762.8719482421875 0 11327.233
763.3690185546875 0 4161.6895
765.3916015625 0 3735.145
765.8914794921875 0 3878.47
766.8997802734375 0 2129.4976
769.877197265625 0 13534.735 y 4
770.3806762695312 0 7796.387
770.883056640625 0 4918.007
772.4437255859375 0 2457.4624
772.8870849609375 0 7805.9053
773.390380859375 0 141147.83 c 12
773.4512329101562 0 28987.375
773.889892578125 0 139466.95
774.392333984375 0 62566.406
774.4530029296875 0 8989.45
774.8898315429688 0 30958.281
775.392578125 0 9223.454
779.4325561523438 0 2111.3606
784.4058837890625 0 2693.0774
786.397216796875 0 2264.7664
789.4466552734375 0 22901.037 y 10
790.4482421875 0 7298.6025
791.9119873046875 0 3191.8247
792.4056396484375 0 2879.3428
792.9081420898438 0 5432.381
793.4065551757812 0 14454.102
793.9105224609375 0 10339.342
794.405517578125 0 5223.122
794.90673828125 0 3342.69
797.4369506835938 0 4876.7603
798.4478149414062 0 249120.48 c 6
798.8922119140625 0 3380.5757
799.4505004882812 0 106139.695
800.4523315429688 0 26204.258
800.9146118164062 0 12243.497
801.4171142578125 0 11172.066
801.9169921875 0 8533.264
802.4183959960938 0 4832.877
803.4087524414062 0 5530.836
805.399658203125 0 2452.0588
807.4025268554688 0 35865.27
807.9031982421875 0 31854.705
808.403076171875 0 17516.83
808.905029296875 0 9829.986
809.403076171875 0 3843.2366 y Water loss 3
809.89697265625 0 3530.5295 y Ammonia loss 3
812.4682006835938 0 2729.8354
813.4146118164062 0 46101.29
813.9140625 0 43270.266 c Water loss 13
814.4127197265625 0 33565.25 c Ammonia loss 13
814.4937744140625 0 3180.1536
814.91259765625 0 15097.668
815.4125366210938 0 11693.3125
815.9174194335938 0 4554.7866
816.4105834960938 0 2232.06
817.397216796875 0 2670.2773
817.90234375 0 2845.829
818.4059448242188 0 71793.086 y 3
818.9058837890625 0 65309.332
819.4073486328125 0 29242.785
819.90771484375 0 13317.777
820.407958984375 0 3794.5598
822.4191284179688 0 79386.69
822.9213256835938 0 101084.945 c 13
823.4219970703125 0 89654.06
823.923828125 0 43572.277
824.4359130859375 0 14781.895
824.9215087890625 0 4877.1826
825.4649047851562 0 4412.2646
830.4635009765625 0 7453.399
831.4654541015625 0 5173.843
832.4140625 0 2701.7146
832.907958984375 0 3624.1235
834.931884765625 0 13664.267
835.4310302734375 0 17144.252
835.9292602539062 0 7406.4224
836.430908203125 0 6941.4224
836.92822265625 0 4741.307
843.9271850585938 0 3372.5823
844.432373046875 0 2963.315
844.9261474609375 0 4838.9775
845.422119140625 0 9580.953 w 2
845.9138793945312 0 9268.828
846.4175415039062 0 3577.4727
849.4180908203125 0 5183.117
856.46533203125 0 4662.492 z Water loss 9
856.933837890625 0 2509.8464
857.4725952148438 0 12748.83 w 9
857.927734375 0 14761.404 c Ammonia loss 14
858.4285888671875 0 15562.21 y Ammonia loss 2
858.9270629882812 0 11924.754 z 2
859.44970703125 0 25329.92 w 9
859.925537109375 0 5065.4565
860.4542236328125 0 10243.973
865.4472045898438 0 6554.5254
865.9349975585938 0 19021.828
866.4388427734375 0 213491.98 c 14
866.9345703125 0 485820 y 2
867.4344482421875 0 381265.47
867.9345703125 0 205261.66
868.0377807617188 0 2961.3706
868.4346923828125 0 76448.3
868.9362182617188 0 22957.236
869.4376831054688 0 5345.915
872.4689331054688 0 4532.5264 y Water loss 9
872.9461059570312 0 3086.1448
873.46630859375 0 23169.572 y Ammonia loss 9
874.475830078125 0 126793.4 z 9
875.479248046875 0 69212.03
876.4812622070312 0 21826.393
877.4832153320312 0 5360.892
879.4500732421875 0 10669.805
879.9501342773438 0 12685.603
880.4486083984375 0 4541.3774
880.9462280273438 0 3520.32
881.4344482421875 0 6416.4893
882.4330444335938 0 2621.8728
883.4659423828125 0 4883.485
884.470458984375 0 3097.5415
887.4349975585938 0 7115.622
887.938232421875 0 5720.3906
888.4512939453125 0 3447.4485
889.488037109375 0 3546.5322
890.4944458007812 0 45039.86 y 9
891.4961547851562 0 17281.533
891.9715576171875 0 7110.058
892.4876098632812 0 9625.34
892.9657592773438 0 5250.323
893.4574584960938 0 3959.8838
893.9365844726562 0 83952.3 w 1
894.4376831054688 0 82312.71
894.93896484375 0 45061.117
895.4390869140625 0 27148.701
895.9393920898438 0 5387.6465
900.9761352539062 0 8977.565
901.4760131835938 0 15094.133
901.9777221679688 0 14166.799
902.47705078125 0 11046.691
902.9835815429688 0 2719.9014
903.4695434570312 0 4170.9614
906.4719848632812 0 5055.4395 z Water loss 1
906.969482421875 0 6196.4507
907.4766235351562 0 5731.8984
908.45849609375 0 2478.1255 c Ammonia loss 15
909.4563598632812 0 2590.8574
913.4795532226562 0 2271.6174
913.9771728515625 0 3097.6436
914.4741821289062 0 11836.748 y Water loss 1
914.9656372070312 0 15435.385 y Ammonia loss 1
915.467529296875 0 48401.33 z 1
915.9672241210938 0 46462.938
916.4694213867188 0 32531.4
916.9625244140625 0 18366.516 c 15
917.4564208984375 0 8548.43
920.4881591796875 0 7853.2324
920.9912719726562 0 3857.2874
921.4886474609375 0 4604.1797
922.4842529296875 0 7629.6304
922.98095703125 0 197797.92
923.4813842773438 0 191230.83 y 1
923.9832153320312 0 141006.23
924.4827880859375 0 72919.65
924.9827880859375 0 27781.938
925.4854736328125 0 8174.913
926.48388671875 0 7399.5483
927.4901733398438 0 241271.61 c 7
928.4931030273438 0 124756.36
928.9791870117188 0 8024.164
929.494384765625 0 41570.688
929.96875 0 13560.411
930.4852905273438 0 21234.81
930.97314453125 0 6054.278
931.4918823242188 0 7407.512
931.9652709960938 0 3510.3306
932.491455078125 0 2489.3376
934.499267578125 0 4622.5537
934.9915161132812 0 7182.036
935.4927978515625 0 5622.5884
935.9830932617188 0 4812.792
936.4807739257812 0 7490.4473
936.9823608398438 0 6272.477
937.470947265625 0 14272.528
937.958984375 0 82734.51
938.4620361328125 0 83925.2
938.96484375 0 59282.82
939.4708862304688 0 27877.232
939.9745483398438 0 9268.615
940.4739379882812 0 3386.5327
941.5166625976562 0 5585.1865
942.498046875 0 22120.447
942.9938354492188 0 27054.992
943.491943359375 0 31332.771
943.9888916015625 0 22877.709
944.48095703125 0 20128.86
944.9816284179688 0 48692.387
945.4822387695312 0 43998.42
945.9848022460938 0 24350.398
946.4736938476562 0 37430.875
946.9718627929688 0 35276.73
947.4718017578125 0 12842.112
947.9725952148438 0 6574.2812
948.4757690429688 0 3824.3635
951.499755859375 0 11577.753
951.9903564453125 0 43930.492
952.4926147460938 0 48984.344
952.9905395507812 0 50511.984
953.4932250976562 0 32008.64
953.9869384765625 0 13757.901
954.4907836914062 0 7238.275
956.489013671875 0 18150.465
956.9866333007812 0 39367.02
957.4874877929688 0 43056.035
957.9859619140625 0 24571.436
958.5148315429688 0 34639.715
958.9869995117188 0 8934.484
959.5276489257812 0 98388.445
959.970947265625 0 2358.7947
960.5301513671875 0 55271.277
961.5316772460938 0 15698.354
962.5339965820312 0 4890.378
965.494140625 0 61159.77
965.9908447265625 0 302769.84
966.4917602539062 0 292998.66
966.991943359375 0 186198.78
967.4921875 0 102493.71
967.99267578125 0 29892.1
968.520263671875 0 13646.959
969.5320434570312 0 7937.293
973.9971923828125 0 219664.56
974.4991455078125 0 370235.3
975.0005493164062 0 301457.6
975.501708984375 0 178010.97
976.0028076171875 0 77582.445
976.5037231445312 0 20960.549
977.0033569335938 0 4786.178
1003.5026245117188 0 3057.0742 z Water loss 8
1004.5123901367188 0 3011.058
1005.51025390625 0 3266.1
1006.514892578125 0 4307.296
1012.5026245117188 0 6503.5894
1013.4937133789062 0 2916.5786
1015.505859375 0 4565.1416
1016.500244140625 0 3292.6455
1021.511962890625 0 28349.703 z 8
1022.515380859375 0 25409.94
1023.51904296875 0 10005.324
1030.5115966796875 0 26060.264
1031.51708984375 0 19983.635
1032.514404296875 0 11006.136
1033.5194091796875 0 4319.6387
1037.53125 0 15044.86 y 8
1038.5316162109375 0 10827.516
1049.5302734375 0 14885.824
1050.533447265625 0 6269.23
1055.502197265625 0 3524.628
1073.5213623046875 0 5039.359
1074.5257568359375 0 255996.25 c 8
1075.5284423828125 0 149509.61
1076.5289306640625 0 57951.758
1077.5281982421875 0 21402.254
1078.5308837890625 0 4931.029
1079.52294921875 0 2185.285
1087.5640869140625 0 6588.0903
1088.559814453125 0 4409.2124
1091.5419921875 0 33484.617 w 7
1092.5430908203125 0 22730.701
1093.54296875 0 9917.88
1094.500732421875 0 6537.203
1095.4931640625 0 2481.0713
1113.552490234375 0 6292.895
1114.54443359375 0 3189.5571
1131.55908203125 0 5836.098
1132.5523681640625 0 9065.886 z Water loss 7
1133.5714111328125 0 2718.984
1134.5626220703125 0 4826.4106
1149.5721435546875 0 2365.7651
1150.553955078125 0 104965.59 z 7
1151.5572509765625 0 76766.43
1152.556640625 0 38299.547
1153.561279296875 0 11473.251
1154.568115234375 0 2373.571
1166.5731201171875 0 7617.9873 y 7
1167.573974609375 0 4929.9224
1175.572998046875 0 120494.82 c 9
1176.57568359375 0 74041.64
1177.5760498046875 0 35900.543
1178.57763671875 0 16046.5205
1179.57763671875 0 4281.329
1207.5701904296875 0 2710.7012
1208.5755615234375 0 3885.7434
1220.579833984375 0 20269.006 w 6
1221.5836181640625 0 14321.709
1222.5897216796875 0 8258.298
1223.5458984375 0 3206.6885
1259.6517333984375 0 8129.8286
1260.6595458984375 0 8031.1416
1261.646728515625 0 2568.264
1279.595458984375 0 83898.13 z 6
1280.5975341796875 0 57612.246
1281.5992431640625 0 31101.562
1282.59912109375 0 8313.91
1284.6558837890625 0 2782.238
1285.64306640625 0 3379.8728 c Water loss 10
1286.65087890625 0 2839.4753 c Ammonia loss 10
1294.5855712890625 0 2308.5586
1295.615478515625 0 27289.82 y 6
1296.6156005859375 0 17148.969
1297.6163330078125 0 9531.239
1298.623046875 0 5113.8086
1302.65771484375 0 10226.301
1303.66455078125 0 23558.03 c 10
1304.6685791015625 0 16595.78
1305.669189453125 0 6550.5073
1306.68359375 0 3128.299
1333.672119140625 0 2708.5095
1334.6610107421875 0 6554.888
1361.669677734375 0 2745.614
1373.7000732421875 0 3605.0684
1374.7069091796875 0 3223.895
1376.682373046875 0 3381.9373
1379.6436767578125 0 3012.4126
1416.698974609375 0 6251.2656
1417.7078857421875 0 47244.645 c 11
1418.71044921875 0 41961.645
1419.7098388671875 0 16716.893
1420.71240234375 0 8052.382
1421.686767578125 0 4450.552
1422.7069091796875 0 2433.398
1435.69677734375 0 63029.63 z 5
1436.7010498046875 0 110349.71
1437.70263671875 0 66518.36
1438.7054443359375 0 34247.473
1439.70654296875 0 9009.72
1450.7076416015625 0 8013.5137
1451.713134765625 0 38494.914 y 5
1452.715576171875 0 24727.52
1453.7178955078125 0 12597.64
1454.7239990234375 0 5918.9487
1502.76220703125 0 6044.229
1522.7294921875 0 5560.3164 z 4
1523.73681640625 0 26930.488
1524.7373046875 0 23919.635
1525.741455078125 0 11642.113
1526.747802734375 0 5702.839
1529.7542724609375 0 2695.8403
1538.747314453125 0 7581.4033 y 4
1539.743896484375 0 7725.889
1544.7655029296875 0 2969.1614
1545.76708984375 0 68916.22 c 12
1546.7706298828125 0 54844.02
1547.769287109375 0 29438.256
1548.7711181640625 0 10612.444
1549.7760009765625 0 4599.593
1601.8270263671875 0 5230.833
1603.8240966796875 0 2493.5422
1614.7996826171875 0 6269.34
1615.8060302734375 0 5519.577
1617.809814453125 0 2684.9624 y Water loss 3
1626.821044921875 0 2961.4587 c Water loss 13
1627.8192138671875 0 7083.165 c Ammonia loss 13
1628.817138671875 0 6529.178
1629.8189697265625 0 5012.0664
1635.79931640625 0 21442 y 3
1636.799072265625 0 19346.24
1637.8037109375 0 11784.6875
1638.81787109375 0 5040.01
1643.830810546875 0 9182.821
1644.8350830078125 0 75811.23 c 13
1645.839111328125 0 66781.61
1646.84033203125 0 39275.777
1647.843994140625 0 16291.826
1648.8463134765625 0 6922.6465
1649.514404296875 0 2486.7898
1670.85498046875 0 3355.5042
1671.856201171875 0 3315.3281
1674.82470703125 0 2863.9287
1687.867919921875 0 2572.3665
1688.856689453125 0 6612.626
1689.873046875 0 5234.258
1715.8572998046875 0 5333.487
1716.8548583984375 0 6062.6626 z 2
1717.8712158203125 0 4307.757
1728.8966064453125 0 7912.8223
1729.8963623046875 0 6991.2554
1730.8880615234375 0 4695.654
1731.868896484375 0 51327.004 c 14
1732.8741455078125 0 68340.58 y 2
1733.87353515625 0 47472.223
1734.8743896484375 0 29123.438
1735.8739013671875 0 12394.619
1736.8795166015625 0 4630.585
1785.9078369140625 0 3009.0571
1786.90869140625 0 4491.3267
1787.9171142578125 0 3022.98
1800.943359375 0 4304.4995
1801.945068359375 0 17454.676
1802.945556640625 0 16474.756
1803.9451904296875 0 9177.153
1804.9517822265625 0 6142.7153
1816.9580078125 0 3055.4915
1828.929443359375 0 7233.8125 y Ammonia loss 1
1829.9251708984375 0 21476.873 z 1
1830.9259033203125 0 77847.484
1831.9278564453125 0 69628.02
1832.92431640625 0 43986.938 c 15
1833.9251708984375 0 19440.361
1834.937255859375 0 6319.0444
1844.951171875 0 20966.549
1845.958984375 0 48810.43 y 1
1846.961669921875 0 52003.082
1847.961181640625 0 34431.574
1848.956787109375 0 17670.342
1849.9630126953125 0 7750.888
1850.8946533203125 0 3031.816
1871.93310546875 0 4963.762
1872.95556640625 0 3618.2944
1877.95068359375 0 2914.672
1884.99609375 0 6849.8247
1885.9932861328125 0 8371.219
1886.972412109375 0 12550.263
1887.964599609375 0 12547.251
1888.9595947265625 0 11597.757
1889.958740234375 0 24535.29
1890.958984375 0 23029.97
1891.963623046875 0 11319.777
1892.9681396484375 0 6842.3857
1902.9822998046875 0 17378.973
1903.9847412109375 0 55293.19
1904.9835205078125 0 46822.65
1905.9835205078125 0 28079.793
1906.983154296875 0 12399.703
1907.9873046875 0 4642.1636
1913.9619140625 0 12212.41
1914.9581298828125 0 16492.36
1915.9580078125 0 15238.379
1916.9571533203125 0 6062.4233
1917.9525146484375 0 3010.519
1919.992431640625 0 9056.802
1921.0045166015625 0 39213.42
1922.0067138671875 0 37279.27
1923.00927734375 0 22014.117
1924.0179443359375 0 9844.34
1924.9912109375 0 4769.0923
1929.975341796875 0 4424.6055
1930.9754638671875 0 55084.832
1931.9752197265625 0 176706.08
1932.9764404296875 0 168872.4
1933.9776611328125 0 106637.31
1934.9815673828125 0 44656.203
1935.9779052734375 0 17278.615
1936.9774169921875 0 5892.0996
1946.983642578125 0 20830.326
1947.989013671875 0 108965.63
1948.9954833984375 0 494190.84
1949.9996337890625 0 469580.56
1951.0013427734375 0 296976.7
1952.002197265625 0 137849.2
1953.003173828125 0 46061.863
1954.005126953125 0 14009.314
2152.074462890625 0 2860.8787
3083.98193359375 0 2207.9036
3084.62353515625 0 2336.242

Spectrum Details

|  |  |
| --- | --- |
| Matched peaks? Matched peaks The total absolute number of peaks matched. Additionally in brackets the total fraction of peaks matched and the total number of peaks is shown. | 94 (15.24% of 617) |
| FDR? FDR The false discovery rate estimated for this peptide. It is calculated by matching all theoretical fragments with a non-integer shift with the raw peaks for this spectrum. This is done with 40 different shifts. The resulting percentage is the average number of annotated peaks over the number of annotated peaks with the correct spectrum. | 0.79% |
| Satellite FDR? Satellite FDR See the FDR for details on its calculation. This satellite ion specific FDR only contains the satellite ions (d/w) for I/L/J positions. | 11.90% |
| PSM Score? PSM Score The PSM Score as given by Hecklib to this annotated spectrum. It is shown with three significant figures. | 681 |

#### Spectrum 4116? Spectrum 4116 The raw spectrum of this peptide as annotated by Hecklib. The fragments are coloured according to ion type (see legend). Any peaks with a star '\*' as text can be hovered over to see the full details, first the ion type second the mass shift type. By hovering over the amino acids in the peptide or ions in the legend the corresponding peaks are highlighted. By toggling the 'Unassigned' label you can turn the background (unassigned) peaks on or off in the plot. By updating the slider in the Ion legend you can update the spectrum to only show the top X% of the peaks with labels. The top X% means any peak that is within X% of the highest intensity. By dragging in the spectrum you can zoom in to a specific part of the spectrum and use 'Zoom Out' to get back to the original zoom level. The annotation of the spectrum is based on the given sequence in the peptides file and is done with different software so inconsistencies are likely. The peaks are annotated based on the given sequence, with 20 ppm tolerance.

Copy Data

##### Spectrum 4116 (TSV)

###### Preview

```
Loading example...
```

*Click on the button to copy the data to your clipboard.*

Mz MinMz MaxIntensity Max

WidthHeightPeptide font sizePeptide stroke widthSpectrum font sizeSpectrum stroke widthCompact peptide

Ion legend

wxyz

abcd

OtherUnassignedIon^Charge^_Position_Show for top:%

TLPPSREEMTKNQVSTJ

06.62e+41.32e+51.99e+52.65e+5

Zoom Out

y^+2^_4_y^+1^_2_y^+1^_2_y^+1^_3_y^+1^_3_c^+1^_4_w^+1^_5_c^+3^_13_c^+1^_5_y^+1^_5_z^+1^_5_y^+1^_5_c^+3^_15_y^+3^_15_z^+3^_15_c^+3^_15_y^+3^_15_c^+2^_10_z^+1^_6_z^+1^_6_c^+2^_11_y^+1^_6_c^+1^_6_c^+2^_12_c^+2^_12_c^+2^_12_z^+2^_12_y^+2^_12_z^+2^_13_y^+2^_13_z^+2^_13_c^+2^_13_y^+2^_13_c^+2^_13_y^+1^_7_w^+2^_14_c^+1^_7_y^+2^_14_c^+2^_14_c^+2^_14_y^+2^_14_c^+2^_14_w^+2^_15_z^+1^_8_w^+1^_8_c^+2^_15_y^+2^_15_z^+2^_15_w^+1^_8_c^+2^_15_y^+2^_15_y^+1^_8_z^+1^_8_y^+1^_8_w^+2^_16_z^+2^_16_c^+2^_16_c^+2^_16_y^+2^_16_y^+2^_16_z^+2^_16_c^+2^_16_y^+2^_16_c^+1^_8_z^+1^_9_z^+1^_9_y^+1^_9_c^+1^_9_c^+1^_9_w^+1^_10_z^+1^_10_y^+1^_10_c^+1^_10_w^+1^_11_z^+1^_11_c^+1^_11_y^+1^_11_c^+1^_11_c^+1^_12_y^+1^_12_z^+1^_12_y^+1^_12_z^+1^_13_c^+1^_13_y^+1^_13_c^+1^_13_c^+1^_14_y^+1^_14_c^+1^_14_z^+1^_15_c^+1^_15_y^+1^_15_y^+1^_16_z^+1^_16_c^+1^_16_y^+1^_16_

0543108716302174

Fragment Matches Table

Show background peaks

| Position | Ion type | Intensity | mz Theoretical | mz Error (Th) | mz Error (ppm) | Charge | Series Number |
| --- | --- | --- | --- | --- | --- | --- | --- |
| - | - | 8159 | 120.1 | - | - | 0 | - |
| - | - | 647.1 | 122.4 | - | - | 0 | - |
| - | - | 671.4 | 125.3 | - | - | 0 | - |
| - | - | 640.3 | 133.6 | - | - | 0 | - |
| - | - | 715.4 | 148.9 | - | - | 0 | - |
| - | - | 837.1 | 155.9 | - | - | 0 | - |
| - | - | 731.1 | 156.1 | - | - | 0 | - |
| - | - | 861.3 | 164 | - | - | 0 | - |
| - | - | 934.9 | 168.4 | - | - | 0 | - |
| - | - | 808.7 | 173.1 | - | - | 0 | - |
| - | - | 4399 | 173.4 | - | - | 0 | - |
| - | - | 9.108E+04 | 187.1 | - | - | 0 | - |
| - | - | 8754 | 188.1 | - | - | 0 | - |
| - | - | 876.8 | 190.8 | - | - | 0 | - |
| 14 | y | 3532 | 201.1 | 0.0002579 | 1.282 | +2 | 4 |
| - | - | 837.9 | 207.9 | - | - | 0 | - |
| - | - | 2550 | 212.1 | - | - | 0 | - |
| 16 | y | 5.948E+04 | 215.1 | 0.0004006 | 1.862 | +1 | 2 |
| - | - | 6340 | 216.1 | - | - | 0 | - |
| - | - | 1082 | 226.1 | - | - | 0 | - |
| 16 | y | 5223 | 233.1 | 0.0003645 | 1.563 | +1 | 2 |
| - | - | 1054 | 234.2 | - | - | 0 | - |
| - | - | 1500 | 256.2 | - | - | 0 | - |
| - | - | 931.6 | 257.2 | - | - | 0 | - |
| - | - | 9990 | 299.2 | - | - | 0 | - |
| - | - | 1435 | 300.2 | - | - | 0 | - |
| 15 | y | 1898 | 302.2 | 0.0005225 | 1.729 | +1 | 3 |
| - | - | 2224 | 312.2 | - | - | 0 | - |
| - | - | 912.4 | 317.8 | - | - | 0 | - |
| 15 | y | 3220 | 320.2 | 0.0003948 | 1.233 | +1 | 3 |
| - | - | 848 | 347.3 | - | - | 0 | - |
| - | - | 1057 | 382.3 | - | - | 0 | - |
| - | - | 1342 | 383.3 | - | - | 0 | - |
| - | - | 1128 | 385.2 | - | - | 0 | - |
| - | - | 3305 | 407.3 | - | - | 0 | - |
| - | - | 2529 | 425.3 | - | - | 0 | - |
| 4 | c | 3682 | 426.3 | 0.0006489 | 1.522 | +1 | 4 |
| - | - | 1444 | 427.3 | - | - | 0 | - |
| - | - | 1378 | 429.1 | - | - | 0 | - |
| - | - | 914.3 | 444.8 | - | - | 0 | - |
| - | - | 2008 | 451.3 | - | - | 0 | - |
| - | - | 4880 | 470.3 | - | - | 0 | - |
| - | - | 1492 | 471.9 | - | - | 0 | - |
| 13 | w | 1235 | 473.3 | 0.001342 | 2.835 | +1 | 5 |
| - | - | 1039 | 491.9 | - | - | 0 | - |
| - | - | 2542 | 494.3 | - | - | 0 | - |
| - | - | 1302 | 495.6 | - | - | 0 | - |
| - | - | 1350 | 499.6 | - | - | 0 | - |
| - | - | 1496 | 500.9 | - | - | 0 | - |
| - | - | 3213 | 501.2 | - | - | 0 | - |
| 13 | c | 936.2 | 509.9 | 0.00655 | 12.84 | +3 | 13 |
| - | - | 1871 | 512.3 | - | - | 0 | - |
| 5 | c | 5.807E+04 | 513.3 | 0.000831 | 1.619 | +1 | 5 |
| - | - | 1.901E+04 | 514.3 | - | - | 0 | - |
| - | - | 2752 | 515.3 | - | - | 0 | - |
| - | - | 1415 | 520.2 | - | - | 0 | - |
| - | - | 1835 | 523.3 | - | - | 0 | - |
| - | - | 1446 | 523.6 | - | - | 0 | - |
| - | - | 1107 | 526.3 | - | - | 0 | - |
| - | - | 2474 | 526.9 | - | - | 0 | - |
| - | - | 2231 | 527.3 | - | - | 0 | - |
| - | - | 5916 | 529.3 | - | - | 0 | - |
| - | - | 4577 | 529.6 | - | - | 0 | - |
| - | - | 1223 | 529.9 | - | - | 0 | - |
| 13 | y | 1874 | 530.3 | 0.008006 | 15.1 | +1 | 5 |
| 13 | z | 6862 | 531.3 | 0.001625 | 3.058 | +1 | 5 |
| - | - | 4954 | 532.3 | - | - | 0 | - |
| - | - | 7025 | 532.6 | - | - | 0 | - |
| - | - | 8761 | 532.9 | - | - | 0 | - |
| - | - | 4135 | 533.3 | - | - | 0 | - |
| - | - | 1224 | 533.3 | - | - | 0 | - |
| - | - | 2747 | 533.6 | - | - | 0 | - |
| - | - | 1.805E+04 | 538.6 | - | - | 0 | - |
| - | - | 1.22E+04 | 538.9 | - | - | 0 | - |
| - | - | 8805 | 539.3 | - | - | 0 | - |
| - | - | 3756 | 539.6 | - | - | 0 | - |
| - | - | 1201 | 544.3 | - | - | 0 | - |
| 13 | y | 4514 | 547.3 | 0.001394 | 2.547 | +1 | 5 |
| - | - | 1188 | 553.3 | - | - | 0 | - |
| - | - | 4019 | 557 | - | - | 0 | - |
| - | - | 3239 | 557.3 | - | - | 0 | - |
| - | - | 3342 | 557.6 | - | - | 0 | - |
| - | - | 2479 | 558 | - | - | 0 | - |
| - | - | 1101 | 564.3 | - | - | 0 | - |
| - | - | 1801 | 570.3 | - | - | 0 | - |
| - | - | 4698 | 571.3 | - | - | 0 | - |
| 15 | c | 1.455E+04 | 572.3 | 0.002333 | 4.076 | +3 | 15 |
| - | - | 1209 | 572.3 | - | - | 0 | - |
| 3 | y | 1.518E+04 | 572.6 | 0.007641 | 13.34 | +3 | 15 |
| 3 | z | 8593 | 572.9 | 0.005927 | 10.35 | +3 | 15 |
| - | - | 4251 | 573.3 | - | - | 0 | - |
| - | - | 3155 | 573.3 | - | - | 0 | - |
| - | - | 2581 | 573.6 | - | - | 0 | - |
| - | - | 1043 | 574 | - | - | 0 | - |
| - | - | 1214 | 574.3 | - | - | 0 | - |
| 15 | c | 1875 | 578 | 0.001743 | 3.016 | +3 | 15 |
| 3 | y | 2.545E+05 | 578.3 | 0.002779 | 4.805 | +3 | 15 |
| - | - | 2.565E+05 | 578.6 | - | - | 0 | - |
| - | - | 1.475E+05 | 579 | - | - | 0 | - |
| 10 | c | 5.89E+04 | 579.3 | 0.009783 | 16.89 | +2 | 10 |
| - | - | 2.229E+04 | 579.6 | - | - | 0 | - |
| - | - | 6221 | 580 | - | - | 0 | - |
| - | - | 3513 | 607.4 | - | - | 0 | - |
| - | - | 970.1 | 615.3 | - | - | 0 | - |
| - | - | 7916 | 625.4 | - | - | 0 | - |
| - | - | 4747 | 626.4 | - | - | 0 | - |
| 12 | z | 1213 | 627.3 | 0.00077 | 1.227 | +1 | 6 |
| - | - | 1124 | 629.4 | - | - | 0 | - |
| - | - | 3493 | 643.7 | - | - | 0 | - |
| - | - | 3171 | 644 | - | - | 0 | - |
| 12 | z | 2452 | 645.3 | 0.0004704 | 0.7289 | +1 | 6 |
| - | - | 6893 | 646.3 | - | - | 0 | - |
| - | - | 1277 | 647.3 | - | - | 0 | - |
| - | - | 1527 | 648.3 | - | - | 0 | - |
| - | - | 2890 | 649.3 | - | - | 0 | - |
| - | - | 2766 | 649.7 | - | - | 0 | - |
| - | - | 2908 | 649.8 | - | - | 0 | - |
| 11 | c | 1.138E+04 | 652.3 | 0.003284 | 5.034 | +2 | 11 |
| - | - | 6921 | 652.8 | - | - | 0 | - |
| - | - | 4129 | 653.3 | - | - | 0 | - |
| 12 | y | 9921 | 661.4 | 0.0008251 | 1.248 | +1 | 6 |
| - | - | 2414 | 662.4 | - | - | 0 | - |
| 6 | c | 9.497E+04 | 669.4 | 0.00116 | 1.733 | +1 | 6 |
| - | - | 3.49E+04 | 670.4 | - | - | 0 | - |
| - | - | 7766 | 671.4 | - | - | 0 | - |
| - | - | 2948 | 672.4 | - | - | 0 | - |
| - | - | 1468 | 673.4 | - | - | 0 | - |
| - | - | 2446 | 686.8 | - | - | 0 | - |
| - | - | 2002 | 695.4 | - | - | 0 | - |
| - | - | 2551 | 699.9 | - | - | 0 | - |
| 12 | c | 2774 | 700.4 | 0.001606 | 2.293 | +2 | 12 |
| 12 | c | 2005 | 700.8 | 0.007973 | 11.38 | +2 | 12 |
| - | - | 8412 | 707.3 | - | - | 0 | - |
| - | - | 6172 | 707.8 | - | - | 0 | - |
| - | - | 3603 | 708.3 | - | - | 0 | - |
| - | - | 1.061E+04 | 708.9 | - | - | 0 | - |
| 12 | c | 6.192E+04 | 709.4 | 0.002999 | 4.228 | +2 | 12 |
| - | - | 4.539E+04 | 709.9 | - | - | 0 | - |
| - | - | 2.1E+04 | 710.4 | - | - | 0 | - |
| - | - | 8916 | 710.9 | - | - | 0 | - |
| - | - | 2372 | 711.4 | - | - | 0 | - |
| - | - | 1690 | 715.4 | - | - | 0 | - |
| 6 | z | 3699 | 718.3 | 0.006326 | 8.806 | +2 | 12 |
| - | - | 6068 | 718.9 | - | - | 0 | - |
| - | - | 3531 | 719.4 | - | - | 0 | - |
| - | - | 1459 | 719.9 | - | - | 0 | - |
| - | - | 1797 | 720.4 | - | - | 0 | - |
| - | - | 3552 | 722.4 | - | - | 0 | - |
| - | - | 2064 | 722.9 | - | - | 0 | - |
| - | - | 1110 | 723.4 | - | - | 0 | - |
| - | - | 2631 | 725.9 | - | - | 0 | - |
| 6 | y | 9361 | 726.4 | 0.004288 | 5.903 | +2 | 12 |
| - | - | 5804 | 726.9 | - | - | 0 | - |
| - | - | 3389 | 727.4 | - | - | 0 | - |
| - | - | 1308 | 727.9 | - | - | 0 | - |
| - | - | 1.227E+04 | 729.4 | - | - | 0 | - |
| - | - | 6360 | 730.4 | - | - | 0 | - |
| - | - | 1876 | 741.9 | - | - | 0 | - |
| - | - | 1737 | 742.4 | - | - | 0 | - |
| - | - | 1837 | 743.9 | - | - | 0 | - |
| - | - | 8660 | 750.9 | - | - | 0 | - |
| - | - | 1.044E+04 | 751.4 | - | - | 0 | - |
| - | - | 3476 | 751.9 | - | - | 0 | - |
| - | - | 2870 | 752.4 | - | - | 0 | - |
| 5 | z | 1461 | 752.9 | 0.00243 | 3.228 | +2 | 13 |
| - | - | 1765 | 754.4 | - | - | 0 | - |
| - | - | 1206 | 755.4 | - | - | 0 | - |
| - | - | 2037 | 758.4 | - | - | 0 | - |
| - | - | 2273 | 759.4 | - | - | 0 | - |
| 5 | y | 1413 | 761.4 | 0.001243 | 1.633 | +2 | 13 |
| 5 | z | 9716 | 761.9 | 0.003434 | 4.508 | +2 | 13 |
| - | - | 1.045E+04 | 762.4 | - | - | 0 | - |
| - | - | 3986 | 762.9 | - | - | 0 | - |
| - | - | 1805 | 763.4 | - | - | 0 | - |
| - | - | 1162 | 763.9 | - | - | 0 | - |
| 13 | c | 2042 | 764.4 | 0.002758 | 3.608 | +2 | 13 |
| - | - | 1563 | 764.9 | - | - | 0 | - |
| - | - | 3520 | 765.4 | - | - | 0 | - |
| - | - | 2041 | 765.9 | - | - | 0 | - |
| 5 | y | 4455 | 769.9 | 0.003777 | 4.906 | +2 | 13 |
| - | - | 7029 | 770.4 | - | - | 0 | - |
| - | - | 3795 | 770.9 | - | - | 0 | - |
| - | - | 4929 | 772.9 | - | - | 0 | - |
| 13 | c | 6.893E+04 | 773.4 | 0.005266 | 6.809 | +2 | 13 |
| - | - | 1.764E+04 | 773.5 | - | - | 0 | - |
| - | - | 7.958E+04 | 773.9 | - | - | 0 | - |
| - | - | 3.161E+04 | 774.4 | - | - | 0 | - |
| - | - | 6164 | 774.5 | - | - | 0 | - |
| - | - | 1.883E+04 | 774.9 | - | - | 0 | - |
| - | - | 4161 | 775.4 | - | - | 0 | - |
| - | - | 1713 | 775.9 | - | - | 0 | - |
| - | - | 2216 | 779.4 | - | - | 0 | - |
| - | - | 1426 | 785.4 | - | - | 0 | - |
| - | - | 1649 | 785.9 | - | - | 0 | - |
| - | - | 1151 | 786.4 | - | - | 0 | - |
| - | - | 1089 | 787.4 | - | - | 0 | - |
| - | - | 1182 | 788.4 | - | - | 0 | - |
| 11 | y | 1.386E+04 | 789.4 | 0.001199 | 1.519 | +1 | 7 |
| - | - | 5024 | 790.4 | - | - | 0 | - |
| - | - | 1860 | 791.9 | - | - | 0 | - |
| - | - | 3938 | 792.9 | - | - | 0 | - |
| - | - | 6435 | 793.4 | - | - | 0 | - |
| - | - | 5940 | 793.9 | - | - | 0 | - |
| - | - | 2109 | 794.4 | - | - | 0 | - |
| - | - | 1961 | 794.9 | - | - | 0 | - |
| - | - | 1176 | 795.4 | - | - | 0 | - |
| 4 | w | 1830 | 796.9 | 0.002584 | 3.242 | +2 | 14 |
| - | - | 1912 | 797.4 | - | - | 0 | - |
| 7 | c | 1.358E+05 | 798.4 | 0.001597 | 2 | +1 | 7 |
| - | - | 3278 | 798.9 | - | - | 0 | - |
| - | - | 5.912E+04 | 799.5 | - | - | 0 | - |
| - | - | 1616 | 799.9 | - | - | 0 | - |
| - | - | 1.392E+04 | 800.5 | - | - | 0 | - |
| - | - | 6436 | 800.9 | - | - | 0 | - |
| - | - | 6363 | 801.4 | - | - | 0 | - |
| - | - | 4740 | 801.9 | - | - | 0 | - |
| - | - | 1529 | 802.4 | - | - | 0 | - |
| - | - | 2.067E+04 | 807.4 | - | - | 0 | - |
| - | - | 2.037E+04 | 807.9 | - | - | 0 | - |
| - | - | 1609 | 808 | - | - | 0 | - |
| - | - | 6156 | 808.4 | - | - | 0 | - |
| - | - | 4781 | 808.9 | - | - | 0 | - |
| 4 | y | 1659 | 809.4 | 0.01197 | 14.79 | +2 | 14 |
| - | - | 2.2E+04 | 813.4 | - | - | 0 | - |
| 14 | c | 2.745E+04 | 813.9 | 8.381E-05 | 0.103 | +2 | 14 |
| 14 | c | 1.763E+04 | 814.4 | 0.006916 | 8.492 | +2 | 14 |
| - | - | 1.075E+04 | 814.9 | - | - | 0 | - |
| - | - | 6661 | 815.4 | - | - | 0 | - |
| - | - | 3005 | 815.9 | - | - | 0 | - |
| - | - | 1384 | 817.4 | - | - | 0 | - |
| - | - | 1972 | 817.9 | - | - | 0 | - |
| 4 | y | 3.885E+04 | 818.4 | 0.004372 | 5.343 | +2 | 14 |
| - | - | 3.634E+04 | 818.9 | - | - | 0 | - |
| - | - | 2.037E+04 | 819.4 | - | - | 0 | - |
| - | - | 7133 | 819.9 | - | - | 0 | - |
| - | - | 4770 | 820.4 | - | - | 0 | - |
| - | - | 4.422E+04 | 822.4 | - | - | 0 | - |
| 14 | c | 6.94E+04 | 822.9 | 0.002004 | 2.435 | +2 | 14 |
| - | - | 4.544E+04 | 823.4 | - | - | 0 | - |
| - | - | 2.81E+04 | 823.9 | - | - | 0 | - |
| - | - | 6716 | 824.4 | - | - | 0 | - |
| - | - | 2781 | 824.9 | - | - | 0 | - |
| - | - | 1302 | 825.5 | - | - | 0 | - |
| - | - | 3905 | 830.5 | - | - | 0 | - |
| - | - | 3066 | 831.5 | - | - | 0 | - |
| - | - | 7995 | 834.9 | - | - | 0 | - |
| - | - | 7693 | 835.4 | - | - | 0 | - |
| - | - | 7779 | 835.9 | - | - | 0 | - |
| - | - | 4919 | 836.4 | - | - | 0 | - |
| - | - | 1533 | 843.9 | - | - | 0 | - |
| - | - | 2051 | 844.4 | - | - | 0 | - |
| - | - | 4743 | 844.9 | - | - | 0 | - |
| 3 | w | 5504 | 845.4 | 0.01081 | 12.79 | +2 | 15 |
| - | - | 5838 | 845.9 | - | - | 0 | - |
| - | - | 2657 | 846.4 | - | - | 0 | - |
| - | - | 1333 | 846.9 | - | - | 0 | - |
| - | - | 2715 | 849.4 | - | - | 0 | - |
| - | - | 1567 | 852.4 | - | - | 0 | - |
| 10 | z | 2092 | 856.5 | 0.007608 | 8.883 | +1 | 8 |
| 10 | w | 8225 | 857.5 | 0.001639 | 1.912 | +1 | 8 |
| 15 | c | 9690 | 857.9 | 0.006832 | 7.964 | +2 | 15 |
| 3 | y | 8415 | 858.4 | 0.01397 | 16.27 | +2 | 15 |
| 3 | z | 8676 | 858.9 | 0.007616 | 8.867 | +2 | 15 |
| 10 | w | 1.386E+04 | 859.5 | 0.003243 | 3.773 | +1 | 8 |
| - | - | 3230 | 859.9 | - | - | 0 | - |
| - | - | 6324 | 860.5 | - | - | 0 | - |
| - | - | 1642 | 861.5 | - | - | 0 | - |
| - | - | 7728 | 865.9 | - | - | 0 | - |
| 15 | c | 1.158E+05 | 866.4 | 0.003506 | 4.047 | +2 | 15 |
| 3 | y | 2.622E+05 | 866.9 | 0.007043 | 8.124 | +2 | 15 |
| - | - | 2.142E+05 | 867.4 | - | - | 0 | - |
| - | - | 1.168E+05 | 867.9 | - | - | 0 | - |
| - | - | 4.79E+04 | 868.4 | - | - | 0 | - |
| - | - | 1.466E+04 | 868.9 | - | - | 0 | - |
| - | - | 2605 | 869.4 | - | - | 0 | - |
| - | - | 1545 | 872.5 | - | - | 0 | - |
| 10 | y | 1.214E+04 | 873.5 | 0.000216 | 0.2473 | +1 | 8 |
| 10 | z | 7.283E+04 | 874.5 | 0.000809 | 0.9252 | +1 | 8 |
| - | - | 3.785E+04 | 875.5 | - | - | 0 | - |
| - | - | 1.221E+04 | 876.5 | - | - | 0 | - |
| - | - | 2936 | 877.5 | - | - | 0 | - |
| - | - | 5206 | 879.5 | - | - | 0 | - |
| - | - | 5760 | 880 | - | - | 0 | - |
| - | - | 2932 | 880.5 | - | - | 0 | - |
| - | - | 2784 | 880.9 | - | - | 0 | - |
| - | - | 1263 | 881.4 | - | - | 0 | - |
| - | - | 2378 | 881.9 | - | - | 0 | - |
| - | - | 3478 | 883.5 | - | - | 0 | - |
| - | - | 2641 | 884.5 | - | - | 0 | - |
| - | - | 1218 | 885.5 | - | - | 0 | - |
| - | - | 1623 | 886 | - | - | 0 | - |
| - | - | 2444 | 887.4 | - | - | 0 | - |
| - | - | 2449 | 887.9 | - | - | 0 | - |
| - | - | 2689 | 889.5 | - | - | 0 | - |
| 10 | y | 2.407E+04 | 890.5 | 0.0006396 | 0.7183 | +1 | 8 |
| - | - | 1.282E+04 | 891.5 | - | - | 0 | - |
| - | - | 5011 | 892 | - | - | 0 | - |
| - | - | 6681 | 892.5 | - | - | 0 | - |
| - | - | 4096 | 893 | - | - | 0 | - |
| - | - | 4967 | 893.5 | - | - | 0 | - |
| 2 | w | 4.341E+04 | 893.9 | 0.004629 | 5.179 | +2 | 16 |
| - | - | 4.983E+04 | 894.4 | - | - | 0 | - |
| - | - | 2.915E+04 | 894.9 | - | - | 0 | - |
| - | - | 1.23E+04 | 895.4 | - | - | 0 | - |
| - | - | 5031 | 895.9 | - | - | 0 | - |
| - | - | 1337 | 899.5 | - | - | 0 | - |
| - | - | 1521 | 900.5 | - | - | 0 | - |
| - | - | 6506 | 901 | - | - | 0 | - |
| - | - | 1.036E+04 | 901.5 | - | - | 0 | - |
| - | - | 8068 | 902 | - | - | 0 | - |
| - | - | 6992 | 902.5 | - | - | 0 | - |
| - | - | 2604 | 903.5 | - | - | 0 | - |
| - | - | 1534 | 906 | - | - | 0 | - |
| 2 | z | 1805 | 906.5 | 0.005901 | 6.51 | +2 | 16 |
| - | - | 3787 | 907 | - | - | 0 | - |
| - | - | 3542 | 907.5 | - | - | 0 | - |
| 16 | c | 1787 | 908 | 0.006495 | 7.153 | +2 | 16 |
| 16 | c | 2008 | 908.4 | 0.01681 | 18.5 | +2 | 16 |
| - | - | 2918 | 909.5 | - | - | 0 | - |
| 2 | y | 5243 | 914.5 | 0.009539 | 10.43 | +2 | 16 |
| 2 | y | 7901 | 915 | 0.01015 | 11.09 | +2 | 16 |
| 2 | z | 2.661E+04 | 915.5 | 0.006722 | 7.343 | +2 | 16 |
| - | - | 2.934E+04 | 916 | - | - | 0 | - |
| - | - | 1.853E+04 | 916.5 | - | - | 0 | - |
| 16 | c | 9714 | 917 | 0.003288 | 3.586 | +2 | 16 |
| - | - | 3253 | 917.5 | - | - | 0 | - |
| - | - | 1806 | 918 | - | - | 0 | - |
| - | - | 5556 | 920.5 | - | - | 0 | - |
| - | - | 2939 | 921 | - | - | 0 | - |
| - | - | 2677 | 921.5 | - | - | 0 | - |
| - | - | 2849 | 922.5 | - | - | 0 | - |
| - | - | 1.143E+05 | 923 | - | - | 0 | - |
| 2 | y | 1.196E+05 | 923.5 | 0.01237 | 13.4 | +2 | 16 |
| - | - | 8.572E+04 | 924 | - | - | 0 | - |
| - | - | 3.427E+04 | 924.5 | - | - | 0 | - |
| - | - | 1.548E+04 | 925 | - | - | 0 | - |
| - | - | 4458 | 925.5 | - | - | 0 | - |
| - | - | 4841 | 926.5 | - | - | 0 | - |
| 8 | c | 1.358E+05 | 927.5 | 0.001301 | 1.403 | +1 | 8 |
| - | - | 7.607E+04 | 928.5 | - | - | 0 | - |
| - | - | 4061 | 929 | - | - | 0 | - |
| - | - | 2.218E+04 | 929.5 | - | - | 0 | - |
| - | - | 6919 | 930 | - | - | 0 | - |
| - | - | 1.008E+04 | 930.5 | - | - | 0 | - |
| - | - | 5594 | 931 | - | - | 0 | - |
| - | - | 3491 | 931.5 | - | - | 0 | - |
| - | - | 1357 | 932 | - | - | 0 | - |
| - | - | 1568 | 933.5 | - | - | 0 | - |
| - | - | 1671 | 934 | - | - | 0 | - |
| - | - | 1743 | 934.5 | - | - | 0 | - |
| - | - | 2655 | 935 | - | - | 0 | - |
| - | - | 2742 | 935.5 | - | - | 0 | - |
| - | - | 2459 | 936 | - | - | 0 | - |
| - | - | 6560 | 936.5 | - | - | 0 | - |
| - | - | 6034 | 937 | - | - | 0 | - |
| - | - | 8692 | 937.5 | - | - | 0 | - |
| - | - | 4.655E+04 | 938 | - | - | 0 | - |
| - | - | 4.93E+04 | 938.5 | - | - | 0 | - |
| - | - | 3.21E+04 | 939 | - | - | 0 | - |
| - | - | 1.664E+04 | 939.5 | - | - | 0 | - |
| - | - | 7595 | 940 | - | - | 0 | - |
| - | - | 2464 | 940.5 | - | - | 0 | - |
| - | - | 2661 | 941.5 | - | - | 0 | - |
| - | - | 9673 | 942.5 | - | - | 0 | - |
| - | - | 1.605E+04 | 943 | - | - | 0 | - |
| - | - | 1.776E+04 | 943.5 | - | - | 0 | - |
| - | - | 1.379E+04 | 944 | - | - | 0 | - |
| - | - | 1.056E+04 | 944.5 | - | - | 0 | - |
| - | - | 2.645E+04 | 945 | - | - | 0 | - |
| - | - | 2.643E+04 | 945.5 | - | - | 0 | - |
| - | - | 1.573E+04 | 946 | - | - | 0 | - |
| - | - | 1.68E+04 | 946.5 | - | - | 0 | - |
| - | - | 1.597E+04 | 947 | - | - | 0 | - |
| - | - | 1.123E+04 | 947.5 | - | - | 0 | - |
| - | - | 4483 | 948 | - | - | 0 | - |
| - | - | 1791 | 948.5 | - | - | 0 | - |
| - | - | 1324 | 950.5 | - | - | 0 | - |
| - | - | 3941 | 951.5 | - | - | 0 | - |
| - | - | 2.433E+04 | 952 | - | - | 0 | - |
| - | - | 2.819E+04 | 952.5 | - | - | 0 | - |
| - | - | 2.554E+04 | 953 | - | - | 0 | - |
| - | - | 1.639E+04 | 953.5 | - | - | 0 | - |
| - | - | 7491 | 954 | - | - | 0 | - |
| - | - | 2911 | 954.5 | - | - | 0 | - |
| - | - | 1.058E+04 | 956.5 | - | - | 0 | - |
| - | - | 1.874E+04 | 957 | - | - | 0 | - |
| - | - | 2.022E+04 | 957.5 | - | - | 0 | - |
| - | - | 1.321E+04 | 958 | - | - | 0 | - |
| - | - | 1.67E+04 | 958.5 | - | - | 0 | - |
| - | - | 4324 | 959 | - | - | 0 | - |
| - | - | 5.735E+04 | 959.5 | - | - | 0 | - |
| - | - | 2.947E+04 | 960.5 | - | - | 0 | - |
| - | - | 9987 | 961.5 | - | - | 0 | - |
| - | - | 1964 | 962.5 | - | - | 0 | - |
| - | - | 1199 | 965 | - | - | 0 | - |
| - | - | 3.646E+04 | 965.5 | - | - | 0 | - |
| - | - | 1.711E+05 | 966 | - | - | 0 | - |
| - | - | 1.845E+05 | 966.5 | - | - | 0 | - |
| - | - | 1.13E+05 | 967 | - | - | 0 | - |
| - | - | 5.243E+04 | 967.5 | - | - | 0 | - |
| - | - | 1.935E+04 | 968 | - | - | 0 | - |
| - | - | 9346 | 968.5 | - | - | 0 | - |
| - | - | 3413 | 969.5 | - | - | 0 | - |
| - | - | 3230 | 973.5 | - | - | 0 | - |
| - | - | 1.233E+05 | 974 | - | - | 0 | - |
| - | - | 2.174E+05 | 974.5 | - | - | 0 | - |
| - | - | 1.851E+05 | 975 | - | - | 0 | - |
| - | - | 1.071E+05 | 975.5 | - | - | 0 | - |
| - | - | 4.689E+04 | 976 | - | - | 0 | - |
| - | - | 1.679E+04 | 976.5 | - | - | 0 | - |
| - | - | 2342 | 977 | - | - | 0 | - |
| - | - | 1459 | 978.5 | - | - | 0 | - |
| 9 | z | 2547 | 1003 | 0.006341 | 6.319 | +1 | 9 |
| - | - | 2006 | 1006 | - | - | 0 | - |
| - | - | 1741 | 1007 | - | - | 0 | - |
| - | - | 1088 | 1013 | - | - | 0 | - |
| - | - | 1304 | 1014 | - | - | 0 | - |
| - | - | 1442 | 1015 | - | - | 0 | - |
| - | - | 1216 | 1016 | - | - | 0 | - |
| - | - | 1482 | 1017 | - | - | 0 | - |
| 9 | z | 1.482E+04 | 1022 | 0.005847 | 5.724 | +1 | 9 |
| - | - | 1.573E+04 | 1023 | - | - | 0 | - |
| - | - | 6673 | 1024 | - | - | 0 | - |
| - | - | 2823 | 1025 | - | - | 0 | - |
| - | - | 1.944E+04 | 1031 | - | - | 0 | - |
| - | - | 1.105E+04 | 1032 | - | - | 0 | - |
| - | - | 5617 | 1033 | - | - | 0 | - |
| - | - | 1610 | 1034 | - | - | 0 | - |
| 9 | y | 7217 | 1038 | 0.005739 | 5.531 | +1 | 9 |
| - | - | 4252 | 1039 | - | - | 0 | - |
| - | - | 2001 | 1040 | - | - | 0 | - |
| - | - | 6890 | 1050 | - | - | 0 | - |
| - | - | 3621 | 1051 | - | - | 0 | - |
| - | - | 2688 | 1052 | - | - | 0 | - |
| - | - | 2080 | 1056 | - | - | 0 | - |
| 9 | c | 1296 | 1057 | 0.0007906 | 0.7483 | +1 | 9 |
| - | - | 4116 | 1074 | - | - | 0 | - |
| 9 | c | 1.44E+05 | 1075 | 0.006339 | 5.9 | +1 | 9 |
| - | - | 8.245E+04 | 1076 | - | - | 0 | - |
| - | - | 3.131E+04 | 1077 | - | - | 0 | - |
| - | - | 1.432E+04 | 1078 | - | - | 0 | - |
| - | - | 1159 | 1079 | - | - | 0 | - |
| - | - | 2386 | 1080 | - | - | 0 | - |
| - | - | 1212 | 1081 | - | - | 0 | - |
| - | - | 3101 | 1088 | - | - | 0 | - |
| - | - | 2789 | 1089 | - | - | 0 | - |
| - | - | 1531 | 1090 | - | - | 0 | - |
| 8 | w | 1.87E+04 | 1092 | 0.006771 | 6.203 | +1 | 10 |
| - | - | 1.056E+04 | 1093 | - | - | 0 | - |
| - | - | 4504 | 1094 | - | - | 0 | - |
| - | - | 3633 | 1094 | - | - | 0 | - |
| - | - | 1976 | 1095 | - | - | 0 | - |
| - | - | 1271 | 1107 | - | - | 0 | - |
| - | - | 3595 | 1114 | - | - | 0 | - |
| - | - | 2964 | 1115 | - | - | 0 | - |
| - | - | 1798 | 1116 | - | - | 0 | - |
| - | - | 4379 | 1132 | - | - | 0 | - |
| - | - | 4064 | 1133 | - | - | 0 | - |
| - | - | 1836 | 1134 | - | - | 0 | - |
| - | - | 2864 | 1135 | - | - | 0 | - |
| - | - | 1903 | 1136 | - | - | 0 | - |
| 8 | z | 5.52E+04 | 1151 | 0.006406 | 5.567 | +1 | 10 |
| - | - | 4.396E+04 | 1152 | - | - | 0 | - |
| - | - | 2.044E+04 | 1153 | - | - | 0 | - |
| - | - | 5897 | 1154 | - | - | 0 | - |
| - | - | 2071 | 1155 | - | - | 0 | - |
| 8 | y | 3730 | 1167 | 0.00587 | 5.032 | +1 | 10 |
| - | - | 3076 | 1168 | - | - | 0 | - |
| - | - | 1988 | 1169 | - | - | 0 | - |
| 10 | c | 7.703E+04 | 1176 | 0.006146 | 5.228 | +1 | 10 |
| - | - | 4.057E+04 | 1177 | - | - | 0 | - |
| - | - | 2.437E+04 | 1178 | - | - | 0 | - |
| - | - | 6741 | 1179 | - | - | 0 | - |
| - | - | 2489 | 1180 | - | - | 0 | - |
| - | - | 1765 | 1181 | - | - | 0 | - |
| - | - | 1731 | 1208 | - | - | 0 | - |
| 7 | w | 1.021E+04 | 1221 | 0.002141 | 1.754 | +1 | 11 |
| - | - | 7110 | 1222 | - | - | 0 | - |
| - | - | 4149 | 1223 | - | - | 0 | - |
| - | - | 2845 | 1224 | - | - | 0 | - |
| - | - | 1266 | 1236 | - | - | 0 | - |
| - | - | 3656 | 1260 | - | - | 0 | - |
| - | - | 4128 | 1261 | - | - | 0 | - |
| - | - | 1795 | 1262 | - | - | 0 | - |
| - | - | 1221 | 1263 | - | - | 0 | - |
| 7 | z | 4.576E+04 | 1280 | 0.005683 | 4.441 | +1 | 11 |
| - | - | 3.855E+04 | 1281 | - | - | 0 | - |
| - | - | 1.57E+04 | 1282 | - | - | 0 | - |
| - | - | 8028 | 1283 | - | - | 0 | - |
| 11 | c | 1878 | 1286 | 0.002558 | 1.99 | +1 | 11 |
| 7 | y | 1.878E+04 | 1296 | 0.005635 | 4.35 | +1 | 11 |
| - | - | 1.205E+04 | 1297 | - | - | 0 | - |
| - | - | 5412 | 1298 | - | - | 0 | - |
| - | - | 1909 | 1299 | - | - | 0 | - |
| - | - | 1463 | 1300 | - | - | 0 | - |
| - | - | 5666 | 1303 | - | - | 0 | - |
| 11 | c | 1.446E+04 | 1304 | 0.004689 | 3.597 | +1 | 11 |
| - | - | 1.087E+04 | 1305 | - | - | 0 | - |
| - | - | 3752 | 1306 | - | - | 0 | - |
| - | - | 1794 | 1307 | - | - | 0 | - |
| - | - | 1449 | 1334 | - | - | 0 | - |
| - | - | 3067 | 1335 | - | - | 0 | - |
| - | - | 1272 | 1336 | - | - | 0 | - |
| - | - | 1557 | 1365 | - | - | 0 | - |
| - | - | 1498 | 1374 | - | - | 0 | - |
| - | - | 1763 | 1375 | - | - | 0 | - |
| - | - | 1849 | 1376 | - | - | 0 | - |
| - | - | 3252 | 1417 | - | - | 0 | - |
| 12 | c | 3.026E+04 | 1418 | 0.002899 | 2.045 | +1 | 12 |
| - | - | 2.377E+04 | 1419 | - | - | 0 | - |
| - | - | 1.027E+04 | 1420 | - | - | 0 | - |
| - | - | 5022 | 1421 | - | - | 0 | - |
| - | - | 4020 | 1422 | - | - | 0 | - |
| 6 | y | 1328 | 1435 | 0.02604 | 18.15 | +1 | 12 |
| - | - | 770.9 | 1436 | - | - | 0 | - |
| 6 | z | 3.696E+04 | 1436 | 0.005035 | 3.507 | +1 | 12 |
| - | - | 5.795E+04 | 1437 | - | - | 0 | - |
| - | - | 3.867E+04 | 1438 | - | - | 0 | - |
| - | - | 1.635E+04 | 1439 | - | - | 0 | - |
| - | - | 5777 | 1440 | - | - | 0 | - |
| - | - | 4996 | 1451 | - | - | 0 | - |
| 6 | y | 1.956E+04 | 1452 | 0.004134 | 2.847 | +1 | 12 |
| - | - | 1.486E+04 | 1453 | - | - | 0 | - |
| - | - | 8596 | 1454 | - | - | 0 | - |
| - | - | 3447 | 1455 | - | - | 0 | - |
| - | - | 2487 | 1503 | - | - | 0 | - |
| - | - | 1778 | 1505 | - | - | 0 | - |
| 5 | z | 3981 | 1523 | 0.01268 | 8.327 | +1 | 13 |
| - | - | 1.53E+04 | 1524 | - | - | 0 | - |
| - | - | 1.128E+04 | 1525 | - | - | 0 | - |
| - | - | 5901 | 1526 | - | - | 0 | - |
| - | - | 2709 | 1527 | - | - | 0 | - |
| 13 | c | 1853 | 1528 | 0.01191 | 7.796 | +1 | 13 |
| 5 | y | 4448 | 1539 | 0.006895 | 4.481 | +1 | 13 |
| - | - | 4080 | 1540 | - | - | 0 | - |
| - | - | 1997 | 1541 | - | - | 0 | - |
| - | - | 1481 | 1542 | - | - | 0 | - |
| - | - | 1636 | 1545 | - | - | 0 | - |
| 13 | c | 3.195E+04 | 1546 | 0.004014 | 2.597 | +1 | 13 |
| - | - | 3.221E+04 | 1547 | - | - | 0 | - |
| - | - | 1.735E+04 | 1548 | - | - | 0 | - |
| - | - | 6290 | 1549 | - | - | 0 | - |
| - | - | 2366 | 1602 | - | - | 0 | - |
| - | - | 3770 | 1603 | - | - | 0 | - |
| - | - | 2057 | 1604 | - | - | 0 | - |
| - | - | 1886 | 1605 | - | - | 0 | - |
| - | - | 2218 | 1615 | - | - | 0 | - |
| - | - | 4097 | 1616 | - | - | 0 | - |
| - | - | 1892 | 1617 | - | - | 0 | - |
| - | - | 1670 | 1618 | - | - | 0 | - |
| 14 | c | 4819 | 1628 | 0.008048 | 4.944 | +1 | 14 |
| - | - | 3565 | 1629 | - | - | 0 | - |
| 4 | y | 1.095E+04 | 1636 | 0.001373 | 0.8391 | +1 | 14 |
| - | - | 9196 | 1637 | - | - | 0 | - |
| - | - | 7368 | 1638 | - | - | 0 | - |
| - | - | 4577 | 1639 | - | - | 0 | - |
| - | - | 1220 | 1640 | - | - | 0 | - |
| - | - | 6689 | 1644 | - | - | 0 | - |
| 14 | c | 4.322E+04 | 1645 | 0.004326 | 2.63 | +1 | 14 |
| - | - | 3.78E+04 | 1646 | - | - | 0 | - |
| - | - | 2.396E+04 | 1647 | - | - | 0 | - |
| - | - | 1.004E+04 | 1648 | - | - | 0 | - |
| - | - | 3533 | 1649 | - | - | 0 | - |
| - | - | 1549 | 1650 | - | - | 0 | - |
| - | - | 2390 | 1672 | - | - | 0 | - |
| - | - | 1810 | 1688 | - | - | 0 | - |
| - | - | 2886 | 1689 | - | - | 0 | - |
| - | - | 2964 | 1690 | - | - | 0 | - |
| - | - | 3771 | 1716 | - | - | 0 | - |
| 3 | z | 3080 | 1717 | 0.02507 | 14.6 | +1 | 15 |
| - | - | 2099 | 1728 | - | - | 0 | - |
| - | - | 4578 | 1729 | - | - | 0 | - |
| - | - | 3916 | 1730 | - | - | 0 | - |
| - | - | 3094 | 1731 | - | - | 0 | - |
| 15 | c | 2.4E+04 | 1732 | 0.006477 | 3.74 | +1 | 15 |
| 3 | y | 3.614E+04 | 1733 | 0.02539 | 14.65 | +1 | 15 |
| - | - | 2.972E+04 | 1734 | - | - | 0 | - |
| - | - | 1.546E+04 | 1735 | - | - | 0 | - |
| - | - | 7195 | 1736 | - | - | 0 | - |
| - | - | 2812 | 1737 | - | - | 0 | - |
| - | - | 1721 | 1786 | - | - | 0 | - |
| - | - | 1408 | 1787 | - | - | 0 | - |
| - | - | 1844 | 1788 | - | - | 0 | - |
| - | - | 1791 | 1801 | - | - | 0 | - |
| - | - | 9983 | 1802 | - | - | 0 | - |
| - | - | 8981 | 1803 | - | - | 0 | - |
| - | - | 5116 | 1804 | - | - | 0 | - |
| - | - | 3170 | 1805 | - | - | 0 | - |
| - | - | 1680 | 1814 | - | - | 0 | - |
| 2 | y | 3894 | 1829 | 0.03184 | 17.41 | +1 | 16 |
| 2 | z | 8675 | 1830 | 0.01047 | 5.719 | +1 | 16 |
| - | - | 3.435E+04 | 1831 | - | - | 0 | - |
| - | - | 3.294E+04 | 1832 | - | - | 0 | - |
| 16 | c | 2.017E+04 | 1833 | 0.01629 | 8.889 | +1 | 16 |
| - | - | 1.024E+04 | 1834 | - | - | 0 | - |
| - | - | 4808 | 1835 | - | - | 0 | - |
| - | - | 1.201E+04 | 1845 | - | - | 0 | - |
| 2 | y | 3.239E+04 | 1846 | 0.02617 | 14.17 | +1 | 16 |
| - | - | 2.628E+04 | 1847 | - | - | 0 | - |
| - | - | 1.828E+04 | 1848 | - | - | 0 | - |
| - | - | 7201 | 1849 | - | - | 0 | - |
| - | - | 3974 | 1850 | - | - | 0 | - |
| - | - | 1275 | 1862 | - | - | 0 | - |
| - | - | 2652 | 1871 | - | - | 0 | - |
| - | - | 2920 | 1872 | - | - | 0 | - |
| - | - | 1785 | 1873 | - | - | 0 | - |
| - | - | 2040 | 1874 | - | - | 0 | - |
| - | - | 2086 | 1875 | - | - | 0 | - |
| - | - | 2434 | 1876 | - | - | 0 | - |
| - | - | 1494 | 1877 | - | - | 0 | - |
| - | - | 3377 | 1885 | - | - | 0 | - |
| - | - | 6226 | 1886 | - | - | 0 | - |
| - | - | 7183 | 1887 | - | - | 0 | - |
| - | - | 5511 | 1888 | - | - | 0 | - |
| - | - | 6614 | 1889 | - | - | 0 | - |
| - | - | 1.391E+04 | 1890 | - | - | 0 | - |
| - | - | 1.037E+04 | 1891 | - | - | 0 | - |
| - | - | 7421 | 1892 | - | - | 0 | - |
| - | - | 2511 | 1893 | - | - | 0 | - |
| - | - | 1325 | 1894 | - | - | 0 | - |
| - | - | 9440 | 1903 | - | - | 0 | - |
| - | - | 2.582E+04 | 1904 | - | - | 0 | - |
| - | - | 2.501E+04 | 1905 | - | - | 0 | - |
| - | - | 1.401E+04 | 1906 | - | - | 0 | - |
| - | - | 8480 | 1907 | - | - | 0 | - |
| - | - | 3390 | 1908 | - | - | 0 | - |
| - | - | 1683 | 1913 | - | - | 0 | - |
| - | - | 4410 | 1914 | - | - | 0 | - |
| - | - | 9983 | 1915 | - | - | 0 | - |
| - | - | 7359 | 1916 | - | - | 0 | - |
| - | - | 3596 | 1917 | - | - | 0 | - |
| - | - | 1788 | 1918 | - | - | 0 | - |
| - | - | 5013 | 1920 | - | - | 0 | - |
| - | - | 1.999E+04 | 1921 | - | - | 0 | - |
| - | - | 2.017E+04 | 1922 | - | - | 0 | - |
| - | - | 1.304E+04 | 1923 | - | - | 0 | - |
| - | - | 5993 | 1924 | - | - | 0 | - |
| - | - | 1873 | 1925 | - | - | 0 | - |
| - | - | 2113 | 1930 | - | - | 0 | - |
| - | - | 2.551E+04 | 1931 | - | - | 0 | - |
| - | - | 8.749E+04 | 1932 | - | - | 0 | - |
| - | - | 8.723E+04 | 1933 | - | - | 0 | - |
| - | - | 5.307E+04 | 1934 | - | - | 0 | - |
| - | - | 2.585E+04 | 1935 | - | - | 0 | - |
| - | - | 7879 | 1936 | - | - | 0 | - |
| - | - | 2735 | 1937 | - | - | 0 | - |
| - | - | 1.171E+04 | 1947 | - | - | 0 | - |
| - | - | 5.625E+04 | 1948 | - | - | 0 | - |
| - | - | 2.564E+05 | 1949 | - | - | 0 | - |
| - | - | 2.503E+05 | 1950 | - | - | 0 | - |
| - | - | 1.64E+05 | 1951 | - | - | 0 | - |
| - | - | 7.51E+04 | 1952 | - | - | 0 | - |
| - | - | 2.565E+04 | 1953 | - | - | 0 | - |
| - | - | 6599 | 1954 | - | - | 0 | - |
| - | - | 1733 | 2152 | - | - | 0 | - |

m/z Charge Intensity FragmentType MassShift Position
120.06586456298828 0 8159.134
122.4383316040039 0 647.132
125.33061218261719 0 671.3798
133.56094360351562 0 640.2713
148.91636657714844 0 715.3752
155.90859985351562 0 837.1445
156.05467224121094 0 731.1042
164.0065155029297 0 861.3409
168.42288208007812 0 934.8795
173.12945556640625 0 808.666
173.43955993652344 0 4399.329
187.14450073242188 0 91080.42
188.14796447753906 0 8754.44
190.76992797851562 0 876.7935
201.12362670898438 0 3532.1973 y Water loss 13
207.94073486328125 0 837.9434
212.1395721435547 0 2550.041
215.13941955566406 0 59478.13 y Water loss 15
216.14273071289062 0 6339.756
226.1307830810547 0 1081.9143
233.1499481201172 0 5223.2593 y 15
234.1539764404297 0 1054.4015
256.16619873046875 0 1500.1652
257.170654296875 0 931.6363
299.1719970703125 0 9990.4375
300.17559814453125 0 1435.164
302.17156982421875 0 1897.9084 y Water loss 14
312.1927795410156 0 2224.0254
317.8163757324219 0 912.36444
320.1820068359375 0 3219.6982 y 14
347.2649841308594 0 848.0126
382.2569580078125 0 1057.3147
383.263427734375 0 1341.8033
385.1700439453125 0 1128.1388
407.2533264160156 0 3305.1555
425.26513671875 0 2528.516
426.27044677734375 0 3681.8486 c 3
427.27288818359375 0 1444.292
429.0906982421875 0 1377.8173
444.83331298828125 0 914.2857
451.2814636230469 0 2008.002
470.2982177734375 0 4879.5864
471.8990478515625 0 1492.074
473.2619323730469 0 1235.2437 w 12
491.9126281738281 0 1039.2734
494.28662109375 0 2542.0889
495.5729675292969 0 1301.7042
499.5782165527344 0 1349.7195
500.9111022949219 0 1496.0133
501.24273681640625 0 3213.0789
509.9292297363281 0 936.2127 c Water loss 12
512.2975463867188 0 1871.3417
513.303955078125 0 58065.203 c 4
514.3070068359375 0 19013.344
515.310302734375 0 2752.4773
520.2326049804688 0 1415.472
523.268798828125 0 1835.3328
523.6021118164062 0 1446.4729
526.2782592773438 0 1107.4465
526.9263916015625 0 2474.3044
527.28515625 0 2231.1755
529.2730712890625 0 5915.632
529.606689453125 0 4576.564
529.9423217773438 0 1222.9609
530.2740478515625 0 1874.4008 y Ammonia loss 12
531.29150390625 0 6862.06 z 12
532.2975463867188 0 4953.915
532.5999145507812 0 7024.879
532.93310546875 0 8761.334
533.2651977539062 0 4135.2407
533.3055419921875 0 1224.3511
533.5988159179688 0 2746.7095
538.6045532226562 0 18052.959
538.938232421875 0 12199.283
539.2717895507812 0 8804.767
539.6043701171875 0 3756.193
544.3043823242188 0 1201.189
547.3099975585938 0 4513.5107 y 12
553.3068237304688 0 1188.4285
556.9578247070312 0 4019.1665
557.2903442382812 0 3238.9526
557.624755859375 0 3342.0508
557.9576416015625 0 2479.3945
564.2850341796875 0 1101.2079
570.3134765625 0 1801.3483
571.3203125 0 4697.6606
572.2864990234375 0 14550.396 c Ammonia loss 14
572.3265991210938 0 1209.1171
572.6198120117188 0 15183.561 y Ammonia loss 2
572.9540405273438 0 8593.004 z 2
573.2874755859375 0 4251.4243
573.3372802734375 0 3154.8638
573.6204223632812 0 2580.9521
573.95068359375 0 1043.411
574.33740234375 0 1213.6519
577.96142578125 0 1874.8118 c 14
578.2904663085938 0 254541.66 y 2
578.6243286132812 0 256504.78
578.9578857421875 0 147462.39
579.2919311523438 0 58900.848 c Water loss 9
579.6253662109375 0 22293.287
579.9593505859375 0 6220.773
607.380615234375 0 3512.8533
615.348388671875 0 970.0695
625.39111328125 0 7916.164
626.395751953125 0 4746.8325
627.3214721679688 0 1213.0972 z Water loss 11
629.3927001953125 0 1124.0886
643.6624755859375 0 3493.0547
643.9945068359375 0 3171.2942
645.3323364257812 0 2451.634 z 11
646.3404541015625 0 6893.355
647.341552734375 0 1276.927
648.3405151367188 0 1527.1989
649.3363037109375 0 2890.0605
649.6691284179688 0 2765.8372
649.849609375 0 2908.2905
652.3381958007812 0 11375.513 c 10
652.840087890625 0 6921.01
653.3412475585938 0 4129.1714
661.3523559570312 0 9920.512 y 11
662.3546752929688 0 2413.6575
669.4053955078125 0 94970.3 c 5
670.4081420898438 0 34900.586
671.410400390625 0 7766.2095
672.4039916992188 0 2947.6516
673.409423828125 0 1468.4528
686.8478393554688 0 2446.3547
695.4208374023438 0 2002.2532
699.8515625 0 2550.8418
700.3494873046875 0 2774.0305 c Water loss 11
700.85107421875 0 2004.8901 c Ammonia loss 11
707.3445434570312 0 8411.504
707.8460693359375 0 6172.4937
708.3455200195312 0 3602.9697
708.8549194335938 0 10606.714
709.359375 0 61916.887 c 11
709.8607788085938 0 45385.48
710.3614501953125 0 21004.35
710.8604736328125 0 8915.556
711.3570556640625 0 2371.939
715.4005126953125 0 1690.2976
718.3560791015625 0 3699.3496 z 5
718.8540649414062 0 6068.4697
719.3557739257812 0 3530.5278
719.859619140625 0 1458.7258
720.3689575195312 0 1796.593
722.3657836914062 0 3551.5747
722.864990234375 0 2064.1042
723.3646850585938 0 1110.0588
725.85693359375 0 2631.3484
726.3634033203125 0 9360.902 y 5
726.8624267578125 0 5803.869
727.3614501953125 0 3389.4177
727.8642578125 0 1307.9385
729.4273071289062 0 12270.332
730.4300537109375 0 6359.915
741.8535766601562 0 1876.2056
742.3606567382812 0 1736.7084
743.8755493164062 0 1836.6073
750.8598022460938 0 8659.979
751.369384765625 0 10440.718
751.8662109375 0 3475.5793
752.366943359375 0 2870.2102
752.8629150390625 0 1461.2318 z Water loss 4
754.4368286132812 0 1764.8976
755.432861328125 0 1205.857
758.3880004882812 0 2036.8809
759.3776245117188 0 2272.662
761.3630981445312 0 1413.4174 y Ammonia loss 4
761.8692016601562 0 9715.738 z 4
762.3709716796875 0 10453.247
762.8716430664062 0 3985.9087
763.3755493164062 0 1805.2817
763.8779296875 0 1162.2897
764.3776245117188 0 2042.4492 c Water loss 12
764.8969116210938 0 1562.629
765.3939208984375 0 3519.5159
765.8976440429688 0 2041.4528
769.87890625 0 4455.1313 y 4
770.3806762695312 0 7029.249
770.8807373046875 0 3794.558
772.8859252929688 0 4928.5293
773.3909301757812 0 68934.34 c 12
773.4515380859375 0 17643.127
773.89013671875 0 79580.305
774.3931274414062 0 31610.145
774.4571533203125 0 6164.078
774.891357421875 0 18827.896
775.437744140625 0 4161.034
775.896240234375 0 1712.8943
779.4199829101562 0 2216.2695
785.3921508789062 0 1426.4677
785.8779296875 0 1648.9098
786.3839721679688 0 1150.85
787.400634765625 0 1088.8817
788.4462890625 0 1181.8954
789.4476928710938 0 13855.291 y 10
790.449462890625 0 5023.6357
791.9234008789062 0 1860.106
792.9117431640625 0 3937.6992
793.4093017578125 0 6435.175
793.9107666015625 0 5939.7505
794.403076171875 0 2109.456
794.9000854492188 0 1961.2365
795.3921508789062 0 1175.8121
796.8829956054688 0 1829.8383 w 3
797.4407348632812 0 1911.6062
798.4484252929688 0 135831.36 c 6
798.8960571289062 0 3277.9321
799.45068359375 0 59124.348
799.8959350585938 0 1615.9772
800.4536743164062 0 13920.548
800.9140014648438 0 6436.2886
801.4164428710938 0 6362.594
801.91796875 0 4739.5483
802.4159545898438 0 1528.7815
807.4019165039062 0 20670.385
807.903564453125 0 20367.928
807.9833984375 0 1609.3878
808.4050903320312 0 6155.9473
808.903564453125 0 4781.499
809.408203125 0 1659.0848 y Water loss 3
813.4144897460938 0 22004.586
813.9146728515625 0 27453.215 c Water loss 13
814.4135131835938 0 17627.875 c Ammonia loss 13
814.9132080078125 0 10747.168
815.4135131835938 0 6660.92
815.9163818359375 0 3005.1897
817.4154663085938 0 1383.9962
817.9032592773438 0 1972.1304
818.4058837890625 0 38853.47 y 3
818.9063110351562 0 36336.293
819.40673828125 0 20367.65
819.9071655273438 0 7132.644
820.40673828125 0 4770.0884
822.4195556640625 0 44224.754
822.921875 0 69403.79 c 13
823.422607421875 0 45435.453
823.9232788085938 0 28103.01
824.4256591796875 0 6716.265
824.9274291992188 0 2780.5566
825.4603881835938 0 1301.983
830.4632568359375 0 3905.2864
831.4668579101562 0 3065.8323
834.9336547851562 0 7995.2026
835.4315795898438 0 7692.8555
835.9326171875 0 7778.981
836.4292602539062 0 4919.189
843.9308471679688 0 1532.5295
844.4244995117188 0 2051.3171
844.9348754882812 0 4743.38
845.4176025390625 0 5504.226 w 2
845.9139404296875 0 5837.8467
846.4151611328125 0 2656.5327
846.9091186523438 0 1333.2926
849.4132080078125 0 2714.8572
852.4158935546875 0 1567.4801
856.457275390625 0 2092.0435 z Water loss 9
857.4710693359375 0 8225.019 w 9
857.929443359375 0 9690.349 c Ammonia loss 14
858.4285888671875 0 8414.575 y Ammonia loss 2
858.9261474609375 0 8675.756 z 2
859.44873046875 0 13864.763 w 9
859.9248046875 0 3230.1895
860.4552612304688 0 6323.5044
861.4532470703125 0 1642.2537
865.9367065429688 0 7728.2666
866.4393920898438 0 115803.04 c 14
866.9349365234375 0 262235.94 y 2
867.4347534179688 0 214246.84
867.9345703125 0 116834.87
868.4352416992188 0 47902.96
868.9352416992188 0 14658.677
869.4380493164062 0 2604.801
872.4613037109375 0 1545.2666
873.4674072265625 0 12138.991 y Ammonia loss 9
874.4762573242188 0 72826.18 z 9
875.47900390625 0 37853.996
876.47998046875 0 12213.833
877.481689453125 0 2935.7168
879.455078125 0 5205.997
879.9536743164062 0 5759.556
880.4505615234375 0 2931.5679
880.9451293945312 0 2783.755
881.4310302734375 0 1262.5656
881.9332885742188 0 2378.1157
883.4737548828125 0 3477.5156
884.4775390625 0 2640.6768
885.4744873046875 0 1217.961
885.9562377929688 0 1623.3419
887.4317016601562 0 2443.9028
887.937744140625 0 2448.7473
889.4918212890625 0 2688.6006
890.4948120117188 0 24072.324 y 9
891.4959106445312 0 12819.606
891.9690551757812 0 5011.33
892.4802856445312 0 6681.25
892.9697265625 0 4095.7056
893.4775390625 0 4967.447
893.9378051757812 0 43405.97 w 1
894.4376831054688 0 49831.168
894.9383544921875 0 29150.754
895.4396362304688 0 12302.962
895.9392700195312 0 5031.0156
899.4736938476562 0 1336.9957
900.4732666015625 0 1520.6204
900.9716796875 0 6506.2954
901.4760131835938 0 10358.934
901.9774780273438 0 8068.2734
902.47509765625 0 6991.8203
903.4646606445312 0 2604.1746
905.9600219726562 0 1534.0142
906.461181640625 0 1804.9329 z Water loss 1
906.9710693359375 0 3787.3425
907.4760131835938 0 3542.093
907.9609375 0 1786.5906 c Water loss 15
908.4632568359375 0 2007.809 c Ammonia loss 15
909.4591064453125 0 2917.5713
914.4741821289062 0 5243.3696 y Water loss 1
914.966796875 0 7901.3413 y Ammonia loss 1
915.46728515625 0 26605.29 z 1
915.966064453125 0 29337.04
916.4697875976562 0 18525.275
916.9630126953125 0 9714.06 c 15
917.4622802734375 0 3252.712
917.9652709960938 0 1806.0754
920.4873046875 0 5556.298
920.9852294921875 0 2939.1611
921.4805908203125 0 2677.4688
922.4804077148438 0 2849.235
922.9816284179688 0 114332.81
923.4822998046875 0 119585.84 y 1
923.9822387695312 0 85715
924.482421875 0 34273.746
924.9841918945312 0 15483.6045
925.4826049804688 0 4457.6953
926.4818725585938 0 4840.5977
927.49072265625 0 135824.05 c 7
928.493408203125 0 76074.734
928.9762573242188 0 4061.176
929.4925537109375 0 22181.223
929.9743041992188 0 6918.7935
930.481689453125 0 10079.43
930.9718627929688 0 5593.621
931.4905395507812 0 3490.962
931.9710083007812 0 1356.7302
933.4983520507812 0 1567.9878
933.9999389648438 0 1670.868
934.4998168945312 0 1743.4541
934.9863891601562 0 2655.0232
935.4835205078125 0 2741.7734
935.979248046875 0 2459.438
936.4808349609375 0 6559.873
936.9757690429688 0 6034.2026
937.47314453125 0 8692.036
937.9598999023438 0 46554.086
938.4617309570312 0 49301.457
938.9656982421875 0 32095.541
939.466552734375 0 16644.34
939.972900390625 0 7594.6436
940.4637451171875 0 2464.4163
941.5160522460938 0 2660.7612
942.497314453125 0 9673.245
942.9945068359375 0 16052.569
943.4935913085938 0 17757.016
943.9888916015625 0 13788.719
944.482177734375 0 10558.57
944.9830322265625 0 26454.055
945.4835815429688 0 26432.057
945.9861450195312 0 15726.284
946.4765625 0 16795.621
946.9738159179688 0 15974.904
947.4710083007812 0 11230.443
947.9743041992188 0 4483.496
948.4766845703125 0 1790.843
950.5125122070312 0 1323.8499
951.4971313476562 0 3941.4558
951.9921875 0 24332.6
952.4927978515625 0 28193.746
952.9911499023438 0 25540.014
953.4927978515625 0 16385.258
953.9879150390625 0 7490.7227
954.4906005859375 0 2910.888
956.4900512695312 0 10584.735
956.985107421875 0 18743.488
957.4886474609375 0 20219.865
957.9849243164062 0 13213.291
958.5130004882812 0 16704.262
958.9862060546875 0 4323.975
959.5284423828125 0 57349.773
960.5298461914062 0 29466.105
961.5331420898438 0 9986.738
962.5315551757812 0 1964.049
964.9903564453125 0 1198.9032
965.4953002929688 0 36460.97
965.9918823242188 0 171051.58
966.4918212890625 0 184489.88
966.9922485351562 0 112995.92
967.4933471679688 0 52429.75
967.9924926757812 0 19348.51
968.5145263671875 0 9345.996
969.5319213867188 0 3412.6074
973.5008544921875 0 3229.9749
973.99755859375 0 123321.79
974.4996337890625 0 217424.34
975.000732421875 0 185093.39
975.50146484375 0 107057.71
976.001708984375 0 46888.445
976.504638671875 0 16786.002
977.00390625 0 2341.667
978.4581909179688 0 1458.5073
1003.501708984375 0 2547.0596 z Water loss 8
1005.5200805664062 0 2006.4696
1006.5113525390625 0 1740.6208
1012.5105590820312 0 1087.6681
1013.5001220703125 0 1303.8159
1014.5072631835938 0 1441.959
1015.5100708007812 0 1215.8453
1016.503662109375 0 1481.9441
1021.5117797851562 0 14824.527 z 8
1022.5151977539062 0 15729.942
1023.518310546875 0 6673.4355
1024.5164794921875 0 2823.3245
1030.512451171875 0 19435.127
1031.5169677734375 0 11049.58
1032.51806640625 0 5617.0083
1033.52392578125 0 1610.424
1037.5303955078125 0 7217.0454 y 8
1038.529541015625 0 4251.703
1039.5306396484375 0 2001.0134
1049.530517578125 0 6890.0767
1050.5362548828125 0 3621.1423
1051.53076171875 0 2687.596
1055.509521484375 0 2080.2297
1056.5101318359375 0 1295.9838 c Water loss 8
1073.5201416015625 0 4115.978
1074.5262451171875 0 143959.08 c 8
1075.529052734375 0 82449.37
1076.5303955078125 0 31311.518
1077.53076171875 0 14324.987
1078.533447265625 0 1158.7806
1079.517333984375 0 2386.4895
1080.5076904296875 0 1212.1841
1087.5643310546875 0 3100.6824
1088.560302734375 0 2788.7012
1089.557373046875 0 1530.5641
1091.5419921875 0 18703.318 w 7
1092.5430908203125 0 10555.6455
1093.5396728515625 0 4503.539
1094.4981689453125 0 3632.825
1095.494140625 0 1976.2096
1106.5357666015625 0 1270.7887
1113.55224609375 0 3595.4072
1114.5474853515625 0 2964.3281
1115.544677734375 0 1798.0588
1131.5606689453125 0 4379.3564
1132.5611572265625 0 4063.9731
1133.5679931640625 0 1835.7013
1134.559814453125 0 2863.958
1135.5509033203125 0 1902.6168
1150.554931640625 0 55201.805 z 7
1151.556396484375 0 43960.742
1152.5557861328125 0 20440.56
1153.5606689453125 0 5896.573
1154.557861328125 0 2070.6272
1166.5731201171875 0 3730.1223 y 7
1167.576171875 0 3076.4226
1168.581298828125 0 1988.2454
1175.57373046875 0 77025.57 c 9
1176.576416015625 0 40571.016
1177.578369140625 0 24369.209
1178.5751953125 0 6741.015
1179.5731201171875 0 2489.1394
1180.5802001953125 0 1764.9208
1207.5777587890625 0 1731.1271
1220.5799560546875 0 10206.463 w 6
1221.5833740234375 0 7110.1523
1222.580078125 0 4149.404
1223.5616455078125 0 2844.553
1235.5897216796875 0 1266.138
1259.653076171875 0 3655.8516
1260.6595458984375 0 4128.1914
1261.65771484375 0 1794.8792
1262.6654052734375 0 1221.4396
1279.5968017578125 0 45758.574 z 6
1280.5986328125 0 38548.72
1281.5980224609375 0 15700.721
1282.5994873046875 0 8027.9585
1285.654541015625 0 1878.4463 c Water loss 10
1295.615478515625 0 18780.8 y 6
1296.615234375 0 12050.205
1297.6253662109375 0 5411.9165
1298.63037109375 0 1909.3575
1299.60498046875 0 1462.651
1302.65966796875 0 5666.3154
1303.667236328125 0 14464.997 c 10
1304.669921875 0 10872.953
1305.672119140625 0 3752.185
1306.6810302734375 0 1794.122
1333.659912109375 0 1449.2163
1334.6722412109375 0 3067.2346
1335.6739501953125 0 1272.4034
1364.6654052734375 0 1557.3016
1373.690673828125 0 1498.008
1374.708740234375 0 1763.0315
1375.710205078125 0 1849.4601
1416.6986083984375 0 3251.7268
1417.7083740234375 0 30256.92 c 11
1418.71142578125 0 23770.93
1419.7105712890625 0 10267.178
1420.7115478515625 0 5021.7905
1421.70458984375 0 4020.499
1434.71044921875 0 1327.5607 y Ammonia loss 5
1435.5391845703125 0 770.9051
1435.697265625 0 36964.492 z 5
1436.702392578125 0 57952.465
1437.703857421875 0 38669.188
1438.7060546875 0 16345.114
1439.7080078125 0 5776.712
1450.7091064453125 0 4996.264
1451.715087890625 0 19561.947 y 5
1452.7147216796875 0 14855.791
1453.7213134765625 0 8596.36
1454.716552734375 0 3446.59
1502.75537109375 0 2487.0366
1504.7467041015625 0 1777.6353
1522.7369384765625 0 3980.6753 z 4
1523.7354736328125 0 15299.589
1524.7408447265625 0 11282.0205
1525.736328125 0 5901.475
1526.7401123046875 0 2708.9219
1527.7415771484375 0 1853.2692 c Water loss 12
1538.7498779296875 0 4448.3496 y 4
1539.75244140625 0 4080.0273
1540.7567138671875 0 1997.288
1541.7730712890625 0 1481.4829
1544.75634765625 0 1635.9739
1545.76806640625 0 31947.896 c 12
1546.7701416015625 0 32206.836
1547.770751953125 0 17354.438
1548.7730712890625 0 6289.8296
1601.8326416015625 0 2366.0046
1602.8253173828125 0 3770.4292
1603.82958984375 0 2057.2131
1604.8226318359375 0 1885.7628
1614.805419921875 0 2218.2441
1615.8065185546875 0 4096.515
1616.8194580078125 0 1891.5747
1617.833740234375 0 1670.0836
1627.81396484375 0 4819.2905 c Ammonia loss 13
1628.8193359375 0 3565.2554
1635.797119140625 0 10945.213 y 3
1636.8016357421875 0 9195.618
1637.802490234375 0 7368.172
1638.809814453125 0 4577.288
1639.806884765625 0 1220.2094
1643.8306884765625 0 6689.454
1644.8367919921875 0 43220.117 c 13
1645.839111328125 0 37804.4
1646.84130859375 0 23961.99
1647.8446044921875 0 10037.234
1648.842041015625 0 3533.0605
1649.8446044921875 0 1548.6542
1671.841796875 0 2390.111
1687.85791015625 0 1810.0159
1688.8607177734375 0 2886.0906
1689.8538818359375 0 2964.4214
1715.8636474609375 0 3771.3176
1716.8548583984375 0 3079.588 z 2
1727.8941650390625 0 2099.2358
1728.8895263671875 0 4578.36
1729.8878173828125 0 3916.0847
1730.8865966796875 0 3093.6606
1731.8709716796875 0 23997.277 c 14
1732.8739013671875 0 36138.816 y 2
1733.87646484375 0 29716.414
1734.873291015625 0 15458.022
1735.86865234375 0 7195.375
1736.8760986328125 0 2812.4897
1785.9178466796875 0 1721.433
1786.90283203125 0 1408.2478
1787.9140625 0 1843.5314
1800.9404296875 0 1790.5974
1801.9481201171875 0 9983.256
1802.94775390625 0 8980.753
1803.9459228515625 0 5115.6885
1804.9515380859375 0 3169.6653
1813.8983154296875 0 1680.4088
1828.9378662109375 0 3893.5554 y Ammonia loss 1
1829.92431640625 0 8675.405 z 1
1830.9263916015625 0 34348.152
1831.9276123046875 0 32942.113
1832.928466796875 0 20171.523 c 15
1833.9210205078125 0 10238.3
1834.9317626953125 0 4808.381
1844.95654296875 0 12008.831
1845.958740234375 0 32394.19 y 1
1846.9622802734375 0 26279.371
1847.963623046875 0 18282.824
1848.9573974609375 0 7201.398
1849.9644775390625 0 3974.2734
1861.9775390625 0 1275.0121
1870.9432373046875 0 2652.294
1871.9327392578125 0 2920.399
1872.9498291015625 0 1784.9819
1873.933837890625 0 2039.6664
1874.9388427734375 0 2086.216
1875.92919921875 0 2433.5613
1876.984375 0 1494.2269
1885.0018310546875 0 3376.674
1885.9984130859375 0 6225.949
1886.9757080078125 0 7182.7124
1887.9713134765625 0 5510.9355
1888.9678955078125 0 6613.806
1889.963134765625 0 13909.464
1890.96484375 0 10371.474
1891.9647216796875 0 7420.6265
1892.9619140625 0 2511.461
1893.9710693359375 0 1324.9467
1902.9844970703125 0 9440.097
1903.984130859375 0 25821.594
1904.9833984375 0 25013.006
1905.9818115234375 0 14007.335
1906.9814453125 0 8480.45
1907.9923095703125 0 3389.5107
1912.9822998046875 0 1682.6571
1913.96875 0 4409.9487
1914.9586181640625 0 9982.757
1915.9554443359375 0 7358.5986
1916.9571533203125 0 3596.153
1917.9710693359375 0 1788.4274
1919.9945068359375 0 5012.8164
1921.00390625 0 19994.957
1922.006591796875 0 20170.125
1923.010498046875 0 13040.945
1924.0040283203125 0 5993.4785
1925.01513671875 0 1872.7502
1929.991455078125 0 2112.7708
1930.9761962890625 0 25505.197
1931.9764404296875 0 87492.54
1932.9764404296875 0 87230.98
1933.97900390625 0 53069.035
1934.9803466796875 0 25853.883
1935.9801025390625 0 7878.617
1936.993896484375 0 2734.8242
1946.990234375 0 11710.586
1947.989990234375 0 56249.67
1948.99658203125 0 256419.2
1950.0001220703125 0 250322.28
1951.0018310546875 0 163967.3
1952.0023193359375 0 75095.195
1953.0048828125 0 25651.021
1954.0035400390625 0 6598.864
2152.064697265625 0 1732.9943

Spectrum Details

|  |  |
| --- | --- |
| Matched peaks? Matched peaks The total absolute number of peaks matched. Additionally in brackets the total fraction of peaks matched and the total number of peaks is shown. | 96 (14.66% of 655) |
| FDR? FDR The false discovery rate estimated for this peptide. It is calculated by matching all theoretical fragments with a non-integer shift with the raw peaks for this spectrum. This is done with 40 different shifts. The resulting percentage is the average number of annotated peaks over the number of annotated peaks with the correct spectrum. | 0.72% |
| Satellite FDR? Satellite FDR See the FDR for details on its calculation. This satellite ion specific FDR only contains the satellite ions (d/w) for I/L/J positions. | 11.90% |
| PSM Score? PSM Score The PSM Score as given by Hecklib to this annotated spectrum. It is shown with three significant figures. | 645 |

#### Spectrum 4342? Spectrum 4342 The raw spectrum of this peptide as annotated by Hecklib. The fragments are coloured according to ion type (see legend). Any peaks with a star '\*' as text can be hovered over to see the full details, first the ion type second the mass shift type. By hovering over the amino acids in the peptide or ions in the legend the corresponding peaks are highlighted. By toggling the 'Unassigned' label you can turn the background (unassigned) peaks on or off in the plot. By updating the slider in the Ion legend you can update the spectrum to only show the top X% of the peaks with labels. The top X% means any peak that is within X% of the highest intensity. By dragging in the spectrum you can zoom in to a specific part of the spectrum and use 'Zoom Out' to get back to the original zoom level. The annotation of the spectrum is based on the given sequence in the peptides file and is done with different software so inconsistencies are likely. The peaks are annotated based on the given sequence, with 20 ppm tolerance.

Copy Data

##### Spectrum 4342 (TSV)

###### Preview

```
Loading example...
```

*Click on the button to copy the data to your clipboard.*

Mz MinMz MaxIntensity Max

WidthHeightPeptide font sizePeptide stroke widthSpectrum font sizeSpectrum stroke widthCompact peptide

Ion legend

wxyz

abcd

OtherUnassignedIon^Charge^_Position_Show for top:%

TLPPSREEMTKNQVSTJ

02.28e+44.57e+46.85e+49.13e+4

Zoom Out

c^+3^_5_y^+2^_4_y^+1^_2_y^+1^_2_y^+1^_3_c^+1^_4_c^+1^_5_z^+1^_5_y^+1^_5_c^+3^_15_y^+3^_15_z^+3^_15_y^+3^_15_c^+2^_10_z^+1^_6_c^+2^_11_y^+1^_6_c^+1^_6_c^+2^_12_c^+2^_12_z^+2^_12_y^+2^_12_z^+2^_13_y^+2^_13_c^+2^_13_y^+1^_7_c^+1^_7_y^+2^_14_c^+2^_14_c^+2^_14_y^+2^_14_c^+2^_14_w^+2^_15_w^+1^_8_c^+2^_15_y^+2^_15_z^+2^_15_w^+1^_8_c^+2^_15_y^+2^_15_y^+1^_8_z^+1^_8_y^+1^_8_w^+2^_16_y^+2^_16_y^+2^_16_z^+2^_16_c^+2^_16_y^+2^_16_c^+1^_8_z^+1^_9_y^+1^_9_c^+1^_9_w^+1^_10_z^+1^_10_z^+1^_10_y^+1^_10_c^+1^_10_w^+1^_11_z^+1^_11_y^+1^_11_c^+1^_11_c^+1^_12_z^+1^_12_y^+1^_12_z^+1^_13_c^+1^_13_y^+1^_13_c^+1^_13_c^+1^_14_c^+1^_14_y^+1^_14_c^+1^_14_y^+1^_15_z^+1^_15_c^+1^_15_y^+1^_15_y^+1^_16_z^+1^_16_c^+1^_16_y^+1^_16_

049398714801974

Fragment Matches Table

Show background peaks

| Position | Ion type | Intensity | mz Theoretical | mz Error (Th) | mz Error (ppm) | Charge | Series Number |
| --- | --- | --- | --- | --- | --- | --- | --- |
| - | - | 2178 | 120.1 | - | - | 0 | - |
| - | - | 388.7 | 127 | - | - | 0 | - |
| - | - | 426.2 | 141.7 | - | - | 0 | - |
| - | - | 387.4 | 142 | - | - | 0 | - |
| - | - | 372.1 | 152.1 | - | - | 0 | - |
| 5 | c | 438.8 | 165.8 | 0.0003454 | 2.084 | +3 | 5 |
| - | - | 531 | 168.1 | - | - | 0 | - |
| - | - | 698.8 | 173.1 | - | - | 0 | - |
| - | - | 2.185E+04 | 187.1 | - | - | 0 | - |
| - | - | 1924 | 188.1 | - | - | 0 | - |
| - | - | 473.2 | 195.7 | - | - | 0 | - |
| 14 | y | 701 | 201.1 | 5.95E-05 | 0.2958 | +2 | 4 |
| - | - | 1657 | 212.1 | - | - | 0 | - |
| 16 | y | 1.392E+04 | 215.1 | 0.0001565 | 0.7275 | +1 | 2 |
| - | - | 1850 | 216.1 | - | - | 0 | - |
| 16 | y | 1104 | 233.1 | 2.883E-05 | 0.1237 | +1 | 2 |
| - | - | 546.1 | 238.2 | - | - | 0 | - |
| - | - | 499.4 | 276.2 | - | - | 0 | - |
| - | - | 583.8 | 298.6 | - | - | 0 | - |
| - | - | 3997 | 299.2 | - | - | 0 | - |
| - | - | 935.8 | 300.2 | - | - | 0 | - |
| - | - | 637.2 | 319.2 | - | - | 0 | - |
| 15 | y | 808.6 | 320.2 | 0.0002728 | 0.8519 | +1 | 3 |
| - | - | 560.8 | 321.9 | - | - | 0 | - |
| - | - | 501 | 332.7 | - | - | 0 | - |
| - | - | 724.9 | 348.7 | - | - | 0 | - |
| - | - | 779.2 | 359 | - | - | 0 | - |
| - | - | 792.7 | 407.3 | - | - | 0 | - |
| - | - | 555.4 | 425 | - | - | 0 | - |
| 4 | c | 641.4 | 426.3 | 0.0007854 | 1.842 | +1 | 4 |
| - | - | 1768 | 429.1 | - | - | 0 | - |
| - | - | 1620 | 470.3 | - | - | 0 | - |
| - | - | 745.2 | 494.3 | - | - | 0 | - |
| - | - | 678.2 | 509.3 | - | - | 0 | - |
| 5 | c | 1.618E+04 | 513.3 | 0.0002206 | 0.4298 | +1 | 5 |
| - | - | 5035 | 514.3 | - | - | 0 | - |
| - | - | 822.8 | 515.3 | - | - | 0 | - |
| - | - | 773.9 | 523.3 | - | - | 0 | - |
| - | - | 629.1 | 526.3 | - | - | 0 | - |
| - | - | 749.7 | 529.3 | - | - | 0 | - |
| - | - | 1366 | 529.6 | - | - | 0 | - |
| - | - | 926.1 | 529.9 | - | - | 0 | - |
| 13 | z | 1497 | 531.3 | 0.0008311 | 1.564 | +1 | 5 |
| - | - | 1881 | 532.3 | - | - | 0 | - |
| - | - | 1364 | 532.6 | - | - | 0 | - |
| - | - | 2322 | 532.9 | - | - | 0 | - |
| - | - | 1004 | 533.3 | - | - | 0 | - |
| - | - | 4815 | 538.6 | - | - | 0 | - |
| - | - | 3607 | 538.9 | - | - | 0 | - |
| - | - | 2232 | 539.3 | - | - | 0 | - |
| - | - | 668.3 | 539.9 | - | - | 0 | - |
| 13 | y | 1079 | 547.3 | 0.0009058 | 1.655 | +1 | 5 |
| - | - | 911.3 | 557 | - | - | 0 | - |
| - | - | 1359 | 557.3 | - | - | 0 | - |
| - | - | 779.5 | 558 | - | - | 0 | - |
| - | - | 1529 | 571.3 | - | - | 0 | - |
| 15 | c | 1831 | 572.3 | 0.003065 | 5.356 | +3 | 15 |
| 3 | y | 2379 | 572.6 | 0.005626 | 9.826 | +3 | 15 |
| 3 | z | 2671 | 572.9 | 0.002876 | 5.019 | +3 | 15 |
| - | - | 1787 | 573.3 | - | - | 0 | - |
| - | - | 762.8 | 573.3 | - | - | 0 | - |
| 3 | y | 5.797E+04 | 578.3 | 0.002046 | 3.538 | +3 | 15 |
| - | - | 5.987E+04 | 578.6 | - | - | 0 | - |
| - | - | 3.767E+04 | 579 | - | - | 0 | - |
| 10 | c | 1.397E+04 | 579.3 | 0.008501 | 14.68 | +2 | 10 |
| - | - | 5256 | 579.6 | - | - | 0 | - |
| - | - | 1155 | 580 | - | - | 0 | - |
| - | - | 1654 | 625.4 | - | - | 0 | - |
| - | - | 1019 | 626.4 | - | - | 0 | - |
| 12 | z | 1120 | 645.3 | 0.0013 | 2.014 | +1 | 6 |
| - | - | 2056 | 646.3 | - | - | 0 | - |
| - | - | 666.6 | 647.3 | - | - | 0 | - |
| - | - | 817.5 | 649.3 | - | - | 0 | - |
| - | - | 1219 | 649.9 | - | - | 0 | - |
| - | - | 884.3 | 650.3 | - | - | 0 | - |
| - | - | 953.9 | 650.8 | - | - | 0 | - |
| 11 | c | 3005 | 652.3 | 0.003467 | 5.315 | +2 | 11 |
| - | - | 2129 | 652.8 | - | - | 0 | - |
| 12 | y | 2465 | 661.4 | 0.0001537 | 0.2324 | +1 | 6 |
| - | - | 728.1 | 662.4 | - | - | 0 | - |
| 6 | c | 2.833E+04 | 669.4 | 0.0001228 | 0.1834 | +1 | 6 |
| - | - | 9530 | 670.4 | - | - | 0 | - |
| - | - | 1835 | 671.4 | - | - | 0 | - |
| - | - | 714.4 | 672.4 | - | - | 0 | - |
| - | - | 734.3 | 693.8 | - | - | 0 | - |
| - | - | 1019 | 699.9 | - | - | 0 | - |
| 12 | c | 916.4 | 700.4 | 0.003132 | 4.472 | +2 | 12 |
| - | - | 3150 | 707.3 | - | - | 0 | - |
| - | - | 928 | 707.8 | - | - | 0 | - |
| - | - | 1054 | 708.3 | - | - | 0 | - |
| - | - | 2306 | 708.9 | - | - | 0 | - |
| 12 | c | 1.962E+04 | 709.4 | 0.00184 | 2.593 | +2 | 12 |
| - | - | 1.518E+04 | 709.9 | - | - | 0 | - |
| - | - | 4793 | 710.4 | - | - | 0 | - |
| - | - | 3256 | 710.9 | - | - | 0 | - |
| - | - | 876.6 | 711.4 | - | - | 0 | - |
| 6 | z | 854.4 | 718.3 | 0.006631 | 9.231 | +2 | 12 |
| - | - | 1008 | 718.9 | - | - | 0 | - |
| 6 | y | 2915 | 726.4 | 0.001663 | 2.29 | +2 | 12 |
| - | - | 2730 | 726.9 | - | - | 0 | - |
| - | - | 1150 | 727.4 | - | - | 0 | - |
| - | - | 3505 | 729.4 | - | - | 0 | - |
| - | - | 2141 | 730.4 | - | - | 0 | - |
| - | - | 1574 | 750.9 | - | - | 0 | - |
| - | - | 1988 | 751.4 | - | - | 0 | - |
| - | - | 2260 | 751.9 | - | - | 0 | - |
| 5 | z | 1744 | 761.9 | 0.002397 | 3.146 | +2 | 13 |
| - | - | 1625 | 762.4 | - | - | 0 | - |
| - | - | 785.2 | 762.9 | - | - | 0 | - |
| - | - | 732.6 | 766.4 | - | - | 0 | - |
| 5 | y | 2306 | 769.9 | 0.001763 | 2.289 | +2 | 13 |
| - | - | 1185 | 770.4 | - | - | 0 | - |
| - | - | 949 | 772.9 | - | - | 0 | - |
| 13 | c | 2.016E+04 | 773.4 | 0.003923 | 5.072 | +2 | 13 |
| - | - | 5585 | 773.5 | - | - | 0 | - |
| - | - | 2.465E+04 | 773.9 | - | - | 0 | - |
| - | - | 1.163E+04 | 774.4 | - | - | 0 | - |
| - | - | 4778 | 774.9 | - | - | 0 | - |
| - | - | 1827 | 775.4 | - | - | 0 | - |
| - | - | 834.3 | 779.4 | - | - | 0 | - |
| - | - | 876.5 | 786.4 | - | - | 0 | - |
| 11 | y | 3571 | 789.4 | 0.0001614 | 0.2044 | +1 | 7 |
| - | - | 2245 | 790.4 | - | - | 0 | - |
| - | - | 980.3 | 791.4 | - | - | 0 | - |
| - | - | 1395 | 792.9 | - | - | 0 | - |
| - | - | 1770 | 793.4 | - | - | 0 | - |
| - | - | 1658 | 793.9 | - | - | 0 | - |
| - | - | 1159 | 794.4 | - | - | 0 | - |
| 7 | c | 4.018E+04 | 798.4 | 0.0004374 | 0.5478 | +1 | 7 |
| - | - | 926.9 | 798.9 | - | - | 0 | - |
| - | - | 1.85E+04 | 799.4 | - | - | 0 | - |
| - | - | 4468 | 800.5 | - | - | 0 | - |
| - | - | 1721 | 800.9 | - | - | 0 | - |
| - | - | 2744 | 801.4 | - | - | 0 | - |
| - | - | 1496 | 801.9 | - | - | 0 | - |
| - | - | 861.5 | 802.4 | - | - | 0 | - |
| - | - | 4248 | 807.4 | - | - | 0 | - |
| - | - | 4656 | 807.9 | - | - | 0 | - |
| - | - | 2340 | 808.4 | - | - | 0 | - |
| - | - | 901.1 | 808.9 | - | - | 0 | - |
| 4 | y | 707.9 | 809.9 | 0.008186 | 10.11 | +2 | 14 |
| - | - | 7472 | 813.4 | - | - | 0 | - |
| 14 | c | 7538 | 813.9 | 0.00193 | 2.372 | +2 | 14 |
| 14 | c | 4601 | 814.4 | 0.005574 | 6.844 | +2 | 14 |
| - | - | 2345 | 814.9 | - | - | 0 | - |
| - | - | 866.5 | 815.4 | - | - | 0 | - |
| 4 | y | 1.223E+04 | 818.4 | 0.002724 | 3.329 | +2 | 14 |
| - | - | 9055 | 818.9 | - | - | 0 | - |
| - | - | 5305 | 819.4 | - | - | 0 | - |
| - | - | 1899 | 819.9 | - | - | 0 | - |
| - | - | 1.183E+04 | 822.4 | - | - | 0 | - |
| 14 | c | 1.876E+04 | 822.9 | 0.0008439 | 1.026 | +2 | 14 |
| - | - | 1.418E+04 | 823.4 | - | - | 0 | - |
| - | - | 8261 | 823.9 | - | - | 0 | - |
| - | - | 3498 | 824.4 | - | - | 0 | - |
| - | - | 1411 | 824.9 | - | - | 0 | - |
| - | - | 815.8 | 825.5 | - | - | 0 | - |
| - | - | 885.3 | 830.5 | - | - | 0 | - |
| - | - | 2290 | 834.9 | - | - | 0 | - |
| - | - | 1799 | 835.4 | - | - | 0 | - |
| - | - | 2189 | 835.9 | - | - | 0 | - |
| - | - | 1239 | 836.4 | - | - | 0 | - |
| - | - | 909.4 | 836.9 | - | - | 0 | - |
| - | - | 797.7 | 844.4 | - | - | 0 | - |
| 3 | w | 1722 | 845.4 | 0.01008 | 11.92 | +2 | 15 |
| - | - | 1832 | 845.9 | - | - | 0 | - |
| - | - | 1085 | 846.4 | - | - | 0 | - |
| - | - | 772.4 | 849.4 | - | - | 0 | - |
| 10 | w | 2179 | 857.5 | 0.001944 | 2.268 | +1 | 8 |
| 15 | c | 3393 | 857.9 | 0.002255 | 2.628 | +2 | 15 |
| 3 | y | 2425 | 858.4 | 0.01489 | 17.34 | +2 | 15 |
| 3 | z | 1503 | 858.9 | 0.006762 | 7.872 | +2 | 15 |
| 10 | w | 4172 | 859.5 | 0.0001752 | 0.2039 | +1 | 8 |
| - | - | 2026 | 860.5 | - | - | 0 | - |
| - | - | 839.8 | 864.9 | - | - | 0 | - |
| - | - | 1820 | 865.9 | - | - | 0 | - |
| - | - | 443.4 | 866.4 | - | - | 0 | - |
| 15 | c | 3.394E+04 | 866.4 | 0.00253 | 2.92 | +2 | 15 |
| 3 | y | 6.851E+04 | 866.9 | 0.005883 | 6.787 | +2 | 15 |
| - | - | 5.656E+04 | 867.4 | - | - | 0 | - |
| - | - | 3.27E+04 | 867.9 | - | - | 0 | - |
| - | - | 1.331E+04 | 868.4 | - | - | 0 | - |
| - | - | 5614 | 868.9 | - | - | 0 | - |
| 10 | y | 3838 | 873.5 | 0.003634 | 4.16 | +1 | 8 |
| 10 | z | 2.045E+04 | 874.5 | 0.0005338 | 0.6104 | +1 | 8 |
| - | - | 1.228E+04 | 875.5 | - | - | 0 | - |
| - | - | 3556 | 876.5 | - | - | 0 | - |
| - | - | 1583 | 877.5 | - | - | 0 | - |
| - | - | 2285 | 879.5 | - | - | 0 | - |
| - | - | 1244 | 879.9 | - | - | 0 | - |
| - | - | 911.5 | 882.4 | - | - | 0 | - |
| - | - | 1521 | 887.4 | - | - | 0 | - |
| 10 | y | 6420 | 890.5 | 0.00113 | 1.269 | +1 | 8 |
| - | - | 3766 | 891.5 | - | - | 0 | - |
| - | - | 1477 | 892 | - | - | 0 | - |
| - | - | 2011 | 892.5 | - | - | 0 | - |
| 2 | w | 1.18E+04 | 893.9 | 0.002737 | 3.062 | +2 | 16 |
| - | - | 1.199E+04 | 894.4 | - | - | 0 | - |
| - | - | 8204 | 894.9 | - | - | 0 | - |
| - | - | 3142 | 895.4 | - | - | 0 | - |
| - | - | 1509 | 895.9 | - | - | 0 | - |
| - | - | 1214 | 901 | - | - | 0 | - |
| - | - | 2073 | 901.5 | - | - | 0 | - |
| - | - | 1809 | 902 | - | - | 0 | - |
| - | - | 1796 | 902.5 | - | - | 0 | - |
| - | - | 912.7 | 907 | - | - | 0 | - |
| - | - | 839.3 | 912.5 | - | - | 0 | - |
| 2 | y | 2172 | 914.5 | 0.005206 | 5.693 | +2 | 16 |
| 2 | y | 1673 | 915 | 0.001967 | 2.15 | +2 | 16 |
| 2 | z | 6953 | 915.5 | 0.002022 | 2.209 | +2 | 16 |
| - | - | 7298 | 916 | - | - | 0 | - |
| - | - | 4247 | 916.5 | - | - | 0 | - |
| 16 | c | 3456 | 917 | 0.0008625 | 0.9406 | +2 | 16 |
| - | - | 899.5 | 917.5 | - | - | 0 | - |
| - | - | 866 | 917.9 | - | - | 0 | - |
| - | - | 1060 | 920.5 | - | - | 0 | - |
| - | - | 3.033E+04 | 923 | - | - | 0 | - |
| 2 | y | 3.546E+04 | 923.5 | 0.0106 | 11.48 | +2 | 16 |
| - | - | 2.321E+04 | 924 | - | - | 0 | - |
| - | - | 1.132E+04 | 924.5 | - | - | 0 | - |
| - | - | 4545 | 925 | - | - | 0 | - |
| - | - | 1996 | 925.5 | - | - | 0 | - |
| - | - | 1565 | 926.5 | - | - | 0 | - |
| 8 | c | 4.045E+04 | 927.5 | 0.0001025 | 0.1105 | +1 | 8 |
| - | - | 970.8 | 928.4 | - | - | 0 | - |
| - | - | 1.889E+04 | 928.5 | - | - | 0 | - |
| - | - | 5950 | 929.5 | - | - | 0 | - |
| - | - | 1600 | 930 | - | - | 0 | - |
| - | - | 2678 | 930.5 | - | - | 0 | - |
| - | - | 2109 | 931 | - | - | 0 | - |
| - | - | 1262 | 931.5 | - | - | 0 | - |
| - | - | 982.6 | 934.5 | - | - | 0 | - |
| - | - | 710.7 | 935 | - | - | 0 | - |
| - | - | 943.1 | 936 | - | - | 0 | - |
| - | - | 851.5 | 936.5 | - | - | 0 | - |
| - | - | 1258 | 937 | - | - | 0 | - |
| - | - | 2053 | 937.5 | - | - | 0 | - |
| - | - | 1.418E+04 | 938 | - | - | 0 | - |
| - | - | 1.573E+04 | 938.5 | - | - | 0 | - |
| - | - | 8974 | 939 | - | - | 0 | - |
| - | - | 6292 | 939.5 | - | - | 0 | - |
| - | - | 2473 | 940 | - | - | 0 | - |
| - | - | 1159 | 940.5 | - | - | 0 | - |
| - | - | 3106 | 942.5 | - | - | 0 | - |
| - | - | 3195 | 943 | - | - | 0 | - |
| - | - | 5678 | 943.5 | - | - | 0 | - |
| - | - | 4542 | 944 | - | - | 0 | - |
| - | - | 3008 | 944.5 | - | - | 0 | - |
| - | - | 7666 | 945 | - | - | 0 | - |
| - | - | 7576 | 945.5 | - | - | 0 | - |
| - | - | 4204 | 946 | - | - | 0 | - |
| - | - | 5422 | 946.5 | - | - | 0 | - |
| - | - | 5962 | 947 | - | - | 0 | - |
| - | - | 3292 | 947.5 | - | - | 0 | - |
| - | - | 2089 | 948 | - | - | 0 | - |
| - | - | 1410 | 951.5 | - | - | 0 | - |
| - | - | 7376 | 952 | - | - | 0 | - |
| - | - | 8181 | 952.5 | - | - | 0 | - |
| - | - | 7697 | 953 | - | - | 0 | - |
| - | - | 6625 | 953.5 | - | - | 0 | - |
| - | - | 2850 | 954 | - | - | 0 | - |
| - | - | 1763 | 954.5 | - | - | 0 | - |
| - | - | 1517 | 955.4 | - | - | 0 | - |
| - | - | 1554 | 955.9 | - | - | 0 | - |
| - | - | 2494 | 956.5 | - | - | 0 | - |
| - | - | 5478 | 957 | - | - | 0 | - |
| - | - | 6332 | 957.5 | - | - | 0 | - |
| - | - | 2723 | 958 | - | - | 0 | - |
| - | - | 6219 | 958.5 | - | - | 0 | - |
| - | - | 898.4 | 959 | - | - | 0 | - |
| - | - | 1.638E+04 | 959.5 | - | - | 0 | - |
| - | - | 9683 | 960.5 | - | - | 0 | - |
| - | - | 4125 | 961.5 | - | - | 0 | - |
| - | - | 914.4 | 964.4 | - | - | 0 | - |
| - | - | 1.255E+04 | 965.5 | - | - | 0 | - |
| - | - | 4.67E+04 | 966 | - | - | 0 | - |
| - | - | 4.889E+04 | 966.5 | - | - | 0 | - |
| - | - | 3.297E+04 | 967 | - | - | 0 | - |
| - | - | 1.561E+04 | 967.5 | - | - | 0 | - |
| - | - | 7140 | 968 | - | - | 0 | - |
| - | - | 2243 | 968.5 | - | - | 0 | - |
| - | - | 1032 | 969.5 | - | - | 0 | - |
| - | - | 2.981E+04 | 974 | - | - | 0 | - |
| - | - | 5.828E+04 | 974.5 | - | - | 0 | - |
| - | - | 4.979E+04 | 975 | - | - | 0 | - |
| - | - | 3.097E+04 | 975.5 | - | - | 0 | - |
| - | - | 1.196E+04 | 976 | - | - | 0 | - |
| - | - | 3730 | 976.5 | - | - | 0 | - |
| - | - | 688.7 | 977 | - | - | 0 | - |
| - | - | 669.4 | 992.4 | - | - | 0 | - |
| - | - | 905.7 | 1013 | - | - | 0 | - |
| - | - | 612.2 | 1016 | - | - | 0 | - |
| 9 | z | 3576 | 1022 | 0.004443 | 4.35 | +1 | 9 |
| - | - | 3611 | 1023 | - | - | 0 | - |
| - | - | 1781 | 1024 | - | - | 0 | - |
| - | - | 765.7 | 1025 | - | - | 0 | - |
| - | - | 5344 | 1031 | - | - | 0 | - |
| - | - | 2220 | 1032 | - | - | 0 | - |
| - | - | 1347 | 1033 | - | - | 0 | - |
| 9 | y | 1837 | 1038 | 0.004884 | 4.707 | +1 | 9 |
| - | - | 1067 | 1039 | - | - | 0 | - |
| - | - | 938.1 | 1040 | - | - | 0 | - |
| - | - | 2340 | 1050 | - | - | 0 | - |
| - | - | 1480 | 1051 | - | - | 0 | - |
| - | - | 880.8 | 1074 | - | - | 0 | - |
| 9 | c | 4.234E+04 | 1075 | 0.004752 | 4.423 | +1 | 9 |
| - | - | 2.558E+04 | 1076 | - | - | 0 | - |
| - | - | 520.9 | 1076 | - | - | 0 | - |
| - | - | 1.092E+04 | 1077 | - | - | 0 | - |
| - | - | 2502 | 1078 | - | - | 0 | - |
| - | - | 1183 | 1088 | - | - | 0 | - |
| - | - | 1128 | 1089 | - | - | 0 | - |
| - | - | 703.1 | 1090 | - | - | 0 | - |
| 8 | w | 5447 | 1092 | 0.005184 | 4.749 | +1 | 10 |
| - | - | 3126 | 1093 | - | - | 0 | - |
| - | - | 1445 | 1094 | - | - | 0 | - |
| - | - | 984.6 | 1094 | - | - | 0 | - |
| - | - | 790.7 | 1114 | - | - | 0 | - |
| - | - | 889.4 | 1115 | - | - | 0 | - |
| - | - | 792.4 | 1132 | - | - | 0 | - |
| 8 | z | 1135 | 1133 | 0.01282 | 11.32 | +1 | 10 |
| 8 | z | 1.537E+04 | 1151 | 0.003964 | 3.446 | +1 | 10 |
| - | - | 1.311E+04 | 1152 | - | - | 0 | - |
| - | - | 6687 | 1153 | - | - | 0 | - |
| - | - | 2516 | 1154 | - | - | 0 | - |
| 8 | y | 795.3 | 1167 | 0.008434 | 7.229 | +1 | 10 |
| - | - | 782.5 | 1168 | - | - | 0 | - |
| 10 | c | 2.346E+04 | 1176 | 0.003827 | 3.255 | +1 | 10 |
| - | - | 1.442E+04 | 1177 | - | - | 0 | - |
| - | - | 5776 | 1178 | - | - | 0 | - |
| - | - | 1119 | 1179 | - | - | 0 | - |
| - | - | 849 | 1180 | - | - | 0 | - |
| 7 | w | 3276 | 1221 | 0.004949 | 4.055 | +1 | 11 |
| - | - | 3361 | 1222 | - | - | 0 | - |
| - | - | 1166 | 1223 | - | - | 0 | - |
| - | - | 945.2 | 1224 | - | - | 0 | - |
| - | - | 1378 | 1260 | - | - | 0 | - |
| - | - | 864.6 | 1261 | - | - | 0 | - |
| 7 | z | 1.326E+04 | 1280 | 0.003852 | 3.01 | +1 | 11 |
| - | - | 1.064E+04 | 1281 | - | - | 0 | - |
| - | - | 4969 | 1282 | - | - | 0 | - |
| - | - | 1805 | 1283 | - | - | 0 | - |
| - | - | 820.8 | 1284 | - | - | 0 | - |
| 7 | y | 5096 | 1296 | 0.002828 | 2.183 | +1 | 11 |
| - | - | 2990 | 1297 | - | - | 0 | - |
| - | - | 2245 | 1298 | - | - | 0 | - |
| - | - | 2433 | 1303 | - | - | 0 | - |
| 11 | c | 5446 | 1304 | 0.002614 | 2.005 | +1 | 11 |
| - | - | 3849 | 1305 | - | - | 0 | - |
| - | - | 732.7 | 1306 | - | - | 0 | - |
| - | - | 1112 | 1335 | - | - | 0 | - |
| - | - | 734 | 1336 | - | - | 0 | - |
| - | - | 926.6 | 1377 | - | - | 0 | - |
| - | - | 1467 | 1417 | - | - | 0 | - |
| 12 | c | 6418 | 1418 | 0.001556 | 1.098 | +1 | 12 |
| - | - | 5583 | 1419 | - | - | 0 | - |
| - | - | 3332 | 1420 | - | - | 0 | - |
| - | - | 2545 | 1421 | - | - | 0 | - |
| - | - | 1061 | 1422 | - | - | 0 | - |
| - | - | 785.3 | 1423 | - | - | 0 | - |
| 6 | z | 1.037E+04 | 1436 | 0.003204 | 2.232 | +1 | 12 |
| - | - | 1.647E+04 | 1437 | - | - | 0 | - |
| - | - | 1.177E+04 | 1438 | - | - | 0 | - |
| - | - | 6541 | 1439 | - | - | 0 | - |
| - | - | 1606 | 1440 | - | - | 0 | - |
| - | - | 1332 | 1451 | - | - | 0 | - |
| 6 | y | 6699 | 1452 | 0.002425 | 1.67 | +1 | 12 |
| - | - | 5351 | 1453 | - | - | 0 | - |
| - | - | 3201 | 1454 | - | - | 0 | - |
| 5 | z | 865.1 | 1523 | 0.00206 | 1.353 | +1 | 13 |
| - | - | 4247 | 1524 | - | - | 0 | - |
| - | - | 3611 | 1525 | - | - | 0 | - |
| - | - | 2133 | 1526 | - | - | 0 | - |
| - | - | 1199 | 1527 | - | - | 0 | - |
| 13 | c | 855.8 | 1529 | 0.02031 | 13.28 | +1 | 13 |
| 5 | y | 1464 | 1539 | 0.002992 | 1.945 | +1 | 13 |
| - | - | 1275 | 1540 | - | - | 0 | - |
| 13 | c | 1.228E+04 | 1546 | 0.001328 | 0.8594 | +1 | 13 |
| - | - | 9823 | 1547 | - | - | 0 | - |
| - | - | 4959 | 1548 | - | - | 0 | - |
| - | - | 2246 | 1549 | - | - | 0 | - |
| - | - | 1301 | 1602 | - | - | 0 | - |
| - | - | 1137 | 1603 | - | - | 0 | - |
| - | - | 1525 | 1615 | - | - | 0 | - |
| - | - | 852.6 | 1617 | - | - | 0 | - |
| 14 | c | 1554 | 1627 | 0.003172 | 1.95 | +1 | 14 |
| 14 | c | 1579 | 1628 | 0.006705 | 4.119 | +1 | 14 |
| - | - | 1348 | 1629 | - | - | 0 | - |
| - | - | 779.8 | 1630 | - | - | 0 | - |
| - | - | 727.7 | 1631 | - | - | 0 | - |
| 4 | y | 3752 | 1636 | 0.004302 | 2.63 | +1 | 14 |
| - | - | 3502 | 1637 | - | - | 0 | - |
| - | - | 2371 | 1638 | - | - | 0 | - |
| - | - | 1413 | 1639 | - | - | 0 | - |
| - | - | 1975 | 1644 | - | - | 0 | - |
| 14 | c | 1.062E+04 | 1645 | 0.002495 | 1.517 | +1 | 14 |
| - | - | 1.274E+04 | 1646 | - | - | 0 | - |
| - | - | 7126 | 1647 | - | - | 0 | - |
| - | - | 3026 | 1648 | - | - | 0 | - |
| - | - | 1461 | 1649 | - | - | 0 | - |
| - | - | 783.4 | 1672 | - | - | 0 | - |
| - | - | 1004 | 1689 | - | - | 0 | - |
| - | - | 731.6 | 1691 | - | - | 0 | - |
| 3 | y | 866.5 | 1716 | 0.0307 | 17.89 | +1 | 15 |
| 3 | z | 711.4 | 1717 | 0.02287 | 13.32 | +1 | 15 |
| - | - | 1164 | 1729 | - | - | 0 | - |
| - | - | 857.2 | 1730 | - | - | 0 | - |
| - | - | 1047 | 1731 | - | - | 0 | - |
| 15 | c | 5756 | 1732 | 0.00196 | 1.132 | +1 | 15 |
| 3 | y | 1.186E+04 | 1733 | 0.02039 | 11.76 | +1 | 15 |
| - | - | 9824 | 1734 | - | - | 0 | - |
| - | - | 5931 | 1735 | - | - | 0 | - |
| - | - | 2578 | 1736 | - | - | 0 | - |
| - | - | 721.1 | 1785 | - | - | 0 | - |
| - | - | 1145 | 1786 | - | - | 0 | - |
| - | - | 2265 | 1802 | - | - | 0 | - |
| - | - | 3247 | 1803 | - | - | 0 | - |
| - | - | 1771 | 1804 | - | - | 0 | - |
| - | - | 1141 | 1805 | - | - | 0 | - |
| - | - | 788.3 | 1814 | - | - | 0 | - |
| 2 | y | 1018 | 1829 | 0.01451 | 7.932 | +1 | 16 |
| 2 | z | 2725 | 1830 | 0.01803 | 9.855 | +1 | 16 |
| - | - | 1.215E+04 | 1831 | - | - | 0 | - |
| - | - | 1.207E+04 | 1832 | - | - | 0 | - |
| 16 | c | 7977 | 1833 | 0.01165 | 6.359 | +1 | 16 |
| - | - | 4370 | 1834 | - | - | 0 | - |
| - | - | 1191 | 1835 | - | - | 0 | - |
| - | - | 4570 | 1845 | - | - | 0 | - |
| 2 | y | 9518 | 1846 | 0.02275 | 12.32 | +1 | 16 |
| - | - | 9624 | 1847 | - | - | 0 | - |
| - | - | 5694 | 1848 | - | - | 0 | - |
| - | - | 3299 | 1849 | - | - | 0 | - |
| - | - | 1120 | 1850 | - | - | 0 | - |
| - | - | 1421 | 1885 | - | - | 0 | - |
| - | - | 2184 | 1886 | - | - | 0 | - |
| - | - | 3138 | 1887 | - | - | 0 | - |
| - | - | 2358 | 1888 | - | - | 0 | - |
| - | - | 1356 | 1889 | - | - | 0 | - |
| - | - | 5255 | 1890 | - | - | 0 | - |
| - | - | 4719 | 1891 | - | - | 0 | - |
| - | - | 3166 | 1892 | - | - | 0 | - |
| - | - | 2873 | 1903 | - | - | 0 | - |
| - | - | 9432 | 1904 | - | - | 0 | - |
| - | - | 9495 | 1905 | - | - | 0 | - |
| - | - | 5311 | 1906 | - | - | 0 | - |
| - | - | 2677 | 1907 | - | - | 0 | - |
| - | - | 762.9 | 1912 | - | - | 0 | - |
| - | - | 1507 | 1914 | - | - | 0 | - |
| - | - | 2477 | 1915 | - | - | 0 | - |
| - | - | 2772 | 1916 | - | - | 0 | - |
| - | - | 1524 | 1917 | - | - | 0 | - |
| - | - | 1060 | 1918 | - | - | 0 | - |
| - | - | 1885 | 1920 | - | - | 0 | - |
| - | - | 7288 | 1921 | - | - | 0 | - |
| - | - | 7651 | 1922 | - | - | 0 | - |
| - | - | 3747 | 1923 | - | - | 0 | - |
| - | - | 2359 | 1924 | - | - | 0 | - |
| - | - | 769.9 | 1925 | - | - | 0 | - |
| - | - | 853.4 | 1930 | - | - | 0 | - |
| - | - | 8793 | 1931 | - | - | 0 | - |
| - | - | 2.941E+04 | 1932 | - | - | 0 | - |
| - | - | 3.101E+04 | 1933 | - | - | 0 | - |
| - | - | 1.899E+04 | 1934 | - | - | 0 | - |
| - | - | 7858 | 1935 | - | - | 0 | - |
| - | - | 4477 | 1936 | - | - | 0 | - |
| - | - | 880.8 | 1937 | - | - | 0 | - |
| - | - | 3073 | 1947 | - | - | 0 | - |
| - | - | 1.871E+04 | 1948 | - | - | 0 | - |
| - | - | 8.567E+04 | 1949 | - | - | 0 | - |
| - | - | 9.042E+04 | 1950 | - | - | 0 | - |
| - | - | 5.59E+04 | 1951 | - | - | 0 | - |
| - | - | 2.391E+04 | 1952 | - | - | 0 | - |
| - | - | 1.032E+04 | 1953 | - | - | 0 | - |
| - | - | 2265 | 1954 | - | - | 0 | - |

m/z Charge Intensity FragmentType MassShift Position
120.06572723388672 0 2178.2183
126.98662567138672 0 388.7463
141.73240661621094 0 426.24017
142.04287719726562 0 387.38074
152.13735961914062 0 372.11258
165.76869201660156 0 438.77988 c Water loss 4
168.12637329101562 0 531.0015
173.12843322753906 0 698.7716
187.14430236816406 0 21850.408
188.14752197265625 0 1924.1395
195.69448852539062 0 473.21887
201.12342834472656 0 701.0034 y Water loss 13
212.13955688476562 0 1657.1361
215.13917541503906 0 13921.789 y Water loss 15
216.14288330078125 0 1849.7982
233.1496124267578 0 1104.3062 y 15
238.15573120117188 0 546.1083
276.16558837890625 0 499.43414
298.62689208984375 0 583.8358
299.17169189453125 0 3996.9907
300.17559814453125 0 935.82416
319.1965637207031 0 637.2035
320.181884765625 0 808.6189 y 14
321.8746032714844 0 560.75525
332.6846618652344 0 500.9951
348.66845703125 0 724.8826
359.0281677246094 0 779.17255
407.2533264160156 0 792.7319
424.9517517089844 0 555.3721
426.2718811035156 0 641.41205 c 3
429.0891418457031 0 1767.9784
470.298828125 0 1619.6952
494.2844543457031 0 745.1673
509.2676696777344 0 678.21075
513.3033447265625 0 16179.596 c 4
514.3062744140625 0 5034.5273
515.3118286132812 0 822.8019
523.2687377929688 0 773.8859
526.280517578125 0 629.13434
529.2758178710938 0 749.67206
529.6067504882812 0 1366.4573
529.9344482421875 0 926.1165
531.2907104492188 0 1496.9453 z 12
532.2960205078125 0 1880.5022
532.5986938476562 0 1363.8618
532.9318237304688 0 2322.0874
533.2661743164062 0 1003.6377
538.6033325195312 0 4814.5405
538.936279296875 0 3607.2341
539.2708129882812 0 2232.3784
539.90380859375 0 668.31555
547.3095092773438 0 1079.2572 y 12
556.9542846679688 0 911.25336
557.2892456054688 0 1358.6816
557.9588012695312 0 779.5215
571.3184204101562 0 1528.7771
572.2872314453125 0 1831.4385 c Ammonia loss 14
572.6177978515625 0 2378.7861 y Ammonia loss 2
572.9509887695312 0 2671.342 z 2
573.2822875976562 0 1786.6421
573.3336181640625 0 762.7533
578.2897338867188 0 57972.457 y 2
578.6233520507812 0 59870.395
578.95703125 0 37666.89
579.2906494140625 0 13969.682 c Water loss 9
579.624755859375 0 5256.1816
579.955322265625 0 1155.1346
625.3905029296875 0 1653.8315
626.3968505859375 0 1018.6692
645.3341064453125 0 1119.5942 z 11
646.3379516601562 0 2056.1501
647.3416748046875 0 666.632
649.310546875 0 817.5005
649.8544311523438 0 1218.9626
650.3486328125 0 884.3409
650.84912109375 0 953.8842
652.33837890625 0 3004.8784 c 10
652.8388671875 0 2129.2886
661.3516845703125 0 2465.3167 y 11
662.3558349609375 0 728.1459
669.4043579101562 0 28330.354 c 5
670.4078369140625 0 9529.796
671.4087524414062 0 1835.0264
672.4034423828125 0 714.4218
693.8402709960938 0 734.27594
699.8500366210938 0 1019.4554
700.3479614257812 0 916.4494 c Water loss 11
707.3438110351562 0 3150.1367
707.8489990234375 0 928.03815
708.3416748046875 0 1053.7203
708.8533935546875 0 2306.4421
709.3582153320312 0 19615.031 c 11
709.8599853515625 0 15176.067
710.3594970703125 0 4793.2837
710.86083984375 0 3255.695
711.3699340820312 0 876.59546
718.3563842773438 0 854.37195 z 5
718.8555908203125 0 1008.45233
726.3607788085938 0 2914.6062 y 5
726.8638916015625 0 2730.067
727.3648071289062 0 1150.3693
729.422607421875 0 3505.1501
730.4260864257812 0 2140.5615
750.8595581054688 0 1574.0875
751.3621826171875 0 1987.9355
751.86669921875 0 2259.8372
761.8681640625 0 1744.1344 z 4
762.3720092773438 0 1624.9255
762.86572265625 0 785.2049
766.3964233398438 0 732.59186
769.8768920898438 0 2306.4456 y 4
770.3760986328125 0 1184.7275
772.884521484375 0 948.9813
773.3895874023438 0 20164.36 c 12
773.4512939453125 0 5585.357
773.8888549804688 0 24649.904
774.3917236328125 0 11625.111
774.886962890625 0 4777.982
775.389892578125 0 1827.0109
779.4200439453125 0 834.3415
786.3981323242188 0 876.5195
789.4466552734375 0 3571.22 y 10
790.4470825195312 0 2244.5776
791.4341430664062 0 980.33264
792.9103393554688 0 1395.4801
793.4051513671875 0 1769.8845
793.9111938476562 0 1658.0175
794.4003295898438 0 1158.9845
798.447265625 0 40180.965 c 6
798.907958984375 0 926.92957
799.449951171875 0 18503.518
800.4512939453125 0 4467.7427
800.9141235351562 0 1721.4747
801.4151611328125 0 2744.2795
801.9204711914062 0 1495.6923
802.4141235351562 0 861.50073
807.4017944335938 0 4247.9404
807.9017944335938 0 4655.583
808.4014282226562 0 2340.2651
808.8975830078125 0 901.0671
809.8964233398438 0 707.8518 y Ammonia loss 3
813.4130859375 0 7472.3857
813.9126586914062 0 7538.3125 c Water loss 13
814.4121704101562 0 4600.5107 c Ammonia loss 13
814.91455078125 0 2344.5198
815.4163818359375 0 866.5047
818.4042358398438 0 12226.396 y 3
818.904296875 0 9055.073
819.4069213867188 0 5305.1797
819.9087524414062 0 1898.8369
822.4183959960938 0 11831.259
822.9207153320312 0 18756.836 c 13
823.4214477539062 0 14179.342
823.92236328125 0 8261.101
824.4242553710938 0 3498.3782
824.9259643554688 0 1410.6509
825.46142578125 0 815.8067
830.4686889648438 0 885.3038
834.93359375 0 2290.4
835.435791015625 0 1798.7284
835.92919921875 0 2189.3186
836.4267578125 0 1239.3865
836.9296875 0 909.40283
844.4282836914062 0 797.66345
845.4168701171875 0 1722.383 w 2
845.9117431640625 0 1832.277
846.41162109375 0 1085.0308
849.417724609375 0 772.3893
857.4707641601562 0 2179.4485 w 9
857.9248657226562 0 3392.8914 c Ammonia loss 14
858.4295043945312 0 2425.2405 y Ammonia loss 2
858.92529296875 0 1503.3154 z 2
859.4521484375 0 4172.17 w 9
860.453857421875 0 2026.3708
864.9404907226562 0 839.8402
865.9319458007812 0 1820.4648
866.360595703125 0 443.41205
866.4384155273438 0 33937.125 c 14
866.9337768554688 0 68506.3 y 2
867.4340209960938 0 56563.89
867.9331665039062 0 32701.297
868.4344482421875 0 13305.517
868.933349609375 0 5614.2593
873.4639892578125 0 3838.2937 y Ammonia loss 9
874.4749145507812 0 20450.037 z 9
875.4760131835938 0 12284.269
876.4782104492188 0 3556.1365
877.4771118164062 0 1583.2306
879.4508666992188 0 2285.3428
879.9481811523438 0 1243.8586
882.43505859375 0 911.5246
887.4361572265625 0 1520.7767
890.4930419921875 0 6420.494 y 9
891.4923095703125 0 3765.6653
891.9715576171875 0 1477.1733
892.4851684570312 0 2010.8835
893.9359130859375 0 11799.045 w 1
894.435791015625 0 11987.5205
894.93505859375 0 8204.484
895.4337158203125 0 3142.47
895.9421997070312 0 1508.8127
900.9789428710938 0 1214.4958
901.4747924804688 0 2073.1445
901.9755859375 0 1808.9827
902.477294921875 0 1796.4724
906.9720458984375 0 912.6747
912.48486328125 0 839.3341
914.4698486328125 0 2171.7805 y Water loss 1
914.9586181640625 0 1672.8608 y Ammonia loss 1
915.4625854492188 0 6953.071 z 1
915.9667358398438 0 7297.552
916.4684448242188 0 4247.0933
916.9588623046875 0 3456.437 c 15
917.45947265625 0 899.5253
917.9480590820312 0 866.0231
920.487548828125 0 1060.3921
922.9801635742188 0 30330.605
923.4805297851562 0 35464.215 y 1
923.9811401367188 0 23206.463
924.4819946289062 0 11323.093
924.9813232421875 0 4545.124
925.4864501953125 0 1995.6982
926.4810180664062 0 1564.5668
927.4893188476562 0 40451.5 c 7
928.3840942382812 0 970.7738
928.491943359375 0 18893.59
929.4925537109375 0 5950.0625
929.970458984375 0 1599.8112
930.481689453125 0 2677.6763
930.9677734375 0 2108.5586
931.4837036132812 0 1262.3711
934.4946899414062 0 982.64984
934.9890747070312 0 710.65894
935.9805297851562 0 943.148
936.4818115234375 0 851.541
936.9833984375 0 1257.8015
937.4716796875 0 2052.9976
937.9603881835938 0 14178.6455
938.4616088867188 0 15726.578
938.9632568359375 0 8973.594
939.4682006835938 0 6292.48
939.966064453125 0 2473.2844
940.4716796875 0 1158.9421
942.4967041015625 0 3105.9805
942.9920043945312 0 3195.0342
943.4903564453125 0 5678.1055
943.9854736328125 0 4542.334
944.480224609375 0 3008.2615
944.9788208007812 0 7666.1436
945.4824829101562 0 7576.371
945.9848022460938 0 4204.2695
946.4725952148438 0 5422.3486
946.9697875976562 0 5961.763
947.4682006835938 0 3292.1575
947.9683227539062 0 2089.4307
951.4996337890625 0 1409.6495
951.9909057617188 0 7375.8687
952.4926147460938 0 8181.0786
952.9892578125 0 7696.9473
953.4913330078125 0 6624.833
953.980712890625 0 2850.014
954.4867553710938 0 1762.6312
955.4288940429688 0 1517.2191
955.9359741210938 0 1553.6178
956.4866943359375 0 2493.772
956.98388671875 0 5477.9478
957.4866333007812 0 6331.9976
957.98583984375 0 2722.866
958.5125122070312 0 6219.3843
958.9758911132812 0 898.4027
959.5265502929688 0 16382.938
960.5286254882812 0 9683.287
961.52734375 0 4124.687
964.4221801757812 0 914.36206
965.4946899414062 0 12546.865
965.990478515625 0 46699.24
966.490234375 0 48888.297
966.9906005859375 0 32965.77
967.4911499023438 0 15606.604
967.9923706054688 0 7140.069
968.5082397460938 0 2242.5823
969.5357055664062 0 1032.1034
973.9965209960938 0 29812.273
974.4979248046875 0 58281.836
974.9989624023438 0 49793.168
975.5001220703125 0 30971.56
976.0011596679688 0 11959.18
976.5009155273438 0 3729.5981
977.0087280273438 0 688.6702
992.4190673828125 0 669.4348
1012.5032958984375 0 905.7156
1015.50830078125 0 612.1877
1021.5103759765625 0 3575.7896 z 8
1022.5152587890625 0 3611.3975
1023.5115966796875 0 1780.7036
1024.508544921875 0 765.65717
1030.5108642578125 0 5344.22
1031.5145263671875 0 2219.9004
1032.5164794921875 0 1346.7982
1037.529541015625 0 1837.086 y 8
1038.52978515625 0 1066.9556
1039.52880859375 0 938.1405
1049.5289306640625 0 2340.383
1050.5255126953125 0 1479.5127
1073.5108642578125 0 880.80237
1074.524658203125 0 42340.406 c 8
1075.5274658203125 0 25580.26
1076.40625 0 520.8826
1076.5283203125 0 10918.958
1077.5269775390625 0 2502.447
1087.561767578125 0 1182.6348
1088.5660400390625 0 1128.1265
1089.55224609375 0 703.0887
1091.5404052734375 0 5446.694 w 7
1092.5428466796875 0 3126.2756
1093.541015625 0 1444.9609
1094.4981689453125 0 984.62146
1113.5443115234375 0 790.70953
1114.5438232421875 0 889.4102
1131.5621337890625 0 792.3613
1132.55078125 0 1135.1635 z Water loss 7
1150.552490234375 0 15366.164 z 7
1151.5543212890625 0 13114.994
1152.5548095703125 0 6687.0264
1153.5567626953125 0 2516.2083
1166.57568359375 0 795.33966 y 7
1167.5711669921875 0 782.5114
1175.5714111328125 0 23460.46 c 9
1176.573974609375 0 14423.594
1177.5740966796875 0 5775.996
1178.5816650390625 0 1119.3234
1179.58203125 0 849.01013
1220.582763671875 0 3275.6113 w 6
1221.5799560546875 0 3360.7302
1222.5811767578125 0 1166.2856
1223.569091796875 0 945.1574
1259.650146484375 0 1377.8545
1260.658447265625 0 864.56555
1279.594970703125 0 13260.958 z 6
1280.59619140625 0 10638.51
1281.592529296875 0 4969.171
1282.59423828125 0 1804.815
1283.596923828125 0 820.82153
1295.6126708984375 0 5095.52 y 6
1296.615966796875 0 2989.9304
1297.6103515625 0 2244.676
1302.6583251953125 0 2433.2993
1303.6651611328125 0 5446.246 c 10
1304.66845703125 0 3849.217
1305.66943359375 0 732.69666
1334.6707763671875 0 1112.2012
1335.6634521484375 0 733.9957
1376.6805419921875 0 926.573
1416.693115234375 0 1467.2557
1417.70703125 0 6417.6167 c 11
1418.7076416015625 0 5582.9785
1419.7091064453125 0 3332.1418
1420.707763671875 0 2545.1064
1421.699462890625 0 1061.2118
1422.6842041015625 0 785.33
1435.6954345703125 0 10372.204 z 5
1436.699951171875 0 16470.494
1437.6995849609375 0 11768.515
1438.7032470703125 0 6540.8896
1439.7003173828125 0 1606.225
1450.7100830078125 0 1332.3951
1451.71337890625 0 6699.2603 y 5
1452.7119140625 0 5351.4683
1453.711181640625 0 3201.271
1522.726318359375 0 865.0858 z 4
1523.72998046875 0 4246.693
1524.7354736328125 0 3611.13
1525.7425537109375 0 2133.299
1526.7427978515625 0 1198.8264
1528.7578125 0 855.8272 c Ammonia loss 12
1538.739990234375 0 1464.0977 y 4
1539.748046875 0 1274.9227
1545.765380859375 0 12284.813 c 12
1546.7681884765625 0 9823.203
1547.7694091796875 0 4958.5894
1548.769287109375 0 2245.5608
1601.820556640625 0 1300.6617
1602.8238525390625 0 1137.2245
1614.798583984375 0 1525.1084
1616.808349609375 0 852.59357
1626.8250732421875 0 1553.8346 c Water loss 13
1627.8126220703125 0 1578.5115 c Ammonia loss 13
1628.8116455078125 0 1347.6256
1629.8016357421875 0 779.7811
1630.815185546875 0 727.68353
1635.800048828125 0 3752.2056 y 3
1636.7919921875 0 3502.347
1637.8087158203125 0 2370.5403
1638.79638671875 0 1412.6355
1643.8309326171875 0 1974.6958
1644.8349609375 0 10624.241 c 13
1645.8358154296875 0 12743.483
1646.836669921875 0 7125.8184
1647.8460693359375 0 3026.5
1648.8497314453125 0 1461.4403
1671.859375 0 783.38684
1688.8544921875 0 1004.45215
1690.8392333984375 0 731.564
1715.8526611328125 0 866.5096 y Ammonia loss 2
1716.8526611328125 0 711.3753 z 2
1728.88916015625 0 1164.3624
1729.90576171875 0 857.1852
1730.872314453125 0 1047.0049
1731.866455078125 0 5756.324 c 14
1732.868896484375 0 11861.279 y 2
1733.8714599609375 0 9823.528
1734.8704833984375 0 5930.8916
1735.8719482421875 0 2577.7524
1784.9630126953125 0 721.0599
1785.9217529296875 0 1145.046
1801.946044921875 0 2264.8901
1802.9461669921875 0 3246.6304
1803.9359130859375 0 1770.9626
1804.93798828125 0 1140.6161
1813.93212890625 0 788.282
1828.9205322265625 0 1017.7759 y Ammonia loss 1
1829.931884765625 0 2725.4292 z 1
1830.9229736328125 0 12149.411
1831.92431640625 0 12070.351
1832.923828125 0 7976.7896 c 15
1833.92041015625 0 4370.0605
1834.9266357421875 0 1191.2322
1844.945556640625 0 4570.137
1845.955322265625 0 9517.818 y 1
1846.955322265625 0 9624.416
1847.960205078125 0 5693.978
1848.9605712890625 0 3298.8198
1849.948486328125 0 1120.3375
1885.02001953125 0 1421.2738
1885.9727783203125 0 2183.9788
1886.9764404296875 0 3137.7415
1887.96728515625 0 2358.2893
1888.95751953125 0 1355.7146
1889.9569091796875 0 5254.6963
1890.95947265625 0 4718.6665
1891.9560546875 0 3166.0916
1902.986083984375 0 2873.111
1903.980712890625 0 9431.686
1904.9788818359375 0 9495.384
1905.9815673828125 0 5310.5044
1906.987060546875 0 2676.5422
1911.863037109375 0 762.85364
1913.9512939453125 0 1507.309
1914.95849609375 0 2476.981
1915.954345703125 0 2772.2734
1916.94189453125 0 1524.4342
1917.947998046875 0 1060.4385
1919.985595703125 0 1885.2606
1921.0028076171875 0 7288.0513
1922.000732421875 0 7650.8105
1923.0023193359375 0 3746.7732
1924.0042724609375 0 2359.3567
1924.996826171875 0 769.9409
1929.9810791015625 0 853.4344
1930.9749755859375 0 8792.598
1931.9732666015625 0 29414.062
1932.9720458984375 0 31013.582
1933.974365234375 0 18994.836
1934.976318359375 0 7858.1885
1935.9727783203125 0 4477.168
1936.978515625 0 880.81976
1946.98046875 0 3073.293
1947.98681640625 0 18710.14
1948.993408203125 0 85665.96
1949.9959716796875 0 90418.6
1950.9970703125 0 55902.84
1951.9970703125 0 23910.004
1952.99755859375 0 10319.598
1954.0045166015625 0 2264.5327

Spectrum Details

|  |  |
| --- | --- |
| Matched peaks? Matched peaks The total absolute number of peaks matched. Additionally in brackets the total fraction of peaks matched and the total number of peaks is shown. | 81 (17.09% of 474) |
| FDR? FDR The false discovery rate estimated for this peptide. It is calculated by matching all theoretical fragments with a non-integer shift with the raw peaks for this spectrum. This is done with 40 different shifts. The resulting percentage is the average number of annotated peaks over the number of annotated peaks with the correct spectrum. | 0.50% |
| Satellite FDR? Satellite FDR See the FDR for details on its calculation. This satellite ion specific FDR only contains the satellite ions (d/w) for I/L/J positions. | 9.52% |
| PSM Score? PSM Score The PSM Score as given by Hecklib to this annotated spectrum. It is shown with three significant figures. | 556 |

#### Spectrum 4919? Spectrum 4919 The raw spectrum of this peptide as annotated by Hecklib. The fragments are coloured according to ion type (see legend). Any peaks with a star '\*' as text can be hovered over to see the full details, first the ion type second the mass shift type. By hovering over the amino acids in the peptide or ions in the legend the corresponding peaks are highlighted. By toggling the 'Unassigned' label you can turn the background (unassigned) peaks on or off in the plot. By updating the slider in the Ion legend you can update the spectrum to only show the top X% of the peaks with labels. The top X% means any peak that is within X% of the highest intensity. By dragging in the spectrum you can zoom in to a specific part of the spectrum and use 'Zoom Out' to get back to the original zoom level. The annotation of the spectrum is based on the given sequence in the peptides file and is done with different software so inconsistencies are likely. The peaks are annotated based on the given sequence, with 20 ppm tolerance.

Copy Data

##### Spectrum 4919 (TSV)

###### Preview

```
Loading example...
```

*Click on the button to copy the data to your clipboard.*

Mz MinMz MaxIntensity Max

WidthHeightPeptide font sizePeptide stroke widthSpectrum font sizeSpectrum stroke widthCompact peptide

Ion legend

wxyz

abcd

OtherUnassignedIon^Charge^_Position_Show for top:%

TLPPSREEMTKNQVSTJ

01.38e+42.76e+44.14e+45.52e+4

Zoom Out

y^+1^_2_y^+1^_3_c^+1^_4_y^+3^_12_c^+1^_5_z^+1^_5_c^+3^_15_y^+2^_10_z^+3^_15_y^+3^_15_y^+1^_6_c^+2^_11_y^+1^_6_c^+1^_6_c^+2^_12_y^+2^_12_y^+2^_13_c^+2^_13_z^+1^_7_y^+1^_7_c^+1^_7_c^+2^_14_c^+2^_14_y^+2^_14_c^+2^_14_w^+2^_15_c^+2^_15_y^+2^_15_z^+2^_15_w^+1^_8_c^+2^_15_y^+2^_15_w^+1^_8_y^+1^_8_z^+1^_8_w^+2^_16_y^+1^_8_z^+2^_16_c^+2^_16_y^+2^_16_c^+1^_8_w^+1^_9_z^+1^_9_y^+1^_9_c^+1^_9_w^+1^_10_z^+1^_10_y^+1^_10_c^+1^_10_w^+1^_11_z^+1^_11_c^+1^_11_y^+1^_11_c^+1^_11_c^+1^_12_z^+1^_12_y^+1^_12_z^+1^_13_y^+1^_13_c^+1^_13_c^+1^_14_y^+1^_14_c^+1^_14_c^+1^_15_y^+1^_15_z^+1^_16_c^+1^_16_y^+1^_16_

0877175426323509

Fragment Matches Table

Show background peaks

| Position | Ion type | Intensity | mz Theoretical | mz Error (Th) | mz Error (ppm) | Charge | Series Number |
| --- | --- | --- | --- | --- | --- | --- | --- |
| - | - | 1068 | 120.1 | - | - | 0 | - |
| - | - | 548.4 | 120.1 | - | - | 0 | - |
| - | - | 387.3 | 135.2 | - | - | 0 | - |
| - | - | 492.5 | 148.9 | - | - | 0 | - |
| - | - | 506.2 | 148.9 | - | - | 0 | - |
| - | - | 1206 | 148.9 | - | - | 0 | - |
| - | - | 1267 | 148.9 | - | - | 0 | - |
| - | - | 1534 | 148.9 | - | - | 0 | - |
| - | - | 3478 | 148.9 | - | - | 0 | - |
| - | - | 5992 | 149 | - | - | 0 | - |
| - | - | 3306 | 149 | - | - | 0 | - |
| - | - | 1422 | 149 | - | - | 0 | - |
| - | - | 1337 | 149 | - | - | 0 | - |
| - | - | 1213 | 149 | - | - | 0 | - |
| - | - | 759.1 | 149 | - | - | 0 | - |
| - | - | 916.3 | 149 | - | - | 0 | - |
| - | - | 782.8 | 149 | - | - | 0 | - |
| - | - | 669.1 | 149 | - | - | 0 | - |
| - | - | 394.1 | 149 | - | - | 0 | - |
| - | - | 463.3 | 149.1 | - | - | 0 | - |
| - | - | 411.9 | 163.6 | - | - | 0 | - |
| - | - | 594 | 166.1 | - | - | 0 | - |
| - | - | 890.7 | 173.4 | - | - | 0 | - |
| - | - | 721.6 | 187.1 | - | - | 0 | - |
| - | - | 1.248E+04 | 187.1 | - | - | 0 | - |
| - | - | 828.9 | 188.1 | - | - | 0 | - |
| - | - | 554 | 215.1 | - | - | 0 | - |
| 16 | y | 7184 | 215.1 | 0.0003318 | 1.542 | +1 | 2 |
| - | - | 759 | 216.1 | - | - | 0 | - |
| - | - | 1087 | 223.1 | - | - | 0 | - |
| - | - | 517.1 | 244.3 | - | - | 0 | - |
| - | - | 579.2 | 250.1 | - | - | 0 | - |
| - | - | 582.5 | 250.6 | - | - | 0 | - |
| - | - | 2694 | 299.2 | - | - | 0 | - |
| 15 | y | 719.8 | 320.2 | 0.0008564 | 2.675 | +1 | 3 |
| - | - | 752.2 | 407.3 | - | - | 0 | - |
| 4 | c | 721.2 | 426.3 | 0.001259 | 2.954 | +1 | 4 |
| - | - | 729.3 | 428.3 | - | - | 0 | - |
| - | - | 1169 | 460.2 | - | - | 0 | - |
| - | - | 1565 | 470.3 | - | - | 0 | - |
| 6 | y | 1678 | 473.2 | 0.0001008 | 0.2131 | +3 | 12 |
| - | - | 2807 | 478.2 | - | - | 0 | - |
| - | - | 1521 | 494.3 | - | - | 0 | - |
| 5 | c | 8869 | 513.3 | 2.354E-05 | 0.04586 | +1 | 5 |
| - | - | 2078 | 514.3 | - | - | 0 | - |
| - | - | 819.6 | 524.3 | - | - | 0 | - |
| - | - | 729.2 | 526.3 | - | - | 0 | - |
| - | - | 844.8 | 527.3 | - | - | 0 | - |
| 13 | z | 1449 | 531.3 | 0.0007701 | 1.449 | +1 | 5 |
| - | - | 702.1 | 532.3 | - | - | 0 | - |
| - | - | 1447 | 533.3 | - | - | 0 | - |
| - | - | 1620 | 533.6 | - | - | 0 | - |
| - | - | 733.8 | 533.9 | - | - | 0 | - |
| 15 | c | 2010 | 567 | 0.0001867 | 0.3293 | +3 | 15 |
| 8 | y | 2159 | 567.3 | 0.008243 | 14.53 | +2 | 10 |
| 3 | z | 1081 | 567.6 | 0.003713 | 6.542 | +3 | 15 |
| 3 | y | 3.433E+04 | 573 | 0.0005955 | 1.039 | +3 | 15 |
| - | - | 3.05E+04 | 573.3 | - | - | 0 | - |
| - | - | 1.53E+04 | 573.6 | - | - | 0 | - |
| - | - | 9148 | 574 | - | - | 0 | - |
| - | - | 2242 | 574.3 | - | - | 0 | - |
| - | - | 1578 | 594.2 | - | - | 0 | - |
| - | - | 1090 | 607.4 | - | - | 0 | - |
| - | - | 903 | 625.4 | - | - | 0 | - |
| - | - | 726.6 | 638.7 | - | - | 0 | - |
| 12 | y | 861.8 | 643.3 | 0.004066 | 6.319 | +1 | 6 |
| - | - | 1.994E+04 | 643.9 | - | - | 0 | - |
| 11 | c | 1.938E+04 | 644.3 | 0.009575 | 14.86 | +2 | 11 |
| - | - | 7123 | 644.9 | - | - | 0 | - |
| - | - | 3026 | 645.4 | - | - | 0 | - |
| - | - | 1061 | 645.9 | - | - | 0 | - |
| - | - | 1543 | 646.3 | - | - | 0 | - |
| 12 | y | 2399 | 661.4 | 0.001128 | 1.706 | +1 | 6 |
| - | - | 896.6 | 665.2 | - | - | 0 | - |
| - | - | 623.4 | 666.5 | - | - | 0 | - |
| 6 | c | 1.306E+04 | 669.4 | 0.0006097 | 0.9108 | +1 | 6 |
| - | - | 4617 | 670.4 | - | - | 0 | - |
| - | - | 725.1 | 671.4 | - | - | 0 | - |
| - | - | 649.7 | 691.4 | - | - | 0 | - |
| - | - | 661.7 | 695.4 | - | - | 0 | - |
| - | - | 576.9 | 696.4 | - | - | 0 | - |
| - | - | 1775 | 699.3 | - | - | 0 | - |
| - | - | 1328 | 699.8 | - | - | 0 | - |
| - | - | 791.6 | 700.3 | - | - | 0 | - |
| 12 | c | 1.059E+04 | 701.4 | 0.0003527 | 0.5029 | +2 | 12 |
| - | - | 6237 | 701.9 | - | - | 0 | - |
| - | - | 3168 | 702.4 | - | - | 0 | - |
| - | - | 945.8 | 702.9 | - | - | 0 | - |
| - | - | 953.2 | 711.4 | - | - | 0 | - |
| - | - | 856.3 | 714.4 | - | - | 0 | - |
| - | - | 727.1 | 717.9 | - | - | 0 | - |
| 6 | y | 1538 | 718.4 | 0.0006308 | 0.8781 | +2 | 12 |
| - | - | 1831 | 718.9 | - | - | 0 | - |
| - | - | 1405 | 729.4 | - | - | 0 | - |
| - | - | 1001 | 730.4 | - | - | 0 | - |
| - | - | 1731 | 742.9 | - | - | 0 | - |
| - | - | 1170 | 743.4 | - | - | 0 | - |
| - | - | 818.4 | 750.9 | - | - | 0 | - |
| - | - | 939.6 | 754.4 | - | - | 0 | - |
| - | - | 723.7 | 757.4 | - | - | 0 | - |
| 5 | y | 905.5 | 761.9 | 0.0006688 | 0.8778 | +2 | 13 |
| - | - | 570.8 | 762.9 | - | - | 0 | - |
| - | - | 746 | 763.4 | - | - | 0 | - |
| - | - | 901.2 | 764.9 | - | - | 0 | - |
| 13 | c | 1.489E+04 | 765.4 | 0.001077 | 1.407 | +2 | 13 |
| - | - | 1.252E+04 | 765.9 | - | - | 0 | - |
| - | - | 7033 | 766.4 | - | - | 0 | - |
| - | - | 2834 | 766.9 | - | - | 0 | - |
| - | - | 970.9 | 767.4 | - | - | 0 | - |
| 11 | z | 5823 | 773.4 | 0.00796 | 10.29 | +1 | 7 |
| - | - | 3374 | 774.4 | - | - | 0 | - |
| - | - | 627.3 | 775.4 | - | - | 0 | - |
| - | - | 1022 | 777.4 | - | - | 0 | - |
| - | - | 976.8 | 785.4 | - | - | 0 | - |
| - | - | 790.2 | 785.9 | - | - | 0 | - |
| - | - | 887.4 | 786.4 | - | - | 0 | - |
| - | - | 913.3 | 786.9 | - | - | 0 | - |
| 11 | y | 2115 | 789.4 | 0.001077 | 1.364 | +1 | 7 |
| - | - | 659.1 | 791 | - | - | 0 | - |
| - | - | 950.3 | 792.9 | - | - | 0 | - |
| - | - | 1436 | 793.4 | - | - | 0 | - |
| - | - | 1184 | 794.4 | - | - | 0 | - |
| 7 | c | 3.036E+04 | 798.4 | 0.0004782 | 0.5989 | +1 | 7 |
| - | - | 1.124E+04 | 799.4 | - | - | 0 | - |
| - | - | 2775 | 799.9 | - | - | 0 | - |
| - | - | 2627 | 800.5 | - | - | 0 | - |
| - | - | 5496 | 805.4 | - | - | 0 | - |
| 14 | c | 5938 | 805.9 | 0.003757 | 4.661 | +2 | 14 |
| 14 | c | 3706 | 806.4 | 0.006799 | 8.431 | +2 | 14 |
| - | - | 1320 | 806.9 | - | - | 0 | - |
| - | - | 840.9 | 809.9 | - | - | 0 | - |
| 4 | y | 5314 | 810.4 | 0.0005666 | 0.6992 | +2 | 14 |
| - | - | 4117 | 810.9 | - | - | 0 | - |
| - | - | 3218 | 811.4 | - | - | 0 | - |
| - | - | 1016 | 811.9 | - | - | 0 | - |
| - | - | 1168 | 812.4 | - | - | 0 | - |
| - | - | 4471 | 814.4 | - | - | 0 | - |
| 14 | c | 9607 | 814.9 | 0.002508 | 3.078 | +2 | 14 |
| - | - | 5651 | 815.4 | - | - | 0 | - |
| - | - | 3923 | 815.9 | - | - | 0 | - |
| - | - | 1525 | 816.4 | - | - | 0 | - |
| - | - | 655.1 | 818.4 | - | - | 0 | - |
| - | - | 1719 | 820.4 | - | - | 0 | - |
| - | - | 703.5 | 820.9 | - | - | 0 | - |
| - | - | 729.2 | 830.5 | - | - | 0 | - |
| - | - | 660.4 | 831.5 | - | - | 0 | - |
| - | - | 713 | 836.4 | - | - | 0 | - |
| 3 | w | 1396 | 837.4 | 0.00587 | 7.009 | +2 | 15 |
| - | - | 1091 | 837.9 | - | - | 0 | - |
| - | - | 1426 | 838.4 | - | - | 0 | - |
| - | - | 1179 | 838.9 | - | - | 0 | - |
| 15 | c | 1757 | 849.9 | 0.0009778 | 1.15 | +2 | 15 |
| 3 | y | 658.7 | 850.4 | 0.003843 | 4.519 | +2 | 15 |
| 3 | z | 993.6 | 850.9 | 0.001168 | 1.373 | +2 | 15 |
| - | - | 769.5 | 851.4 | - | - | 0 | - |
| 10 | w | 1555 | 857.5 | 0.003348 | 3.905 | +1 | 8 |
| 15 | c | 1.748E+04 | 858.4 | 0.0001507 | 0.1756 | +2 | 15 |
| 3 | y | 4.641E+04 | 858.9 | 0.003752 | 4.368 | +2 | 15 |
| 10 | w | 3.546E+04 | 859.5 | 0.01453 | 16.91 | +1 | 8 |
| - | - | 1.761E+04 | 859.9 | - | - | 0 | - |
| - | - | 5321 | 860.4 | - | - | 0 | - |
| - | - | 1633 | 860.9 | - | - | 0 | - |
| - | - | 1099 | 871.4 | - | - | 0 | - |
| - | - | 1598 | 872 | - | - | 0 | - |
| 10 | y | 1080 | 873.5 | 0.00131 | 1.5 | +1 | 8 |
| 10 | z | 1.32E+04 | 874.5 | 0.001144 | 1.308 | +1 | 8 |
| - | - | 6216 | 875.5 | - | - | 0 | - |
| - | - | 1636 | 876.5 | - | - | 0 | - |
| - | - | 842.4 | 885 | - | - | 0 | - |
| 2 | w | 8089 | 885.9 | 0.001286 | 1.452 | +2 | 16 |
| - | - | 8433 | 886.4 | - | - | 0 | - |
| - | - | 5298 | 886.9 | - | - | 0 | - |
| - | - | 2596 | 887.4 | - | - | 0 | - |
| 10 | y | 2486 | 890.5 | 0.001558 | 1.749 | +1 | 8 |
| - | - | 945.1 | 891.5 | - | - | 0 | - |
| - | - | 952.2 | 893 | - | - | 0 | - |
| - | - | 1260 | 893.5 | - | - | 0 | - |
| - | - | 1014 | 894 | - | - | 0 | - |
| 2 | z | 5813 | 907.5 | 0.003614 | 3.982 | +2 | 16 |
| - | - | 4578 | 908 | - | - | 0 | - |
| - | - | 2186 | 908.5 | - | - | 0 | - |
| 16 | c | 2500 | 909 | 0.008237 | 9.062 | +2 | 16 |
| - | - | 701.2 | 910 | - | - | 0 | - |
| - | - | 826.1 | 913.5 | - | - | 0 | - |
| - | - | 1156 | 914 | - | - | 0 | - |
| - | - | 1.697E+04 | 915 | - | - | 0 | - |
| 2 | y | 1.905E+04 | 915.5 | 0.008412 | 9.189 | +2 | 16 |
| - | - | 1.237E+04 | 916 | - | - | 0 | - |
| - | - | 6043 | 916.5 | - | - | 0 | - |
| - | - | 1436 | 917 | - | - | 0 | - |
| - | - | 1501 | 917.5 | - | - | 0 | - |
| - | - | 1176 | 919.4 | - | - | 0 | - |
| - | - | 972.2 | 919.9 | - | - | 0 | - |
| - | - | 920.3 | 921.5 | - | - | 0 | - |
| - | - | 1004 | 922 | - | - | 0 | - |
| - | - | 2266 | 922.5 | - | - | 0 | - |
| - | - | 1233 | 923 | - | - | 0 | - |
| - | - | 1262 | 923.5 | - | - | 0 | - |
| 8 | c | 2.299E+04 | 927.5 | 0.001079 | 1.163 | +1 | 8 |
| - | - | 1.258E+04 | 928.5 | - | - | 0 | - |
| - | - | 1208 | 929 | - | - | 0 | - |
| - | - | 5592 | 929.5 | - | - | 0 | - |
| - | - | 1.082E+04 | 930 | - | - | 0 | - |
| - | - | 1.128E+04 | 930.5 | - | - | 0 | - |
| - | - | 7188 | 931 | - | - | 0 | - |
| - | - | 3267 | 931.5 | - | - | 0 | - |
| - | - | 1109 | 932 | - | - | 0 | - |
| - | - | 1060 | 934.5 | - | - | 0 | - |
| - | - | 959.5 | 935 | - | - | 0 | - |
| - | - | 891.2 | 935.5 | - | - | 0 | - |
| - | - | 1723 | 936 | - | - | 0 | - |
| - | - | 1704 | 936.5 | - | - | 0 | - |
| - | - | 4381 | 937 | - | - | 0 | - |
| - | - | 4122 | 937.5 | - | - | 0 | - |
| - | - | 2613 | 938 | - | - | 0 | - |
| - | - | 3574 | 938.5 | - | - | 0 | - |
| - | - | 2765 | 939 | - | - | 0 | - |
| - | - | 2324 | 939.5 | - | - | 0 | - |
| - | - | 894.5 | 943.5 | - | - | 0 | - |
| - | - | 6250 | 944 | - | - | 0 | - |
| 9 | w | 1.121E+04 | 944.5 | 0.004615 | 4.886 | +1 | 9 |
| - | - | 4572 | 945 | - | - | 0 | - |
| - | - | 5740 | 945.5 | - | - | 0 | - |
| - | - | 836.2 | 946 | - | - | 0 | - |
| - | - | 1443 | 946.5 | - | - | 0 | - |
| - | - | 1210 | 948.5 | - | - | 0 | - |
| - | - | 2933 | 949 | - | - | 0 | - |
| - | - | 2539 | 949.5 | - | - | 0 | - |
| - | - | 1953 | 950 | - | - | 0 | - |
| - | - | 1229 | 950.5 | - | - | 0 | - |
| - | - | 703.6 | 951 | - | - | 0 | - |
| - | - | 937.5 | 952.4 | - | - | 0 | - |
| - | - | 4253 | 957.5 | - | - | 0 | - |
| - | - | 2.093E+04 | 958 | - | - | 0 | - |
| - | - | 2.211E+04 | 958.5 | - | - | 0 | - |
| - | - | 9995 | 959 | - | - | 0 | - |
| - | - | 5414 | 959.5 | - | - | 0 | - |
| - | - | 1654 | 960 | - | - | 0 | - |
| - | - | 653.4 | 960.5 | - | - | 0 | - |
| - | - | 675.1 | 963.9 | - | - | 0 | - |
| - | - | 1.896E+04 | 966 | - | - | 0 | - |
| - | - | 3.647E+04 | 966.5 | - | - | 0 | - |
| - | - | 2.547E+04 | 967 | - | - | 0 | - |
| - | - | 1.29E+04 | 967.5 | - | - | 0 | - |
| - | - | 5661 | 968 | - | - | 0 | - |
| - | - | 1.12E+04 | 968.4 | - | - | 0 | - |
| - | - | 1.739E+04 | 969.4 | - | - | 0 | - |
| - | - | 8137 | 970.4 | - | - | 0 | - |
| - | - | 3486 | 971.4 | - | - | 0 | - |
| - | - | 810.2 | 972.5 | - | - | 0 | - |
| - | - | 992 | 976.1 | - | - | 0 | - |
| - | - | 873.1 | 976.4 | - | - | 0 | - |
| - | - | 926.9 | 977.4 | - | - | 0 | - |
| - | - | 952.5 | 978.4 | - | - | 0 | - |
| 9 | z | 9609 | 1006 | 0.0007352 | 0.7311 | +1 | 9 |
| - | - | 6075 | 1007 | - | - | 0 | - |
| - | - | 1852 | 1008 | - | - | 0 | - |
| - | - | 1533 | 1015 | - | - | 0 | - |
| - | - | 1538 | 1016 | - | - | 0 | - |
| 9 | y | 1777 | 1022 | 0.0001331 | 0.1303 | +1 | 9 |
| - | - | 832.7 | 1027 | - | - | 0 | - |
| - | - | 1146 | 1034 | - | - | 0 | - |
| - | - | 858.9 | 1057 | - | - | 0 | - |
| 9 | c | 2.668E+04 | 1059 | 0.0009753 | 0.9214 | +1 | 9 |
| - | - | 1.599E+04 | 1060 | - | - | 0 | - |
| - | - | 7458 | 1061 | - | - | 0 | - |
| - | - | 2154 | 1062 | - | - | 0 | - |
| - | - | 1724 | 1068 | - | - | 0 | - |
| - | - | 1224 | 1068 | - | - | 0 | - |
| - | - | 1460 | 1069 | - | - | 0 | - |
| - | - | 752.1 | 1069 | - | - | 0 | - |
| - | - | 1054 | 1074 | - | - | 0 | - |
| - | - | 945 | 1074 | - | - | 0 | - |
| - | - | 1331 | 1074 | - | - | 0 | - |
| - | - | 872.7 | 1074 | - | - | 0 | - |
| - | - | 1492 | 1075 | - | - | 0 | - |
| 8 | w | 2798 | 1076 | 0.0005439 | 0.5057 | +1 | 10 |
| - | - | 2256 | 1077 | - | - | 0 | - |
| - | - | 826.3 | 1080 | - | - | 0 | - |
| - | - | 919.8 | 1088 | - | - | 0 | - |
| - | - | 884.1 | 1106 | - | - | 0 | - |
| - | - | 901.9 | 1107 | - | - | 0 | - |
| - | - | 784.8 | 1118 | - | - | 0 | - |
| - | - | 6832 | 1126 | - | - | 0 | - |
| - | - | 8880 | 1126 | - | - | 0 | - |
| - | - | 5586 | 1127 | - | - | 0 | - |
| - | - | 3080 | 1128 | - | - | 0 | - |
| - | - | 1080 | 1128 | - | - | 0 | - |
| - | - | 731.3 | 1129 | - | - | 0 | - |
| 8 | z | 1.523E+04 | 1135 | 0.0009088 | 0.801 | +1 | 10 |
| - | - | 8354 | 1136 | - | - | 0 | - |
| - | - | 3992 | 1137 | - | - | 0 | - |
| - | - | 1203 | 1138 | - | - | 0 | - |
| 8 | y | 1565 | 1151 | 0.00474 | 4.12 | +1 | 10 |
| - | - | 8208 | 1153 | - | - | 0 | - |
| - | - | 4043 | 1154 | - | - | 0 | - |
| - | - | 2194 | 1155 | - | - | 0 | - |
| 10 | c | 9044 | 1160 | 0.0001918 | 0.1654 | +1 | 10 |
| - | - | 6477 | 1161 | - | - | 0 | - |
| - | - | 2387 | 1162 | - | - | 0 | - |
| - | - | 769.9 | 1163 | - | - | 0 | - |
| 7 | w | 1027 | 1205 | 0.008499 | 7.055 | +1 | 11 |
| - | - | 712.2 | 1206 | - | - | 0 | - |
| - | - | 983.2 | 1244 | - | - | 0 | - |
| - | - | 1072 | 1245 | - | - | 0 | - |
| 7 | z | 7000 | 1264 | 0.002242 | 1.774 | +1 | 11 |
| - | - | 5783 | 1265 | - | - | 0 | - |
| - | - | 2094 | 1266 | - | - | 0 | - |
| 11 | c | 1547 | 1271 | 0.01855 | 14.6 | +1 | 11 |
| - | - | 834.7 | 1272 | - | - | 0 | - |
| - | - | 1009 | 1273 | - | - | 0 | - |
| - | - | 1259 | 1274 | - | - | 0 | - |
| 7 | y | 3165 | 1280 | 0.004487 | 3.506 | +1 | 11 |
| - | - | 2418 | 1281 | - | - | 0 | - |
| - | - | 806.7 | 1286 | - | - | 0 | - |
| - | - | 2621 | 1287 | - | - | 0 | - |
| 11 | c | 1.646E+04 | 1288 | 0.02215 | 17.21 | +1 | 11 |
| - | - | 710.7 | 1289 | - | - | 0 | - |
| - | - | 1.368E+04 | 1289 | - | - | 0 | - |
| - | - | 5790 | 1290 | - | - | 0 | - |
| - | - | 2161 | 1291 | - | - | 0 | - |
| - | - | 1280 | 1291 | - | - | 0 | - |
| - | - | 982 | 1292 | - | - | 0 | - |
| - | - | 2114 | 1308 | - | - | 0 | - |
| - | - | 813.9 | 1401 | - | - | 0 | - |
| 12 | c | 4063 | 1402 | 0.006124 | 4.369 | +1 | 12 |
| - | - | 3881 | 1403 | - | - | 0 | - |
| - | - | 1776 | 1404 | - | - | 0 | - |
| - | - | 824.5 | 1405 | - | - | 0 | - |
| 6 | z | 5237 | 1420 | 0.002279 | 1.605 | +1 | 12 |
| - | - | 1.166E+04 | 1421 | - | - | 0 | - |
| - | - | 6267 | 1422 | - | - | 0 | - |
| - | - | 2730 | 1423 | - | - | 0 | - |
| - | - | 1548 | 1435 | - | - | 0 | - |
| 6 | y | 5713 | 1436 | 0.002815 | 1.96 | +1 | 12 |
| - | - | 3217 | 1437 | - | - | 0 | - |
| - | - | 2185 | 1438 | - | - | 0 | - |
| - | - | 1412 | 1468 | - | - | 0 | - |
| 5 | z | 911.4 | 1507 | 0.004644 | 3.082 | +1 | 13 |
| - | - | 2684 | 1508 | - | - | 0 | - |
| - | - | 2216 | 1509 | - | - | 0 | - |
| 5 | y | 1115 | 1523 | 0.01336 | 8.773 | +1 | 13 |
| - | - | 848.8 | 1524 | - | - | 0 | - |
| 13 | c | 7653 | 1530 | 0.001836 | 1.2 | +1 | 13 |
| - | - | 4574 | 1531 | - | - | 0 | - |
| - | - | 3296 | 1532 | - | - | 0 | - |
| - | - | 861.7 | 1533 | - | - | 0 | - |
| - | - | 792.9 | 1558 | - | - | 0 | - |
| - | - | 1415 | 1581 | - | - | 0 | - |
| - | - | 1183 | 1582 | - | - | 0 | - |
| - | - | 1065 | 1583 | - | - | 0 | - |
| - | - | 857.5 | 1587 | - | - | 0 | - |
| - | - | 1000 | 1597 | - | - | 0 | - |
| - | - | 1408 | 1599 | - | - | 0 | - |
| - | - | 817.5 | 1601 | - | - | 0 | - |
| - | - | 1065 | 1602 | - | - | 0 | - |
| - | - | 853.3 | 1603 | - | - | 0 | - |
| - | - | 1325 | 1611 | - | - | 0 | - |
| - | - | 1483 | 1611 | - | - | 0 | - |
| 14 | c | 797.1 | 1612 | 0.02209 | 13.71 | +1 | 14 |
| 4 | y | 1827 | 1620 | 0.008261 | 5.1 | +1 | 14 |
| - | - | 2188 | 1621 | - | - | 0 | - |
| - | - | 1360 | 1622 | - | - | 0 | - |
| - | - | 1458 | 1628 | - | - | 0 | - |
| 14 | c | 9578 | 1629 | 0.002989 | 1.835 | +1 | 14 |
| - | - | 8461 | 1630 | - | - | 0 | - |
| - | - | 4607 | 1631 | - | - | 0 | - |
| - | - | 998.3 | 1632 | - | - | 0 | - |
| - | - | 1165 | 1633 | - | - | 0 | - |
| - | - | 1033 | 1673 | - | - | 0 | - |
| - | - | 1413 | 1713 | - | - | 0 | - |
| - | - | 950.6 | 1714 | - | - | 0 | - |
| 15 | c | 4826 | 1716 | 0.0009596 | 0.5592 | +1 | 15 |
| 3 | y | 6819 | 1717 | 0.01844 | 10.74 | +1 | 15 |
| - | - | 5652 | 1718 | - | - | 0 | - |
| - | - | 2510 | 1719 | - | - | 0 | - |
| - | - | 1629 | 1720 | - | - | 0 | - |
| - | - | 1086 | 1786 | - | - | 0 | - |
| - | - | 1962 | 1787 | - | - | 0 | - |
| - | - | 1166 | 1788 | - | - | 0 | - |
| 2 | z | 1655 | 1814 | 0.00425 | 2.343 | +1 | 16 |
| - | - | 7365 | 1815 | - | - | 0 | - |
| - | - | 6493 | 1816 | - | - | 0 | - |
| 16 | c | 4342 | 1817 | 0.005805 | 3.195 | +1 | 16 |
| - | - | 1748 | 1818 | - | - | 0 | - |
| - | - | 894.6 | 1819 | - | - | 0 | - |
| - | - | 2272 | 1829 | - | - | 0 | - |
| 2 | y | 5348 | 1830 | 0.02019 | 11.04 | +1 | 16 |
| - | - | 5726 | 1831 | - | - | 0 | - |
| - | - | 3248 | 1832 | - | - | 0 | - |
| - | - | 1385 | 1833 | - | - | 0 | - |
| - | - | 1013 | 1834 | - | - | 0 | - |
| - | - | 881.9 | 1857 | - | - | 0 | - |
| - | - | 2883 | 1874 | - | - | 0 | - |
| - | - | 2260 | 1875 | - | - | 0 | - |
| - | - | 2202 | 1876 | - | - | 0 | - |
| - | - | 1881 | 1887 | - | - | 0 | - |
| - | - | 5777 | 1888 | - | - | 0 | - |
| - | - | 5899 | 1889 | - | - | 0 | - |
| - | - | 4390 | 1890 | - | - | 0 | - |
| - | - | 1349 | 1891 | - | - | 0 | - |
| - | - | 1733 | 1899 | - | - | 0 | - |
| - | - | 1182 | 1900 | - | - | 0 | - |
| - | - | 933.6 | 1901 | - | - | 0 | - |
| - | - | 839.8 | 1904 | - | - | 0 | - |
| - | - | 4681 | 1905 | - | - | 0 | - |
| - | - | 4356 | 1906 | - | - | 0 | - |
| - | - | 2121 | 1907 | - | - | 0 | - |
| - | - | 1133 | 1908 | - | - | 0 | - |
| - | - | 3879 | 1915 | - | - | 0 | - |
| - | - | 1.803E+04 | 1916 | - | - | 0 | - |
| - | - | 1.753E+04 | 1917 | - | - | 0 | - |
| - | - | 670.4 | 1917 | - | - | 0 | - |
| - | - | 1.188E+04 | 1918 | - | - | 0 | - |
| - | - | 3800 | 1919 | - | - | 0 | - |
| - | - | 1041 | 1920 | - | - | 0 | - |
| - | - | 935 | 1929 | - | - | 0 | - |
| - | - | 998.2 | 1930 | - | - | 0 | - |
| - | - | 2844 | 1931 | - | - | 0 | - |
| - | - | 1.247E+04 | 1932 | - | - | 0 | - |
| - | - | 5.461E+04 | 1933 | - | - | 0 | - |
| - | - | 4.99E+04 | 1934 | - | - | 0 | - |
| - | - | 2.857E+04 | 1935 | - | - | 0 | - |
| - | - | 1.281E+04 | 1936 | - | - | 0 | - |
| - | - | 2952 | 1937 | - | - | 0 | - |
| - | - | 718.5 | 2047 | - | - | 0 | - |
| - | - | 1062 | 2236 | - | - | 0 | - |
| - | - | 895 | 2251 | - | - | 0 | - |
| - | - | 2259 | 2252 | - | - | 0 | - |
| - | - | 2644 | 2253 | - | - | 0 | - |
| - | - | 1990 | 2254 | - | - | 0 | - |
| - | - | 1061 | 2255 | - | - | 0 | - |
| - | - | 700.5 | 2922 | - | - | 0 | - |
| - | - | 843.5 | 3082 | - | - | 0 | - |
| - | - | 687.7 | 3474 | - | - | 0 | - |

m/z Charge Intensity FragmentType MassShift Position
120.06546783447266 0 1067.7754
120.08067321777344 0 548.4049
135.1719207763672 0 387.27075
148.8673858642578 0 492.51297
148.91036987304688 0 506.20898
148.91770935058594 0 1206.3124
148.92495727539062 0 1266.751
148.9320526123047 0 1533.8721
148.9398651123047 0 3477.7148
148.9563446044922 0 5992.204
148.9641571044922 0 3306.4565
148.97164916992188 0 1422.4451
148.97882080078125 0 1336.5397
148.98605346679688 0 1213.12
148.99302673339844 0 759.07556
149.00108337402344 0 916.2686
149.00799560546875 0 782.8075
149.01513671875 0 669.07007
149.02955627441406 0 394.11472
149.0725555419922 0 463.2765
163.61888122558594 0 411.94415
166.08612060546875 0 593.9779
173.44960021972656 0 890.6837
187.13577270507812 0 721.62177
187.14376831054688 0 12477.019
188.14752197265625 0 828.9432
215.12928771972656 0 553.99097
215.13868713378906 0 7183.8022 y Water loss 15
216.14276123046875 0 759.03925
223.10792541503906 0 1087.2124
244.2624969482422 0 517.1038
250.07403564453125 0 579.16815
250.5540008544922 0 582.4503
299.17095947265625 0 2694.4395
320.1807556152344 0 719.7913 y 14
407.2530212402344 0 752.2071
426.26983642578125 0 721.1508 c 3
428.274658203125 0 729.3264
460.1819152832031 0 1169.0232
470.2953186035156 0 1564.7064
473.2415466308594 0 1677.6973 y Water loss 5
478.1930847167969 0 2806.9175
494.2835998535156 0 1520.9098
513.3031005859375 0 8869.458 c 4
514.3056030273438 0 2078.0986
524.27392578125 0 819.61346
526.27197265625 0 729.1951
527.275146484375 0 844.849
531.2906494140625 0 1449.284 z 12
532.2962036132812 0 702.0709
533.2719116210938 0 1447.2211
533.6035766601562 0 1619.6888
533.9402465820312 0 733.84924
566.9539794921875 0 2010.1748 c Ammonia loss 14
567.2872314453125 0 2158.558 y Ammonia loss 7
567.621826171875 0 1080.9579 z 2
572.9570922851562 0 34329.656 y 2
573.2913818359375 0 30504.914
573.6253662109375 0 15299.993
573.9583129882812 0 9148.401
574.2931518554688 0 2241.9514
594.2072143554688 0 1577.5082
607.3810424804688 0 1089.6238
625.3887939453125 0 902.9618
638.6624755859375 0 726.642
643.3450317382812 0 861.81134 y Water loss 11
643.850830078125 0 19942.111
644.3494873046875 0 19382.166 c 10
644.8524780273438 0 7123.0415
645.3705444335938 0 3025.8564
645.8667602539062 0 1061.1385
646.3397827148438 0 1543.4736
661.3504028320312 0 2398.953 y 11
665.2439575195312 0 896.5575
666.5000610351562 0 623.4201
669.4036254882812 0 13060.387 c 5
670.4058837890625 0 4617.067
671.411865234375 0 725.0562
691.420166015625 0 649.7006
695.4252319335938 0 661.73346
696.4218139648438 0 576.94696
699.3447265625 0 1775.4313
699.8445434570312 0 1327.7522
700.3408203125 0 791.64374
701.3610229492188 0 10590.884 c 11
701.8619995117188 0 6237.258
702.364990234375 0 3167.8787
702.86572265625 0 945.7982
711.3587036132812 0 953.1614
714.3687744140625 0 856.26483
717.8602294921875 0 727.1325
718.36474609375 0 1537.7734 y 5
718.8634643554688 0 1831.0858
729.4288940429688 0 1404.8718
730.4312133789062 0 1001.3897
742.8627319335938 0 1730.798
743.368408203125 0 1169.9186
750.8757934570312 0 818.4098
754.3719482421875 0 939.5617
757.4005126953125 0 723.6785
761.8807983398438 0 905.5088 y 4
762.8779296875 0 570.7604
763.3777465820312 0 746.0301
764.8897094726562 0 901.24255
765.3895874023438 0 14891.025 c 12
765.891357421875 0 12516.77
766.3916015625 0 7033.068
766.8931274414062 0 2834.3364
767.3954467773438 0 970.88513
773.4357299804688 0 5822.929 z 10
774.4363403320312 0 3374.3022
775.4432373046875 0 627.3244
777.4077758789062 0 1022.13135
785.41162109375 0 976.8047
785.9024658203125 0 790.15283
786.4091186523438 0 887.4053
786.8895263671875 0 913.29816
789.4475708007812 0 2115.2092 y 10
791.0338134765625 0 659.06683
792.9180908203125 0 950.2803
793.418701171875 0 1436.0117
794.4254760742188 0 1184.2179
798.4463500976562 0 30355.385 c 6
799.448974609375 0 11241.26
799.9053955078125 0 2775.1147
800.4523315429688 0 2627.0676
805.41552734375 0 5496.081
805.9158325195312 0 5938.383 c Water loss 13
806.4183959960938 0 3706.044 c Ammonia loss 13
806.9178466796875 0 1320.4443
809.9020385742188 0 840.8765
810.4059448242188 0 5313.7334 y 3
810.9063720703125 0 4117.249
811.4082641601562 0 3218.059
811.9025268554688 0 1016.16205
812.4080200195312 0 1168.1273
814.4205932617188 0 4470.7153
814.92236328125 0 9606.934 c 13
815.4234008789062 0 5651.2217
815.9259033203125 0 3923.1653
816.423095703125 0 1524.6837
818.4102172851562 0 655.0587
820.3772583007812 0 1719.1731
820.8751831054688 0 703.54785
830.4672241210938 0 729.22473
831.4652709960938 0 660.43005
836.4228515625 0 713.04626
837.4176635742188 0 1396.11 w 2
837.9183959960938 0 1090.532
838.41015625 0 1425.5874
838.9149169921875 0 1178.9312
849.9285888671875 0 1756.6445 c Ammonia loss 14
850.4234619140625 0 658.68054 y Ammonia loss 2
850.92236328125 0 993.5992 z 2
851.4255981445312 0 769.50055
857.4693603515625 0 1554.9517 w 9
858.4407348632812 0 17482.885 c 14
858.9366455078125 0 46406.63 y 2
859.4374389648438 0 35457.816 w 9
859.93603515625 0 17607.826
860.4388427734375 0 5320.8066
860.9329833984375 0 1633.1771
871.4495239257812 0 1099.3912
871.9541015625 0 1598.455
873.4689331054688 0 1080.309 y Ammonia loss 9
874.4743041992188 0 13197.192 z 9
875.4772338867188 0 6215.823
876.4819946289062 0 1635.8278
884.9638671875 0 842.409
885.9368896484375 0 8088.627 w 1
886.4387817382812 0 8433.302
886.9383544921875 0 5298.2275
887.4430541992188 0 2595.8345
890.4926147460938 0 2486.3674 y 9
891.4947509765625 0 945.1371
892.9742431640625 0 952.221
893.4808959960938 0 1259.5182
893.9624633789062 0 1013.7976
907.4691772460938 0 5813.0938 z 1
907.9703979492188 0 4578.4097
908.4667358398438 0 2186.2527
908.9729614257812 0 2500.377 c 15
909.9583740234375 0 701.21313
913.485107421875 0 826.09503
913.987548828125 0 1156.0013
914.982177734375 0 16971.932
915.4833374023438 0 19053.57 y 1
915.9834594726562 0 12370.855
916.4845581054688 0 6043.3965
916.9808959960938 0 1435.9636
917.4853515625 0 1501.413
919.43603515625 0 1176.2499
919.9395751953125 0 972.2409
921.4772338867188 0 920.3069
921.971923828125 0 1003.57355
922.4735717773438 0 2266.0776
922.97998046875 0 1232.9491
923.4765625 0 1262.0275
927.4883422851562 0 22991.602 c 7
928.4899291992188 0 12584.527
928.9767456054688 0 1208.3547
929.4907836914062 0 5592.2124
929.9625854492188 0 10822.806
930.4614868164062 0 11280.734
930.9635620117188 0 7187.598
931.4625244140625 0 3267.1135
931.968994140625 0 1108.983
934.4774169921875 0 1060.0817
934.9893188476562 0 959.5213
935.4923706054688 0 891.2151
935.993408203125 0 1722.8627
936.4832153320312 0 1703.9305
936.9833984375 0 4380.5293
937.4816284179688 0 4122.0615
937.9877319335938 0 2613.1528
938.473388671875 0 3573.7737
938.9725952148438 0 2765.2498
939.473388671875 0 2323.5437
943.5031127929688 0 894.54535
943.9901123046875 0 6249.8896
944.5001220703125 0 11206.177 w 8
944.9907836914062 0 4572.4683
945.4967041015625 0 5740.098
945.9923095703125 0 836.1764
946.5065307617188 0 1443.214
948.4986572265625 0 1210.0413
948.987060546875 0 2933.1763
949.482421875 0 2539.1543
949.9884033203125 0 1952.591
950.4828491210938 0 1228.9735
950.9931030273438 0 703.5882
952.4153442382812 0 937.51526
957.4959106445312 0 4252.5938
957.9910888671875 0 20928.342
958.4918823242188 0 22112.846
958.9904174804688 0 9994.635
959.4942016601562 0 5413.846
959.9891357421875 0 1653.6191
960.5161743164062 0 653.37524
963.930419921875 0 675.12524
965.9989013671875 0 18963.775
966.50048828125 0 36470.645
967.0020141601562 0 25469.393
967.5035400390625 0 12901.403
968.0040283203125 0 5660.6978
968.4370727539062 0 11200.011
969.44384765625 0 17389.816
970.4459228515625 0 8136.7886
971.448486328125 0 3486.357
972.4598388671875 0 810.2041
976.1047973632812 0 991.99945
976.43798828125 0 873.10266
977.432861328125 0 926.9242
978.43994140625 0 952.50104
1005.5151977539062 0 9608.651 z 8
1006.5181884765625 0 6075.3027
1007.5164794921875 0 1851.7253
1014.51513671875 0 1532.7238
1015.5122680664062 0 1537.8462
1021.5347900390625 0 1777.0974 y 8
1027.4432373046875 0 832.74817
1033.5333251953125 0 1145.5586
1057.4749755859375 0 858.9107
1058.5289306640625 0 26680.736 c 8
1059.53173828125 0 15989.99
1060.532470703125 0 7457.9414
1061.5357666015625 0 2153.787
1068.138671875 0 1724.2666
1068.4725341796875 0 1224.1384
1068.8104248046875 0 1460.0165
1069.137939453125 0 752.0831
1073.5401611328125 0 1053.8578
1073.8138427734375 0 944.9663
1074.1466064453125 0 1330.9952
1074.4781494140625 0 872.72546
1074.81298828125 0 1492.0819
1075.544677734375 0 2798.4272 w 7
1076.544921875 0 2255.8699
1080.4990234375 0 826.2744
1087.5638427734375 0 919.7986
1106.4742431640625 0 884.12646
1107.47705078125 0 901.9111
1117.9913330078125 0 784.83105
1125.99560546875 0 6832.3916
1126.49755859375 0 8879.946
1126.9991455078125 0 5585.6074
1127.5018310546875 0 3080.3845
1128.0096435546875 0 1080.1642
1128.503662109375 0 731.26373
1134.5576171875 0 15226.633 z 7
1135.56103515625 0 8353.977
1136.5628662109375 0 3991.9082
1137.565185546875 0 1203.3164
1150.572509765625 0 1565.1913 y 7
1152.63427734375 0 8207.588
1153.6387939453125 0 4043.4854
1154.6397705078125 0 2194.3354
1159.577392578125 0 9043.587 c 9
1160.5811767578125 0 6477.4414
1161.5838623046875 0 2386.8398
1162.572998046875 0 769.90497
1204.5963134765625 0 1026.5504 w 6
1205.5897216796875 0 712.2326
1243.6619873046875 0 983.2322
1244.672119140625 0 1072.4664
1263.598876953125 0 7000.117 z 6
1264.6024169921875 0 5783.249
1265.6029052734375 0 2093.7708
1270.66455078125 0 1546.7886 c Ammonia loss 10
1271.668701171875 0 834.7391
1272.5989990234375 0 1009.21277
1273.594970703125 0 1259.4202
1279.6153564453125 0 3164.566 y 6
1280.6207275390625 0 2417.5476
1285.630615234375 0 806.70374
1286.678955078125 0 2620.9385
1287.6947021484375 0 16460.76 c 10
1288.53564453125 0 710.72174
1288.7000732421875 0 13682.021
1289.7027587890625 0 5789.8955
1290.60693359375 0 2161.4211
1290.7506103515625 0 1279.6948
1291.612060546875 0 982.0132
1307.5958251953125 0 2113.8967
1400.698974609375 0 813.93945
1401.7093505859375 0 4063.1887 c 11
1402.7127685546875 0 3881.3945
1403.7032470703125 0 1775.6628
1404.688232421875 0 824.4658
1419.699951171875 0 5236.525 z 5
1420.70556640625 0 11661.015
1421.7071533203125 0 6266.6177
1422.712890625 0 2729.8967
1434.7158203125 0 1547.661
1435.7181396484375 0 5712.8086 y 5
1436.7220458984375 0 3216.5295
1437.722900390625 0 2184.995
1467.615478515625 0 1411.6216
1506.7296142578125 0 911.43756 z 4
1507.735595703125 0 2684.042
1508.7398681640625 0 2215.638
1522.7396240234375 0 1115.2112 y 4
1523.76123046875 0 848.7603
1529.772216796875 0 7652.9365 c 12
1530.7734375 0 4574.07
1531.7786865234375 0 3296.2292
1532.7806396484375 0 861.7237
1558.1864013671875 0 792.85803
1580.7017822265625 0 1415.2859
1581.6962890625 0 1182.7448
1582.708984375 0 1065.2299
1586.8267822265625 0 857.48486
1596.7193603515625 0 1000.04004
1598.79052734375 0 1408.0646
1600.8172607421875 0 817.4718
1602.2000732421875 0 1064.6349
1602.7186279296875 0 853.26636
1610.715087890625 0 1325.011
1611.2127685546875 0 1483.1786
1611.7938232421875 0 797.12616 c Ammonia loss 13
1619.7974853515625 0 1827.2491 y 3
1620.8070068359375 0 2188.3367
1621.804931640625 0 1360.4774
1627.83056640625 0 1457.8429
1628.8394775390625 0 9578.286 c 13
1629.842041015625 0 8461.48
1630.843017578125 0 4606.578
1631.8529052734375 0 998.3122
1632.8511962890625 0 1165.109
1672.8775634765625 0 1032.8151
1712.8929443359375 0 1413.2739
1713.8992919921875 0 950.6246
1715.87353515625 0 4825.5205 c 14
1716.876953125 0 6818.628 y 2
1717.8753662109375 0 5652.4106
1718.8748779296875 0 2510.014
1719.8812255859375 0 1628.9341
1785.9635009765625 0 1086.0713
1786.955810546875 0 1962.2136
1787.95166015625 0 1166.37
1813.9281005859375 0 1655.4517 z 1
1814.932861328125 0 7365.2407
1815.9287109375 0 6492.954
1816.927978515625 0 4342.1714 c 15
1817.9239501953125 0 1747.844
1818.931396484375 0 894.5795
1828.9473876953125 0 2271.6807
1829.9627685546875 0 5348.3 y 1
1830.9654541015625 0 5726.236
1831.9677734375 0 3248.3076
1832.9630126953125 0 1384.8584
1833.95947265625 0 1013.3504
1856.94580078125 0 881.8954
1873.957275390625 0 2883.1807
1874.962158203125 0 2259.703
1875.9703369140625 0 2201.6675
1886.979736328125 0 1880.6997
1887.9859619140625 0 5776.6484
1888.985595703125 0 5898.9277
1889.9857177734375 0 4390.0327
1890.9842529296875 0 1348.9475
1898.964599609375 0 1733.0518
1899.962158203125 0 1182.4652
1900.9510498046875 0 933.60284
1904.0018310546875 0 839.811
1905.0074462890625 0 4681.1743
1906.0069580078125 0 4356.0176
1907.011962890625 0 2120.5981
1908.0054931640625 0 1132.9227
1914.9705810546875 0 3879.221
1915.9774169921875 0 18033.855
1916.981201171875 0 17530.05
1917.2930908203125 0 670.42865
1917.9820556640625 0 11881.204
1918.98193359375 0 3800.1982
1919.9620361328125 0 1041.015
1928.8931884765625 0 935.00165
1929.9268798828125 0 998.2464
1930.9853515625 0 2844.001
1931.995849609375 0 12471.53
1933.0030517578125 0 54609.07
1934.005615234375 0 49902.734
1935.0069580078125 0 28566.984
1936.0067138671875 0 12808.871
1937.008544921875 0 2952.4146
2046.9439697265625 0 718.5162
2235.9599609375 0 1062.03
2250.9638671875 0 894.95874
2251.993896484375 0 2258.8496
2252.994140625 0 2644.246
2254.00634765625 0 1989.817
2254.992919921875 0 1060.6724
2922.013427734375 0 700.54065
3082.0859375 0 843.51764
3474.21533203125 0 687.708

Spectrum Details

|  |  |
| --- | --- |
| Matched peaks? Matched peaks The total absolute number of peaks matched. Additionally in brackets the total fraction of peaks matched and the total number of peaks is shown. | 68 (15.63% of 435) |
| FDR? FDR The false discovery rate estimated for this peptide. It is calculated by matching all theoretical fragments with a non-integer shift with the raw peaks for this spectrum. This is done with 40 different shifts. The resulting percentage is the average number of annotated peaks over the number of annotated peaks with the correct spectrum. | 1.86% |
| Satellite FDR? Satellite FDR See the FDR for details on its calculation. This satellite ion specific FDR only contains the satellite ions (d/w) for I/L/J positions. | 7.14% |
| PSM Score? PSM Score The PSM Score as given by Hecklib to this annotated spectrum. It is shown with three significant figures. | 455 |

#### Reverse Lookup? Reverse Lookup All places where this read could be placed.

| Group | Segment | Template | Template Part | Read Part | Score | Unique |
| --- | --- | --- | --- | --- | --- | --- |
| Homo sapiens Heavy Chain | IGHC | IGHG3 | [279..296] | [0..17] | 118 | False |
| Homo sapiens Heavy Chain | IGHC | IGHG2 | [228..245] | [0..17] | 118 | False |
| Homo sapiens Heavy Chain | IGHC | IGHG4 | [229..246] | [0..17] | 109 | False |

| Recombined | Template Part | Read Part | Score | Unique |
| --- | --- | --- | --- | --- |
| REC-0-1 | [357..374] | [0..17] | 104 | True |

#### Meta Information from Multiple reads

##### Number of combined reads

7

##### Intensity

0.8403

##### TotalArea

3.087E+09

##### Changes to the peptide sequence

TLPPSREEMTKNQVSTJ

L→JNo support for either Leucine or Isoleucine based on side chain ions (Position: 17)

#### Positional Score

Copy Data

##### Positional Score (TSV)

###### Preview

```
Loading example...
```

*Click on the button to copy the data to your clipboard.*

10012345678910111213141516

Label Value
"0" 0.684
"1" 0.64
"2" 0.667
"3" 0.699
"4" 0.711
"5" 0.714
"6" 0.714
"7" 0.714
"8" 0.714
"9" 0.707
"10" 0.693
"11" 0.671
"12" 0.653
"13" 0.66
"14" 0.687
"15" 0.709
"16" 0.713

#### Meta Information from PEAKS

##### Scan Identifier

F3:3817

##### Original sequence

T

L

P

P

S

R

E

E

M

+15.99

T

K

N

Q

V

S

T

L

##### Posttranslational Modifications

Oxidation (M)

##### Source File

D:\separate\_stitch\_analyses\xle-disambiguation\raw\20210323\_F1\_UM1\_Peng0013\_SA\_F59\_ingel\_3ug\_chymo.raw

##### Fraction

3

##### Scan Feature

F3:10186

##### De Novo Score

97

##### ConfidenceScore

97

### m/z

649.6663

##### Mass

1945.9778

##### Charge

3

##### Retention Time

21

##### Predicted Retention Time

-

##### Area

5.095E+08

##### Fragmentation mode

ETHCD

##### Originating file

01 D:\separate\_stitch\_analyses\xle-disambiguation\20210325\_F59\_3ug\_DENOVO\_12.csv

#### Meta Information from PEAKS

##### Scan Identifier

F3:4052

##### Original sequence

T

L

P

P

S

R

E

E

M

+15.99

T

K

N

Q

V

S

T

L

##### Posttranslational Modifications

Oxidation (M)

##### Source File

D:\separate\_stitch\_analyses\xle-disambiguation\raw\20210323\_F1\_UM1\_Peng0013\_SA\_F59\_ingel\_3ug\_chymo.raw

##### Fraction

3

##### Scan Feature

F3:10186

##### De Novo Score

97

##### ConfidenceScore

97

### m/z

649.6663

##### Mass

1945.9778

##### Charge

3

##### Retention Time

21

##### Predicted Retention Time

-

##### Area

5.095E+08

##### Fragmentation mode

ETHCD

##### Originating file

01 D:\separate\_stitch\_analyses\xle-disambiguation\20210325\_F59\_3ug\_DENOVO\_12.csv

#### Meta Information from PEAKS

##### Scan Identifier

F3:4280

##### Original sequence

T

L

P

P

S

R

E

E

M

+15.99

T

K

N

Q

V

S

T

L

##### Posttranslational Modifications

Oxidation (M)

##### Source File

D:\separate\_stitch\_analyses\xle-disambiguation\raw\20210323\_F1\_UM1\_Peng0013\_SA\_F59\_ingel\_3ug\_chymo.raw

##### Fraction

3

##### Scan Feature

F3:10186

##### De Novo Score

97

##### ConfidenceScore

97

### m/z

649.6663

##### Mass

1945.9778

##### Charge

3

##### Retention Time

21

##### Predicted Retention Time

-

##### Area

5.095E+08

##### Fragmentation mode

ETHCD

##### Originating file

01 D:\separate\_stitch\_analyses\xle-disambiguation\20210325\_F59\_3ug\_DENOVO\_12.csv

#### Meta Information from PEAKS

##### Scan Identifier

F3:3951

##### Original sequence

T

L

P

P

S

R

E

E

M

+15.99

T

K

N

Q

V

S

T

L

##### Posttranslational Modifications

Oxidation (M)

##### Source File

D:\separate\_stitch\_analyses\xle-disambiguation\raw\20210323\_F1\_UM1\_Peng0013\_SA\_F59\_ingel\_3ug\_chymo.raw

##### Fraction

3

##### Scan Feature

F3:10186

##### De Novo Score

97

##### ConfidenceScore

97

### m/z

649.6663

##### Mass

1945.9778

##### Charge

3

##### Retention Time

21

##### Predicted Retention Time

-

##### Area

5.095E+08

##### Fragmentation mode

ETHCD

##### Originating file

01 D:\separate\_stitch\_analyses\xle-disambiguation\20210325\_F59\_3ug\_DENOVO\_12.csv

#### Meta Information from PEAKS

##### Scan Identifier

F3:4116

##### Original sequence

T

L

P

P

S

R

E

E

M

+15.99

T

K

N

Q

V

S

T

L

##### Posttranslational Modifications

Oxidation (M)

##### Source File

D:\separate\_stitch\_analyses\xle-disambiguation\raw\20210323\_F1\_UM1\_Peng0013\_SA\_F59\_ingel\_3ug\_chymo.raw

##### Fraction

3

##### Scan Feature

F3:10186

##### De Novo Score

96

##### ConfidenceScore

96

### m/z

649.6663

##### Mass

1945.9778

##### Charge

3

##### Retention Time

21

##### Predicted Retention Time

-

##### Area

5.095E+08

##### Fragmentation mode

ETHCD

##### Originating file

01 D:\separate\_stitch\_analyses\xle-disambiguation\20210325\_F59\_3ug\_DENOVO\_12.csv

#### Meta Information from PEAKS

##### Scan Identifier

F3:4342

##### Original sequence

T

L

P

P

S

R

E

E

M

+15.99

T

K

N

Q

V

S

T

L

##### Posttranslational Modifications

Oxidation (M)

##### Source File

D:\separate\_stitch\_analyses\xle-disambiguation\raw\20210323\_F1\_UM1\_Peng0013\_SA\_F59\_ingel\_3ug\_chymo.raw

##### Fraction

3

##### Scan Feature

F3:10186

##### De Novo Score

96

##### ConfidenceScore

96

### m/z

649.6663

##### Mass

1945.9778

##### Charge

3

##### Retention Time

21

##### Predicted Retention Time

-

##### Area

5.095E+08

##### Fragmentation mode

ETHCD

##### Originating file

01 D:\separate\_stitch\_analyses\xle-disambiguation\20210325\_F59\_3ug\_DENOVO\_12.csv

#### Meta Information from PEAKS

##### Scan Identifier

F3:4919

##### Original sequence

T

L

P

P

S

R

E

E

M

T

K

N

Q

V

S

T

L

##### Posttranslational Modifications

##### Source File

D:\separate\_stitch\_analyses\xle-disambiguation\raw\20210323\_F1\_UM1\_Peng0013\_SA\_F59\_ingel\_3ug\_chymo.raw

##### Fraction

3

##### Scan Feature

F3:9987

##### De Novo Score

95

##### ConfidenceScore

95

### m/z

644.3353

##### Mass

1929.9829

##### Charge

3

##### Retention Time

27.52

##### Predicted Retention Time

-

##### Area

3.011E+07

##### Parts Per Million

0.5

##### Fragmentation mode

ETHCD

##### Originating file

01 D:\separate\_stitch\_analyses\xle-disambiguation\20210325\_F59\_3ug\_DENOVO\_12.csv
