## Supplementary material for "A handle on mass coincidence errors in *de novo* sequencing of antibodies by bottom-up proteomics": Combined_097.html

Details Combined\_097 | Stitch OverviewUndefined

### Read Combined\_097

#### Sequence (length=7)

GTJMISR

#### Spectrum 3929? Spectrum 3929 The raw spectrum of this peptide as annotated by Hecklib. The fragments are coloured according to ion type (see legend). Any peaks with a star '\*' as text can be hovered over to see the full details, first the ion type second the mass shift type. By hovering over the amino acids in the peptide or ions in the legend the corresponding peaks are highlighted. By toggling the 'Unassigned' label you can turn the background (unassigned) peaks on or off in the plot. By updating the slider in the Ion legend you can update the spectrum to only show the top X% of the peaks with labels. The top X% means any peak that is within X% of the highest intensity. By dragging in the spectrum you can zoom in to a specific part of the spectrum and use 'Zoom Out' to get back to the original zoom level. The annotation of the spectrum is based on the given sequence in the peptides file and is done with different software so inconsistencies are likely. The peaks are annotated based on the given sequence, with 20 ppm tolerance.

Copy Data

##### Spectrum 3929 (TSV)

###### Preview

```
Loading example...
```

*Click on the button to copy the data to your clipboard.*

Mz MinMz MaxIntensity Max

WidthHeightPeptide font sizePeptide stroke widthSpectrum font sizeSpectrum stroke widthCompact peptide

Ion legend

wxyz

abcd

OtherUnassignedIon^Charge^_Position_Show for top:%

GTJMISR

02.20e+54.39e+56.59e+58.79e+5

Zoom Out

y^+1^_1_y^+1^_1_y^+2^_3_b^+1^_2_y^+1^_2_y^+1^_2_y^+2^_4_y^+2^_4_y^+2^_4_y^+1^_2_a^+1^_3_a^+1^_3_y^+2^_5_b^+1^_3_y^+2^_5_b^+1^_3_y^+1^_3_y^+1^_3_y^+2^_6_y^+2^_6_y^+1^_3_\*\*b^+1^_4_b^+1^_4_y^+1^_4_y^+1^_4_y^+1^_4_b^+1^_5_b^+1^_5_y^+1^_5_y^+1^_5_y^+1^_5_b^+1^_6_y^+1^_6_y^+1^_6_

0703140621092812

Fragment Matches Table

Show background peaks

| Position | Ion type | Intensity | mz Theoretical | mz Error (Th) | mz Error (ppm) | Charge | Series Number |
| --- | --- | --- | --- | --- | --- | --- | --- |
| - | - | 694.9 | 120.1 | - | - | 0 | - |
| - | - | 1084 | 122.1 | - | - | 0 | - |
| - | - | 1.686E+04 | 124.1 | - | - | 0 | - |
| - | - | 7186 | 125.1 | - | - | 0 | - |
| - | - | 1155 | 125.1 | - | - | 0 | - |
| - | - | 1372 | 125.1 | - | - | 0 | - |
| - | - | 1.519E+05 | 126.1 | - | - | 0 | - |
| - | - | 1124 | 126.1 | - | - | 0 | - |
| - | - | 2952 | 126.1 | - | - | 0 | - |
| - | - | 5840 | 127.1 | - | - | 0 | - |
| - | - | 8107 | 127.1 | - | - | 0 | - |
| - | - | 2262 | 127.1 | - | - | 0 | - |
| - | - | 5.78E+04 | 129.1 | - | - | 0 | - |
| - | - | 769.2 | 129.1 | - | - | 0 | - |
| - | - | 4358 | 129.1 | - | - | 0 | - |
| - | - | 1619 | 130 | - | - | 0 | - |
| - | - | 8214 | 130.1 | - | - | 0 | - |
| - | - | 5.141E+04 | 130.1 | - | - | 0 | - |
| - | - | 2314 | 130.1 | - | - | 0 | - |
| - | - | 1.051E+04 | 130.1 | - | - | 0 | - |
| - | - | 1240 | 131.1 | - | - | 0 | - |
| - | - | 4249 | 133.1 | - | - | 0 | - |
| - | - | 846.5 | 134 | - | - | 0 | - |
| - | - | 5300 | 136 | - | - | 0 | - |
| - | - | 963.8 | 137.1 | - | - | 0 | - |
| - | - | 3403 | 138.1 | - | - | 0 | - |
| - | - | 3364 | 139.1 | - | - | 0 | - |
| - | - | 934.1 | 139.1 | - | - | 0 | - |
| - | - | 1426 | 140.1 | - | - | 0 | - |
| - | - | 3062 | 141.1 | - | - | 0 | - |
| - | - | 5679 | 141.1 | - | - | 0 | - |
| - | - | 1.769E+04 | 141.1 | - | - | 0 | - |
| - | - | 4712 | 142.1 | - | - | 0 | - |
| - | - | 6957 | 142.1 | - | - | 0 | - |
| - | - | 1306 | 142.1 | - | - | 0 | - |
| - | - | 1526 | 143 | - | - | 0 | - |
| - | - | 8657 | 143.1 | - | - | 0 | - |
| - | - | 8732 | 143.1 | - | - | 0 | - |
| - | - | 5.073E+05 | 144.1 | - | - | 0 | - |
| - | - | 892.3 | 144.1 | - | - | 0 | - |
| - | - | 2722 | 145.1 | - | - | 0 | - |
| - | - | 3.115E+04 | 145.1 | - | - | 0 | - |
| - | - | 2400 | 146.1 | - | - | 0 | - |
| - | - | 2583 | 148 | - | - | 0 | - |
| - | - | 865.2 | 149 | - | - | 0 | - |
| - | - | 1438 | 150.1 | - | - | 0 | - |
| - | - | 871.9 | 151.1 | - | - | 0 | - |
| - | - | 1248 | 151.1 | - | - | 0 | - |
| - | - | 9911 | 152.1 | - | - | 0 | - |
| - | - | 4.083E+04 | 153.1 | - | - | 0 | - |
| - | - | 1994 | 153.1 | - | - | 0 | - |
| - | - | 956.6 | 153.1 | - | - | 0 | - |
| - | - | 1.93E+04 | 154.1 | - | - | 0 | - |
| - | - | 3311 | 154.1 | - | - | 0 | - |
| - | - | 1682 | 154.1 | - | - | 0 | - |
| - | - | 1436 | 154.1 | - | - | 0 | - |
| - | - | 961.7 | 155 | - | - | 0 | - |
| - | - | 832.7 | 155.1 | - | - | 0 | - |
| - | - | 3944 | 155.1 | - | - | 0 | - |
| - | - | 2986 | 155.1 | - | - | 0 | - |
| - | - | 1367 | 156.1 | - | - | 0 | - |
| - | - | 2315 | 157.1 | - | - | 0 | - |
| - | - | 5721 | 157.1 | - | - | 0 | - |
| - | - | 1.277E+04 | 157.1 | - | - | 0 | - |
| - | - | 3374 | 157.1 | - | - | 0 | - |
| 7 | y | 1.088E+05 | 158.1 | 0.0004161 | 2.632 | +1 | 1 |
| - | - | 911.6 | 163.6 | - | - | 0 | - |
| - | - | 1225 | 164 | - | - | 0 | - |
| - | - | 5387 | 166.1 | - | - | 0 | - |
| - | - | 2262 | 166.1 | - | - | 0 | - |
| - | - | 1007 | 167.1 | - | - | 0 | - |
| - | - | 3736 | 167.1 | - | - | 0 | - |
| - | - | 2059 | 169.1 | - | - | 0 | - |
| - | - | 1.415E+05 | 169.1 | - | - | 0 | - |
| - | - | 4.392E+04 | 170.1 | - | - | 0 | - |
| - | - | 3199 | 170.1 | - | - | 0 | - |
| - | - | 1090 | 170.1 | - | - | 0 | - |
| - | - | 1.198E+04 | 170.1 | - | - | 0 | - |
| - | - | 8.816E+04 | 171.1 | - | - | 0 | - |
| - | - | 2420 | 171.1 | - | - | 0 | - |
| - | - | 2134 | 171.1 | - | - | 0 | - |
| - | - | 1156 | 171.1 | - | - | 0 | - |
| - | - | 7264 | 172.1 | - | - | 0 | - |
| - | - | 5518 | 172.1 | - | - | 0 | - |
| - | - | 1033 | 172.1 | - | - | 0 | - |
| - | - | 3844 | 173.1 | - | - | 0 | - |
| - | - | 1980 | 173.1 | - | - | 0 | - |
| - | - | 1.309E+04 | 173.1 | - | - | 0 | - |
| - | - | 2896 | 173.4 | - | - | 0 | - |
| 7 | y | 3.126E+05 | 175.1 | 0.0004631 | 2.645 | +1 | 1 |
| - | - | 792.1 | 175.6 | - | - | 0 | - |
| - | - | 1957 | 176.1 | - | - | 0 | - |
| - | - | 1.711E+04 | 176.1 | - | - | 0 | - |
| - | - | 1669 | 177.1 | - | - | 0 | - |
| - | - | 1381 | 178.1 | - | - | 0 | - |
| 5 | y | 7354 | 179.1 | 0.002393 | 13.36 | +2 | 3 |
| - | - | 1.641E+05 | 180.1 | - | - | 0 | - |
| - | - | 9026 | 181.1 | - | - | 0 | - |
| - | - | 2884 | 181.1 | - | - | 0 | - |
| - | - | 1.538E+04 | 181.1 | - | - | 0 | - |
| - | - | 1053 | 181.1 | - | - | 0 | - |
| - | - | 2458 | 182 | - | - | 0 | - |
| - | - | 3749 | 182.1 | - | - | 0 | - |
| - | - | 1054 | 183.1 | - | - | 0 | - |
| - | - | 4005 | 183.1 | - | - | 0 | - |
| - | - | 838.1 | 183.9 | - | - | 0 | - |
| - | - | 6274 | 184.1 | - | - | 0 | - |
| - | - | 6402 | 185.1 | - | - | 0 | - |
| - | - | 1907 | 185.1 | - | - | 0 | - |
| - | - | 1.866E+04 | 185.1 | - | - | 0 | - |
| - | - | 874.9 | 186.1 | - | - | 0 | - |
| - | - | 1.449E+05 | 187.1 | - | - | 0 | - |
| - | - | 1.342E+04 | 188.1 | - | - | 0 | - |
| - | - | 5.589E+05 | 189.1 | - | - | 0 | - |
| - | - | 3.7E+04 | 190.1 | - | - | 0 | - |
| - | - | 3689 | 191.1 | - | - | 0 | - |
| - | - | 1392 | 191.1 | - | - | 0 | - |
| - | - | 1410 | 193.1 | - | - | 0 | - |
| - | - | 3139 | 193.1 | - | - | 0 | - |
| - | - | 978.7 | 195.1 | - | - | 0 | - |
| - | - | 7654 | 195.1 | - | - | 0 | - |
| - | - | 1548 | 195.1 | - | - | 0 | - |
| - | - | 1893 | 196.1 | - | - | 0 | - |
| - | - | 1.587E+05 | 197.1 | - | - | 0 | - |
| - | - | 2.236E+04 | 198.1 | - | - | 0 | - |
| - | - | 1.68E+04 | 198.1 | - | - | 0 | - |
| - | - | 1.869E+05 | 199.1 | - | - | 0 | - |
| - | - | 4677 | 199.1 | - | - | 0 | - |
| - | - | 957.5 | 199.1 | - | - | 0 | - |
| - | - | 1529 | 199.1 | - | - | 0 | - |
| - | - | 3340 | 200.1 | - | - | 0 | - |
| - | - | 1.401E+04 | 200.1 | - | - | 0 | - |
| - | - | 3734 | 200.1 | - | - | 0 | - |
| - | - | 968 | 200.1 | - | - | 0 | - |
| - | - | 1020 | 200.4 | - | - | 0 | - |
| - | - | 1381 | 201.1 | - | - | 0 | - |
| - | - | 1751 | 201.1 | - | - | 0 | - |
| - | - | 2.284E+04 | 201.1 | - | - | 0 | - |
| - | - | 3027 | 201.1 | - | - | 0 | - |
| - | - | 2.889E+04 | 202.1 | - | - | 0 | - |
| - | - | 2026 | 202.1 | - | - | 0 | - |
| - | - | 1.17E+04 | 203.1 | - | - | 0 | - |
| - | - | 1632 | 203.1 | - | - | 0 | - |
| - | - | 1870 | 206.1 | - | - | 0 | - |
| - | - | 1024 | 206.3 | - | - | 0 | - |
| - | - | 939.8 | 206.5 | - | - | 0 | - |
| - | - | 7665 | 207.1 | - | - | 0 | - |
| - | - | 1470 | 208.1 | - | - | 0 | - |
| - | - | 2235 | 209.1 | - | - | 0 | - |
| - | - | 3020 | 209.1 | - | - | 0 | - |
| - | - | 1811 | 209.1 | - | - | 0 | - |
| - | - | 9601 | 210.1 | - | - | 0 | - |
| - | - | 1097 | 210.1 | - | - | 0 | - |
| - | - | 4793 | 211.1 | - | - | 0 | - |
| - | - | 4100 | 211.1 | - | - | 0 | - |
| - | - | 2876 | 212.1 | - | - | 0 | - |
| - | - | 1789 | 213.1 | - | - | 0 | - |
| - | - | 959.6 | 213.1 | - | - | 0 | - |
| - | - | 1842 | 214.2 | - | - | 0 | - |
| - | - | 1089 | 214.6 | - | - | 0 | - |
| - | - | 3186 | 215.1 | - | - | 0 | - |
| - | - | 3493 | 215.1 | - | - | 0 | - |
| - | - | 3.391E+04 | 215.1 | - | - | 0 | - |
| - | - | 5731 | 216.1 | - | - | 0 | - |
| - | - | 4076 | 216.1 | - | - | 0 | - |
| 2 | b | 3.169E+05 | 217.1 | 0.004082 | 18.8 | +1 | 2 |
| - | - | 2.474E+04 | 218.1 | - | - | 0 | - |
| - | - | 770.1 | 218.1 | - | - | 0 | - |
| - | - | 2030 | 219.1 | - | - | 0 | - |
| - | - | 2103 | 220.1 | - | - | 0 | - |
| - | - | 1.108E+04 | 220.1 | - | - | 0 | - |
| - | - | 1258 | 220.1 | - | - | 0 | - |
| - | - | 3948 | 221.1 | - | - | 0 | - |
| - | - | 2185 | 222.1 | - | - | 0 | - |
| - | - | 3654 | 223.1 | - | - | 0 | - |
| - | - | 2704 | 223.1 | - | - | 0 | - |
| - | - | 1104 | 224.1 | - | - | 0 | - |
| - | - | 2783 | 224.1 | - | - | 0 | - |
| - | - | 3.307E+04 | 225.1 | - | - | 0 | - |
| - | - | 4609 | 226.1 | - | - | 0 | - |
| - | - | 2.475E+04 | 227.1 | - | - | 0 | - |
| - | - | 1772 | 227.1 | - | - | 0 | - |
| - | - | 9498 | 228.1 | - | - | 0 | - |
| - | - | 2417 | 228.1 | - | - | 0 | - |
| - | - | 2492 | 228.1 | - | - | 0 | - |
| - | - | 973.6 | 228.6 | - | - | 0 | - |
| - | - | 1503 | 229 | - | - | 0 | - |
| - | - | 5845 | 229.1 | - | - | 0 | - |
| - | - | 5.104E+04 | 229.6 | - | - | 0 | - |
| - | - | 1.005E+04 | 230.1 | - | - | 0 | - |
| - | - | 9182 | 230.1 | - | - | 0 | - |
| - | - | 1222 | 230.6 | - | - | 0 | - |
| - | - | 5055 | 232.1 | - | - | 0 | - |
| - | - | 1910 | 233.1 | - | - | 0 | - |
| - | - | 4.327E+04 | 233.1 | - | - | 0 | - |
| - | - | 4154 | 234.1 | - | - | 0 | - |
| - | - | 1108 | 235.1 | - | - | 0 | - |
| - | - | 3212 | 236.1 | - | - | 0 | - |
| - | - | 1324 | 237.1 | - | - | 0 | - |
| - | - | 1.484E+04 | 238.2 | - | - | 0 | - |
| - | - | 6309 | 239.1 | - | - | 0 | - |
| - | - | 1385 | 239.2 | - | - | 0 | - |
| - | - | 1.069E+04 | 240.1 | - | - | 0 | - |
| - | - | 1513 | 240.2 | - | - | 0 | - |
| - | - | 1938 | 241.1 | - | - | 0 | - |
| - | - | 1005 | 242.6 | - | - | 0 | - |
| - | - | 1777 | 243.1 | - | - | 0 | - |
| - | - | 6328 | 243.6 | - | - | 0 | - |
| - | - | 2.137E+04 | 244.1 | - | - | 0 | - |
| 6 | y | 2.347E+04 | 244.1 | 0.0003617 | 1.482 | +1 | 2 |
| 6 | y | 2.06E+05 | 245.1 | 0.000538 | 2.195 | +1 | 2 |
| - | - | 1073 | 246.1 | - | - | 0 | - |
| - | - | 2.098E+04 | 246.1 | - | - | 0 | - |
| - | - | 1806 | 247.1 | - | - | 0 | - |
| - | - | 1.037E+04 | 249.1 | - | - | 0 | - |
| - | - | 2189 | 250.2 | - | - | 0 | - |
| - | - | 7461 | 251.1 | - | - | 0 | - |
| - | - | 1109 | 251.1 | - | - | 0 | - |
| - | - | 1166 | 251.2 | - | - | 0 | - |
| - | - | 2347 | 252.1 | - | - | 0 | - |
| 4 | y | 903 | 252.6 | 0.002028 | 8.026 | +2 | 4 |
| 4 | y | 3671 | 253.1 | 0.002497 | 9.866 | +2 | 4 |
| - | - | 3634 | 254.2 | - | - | 0 | - |
| - | - | 2957 | 255.1 | - | - | 0 | - |
| - | - | 2825 | 256.1 | - | - | 0 | - |
| - | - | 2682 | 256.2 | - | - | 0 | - |
| - | - | 2587 | 256.6 | - | - | 0 | - |
| - | - | 1055 | 257.6 | - | - | 0 | - |
| - | - | 3676 | 258.1 | - | - | 0 | - |
| - | - | 967.9 | 260.1 | - | - | 0 | - |
| - | - | 1306 | 261.1 | - | - | 0 | - |
| - | - | 8.331E+04 | 261.1 | - | - | 0 | - |
| 4 | y | 3.321E+04 | 261.6 | 0.002818 | 10.77 | +2 | 4 |
| 6 | y | 3.91E+05 | 262.2 | 0.0004171 | 1.591 | +1 | 2 |
| - | - | 1315 | 262.6 | - | - | 0 | - |
| - | - | 4717 | 263.1 | - | - | 0 | - |
| - | - | 3759 | 263.1 | - | - | 0 | - |
| - | - | 3.815E+04 | 263.2 | - | - | 0 | - |
| - | - | 3530 | 264.1 | - | - | 0 | - |
| - | - | 3179 | 264.2 | - | - | 0 | - |
| - | - | 2098 | 265.2 | - | - | 0 | - |
| - | - | 1.903E+04 | 266.2 | - | - | 0 | - |
| - | - | 2.125E+04 | 267.1 | - | - | 0 | - |
| - | - | 2761 | 267.2 | - | - | 0 | - |
| - | - | 7868 | 268.1 | - | - | 0 | - |
| - | - | 4111 | 268.7 | - | - | 0 | - |
| - | - | 999.1 | 269.1 | - | - | 0 | - |
| - | - | 6602 | 270.1 | - | - | 0 | - |
| - | - | 2107 | 271.2 | - | - | 0 | - |
| - | - | 7151 | 272.1 | - | - | 0 | - |
| - | - | 6645 | 277.1 | - | - | 0 | - |
| - | - | 1907 | 277.2 | - | - | 0 | - |
| - | - | 1678 | 277.7 | - | - | 0 | - |
| - | - | 990.9 | 278.1 | - | - | 0 | - |
| - | - | 2054 | 278.2 | - | - | 0 | - |
| - | - | 7535 | 280.2 | - | - | 0 | - |
| - | - | 1333 | 281.2 | - | - | 0 | - |
| - | - | 1226 | 282.1 | - | - | 0 | - |
| - | - | 1440 | 282.1 | - | - | 0 | - |
| - | - | 7471 | 282.2 | - | - | 0 | - |
| 3 | a | 1.641E+05 | 284.2 | 0.004023 | 14.16 | +1 | 3 |
| - | - | 3423 | 285.1 | - | - | 0 | - |
| - | - | 2.034E+04 | 285.2 | - | - | 0 | - |
| - | - | 4427 | 286.1 | - | - | 0 | - |
| - | - | 1.268E+05 | 286.2 | - | - | 0 | - |
| - | - | 3.231E+04 | 286.7 | - | - | 0 | - |
| - | - | 1601 | 287.1 | - | - | 0 | - |
| - | - | 5809 | 287.2 | - | - | 0 | - |
| - | - | 1664 | 288.1 | - | - | 0 | - |
| - | - | 1817 | 290.1 | - | - | 0 | - |
| - | - | 1408 | 291.2 | - | - | 0 | - |
| - | - | 1856 | 292.2 | - | - | 0 | - |
| - | - | 1142 | 293.2 | - | - | 0 | - |
| - | - | 2047 | 293.9 | - | - | 0 | - |
| - | - | 7687 | 294.1 | - | - | 0 | - |
| - | - | 6926 | 295.1 | - | - | 0 | - |
| - | - | 1336 | 295.1 | - | - | 0 | - |
| - | - | 1944 | 296.2 | - | - | 0 | - |
| - | - | 1445 | 297.1 | - | - | 0 | - |
| - | - | 1898 | 297.2 | - | - | 0 | - |
| - | - | 1432 | 297.2 | - | - | 0 | - |
| - | - | 3895 | 298.2 | - | - | 0 | - |
| - | - | 1.073E+04 | 299.2 | - | - | 0 | - |
| - | - | 5175 | 299.2 | - | - | 0 | - |
| - | - | 2648 | 299.7 | - | - | 0 | - |
| - | - | 1581 | 299.7 | - | - | 0 | - |
| 3 | a | 1.068E+04 | 302.2 | 0.003754 | 12.42 | +1 | 3 |
| - | - | 2580 | 303.2 | - | - | 0 | - |
| - | - | 1612 | 305.7 | - | - | 0 | - |
| - | - | 1.097E+04 | 308.2 | - | - | 0 | - |
| - | - | 2099 | 308.2 | - | - | 0 | - |
| 3 | y | 1824 | 309.2 | 0.00504 | 16.3 | +2 | 5 |
| - | - | 1011 | 309.2 | - | - | 0 | - |
| - | - | 1262 | 309.7 | - | - | 0 | - |
| - | - | 4925 | 310.2 | - | - | 0 | - |
| - | - | 1.179E+04 | 310.2 | - | - | 0 | - |
| - | - | 1429 | 311.2 | - | - | 0 | - |
| - | - | 3369 | 311.9 | - | - | 0 | - |
| 3 | b | 2.517E+05 | 312.2 | 0.003912 | 12.53 | +1 | 3 |
| - | - | 3.944E+04 | 313.2 | - | - | 0 | - |
| - | - | 5017 | 314.2 | - | - | 0 | - |
| - | - | 2038 | 314.7 | - | - | 0 | - |
| - | - | 3256 | 315.2 | - | - | 0 | - |
| - | - | 2944 | 317.2 | - | - | 0 | - |
| 3 | y | 2.869E+05 | 318.2 | 0.003114 | 9.788 | +2 | 5 |
| - | - | 8.842E+04 | 318.7 | - | - | 0 | - |
| - | - | 2.863E+04 | 319.2 | - | - | 0 | - |
| - | - | 1.23E+04 | 319.7 | - | - | 0 | - |
| - | - | 1407 | 320.2 | - | - | 0 | - |
| - | - | 2128 | 320.7 | - | - | 0 | - |
| - | - | 1216 | 322.2 | - | - | 0 | - |
| - | - | 1751 | 323.2 | - | - | 0 | - |
| - | - | 1504 | 324.7 | - | - | 0 | - |
| - | - | 3296 | 325.2 | - | - | 0 | - |
| - | - | 3466 | 326.1 | - | - | 0 | - |
| - | - | 7596 | 327.7 | - | - | 0 | - |
| - | - | 1079 | 328.1 | - | - | 0 | - |
| - | - | 5334 | 328.2 | - | - | 0 | - |
| - | - | 3288 | 328.2 | - | - | 0 | - |
| 3 | b | 7.528E+04 | 330.2 | 0.003704 | 11.22 | +1 | 3 |
| - | - | 1.171E+04 | 331.2 | - | - | 0 | - |
| - | - | 9772 | 331.2 | - | - | 0 | - |
| - | - | 2807 | 331.7 | - | - | 0 | - |
| - | - | 9024 | 332.7 | - | - | 0 | - |
| - | - | 4671 | 333.2 | - | - | 0 | - |
| - | - | 1926 | 333.2 | - | - | 0 | - |
| - | - | 2.506E+04 | 336.7 | - | - | 0 | - |
| - | - | 1992 | 337.2 | - | - | 0 | - |
| - | - | 6865 | 337.2 | - | - | 0 | - |
| - | - | 1.366E+04 | 340.2 | - | - | 0 | - |
| - | - | 1810 | 340.2 | - | - | 0 | - |
| - | - | 1520 | 341.2 | - | - | 0 | - |
| - | - | 2702 | 341.2 | - | - | 0 | - |
| - | - | 2.331E+04 | 341.7 | - | - | 0 | - |
| - | - | 5697 | 342.2 | - | - | 0 | - |
| - | - | 1861 | 342.7 | - | - | 0 | - |
| - | - | 1424 | 343.7 | - | - | 0 | - |
| - | - | 5143 | 344.2 | - | - | 0 | - |
| - | - | 1482 | 344.2 | - | - | 0 | - |
| - | - | 1479 | 345.2 | - | - | 0 | - |
| - | - | 4453 | 345.2 | - | - | 0 | - |
| - | - | 1977 | 345.7 | - | - | 0 | - |
| - | - | 3732 | 346.1 | - | - | 0 | - |
| - | - | 1.083E+04 | 346.1 | - | - | 0 | - |
| - | - | 1181 | 346.7 | - | - | 0 | - |
| - | - | 1384 | 347.1 | - | - | 0 | - |
| - | - | 5700 | 348.2 | - | - | 0 | - |
| - | - | 2323 | 349.2 | - | - | 0 | - |
| - | - | 2079 | 349.2 | - | - | 0 | - |
| - | - | 2153 | 351.2 | - | - | 0 | - |
| - | - | 2508 | 351.7 | - | - | 0 | - |
| - | - | 1387 | 352.2 | - | - | 0 | - |
| - | - | 2671 | 354.1 | - | - | 0 | - |
| - | - | 2207 | 354.2 | - | - | 0 | - |
| - | - | 4971 | 354.7 | - | - | 0 | - |
| - | - | 2510 | 354.9 | - | - | 0 | - |
| - | - | 2426 | 355.2 | - | - | 0 | - |
| - | - | 1508 | 355.2 | - | - | 0 | - |
| - | - | 8905 | 355.7 | - | - | 0 | - |
| - | - | 3257 | 356.2 | - | - | 0 | - |
| 5 | y | 4.19E+04 | 357.2 | 0.0005262 | 1.473 | +1 | 3 |
| 5 | y | 4.156E+04 | 358.2 | 0.0004279 | 1.195 | +1 | 3 |
| - | - | 4292 | 358.2 | - | - | 0 | - |
| - | - | 3929 | 358.7 | - | - | 0 | - |
| - | - | 6775 | 359.2 | - | - | 0 | - |
| 2 | y | 7901 | 359.7 | 0.002746 | 7.634 | +2 | 6 |
| - | - | 1599 | 360.2 | - | - | 0 | - |
| - | - | 5571 | 362.2 | - | - | 0 | - |
| - | - | 2.981E+04 | 362.2 | - | - | 0 | - |
| - | - | 9070 | 362.7 | - | - | 0 | - |
| - | - | 6615 | 363.2 | - | - | 0 | - |
| - | - | 1322 | 363.7 | - | - | 0 | - |
| - | - | 2559 | 364.1 | - | - | 0 | - |
| - | - | 1.237E+04 | 364.7 | - | - | 0 | - |
| - | - | 3303 | 365.2 | - | - | 0 | - |
| - | - | 1248 | 365.7 | - | - | 0 | - |
| - | - | 5673 | 367.2 | - | - | 0 | - |
| - | - | 1.083E+04 | 367.7 | - | - | 0 | - |
| - | - | 7939 | 368.2 | - | - | 0 | - |
| 2 | y | 3.494E+04 | 368.7 | 0.003048 | 8.267 | +2 | 6 |
| - | - | 1.259E+04 | 369.2 | - | - | 0 | - |
| - | - | 3345 | 369.7 | - | - | 0 | - |
| - | - | 2.721E+04 | 372.2 | - | - | 0 | - |
| - | - | 1452 | 372.2 | - | - | 0 | - |
| - | - | 3692 | 373.2 | - | - | 0 | - |
| - | - | 5909 | 373.2 | - | - | 0 | - |
| - | - | 1.71E+04 | 373.7 | - | - | 0 | - |
| - | - | 1347 | 374.2 | - | - | 0 | - |
| - | - | 6081 | 374.2 | - | - | 0 | - |
| - | - | 1641 | 374.7 | - | - | 0 | - |
| 5 | y | 7.055E+05 | 375.2 | 0.0004901 | 1.306 | +1 | 3 |
| - | - | 1.066E+04 | 376.2 | - | - | 0 | - |
| - | - | 1.152E+05 | 376.2 | - | - | 0 | - |
| - | - | 7129 | 376.7 | - | - | 0 | - |
| - | - | 1601 | 377.2 | - | - | 0 | - |
| - | - | 1.624E+04 | 377.2 | - | - | 0 | - |
| - | - | 1499 | 377.7 | - | - | 0 | - |
| - | - | 3036 | 379.2 | - | - | 0 | - |
| - | - | 2561 | 380.2 | - | - | 0 | - |
| - | - | 1256 | 381.2 | - | - | 0 | - |
| - | - | 2317 | 384.2 | - | - | 0 | - |
| - | - | 1.146E+05 | 385.2 | - | - | 0 | - |
| - | - | 2.216E+04 | 385.7 | - | - | 0 | - |
| - | - | 1.483E+04 | 386.2 | - | - | 0 | - |
| - | - | 5614 | 386.7 | - | - | 0 | - |
| - | - | 2093 | 387.2 | - | - | 0 | - |
| - | - | 1784 | 387.7 | - | - | 0 | - |
| - | - | 7088 | 390.7 | - | - | 0 | - |
| - | - | 2822 | 391.2 | - | - | 0 | - |
| - | - | 1084 | 391.7 | - | - | 0 | - |
| - | - | 1.033E+05 | 394.2 | - | - | 0 | - |
| - | - | 4.367E+04 | 394.7 | - | - | 0 | - |
| - | - | 2117 | 395.2 | - | - | 0 | - |
| - | - | 9459 | 395.2 | - | - | 0 | - |
| - | - | 1526 | 395.7 | - | - | 0 | - |
| - | - | 2578 | 396.2 | - | - | 0 | - |
| - | - | 4826 | 397.2 | - | - | 0 | - |
| - | - | 1202 | 398.1 | - | - | 0 | - |
| - | - | 4259 | 399.2 | - | - | 0 | - |
| - | - | 1.786E+04 | 399.7 | - | - | 0 | - |
| - | - | 2019 | 400.2 | - | - | 0 | - |
| - | - | 6263 | 400.2 | - | - | 0 | - |
| - | - | 2071 | 400.7 | - | - | 0 | - |
| - | - | 8656 | 401.2 | - | - | 0 | - |
| - | - | 4.971E+04 | 402.2 | - | - | 0 | - |
| - | - | 7286 | 403.2 | - | - | 0 | - |
| - | - | 1324 | 404.2 | - | - | 0 | - |
| - | - | 2367 | 405.2 | - | - | 0 | - |
| - | - | 3920 | 406.2 | - | - | 0 | - |
| - | - | 8108 | 408.2 | - | - | 0 | - |
| - | - | 7893 | 408.7 | - | - | 0 | - |
| - | - | 2487 | 409.2 | - | - | 0 | - |
| - | - | 2041 | 409.7 | - | - | 0 | - |
| - | - | 1142 | 411.2 | - | - | 0 | - |
| - | - | 1048 | 412.3 | - | - | 0 | - |
| - | - | 6214 | 413.2 | - | - | 0 | - |
| - | - | 1091 | 414.2 | - | - | 0 | - |
| - | - | 1555 | 414.2 | - | - | 0 | - |
| - | - | 1432 | 415.2 | - | - | 0 | - |
| - | - | 1385 | 415.3 | - | - | 0 | - |
| - | - | 1412 | 416.1 | - | - | 0 | - |
| - | - | 6530 | 416.2 | - | - | 0 | - |
| - | - | 2827 | 416.3 | - | - | 0 | - |
| 0 | Precursor | 1.536E+05 | 417.2 | 0.0008385 | 2.01 | +2 | -1 |
| - | - | 6.335E+04 | 417.7 | - | - | 0 | - |
| - | - | 2.802E+04 | 418.2 | - | - | 0 | - |
| - | - | 1921 | 418.7 | - | - | 0 | - |
| - | - | 1780 | 421.2 | - | - | 0 | - |
| - | - | 1938 | 422.2 | - | - | 0 | - |
| - | - | 1447 | 422.3 | - | - | 0 | - |
| - | - | 2.149E+04 | 423.2 | - | - | 0 | - |
| - | - | 7259 | 424.2 | - | - | 0 | - |
| - | - | 6008 | 424.2 | - | - | 0 | - |
| - | - | 2791 | 425.2 | - | - | 0 | - |
| 0 | Precursor | 1.814E+04 | 426.2 | 0.001141 | 2.677 | +2 | -1 |
| - | - | 6919 | 426.7 | - | - | 0 | - |
| - | - | 2706 | 427.2 | - | - | 0 | - |
| - | - | 3774 | 428.3 | - | - | 0 | - |
| - | - | 1.055E+04 | 430.2 | - | - | 0 | - |
| - | - | 1.423E+05 | 431.2 | - | - | 0 | - |
| - | - | 3.2E+04 | 432.3 | - | - | 0 | - |
| - | - | 1841 | 433.2 | - | - | 0 | - |
| - | - | 5064 | 433.3 | - | - | 0 | - |
| - | - | 2263 | 439.2 | - | - | 0 | - |
| - | - | 2.989E+04 | 440.3 | - | - | 0 | - |
| - | - | 3182 | 441.2 | - | - | 0 | - |
| - | - | 8.258E+04 | 441.2 | - | - | 0 | - |
| - | - | 2907 | 442.2 | - | - | 0 | - |
| - | - | 1.855E+04 | 442.2 | - | - | 0 | - |
| - | - | 1248 | 443.2 | - | - | 0 | - |
| - | - | 3641 | 443.3 | - | - | 0 | - |
| - | - | 5674 | 444.3 | - | - | 0 | - |
| - | - | 4350 | 456.3 | - | - | 0 | - |
| - | - | 8.186E+05 | 458.3 | - | - | 0 | - |
| 4 | b | 4.399E+04 | 459.2 | 0.001248 | 2.717 | +1 | 4 |
| - | - | 1.769E+05 | 459.3 | - | - | 0 | - |
| - | - | 7891 | 460.2 | - | - | 0 | - |
| - | - | 2.673E+04 | 460.3 | - | - | 0 | - |
| - | - | 1678 | 461.2 | - | - | 0 | - |
| - | - | 1558 | 461.2 | - | - | 0 | - |
| - | - | 1143 | 461.3 | - | - | 0 | - |
| - | - | 1648 | 462.2 | - | - | 0 | - |
| - | - | 1394 | 462.3 | - | - | 0 | - |
| - | - | 1.023E+04 | 468.3 | - | - | 0 | - |
| - | - | 1823 | 469.3 | - | - | 0 | - |
| - | - | 1459 | 476.2 | - | - | 0 | - |
| - | - | 1562 | 476.3 | - | - | 0 | - |
| 4 | b | 6.755E+04 | 477.2 | 0.001212 | 2.539 | +1 | 4 |
| - | - | 1.512E+04 | 478.2 | - | - | 0 | - |
| - | - | 4404 | 479.2 | - | - | 0 | - |
| - | - | 1433 | 480.3 | - | - | 0 | - |
| - | - | 2398 | 483.3 | - | - | 0 | - |
| - | - | 8796 | 484.3 | - | - | 0 | - |
| - | - | 2824 | 485.3 | - | - | 0 | - |
| - | - | 1.926E+04 | 486.3 | - | - | 0 | - |
| - | - | 1748 | 487.2 | - | - | 0 | - |
| - | - | 5352 | 487.3 | - | - | 0 | - |
| - | - | 1.154E+04 | 488.3 | - | - | 0 | - |
| - | - | 3232 | 489.3 | - | - | 0 | - |
| - | - | 3426 | 492.3 | - | - | 0 | - |
| - | - | 996.2 | 497.3 | - | - | 0 | - |
| - | - | 3880 | 498.3 | - | - | 0 | - |
| - | - | 5.953E+04 | 501.3 | - | - | 0 | - |
| - | - | 1.396E+04 | 502.3 | - | - | 0 | - |
| - | - | 2494 | 503.3 | - | - | 0 | - |
| 4 | y | 2.083E+04 | 504.3 | 0.00535 | 10.61 | +1 | 4 |
| 4 | y | 1.02E+04 | 505.2 | 0.00751 | 14.87 | +1 | 4 |
| - | - | 2599 | 506.2 | - | - | 0 | - |
| - | - | 1.637E+04 | 507.2 | - | - | 0 | - |
| - | - | 2889 | 508.3 | - | - | 0 | - |
| - | - | 1159 | 509.2 | - | - | 0 | - |
| - | - | 2.008E+04 | 515.3 | - | - | 0 | - |
| - | - | 6042 | 516.3 | - | - | 0 | - |
| 4 | y | 8.702E+05 | 522.3 | 0.005589 | 10.7 | +1 | 4 |
| - | - | 2.05E+05 | 523.3 | - | - | 0 | - |
| - | - | 5.802E+04 | 524.3 | - | - | 0 | - |
| - | - | 4849 | 525.3 | - | - | 0 | - |
| - | - | 3448 | 526.3 | - | - | 0 | - |
| - | - | 1078 | 527.3 | - | - | 0 | - |
| - | - | 9936 | 528.3 | - | - | 0 | - |
| - | - | 5197 | 529.3 | - | - | 0 | - |
| - | - | 8988 | 532.3 | - | - | 0 | - |
| - | - | 1829 | 533.3 | - | - | 0 | - |
| - | - | 2562 | 536.3 | - | - | 0 | - |
| - | - | 3766 | 537.3 | - | - | 0 | - |
| - | - | 1714 | 541.3 | - | - | 0 | - |
| - | - | 9255 | 543.3 | - | - | 0 | - |
| - | - | 3349 | 544.3 | - | - | 0 | - |
| - | - | 1.953E+04 | 544.3 | - | - | 0 | - |
| - | - | 5651 | 545.3 | - | - | 0 | - |
| - | - | 2.186E+04 | 553.3 | - | - | 0 | - |
| - | - | 1.809E+04 | 554.3 | - | - | 0 | - |
| - | - | 5043 | 555.3 | - | - | 0 | - |
| - | - | 1378 | 556.3 | - | - | 0 | - |
| - | - | 1316 | 557.3 | - | - | 0 | - |
| - | - | 4008 | 562.3 | - | - | 0 | - |
| - | - | 5.873E+05 | 571.4 | - | - | 0 | - |
| 5 | b | 4725 | 572.3 | 8.565E-06 | 0.01497 | +1 | 5 |
| - | - | 1.68E+05 | 572.4 | - | - | 0 | - |
| - | - | 3.067E+04 | 573.4 | - | - | 0 | - |
| - | - | 1791 | 574.4 | - | - | 0 | - |
| - | - | 5132 | 581.3 | - | - | 0 | - |
| - | - | 2059 | 582.3 | - | - | 0 | - |
| 5 | b | 5762 | 590.3 | 0.001468 | 2.487 | +1 | 5 |
| - | - | 1534 | 591.3 | - | - | 0 | - |
| - | - | 1952 | 595.3 | - | - | 0 | - |
| - | - | 2697 | 597.3 | - | - | 0 | - |
| - | - | 1645 | 598.3 | - | - | 0 | - |
| - | - | 1044 | 602.3 | - | - | 0 | - |
| - | - | 1163 | 610.3 | - | - | 0 | - |
| - | - | 2475 | 613.3 | - | - | 0 | - |
| - | - | 1104 | 614.3 | - | - | 0 | - |
| 3 | y | 8315 | 617.3 | 0.005088 | 8.241 | +1 | 5 |
| 3 | y | 5533 | 618.3 | 0.009048 | 14.63 | +1 | 5 |
| - | - | 1.08E+04 | 620.3 | - | - | 0 | - |
| - | - | 3062 | 621.3 | - | - | 0 | - |
| - | - | 1269 | 622.3 | - | - | 0 | - |
| - | - | 1232 | 626.3 | - | - | 0 | - |
| 3 | y | 5.614E+05 | 635.3 | 0.005448 | 8.575 | +1 | 5 |
| - | - | 1.649E+05 | 636.4 | - | - | 0 | - |
| - | - | 5.531E+04 | 637.4 | - | - | 0 | - |
| - | - | 1.594E+04 | 638.4 | - | - | 0 | - |
| - | - | 4297 | 639.4 | - | - | 0 | - |
| - | - | 3846 | 645.3 | - | - | 0 | - |
| - | - | 1836 | 646.3 | - | - | 0 | - |
| - | - | 4217 | 654.4 | - | - | 0 | - |
| - | - | 1276 | 656.3 | - | - | 0 | - |
| - | - | 1480 | 661.3 | - | - | 0 | - |
| - | - | 4216 | 664.4 | - | - | 0 | - |
| - | - | 1577 | 665.4 | - | - | 0 | - |
| - | - | 1.28E+04 | 672.4 | - | - | 0 | - |
| - | - | 4320 | 673.4 | - | - | 0 | - |
| 6 | b | 1910 | 677.3 | 0.00224 | 3.308 | +1 | 6 |
| - | - | 4.666E+04 | 682.4 | - | - | 0 | - |
| - | - | 1.731E+04 | 683.4 | - | - | 0 | - |
| - | - | 3928 | 684.4 | - | - | 0 | - |
| - | - | 3141 | 702.4 | - | - | 0 | - |
| - | - | 1217 | 703.4 | - | - | 0 | - |
| 2 | y | 3338 | 718.4 | 0.006787 | 9.447 | +1 | 6 |
| - | - | 4261 | 728.4 | - | - | 0 | - |
| - | - | 1406 | 729.4 | - | - | 0 | - |
| 2 | y | 9750 | 736.4 | 0.005194 | 7.054 | +1 | 6 |
| - | - | 3815 | 737.4 | - | - | 0 | - |
| - | - | 8.072E+04 | 746.4 | - | - | 0 | - |
| - | - | 2.794E+04 | 747.4 | - | - | 0 | - |
| - | - | 9835 | 748.4 | - | - | 0 | - |
| - | - | 1212 | 1302 | - | - | 0 | - |
| - | - | 1163 | 2784 | - | - | 0 | - |

m/z Charge Intensity FragmentType MassShift Position
120.08157348632812 0 694.9088
122.09652709960938 0 1084.433
124.11244201660156 0 16857.152
125.07137298583984 0 7185.6284
125.10750579833984 0 1154.995
125.11579132080078 0 1372.1239
126.05537414550781 0 151898.98
126.09181213378906 0 1123.7922
126.12814331054688 0 2951.7021
127.0506362915039 0 5840.0894
127.05867767333984 0 8106.947
127.08692169189453 0 2262.1636
129.06625366210938 0 57799.992
129.10289001464844 0 769.1885
129.11387634277344 0 4357.511
130.03257751464844 0 1618.6809
130.05029296875 0 8213.871
130.0614776611328 0 51410.85
130.06985473632812 0 2314.365
130.09786987304688 0 10510.033
131.12974548339844 0 1239.7461
133.09751892089844 0 4249.1953
134.04507446289062 0 846.481
136.0397186279297 0 5300.238
137.071044921875 0 963.76184
138.0553436279297 0 3403.3599
139.05072021484375 0 3364.4282
139.08685302734375 0 934.12994
140.08209228515625 0 1425.9398
141.0663604736328 0 3062.321
141.10264587402344 0 5679.3228
141.1390380859375 0 17688.947
142.11053466796875 0 4712.075
142.12306213378906 0 6956.7427
142.1426544189453 0 1306.2972
143.04542541503906 0 1525.5892
143.08197021484375 0 8656.72
143.11831665039062 0 8732.459
144.06597900390625 0 507296.34
144.1215057373047 0 892.3479
145.06243896484375 0 2721.82
145.0693359375 0 31153.014
146.0702362060547 0 2399.7593
148.04293823242188 0 2582.6565
149.02711486816406 0 865.19604
150.0913848876953 0 1437.6094
151.0946502685547 0 871.86237
151.12289428710938 0 1247.9519
152.1073760986328 0 9911.296
153.06626892089844 0 40832.56
153.1025390625 0 1994.2695
153.11094665527344 0 956.6097
154.0502471923828 0 19302.594
154.069580078125 0 3310.8604
154.09803771972656 0 1681.6981
154.12298583984375 0 1436.2833
155.0459747314453 0 961.7338
155.0529327392578 0 832.6536
155.08187866210938 0 3944.1091
155.11819458007812 0 2985.7368
156.11358642578125 0 1367.0542
157.06085205078125 0 2315.008
157.09762573242188 0 5721.4995
157.10874938964844 0 12774.894
157.1339111328125 0 3374.0652
158.0928192138672 0 108774.59 y Ammonia loss 6
163.57620239257812 0 911.5997
164.0348358154297 0 1225.4349
166.05352783203125 0 5387.131
166.09776306152344 0 2261.7402
167.08181762695312 0 1007.4287
167.1182861328125 0 3736.2632
169.0978240966797 0 2058.6865
169.13397216796875 0 141514.62
170.0928192138672 0 43924.25
170.10556030273438 0 3198.856
170.13040161132812 0 1089.5194
170.13729858398438 0 11984.094
171.0768280029297 0 88158.4
171.0955047607422 0 2419.729
171.11338806152344 0 2134.017
171.14947509765625 0 1155.8647
172.06085205078125 0 7263.824
172.08021545410156 0 5518.1143
172.1083984375 0 1032.6073
173.05613708496094 0 3843.7483
173.0923309326172 0 1979.7554
173.12881469726562 0 13088.116
173.4392547607422 0 2895.5476
175.11941528320312 0 312555.47 y 6
175.61654663085938 0 792.0503
176.1160430908203 0 1957.2373
176.1228485107422 0 17109.504
177.1024627685547 0 1669.3195
178.09780883789062 0 1381.4698
179.11827087402344 0 7354.188 y Water loss 4
180.10235595703125 0 164145.17
181.0611572265625 0 9026.027
181.09727478027344 0 2884.423
181.10589599609375 0 15378.521
181.13368225097656 0 1052.9275
182.04495239257812 0 2458.4822
182.0928192138672 0 3748.6323
183.0764923095703 0 1053.8755
183.1132049560547 0 4005.2686
183.8920135498047 0 838.1256
184.10845947265625 0 6273.6094
185.0925750732422 0 6401.5913
185.1029052734375 0 1906.8635
185.14019775390625 0 18655.941
186.09634399414062 0 874.9081
187.14456176757812 0 144894.81
188.14794921875 0 13420.562
189.0874786376953 0 558893.4
190.09078979492188 0 37004.617
191.09197998046875 0 3688.8193
191.11900329589844 0 1391.698
193.097412109375 0 1409.9506
193.11387634277344 0 3138.8752
195.07655334472656 0 978.6683
195.1131591796875 0 7654.3174
195.14926147460938 0 1547.8403
196.14511108398438 0 1892.8444
197.12884521484375 0 158698.16
198.08767700195312 0 22364.148
198.13233947753906 0 16799.334
199.07171630859375 0 186864.33
199.11929321289062 0 4677.334
199.13487243652344 0 957.4939
199.14389038085938 0 1529.3785
200.05580139160156 0 3339.5654
200.0751190185547 0 14008.932
200.10325622558594 0 3733.9558
200.14105224609375 0 968.0004
200.36770629882812 0 1020.1193
201.0767364501953 0 1380.6129
201.09873962402344 0 1750.704
201.12376403808594 0 22841.963
201.1349334716797 0 3027.455
202.1189727783203 0 28887.7
202.143310546875 0 2026.1085
203.10301208496094 0 11695.89
203.1215057373047 0 1631.8811
206.12916564941406 0 1870.2402
206.2869415283203 0 1024.4768
206.45277404785156 0 939.8138
207.11260986328125 0 7665.3784
208.14527893066406 0 1470.2991
209.09249877929688 0 2235.033
209.10385131835938 0 3019.7842
209.1283416748047 0 1811.498
210.08763122558594 0 9600.96
210.1234893798828 0 1096.7032
211.10794067382812 0 4792.749
211.1444854736328 0 4099.9824
212.10324096679688 0 2875.5793
213.08787536621094 0 1789.2513
213.12400817871094 0 959.6345
214.1552276611328 0 1842.0535
214.6331329345703 0 1088.5822
215.11328125 0 3185.6177
215.12156677246094 0 3492.756
215.1394805908203 0 33907.09
216.09823608398438 0 5731.083
216.14315795898438 0 4075.8364
217.08233642578125 0 316874.44 b 1
218.08572387695312 0 24735.916
218.10592651367188 0 770.08765
219.08680725097656 0 2030.3668
220.11859130859375 0 2103.2725
220.12969970703125 0 11081.271
220.14149475097656 0 1257.9435
221.12828063964844 0 3948.3313
222.12403869628906 0 2184.9949
223.10826110839844 0 3654.2244
223.14462280273438 0 2703.7468
224.10362243652344 0 1103.9803
224.1397247314453 0 2782.8909
225.1238250732422 0 33070.453
226.12864685058594 0 4609.077
227.1143035888672 0 24749.197
227.12686157226562 0 1771.6396
228.09849548339844 0 9497.925
228.11831665039062 0 2417.247
228.13455200195312 0 2492.1665
228.6326446533203 0 973.60077
229.00146484375 0 1502.7148
229.13015747070312 0 5845.0825
229.6401824951172 0 51039.63
230.12527465820312 0 10051.796
230.14170837402344 0 9182.413
230.64329528808594 0 1222.0745
232.140869140625 0 5054.989
233.09239196777344 0 1910.4968
233.13221740722656 0 43274.71
234.135498046875 0 4154.1523
235.12852478027344 0 1108.1566
236.14002990722656 0 3211.8052
237.12399291992188 0 1324.3673
238.15545654296875 0 14840.652
239.13949584960938 0 6309.262
239.15834045410156 0 1385.0978
240.1346893310547 0 10691.12
240.17135620117188 0 1513.2634
241.12969970703125 0 1937.7402
242.64854431152344 0 1004.7552
243.1179962158203 0 1776.6849
243.6377716064453 0 6328.3657
244.1007080078125 0 21369.986
244.14077758789062 0 23472.055 y Water loss 5
245.12496948242188 0 205989.88 y Ammonia loss 5
246.10792541503906 0 1073.1199
246.12998962402344 0 20976.19
247.13133239746094 0 1805.5875
249.1237335205078 0 10371.348
250.1557159423828 0 2188.854
251.10304260253906 0 7460.536
251.14195251464844 0 1109.0355
251.16018676757812 0 1166.2668
252.1343231201172 0 2346.5803
252.63314819335938 0 902.9551 y Water loss 3
253.12562561035156 0 3671.2085 y Ammonia loss 3
254.1505126953125 0 3634.3853
255.10955810546875 0 2957.0076
256.12969970703125 0 2825.284
256.1659240722656 0 2682.275
256.6451416015625 0 2586.638
257.61602783203125 0 1055.1265
258.1459045410156 0 3676.0747
260.13519287109375 0 967.89435
261.1044921875 0 1306.2708
261.12713623046875 0 83311.36
261.63922119140625 0 33214.434 y 3
262.1513977050781 0 391037.16 y 5
262.6385803222656 0 1315.3313
263.070068359375 0 4716.5537
263.12225341796875 0 3758.507
263.1545715332031 0 38145.17
264.1344299316406 0 3530.0574
264.15594482421875 0 3179.0024
265.19244384765625 0 2097.9556
266.15020751953125 0 19033.604
267.13421630859375 0 21251.443
267.152587890625 0 2761.11
268.1299133300781 0 7868.255
268.6631774902344 0 4110.687
269.1334533691406 0 999.1403
270.1200256347656 0 6601.653
271.1765441894531 0 2107.1172
272.1357116699219 0 7151.1387
277.1195373535156 0 6645.2793
277.1774597167969 0 1907.0836
277.66754150390625 0 1678.0345
278.12237548828125 0 990.93524
278.171142578125 0 2053.9294
280.1660461425781 0 7535.096
281.1692810058594 0 1332.7744
282.10968017578125 0 1226.3748
282.14508056640625 0 1440.2617
282.218017578125 0 7470.6196
284.1609802246094 0 164062.3 a Water loss 2
285.14501953125 0 3422.9087
285.1642150878906 0 20343.836
286.141357421875 0 4426.5938
286.18231201171875 0 126806.9
286.68389892578125 0 32307.47
287.1069030761719 0 1601.1658
287.1847839355469 0 5809.385
288.1313171386719 0 1664.4432
290.14990234375 0 1816.9995
291.1752624511719 0 1407.679
292.2031555175781 0 1855.6211
293.1882019042969 0 1141.7473
293.93548583984375 0 2047.2838
294.1452331542969 0 7686.9556
295.1296081542969 0 6925.96
295.14703369140625 0 1335.6576
296.16094970703125 0 1944.3558
297.0587463378906 0 1445.3601
297.17486572265625 0 1898.3229
297.1922912597656 0 1432.0405
298.1767272949219 0 3894.512
299.1717834472656 0 10734.407
299.1904296875 0 5174.7837
299.6734924316406 0 2648.119
299.6913146972656 0 1580.5859
302.17181396484375 0 10684.872 a 2
303.17535400390625 0 2580.475
305.68719482421875 0 1611.8347
308.1613464355469 0 10971.876
308.1978759765625 0 2099.291
309.1781921386719 0 1824.0288 y Water loss 2
309.20404052734375 0 1010.50616
309.67974853515625 0 1262.2787
310.1870422363281 0 4925.107
310.21319580078125 0 11794.878
311.2181396484375 0 1428.7089
311.9456481933594 0 3369.2798
312.156005859375 0 251697.92 b Water loss 2
313.1591491699219 0 39437.645
314.1612854003906 0 5016.938
314.69378662109375 0 2037.7998
315.2023010253906 0 3256.2034
317.1571044921875 0 2943.77
318.1815490722656 0 286948.94 y 2
318.682861328125 0 88421.48
319.1823425292969 0 28627.941
319.6851501464844 0 12303.769
320.18621826171875 0 1407.063
320.6682434082031 0 2127.6377
322.1882019042969 0 1215.625
323.17266845703125 0 1750.5099
324.6755065917969 0 1504.485
325.1876220703125 0 3296.1758
326.1477355957031 0 3465.5764
327.700927734375 0 7596.1016
328.1415100097656 0 1079.4382
328.1625671386719 0 5334.499
328.202880859375 0 3288.2236
330.1667785644531 0 75283.38 b 2
331.16973876953125 0 11709.211
331.1895751953125 0 9771.946
331.6906433105469 0 2807.1765
332.693115234375 0 9023.767
333.1940002441406 0 4671.4233
333.2143859863281 0 1926.2793
336.70635986328125 0 25057.688
337.1866760253906 0 1991.6495
337.20794677734375 0 6864.775
340.1983642578125 0 13658.076
340.2197265625 0 1810.1272
341.1817932128906 0 1520.0164
341.2023620605469 0 2701.978
341.69842529296875 0 23307.363
342.1997375488281 0 5697.1274
342.7018737792969 0 1861.1118
343.6712341308594 0 1423.9641
344.1650085449219 0 5142.758
344.1919860839844 0 1481.7136
345.1923522949219 0 1479.3823
345.2252502441406 0 4453.1274
345.6819152832031 0 1976.7025
346.1071472167969 0 3731.6787
346.14373779296875 0 10832.192
346.68988037109375 0 1181.2218
347.1479187011719 0 1383.8909
348.1590270996094 0 5700.076
349.1613464355469 0 2322.9307
349.1855773925781 0 2078.7935
351.1658935546875 0 2153.4666
351.68414306640625 0 2507.5442
352.1845397949219 0 1386.5475
354.1477355957031 0 2670.711
354.1971130371094 0 2206.5203
354.6883239746094 0 4970.779
354.8859558105469 0 2509.8335
355.1889343261719 0 2425.6084
355.21075439453125 0 1507.9421
355.6952819824219 0 8904.909
356.1971130371094 0 3257.2847
357.2250061035156 0 41896.23 y Water loss 4
358.20892333984375 0 41558.85 y Ammonia loss 4
358.2272644042969 0 4292.1636
358.6907043457031 0 3929.1157
359.2128601074219 0 6775.2876
359.6997375488281 0 7901.4604 y Water loss 1
360.1996154785156 0 1598.9065
362.1744384765625 0 5571.245
362.2115478515625 0 29805.021
362.71240234375 0 9070.458
363.2017517089844 0 6615.0986
363.7003173828125 0 1322.3611
364.1169128417969 0 2559.446
364.6921081542969 0 12371.6875
365.1922302246094 0 3302.9426
365.6930847167969 0 1247.6663
367.2049865722656 0 5673.18
367.6965026855469 0 10831.039
368.19482421875 0 7939.1533
368.705322265625 0 34941.96 y 1
369.2064514160156 0 12586.968
369.7064208984375 0 3344.5193
372.1593322753906 0 27209.764
372.20233154296875 0 1452.417
373.1621398925781 0 3692.1284
373.21942138671875 0 5908.804
373.6972961425781 0 17097.512
374.15447998046875 0 1346.8916
374.21051025390625 0 6081.303
374.6982727050781 0 1641.4146
375.23553466796875 0 705500.6 y 4
376.209228515625 0 10655.057
376.2384033203125 0 115192.93
376.7031555175781 0 7128.827
377.2119445800781 0 1601.0259
377.24066162109375 0 16241.265
377.6804504394531 0 1499.3561
379.2345275878906 0 3035.7283
380.2284851074219 0 2560.7463
381.21221923828125 0 1256.0756
384.19873046875 0 2317.3616
385.21673583984375 0 114629.734
385.7152404785156 0 22156.309
386.22027587890625 0 14829.169
386.68853759765625 0 5614.0107
387.2001647949219 0 2093.1665
387.69158935546875 0 1784.3564
390.6893615722656 0 7087.5713
391.19073486328125 0 2821.6697
391.6895751953125 0 1084.1351
394.2178649902344 0 103263.08
394.71917724609375 0 43673.76
395.19207763671875 0 2116.9312
395.2200927734375 0 9458.515
395.72247314453125 0 1525.679
396.1910705566406 0 2578.2585
397.24542236328125 0 4825.8896
398.1397399902344 0 1202.0144
399.2015686035156 0 4258.6353
399.6951904296875 0 17861.346
400.153564453125 0 2019.0034
400.19659423828125 0 6263.1123
400.696044921875 0 2070.679
401.2152099609375 0 8655.653
402.2101745605469 0 49709.26
403.2127990722656 0 7285.997
404.2091979980469 0 1324.3468
405.22589111328125 0 2366.5405
406.208984375 0 3920.3743
408.208984375 0 8107.771
408.7019348144531 0 7893.248
409.2035827636719 0 2487.1978
409.7015075683594 0 2040.7632
411.2353210449219 0 1142.0367
412.26715087890625 0 1047.7706
413.2065124511719 0 6214.0864
414.2115783691406 0 1090.9155
414.24456787109375 0 1554.9644
415.16741943359375 0 1431.9335
415.2564392089844 0 1384.8063
416.1495056152344 0 1412.4628
416.2259521484375 0 6530.191
416.2514953613281 0 2826.7527
417.21356201171875 0 153632.78 Precursor Water loss
417.71484375 0 63346.82
418.2147216796875 0 28019.23
418.7129211425781 0 1920.641
421.245849609375 0 1779.9661
422.20330810546875 0 1938.4795
422.2525939941406 0 1446.9022
423.23565673828125 0 21485.377
424.15380859375 0 7258.728
424.21966552734375 0 6008.5
425.21728515625 0 2791.0156
426.2191467285156 0 18144.758 Precursor
426.7202453613281 0 6919.1465
427.2191467285156 0 2706.233
428.2614440917969 0 3774.1997
430.2417297363281 0 10550.274
431.2493896484375 0 142270.72
432.2532043457031 0 32004.188
433.1752014160156 0 1841.0754
433.257080078125 0 5064.252
439.231201171875 0 2263.2249
440.2622985839844 0 29893.541
441.1819763183594 0 3181.9448
441.2463073730469 0 82577.664
442.1649475097656 0 2906.788
442.2494201660156 0 18554.041
443.1697998046875 0 1248.2198
443.2507019042969 0 3640.7197
444.2575988769531 0 5674.1274
456.2571105957031 0 4349.6875
458.2729797363281 0 818566.44
459.191650390625 0 43985.504 b Water loss 3
459.2756652832031 0 176920.02
460.19439697265625 0 7890.9043
460.27801513671875 0 26730.006
461.1874084472656 0 1677.7762
461.2435302734375 0 1558.2058
461.2817077636719 0 1142.5555
462.2389831542969 0 1647.5206
462.2729187011719 0 1394.1653
468.2572021484375 0 10225.495
469.261474609375 0 1823.4249
476.2282409667969 0 1459.1014
476.2856140136719 0 1562.1412
477.2021789550781 0 67551.04 b 3
478.20513916015625 0 15121.504
479.20135498046875 0 4403.5806
480.28460693359375 0 1433.0538
483.267822265625 0 2398.052
484.2521667480469 0 8795.952
485.2507019042969 0 2824.2615
486.267578125 0 19258.889
487.2330322265625 0 1747.7408
487.2694396972656 0 5351.774
488.2711181640625 0 11536.57
489.27532958984375 0 3231.9775
492.260498046875 0 3425.78
497.29168701171875 0 996.21094
498.2939758300781 0 3879.668
501.2787780761719 0 59530.496
502.28125 0 13958.607
503.2821350097656 0 2494.2354
504.26031494140625 0 20833.36 y Water loss 3
505.2464904785156 0 10198.755 y Ammonia loss 3
506.2495422363281 0 2599.4624
507.2477722167969 0 16368.278
508.250244140625 0 2888.9714
509.2433776855469 0 1159.4604
515.2942504882812 0 20083.096
516.2979736328125 0 6041.5156
522.2711181640625 0 870169.44 y 3
523.273681640625 0 204966.25
524.2711791992188 0 58018.89
525.2725219726562 0 4848.9663
526.2870483398438 0 3448.1033
527.2926635742188 0 1077.7578
528.314453125 0 9936.295
529.3119506835938 0 5197.024
532.255615234375 0 8987.529
533.2593383789062 0 1829.4946
536.3189697265625 0 2562.3855
537.3047485351562 0 3765.8862
541.34326171875 0 1713.9622
543.3258666992188 0 9254.578
544.2820434570312 0 3348.9514
544.3328247070312 0 19529.375
545.3350219726562 0 5650.772
553.3460083007812 0 21855.203
554.3320922851562 0 18094.832
555.3344116210938 0 5042.538
556.3367919921875 0 1378.2117
557.3427124023438 0 1315.972
562.2907104492188 0 4008.1675
571.3568725585938 0 587257.94
572.2744750976562 0 4724.7046 b Water loss 4
572.3594360351562 0 167976.34
573.362060546875 0 30668.68
574.3680419921875 0 1790.7578
581.341064453125 0 5131.895
582.3435668945312 0 2059.3054
590.2864990234375 0 5762.076 b 4
591.2893676757812 0 1534.1157
595.3110961914062 0 1952.2292
597.333740234375 0 2697.2063
598.3329467773438 0 1644.8152
602.3231201171875 0 1043.9952
610.3240966796875 0 1163.2085
613.3179321289062 0 2475.2092
614.3228149414062 0 1104.3506
617.3441162109375 0 8315.427 y Water loss 2
618.3320922851562 0 5532.8843 y Ammonia loss 2
620.3328247070312 0 10798.92
621.335693359375 0 3062.3767
622.3414306640625 0 1268.8156
626.3280029296875 0 1232.23
635.3550415039062 0 561415.94 y 2
636.3577270507812 0 164918.05
637.3564453125 0 55313.62
638.3607788085938 0 15936.785
639.365234375 0 4297.3804
645.3391723632812 0 3846.383
646.3425903320312 0 1836.0056
654.3922119140625 0 4216.5845
656.3367919921875 0 1276.2659
661.329345703125 0 1479.94
664.3782348632812 0 4215.832
665.382080078125 0 1576.5012
672.40380859375 0 12796.908
673.4071655273438 0 4320.1807
677.3148193359375 0 1910.4917 b 5
682.3885498046875 0 46657.9
683.3912353515625 0 17309.473
684.3927612304688 0 3927.8674
702.3592529296875 0 3141.1672
703.3703002929688 0 1216.9766
718.3934936523438 0 3338.38 y Water loss 1
728.3776245117188 0 4260.7896
729.3777465820312 0 1406.3657
736.4024658203125 0 9749.502 y 1
737.4049682617188 0 3814.6814
746.3867797851562 0 80723.516
747.3897705078125 0 27940.246
748.3895874023438 0 9835.385
1301.748779296875 0 1211.6702
2784.035400390625 0 1163.4999

Spectrum Details

|  |  |
| --- | --- |
| Matched peaks? Matched peaks The total absolute number of peaks matched. Additionally in brackets the total fraction of peaks matched and the total number of peaks is shown. | 36 (6.12% of 588) |
| FDR? FDR The false discovery rate estimated for this peptide. It is calculated by matching all theoretical fragments with a non-integer shift with the raw peaks for this spectrum. This is done with 40 different shifts. The resulting percentage is the average number of annotated peaks over the number of annotated peaks with the correct spectrum. | 0.40% |
| Satellite FDR? Satellite FDR See the FDR for details on its calculation. This satellite ion specific FDR only contains the satellite ions (d/w) for I/L/J positions. | - |
| PSM Score? PSM Score The PSM Score as given by Hecklib to this annotated spectrum. It is shown with three significant figures. | 448 |

#### Spectrum 3876? Spectrum 3876 The raw spectrum of this peptide as annotated by Hecklib. The fragments are coloured according to ion type (see legend). Any peaks with a star '\*' as text can be hovered over to see the full details, first the ion type second the mass shift type. By hovering over the amino acids in the peptide or ions in the legend the corresponding peaks are highlighted. By toggling the 'Unassigned' label you can turn the background (unassigned) peaks on or off in the plot. By updating the slider in the Ion legend you can update the spectrum to only show the top X% of the peaks with labels. The top X% means any peak that is within X% of the highest intensity. By dragging in the spectrum you can zoom in to a specific part of the spectrum and use 'Zoom Out' to get back to the original zoom level. The annotation of the spectrum is based on the given sequence in the peptides file and is done with different software so inconsistencies are likely. The peaks are annotated based on the given sequence, with 20 ppm tolerance.

Copy Data

##### Spectrum 3876 (TSV)

###### Preview

```
Loading example...
```

*Click on the button to copy the data to your clipboard.*

Mz MinMz MaxIntensity Max

WidthHeightPeptide font sizePeptide stroke widthSpectrum font sizeSpectrum stroke widthCompact peptide

Ion legend

wxyz

abcd

OtherUnassignedIon^Charge^_Position_Show for top:%

GTJMISR

07.27e+51.45e+62.18e+62.91e+6

Zoom Out

y^+1^_1_z^+1^_1_y^+1^_1_c^+1^_2_w^+1^_2_y^+1^_2_y^+1^_2_z^+1^_2_y^+2^_4_y^+1^_2_y^+2^_5_y^+2^_5_w^+1^_3_w^+1^_3_y^+1^_3_y^+1^_3_z^+1^_3_y^+2^_6_y^+2^_6_y^+1^_3_y^+1^_4_y^+1^_4_z^+1^_4_y^+1^_4_w^+1^_5_c^+1^_5_z^+1^_5_y^+1^_5_c^+1^_6_w^+1^_6_w^+1^_6_y^+1^_6_z^+1^_6_

0790158023703160

Fragment Matches Table

Show background peaks

| Position | Ion type | Intensity | mz Theoretical | mz Error (Th) | mz Error (ppm) | Charge | Series Number |
| --- | --- | --- | --- | --- | --- | --- | --- |
| - | - | 6717 | 126.1 | - | - | 0 | - |
| - | - | 4918 | 129.1 | - | - | 0 | - |
| - | - | 8.565E+04 | 144.1 | - | - | 0 | - |
| - | - | 4450 | 145.1 | - | - | 0 | - |
| - | - | 3483 | 153.1 | - | - | 0 | - |
| - | - | 2294 | 154.1 | - | - | 0 | - |
| 7 | y | 7602 | 158.1 | 0.0003856 | 2.439 | +1 | 1 |
| 7 | z | 1.788E+05 | 159.1 | 0.0003578 | 2.249 | +1 | 1 |
| - | - | 3.078E+05 | 160.1 | - | - | 0 | - |
| - | - | 2624 | 161.1 | - | - | 0 | - |
| - | - | 1.613E+04 | 161.1 | - | - | 0 | - |
| - | - | 2673 | 166.1 | - | - | 0 | - |
| - | - | 4559 | 169.1 | - | - | 0 | - |
| - | - | 9959 | 170.1 | - | - | 0 | - |
| - | - | 2.724E+04 | 171.1 | - | - | 0 | - |
| - | - | 7071 | 173.4 | - | - | 0 | - |
| - | - | 4448 | 174.1 | - | - | 0 | - |
| 7 | y | 7.543E+04 | 175.1 | 0.0003258 | 1.86 | +1 | 1 |
| - | - | 4167 | 176.1 | - | - | 0 | - |
| - | - | 3.464E+04 | 180.1 | - | - | 0 | - |
| - | - | 3284 | 181.1 | - | - | 0 | - |
| - | - | 8820 | 185.1 | - | - | 0 | - |
| - | - | 3.278E+04 | 187.1 | - | - | 0 | - |
| - | - | 2932 | 188.1 | - | - | 0 | - |
| - | - | 3533 | 188.1 | - | - | 0 | - |
| - | - | 3.192E+05 | 189.1 | - | - | 0 | - |
| - | - | 2.318E+04 | 190.1 | - | - | 0 | - |
| - | - | 2.147E+04 | 197.1 | - | - | 0 | - |
| - | - | 6.897E+04 | 199.1 | - | - | 0 | - |
| - | - | 5298 | 200.1 | - | - | 0 | - |
| - | - | 3.215E+04 | 200.1 | - | - | 0 | - |
| - | - | 4201 | 201.1 | - | - | 0 | - |
| - | - | 9064 | 202.1 | - | - | 0 | - |
| - | - | 2982 | 203.1 | - | - | 0 | - |
| - | - | 8586 | 203.2 | - | - | 0 | - |
| - | - | 2.182E+04 | 215.1 | - | - | 0 | - |
| 2 | c | 2948 | 216.1 | 0.003892 | 18.01 | +1 | 2 |
| - | - | 2.731E+05 | 217.1 | - | - | 0 | - |
| - | - | 2530 | 218.1 | - | - | 0 | - |
| - | - | 2.493E+04 | 218.1 | - | - | 0 | - |
| - | - | 2310 | 219.1 | - | - | 0 | - |
| - | - | 8463 | 225.1 | - | - | 0 | - |
| - | - | 2223 | 226.1 | - | - | 0 | - |
| 6 | w | 2930 | 229.1 | 7.604E-05 | 0.3319 | +1 | 2 |
| - | - | 1.482E+04 | 229.6 | - | - | 0 | - |
| - | - | 3988 | 230.1 | - | - | 0 | - |
| - | - | 4815 | 230.1 | - | - | 0 | - |
| - | - | 2316 | 230.1 | - | - | 0 | - |
| - | - | 2072 | 232.1 | - | - | 0 | - |
| - | - | 2674 | 232.7 | - | - | 0 | - |
| - | - | 9425 | 233.1 | - | - | 0 | - |
| - | - | 3902 | 240.1 | - | - | 0 | - |
| - | - | 1.281E+04 | 244.1 | - | - | 0 | - |
| - | - | 2.239E+04 | 244.1 | - | - | 0 | - |
| 6 | y | 2916 | 244.1 | 0.0001786 | 0.7315 | +1 | 2 |
| 6 | y | 1.718E+05 | 245.1 | 0.0003854 | 1.572 | +1 | 2 |
| 6 | z | 9.075E+04 | 246.1 | 0.0001154 | 0.4688 | +1 | 2 |
| - | - | 5.453E+05 | 247.1 | - | - | 0 | - |
| - | - | 5.004E+04 | 248.1 | - | - | 0 | - |
| - | - | 4836 | 249.1 | - | - | 0 | - |
| - | - | 2590 | 251.1 | - | - | 0 | - |
| - | - | 4.124E+04 | 261.1 | - | - | 0 | - |
| - | - | 3.207E+04 | 261.1 | - | - | 0 | - |
| 4 | y | 3.567E+04 | 261.6 | 0.002635 | 10.07 | +2 | 4 |
| 6 | y | 3.129E+05 | 262.2 | 0.0002645 | 1.009 | +1 | 2 |
| - | - | 3924 | 262.6 | - | - | 0 | - |
| - | - | 3285 | 263.1 | - | - | 0 | - |
| - | - | 2.555E+04 | 263.2 | - | - | 0 | - |
| - | - | 7277 | 266.2 | - | - | 0 | - |
| - | - | 3326 | 267.1 | - | - | 0 | - |
| - | - | 2401 | 268.1 | - | - | 0 | - |
| - | - | 1.574E+04 | 270.1 | - | - | 0 | - |
| - | - | 8.254E+04 | 284.2 | - | - | 0 | - |
| - | - | 4545 | 284.2 | - | - | 0 | - |
| - | - | 3049 | 285.1 | - | - | 0 | - |
| - | - | 7804 | 285.2 | - | - | 0 | - |
| - | - | 5.511E+04 | 286.2 | - | - | 0 | - |
| - | - | 9899 | 286.7 | - | - | 0 | - |
| - | - | 3347 | 287.2 | - | - | 0 | - |
| - | - | 3672 | 288.1 | - | - | 0 | - |
| - | - | 4719 | 289.1 | - | - | 0 | - |
| - | - | 3.656E+04 | 290.1 | - | - | 0 | - |
| - | - | 5124 | 294.1 | - | - | 0 | - |
| - | - | 3499 | 295.1 | - | - | 0 | - |
| - | - | 5618 | 299.2 | - | - | 0 | - |
| - | - | 2505 | 300.1 | - | - | 0 | - |
| - | - | 7009 | 302.2 | - | - | 0 | - |
| 3 | y | 3645 | 309.2 | 0.002873 | 9.292 | +2 | 5 |
| - | - | 2.013E+05 | 312.2 | - | - | 0 | - |
| - | - | 2.589E+04 | 313.2 | - | - | 0 | - |
| 3 | y | 3.649E+05 | 318.2 | 0.002962 | 9.308 | +2 | 5 |
| - | - | 1.047E+05 | 318.7 | - | - | 0 | - |
| - | - | 3.326E+04 | 319.2 | - | - | 0 | - |
| - | - | 3718 | 319.7 | - | - | 0 | - |
| 5 | w | 1.076E+05 | 330.2 | 0.0003154 | 0.9554 | +1 | 3 |
| - | - | 6936 | 331.2 | - | - | 0 | - |
| - | - | 1.299E+04 | 331.2 | - | - | 0 | - |
| - | - | 2504 | 332.5 | - | - | 0 | - |
| - | - | 1.494E+04 | 336.7 | - | - | 0 | - |
| - | - | 3787 | 337.2 | - | - | 0 | - |
| - | - | 9695 | 341.7 | - | - | 0 | - |
| 5 | w | 1.162E+04 | 344.2 | 0.0008497 | 2.469 | +1 | 3 |
| - | - | 9132 | 346.1 | - | - | 0 | - |
| - | - | 4317 | 354.7 | - | - | 0 | - |
| - | - | 3069 | 355.7 | - | - | 0 | - |
| 5 | y | 9519 | 357.2 | 0.0004957 | 1.388 | +1 | 3 |
| 5 | y | 8075 | 358.2 | 0.0002753 | 0.7685 | +1 | 3 |
| 5 | z | 4.677E+05 | 359.2 | 0.0002322 | 0.6465 | +1 | 3 |
| 2 | y | 1.109E+04 | 359.7 | 0.00238 | 6.616 | +2 | 6 |
| - | - | 7534 | 360.2 | - | - | 0 | - |
| - | - | 1.827E+05 | 360.2 | - | - | 0 | - |
| - | - | 2.803E+04 | 361.2 | - | - | 0 | - |
| - | - | 2861 | 362.2 | - | - | 0 | - |
| - | - | 3519 | 362.2 | - | - | 0 | - |
| - | - | 5061 | 363.2 | - | - | 0 | - |
| - | - | 3187 | 364.1 | - | - | 0 | - |
| - | - | 8427 | 364.7 | - | - | 0 | - |
| - | - | 5403 | 365.2 | - | - | 0 | - |
| - | - | 4136 | 367.7 | - | - | 0 | - |
| 2 | y | 6.583E+04 | 368.7 | 0.002499 | 6.778 | +2 | 6 |
| - | - | 1.994E+04 | 369.2 | - | - | 0 | - |
| - | - | 6842 | 369.7 | - | - | 0 | - |
| - | - | 1.789E+04 | 372.2 | - | - | 0 | - |
| - | - | 4901 | 373.2 | - | - | 0 | - |
| - | - | 1.887E+04 | 373.7 | - | - | 0 | - |
| - | - | 5571 | 374.2 | - | - | 0 | - |
| - | - | 3.673E+04 | 374.2 | - | - | 0 | - |
| 5 | y | 4.762E+05 | 375.2 | 0.0001544 | 0.4115 | +1 | 3 |
| - | - | 4001 | 376.2 | - | - | 0 | - |
| - | - | 5323 | 376.2 | - | - | 0 | - |
| - | - | 7.991E+04 | 376.2 | - | - | 0 | - |
| - | - | 3798 | 376.7 | - | - | 0 | - |
| - | - | 1.263E+04 | 377.2 | - | - | 0 | - |
| - | - | 7.046E+04 | 385.2 | - | - | 0 | - |
| - | - | 2.738E+04 | 385.7 | - | - | 0 | - |
| - | - | 6228 | 386.2 | - | - | 0 | - |
| - | - | 7460 | 386.7 | - | - | 0 | - |
| - | - | 4714 | 387.2 | - | - | 0 | - |
| - | - | 2.581E+04 | 388.2 | - | - | 0 | - |
| - | - | 4111 | 389.2 | - | - | 0 | - |
| - | - | 7366 | 390.7 | - | - | 0 | - |
| - | - | 1.211E+05 | 394.2 | - | - | 0 | - |
| - | - | 4.553E+04 | 394.7 | - | - | 0 | - |
| - | - | 1.334E+04 | 395.2 | - | - | 0 | - |
| - | - | 3927 | 399.2 | - | - | 0 | - |
| - | - | 2.252E+04 | 399.7 | - | - | 0 | - |
| - | - | 9360 | 400.2 | - | - | 0 | - |
| - | - | 1.684E+04 | 401.2 | - | - | 0 | - |
| - | - | 9214 | 408.2 | - | - | 0 | - |
| - | - | 9738 | 408.7 | - | - | 0 | - |
| - | - | 3678 | 409.2 | - | - | 0 | - |
| - | - | 3560 | 409.7 | - | - | 0 | - |
| - | - | 5860 | 413.2 | - | - | 0 | - |
| - | - | 2518 | 415.2 | - | - | 0 | - |
| - | - | 4.489E+04 | 415.2 | - | - | 0 | - |
| - | - | 3.771E+04 | 416.2 | - | - | 0 | - |
| - | - | 2.304E+05 | 417.2 | - | - | 0 | - |
| - | - | 9.206E+04 | 417.7 | - | - | 0 | - |
| - | - | 2.937E+04 | 418.2 | - | - | 0 | - |
| - | - | 2846 | 418.7 | - | - | 0 | - |
| - | - | 9371 | 424.2 | - | - | 0 | - |
| - | - | 4.694E+04 | 426.2 | - | - | 0 | - |
| - | - | 1.623E+04 | 426.7 | - | - | 0 | - |
| - | - | 7739 | 427.2 | - | - | 0 | - |
| - | - | 1.063E+04 | 429.2 | - | - | 0 | - |
| - | - | 2938 | 430.2 | - | - | 0 | - |
| - | - | 1.39E+04 | 431.2 | - | - | 0 | - |
| - | - | 9099 | 441.2 | - | - | 0 | - |
| - | - | 4503 | 442.2 | - | - | 0 | - |
| - | - | 3.739E+04 | 442.3 | - | - | 0 | - |
| - | - | 1.225E+05 | 443.3 | - | - | 0 | - |
| - | - | 1.174E+05 | 444.3 | - | - | 0 | - |
| - | - | 2.22E+04 | 445.3 | - | - | 0 | - |
| - | - | 4151 | 446.3 | - | - | 0 | - |
| - | - | 5737 | 450.2 | - | - | 0 | - |
| - | - | 1.865E+05 | 458.3 | - | - | 0 | - |
| - | - | 4.529E+04 | 459.2 | - | - | 0 | - |
| - | - | 4.372E+04 | 459.3 | - | - | 0 | - |
| - | - | 8351 | 460.2 | - | - | 0 | - |
| - | - | 5003 | 460.3 | - | - | 0 | - |
| - | - | 8.638E+04 | 477.2 | - | - | 0 | - |
| - | - | 1.616E+04 | 478.2 | - | - | 0 | - |
| - | - | 5133 | 479.2 | - | - | 0 | - |
| 4 | y | 9216 | 504.3 | 0.004771 | 9.461 | +1 | 4 |
| 4 | y | 6511 | 505.2 | 0.007144 | 14.14 | +1 | 4 |
| 4 | z | 1.954E+04 | 506.2 | 0.005453 | 10.77 | +1 | 4 |
| - | - | 1.146E+04 | 507.3 | - | - | 0 | - |
| - | - | 2925 | 508.3 | - | - | 0 | - |
| - | - | 3719 | 521.3 | - | - | 0 | - |
| 4 | y | 8.161E+05 | 522.3 | 0.005162 | 9.883 | +1 | 4 |
| - | - | 1.948E+05 | 523.3 | - | - | 0 | - |
| - | - | 4.999E+04 | 524.3 | - | - | 0 | - |
| - | - | 4031 | 525.3 | - | - | 0 | - |
| - | - | 4420 | 526.3 | - | - | 0 | - |
| - | - | 2352 | 529.3 | - | - | 0 | - |
| - | - | 5322 | 532.3 | - | - | 0 | - |
| - | - | 5331 | 544.3 | - | - | 0 | - |
| - | - | 2774 | 553.4 | - | - | 0 | - |
| - | - | 7639 | 556.3 | - | - | 0 | - |
| - | - | 2926 | 557.3 | - | - | 0 | - |
| - | - | 6413 | 562.3 | - | - | 0 | - |
| - | - | 2796 | 563.3 | - | - | 0 | - |
| - | - | 1.756E+05 | 571.4 | - | - | 0 | - |
| - | - | 6336 | 572.3 | - | - | 0 | - |
| - | - | 4.883E+04 | 572.4 | - | - | 0 | - |
| - | - | 9541 | 573.4 | - | - | 0 | - |
| 3 | w | 1.302E+05 | 576.3 | 0.004851 | 8.418 | +1 | 5 |
| - | - | 3.338E+04 | 577.3 | - | - | 0 | - |
| - | - | 1.133E+04 | 578.3 | - | - | 0 | - |
| - | - | 1.906E+04 | 590.3 | - | - | 0 | - |
| - | - | 3501 | 591.3 | - | - | 0 | - |
| 5 | c | 1.078E+04 | 607.3 | 0.001164 | 1.916 | +1 | 5 |
| - | - | 3440 | 608.3 | - | - | 0 | - |
| - | - | 2750 | 613.3 | - | - | 0 | - |
| 3 | z | 5.08E+04 | 619.3 | 0.004885 | 7.888 | +1 | 5 |
| - | - | 1.778E+04 | 620.3 | - | - | 0 | - |
| - | - | 8766 | 621.3 | - | - | 0 | - |
| - | - | 3444 | 622.3 | - | - | 0 | - |
| - | - | 3681 | 630.4 | - | - | 0 | - |
| 3 | y | 5.573E+05 | 635.3 | 0.004899 | 7.711 | +1 | 5 |
| - | - | 1.771E+05 | 636.4 | - | - | 0 | - |
| - | - | 5.265E+04 | 637.4 | - | - | 0 | - |
| - | - | 4757 | 638.4 | - | - | 0 | - |
| - | - | 3852 | 643.4 | - | - | 0 | - |
| - | - | 4949 | 645.3 | - | - | 0 | - |
| - | - | 2677 | 656.4 | - | - | 0 | - |
| - | - | 5847 | 657.4 | - | - | 0 | - |
| - | - | 1.397E+04 | 658.4 | - | - | 0 | - |
| - | - | 4787 | 659.3 | - | - | 0 | - |
| - | - | 4755 | 659.4 | - | - | 0 | - |
| - | - | 3.984E+04 | 662.3 | - | - | 0 | - |
| - | - | 1.622E+04 | 663.3 | - | - | 0 | - |
| - | - | 8.704E+04 | 664.3 | - | - | 0 | - |
| - | - | 2.816E+04 | 665.3 | - | - | 0 | - |
| - | - | 7676 | 666.3 | - | - | 0 | - |
| - | - | 7952 | 673.4 | - | - | 0 | - |
| - | - | 4455 | 674.4 | - | - | 0 | - |
| - | - | 4101 | 675.4 | - | - | 0 | - |
| - | - | 2964 | 676.4 | - | - | 0 | - |
| - | - | 3.076E+04 | 677.3 | - | - | 0 | - |
| - | - | 1.572E+04 | 677.4 | - | - | 0 | - |
| - | - | 8277 | 678.3 | - | - | 0 | - |
| - | - | 6239 | 678.4 | - | - | 0 | - |
| - | - | 8447 | 682.4 | - | - | 0 | - |
| - | - | 2991 | 683.4 | - | - | 0 | - |
| - | - | 1.611E+04 | 693.3 | - | - | 0 | - |
| 6 | c | 9.897E+04 | 694.3 | 0.0003244 | 0.4672 | +1 | 6 |
| - | - | 3.604E+04 | 695.3 | - | - | 0 | - |
| - | - | 8954 | 696.3 | - | - | 0 | - |
| - | - | 2.482E+04 | 701.4 | - | - | 0 | - |
| - | - | 8939 | 702.4 | - | - | 0 | - |
| 2 | w | 1.388E+04 | 703.4 | 0.001878 | 2.669 | +1 | 6 |
| - | - | 1.44E+04 | 704.4 | - | - | 0 | - |
| 2 | w | 2.907E+04 | 705.4 | 0.008819 | 12.5 | +1 | 6 |
| - | - | 9287 | 706.4 | - | - | 0 | - |
| - | - | 7423 | 717.4 | - | - | 0 | - |
| - | - | 8.624E+04 | 718.4 | - | - | 0 | - |
| 2 | y | 2.856E+05 | 719.4 | 0.004339 | 6.031 | +1 | 6 |
| 2 | z | 3.873E+05 | 720.4 | 0.003594 | 4.988 | +1 | 6 |
| - | - | 1.739E+05 | 721.4 | - | - | 0 | - |
| - | - | 6.443E+04 | 722.4 | - | - | 0 | - |
| - | - | 1.654E+04 | 723.4 | - | - | 0 | - |
| - | - | 2743 | 727.4 | - | - | 0 | - |
| - | - | 3104 | 728.4 | - | - | 0 | - |
| - | - | 5.573E+04 | 728.4 | - | - | 0 | - |
| - | - | 6136 | 729.4 | - | - | 0 | - |
| - | - | 3.892E+04 | 729.4 | - | - | 0 | - |
| - | - | 1.393E+04 | 730.4 | - | - | 0 | - |
| - | - | 6738 | 732.4 | - | - | 0 | - |
| - | - | 3171 | 733.4 | - | - | 0 | - |
| - | - | 1.851E+05 | 735.4 | - | - | 0 | - |
| - | - | 5.766E+04 | 736.4 | - | - | 0 | - |
| - | - | 2.132E+04 | 737.4 | - | - | 0 | - |
| - | - | 3854 | 743.4 | - | - | 0 | - |
| - | - | 1.219E+04 | 745.4 | - | - | 0 | - |
| - | - | 8.124E+04 | 746.4 | - | - | 0 | - |
| - | - | 4.147E+04 | 747.4 | - | - | 0 | - |
| - | - | 4.896E+04 | 748.4 | - | - | 0 | - |
| - | - | 1.712E+04 | 749.4 | - | - | 0 | - |
| - | - | 3521 | 750.4 | - | - | 0 | - |
| - | - | 2.88E+04 | 751.3 | - | - | 0 | - |
| - | - | 1.116E+04 | 752.3 | - | - | 0 | - |
| - | - | 2726 | 753.3 | - | - | 0 | - |
| - | - | 3071 | 756.4 | - | - | 0 | - |
| - | - | 4081 | 758.4 | - | - | 0 | - |
| - | - | 3.772E+04 | 760.4 | - | - | 0 | - |
| - | - | 5216 | 761.3 | - | - | 0 | - |
| - | - | 1.203E+04 | 761.4 | - | - | 0 | - |
| - | - | 9635 | 762.4 | - | - | 0 | - |
| - | - | 8460 | 763.4 | - | - | 0 | - |
| - | - | 4187 | 764.4 | - | - | 0 | - |
| - | - | 2.275E+04 | 766.4 | - | - | 0 | - |
| - | - | 6975 | 767.4 | - | - | 0 | - |
| - | - | 5.452E+04 | 772.4 | - | - | 0 | - |
| - | - | 3441 | 773.3 | - | - | 0 | - |
| - | - | 1.268E+05 | 773.4 | - | - | 0 | - |
| - | - | 4.749E+04 | 774.4 | - | - | 0 | - |
| - | - | 8935 | 775.4 | - | - | 0 | - |
| - | - | 3.831E+04 | 776.4 | - | - | 0 | - |
| - | - | 1.408E+04 | 777.4 | - | - | 0 | - |
| - | - | 1.155E+05 | 778.4 | - | - | 0 | - |
| - | - | 1.518E+05 | 779.4 | - | - | 0 | - |
| - | - | 5.116E+04 | 780.4 | - | - | 0 | - |
| - | - | 1.704E+04 | 780.4 | - | - | 0 | - |
| - | - | 1.387E+04 | 781.3 | - | - | 0 | - |
| - | - | 6430 | 781.4 | - | - | 0 | - |
| - | - | 5.181E+04 | 788.4 | - | - | 0 | - |
| - | - | 3.385E+04 | 789.4 | - | - | 0 | - |
| - | - | 1.16E+05 | 790.4 | - | - | 0 | - |
| - | - | 2.002E+06 | 791.4 | - | - | 0 | - |
| - | - | 1.299E+06 | 792.4 | - | - | 0 | - |
| - | - | 4.602E+05 | 793.4 | - | - | 0 | - |
| - | - | 1.09E+05 | 794.4 | - | - | 0 | - |
| - | - | 1.405E+04 | 795.4 | - | - | 0 | - |
| - | - | 1.306E+05 | 796.4 | - | - | 0 | - |
| - | - | 4.659E+04 | 797.4 | - | - | 0 | - |
| - | - | 1.873E+04 | 798.4 | - | - | 0 | - |
| - | - | 4402 | 806.4 | - | - | 0 | - |
| - | - | 2.88E+04 | 806.4 | - | - | 0 | - |
| - | - | 1.614E+04 | 807.4 | - | - | 0 | - |
| - | - | 1.999E+04 | 808.4 | - | - | 0 | - |
| - | - | 5.669E+05 | 809.4 | - | - | 0 | - |
| - | - | 2.255E+05 | 810.4 | - | - | 0 | - |
| - | - | 7.657E+04 | 811.4 | - | - | 0 | - |
| - | - | 4762 | 812.4 | - | - | 0 | - |
| - | - | 1.011E+05 | 817.4 | - | - | 0 | - |
| - | - | 4.882E+04 | 818.4 | - | - | 0 | - |
| - | - | 1.522E+04 | 819.4 | - | - | 0 | - |
| - | - | 1.891E+04 | 820.4 | - | - | 0 | - |
| - | - | 7027 | 821.4 | - | - | 0 | - |
| - | - | 3900 | 822.4 | - | - | 0 | - |
| - | - | 2.424E+04 | 823.4 | - | - | 0 | - |
| - | - | 4.39E+04 | 824.4 | - | - | 0 | - |
| - | - | 1.222E+04 | 825.4 | - | - | 0 | - |
| - | - | 3619 | 826.4 | - | - | 0 | - |
| - | - | 6.887E+04 | 834.4 | - | - | 0 | - |
| - | - | 2.026E+06 | 835.4 | - | - | 0 | - |
| - | - | 8.015E+05 | 836.4 | - | - | 0 | - |
| - | - | 3.63E+05 | 837.4 | - | - | 0 | - |
| - | - | 5.618E+04 | 838.4 | - | - | 0 | - |
| - | - | 7338 | 839.4 | - | - | 0 | - |
| - | - | 8.544E+05 | 851.4 | - | - | 0 | - |
| - | - | 2.879E+06 | 852.4 | - | - | 0 | - |
| - | - | 1.126E+06 | 853.4 | - | - | 0 | - |
| - | - | 3.971E+05 | 854.4 | - | - | 0 | - |
| - | - | 3.72E+04 | 855.4 | - | - | 0 | - |
| - | - | 3780 | 867.4 | - | - | 0 | - |
| - | - | 4134 | 884.4 | - | - | 0 | - |
| - | - | 3623 | 885.4 | - | - | 0 | - |
| - | - | 2900 | 1062 | - | - | 0 | - |
| - | - | 2796 | 1103 | - | - | 0 | - |
| - | - | 3014 | 1156 | - | - | 0 | - |
| - | - | 2858 | 1245 | - | - | 0 | - |
| - | - | 2622 | 1385 | - | - | 0 | - |
| - | - | 3165 | 1857 | - | - | 0 | - |
| - | - | 2932 | 1907 | - | - | 0 | - |
| - | - | 2916 | 3129 | - | - | 0 | - |

m/z Charge Intensity FragmentType MassShift Position
126.05525970458984 0 6716.698
129.06602478027344 0 4918.268
144.06585693359375 0 85650.07
145.0691680908203 0 4450.08
153.0659637451172 0 3482.8591
154.05020141601562 0 2293.8674
158.09278869628906 0 7602.161 y Ammonia loss 6
159.1005859375 0 178821.72 z 6
160.1083984375 0 307797.38
161.1053924560547 0 2624.2024
161.1117706298828 0 16129.063
166.0532989501953 0 2673.0784
169.13365173339844 0 4558.8203
170.09254455566406 0 9959.388
171.07679748535156 0 27244.688
173.4387969970703 0 7070.6904
174.1111297607422 0 4448.134
175.11927795410156 0 75434.73 y 6
176.1228790283203 0 4166.9507
180.10223388671875 0 34638.598
181.1061248779297 0 3284.157
185.14007568359375 0 8820.0205
187.14443969726562 0 32778.543
188.09205627441406 0 2932.4858
188.14756774902344 0 3532.8662
189.0873260498047 0 319234.06
190.09078979492188 0 23175.418
197.12884521484375 0 21465.078
199.0716094970703 0 68969.08
200.07504272460938 0 5297.523
200.12698364257812 0 32151.684
201.1351318359375 0 4200.874
202.14247131347656 0 9064.19
203.1142578125 0 2981.7637
203.15036010742188 0 8586.217
215.13925170898438 0 21824.502
216.0985107421875 0 2947.6567 c Water loss 1
217.08218383789062 0 273070.84
218.06854248046875 0 2529.5159
218.08554077148438 0 24934.6
219.08665466308594 0 2309.9155
225.12380981445312 0 8463.343
226.12939453125 0 2223.2415
229.1295928955078 0 2930.1797 w 5
229.63998413085938 0 14820.502
230.1141357421875 0 3988.0872
230.12542724609375 0 4815.1313
230.14093017578125 0 2316.2695
232.11483764648438 0 2072.0398
232.66429138183594 0 2674.0298
233.13221740722656 0 9424.786
240.1345672607422 0 3902.2742
244.10084533691406 0 12810.11
244.11695861816406 0 22392.518
244.14059448242188 0 2915.7861 y Water loss 5
245.12481689453125 0 171758.95 y Ammonia loss 5
246.13214111328125 0 90751.805 z 5
247.14044189453125 0 545336.6
248.14370727539062 0 50040.656
249.1450653076172 0 4835.979
251.1021728515625 0 2589.5903
261.1269836425781 0 41243.176
261.1434326171875 0 32065.863
261.6390380859375 0 35668.46 y 3
262.1512451171875 0 312855.2 y 5
262.63739013671875 0 3924.357
263.0709533691406 0 3285.0996
263.15411376953125 0 25547.049
266.1501770019531 0 7277.245
267.1330871582031 0 3326.3037
268.12908935546875 0 2401.0508
270.1199645996094 0 15744.45
284.1609191894531 0 82535.21
284.1779479980469 0 4544.5786
285.1453857421875 0 3048.733
285.1644592285156 0 7803.8496
286.1820983886719 0 55112.223
286.6839294433594 0 9898.955
287.18414306640625 0 3346.946
288.13018798828125 0 3671.6113
289.1389465332031 0 4719.3
290.14630126953125 0 36561.258
294.1446228027344 0 5123.8857
295.1295471191406 0 3499.476
299.17181396484375 0 5618.041
300.1246643066406 0 2505.1519
302.1709289550781 0 7008.792
309.176025390625 0 3644.8674 y Water loss 2
312.15582275390625 0 201347.81
313.15875244140625 0 25889.184
318.181396484375 0 364888.8 y 2
318.6827392578125 0 104740.805
319.18243408203125 0 33259.258
319.6842041015625 0 3717.5056
330.1768798828125 0 107633.52 w 4
331.16766357421875 0 6935.88
331.1855773925781 0 12991.542
332.5271301269531 0 2504.2883
336.7062683105469 0 14939.342
337.2071228027344 0 3787.1055
341.69854736328125 0 9694.568
344.1936950683594 0 11619.418 w 4
346.1439208984375 0 9131.558
354.6871643066406 0 4317.357
355.69256591796875 0 3068.8381
357.2249755859375 0 9519.307 y Water loss 4
358.2087707519531 0 8074.5684 y Ammonia loss 4
359.216552734375 0 467683.72 z 4
359.6993713378906 0 11086.885 y Water loss 1
360.1753234863281 0 7533.857
360.2228088378906 0 182709.84
361.2257080078125 0 28025.902
362.1744689941406 0 2860.54
362.2080383300781 0 3518.7427
363.19940185546875 0 5061.3765
364.11968994140625 0 3187.2883
364.6910705566406 0 8426.868
365.1918640136719 0 5402.8413
367.6962585449219 0 4135.7095
368.70477294921875 0 65834.766 y 1
369.2059631347656 0 19939.25
369.70501708984375 0 6841.885
372.1588439941406 0 17885.295
373.2201232910156 0 4901.49
373.6971740722656 0 18867.027
374.2002868652344 0 5570.6235
374.227294921875 0 36726.785
375.2351989746094 0 476236.34 y 4
376.1549072265625 0 4001.3457
376.20916748046875 0 5323.3657
376.2381286621094 0 79905.13
376.70758056640625 0 3797.9614
377.2411804199219 0 12627.299
385.21392822265625 0 70455.516
385.715087890625 0 27381.447
386.2156066894531 0 6228.373
386.6876525878906 0 7459.9253
387.1993713378906 0 4714.3174
388.20660400390625 0 25809.932
389.2071533203125 0 4111.0674
390.6891174316406 0 7365.933
394.2177734375 0 121066.67
394.71905517578125 0 45531.613
395.2201232910156 0 13340.331
399.2028503417969 0 3927.4128
399.6949462890625 0 22524.4
400.19744873046875 0 9359.782
401.2146301269531 0 16840.037
408.2082214355469 0 9214.436
408.7027282714844 0 9738.499
409.20111083984375 0 3677.6
409.70135498046875 0 3559.6304
413.2041015625 0 5859.97
415.1621398925781 0 2517.864
415.2304992675781 0 44889.547
416.2373352050781 0 37706.51
417.21331787109375 0 230426.03
417.71453857421875 0 92059.54
418.2147521972656 0 29373.576
418.7131652832031 0 2845.673
424.1544189453125 0 9370.841
426.21875 0 46939.887
426.71990966796875 0 16233.422
427.2196350097656 0 7739.443
429.2466125488281 0 10634.121
430.24761962890625 0 2937.575
431.24932861328125 0 13896.491
441.2459716796875 0 9099.153
442.1631164550781 0 4502.766
442.2538757324219 0 37391.12
443.2615051269531 0 122461.88
444.2687072753906 0 117395.31
445.2724914550781 0 22202.867
446.27374267578125 0 4151.0054
450.18896484375 0 5736.7544
458.2726135253906 0 186473.03
459.1912841796875 0 45285.137
459.2752685546875 0 43722.3
460.1938171386719 0 8351.384
460.2772521972656 0 5003.1943
477.2032165527344 0 86382.03
478.2065734863281 0 16162.663
479.20660400390625 0 5133.0723
504.2597351074219 0 9216.03 y Water loss 3
505.2461242675781 0 6510.7573 y Ammonia loss 3
506.25225830078125 0 19539.154 z 3
507.25390625 0 11461.431
508.2658996582031 0 2924.6628
521.262451171875 0 3719.4387
522.2706909179688 0 816099.44 y 3
523.2733154296875 0 194836
524.2713012695312 0 49994.664
525.273681640625 0 4031.3877
526.2847290039062 0 4420.1577
529.3189086914062 0 2351.5886
532.253173828125 0 5321.551
544.3344116210938 0 5331.3477
553.3523559570312 0 2773.9856
556.3435668945312 0 7638.5703
557.3489990234375 0 2925.9326
562.2894287109375 0 6413.4673
563.2985229492188 0 2795.7117
571.3563232421875 0 175564.73
572.274169921875 0 6336.378
572.3589477539062 0 48832.305
573.36181640625 0 9540.632
576.2809448242188 0 130201.96 w 2
577.2837524414062 0 33378.082
578.2819213867188 0 11329.489
590.2855224609375 0 19060.32
591.288330078125 0 3500.8916
607.312744140625 0 10776.584 c 4
608.315185546875 0 3439.701
613.3198852539062 0 2749.5867
619.3357543945312 0 50798.906 z 2
620.3370361328125 0 17784.975
621.3413696289062 0 8765.559
622.3495483398438 0 3444.3635
630.3687744140625 0 3680.8057
635.3544921875 0 557279.75 y 2
636.3572998046875 0 177095.03
637.3563232421875 0 52650.566
638.3541259765625 0 4756.8853
643.3760986328125 0 3851.9868
645.3405151367188 0 4948.8115
656.3814697265625 0 2676.9207
657.3928833007812 0 5847.1733
658.39990234375 0 13969.757
659.3075561523438 0 4786.581
659.4029541015625 0 4754.7036
662.305419921875 0 39841.484
663.3079833984375 0 16215.477
664.3204345703125 0 87035.59
665.3239135742188 0 28162.422
666.3211059570312 0 7675.802
673.3764038085938 0 7952.2227
674.3755493164062 0 4455.2188
675.3726196289062 0 4101.244
676.3838500976562 0 2963.8228
677.3240966796875 0 30759.639
677.3795166015625 0 15723.29
678.3279418945312 0 8276.924
678.3831787109375 0 6239.4473
682.3871459960938 0 8447.35
683.3945922851562 0 2991.0771
693.3353271484375 0 16110.303
694.3439331054688 0 98972.92 c 5
695.3458251953125 0 36044.953
696.3463745117188 0 8953.554
701.4057006835938 0 24821.99
702.40673828125 0 8939.439
703.377685546875 0 13877.109 w 1
704.374755859375 0 14401.17
705.3638916015625 0 29065.37 w 1
706.3701782226562 0 9287.113
717.3546142578125 0 7423.4976
718.3676147460938 0 86236.91
719.3750610351562 0 285575.75 y Ammonia loss 1
720.3821411132812 0 387293.12 z 1
721.3856811523438 0 173917.8
722.3904418945312 0 64429.21
723.395751953125 0 16540.312
727.4136962890625 0 2742.8325
728.3740844726562 0 3104.0435
728.4307250976562 0 55731.58
729.3616943359375 0 6135.858
729.4338989257812 0 38920.62
730.4351196289062 0 13930.039
732.3742065429688 0 6737.8325
733.3817749023438 0 3170.554
735.3576049804688 0 185090.36
736.3607177734375 0 57662.977
737.3580932617188 0 21319.793
743.4496459960938 0 3853.704
745.3958740234375 0 12188.909
746.3870849609375 0 81237.77
747.3922119140625 0 41473.016
748.4116821289062 0 48956.32
749.4173583984375 0 17120.78
750.3636474609375 0 3520.9373
751.3419799804688 0 28796.074
752.3475341796875 0 11155.16
753.3438720703125 0 2726.3315
756.3829956054688 0 3071.499
758.3920288085938 0 4080.6042
760.4074096679688 0 37718.758
761.3353881835938 0 5215.894
761.40869140625 0 12031.781
762.4208374023438 0 9634.892
763.3909301757812 0 8459.916
764.3851928710938 0 4187.3164
766.4112548828125 0 22748.25
767.412109375 0 6975.218
772.4198608398438 0 54518.574
773.288330078125 0 3440.9058
773.4137573242188 0 126833.74
774.4154052734375 0 47494.973
775.41455078125 0 8934.78
776.3967895507812 0 38312.812
777.3984375 0 14079.694
778.3650512695312 0 115546.06
779.3511352539062 0 151785.08
780.352783203125 0 51159.1
780.4178466796875 0 17038.611
781.3485107421875 0 13871.057
781.4197998046875 0 6429.8516
788.43798828125 0 51807.5
789.443603515625 0 33846.426
790.4158935546875 0 115969.1
791.4205322265625 0 2001721.1
792.4196166992188 0 1299047.1
793.4175415039062 0 460168.38
794.41357421875 0 109022.95
795.39794921875 0 14046.909
796.3743286132812 0 130600.27
797.377197265625 0 46589.242
798.3779907226562 0 18733.594
806.3602905273438 0 4402.2153
806.431396484375 0 28798.998
807.4346923828125 0 16135.198
808.4359130859375 0 19990.75
809.40673828125 0 566882.9
810.4096069335938 0 225462.05
811.4091796875 0 76567.34
812.405029296875 0 4761.8926
817.3995971679688 0 101053.79
818.4026489257812 0 48816.633
819.4024658203125 0 15219.612
820.3887329101562 0 18909.738
821.38916015625 0 7026.784
822.426513671875 0 3900.1511
823.408203125 0 24244.135
824.4388427734375 0 43899.457
825.4405517578125 0 12217.84
826.4431762695312 0 3618.83
834.424072265625 0 68874.555
835.4155883789062 0 2025827.9
836.4181518554688 0 801465.2
837.416748046875 0 363045.12
838.4180908203125 0 56177.133
839.4176635742188 0 7337.839
851.4285888671875 0 854370
852.4359130859375 0 2879088
853.4380493164062 0 1125854
854.4388427734375 0 397079
855.4404907226562 0 37196.22
867.397705078125 0 3780.4937
884.42431640625 0 4133.5566
885.4332885742188 0 3623.0273
1061.6588134765625 0 2900.3237
1103.3717041015625 0 2795.6765
1156.032470703125 0 3013.9404
1244.7659912109375 0 2857.6226
1385.3475341796875 0 2622.3977
1856.98974609375 0 3164.5625
1907.3541259765625 0 2931.8762
3128.949462890625 0 2916.4387

Spectrum Details

|  |  |
| --- | --- |
| Matched peaks? Matched peaks The total absolute number of peaks matched. Additionally in brackets the total fraction of peaks matched and the total number of peaks is shown. | 33 (9.24% of 357) |
| FDR? FDR The false discovery rate estimated for this peptide. It is calculated by matching all theoretical fragments with a non-integer shift with the raw peaks for this spectrum. This is done with 40 different shifts. The resulting percentage is the average number of annotated peaks over the number of annotated peaks with the correct spectrum. | 0.00% |
| Satellite FDR? Satellite FDR See the FDR for details on its calculation. This satellite ion specific FDR only contains the satellite ions (d/w) for I/L/J positions. | 0.00% |
| PSM Score? PSM Score The PSM Score as given by Hecklib to this annotated spectrum. It is shown with three significant figures. | 318 |

#### Spectrum 3823? Spectrum 3823 The raw spectrum of this peptide as annotated by Hecklib. The fragments are coloured according to ion type (see legend). Any peaks with a star '\*' as text can be hovered over to see the full details, first the ion type second the mass shift type. By hovering over the amino acids in the peptide or ions in the legend the corresponding peaks are highlighted. By toggling the 'Unassigned' label you can turn the background (unassigned) peaks on or off in the plot. By updating the slider in the Ion legend you can update the spectrum to only show the top X% of the peaks with labels. The top X% means any peak that is within X% of the highest intensity. By dragging in the spectrum you can zoom in to a specific part of the spectrum and use 'Zoom Out' to get back to the original zoom level. The annotation of the spectrum is based on the given sequence in the peptides file and is done with different software so inconsistencies are likely. The peaks are annotated based on the given sequence, with 20 ppm tolerance.

Copy Data

##### Spectrum 3823 (TSV)

###### Preview

```
Loading example...
```

*Click on the button to copy the data to your clipboard.*

Mz MinMz MaxIntensity Max

WidthHeightPeptide font sizePeptide stroke widthSpectrum font sizeSpectrum stroke widthCompact peptide

Ion legend

wxyz

abcd

OtherUnassignedIon^Charge^_Position_Show for top:%

GTJMISR

06.58e+61.32e+71.97e+72.63e+7

Zoom Out

y^+1^_1_z^+1^_1_y^+1^_1_y^+1^_2_y^+1^_2_z^+1^_2_y^+2^_4_y^+1^_2_y^+2^_5_w^+1^_3_w^+1^_3_z^+2^_6_y^+1^_3_y^+1^_3_z^+1^_3_y^+2^_6_y^+2^_6_y^+1^_3_y^+1^_4_y^+1^_4_z^+1^_4_y^+1^_4_w^+1^_5_c^+1^_5_y^+1^_5_y^+1^_5_z^+1^_5_y^+1^_5_c^+1^_6_w^+1^_6_w^+1^_6_y^+1^_6_z^+1^_6_

0779155823373116

Fragment Matches Table

Show background peaks

| Position | Ion type | Intensity | mz Theoretical | mz Error (Th) | mz Error (ppm) | Charge | Series Number |
| --- | --- | --- | --- | --- | --- | --- | --- |
| - | - | 6.734E+04 | 126.1 | - | - | 0 | - |
| - | - | 6.276E+04 | 129.1 | - | - | 0 | - |
| - | - | 8.064E+05 | 144.1 | - | - | 0 | - |
| - | - | 4.913E+04 | 145.1 | - | - | 0 | - |
| - | - | 1.994E+04 | 149 | - | - | 0 | - |
| - | - | 4.825E+04 | 153.1 | - | - | 0 | - |
| - | - | 1.392E+04 | 156.7 | - | - | 0 | - |
| 7 | y | 4.433E+04 | 158.1 | 0.0002788 | 1.764 | +1 | 1 |
| - | - | 3.579E+04 | 159.1 | - | - | 0 | - |
| 7 | z | 1.683E+06 | 159.1 | 0.0004341 | 2.729 | +1 | 1 |
| - | - | 2.727E+06 | 160.1 | - | - | 0 | - |
| - | - | 2.098E+04 | 161.1 | - | - | 0 | - |
| - | - | 1.42E+05 | 161.1 | - | - | 0 | - |
| - | - | 2.014E+04 | 166.1 | - | - | 0 | - |
| - | - | 2.056E+04 | 168.1 | - | - | 0 | - |
| - | - | 6.039E+04 | 169.1 | - | - | 0 | - |
| - | - | 9.121E+04 | 170.1 | - | - | 0 | - |
| - | - | 2.903E+05 | 171.1 | - | - | 0 | - |
| - | - | 3.048E+04 | 172.1 | - | - | 0 | - |
| - | - | 2.482E+04 | 172.1 | - | - | 0 | - |
| - | - | 1.764E+04 | 173.1 | - | - | 0 | - |
| - | - | 1.631E+04 | 173.4 | - | - | 0 | - |
| - | - | 5.693E+04 | 174.1 | - | - | 0 | - |
| 7 | y | 7.874E+05 | 175.1 | 0.0003868 | 2.209 | +1 | 1 |
| - | - | 2.024E+04 | 176.1 | - | - | 0 | - |
| - | - | 3.694E+05 | 180.1 | - | - | 0 | - |
| - | - | 1.575E+04 | 183.1 | - | - | 0 | - |
| - | - | 8.763E+04 | 185.1 | - | - | 0 | - |
| - | - | 3.551E+05 | 187.1 | - | - | 0 | - |
| - | - | 2.233E+04 | 188.1 | - | - | 0 | - |
| - | - | 3.663E+04 | 188.1 | - | - | 0 | - |
| - | - | 3.161E+06 | 189.1 | - | - | 0 | - |
| - | - | 1.854E+05 | 190.1 | - | - | 0 | - |
| - | - | 1.738E+05 | 197.1 | - | - | 0 | - |
| - | - | 6.623E+05 | 199.1 | - | - | 0 | - |
| - | - | 5.445E+04 | 200.1 | - | - | 0 | - |
| - | - | 3.13E+05 | 200.1 | - | - | 0 | - |
| - | - | 5.904E+04 | 201.1 | - | - | 0 | - |
| - | - | 9.282E+04 | 202.1 | - | - | 0 | - |
| - | - | 3.813E+04 | 203.1 | - | - | 0 | - |
| - | - | 5.993E+04 | 203.2 | - | - | 0 | - |
| - | - | 3.55E+04 | 207.1 | - | - | 0 | - |
| - | - | 2.367E+05 | 215.1 | - | - | 0 | - |
| - | - | 2.814E+06 | 217.1 | - | - | 0 | - |
| - | - | 2.192E+05 | 218.1 | - | - | 0 | - |
| - | - | 2.23E+04 | 219.1 | - | - | 0 | - |
| - | - | 1.844E+04 | 222.4 | - | - | 0 | - |
| - | - | 1.761E+04 | 225.1 | - | - | 0 | - |
| - | - | 8.516E+04 | 225.1 | - | - | 0 | - |
| - | - | 1.285E+05 | 229.6 | - | - | 0 | - |
| - | - | 1.94E+04 | 230.1 | - | - | 0 | - |
| - | - | 4.134E+04 | 230.1 | - | - | 0 | - |
| - | - | 4.16E+04 | 233.1 | - | - | 0 | - |
| - | - | 1.651E+04 | 233.2 | - | - | 0 | - |
| - | - | 1.743E+04 | 234.9 | - | - | 0 | - |
| - | - | 2.284E+04 | 240.1 | - | - | 0 | - |
| - | - | 8.994E+04 | 244.1 | - | - | 0 | - |
| - | - | 2.005E+05 | 244.1 | - | - | 0 | - |
| 6 | y | 2.042E+04 | 244.1 | 0.0002792 | 1.143 | +1 | 2 |
| 6 | y | 1.603E+06 | 245.1 | 0.000538 | 2.195 | +1 | 2 |
| 6 | z | 8.149E+05 | 246.1 | 9.823E-05 | 0.3991 | +1 | 2 |
| - | - | 4.985E+06 | 247.1 | - | - | 0 | - |
| - | - | 4.917E+05 | 248.1 | - | - | 0 | - |
| - | - | 1.962E+04 | 249.1 | - | - | 0 | - |
| - | - | 3.936E+04 | 249.1 | - | - | 0 | - |
| - | - | 2.811E+04 | 258.1 | - | - | 0 | - |
| - | - | 4.224E+05 | 261.1 | - | - | 0 | - |
| - | - | 2.988E+05 | 261.1 | - | - | 0 | - |
| 4 | y | 2.893E+05 | 261.6 | 0.002879 | 11.01 | +2 | 4 |
| 6 | y | 2.884E+06 | 262.2 | 0.0003866 | 1.475 | +1 | 2 |
| - | - | 2.974E+05 | 263.2 | - | - | 0 | - |
| - | - | 5.323E+04 | 264.2 | - | - | 0 | - |
| - | - | 3.944E+04 | 266.1 | - | - | 0 | - |
| - | - | 6.979E+04 | 267.1 | - | - | 0 | - |
| - | - | 2.097E+04 | 267.3 | - | - | 0 | - |
| - | - | 1.4E+05 | 270.1 | - | - | 0 | - |
| - | - | 1.895E+04 | 271.1 | - | - | 0 | - |
| - | - | 2.64E+04 | 275.1 | - | - | 0 | - |
| - | - | 2.143E+04 | 277.1 | - | - | 0 | - |
| - | - | 7.312E+05 | 284.2 | - | - | 0 | - |
| - | - | 3.696E+04 | 285.1 | - | - | 0 | - |
| - | - | 7.482E+04 | 285.2 | - | - | 0 | - |
| - | - | 5.389E+04 | 286.1 | - | - | 0 | - |
| - | - | 6.218E+05 | 286.2 | - | - | 0 | - |
| - | - | 1.477E+05 | 286.7 | - | - | 0 | - |
| - | - | 3.445E+04 | 288.1 | - | - | 0 | - |
| - | - | 4.56E+04 | 289.1 | - | - | 0 | - |
| - | - | 3.549E+05 | 290.1 | - | - | 0 | - |
| - | - | 4.377E+04 | 291.1 | - | - | 0 | - |
| - | - | 6.121E+04 | 294.1 | - | - | 0 | - |
| - | - | 2.881E+04 | 295.1 | - | - | 0 | - |
| - | - | 5.764E+04 | 299.2 | - | - | 0 | - |
| - | - | 2.153E+04 | 299.2 | - | - | 0 | - |
| - | - | 2.094E+04 | 299.7 | - | - | 0 | - |
| - | - | 1.818E+04 | 302.1 | - | - | 0 | - |
| - | - | 1.002E+05 | 302.2 | - | - | 0 | - |
| - | - | 1.975E+06 | 312.2 | - | - | 0 | - |
| - | - | 2.548E+05 | 313.2 | - | - | 0 | - |
| - | - | 3.543E+04 | 314.2 | - | - | 0 | - |
| 3 | y | 3.723E+06 | 318.2 | 0.003145 | 9.884 | +2 | 5 |
| - | - | 1.083E+06 | 318.7 | - | - | 0 | - |
| - | - | 3.767E+05 | 319.2 | - | - | 0 | - |
| - | - | 7.308E+04 | 319.7 | - | - | 0 | - |
| - | - | 3.369E+04 | 327.7 | - | - | 0 | - |
| 5 | w | 1.098E+06 | 330.2 | 7.13E-05 | 0.216 | +1 | 3 |
| - | - | 1.333E+05 | 331.2 | - | - | 0 | - |
| - | - | 4.203E+04 | 332.7 | - | - | 0 | - |
| - | - | 1.556E+05 | 336.7 | - | - | 0 | - |
| - | - | 5.3E+04 | 337.2 | - | - | 0 | - |
| - | - | 9.886E+04 | 341.7 | - | - | 0 | - |
| 5 | w | 1.202E+05 | 344.2 | 0.0003919 | 1.139 | +1 | 3 |
| - | - | 9.455E+04 | 346.1 | - | - | 0 | - |
| - | - | 2.264E+04 | 347.1 | - | - | 0 | - |
| 2 | z | 5.862E+04 | 351.7 | 0.003181 | 9.046 | +2 | 6 |
| - | - | 3.765E+04 | 354.7 | - | - | 0 | - |
| 5 | y | 7.067E+04 | 357.2 | 0.0007399 | 2.071 | +1 | 3 |
| 5 | y | 9.202E+04 | 358.2 | 0.0005194 | 1.45 | +1 | 3 |
| 5 | z | 4.25E+06 | 359.2 | 0.0004154 | 1.156 | +1 | 3 |
| 2 | y | 9.166E+04 | 359.7 | 0.003204 | 8.906 | +2 | 6 |
| - | - | 5.166E+04 | 360.2 | - | - | 0 | - |
| - | - | 1.666E+06 | 360.2 | - | - | 0 | - |
| - | - | 3.113E+05 | 361.2 | - | - | 0 | - |
| - | - | 3.737E+04 | 362.2 | - | - | 0 | - |
| - | - | 7.574E+04 | 362.2 | - | - | 0 | - |
| - | - | 4.087E+04 | 362.7 | - | - | 0 | - |
| - | - | 3.238E+04 | 363.2 | - | - | 0 | - |
| - | - | 1.158E+05 | 364.7 | - | - | 0 | - |
| - | - | 3.458E+04 | 365.2 | - | - | 0 | - |
| - | - | 2.672E+04 | 367.2 | - | - | 0 | - |
| - | - | 4.359E+04 | 367.7 | - | - | 0 | - |
| 2 | y | 5.641E+05 | 368.7 | 0.002835 | 7.688 | +2 | 6 |
| - | - | 1.907E+05 | 369.2 | - | - | 0 | - |
| - | - | 4.62E+04 | 369.7 | - | - | 0 | - |
| - | - | 1.644E+05 | 372.2 | - | - | 0 | - |
| - | - | 2.447E+04 | 373.2 | - | - | 0 | - |
| - | - | 1.899E+05 | 373.7 | - | - | 0 | - |
| - | - | 4.289E+04 | 374.2 | - | - | 0 | - |
| - | - | 3.31E+05 | 374.2 | - | - | 0 | - |
| 5 | y | 4.489E+06 | 375.2 | 0.0004291 | 1.143 | +1 | 3 |
| - | - | 3.884E+04 | 376.2 | - | - | 0 | - |
| - | - | 4.761E+04 | 376.2 | - | - | 0 | - |
| - | - | 7.651E+05 | 376.2 | - | - | 0 | - |
| - | - | 2.537E+04 | 376.7 | - | - | 0 | - |
| - | - | 9.958E+04 | 377.2 | - | - | 0 | - |
| - | - | 7.673E+05 | 385.2 | - | - | 0 | - |
| - | - | 2.431E+05 | 385.7 | - | - | 0 | - |
| - | - | 2.883E+04 | 386.2 | - | - | 0 | - |
| - | - | 5.879E+04 | 386.2 | - | - | 0 | - |
| - | - | 5.547E+04 | 386.7 | - | - | 0 | - |
| - | - | 6.063E+04 | 387.2 | - | - | 0 | - |
| - | - | 2.259E+04 | 387.7 | - | - | 0 | - |
| - | - | 2.507E+05 | 388.2 | - | - | 0 | - |
| - | - | 4.866E+04 | 389.2 | - | - | 0 | - |
| - | - | 7.65E+04 | 390.7 | - | - | 0 | - |
| - | - | 1.105E+06 | 394.2 | - | - | 0 | - |
| - | - | 4.796E+05 | 394.7 | - | - | 0 | - |
| - | - | 1.335E+05 | 395.2 | - | - | 0 | - |
| - | - | 3.204E+04 | 399.2 | - | - | 0 | - |
| - | - | 1.734E+05 | 399.7 | - | - | 0 | - |
| - | - | 7.141E+04 | 400.2 | - | - | 0 | - |
| - | - | 1.472E+05 | 401.2 | - | - | 0 | - |
| - | - | 8.403E+04 | 408.2 | - | - | 0 | - |
| - | - | 7.902E+04 | 408.7 | - | - | 0 | - |
| - | - | 3.916E+04 | 409.2 | - | - | 0 | - |
| - | - | 2.532E+04 | 413.2 | - | - | 0 | - |
| - | - | 4.23E+05 | 415.2 | - | - | 0 | - |
| - | - | 4.042E+05 | 416.2 | - | - | 0 | - |
| - | - | 2.386E+06 | 417.2 | - | - | 0 | - |
| - | - | 9.705E+05 | 417.7 | - | - | 0 | - |
| - | - | 3.244E+05 | 418.2 | - | - | 0 | - |
| - | - | 7.916E+04 | 424.2 | - | - | 0 | - |
| - | - | 4.073E+05 | 426.2 | - | - | 0 | - |
| - | - | 2.239E+04 | 426.3 | - | - | 0 | - |
| - | - | 1.636E+05 | 426.7 | - | - | 0 | - |
| - | - | 8.979E+04 | 427.2 | - | - | 0 | - |
| - | - | 1.093E+05 | 429.2 | - | - | 0 | - |
| - | - | 2.663E+04 | 430.2 | - | - | 0 | - |
| - | - | 1.578E+05 | 431.2 | - | - | 0 | - |
| - | - | 2.847E+04 | 432.3 | - | - | 0 | - |
| - | - | 3.315E+04 | 441.2 | - | - | 0 | - |
| - | - | 1.029E+05 | 441.2 | - | - | 0 | - |
| - | - | 2.533E+04 | 442.2 | - | - | 0 | - |
| - | - | 3.611E+05 | 442.3 | - | - | 0 | - |
| - | - | 1.058E+06 | 443.3 | - | - | 0 | - |
| - | - | 1.099E+06 | 444.3 | - | - | 0 | - |
| - | - | 1.927E+05 | 445.3 | - | - | 0 | - |
| - | - | 3.308E+04 | 446.3 | - | - | 0 | - |
| - | - | 3.667E+04 | 450.2 | - | - | 0 | - |
| - | - | 2.007E+06 | 458.3 | - | - | 0 | - |
| - | - | 4.914E+05 | 459.2 | - | - | 0 | - |
| - | - | 4.194E+05 | 459.3 | - | - | 0 | - |
| - | - | 8.608E+04 | 460.2 | - | - | 0 | - |
| - | - | 5.681E+04 | 460.3 | - | - | 0 | - |
| - | - | 2.364E+04 | 461.2 | - | - | 0 | - |
| - | - | 8.707E+05 | 477.2 | - | - | 0 | - |
| - | - | 2.068E+05 | 478.2 | - | - | 0 | - |
| - | - | 6.422E+04 | 479.2 | - | - | 0 | - |
| - | - | 2.888E+04 | 498.3 | - | - | 0 | - |
| 4 | y | 1.174E+05 | 504.3 | 0.00532 | 10.55 | +1 | 4 |
| 4 | y | 3.809E+04 | 505.2 | 0.002872 | 5.684 | +1 | 4 |
| 4 | z | 1.661E+05 | 506.2 | 0.005362 | 10.59 | +1 | 4 |
| - | - | 7.414E+04 | 507.3 | - | - | 0 | - |
| - | - | 2.439E+04 | 508.3 | - | - | 0 | - |
| 4 | y | 7.788E+06 | 522.3 | 0.005528 | 10.58 | +1 | 4 |
| - | - | 1.894E+06 | 523.3 | - | - | 0 | - |
| - | - | 5.943E+05 | 524.3 | - | - | 0 | - |
| - | - | 2.825E+04 | 525.3 | - | - | 0 | - |
| - | - | 4.143E+04 | 526.3 | - | - | 0 | - |
| - | - | 5.363E+04 | 532.3 | - | - | 0 | - |
| - | - | 2.253E+04 | 542.3 | - | - | 0 | - |
| - | - | 4.704E+04 | 544.3 | - | - | 0 | - |
| - | - | 2.164E+04 | 545.3 | - | - | 0 | - |
| - | - | 2.983E+04 | 555.3 | - | - | 0 | - |
| - | - | 6.465E+04 | 556.3 | - | - | 0 | - |
| - | - | 2.843E+04 | 557.3 | - | - | 0 | - |
| - | - | 6.011E+04 | 562.3 | - | - | 0 | - |
| - | - | 3.429E+04 | 563.3 | - | - | 0 | - |
| - | - | 2.454E+04 | 564.3 | - | - | 0 | - |
| - | - | 1.751E+06 | 571.4 | - | - | 0 | - |
| - | - | 6.402E+04 | 572.3 | - | - | 0 | - |
| - | - | 4.701E+05 | 572.4 | - | - | 0 | - |
| - | - | 8.757E+04 | 573.4 | - | - | 0 | - |
| 3 | w | 1.213E+06 | 576.3 | 0.0054 | 9.371 | +1 | 5 |
| - | - | 3.352E+05 | 577.3 | - | - | 0 | - |
| - | - | 8.416E+04 | 578.3 | - | - | 0 | - |
| - | - | 1.494E+05 | 590.3 | - | - | 0 | - |
| - | - | 4.305E+04 | 591.3 | - | - | 0 | - |
| - | - | 2.628E+04 | 602.3 | - | - | 0 | - |
| - | - | 2.385E+04 | 606.3 | - | - | 0 | - |
| 5 | c | 8.015E+04 | 607.3 | 5.688E-05 | 0.09366 | +1 | 5 |
| 3 | y | 5.474E+04 | 617.3 | 0.006369 | 10.32 | +1 | 5 |
| 3 | y | 4.605E+04 | 618.3 | 0.008804 | 14.24 | +1 | 5 |
| 3 | z | 3.507E+05 | 619.3 | 0.005313 | 8.578 | +1 | 5 |
| - | - | 1.687E+05 | 620.3 | - | - | 0 | - |
| - | - | 7.56E+04 | 621.3 | - | - | 0 | - |
| - | - | 5.982E+04 | 622.3 | - | - | 0 | - |
| - | - | 2.637E+04 | 630.4 | - | - | 0 | - |
| - | - | 2.213E+04 | 634.4 | - | - | 0 | - |
| 3 | y | 5.528E+06 | 635.3 | 0.005448 | 8.575 | +1 | 5 |
| - | - | 1.695E+06 | 636.4 | - | - | 0 | - |
| - | - | 5.188E+05 | 637.4 | - | - | 0 | - |
| - | - | 4.731E+04 | 638.4 | - | - | 0 | - |
| - | - | 3.76E+04 | 645.3 | - | - | 0 | - |
| - | - | 2.318E+04 | 647.3 | - | - | 0 | - |
| - | - | 5.538E+04 | 657.4 | - | - | 0 | - |
| - | - | 1.367E+05 | 658.4 | - | - | 0 | - |
| - | - | 4.089E+04 | 659.3 | - | - | 0 | - |
| - | - | 5.838E+04 | 659.4 | - | - | 0 | - |
| - | - | 3.278E+05 | 662.3 | - | - | 0 | - |
| - | - | 2.32E+04 | 662.4 | - | - | 0 | - |
| - | - | 1.168E+05 | 663.3 | - | - | 0 | - |
| - | - | 8.083E+05 | 664.3 | - | - | 0 | - |
| - | - | 2.413E+05 | 665.3 | - | - | 0 | - |
| - | - | 6.032E+04 | 666.3 | - | - | 0 | - |
| - | - | 6.734E+04 | 673.4 | - | - | 0 | - |
| - | - | 4.183E+04 | 674.4 | - | - | 0 | - |
| - | - | 5.129E+04 | 675.4 | - | - | 0 | - |
| - | - | 4.237E+04 | 676.4 | - | - | 0 | - |
| - | - | 2.388E+05 | 677.3 | - | - | 0 | - |
| - | - | 1.699E+05 | 677.4 | - | - | 0 | - |
| - | - | 7.534E+04 | 678.3 | - | - | 0 | - |
| - | - | 6.157E+04 | 678.4 | - | - | 0 | - |
| - | - | 2.932E+04 | 679.3 | - | - | 0 | - |
| - | - | 7.973E+04 | 682.4 | - | - | 0 | - |
| - | - | 4.976E+04 | 683.4 | - | - | 0 | - |
| - | - | 2.41E+04 | 686.4 | - | - | 0 | - |
| - | - | 2.607E+04 | 692.4 | - | - | 0 | - |
| - | - | 1.189E+05 | 693.3 | - | - | 0 | - |
| 6 | c | 8.509E+05 | 694.3 | 0.0004464 | 0.643 | +1 | 6 |
| - | - | 2.919E+05 | 695.3 | - | - | 0 | - |
| - | - | 8.516E+04 | 696.3 | - | - | 0 | - |
| - | - | 1.972E+05 | 701.4 | - | - | 0 | - |
| - | - | 7.96E+04 | 702.4 | - | - | 0 | - |
| 2 | w | 1.277E+05 | 703.4 | 0.004563 | 6.488 | +1 | 6 |
| - | - | 1.093E+05 | 704.4 | - | - | 0 | - |
| 2 | w | 2.84E+05 | 705.4 | 0.009246 | 13.11 | +1 | 6 |
| - | - | 9.016E+04 | 706.4 | - | - | 0 | - |
| - | - | 2.633E+04 | 707.4 | - | - | 0 | - |
| - | - | 6.461E+04 | 717.4 | - | - | 0 | - |
| - | - | 8.09E+05 | 718.4 | - | - | 0 | - |
| 2 | y | 2.481E+06 | 719.4 | 0.004766 | 6.625 | +1 | 6 |
| 2 | z | 3.614E+06 | 720.4 | 0.004082 | 5.666 | +1 | 6 |
| - | - | 1.608E+06 | 721.4 | - | - | 0 | - |
| - | - | 5.909E+05 | 722.4 | - | - | 0 | - |
| - | - | 1.03E+05 | 723.4 | - | - | 0 | - |
| - | - | 2.71E+04 | 724.4 | - | - | 0 | - |
| - | - | 4.06E+04 | 728.4 | - | - | 0 | - |
| - | - | 5.599E+05 | 728.4 | - | - | 0 | - |
| - | - | 3.84E+04 | 729.4 | - | - | 0 | - |
| - | - | 4.351E+05 | 729.4 | - | - | 0 | - |
| - | - | 9.809E+04 | 730.4 | - | - | 0 | - |
| - | - | 9.601E+04 | 732.4 | - | - | 0 | - |
| - | - | 2.381E+04 | 734.4 | - | - | 0 | - |
| - | - | 1.827E+06 | 735.4 | - | - | 0 | - |
| - | - | 5.397E+05 | 736.4 | - | - | 0 | - |
| - | - | 1.722E+05 | 737.4 | - | - | 0 | - |
| - | - | 4.151E+04 | 744.4 | - | - | 0 | - |
| - | - | 7.884E+04 | 745.4 | - | - | 0 | - |
| - | - | 8.997E+05 | 746.4 | - | - | 0 | - |
| - | - | 4.485E+05 | 747.4 | - | - | 0 | - |
| - | - | 4.086E+05 | 748.4 | - | - | 0 | - |
| - | - | 1.589E+05 | 749.4 | - | - | 0 | - |
| - | - | 3.351E+04 | 750.4 | - | - | 0 | - |
| - | - | 2.243E+05 | 751.3 | - | - | 0 | - |
| - | - | 1.288E+05 | 752.3 | - | - | 0 | - |
| - | - | 2.822E+04 | 756.4 | - | - | 0 | - |
| - | - | 3.576E+05 | 760.4 | - | - | 0 | - |
| - | - | 6.915E+04 | 761.3 | - | - | 0 | - |
| - | - | 1.676E+05 | 761.4 | - | - | 0 | - |
| - | - | 5.427E+04 | 762.4 | - | - | 0 | - |
| - | - | 7.495E+04 | 762.4 | - | - | 0 | - |
| - | - | 8.453E+04 | 763.4 | - | - | 0 | - |
| - | - | 2.733E+04 | 764.4 | - | - | 0 | - |
| - | - | 4.396E+04 | 765.4 | - | - | 0 | - |
| - | - | 2.099E+05 | 766.4 | - | - | 0 | - |
| - | - | 8.192E+04 | 767.4 | - | - | 0 | - |
| - | - | 4.341E+05 | 772.4 | - | - | 0 | - |
| - | - | 1.14E+06 | 773.4 | - | - | 0 | - |
| - | - | 4.86E+05 | 774.4 | - | - | 0 | - |
| - | - | 1.593E+05 | 775.4 | - | - | 0 | - |
| - | - | 4.014E+05 | 776.4 | - | - | 0 | - |
| - | - | 1.299E+05 | 777.4 | - | - | 0 | - |
| - | - | 9.466E+05 | 778.4 | - | - | 0 | - |
| - | - | 1.345E+06 | 779.4 | - | - | 0 | - |
| - | - | 4.549E+05 | 780.4 | - | - | 0 | - |
| - | - | 1.529E+05 | 780.4 | - | - | 0 | - |
| - | - | 1.475E+05 | 781.3 | - | - | 0 | - |
| - | - | 5.827E+04 | 781.4 | - | - | 0 | - |
| - | - | 4.612E+04 | 782.4 | - | - | 0 | - |
| - | - | 4.369E+04 | 787.4 | - | - | 0 | - |
| - | - | 2.578E+04 | 788.4 | - | - | 0 | - |
| - | - | 5.445E+05 | 788.4 | - | - | 0 | - |
| - | - | 2.54E+05 | 789.4 | - | - | 0 | - |
| - | - | 1.001E+06 | 790.4 | - | - | 0 | - |
| - | - | 1.823E+07 | 791.4 | - | - | 0 | - |
| - | - | 1.182E+07 | 792.4 | - | - | 0 | - |
| - | - | 4.339E+06 | 793.4 | - | - | 0 | - |
| - | - | 9.207E+05 | 794.4 | - | - | 0 | - |
| - | - | 1.607E+05 | 795.4 | - | - | 0 | - |
| - | - | 1.119E+06 | 796.4 | - | - | 0 | - |
| - | - | 4.752E+05 | 797.4 | - | - | 0 | - |
| - | - | 1.205E+05 | 798.4 | - | - | 0 | - |
| - | - | 4.946E+04 | 806.4 | - | - | 0 | - |
| - | - | 2.591E+05 | 806.4 | - | - | 0 | - |
| - | - | 1.334E+05 | 807.4 | - | - | 0 | - |
| - | - | 1.953E+05 | 808.4 | - | - | 0 | - |
| - | - | 5.559E+06 | 809.4 | - | - | 0 | - |
| - | - | 2.004E+06 | 810.4 | - | - | 0 | - |
| - | - | 8.096E+05 | 811.4 | - | - | 0 | - |
| - | - | 6.194E+04 | 812.4 | - | - | 0 | - |
| - | - | 9.477E+05 | 817.4 | - | - | 0 | - |
| - | - | 4.055E+05 | 818.4 | - | - | 0 | - |
| - | - | 1.404E+05 | 819.4 | - | - | 0 | - |
| - | - | 1.719E+05 | 820.4 | - | - | 0 | - |
| - | - | 6.766E+04 | 821.4 | - | - | 0 | - |
| - | - | 4.653E+04 | 822.4 | - | - | 0 | - |
| - | - | 2.928E+05 | 823.4 | - | - | 0 | - |
| - | - | 4.091E+05 | 824.4 | - | - | 0 | - |
| - | - | 1.485E+05 | 825.4 | - | - | 0 | - |
| - | - | 6.335E+04 | 826.4 | - | - | 0 | - |
| - | - | 3.936E+04 | 833.4 | - | - | 0 | - |
| - | - | 6.635E+05 | 834.4 | - | - | 0 | - |
| - | - | 1.859E+07 | 835.4 | - | - | 0 | - |
| - | - | 7.417E+06 | 836.4 | - | - | 0 | - |
| - | - | 3.344E+06 | 837.4 | - | - | 0 | - |
| - | - | 4.82E+05 | 838.4 | - | - | 0 | - |
| - | - | 1.215E+05 | 839.4 | - | - | 0 | - |
| - | - | 7.782E+06 | 851.4 | - | - | 0 | - |
| - | - | 2.605E+07 | 852.4 | - | - | 0 | - |
| - | - | 1.073E+07 | 853.4 | - | - | 0 | - |
| - | - | 3.632E+06 | 854.4 | - | - | 0 | - |
| - | - | 2.561E+05 | 855.4 | - | - | 0 | - |
| - | - | 3.95E+04 | 867.4 | - | - | 0 | - |
| - | - | 5.266E+04 | 884.4 | - | - | 0 | - |
| - | - | 2.274E+04 | 1970 | - | - | 0 | - |
| - | - | 2.452E+04 | 2815 | - | - | 0 | - |
| - | - | 2.346E+04 | 2917 | - | - | 0 | - |
| - | - | 3.754E+04 | 3085 | - | - | 0 | - |

m/z Charge Intensity FragmentType MassShift Position
126.0554428100586 0 67341.77
129.06622314453125 0 62763.36
144.06593322753906 0 806440.3
145.0695037841797 0 49129.95
148.95538330078125 0 19942.246
153.06622314453125 0 48247.74
156.73406982421875 0 13923.567
158.09268188476562 0 44330.695 y Ammonia loss 6
159.09288024902344 0 35788.5
159.1006622314453 0 1683081.2 z 6
160.10848999023438 0 2726549
161.1056671142578 0 20978.2
161.1117706298828 0 142012
166.053955078125 0 20138.953
168.11373901367188 0 20555.64
169.13404846191406 0 60389.72
170.09295654296875 0 91205.766
171.0768585205078 0 290294.72
172.06069946289062 0 30478.975
172.08029174804688 0 24815.46
173.05599975585938 0 17644.008
173.43771362304688 0 16305.197
174.11151123046875 0 56931.316
175.1193389892578 0 787433.7 y 6
176.12301635742188 0 20239.78
180.10232543945312 0 369413.5
183.10430908203125 0 15745.75
185.1400909423828 0 87631.73
187.1445770263672 0 355106.22
188.09173583984375 0 22327.982
188.14825439453125 0 36629.305
189.0874786376953 0 3161173
190.0910186767578 0 185420.16
197.1288299560547 0 173766.72
199.07171630859375 0 662270.75
200.07496643066406 0 54454.97
200.1271514892578 0 313036.38
201.13482666015625 0 59039.008
202.14271545410156 0 92816.56
203.11380004882812 0 38128.52
203.15061950683594 0 59933.08
207.11151123046875 0 35495.652
215.13941955566406 0 236749.72
217.0823211669922 0 2813857
218.08570861816406 0 219189.22
219.0876922607422 0 22299.229
222.41558837890625 0 18435.457
225.079833984375 0 17613.809
225.12384033203125 0 85163.76
229.6402130126953 0 128460.695
230.113525390625 0 19400.51
230.1249542236328 0 41339.844
233.1324920654297 0 41604.61
233.2041778564453 0 16507.889
234.87451171875 0 17425.295
240.13406372070312 0 22838.574
244.10072326660156 0 89940.8
244.11737060546875 0 200509.9
244.14013671875 0 20422.266 y Water loss 5
245.12496948242188 0 1602966.9 y Ammonia loss 5
246.13235473632812 0 814909.3 z 5
247.1405792236328 0 4984725
248.14378356933594 0 491666.12
249.12429809570312 0 19623.883
249.14501953125 0 39358.867
258.144775390625 0 28111.604
261.1270751953125 0 422449.25
261.1436462402344 0 298814.22
261.6392822265625 0 289304.38 y 3
262.1513671875 0 2883535.8 y 5
263.1543884277344 0 297404.2
264.15643310546875 0 53226.703
266.14990234375 0 39439.965
267.13458251953125 0 69786
267.2811584472656 0 20966.1
270.1201477050781 0 140010.48
271.1234130859375 0 18945.309
275.1360168457031 0 26402.143
277.1187744140625 0 21429.088
284.16107177734375 0 731210.3
285.1451110839844 0 36962.824
285.1640319824219 0 74821.82
286.1407165527344 0 53892.75
286.1823425292969 0 621781.4
286.6838684082031 0 147720.27
288.13018798828125 0 34450.945
289.13897705078125 0 45602.754
290.1464538574219 0 354925.4
291.1493835449219 0 43768.113
294.1452941894531 0 61212.96
295.1290283203125 0 28812.482
299.171875 0 57642.977
299.18951416015625 0 21525.055
299.6757507324219 0 20939.871
302.12164306640625 0 18179.885
302.1719970703125 0 100181.19
312.1560363769531 0 1974663.6
313.1591491699219 0 254836.94
314.1621398925781 0 35427.867
318.18157958984375 0 3723431.2 y 2
318.6829833984375 0 1082509.6
319.1822509765625 0 376748.78
319.68377685546875 0 73076.73
327.70025634765625 0 33693.75
330.1771240234375 0 1098448.2 w 4
331.1829528808594 0 133313.86
332.69390869140625 0 42025.566
336.7062072753906 0 155632.36
337.2075500488281 0 53002.145
341.6982421875 0 98862.99
344.1932373046875 0 120240.16 w 4
346.1439514160156 0 94553.625
347.1462097167969 0 22638.43
351.6844482421875 0 58616.52 z Water loss 1
354.68878173828125 0 37649.16
357.2252197265625 0 70672.23 y Water loss 4
358.2090148925781 0 92022.61 y Ammonia loss 4
359.21673583984375 0 4250284.5 z 4
359.7001953125 0 91658.266 y Water loss 1
360.17572021484375 0 51663.855
360.22308349609375 0 1665795.4
361.2257385253906 0 311255
362.17529296875 0 37367.957
362.2112731933594 0 75736.1
362.7133483886719 0 40872.43
363.198974609375 0 32376.91
364.6917419433594 0 115755.805
365.1920166015625 0 34578.914
367.1981506347656 0 26720.754
367.6964111328125 0 43589.336
368.7051086425781 0 564112.06 y 1
369.206298828125 0 190717.42
369.7049255371094 0 46202.715
372.1591491699219 0 164358.45
373.2198181152344 0 24467.295
373.6969299316406 0 189944.5
374.2030944824219 0 42886.598
374.2278747558594 0 331041.78
375.2354736328125 0 4489031 y 4
376.15460205078125 0 38837.71
376.2102966308594 0 47611.266
376.23858642578125 0 765051.56
376.7001647949219 0 25370.67
377.2398681640625 0 99576.445
385.21441650390625 0 767306.5
385.7155456542969 0 243149.19
386.19287109375 0 28833.748
386.2181396484375 0 58794.836
386.6878356933594 0 55470.504
387.1985778808594 0 60627.59
387.6925354003906 0 22585.012
388.2068786621094 0 250679.33
389.21026611328125 0 48660.516
390.6890869140625 0 76501.99
394.21826171875 0 1105318.8
394.71923828125 0 479581.25
395.220703125 0 133548.67
399.20196533203125 0 32035.61
399.69488525390625 0 173404.88
400.1970520019531 0 71407.41
401.2148132324219 0 147153.98
408.2080993652344 0 84034.305
408.70428466796875 0 79021.734
409.201416015625 0 39156.867
413.2044372558594 0 25320.826
415.2307434082031 0 423047.8
416.2374572753906 0 404161.03
417.21356201171875 0 2386430.5
417.7148132324219 0 970516.1
418.214599609375 0 324393.8
424.154541015625 0 79162.914
426.2190856933594 0 407253.94
426.25433349609375 0 22390.098
426.72015380859375 0 163582.14
427.2196960449219 0 89793
429.24658203125 0 109252.805
430.24609375 0 26632.693
431.2496643066406 0 157777.1
432.2522277832031 0 28467.652
441.18231201171875 0 33148.77
441.2456970214844 0 102923.914
442.1672668457031 0 25328.02
442.254150390625 0 361111.22
443.2618713378906 0 1057503.4
444.26904296875 0 1098525
445.27276611328125 0 192708.78
446.2745666503906 0 33082.86
450.1895751953125 0 36673.406
458.2729187011719 0 2006956.2
459.1915283203125 0 491399.47
459.2757263183594 0 419400.22
460.1938781738281 0 86082.93
460.2774353027344 0 56812.203
461.18841552734375 0 23635.059
477.2033386230469 0 870727.9
478.206298828125 0 206836.5
479.2037048339844 0 64216.727
498.29229736328125 0 28877.725
504.2602844238281 0 117369.91 y Water loss 3
505.2418518066406 0 38089.203 y Ammonia loss 3
506.2521667480469 0 166123.19 z 3
507.25518798828125 0 74142.64
508.2742919921875 0 24394.182
522.2710571289062 0 7787995.5 y 3
523.273681640625 0 1894334.2
524.2711181640625 0 594258.7
525.2716064453125 0 28249.086
526.287353515625 0 41434.52
532.2562866210938 0 53632.71
542.3285522460938 0 22530.22
544.3326416015625 0 47038.67
545.2634887695312 0 21641.549
555.3347778320312 0 29834.404
556.3464965820312 0 64653.734
557.3489379882812 0 28433.348
562.291015625 0 60113.914
563.2943115234375 0 34293.418
564.3014526367188 0 24539.117
571.3568115234375 0 1750970.1
572.2756958007812 0 64022.36
572.3597412109375 0 470094.1
573.361572265625 0 87570.52
576.281494140625 0 1212620.5 w 2
577.2838745117188 0 335204.5
578.2850341796875 0 84159.91
590.285888671875 0 149388.1
591.2861938476562 0 43046.438
602.3309936523438 0 26283.096
606.2794799804688 0 23848.68
607.3115234375 0 80154.125 c 4
617.3453979492188 0 54737.016 y Water loss 2
618.3318481445312 0 46045.48 y Ammonia loss 2
619.336181640625 0 350696.38 z 2
620.3375854492188 0 168700.11
621.3427734375 0 75595.46
622.3458251953125 0 59820.64
630.371826171875 0 26366.965
634.3551025390625 0 22128.693
635.3550415039062 0 5527795 y 2
636.35791015625 0 1694866.9
637.356201171875 0 518838.1
638.36083984375 0 47308.957
645.3399658203125 0 37595.043
647.3045043945312 0 23176.633
657.3929443359375 0 55376.11
658.4013671875 0 136668.28
659.3092041015625 0 40889.047
659.4026489257812 0 58379.99
662.3056030273438 0 327828.88
662.3577270507812 0 23198.354
663.3097534179688 0 116826.1
664.3206787109375 0 808264.4
665.3247680664062 0 241320.34
666.3214111328125 0 60315.523
673.376708984375 0 67343.84
674.3764038085938 0 41832.633
675.3682250976562 0 51289.066
676.387451171875 0 42366.72
677.3250732421875 0 238785.56
677.3810424804688 0 169937.31
678.328369140625 0 75339.07
678.3829956054688 0 61574.742
679.3242797851562 0 29315.197
682.3897094726562 0 79732.97
683.390625 0 49758.39
686.3804321289062 0 24103.385
692.356201171875 0 26067.398
693.3367919921875 0 118887.43
694.3440551757812 0 850928.94 c 5
695.3472290039062 0 291945.47
696.346923828125 0 85158.89
701.4074096679688 0 197174.73
702.408935546875 0 79603.43
703.38037109375 0 127745.04 w 1
704.3739624023438 0 109309.72
705.3643188476562 0 283952.44 w 1
706.3706665039062 0 90158.414
707.3744506835938 0 26328.01
717.3582153320312 0 64605.027
718.3688354492188 0 809022.44
719.37548828125 0 2481226.5 y Ammonia loss 1
720.3826293945312 0 3614477 z 1
721.385986328125 0 1607646
722.3909301757812 0 590894.5
723.39697265625 0 103039.66
724.4071044921875 0 27098.188
728.3656616210938 0 40599.938
728.4306030273438 0 559868.3
729.3638305664062 0 38402.348
729.4347534179688 0 435109.84
730.437255859375 0 98093.08
732.3786010742188 0 96006.77
734.3838500976562 0 23812.076
735.3580322265625 0 1827136.5
736.3613891601562 0 539672.2
737.359619140625 0 172182.47
744.4486694335938 0 41512.92
745.3984375 0 78839.88
746.3873901367188 0 899698.75
747.3936767578125 0 448536.3
748.4119262695312 0 408633.62
749.4181518554688 0 158856.25
750.4276733398438 0 33507.758
751.3421630859375 0 224334
752.3450317382812 0 128830.9
756.3828735351562 0 28223.945
760.406982421875 0 357612.44
761.3370971679688 0 69153.64
761.4105834960938 0 167604.67
762.3773803710938 0 54274.125
762.4236450195312 0 74950.77
763.3909301757812 0 84525.625
764.3934936523438 0 27331.666
765.399658203125 0 43955.355
766.4119873046875 0 209924.72
767.4161987304688 0 81917.44
772.4203491210938 0 434073.22
773.4135131835938 0 1140021.1
774.4166870117188 0 485996.4
775.4129638671875 0 159347.9
776.3970336914062 0 401440.25
777.3997192382812 0 129866.49
778.3662109375 0 946636.56
779.35205078125 0 1345305.5
780.35302734375 0 454890.3
780.4183959960938 0 152906.11
781.3489379882812 0 147549.06
781.420654296875 0 58270.47
782.421630859375 0 46121.586
787.4317016601562 0 43685.4
788.3567504882812 0 25782.732
788.4384765625 0 544486.4
789.443359375 0 253996.72
790.418212890625 0 1001054.1
791.4212036132812 0 18231834
792.4202270507812 0 11819534
793.4180297851562 0 4339493.5
794.413818359375 0 920657.94
795.3967895507812 0 160737.98
796.374755859375 0 1119078.2
797.3777465820312 0 475157.53
798.3755493164062 0 120510.42
806.3756103515625 0 49458.42
806.4315185546875 0 259132.55
807.4349365234375 0 133355.47
808.43798828125 0 195280.44
809.4072265625 0 5559105.5
810.409912109375 0 2004494
811.4091796875 0 809596
812.4137573242188 0 61939.395
817.3994750976562 0 947736.06
818.4027099609375 0 405494.2
819.4029541015625 0 140380.7
820.3897705078125 0 171850.6
821.3901977539062 0 67660.63
822.4276733398438 0 46533.305
823.4102172851562 0 292816.9
824.4385375976562 0 409075.25
825.4442749023438 0 148530.66
826.4435424804688 0 63345.645
833.4198608398438 0 39360.52
834.4248657226562 0 663510.94
835.4161376953125 0 18594606
836.4183349609375 0 7417181.5
837.4173583984375 0 3343947
838.4171142578125 0 482041.28
839.4161376953125 0 121501.68
851.4293212890625 0 7782140.5
852.4365234375 0 26051282
853.4389038085938 0 10727519
854.4396362304688 0 3632405.8
855.4396362304688 0 256065.52
867.3978881835938 0 39503.477
884.425048828125 0 52659.438
1969.83349609375 0 22744.475
2814.58349609375 0 24520.926
2916.8525390625 0 23460.152
3084.677001953125 0 37535.145

Spectrum Details

|  |  |
| --- | --- |
| Matched peaks? Matched peaks The total absolute number of peaks matched. Additionally in brackets the total fraction of peaks matched and the total number of peaks is shown. | 33 (8.73% of 378) |
| FDR? FDR The false discovery rate estimated for this peptide. It is calculated by matching all theoretical fragments with a non-integer shift with the raw peaks for this spectrum. This is done with 40 different shifts. The resulting percentage is the average number of annotated peaks over the number of annotated peaks with the correct spectrum. | 0.07% |
| Satellite FDR? Satellite FDR See the FDR for details on its calculation. This satellite ion specific FDR only contains the satellite ions (d/w) for I/L/J positions. | 0.00% |
| PSM Score? PSM Score The PSM Score as given by Hecklib to this annotated spectrum. It is shown with three significant figures. | 338 |

#### Reverse Lookup? Reverse Lookup All places where this read could be placed.

| Group | Segment | Template | Template Part | Read Part | Score | Unique |
| --- | --- | --- | --- | --- | --- | --- |
| Homo sapiens Heavy Chain | IGHC | IGHG1 | [132..138] | [0..7] | 47 | False |
| Homo sapiens Heavy Chain | IGHC | IGHG3 | [179..185] | [0..7] | 47 | False |
| Homo sapiens Heavy Chain | IGHC | IGHG2 | [128..134] | [0..7] | 47 | False |
| Homo sapiens Heavy Chain | IGHC | IGHG4 | [129..135] | [0..7] | 47 | False |

| Recombined | Template Part | Read Part | Score | Unique |
| --- | --- | --- | --- | --- |
| REC-0-1 | [255..261] | [0..7] | 47 | True |

#### Meta Information from Multiple reads

##### Number of combined reads

3

##### Intensity

0.93

##### TotalArea

3.895E+09

##### Changes to the peptide sequence

GTJMISR

J→ISupport for Isoleucine based on side chain ions (2 for I 0 for L) (Position: 5)

L→JNo support for either Leucine or Isoleucine based on side chain ions (Position: 5)

L→JNo support for either Leucine or Isoleucine based on side chain ions (Position: 3)

#### Positional Score

Copy Data

##### Positional Score (TSV)

###### Preview

```
Loading example...
```

*Click on the button to copy the data to your clipboard.*

100123456

Label Value
"0" 0.293
"1" 0.3
"2" 0.327
"3" 0.323
"4" 0.32
"5" 0.327
"6" 0.33

#### Meta Information from PEAKS

##### Scan Identifier

F4:3929

##### Original sequence

G

+58.01

T

L

M

+15.99

L

S

R

##### Posttranslational Modifications

Carboxymethyl (KW X@N-term); Oxidation (M)

##### Source File

D:\separate\_stitch\_analyses\xle-disambiguation\raw\20210323\_F1\_UM1\_Peng0013\_SA\_F59\_ingel\_3ug\_tryp.raw

##### Fraction

4

##### Scan Feature

F4:1263

##### De Novo Score

98

##### ConfidenceScore

98

### m/z

426.2188

##### Mass

850.4219

##### Charge

2

##### Retention Time

20.29

##### Predicted Retention Time

18.90

##### Area

1.298E+09

##### Parts Per Million

1.5

##### Fragmentation mode

HCD

##### Originating file

01 D:\separate\_stitch\_analyses\xle-disambiguation\20210325\_F59\_3ug\_DENOVO\_12.csv

#### Meta Information from PEAKS

##### Scan Identifier

F4:3876

##### Original sequence

G

+58.01

T

L

M

+15.99

L

S

R

##### Posttranslational Modifications

Carboxymethyl (KW X@N-term); Oxidation (M)

##### Source File

D:\separate\_stitch\_analyses\xle-disambiguation\raw\20210323\_F1\_UM1\_Peng0013\_SA\_F59\_ingel\_3ug\_tryp.raw

##### Fraction

4

##### Scan Feature

F4:1263

##### De Novo Score

97

##### ConfidenceScore

95

### m/z

426.2188

##### Mass

850.4219

##### Charge

2

##### Retention Time

20.29

##### Predicted Retention Time

18.90

##### Area

1.298E+09

##### Parts Per Million

1.5

##### Fragmentation mode

ETHCD

##### Originating file

01 D:\separate\_stitch\_analyses\xle-disambiguation\20210325\_F59\_3ug\_DENOVO\_12.csv

#### Meta Information from PEAKS

##### Scan Identifier

F4:3823

##### Original sequence

G

+58.01

T

L

M

+15.99

L

S

R

##### Posttranslational Modifications

Carboxymethyl (KW X@N-term); Oxidation (M)

##### Source File

D:\separate\_stitch\_analyses\xle-disambiguation\raw\20210323\_F1\_UM1\_Peng0013\_SA\_F59\_ingel\_3ug\_tryp.raw

##### Fraction

4

##### Scan Feature

F4:1263

##### De Novo Score

97

##### ConfidenceScore

95

### m/z

426.2188

##### Mass

850.4219

##### Charge

2

##### Retention Time

20.29

##### Predicted Retention Time

18.90

##### Area

1.298E+09

##### Parts Per Million

1.5

##### Fragmentation mode

ETHCD

##### Originating file

01 D:\separate\_stitch\_analyses\xle-disambiguation\20210325\_F59\_3ug\_DENOVO\_12.csv
